# ProtRAP-LM: Fast and accurate protein relative accessibility prediction and membrane protein screening through protein language model embeddings

**DOI:** 10.1101/2025.01.20.633985

**Authors:** Lei Wang, Kai Kang, Chen Song

## Abstract

Membrane proteins play pivotal roles in cellular signaling and transport, making them prime targets for therapeutic intervention. Therefore, expeditious screening and accurate property prediction of these proteins is crucial. Recently, we proposed a new metric, the membrane contact probability (MCP), to characterize the membrane-contacting features of membrane proteins and further refine the prediction of the relative accessibility of proteins (ProtRAP). However, these models relied on evolutionary information in the form of multiple sequence alignments (MSAs), which hindered rapid predictions. In this study, we present a novel transformer-based model, ProtRAP-LM, utilizing language model (LM) embeddings as input features, to quickly and accurately predict MCP and relative accessibility for each residue of a given protein sequence. ProtRAP-LM can achieve accurate predictions for entire proteomes within hours, demonstrating superior performance compared to previous MSA-based models, with a speedup of over 300 times. This empowers us to furnish more thorough annotations of membrane protein sequences on a proteome-wide scale, particularly for single-pass transmembrane proteins, membrane-anchored proteins, and β-sheet-containing membrane proteins, which have long been a challenge in the field. In the end, we provide a comprehensive list of membrane proteins for 48 living organisms, offering a rich resource for investigating the structure and function of these essential biomolecules in the future.

## 1. Introduction

Membrane proteins constitute a large subgroup within proteomes, comprising ~20-30% of all proteins within human proteome. ^1–4^ They play a crucial role in diverse essential cellular functions, including cell communication, transport, enzymatic activities, and cell structure maintaining. ^5–7^ Moreover, membrane proteins attract significant biomedical interest and serve as the targets of over 50% of known drugs. ^4,8^ Despite the important biological roles, their embeddedness in lipid layers makes it difficult to obtain their high-resolution structures and dictates the need to develop reliable and efficient computational methods for the studies of membrane protein structure and function. ^9,10^ The membrane proteins can be grouped into three categories based on their spatial relationships with the lipid bilayers: transmembrane proteins, membrane-anchored proteins, and peripheral membrane proteins. ^11,12^ Existing prediction methods have primarily focused on the studies of α-helical and β-barrel transmembrane proteins, ^13–15^ which may ignore a considerable proportion of the other vital proteins present in the cell membranes, such as membrane-anchored proteins and peripheral membrane proteins. ^16–18^ Therefore, a more comprehensive computational model for membrane protein identification across the entire proteomes is needed.

In our recent work, we proposed a novel characteristic and predictive quantity called the membrane contact probability (MCP), which describes the likelihood of direct contact between the protein amino acids and the hydrophobic acyl chains of lipid molecules. ^19^ We also trained an MCP predictor, which performed well for identifying both α-helical and β-barrel transmembrane proteins, as well as for membrane-anchored proteins that interact directly with the lipid bilayers. ^19^ Building upon MCP, we further developed ProtRAP (protein relative accessibility predictor), to predict the relative accessibility of residues from a given protein sequence. ^20^ However, both the MCP predictor and ProtRAP require multiple sequence alignments against the protein sequence database and generating a position-specific scoring matrix (PSSM) as input. This is a common trait among many property predictors, ^21–23^ and it is also the major time-consuming step that limits the analysis scale.

Language models (LMs) have demonstrated exceptional performance in natural language processing ^24^ and computer vision. ^25^ And protein LMs trained on unsupervised protein sequences such as ProtTrans, ^26^ ESM-1b, ^27^ ProGen2, ^28^ recently developed ESM-2^29^ and ESM-3^30^ are able to capture protein structural and functional properties. By utilizing information-rich representations of protein sequences generated from these protein LMs, several new predictors were developed, which no longer require MSAs and achieve comparable accuracy with a substantial increase in calculation speed. ^15,31–33^

In this work, we combined the prediction targets of our two previous models (MCP and ProtRAP) ^19,20^ and further developed a novel transformer-based model for protein relative accessibility prediction through protein language model embeddings (ProtRAP-LM). This new model enables fast and highly accurate predictions of the relative accessible surface area (RASA) and membrane contact probability (MCP) from a given protein sequence (Fig. 1A-1B). Furthermore, it provides the relative lipid accessibility (RLA), relative solvent accessibility (RSA), and relative buried surface area (RBSA) of each protein residue (Fig. 1C). Notably, ProtRAP-LM is over 300 times faster than the MSA-based MCP predictor on the MemProtMD_2022 dataset presented in this paper (described in the ‘Methods’ section), thus enabling the screening and analysis of proteome-scale membrane proteins within hours (Fig. S1). The implementation of ProtRAP-LM significantly improved our ability to identify and uncover likely membrane proteins, providing a more comprehensive picture of their structures and functions.

**Figure 1:**
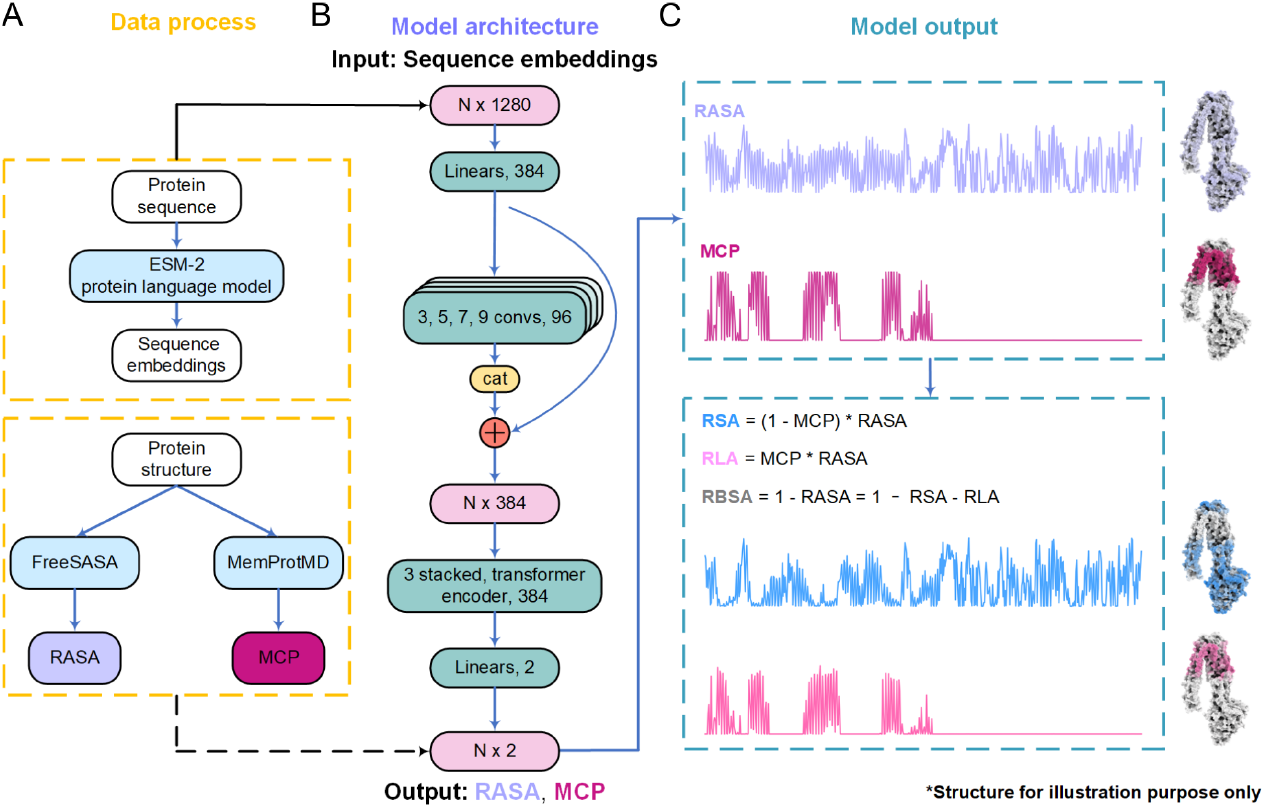
Framework of the ProtRAP-LM. (A) Data processing. For a protein sequence, the embeddings were generated using the ESM-2 model, ^29^ which served as the input for our model. Protein structures were analyzed to get the values of the RASA using FreeSASA ^73^ and the MCP from the MemProtMD database, ^70,71^ which were utilized as the labels for our datasets. (B) Model architecture. The model mainly contains one-dimension multiscale convolution layers and transformer encoder modules. (C) Model output. The model can predict the RASA and the MCP of each residue directly, which were further used to calculate the RLA, RSA, and RBSA for the additional outputs.

## 2. Results

### 2.1. Overall performance of the ProtRAP-LM predictor

ProtRAP-LM can predict the RASA and MCP of each residue directly, and provide the predicted relative accessibility (RLA, RSA, and RBSA) in one run for a given sequence. With the refined model, the 10-fold cross-validation of the ProtRAP-LM model showed low standard deviations (Table S1). ProtRAP-LM achieved very satisfactory performance on both the residue level (Tables 1 and S2) and the protein level (Fig. S2) on the MemProtMD_2022 dataset with the overall prediction Pearson correlation coefficients (PCCs) on the residue level of 0.915, 0.782, 0.863, and 0.803 for the MCP, RASA, RLA, and RSA, respectively. Of all the state-of-the-art protein relative accessibility predictors, ProtRAP-LM achieved the best performance (Table 1). Particularly, compared with NetSurP 3.0, ^31^ which focused on the prediction of RASA, ProtRAP-LM demonstrated a 0.089 PCC improvement of RASA on the residue level (Table 1), and a 0.055 PCC improvement on the protein level (Fig. S2).

**Table 1.** Comparison of the membrane contact probability (MCP) and protein relative accessible surface area (RASA) prediction performance using the MemProtMD_2022 dataset with different methods (on the residue level).

|  | Evaluation | ProtRAP-LM | MCP predictor | ProtRAP | NetSurfP-3.0 | NetSurfP-2.0 |
| --- | --- | --- | --- | --- | --- | --- |
| MCP | PCC | **0.9152** | 0.8636 | - | - | - |
|  | MAE | **0.0423** | 0.0600 | - | - | - |
| RASA | PCC | **0.7816** | - | 0.7137 | 0.6927 | 0.6800 |
|  | MAE | **0.1170** | - | 0.1333 | 0.1384 | 0.1390 |

Compared to our previous work, on the residue level, the MCP prediction was improved by 0.053 of PCC compared to the MSA-based MCP predictor, ^19^ as shown in Table 1. And the predictions of RASA, RLA, and RSA were improved by 0.068, 0.023, and 0.056 of PCCs compared to our previous MSA-based ProtRAP, ^20^ as shown in Tables 1 and S2.

More importantly, according to the runtime analysis of the ProtRAP-LM and the MSA-based MCP predictor, the average runtimes of ProtRAP-LM and MCP predictor were 0.10 s and 32.75 s per sequence for the MemProtMD_2022 dataset, respectively. Therefore, ProtRAP-LM achieved an over 300-fold speed-up versus the MSA-based MCP predictor (Fig. S1A), which enables us to conduct quick and accurate proteome-wide prediction of membrane proteins.

### 2.2 Proteome-wide analysis of membrane proteins

In our previous study, we developed an empirical approach for membrane protein identification based on MCP prediction. ^19^ Specifically, if there exist more than or equal to three high-MCP (MCP ≥ 0.5) predictions in ten successive amino acids of a protein, we consider the protein highly likely to direct contact with the hydrophobic acyl chains of lipid molecules and therefore a potentially membrane-contacting protein, otherwise a non-membrane protein. To evaluate the membrane protein identification performance of our ProtRAP-LM model, we constructed three large datasets (please refer to the section of ‘Methods’ for details) and compared the prediction performance with the state-of-the-art transmembrane protein predictor TMbed. ^15^ TMbed is a powerful tool developed by Bernhofer and Rost, which combined language model embeddings with the convolutional neural network to classify each residue into four different classes, including transmembrane helix, transmembrane strand, signal peptide, or other.

Therefore, TMbed can be utilized as a high-throughput filter for transmembrane protein detection and transmembrane span prediction. According to the analysis results presented in Tables 2 and S3, our criteria for membrane protein identification demonstrated high reliability and outperform TMbed ^15^ on the recall and false positive rate (FPR) for the UniProt_Mem dataset and UniProt_Sol dataset presented in this paper, which respectively comprise the reviewed membrane and soluble protein sequences from the UniProtKB database, as described in the ‘Methods’ section. For the PDBTM dataset, which consists of transmembrane proteins with known structures, ProtRAP-LM achieved comparable results to TMbed in terms of recall (Table 2). Given the satisfactory performance of ProtRAP-LM (Table 2), we implemented a protocol, as illustrated in Fig. 2A, to conduct a comprehensive membrane protein analysis and estimate the membrane protein fractions within 48 representative organism proteomes (Table S4), for which AlphaFold built the protein structure database ^34^ (https://alphafold.ebi.ac.uk/).

**Table 2.**
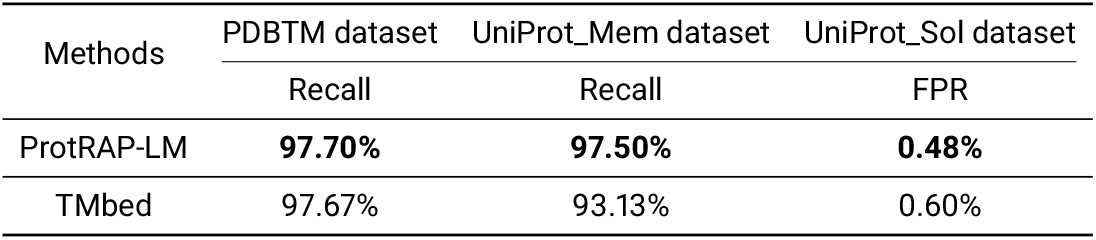
Comparison of the membrane protein identification performance of ProtRAP-LM with the results obtained from TMbed across the three large datasets.

**Figure 2:**
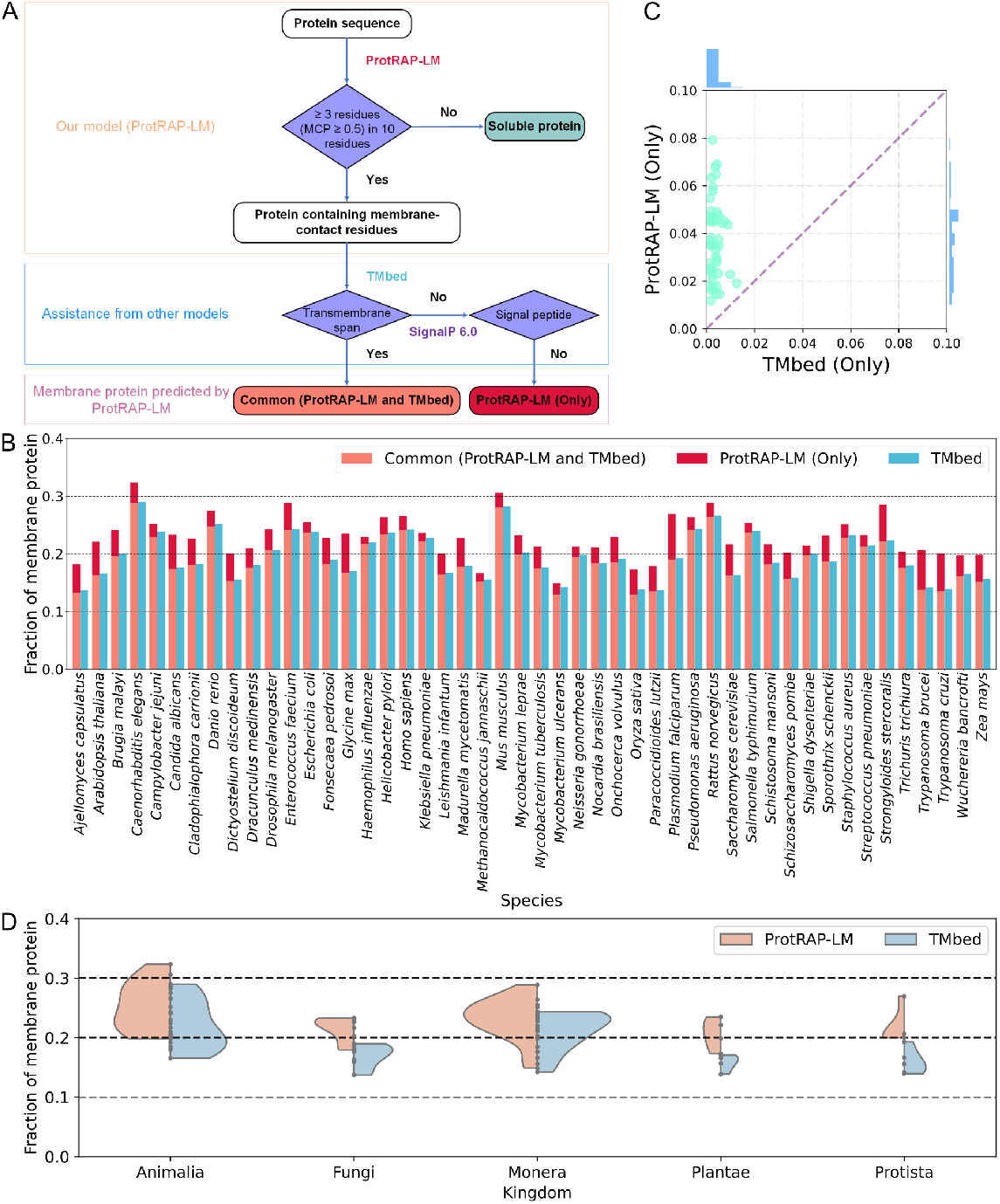
Proteome-wide prediction of membrane proteins. (A) Our protocol of membrane protein identification. First, with ProtRAP-LM, we can identify the proteins containing high-MCP residues as potentially membrane-contacting proteins. Second, with the assistance of the existing tools TMbed ^15^ and SignalP 6.0, ^32^ we can obtain the common subset of transmembrane proteins between ProtRAP-LM and TMbed (termed ‘Common (ProtRAP-LM and TMbed)’) (salmon) and the other part of membrane proteins only identified by ProtRAP-LM (termed ‘ProtRAP-LM (Only)’) (crimson). Finally, we can get the membrane proteins predicted by ProtRAP-LM containing two parts (salmon+crimson). (B) The membrane protein fractions identified within different proteomes. As can be seen, the ProtRAP-LM (salmon+crimson) can predict higher membrane protein fractions compared to the fractions predicted by TMbed (teal) within each proteome. (C) Comparison of the likely membrane protein fractions only identified by ProtRAP-LM (y-axis) and TMbed (x-axis) out of the entire proteome within 48 different species. The dashed line is the function *y = x*. Each point represents a proteome. (D) The predicted membrane protein fractions within five different biological kingdoms.

First, we identified the potentially membrane-contacting proteins as described above using ProtRAP-LM. Then, we utilized two other existing predictors, TMbed ^15^ and SignalP 6.0, ^32^ to further classify the identified membrane proteins.

With the prediction results of TMbed, we can obtain the likely membrane proteins with at least one transmembrane span. The majority of these TMbed-identified membrane proteins form a common part with the ProtRAP-LM-identified membrane proteins (termed ‘Common (ProtRAP-LM and TMbed)’). Then, we used SignalP 6.0^32^ to identify the likely signal peptides in secreted proteins, which were not considered membrane proteins in this study. With the above procedure, we can rule out the membrane proteins that can be identified by existing state-of-the-art tools and find potentially *unknown* membrane proteins (termed ‘ProtRAP-LM (Only)’).

Notably, we compared the TMbed-identified membrane proteins to our ProtRAP-LM-identified ones, and the results showed that the ProtRAP-LM predictions gave good coverage of the TMbed results. The membrane proteins identified by TMbed but not by ProtRAP-LM (termed ‘TMbed (Only)’) account for only a very small part of all the likely membrane proteins within each proteome (Fig. 2B-2C). For instance, for the human proteome, our ProtRAP prediction covered 99.4% of the transmembrane proteins identified by TMbed. This indicates that the ProtRAP-LM predictor can accurately identify the likely transmembrane proteins, on par with the TMbed predictor.

Notable variations in the membrane protein fractions can also be observed among different species (Fig. 2B). Further, we discussed the membrane protein fractions within 48 species according to the division of the five biological kingdoms (Fig. 2D). In both the ProtRAP-LM and TMbed predictions, the predicted membrane proteins constitute ~20–30% within the proteomes of most bacteria and animals (Fig. S3A-S3B), which is consistent with the conclusions of previous assessments. ^2^ In contrast, membrane proteins appear to represent a relatively small portion of the proteome in the Fungi and Plantae kingdoms. Our method estimated this fraction to be around 20% (Fig. S3A), whereas only approximately 10-20% of transmembrane proteins were identified based on the results from TMbed (Fig. S3B).

More interestingly, our findings reveal that ProtRAP-LM has the ability to predict a greater number of membrane proteins than TMbed (Figs. 2B-2D and S3C-S3D). For instance, ProtRAP-LM predicted that about 26.5% of human proteins are likely membrane proteins, compared to 24.2% by TMbed. This is reasonable as ProtRAP-LM was designed to identify not only membrane-spanning proteins but also membrane-anchoring proteins, which will be further discussed below.

### 2.3. ProtRAP-LM can better identify single-pass transmembrane proteins and membrane-anchored proteins

The ‘ProtRAP-LM (Only)’ proteins are the most interesting ones to explore, as they may represent the *unknown* membrane proteins. To further extract the characteristics of the likely membrane proteins only identified by ProtRAP-LM, we conduct a more in-depth study of the ‘ProtRAP-LM (Only)’ proteins with the following two steps.

In the first step, we analyzed the MCP predictions for the ‘ProtRAP-LM (Only)’ protein sequences and found that the amino acids with high MCP values are predominantly located at both ends of the proteins (Figs. 3A and S4A). This led us to speculate that our ProtRAP-LM model can better identify single-pass transmembrane proteins or tail-anchored membrane proteins. Therefore, we evaluated the performance of our ProtRAP-LM model on the Membranome database composed of single-pass transmembrane proteins from six evolutionarily distant organisms, ^35^ as shown in Fig. 3B. Indeed, it appears that the distribution of MCP predictions for the ‘ProtRAP-LM (Only)’ proteins within the 48 proteomes is similar to the results of the database only composed of single-pass transmembrane proteins (Fig. S4B), with the high-MCP predictions mainly distributed at the termini of the proteins. Therefore, our model has the ability to identify a significantly higher number of single-pass transmembrane proteins in various organisms compared to TMbed. In some cases, the additional fraction of correctly identified membrane proteins to total single-pass transmembrane proteins can reach up to 30% for certain proteomes (Fig. 3C). Our ProtRAP-LM predictor may offer a more accurate approach to single-pass transmembrane protein prediction, as it focuses on identifying patterns in membrane-contract residues rather than relying on transmembrane span information, which can be challenging to resolve experimentally and predict computationally. ^36^

**Figure 3:**
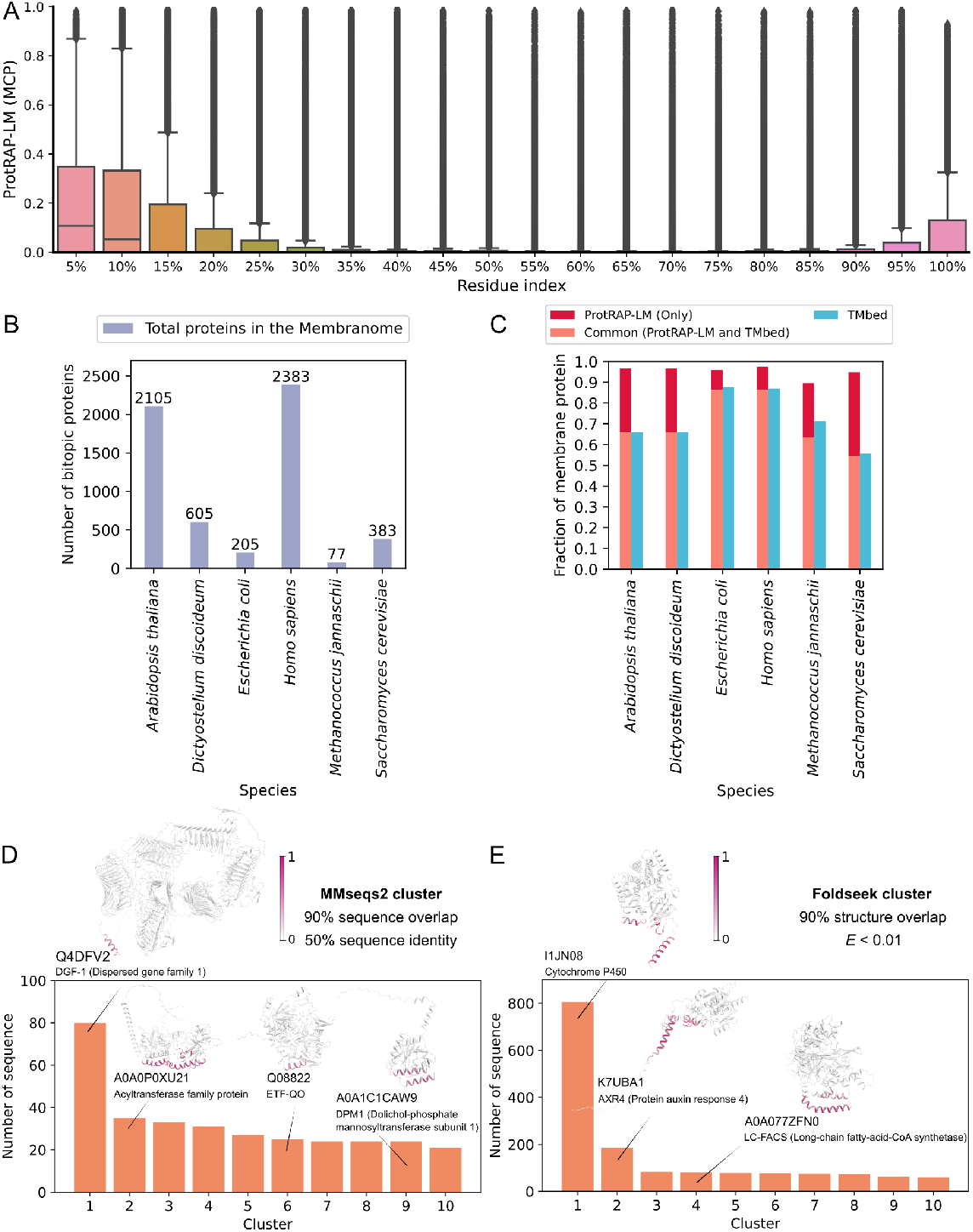
Analysis of the ‘ProtRAP-LM (Only)’ proteins within 48 proteomes. (A) The distribution of MCP predictions in the ‘ProtRAP-LM (Only)’ protein sequences within 48 proteomes. Each protein sequence was divided into 20 equal parts, and the average of MCP predictions in each part was calculated. (B) The number of single-pass transmembrane (bitopic) proteins from six organisms in the Membranome database. ^35^ (C) The membrane protein fractions identified by ProtRAP-LM containing the parts of ‘Common (ProtRAP-LM and TMbed)’ (salmon) and ‘ProtRAP-LM (Only)’ (crimson) are higher than the fractions predicted by TMbed (teal) within each proteome. (D) The top 10 sequential-alignment-based cluster representatives within the ‘ProtRAP-LM (Only)’ proteins. (E) The top 10 structural-alignment-based cluster representatives within the ‘ProtRAP-LM (Only)’ proteins. The representative structures in the (D) and (E) predicted by AlphaFold2 were colored according to the predicted MCP by ProtRAP-LM, visualized as cartoon.

In the second step, we try to cluster the ‘ProtRAP-LM (Only)’ proteins to obtain their representative features. We employed two clustering methods, namely MMseqs2^37,38^ and Fold-seek, ^38,39^ to cluster the ‘ProtRAP-LM (Only)’ proteins based on sequence identity and structural similarity, ^38^ respectively. To cluster the ‘ProtRAP-LM (Only)’ sequences within 48 proteomes, we utilized the MMseqs2 clustering module, employing a criterion of at least 50% sequence identity and 90% sequence alignment overlap. This allowed us to group similar sequences and identify the top 10 cluster representatives based on their shared sequence characteristics (Fig. 3D). The largest sequential-alignment-based cluster representative identified through the MMseqs2 clustering is the dispersed gene family protein 1 (DGF-1), a likely single-pass transmembrane protein according to the MCP-colored positions in the AlphaFold2-predicted structure (Fig. 3D). DGF-1 is known to be a highly abundant protein family in the *Trypanosoma cruzi* and is speculated to localize in the membrane. ^40^

Similarly, by applying the Foldseek clustering method, we clustered the AlphaFold2-predicted structures of the ‘ProtRAP-LM (Only)’ proteins within 48 proteomes. The clustering was performed using an *E*-value threshold of 0.01 and a structural alignment overlap of 90%. Through this process, we identified the top 10 cluster representatives based on their structural similarity. These representatives provide insights into the most prominent and representative structural motifs within the ‘ProtRAP-LM (Only)’ proteins, aiding in the understanding of the overall structural landscape. The largest structural-alignment-based cluster representative obtained from the Foldseek clustering analysis is the cytochrome P450 protein (Fig. 3E). The cytochrome P450 proteins encompass a diverse family of enzymes involved in the various metabolic processes, including drug metabolism and synthesis of important molecules, and some isoforms of the cytochrome P450 protein family were known to be membrane-attached proteins, with their N-terminal helix facilitating their attachment to the membrane. ^41^ Despite exhibiting low sequence identity, these isoforms share a common structural fold, indicating a conserved structural motif that underlies their functional similarities. ^42^

The clustered representatives of ‘ProtRAP-LM (Only)’ proteins shown in the above second-step analysis revealed that the amino acid residues with high MCP values of some representatives (UniProt IDs: Q4DFV2, I1JN08, etc.) are predominantly located at the terminal regions, which were speculated to be likely single-pass transmembrane proteins (Figs. 3D-3E and S5). In addition, our analysis also revealed cluster representatives of likely membrane-anchored proteins (UniProt IDs: A0A0P0XU21, Q08822, etc.) based on the MCP-colored positions of the AlphaFold2-predicted structures. For instance, electron transfer flavoprotein-ubiquinone oxidoreductase (ETF-QO, UniProt ID: Q08822), a known membrane-anchored protein located in the inner mitochondrial membrane, shows consistency between the high-MCP region and the α9-helix in the putative membrane-binding surface as described in previous literature ^43,44^ (Fig. 3D). The ability to identify membrane-anchored proteins is unique to our method compared to the existing transmembrane span-based membrane protein predictors.

### 2.4. ProtRAP-LM can identify more β-sheet-containing membrane proteins

Based on the analysis of protein secondary structures using DSSP ^45^ on AlphaFold2-predicted structures, we identified proteins containing high-MCP β-sheets using ProtRAP-LM. Specifically, a β-sheet is considered a high-MCP sheet if it contains three or more residues with MCP ≥ 0.5, indicating a strong likelihood of membrane contact. For comparison, we also identified transmembrane-strand-containing proteins within the TMbed-identified membrane proteins.

Similarly to above, we classified the proteins that contain at least one high-MCP β-sheet or transmembrane strand into three categories, which were labeled as ‘Common (ProtRAP-LM and TMbed)’ (915 proteins), ‘ProtRAP-LM (Only)’ (244 proteins), and ‘TMbed (Only)’ (83 proteins). In 28 out of 48 protein proteomes, ProtRAP-LM identified more β-sheet-containing membrane proteins than TMbed (Fig. S6A), with most of these species belonging to the Plan-tae and Protista kingdoms. These 28 proteomes contained 479 ‘Common (ProtRAP-LM and TMbed)’ proteins, 230 ‘ProtRAP-LM (Only)’ proteins, and 39 ‘TMbed (Only)’ proteins. From another eight out of the 48 proteomes, we identified the same amount of β-sheet-containing membrane proteins, with a total number of 242 ‘Common (ProtRAP-LM and TMbed)’ proteins, with only one ‘ProtRAP-LM (Only)’ and one ‘TMbed (Only)’ protein. In the remaining 12 proteomes where TMbed predicted more membrane proteins, we found 194 ‘Common (ProtRAP-LM and TMbed)’ proteins, 13 ‘ProtRAP-LM (Only)’ proteins, and 43 ‘TMbed (Only)’ proteins. For the ‘TMbed (Only)’ proteins not detected by our method, secondary structure identification may be an influencing factor (Fig. S7). For example, in the *Danio rerio* proteome, ProtRAP-LM and TMbed both accurately identified similar transmembrane segments for the two ‘Common (ProtRAP-LM and TMbed)’ proteins (Fig. S7A-S7D), and both methods predicted a similar region for the one ‘TMbed (Only)’ protein (Aerolysin-like protein Aep1, UniProt ID: Q5CZR5). However, ProtRAP-LM did not identify it as containing a high-MCP β-sheet, since these regions do not form a continuous β-sheet according to AlphaFold2 predictions. The identified region of this protein may form a β-barrel through a conformational change, ^46^ probably indicating a false negative of our method due to the possible alternative conformation not obtained in AlphaFold2 predictions. Nonetheless, this does not undermine the ability of our method to identify it as a membrane protein. In another case, the ‘TMbed (Only)’ protein from *Staphylococcus aureus* (Uncharacterized protein, UniProt ID: Q2FW43), was predicted by both ProtRAP-LM and TMbed to have a similar membrane-contacting region, but whether this region will form a β-sheet remains uncertain (Fig. S7G-S7H).

Thus, both ‘ProtRAP-LM (Only)’ and ‘TMbed (Only)’ proteins warrant further investigation. However, given the challenges in accurately determining the correct secondary structures for some proteins, and the fact that the number of ‘ProtRAP-LM (Only)’ proteins is nearly three times that of ‘TMbed (Only)’ proteins, our discussion will focus primarily on the ‘ProtRAP-LM (Only)’ proteins.

Similarly to the above section, we can obtain the cluster representatives of these ‘ProtRAP-LM (Only)’ β-sheet-containing membrane proteins (Fig. S6B-S6C). Among them, two specific types of proteins deserve the most attention. One type of these proteins possesses a likely lipid-binding YAB/SYLF domain (Fig. S6B-S6C), suggesting its potential localization on the cell membrane. ^47,48^ And the other type contains massive β-strands (Fig. S6B). In Fig. 4A, we showed all these likely membrane proteins containing massive high-MCP sheets or transmembrane strands (n > 6) from different proteomes, which only appeared within the ProtRAP-LM predictions.

**Figure 4:**
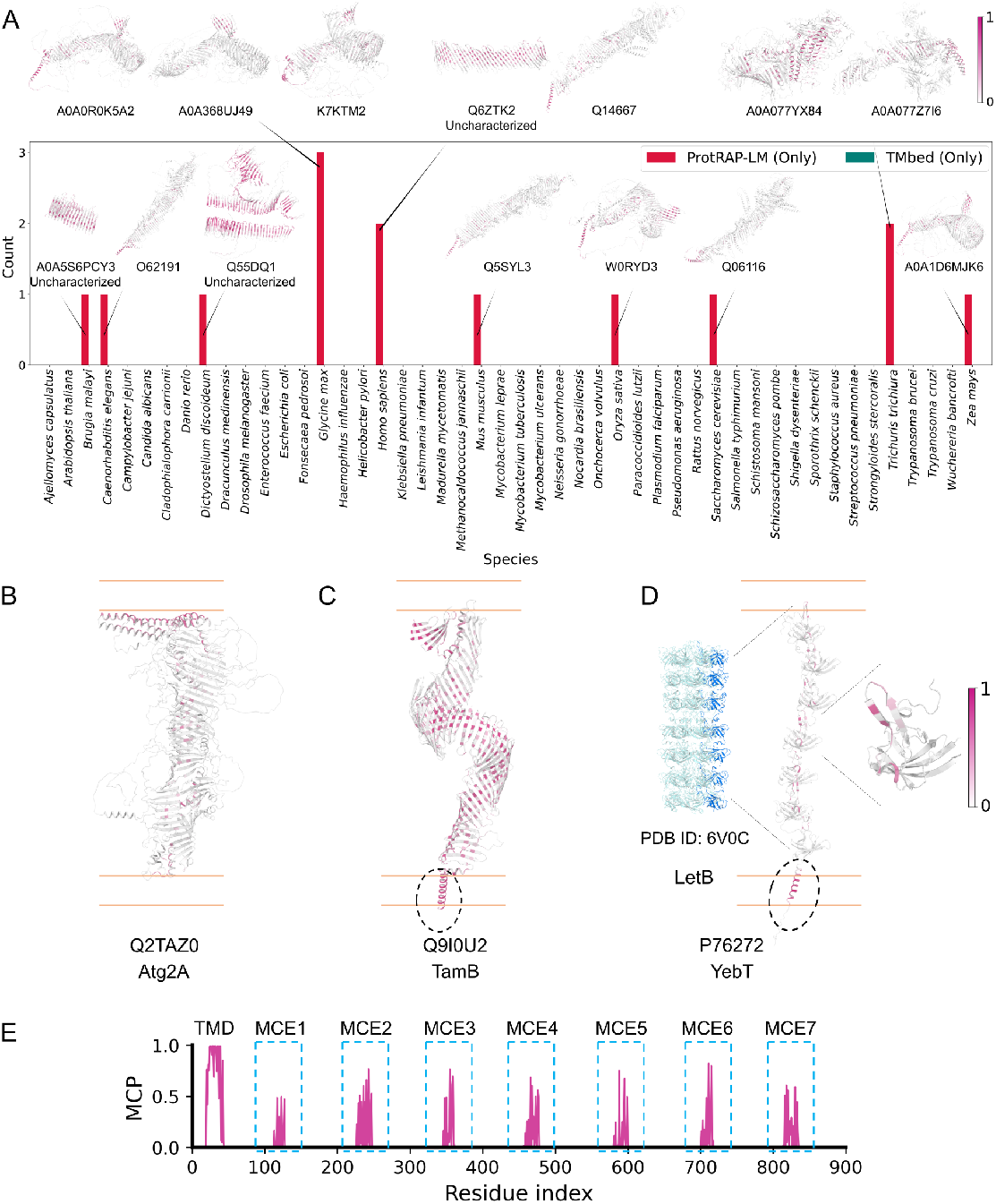
The identified β-sheet-containing membrane proteins with a significant number (n > 6) of β-strands. (A) The β-sheet-containing membrane proteins containing massive β-strands within 48 proteomes. The structures of this part predicted by AlphaFold2^34,63^ were colored according to the predicted MCP by ProtRAP-LM, represented as cartoon. Three known intermembrane lipid transfer proteins of (B) Atg2A, ^50^ (C) TamB ^51,52^ and, (D) YebT ^53^ that direct lipid transfer between membranes were shown. The full-length protein structures predicted by AlphaFold2 and the solved protein structure (PDB ID: 6V0C) were colored according to the predicted MCP by ProtRAP-LM, represented as cartoon. TMbed can only identify the trans-membrane region of the protein TamB and YebT, represented by a black dotted circle in the (C) and (D). The orange parallel lines in the (B-D) indicate the putative positions of the membrane. (E) The predicted MCP of the full-length protein YebT by ProtRAP-LM (warmpink). Our ProtRAP-LM model can effectively identify the membrane-contacting residues located in both the transmembrane domain (TMD) and each mammalian cell entry (MCE) subunit.

There are a couple of noteworthy discoveries here. First, we found there exist structural outliers (bigger circles) that consist of many high-MCP β-strands (UniProt IDs: Q6ZTK2, A0A5S6PCY3, and Q55DQ1), which appear to be functionally dark proteins. ^49^ Our predictions illustrate the possibility that these proteins may be localized in the membrane environments. Next, among these proteins, we found an interesting class of proteins, bridge-like lipid transfer proteins that direct lipid transfer between membranes (UniProt IDs: Q14667, Q5SYL3, etc.). Based on this finding, we evaluated three classes of known intermembrane lipid transfer proteins, including mammalian Atg2A, ^50^ TamB, ^51,52^ and YebT ^53^ in Gram-negative bacteria (Fig. 4B-4D). In these three proteins, we identified high-MCP amino acids in the intermembrane regions (Fig. 4B-4D). According to the predicted MCP of the full-length protein YebT by ProtRAP-LM (Fig. 4E), we found our ProtRAP-LM model can identify the specific lipid-contacting residues located in the transmembrane domain and pore-lining loops (PPLs) of each mammalian cell entry (MCE) subunit. It was shown that the intact hydrophobic tunnel lining formed by the PLLs is essential for the function of lipid transfer. ^53,54^ Therefore, our ProtRAP-LM model may be able to predict intermembrane lipid transfer proteins and identify potential lipid-binding sites and lipid-translocating pathways within their intermembrane domains. Considering that the training data of our ProtRAP-LM predictor does not contain any intermembrane lipid transfer proteins, we speculate that the ProtRAP-LM model has learned the characteristics of lipid-interacting and lipid-binding sites, enabling it to effectively predict lipid-interacting sites within novel membrane proteins such as the intermembrane lipid transfer proteins. Such predictions are challenging to achieve with the other existing methods.

### 2.5. Screening unknown membrane proteins within human proteome

Given the advantages of ProtRAP-LM in identifying the single-pass transmembrane proteins, the membrane-anchored proteins, and the β-sheet-containing membrane proteins, we explored whether there exist previously *unknown* membrane proteins in the human proteome. The human proteome is of particular interest due to its significance in biomedical research, the availability of comprehensive annotation information, and its relevance to human health.

First, we compared the potentially membrane-contacting proteins identified by ProtRAP-LM with known membrane proteins, including the annotated membrane proteins in the public databases and likely membrane proteins identified by other existing tools. The sequence lists in this section are available in the Supplementary File S1. Specifically, the process was divided into the following four steps (Fig. 5).

**Figure 5:**
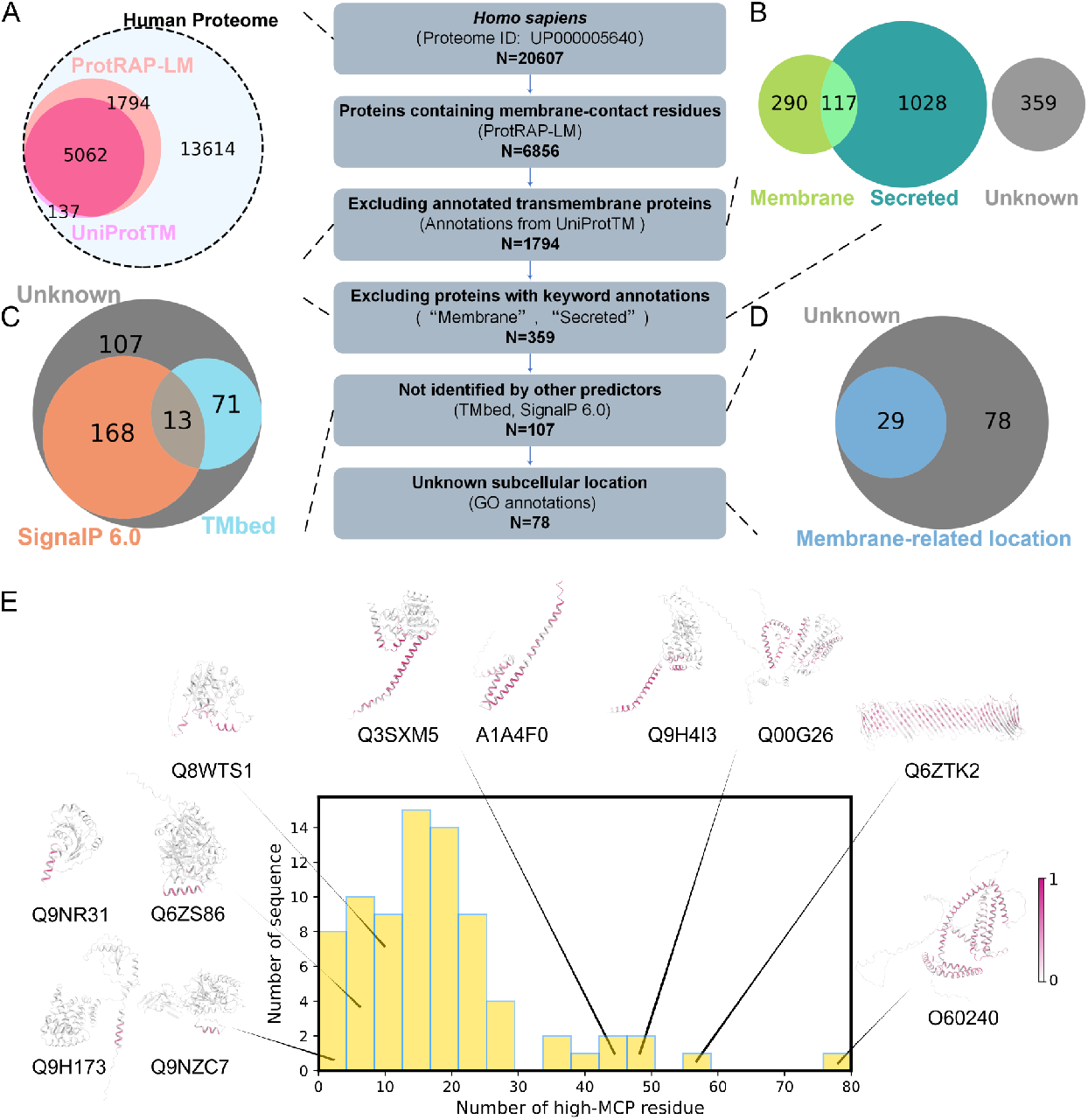
Membrane protein screening within human proteome. (A) Comparison of the potentially membrane-contacting proteins identified by ProtRAP-LM (salmon) with the annotated transmembrane proteins in the UniProt (purple) within the whole human proteome (Alice blue). (B) Comparison of the remaining ProtRAP-LM-identified proteins (excluding the annotated transmembrane proteins) with the protein lists with keyword annotations of “Membrane [SL-0039]” (yellow-green) and “Secreted [SL-0243]” (dark cyan) in the UniProt. ^55^ The proteins without above annotated information of membrane-related subcellular location in the UniProt were shown in gray. (C) Some of the remaining proteins can be identified by TMbed ^15^ with transmembrane span (teal) and SignalP 6.0^32^ with signal peptide (orange). The unidentified proteins were shown in gray. (D) The 29 proteins with membrane-related subcellular location terms from GO annotations (dodgerblue) and the rest for the unknown proteins (gray). The detailed information on these 29 proteins is presented in Table S5. (E) The amount of high-MCP ≥ (MCP 0.5) residues within 78 likely membrane proteins identified by ProtRAP-LM. The representative structures predicted by AlphaFold2 were colored according to the predicted MCP by ProtRAP-LM, represented by cartoon. The UniProt ID of each protein sequence is displayed below the structure diagram. Table S6 provides a comprehensive view of the detailed information for these 78 proteins.

1. Comparing all the ProtRAP-LM-identified proteins to the reviewed sequences with the keyword “Transmembrane [KW-0812]” that were released on 02 June 2021 in UniProtKB database. ^55^ In this step, we found that ProtRAP-LM identified the vast majority of transmembrane proteins annotated in the UniProt (UniProtTM), up to 97.4%, as shown in Fig. 5A. Apart from these annotated membrane proteins, ProtRAP-LM identified 1,794 extra likely membrane proteins.
2. Comparing the 1,794 extra likely membrane proteins against the reviewed sequences with the keyword annotations of “Membrane [SL-0039]” for membrane proteins (released on 02 June 2021) and “Secreted [SL-0243]” for secreted proteins (released on 02 June 2021) in UniProtKB database, ^55^ respectively. We found that a considerable portion of the 1,794 proteins are localized on the membrane (N=407) or classified as secreted proteins with a signal peptide (N=1145) (Fig. 5B). There were still 359 likely membrane proteins left after removing the membrane proteins or secreted proteins previously known in the above databases.
3. Analyzing the remaining 359 proteins in the list with the existing tools, TMbed ^15^ and SignalP 6.0. ^32^ In this step, some of the proteins were identified by TMbed ^15^ and SignalP 6.0, ^32^ which were considered as known membrane proteins or secreted proteins in this study. Interestingly, there were still 107 proteins unidentified by the above tools (Fig. 5C).
4. Searching the GO annotations of cellular localization included in the UniProtKB entry view. ^56^ After this step, 29 out of the 107 remaining proteins were shown to be likely associated or attached to the biological membranes (Fig. 5D and Table S5), and there are still 78 proteins (Fig. 5D and Table S6) without sufficient evidence showing that they are membrane proteins or localized in the membrane.

These 78 previously unknown and potential human membrane proteins, which have not been discovered in the public databases nor identified by the existing tools, were listed in Table S6. We have not found strong evidence that these proteins are indeed membrane proteins, but we believe they are worth in-depth investigations in the future, given the critical importance of membrane proteins in biological functions and medical potentials.

## 3. Discussion

The precise identification of membrane proteins within a proteome is a critical endeavor, as it bears immense potential for drug target discovery and drug design. Over time, a multitude of methods have been developed for membrane protein topology prediction and membrane protein classification from a given sequence, ^2,15,57,58^ some of which were applied for proteome-wide membrane protein identification. ^2,15^ Among these methods, a significant body of research has been dedicated to studying the human proteome. ^59–61^ As the number of known β-barrel transmembrane proteins is much smaller than that of α-helical transmembrane proteins, ^13^ most prediction methods focused on α-helical transmembrane proteins, while only a few predictors have been specifically designed to predict both types of transmembrane pro-_teins._ 10,14,15

In our previous work, ^19^ we found that the membrane contact probability (MCP) predictor performed well for both α-helical and β-barrel transmembrane proteins. Additionally, the MCP predictor also showed potential in identifying membrane-anchored proteins. ^19^ This was not totally unexpected, as the MCP predictor does not rely on the transmembrane topology prediction. Rather, it aims to predict the residues that directly interact with membranes. Therefore, the MCP predictor may be able to identify a greater number of membrane proteins compared to the transmembrane-span based methods such as TMbed (Figs. 2 and S3). Indeed, according to the results presented above, our initial speculation was well-supported, showing that ProtRAP-LM can identify more single-pass transmembrane proteins, membrane-anchored proteins, and β-sheet-containing membrane proteins. In fact, among the ProtRAP-LM-identified proteins within the human proteome, a substantial number of proteins were membrane-anchored (Fig. 5B). Supportively, the MCP-colored locations of the AlphaFold2-predicted structures also suggest the presence of the likely membrane-anchored proteins within the ProtRAP-LM-identified proteins (Fig. 3D-3E). Based on this observation, we speculated that the MCP-based method possesses certain advantages in the identification of membrane proteins compared to the methods solely based on the prediction of transmembrane regions. Our method appears to learn more universal features shared across all types of experimentally solved transmembrane proteins. One challenge in proteome-wide predictions of transmembrane proteins is the confusion between signal peptides and N-terminal transmembrane regions, leading to inaccurate predictions and an overestimation of transmembrane protein fractions. ^62^ To address this issue, we utilized the state-of-the-art predictor SignalP 6.0^32^ to rule out the secreted proteins with a signal peptide (Fig. 2A).

To thoroughly evaluate the performance of our model in proteome-wide predictions and explore unknown membrane proteins, we specifically focus on the human proteome, which benefits from the availability of extensive annotated information. ^4^ Firstly, we conducted a comparative analysis by juxtaposing our results with other high-quality databases of human membrane proteins (Figs. 5 and S8). ProtRAP-LM-identified membrane proteins provided comprehensive coverage of the annotated membrane proteins provided by UniProt (Fig. 5). Specifically, most ProtRAP-LM-identified membrane proteins (up to 94.8%) were already annotated as membrane proteins or secreted proteins with a signal peptide, indicating that most of the membrane proteins have been clearly identified. Recent advancements in protein structure prediction methods, most notably AlphaFold2, ^34,63^ have enabled the identification of transmembrane proteins using high-quality structural models. ^64,65^ These advancements have significantly reduced false positives in data annotation compared to previous methods. Interestingly, in the comparison, our predictions demonstrated a higher coverage for the AlphaFold2-based human transmembrane protein databases compared to both the protein entries of the UniProtTM^66^ and the human transmembrane proteome (HTP) database, ^61^ as shown in Fig. S8A-S8D. Notably, our method achieved coverage of 98.9% and 98.7% for the protein entries in the AlphaFold2-based databases AFTM and TmAlphaFold, respectively. However, it is important to note that AlphaFold2-based approaches also have limitations, and the accuracy still relies on the quality of the predicted structures. For instance, the AFTM may fail to identify transmembrane regions in certain single-pass transmembrane proteins. ^64^ In contrast, our method demonstrated the capability to identify 96.9% of all the known single-pass transmembrane proteins within the human proteome (Fig. S8E), whereas the AFTM procedure missed ^64^ ~20% of single-pass transmembrane compared to the data from UniProt. Combined with the results shown in Fig. 3, it is evident that our method exhibits a remarkable ability to identify single-pass transmembrane proteins, despite the limited availability of experimentally solved structures as a reference.

Our findings also demonstrated the potential of our model in effectively identifying unique β-sheet-containing membrane proteins including intermembrane lipid transfer proteins (Figs. 4 and S6). Among them, the identification of intermembrane lipid transfer proteins by our method came as a surprising discovery that exceeded our expectations. These proteins are entirely independent of our training set, and their limited structural information at this stage adds to the intrigue. However, most of our structural information about this protein type comes from AlphaFold2 predictions, which may be prone to introducing errors (Fig. S7). Therefore, we suggest conducting further experimental research on these proteins, which may hold immense potential to offer valuable insights into the structure and function of this protein type.

Interestingly, through the utilization of the annotations provided by UniProt, the assistance of two existing tools, and the inclusion of subcellular location information provided by GO annotations (Fig. 5A-5D and Table S5), we have identified 78 potentially human membrane proteins previously unknown (Fig. 5E and Table S6). Manual literature inspection on these proteins revealed that some of them could indeed be true membrane proteins, based on available experimental evidence. For example, the first and third most probable membrane proteins correspond to UniProt IDs O60240 and Q00G26 (Table S6), which represent the proteins PLIN1 and PLIN5, respectively. They are both lipid droplet-associated proteins, and serve as a physical and metabolic linkage between the lipid droplet and the endoplasmic reticulum, and between the lipid droplet and mitochondria, respectively. ^67,68^ Recent evidence suggests that PLIN1 is an integral membrane protein dually localizing to the endoplasmic reticulum and lipid droplet. ^69^ While PLIN1 does not contain any TMHMM-predicted and TMbed-predicted transmembrane segments, our ProtRAP-LM prediction reveals high-MCP residues in this protein (Fig. 5E), which is consistent with the experimental findings. This makes the 78 potentially new membrane proteins highly intriguing, which may lead to discoveries of new human membrane proteins with novel functions. We hope these findings will stimulate experimental interest in this direction.

In summary, our study presents a novel transformer- and protein language model-based protein relative accessibility predictor, ProtRAP-LM, which exhibits much improved accuracy and speed. In proteome analysis, we demonstrated that our method offers significant advantages in identifying single-pass transmembrane proteins, membrane-anchored proteins, and β-sheet-containing membrane proteins, when compared to other existing tools and transmembrane protein databases. Therefore, our predictor holds great potential for high throughput membrane protein analysis and membrane protein design, enabling more comprehensive investigations on membrane protein properties and functions.

We provide a more comprehensive list of likely membrane proteins of 48 living organisms identified by our ProtRAP-LM predictor for future reference (Supplementary File S2). Particularly, we propose 78 potential human membrane proteins previously unknown. These functionally dark (membrane) proteins should be further explored in the future.

## 4. Methods

### 4.1. Datasets

Same as the older version of MSA-based ProtRAP, we used the 5650 structures of membrane proteins (released before 2022) with membrane contact probability (MCP) recorded from the MemProtMD database, ^70,71^ and split them into single-chain sequences. We removed the sequences with over 1000 residues, and filtered the rest with over 40% identity by BlastP (version 2.9.0+). ^72^ Finally, 1362 chains of membrane proteins were left. Based on the ablation experiments in our previous work, ^20^ 7440 chains of soluble proteins with lower sequence identity (<25%) were also considered in our dataset. The MCP of each soluble protein residue was considered to be 0. The RASA of each residue was calculated using FreeSASA with a probe radius of 1.4 Å. ^73^ With the RASA and MCP of each residue (Fig. 1A), the RLA, RSA, and RBSA can be calculated (Fig. 1C), which were then used to label the dataset. The above data of membrane proteins and soluble proteins were randomly split into 10 sub-datasets separately for the training and 10-fold cross-validation. Please refer to the previous publications for more details. ^20^

For further evaluation and testing compared to other predictors, we constructed a membrane protein test set with less than 40% sequence identity against the training set, which was composed of the 184 protein sequences released between 01 January 2022 and 31 December 2022 from the MemProtMD database (termed ‘MemProtMD_2022 dataset’). The above method was then used to get the corresponding labels for each protein sequence in the Mem-ProtMD_2022 dataset.

To validate the generalization ability and the performance of membrane protein identification, we constructed three large datasets. For the large dataset 1 (termed ‘PDBTM dataset’), we downloaded the 29795 transmembrane protein sequences with known structures from the widely used transmembrane protein database PDBTM ^74^ (released on 10 November 2022) at http://pdbtm.enzim.hu/. For the large dataset 2 (termed ‘UniProt_Mem dataset’), we collected the 80634 reviewed sequences in the UniProtKB database ^55^ (released on 12 October 2022) with the keyword “Transmembrane [KW-0812]”. The UniProt_Mem dataset consists of the proteins with at least one helix or β-strand embedded in a membrane according to the annotations of UniProt. ^55,66,75^ The PDBTM dataset and UniProt_Mem dataset were regarded as the ground truth of the transmembrane protein databases here. For the large dataset 3 (termed ‘UniProt_Sol dataset’), we constructed a non-membrane protein dataset, which contains the 371069 reviewed sequences in the UniProtKB database ^55^ (released on 14 December 2022) without any keyword about membrane protein or secreted protein. The keywords we utilized include the general keywords “Transmembrane [KW-0812]”, “Membrane [KW-0472]”, subcellular location keywords “Membrane [SL-0162]”, secreted protein related keywords “Secreted [SL-0243]” and “Signal”, as well as plain text keyword “Lipid”. The UniProt_Sol dataset was used as the ground truth of the non-membrane protein database.

The datasets used in this work are available in the Supplementary File S3.

### 4.2. Input features

We generated the embeddings for each sequence using the transformer-based protein language model Evolutionary Scale Modeling 2 with 650M parameters (ESM2_650M) at https://github.com/facebookresearch/esm. ^29^ The encoder model converts a protein sequence into an embedding tensor that represents each residue in the protein by a 1280-dimension vector containing global and local context information.

### 4.3. The ProtRAP-LM architecture

ProtRAP-LM mainly contains one-dimension multiscale convolution layers and stacked transformer encoder modules. The architecture of ProtRAP-LM is shown in Fig. 1B. The input of our model is a 1280-dimension tensor from the ESM-2 model. First, the embeddings were sent to the linear layers to scale down. Then, we used one-dimension multiscale convolution layers to capture local information. The sizes of the convolution kernel were set as 3, 5, 7, and 9, respectively.

Then, the outputs of the convolution layers were concatenated together and added with the inputs of the convolution layers. The results were sent to the transformer encoder to capture more complex dependencies in the input sequence. The transformer encoder in the model contains three layers with four attention heads. Finally, the outputs of the transformer encoder were sent to the linear layers. The outputs of the last layer were processed by the Sigmoid function for the prediction of RASA and MCP, which were further used to calculate the RLA, RSA, and RBSA of each residue for the additional outputs (Fig. 1C). We applied dropout and *L*_2_ regularization to avoid overfitting. Our code was implemented with PyTorch (https://pytorch.org/).

### 4.4. Parameter settings

As introduced before, our dataset was randomly divided into 10 sub-datasets. We used nine of them for training and the remaining one for validation. Hence, 10 different combinations of training and validation sets were generated, and the hyperparameters were determined by the first fold and applied to the remaining nine folds. The predictions for the testing process in this paper were the average results of the ten models.

The batch size of each training epoch was set to 32. The AMSGrad optimizer was used with a basic learning rate of 1 × 10^−4^. When training the models, we employed 10 epochs for the warm-up training and 20 epochs for the normal training. During the warm-up training, the learning rate increased linearly from 2 × 10^−6^ to the basic learning rate. During the normal training, the learning rate decreased exponentially, each epoch dropping to 0.9 of the previous epochs. The loss function was set as the mean square error between the prediction and ground truth (obtained from the MemProtMD database ^70,71^ and the results of FreeSASA ^73^). A similar training process achieved satisfactory results in our previous work. ^20^

### 4.5. Performance evaluation

The Pearson correlation coefficient (PCC) and mean absolute error (MAE) were used to analyze the regression performances of our model, and 10-fold cross-validation was performed to evaluate the generalization and stability of the model with the mean value and standard deviation calculated (Table S1). Each fold was trained with the same training conditions and hyperparameters.

In the runtime analysis, the testing process was conducted on a workstation containing a ten-core (20-thread) Intel Xeon W-2255 CPU and an RTX 3080 Nvidia GPU with 10 GB of GPU memory.

To evaluate the generalization ability and the membrane protein identification performance of our model, recall and false positive rate (FPR) were used (Tables 2 and S3). In this paper, the recall was defined as the percentage of membrane proteins correctly identified out of the total annotated membrane proteins in the PDBTM dataset and UniProt_Mem dataset. The false positive rate is the proportion of all non-membrane proteins that are incorrectly classified as membrane proteins out of the total non-membrane proteins in the UniProt_Sol dataset.

### 4.6. Comparing ProtRAP-LM to other approaches

We used the MemProtMD_2022 dataset as the test set to compare the performance of our ProtRAP-LM model with the MSA-based MCP predictor^19^ and ProtRAP, ^20^ running locally, and other state-of-the-art predictors, NetSurfP-2.0^23^ and NetSurfP-3.0, ^31^ using the official server at https://services.healthtech.dtu.dk/services/NetSurfP-2.0/ and https://services.healthtech.dtu.dk/services/NetSurfP-3.0/.

For the membrane protein identification within entire proteomes, we compared the predicted membrane protein fractions with the state-of-the-art transmembrane protein predictor TMbed, ^15^ which was installed and run locally.

## 5. Data and code availability

An online computation server of ProtRAP-LM is available for protein relative accessibility (RASA, RLA, RSA, and RBSA) and membrane contact probability (MCP) predictions at http://www.songlab.cn/ProtRAP-LM/home/.

The source code and its corresponding materials are available at https://github.com/ComputBiophys/ProtRAP-LM. It is available under the permissive GPL-3.0 Licence.

## 6. Acknowledgments

This work was supported by the National Key R&D Program of China (2024YFA0916800) and the Science Fund for Creative Research Groups of the National Natural Science Foundation of China (T2321001). W.L. was supported by the Postdoctoral Fellowship of Peking-Tsinghua Center for Life Sciences. Part of the computations were performed on the Computing Platform of the Center for Life Sciences at Peking University.

## 7. Author contributions

C.S. designed and supervised the project. L.W. conducted the research, including the proteome-wide analysis of membrane proteins. K.K. trained the ProtRAP-LM predictor. All the authors participated in the writing of the manuscript.

## 8. Competing interests

The authors declare no competing interests.

## Notes

### Competing Interest Statement

The authors have declared no competing interest.

## References

[1] Wallin, E.; Heijne, G. V. Genome-wide analysis of integral membrane proteins from eubacterial, archaean, and eukaryotic organisms. Protein Sci. 1998, 7, 1038–.

[2] Krogh, A.; Larsson, B.; Von Heijne, G.; Sonnhammer, E. L. Predicting transmembrane protein topology with a hidden Markov model: Application to complete genomes. J. Mol. Biol. 2001, 305, 580–.

[3] Wu, C. C.; MacCoss, M. J.; Howell, K. E.; Yates III, J. R. A method for the comprehensive proteomic analysis of membrane proteins. Nat. Biotechnol. 2003, 21, 538–.

[4] Uhlén, M.; Fagerberg, L.; Hallström, B. M.; Lindskog, C.; Oksvold, P.; Mardinoglu, A.; Sivertsson, Å.; Kampf, C.; Sjöstedt, E.; Asplund, A.; others Tissue-based map of the human proteome. Science 2015, 347, 1260419.

[5] Tan, S.; Tan, H. T.; Chung, M. C. Membrane proteins and membrane proteomics. Proteomics 2008, 8, 3932–.

[6] Ryu, H.; Fuwad, A.; Yoon, S.; Jang, H.; Lee, J. C.; Kim, S. M.; Jeon, T.-J. Biomimetic membranes with transmembrane proteins: State-of-the-art in transmembrane protein applications. Int. J. Mol. Sci. 2019, 20, 1437.

[7] Li, F.; Egea, P. F.; Vecchio, A. J.; Asial, I.; Gupta, M.; Paulino, J.; Bajaj, R.; Dickinson, M. S.; Ferguson-Miller, S.; Monk, B. C.; others Highlighting membrane protein structure and function: A celebration of the Protein Data Bank. J. Biol. Chem. 2021, 296.

[8] Overington, J. P.; Al-Lazikani, B.; Hopkins, A. L. How many drug targets are there? Nat. Rev. Drug Discov. 2006, 5, 996–.

[9] Almeida, J. G.; Preto, A. J.; Koukos, P. I.; Bonvin, A. M.; Moreira, I. S. Membrane proteins structures: A review on computational modeling tools. Biochim. Biophys. Acta. - Biomembr. 2017, 1859, 2039–.

[10] Sun, J.; Kulandaisamy, A.; Liu, J.; Hu, K.; Gromiha, M. M.; Zhang, Y. Machine learning in computational modelling of membrane protein sequences and structures: From methodologies to applications. Comput. Struct. Biotechnol. J. 2023, 21, 1226–.

[11] Chou, K.-C.; Elrod, D. W. Prediction of membrane protein types and subcellular locations. Proteins 1999, 34, 153–.

[12] Chou, K.-C.; Cai, Y.-D. Prediction of membrane protein types by incorporating amphipathic effects. Journal of chemical information and modeling 2005, 45, 413–.

[13] Punta, M.; Forrest, L. R.; Bigelow, H.; Kernytsky, A.; Liu, J.; Rost, B. Membrane protein prediction methods. Methods 2007, 41, 474–.

[14] Hallgren, J.; Tsirigos, K. D.; Pedersen, M. D.; Armenteros, J. J. A.; Marcatili, P.; Nielsen, H.; Krogh, A.; Winther, O. DeepTMHMM predicts alpha and beta transmembrane proteins using deep neural networks. bioRxiv 2022,

[15] Bernhofer, M.; Rost, B. TMbed: Transmembrane proteins predicted through language model embeddings. BMC Bioinformatics 2022, 23, 326.

[16] Sudhahar, C.; Haney, R.; Xue, Y.; Stahelin, R. Cellular membranes and lipid-binding domains as attractive targets for drug development. Curr. Drug Targets 2008, 9, 613–.

[17] Pataki, C. I.; Rodrigues, J.; Zhang, L.; Qian, J.; Efron, B.; Hastie, T.; Elias, J. E.; Levitt, M.; Kopito, R. R. Proteomic analysis of monolayer-integrated proteins on lipid droplets identifies amphipathic interfacial α-helical membrane anchors. Proc. Natl. Acad. Sci. U.S.A. 2018, 115, E8172–E8180.

[18] Boes, D. M.; Godoy-Hernandez, A.; McMillan, D. G. Peripheral membrane proteins: Promising therapeutic targets across domains of life. Membranes 2021, 11, 346.

[19] Wang, L.; Zhang, J.; Wang, D.; Song, C. Membrane contact probability: An essential and predictive character for the structural and functional studies of membrane proteins. PLoS Comput. Biol. 2022, 18, e1009972.

[20] Kang, K.; Wang, L.; Song, C. ProtRAP: Predicting lipid accessibility together with solvent accessibility of proteins in one run. J. Chem. Inf. Model 2023, 63, 1065–.

[21] Wang, S.; Li, W.; Liu, S.; Xu, J. RaptorX-Property: A web server for protein structure property prediction. Nucleic Acids Res. 2016, 44, W430–W435.

[22] Wang, S.; Sun, S.; Li, Z.; Zhang, R.; Xu, J. Accurate de novo prediction of protein contact map by ultra-deep learning model. PLoS computational biology 2017, 13, e1005324.

[23] Klausen, M. S.; Jespersen, M. C.; Nielsen, H.; Jensen, K. K.; Jurtz, V. I.; Soenderby, C. K.; Sommer, M. O. A.; Winther, O.; Nielsen, M.; Petersen, B.; others NetSurfP-2.0: Improved prediction of protein structural features by integrated deep learning. Proteins 2019, 87, 527–.

[24] Khurana, D.; Koli, A.; Khatter, K.; Singh, S. Natural language processing: State of the art, current trends and challenges. Multimed. Tools. Appl. 2023, 82, 3744–.

[25] El-Komy, A.; Shahin, O. R.; Abd El-Aziz, R. M.; Taloba, A. I. Integration of computer vision and natural language processing in multimedia robotics application. Inf. Sci 2022, 7.

[26] Elnaggar, A.; Heinzinger, M.; Dallago, C.; Rehawi, G.; Wang, Y.; Jones, L.; Gibbs, T.; Feher, T.; Angerer, C.; Steinegger, M.; Bhowmik, D.; Rost, B. ProtTrans: Toward Understanding the Language of Life Through Self-Supervised Learning. IEEE Trans. Pattern Anal. Mach. Intell. 2022, 44, 7127–.

[27] Meier, J.; Rao, R.; Verkuil, R.; Liu, J.; Sercu, T.; Rives, A. Language models enable zero-shot prediction of the effects of mutations on protein function. Adv. Neural Inf. Process. 2021, 34, 29303–.

[28] Nijkamp, E.; Ruffolo, J. A.; Weinstein, E. N.; Naik, N.; Madani, A. Progen2: exploring the boundaries of protein language models. Cell Syst. 2023, 14, 978–.

[29] Lin, Z.; Akin, H.; Rao, R.; Hie, B.; Zhu, Z.; Lu, W.; Smetanin, N.; Verkuil, R.; Kabeli, O.; Shmueli, Y.; others Evolutionary-scale prediction of atomic-level protein structure with a language model. Science 2023, 379, 1130–.

[30] Hayes, T.; Rao, R.; Akin, H.; Sofroniew, N. J.; Oktay, D.; Lin, Z.; Verkuil, R.; Tran, V. Q.; Deaton, J.; Wiggert, M.; others Simulating 500 million years of evolution with a language model. Science 2025, eads0018.

[31] Høie, M. H.; Kiehl, E. N.; Petersen, B.; Nielsen, M.; Winther, O.; Nielsen, H.; Hallgren, J.; Marcatili, P. NetSurfP-3.0: Accurate and fast prediction of protein structural features by protein language models and deep learning. Nucleic Acids Res. 2022, 50, W510–W515.

[32] Teufel, F.; Almagro Armenteros, J. J.; Johansen, A. R.; Gíslason, M. H.; Pihl, S. I.; Tsirigos, K. D.; Winther, O.; Brunak, S.; von Heijne, G.; Nielsen, H. SignalP 6.0 predicts all five types of signal peptides using protein language models. Nat. Biotechnol. 2022, 40, 1025–.

[33] Gu, Z.; Luo, X.; Chen, J.; Deng, M.; Lai, L. Hierarchical graph transformer with contrastive learning for protein function prediction. Bioinformatics 2023, 39, btad410.

[34] Tunyasuvunakool, K.; Adler, J.; Wu, Z.; Green, T.; Zielinski, M.; Žídek, A.; Bridgland, A.; Cowie, A.; Meyer, C.; Laydon, A.; others Highly accurate protein structure prediction for the human proteome. Nature 2021, 596, 596–.

[35] Lomize, A. L.; Schnitzer, K. A.; Todd, S. C.; Cherepanov, S.; Outeiral, C.; Deane, C. M.; Pogozheva, I. D. Membranome 3.0: Database of single-pass membrane proteins with AlphaFold models. Protein Sci. 2022, 31, e4318.

[36] Bugge, K.; Lindorff-Larsen, K.; Kragelund, B. B. Understanding single-pass transmembrane receptor signaling from a structural viewpoint—what are we missing? FEBS J. 2016, 283, 4451–.

[37] Steinegger, M.; Söding, J. MMseqs2 enables sensitive protein sequence searching for the analysis of massive data sets. Nat. Biotechnol. 2017, 35, 1028–.

[38] Barrio-Hernandez, I.; Yeo, J.; Jänes, J.; Mirdita, M.; Gilchrist, C. L.; Wein, T.; Varadi, M.; Velankar, S.; Beltrao, P.; Steinegger, M. Clustering predicted structures at the scale of the known protein universe. Nature 2023, 622, 645–.

[39] van Kempen, M.; Kim, S. S.; Tumescheit, C.; Mirdita, M.; Lee, J.; Gilchrist, C. L.; Söding, J.; Steinegger, M. Fast and accurate protein structure search with Foldseek. Nat. Biotechnol. 2023, 1–4.

[40] Lander, N.; Bernal, C.; Diez, N.; Anez, N.; Docampo, R.; Ramírez, J. L. Localization and developmental regulation of a dispersed gene family 1 protein in Trypanosoma cruzi. Infect. Immun. 2010, 78, 240–.

[41] Murakami, K.; Mihara, K.; Omura, T. The transmembrane region of microsomal cytochrome P450 identified as the endoplasmic reticulum retention signal. J. Biochem. 1994, 116, 175–.

[42] Dorner, M. E.; McMunn, R. D.; Bartholow, T. G.; Calhoon, B. E.; Conlon, M. R.; Dulli, J. M.; Fehling, S. C.; Fisher, C. R.; Hodgson, S. W.; Keenan, S. W.; others Comparison of intrinsic dynamics of cytochrome p450 proteins using normal mode analysis. Protein Sci. 2015, 24, 1507–.

[43] Zhang, J.; Frerman, F. E.; Kim, J.-J. P. Structure of electron transfer flavoprotein-ubiquinone oxidoreductase and electron transfer to the mitochondrial ubiquinone pool. Proc. Natl. Acad. Sci. U. S. A. 2006, 103, 16217–.

[44] Watmough, N. J.; Frerman, F. E. The electron transfer flavoprotein: ubiquinone oxidoreductases. Biochim. Biophys. Acta - Bioenerg. 2010, 1797, 1916–.

[45] Kabsch, W.; Sander, C. Dictionary of protein secondary structure: Pattern recognition of hydrogen-bonded and geometrical features. Biopolymers 1983, 22, 2637–.

[46] Jia, N.; Liu, N.; Cheng, W.; Jiang, Y.-L.; Sun, H.; Chen, L.-L.; Peng, J.; Zhang, Y.; Ding, Y.-H.; Zhang, Z.-H.; others Structural basis for receptor recognition and pore formation of a zebrafish aerolysin-like protein. EMBO Rep. 2016, 17, 248–.

[47] Hasegawa, J.; Tokuda, E.; Tenno, T.; Tsujita, K.; Sawai, H.; Hiroaki, H.; Takenawa, T.; Itoh, T. SH3YL1 regulates dorsal ruffle formation by a novel phosphoinositide-binding domain. J. Cell Biol. 2011, 193, 916–.

[48] Quilici, G.; Berardi, A.; Fabris, C.; Ghitti, M.; Punta, M.; Gourlay, L. J.; Bolognesi, M.; Musco, G. Solution structure of the BPSL1445 protein of Burkholderia pseudomallei reveals the SYLF domain three-dimensional fold. ACS Chem. Biol. 2021, 17, 239–.

[49] Durairaj, J.; Waterhouse, A. M.; Mets, T.; Brodiazhenko, T.; Abdullah, M.; Studer, G.; Tauriello, G.; Akdel, M.; Andreeva, A.; Bateman, A.; others Uncovering new families and folds in the natural protein universe. Nature 2023, 622, 653–.

[50] Maeda, S.; Otomo, C.; Otomo, T. The autophagic membrane tether ATG2A transfers lipids between membranes. elife 2019, 8, e45777.

[51] Ruiz, N.; Davis, R. M.; Kumar, S. YhdP, TamB, and YdbH are redundant but essential for growth and lipid homeostasis of the Gram-negative outer membrane. MBio 2021, 12, e02714–21.

[52] Douglass, M. V.; McLean, A. B.; Trent, M. S. Absence of YhdP, TamB, and YdbH leads to defects in glycerophospholipid transport and cell morphology in Gram-negative bacteria. PLoS Genet. 2022, 18, e1010096.

[53] Isom, G. L.; Coudray, N.; MacRae, M. R.; McManus, C. T.; Ekiert, D. C.; Bhabha, G. LetB structure reveals a tunnel for lipid transport across the bacterial envelope. Cell 2020, 181, 664–.

[54] Liu, C.; Ma, J.; Wang, J.; Wang, H.; Zhang, L. Cryo-EM structure of a bacterial lipid transporter YebT. J. Mol. Biol. 2020, 432, 1019–.

[55] Consortium, T. U. UniProt: A worldwide hub of protein knowledge. Nucleic Acids Res. 2019, 47, D506–D515.

[56] Dimmer, E. C.; Huntley, R. P.; Alam-Faruque, Y.; Sawford, T.; O’Donovan, C.; Martin, M. J.; Bely, B.; Browne, P.; Mun Chan, W.; Eberhardt, R.; others The UniProt-GO annotation database in 2011. Nucleic Acids Res. 2012, 40, D565–D570.

[57] Viklund, H.; Elofsson, A. OCTOPUS: Improving topology prediction by two-track ANN-based preference scores and an extended topological grammar. Bioinformatics 2008, 24, 1668–.

[58] Bernsel, A.; Viklund, H.; Hennerdal, A.; Elofsson, A. TOPCONS: Consensus prediction of membrane protein topology. Nucleic Acids Res. 2009, 37, W465–W468.

[59] Almén, M. S.; Nordström, K. J.; Fredriksson, R.; Schiöth, H. B. Mapping the human membrane proteome: A majority of the human membrane proteins can be classified according to function and evolutionary origin. BMC Biol. 2009, 7, 14–.

[60] Fagerberg, L.; Jonasson, K.; von Heijne, G.; Uhlén, M.; Berglund, L. Prediction of the human membrane proteome. Proteomics 2010, 10, 1149–.

[61] Dobson, L.; Reményi, I.; Tusnády, G. E. The human transmembrane proteome. Biol. Direct 2015, 10, 18–.

[62] Tsirigos, K. D.; Peters, C.; Shu, N.; Käll, L.; Elofsson, A. The TOPCONS web server for consensus prediction of membrane protein topology and signal peptides. Nucleic Acids Res. 2015, 43, W401–W407.

[63] Jumper, J.; Evans, R.; Pritzel, A.; Green, T.; Figurnov, M.; Ronneberger, O.; Tunyasuvunakool, K.; Bates, R.; Žídek, A.; Potapenko, A.; others Highly accurate protein structure prediction with AlphaFold. Nature 2021, 596, 589–.

[64] Pei, J.; Cong, Q. AFTM: A database of transmembrane regions in the human proteome predicted by AlphaFold. Database 2023, 2023, baad008.

[65] Dobson, L.; Szekeres, L. I.; Gerdán, C.; Langó, T.; Zeke, A.; Tusnády, G. E. TmAlphaFold database: Membrane localization and evaluation of AlphaFold2 predicted alpha-helical transmembrane protein structures. Nucleic Acids Res. 2023, 51, D517–D522.

[66] Consortium, T. U. UniProt: The universal protein knowledgebase in 2021. Nucleic Acids Res. 2021, 49, D480–D489.

[67] Sztalryd, C.; Brasaemle, D. L. The perilipin family of lipid droplet proteins: Gatekeepers of intracellular lipolysis. Biochim. Biophys. Acta - Mol. Cell Biol. Lipids 2017, 1862, 1232–.

[68] Najt, C. P.; Devarajan, M.; Mashek, D. G. Perilipins at a glance. J. Cell Sci. 2022, 135, jcs259501.

[69] Majchrzak, M.; Stojanović, O.; Ajjaji, D.; M’barek, K. B.; Omrane, M.; Thiam, A. R.; Klemm, R. W. Perilipin membrane integration determines lipid droplet heterogeneity in differentiating adipocytes. Cell Rep. 2024, 43.

[70] Stansfeld, P. J.; Goose, J. E.; Caffrey, M.; Carpenter, E. P.; Parker, J. L.; Newstead, S.; San-som, M. S. MemProtMD: Automated insertion of membrane protein structures into explicit lipid membranes. Structure 2015, 23, 1361–.

[71] Newport, T. D.; Sansom, M. S. P.; Stansfeld, P. J. The MemProtMD database: A resource for membrane-embedded protein structures and their lipid interactions. Nucleic Acids Res. 2019, 47, D390–D397.

[72] Altschul, S. F.; Gish, W.; Miller, W.; Myers, E. W.; Lipman, D. J. Basic local alignment search tool. J. Mol. Biol. 1990, 215, 410–.

[73] Mitternacht, S. FreeSASA: An open source C library for solvent accessible surface area calculations. F1000Res. 2016, 5.

[74] Kozma, D.; Simon, I.; Tusnady, G. E. PDBTM: Protein Data Bank of transmembrane proteins after 8 years. Nucleic Acids Res. 2012, 41, D524–D529.

[75] Consortium, T. U. UniProt: The universal protein knowledgebase in 2023. Nucleic Acids Res. 2023, 51, D523–D531.

