## Supplementary Information for "ProtRAP-LM: Fast and accurate protein relative accessibility prediction and membrane protein screening through protein language model embeddings"

### 1 Supplementary Tables

**Table S1:** Overall prediction performance of ProtRAP-LM in the 10-fold cross-validation (on the residue level).

| | Evaluation | Mean value | Standard deviation |
|---|---|---|---|
| MCP | PCC | 0.9012 | 0.0066 |
| | MAE | 0.0139 | 0.0013 |
| RASA | PCC | 0.7969 | 0.0031 |
| | MAE | 0.1129 | 0.0009 |
| RLA | PCC | 0.8478 | 0.0078 |
| | MAE | 0.0072 | 0.0005 |
| RSA | PCC | 0.8043 | 0.0029 |
| | MAE | 0.1095 | 0.0008 |

**Table S2:** Comparison of the protein relative accessibility (RLA and RSA) prediction performance using the MemProtMD_2022 dataset among different methods (on the residue level).

| | Evaluation | ProtRAP-LM | ProtRAP | NetSurfP-3.0 | NetSurfP-2.0 |
|---|---|---|---|---|---|
| RLA | PCC | **0.8634** | 0.8407 | - | - |
| | MAE | **0.0240** | 0.0282 | - | - |
| RSA | PCC | **0.8031** | 0.7468 | 0.6827 | 0.6051 |
| | MAE | **0.1055** | 0.1202 | 0.1451 | 0.1592 |

**Table S3:** Comparative analysis of the membrane protein identification performance using ProtRAP-LM with the varying parameters, specifically focusing on the amount of high-MCP (MCP $\geq$ 0.5) predictions in ten successive amino acids of a protein.

| Parameter | PDBTM dataset | UniProt_Mem dataset | UniProt_Sol dataset |
|---|---|---|---|
| | Recall | Recall | FPR |
| 1 | 98.13% | 97.89% | 0.66% |
| 2 | 97.87% | 97.72% | 0.52% |
| 3 | 97.70% | 97.50% | 0.48% |
| 4 | 96.33% | 97.25% | 0.46% |
| 5 | 94.62% | 96.73% | 0.41% |
| 6 | 91.74% | 95.81% | 0.36% |

**Table S4:** The 48 proteomes deposited in the AlphaFold Database (version 4).

| Species | Common Name | Proteome ID | Protein number |
|---|---|---|---|
| *Ajellomyces capsulatus* | *Ajellomyces capsulatus* | UP000001631 | 9,214 |
| *Arabidopsis thaliana* | *Arabidopsis* | UP000006548 | 27,487 |
| *Brugia malayi* | *Brugia malayi* | UP000006672 | 11,497 |
| *Caenorhabditis elegans* | *Nematode worm* | UP000001940 | 19,838 |
| *Campylobacter jejuni* | *C. jejuni* | UP000000799 | 1,623 |
| *Candida albicans* | *C. albicans* | UP000000559 | 6,035 |
| *Cladophialophora carrionii* | *Cladophialophora carrionii* | UP000094526 | 11,181 |
| *Danio rerio* | *Zebrafish* | UP000000437 | 20,358 |
| *Dictyostelium discoideum* | *Dictyostelium* | UP000002195 | 12,727 |
| *Dracunculus medinensis* | *Dracunculus medinensis* | UP000274756 | 10,868 |
| *Drosophila melanogaster* | *Fruit fly* | UP000000803 | 13,821 |
| *Enterococcus faecium* | *Enterococcus faecium* | UP000325664 | 2,823 |
| *Escherichia coli* | *E. coli* | UP000000625 | 4,402 |
| *Fonsecaea pedrosoi* | *Fonsecaea pedrosoi* | UP000053029 | 12,525 |
| *Glycine max* | *Soybean* | UP000008827 | 55,855 |
| *Haemophilus influenzae* | *H. influenzae* | UP000000579 | 1,704 |
| *Helicobacter pylori* | *H. pylori* | UP000000429 | 1,554 |
| *Homo sapiens* | *Human* | UP000005640 | 20,607 |
| *Klebsiella pneumoniae* | *K. pneumoniae* | UP000007841 | 5,728 |
| *Leishmania infantum* | *L. infantum* | UP000008153 | 8,045 |
| *Madurella mycetomatis* | *Madurella mycetomatis* | UP000078237 | 9,733 |
| *Methanocaldococcus jannaschii* | *M. jannaschii* | UP000000805 | 1,787 |
| *Mus musculus* | *Mouse* | UP000000589 | 21,985 |
| *Mycobacterium leprae* | *Mycobacterium leprae* | UP000000806 | 1,603 |
| *Mycobacterium tuberculosis* | *M. tuberculosis* | UP000001584 | 3,995 |
| *Mycobacterium ulcerans* | *Mycobacterium ulcerans* | UP000020681 | 9,033 |
| *Neisseria gonorrhoeae* | *N. gonorrhoeae* | UP000000535 | 2,106 |
| *Nocardia brasiliensis* | *Nocardia brasiliensis* | UP000006304 | 8,414 |
| *Onchocerca volvulus* | *Onchocerca volvulus* | UP000024404 | 12,111 |
| *Oryza sativa* | *Asian rice* | UP000059680 | 43,673 |
| *Paracoccidioides lutzii* | *Paracoccidioides lutzii* | UP000002059 | 8,811 |
| *Plasmodium falciparum* | *P. falciparum* | UP000001450 | 5,372 |
| *Pseudomonas aeruginosa* | *P. aeruginosa* | UP000002438 | 5,564 |
| *Rattus norvegicus* | *Rat* | UP000002494 | 22,859 |
| *Saccharomyces cerevisiae* | *Budding yeast* | UP000002311 | 6,060 |
| *Salmonella typhimurium* | *S. typhimurium* | UP000001014 | 4,533 |

Continued on next page

| Species | Common Name | Proteome ID | Protein number |
|---|---|---|---|
| *Schistosoma mansoni* | *Schistosoma mansoni* | UP000008854 | 10,026 |
| *Schizosaccharomyces pombe* | *Fission yeast* | UP000002485 | 5,122 |
| *Shigella dysenteriae* | *S. dysenteriae* | UP000002716 | 3,897 |
| *Sporothrix schenckii* | *Sporothrix schenckii* | UP000018087 | 8,673 |
| *Staphylococcus aureus* | *S. aureus* | UP000008816 | 2,889 |
| *Streptococcus pneumoniae* | *S. pneumoniae* | UP000000586 | 2,030 |
| *Strongyloides stercoralis* | *Strongyloides stercoralis* | UP000035681 | 12,825 |
| *Trichuris trichiura* | *Trichuris trichiura* | UP000030665 | 9,624 |
| *Trypanosoma brucei* | *Trypanosoma brucei* | UP000008524 | 8,561 |
| *Trypanosoma cruzi* | *T. cruzi* | UP000002296 | 19,242 |
| *Wuchereria bancrofti* | *Wuchereria bancrofti* | UP000270924 | 13,000 |
| *Zea mays* | *Maize* | UP000007305 | 39,208 |

**Table S5:** The membrane-related subcellular location terms from the GO annotations of UniProtKB entries for the 29 likely membrane proteins in Fig. 5D.

| Number | UniProt ID | The membrane-related subcellular location terms |
|---|---|---|
| 1 | P36551 | Mitochondrial inner membrane |
| 2 | Q9Y4L1 | Membrane |
| 3 | Q8IVK1 | Plasma membrane |
| 4 | Q9NRG9 | Membrane, nuclear membrane |
| 5 | Q96HL8 | Ruffle membrane |
| 6 | Q8N2G8 | Membrane, nuclear envelope |
| 7 | Q16795 | Mitochondrial membrane |
| 8 | Q14249 | Mitochondrial inner membrane |
| 9 | Q9UF12 | Mitochondrial inner membrane |
| 10 | Q8N0X7 | Mitochondrial outer membrane, plasma membrane |
| 11 | Q07973 | Mitochondrial inner membrane |
| 12 | Q9NQZ5 | Mitochondrial outer membrane |
| 13 | Q8IXB1 | Membrane |
| 14 | Q8NBX0 | Membrane |
| 15 | P07686 | Membrane |
| 16 | Q9Y3E5 | Membrane |
| 17 | O60762 | Endoplasmic reticulum membrane, membrane |
| 18 | Q96HE9 | Membrane |
| 19 | P43304 | Mitochondrial inner membrane |
| 20 | Q8IYU8 | Mitochondrial inner membrane |
| 21 | O43272 | Mitochondrial inner membrane |
| 22 | Q9Y4P3 | Endoplasmic reticulum membrane |
| 23 | Q5THJ4 | Extrinsic component of membrane |
| 24 | Q8WWC4 | Mitochondrial inner membrane |
| 25 | Q8IXM3 | Mitochondrial inner membrane |
| 26 | Q7Z6Z6 | Membrane |
| 27 | Q8TB40 | Endoplasmic reticulum membrane |
| 28 | Q9HBH5 | Endoplasmic reticulum membrane, lysosomal membrane, membrane |
| 29 | O43610 | Membrane |

**Table S6:** The information of the 78 previously unknown and potential human membrane proteins in Fig. 5D that have not been observed in the public datasets nor identified by other existing tools. The sequences identified by the MSA-based MCP predictor were indicated in bold font in the column one.

| UniProt ID | Amount of high-MCP residues | Mean pLDDT (MCP $\geq$ 0.5) | Length | Mean pLDDT |
|---|---|---|---|---|
| Q9H173 | 3 | 71.56 | 461 | 81.82 |
| Q9NZC7 | 3 | 41.41 | 414 | 85.59 |
| **Q7RTY3** | 3 | 43.88 | 260 | 83.19 |
| O75920 | 3 | 87.20 | 110 | 62.39 |
| A0A1B0GVS7 | 4 | 61.51 | 189 | 54.06 |
| A6NIN4 | 4 | 77.78 | 190 | 69.30 |
| Q96T59 | 4 | 79.65 | 188 | 51.31 |
| Q9UHT4 | 4 | 87.10 | 67 | 65.45 |
| Q6ZS86 | 5 | 96.00 | 529 | 94.12 |
| Q9NRG7 | 6 | 95.01 | 293 | 97.25 |
| **P83111** | 7 | 36.05 | 547 | 79.17 |
| Q96LJ7 | 7 | 80.80 | 313 | 91.61 |
| **P15586** | 7 | 45.61 | 552 | 89.38 |
| Q9NR28 | 7 | 53.22 | 239 | 82.48 |
| Q6S5H5 | 7 | 35.81 | 508 | 65.07 |
| Q8IXL9 | 7 | 97.87 | 164 | 88.85 |
| **Q6NUM6** | 8 | 79.49 | 668 | 75.65 |
| **Q9NR31** | 8 | 69.37 | 198 | 86.11 |
| **Q96DB5** | 9 | 74.14 | 314 | 83.63 |
| Q8WTS1 | 9 | 79.79 | 349 | 87.61 |
| P55327 | 9 | 68.33 | 224 | 67.15 |
| P0C2W7 | 10 | 85.73 | 299 | 51.87 |
| Q8WYQ3 | 10 | 50.92 | 142 | 61.08 |
| Q5T1J5 | 10 | 52.65 | 151 | 60.79 |
| Q96J77 | 10 | 68.65 | 140 | 71.01 |
| Q96EZ4 | 10 | 80.34 | 313 | 30.99 |
| Q9H078 | 12 | 40.15 | 707 | 72.60 |
| Q9UKA2 | 13 | 44.20 | 621 | 86.73 |
| **Q8IZ16** | 13 | 82.56 | 206 | 61.74 |
| Q9Y6H1 | 13 | 53.29 | 151 | 62.10 |
| **Q8IZ81** | 13 | 74.37 | 293 | 94.61 |
| P51861 | 13 | 72.80 | 262 | 49.20 |
| **Q8N128** | 14 | 82.28 | 213 | 64.26 |
| **Q99675** | 14 | 90.04 | 332 | 80.25 |
| **P0C7X4** | 14 | 41.79 | 201 | 76.65 |

Continued on next page

| UniProt ID | Amount of high-MCP residues | Mean pLDDT (MCP $\geq$ 0.5) | Length | Mean pLDDT |
|---|---|---|---|---|
| A6PVY3 | 15 | 75.11 | 158 | 63.31 |
| O76070 | 15 | 88.72 | 127 | 77.81 |
| **Q9UG22** | 16 | 90.76 | 337 | 88.92 |
| **P51687** | 16 | 44.77 | 545 | 85.09 |
| **Q96BQ5** | 16 | 66.81 | 260 | 87.08 |
| Q9NRI7 | 16 | 90.78 | 21 | 88.65 |
| **Q9NPH0** | 16 | 46.40 | 428 | 90.83 |
| Q96AQ6 | 17 | 84.98 | 731 | 57.33 |
| Q96EG1 | 17 | 37.95 | 525 | 91.15 |
| **Q96MZ0** | 18 | 65.70 | 367 | 78.11 |
| Q5VYY1 | 18 | 90.11 | 191 | 94.46 |
| **Q8N336** | 18 | 87.57 | 334 | 91.45 |
| Q9H6V9 | 18 | 90.39 | 325 | 90.41 |
| O43399 | 18 | 72.43 | 206 | 68.76 |
| Q9Y2L9 | 18 | 85.42 | 728 | 63.44 |
| Q8IW45 | 20 | 46.54 | 347 | 88.77 |
| **Q9NZF1** | 20 | 95.43 | 115 | 89.73 |
| **Q8N8L6** | 21 | 51.46 | 244 | 77.99 |
| Q8WWH4 | 21 | 38.81 | 475 | 67.88 |
| Q9Y2W6 | 21 | 77.79 | 561 | 74.09 |
| Q69YL0 | 21 | 72.91 | 99 | 67.63 |
| **Q96CS3** | 22 | 74.99 | 445 | 85.13 |
| Q9H0P0 | 22 | 67.09 | 336 | 91.76 |
| **Q9BYD5** | 22 | 87.89 | 112 | 85.06 |
| **Q9H1A3** | 24 | 60.21 | 318 | 82.90 |
| **Q8NBQ5** | 24 | 90.32 | 300 | 93.28 |
| O15033 | 24 | 74.60 | 823 | 83.35 |
| Q8NEX9 | 25 | 80.82 | 313 | 93.31 |
| **A1L4L8** | 25 | 93.83 | 177 | 72.23 |
| **Q7Z5P4** | 25 | 92.31 | 300 | 93.11 |
| **Q9NV66** | 26 | 80.29 | 732 | 80.73 |
| Q6NXR0 | 27 | 56.56 | 463 | 69.33 |
| A0A494C176 | 27 | 78.29 | 62 | 72.40 |
| **Q8WZA9** | 28 | 49.71 | 623 | 66.76 |
| Q16143 | 34 | 66.60 | 134 | 60.56 |
| **Q5VUY0** | 34 | 92.79 | 407 | 94.82 |
| **Q9H3Z7** | 38 | 59.96 | 469 | 85.83 |
| **A1A4F0** | 45 | 71.71 | 135 | 66.24 |
| **Q3SXM5** | 45 | 87.13 | 330 | 92.03 |

Continued on next page

| UniProt ID | Amount of high-MCP residues | Mean pLDDT (MCP ≥ 0.5) | Length | Mean pLDDT |
|---|---|---|---|---|
| **Q9H4I3** | 47 | 85.50 | 376 | 80.35 |
| Q00G26 | 48 | 66.33 | 463 | 67.99 |
| Q6ZTK2 | 58 | 93.83 | 550 | 87.66 |
| O60240 | 78 | 64.17 | 522 | 53.46 |

#### 2 Supplementary Figures

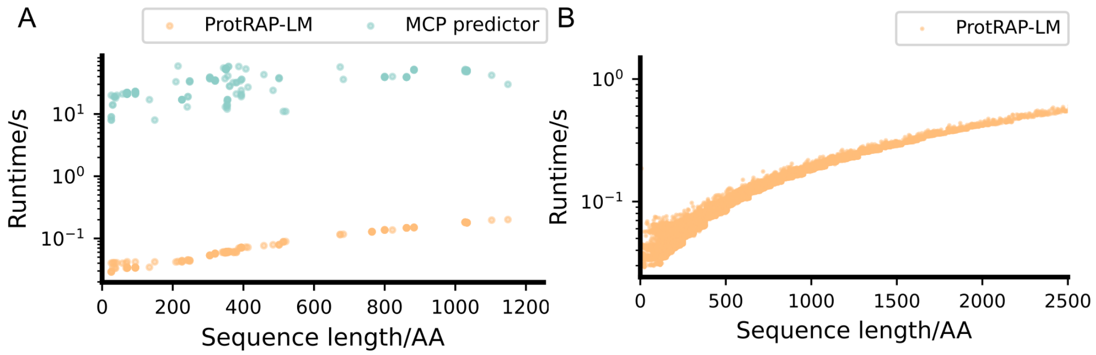


**Figure S1: Runtime analysis.** (A) Runtime analysis of the ProtRAP-LM and the MSA-based MCP predictor using the MemProtMD_2022 dataset (N=184). The x-axis presents the sequence length calculated in amino acids (AA). A logarithmic scale was used in y-axis to present the time usage in seconds (s). Obviously, the ProtRAP-LM is two orders of magnitude faster than the MSA-based MCP predictor. The average time of ProtRAP-LM and the MSA-based MCP predictor was 0.10 s and 32.75 s per sequence, respectively. (B) Runtime in seconds (s) for analyzing the human proteome (N=20607) using the ProtRAP-LM model. The runtime was 36.4 minutes for the whole human proteome.

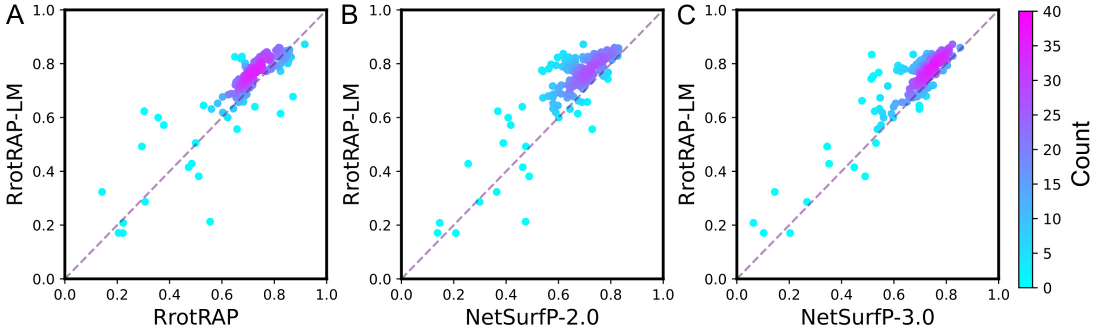


**Figure S2: Comparison of the relative accessible surface area (RASA) prediction performance of PCCs among different methods (on the protein level).** The comparative performance of (A) ProtRAP-LM and ProtRAP, (B) ProtRAP-LM and NetSurfP-2.0, and (C) ProtRAP-LM and NetSurfP-3.0 was illustrated. The overall prediction PCCs of ProtRAP-LM, ProtRAP, NetSurfP-2.0, and NetSurfP-3.0 were 0.729, 0.694, 0.661, and 0.674, respectively. The y-axis depicts the PCC values of the ProtRAP-LM, while the x-axis represents the PCC values of other predictors on the MemProtMD_2022 dataset. The dashed line represents the function $y = x$. Each point represents a test protein, colored by the density of local points.

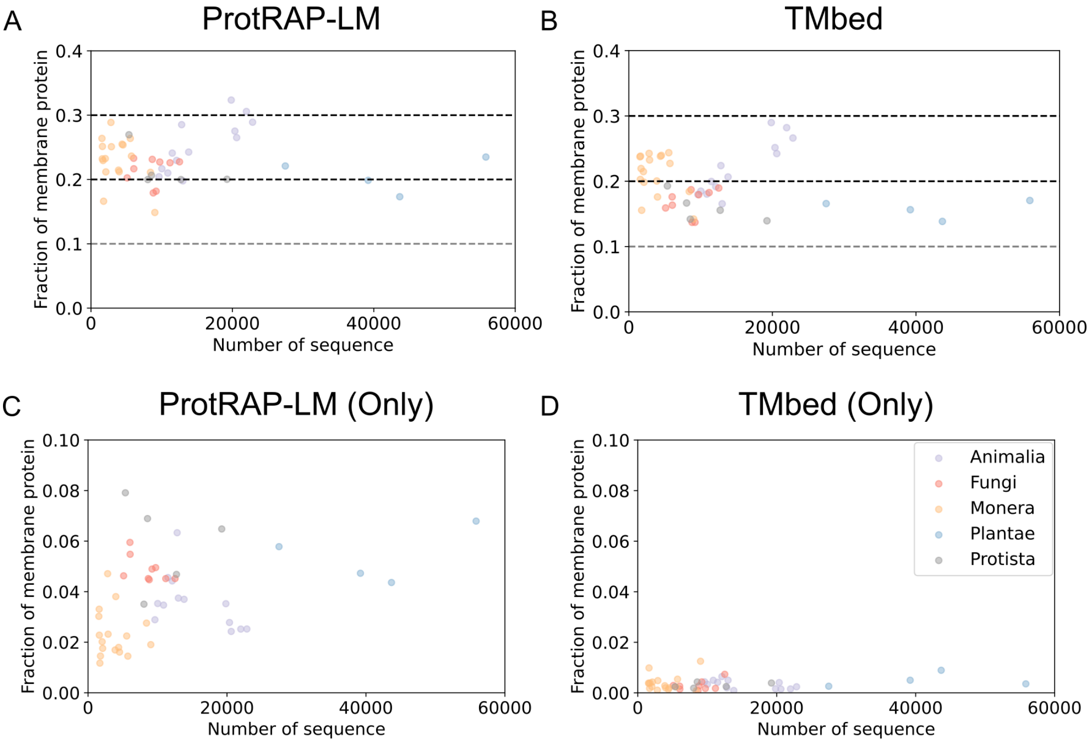


**Figure S3: The predicted membrane protein fractions across the five different biological kingdoms.** The predicted membrane protein fractions of (A) ProtRAP-LM-identified, (B) TMbed-identified, (C) 'ProtRAP-LM (Only)', and (D) 'TMbed (Only)' were shown. The x-axis depicts the number of protein-coding genes within each proteome, while the y-axis represents the fraction of membrane proteins present in each proteome.

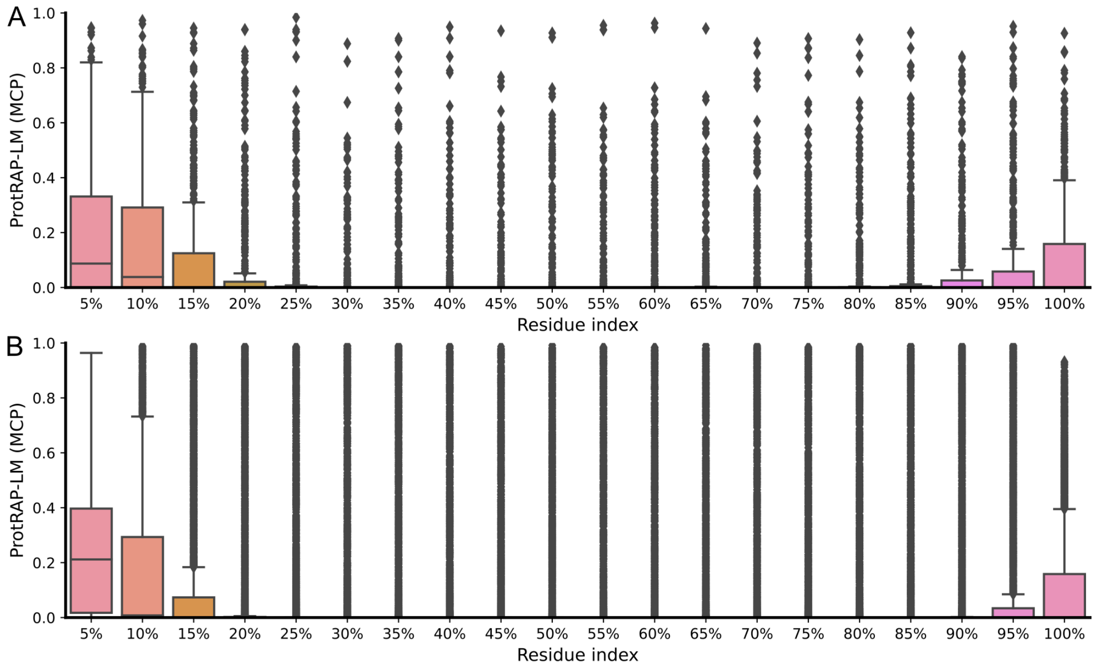


**Figure S4: The distribution of MCP predictions across the analyzed sequences.** The distribution of MCP predictions in the sequences of (A) the 'ProtRAP-LM (Only)' proteins within human proteome and (B) single-pass transmembrane proteins from six evolutionarily distant organisms in the Membranome database[1] was shown. In this analysis, we first divided each protein sequence into 20 equal parts and calculate the average value of MCP predictions in each part. Then, we visualized the average MCP distribution of protein sequences across different datasets here.

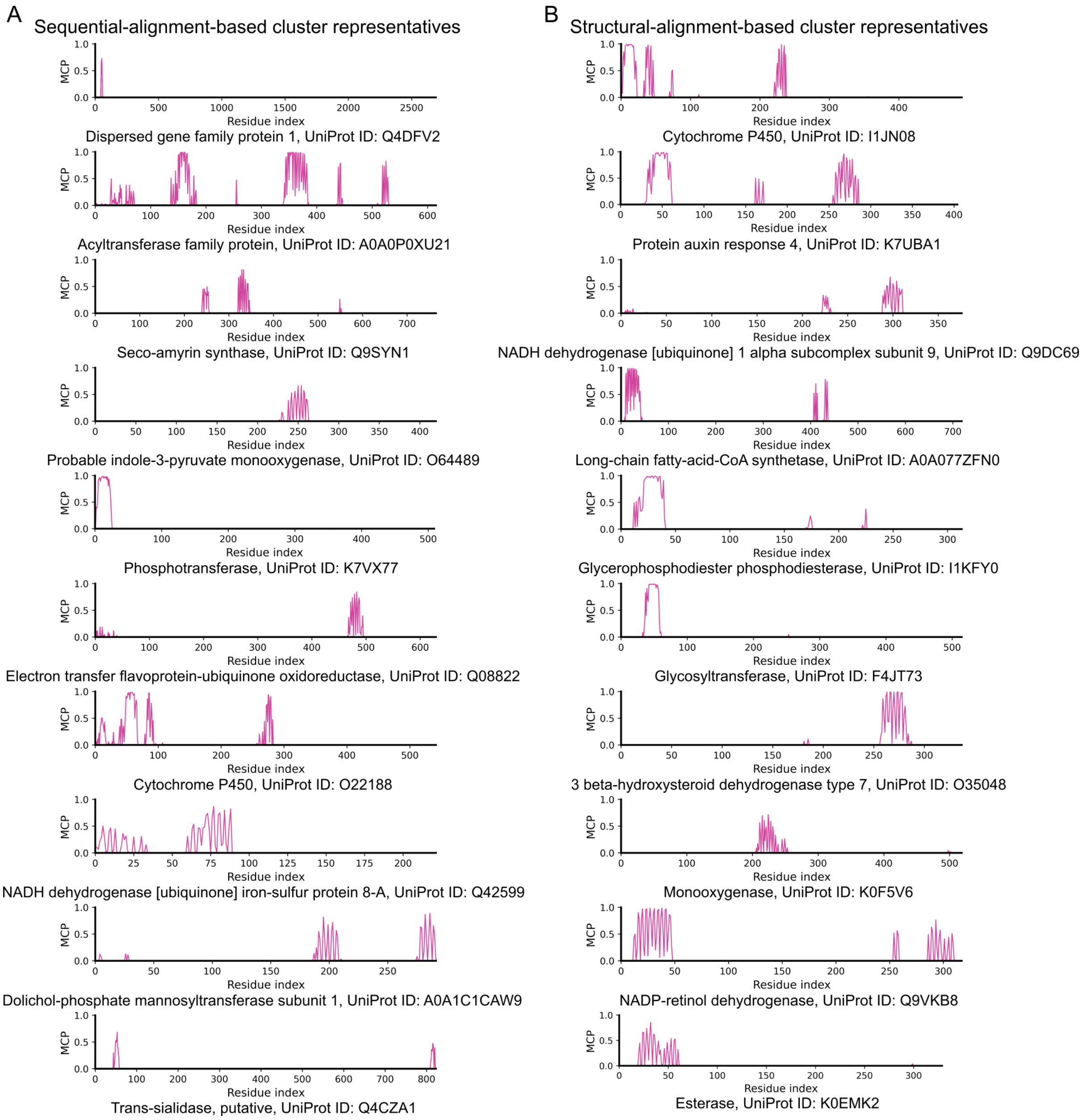


**Figure S5: The predicted MCP of the cluster representatives within the 'ProtRAP-LM (Only)' proteins by ProtRAP-LM.** The predicted MCP of the top 10 representatives clustered by (A) sequential alignment and (B) structural alignment within the 'ProtRAP-LM (Only)' proteins.

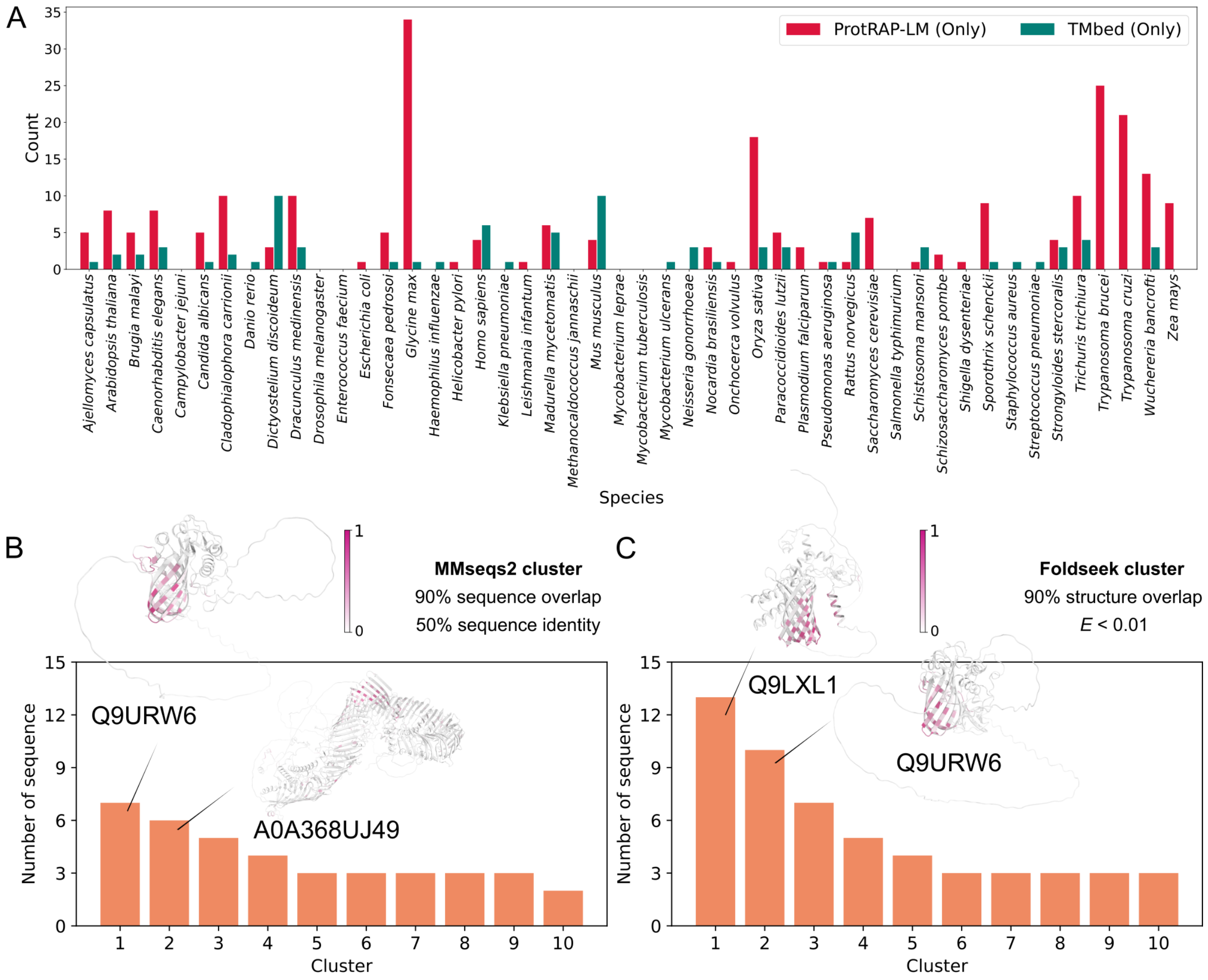


**Figure S6: Analysis of the likely β-sheet-containing membrane proteins of the 'ProtRAP-LM (Only)' and 'TMbed (Only)' parts within 48 proteomes.** (A) The number of the likely β-sheet-containing membrane proteins identified within different proteomes. (B) The top 10 sequential-alignment-based cluster representatives within the 'ProtRAP-LM (Only)' β-sheet-containing membrane proteins. (C) The top 10 structural-alignment-based cluster representatives within the 'ProtRAP-LM (Only)' β-sheet-containing membrane proteins. The structures in the (B) and (C) predicted by AlphaFold2 were colored according to the predicted MCP by ProtRAP-LM, represented as cartoon, and the top two cluster representative structures are shown.

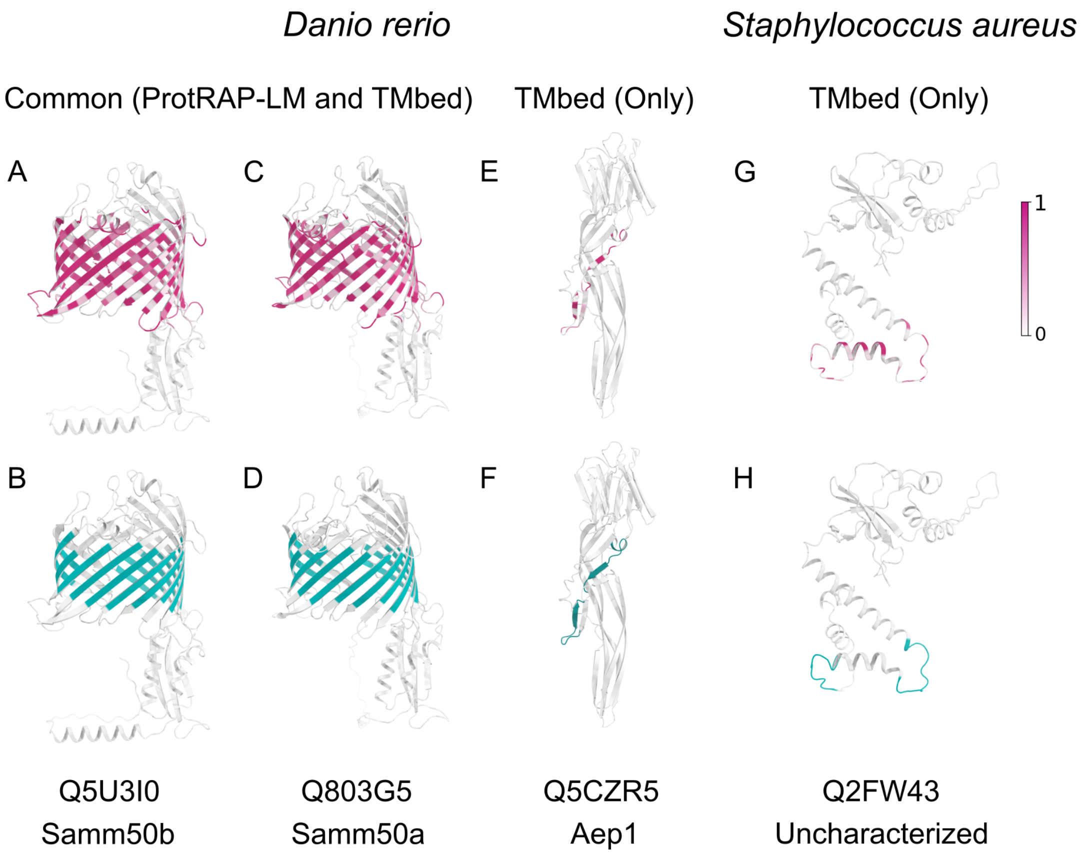


**Figure S7: The transmembrane-strand-containing proteins identified by TMbed within the proteomes of *Danio rerio* and *Staphylococcus aureus*.** The structures predicted by AlphaFold2 were colored according to the predicted MCP values by ProtRAP-LM (upper) and the positions of transmembrane strands predicted by TMbed (lower). (A-D) The only two 'Common (ProtRAP-LM and TMbed)' proteins of *Danio rerio* are the β-sheet-containing proteins recognized by both ProtRAP-LM and TMbed simultaneously. (E-F) The only one 'TMbed (Only)' protein in the *Danio rerio* proteome. However, both ProtRAP-LM and TMbed can recognize similar regions, but only TMbed classified these regions as strands, while ProtRAP-LM, based on AlphaFold2 secondary structure predictions, did not recognize these regions as strands. (G-H) Similar to (E-F), but for the only one 'TMbed (Only)' protein from *Staphylococcus aureus*, a Gram-positive bacterium that does not contain any β-barrel transmembrane protein or 'Common (ProtRAP-LM and TMbed)' protein.

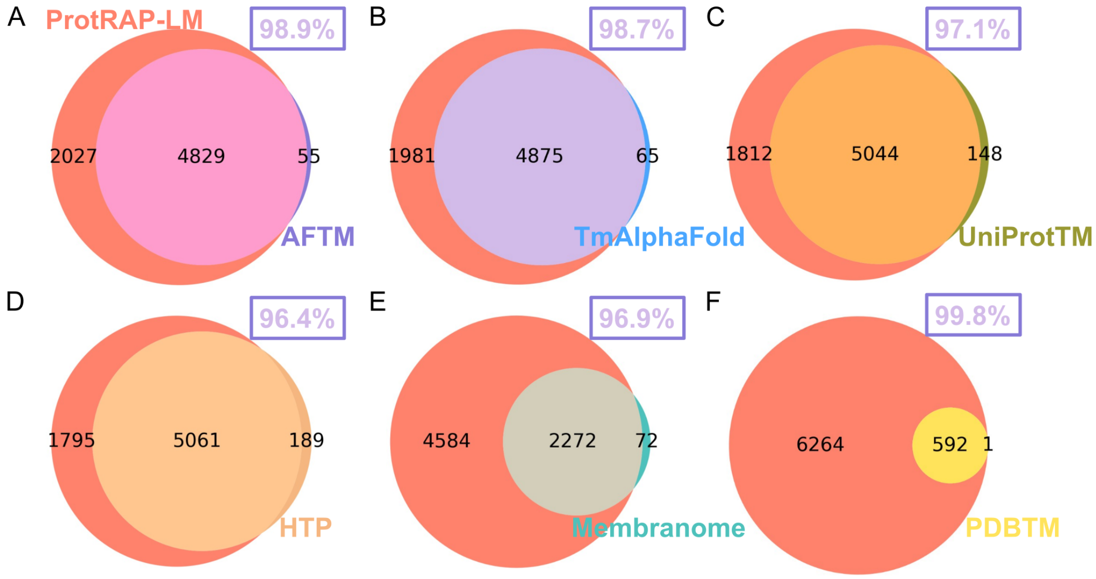


**Figure S8: Comparison of the potentially membrane-contacting proteins predicted by ProtRAP-LM with the public human transmembrane databases.** The public human transmembrane databases are (A) AFTM,[2] (B) TmAlphafold,[3] (C) UniProtTM,[4] (D) HTP,[5] (E) Membranome,[1] and (F) PDBTM.[6] The protein lists of AFTM, TmAlphafold, UniProtTM, HTP, Membranome, and PDBTM were provided by the online database of AFTM (http://conglab.swmed.edu/AFTM). The protein list of UniProtTM from the AFTM database was the reviewed UniProt entries with annotated transmembrane protein spans released on 23 February 2022, which is why the protein list of transmembrane proteins slightly differently in Fig. 5A than in here. The data in the upper right corner of each panel indicates the proportion of the common entries out of all entries within each public human transmembrane database, showing that the ProtRAP-LM-identified membrane proteins cover most of the entries in the databases.

#### References


[1] Lomize, A. L.; Schnitzer, K. A.; Todd, S. C.; Cherepanov, S.; Outeiral, C.; Deane, C. M.; Pogozheva, I. D. Membranome 3.0: Database of single-pass membrane proteins with AlphaFold models. *Protein Sci.* **2022**, *31*, e4318.

[2] Pei, J.; Cong, Q. AFTM: A database of transmembrane regions in the human proteome predicted by AlphaFold. *Database* **2023**, *2023*, baad008.

[3] Dobson, L.; Szekeres, L. I.; Gerdán, C.; Langó, T.; Zeke, A.; Tusnády, G. E. TmAlphaFold database: Membrane localization and evaluation of AlphaFold2 predicted alpha-helical transmembrane protein structures. *Nucleic Acids Res.* **2023**, *51*, D517–D522.

[4] Consortium, T. U. UniProt: The universal protein knowledgebase in 2021. *Nucleic Acids Res.* **2021**, *49*, D480–D489.

[5] Dobson, L.; Reményi, I.; Tusnády, G. E. The human transmembrane proteome. *Biol. Direct* **2015**, *10*, 1–18.

[6] Kozma, D.; Simon, I.; Tusnady, G. E. PDBTM: Protein Data Bank of transmembrane proteins after 8 years. *Nucleic Acids Res.* **2012**, *41*, D524–D529.
