## Supplementary material for "ProtRAP-LM: Fast and accurate protein relative accessibility prediction and membrane protein screening through protein language model embeddings": File S1

### The sequences in the section of “Screening unknown membrane proteins within human proteome”

#### 1. The sequences shown in Fig. 5A

##### 1.1 The potentially membrane-contacting proteins identified by ProtRAP-LM (6856 sequences)

A0A075B6K0 A0A075B6K4 A0A075B6K5 A0A075B6L2 A0A075B6N3 A0A075B6R0 A0A075B6R2
A0A075B6T8 A0A075B6U4 A0A075B6U6 A0A075B6X5 A0A075B734 A0A075B7B6 A0A087WSY4
A0A087WT02 A0A087WT03 A0A087WTH1 A0A087WTH5 A0A087WU88 A0A087WW49 A0A087X1C5
A0A087X1L8 A0A096LNH4 A0A096LNP1 A0A096LP01 A0A096LPK9 A0A0A0MS00 A0A0A0MS02
A0A0A6YYC5 A0A0A6YYJ7 A0A0A6YYK1 A0A0A6YYK6 A0A0A6YYK7 A0A0B4J1T7 A0A0B4J1U4
A0A0B4J1U7 A0A0B4J237 A0A0B4J238 A0A0B4J240 A0A0B4J244 A0A0B4J248 A0A0B4J262
A0A0B4J263 A0A0B4J264 A0A0B4J274 A0A0B4J277 A0A0B4J280 A0A0B4J2F0 A0A0C4DH27
A0A0C4DH28 A0A0C4DH34 A0A0C4DH41 A0A0D9SF12 A0A0G2JJF7 A0A0G2JKD1 A0A0G2JLG4
A0A0G2JLY3 A0A0G2JNF4 A0A0G2JNH3 A0A0J9YWU9 A0A0J9YXV3 A0A0K0K1B3 A0A0K2S4Q6
A0A0U1RQS6 A0A0U1RRA0 A0A0U1RRN3 A0A0X1KG70 A0A126GWB0 A0A126GWI2 A0A140T8X8
A0A191URJ7 A0A1B0GTB2 A0A1B0GTC6 A0A1B0GTE1 A0A1B0GTG8 A0A1B0GTI8 A0A1B0GTL2
A0A1B0GTQ1 A0A1B0GTQ4 A0A1B0GTR0 A0A1B0GTU2 A0A1B0GTW7 A0A1B0GTY4 A0A1B0GU29
A0A1B0GUA5 A0A1B0GUA7 A0A1B0GUW6 A0A1B0GUW7 A0A1B0GUY1 A0A1B0GUZ9 A0A1B0GV85
A0A1B0GV90 A0A1B0GVD1 A0A1B0GVG4 A0A1B0GVH4 A0A1B0GVN3 A0A1B0GVQ0 A0A1B0GVS7
A0A1B0GVT2 A0A1B0GVV1 A0A1B0GVX0 A0A1B0GVY4 A0A1B0GVZ9 A0A1B0GW54 A0A1B0GW64
A0A1B0GWB2 A0A1B0GWG4 A0A1B0GWH6 A0A1B0GX56 A0A1W2PN81 A0A1W2PP97 A0A1W2PQM1
A0A1W2PQU2 A0A1W2PS18 A0A286YEU6 A0A286YF18 A0A286YF58 A0A286YFK9 A0A2R8Y4L6
A0A2R8Y4M2 A0A2R8Y4M4 A0A2R8Y550 A0A2R8Y7Y5 A0A2R8YCJ5 A0A2R8YE69 A0A2R8YED5
A0A2R8YEG4 A0A2R8YEH3 A0A2R8YEV3 A0A2U3TZM8 A0A3B3IT45 A0A494BZU4 A0A494C0I6
A0A494C103 A0A494C176 A0A494C1I1 A0A4W9AIG4 A0A5B9 A0A5F9ZH02 A0A5K1VDZ0
A0A6I8PU40 A0A7I2V2S6 A0A7I2V3R4 A0A8I5KY86 A0A8Q3SIG1 A0A8Q3SIZ7 A0A8Q3WLD3
A0AV02 A0AVI2 A0AVI4 A0FGR8 A0FGR9 A0JD32 A0JD37 A0PJK1 A0PJW6 A0PJX4 A0PJX8
A0PJZ3 A0PK00 A0PK05 A0PK11 A0ZSE6 A1A4F0 A1A5B4 A1A5C7 A1E959 A1L0T0 A1L157
A1L1A6 A1L3X0 A1L453 A1L4H1 A1L4L8 A2A2V5 A2A2Y4 A2RRL7 A2RU14 A2RU48 A2RU67
A2RUG3 A2RUT3 A2RUU4 A2VDJ0 A3KFT3 A3KN74 A4D0S4 A4D0T7 A4D0V7 A4D1S0 A4D1T9
A4D256 A4D2G3 A4D2H0 A4FU28 A4IF30 A5D6W6 A5D8T8 A5PLL7 A5X5Y0 A6BM72 A6H8M9
A6NC51 A6NCI5 A6NCL2 A6NCQ9 A6NCV1 A6ND01 A6ND48 A6NDA9 A6NDD5 A6NDH6 A6NDL8
A6NDP7 A6NDV4 A6NEH6 A6NET4 A6NF34 A6NF89 A6NFA1 A6NFC5 A6NFC9 A6NFE2 A6NFR6
A6NFU0 A6NFX1 A6NFY4 A6NFZ4 A6NG13 A6NGA9 A6NGB0 A6NGB7 A6NGC4 A6NGN9 A6NGU5
A6NGY5 A6NGZ8 A6NH00 A6NH11 A6NH21 A6NH52 A6NHA9 A6NHG9 A6NHM9 A6NHN0 A6NHN6
A6NHS7 A6NI61 A6NI73 A6NIE9 A6NIJ9 A6NIM6 A6NIN4 A6NJU9 A6NJW4 A6NJW9 A6NJY1
A6NJY4 A6NJZ3 A6NK97 A6NKB5 A6NKF7 A6NKK0 A6NKL6 A6NKP2 A6NKQ9 A6NKW6 A6NKX4
A6NL05 A6NL08 A6NL26 A6NL88 A6NL99 A6NLU5 A6NLX4 A6NM03 A6NM10 A6NM11 A6NM45
A6NM62 A6NM76 A6NMB1 A6NMD0 A6NML5 A6NMS3 A6NMS7 A6NMU1 A6NMZ5 A6NN92 A6NNB3
A6NNC1 A6NND4 A6NNE9 A6NNL5 A6NNN8 A6NNS2 A6PVL3 A6PVY3 A7MBM2 A8CG34 A8K4G0

A8K7I4 A8MPY1 A8MRT5 A8MTI9 A8MTT3 A8MTW9 A8MUP6 A8MV23 A8MV81 A8MVS5 A8MVW0
A8MVW5 A8MVZ5 A8MWK0 A8MWL6 A8MWL7 A8MWV9 A8MWY0 A8MXE2 A8MXK1 A8MXU0 A8MXV6
A8MYB1 A8MYU2 A8MZ97 A8MZH6 A9Z1Z3 B0FP48 B0YJ81 B1AKI9 B2RN74 B2RNN3 B2RUY7
B2RUZ4 B2RXF0 B3GLJ2 B3SHH9 B4DJY2 B4DS77 B4DYI2 B6A8C7 B6SEH8 B6SEH9 B7U540
B7Z8K6 B7ZAQ6 B8ZZ34 B9EJG8 C9JDP6 C9JG80 C9JH25 C9JI98 C9JL84 C9JQL5 C9JUS6
C9JVW0 C9JXX5 D3DTV9 D3W0D1 E0CX11 E2RYF6 E2RYF7 E5RHQ5 E5RIL1 E7ERA6 E9PQ53
E9PQX1 F2Z333 F5H4A9 F8W0I5 F8WCM5 G3V0H7 H0YL14 H3BQJ8 H3BR10 H3BS89 H3BTG2
H3BV60 H7C241 H7C350 I3L273 I3L3R5 K7EJ46 K9M1U5 M0QZC1 M5A8F1 O00115 O00124
O00144 O00155 O00168 O00180 O00187 O00198 O00206 O00217 O00219 O00220 O00222
O00230 O00237 O00238 O00241 O00253 O00254 O00258 O00264 O00270 O00292 O00322
O00337 O00339 O00341 O00391 O00398 O00400 O00421 O00445 O00451 O00453 O00461
O00462 O00468 O00469 O00476 O00478 O00481 O00483 O00501 O00519 O00526 O00533
O00548 O00555 O00559 O00574 O00584 O00587 O00590 O00591 O00592 O00602 O00622
O00623 O00624 O00631 O00744 O00748 O00754 O00767 O14493 O14494 O14495 O14498
O14511 O14514 O14520 O14521 O14522 O14523 O14524 O14525 O14548 O14569 O14581
O14609 O14626 O14638 O14649 O14653 O14656 O14657 O14662 O14668 O14669 O14672
O14678 O14681 O14683 O14684 O14718 O14735 O14756 O14763 O14764 O14773 O14786
O14788 O14791 O14792 O14793 O14798 O14804 O14817 O14828 O14836 O14842 O14843
O14863 O14880 O14894 O14904 O14905 O14917 O14925 O14931 O14944 O14949 O14957
O14958 O14960 O14967 O14975 O14983 O15031 O15033 O15072 O15079 O15118 O15120
O15121 O15123 O15126 O15127 O15130 O15146 O15155 O15165 O15173 O15197 O15218
O15229 O15232 O15239 O15240 O15243 O15244 O15245 O15258 O15260 O15263 O15269
O15270 O15303 O15321 O15335 O15342 O15354 O15374 O15375 O15389 O15393 O15394
O15399 O15400 O15403 O15427 O15431 O15432 O15438 O15439 O15440 O15442 O15455
O15460 O15466 O15482 O15503 O15529 O15533 O15537 O15547 O15551 O15552 O15554
O43155 O43157 O43169 O43173 O43174 O43184 O43193 O43194 O43240 O43246 O43272
O43278 O43280 O43286 O43291 O43292 O43300 O43306 O43315 O43323 O43399 O43405
O43424 O43451 O43462 O43464 O43490 O43493 O43497 O43505 O43506 O43508 O43511
O43520 O43525 O43526 O43529 O43555 O43556 O43557 O43561 O43567 O43570 O43581
O43597 O43603 O43609 O43610 O43612 O43613 O43614 O43615 O43653 O43657 O43674
O43676 O43677 O43688 O43692 O43699 O43731 O43736 O43749 O43752 O43759 O43760
O43761 O43772 O43808 O43819 O43820 O43825 O43826 O43827 O43852 O43854 O43861
O43866 O43868 O43869 O43889 O43895 O43908 O43909 O43914 O43916 O43921 O43927
O43934 O60235 O60238 O60240 O60241 O60242 O60243 O60245 O60259 O60266 O60279
O60309 O60312 O60313 O60320 O60330 O60337 O60353 O60359 O60383 O60391 O60397
O60403 O60404 O60412 O60423 O60427 O60431 O60449 O60462 O60469 O60476 O60478
O60486 O60487 O60488 O60499 O60500 O60503 O60507 O60512 O60513 O60565 O60568
O60575 O60602 O60603 O60609 O60613 O60635 O60636 O60637 O60656 O60664 O60667
O60669 O60676 O60683 O60687 O60704 O60706 O60721 O60725 O60741 O60755 O60762
O60774 O60779 O60830 O60831 O60840 O60844 O60858 O60883 O60888 O60894 O60895
O60896 O60906 O60909 O60911 O60928 O60931 O60938 O60939 O75015 O75019 O75022
O75023 O75027 O75051 O75054 O75056 O75063 O75069 O75071 O75072 O75074 O75077
O75078 O75084 O75093 O75094 O75095 O75096 O75106 O75110 O75121 O75129 O75144
O75185 O75192 O75197 O75200 O75204 O75264 O75298 O75309 O75310 O75311 O75324

O75325 O75326 O75339 O75352 O75354 O75355 O75356 O75379 O75381 O75387 O75388
O75396 O75425 O75427 O75438 O75452 O75460 O75462 O75473 O75477 O75487 O75493
O75503 O75508 O75509 O75556 O75578 O75581 O75594 O75596 O75610 O75629 O75631
O75636 O75711 O75712 O75715 O75718 O75746 O75751 O75752 O75762 O75783 O75787
O75795 O75829 O75841 O75844 O75845 O75871 O75880 O75881 O75882 O75888 O75899
O75900 O75907 O75908 O75911 O75915 O75920 O75923 O75951 O75954 O75955 O75964
O75973 O75976 O76000 O76001 O76002 O76024 O76036 O76061 O76062 O76070 O76076
O76082 O76090 O76095 O76099 O76100 O94759 O94766 O94769 O94772 O94777 O94778
O94779 O94823 O94826 O94856 O94876 O94886 O94898 O94901 O94905 O94910 O94911
O94919 O94923 O94933 O94956 O94966 O94985 O94991 O95006 O95007 O95013 O95047
O95069 O95070 O95084 O95136 O95139 O95140 O95150 O95156 O95157 O95158 O95159
O95167 O95168 O95169 O95178 O95180 O95183 O95185 O95196 O95197 O95202 O95206
O95210 O95214 O95221 O95222 O95236 O95237 O95249 O95255 O95256 O95258 O95259
O95264 O95274 O95279 O95292 O95297 O95298 O95302 O95342 O95371 O95377 O95389
O95390 O95393 O95395 O95399 O95406 O95407 O95415 O95424 O95427 O95428 O95436
O95450 O95452 O95460 O95461 O95470 O95471 O95473 O95476 O95477 O95479 O95484
O95490 O95497 O95498 O95500 O95502 O95528 O95562 O95563 O95573 O95622 O95631
O95633 O95665 O95672 O95674 O95711 O95715 O95727 O95750 O95754 O95772 O95800
O95803 O95807 O95813 O95831 O95832 O95838 O95841 O95847 O95857 O95858 O95859
O95864 O95866 O95867 O95868 O95870 O95873 O95881 O95897 O95907 O95918 O95925
O95944 O95965 O95967 O95968 O95969 O95971 O95976 O95977 O95980 O95992 O95994
O96000 O96002 O96005 O96008 O96009 O96011 O96014 O96024 P00156 P00167 P00387
P00395 P00403 P00414 P00450 P00533 P00709 P00734 P00736 P00738 P00746 P00747
P00748 P00749 P00750 P00797 P00846 P00995 P01009 P01011 P01019 P01031 P01034
P01036 P01037 P01127 P01130 P01133 P01135 P01137 P01138 P01148 P01160 P01189
P01210 P01213 P01215 P01222 P01225 P01229 P01258 P01275 P01282 P01286 P01298
P01303 P01308 P01344 P01350 P01374 P01375 P01574 P01579 P01588 P01589 P01591
P01715 P01717 P01730 P01732 P01737 P01814 P01824 P01825 P01833 P01848 P01850
P01889 P01893 P01903 P01906 P01909 P01911 P01920 P02452 P02458 P02461 P02462
P02649 P02652 P02654 P02655 P02656 P02671 P02675 P02679 P02708 P02724 P02730
P02741 P02745 P02746 P02747 P02749 P02753 P02765 P02766 P02768 P02774 P02775
P02776 P02778 P02786 P02787 P02788 P02790 P02808 P02810 P02812 P02814 P02818
P03886 P03891 P03897 P03901 P03905 P03915 P03923 P03928 P03950 P03952 P03956
P03971 P03979 P03986 P03999 P04000 P04001 P04004 P04035 P04054 P04062 P04066
P04085 P04090 P04114 P04118 P04141 P04155 P04156 P04180 P04196 P04201 P04216
P04233 P04234 P04275 P04278 P04279 P04280 P04439 P04440 P04626 P04628 P04629
P04746 P04798 P04808 P04839 P04843 P04844 P04920 P04921 P05019 P05023 P05026
P05060 P05067 P05090 P05093 P05106 P05107 P05111 P05112 P05113 P05121 P05141
P05154 P05155 P05156 P05164 P05177 P05181 P05186 P05187 P05305 P05362 P05408
P05452 P05496 P05538 P05543 P05546 P05556 P05814 P05981 P05997 P06028 P06126
P06127 P06133 P06213 P06276 P06280 P06307 P06331 P06340 P06396 P06729 P06731
P06734 P06756 P06850 P06858 P06865 P06870 P07093 P07098 P07099 P07202 P07204
P07225 P07237 P07288 P07306 P07307 P07333 P07339 P07357 P07358 P07359 P07477
P07478 P07492 P07498 P07510 P07550 P07585 P07602 P07686 P07711 P07766 P07858

P07911 P07942 P07949 P07988 P07996 P08034 P08069 P08100 P08118 P08123 P08138
P08172 P08173 P08174 P08183 P08185 P08195 P08217 P08218 P08246 P08247 P08253
P08254 P08294 P08311 P08473 P08476 P08493 P08514 P08519 P08571 P08574 P08575
P08581 P08582 P08588 P08620 P08637 P08648 P08684 P08686 P08700 P08709 P08833
P08842 P08861 P08887 P08908 P08910 P08912 P08913 P08922 P08949 P08962 P08F94
P09093 P09131 P09228 P09237 P09238 P09326 P09341 P09486 P09529 P09544 P09564
P09601 P09603 P09619 P09668 P09669 P09681 P09683 P09693 P09758 P09848 P09871
P09912 P09923 P09958 P0C091 P0C0P6 P0C2L3 P0C2S0 P0C2W7 P0C604 P0C617 P0C623
P0C626 P0C628 P0C629 P0C645 P0C646 P0C672 P0C6S8 P0C6T2 P0C7L1 P0C7M8 P0C7N1
P0C7N4 P0C7N5 P0C7N8 P0C7P4 P0C7Q5 P0C7Q6 P0C7T2 P0C7T3 P0C7T8 P0C7U0 P0C7U3
P0C7V7 P0C7X4 P0C851 P0C862 P0C874 P0C8F1 P0CF51 P0CG01 P0CG08 P0CG36 P0CG37
P0CG41 P0CK96 P0CK97 P0CW18 P0DI80 P0DJ07 P0DJ93 P0DJD7 P0DJD8 P0DJD9 P0DJI8
P0DJI9 P0DKB5 P0DKB6 P0DKV0 P0DKX4 P0DL12 P0DMC3 P0DML3 P0DMQ5 P0DMR2 P0DMS8
P0DMS9 P0DMT0 P0DMU2 P0DN25 P0DN77 P0DN78 P0DN80 P0DN81 P0DN82 P0DN84 P0DN86
P0DN87 P0DP06 P0DP07 P0DP42 P0DP57 P0DP58 P0DP72 P0DP73 P0DP74 P0DPA2 P0DPD6
P0DPD8 P0DPE3 P0DPK3 P0DQD5 P0DSN6 P0DTE0 P0DTE4 P0DTE5 P0DTE7 P0DTE8 P0DTF9
P0DTL5 P0DTU3 P0DTU4 P0DUB6 P10082 P10092 P10124 P10144 P10145 P10153 P10163
P10176 P10253 P10321 P10323 P10415 P10451 P10586 P10600 P10619 P10620 P10632
P10635 P10643 P10645 P10646 P10696 P10720 P10721 P10747 P10767 P10909 P10912
P10915 P10966 P10997 P11021 P11047 P11049 P11117 P11150 P11166 P11168 P11169
P11215 P11226 P11229 P11230 P11279 P11362 P11487 P11509 P11511 P11597 P11678
P11684 P11686 P11712 P11717 P11836 P11912 P12034 P12074 P12109 P12110 P12111
P12235 P12236 P12272 P12273 P12314 P12318 P12319 P12532 P12643 P12645 P12724
P12821 P12830 P12838 P12872 P13073 P13164 P13224 P13232 P13284 P13385 P13473
P13497 P13498 P13521 P13569 P13584 P13591 P13598 P13612 P13637 P13667 P13671
P13674 P13686 P13688 P13726 P13727 P13747 P13762 P13765 P13866 P13942 P13945
P13987 P14060 P14091 P14138 P14151 P14207 P14209 P14222 P14314 P14384 P14406
P14410 P14415 P14416 P14543 P14555 P14616 P14625 P14672 P14679 P14770 P14778
P14780 P14784 P14867 P15085 P15086 P15088 P15144 P15151 P15169 P15248 P15260
P15289 P15291 P15309 P15328 P15382 P15391 P15421 P15502 P15509 P15514 P15515
P15516 P15529 P15586 P15692 P15812 P15813 P15814 P15848 P15907 P15941 P15954
P16035 P16066 P16070 P16109 P16144 P16150 P16233 P16234 P16260 P16278 P16284
P16389 P16410 P16422 P16435 P16442 P16444 P16471 P16473 P16562 P16581 P16615
P16662 P16671 P16860 P16870 P16871 P17050 P17152 P17181 P17213 P17301 P17302
P17342 P17405 P17538 P17540 P17643 P17658 P17693 P17706 P17787 P17813 P17900
P17927 P17936 P17948 P18031 P18065 P18075 P18084 P18089 P18405 P18428 P18433
P18505 P18507 P18509 P18564 P18577 P18627 P18825 P18827 P18850 P19021 P19022
P19075 P19224 P19235 P19256 P19320 P19397 P19438 P19440 P19526 P19634 P19801
P19823 P19827 P19835 P19875 P19876 P19883 P19957 P19961 P20020 P20023 P20036
P20061 P20062 P20138 P20142 P20151 P20155 P20160 P20231 P20273 P20292 P20309
P20333 P20366 P20382 P20396 P20594 P20645 P20648 P20701 P20702 P20774 P20783
P20800 P20809 P20813 P20815 P20827 P20849 P20851 P20853 P20908 P20916 P20933
P20963 P21128 P21145 P21217 P21246 P21397 P21439 P21452 P21453 P21462 P21554
P21579 P21583 P21589 P21709 P21728 P21730 P21731 P21741 P21754 P21757 P21796

P21802 P21815 P21817 P21854 P21860 P21917 P21918 P21926 P21964 P22001 P22004
P22079 P22083 P22223 P22301 P22303 P22304 P22309 P22310 P22352 P22362 P22413
P22455 P22459 P22460 P22466 P22607 P22680 P22692 P22732 P22748 P22749 P22760
P22792 P22794 P22888 P22894 P22897 P23141 P23142 P23219 P23229 P23276 P23280
P23284 P23327 P23352 P23415 P23416 P23435 P23467 P23468 P23469 P23470 P23471
P23510 P23515 P23634 P23763 P23942 P23945 P23946 P23975 P24001 P24043 P24046
P24071 P24158 P24310 P24311 P24347 P24387 P24390 P24394 P24462 P24530 P24539
P24557 P24592 P24593 P24855 P24903 P25021 P25024 P25025 P25063 P25067 P25089
P25090 P25092 P25100 P25101 P25103 P25105 P25106 P25116 P25189 P25311 P25391
P25445 P25774 P25874 P25929 P25940 P25942 P26006 P26010 P26012 P26022 P26436
P26439 P26572 P26678 P26715 P26717 P26718 P26842 P26885 P26927 P26951 P26992
P27037 P27105 P27169 P27338 P27449 P27469 P27487 P27539 P27544 P27658 P27701
P27797 P27824 P27918 P27930 P28039 P28067 P28068 P28221 P28222 P28223 P28288
P28300 P28325 P28328 P28335 P28336 P28472 P28476 P28566 P28827 P28845 P28906
P28907 P28908 P29016 P29017 P29033 P29120 P29122 P29274 P29275 P29279 P29317
P29320 P29322 P29323 P29371 P29376 P29400 P29965 P29972 P29973 P30040 P30101
P30203 P30273 P30301 P30408 P30411 P30511 P30518 P30519 P30530 P30531 P30532
P30533 P30536 P30542 P30550 P30556 P30559 P30825 P30872 P30874 P30926 P30939
P30953 P30954 P30968 P30988 P30989 P30990 P31213 P31358 P31391 P31415 P31431
P31512 P31513 P31639 P31641 P31644 P31645 P31785 P31994 P31995 P31997 P32004
P32189 P32238 P32239 P32241 P32245 P32246 P32247 P32248 P32249 P32297 P32302
P32418 P32745 P32856 P32926 P32927 P32942 P32970 P32971 P33032 P33121 P33151
P33260 P33261 P33527 P33681 P33897 P33908 P33947 P34059 P34130 P34741 P34810
P34820 P34903 P34910 P34925 P34969 P34972 P34981 P34982 P34995 P34998 P35052
P35070 P35212 P35232 P35247 P35318 P35346 P35348 P35354 P35367 P35368 P35372
P35408 P35410 P35414 P35442 P35443 P35462 P35475 P35498 P35499 P35503 P35504
P35523 P35542 P35575 P35590 P35610 P35613 P35625 P35670 P35858 P35916 P35968
P36021 P36222 P36268 P36269 P36382 P36383 P36537 P36544 P36551 P36888 P36894
P36896 P36897 P36941 P36955 P36956 P37023 P37058 P37059 P37088 P37173 P37287
P37288 P37840 P38435 P38484 P38567 P38570 P38571 P39060 P39086 P39210 P39656
P39877 P39900 P40126 P40145 P40189 P40197 P40198 P40199 P40200 P40238 P40259
P40305 P40313 P40879 P40967 P41143 P41145 P41146 P41180 P41181 P41217 P41221
P41231 P41271 P41273 P41439 P41440 P41586 P41587 P41594 P41595 P41597 P41732
P41968 P42081 P42127 P42261 P42262 P42263 P42658 P42701 P42702 P42785 P42857
P42892 P43003 P43004 P43005 P43007 P43026 P43088 P43115 P43116 P43119 P43121
P43146 P43220 P43234 P43235 P43251 P43304 P43307 P43308 P43489 P43626 P43627
P43628 P43629 P43630 P43631 P43632 P43652 P43657 P43681 P45452 P45844 P45877
P45880 P46059 P46089 P46091 P46092 P46093 P46094 P46095 P46098 P46531 P46663
P46695 P46721 P46977 P47211 P47710 P47775 P47804 P47869 P47870 P47871 P47872
P47881 P47883 P47884 P47887 P47888 P47890 P47893 P47898 P47900 P47901 P47972
P47985 P47992 P48023 P48029 P48039 P48048 P48050 P48051 P48052 P48058 P48060
P48065 P48066 P48067 P48145 P48146 P48165 P48167 P48169 P48201 P48230 P48304
P48307 P48357 P48449 P48509 P48544 P48546 P48547 P48549 P48551 P48645 P48651
P48664 P48723 P48745 P48751 P48764 P48960 P48995 P49019 P49069 P49146 P49184

P49190 P49223 P49238 P49257 P49279 P49281 P49286 P49326 P49447 P49641 P49682
P49683 P49685 P49746 P49747 P49755 P49765 P49767 P49768 P49771 P49788 P49810
P49863 P49895 P49908 P49913 P49961 P50052 P50281 P50391 P50402 P50406 P50416
P50443 P50454 P50591 P50876 P50895 P50897 P50993 P51124 P51164 P51168 P51170
P51172 P51460 P51511 P51512 P51571 P51572 P51575 P51582 P51589 P51636 P51648
P51654 P51674 P51677 P51679 P51681 P51684 P51685 P51686 P51687 P51688 P51689
P51690 P51693 P51787 P51788 P51790 P51793 P51795 P51797 P51798 P51800 P51801
P51805 P51809 P51810 P51811 P51828 P51841 P51861 P51884 P51888 P51993 P52429
P52569 P52797 P52798 P52799 P52803 P52823 P52848 P52849 P52961 P53007 P53634
P53701 P53708 P53794 P53801 P53816 P53985 P54107 P54108 P54219 P54289 P54315
P54317 P54707 P54709 P54710 P54753 P54756 P54760 P54762 P54764 P54793 P54802
P54803 P54826 P54829 P54849 P54851 P54852 P54855 P55000 P55001 P55011 P55017
P55056 P55058 P55061 P55064 P55073 P55075 P55082 P55083 P55085 P55087 P55089
P55103 P55107 P55145 P55157 P55259 P55268 P55283 P55285 P55286 P55287 P55289
P55290 P55291 P55327 P55344 P55774 P55808 P55851 P55899 P55916 P56134 P56159
P56180 P56199 P56202 P56373 P56378 P56385 P56539 P56557 P56589 P56696 P56703
P56704 P56730 P56746 P56747 P56748 P56749 P56750 P56817 P56851 P56856 P56880
P56937 P56962 P56975 P57054 P57057 P57087 P57088 P57103 P57105 P57679 P57727
P57738 P57739 P57773 P57789 P58062 P58166 P58170 P58173 P58180 P58181 P58182
P58215 P58294 P58335 P58417 P58418 P58499 P58511 P58549 P58550 P58658 P58743
P58872 P59025 P59533 P59534 P59535 P59536 P59537 P59538 P59539 P59540 P59541
P59542 P59543 P59544 P59551 P59646 P59665 P59666 P59773 P59796 P59826 P59827
P59861 P59901 P59922 P60022 P60033 P60059 P60201 P60468 P60507 P60508 P60509
P60568 P60602 P60606 P60608 P60827 P60852 P60893 P60985 P61009 P61073 P61165
P61266 P61278 P61366 P61550 P61565 P61619 P61626 P61647 P61769 P61803 P61812
P61916 P62079 P62341 P62952 P62955 P63027 P63135 P63252 P67812 P69849 P78310
P78324 P78325 P78329 P78333 P78334 P78348 P78357 P78363 P78369 P78380 P78381
P78382 P78383 P78410 P78423 P78504 P78508 P78536 P78539 P78552 P78562 P79483
P80303 P80365 P80370 P80748 P81172 P81277 P81408 P81534 P81605 P82251 P82279
P83105 P83110 P83111 P83859 P84157 P98066 P98073 P98095 P98153 P98155 P98161
P98164 P98172 P98173 P98187 P98194 P98196 P98198 Q00325 Q00604 Q00765 Q00973
Q00975 Q00G26 Q00LT1 Q01113 Q01118 Q01151 Q01344 Q01362 Q01453 Q01459 Q01523
Q01524 Q01628 Q01629 Q01638 Q01650 Q01668 Q01718 Q01726 Q01740 Q01814 Q01955
Q01959 Q01973 Q01974 Q02083 Q02094 Q02127 Q02161 Q02221 Q02223 Q02246 Q02297
Q02318 Q02325 Q02338 Q02383 Q02388 Q02413 Q02487 Q02505 Q02643 Q02742 Q02747
Q02763 Q02809 Q02818 Q02846 Q02928 Q02978 Q03135 Q03167 Q03395 Q03403 Q03405
Q03431 Q03518 Q03519 Q03692 Q03721 Q04118 Q04609 Q04656 Q04671 Q04721 Q04756
Q04771 Q04844 Q04900 Q04912 Q04941 Q05586 Q05901 Q05940 Q05996 Q06033 Q06055
Q06136 Q06141 Q06418 Q06432 Q06481 Q06495 Q06643 Q06828 Q07001 Q07011 Q07065
Q07075 Q07108 Q07326 Q07444 Q07507 Q07654 Q07699 Q07812 Q07817 Q07820 Q07837
Q07954 Q07973 Q08174 Q08334 Q08345 Q08357 Q08380 Q08397 Q08431 Q08462 Q08477
Q08554 Q08648 Q08708 Q08722 Q08828 Q08830 Q08AI6 Q08ET2 Q09327 Q09328 Q09428
Q09470 Q0D2K0 Q0GE19 Q0P5P2 Q0P670 Q0P6D2 Q0P6H9 Q0VAF6 Q0VAQ4 Q0VDE8 Q0VDI3
Q10469 Q10471 Q10472 Q10588 Q10589 Q10981 Q11128 Q11130 Q11201 Q11203 Q11206

Q12767 Q12770 Q12772 Q12791 Q12794 Q12797 Q12805 Q12809 Q12836 Q12841 Q12846
Q12860 Q12864 Q12866 Q12879 Q12884 Q12887 Q12889 Q12891 Q12893 Q12907 Q12908
Q12912 Q12913 Q12918 Q12981 Q12983 Q12999 Q13002 Q13003 Q13018 Q13021 Q13061
Q13087 Q13093 Q13113 Q13145 Q13162 Q13183 Q13190 Q13201 Q13214 Q13217 Q13219
Q13224 Q13241 Q13253 Q13255 Q13258 Q13261 Q13275 Q13277 Q13286 Q13291 Q13296
Q13304 Q13308 Q13316 Q13323 Q13324 Q13326 Q13332 Q13336 Q13349 Q13361 Q13370
Q13410 Q13421 Q13423 Q13433 Q13438 Q13443 Q13444 Q13445 Q13449 Q13454 Q13467
Q13477 Q13478 Q13488 Q13491 Q13505 Q13507 Q13508 Q13510 Q13519 Q13520 Q13530
Q13563 Q13571 Q13585 Q13586 Q13591 Q13606 Q13607 Q13609 Q13621 Q13634 Q13635
Q13639 Q13641 Q13651 Q13683 Q13698 Q13705 Q13724 Q13733 Q13740 Q13751 Q13790
Q13797 Q13873 Q13936 Q14002 Q14003 Q14028 Q14031 Q14050 Q14055 Q14108 Q14114
Q14118 Q14126 Q14154 Q14162 Q14165 Q14210 Q14213 Q14242 Q14246 Q14249 Q14257
Q14264 Q14318 Q14330 Q14332 Q14392 Q14393 Q14406 Q14409 Q14410 Q14416 Q14432
Q14435 Q14439 Q14442 Q14500 Q14507 Q14508 Q14512 Q14515 Q14517 Q14520 Q14524
Q14534 Q14542 Q14554 Q14563 Q14571 Q14573 Q14574 Q14623 Q14624 Q14626 Q14627
Q14641 Q14643 Q14654 Q14656 Q14667 Q14696 Q14697 Q14703 Q14714 Q14721 Q14728
Q14739 Q14761 Q14766 Q14773 Q14802 Q14831 Q14832 Q14833 Q14849 Q14916 Q14940
Q14943 Q14952 Q14953 Q14954 Q14956 Q14957 Q14973 Q14982 Q14BN4 Q14C87 Q14CN2
Q14CX5 Q14CZ8 Q14DG7 Q15005 Q15011 Q15012 Q15035 Q15041 Q15043 Q15049 Q15053
Q15063 Q15070 Q15077 Q15084 Q15109 Q15113 Q15116 Q15125 Q15155 Q15165 Q15166
Q15195 Q15198 Q15223 Q15256 Q15262 Q15293 Q15303 Q15363 Q15375 Q15388 Q15389
Q15391 Q15392 Q15399 Q15413 Q15465 Q15517 Q15526 Q15546 Q15582 Q15612 Q15615
Q15617 Q15619 Q15620 Q15622 Q15629 Q15661 Q15722 Q15726 Q15738 Q15743 Q15758
Q15760 Q15761 Q15762 Q15768 Q15782 Q15800 Q15818 Q15822 Q15825 Q15828 Q15836
Q15842 Q15846 Q15848 Q15849 Q15858 Q15878 Q15884 Q15904 Q16099 Q16134 Q16143
Q16270 Q16280 Q16281 Q16288 Q16322 Q16348 Q16378 Q16394 Q16445 Q16478 Q16515
Q16517 Q16538 Q16549 Q16553 Q16558 Q16563 Q16568 Q16570 Q16572 Q16581 Q16585
Q16586 Q16602 Q16609 Q16611 Q16617 Q16620 Q16623 Q16625 Q16627 Q16635 Q16647
Q16651 Q16653 Q16655 Q16661 Q16671 Q16678 Q16696 Q16706 Q16720 Q16739 Q16769
Q16790 Q16795 Q16799 Q16819 Q16820 Q16821 Q16827 Q16832 Q16842 Q16849 Q16850
Q16853 Q16873 Q16880 Q16891 Q17R55 Q17RF5 Q17RQ9 Q17RR3 Q17RW2 Q17RY6 Q19T08
Q1EHB4 Q1HG43 Q1HG44 Q1W4C9 Q1ZYL8 Q24JP5 Q24JQ0 Q29980 Q29983 Q2HXU8 Q2I0M4
Q2I0M5 Q2KHT4 Q2M2E3 Q2M2H8 Q2M2W7 Q2M385 Q2M3C6 Q2M3G0 Q2M3M2 Q2M3R5 Q2M3T9
Q2MJR0 Q2MKA7 Q2PZI1 Q2QL34 Q2T9K0 Q2TAA5 Q2TAP0 Q2TAZ0 Q2TBF2 Q2UY09 Q2VWP7
Q2VYF4 Q2WGJ8 Q2WGJ9 Q2Y0W8 Q30154 Q30201 Q30KP8 Q30KP9 Q30KQ1 Q30KQ7 Q30KQ9
Q30KR1 Q32M45 Q32P28 Q32ZL2 Q330K2 Q3B7T3 Q3C1V0 Q3KNS1 Q3KNT9 Q3KNW5 Q3KP22
Q3KPI0 Q3KQZ1 Q3KR37 Q3MIP1 Q3MIR4 Q3MIW9 Q3MIX3 Q3MUY2 Q3SXM5 Q3SXP7 Q3SXY7
Q3SY17 Q3SY77 Q3SYC2 Q3T906 Q3V5L5 Q3YBM2 Q3ZAQ7 Q3ZCQ3 Q3ZCQ8 Q401N2 Q494W8
Q495A1 Q495M3 Q495N2 Q495T6 Q495W5 Q496F6 Q496H8 Q496J9 Q49A17 Q49AH0 Q49SQ1
Q4G0G5 Q4G0I0 Q4G0M1 Q4G0N0 Q4G0N8 Q4G0T1 Q4G148 Q4G1C9 Q4KMG0 Q4KMG9 Q4KMQ2
Q4KMZ8 Q4LDR2 Q4QY38 Q4U2R8 Q4V9L6 Q4VC39 Q4VNC0 Q4VNC1 Q4VXA5 Q4VXF1 Q4W5P6
Q4ZHG4 Q4ZIN3 Q4ZJI4 Q504Y0 Q504Y2 Q50LG9 Q52LC2 Q53EL9 Q53EP0 Q53EU6 Q53F39
Q53FP2 Q53FV1 Q53GD3 Q53GQ0 Q53H12 Q53H76 Q53HI1 Q53R12 Q53RD9 Q53RY4 Q53S58
Q53TN4 Q567V2 Q56VL3 Q587I9 Q58DX5 Q58EX2 Q58HT5 Q5BIV9 Q5BJD5 Q5BJF2 Q5BJH2

Q5BJH7 Q5BKT4 Q5BKX6 Q5BLP8 Q5BVD1 Q5DID0 Q5DT21 Q5DX21 Q5EB52 Q5FWE3 Q5FYA8
Q5FYB0 Q5FYB1 Q5GAN6 Q5GH70 Q5GH72 Q5GH73 Q5GH76 Q5GH77 Q5H8A3 Q5H8A4 Q5H943
Q5H9E4 Q5H9R4 Q5HYA8 Q5HYJ1 Q5HYL7 Q5I7T1 Q5IJ48 Q5J8M3 Q5J8X5 Q5JPE7 Q5JQD4
Q5JQS5 Q5JRA6 Q5JRM2 Q5JRS4 Q5JRV8 Q5JS37 Q5JTB6 Q5JTV8 Q5JU69 Q5JUK3 Q5JW98
Q5JX69 Q5JX71 Q5JXA9 Q5JXM2 Q5JXX7 Q5JZY3 Q5K4E3 Q5K4L6 Q5KU26 Q5M7Z0 Q5M8T2
Q5MY95 Q5NDL2 Q5NUL3 Q5PT55 Q5QFB9 Q5QGT7 Q5QGZ9 Q5QJU3 Q5R387 Q5R3F8 Q5R3K3
Q5RGS3 Q5RI15 Q5SGD2 Q5SNT2 Q5SQ64 Q5SR56 Q5SRD1 Q5SRI9 Q5SRN2 Q5SSG8 Q5STR5
Q5SV17 Q5SVS4 Q5SWH9 Q5SWX8 Q5SY80 Q5SZI1 Q5SZK8 Q5T0T0 Q5T197 Q5T1A1 Q5T1J5
Q5T1Q4 Q5T1S8 Q5T292 Q5T2D2 Q5T2N8 Q5T3F8 Q5T3U5 Q5T442 Q5T4B2 Q5T4D3 Q5T4F4
Q5T4F7 Q5T4T1 Q5T4W7 Q5T601 Q5T6X4 Q5T6X5 Q5T700 Q5T7M4 Q5T7M9 Q5T7P6 Q5T7P8
Q5T7R7 Q5T848 Q5T8D3 Q5T9A4 Q5T9G4 Q5T9L3 Q5T9Z0 Q5TAH2 Q5TAT6 Q5TCH4 Q5TEA6
Q5TF21 Q5TF39 Q5TGI0 Q5TGU0 Q5TGY1 Q5TGZ0 Q5THJ4 Q5TZ20 Q5TZJ5 Q5U3C3 Q5U4P2
Q5UAW9 Q5UCC4 Q5VSG8 Q5VST6 Q5VT66 Q5VT99 Q5VTJ3 Q5VTY9 Q5VU36 Q5VU65 Q5VU97
Q5VUB5 Q5VUD6 Q5VUY0 Q5VUY2 Q5VV42 Q5VV43 Q5VV63 Q5VVB8 Q5VVP1 Q5VW38 Q5VWC8
Q5VWK5 Q5VWW1 Q5VX71 Q5VXJ0 Q5VXM1 Q5VXT5 Q5VXU1 Q5VXU3 Q5VY43 Q5VY80 Q5VYJ5
Q5VYP0 Q5VYY1 Q5VYY2 Q5VZ72 Q5VZI3 Q5VZR4 Q5VZY2 Q5W0B7 Q5W0Z9 Q5XG92 Q5XG99
Q5XKP0 Q5XXA6 Q5ZPR3 Q629K1 Q63HM2 Q63HQ2 Q63ZE4 Q641Q3 Q643R3 Q658N2 Q658P3
Q66K66 Q66K79 Q674R7 Q685J3 Q687X5 Q68BL7 Q68BL8 Q68CJ9 Q68CP4 Q68CQ7 Q68CR1
Q68CR7 Q68D42 Q68D85 Q68DH5 Q68DV7 Q68G75 Q695T7 Q69YG0 Q69YL0 Q69YU5 Q69YW2
Q69YZ2 Q6AI14 Q6AZY7 Q6DD88 Q6DKI7 Q6DN12 Q6DN14 Q6DN72 Q6DWJ6 Q6E0U4 Q6E213
Q6EBC2 Q6EIG7 Q6EMK4 Q6GMR7 Q6GPH6 Q6GPI1 Q6GTS8 Q6GTX8 Q6GV28 Q6H3X3 Q6H9L7
Q6HA08 Q6IA17 Q6IAN0 Q6IC98 Q6ICH7 Q6ICI0 Q6ICL7 Q6IE38 Q6IEE7 Q6IEE8 Q6IEU7
Q6IEV9 Q6IEY1 Q6IEZ7 Q6IF00 Q6IF36 Q6IF42 Q6IF63 Q6IF82 Q6IF99 Q6IFG1 Q6IFH4
Q6IFN5 Q6IS24 Q6ISU1 Q6IWH7 Q6J4K2 Q6J9G0 Q6JVE6 Q6KCM7 Q6KF10 Q6L9W6 Q6MZM0
Q6MZM9 Q6MZW2 Q6N022 Q6N075 Q6NSJ0 Q6NSJ5 Q6NT16 Q6NT52 Q6NT55 Q6NTF9 Q6NUI6
Q6NUJ1 Q6NUJ2 Q6NUK1 Q6NUK4 Q6NUM6 Q6NUM9 Q6NUQ4 Q6NUS6 Q6NUS8 Q6NUT2 Q6NUT3
Q6NV75 Q6NVV3 Q6NW40 Q6NXN4 Q6NXR0 Q6NXT4 Q6NXT6 Q6NZ63 Q6P093 Q6P0A1 Q6P1A2
Q6P1J6 Q6P1K1 Q6P1M0 Q6P1Q0 Q6P1S2 Q6P2H8 Q6P499 Q6P4A7 Q6P4A8 Q6P4E1 Q6P4F1
Q6P4H8 Q6P4Q7 Q6P531 Q6P5S2 Q6P5S7 Q6P5W5 Q6P5X7 Q6P7N7 Q6P988 Q6P995 Q6P9A2
Q6P9F7 Q6P9G4 Q6PCB0 Q6PCB6 Q6PCB7 Q6PCB8 Q6PDA7 Q6PEW0 Q6PEX7 Q6PEY0 Q6PEY1
Q6PEZ8 Q6PHW0 Q6PI25 Q6PI73 Q6PI78 Q6PIS1 Q6PIU1 Q6PIU2 Q6PIV7 Q6PIZ9 Q6PJF5
Q6PJG9 Q6PJW8 Q6PK18 Q6PKC3 Q6PL45 Q6PML9 Q6PP77 Q6PRD1 Q6PXP3 Q6Q0C1 Q6Q4G3
Q6Q788 Q6Q8B3 Q6QAJ8 Q6QHC5 Q6QNK2 Q6RW13 Q6S5H5 Q6T423 Q6T4P5 Q6TCH4 Q6TCH7
Q6U736 Q6U841 Q6UDR6 Q6UE05 Q6UQ28 Q6UVK1 Q6UVM3 Q6UVW9 Q6UVY6 Q6UW01 Q6UW02
Q6UW10 Q6UW15 Q6UW32 Q6UW49 Q6UW56 Q6UW60 Q6UW63 Q6UW68 Q6UW78 Q6UW88 Q6UWB1
Q6UWB4 Q6UWD8 Q6UWE3 Q6UWF3 Q6UWF7 Q6UWF9 Q6UWH4 Q6UWH6 Q6UWI2 Q6UWI4 Q6UWJ1
Q6UWJ8 Q6UWK7 Q6UWL2 Q6UWL6 Q6UWM5 Q6UWM7 Q6UWM9 Q6UWN0 Q6UWN5 Q6UWN8 Q6UWP7
Q6UWP8 Q6UWQ7 Q6UWR7 Q6UWS5 Q6UWT2 Q6UWT4 Q6UWU2 Q6UWU4 Q6UWV2 Q6UWV6 Q6UWV7
Q6UWW9 Q6UWY0 Q6UWY2 Q6UWY5 Q6UX01 Q6UX06 Q6UX07 Q6UX15 Q6UX27 Q6UX34 Q6UX39
Q6UX40 Q6UX41 Q6UX46 Q6UX53 Q6UX65 Q6UX68 Q6UX71 Q6UX72 Q6UX73 Q6UX82 Q6UX98
Q6UXA7 Q6UXB1 Q6UXB2 Q6UXB3 Q6UXB4 Q6UXB8 Q6UXC1 Q6UXD1 Q6UXD5 Q6UXD7 Q6UXE8
Q6UXF1 Q6UXF7 Q6UXG2 Q6UXG3 Q6UXG8 Q6UXH0 Q6UXH1 Q6UXH8 Q6UXI7 Q6UXI9 Q6UXK2
Q6UXK5 Q6UXL0 Q6UXM1 Q6UXN2 Q6UXN7 Q6UXN8 Q6UXP3 Q6UXQ4 Q6UXS0 Q6UXT8 Q6UXT9
Q6UXU4 Q6UXU6 Q6UXV0 Q6UXV1 Q6UXV4 Q6UXX9 Q6UXY8 Q6UXZ0 Q6UXZ3 Q6UXZ4 Q6UY09

Q6UY11 Q6UY13 Q6UY18 Q6UY27 Q6V0I7 Q6V0L0 Q6V1P9 Q6VVX0 Q6W3E5 Q6W5P4 Q6X4U4
Q6X784 Q6XPS3 Q6XR72 Q6XYQ8 Q6XZB0 Q6Y1H2 Q6Y288 Q6Y2X3 Q6YBV0 Q6YHK3 Q6YI46
Q6ZMB0 Q6ZMB5 Q6ZMC9 Q6ZMD2 Q6ZMG9 Q6ZMH5 Q6ZMI3 Q6ZMJ2 Q6ZMM2 Q6ZMQ8 Q6ZMR5
Q6ZMZ0 Q6ZMZ3 Q6ZN44 Q6ZN68 Q6ZNA5 Q6ZNB6 Q6ZNB7 Q6ZNC8 Q6ZNF0 Q6ZNI0 Q6ZNR0
Q6ZP29 Q6ZP80 Q6ZPD8 Q6ZPD9 Q6ZQN7 Q6ZQQ2 Q6ZRH7 Q6ZRP7 Q6ZRR5 Q6ZS10 Q6ZS82
Q6ZS86 Q6ZSA7 Q6ZSJ9 Q6ZSM3 Q6ZSS7 Q6ZSY5 Q6ZT21 Q6ZT89 Q6ZTK2 Q6ZTQ4 Q6ZU45
Q6ZUB0 Q6ZUB1 Q6ZUK4 Q6ZUX7 Q6ZV29 Q6ZVE7 Q6ZVK1 Q6ZVL6 Q6ZVN8 Q6ZVX9 Q6ZW05
Q6ZWJ8 Q6ZWK4 Q6ZWK6 Q6ZWL3 Q6ZWT7 Q6ZXV5 Q709C8 Q70CQ3 Q70HW3 Q70JA7 Q70SY1
Q70UQ0 Q70Z44 Q71H61 Q71RC9 Q71RG4 Q71RH2 Q71RS6 Q75T13 Q75V66 Q765I0 Q76EJ3
Q76KP1 Q76M96 Q76MJ5 Q7KYR7 Q7KZN9 Q7L0J3 Q7L0L9 Q7L0X0 Q7L1I2 Q7L1S5 Q7L1W4
Q7L211 Q7L311 Q7L4E1 Q7L4S7 Q7L5A8 Q7L5L3 Q7L5N7 Q7L8C5 Q7L985 Q7LBE3 Q7LFX5
Q7LGA3 Q7LGC8 Q7RTM1 Q7RTP0 Q7RTR8 Q7RTS5 Q7RTS6 Q7RTS9 Q7RTT9 Q7RTW8 Q7RTX0
Q7RTX1 Q7RTX7 Q7RTX9 Q7RTY0 Q7RTY1 Q7RTY3 Q7RTY7 Q7RTY8 Q7RTY9 Q7RTZ1 Q7Z2D5
Q7Z2H8 Q7Z2K6 Q7Z2Q7 Q7Z2W7 Q7Z388 Q7Z3B0 Q7Z3B1 Q7Z3C6 Q7Z3D4 Q7Z3F1 Q7Z3Q1
Q7Z3S7 Q7Z3T1 Q7Z402 Q7Z403 Q7Z404 Q7Z407 Q7Z408 Q7Z410 Q7Z412 Q7Z418 Q7Z419
Q7Z429 Q7Z434 Q7Z442 Q7Z443 Q7Z449 Q7Z4F1 Q7Z4H4 Q7Z4J2 Q7Z4L0 Q7Z4N2 Q7Z4N8
Q7Z4P5 Q7Z4R8 Q7Z4T8 Q7Z4Y8 Q7Z553 Q7Z5A4 Q7Z5A7 Q7Z5A8 Q7Z5A9 Q7Z5B4 Q7Z5G4
Q7Z5H4 Q7Z5H5 Q7Z5J1 Q7Z5L0 Q7Z5L3 Q7Z5L4 Q7Z5L7 Q7Z5M5 Q7Z5N4 Q7Z5P4 Q7Z5S9
Q7Z5Y6 Q7Z601 Q7Z602 Q7Z692 Q7Z695 Q7Z698 Q7Z699 Q7Z6A9 Q7Z6J6 Q7Z6L0 Q7Z6M3
Q7Z6W1 Q7Z6Z6 Q7Z769 Q7Z7B1 Q7Z7B7 Q7Z7B8 Q7Z7D3 Q7Z7G0 Q7Z7G8 Q7Z7H5 Q7Z7J7
Q7Z7M0 Q7Z7M1 Q7Z7M8 Q7Z7M9 Q7Z7N9 Q86SF2 Q86SG7 Q86SH4 Q86SJ2 Q86SJ6 Q86SK9
Q86SM5 Q86SM8 Q86SP6 Q86SQ3 Q86SQ4 Q86SQ6 Q86SR1 Q86SS6 Q86SU0 Q86T03 Q86T13
Q86T20 Q86T26 Q86T96 Q86TD4 Q86TE4 Q86TG1 Q86TH1 Q86TL2 Q86TM6 Q86TW2 Q86TY3
Q86U02 Q86UB9 Q86UD1 Q86UD3 Q86UD5 Q86UE4 Q86UE6 Q86UF1 Q86UF2 Q86UG4 Q86UK0
Q86UK5 Q86UL3 Q86UN2 Q86UN3 Q86UP0 Q86UP2 Q86UP6 Q86UP9 Q86UQ4 Q86UQ5 Q86UU9
Q86UW1 Q86UW2 Q86UW8 Q86UX2 Q86V24 Q86V35 Q86V40 Q86V85 Q86VB7 Q86VD7 Q86VD9
Q86VE9 Q86VF5 Q86VH4 Q86VH5 Q86VI4 Q86VL8 Q86VR2 Q86VR7 Q86VR8 Q86VU5 Q86VW1
Q86VY9 Q86VZ1 Q86VZ4 Q86VZ5 Q86W10 Q86W33 Q86W47 Q86W74 Q86WA9 Q86WB7 Q86WC4
Q86WD7 Q86WI0 Q86WK6 Q86WK7 Q86WK9 Q86WS5 Q86WV6 Q86X19 Q86X29 Q86X52 Q86XE3
Q86XJ0 Q86XK7 Q86XL3 Q86XM0 Q86XP6 Q86XQ3 Q86XR5 Q86XS5 Q86XS8 Q86XT9 Q86XX4
Q86Y07 Q86Y22 Q86Y34 Q86Y38 Q86Y39 Q86Y78 Q86Y82 Q86YB8 Q86YC3 Q86YD3 Q86YD5
Q86YJ5 Q86YL7 Q86YN1 Q86YQ2 Q86YT5 Q86YT9 Q86YW5 Q86YW7 Q86Z14 Q86Z23 Q8IU57
Q8IU68 Q8IU80 Q8IU89 Q8IU99 Q8IUA0 Q8IUA7 Q8IUB3 Q8IUB5 Q8IUC8 Q8IUH2 Q8IUH4
Q8IUH5 Q8IUH8 Q8IUK5 Q8IUK8 Q8IUL8 Q8IUN9 Q8IUR5 Q8IUS5 Q8IUW5 Q8IUX1 Q8IUX7
Q8IUX8 Q8IUY3 Q8IV01 Q8IV08 Q8IV16 Q8IV31 Q8IV45 Q8IV77 Q8IVB4 Q8IVJ1 Q8IVJ8
Q8IVK1 Q8IVL5 Q8IVL6 Q8IVL8 Q8IVM8 Q8IVN8 Q8IVP5 Q8IVQ6 Q8IVU1 Q8IVV8 Q8IVW8
Q8IVY1 Q8IW00 Q8IW45 Q8IW52 Q8IW70 Q8IW75 Q8IW92 Q8IWA4 Q8IWA5 Q8IWB1 Q8IWB4
Q8IWB9 Q8IWD5 Q8IWF2 Q8IWK6 Q8IWL1 Q8IWL2 Q8IWR1 Q8IWT1 Q8IWT6 Q8IWU2 Q8IWU4
Q8IWU5 Q8IWU6 Q8IWV1 Q8IWV2 Q8IWX5 Q8IWY4 Q8IX05 Q8IX19 Q8IX30 Q8IX94 Q8IXA5
Q8IXB1 Q8IXB3 Q8IXE1 Q8IXF9 Q8IXH8 Q8IXI1 Q8IXI2 Q8IXK2 Q8IXL6 Q8IXL7 Q8IXL9
Q8IXM3 Q8IXM6 Q8IXU6 Q8IY17 Q8IY26 Q8IY34 Q8IY49 Q8IY50 Q8IY95 Q8IYJ0 Q8IYK4
Q8IYL9 Q8IYP2 Q8IYP9 Q8IYR6 Q8IYS0 Q8IYS2 Q8IYU8 Q8IYV9 Q8IZ08 Q8IZ16 Q8IZ52
Q8IZ57 Q8IZ81 Q8IZ96 Q8IZA0 Q8IZD6 Q8IZF0 Q8IZF2 Q8IZF3 Q8IZF4 Q8IZF5 Q8IZF6
Q8IZF7 Q8IZJ1 Q8IZJ3 Q8IZK6 Q8IZM9 Q8IZN3 Q8IZN7 Q8IZP7 Q8IZP9 Q8IZR5 Q8IZS8

Q8IZT8 Q8IZU8 Q8IZU9 Q8IZV2 Q8IZV5 Q8IZY2 Q8J025 Q8MH63 Q8N0U2 Q8N0U8 Q8N0V4
Q8N0V5 Q8N0W4 Q8N0W7 Q8N0X7 Q8N0Y3 Q8N0Y5 Q8N0Z9 Q8N109 Q8N111 Q8N112 Q8N114
Q8N118 Q8N119 Q8N126 Q8N127 Q8N128 Q8N129 Q8N130 Q8N131 Q8N135 Q8N138 Q8N139
Q8N144 Q8N145 Q8N146 Q8N148 Q8N149 Q8N158 Q8N162 Q8N1C3 Q8N1E2 Q8N1L4 Q8N1M1
Q8N1N0 Q8N1N2 Q8N1S5 Q8N205 Q8N271 Q8N292 Q8N2A8 Q8N2E2 Q8N2F6 Q8N2G4 Q8N2G8
Q8N2H4 Q8N2K0 Q8N2K1 Q8N2M4 Q8N2Q7 Q8N2S1 Q8N2U0 Q8N2U9 Q8N307 Q8N323 Q8N326
Q8N336 Q8N349 Q8N350 Q8N357 Q8N370 Q8N386 Q8N387 Q8N394 Q8N3F9 Q8N3G9 Q8N3H0
Q8N3J6 Q8N3S3 Q8N3T1 Q8N3T6 Q8N3Y3 Q8N3Y7 Q8N3Z0 Q8N413 Q8N423 Q8N428 Q8N434
Q8N436 Q8N441 Q8N468 Q8N474 Q8N475 Q8N490 Q8N4A0 Q8N4C9 Q8N4F0 Q8N4F4 Q8N4F7
Q8N4H5 Q8N4K4 Q8N4L1 Q8N4L2 Q8N4L4 Q8N4M1 Q8N4S7 Q8N4S9 Q8N4T0 Q8N4V1 Q8N4V2
Q8N4W6 Q8N511 Q8N539 Q8N565 Q8N5B7 Q8N5C1 Q8N5D6 Q8N5G0 Q8N5G2 Q8N5I4 Q8N5K1
Q8N5M9 Q8N5S1 Q8N5U1 Q8N5W8 Q8N5Y8 Q8N608 Q8N609 Q8N614 Q8N628 Q8N661 Q8N682
Q8N688 Q8N690 Q8N695 Q8N697 Q8N699 Q8N6C5 Q8N6D2 Q8N6F1 Q8N6G5 Q8N6G6 Q8N6I4
Q8N6K0 Q8N6L0 Q8N6L1 Q8N6L7 Q8N6M3 Q8N6P7 Q8N6Q1 Q8N6Q3 Q8N6R1 Q8N6S5 Q8N6U8
Q8N6Y1 Q8N6Y2 Q8N729 Q8N743 Q8N755 Q8N766 Q8N7C0 Q8N7C4 Q8N7C7 Q8N7P1 Q8N7P3
Q8N7S2 Q8N7X8 Q8N808 Q8N816 Q8N8D7 Q8N8F6 Q8N8F7 Q8N8J7 Q8N8L6 Q8N8N0 Q8N8Q1
Q8N8Q8 Q8N8Q9 Q8N8R3 Q8N8Z6 Q8N907 Q8N912 Q8N966 Q8N967 Q8N9A8 Q8N9F0 Q8N9F7
Q8N9I0 Q8N9I5 Q8NA29 Q8NA58 Q8NAC3 Q8NAN2 Q8NAT1 Q8NAU1 Q8NB49 Q8NB59 Q8NBD8
Q8NBI2 Q8NBI3 Q8NBI5 Q8NBI6 Q8NBJ4 Q8NBJ5 Q8NBJ7 Q8NBJ9 Q8NBK3 Q8NBL1 Q8NBL3
Q8NBM4 Q8NBM8 Q8NBN3 Q8NBN7 Q8NBP0 Q8NBP5 Q8NBP7 Q8NBQ5 Q8NBQ7 Q8NBR0 Q8NBS3
Q8NBS9 Q8NBT3 Q8NBU5 Q8NBV4 Q8NBV8 Q8NBW4 Q8NBX0 Q8NBZ7 Q8NC01 Q8NC24 Q8NC42
Q8NC44 Q8NC54 Q8NC56 Q8NC67 Q8NCC3 Q8NCC5 Q8NCF0 Q8NCG5 Q8NCG7 Q8NCH0 Q8NCK7
Q8NCL4 Q8NCL8 Q8NCL9 Q8NCM2 Q8NCQ3 Q8NCR0 Q8NCR9 Q8NCS4 Q8NCS7 Q8NCU8 Q8NCW0
Q8NCW6 Q8ND94 Q8NDV1 Q8NDV2 Q8NDX2 Q8NDX9 Q8NDY8 Q8NDZ4 Q8NDZ6 Q8NE00 Q8NE01
Q8NE79 Q8NE86 Q8NEA5 Q8NEB5 Q8NEB7 Q8NEC5 Q8NEN9 Q8NEQ5 Q8NER1 Q8NER5 Q8NES3
Q8NES8 Q8NET1 Q8NET5 Q8NET6 Q8NET8 Q8NEW0 Q8NEW7 Q8NEX5 Q8NEX9 Q8NF37 Q8NF86
Q8NFB2 Q8NFF2 Q8NFJ5 Q8NFJ6 Q8NFK1 Q8NFL0 Q8NFM4 Q8NFM7 Q8NFN8 Q8NFP4 Q8NFQ5
Q8NFQ8 Q8NFR3 Q8NFR9 Q8NFT2 Q8NFT8 Q8NFU0 Q8NFU1 Q8NFU4 Q8NFY4 Q8NFZ3 Q8NFZ4
Q8NFZ6 Q8NFZ8 Q8NG04 Q8NG11 Q8NG35 Q8NG41 Q8NG75 Q8NG76 Q8NG77 Q8NG78 Q8NG80
Q8NG81 Q8NG83 Q8NG84 Q8NG85 Q8NG92 Q8NG94 Q8NG95 Q8NG97 Q8NG98 Q8NG99 Q8NGA0
Q8NGA1 Q8NGA2 Q8NGA4 Q8NGA5 Q8NGA6 Q8NGA8 Q8NGB2 Q8NGB4 Q8NGB6 Q8NGB8 Q8NGB9
Q8NGC0 Q8NGC1 Q8NGC2 Q8NGC3 Q8NGC4 Q8NGC5 Q8NGC6 Q8NGC7 Q8NGC8 Q8NGC9 Q8NGD0
Q8NGD1 Q8NGD2 Q8NGD3 Q8NGD4 Q8NGD5 Q8NGE0 Q8NGE1 Q8NGE2 Q8NGE3 Q8NGE5 Q8NGE7
Q8NGE8 Q8NGE9 Q8NGF0 Q8NGF1 Q8NGF3 Q8NGF4 Q8NGF6 Q8NGF7 Q8NGF8 Q8NGF9 Q8NGG0
Q8NGG1 Q8NGG2 Q8NGG3 Q8NGG4 Q8NGG5 Q8NGG6 Q8NGG7 Q8NGG8 Q8NGH3 Q8NGH5 Q8NGH6
Q8NGH7 Q8NGH8 Q8NGH9 Q8NGI0 Q8NGI1 Q8NGI2 Q8NGI3 Q8NGI4 Q8NGI6 Q8NGI7 Q8NGI8
Q8NGI9 Q8NGJ0 Q8NGJ1 Q8NGJ2 Q8NGJ3 Q8NGJ4 Q8NGJ5 Q8NGJ6 Q8NGJ7 Q8NGJ8 Q8NGJ9
Q8NGK0 Q8NGK1 Q8NGK2 Q8NGK3 Q8NGK4 Q8NGK5 Q8NGK6 Q8NGK9 Q8NGL0 Q8NGL1 Q8NGL2
Q8NGL3 Q8NGL4 Q8NGL6 Q8NGL7 Q8NGL9 Q8NGM1 Q8NGM8 Q8NGM9 Q8NGN0 Q8NGN1 Q8NGN2
Q8NGN3 Q8NGN4 Q8NGN5 Q8NGN6 Q8NGN7 Q8NGN8 Q8NGP0 Q8NGP2 Q8NGP3 Q8NGP4 Q8NGP6
Q8NGP8 Q8NGP9 Q8NGQ1 Q8NGQ2 Q8NGQ3 Q8NGQ4 Q8NGQ5 Q8NGQ6 Q8NGR1 Q8NGR2 Q8NGR3
Q8NGR4 Q8NGR5 Q8NGR6 Q8NGR8 Q8NGR9 Q8NGS0 Q8NGS1 Q8NGS2 Q8NGS3 Q8NGS4 Q8NGS5
Q8NGS6 Q8NGS7 Q8NGS8 Q8NGS9 Q8NGT0 Q8NGT1 Q8NGT2 Q8NGT5 Q8NGT7 Q8NGT9 Q8NGU1
Q8NGU2 Q8NGU4 Q8NGU9 Q8NGV0 Q8NGV5 Q8NGV6 Q8NGV7 Q8NGW1 Q8NGW6 Q8NGX0 Q8NGX1

Q8NGX2 Q8NGX3 Q8NGX5 Q8NGX6 Q8NGX8 Q8NGX9 Q8NGY0 Q8NGY1 Q8NGY2 Q8NGY3 Q8NGY5

Q8NGY6 Q8NGY7 Q8NGY9 Q8NGZ0 Q8NGZ2 Q8NGZ3 Q8NGZ4 Q8NGZ5 Q8NGZ6 Q8NGZ9 Q8NH00

Q8NH01 Q8NH02 Q8NH03 Q8NH04 Q8NH05 Q8NH06 Q8NH07 Q8NH08 Q8NH09 Q8NH10 Q8NH16

Q8NH18 Q8NH19 Q8NH21 Q8NH37 Q8NH40 Q8NH41 Q8NH42 Q8NH43 Q8NH48 Q8NH49 Q8NH50

Q8NH51 Q8NH53 Q8NH54 Q8NH55 Q8NH56 Q8NH57 Q8NH59 Q8NH60 Q8NH61 Q8NH63 Q8NH64

Q8NH67 Q8NH69 Q8NH70 Q8NH72 Q8NH73 Q8NH74 Q8NH76 Q8NH79 Q8NH80 Q8NH81 Q8NH83

Q8NH85 Q8NH87 Q8NH89 Q8NH90 Q8NH92 Q8NH93 Q8NH94 Q8NH95 Q8NHA4 Q8NHA6 Q8NHA8

Q8NHB1 Q8NHB7 Q8NHB8 Q8NHC4 Q8NHC5 Q8NHC6 Q8NHC7 Q8NHC8 Q8NHE4 Q8NHH9 Q8NHJ6

Q8NHK3 Q8NHL6 Q8NHM4 Q8NHP6 Q8NHP8 Q8NHS1 Q8NHS3 Q8NHU3 Q8NHV5 Q8NHW6 Q8NHX9

Q8NHY0 Q8NI17 Q8NI22 Q8NI32 Q8NI60 Q8NI99 Q8TAA1 Q8TAA9 Q8TAB3 Q8TAC9 Q8TAD2

Q8TAD4 Q8TAE7 Q8TAF8 Q8TAG5 Q8TAL6 Q8TAQ9 Q8TAT2 Q8TAV3 Q8TAV4 Q8TAX7 Q8TAZ6

Q8TB36 Q8TB40 Q8TB61 Q8TB68 Q8TB73 Q8TB96 Q8TBA6 Q8TBB6 Q8TBE1 Q8TBE3 Q8TBE7

Q8TBF5 Q8TBG9 Q8TBJ4 Q8TBM7 Q8TBM8 Q8TBP5 Q8TBP6 Q8TBQ9 Q8TBR7 Q8TC12 Q8TC26

Q8TC27 Q8TC36 Q8TC41 Q8TCB6 Q8TCC7 Q8TCD1 Q8TCG5 Q8TCJ2 Q8TCQ1 Q8TCT6 Q8TCT7

Q8TCT8 Q8TCT9 Q8TCU3 Q8TCU5 Q8TCV5 Q8TCW7 Q8TCW9 Q8TCY5 Q8TCZ2 Q8TD06 Q8TD07

Q8TD20 Q8TD22 Q8TD33 Q8TD43 Q8TD46 Q8TD84 Q8TDB4 Q8TDB8 Q8TDD5 Q8TDF5 Q8TDI7

Q8TDI8 Q8TDM5 Q8TDN1 Q8TDN2 Q8TDN7 Q8TDQ0 Q8TDQ1 Q8TDS4 Q8TDS5 Q8TDS7 Q8TDT2

Q8TDU5 Q8TDU6 Q8TDU9 Q8TDV0 Q8TDV2 Q8TDV5 Q8TDW0 Q8TDW4 Q8TDW7 Q8TDX6 Q8TDX9

Q8TDY8 Q8TE23 Q8TE54 Q8TE56 Q8TE57 Q8TE58 Q8TE99 Q8TEB7 Q8TEB9 Q8TED1 Q8TED4

Q8TEF2 Q8TEM1 Q8TEQ8 Q8TER0 Q8TEU8 Q8TEY5 Q8TEZ7 Q8TF08 Q8TF62 Q8TF66 Q8TF71

Q8WTQ1 Q8WTR4 Q8WTR8 Q8WTS1 Q8WTT0 Q8WTU2 Q8WTV0 Q8WTX9 Q8WU17 Q8WU66 Q8WU67

Q8WUA8 Q8WUD6 Q8WUG5 Q8WUH6 Q8WUJ1 Q8WUK0 Q8WUM9 Q8WUS8 Q8WUT4 Q8WUT9 Q8WUU8

Q8WUX1 Q8WUY1 Q8WUY8 Q8WV15 Q8WV19 Q8WV48 Q8WV83 Q8WVC6 Q8WVE6 Q8WVE7 Q8WVF2

Q8WVI0 Q8WVN6 Q8WVP7 Q8WVQ1 Q8WVV5 Q8WVX3 Q8WVX9 Q8WVZ1 Q8WVZ7 Q8WW34 Q8WW43

Q8WW52 Q8WW62 Q8WWA0 Q8WWA1 Q8WWB7 Q8WWC4 Q8WWF1 Q8WWF3 Q8WWF5 Q8WWG1 Q8WWG9

Q8WWH4 Q8WWI5 Q8WWP7 Q8WWQ2 Q8WWQ8 Q8WWT9 Q8WWU7 Q8WWV6 Q8WWX8 Q8WWX9 Q8WWY7

Q8WWY8 Q8WWZ4 Q8WWZ7 Q8WX69 Q8WX77 Q8WXA2 Q8WXA8 Q8WXD0 Q8WXD2 Q8WXF3 Q8WXF7

Q8WXH2 Q8WXI8 Q8WXQ8 Q8WXS4 Q8WXS5 Q8WXS8 Q8WY07 Q8WY21 Q8WY22 Q8WY98 Q8WYK1

Q8WYQ3 Q8WZ04 Q8WZ55 Q8WZ59 Q8WZ71 Q8WZ75 Q8WZ79 Q8WZ84 Q8WZ92 Q8WZ94 Q8WZA1

Q8WZA6 Q8WZA9 Q902F9 Q92185 Q92186 Q92187 Q92478 Q92482 Q92484 Q92485 Q92504

Q92508 Q92520 Q92521 Q92523 Q92535 Q92536 Q92537 Q92542 Q92543 Q92544 Q92545

Q92563 Q92575 Q92581 Q92604 Q92611 Q92617 Q92626 Q92629 Q92633 Q92637 Q92643

Q92667 Q92673 Q92685 Q92692 Q92729 Q92736 Q92743 Q92765 Q92781 Q92791 Q92806

Q92813 Q92819 Q92820 Q92823 Q92824 Q92832 Q92838 Q92839 Q92843 Q92847 Q92854

Q92859 Q92874 Q92876 Q92887 Q92896 Q92903 Q92911 Q92932 Q92935 Q92952 Q92953

Q92954 Q92956 Q92959 Q92968 Q92982 Q93033 Q93038 Q93050 Q93063 Q93070 Q93084

Q93086 Q93097 Q93098 Q95460 Q969D9 Q969E1 Q969E2 Q969E3 Q969F0 Q969F8 Q969H8

Q969I6 Q969K7 Q969L2 Q969M1 Q969M2 Q969M3 Q969N2 Q969N4 Q969P0 Q969S0 Q969S6

Q969V1 Q969V3 Q969V5 Q969W0 Q969W1 Q969W9 Q969X1 Q969X2 Q969X5 Q969Y0 Q969Z3

Q969Z4 Q96A11 Q96A25 Q96A26 Q96A28 Q96A29 Q96A33 Q96A46 Q96A54 Q96A57 Q96A59

Q96A83 Q96A84 Q96A98 Q96AA3 Q96AD5 Q96AG3 Q96AG4 Q96AJ9 Q96AM1 Q96AN5 Q96AP7

Q96AQ2 Q96AQ6 Q96AQ8 Q96AW1 Q96B21 Q96B33 Q96B42 Q96B49 Q96B77 Q96B86 Q96B96

Q96BA8 Q96BD0 Q96BF3 Q96BI1 Q96BI3 Q96BM0 Q96BQ1 Q96BQ5 Q96BY7 Q96BY9 Q96BZ4

Q96C03 Q96CC6 Q96CE8 Q96CG8 Q96CH1 Q96CP6 Q96CP7 Q96CQ1 Q96CS3 Q96CW9 Q96D05

Q96D15 Q96D31 Q96D42 Q96D53 Q96D59 Q96D96 Q96DA0 Q96DA6 Q96DB5 Q96DB9 Q96DD7
Q96DL1 Q96DN0 Q96DN2 Q96DR5 Q96DR8 Q96DS6 Q96DU3 Q96DW6 Q96DX8 Q96DZ1 Q96DZ7
Q96DZ9 Q96E16 Q96E22 Q96E52 Q96E93 Q96EC8 Q96EE4 Q96EG1 Q96EP9 Q96ER9 Q96ES6
Q96ET8 Q96EU7 Q96EX1 Q96EX2 Q96EZ4 Q96F05 Q96F15 Q96F25 Q96F46 Q96F81 Q96FE5
Q96FE7 Q96FL8 Q96FL9 Q96FM1 Q96FT7 Q96FV3 Q96FX8 Q96FZ5 Q96G23 Q96G27 Q96G30
Q96G79 Q96G91 Q96G97 Q96GC9 Q96GE9 Q96GF1 Q96GL9 Q96GM1 Q96GP6 Q96GQ5 Q96GR4
Q96GS6 Q96GW7 Q96GX1 Q96GZ6 Q96H15 Q96H72 Q96H78 Q96H96 Q96HA1 Q96HA4 Q96HA9
Q96HD1 Q96HE7 Q96HE8 Q96HE9 Q96HH4 Q96HH6 Q96HH9 Q96HJ5 Q96HL8 Q96HP8 Q96HR9
Q96HS1 Q96HV5 Q96HY6 Q96I36 Q96I45 Q96I82 Q96ID5 Q96IK0 Q96IL0 Q96IQ7 Q96IV6
Q96IW7 Q96IX5 Q96IY4 Q96IZ2 Q96J42 Q96J65 Q96J66 Q96J77 Q96J84 Q96J86 Q96JA1
Q96JA4 Q96JB6 Q96JF0 Q96JJ6 Q96JJ7 Q96JK4 Q96JN2 Q96JP9 Q96JQ0 Q96JQ2 Q96JQ5
Q96JT2 Q96JW4 Q96JX3 Q96K12 Q96K19 Q96K37 Q96K49 Q96K78 Q96KA5 Q96KC8 Q96KF7
Q96KG7 Q96KJ9 Q96KK3 Q96KK4 Q96KN2 Q96KN9 Q96KR4 Q96KR6 Q96KT7 Q96KV6 Q96KW9
Q96KX0 Q96L08 Q96L11 Q96L12 Q96L42 Q96L58 Q96LA5 Q96LA6 Q96LA9 Q96LB0 Q96LB1
Q96LB2 Q96LB8 Q96LC7 Q96LD1 Q96LJ7 Q96LL3 Q96LL9 Q96LR4 Q96LR9 Q96LT4 Q96LU5
Q96LU7 Q96LZ7 Q96M19 Q96MC6 Q96MG2 Q96MH6 Q96MK3 Q96MM7 Q96MS0 Q96MT1 Q96MU8
Q96MV1 Q96MV8 Q96MX0 Q96MZ0 Q96N19 Q96N66 Q96N68 Q96N87 Q96NA8 Q96NB2 Q96ND0
Q96NI6 Q96NL1 Q96NR3 Q96NR8 Q96NT5 Q96NU0 Q96NY8 Q96NZ8 Q96NZ9 Q96P31 Q96P56
Q96P65 Q96P66 Q96P67 Q96P68 Q96P69 Q96P88 Q96PB1 Q96PB8 Q96PC5 Q96PD2 Q96PD5
Q96PD6 Q96PD7 Q96PE1 Q96PE5 Q96PG1 Q96PG2 Q96PH1 Q96PJ5 Q96PL1 Q96PL2 Q96PL5
Q96PQ0 Q96PQ1 Q96PR1 Q96PS8 Q96PX8 Q96PZ7 Q96Q04 Q96Q45 Q96Q80 Q96Q91 Q96QD8
Q96QE2 Q96QE4 Q96QI5 Q96QK8 Q96QR1 Q96QS1 Q96QT4 Q96QU1 Q96QV1 Q96QZ0 Q96R08
Q96R09 Q96R27 Q96R28 Q96R30 Q96R45 Q96R47 Q96R48 Q96R54 Q96R67 Q96R69 Q96R72
Q96R84 Q96RA2 Q96RB7 Q96RC9 Q96RD0 Q96RD1 Q96RD2 Q96RD3 Q96RD6 Q96RD7 Q96RD9
Q96RI0 Q96RI8 Q96RI9 Q96RJ0 Q96RJ3 Q96RL6 Q96RL7 Q96RN1 Q96RP3 Q96RP7 Q96RP8
Q96RQ1 Q96RQ9 Q96RT6 Q96RV3 Q96S06 Q96S16 Q96S37 Q96S52 Q96S66 Q96S86 Q96S97
Q96SA4 Q96SE0 Q96SJ8 Q96SK2 Q96SL1 Q96SL4 Q96SM3 Q96SN7 Q96SQ9 Q96T52 Q96T53
Q96T54 Q96T55 Q96T59 Q96T83 Q96T91 Q96TA0 Q96TA2 Q96TC7 Q99062 Q99075 Q99217
Q99218 Q99250 Q99435 Q99437 Q99442 Q99445 Q99463 Q99466 Q99467 Q99470 Q99500
Q99518 Q99519 Q99523 Q99527 Q99538 Q99541 Q99542 Q99571 Q99572 Q99574 Q99595
Q99623 Q99624 Q99640 Q99643 Q99645 Q99650 Q99665 Q99674 Q99675 Q99677 Q99678
Q99679 Q99680 Q99705 Q99706 Q99712 Q99720 Q99726 Q99727 Q99731 Q99732 Q99735
Q99748 Q99758 Q99788 Q99795 Q99805 Q99807 Q99808 Q99835 Q99884 Q99895 Q99928
Q99935 Q99941 Q99942 Q99943 Q99944 Q99946 Q99954 Q99965 Q99969 Q99972 Q99983
Q99988 Q99999 Q9BPV8 Q9BPW4 Q9BPW9 Q9BPX6 Q9BQ08 Q9BQ16 Q9BQ31 Q9BQ49 Q9BQ51
Q9BQA9 Q9BQB4 Q9BQB6 Q9BQD7 Q9BQE4 Q9BQE5 Q9BQG1 Q9BQI4 Q9BQJ4 Q9BQP9 Q9BQQ7
Q9BQR3 Q9BQS2 Q9BQS7 Q9BQT8 Q9BQT9 Q9BR26 Q9BR39 Q9BRB3 Q9BRI3 Q9BRK0 Q9BRK3
Q9BRK5 Q9BRL7 Q9BRN9 Q9BRQ5 Q9BRR3 Q9BRR6 Q9BRV3 Q9BRX8 Q9BRY0 Q9BS86 Q9BS91
Q9BSA4 Q9BSA9 Q9BSE2 Q9BSE4 Q9BSF4 Q9BSG0 Q9BSG5 Q9BSJ5 Q9BSJ8 Q9BSK0 Q9BSK2
Q9BSN7 Q9BSR8 Q9BT09 Q9BT22 Q9BT56 Q9BT67 Q9BT76 Q9BT88 Q9BTD3 Q9BTN0 Q9BTV4
Q9BTX1 Q9BTX3 Q9BTY2 Q9BU23 Q9BU79 Q9BUB7 Q9BUD6 Q9BUF7 Q9BUJ0 Q9BUM1 Q9BUN1
Q9BUN8 Q9BUR5 Q9BUV8 Q9BV10 Q9BV23 Q9BV35 Q9BV40 Q9BV81 Q9BV87 Q9BV94 Q9BVA6
Q9BVC6 Q9BVG9 Q9BVH7 Q9BVK2 Q9BVK6 Q9BVK8 Q9BVT8 Q9BVV7 Q9BVV8 Q9BVW6 Q9BVX2
Q9BW60 Q9BW72 Q9BWH2 Q9BWL3 Q9BWM7 Q9BWQ6 Q9BWQ8 Q9BWS9 Q9BWV1 Q9BWV2 Q9BWW8

Q9BWW9 Q9BX59 Q9BX67 Q9BX73 Q9BX74 Q9BX79 Q9BX84 Q9BX93 Q9BX95 Q9BX97 Q9BXA5
Q9BXB1 Q9BXC0 Q9BXC1 Q9BXE9 Q9BXI2 Q9BXI9 Q9BXJ0 Q9BXJ1 Q9BXJ2 Q9BXJ3 Q9BXJ4
Q9BXJ5 Q9BXJ7 Q9BXJ8 Q9BXK5 Q9BXM7 Q9BXN1 Q9BXN2 Q9BXP2 Q9BXQ6 Q9BXR5 Q9BXS0
Q9BXS4 Q9BXS9 Q9BXT2 Q9BXU9 Q9BXX0 Q9BXY4 Q9BY07 Q9BY08 Q9BY10 Q9BY14 Q9BY15
Q9BY19 Q9BY21 Q9BY50 Q9BY64 Q9BY67 Q9BY71 Q9BY76 Q9BY78 Q9BY79 Q9BYC5 Q9BYD5
Q9BYE2 Q9BYE9 Q9BYF1 Q9BYG0 Q9BYH1 Q9BYJ0 Q9BYT1 Q9BYT9 Q9BYW1 Q9BYZ8 Q9BZ11
Q9BZ76 Q9BZA7 Q9BZA8 Q9BZC5 Q9BZC7 Q9BZD2 Q9BZD6 Q9BZD7 Q9BZF1 Q9BZG2 Q9BZJ3
Q9BZJ4 Q9BZJ6 Q9BZJ7 Q9BZJ8 Q9BZL3 Q9BZM1 Q9BZM2 Q9BZM4 Q9BZM5 Q9BZM6 Q9BZP6
Q9BZQ6 Q9BZR6 Q9BZV2 Q9BZV3 Q9BZW2 Q9BZW4 Q9BZW5 Q9BZW8 Q9BZZ2 Q9C002 Q9C004
Q9C0A0 Q9C0B5 Q9C0B6 Q9C0C4 Q9C0D9 Q9C0E8 Q9C0H2 Q9C0I4 Q9C0J1 Q9C0K1 Q9GIP4
Q9GZK3 Q9GZK4 Q9GZK6 Q9GZK7 Q9GZM5 Q9GZM6 Q9GZM7 Q9GZN0 Q9GZN6 Q9GZP0 Q9GZP1
Q9GZP7 Q9GZP9 Q9GZQ4 Q9GZQ6 Q9GZR5 Q9GZS9 Q9GZT6 Q9GZU1 Q9GZU3 Q9GZU5 Q9GZV3
Q9GZV9 Q9GZW8 Q9GZX3 Q9GZX9 Q9GZY4 Q9GZY6 Q9GZY8 Q9GZZ6 Q9GZZ7 Q9GZZ8 Q9H013
Q9H015 Q9H061 Q9H078 Q9H0B8 Q9H0C2 Q9H0C3 Q9H0P0 Q9H0Q3 Q9H0R3 Q9H0U3 Q9H0V1
Q9H0V9 Q9H0X4 Q9H0X9 Q9H114 Q9H156 Q9H158 Q9H159 Q9H172 Q9H173 Q9H195 Q9H1A3
Q9H1B5 Q9H1C0 Q9H1C3 Q9H1C4 Q9H1C7 Q9H1D0 Q9H1E5 Q9H1F0 Q9H1K4 Q9H1N7 Q9H1U4
Q9H1U9 Q9H1V8 Q9H1X3 Q9H1Y3 Q9H1Z8 Q9H1Z9 Q9H205 Q9H207 Q9H208 Q9H209 Q9H210
Q9H221 Q9H222 Q9H228 Q9H237 Q9H239 Q9H244 Q9H251 Q9H252 Q9H255 Q9H295 Q9H2A7
Q9H2A9 Q9H2B2 Q9H2B4 Q9H2C2 Q9H2C5 Q9H2C8 Q9H2D1 Q9H2E6 Q9H2F3 Q9H2H9 Q9H2J7
Q9H2L4 Q9H2R5 Q9H2S1 Q9H2S6 Q9H2U9 Q9H2V7 Q9H2W1 Q9H2X0 Q9H2X3 Q9H2X8 Q9H2X9
Q9H2Y9 Q9H300 Q9H305 Q9H306 Q9H310 Q9H313 Q9H324 Q9H330 Q9H339 Q9H340 Q9H341
Q9H342 Q9H343 Q9H344 Q9H346 Q9H3G5 Q9H3H5 Q9H3K2 Q9H3M0 Q9H3N1 Q9H3N8 Q9H3Q3
Q9H3R1 Q9H3R2 Q9H3S1 Q9H3S3 Q9H3S5 Q9H3T2 Q9H3T3 Q9H3U5 Q9H3U7 Q9H3V2 Q9H3W5
Q9H3Z4 Q9H3Z7 Q9H400 Q9H427 Q9H461 Q9H488 Q9H490 Q9H497 Q9H4A9 Q9H4B8 Q9H4D0
Q9H4F1 Q9H4F8 Q9H4I3 Q9H4I9 Q9H553 Q9H598 Q9H5I5 Q9H5J4 Q9H5K3 Q9H5V8 Q9H5Y7
Q9H665 Q9H6A9 Q9H6B4 Q9H6B9 Q9H6D3 Q9H6D8 Q9H6E4 Q9H6F2 Q9H6H4 Q9H6K4 Q9H6L2
Q9H6L5 Q9H6R6 Q9H6U8 Q9H6V9 Q9H6X2 Q9H6X4 Q9H6Y7 Q9H720 Q9H741 Q9H756 Q9H772
Q9H7F0 Q9H7F4 Q9H7M9 Q9H7T0 Q9H7V2 Q9H7X0 Q9H7X2 Q9H7Y0 Q9H7Z7 Q9H813 Q9H819
Q9H841 Q9H8H3 Q9H8J5 Q9H8M5 Q9H8M9 Q9H8P0 Q9H8X9 Q9H902 Q9H920 Q9H936 Q9H9B4
Q9H9K5 Q9H9P2 Q9H9S3 Q9H9S5 Q9H9V4 Q9HA72 Q9HA82 Q9HAB3 Q9HAR2 Q9HAS3 Q9HAT1
Q9HAT2 Q9HAV5 Q9HAW7 Q9HAW8 Q9HAW9 Q9HB03 Q9HB14 Q9HB15 Q9HB29 Q9HB40 Q9HB55
Q9HB63 Q9HB66 Q9HB89 Q9HBA0 Q9HBB8 Q9HBE4 Q9HBE5 Q9HBG4 Q9HBG7 Q9HBH5 Q9HBI6
Q9HBJ8 Q9HBL6 Q9HBL7 Q9HBM0 Q9HBR0 Q9HBT6 Q9HBU9 Q9HBV1 Q9HBV2 Q9HBW0 Q9HBW1
Q9HBW9 Q9HBX8 Q9HBX9 Q9HBY0 Q9HC07 Q9HC10 Q9HC21 Q9HC23 Q9HC24 Q9HC56 Q9HC57
Q9HC58 Q9HC73 Q9HC97 Q9HCB6 Q9HCC8 Q9HCE9 Q9HCF6 Q9HCJ1 Q9HCJ2 Q9HCK4 Q9HCL0
Q9HCL2 Q9HCM2 Q9HCM3 Q9HCN3 Q9HCN6 Q9HCN8 Q9HCP6 Q9HCQ5 Q9HCQ7 Q9HCS2 Q9HCT0
Q9HCU0 Q9HCU4 Q9HCU5 Q9HCX4 Q9HD20 Q9HD23 Q9HD36 Q9HD43 Q9HD45 Q9HD89 Q9HDC5
Q9HDC9 Q9N2J8 Q9N2K0 Q9NNX6 Q9NNZ3 Q9NP55 Q9NP58 Q9NP59 Q9NP60 Q9NP70 Q9NP78
Q9NP84 Q9NP85 Q9NP91 Q9NP94 Q9NP99 Q9NPA0 Q9NPA1 Q9NPA2 Q9NPB0 Q9NPB9 Q9NPC1
Q9NPC2 Q9NPC4 Q9NPD5 Q9NPD7 Q9NPE6 Q9NPF0 Q9NPF2 Q9NPF7 Q9NPG1 Q9NPG4 Q9NPG8
Q9NPH0 Q9NPH3 Q9NPH5 Q9NPH9 Q9NPI0 Q9NPI9 Q9NPL8 Q9NPR2 Q9NPR9 Q9NPU4 Q9NPY3
Q9NPZ5 Q9NQ11 Q9NQ25 Q9NQ30 Q9NQ34 Q9NQ36 Q9NQ38 Q9NQ40 Q9NQ60 Q9NQ76 Q9NQ79
Q9NQ84 Q9NQ90 Q9NQA5 Q9NQC3 Q9NQG1 Q9NQG6 Q9NQN1 Q9NQQ7 Q9NQR9 Q9NQS3 Q9NQS5
Q9NQW8 Q9NQX5 Q9NQX7 Q9NQZ5 Q9NQZ7 Q9NR16 Q9NR23 Q9NR28 Q9NR31 Q9NR34 Q9NR61

Q9NR63 Q9NR71 Q9NR77 Q9NR82 Q9NR96 Q9NR97 Q9NRA1 Q9NRA2 Q9NRB3 Q9NRC1 Q9NRC9
Q9NRD8 Q9NRD9 Q9NRE1 Q9NRG7 Q9NRG9 Q9NRI6 Q9NRI7 Q9NRJ7 Q9NRK6 Q9NRM0 Q9NRM1
Q9NRM6 Q9NRN5 Q9NRP0 Q9NRQ5 Q9NRR1 Q9NRR2 Q9NRS4 Q9NRU3 Q9NRX3 Q9NRX5 Q9NRX6
Q9NRZ5 Q9NRZ7 Q9NS00 Q9NS15 Q9NS40 Q9NS62 Q9NS64 Q9NS66 Q9NS67 Q9NS68 Q9NS69
Q9NS71 Q9NS75 Q9NS82 Q9NS84 Q9NS93 Q9NSA0 Q9NSA1 Q9NSA2 Q9NSC7 Q9NSD5 Q9NSD7
Q9NSI5 Q9NSK7 Q9NST1 Q9NSU2 Q9NT22 Q9NT68 Q9NT99 Q9NTG1 Q9NTI2 Q9NTJ5 Q9NTN3
Q9NTN9 Q9NTQ9 Q9NTU7 Q9NU53 Q9NUB4 Q9NUD9 Q9NUE0 Q9NUH8 Q9NUM3 Q9NUM4 Q9NUN5
Q9NUN7 Q9NUQ2 Q9NUR3 Q9NUT2 Q9NUU6 Q9NUV7 Q9NV12 Q9NV29 Q9NV58 Q9NV64 Q9NV66
Q9NV92 Q9NV96 Q9NVA4 Q9NVC3 Q9NVH0 Q9NVH1 Q9NVI7 Q9NVM1 Q9NVV0 Q9NVV5 Q9NW15
Q9NW97 Q9NWC5 Q9NWD8 Q9NWF4 Q9NWH2 Q9NWM8 Q9NWQ8 Q9NWR8 Q9NWS8 Q9NWW5 Q9NWW9
Q9NX00 Q9NX14 Q9NX40 Q9NX47 Q9NX52 Q9NX61 Q9NX62 Q9NX76 Q9NX77 Q9NX78 Q9NX94
Q9NX95 Q9NXB9 Q9NXE4 Q9NXF8 Q9NXG6 Q9NXH8 Q9NXI6 Q9NXJ0 Q9NXK6 Q9NXL6 Q9NXS2
Q9NXW2 Q9NY15 Q9NY25 Q9NY26 Q9NY28 Q9NY35 Q9NY37 Q9NY46 Q9NY47 Q9NY59 Q9NY64
Q9NY72 Q9NY91 Q9NY97 Q9NYB5 Q9NYG2 Q9NYG8 Q9NYJ7 Q9NYK1 Q9NYL4 Q9NYL5 Q9NYM4
Q9NYM9 Q9NYP7 Q9NYP8 Q9NYQ6 Q9NYQ7 Q9NYQ8 Q9NYV7 Q9NYV8 Q9NYV9 Q9NYW0 Q9NYW1
Q9NYW2 Q9NYW3 Q9NYW4 Q9NYW5 Q9NYW6 Q9NYW7 Q9NYX4 Q9NYZ1 Q9NYZ2 Q9NYZ4 Q9NZ01
Q9NZ08 Q9NZ42 Q9NZ43 Q9NZ45 Q9NZ53 Q9NZ94 Q9NZC2 Q9NZC3 Q9NZC7 Q9NZD1 Q9NZF1
Q9NZG7 Q9NZH0 Q9NZJ5 Q9NZJ7 Q9NZK5 Q9NZK7 Q9NZM1 Q9NZM6 Q9NZN1 Q9NZP0 Q9NZP2
Q9NZP5 Q9NZP8 Q9NZQ7 Q9NZQ8 Q9NZR2 Q9NZS2 Q9NZS9 Q9NZU0 Q9NZU1 Q9NZV1 Q9NZV5
Q9NZV8 Q9NZW4 Q9P003 Q9P035 Q9P055 Q9P0B6 Q9P0I2 Q9P0J0 Q9P0K1 Q9P0K9 Q9P0L0
Q9P0L9 Q9P0M4 Q9P0N5 Q9P0N8 Q9P0S2 Q9P0S3 Q9P0S9 Q9P0T7 Q9P0U1 Q9P0V8 Q9P0X4
Q9P109 Q9P121 Q9P126 Q9P1C3 Q9P1P4 Q9P1P5 Q9P1Q5 Q9P1W3 Q9P1W8 Q9P1Z3 Q9P232
Q9P241 Q9P244 Q9P246 Q9P273 Q9P283 Q9P291 Q9P296 Q9P298 Q9P2B2 Q9P2C4 Q9P2E5
Q9P2E7 Q9P2E8 Q9P2E9 Q9P2J2 Q9P2K9 Q9P2S2 Q9P2U7 Q9P2U8 Q9P2V4 Q9P2W7 Q9P2W9
Q9P2X0 Q9UBC7 Q9UBD3 Q9UBD6 Q9UBG0 Q9UBH6 Q9UBI4 Q9UBJ2 Q9UBK5 Q9UBL9 Q9UBM1
Q9UBM4 Q9UBM7 Q9UBM8 Q9UBN1 Q9UBN4 Q9UBN6 Q9UBP0 Q9UBP4 Q9UBQ6 Q9UBR2 Q9UBR5
Q9UBS3 Q9UBS5 Q9UBS9 Q9UBT3 Q9UBU2 Q9UBU3 Q9UBU6 Q9UBV2 Q9UBV4 Q9UBV7 Q9UBX1
Q9UBX3 Q9UBX5 Q9UBX7 Q9UBX8 Q9UBY0 Q9UBY5 Q9UBY8 Q9UDW1 Q9UDX5 Q9UEF7 Q9UEU0
Q9UEW3 Q9UF02 Q9UF12 Q9UF33 Q9UF47 Q9UFP1 Q9UG22 Q9UG56 Q9UGF5 Q9UGF6 Q9UGF7
Q9UGH3 Q9UGI6 Q9UGM1 Q9UGN4 Q9UGP8 Q9UGQ2 Q9UGQ3 Q9UGT4 Q9UH62 Q9UH99 Q9UHC3
Q9UHC6 Q9UHC9 Q9UHE5 Q9UHE8 Q9UHF0 Q9UHF1 Q9UHF3 Q9UHF4 Q9UHF5 Q9UHG2 Q9UHG3
Q9UHI5 Q9UHI7 Q9UHI8 Q9UHJ9 Q9UHL4 Q9UHM6 Q9UHN6 Q9UHP7 Q9UHQ4 Q9UHQ9 Q9UHT4
Q9UHW9 Q9UHX3 Q9UI14 Q9UI33 Q9UI38 Q9UI40 Q9UI42 Q9UIB8 Q9UIG4 Q9UIG8 Q9UIJ5
Q9UIK5 Q9UIQ6 Q9UIW2 Q9UIX4 Q9UJ14 Q9UJ37 Q9UJ42 Q9UJ71 Q9UJ90 Q9UJ96 Q9UJ99
Q9UJA2 Q9UJA9 Q9UJG1 Q9UJH8 Q9UJJ9 Q9UJQ1 Q9UJS0 Q9UJZ1 Q9UK00 Q9UK05 Q9UK17
Q9UK23 Q9UK28 Q9UK55 Q9UK85 Q9UKA2 Q9UKB5 Q9UKF2 Q9UKF5 Q9UKG4 Q9UKJ0 Q9UKJ1
Q9UKJ5 Q9UKJ8 Q9UKL2 Q9UKL4 Q9UKM7 Q9UKP4 Q9UKP6 Q9UKQ2 Q9UKR0 Q9UKR3 Q9UKR5
Q9UKR8 Q9UKU0 Q9UKU6 Q9UKU9 Q9UKV5 Q9UKX5 Q9UKY0 Q9UKY3 Q9UKY4 Q9UKZ4 Q9UKZ9
Q9UL01 Q9UL19 Q9UL51 Q9UL52 Q9UL54 Q9UL62 Q9ULB1 Q9ULB4 Q9ULB5 Q9ULC0 Q9ULC5
Q9ULC8 Q9ULD8 Q9ULF5 Q9ULG6 Q9ULH0 Q9ULH4 Q9ULI3 Q9ULK0 Q9ULK5 Q9ULK6 Q9ULL4
Q9ULQ1 Q9ULS5 Q9ULS6 Q9ULT6 Q9ULV1 Q9ULW2 Q9ULX5 Q9ULX7 Q9ULY5 Q9ULZ1 Q9ULZ9
Q9UM00 Q9UM01 Q9UM21 Q9UM22 Q9UM44 Q9UM47 Q9UM73 Q9UMD9 Q9UMF0 Q9UMR5 Q9UMR7
Q9UMS5 Q9UMX3 Q9UMX5 Q9UMX9 Q9UMZ3 Q9UN42 Q9UN66 Q9UN67 Q9UN70 Q9UN71 Q9UN72
Q9UN73 Q9UN74 Q9UN75 Q9UN76 Q9UN88 Q9UNA0 Q9UNA3 Q9UNE0 Q9UNG2 Q9UNK0 Q9UNK4

Q9UNL2 Q9UNN8 Q9UNP4 Q9UNQ0 Q9UNU6 Q9UNW1 Q9UNW8 Q9UNX9 Q9UP38 Q9UP52 Q9UP95 Q9UPC5 Q9UPI3 Q9UPQ8 Q9UPR5 Q9UPU3 Q9UPX0 Q9UPX6 Q9UPY5 Q9UPZ6 Q9UQ05 Q9UQ52 Q9UQ53 Q9UQ90 Q9UQC9 Q9UQD0 Q9UQF0 Q9UQP3 Q9UQQ1 Q9UQV4 Q9Y210 Q9Y219 Q9Y225 Q9Y226 Q9Y227 Q9Y228 Q9Y231 Q9Y240 Q9Y241 Q9Y251 Q9Y256 Q9Y257 Q9Y258 Q9Y264 Q9Y267 Q9Y271 Q9Y274 Q9Y275 Q9Y276 Q9Y277 Q9Y278 Q9Y279 Q9Y282 Q9Y284 Q9Y286 Q9Y287 Q9Y289 Q9Y2A9 Q9Y2B0 Q9Y2B1 Q9Y2B2 Q9Y2C2 Q9Y2C3 Q9Y2C4 Q9Y2C5 Q9Y2C9 Q9Y2D2 Q9Y2E5 Q9Y2E8 Q9Y2G1 Q9Y2G3 Q9Y2G5 Q9Y2G8 Q9Y2H6 Q9Y2I2 Q9Y2J2 Q9Y2L9 Q9Y2P4 Q9Y2P5 Q9Y2Q0 Q9Y2R0 Q9Y2T5 Q9Y2T6 Q9Y2U2 Q9Y2U8 Q9Y2W3 Q9Y2W6 Q9Y2Y6 Q9Y2Y8 Q9Y2Z9 Q9Y320 Q9Y328 Q9Y334 Q9Y336 Q9Y337 Q9Y342 Q9Y345 Q9Y385 Q9Y394 Q9Y397 Q9Y3A0 Q9Y3A6 Q9Y3B3 Q9Y3D6 Q9Y3E0 Q9Y3E5 Q9Y3N9 Q9Y3P4 Q9Y3P8 Q9Y3Q0 Q9Y3Q3 Q9Y3Q4 Q9Y3Q7 Q9Y426 Q9Y487 Q9Y493 Q9Y4A9 Q9Y4C0 Q9Y4C5 Q9Y4D2 Q9Y4D7 Q9Y4K0 Q9Y4L1 Q9Y4P3 Q9Y4W6 Q9Y512 Q9Y519 Q9Y548 Q9Y561 Q9Y581 Q9Y584 Q9Y585 Q9Y5C1 Q9Y5E1 Q9Y5E2 Q9Y5E3 Q9Y5E4 Q9Y5E5 Q9Y5E6 Q9Y5E7 Q9Y5E8 Q9Y5E9 Q9Y5F0 Q9Y5F1 Q9Y5F2 Q9Y5F3 Q9Y5F6 Q9Y5F7 Q9Y5F8 Q9Y5F9 Q9Y5G0 Q9Y5G1 Q9Y5G2 Q9Y5G3 Q9Y5G4 Q9Y5G5 Q9Y5G6 Q9Y5G7 Q9Y5G8 Q9Y5G9 Q9Y5H0 Q9Y5H1 Q9Y5H2 Q9Y5H3 Q9Y5H4 Q9Y5H5 Q9Y5H6 Q9Y5H7 Q9Y5H8 Q9Y5H9 Q9Y5I0 Q9Y5I1 Q9Y5I2 Q9Y5I3 Q9Y5I4 Q9Y5I7 Q9Y5L2 Q9Y5L3 Q9Y5M8 Q9Y5N1 Q9Y5P0 Q9Y5P1 Q9Y5Q0 Q9Y5Q5 Q9Y5Q6 Q9Y5R2 Q9Y5S1 Q9Y5S8 Q9Y5T4 Q9Y5U4 Q9Y5U5 Q9Y5U8 Q9Y5U9 Q9Y5W5 Q9Y5W7 Q9Y5W8 Q9Y5X5 Q9Y5X9 Q9Y5Y0 Q9Y5Y3 Q9Y5Y4 Q9Y5Y5 Q9Y5Y6 Q9Y5Y7 Q9Y5Y9 Q9Y5Z0 Q9Y5Z6 Q9Y5Z9 Q9Y619 Q9Y624 Q9Y625 Q9Y639 Q9Y644 Q9Y646 Q9Y653 Q9Y661 Q9Y662 Q9Y663 Q9Y666 Q9Y672 Q9Y673 Q9Y679 Q9Y680 Q9Y691 Q9Y693 Q9Y694 Q9Y698 Q9Y6A1 Q9Y6A2 Q9Y6A9 Q9Y6B6 Q9Y6C5 Q9Y6C9 Q9Y6D0 Q9Y6F6 Q9Y6F9 Q9Y6G1 Q9Y6H1 Q9Y6H6 Q9Y6H8 Q9Y6I8 Q9Y6I9 Q9Y6J6 Q9Y6K0 Q9Y6L6 Q9Y6L7 Q9Y6M0 Q9Y6M5 Q9Y6M7 Q9Y6N1 Q9Y6N6 Q9Y6N7 Q9Y6N8 Q9Y6Q6 Q9Y6R1 Q9Y6U7 Q9Y6W8 Q9Y6X1 Q9Y6X5 Q9Y6Y9 U3KPV4 V9GZ13 W5XKT8

#### 1.2 The annotated transmembrane proteins in the UniProt (5199 sequences)

A0A075B734 A0A087WTH1 A0A087WTH5 A0A087X1C5 A0A096LP01 A0A096LPK9 A0A0B4J2F0 A0A0D9SF12 A0A0K2S4Q6 A0A0U1RQS6 A0A0U1RRA0 A0A0U1RRN3 A0A0X1KG70 A0A1B0GTB2 A0A1B0GTI8 A0A1B0GTK4 A0A1B0GTQ4 A0A1B0GTU2 A0A1B0GTW7 A0A1B0GTY4 A0A1B0GU29 A0A1B0GUA5 A0A1B0GUA7 A0A1B0GUW6 A0A1B0GUW7 A0A1B0GV85 A0A1B0GV90 A0A1B0GVN3 A0A1B0GVQ0 A0A1B0GVT2 A0A1B0GVV1 A0A1B0GVY4 A0A1B0GVZ9 A0A1B0GW54 A0A1B0GW64 A0A1B0GWB2 A0A1B0GWG4 A0A1W2PS18 A0A286YF18 A0A286YF58 A0A286YFK9 A0A2R8YCJ5 A0A494BZU4 A0A5B9 A0A5F9ZH02 A0A6I8PU40 A0AV02 A0AVI2 A0AVI4 A0FGR8 A0FGR9 A0PJK1 A0PJW6 A0PJX4 A0PJX8 A0PJZ3 A0PK00 A0PK05 A0PK11 A0ZSE6 A1A5B4 A1A5C7 A1KXE4 A1L0T0 A1L157 A1L1A6 A1L3X0 A2A2V5 A2A2Y4 A2RRL7 A2RU14 A2RU48 A2RU67 A2RUG3 A2RUT3 A2VDJ0 A3KFT3 A4D0T7 A4D1S0 A4D256 A4D2G3 A4D2H0 A4FU28 A4IF30 A5D6W6 A5PLK6 A5PLL7 A5X5Y0 A6BM72 A6H8M9 A6NC51 A6NC97 A6NCI5 A6NCQ9 A6NCV1 A6ND48 A6NDA9 A6NDD5 A6NDH6 A6NDL8 A6NDP7 A6NDV4 A6NDX4 A6NEH6 A6NET4 A6NF34 A6NF89 A6NFA0 A6NFA1 A6NFC5 A6NFC9 A6NFE2 A6NFR6 A6NFU0 A6NFX1 A6NFY4 A6NG13 A6NGA9 A6NGB0 A6NGB7 A6NGC4 A6NGU5 A6NGY5 A6NGZ8 A6NH00 A6NH21 A6NH52 A6NHA9 A6NHG9 A6NHS7 A6NI61 A6NI73 A6NIJ9 A6NIM6 A6NJU9 A6NJW4 A6NJW9 A6NJY1 A6NJY4 A6NJZ3 A6NK97 A6NKB5 A6NKF7 A6NKG5 A6NKK0 A6NKL6 A6NKW6 A6NKX4 A6NL05 A6NL08

A6NL26 A6NL88 A6NL99 A6NLE4 A6NLU5 A6NLX4 A6NM03 A6NM10 A6NM11 A6NM45 A6NM62
A6NM76 A6NMB1 A6NMD0 A6NML5 A6NMS3 A6NMS7 A6NMU1 A6NMZ5 A6NN92 A6NNB3 A6NNC1
A6NND4 A6NNE9 A6NNN8 A6PVL3 A6QL63 A7MBM2 A8CG34 A8K4G0 A8MPY1 A8MRT5 A8MTT3
A8MUP6 A8MV81 A8MVS5 A8MVW0 A8MVW5 A8MVZ5 A8MWK0 A8MWL6 A8MWL7 A8MWV9 A8MWY0
A8MXE2 A8MXK1 A8MXV6 A8MYB1 A8MYU2 A8MZ97 A9Z1Z3 B0FP48 B0L3A2 B0YJ81 B1AL88
B2RN74 B2RTY4 B2RUZ4 B2RXF0 B3SHH9 B4DJY2 B4DS77 B4DYI2 B6A8C7 B6SEH8 B6SEH9
B7U540 B7Z8K6 B7ZAQ6 B8ZZ34 B9EJG8 C9JDP6 C9JG80 C9JH25 C9JI98 C9JQI7 C9JQL5
C9JVW0 D3W0D1 E0CX11 E2RYF6 E5RHQ5 E5RIL1 E7ERA6 E9PQ53 E9PQX1 F2Z333 F5H4A9
F8W0I5 G3V0H7 H0YL14 H3BR10 H3BS89 H3BV60 H7C241 H7C350 I3L273 K7EJ46 M0QZC1
O00124 O00144 O00155 O00168 O00180 O00198 O00206 O00219 O00220 O00222 O00237
O00238 O00241 O00254 O00258 O00264 O00270 O00299 O00322 O00337 O00341 O00391
O00398 O00400 O00421 O00445 O00453 O00461 O00468 O00476 O00478 O00481 O00483
O00501 O00519 O00526 O00533 O00548 O00555 O00559 O00574 O00587 O00590 O00591
O00592 O00623 O00624 O00631 O00767 O14493 O14494 O14495 O14511 O14514 O14520
O14521 O14522 O14523 O14524 O14525 O14569 O14581 O14609 O14626 O14638 O14649
O14653 O14662 O14668 O14669 O14672 O14678 O14681 O14683 O14684 O14718 O14735
O14763 O14764 O14786 O14788 O14804 O14817 O14828 O14836 O14842 O14843 O14863
O14880 O14894 O14917 O14925 O14931 O14944 O14949 O14957 O14967 O14975 O14983
O15031 O15079 O15118 O15120 O15121 O15126 O15127 O15146 O15155 O15162 O15165
O15173 O15197 O15218 O15229 O15239 O15243 O15244 O15245 O15247 O15258 O15260
O15269 O15270 O15303 O15321 O15342 O15354 O15374 O15375 O15389 O15393 O15394
O15399 O15400 O15403 O15427 O15431 O15432 O15438 O15439 O15440 O15455 O15466
O15482 O15503 O15529 O15533 O15547 O15551 O15552 O15554 O43155 O43157 O43169
O43173 O43184 O43193 O43194 O43246 O43286 O43291 O43292 O43300 O43306 O43315
O43424 O43451 O43462 O43464 O43490 O43493 O43497 O43505 O43506 O43508 O43511
O43520 O43525 O43526 O43529 O43556 O43557 O43561 O43567 O43570 O43581 O43603
O43613 O43614 O43657 O43674 O43676 O43677 O43688 O43699 O43731 O43736 O43749
O43752 O43759 O43760 O43761 O43772 O43808 O43819 O43825 O43826 O43861 O43868
O43869 O43889 O43908 O43909 O43914 O43916 O43934 O60235 O60238 O60241 O60242
O60243 O60245 O60266 O60279 O60309 O60312 O60313 O60320 O60330 O60337 O60353
O60359 O60391 O60403 O60404 O60412 O60423 O60427 O60431 O60443 O60449 O60462
O60469 O60476 O60478 O60486 O60487 O60488 O60499 O60500 O60503 O60507 O60512
O60513 O60602 O60603 O60635 O60636 O60637 O60656 O60667 O60669 O60704 O60706
O60721 O60725 O60733 O60741 O60755 O60774 O60779 O60830 O60831 O60840 O60858
O60883 O60894 O60895 O60896 O60906 O60909 O60928 O60931 O60939 O75019 O75022
O75023 O75027 O75051 O75054 O75056 O75063 O75069 O75072 O75074 O75077 O75078
O75084 O75096 O75110 O75121 O75129 O75144 O75185 O75192 O75197 O75204 O75264
O75298 O75309 O75310 O75311 O75324 O75325 O75352 O75354 O75355 O75379 O75387
O75388 O75396 O75425 O75438 O75445 O75452 O75460 O75473 O75477 O75503 O75508
O75509 O75578 O75581 O75631 O75712 O75746 O75751 O75752 O75762 O75783 O75787
O75795 O75829 O75841 O75844 O75845 O75871 O75880 O75881 O75882 O75899 O75900
O75907 O75908 O75911 O75915 O75923 O75949 O75954 O75976 O76000 O76001 O76002
O76024 O76036 O76062 O76082 O76090 O76095 O76099 O76100 O94759 O94766 O94777
O94778 O94823 O94826 O94856 O94876 O94886 O94898 O94901 O94905 O94910 O94911

O94923 O94933 O94956 O94966 O94985 O94991 O95006 O95007 O95013 O95047 O95069
O95070 O95136 O95139 O95140 O95150 O95159 O95167 O95168 O95169 O95180 O95183
O95185 O95196 O95197 O95202 O95206 O95210 O95214 O95221 O95222 O95237 O95249
O95255 O95256 O95258 O95259 O95264 O95279 O95292 O95297 O95298 O95342 O95371
O95377 O95395 O95406 O95415 O95427 O95436 O95452 O95461 O95470 O95471 O95473
O95476 O95477 O95484 O95490 O95500 O95502 O95528 O95562 O95563 O95573 O95622
O95665 O95672 O95674 O95727 O95754 O95772 O95800 O95803 O95807 O95832 O95833
O95838 O95847 O95857 O95858 O95859 O95864 O95866 O95870 O95907 O95918 O95944
O95976 O95977 O95992 O96002 O96005 O96008 O96011 O96024 P00156 P00167 P00395
P00403 P00414 P00533 P00846 P01130 P01133 P01135 P01375 P01589 P01730 P01732
P01833 P01848 P01850 P01871 P01880 P01889 P01893 P01903 P01906 P01909 P01911
P01920 P02708 P02724 P02730 P02748 P02786 P03372 P03886 P03891 P03897 P03901
P03905 P03915 P03923 P03928 P03986 P03999 P04000 P04001 P04035 P04201 P04233
P04234 P04439 P04440 P04626 P04629 P04839 P04843 P04844 P04920 P04921 P05023
P05026 P05067 P05106 P05107 P05141 P05187 P05362 P05496 P05538 P05556 P05981
P06028 P06126 P06127 P06133 P06213 P06340 P06729 P06734 P06756 P07099 P07202
P07204 P07306 P07307 P07333 P07357 P07359 P07510 P07550 P07766 P07949 P08034
P08069 P08100 P08138 P08172 P08173 P08183 P08195 P08247 P08473 P08514 P08574
P08575 P08581 P08588 P08637 P08648 P08684 P08842 P08887 P08908 P08910 P08912
P08913 P08922 P08962 P08F94 P09131 P09172 P09564 P09601 P09603 P09619 P09669
P09693 P09758 P09848 P09912 P09958 P0C2L3 P0C2S0 P0C604 P0C617 P0C623 P0C626
P0C628 P0C629 P0C645 P0C646 P0C672 P0C6S8 P0C6T2 P0C7M8 P0C7N1 P0C7N4 P0C7N5
P0C7N8 P0C7P4 P0C7Q5 P0C7Q6 P0C7T2 P0C7T3 P0C7T8 P0C7U0 P0C7U3 P0C7U9 P0C7V7
P0C851 P0C874 P0CF51 P0CG08 P0CG41 P0CK96 P0CK97 P0DI80 P0DJ07 P0DJ93 P0DKB5
P0DKB6 P0DKV0 P0DKX4 P0DL12 P0DMQ5 P0DMS8 P0DMS9 P0DMT0 P0DMU2 P0DN25 P0DN77
P0DN78 P0DN80 P0DN81 P0DN82 P0DN84 P0DP42 P0DP72 P0DPA2 P0DPD6 P0DPD8 P0DPE3
P0DQD5 P0DSN6 P0DTE0 P0DTE4 P0DTE5 P0DTF9 P0DTL5 P0DTU3 P0DTU4 P10176 P10321
P10415 P10586 P10620 P10721 P10747 P10912 P10966 P11049 P11117 P11166 P11168
P11169 P11215 P11229 P11230 P11279 P11362 P11511 P11717 P11836 P11912 P12074
P12235 P12236 P12314 P12318 P12319 P12821 P12830 P13073 P13164 P13224 P13473
P13569 P13591 P13598 P13612 P13637 P13688 P13726 P13747 P13762 P13765 P13866
P13945 P14060 P14151 P14209 P14222 P14406 P14410 P14415 P14416 P14616 P14672
P14679 P14770 P14778 P14784 P14867 P15144 P15151 P15260 P15291 P15382 P15391
P15421 P15509 P15514 P15529 P15812 P15813 P15907 P15941 P15954 P16066 P16070
P16109 P16144 P16150 P16234 P16260 P16284 P16389 P16410 P16422 P16435 P16442
P16471 P16473 P16581 P16615 P16662 P16671 P16871 P17152 P17181 P17301 P17302
P17342 P17643 P17658 P17693 P17787 P17813 P17861 P17927 P17948 P18084 P18089
P18405 P18433 P18505 P18507 P18564 P18577 P18627 P18825 P18827 P18850 P19021
P19022 P19075 P19224 P19235 P19256 P19320 P19397 P19438 P19440 P19526 P19634
P20020 P20023 P20036 P20138 P20273 P20292 P20309 P20333 P20594 P20645 P20648
P20701 P20702 P20916 P20963 P21145 P21217 P21397 P21439 P21452 P21453 P21462
P21554 P21579 P21583 P21709 P21728 P21730 P21731 P21754 P21757 P21796 P21802
P21817 P21854 P21860 P21917 P21918 P21926 P21964 P22001 P22083 P22223 P22309
P22310 P22413 P22455 P22459 P22460 P22607 P22680 P22732 P22760 P22794 P22888

P22897 P23229 P23276 P23415 P23416 P23467 P23468 P23469 P23470 P23471 P23510
P23634 P23763 P23942 P23945 P23975 P24046 P24071 P24310 P24311 P24390 P24394
P24530 P24557 P25021 P25024 P25025 P25089 P25090 P25092 P25100 P25101 P25103
P25105 P25106 P25116 P25189 P25445 P25874 P25929 P25942 P26006 P26010 P26012
P26439 P26572 P26678 P26715 P26717 P26718 P26842 P26951 P27037 P27338 P27449
P27487 P27544 P27701 P27824 P27930 P28067 P28068 P28221 P28222 P28223 P28288
P28328 P28335 P28336 P28472 P28476 P28566 P28827 P28845 P28906 P28907 P28908
P29016 P29017 P29033 P29274 P29275 P29317 P29320 P29322 P29323 P29371 P29376
P29965 P29972 P29973 P30203 P30273 P30301 P30408 P30411 P30511 P30518 P30519
P30530 P30531 P30532 P30536 P30542 P30550 P30556 P30559 P30825 P30872 P30874
P30926 P30939 P30953 P30954 P30968 P30988 P30989 P31213 P31391 P31431 P31512
P31513 P31639 P31641 P31644 P31645 P31785 P31994 P31995 P32004 P32238 P32239
P32241 P32245 P32246 P32247 P32248 P32249 P32297 P32302 P32418 P32745 P32856
P32926 P32927 P32942 P32970 P32971 P33032 P33121 P33151 P33527 P33681 P33897
P33908 P33947 P34741 P34810 P34903 P34910 P34925 P34969 P34972 P34981 P34982
P34995 P34998 P35070 P35212 P35346 P35348 P35367 P35368 P35372 P35408 P35410
P35414 P35462 P35498 P35499 P35503 P35504 P35523 P35575 P35590 P35610 P35613
P35670 P35916 P35968 P36021 P36269 P36382 P36383 P36537 P36544 P36888 P36894
P36896 P36897 P36941 P36956 P37023 P37059 P37088 P37173 P37268 P37287 P37288
P38435 P38484 P38570 P39086 P39210 P39656 P40126 P40145 P40189 P40197 P40198
P40200 P40238 P40259 P40305 P40879 P40967 P41143 P41145 P41146 P41180 P41181
P41217 P41231 P41273 P41440 P41586 P41587 P41594 P41595 P41597 P41732 P41968
P42081 P42261 P42262 P42263 P42658 P42701 P42702 P42857 P42892 P43003 P43004
P43005 P43007 P43088 P43115 P43116 P43119 P43121 P43146 P43220 P43307 P43308
P43489 P43626 P43627 P43628 P43629 P43630 P43631 P43632 P43657 P43681 P45844
P45880 P46059 P46089 P46091 P46092 P46093 P46094 P46095 P46098 P46531 P46663
P46695 P46721 P46977 P47211 P47775 P47804 P47869 P47870 P47871 P47872 P47881
P47883 P47884 P47887 P47888 P47890 P47893 P47898 P47900 P47901 P47985 P48023
P48029 P48039 P48048 P48050 P48051 P48058 P48060 P48065 P48066 P48067 P48145
P48146 P48165 P48167 P48169 P48201 P48230 P48357 P48509 P48544 P48546 P48547
P48549 P48551 P48651 P48664 P48751 P48764 P48960 P48995 P49019 P49069 P49146
P49190 P49238 P49257 P49279 P49281 P49286 P49326 P49447 P49641 P49682 P49683
P49685 P49755 P49768 P49771 P49788 P49810 P49895 P49961 P50052 P50281 P50391
P50402 P50406 P50416 P50443 P50591 P50851 P50876 P50895 P50993 P51164 P51168
P51170 P51172 P51511 P51512 P51571 P51572 P51575 P51582 P51648 P51674 P51677
P51679 P51681 P51684 P51685 P51686 P51693 P51787 P51788 P51790 P51793 P51795
P51797 P51798 P51800 P51801 P51805 P51809 P51810 P51811 P51828 P51841 P51993
P52429 P52569 P52799 P52848 P52849 P53007 P53708 P53794 P53801 P53816 P53985
P54219 P54289 P54707 P54709 P54710 P54753 P54756 P54760 P54762 P54764 P54829
P54849 P54851 P54852 P54855 P55011 P55017 P55061 P55064 P55073 P55082 P55085
P55087 P55160 P55283 P55285 P55286 P55287 P55289 P55291 P55344 P55808 P55851
P55899 P55916 P56134 P56180 P56199 P56373 P56378 P56557 P56589 P56696 P56746
P56747 P56748 P56749 P56750 P56817 P56856 P56880 P56937 P56962 P56975 P57054
P57057 P57087 P57088 P57103 P57105 P57679 P57727 P57738 P57739 P57764 P57773

P57789 P58170 P58173 P58180 P58181 P58182 P58335 P58418 P58511 P58549 P58550
P58658 P58743 P58872 P59025 P59533 P59534 P59535 P59536 P59537 P59538 P59539
P59540 P59541 P59542 P59543 P59544 P59551 P59646 P59773 P59901 P59922 P60033
P60059 P60201 P60468 P60507 P60508 P60509 P60602 P60606 P60852 P60893 P61009
P61073 P61165 P61266 P61550 P61565 P61619 P61647 P61803 P62079 P62341 P62952
P62955 P63027 P63252 P67812 P69849 P78310 P78324 P78325 P78334 P78348 P78357
P78363 P78369 P78380 P78381 P78382 P78383 P78410 P78423 P78504 P78508 P78536
P78552 P78562 P79483 P80370 P81408 P81605 P82251 P82279 P84157 P98073 P98153
P98155 P98161 P98164 P98172 P98187 P98194 P98196 P98198 Q00325 Q00765 Q00973
Q00975 Q01113 Q01118 Q01151 Q01344 Q01362 Q01453 Q01628 Q01629 Q01638 Q01650
Q01668 Q01718 Q01726 Q01740 Q01814 Q01959 Q01973 Q01974 Q02094 Q02127 Q02161
Q02221 Q02223 Q02297 Q02413 Q02487 Q02505 Q02643 Q02742 Q02763 Q02846 Q02978
Q03001 Q03167 Q03395 Q03431 Q03518 Q03519 Q03721 Q04609 Q04656 Q04671 Q04721
Q04771 Q04844 Q04900 Q04912 Q04941 Q05586 Q05901 Q05940 Q05996 Q06055 Q06136
Q06418 Q06432 Q06481 Q06495 Q06643 Q07001 Q07011 Q07065 Q07075 Q07108 Q07326
Q07444 Q07699 Q07812 Q07817 Q07820 Q07837 Q07954 Q08174 Q08334 Q08345 Q08357
Q08462 Q08477 Q08554 Q08708 Q08722 Q08828 Q08AI6 Q08ET2 Q09013 Q09327 Q09328
Q09428 Q09470 Q0D2K0 Q0GE19 Q0P670 Q0P6D2 Q0P6H9 Q0VAQ4 Q0VDE8 Q0VDI3 Q10469
Q10471 Q10472 Q10589 Q10981 Q11128 Q11130 Q11201 Q11203 Q11206 Q12767 Q12770
Q12772 Q12791 Q12797 Q12809 Q12836 Q12846 Q12864 Q12866 Q12879 Q12884 Q12887
Q12893 Q12907 Q12908 Q12912 Q12913 Q12918 Q12981 Q12983 Q12999 Q13002 Q13003
Q13018 Q13021 Q13061 Q13113 Q13145 Q13183 Q13190 Q13224 Q13241 Q13255 Q13258
Q13261 Q13277 Q13286 Q13291 Q13304 Q13308 Q13323 Q13324 Q13326 Q13332 Q13336
Q13349 Q13370 Q13410 Q13423 Q13433 Q13443 Q13444 Q13445 Q13454 Q13467 Q13477
Q13478 Q13488 Q13491 Q13505 Q13507 Q13520 Q13530 Q13563 Q13571 Q13585 Q13586
Q13591 Q13606 Q13607 Q13621 Q13634 Q13635 Q13639 Q13641 Q13651 Q13683 Q13698
Q13705 Q13724 Q13733 Q13740 Q13797 Q13873 Q13936 Q14003 Q14028 Q14108 Q14114
Q14118 Q14126 Q14162 Q14165 Q14242 Q14246 Q14318 Q14330 Q14332 Q14392 Q14410
Q14416 Q14432 Q14435 Q14439 Q14494 Q14500 Q14517 Q14524 Q14542 Q14571 Q14573
Q14574 Q14626 Q14627 Q14643 Q14654 Q14656 Q14703 Q14714 Q14721 Q14728 Q14739
Q14761 Q14773 Q14789 Q147U7 Q14802 Q14831 Q14832 Q14833 Q14849 Q14916 Q14940
Q14943 Q14952 Q14953 Q14954 Q14956 Q14957 Q14973 Q14BN4 Q14C87 Q14CN2 Q14CX5
Q14CZ8 Q14D33 Q14DG7 Q15005 Q15011 Q15012 Q15035 Q15041 Q15043 Q15049 Q15053
Q15070 Q15077 Q15109 Q15116 Q15125 Q15155 Q15223 Q15256 Q15262 Q15303 Q15363
Q15375 Q15388 Q15391 Q15392 Q15399 Q15413 Q15526 Q15546 Q15612 Q15615 Q15617
Q15619 Q15620 Q15622 Q15629 Q15722 Q15738 Q15743 Q15758 Q15760 Q15761 Q15762
Q15768 Q15800 Q15822 Q15825 Q15836 Q15842 Q15849 Q15858 Q15878 Q15884 Q15904
Q16099 Q16280 Q16281 Q16288 Q16322 Q16348 Q16394 Q16445 Q16478 Q16515 Q16538
Q16549 Q16558 Q16563 Q16570 Q16572 Q16581 Q16585 Q16586 Q16602 Q16611 Q16617
Q16620 Q16623 Q16625 Q16647 Q16651 Q16653 Q16655 Q16671 Q16706 Q16720 Q16739
Q16790 Q16799 Q16819 Q16820 Q16821 Q16827 Q16832 Q16842 Q16849 Q16850 Q16853
Q16873 Q16880 Q16891 Q17R55 Q17RQ9 Q19T08 Q1AE95 Q1EHB4 Q1HG43 Q1HG44 Q24JP5
Q24JQ0 Q29980 Q29983 Q2HXU8 Q2I0M4 Q2KHT4 Q2LD37 Q2M2E3 Q2M2H8 Q2M385 Q2M3C6
Q2M3G0 Q2M3M2 Q2M3R5 Q2M3T9 Q2PZI1 Q2QL34 Q2T9K0 Q2TAA5 Q2TBF2 Q2VWP7 Q2VYF4

Q2WGJ8 Q2WGJ9 Q2Y0W8 Q30154 Q30201 Q32M45 Q32ZL2 Q3B7S5 Q3B7T3 Q3C1V0 Q3KNS1
Q3KNT9 Q3KNW5 Q3KP22 Q3KPI0 Q3KQZ1 Q3KR37 Q3MIP1 Q3MIR4 Q3MIW9 Q3MIX3 Q3MUY2
Q3SXP7 Q3SXY7 Q3SY17 Q3SY77 Q3SYC2 Q3T906 Q3V5L5 Q3YBM2 Q3ZAQ7 Q3ZCQ3 Q3ZCQ8
Q401N2 Q494W8 Q495A1 Q495M3 Q495N2 Q495T6 Q495W5 Q496F6 Q496J9 Q49A17 Q49SQ1
Q4G0I0 Q4G0N0 Q4G0N8 Q4G0T1 Q4G148 Q4G1C9 Q4KMG0 Q4KMG9 Q4KMQ2 Q4KMZ8 Q4LDR2
Q4U2R8 Q4V9L6 Q4VC39 Q4VNC0 Q4VNC1 Q4VXA5 Q4VXF1 Q4W5P6 Q4ZG55 Q4ZIN3 Q4ZJI4
Q504Y0 Q50LG9 Q52LC2 Q53EL9 Q53EP0 Q53EU6 Q53F39 Q53FP2 Q53FV1 Q53GD3 Q53GQ0
Q53HI1 Q53R12 Q53RT3 Q53RY4 Q53S58 Q53TN4 Q567V2 Q587I9 Q58DX5 Q58EX2 Q58HT5
Q5BJD5 Q5BJF2 Q5BJH2 Q5BJH7 Q5BKT4 Q5BKX6 Q5BVD1 Q5DID0 Q5DX21 Q5EB52 Q5FWE3
Q5FYA8 Q5GH70 Q5GH72 Q5GH73 Q5GH76 Q5GH77 Q5H8A4 Q5H943 Q5H9E4 Q5H9R4 Q5H9S7
Q5HYA8 Q5HYJ1 Q5HYL7 Q5I7T1 Q5IJ48 Q5J8M3 Q5J8X5 Q5JPE7 Q5JQS5 Q5JRA6 Q5JRM2
Q5JRS4 Q5JRV8 Q5JTH9 Q5JTV8 Q5JUK3 Q5JW98 Q5JX69 Q5JX71 Q5JXA9 Q5JXX7 Q5JZY3
Q5K4L6 Q5KU26 Q5M7Z0 Q5M8T2 Q5MY95 Q5NUL3 Q5PT55 Q5QFB9 Q5QGT7 Q5QGZ9 Q5QJU3
Q5R3F8 Q5R3K3 Q5RGS3 Q5RI15 Q5SGD2 Q5SNT2 Q5SQ64 Q5SR56 Q5SRD1 Q5SRI9 Q5SRN2
Q5SSG8 Q5STR5 Q5SV17 Q5SVS4 Q5SWH9 Q5SWX8 Q5SY80 Q5SZI1 Q5SZK8 Q5T0T0 Q5T197
Q5T1A1 Q5T1Q4 Q5T1S8 Q5T292 Q5T2D2 Q5T2E6 Q5T3F8 Q5T3U5 Q5T442 Q5T4D3 Q5T4F4
Q5T4S7 Q5T4T1 Q5T601 Q5T6L9 Q5T6X4 Q5T6X5 Q5T700 Q5T7M9 Q5T7P6 Q5T7P8 Q5T7R7
Q5T848 Q5T8D3 Q5T9L3 Q5T9Z0 Q5TAH2 Q5TAT6 Q5TEA6 Q5TF21 Q5TF39 Q5TGI0 Q5TGU0
Q5TGY1 Q5TGZ0 Q5TH69 Q5TZ20 Q5TZJ5 Q5U3C3 Q5U4P2 Q5UAW9 Q5UCC4 Q5VSG8 Q5VT66
Q5VT99 Q5VTJ3 Q5VTT2 Q5VTY9 Q5VU36 Q5VU65 Q5VU97 Q5VUB5 Q5VUD6 Q5VUY2 Q5VV42
Q5VV43 Q5VV63 Q5VVB8 Q5VVP1 Q5VW36 Q5VW38 Q5VWC8 Q5VWK5 Q5VX71 Q5VXT5 Q5VXU1
Q5VY43 Q5VYJ5 Q5VYP0 Q5VZ72 Q5VZI3 Q5VZR4 Q5VZY2 Q5W0B7 Q5W0N0 Q5W0Z9 Q5XG99
Q5XKP0 Q5XXA6 Q5ZPR3 Q629K1 Q63HM2 Q63ZE4 Q643R3 Q658N2 Q658P3 Q66K14 Q66K66
Q674R7 Q685J3 Q687X5 Q68CJ9 Q68CP4 Q68CQ1 Q68CQ7 Q68CR1 Q68CR7 Q68D42 Q68D85
Q68DH5 Q68DV7 Q68G75 Q695T7 Q69YG0 Q69YU5 Q69YW2 Q69YZ2 Q6AI14 Q6AZY7 Q6DD88
Q6DKI7 Q6DN12 Q6DN14 Q6DN72 Q6DWJ6 Q6E213 Q6EIG7 Q6EMK4 Q6GMR7 Q6GPH6 Q6GTX8
Q6GV28 Q6H3X3 Q6IA17 Q6IAN0 Q6IC98 Q6ICH7 Q6ICI0 Q6ICL7 Q6IEE7 Q6IEE8 Q6IEU7
Q6IEV9 Q6IEY1 Q6IEZ7 Q6IF00 Q6IF36 Q6IF42 Q6IF63 Q6IF82 Q6IF99 Q6IFG1 Q6IFH4
Q6IFN5 Q6IS24 Q6ISU1 Q6IWH7 Q6J4K2 Q6J9G0 Q6KCM7 Q6L9W6 Q6MZM0 Q6N022 Q6N075
Q6NSJ0 Q6NSJ5 Q6NT16 Q6NT55 Q6NTF9 Q6NUI2 Q6NUJ2 Q6NUK1 Q6NUK4 Q6NUQ4 Q6NUS6
Q6NUS8 Q6NUT2 Q6NUT3 Q6NV75 Q6NVV3 Q6NXN4 Q6NXT4 Q6NXT6 Q6NZ63 Q6P179 Q6P1A2
Q6P1J6 Q6P1K1 Q6P1M0 Q6P1Q0 Q6P1S2 Q6P2H8 Q6P499 Q6P4A7 Q6P4E1 Q6P4F1 Q6P4H8
Q6P4Q7 Q6P531 Q6P5S7 Q6P5W5 Q6P5X7 Q6P7N7 Q6P995 Q6P9A2 Q6P9B9 Q6P9F7 Q6P9G4
Q6PCB7 Q6PCB8 Q6PEX7 Q6PEY0 Q6PEY1 Q6PHW0 Q6PI25 Q6PI73 Q6PI78 Q6PIS1 Q6PIU1
Q6PIU2 Q6PIV7 Q6PIZ9 Q6PJF5 Q6PJG9 Q6PJW8 Q6PK18 Q6PKC3 Q6PL45 Q6PML9 Q6PP77
Q6PRD1 Q6PXP3 Q6Q0C1 Q6Q4G3 Q6Q8B3 Q6QAJ8 Q6QHC5 Q6QNK2 Q6RW13 Q6T423 Q6T4P5
Q6TCH4 Q6TCH7 Q6U736 Q6U841 Q6UE05 Q6UVK1 Q6UVM3 Q6UVW9 Q6UVY6 Q6UW02 Q6UW56
Q6UW60 Q6UW68 Q6UW78 Q6UW88 Q6UWB1 Q6UWD8 Q6UWF3 Q6UWF5 Q6UWH4 Q6UWH6 Q6UWI2
Q6UWI4 Q6UWJ1 Q6UWJ8 Q6UWL2 Q6UWL6 Q6UWM7 Q6UWM9 Q6UWP7 Q6UWU4 Q6UWV2 Q6UWV6
Q6UWV7 Q6UWW9 Q6UX01 Q6UX15 Q6UX27 Q6UX34 Q6UX40 Q6UX41 Q6UX65 Q6UX68 Q6UX71
Q6UX72 Q6UX98 Q6UXB4 Q6UXC1 Q6UXD1 Q6UXD5 Q6UXD7 Q6UXE8 Q6UXF1 Q6UXG2 Q6UXG3
Q6UXG8 Q6UXK2 Q6UXK5 Q6UXL0 Q6UXM1 Q6UXN7 Q6UXN8 Q6UXP3 Q6UXU4 Q6UXU6 Q6UXV0
Q6UXV1 Q6UXV4 Q6UXY8 Q6UXZ0 Q6UXZ3 Q6UXZ4 Q6UY09 Q6UY11 Q6UY18 Q6V0I7 Q6V0L0
Q6V1P9 Q6W3E5 Q6W5P4 Q6XPS3 Q6XR72 Q6XYQ8 Q6Y1H2 Q6Y288 Q6Y2X3 Q6YBV0 Q6YI46

Q6ZMB0 Q6ZMB5 Q6ZMC9 Q6ZMD2 Q6ZMG9 Q6ZMH5 Q6ZMI3 Q6ZMJ2 Q6ZMQ8 Q6ZMR5 Q6ZMZ0
Q6ZMZ3 Q6ZN44 Q6ZN68 Q6ZNA5 Q6ZNB6 Q6ZNB7 Q6ZNC8 Q6ZNI0 Q6ZNR0 Q6ZP29 Q6ZP80
Q6ZPB5 Q6ZPD8 Q6ZPD9 Q6ZQN7 Q6ZQQ2 Q6ZRH7 Q6ZRP7 Q6ZRR5 Q6ZS10 Q6ZS62 Q6ZS82
Q6ZSA7 Q6ZSJ9 Q6ZSM3 Q6ZSS7 Q6ZSY5 Q6ZT12 Q6ZT21 Q6ZT89 Q6ZTQ4 Q6ZU64 Q6ZU69
Q6ZUB0 Q6ZUB1 Q6ZUK4 Q6ZUT9 Q6ZUX7 Q6ZV29 Q6ZVE7 Q6ZVK1 Q6ZVL6 Q6ZVX9 Q6ZW05
Q6ZWK4 Q6ZWK6 Q6ZWL3 Q6ZWT7 Q6ZXV5 Q70CQ3 Q70HW3 Q70JA7 Q70SY1 Q70UQ0 Q70Z44
Q71H61 Q71RC9 Q71RG4 Q71RH2 Q71RS6 Q75NE6 Q75T13 Q75V66 Q76EJ3 Q76KP1 Q76MJ5
Q7KYR7 Q7KZN9 Q7L0J3 Q7L0L9 Q7L0X0 Q7L1I2 Q7L1S5 Q7L1W4 Q7L211 Q7L311 Q7L4E1
Q7L4S7 Q7L5A8 Q7L5L3 Q7L5N7 Q7L8C5 Q7L985 Q7LBE3 Q7LFX5 Q7LGA3 Q7LGC8 Q7RTM1
Q7RTP0 Q7RTR8 Q7RTS5 Q7RTS6 Q7RTT9 Q7RTX0 Q7RTX1 Q7RTX7 Q7RTX9 Q7RTY0 Q7RTY1
Q7RTY8 Q7Z2D5 Q7Z2H8 Q7Z2K6 Q7Z2Q7 Q7Z2W7 Q7Z388 Q7Z3B0 Q7Z3C6 Q7Z3D4 Q7Z3F1
Q7Z3J2 Q7Z3Q1 Q7Z3S7 Q7Z3T1 Q7Z402 Q7Z403 Q7Z404 Q7Z407 Q7Z408 Q7Z410 Q7Z412
Q7Z418 Q7Z419 Q7Z429 Q7Z434 Q7Z442 Q7Z443 Q7Z449 Q7Z4F1 Q7Z4J2 Q7Z4L0 Q7Z4N2
Q7Z4T8 Q7Z5B4 Q7Z5H4 Q7Z5H5 Q7Z5J8 Q7Z5M5 Q7Z5N4 Q7Z5S9 Q7Z601 Q7Z602 Q7Z692
Q7Z695 Q7Z6A9 Q7Z6J6 Q7Z6L0 Q7Z6M3 Q7Z6W1 Q7Z769 Q7Z7B1 Q7Z7D3 Q7Z7H5 Q7Z7J7
Q7Z7M0 Q7Z7M1 Q7Z7M8 Q7Z7M9 Q7Z7N9 Q86SF2 Q86SJ2 Q86SJ6 Q86SK9 Q86SM5 Q86SM8
Q86SP6 Q86SQ3 Q86SQ4 Q86SQ6 Q86SR1 Q86SS6 Q86SU0 Q86T03 Q86T13 Q86T20 Q86T26
Q86T96 Q86TG1 Q86TL2 Q86TM6 Q86TY3 Q86U02 Q86UB9 Q86UD3 Q86UD5 Q86UE4 Q86UE6
Q86UF1 Q86UF2 Q86UG4 Q86UK0 Q86UK5 Q86UL3 Q86UP0 Q86UP2 Q86UP6 Q86UP9 Q86UQ4
Q86UQ5 Q86UW1 Q86UW2 Q86V24 Q86V35 Q86V40 Q86V85 Q86VB7 Q86VD7 Q86VD9 Q86VE9
Q86VF5 Q86VH4 Q86VH5 Q86VI4 Q86VL8 Q86VR2 Q86VR7 Q86VU5 Q86VW1 Q86VY9 Q86VZ1
Q86VZ4 Q86VZ5 Q86W10 Q86W33 Q86W47 Q86W74 Q86WA9 Q86WB7 Q86WC4 Q86WI0 Q86WI1
Q86WK6 Q86WK7 Q86WK9 Q86WR0 Q86WS5 Q86WV6 Q86X19 Q86X29 Q86X52 Q86XE3 Q86XJ0
Q86XK7 Q86XL3 Q86XM0 Q86XQ3 Q86XR5 Q86XS8 Q86XT9 Q86XX4 Q86Y07 Q86Y22 Q86Y34
Q86Y38 Q86Y39 Q86Y82 Q86YA3 Q86YC3 Q86YD3 Q86YD5 Q86YJ5 Q86YL7 Q86YN1 Q86YT5
Q86YT9 Q86YW5 Q86Z14 Q8IU57 Q8IU68 Q8IU80 Q8IU89 Q8IU99 Q8IUA7 Q8IUC8 Q8IUH4
Q8IUH5 Q8IUH8 Q8IUK5 Q8IUN9 Q8IUR5 Q8IUS5 Q8IUW5 Q8IUX1 Q8IUY3 Q8IV01 Q8IV08
Q8IV31 Q8IV45 Q8IV77 Q8IVB4 Q8IVJ1 Q8IVJ8 Q8IVM8 Q8IVP5 Q8IVQ6 Q8IVU1 Q8IVV8
Q8IVW8 Q8IVY1 Q8IW00 Q8IW52 Q8IW70 Q8IWA4 Q8IWA5 Q8IWB1 Q8IWB4 Q8IWB9 Q8IWD5
Q8IWK6 Q8IWR1 Q8IWT1 Q8IWT6 Q8IWU2 Q8IWU4 Q8IWV1 Q8IWX5 Q8IWY9 Q8IX05 Q8IX19
Q8IX94 Q8IXA5 Q8IXB3 Q8IXE1 Q8IXF9 Q8IXH8 Q8IXI1 Q8IXI2 Q8IXK2 Q8IXL6 Q8IXM6
Q8IXU6 Q8IXX5 Q8IY17 Q8IY26 Q8IY34 Q8IY49 Q8IY50 Q8IY95 Q8IYJ0 Q8IYL9 Q8IYP9
Q8IYR6 Q8IYS0 Q8IYS2 Q8IYS5 Q8IYV9 Q8IZ08 Q8IZ52 Q8IZ57 Q8IZ96 Q8IZA0 Q8IZD6
Q8IZF0 Q8IZF2 Q8IZF3 Q8IZF4 Q8IZF5 Q8IZF6 Q8IZF7 Q8IZJ1 Q8IZK6 Q8IZM9 Q8IZN3
Q8IZP7 Q8IZP9 Q8IZR5 Q8IZS7 Q8IZS8 Q8IZT8 Q8IZU8 Q8IZU9 Q8IZV2 Q8IZV5 Q8IZY2
Q8J025 Q8MH63 Q8N0U2 Q8N0U8 Q8N0V5 Q8N0W4 Q8N0W7 Q8N0Y3 Q8N0Y5 Q8N0Z9 Q8N109
Q8N111 Q8N112 Q8N114 Q8N118 Q8N126 Q8N127 Q8N130 Q8N131 Q8N138 Q8N139 Q8N144
Q8N146 Q8N148 Q8N149 Q8N162 Q8N1C3 Q8N1L4 Q8N1M1 Q8N1N0 Q8N1N2 Q8N1S5 Q8N1Y9
Q8N201 Q8N205 Q8N271 Q8N292 Q8N2A8 Q8N2C7 Q8N2F6 Q8N2H4 Q8N2K0 Q8N2K1 Q8N2M4
Q8N2Q7 Q8N2U0 Q8N2U9 Q8N326 Q8N349 Q8N350 Q8N357 Q8N370 Q8N386 Q8N387 Q8N394
Q8N3F9 Q8N3G9 Q8N3J6 Q8N3S3 Q8N3T1 Q8N3T6 Q8N3Y3 Q8N3Y7 Q8N413 Q8N423 Q8N428
Q8N434 Q8N441 Q8N468 Q8N4A0 Q8N4C9 Q8N4F4 Q8N4F7 Q8N4H5 Q8N4K4 Q8N4L1 Q8N4L2
Q8N4L4 Q8N4M1 Q8N4S7 Q8N4S9 Q8N4V1 Q8N4V2 Q8N4W6 Q8N511 Q8N539 Q8N5B7 Q8N5C1
Q8N5D6 Q8N5G0 Q8N5G2 Q8N5K1 Q8N5M9 Q8N5S1 Q8N5U1 Q8N5Y8 Q8N608 Q8N609 Q8N614

Q8N628 Q8N661 Q8N682 Q8N695 Q8N697 Q8N6C5 Q8N6D2 Q8N6F1 Q8N6G5 Q8N6I4 Q8N6K0
Q8N6L0 Q8N6L1 Q8N6L7 Q8N6M3 Q8N6P7 Q8N6Q1 Q8N6R1 Q8N6S5 Q8N6U8 Q8N6Y1 Q8N743
Q8N755 Q8N766 Q8N7C0 Q8N7C4 Q8N7C7 Q8N7P1 Q8N7P3 Q8N7S6 Q8N7X8 Q8N808 Q8N816
Q8N8D7 Q8N8F6 Q8N8F7 Q8N8J7 Q8N8N0 Q8N8Q1 Q8N8Q8 Q8N8Q9 Q8N8R3 Q8N8V8 Q8N8Z6
Q8N912 Q8N966 Q8N967 Q8N9A8 Q8N9F0 Q8N9F7 Q8N9I0 Q8N9I5 Q8N9M5 Q8N9R8 Q8N9W7
Q8N9X5 Q8NA29 Q8NA58 Q8NAC3 Q8NAN2 Q8NAT1 Q8NAU1 Q8NB49 Q8NB59 Q8NBD8 Q8NBF6
Q8NBI2 Q8NBI5 Q8NBI6 Q8NBJ4 Q8NBJ9 Q8NBL3 Q8NBM4 Q8NBN3 Q8NBP5 Q8NBQ7 Q8NBR0
Q8NBS3 Q8NBT3 Q8NBU5 Q8NBV4 Q8NBV8 Q8NBW4 Q8NBZ7 Q8NC01 Q8NC24 Q8NC42 Q8NC44
Q8NC54 Q8NC56 Q8NC67 Q8NCC5 Q8NCG5 Q8NCG7 Q8NCH0 Q8NCK7 Q8NCL4 Q8NCL8 Q8NCL9
Q8NCM2 Q8NCQ3 Q8NCR0 Q8NCR9 Q8NCS4 Q8NCS7 Q8NCU8 Q8NCW0 Q8NCW6 Q8ND61 Q8ND94
Q8NDB6 Q8NDH2 Q8NDV1 Q8NDV2 Q8NDX2 Q8NDY8 Q8NDZ6 Q8NE00 Q8NE01 Q8NE79 Q8NE86
Q8NEA5 Q8NEB5 Q8NEC5 Q8NEN9 Q8NEQ5 Q8NER1 Q8NER5 Q8NES3 Q8NET5 Q8NET6 Q8NET8
Q8NEW0 Q8NEW7 Q8NF37 Q8NF91 Q8NFB2 Q8NFF2 Q8NFJ5 Q8NFJ6 Q8NFK1 Q8NFL0 Q8NFM4
Q8NFM7 Q8NFN8 Q8NFQ8 Q8NFR3 Q8NFR9 Q8NFT2 Q8NFT8 Q8NFU0 Q8NFU1 Q8NFY4 Q8NFZ3
Q8NFZ4 Q8NFZ6 Q8NFZ8 Q8NG04 Q8NG11 Q8NG75 Q8NG76 Q8NG77 Q8NG78 Q8NG80 Q8NG81
Q8NG83 Q8NG84 Q8NG85 Q8NG92 Q8NG94 Q8NG95 Q8NG97 Q8NG98 Q8NG99 Q8NGA0 Q8NGA1
Q8NGA2 Q8NGA4 Q8NGA5 Q8NGA6 Q8NGA8 Q8NGB2 Q8NGB4 Q8NGB6 Q8NGB8 Q8NGB9 Q8NGC0
Q8NGC1 Q8NGC2 Q8NGC3 Q8NGC4 Q8NGC5 Q8NGC6 Q8NGC7 Q8NGC8 Q8NGC9 Q8NGD0 Q8NGD1
Q8NGD2 Q8NGD3 Q8NGD4 Q8NGD5 Q8NGE0 Q8NGE1 Q8NGE2 Q8NGE3 Q8NGE5 Q8NGE7 Q8NGE8
Q8NGE9 Q8NGF0 Q8NGF1 Q8NGF3 Q8NGF4 Q8NGF6 Q8NGF7 Q8NGF8 Q8NGF9 Q8NGG0 Q8NGG1
Q8NGG2 Q8NGG3 Q8NGG4 Q8NGG5 Q8NGG6 Q8NGG7 Q8NGG8 Q8NGH3 Q8NGH5 Q8NGH6 Q8NGH7
Q8NGH8 Q8NGH9 Q8NGI0 Q8NGI1 Q8NGI2 Q8NGI3 Q8NGI4 Q8NGI6 Q8NGI7 Q8NGI8 Q8NGI9
Q8NGJ0 Q8NGJ1 Q8NGJ2 Q8NGJ3 Q8NGJ4 Q8NGJ5 Q8NGJ6 Q8NGJ7 Q8NGJ8 Q8NGJ9 Q8NGK0
Q8NGK1 Q8NGK2 Q8NGK3 Q8NGK4 Q8NGK5 Q8NGK6 Q8NGK9 Q8NGL0 Q8NGL1 Q8NGL2 Q8NGL3
Q8NGL4 Q8NGL6 Q8NGL7 Q8NGL9 Q8NGM1 Q8NGM8 Q8NGM9 Q8NGN0 Q8NGN1 Q8NGN2 Q8NGN3
Q8NGN4 Q8NGN5 Q8NGN6 Q8NGN7 Q8NGN8 Q8NGP0 Q8NGP2 Q8NGP3 Q8NGP4 Q8NGP6 Q8NGP8
Q8NGP9 Q8NGQ1 Q8NGQ2 Q8NGQ3 Q8NGQ4 Q8NGQ5 Q8NGQ6 Q8NGR1 Q8NGR2 Q8NGR3 Q8NGR4
Q8NGR5 Q8NGR6 Q8NGR8 Q8NGR9 Q8NGS0 Q8NGS1 Q8NGS2 Q8NGS3 Q8NGS4 Q8NGS5 Q8NGS6
Q8NGS7 Q8NGS8 Q8NGS9 Q8NGT0 Q8NGT1 Q8NGT2 Q8NGT5 Q8NGT7 Q8NGT9 Q8NGU1 Q8NGU2
Q8NGU4 Q8NGU9 Q8NGV0 Q8NGV5 Q8NGV6 Q8NGV7 Q8NGW1 Q8NGW6 Q8NGX0 Q8NGX1 Q8NGX2
Q8NGX3 Q8NGX5 Q8NGX6 Q8NGX8 Q8NGX9 Q8NGY0 Q8NGY1 Q8NGY2 Q8NGY3 Q8NGY5 Q8NGY6
Q8NGY7 Q8NGY9 Q8NGZ0 Q8NGZ2 Q8NGZ3 Q8NGZ4 Q8NGZ5 Q8NGZ6 Q8NGZ9 Q8NH00 Q8NH01
Q8NH02 Q8NH03 Q8NH04 Q8NH05 Q8NH06 Q8NH07 Q8NH08 Q8NH09 Q8NH10 Q8NH16 Q8NH18
Q8NH19 Q8NH21 Q8NH37 Q8NH40 Q8NH41 Q8NH42 Q8NH43 Q8NH48 Q8NH49 Q8NH50 Q8NH51
Q8NH53 Q8NH54 Q8NH55 Q8NH56 Q8NH57 Q8NH59 Q8NH60 Q8NH61 Q8NH63 Q8NH64 Q8NH67
Q8NH69 Q8NH70 Q8NH72 Q8NH73 Q8NH74 Q8NH76 Q8NH79 Q8NH80 Q8NH81 Q8NH83 Q8NH85
Q8NH87 Q8NH89 Q8NH90 Q8NH92 Q8NH93 Q8NH94 Q8NH95 Q8NHA4 Q8NHA6 Q8NHA8 Q8NHB1
Q8NHB7 Q8NHB8 Q8NHC4 Q8NHC5 Q8NHC6 Q8NHC7 Q8NHC8 Q8NHE4 Q8NHH9 Q8NHJ6 Q8NHK3
Q8NHL6 Q8NHP6 Q8NHS1 Q8NHS3 Q8NHU3 Q8NHV5 Q8NHX9 Q8NHY0 Q8NI17 Q8NI28 Q8NI60
Q8TAA9 Q8TAB3 Q8TAC9 Q8TAD4 Q8TAE7 Q8TAF8 Q8TAQ9 Q8TAV4 Q8TAX9 Q8TAZ6 Q8TB36
Q8TB61 Q8TB68 Q8TB96 Q8TBA6 Q8TBB6 Q8TBE1 Q8TBE3 Q8TBE7 Q8TBF5 Q8TBG9 Q8TBJ4
Q8TBM7 Q8TBM8 Q8TBP5 Q8TBP6 Q8TBQ9 Q8TBR7 Q8TC12 Q8TC26 Q8TC27 Q8TC36 Q8TC41
Q8TCB6 Q8TCC7 Q8TCG1 Q8TCG5 Q8TCJ2 Q8TCP9 Q8TCQ1 Q8TCT6 Q8TCT7 Q8TCT8 Q8TCT9
Q8TCU3 Q8TCU5 Q8TCW7 Q8TCW9 Q8TCY5 Q8TCZ2 Q8TD07 Q8TD20 Q8TD22 Q8TD43 Q8TD46

Q8TD84 Q8TDB4 Q8TDB8 Q8TDD5 Q8TDF5 Q8TDI7 Q8TDI8 Q8TDN1 Q8TDN2 Q8TDN7 Q8TDQ0
Q8TDQ1 Q8TDS4 Q8TDS5 Q8TDS7 Q8TDT2 Q8TDU5 Q8TDU6 Q8TDU9 Q8TDV0 Q8TDV2 Q8TDV5
Q8TDW0 Q8TDW4 Q8TDW7 Q8TDX6 Q8TDX9 Q8TDY8 Q8TE23 Q8TE54 Q8TE99 Q8TEB7 Q8TEB9
Q8TED1 Q8TED4 Q8TEF2 Q8TEM1 Q8TEQ8 Q8TEY5 Q8TEZ7 Q8TF08 Q8TF62 Q8TF66 Q8TF71
Q8WTR4 Q8WTT0 Q8WTV0 Q8WTX9 Q8WU17 Q8WU67 Q8WUD6 Q8WUG5 Q8WUH6 Q8WUM9 Q8WUS8
Q8WUT4 Q8WUT9 Q8WUU8 Q8WUX1 Q8WUY8 Q8WV15 Q8WV19 Q8WV48 Q8WV83 Q8WVE6 Q8WVE7
Q8WVI0 Q8WVN6 Q8WVP7 Q8WVQ1 Q8WVV5 Q8WVX3 Q8WVX9 Q8WVZ1 Q8WVZ7 Q8WW34 Q8WW43
Q8WW52 Q8WW62 Q8WWA1 Q8WWB7 Q8WWF3 Q8WWF5 Q8WWG1 Q8WWG9 Q8WWI5 Q8WWP7 Q8WWQ8
Q8WWT9 Q8WWU5 Q8WWV6 Q8WWX8 Q8WWZ4 Q8WWZ7 Q8WXA8 Q8WXD0 Q8WXF7 Q8WXG9 Q8WXH0
Q8WXH2 Q8WXI7 Q8WXI8 Q8WXS4 Q8WXS5 Q8WY07 Q8WY21 Q8WY22 Q8WY98 Q8WYK1 Q8WZ04
Q8WZ55 Q8WZ59 Q8WZ71 Q8WZ84 Q8WZ92 Q8WZ94 Q8WZA1 Q8WZA6 Q902F9 Q92185 Q92186
Q92187 Q92478 Q92482 Q92504 Q92508 Q92521 Q92523 Q92535 Q92536 Q92537 Q92542
Q92544 Q92545 Q92581 Q92604 Q92611 Q92617 Q92629 Q92633 Q92637 Q92643 Q92667
Q92673 Q92685 Q92692 Q92729 Q92736 Q92781 Q92806 Q92813 Q92819 Q92823 Q92824
Q92838 Q92839 Q92847 Q92854 Q92859 Q92887 Q92896 Q92903 Q92911 Q92932 Q92935
Q92952 Q92953 Q92956 Q92959 Q92968 Q92982 Q93033 Q93038 Q93050 Q93063 Q93084
Q93086 Q95460 Q969E2 Q969F0 Q969F8 Q969I6 Q969K7 Q969L2 Q969M1 Q969M2 Q969M3
Q969N2 Q969N4 Q969P0 Q969S0 Q969S6 Q969V1 Q969V3 Q969V5 Q969W0 Q969W1 Q969W9
Q969X1 Q969X2 Q969X5 Q969Z4 Q96A11 Q96A25 Q96A26 Q96A28 Q96A29 Q96A33 Q96A46
Q96A54 Q96A57 Q96A59 Q96AA3 Q96AD5 Q96AG3 Q96AG4 Q96AJ9 Q96AM1 Q96AN5 Q96AP7
Q96AQ2 Q96AQ8 Q96AW1 Q96B21 Q96B33 Q96B42 Q96B77 Q96B96 Q96BA8 Q96BD0 Q96BF3
Q96BI1 Q96BI3 Q96BM0 Q96BY9 Q96BZ4 Q96BZ9 Q96C03 Q96CC6 Q96CE8 Q96CH1 Q96CP6
Q96CP7 Q96CQ1 Q96CU9 Q96D05 Q96D31 Q96D42 Q96D53 Q96D59 Q96D96 Q96DA6 Q96DB9
Q96DC7 Q96DD7 Q96DL1 Q96DS6 Q96DU3 Q96DW6 Q96DX8 Q96DZ7 Q96DZ9 Q96E16 Q96E22
Q96E52 Q96E93 Q96EC8 Q96EP9 Q96ER9 Q96ES6 Q96ET8 Q96EU7 Q96EX1 Q96EX2 Q96F05
Q96F15 Q96F25 Q96F46 Q96F81 Q96FB5 Q96FE5 Q96FE7 Q96FL8 Q96FL9 Q96FM1 Q96FT7
Q96FV3 Q96FX8 Q96FZ5 Q96G23 Q96G30 Q96G79 Q96G91 Q96G97 Q96GC9 Q96GE9 Q96GF1
Q96GL9 Q96GM1 Q96GP6 Q96GQ5 Q96GR4 Q96GX1 Q96GZ6 Q96H15 Q96H72 Q96H78 Q96H96
Q96HA1 Q96HA4 Q96HA9 Q96HD1 Q96HE8 Q96HG1 Q96HH4 Q96HH6 Q96HJ5 Q96HP8 Q96HR9
Q96HS1 Q96HV5 Q96I36 Q96I45 Q96IK0 Q96IQ7 Q96IV6 Q96IW7 Q96IX5 Q96IZ2 Q96J42
Q96J65 Q96J66 Q96J84 Q96J86 Q96JA1 Q96JA4 Q96JF0 Q96JJ6 Q96JJ7 Q96JN2 Q96JP9
Q96JQ0 Q96JQ2 Q96JQ5 Q96JT2 Q96JW4 Q96JX3 Q96K12 Q96K19 Q96K37 Q96K49 Q96K78
Q96KA5 Q96KC8 Q96KF7 Q96KG7 Q96KJ4 Q96KJ9 Q96KK3 Q96KK4 Q96KN9 Q96KR6 Q96KT7
Q96KV6 Q96L08 Q96L42 Q96L58 Q96LA5 Q96LA6 Q96LA9 Q96LB0 Q96LB1 Q96LB2 Q96LC7
Q96LD1 Q96LL3 Q96LL9 Q96LR9 Q96LT4 Q96LU7 Q96LW7 Q96LZ7 Q96M19 Q96MC6 Q96MH6
Q96MM7 Q96MS0 Q96MT1 Q96MU8 Q96MV1 Q96MV8 Q96MX0 Q96N19 Q96N35 Q96N66 Q96N68
Q96N87 Q96NA8 Q96NB2 Q96ND0 Q96NI6 Q96NL1 Q96NR3 Q96NT5 Q96NU0 Q96NY7 Q96NY8
Q96P31 Q96P56 Q96P65 Q96P66 Q96P67 Q96P68 Q96P69 Q96P88 Q96PB1 Q96PB8 Q96PC5
Q96PD2 Q96PD6 Q96PD7 Q96PE1 Q96PE5 Q96PG1 Q96PG2 Q96PH1 Q96PJ5 Q96PL5 Q96PQ0
Q96PQ1 Q96PR1 Q96PS6 Q96PS8 Q96PX8 Q96PZ7 Q96Q04 Q96Q45 Q96Q80 Q96Q91 Q96QA5
Q96QD8 Q96QE2 Q96QE4 Q96QI5 Q96QK8 Q96QS1 Q96QT4 Q96QU1 Q96QZ0 Q96R08 Q96R09
Q96R27 Q96R28 Q96R30 Q96R45 Q96R47 Q96R48 Q96R54 Q96R67 Q96R69 Q96R72 Q96R84
Q96RA2 Q96RB7 Q96RC9 Q96RD0 Q96RD1 Q96RD2 Q96RD3 Q96RD6 Q96RD7 Q96RD9 Q96RI0
Q96RI8 Q96RI9 Q96RJ0 Q96RJ3 Q96RL6 Q96RN1 Q96RP7 Q96RP8 Q96RQ1 Q96RT6 Q96RV3

Q96S06 Q96S37 Q96S52 Q96S66 Q96S97 Q96SA4 Q96SE0 Q96SJ8 Q96SK2 Q96SL1 Q96SN7
Q96T52 Q96T53 Q96T54 Q96T55 Q96T83 Q96TA0 Q96TA2 Q96TC7 Q99062 Q99075 Q99102
Q99250 Q99437 Q99442 Q99463 Q99466 Q99467 Q99500 Q99518 Q99523 Q99527 Q99571
Q99572 Q99595 Q99624 Q99643 Q99650 Q99665 Q99677 Q99678 Q99679 Q99680 Q99705
Q99706 Q99712 Q99720 Q99726 Q99735 Q99758 Q99788 Q99795 Q99805 Q99808 Q99835
Q99884 Q99928 Q99941 Q99942 Q99943 Q99946 Q99965 Q99999 Q9BPV8 Q9BPX6 Q9BQ31
Q9BQ49 Q9BQ51 Q9BQA9 Q9BQB6 Q9BQD7 Q9BQE4 Q9BQG1 Q9BQI7 Q9BQJ4 Q9BQQ7 Q9BQS2
Q9BQS7 Q9BQT8 Q9BQT9 Q9BR10 Q9BR26 Q9BR39 Q9BRB3 Q9BRI3 Q9BRK0 Q9BRK3 Q9BRL7
Q9BRN9 Q9BRQ5 Q9BRQ8 Q9BRR3 Q9BRV3 Q9BRY0 Q9BS91 Q9BSA4 Q9BSA9 Q9BSE2 Q9BSE4
Q9BSF4 Q9BSJ5 Q9BSJ8 Q9BSK0 Q9BSK2 Q9BSN7 Q9BSR8 Q9BT22 Q9BT67 Q9BT76 Q9BT88
Q9BTD3 Q9BTN0 Q9BTV4 Q9BTX1 Q9BTX3 Q9BU23 Q9BU79 Q9BUB7 Q9BUF7 Q9BUJ0 Q9BUM1
Q9BUN8 Q9BUR5 Q9BUV8 Q9BV10 Q9BV23 Q9BV35 Q9BV40 Q9BV81 Q9BV87 Q9BVA6 Q9BVC6
Q9BVG9 Q9BVH7 Q9BVI4 Q9BVK2 Q9BVK6 Q9BVK8 Q9BVT8 Q9BVV7 Q9BVV8 Q9BVW6 Q9BVX2
Q9BW60 Q9BW72 Q9BWH2 Q9BWL3 Q9BWM7 Q9BWQ6 Q9BWQ8 Q9BWV1 Q9BWV2 Q9BX59 Q9BX67
Q9BX73 Q9BX74 Q9BX79 Q9BX84 Q9BX95 Q9BX97 Q9BXA5 Q9BXB1 Q9BXC0 Q9BXC1 Q9BXE9
Q9BXI2 Q9BXJ7 Q9BXJ8 Q9BXK5 Q9BXM7 Q9BXN2 Q9BXP2 Q9BXR5 Q9BXS0 Q9BXS4 Q9BXS9
Q9BXT2 Q9BXU9 Q9BY07 Q9BY08 Q9BY10 Q9BY15 Q9BY19 Q9BY21 Q9BY50 Q9BY64 Q9BY67
Q9BY71 Q9BY78 Q9BY79 Q9BYC5 Q9BYE2 Q9BYE9 Q9BYF1 Q9BYG0 Q9BYG8 Q9BYH1 Q9BYT1
Q9BYT9 Q9BYW1 Q9BZ11 Q9BZ76 Q9BZ97 Q9BZA7 Q9BZA8 Q9BZC7 Q9BZD2 Q9BZD6 Q9BZD7
Q9BZF1 Q9BZG2 Q9BZJ4 Q9BZJ6 Q9BZJ7 Q9BZJ8 Q9BZL3 Q9BZV2 Q9BZV3 Q9BZW2 Q9BZW4
Q9BZW5 Q9BZW8 Q9BZZ2 Q9C091 Q9C0A0 Q9C0B5 Q9C0B7 Q9C0C4 Q9C0D9 Q9C0E8 Q9C0H2
Q9C0I4 Q9C0J1 Q9C0K1 Q9GIP4 Q9GZK3 Q9GZK4 Q9GZK6 Q9GZK7 Q9GZM5 Q9GZM6 Q9GZN0
Q9GZN6 Q9GZP1 Q9GZP7 Q9GZP9 Q9GZQ4 Q9GZQ6 Q9GZR5 Q9GZS9 Q9GZT6 Q9GZU1 Q9GZU3
Q9GZV3 Q9GZW8 Q9GZX3 Q9GZY4 Q9GZY6 Q9GZY8 Q9GZZ6 Q9H013 Q9H015 Q9H061 Q9H0A3
Q9H0C2 Q9H0C3 Q9H0H0 Q9H0Q3 Q9H0R3 Q9H0U3 Q9H0V1 Q9H0V9 Q9H0X4 Q9H0X9 Q9H156
Q9H158 Q9H159 Q9H172 Q9H195 Q9H1B5 Q9H1C0 Q9H1C3 Q9H1C4 Q9H1C7 Q9H1D0 Q9H1E5
Q9H1K4 Q9H1N7 Q9H1U4 Q9H1U9 Q9H1V8 Q9H1X3 Q9H1Y3 Q9H1Z9 Q9H205 Q9H207 Q9H208
Q9H209 Q9H210 Q9H221 Q9H222 Q9H228 Q9H237 Q9H244 Q9H251 Q9H252 Q9H255 Q9H295
Q9H2A7 Q9H2A9 Q9H2B2 Q9H2B4 Q9H2C2 Q9H2C5 Q9H2C8 Q9H2D1 Q9H2E6 Q9H2F3 Q9H2H9
Q9H2J7 Q9H2L4 Q9H2S1 Q9H2S6 Q9H2U9 Q9H2V7 Q9H2W1 Q9H2X3 Q9H2X8 Q9H2X9 Q9H2Y9
Q9H300 Q9H310 Q9H313 Q9H330 Q9H339 Q9H340 Q9H341 Q9H342 Q9H343 Q9H344 Q9H346
Q9H354 Q9H3H5 Q9H3K2 Q9H3M0 Q9H3N1 Q9H3N8 Q9H3Q3 Q9H3R1 Q9H3R2 Q9H3S1 Q9H3S3
Q9H3S5 Q9H3T2 Q9H3T3 Q9H3U5 Q9H3V2 Q9H3W5 Q9H400 Q9H427 Q9H461 Q9H490 Q9H4D0
Q9H4F1 Q9H4I9 Q9H553 Q9H598 Q9H5I5 Q9H5J4 Q9H5K3 Q9H5V8 Q9H5Y7 Q9H665 Q9H6A9
Q9H6B4 Q9H6B9 Q9H6D3 Q9H6D8 Q9H6F2 Q9H6H4 Q9H6L2 Q9H6L5 Q9H6R6 Q9H6U8 Q9H6X2
Q9H6X4 Q9H6Y7 Q9H720 Q9H741 Q9H756 Q9H799 Q9H7F0 Q9H7F4 Q9H7M9 Q9H7T0 Q9H7V2
Q9H7X2 Q9H7Z7 Q9H813 Q9H819 Q9H841 Q9H8J5 Q9H8M5 Q9H8M9 Q9H8P0 Q9H8X9 Q9H902
Q9H920 Q9H936 Q9H9B4 Q9H9K5 Q9H9P2 Q9H9S3 Q9H9S5 Q9H9V4 Q9HA72 Q9HA82 Q9HAB3
Q9HAR2 Q9HAS3 Q9HAT1 Q9HAV5 Q9HAW7 Q9HAW8 Q9HAW9 Q9HB03 Q9HB14 Q9HB15 Q9HB29
Q9HB89 Q9HBA0 Q9HBB8 Q9HBE5 Q9HBG4 Q9HBG7 Q9HBI6 Q9HBJ8 Q9HBL6 Q9HBL7 Q9HBM0
Q9HBR0 Q9HBT6 Q9HBU9 Q9HBV1 Q9HBV2 Q9HBW0 Q9HBW1 Q9HBW9 Q9HBX8 Q9HBX9 Q9HBY0
Q9HC07 Q9HC10 Q9HC21 Q9HC24 Q9HC56 Q9HC58 Q9HC73 Q9HC97 Q9HCC8 Q9HCE9 Q9HCF6
Q9HCJ1 Q9HCJ2 Q9HCK4 Q9HCL0 Q9HCL2 Q9HCM2 Q9HCM3 Q9HCN3 Q9HCN6 Q9HCP6 Q9HCQ5
Q9HCS2 Q9HCU0 Q9HCU4 Q9HCU5 Q9HCX4 Q9HD20 Q9HD23 Q9HD36 Q9HD43 Q9HD45 Q9HD87

Q9HDC5 Q9HDC9 Q9HDD0 Q9N2J8 Q9N2K0 Q9NNX6 Q9NNZ3 Q9NP58 Q9NP59 Q9NP60 Q9NP78
Q9NP80 Q9NP84 Q9NP91 Q9NP94 Q9NP99 Q9NPA0 Q9NPA1 Q9NPB0 Q9NPB9 Q9NPC1 Q9NPC2
Q9NPC4 Q9NPD5 Q9NPE6 Q9NPF0 Q9NPF2 Q9NPG1 Q9NPG4 Q9NPG8 Q9NPH3 Q9NPH5 Q9NPI0
Q9NPI9 Q9NPL8 Q9NPR2 Q9NPR9 Q9NPU4 Q9NPY3 Q9NPZ5 Q9NQ11 Q9NQ25 Q9NQ34 Q9NQ40
Q9NQ60 Q9NQ84 Q9NQ90 Q9NQA5 Q9NQC3 Q9NQG1 Q9NQG6 Q9NQN1 Q9NQQ7 Q9NQR9 Q9NQS3
Q9NQS5 Q9NQW8 Q9NQX5 Q9NQX7 Q9NQZ7 Q9NR16 Q9NR34 Q9NR61 Q9NR71 Q9NR77 Q9NR82
Q9NR96 Q9NR97 Q9NRA2 Q9NRB3 Q9NRC1 Q9NRD8 Q9NRD9 Q9NRJ7 Q9NRK6 Q9NRM0 Q9NRM6
Q9NRP0 Q9NRQ2 Q9NRQ5 Q9NRR2 Q9NRS4 Q9NRU3 Q9NRX5 Q9NRX6 Q9NRY6 Q9NRY7 Q9NRZ5
Q9NRZ7 Q9NS00 Q9NS40 Q9NS62 Q9NS64 Q9NS66 Q9NS67 Q9NS68 Q9NS69 Q9NS75 Q9NS82
Q9NS84 Q9NS93 Q9NSA0 Q9NSA2 Q9NSC7 Q9NSD5 Q9NSD7 Q9NSI5 Q9NSK7 Q9NST1 Q9NT68
Q9NT99 Q9NTG1 Q9NTI2 Q9NTJ5 Q9NTN3 Q9NTN9 Q9NTQ9 Q9NU53 Q9NUB4 Q9NUD9 Q9NUE0
Q9NUH8 Q9NUM3 Q9NUM4 Q9NUN5 Q9NUN7 Q9NUQ2 Q9NUR3 Q9NUT2 Q9NUV7 Q9NV12 Q9NV29
Q9NV58 Q9NV64 Q9NV92 Q9NV96 Q9NVA4 Q9NVC3 Q9NVH0 Q9NVI7 Q9NVM1 Q9NVV0 Q9NVV5
Q9NW15 Q9NW97 Q9NWC5 Q9NWD8 Q9NWF4 Q9NWH2 Q9NWQ8 Q9NWR8 Q9NWS6 Q9NWW5 Q9NWW9
Q9NX00 Q9NX14 Q9NX47 Q9NX52 Q9NX61 Q9NX62 Q9NX76 Q9NX77 Q9NX78 Q9NX94 Q9NX95
Q9NXB9 Q9NXE4 Q9NXF8 Q9NXG6 Q9NXH8 Q9NXI6 Q9NXJ0 Q9NXK6 Q9NXL6 Q9NXS2 Q9NXW2
Q9NY15 Q9NY25 Q9NY26 Q9NY28 Q9NY35 Q9NY37 Q9NY46 Q9NY47 Q9NY64 Q9NY72 Q9NY91
Q9NY97 Q9NYB5 Q9NYG2 Q9NYG8 Q9NYJ7 Q9NYK1 Q9NYL4 Q9NYL5 Q9NYM4 Q9NYM9 Q9NYP7
Q9NYQ6 Q9NYQ7 Q9NYQ8 Q9NYR8 Q9NYV7 Q9NYV8 Q9NYV9 Q9NYW0 Q9NYW1 Q9NYW2 Q9NYW3
Q9NYW4 Q9NYW5 Q9NYW6 Q9NYW7 Q9NYX4 Q9NYZ1 Q9NYZ2 Q9NYZ4 Q9NZ01 Q9NZ08 Q9NZ42
Q9NZ43 Q9NZ45 Q9NZ53 Q9NZ94 Q9NZA1 Q9NZC2 Q9NZC3 Q9NZD1 Q9NZG7 Q9NZH0 Q9NZJ5
Q9NZJ7 Q9NZM1 Q9NZM6 Q9NZN1 Q9NZP0 Q9NZP2 Q9NZP5 Q9NZQ7 Q9NZQ8 Q9NZR2 Q9NZS2
Q9NZS9 Q9NZU0 Q9NZU1 Q9NZV1 Q9NZV8 Q9P003 Q9P035 Q9P055 Q9P0B6 Q9P0I2 Q9P0J0
Q9P0K1 Q9P0K9 Q9P0L0 Q9P0L9 Q9P0N5 Q9P0N8 Q9P0S2 Q9P0S3 Q9P0S9 Q9P0T7 Q9P0U1
Q9P0V8 Q9P0X4 Q9P109 Q9P126 Q9P1P4 Q9P1P5 Q9P1Q5 Q9P1W3 Q9P1W8 Q9P1Z3 Q9P241
Q9P244 Q9P246 Q9P273 Q9P283 Q9P291 Q9P296 Q9P298 Q9P2B2 Q9P2C4 Q9P2D8 Q9P2E5
Q9P2E7 Q9P2E8 Q9P2E9 Q9P2J2 Q9P2K9 Q9P2P1 Q9P2S2 Q9P2U7 Q9P2U8 Q9P2V4 Q9P2W7
Q9P2W9 Q9P2X0 Q9UBD6 Q9UBG0 Q9UBH6 Q9UBI4 Q9UBJ2 Q9UBK5 Q9UBL9 Q9UBM1 Q9UBM7
Q9UBM8 Q9UBN1 Q9UBN4 Q9UBN6 Q9UBQ6 Q9UBR5 Q9UBS5 Q9UBS9 Q9UBU6 Q9UBV2 Q9UBV7
Q9UBX3 Q9UBX8 Q9UBY0 Q9UBY5 Q9UBY8 Q9UDW1 Q9UDX5 Q9UEF7 Q9UEU0 Q9UEW3 Q9UF02
Q9UF33 Q9UG56 Q9UGF5 Q9UGF6 Q9UGF7 Q9UGH3 Q9UGI6 Q9UGM1 Q9UGN4 Q9UGP8 Q9UGQ2
Q9UGQ3 Q9UGT4 Q9UH62 Q9UH99 Q9UHC3 Q9UHC6 Q9UHC9 Q9UHE5 Q9UHE8 Q9UHF3 Q9UHF4
Q9UHI5 Q9UHI7 Q9UHJ9 Q9UHM6 Q9UHN6 Q9UHP7 Q9UHQ4 Q9UHQ9 Q9UHW9 Q9UHX3 Q9UI14
Q9UI33 Q9UI40 Q9UIB8 Q9UIG8 Q9UIJ5 Q9UIK5 Q9UIQ6 Q9UIR0 Q9UIW2 Q9UIX4 Q9UJ14
Q9UJ37 Q9UJ42 Q9UJ71 Q9UJ90 Q9UJ96 Q9UJ99 Q9UJA2 Q9UJA9 Q9UJG1 Q9UJQ1 Q9UJS0
Q9UK00 Q9UK17 Q9UK23 Q9UK28 Q9UKB5 Q9UKF2 Q9UKF5 Q9UKG4 Q9UKJ0 Q9UKJ1 Q9UKJ8
Q9UKL2 Q9UKL4 Q9UKM7 Q9UKN1 Q9UKP6 Q9UKQ2 Q9UKR5 Q9UKR8 Q9UKU0 Q9UKU6 Q9UKV5
Q9UKX5 Q9UKY4 Q9UKZ4 Q9UL01 Q9UL19 Q9UL51 Q9UL52 Q9UL54 Q9UL62 Q9ULB1 Q9ULB4
Q9ULB5 Q9ULC0 Q9ULC5 Q9ULC8 Q9ULD8 Q9ULF5 Q9ULG6 Q9ULH0 Q9ULH4 Q9ULI3 Q9ULK0
Q9ULK5 Q9ULK6 Q9ULL4 Q9ULQ1 Q9ULS5 Q9ULS6 Q9ULT6 Q9ULV1 Q9ULW2 Q9ULX5 Q9ULX7
Q9ULY5 Q9UM00 Q9UM01 Q9UM21 Q9UM44 Q9UM47 Q9UM73 Q9UMD9 Q9UMF0 Q9UMR7 Q9UMS5
Q9UMX3 Q9UMX9 Q9UMZ3 Q9UN42 Q9UN66 Q9UN67 Q9UN70 Q9UN71 Q9UN72 Q9UN73 Q9UN74
Q9UN75 Q9UN76 Q9UN88 Q9UNA3 Q9UNE0 Q9UNG2 Q9UNK0 Q9UNL2 Q9UNN8 Q9UNP4 Q9UNQ0
Q9UNU6 Q9UNW8 Q9UNX9 Q9UP38 Q9UP52 Q9UP95 Q9UPC5 Q9UPI3 Q9UPQ8 Q9UPR5 Q9UPU3

Q9UPX0 Q9UPX6 Q9UPY5 Q9UPZ6 Q9UQ05 Q9UQ53 Q9UQ90 Q9UQC9 Q9UQD0 Q9UQF0 Q9UQQ1
Q9UQV4 Q9Y210 Q9Y219 Q9Y225 Q9Y226 Q9Y227 Q9Y228 Q9Y231 Q9Y241 Q9Y256 Q9Y257
Q9Y267 Q9Y271 Q9Y274 Q9Y275 Q9Y276 Q9Y277 Q9Y278 Q9Y279 Q9Y282 Q9Y284 Q9Y286
Q9Y287 Q9Y289 Q9Y2A7 Q9Y2A9 Q9Y2B1 Q9Y2B2 Q9Y2C2 Q9Y2C3 Q9Y2C5 Q9Y2C9 Q9Y2D2
Q9Y2E8 Q9Y2G1 Q9Y2G3 Q9Y2G8 Q9Y2H6 Q9Y2P4 Q9Y2P5 Q9Y2Q0 Q9Y2R0 Q9Y2T5 Q9Y2T6
Q9Y2U2 Q9Y2U8 Q9Y2W3 Q9Y2Y6 Q9Y320 Q9Y328 Q9Y336 Q9Y342 Q9Y345 Q9Y385 Q9Y397
Q9Y3A6 Q9Y3B3 Q9Y3D6 Q9Y3E0 Q9Y3N9 Q9Y3P4 Q9Y3P8 Q9Y3Q0 Q9Y3Q3 Q9Y3Q4 Q9Y3Q7
Q9Y426 Q9Y442 Q9Y487 Q9Y493 Q9Y4A9 Q9Y4C0 Q9Y4C5 Q9Y4D2 Q9Y4D7 Q9Y4D8 Q9Y4W6
Q9Y512 Q9Y519 Q9Y548 Q9Y561 Q9Y584 Q9Y585 Q9Y5E1 Q9Y5E2 Q9Y5E3 Q9Y5E4 Q9Y5E5
Q9Y5E6 Q9Y5E7 Q9Y5E8 Q9Y5E9 Q9Y5F0 Q9Y5F1 Q9Y5F2 Q9Y5F3 Q9Y5F6 Q9Y5F7 Q9Y5F8
Q9Y5F9 Q9Y5G0 Q9Y5G1 Q9Y5G2 Q9Y5G3 Q9Y5G4 Q9Y5G5 Q9Y5G6 Q9Y5G7 Q9Y5G8 Q9Y5G9
Q9Y5H0 Q9Y5H1 Q9Y5H2 Q9Y5H3 Q9Y5H4 Q9Y5H5 Q9Y5H6 Q9Y5H7 Q9Y5H8 Q9Y5H9 Q9Y5I0
Q9Y5I1 Q9Y5I2 Q9Y5I3 Q9Y5I4 Q9Y5I7 Q9Y5L2 Q9Y5L3 Q9Y5M8 Q9Y5N1 Q9Y5P0 Q9Y5P1
Q9Y5Q0 Q9Y5Q5 Q9Y5R2 Q9Y5S1 Q9Y5S8 Q9Y5T4 Q9Y5U4 Q9Y5U5 Q9Y5U8 Q9Y5U9 Q9Y5W7
Q9Y5X5 Q9Y5Y0 Q9Y5Y3 Q9Y5Y4 Q9Y5Y5 Q9Y5Y6 Q9Y5Y7 Q9Y5Y9 Q9Y5Z0 Q9Y5Z6 Q9Y5Z9
Q9Y619 Q9Y624 Q9Y639 Q9Y644 Q9Y653 Q9Y661 Q9Y662 Q9Y663 Q9Y666 Q9Y672 Q9Y673
Q9Y691 Q9Y693 Q9Y694 Q9Y696 Q9Y698 Q9Y6A1 Q9Y6A2 Q9Y6A9 Q9Y6C5 Q9Y6C9 Q9Y6D0
Q9Y6F6 Q9Y6G1 Q9Y6H6 Q9Y6H8 Q9Y6I8 Q9Y6I9 Q9Y6J6 Q9Y6K0 Q9Y6L6 Q9Y6M5 Q9Y6M7
Q9Y6N1 Q9Y6N7 Q9Y6N8 Q9Y6Q6 Q9Y6R1 Q9Y6U7 Q9Y6W8 Q9Y6X1 Q9Y6X5 U3KPV4 W5XKT8

#### 2. The sequences shown in Fig. 5B

##### 2.1 The proteins with keyword of "Membrane" (407 sequences)

A0A075B6K0 A0A075B6K4 A0A075B6K5 A0A075B6L2 A0A075B6N3 A0A075B6R0 A0A075B6R2 A0A075B6T8 A0A075B6U4 A0A075B6X5 A0A087WSY4 A0A087WT02 A0A087WT03 A0A0A0MS00 A0A0A0MS02 A0A0A6YYC5 A0A0A6YYJ7 A0A0A6YYK1 A0A0A6YYK6 A0A0A6YYK7 A0A0B4J1U4 A0A0B4J1U7 A0A0B4J237 A0A0B4J238 A0A0B4J240 A0A0B4J244 A0A0B4J248 A0A0B4J262 A0A0B4J263 A0A0B4J264 A0A0B4J274 A0A0B4J277 A0A0B4J280 A0A0C4DH27 A0A0C4DH28 A0A0C4DH34 A0A0C4DH41 A0A0K0K1B3 A0A1B0GVX0 A0A1B0GX56 A0JD32 A0JD37 A6ND01 A6NNS2 A8K7I4 H3BQJ8 O00217 O00451 O00469 O00602 O14548 O14656 O14657 O14756 O14798 O15537 O43174 O43280 O43323 O43597 O43609 O43615 O43653 O43820 O43852 O43895 O43921 O60397 O60568 O60609 O60664 O60683 O75015 O75106 O75326 O75381 O75487 O75955 O75964 O94772 O94779 O95178 O95274 O95497 O95498 O95831 O95867 O95868 O95897 O95971 O95980 O96000 P00387 P00797 P01374 P01715 P01717 P01737 P01814 P01824 P01825 P03979 P04062 P04156 P04216 P04798 P05093 P05177 P05181 P05186 P06331 P06731 P06858 P07237 P07711 P07858 P07911 P08174 P08253 P08311 P08571 P08582 P08686 P09326 P09923 P0CG37 P0DP06 P0DP07 P0DP58 P10253 P10632 P10635 P10646 P10696 P10909 P11509 P11712 P12110 P12532 P13385 P13498 P13584 P13987 P14207 P14384 P14555 P15309 P15328 P16444 P16870 P17213 P17540 P17706 P18031 P18428 P20160 P20813 P20815 P20827 P20853 P21589 P22303 P22748 P23219 P23352 P23435 P23515 P24158 P24462 P24539 P24903 P25063 P26885 P26992 P27105 P29122 P31358 P31415 P31997 P32189 P33260 P33261 P35052 P35232 P35354 P37840 P38567 P39877 P40199 P43026 P43235 P48449 P51589 P51636 P51654 P52797 P52798 P52803 P52961 P53701 P54317 P54826 P55259 P55290 P56159 P56385 P56539 P60022 P78329 P78333 P80303 P80748 Q00LT1 Q02083 Q02246 Q02318 Q02338 Q02809 Q02818 Q02928 Q03135 Q03405 Q08431 Q0VAF6 Q10588 Q12860 Q12891 Q13421 Q13449 Q13508 Q14002 Q14154 Q14210 Q14409 Q14512 Q14534 Q14623 Q14982 Q15084 Q15165 Q15465 Q16134 Q16553 Q16635 Q16678 Q16696 Q17RY6 Q2MJR0 Q2TAP0 Q2TAZ0 Q330K2 Q496H8 Q53H12 Q5BIV9 Q5T9A4 Q5T9G4 Q5TCH4 Q5VST6 Q5VXU3 Q5VY80 Q6HA08 Q6NUM9 Q6NW40 Q6PCB6 Q6UQ28 Q6UWB4 Q6UWM5 Q6UWN0 Q6UWN5 Q6UWR7 Q6UX46 Q6UX53 Q6UX82 Q6UXB3 Q6UXT8 Q6VVX0 Q6XZB0 Q6YHK3 Q6ZVN8 Q709C8 Q7RTS9 Q7RTW8 Q7RTY9 Q7Z3B1 Q7Z4Y8 Q7Z553 Q7Z5A4 Q7Z5G4 Q7Z5L4 Q7Z698 Q7Z699 Q7Z7G8 Q86TD4 Q86UN2 Q86UN3 Q86WD7 Q86Y78 Q86YB8 Q8IV16 Q8IVL8 Q8IWV2 Q8IWY4 Q8IZJ3 Q8N158 Q8N2G4 Q8N307 Q8N490 Q8N565 Q8N6Q3 Q8N7S2 Q8NBN7 Q8NCC3 Q8NFP4 Q8NI32 Q8TAV3 Q8TDM5 Q8WUK0 Q8WWA0 Q8WWY8 Q8WXD2 Q92485 Q92543 Q92575 Q92743 Q92843 Q93070 Q969Z3 Q96B49 Q96B86 Q96BY7 Q96CW9 Q96DR8 Q96GS6 Q96GW7 Q96HE7 Q96HY6 Q96ID5 Q96IL0 Q96LU5 Q96MG2 Q96NR8 Q96PD5 Q96PL2 Q96QV1 Q96RL7 Q96SQ9 Q99445 Q99519 Q99541 Q99623 Q99640 Q99732 Q99807 Q99972 Q9BPW9 Q9BRK5 Q9BS86 Q9BSG5 Q9BY14 Q9BZM2 Q9BZM4 Q9BZM5 Q9BZM6 Q9BZR6 Q9C004 Q9GZZ7 Q9H1Z8 Q9H305 Q9H306 Q9H3Z4 Q9H4A9 Q9H4B8 Q9H7X0 Q9H8H3 Q9HB55 Q9NP85 Q9NPA2 Q9NPD7 Q9NR63 Q9NRA1 Q9NSU2 Q9NUU6 Q9NVH1 Q9NY59 Q9NZV5 Q9P121 Q9P232 Q9UBP0 Q9UF47 Q9UFP1 Q9UJZ1 Q9UKJ5 Q9UKY0 Q9ULZ9 Q9UQ52 Q9Y251 Q9Y2C4 Q9Y2I2 Q9Y2J2 Q9Y2Z9 Q9Y394 Q9Y3A0 Q9Y5W8 Q9Y625 Q9Y679 Q9Y6B6 Q9Y6M0

#### 2.2 The proteins with keyword of "Secreted" (1145 sequences)

A0A075B6K0 A0A075B6K4 A0A075B6K5 A0A075B6R2 A0A087WSY4 A0A096LNP1 A0A0A0MS00

A0A0B4J1U7 A0A0C4DH34 A0A0C4DH41 A0A1B0GTR0 A0A1B0GVH4 A1E959 A1L453 A1L4H1

A2RUU4 A4D0S4 A4D1T9 A5D8T8 A6NFZ4 A6NGN9 A6NHN0 A6NHN6 A6NIE9 A6NKQ9 A6NNL5

A8K7I4 A8MTI9 A8MTW9 A8MV23 A8MXU0 A8MZH6 B1AKI9 B2RNN3 B2RUY7 B3GLJ2 C9JL84

C9JUS6 C9JXX5 E2RYF7 I3L3R5 K9M1U5 M5A8F1 O00187 O00230 O00253 O00292 O00339

O00584 O00602 O00622 O00744 O14498 O14791 O14793 O14904 O14905 O14960 O15072

O15123 O15130 O15232 O15240 O15263 O15335 O15537 O43240 O43278 O43323 O43405

O43555 O43692 O43820 O43827 O43852 O43854 O43866 O43927 O60259 O60383 O60565

O60568 O60575 O60676 O60687 O60844 O60938 O75015 O75093 O75094 O75095 O75200

O75339 O75356 O75462 O75487 O75493 O75556 O75594 O75596 O75610 O75629 O75636

O75711 O75715 O75718 O75888 O75951 O75973 O76061 O76076 O94769 O94919 O95084

O95156 O95157 O95158 O95389 O95390 O95393 O95399 O95407 O95428 O95450 O95460

O95631 O95633 O95711 O95715 O95750 O95813 O95841 O95897 O95925 O95965 O95967

O95968 O95969 O95971 O95994 O96009 O96014 P00450 P00709 P00734 P00736 P00738

P00746 P00747 P00748 P00749 P00750 P00797 P00995 P01009 P01011 P01019 P01031

P01034 P01036 P01037 P01127 P01137 P01138 P01148 P01160 P01189 P01210 P01213

P01215 P01222 P01225 P01229 P01258 P01275 P01282 P01286 P01298 P01303 P01308

P01344 P01350 P01374 P01574 P01579 P01588 P01591 P01715 P01717 P01814 P01824

P01825 P02452 P02458 P02461 P02462 P02649 P02652 P02654 P02655 P02656 P02671

P02675 P02679 P02741 P02745 P02746 P02747 P02749 P02753 P02765 P02766 P02768

P02774 P02775 P02776 P02778 P02787 P02788 P02790 P02808 P02810 P02812 P02814

P02818 P03950 P03952 P03956 P03971 P04004 P04054 P04085 P04090 P04114 P04118

P04141 P04155 P04180 P04196 P04275 P04278 P04279 P04280 P04628 P04746 P04808

P05019 P05060 P05090 P05111 P05112 P05113 P05121 P05154 P05155 P05156 P05305

P05408 P05452 P05543 P05814 P05997 P06276 P06307 P06331 P06396 P06850 P06858

P07093 P07098 P07225 P07288 P07339 P07358 P07477 P07478 P07492 P07498 P07585

P07602 P07711 P07858 P07911 P07942 P07988 P07996 P08118 P08123 P08174 P08185

P08217 P08218 P08253 P08254 P08294 P08311 P08476 P08493 P08571 P08620 P08700

P08709 P08833 P08949 P09228 P09237 P09238 P09326 P09341 P09486 P09529 P09544

P09681 P09683 P0C091 P0C0P6 P0C7L1 P0C862 P0C8F1 P0CG01 P0CG36 P0CG37 P0DJD7
P0DJD8 P0DJD9 P0DJI8 P0DJI9 P0DMC3 P0DML3 P0DMR2 P0DN86 P0DN87 P0DP06 P0DP07
P0DP57 P0DP73 P0DP74 P0DPK3 P0DTE7 P0DTE8 P0DUB6 P10082 P10092 P10124 P10144
P10145 P10163 P10451 P10600 P10643 P10645 P10646 P10720 P10767 P10909 P10915
P10997 P11047 P11150 P11226 P11487 P11597 P11684 P11686 P12034 P12109 P12110
P12111 P12272 P12273 P12643 P12645 P12724 P12838 P12872 P13232 P13284 P13385
P13497 P13521 P13671 P13727 P13942 P13987 P14138 P14207 P14543 P14555 P14780
P15085 P15086 P15169 P15248 P15309 P15328 P15502 P15515 P15516 P15692 P15814
P16035 P16233 P16562 P16860 P16870 P17213 P17405 P17538 P17936 P18065 P18075
P18428 P18509 P19801 P19823 P19827 P19835 P19875 P19876 P19883 P19957 P19961
P20061 P20062 P20142 P20155 P20231 P20366 P20382 P20396 P20774 P20783 P20800
P20809 P20827 P20849 P20851 P20908 P21128 P21246 P21741 P21815 P22004 P22079
P22301 P22303 P22352 P22362 P22466 P22692 P22749 P22792 P22894 P23142 P23280
P23352 P23435 P23946 P24001 P24043 P24158 P24347 P24387 P24592 P24593 P24855
P25067 P25311 P25391 P25774 P25940 P26022 P26927 P27169 P27539 P27658 P27797
P27918 P28039 P28300 P28325 P29122 P29279 P29400 P30990 P34130 P34820 P35052
P35247 P35318 P35443 P35542 P35625 P35858 P36222 P36955 P37840 P39060 P39877
P39900 P41221 P41271 P41439 P42127 P43026 P43235 P43251 P43652 P45452 P47710
P47972 P47992 P48052 P48304 P48307 P48645 P48745 P49223 P49747 P49765 P49767
P49863 P49908 P49913 P50897 P51124 P51460 P51884 P51888 P52798 P52823 P54108
P54315 P54317 P54793 P55000 P55001 P55056 P55058 P55075 P55083 P55089 P55103
P55107 P55145 P55259 P55268 P55774 P56703 P56704 P56730 P56851 P58062 P58166
P58215 P58294 P58417 P58499 P59665 P59666 P59796 P59826 P59827 P59861 P60022
P60568 P60827 P60985 P61278 P61366 P61626 P61769 P61812 P61916 P78333 P80303
P80748 P81172 P81277 P81534 P83105 P83110 P83859 P98066 P98095 P98173 Q00604
Q01523 Q01524 Q01955 Q02325 Q02383 Q02388 Q02747 Q02818 Q03403 Q03405 Q03692
Q04118 Q04756 Q06033 Q06141 Q06828 Q07507 Q07654 Q08380 Q08397 Q08431 Q08648
Q08830 Q0P5P2 Q12794 Q12805 Q12841 Q13093 Q13201 Q13214 Q13219 Q13253 Q13275
Q13296 Q13316 Q13361 Q13421 Q13510 Q13519 Q13609 Q13751 Q13790 Q14031 Q14050
Q14055 Q14213 Q14393 Q14406 Q14507 Q14508 Q14512 Q14515 Q14520 Q14563 Q14623

Q14624 Q14641 Q14667 Q14766 Q15063 Q15113 Q15166 Q15195 Q15198 Q15389 Q15465

Q15517 Q15582 Q15661 Q15726 Q15782 Q15828 Q15846 Q15848 Q16270 Q16378 Q16568

Q16609 Q16627 Q16661 Q16769 Q17RF5 Q17RR3 Q17RW2 Q17RY6 Q1W4C9 Q1ZYL8 Q2I0M5

Q2MKA7 Q2UY09 Q30KP8 Q30KP9 Q30KQ1 Q30KQ7 Q30KQ9 Q30KR1 Q32P28 Q49AH0 Q4G0G5

Q4G0M1 Q4QY38 Q4ZHG4 Q504Y2 Q53H76 Q53RD9 Q5BLP8 Q5DT21 Q5FYB0 Q5FYB1 Q5GAN6

Q5H8A3 Q5JQD4 Q5JS37 Q5JTB6 Q5JXM2 Q5K4E3 Q5R387 Q5T4F7 Q5T4W7 Q5T7M4 Q5VWW1

Q5VXJ0 Q5VXM1 Q5VYY2 Q5XG92 Q63HQ2 Q641Q3 Q66K79 Q68BL7 Q68BL8 Q6E0U4 Q6EBC2

Q6GPI1 Q6GTS8 Q6H9L7 Q6IE38 Q6JVE6 Q6KF10 Q6MZM9 Q6MZW2 Q6NT52 Q6NUI6 Q6NUJ1

Q6P093 Q6P0A1 Q6P5S2 Q6P988 Q6PCB0 Q6PDA7 Q6PEW0 Q6PEZ8 Q6Q788 Q6UDR6 Q6UW01

Q6UW10 Q6UW15 Q6UW32 Q6UWE3 Q6UWF7 Q6UWF9 Q6UWK7 Q6UWN8 Q6UWP8 Q6UWQ7 Q6UWT2

Q6UWT4 Q6UWU2 Q6UWY0 Q6UWY2 Q6UWY5 Q6UX06 Q6UX07 Q6UX39 Q6UX46 Q6UX73 Q6UX82

Q6UXA7 Q6UXB1 Q6UXB2 Q6UXB8 Q6UXF7 Q6UXH0 Q6UXH8 Q6UXI7 Q6UXI9 Q6UXN2 Q6UXQ4

Q6UXS0 Q6UXT8 Q6UXT9 Q6UXX9 Q6UY13 Q6UY27 Q6X4U4 Q6X784 Q6XZB0 Q6ZMM2 Q6ZNF0

Q6ZWJ8 Q765I0 Q76M96 Q7RTW8 Q7RTY7 Q7RTZ1 Q7Z4H4 Q7Z4P5 Q7Z4R8 Q7Z5A7 Q7Z5A8

Q7Z5A9 Q7Z5J1 Q7Z5L0 Q7Z5L3 Q7Z5L7 Q7Z5Y6 Q7Z7B7 Q7Z7B8 Q7Z7G0 Q86SG7 Q86SH4

Q86TE4 Q86TH1 Q86TW2 Q86UU9 Q86UW8 Q86UX2 Q86VR8 Q86WD7 Q86XP6 Q86XS5 Q86Y78

Q86YQ2 Q86YW7 Q86Z23 Q8IUA0 Q8IUB3 Q8IUB5 Q8IUH2 Q8IUK8 Q8IUL8 Q8IUX7 Q8IUX8

Q8IVN8 Q8IW75 Q8IW92 Q8IWL1 Q8IWL2 Q8IWV2 Q8IWY4 Q8IX30 Q8IYP2 Q8IZJ3 Q8IZN7

Q8N0V4 Q8N119 Q8N129 Q8N135 Q8N145 Q8N158 Q8N1E2 Q8N2E2 Q8N2S1 Q8N307 Q8N323

Q8N3Z0 Q8N436 Q8N474 Q8N475 Q8N4F0 Q8N4T0 Q8N5I4 Q8N5W8 Q8N688 Q8N690 Q8N6G6

Q8N6Q3 Q8N6Y2 Q8N729 Q8N907 Q8NBI3 Q8NBM8 Q8NBP7 Q8NCC3 Q8NCF0 Q8NDX9 Q8NDZ4

Q8NEB7 Q8NES8 Q8NET1 Q8NEX5 Q8NF86 Q8NFQ5 Q8NFU4 Q8NG35 Q8NG41 Q8NHM4 Q8NHW6

Q8NI99 Q8TAA1 Q8TAD2 Q8TAG5 Q8TAL6 Q8TAT2 Q8TAX7 Q8TB73 Q8TCV5 Q8TD33 Q8TE56

Q8TE57 Q8TE58 Q8TER0 Q8TEU8 Q8WTQ1 Q8WTR8 Q8WTU2 Q8WU66 Q8WUA8 Q8WUJ1 Q8WUY1

Q8WVF2 Q8WWA0 Q8WWF1 Q8WWQ2 Q8WWU7 Q8WWY7 Q8WWY8 Q8WX77 Q8WXA2 Q8WXD2 Q8WXF3

Q8WXQ8 Q8WXS8 Q92484 Q92485 Q92520 Q92563 Q92626 Q92743 Q92765 Q92820 Q92832

Q92874 Q92876 Q92954 Q93097 Q93098 Q969D9 Q969E1 Q969E3 Q969H8 Q969Y0 Q96A83

Q96A84 Q96A98 Q96BQ1 Q96CG8 Q96DA0 Q96DN2 Q96DR5 Q96DR8 Q96EE4 Q96GW7 Q96HE7

Q96I82 Q96IY4 Q96JB6 Q96JK4 Q96KN2 Q96KW9 Q96KX0 Q96L11 Q96LB8 Q96LR4 Q96MK3

Q96NZ8 Q96NZ9 Q96PD5 Q96PL1 Q96PL2 Q96QR1 Q96QV1 Q96RP3 Q96RQ9 Q96S86 Q96SL4

Q96SM3 Q96T91 Q99217 Q99218 Q99435 Q99470 Q99542 Q99574 Q99645 Q99674 Q99727

Q99731 Q99748 Q99935 Q99944 Q99954 Q99969 Q99972 Q99983 Q99988 Q9BPW4 Q9BQ08

Q9BQ16 Q9BQB4 Q9BQI4 Q9BQP9 Q9BQR3 Q9BRR6 Q9BRX8 Q9BS86 Q9BSG0 Q9BSG5 Q9BT56

Q9BTY2 Q9BUD6 Q9BUN1 Q9BWS9 Q9BX93 Q9BXI9 Q9BXJ0 Q9BXJ1 Q9BXJ2 Q9BXJ3 Q9BXJ4

Q9BXJ5 Q9BXN1 Q9BXX0 Q9BXY4 Q9BY14 Q9BY76 Q9BYJ0 Q9BYZ8 Q9BZJ3 Q9BZM1 Q9BZM2

Q9BZM5 Q9BZP6 Q9C0B6 Q9GZM7 Q9GZP0 Q9GZU5 Q9GZV9 Q9GZX9 Q9GZZ7 Q9GZZ8 Q9H0B8

Q9H114 Q9H1F0 Q9H1Z8 Q9H239 Q9H2R5 Q9H2X0 Q9H324 Q9H3U7 Q9H4F8 Q9H6E4 Q9H772

Q9H7Y0 Q9HAT2 Q9HB40 Q9HB63 Q9HBE4 Q9HC23 Q9HC57 Q9HCB6 Q9HCQ7 Q9HCT0 Q9HD89

Q9NP55 Q9NP70 Q9NPA2 Q9NPF7 Q9NPH9 Q9NQ30 Q9NQ36 Q9NQ38 Q9NQ76 Q9NQ79 Q9NR23

Q9NRA1 Q9NRC9 Q9NRE1 Q9NRI6 Q9NRM1 Q9NRN5 Q9NRR1 Q9NS15 Q9NS71 Q9NSA1 Q9NT22

Q9NTU7 Q9NZK5 Q9NZK7 Q9NZP8 Q9NZW4 Q9P0M4 Q9P1C3 Q9UBC7 Q9UBD3 Q9UBM4 Q9UBP4

Q9UBT3 Q9UBU2 Q9UBU3 Q9UBV4 Q9UBX5 Q9UBX7 Q9UFP1 Q9UHF0 Q9UHF1 Q9UHF5 Q9UHG2

Q9UHI8 Q9UHL4 Q9UI42 Q9UIG4 Q9UJH8 Q9UJJ9 Q9UK05 Q9UK55 Q9UK85 Q9UKP4 Q9UKR0

Q9UKR3 Q9UKU9 Q9UKY3 Q9UKZ9 Q9ULZ1 Q9ULZ9 Q9UM22 Q9UMX5 Q9UNA0 Q9UNK4 Q9UQP3

Q9Y240 Q9Y251 Q9Y258 Q9Y264 Q9Y2E5 Q9Y334 Q9Y337 Q9Y4K0 Q9Y581 Q9Y5C1 Q9Y5Q6

Q9Y5W5 Q9Y5X9 Q9Y625 Q9Y646 Q9Y6F9 Q9Y6L7 Q9Y6N6 Q9Y6Y9

#### 3. The sequences shown in Fig. 5C

##### 3.1 The proteins identified by TMbed (84 sequences)

A0A087WU88 A0A087X1L8 A0A096LNH4 A0A0G2JJF7 A0A0G2JKD1 A0A0G2JLG4 A0A0G2JNF4

A0A0G2JNH3 A0A126GWB0 A0A126GWI2 A0A140T8X8 A0A191URJ7 A0A1B0GTG8 A0A1B0GTQ1

A0A1B0GUZ9 A0A1B0GVG4 A0A1B0GWH6 A0A1W2PN81 A0A1W2PQM1 A0A1W2PQU2 A0A286YEU6

A0A2R8Y4L6 A0A2R8Y4M2 A0A2R8Y4M4 A0A2R8Y550 A0A2R8Y7Y5 A0A2R8YE69 A0A2R8YED5

A0A2R8YEG4 A0A2R8YEH3 A0A2R8YEV3 A0A2U3TZM8 A0A3B3IT45 A0A494C0I6 A0A494C1I1

A0A4W9AIG4 A0A5K1VDZ0 A0A7I2V2S6 A0A7I2V3R4 A0A8Q3SIZ7 A4D0V7 A6NH11 A6NKP2

H3BTG2 O60888 O75071 O75427 O95236 O95424 O95873 P27469 P36268 P37058 P48723

P51689 P51690 P60608 P63135 P80365 Q14442 Q16517 Q56VL3 Q5T2N8 Q6UWS5 Q8N699

Q8TCD1 Q8WVC6 Q8WX69 Q8WZ75 Q96G27 Q96HH9 Q96KR4 Q9BQE5 Q9BWW8 Q9BWW9 Q9BXQ6

Q9BZC5 Q9C002 Q9H6K4 Q9HB66 Q9NRX3 Q9NWS8 Q9NX40 V9GZ13

##### 3.2 The proteins identified by SignalP 6.0 (181 sequences)

A0A075B6U6 A0A075B7B6 A0A087WW49 A0A087X1L8 A0A0B4J1T7 A0A0G2JJF7 A0A0G2JKD1

A0A0G2JLY3 A0A0G2JNF4 A0A0J9YWU9 A0A0J9YXV3 A0A140T8X8 A0A191URJ7 A0A1B0GTC6

A0A1B0GTE1 A0A1B0GTL2 A0A1B0GUY1 A0A1B0GVD1 A0A1W2PN81 A0A1W2PP97 A0A1W2PQM1

A0A2R8Y7Y5 A0A494C103 A0A5K1VDZ0 A0A8I5KY86 A0A8Q3SIG1 A0A8Q3WLD3 A3KN74

A6NCL2 A6NHM9 D3DTV9 F8WCM5 O00115 O00462 O00748 O00754 O14773 O14792 O14958

O15442 O15460 O43612 O60613 O60911 O95302 O95479 O95881 P04066 P05164 P05546

P06280 P06865 P06870 P08246 P08519 P08861 P09093 P09668 P09871 P0CW18 P10153

P10323 P10619 P11021 P11678 P13667 P13674 P13686 P14091 P14314 P14625 P15088

P15289 P15848 P16278 P17050 P17900 P20151 P20933 P22304 P23141 P23284 P23327

P26436 P29120 P30040 P30101 P30533 P34059 P35442 P35475 P38571 P40313 P42785

P43234 P45877 P49184 P49746 P50454 P51688 P51689 P51690 P53634 P54107 P54802

P54803 P55157 P56202 P78539 Q01459 Q12889 Q13087 Q13162 Q13217 Q13438 Q14257

Q14264 Q14554 Q14696 Q14697 Q15293 Q15818 Q2M2W7 Q5JU69 Q5NDL2 Q5T4B2 Q6P4A8

Q6UW49 Q6UW63 Q6UXH1 Q6ZU45 Q7Z4N8 Q86UD1 Q8IVL5 Q8IVL6 Q8IWF2 Q8IWU5 Q8IWU6

Q8IXL7 Q8IYK4 Q8N3H0 Q8NBJ5 Q8NBJ7 Q8NBK3 Q8NBL1 Q8NBP0 Q8NBS9 Q8NHP8 Q8NI22

Q8TD06 Q8WWX9 Q8WZ75 Q8WZ79 Q92791 Q96D15 Q96DN0 Q96DZ1 Q96L12 Q96S16 Q99538

Q99895 Q9BT09 Q9BV94 Q9BZQ6 Q9H3G5 Q9H488 Q9H497 Q9HCN8 Q9NWM8 Q9NYP8 Q9UBR2

Q9UBS3 Q9UBX1 Q9UHG3 Q9UI38 Q9UMR5 Q9UNW1 Q9Y2B0 Q9Y2G5 Q9Y2Y8 Q9Y680

#### 4. The sequences shown in Fig. 5D

##### 4.1 The proteins with membrane-related subcellular location terms from GO annotations (29 sequences)

O43272 O43610 O60762 P07686 P36551 P43304 Q07973 Q14249 Q16795 Q5THJ4 Q7Z6Z6 Q8IVK1 Q8IXB1 Q8IXM3 Q8IYU8 Q8N0X7 Q8N2G8 Q8NBX0 Q8TB40 Q8WWC4 Q96HE9 Q96HL8 Q9HBH5 Q9NQZ5 Q9NRG9 Q9UF12 Q9Y3E5 Q9Y4L1 Q9Y4P3

##### 4.2 The potentially new membrane proteins (78 sequences)

A0A1B0GVS7 A0A494C176 A1A4F0 A1L4L8 A6NIN4 A6PVY3 O15033 O43399 O60240 O75920 O76070 P0C2W7 P0C7X4 P15586 P51687 P51861 P55327 P83111 Q00G26 Q16143 Q3SXM5 Q5T1J5 Q5VUY0 Q5VYY1 Q69YL0 Q6NUM6 Q6NXR0 Q6S5H5 Q6ZS86 Q6ZTK2 Q7RTY3 Q7Z5P4 Q8IW45 Q8IXL9 Q8IZ16 Q8IZ81 Q8N128 Q8N336 Q8N8L6 Q8NBQ5 Q8NEX9 Q8WTS1 Q8WWH4 Q8WYQ3 Q8WZA9 Q96AQ6 Q96BQ5 Q96CS3 Q96DB5 Q96EG1 Q96EZ4 Q96J77 Q96LJ7 Q96MZ0 Q96T59 Q99675 Q9BYD5 Q9H078 Q9H0P0 Q9H173 Q9H1A3 Q9H3Z7 Q9H4I3 Q9H6V9 Q9NPH0 Q9NR28 Q9NR31 Q9NRG7 Q9NRI7 Q9NV66 Q9NZC7 Q9NZF1 Q9UG22 Q9UHT4 Q9UKA2 Q9Y2L9 Q9Y2W6 Q9Y6H1
