## Supplementary material for "ProtRAP-LM: Fast and accurate protein relative accessibility prediction and membrane protein screening through protein language model embeddings": File S3

### Datasets Used in the Work

#### 1. The datasets for the ProtRAP-LM model

##### 1.1 The training set & validation set (1362 membrane protein sequences)

1A0SA 1A91A 1AY2A 1DX7A 1E54A 1E7PC 1EHKB 1FFTA 1FFTB 1FW2A 1FX8A 1H6SA
1HGVA 1IFMA 1IFPA 1IIJA 1K24A 1KF6A 1KF6C 1KMOA 1KQFB 1KQFC 1KYOB 1KZUA
1KZUB 1L7VC 1L9BD 1LGHB 1M56B 1MHSA 1MM4A 1MPRA 1NQEA 1OQWA 1P49A 1P4TA
1PW4A 1Q16B 1Q16C 1Q90B 1Q90D 1Q90E 1Q90G 1Q90H 1Q90I 1QFFA 1QLEC 1QLED
1RH5C 1SPFA 1T16A 1TLWA 1UUNA 1UYNA 1VF5E 1XIOA 1XKWA 1XL4A 1XMEC 1YC9A
1ZZAA 2A65A 2A9HE 2B2FA 2BL2A 2BS2A 2BS2B 2C0XA 2ERVA 2F1CA 2F93A 2F95B
2FYNC 2HDFA 2HDIB 2HI7A 2IBZH 2IBZI 2J5DA 2JLNA 2K0LA 2K1KA 2K73A 2K9YA
2KLUA 2KNCA 2KS9A 2KS9B 2KSDA 2KSEA 2KSFA 2KV5A 2L16A 2L2TA 2L35A 2L5BA
2L6WA 2L8SA 2L9UA 2LATA 2LBGA 2LCKA 2LHFA 2LIFA 2LMEA 2LOQA 2LORA 2LOSA
2LOWA 2LS3A 2LX0A 2LZLA 2LZRA 2M3EA 2M59A 2M67A 2MC7A 2MFRA 2MGYA 2MI2A
2MJOA 2MLHA 2MMUA 2MOFA 2MPNA 2MXBA 2N1PA 2N28A 2N2AA 2N4XA 2N5SA 2N7IA
2N90A 2NA6A 2NA7A 2NA8A 2NMRA 2NQ2A 2NQ2B 2NR9A 2NRGA 2NUUG 2O4VA 2OARA
2OAUA 2ONKI 2PORA 2QFIA 2RDDB 2RQ2A 2VOYA 2VPWA 2VPWB 2VPWC 2W1PA 2WITA
2WJQA 2WSEC 2WSXA 2WW9A 2WW9C 2WW9H 2WW9I 2WWBC 2X27A 2X2VA 2XQ2A 2XZBB
2Y5YA 2Y69E 2YEVA 2YEVB 2YEVC 2YNKA 2YVXA 2Z73A 2ZJSB 2ZT9D 2ZT9E 2ZT9F
2ZT9H 2ZXEB 2ZXEC 3A7KA 3ANZA 3AYFA 3B4RA 3B5DA 3B9WA 3BO0E 3C02A 3CSLA
3CSLC 3CX5F 3D31A 3D31C 3DINA 3DIND 3DWOA 3DZMA 3EFMA 3EH3A 3EMOA 3FHHA
3FIDA 3GI8A 3HD6A 3HD7A 3HD7B 3HD7C 3HD7D 3J2PB 3J45F 3J46G 3J9TJ 3J9TK
3J9TL 3J9VE 3JC2C 3JTYA 3K3FA 3KP9A 3KVNA 3LLQA 3M71A 3MK7A 3MK7B 3MK7C
3MP7B 3NE2A 3NE5A 3NJTA 3NOCD 3O0RC 3O7PA 3ORGA 3P5NA 3PIKA 3PJZA 3PL9A
3PQRB 3PUVB 3Q7KA 3QF4A 3QNQA 3QQ2A 3QRAA 3RCEA 3RFUA 3RKOA 3RKOF 3RLBA
3S0XA 3SLJA 3SOKA 3SY9A 3SYBA 3TUIC 3UDCA 3UKMA 3UQ4A 3UX4A 3V5SA 3V89A
3V8XB 3VMQA 3VR8C 3VR8D 3VVRB 3VVSB 3W5AC 3W9TA 3WAJA 3WBNB 3WDOA 3WMFA
3WMGB 3WMMA 3WMMA 3WMMB 3WO7A 3X2RA 3X3BA 3ZD0A 3ZE3A 4A2NA 4AIQA 4AQ5A
4ATVA 4AUIA 4AYWA 4B4AA 4BEMA 4BGNA 4BOIE 4BPMA 4BUMA 4BWZA 4C00A 4CADC
4CDIB 4CG6D 4CHVA 4CU4B 4CZ8A 4CZBA 4D5BA 4DBLA 4DBLE 4DCBA 4DCBB 4DJIA
4DVEA 4DXWA 4E1TA 4EPAA 4EV6A 4EZCA 4F35A 4FC4A 4FQEA 4FSOA 4FZ0D 4G7VC
4GBYA 4GD3A 4GD3B 4GD3I 4GEYA 4GX0A 4H56A 4HEAF 4HEAG 4HEAH 4HEAI 4HKRA
4HUKA 4HUQB 4HUQC 4HW9A 4HYCA 4HYJA 4HZUD 4IL3A 4IN7A 4J05A 4J72A 4J7CI
4JBWB 4JBWE 4K1CA 4K3BA 4KJRA 4KNFA 4KPPA 4KYTB 4LDSA 4M58A 4M64A 4MLBA
4MNDA 4MT4A 4N7XA 4NJNA 4NTWB 4NTWC 4O6MA 4O6YA 4O9PA 4O9UB 4O9UE 4OD4A
4OH3A 4OJ2A 4P00A 4P6JA 4P6VA 4P6VB 4P6VC 4P6VD 4P6VE 4P6VF 4P79A 4PGRA
4PHZE 4PL0A 4PX7A 4Q2EA 4Q65A 4QNDA 4QTNA 4QUVA 4R1IA 4RDRA 4RFSC 4RI2A
4RL8A 4RLBA 4RLCA 4RNGA 4RP8A 4RY2A 4RYIA 4TKRA 4TQ3A 4TQUA 4TQUB 4TQUC
4TQUD 4TSYA 4U4TA 4U5BE 4UC1A 4UMVA 4US3A 4V1FA 4W6VA 4WD7A 4WMZA 4X5MA
4XE5C 4XNKA 4XNVA 4XT1A 4XT1B 4XU4A 4XYDB 4Y25A 4Y7JA 4YBQA 4YMSC 4YXDA
4YXDB 4YXDC 4YZFA 4Z3NA 4Z7FA 4ZGVA 4ZR0A 4ZYOA 5A1SA 5A40A 5A63A 5A63B
5A63C 5AEZA 5AWWC 5AWZA 5AYMA 5AZBA 5AZOA 5AZSA 5B57A 5B57C 5B58E 5BUTA

5C6OA 5CBFA 5CTGA 5D0QB 5D0QC 5D3MB 5D3MC 5D3MD 5DA0A 5DIRA 5DJQM 5DL5A
5DL7A 5DL8A 5DN6J 5DN6K 5DN6L 5DN6M 5DO7A 5DOQA 5DOQB 5DOQC 5DWYA 5E1JA
5EC2C 5EDLA 5EIKA 5EKEA 5EULB 5EULC 5EZMA 5F1CA 5FGNA 5FGZA 5FL7G 5FP1A
5FQ6A 5FQ6B 5FQ6E 5FQ7G 5FR8A 5GAEC 5GAED 5GAEE 5GAEF 5GAEG 5GAEH 5GAEI
5GASM 5GASO 5GPJA 5GUFA 5H1QA 5H2FA 5HI9A 5HK1A 5I20A 5I32A 5I6CA 5I9KA
5IJIA 5ITCA 5IV8B 5IV9A 5IVAA 5IVAB 5IWSA 5J4IA 5JKIA 5JNQA 5JRWA 5JSIA
5K7LA 5KBNA 5KC0A 5KKZB 5KSDA 5L22A 5L24A 5L25A 5L75C 5LBJA 5LDVA 5LILA
5LQXL 5LQXM 5LWEA 5LXGA 5MDOA 5MG3B 5MG3C 5MG3D 5MG3E 5MG3F 5MLZA 5MRWA
5N6MA 5N9YA 5NAOA 5NUQG 5NV9A 5O0TA 5O65A 5O8OA 5OC0A 5OC9A 5ODWA 5ODWC
5OEKA 5OGEA 5OKDH 5ONUA 5OQKA 5OQTA 5OYBA 5OYKA 5SV9A 5SVJA 5SY1B 5T77A
5TCXA 5TSAA 5TTPA 5U1DB 5U1DC 5UIWB 5UOWD 5V2SA 5V4SA 5V7PA 5V8KA 5V8KB
5VA2A 5VKVA 5W63A 5WPQA 5WUCA 5WUFA 5X3XA 5X3XC 5X41D 5X5YC 5X5YD 5XAMA
5XJ6A 5XLSA 5XNNC 5XNND 5XTED 5XTEF 5XTEG 5XU1C 5XW6A 5Y78A 5Y83A 5YCKA
5YHLA 5YQZB 5YS3A 5Z1FA 5Z1LA 5Z62D 5Z62G 5Z62I 5ZAZA 5ZBGA 5ZBQA 5ZKPA
5ZLEA 6A2JA 6A90B 6A93A 6AB9A 6AWFB 6AWFD 6B3JB 6B85A 6B87A 6BH8A 6BHPA
6BMSA 6BPMA 6BTMA 6BTMB 6BTMC 6BTMD 6BTME 6BTMF 6BUGA 6BUIA 6BW6A 6C08C
6C14A 6C14B 6C26F 6C3IA 6C4XA 6C6LD 6C70A 6C96A 6CB2A 6CFWA 6CFWB 6CFWC
6CFWD 6CFWE 6CFWF 6CFWG 6CFWH 6CFWI 6CJQA 6CNMA 6COYA 6CP7K 6CP7L 6CP7O
6CSEC 6CVLE 6CXHB 6CXHC 6D0JA 6D9ZA 6DDEE 6DT0A 6DU8A 6DZ7A 6E14A 6E14B
6E14C 6E14D 6E15A 6E2GA 6E3YA 6E4VA 6E8JA 6E9NA 6EBUA 6EDQA 6EI3A 6EIDA
6EU6A 6EUQA 6EUSA 6EYUA 6EZNA 6EZNE 6EZNH 6F0KF 6F0KG 6F2DG 6F2GA 6F46A
6FFVA 6FKFJ 6FKHA 6FKIF 6FOKA 6FOSA 6FOSC 6FV6A 6G1KA 6G79D 6GCIA 6GIEA
6GS1A 6GZ9A 6H2FA 6H3IB 6H3IC 6H5AA 6H7DA 6H7FA 6HCYA 6HD9B 6HJAA 6HJRA
6HJRB 6HRAC 6HRAD 6HUGC 6HUMA 6HUMB 6HUMD 6I1PB 6I53A 6I8WA 6I97C 6IBBA
6IDFE 6IDPA 6IGKB 6IGLB 6IGZE 6IGZG 6IJJC 6IJOG 6ITCE 6IU3A 6IVAA 6IVAB
6IYCE 6IZ3A 6J54I 6J8EC 6JBHA 6JBHC 6JI1A 6JP8A 6JPAD 6JXRA 6JXRC 6JXRD
6JXRE 6K1HA 6K1HB 6K41D 6K61B 6K61C 6K61E 6K61I 6K7YI 6KHJF 6KHJG 6KHJI
6KI1A 6KKIA 6KO5A 6KOBC 6KOEB 6KSWA 6L3TA 6L85A 6L95A 6LCPB 6LEOA 6LFMD
6LFMF 6LI3A 6LI9A 6LKNB 6LM0A 6LMVA 6LO8A 6LO8B 6LO8C 6LO8D 6LO8E 6LO8F
6LO8H 6LODA 6LODB 6LODD 6LODE 6LODF 6LRYA 6LRYB 6LU1I 6LUMA 6LUMB 6LUMC
6LUME 6LVFA 6LY9A 6LY9O 6LYGA 6LYQB 6M17B 6M17C 6M1HB 6M20A 6M32A 6M32D
6M32E 6M32F 6M49A 6M49B 6M4NA 6M4NB 6M4NC 6M4ND 6M96B 6MCTA 6MEOA 6MGVA
6MHOA 6MHUB 6MHZA 6MI7E 6MPNA 6MQUA 6N1GA 6N3QF 6N4BD 6N4IA 6N4IE 6NBFB
6NF4A 6NHWA 6NIYF 6NK5A 6NK5B 6NK5C 6NK7M 6NT4B 6NT8A 6NWDA 6NZWA 6O20C
6O58B 6O7UA 6O7UB 6O7UF 6OFRA 6OH2A 6OIJA 6OIJB 6OMMA 6OQRA 6OQRF 6OQRG
6OQRT 6OSAA 6OV2A 6P25A 6P25B 6P46A 6P9XE 6P9YE 6P9YF 6PB0B 6PE4F 6PE4P
6PISA 6PL5B 6PVRA 6PW4A 6Q53A 6QIMA 6QUMD 6QUMG 6QUMH 6QUMJ 6R2QA 6R2QB
6R2QC 6R6BA 6R6BF 6R6BG 6RAFA 6RAFB 6RBGA 6RD7A 6RD7F 6RD7G 6RD7R 6RDPN
6REUK 6RKOA 6RKOC 6RQFC 6RQFE 6RQFG 6RV2A 6RX4A 6RX4C 6RYOA 6RZ6A 6S3LG
6S3RA 6S3SF 6S7OC 6S7OD 6S7OH 6S7TA 6S7TE 6S7TF 6S7TH 6S7TI 6S7VA 6SEMA
6SGZE 6SH3A 6SL5I 6SLIA 6SLIC 6SLJA 6SLJE 6SLNE 6SM3C 6SMLC 6SMQC 6SNHA
6SO5A 6SO5C 6SO5E 6SSRA 6SU3A 6T1ZA 6T6VA 6TD6B 6TDXA 6TDXB 6TDXC 6TDXD
6TDXN 6TEJA 6TJVD 6TJVF 6TMHA 6TMHB 6TMHE 6TMHH 6TMHI 6TMHJ 6TMHK 6TMHL
6TQEA 6U0OA 6U0OB 6U1SA 6U5RA 6UCUB 6UCUE 6UKJA 6UMMB 6UMMD 6UMME 6UUSE

6UVAF 6UWMA 6UWZF 6UZZA 6V01B 6V05F 6V1QA 6V22B 6V2JA 6V4AA 6V4DA 6V4KE

6V78A 6V9ZB 6VAIA 6VD7A 6VEJD 6VGCA 6VJAA 6VJYB 6VJZA 6VK0A 6VK1A 6VMBJ

6VMBK 6VMSC 6VOHJ 6VQCA 6VQHB 6VQHC 6VQHF 6VQHI 6VQQA 6VWKK 6VX5A 6VXMA

6W2XB 6W4SA 6W5VB 6W6MA 6W6OB 6W8NA 6WB9A 6WBVA 6WHGA 6WHRA 6WILA 6WIMA

6WJ6A 6WJ6D 6WJ6E 6WJ6F 6WJ6G 6WJ6H 6WJ6I 6WK5A 6WLWM 6WLWN 6WLWO 6WLWP

6WM0A 6WR4A 6WU0A 6WU1A 6WU6B 6WU6C 6WVGA 6WW7A 6WW7B 6WW7C 6WW7F 6WW7G

6WWZB 6WXHA 6WXHD 6WZGA 6X4SA 6X9ZA 6XBJA 6XCRA 6XDCA 6XJHA 6XJHC 6XJVJ

6XXDA 6XXEA 6Y1ZA 6YB1A 6YKMA 6YMYA 6YMYB 6YMYC 6YMYD 6YMYE 6YMYG 6YMYH

6YMYI 6YNWA 6YNWL 6YNWM 6YS8A 6YS8C 6YSFA 6YSFC 6YSLA 6YSLB 6YV7A 6Z0FA

6Z0GA 6Z16A 6Z16B 6Z16C 6Z16D 6Z16E 6Z16F 6Z16G 6Z1RJ 6Z34A 6Z3YA 6Z5UE

6Z5UJ 6Z8AA 6Z9AA 6ZA9A 6ZA9I 6ZBVA 6ZDBA 6ZG3C 6ZHHA 6ZMRL 6ZMRO 6ZMRP

6ZPLC 6ZQJA 6ZQJB 6ZXAA 6ZXAB 6ZXRA 6ZY3G 7ABWA 7AD3A 7AD3C 7AD3E 7AD3F

7AD3G 7ADOE 7ADPD 7ADPG 7ADPH 7AHCA 7AHDA 7AJQA 7AJQF 7AKXA 7ARHA 7ARJC

7ARMF 7ATBA 7AUEA 7B0OA 7B9FA 7B9FB 7B9FE 7BB6A 7BC6A 7BMHA 7BP3A 7BPUA

7BR3A 7BRMJ 7BSBA 7BVEC 7C77A 7C77B 7C7SA 7C83A 7CADC 7CADD 7CAFA 7CALA

7CCSB 7CFFA 7CFTD 7CG4A 7CH1A 7CH8A 7CH8G 7CH9K 7CJ3A 7CK6A 7CK6C 7CMHA

7CP9A 7CP9C 7CP9E 7CQ5A 7CQ5C 7CR0A 7CU3B 7CUBD 7CX2A 7CYEA 7CYFD 7CZ5C

7CZ5D 7CZBA 7D06C 7D0IA 7D3EB 7D3UA 7D3UB 7D3UC 7D3UD 7D3UE 7D3UF 7D5PA

7D68E 7D68F 7D6VA 7D6VB 7D6VC 7D76D 7D7ED 7DB6E 7DEGA 7DEGB 7DEGC 7DH5E

7DL9A 7DQKA 7DRTA 7DSXA 7DSXB 7DTYB 7DWQF 7DXAB 7DXHA 7DYRC 7E1VA 7E1VC

7E1VE 7E1VF 7E4HC 7E4IA 7E4IB 7E4IE 7E4IF 7E8EA 7E8EC 7EJ1A 7EJ1B 7EKIA

7EKLA 7EN0A 7EW1E 7F0JA 7F16A 7F16B 7F4DF 7F56A 7F56E 7F8VE 7F8WE 7F9YC

7FIGF 7FIHD 7FIHF 7JGAG 7JGAH 7JGAJ 7JGAK 7JGBA 7JHJE 7JKCA 7JNAA 7JRGB

7JRGD 7JRGG 7JRGH 7JRGI 7JROD 7JROE 7JROF 7JROG 7JROH 7JROI 7JSJA 7JV5A

7K3GA 7K7AA 7KAID 7KAKA 7KAKB 7KAKD 7KAKE 7KAKF 7KALG 7KB5E 7KC4A 7KC4B

7KI1F 7KJRC 7KN0B 7KP4B 7KRAA 7KRAB 7KRAC 7KRAF 7KRAG 7KTXD 7KTXE 7KY7B

7KYOA 7KYPB 7L0PE 7L1UE 7L2ME 7LC3C 7LC8A 7LD3B 7LFTA 7LGUA 7LHGA 7LHGB

7LHGC 7LHIB 7LHIC 7LHVA 7LICA 7LOIA 7MBZA 7MDYA 7MHZB 7MIXB 7MJSC 7MLUK

7MPAA 7MTRA 7N6QA 7NCSE 7NH9A 7NHRA 7NKZA 7NKZB 7NP3A 7NS1A 7NSTE 7NSUD

7NYRC 7NYRD 7NYRE 7NYVG 7O3HA 7O3HB 7O3HC 7O3HG 7O7FC 7OCAC 7OFOA 7OJ8A

7OQZA 7OZ1A 7P14A 7P34A 7P4IA 7P5CA 7PHPA 7PHQC 7R87B 7RH6D 7RH7G 7RH7H

7RJAA 7RJAD 7RJBH 7RNNB 7RTMA 7RTTA

#### 1.2 The training set & validation set (7440 soluble protein sequences)

12ASA 1A1XA 1A2XB 1A32A 1A62A 1A76A 1A8LA 1A9XB 1AD6A 1AFRA 1AH7A 1AHSA

1AL3A 1ALYA 1AM7A 1AOLA 1ATZA 1AYOA 1B33N 1B35A 1B35B 1B35C 1B3UA 1B5EA

1B63A 1B67A 1B77A 1B9WA 1BAZA 1BBPA 1BF2A 1BG2A 1BGFA 1BHEA 1BJAA 1BKBA

1BKRA 1BM8A 1BNDA 1BTKB 1BXYA 1BYFA 1BYIA 1C0PA 1C1KA 1C1XA 1C5EA 1C7KA

1C8KA 1C8NA 1C8UA 1C8ZA 1CAUB 1CB8A 1CC8A 1CCWA 1CCWB 1CFBA 1CFRA 1CHDA

1CHMA 1CI9A 1CJCA 1CQ3A 1CV8A 1CVRA 1CY5A 1D2NA 1D2OA 1D2SA 1D2ZB 1D3BA

1D4AA 1D4VB 1D5TA 1D9CA 1DBTA 1DBWA 1DE6A 1DEVB 1DGWA 1DJ0A 1DJ8A 1DP4A

1DP5B 1DQGA 1DQPA 1DS1A 1DUNA 1DVOA 1DYNA 1DYOA 1DYPA 1DZFA 1E20A 1E3JA

1E58A 1E5KA 1E6UA 1E7LA 1EAQA 1EAYC 1EAZA 1EB6A 1EERA 1EF1C 1EFDN 1EGPA

1EI5A 1EI9A 1EJ8A 1EJFA 1EK6A 1EL6A 1ELKA 1ELUA 1EM9A 1EOKA 1EPAA 1EQ2D

1ES5A 1ES9A 1ESCA 1EUWA 1EVLA 1EW4A 1EWFA 1EZGA 1EZJA 1EZWA 1F08A 1F1MA
1F3UA 1F3VA 1F46A 1F86A 1FC3A 1FCYA 1FD3A 1FG7A 1FJRA 1FLTX 1FN9A 1FP3A
1FPOA 1FS1A 1FSGA 1FT5A 1FTRA 1FX2A 1FXKB 1FXKC 1G2RA 1G3KA 1G5BA 1G61A
1G66A 1G6CA 1G6GA 1G6XA 1G8AA 1G8EA 1G8MA 1GAKA 1GCIA 1GHEA 1GJWA 1GK8I
1GK9A 1GK9B 1GKMA 1GMUA 1GMXA 1GNLA 1GNYA 1GO3F 1GP0A 1GPLA 1GPRA 1GQEA
1GQIA 1GS9A 1GSAA 1GU2A 1GU7A 1GUIA 1GUQA 1GUTA 1GUXB 1GVEA 1GVPA 1GWEA
1GWMA 1GXMA 1GXYA 1GZSB 1H12A 1H16A 1H2VZ 1H3OB 1H4XA 1H6GA 1H7CA 1H8EH
1H8PA 1H8UA 1H97A 1HDFA 1HDHA 1HDOA 1HE1A 1HF2B 1HH8A 1HI9A 1HQ0A 1HQZ1
1HRUA 1HTRP 1HULA 1HW7A 1HX6A 1HXIA 1HXNA 1HXRB 1HYOA 1HYPA 1HZ4A 1HZ6A
1HZTA 1I1QA 1I1RB 1I1WA 1I24A 1I27A 1I2TA 1I4UA 1I58A 1I5RA 1I60A 1I7NA
1I8AA 1I8NA 1IA9B 1ID0A 1IDPA 1IFGA 1IFQA 1IFRA 1IG0A 1IGQA 1IKOP 1IKTA
1ILKA 1INLC 1INPA 1IO0A 1IQ4A 1IQZA 1IRQA 1IS3A 1ISUA 1IT2A 1IUGA 1IX9A
1IXHA 1IZCA 1J0PA 1J1TA 1J24A 1J27A 1J3WA 1J5UA 1J6RA 1J77A 1J7XA 1J83A
1J98A 1JADA 1JB3A 1JBEA 1JCDA 1JCLA 1JEOA 1JETA 1JF3A 1JFBA 1JH6A 1JHJA
1JHSA 1JKEA 1JLJA 1JLYA 1JMKO 1JMVA 1JNRA 1JO0A 1JOGA 1JOSA 1JOVA 1JR2A
1JS1X 1JSUC 1JU2A 1JUVA 1JX6A 1JYAA 1JYHA 1JYOA 1K2XA 1K3RA 1K4IA 1K5CA
1K68A 1K6KA 1K77A 1K7CA 1K7JA 1K8KD 1K8KE 1K8KF 1K8UA 1KA1A 1KAEA 1KAFA
1KCMA 1KDGA 1KHYD 1KKOA 1KMTA 1KOEA 1KPGA 1KPTA 1KQ3A 1KQ6A 1KQPA 1KT6A
1KTGA 1KVEB 1KW2A 1KWFA 1KYFA 1KYQB 1L0SA 1L2WI 1L7AA 1L9LA 1L9XA 1LBVA
1LC5A 1LDDA 1LF6A 1LFPA 1LFWA 1LJ8A 1LK2B 1LKIA 1LMIA 1LN1A 1LNIA 1LO7A
1LPBA 1LQ9A 1LQAA 1LQVA 1LR0A 1LSHB 1LTSC 1LXJA 1M15A 1M1HA 1M1QA 1M1SA
1M2DA 1M2XA 1M4LA 1M4RA 1M5WA 1M6SA 1M6UA 1M7GA 1M9ZA 1MAIA 1MBMA 1MIXA
1MJ5A 1MJNA 1MK4A 1MKFA 1MN8A 1MO9A 1MPGA 1MSCA 1MT6A 1MTZA 1MUWA 1MW5A
1MWQA 1MXRA 1MY7A 1N08A 1N12A 1N13A 1N13B 1N1EA 1N2DC 1N57A 1N62A 1N62B
1N62C 1N67A 1N69A 1N7KA 1N7SC 1N7UA 1N7ZA 1N81A 1N8VA 1NBWB 1NC7A 1NEPA
1NEZG 1NFPA 1NG6A 1NH1A 1NIJA 1NKDA 1NKGA 1NKIA 1NLQA 1NMOA 1NN3A 1NNWA
1NO1A 1NR0A 1NSZA 1NTVA 1NU7D 1NUYA 1NWZA 1NXMA 1NXUA 1NYCA 1NYTA 1NZ0A
1NZEA 1O1YA 1O22A 1O26A 1O4WA 1O6DA 1O7IA 1O7JA 1O7ZA 1O94A 1O9GA 1O9YA
1OA8A 1OAAA 1OAIA 1OAOC 1OC0B 1OC7A 1OCKA 1OCYA 1OD3A 1ODMA 1OEWA 1OFLA
1OFUX 1OGQA 1OH4A 1OHUB 1OIOA 1OJHA 1OK0A 1OKSA 1OMSA 1ON0A 1OO0A 1OOEA
1OQ1A 1OQJA 1OR4A 1ORUA 1OSYA 1OTKA 1OU9A 1OUWC 1OW4A 1P1JA 1P1MA 1P35A
1P3CA 1P3QR 1P57A 1P5VB 1P6OA 1P6XA 1P90A 1P9GA 1PAQA 1PBWA 1PBYC 1PDKB
1PDOA 1PI1A 1PKHA 1PMHX 1PN4A 1PP0B 1PQZA 1PSRA 1PU6A 1PUCA 1PV5A 1PYMA
1PYOB 1PYTA 1PZ4A 1Q19A 1Q1FA 1Q2HA 1Q2LA 1Q5YA 1Q5ZA 1Q6OA 1Q6ZA 1Q7LA
1Q7LB 1Q88B 1Q8BA 1QAZA 1QB2A 1QC7A 1QD1B 1QEXA 1QFHA 1QFTA 1QHDA 1QKRA
1QKSA 1QLWA 1QNRA 1QQ5A 1QQFA 1QQRA 1QR0A 1QSTA 1QW2A 1QW9A 1QWIA 1QWOA
1QYNA 1QZ7B 1QZ9A 1QZMA 1R0DA 1R44A 1R45A 1R4VA 1R4XA 1R5ZA 1R6DA 1R6JA
1R6UA 1R6XA 1R7JA 1R7LA 1R89A 1R9LA 1RA0A 1RCQA 1RFYA 1RG8A 1RHCA 1RIFA
1RJDA 1RK6A 1RK8C 1RKIA 1RKTA 1RKUA 1RL0A 1RLMA 1RMGA 1RO2A 1RO5A 1RO7A
1ROCA 1RP4A 1RQPA 1RTQA 1RTTA 1RTWA 1RWHA 1RWZA 1RXQA 1RY9A 1RYIA 1RYOA
1RYPL 1S0PA 1S12A 1S1DA 1S2OA 1S2XA 1S35A 1S3CA 1S4CB 1S5UA 1S67L 1S7HA
1S7ZA 1S98A 1S9RA 1SAUA 1SBXA 1SBYA 1SD4A 1SEDA 1SEFA 1SEIA 1SENA 1SFPA
1SFSA 1SG4A 1SGMA 1SHUX 1SI7A 1SKZA 1SMXA 1SQGA 1SQHA 1SQWA 1SR4A 1SR8A
1ST0A 1ST9A 1STMA 1STZA 1SULB 1SUMB 1SVVA 1SYXB 1SZ9A 1SZHA 1T07A 1T0BA

1T0FA 1T1IA 1T1UA 1T2DA 1T4BA 1T57A 1T61A 1T6CA 1T6UA 1T77A 1T82A 1T8KA
1T92A 1TBFA 1TE5A 1TF1A 1TFEA 1TFKB 1TH7A 1TIIC 1TIQA 1TJLA 1TKEA 1TKIA
1TKJA 1TQGA 1TQYB 1TS9A 1TT8A 1TTWA 1TU1A 1TU3F 1TU9A 1TUAA 1TUKA 1TUVA
1TUWA 1TV8A 1TVGA 1TWUA 1U07A 1U14A 1U2HA 1U2KA 1U2MC 1U53A 1U5UA 1U60A
1U6TA 1U7IA 1U84A 1U9KA 1UA7A 1UCDA 1UCRA 1UCSA 1UEBA 1UEKA 1UF5A 1UFIA
1UFOA 1UFYA 1UGIA 1UIXA 1UKFA 1UM0A 1UNQA 1UPTB 1URQA 1US0A 1USCA 1USGA
1USOA 1UTGA 1UTYA 1UUJA 1UW4B 1UWKA 1UZ3A 1V05A 1V2ZA 1V30A 1V33A 1V4EA
1V4PA 1V58A 1V5IB 1V5VA 1V6PA 1V6ZA 1V74A 1V77A 1V7ZA 1V8HA 1V96B 1V9FA
1VAJA 1VBKA 1VBWA 1VCCA 1VD6A 1VF6C 1VF7A 1VH5A 1VHNA 1VHQA 1VI6A 1VJ0A
1VJFA 1VJGA 1VJQA 1VK1A 1VKEB 1VKHB 1VKKA 1VL1A 1VL7A 1VLJA 1VLYA 1VMHA
1VMOA 1VPMA 1VPQA 1VPRA 1VQRC 1VQSA 1VR7A 1VRMA 1VSRA 1VYBA 1VYRA 1VZJA
1VZYA 1W07B 1W0NA 1W1HA 1W23A 1W2WA 1W2WB 1W41A 1W4SA 1W5RA 1W66A 1W6SA
1W6SB 1W85C 1W8IA 1W99A 1WB4A 1WC9A 1WCWA 1WDDA 1WDJA 1WEHA 1WERA 1WF3A
1WHIA 1WHSA 1WHZA 1WKAA 1WKQA 1WL8A 1WLGA 1WLUA 1WLZA 1WMGA 1WMHB 1WMIA
1WMWA 1WN2A 1WNAA 1WOLA 1WPBA 1WQ6A 1WQJB 1WQLB 1WRDA 1WTJA 1WU3I 1WURA
1WV3A 1WVFA 1WVQA 1WWIA 1WWPA 1WZ3A 1WZDA 1WZZA 1X19A 1X3LA 1X54A 1X6IA
1X6MA 1X6ZA 1X7DA 1X7YB 1X8QA 1X91A 1X9IA 1X9UA 1X9ZA 1XAKA 1XAUA 1XAWA
1XBIA 1XCRA 1XD3A 1XDNA 1XFFA 1XFKA 1XG0A 1XG0D 1XG5C 1XHNA 1XK5A 1XK7A
1XKNA 1XKPB 1XKPC 1XKRA 1XL3B 1XL3C 1XLQA 1XMKA 1XMTA 1XQOA 1XQRA 1XR4A
1XRUA 1XSVA 1XSZA 1XTEA 1XU1A 1XUBA 1XUVA 1XV5A 1XW3A 1XXLA 1Y0HA 1Y2OA
1Y43B 1Y60A 1Y6XA 1Y7PB 1Y88A 1Y8AA 1Y93A 1Y9LA 1YA5T 1YACA 1YB3A 1YCDA
1YDXA 1YF2A 1YFQA 1YG9A 1YGAA 1YGEA 1YGTA 1YHTA 1YIXA 1YIZA 1YK3A 1YLLC
1YLMA 1YLOA 1YLXA 1YN3A 1YNHA 1YNSA 1YOCA 1YOZB 1YP0A 1YPYA 1YQHA 1YQSA
1YRBA 1YREA 1YRKA 1YTLA 1YU0A 1YWQA 1YX1A 1YYQA 1YZVA 1YZYA 1Z0SA 1Z1SA
1Z2NX 1Z2UA 1Z3EB 1Z5GA 1Z6NA 1Z6OA 1Z7XW 1ZAVA 1ZB1A 1ZBA2 1ZBPA 1ZBSA
1ZC3B 1ZCEA 1ZD0A 1ZE3H 1ZEQX 1ZGHA 1ZHSA 1ZHVA 1ZI8A 1ZJCA 1ZJRA 1ZK4A
1ZKGA 1ZL0A 1ZO2A 1ZPSA 1ZR6A 1ZSQA 1ZT3A 1ZUUA 1ZV1A 1ZVFA 1ZVTB 1ZX8A
1ZXXA 1ZYNA 1ZZKA 2A0BA 2A0MA 2A14A 2A1KA 2A1VA 2A26A 2A2MA 2A40B 2A4NA
2A5HA 2A5ZA 2A6BA 2A6ZA 2A7KB 2A9UA 2AAMA 2AB5A 2ABWB 2ABZC 2AEGA 2AEUA
2AIBA 2AJ7A 2AKAA 2AKAB 2AKZA 2AMLA 2AOTA 2APCA 2APJA 2AQ6A 2ARCA 2ASFA
2ASKA 2ATMA 2AU3A 2AVWA 2AXQA 2AXWA 2AYDA 2AZ4A 2AZJA 2B0AA 2B0TA 2B0VA
2B1YA 2B3YA 2B4HA 2B56A 2B5WA 2B67A 2B8MA 2B8TA 2B97A 2B9WA 2BA2A 2BAYA
2BBDA 2BCMA 2BDRA 2BEZF 2BHZA 2BI0A 2BJIA 2BJQA 2BK9A 2BKFA 2BKMA 2BKRA
2BKXA 2BKYA 2BL8B 2BMOA 2BMOB 2BNLC 2BSJA 2BT9A 2BW0A 2BW3B 2BWRA 2BYKA
2BZ1A 2C0DA 2C0GA 2C1LA 2C1MB 2C2UA 2C3GA 2C3VA 2C5AA 2C5JB 2C5QA 2C71A
2C7NA 2C8SA 2C92A 2CARA 2CAYA 2CB8A 2CBOA 2CC6A 2CCQA 2CDPA 2CE2X 2CE8A
2CG7A 2CI1A 2CIUA 2CIWA 2CJ4A 2CJTC 2CKKA 2CLBA 2CLSA 2CMGA 2CO3A 2COVD
2CW9A 2CWSA 2CX7A 2CXHA 2CXIA 2CXKA 2CXNA 2CXYA 2CY5A 2CYJA 2CZCA 2CZLA
2CZVC 2D0OA 2D1LA 2D1SA 2D27A 2D3DA 2D4PA 2D58A 2D5WA 2D68B 2D73A 2D7VA
2DC4A 2DDRC 2DDXA 2DDZA 2DG1A 2DHOA 2DKJA 2DKOA 2DKOB 2DLBA 2DM9A 2DP9A
2DRVB 2DSKA 2DT4A 2DT8A 2DTCA 2DTJA 2DURA 2DWUA 2DXAA 2DXQA 2DY1A 2DYJA
2E11A 2E12A 2E1HB 2E1NA 2E2DC 2E2OA 2E3PA 2E4MC 2E4TA 2E56A 2E5FA 2E5YA
2E8BA 2E8EA 2E9XA 2E9XB 2EABA 2EBEA 2EBFX 2ECEA 2ECUA 2ED6A 2EFEA 2EFVA
2EGJA 2EGVA 2EHPA 2EHWA 2EHZA 2EI9A 2EIYB 2EJQA 2EJXA 2EKDA 2EKLA 2ELCA

2ENDA 2EPOA 2ERFA 2ES4E 2ETSA 2ETVA 2EV1A 2EW0A 2EWCA 2EX2A 2EY4C 2EYUA

2F01A 2F1FA 2F1NA 2F1SA 2F20A 2F22A 2F23A 2F2HA 2F46A 2F48A 2F4IA 2F4MA

2F5GA 2F5TX 2F5VA 2F60K 2F6MB 2F9HA 2FA1A 2FAOA 2FB0A 2FB6A 2FBAA 2FBIA

2FBYA 2FCJA 2FCLA 2FCUA 2FDBM 2FE1A 2FE7B 2FEAA 2FFGA 2FGEA 2FHPA 2FHZA

2FHZB 2FI1A 2FIPA 2FIUA 2FIWA 2FJ8A 2FJI1 2FJRA 2FL4A 2FMAA 2FNOA 2FNUA

2FOTC 2FP1A 2FPHX 2FQ3A 2FR5A 2FREA 2FSDA 2FT0A 2FTRA 2FUEA 2FULB 2FVYA

2FW5A 2FWHA 2FX5A 2FYZA 2FZ4A 2FZPA 2G0WA 2G19A 2G30A 2G3RA 2G3WA 2G5CB

2G5GX 2G7GA 2G7OA 2G7SA 2G8LA 2GA1A 2GAGB 2GAGC 2GAXA 2GB4A 2GBBA 2GDMA

2GDQA 2GECA 2GENA 2GEYA 2GF3A 2GF4A 2GFFA 2GFQA 2GGCA 2GH0A 2GHSA 2GHTA

2GIAB 2GK4A 2GKEA 2GKGA 2GKPA 2GLZA 2GMYA 2GNOA 2GOMA 2GPIA 2GQ0A 2GQTA

2GQWA 2GRRB 2GS5A 2GSCB 2GSOA 2GSVA 2GT1A 2GT4A 2GU3A 2GU9A 2GUDA 2GUIA

2GUKA 2GUPA 2GUZB 2GWFA 2GWMA 2GXGA 2GXQA 2GYQB 2GZ4A 2H1TA 2H1VA 2H21B

2H30A 2H5CA 2H62C 2H6FA 2H8EA 2H8GA 2H8LA 2H9AA 2H9AB 2HA8A 2HALA 2HBAA

2HBJA 2HBOA 2HBVA 2HCJB 2HD9A 2HDOA 2HDZA 2HEUA 2HEWF 2HF9A 2HFSA 2HGSA

2HINA 2HKJA 2HKVA 2HNGA 2HNHA 2HNUA 2HOXA 2HP0A 2HQ7A 2HQLA 2HQTB 2HQYA

2HS3A 2HSBA 2HSNA 2HTDA 2HUHA 2HUJA 2HW2A 2HX5A 2HXRA 2HXTA 2HY5A 2HY5B

2HZCA 2HZGA 2I00A 2I2CA 2I2OA 2I39A 2I3DA 2I44A 2I49A 2I4LA 2I53A 2I5IA

2I5UA 2I5VO 2I6HA 2I74A 2I79A 2I7GA 2I7HA 2I7RA 2I8DA 2I8EA 2I9CA 2I9DA

2I9IA 2I9XA 2IA7A 2IAYA 2IAZB 2IB0A 2IBDA 2IBGA 2IBLA 2IC6A 2ICCA 2ICGA

2ICWG 2ID4A 2IDLA 2IECA 2IF6A 2IFCA 2IG6A 2IG8A 2IGPA 2IHUA 2II2A 2IIHA

2IJES 2IJQA 2IJXA 2IKSA 2IMFA 2IMJA 2IMQX 2IN9A 2INWA 2IP6A 2IQJA 2IRPA

2ISBA 2ISKA 2IT9B 2IU4A 2IUMA 2IVFC 2IVNA 2IW1A 2IWRA 2IX2B 2IXAA 2IXSA

2IYVA 2IZXA 2J1VA 2J2JA 2J2ZB 2J3TC 2J3WA 2J3WB 2J43A 2J45A 2J4JF 2J4OA

2J5GA 2J6BA 2J6PA 2J6VA 2J6YA 2J7QA 2J85A 2J8KA 2J92A 2J9OA 2J9WA 2JA2A

2JBWA 2JCBA 2JCDA 2JDAA 2JDII 2JDJA 2JE6B 2JE8A 2JEKA 2JFKA 2JFRA 2JGPA

2JHFA 2JISB 2MHRA 2NLVA 2NMLA 2NN5A 2NNUA 2NO7A 2NOGA 2NP9A 2NQWA 2NR5C

2NR7A 2NS9A 2NSAA 2NSCA 2NSFA 2NSNA 2NSZA 2NTKA 2NTPA 2NUHA 2NUJA 2NV9B

2NVOA 2NW8A 2NWIA 2NX2A 2NX4A 2NXPA 2NZXC 2O0JA 2O0MA 2O0QA 2O14A 2O1KA

2O1MA 2O1QA 2O2GA 2O2XA 2O34A 2O38A 2O3BB 2O4TA 2O57A 2O5HB 2O5NB 2O62A

2O6KA 2O6PA 2O71A 2O7IA 2O8PA 2O90A 2O9SA 2O9UX 2OAFA 2OB3A 2OBDA 2OBLA

2OCTA 2OD0A 2OD4A 2OD6A 2ODFA 2ODKA 2ODLA 2OEBA 2OF3A 2OFCA 2OG4A 2OH1A

2OITA 2OIWA 2OIZA 2OIZD 2OJHA 2OKMA 2OKQA 2OKTA 2OKUA 2OLNA 2OLTA 2OM6A

2OMKA 2ONFA 2OOJB 2OP5A 2OPCA 2OPEA 2OPLA 2OQBA 2OQMA 2ORYB 2OSXA 2OTMA

2OU3A 2OU5A 2OU6A 2OUIA 2OV0A 2OV9A 2OVGA 2OVJA 2OWLA 2OWNA 2OX6A 2OX7A

2OXGB 2OXLA 2OYOA 2OZEA 2OZGA 2OZHA 2OZNA 2OZTA 2P0HA 2P0MA 2P0NA 2P0SB

2P0TA 2P11B 2P12A 2P14A 2P17A 2P2EA 2P2SA 2P38A 2P3HA 2P3PA 2P51A 2P58A

2P58C 2P65A 2P6WA 2P8IA 2P90A 2P97A 2P9HA 2P9XA 2PA7A 2PBLA 2PC1A 2PD0A

2PD1A 2PFMA 2PGFA 2PGNA 2PHNA 2PHPA 2PIMA 2PJSA 2PKDA 2PKFA 2PLRA 2PN2A

2PNEA 2POCB 2POFA 2PPXA 2PQRD 2PQVA 2PQXA 2PR7A 2PRVA 2PTTA 2PU3A 2PUZA

2PV2A 2PV4A 2PVBA 2PWWA 2PXXA 2PYWA 2PYXA 2Q03A 2Q0OC 2Q0TA 2Q0XA 2Q14A

2Q28A 2Q2FA 2Q35A 2Q3QA 2Q3TA 2Q3WA 2Q4WA 2Q4XA 2Q4ZA 2Q58A 2Q5CA 2Q5WE

2Q7AA 2Q7SB 2Q88A 2Q8OA 2Q8PA 2Q9KA 2Q9RA 2QAPA 2QB7A 2QBUA 2QCUA 2QDLA

2QE8A 2QE9B 2QEEA 2QEUA 2QEVA 2QF4A 2QFAB 2QFAC 2QFEA 2QGYA 2QH9A 2QHQA

2QIHA 2QJLA 2QJVA 2QJWA 2QJZA 2QK1A 2QL8A 2QLCA 2QLWA 2QMLA 2QMQA 2QN6B

2QNEA 2QNGA 2QNLA 2QNUA 2QPVA 2QPXA 2QQYA 2QQZA 2QRDD 2QRLA 2QRYA 2QSBA
2QSVA 2QSWA 2QTDA 2QTIA 2QTRA 2QU7A 2QUDA 2QV5A 2QV6A 2QVGA 2QW5A 2QWUA
2QYAA 2QYBA 2QYCA 2QYUA 2QZIA 2QZQA 2R01A 2R0XA 2R16A 2R1FA 2R25A 2R2AA
2R2ZA 2R31A 2R4FA 2R4IA 2R51A 2R58A 2R5OA 2R5UA 2R6JA 2R6VA 2R78A 2R7DA
2R87A 2R9FA 2RA1A 2RA9A 2RB7A 2RBBA 2RBCA 2RBDA 2RBKA 2RCIA 2RD9A 2RDCA
2RDEA 2RDGA 2RDMA 2RDQA 2REEA 2RETA 2RFQA 2RG4A 2RG8A 2RGQA 2RH0A 2RH3A
2RHFA 2RHWA 2RIQA 2RJ2A 2RJIA 2RKLA 2RL8B 2RLDB 2SAKA 2SASA 2SCPA 2SPCA
2TNFA 2TRCP 2UU8A 2UUZA 2UV4A 2UVKB 2UVPA 2UYTA 2V0OA 2V1MA 2V1YA 2V1YB
2V2GA 2V33A 2V3GA 2V3IA 2V4XA 2V5CA 2V6KA 2V6VA 2V76B 2V78A 2V79A 2V82A
2V84A 2V89A 2V8FA 2V8TA 2V94A 2V9LA 2V9VA 2VB1A 2VBUA 2VECA 2VEZA 2VFOA
2VFRA 2VFXE 2VGAA 2VH3A 2VHAA 2VJQA 2VK2A 2VK8A 2VLGC 2VLIB 2VM9A 2VNGA
2VOVA 2VPNB 2VQ2A 2VQGA 2VRSA 2VS0A 2VSYA 2VVEA 2VVWA 2VW8A 2VWSA 2VX8A
2VXNA 2VXTI 2VZCA 2VZPA 2VZZA 2W0GA 2W1JA 2W1RA 2W1VA 2W20A 2W2XC 2W31A
2W39A 2W3GA 2W3NA 2W3XA 2W40A 2W5EA 2W5QA 2W70A 2W72B 2W7AA 2W7ZB 2W8XA
2W91A 2WAGA 2WAWA 2WBMA 2WBNA 2WBWB 2WCRA 2WDCA 2WDSA 2WE5A 2WEWA 2WFIA
2WFKA 2WFOA 2WH6A 2WI8A 2WJ5A 2WJ6A 2WL8A 2WLRA 2WLVA 2WMMA 2WMYA 2WOLA
2WOYA 2WP7A 2WPVA 2WPVB 2WQKA 2WSHA 2WTGA 2WTMA 2WVBB 2WVQA 2WW5A 2WWEA
2WWXB 2WY3B 2WY4A 2WY8Q 2WYAA 2WZLA 2WZOA 2X1CA 2X32A 2X3JA 2X3MA 2X46A
2X49A 2X4DA 2X4LA 2X5FA 2X5XA 2X5YA 2X65A 2X9JA 2X9ZA 2XC8A 2XCIA 2XCMC
2XCME 2XDWA 2XDYA 2XE4A 2XF3B 2XFGB 2XFRA 2XFVA 2XFWA 2XHFA 2XHGA 2XI3A
2XIJA 2XIOA 2XJ4A 2XJPA 2XKIA 2XLAA 2XM5A 2XODA 2XOLA 2XOMA 2XP1A 2XPPA
2XQHA 2XQQA 2XQXA 2XSGA 2XT2A 2XTMA 2XTSA 2XTSB 2XU2A 2XU3A 2XU8A 2XUVA
2XVOA 2XVSA 2XVYA 2XW6A 2XWLA 2XWSA 2XWVA 2XXNA 2XXPA 2XZ2A 2XZ8A 2XZ9A
2Y0OA 2Y1BA 2Y27A 2Y4RA 2Y4XA 2Y5PA 2Y6XA 2Y7EA 2Y7PA 2Y88A 2Y8FA 2Y9UA
2Y9WA 2YAYA 2YCEA 2YFAA 2YFOA 2YFUA 2YGGA 2YGNA 2YH5A 2YHFA 2YHGA 2YILA
2YIZA 2YJLA 2YKZA 2YMVA 2YNMC 2YVIA 2YVTA 2YXHA 2YXOA 2YXXA 2YYJA 2YYOA
2YYSA 2YZTA 2Z0AA 2Z0JA 2Z0RA 2Z0VA 2Z0XA 2Z1DA 2Z26A 2Z2NA 2Z30B 2Z3HA
2Z3OA 2Z3QC 2Z3VA 2Z51A 2Z5WA 2Z6OA 2Z6RA 2Z72A 2Z7BA 2Z8FA 2Z8XA 2Z9WA
2ZAYA 2ZCAA 2ZD7A 2ZDJA 2ZDPA 2ZDSA 2ZE3A 2ZE7A 2ZEXA 2ZGYA 2ZHJA 2ZK9X
2ZKZA 2ZNLB 2ZNRA 2ZOGA 2ZOSB 2ZPDA 2ZPTX 2ZRRA 2ZS0C 2ZSJA 2ZUSA 2ZUYA
2ZVCA 2ZW5A 2ZXCA 2ZY4B 2ZYHA 2ZYJA 2ZYZA 2ZYZB 2ZZJA 3A02A 3A09A 3A0OA
3A0YB 3A1GB 3A1SA 3A2ZA 3A35A 3A4CA 3A4RA 3A57A 3A5FA 3A5PA 3A6RA 3A72A
3A8GA 3A8GB 3A8UX 3A9SA 3AA0B 3ABDX 3ABHA 3ACHA 3ACXA 3ADRA 3AEIA 3AEKB
3AGCA 3AGNA 3AGYA 3AHCA 3AHNA 3AIAA 3AJ4A 3AJFA 3AJVA 3AK8A 3AKBA 3ALMA
3AMLA 3AMRA 3ANOB 3AONA 3AOTA 3APTA 3AQBA 3AU4B 3AXBA 3AXJA 3AXSA 3AYHA
3AZOA 3B0CW 3B0FA 3B0GA 3B0TA 3B0XA 3B21A 3B33A 3B34A 3B42A 3B4QB 3B4UA
3B5EA 3B5MA 3B64A 3B79A 3B7CA 3B8FA 3B8ZA 3BA3A 3BB6A 3BBLA 3BBZA 3BC1B
3BC9A 3BCBA 3BCYA 3BDVA 3BE3A 3BE6A 3BEDA 3BEIA 3BEMA 3BESR 3BEYA 3BF4A
3BF5A 3BF7A 3BFMA 3BFOA 3BGUA 3BGYA 3BH0A 3BH7B 3BHDB 3BHNA 3BJAA 3BJEA
3BJNA 3BJOA 3BKXA 3BLNA 3BLZA 3BM7A 3BMAA 3BMZA 3BN3B 3BN7A 3BNKA 3BODA
3BOEA 3BOSA 3BPKB 3BPQD 3BQPA 3BQWA 3BQXA 3BRCA 3BROA 3BRQB 3BRUA 3BRVA
3BS5A 3BUUA 3BUXB 3BV8A 3BWHA 3BWZA 3BY8A 3BYQA 3C0FB 3C18A 3C1YA 3C2QB
3C2UA 3C3YA 3C4RA 3C5EA 3C5XC 3C70A 3C7TA 3C7XA 3C8CA 3C8FA 3C8IA 3C8LA
3C8OA 3C8WA 3C8YA 3C97A 3C9AA 3C9PA 3C9UA 3CA8A 3CAIA 3CAXA 3CC1A 3CCDA

3CCGA 3CCYA 3CE9A 3CEAA 3CEBA 3CECA 3CETB 3CEXA 3CF4A 3CGXA 3CHJA 3CHMA

3CI3A 3CIHA 3CIJA 3CIMA 3CINA 3CJIA 3CJLA 3CJMA 3CJPA 3CJSB 3CK1A 3CK2A

3CK6A 3CK9B 3CKMA 3CL5A 3CL6A 3CLWA 3CM3A 3CMBA 3CNBA 3CNHA 3CNIA 3CNUA

3CNYA 3COVA 3CP7A 3CPXA 3CQ1A 3CQRA 3CQXC 3CR3A 3CRKC 3CRYA 3CS5A 3CT6A

3CU2A 3CUZA 3CVEA 3CVGC 3CVJA 3CWNA 3CWRA 3CXNA 3CZ1A 3CZ7A 3CZXA 3D03A

3D0FA 3D0JA 3D0WA 3D19A 3D1PA 3D2YA 3D34A 3D3BA 3D3BJ 3D3OA 3D3RA 3D3YA

3D4EA 3D5LA 3D5PA 3D5YA 3D79A 3D7AA 3D7JA 3D7RA 3D9NA 3D9RA 3D9XA 3DA0A

3DA5A 3DA8A 3DAOA 3DB0B 3DBYA 3DD7A 3DDDA 3DDEA 3DEEA 3DELB 3DF6A 3DF8A

3DFFA 3DFUA 3DGBA 3DGPB 3DHAA 3DHXA 3DI5A 3DK9A 3DKRA 3DLCA 3DLIA 3DM8A

3DMEA 3DMWA 3DNJA 3DNPA 3DNSA 3DO8B 3DQPA 3DQYA 3DR5A 3DRFA 3DS8A 3DSBA

3DT5A 3DTDB 3DTNB 3DTZD 3DUPA 3DX5A 3DXEA 3DXYA 3DYJA 3DZ1A 3E05B 3E0IA

3E0RA 3E0XA 3E18A 3E1EA 3E1IA 3E48A 3E4GA 3E4VA 3E4WA 3E56A 3E59A 3E5XA

3E5ZA 3E6MD 3E6QA 3E7JA 3E8MA 3E8OA 3E8TA 3E96A 3E98A 3E9TA 3E9VA 3EA6A

3EAFA 3EBBA 3EBTA 3ED5A 3EDNA 3EDOA 3EDVA 3EEHA 3EF8A 3EFPA 3EHCA 3EI3B

3EJFA 3EJKA 3EJVA 3EK6A 3EMFA 3EMIA 3EMXA 3EN0A 3EN8A 3ENPB 3EO4B 3EO6A

3EOIA 3EPWA 3EQZA 3ER6A 3ER7A 3ERMA 3ESLB 3ESMA 3ETOA 3ETZA 3EUDA 3EUNA

3EUOA 3EURA 3EVYA 3EXNA 3EXQA 3EXZA 3EY5A 3EYEA 3EZIA 3EZSA 3F0HA 3F14A

3F1IC 3F1IH 3F1LA 3F1TA 3F1ZA 3F2EA 3F2ZA 3F3KB 3F40A 3F42A 3F43A 3F4AA

3F4LA 3F4MA 3F67A 3F6OA 3F6YA 3F7CA 3F7EA 3F7SA 3F7WA 3F7XA 3F8XA 3F9TA

3F9UA 3F9XA 3FAVB 3FAYA 3FB9A 3FBGA 3FCNA 3FD3A 3FD4A 3FDSC 3FEFA 3FETA

3FF1A 3FF2A 3FFDP 3FFMA 3FFRA 3FFVA 3FFYA 3FG9A 3FGRB 3FGTA 3FGVA 3FGYA

3FH1A 3FH3A 3FHVA 3FI7A 3FILA 3FJ1A 3FJ2A 3FKAA 3FKCA 3FKEA 3FKJA 3FKQA

3FM8A 3FMCA 3FN2A 3FNCA 3FO5A 3FO8D 3FPNA 3FPNB 3FPZA 3FQMA 3FRQA 3FS9A

3FSGA 3FT1A 3FTDA 3FVWB 3FVZA 3FW2A 3FWNA 3FXAA 3FXHA 3FYBA 3FYRA 3FZ4A

3FZXA 3G0KA 3G0MA 3G13B 3G16A 3G1PA 3G21A 3G2BA 3G2EB 3G36A 3G3RA 3G3SA

3G46A 3G48A 3G5TA 3G67A 3G6KB 3G74B 3G7NB 3G7PA 3G7RB 3G85A 3G8BA 3G8RA

3G8YA 3G91A 3G98A 3G9GA 3GA3A 3GAEA 3GBVA 3GBWA 3GC2A 3GCGB 3GD6A 3GDWA

3GDZA 3GE2A 3GE3A 3GE3B 3GE3E 3GF6A 3GF8A 3GFAA 3GFPA 3GG7A 3GGDA 3GGHA

3GGYA 3GI7A 3GIUA 3GIWA 3GJYA 3GKEA 3GKRA 3GMFA 3GMGA 3GMXA 3GN6C 3GNEA

3GO5A 3GOEA 3GONA 3GOSB 3GOZA 3GPVA 3GQHA 3GR0A 3GR3A 3GR5A 3GRAA 3GRCA

3GRDA 3GRHA 3GT5A 3GU3A 3GUYG 3GV0A 3GW4A 3GWLA 3GWNA 3GWRA 3GWYB 3GXVA

3GY9A 3GYCA 3GYDA 3GZAA 3GZBA 3GZHA 3GZRA 3H09B 3H0LB 3H0NA 3H0OA 3H1NA

3H35A 3H36A 3H3HB 3H3IA 3H3LA 3H4XA 3H51A 3H5JA 3H5LA 3H6JA 3H6PA 3H6PC

3H6QA 3H6RA 3H79A 3H7HA 3H7IA 3H87C 3H8DA 3H8GA 3H9WA 3HA2A 3HA4A 3HA9A

3HBNA 3HBZA 3HDGA 3HF5A 3HFIA 3HFTA 3HG9A 3HHFA 3HI2B 3HIEC 3HJ4A 3HJ7A

3HJEA 3HK4A 3HL4A 3HLXA 3HLZA 3HM4A 3HMSA 3HN0A 3HN5A 3HOIA 3HP0A 3HPCX

3HPXA 3HPYA 3HR0A 3HRDC 3HRGA 3HRPA 3HRZB 3HS3A 3HSAA 3HSHA 3HTKA 3HTKB

3HTRA 3HTUA 3HUGB 3HUHA 3HUTA 3HVAA 3HVWA 3HWUA 3HX3A 3HX8A 3HY0A 3HYNA

3HYOA 3HZPA 3I09A 3I0ZA 3I10A 3I1AA 3I2KA 3I38A 3I3VA 3I3WA 3I45A 3I4GA

3I4ZB 3I57A 3I5QA 3I7AA 3I7DA 3I7MA 3I83A 3I94A 3IACA 3IALB 3IB5A 3IC9A

3ICUA 3IE7A 3IECE 3IEEA 3IEZA 3IFEA 3IFUA 3IG5A 3IG9A 3IGHX 3IGSA 3IHMA

3IHUA 3IHVA 3II2A 3IISM 3IJ6A 3IJLA 3IJMA 3IKBA 3IL0A 3ILSA 3ILWA 3IM1A

3IM4C 3IMKA 3IMNA 3IN6A 3IP0A 3IP3A 3IP4C 3IPFA 3IPJA 3IQTA 3IQUA 3IR4A

3IRAA 3IRBA 3IRSA 3IRUA 3ISAA 3ISQA 3ISXA 3IT3A 3IT4A 3IT4B 3IT5A 3ITFA

3ITWA 3IU6A 3IUKA 3IUOA 3IUSA 3IUUA 3IUWA 3IV4A 3IVLA 3IVVA 3IX3A 3IXSA
3JQHA 3JR1A 3JRVB 3JS8A 3JSYA 3JSZA 3JTNA 3JTWA 3JTZA 3JU1A 3JU4A 3JU7A
3JUMA 3JV1A 3JVOA 3JXOA 3JYBA 3JYOA 3JZ0A 3JZLA 3K0ZA 3K11A 3K1HA 3K1TA
3K29A 3K2OA 3K3CA 3K3SA 3K4OA 3K5JA 3K63A 3K67A 3K6GD 3K6OA 3K6QA 3K7CA
3K7DA 3K8JA 3K93A 3K94A 3KB2A 3KBQB 3KBYA 3KD4A 3KD6A 3KDFA 3KDWA 3KE3A
3KE7A 3KEOB 3KEWA 3KF6B 3KF8A 3KF8B 3KFFA 3KFOA 3KGWA 3KGYA 3KGZA 3KHWA
3KIZA 3KJHA 3KKFA 3KKGA 3KKZA 3KL7A 3KLJA 3KM3A 3KM5A 3KMIA 3KNVA 3KORA
3KOSA 3KRAB 3KREA 3KS6A 3KS7A 3KTCA 3KUVA 3KVPA 3KWEA 3KWKA 3KWLA 3KWOA
3KWRA 3KWSA 3KXRA 3KYEA 3KYFA 3KYJA 3KZ3A 3KZPA 3KZTA 3L0AA 3L0GA 3L0QA
3L1WA 3L22A 3L34A 3L39A 3L46A 3L49A 3L51B 3L5AA 3L6DA 3L6IA 3L7HA 3L7ZC
3L8WA 3L9AX 3L9BA 3LAEA 3LB2A 3LD7A 3LEDA 3LETA 3LF9A 3LFJA 3LFKA 3LGBB
3LGDA 3LHIA 3LHLA 3LHNA 3LHOA 3LHSA 3LIDA 3LK2T 3LKBA 3LKDB 3LKEA 3LL3A
3LLUA 3LM2A 3LM4A 3LMFA 3LMOA 3LMZA 3LOFA 3LOGA 3LOIA 3LOPA 3LQBA 3LQCB
3LQKA 3LQMA 3LR2A 3LRTA 3LS9A 3LUAA 3LUBA 3LUQB 3LURA 3LUYA 3LVJC 3LVYA
3LW3A 3LW6A 3LX3A 3LXQA 3LXRF 3LXZA 3LY1A 3LYDA 3LYGA 3LYHA 3LYWA 3LZ6A
3LZKA 3M0FA 3M0MA 3M0ZA 3M1UB 3M2TA 3M33A 3M5BA 3M5QA 3M66A 3M6ZA 3M7AA
3M84A 3M86A 3M8JA 3M92A 3M9LA 3M9QA 3MADA 3MAOA 3MB2B 3MBRX 3MC3A 3MCBB
3MCWA 3MCXA 3MCZA 3MDKA 3MDQA 3MDUA 3ME0B 3ME7A 3MEMA 3MEUA 3MEZA 3MFBA
3MFNA 3MGAA 3MGDA 3MHSB 3MI9C 3MILA 3MIZA 3MJFA 3MJVA 3MKHA 3MMGA 3MMHA
3MN2A 3MN7S 3MQ2A 3MQDA 3MQQA 3MQZA 3MREA 3MSWA 3MSXB 3MTQA 3MTRA 3MTWA
3MU7A 3MUDA 3MVPA 3MVUA 3MW6A 3MWCA 3MWXA 3MWZA 3MX1A 3MXNA 3MYXA 3MZ0A
3MZOA 3N01A 3N08A 3N0XA 3N10A 3N17A 3N1EA 3N2OA 3N3MA 3N40P 3N4JA 3N5BB
3N5LA 3N6TA 3N72B 3N8BA 3N91A 3N98A 3NA5A 3NA6A 3NB0A 3NBCA 3NBMA 3NCEA
3NCEB 3NDCA 3NDQA 3NECA 3NEHA 3NFIA 3NGGA 3NGVA 3NGWA 3NHEA 3NI0A 3NIRA
3NJCA 3NJEB 3NJNA 3NK7B 3NKGA 3NKZA 3NL9A 3NNBA 3NO0A 3NO2A 3NO3A 3NO4A
3NO6A 3NO7A 3NOJA 3NOQA 3NPDA 3NPHB 3NQBA 3NREA 3NRLA 3NRWA 3NSWA 3NTVA
3NUFA 3NV0A 3NV9A 3NVNA 3NVSA 3NVWB 3NVWC 3NW4A 3NYCA 3NZNA 3O0GD 3O0LA
3O0YB 3O10A 3O12A 3O14A 3O1NA 3O2EA 3O3MA 3O3MB 3O3VA 3O4PA 3O5YA 3O7HA
3O94A 3O9ZA 3OA4A 3OA5A 3OAJA 3OAMA 3OC8A 3OCUA 3OD3A 3ODSA 3OE3C 3OFHA
3OFKA 3OG6B 3OGNA 3OHEA 3OHGA 3OISA 3OKQA 3OKXB 3OLCX 3OLOA 3OMDA 3ON4A
3ON7B 3ON9A 3ONHA 3ONJA 3ONPA 3ONQA 3OO8A 3OOSA 3OOVA 3OPFB 3OQ2A 3OQPA
3OR1A 3OR1B 3ORHA 3ORSA 3ORUA 3OS4A 3OSEA 3OSTA 3OT2A 3OTIA 3OTKA 3OTNA
3OULA 3OV8A 3OV9A 3OVPA 3OWCA 3OXPA 3OY2A 3OYVA 3OZ2A 3OZPA 3P06A 3P0FA
3P0KA 3P0YA 3P2TA 3P42A 3P4GA 3P4HA 3P51A 3P6LA 3P8AB 3P8BB 3PA8A 3PB6X
3PC7A 3PD7A 3PDDA 3PDYA 3PE6A 3PETA 3PF0A 3PF7A 3PFEA 3PFGA 3PFOA 3PG7A
3PH1A 3PICA 3PIJA 3PIWA 3PJ0A 3PJVD 3PKZA 3PL0A 3PLUA 3PLWA 3PM2A 3PMOA
3PMSA 3PN3A 3PNNA 3PNXA 3POPA 3PP5A 3PPLA 3PQHA 3PSHA 3PSMA 3PT8B 3PU5A
3PU9A 3PUCA 3PVHA 3PVTA 3PYCA 3PYWA 3Q1NA 3Q1TB 3Q20A 3Q2BA 3Q2KA 3Q3EA
3Q46A 3Q4OA 3Q63A 3Q64A 3Q6AB 3Q7RA 3Q8DA 3Q9LA 3QBMA 3QC0A 3QC7A 3QD7X
3QF7C 3QFMA 3QFSA 3QGUA 3QHEB 3QHPA 3QHQA 3QHTA 3QKXB 3QL9A 3QOOA 3QORA
3QP4A 3QPAA 3QQAA 3QR7A 3QRLA 3QS2A 3QSZA 3QTAA 3QTHA 3QU3A 3QU5A 3QVFA
3QVPA 3QVSA 3QW9A 3QWBA 3QWGA 3QWNA 3QWUA 3QX1A 3QXBA 3QXCA 3QY3A 3QY7A
3QY9B 3QZBA 3QZMA 3QZRA 3R0AA 3R15A 3R2QA 3R2UA 3R5TA 3R62A 3R6DA 3R72A
3R87A 3R8JA 3R90A 3R9FA 3R9MA 3R9VA 3RAGA 3RBYA 3RC1A 3RENA 3RF0A 3RF1A

3RF3A 3RGAA 3RGCB 3RH0A 3RHGA 3RHTA 3RHZA 3RIOA 3RJ2X 3RJTA 3RJUA 3RJVA
3RKCA 3RKGA 3RKLA 3RLGA 3RLKA 3RNRB 3RNVA 3RO3A 3ROBA 3ROFA 3ROTA 3RPCA
3RPDA 3RPEB 3RPFA 3RPJB 3RPNA 3RPZA 3RQ1A 3RQ4A 3RQ9A 3RQTA 3RRIA 3RRLA
3RRLB 3RRSA 3RS1A 3RT2A 3RT3C 3RTLA 3RU8X 3RUIA 3RUVA 3RWNA 3RWXA 3RX6A
3RX9A 3RXYA 3RZNA 3S0AA 3S1SA 3S2JA 3S2RA 3S2WF 3S44A 3S4EA 3S5BB 3S5QA
3S64A 3S6EA 3S6FA 3S84A 3S8SA 3S93A 3S9JA 3S9XA 3SAOB 3SBMA 3SBUA 3SCYA
3SD7A 3SEEA 3SFVB 3SGGA 3SHGA 3SHGB 3SHPA 3SI5A 3SIHA 3SJ5A 3SJHB 3SK2A
3SKQA 3SL9C 3SMPB 3SO6A 3SOJA 3SONA 3SS7X 3SSOA 3ST1A 3ST7A 3SU6A 3SUKA
3SUMA 3SXKA 3SXNA 3SXUB 3SY1A 3SZ3A 3SZYA 3T0YB 3T2CA 3T33A 3T3LA 3T47A
3T4RA 3T4SA 3T5BA 3T5XA 3T5XB 3T6AA 3T6OA 3T6QC 3T6SA 3T7AA 3T7FA 3T7ZA
3T8KA 3T92A 3T94A 3T9OB 3TB6A 3TBDA 3TBMA 3TC1B 3TC8A 3TCVA 3TDQA 3TDUA
3TE8A 3TEKA 3TEQA 3TEUA 3TFFA 3TFJA 3TG0A 3TG2A 3TG9B 3THGA 3THIA 3TJ7A
3TJ8A 3TJYA 3TL4X 3TLQA 3TM8A 3TO7A 3TOSA 3TOWA 3TSOC 3TT9A 3TUFA 3TUOA
3TUTA 3TVAA 3TVJA 3TVQA 3TXSA 3TY1A 3TYSA 3U01A 3U0CA 3U0HA 3U0OA 3U12A
3U1DA 3U28B 3U28C 3U2AA 3U2GA 3U2UA 3U3GA 3U3ZA 3U4TA 3U4VA 3U52C 3U64A
3U65B 3U6GA 3U7IA 3U7ZA 3U97A 3U9JA 3U9RB 3U9WA 3UAFA 3UB1B 3UB6A 3UC2A
3UCEA 3UD1A 3UF6A 3UFEA 3UFIA 3UI4A 3UIDA 3UIPD 3UJCA 3ULJA 3ULQA 3ULTA
3UMHA 3UMVA 3UP3A 3UPLA 3UPSA 3UQCA 3UR8A 3URRA 3URZB 3US3A 3USHA 3UT4A
3UUEA 3UULA 3UUWA 3UV0A 3UV1A 3UWPA 3UX3A 3UXJA 3V1AA 3V3LA 3V46A 3V4KA
3V4OA 3V5CA 3V67A 3V68A 3V6IB 3V75A 3V7BA 3V93B 3VA4A 3VA9A 3VC1A 3VC8A
3VDMA 3VENA 3VG7A 3VG8A 3VGLA 3VIIA 3VIQA 3VJ9A 3VJFA 3VK5A 3VKIA 3VL1A
3VLAA 3VLDA 3VMKA 3VMNA 3VN5A 3VOMA 3VORA 3VP5A 3VPBA 3VPZA 3VQJA 3VR0A
3VRDA 3VRDB 3VS8A 3VSEA 3VSJB 3VSVA 3VUPA 3VV1A 3VWCA 3VYGB 3VYWB 3VZ9B
3VZ9D 3VZXA 3W06A 3W0KA 3W0OA 3W19C 3W1EA 3W1OA 3W3WB 3W42A 3W4SA 3W5SA
3W6BA 3W6SA 3W7TA 3W81A 3W9SA 3WA2X 3WASA 3WBZC 3WDCA 3WDNA 3WDQA 3WEUB
3WFIA 3WG3A 3WISA 3WJPA 3WJTA 3WK3A 3WKGA 3WLIA 3WMVA 3WNDA 3WNZA 3WOEA
3WOZA 3WQCA 3WRBA 3WRYB 3WSGA 3WT0A 3WTDB 3WU4A 3WUCA 3WURA 3WUZA 3WV7A
3WVAA 3WVQA 3WVTA 3WWCA 3WWVA 3WX7A 3WXMB 3WYGD 3WZ3A 3X0IA 3X0TA 3X0VA
3X27A 3X2MA 3X37A 3X37B 3X38A 3ZBDA 3ZBGA 3ZBOA 3ZCNA 3ZDMA 3ZDOA 3ZDSA
3ZEDB 3ZEDD 3ZEUA 3ZGHA 3ZH5B 3ZH9B 3ZHFA 3ZIEA 3ZILA 3ZIUB 3ZJAA 3ZL8A
3ZLEA 3ZN4A 3ZNUA 3ZNVA 3ZOQB 3ZPNA 3ZPPA 3ZQSA 3ZR8X 3ZRDA 3ZRIA 3ZRXA
3ZSJA 3ZSUA 3ZT9A 3ZTHA 3ZUCA 3ZUIA 3ZVSA 3ZXKA 3ZXNB 3ZYLA 3ZYTA 3ZZLA
3ZZOA 3ZZPA 3ZZYA 4A02A 4A1RA 4A29A 4A37A 4A3ZA 4A56A 4A57A 4A5KA 4A5NA
4A69C 4A6QA 4A7UA 4A9VA 4ABMA 4ABXA 4AC1X 4ACJA 4ADNA 4AF1A 4AFLA 4AFMA
4AGHA 4AIVA 4AK2A 4AK5A 4ALZA 4ANBA 4ANNA 4APOA 4AQLA 4AQOA 4AQRD 4AR9A
4ARLB 4AS2A 4ASMB 4AT7A 4ATEA 4ATGA 4AU1A 4AUKA 4AVSA 4AXOA 4AY7A 4AYOA
4AZ6A 4B0ZA 4B1MA 4B28A 4B2FA 4B2OA 4B4UA 4B4YA 4B5ZA 4B6GA 4B6MA 4B6WA
4B89A 4B9GA 4BC3A 4BE3A 4BEUA 4BFCA 4BFOA 4BGBA 4BHRB 4BHUA 4BI3A 4BJ0A
4BJJA 4BJSA 4BJZA 4BK0A 4BK7A 4BKRA 4BLPB 4BLUA 4BN4A 4BNDA 4BOEA 4BOJA
4BOPA 4BOQA 4BOUA 4BPFA 4BPSA 4BPZA 4BQ2A 4BQNA 4BRCA 4BT7A 4BVXA 4BVXB
4BWCA 4BWQB 4BWRA 4BWVA 4BYZA 4C0FA 4C0XA 4C12A 4C1OA 4C24A 4C2LA 4C2VC
4C3SA 4C5KD 4C5WA 4C6AA 4C6RA 4C6SA 4C7NB 4C9RA 4CAHB 4CAYB 4CCOB 4CCSA
4CCVA 4CCWA 4CD8A 4CGSA 4CGUB 4CHDA 4CHGA 4CHIA 4CHMA 4CIHA 4CIIA 4CIJA
4CJ0A 4CJNA 4CK4A 4CKMB 4CLQA 4CMRA 4CMXA 4CNGA 4CNIC 4CO8A 4COGA 4CP6A

4CQIA 4CRUB 4CS4A 4CSHA 4CT0B 4CT7A 4CU5A 4CUAA 4CVDA 4CVRA 4CW4A 4CZXA
4D05A 4D06A 4D0KB 4D0NB 4D0PA 4D0QA 4D2MA 4D53A 4D5AA 4D5TA 4D6GA 4D70A
4D7JA 4D8BA 4D97A 4DCAA 4DELA 4DEVA 4DGFA 4DHXA 4DI9A 4DIDB 4DIXA 4DJBA
4DKCA 4DKWA 4DMVA 4DN1A 4DN7A 4DN9A 4DNNA 4DNYA 4DO4A 4DOOA 4DOVA 4DP1X
4DQ6A 4DQAA 4DQJA 4DSDA 4DT5A 4DTHA 4DV8A 4DVKA 4DWEA 4DWRA 4DZZA 4E0AA
4E15A 4E1OA 4E2TA 4E2XA 4E3YA 4E40A 4E4RA 4E57A 4E5VA 4E5XG 4E6FA 4E6SA
4E8OA 4EACA 4EADA 4EAEB 4EBBA 4EBGA 4EBJA 4ECFA 4EDHA 4EE6A 4EEKA 4EFOA
4EFPA 4EG9A 4EGUA 4EHSA 4EI7A 4EICA 4EIUA 4EJRA 4EL4A 4EMKA 4EMNA 4EO0A
4EO1A 4EOJB 4EP4A 4EQ3A 4EQ9A 4EQAC 4EQPA 4EQQA 4EQSA 4ERNA 4ES6A 4ES8A
4ESMA 4ESNA 4ESQA 4ESWA 4ETXA 4EU9A 4EUNA 4EUOA 4EUUA 4EV1A 4EVFA 4EVUB
4EVWA 4EW5A 4EW7A 4EXJB 4EXOA 4EXRA 4EYSA 4EYYR 4EYZA 4EZBA 4EZGA 4EZIA
4F03A 4F06A 4F0QA 4F0WA 4F1JA 4F1VA 4F2FA 4F2LA 4F3VA 4F4FA 4F55A 4F67A
4F7GB 4F7HA 4F8CA 4F8LA 4F98A 4F9ZA 4FBCA 4FBJA 4FBWD 4FC3E 4FC9A 4FCHA
4FFUA 4FHRA 4FIBA 4FK9A 4FKEA 4FKZA 4FLBA 4FLEA 4FM3A 4FMAA 4FN7A 4FOKA
4FP5D 4FPPA 4FPRA 4FQGA 4FQNA 4FR9A 4FSDA 4FTDA 4FTFA 4FTXA 4FYYB 4FZLA
4FZPA 4G08A 4G0XA 4G1QA 4G26A 4G2AA 4G2CA 4G2NA 4G2SA 4G3VA 4G4KA 4G67A
4G6IA 4G6TA 4G6UA 4G7NA 4G7XA 4G7XB 4G9EA 4G9QA 4G9SA 4G9SB 4GA2A 4GAKA
4GB5A 4GBFA 4GBMA 4GBSB 4GC3A 4GCIA 4GEHD 4GEIA 4GEKA 4GF3B 4GIMA 4GIOA
4GIPA 4GJ4A 4GKHA 4GM6A 4GMQA 4GMUA 4GN1A 4GN5A 4GNRA 4GOQA 4GQOA 4GQZB
4GS1A 4GS5A 4GT6A 4GT8A 4GUCA 4GVBA 4GVFA 4GVQA 4GWBA 4GXBB 4GXTA 4GXWA
4GYTA 4GZCA 4GZKA 4H08A 4H0CA 4H0FA 4H14A 4H2DA 4H2HA 4H2WA 4H2WC 4H31A
4H3UA 4H3WA 4H3ZB 4H41C 4H4DA 4H59A 4H5BA 4H5IA 4H6CA 4H6QA 4H7WA 4H86A
4H8EA 4HA6A 4HAHA 4HATB 4HC5A 4HCEA 4HCJA 4HCSA 4HDJA 4HDQA 4HE6A 4HEHA
4HEIA 4HESA 4HFKB 4HFSA 4HFVA 4HGUA 4HH3A 4HHVA 4HI0B 4HI6A 4HJ1A 4HJIA
4HL4A 4HMSA 4HN7A 4HNLA 4HNOA 4HPMA 4HR3A 4HROA 4HRVB 4HS1A 4HS2A 4HSPA
4HSQA 4HSTB 4HSUB 4HTGA 4HVTA 4HW6A 4HWHB 4HWMA 4HWVA 4HY4A 4HYLA 4HYZA
4HZ9B 4I0WA 4I0XA 4I0XB 4I17A 4I1KA 4I1SB 4I3YA 4I4CA 4I4OA 4I6RA 4I6VA
4I6YA 4I86A 4I8HA 4I8IA 4I90A 4I93A 4IABA 4IAUA 4IC1D 4IC4A 4IC9A 4ICVA
4ID2A 4ID9A 4IEJA 4IF5A 4IGBC 4IGIA 4IGVA 4IHUA 4IILA 4IIYA 4IJNA 4IL7A
4ILOA 4IM9A 4IN0A 4INWA 4IPAA 4IPIA 4IPUA 4IRFA 4IRVE 4IU3B 4IULB 4IUMA
4IUWA 4IWBA 4IX3A 4IX9A 4IYJA 4J0DA 4J27A 4J2CA 4J32B 4J42A 4J4ZA 4J5RA
4J6OA 4J7BC 4J7NA 4J7OA 4J8CA 4J8LA 4J8PA 4J8SA 4JAQA 4JB3A 4JBBA 4JBDA
4JBEA 4JCCA 4JDEA 4JDNA 4JE3A 4JEJA 4JEMA 4JF8A 4JG2A 4JGLA 4JGPA 4JHCA
4JHTA 4JHYA 4JIMA 4JIUA 4JJ9A 4JK8A 4JL5A 4JLEA 4JM1A 4JN7A 4JNUA 4JOIC
4JOQA 4JOXA 4JP6A 4JPKA 4JPNA 4JQRA 4JS1A 4JWJA 4JXRA 4JYJA 4JYKA 4JZ5A
4K0DA 4K0NA 4K12A 4K12B 4K2JA 4K35A 4K37B 4K3ZA 4K6NA 4K70A 4K7BA 4K7CA
4K7JA 4K84A 4K8WA 4K90A 4KALA 4KBXA 4KC5A 4KCFA 4KDWA 4KEMA 4KF9A 4KFUA
4KG0A 4KGDA 4KH8A 4KM6A 4KMCA 4KNKA 4KNVA 4KOQA 4KP1A 4KP3C 4KQ7A 4KQ9A
4KQIA 4KQPA 4KRDB 4KRGA 4KSNA 4KT6A 4KT6B 4KTBA 4KTPA 4KTWA 4KU0A 4KU0D
4KUNA 4KV2A 4KWDA 4KYQA 4L07A 4L0JA 4L0NA 4L1JA 4L1PB 4L2HA 4L2IA 4L2IB
4L2WA 4L3UA 4L4EA 4L4WB 4L57A 4L5GA 4L7AA 4L7MA 4L8AA 4L8JA 4L8KA 4L8PA
4L9HA 4L9PB 4L9UA 4LA6A 4LANA 4LB8A 4LBAA 4LD1A 4LD3B 4LE3A 4LEBA 4LERA
4LESA 4LF0A 4LFGA 4LG3A 4LGJA 4LH6A 4LHPA 4LHSA 4LIRA 4LIZA 4LJIA 4LJOA
4LJSA 4LMIA 4LMOA 4LMYA 4LO6A 4LPQA 4LQBA 4LQEA 4LQXA 4LQZA 4LRTA 4LRTB

4LTTA 4LTYA 4LUKA 4LUQC 4LVNP 4LWSA 4LXQA 4LY4A 4M0NA 4M0QA 4M1AA 4M1HA

4M1XA 4M2MA 4M3LA 4M51A 4M6VA 4M73A 4M7OA 4M7RA 4M7XA 4M82A 4M83A 4M8DA

4M8KA 4MAKA 4MAMA 4MAXA 4MB0A 4MB7A 4MCJB 4MCWA 4ME2A 4MEJA 4MF5A 4MF9A

4MGQA 4MI8A 4MJDA 4MKMA 4MKOA 4MLMA 4MLZA 4MM2A 4MMWA 4MNOA 4MP8A 4MPTA

4MQ3A 4MQDA 4MQVA 4MT2A 4MT8A 4MTHA 4MTUA 4MUDA 4MUQA 4MVEA 4MYDA 4MZAA

4MZCA 4MZJA 4MZYA 4N02A 4N03A 4N06A 4N0RA 4N1IA 4N2PA 4N2XA 4N3YB 4N4UA

4N6OB 4N6QA 4N7TA 4N8NA 4NARA 4NBPA 4NC6A 4NC7A 4NDOA 4NDSA 4NFUA 4NG0A

4NG2E 4NHEA 4NJ6A 4NJCA 4NK6A 4NL9C 4NLHA 4NMWA 4NMYA 4NN5B 4NNOA 4NOAA

4NOCC 4NOGA 4NOHA 4NOOB 4NQGA 4NQID 4NQJA 4NQWB 4NRHB 4NRTA 4NSVA 4NT1A

4NTDA 4NTKA 4NTQA 4NTQB 4NURA 4NUTB 4NV0A 4NV4A 4NX1B 4NXIB 4NXYA 4NYHA

4NYQA 4NZGA 4NZKA 4NZRM 4O0AA 4O1EA 4O2HA 4O4BB 4O5FA 4O6UA 4O7HA 4O7JA

4O7KA 4O7QA 4O87B 4O8SA 4O8WA 4OA3A 4OBMA 4OBXA 4OCVA 4OD6A 4OD7A 4OD8A

4OE9A 4OEVA 4OF6A 4OG4A 4OHCA 4OHJA 4OI4A 4OI4B 4OIEA 4OJ5A 4OJ8B 4OJJA

4OK7A 4OKEA 4OKZA 4OLSA 4OM8A 4OMFB 4OO3A 4OPCA 4OPWA 4OQ2A 4OQPA 4ORZB

4OT1A 4OTNA 4OU9A 4OVJA 4OVXA 4OVYA 4OWTA 4OWTC 4OX0A 4OYBA 4OYUA 4OZKA

4OZUA 4P1MA 4P1ND 4P1ZA 4P3FB 4P3HA 4P5PA 4P6IC 4P7QA 4P7XA 4P82A 4P98A

4PAGA 4PCAA 4PE3A 4PE6A 4PEDA 4PF3A 4PH2A 4PH8A 4PHJA 4PHRA 4PIBB 4PIOA

4PJ2A 4PK9A 4PKFB 4PKLA 4PMOA 4POWA 4PP4A 4PQ9A 4PQGA 4PQHA 4PQZA 4PS6A

4PSNA 4PSRA 4PU5A 4PUXA 4PVKA 4PW0A 4PWSA 4PWUA 4PXYA 4PZ0A 4PZ1A 4PZ7A

4Q0YA 4Q1QA 4Q1VA 4Q29A 4Q2SA 4Q3KA 4Q4W1 4Q4W2 4Q4W3 4Q51A 4Q53A 4Q5KA

4Q5WA 4Q63A 4Q7FA 4Q7OA 4Q82A 4Q86A 4Q8MA 4Q98A 4QA8A 4QAFA 4QANA 4QASA

4QBOA 4QBSA 4QC6A 4QDCA 4QE0A 4QEKA 4QFTA 4QFUG 4QGPA 4QHQA 4QHWA 4QI3A

4QJVA 4QJVB 4QLPA 4QLPB 4QM9A 4QMDA 4QMHA 4QN8A 4QP5A 4QPNA 4QPOA 4QPVA

4QPWA 4QREA 4QRKA 4QRLA 4QRNA 4QTCA 4QTPA 4QTQA 4QURA 4QXBA 4QXBB 4QXZA

4QY7A 4R03A 4R0MA 4R12A 4R16A 4R1DB 4R1JA 4R1QA 4R1SA 4R29B 4R2FA 4R2KA

4R3NA 4R4KA 4R5RA 4R5ZA 4R60A 4R6HA 4R6YA 4R75A 4R78A 4R7FA 4R80A 4R9OA

4R9PA 4RA9A 4RAJA 4RAYA 4RBRA 4RD4A 4RD8A 4REKA 4REOA 4REPA 4REXA 4RFUA

4RGIA 4RGPA 4RHOA 4RHZB 4RI1A 4RJ2A 4RK5A 4RKQA 4RKSA 4RL3A 4RLEA 4RLZA

4RP3A 4RPCA 4RPMA 4RPTA 4RRIA 4RS2A 4RS7A 4RSWA 4RT0A 4RTIA 4RU1A 4RU3A

4RU4A 4RUQA 4RV5A 4RW0A 4RXMA 4RXTA 4RXVA 4RXXA 4RY1A 4RYAA 4RZ9A 4RZLA

4S0UC 4S12A 4S1HA 4S1PA 4S1VA 4S28A 4S37A 4S39A 4S3IA 4SGBI 4TJVA 4TKCA

4TLAE 4TM3A 4TMXA 4TO7A 4TPSA 4TPSB 4TPWA 4TR6A 4TRHA 4TRTA 4TSDA 4TVEA

4TVVB 4TX5A 4TXRA 4TXRC 4TYZA 4TZ1A 4TZQC 4U0NA 4U12A 4U19A 4U1EB 4U1EG

4U5HA 4U5RA 4U7WB 4U8FA 4U8PA 4U98A 4UA6A 4UASA 4UC8A 4UDXX 4UE0A 4UEYA

4UF7A 4UFQA 4UFRB 4UHCA 4UHVA 4UIQA 4UJ7A 4UNMB 4UOBA 4UOSA 4UP3A 4UQWA

4UQXA 4UQZB 4USKA 4UU3A 4UULA 4UW9A 4UYIA 4UYPA 4UYTA 4UZSA 4UZYB 4UZZA

4V12A 4V17A 4V1KA 4V1TC 4V23A 4V2KA 4W4KA 4W4KB 4W4LC 4W5XA 4W64A 4W66A

4W6YA 4W78B 4W79A 4W7LA 4W7WA 4W7ZA 4W8HA 4W8KB 4W8OA 4W8PA 4W9WA 4WA0A

4WBEA 4WBJA 4WCJA 4WCKA 4WEEA 4WFWA 4WH5A 4WH9A 4WHIA 4WHNA 4WHSA 4WIAA

4WIDA 4WJKA 4WJTA 4WKAA 4WKSC 4WKZB 4WLRB 4WMAA 4WN5B 4WNDB 4WPGA 4WPKA

4WPMA 4WPYA 4WQKA 4WRIA 4WSFA 4WT3A 4WU0A 4WUIA 4WUVA 4WVRA 4WVZA 4WWFA

4WWHA 4WX0A 4WX4A 4WXAA 4WXMA 4WYHA 4WZXA 4X1ZA 4X5PA 4X7GA 4X7RA 4X7YA

4X84A 4X86B 4X8YA 4X9RA 4X9TA 4X9UB 4X9ZA 4XALA 4XAXA 4XB4B 4XB6A 4XB6B

4XB6C 4XBAA 4XCAA 4XCPA 4XDXA 4XE7A 4XEAA 4XEKA 4XEZA 4XFJB 4XFKA 4XFRA

4XHTA 4XJYB 4XKZA 4XLGB 4XLXA 4XMQA 4XO1A 4XOMA 4XOTA 4XPXA 4XQCA 4XRIA
4XRTA 4XT0A 4XT4A 4XURA 4XUWA 4XWHA 4XXHA 4XXIA 4XY5A 4XYPA 4XZ7B 4Y0HA
4Y1BA 4Y2FA 4Y5JA 4Y68A 4Y6CA 4Y7DA 4Y7MC 4Y7SA 4Y99B 4Y9IA 4Y9MA 4Y9VA
4YAAA 4YAPA 4YARA 4YB8A 4YB8B 4YC0A 4YCAB 4YDPA 4YDRA 4YE7A 4YECA 4YECB
4YEPA 4YFBA 4YGTA 4YH8B 4YHVA 4YI8A 4YJIA 4YK2A 4YMYA 4YNXA 4YORA 4YP6A
4YPCA 4YSLA 4YTBA 4YTDA 4YTWA 4YTWB 4YUCA 4YUDA 4YWZA 4YX1A 4YXPB 4YY2A
4YZ0A 4YZGA 4Z02A 4Z04A 4Z0YA 4Z24A 4Z29A 4Z2BA 4Z2ZB 4Z39A 4Z3GA 4Z3TA
4Z3XA 4Z3XF 4Z48A 4Z4AA 4Z67A 4Z8EA 4Z8TB 4Z8UA 4ZA9A 4ZAVA 4ZB3A 4ZBHA
4ZBOA 4ZC3A 4ZCDB 4ZCEA 4ZDSA 4ZE8A 4ZEVA 4ZFLA 4ZFVA 4ZGFA 4ZH5A 4ZHBA
4ZI3C 4ZJHA 4ZJUA 4ZKQA 4ZLFA 4ZMHA 4ZMIA 4ZMKA 4ZNMA 4ZO2A 4ZOSA 4ZOXA
4ZPCA 4ZQAA 4ZQXA 4ZR8B 4ZRLB 4ZRSA 4ZSIA 4ZTYA 4ZURA 4ZV0B 4ZV5A 4ZVCA
4ZVFA 4ZW2A 4ZWNA 4ZWQA 4ZWVA 4ZX2A 4ZY7A 4ZY9A 4ZZFA 5A04A 5A07A 5A0LA
5A0NA 5A10A 5A12A 5A1NA 5A35A 5A3VA 5A4AA 5A5HB 5A5XA 5A5YA 5A61A 5A62A
5A6MA 5A6SB 5A7ZA 5A8CA 5A8JA 5A8SA 5A99A 5AB5A 5ABSA 5ABYB 5AE0A 5AEGB
5AEJA 5AGIA 5AIGA 5AIZA 5AJSA 5AL6A 5AMTA 5AN6A 5ANRC 5AOGA 5AOTA 5AOZA
5APAA 5APGA 5AQ5A 5AQCA 5AQMB 5AWSA 5AX2A 5AYSC 5AZWA 5AZXA 5B08A 5B0HA
5B19A 5B1QA 5B21A 5B3DA 5B3GA 5B3PA 5B3TA 5B46B 5B4BA 5B4ZA 5B5IA 5B5LA
5B6CA 5B7WA 5B89A 5B8DA 5BJXA 5BK9A 5BMNA 5BMOA 5BMQA 5BMTA 5BNZA 5BO6A
5BOBB 5BOWA 5BP7A 5BP8A 5BPZA 5BQ5A 5BR4A 5BRKB 5BS1D 5BSEA 5BT9A 5BTYA
5BU1A 5BV8A 5BXAA 5BXGA 5BXRA 5BY4A 5BY5A 5BY8A 5BY8B 5C17A 5C2MA 5C2UA
5C3QA 5C40A 5C50A 5C50B 5C54A 5C5GA 5C5ZA 5C6KA 5C79A 5C86A 5C8AA 5C8HA
5C8ZA 5C9FA 5CAJA 5CDKA 5CE7A 5CEGB 5CGEB 5CGZA 5CH8A 5CHLA 5CJ3A 5CKLA
5CL2A 5CN2A 5COFA 5COSA 5COWA 5COZA 5CPHA 5CR4A 5CR8A 5CSRA 5CTDC 5CTMA
5CTVA 5CWGA 5CWHA 5CWLA 5CX8A 5CXXA 5CYAB 5CYWB 5CYZC 5CZCA 5CZYA 5D04C
5D16A 5D22A 5D2KA 5D2MG 5D3KA 5D3UA 5D4VA 5D5YA 5D6EA 5D6JA 5D6OA 5D78A
5D7WA 5D8VA 5D9GA 5D9RA 5DBLA 5DCYA 5DE3A 5DGQA 5DHDA 5DICA 5DILA 5DJEA
5DKOA 5DLEA 5DLYA 5DNKA 5DOBB 5DOCB 5DOFD 5DP2A 5DPOA 5DS2A 5DVIA 5DWDA
5DYQA 5DZEA 5DZOA 5E0JA 5E0UA 5E1ZB 5E2CA 5E3EA 5E3EB 5E3PA 5E4BA 5E5YA
5E6XA 5E8JA 5E9PA 5EB9A 5ECDA 5ECKA 5EDFA 5EFVB 5EHAA 5EHIA 5EJ8A 5EJYA
5EK5A 5EL3A 5EL9A 5EMBA 5EMIA 5EMJB 5EMXA 5EO4A 5EO6A 5EP2A 5EP6A 5EPAA
5EQVA 5EQWA 5ER9A 5EU0B 5EUVA 5EVHA 5EWOA 5EWUA 5EXEA 5EXEC 5EXJA 5EYIA
5EYNA 5EZQA 5EZUA 5F1BA 5F2KA 5F3OA 5F47A 5F4CA 5F4WA 5F5OA 5F5SB 5F6RA
5F74B 5FAAA 5FAFA 5FAIA 5FAVA 5FBFA 5FCEA 5FCNA 5FEBA 5FEWA 5FFGB 5FFSA
5FHKD 5FI3A 5FIAB 5FIDA 5FIIA 5FISA 5FJZA 5FLGB 5FLHA 5FLWA 5FLYA 5FMOL
5FMOS 5FMUA 5FN7A 5FNOA 5FNPA 5FPZA 5FQ4B 5FQEA 5FRDA 5FRYA 5FTWA 5FU5A
5FUHB 5FUSA 5FVKA 5FWZA 5FZPA 5G0AA 5G0ZA 5G1SG 5G1XB 5G38A 5G4AA 5G4KA
5G51A 5G59A 5G5OB 5GGNA 5GJHA 5GJIA 5GK1A 5GKXA 5GMDA 5GMTA 5GN2A 5GNGA
5GOOA 5GP4A 5GPYB 5GQIA 5GS7A 5GSMA 5GT5A 5GTQA 5GTUA 5GUQA 5GV0A 5GV3B
5GV8A 5GVAA 5GW9A 5GXEA 5GYQA 5GYYA 5GYYG 5GZ3A 5GZAA 5GZKA 5GZTA 5GZXA
5H0JA 5H0MA 5H0QA 5H1XA 5H3VA 5H3XA 5H4EA 5H4SA 5H5FA 5H62B 5H65B 5H66B
5H66C 5H6TA 5H6XA 5H7EA 5H7TA 5H92A 5H9CA 5H9IA 5H9MA 5H9NA 5HALA 5HB6A
5HB7A 5HCAA 5HCIA 5HD9A 5HDAA 5HDKD 5HDWA 5HE9A 5HEEA 5HFSA 5HGJA 5HHEA
5HHJA 5HI8A 5HJ1A 5HJ9A 5HJFA 5HKPC 5HKQA 5HKQI 5HL8A 5HLJA 5HMLA 5HNOB
5HOBA 5HOEA 5HONA 5HQHA 5HRAA 5HT6B 5HTLA 5HTXA 5HUBA 5HWAA 5HWNA 5HXKA

5I0QA 5I3EB 5I45A 5I4CA 5I4MA 5I4QB 5I5NA 5I95A 5I9JA 5IATA 5IDBA 5IG0A
5IG6A 5IGIA 5IGOA 5IHFB 5IHWA 5II8A 5IJJA 5IK4A 5IL2A 5IL3A 5ILBA 5IMAA
5IMUA 5INBB 5INRA 5IO9A 5IP4D 5IQJA 5IR2A 5IRCA 5ISVA 5ITZD 5IU1A 5IUFA
5IW9A 5IWBA 5IWHA 5IX8A 5IXGA 5IXHA 5IXPA 5IZAA 5IZEA 5IZWA 5J1GA 5J1JA
5J1NA 5J1SA 5J1SB 5J3TA 5J41A 5J4AA 5J4AB 5J4FA 5J4OA 5J4UA 5J5LA 5J81A
5J90A 5JAZA 5JBNA 5JBRA 5JBXA 5JCAS 5JDAA 5JDKA 5JEDA 5JELA 5JENA 5JGKA
5JH8A 5JIWA 5JJ2A 5JKJA 5JKQA 5JLVC 5JMBA 5JO8A 5JODA 5JP1A 5JRBA 5JRHB
5JRTA 5JS4A 5JSOA 5JUGA 5JVIE 5JW8A 5JW9A 5JY4A 5K0AA 5K2LA 5K2XA 5K34A
5K3QA 5K4CA 5K8CA 5K8JA 5K8JB 5K8PA 5K9GA 5KARA 5KAXA 5KAYA 5KB6A 5KC1A
5KCIA 5KCMA 5KD5A 5KDSA 5KDWA 5KEFA 5KEWA 5KEWB 5KF9A 5KHLB 5KHTA 5KJVA
5KO3A 5KO5A 5KTCA 5KTNA 5KU5A 5KUXA 5KVBA 5KVCA 5KVSA 5KWBA 5KY2A 5L09B
5L0VA 5L20A 5L37A 5L44A 5L4LA 5L73A 5L74A 5L87A 5L8XB 5LALA 5LCYA 5LE5E
5LE5H 5LE5I 5LE5J 5LE5L 5LE5M 5LJWA 5LLTA 5LOSA 5LOUB 5LP9A 5LQ7A 5LS4A
5LS7B 5LSFA 5LSFB 5LSFC 5LSID 5LSLE 5LT5A 5LTEA 5LTJA 5LU5A 5LUNC 5LUSA
5LW3A 5LW6A 5LXFA 5LXXA 5LY3A 5LY8A 5LZGA 5M0WA 5M0YB 5M17A 5M1MA 5M1PA
5M1XD 5M26A 5M26B 5M2PA 5M2YA 5M3IA 5M45A 5M45B 5M5EC 5M5ZA 5M6QA 5M72A
5M72B 5M7YA 5M86A 5M89A 5M8BA 5M97A 5MALA 5MAOA 5MAWE 5MDIA 5MDUA 5MEBA
5MFOA 5MGWA 5MH6A 5MIYA 5MJRA 5MK2B 5ML3B 5MLDA 5MLTA 5MPTA 5MQ8A 5MQPA
5MR1A 5MR3H 5MRIA 5MS2A 5MSMB 5MSMC 5MU3A 5MU3B 5MU3C 5MU9A 5MUNA 5MUXA
5MUZA 5MV0A 5MVXA 5MZWB 5N0OB 5N2CA 5N2PA 5N3JA 5N3UB 5N40A 5N41A 5N5PA
5N86A 5N8AX 5N9MA 5NB4A 5NCJA 5NCWB 5NCXA 5NFJA 5NFMA 5NGWA 5NHUJ 5NJ9A
5NJ9B 5NJOA 5NKGA 5NL6A 5NMOB 5NNYA 5NODA 5NOHA 5NRHA 5NRMA 5NRRB 5NSAA
5NSLA 5NT7A 5NWPA 5NXFA 5NXKA 5NYOA 5O0JA 5O15A 5O29A 5O2DA 5O2XA 5O33B
5O37A 5O3NA 5O45A 5O6HA 5O75A 5O99A 5O9EA 5O9FA 5O9JA 5O9XA 5OBYA 5ODQB
5OE8A 5OEIA 5OEMA 5OH5A 5OHQA 5OI0A 5OIXA 5OK6A 5OL9A 5OLPA 5OLRA 5OMTA
5ONKA 5OOOA 5OPFA 5OPZA 5OQ3A 5OUOA 5OVOA 5P9VA 5Q22A 5QI0A 5QIVA 5QOQA
5R7XA 5R8QA 5RW1A 5SBFA 5SBQA 5SUZA 5SV2A 5SV5A 5SVHB 5SWCA 5SWUA 5SY4A
5SY8O 5SZCA 5T0FB 5T1IA 5T38A 5T3BA 5T3EA 5T3PA 5T51B 5T59C 5T5IB 5T5IC
5T6JA 5T6JB 5T7AA 5T86A 5T86I 5T95A 5T9YA 5TA0A 5TCBA 5TD6A 5TDAA 5TEDA
5TF3A 5TFPA 5TFQA 5TGFA 5TGQA 5THKA 5TIFA 5TIPA 5TJZA 5TK8A 5TKWA 5TNVA
5TOQA 5TPHA 5TPIA 5TQBB 5TQIA 5TRQA 5TS9A 5TSEA 5TSQA 5TT6A 5TTYA 5TUIA
5TUUA 5TVMB 5TVOB 5TWBA 5TZDA 5TZJA 5TZMA 5TZPA 5U22A 5U3AA 5U4HA 5U55A
5U5NA 5U69A 5U7AA 5U81A 5U9JA 5UAMA 5UB3A 5UBPA 5UBPB 5UE1A 5UEBA 5UEJA
5UFNA 5UFTA 5UFYA 5UGRA 5UH7A 5UJ1A 5UJCA 5UKIA 5UKVA 5ULBA 5UM2A 5UMHA
5UMRA 5UMSA 5UMVA 5UOFA 5UPBA 5UPKA 5UQ6A 5UQSA 5UQZA 5USWA 5UT3A 5UUJA
5UUKA 5UVDA 5UVRA 5UWCI 5UWZA 5UX1A 5UXMA 5UZGA 5UZXA 5V01A 5V1TA 5V1YA
5V2OA 5V4RA 5V4TA 5V50A 5V5HA 5V5TA 5V6JA 5V77A 5V8EA 5V8SA 5V8WD 5VACA
5VFBA 5VG7A 5VGBB 5VGLA 5VHGA 5VIPD 5VIQA 5VJ2A 5VJIA 5VKYA 5VMDA 5VN4A
5VNYA 5VOGA 5VPSA 5VPUA 5VVIA 5VW4A 5VX1A 5VXFA 5VXVA 5VYQA 5VYRB 5VZ4B
5W0HA 5W15A 5W1AB 5W2FA 5W3XA 5W4AA 5W4NA 5W53A 5W5CE 5W6JB 5W6YA 5W7BA
5W7CC 5W7WA 5W82A 5W8EA 5WASA 5WASB 5WB4A 5WC6A 5WECA 5WEEA 5WF2A 5WFBA
5WFYA 5WJCB 5WJPA 5WK0A 5WKRA 5WKWA 5WL1A 5WM2A 5WMKA 5WQJB 5WRIA 5WS7A
5WSFA 5WSTA 5WUJB 5WWDA 5WWLM 5WWOC 5WXXA 5WZFA 5X1EB 5X1ED 5X1JA 5X2EA
5X3DA 5X3GA 5X42B 5X42C 5X4KA 5X4RA 5X5MA 5X62B 5X6SA 5X6VG 5X7LA 5X9CA

5X9IA 5X9LA 5XA5A 5XA5B 5XAQA 5XAUC 5XB6A 5XB7F 5XBCA 5XBIB 5XBJA 5XBNA
5XC5A 5XCCA 5XDCA 5XDHA 5XEYA 5XFMA 5XFTA 5XGAA 5XGTA 5XISA 5XJ1A 5XJGB
5XK6A 5XLJA 5XLLA 5XLNB 5XOMA 5XOYB 5XP0A 5XPVA 5XRLA 5XSJL 5XSWA 5XTKA
5XV0A 5XVJA 5XVRA 5XVTA 5XYFB 5Y11C 5Y1FA 5Y24B 5Y27A 5Y4RC 5Y4UA 5Y4ZA
5Y5RD 5Y6GA 5Y6HA 5Y7WA 5Y9QA 5Y9WC 5Y9XA 5Y9ZA 5YA6A 5YADA 5YB7A 5YCEA
5YDDA 5YDEA 5YDNA 5YDRB 5YDVA 5YFCA 5YGEA 5YH1A 5YH4A 5YHRA 5YHUA 5YI6A
5YJ6A 5YJCA 5YKRA 5YKUA 5YKZA 5YLBA 5YNVB 5YNXA 5YQ0C 5YQJA 5YQWA 5YR0B
5YRVA 5YRVB 5YRVC 5YSCA 5YSIA 5YSNA 5YSNB 5YSQB 5YVKA 5YVNA 5YVQB 5YZCA
5YZOA 5YZPA 5Z0DA 5Z0UA 5Z1AA 5Z1GA 5Z1NA 5Z1VA 5Z28A 5Z2HA 5Z38A 5Z3EA
5Z3KB 5Z3RA 5Z42A 5Z4GA 5Z51A 5Z5OB 5Z6DA 5Z8AA 5Z8OA 5Z99A 5Z9ZA 5ZB2A
5ZBFA 5ZBTA 5ZBYA 5ZCTA 5ZDMA 5ZGXA 5ZHOA 5ZHZA 5ZI2A 5ZIKA 5ZIQA 5ZKEA
5ZKKA 5ZLAB 5ZN9B 5ZNGC 5ZNTA 5ZRTA 5ZRYA 5ZT3A 5ZTCA 5ZW7A 5ZWBA 5ZWLG
5ZWRA 5ZWUA 5ZYFA 5ZZAA 5ZZAP 6A02A 6A0EA 6A56A 6A5DA 6A5HA 6A6FA 6A7VD
6A8MA 6AC0A 6AD3A 6AE8C 6AE9A 6AEFA 6AEPA 6AHPA 6AHTA 6AJPA 6AKMA 6AL9A
6AM3A 6ANSA 6ANWA 6ANZA 6AO3A 6AO7A 6AONA 6APPB 6AQEA 6AR7A 6ARIA 6AS3A
6ASKA 6AT0A 6AT4A 6ATGB 6AU8C 6AVXA 6AZ5A 6B12A 6B1KA 6B29A 6B2VA 6B4EA
6B4HA 6B4HB 6B6UB 6B7MA 6B9MD 6B9RA 6B9XA 6B9XC 6B9XE 6BA9A 6BBWA 6BCDA
6BDEA 6BDUA 6BE0A 6BEVA 6BJTB 6BL5A 6BLKB 6BM5A 6BO0A 6BR8A 6BS9A 6BSCA
6BSCB 6BSUA 6BTCA 6BUMA 6BW9B 6BWGA 6BWRA 6BXOA 6BXRA 6BZFA 6C0YA 6C1WA
6C1XA 6C29A 6C30A 6C33A 6C3CA 6C4QA 6C4VA 6C62C 6C72A 6C74A 6C8CA 6C8RA
6C90B 6C9XA 6CB7A 6CBUA 6CCDA 6CD9A 6CF9A 6CGKA 6CGOA 6CH1A 6CHXA 6CJ6A
6CK0A 6CKAA 6CKMA 6CNGA 6CNWA 6COJA 6COJB 6CP8A 6CPBA 6CPDA 6CR0A 6CTZA
6CUMA 6CWMA 6CXBA 6CXGA 6D0GA 6D0HA 6D0HB 6D0TA 6D3VA 6D4KA 6D6YA 6D7RA
6D7YA 6D7YB 6D97A 6D9FA 6D9NA 6DA0A 6DAOA 6DB1A 6DBPA 6DCJA 6DCMA 6DETA
6DEWA 6DEXB 6DF3C 6DF3H 6DG4A 6DGAA 6DGMA 6DHXA 6DKQB 6DLMA 6DNMA 6DNQE
6DNVA 6DQHA 6DQPA 6DS9A 6DT3A 6DXSA 6DXYA 6DYMA 6E0KA 6E1FA 6E1XA 6E1ZA
6E28C 6E3AA 6E4DA 6E4LA 6E55A 6E60A 6E6OA 6E6YA 6E7EA 6E85A 6E8MA 6EA6A
6EACA 6ECVA 6EDVA 6EF7A 6EHIA 6EIOA 6EKBA 6EKZA 6ELMA 6ELVA 6ENNA 6ENOA
6EPGD 6EPZA 6ER1A 6ER4A 6ES9A 6ESZC 6ET0B 6ET6A 6ETCX 6ETLA 6EU4A 6EUNA
6EVNA 6EVUA 6EWHA 6EWLA 6EWMA 6EWZA 6EX6A 6EXAB 6EXPA 6EXXA 6EY6A 6F03A
6F1SA 6F3PA 6F43A 6F45D 6F4JA 6F4QA 6F5XA 6F5ZC 6F7DA 6F8AA 6F9OA 6FANA
6FC0B 6FC1B 6FCTA 6FDFA 6FDGA 6FDKA 6FF1A 6FFAA 6FGEA 6FIGA 6FIYA 6FJLA
6FJNA 6FKGA 6FM8A 6FMBA 6FMCA 6FMEA 6FOHB 6FOPA 6FP5A 6FPGB 6FSNA 6FTOA
6FTOC 6FU4A 6FU9A 6FU9B 6FVIA 6FW8A 6FWYA 6FX6A 6FXDA 6FXQA 6FY5A 6FYRA
6FYWA 6FZ6A 6FZZA 6G1BB 6G1CB 6G1IA 6G21A 6G43A 6G49A 6G4JA 6G6KA 6G7BA
6G7NA 6G86B 6G8YA 6GAJA 6GBIA 6GBOA 6GCBA 6GDJA 6GEHA 6GEUA 6GF6A 6GG7A
6GG7C 6GGCA 6GGRA 6GGYA 6GH8B 6GHTA 6GHUA 6GI4B 6GITA 6GKIA 6GKWA 6GMOA
6GN5A 6GNCA 6GNYA 6GPAA 6GPKA 6GQ4A 6GQZA 6GRFB 6GRH1 6GRHC 6GRHD 6GT9A
6GU1A 6GUMB 6GUOD 6GV5A 6GVDA 6GWJB 6GX2A 6GXRA 6GXSA 6GY5A 6GZ8A 6GZJB
6H0CA 6H10A 6H17A 6H20A 6H24A 6H3SA 6H3WA 6H3XA 6H40A 6H4EA 6H4LA 6H6OA
6H8FA 6H8GA 6H8OA 6H96A 6HAVA 6HB1A 6HBBA 6HBVA 6HC0B 6HCWA 6HEMA 6HFMA
6HFQA 6HFZC 6HGMB 6HGOA 6HHKA 6HHMA 6HIKL 6HIUA 6HJ6A 6HK9A 6HLMA 6HMVB
6HN1A 6HNIA 6HPHA 6HQ9A 6HQCA 6HQZA 6HSAA 6HT0A 6HTOB 6HU8A 6HURA 6HX0A
6HXAA 6HXMA 6HXPA 6HYYA 6HZNA 6I01B 6I03A 6I05A 6I18A 6I1AA 6I31A 6I35B

6I3BA 6I4EG 6I5OA 6I5RA 6I6MB 6I86A 6I8MA 6I9AA 6I9WA 6IAHA 6IBEA 6IBGA
6IBHA 6IC9A 6ICNA 6ID6A 6IE1A 6IEJA 6IF2A 6IF4A 6IF6A 6IGGA 6IH0A 6IHGA
6II6A 6IIPA 6IIXA 6IJBB 6IJEA 6IL9A 6ILUA 6IMVA 6IN7A 6INRA 6IPBA 6IPWA
6IQCA 6IQEA 6ISTA 6ITAA 6ITWA 6IUBA 6IUFA 6IUHA 6IUXA 6IWDB 6IWVA 6IX1A
6IX4B 6IX8A 6IY4I 6IY9A 6IYYA 6IZBA 6J0PA 6J27A 6J2ZA 6J4PA 6J4PB 6J67A
6J6AA 6J93A 6J98A 6J9LA 6J9LE 6J9MA 6JALA 6JAUB 6JC0A 6JC4A 6JEBA 6JFKA
6JHWB 6JIEA 6JILA 6JKKA 6JL9A 6JLCA 6JLEA 6JLEE 6JLIA 6JM5A 6JMKA 6JNJA
6JP4A 6JPTA 6JQ9A 6JQFA 6JQWA 6JSAA 6JU1A 6JV0A 6JV7A 6JWFA 6JWMA 6JYUA
6JZ2B 6K06A 6K2FA 6K2HA 6K39A 6K6LB 6K7CA 6K8EA 6K93A 6KA3A 6KBBE 6KCWA
6KETA 6KEWA 6KFNA 6KFSA 6KHLA 6KHUA 6KI2A 6KIAA 6KIIA 6KISA 6KJKA 6KKOC
6KLZA 6KM7A 6KMEA 6KMLA 6KNLA 6KO8A 6KP0B 6KPLA 6KSRA 6KSYA 6KTAA 6KU3A
6KWZA 6KXDB 6KXTA 6KY9A 6KZJB 6L0OA 6L0RA 6L0TB 6L1MA 6L1PA 6L2WB 6L3MA
6L4PB 6L5HA 6L6GA 6L7QA 6L7RB 6L7XA 6L8GB 6L8SA 6L8XA 6LACA 6LCEA 6LDLA
6LE1A 6LJ9A 6LJHA 6LKIB 6LKKA 6LL8A 6LLAA 6LN3A 6LNLA 6LNMB 6LPHA 6LPNA
6LR3A 6LU2A 6LUJA 6LXSA 6LXUA 6M2IA 6M3HA 6M4BA 6M64A 6M6UB 6M76A 6M7AC
6M8NA 6M9KD 6MAGA 6MBHA 6MEZA 6MFUA 6MGPB 6MICA 6MJBB 6MJOC 6MM2A 6MOSA
6MPRA 6MQ5A 6MQ7A 6MRRA 6MRSA 6MTZB 6MU0A 6MUQA 6MYDA 6N0SA 6N1BA 6N1FD
6N1LA 6N2CA 6N2NA 6N4LA 6N63A 6N6RB 6N7OA 6N9HA 6N9MA 6NAUA 6NBGA 6NCHA
6ND7A 6NDSA 6NDTB 6NE6A 6NEPB 6NEYA 6NFFA 6NIEA 6NIOA 6NJYA 6NK8B 6NKHA
6NL2A 6NLPA 6NLQA 6NMGB 6NMOA 6NNWA 6NOPA 6NOZA 6NP3A 6NQ6B 6NQCA 6NQWA
6NR6A 6NRXA 6NSVA 6NTRA 6NU8A 6NWXA 6NX3A 6NY9B 6NYRA 6NZ4A 6NZSA 6O1WA
6O2VA 6O3XA 6O43A 6O4OA 6O5KA 6O5WA 6O6YA 6OC0A 6OD1B 6OD2A 6OFQA 6OG4C
6OGHA 6OH6A 6OJ1A 6OJFA 6OJLA 6OJMB 6OK1B 6OKHA 6OMPA 6ON1A 6ONBA 6OP8B
6OQAC 6ORMA 6OS6A 6OUXA 6OVTA 6OYCB 6OZ1A 6OZEA 6OZUA 6P0FA 6P0QA 6P1BA
6P1EA 6P20D 6P22D 6P29A 6P2DA 6P2KA 6P2LA 6P3HA 6P5HA 6P5TA 6P61A 6P78A
6P7LA 6P7RA 6P80A 6P8JA 6P8PA 6PBMA 6PCDD 6PCFA 6PCZA 6PD2A 6PDKA 6PEUA
6PFQA 6PI6A 6PIRB 6PLJA 6PNVA 6PNYA 6POLB 6PQKB 6PRHA 6PSEA 6PT4A 6PT8A
6PUQA 6PVJA 6PWGA 6PWSA 6PX4A 6PX4B 6PYKA 6PZDA 6PZJA 6Q00A 6Q00B 6Q0MA
6Q10A 6Q1MA 6Q3PB 6Q3PC 6Q3VA 6Q43A 6Q43B 6Q43C 6Q56A 6Q5AA 6Q9CB 6Q9LA
6QAIA 6QAZA 6QEQA 6QFSA 6QGRA 6QHJA 6QJ6A 6QJAA 6QLCA 6QNNB 6QO0A 6QO9A
6QPKB 6QPSA 6QQ4A 6QSPB 6QTAB 6QUBA 6QV7A 6QVFA 6QVPA 6QW5A 6QWOA 6QXRA
6QXUA 6QYYA 6R01A 6R09A 6R0NB 6R0RA 6R17A 6R1DA 6R1MB 6R2WL 6R2WT 6R3RA
6R45A 6R4ZA 6R5JA 6R5WA 6R6KA 6R6MA 6R7VA 6RCGA 6RFGA 6RG2B 6RGNA 6RI6A
6RIVA 6RJXA 6RK0A 6RMVB 6RNIA 6RO0D 6RO6A 6RP3A 6RRAA 6RRVA 6RS4A 6RSNA
6RSWB 6RVQA 6RVUA 6RWCA 6RX3A 6RXAA 6RY0A 6RY3A 6RYKA 6RZ0A 6RZOA 6S07A
6S0FA 6S2MA 6S2RA 6S3FA 6S3YA 6S5MA 6S5WA 6S5ZA 6S95A 6S9TA 6SAOA 6SBCA
6SCQA 6SD8A 6SE1A 6SERA 6SF3A 6SF4A 6SGFA 6SH6B 6SI6A 6SIBA 6SIDA 6SIUA
6SJ3A 6SJ8A 6SJAB 6SJQA 6SK0A 6SKHA 6SLDA 6SLFA 6SLLA 6SM4A 6SMTA 6SMZA
6SPOA 6SQPA 6SREA 6SRHB 6SSDA 6ST4A 6STLA 6SU5A 6SUKA 6SUNA 6SV1A 6SVKA
6SW2A 6SWIA 6SXTA 6SYVA 6T02B 6T0YA 6T2TA 6T3OA 6T40D 6T6EA 6T6HA 6T84A
6T85A 6T8IA 6T9QA 6TACA 6TB5A 6TBIA 6TCBA 6TCCA 6TD9A 6TEQA 6TFXA 6TG6A
6TGJA 6TGSA 6TGUA 6THOA 6TITA 6TJ0A 6TJ4A 6TJ7A 6TJRA 6TKUA 6TL1A 6TL7A
6TM3A 6TM6A 6TMEA 6TMEC 6TQDA 6TQPA 6TQRA 6TQTA 6TR4A 6TRJA 6TRKA 6TS3A
6TTOA 6TUNC 6TV0A 6TV2A 6TVEP 6TWKA 6TYJA 6TYKA 6TYWA 6TYYA 6TZCB 6TZNA

6TZXA 6U08A 6U08F 6U1RA 6U1VA 6U2DA 6U3UC 6U3WA 6U4ZA 6U54B 6U66A 6U7BB
6U83A 6U9IB 6UAQA 6UB8A 6UBDA 6UBLA 6UBOA 6UD7D 6UDVA 6UEXA 6UIDA 6UIOA
6ULLA 6ULOA 6UN8A 6UO3A 6UOFA 6UPQA 6UPUA 6UQJA 6UQVA 6UT9A 6UTCA 6UUFA
6UUIC 6UUYA 6UW5B 6UWOA 6UXCB 6UXUA 6V04A 6V0MA 6V1CA 6V4VA 6V67A 6V7UA
6VANA 6VDAA 6VE6A 6VEKI 6VH5A 6VH6A 6VIGA 6VJRA 6VK5D 6VK6A 6VK6B 6VK6C
6VKZA 6VLIA 6VMEB 6VMEC 6VNAB 6VP4A 6VPQA 6VTBA 6VVFA 6VVNA 6VYDA 6VZ0A
6VZDA 6VZFA 6W0VA 6W0VB 6W1IA 6W2GA 6W3WA 6W4QA 6W54A 6W6BA 6W6ZA 6W92A
6W9RA 6WAOA 6WB7A 6WBPA 6WESA 6WFNA 6WFVA 6WG4B 6WGMA 6WI5A 6WINA 6WISA
6WJOA 6WKCA 6WL5A 6WN2A 6WN9A 6WQYA 6WRTA 6WSMA 6WW3A 6WY9A 6WY9B 6X1GA
6X1XA 6X6BA 6X6FA 6X6XA 6X7HA 6X7JA 6X7NA 6X7QA 6X84A 6X9NA 6XAWA 6XAWB
6XAZB 6XB3A 6XB4A 6XB6A 6XCQA 6XFJA 6XHHA 6XIPA 6XIPB 6XJ6A 6XKGB 6XL7A
6XM7A 6XNUB 6XODB 6XOJA 6XP8A 6XRBA 6XRUA 6XRUB 6XRWA 6XUBA 6XVMC 6XWVE
6XXJA 6XYAB 6XYZA 6XZ4A 6XZLA 6XZUB 6Y04A 6Y1XA 6Y1YA 6Y3AA 6Y4EA 6Y4JA
6Y4LA 6Y56A 6Y5UA 6YAVA 6YB5A 6YB5C 6YB7A 6YDRA 6YGAA 6YGAC 6YGUA 6YGUB
6YHVA 6YIGA 6YIIA 6YIPA 6YIZA 6YLDA 6YN1J 6YOOA 6YP6A 6YSPA 6YTZA 6YUQA
6YV5A 6YVGA 6YWNA 6YX3A 6YX5B 6YXSA 6YYLA 6Z01A 6Z0MB 6Z1KA 6Z2LA 6Z2TA
6Z3JC 6Z5YA 6Z68A 6Z6EA 6Z96A 6Z9KA 6Z9UB 6ZA0A 6ZAYC 6ZBKB 6ZC2A 6ZCOA
6ZEGC 6ZGQA 6ZH1A 6ZM8A 6ZMPA 6ZNVA 6ZP9A 6ZPAA 6ZPEA 6ZPJA 6ZRNC 6ZRWA
6ZSIC 6ZSYA 6ZT4A 6ZTUA 6ZUSA 6ZXQA 6ZZJA 7A03A 7A0HA 7A0QA 7A3HA 7A5MA
7A66A 7A76A 7A8RA 7AAHA 7AALA 7AB3C 7ACOA 7ACWA 7ACWB 7AD5A 7ADRA 7ADRB
7ADRC 7AE6A 7AEDA 7AEXA 7AFWA 7AGWA 7AJOA 7AK7C 7AL1A 7AL7A 7ALJA 7AMKA
7AN7A 7ANIB 7AO3A 7AOOA 7APEA 7AQXA 7AS1A 7ASGA 7ASIA 7ASVA 7AT0A 7ATRA
7AVKA 7AWKA 7B0EA 7B0XC 7B1FA 7B1SA 7B1SB 7B1SC 7B29A 7B3AA 7B5WA 7B67A
7B7PA 7B7ZA 7BAFA 7BBLA 7BBLB 7BBLE 7BBXA 7BIUA 7BIZA 7BJ9A 7BJSA 7BJSC
7BLKA 7BLLA 7BMKA 7BNHA 7BQGA 7BQIA 7BR0A 7BR1A 7BR2A 7BRDA 7BSXA 7BU0A
7BUGA 7BV7C 7BVJA 7BVTA 7BWFD 7BX9A 7BXDA 7BXXA 7BY3B 7BYJA 7BYWA 7BZKA
7C0DA 7C1IA 7C1UA 7C2BB 7C2FB 7C42A 7C6CA 7C70A 7C7IC 7C8NA 7CCBA 7CCEA
7CDYA 7CFBB 7CHQA 7CHRA 7CIKA 7CIZC 7CIZD 7CJ7A 7CL2A 7CLUA 7CLYA 7CN7C
7CNBA 7CNPA 7CNWB 7CPRA 7CQ3A 7CQHA 7CQNA 7CRAA 7CRVA 7CSLC 7CT3A 7CT6A
7CTMA 7CTQA 7CU9A 7CUPA 7CUSA 7CX5A 7CXZA 7CZGB 7CZJA 7D1TU 7D2GA 7D2SB
7D48A 7D53A 7D55A 7D5MA 7D78A 7D8DA 7D8QA 7DA0B 7DA0C 7DA9A 7DB5A 7DBLA
7DBUA 7DC4A 7DCNA 7DE2A 7DE7A 7DEDA 7DFSA 7DG2C 7DH7A 7DHGB 7DHPA 7DIEA
7DKAA 7DKLA 7DL3A 7DM0A 7DMAA 7DMAB 7DMSA 7DNNA 7DNNB 7DOBA 7DOQA 7DPYA
7DQGA 7DQTA 7DRGB 7DRYA 7DS2C 7DS6A 7DTLA 7DU0A 7DU7A 7DWCA 7DZ9A 7E04A
7E0MA 7E1GA 7E2KA 7E2SA 7E38A 7E4NA 7E4VA 7E5BC 7E5CA 7E5TA 7E72E 7E8JA
7EAMA 7EBAA 7EBLA 7EC0D 7EC1A 7ECQA 7ECRA 7EDPB 7EE5A 7EF0A 7EF6A 7EGSA
7EGSB 7EHGA 7EHTA 7EHUA 7EIJA 7EJGC 7EJWC 7EK6A 7EMEA 7EN7A 7ENMA 7EP3A
7EPNB 7EQXA 7ERQA 7ES4A 7ESKA 7ESRA 7ESWA 7ET5A 7ETKA 7EUNA 7EVCB 7EXBB
7EXMA 7EYLA 7EZIA 7F0AA 7F0YA 7F22A 7F2YA 7F6QA 7F82A 7FDSB 7FE1A 7FE4A
7FFPA 7FH3A 7FH8A 7FHMA 7JFLC 7JGUA 7JIDA 7JJAA 7JJKA 7JKAA 7JL6A 7JOOC
7JP2A 7JRKA 7JRWA 7JT8I 7JTOL 7JUMA 7JV7B 7JW2A 7JW9A 7JX6A 7JXRA 7JXUA
7JZKB 7JZSA 7K34A 7K3TA 7K44A 7K7PB 7K7VA 7K9CA 7KAWC 7KBGA 7KC8A 7KCJA
7KD7A 7KDYB 7KFRA 7KHSA 7KIHA 7KJGA 7KJRC 7KLDB 7KLJA 7KOMA 7KPJE 7KPOA
7KQ5A 7KQ7B 7KQQA 7KQUB 7KR0A 7KRIA 7KS4B 7KZWA 7L00A 7L0AB 7L1IA 7L4AA

7L6GA 7L6LA 7L71A 7L78A 7LC5A 7LCZA 7LD8A 7LD9A 7LD9B 7LDAA 7LDQA 7LFAA
7LGAA 7LGCA 7LJIA 7LJOA 7LJPA 7LKLA 7LKLB 7LKNA 7LKNC 7LPZA 7LQ2A 7LQXA
7LRZA 7LSVA 7LT1A 7LVMA 7LWBD 7LWEA 7M4MA 7M5FA 7M5FC 7M8HA 7MC5A 7ME5A
7MELA 7MF4A 7MFOA 7MGVA 7MIWA 7MODA 7MPDA 7MPRA 7MPZA 7MQ3A 7MQQA 7MS0A
7MS7A 7N7HA 7NA9D 7NAJA 7NAZA 7NBIA 7NBVA 7NCEA 7NCYB 7NDHA 7NDYG 7NE2A
7NGEA 7NIYA 7NK7A 7NLCA 7NLOA 7NNAA 7NNIA 7NQEA 7NTEA 7NUUA 7NUVA 7NX0A
7NYQA 7NZ9A 7NZAA 7NZJA 7O39A 7O4FA 7O4PA 7O5YA 7O91A 7O97A 7OBVA 7OBVB
7OC4A 7OC9A 7OCXC 7ODXA 7ODYC 7OECA 7OEXA 7OFVA 7OGWA 7OL3A 7OM2B 7OOFA
7OUZA 7OWCB 7OYWA 7P18A 7P1QA 7P1WA 7P6SA 7P8MC 7PB9A 7PD1A 7PHYA 7PJDA
7PLPA 7PLQB 7POKA 7PPAA 7PQFA 7PSYA 7PT1A 7PTBA 7PVHA 7PYTB 7Q06A 7Q1GA
7Q47A 7Q4IA 7QBPA 7QDVA 7QGSB 7QIHA 7QIHB 7QZQA 7R09A 7R1ZA 7R25A 7R4UA
7R5YA 7R5ZA 7R65A 7R7BA 7R84A 7RC2A 7REFA 7RFTA 7RG8A 7RG8B 7RH8A 7RLMA
7RM7A 7RPYA 7RQGA 7RTYA 7RWGA 7RWKA 7RZBA 7S0MA 7S13C 7S2SD 7S3WA 7S56A
7S5HA 7S5NA 7S5OA 7S6FA 7S7YA 7SCIA 7SCXB 7SF6A 7SGPA 7SKLB 7SLVA 7SMJA
7SNEA 7SR1A 7SULC 7SZYA 7T26A 7T7ZA 7T88A 7T9WA 7TB6A 7TCSA 7TCXC 7TEMA
7TFMA 7THHA 7TI7A 7TJ1A 7TMGA 7TOIA 7TT9A 7TVCA 7TWCA 7TXNA 7TXSA 7TZHB
7U1YA 7UW7A 7V09A 7V4OA 7V5ZA 7V8ED 7V9OA 7VI7A 7VJOA 7VLKG 7VLKI 7VNIA
7VOBC 7VRCB 7VTDA 7VUXA 7VW0A 7VWKA 7W26A 7W5GA 7W5SA 7W83A 7WABA 7WJCA
7WKHA 7WRKA 7X4EA 7X8VA 7XGCA 7YXVA 7Z18D 7Z2TA 7Z5GA 7ZCVA 7ZGHA 8ABPA

##### 1.3 The MemProtMD_2022 dataset (184 sequences)

7TDMA 7TDMB 7TE9A 7TE9B 7TEBA 7TEBB 7TEEA 7TEEB 7TEQA 7TEQB 7TERA 7TERB
7TERF 7TESA 7TESB 7TETA 7TETB 7TJ8A 7TJ9A 7TNJA 7TNKA 7TNLA 7TNMA 7TNNA
7TNOA 7TNPA 7TPJA 7TT0A 7TT0B 7TT0C 7TT0D 7TT0E 7TT2A 7TT2B 7TT2C 7TT2D
7TT2E 7TT2F 7TT3A 7TT3B 7TT3C 7TT3D 7TT3E 7TT3F 7TT5A 7TT5B 7TT5C 7TT5D
7TT5E 7TT6A 7TT6B 7TT6C 7TT6D 7TT6E 7TT7A 7TT7B 7TT7C 7TT7D 7TT7E 7TYIA
7TYIB 7TYIC 7TYID 7TYIE 7TYLA 7TYLB 7TYLC 7TYLE 7TYLF 7TYOA 7TYOB 7TYOC
7TYOD 7TYOE 7U2LA 7U2LB 7U2LC 7U2LD 7UDVA 7UDWA 7UDXA 7UDYA 7UDZA 7UHFA
7UHFB 7UHFC 7WJIA 7WJIB 7WJIC 7WJID 7WJIE 7WLIA 7WLJA 7WLKA 7WLLA 7WQ3A
7WQ3B 7WQ3C 7WQ3D 7WQ3F 7WQ4A 7WQ4B 7WQ4C 7WQ4D 7WQ4F 7WSNA 7WSVA 7WU2A
7WU2B 7WU2C 7WU2E 7WU3A 7WU3B 7WU3C 7WU3E 7WU4A 7WU4B 7WU4C 7WU4D 7WUIA
7WUIB 7WUIC 7WUIE 7WUQA 7WUQB 7WUQC 7WUQE 7WVUA 7WVUB 7WVUC 7WVUD 7WVVB
7WVVC 7WVVD 7WVVE 7WVWB 7WVWC 7WVWD 7WVWE 7WVYA 7WVYB 7WVYC 7WVYD 7WVYE
7WYUA 7WYUB 7WYUC 7WYVA 7WYVB 7WYVC 7WYWA 7WYWB 7WYWC 7WYXA 7WYXB 7WYXC
7WYYA 7WYYB 7WYYC 7WYZA 7WYZB 7WYZC 7X9BA 7X9BB 7X9BC 7X9BD 7X9BE 7X9CA
7X9CB 7X9CC 7X9CD 7X9CE 7XJHA 7XJHB 7XJHC 7XJHE 7XJIA 7XJIB 7XJID 7XJIE
7Z0AA 7Z0CA 7Z0DA 7Z0EA

#### 2. The datasets for validating the membrane protein identification performance

##### 2.1 The PDBTM dataset (29795 sequences)

1A0SP 1A0SQ 1A0SR 1A0TP 1A0TQ 1A0TR 1A11A 1A91A 1AF6A 1AF6B 1AF6C 1AFOA
1AFOB 1AIGH 1AIGL 1AIGM 1AIJH 1AIJL 1AIJM 1AMTA 1AP9A 1AP9B 1AP9C 1AR1A
1AR1B 1AT9A 1AT9B 1AT9C 1ATYA 1B9UA 1BA4A 1BACA 1BADA 1BCCC 1BCCD 1BCCE
1BCCG 1BCCJ 1BCCM 1BCCN 1BCCP 1BCCR 1BCCU 1BCTA 1BE3C 1BE3D 1BE3E 1BE3G
1BE3J 1BE3K 1BGYC 1BGYD 1BGYE 1BGYG 1BGYJ 1BGYK 1BGYO 1BGYP 1BGYQ 1BGYS
1BGYV 1BGYW 1BH3A 1BH3B 1BH3C 1BHAA 1BHBA 1BL1A 1BL8A 1BL8B 1BL8C 1BL8D
1BM1A 1BM1B 1BM1C 1BNXA 1BOJA 1BOKA 1BRDA 1BRDB 1BRDC 1BRRA 1BRRB 1BRRC
1BRXA 1BRXB 1BRXC 1BT9A 1BT9B 1BT9C 1BTRA 1BTTA 1BXWA 1BY3A 1BY5A 1BZKA
1C0VA 1C17A 1C17B 1C17C 1C17D 1C17E 1C17F 1C17G 1C17H 1C17I 1C17J 1C17K
1C17L 1C17M 1C3WA 1C3WB 1C3WC 1C51A 1C51B 1C51F 1C51G 1C51H 1C51K 1C51L
1C51N 1C51P 1C51Q 1C51R 1C51S 1C51W 1C51X 1C51Y 1C8RA 1C8RB 1C8RC 1C8SA
1C8SB 1C8SC 1C99A 1CLTL 1CLTM 1CWQA 1CWQC 1CWQE 1DDD1 1DDD2 1DDD3 1DDD4
1DDD5 1DDD6 1DDD7 1DHLA 1DOPA 1DOPD 1DS8H 1DS8L 1DS8M 1DV3H 1DV3L 1DV3M
1DV6H 1DV6L 1DV6M 1DX7A 1DXRH 1DXRL 1DXRM 1DXZA 1DZEA 1DZEC 1DZED 1E0PA
1E0PB 1E0PC 1E12A 1E12C 1E12D 1E14H 1E14L 1E14M 1E54A 1E6DH 1E6DL 1E6DM
1E7PC 1E7PF 1EHKA 1EHKB 1EHKC 1EK9A 1EK9B 1EK9C 1EQ8A 1ET2S 1ET3S 1EYSH
1EYSL 1EYSM 1EZVC 1EZVD 1EZVE 1EZVG 1EZVI 1EZVL 1EZVM 1EZVN 1EZVR 1EZVS
1F4ZA 1F4ZB 1F4ZC 1F50A 1F50B 1F50C 1F6GA 1F6GB 1F6GC 1F6GD 1F6NH 1F6NL
1F6NM 1F88A 1FBBA 1FBBB 1FBBC 1FBKA 1FBKB 1FBKC 1FCPA 1FDFA 1FDMA 1FE1A
1FE1B 1FE1C 1FE1D 1FE1E 1FE1F 1FE1G 1FE1J 1FE1K 1FE1L 1FE1M 1FE1N 1FE1O
1FE1P 1FEPA 1FFTA 1FFTB 1FFTC 1FFTD 1FI1A 1FJKA 1FJPA 1FNPH 1FNPL 1FNPM
1FNQH 1FNQL 1FNQM 1FQUA 1FQYA 1FQYB 1FQYC 1FQYD 1FW2A 1FW3A 1FW3B 1FX8A
1FX8B 1FX8C 1FX8D 1G90A 1GFMA 1GFMB 1GFMC 1GFNA 1GFNB 1GFNC 1GFOA 1GFOB
1GFOC 1GFPA 1GFPB 1GFPC 1GFQA 1GFQB 1GFQC 1GU8A 1GUEA 1GZMA 1H2SA 1H2SB
1H2SC 1H2SD 1H68A 1H6IA 1H6S1 1HXTA 1HXTB 1HXTC 1HXUA 1HXUB 1HXUC 1HXXA
1HXXB 1HXXC 1HZXA 1I15A 1I15B 1I15C 1I15D 1I15E 1I15F 1I15G 1I78A 1IGKA
1IGKB 1IGKC 1IGKD 1IGKE 1IGKF 1IGKG 1IH5A 1IH5B 1IH5C 1IH5D 1IIJA 1IIVA
1IIVB 1IIVC 1IJDA 1IJDB 1IJDC 1IJDD 1IJDE 1IJDF 1IJDG 1IJDH 1IJDI 1IJDJ
1IJDK 1IJDL 1IJDM 1IJDN 1IJDP 1IJDQ 1IJDR 1IJDS 1IJPA 1ILDA 1ILXA 1ILXB
1ILXC 1ILXD 1ILXE 1ILXF 1ILXG 1ILXJ 1ILXK 1ILXL 1ILXM 1ILXN 1ILXO 1ILXP
1ILZA 1IM0A 1IW6A 1IW6C 1IW6D 1IW9A 1IW9C 1IW9D 1IWGA 1IWGB 1IWGC 1IWOA
1IXFA 1IXFC 1IXFD 1J4NA 1J4NB 1J4NC 1J4ND 1J7FA 1J7FB 1J7FC 1J7FD 1J7FE
1J7FF 1J7FG 1J7FH 1J7FI 1J7FJ 1J7FK 1J7FL 1J95A 1J95B 1J95C 1J95D 1JB0A
1JB0B 1JB0F 1JB0I 1JB0J 1JB0K 1JB0L 1JB0M 1JB0X 1JDMA 1JFPA 1JGJA 1JGWH
1JGWL 1JGWM 1JGXH 1JGXL 1JGXM 1JGYH 1JGYL 1JGYM 1JGZH 1JGZL 1JGZM 1JH0H
1JH0L 1JH0M 1JO5A 1JQ1A 1JQ1B 1JQ1C 1JQ1D 1JQ2A 1JQ2B 1JQ2C 1JQ2D 1JV6A
1JV7A 1JVMA 1JVMB 1JVMC 1JVMD 1K24A 1K4CC 1K4CF 1K4CI 1K4CL 1K4DC 1K4DF
1K4DI 1K4DL 1K6LH 1K6LL 1K6LM 1K6NH 1K6NL 1K6NM 1K9NA 1K9NB 1K9NC 1K9ND
1K9NE 1KADA 1KB9C 1KB9D 1KB9E 1KB9H 1KB9I 1KBYH 1KBYL 1KBYM 1KCHA 1KCHB
1KCHC 1KCHD 1KCHE 1KF6C 1KF6D 1KF6O 1KF6P 1KFYC 1KFYD 1KFYO 1KFYP 1KG8A

1KG8B 1KG8C 1KG9A 1KG9B 1KG9C 1KGBA 1KGBB 1KGBC 1KJUA 1KMEA 1KMOA 1KMPA
1KP1A 1KPKA 1KPKB 1KPLA 1KPLB 1KPNA 1KPWA 1KPXA 1KQFB 1KQFC 1KQFE 1KQFF
1KQFH 1KQFI 1KQGB 1KQGC 1KQGE 1KQGF 1KQGH 1KQGI 1KYKA 1KYKB 1KYKC 1KYKD
1KYKE 1KYLA 1KYLB 1KYLC 1KYLD 1KYLE 1KYOC 1KYOD 1KYOE 1KYOH 1KYOI 1KYON
1KYOO 1KYOP 1KYOS 1KYOT 1KZUA 1KZUB 1KZUC 1KZUD 1KZUE 1KZUF 1KZUG 1KZUH
1KZUI 1KZUJ 1KZUK 1KZUL 1KZUM 1KZUN 1KZUP 1KZUQ 1KZUR 1KZUS 1L0LC 1L0LD
1L0LE 1L0LG 1L0LJ 1L0LK 1L0MA 1L0NC 1L0ND 1L0NE 1L0NG 1L0NJ 1L0NK 1L0VC
1L0VD 1L0VO 1L0VP 1L6TA 1L7VA 1L7VB 1L9BH 1L9BL 1L9BM 1L9HA 1L9JH 1L9JL
1L9JM 1LBNA 1LDAA 1LDAB 1LDAC 1LDAD 1LDFA 1LDFB 1LDFC 1LDFD 1LDIA 1LDIB
1LDIC 1LDID 1LGHA 1LGHB 1LGHD 1LGHE 1LGHM 1LGHN 1LGHP 1LGHQ 1LGHR 1LGHS
1LGHT 1LGHU 1LGHV 1LGHW 1LGHX 1LGHY 1LLKA 1LLYA 1LN6A 1LNQA 1LNQB 1LNQC
1LNQD 1LVIA 1M0KA 1M0KB 1M0KC 1M0LA 1M0LB 1M0LC 1M0MA 1M0MB 1M0MC 1M2UA
1M3XH 1M3XL 1M3XM 1M56A 1M56B 1M56C 1M56D 1M57A 1M57B 1M57C 1M57D 1MALA
1MALB 1MALC 1MGYA 1MHSA 1MHSB 1MM4A 1MM5A 1MMH1 1MMH2 1MMH3 1MMH4 1MMH5
1MMH6 1MMH7 1MOTA 1MPFA 1MPFB 1MPFC 1MPMA 1MPMB 1MPMC 1MPNA 1MPNB 1MPNC
1MPOA 1MPOB 1MPOC 1MPQA 1MPQB 1MPQC 1MPRA 1MPRB 1MPRC 1MPSH 1MPSL 1MPSM
1MSRA 1MSRB 1N3MA 1N3MC 1N7LA 1ND8A 1NE0A 1NEKC 1NEKD 1NEKG 1NEKH 1NEKK
1NEKL 1NENC 1NEND 1NENG 1NENH 1NENK 1NENL 1NKZA 1NKZB 1NKZC 1NKZD 1NKZE
1NKZF 1NKZG 1NKZH 1NKZI 1NKZJ 1NKZK 1NKZL 1NKZM 1NKZN 1NKZP 1NKZQ 1NKZR
1NKZS 1NQEA 1NQFA 1NQGA 1NQHA 1NTKC 1NTKD 1NTKE 1NTKG 1NTKJ 1NTKK 1NTMC
1NTMD 1NTME 1NTMG 1NTMJ 1NTMK 1NTZC 1NTZD 1NTZE 1NTZG 1NTZJ 1NTZK 1NU1C
1NU1D 1NU1E 1NU1G 1NU1J 1NU1K 1NW0A 1NW0B 1O0AA 1O0AB 1O0AC 1O5WA 1OCCA
1OCCB 1OCCC 1OCCD 1OCCG 1OCCI 1OCCJ 1OCCK 1OCCL 1OCCM 1OCCN 1OCCO 1OCCP
1OCCQ 1OCCT 1OCCV 1OCCW 1OCCX 1OCCY 1OCCZ 1OCOA 1OCOB 1OCOC 1OCOD 1OCOG
1OCOI 1OCOJ 1OCOK 1OCOL 1OCOM 1OCON 1OCOO 1OCOP 1OCOQ 1OCOT 1OCOV 1OCOW
1OCOX 1OCOY 1OCOZ 1OCRA 1OCRB 1OCRC 1OCRD 1OCRG 1OCRI 1OCRJ 1OCRK 1OCRL
1OCRM 1OCRN 1OCRO 1OCRP 1OCRQ 1OCRT 1OCRV 1OCRW 1OCRX 1OCRY 1OCRZ 1OCZA
1OCZB 1OCZC 1OCZD 1OCZG 1OCZI 1OCZJ 1OCZK 1OCZL 1OCZM 1OCZN 1OCZO 1OCZP
1OCZQ 1OCZT 1OCZV 1OCZW 1OCZX 1OCZY 1OCZZ 1OEAA 1OEDA 1OEDB 1OEDC 1OEDD
1OEDE 1OGVH 1OGVL 1OGVM 1OH2P 1OH2Q 1OH2R 1OJ9A 1OJAA 1OJCA 1OKCA 1OPFA
1OPFB 1OPFC 1OPNA 1ORMA 1ORQC 1ORQD 1ORQE 1ORQF 1ORSC 1OSMA 1OSMB 1OSMC
1OTSA 1OTSB 1OTTA 1OTTB 1OTUA 1OTUB 1OV0A 1OV1A 1OY6A 1OY6B 1OY6C 1OY8A
1OY8B 1OY8C 1OY9A 1OY9B 1OY9C 1OYDA 1OYDB 1OYDC 1OYEA 1OYEB 1OYEC 1OZ5A
1OZCA 1P49A 1P4TA 1P4TB 1P4TC 1P7BA 1P7BB 1P7BC 1P7BD 1P84C 1P84D 1P84E
1P84H 1P84I 1P84N 1P84O 1P84P 1P84S 1P84T 1P8HA 1P8HB 1P8HC 1P8IA 1P8IB
1P8IC 1P8UA 1P8UB 1P8UC 1PB2A 1PB4C 1PB4D 1PCRH 1PCRL 1PCRM 1PHOA 1PHOB
1PHOC 1PLNA 1PLNB 1PLNC 1PLND 1PLNE 1PNZA 1PO0A 1PO3A 1PP9C 1PP9D 1PP9E
1PP9G 1PP9J 1PP9P 1PP9Q 1PP9R 1PP9T 1PP9W 1PPJC 1PPJD 1PPJE 1PPJG 1PPJJ
1PPJP 1PPJQ 1PPJR 1PPJT 1PPJW 1PRCH 1PRCL 1PRCM 1PRNA 1PRNB 1PRNC 1PSLA
1PSLB 1PSLC 1PSLD 1PSLE 1PSSH 1PSSL 1PSSM 1PSTH 1PSTL 1PSTM 1PV6A 1PV7A
1PW4A 1PXRA 1PXRC 1PXRD 1PXSA 1PXSC 1PXSD 1PY6A 1PY6C 1PY6D 1PYH1 1PYH2
1PYH3 1PYH4 1PYH5 1PYH6 1PYH7 1PYH8 1PYHA 1PYHB 1PYHC 1PYHD 1PYHE 1PYHF
1PYHG 1PYHH 1PYHI 1PYHJ 1PYHK 1PYHL 1PYHM 1PYHN 1PYHO 1PYHP 1PYHQ 1PYHR
1PYHS 1PYHT 1PYHU 1PYHV 1PYHW 1PYHX 1PYHY 1PYHZ 1Q16C 1Q16F 1Q5IA 1Q5IC

1Q5ID 1Q5JA 1Q5JC 1Q5JD 1Q90A 1Q90B 1Q90D 1Q90E 1Q90F 1Q90G 1Q90I 1Q90J
1Q90K 1Q90L 1Q90M 1Q90N 1Q90O 1Q90P 1Q90Q 1Q90R 1Q9FA 1Q9GA 1QCRC 1QCRD
1QCRE 1QCRG 1QCRJ 1QCRK 1QD5A 1QD6C 1QD6D 1QFFA 1QFGA 1QHJA 1QHJB 1QHJC
1QJ8A 1QJ9A 1QJPA 1QJQA 1QKCA 1QKOA 1QKOB 1QKOC 1QKPA 1QKPB 1QKPC 1QLBC
1QLBF 1QLEA 1QLEB 1QLEC 1QLED 1QM8A 1QM8C 1QM8D 1QO1K 1QO1L 1QO1M 1QO1N
1QO1O 1QO1P 1QO1Q 1QO1R 1QO1S 1QO1T 1QOVH 1QOVL 1QOVM 1R2CH 1R2CL 1R2CM
1R2NA 1R3IC 1R3ID 1R3IG 1R3IK 1R3JC 1R3JF 1R3JI 1R3JL 1R3KC 1R3KF 1R3KI
1R3KL 1R3LC 1R3LF 1R3LI 1R3LL 1R84A 1RC2B 1RG5H 1RG5L 1RG5M 1RGNH 1RGNL
1RGNM 1RH5A 1RH5B 1RH5C 1RHZA 1RHZB 1RHZC 1RKLA 1RQKH 1RQKL 1RQKM 1RVJH
1RVJL 1RVJM 1RWTB 1RWTF 1RWTG 1RY5H 1RY5L 1RY5M 1RZHH 1RZHL 1RZHM 1RZZH
1RZZL 1RZZM 1S00H 1S00L 1S00M 1S2QA 1S2YA 1S33A 1S33B 1S33C 1S33D 1S3BA
1S3EA 1S51A 1S51C 1S51E 1S52A 1S53A 1S53C 1S53E 1S54A 1S54C 1S54E 1S5HC
1S5HF 1S5HI 1S5HL 1S5LA 1S5LA 1S5LB 1S5LB 1S5LC 1S5LC 1S5LD 1S5LD 1S5LE
1S5LE 1S5LF 1S5LF 1S5LH 1S5LH 1S5LI 1S5LI 1S5LJ 1S5LJ 1S5LK 1S5LK 1S5LL
1S5LL 1S5LM 1S5LM 1S5LN 1S5LN 1S5LT 1S5LT 1S5LX 1S5LX 1S5LZ 1S5LZ 1S6EA
1S8JA 1S8LA 1SIWC 1SIWF 1SKHA 1SORA 1SORB 1SORC 1SORD 1SQBC 1SQBD 1SQBE
1SQBG 1SQBJ 1SQBK 1SQPC 1SQPD 1SQPE 1SQPG 1SQPJ 1SQPK 1SQQC 1SQQD 1SQQE
1SQQG 1SQQJ 1SQQK 1SQVC 1SQVD 1SQVE 1SQVG 1SQVJ 1SQVK 1SQXC 1SQXD 1SQXE
1SQXG 1SQXJ 1SQXK 1SR1A 1SU4A 1SUKA 1T16A 1T1LA 1T1LB 1T5SA 1T5TA 1T78A
1T9TA 1T9TB 1T9TC 1T9UA 1T9UB 1T9UC 1T9VA 1T9VB 1T9VC 1T9WA 1T9WB 1T9WC
1T9XA 1T9XB 1T9XC 1T9YA 1T9YB 1T9YC 1THQA 1TLWA 1TLYA 1TLZA 1TN0A 1TN0C
1TN0D 1TN5A 1TN5C 1TN5D 1TQQA 1TQQB 1TQQC 1TXHA 1TXHB 1TXHC 1TXHD 1TXHE
1TXHF 1U19A 1U5NA 1U77A 1U77B 1U77C 1U7CA 1U7CB 1U7CC 1U7GA 1U7GB 1U7GC
1UAZA 1UAZC 1UAZF 1UCQA 1UCQC 1UCQD 1UFDA 1UFDB 1UFDC 1UFDD 1UJWA 1UMXH
1UMXL 1UMXM 1UPEA 1UUNA 1UUNB 1UUNC 1UUND 1UUNE 1UUNF 1UUNG 1UUNH 1UV3A
1UXFA 1UYNX 1UYOX 1V36A 1V36B 1V36C 1V36D 1V36E 1V36F 1V36G 1V54A 1V54B
1V54C 1V54D 1V54G 1V54I 1V54J 1V54K 1V54L 1V54M 1V54N 1V54O 1V54P 1V54Q
1V54T 1V54V 1V54W 1V54X 1V54Y 1V54Z 1V55A 1V55B 1V55C 1V55D 1V55G 1V55I
1V55J 1V55K 1V55L 1V55M 1V55N 1V55O 1V55P 1V55Q 1V55T 1V55V 1V55W 1V55X
1V55Y 1V55Z 1VCRA 1VF5A 1VF5B 1VF5C 1VF5D 1VF5E 1VF5F 1VF5G 1VF5H 1VF5N
1VF5O 1VF5P 1VF5Q 1VF5R 1VF5S 1VF5T 1VF5U 1VFPA 1VGOA 1VGOC 1VGOF 1VJMA
1VJMB 1VJMC 1VRNH 1VRNL 1VRNM 1VRYA 1VZ1A 1W5CA 1W5CB 1W5CC 1W5CD 1W5CE
1W5CF 1W5CG 1W5CH 1W5CI 1W5CJ 1W5CK 1W5CL 1W5CX 1W5CY 1WAZA 1WP1A 1WP1C
1WP1D 1WPGA 1WRGA 1WU0A 1X0I1 1X0K1 1X0SA 1X0SC 1X0SD 1XFHA 1XFHB 1XFHC
1XIOA 1XIOB 1XJIA 1XKHA 1XKWA 1XL4A 1XL4B 1XL4C 1XL4D 1XL6A 1XL6B 1XL6C
1XL6D 1XMEA 1XMEB 1XMEC 1XNUA 1XNUB 1XNUC 1XNUD 1XNUE 1XP5A 1XQEA 1XQEB
1XQEC 1XQFA 1XQFB 1XQFC 1XRDA 1Y2LA 1Y36A 1Y4ZC 1Y4ZF 1Y5DA 1Y5IC 1Y5IF
1Y5LC 1Y5LF 1Y5NC 1Y5NF 1Y8SA 1Y9CA 1YC9A 1YC9B 1YC9C 1YCEA 1YCEB 1YCEC
1YCED 1YCEE 1YCEF 1YCEG 1YCEH 1YCEI 1YCEJ 1YCEK 1YF6H 1YF6L 1YF6M 1YG1A
1YG7A 1YMGA 1YMGB 1YMGC 1YMGD 1YO91 1YO92 1YO93 1YO94 1YO9A 1YO9B 1YO9F
1YO9G 1YO9H 1YO9I 1YO9J 1YO9K 1YO9L 1YQ3C 1YQ3D 1YQ4C 1YQ4D 1YSTH 1YSTL
1YSTM 1Z65A 1Z8EA 1Z98A 1Z98B 1Z98C 1Z98D 1Z9JA 1Z9JB 1Z9JC 1Z9KA 1Z9KB
1Z9KC 1ZC7A 1ZCDA 1ZCDC 1ZLLA 1ZLLB 1ZLLC 1ZLLD 1ZLLE 1ZOYC 1ZOYD 1ZP0C
1ZP0D 1ZRTC 1ZRTD 1ZRTE 1ZRTP 1ZRTQ 1ZRTR 1ZTIA 1ZV0B 1ZWIC 1ZWID 1ZWIE

1ZWIF 1ZYYA 1ZYYB 1ZZAA 2A06C 2A06D 2A06E 2A06G 2A06P 2A06Q 2A06R 2A06T
2A06W 2A0DA 2A65A 2A65B 2A95B 2A9HA 2A9HB 2A9HC 2A9HD 2ABMA 2ABMB 2ABMC
2ABMD 2AC6A 2ACZC 2ACZD 2ACZG 2ACZH 2ACZK 2ACZL 2AD0C 2AD0D 2AFLA 2AFLB
2AFLC 2AFLD 2AGVA 2AH3A 2AHYA 2AHYB 2AHYC 2AHYD 2AHZA 2AHZB 2AHZC 2AHZD
2AKHA 2AKHB 2AKHC 2AKHX 2AKHY 2AKHZ 2AKIA 2AKIB 2AKIC 2AKIX 2AKIY 2AKIZ
2AMKA 2ASGA 2ASGB 2ASGC 2ASGD 2ASGE 2AT9A 2AT9B 2AT9C 2ATKC 2ATKD 2ATKE
2ATKF 2AUIA 2AULA 2AVLA 2AXTA 2AXTA 2AXTB 2AXTB 2AXTC 2AXTC 2AXTD 2AXTD
2AXTE 2AXTE 2AXTF 2AXTF 2AXTH 2AXTH 2AXTI 2AXTI 2AXTJ 2AXTJ 2AXTK 2AXTK
2AXTL 2AXTL 2AXTM 2AXTM 2AXTT 2AXTT 2AXTX 2AXTX 2AXTZ 2AXTZ 2B0XA 2B2FA
2B2FB 2B2FC 2B2HA 2B2HB 2B2HC 2B2IA 2B2IB 2B2IC 2B2JA 2B2JB 2B2JC 2B5FA
2B5FB 2B5FC 2B5FD 2B6OA 2B6OI 2B6OJ 2B6OK 2B6PA 2B6PB 2B6PC 2B6PD 2B6QA
2B6RA 2B6SA 2B6UA 2B6VA 2B76C 2B76D 2B76O 2B76P 2BBJA 2BBJB 2BBJD 2BBJE
2BBJF 2BCCC 2BCCD 2BCCE 2BCCG 2BCCJ 2BG9A 2BG9B 2BG9C 2BG9D 2BG9E 2BHWA
2BHWB 2BHWC 2BL2A 2BL2B 2BL2C 2BL2D 2BL2E 2BL2F 2BL2G 2BL2H 2BL2I 2BL2J
2BMNA 2BNPA 2BNPB 2BNPC 2BNSA 2BNSB 2BNSC 2BOBC 2BOBD 2BOBE 2BOBF 2BOCC
2BOCD 2BOCE 2BOCF 2BOZH 2BOZL 2BOZM 2BRDA 2BRDB 2BRDC 2BS2C 2BS2F 2BS3C
2BS3F 2BS4C 2BS4F 2BY4A 2C32A 2C32B 2C32C 2C32D 2C3EA 2C64A 2C65A 2C66A
2C67A 2C70A 2C72A 2C73A 2C75A 2C76A 2C88A 2C8KA 2C8LA 2C9MA 2CFPA 2CFQA
2CPBA 2CPSA 2CYDA 2CYDB 2CYDC 2CYDD 2CYDE 2CYDF 2CYDG 2CYDH 2CYDI 2CYDJ
2D2CA 2D2CB 2D2CC 2D2CD 2D2CE 2D2CF 2D2CG 2D2CH 2D2CN 2D2CO 2D2CP 2D2CQ
2D2CR 2D2CS 2D2CT 2D2CU 2D57A 2D57B 2D57C 2D57D 2DB4A 2DB4B 2DB4C 2DB4D
2DB4E 2DB4F 2DB4G 2DB4H 2DB4I 2DB4J 2DF9A 2DHHA 2DHHB 2DHHC 2DHLA 2DQSA
2DR6A 2DR6B 2DR6C 2DRDA 2DRDB 2DRDC 2DW3A 2DWDC 2DWDD 2DWDE 2DWDF 2DWEC
2DWED 2DWEE 2DWEF 2DYRA 2DYRB 2DYRC 2DYRD 2DYRG 2DYRI 2DYRJ 2DYRK 2DYRL
2DYRM 2DYRN 2DYRO 2DYRP 2DYRQ 2DYRT 2DYRV 2DYRW 2DYRX 2DYRY 2DYRZ 2DYSA
2DYSB 2DYSC 2DYSD 2DYSG 2DYSI 2DYSJ 2DYSK 2DYSL 2DYSM 2DYSN 2DYSO 2DYSP
2DYSQ 2DYST 2DYSV 2DYSW 2DYSX 2DYSY 2DYSZ 2E74A 2E74B 2E74C 2E74D 2E74E
2E74F 2E74G 2E74H 2E74I 2E74J 2E74K 2E74L 2E74M 2E74N 2E74P 2E74Q 2E75A
2E75B 2E75C 2E75D 2E75E 2E75F 2E75G 2E75H 2E75I 2E75J 2E75K 2E75L 2E75M
2E75N 2E75P 2E75Q 2E76A 2E76B 2E76C 2E76D 2E76E 2E76F 2E76G 2E76H 2E76I
2E76J 2E76K 2E76L 2E76M 2E76N 2E76P 2E76Q 2EARA 2EATA 2EAUA 2EI4A 2EI4B
2EI4C 2EIJA 2EIJB 2EIJC 2EIJD 2EIJG 2EIJI 2EIJJ 2EIJK 2EIJL 2EIJM 2EIJN
2EIJO 2EIJP 2EIJQ 2EIJT 2EIJV 2EIJW 2EIJX 2EIJY 2EIJZ 2EIKA 2EIKB 2EIKC
2EIKD 2EIKG 2EIKI 2EIKJ 2EIKK 2EIKL 2EIKM 2EIKN 2EIKO 2EIKP 2EIKQ 2EIKT
2EIKV 2EIKW 2EIKX 2EIKY 2EIKZ 2EILA 2EILB 2EILC 2EILD 2EILG 2EILI 2EILJ
2EILK 2EILL 2EILM 2EILN 2EILO 2EILP 2EILQ 2EILT 2EILV 2EILW 2EILX 2EILY
2EILZ 2EIMA 2EIMB 2EIMC 2EIMD 2EIMG 2EIMI 2EIMJ 2EIMK 2EIML 2EIMM 2EIMN
2EIMO 2EIMP 2EIMQ 2EIMT 2EIMV 2EIMW 2EIMX 2EIMY 2EIMZ 2EINA 2EINB 2EINC
2EIND 2EING 2EINI 2EINJ 2EINK 2EINL 2EINM 2EINN 2EINO 2EINP 2EINQ 2EINT
2EINV 2EINW 2EINX 2EINY 2EINZ 2ERVA 2ERVB 2EVUA 2EVUB 2EVUC 2EVUD 2EXWA
2EXWB 2EXYA 2EXYB 2EZ0A 2EZ0B 2F1CX 2F1TA 2F1UA 2F1UB 2F1UC 2F1UD 2F1UE
2F1UF 2F1UG 2F1VA 2F1VB 2F2BA 2F2BB 2F2BC 2F2BD 2F75A 2F93A 2F93B 2F93C
2F93D 2F95A 2F95B 2F95C 2F95D 2FBWC 2FBWD 2FBWQ 2FCPA 2FECA 2FECB 2FEDA
2FEDB 2FEEA 2FEEB 2FF9A 2FGQA 2FGQB 2FGQX 2FGRA 2FGRC 2FGRE 2FKWA 2FKWB

2FKWC 2FKWD 2FKWE 2FKWF 2FKWG 2FKWH 2FKWI 2FKWJ 2FKWK 2FKWL 2FKWM 2FKWN
2FKWO 2FKWP 2FKWR 2FKWS 2FYNA 2FYNB 2FYNC 2FYND 2FYNE 2FYNF 2FYUC 2FYUD
2FYUE 2FYUG 2FYUJ 2FYUK 2G1XA 2G2AA 2G87A 2GE4A 2GFPA 2GFZA 2GIFA 2GIFB
2GIFC 2GMRH 2GMRL 2GMRM 2GNUH 2GNUL 2GNUM 2GR7A 2GR7B 2GR7C 2GR7D 2GR7E
2GR7F 2GR8A 2GR8C 2GR8D 2GRXA 2GSKA 2GSMA 2GSMB 2GUFA 2H2PA 2H2PB 2H2SA
2H2SB 2H3OA 2H88C 2H88D 2H89C 2H89D 2H8AA 2H8AB 2H8AC 2H8PC 2H8PD 2H8PE
2H8PF 2H8PG 2H8PH 2H8PI 2H8PJ 2HACA 2HACB 2HDFA 2HDIA 2HE6 2HFEC 2HFED 2HFEE
2HFEF 2HFEG 2HFEH 2HFEI 2HFEJ 2HG3H 2HG3L 2HG3M 2HG5C 2HG5D 2HG5E 2HG5F
2HG5G 2HG5H 2HG5I 2HG5J 2HG9H 2HG9L 2HG9M 2HH1H 2HH1L 2HH1M 2HHKH 2HHKL
2HHKM 2HI7B 2HITH 2HITL 2HITM 2HJ6H 2HJ6L 2HJ6M 2HJFC 2HJFD 2HJFE 2HJFF
2HLFA 2HLFB 2HN2A 2HN2B 2HN2C 2HN2D 2HN2E 2HPYA 2HQCA 2HQCB 2HQCC 2HQDA
2HQDB 2HQDC 2HQFA 2HQFB 2HQFC 2HQGA 2HQGB 2HQGC 2HRTA 2HRTB 2HRTC 2HT2A
2HT2B 2HT3A 2HT3B 2HT4A 2HT4B 2HTGA 2HTKA 2HTKB 2HTLA 2HTLB 2HVJC 2HVJD
2HVJE 2HVJF 2HVKC 2HVKD 2HVKE 2HVKF 2HYDA 2HYDB 2HYNA 2HYNB 2HYNC 2HYND
2HYNE 2I1XA 2I1XB 2I1XC 2I20A 2I20B 2I20C 2I21A 2I21B 2I21C 2I35A 2I36A
2I36B 2I37A 2I37B 2I5NH 2I5NL 2I5NM 2I6WA 2I6WB 2I6WC 2IAHA 2IBZC 2IBZD
2IBZE 2IBZG 2IBZI 2IC8A 2IH1C 2IH1D 2IH1E 2IH1F 2IH3C 2IH3D 2IH3E 2IH3F 2IIL
2IK3 2IK5 2IQ4 2IQKA 2IQKC 2IQLA 2IQMA 2IQNA 2IQOA 2IQPA 2IQRA 2IQSA 2IQUA
2IQVA 2IQWA 2IRVA 2IRVC 2IRVD 2ITAA 2ITCC 2ITCD 2ITCE 2ITCF 2ITDC 2ITDD
2ITDE 2ITDF 2IUBA 2IUBB 2IUBC 2IUBD 2IUBE 2IWVA 2IWWA 2J1HA 2J1IA 2J1IB
2J1IC 2J1ID 2J1NA 2J1NB 2J1NC 2J4UP 2J4UQ 2J4UR 2J4YA 2J58A 2J58B 2J58C
2J58D 2J58E 2J58F 2J58G 2J58H 2J7AC 2J7AF 2J8CH 2J8CL 2J8CM 2J8DH 2J8DL
2J8DM 2J8SA 2J8SB 2J8SC 2JAFA 2JAFB 2JAFC 2JAGA 2JAGB 2JAGC 2JBLH 2JBLL
2JBLM 2JIYH 2JIYL 2JIYM 2JJ0H 2JJ0L 2JJ0M 2JK4A 2JK5C 2JK5D 2JK5E 2JK5F
2JLNA 2JMMA 2JO1A 2JP3A 2JQYA 2JTWA 2JWAA 2JWAB 2K0LA 2K1AA 2K1KA 2K1KB
2K1LA 2K1LB 2K3CA 2K4TA 2K58B 2K59B 2K73A 2K74A 2K9JA 2K9JB 2K9PA 2K9YA
2KA1A 2KA1B 2KA2A 2KA2B 2KB7P 2KBVA 2KDCA 2KDCB 2KDCC 2KI9A 2KIHA 2KIHB
2KIHC 2KIHD 2KLUA 2KNCA 2KOGA 2KPEA 2KPEB 2KPFA 2KPNA 2KS1A 2KS1B 2KS9A
2KSAA 2KSBA 2KSDA 2KSEA 2KSFA 2KSRA 2KSYA 2KWXA 2KWXB 2KWXC 2KWXD 2KYHA
2KYVA 2KYVB 2KYVC 2KYVD 2KYVE 2L0JA 2L0JB 2L0JC 2L0JD 2L0LA 2L0MA 2L0NA
2L0OA 2L2TA 2L2TB 2L34A 2L34B 2L35A 2L35B 2L6WA 2L6XA 2L8SA 2L91A 2L9UA
2LATA 2LBGA 2LCKA 2LCXA 2LCXB 2LEGB 2LHFA 2LJ2A 2LKGA 2LKHA 2LLMA 2LLYA
2LM2A 2LMEA 2LMEB 2LMEC 2LNLA 2LOHA 2LOHB 2LOMA 2LOOA 2LOPA 2LOQA 2LORA
2LOSA 2LOTA 2LOUA 2LOVA 2LOWA 2LP1A 2LPFA 2LS2A 2LS3A 2LS4A 2LTQA 2LX0A
2LZ3A 2LZ3B 2LZ4A 2LZ4B 2LZLA 2M06A 2M07A 2M0BA 2M0QA 2M20A 2M20B 2M3BA
2M3BB 2M3BC 2M3BD 2M3BE 2M3EA 2M3GA 2M3GB 2M3GC 2M59A 2M59B 2M67A 2M6BA
2M6IA 2M6IB 2M6IC 2M6ID 2M6IE 2M7GA 2M7XA 2MAFA 2MAWA 2MC7A 2META 2METB
2METC 2MEUA 2MEUB 2MFRA 2MGYA 2MHLA 2MICA 2MICB 2MJOA 2MK9A 2MKAA 2MKAB
2MKVA 2MLHA 2MMUA 2MN6B 2MN7A 2MNHA 2MOFA 2MOMA 2MOMB 2MOMC 2MOZA 2MPNA
2MPNB 2MPRA 2MPRB 2MPRC 2MV6A 2MXBA 2N02A 2N2AA 2N2AB 2N2LA 2N2MA 2N4XA
2N5SA 2N6LA 2N6PA 2N7IA 2N7QA 2N7RA 2N90A 2N90B 2N9YA 2N9YB 2NA6A 2NA6B
2NA6C 2NA7A 2NA7B 2NA7C 2NA8A 2NA9A 2NDJA 2NLJC 2NLJD 2NLJE 2NLJF 2NMRA
2NMRB 2NMRC 2NOPA 2NOPB 2NOPC 2NOWA 2NOWB 2NOWC 2NPCA 2NPCB 2NPCC 2NPDA
2NPDB 2NPDC 2NPEA 2NPEB 2NPEC 2NPGA 2NPGB 2NPGC 2NPJA 2NPJB 2NPJC 2NPKA

2NPKB 2NPKC 2NQ2A 2NQ2B 2NR1A 2NR9A 2NRFA 2NRFC 2NRFD 2NRGA 2NS1A 2NS1C
2NS1E 2NTUA 2NTUB 2NTUC 2NTWA 2NTWB 2NTWC 2NUUA 2NUUB 2NUUC 2NWLA 2NWLB
2NWLC 2NWWA 2NWWB 2NWWC 2NWXA 2NWXB 2NWXC 2O011 2O012 2O013 2O014 2O01A
2O01B 2O01F 2O01G 2O01H 2O01I 2O01J 2O01L 2O4VA 2O4VB 2O4VC 2O5PA 2O7LA
2O9DA 2O9EA 2O9EB 2O9EC 2O9ED 2O9FA 2O9GA 2O9GB 2O9GC 2O9GD 2O9JA 2OA0A
2OARA 2OARB 2OARC 2OARD 2OARE 2OAUA 2OAUB 2OAUC 2OAUD 2OAUE 2OAUF 2OAUG
2OCCA 2OCCB 2OCCC 2OCCD 2OCCG 2OCCI 2OCCJ 2OCCK 2OCCL 2OCCM 2OCCN 2OCCO
2OCCP 2OCCQ 2OCCT 2OCCV 2OCCW 2OCCX 2OCCY 2OCCZ 2ODJA 2OMFA 2OMFB 2OMFC
2ONJA 2ONJB 2ONKC 2ONKD 2P7TC 2P7TD 2P7TE 2P7TF 2PEDA 2PNOA 2PNOB 2PNOC
2PORA 2PORB 2PORC 2PPSA 2PPSB 2PPSF 2PPSK 2PPSL 2PRCH 2PRCL 2PRCM 2PRNA
2PRNB 2PRNC 2Q67A 2Q67B 2Q67C 2Q67D 2Q68A 2Q68B 2Q68C 2Q68D 2Q69A 2Q69B
2Q69C 2Q69D 2Q6AA 2Q6AB 2Q6AC 2Q6AD 2Q6HA 2Q72A 2Q72B 2Q7MA 2Q7MB 2Q7MC
2Q7RA 2Q7RB 2Q7RC 2QB4A 2QB4B 2QEIA 2QEIB 2QFIA 2QI9A 2QI9B 2QJKA 2QJKB
2QJKC 2QJKD 2QJKE 2QJKF 2QJPA 2QJPB 2QJPC 2QJPD 2QJPE 2QJPF 2QJUA 2QJYA
2QJYB 2QJYC 2QJYD 2QJYE 2QJYF 2QKSA 2QKSC 2QKSD 2QKSE 2QOMA 2QPDA 2QPDB
2QPDC 2QPEA 2QPEB 2QPEC 2QTKA 2QTOA 2QTOB 2QTOC 2QTOD 2QTSA 2QTSB 2QTSC
2R4LB 2R4NA 2R4OA 2R4PA 2R4RA 2R4SA 2R6GF 2R6GG 2R88A 2R89A 2R8AA 2R9HA
2R9HB 2R9RB 2R9RC 2R9RD 2R9RE 2RCRH 2RCRL 2RCRM 2RDDA 2RDDB 2RDDC 2RDDD
2RDDE 2RDDF 2RH1A 2RMZA 2RN0A 2UUHA 2UUHB 2UUHC 2UUIA 2UUIB 2UUIC 2UWSH
2UWSL 2UWSM 2UWTH 2UWTL 2UWTM 2UWUH 2UWUL 2UWUM 2UWVH 2UWVL 2UWVM 2UWWH
2UWWL 2UWWM 2UX3H 2UX3L 2UX3M 2UX4H 2UX4L 2UX4M 2UX5H 2UX5L 2UX5M 2UXJH
2UXJL 2UXJM 2UXKH 2UXKL 2UXKM 2UXLH 2UXLL 2UXLM 2UXMH 2UXML 2UXMM 2V50A
2V50B 2V50C 2V5ZA 2V60A 2V61A 2V8NA 2VDDA 2VDDB 2VDDC 2VDEA 2VDEB 2VDEC
2VDFA 2VL0A 2VL0B 2VL0C 2VL0D 2VL0E 2VOYB 2VOYC 2VOYD 2VOYE 2VOYG 2VOYH
2VOYK 2VOYL 2VPWC 2VPWG 2VPXC 2VPXG 2VPYC 2VPYG 2VPZC 2VPZG 2VQIA 2VR0C
2VR0F 2VRLA 2VRMA 2VT4A 2VV5A 2VV5B 2VV5C 2VV5D 2VV5E 2VV5F 2VV5G 2W0FC
2W0FD 2W0FE 2W0FF 2W16A 2W1BA 2W1BB 2W1BC 2W1PA 2W1PB 2W1PC 2W1PD 2W2EA
2W2EB 2W2EC 2W2ED 2W5JA 2W5JB 2W5JC 2W5JD 2W5JE 2W5JF 2W5JG 2W5JH 2W5JI
2W5JJ 2W5JK 2W5JL 2W5JM 2W5JV 2W6TA 2W6UA 2W75A 2W76A 2W77A 2W78A 2WDQC
2WDQD 2WDQG 2WDQH 2WDRC 2WDRD 2WDRG 2WDRH 2WDVC 2WDVD 2WDVG 2WDVH 2WDVK
2WDVL 2WGMA 2WGMB 2WGMC 2WGMD 2WGME 2WGMF 2WGMG 2WGMH 2WGMI 2WGMJ 2WGMK
2WIEA 2WIEB 2WIEC 2WIED 2WIEE 2WIEF 2WIEG 2WIEH 2WIEI 2WIEJ 2WIEK 2WIEL
2WIEM 2WIEN 2WIEP 2WITA 2WITB 2WITC 2WJKA 2WJKB 2WJKC 2WJLA 2WJLB 2WJLC
2WJMH 2WJML 2WJMM 2WJNH 2WJNL 2WJNM 2WJQA 2WJRA 2WLHA 2WLHB 2WLHC 2WLHD
2WLIA 2WLIB 2WLID 2WLIE 2WLJA 2WLJB 2WLJC 2WLJD 2WLKA 2WLKB 2WLKC 2WLKD
2WLLA 2WLLB 2WLLC 2WLLD 2WLMA 2WLMB 2WLMC 2WLMD 2WLNA 2WLNB 2WLNC 2WLND
2WLOA 2WLOB 2WOXA 2WP9C 2WP9D 2WP9G 2WP9H 2WPDJ 2WPDK 2WPDL 2WPDM 2WPDN
2WPDO 2WPDP 2WPDQ 2WPDR 2WPDS 2WQYC 2WQYD 2WS3C 2WS3D 2WS3G 2WS3H 2WS3K
2WS3L 2WSC1 2WSC2 2WSC3 2WSC4 2WSCA 2WSCB 2WSCF 2WSCG 2WSCH 2WSCI 2WSCJ
2WSCK 2WSCL 2WSCR 2WSE1 2WSE2 2WSE3 2WSE4 2WSEA 2WSEB 2WSEF 2WSEG 2WSEH
2WSEI 2WSEJ 2WSEK 2WSF1 2WSF2 2WSF3 2WSF4 2WSFA 2WSFB 2WSFF 2WSFG 2WSFH
2WSFI 2WSFJ 2WSFK 2WSWA 2WSWB 2WSWC 2WSXA 2WSXB 2WSXC 2WU2C 2WU2D 2WU2G
2WU2H 2WU2K 2WU2L 2WU5C 2WU5D 2WU5G 2WU5H 2WU5K 2WU5L 2WVPA 2WX5H 2WX5L
2WX5M 2X27X 2X2VA 2X2VB 2X2VC 2X2VD 2X2VE 2X2VF 2X2VG 2X2VH 2X2VI 2X2VJ

2X2VK 2X2VL 2X2VM 2X4MA 2X55A 2X56A 2X5UH 2X5UL 2X5UM 2X5VH 2X5VL 2X5VM
2X6AA 2X6AB 2X6AC 2X6AD 2X6BA 2X6BB 2X6BC 2X6BD 2X6CA 2X6CB 2X6CC 2X6CD
2X72A 2X72D 2X79A 2X9KA 2XE1A 2XE1B 2XE1C 2XE2A 2XE2B 2XE2C 2XE3A 2XE3C
2XE3D 2XE5A 2XE5B 2XE5C 2XFNA 2XFOA 2XFPA 2XFQA 2XFUA 2XG6A 2XG6B 2XG6C
2XMNA 2XMNB 2XMNC 2XNDJ 2XNDK 2XNDL 2XNDM 2XNDN 2XNDO 2XNDP 2XNDQ 2XOKK
2XOKL 2XOKM 2XOKN 2XOKO 2XOKP 2XOKQ 2XOKR 2XOKS 2XOKT 2XOVA 2XOWA 2XQ2A
2XQ2B 2XQ3A 2XQ3B 2XQ3C 2XQ3D 2XQ3E 2XQ4A 2XQ4B 2XQ4C 2XQ4D 2XQ4E 2XQ5A
2XQ5B 2XQ5C 2XQ5D 2XQ5E 2XQ6A 2XQ6B 2XQ6C 2XQ6D 2XQ6E 2XQ7A 2XQ7B 2XQ7C
2XQ7D 2XQ7E 2XQ8A 2XQ8B 2XQ8C 2XQ8D 2XQ8E 2XQ9A 2XQ9B 2XQ9C 2XQ9D 2XQ9E
2XQAA 2XQAB 2XQAC 2XQAD 2XQAE 2XQSA 2XQSB 2XQSC 2XQSD 2XQSE 2XQSF 2XQSG
2XQSH 2XQSI 2XQSJ 2XQSK 2XQSL 2XQSM 2XQSN 2XQSP 2XQTA 2XQTB 2XQTC 2XQTD
2XQTE 2XQTF 2XQTG 2XQTH 2XQTI 2XQTJ 2XQTK 2XQTL 2XQTM 2XQTN 2XQTP 2XQUA
2XQUB 2XQUC 2XQUD 2XQUE 2XQUF 2XQUG 2XQUH 2XQUI 2XQUJ 2XQUK 2XQUL 2XQUM
2XQUN 2XQUP 2XTUA 2XTVA 2XUTA 2XZBA 2XZBB 2Y00A 2Y01A 2Y02A 2Y03A 2Y04A
2Y0HA 2Y0KA 2Y0LA 2Y2XA 2Y5YA 2Y69A 2Y69B 2Y69C 2Y69D 2Y69G 2Y69I 2Y69J
2Y69K 2Y69L 2Y69M 2Y69N 2Y69O 2Y69P 2Y69Q 2Y69T 2Y69V 2Y69W 2Y69X 2Y69Y
2Y69Z 2YBBC 2YBBC 2YBBD 2YBBD 2YBBE 2YBBE 2YBBG 2YBBG 2YBBJ 2YBBJ 2YBBK
2YBBK 2YBBL 2YBBM 2YBBM 2YBBN 2YBBN 2YBBO 2YBBO 2YBBP 2YBBR 2YBBT 2YBBU
2YBBV 2YBBW 2YBBX 2YCWA 2YCXA 2YCYA 2YCZA 2YDOA 2YDVA 2YEVA 2YEVB 2YEVC
2YFYA 2YIUA 2YIUB 2YIUC 2YIUD 2YIUE 2YIUF 2YKSA 2YKSB 2YKSC 2YKSD 2YKSE
2YN6A 2YN6B 2YN6C 2YN6D 2YN6E 2YN9A 2YN9B 2YNKA 2YOEA 2YOEB 2YOEC 2YOED
2YOEE 2YPTA 2YSUA 2YVXA 2YVXB 2YXQA 2YXQB 2YXQC 2YXRA 2YXRB 2YXRC 2Z55A
2Z55I 2Z55J 2Z5XA 2Z5YA 2Z73A 2ZBDA 2ZBEA 2ZBFA 2ZBGA 2ZD9A 2ZD9B 2ZD9C
2ZD9D 2ZFEA 2ZFEC 2ZFED 2ZFGA 2ZFGB 2ZFGC 2ZIYA 2ZJSE 2ZJSY 2ZLDB 2ZLDE
2ZLDG 2ZLED 2ZLEN 2ZLEP 2ZQPE 2ZQPY 2ZT9A 2ZT9B 2ZT9C 2ZT9D 2ZT9E 2ZT9F
2ZT9G 2ZT9H 2ZT9I 2ZT9J 2ZT9K 2ZT9L 2ZT9M 2ZT9N 2ZT9P 2ZT9Q 2ZUPB 2ZUQA
2ZW3A 2ZW3B 2ZW3C 2ZW3D 2ZW3E 2ZW3F 2ZXEA 2ZXEB 2ZXEG 2ZXWA 2ZXWB 2ZXWC
2ZXWD 2ZXWG 2ZXWI 2ZXWJ 2ZXWK 2ZXWL 2ZXWM 2ZXWN 2ZXWO 2ZXWP 2ZXWQ 2ZXWT
2ZXWV 2ZXWW 2ZXWX 2ZXWY 2ZXWZ 2ZY9A 2ZY9B 2ZZLA 2ZZLC 2ZZLD 3A0BA 3A0BA
3A0BB 3A0BB 3A0BC 3A0BC 3A0BD 3A0BD 3A0BE 3A0BE 3A0BF 3A0BF 3A0BH 3A0BH
3A0BI 3A0BI 3A0BJ 3A0BJ 3A0BK 3A0BK 3A0BL 3A0BL 3A0BM 3A0BM 3A0BN 3A0BN
3A0BT 3A0BT 3A0BX 3A0BX 3A0BY 3A0BY 3A0BZ 3A0BZ 3A0HA 3A0HA 3A0HB 3A0HB
3A0HC 3A0HC 3A0HD 3A0HD 3A0HE 3A0HE 3A0HF 3A0HF 3A0HH 3A0HH 3A0HI 3A0HI
3A0HJ 3A0HJ 3A0HK 3A0HK 3A0HL 3A0HL 3A0HM 3A0HM 3A0HN 3A0HN 3A0HT 3A0HT
3A0HX 3A0HX 3A0HY 3A0HY 3A0HZ 3A0HZ 3A2SB 3A2SC 3A2SX 3A3YA 3A3YB 3A3YG
3A7KA 3A7KB 3A7KD 3ABKA 3ABKB 3ABKC 3ABKD 3ABKG 3ABKI 3ABKJ 3ABKK 3ABKL
3ABKM 3ABKN 3ABKO 3ABKP 3ABKQ 3ABKT 3ABKV 3ABKW 3ABKX 3ABKY 3ABKZ 3ABLA
3ABLB 3ABLC 3ABLD 3ABLG 3ABLI 3ABLJ 3ABLK 3ABLL 3ABLM 3ABLN 3ABLO 3ABLP
3ABLQ 3ABLT 3ABLV 3ABLW 3ABLX 3ABLY 3ABLZ 3ABMA 3ABMB 3ABMC 3ABMD 3ABMG
3ABMI 3ABMJ 3ABMK 3ABML 3ABMM 3ABMN 3ABMO 3ABMP 3ABMQ 3ABMT 3ABMV 3ABMW
3ABMX 3ABMY 3ABMZ 3ABVC 3ABVD 3ABWA 3ABWB 3ABWD 3AE1C 3AE1D 3AE2C 3AE2D
3AE3C 3AE3D 3AE4C 3AE4D 3AE5C 3AE5D 3AE6C 3AE6D 3AE7C 3AE7D 3AE8C 3AE8D
3AE9C 3AE9D 3AEAC 3AEAD 3AEBC 3AEBD 3AECC 3AECD 3AEDC 3AEDD 3AEEC 3AEED
3AEFC 3AEFD 3AEGC 3AEGD 3AEHA 3AEHB 3AG1A 3AG1B 3AG1C 3AG1D 3AG1G 3AG1I

3AG1J 3AG1K 3AG1L 3AG1M 3AG1N 3AG1O 3AG1P 3AG1Q 3AG1T 3AG1V 3AG1W 3AG1X
3AG1Y 3AG1Z 3AG2A 3AG2B 3AG2C 3AG2D 3AG2G 3AG2I 3AG2J 3AG2K 3AG2L 3AG2M
3AG2N 3AG2O 3AG2P 3AG2Q 3AG2T 3AG2V 3AG2W 3AG2X 3AG2Y 3AG2Z 3AG3A 3AG3B
3AG3C 3AG3D 3AG3G 3AG3I 3AG3J 3AG3K 3AG3L 3AG3M 3AG3N 3AG3O 3AG3P 3AG3Q
3AG3T 3AG3V 3AG3W 3AG3X 3AG3Y 3AG3Z 3AG4A 3AG4B 3AG4C 3AG4D 3AG4G 3AG4I
3AG4J 3AG4K 3AG4L 3AG4M 3AG4N 3AG4O 3AG4P 3AG4Q 3AG4T 3AG4V 3AG4W 3AG4X
3AG4Y 3AG4Z 3AM6A 3AM6B 3AM6C 3AM6D 3ANZA 3ANZB 3ANZC 3ANZD 3ANZE 3ANZF
3ANZG 3AOAA 3AOAB 3AOAC 3AOBA 3AOBB 3AOBC 3AOCA 3AOCB 3AOCC 3AODA 3AODB
3AODC 3AOUA 3AOUB 3AOUC 3AOUD 3AOUE 3AOUF 3AOUG 3AOUH 3AOUI 3AOUJ 3AQPA
3AR2A 3AR3A 3AR4A 3AR5A 3AR6A 3AR7A 3AR8A 3AR9A 3ASNA 3ASNB 3ASNC 3ASND
3ASNG 3ASNI 3ASNJ 3ASNK 3ASNL 3ASNM 3ASNN 3ASNO 3ASNP 3ASNQ 3ASNT 3ASNV
3ASNW 3ASNX 3ASNY 3ASNZ 3ASOA 3ASOB 3ASOC 3ASOD 3ASOG 3ASOI 3ASOJ 3ASOK
3ASOL 3ASOM 3ASON 3ASOO 3ASOP 3ASOQ 3ASOT 3ASOV 3ASOW 3ASOX 3ASOY 3ASOZ
3AYFA 3AYGA 3AYMA 3AYNA 3B29A 3B29B 3B29C 3B44A 3B45A 3B4RA 3B5DA 3B5WA
3B5WB 3B5XA 3B5XB 3B5YA 3B5YB 3B5ZA 3B5ZB 3B60A 3B60B 3B61A 3B8EC 3B8ED
3B8EH 3B9BA 3B9RA 3B9WA 3B9WB 3B9WC 3B9YA 3B9YB 3B9YC 3B9ZA 3B9ZB 3B9ZC
3BA6A 3BCCC 3BCCD 3BCCE 3BCCG 3BCCJ 3BCCM 3BCCN 3BCCP 3BCCR 3BCCU 3BEHA
3BEHB 3BEHC 3BEHD 3BHSA 3BHSB 3BHSC 3BRYA 3BRZA 3BS0A 3BVDA 3BVDB 3BVDC
3C02A 3C02B 3C02C 3C02D 3C1GA 3C1GB 3C1GC 3C1HA 3C1HB 3C1HC 3C1IA 3C1IB
3C1IC 3C1JA 3C1JB 3C1JC 3C9LA 3C9MA 3CAPA 3CAPB 3CHXA 3CHXB 3CHXC 3CHXE
3CHXF 3CHXG 3CHXI 3CHXJ 3CHXK 3CHXM 3CHXN 3CHXO 3CIRC 3CIRD 3CIRO 3CIRP
3CLLA 3CLLB 3CLLC 3CLLD 3CN5A 3CN5B 3CN5C 3CN5D 3CN6A 3CN6C 3CN6D 3CN6E
3COCA 3COCC 3COCD 3CODA 3CSLA 3CSNA 3CWBC 3CWBD 3CWBE 3CWBG 3CWBJ 3CWBP
3CWBQ 3CWBR 3CWBT 3CWBW 3CX5C 3CX5D 3CX5E 3CX5H 3CX5I 3CX5N 3CX5O 3CX5P
3CX5S 3CX5T 3CXHC 3CXHD 3CXHE 3CXHH 3CXHI 3CXHN 3CXHO 3CXHP 3CXHS 3CXHT
3D31C 3D31D 3D38H 3D38L 3D38M 3D4SA 3D9BA 3D9BB 3D9BC 3D9SA 3D9SB 3D9SC
3D9SD 3DDLA 3DDRB 3DETA 3DETB 3DH4A 3DH4B 3DHLA 3DHWA 3DHWB 3DINC 3DIND
3DINE 3DL8C 3DL8E 3DL8G 3DQBA 3DSYH 3DSYL 3DSYM 3DTAH 3DTAL 3DTAM 3DTRH
3DTRL 3DTRM 3DTSH 3DTSL 3DTSM 3DTUA 3DTUB 3DU2H 3DU2L 3DU2M 3DU3H 3DU3L
3DU3M 3DUQH 3DUQL 3DUQM 3DWNA 3DWOX 3DWWA 3DWWB 3DWWC 3DZMA 3E83A 3E83C
3E83D 3E83E 3E86A 3E86C 3E86D 3E86E 3E89A 3E89C 3E89D 3E89E 3E8BA 3E8BC
3E8BD 3E8BE 3E8FA 3E8FC 3E8FD 3E8FE 3E8GA 3E8GC 3E8GD 3E8GE 3E8HA 3E8HC
3E8HD 3E8HE 3E9JC 3EAMA 3EAMB 3EAMC 3EAMD 3EAME 3EFFK 3EFFL 3EFFM 3EFFN
3EFMA 3EGWC 3EGWF 3EH3A 3EH3B 3EH3C 3EH4A 3EH4B 3EH4C 3EH5A 3EH5B 3EH5C
3EHBA 3EHBB 3EHZA 3EHZB 3EHZC 3EHZD 3EHZE 3EI0A 3EI0B 3EI0C 3EI0D 3EI0E
3EJYA 3EJYB 3EJZA 3EJZB 3EMLA 3EMNX 3EMOA 3EMOB 3EMOC 3F3AA 3F3AB 3F3CA
3F3CB 3F3DA 3F3DB 3F3EA 3F3EB 3F48A 3F48B 3F4IA 3F4IB 3F4JA 3F4JB 3F5WC
3F5WD 3F5WE 3F5WF 3F7VC 3F7VD 3F7VE 3F7VF 3F7YC 3F7YD 3F7YE 3F7YF 3FB5C
3FB5D 3FB5E 3FB5F 3FB6C 3FB6D 3FB6E 3FB6F 3FB7C 3FB7D 3FB7E 3FB7F 3FB8C
3FB8E 3FB8F 3FB8G 3FGOA 3FH6F 3FH6G 3FHHA 3FI1A 3FI1B 3FIDA 3FIPA 3FPBA
3FPSA 3FWLA 3FYEA 3FYEB 3FYIA 3FYIB 3FYXA 3FYXB 3FYXC 3G5UA 3G60A 3G61A
3G7FH 3G7FL 3G7FM 3GB7C 3GB7D 3GB7E 3GB7F 3GD8A 3GD8B 3GD8C 3GD8D 3GI8C
3GI9C 3GIAA 3GJCA 3GJCB 3GJDA 3GJDB 3GP6A 3GWUA 3GWUB 3GWVA 3GWVB 3GWWA
3GWWB 3H1HC 3H1HD 3H1HE 3H1HG 3H1HJ 3H1HP 3H1HQ 3H1HR 3H1HT 3H1HW 3H1IC

3H1ID 3H1IE 3H1IG 3H1IJ 3H1IP 3H1IQ 3H1IR 3H1IT 3H1IW 3H1JC 3H1JD 3H1JE
3H1JG 3H1JJ 3H1JP 3H1JQ 3H1JR 3H1JT 3H1JW 3H1KC 3H1KD 3H1KE 3H1KG 3H1KJ
3H1KP 3H1KQ 3H1KR 3H1KT 3H1KW 3H1LC 3H1LD 3H1LE 3H1LG 3H1LJ 3H1LP 3H1LQ
3H1LR 3H1LT 3H1LW 3H90A 3H9VA 3H9VD 3HANA 3HAOA 3HAOC 3HAOD 3HAPA 3HAQA
3HARA 3HASA 3HB3A 3HB3B 3HD6A 3HD6B 3HD6C 3HD7A 3HD7B 3HFXA 3HFXB 3HFXC
3HIGA 3HIIA 3HKKA 3HKKB 3HKKC 3HPLC 3HPLD 3HPLE 3HPLF 3HW9A 3HWBA 3HWBC
3HWBD 3I4DH 3I4DL 3I4DM 3I5DA 3I5DB 3I5DC 3IFXA 3IFXB 3IFXC 3IFXD 3IGAC
3IGAD 3IGAE 3IGAF 3IJ4A 3IJ4B 3IJ4C 3IPDA 3IPDB 3IR5C 3IR5F 3IR6C 3IR6F
3IR7C 3IR7H 3IXZA 3IXZB 3IYZA 3IYZB 3IYZC 3IYZD 3IZ1A 3IZ1B 3IZ1C 3IZ1D
3IZ1E 3IZ1F 3IZ2A 3IZ2B 3IZ2C 3IZ2D 3IZ2E 3IZ2F 3J08A 3J08B 3J09A 3J09B
3J1ZP 3J1ZQ 3J41A 3J41B 3J41C 3J41D 3J5PA 3J5PB 3J5PC 3J5PD 3J5QB 3J5QD
3J5QE 3J5QG 3J5RA 3J5RB 3J5RC 3J5RD 3J7TA 3J9JA 3J9JB 3J9JC 3J9JD 3J9PA
3J9PB 3J9PC 3J9PD 3J9TA 3J9TR 3J9TS 3J9TT 3J9TU 3J9TV 3J9TW 3J9TX 3J9TY
3J9TZ 3J9UA 3J9UR 3J9US 3J9UT 3J9UU 3J9UV 3J9UW 3J9UX 3J9UY 3J9UZ 3J9VA
3J9VR 3J9VS 3J9VT 3J9VU 3J9VV 3J9VW 3J9VX 3J9VY 3J9VZ 3JACA 3JACB 3JACC
3JADA 3JADB 3JADC 3JADD 3JADE 3JAEA 3JAEB 3JAEC 3JAED 3JAEE 3JAFA 3JAFB
3JAFC 3JAFD 3JAFE 3JAVA 3JAVB 3JAVC 3JAVD 3JBRA 3JBRE 3JC21 3JC22 3JC23
3JCFA 3JCFB 3JCFC 3JCFD 3JCFE 3JCGA 3JCGB 3JCGC 3JCGD 3JCGE 3JCHA 3JCHB
3JCHC 3JCHD 3JCHE 3JCUA 3JCUA 3JCUB 3JCUB 3JCUC 3JCUC 3JCUD 3JCUD 3JCUE
3JCUE 3JCUF 3JCUF 3JCUG 3JCUG 3JCUH 3JCUH 3JCUI 3JCUI 3JCUJ 3JCUJ 3JCUK
3JCUK 3JCUL 3JCUL 3JCUM 3JCUM 3JCUN 3JCUN 3JCUR 3JCUR 3JCUS 3JCUS 3JCUT
3JCUT 3JCUW 3JCUW 3JCUX 3JCUX 3JCUY 3JCUY 3JCUZ 3JCUZ 3JD8A 3JTYA 3JYCA
3JYCB 3JYCC 3JYCD 3K03A 3K03C 3K03D 3K03E 3K04A 3K04C 3K04D 3K04E 3K06A
3K06C 3K06D 3K06E 3K07A 3K07B 3K07C 3K08A 3K08C 3K08D 3K08E 3K0DA 3K0DC
3K0DD 3K0DE 3K0GA 3K0GC 3K0GD 3K0GE 3K0IA 3K0IB 3K0IC 3K19A 3K19B 3K19C
3K1BA 3K1BB 3K1BC 3K3FA 3K3FB 3K3FC 3K3GA 3K3GB 3K3GC 3KBCA 3KBCB 3KBCC
3KCUA 3KCUB 3KCUC 3KCUD 3KCUE 3KCVA 3KCVB 3KCVC 3KCVD 3KCVE 3KDPC 3KDPD
3KDPH 3KG2A 3KG2B 3KG2C 3KG2D 3KJ6A 3KLYA 3KLYB 3KLYC 3KLYD 3KLYE 3KLZA
3KLZB 3KLZC 3KLZD 3KLZE 3KP9A 3KSOA 3KSOB 3KSOC 3KSSA 3KSSB 3KSSC 3KVNX
3KZIA 3KZIB 3KZIC 3KZID 3KZIE 3KZIF 3KZIH 3KZII 3KZIJ 3KZIK 3KZIL 3KZIM
3KZIT 3KZIX 3KZIY 3KZIZ 3L1LA 3L70C 3L70D 3L70E 3L70G 3L70J 3L70P 3L70Q
3L70R 3L70T 3L70W 3L71C 3L71D 3L71E 3L71G 3L71J 3L71P 3L71Q 3L71R 3L71T
3L71W 3L72C 3L72D 3L72E 3L72G 3L72J 3L72P 3L72Q 3L72R 3L72T 3L72W 3L73C
3L73D 3L73E 3L73G 3L73J 3L73P 3L73Q 3L73R 3L73T 3L73W 3L74C 3L74D 3L74E
3L74G 3L74J 3L74P 3L74Q 3L74R 3L74T 3L74W 3L75C 3L75D 3L75E 3L75G 3L75J
3L75P 3L75Q 3L75R 3L75T 3L75W 3LDCA 3LDCB 3LDCC 3LDCD 3LDDA 3LDDB 3LDDC
3LDDD 3LDEA 3LDEB 3LDEC 3LDED 3LEOA 3LEOB 3LEOC 3LLQA 3LNMB 3LNME 3LNMF
3LNMG 3LRBA 3LRBB 3LRCA 3LRCB 3LSVA 3LSVB 3LSVC 3LSVD 3LSVE 3LUTB 3LUTD
3LUTF 3LUTH 3LW51 3LW52 3LW53 3LW54 3LW5A 3LW5B 3LW5F 3LW5G 3LW5H 3LW5I
3LW5J 3LW5K 3LW5L 3M2LA 3M2LB 3M2LC 3M2LD 3M2LE 3M2LF 3M2LG 3M3RA 3M3RB
3M3RC 3M3RD 3M3RE 3M3RF 3M3RG 3M4DA 3M4DB 3M4DC 3M4DD 3M4DE 3M4DF 3M4DG
3M4EA 3M4EB 3M4EC 3M4ED 3M4EE 3M4EF 3M4EG 3M6EA 3M6EB 3M6EC 3M71A 3M71B
3M71C 3M72A 3M72B 3M72C 3M73A 3M73B 3M73C 3M74A 3M74B 3M74C 3M75A 3M75B
3M75C 3M76A 3M76B 3M76C 3M77A 3M77B 3M77C 3M78A 3M78B 3M78C 3M7BA 3M7BB

3M7BC 3M7CA 3M7CB 3M7CC 3M7EA 3M7EB 3M7EC 3M7LA 3M7LB 3M7LC 3M8BA 3M8DA
3M9CL 3M9CM 3M9CN 3M9CR 3M9IA 3M9SH 3M9SL 3M9SM 3M9SN 3M9SR 3MBVA 3MBVB
3MBVC 3MK7A 3MK7B 3MK7C 3MK7U 3MKTA 3MKUA 3MP7A 3MP7B 3MPNA 3MPNB 3MPQA
3MPQB 3MRAA 3N23A 3N23B 3N23G 3N5KA 3N8GA 3NALA 3NAMA 3NANA 3NCYA 3NCYB
3ND0A 3ND0B 3NE2A 3NE2B 3NE2C 3NE2D 3NE5A 3NE5H 3NE5I 3NJTA 3NK5A 3NKAA
3NKAC 3NKAD 3NKAE 3NKCA 3NKCC 3NKCD 3NKCE 3NMOA 3NOCA 3NOCB 3NOCC 3NOGA
3NOGB 3NOGC 3NS0A 3NS0B 3NS0C 3NSBA 3NSBB 3NSBC 3NSGA 3NSGB 3NSGC 3NY8A
3NY9A 3NYAA 3NYMA 3O0EB 3O0RB 3O0RC 3O44A 3O44B 3O44C 3O44D 3O44E 3O44F
3O44G 3O7PA 3O7QA 3OAXA 3OB6A 3OB6B 3ODJA 3ODUA 3ODUB 3OE0A 3OE0B 3OE6A
3OE6B 3OE8B 3OE8C 3OE9A 3OE9B 3OGCC 3OGCD 3OGCE 3OGCF 3OHNA 3OM3A 3OM3B
3OMAA 3OMAB 3OMIA 3OMIB 3OMNA 3OMNB 3OR6C 3OR6D 3OR6E 3OR6F 3OR7C 3OR7D
3OR7E 3OR7F 3ORGA 3ORGD 3OUFA 3OUFC 3OUFD 3OUFE 3OUSA 3OUSB 3OUSC 3OUSD
3P03A 3P03B 3P03C 3P4PC 3P4PD 3P4PO 3P4PP 3P4QC 3P4QD 3P4QO 3P4QP 3P4RC
3P4RD 3P4RO 3P4RP 3P4SC 3P4SD 3P4SO 3P4SP 3P4WA 3P4WB 3P4WC 3P4WD 3P4WE
3P50A 3P50B 3P50C 3P50D 3P50E 3P5NA 3PBLA 3PCQA 3PCQB 3PCQF 3PCQI 3PCQJ
3PCQK 3PCQL 3PCQM 3PCQX 3PCVA 3PCVB 3PCVC 3PDSA 3PF1A 3PGRA 3PGSA 3PGUA
3PIKA 3PIKB 3PIKC 3PJSK 3PJSL 3PJSM 3PJSN 3PJZA 3PJZB 3PL9A 3PO7A 3POQA
3POQB 3POQC 3PORA 3PORB 3PORC 3POUA 3POUB 3POUC 3POXA 3POXB 3POXC 3PQRA
3PRCH 3PRCL 3PRCM 3PRNA 3PRNB 3PRNC 3PUVF 3PUVG 3PUWF 3PUWG 3PUXF 3PUXG
3PUYF 3PUYG 3PUZF 3PUZG 3PV0F 3PV0G 3PWHA 3PXOA 3Q17A 3Q17B 3Q7KA 3Q7KB
3Q7KC 3Q7KD 3Q7KE 3QAKA 3QAPA 3QBGA 3QBGB 3QBGD 3QBIA 3QBIB 3QBID 3QBKA
3QBKB 3QBKD 3QBLA 3QBLB 3QBLD 3QDCA 3QE7A 3QE7B 3QF4A 3QF4B 3QJQA 3QJQB
3QJQC 3QJRA 3QJRB 3QJRC 3QJSA 3QJSB 3QJSC 3QJTA 3QJTB 3QJTC 3QJUA 3QJUB
3QJUC 3QJVA 3QJVB 3QJVC 3QLBA 3QLBB 3QNQA 3QNQB 3QQ2A 3QRAA 3QRCA 3QS4A
3QS4B 3QS5A 3QS5B 3QS6A 3QS6B 3R65A 3R65B 3R65C 3R65D 3RBHA 3RCEA 3REYA
3RFMA 3RFUA 3RFZB 3RGBA 3RGBB 3RGBC 3RGBE 3RGBF 3RGBG 3RGBI 3RGBJ 3RGBK
3RGMA 3RGNA 3RHWA 3RHWB 3RHWC 3RHWD 3RHWE 3RI5A 3RI5B 3RI5C 3RI5D 3RI5E
3RIAA 3RIAB 3RIAC 3RIAD 3RIAE 3RIFA 3RIFB 3RIFC 3RIFD 3RIFE 3RKOA 3RKOJ
3RKOK 3RKOL 3RKOM 3RKON 3RLBA 3RLFF 3RLFG 3RQUA 3RQUB 3RQUC 3RQUD 3RQUE
3RQWA 3RQWB 3RQWC 3RQWD 3RQWE 3RVYA 3RVYB 3RVYC 3RVYD 3RVZA 3RVZB 3RVZC
3RVZD 3RW0A 3RW0B 3RW0C 3RW0D 3RZEA 3S0XA 3S0XB 3S33A 3S33B 3S33C 3S38A
3S38B 3S38C 3S39A 3S39B 3S39C 3S3AA 3S3AB 3S3AC 3S3BA 3S3BB 3S3BC 3S3CA
3S3CB 3S3CC 3S3DA 3S3DB 3S3DC 3S3WA 3S3WB 3S3WC 3S3XA 3S3XB 3S3XC 3S8FA
3S8FB 3S8FC 3S8GA 3S8GB 3S8GC 3SFDC 3SFDD 3SFEC 3SFED 3SLJA 3SLOA 3SLTA
3SN6R 3SPCA 3SPCB 3SPCC 3SPCD 3SPGA 3SPGB 3SPGC 3SPGD 3SPHA 3SPHB 3SPHC
3SPHD 3SPIA 3SPIB 3SPIC 3SPID 3SPJA 3SPJB 3SPJC 3SPJD 3STLC 3STLD 3STLE
3STLF 3STZC 3STZD 3STZE 3STZF 3SY7A 3SY9A 3SYAA 3SYAB 3SYAC 3SYAD 3SYBA
3SYCA 3SYCB 3SYCC 3SYCD 3SYOA 3SYOB 3SYOC 3SYOD 3SYPA 3SYPB 3SYPC 3SYPD
3SYQA 3SYQB 3SYQC 3SYQD 3SYSA 3SZDB 3SZVA 3T0SA 3T1CA 3T1CC 3T1CD 3T1CE
3T20A 3T24A 3T2MA 3T2MC 3T2MD 3T2ME 3T45A 3T45B 3T45C 3T4DB 3T4DF 3T4DG
3T4DH 3T4ZA 3T4ZC 3T4ZD 3T4ZE 3T51A 3T51F 3T51I 3T53A 3T53F 3T53I 3T56A
3T56D 3T56G 3T6DH 3T6DL 3T6DM 3T6EH 3T6EL 3T6EM 3T9NA 3T9NB 3T9NC 3T9ND
3T9NE 3T9NF 3T9NG 3TCUA 3TCUC 3TCUD 3TCUE 3TDOA 3TDOB 3TDOC 3TDOD 3TDOE
3TDPA 3TDPB 3TDPC 3TDPD 3TDPE 3TDRA 3TDRB 3TDRC 3TDRD 3TDRE 3TDSA 3TDSB

3TDSC 3TDSD 3TDSE 3TDXA 3TDXB 3TDXC 3TDXD 3TDXE 3TE0A 3TE0B 3TE0C 3TE0D
3TE0E 3TE1A 3TE1B 3TE1C 3TE1D 3TE1E 3TE2A 3TE2B 3TE2C 3TE2D 3TE2E 3TETA
3TETC 3TETD 3TETE 3TGUC 3TGUD 3TGUE 3TGUG 3TGUJ 3TGUP 3TGUQ 3TGUR 3TGUT
3TGUW 3TIJA 3TIJB 3TIJC 3TLMA 3TLSA 3TLSB 3TLSC 3TLSD 3TLSE 3TLTA 3TLTB
3TLTC 3TLTD 3TLTE 3TLUA 3TLUB 3TLUC 3TLUD 3TLUE 3TLVA 3TLVB 3TLVC 3TLVD
3TLVE 3TLWA 3TLWB 3TLWC 3TLWD 3TLWE 3TT1A 3TT1B 3TT3A 3TT3B 3TU0A 3TU0B
3TUIA 3TUIB 3TUJA 3TUJB 3TUZA 3TUZB 3TX3A 3TXTA 3TXTB 3TXTC 3U2FA 3U2FB
3U2FC 3U2FD 3U2FE 3U2FK 3U2FL 3U2FM 3U2FN 3U2FO 3U2YA 3U2YB 3U2YC 3U2YD
3U2YE 3U2YK 3U2YL 3U2YM 3U2YN 3U2YO 3U32A 3U32B 3U32C 3U32D 3U32E 3U32K
3U32L 3U32M 3U32N 3U32O 3UBBA 3UBBB 3UBBC 3UD0A 3UD0B 3UD0C 3UD0D 3UD0E
3UD0K 3UD0L 3UD0M 3UD0N 3UD0O 3UDCA 3UDCB 3UDCC 3UDCD 3UDCE 3UDCF 3UDCG
3UG9A 3UG9B 3UKMA 3UKMB 3UM7A 3UM7B 3UONA 3UPGA 3UPGB 3UPGC 3UQ4A 3UQ4B
3UQ4C 3UQ4D 3UQ4E 3UQ5A 3UQ5B 3UQ5C 3UQ5D 3UQ5E 3UQ7A 3UQ7B 3UQ7C 3UQ7D
3UQ7E 3USGA 3USGB 3USIA 3USJA 3USKA 3USKB 3USLA 3USLB 3USMA 3USMB 3USOA
3USOB 3USPA 3USPB 3UTVA 3UTWA 3UTXA 3UTYA 3UU2A 3UU2B 3UU2C 3UU3A 3UU3B
3UU3C 3UU3D 3UU3E 3UU4A 3UU4B 3UU4C 3UU4D 3UU4E 3UU5A 3UU5B 3UU5C 3UU5D
3UU5E 3UU6A 3UU6B 3UU6C 3UU6D 3UU6E 3UU8A 3UU8B 3UU8C 3UU8D 3UU8E 3UUBA
3UUBB 3UUBC 3UUBD 3UUBE 3UX4A 3UX4B 3UX4C 3UZAA 3UZCA 3V2WA 3V2YA 3V3CA
3V3CB 3V3CC 3V3CD 3V3CE 3V3CF 3V3CG 3V3CH 3V3CI 3V3CJ 3V3CK 3V3CL 3V3CM
3V3CN 3V3YH 3V3YL 3V3YM 3V3ZH 3V3ZL 3V3ZM 3V5SA 3V5UA 3V89A 3V8FA 3V8FB
3V8FC 3V8GA 3V8GB 3V8GC 3V8XA 3VG9A 3VGAA 3VHZA 3VHZC 3VHZD 3VI0A 3VI0C
3VI0D 3VMAA 3VMQA 3VMRA 3VMSA 3VMTA 3VOUA 3VOUB 3VOUC 3VOUD 3VR8C 3VR8D
3VR9C 3VR9D 3VRAC 3VRAD 3VRBC 3VRBD 3VVKA 3VVKB 3VVKC 3VVNA 3VVOA 3VVPA
3VVRA 3VVSA 3VW7A 3VY8A 3VY8B 3VY8X 3VY9A 3VY9B 3VY9X 3VZTA 3VZTB 3VZTX
3VZUA 3VZUB 3VZUX 3VZWA 3VZWB 3VZWX 3W4TA 3W5AA 3W5AC 3W5BA 3W5CA 3W5DA
3W9HA 3W9HB 3W9HC 3W9IA 3W9IB 3W9IC 3W9JA 3W9JB 3W9JC 3W9TA 3W9TB 3W9TC
3W9TD 3W9TE 3W9TF 3W9TG 3WAJA 3WAKA 3WBNA 3WDOA 3WFBB 3WFBC 3WFCB 3WFCC
3WFDB 3WFDC 3WFEB 3WFEC 3WG7A 3WG7B 3WG7C 3WG7D 3WG7G 3WG7I 3WG7J 3WG7K
3WG7L 3WG7M 3WG7N 3WG7O 3WG7P 3WG7Q 3WG7T 3WG7V 3WG7W 3WG7X 3WG7Y 3WG7Z
3WGUC 3WGUD 3WGUE 3WGVC 3WGVD 3WGVE 3WI4A 3WI4B 3WI4C 3WI5A 3WI5B 3WI5C
3WKVA 3WKVB 3WKVC 3WMEA 3WMEB 3WMFA 3WMFB 3WMGA 3WMM0 3WMM1 3WMM2 3WMM3
3WMM4 3WMM5 3WMM6 3WMM7 3WMM8 3WMM9 3WMMA 3WMMB 3WMMD 3WMME 3WMMF 3WMMG
3WMMH 3WMMI 3WMMJ 3WMMK 3WMML 3WMMM 3WMMN 3WMMO 3WMMP 3WMMR 3WMMS 3WMMT
3WMMU 3WMMV 3WMMW 3WMMX 3WMMY 3WMMZ 3WO6A 3WO7A 3WQJA 3WQJB 3WQJC 3WU2A
3WU2A 3WU2B 3WU2B 3WU2C 3WU2C 3WU2D 3WU2D 3WU2E 3WU2E 3WU2F 3WU2F 3WU2H
3WU2H 3WU2I 3WU2I 3WU2J 3WU2J 3WU2K 3WU2K 3WU2L 3WU2L 3WU2M 3WU2M 3WU2T
3WU2T 3WU2X 3WU2X 3WU2Y 3WU2Y 3WU2Z 3WU2Z 3WVFA 3X29A 3X2QA 3X2QB 3X2QC
3X2QD 3X2QG 3X2QI 3X2QJ 3X2QK 3X2QL 3X2QM 3X2QN 3X2QO 3X2QP 3X2QQ 3X2QT
3X2QV 3X2QW 3X2QX 3X2QY 3X2QZ 3X2RA 3X2RB 3X2RC 3X2RD 3X2RE 3X2RF 3X2RG
3X2RH 3X2RI 3X3BA 3X3BB 3X3CA 3X3CB 3ZDQA 3ZDQB 3ZE3A 3ZE3B 3ZE3C 3ZE4A
3ZE4B 3ZE4C 3ZE5A 3ZE5B 3ZE5C 3ZEBA 3ZEVA 3ZJZA 3ZJZB 3ZJZE 3ZJZF 3ZK1A
3ZK1B 3ZK1C 3ZK1D 3ZK1E 3ZK1F 3ZK1G 3ZK1H 3ZK1I 3ZK1J 3ZK1K 3ZK2A 3ZK2B
3ZK2C 3ZK2D 3ZK2E 3ZK2F 3ZK2G 3ZK2H 3ZK2I 3ZK2J 3ZK2K 3ZKRA 3ZKRB 3ZKRC
3ZKRD 3ZKRE 3ZMHA 3ZMIA 3ZMJA 3ZO6A 3ZO6B 3ZO6C 3ZO6D 3ZO6E 3ZO6F 3ZO6H

3ZO6I 3ZO6J 3ZO6K 3ZO6L 3ZO6M 3ZOJA 3ZOJB 3ZOJC 3ZOJD 3ZOTA 3ZPQA 3ZPRA

3ZRSA 3ZRSB 3ZRSC 3ZRSD 3ZRYJ 3ZRYK 3ZUMH 3ZUML 3ZUMM 3ZUWH 3ZUWL 3ZUWM

3ZUXA 3ZUYA 4A01A 4A01B 4A2NB 4A4MA 4A4MD 4A79A 4A7AA 4A82A 4A82B 4A97A

4A97B 4A97C 4A97D 4A97E 4A98A 4A98B 4A98C 4A98D 4A98E 4AC5H 4AC5L 4AC5M

4AFKA 4AGEA 4AGEB 4AGEC 4AGED 4AGEE 4AGEF 4AGEG 4AGFA 4AGFB 4AGFC 4AGFD

4AGFE 4AGFF 4AGFG 4AINA 4AINB 4AINC 4AIPA 4AIPB 4AIPC 4AIQA 4AL0A 4AL0B

4AL0C 4AL1A 4AL1B 4AL1C 4AMIA 4AMJA 4APSA 4AQ5A 4AQ5B 4AQ5C 4AQ5D 4AQ5E

4AQ9A 4AQ9B 4AQ9C 4AQ9D 4AQ9E 4ATVA 4ATVB 4AU5A 4AU5B 4AV3A 4AV3B 4AV6A

4AV6B 4AW6A 4AYTA 4AYTB 4AYWA 4AYWB 4AYXA 4AYXB 4AZLA 4B2QJ 4B2QK 4B2QL

4B2QM 4B2QN 4B2QO 4B2QP 4B2QQ 4B2QR 4B2QS 4B4AA 4B61A 4B7OA 4BBJA 4BEMA

4BEMB 4BEMC 4BEMD 4BEME 4BEMF 4BEMG 4BEMH 4BEMI 4BEMJ 4BEVA 4BEWA 4BEYA

4BEZA 4BEZC 4BGNA 4BGNC 4BGND 4BGNE 4BOGA 4BOGB 4BOGC 4BOGD 4BOGE 4BOIA

4BOIB 4BOIC 4BOID 4BOIE 4BONA 4BONB 4BONC 4BOND 4BONE 4BOOA 4BOOB 4BOOC

4BOOD 4BOOE 4BORA 4BORB 4BORC 4BORD 4BORE 4BOTA 4BOTB 4BOTC 4BOTD 4BOTE

4BPDA 4BPDB 4BPDC 4BPMA 4BPMB 4BPMC 4BPQA 4BPQB 4BRBA 4BRBB 4BRBC 4BRRA

4BRRB 4BRRC 4BSKC 4BUMX 4BUOA 4BV0A 4BVNA 4BW5A 4BW5B 4BWBA 4BWZA 4BYGA

4C00A 4C48A 4C48C 4C48D 4C48F 4C48G 4C48I 4C4VA 4C4VB 4C69X 4C7RA 4C7RB

4C7RC 4C9GA 4C9HA 4C9JA 4C9QA 4C9QB 4CADC 4CASB 4CASC 4CASD 4CBCA 4CBCB

4CBCG 4CBCH 4CBJA 4CBJB 4CBJC 4CBJD 4CBJE 4CBJF 4CBJG 4CBJH 4CBJI 4CBJJ

4CBJK 4CBJL 4CBJM 4CBKA 4CBKB 4CBKC 4CBKD 4CBKE 4CBKF 4CBKG 4CBKH 4CBKI

4CBKJ 4CBKK 4CBKL 4CBKM 4CDIA 4CDIB 4CDIC 4CDID 4CDIE 4CDIF 4CG5A 4CG5B

4CG5C 4CG6A 4CG6B 4CG6C 4CG6D 4CG7A 4CG7B 4CG7C 4CHVA 4CHVB 4CHVC 4CHVD

4CHWA 4CHWB 4CHWC 4CHWD 4CJZA 4CJZB 4CJZC 4CK0A 4CK0B 4CK0C 4COFA 4COFB

4COFC 4COFD 4COFE 4CRTA 4CSKA 4CSKB 4CSKC 4CSKD 4CTDA 4CU4A 4CY4A 4CY4B

4CY4C 4CY4D 4CY4E 4CZ8A 4CZ8B 4CZ9A 4CZ9B 4CZAA 4CZAB 4CZBA 4CZBB 4D0AA

4D0AB 4D1AA 4D1BA 4D1CA 4D1DA 4D2BA 4D2CA 4D2DA 4D2EA 4D2EB 4D2EC 4D51A

4D5BA 4D5DA 4D5UA 4D5UB 4D5UC 4D64A 4D64B 4D64C 4D65D 4D65E 4D65F 4D6TC

4D6TD 4D6TE 4D6TG 4D6TJ 4D6TP 4D6TQ 4D6TR 4D6TT 4D6TW 4D6UC 4D6UD 4D6UE

4D6UG 4D6UJ 4D6UP 4D6UQ 4D6UR 4D6UT 4D6UW 4DAJA 4DBLA 4DBLB 4DCBA 4DJHA

4DJHB 4DJIA 4DJKA 4DKLA 4DKLB 4DNRA 4DNRF 4DNRI 4DNTA 4DNTF 4DNTI 4DOJA

4DOJB 4DOJC 4DOPA 4DOPF 4DOPI 4DVEA 4DW0A 4DW0C 4DW0D 4DW1A 4DW1C 4DW1D

4DX5A 4DX5B 4DX5C 4DX6A 4DX6B 4DX6C 4DX7A 4DX7B 4DX7C 4DXWA 4DXWB 4DXWC

4DXWD 4E1SA 4E1TA 4EA3A 4EEBA 4EEBB 4EEBC 4EEBD 4EEBE 4EEDA 4EEDB 4EEDC

4EEDD 4EEDE 4EIYA 4EJ4A 4EKWA 4EKWB 4EKWE 4EKWF 4ENEA 4ENEB 4EPAA 4EV6A

4EV6B 4EV6C 4EV6D 4EV6E 4EZCA 4EZCB 4EZCC 4EZDA 4EZDB 4EZDC 4F35A 4F35B

4F4CA 4F4LA 4F4LB 4F4LC 4F4LD 4F4SK 4F4SL 4F4SM 4F4SN 4F4SO 4F4SP 4F4SQ

4F4SR 4F4SS 4F4ST 4F8HA 4F8HB 4F8HC 4F8HD 4F8HE 4FA7A 4FA7B 4FA7C 4FAAA

4FAAB 4FAAC 4FBYA 4FBYA 4FBYB 4FBYB 4FBYC 4FBYC 4FBYD 4FBYD 4FBYE 4FBYE

4FBYF 4FBYG 4FBYG 4FBYH 4FBYI 4FBYJ 4FBYJ 4FBYK 4FBYK 4FBYL 4FBYL 4FBYM

4FBYM 4FBYN 4FBYP 4FBYQ 4FBYR 4FBYS 4FBYT 4FBYW 4FBYX 4FBYY 4FBYY 4FBYZ

4FBZA 4FBZB 4FBZC 4FC4A 4FC4B 4FC4C 4FC4D 4FC4E 4FE1A 4FE1B 4FE1F 4FE1I

4FE1J 4FE1K 4FE1L 4FE1M 4FE1X 4FG6A 4FG6B 4FI3A 4FI3B 4FMSB 4FOZA 4FPDA

4FPDB 4FPDC 4FQEA 4FRTA 4FRXA 4FSOA 4FSPA 4FT6A 4FUVA 4FUVB 4FXZA 4FXZB

4FY0A 4FY0B 4FZ0A 4FZ0B 4FZ0C 4FZ1A 4FZ1B 4FZ1E 4G1UA 4G1UB 4G70A 4G70B

4G70C 4G71A 4G71B 4G71C 4G72A 4G72B 4G72C 4G7QA 4G7QB 4G7QC 4G7RA 4G7RB
4G7RC 4G7SA 4G7SB 4G7SC 4G7VS 4G7YS 4G80S 4G80T 4GBRA 4GBYA 4GBZA 4GC0A
4GCPA 4GCQA 4GCSA 4GD3A 4GD3B 4GD3Q 4GD3R 4GD3S 4GD3T 4GEYA 4GF4A 4GP4A
4GP4B 4GP4C 4GP5A 4GP5B 4GP5C 4GP8A 4GP8B 4GP8C 4GPOA 4GPOB 4GRVA 4GX0A
4GX0B 4GX0F 4GX0G 4GX1A 4GX1B 4GX1E 4GX1F 4GX2A 4GX2B 4GX2E 4GX2F 4GX5A
4GX5B 4GX5E 4GX5F 4GYCA 4GYCB 4GYCC 4GYCD 4H0LA 4H0LB 4H0LC 4H0LD 4H0LE
4H0LF 4H0LG 4H0LH 4H0LI 4H0LJ 4H0LK 4H0LL 4H0LM 4H0LN 4H0LO 4H0LP 4H13A
4H13B 4H13C 4H13D 4H13E 4H13F 4H13G 4H13H 4H13I 4H13J 4H13K 4H13L 4H13M
4H13N 4H13O 4H13P 4H1DA 4H1DB 4H1DC 4H1WA 4H1WB 4H33A 4H33B 4H33C 4H33D
4H37A 4H37B 4H37C 4H37D 4H44A 4H44B 4H44C 4H44D 4H44E 4H44F 4H44G 4H44H
4H44I 4H44J 4H44K 4H44L 4H44M 4H44N 4H44O 4H44P 4H99H 4H99L 4H99M 4H9LH
4H9LL 4H9LM 4HBHH 4HBHL 4HBHM 4HBJH 4HBJL 4HBJM 4HE8B 4HE8C 4HE8D 4HE8E
4HE8F 4HE8G 4HE8I 4HEAA 4HEAH 4HEAJ 4HEAK 4HEAL 4HEAM 4HEAN 4HFBA 4HFBB
4HFBC 4HFBD 4HFBE 4HFCA 4HFCB 4HFCC 4HFCD 4HFCE 4HFDA 4HFDB 4HFDC 4HFDD
4HFDE 4HFEA 4HFEB 4HFEC 4HFED 4HFEE 4HFHA 4HFHB 4HFHC 4HFHD 4HFHE 4HFIA
4HFIB 4HFIC 4HFID 4HFIE 4HG6A 4HG6B 4HKRA 4HKRB 4HKRC 4HKRD 4HKRE 4HKRF
4HKSA 4HKSB 4HKSC 4HKSD 4HKSE 4HKSF 4HMKA 4HMKB 4HODA 4HODB 4HQJC 4HQJD
4HQJE 4HTSA 4HTTA 4HUKA 4HULA 4HUMA 4HUNA 4HUQS 4HUQT 4HW9A 4HW9B 4HW9C
4HW9D 4HW9E 4HW9F 4HW9G 4HWAA 4HWAB 4HWAC 4HWAD 4HWAE 4HWAF 4HWAG 4HWLA
4HYCA 4HYCB 4HYCC 4HYCD 4HYDA 4HYDB 4HYDC 4HYDD 4HYGA 4HYJA 4HYOA 4HYOB
4HYOC 4HYOD 4HYTA 4HYTB 4HYTG 4HYXA 4HZ3A 4HZ3B 4HZ3C 4HZ3D 4HZUS 4HZUT
4I0UA 4I0UB 4I0UC 4I0UD 4I0UE 4I7ZA 4I7ZB 4I7ZC 4I7ZD 4I7ZE 4I7ZF 4I7ZG
4I7ZH 4I7ZI 4I7ZJ 4I7ZK 4I7ZL 4I7ZM 4I7ZN 4I7ZO 4I7ZP 4I9WA 4I9WB 4IA4A
4IA4B 4IA4C 4IA4D 4IAQA 4IARA 4IB4A 4IFFA 4IFFB 4IKVA 4IKWA 4IKXA 4IKYA
4IKZA 4IL3A 4IL4A 4IL4B 4IL4C 4IL4D 4IL4E 4IL6A 4IL6A 4IL6B 4IL6B 4IL6C
4IL6C 4IL6D 4IL6D 4IL6E 4IL6E 4IL6F 4IL6F 4IL6H 4IL6H 4IL6I 4IL6I 4IL6J
4IL6J 4IL6K 4IL6K 4IL6L 4IL6L 4IL6M 4IL6M 4IL6R 4IL6T 4IL6T 4IL6X 4IL6X
4IL6Y 4IL6Y 4IL6Z 4IL6Z 4IL9A 4IL9B 4IL9C 4IL9D 4IL9E 4ILAA 4ILAB 4ILAC
4ILAD 4ILAE 4ILBA 4ILBB 4ILBC 4ILBD 4ILBE 4ILCA 4ILCB 4ILCC 4ILCD 4ILCE
4IN5H 4IN5L 4IN5M 4IN6H 4IN6L 4IN6M 4IN7H 4IN7L 4IN7M 4IREA 4IREB 4IREC
4IRED 4IREE 4IU8A 4IU9B 4IXQA 4IXQA 4IXQB 4IXQB 4IXQC 4IXQC 4IXQD 4IXQD
4IXQE 4IXQE 4IXQF 4IXQF 4IXQG 4IXQG 4IXQH 4IXQH 4IXQI 4IXQI 4IXQJ 4IXQJ
4IXQK 4IXQK 4IXQL 4IXQL 4IXQM 4IXQM 4IXQT 4IXQT 4IXQX 4IXQX 4IXQY 4IXQY
4IXQZ 4IXQZ 4IXRA 4IXRA 4IXRB 4IXRB 4IXRC 4IXRC 4IXRD 4IXRD 4IXRE 4IXRE
4IXRF 4IXRF 4IXRG 4IXRG 4IXRH 4IXRH 4IXRI 4IXRI 4IXRJ 4IXRJ 4IXRK 4IXRK
4IXRL 4IXRL 4IXRM 4IXRM 4IXRT 4IXRT 4IXRX 4IXRX 4IXRY 4IXRY 4IXRZ 4IXRZ
4IZMA 4IZMB 4IZMC 4J2TA 4J4QA 4J72A 4J72B 4J7CK 4J7CL 4J7TA 4J7TB 4J7TC
4J7YA 4J7YB 4J7YC 4J9UA 4J9UB 4JA3A 4JA4A 4JBWF 4JBWG 4JC6A 4JC6B 4JC6C
4JC6D 4JC7A 4JC7B 4JC7C 4JCZA 4JCZB 4JCZC 4JFBA 4JFBB 4JFBC 4JKVA 4JKVB
4JQ6A 4JQ6B 4JQ6C 4JQ6D 4JQ6E 4JQ6F 4JR8A 4JR8B 4JR8C 4JR9A 4JREA 4JRZA
4JRZB 4JRZC 4JTAB 4JTAD 4JTAF 4JTAH 4JTCH 4JTCL 4JTCN 4JTCP 4JTDB 4K0EA
4K0EB 4K0EC 4K0JA 4K0JB 4K0JC 4K1CA 4K3BA 4K3CA 4K5YA 4K7QA 4K7QB 4K7QC
4K7RA 4K7RB 4K7RC 4KFMA 4KFMC 4KFMF 4KFMJ 4KHZF 4KHZG 4KI0F 4KI0G 4KJPA
4KJPB 4KJQA 4KJQB 4KJRA 4KJRC 4KJRD 4KJSA 4KJSC 4KJSD 4KJWA 4KJWB 4KK5A

4KK5B 4KK6A 4KK6B 4KK8A 4KK8B 4KK9A 4KK9B 4KKAA 4KKAB 4KKBA 4KKBB 4KKCA
4KKCB 4KKLA 4KKLB 4KLYA 4KLYB 4KLYC 4KLYD 4KLYE 4KNFA 4KNFB 4KNFC 4KNFD
4KNFE 4KPPA 4KR4A 4KR4B 4KR4C 4KR8A 4KR8B 4KR8C 4KRAA 4KRAB 4KRAC 4KSBA
4KSCA 4KSDA 4KT0A 4KT0B 4KT0F 4KT0J 4KT0K 4KT0M 4KX6C 4KX6D 4KX6O 4KX6P
4KY0A 4KY0B 4KY0C 4KYTA 4KYTB 4L35A 4L35B 4L35C 4L6RA 4L6VA 4L6VB 4L6VF
4L6VK 4L6VL 4L6VM 4LBEC 4LBED 4LBEE 4LBEF 4LCUC 4LCUD 4LCUE 4LCUF 4LCZA
4LCZB 4LCZC 4LDEA 4LDLA 4LDOA 4LDSA 4LEPA 4LEPB 4LLHA 4LLHB 4LLHC 4LMJA
4LMJB 4LMJC 4LMJD 4LMJE 4LMKA 4LMKB 4LMKC 4LMKD 4LMKE 4LMLA 4LMLB 4LMLC
4LMLD 4LMLE 4LOUA 4LOUB 4LP8A 4LP8B 4LP8C 4LP8D 4LSEA 4LSEB 4LSEC 4LSFA
4LSFC 4LSFD 4LSGA 4LSHA 4LSHC 4LSHD 4LSIA 4LSIB 4LSIC 4LTOA 4LTOB 4LTOC
4LTOD 4LTPA 4LTPB 4LTPC 4LTPD 4LTQA 4LTQB 4LTQC 4LTQD 4LTRA 4LTRB 4LTRC
4LTRD 4LWYH 4LWYL 4LWYM 4LXJA 4LZ6A 4LZ9A 4M1MA 4M2SA 4M2TA 4M48A 4M58A
4M5BA 4M5CA 4M64B 4M8JA 4M8JB 4M8JC 4MBSA 4MD1A 4MD1B 4MD1C 4MD2A 4MD2B
4MD2C 4MEEA 4MESA 4MHWA 4MJNA 4MJNB 4MJNC 4MJND 4MJNE 4MJNF 4MJNG 4MJNH
4MJNI 4MJNJ 4MJNK 4MJNL 4MJNM 4MJNN 4MLBA 4MM4A 4MM4B 4MM5A 4MM6A 4MM7A
4MM8A 4MM8B 4MM9A 4MMAA 4MMBA 4MMCA 4MMDA 4MMDB 4MMEA 4MMEB 4MMFA 4MMFB
4MNDA 4MNDB 4MQSA 4MQTA 4MQXA 4MQXB 4MRNA 4MRNB 4MRPA 4MRPB 4MRRA 4MRRB
4MRSA 4MRSB 4MRVA 4MRVB 4MSWC 4MSWD 4MSWE 4MSWF 4MT0A 4MT0B 4MT0C 4MT1A
4MT1B 4MT1C 4MT4A 4MT4B 4MT4C 4MUUA 4MYCA 4MYCB 4MYHA 4MYHB 4N4DA 4N4RA
4N4WA 4N4YA 4N4YB 4N4YC 4N6HA 4N74A 4N75A 4N7KH 4N7KL 4N7KM 4N7LH 4N7LL
4N7LM 4N7WA 4N7WB 4N7XA 4NABA 4NC3A 4NEFA 4NEFB 4NEFC 4NEFD 4NH2A 4NH2B
4NH2C 4NH2D 4NH2E 4NH2F 4NJNA 4NJPA 4NPPA 4NPPB 4NPPC 4NPPD 4NPPE 4NTAA
4NTAB 4NTAC 4NTBA 4NTBB 4NTBC 4NTFA 4NTFB 4NTFC 4NTJA 4NTWA 4NTWD 4NTWG
4NTXA 4NTXD 4NTXG 4NTYA 4NTYD 4NTYG 4NV2A 4NV5A 4NV6A 4NYKA 4NYKB 4NYKC
4O6MA 4O6MB 4O6NA 4O6NB 4O6YA 4O6YB 4O79A 4O79B 4O7GA 4O7GB 4O93A 4O93B
4O93C 4O93D 4O9PA 4O9PB 4O9PC 4O9PD 4O9RA 4O9TA 4O9TB 4O9TC 4O9TD 4O9UA
4O9UB 4O9UC 4O9UD 4OAAA 4OD4A 4OD5A 4OGQA 4OGQB 4OGQC 4OGQD 4OGQE 4OGQF
4OGQG 4OGQH 4OGQI 4OGQJ 4OGQK 4OGQL 4OGQM 4OGQN 4OGQO 4OGQP 4OH3A 4OH3B
4OJ2A 4OJ2B 4OJ2C 4OJ2X 4OO9A 4OR2A 4OR2B 4OV0A 4OV0B 4OV0C 4OXSA 4OXSB
4OXSE 4OXSF 4OYEA 4OYEB 4OYEC 4OYFA 4OYFB 4OYFC 4P00A 4P00B 4P02A 4P02B
4P19A 4P19B 4P19C 4P1AA 4P1AB 4P1AC 4P2ZA 4P2ZB 4P2ZE 4P2ZF 4P30A 4P30B
4P30E 4P30F 4P3JA 4P3JB 4P3JC 4P6HA 4P6HB 4P6HC 4P6JA 4P6JB 4P6KA 4P6KB
4P6LA 4P6LB 4P6VB 4P6VC 4P6VD 4P6VE 4P6VF 4P79A 4P9OA 4P9OB 4P9OE 4P9OF
4P9PA 4P9PB 4P9PE 4P9PF 4PA3A 4PA3B 4PA3E 4PA3F 4PA4A 4PA4B 4PA4E 4PA4F
4PA6A 4PA6B 4PA6E 4PA6F 4PA7A 4PA7B 4PA7E 4PA7F 4PA9A 4PA9B 4PA9E 4PA9F
4PB1A 4PB1B 4PB1C 4PB2A 4PB2B 4PB2C 4PBUA 4PBUA 4PBUB 4PBUB 4PBUC 4PBUC
4PBUD 4PBUD 4PBUE 4PBUE 4PBUF 4PBUF 4PBUH 4PBUH 4PBUI 4PBUI 4PBUJ 4PBUJ
4PBUK 4PBUK 4PBUL 4PBUL 4PBUM 4PBUM 4PBUT 4PBUT 4PBUX 4PBUX 4PBUY 4PBUY
4PBUZ 4PBUZ 4PD4C 4PD4D 4PD4E 4PD4H 4PD4I 4PD5A 4PD5B 4PD5C 4PD6A 4PD6B
4PD6C 4PD7A 4PD7B 4PD7C 4PD8A 4PD8B 4PD8C 4PD9A 4PD9B 4PD9C 4PDAA 4PDAB
4PDAC 4PDLA 4PDLC 4PDLD 4PDLE 4PDMA 4PDMC 4PDMD 4PDME 4PDRB 4PDRF 4PDRG
4PDRH 4PDVA 4PDVC 4PDVD 4PDVE 4PE5A 4PE5B 4PE5C 4PE5D 4PGRA 4PGSA 4PGUA
4PGVA 4PGWA 4PHUA 4PHZA 4PHZB 4PHZC 4PHZD 4PHZE 4PHZF 4PHZG 4PHZH 4PHZI
4PHZJ 4PHZK 4PHZN 4PI0A 4PI0B 4PI0C 4PI0D 4PI0E 4PI0F 4PI0G 4PI0H 4PI0I

4PI0J 4PI0K 4PI0N 4PI2A 4PI2B 4PI2C 4PI2D 4PI2E 4PI2F 4PI2G 4PI2H 4PI2I
4PI2J 4PI2K 4PI2N 4PIRA 4PIRB 4PJ0A 4PJ0A 4PJ0B 4PJ0B 4PJ0C 4PJ0C 4PJ0D
4PJ0D 4PJ0E 4PJ0E 4PJ0F 4PJ0F 4PJ0H 4PJ0H 4PJ0I 4PJ0I 4PJ0J 4PJ0J 4PJ0K
4PJ0K 4PJ0L 4PJ0L 4PJ0M 4PJ0M 4PJ0R 4PJ0R 4PJ0T 4PJ0T 4PJ0X 4PJ0X 4PJ0Y
4PJ0Y 4PJ0Z 4PJ0Z 4PL0A 4PL0B 4POPA 4POVA 4PR7A 4PV1A 4PV1B 4PV1C 4PV1D
4PV1E 4PV1F 4PV1G 4PV1H 4PV1I 4PV1J 4PV1K 4PV1L 4PV1M 4PV1N 4PV1O 4PV1P
4PX7A 4PXFA 4PXFE 4PXKA 4PXKB 4PXKC 4PXZA 4PY0A 4PYPA 4Q2EA 4Q2EB 4Q2GA
4Q2GB 4Q35A 4Q4AA 4Q4AB 4Q4HA 4Q4HB 4Q4JA 4Q4JB 4Q65A 4Q7CA 4Q7CB 4Q9HA
4Q9IA 4Q9JA 4Q9KA 4Q9LA 4QE7A 4QE7B 4QE7C 4QE7D 4QE9A 4QE9B 4QE9C 4QE9D
4QH1A 4QH1B 4QH1C 4QH1D 4QH1E 4QH4A 4QH4B 4QH4C 4QH4D 4QH4E 4QH5A 4QH5B
4QH5C 4QH5D 4QH5E 4QI1A 4QI1B 4QI1C 4QIDA 4QIMA 4QINA 4QINB 4QIQA 4QKXA
4QKYA 4QL0A 4QNCA 4QNCB 4QNDA 4QNDB 4QNZA 4QO0A 4QO0B 4QO2A 4QRYA 4QRYB
4QRYC 4QTNA 4QTNB 4QTNC 4QUVA 4R0CA 4R0CB 4R1IA 4R1IB 4R50A 4R50C 4R50D
4R50E 4R6ZA 4R6ZC 4R6ZD 4R6ZE 4R7CA 4R7CB 4R7CE 4R7CF 4R8CA 4R8CC 4R8CD
4R8CE 4R9UA 4R9UB 4RAIA 4RAIC 4RAID 4RAIE 4RCRH 4RCRL 4RCRM 4RDQA 4RDQB
4RDQC 4RDQD 4RDQE 4RDRA 4RDTA 4RESA 4RESB 4RESG 4RETA 4RETB 4RETG 4RFSS
4RFST 4RHBA 4RI2A 4RI2B 4RI3A 4RI3B 4RJWA 4RJWB 4RJWC 4RJXA 4RJXB 4RJXC
4RL8A 4RL9A 4RLBA 4RLBB 4RLCA 4RNGA 4RNGC 4RO2A 4RO2B 4RO2E 4RO2F 4RP8A
4RP8C 4RP9A 4RP9B 4RUEA 4RUEB 4RUFA 4RUFB 4RVWA 4RVYA 4RVYA 4RVYB 4RVYB
4RVYC 4RVYC 4RVYD 4RVYD 4RVYE 4RVYE 4RVYF 4RVYF 4RVYH 4RVYH 4RVYI 4RVYI
4RVYJ 4RVYJ 4RVYK 4RVYK 4RVYL 4RVYL 4RVYM 4RVYM 4RVYT 4RVYT 4RVYX 4RVYX
4RVYY 4RVYY 4RVYZ 4RVYZ 4RWAA 4RWAB 4RWDA 4RWDB 4RY2A 4RY2B 4RYIA 4RYIB
4RYJA 4RYJB 4RYMA 4RYNA 4RYOA 4RYQA 4RYRA 4S0FA 4S0FB 4S0VA 4TKQA 4TKRA
4TL3A 4TL3B 4TLLA 4TLLB 4TLLC 4TLLD 4TLMA 4TLMB 4TLMC 4TLMD 4TNHA 4TNHA
4TNHB 4TNHB 4TNHC 4TNHC 4TNHD 4TNHD 4TNHE 4TNHE 4TNHF 4TNHF 4TNHG 4TNHG
4TNHH 4TNHH 4TNHI 4TNHI 4TNHJ 4TNHJ 4TNHK 4TNHK 4TNHL 4TNHL 4TNHM 4TNHM
4TNHT 4TNHT 4TNHX 4TNHX 4TNHY 4TNHY 4TNHZ 4TNHZ 4TNIA 4TNIA 4TNIB 4TNIB
4TNIC 4TNIC 4TNID 4TNID 4TNIE 4TNIE 4TNIF 4TNIF 4TNIG 4TNIG 4TNIH 4TNIH
4TNII 4TNII 4TNIJ 4TNIJ 4TNIK 4TNIK 4TNIL 4TNIL 4TNIM 4TNIM 4TNIT 4TNIT
4TNIX 4TNIX 4TNIY 4TNIY 4TNIZ 4TNIZ 4TNJA 4TNJA 4TNJB 4TNJB 4TNJC 4TNJC
4TNJD 4TNJD 4TNJE 4TNJE 4TNJF 4TNJF 4TNJG 4TNJG 4TNJH 4TNJH 4TNJI 4TNJI
4TNJJ 4TNJJ 4TNJK 4TNJK 4TNJL 4TNJL 4TNJM 4TNJM 4TNJT 4TNJT 4TNJX 4TNJX
4TNJY 4TNJY 4TNJZ 4TNJZ 4TNKA 4TNKA 4TNKB 4TNKB 4TNKC 4TNKC 4TNKD 4TNKD
4TNKE 4TNKE 4TNKF 4TNKF 4TNKG 4TNKG 4TNKH 4TNKH 4TNKI 4TNKI 4TNKJ 4TNKJ
4TNKK 4TNKK 4TNKL 4TNKL 4TNKM 4TNKM 4TNKT 4TNKT 4TNKX 4TNKX 4TNKY 4TNKY
4TNKZ 4TNKZ 4TNVA 4TNVB 4TNVC 4TNVD 4TNVE 4TNWA 4TNWB 4TNWC 4TNWD 4TNWE
4TPGA 4TPGB 4TPHA 4TPHB 4TPJA 4TPJB 4TQ3A 4TQ4A 4TQ5A 4TQ6A 4TQQH 4TQQL
4TQQM 4TQUM 4TQUN 4TQVA 4TQVB 4TSYA 4TSYB 4TSYC 4TSYD 4TSYE 4TSYF 4TSYG
4TSYH 4TWDA 4TWDB 4TWDC 4TWDD 4TWDE 4TWFA 4TWFB 4TWFC 4TWFD 4TWFE 4TWHA
4TWHB 4TWHC 4TWHD 4TWHE 4TWKA 4TWKB 4U14A 4U15A 4U16A 4U1WA 4U1WB 4U1WC
4U1WD 4U1XA 4U1XB 4U1XC 4U1XD 4U1YA 4U1YB 4U1YC 4U1YD 4U2PA 4U2PB 4U2PC
4U2PD 4U2QA 4U2QB 4U2QC 4U2QD 4U3FC 4U3FD 4U3FE 4U3FG 4U3FJ 4U3FP 4U3FQ
4U3FR 4U3FT 4U3FW 4U4FA 4U4FB 4U4FC 4U4FD 4U4GA 4U4GB 4U4GC 4U4GD 4U4TA
4U4VA 4U4WA 4U4WB 4U5BA 4U5BB 4U5BC 4U5BD 4U5CA 4U5CB 4U5CC 4U5CD 4U5DA

4U5DB 4U5DC 4U5DD 4U5EA 4U5EB 4U5EC 4U5ED 4U5FA 4U5FB 4U5FC 4U5FD 4U8VA
4U8VB 4U8YA 4U8YB 4U95A 4U95B 4U96A 4U96B 4U96C 4U9LA 4U9LB 4U9NA 4U9NB
4UB6A 4UB6A 4UB6B 4UB6B 4UB6C 4UB6C 4UB6D 4UB6D 4UB6E 4UB6E 4UB6F 4UB6F
4UB6H 4UB6H 4UB6I 4UB6I 4UB6J 4UB6J 4UB6K 4UB6K 4UB6L 4UB6L 4UB6M 4UB6M
4UB6R 4UB6T 4UB6T 4UB6X 4UB6X 4UB6Y 4UB6Y 4UB6Z 4UB6Z 4UB8A 4UB8A 4UB8B
4UB8B 4UB8C 4UB8C 4UB8D 4UB8D 4UB8E 4UB8E 4UB8F 4UB8F 4UB8H 4UB8H 4UB8I
4UB8I 4UB8J 4UB8J 4UB8K 4UB8K 4UB8L 4UB8L 4UB8M 4UB8M 4UB8R 4UB8T 4UB8T
4UB8X 4UB8X 4UB8Y 4UB8Y 4UB8Z 4UB8Z 4UC1A 4UC1B 4UC2A 4UC2B 4UC3A 4UC3B
4UG2A 4UHRA 4UISA 4UISB 4UISC 4UISD 4UMVA 4UMWA 4UP6A 4UP6B 4UP6C 4UQJA
4UQJB 4UQJC 4UQJD 4UQQA 4UQQB 4UQQC 4UQQD 4US3A 4US4A 4UU0A 4UU1A 4UUJC
4UUJD 4UUJE 4UUJF 4UV3A 4UV3B 4UV3C 4UV3D 4UV3E 4UV3F 4UV3G 4UV3H 4UV3I
4UVMA 4UWAA 4UWAB 4UWAC 4UWAD 4UWEA 4UWEB 4UWEC 4UWED 4UX1A 4UX1B 4UX2A
4UX2B 4UXWA 4UXWB 4UXWC 4UXXA 4UXXB 4UXXC 4UXZA 4UXZB 4UXZC 4UYOA 4UYOB
4UYOC 4V1FA 4V1FB 4V1FC 4V1FD 4V1FE 4V1FF 4V1FG 4V1FH 4V1FI 4V1GA 4V1GB
4V1GC 4V1GD 4V1GE 4V1GF 4V1GG 4V1GH 4V1GI 4V1HA 4V1HB 4V1HC 4V1HD 4V1HE
4V1HF 4V1HG 4V1HH 4V1HI 4V3GA 4V3HA 4W6VA 4WABA 4WABB 4WABC 4WAVA 4WD7A
4WD7B 4WD7C 4WD7D 4WD7E 4WD8A 4WD8B 4WD8C 4WD8D 4WD8E 4WFEA 4WFEB 4WFFA
4WFFB 4WFGA 4WFGB 4WFHA 4WFHB 4WGVA 4WGWA 4WIBA 4WIBB 4WISA 4WISB 4WITA
4WITB 4WMZA 4WO1A 4WOLA 4WOLB 4WW3A 4X1HA 4X2SA 4X2SB 4X2SC 4X31A 4X31B
4X31C 4X32A 4X32B 4X32C 4X5MB 4X5MC 4X5NA 4X5NB 4X5TA 4X5TB 4X5TC 4X5TD
4X5TE 4X88A 4X88B 4X88E 4X88F 4X89A 4X89B 4X89E 4X89F 4X8AA 4X8AB 4X8AE
4X8AF 4XDJA 4XDJB 4XDLA 4XDLB 4XE5A 4XE5B 4XE5G 4XEEA 4XESA 4XIGM 4XIGN
4XK81 4XK82 4XK84 4XK8A 4XK8B 4XK8F 4XK8G 4XK8H 4XK8I 4XK8J 4XK8K 4XK8L
4XNIA 4XNJA 4XNKA 4XNLA 4XNUA 4XNVA 4XNWA 4XNXA 4XOUA 4XP1A 4XP4A 4XP5A
4XP6A 4XP9C 4XPAA 4XPBA 4XPFA 4XPGA 4XPHA 4XPTA 4XTCM 4XTCN 4XTLA 4XTNA
4XTNB 4XTNC 4XTND 4XTNE 4XTOA 4XTOB 4XTOC 4XTOD 4XTOE 4XU4A 4XU4B 4XU4C
4XU5A 4XU5B 4XU5C 4XU6A 4XU6B 4XU6C 4XWKA 4XXJA 4XXJB 4XXJC 4XYDA 4XYDB
4Y1KA 4Y1KB 4Y1KC 4Y25A 4Y281 4Y282 4Y283 4Y284 4Y28A 4Y28B 4Y28F 4Y28G
4Y28H 4Y28I 4Y28J 4Y28K 4Y28L 4Y3UA 4Y3UB 4Y3UC 4Y6KA 4Y7JA 4Y7JB 4Y7JC
4Y7JD 4Y7JE 4Y7KA 4Y7KB 4Y7KC 4Y7KD 4Y7KE 4Y9HA 4YAYA 4YB9D 4YBQA 4YCLA
4YCMA 4YCNA 4YCRA 4YCRB 4YCRC 4YEUA 4YEUB 4YEUC 4YEUD 4YEUE 4YK5A 4YK5B
4YK5C 4YL0A 4YL0B 4YL0C 4YL1A 4YL1B 4YL1C 4YL3A 4YL3B 4YL3C 4YMKA 4YMKD
4YMSC 4YMSD 4YMTB 4YMTC 4YMUC 4YMUD 4YMVC 4YMVD 4YMWC 4YMWD 4YSXC 4YSXD
4YSYC 4YSYD 4YSZC 4YSZD 4YT0C 4YT0D 4YTMC 4YTMD 4YTNC 4YTND 4YTPC 4YTPD
4YXDC 4YXDD 4YZFA 4YZFB 4YZIA 4YZIB 4Z34A 4Z35A 4Z36A 4Z3NA 4Z3PA 4Z7FA
4Z90A 4Z90B 4Z90C 4Z90D 4Z90E 4Z91A 4Z91B 4Z91C 4Z91D 4Z91E 4ZBMA 4ZBMC
4ZBMD 4ZBME 4ZGVA 4ZITA 4ZITB 4ZITC 4ZIVA 4ZIVB 4ZIVC 4ZIWA 4ZIWB 4ZIWC
4ZJ8A 4ZJCA 4ZJLA 4ZJLB 4ZJLC 4ZJOA 4ZJOB 4ZJOC 4ZJQA 4ZJQB 4ZJQC 4ZLJA
4ZLJB 4ZLJC 4ZLLA 4ZLLB 4ZLLC 4ZLNA 4ZLNB 4ZLNC 4ZOWA 4ZP0A 4ZP2A 4ZR0A
4ZR1A 4ZUDA 4ZW9A 4ZWBA 4ZWCA 4ZWJA 4ZYOA 4ZYRA 4ZZBA 4ZZBB 4ZZBC 4ZZBD
4ZZBE 4ZZCA 4ZZCB 4ZZCC 4ZZCD 4ZZCE 5A1SA 5A1SB 5A2NA 5A2NB 5A2OA 5A2OB
5A3QA 5A3RA 5A3SA 5A40A 5A40B 5A43A 5A43B 5A44A 5A44B 5A44C 5A45A 5A45B
5A45C 5A63A 5A63B 5A63C 5A63D 5A6EA 5A6EB 5A6GA 5A6GB 5A6UA 5A6UB 5A6UG
5A8EA 5ABBA 5ABBB 5ABBZ 5AEXA 5AEXB 5AEXC 5AEZA 5AEZB 5AEZC 5AF1A 5AF1C

5AF1D 5AH3A 5AH3B 5AH3C 5AHYA 5AHYB 5AHYC 5AHZA 5AHZB 5AHZC 5AIDA 5AIDB
5AIDC 5AJIA 5AJIB 5AJIC 5AJID 5AJIE 5AJIF 5AJIG 5AN8A 5AN8B 5AN8C 5AN8D
5ARAJ 5ARAK 5ARAL 5ARAM 5ARAN 5ARAO 5ARAP 5ARAQ 5ARAT 5ARAW 5AREJ 5AREK
5AREL 5AREM 5AREN 5AREO 5AREP 5AREQ 5ARET 5AREW 5ARHJ 5ARHK 5ARHL 5ARHM
5ARHN 5ARHO 5ARHP 5ARHQ 5ARHT 5ARHW 5ARIJ 5ARIK 5ARIL 5ARIM 5ARIN 5ARIO
5ARIP 5ARIQ 5ARIT 5ARIW 5AVQA 5AVQB 5AVQG 5AVRA 5AVRB 5AVRG 5AVSA 5AVSB
5AVSG 5AVTA 5AVTB 5AVTG 5AVUA 5AVUB 5AVUG 5AVVA 5AVVB 5AVVG 5AVWA 5AVWB
5AVWG 5AVXA 5AVXB 5AVXG 5AVYA 5AVYB 5AVYG 5AVZA 5AVZB 5AVZG 5AW0A 5AW0B
5AW0G 5AW1A 5AW1B 5AW1G 5AW2A 5AW2B 5AW2G 5AW3A 5AW3B 5AW3G 5AW4A 5AW4B
5AW4G 5AW5A 5AW5B 5AW5G 5AW6A 5AW6B 5AW6G 5AW7A 5AW7B 5AW7G 5AW8A 5AW8B
5AW8G 5AW9A 5AW9B 5AW9G 5AWWE 5AWWG 5AWWY 5AWZA 5AX0A 5AX1A 5AYMA 5AYNA
5AYOA 5AYWA 5AZBA 5AZCA 5AZDA 5AZDB 5AZDC 5AZDD 5B0WA 5B0WB 5B0WD 5B1AA
5B1AB 5B1AC 5B1AD 5B1AG 5B1AI 5B1AJ 5B1AK 5B1AL 5B1AM 5B1AN 5B1AO 5B1AP
5B1AQ 5B1AT 5B1AV 5B1AW 5B1AX 5B1AY 5B1AZ 5B1BA 5B1BB 5B1BC 5B1BD 5B1BG
5B1BI 5B1BJ 5B1BK 5B1BL 5B1BM 5B1BN 5B1BO 5B1BP 5B1BQ 5B1BT 5B1BV 5B1BW
5B1BX 5B1BY 5B1BZ 5B2GA 5B2NA 5B34A 5B35A 5B3SA 5B3SB 5B3SC 5B3SD 5B3SG
5B3SI 5B3SJ 5B3SK 5B3SL 5B3SM 5B3SN 5B3SO 5B3SP 5B3SQ 5B3ST 5B3SV 5B3SW
5B3SX 5B3SY 5B3SZ 5B57A 5B57B 5B58A 5B58B 5B5EA 5B5EA 5B5EB 5B5EB 5B5EC
5B5EC 5B5ED 5B5ED 5B5EE 5B5EE 5B5EF 5B5EF 5B5EH 5B5EH 5B5EI 5B5EI 5B5EJ
5B5EJ 5B5EK 5B5EK 5B5EL 5B5EL 5B5EM 5B5EM 5B5ET 5B5ET 5B5EX 5B5EX 5B5EY
5B5EY 5B5EZ 5B5EZ 5B66A 5B66A 5B66B 5B66B 5B66C 5B66C 5B66D 5B66D 5B66E
5B66E 5B66F 5B66F 5B66H 5B66H 5B66I 5B66I 5B66J 5B66J 5B66K 5B66K 5B66L
5B66L 5B66M 5B66M 5B66T 5B66T 5B66X 5B66X 5B66Y 5B66Y 5B66Z 5B66Z 5B6VA
5B6VB 5B6VC 5B6WA 5B6WB 5B6WC 5B6XA 5B6XB 5B6XC 5B6YA 5B6YB 5B6YC 5B6ZA
5B6ZB 5B6ZC 5BKFA 5BKFB 5BKFC 5BKFD 5BKFE 5BKGA 5BKGB 5BKGC 5BKGD 5BKGE
5BKIA 5BKIC 5BKIE 5BKIG 5BKJA 5BKJC 5BKJE 5BKJG 5BKKA 5BKKC 5BKKE 5BKKG
5BN2A 5BN2B 5BN2C 5BN2D 5BPSA 5BPSB 5BPSC 5BPSD 5BPSE 5BPSF 5BPSG 5BPSH
5BPSI 5BPSJ 5BQ6A 5BQ6B 5BQ6C 5BQ6D 5BQ6E 5BQ6F 5BQ6G 5BQ6H 5BQ6I 5BQ6J
5BQAA 5BQAB 5BQAC 5BQAD 5BQAE 5BQAF 5BQAG 5BQAH 5BQAI 5BQAJ 5BQGA 5BQGB
5BQHA 5BQHB 5BQIA 5BQIB 5BQJA 5BQJB 5BQJC 5BQJD 5BQJE 5BQJF 5BQJG 5BQJH
5BQJI 5BQJJ 5BR2A 5BR2B 5BR2C 5BR5A 5BR5B 5BR5C 5BUTI 5BUTJ 5BZ2A 5BZ2B
5BZ3A 5BZ3B 5BZBA 5BZBB 5BZBE 5BZBF 5C1MA 5C2TC 5C2TD 5C3JC 5C3JD 5C58A
5C5XA 5C5XB 5C5XC 5C5XD 5C65A 5C6NA 5C6OA 5C6PA 5C73C 5C73K 5C76A 5C76D
5C78A 5C78D 5C8JI 5CBFA 5CBFB 5CBFD 5CBFE 5CBGA 5CBGB 5CBGD 5CBGE 5CBHA
5CBHB 5CBHD 5CBHE 5CFBA 5CFBB 5CFBC 5CFBD 5CFBE 5CFYD 5CFYE 5CFYF 5CGCA
5CGDA 5CH4E 5CH4G 5CH4Y 5CKRA 5CKRB 5CTGA 5CTGB 5CTGC 5CTHA 5CTHB 5CTHC
5CXVA 5D0OA 5D0QA 5D0YA 5D0YB 5D3MD 5D56A 5D56B 5D56C 5D57A 5D57B 5D57C
5D58A 5D59A 5D5AA 5D5BA 5D5DA 5D6IA 5D6IB 5D6IC 5D6KA 5D6LA 5D91A 5D92A
5D92D 5DA0A 5DGYA 5DHGA 5DHHA 5DIRA 5DJQA 5DJQB 5DJQC 5DJQN 5DL5A 5DL6A
5DL7A 5DL8B 5DN6J 5DN6K 5DN6L 5DN6M 5DN6N 5DN6O 5DN6P 5DN6Q 5DN6R 5DN6S
5DN6T 5DN6U 5DN6X 5DN6Y 5DO7A 5DO7B 5DOQA 5DOQB 5DOQC 5DQQA 5DQQB 5DSGA
5DUOA 5DUOB 5DWKA 5DWKB 5DWYA 5DWYB 5DWYC 5DYEA 5DYEB 5DYEC 5DYED 5DYSA
5E1AC 5E1AD 5E1AE 5E1AF 5E1JA 5E1JB 5E79A 5E79A 5E79B 5E79B 5E79C 5E79C
5E79D 5E79D 5E79E 5E79E 5E79F 5E79F 5E79H 5E79H 5E79I 5E79I 5E79J 5E79J

5E79K 5E79K 5E79L 5E79L 5E79M 5E79M 5E79T 5E79T 5E79X 5E79X 5E79Y 5E79Y
5E79Z 5E79Z 5E7CA 5E7CA 5E7CB 5E7CB 5E7CC 5E7CC 5E7CD 5E7CD 5E7CE 5E7CE
5E7CF 5E7CF 5E7CH 5E7CH 5E7CI 5E7CI 5E7CJ 5E7CJ 5E7CK 5E7CK 5E7CL 5E7CL
5E7CM 5E7CM 5E7CT 5E7CT 5E7CX 5E7CX 5E7CY 5E7CY 5E7CZ 5E7CZ 5E9SA 5E9SB
5E9SC 5EBLC 5EBLD 5EBLE 5EBLF 5EBMC 5EBMD 5EBME 5EBMF 5EBWC 5EBWD 5EBWE
5EBWF 5EC1C 5EC1D 5EC1E 5EC1F 5EC2C 5EC2D 5EC2E 5EC2F 5EC5A 5EC5B 5EC5C
5EC5D 5EC5E 5EC5F 5EC5G 5EC5H 5EC5I 5EDLA 5EE7A 5EG1A 5EG1B 5EGIA 5EGIB
5EGIC 5EH4A 5EH4B 5EH6A 5EIKA 5EIKB 5EIKC 5EIYA 5EIYB 5EJ1A 5EJ1B 5EJZA
5EJZB 5EK0A 5EK0B 5EK0C 5EK0D 5EKEA 5EKEB 5EKEC 5EKED 5EKPA 5EKPB 5EKPC
5EKPD 5EKQA 5EN0A 5EQGA 5EQHA 5EQIA 5ER7A 5ER7B 5ER7C 5ER7D 5ER7E 5ER7F
5ERAA 5ERAB 5ERAC 5ERAD 5ERAE 5ERAF 5ETZA 5ETZB 5ETZD 5EULA 5EULE 5EULY
5EZMA 5F15A 5F1CA 5F1CB 5F1CC 5F5BA 5F5DA 5F5GA 5F5JA 5F5KA 5F8UA 5F8UB
5FGNA 5FGZA 5FIJJ 5FIJK 5FIJL 5FIJM 5FIJN 5FIJO 5FIJP 5FIJQ 5FIJT 5FIJW
5FIKJ 5FIKK 5FIKL 5FIKM 5FIKN 5FIKO 5FIKP 5FIKQ 5FIKT 5FIKW 5FILJ 5FILK
5FILL 5FILM 5FILN 5FILO 5FILP 5FILQ 5FILT 5FILW 5FJGA 5FL7K 5FL7L 5FL7M
5FL7N 5FL7O 5FL7P 5FL7Q 5FL7R 5FL7S 5FL7T 5FMWA 5FMWB 5FMWC 5FMWD 5FMWE
5FMWF 5FMWG 5FMWH 5FMWI 5FMWJ 5FMWK 5FMWL 5FMWM 5FMWN 5FMWO 5FMWP 5FMWQ
5FMWR 5FMWS 5FMWT 5FMWU 5FMWV 5FN2A 5FN2B 5FN2C 5FN2D 5FN5A 5FN5B 5FN5C
5FN5D 5FOKA 5FP1A 5FQ6B 5FQ6I 5FQ7B 5FQ7D 5FQ8B 5FQ8D 5FR8A 5FUFA 5FUFB
5FUFC 5FVNA 5FVNB 5FVNC 5FXHA 5FXHB 5FXHC 5FXHD 5FXJA 5FXJB 5FXJC 5FXJD
5FXKA 5FXKB 5FXKC 5FXKD 5G26A 5G28A 5G2AA 5G2CA 5G2DA 5G36A 5G36B 5G36C
5G53A 5G54A 5GARN 5GARO 5GARP 5GARQ 5GARR 5GARS 5GART 5GARU 5GARV 5GARW
5GARX 5GARY 5GARZ 5GASN 5GASO 5GASP 5GASQ 5GASR 5GASS 5GAST 5GASU 5GASV
5GASW 5GASX 5GASY 5GASZ 5GJVA 5GJVE 5GJWA 5GJWE 5GKOA 5GKOB 5GLHA 5GLIA
5GMYA 5GPJA 5GPJB 5GTHA 5GTHA 5GTHB 5GTHB 5GTHC 5GTHC 5GTHD 5GTHD 5GTHE
5GTHE 5GTHF 5GTHF 5GTHH 5GTHH 5GTHI 5GTHI 5GTHJ 5GTHJ 5GTHK 5GTHK 5GTHL
5GTHL 5GTHM 5GTHM 5GTHR 5GTHT 5GTHT 5GTHX 5GTHX 5GTHY 5GTHY 5GTHZ 5GTHZ
5GTIA 5GTIA 5GTIB 5GTIB 5GTIC 5GTIC 5GTID 5GTID 5GTIE 5GTIE 5GTIF 5GTIF
5GTIH 5GTIH 5GTII 5GTII 5GTIJ 5GTIJ 5GTIK 5GTIK 5GTIL 5GTIL 5GTIM 5GTIM
5GTIR 5GTIT 5GTIT 5GTIX 5GTIX 5GTIY 5GTIY 5GTIZ 5GTIZ 5GUFA 5GUFB 5GUWA
5GUWB 5GUXB 5GUXC 5GXBA 5H1QA 5H1QB 5H1QC 5H1QD 5H1QE 5H1QF 5H1QG 5H1QH
5H1RA 5H1RB 5H1RC 5H1RD 5H1RE 5H1RF 5H1RG 5H1RH 5H2FA 5H2FA 5H2FB 5H2FB
5H2FC 5H2FC 5H2FD 5H2FD 5H2FE 5H2FE 5H2FF 5H2FF 5H2FH 5H2FH 5H2FI 5H2FI
5H2FJ 5H2FJ 5H2FK 5H2FK 5H2FL 5H2FL 5H2FT 5H2FT 5H2FX 5H2FX 5H2FY 5H2FY
5H2FZ 5H2FZ 5H2HA 5H2HB 5H2HC 5H2IA 5H2IB 5H2IC 5H2JA 5H2JB 5H2JC 5H2KA
5H2KB 5H2KC 5H2LA 5H2LB 5H2LC 5H2MA 5H2MB 5H2MC 5H2NA 5H2NB 5H2NC 5H2OA
5H2OB 5H2OC 5H2PA 5H2PB 5H2PC 5H35C 5H35D 5H35E 5H36A 5H36B 5H36C 5H3OA
5H3OB 5H3OC 5H3OD 5HCJA 5HCJB 5HCJC 5HCJD 5HCJE 5HCMA 5HCMB 5HCMC 5HCMD
5HCME 5HD8A 5HD8B 5HEGA 5HEGB 5HEGC 5HEGD 5HEGE 5HEHA 5HEHB 5HEHC 5HEHD
5HEHE 5HEJA 5HEJB 5HEJC 5HEJD 5HEJE 5HEOA 5HEOB 5HEOC 5HEOD 5HEOE 5HEUA
5HEUB 5HEUC 5HEUD 5HEUE 5HEWA 5HEWB 5HEWC 5HEWD 5HEWE 5HI9A 5HI9B 5HI9C
5HI9D 5HJ8A 5HJ8B 5HJ8C 5HJ8D 5HK1A 5HK2A 5HK6A 5HK6B 5HK6C 5HK6D 5HK7A
5HK7B 5HK7C 5HK7D 5HKDA 5HKDB 5HKDC 5HKDD 5HL9A 5HLAA 5HLBA 5HLDA 5HS1A
5HV9A 5HV9B 5HV9C 5HVDA 5HVDB 5HVDC 5HVDD 5HVXA 5HVXB 5HVXC 5HVXD 5HWXA

5HWYA 5HXCA 5HXEA 5HXHA 5HXRA 5HXSA 5HYAA 5I1MV 5I20A 5I32A 5I32B 5I32C
5I32D 5I6CA 5I6CB 5I6XA 5I6ZA 5I71A 5I73A 5I74A 5I75A 5I9KA 5I9KB 5I9KC
5IA9A 5IA9B 5IA9C 5ID3A 5ID3B 5ID3C 5ID3D 5ID3E 5IDEA 5IDEB 5IDEC 5IDED
5IDFA 5IDFB 5IDFC 5IDFD 5IJIA 5IJIB 5IOFA 5IOUA 5IOUB 5IOUC 5IOUD 5IOVA
5IOVB 5IOVC 5IOVD 5IR6A 5IR6B 5IR6C 5IRXA 5IRXB 5IRXC 5IRXD 5IRZB 5IRZC
5IRZD 5IRZE 5IS0B 5IS0C 5IS0D 5IS0E 5ITCA 5ITCB 5ITCC 5ITEA 5ITEB 5ITEC
5IU4A 5IU7A 5IU8A 5IUAA 5IUBA 5IUXA 5IUXB 5IUXC 5IUXD 5IUXE 5IUYA 5IUYB
5IUYC 5IV8A 5IV8B 5IV9A 5IV9B 5IVAA 5IVAB 5IWKA 5IWKB 5IWKC 5IWKD 5IWNA
5IWNB 5IWNC 5IWND 5IWOA 5IWOB 5IWOC 5IWOD 5IWPA 5IWPB 5IWPC 5IWPD 5IWRA
5IWRB 5IWRC 5IWRD 5IWSA 5IWSB 5IWTA 5IWTB 5IWTC 5IWTD 5IXMA 5IXMB 5IY5A
5IY5B 5IY5C 5IY5D 5IY5G 5IY5I 5IY5J 5IY5K 5IY5L 5IY5M 5IY5N 5IY5O 5IY5P
5IY5Q 5IY5T 5IY5V 5IY5W 5IY5X 5IY5Y 5IY5Z 5IYUA 5J0ZA 5J0ZB 5J0ZC 5J0ZD
5J0ZE 5J4IA 5J4IB 5J4NA 5J4NB 5J7AA 5J7AB 5J7AC 5J9PC 5J9PD 5J9PE 5J9PF
5JAEA 5JAEB 5JAFA 5JAGA 5JDFA 5JDGA 5JDHA 5JDLA 5JDMA 5JDNA 5JDPA 5JDQA
5JEFA 5JEFB 5JEQA 5JEQB 5JGPA 5JGPB 5JJEA 5JJEB 5JJEC 5JJED 5JJJA 5JJJB
5JJJC 5JJJD 5JJNA 5JJNB 5JJNC 5JJND 5JKIA 5JLCA 5JMNA 5JMNB 5JMNC 5JNQA
5JNQB 5JQHA 5JR0A 5JRFA 5JRWA 5JRWB 5JRWC 5JRWD 5JRWE 5JSIA 5JSZD 5JTBA
5JTGA 5JTGB 5JTGC 5JTGD 5JTGE 5JWYA 5K0IA 5K0IB 5K0IC 5K2AA 5K2BA 5K2CA
5K2DA 5K47A 5K47B 5K7LA 5K7LC 5K7LE 5K7LG 5KAFA 5KAFA 5KAFB 5KAFB 5KAFC
5KAFC 5KAFD 5KAFD 5KAFE 5KAFE 5KAFF 5KAFF 5KAFH 5KAFH 5KAFI 5KAFI 5KAFJ
5KAFJ 5KAFK 5KAFK 5KAFL 5KAFL 5KAFM 5KAFM 5KAFR 5KAFR 5KAFT 5KAFT 5KAFX
5KAFX 5KAFY 5KAFY 5KAFZ 5KAFZ 5KAIA 5KAIA 5KAIB 5KAIB 5KAIC 5KAIC 5KAID
5KAID 5KAIE 5KAIE 5KAIF 5KAIF 5KAIH 5KAIH 5KAII 5KAII 5KAIJ 5KAIJ 5KAIK
5KAIK 5KAIL 5KAIL 5KAIM 5KAIM 5KAIR 5KAIR 5KAIT 5KAIT 5KAIX 5KAIX 5KAIY
5KAIY 5KAIZ 5KAIZ 5KBNB 5KBSA 5KBSB 5KBSC 5KBSD 5KBTA 5KBTB 5KBTC 5KBTD
5KBUA 5KBUB 5KBUC 5KBUD 5KBVA 5KBVB 5KBVC 5KBVD 5KBWA 5KC0A 5KC4A 5KHNA
5KHNB 5KHSA 5KHSB 5KK2A 5KK2B 5KK2C 5KK2D 5KK2E 5KK2F 5KK2G 5KK2H 5KKHA
5KKHB 5KKHC 5KKZA 5KKZB 5KKZC 5KKZE 5KKZF 5KKZG 5KLIA 5KLIB 5KLIC 5KLIE
5KLIF 5KLIG 5KLVC 5KLVD 5KLVE 5KLVG 5KLVJ 5KLVK 5KMDA 5KMDB 5KMDC 5KMDD
5KO2A 5KOMB 5KOYA 5KPDA 5KPIA 5KPJA 5KSDA 5KTEA 5KTFA 5KUFA 5KUFB 5KUFC
5KUFD 5KUKA 5KUKB 5KUKC 5KUKD 5KUMA 5KUMB 5KUMC 5KUMD 5KW2A 5KXIA 5KXIB
5KXIC 5KXID 5KXIE 5KYHA 5KZOA 5L1BA 5L1BB 5L1BC 5L1BD 5L1EA 5L1EB 5L1EC
5L1ED 5L1FA 5L1FB 5L1FC 5L1FD 5L1GA 5L1GB 5L1GC 5L1GD 5L1HA 5L1HB 5L1HC
5L1HD 5L22A 5L22B 5L24A 5L24B 5L24C 5L25A 5L25B 5L26A 5L26B 5L26C 5L27A
5L27B 5L27C 5L2AA 5L2AB 5L2AC 5L2BA 5L2BB 5L2BC 5L47A 5L47B 5L47C 5L47D
5L47E 5L4EA 5L4EB 5L4EC 5L4ED 5L4EE 5L4HA 5L4HB 5L4HC 5L4HD 5L4HE 5L75F
5L75G 5L7DA 5L7IA 5L8R1 5L8R2 5L8R3 5L8R4 5L8RA 5L8RB 5L8RF 5L8RG 5L8RH
5L8RI 5L8RJ 5L8RK 5L8RL 5LC5A 5LC5A 5LC5B 5LC5C 5LC5D 5LC5F 5LC5G 5LC5H
5LC5H 5LC5I 5LC5J 5LC5J 5LC5K 5LC5K 5LC5L 5LC5L 5LC5M 5LC5M 5LC5N 5LC5Y
5LC5Z 5LDTA 5LDTB 5LDTC 5LDVA 5LDVB 5LDVC 5LDWA 5LDWA 5LDWB 5LDWC 5LDWD
5LDWF 5LDWG 5LDWH 5LDWH 5LDWI 5LDWJ 5LDWJ 5LDWK 5LDWK 5LDWL 5LDWL 5LDWM
5LDWM 5LDWN 5LDWY 5LDWZ 5LDXA 5LDXA 5LDXB 5LDXC 5LDXD 5LDXF 5LDXG 5LDXH
5LDXH 5LDXI 5LDXJ 5LDXJ 5LDXK 5LDXK 5LDXL 5LDXL 5LDXM 5LDXM 5LDXN 5LDXY
5LDXZ 5LEVA 5LG3A 5LG3B 5LG3C 5LG3D 5LG3E 5LIDA 5LIDB 5LIDC 5LIDD 5LIDE

5LILA 5LILB 5LJ6A 5LJ6B 5LJ7A 5LJ7B 5LJOA 5LKIA 5LKIB 5LKIC 5LKID 5LKIE
5LNKA 5LNKH 5LNKJ 5LNKK 5LNKL 5LNKM 5LNKM 5LNKN 5LNKN 5LNKO 5LNKP 5LNKQ
5LNKR 5LNKU 5LNKV 5LNKV 5LNKW 5LNKW 5LNKX 5LNKY 5LNKZ 5LQ3A 5LQ3B 5LQ3C
5LQX1 5LQX4 5LQXK 5LQXL 5LQXM 5LQXN 5LQXO 5LQXP 5LQXQ 5LQXR 5LQXS 5LQXT
5LQXV 5LQXY 5LQY1 5LQY4 5LQYK 5LQYL 5LQYM 5LQYN 5LQYO 5LQYP 5LQYQ 5LQYR
5LQYS 5LQYT 5LQYV 5LQYY 5LQZ1 5LQZ3 5LQZ4 5LQZK 5LQZL 5LQZM 5LQZN 5LQZO
5LQZP 5LQZQ 5LQZR 5LQZS 5LQZT 5LQZV 5LQZY 5LQZZ 5LRIH 5LRIL 5LRIM 5LSEH
5LSEL 5LSEM 5LV6A 5LV6B 5LWEA 5LWEB 5LWYA 5LX9A 5LXAA 5LXGA 5LZQA 5LZQB
5LZRA 5LZRB 5M5HA 5M7JB 5M7JC 5M7JD 5M7KB 5M7KC 5M7KD 5M7LB 5M7LC 5M7LD
5M87A 5M8AA 5M8JA 5M8KA 5M94A 5M94C 5M95A 5M95C 5M9BA 5MDOA 5MDOB 5MDOC
5MDPA 5MDPB 5MDPC 5MDQA 5MDQB 5MDQC 5MDRA 5MDRB 5MDRC 5MDSA 5MDSB 5MDSC
5MDX1 5MDX2 5MDX3 5MDX4 5MDX5 5MDX6 5MDX7 5MDX8 5MDXA 5MDXA 5MDXB 5MDXB
5MDXC 5MDXC 5MDXD 5MDXD 5MDXE 5MDXE 5MDXF 5MDXF 5MDXG 5MDXG 5MDXH 5MDXH
5MDXI 5MDXI 5MDXK 5MDXK 5MDXL 5MDXL 5MDXM 5MDXM 5MDXN 5MDXN 5MDXR 5MDXR
5MDXS 5MDXS 5MDXT 5MDXT 5MDXW 5MDXW 5MDXX 5MDXX 5MDXY 5MDXY 5MDXZ 5MDXZ
5MG3C 5MG3D 5MG3E 5MG3F 5MG3G 5MG3Y 5MKEA 5MKEB 5MKEC 5MKED 5MKFA 5MKFB
5MKFC 5MKFD 5MKKA 5MKKB 5MLZA 5MM0A 5MM1A 5MMTA 5MPMA 5MRLA 5MRWA 5MRWB
5MRWC 5MRWD 5MT6A 5MT7A 5MT8A 5MTFA 5MTFB 5MTFE 5MUOA 5MUOB 5MUOC 5MUOD
5MUOE 5MURA 5MURB 5MURC 5MURD 5MURE 5MVMA 5MVMB 5MVMC 5MVMD 5MVME 5MVNA
5MVNB 5MVNC 5MVND 5MVNE 5MWVA 5MX2A 5MX2A 5MX2B 5MX2B 5MX2C 5MX2C 5MX2D
5MX2D 5MX2E 5MX2E 5MX2F 5MX2F 5MX2H 5MX2H 5MX2I 5MX2I 5MX2J 5MX2J 5MX2K
5MX2K 5MX2L 5MX2L 5MX2M 5MX2M 5MX2R 5MX2R 5MX2T 5MX2T 5MX2X 5MX2X 5MX2Y
5MX2Y 5MX2Z 5MX2Z 5MZJA 5MZPA 5MZQA 5MZQB 5MZQC 5MZQD 5MZQE 5MZRA 5MZRB
5MZRC 5MZRD 5MZRE 5MZSA 5MZTA 5MZTB 5MZTC 5MZTD 5MZTE 5N2RA 5N2SA 5N6HA
5N6LA 5N6MA 5N9YA 5N9YB 5N9YC 5N9YD 5N9YE 5NAMA 5NAOA 5NBDA 5NBDB 5NC3A
5NC4A 5NC5A 5NC5B 5NC5G 5NCQA 5NDAA 5NDCA 5NDCB 5NDCC 5NDDA 5NDZA 5NECA
5NECB 5NF8A 5NF8B 5NG5C 5NG5F 5NG5I 5NG5J 5NG5K 5NG5L 5NG5M 5NG5N 5NG5O
5NIKA 5NIKB 5NIKC 5NIKJ 5NIKK 5NILA 5NILB 5NILC 5NILJ 5NILK 5NJ3A 5NJ3B
5NJ4H 5NJ4L 5NJ4M 5NJ6A 5NJGA 5NJGB 5NJYA 5NJYB 5NJYC 5NJYD 5NJYE 5NKJA
5NKJB 5NKJC 5NKJD 5NKJE 5NKQA 5NKQB 5NKQC 5NKQD 5NL2A 5NL2B 5NLXA 5NM2A
5NM4A 5NMIC 5NMID 5NMIE 5NMIG 5NMIJ 5NMIK 5NMIP 5NMIQ 5NMIR 5NMIT 5NMIW
5NMIX 5NR2A 5NSWA 5NSWB 5NSWC 5NUOA 5NUOC 5NUOE 5NUPA 5NUPB 5NUPC 5NUQA
5NUQC 5NUQE 5NV9A 5NVAA 5NX2A 5NXNA 5NXNB 5NXNC 5NXRA 5NXRC 5NXUA 5NXUB
5NXUC 5O0TA 5O31A 5O31A 5O31B 5O31C 5O31D 5O31F 5O31G 5O31H 5O31H 5O31I
5O31J 5O31J 5O31K 5O31K 5O31L 5O31L 5O31M 5O31M 5O31N 5O31Y 5O31Z 5O4CH
5O4CL 5O4CM 5O5EA 5O5EB 5O64H 5O64L 5O64M 5O65A 5O65B 5O65C 5O66A 5O66B
5O66C 5O66J 5O66K 5O66L 5O66M 5O66N 5O66O 5O67A 5O67B 5O67C 5O68A 5O68B
5O68C 5O77A 5O77B 5O77C 5O78A 5O78B 5O78C 5O79A 5O79B 5O79C 5O8FA 5O8FB
5O8FC 5O8FD 5O8FE 5O8OA 5O9CC 5O9CE 5O9CF 5O9HA 5O9HB 5OB4A 5OC0A 5OC9A
5OC9B 5OCHA 5OCHB 5ODWA 5ODWB 5OEKA 5OFPA 5OFPB 5OFRA 5OFRB 5OGEA 5OGKA
5OGKB 5OGLA 5OHDA 5OJMA 5OJMB 5OJMC 5OJMD 5OJME 5OKDC 5OKDD 5OKDE 5OKDG
5OKDJ 5OLOA 5ONUA 5ONUB 5ONUC 5OONA 5OQKA 5OQTA 5OQTC 5OR1A 5OSAA 5OSAB
5OSAC 5OSAD 5OSAE 5OSBA 5OSBB 5OSBC 5OSBD 5OSBE 5OSCA 5OSCB 5OSCC 5OSCD
5OSCE 5OUTA 5OXKA 5OXLA 5OXMA 5OXNA 5OXOA 5OXPA 5OXQA 5OY0A 5OY0A 5OY0B

5OY0B 5OY0F 5OY0F 5OY0I 5OY0I 5OY0J 5OY0J 5OY0K 5OY0K 5OY0L 5OY0L 5OY0M
5OY0M 5OYBA 5OYBB 5OYGA 5OYGB 5OYKA 5PRCH 5PRCL 5PRCM 5PRNA 5PRNB 5PRNC
5SV0A 5SV0B 5SV0C 5SV0D 5SV0F 5SV1A 5SV1B 5SV1C 5SV1D 5SV1E 5SV1Y 5SV9A
5SV9B 5SVJA 5SVJB 5SVJC 5SVKA 5SVKC 5SVKD 5SVLA 5SVLC 5SVLD 5SVMA 5SVMC
5SVMD 5SVPA 5SVPC 5SVPD 5SVQA 5SVQB 5SVQC 5SVRA 5SVRB 5SVRC 5SVSA 5SVSB
5SVSC 5SVTA 5SVTB 5SVTC 5SXUA 5SXUB 5SXUC 5SXUD 5SXUE 5SXVA 5SXVB 5SXVC
5SXVD 5SXVE 5SY1A 5SY1B 5SYTA 5T04A 5T0OA 5T0OB 5T0OC 5T1AA 5T36A 5T36B
5T36C 5T37A 5T37B 5T37C 5T3RD 5T4DA 5T4DB 5T4DC 5T4DD 5T4OI 5T4OJ 5T4OK
5T4OM 5T4ON 5T4OO 5T4OP 5T4OQ 5T4OR 5T4OS 5T4OT 5T4OU 5T4OV 5T4PI 5T4PJ
5T4PK 5T4PM 5T4PN 5T4PO 5T4PP 5T4PQ 5T4PR 5T4PS 5T4PT 5T4PU 5T4PV 5T4QI
5T4QJ 5T4QK 5T4QM 5T4QN 5T4QO 5T4QP 5T4QQ 5T4QR 5T4QS 5T4QT 5T4QU 5T4QV
5T4YC 5T4YD 5T5NA 5T5NB 5T5NC 5T5ND 5T5NE 5T77A 5TCXA 5TE3A 5TE3B 5TE5A
5TGZA 5TINA 5TINB 5TINC 5TIND 5TINE 5TIOA 5TIOB 5TIOC 5TIOD 5TIOE 5TISA
5TISA 5TISB 5TISB 5TISC 5TISC 5TISD 5TISD 5TISE 5TISE 5TISF 5TISF 5TISH
5TISH 5TISI 5TISI 5TISJ 5TISJ 5TISK 5TISK 5TISL 5TISL 5TISM 5TISM 5TISR
5TISR 5TIST 5TIST 5TISX 5TISX 5TISY 5TISY 5TISZ 5TISZ 5TJ5A 5TJ5B 5TJ5D
5TJ5E 5TJ5F 5TJ5G 5TJ5H 5TJ5I 5TJ5J 5TJ5L 5TJ5M 5TJ5N 5TJ5O 5TJ6A 5TJ6B
5TJ6C 5TJ6D 5TJIA 5TJIB 5TJIC 5TJID 5TL9A 5TL9B 5TL9C 5TQQA 5TQQB 5TR1A
5TR1B 5TSAA 5TSBA 5TSJN 5TSJO 5TSJP 5TSJQ 5TSJR 5TSJS 5TSJT 5TSJU 5TSJV
5TSJW 5TSJX 5TSJY 5TSJZ 5TTPA 5TTPB 5TUAA 5TUAB 5TUDA 5TUDD 5TV4A 5TV4B
5TVNA 5TWVA 5TWVB 5TWVC 5TWVD 5TWVE 5TWVF 5TWVG 5TWVH 5TZRA 5TZYA 5U09A
5U1LA 5U1LB 5U1LC 5U1UA 5U1UB 5U1UC 5U1VA 5U1VB 5U1VC 5U1WA 5U1WB 5U1WC
5U1XA 5U1XB 5U1XC 5U1YA 5U1YB 5U1YC 5U2HA 5U2HC 5U2HD 5U6OA 5U6OB 5U6OC
5U6OD 5U6PA 5U6PB 5U6PC 5U6PD 5U70A 5U70B 5U70C 5U70D 5U73A 5U74A 5U76A
5U76B 5U76C 5U76D 5U9WA 5U9WB 5U9WC 5UAKA 5UARA 5UENA 5UENB 5UHQA 5UHQB
5UHSA 5UHSB 5UIGA 5UIWA 5UJ9A 5UJAA 5UK6A 5UK6B 5UK6C 5UL0A 5UL7A 5UL7B
5UL9A 5UL9B 5ULDA 5ULDB 5ULEA 5ULEB 5UN1A 5UN1B 5UN1C 5UN1D 5UNFA 5UNGB
5UNHA 5UNIA 5UNIB 5UNIC 5UNID 5UOWA 5UOWB 5UOWC 5UOWD 5UP2A 5UP2B 5UP2C
5UP2D 5UVIA 5UZ7R 5V2CA 5V2CA 5V2CB 5V2CB 5V2CC 5V2CC 5V2CD 5V2CD 5V2CE
5V2CE 5V2CF 5V2CF 5V2CH 5V2CH 5V2CI 5V2CI 5V2CJ 5V2CJ 5V2CK 5V2CK 5V2CL
5V2CL 5V2CM 5V2CM 5V2CT 5V2CT 5V2CX 5V2CX 5V2CY 5V2CY 5V2CZ 5V2CZ 5V33H
5V33L 5V33M 5V4SA 5V4SB 5V4SC 5V4SD 5V54A 5V56A 5V57A 5V5SA 5V5SB 5V5SC
5V5SJ 5V5SK 5V5SL 5V5ZA 5V6NA 5V6NB 5V6NC 5V6ND 5V6NE 5V6OA 5V6OB 5V6OC
5V6OD 5V6OE 5V6PA 5V6PB 5V7PA 5V8KA 5V8KB 5V8KC 5V8KD 5VA1A 5VA1B 5VA1C
5VA1D 5VA2A 5VA2B 5VA2C 5VA2D 5VA3A 5VA3B 5VA3C 5VA3D 5VAIR 5VB2A 5VB2B
5VB2C 5VB2D 5VB8A 5VB8B 5VB8C 5VB8D 5VBLB 5VDHA 5VDHB 5VDHC 5VDHD 5VDHE
5VDIA 5VDIB 5VDIC 5VDID 5VDIE 5VEWA 5VEXA 5VG9A 5VHWA 5VHWB 5VHWC 5VHWD
5VHXA 5VHXB 5VHXC 5VHXD 5VHXE 5VHYA 5VHYB 5VHYC 5VHYD 5VHYE 5VHYF 5VHZA
5VHZB 5VHZC 5VHZD 5VHZE 5VHZF 5VJ8A 5VK6C 5VK6F 5VK6I 5VK6L 5VKEC 5VKEF
5VKEI 5VKEL 5VKHC 5VKHF 5VKHI 5VKHL 5VKQA 5VKQB 5VKQC 5VKQD 5VKVA 5VMSA
5VMSC 5VMSE 5VMSG 5VN7A 5VN7B 5VN9A 5VOTA 5VOTB 5VOTC 5VOTD 5VOTE 5VOTF
5VOTG 5VOTH 5VOUA 5VOUB 5VOUC 5VOUD 5VOUE 5VOUF 5VOUG 5VOUH 5VOVA 5VOVB
5VOVC 5VOVD 5VOVE 5VOVF 5VOVG 5VOVH 5VOXA 5VOXB 5VOXC 5VOXD 5VOXR 5VOXS
5VOXT 5VOXU 5VOXV 5VOXW 5VOXX 5VOXY 5VOXZ 5VOYA 5VOYB 5VOYC 5VOYD 5VOYR

5VOYS 5VOYT 5VOYU 5VOYV 5VOYW 5VOYX 5VOYY 5VOYZ 5VOZA 5VOZB 5VOZC 5VOZD

5VOZR 5VOZS 5VOZT 5VOZU 5VOZV 5VOZW 5VOZX 5VOZY 5VOZZ 5VPNC 5VPND 5VPNG

5VPNH 5VRAA 5VREA 5VREB 5VREC 5VRED 5VRFA 5VRFB 5VRGA 5VRHA 5W0PA 5W3SA

5W3SB 5W3SC 5W3SD 5W68A 5W68B 5W68C 5W68D 5W68E 5W68F 5W68G 5W68H 5W68I

5W68J 5W68K 5W68L 5W68M 5W68N 5W68O 5W81A 5W97A 5W97A 5W97B 5W97B 5W97C

5W97C 5W97D 5W97D 5W97G 5W97G 5W97I 5W97I 5W97J 5W97J 5W97K 5W97K 5W97L

5W97L 5W97M 5W97M 5WAUA 5WAUA 5WAUB 5WAUB 5WAUC 5WAUC 5WAUD 5WAUD 5WAUG

5WAUG 5WAUI 5WAUI 5WAUJ 5WAUJ 5WAUK 5WAUK 5WAUL 5WAUL 5WAUM 5WAUM 5WEHA

5WEHB 5WEHC 5WEHD 5WEKA 5WEKB 5WEKC 5WEKD 5WELA 5WELB 5WEMA 5WEMB 5WEMC

5WEMD 5WENA 5WENB 5WEOA 5WEOB 5WEOC 5WEOD 5WF5A 5WF6A 5WIEB 5WIED 5WIEF

5WIEJ 5WIUA 5WIVA 5WJ5A 5WJ5B 5WJ5C 5WJ5D 5WJ9A 5WJ9B 5WJ9C 5WJ9D 5WKTA

5WKUA 5WKUB 5WKUC 5WKVA 5WKVB 5WKVC 5WKXA 5WKXB 5WKXC 5WKYA 5WKYB 5WKYC

5WLNA 5WLNB 5WLNC 5WLND 5WLNE 5WLNF 5WLNG 5WLNH 5WLNI 5WLNJ 5WLNK 5WLNL

5WLNM 5WLNN 5WLNO 5WO6A 5WO7A 5WO8A 5WO9A 5WP6A 5WP6B 5WP6C 5WP6D 5WPQA

5WPQB 5WPQC 5WPQD 5WPTA 5WPTB 5WPTC 5WPTD 5WPVA 5WPVB 5WPVC 5WPVD 5WQ7A

5WQ7B 5WQ7C 5WQ7D 5WQ7E 5WQ7F 5WQ7G 5WQ7H 5WQ7I 5WQ7J 5WQ7K 5WQ7L 5WQ7M

5WQ7N 5WQ7O 5WQ8A 5WQ8B 5WQ8C 5WQ8D 5WQ8E 5WQ8F 5WQ8G 5WQ8H 5WQ8I 5WQ8J

5WQ8K 5WQ8L 5WQ8M 5WQ8N 5WQ8O 5WQ9A 5WQ9B 5WQ9C 5WQ9D 5WQ9E 5WQ9F 5WQ9G

5WQ9H 5WQ9I 5WQ9J 5WQ9K 5WQ9L 5WQ9M 5WQ9N 5WQ9O 5WQCA 5WS3A 5WS4A 5WS4B

5WS5A 5WS5A 5WS5B 5WS5B 5WS5C 5WS5C 5WS5D 5WS5D 5WS5E 5WS5E 5WS5F 5WS5F

5WS5H 5WS5H 5WS5I 5WS5I 5WS5J 5WS5J 5WS5K 5WS5K 5WS5L 5WS5L 5WS5M 5WS5M

5WS5R 5WS5T 5WS5T 5WS5X 5WS5X 5WS5Y 5WS5Y 5WS5Z 5WS5Z 5WS6A 5WS6A 5WS6B

5WS6B 5WS6C 5WS6C 5WS6D 5WS6D 5WS6E 5WS6E 5WS6F 5WS6F 5WS6H 5WS6H 5WS6I

5WS6I 5WS6J 5WS6J 5WS6K 5WS6K 5WS6L 5WS6L 5WS6M 5WS6M 5WS6R 5WS6T 5WS6T

5WS6X 5WS6X 5WS6Y 5WS6Y 5WS6Z 5WS6Z 5WTRA 5WTRB 5WTRC 5WUAA 5WUAB 5WUAC

5WUAD 5WUAE 5WUAF 5WUAG 5WUAH 5WUCA 5WUCB 5WUCC 5WUDA 5WUDB 5WUDC 5WUEA

5WUEB 5WUEC 5WUFA 5WUFB 5WUFC 5WZYA 5WZYB 5WZYC 5X0MA 5X19A 5X19B 5X19C

5X19D 5X19G 5X19I 5X19J 5X19K 5X19L 5X19M 5X19N 5X19O 5X19P 5X19Q 5X19T

5X19V 5X19W 5X19X 5X19Y 5X19Z 5X1BA 5X1BB 5X1BC 5X1BD 5X1BG 5X1BI 5X1BJ

5X1BK 5X1BL 5X1BM 5X1BN 5X1BO 5X1BP 5X1BQ 5X1BT 5X1BV 5X1BW 5X1BX 5X1BY

5X1BZ 5X1FA 5X1FB 5X1FC 5X1FD 5X1FG 5X1FI 5X1FJ 5X1FK 5X1FL 5X1FM 5X1FN

5X1FO 5X1FP 5X1FQ 5X1FT 5X1FV 5X1FW 5X1FX 5X1FY 5X1FZ 5X33A 5X3XM 5X3XQ

5X5YF 5X5YG 5X7DA 5X87A 5X87B 5X87C 5X87D 5X87E 5X93A 5X9HA 5X9HB 5X9RA

5X9RB 5XA7A 5XA8A 5XA9A 5XAAA 5XABA 5XAMA 5XANA 5XANB 5XAPA 5XARA 5XARB

5XASA 5XASB 5XATA 5XDNA 5XDOA 5XDQA 5XDQB 5XDQC 5XDQD 5XDQG 5XDQI 5XDQJ

5XDQK 5XDQL 5XDQM 5XDQN 5XDQO 5XDQP 5XDQQ 5XDQT 5XDQV 5XDQW 5XDQX 5XDQY

5XDQZ 5XDXA 5XDXB 5XDXC 5XDXD 5XDXG 5XDXI 5XDXJ 5XDXK 5XDXL 5XDXM 5XDXN

5XDXO 5XDXP 5XDXQ 5XDXT 5XDXV 5XDXW 5XDXX 5XDXY 5XDXZ 5XEFA 5XEZA 5XF1A

5XHQA 5XJ5A 5XJ6A 5XJ7A 5XJ8A 5XJ9A 5XJJA 5XJMA 5XJYA 5XLSA 5XMJC 5XMJO

5XNL1 5XNL2 5XNL3 5XNL4 5XNL5 5XNL6 5XNL7 5XNL8 5XNLA 5XNLA 5XNLB 5XNLB

5XNLC 5XNLC 5XNLD 5XNLD 5XNLE 5XNLE 5XNLF 5XNLF 5XNLG 5XNLG 5XNLH 5XNLH

5XNLI 5XNLI 5XNLJ 5XNLJ 5XNLK 5XNLK 5XNLL 5XNLL 5XNLM 5XNLM 5XNLN 5XNLN

5XNLR 5XNLR 5XNLS 5XNLS 5XNLT 5XNLT 5XNLW 5XNLW 5XNLX 5XNLX 5XNLY 5XNLY

5XNLZ 5XNLZ 5XNM1 5XNM2 5XNM3 5XNM4 5XNM5 5XNM6 5XNM7 5XNM8 5XNMA 5XNMA

5XNMB 5XNMB 5XNMC 5XNMC 5XNMD 5XNMD 5XNME 5XNME 5XNMF 5XNMF 5XNMG 5XNMG
5XNMH 5XNMH 5XNMI 5XNMI 5XNMJ 5XNMJ 5XNMK 5XNMK 5XNML 5XNML 5XNMM 5XNMM
5XNMN 5XNMN 5XNMR 5XNMR 5XNMS 5XNMS 5XNMT 5XNMT 5XNMW 5XNMW 5XNMX 5XNMX
5XNMY 5XNMY 5XNMZ 5XNMZ 5XNN1 5XNN2 5XNN3 5XNN4 5XNO1 5XNO2 5XNO3 5XNO4
5XPDA 5XPRA 5XR8A 5XRAA 5XSYA 5XSYB 5XSZA 5XTCA 5XTCB 5XTCC 5XTCE 5XTCF
5XTCG 5XTCI 5XTCJ 5XTCK 5XTCL 5XTCM 5XTCN 5XTCO 5XTCR 5XTCS 5XTCS 5XTCU
5XTCV 5XTCW 5XTCY 5XTCZ 5XTDA 5XTDB 5XTDC 5XTDE 5XTDF 5XTDG 5XTDI 5XTDJ
5XTDK 5XTDL 5XTDM 5XTDN 5XTDO 5XTDR 5XTDS 5XTDS 5XTDU 5XTDV 5XTDW 5XTDY
5XTDZ 5XTEA 5XTEC 5XTED 5XTEG 5XTEH 5XTEJ 5XTEN 5XTEP 5XTEQ 5XTET 5XTEU
5XTEV 5XU1M 5XU1S 5XW6A 5XW6B 5XW6C 5Y0BA 5Y0BB 5Y4OA 5Y4OB 5Y4OC 5Y4OD
5Y4OE 5Y4OF 5Y4OG 5Y50A 5Y5SA 5Y5SB 5Y5SD 5Y5SE 5Y5SF 5Y5SH 5Y5SL 5Y5SM
5Y5XN 5Y5XO 5Y5XP 5Y5XQ 5Y5XR 5Y5XS 5Y5XT 5Y5XU 5Y5XV 5Y5XW 5Y5XX 5Y5XY
5Y5XZ 5Y5ZN 5Y5ZO 5Y5ZP 5Y5ZQ 5Y5ZR 5Y5ZS 5Y5ZT 5Y5ZU 5Y5ZV 5Y5ZW 5Y5ZX
5Y5ZY 5Y5ZZ 5Y60N 5Y60O 5Y60P 5Y60Q 5Y60R 5Y60S 5Y60T 5Y60U 5Y60V 5Y60W
5Y60X 5Y60Y 5Y60Z 5Y78A 5Y78B 5Y79A 5Y79B 5Y83A 5YC8A 5YCKA 5YDZA 5YDZB
5YDZC 5YDZD 5YE1A 5YE1B 5YE1C 5YE1D 5YE2A 5YE2B 5YE2C 5YE2D 5YE5A 5YE5B
5YE5C 5YE5D 5YHFA 5YHLA 5YILA 5YILB 5YILC 5YKEA 5YKEB 5YKEC 5YKED 5YKEE
5YKEF 5YKEG 5YKEH 5YKFA 5YKFB 5YKFC 5YKFD 5YKFE 5YKFF 5YKFG 5YKFH 5YKGA
5YKGB 5YKGC 5YKGD 5YKGE 5YKGF 5YKGG 5YKGH 5YLUA 5YLUB 5YLVA 5YLVB 5YQ70
5YQ71 5YQ72 5YQ73 5YQ74 5YQ75 5YQ76 5YQ77 5YQ78 5YQ79 5YQ7A 5YQ7B 5YQ7C
5YQ7D 5YQ7E 5YQ7F 5YQ7G 5YQ7H 5YQ7I 5YQ7J 5YQ7K 5YQ7L 5YQ7M 5YQ7N 5YQ7O
5YQ7P 5YQ7Q 5YQ7R 5YQ7S 5YQ7T 5YQ7U 5YQ7V 5YQ7W 5YQ7X 5YQ7Y 5YS3A 5YS3B
5YS3C 5YS3D 5YS3E 5YS3F 5YS8A 5YS8B 5YS8C 5YS8D 5YS8E 5YS8F 5YUAA 5YUAB
5YUAC 5YUAD 5YUBA 5YUBB 5YUBC 5YUBD 5YUCA 5YUCB 5YUCC 5YUCD 5YVEA 5YVEB
5YVEC 5YW7B 5YW8A 5YW8B 5YW8C 5YW8D 5YW8E 5YW8F 5YW8G 5YW8H 5YW9A 5YW9B
5YW9C 5YW9D 5YW9E 5YW9F 5YW9G 5YW9H 5YWAA 5YWAB 5YWAC 5YWAD 5YWAE 5YWAF
5YWAG 5YWAH 5YWBA 5YWBB 5YWBC 5YWBD 5YWBE 5YWBF 5YWBG 5YWBH 5YWCA 5YWCB
5YWCC 5YWCD 5YWCE 5YWCF 5YWCG 5YWCH 5YWDB 5YWYA 5YX9A 5YX9B 5YX9C 5YX9D
5Z10A 5Z1FA 5Z1FB 5Z1WA 5Z1WB 5Z1WC 5Z1WD 5Z62C 5Z62G 5Z62J 5Z84A 5Z84B
5Z84C 5Z84D 5Z84G 5Z84I 5Z84J 5Z84K 5Z84L 5Z84M 5Z84N 5Z84O 5Z84P 5Z84Q
5Z84T 5Z84V 5Z84W 5Z84X 5Z84Y 5Z84Z 5Z85A 5Z85B 5Z85C 5Z85D 5Z85G 5Z85I
5Z85J 5Z85K 5Z85L 5Z85M 5Z85N 5Z85O 5Z85P 5Z85Q 5Z85T 5Z85V 5Z85W 5Z85X
5Z85Y 5Z85Z 5Z86A 5Z86B 5Z86C 5Z86D 5Z86G 5Z86I 5Z86J 5Z86K 5Z86L 5Z86M
5Z86N 5Z86O 5Z86P 5Z86Q 5Z86T 5Z86V 5Z86W 5Z86X 5Z86Y 5Z86Z 5Z96A 5Z96B
5Z96C 5Z96D 5ZAZA 5ZBGA 5ZBGB 5ZBGC 5ZBGD 5ZBHA 5ZBQA 5ZCOA 5ZCOB 5ZCOC
5ZCOD 5ZCOG 5ZCOI 5ZCOJ 5ZCOK 5ZCOL 5ZCOM 5ZCON 5ZCOO 5ZCOP 5ZCOQ 5ZCOT
5ZCOV 5ZCOW 5ZCOX 5ZCOY 5ZCOZ 5ZCPA 5ZCPB 5ZCPC 5ZCPD 5ZCPG 5ZCPI 5ZCPJ
5ZCPK 5ZCPL 5ZCPM 5ZCPN 5ZCPO 5ZCPP 5ZCPQ 5ZCPT 5ZCPV 5ZCPW 5ZCPX 5ZCPY
5ZCPZ 5ZCQA 5ZCQB 5ZCQC 5ZCQD 5ZCQG 5ZCQI 5ZCQJ 5ZCQK 5ZCQL 5ZCQM 5ZCQN
5ZCQO 5ZCQP 5ZCQQ 5ZCQT 5ZCQV 5ZCQW 5ZCQX 5ZCQY 5ZCQZ 5ZDHA 5ZDHB 5ZDHC
5ZDHD 5ZDHE 5ZDHF 5ZDHG 5ZDHH 5ZDHI 5ZDHJ 5ZDHK 5ZDHL 5ZDHM 5ZDHN 5ZDHO
5ZFPA 5ZFPB 5ZFPC 5ZFPD 5ZFPE 5ZFPF 5ZFUA 5ZFUB 5ZFUC 5ZFUD 5ZFUE 5ZFUF
5ZFUG 5ZFUH 5ZFUI 5ZFVA 5ZFVB 5ZFVC 5ZFVD 5ZFVE 5ZFVF 5ZGB1 5ZGB2 5ZGB3
5ZGB4 5ZGB5 5ZGBA 5ZGBB 5ZGBF 5ZGBI 5ZGBJ 5ZGBK 5ZGBL 5ZGBM 5ZGBO 5ZGGA

5ZGH1 5ZGH2 5ZGH3 5ZGHA 5ZGHB 5ZGHF 5ZGHI 5ZGHJ 5ZGHK 5ZGHL 5ZGHM 5ZGHO

5ZIHA 5ZIHB 5ZILA 5ZILB 5ZIMA 5ZIMB 5ZINA 5ZINB 5ZJI1 5ZJI2 5ZJI3 5ZJI4

5ZJIA 5ZJIB 5ZJIF 5ZJIG 5ZJIH 5ZJII 5ZJIJ 5ZJIK 5ZJIL 5ZJIO 5ZJIX 5ZJIY

5ZJIZ 5ZK3A 5ZK8A 5ZKBA 5ZKCA 5ZKPA 5ZLEA 5ZLEB 5ZLGA 5ZLGB 5ZMVA 5ZMWA

5ZOVA 5ZOVB 5ZSUA 5ZSUB 5ZSUC 5ZSUD 5ZSUE 5ZSUF 5ZTFA 5ZTKA 5ZTLA 5ZUGA

5ZUGB 5ZUGC 5ZUGD 5ZUGE 5ZUGF 5ZX5A 5ZX5B 5ZX5C 5ZX5D 5ZZNA 5ZZNA 5ZZNB

5ZZNB 5ZZNC 5ZZNC 5ZZND 5ZZND 5ZZNE 5ZZNE 5ZZNF 5ZZNF 5ZZNH 5ZZNH 5ZZNI

5ZZNI 5ZZNJ 5ZZNJ 5ZZNK 5ZZNK 5ZZNL 5ZZNL 5ZZNM 5ZZNM 5ZZNT 5ZZNT 5ZZNX

5ZZNX 5ZZNY 5ZZNY 5ZZNZ 5ZZNZ 6A2JA 6A2WA 6A69A 6A69B 6A6MA 6A6MB 6A6NA

6A6NB 6A70A 6A70B 6A70F 6A70G 6A90A 6A91A 6A93A 6A94A 6A95A 6A96A 6A96B

6A96C 6A96D 6A96E 6AB9A 6ABAA 6AD7A 6AD7B 6AD8A 6AD8B 6ADAA 6ADAB 6ADBA

6ADBB 6ADCA 6ADCB 6ADQA 6ADQB 6ADQC 6ADQD 6ADQE 6ADQF 6ADQG 6ADQH 6ADQI

6ADQM 6ADQN 6ADQO 6ADQP 6ADQQ 6ADQR 6ADQS 6ADQT 6ADQU 6AEIA 6AEIB 6AEIC

6AEID 6AFSA 6AFSB 6AFTA 6AFTB 6AFUA 6AFUB 6AFVA 6AFVB 6AFWA 6AFWB 6AFXA

6AFXB 6AFYA 6AFYB 6AFZA 6AFZB 6AGFA 6AGFB 6AH4A 6AH4B 6AH5A 6AH5B 6AK3A

6AK3B 6AKEA 6AKFA 6AKGA 6AKIA 6AKIB 6AKIC 6AKID 6AKIE 6AKIF 6AKXA 6AKXB

6AKYA 6AL2A 6ALEB 6AQFA 6AVEA 6AVEB 6AVEC 6AWFC 6AWFD 6AWNA 6AWOA 6AWPA

6AWQA 6AYEA 6AYEB 6AYEC 6AYED 6AYFA 6AYFB 6AYFC 6AYFD 6AYGA 6AYGB 6AYGC

6AYGD 6B21A 6B21B 6B21C 6B24A 6B24B 6B2AA 6B2AB 6B2BA 6B2BB 6B2DA 6B2DB

6B2ZB 6B2ZC 6B2ZD 6B2ZE 6B2ZF 6B2ZG 6B2ZH 6B2ZI 6B2ZJ 6B2ZK 6B2ZL 6B2ZM

6B2ZN 6B2ZQ 6B2ZS 6B2ZT 6B3JR 6B5VA 6B5VB 6B5VC 6B5VD 6B73A 6B85D 6B85E

6B85F 6B85J 6B87A 6B87E 6BAAA 6BAAB 6BAAC 6BAAD 6BAAE 6BAAF 6BAAG 6BAAH

6BAJA 6BAJB 6BAJC 6BARA 6BASA 6BATA 6BATB 6BATC 6BAUA 6BAUB 6BAUC 6BAVA

6BAVB 6BAVC 6BBFA 6BBFB 6BBFC 6BBFD 6BBFE 6BBFF 6BBGA 6BBGB 6BBGC 6BBGD

6BBGE 6BBGF 6BBHA 6BBHB 6BBHC 6BBHD 6BBHE 6BBHF 6BBIA 6BBIB 6BBIC 6BBID

6BBIE 6BBIF 6BBJA 6BBJB 6BBJC 6BBJD 6BCJA 6BCJB 6BCJC 6BCJD 6BCLA 6BCLB

6BCLC 6BCLD 6BCOA 6BCOB 6BCOC 6BCOD 6BCQA 6BCQB 6BCQC 6BCQD 6BD4A 6BE1A

6BE1B 6BE1C 6BE1D 6BE1E 6BGIA 6BGIB 6BGJA 6BGJB 6BH8A 6BHPA 6BHPB 6BHPC

6BHUA 6BL6A 6BL6C 6BMIA 6BMIB 6BMIC 6BMLA 6BMMA 6BMNA 6BMSA 6BO4A 6BO4B

6BO4C 6BO4D 6BO5A 6BO5B 6BO5C 6BO5D 6BO8A 6BO8B 6BO8C 6BO8D 6BO9A 6BO9B

6BO9C 6BO9D 6BOAA 6BOAB 6BOAC 6BOAD 6BOBA 6BOBB 6BOBC 6BOBD 6BPLA 6BPLB

6BPMA 6BPMC 6BPNA 6BPOA 6BPPA 6BPPB 6BPQA 6BPQB 6BPQC 6BPQD 6BPZA 6BPZB

6BPZC 6BQGA 6BQHA 6BQNA 6BQNB 6BQNC 6BQOA 6BQRA 6BQRB 6BQRC 6BQRD 6BQVA

6BQVB 6BQVC 6BQVD 6BTMA 6BTMC 6BTMD 6BTMF 6BTXA 6BU5A 6BUGC 6BUHC 6BUIC

6BVGA 6BVGB 6BW5A 6BW5B 6BW6A 6BW6B 6BWDA 6BWDB 6BWDC 6BWDD 6BWFA 6BWFB

6BWFC 6BWFD 6BWIA 6BWIB 6BWIC 6BWID 6BWJA 6BWJB 6BWJC 6BWJD 6BWMA 6BWMB

6BWMC 6BWMD 6BX4A 6BX5A 6BY2C 6BY3C 6BYOA 6C08C 6C0VA 6C14A 6C14B 6C14C

6C14D 6C1EA 6C1EB 6C1EC 6C1ED 6C1KA 6C1KB 6C1KC 6C1KD 6C1MA 6C1MB 6C1MC

6C1MD 6C1PA 6C1PB 6C1PC 6C1PD 6C1QB 6C1RB 6C261 6C262 6C263 6C264 6C265

6C26A 6C26B 6C26C 6C3IA 6C3OA 6C3OB 6C3OC 6C3OD 6C3OE 6C3OF 6C3OG 6C3OH

6C3PA 6C3PB 6C3PC 6C3PD 6C3PE 6C3PF 6C3PG 6C3PH 6C4XA 6C4XB 6C4XC 6C4XD

6C4XE 6C4XF 6C4XG 6C4XH 6C5WA 6C5WB 6C6LA 6C6LC 6C6LD 6C6LE 6C6LF 6C6LG

6C6LH 6C6LI 6C6LJ 6C6LK 6C6LL 6C6LM 6C6LN 6C6LO 6C70A 6C70B 6C8FA 6C8FB

6C8FC 6C8FD 6C8GA 6C8GB 6C8GC 6C8GD 6C8HA 6C8HB 6C8HC 6C8HD 6C96A 6C96B

6C9AA 6C9AB 6C9WA 6CAAA 6CAAB 6CB2A 6CB8A 6CB8A 6CB8B 6CB8C 6CB8D 6CB8E
6CB8F 6CB8G 6CB8H 6CB8I 6CB8J 6CB8K 6CB8L 6CB8M 6CB8N 6CB8O 6CB8P 6CB8Q
6CB8R 6CB8S 6CB8T 6CB8U 6CB8V 6CB8W 6CB8X 6CB8Y 6CB8Z 6CC4A 6CDUA 6CDUB
6CDUC 6CDUD 6CDUE 6CFWA 6CFWB 6CFWC 6CFWD 6CFWE 6CFWF 6CFWG 6CFWH 6CFWI
6CFWM 6CI0A 6CI0B 6CJQA 6CJQB 6CJQC 6CJQD 6CJTA 6CJTB 6CJTC 6CJTD 6CJUA
6CJUB 6CJUC 6CJUD 6CM4A 6CMCA 6CMCB 6CMCC 6CMOR 6CNJA 6CNJB 6CNJC 6CNJD
6CNJE 6CNKA 6CNKB 6CNKC 6CNKD 6CNKE 6CNMA 6CNMB 6CNMC 6CNMD 6CNNA 6CNNB
6CNNC 6CNND 6CNOA 6CNOB 6CNOC 6CNOD 6CO7A 6CO7B 6CO7C 6CO7D 6COYA 6COYB
6CP38 6CP3J 6CP3K 6CP3L 6CP3U 6CP3X 6CP3Z 6CP58 6CP5J 6CP5K 6CP5L 6CP5M
6CP5N 6CP5O 6CP5P 6CP5Q 6CP5R 6CP5S 6CP5T 6CP5U 6CP5X 6CP5Z 6CP68 6CP6J
6CP6K 6CP6L 6CP6U 6CP6X 6CP6Z 6CP78 6CP7J 6CP7K 6CP7L 6CP7M 6CP7N 6CP7O
6CP7P 6CP7Q 6CP7R 6CP7S 6CP7T 6CP7U 6CP7X 6CP7Z 6CPVA 6CPVC 6CPVD 6CPVE
6CQ6A 6CQ6B 6CQ8A 6CQ8B 6CQ9A 6CQ9B 6CSEM 6CSFM 6CSMA 6CSMB 6CSNA 6CSNB
6CSOA 6CSOB 6CSXA 6CSXB 6CSXC 6CTDA 6CTDB 6CTDC 6CTDD 6CTDE 6CTFA 6CTFB
6CTFC 6CUDA 6CUDB 6CUDC 6CUDD 6CVLA 6CVLB 6CX0A 6CX0B 6CXHA 6CXHB 6CXHC
6CXHD 6CXHE 6CXHF 6CXHG 6CXHH 6CZQB 6D0JA 6D0JB 6D0KA 6D0KB 6D0NA 6D0NB
6D1SA 6D1SB 6D1SC 6D1SD 6D1SE 6D1WA 6D1WB 6D1WC 6D1WD 6D32A 6D35A 6D3RA
6D3SA 6D6TA 6D6TB 6D6TC 6D6TD 6D6TE 6D6UA 6D6UB 6D6UC 6D6UD 6D6UE 6D73A
6D73B 6D73C 6D73D 6D79A 6D7OA 6D7OB 6D7OC 6D7OD 6D7PA 6D7PB 6D7PC 6D7PD
6D7QA 6D7QB 6D7QC 6D7QD 6D7SA 6D7SB 6D7SC 6D7SD 6D7TA 6D7TB 6D7TC 6D7TD
6D7VA 6D7VB 6D7VC 6D7VD 6D7WA 6D7WB 6D7WC 6D7WD 6D7XA 6D7XB 6D7XC 6D7XD
6D80A 6D80B 6D80C 6D80D 6D91A 6D9HR 6D9WA 6D9ZF 6DD9A 6DDER 6DDFR 6DG7A
6DG7B 6DG7C 6DG7D 6DG7E 6DG8A 6DG8B 6DG8C 6DG8D 6DG8E 6DJBA 6DJBB 6DJBC
6DJBD 6DJBE 6DJBF 6DJRA 6DJRB 6DJRC 6DJRD 6DJSA 6DJSB 6DJSC 6DJSD 6DJZA
6DJZB 6DJZC 6DK0A 6DK0B 6DK0C 6DK1A 6DK1B 6DK1C 6DLWA 6DLWB 6DLWC 6DLWD
6DLWE 6DLWF 6DLWG 6DLWH 6DLWI 6DLWJ 6DLWK 6DLWL 6DLWM 6DLWN 6DLWO 6DLWP
6DLWQ 6DLWR 6DLWS 6DLWT 6DLWU 6DLWV 6DLZA 6DLZB 6DLZC 6DLZD 6DM0A 6DM0B
6DM0C 6DM0D 6DM1A 6DM1B 6DM1C 6DM1D 6DMBA 6DMOA 6DMRA 6DMRB 6DMRC 6DMRD
6DMUA 6DMUB 6DMUC 6DMUD 6DMWA 6DMWB 6DMWC 6DMWD 6DMYA 6DNFA 6DNFB 6DNFC
6DNFD 6DO1A 6DO1B 6DRJA 6DRJB 6DRJC 6DRJD 6DRKA 6DRKB 6DRKC 6DRKD 6DRXA
6DRYA 6DRZA 6DS0A 6DT0A 6DT0B 6DT0C 6DT0D 6DU8A 6DU8B 6DU8C 6DU8D 6DVWA
6DVWB 6DVWC 6DVWD 6DVYA 6DVYB 6DVYC 6DVYD 6DVZA 6DVZB 6DVZC 6DVZD 6DW0A
6DW0B 6DW0C 6DW0D 6DW0E 6DZ1A 6DZ1C 6DZ1D 6DZ1E 6DZ7A 6DZ7B 6DZVA 6DZWA
6DZYA 6DZZA 6E0HA 6E0HB 6E10A 6E10B 6E10C 6E10D 6E10E 6E10F 6E10G 6E14D
6E15D 6E1HA 6E1HB 6E1KA 6E1KB 6E1MA 6E1MB 6E1NA 6E1NB 6E1OA 6E1OB 6E1PA
6E1PB 6E2FA 6E2FB 6E2FC 6E2FD 6E2GA 6E2GB 6E2GC 6E2GD 6E3YE 6E3YR 6E4VA
6E59A 6E67A 6E7PA 6E7PB 6E7PC 6E7PD 6E7YA 6E7YB 6E7YC 6E7YD 6E7ZA 6E7ZB
6E7ZC 6E7ZD 6E8JA 6E9CA 6E9NA 6E9OB 6EBAA 6EBKB 6EBKD 6EBKF 6EBKH 6EBMB
6EBMD 6EBMF 6EBMH 6EBUA 6EDQA 6EDQB 6EHBA 6EHBB 6EHBC 6EHCA 6EHDA 6EHEA
6EHFA 6EHFB 6EHFC 6EI3A 6EIAA 6EIDA 6EIDC 6EIGA 6EIGC 6EJ6A 6EJ6B 6EJ6C
6EJHA 6EJHB 6EJHC 6EL1K 6EL1L 6EL1M 6EL1N 6EL1O 6EL1P 6EL1Q 6EL1R 6EL1S
6EL1T 6EMXA 6EMXB 6EMXC 6EMXD 6EMXE 6ENEA 6ENEB 6ENEC 6EO1A 6EO1B 6EO1C
6EO1D 6ET51 6ET52 6ET53 6ET54 6ET55 6ET56 6ET57 6ET5A 6ET5B 6ET5C 6ET5D
6ET5E 6ET5F 6ET5F 6ET5G 6ET5G 6ET5H 6ET5H 6ET5I 6ET5I 6ET5J 6ET5K 6ET5K

6ET5L 6ET5L 6ET5M 6ET5M 6ET5N 6ET5N 6ET5O 6ET5O 6ET5P 6ET5P 6ET5Q 6ET5Q
6ET5R 6ET5R 6ET5S 6ET5S 6ET5T 6ET5T 6ET5U 6ET5U 6ET5V 6ET5V 6ET5W 6ET5W
6ET5X 6ET5X 6ET5Y 6ET5Y 6ET5Z 6ET5Z 6ETIA 6ETIB 6EU6A 6EU6B 6EU6C 6EUQA
6EUSA 6EUSB 6EUSC 6EXSA 6EYUA 6EYUB 6EYUC 6EZNA 6EZNB 6EZNC 6EZND 6EZNE
6EZNF 6EZNG 6EZNH 6F0IA 6F0IB 6F0IC 6F0ID 6F0IE 6F0JA 6F0JB 6F0JC 6F0JD
6F0JE 6F0KA 6F0KC 6F0KD 6F0KF 6F0MA 6F0MB 6F0MC 6F0MD 6F0ME 6F0NA 6F0NB
6F0NC 6F0ND 6F0NE 6F0RA 6F0RB 6F0RC 6F0RD 6F0RE 6F0UA 6F0UB 6F0UC 6F0UD
6F0UE 6F0VA 6F0VB 6F0VC 6F0VD 6F0VE 6F0ZA 6F0ZB 6F0ZC 6F0ZD 6F0ZE 6F10A
6F10B 6F10C 6F10D 6F10E 6F11A 6F11B 6F11C 6F11D 6F11E 6F12A 6F12B 6F12C
6F12D 6F12E 6F13A 6F13B 6F13C 6F13D 6F13E 6F15A 6F15B 6F15C 6F15D 6F15E
6F16A 6F16B 6F16C 6F16D 6F16E 6F2DF 6F2DG 6F2DH 6F2DI 6F2DJ 6F2GA 6F2WA
6F34A 6F34C 6F36A 6F36B 6F36C 6F36D 6F36E 6F36F 6F36G 6F36H 6F36I 6F36J
6F36M 6F36N 6F46A 6F7AA 6F7AB 6F7AC 6F7AD 6F7AE 6F7HA 6F7HB 6F7HC 6F7HD
6FEQA 6FEQB 6FFCA 6FFCB 6FFVA 6FHZA 6FIZA 6FIZB 6FIZC 6FIZD 6FJ3A 6FK6A
6FK7A 6FK7C 6FK8A 6FK9A 6FKAA 6FKBA 6FKCA 6FKDA 6FKFA 6FKFB 6FKFG 6FKFH
6FKFI 6FKFJ 6FKFK 6FKFL 6FKFM 6FKFN 6FKFO 6FKFP 6FKFP 6FKFQ 6FKFR 6FKFS
6FKFT 6FKHA 6FKHB 6FKHG 6FKHH 6FKHI 6FKHJ 6FKHK 6FKHL 6FKHM 6FKHN 6FKHO
6FKHP 6FKHP 6FKHQ 6FKHR 6FKHS 6FKHT 6FKIA 6FKIB 6FKIG 6FKIH 6FKII 6FKIJ
6FKIK 6FKIL 6FKIM 6FKIN 6FKIO 6FKIP 6FKIP 6FKIQ 6FKIR 6FKIS 6FKIT 6FL9A
6FL9B 6FL9C 6FL9D 6FL9E 6FLIA 6FLIB 6FLIC 6FLID 6FLIE 6FM9A 6FM9B 6FMRA
6FMSA 6FMTA 6FMVA 6FMWA 6FMXA 6FMYA 6FN1A 6FN4A 6FNPD 6FO0C 6FO0D 6FO0E
6FO0G 6FO0J 6FO0P 6FO0Q 6FO0R 6FO0T 6FO0W 6FO2C 6FO2D 6FO2E 6FO2G 6FO2J
6FO2P 6FO2Q 6FO2R 6FO2T 6FO2W 6FO6C 6FO6D 6FO6E 6FO6G 6FO6J 6FO6P 6FO6Q
6FO6R 6FO6T 6FO6W 6FOKA 6FOKB 6FOMA 6FOS2 6FOS4 6FOSA 6FOSB 6FOSF 6FOSI
6FOSJ 6FOSK 6FOSL 6FOSM 6FOSO 6FSUA 6FSUB 6FUFA 6FV6A 6FV7A 6FV7B 6FV8A
6FV8B 6FVQA 6FVQB 6FVQC 6FVQD 6FVQE 6FVRA 6FVRB 6FVRC 6FVRD 6FVRE 6FVSA
6FVSB 6FVSC 6FVSD 6FVSE 6FVZA 6FW0A 6FWCA 6FWFA 6FWZA 6FWZB 6G2JA 6G2JA
6G2JB 6G2JC 6G2JD 6G2JF 6G2JG 6G2JH 6G2JH 6G2JI 6G2JJ 6G2JJ 6G2JK 6G2JK
6G2JL 6G2JL 6G2JM 6G2JM 6G2JN 6G2JY 6G2JZ 6G6UA 6G72A 6G72A 6G72B 6G72C
6G72D 6G72F 6G72G 6G72H 6G72H 6G72I 6G72J 6G72J 6G72K 6G72K 6G72L 6G72L
6G72M 6G72M 6G72N 6G72Y 6G72Z 6G73A 6G79S 6G7HA 6G7HB 6G7HC 6G7IA 6G7IB
6G7IC 6G7JA 6G7JB 6G7JC 6G7KA 6G7KB 6G7KC 6G7LA 6G7LB 6G7LC 6G7OA 6G8ZA
6G8ZB 6G8ZC 6G8ZD 6G8ZE 6G8ZF 6G94A 6G94B 6G94Q 6G94R 6G94S 6G94T 6G9LA
6G9LB 6G9LC 6G9LD 6G9LE 6G9LF 6G9OA 6G9OB 6G9OC 6G9OD 6G9OE 6G9OF 6G9XA
6GA1A 6GA1B 6GA1C 6GA2A 6GA2B 6GA2C 6GA3A 6GA3B 6GA3C 6GA4A 6GA4B 6GA4C
6GA5A 6GA5B 6GA5C 6GA6A 6GA6B 6GA6C 6GA7A 6GA7B 6GA7C 6GA8A 6GA8B 6GA8C
6GA9A 6GA9B 6GA9C 6GAAA 6GAAB 6GAAC 6GABA 6GABB 6GABC 6GACA 6GACB 6GACC
6GADA 6GADB 6GADC 6GAEA 6GAEB 6GAEC 6GAFA 6GAFB 6GAFC 6GAGA 6GAGB 6GAGC
6GAHA 6GAHB 6GAHC 6GAIA 6GAIB 6GAIC 6GCIA 6GCS1 6GCS2 6GCS3 6GCS4 6GCS5
6GCS6 6GCSA 6GCSB 6GCSC 6GCSD 6GCSE 6GCSG 6GCSI 6GCSJ 6GCSJ 6GCSL 6GCSN
6GCSS 6GCSW 6GCSX 6GCTA 6GCTB 6GCTC 6GDGA 6GDIA 6GHJA 6GIEA 6GIGA 6GPSA
6GPXA 6GPXB 6GRJA 6GRJB 6GRJC 6GRJD 6GRJE 6GRJF 6GRJG 6GRJH 6GRJI 6GRJJ
6GS1A 6GS4A 6GS7A 6GT3A 6GUXA 6GUYA 6GUZA 6GV1A 6GWHA 6GXCA 6GY6A 6GY6B
6GY6C 6GY6D 6GY6E 6GY6F 6GY6G 6GY6H 6GY6I 6GY6J 6GY6K 6GY6L 6GY6M 6GY6N

6GY6O 6GY6P 6GY6Q 6GY6R 6GY6S 6GY6T 6GY6U 6GY6V 6GY6W 6GY6X 6GY6Y 6GY6Z
6GYHA 6GYHB 6GYHC 6GYNA 6GYNB 6GYNC 6GYND 6GYOA 6GYOB 6GYOC 6GYOD 6GZ9A
6H04B 6H04C 6H04F 6H04G 6H04H 6H04I 6H04J 6H04K 6H04L 6H04M 6H04N 6H04O
6H04P 6H04Q 6H04R 6H04S 6H04T 6H04U 6H04V 6H04W 6H04X 6H2FA 6H2FB 6H2FC
6H2FD 6H2FE 6H2FF 6H2FG 6H2FH 6H2FI 6H2FJ 6H3IA 6H3IF 6H53A 6H53B 6H59A
6H59B 6H5AA 6H5AB 6H7DA 6H7FA 6H7FB 6H7FC 6H7JA 6H7LA 6H7MA 6H7NA 6H7OA
6H7UA 6H7VA 6H7VB 6H7VC 6HAWC 6HAWD 6HAWE 6HAWG 6HAWJ 6HBUA 6HBUB 6HCGA
6HCGB 6HCGC 6HCGD 6HCGE 6HCGF 6HCGG 6HCGH 6HCGI 6HCGJ 6HCGK 6HCGL 6HCGM
6HCGN 6HCGO 6HCLA 6HCOA 6HCOB 6HCPA 6HCPB 6HCPC 6HCYA 6HCYC 6HD1A 6HD1B
6HD1C 6HD8B 6HD8C 6HD8D 6HD8E 6HD9B 6HD9D 6HD9E 6HD9F 6HDAB 6HDAC 6HDAD
6HDAE 6HDBB 6HDBC 6HDBD 6HDBE 6HDCB 6HDCC 6HDCD 6HDCE 6HEFA 6HFBA 6HIJA
6HIJB 6HINA 6HINB 6HINC 6HIND 6HINE 6HIOA 6HIOB 6HIOC 6HIOD 6HIOE 6HIQA
6HIQB 6HIQC 6HIQD 6HIQE 6HISA 6HISB 6HISC 6HISD 6HISE 6HJ3A 6HJ3B 6HJ3C
6HJ3D 6HJ3E 6HJAA 6HJAB 6HJAC 6HJAD 6HJAE 6HJBA 6HJBB 6HJBC 6HJBD 6HJBE
6HJIA 6HJIB 6HJIC 6HJID 6HJIE 6HJXA 6HJXB 6HJXC 6HJXD 6HJXE 6HJYA 6HJYB
6HJYC 6HJYD 6HJYE 6HJZA 6HJZB 6HJZC 6HJZD 6HJZE 6HK0A 6HK0B 6HK0C 6HK0D
6HK0E 6HLLA 6HLOA 6HLPA 6HPPA 6HPPB 6HPPC 6HPPD 6HPPE 6HQBA 6HQBB 6HQBF
6HQBI 6HQBJ 6HQBK 6HQBL 6HQBM 6HRAA 6HRAB 6HRAC 6HRAD 6HRBA 6HRBB 6HRBC
6HRBD 6HRCA 6HRCB 6HU9A 6HU9B 6HU9C 6HU9C 6HU9D 6HU9E 6HU9E 6HU9G 6HU9H
6HU9H 6HU9I 6HU9I 6HU9J 6HU9K 6HU9L 6HU9M 6HU9N 6HU9N 6HU9O 6HU9O 6HU9P
6HU9Q 6HU9S 6HU9S 6HU9T 6HU9T 6HU9U 6HU9U 6HU9W 6HU9X 6HUGA 6HUGB 6HUGC
6HUGD 6HUGE 6HUJA 6HUJB 6HUJC 6HUJD 6HUJE 6HUKA 6HUKB 6HUKC 6HUKD 6HUKE
6HUMA 6HUMB 6HUMC 6HUMD 6HUME 6HUMF 6HUMG 6HUML 6HUMP 6HUMQ 6HUOA 6HUOB
6HUOC 6HUOD 6HUOE 6HUPA 6HUPB 6HUPC 6HUPD 6HUPE 6HWHA 6HWHB 6HWHB 6HWHC
6HWHD 6HWHG 6HWHH 6HWHK 6HWHL 6HWHM 6HWHN 6HWHP 6HWHQ 6HWHR 6HWHS 6HWHV
6HWHW 6HWHX 6HWHY 6HWHZ 6HXBA 6HY5A 6HY5B 6HY9A 6HY9B 6HY9C 6HY9D 6HY9E
6HYAA 6HYAB 6HYAC 6HYAD 6HYAE 6HYRA 6HYRB 6HYRC 6HYRD 6HYRE 6HYVA 6HYVB
6HYVC 6HYVD 6HYVE 6HYWA 6HYWB 6HYWC 6HYWD 6HYWE 6HYXA 6HYXB 6HYXC 6HYXD
6HYXE 6HYZA 6HYZB 6HYZC 6HYZD 6HYZE 6HZ0A 6HZ0B 6HZ0C 6HZ0D 6HZ0E 6HZ1A
6HZ1B 6HZ1C 6HZ1D 6HZ1E 6HZ3A 6HZ3B 6HZ3C 6HZ3D 6HZ3E 6HZMA 6HZMB 6HZPA
6HZWA 6HZWB 6HZWC 6HZWD 6HZWE 6I08A 6I08B 6I08C 6I08D 6I08E 6I0DA 6I0DH
6I0DJ 6I0DK 6I0DL 6I0DM 6I0DN 6I1PA 6I1PH 6I1PJ 6I1PK 6I1PL 6I1PM 6I1PN
6I1RA 6I1XA 6I1XB 6I1XC 6I1XD 6I1XE 6I1XF 6I1XG 6I1XH 6I1XI 6I1XJ 6I1XK
6I1XL 6I1XM 6I1XN 6I1XO 6I1YA 6I1YB 6I1YC 6I1YD 6I1YE 6I1YF 6I1YG 6I1YH
6I1YI 6I1YJ 6I1YK 6I1YL 6I1YM 6I1YN 6I1YO 6I1ZA 6I1ZB 6I2JA 6I53A 6I53B
6I53C 6I53D 6I53E 6I6BA 6I6HA 6I6JA 6I8WA 6I8WB 6I96A 6I97A 6I98A 6I9DA
6I9DB 6I9DC 6I9DD 6I9KA 6IAXA 6IAXB 6IAXC 6IAXD 6IBBA 6IBLA 6IC4A 6IC4B
6IC4C 6IC4D 6IC4E 6IC4F 6IC4G 6IC4H 6IDFA 6IDFB 6IDFC 6IDFD 6IDFE 6IDPA
6IDRA 6IDSA 6IEDA 6IGKA 6IGLA 6IGZ0 6IGZ1 6IGZ2 6IGZ3 6IGZ4 6IGZ5 6IGZ6
6IGZ7 6IGZ8 6IGZ9 6IGZA 6IGZB 6IGZF 6IGZG 6IGZH 6IGZI 6IGZJ 6IGZK 6IGZL
6IGZM 6IIAA 6IIAB 6IIUA 6IIVA 6IJJ1 6IJJ3 6IJJ4 6IJJ5 6IJJ6 6IJJA 6IJJB
6IJJF 6IJJI 6IJJJ 6IJJK 6IJJL 6IJZA 6IJZB 6IOKE 6IOKF 6IOKG 6IOLE 6IOLF
6IOLG 6IQLA 6IRAA 6IRAB 6IRAC 6IRAD 6IRFA 6IRFB 6IRFC 6IRFD 6IRGA 6IRGB
6IRGC 6IRGD 6IRHA 6IRHB 6IRHC 6IRHD 6IRSA 6IRSB 6IRTA 6IRTB 6IS6A 6IS6B

6ITCB 6ITCE 6ITCY 6ITHA 6IU3A 6IU3B 6IU4A 6IU4B 6IV0A 6IV0B 6IV0C 6IV0D

6IV0E 6IV1A 6IV1B 6IV1C 6IV1D 6IV1E 6IV2A 6IV2B 6IV2C 6IV2D 6IV2E 6IV3A

6IV3B 6IV3C 6IV3D 6IV3E 6IV4A 6IV4B 6IV4C 6IV4D 6IV4E 6IVAA 6IVAB 6IVAC

6IVAD 6IVJA 6IVJB 6IVJC 6IVJD 6IVJE 6IVKA 6IVKB 6IVKC 6IVKD 6IVKE 6IVLA

6IVLB 6IVLC 6IVLD 6IVLE 6IVMA 6IVMB 6IVMC 6IVMD 6IVME 6IVNA 6IVNB 6IVNC

6IVND 6IVNE 6IVOA 6IVOB 6IVOC 6IVOD 6IVOE 6IVPA 6IVPB 6IVPC 6IVPD 6IVPE

6IVQA 6IVQB 6IVQC 6IVQD 6IVQE 6IVRA 6IVRB 6IVRC 6IVRD 6IVRE 6IVWA 6IVWB

6IVWC 6IVWD 6IVWE 6IWWA 6IWWB 6IWWC 6IWWD 6IWWE 6IWWF 6IYCA 6IYCB 6IYCC

6IYCD 6IYCE 6IYUA 6IYUB 6IYUC 6IYXA 6IYXB 6IYXC 6IYZA 6IYZB 6IYZC 6IZ0A

6IZ0B 6IZ0C 6IZ1A 6IZ1B 6IZ1C 6IZ3A 6IZ3B 6IZ3C 6IZ4A 6IZ4B 6IZ4C 6IZ5A

6IZ5B 6IZ5C 6IZ6A 6IZ6B 6IZ6C 6IZFA 6IZFB 6IZFC 6J5I8 6J5IA 6J5IB 6J5IF

6J5II 6J5IK 6J5IK 6J5IL 6J5IM 6J5IN 6J5IO 6J5IP 6J5IQ 6J5IR 6J5IU 6J5J8

6J5JA 6J5JB 6J5JF 6J5JI 6J5JK 6J5JK 6J5JL 6J5JM 6J5JN 6J5JO 6J5JP 6J5JQ

6J5JR 6J5JU 6J5TC 6J5TF 6J5TG 6J5TL 6J5TO 6J8EA 6J8GA 6J8GB 6J8HA 6J8HB

6J8IA 6J8IB 6J8JA 6J8JB 6J8MA 6J8MB 6J8MC 6J8MD 6J8MG 6J8MI 6J8MJ 6J8MK

6J8ML 6J8MM 6J8MN 6J8MO 6J8MP 6J8MQ 6J8MT 6J8MV 6J8MW 6J8MX 6J8MY 6J8MZ

6JB1A 6JB1B 6JB1C 6JB1D 6JB3B 6JBHC 6JBHD 6JBJA 6JBJB 6JFYA 6JFYB 6JFYC

6JFYD 6JFZA 6JFZB 6JFZC 6JFZD 6JI1A 6JI1B 6JI1C 6JI1D 6JJUA 6JKCA 6JKCB

6JKCC 6JKCD 6JKDA 6JKDB 6JKDC 6JKDD 6JLFA 6JLFB 6JLFC 6JLFD 6JLFE 6JLJA

6JLJA 6JLJB 6JLJB 6JLJC 6JLJC 6JLJD 6JLJD 6JLJE 6JLJE 6JLJF 6JLJF 6JLJH

6JLJH 6JLJI 6JLJI 6JLJJ 6JLJJ 6JLJK 6JLJK 6JLJL 6JLJL 6JLJM 6JLJM 6JLJR

6JLJT 6JLJT 6JLJX 6JLJX 6JLJY 6JLJY 6JLJZ 6JLJZ 6JLKA 6JLKA 6JLKB 6JLKB

6JLKC 6JLKC 6JLKD 6JLKD 6JLKE 6JLKE 6JLKF 6JLKF 6JLKH 6JLKH 6JLKI 6JLKI

6JLKJ 6JLKJ 6JLKK 6JLKK 6JLKL 6JLKL 6JLKM 6JLKM 6JLKR 6JLKT 6JLKT 6JLKX

6JLKX 6JLKY 6JLKY 6JLKZ 6JLKZ 6JLLA 6JLLA 6JLLB 6JLLB 6JLLC 6JLLC 6JLLD

6JLLD 6JLLE 6JLLE 6JLLF 6JLLF 6JLLH 6JLLH 6JLLI 6JLLI 6JLLJ 6JLLJ 6JLLK

6JLLK 6JLLL 6JLLL 6JLLM 6JLLM 6JLLR 6JLLT 6JLLT 6JLLX 6JLLX 6JLLY 6JLLY

6JLLZ 6JLLZ 6JLMA 6JLMA 6JLMB 6JLMB 6JLMC 6JLMC 6JLMD 6JLMD 6JLME 6JLME

6JLMF 6JLMF 6JLMH 6JLMH 6JLMI 6JLMI 6JLMJ 6JLMJ 6JLMK 6JLMK 6JLML 6JLML

6JLMM 6JLMM 6JLMR 6JLMT 6JLMT 6JLMX 6JLMX 6JLMY 6JLMY 6JLMZ 6JLMZ 6JLNA

6JLNA 6JLNB 6JLNB 6JLNC 6JLNC 6JLND 6JLND 6JLNE 6JLNE 6JLNF 6JLNF 6JLNH

6JLNH 6JLNI 6JLNI 6JLNJ 6JLNJ 6JLNK 6JLNK 6JLNL 6JLNL 6JLNM 6JLNM 6JLNR

6JLNT 6JLNT 6JLNX 6JLNX 6JLNY 6JLNY 6JLNZ 6JLNZ 6JLOA 6JLOA 6JLOB 6JLOB

6JLOC 6JLOC 6JLOD 6JLOD 6JLOE 6JLOE 6JLOF 6JLOF 6JLOH 6JLOH 6JLOI 6JLOI

6JLOJ 6JLOJ 6JLOK 6JLOK 6JLOL 6JLOL 6JLOM 6JLOM 6JLOR 6JLOT 6JLOT 6JLOX

6JLOX 6JLOY 6JLOY 6JLOZ 6JLOZ 6JLPA 6JLPA 6JLPB 6JLPB 6JLPC 6JLPC 6JLPD

6JLPD 6JLPE 6JLPE 6JLPF 6JLPF 6JLPH 6JLPH 6JLPI 6JLPI 6JLPJ 6JLPJ 6JLPK

6JLPK 6JLPL 6JLPL 6JLPM 6JLPM 6JLPR 6JLPT 6JLPT 6JLPX 6JLPX 6JLPY 6JLPY

6JLPZ 6JLPZ 6JMQA 6JMQB 6JNFA 6JNFB 6JNFC 6JNFD 6JNFE 6JO51 6JO52 6JO53

6JO54 6JO55 6JO56 6JO57 6JO58 6JO59 6JO5A 6JO5B 6JO5F 6JO5G 6JO5I 6JO5J

6JO5K 6JO5L 6JO5Z 6JO61 6JO63 6JO64 6JO65 6JO66 6JO67 6JO68 6JO6A 6JO6B

6JO6F 6JO6G 6JO6I 6JO6J 6JO6K 6JO6L 6JODA 6JP5A 6JP5E 6JP8A 6JP8E 6JPAA

6JPAE 6JPBA 6JPBE 6JPFA 6JPFB 6JUHA 6JUHB 6JUHC 6JUHD 6JUWA 6JUWB 6JUWC

6JUWD 6JUWG 6JUWI 6JUWJ 6JUWK 6JUWL 6JUWM 6JUWN 6JUWO 6JUWP 6JUWQ 6JUWT

6JUWV 6JUWW 6JUWX 6JUWY 6JUWZ 6JXHA 6JXHB 6JXIA 6JXIB 6JXJA 6JXJB 6JXKA
6JXKB 6JXRA 6JXRB 6JXRD 6JXRE 6JXRF 6JXRG 6JXRM 6JXRN 6JY3A 6JY3B 6JY3C
6JY3D 6JY3G 6JY3I 6JY3J 6JY3K 6JY3L 6JY3M 6JY4A 6JY4B 6JY4C 6JY4D 6JY4G
6JY4I 6JY4J 6JY4K 6JY4L 6JY4M 6JY6A 6JY7A 6JY8A 6JY9A 6JYAA 6JYBA 6JYCA
6JYDA 6JYEA 6JYFA 6JZHA 6JZOA 6JZOB 6JZOC 6JZOD 6K1HE 6K1HF 6K1HY 6K1HZ
6K1QA 6K4JA 6K5AA 6K5AB 6K5DA 6K5DB 6K5FA 6K5FB 6K5IA 6K5IB 6K61A 6K61A
6K61B 6K61B 6K61F 6K61F 6K61I 6K61I 6K61J 6K61J 6K61K 6K61K 6K61L 6K61L
6K61M 6K61M 6K61X 6K61X 6K6IA 6K6IB 6K6IC 6K6JA 6K6JB 6K6JC 6K6KA 6K6KB
6K6KC 6K7GA 6K7GC 6K7HA 6K7HC 6K7IA 6K7IC 6K7JA 6K7JC 6K7KA 6K7KC 6K7LA
6K7LC 6K7MA 6K7MC 6K7NA 6K7NC 6K7XA 6K7XB 6K7XC 6K7XD 6K7XE 6K7XF 6K7XG
6K7XH 6K7YA 6K7YB 6K7YC 6K7YD 6K7YE 6K7YF 6K7YG 6K7YH 6KE5A 6KE5B 6KE5C
6KE5D 6KEBA 6KEBB 6KEBC 6KEBD 6KFFA 6KFFB 6KFFC 6KFFD 6KFFE 6KFFF 6KFFG
6KFFH 6KFGA 6KFGB 6KFGC 6KFGD 6KFGE 6KFGF 6KFGG 6KFGH 6KFHA 6KFHB 6KFHC
6KFHD 6KFHE 6KFHF 6KFHG 6KFHH 6KFQA 6KG7A 6KHIA 6KHIB 6KHIC 6KHID 6KHIE
6KHIF 6KHIG 6KHIL 6KHIP 6KHIQ 6KHJA 6KHJB 6KHJC 6KHJD 6KHJE 6KHJF 6KHJG
6KHJL 6KHJP 6KHJQ 6KI1A 6KJVA 6KK1A 6KK7A 6KKIA 6KKJB 6KKKA 6KKLA 6KKRA
6KKRB 6KKTA 6KKTB 6KKUA 6KKUB 6KLSA 6KLSB 6KLSC 6KLSD 6KLSE 6KLSF 6KLVA
6KLVB 6KLVC 6KLVD 6KLVE 6KLVF 6KO5A 6KOBA 6KOBB 6KOBC 6KOBD 6KOCA 6KOCB
6KOCC 6KOCD 6KOEA 6KOEB 6KOEC 6KOED 6KP6A 6KPCA 6KPFR 6KPGR 6KQIA 6KR8A
6KRZA 6KS0A 6KS1A 6KSPA 6KSPB 6KSPC 6KSPD 6KSSA 6KSSB 6KSSC 6KSSD 6KSWA
6KSWB 6KSWC 6KUWA 6KUWB 6KUXA 6KUYA 6KXWA 6KXWB 6KXWC 6KXWD 6KZMA 6KZMB
6KZMC 6KZMD 6KZOA 6KZPA 6L1XA 6L352 6L353 6L355 6L356 6L35A 6L35B 6L35F
6L35G 6L35H 6L35I 6L35J 6L35K 6L35L 6L35M 6L3HA 6L3HB 6L3TA 6L3TB 6L3TC
6L3TD 6L3TE 6L3TF 6L3UA 6L3UB 6L3UC 6L3UD 6L3UE 6L3UF 6L3VA 6L3VB 6L3VC
6L3VD 6L3VE 6L3VF 6L47A 6L47B 6L48A 6L48B 6L6FA 6L6FB 6L6FC 6L6FD 6L6IA
6L6IB 6L6IC 6L6NA 6L6NB 6L6NC 6L6PA 6L6PB 6L6PC 6L7OA 6L7OB 6L7OC 6L7OD
6L7OE 6L7OF 6L7OG 6L7OL 6L7OP 6L7OQ 6L7PA 6L7PB 6L7PC 6L7PD 6L7PE 6L7PF
6L7PG 6L7PL 6L7PP 6L7PQ 6L85A 6L85B 6L95A 6LARA 6LARB 6LARC 6LARD 6LARE
6LARG 6LARH 6LARI 6LBHA 6LBHB 6LCPA 6LCPB 6LCRA 6LCRB 6LEOA 6LEPA 6LFLA
6LFMR 6LFOR 6LGPA 6LGPB 6LGPC 6LGPD 6LGVA 6LGYA 6LGZA 6LH0A 6LH1A 6LI0A
6LI1A 6LI2A 6LI3R 6LI9A 6LI9B 6LI9C 6LI9D 6LIDA 6LIDB 6LIDC 6LIDD 6LKDA
6LKDB 6LKEA 6LKEB 6LKNA 6LKNC 6LLEA 6LLYA 6LM0A 6LM0B 6LM0C 6LM1A 6LM1B
6LM1C 6LMKR 6LMLR 6LMTA 6LMTB 6LMTC 6LMTD 6LMTE 6LMTF 6LMTG 6LMTH 6LMUA
6LMUB 6LMUC 6LMUD 6LMUE 6LMUF 6LMUG 6LMUH 6LMUI 6LMUJ 6LMUK 6LMVA 6LMVB
6LMVC 6LMVD 6LMVE 6LMVF 6LMVG 6LMVH 6LMVI 6LMWA 6LMWB 6LMWC 6LMWD 6LMWE
6LMWF 6LMWG 6LMWH 6LMXA 6LMXB 6LMXC 6LMXD 6LMXE 6LMXF 6LMXG 6LMXH 6LMXI
6LN2A 6LN5A 6LN6A 6LN7A 6LN8A 6LN9A 6LO8A 6LO8B 6LO8C 6LO8D 6LODA 6LODC
6LODD 6LODF 6LOEA 6LOEC 6LOED 6LOEF 6LOMA 6LOMB 6LOMC 6LOMD 6LOME 6LOMF
6LOMG 6LOMH 6LOMI 6LOMJ 6LPBR 6LPJA 6LPKA 6LPLA 6LQAB 6LQGA 6LQGB 6LQGC
6LQGD 6LQIA 6LR0U 6LR4A 6LR4B 6LR4C 6LR4D 6LRYA 6LSCA 6LSCB 6LTNA 6LTNB
6LTNC 6LTND 6LTNE 6LTNF 6LTNG 6LTOA 6LTOB 6LTOC 6LTOD 6LTOE 6LTOF 6LTOG
6LU1A 6LU1B 6LU1F 6LU1I 6LU1L 6LU1M 6LULA 6LULB 6LUMC 6LUMD 6LUME 6LUMG
6LUMH 6LUMI 6LUQA 6LVFA 6LVFB 6LW5A 6LY5A 6LY5A 6LY5B 6LY5B 6LY5C 6LY5D
6LY5E 6LY5F 6LY5F 6LY5G 6LY5H 6LY5H 6LY5I 6LY5J 6LY5J 6LY5K 6LY5L 6LY5M

6LY5N 6LY5O 6LY5P 6LY5Q 6LY5R 6LY5S 6LY5V 6LY5W 6LY5X 6LYGA 6LYGB 6LYGC
6LYGD 6LYGE 6LYGF 6LYGG 6LYGH 6LYPA 6LYPB 6LYPC 6LYPD 6LYPE 6LYPF 6LYPG
6LYUA 6LYYA 6LYYB 6LZ0A 6LZ0B 6M02A 6M02B 6M02C 6M02D 6M02E 6M02F 6M02G
6M04A 6M04B 6M04C 6M04D 6M04E 6M04F 6M0FA 6M0RA 6M0RC 6M0RD 6M0RE 6M0RF
6M0RG 6M0RH 6M0RI 6M0RJ 6M0RK 6M0RL 6M0RM 6M0RN 6M0RO 6M0SA 6M0SC 6M0SD
6M0SE 6M0SF 6M0SG 6M0SH 6M0SI 6M0SJ 6M0SK 6M0SL 6M0SM 6M0SN 6M0SO 6M0ZA
6M18A 6M18B 6M18C 6M18D 6M1DA 6M1DB 6M1DC 6M1DD 6M1HA 6M1IA 6M1YA 6M1YB
6M20A 6M20B 6M22A 6M22B 6M23A 6M23B 6M2LA 6M2RA 6M31A 6M31B 6M32A 6M32A
6M34A 6M34B 6M38A 6M3ZA 6M47A 6M49A 6M49B 6M4NA 6M4NC 6M4ND 6M4NE 6M4NG
6M4NH 6M4OA 6M4OB 6M4OE 6M66A 6M66B 6M66C 6M66D 6M66E 6M66F 6M66G 6M67A
6M67B 6M67C 6M67D 6M67E 6M67F 6M67G 6M68A 6M68B 6M68C 6M68D 6M68E 6M68F
6M68G 6M6ZA 6M6ZB 6M6ZC 6M6ZD 6M84A 6M84B 6M84C 6M84D 6M85A 6M85B 6M85C
6M85D 6M86A 6M86B 6M86C 6M86D 6M96B 6M96D 6M97A 6M97B 6M97C 6M98A 6M98B
6M98C 6M9TA 6MCTA 6MCTB 6MCTC 6MCTD 6MCTE 6ME2A 6ME3A 6ME4A 6ME5A 6ME6A
6ME7A 6ME8A 6ME9A 6MEMA 6MEMB 6MEMC 6MEMD 6MEME 6MEMF 6MEMG 6MEMH 6MEMI
6MEMJ 6MEMK 6MEML 6MEMM 6MEMO 6MEMR 6MEMS 6MEMT 6MEMU 6MEMV 6MEMW 6MEMX
6MF8A 6MG8A 6MGVA 6MGVB 6MGWA 6MGWB 6MH8A 6MHOA 6MHOB 6MHOC 6MHOD 6MHQG
6MHQH 6MHQI 6MHQJ 6MHQK 6MHQL 6MHSA 6MHSB 6MHSC 6MHSD 6MHUF 6MHUG 6MHVA
6MHVB 6MHVC 6MHVD 6MHWA 6MHWB 6MHWC 6MHWD 6MHXA 6MHXB 6MHXC 6MHXD 6MHYG
6MHYH 6MHYI 6MHYJ 6MHYK 6MHYL 6MHZF 6MHZG 6MI7C 6MI7F 6MI7G 6MI8F 6MI8G
6MIEA 6MITC 6MITF 6MITG 6MIXA 6MIXB 6MIXC 6MIXD 6MIZA 6MIZB 6MIZC 6MIZD
6MJ2A 6MJ2B 6MJ2C 6MJ2D 6MJPC 6MJPF 6MJPG 6MM9A 6MM9B 6MM9C 6MM9D 6MMAA
6MMAB 6MMAC 6MMAD 6MMBA 6MMBB 6MMBC 6MMBD 6MMGA 6MMGB 6MMGC 6MMGD 6MMHA
6MMHB 6MMHC 6MMHD 6MMIA 6MMIB 6MMIC 6MMID 6MMJA 6MMJB 6MMJC 6MMJD 6MMKA
6MMKB 6MMKC 6MMKD 6MMLA 6MMLB 6MMLC 6MMLD 6MMMA 6MMMB 6MMMC 6MMMD 6MMNA
6MMNB 6MMNC 6MMND 6MMPA 6MMPB 6MMPC 6MMPD 6MMRA 6MMRB 6MMRC 6MMRD 6MMSA
6MMSB 6MMSC 6MMSD 6MMTA 6MMTB 6MMTC 6MMTD 6MMUA 6MMUB 6MMUC 6MMUD 6MMWA
6MMWB 6MMWC 6MMWD 6MMXA 6MMXB 6MMXC 6MMXD 6MPWA 6MPWB 6MPWC 6MPWD 6MPWE
6MQ2A 6MQ2B 6MQ2C 6MQ2D 6MQ2E 6MQUA 6MQUB 6MQUC 6MQUD 6MQUE 6MRTA 6MRTB
6MRTC 6MRTD 6MRTE 6MRTF 6MRTG 6MRTH 6MRTI 6MRTJ 6MRTK 6MRTL 6MRUA 6MRUB
6MRUC 6MRUD 6MRUE 6MRUF 6MRUG 6MRUH 6MRUI 6MRUJ 6MRUK 6MRUL 6MRUM 6MRWA
6MRWB 6MRWC 6MRWD 6MRWE 6MRWF 6MRWG 6MRWH 6MRWI 6MRWJ 6MRWK 6MRWL 6MRWM
6MRWN 6MSMA 6MU1A 6MU2A 6MU2B 6MU2C 6MU2D 6MVVA 6MVVB 6MVVC 6MVVD 6MVWA
6MVWB 6MVWC 6MVWD 6MVXA 6MVXB 6MVXC 6MVXD 6MVYA 6MVYC 6MVYD 6MVYE 6MWAA
6MWAB 6MWAC 6MWAD 6MWBA 6MWBB 6MWBC 6MWBD 6MWDA 6MWDB 6MWDC 6MWDD 6MWGA
6MWGB 6MWGC 6MWGD 6MXTA 6MYOC 6MYOD 6MYPC 6MYPD 6MYQC 6MYQD 6MYRC 6MYRD
6MYSC 6MYSD 6MYTC 6MYTD 6MYUC 6MYUD 6N1GA 6N1GC 6N23A 6N23B 6N23C 6N23D
6N23E 6N24A 6N24B 6N24C 6N24D 6N24E 6N25A 6N25B 6N25C 6N25D 6N25E 6N26A
6N26B 6N26C 6N26D 6N26E 6N27A 6N27B 6N27C 6N27D 6N27E 6N28A 6N28B 6N28C
6N28D 6N28E 6N38D 6N38E 6N38F 6N3IA 6N3QA 6N3QB 6N3QC 6N3QD 6N40A 6N48A
6N4BR 6N4IA 6N4IB 6N4IC 6N4ID 6N4QA 6N4QB 6N4QC 6N4QD 6N4RA 6N4RB 6N4RC
6N4RD 6N51A 6N51B 6N52A 6N52B 6N7GA 6N7GB 6N7HA 6N7HB 6N7KA 6N7KB 6NBFR
6NBHR 6NBIR 6NBQA 6NBQB 6NBQC 6NBQD 6NBQE 6NBQF 6NBQG 6NBQL 6NBQP 6NBXA
6NBXB 6NBXC 6NBXD 6NBXE 6NBXF 6NBXG 6NBXL 6NBXP 6NBXQ 6NBYA 6NBYB 6NBYC

6NBYD 6NBYE 6NBYF 6NBYG 6NBYL 6NBYP 6NBYQ 6NC6A 6NC7A 6NC8A 6NC9A 6ND0A
6ND0B 6ND0C 6ND0D 6ND1A 6ND1B 6ND1C 6ND1D 6ND1E 6NF4A 6NF4B 6NF6A 6NF6B
6NFUC 6NFUD 6NFUE 6NFUF 6NFVC 6NFVD 6NFVE 6NFVF 6NHGC 6NHGD 6NHGE 6NHGG
6NHGJ 6NHGK 6NHHA 6NHHB 6NHHC 6NHHE 6NHHF 6NHHG 6NHWA 6NHWB 6NHWC 6NHYA
6NHYB 6NHYC 6NI3R 6NINA 6NINB 6NINC 6NINE 6NINF 6NING 6NIYR 6NKNA 6NKNB
6NKNC 6NKND 6NKNG 6NKNI 6NKNJ 6NKNK 6NKNL 6NKNM 6NKNN 6NKNO 6NKNP 6NKNQ
6NKNT 6NKNV 6NKNW 6NKNX 6NKNY 6NKNZ 6NLEA 6NMFA 6NMFB 6NMFC 6NMFD 6NMFG
6NMFI 6NMFJ 6NMFK 6NMFL 6NMFM 6NMFN 6NMFO 6NMFP 6NMFQ 6NMFT 6NMFV 6NMFW
6NMFX 6NMFY 6NMFZ 6NMPA 6NMPB 6NMPC 6NMPD 6NMPG 6NMPI 6NMPJ 6NMPK 6NMPL
6NMPM 6NMPN 6NMPO 6NMPP 6NMPQ 6NMPT 6NMPV 6NMPW 6NMPX 6NMPY 6NMPZ 6NP0A
6NP0B 6NP0C 6NP0D 6NP0E 6NPHA 6NPHB 6NPKA 6NPKB 6NPLA 6NPLB 6NQ0A 6NQ0B
6NQ1A 6NQ1B 6NQ2A 6NQ2B 6NQ7A 6NQ8A 6NQ9A 6NR2A 6NR2B 6NR2C 6NR2D 6NR3A
6NR3B 6NR3C 6NR3D 6NR4A 6NR4B 6NR4C 6NR4D 6NSJA 6NSJB 6NSJC 6NSJD 6NSJE
6NSJF 6NSKA 6NSKB 6NSKC 6NSKD 6NSKE 6NSKF 6NT3A 6NT4A 6NT5A 6NT5B 6NT6A
6NT6B 6NT7A 6NT7B 6NT8A 6NT8B 6NWDA 6NWDB 6NWEA 6NWRA 6NZIA 6NZIB 6NZIC
6NZWA 6NZWB 6NZWC 6NZWD 6NZWE 6NZWF 6NZZA 6NZZB 6NZZC 6NZZD 6NZZE 6NZZF
6O00A 6O00B 6O00C 6O00D 6O00E 6O00F 6O1NA 6O1NB 6O1NC 6O1ND 6O1PA 6O1PB
6O1PC 6O1PD 6O1UA 6O1UB 6O1UC 6O1UD 6O1VA 6O20A 6O20B 6O20C 6O20D 6O2PA
6O30A 6O30C 6O3CA 6O58I 6O58J 6O58K 6O58L 6O58M 6O58N 6O58O 6O58P 6O6AA
6O6AB 6O6AC 6O6AD 6O6RA 6O6RB 6O6RC 6O6RD 6O72A 6O72B 6O72C 6O72D 6O77A
6O77B 6O77C 6O77D 6O7TA 6O7TB 6O7TC 6O7TE 6O7TF 6O7TG 6O7TH 6O7TI 6O7TJ
6O7TK 6O7TL 6O7TM 6O7TN 6O7TO 6O7UA 6O7UB 6O7UC 6O7UE 6O7UF 6O7UG 6O7UH
6O7UI 6O7UJ 6O7UK 6O7UL 6O7UM 6O7UN 6O7UO 6O7VA 6O7VB 6O7VC 6O7VE 6O7VF
6O7VG 6O7VH 6O7VI 6O7VJ 6O7VK 6O7VL 6O7VM 6O7VN 6O7VO 6O7WA 6O7WB 6O7WC
6O7WE 6O7WG 6O7WH 6O7WI 6O7WJ 6O7WK 6O7WL 6O7WM 6O7WN 6O7WO 6O7XA 6O7XB
6O7XC 6O7XE 6O7XF 6O7XG 6O7XH 6O7XI 6O7XJ 6O7XK 6O7XL 6O7XM 6O7XN 6O7XO
6O84A 6O84B 6O9GA 6O9GB 6O9GC 6O9GD 6O9TA 6O9TB 6O9TC 6O9TD 6O9UA 6O9UB
6O9UC 6O9UD 6O9VA 6O9VB 6O9VC 6O9VD 6OB6A 6OB7A 6OBAA 6OCEA 6OCEB 6OEUA
6OEVA 6OFJA 6OFJB 6OFRA 6OH2A 6OH3A 6OH4A 6OHTA 6OHTB 6OHUA 6OHUB 6OIHC
6OIHD 6OIJR 6OIKR 6OKZC 6OKZD 6OL0C 6OL0D 6OL1C 6OL1D 6OL9A 6OLYA 6OLYB
6OLYE 6OLYF 6OMMR 6OO3A 6OO3B 6OO3C 6OO3D 6OO4A 6OO4B 6OO4C 6OO4D 6OO5A
6OO5B 6OO5C 6OO5D 6OO7A 6OO7B 6OO7C 6OO7D 6OOMA 6OOPA 6OOQA 6OQHA 6OQRA
6OQRI 6OQRJ 6OQRL 6OQRM 6OQRN 6OQRO 6OQRP 6OQRQ 6OQRR 6OQRS 6OQRX 6OQRY
6OQSA 6OQSI 6OQSJ 6OQSL 6OQSM 6OQSN 6OQSO 6OQSP 6OQSQ 6OQSR 6OQSS 6OQSX
6OQSY 6OQTA 6OQTI 6OQTJ 6OQTL 6OQTM 6OQTN 6OQTO 6OQTP 6OQTQ 6OQTR 6OQTS
6OQTX 6OQTY 6OQUA 6OQUI 6OQUJ 6OQUL 6OQUM 6OQUN 6OQUO 6OQUP 6OQUQ 6OQUR
6OQUS 6OQUX 6OQUY 6OQVA 6OQVI 6OQVJ 6OQVX 6OQWA 6OQWI 6OQWJ 6OQWL 6OQWM
6OQWN 6OQWO 6OQWP 6OQWQ 6OQWR 6OQWS 6OQWX 6OQWY 6OR2A 6OS0A 6OS1A 6OS2A
6OS9R 6OSAR 6OT0R 6OT2A 6OT2B 6OT2C 6OT2D 6OT5A 6OT5B 6OT5C 6OT5D 6OV2A
6OV3A 6OWSA 6OWSB 6OWSC 6OY3A 6OY3B 6OY9R 6OYAR 6OYHA 6OYHB 6OYZA 6OYZB
6OZ6A 6OZ6B 6P25A 6P25B 6P2JA 6P2JB 6P2PA 6P2PB 6P2RA 6P2RB 6P46A 6P46B
6P47A 6P47B 6P48A 6P48B 6P49A 6P49B 6P6IA 6P6IB 6P6JA 6P6JB 6P6WA 6P6WB
6P6WC 6P6WD 6P6XA 6P6XB 6P6XC 6P6XD 6P6YA 6P6YB 6P6YC 6P6YD 6P9XR 6P9YR
6PA0C 6PA0D 6PA0E 6PA0F 6PAMA 6PAMB 6PANA 6PANB 6PAOA 6PAOB 6PAQA 6PAQB

6PARA 6PARB 6PB0R 6PB1P 6PBEA 6PBEB 6PBEC 6PBED 6PBFA 6PBFB 6PBFC 6PBFD
6PBXA 6PBXC 6PBXE 6PBXG 6PBYA 6PBYC 6PBYE 6PBYG 6PE4A 6PE4B 6PE4E 6PE4F
6PE4G 6PE4H 6PE4I 6PE4J 6PE4K 6PE4L 6PE4M 6PE4N 6PE4O 6PE4P 6PE5A 6PE5B
6PE5E 6PE5F 6PE5G 6PE5H 6PE5I 6PE5J 6PE5K 6PE5L 6PE5M 6PE5N 6PE5O 6PE5P
6PELA 6PEQA 6PEQB 6PEQC 6PEQD 6PEQE 6PEQF 6PEQG 6PEQH 6PFYA 6PFYB 6PFYD
6PFYE 6PFYF 6PFYF 6PFYG 6PFYG 6PFYH 6PFYH 6PFYI 6PFYI 6PFYJ 6PFYJ 6PFYK
6PFYL 6PFYM 6PFYQ 6PFYR 6PFYS 6PFYT 6PFYU 6PFYV 6PFYW 6PFYX 6PFYY 6PFYZ
6PGIA 6PGKA 6PGKB 6PGKD 6PGKE 6PGKF 6PGKF 6PGKG 6PGKG 6PGKH 6PGKH 6PGKI
6PGKI 6PGKJ 6PGKJ 6PGKK 6PGKL 6PGKM 6PGKQ 6PGKR 6PGKS 6PGKT 6PGKU 6PGKV
6PGKW 6PGKX 6PGKY 6PGKZ 6PGSA 6PH7A 6PH7C 6PISA 6PISB 6PJ4A 6PJ5A 6PJ7A
6PJ8A 6PJ9A 6PJAA 6PJPA 6PJQA 6PJRA 6PJUA 6PKVA 6PKVB 6PKVC 6PKVD 6PKWA
6PKWB 6PKWC 6PKWD 6PKXA 6PKXB 6PKXC 6PKXD 6PL5A 6PL5B 6PL6A 6PL6B 6PLOA
6PLOB 6PLOC 6PLOD 6PLOE 6PLPA 6PLPB 6PLPC 6PLPD 6PLPE 6PLQA 6PLQB 6PLQC
6PLQD 6PLQE 6PLRA 6PLRB 6PLRC 6PLRD 6PLRE 6PLSA 6PLSB 6PLSC 6PLSD 6PLSE
6PLTA 6PLTB 6PLTC 6PLTD 6PLTE 6PLUA 6PLUB 6PLUC 6PLUD 6PLUE 6PLVA 6PLVB
6PLVC 6PLVD 6PLVE 6PLWA 6PLWB 6PLWC 6PLWD 6PLWE 6PLXA 6PLXB 6PLXC 6PLXD
6PLXE 6PLYA 6PLYB 6PLYC 6PLYD 6PLYE 6PLZA 6PLZB 6PLZC 6PLZD 6PLZE 6PM0A
6PM0B 6PM0C 6PM0D 6PM0E 6PM1A 6PM1B 6PM1C 6PM1D 6PM1E 6PM2A 6PM2B 6PM2C
6PM2D 6PM2E 6PM3A 6PM3B 6PM3C 6PM3D 6PM3E 6PM4A 6PM4B 6PM4C 6PM4D 6PM4E
6PM5A 6PM5B 6PM5C 6PM5D 6PM5E 6PM6A 6PM6B 6PM6C 6PM6D 6PM6E 6PNJA 6PNJB
6PNJF 6PNJG 6PNJH 6PNJI 6PNJJ 6PNJK 6PNJL 6PNJM 6PNJQ 6PNJR 6PNJS 6PNJT
6PNJU 6PNJV 6PNJW 6PNJX 6POJA 6PQOA 6PQOB 6PQOC 6PQOD 6PQPA 6PQPB 6PQPC
6PQPD 6PQQA 6PQQB 6PQQC 6PQQD 6PQVA 6PQVI 6PQVJ 6PQVL 6PQVM 6PQVN 6PQVO
6PQVP 6PQVQ 6PQVR 6PQVS 6PQVX 6PQVY 6PRCH 6PRCL 6PRCM 6PRNA 6PRNB 6PRNC
6PRZA 6PS0A 6PS1A 6PS2A 6PS3A 6PS4A 6PS5A 6PS6A 6PS7A 6PS8A 6PSXA 6PSXE
6PSYA 6PSYE 6PT0R 6PT2A 6PT3A 6PUOA 6PUOB 6PUOC 6PUOD 6PURA 6PURB 6PURC
6PURD 6PUSA 6PUSB 6PUSC 6PUSD 6PUUA 6PUUB 6PUUC 6PUUD 6PV7A 6PV7B 6PV7C
6PV7D 6PV7E 6PV8A 6PV8B 6PV8C 6PV8D 6PV8E 6PVLA 6PVLB 6PVLC 6PVLD 6PVMA
6PVMB 6PVMC 6PVMD 6PVNA 6PVNB 6PVNC 6PVND 6PVOA 6PVOB 6PVOC 6PVOD 6PVPA
6PVPB 6PVPC 6PVPD 6PVQA 6PVQB 6PVQC 6PVQD 6PW0A 6PW0B 6PW1A 6PW1B 6PW4A
6PW4B 6PW4C 6PW4D 6PW5A 6PW5B 6PW5C 6PW5D 6PWCR 6PWNA 6PWNB 6PWNC 6PWND
6PWNE 6PWNF 6PWNG 6PWOA 6PWOB 6PWOC 6PWOD 6PWOE 6PWOF 6PWOG 6PWPA 6PWPB
6PWPC 6PWPD 6PWPE 6PWPF 6PWPG 6PXDA 6PXDB 6PXDC 6PXDD 6PXDE 6PZ9C 6PZBG
6PZCH 6PZIG 6PZTA 6PZTB 6Q3AA 6Q4NA 6Q4NB 6Q4NC 6Q4OA 6Q4OB 6Q4OC 6Q4PA
6Q4PB 6Q4PC 6Q53A 6Q53B 6Q53D 6Q53E 6Q5EA 6Q81A 6Q8OA 6Q8OH 6Q8OJ 6Q8OK
6Q8OL 6Q8OM 6Q8ON 6Q8WA 6Q8WH 6Q8WJ 6Q8WK 6Q8WL 6Q8WM 6Q8WN 6Q8XA 6Q8XH
6Q8XJ 6Q8XK 6Q8XL 6Q8XM 6Q8XN 6QAMA 6QCYA 6QCYB 6QCYC 6QCYD 6QCZA 6QCZB
6QCZC 6QCZD 6QD0A 6QD0B 6QD0C 6QD0D 6QD1A 6QD1B 6QD1C 6QD1D 6QD2A 6QD2B
6QD2C 6QD2D 6QD3A 6QD3B 6QD3C 6QD3D 6QD4A 6QD4B 6QD4C 6QD4D 6QD5A 6QD5B
6QD5C 6QEEA 6QEXA 6QF5A 6QF5B 6QF5C 6QF5D 6QFAA 6QFAB 6QFAC 6QFAD 6QFAE
6QGXA 6QGYA 6QIMA 6QIMB 6QIMC 6QIMD 6QKCA 6QKCB 6QKCC 6QKCD 6QKCI 6QKCJ
6QKZA 6QKZB 6QKZC 6QKZD 6QKZI 6QKZJ 6QM4A 6QM4B 6QM5A 6QM5B 6QM6A 6QM6B
6QM9A 6QM9B 6QMAA 6QMAB 6QMBA 6QMBB 6QNOR 6QP6A 6QP6B 6QPBA 6QPBB 6QPCA
6QPCB 6QPH1 6QPH2 6QPH3 6QPH4 6QPHA 6QPHB 6QPHF 6QPHJ 6QQ5A 6QQ5B 6QQ6A

6QQ6B 6QSKA 6QSKE 6QTIA 6QTIB 6QU1A 6QUEA 6QUEB 6QUMN 6QUMO 6QUMP 6QUMQ
6QUMR 6QUMS 6QUMT 6QUMU 6QUMV 6QUMW 6QUMX 6QUMY 6QUMZ 6QUZA 6QUZB 6QV0A
6QV0B 6QV1A 6QV1B 6QV2A 6QV2B 6QV6A 6QV6B 6QVBA 6QVBB 6QVCA 6QVCB 6QVDA
6QVDB 6QVUA 6QVUB 6QXAA 6QXAB 6QZIA 6QZIB 6QZIC 6QZID 6QZJA 6QZJB 6QZJC
6QZJD 6R0WN 6R0WO 6R0WP 6R0WQ 6R0WR 6R0WS 6R0WT 6R0WU 6R0WV 6R0WW 6R0WX
6R0WY 6R0WZ 6R0YN 6R0YO 6R0YP 6R0YQ 6R0YR 6R0YS 6R0YT 6R0YU 6R0YV 6R0YW
6R0YX 6R0YY 6R0YZ 6R0ZN 6R0ZO 6R0ZP 6R0ZQ 6R0ZR 6R0ZS 6R0ZT 6R0ZU 6R0ZV
6R0ZW 6R0ZX 6R0ZY 6R0ZZ 6R10N 6R10O 6R10P 6R10Q 6R10R 6R10S 6R10T 6R10U
6R10V 6R10W 6R10X 6R10Y 6R10Z 6R1FA 6R2QB 6R3QA 6R4LA 6R4OA 6R65A 6R65B
6R69E 6R69F 6R69G 6R69H 6R69I 6R69J 6R6BD 6R6BE 6R6BF 6R6BG 6R6BH 6R6BI
6R6BJ 6R72A 6R72B 6R7DA 6R7DB 6R7DC 6R7LE 6R7LG 6R7LY 6R7RA 6R7RB 6R7RC
6R7XA 6R7XB 6R7YA 6R7YB 6R7ZA 6R7ZB 6R81A 6R81B 6RAGA 6RAGB 6RAHA 6RAHB
6RAIA 6RAIB 6RAJA 6RAJB 6RAKA 6RAKB 6RALA 6RALB 6RAMA 6RAMB 6RANA 6RANB
6RB2A 6RBGA 6RBGB 6RBGC 6RBGD 6RBGE 6RBGF 6RBGG 6RBGH 6RCKA 6RCKB 6RCKC
6RCPA 6RCPB 6RCPF 6RD3A 6RD3B 6RD3C 6RD40 6RD43 6RD45 6RD46 6RD48 6RD49
6RD4A 6RD4B 6RD4C 6RD4D 6RD4E 6RD4F 6RD4G 6RD4H 6RD4I 6RD4J 6RD4M 6RD70
6RD73 6RD75 6RD76 6RD78 6RD79 6RD7A 6RD7B 6RD7C 6RD7D 6RD7E 6RD7F 6RD7G
6RD7H 6RD7I 6RD7J 6RD7M 6RD80 6RD83 6RD85 6RD86 6RD88 6RD89 6RD8A 6RD8B
6RD8C 6RD8D 6RD8E 6RD8F 6RD8G 6RD8H 6RD8I 6RD8J 6RD8M 6RD90 6RD93 6RD95
6RD96 6RD98 6RD99 6RD9A 6RD9B 6RD9C 6RD9D 6RD9E 6RD9F 6RD9G 6RD9H 6RD9I
6RD9J 6RD9M 6RDA0 6RDA5 6RDA6 6RDA8 6RDA9 6RDAM 6RDBA 6RDBB 6RDBC 6RDBD
6RDBE 6RDBF 6RDBG 6RDBH 6RDBI 6RDBJ 6RDC0 6RDC3 6RDC5 6RDC6 6RDC8 6RDC9
6RDCA 6RDCB 6RDCC 6RDCD 6RDCE 6RDCF 6RDCG 6RDCH 6RDCI 6RDCJ 6RDCM 6RDD0
6RDD5 6RDD6 6RDD8 6RDD9 6RDDM 6RDEA 6RDEB 6RDEC 6RDED 6RDEE 6RDEF 6RDEG
6RDEH 6RDEI 6RDEJ 6RDF0 6RDF5 6RDF6 6RDF8 6RDF9 6RDFM 6RDGA 6RDGB 6RDGC
6RDGD 6RDGE 6RDGF 6RDGG 6RDGH 6RDGI 6RDGJ 6RDH0 6RDH3 6RDH5 6RDH6 6RDH8
6RDH9 6RDHA 6RDHB 6RDHC 6RDHD 6RDHE 6RDHF 6RDHG 6RDHH 6RDHI 6RDHJ 6RDHM
6RDI0 6RDI3 6RDI5 6RDI6 6RDI8 6RDI9 6RDIA 6RDIB 6RDIC 6RDID 6RDIE 6RDIF
6RDIG 6RDIH 6RDII 6RDIJ 6RDIM 6RDJA 6RDJB 6RDJC 6RDJD 6RDJE 6RDJF 6RDJG
6RDJH 6RDJI 6RDJJ 6RDK0 6RDK3 6RDK5 6RDK6 6RDK8 6RDK9 6RDKA 6RDKB 6RDKC
6RDKD 6RDKE 6RDKF 6RDKG 6RDKH 6RDKI 6RDKJ 6RDL0 6RDL3 6RDL5 6RDL6 6RDL8
6RDL9 6RDLA 6RDLB 6RDLC 6RDLD 6RDLE 6RDLF 6RDLG 6RDLH 6RDLI 6RDLJ 6RDMA
6RDMB 6RDMC 6RDMD 6RDME 6RDMF 6RDMG 6RDMH 6RDMI 6RDMJ 6RDN0 6RDN3 6RDN5
6RDN6 6RDN8 6RDN9 6RDNA 6RDNB 6RDNC 6RDND 6RDNE 6RDNF 6RDNG 6RDNH 6RDNI
6RDNJ 6RDNM 6RDO0 6RDO3 6RDO5 6RDO6 6RDO8 6RDO9 6RDOA 6RDOB 6RDOC 6RDOD
6RDOE 6RDOF 6RDOG 6RDOH 6RDOI 6RDOJ 6RDOM 6RDPA 6RDPB 6RDPC 6RDPD 6RDPE
6RDPF 6RDPG 6RDPH 6RDPI 6RDPJ 6RDQ0 6RDQ3 6RDQ5 6RDQ6 6RDQ8 6RDQ9 6RDQA
6RDQB 6RDQC 6RDQD 6RDQE 6RDQF 6RDQG 6RDQH 6RDQI 6RDQJ 6RDQM 6RDR0 6RDR3
6RDR5 6RDR6 6RDR8 6RDR9 6RDRA 6RDRB 6RDRC 6RDRD 6RDRE 6RDRF 6RDRG 6RDRH
6RDRI 6RDRJ 6RDRM 6RDSA 6RDSB 6RDSC 6RDSD 6RDSE 6RDSF 6RDSG 6RDSH 6RDSI
6RDSJ 6RDT0 6RDT3 6RDT5 6RDT6 6RDT8 6RDT9 6RDTA 6RDTB 6RDTC 6RDTD 6RDTE
6RDTF 6RDTG 6RDTH 6RDTI 6RDTJ 6RDTM 6RDU0 6RDU3 6RDU5 6RDU6 6RDU8 6RDU9
6RDUA 6RDUB 6RDUC 6RDUD 6RDUE 6RDUF 6RDUG 6RDUH 6RDUI 6RDUJ 6RDUM 6RDVA
6RDVB 6RDVC 6RDVD 6RDVE 6RDVF 6RDVG 6RDVH 6RDVI 6RDVJ 6RDW0 6RDW3 6RDW5

6RDW6 6RDW8 6RDW9 6RDWA 6RDWB 6RDWC 6RDWD 6RDWE 6RDWF 6RDWG 6RDWH 6RDWI
6RDWJ 6RDWM 6RDX0 6RDX3 6RDX5 6RDX6 6RDX8 6RDX9 6RDXA 6RDXB 6RDXC 6RDXD
6RDXE 6RDXF 6RDXG 6RDXH 6RDXI 6RDXJ 6RDXM 6RDYA 6RDYB 6RDYC 6RDYD 6RDYE
6RDYF 6RDYG 6RDYH 6RDYI 6RDYJ 6RDZ0 6RDZ3 6RDZ5 6RDZ6 6RDZ8 6RDZ9 6RDZA
6RDZB 6RDZC 6RDZD 6RDZE 6RDZF 6RDZG 6RDZH 6RDZI 6RDZJ 6RDZM 6RE00 6RE03
6RE05 6RE06 6RE08 6RE09 6RE0A 6RE0B 6RE0C 6RE0D 6RE0E 6RE0F 6RE0G 6RE0H
6RE0I 6RE0J 6RE0M 6RE1A 6RE1B 6RE1C 6RE1D 6RE1E 6RE1F 6RE1G 6RE1H 6RE1I
6RE1J 6RE20 6RE23 6RE25 6RE26 6RE28 6RE29 6RE2A 6RE2B 6RE2C 6RE2D 6RE2E
6RE2F 6RE2G 6RE2H 6RE2I 6RE2J 6RE2M 6RE30 6RE33 6RE35 6RE36 6RE38 6RE39
6RE3A 6RE3B 6RE3C 6RE3D 6RE3E 6RE3F 6RE3G 6RE3H 6RE3I 6RE3J 6RE3M 6RE4A
6RE4B 6RE4C 6RE4D 6RE4E 6RE4F 6RE4G 6RE4H 6RE4I 6RE4J 6RE50 6RE53 6RE55
6RE56 6RE58 6RE59 6RE5A 6RE5B 6RE5C 6RE5D 6RE5E 6RE5F 6RE5G 6RE5H 6RE5I
6RE5J 6RE5M 6RE60 6RE63 6RE65 6RE66 6RE68 6RE69 6RE6A 6RE6B 6RE6C 6RE6D
6RE6E 6RE6F 6RE6G 6RE6H 6RE6I 6RE6J 6RE6M 6RE7A 6RE7B 6RE7C 6RE7D 6RE7E
6RE7F 6RE7G 6RE7H 6RE7I 6RE7J 6RE80 6RE83 6RE85 6RE86 6RE88 6RE89 6RE8A
6RE8B 6RE8C 6RE8D 6RE8E 6RE8F 6RE8G 6RE8H 6RE8I 6RE8J 6RE8M 6RE90 6RE93
6RE95 6RE96 6RE98 6RE99 6RE9A 6RE9B 6RE9C 6RE9D 6RE9E 6RE9F 6RE9G 6RE9H
6RE9I 6RE9J 6RE9M 6REAA 6REAB 6REAC 6READ 6REAE 6REAF 6REAG 6REAH 6REAI
6REAJ 6REB0 6REB3 6REB5 6REB6 6REB8 6REB9 6REBA 6REBB 6REBC 6REBD 6REBE
6REBF 6REBG 6REBH 6REBI 6REBJ 6REBM 6REC0 6REC3 6REC5 6REC6 6REC8 6REC9
6RECA 6RECB 6RECC 6RECD 6RECE 6RECF 6RECG 6RECH 6RECI 6RECJ 6RECM 6REDA
6REDB 6REDC 6REDD 6REDE 6REDF 6REDG 6REDH 6REDI 6REDJ 6REE0 6REE3 6REE5
6REE6 6REE8 6REE9 6REEA 6REEB 6REEC 6REED 6REEE 6REEF 6REEG 6REEH 6REEI
6REEJ 6REEM 6REF0 6REF3 6REF5 6REF6 6REF8 6REF9 6REFA 6REFB 6REFC 6REFD
6REFE 6REFF 6REFG 6REFH 6REFI 6REFJ 6REFM 6REP0 6REP3 6REP5 6REP6 6REP8
6REP9 6REPA 6REPB 6REPC 6REPD 6REPE 6REPF 6REPG 6REPH 6REPI 6REPJ 6REPM
6RERA 6RERB 6RERC 6RERD 6RERE 6RERF 6RERG 6RERH 6RERI 6RERJ 6RES0 6RES3
6RES5 6RES6 6RES8 6RES9 6RESA 6RESB 6RESC 6RESD 6RESE 6RESF 6RESG 6RESH
6RESI 6RESJ 6RESM 6RET0 6RET3 6RET5 6RET6 6RET8 6RET9 6RETA 6RETB 6RETC
6RETD 6RETE 6RETF 6RETG 6RETH 6RETI 6RETJ 6RETM 6REUA 6REUB 6REUC 6REUD
6REUE 6REUF 6REUG 6REUH 6REUI 6REUJ 6REWA 6REWB 6REWC 6REWD 6REWE 6REXA
6REXB 6REXC 6REXD 6REXE 6REZA 6REZB 6REZC 6REZD 6REZE 6RF0A 6RF0B 6RF0C
6RF0D 6RF0E 6RF1A 6RF1B 6RF1C 6RF1D 6RF1E 6RF3A 6RF3B 6RF3C 6RF3D 6RF3E
6RF4A 6RF4B 6RF4C 6RF4D 6RF4E 6RF5A 6RF6A 6RF7A 6RF9A 6RFAA 6RFBA 6RFCA
6RFQ1 6RFQ2 6RFQ3 6RFQ4 6RFQ5 6RFQ6 6RFQA 6RFQB 6RFQD 6RFQE 6RFQG 6RFQI
6RFQJ 6RFQJ 6RFQL 6RFQN 6RFQS 6RFQW 6RFQX 6RFR1 6RFR2 6RFR3 6RFR4 6RFR5
6RFR6 6RFRA 6RFRB 6RFRC 6RFRD 6RFRE 6RFRG 6RFRI 6RFRJ 6RFRJ 6RFRL 6RFRN
6RFRS 6RFRW 6RFRX 6RFS1 6RFS2 6RFS3 6RFS4 6RFS5 6RFS6 6RFSA 6RFSB 6RFSC
6RFSD 6RFSE 6RFSG 6RFSI 6RFSJ 6RFSJ 6RFSL 6RFSN 6RFSS 6RFSW 6RFSX 6RHZ1
6RHZ2 6RHZ3 6RHZ4 6RHZA 6RHZB 6RHZF 6RHZJ 6RKBA 6RKOA 6RKOB 6RKOH 6RKOX
6RKPA 6RLDA 6RLDB 6RLDC 6RLDD 6RLDE 6RLDF 6RLDG 6RLEA 6RMGA 6RMKA 6RMKB
6RMKC 6RNJA 6RNJB 6RNJC 6RNKA 6ROHA 6ROHC 6ROIA 6ROIC 6ROJA 6ROJC 6RPHA
6RPHB 6RQFA 6RQFB 6RQFC 6RQFD 6RQFE 6RQFF 6RQFG 6RQFH 6RQFI 6RQFJ 6RQFK
6RQFL 6RQFM 6RQFN 6RQFO 6RQFP 6RQOA 6RQOB 6RQOC 6RQPA 6RQPB 6RQPC 6RTCA

6RTCB 6RTFA 6RTFB 6RUQA 6RV2A 6RV2B 6RV3A 6RV3B 6RV4A 6RV4B 6RVDA 6RVDB

6RW3A 6RW3B 6RX4A 6RX4B 6RX4C 6RX4D 6RYOA 6RYPA 6RZ4A 6RZ5A 6RZ6A 6RZ7A

6RZ8A 6RZ9A 6S0LA 6S0QA 6S3KA 6S3LG 6S3LH 6S3LI 6S3LJ 6S3LK 6S3QA 6S3QB

6S3QC 6S3RC 6S3RD 6S3RE 6S3RF 6S3RG 6S3RH 6S3RI 6S3RJ 6S3RK 6S3SE 6S3SF

6S3SG 6S3SH 6S3SI 6S3SJ 6S4MA 6S59A 6S59B 6S63A 6S6CA 6S7PA 6S7TA 6S7TB

6S7TC 6S7TD 6S7TE 6S7TF 6S7TG 6S7TH 6S7TI 6S7VA 6S8GF 6S8GG 6S8HF 6S8HG

6S8NC 6S8NF 6S8NG 6SB5A 6SB5B 6SB5C 6SB5D 6SB5E 6SB5F 6SB5G 6SB5H 6SB5I

6SB5J 6SB5K 6SB5L 6SB5M 6SB5N 6SB5O 6SB5P 6SE3B 6SE3D 6SEMA 6SEMD 6SGRA

6SGRB 6SGRC 6SGRF 6SGRG 6SGRH 6SGSA 6SGSB 6SGSC 6SGSF 6SGSG 6SGSH 6SGTA

6SGTB 6SGTC 6SGUA 6SGUB 6SGUC 6SGWA 6SGWB 6SGWC 6SGWD 6SGWE 6SGWG 6SGWH

6SGWI 6SGXA 6SGXB 6SGXC 6SGXG 6SGZD 6SGZE 6SGZH 6SGZI 6SH3A 6SH3B 6SH3C

6SH3D 6SH3E 6SH3F 6SH3G 6SH4A 6SH4B 6SH4C 6SH4D 6SH4E 6SH4F 6SH4G 6SHKA

6SL51 6SL52 6SL53 6SL54 6SL55 6SL56 6SL5A 6SL5B 6SL5F 6SL5G 6SL5H 6SL5I

6SL5J 6SL5K 6SL5L 6SL5O 6SLIG 6SLIJ 6SNHX 6SNIX 6SO5C 6SO5D 6SO5E 6SO5F

6SO8A 6SOAA 6SOCA 6SOHA 6SP2A 6SP2B 6SP2C 6SP2D 6SP2E 6SP2F 6SSIA 6SSIB

6SSPA 6SSPB 6SSPC 6SSPD 6SSPE 6SSRA 6SSRB 6SSRC 6SSSA 6SSSB 6SSUA 6SSUB

6SSUC 6SSWA 6SSWB 6SSWC 6SU3A 6SU3X 6SU4A 6SU4X 6SWRB 6SWRE 6SX5A 6SX5B

6SX5C 6SX5D 6SX7A 6SX7B 6SX7C 6SX7D 6SXCA 6SXCB 6SXCC 6SXCD 6SXEA 6SXEB

6SXEC 6SXED 6SXFA 6SXFC 6SXFD 6SXFE 6SXGA 6SXGB 6SXGC 6SXGD 6T0BA 6T0BB

6T0BC 6T0BC 6T0BD 6T0BE 6T0BE 6T0BG 6T0BH 6T0BH 6T0BI 6T0BI 6T0BJ 6T0BK

6T0BL 6T0BM 6T0BN 6T0BN 6T0BO 6T0BO 6T0BP 6T0BP 6T0BR 6T0BS 6T0BT 6T0BT

6T0BU 6T0BU 6T0BV 6T0BX 6T0BY 6T0BZ 6T15A 6T15B 6T15C 6T15C 6T15D 6T15E

6T15E 6T15G 6T15H 6T15H 6T15I 6T15I 6T15J 6T15K 6T15L 6T15M 6T15N 6T15O

6T15P 6T15S 6T15T 6T15U 6T1WA 6T1WB 6T1ZA 6T6VA 6T7SJ 6T7SK 6T7SL 6T9NA

6T9NB 6T9NC 6T9ND 6T9OA 6T9OB 6T9OC 6T9OD 6TBUA 6TD6A 6TDXO 6TDXP 6TDXQ

6TDXR 6TDXS 6TDXT 6TDXU 6TDXV 6TDXW 6TDXX 6TDYO 6TDYP 6TDYQ 6TDYR 6TDYS

6TDYT 6TDYU 6TDYV 6TDYW 6TDYX 6TDZO 6TDZP 6TDZQ 6TDZR 6TDZS 6TDZT 6TDZU

6TDZV 6TDZW 6TDZX 6TE0O 6TE0P 6TE0Q 6TE0R 6TE0S 6TE0T 6TE0U 6TE0V 6TE0W

6TE0X 6TEJA 6TEJB 6THAA 6TIRA 6TJVA 6TJVB 6TJVC 6TJVD 6TJVE 6TJVF 6TJVG

6TJVL 6TK1A 6TK2A 6TK3A 6TK4A 6TK5A 6TK6A 6TK7A 6TKOA 6TL2A 6TMHH 6TMHI

6TMHJ 6TMHK 6TMHL 6TMHM 6TMHN 6TMHO 6TMHP 6TMHQ 6TO7A 6TODA 6TOSA 6TOTA

6TP3A 6TP4A 6TP6A 6TPGA 6TPJA 6TPKA 6TPNA 6TQ4A 6TQ6A 6TQ7A 6TQ9A 6TQEA

6TQEB 6TQFA 6TQFB 6TQJA 6TQJB 6TQJC 6TQJD 6TQJE 6TQJF 6TQJG 6TQJH 6TQJI

6TQJJ 6TQJK 6TQJL 6TQJM 6TQJN 6TRAA 6TRAB 6TRAF 6TRAI 6TRAJ 6TRAK 6TRAL

6TRAM 6TRAX 6TRC0 6TRC1 6TRC2 6TRC6 6TRC7 6TRC8 6TRC9 6TRCA 6TRCA 6TRCB

6TRCB 6TRCF 6TRCF 6TRCI 6TRCI 6TRCJ 6TRCJ 6TRCK 6TRCK 6TRCL 6TRCL 6TRCM

6TRCM 6TRCX 6TRCX 6TRCY 6TRCZ 6TRD0 6TRD1 6TRD2 6TRD6 6TRD7 6TRD8 6TRD9

6TRDA 6TRDA 6TRDB 6TRDB 6TRDF 6TRDF 6TRDI 6TRDI 6TRDJ 6TRDJ 6TRDK 6TRDK

6TRDL 6TRDL 6TRDM 6TRDM 6TRDX 6TRDX 6TRDY 6TRDZ 6TT71 6TT72 6TT7K 6TT7N

6TT7O 6TT7P 6TT7Q 6TT7R 6TYIA 6TYIB 6TYIC 6TYID 6TYIE 6TYIY 6TYIZ 6TZKA

6U0OA 6U0OB 6U1NR 6U1SA 6U1SB 6U1SD 6U1SF 6U1SH 6U1SJ 6U1SL 6U1SN 6U5RA

6U5RC 6U5RE 6U5RG 6U6DA 6U6DC 6U6DE 6U6DG 6U84A 6U84B 6U86A 6U86B 6U86C

6U86D 6U88A 6U88B 6U88C 6U88D 6U8AA 6U8AB 6U8AC 6U8AD 6U8YA 6U8YA 6U8YB

6U8YB 6U8YC 6U8YC 6U8YD 6U8YD 6U8YE 6U8YE 6U8YG 6U8YG 6U8YH 6U8YH 6U8YM

6U8YM 6U8YX 6U8YX 6U9PA 6U9PB 6U9PC 6U9PD 6U9TA 6U9TB 6U9TC 6U9TD 6U9VA
6U9VB 6U9VC 6U9WA 6U9WB 6U9WC 6U9YA 6U9YB 6U9YC 6U9YD 6U9ZA 6U9ZB 6U9ZC
6U9ZD 6UBSA 6UBSB 6UBSC 6UBSD 6UBSE 6UBTA 6UBTB 6UBTC 6UBTD 6UBTE 6UCBA
6UCBB 6UCBC 6UCBD 6UCBE 6UCBF 6UCBG 6UCBH 6UD3A 6UD3B 6UD3C 6UD3D 6UD3E
6UD4A 6UD4B 6UD4C 6UD4D 6UD4E 6UD4F 6UD4G 6UD4H 6UD8A 6UD8B 6UD8C 6UD8D
6UD8E 6UD8F 6UD8G 6UD8H 6UFEA 6UFEC 6UFED 6UFEE 6UH3A 6UH3B 6UIVA 6UIVB
6UIVC 6UIVD 6UIVE 6UIVF 6UIVG 6UIVH 6UIVI 6UIVJ 6UIVK 6UIWA 6UIWB 6UIWC
6UIWD 6UIWE 6UIWF 6UIWG 6UIWH 6UIWI 6UIWJ 6UIWK 6UIXA 6UIXB 6UIXC 6UIXD
6UIXE 6UIXF 6UIXG 6UIXH 6UIXI 6UIXJ 6UIXK 6UJNA 6UJPA 6UJRA 6UJSA 6UJTA
6UJWA 6UKJA 6UKNA 6UMMA 6UMMB 6UMMC 6UMMD 6UMMF 6UMMG 6UMMH 6UMMI 6UO8A
6UO8B 6UO9A 6UO9B 6UOAA 6UOAB 6UOXA 6UP7R 6UQFA 6UQFB 6UQFC 6UQFD 6UQGA
6UQGB 6UQGC 6UQGD 6UQKA 6UQKB 6UQKC 6UQKD 6UR8A 6UR8B 6UR8C 6UR8D 6UR8E
6URTA 6URTB 6URTC 6URTD 6URTE 6URTF 6URTG 6UUSE 6UUSR 6UVRA 6UVRB 6UVRC
6UVRD 6UVRH 6UVRI 6UVSA 6UVSB 6UVSC 6UVSD 6UVSH 6UVSI 6UVTA 6UVTB 6UVTC
6UVTD 6UVTH 6UVTI 6UW4A 6UW4B 6UW4C 6UW4D 6UW6A 6UW6B 6UW6C 6UW6D 6UW8A
6UW8B 6UW8C 6UW8D 6UW9A 6UW9B 6UW9C 6UW9D 6UWFA 6UWLA 6UWMA 6UWMB 6UWMC
6UWMD 6UWZA 6UWZB 6UWZC 6UWZD 6UWZE 6UY0A 6UZ2A 6UZ2B 6UZ8A 6UZ8B 6UZ8C
6UZ8D 6UZAA 6UZAB 6UZAC 6UZAD 6UZBA 6UZBB 6UZBC 6UZBD 6UZBE 6UZBF 6UZBG
6UZDA 6UZDB 6UZDC 6UZDD 6UZDE 6UZDF 6UZDG 6UZEA 6UZEB 6UZEC 6UZED 6UZEE
6UZEF 6UZEG 6UZHA 6UZHB 6UZHC 6UZHD 6UZHE 6UZHF 6UZHG 6UZLA 6UZLB 6UZV0
6UZV1 6UZV2 6UZV6 6UZV7 6UZV8 6UZV9 6UZVA 6UZVA 6UZVB 6UZVB 6UZVF 6UZVF
6UZVH 6UZVI 6UZVI 6UZVJ 6UZVJ 6UZVK 6UZVK 6UZVL 6UZVL 6UZVM 6UZVM 6UZYA
6UZYB 6UZYC 6UZYD 6UZYE 6UZYF 6UZYG 6UZZA 6UZZC 6UZZE 6UZZG 6V00A 6V00C
6V00D 6V00F 6V00G 6V00I 6V00J 6V00L 6V01A 6V01C 6V01D 6V01F 6V01G 6V01I
6V01J 6V01L 6V03A 6V03B 6V03C 6V03D 6V03E 6V05A 6V05F 6V0BA 6V0BB 6V0BC
6V0BD 6V0BE 6V1QA 6V1QB 6V1XA 6V1XB 6V1XC 6V1XD 6V1YA 6V1YB 6V1YC 6V1YD
6V22A 6V22B 6V22C 6V22D 6V22E 6V22F 6V22G 6V22H 6V2JA 6V2JB 6V35A 6V35B
6V35C 6V35D 6V35E 6V35F 6V35G 6V35H 6V36A 6V36B 6V37A 6V37B 6V38A 6V38B
6V38C 6V38D 6V3CA 6V3CB 6V3FA 6V3GA 6V3GB 6V3GC 6V3GD 6V3HA 6V3IA 6V3IB
6V4AA 6V4AB 6V4AC 6V4AD 6V4AE 6V4DA 6V4JA 6V4JB 6V4KA 6V4LA 6V4SA 6V4SB
6V4SC 6V4SD 6V4SE 6V6DA 6V6DB 6V6DC 6V6DD 6V6DE 6V6DF 6V6DG 6V8GA 6V8GB
6V8GC 6V8QA 6V8YA 6V8YB 6V8YC 6V8YD 6V9SA 6V9VA 6V9VB 6V9VC 6V9VD 6V9WA
6V9WB 6V9WC 6V9WD 6V9XA 6V9XB 6V9XC 6V9XD 6V9YA 6V9YB 6V9YC 6V9YD 6V9ZA
6V9ZB 6VAIA 6VAIB 6VAIC 6VAID 6VAIE 6VAIF 6VAIG 6VAIH 6VAII 6VAIJ 6VAIK
6VAKA 6VAKB 6VAKC 6VAKD 6VAKE 6VAKF 6VAKG 6VAKH 6VAKI 6VAKJ 6VAKK 6VALA
6VALB 6VALC 6VALD 6VALE 6VALF 6VALG 6VALH 6VALI 6VALJ 6VALK 6VBGA 6VCBR
6VD7A 6VD7B 6VD7C 6VD7D 6VD7E 6VD7F 6VD7G 6VEJA 6VEJB 6VEJC 6VGCA 6VGCB
6VGCC 6VGIA 6VGIE 6VGIF 6VI4A 6VJ8A 6VJ9A 6VJAC 6VJAD 6VJMA 6VJMB 6VJYB
6VJZB 6VJZC 6VK0B 6VK0C 6VK0D 6VK1B 6VKSA 6VKSB 6VKSC 6VKTA 6VKTB 6VKTC
6VL4A 6VL4B 6VL4C 6VM0A 6VM0B 6VM0C 6VM0D 6VM0E 6VM1A 6VM1J 6VM1M 6VM1N
6VM1O 6VM1P 6VM1Q 6VM1R 6VM1S 6VM1T 6VM1U 6VM1V 6VM1W 6VM1X 6VM1Y 6VM1Z
6VM2A 6VM2B 6VM2C 6VM2D 6VM2E 6VM3A 6VM3B 6VM3C 6VM3D 6VM3E 6VM4A 6VM4I
6VM4J 6VM4M 6VM4N 6VM4O 6VM4P 6VM4Q 6VM4R 6VM4S 6VM4T 6VM4U 6VM4V 6VM4W
6VM4X 6VM4Y 6VM4Z 6VMBA 6VMBI 6VMBJ 6VMBM 6VMBN 6VMBO 6VMBP 6VMBQ 6VMBR

6VMBS 6VMBT 6VMBU 6VMBV 6VMBW 6VMBX 6VMBY 6VMBZ 6VMGA 6VMGJ 6VMGM 6VMGN

6VMGO 6VMGP 6VMGQ 6VMGR 6VMGS 6VMGT 6VMGU 6VMGV 6VMGW 6VMGX 6VMGY 6VMGZ

6VMSR 6VN7R 6VOFA 6VOFI 6VOFJ 6VOFM 6VOFN 6VOFO 6VOFP 6VOFQ 6VOFR 6VOFS

6VOFT 6VOFU 6VOFV 6VOFW 6VOFX 6VOFY 6VOFZ 6VOHA 6VOHI 6VOHJ 6VOHM 6VOHN

6VOHO 6VOHP 6VOHQ 6VOHR 6VOHS 6VOHT 6VOHU 6VOHV 6VOHW 6VOHX 6VOHY 6VOHZ

6VOJA 6VOJI 6VOJJ 6VOJM 6VOJN 6VOJO 6VOJP 6VOJQ 6VOJR 6VOJS 6VOJT 6VOJU

6VOJV 6VOJW 6VOJX 6VOJY 6VOJZ 6VOLA 6VOLI 6VOLJ 6VOLM 6VOLN 6VOLO 6VOLP

6VOLQ 6VOLR 6VOLS 6VOLT 6VOLU 6VOLV 6VOLW 6VOLX 6VOLY 6VOLZ 6VONA 6VONI

6VONJ 6VONM 6VONN 6VONO 6VONP 6VONQ 6VONR 6VONS 6VONT 6VONU 6VONV 6VONW

6VONX 6VONY 6VONZ 6VP0C 6VP0E 6VPV0 6VPV1 6VPV2 6VPV6 6VPV7 6VPV8 6VPV9

6VPVA 6VPVA 6VPVB 6VPVB 6VPVF 6VPVF 6VPVI 6VPVI 6VPVJ 6VPVJ 6VPVK 6VPVK

6VPVL 6VPVL 6VPVM 6VPVM 6VPVZ 6VQ8A 6VQ8B 6VQ8C 6VQ8E 6VQ8F 6VQ8G 6VQ8H

6VQ8I 6VQ8J 6VQ8K 6VQ8L 6VQ8M 6VQ8N 6VQ8O 6VQ8P 6VQCA 6VQCB 6VQCC 6VQCE

6VQCF 6VQCG 6VQCH 6VQCI 6VQCJ 6VQCK 6VQCL 6VQCM 6VQCN 6VQCO 6VQCP 6VQGA

6VQGB 6VQGC 6VQGE 6VQGF 6VQGG 6VQGH 6VQGI 6VQGJ 6VQGK 6VQGL 6VQGM 6VQGN

6VQGO 6VQGP 6VQHA 6VQHB 6VQHC 6VQHE 6VQHF 6VQHG 6VQHH 6VQHI 6VQHJ 6VQHK

6VQHL 6VQHM 6VQHN 6VQHO 6VQHP 6VQQA 6VQQB 6VQQC 6VQQD 6VQQE 6VQRA 6VQRB

6VQRC 6VQRD 6VQRE 6VQTA 6VQTB 6VQUA 6VQUB 6VRHA 6VRKA 6VRLA 6VRZA 6VS0A

6VS1A 6VS2A 6VTKA 6VTKB 6VTKC 6VTLA 6VTLB 6VTLC 6VUMA 6VUMB 6VWKA 6VWKI

6VWKJ 6VWKL 6VWKM 6VWKN 6VWKO 6VWKP 6VWKQ 6VWKR 6VWKS 6VWKX 6VWKY 6VWXA

6VWXD 6VWXE 6VWXF 6VX3A 6VX3B 6VX3C 6VX3D 6VX5A 6VX5B 6VX5C 6VX5D 6VX5E

6VX6A 6VX6B 6VX6C 6VX6D 6VX6E 6VX7A 6VX7B 6VX7C 6VX7D 6VX7E 6VX8A 6VX8B

6VX8C 6VX8D 6VX8E 6VX9A 6VX9B 6VX9C 6VX9D 6VX9E 6VXFA 6VXFB 6VXHA 6VXHB

6VXIA 6VXIB 6VXJA 6VXJB 6VXMA 6VXMB 6VXMC 6VXMD 6VXME 6VXMF 6VXMG 6VXNA

6VXNB 6VXNC 6VXND 6VXNE 6VXNF 6VXNG 6VXOA 6VXOC 6VXOD 6VXOE 6VXPA 6VXPB

6VXPC 6VXPD 6VXPE 6VXPF 6VXPG 6VXZA 6VXZB 6VXZC 6VXZD 6VY0A 6VY0B 6VY0C

6VY0D 6VYHA 6VYIA 6VYIB 6VYKA 6VYKB 6VYKC 6VYKD 6VYKE 6VYKF 6VYKG 6VYLA

6VYLB 6VYLC 6VYLD 6VYLE 6VYLF 6VYLG 6VYMA 6VYMB 6VYMC 6VYMD 6VYME 6VYMF

6VYMG 6VZ1A 6VZ1B 6W0AC 6W0AD 6W0AE 6W0AF 6W0BC 6W0BD 6W0BE 6W0BF 6W0CC

6W0CD 6W0CE 6W0CF 6W0DC 6W0DD 6W0DE 6W0DF 6W0EC 6W0ED 6W0EE 6W0EF 6W0FC

6W0FD 6W0FE 6W0FF 6W0GC 6W0GD 6W0GE 6W0GF 6W0HC 6W0HD 6W0HE 6W0HF 6W0IC

6W0ID 6W0IE 6W0IF 6W0JC 6W0JD 6W0JE 6W0JF 6W1JA 6W1JB 6W1JC 6W1JD 6W1JE

6W1MA 6W1MB 6W1MC 6W1MD 6W1ME 6W1YA 6W1YB 6W1YC 6W1YD 6W1YE 6W25A 6W2BA

6W2CA 6W2XA 6W2XB 6W2YA 6W2YB 6W4SF 6W5RA 6W5SA 6W5TA 6W5UA 6W5VA 6W6OA

6W6OD 6W6OF 6W6OH 6W7BA 6W7BB 6W7CA 6W7CB 6W7DA 6W7DB 6W7EA 6W7EB 6W82A

6W82B 6W83A 6W83B 6W84A 6W84B 6W85A 6W85B 6W86A 6W86B 6W87A 6W87B 6W88A

6W88B 6W8AA 6W8AB 6W8CA 6W8CB 6W8FA 6W8FB 6W8NA 6W8NB 6W8OA 6W8OB 6W8PA

6W8PB 6W98A 6W9YA 6W9ZA 6W9ZB 6WA0A 6WA0B 6WA0C 6WADA 6WB8A 6WB8B 6WB8C

6WB8D 6WB91 6WB93 6WB94 6WB95 6WB96 6WBFA 6WBFB 6WBGA 6WBGB 6WBIA 6WBIB

6WBIC 6WBID 6WBIE 6WBIF 6WBIG 6WBKA 6WBKB 6WBKC 6WBKD 6WBKE 6WBKF 6WBKG

6WBLA 6WBLB 6WBLC 6WBLD 6WBLE 6WBLF 6WBLG 6WBMA 6WBMB 6WBNA 6WBNB 6WBVA

6WBXA 6WBYA 6WC9A 6WC9B 6WCAA 6WCAB 6WCBA 6WCBB 6WCCA 6WCCB 6WDNC 6WDND

6WDNE 6WDNF 6WDNG 6WDNH 6WDNI 6WDNJ 6WEJA 6WEJB 6WEJC 6WEJD 6WEKA 6WEKB

6WEKC 6WEKD 6WELA 6WELB 6WELC 6WELD 6WF2A 6WGTA 6WGTB 6WGTC 6WH4A 6WH4B

6WH4C 6WHAA 6WHCR 6WHGA 6WHGB 6WHGC 6WHGD 6WHRA 6WHRB 6WHRC 6WHRD 6WHSA
6WHSB 6WHSC 6WHSD 6WHTA 6WHTB 6WHTC 6WHTD 6WHUA 6WHUB 6WHUC 6WHUD 6WHVA
6WHVB 6WHVC 6WHVD 6WHWA 6WHWB 6WHWC 6WHWD 6WHXA 6WHXB 6WHXC 6WHXD 6WHYA
6WHYB 6WHYC 6WHYD 6WI0A 6WI0B 6WI0C 6WI0D 6WI1A 6WI1B 6WI1C 6WI1D 6WI9R
6WIKA 6WIVA 6WIVB 6WJ5A 6WJ5B 6WJ5C 6WJ5D 6WJ6A 6WJ6B 6WJ6C 6WJ6D 6WJ6E
6WJ6F 6WJ6H 6WJ6I 6WJ6K 6WJ6L 6WJ6M 6WJ6T 6WJ6X 6WJCA 6WK9A 6WKNA 6WKNB
6WKNC 6WKND 6WLBA 6WLBC 6WLVA 6WLVB 6WLW0 6WLW1 6WLW2 6WLW3 6WLW4 6WLW5
6WLW6 6WLW7 6WLW8 6WLW9 6WLWR 6WLWS 6WLWT 6WLWU 6WLWV 6WM0A 6WM0B 6WM20
6WM21 6WM22 6WM23 6WM24 6WM25 6WM26 6WM27 6WM28 6WM29 6WM2R 6WM2S 6WM2T
6WM2U 6WM2V 6WM40 6WM41 6WM42 6WM4R 6WM4S 6WM4T 6WM4U 6WM4V 6WM5A 6WM5C
6WMVA 6WMVC 6WNQA 6WNQI 6WNQJ 6WNQL 6WNQM 6WNQN 6WNQO 6WNQP 6WNQQ 6WNQR
6WNQS 6WNQX 6WNQY 6WNRA 6WNRI 6WNRJ 6WNRL 6WNRM 6WNRN 6WNRO 6WNRP 6WNRQ
6WNRR 6WNRS 6WNRX 6WNRY 6WP8A 6WP8B 6WP8C 6WPWR 6WQAA 6WQZA 6WQZB 6WQZC
6WR4A 6WR4B 6WR4C 6WTD8 6WTDJ 6WTDK 6WTDL 6WTDM 6WTDN 6WTDO 6WTDP 6WTDQ
6WTDR 6WTDS 6WTDT 6WTDU 6WTDX 6WTDZ 6WTIA 6WTIB 6WTIC 6WTID 6WTWA 6WTWB
6WTXC 6WTXD 6WU0B 6WU1A 6WU1B 6WU2A 6WU2B 6WU3A 6WU3C 6WU4A 6WU4B 6WU6C
6WU6D 6WU6G 6WU6H 6WU6K 6WU6L 6WUJB 6WULB 6WULC 6WUMB 6WUMC 6WUNB 6WUND
6WV3A 6WV4A 6WV5A 6WV6A 6WV7A 6WV9A 6WVAA 6WVDA 6WVEA 6WVFA 6WVGA 6WVHA
6WW2R 6WW5A 6WW5B 6WW7A 6WW7C 6WW7E 6WW7F 6WWZR 6WXRA 6WXRB 6WXUA 6WXUB
6WXUC 6WXUD 6WXVA 6WXVB 6WYJA 6WYKA 6WYLA 6WYLB 6WYLC 6WZBA 6WZBB 6WZBC
6WZGR 6X01A 6X01B 6X01C 6X0OA 6X12A 6X13A 6X14A 6X14B 6X14C 6X15A 6X15B
6X15C 6X16A 6X16B 6X16C 6X17A 6X17B 6X17C 6X18R 6X19R 6X1AR 6X2JA 6X2JB
6X2JC 6X2JD 6X2LA 6X2LB 6X2LC 6X2ZA 6X3EA 6X3EB 6X3EC 6X3FA 6X3FB 6X3FC
6X3SA 6X3SB 6X3SC 6X3SD 6X3SE 6X3TA 6X3TB 6X3TC 6X3TD 6X3TE 6X3UA 6X3UB
6X3UC 6X3UD 6X3UE 6X3VA 6X3VB 6X3VC 6X3VD 6X3VE 6X3WA 6X3WB 6X3WC 6X3WD
6X3WE 6X3XA 6X3XB 6X3XC 6X3XD 6X3XE 6X3ZA 6X3ZB 6X3ZC 6X3ZD 6X3ZE 6X40A
6X40B 6X40C 6X40D 6X40E 6X4SA 6X4SB 6X4SC 6X4SD 6X4SE 6X4SF 6X4SG 6X4SH
6X7IA 6X9HA 6X9HB 6X9HC 6X9ZA 6XBKR 6XBOA 6XCRA 6XCUA 6XE6A 6XEUA 6XEUD
6XEUG 6XEUJ 6XEVA 6XEVE 6XEVI 6XEVM 6XI0C 6XI0D 6XI0E 6XI0P 6XI0Q 6XI0R
6XISA 6XISB 6XISC 6XISD 6XITA 6XITB 6XITC 6XITD 6XIWA 6XJHA 6XJHB 6XJIA
6XJIB 6XJVA 6XJVB 6XJVC 6XJVD 6XJVE 6XJVF 6XJVG 6XJVH 6XJXA 6XJXB 6XJXC
6XJXD 6XJXE 6XJXF 6XJXG 6XJXH 6XKTC 6XKTD 6XKTE 6XKTP 6XKTQ 6XKTR 6XKUC
6XKUD 6XKUE 6XKUP 6XKUQ 6XKVC 6XKVD 6XKVE 6XKVP 6XKVQ 6XKVR 6XKWC 6XKWD
6XKWE 6XKWH 6XKWN 6XKWO 6XKWP 6XKWP 6XKWQ 6XKWR 6XKWY 6XKXC 6XKXD 6XKXE
6XKXN 6XKXO 6XKXP 6XKXP 6XKXQ 6XKXR 6XKYA 6XKZC 6XKZD 6XKZE 6XKZN 6XKZO
6XKZP 6XKZP 6XKZQ 6XKZR 6XL3A 6XL3C 6XL3D 6XLPA 6XMPA 6XMQA 6XMSA 6XMTA
6XMUA 6XMUB 6XOXR 6XPDA 6XPDB 6XPEA 6XPEB 6XPFA 6XPFB 6XROA 6XRPA 6XSRA
6XSRB 6XSRC 6XSRD 6XVFC 6XVFD 6XVFE 6XVFG 6XVFJ 6XVFN 6XVFO 6XVFQ 6XVFT
6XWMA 6XWMB 6XWNA 6XWOA 6XWPA 6XWQA 6XWQB 6XWQC 6XWRA 6XYNA 6XYNB 6XYTA
6XYTB 6XYTC 6XYTD 6XYTE 6Y11A 6Y11H 6Y11J 6Y11K 6Y11L 6Y11M 6Y11N 6Y1ZA
6Y59A 6Y59B 6Y59C 6Y59D 6Y59E 6Y5AA 6Y5AB 6Y5AC 6Y5AD 6Y5AE 6Y5BA 6Y5BB
6Y5BC 6Y5BD 6Y5BE 6Y5RA 6Y5RB 6Y5VA 6Y5VB 6Y791 6Y792 6Y793 6Y794 6Y795
6Y796 6Y79A 6Y79B 6Y79D 6Y79E 6Y79G 6Y79I 6Y79J 6Y79J 6Y79L 6Y79N 6Y79S
6Y79W 6Y79X 6Y7FA 6Y7VA 6Y86A 6Y90A 6Y90B 6Y92A 6Y92B 6Y97A 6Y97B 6Y9AA

| | | | | | | | | | | | |
|---|---|---|---|---|---|---|---|---|---|---|---|
| 6Y9AB | 6Y9BA | 6Y9BC | 6YAAA | 6YAC1 | 6YAC2 | 6YAC4 | 6YACA | 6YACB | 6YACF | 6YACG | 6YACH |
| 6YACI | 6YACJ | 6YACK | 6YACL | 6YB1A | 6YB1B | 6YB1C | 6YB1D | 6YB1E | 6YB1F | 6YB1G | 6YB1H |
| 6YBYA | 6YBZA | 6YBZB | 6YBZC | 6YBZD | 6YBZE | 6YC0A | 6YC0B | 6YC0C | 6YC0D | 6YC0E | 6YC1A |
| 6YC1B | 6YC1C | 6YC1D | 6YC1E | 6YC2A | 6YC2B | 6YC2C | 6YC2D | 6YC2E | 6YC3A | 6YC3B | 6YC3C |
| 6YC3D | 6YC3E | 6YC4A | 6YC4B | 6YC4C | 6YC4D | 6YC4E | 6YE4A | 6YE4B | 6YE4C | 6YE4D | 6YE4E |
| 6YEZ1 | 6YEZ2 | 6YEZ4 | 6YEZA | 6YEZB | 6YEZF | 6YEZG | 6YEZH | 6YEZI | 6YEZJ | 6YEZK | 6YEZL |
| 6YHFA | 6YHIA | 6YHOA | 6YHPA | 6YHXA | 6YJ4A | 6YJ4A | 6YJ4B | 6YJ4D | 6YJ4G | 6YJ4H | 6YJ4H |
| 6YJ4I | 6YJ4J | 6YJ4J | 6YJ4K | 6YJ4K | 6YJ4L | 6YJ4L | 6YJ4M | 6YJ4M | 6YJ4N | 6YJ4O | 6YJ4Y |
| 6YJ4Z | 6YKMA | 6YKMB | 6YKMC | 6YKMD | 6YKME | 6YKMF | 6YKMG | 6YKPA | 6YKPB | 6YKPC | 6YKPD |
| 6YKPE | 6YKPF | 6YKPG | 6YKRA | 6YKRB | 6YKRC | 6YKRD | 6YKRE | 6YKRF | 6YKRG | 6YMXA | 6YMXB |
| 6YMXC | 6YMXC | 6YMXD | 6YMXE | 6YMXE | 6YMXG | 6YMXH | 6YMXH | 6YMXI | 6YMXI | 6YMXK | 6YMXM |
| 6YMXN | 6YMXO | 6YMXP | 6YMXS | 6YMXT | 6YMXU | 6YMYA | 6YMYB | 6YMYC | 6YMYE | 6YMYG | 6YMYH |
| 6YMYI | 6YMYK | 6YMYM | 6YNWH | 6YNWI | 6YNWJ | 6YNWK | 6YNWL | 6YNWM | 6YNWN | 6YNWO | 6YNWP |
| 6YNWQ | 6YOFA | 6YOGA | 6YP7A | 6YP7A | 6YP7B | 6YP7B | 6YP7C | 6YP7C | 6YP7D | 6YP7D | 6YP7E |
| 6YP7E | 6YP7F | 6YP7F | 6YP7G | 6YP7G | 6YP7H | 6YP7H | 6YP7I | 6YP7I | 6YP7J | 6YP7J | 6YP7K |
| 6YP7K | 6YP7L | 6YP7L | 6YP7M | 6YP7M | 6YP7N | 6YP7N | 6YP7R | 6YP7R | 6YP7S | 6YP7S | 6YP7T |
| 6YP7T | 6YP7W | 6YP7W | 6YP7X | 6YP7X | 6YP7Y | 6YP7Y | 6YP7Z | 6YP7Z | 6YS8A | 6YS8B | 6YS8C |
| 6YS8D | 6YS8E | 6YS8F | 6YS8G | 6YSFA | 6YSFB | 6YSFC | 6YSFD | 6YSFE | 6YSFF | 6YSFG | 6YSLA |
| 6YSLB | 6YSLC | 6YSLD | 6YSLE | 6YSLF | 6YSLG | 6YSNA | 6YSNB | 6YSNC | 6YSND | 6YSOA | 6YSOB |
| 6YT2A | 6YT4A | 6YT4B | 6YT4C | 6YT4D | 6YT4E | 6YTKA | 6YTKB | 6YTKC | 6YTKD | 6YTKE | 6YTKF |
| 6YTKG | 6YTKH | 6YTKI | 6YTKJ | 6YTLA | 6YTLB | 6YTLC | 6YTLD | 6YTLE | 6YTLF | 6YTLG | 6YTLH |
| 6YTLI | 6YTLJ | 6YTLK | 6YTOA | 6YTOB | 6YTOC | 6YTOD | 6YTOE | 6YTOF | 6YTOG | 6YTOH | 6YTOI |
| 6YTOJ | 6YTQA | 6YTQB | 6YTQC | 6YTQD | 6YTQE | 6YTQF | 6YTQG | 6YTQH | 6YTQI | 6YTQJ | 6YTQK |
| 6YTVA | 6YTVB | 6YTVC | 6YTVD | 6YTVE | 6YTVF | 6YTVG | 6YTVH | 6YTVI | 6YTVJ | 6YTXA | 6YTXB |
| 6YTXC | 6YTXD | 6YTXE | 6YTXF | 6YTXG | 6YTXH | 6YTXI | 6YTXJ | 6YTXK | 6YU2A | 6YU2B | 6YU3A |
| 6YU4A | 6YU5A | 6YU6A | 6YU7A | 6YUEA | 6YUEB | 6YUPA | 6YUPC | 6YUPD | 6YUPE | 6YUZA | 6YUZC |
| 6YV1A | 6YX9A | 6YXDA | 6YXFA | 6YXGA | 6YXHA | 6YXR1 | 6YXR2 | 6YXR3 | 6YXR4 | 6YXRA | 6YXRB |
| 6YXRF | 6YXRJ | 6YZ0A | 6YZ2A | 6YZ2B | 6YZ2C | 6YZ2D | 6Z02H | 6Z02L | 6Z02M | 6Z0FA | 6Z0FB |
| 6Z0GA | 6Z0HA | 6Z0IA | 6Z10A | 6Z12A | 6Z12B | 6Z12C | 6Z16A | 6Z16A | 6Z16B | 6Z16B | 6Z16C |
| 6Z16C | 6Z16D | 6Z16D | 6Z16E | 6Z16E | 6Z16F | 6Z16F | 6Z16G | 6Z16G | 6Z1JH | 6Z1JL | 6Z1JM |
| 6Z1RK | 6Z1RL | 6Z1RM | 6Z1RN | 6Z1RO | 6Z1RP | 6Z1RQ | 6Z1RR | 6Z1UK | 6Z1UL | 6Z1UM | 6Z1UN |
| 6Z1UO | 6Z1UP | 6Z1UQ | 6Z1UR | 6Z27H | 6Z27L | 6Z27M | 6Z38A | 6Z3TA | 6Z3TB | 6Z3TC | 6Z3WA |
| 6Z3WC | 6Z3WD | 6Z3WE | 6Z3WF | 6Z3WG | 6Z3YA | 6Z3YB | 6Z3ZA | 6Z3ZB | 6Z5R0 | 6Z5R1 | 6Z5R2 |
| 6Z5R3 | 6Z5R4 | 6Z5R5 | 6Z5R6 | 6Z5R7 | 6Z5R8 | 6Z5R9 | 6Z5RA | 6Z5RB | 6Z5RC | 6Z5RD | 6Z5RE |
| 6Z5RF | 6Z5RG | 6Z5RH | 6Z5RI | 6Z5RJ | 6Z5RK | 6Z5RL | 6Z5RM | 6Z5RN | 6Z5RO | 6Z5RP | 6Z5RQ |
| 6Z5RR | 6Z5RS | 6Z5RT | 6Z5RU | 6Z5RV | 6Z5RX | 6Z5RY | 6Z5RZ | 6Z5S1 | 6Z5S2 | 6Z5S3 | 6Z5S4 |
| 6Z5S5 | 6Z5S6 | 6Z5SA | 6Z5SB | 6Z5SC | 6Z5SD | 6Z5SE | 6Z5SF | 6Z5SG | 6Z5SH | 6Z5SI | 6Z5SJ |
| 6Z5SK | 6Z5SL | 6Z5SM | 6Z5SN | 6Z5SO | 6Z5SP | 6Z5SQ | 6Z5SR | 6Z5SS | 6Z5ST | 6Z5SU | 6Z5SV |
| 6Z5SW | 6Z5SX | 6Z5SY | 6Z5SZ | 6Z5UA | 6Z5UB | 6Z5UE | 6Z5UG | 6Z70A | 6Z70B | 6Z71A | 6Z71B |
| 6Z8CA | 6Z8CB | 6Z8CC | 6Z8CD | 6ZA92 | 6ZA9K | 6ZA9N | 6ZA9O | 6ZA9P | 6ZA9Q | 6ZA9R | 6ZBBK |
| 6ZBBL | 6ZBBM | 6ZBBN | 6ZBBO | 6ZBBP | 6ZBBQ | 6ZBBR | 6ZBVA | 6ZDBA | 6ZDBB | 6ZDRA | 6ZDVA |
| 6ZFSC | 6ZFSD | 6ZFSE | 6ZFSG | 6ZFSJ | 6ZFTC | 6ZFTD | 6ZFTE | 6ZFTG | 6ZFTJ | 6ZFUC | 6ZFUD |
| 6ZFUE | 6ZFUG | 6ZFUJ | 6ZG3H | 6ZG3I | 6ZG4A | 6ZG7K | 6ZG7L | 6ZG7M | 6ZG7N | 6ZG7O | 6ZG7P |
| 6ZG7Q | 6ZG7R | 6ZG8K | 6ZG8L | 6ZG8M | 6ZG8N | 6ZG8O | 6ZG8P | 6ZG8Q | 6ZG8R | 6ZG9A | 6ZGBA |

6ZGBB 6ZGBC 6ZGDA 6ZGDB 6ZGDC 6ZGDD 6ZGDE 6ZGJA 6ZGJB 6ZGJC 6ZGJD 6ZGJE
6ZGKA 6ZGKB 6ZGKC 6ZGKD 6ZGKE 6ZGRA 6ZGSA 6ZGTA 6ZGUA 6ZHFA 6ZHGA 6ZHHA
6ZHVA 6ZHVB 6ZHWH 6ZHWL 6ZHWM 6ZI4H 6ZI4L 6ZI4M 6ZI5H 6ZI5L 6ZI5M 6ZI6H
6ZI6L 6ZI6M 6ZI9H 6ZI9L 6ZI9M 6ZIAH 6ZIAL 6ZIAM 6ZIDH 6ZIDL 6ZIDM 6ZIKK
6ZIKL 6ZIKM 6ZIKN 6ZIKO 6ZIKP 6ZIKQ 6ZIKR 6ZKAA 6ZKAH 6ZKAJ 6ZKAK 6ZKAL
6ZKAM 6ZKAM 6ZKAN 6ZKAN 6ZKAO 6ZKAP 6ZKAQ 6ZKAR 6ZKAU 6ZKAV 6ZKAV 6ZKAW
6ZKAW 6ZKAX 6ZKAY 6ZKAZ 6ZKBA 6ZKBH 6ZKBJ 6ZKBK 6ZKBL 6ZKBM 6ZKBM 6ZKBN
6ZKBN 6ZKBO 6ZKBP 6ZKBQ 6ZKBR 6ZKBU 6ZKBV 6ZKBV 6ZKBW 6ZKBW 6ZKBX 6ZKBY
6ZKBZ 6ZKCA 6ZKCH 6ZKCJ 6ZKCK 6ZKCL 6ZKCM 6ZKCM 6ZKCN 6ZKCN 6ZKCO 6ZKCP
6ZKCQ 6ZKCR 6ZKCU 6ZKCV 6ZKCV 6ZKCW 6ZKCW 6ZKCX 6ZKCY 6ZKCZ 6ZKDA 6ZKDH
6ZKDJ 6ZKDK 6ZKDL 6ZKDM 6ZKDM 6ZKDN 6ZKDN 6ZKDO 6ZKDP 6ZKDQ 6ZKDR 6ZKDU
6ZKDV 6ZKDV 6ZKDW 6ZKDW 6ZKDX 6ZKDY 6ZKDZ 6ZKEA 6ZKEH 6ZKEJ 6ZKEK 6ZKEL
6ZKEM 6ZKEM 6ZKEN 6ZKEN 6ZKEO 6ZKEP 6ZKEQ 6ZKER 6ZKEU 6ZKEV 6ZKEV 6ZKEW
6ZKEW 6ZKEX 6ZKEY 6ZKEZ 6ZKFA 6ZKFH 6ZKFJ 6ZKFK 6ZKFL 6ZKFM 6ZKFM 6ZKFN
6ZKFN 6ZKFO 6ZKFP 6ZKFQ 6ZKFR 6ZKFU 6ZKFV 6ZKFV 6ZKFW 6ZKFW 6ZKFX 6ZKFY
6ZKFZ 6ZKGA 6ZKGH 6ZKGJ 6ZKGK 6ZKGL 6ZKGM 6ZKGM 6ZKGN 6ZKGN 6ZKGO 6ZKGP
6ZKGQ 6ZKGR 6ZKGU 6ZKGV 6ZKGV 6ZKGW 6ZKGW 6ZKGX 6ZKGY 6ZKGZ 6ZKHA 6ZKHH
6ZKHJ 6ZKHK 6ZKHL 6ZKHM 6ZKHM 6ZKHN 6ZKHN 6ZKHO 6ZKHP 6ZKHQ 6ZKHR 6ZKHU
6ZKHV 6ZKHV 6ZKHW 6ZKHW 6ZKHX 6ZKHY 6ZKHZ 6ZKIA 6ZKIH 6ZKIJ 6ZKIK 6ZKIL
6ZKIM 6ZKIM 6ZKIN 6ZKIN 6ZKIO 6ZKIP 6ZKIQ 6ZKIR 6ZKIU 6ZKIV 6ZKIV 6ZKIW
6ZKIW 6ZKIX 6ZKIY 6ZKIZ 6ZKJA 6ZKJH 6ZKJJ 6ZKJK 6ZKJL 6ZKJM 6ZKJM 6ZKJN
6ZKJN 6ZKJO 6ZKJP 6ZKJQ 6ZKJR 6ZKJU 6ZKJV 6ZKJV 6ZKJW 6ZKJW 6ZKJX 6ZKJY
6ZKJZ 6ZKKA 6ZKKH 6ZKKJ 6ZKKK 6ZKKL 6ZKKM 6ZKKM 6ZKKN 6ZKKN 6ZKKO 6ZKKP
6ZKKQ 6ZKKR 6ZKKU 6ZKKV 6ZKKV 6ZKKW 6ZKKW 6ZKKX 6ZKKY 6ZKKZ 6ZKLA 6ZKLH
6ZKLJ 6ZKLK 6ZKLL 6ZKLM 6ZKLM 6ZKLN 6ZKLN 6ZKLO 6ZKLP 6ZKLQ 6ZKLR 6ZKLU
6ZKLV 6ZKLV 6ZKLW 6ZKLW 6ZKLX 6ZKLY 6ZKLZ 6ZKMA 6ZKMH 6ZKMJ 6ZKMK 6ZKML
6ZKMM 6ZKMM 6ZKMN 6ZKMN 6ZKMO 6ZKMP 6ZKMQ 6ZKMR 6ZKMV 6ZKMW 6ZKMW 6ZKMX
6ZKMY 6ZKMZ 6ZKNA 6ZKNH 6ZKNJ 6ZKNK 6ZKNL 6ZKNM 6ZKNM 6ZKNN 6ZKNN 6ZKNO
6ZKNP 6ZKNQ 6ZKNR 6ZKNV 6ZKNW 6ZKNW 6ZKNX 6ZKNY 6ZKNZ 6ZKOA 6ZKOH 6ZKOJ
6ZKOK 6ZKOL 6ZKOM 6ZKOM 6ZKON 6ZKON 6ZKOO 6ZKOP 6ZKOQ 6ZKOR 6ZKOU 6ZKOV
6ZKOV 6ZKOW 6ZKOW 6ZKOX 6ZKOY 6ZKOZ 6ZKPA 6ZKPH 6ZKPJ 6ZKPK 6ZKPL 6ZKPM
6ZKPM 6ZKPN 6ZKPN 6ZKPO 6ZKPP 6ZKPQ 6ZKPR 6ZKPU 6ZKPV 6ZKPV 6ZKPW 6ZKPW
6ZKPX 6ZKPY 6ZKPZ 6ZKQA 6ZKQH 6ZKQJ 6ZKQK 6ZKQL 6ZKQM 6ZKQM 6ZKQN 6ZKQN
6ZKQO 6ZKQP 6ZKQQ 6ZKQR 6ZKQU 6ZKQV 6ZKQV 6ZKQW 6ZKQW 6ZKQX 6ZKQY 6ZKQZ
6ZKRA 6ZKRH 6ZKRJ 6ZKRK 6ZKRL 6ZKRM 6ZKRM 6ZKRN 6ZKRN 6ZKRO 6ZKRP 6ZKRQ
6ZKRR 6ZKRU 6ZKRV 6ZKRV 6ZKRW 6ZKRW 6ZKRX 6ZKRY 6ZKRZ 6ZKSA 6ZKSH 6ZKSJ
6ZKSK 6ZKSL 6ZKSM 6ZKSM 6ZKSN 6ZKSN 6ZKSO 6ZKSP 6ZKSQ 6ZKSR 6ZKSU 6ZKSV
6ZKSV 6ZKSW 6ZKSW 6ZKSX 6ZKSY 6ZKSZ 6ZKTA 6ZKTH 6ZKTJ 6ZKTK 6ZKTL 6ZKTM
6ZKTM 6ZKTN 6ZKTN 6ZKTO 6ZKTP 6ZKTQ 6ZKTR 6ZKTU 6ZKTV 6ZKTV 6ZKTW 6ZKTW
6ZKTX 6ZKTY 6ZKTZ 6ZKUA 6ZKUH 6ZKUJ 6ZKUK 6ZKUL 6ZKUM 6ZKUM 6ZKUN 6ZKUN
6ZKUO 6ZKUP 6ZKUQ 6ZKUR 6ZKUU 6ZKUV 6ZKUW 6ZKUW 6ZKUX 6ZKUY 6ZKUZ 6ZKVA
6ZKVH 6ZKVJ 6ZKVK 6ZKVL 6ZKVM 6ZKVM 6ZKVN 6ZKVN 6ZKVO 6ZKVP 6ZKVQ 6ZKVR
6ZKVU 6ZKVV 6ZKVW 6ZKVW 6ZKVX 6ZKVY 6ZKVZ 6ZL4A 6ZL4B 6ZL4C 6ZLHA 6ZLHB
6ZLHC 6ZMQA 6ZO5A 6ZO5B 6ZO5C 6ZO6A 6ZO6B 6ZO6C 6ZO7A 6ZO7B 6ZO7C 6ZO8A

6ZO8B 6ZO8C 6ZO9A 6ZO9B 6ZO9C 6ZOAA 6ZOAB 6ZOAC 6ZOBA 6ZOBB 6ZOBC 6ZOCA
6ZOCB 6ZOCC 6ZODA 6ZODB 6ZODC 6ZOEA 6ZOEC 6ZOEE 6ZOFA 6ZOFB 6ZOFC 6ZOGA
6ZOGB 6ZOGC 6ZOHA 6ZOHB 6ZOHC 6ZOO1 6ZOO2 6ZOO4 6ZOOA 6ZOOB 6ZOOF 6ZOOG
6ZOOH 6ZOOI 6ZOOJ 6ZOOK 6ZOOL 6ZPLA 6ZPLB 6ZPO8 6ZPOA 6ZPOB 6ZPOE 6ZPOF
6ZPOG 6ZPOJ 6ZPOK 6ZPOK 6ZPOL 6ZPOM 6ZPON 6ZPOO 6ZPOP 6ZPOQ 6ZPOR 6ZQMK
6ZQML 6ZQMM 6ZQMN 6ZQMO 6ZQMP 6ZQMQ 6ZQMR 6ZQN8 6ZQNA 6ZQNB 6ZQNF 6ZQNG
6ZQNJ 6ZQNK 6ZQNK 6ZQNL 6ZQNM 6ZQNN 6ZQNO 6ZQNP 6ZQNQ 6ZQNR 6ZR2A 6ZR2A
6ZR2B 6ZR2C 6ZR2D 6ZR2F 6ZR2G 6ZR2H 6ZR2H 6ZR2I 6ZR2J 6ZR2J 6ZR2K 6ZR2K
6ZR2L 6ZR2L 6ZR2M 6ZR2M 6ZR2N 6ZR2Y 6ZR2Z 6ZREB 6ZREE 6ZREF 6ZTQA 6ZTQA
6ZTQB 6ZTQC 6ZTQD 6ZTQF 6ZTQG 6ZTQH 6ZTQH 6ZTQI 6ZTQJ 6ZTQJ 6ZTQK 6ZTQK
6ZTQL 6ZTQL 6ZTQM 6ZTQM 6ZTQN 6ZTQY 6ZTQZ 6ZXAA 6ZXAB 6ZXAC 6ZXAD 6ZXAE
6ZXAF 6ZXAG 6ZXAH 6ZXAI 6ZXAJ 6ZXAK 6ZXAL 6ZXAM 6ZXAN 6ZXRA 6ZXS1 6ZXS2
6ZXS4 6ZXSA 6ZXSB 6ZXSF 6ZXSG 6ZXSH 6ZXSI 6ZXSJ 6ZXSK 6ZXSL 6ZY2A 6ZY2D
6ZY2E 6ZY2H 6ZY2I 6ZY2J 6ZY2K 6ZY2L 6ZY3A 6ZY3D 6ZY3E 6ZY3H 6ZY3I 6ZY3J
6ZY3K 6ZY3L 6ZY4A 6ZY4E 6ZY4H 6ZY4I 6ZY4J 6ZY4L 6ZY9A 6ZY9E 6ZY9H 6ZY9I
6ZY9J 6ZY9K 6ZY9L 6ZYDA 6ZYDB 6ZYDC 6ZYDD 6ZYDE 6ZYDF 6ZYDG 6ZYEA 6ZYEB
6ZYEC 6ZYED 6ZYEE 6ZYEF 6ZYEG 6ZZX1 6ZZX2 6ZZX3 6ZZX4 6ZZX5 6ZZX6 6ZZX7
6ZZX8 6ZZX9 6ZZXA 6ZZXA 6ZZXB 6ZZXF 6ZZXG 6ZZXH 6ZZXI 6ZZXJ 6ZZXK 6ZZXL
6ZZXM 6ZZXO 6ZZY1 6ZZY2 6ZZY3 6ZZY4 6ZZY5 6ZZY6 6ZZY7 6ZZY8 6ZZY9 6ZZYA
6ZZYA 6ZZYB 6ZZYF 6ZZYG 6ZZYH 6ZZYI 6ZZYJ 6ZZYK 6ZZYL 6ZZYM 7A0WA 7A0WB
7A0XA 7A0XB 7A0YA 7A0YB 7A46A 7A46B 7A46C 7A46D 7A46E 7A46F 7A46G 7A4P1
7A4P3 7A4P4 7A4P5 7A4P6 7A4P7 7A4P8 7A4PA 7A4PA 7A4PB 7A4PF 7A4PG 7A4PI
7A4PJ 7A4PK 7A4PL 7A4PM 7A5VA 7A5VD 7A5VG 7A5VJ 7A5VM 7A65A 7A69A 7A6CA
7A6EA 7A6FA 7AA5A 7AA5B 7AA5C 7AA5D 7AAQA 7AARA 7ABWA 7AC6A 7ACIA 7AD3A
7AD3B 7ADIA 7ADIB 7ADIC 7ADID 7ADPA 7ADPC 7ADPE 7ADPF 7AFTA 7AFTB 7AFTC
7AFTD 7AFTG 7AHCA 7AHCB 7AHDA 7AHDB 7AHEA 7AHEB 7AHHA 7AHHB 7AHKA 7AHKB
7AHKC 7AHLA 7AHLB 7AHLC 7AHLD 7AHLE 7AHLF 7AHLG 7AINA 7AINB 7AIOA 7AIOB
7AIPA 7AIPB 7AIQA 7AIQB 7AIRA 7AIRB 7AJQA 7AJQB 7AJQC 7AJQD 7AJQE 7AJQG
7AK5B 7AK5C 7AK5D 7AK5F 7AK5G 7AK5H 7AK5H 7AK5I 7AK5J 7AK5J 7AK5K 7AK5K
7AK5L 7AK5L 7AK5M 7AK5M 7AK5N 7AK5Y 7AK5Z 7AK6A 7AK6A 7AK6B 7AK6C 7AK6D
7AK6F 7AK6G 7AK6H 7AK6H 7AK6I 7AK6J 7AK6J 7AK6K 7AK6K 7AK6L 7AK6L 7AK6M
7AK6M 7AK6N 7AK6Y 7AK6Z 7AKZA 7AKZC 7AKZD 7AQOC 7AQQA 7AQQA 7AQQB 7AQQD
7AQQF 7AQQH 7AQQJ 7AQQK 7AQQM 7AQQN 7AQQU 7AQQV 7AQQZ 7AQWC 7AQWG 7AQWJ
7AQWK 7AQWL 7AQWL 7AQWM 7AQWM 7AR7A 7AR7A 7AR7B 7AR7C 7AR7D 7AR7F 7AR7G
7AR7H 7AR7J 7AR7J 7AR7K 7AR7K 7AR7L 7AR7L 7AR7M 7AR7M 7AR7N 7AR7U 7AR7V
7AR7Z 7AR8A 7AR8A 7AR8B 7AR8C 7AR8D 7AR8F 7AR8G 7AR8H 7AR8J 7AR8J 7AR8K
7AR8L 7AR8L 7AR8M 7AR8M 7AR8N 7AR8U 7AR8V 7AR8Z 7AR9A 7AR9A 7AR9B 7AR9C
7AR9D 7AR9F 7AR9G 7AR9H 7AR9H 7AR9J 7AR9J 7AR9K 7AR9K 7AR9L 7AR9L 7AR9M
7AR9M 7AR9N 7AR9S 7AR9T 7AR9U 7AR9W 7AR9Y 7AR9Z 7ARBA 7ARBA 7ARBB 7ARBC
7ARBD 7ARBF 7ARBG 7ARBH 7ARBJ 7ARBJ 7ARBK 7ARBL 7ARBL 7ARBM 7ARBM 7ARBN
7ARBU 7ARBV 7ARBZ 7ARDA 7ARDA 7ARDB 7ARDC 7ARDD 7ARDF 7ARDG 7ARDH 7ARDH
7ARDJ 7ARDJ 7ARDK 7ARDK 7ARDL 7ARDL 7ARDM 7ARDM 7ARDN 7ARDS 7ARDT 7ARDU
7ARDW 7ARDY 7ARDZ 7ARHC 7ARHE 7ARIC 7ARIE 7ARJC 7ARJE 7ARKC 7ARKE 7ARLC
7ARLE 7ARMC 7ARME 7AROA 7ASYA 7AT7A 7ATBA 7ATEA 7ATEB 7ATEC 7ATED 7ATNA

7ATNB 7ATNC 7ATND 7AU3A 7AU3B 7AU3C 7AU3D 7AU6A 7AU6B 7AU6C 7AU6D 7AUER
7AVNA 7AVOA 7AVOB 7AVPA 7AWLA 7AWLB 7AWLC 7B00A 7B00B 7B03A 7B03B 7B03C
7B03D 7B03E 7B0KA 7B0OA 7B0OB 7B0PA 7B0PB 7B0QA 7B0RA 7B1KA 7B1KB 7B1LA
7B1LB 7B1NA 7B1NB 7B1UA 7B3JA 7B3KA 7B4LA 7B4LB 7B4MA 7B4MB 7B5CA 7B5CB
7B5DA 7B5DB 7B5EA 7B5EB 7B5PA 7B5PB 7B5PC 7B7OA 7B8PA 7B8PB 7B8PC 7B8QA
7B8QB 7B8QC 7B93A 7B93A 7B93B 7B93C 7B93D 7B93F 7B93G 7B93H 7B93H 7B93I
7B93J 7B93J 7B93K 7B93K 7B93L 7B93L 7B93M 7B93M 7B93N 7B93Y 7B93Z 7B9FB
7B9FD 7B9FX 7B9S1 7B9S2 7B9S3 7B9S5 7B9SA 7B9SB 7B9SC 7B9SD 7B9SF 7B9SG
7B9SI 7B9SJ 7B9SK 7B9SL 7B9SO 7B9SP 7B9SQ 7B9SS 7B9ST 7B9SV 7B9SW 7B9SX
7B9SZ 7BB3A 7BB3B 7BB6A 7BB7A 7BC6A 7BC7A 7BCWA 7BCWB 7BG4A 7BG4B 7BGHA
7BGI1 7BGI2 7BGI3 7BGI4 7BGI5 7BGI6 7BGI7 7BGI8 7BGI9 7BGIA 7BGIB 7BGIF
7BGIG 7BGII 7BGIJ 7BGIK 7BGIL 7BGYA 7BGYB 7BGYC 7BGYD 7BH1A 7BH1B 7BH1C
7BH1D 7BH2A 7BH2B 7BH2C 7BH2D 7BLX1 7BLX3 7BLX4 7BLX5 7BLX6 7BLX7 7BLX8
7BLXA 7BLXB 7BLXF 7BLXG 7BLXI 7BLXJ 7BLXK 7BLXL 7BLZ1 7BLZ2 7BLZ3 7BLZA
7BLZB 7BLZF 7BLZI 7BLZJ 7BLZK 7BLZL 7BLZM 7BLZO 7BMHA 7BMHB 7BNQA 7BP3A
7BP3B 7BPUA 7BR3A 7BSPA 7BSPC 7BSQA 7BSQC 7BSSA 7BSSC 7BSUA 7BSUC 7BSVA
7BSVC 7BSWA 7BSWC 7BT2A 7BTSA 7BU6A 7BU7A 7BVCA 7BVCB 7BVEA 7BVEB 7BVFA
7BVFB 7BVGA 7BVGB 7BVHA 7BVHB 7BVQA 7BW0R 7BW1A 7BWRA 7BWRB 7BX8A 7BX8B
7BXUA 7BXUB 7BXUC 7BXUD 7BYLA 7BYLC 7BYLE 7BYLG 7BYMA 7BYMC 7BYME 7BYMG
7BYNA 7BYNC 7BYNE 7BYNG 7BZ2R 7C2ER 7C2MA 7C2NA 7C520 7C521 7C522 7C523
7C524 7C525 7C526 7C527 7C528 7C529 7C52A 7C52B 7C52D 7C52E 7C52F 7C52G
7C52H 7C52I 7C52J 7C52K 7C52L 7C52M 7C52N 7C52O 7C52P 7C52Q 7C52R 7C52S
7C52T 7C52U 7C52V 7C52W 7C52X 7C52Y 7C52Z 7C61A 7C6AA 7C76A 7C76B 7C77A
7C77B 7C7QA 7C7QB 7C7SA 7C7SB 7C83A 7C9IA 7C9IB 7C9IC 7C9ID 7C9R0 7C9R1
7C9R2 7C9R3 7C9R4 7C9R5 7C9R6 7C9R7 7C9R8 7C9R9 7C9RA 7C9RB 7C9RD 7C9RE
7C9RF 7C9RG 7C9RH 7C9RI 7C9RJ 7C9RK 7C9RL 7C9RM 7C9RN 7C9RO 7C9RP 7C9RQ
7C9RR 7C9RS 7C9RT 7C9RU 7C9RV 7C9RW 7C9RX 7C9RY 7C9RZ 7CA3A 7CA3B 7CA5A
7CA5B 7CADA 7CADB 7CAEA 7CAEB 7CAFA 7CAFB 7CAGA 7CAGB 7CALA 7CALB 7CCSA
7CCSB 7CFFA 7CFFB 7CFGA 7CFGB 7CFMR 7CFNR 7CFSA 7CFSB 7CFSC 7CFTA 7CFTB
7CFTC 7CGEA 7CGED 7CGEG 7CGNA 7CGND 7CGNG 7CGNH 7CH0A 7CH0D 7CH0G 7CH1A
7CH1B 7CH6A 7CH6B 7CH7A 7CH7B 7CH8A 7CH8B 7CH8C 7CH8D 7CH8E 7CH8F 7CH8G
7CH8H 7CH9A 7CH9B 7CH9C 7CH9D 7CH9E 7CH9F 7CH9G 7CH9H 7CHAA 7CHAB 7CHAG
7CHAH 7CJ3A 7CJ3B 7CJIA 7CJIA 7CJIB 7CJIB 7CJIC 7CJIC 7CJID 7CJID 7CJIE
7CJIE 7CJIF 7CJIF 7CJIH 7CJIH 7CJII 7CJII 7CJIJ 7CJIJ 7CJIK 7CJIK 7CJIL
7CJIL 7CJIM 7CJIM 7CJIR 7CJIT 7CJIT 7CJIX 7CJIX 7CJIY 7CJIY 7CJIZ 7CJIZ
7CJJA 7CJJA 7CJJB 7CJJB 7CJJC 7CJJC 7CJJD 7CJJD 7CJJE 7CJJE 7CJJF 7CJJF
7CJJH 7CJJH 7CJJI 7CJJI 7CJJJ 7CJJJ 7CJJK 7CJJK 7CJJL 7CJJL 7CJJM 7CJJM
7CJJR 7CJJT 7CJJT 7CJJX 7CJJX 7CJJY 7CJJY 7CJJZ 7CJJZ 7CJSA 7CJSB 7CJSC
7CJSD 7CKOA 7CKOB 7CKRA 7CKRB 7CKWR 7CKXR 7CKYR 7CKZR 7CLJA 7CLJB 7CLVA
7CLVB 7CM3A 7CMHA 7CMHB 7CMIA 7CMIB 7CMUR 7CMVR 7CN0A 7CN0B 7CN0C 7CN0D
7CN1A 7CN1B 7CN1C 7CN1D 7COHA 7COHB 7COHC 7COHD 7COHG 7COHI 7COHJ 7COHK
7COHL 7COHM 7COHN 7COHO 7COHP 7COHQ 7COHT 7COHV 7COHW 7COHX 7COHY 7COHZ
7COUA 7COUA 7COUB 7COUB 7COUC 7COUC 7COUD 7COUD 7COUE 7COUE 7COUF 7COUF
7COUH 7COUH 7COUI 7COUI 7COUJ 7COUJ 7COUK 7COUK 7COUL 7COUL 7COUM 7COUM

7COUR 7COUT 7COUT 7COUX 7COUX 7COUY 7COUY 7COUZ 7COUZ 7CP5A 7CP5B 7CP5C
7CP5D 7CP5G 7CP5I 7CP5J 7CP5K 7CP5L 7CP5M 7CP5N 7CP5O 7CP5P 7CP5Q 7CP5T
7CP5V 7CP5W 7CP5X 7CP5Y 7CP5Z 7CP9A 7CP9B 7CP9C 7CP9D 7CP9E 7CP9F 7CP9G
7CP9H 7CP9I 7CP9J 7CQ5A 7CQ5B 7CQ5C 7CQ5D 7CQ6A 7CQ6B 7CQ6C 7CQ6D 7CQ7A
7CQ7B 7CQ7C 7CQ7D 7CQIA 7CQIC 7CQIE 7CQKA 7CQKC 7CQKE 7CQMA 7CR0A 7CR0B
7CR0C 7CR0D 7CR1A 7CR1B 7CR1C 7CR1D 7CR2A 7CR2B 7CR2C 7CR2D 7CR3A 7CR3B
7CR3D 7CR3G 7CR4A 7CR4B 7CR4D 7CR4F 7CR7A 7CR7B 7CR7D 7CR7G 7CRHR 7CRIA
7CRJA 7CRKA 7CRLA 7CRSA 7CRTA 7CRXA 7CRYA 7CRZA 7CU3A 7CUBA 7CUBB 7CUBC
7CUBD 7CUQA 7CUQB 7CUQC 7CUQD 7CUWA 7CUWB 7CUWC 7CUWD 7CVSA 7CVSB 7CVTA
7CVTB 7CX2R 7CX3R 7CX4R 7CXRA 7CXRB 7CYEA 7CYEB 7CYEC 7CYFA 7CYFB 7CYFC
7CYGA 7CYGB 7CYKA 7CYKB 7CYNB 7CYND 7CZ5R 7CZ9A 7CZ9B 7CZ9C 7CZBA 7CZBB
7CZBC 7CZBD 7CZLA 7CZLA 7CZLB 7CZLB 7CZLC 7CZLC 7CZLD 7CZLD 7CZLE 7CZLE
7CZLF 7CZLF 7CZLH 7CZLH 7CZLI 7CZLI 7CZLK 7CZLK 7CZLL 7CZLL 7CZLM 7CZLM
7CZLT 7CZLT 7CZLX 7CZLX 7CZLY 7CZLY 7CZLZ 7CZLZ 7D06A 7D06D 7D06G 7D06H
7D06I 7D06J 7D06K 7D06L 7D08A 7D08D 7D08G 7D08H 7D08I 7D08J 7D08K 7D08L
7D09A 7D09D 7D09G 7D09H 7D09I 7D09J 7D09K 7D09L 7D0AA 7D0AD 7D0AG 7D0AH
7D0AI 7D0AJ 7D0AK 7D0AL 7D0IB 7D0ID 7D0IF 7D0J1 7D0J2 7D0J3 7D0J4 7D0J5
7D0J6 7D0J7 7D0J8 7D0J9 7D0JA 7D0JA 7D0JB 7D0JF 7D0JG 7D0JH 7D0JI 7D0JJ
7D0JK 7D0JL 7D0JO 7D0JP 7D0JQ 7D0JR 7D0JS 7D0JT 7D0JU 7D10A 7D10B 7D14A
7D14B 7D1TA 7D1TA 7D1TB 7D1TB 7D1TC 7D1TC 7D1TD 7D1TD 7D1TE 7D1TE 7D1TF
7D1TF 7D1TH 7D1TH 7D1TI 7D1TI 7D1TJ 7D1TJ 7D1TK 7D1TK 7D1TL 7D1TL 7D1TM
7D1TM 7D1TR 7D1TR 7D1TT 7D1TT 7D1TX 7D1TX 7D1TY 7D1TY 7D1TZ 7D1TZ 7D1UA
7D1UA 7D1UB 7D1UB 7D1UC 7D1UC 7D1UD 7D1UD 7D1UE 7D1UE 7D1UF 7D1UF 7D1UH
7D1UH 7D1UI 7D1UI 7D1UJ 7D1UJ 7D1UK 7D1UK 7D1UL 7D1UL 7D1UM 7D1UM 7D1UR
7D1UR 7D1UT 7D1UT 7D1UX 7D1UX 7D1UY 7D1UY 7D1UZ 7D1UZ 7D2KA 7D3EA 7D3EB
7D3EC 7D3ED 7D3FA 7D3FB 7D3FC 7D3FD 7D3SR 7D3UA 7D3UC 7D3UD 7D3UE 7D3UF
7D3UG 7D4PA 7D4PB 7D4PC 7D4PD 7D4QA 7D4QB 7D4QC 7D4QD 7D5IA 7D5IB 7D5KA
7D5KB 7D5KC 7D5PA 7D5QA 7D5WA 7D5WB 7D5WC 7D5WD 7D5WG 7D5WI 7D5WJ 7D5WK
7D5WL 7D5WM 7D5WN 7D5WO 7D5WP 7D5WQ 7D5WT 7D5WV 7D5WW 7D5WX 7D5WY 7D5WZ
7D5XA 7D5XB 7D5XC 7D5XD 7D5XG 7D5XI 7D5XJ 7D5XK 7D5XL 7D5XM 7D5XN 7D5XO
7D5XP 7D5XQ 7D5XT 7D5XV 7D5XW 7D5XX 7D5XY 7D5XZ 7D60A 7D60B 7D60C 7D60D
7D60E 7D60F 7D60G 7D60H 7D60I 7D60J 7D60K 7D61A 7D61B 7D61C 7D61D 7D61E
7D61F 7D61G 7D61H 7D61I 7D61J 7D61K 7D65A 7D65B 7D65C 7D65D 7D65E 7D65F
7D65G 7D65H 7D65I 7D65J 7D65K 7D68R 7D6VC 7D6XC 7D76R 7D77R 7D7EA 7D7EB
7D7EC 7D7ED 7D7FA 7D7FB 7D7FC 7D7FD 7D7NA 7D7NB 7D7QA 7D7QB 7D7RA 7D7RB
7D8XA 7D8XB 7D8XC 7D8XD 7D8ZA 7D8ZB 7D90A 7D90B 7D91A 7D91B 7D91G 7D92A
7D92B 7D92G 7D99A 7D99B 7DA5A 7DA5B 7DB6D 7DCEB 7DCEX 7DCVA 7DD5A 7DD5B
7DD6A 7DD6B 7DD7A 7DD7B 7DDFA 7DDFB 7DDFE 7DDFG 7DDQT 7DDQT 7DDZA 7DEGA
7DEGB 7DEGC 7DEGD 7DEGE 7DEGF 7DF8A 7DFLR 7DFPA 7DFWA 7DFZA 7DGDA 7DGDB
7DGEA 7DGEB 7DGZ1 7DGZ2 7DGZ3 7DGZ4 7DGZ5 7DGZ6 7DGZ7 7DGZA 7DGZB 7DGZC
7DGZE 7DGZG 7DGZH 7DGZI 7DGZJ 7DGZJ 7DGZL 7DGZT 7DGZU 7DGZV 7DGZX 7DGZY
7DH01 7DH02 7DH03 7DH04 7DH05 7DH06 7DH07 7DH0A 7DH0B 7DH0C 7DH0E 7DH0H
7DH0I 7DH0J 7DH0J 7DH0L 7DH0T 7DH0V 7DH0X 7DH0Y 7DH5R 7DHIR 7DHRR 7DIIA
7DIIB 7DIXA 7DIXB 7DJ1A 7DJ1B 7DJ2A 7DJ2B 7DJCA 7DKZ1 7DKZ2 7DKZ3 7DKZ4

7DKZA 7DKZB 7DKZF 7DKZG 7DKZH 7DKZI 7DKZJ 7DKZK 7DKZL 7DL9A 7DLAA 7DLUA
7DLUB 7DLUC 7DLUD 7DLUE 7DLUF 7DLUG 7DNYA 7DNYB 7DNZA 7DNZB 7DQKA 7DQVA
7DQVB 7DR0A 7DR0B 7DR0F 7DR0I 7DR0J 7DR0K 7DR0L 7DR0M 7DR1A 7DR1B 7DR1F
7DR1I 7DR1J 7DR1L 7DR1M 7DRJA 7DRJB 7DRKA 7DRKB 7DRTB 7DRXA 7DRXB 7DSCA
7DSCB 7DSCC 7DSCD 7DSCE 7DSCF 7DSCG 7DSCH 7DSDA 7DSDB 7DSDC 7DSDD 7DSDE
7DSDF 7DSDG 7DSEA 7DSEB 7DSEC 7DSED 7DSEE 7DSEF 7DSEG 7DSHA 7DSHB 7DSIA
7DSIB 7DSKA 7DSKB 7DSLA 7DSLB 7DSNA 7DSNB 7DSQA 7DSQB 7DSWA 7DSWB 7DSXA
7DSXB 7DTCA 7DTDA 7DTTA 7DTTB 7DTUA 7DTUB 7DTVA 7DTVB 7DTWA 7DTWB 7DTYR
7DUQR 7DURR 7DUWA 7DUWB 7DV5U 7DVMA 7DVMB 7DVMC 7DW9R 7DWBA 7DWBB 7DWBC
7DWBD 7DWBE 7DWBF 7DWBG 7DWQA 7DWQB 7DWQF 7DWQJ 7DWQL 7DWQM 7DWQW 7DXBA
7DXBB 7DXBC 7DXBD 7DXCA 7DXCB 7DXCC 7DXCD 7DXDA 7DXDB 7DXDC 7DXDD 7DXEA
7DXEB 7DXEC 7DXED 7DXFA 7DXFB 7DXFC 7DXFD 7DXGA 7DXGB 7DXGC 7DXGD 7DXHA
7DXHB 7DXHB 7DXHC 7DXHC 7DXHD 7DXHE 7DXHF 7DXHH 7DXHI 7DXHK 7DXHL 7DXHM
7DXHT 7DXHX 7DXHY 7DXHZ 7DY8H 7DYRA 7DYRB 7DYRC 7DYRD 7DYRE 7DYRF 7DYRG
7DYRY 7DYRZ 7DYSA 7DYSB 7DYSC 7DYSD 7DZ71 7DZ72 7DZ73 7DZ74 7DZ75 7DZ76
7DZ77 7DZ78 7DZ79 7DZ7A 7DZ7A 7DZ7B 7DZ7F 7DZ7G 7DZ7H 7DZ7I 7DZ7J 7DZ7K
7DZ7L 7DZ7O 7DZ7U 7DZ7V 7DZ7W 7DZ7X 7DZ7Y 7DZ7Z 7DZ81 7DZ82 7DZ83 7DZ84
7DZ85 7DZ86 7DZ87 7DZ88 7DZ89 7DZ8A 7DZ8A 7DZ8B 7DZ8F 7DZ8G 7DZ8H 7DZ8I
7DZ8J 7DZ8K 7DZ8L 7DZ8O 7DZ8U 7DZ8V 7DZ8W 7DZ8X 7DZ8Y 7DZ8Z 7E0HX 7E0HY
7E0HZ 7E0IU 7E0IV 7E0IW 7E0JX 7E0JY 7E0JZ 7E0KU 7E0KV 7E0KW 7E14R 7E1AU
7E1VA 7E1VB 7E1VC 7E1VD 7E1VE 7E1VF 7E1VG 7E1VH 7E1VI 7E1VM 7E1VN 7E1VO
7E1VP 7E1VQ 7E1VR 7E1VS 7E1VT 7E1VU 7E1WA 7E1WB 7E1WC 7E1WD 7E1WE 7E1WF
7E1WG 7E1WH 7E1WI 7E1WM 7E1WN 7E1WO 7E1WP 7E1WQ 7E1WR 7E1WS 7E1WT 7E1WU
7E1XA 7E1XB 7E1XC 7E1XD 7E1XE 7E1XF 7E1XG 7E1XH 7E1XI 7E1XM 7E1XN 7E1XO
7E1XP 7E1XQ 7E1XR 7E1XS 7E1XT 7E1XU 7E1ZA 7E1ZB 7E1ZC 7E20A 7E20B 7E20C
7E21A 7E21B 7E21C 7E26A 7E26B 7E26C 7E26D 7E26E 7E27A 7E27B 7E27C 7E27D
7E27E 7E2GD 7E2HD 7E2HE 7E2ID 7E2XR 7E2YR 7E2ZR 7E32R 7E33R 7E4GA 7E4GB
7E4GC 7E4IA 7E4ID 7E4IE 7E4IF 7E4TA 7E4TB 7E4TC 7E4TD 7E6TA 7E6TB 7E6UA
7E6UC 7E6XA 7E6XB 7E6YA 7E6YB 7E6ZA 7E6ZB 7E70A 7E70B 7E71A 7E71B 7E7IA
7E7OA 7E7QA 7E7SA 7E7ZA 7E7ZB 7E7ZC 7E7ZD 7E84A 7E84B 7E84D 7E84G 7E87A
7E87B 7E87C 7E87D 7E87E 7E87F 7E87I 7E87J 7E8BA 7E8BB 7E8BC 7E8BD 7E8BI
7E8BJ 7E8BK 7E8BL 7E8EA 7E8EB 7E8EC 7E8ED 7E8EI 7E8EJ 7E8EK 7E8EL 7E8HA
7E8HB 7E8HC 7E8HD 7E8HI 7E8HJ 7E8HK 7E8HL 7E9GR 7E9GS 7E9HR 7E9HS 7E9SA
7EDAA 7EDAB 7EDAD 7EDAE 7EDAF 7EDAH 7EDAI 7EDAJ 7EDAK 7EDAL 7EDAM 7EDAR
7EDAT 7EDAX 7EDAY 7EDAZ 7EFLA 7EFLB 7EFMA 7EFMB 7EFNA 7EFNB 7EFOF 7EFOG
7EGKA 7EGKC 7EGKE 7EGLA 7EGLC 7EGLE 7EHLA 7EHLB 7EIBA 7EJ0R 7EJ1B 7EJ1D
7EJ1F 7EJ1H 7EJ2B 7EJ2D 7EJ2F 7EJ2H 7EJ8R 7EJAR 7EJKR 7EJXR 7EK1A 7EK1B
7EK1C 7EK1D 7EK1E 7EK2A 7EK2B 7EK2C 7EK2D 7EK2E 7EK3A 7EK3B 7EK3C 7EK3D
7EK3E 7EKIA 7EKIB 7EKIC 7EKID 7EKIE 7EKLA 7EKLB 7EKPA 7EKPB 7EKPC 7EKPD
7EKPE 7EKTA 7EKTB 7EKTC 7EKTD 7EKTE 7EN0A 7EN0B 7EN0C 7EO2A 7EO4A 7EOQA
7EOQB 7EOQC 7EOQD 7EORA 7EORB 7EORC 7EORD 7EOSA 7EOSB 7EOSC 7EOSD 7EOTA
7EOTB 7EOTC 7EOTD 7EOUA 7EOUB 7EOUC 7EOUD 7EPAA 7EPAB 7EPBA 7EPBB 7EPCA
7EPCB 7EPDA 7EPDB 7EPEA 7EPFA 7EPTR 7EPZA 7EPZB 7EQDA 7EQDB 7EQDD 7EQDE
7EQDH 7EQDL 7EQDM 7EQMA 7EQMB 7EQMC 7EQRA 7EQRB 7EQRC 7ESGA 7ET1A 7ET1B

7ETWA 7ETWB 7EU30 7EU39 7EU3A 7EU3B 7EU3C 7EU3D 7EU3E 7EU3F 7EU3G 7EU3L
7EU7A 7EU7B 7EU7C 7EU7D 7EU8A 7EU8B 7EU8C 7EU8D 7EUOR 7EV7A 7EV7B 7EV7C
7EV7D 7EV7G 7EV7I 7EV7J 7EV7K 7EV7L 7EV7M 7EV7N 7EV7O 7EV7P 7EV7Q 7EV7T
7EV7V 7EV7W 7EV7X 7EV7Y 7EV7Z 7EV9A 7EV9B 7EV9C 7EV9E 7EV9F 7EV9G 7EV9I
7EV9J 7EV9K 7EVMR 7EVYD 7EVZD 7EW0D 7EW1A 7EW2R 7EW3R 7EW4R 7EW61 7EW62
7EW63 7EW65 7EW6A 7EW6B 7EW6F 7EW6H 7EW6I 7EW6J 7EW6K 7EW6L 7EW7D 7EWK1
7EWK3 7EWK4 7EWK6 7EWKA 7EWKB 7EWKF 7EWKH 7EWKI 7EWKJ 7EWKK 7EWKL 7EWLA
7EWLB 7EWPA 7EWPB 7EWRA 7EWRB 7EWTA 7EXDR 7EZCA 7EZHD 7EZKD 7EZMD 7F0JB
7F0JC 7F0L1 7F0L2 7F0L3 7F0L4 7F0L5 7F0L6 7F0L7 7F0L8 7F0LA 7F0LB 7F0LD
7F0LE 7F0LF 7F0LG 7F0LH 7F0LI 7F0LJ 7F0LK 7F0LL 7F0LM 7F0LN 7F0LO 7F0LP
7F0LQ 7F0LR 7F0LS 7F0LT 7F0LU 7F0LV 7F0LW 7F0LX 7F0LY 7F0LZ 7F0TF 7F16R
7F1OF 7F1QR 7F1RR 7F1SR 7F1TA 7F1ZF 7F23F 7F24F 7F2OR 7F3FA 7F3FB 7F3FD
7F3FG 7F3TA 7F3TB 7F3UA 7F3UB 7F3XA 7F3XB 7F40A 7F40B 7F47A 7F47B 7F4DR
7F4FR 7F4HR 7F4IR 7F53R 7F54R 7F55R 7F56A 7F56B 7F57A 7F57B 7F57C 7F57D
7F57E 7F58R 7F59A 7F59B 7F59C 7F59D 7F59E 7F5AA 7F5AB 7F5AC 7F5AD 7F5AE
7F5AF 7F5BA 7F5BB 7F5BC 7F5BD 7F5BE 7F61A 7F6HA 7F6IA 7F6VA 7F6VB 7F73A
7F73B 7F7FA 7F7FB 7F83A 7F8JA 7F8JB 7F8JC 7F8JD 7F8JE 7F8JF 7F8JG 7F8NA
7F8NB 7F8NC 7F8ND 7F8NE 7F8NF 7F8NG 7F8OA 7F8OB 7F8OC 7F8OD 7F8OE 7F8OF
7F8OG 7F8VR 7F8WR 7F92A 7F92B 7F92C 7F92D 7F92E 7F92F 7F93A 7F93B 7F93C
7F93D 7F93E 7F93F 7F94A 7F94B 7F94C 7F94D 7F94E 7F94F 7F9YR 7F9ZR 7FC9A
7FC9B 7FCIA 7FD8A 7FD8B 7FD9A 7FD9B 7FDAA 7FDAB 7FDAC 7FDAD 7FDAE 7FDAF
7FDAQ 7FDAT 7FDAU 7FDAV 7FDAW 7FDAX 7FDAY 7FDAZ 7FDBA 7FDBB 7FDBC 7FDBD
7FDBE 7FDBF 7FDBQ 7FDBT 7FDBU 7FDBV 7FDBW 7FDBX 7FDBY 7FDBZ 7FDCA 7FDCB
7FDCC 7FDCD 7FDCE 7FDCF 7FDCQ 7FDCT 7FDCU 7FDCV 7FDCW 7FDCX 7FDCY 7FDCZ
7FDVA 7FDVD 7FEEA 7FH1A 7FH1B 7FIFA 7FIGR 7FIHR 7FIIR 7FIJR 7FIMR 7FINR
7FIYR 7FJDA 7FJDB 7FJDD 7FJDE 7FJDF 7FJDG 7FJDM 7FJDN 7FJEA 7FJEB 7FJED
7FJEE 7FJEF 7FJEG 7FJEM 7FJEN 7FJFA 7FJFB 7FJFD 7FJFE 7FJFF 7FJFG 7FJFM
7FJFN 7JG51 7JG52 7JG53 7JG54 7JG55 7JG56 7JG57 7JG58 7JG59 7JG5A 7JG5B
7JG5D 7JG61 7JG62 7JG63 7JG64 7JG65 7JG66 7JG67 7JG68 7JG69 7JG71 7JG72
7JG73 7JG74 7JG75 7JG76 7JG77 7JG78 7JG79 7JG81 7JG82 7JG83 7JG84 7JG85
7JG86 7JG87 7JG88 7JG89 7JG91 7JG92 7JG93 7JG94 7JG95 7JG96 7JG97 7JG98
7JG99 7JGA1 7JGA2 7JGA3 7JGA4 7JGA5 7JGA6 7JGA7 7JGA8 7JGA9 7JGAA 7JGAB
7JGAD 7JGB1 7JGB2 7JGBA 7JGBB 7JGBD 7JGC1 7JGC2 7JGCA 7JGCB 7JGCD 7JI3A
7JI3B 7JI3C 7JICA 7JICB 7JJOR 7JJPA 7JJPB 7JJPC 7JJPD 7JJPE 7JJPF 7JK8A
7JKCA 7JKCB 7JKCC 7JKCD 7JKCE 7JKCF 7JLOA 7JLOB 7JLOC 7JLPA 7JLPB 7JLPC
7JLQA 7JLQB 7JLQC 7JLWA 7JLWB 7JLWC 7JLWD 7JLWE 7JLWF 7JM6A 7JM6B 7JM7A
7JM7B 7JM7C 7JM7D 7JM9A 7JM9B 7JM9C 7JM9D 7JM9E 7JM9F 7JMCA 7JMCB 7JMCC
7JMCD 7JMCE 7JMCF 7JMDA 7JMDB 7JMDC 7JMDD 7JMDE 7JMDF 7JN0A 7JN0B 7JN0C
7JN0D 7JN0E 7JN0F 7JN1A 7JN1B 7JN1C 7JN1D 7JN1E 7JN1F 7JNAA 7JNAB 7JNAC
7JNCA 7JNCB 7JNCC 7JNIA 7JPKA 7JPKE 7JPLA 7JPLE 7JPVA 7JPVE 7JPWA 7JPWE
7JPXA 7JPXE 7JR7A 7JR7B 7JRGC 7JRGD 7JRGE 7JRGG 7JRGJ 7JRGK 7JRGO 7JRGP
7JRGQ 7JRGS 7JRGV 7JRGW 7JROA 7JROB 7JROC 7JROD 7JROF 7JROH 7JROI 7JROJ
7JRPA 7JRPB 7JRPC 7JRPC 7JRPD 7JRPD 7JRPE 7JRPF 7JRPG 7JRPH 7JRPI 7JRPJ
7JRPJ 7JRPK 7JRPO 7JRPP 7JRPQ 7JRPS 7JRPV 7JRPW 7JSJA 7JSJB 7JSKA 7JSKB

7JUPA 7JUPB 7JUPC 7JUPD 7JV5R 7JVPR 7JVQR 7JVRR 7JZ2C 7JZ2D 7JZ2G 7JZ2H
7JZ2K 7JZ2L 7K0IC 7K0IF 7K0JC 7K0KC 7K0LC 7K0MA 7K0MC 7K0MD 7K0ME 7K0MG
7K0MH 7K0NA 7K0NC 7K0ND 7K0NE 7K0NG 7K0NH 7K0OA 7K0OC 7K0OD 7K0OE 7K0OG
7K0OH 7K0PA 7K0PC 7K0PD 7K0PE 7K0PG 7K0PH 7K0QC 7K0QD 7K15A 7K18A 7K2TB
7K2TD 7K48A 7K48B 7K48C 7K48D 7K4AA 7K4AB 7K4AC 7K4AD 7K4BA 7K4BB 7K4BC
7K4BD 7K4CA 7K4CB 7K4CC 7K4CD 7K4DA 7K4DB 7K4DC 7K4DD 7K4EA 7K4EB 7K4EC
7K4ED 7K4FA 7K4FB 7K4FC 7K4FD 7K65A 7K7AA 7K7AB 7K7AC 7K7MA 7K8AA 7K8BA
7K8CA 7K8DA 7KAHA 7KAHB 7KAHC 7KAHD 7KAIA 7KAIB 7KAIC 7KAID 7KAIG 7KAJA
7KAJB 7KAJC 7KAJD 7KAJG 7KAKA 7KAKB 7KAKC 7KAKD 7KAKE 7KALA 7KALB 7KALC
7KALD 7KALE 7KALG 7KAMA 7KAMB 7KAMC 7KAMD 7KAME 7KAMG 7KANA 7KANB 7KANC
7KAND 7KANE 7KAOA 7KAOB 7KAOC 7KAOD 7KAPA 7KAPB 7KAPC 7KAPD 7KAPG 7KAQA
7KAQB 7KAQC 7KAQD 7KAQG 7KARA 7KARB 7KARC 7KARD 7KASA 7KASB 7KASC 7KASD
7KASG 7KATA 7KATB 7KATC 7KATD 7KAUA 7KAUB 7KAUC 7KAUD 7KAUG 7KB5A 7KB5B
7KB5C 7KB5D 7KC4B 7KF5A 7KF5B 7KF5C 7KF6A 7KF6B 7KF6C 7KF7A 7KF7B 7KF7C
7KF8A 7KF8B 7KF8C 7KGDA 7KGDB 7KGDC 7KGEA 7KGEB 7KGEC 7KGFA 7KGFB 7KGFC
7KGGA 7KGGB 7KGGC 7KGHA 7KGHB 7KGHC 7KGIA 7KGIB 7KGIC 7KGVA 7KH0R 7KHMA
7KHMB 7KI0R 7KI1R 7KK8B 7KK9B 7KKAB 7KKBB 7KKRB 7KN0A 7KN0B 7KNTE 7KNTR
7KNUE 7KNUR 7KOOA 7KOOB 7KOQA 7KOQB 7KOXA 7KOXB 7KP4A 7KR5A 7KR5B 7KR5C
7KR5D 7KR5E 7KR5F 7KRAA 7KRAC 7KRAD 7KRAE 7KRAF 7KRAM 7KRAN 7KS0A 7KS0B
7KS0C 7KS0D 7KS3A 7KS3B 7KS3C 7KS3D 7KSQ1 7KSQ2 7KSQ3 7KSQ4 7KSQA 7KSQB
7KSQF 7KSQG 7KSQH 7KSQI 7KSQJ 7KSQK 7KSQL 7KSQM 7KSQO 7KTXA 7KTXC 7KTXD
7KTXE 7KTXF 7KTXM 7KTXN 7KU5O 7KUX1 7KUX2 7KUX3 7KUX4 7KUXA 7KUXB 7KUXF
7KUXG 7KUXH 7KUXI 7KUXJ 7KUXK 7KUXL 7KUXM 7KUYA 7KUYB 7KUYC 7KUYD 7KUYE
7KXRA 7KXRB 7KXRC 7KXRD 7KXRE 7KXRF 7KXRG 7KY5A 7KY5B 7KY6A 7KY6B 7KY7A
7KY7B 7KY8A 7KY8B 7KY9A 7KY9B 7KYAA 7KYAB 7KYBA 7KYBB 7KYCA 7KYCB 7KYPB
7KYPD 7KZXA 7KZXB 7KZZA 7KZZB 7L0PC 7L0QC 7L0RC 7L0SC 7L16A 7L17A 7L1EA
7L1EB 7L1UR 7L1VR 7L2HA 7L2HB 7L2HC 7L2HD 7L2IA 7L2IB 7L2IC 7L2ID 7L2JA
7L2JB 7L2JC 7L2JD 7L2KA 7L2KB 7L2KC 7L2KD 7L2LA 7L2LB 7L2LC 7L2LD 7L2MA
7L2MB 7L2MC 7L2MD 7L2NA 7L2NB 7L2NC 7L2ND 7L2OA 7L2OB 7L2OC 7L2OD 7L2PA
7L2PB 7L2PC 7L2PD 7L2RA 7L2RB 7L2RC 7L2RD 7L2SA 7L2SB 7L2SC 7L2SD 7L2TA
7L2TB 7L2TC 7L2TD 7L2UA 7L2UB 7L2UC 7L2UD 7L2VA 7L2VB 7L2VC 7L2VD 7L2WA
7L2WB 7L2WC 7L2WD 7L2XA 7L2XB 7L2XC 7L2XD 7L31A 7L31B 7L31C 7L31D 7L31E
7L6QA 7L6QB 7L6QC 7L6QD 7L6QE 7L6UA 7L6UB 7L6UC 7L6UD 7L6UE 7LB8B 7LC3A
7LC3B 7LC3C 7LC3D 7LC6A 7LC6B 7LC6C 7LC6D 7LCIR 7LCJR 7LCKR 7LD3R 7LD4R
7LEPA 7LEPB 7LEPC 7LEPE 7LEPF 7LEPG 7LEPH 7LF6A 7LF6B 7LFTA 7LFTB 7LFTC
7LFTD 7LFWA 7LFWB 7LFWC 7LFWD 7LFXA 7LFXB 7LFXC 7LFXD 7LFYA 7LFYB 7LFYC
7LFYD 7LG1A 7LG1B 7LG1C 7LG1D 7LGUA 7LGUB 7LGWA 7LGWB 7LH2A 7LH2B 7LH3A
7LH3B 7LHEA 7LHEB 7LHEC 7LHED 7LHFA 7LHFB 7LHFC 7LHFD 7LHIC 7LHVA 7LHVB
7LI6A 7LI7A 7LI8A 7LI9A 7LIAA 7LICA 7LICB 7LIDA 7LIDB 7LIGA 7LIGB 7LJ4A
7LJ4B 7LJ5A 7LJ5B 7LJAA 7LJAB 7LJBA 7LJBB 7LJCR 7LJDR 7LKPA 7LKZA 7LLLR
7LLYR 7LO7Z 7LO8Z 7LP9A 7LP9B 7LP9C 7LP9D 7LPAA 7LPAB 7LPAC 7LPAD 7LPBA
7LPBB 7LPBC 7LPBD 7LPCA 7LPCB 7LPCC 7LPCD 7LPDA 7LPDB 7LPDC 7LPDD 7LPEA
7LPEB 7LPEC 7LPED 7LQJA 7LQJB 7LQKA 7LQKB 7LQLA 7LQLB 7LQYA 7LQYB 7LQYC
7LQYD 7LQZA 7LQZB 7LQZC 7LQZD 7LR0A 7LR0B 7LR0C 7LR0D 7LVTA 7LVTB 7LWDA

7LX0A 7LX0A 7LX0B 7LX0B 7LX0F 7LX0F 7LX0G 7LX0H 7LX0I 7LX0I 7LX0J 7LX0J
7LX0K 7LX0K 7LX0L 7LX0L 7LX0M 7LX0M 7LX0Q 7LX0R 7LX0S 7LX0T 7LX0U 7LX0V
7LX0W 7LX0X 7LX0X 7LZHA 7LZHB 7LZHC 7LZHD 7LZIA 7LZIB 7LZIC 7LZID 7M17A
7M17B 7M19A 7M19B 7M1PA 7M1QA 7M1TA 7M1TB 7M1UA 7M1UB 7M2HC 7M2HI 7M2HL
7M2HO 7M2IC 7M2IF 7M2II 7M2IL 7M2JC 7M2JF 7M2JI 7M2JL 7M33C 7M33D 7M3EA
7M3EB 7M3FA 7M3FB 7M3GA 7M3GB 7M3JA 7M3JB 7M4PA 7M4PB 7M4QA 7M4QB 7M5VA
7M5XA 7M5YA 7M68A 7M69A 7M6MA 7M6MB 7M6MC 7M6MD 7M6ME 7M6NA 7M6NB 7M6NC
7M6ND 7M6NE 7M6OA 7M6OB 7M6OC 7M6OD 7M6OE 7M6PA 7M6PB 7M6PC 7M6PD 7M6PE
7M6QA 7M6QB 7M6QC 7M6QD 7M6QE 7M6RA 7M6RB 7M6RC 7M6RD 7M6RE 7M6SA 7M6SB
7M6SC 7M6SD 7M6SE 7M75A 7M75B 7M75F 7M75I 7M75J 7M75K 7M75L 7M75M 7M75X
7M76A 7M76B 7M76F 7M76I 7M76J 7M76K 7M76L 7M76M 7M76X 7M78A 7M78B 7M78F
7M78I 7M78J 7M78K 7M78L 7M78M 7M78X 7M8WA 7M93A 7M93B 7MBPA 7MBPB 7MBPC
7MBPD 7MBQA 7MBQB 7MBQC 7MBQD 7MBRA 7MBRB 7MBRC 7MBRD 7MBSA 7MBSB 7MBSC
7MBSD 7MBTA 7MBTB 7MBTC 7MBTD 7MBUA 7MBUB 7MBUC 7MBUD 7MBVA 7MBVB 7MBVC
7MBVD 7MBXR 7MBYR 7MBZA 7MBZB 7MC0A 7MC0B 7MDCA 7MDEA 7MDEB 7MDFA 7MDFB
7MDJC 7MDJD 7MDJE 7MDJF 7MDXA 7MDXB 7MDYA 7MDYB 7META 7METB 7MEWA 7MEWB
7MFIA 7MFIB 7MGLA 7MGLB 7MGLC 7MGLD 7MGWA 7MGXA 7MH3H 7MH3L 7MH3M 7MH4H
7MH4L 7MH4M 7MH5H 7MH5L 7MH5M 7MH6A 7MH6B 7MH8H 7MH8L 7MH8M 7MH9H 7MH9L
7MH9M 7MHAH 7MHAL 7MHAM 7MHRC 7MHRD 7MHRE 7MHRF 7MHXC 7MHXD 7MHXE 7MHXF
7MHYA 7MHZA 7MIJA 7MIJB 7MIJC 7MIJD 7MIKA 7MIKB 7MIKC 7MIKD 7MILA 7MILB
7MILC 7MILD 7MIMA 7MIMB 7MIMC 7MIMD 7MINA 7MINB 7MINC 7MIND 7MIOA 7MIOB
7MIOC 7MIOD 7MITA 7MITB 7MITC 7MITD 7MITE 7MIXA 7MIYA 7MJOA 7MJOB 7MJOC
7MJOD 7MJOE 7MJOG 7MJPA 7MJPB 7MJPC 7MJPD 7MJPE 7MJQA 7MJQB 7MJQC 7MJQD
7MJQE 7MJQG 7MJSX 7MJTC 7MJTD 7MJTE 7MJTF 7MK6C 7MK6D 7MK6E 7MK6F 7MLUA
7MLUB 7MLUC 7MLUD 7MLUE 7MPAA 7MPEA 7MT8R 7MT9R 7MTAR 7MTBR 7MTQA 7MTQB
7MTRA 7MTRB 7MTSA 7MTSB 7MUBC 7MUBD 7MUBE 7MUBF 7MXOA 7MXOB 7MXYA 7MXYB
7MXYC 7MXYD 7MXYE 7MYZC 7MZ5A 7MZ5B 7MZ5C 7MZ5D 7MZ6A 7MZ6B 7MZ6C 7MZ6D
7MZ7A 7MZ7B 7MZ7C 7MZ7D 7MZ9A 7MZ9B 7MZ9C 7MZ9D 7MZAA 7MZAB 7MZAC 7MZAD
7MZBA 7MZBB 7MZBC 7MZBD 7MZCA 7MZCB 7MZCC 7MZCD 7MZDA 7MZDB 7MZDC 7MZDD
7MZEA 7MZEB 7MZEC 7MZED 7N0KA 7N0KB 7N0LA 7N0LB 7N0MA 7N0MB 7N0MC 7N0MD
7N0NA 7N0NB 7N0NC 7N0ND 7N15A 7N15B 7N15C 7N15D 7N16A 7N16B 7N16C 7N16D
7N17A 7N17B 7N17C 7N17D 7N3NA 7N3NB 7N4TA 7N4TB 7N4TC 7N4TD 7N4TE 7N4TF
7N4TG 7N4UA 7N4UB 7N4VA 7N4VB 7N4XA 7N58A 7N58B 7N59A 7N59B 7N5AA 7N5AB
7N5BA 7N5BB 7N6BA 7N6QA 7N6QB 7N6RA 7N6RB 7N70A 7N72A 7N73A 7N74A 7N75A
7N76A 7N77A 7N78A 7N7PA 7N7PB 7N8OA 7N8OA 7N8OB 7N8OB 7N8OC 7N8OC 7N8OD
7N8OD 7N8OE 7N8OE 7N8OF 7N8OF 7N8OH 7N8OH 7N8OI 7N8OI 7N8OJ 7N8OJ 7N8OK
7N8OK 7N8OL 7N8OL 7N8OM 7N8OM 7N8OR 7N8OR 7N8OT 7N8OT 7N8OX 7N8OX 7N8OY
7N8OY 7N8OZ 7N8OZ 7N98A 7N9KA 7N9KB 7N9KC 7N9KD 7N9LA 7N9LB 7N9LC 7N9LD
7N9ZF 7N9ZG 7N9ZH 7N9ZI 7NA7R 7NA8R 7NB6A 7NB6B 7NEQA 7NEQB 7NEZA 7NEZB
7NF6A 7NF6B 7NF8A 7NF8B 7NF8E 7NF8F 7NFDA 7NFDB 7NG9A 7NG9B 7NG9C 7NGBA
7NGBB 7NGHA 7NGHB 7NGHC 7NH9A 7NH9B 7NH9C 7NH9D 7NH9E 7NHOA 7NHOB 7NHOC
7NHOD 7NHOE 7NHOF 7NHOH 7NHOI 7NHOK 7NHOL 7NHOM 7NHOT 7NHOX 7NHOY 7NHOZ
7NHP3 7NHPA 7NHPB 7NHPC 7NHPD 7NHPE 7NHPF 7NHPH 7NHPI 7NHPK 7NHPL 7NHPM
7NHPT 7NHPX 7NHPY 7NHPZ 7NHQ3 7NHQA 7NHQB 7NHQC 7NHQD 7NHQE 7NHQF 7NHQH

7NHQI 7NHQK 7NHQL 7NHQM 7NHQT 7NHQX 7NHQY 7NHQZ 7NHRA 7NHRB 7NHRC 7NHRD
7NHRE 7NHRF 7NHRG 7NHRH 7NHSA 7NHSB 7NHSC 7NHSD 7NHSE 7NHSF 7NHSG 7NHSH
7NI2A 7NI2B 7NI2C 7NI2D 7NI2E 7NI2F 7NI2G 7NI2H 7NIBA 7NIBB 7NIBC 7NIBD
7NIBE 7NIBF 7NIBG 7NIBH 7NIHA 7NIHB 7NIHC 7NIHD 7NIHE 7NIHF 7NIHG 7NIHH
7NIIA 7NIIB 7NIIC 7NIID 7NIIE 7NIIF 7NIIG 7NIIH 7NIVA 7NIWA 7NJKA 7NJKB
7NJKD 7NJKL 7NJKM 7NJKN 7NJKO 7NJKP 7NJKQ 7NJKR 7NJKS 7NJKT 7NJLA 7NJLB
7NJLD 7NJLL 7NJLM 7NJLN 7NJLO 7NJLP 7NJLQ 7NJLR 7NJLS 7NJLT 7NJMA 7NJMB
7NJMD 7NJML 7NJMM 7NJMN 7NJMO 7NJMP 7NJMQ 7NJMR 7NJMS 7NJMT 7NJNA 7NJNB
7NJND 7NJNL 7NJNM 7NJNN 7NJNO 7NJNP 7NJNQ 7NJNR 7NJNS 7NJNT 7NJOA 7NJOB
7NJOD 7NJOL 7NJOM 7NJON 7NJOO 7NJOP 7NJOQ 7NJOR 7NJOS 7NJOT 7NJPA 7NJPB
7NJPD 7NJPL 7NJPM 7NJPN 7NJPO 7NJPP 7NJPQ 7NJPR 7NJPS 7NJPT 7NJQA 7NJQB
7NJQD 7NJQL 7NJQM 7NJQN 7NJQO 7NJQP 7NJQQ 7NJQR 7NJQS 7NJQT 7NJRA 7NJRB
7NJRD 7NJRL 7NJRM 7NJRN 7NJRO 7NJRP 7NJRQ 7NJRR 7NJRS 7NJRT 7NJSA 7NJSB
7NJSD 7NJSL 7NJSM 7NJSN 7NJSO 7NJSP 7NJSQ 7NJSR 7NJSS 7NJST 7NJTA 7NJTB
7NJTD 7NJTL 7NJTM 7NJTN 7NJUA 7NJUB 7NJUD 7NJUL 7NJUM 7NJUN 7NJUO 7NJUP
7NJUQ 7NJUR 7NJUS 7NJUT 7NJVA 7NJVB 7NJVD 7NJVL 7NJVM 7NJVN 7NJVO 7NJVP
7NJVQ 7NJVR 7NJVS 7NJVT 7NJWA 7NJWB 7NJWD 7NJWL 7NJWM 7NJWN 7NJWO 7NJWP
7NJWQ 7NJWR 7NJWS 7NJWT 7NJXA 7NJXB 7NJXD 7NJXL 7NJXM 7NJXN 7NJXO 7NJXP
7NJXQ 7NJXR 7NJXS 7NJXT 7NJYA 7NJYB 7NJYD 7NJYL 7NJYM 7NJYN 7NJYO 7NJYP
7NJYQ 7NJYR 7NJYS 7NJYT 7NK9A 7NK9B 7NK9D 7NK9L 7NK9M 7NK9N 7NK9O 7NK9P
7NK9Q 7NK9R 7NK9S 7NK9T 7NKBL 7NKBM 7NKBN 7NKBO 7NKBP 7NKBQ 7NKBR 7NKBS
7NKBT 7NKKL 7NKKM 7NKKN 7NKKO 7NKKP 7NKKQ 7NKKR 7NKKS 7NKKT 7NKNL 7NKNM
7NKNN 7NKNO 7NKNP 7NKNQ 7NKNR 7NKNS 7NKNT 7NKPA 7NKPB 7NKPD 7NKPL 7NKPM
7NKPN 7NKPO 7NKPP 7NKPQ 7NKPR 7NKPS 7NKPT 7NKZA 7NKZB 7NL4A 7NL4B 7NL4C
7NL4D 7NL9A 7NL9B 7NL9D 7NL9L 7NL9M 7NL9N 7NL9O 7NL9P 7NL9Q 7NL9R 7NL9S
7NL9T 7NMNA 7NMNB 7NMNC 7NMND 7NNLA 7NNLB 7NNLC 7NNLD 7NNPA 7NNPB 7NNPC
7NNPD 7NNTC 7NNTD 7NNUC 7NNUD 7NP3A 7NP3B 7NP3C 7NP3D 7NP4A 7NP4B 7NP4C
7NP4D 7NPWA 7NPWB 7NPWC 7NQKA 7NS1A 7NSGA 7NSGB 7NSGC 7NVHA 7NXFA 7NY1A
7NY1B 7NY1C 7NY1D 7NY1E 7NY1F 7NYHA 7NYHH 7NYHJ 7NYHK 7NYHL 7NYHM 7NYHN
7NYRA 7NYRH 7NYRJ 7NYRK 7NYRL 7NYRM 7NYRN 7NYUA 7NYUH 7NYUJ 7NYUK 7NYUL
7NYUM 7NYUN 7NYVA 7NYVH 7NYVJ 7NYVK 7NYVL 7NYVM 7NYVN 7O011 7O012 7O013
7O014 7O015 7O016 7O017 7O018 7O019 7O01A 7O01A 7O01B 7O01B 7O01F 7O01F
7O01G 7O01G 7O01I 7O01I 7O01J 7O01J 7O01K 7O01K 7O01L 7O01L 7O01Q 7O01R
7O01S 7O01T 7O01U 7O01V 7O0YD 7O0YE 7O10D 7O10E 7O11D 7O11E 7O12D 7O12E
7O14D 7O14E 7O15D 7O15E 7O16D 7O16E 7O17D 7O17E 7O1VA 7O1VB 7O1VK 7O1VM
7O37A 7O37B 7O37C 7O37C 7O37D 7O37D 7O37E 7O37G 7O37G 7O37I 7O37I 7O37J
7O37K 7O37K 7O37L 7O37M 7O37N 7O37O 7O37P 7O37R 7O37U 7O37V 7O3CA 7O3CB
7O3CC 7O3CC 7O3CD 7O3CD 7O3CE 7O3CG 7O3CG 7O3CI 7O3CI 7O3CJ 7O3CK 7O3CK
7O3CL 7O3CM 7O3CN 7O3CO 7O3CP 7O3CR 7O3CU 7O3CV 7O3EA 7O3EB 7O3EC 7O3EC
7O3ED 7O3ED 7O3EG 7O3EG 7O3EI 7O3EI 7O3EK 7O3EL 7O3EM 7O3EN 7O3EO 7O3ER
7O3HC 7O3HD 7O3HE 7O3HG 7O3HJ 7O3HK 7O3HN 7O3HO 7O3HP 7O3HR 7O3HU 7O3HV
7O3JA 7O3JB 7O3JD 7O3JE 7O3JG 7O3JH 7O3JJ 7O3JK 7O3JM 7O3JN 7O3JP 7O3JS
7O3JV 7O3JY 7O3LA 7O3LB 7O3LC 7O3MA 7O3MB 7O3MC 7O3NA 7O3NB 7O3NC 7O3VF
7O3VG 7O3VJ 7O41B 7O41C 7O41C 7O41D 7O41D 7O41E 7O41E 7O41F 7O41G 7O41H

7O41I 7O41J 7O41J 7O41K 7O41L 7O41M 7O41P 7O41Q 7O41R 7O41S 7O41V 7O41W
7O41X 7O41Y 7O4HA 7O4HB 7O4HC 7O4HD 7O6Y1 7O6Y2 7O6Y3 7O6Y4 7O6Y5 7O6Y6
7O6YA 7O6YB 7O6YC 7O6YD 7O6YE 7O6YG 7O6YI 7O6YJ 7O6YJ 7O6YL 7O6YN 7O6YS
7O6YW 7O6YX 7O711 7O712 7O713 7O714 7O715 7O716 7O71A 7O71B 7O71C 7O71D
7O71E 7O71G 7O71I 7O71J 7O71J 7O71L 7O71N 7O71S 7O71W 7O71X 7O7FC 7O82A
7O82B 7O8FA 7O8GA 7O8HA 7O8IA 7O8JA 7O8KA 7O8LA 7O8MA 7O8NA 7O8OA 7O8PA
7O8QA 7O8RA 7O8SA 7O8TA 7O8UA 7O8VA 7O8YA 7O8ZA 7O9WA 7OCAA 7OCAB 7OCAC
7OCAD 7OCAE 7OCAG 7OCAI 7OCAJ 7OCDA 7OCDB 7OCDC 7OCDD 7OCDI 7OCDJ 7OCEA
7OCEB 7OCEC 7OCED 7OCEE 7OCEG 7OCEI 7OCEJ 7OCFA 7OCFB 7OCFC 7OCFD 7OCFE
7OCFG 7OCFI 7OCFJ 7OCIA 7OCIB 7OCIC 7OCID 7OCIE 7OCIF 7OCIG 7OCIH 7OCII
7OD5H 7OD5L 7OD5M 7OFMA 7OFOA 7OH4A 7OH4C 7OH5A 7OH5C 7OH6A 7OH6C 7OH7A
7OH7C 7OIUD 7OIUE 7OIUF 7OJ8A 7OJ8B 7OJHA 7OJHB 7OJIA 7OJIB 7OK9E 7OMMA
7ONJA 7ONJB 7ONJC 7ONJD 7ONJE 7ONJF 7ONJG 7ONLA 7ONLB 7ONLC 7ONLD 7ONLE
7ONLF 7ONLG 7OO0A 7OO0B 7OO0C 7OO0D 7OO0E 7OO0F 7OO0G 7OO6A 7OO6B 7OO6C
7OO6D 7OO6E 7OO6F 7OO6G 7OO8A 7OO8B 7OO8C 7OO8D 7OO8E 7OO8F 7OO8G 7OOAA
7OOAB 7OOAC 7OOAD 7OOAE 7OOAF 7OOAG 7OORA 7OORB 7OORC 7OORD 7OOUA 7OOUB
7OOUC 7OOUD 7OP1A 7OP3A 7OP5A 7OP8A 7OPHA 7OPHB 7OPHC 7OPHD 7OQ1A 7OQ1B
7OQ1C 7OQ1D 7OQ2A 7OQ2B 7OQ2C 7OQ2D 7OQZA 7OQZB 7OQZC 7OQZD 7OR0A 7OR0B
7OR0C 7OR0D 7OR1A 7OR1B 7OR1C 7OR1D 7OS4B 7OSEA 7OSEB 7OSEC 7OSED 7OSEE
7OSEF 7OSFD 7OSFE 7OSGD 7OSGE 7OSHD 7OSHE 7OSID 7OSIE 7OSJD 7OSJE 7OT9A
7OT9B 7OT9C 7OT9D 7OT9E 7OTGA 7OTIA 7OUI1 7OUI2 7OUI3 7OUI4 7OUI5 7OUI6
7OUI7 7OUI8 7OUIA 7OUIA 7OUIB 7OUIB 7OUIC 7OUIC 7OUID 7OUID 7OUIE 7OUIE
7OUIF 7OUIF 7OUIG 7OUIG 7OUIH 7OUIH 7OUII 7OUII 7OUIK 7OUIK 7OUIL 7OUIL
7OUIM 7OUIM 7OUIN 7OUIN 7OUIR 7OUIR 7OUIS 7OUIS 7OUIT 7OUIT 7OUIW 7OUIW
7OUIX 7OUIX 7OUIY 7OUIY 7OUIZ 7OUIZ 7OUKA 7OUKB 7OUKC 7OULA 7OULB 7OULC
7OUMA 7OUMB 7OUMC 7OW8A 7OW8B 7OXEA 7OY2B 7OY2C 7OY2X 7OY81 7OY82 7OY83
7OY84 7OY85 7OY86 7OY87 7OY88 7OY89 7OY8A 7OY8D 7OY8D 7OY8E 7OY8F 7OY8G
7OY8H 7OY8I 7OY8J 7OY8K 7OY8L 7OY8M 7OY8M 7OY8N 7OY8N 7OY8O 7OY8Q 7OY8R
7OY8S 7OY8T 7OY8U 7OY8V 7OY8W 7OY8X 7OY8Y 7OY8Z 7OYEA 7OZ1A 7OZ1B 7P03A
7P04A 7P05A 7P06A 7P0QH 7P0QL 7P0QM 7P14A 7P16A 7P17H 7P17L 7P17M 7P1CA
7P1IA 7P1IB 7P1JA 7P1JB 7P1KA 7P1KB 7P2CH 7P2CL 7P2CM 7P2PA 7P2PB 7P2PC
7P2PD 7P2QA 7P2QB 7P2QC 7P2QD 7P2YA 7P2YB 7P2YG 7P2YH 7P2YJ 7P2YK 7P2YL
7P2YO 7P2YP 7P2YP 7P2YQ 7P2YR 7P2YS 7P34A 7P34B 7P3NA 7P3NB 7P3NG 7P3NH
7P3RA 7P3RB 7P3WA 7P3WB 7P3WG 7P3WH 7P3WJ 7P3WK 7P3WL 7P3WO 7P3WP 7P3WP
7P3WQ 7P3WR 7P3WS 7P4FA 7P4IA 7P4IB 7P4IC 7P54A 7P54B 7P5CA 7P5CB 7P5JA
7P5JB 7P5MA 7P5MB 7P5VA 7P5VB 7P5VC 7P5VD 7P5VE 7P5VF 7P5WA 7P5WB 7P5WC
7P5WD 7P5WE 7P5WF 7P5YA 7P5YB 7P5YC 7P5YD 7P5YE 7P5YF 7P60A 7P60B 7P60C
7P60D 7P60E 7P60F 7P61A 7P61H 7P61J 7P61K 7P61L 7P61M 7P61N 7P62A 7P62H
7P62J 7P62K 7P62L 7P62M 7P62N 7P63A 7P63H 7P63J 7P63K 7P63L 7P63M 7P63N
7P64A 7P64H 7P64J 7P64K 7P64L 7P64M 7P64N 7P69A 7P69H 7P69J 7P69K 7P69L
7P69M 7P69N 7P6KA 7P6KB 7P6KC 7P6KD 7P6KE 7P6KF 7P7CA 7P7CH 7P7CJ 7P7CK
7P7CL 7P7CM 7P7CN 7P7EA 7P7EH 7P7EJ 7P7EK 7P7EL 7P7EM 7P7EN 7P7JA 7P7JH
7P7JJ 7P7JK 7P7JL 7P7JM 7P7JN 7P7KA 7P7KH 7P7KJ 7P7KK 7P7KL 7P7KM 7P7KN
7P7LA 7P7LH 7P7LJ 7P7LK 7P7LL 7P7LM 7P7LN 7P7MA 7P7MH 7P7MJ 7P7MK 7P7ML

7P7MM 7P7MN 7P9UA 7P9UB 7P9VA 7P9VB 7PA0A 7PA0B 7PA0C 7PA0D 7PAFA 7PAFD
7PAGA 7PD4A 7PD8A 7PDEA 7PDGA 7PEMA 7PEMC 7PG8C 7PG8E 7PG8H 7PG8K 7PGBC
7PGBD 7PGBL 7PGBO 7PGBW 7PGEB 7PGFC 7PGFD 7PGGA 7PGGB 7PGGC 7PGGD 7PGHA
7PGHD 7PGHE 7PGHF 7PGIA 7PGIB 7PGIC 7PGID 7PH2A 7PH2B 7PH3A 7PH3B 7PH4A
7PH4B 7PH7A 7PH7B 7PH8A 7PHHA 7PHIA 7PHIB 7PHIC 7PHID 7PHKA 7PHKB 7PHKC
7PHKD 7PHLA 7PHLB 7PHLC 7PHLD 7PHPA 7PHQB 7PHQD 7PHRA 7PHRB 7PHRC 7PHRD
7PHRE 7PHRE 7PHRZ 7PHRZ 7PHTA 7PI0A 7PI0A 7PI0B 7PI0B 7PI0C 7PI0C 7PI0D
7PI0D 7PI0E 7PI0E 7PI0F 7PI0F 7PI0G 7PI0G 7PI0H 7PI0H 7PI0I 7PI0I 7PI0J
7PI0J 7PI0K 7PI0K 7PI0L 7PI0L 7PI0M 7PI0M 7PI0N 7PI0N 7PI0R 7PI0R 7PI0S
7PI0S 7PI0T 7PI0T 7PI0V 7PI0V 7PI0W 7PI0W 7PI0X 7PI0X 7PI0Y 7PI0Y 7PI0Z
7PI0Z 7PIJB 7PIUR 7PIVR 7PL4A 7PL9A 7PL9B 7PMWA 7PMXA 7PMYA 7PN1A 7PNCA
7POWA 7POWB 7PQGA 7PQQA 7PQTA 7PQTB 7PQUA 7PQUB 7PQUC 7PQUD 7PQXA 7PQXB
7PR1A 7PR1B 7PRCH 7PRCL 7PRCM 7PRNA 7PRNB 7PRNC 7PROA 7PROB 7PRUA 7PRUB
7PSDA 7PSDB 7PSLA 7PSLB 7PSMA 7PSMB 7PSNA 7PSNB 7PX4A 7PXEA 7PXEB 7PXEC
7PXED 7PXFA 7PXFB 7PXFC 7PXFD 7PXGA 7PXGB 7PXGC 7PXGD 7PXHA 7PXHB 7PXHC
7PXHD 7PY4A 7PY4B 7PYRA 7PZFA 7PZFB 7PZFC 7Q0LA 7Q0MA 7Q1UA 7Q21A 7Q21A
7Q21B 7Q21B 7Q21C 7Q21C 7Q21D 7Q21D 7Q21E 7Q21E 7Q21F 7Q21F 7Q21G 7Q21G
7Q21H 7Q21H 7Q21K 7Q21K 7Q21Y 7Q21Y 7Q35A 7Q36A 7Q37A 7Q38A 7Q3TA 7Q3TB
7Q3TC 7Q6ZA 7QA8A 7QA8B 7QAMA 7QAOA 7QAPA 7QB9A 7QB9B 7QBAD 7QBAE 7QBCA
7QBCB 7QBIA 7QBIB 7QBZA 7QC0A 7QCOA 7QCOA 7QCOB 7QCOB 7QCOE 7QCOE 7QCOF
7QCOF 7QCOG 7QCOG 7QCOI 7QCOI 7QCOJ 7QCOJ 7QCOK 7QCOK 7QCOL 7QCOL 7QCOM
7QCOM 7QCOO 7QCOO 7QCOP 7QCOP 7QCOQ 7QCOQ 7QCOR 7QCOR 7QCOS 7QCOS 7QCOT
7QCOT 7QE5A 7QEOA 7QEOB 7QEQA 7QEQB 7QEQC 7QEQD 7QEQE 7QEQF 7QERA 7QERB
7QERC 7QERD 7QERE 7QERF 7QESA 7QESB 7QETA 7QETB 7QETC 7QETD 7QETE 7QETF
7QEUA 7QEUB 7QEVA 7QEVB 7QEWG 7QEWH 7QEWI 7QEWJ 7QEWK 7QEWL 7QEYG 7QEYH
7QEYI 7QEYJ 7QEYK 7QEYL 7QHBA 7QHBB 7QHBC 7QHBD 7QHBI 7QHBJ 7QHHA 7QHHB
7QHHC 7QHHD 7QHHI 7QHHJ 7QHMA 7QHMB 7QHMC 7QHMD 7QHME 7QHMF 7QHMG 7QHMI
7QHMJ 7QHML 7QHMN 7QHMO 7QHMP 7QHMQ 7QHMR 7QHMS 7QHMT 7QHMV 7QHMW 7QHMY
7QHOA 7QHOB 7QHOC 7QHOD 7QHOE 7QHOF 7QHOG 7QHOI 7QHOJ 7QHOL 7QHON 7QHOO
7QHOP 7QHOQ 7QHOR 7QHOS 7QHOT 7QHOV 7QHOW 7QHOY 7QI2A 7QIAA 7QICA 7QJIA
7QJJA 7QKOA 7QKOB 7QKOC 7QKOD 7QKOE 7QKRA 7QKSA 7QL5A 7QL5B 7QL5C 7QL5D
7QL5E 7QL6A 7QL6B 7QL6C 7QL6D 7QL6E 7QN5A 7QN5B 7QN5C 7QN5D 7QN5E 7QN6A
7QN6B 7QN6C 7QN6D 7QN6E 7QN7A 7QN7B 7QN7C 7QN7D 7QN7E 7QN8A 7QN8B 7QN8C
7QN8D 7QN8E 7QN9A 7QN9B 7QN9C 7QN9D 7QN9E 7QNAA 7QNAB 7QNAC 7QNAD 7QNAE
7QNBA 7QNBB 7QNBC 7QNBD 7QNBE 7QNCA 7QNCB 7QNCC 7QNCD 7QNCE 7QNDA 7QNDB
7QNDC 7QNDD 7QNDE 7QNEA 7QNEB 7QNEC 7QNED 7QNEE 7QOAA 7QOOQ 7QOOU 7QP9A
7QP9B 7QPAA 7QPAB 7QPCA 7QPCB 7QSDA 7QSDA 7QSDB 7QSDC 7QSDD 7QSDF 7QSDG
7QSDH 7QSDH 7QSDI 7QSDJ 7QSDJ 7QSDK 7QSDK 7QSDL 7QSDL 7QSDM 7QSDM 7QSDN
7QSDY 7QSDZ 7QSKA 7QSKA 7QSKB 7QSKC 7QSKD 7QSKF 7QSKG 7QSKH 7QSKH 7QSKI
7QSKJ 7QSKJ 7QSKK 7QSKK 7QSKL 7QSKL 7QSKM 7QSKM 7QSKN 7QSKY 7QSKZ 7QSLA
7QSLA 7QSLB 7QSLC 7QSLD 7QSLF 7QSLG 7QSLH 7QSLH 7QSLI 7QSLJ 7QSLJ 7QSLK
7QSLK 7QSLL 7QSLL 7QSLM 7QSLM 7QSLN 7QSLY 7QSLZ 7QSMA 7QSMA 7QSMB 7QSMC
7QSMD 7QSMF 7QSMG 7QSMH 7QSMH 7QSMI 7QSMJ 7QSMJ 7QSMK 7QSMK 7QSML 7QSML
7QSMM 7QSMM 7QSMN 7QSMY 7QSMZ 7QSNA 7QSNA 7QSNB 7QSNC 7QSND 7QSNF 7QSNG

7QSNH 7QSNH 7QSNI 7QSNJ 7QSNJ 7QSNK 7QSNK 7QSNL 7QSNL 7QSNM 7QSNM 7QSNN
7QSNY 7QSNZ 7QSOA 7QSOA 7QSOB 7QSOC 7QSOD 7QSOF 7QSOG 7QSOH 7QSOH 7QSOI
7QSOJ 7QSOJ 7QSOK 7QSOK 7QSOL 7QSOL 7QSOM 7QSOM 7QSON 7QSOZ 7QVMR 7R0CA
7R0GA 7R0HA 7R0IA 7R0WA 7R0WB 7R0WC 7R0WD 7R0WE 7R0WF 7R0WG 7R0WH 7R0WI
7R0WJ 7R0WK 7R0WL 7R0WM 7R0WN 7R0WO 7R0WP 7R1VA 7R1WA 7R3VC 7R3VD 7R3VE
7R3VG 7R3VJ 7R87A 7R87B 7R88A 7R88B 7R89A 7R89B 7R8AA 7R8AB 7R8BA 7R8BB
7R8CA 7R8CB 7R8DA 7R8DB 7R8EA 7R8EB 7RA3R 7RANA 7RASA 7RASB 7RASC 7RASD
7RAUA 7RAUB 7RAUC 7RAUD 7RAZA 7RAZB 7RAZC 7RAZD 7RAZE 7RAZF 7RAZG 7RBTR
7RCPA 7RCPB 7RCPC 7RCVA 7RCVA 7RCVB 7RCVB 7RCVC 7RCVC 7RCVD 7RCVD 7RCVE
7RCVE 7RCVF 7RCVF 7RCVH 7RCVH 7RCVI 7RCVI 7RCVJ 7RCVJ 7RCVK 7RCVK 7RCVL
7RCVL 7RCVM 7RCVM 7RCVR 7RCVR 7RCVT 7RCVT 7RCVX 7RCVX 7RCVY 7RCVY 7RCVZ
7RCVZ 7RD6A 7RD7A 7RD8A 7RG9R 7RGPR 7RH5A 7RH5E 7RH5F 7RH5I 7RH5J 7RH5K
7RH5L 7RH5M 7RH5O 7RH5P 7RH5Q 7RH5R 7RH5S 7RH5T 7RH5U 7RH5X 7RH5Y 7RH5Z
7RH6A 7RH6E 7RH6F 7RH6I 7RH6J 7RH6K 7RH6L 7RH6M 7RH6O 7RH6P 7RH6Q 7RH6R
7RH6S 7RH6T 7RH6U 7RH6X 7RH6Y 7RH6Z 7RH7A 7RH7E 7RH7F 7RH7I 7RH7J 7RH7K
7RH7L 7RH7M 7RH7O 7RH7P 7RH7Q 7RH7R 7RH7S 7RH7T 7RH7U 7RH7X 7RH7Y 7RH7Z
7RH9A 7RH9B 7RH9C 7RH9D 7RHGA 7RHGB 7RHGC 7RHGD 7RHHA 7RHHB 7RHHC 7RHHD
7RHIA 7RHIB 7RHIC 7RHID 7RHJA 7RHJB 7RHJC 7RHJD 7RHKA 7RHKB 7RHKC 7RHKD
7RHLA 7RHLB 7RHLC 7RHLD 7RHQA 7RHRA 7RHSA 7RHSB 7RHSC 7RHSD 7RI4A 7RI4G
7RI5A 7RI6A 7RI7A 7RI9A 7RITA 7RITB 7RJ5A 7RJ5G 7RJAC 7RJAD 7RJAF 7RJAI
7RJAK 7RJAM 7RJAN 7RJAP 7RJAS 7RJAT 7RJBD 7RJBF 7RJBI 7RJBK 7RJBM 7RJCD
7RJCF 7RJCI 7RJCK 7RJCM 7RJDD 7RJDF 7RJDI 7RJDK 7RJDM 7RJEC 7RJED 7RJEF
7RJEI 7RJEK 7RJEM 7RJEN 7RJEO 7RJER 7RJET 7RJTA 7RJTB 7RJTC 7RJTD 7RK6A
7RK6B 7RK6C 7RK6D 7RM5A 7RMGR 7RMHR 7RMIR 7RNND 7ROQA 7RP0C 7RP0F 7RP0I
7RP0L 7RPHA 7RPIA 7RPJA 7RPKA 7RPMA 7RPMB 7RPMC 7RPMD 7RPME 7RQUA 7RQUB
7RQUC 7RQUD 7RQVA 7RQVB 7RQVC 7RQVD 7RQWA 7RQWB 7RQWC 7RQWD 7RQXA 7RQXB
7RQXC 7RQXD 7RQYA 7RQYB 7RQYC 7RQYD 7RQZA 7RQZB 7RQZC 7RQZD 7RR6A 7RR6B
7RR6C 7RR7A 7RR7B 7RR7C 7RR8A 7RR8B 7RR8C 7RR9A 7RR9B 7RRAA 7RRAB 7RSLA
7RSLB 7RSLC 7RSLD 7RSLE 7RSLF 7RSLG 7RSLH 7RSLI 7RSLJ 7RTBR 7RTMA 7RTMB
7RTTA 7RTTB 7RTUA 7RTVA 7RTWA 7RTWB 7RU6A 7RUGA 7RWJA 7RWJB 7RX2A 7RX2B
7RX3A 7RX3B 7RXAA 7RXAB 7RXBA 7RXBB 7RXCK 7RXGA 7RXGB 7RXHA 7RY3A 7RY3B
7RY3C 7RYCO 7RYXA 7RYYA 7RYYB 7RYYC 7RYYD 7RYZA 7RYZB 7RYZC 7RYZD 7RZ4A
7RZ4B 7RZ4C 7RZ4D 7RZ5A 7RZ5B 7RZ5C 7RZ5D 7RZ6A 7RZ6B 7RZ6C 7RZ6D 7RZ7A
7RZ7B 7RZ7C 7RZ7D 7RZ8A 7RZ8B 7RZ8C 7RZ8D 7RZ9A 7RZ9B 7RZ9C 7RZ9D 7RZAA
7RZAB 7RZAC 7RZAD 7S0FR 7S0GR 7S1MR 7S1XA 7S1XB 7S1YA 7S1YB 7S1ZA 7S1ZB
7S24A 7S3DA 7S3DA 7S3DB 7S3DB 7S3DF 7S3DF 7S3DG 7S3DH 7S3DI 7S3DI 7S3DJ
7S3DJ 7S3DK 7S3DK 7S3DL 7S3DL 7S3DM 7S3DM 7S3DQ 7S3DR 7S3DS 7S3DT 7S3DU
7S3DV 7S3IR 7S4HA 7S4HB 7S4HC 7S4HE 7S4HF 7S4HG 7S4HI 7S4HJ 7S4HK 7S4IA
7S4IB 7S4IC 7S4IE 7S4IF 7S4IG 7S4II 7S4IJ 7S4IK 7S4JA 7S4JB 7S4JC 7S4JE
7S4JF 7S4JG 7S4JI 7S4JJ 7S4JK 7S4KA 7S4KB 7S4KC 7S4KE 7S4KF 7S4KG 7S4KI
7S4KJ 7S4KK 7S4LA 7S4LB 7S4LC 7S4LD 7S4LE 7S4LF 7S4LG 7S4LH 7S4LI 7S4MA
7S4MB 7S4MC 7S4MD 7S4ME 7S4MF 7S4MG 7S4MH 7S4MI 7S4MJ 7S4MK 7S4MN 7S5TA
7S5TB 7S5TC 7S5TD 7S5VE 7S5XA 7S5XB 7S5XC 7S5XD 7S5XE 7S5YA 7S5YB 7S5YC
7S5YD 7S5YE 7S5ZA 7S5ZB 7S5ZC 7S5ZD 7S5ZE 7S60A 7S60B 7S60C 7S60D 7S60E

7S61A 7S61B 7S61C 7S61D 7S61E 7S88A 7S88B 7S89A 7S89B 7S89C 7S89D 7S8BA
7S8BB 7S8BC 7S8BD 7S8CA 7S8CB 7S8CC 7S8CD 7S8LR 7S8NR 7S8OR 7S8UA 7S8XA
7S8XB 7S9AA 7S9AB 7S9BA 7S9BB 7S9CA 7S9CB 7S9DA 7S9DB 7S9EA 7S9EB 7S9YA
7S9ZA 7SA3A 7SA3B 7SA3C 7SA3D 7SA3E 7SA3F 7SA3I 7SA3K 7SA3N 7SAAA 7SAAB
7SAAC 7SAAD 7SABA 7SABB 7SABC 7SABD 7SACA 7SACB 7SACC 7SACD 7SADA 7SADB
7SADC 7SADD 7SATA 7SATB 7SATC 7SATD 7SATE 7SATF 7SATG 7SAUA 7SAUB 7SAUC
7SAUD 7SAUE 7SAUF 7SAUG 7SAXA 7SAXB 7SAXC 7SAXD 7SAXE 7SAXF 7SAXG 7SAZA
7SAZB 7SAZC 7SAZD 7SAZE 7SAZF 7SAZG 7SBFR 7SCGD 7SELA 7SELB 7SF7A 7SF8A
7SFJA 7SFJB 7SFJC 7SFKA 7SFKB 7SFKC 7SFLA 7SFLB 7SFQA 7SFQB 7SGRA 7SGRB
7SGRC 7SGRD 7SGRE 7SGRF 7SGRG 7SGRH 7SGRI 7SGRJ 7SGRK 7SGRL 7SHEA 7SHEB
7SHFA 7SHFB 7SHMA 7SHMB 7SHNA 7SHNB 7SHSA 7SHSB 7SHSC 7SI3A 7SI6A 7SI7A
7SIIA 7SIIB 7SIIC 7SIID 7SILA 7SILB 7SIMA 7SIMB 7SINA 7SINB 7SIPA 7SIPB
7SIPC 7SIPD 7SITB 7SITF 7SITH 7SITJ 7SIZB 7SIZF 7SIZH 7SIZJ 7SJ1A 7SJ1B
7SJ1C 7SJ1D 7SK0A 7SK0B 7SK1A 7SK1B 7SK2A 7SK3A 7SK4A 7SK5A 7SK6A 7SK7A
7SK8A 7SK9A 7SL8A 7SL9A 7SLAA 7SMMA 7SMMB 7SMMC 7SMMD 7SMME 7SMPA 7SMPB
7SMQA 7SMQB 7SMQC 7SMQD 7SMQE 7SMRA 7SMRB 7SMRC 7SMRD 7SMRE 7SMSA 7SMSB
7SMSC 7SMSD 7SMSE 7SMTA 7SMTB 7SMTC 7SMTD 7SMTE 7SP5A 7SPSA 7SPSB 7SPTA
7SQ6A 7SQ6B 7SQ7A 7SQ7B 7SQ7C 7SQ7D 7SQ8A 7SQ8B 7SQ8C 7SQ8D 7SQ9A 7SQ9B
7SQ9C 7SQ9D 7SQGA 7SQGB 7SQOR 7SQWC 7SR8R 7SRQR 7SRRR 7SRSR 7SSUA 7SSUB
7SSXA 7SSXB 7SSXC 7SSXD 7SSYA 7SSYB 7SSYC 7SSYD 7SSZA 7SSZB 7SSZC 7SSZD
7STLA 7STLB 7STMA 7STMB 7STNA 7STNB 7STOA 7STOB 7SUNA 7SUNB 7SUSA 7SV7A
7SV9A 7SV9B 7SVDA 7SVRA 7SVXA 7SVXB 7SWJA 7SWJB 7SWJC 7SWJD 7SX3A 7SX4A
7SZTA 7SZTB 7T00A 7T00B 7T10R 7T11R 7T32A 7T37A 7T37B 7T37C 7T37D 7T38A
7T38B 7T38C 7T38D 7T3UA 7T3UB 7T4EA 7T4EB 7T4EC 7T4ED 7T4EE 7T4EF 7T4EG
7T4EH 7T4OA 7T4OB 7T4OC 7T4OE 7T4OF 7T4OG 7T4OI 7T4OJ 7T4OK 7T4PA 7T4PB
7T4PC 7T4PE 7T4PF 7T4PG 7T4PI 7T4PJ 7T4PK 7T4XA 7T4XB 7T4XC 7T4XD 7T54A
7T54B 7T55A 7T55B 7T56A 7T56B 7T57A 7T57B 7T6BR 7T6DA 7T6DB 7T6DC 7T6DD
7T6JA 7T6JB 7T6KA 7T6KB 7T6KC 7T6KD 7T6LA 7T6LB 7T6LC 7T6LD 7T6MA 7T6MB
7T6MC 7T6MD 7T6NA 7T6NB 7T6NC 7T6ND 7T6OA 7T6OB 7T6OC 7T6OD 7T6PA 7T6PB
7T6PC 7T6PD 7T6QA 7T6QB 7T6QC 7T6QD 7T6RA 7T6RB 7T6RC 7T6RD 7T6SR 7T6TR
7T6UR 7T6VR 7T92A 7T92B 7T92C 7T9FA 7T9FB 7T9GA 7T9GY 7T9IR 7T9MR 7TAOA
7TAOC 7TAOD 7TAOE 7TAOF 7TAOG 7TAOH 7TAOI 7TAOJ 7TAOK 7TAOL 7TAOM 7TAON
7TAOO 7TAPA 7TAPC 7TAPD 7TAPE 7TAPF 7TAPG 7TAPH 7TAPI 7TAPJ 7TAPK 7TAPL
7TAPM 7TAPN 7TAPO 7TBGA 7TBGB 7TBWA 7TBYA 7TBZA 7TC0A 7TCGA 7TCHA 7TCIA
7TCIC 7TCIE 7TCIG 7TCPA 7TCPC 7TCPE 7TCPG 7TD0R 7TD1R 7TD2R 7TD3R 7TD4R
7TD61 7TDDA 7TDDB 7TDEA 7TDEB 7TDFA 7TDFB 7TDMA 7TDNA 7TE9A 7TE9B 7TE9C
7TE9D 7TEBA 7TEBB 7TEBC 7TEBD 7TEEA 7TEEB 7TEEC 7TEED 7TEQA 7TEQB 7TEQC
7TEQD 7TERA 7TERB 7TERC 7TERD 7TESA 7TESB 7TESC 7TESD 7TETA 7TETB 7TETC
7TETD 7TJ5A 7TJ5B 7TJ5C 7TJ5D 7TJ6A 7TJ6B 7TJ6C 7TJ6D 7TJ8A 7TJ8B 7TJ9A
7TJ9B 7TJZ0 7TJZ1 7TJZ2 7TJZ3 7TJZ4 7TJZ5 7TJZ6 7TJZ7 7TJZ8 7TJZ9 7TJZT
7TJZU 7TJZW 7TJZY 7TJZZ 7TK10 7TK11 7TK12 7TK13 7TK14 7TK15 7TK16 7TK17
7TK18 7TK19 7TK1T 7TK1U 7TK1W 7TK1Y 7TK1Z 7TK20 7TK21 7TK22 7TK23 7TK24
7TK25 7TK26 7TK27 7TK28 7TK29 7TK2T 7TK2U 7TK2W 7TK2Y 7TK2Z 7TK40 7TK41
7TK42 7TK43 7TK44 7TK45 7TK46 7TK47 7TK48 7TK49 7TK4T 7TK4U 7TK4W 7TK4Y

7TK4Z 7TK50 7TK51 7TK52 7TK53 7TK54 7TK55 7TK56 7TK57 7TK58 7TK59 7TK5T

7TK5U 7TK5W 7TK5Y 7TK5Z 7TK70 7TK71 7TK72 7TK73 7TK74 7TK75 7TK76 7TK77

7TK78 7TK79 7TK7T 7TK7U 7TK7W 7TK7Y 7TK7Z 7TK80 7TK81 7TK82 7TK83 7TK84

7TK85 7TK86 7TK87 7TK88 7TK89 7TK8T 7TK8U 7TK8W 7TK8Y 7TK8Z 7TK90 7TK91

7TK92 7TK93 7TK94 7TK95 7TK96 7TK97 7TK98 7TK99 7TK9T 7TK9U 7TK9W 7TK9Y

7TK9Z 7TKA0 7TKA1 7TKA2 7TKA3 7TKA4 7TKA5 7TKA6 7TKA7 7TKA8 7TKA9 7TKAT

7TKAU 7TKAW 7TKAY 7TKAZ 7TKB0 7TKB1 7TKB2 7TKB3 7TKB4 7TKB5 7TKB6 7TKB7

7TKB8 7TKB9 7TKBT 7TKBU 7TKBW 7TKBY 7TKBZ 7TKC0 7TKC1 7TKC2 7TKC3 7TKC4

7TKC5 7TKC6 7TKC7 7TKC8 7TKC9 7TKCT 7TKCU 7TKCW 7TKCY 7TKCZ 7TKD0 7TKD1

7TKD2 7TKD3 7TKD4 7TKD5 7TKD6 7TKD7 7TKD8 7TKD9 7TKDT 7TKDU 7TKDW 7TKDY

7TKDZ 7TKE0 7TKE1 7TKE2 7TKE3 7TKE4 7TKE5 7TKE6 7TKE7 7TKE8 7TKE9 7TKET

7TKEU 7TKEW 7TKEY 7TKEZ 7TKF0 7TKF1 7TKF2 7TKF3 7TKF4 7TKF5 7TKF6 7TKF7

7TKF8 7TKF9 7TKFT 7TKFU 7TKFW 7TKFY 7TKFZ 7TKG0 7TKG1 7TKG2 7TKG3 7TKG4

7TKG5 7TKG6 7TKG7 7TKG8 7TKG9 7TKGT 7TKGU 7TKGW 7TKGY 7TKGZ 7TKH0 7TKH1

7TKH2 7TKH3 7TKH4 7TKH5 7TKH6 7TKH7 7TKH8 7TKH9 7TKHT 7TKHU 7TKHW 7TKHY

7TKHZ 7TKI0 7TKI1 7TKI2 7TKI3 7TKI4 7TKI5 7TKI6 7TKI7 7TKI8 7TKI9 7TKIT

7TKIU 7TKIW 7TKIY 7TKIZ 7TKJ0 7TKJ1 7TKJ2 7TKJ3 7TKJ4 7TKJ5 7TKJ6 7TKJ7

7TKJ8 7TKJ9 7TKJT 7TKJU 7TKJW 7TKJY 7TKJZ 7TKK0 7TKK1 7TKK2 7TKK3 7TKK4

7TKK5 7TKK6 7TKK7 7TKK8 7TKK9 7TKKT 7TKKU 7TKKW 7TKKY 7TKKZ 7TKL0 7TKL1

7TKL2 7TKL3 7TKL4 7TKL5 7TKL6 7TKL7 7TKL8 7TKL9 7TKLT 7TKLU 7TKLW 7TKLY

7TKLZ 7TKN0 7TKN1 7TKN2 7TKN3 7TKN4 7TKN5 7TKN6 7TKN7 7TKN8 7TKN9 7TKNT

7TKNU 7TKNW 7TKNY 7TKNZ 7TKO0 7TKO1 7TKO2 7TKO3 7TKO4 7TKO5 7TKO6 7TKO7

7TKO8 7TKO9 7TKOT 7TKOU 7TKOW 7TKOY 7TKOZ 7TKP0 7TKP1 7TKP2 7TKP3 7TKP4

7TKP5 7TKP6 7TKP7 7TKP8 7TKP9 7TKPT 7TKPU 7TKPW 7TKPY 7TKPZ 7TKR0 7TKR1

7TKR2 7TKR3 7TKR4 7TKR5 7TKR6 7TKR7 7TKR8 7TKR9 7TKRT 7TKRU 7TKRW 7TKRY

7TKRZ 7TKS0 7TKS1 7TKS2 7TKS3 7TKS4 7TKS5 7TKS6 7TKS7 7TKS8 7TKS9 7TKST

7TKSU 7TKSW 7TKSY 7TKSZ 7TKTA 7TKTB 7TKTC 7TKTD 7TMRA 7TMRB 7TMRC 7TMRE

7TMRF 7TMRG 7TMRH 7TMRI 7TMRJ 7TMRK 7TMRL 7TMRM 7TMRN 7TMRO 7TNJA 7TNKA

7TNKB 7TNKC 7TNKD 7TNLA 7TNLB 7TNLC 7TNLD 7TNMA 7TNMB 7TNMC 7TNMD 7TNNA

7TNNB 7TNNC 7TNND 7TNOA 7TNOB 7TNOC 7TNOD 7TNPA 7TNPB 7TNPC 7TNPD 7TPGB

7TPJB 7TSZA 7TSZP 7TT0A 7TT0P 7TT1A 7TT1P 7TT2A 7TT2P 7TT3A 7TT3P 7TT4A

7TT4P 7TT5A 7TT5P 7TT6A 7TT6P 7TT7A 7TT7P 7TTCA 7TTCP 7TTHA 7TTHB 7TTIA

7TVZA 7TVZB 7TW0A 7TW0B 7TW1A 7TW1B 7TW3A 7TW3B 7TW5A 7TW5B 7TW6A 7TW6B

7TX6A 7TX7A 7TYIR 7TYLR 7TYNR 7TYOR 7U2LD 7U55A 7U55B 7U8GA 7U8GB 7UDVA

7UDVB 7UDVC 7UDVD 7UDVE 7UDWA 7UDWB 7UDWC 7UDWD 7UDWE 7UDXA 7UDXB 7UDXC

7UDXD 7UDXE 7UDYA 7UDYB 7UDYC 7UDYD 7UDYE 7UDZA 7UDZB 7UDZC 7UDZD 7UDZE

7UEAA 7UEAA 7UEAC 7UEAC 7UGGA 7UGGB 7UGGC 7UGGD 7UHFA 7UHGA 7UK5A 7UK5B

7UK5C 7UK5D 7UKCA 7UKCB 7UKCC 7UKCD 7UKDA 7UKDB 7UKDC 7UKDD 7UKEA 7UKEB

7UKEC 7UKED 7UKFA 7UKFB 7UKFC 7UKFD 7UKGA 7UKGB 7UKGC 7UKGD 7UKGI 7UKGJ

7UKGK 7UKGL 7UL2R 7UL3A 7UL4A 7UL5A 7UL6A 7UL6B 7UM4A 7UM5A 7UM6A 7UM7A

7UNLA 7UNLB 7UNMA 7UNMB 7URAA 7URCA 7URDA 7UREA 7URFA 7USWA 7USWB 7USWD

7USWF 7USXA 7USXB 7USXD 7USXF 7USYA 7USYB 7USYD 7USYF 7UTZR 7UZ3B 7UZ3C

7UZ3D 7UZ3E 7UZEA 7UZEB 7UZEC 7UZED 7UZQK 7UZQL 7UZQQ 7V07B 7V07C 7V07D

7V07E 7V0SK 7V0SL 7V0SQ 7V19B 7V19C 7V19D 7V19E 7V2CA 7V2CB 7V2CC 7V2CE

7V2CF 7V2CG 7V2CI 7V2CJ 7V2CK 7V2CL 7V2CM 7V2CN 7V2CO 7V2CR 7V2CS 7V2CS
7V2CU 7V2CV 7V2CW 7V2CY 7V2CZ 7V2DA 7V2DB 7V2DC 7V2DE 7V2DF 7V2DG 7V2DI
7V2DJ 7V2DK 7V2DL 7V2DM 7V2DN 7V2DO 7V2DR 7V2DS 7V2DS 7V2DU 7V2DV 7V2DW
7V2DY 7V2DZ 7V2EA 7V2EB 7V2EC 7V2EE 7V2EF 7V2EG 7V2EI 7V2EJ 7V2EK 7V2EL
7V2EM 7V2EN 7V2EO 7V2ER 7V2ES 7V2ES 7V2EU 7V2EV 7V2EW 7V2EY 7V2EZ 7V2FA
7V2FB 7V2FC 7V2FE 7V2FF 7V2FG 7V2FI 7V2FJ 7V2FK 7V2FL 7V2FM 7V2FN 7V2FO
7V2FR 7V2FS 7V2FS 7V2FU 7V2FV 7V2FW 7V2FY 7V2FZ 7V2HA 7V2HB 7V2HC 7V2HE
7V2HF 7V2HG 7V2HI 7V2HJ 7V2HK 7V2HL 7V2HM 7V2HN 7V2HO 7V2HR 7V2HS 7V2HS
7V2HU 7V2HV 7V2HW 7V2HY 7V2HZ 7V2KA 7V2KB 7V2KC 7V2KE 7V2KF 7V2KG 7V2KI
7V2KJ 7V2KK 7V2KL 7V2KM 7V2KN 7V2KO 7V2KR 7V2KS 7V2KS 7V2KU 7V2KV 7V2KW
7V2KY 7V2KZ 7V2RA 7V2RB 7V2RC 7V2RE 7V2RF 7V2RG 7V2RI 7V2RJ 7V2RK 7V2RL
7V2RM 7V2RN 7V2RO 7V2RR 7V2RS 7V2RS 7V2RU 7V2RV 7V2RW 7V2RY 7V2RZ 7V30A
7V30B 7V30C 7V30D 7V30E 7V30F 7V30G 7V30I 7V30J 7V30K 7V30L 7V30M 7V30N
7V30O 7V30R 7V30S 7V30S 7V30U 7V30V 7V30W 7V30Y 7V30Z 7V31A 7V31B 7V31C
7V31E 7V31F 7V31G 7V31I 7V31J 7V31K 7V31L 7V31M 7V31N 7V31O 7V31R 7V31S
7V31S 7V31U 7V31V 7V31W 7V31Y 7V31Z 7V32A 7V32B 7V32C 7V32E 7V32F 7V32G
7V32I 7V32J 7V32K 7V32L 7V32M 7V32N 7V32O 7V32R 7V32S 7V32S 7V32U 7V32V
7V32W 7V32Y 7V32Z 7V33A 7V33B 7V33C 7V33E 7V33F 7V33G 7V33I 7V33J 7V33K
7V33L 7V33M 7V33N 7V33O 7V33R 7V33S 7V33S 7V33U 7V33V 7V33W 7V33Y 7V33Z
7V35R 7V3MA 7V3MB 7V3MC 7V3MD 7V3ME 7V3MF 7V3MG 7V3MI 7V3MJ 7V3MK 7V3ML
7V3MM 7V3MN 7V3MO 7V3MR 7V3MS 7V3MS 7V3MU 7V3MV 7V3MW 7V3MY 7V3MZ 7V3ZA
7V5CA 7V5CB 7V5DA 7V5DB 7V68R 7V69R 7V6AR 7V6YA 7V6ZA 7V73A 7V73B 7V74A
7V74B 7V75A 7V75B 7V8IC 7V8IE 7V8LC 7V8LE 7V8MC 7V8ME 7V9LR 7V9MR 7VABR
7VADA 7VAEA 7VAFA 7VAGA 7VBHR 7VBIR 7VBLA 7VBLB 7VBLC 7VBLE 7VBLF 7VBLG
7VBLI 7VBLJ 7VBLK 7VBLL 7VBLM 7VBLN 7VBLO 7VBLR 7VBLS 7VBLS 7VBLU 7VBLV
7VBLW 7VBLY 7VBLZ 7VBPA 7VBPB 7VBPC 7VBPE 7VBPF 7VBPG 7VBPI 7VBPJ 7VBPK
7VBPL 7VBPM 7VBPN 7VBPO 7VBPR 7VBPS 7VBPS 7VBPU 7VBPV 7VBPW 7VBPY 7VBPZ
7VBYA 7VBYB 7VBYC 7VBYD 7VBYE 7VBYF 7VBYG 7VBYH 7VBYI 7VBYJ 7VC0A 7VC0B
7VC0C 7VC0E 7VC0F 7VC0G 7VC0I 7VC0J 7VC0K 7VC0L 7VC0M 7VC0N 7VC0O 7VC0R
7VC0S 7VC0S 7VC0U 7VC0V 7VC0W 7VC0Y 7VC0Z 7VC4A 7VC4B 7VC4C 7VC4D 7VC4E
7VC4F 7VC4G 7VC4H 7VC4I 7VC4J 7VD2B 7VD2I 7VDDB 7VDDI 7VDHR 7VDLR 7VDMR
7VFIA 7VFIB 7VFSA 7VFUA 7VFVA 7VFWA 7VFXR 7VGFA 7VGFB 7VGHA 7VGHB 7VGIA
7VGIB 7VGJA 7VGJB 7VGTA 7VGUA 7VGVA 7VGXR 7VGYA 7VGZB 7VH0A 7VH5A 7VH5B
7VH5C 7VH5D 7VH5E 7VH5F 7VH6A 7VH6B 7VH6C 7VH6D 7VH6E 7VH6F 7VHPE 7VHPF
7VHPG 7VHPG 7VHPH 7VHPH 7VHPI 7VHPI 7VHPJ 7VHPK 7VHPL 7VHPM 7VHPN 7VHPN
7VHPO 7VHPO 7VHPP 7VHPP 7VHPQ 7VHPQ 7VHPR 7VHPR 7VHPS 7VHPU 7VHQE 7VHQF
7VHQG 7VHQH 7VHQI 7VHQU 7VIEF 7VIFF 7VIGF 7VIHF 7VKTA 7VL8R 7VL9R 7VLAR
7VLRA 7VLSA 7VLTA 7VLUA 7VLXA 7VLXB 7VLXC 7VLXD 7VLXY 7VLXZ 7VLYA 7VLYC
7VLYD 7VLYE 7VLYF 7VLYG 7VLYH 7VLYY 7VLYZ 7VM2A 7VM2B 7VM2C 7VM2D 7VNM0
7VNM1 7VNM2 7VNM7 7VNM8 7VNM9 7VNMA 7VNMB 7VNMC 7VNMD 7VNME 7VNMF 7VNMG
7VNMH 7VNMI 7VNMJ 7VNMK 7VNML 7VNMM 7VNMN 7VNMO 7VNMP 7VNMQ 7VNMR 7VNMS
7VNMT 7VNMU 7VNMV 7VNMW 7VNMX 7VNPA 7VNPC 7VNPE 7VNPG 7VNQA 7VNQC 7VNQE
7VNQG 7VNRA 7VNRC 7VNRE 7VNRG 7VNY0 7VNY1 7VNY2 7VNY3 7VNY7 7VNY8 7VNY9
7VNYA 7VNYB 7VNYC 7VNYD 7VNYE 7VNYF 7VNYG 7VNYH 7VNYI 7VNYJ 7VNYK 7VNYL

7VNYM 7VNYN 7VNYO 7VNYP 7VNYQ 7VNYR 7VNYS 7VNYT 7VNYU 7VNYV 7VNYW 7VNYX
7VNYY 7VNYZ 7VODA 7VOEA 7VOJA 7VOJB 7VOY0 7VOY1 7VOY2 7VOY3 7VOY4 7VOY5
7VOY6 7VOY7 7VOY8 7VOY9 7VOYA 7VOYB 7VOYC 7VOYD 7VOYE 7VOYF 7VOYG 7VOYH
7VOYI 7VOYJ 7VOYK 7VOYL 7VOYM 7VOYN 7VOYO 7VOYP 7VOYQ 7VOYR 7VOYS 7VOYT
7VOYT 7VOYU 7VOYV 7VOYW 7VOYX 7VOYY 7VOYZ 7VPIA 7VPJA 7VPKA 7VPLA 7VQ1A
7VQ1B 7VQ1C 7VQ1D 7VQ2A 7VQ2B 7VQ2C 7VQ2D 7VQ3A 7VQ3B 7VQ4A 7VQ4B 7VQ5A
7VQ5B 7VQ7A 7VQ7B 7VQXR 7VR1A 7VR1B 7VR5A 7VR5B 7VR7A 7VR7B 7VR7C 7VR8A
7VR8B 7VR8C 7VRJ0 7VRJ1 7VRJ2 7VRJ3 7VRJ4 7VRJ5 7VRJ6 7VRJ7 7VRJ8 7VRJ9
7VRJA 7VRJB 7VRJD 7VRJE 7VRJF 7VRJG 7VRJH 7VRJI 7VRJJ 7VRJK 7VRJL 7VRJM
7VRJN 7VRJO 7VRJP 7VRJQ 7VRJR 7VRJS 7VRJT 7VRJU 7VRJV 7VRJW 7VRJX 7VRJY
7VRJZ 7VSGA 7VSGB 7VSHA 7VSHC 7VSIA 7VSIB 7VSOA 7VU5A 7VU5B 7VUGR 7VUYR
7VUZR 7VV0R 7VV3R 7VV4R 7VV5R 7VV6R 7VVJR 7VVKR 7VVLR 7VVMR 7VVNR 7VVOR
7VWCA 7VWCB 7VWLA 7VWLB 7VWLC 7VWLD 7VWLE 7VWLF 7VWLG 7VWLI 7VWLJ 7VWLK
7VWLL 7VWLM 7VWLN 7VWLO 7VWLR 7VWLS 7VWLS 7VWLU 7VWLV 7VWLW 7VWLY 7VWLZ
7VX8A 7VX8B 7VY3A 7VY3B 7VY3D 7VY3E 7VY3H 7VY3L 7VY3M 7VY3X 7VZBA 7VZBB
7W01A 7W02A 7W0LQ 7W0LR 7W0MR 7W0NQ 7W0NR 7W0OR 7W0PR 7W2BA 7W2BC 7W2CA
7W2CC 7W2EA 7W2EB 7W2ZR 7W47A 7W47B 7W48A 7W48B 7W49A 7W49B 7W4AA 7W4AB
7W4OA 7W4OB 7W4OC 7W4OD 7W4OE 7W4OF 7W4OG 7W4OH 7W4PA 7W4PB 7W4PC 7W4PD
7W4PE 7W4PF 7W4PG 7W4PH 7W53R 7W55R 7W56R 7W57R 7W6KA 7W6KB 7W6PR 7W6XA
7W6YA 7W6ZA 7W72A 7W72K 7W72S 7W72T 7W72U 7W74A 7W77B 7W77D 7W78B 7W78D
7W79B 7W79D 7W7AB 7W7AD 7W7BB 7W7BD 7W7CB 7W7CD 7W7DB 7W7DD 7W7ER 7W7FB
7W7FD 7W7GC 7W7RA 7W7SA 7W7SB 7W7SC 7W7SD 7W9KA 7W9KB 7W9LA 7W9LB 7W9MA
7W9MB 7W9PA 7W9PB 7W9TA 7W9TB 7W9WA 7W9WB 7W9WC 7WAGA 7WAWA 7WAXA 7WBJR
7WC4A 7WC5A 7WC6A 7WC7A 7WC8A 7WC9A 7WDBA 7WE4A 7WELA 7WEMA 7WEMB 7WEMC
7WEMD 7WEME 7WEMF 7WEMG 7WEMH 7WEMI 7WEMJ 7WF3B 7WF3D 7WF3F 7WF3H 7WF4B
7WF4D 7WF4F 7WF4H 7WF7A 7WFFA 7WFFB 7WFFC 7WFFD 7WFFD 7WFFE 7WFFE 7WFFF
7WFFG 7WFRA 7WFWA 7WGDA 7WGTA 7WHVA 7WHVB 7WHWA 7WHWB 7WI3A 7WI3A 7WI3B
7WI3B 7WI3C 7WI3C 7WI3D 7WI3D 7WI3E 7WI3E 7WI3F 7WI3F 7WI3G 7WI3G 7WI3H
7WI3H 7WI3I 7WI3I 7WI3J 7WI3J 7WI3K 7WI3K 7WI3L 7WI3L 7WI3M 7WI3M 7WI3N
7WI3N 7WI3O 7WI3O 7WI3P 7WI3P 7WI3Q 7WI3Q 7WI3R 7WI3R 7WI3S 7WI3S 7WI3T
7WI3T 7WI3U 7WI3U 7WI3V 7WI3V 7WI3W 7WI3X 7WI3Y 7WI3Z 7WI6A 7WI6B 7WI8A
7WI8B 7WICR 7WIGR 7WIHA 7WIHB 7WJ5R 7WJIC 7WJMA 7WJMB 7WJNA 7WJNB 7WJOA
7WJOB 7WKDR 7WKSA 7WKSB 7WKWA 7WKWB 7WLDG 7WLDK 7WLDS 7WLDT 7WLDU 7WLIA
7WLJA 7WLKA 7WLLA 7WLTA 7WLUA 7WLWA 7WNQA 7WNQB 7WNQC 7WNXA 7WQ3R 7WQ4R
7WRAA 7WRAB 7WRAC 7WRAD 7WRBA 7WRBB 7WRBC 7WRBD 7WRCA 7WRCB 7WRCC 7WRCD
7WRDA 7WRDB 7WRDC 7WRDD 7WREA 7WREB 7WREC 7WRED 7WRFA 7WRFB 7WRFC 7WRFD
7WSIA 7WSMA 7WSNA 7WSOA 7WSOB 7WSOC 7WSOK 7WSVA 7WSVB 7WSVC 7WSVD 7WSVE
7WSVF 7WSVG 7WU2R 7WU3R 7WU4R 7WU5R 7WU9R 7WUIR 7WUJE 7WUQR 7WV9R 7WVUR
7WVVR 7WVWR 7WVXR 7WVYR 7WWBA 7WWBB 7WXUR 7WXWR 7WY0R 7WY5R 7WY8R 7WYBR
7WYSA 7WYSB 7WYSE 7WYSG 7WYTA 7WYTB 7WYTE 7WYTG 7WYUA 7WYUB 7WYUG 7WYVC
7WYVD 7WYVE 7WYWA 7WYWB 7WYWG 7WYXA 7WYXB 7WYXG 7WYYA 7WYYB 7WYYG 7WYZA
7WYZB 7WYZG 7WZ0A 7WZ0B 7WZ0G 7WZ4R 7WZ7R 7X05A 7X05B 7X06A 7X06B 7X1TA
7X1UA 7X20A 7X20B 7X21A 7X21B 7X22A 7X22B 7X23A 7X23B 7X24A 7X24B 7X2CF
7X2DF 7X2FF 7X2UA 7X2UB 7X2VR 7X5HR 7X5VA 7X5VB 7X5VC 7X5VD 7X8RR 7X8SR

7X9AR 7X9BR 7X9CR 7X9YR 7XA3R 7XATA 7XAUA 7XAVA 7XBDA 7XBWR 7XBXR 7XEMA
7XEMB 7XEMC 7XEMD 7XEOA 7XEOB 7XEOC 7XEOD 7XERA 7XERB 7XERC 7XERD 7XEUA
7XEUB 7XEUC 7XEUD 7XEVA 7XEVB 7XEVC 7XEVD 7XEWA 7XEWB 7XEWC 7XEWD 7XJHR
7XJIR 7XJKF 7XJLF 7XK3B 7XK3C 7XK3D 7XK3E 7XK3F 7XK4B 7XK4C 7XK4D 7XK4E
7XK4F 7XK5B 7XK5C 7XK5D 7XK5E 7XK5F 7XK6B 7XK6C 7XK6D 7XK6E 7XK6F 7XK7B
7XK7C 7XK7D 7XK7E 7XK7F 7XKDR 7XKER 7XKFR 7XMRR 7XMSR 7XMTR 7XN9A 7XNAA
7XNOA 7XNOC 7XNOD 7XNOE 7XNOG 7XNOH 7XNOI 7XNOY 7XNOZ 7XOUR 7XOVR 7XOWR
7XOXR 7XP4R 7XP5R 7XP6R 7XPZA 7XQ0A 7XQ1A 7XQ2A 7XR4A 7XR4B 7XR4C 7XR6A
7XR6B 7XR6C 7XT6A 7XT6B 7XT6C 7XT6D 7XT8R 7XT9R 7XTAR 7XTBR 7XTCR 7XTQR
7XVEA 7XVEB 7XVFA 7XVFB 7XW5R 7XW6R 7XW7R 7XXBA 7XXBB 7XZ5R 7XZ6R 7Y12R
7Y13R 7Y14R 7Y15R 7Y24E 7Y26E 7Y27E 7Y44A 7Y44B 7Y44C 7Y44D 7Y44G 7Y44I
7Y44J 7Y44K 7Y44L 7Y44M 7Y44N 7Y44O 7Y44P 7Y44Q 7Y44T 7Y44V 7Y44W 7Y44X
7Y44Y 7Y44Z 7Y45C 7Y45D 7Y45E 7Y46C 7Y46D 7Y46E 7Y89A 7Y9TA 7Y9TB 7Y9UA
7Y9UB 7Y9VA 7Y9VB 7YEPA 7YEPB 7YEPC 7YEPD 7YL9H 7YL9I 7YL9J 7YL9K 7YL9L
7YL9M 7YL9N 7YP7R 7YPYA 7YPYB 7YPYC 7YPYD 7YPYG 7YPYI 7YPYJ 7YPYK 7YPYL
7YPYM 7YPYN 7YPYO 7YPYP 7YPYQ 7YPYT 7YPYV 7YPYW 7YPYX 7YPYY 7YPYZ 7YR5A
7YR5B 7YU3R 7YU4A 7YU5R 7YU6R 7YU7R 7YU8R 7YXAA 7YZIA 7YZIB 7YZKA 7YZKB
7YZYA 7YZYB 7YZYC 7YZYE 7YZYF 7YZYG 7YZYI 7YZYJ 7YZYK 7Z09A 7Z0AA 7Z0CA
7Z0DA 7Z0EP 7Z0SC 7Z0SD 7Z0TC 7Z0TD 7Z10A 7Z10B 7Z10C 7Z10E 7Z10G 7Z10H
7Z10I 7Z14A 7Z14B 7Z14C 7Z14D 7Z14E 7Z6MA 7Z6NA 7Z6NB 7Z7RA 7Z7RH 7Z7RJ
7Z7RK 7Z7RL 7Z7RM 7Z7RN 7Z7SA 7Z7SH 7Z7SJ 7Z7SK 7Z7SL 7Z7SM 7Z7SN 7Z7TA
7Z7TH 7Z7TJ 7Z7TK 7Z7TL 7Z7TM 7Z7TN 7Z7VA 7Z7VH 7Z7VJ 7Z7VK 7Z7VL 7Z7VM
7Z7VN 7Z80A 7Z80H 7Z80J 7Z80K 7Z80L 7Z80M 7Z80N 7Z83A 7Z83H 7Z83J 7Z83K
7Z83L 7Z83M 7Z83N 7Z84A 7Z84H 7Z84J 7Z84K 7Z84L 7Z84M 7Z84N 7ZC2A 7ZC5A
7ZC5H 7ZC5J 7ZC5K 7ZC5L 7ZC5M 7ZC5N 7ZC6A 7ZC6B 7ZC6D 7ZC6E 7ZC6G 7ZCIA
7ZCIH 7ZCIJ 7ZCIK 7ZCIL 7ZCIM 7ZCIN 7ZCUA 7ZCUB 7ZCUC 7ZCUD 7ZCUE 7ZCUF
7ZCUG 7ZCUH 7ZCUI 7ZCUJ 7ZCUK 7ZCUL 7ZCUM 7ZCUN 7ZCUO 7ZCUP 7ZCUQ 7ZCUR
7ZD6A 7ZD6H 7ZD6J 7ZD6K 7ZD6L 7ZD6M 7ZD6M 7ZD6N 7ZD6N 7ZD6O 7ZD6P 7ZD6Q
7ZD6R 7ZD6U 7ZD6V 7ZD6V 7ZD6W 7ZD6W 7ZD6X 7ZD6Y 7ZD6Z 7ZDHA 7ZDHH 7ZDHJ
7ZDHK 7ZDHL 7ZDHM 7ZDHM 7ZDHN 7ZDHN 7ZDHO 7ZDHP 7ZDHQ 7ZDHR 7ZDHU 7ZDHV
7ZDHV 7ZDHW 7ZDHW 7ZDHX 7ZDHY 7ZDHZ 7ZDIA 7ZDIB 7ZDIC 7ZDID 7ZDJA 7ZDJH
7ZDJJ 7ZDJK 7ZDJL 7ZDJM 7ZDJM 7ZDJN 7ZDJN 7ZDJO 7ZDJP 7ZDJQ 7ZDJR 7ZDJU
7ZDJV 7ZDJV 7ZDJW 7ZDJW 7ZDJX 7ZDJY 7ZDJZ 7ZDMA 7ZDMH 7ZDMJ 7ZDMK 7ZDML
7ZDMM 7ZDMM 7ZDMN 7ZDMN 7ZDMO 7ZDMP 7ZDMQ 7ZDMR 7ZDMU 7ZDMV 7ZDMV 7ZDMW
7ZDMW 7ZDMX 7ZDMY 7ZDMZ 7ZDPA 7ZDPH 7ZDPJ 7ZDPK 7ZDPL 7ZDPM 7ZDPM 7ZDPN
7ZDPN 7ZDPO 7ZDPP 7ZDPQ 7ZDPR 7ZDPU 7ZDPV 7ZDPV 7ZDPW 7ZDPW 7ZDPX 7ZDPY
7ZDPZ 7ZDZA 7ZDZB 7ZDZC 7ZDZD 7ZE3A 7ZE3B 7ZE3C 7ZE3D 7ZE3E 7ZE3F 7ZE3G
7ZE3H 7ZE3I 7ZE3J 7ZE3K 7ZE3L 7ZE3M 7ZE3N 7ZE3O 7ZE3P 7ZE3Q 7ZE3R 7ZE8A
7ZE8B 7ZE8C 7ZE8D 7ZE8E 7ZE8F 7ZE8G 7ZE8H 7ZE8I 7ZE8J 7ZE8K 7ZE8L 7ZE8M
7ZE8N 7ZE8O 7ZE8P 7ZE8Q 7ZE8R 7ZEBA 7ZEBH 7ZEBJ 7ZEBK 7ZEBL 7ZEBM 7ZEBM
7ZEBN 7ZEBN 7ZEBO 7ZEBP 7ZEBQ 7ZEBR 7ZEBU 7ZEBV 7ZEBV 7ZEBW 7ZEBW 7ZEBX
7ZEBY 7ZEBZ 7ZGOA 7ZGOB 7ZI0A 7ZK1A 7ZK3A 7ZK3B 7ZKWA 7ZKWB 7ZKZA 7ZNQY
7ZNQY 7ZXYA 7ZXYB 7ZXYC 7ZXYD 7ZXYE 7ZXYF 7ZXYG 7ZXYH 7ZXYI 7ZXYJ 7ZXYK
7ZXYL 7ZXYM 7ZXYN 7ZXYO 7ZXYP 7ZYIA 8A8CA 8AP9O 8AP9P 8AP9Q 8AP9R 8AP9S

8AP9T 8AP9U 8AP9V 8AP9W 8AP9X 8CRQB 8CRQC 8CRQD 8CRQE 8CRRB 8CRRC 8CRRD
8CRRE 8CRTB 8CRTC 8CRTD 8CRTE 8CRTK 8CRTL 8CRTP 8CRTQ 8CSXK 8CSXL 8CSXQ
8CT2B 8CT2D 8CT3B 8CT3C 8CT3D 8CT3E 8CTED 8CTEK 8CTEL 8CTEM 8CTEN 8CTEO
8CTEP 8CTEQ 8CTER 8CTES 8CTET 8CTJA 8CTKA 8CTKB 8CU6A 8CU7A 8CUBA 8CUBB
8CXOA 8D1EA 8D1EB 8D1EC 8D1ED 8D1EE 8D1FA 8D1FB 8D1FC 8D1FD 8D1FE 8D1GA
8D1GB 8D1GC 8D1GD 8D1GE 8D1HA 8D1HB 8D1HC 8D1HD 8D1HE 8D1IA 8D1IB 8D1IC
8D1ID 8D1IE 8D1JA 8D1JB 8D1JC 8D1JD 8D1JE 8D1KA 8D1KB 8D1KC 8D1KD 8D1KE
8D1LA 8D1LB 8D1LC 8D1LD 8D1LE 8D1MA 8D1MB 8D1MC 8D1MD 8D1ME 8D1NA 8D1NB
8D1NC 8D1ND 8D1NE 8D1OA 8D1OB 8D1OC 8D1OD 8D1OE 8D3FA 8D3UA 8D3UB 8D3UG
8D3VA 8D3VB 8D3VG 8D3WA 8D3WB 8D3WG 8D3XA 8D3XB 8D3XG 8D3YA 8D3YB 8D3YG
8D4TM 8D4TN 8D4TO 8D4TP 8D4TQ 8D4TT 8D4TV 8D4TW 8D4TX 8D4TY 8DCKA 8DCKB
8DCKD 8DCKE 8DCKF 8DCKH 8DCKI 8DCKJ 8DCKL 8DCRR 8DCSR 8DD2A 8DD2B 8DD2C
8DD2D 8DD2E 8DEPA 8DEUA 8DEUB 8DEUC 8DEVA 8DEVB 8DEVC 8DEWA 8DEWB 8DEWC
8DFLA 8DFLB 8DFLC 8DFLD 8DH6A 8DH6B 8DH6C 8DH6E 8DH6G 8DH6H 8DH6I 8DJKA
8DJKB 8DJMA 8DJMB 8DKEP 8DKIP 8DKMP 8DKUB 8DKUD 8DKWP 8DKXP 8DL0B 8DL0D
8DN8B 8DN8D 8DNCB 8DNCD 8DNEB 8DNED 8DOUB 8DOUD 8DPFA 8DPGA 8DPHA 8DPIA
8DU3A 8E4LA 8E4LB 8E4LC 8E4LD 8E4MA 8E4MB 8E4MC 8E4MD 8E4NA 8E4NB 8E4NC
8E4ND 8E4OA 8E4OB 8E4OC 8E4OD 8E4PA 8E4PB 8E4PC 8E4PD 8E4QA 8E4QB 8E4QC
8E4QD 8E9GA 8E9GH 8E9GJ 8E9GK 8E9GL 8E9GM 8E9GN 8E9HA 8E9HH 8E9HJ 8E9HK
8E9HL 8E9HM 8E9HN 8E9IA 8E9IH 8E9IJ 8E9IK 8E9IL 8E9IM 8E9IN 8EIGA 8EIOA
8EIQA 8EJ1A 8GOEA 8GOFA 8GQYA 8GQYB 8GQYC 8GQYD 8GQYE 8GVMA 8GVMB 8GVMC
8GVMD 8GVMG 8GVMI 8GVMJ 8GVMK 8GVML 8GVMM 8GVMN 8GVMO 8GVMP 8GVMQ 8GVMT
8GVMV 8GVMW 8GVMX 8GVMY 8GVMZ 8PRNA 8PRNB 8PRNC

#### 2.2 The UniProt_Mem dataset (80634 sequences)

A0A017SFB8 A0A023I4C8 A0A023PXB5 A0A023PXB9 A0A023PXC2 A0A023PXC7 A0A023PXD5
A0A023PXD9 A0A023PXF8 A0A023PXG3 A0A023PXG7 A0A023PXH4 A0A023PXH6 A0A023PXH9
A0A023PXI0 A0A023PXJ3 A0A023PXJ7 A0A023PXK2 A0A023PXK7 A0A023PXL1 A0A023PXL7
A0A023PXM7 A0A023PXN9 A0A023PXQ4 A0A023PXQ9 A0A023PYC6 A0A023PYD0 A0A023PYD7
A0A023PYD9 A0A023PYE4 A0A023PYG1 A0A023PYI5 A0A023PYJ0 A0A023PYJ7 A0A023PYK2
A0A023PZ94 A0A023PZ99 A0A023PZB8 A0A023PZD0 A0A023PZE1 A0A023PZE2 A0A023PZE6
A0A023PZF3 A0A023PZF9 A0A023PZG0 A0A023PZG9 A0A023PZH1 A0A023PZH4 A0A023PZH5
A0A023PZI4 A0A023PZJ3 A0A023PZK9 A0A023PZL2 A0A023PZM3 A0A024B7W1 A0A024FA41
A0A026W182 A0A059J0G5 A0A059JJ46 A0A059JK44 A0A060A682 A0A060L102 A0A060L4I9
A0A061ACU2 A0A061I403 A0A067XMT5 A0A068A9T2 A0A068AA78 A0A068ACU3 A0A068Q5V6
A0A068Q605 A0A068Q609 A0A068Q6L2 A0A068Q721 A0A072U307 A0A072VHJ1 A0A072VMJ3
A0A075B734 A0A075F7E9 A0A075F932 A0A075FBT3 A0A075HNX4 A0A075TMP8 A0A075TRA9
A0A075TRL0 A0A075TRL5 A0A077K8G3 A0A077K9K6 A0A077YBL0 A0A078BQP2 A0A078H868
A0A084AFH0 A0A084API1 A0A084B9Z2 A0A084B9Z5 A0A084R1J1 A0A084R1M7 A0A086F3E3
A0A087WTH1 A0A087WTH5 A0A087X1C5 A0A089FNE5 A0A089FRP6 A0A095C325 A0A096LP01
A0A096LPK9 A0A097ZPD8 A0A097ZPE3 A0A097ZPE6 A0A097ZPF7 A0A098D1J7 A0A098DDI4
A0A098DJ84 A0A098DME4 A0A0A1C3I2 A0A0A1C930 A0A0A1GNF2 A0A0A1H7M6 A0A0A2IBP6
A0A0A2K5R6 A0A0B2XGM8 A0A0B4J1G0 A0A0B4J2F0 A0A0B4K692 A0A0B4K753 A0A0B4KEE4
A0A0B4KZX8 A0A0B4L1W8 A0A0B4LFY9 A0A0B4ZTQ2 A0A0B5A051 A0A0B5A886 A0A0B5EMG9

A0A0B5GR44 A0A0B5KYT4 A0A0B5L778 A0A0B5LB52 A0A0B7P9G0 A0A0C1BV72 A0A0C1C354
A0A0C1E1L9 A0A0C2W6G6 A0A0C4G489 A0A0C5PHQ7 A0A0C5PRW9 A0A0C5Q309 A0A0C5Q4Y6
A0A0C5QRZ2 A0A0C6DUU3 A0A0D1BUH6 A0A0D1C6P2 A0A0D1CVX2 A0A0D1CZ63 A0A0D1DMJ7
A0A0D1DT62 A0A0D1DT68 A0A0D1DYJ6 A0A0D1E6B3 A0A0D1E6R6 A0A0D2YFZ8 A0A0D3MU35
A0A0D3MU50 A0A0D9SF12 A0A0E0SNE8 A0A0E3D8K9 A0A0E3D8L0 A0A0E3D8L1 A0A0E3D8L3
A0A0E3D8L5 A0A0E3D8L6 A0A0E3D8M1 A0A0E3D8M2 A0A0E3D8M3 A0A0E3D8M6 A0A0E3D8N1
A0A0E3D8N7 A0A0E3D8P0 A0A0E3D8P5 A0A0E3D8P8 A0A0E3D8Q2 A0A0E3NFS5 A0A0E4AZP4
A0A0E4BZH1 A0A0F6B506 A0A0F6BAS6 A0A0F6P9C0 A0A0F7GG06 A0A0F7TN11 A0A0F7TN60
A0A0F7TSZ5 A0A0F7TXA0 A0A0F7TXA1 A0A0F7TZE0 A0A0F7U0Z9 A0A0F9XXG3 A0A0G2K1Q8
A0A0G2K309 A0A0G2K5L2 A0A0G2KBC9 A0A0G2KQY6 A0A0G2KTI4 A0A0G2RKY1 A0A0H2V871
A0A0H2V8B5 A0A0H2VAP9 A0A0H2VCA1 A0A0H2VDI7 A0A0H2VG78 A0A0H2WHF1 A0A0H2XEK8
A0A0H2XFP1 A0A0H2XG66 A0A0H2ZC68 A0A0H2ZFV0 A0A0H2ZGW7 A0A0H2ZH12 A0A0H2ZHZ4
A0A0H2ZI93 A0A0H2ZLQ1 A0A0H2ZMF9 A0A0H2ZNG3 A0A0H2ZQ75 A0A0H3AJF5 A0A0H3AKU6
A0A0H3C677 A0A0H3CC47 A0A0H3CMG2 A0A0H3FQN0 A0A0H3FRB6 A0A0H3G4K1 A0A0H3G586
A0A0H3GCG4 A0A0H3GDH9 A0A0H3GFN0 A0A0H3GGE2 A0A0H3GGY3 A0A0H3GM48 A0A0H3H546
A0A0H3H7Y9 A0A0H3HA88 A0A0H3HDD6 A0A0H3HIJ5 A0A0H3JML2 A0A0H3JTK0 A0A0H3JU73
A0A0H3K104 A0A0H3M5A8 A0A0H3M5L9 A0A0H3MAA9 A0A0H3MD02 A0A0H3MFZ3 A0A0H3MGR4
A0A0H3N9T8 A0A0H3NBQ0 A0A0H3NIL4 A0A0H3NJI9 A0A0K1IRS6 A0A0K2S4Q6 A0A0K3AWM6
A0A0L7K812 A0A0M3R8G1 A0A0M3STV6 A0A0M3STX4 A0A0M4FCN7 A0A0M4FLW6 A0A0M4FTF3
A0A0M4KRN2 A0A0M9EQT6 A0A0N0DCA4 A0A0N0DCA8 A0A0N7D7C9 A0A0N7F297 A0A0N9HKQ7
A0A0N9HT29 A0A0N9HTU1 A0A0P0VIP0 A0A0P0WL81 A0A0P0WY03 A0A0P0X004 A0A0P0XII1
A0A0P0XL10 A0A0P0XXR9 A0A0P0ZEA9 A0A0P1AM89 A0A0P1ATX0 A0A0R0HPY5 A0A0R4IDP3
A0A0R4IDX9 A0A0R4IES7 A0A0R4IF99 A0A0R4IGV4 A0A0R4IM31 A0A0R4IMY7 A0A0R4IP11
A0A0R4IQZ2 A0A0R4IY06 A0A0R6L508 A0A0R8YXT5 A0A0S1TP26 A0A0S1TPC7 A0A0S1TQ04
A0A0S2DN74 A0A0S2DNG5 A0A0S2IHL2 A0A0S2II38 A0A0S6XH62 A0A0U1LQE1 A0A0U1LQF6
A0A0U1LR74 A0A0U1LSP7 A0A0U1QT59 A0A0U1RPR8 A0A0U1RQ45 A0A0U1RQS6 A0A0U1RRA0
A0A0U1RRN3 A0A0U2IR85 A0A0U2U8U5 A0A0U2UXG3 A0A0U2V7I8 A0A0U4EBH5 A0A0U4ZPJ7
A0A0U5CJM3 A0A0U5CJT8 A0A0U5CJU6 A0A0U5CJV1 A0A0U5GIU9 A0A0U5GJZ5 A0A0U5GRB4
A0A0X1KG70 A0A0Y0GRS3 A0A125QXJ1 A0A125QZE2 A0A131KVX9 A0A131MCZ8 A0A140FAN3
A0A140H545 A0A140H546 A0A140IL90 A0A140IL91 A0A140IL92 A0A140JWM8 A0A140JWS3
A0A140JWS9 A0A140JWT0 A0A140JWT1 A0A140JWT2 A0A140JWT3 A0A140JWT4 A0A140JWT6
A0A140JWT8 A0A140JWT9 A0A140JWU1 A0A140LFM6 A0A140LHF2 A0A140LHV9 A0A140NFA3
A0A142I5B9 A0A142ZC57 A0A144KPK9 A0A144YEA5 A0A150XSR0 A0A159AKG3 A0A160PAH3
A0A161CLJ6 A0A161GJD5 A0A161IUT7 A0A165U5Z9 A0A166YZU9 A0A166Z003 A0A167LUR6
A0A167LUS5 A0A172M476 A0A172M477 A0A178WF56 A0A179H0I3 A0A179H0I7 A0A179H0T5
A0A179HJU2 A0A182BSS3 A0A182BSS5 A0A193AUK9 A0A193PS58 A0A1B0GQX2 A0A1B0GQX3
A0A1B0GRQ0 A0A1B0GSN8 A0A1B0GST9 A0A1B0GSZ0 A0A1B0GTB2 A0A1B0GTI8 A0A1B0GTK4
A0A1B0GTQ4 A0A1B0GTU2 A0A1B0GTW7 A0A1B0GTY4 A0A1B0GU29 A0A1B0GUA5 A0A1B0GUA7
A0A1B0GUW6 A0A1B0GUW7 A0A1B0GV85 A0A1B0GV90 A0A1B0GVN3 A0A1B0GVQ0 A0A1B0GVT2
A0A1B0GVV1 A0A1B0GVY4 A0A1B0GVZ9 A0A1B0GW54 A0A1B0GW64 A0A1B0GWB2 A0A1B0GWG4
A0A1B2CTA8 A0A1B2CTB0 A0A1B4XBH0 A0A1B4XBH1 A0A1B4XBH8 A0A1B4XBI1 A0A1B4XBI3
A0A1B4XBI5 A0A1B4XBJ9 A0A1B5L6A5 A0A1B5L780 A0A1B7YCK2 A0A1B7YCL6 A0A1B7YCX1
A0A1C5G2D0 A0A1C7E424 A0A1D5NSK0 A0A1D5NY17 A0A1D5PXA5 A0A1D6F9Y9 A0A1D6GQ67
A0A1D6HSP4 A0A1D6L709 A0A1D6N272 A0A1D8PCB9 A0A1D8PCL1 A0A1D8PD39 A0A1D8PH98

A0A1D8PI71 A0A1D8PI78 A0A1D8PIC7 A0A1D8PJ25 A0A1D8PK89 A0A1D8PMB1 A0A1D8PN12
A0A1D8PN14 A0A1D8PN88 A0A1D8PNP3 A0A1D8PPG4 A0A1D8PPI5 A0A1D8PQG0 A0A1D8QMD1
A0A1D8QMD2 A0A1D8QMG4 A0A1E1FFK8 A0A1E1FFL4 A0A1E1FFL6 A0A1E1FFM3 A0A1E1FFM8
A0A1E1FFM9 A0A1E1FFN3 A0A1E1FFN4 A0A1E1FFP5 A0A1F4 A0A1I9LN01 A0A1I9Q5Z0
A0A1I9R3Y6 A0A1J0M738 A0A1L8F5J9 A0A1L8HYT7 A0A1L8I316 A0A1L9UQW4 A0A1L9WLE2
A0A1L9WN31 A0A1L9WN72 A0A1L9WQP6 A0A1L9WQV4 A0A1L9WUS5 A0A1L9WUV2 A0A1L9WVI3
A0A1N7SYS3 A0A1P7Y0A2 A0A1P7Y0A8 A0A1P7Y0D0 A0A1P7Y0D4 A0A1P8ATR9 A0A1R3RGJ7
A0A1S3ZP85 A0A1S4GYH6 A0A1S4H2E2 A0A1S6M251 A0A1S7LCW6 A0A1T4LJG1 A0A1U8QFC3
A0A1U8QG99 A0A1U8QHS4 A0A1U8QKX8 A0A1U8QT10 A0A1U8QTJ9 A0A1U8QW16 A0A1U8QYH7
A0A1U8QYW8 A0A1U9YHZ8 A0A1U9YHZ9 A0A1U9YI05 A0A1U9YI12 A0A1V0HUU2 A0A1V0QSA9
A0A1V0QSB7 A0A1V0QSE4 A0A1V0QSE7 A0A1V0QSE9 A0A1V0QSF1 A0A1V1FNM9 A0A1V1FNZ5
A0A1V1GB10 A0A1V6NWJ0 A0A1V6NWP3 A0A1V6NYL5 A0A1V6NZ11 A0A1V6PB34 A0A1V6PBC8
A0A1V6PBE7 A0A1W2PS18 A0A1W5SP56 A0A1W5T1T6 A0A1W5T1U4 A0A1W5T1Y3 A0A1W5T1Y6
A0A1W5T3J9 A0A1W5X0D5 A0A1X9Q9F0 A0A1X9Q9H1 A0A1X9QDU4 A0A1X9QDU5 A0A1Y0BRF0
A0A1Y0BRF2 A0A1Y0BRF5 A0A1Y0BRF7 A0A1Y0BRF9 A0A1Y1C7S2 A0A1Z2R986 A0A1Z3GBS4
A0A218N030 A0A218N031 A0A218N033 A0A218N034 A0A218N035 A0A218N036 A0A254TZW7
A0A286LF00 A0A286LF01 A0A286YEC0 A0A286YF18 A0A286YF58 A0A286YFK9 A0A287A2K5
A0A291NUG3 A0A291NUI4 A0A291NUI5 A0A291PQF1 A0A291PQG3 A0A291PQH4 A0A2A5JY22
A0A2A5K1W4 A0A2A5K485 A0A2A5K5H4 A0A2G5I8H1 A0A2G5ID46 A0A2H3CNS9 A0A2H3CNY6
A0A2H3CSA7 A0A2H3CZX2 A0A2H3CZZ2 A0A2H3E985 A0A2H4DGV8 A0A2H5AIX6 A0A2H5AIX7
A0A2H5AIZ9 A0A2H5AJ00 A0A2I0C265 A0A2I1BSU0 A0A2I1BSV1 A0A2I1BSW0 A0A2I1BSW8
A0A2I1BT01 A0A2I1BT09 A0A2I1C3T4 A0A2I1C3U4 A0A2I1C3V3 A0A2I2F271 A0A2I2F284
A0A2I6PIZ2 A0A2I6PIZ9 A0A2I6PJ00 A0A2I6PJ07 A0A2I6PJ08 A0A2I6PJ12 A0A2I6PJ13
A0A2K1X4I9 A0A2K1YKA4 A0A2K3DMP5 A0A2K4Z9J5 A0A2K4Z9K4 A0A2K9RFY0 A0A2K9RG08
A0A2L0P0K0 A0A2L0P0L8 A0A2L0VBG2 A0A2P1DP74 A0A2P1DP75 A0A2P1DP87 A0A2P1DP94
A0A2P1DPA5 A0A2R4SBW0 A0A2R8Q3S9 A0A2R8QF68 A0A2R8RY99 A0A2R8VHF7 A0A2R8YCJ5
A0A2R9TD79 A0A2R9YJI3 A0A2S4MYF8 A0A2S4N3N0 A0A2T4VDM4 A0A2U1KZS6 A0A2U1LIM9
A0A2U8U2K9 A0A2U8U2L0 A0A2U8U2L6 A0A2U8U2M7 A0A2U8U2M8 A0A2V5GRP3 A0A2V5GZ95
A0A2Z4HPY0 A0A2Z4HPY4 A0A2Z4HPZ0 A0A2Z4HPZ5 A0A2Z4HPZ7 A0A2Z4HQ22 A0A2Z5D850
A0A2Z5D852 A0A2Z5D854 A0A2Z5TMB8 A0A2Z5TTA9 A0A2Z5XAK5 A0A2Z6FZ19 A0A2Z6FZ20
A0A315 A0A318 A0A319 A0A321 A0A322 A0A323 A0A328 A0A329 A0A330 A0A331 A0A332
A0A334 A0A335 A0A340 A0A343URW6 A0A343URW7 A0A345BJN8 A0A345BJP1 A0A345BJP3
A0A345BJP8 A0A345BJP9 A0A345BJQ0 A0A345 A0A346B546 A0A346 A0A347 A0A348AXX9
A0A348 A0A349 A0A350 A0A351 A0A352 A0A353 A0A354 A0A355 A0A361 A0A362 A0A363
A0A364 A0A365 A0A366 A0A382 A0A384 A0A384JQH2 A0A384XG60 A0A385 A0A386KZI3
A0A387 A0A388 A0A390 A0A393 A0A396GS61 A0A396HQY2 A0A3B1DZW5 A0A3B1EFP7
A0A3B1EFQ0 A0A3B7TNF2 A0A3F2YED3 A0A3G1DIJ8 A0A3G1DIQ9 A0A3G1DJE2 A0A3G1DJE8
A0A3G1DJF1 A0A3G1DJG1 A0A3G1DJH6 A0A3G1DJL2 A0A3G9GNJ4 A0A3G9GNK2 A0A3G9GUR6
A0A3G9GX61 A0A3G9H2R5 A0A3G9H8P0 A0A3G9H9H1 A0A3G9HB50 A0A3G9HRV8 A0A3G9JTR4
A0A3G9JYH7 A0A3G9K3N1 A0A3G9K5C8 A0A3L7I2I8 A0A3Q0NBH7 A0A3Q1LRJ2 A0A3Q2HW92
A0A3Q2IDB4 A0A3Q7HBJ5 A0A3Q7HS74 A0A3Q9FEJ4 A0A3Q9R4N5 A0A3R0A696 A0A3S5HYN5
A0A3S9NM20 A0A3T0QHT2 A0A401ETL2 A0A411KUQ1 A0A411KUQ2 A0A411KUQ5 A0A411KUU9
A0A411KUX1 A0A411KZY6 A0A411PQP0 A0A411PQQ5 A0A443HJZ5 A0A443HK79 A0A452G813
A0A455LLV4 A0A455LLV8 A0A455LLW0 A0A455LLW3 A0A455LLW7 A0A455LLW9 A0A455LM21

A0A455LM26 A0A455LM27 A0A455LRX2 A0A455M2Z1 A0A455M7R7 A0A455R413 A0A455R4X0
A0A455R4Z0 A0A455R624 A0A455RAK9 A0A455ZJC3 A0A481NR20 A0A481WNM6 A0A481WQD3
A0A481WQK1 A0A482APM8 A0A482APN3 A0A482AQK7 A0A482AQV2 A0A482AR57 A0A482ARC8
A0A482ARE1 A0A482ARN9 A0A482ARP7 A0A482AS24 A0A482ASA5 A0A482AT42 A0A482ATU4
A0A482AU60 A0A482AU66 A0A482NAF7 A0A482NAL3 A0A482NB13 A0A482ND39 A0A484FVL2
A0A484FXR4 A0A494BZU4 A0A4C3GMC1 A0A4D6Q414 A0A4D6Q415 A0A4P7TKC8 A0A4P7TME1
A0A4P7TN82 A0A4P7TSF2 A0A4P8DJC8 A0A4P8DJE5 A0A4P8DK01 A0A4P8DK16 A0A4P8GFD0
A0A4P8GG95 A0A4P8W744 A0A4P8W7F5 A0A4Q4NJ90 A0A4Q4NMP3 A0A4Q9KQH5 A0A4R2UGS4
A0A4V1E8I5 A0A4Y1YTM1 A0A4Z3 A0A509ADH4 A0A509AE54 A0A509AI82 A0A509AK04
A0A509AN59 A0A509APX1 A0A515MEM9 A0A515MEN7 A0A517FNB9 A0A517FNC4 A0A517FNC5
A0A517FNC6 A0A517FNC7 A0A517FNC9 A0A517FND0 A0A517FND1 A0A517FND3 A0A5A4DV62
A0A5B8YU71 A0A5B8YU73 A0A5B9 A0A5C1RGE8 A0A5C1RGL7 A0A5F4BST2 A0A5F9ZH02
A0A5S6RB44 A0A6G9KH50 A0A6I8MX38 A0A6I8PU40 A0A6J2ATK2 A0A6J4B6H5 A0A6J4BC30
A0A6L8Q027 A0A6M3QG69 A0A6S6AA17 A0A6S6QI82 A0A6S6QLX9 A0A6S6QNE4 A0A6S6QP77
A0A6S6QPY4 A0A7D5JNZ7 A0A7J6K338 A0A7L8UVC5 A0A7L8UVD5 A0A804KTT0 A0A8B7HA97
A0A8F4PNE5 A0A8F4SN83 A0AAS4 A0AEM5 A0AGM5 A0AGW6 A0AH81 A0AHP2 A0AI87
A0AIA7 A0AIK1 A0AIM7 A0AJ17 A0AJ43 A0AJN0 A0AJU9 A0AKD7 A0AKG3 A0AKG4 A0AKH0
A0AKX7 A0AL16 A0ALG6 A0ALI7 A0ALL7 A0ALL9 A0AM15 A0AM16 A0AM17 A0AQ71 A0AV02
A0AVI2 A0AVI4 A0AW23 A0AWV9 A0AZ27 A0B212 A0B5W6 A0B8A0 A0B8Y7 A0B9M2 A0B9N5
A0BKG2 A0E2L1 A0ELI2 A0EYN2 A0FGR8 A0FGR9 A0FKI7 A0JJT8 A0JJT9 A0JJX5 A0JM12
A0JM20 A0JM41 A0JMD4 A0JMG9 A0JMH0 A0JMW6 A0JMY5 A0JN69 A0JN87 A0JNA2 A0JNC1
A0JNC3 A0JNC4 A0JND9 A0JNG0 A0JNI9 A0JNK3 A0JNK6 A0JNM1 A0JP80 A0JPA0 A0JPA1
A0JPB1 A0JPB5 A0JPE1 A0JPH4 A0JPN2 A0JPQ8 A0JR38 A0JV91 A0JWQ8 A0JWX7 A0JY68
A0JY70 A0JYK8 A0JZI3 A0K2X9 A0K3W3 A0K3W4 A0K483 A0K4E0 A0K4Q5 A0K593 A0K5C2
A0K5J7 A0K6A8 A0K6E4 A0K7S3 A0K834 A0K875 A0K913 A0K916 A0K923 A0K960 A0K9N3
A0K9T6 A0K9X9 A0KAR0 A0KAU6 A0KBJ5 A0KBN3 A0KDE6 A0KEF8 A0KEG0 A0KEP9 A0KG31
A0KG33 A0KG40 A0KG66 A0KGB3 A0KGK4 A0KGY2 A0KGY3 A0KGY4 A0KGY7 A0KHC9 A0KHD0
A0KHF6 A0KI35 A0KIB2 A0KID0 A0KJ55 A0KJ58 A0KJ61 A0KJ67 A0KJR8 A0KK53 A0KKX4
A0KL02 A0KL08 A0KL09 A0KL13 A0KL61 A0KLJ2 A0KLJ5 A0KME5 A0KMJ3 A0KMY1 A0KMZ3
A0KNB2 A0KNW5 A0KNW8 A0KQI1 A0KQY2 A0KQY3 A0KQY4 A0KQZ7 A0KR32 A0KRQ2 A0KRY5
A0KSE2 A0KSJ1 A0KSW3 A0KTG6 A0KTQ7 A0KU83 A0KV41 A0KVC0 A0KVI3 A0KVP7 A0KVP8
A0KWF0 A0KWW4 A0KX79 A0KX82 A0KX84 A0KY72 A0KZP4 A0KZP6 A0KZT4 A0KZT7 A0L0D2
A0L0V9 A0L190 A0L1M6 A0L1M7 A0L1M8 A0L1P5 A0L2E7 A0L2I3 A0L2T2 A0L2T3 A0L2T4
A0L4S0 A0L5H1 A0L5J3 A0L5N4 A0L833 A0L835 A0L889 A0L9F4 A0L9X3 A0L9Y5 A0LCK8
A0LCK9 A0LD93 A0LDR4 A0LDR7 A0LDS0 A0LDW6 A0LDW8 A0LE49 A0LEQ2 A0LEQ5 A0LEQ8
A0LEU7 A0LFC9 A0LG71 A0LHM9 A0LHQ9 A0LIM7 A0LIN9 A0LIP0 A0LJ91 A0LJL8 A0LJM1
A0LJM4 A0LJM6 A0LK67 A0LLE6 A0LLF4 A0LLG2 A0LLH3 A0LLI5 A0LM36 A0LN68 A0LNN5
A0LP92 A0LR74 A0LRI8 A0LRJ1 A0LRJ4 A0LSL0 A0LSL2 A0LSX9 A0LTM0 A0LTZ0 A0LV02
A0LXT1 A0M0N2 A0M1U3 A0M2K0 A0M530 A0M5T6 A0M6G6 A0M8R7 A0M8S0 A0M8S8 A0M8T1
A0M8U1 A0MCU0 A0MD28 A0MD30 A0MD31 A0MD32 A0MD33 A0MD34 A0MD35 A0MFS9 A0MGZ5
A0MGZ7 A0MSX9 A0N0X6 A0NGI1 A0PC02 A0PH66 A0PJ25 A0PJ29 A0PJK1 A0PJW6 A0PJX4
A0PJX8 A0PJZ3 A0PK00 A0PK05 A0PK11 A0PK84 A0PKC4 A0PLW0 A0PNM3 A0PNU3 A0PP57
A0PPP7 A0PQS8 A0PQW1 A0PR45 A0PR48 A0PR79 A0PTJ0 A0PTP3 A0PUK3 A0PUK4 A0PUK5
A0PUK6 A0PVW0 A0PWB0 A0PWB3 A0PXM8 A0PY40 A0PZ39 A0PZT1 A0Q060 A0Q072 A0Q0K0

A0Q1J7 A0Q1L0 A0Q200 A0Q210 A0Q2Z8 A0Q2Z9 A0Q3T6 A0Q419 A0Q4E4 A0Q4N6 A0Q4X3
A0Q528 A0Q5C1 A0Q6X6 A0Q7B3 A0Q8E3 A0Q8G5 A0Q8L0 A0Q8N8 A0Q8S8 A0QBR0 A0QBY3
A0QCH3 A0QCX4 A0QCX5 A0QEZ8 A0QF49 A0QFC0 A0QGK1 A0QHC4 A0QHG7 A0QHM5 A0QHX0
A0QIR4 A0QJV4 A0QJV7 A0QK47 A0QNF5 A0QNG1 A0QNG3 A0QNG6 A0QNJ0 A0QNL0 A0QNL1
A0QP18 A0QP27 A0QPD4 A0QPU4 A0QQ39 A0QQ40 A0QQ46 A0QQ48 A0QQ68 A0QQF4 A0QQH5
A0QR29 A0QRJ0 A0QSC1 A0QTK3 A0QTT7 A0QU26 A0QU29 A0QVH8 A0QW18 A0QW28 A0QWU7
A0QWY2 A0QWZ7 A0QX51 A0QXC5 A0QYL8 A0QYV1 A0QZ13 A0QZ40 A0QZY5 A0R014 A0R019
A0R022 A0R036 A0R065 A0R203 A0R204 A0R211 A0R2D1 A0R2K8 A0R395 A0R3A7 A0R3I3
A0R3I7 A0R3Y2 A0R541 A0R5K5 A0R611 A0R619 A0R626 A0R627 A0R6H7 A0R6H8 A0R7G2
A0R8Y3 A0R9W6 A0RA12 A0RA13 A0RAB3 A0RBJ8 A0RBT0 A0RDM5 A0REM5 A0RET0 A0REW2
A0RGF6 A0RHF0 A0RHT4 A0RHW3 A0RHW4 A0RIM4 A0RJJ3 A0RJP1 A0RJR6 A0RL87 A0RL99
A0RLA0 A0RLB0 A0RMD8 A0RME1 A0RMW2 A0RN51 A0RNB3 A0RNW2 A0RP01 A0RP13 A0RPI7
A0RPZ4 A0RQA9 A0RQF4 A0RQL9 A0RQU1 A0RRL1 A0RRP1 A0RUY7 A0RW14 A0RZ28 A0RZH5
A0ST42 A0SYQ0 A0T097 A0T098 A0T099 A0T0A0 A0T0A1 A0T0A2 A0T0A3 A0T0A4 A0T0A5
A0T0A6 A0T0A7 A0T0A8 A0T0A9 A0T0B0 A0T0B1 A0T0B4 A0T0B7 A0T0B8 A0T0C4 A0T0C9
A0T0E7 A0T0E8 A0T0E9 A0T0G0 A0T0G8 A0T0G9 A0T0L7 A0T0L8 A0T0L9 A0T0M0 A0T0M2
A0T0M6 A0T0M7 A0T0M8 A0T0M9 A0T0N4 A0T0P0 A0T0P1 A0T0P2 A0T0P6 A0T0P7 A0T0P8
A0T0R9 A0T0S1 A0T0S5 A0T0S6 A0T0S7 A0T0S8 A0T0S9 A0T0T0 A0T0T2 A0T0T6 A0T0T7
A0T0T9 A0T0U0 A0T0U1 A0T0U2 A0T0U3 A0T0U4 A0T0U5 A0T0U9 A0T0V1 A0T0V3 A0T0W1
A0T0W2 A0T2N3 A0ZSE6 A0ZT23 A0ZZ15 A0ZZ18 A0ZZ19 A0ZZ21 A0ZZ22 A0ZZ23 A0ZZ28
A0ZZ29 A0ZZ30 A0ZZ31 A0ZZ32 A0ZZ34 A0ZZ35 A0ZZ40 A0ZZ45 A0ZZ46 A0ZZ47 A0ZZ48
A0ZZ49 A0ZZ50 A0ZZ51 A0ZZ52 A0ZZ53 A0ZZ54 A0ZZ55 A0ZZ61 A0ZZ62 A0ZZ63 A0ZZ64
A0ZZ65 A0ZZ66 A0ZZ82 A0ZZ84 A0ZZ85 A0ZZ87 A0ZZ88 A0ZZ90 A0ZZ93 A1A0M3 A1A0Y8
A1A139 A1A1J0 A1A1M3 A1A2F2 A1A2I0 A1A3C9 A1A3D0 A1A3E3 A1A3M0 A1A442 A1A4H9
A1A4I1 A1A4J6 A1A4J8 A1A4K1 A1A4M2 A1A4N1 A1A4P6 A1A4P9 A1A5B4 A1A5C7 A1A5G6
A1A5S3 A1A5V7 A1A5Z0 A1A652 A1A654 A1A696 A1A697 A1A698 A1A699 A1A771 A1A775
A1A790 A1A7D2 A1A7K1 A1A7M1 A1A897 A1A8M2 A1A8W1 A1A970 A1A971 A1A986 A1A9B7
A1A9E1 A1A9U5 A1A9U9 A1A9W0 A1AAA8 A1AAA9 A1AAB4 A1AAH3 A1AAH9 A1AAI0 A1AAT4
A1AAV3 A1ABA7 A1ABA8 A1ABC7 A1ABE4 A1ABE5 A1ABH2 A1ABH5 A1ABH7 A1ABK4 A1ABP7
A1ABX2 A1ABY8 A1ABY9 A1ABZ5 A1AC64 A1ACB7 A1ACT0 A1ACT1 A1ACT2 A1ACX1 A1AD02
A1AD51 A1ADA6 A1ADA9 A1ADB0 A1ADC4 A1ADC7 A1ADD0 A1ADD6 A1ADE4 A1ADR5 A1ADR7
A1ADS9 A1ADT0 A1AE54 A1AEA8 A1AEU2 A1AF40 A1AF46 A1AF47 A1AFR5 A1AFW3 A1AFW5
A1AFX9 A1AFY1 A1AG06 A1AG28 A1AGA9 A1AGD3 A1AGD4 A1AGD5 A1AGI2 A1AGN6 A1AGP8
A1AGU7 A1AGY2 A1AGY3 A1AH31 A1AH70 A1AHK2 A1AHM0 A1AHN9 A1AHP3 A1AHP9 A1AHQ2
A1AHR8 A1AHS7 A1AHW3 A1AI24 A1AI26 A1AI63 A1AI84 A1AI91 A1AIE8 A1AIK9 A1AIL4
A1AIL7 A1AIQ8 A1AIS9 A1AJ59 A1AJ60 A1AJL2 A1AJN6 A1AK41 A1ALL0 A1ALL1 A1ALL3
A1ALM1 A1ALQ2 A1AM94 A1AMA7 A1AMC9 A1ANC7 A1ANE2 A1AP45 A1AP46 A1APN2 A1ASH1
A1AT11 A1ATB4 A1AU64 A1AU84 A1AUY8 A1AV04 A1AV44 A1AV76 A1AVR9 A1AVS2 A1AVS5
A1AVW1 A1AVY0 A1AW91 A1AWU4 A1AXL0 A1AXT7 A1AXU6 A1AXU8 A1AZA9 A1AZK6 A1AZW2
A1B057 A1B0E4 A1B340 A1B348 A1B3B4 A1B479 A1B482 A1B487 A1B4B3 A1B616 A1B619
A1B677 A1B8C2 A1B946 A1BA38 A1BA40 A1BAK9 A1BBD2 A1BCE9 A1BE06 A1BE50 A1BE93
A1BED5 A1BEZ9 A1BF32 A1BF36 A1BFW8 A1BGS5 A1BGZ7 A1BHB7 A1BJH0 A1BJJ2 A1BJW4
A1BJW6 A1BJY1 A1BJZ7 A1C3L4 A1C3U0 A1C5W7 A1C6D6 A1C6J8 A1C7E9 A1C7M3 A1C7T5
A1C8C2 A1C8C8 A1C8D8 A1C8Z3 A1C9L6 A1C9R4 A1CAI0 A1CAL7 A1CB06 A1CBE6 A1CBT0

A1CDT3 A1CFK8 A1CFL0 A1CFL5 A1CFL6 A1CFM0 A1CGU5 A1CHB5 A1CHC5 A1CHU1 A1CI53
A1CIF6 A1CIK0 A1CIM4 A1CIM7 A1CJQ1 A1CJW1 A1CKG4 A1CL29 A1CLY9 A1CM86 A1CMH6
A1CMM2 A1CNW8 A1CPP3 A1CPR8 A1CPX0 A1CQL5 A1CR68 A1CRG6 A1CRK9 A1CRW4 A1CTI3
A1CU77 A1CUH7 A1CW41 A1CW44 A1CW63 A1CWA4 A1CWM3 A1CWW9 A1CXG2 A1CXH2 A1CXY5
A1CZJ3 A1D1S7 A1D2K1 A1D2R3 A1D3I1 A1D3P4 A1D432 A1D4C8 A1D4H0 A1D4S1 A1D6D8
A1D708 A1D7K7 A1D7R6 A1D9Y2 A1DA52 A1DA60 A1DA63 A1DA65 A1DAY6 A1DB12 A1DCU0
A1DDI7 A1DDZ3 A1DEC4 A1DEV5 A1DEZ0 A1DG37 A1DH10 A1DH66 A1DHW1 A1DI57 A1DI95
A1DIF7 A1DLB6 A1DLJ5 A1DLT0 A1DMX4 A1DN29 A1DNW0 A1DP58 A1DWM3 A1E9H9 A1E9I0
A1E9I5 A1E9I6 A1E9I7 A1E9J0 A1E9J1 A1E9J6 A1E9K2 A1E9K3 A1E9K8 A1E9K9 A1E9L0
A1E9L8 A1E9N9 A1E9Q4 A1E9Q7 A1E9Q8 A1E9Q9 A1E9R0 A1E9R1 A1E9R2 A1E9R3 A1E9R8
A1E9R9 A1E9S0 A1E9S3 A1E9S9 A1E9T4 A1E9T5 A1E9T6 A1E9T7 A1E9T8 A1E9T9 A1E9U0
A1E9U1 A1E9U2 A1E9U3 A1E9U4 A1E9V0 A1E9V1 A1E9V2 A1E9V3 A1E9V4 A1E9V5 A1E9X3
A1E9X4 A1E9X6 A1E9X7 A1E9X9 A1E9Y8 A1E9Z1 A1E9Z2 A1E9Z3 A1E9Z4 A1E9Z5 A1E9Z6
A1E9Z7 A1EA02 A1EA03 A1EA04 A1EA07 A1EA08 A1EA13 A1EA18 A1EA19 A1EA20 A1EA21
A1EA22 A1EA23 A1EA24 A1EA25 A1EA26 A1EA27 A1EA28 A1EA34 A1EA35 A1EA36 A1EA37
A1EA38 A1EA39 A1EA56 A1EA58 A1EA59 A1EA61 A1EA62 A1EA64 A1EC31 A1INM7 A1IPD7
A1IPX1 A1ITY7 A1IVY7 A1JHN0 A1JHR6 A1JHU3 A1JHZ2 A1JID8 A1JID9 A1JIF4 A1JIF6
A1JIK1 A1JIM8 A1JIN7 A1JIQ1 A1JJ53 A1JJ54 A1JJC1 A1JJD7 A1JJE2 A1JJF0 A1JJJ0
A1JJT3 A1JKS1 A1JKW8 A1JKW9 A1JL11 A1JL14 A1JLC3 A1JLD6 A1JLG0 A1JLH7 A1JLI8
A1JLL1 A1JLU9 A1JM45 A1JM66 A1JM73 A1JM85 A1JM88 A1JM96 A1JMG4 A1JMQ7 A1JMY5
A1JN04 A1JNE0 A1JNH2 A1JNP6 A1JNZ2 A1JNZ3 A1JNZ5 A1JPA1 A1JPD0 A1JPE9 A1JPF0
A1JPL6 A1JPL9 A1JPM4 A1JPP5 A1JPP8 A1JPQ9 A1JQ02 A1JQ17 A1JQ40 A1JQP8 A1JQQ3
A1JQS2 A1JQS4 A1JQW4 A1JQW7 A1JQY4 A1JR18 A1JR77 A1JRE5 A1JRE8 A1JRH8 A1JRH9
A1JRI1 A1JRI8 A1JRJ8 A1JRQ2 A1JRU3 A1JRU8 A1JRV7 A1JRZ6 A1JSB0 A1JSF0 A1JSP6
A1JSS6 A1JT67 A1JTD1 A1JTD2 A1JTD4 A1JTG3 A1JTR6 A1JTY1 A1JU76 A1JUB7 A1K1R6
A1K1R7 A1K1R8 A1K2P1 A1K2S9 A1K323 A1K338 A1K3B8 A1K3C4 A1K3U3 A1K4R6 A1K5B5
A1K5B8 A1K5C1 A1K668 A1K6B2 A1K6C7 A1K7F5 A1K7T4 A1K8W8 A1KAC6 A1KAQ4 A1KAU9
A1KAV0 A1KAY2 A1KC82 A1KCJ1 A1KEN2 A1KF14 A1KFR6 A1KG40 A1KH31 A1KH32 A1KH33
A1KHB8 A1KHC1 A1KHG8 A1KHH0 A1KI12 A1KI93 A1KI94 A1KI95 A1KIJ9 A1KIP0 A1KIS1
A1KIW9 A1KIY4 A1KJ30 A1KJ57 A1KKE0 A1KKH9 A1KKJ9 A1KKQ0 A1KKY1 A1KLZ1 A1KML4
A1KNE8 A1KNF1 A1KNI8 A1KQF8 A1KRK6 A1KRP4 A1KRT0 A1KRT3 A1KRZ8 A1KS63 A1KSD3
A1KSH3 A1KSH4 A1KSH5 A1KSK9 A1KSZ7 A1KT72 A1KTU8 A1KTV1 A1KTV8 A1KUB7 A1KV57
A1KVL7 A1KVM7 A1KVS7 A1KVV6 A1KW15 A1KWK2 A1KXC4 A1KXE4 A1L0T0 A1L157 A1L1A6
A1L1C2 A1L1J9 A1L1L2 A1L1P9 A1L1W4 A1L1W9 A1L224 A1L244 A1L272 A1L2F6 A1L2I9
A1L2P2 A1L2R7 A1L2S8 A1L314 A1L3G4 A1L3G9 A1L3I3 A1L3M3 A1L3X0 A1L4X0 A1L504
A1L515 A1QRH1 A1QUS1 A1QZ99 A1QZM8 A1QZQ6 A1R4H0 A1R5F5 A1R6I0 A1R6P8 A1R7V6
A1R7V8 A1R883 A1R944 A1RC71 A1RE49 A1RE84 A1REE1 A1REF8 A1REM8 A1REZ3 A1RFH5
A1RFK8 A1RG29 A1RGC9 A1RGP2 A1RH70 A1RHF4 A1RI59 A1RID4 A1RII6 A1RIL6 A1RJC2
A1RJJ3 A1RJZ2 A1RJZ5 A1RJZ7 A1RKH2 A1RKH3 A1RLQ3 A1RMD9 A1RMH4 A1RMH7 A1RN03
A1RNR8 A1RP80 A1RP81 A1RP82 A1RQB4 A1RQB5 A1RQB6 A1RQE9 A1RQS3 A1RRA6 A1RSF4
A1RT82 A1RUA0 A1RVP5 A1RXB2 A1RZ10 A1S066 A1S1G5 A1S246 A1S2F6 A1S2M4 A1S3J2
A1S3L4 A1S3T2 A1S3W9 A1S426 A1S4D8 A1S5R6 A1S6H4 A1S6M9 A1S6N2 A1S6N3 A1S6N4
A1S6X4 A1S6X5 A1S7I6 A1S7P9 A1SAK0 A1SAK1 A1SB50 A1SBD7 A1SBU4 A1SBU5 A1SBU6
A1SDK4 A1SE34 A1SE37 A1SE40 A1SGV9 A1SHI5 A1SHI6 A1SHI7 A1SHU2 A1SJN0 A1SJR1

A1SJZ1 A1SL83 A1SRC7 A1SRG6 A1SRH4 A1SRS0 A1SRS1 A1SS94 A1SS95 A1SSH6 A1SSQ9
A1SSX2 A1SSX5 A1SSY6 A1SSY7 A1STS8 A1SU12 A1SU16 A1SUM3 A1SVD8 A1SY55 A1SZ07
A1SZP2 A1T087 A1T0A0 A1T0M9 A1T0Z3 A1T0Z5 A1T2V5 A1T3T1 A1T557 A1T636 A1T696
A1T699 A1T7Y9 A1T8M8 A1T8Q8 A1TAC7 A1TAK2 A1TAN7 A1TAS8 A1TAX1 A1TB16 A1TD58
A1TD59 A1TDL8 A1TED2 A1TEL6 A1TEV1 A1TJ24 A1TJ35 A1TJ36 A1TJ37 A1TJR2 A1TKC8
A1TKT8 A1TL12 A1TL13 A1TLL6 A1TLM3 A1TLM6 A1TMB0 A1TNT7 A1TQ84 A1TR25 A1TRC8
A1TSC2 A1TSW8 A1TTA2 A1TTF7 A1TTX1 A1TU04 A1TV68 A1TVU0 A1TWQ8 A1TXV0 A1TZ60
A1TZ62 A1TZ65 A1TZE0 A1TZT8 A1U0A9 A1U1Y9 A1U218 A1U277 A1U3F9 A1U3G1 A1U4Q6
A1U669 A1U672 A1U673 A1U6P5 A1U6R3 A1U6U1 A1U7H0 A1U7H8 A1U7I0 A1U7J4 A1U8T1
A1UA55 A1UAZ4 A1UCT7 A1UD94 A1UD97 A1UF43 A1UFQ0 A1UFT9 A1UFV8 A1UFZ7 A1UHB3
A1UHV6 A1UI57 A1UIA4 A1UJY7 A1UJY8 A1UJY9 A1UK99 A1UKX6 A1UL69 A1ULE2 A1UR06
A1URA3 A1URL2 A1URT7 A1URU2 A1URU4 A1URU5 A1URX1 A1URZ6 A1USH8 A1USS5 A1USX4
A1USX7 A1USY0 A1UT26 A1UTC8 A1UTG9 A1UUG9 A1UZV8 A1V002 A1V0S1 A1V1I5 A1V1J1
A1V1V6 A1V2H2 A1V2L6 A1V2M6 A1V303 A1V4I6 A1V4U9 A1V513 A1V5Y8 A1V6V3 A1V750
A1V794 A1V7D5 A1V8I0 A1V8T5 A1V9G1 A1VAL2 A1VAZ2 A1VBE5 A1VCU2 A1VD51 A1VDC9
A1VDN6 A1VER7 A1VF66 A1VF67 A1VF70 A1VFJ1 A1VHP3 A1VI80 A1VIU8 A1VJ21 A1VJS8
A1VJY2 A1VK09 A1VK47 A1VK48 A1VK53 A1VKM6 A1VKQ9 A1VKX1 A1VLZ3 A1VM60 A1VM67
A1VM73 A1VMI3 A1VNB6 A1VR95 A1VST9 A1VSW7 A1VTC0 A1VTC2 A1VTR6 A1VTR7 A1VWH8
A1VXQ0 A1VXT4 A1VY78 A1VY81 A1VYC7 A1VYF0 A1VYM9 A1VYW8 A1VZK8 A1VZM0 A1W084
A1W0G0 A1W0I8 A1W0J3 A1W1H1 A1W1Q6 A1W1Q7 A1W1Z0 A1W2T1 A1W2T2 A1W2T3 A1W307
A1W3X2 A1W450 A1W451 A1W457 A1W4D7 A1W4M2 A1W4M9 A1W560 A1W5M0 A1W606 A1W6J1
A1W7N8 A1W857 A1W8C0 A1WBJ9 A1WBL5 A1WC09 A1WC76 A1WCI8 A1WCP0 A1WES5 A1WF52
A1WF53 A1WF54 A1WFS5 A1WFS6 A1WG92 A1WH77 A1WHB4 A1WHP5 A1WIU9 A1WIY6 A1WJU9
A1WK80 A1WKC0 A1WLN7 A1WLP4 A1WMU7 A1WPD7 A1WR47 A1WRK8 A1WTR8 A1WUX4 A1WUY5
A1WVM9 A1WW03 A1WWE3 A1WWS3 A1WXV4 A1WXW0 A1WYI3 A1WYP1 A1WYU4 A1WYU6 A1WZI0
A1WZT5 A1WZT7 A1X150 A1XFV1 A1XFW0 A1XFW7 A1XG02 A1XG05 A1XG07 A1XG08 A1XG10
A1XG13 A1XGB4 A1XGM2 A1XGM4 A1XGP3 A1XGT3 A1XGT6 A1XGT8 A1XGT9 A1XGU1 A1XGU4
A1XI29 A1XI30 A1XIP9 A1XJK0 A1XQR6 A1XQR7 A1XQS1 A1XQS2 A1XQS5 A1XQT2 A1XQX0
A1XQX1 A1XQX2 A1XQX3 A1XQX8 A1XQY0 A1XQY1 A1XQY3 A1YG32 A1YGR5 A1YGR6 A1YIY0
A1YJE2 A1YJE3 A1YLR8 A1Z0Q5 A1Z6H7 A1Z6L1 A1Z746 A1Z7G7 A1Z7U3 A1Z8N1 A1Z8R8
A1ZAG4 A1ZAI3 A1ZAI5 A1ZAX0 A1ZBD6 A2A259 A2A2V5 A2A2Y4 A2A3V2 A2A559 A2A699
A2A6C4 A2A7Q9 A2A7V7 A2A7Z8 A2A863 A2A8L5 A2A8T7 A2A8U2 A2A9G7 A2A9Q0 A2AAE1
A2ADF7 A2ADU8 A2ADU9 A2AE42 A2AF53 A2AFS3 A2AGA4 A2AGL3 A2AHL1 A2AIR5 A2AJ15
A2AJ88 A2AJA7 A2AJB2 A2AJN7 A2AJQ3 A2AJX4 A2AKG8 A2AKM2 A2AKQ0 A2ALI5 A2ALW5
A2AN08 A2ANU3 A2APA5 A2APT9 A2APX8 A2AR95 A2ARA8 A2ARI4 A2ARJ3 A2ARP9 A2ARV4
A2ASA8 A2ASI5 A2ASQ1 A2ASZ8 A2AUQ7 A2AV25 A2AVR2 A2AVZ9 A2AWH2 A2BD92 A2BDP1
A2BDX4 A2BFP5 A2BGL3 A2BGS3 A2BGT0 A2BIE7 A2BIU7 A2BM25 A2BN92 A2BNK1 A2BNU4
A2BNV5 A2BP21 A2BP50 A2BP51 A2BP52 A2BP71 A2BP94 A2BP97 A2BP98 A2BP99 A2BPA0
A2BPB5 A2BPB8 A2BPC6 A2BPC7 A2BPF4 A2BPR7 A2BPT0 A2BPU3 A2BPU4 A2BPU5 A2BQ51
A2BQM6 A2BQQ0 A2BQX7 A2BRN6 A2BRU6 A2BS52 A2BS56 A2BSP9 A2BT27 A2BT28 A2BT29
A2BT30 A2BT34 A2BT95 A2BT99 A2BTA0 A2BU19 A2BUC6 A2BUD7 A2BUK1 A2BUN2 A2BUN3
A2BUN4 A2BUQ3 A2BUS5 A2BUS8 A2BUS9 A2BUT0 A2BUT1 A2BUU7 A2BUV5 A2BUV6 A2BV95
A2BVA8 A2BVC2 A2BVC3 A2BVC4 A2BVD0 A2BVK2 A2BVN3 A2BW66 A2BWT5 A2BX44 A2BXA4
A2BXL5 A2BXL9 A2BY34 A2BYH8 A2BYH9 A2BYI0 A2BYI1 A2BYI5 A2BYP5 A2BYP8 A2BYP9

A2BZ16 A2BZ97 A2BZN0 A2BZX6 A2BZY7 A2C057 A2C084 A2C085 A2C0A5 A2C0C8 A2C0D1
A2C0D2 A2C0D3 A2C0F8 A2C0G1 A2C0H1 A2C0H2 A2C0Q4 A2C0R7 A2C0R8 A2C0R9 A2C0S5
A2C127 A2C1K1 A2C1W8 A2C210 A2C302 A2C3M5 A2C3U9 A2C3V3 A2C4A0 A2C4J7 A2C4J8
A2C4J9 A2C4K0 A2C4K5 A2C4V3 A2C4V4 A2C666 A2C6G3 A2C6Q1 A2C6W3 A2C6X0 A2C6X1
A2C6X2 A2C6X3 A2C7C1 A2C7D7 A2C7F4 A2C7F5 A2C7F6 A2C7G0 A2C7Y6 A2C7Z1 A2C9P7
A2C9Q5 A2C9Y5 A2CAB5 A2CAP6 A2CBQ9 A2CBR0 A2CBT5 A2CBT8 A2CC72 A2CC73 A2CCI3
A2CCI4 A2CCI6 A2CCL0 A2CCL2 A2CCP7 A2CCQ0 A2CCQ1 A2CCQ2 A2CCQ3 A2CD41 A2CD61
A2CEQ0 A2CER7 A2CEX1 A2CI37 A2CI41 A2CI71 A2CIR5 A2DSR8 A2DZE8 A2E3C6 A2EI35
A2EK20 A2EK80 A2ELE6 A2FJ32 A2IBY8 A2PZE5 A2Q898 A2Q8F7 A2Q8K8 A2Q9F0 A2Q9L8
A2Q9P2 A2Q9P5 A2QBE9 A2QCH3 A2QCV4 A2QEK7 A2QFJ9 A2QGD9 A2QGH7 A2QGW1 A2QHH4
A2QHQ3 A2QI68 A2QI79 A2QJ14 A2QJH4 A2QK68 A2QLJ9 A2QLK4 A2QM49 A2QMB8 A2QN74
A2QPL8 A2QPW5 A2QQV6 A2QR20 A2QS05 A2QTE7 A2QTE8 A2QTW5 A2QTX8 A2QX52 A2R065
A2R350 A2R3F3 A2R3N6 A2R4N0 A2R5A0 A2R616 A2R6G9 A2R920 A2R9P4 A2RA94 A2RAN5
A2RCI8 A2RD43 A2RD66 A2RDW5 A2REZ8 A2RF66 A2RFC6 A2RFC7 A2RFE5 A2RFS5 A2RG98
A2RGB7 A2RGC2 A2RI03 A2RI45 A2RI47 A2RI75 A2RI76 A2RI86 A2RI87 A2RI97 A2RIH3
A2RIQ0 A2RIT7 A2RIU1 A2RJ04 A2RJB8 A2RJJ9 A2RJV0 A2RKA5 A2RKA6 A2RKG9 A2RKV5
A2RL65 A2RLE8 A2RLT1 A2RM05 A2RMA8 A2RMJ9 A2RMP5 A2RN36 A2RNI1 A2RNI5 A2RNQ7
A2RNT9 A2RNU8 A2RNZ5 A2RNZ6 A2RRL7 A2RRT9 A2RRU4 A2RSQ0 A2RST1 A2RT91 A2RTF1
A2RU14 A2RU48 A2RU67 A2RUG3 A2RUT3 A2RUV0 A2RUW0 A2RUW7 A2RV66 A2RV80 A2RVP7
A2RVQ4 A2RVT6 A2RVU1 A2S1P8 A2S1Q0 A2S2B6 A2S3S4 A2S449 A2S459 A2S4A3 A2S502
A2S5U8 A2S603 A2S646 A2S6K2 A2S759 A2S8D6 A2S8H2 A2S8L4 A2S9B4 A2S9C1 A2S9N6
A2SAC9 A2SBC7 A2SBI5 A2SC47 A2SC65 A2SC66 A2SCF8 A2SCY2 A2SE14 A2SE15 A2SE21
A2SEL8 A2SF35 A2SFL2 A2SFN3 A2SFN6 A2SFP5 A2SGC9 A2SGN6 A2SHH9 A2SIS1 A2SIS3
A2SJW8 A2SKB1 A2SKN7 A2SMA7 A2SPZ1 A2SQ88 A2SQF0 A2SRL0 A2SSD6 A2SSE8 A2SWM2
A2T195 A2T1U1 A2T309 A2T314 A2T316 A2T318 A2T319 A2T320 A2T325 A2T327 A2T329
A2T330 A2T332 A2T338 A2T344 A2T345 A2T346 A2T347 A2T348 A2T351 A2T352 A2T353
A2T359 A2T361 A2T379 A2T383 A2T385 A2T386 A2T388 A2T3Q0 A2TGX5 A2TJ54 A2V7M9
A2VCW5 A2VD01 A2VD98 A2VDC5 A2VDC7 A2VDH3 A2VDJ0 A2VDK9 A2VDL4 A2VDL6 A2VDL9
A2VDN0 A2VDP5 A2VDT6 A2VDV9 A2VDX9 A2VDZ9 A2VE04 A2VE10 A2VE13 A2VE15 A2VE31
A2VE53 A2VE54 A2VE55 A2VE58 A2VE61 A2VE67 A2VEI2 A2VEY9 A2WLC1 A2WLP4 A2WMK1
A2WMK7 A2WNZ9 A2WR31 A2WSD3 A2WSD8 A2WSH0 A2WV32 A2WX64 A2WXD3 A2WXD9 A2WY50
A2WYG9 A2WYI4 A2X2I0 A2X345 A2X3S3 A2X5B4 A2X6E6 A2X8A7 A2X8M8 A2X8W3 A2X933
A2XAY1 A2XCA0 A2XCD4 A2XCJ1 A2XDK8 A2XFP3 A2XFQ8 A2XFT5 A2XFT6 A2XGM7 A2XJ35
A2XL05 A2XL32 A2XLM6 A2XN66 A2XNT2 A2XQD3 A2XSY1 A2XU91 A2XUA1 A2XV81 A2XVC2
A2XWN6 A2XX57 A2XX73 A2XZ94 A2Y075 A2Y0X2 A2Y2B7 A2Y4G9 A2Y6Z7 A2Y7D9 A2Y8U6
A2YA15 A2YAZ1 A2YCI3 A2YD22 A2YFN7 A2YGP9 A2YH25 A2YHR9 A2YMH5 A2YMP7 A2YQB8
A2YU42 A2YVG8 A2YXC7 A2YXI1 A2YZ24 A2Z1C8 A2Z1F5 A2Z212 A2Z669 A2Z6C5 A2Z9A6
A2Z9W7 A2ZAK8 A2ZBW5 A2ZE50 A2ZF66 A2ZHX7 A2ZI32 A2ZI41 A2ZIM4 A2ZMM4 A2ZN77
A2ZU80 A2ZVG7 A2ZX97 A3A2W2 A3A4E0 A3A871 A3A8Q4 A3A9H6 A3AB67 A3AWA4 A3AYR1
A3BF39 A3BI11 A3BV82 A3BWJ9 A3BXL8 A3CK84 A3CL57 A3CL70 A3CLI0 A3CLJ7 A3CM09
A3CM10 A3CM55 A3CMA0 A3CN81 A3CP32 A3CPX8 A3CPZ6 A3CQ79 A3CRI4 A3CU85 A3CVP0
A3CW74 A3CX65 A3CYW8 A3CZ45 A3CZD5 A3CZL8 A3CZR1 A3CZU7 A3D0P2 A3D139 A3D1G6
A3D1Q5 A3D1Q8 A3D1U4 A3D2H6 A3D3H1 A3D3U0 A3D4J3 A3D5N7 A3D7X7 A3D839 A3D9F4
A3D9F5 A3D9F6 A3DAR8 A3DAR9 A3DAS0 A3DAS8 A3DBH1 A3DC20 A3DC27 A3DC75 A3DCG3

A3DDR3 A3DDW4 A3DE29 A3DEE4 A3DEX6 A3DFC2 A3DH97 A3DHI6 A3DHS8 A3DIE5 A3DIM5
A3DJP2 A3DK68 A3DM64 A3DRP0 A3DRP1 A3DRP3 A3EX94 A3EX98 A3EX99 A3EXA0 A3EXD0
A3EXD4 A3EXD5 A3EXD6 A3EXG6 A3EXG7 A3EXG8 A3EXG9 A3F5L2 A3F5L3 A3FIN4 A3FMB2
A3FPG8 A3GF86 A3GFQ7 A3GGS6 A3GI61 A3KF10 A3KFT3 A3KFU9 A3KGK3 A3KGT9 A3KMY4
A3KMZ6 A3KN25 A3KN28 A3KN46 A3KN95 A3KNA7 A3KNB6 A3KNK1 A3KNM5 A3KNN3 A3KNS9
A3KNW0 A3KP48 A3KPA0 A3KPL7 A3KPP4 A3KPQ7 A3KPR5 A3LMX9 A3LN69 A3LN78 A3LNK7
A3LP48 A3LPS1 A3LPS6 A3LQD9 A3LQS0 A3LQW6 A3LRT4 A3LSW7 A3LT10 A3LT66 A3LTB7
A3LU17 A3LU53 A3LV34 A3LVL1 A3LVX1 A3LW83 A3LW86 A3LWK1 A3LWM9 A3LWX1 A3LXS1
A3LXZ3 A3LYZ4 A3LZS3 A3M019 A3M0D8 A3M0S2 A3M137 A3M139 A3M140 A3M1Q4 A3M1X1
A3M1X6 A3M1X7 A3M2P8 A3M2Q4 A3M2Q7 A3M3C7 A3M3H0 A3M4X1 A3M596 A3M5Z2 A3M6K4
A3M6Q3 A3M830 A3M8D6 A3M8J6 A3M8K2 A3M9P9 A3MA10 A3MCG6 A3MCG8 A3MI57 A3MIA1
A3MIB1 A3MIN7 A3MK60 A3MKH4 A3MKN7 A3MLQ2 A3MMF8 A3MMS6 A3MMT3 A3MN08 A3MNP7
A3MNU1 A3MPC2 A3MPT9 A3MQ00 A3MQ44 A3MQJ5 A3MR60 A3MS20 A3MT10 A3MVF0 A3MWD7
A3MXH2 A3MXJ5 A3MXY4 A3MY87 A3MYM6 A3MYM7 A3MYN7 A3MYP1 A3MZ39 A3MZ40 A3MZ84
A3N0C9 A3N0G1 A3N0I2 A3N0X4 A3N0Y0 A3N147 A3N1V3 A3N1Y9 A3N207 A3N216 A3N223
A3N2H4 A3N2P0 A3N2U8 A3N2U9 A3N2V0 A3N3I9 A3N3N9 A3N3P2 A3N3S2 A3N3S3 A3N3S4
A3N477 A3N4D7 A3N588 A3N5D1 A3N5V1 A3N6P6 A3N6V2 A3N7H1 A3N7L7 A3N7M4 A3N7M7
A3N836 A3NA39 A3NAG3 A3NAN4 A3NCH2 A3NCV3 A3NDA6 A3NDW7 A3NE88 A3NF44 A3NLD5
A3NLD7 A3NP37 A3NPX1 A3NQ26 A3NQY8 A3NR29 A3NRJ7 A3NSD0 A3NSI9 A3NT59 A3NTA6
A3NTB3 A3NTB6 A3NTU6 A3NTZ5 A3NVV6 A3NW81 A3NWF4 A3NXZ6 A3NYA3 A3NYA9 A3NYL8
A3NZ16 A3NZL8 A3P021 A3P022 A3P0Z4 A3P6Z7 A3P6Z9 A3P9L2 A3PAC1 A3PAL7 A3PAM8
A3PAU3 A3PAX2 A3PAX4 A3PAX5 A3PAZ3 A3PB16 A3PB19 A3PB20 A3PB21 A3PB22 A3PB37
A3PB40 A3PB49 A3PB50 A3PB74 A3PBF7 A3PBH0 A3PBI3 A3PBI4 A3PBI5 A3PBJ1 A3PBU4
A3PCE8 A3PCH3 A3PCQ0 A3PDG2 A3PDZ2 A3PDZ6 A3PEG1 A3PEU1 A3PEU2 A3PEU3 A3PEU4
A3PEU8 A3PF09 A3PF13 A3PF14 A3PFC1 A3PFJ8 A3PG85 A3PGP0 A3PGT6 A3PGX5 A3PGX7
A3PHS2 A3PIX8 A3PIY2 A3PIZ4 A3PK35 A3PKI6 A3PLU3 A3PMB5 A3PN50 A3PN82 A3PN83
A3PN84 A3PN85 A3PNB1 A3PNJ7 A3PNW1 A3PQK0 A3PRP3 A3PRP7 A3PS63 A3PSE8 A3PTR0
A3PUK0 A3PWJ3 A3PWR1 A3PWR4 A3PYK5 A3PZB2 A3PZE9 A3PZG8 A3Q0V0 A3Q1C2 A3Q1M1
A3Q1R8 A3Q3B4 A3Q3B5 A3Q3B6 A3Q3Q3 A3Q537 A3Q5E2 A3Q5L8 A3Q5U1 A3Q941 A3Q9B0
A3Q9U4 A3QBM4 A3QBM7 A3QBR5 A3QBX3 A3QCX7 A3QDP2 A3QEI9 A3QEN4 A3QEN7 A3QEN8
A3QEN9 A3QEZ3 A3QEZ4 A3QF78 A3QFJ6 A3QGZ6 A3QIE3 A3QIE4 A3QIE5 A3QIF2 A3QIM4
A3QIW3 A3QJR4 A3QJR5 A3QJR6 A3QJT1 A3QJU3 A3QK16 A3QM97 A3QMC6 A3QNZ8 A3QNZ9
A3QP00 A3QP01 A3QP07 A3QP08 A3QP09 A3QU28 A3QUP1 A3R044 A3RFZ7 A3RM22 A4D0T7
A4D1S0 A4D256 A4D2G3 A4D2H0 A4D7R9 A4D9R2 A4D9R3 A4DA06 A4DA85 A4F267 A4F4L0
A4F6L1 A4F8J0 A4F8Q9 A4F9B0 A4FBP2 A4FBZ2 A4FCM4 A4FFN9 A4FL81 A4FLW3 A4FN31
A4FPS8 A4FPT1 A4FPT4 A4FU28 A4FUB8 A4FUD4 A4FUN9 A4FUQ5 A4FUW8 A4FUY1 A4FUY9
A4FUZ5 A4FV27 A4FV45 A4FV48 A4FV52 A4FV75 A4FV93 A4FVP6 A4FVW2 A4FVW3 A4FVW5
A4FW42 A4FX27 A4FXY4 A4G0M1 A4G0P3 A4G1F7 A4G1N1 A4G1Y9 A4G2A5 A4G5J7 A4G5J9
A4G5V6 A4G631 A4G637 A4G6P0 A4G729 A4G736 A4G8J0 A4G8N2 A4G8U1 A4G936 A4G9H5
A4G9I1 A4G9I2 A4G9Q2 A4GA77 A4GAH3 A4GAN3 A4GBX7 A4GBX8 A4GBY0 A4GCH5 A4GCH6
A4GCI6 A4GCI7 A4GCI9 A4GCJ7 A4GCJ8 A4GCK0 A4GCK8 A4GCK9 A4GCL1 A4GCL9 A4GCM0
A4GCM2 A4GG66 A4GG92 A4GG97 A4GG98 A4GGA1 A4GGA2 A4GGA3 A4GGA4 A4GGA9 A4GGB0
A4GGB1 A4GGB3 A4GGB7 A4GGB8 A4GGB9 A4GGC0 A4GGC1 A4GGC4 A4GGC5 A4GGC6 A4GGD1
A4GGD3 A4GGE3 A4GGE5 A4GGE6 A4GGE8 A4GGE9 A4GGF1 A4GGF4 A4GRC6 A4GT88 A4GVD1

A4GYN9 A4GYP2 A4GYP4 A4GYP5 A4GYP6 A4GYQ1 A4GYQ2 A4GYQ4 A4GYQ5 A4GYQ9 A4GYR0
A4GYR5 A4GYS2 A4GYS3 A4GYS4 A4GYS5 A4GYS6 A4GYS9 A4GYT0 A4GYT1 A4GYT7 A4GYT9
A4GYW4 A4GYW5 A4GYW6 A4GYW8 A4GYW9 A4GYX1 A4GYX4 A4HK17 A4I7K1 A4IF30 A4IF75
A4IF94 A4IFD7 A4IFE9 A4IFG4 A4IFH6 A4IFJ1 A4IFK2 A4IFL0 A4IFL1 A4IFL2 A4IFM1
A4IFN5 A4IFP3 A4IFW2 A4IG32 A4IG61 A4IG66 A4IGD2 A4IGI5 A4IGL3 A4IGY6 A4IH46
A4IH88 A4IHB9 A4IHD1 A4IHH4 A4IHK6 A4IHW3 A4IHY6 A4IHZ3 A4II32 A4II83 A4II98
A4IIC3 A4IIC5 A4IID1 A4IIF2 A4IIS8 A4IIU3 A4IIV4 A4IIW9 A4IIY1 A4IJ20 A4IJS3
A4IK45 A4IKE7 A4IKL9 A4IKS8 A4IKY2 A4IL76 A4ILH7 A4ILW9 A4ILX0 A4IM06 A4IM26
A4IMH1 A4IMP7 A4IQD8 A4IQF5 A4IRQ2 A4IRU1 A4ISS2 A4ITI0 A4ITI3 A4ITI7 A4ITJ3
A4ITJ4 A4ITK4 A4ITV2 A4IVT5 A4IVV7 A4IVZ9 A4IW20 A4IXE9 A4IYU3 A4IZH0 A4IZX0
A4J015 A4J2A8 A4J3H9 A4J3P2 A4J550 A4J5G9 A4J5T7 A4J650 A4J660 A4J804 A4J832
A4J9A3 A4J9A5 A4J9B4 A4JA31 A4JAX4 A4JB91 A4JBE7 A4JBP9 A4JC72 A4JD93 A4JEM7
A4JEZ2 A4JF27 A4JGC0 A4JGC3 A4JGD0 A4JGG7 A4JH33 A4JH88 A4JI25 A4JI61 A4JJ20
A4JJ46 A4JM56 A4JQW5 A4K143 A4K144 A4K146 A4K2M4 A4K2N5 A4K2N8 A4K2P6 A4K2Q6
A4K2R3 A4K2S2 A4K2T1 A4K2V2 A4K2W1 A4K2W6 A4K2X4 A4K2Y2 A4K526 A4KDP0 A4KDP1
A4KWA1 A4KWA5 A4KWA6 A4KWA8 A4KX73 A4KX74 A4KXB3 A4L9J0 A4Q9X1 A4QC57 A4QCC1
A4QD87 A4QDG9 A4QE56 A4QEA1 A4QEE9 A4QFF5 A4QFM6 A4QFR6 A4QH18 A4QI94 A4QJ96
A4QJ99 A4QJA1 A4QJA2 A4QJA3 A4QJA8 A4QJA9 A4QJB0 A4QJB1 A4QJB4 A4QJB5 A4QJC0
A4QJC6 A4QJC7 A4QJC8 A4QJC9 A4QJD0 A4QJD3 A4QJD4 A4QJD5 A4QJE1 A4QJE3 A4QJG3
A4QJG5 A4QJG6 A4QJG8 A4QJG9 A4QJH1 A4QJH4 A4QJI0 A4QJI3 A4QJI5 A4QJI6 A4QJI7
A4QJJ2 A4QJJ3 A4QJJ4 A4QJJ5 A4QJJ8 A4QJJ9 A4QJK4 A4QJL0 A4QJL1 A4QJL2 A4QJL3
A4QJL4 A4QJL7 A4QJL8 A4QJL9 A4QJM5 A4QJM7 A4QJP7 A4QJP9 A4QJQ0 A4QJQ2 A4QJQ3
A4QJQ5 A4QJQ8 A4QJR4 A4QJR9 A4QJS0 A4QJS1 A4QJS6 A4QJS7 A4QJS8 A4QJS9 A4QJT2
A4QJT3 A4QJT8 A4QJU4 A4QJU5 A4QJU6 A4QJU9 A4QJV0 A4QJV1 A4QJV7 A4QJV9 A4QJX9
A4QJY1 A4QJY2 A4QJY4 A4QJY5 A4QJY7 A4QJZ0 A4QJZ6 A4QJZ7 A4QJZ8 A4QJZ9 A4QK02
A4QK04 A4QK05 A4QK06 A4QK11 A4QK12 A4QK13 A4QK14 A4QK17 A4QK18 A4QK23 A4QK29
A4QK30 A4QK31 A4QK32 A4QK33 A4QK36 A4QK37 A4QK38 A4QK44 A4QK46 A4QK67 A4QK69
A4QK70 A4QK72 A4QK73 A4QK75 A4QK78 A4QK85 A4QK89 A4QK91 A4QK92 A4QK93 A4QK98
A4QK99 A4QKA0 A4QKA1 A4QKA4 A4QKA5 A4QKB0 A4QKB6 A4QKB7 A4QKB8 A4QKB9 A4QKC0
A4QKC3 A4QKC4 A4QKC5 A4QKD1 A4QKD3 A4QKF4 A4QKF6 A4QKF7 A4QKF9 A4QKG0 A4QKG2
A4QKG5 A4QKH2 A4QKH6 A4QKH8 A4QKH9 A4QKI0 A4QKI5 A4QKI6 A4QKI7 A4QKI8 A4QKJ1
A4QKJ2 A4QKJ7 A4QKK3 A4QKK4 A4QKK5 A4QKK6 A4QKK7 A4QKL0 A4QKL1 A4QKL2 A4QKL8
A4QKM0 A4QKP2 A4QKP4 A4QKP5 A4QKP7 A4QKP8 A4QKQ0 A4QKQ3 A4QKR1 A4QKR5 A4QKR7
A4QKR8 A4QKR9 A4QKS4 A4QKS5 A4QKS6 A4QKS7 A4QKT0 A4QKT1 A4QKT6 A4QKU2 A4QKU3
A4QKU4 A4QKU5 A4QKU6 A4QKU9 A4QKV0 A4QKV1 A4QKV7 A4QKV9 A4QKY1 A4QKY3 A4QKY4
A4QKY6 A4QKY7 A4QKY9 A4QKZ2 A4QL00 A4QL03 A4QL05 A4QL06 A4QL07 A4QL12 A4QL13
A4QL14 A4QL15 A4QL18 A4QL19 A4QL24 A4QL30 A4QL31 A4QL32 A4QL33 A4QL34 A4QL37
A4QL38 A4QL39 A4QL45 A4QL47 A4QL68 A4QL70 A4QL71 A4QL73 A4QL74 A4QL76 A4QL79
A4QL86 A4QL90 A4QL92 A4QL93 A4QL94 A4QL99 A4QLA0 A4QLA1 A4QLA2 A4QLA5 A4QLA6
A4QLB1 A4QLB7 A4QLB8 A4QLB9 A4QLC0 A4QLC1 A4QLC4 A4QLC5 A4QLC6 A4QLD2 A4QLD4
A4QLF6 A4QLF8 A4QLF9 A4QLG1 A4QLG2 A4QLG4 A4QLG7 A4QLH5 A4QLH8 A4QLI0 A4QLI1
A4QLI2 A4QLI7 A4QLI8 A4QLI9 A4QLJ0 A4QLJ3 A4QLJ4 A4QLJ9 A4QLK5 A4QLK6 A4QLK7
A4QLK8 A4QLK9 A4QLL2 A4QLL3 A4QLL4 A4QLM0 A4QLM2 A4QLP4 A4QLP6 A4QLP7 A4QLP9
A4QLQ0 A4QLQ2 A4QLQ5 A4QLR3 A4QLR7 A4QLR9 A4QLS0 A4QLS1 A4QLS6 A4QLS7 A4QLS8

A4QLS9 A4QLT2 A4QLT3 A4QLT8 A4QLU4 A4QLU5 A4QLU6 A4QLU7 A4QLU8 A4QLV1 A4QLV2
A4QLV3 A4QLV9 A4QLW1 A4QLY2 A4QLY4 A4QLY5 A4QLY7 A4QLY8 A4QLZ0 A4QLZ3 A4QM02
A4QM18 A4QM25 A4QM32 A4QN56 A4QNF3 A4QNG1 A4QNL6 A4QNN3 A4QNX2 A4QNX3 A4QP81
A4QPC6 A4QR21 A4QT54 A4QTE3 A4QTI8 A4QVZ8 A4QYQ5 A4R0J5 A4R1D5 A4R254 A4R2N5
A4R8V2 A4R935 A4RC22 A4RE47 A4RE85 A4RF05 A4RF23 A4RGA1 A4RHN5 A4RHX3 A4RI25
A4RM13 A4RM19 A4RME3 A4RMK0 A4RPU0 A4S8B7 A4SDF5 A4SDK1 A4SE48 A4SEC9 A4SEF1
A4SEH5 A4SF48 A4SF67 A4SG03 A4SGR1 A4SGV1 A4SGZ0 A4SH05 A4SH20 A4SHM8 A4SI55
A4SI95 A4SIW6 A4SIW8 A4SJ99 A4SJA2 A4SJT9 A4SK68 A4SKP9 A4SLN5 A4SLN8 A4SLP4
A4SLX1 A4SN46 A4SN74 A4SN80 A4SN81 A4SN85 A4SNG2 A4SNP5 A4SNP8 A4SPR3 A4SPT0
A4SPY8 A4SQI9 A4SQL6 A4SQL7 A4SQW8 A4SQX1 A4SQX2 A4SQX3 A4SR87 A4ST36 A4STB5
A4STK7 A4STK9 A4STL2 A4STP7 A4STP8 A4STP9 A4STS5 A4SUT0 A4SV24 A4SV71 A4SW33
A4SWH9 A4SXF0 A4SXH5 A4SXQ3 A4SYW3 A4SZ14 A4SZG7 A4SZG8 A4SZG9 A4SZT5 A4T079
A4T0I1 A4T0N2 A4T134 A4T190 A4T8J9 A4T8K8 A4T8K9 A4T8W2 A4T9I4 A4T9U2 A4TB05
A4TB40 A4TB90 A4TBF2 A4TBJ8 A4TC38 A4TCS6 A4TDA1 A4TDB2 A4TFF8 A4TGM8 A4TGR2
A4TGV3 A4TGX5 A4TH19 A4TH40 A4TH46 A4THC5 A4THE8 A4THE9 A4THF0 A4THK0 A4THL5
A4THM4 A4THT2 A4THT4 A4TI93 A4TIF8 A4TIM3 A4TIM6 A4TIM7 A4TIM8 A4TIP9 A4TIU0
A4TJ27 A4TJ33 A4TJ73 A4TJ76 A4TJ85 A4TJC7 A4TJC8 A4TJI9 A4TJJ0 A4TJL9 A4TK40
A4TKE6 A4TKE7 A4TKN5 A4TL05 A4TL06 A4TL83 A4TLC6 A4TLD2 A4TLD3 A4TLG5 A4TM24
A4TM27 A4TM30 A4TM36 A4TM43 A4TM91 A4TMF6 A4TMG5 A4TMJ0 A4TMM5 A4TMM7 A4TMR4
A4TMT8 A4TN28 A4TN83 A4TPF3 A4TPL2 A4TPL3 A4TPL4 A4TPW7 A4TPZ6 A4TQ86 A4TQF2
A4TQF7 A4TQJ7 A4TQL6 A4TQL7 A4TQX7 A4TR01 A4TR35 A4TR37 A4TRB5 A4TRG0 A4TRQ4
A4TRR6 A4TRS5 A4TRU8 A4TS08 A4TSA8 A4TSF8 A4TSH8 A4TSI7 A4TSI8 A4TSI9 A4U547
A4U6V2 A4U6V3 A4U6V5 A4U7A6 A4U7A7 A4U7A9 A4UHQ1 A4UHQ6 A4UVI1 A4VCH0 A4VCL2
A4VGD9 A4VGL8 A4VGN3 A4VGY3 A4VI46 A4VIH5 A4VJ38 A4VJ44 A4VKL4 A4VKM9 A4VKR4
A4VL35 A4VLM0 A4VLU8 A4VMI8 A4VMK0 A4VMV0 A4VMV3 A4VNB9 A4VNS0 A4VRA1 A4VRK8
A4VS42 A4VS66 A4VS68 A4VS82 A4VTD2 A4VTE9 A4VVK3 A4VWE0 A4VWI6 A4VWT2 A4VX34
A4VX68 A4VXK6 A4VZK9 A4VZM7 A4W1W1 A4W2P4 A4W2T8 A4W334 A4W3D8 A4W3H1 A4W3V2
A4W485 A4W4S5 A4W5E5 A4W5F0 A4W5F3 A4W5I3 A4W5M3 A4W5P8 A4W5P9 A4W678 A4W693
A4W6D7 A4W6E2 A4W6F3 A4W6J0 A4W6S2 A4W799 A4W811 A4W859 A4W860 A4W861 A4W8L3
A4W8S1 A4W964 A4W978 A4W9G6 A4W9T2 A4W9W3 A4WA57 A4WA58 A4WA77 A4WAE6 A4WAL9
A4WAM0 A4WAM1 A4WAM4 A4WAT9 A4WAU8 A4WB73 A4WB74 A4WB82 A4WB84 A4WBF0 A4WBF5
A4WBH8 A4WBH9 A4WBI4 A4WBI8 A4WCC1 A4WCC2 A4WCC3 A4WCH9 A4WCQ5 A4WCQ8 A4WCR1
A4WCR7 A4WCS4 A4WD10 A4WD24 A4WD52 A4WD94 A4WDV3 A4WE04 A4WE10 A4WE11 A4WE68
A4WEE1 A4WEI1 A4WEI9 A4WEJ1 A4WER2 A4WER3 A4WES2 A4WES8 A4WEU1 A4WF55 A4WF56
A4WF68 A4WF99 A4WFE4 A4WFG6 A4WFK8 A4WFT3 A4WFV0 A4WFY3 A4WG45 A4WG79 A4WG89
A4WGB7 A4WGE0 A4WGE9 A4WGF0 A4WGF1 A4WGH2 A4WIG5 A4WKY2 A4WN93 A4WNY6 A4WNY7
A4WNY8 A4WNY9 A4WP17 A4WP56 A4WQ57 A4WQ59 A4WQD0 A4WQY8 A4WRW9 A4WT80 A4WU21
A4WVB7 A4WWA0 A4WZ55 A4X1Z0 A4X422 A4X4U3 A4X5I0 A4X6R9 A4X752 A4X9F8 A4X9R8
A4XAW6 A4XB61 A4XC28 A4XHU5 A4XI01 A4XKX4 A4XKX5 A4XLD4 A4XM19 A4XND8 A4XNK8
A4XPM2 A4XPM5 A4XPR4 A4XQS1 A4XQV6 A4XRP6 A4XS49 A4XS50 A4XS53 A4XSP4 A4XSP7
A4XT57 A4XUB9 A4XV02 A4XV08 A4XV11 A4XV14 A4XVB4 A4XVY6 A4XW26 A4XW98 A4XWM7
A4Y001 A4Y045 A4Y0P0 A4Y191 A4Y193 A4Y1A0 A4Y2A0 A4Y2G7 A4Y2N3 A4Y2Q1 A4Y2Q2
A4Y2Q3 A4Y370 A4Y3Y9 A4Y4F4 A4Y4F7 A4Y4J2 A4Y517 A4Y624 A4Y6I5 A4Y6I7 A4Y6J0
A4Y6Z6 A4Y768 A4Y7X2 A4Y803 A4Y854 A4Y9P2 A4YA01 A4YA10 A4YAR8 A4YAV1 A4YBV6

A4YC13 A4YC85 A4YCI2 A4YCI3 A4YCI4 A4YCM2 A4YDT1 A4YH85 A4YHQ9 A4YI21 A4YI90
A4YJ27 A4YJM2 A4YKA9 A4YN81 A4YQF7 A4YQU0 A4YRM0 A4YRS1 A4YTR6 A4YV72 A4YV73
A4YVK0 A4YVK3 A4YVQ0 A4YWD8 A4YWG6 A4YWY8 A4YWY9 A4YZG3 A4YZK6 A4Z016 A4Z2B4
A4Z2B6 A4Z2B7 A4Z2D0 A4Z2D2 A4ZU95 A4ZUA7 A4ZUB1 A4ZUB3 A4ZUB4 A4ZUB5 A4ZUB9
A4ZUC0 A4ZUC3 A4ZUC4 A4ZUC6 A4ZUC7 A4ZUC8 A4ZUC9 A4ZUD0 A4ZUD1 A4ZUD5 A4ZYQ5
A5A2P0 A5A4K9 A5A4L1 A5A605 A5A615 A5A616 A5A618 A5A624 A5A627 A5A630 A5A6H8
A5A6J8 A5A6L6 A5A6N6 A5A6S6 A5A761 A5A8K9 A5AB61 A5ABG1 A5BFI4 A5CDC5 A5CDC6
A5CF31 A5CF77 A5CFG7 A5CFN5 A5CM04 A5CML5 A5CQ53 A5CQ56 A5CQ95 A5CRN8 A5CRS6
A5CS54 A5CTB2 A5CUP8 A5CVF9 A5CVI2 A5CWG8 A5CWY7 A5CXA6 A5CXC5 A5CXG4 A5CXV6
A5CXZ0 A5CYC0 A5CYE6 A5D0V1 A5D148 A5D176 A5D178 A5D1P8 A5D2F6 A5D2Q9 A5D6V4
A5D6W6 A5D787 A5D7B2 A5D7B5 A5D7B7 A5D7C3 A5D7C9 A5D7E2 A5D7E9 A5D7H1 A5D7H3
A5D7I4 A5D7K7 A5D7K8 A5D7L5 A5D7M7 A5D7N3 A5D7T4 A5D7U1 A5D7U4 A5D7V5 A5D7V7
A5D8P8 A5D9A7 A5D9S8 A5D9W9 A5DAI1 A5DAT0 A5DB26 A5DC90 A5DCL4 A5DDN8 A5DE01
A5DE11 A5DEJ6 A5DEQ7 A5DEU9 A5DEX5 A5DFM9 A5DGH6 A5DGW8 A5DGY3 A5DH87 A5DHA3
A5DHC2 A5DI86 A5DIN8 A5DIR3 A5DIS9 A5DIZ8 A5DJ60 A5DJQ5 A5DJS9 A5DJU3 A5DL79
A5DLK4 A5DLM7 A5DM93 A5DN48 A5DND6 A5DNZ8 A5DQ25 A5DQE4 A5DR93 A5DRE8 A5DS09
A5DS78 A5DSD2 A5DSF0 A5DSM4 A5DSM9 A5DTG8 A5DTQ6 A5DU71 A5DUA8 A5DUG6 A5DUL5
A5DV09 A5DV96 A5DVB1 A5DVH1 A5DX39 A5DYV2 A5DZ28 A5DZ34 A5DZQ4 A5E0R1 A5E1P1
A5E1Q5 A5E1V8 A5E212 A5E240 A5E2M7 A5E3T2 A5E4Z8 A5E567 A5E5C5 A5E5Y6 A5E708
A5E751 A5E7J0 A5E7U2 A5E8F3 A5E8M5 A5E919 A5EA98 A5EAA0 A5EAB1 A5EAB2 A5EAB4
A5EBW8 A5EG97 A5EHA3 A5EHG1 A5EHG2 A5EJW1 A5EJW2 A5EK88 A5EK91 A5EKD5 A5EKR0
A5EMH9 A5EPH1 A5EPK7 A5EQ10 A5ESV8 A5ESW9 A5EUT7 A5EUU4 A5EVU2 A5EVU3 A5EW93
A5EWR9 A5EWT1 A5EX03 A5EX40 A5EXJ9 A5EXK1 A5EY04 A5EY06 A5EYU0 A5F037 A5F0D5
A5F0D8 A5F120 A5F137 A5F1Q0 A5F1U5 A5F1V9 A5F296 A5F2H8 A5F2R1 A5F2R3 A5F2S7
A5F2S8 A5F2W1 A5F2W2 A5F382 A5F385 A5F396 A5F419 A5F473 A5F475 A5F484 A5F4D3
A5F4E9 A5F4G3 A5F4H4 A5F4P6 A5F4Q8 A5F4U3 A5F4Z1 A5F5N2 A5F5X0 A5F5Y4 A5F5Y5
A5F5Y6 A5F5Y7 A5F661 A5F6K5 A5F6Z3 A5F7I4 A5F7R1 A5F8Y5 A5F8Z4 A5F934 A5F9D1
A5F9D4 A5F9G0 A5FA69 A5FAI5 A5FBK7 A5FIF7 A5FIF9 A5FIY0 A5FJD0 A5FKC3 A5FKJ7
A5FKK0 A5FKK7 A5FL32 A5FL65 A5FLP0 A5FMG4 A5FNQ7 A5FPE4 A5FPF2 A5FPL0 A5FPV4
A5FPV7 A5FQH9 A5FQU1 A5FQX9 A5FRQ1 A5FRR4 A5FT18 A5FT52 A5FTA4 A5FUK7 A5FUT7
A5FUZ5 A5FVI7 A5FVI8 A5FVJ0 A5FVL8 A5FWG2 A5FX21 A5FXI8 A5FXJ1 A5FXK0 A5FXV3
A5FXV4 A5G0F8 A5G0I2 A5G2G1 A5G348 A5G6V8 A5G9A9 A5G9B9 A5G9C3 A5G9C4 A5G9D4
A5G9I7 A5G9V4 A5GAD3 A5GAG0 A5GAG2 A5GAQ7 A5GAT3 A5GBD1 A5GCT1 A5GCZ5 A5GD03
A5GD16 A5GFS8 A5GFW8 A5GIA7 A5GIA8 A5GIB0 A5GIB3 A5GIE8 A5GIH1 A5GIH3 A5GIH4
A5GIM7 A5GJ28 A5GJ31 A5GJ44 A5GJH8 A5GJP1 A5GJU5 A5GKT4 A5GL22 A5GL35 A5GLB4
A5GLJ4 A5GM56 A5GMA2 A5GMA7 A5GMA8 A5GMV6 A5GMW1 A5GMW2 A5GMY0 A5GMZ5 A5GND0
A5GND2 A5GND3 A5GNE2 A5GNP1 A5GNW2 A5GP41 A5GP52 A5GPT3 A5GQF1 A5GQF2 A5GQF4
A5GQF7 A5GQH5 A5GQK3 A5GQQ0 A5GQT7 A5GR11 A5GR14 A5GR39 A5GR40 A5GRR8 A5GRU7
A5GRU8 A5GRV3 A5GRX8 A5GTC7 A5GTF4 A5GTI3 A5GTV1 A5GTY4 A5GTY5 A5GU84 A5GUB8
A5GUE6 A5GUN3 A5GUN8 A5GV74 A5GV75 A5GV76 A5GV77 A5GV87 A5GWF4 A5GWH9 A5GWI1
A5GWI2 A5GX19 A5GZW8 A5H9N6 A5HJM1 A5HUI5 A5HXQ7 A5HY47 A5HY48 A5HYT8 A5HZD5
A5I013 A5I1C7 A5I1U0 A5I4H8 A5I6G4 A5I6T8 A5IA60 A5IAV9 A5IB85 A5IBE5 A5IBS2
A5IDR5 A5IEF8 A5IEQ9 A5IES9 A5IF79 A5IF82 A5IFJ6 A5IG55 A5IGR4 A5IH85 A5IHI2
A5IHV3 A5IHV6 A5II42 A5II94 A5III7 A5IIK4 A5IIP0 A5IKI6 A5IKR3 A5IL48 A5IL85

A5ILW7 A5ILW8 A5IMD8 A5INR0 A5IP58 A5IPD1 A5IPD2 A5IQH5 A5IQH6 A5IQH7 A5IQH8
A5IQH9 A5IQI0 A5IQI1 A5IQL3 A5IQN7 A5IQW5 A5IRC4 A5IRC5 A5IRC6 A5IRC7 A5IRC8
A5IRC9 A5IRD0 A5IRE7 A5IRZ2 A5IS02 A5IS03 A5IS68 A5IS81 A5ISH0 A5ISN0 A5ISN6
A5ISW6 A5ISW7 A5ITA1 A5ITP0 A5IU11 A5IU40 A5IU69 A5IU79 A5IUI3 A5IUN6 A5IUQ2
A5IUQ3 A5IUQ4 A5IUV7 A5IVC1 A5IVD7 A5IVE3 A5IVG9 A5IVW5 A5IVW8 A5IVY3 A5IWB2
A5IYE0 A5IYE1 A5IYF1 A5JYQ9 A5JYX8 A5K168 A5K302 A5LFX5 A5LGM7 A5LGW7 A5N0R3
A5N2J8 A5N466 A5N4S9 A5N7E7 A5N857 A5N884 A5PF08 A5PHT0 A5PJ65 A5PJ82 A5PJA8
A5PJC7 A5PJF4 A5PJK1 A5PJL1 A5PJM4 A5PJQ2 A5PJS2 A5PJW2 A5PJX7 A5PJY4 A5PJZ1
A5PJZ2 A5PK13 A5PK14 A5PK27 A5PK40 A5PLA0 A5PLC9 A5PLE2 A5PLE7 A5PLF5 A5PLH4
A5PLK6 A5PLL7 A5PMW0 A5PMX1 A5PMY6 A5PN38 A5PN43 A5TY81 A5TY85 A5TYD5 A5TZH0
A5TZU3 A5U124 A5U127 A5U173 A5U175 A5U1S2 A5U204 A5U205 A5U206 A5U2B2 A5U2F4
A5U2I5 A5U2R1 A5U2R2 A5U2X0 A5U2Z7 A5U4C0 A5U4G2 A5U4I3 A5U4N6 A5U4X2 A5U5Z1
A5U7B6 A5U7G8 A5U7H1 A5U7L0 A5U8T5 A5U9F2 A5U9Y4 A5UA05 A5UA06 A5UA07 A5UAL3
A5UAM4 A5UAR0 A5UAW7 A5UAX2 A5UAX4 A5UAX5 A5UAX6 A5UBA3 A5UBI7 A5UBI8 A5UBI9
A5UBJ2 A5UBU9 A5UCD6 A5UCQ5 A5UCX1 A5UD02 A5UD03 A5UD12 A5UD33 A5UD72 A5UDH1
A5UDI9 A5UDP2 A5UDP3 A5UDP8 A5UE05 A5UE58 A5UEB0 A5UEC3 A5UEG3 A5UF21 A5UFC0
A5UFC3 A5UFC5 A5UFK5 A5UFL3 A5UFU0 A5UFW9 A5UFX1 A5UFX2 A5UFX3 A5UG30 A5UG86
A5UG94 A5UGZ3 A5UGZ4 A5UGZ5 A5UH14 A5UH89 A5UHA1 A5UHG2 A5UHK2 A5UHX7 A5UHY2
A5UI29 A5UID8 A5UIJ7 A5UIL9 A5UIM7 A5UIM8 A5UIQ9 A5UJ57 A5UKV2 A5ULR8 A5UMH8
A5UPV7 A5UQ59 A5UQN7 A5UQS9 A5UVF9 A5UVR8 A5UW67 A5UXK0 A5UZ79 A5UZW0 A5V212
A5V2F7 A5V3C5 A5V412 A5V5N0 A5V7M2 A5V7N3 A5V804 A5V8S7 A5VAC0 A5VAM7 A5VAN3
A5VBY5 A5VCW2 A5VD56 A5VDA9 A5VDC9 A5VEV5 A5VEV8 A5VF68 A5VI38 A5VIB3 A5VIG5
A5VIQ5 A5VIQ7 A5VJ31 A5VJM1 A5VK34 A5VM74 A5VN85 A5VNQ9 A5VNW2 A5VNW4 A5VNW5
A5VNX4 A5VP40 A5VP42 A5VPF4 A5VPH9 A5VPX0 A5VPZ0 A5VPZ3 A5VQ44 A5VQH6 A5VRD1
A5VRI0 A5VRQ8 A5VS16 A5VSF5 A5VSR0 A5VU89 A5VU90 A5VUT3 A5VV34 A5VVQ3 A5VVU6
A5VW32 A5VWP2 A5VWR8 A5VXI5 A5VYB7 A5VYP1 A5VZL9 A5VZR9 A5VZT0 A5W0A9 A5W0P1
A5W0T9 A5W155 A5W183 A5W186 A5W192 A5W1F0 A5W1H5 A5W1K7 A5W2C8 A5W382 A5W499
A5W5G7 A5W758 A5W7Q2 A5W8Q3 A5W917 A5W949 A5WA47 A5WA59 A5WA99 A5WAE0 A5WAK7
A5WB34 A5WB80 A5WBA7 A5WBA8 A5WBA9 A5WBB7 A5WBC5 A5WBG4 A5WBH0 A5WBL5 A5WBQ6
A5WBV5 A5WBV7 A5WCC8 A5WCD3 A5WCE5 A5WCV5 A5WD44 A5WG39 A5WG42 A5WG48 A5WHG7
A5WHK1 A5WHL4 A5WM93 A5WMW1 A5WV69 A5WVU6 A5WVU9 A5WVX9 A5WWA0 A5X5Y0 A5Z1X6
A5Z2X3 A5Z2X5 A6BM11 A6BM12 A6BM17 A6BM20 A6BM21 A6BM22 A6BM24 A6BM27 A6BM28
A6BM29 A6BM30 A6BM34 A6BM35 A6BM37 A6BM38 A6BM63 A6BM72 A6GXA1 A6GYI6 A6GYR0
A6H039 A6H084 A6H086 A6H0G8 A6H0J1 A6H198 A6H1E4 A6H1Q0 A6H1Q3 A6H1Q6 A6H1R4
A6H239 A6H2D9 A6H2G3 A6H4Q2 A6H4Q3 A6H4Q4 A6H4Q5 A6H4Q6 A6H4Q7 A6H4Q8 A6H5E4
A6H5E9 A6H5F0 A6H5F2 A6H5F3 A6H5F5 A6H5G0 A6H5G2 A6H5G6 A6H5G7 A6H5H1 A6H5H9
A6H5I8 A6H5J0 A6H5J1 A6H5J2 A6H5J3 A6H5J7 A6H5J8 A6H5K2 A6H5K7 A6H5K9 A6H5N8
A6H5P1 A6H5P2 A6H5P4 A6H5P5 A6H5P7 A6H666 A6H770 A6H773 A6H784 A6H789 A6H793
A6H7A0 A6H7B0 A6H7B1 A6H7B8 A6H7C2 A6H7F9 A6H8H5 A6H8M9 A6KZJ8 A6L074 A6L162
A6L168 A6L171 A6L227 A6L3M0 A6L4M1 A6L4M2 A6L5D0 A6L5D2 A6L5L6 A6L685 A6L6E5
A6L6X8 A6L743 A6L898 A6L8E9 A6L8N7 A6L9D2 A6LA91 A6LAI2 A6LD25 A6LD62 A6LDT3
A6LEL8 A6LEU6 A6LIU6 A6LJF8 A6LJI3 A6LJR5 A6LKX5 A6LL73 A6LLB7 A6LM64 A6LQH1
A6LQH2 A6LSR9 A6LSW5 A6LTE4 A6LTS6 A6LVG3 A6LXJ5 A6LXP5 A6LXQ5 A6M387 A6MGW7
A6MHQ4 A6MM16 A6MM20 A6MM22 A6MM23 A6MM24 A6MM29 A6MM30 A6MM31 A6MM32 A6MM35

A6MM41 A6MM47 A6MM48 A6MM49 A6MM50 A6MM51 A6MM54 A6MM55 A6MM56 A6MM62 A6MM64
A6MM84 A6MM86 A6MM87 A6MM89 A6MM90 A6MM95 A6MMA2 A6MMA6 A6MMA8 A6MMA9 A6MMB0
A6MMB5 A6MMB6 A6MMB7 A6MMB8 A6MMC1 A6MMC7 A6MMD3 A6MMD4 A6MMD5 A6MMD6 A6MMD7
A6MME0 A6MME1 A6MME2 A6MME8 A6MMF0 A6MMG9 A6MMH4 A6MMH5 A6MMH7 A6MMH8 A6MMI0
A6MMI8 A6MMJ1 A6MMJ3 A6MMJ4 A6MMJ5 A6MMK0 A6MMK1 A6MMK2 A6MMK3 A6MMK6 A6MML2
A6MML8 A6MML9 A6MMM0 A6MMM1 A6MMM2 A6MMM5 A6MMM6 A6MMM7 A6MMN3 A6MMN5 A6MMQ5
A6MMR0 A6MMR1 A6MMR3 A6MMR4 A6MMR6 A6MMS4 A6MMS8 A6MMT0 A6MMT1 A6MMT2 A6MMT7
A6MMT8 A6MMT9 A6MMU0 A6MMU3 A6MMU9 A6MMV5 A6MMV6 A6MMV7 A6MMV8 A6MMV9 A6MMW2
A6MMW3 A6MMW4 A6MMX0 A6MMX2 A6MMZ2 A6MMZ4 A6MMZ5 A6MMZ7 A6MMZ8 A6MN00 A6MN03
A6MVL7 A6MVS2 A6MVT2 A6MVU6 A6MVU9 A6MVV2 A6MVV3 A6MVV6 A6MVV7 A6MVV9 A6MVW0
A6MVW6 A6MVW7 A6MVW8 A6MVW9 A6MVY3 A6MVY7 A6MVZ4 A6MVZ6 A6MW33 A6N3Q4 A6NC51
A6NC97 A6NCI5 A6NCQ9 A6NCV1 A6ND48 A6NDA9 A6NDD5 A6NDH6 A6NDL8 A6NDP7 A6NDV4
A6NDX4 A6NEH6 A6NET4 A6NF34 A6NF89 A6NFA0 A6NFA1 A6NFC5 A6NFC9 A6NFE2 A6NFR6
A6NFU0 A6NFX1 A6NFY4 A6NG13 A6NGA9 A6NGB0 A6NGB7 A6NGC4 A6NGU5 A6NGY5 A6NGZ8
A6NH00 A6NH21 A6NH52 A6NHA9 A6NHG9 A6NHS7 A6NI61 A6NI73 A6NIJ9 A6NIM6 A6NJU9
A6NJW4 A6NJW9 A6NJY1 A6NJY4 A6NJZ3 A6NK97 A6NKB5 A6NKF7 A6NKG5 A6NKK0 A6NKL6
A6NKW6 A6NKX4 A6NL05 A6NL08 A6NL26 A6NL88 A6NL99 A6NLE4 A6NLU5 A6NLX4 A6NM03
A6NM10 A6NM11 A6NM45 A6NM62 A6NM76 A6NMB1 A6NMD0 A6NML5 A6NMS3 A6NMS7 A6NMU1
A6NMZ5 A6NN92 A6NNB3 A6NNC1 A6NND4 A6NNE9 A6NNN8 A6P320 A6PVL3 A6PWV3 A6Q0Z0
A6Q1Q1 A6Q1Q4 A6Q1R7 A6Q218 A6Q3A6 A6Q3V4 A6Q4C4 A6Q4S3 A6Q544 A6Q6B7 A6Q6G8
A6Q7D4 A6Q7V0 A6Q913 A6Q9S3 A6QA78 A6QAM3 A6QB63 A6QBH1 A6QBN8 A6QCC3 A6QCE0
A6QCE9 A6QCF2 A6QD58 A6QDH3 A6QDN6 A6QDN7 A6QET3 A6QET4 A6QET5 A6QET6 A6QET7
A6QET8 A6QET9 A6QEW9 A6QEZ2 A6QF69 A6QFF6 A6QFF7 A6QFF8 A6QFF9 A6QFG0 A6QFG1
A6QFG2 A6QFI0 A6QFW1 A6QFX0 A6QFX1 A6QG35 A6QG82 A6QG97 A6QGC0 A6QGI5 A6QGQ0
A6QGQ6 A6QGY4 A6QGY6 A6QH29 A6QHB6 A6QHQ1 A6QI28 A6QI56 A6QID0 A6QID9 A6QIN3
A6QIS0 A6QIS1 A6QIS2 A6QIT4 A6QIV1 A6QIV2 A6QIV3 A6QJ09 A6QJI0 A6QJJ7 A6QJK2
A6QJK4 A6QJM8 A6QK27 A6QK30 A6QK47 A6QKE2 A6QKH3 A6QL63 A6QL84 A6QL88 A6QL92
A6QL94 A6QLC6 A6QLE6 A6QLE7 A6QLF8 A6QLI1 A6QLK4 A6QLM0 A6QLP7 A6QLU6 A6QLW8
A6QLZ1 A6QM03 A6QM06 A6QNK1 A6QNL6 A6QNL7 A6QNW6 A6QNY1 A6QNZ8 A6QP01 A6QP55
A6QP70 A6QP72 A6QP74 A6QP79 A6QP84 A6QPA0 A6QPB3 A6QPF4 A6QPF8 A6QPI1 A6QPI4
A6QPI6 A6QPL2 A6QPT7 A6QPY0 A6QQ24 A6QQ59 A6QQ85 A6QQ93 A6QQC6 A6QQL0 A6QQP7
A6QQT9 A6QQU6 A6QQX9 A6QQZ0 A6QR09 A6QR22 A6QRX6 A6QS12 A6QTM4 A6QTW5 A6QU86
A6QWA0 A6QWZ2 A6QX24 A6QX86 A6QYP4 A6R1D5 A6R1T7 A6R2K7 A6R3V7 A6R620 A6RAY2
A6RBB3 A6RBC5 A6RBI0 A6RBW9 A6RBY1 A6REI4 A6RF73 A6RH42 A6RIB9 A6RJQ7 A6RJQ8
A6RN41 A6RP27 A6RRF7 A6RUY7 A6S3E0 A6S3R7 A6S3W1 A6S544 A6S8A1 A6S8E0 A6S950
A6S9E2 A6SAY2 A6SEV8 A6SFL7 A6SG70 A6SI59 A6SL49 A6SL61 A6SSM3 A6SSX6 A6SU72
A6SUU0 A6SV70 A6SVP3 A6SW70 A6SWI5 A6SWX0 A6SXG4 A6SXH1 A6SY12 A6SY89 A6SZ14
A6SZ16 A6SZI9 A6T022 A6T273 A6T2A4 A6T2G1 A6T2T7 A6T365 A6T372 A6T3X6 A6T474
A6T476 A6T4D7 A6T4F6 A6T4G1 A6T4H1 A6T4I3 A6T4N0 A6T4V9 A6T4X9 A6T515 A6T522
A6T525 A6T526 A6T5H1 A6T5L4 A6T688 A6T6D7 A6T6D8 A6T6D9 A6T6E2 A6T6U3 A6T7C6
A6T7D8 A6T7N0 A6T7P0 A6T7V7 A6T7V8 A6T814 A6T884 A6T8S5 A6T8S6 A6T8U5 A6T8Y8
A6TA11 A6TA84 A6TAH6 A6TAJ7 A6TAU9 A6TAW2 A6TAW7 A6TAZ1 A6TAZ2 A6TB00 A6TBB1
A6TBH4 A6TBH5 A6TBH6 A6TBM9 A6TBW2 A6TBW5 A6TBW8 A6TBX4 A6TBY2 A6TC38 A6TC48
A6TCD5 A6TD40 A6TDB4 A6TDC6 A6TDG4 A6TDH1 A6TDH2 A6TDS8 A6TDZ7 A6TE36 A6TE38

A6TEB6 A6TEB7 A6TEC8 A6TED5 A6TES0 A6TEU4 A6TEY9 A6TF43 A6TF94 A6TF95 A6TF96
A6TF99 A6TFB2 A6TFD1 A6TFY7 A6TG08 A6TG19 A6TG40 A6TG41 A6TG42 A6TG51 A6TG76
A6TGB1 A6TGB7 A6TGL5 A6TGU1 A6TGU6 A6TGU9 A6TGZ7 A6TH46 A6TH70 A6TH71 A6THX5
A6THZ2 A6TJ58 A6TJ97 A6TK61 A6TLV6 A6TLZ3 A6TP80 A6TS38 A6TS64 A6TSK9 A6TSZ1
A6TU75 A6TW45 A6TWP7 A6TXG9 A6TXX3 A6TY47 A6TY48 A6TZ99 A6TZA0 A6TZA1 A6TZA2
A6TZA3 A6TZA4 A6TZA5 A6TZD7 A6TZG1 A6TZP0 A6U053 A6U054 A6U055 A6U056 A6U057
A6U058 A6U059 A6U076 A6U0S3 A6U0T3 A6U0T4 A6U102 A6U115 A6U1A5 A6U1G3 A6U1G8
A6U1H4 A6U1Q5 A6U1Q6 A6U245 A6U2I4 A6U2U9 A6U2X8 A6U307 A6U317 A6U3C2 A6U3H6
A6U3J2 A6U3J3 A6U3J4 A6U3P7 A6U465 A6U483 A6U489 A6U4B6 A6U4R9 A6U4S2 A6U4T8
A6U570 A6U5K9 A6U5U5 A6U6B9 A6U6M1 A6U6M4 A6U6M6 A6U6M7 A6U6U6 A6U6U8 A6U760
A6U785 A6U7T3 A6U7X5 A6U809 A6U8N5 A6U9P2 A6U9W5 A6UB88 A6UBQ4 A6UE35 A6UE36
A6UED6 A6UFC2 A6UFL3 A6UGS9 A6UH53 A6ULU4 A6UM20 A6UMS8 A6UNS5 A6UPF1 A6UPS4
A6UQC5 A6UQK6 A6UQK8 A6UQK9 A6UQV8 A6UQY1 A6UUW7 A6UV73 A6UVU6 A6UWB6 A6UWH3
A6UWH6 A6UWH7 A6UWS5 A6UWW7 A6UXP9 A6UXQ5 A6UYE4 A6UZ84 A6UZY0 A6V0C5 A6V1N6
A6V1N7 A6V1N8 A6V1P1 A6V1T4 A6V1T6 A6V1T9 A6V2E4 A6V2M1 A6V398 A6V3A0 A6V3A1
A6V3A2 A6V3R0 A6V4D9 A6V4E2 A6V4E9 A6V6C0 A6V6H6 A6V6L2 A6V6S8 A6V701 A6V7X4
A6V817 A6V8N8 A6V8S8 A6V915 A6VA22 A6VA56 A6VAD3 A6VB55 A6VB88 A6VBU6 A6VBW1
A6VC02 A6VCE4 A6VDE2 A6VDI8 A6VDJ1 A6VDK0 A6VEG6 A6VEV6 A6VF36 A6VF37 A6VF38
A6VF45 A6VFP2 A6VG87 A6VH67 A6VHE8 A6VHF0 A6VHF1 A6VHF2 A6VHU1 A6VHW4 A6VKA8
A6VKX7 A6VL34 A6VL53 A6VL61 A6VL62 A6VL63 A6VL73 A6VLY0 A6VLY1 A6VM40 A6VMM0
A6VMQ5 A6VMW8 A6VMX8 A6VNG9 A6VQ43 A6VQC0 A6VQH9 A6VQI5 A6VQI8 A6VQN6 A6VQU4
A6VQU5 A6VQY2 A6VR17 A6VR61 A6VRF6 A6VRW7 A6VT98 A6VUD9 A6VW03 A6VW13 A6VWQ5
A6VYK1 A6W0S1 A6W0T5 A6W1R8 A6W3L5 A6W3T2 A6W3T4 A6W3V1 A6W4F5 A6W5R0 A6W6R5
A6W6R7 A6W792 A6W7G5 A6W915 A6W9V1 A6WC46 A6WCN9 A6WCX7 A6WHV6 A6WI45 A6WIC8
A6WIE5 A6WIE6 A6WIE7 A6WJS5 A6WJZ2 A6WKD5 A6WKK5 A6WKK8 A6WKP5 A6WL74 A6WM74
A6WMM7 A6WN13 A6WN15 A6WN18 A6WNG2 A6WPJ1 A6WPS6 A6WRX3 A6WSR0 A6WT30 A6WT76
A6WU15 A6WU49 A6WUJ4 A6WUJ5 A6WUJ6 A6WUK4 A6WUW2 A6WVY7 A6WW77 A6WW79 A6WW80
A6WW92 A6WWG0 A6WWG2 A6WX58 A6WXE1 A6WXV6 A6WYA8 A6WYK2 A6WYN0 A6WZF1 A6WZQ3
A6WZX3 A6X053 A6X0T1 A6X1F5 A6X1H1 A6X1M3 A6X1M6 A6X1P0 A6X1W1 A6X1X0 A6X1X3
A6X2C8 A6X4N2 A6X514 A6X570 A6X5H8 A6X5I0 A6X5T5 A6X7E7 A6X919 A6X969 A6X993
A6XA80 A6XIP3 A6XKM2 A6YCJ2 A6YG65 A6YG66 A6YG68 A6YG69 A6YG74 A6YG76 A6YG77
A6YG79 A6YG80 A6YG86 A6YG87 A6YG90 A6YG99 A6YGB2 A6YGB8 A6YIH8 A6ZIQ8 A6ZKT1
A6ZKY8 A6ZL18 A6ZLA1 A6ZLG8 A6ZLK2 A6ZLZ9 A6ZM32 A6ZM65 A6ZMC4 A6ZMD0 A6ZMI6
A6ZMK7 A6ZML5 A6ZMM2 A6ZMR2 A6ZN14 A6ZN18 A6ZN61 A6ZNF7 A6ZNQ4 A6ZNZ1 A6ZP10
A6ZPQ9 A6ZQE4 A6ZQF6 A6ZQG3 A6ZQL9 A6ZQU6 A6ZQY8 A6ZR39 A6ZR50 A6ZR73 A6ZRG6
A6ZRQ9 A6ZRW3 A6ZRW8 A6ZRY0 A6ZS21 A6ZSP9 A6ZT02 A6ZT59 A6ZTA1 A6ZTE8 A6ZTF2
A6ZTG5 A6ZTP2 A6ZTW6 A6ZTX2 A6ZU25 A6ZU94 A6ZUA4 A6ZUB9 A6ZUG8 A6ZUI6 A6ZUP4
A6ZUX3 A6ZV78 A6ZV87 A6ZVB8 A6ZVD0 A6ZVG0 A6ZVM6 A6ZVU2 A6ZW78 A6ZWF1 A6ZX66
A6ZX88 A6ZXH8 A6ZXL1 A6ZXN8 A6ZY50 A6ZYZ4 A6ZZH2 A6ZZJ6 A6ZZS6 A6ZZV7 A6ZZX0
A6ZZY0 A7A047 A7A060 A7A063 A7A082 A7A0F9 A7A0K7 A7A0K8 A7A0Q0 A7A1N4 A7A285
A7AT07 A7E1Z1 A7E2I7 A7E2M3 A7E2Z9 A7E3C4 A7E3U5 A7E3W5 A7E558 A7E559 A7E716
A7E7N1 A7ECE0 A7EEA6 A7EF54 A7EFY0 A7EJE5 A7EKT5 A7EMV1 A7EQ00 A7EQZ1 A7ER02
A7ERA6 A7ERK1 A7EWF5 A7EX99 A7EXE6 A7EYY7 A7F1F4 A7F1G7 A7F2V9 A7F4S1 A7F5K4
A7F679 A7F6C1 A7F8T1 A7F9L8 A7F9Y3 A7FBZ7 A7FCN2 A7FCQ3 A7FD14 A7FD61 A7FDE2

A7FDG9 A7FDH5 A7FDQ9 A7FDT4 A7FDT5 A7FDT6 A7FDY6 A7FE10 A7FE72 A7FE74 A7FF08
A7FF76 A7FFD1 A7FFD2 A7FFD9 A7FFH7 A7FFQ7 A7FFQ9 A7FFR1 A7FFY8 A7FG15 A7FG16
A7FG53 A7FG55 A7FG99 A7FGC1 A7FGC8 A7FGJ2 A7FGP5 A7FGQ5 A7FGR0 A7FGR3 A7FGR6
A7FGZ9 A7FHC3 A7FHG9 A7FHH6 A7FHH7 A7FHH8 A7FHJ7 A7FHN8 A7FHP6 A7FHW4 A7FHZ1
A7FHZ5 A7FHZ7 A7FI39 A7FI41 A7FI50 A7FI95 A7FIB4 A7FIB5 A7FIE5 A7FJB2 A7FJB3
A7FJK1 A7FJX9 A7FK36 A7FLD4 A7FLJ3 A7FLJ5 A7FLX9 A7FM08 A7FM69 A7FMD6 A7FME1
A7FMH9 A7FMJ8 A7FMJ9 A7FMZ5 A7FN78 A7FN99 A7FNG3 A7FNK6 A7FNS4 A7FNW6 A7FP02
A7FP30 A7FPA6 A7FPE4 A7FPE5 A7FPE6 A7FQ49 A7FQH4 A7FQH5 A7FRK3 A7FS72 A7FTG8
A7FVX8 A7FY32 A7FYF9 A7G9C1 A7G9Q4 A7G9Q5 A7GAF5 A7GAZ7 A7GBL1 A7GCY0 A7GFZ1
A7GHV1 A7GIE9 A7GL07 A7GLG4 A7GLG5 A7GLN1 A7GNP0 A7GP96 A7GQJ0 A7GQU4 A7GR09
A7GR22 A7GRA8 A7GRL8 A7GRN4 A7GRN9 A7GRX3 A7GRX4 A7GTP0 A7GTT0 A7GTU7 A7GUU4
A7GV44 A7GV47 A7GV51 A7GV60 A7GV61 A7GVI2 A7GW20 A7GW60 A7GW63 A7GWQ7 A7GWS4
A7GX83 A7GXI2 A7GY23 A7GYJ7 A7GYS2 A7GZM4 A7GZR7 A7H021 A7H0X7 A7H1P4 A7H1R9
A7H2G9 A7H2H4 A7H2J8 A7H2R0 A7H3B5 A7H4X1 A7H513 A7H515 A7H5S4 A7H5Z9 A7H600
A7H688 A7H6N5 A7H8V9 A7H902 A7H9U1 A7H9V2 A7HD40 A7HH11 A7HH64 A7HIH5 A7HIH8
A7HIW7 A7HIW9 A7HIY8 A7HJT4 A7HJW1 A7HK75 A7HLH6 A7HMK7 A7HMT2 A7HP46 A7HQW1
A7HQY4 A7HQY5 A7HQY7 A7HR02 A7HRN0 A7HRV5 A7HS58 A7HSR4 A7HVB1 A7HVU4 A7HXA1
A7HY36 A7HY39 A7HY42 A7HZF9 A7HZV5 A7HZV8 A7HZW1 A7I0F9 A7I0J2 A7I137 A7I173
A7I1B2 A7I1I1 A7I1M8 A7I227 A7I2B9 A7I2Y9 A7I339 A7I3M7 A7I3T2 A7I5F0 A7I5W2
A7I5W4 A7I6E8 A7I6J9 A7I9E7 A7IB66 A7IBA0 A7IBQ1 A7IF35 A7IFV8 A7IG27 A7IGE9
A7IGS7 A7IGS8 A7IGT0 A7IKC3 A7IMS5 A7ING5 A7INN4 A7IP97 A7IPB8 A7KAI7 A7KAJ9
A7KAK4 A7KAK7 A7KAM0 A7KAM5 A7KAM9 A7KAN0 A7KAU2 A7KAU3 A7KTC5 A7KVC2 A7L035
A7L036 A7L9Z8 A7LB59 A7LB60 A7LCJ3 A7M8Y6 A7M8Y8 A7M8Z0 A7M8Z1 A7M8Z4 A7M8Z5
A7M8Z6 A7M8Z7 A7M908 A7M910 A7M911 A7M915 A7M916 A7M917 A7M925 A7M927 A7M944
A7M950 A7M952 A7M953 A7M954 A7M959 A7M960 A7M961 A7M962 A7M969 A7M971 A7M972
A7M974 A7M975 A7M981 A7M983 A7M989 A7M991 A7M993 A7M9A9 A7M9B2 A7MB05 A7MB46
A7MB63 A7MB64 A7MB71 A7MB75 A7MBB3 A7MBC7 A7MBD8 A7MBE0 A7MBI7 A7MBJ4 A7MBM2
A7MBS7 A7MC48 A7MC64 A7ME23 A7ME74 A7MEI3 A7MEJ4 A7MEJ5 A7MF23 A7MFC3 A7MFG8
A7MG31 A7MGB5 A7MGE5 A7MGR4 A7MGT7 A7MGV6 A7MH22 A7MH29 A7MH88 A7MHI9 A7MHJ0
A7MHJ1 A7MHT8 A7MI12 A7MIA0 A7MIF2 A7MIL1 A7MIM2 A7MIS1 A7MIT0 A7MJ56 A7MJB1
A7MJB3 A7MJD1 A7MJD8 A7MJT5 A7MJT8 A7MKD6 A7MKF9 A7MKG5 A7MKM3 A7MKP4 A7MKW3
A7ML09 A7ML63 A7ML93 A7MLI3 A7MMB3 A7MMC7 A7MMH0 A7MMH2 A7MML0 A7MML1 A7MML2
A7MMM2 A7MMV3 A7MMV9 A7MMW0 A7MMX3 A7MN02 A7MN87 A7MNA4 A7MNC8 A7MNK6 A7MNL7
A7MNM2 A7MNQ0 A7MNQ2 A7MNR0 A7MP48 A7MPB4 A7MPB7 A7MPE5 A7MPN7 A7MPP0 A7MQ68
A7MQ82 A7MQD5 A7MQI7 A7MQL5 A7MQU8 A7MQV0 A7MR30 A7MR36 A7MR37 A7MRY4 A7MRY6
A7MS55 A7MSH9 A7MT68 A7MTD7 A7MTT1 A7MTU7 A7MTX4 A7MTY8 A7MVC5 A7MVD8 A7MVI0
A7MWL2 A7MWQ0 A7MWQ3 A7MX22 A7MXL0 A7MXY8 A7MYD3 A7MYQ5 A7MZ43 A7MZ44 A7MZB7
A7MZT6 A7N0X0 A7N0X9 A7N0Y5 A7N0Y8 A7N142 A7N2M2 A7N2U2 A7N2U5 A7N307 A7N5Q9
A7N6E0 A7N6I9 A7N6K9 A7N6S2 A7N9B9 A7N9H7 A7N9L6 A7N9N4 A7NA50 A7NB79 A7NBL2
A7NDX5 A7NEH8 A7NEJ7 A7NEK0 A7NEK3 A7NER4 A7NHM5 A7NIR3 A7NL04 A7NL15 A7NPD6
A7NPD9 A7NRY4 A7NW78 A7NW79 A7P0P3 A7P756 A7PA04 A7PHN8 A7PJ32 A7PMY7 A7PP95
A7PTY8 A7PZL3 A7Q6G6 A7QBZ2 A7QC16 A7QF77 A7R333 A7R385 A7RM45 A7RX69 A7RYM6
A7RYM7 A7S1L6 A7S4N4 A7S641 A7S6Y0 A7S7Z9 A7SIM4 A7SLZ2 A7SVT1 A7SXK3 A7T1N0
A7TDM0 A7TDX5 A7TE03 A7TE94 A7TES0 A7TES5 A7TEV4 A7TFA9 A7TFH0 A7TFH9 A7TFK6

A7TFL9 A7TFU8 A7TG13 A7TGB4 A7TGK3 A7TH22 A7TH24 A7THL9 A7THM1 A7THN6 A7THS1
A7TI28 A7TIN0 A7TIQ0 A7TJY3 A7TKT6 A7TKW8 A7TLA7 A7TLB3 A7TLX5 A7TM20 A7TM72
A7TMB7 A7TMJ2 A7TML0 A7TMN2 A7TMU9 A7TN96 A7TNI7 A7TNL0 A7TNL7 A7TP26 A7TP86
A7TQD5 A7TQE6 A7TQM5 A7TR50 A7TR80 A7TRN7 A7TRZ7 A7TS55 A7TSA7 A7TSS9 A7TSZ6
A7TT43 A7TZE6 A7TZF0 A7TZF3 A7TZG1 A7TZG3 A7VL23 A7VMU4 A7WNB3 A7WWQ7 A7WXI2
A7WXS6 A7WXS7 A7WZ76 A7WZ77 A7WZ78 A7WZ79 A7WZ80 A7WZ82 A7WZ83 A7WZC5 A7WZF5
A7WZQ9 A7X0F5 A7X0F7 A7X0F8 A7X0F9 A7X0G0 A7X0G1 A7X0G4 A7X0I5 A7X111 A7X127
A7X128 A7X154 A7X1C2 A7X1E3 A7X1S5 A7X1Z8 A7X204 A7X211 A7X2C3 A7X2C6 A7X2X2
A7X3E7 A7X3V5 A7X3Z2 A7X430 A7X441 A7X4K2 A7X4S6 A7X4U9 A7X4V0 A7X4V1 A7X534
A7X5U9 A7X5X6 A7X5Y6 A7X618 A7X6P1 A7X6P6 A7X6S1 A7X777 A7XDQ9 A7XUX6 A7XUY5
A7XUZ6 A7XV04 A7XV07 A7XV14 A7XY94 A7XZ53 A7Y2W8 A7Y2X0 A7Y395 A7Y3A3 A7Y3A6
A7Y3A9 A7Y3B0 A7Y3C0 A7Y3C1 A7Y3C4 A7Y3C5 A7Y3E3 A7Y3F2 A7Y3F4 A7Y3F5 A7Y3F7
A7Y3F8 A7Y3G4 A7Y3G5 A7Y3H4 A7Y3H6 A7Y3K2 A7Y3K7 A7Y3L0 A7Y3L2 A7Y3L4 A7Y3L7
A7YD35 A7YW81 A7YWC4 A7YWH9 A7YWM1 A7YWN2 A7YWP2 A7YY44 A7YY49 A7YY55 A7Z064
A7Z070 A7Z1S6 A7Z241 A7Z2K2 A7Z2P1 A7Z2Z0 A7Z380 A7Z3W3 A7Z4B0 A7Z4B1 A7Z4E2
A7Z4G5 A7Z4W7 A7Z558 A7Z575 A7Z6Z1 A7Z7K2 A7Z7N6 A7Z830 A7Z8R4 A7Z972 A7Z9K9
A7Z9Q4 A7Z9Q5 A7Z9Q6 A7Z9R3 A7ZA61 A7ZB16 A7ZBE9 A7ZBF3 A7ZBN0 A7ZBZ7 A7ZC33
A7ZC73 A7ZCB7 A7ZCL2 A7ZD62 A7ZDR5 A7ZER3 A7ZF22 A7ZFX1 A7ZHA6 A7ZHB6 A7ZHD1
A7ZHE0 A7ZHH8 A7ZHP7 A7ZHR7 A7ZI09 A7ZI10 A7ZII5 A7ZIX5 A7ZJ12 A7ZJ79 A7ZJ80
A7ZJ81 A7ZJR5 A7ZJW9 A7ZKF0 A7ZKF6 A7ZKH0 A7ZKV7 A7ZKW2 A7ZL20 A7ZL35 A7ZL36
A7ZLF1 A7ZLH2 A7ZLY4 A7ZLY5 A7ZM41 A7ZM51 A7ZM58 A7ZM59 A7ZM87 A7ZM90 A7ZM92
A7ZMC9 A7ZMH9 A7ZMQ9 A7ZMR9 A7ZMU2 A7ZMU3 A7ZMU8 A7ZMV2 A7ZN93 A7ZNF6 A7ZNP8
A7ZNP9 A7ZNQ0 A7ZNU4 A7ZNW3 A7ZP19 A7ZP72 A7ZP75 A7ZP76 A7ZP77 A7ZP90 A7ZP93
A7ZP96 A7ZPA2 A7ZPA9 A7ZPJ9 A7ZPK2 A7ZPL3 A7ZPL4 A7ZPR6 A7ZPV9 A7ZQ23 A7ZQJ2
A7ZQT4 A7ZQU1 A7ZQU2 A7ZRN9 A7ZRS2 A7ZRT8 A7ZRU0 A7ZS06 A7ZSA7 A7ZSB7 A7ZSD4
A7ZSD5 A7ZSH9 A7ZSM6 A7ZSN9 A7ZSV4 A7ZT30 A7ZT69 A7ZTN8 A7ZTR1 A7ZTR5 A7ZTS1
A7ZTS5 A7ZTU8 A7ZTU9 A7ZTV0 A7ZTV8 A7ZTZ7 A7ZU42 A7ZU91 A7ZUB9 A7ZUC3 A7ZUC8
A7ZUD2 A7ZUI6 A7ZUQ1 A7ZUQ8 A7ZUR2 A7ZUU1 A7ZV24 A7ZV25 A7ZVP9 A7ZVR8 A7ZVW6
A7ZVX5 A7ZVZ0 A7ZVZ9 A7ZW39 A7ZWA3 A7ZWC3 A7ZX84 A7ZXL8 A7ZXQ1 A7ZXV7 A7ZXV8
A7ZXV9 A7ZYF0 A7ZYK2 A7ZZ03 A7ZZ10 A7ZZ23 A7ZZC0 A7ZZC5 A7ZZI5 A7ZZJ0 A7ZZJ1
A7ZZR4 A7ZZT5 A8A067 A8A068 A8A081 A8A082 A8A0C2 A8A0D3 A8A0E0 A8A0E1 A8A0H0
A8A0H3 A8A0H5 A8A0L0 A8A0Q3 A8A0Y2 A8A0Z3 A8A117 A8A118 A8A123 A8A127 A8A1H1
A8A1K6 A8A1U7 A8A1U8 A8A1U9 A8A1X1 A8A1Z0 A8A203 A8A259 A8A2C1 A8A2C4 A8A2C5
A8A2C6 A8A2E5 A8A2E8 A8A2F1 A8A2F7 A8A2G3 A8A2P3 A8A2P6 A8A2Q8 A8A2Q9 A8A2W4
A8A322 A8A390 A8A3M7 A8A3W1 A8A3W9 A8A3X0 A8A4G1 A8A4J6 A8A4L2 A8A4L4 A8A4S9
A8A523 A8A534 A8A549 A8A550 A8A595 A8A5F8 A8A5H1 A8A5N2 A8A5V1 A8A5Y8 A8A6E4
A8A6G7 A8A6H2 A8A6H8 A8A6I1 A8A6J9 A8A6K0 A8A6K1 A8A6L0 A8A6P9 A8A6U2 A8A6Y7
A8A711 A8A714 A8A719 A8A723 A8A774 A8A7D0 A8A7D6 A8A7E0 A8A7G7 A8A7Q0 A8A7Q1
A8A888 A8A8A5 A8A9J7 A8AAZ9 A8ABD7 A8ABZ0 A8ACJ7 A8ACL7 A8ACM0 A8ACP2 A8ACP3
A8ACP4 A8ACQ3 A8ACU6 A8ACY4 A8AD70 A8ADB5 A8ADI0 A8ADI1 A8ADJ8 A8ADU3 A8ADV1
A8ADV7 A8ADW0 A8ADW3 A8AE22 A8AE89 A8AEE4 A8AEE5 A8AEE6 A8AEP3 A8AEQ5 A8AER0
A8AFM2 A8AFM8 A8AFN4 A8AFN6 A8AFR5 A8AG50 A8AG55 A8AG56 A8AGB5 A8AGD8 A8AGM8
A8AGR7 A8AGR9 A8AGU5 A8AGW0 A8AGX4 A8AGX5 A8AH08 A8AH11 A8AH13 A8AH15 A8AH48
A8AHA4 A8AHH7 A8AHJ0 A8AI11 A8AI28 A8AI31 A8AII1 A8AIS0 A8AJB3 A8AJB7 A8AJH7

A8AJL6 A8AK26 A8AL09 A8AL14 A8AL22 A8AL54 A8ALB1 A8ALD3 A8ALK9 A8ALQ4 A8ALR3
A8ALT5 A8ALU0 A8ALY5 A8AM03 A8AM07 A8AMP4 A8AMP5 A8AMR4 A8AN31 A8AN88 A8AN92
A8ANM6 A8ANW2 A8AP43 A8AP55 A8AP56 A8AP76 A8APU0 A8APU2 A8APY4 A8AQ08 A8AQ99
A8AQB4 A8AQD4 A8AQD5 A8AQD6 A8AQF0 A8AQI5 A8AQP0 A8AQR8 A8AQX4 A8AR73 A8ARA1
A8AVH5 A8AVK0 A8AVT2 A8AVU9 A8AW16 A8AW76 A8AWV0 A8AY51 A8AYG5 A8AYG6 A8AYY7
A8AZK5 A8C7R4 A8C8J4 A8C8J5 A8C8J7 A8C8W3 A8C8W4 A8C8W6 A8CDR5 A8CG34 A8DMN5
A8DQI7 A8DQI8 A8DQI9 A8DQK7 A8DQK8 A8DQK9 A8DYE2 A8DYQ1 A8DZE7 A8DZH4 A8E0Y8
A8E5V9 A8E639 A8E7D3 A8E7G5 A8ERK3 A8ERK6 A8ERN6 A8ET78 A8EUA0 A8EUK5 A8EV74
A8EV91 A8EVL5 A8EVM2 A8EW04 A8EWC4 A8EWC8 A8EX89 A8EX90 A8EXB4 A8EXB6 A8EXI0
A8EXI2 A8EY08 A8EY83 A8EY96 A8EZ20 A8EZ38 A8EZ97 A8EZU9 A8EZZ9 A8F0C9 A8F0F4
A8F0L9 A8F0M1 A8F119 A8F1B6 A8F1G3 A8F278 A8F2N4 A8F302 A8F3J8 A8F7F7 A8F7S3
A8F8V8 A8FAL9 A8FB95 A8FBJ4 A8FBS5 A8FCF7 A8FCU5 A8FCU6 A8FCX8 A8FD10 A8FDG4
A8FDQ3 A8FFC3 A8FG51 A8FG97 A8FHE3 A8FI71 A8FIB6 A8FIB7 A8FIB8 A8FIC5 A8FJA6
A8FJX5 A8FK06 A8FKE6 A8FKE9 A8FKJ4 A8FKM0 A8FKU3 A8FLY5 A8FMF0 A8FMN2 A8FMQ9
A8FMR4 A8FNN0 A8FNW6 A8FNW7 A8FP42 A8FPN0 A8FPZ2 A8FQ97 A8FQ99 A8FQF8 A8FRQ8
A8FRQ9 A8FRR0 A8FRR3 A8FS65 A8FS68 A8FSC4 A8FSI3 A8FTV8 A8FU25 A8FUX7 A8FUY0
A8FV66 A8FW97 A8FX04 A8FYV9 A8G002 A8G0T0 A8G0T1 A8G1C2 A8G1W9 A8G1X0 A8G1X1
A8G262 A8G2F5 A8G2G6 A8G2R0 A8G2R2 A8G2R3 A8G2T2 A8G2V5 A8G2V8 A8G2V9 A8G2W0
A8G2W1 A8G2X5 A8G2X8 A8G2Y6 A8G2Y7 A8G311 A8G3E9 A8G3G2 A8G3H5 A8G3H6 A8G3H7
A8G4E3 A8G4L8 A8G5C7 A8G5I7 A8G5N5 A8G5U6 A8G6D8 A8G6V3 A8G6V4 A8G6V5 A8G6V6
A8G6W0 A8G719 A8G7L4 A8G7M2 A8G7M3 A8G7M4 A8G7P8 A8G7S0 A8G823 A8G841 A8G8C0
A8G8T4 A8G8T5 A8G9L0 A8G9L5 A8G9S4 A8G9Z1 A8GAC2 A8GAC3 A8GAC4 A8GAQ0 A8GB06
A8GB60 A8GB61 A8GB62 A8GC34 A8GCA3 A8GCZ5 A8GDN1 A8GDR2 A8GDR6 A8GDR9 A8GDS0
A8GDS1 A8GDZ4 A8GE02 A8GE50 A8GEE7 A8GF29 A8GF86 A8GF88 A8GFA0 A8GFG0 A8GFH6
A8GFH7 A8GFK7 A8GFM7 A8GFM8 A8GH04 A8GH07 A8GH10 A8GH16 A8GH23 A8GH95 A8GHF3
A8GHL1 A8GHQ9 A8GHR0 A8GHR1 A8GHW6 A8GID8 A8GIH8 A8GII5 A8GIT2 A8GJU8 A8GJV0
A8GJX3 A8GJY0 A8GK45 A8GK46 A8GK69 A8GKB7 A8GKC3 A8GKC9 A8GKG9 A8GKP6 A8GKU2
A8GLB1 A8GLI0 A8GLV8 A8GLV9 A8GLY7 A8GLY9 A8GM76 A8GM78 A8GMQ9 A8GN36 A8GNA3
A8GNC3 A8GP56 A8GP69 A8GPQ1 A8GPY7 A8GQA2 A8GQF6 A8GQK5 A8GQK8 A8GQT5 A8GQT7
A8GRD0 A8GRQ1 A8GRX4 A8GSW3 A8GSX5 A8GTK4 A8GTS1 A8GU51 A8GUC7 A8GUC9 A8GUJ2
A8GUJ3 A8GVJ0 A8GVM4 A8GVN8 A8GWM8 A8GXM1 A8GY33 A8GYC5 A8GYD9 A8GYE1 A8GZ02
A8GZR0 A8H157 A8H1B5 A8H1H4 A8H1S6 A8H2K2 A8H328 A8H375 A8H3J1 A8H4R3 A8H536
A8H539 A8H541 A8H5C2 A8H779 A8H806 A8H968 A8H969 A8H970 A8H987 A8H9I2 A8H9S9
A8HAG7 A8HAG8 A8HAG9 A8HAI0 A8HPM2 A8HPM5 A8HR72 A8HSH8 A8HT64 A8HT70 A8HT73
A8HU57 A8HY21 A8HYJ1 A8HZ17 A8HZ57 A8I126 A8I1B9 A8I246 A8I413 A8I421 A8I499
A8I4I6 A8I6P9 A8I7Y5 A8ID91 A8IDL3 A8IDZ2 A8IHV3 A8IJF8 A8ILJ2 A8IW99 A8J0J0
A8J4S9 A8J6J0 A8J785 A8JFU2 A8K4G0 A8KB87 A8KBG2 A8KBL5 A8KYS6 A8L3V9 A8L3W1
A8LC88 A8LC91 A8LHT6 A8LIU9 A8LK55 A8LKH7 A8LKH8 A8LKI0 A8LL79 A8LLI0 A8LMA0
A8LMF5 A8LN45 A8LQR9 A8LS59 A8LSB0 A8LVS8 A8LW09 A8LX83 A8M1G0 A8M2B2 A8M2J7
A8M393 A8M3D7 A8M609 A8M612 A8M7V7 A8MC59 A8MEV0 A8MFC3 A8MFV5 A8MG60 A8MGV3
A8MGZ0 A8MH33 A8MIW2 A8MJW3 A8MJW4 A8MLK3 A8MPR5 A8MPY1 A8MQA4 A8MQR0 A8MQY1
A8MR93 A8MRC7 A8MRE9 A8MRT5 A8MRV9 A8MRY9 A8MS09 A8MTT3 A8MUP6 A8MV81 A8MVS5
A8MVW0 A8MVW5 A8MVZ5 A8MWK0 A8MWL6 A8MWL7 A8MWV9 A8MWY0 A8MXE2 A8MXK1 A8MXV6
A8MYB1 A8MYU2 A8MZ97 A8N1W8 A8N1Z1 A8N513 A8N5E5 A8N5E6 A8N6B4 A8N7T9 A8N9T6

A8NCK4 A8NCK6 A8NWS6 A8NX72 A8NYP0 A8NYS9 A8P006 A8PE82 A8PRN6 A8PRS6 A8PTK5
A8PTV6 A8PYF7 A8Q1U2 A8QAN4 A8QD14 A8R7E6 A8SE40 A8SE55 A8SE63 A8SE66 A8SE70
A8SE90 A8SE94 A8SE98 A8SE99 A8SEA8 A8SEB4 A8SEB5 A8SEB6 A8SEB7 A8SEC2 A8SEC3
A8SEC9 A8SEF0 A8SEF3 A8SEF4 A8SEF7 A8SEF8 A8SEG5 A8W3A5 A8W3A8 A8W3B0 A8W3B1
A8W3B2 A8W3B7 A8W3B8 A8W3B9 A8W3C0 A8W3C9 A8W3D0 A8W3D1 A8W3D2 A8W3D3 A8W3E0
A8W3E1 A8W3E7 A8W3E9 A8W3G6 A8W3G9 A8W3H2 A8W3H4 A8W3H6 A8W3H7 A8W3I0 A8W3I1
A8W3I2 A8W3I3 A8W3J4 A8W3J6 A8W3J7 A8W3J8 A8W3K2 A8W3K3 A8W3L1 A8W3M6 A8W649
A8WCG0 A8WFR0 A8WFS8 A8WG88 A8WGA3 A8WGF7 A8WGS4 A8WGT1 A8WGU8 A8WH34 A8WH72
A8WH74 A8WH75 A8WHP3 A8WHP9 A8WHR0 A8WLF6 A8WMY3 A8WN56 A8WPG9 A8WQK3 A8WQT4
A8WT26 A8WUE9 A8WUP2 A8WUS5 A8WVD0 A8WVX4 A8WWR3 A8WXS4 A8WZ09 A8WZU4 A8X0L4
A8X181 A8X2R2 A8X3A7 A8X481 A8X482 A8X4W9 A8X765 A8X8I4 A8X8R3 A8XA40 A8XC92
A8XDX2 A8XF54 A8XI14 A8XK26 A8XKF2 A8XLW0 A8XMW6 A8XNX8 A8XP14 A8XP97 A8XPY4
A8XQC7 A8XST1 A8XWD1 A8Y1P7 A8Y236 A8Y2C9 A8Y2U2 A8Y3H3 A8Y3M2 A8Y3V5 A8Y432
A8Y4B2 A8Y999 A8Y9A0 A8Y9A1 A8Y9A5 A8Y9A6 A8Y9B2 A8Y9B4 A8Y9D4 A8Y9D6 A8Y9D7
A8Y9D9 A8Y9E0 A8Y9F0 A8Y9F4 A8Y9F5 A8Y9F6 A8Y9F8 A8Y9F9 A8Y9G4 A8Y9G5 A8Y9G6
A8Y9H5 A8Y9I0 A8Y9I1 A8YTR2 A8YUB8 A8YUJ6 A8YUJ7 A8YUN7 A8YUY6 A8YUY9 A8YVS5
A8YWL7 A8YXN4 A8YXZ8 A8YY16 A8YY46 A8YY74 A8YY75 A8YY76 A8YYD4 A8YYU2 A8YZ02
A8YZV4 A8Z037 A8Z053 A8Z054 A8Z055 A8Z056 A8Z057 A8Z058 A8Z059 A8Z075 A8Z0F5
A8Z0M5 A8Z0M6 A8Z144 A8Z145 A8Z146 A8Z147 A8Z148 A8Z149 A8Z150 A8Z182 A8Z1Q0
A8Z1Q1 A8Z1V7 A8Z223 A8Z2N4 A8Z2R6 A8Z2S6 A8Z3D6 A8Z3D9 A8Z3F8 A8Z3M3 A8Z3N3
A8Z414 A8Z415 A8Z4B2 A8Z4U0 A8Z4X8 A8Z4X9 A8Z4Y0 A8Z529 A8Z547 A8Z553 A8Z577
A8Z5I6 A8Z5R3 A8Z6E7 A8Z6N6 A8ZNS1 A8ZNS3 A8ZNZ4 A8ZRR1 A8ZU97 A8ZUM7 A8ZUM8
A8ZVQ5 A8ZVV7 A8ZXM2 A8ZXW6 A8ZZT5 A9A506 A9A677 A9A8W5 A9A8Y7 A9A9I2 A9AAG5
A9AC67 A9ACA0 A9AE12 A9AEI1 A9AFZ4 A9AFZ7 A9AGE2 A9AGJ3 A9AGN5 A9AHH5 A9AI31
A9AIS5 A9AJ19 A9AJG0 A9AJK8 A9AJP4 A9ATU4 A9AVV0 A9AXU9 A9AXV2 A9AYY2 A9B477
A9B488 A9B6A7 A9B6Y0 A9B6Y3 A9B7R3 A9B815 A9B9C4 A9BAB0 A9BAR6 A9BB51 A9BBE7
A9BBV0 A9BCE1 A9BCE2 A9BCE3 A9BCE4 A9BCE8 A9BCK8 A9BCL5 A9BD08 A9BDM2 A9BDM3
A9BDM4 A9BDU1 A9BDU4 A9BDU5 A9BDU6 A9BDU7 A9BDX0 A9BDY2 A9BDY3 A9BE82 A9BE84
A9BE89 A9BEK7 A9BEV4 A9BFL9 A9BFX7 A9BFX8 A9BG78 A9BG79 A9BGS8 A9BHD3 A9BJK3
A9BMQ0 A9BNB4 A9BP42 A9BPU2 A9BPU3 A9BUK3 A9BX57 A9BXA7 A9BZG1 A9BZN0 A9C190
A9C3P0 A9CAZ8 A9CB25 A9CBA0 A9CIB0 A9CIM3 A9CJA4 A9CJX3 A9CK01 A9CK03 A9CMA6
A9EX55 A9EX69 A9EXK6 A9EYV6 A9F575 A9F773 A9F781 A9FDV9 A9FGS5 A9FGS9 A9FI43
A9FXA7 A9GAW6 A9GC83 A9GEU3 A9GHS2 A9GIP6 A9GM39 A9GMF2 A9GMG8 A9GQ89 A9GRC9
A9GV45 A9H0I1 A9H2L4 A9H4G8 A9H8F8 A9HBY2 A9HDM4 A9HDM8 A9HDN1 A9HEV3 A9HI81
A9HKH7 A9HN78 A9HNA0 A9HRS3 A9HVV4 A9HW97 A9HWB1 A9HWL4 A9HXK3 A9HY34 A9HY36
A9I4U2 A9I7J3 A9II17 A9II25 A9IIP9 A9IJB7 A9IJC7 A9IL10 A9IMX7 A9IP09 A9IQ78
A9IQG4 A9IQH9 A9IQI5 A9IQI8 A9IQQ6 A9IR17 A9IRA0 A9IS13 A9IT56 A9ITD6 A9IU40
A9IU75 A9IUM6 A9IUN6 A9IVC8 A9IWA8 A9IZD8 A9JLI2 A9JM73 A9JPE0 A9JPE2 A9JPE3
A9JPE4 A9JRA0 A9JSM3 A9JSP6 A9JTG4 A9JTJ0 A9KBG1 A9KBG3 A9KBK4 A9KBL2 A9KBL5
A9KBT1 A9KCC9 A9KEI7 A9KES2 A9KF31 A9KF59 A9KFT7 A9KFZ1 A9KG14 A9KG69 A9KGW7
A9KIG5 A9KK96 A9KK97 A9KMP0 A9KMU8 A9KNK1 A9KP98 A9KRD5 A9KUH2 A9KUX8 A9KV13
A9KWC9 A9KX10 A9KX11 A9KX12 A9KX20 A9KXA4 A9KY26 A9KY79 A9KYL4 A9KYL5 A9KYL6
A9KZ13 A9L0J6 A9L0J8 A9L0K1 A9L0U1 A9L270 A9L2N8 A9L382 A9L3D9 A9L4L5 A9L4Q8
A9L4U8 A9L578 A9L5I4 A9L5I7 A9L5M4 A9L5Z2 A9L8T6 A9L976 A9L980 A9L982 A9L983

A9L984 A9L989 A9L990 A9L991 A9L992 A9L9A1 A9L9A7 A9L9A9 A9L9B1 A9L9B5 A9L9B6
A9L9C2 A9L9C4 A9L9E2 A9L9E4 A9L9E6 A9L9E7 A9L9E9 A9L9F0 A9LIW2 A9LYA6 A9LYB2
A9LYC6 A9LYE7 A9LYE9 A9LYF0 A9LYF2 A9LYF3 A9LYF8 A9LYG9 A9LYI1 A9LYY1 A9LYY6
A9LYZ3 A9LZL5 A9M119 A9M152 A9M1J8 A9M1Y8 A9M2A6 A9M2A7 A9M2A8 A9M2E2 A9M2H7
A9M2I7 A9M2N3 A9M2Z3 A9M390 A9M3D4 A9M3Q1 A9M4B4 A9M640 A9M693 A9M6J4 A9M794
A9M851 A9M8A1 A9M8F8 A9M8G0 A9M8G1 A9M8H1 A9M8S1 A9M8Z3 A9M8Z5 A9M9N4 A9MA14
A9MA15 A9MAG4 A9MAI6 A9MAI9 A9MAN9 A9MAQ5 A9MB47 A9MBP1 A9MC22 A9MCR7 A9MCW8
A9MEG7 A9MEJ6 A9MF27 A9MFB6 A9MFI4 A9MFR0 A9MG10 A9MGK8 A9MGZ7 A9MH10 A9MH90
A9MHA2 A9MHL4 A9MHQ7 A9MHY5 A9MI48 A9MI51 A9MIA4 A9MIF9 A9MIN5 A9MJ16 A9MJ53
A9MJ97 A9MJA3 A9MJA6 A9MJA9 A9MJC2 A9MJC5 A9MJQ3 A9MJR3 A9MJR4 A9MJR5 A9MJT5
A9MJT7 A9MJX2 A9MK88 A9MKE5 A9MKE7 A9MKS7 A9MKW9 A9MKX0 A9MLJ4 A9MLR0 A9MLS2
A9MLS7 A9MM32 A9MM62 A9MMA7 A9MMF2 A9MN27 A9MN76 A9MNI0 A9MNI4 A9MNJ0 A9MNV6
A9MNW8 A9MNW9 A9MNX0 A9MNY7 A9MNZ9 A9MP57 A9MPD5 A9MPE0 A9MPE1 A9MPI4 A9MPK6
A9MPR1 A9MPT8 A9MPV6 A9MPV8 A9MPX1 A9MQ55 A9MQC5 A9MQG6 A9MQH4 A9MQT3 A9MR45
A9MR50 A9MRK2 A9MRL9 A9MRN4 A9MRQ1 A9MRQ2 A9MRT8 A9MRT9 A9MRW5 A9MRW9 A9MRX4
A9MRX7 A9MS81 A9MSN4 A9MT17 A9MT38 A9MT86 A9MT98 A9MUD9 A9MUE0 A9MUE1 A9MUP5
A9MV69 A9MV77 A9MVT2 A9MVT3 A9MVW2 A9MVW9 A9MWE8 A9MWJ2 A9MWP9 A9MWQ4 A9MWQ5
A9MWW8 A9MWZ0 A9MX83 A9MX86 A9MXB0 A9MXB1 A9MXB2 A9MXC4 A9MXG6 A9MXN2 A9MY99
A9MYF8 A9MYH6 A9MYK0 A9MYL9 A9MYZ7 A9MYZ8 A9MZA1 A9MZD0 A9MZE6 A9MZF0 A9MZG2
A9MZG3 A9MZG7 A9MZL6 A9MZW8 A9MZZ2 A9MZZ3 A9N023 A9N026 A9N028 A9N0Q1 A9N0S6
A9N0Y8 A9N1K2 A9N1K8 A9N1L3 A9N1R2 A9N200 A9N235 A9N2A7 A9N2D4 A9N2L8 A9N2Z8
A9N307 A9N361 A9N3H7 A9N3H8 A9N3J5 A9N3Q1 A9N409 A9N410 A9N4B4 A9N4C4 A9N4W6
A9N586 A9N589 A9N592 A9N5A8 A9N5A9 A9N5B0 A9N5B3 A9N5P5 A9N5Q9 A9N5R5 A9N5W7
A9N5Y5 A9N608 A9N628 A9N6L5 A9N776 A9N7A9 A9N7B4 A9N7D2 A9N7H5 A9N7L0 A9N7L1
A9N7L2 A9N7X0 A9N832 A9N862 A9N863 A9N864 A9N877 A9N8B5 A9N8W2 A9N8W5 A9N8X3
A9N982 A9N9B0 A9N9X0 A9NA35 A9NBA5 A9NBB4 A9NBC4 A9NBC6 A9NBM6 A9NCT0 A9ND08
A9NDU9 A9NE17 A9NF64 A9NGL9 A9NGW6 A9NGW7 A9NMM6 A9NN43 A9NQL1 A9NTU2 A9NYX5
A9P0A6 A9P1V1 A9Q1L1 A9QC36 A9QC58 A9QC74 A9QC80 A9QC93 A9QC94 A9QCA0 A9QNE7
A9QPJ1 A9QYG6 A9QYL1 A9QYP6 A9QYR5 A9QYW1 A9QYW2 A9QYZ2 A9QZ74 A9QZC0 A9QZH1
A9QZJ2 A9QZM5 A9QZU7 A9QZU8 A9R007 A9R013 A9R053 A9R075 A9R076 A9R089 A9R090
A9R091 A9R094 A9R099 A9R0E2 A9R120 A9R127 A9R1E4 A9R1R0 A9R1V8 A9R1V9 A9R1W0
A9R1Y3 A9R2H3 A9R2H5 A9R2R4 A9R2S0 A9R2S1 A9R2Y3 A9R388 A9R3X3 A9R3X4 A9R429
A9R473 A9R4E0 A9R4H0 A9R4J5 A9R4U7 A9R535 A9R563 A9R5H1 A9R5U3 A9R5U4 A9R5U5
A9R5Y6 A9R5Y7 A9R654 A9R6A2 A9R6K9 A9R6L2 A9R6L5 A9R6M0 A9R6M8 A9R7E0 A9R7X6
A9R803 A9R8J0 A9R8K8 A9R8U3 A9R8U8 A9R923 A9R9A1 A9R9A2 A9R9D6 A9RA88 A9RAG1
A9RAG2 A9RAG3 A9RAG4 A9RAG5 A9RAG6 A9RAG7 A9RAG9 A9RAH0 A9RAH1 A9RAH2 A9RAH3
A9RAH4 A9RAH5 A9RAH6 A9RAH7 A9RAH8 A9RGD8 A9RJH1 A9RKK4 A9RLK6 A9RZ57 A9S1T8
A9S848 A9SB31 A9SEY7 A9SG36 A9SHQ9 A9SIZ6 A9STS7 A9STU1 A9SU70 A9SV59 A9SV60
A9SY64 A9SY65 A9T836 A9TKY8 A9TXT1 A9TZ63 A9UKA2 A9UL59 A9ULC7 A9ULE9 A9ULX8
A9UUB8 A9UWF0 A9UY97 A9VFM0 A9VFM1 A9VFM2 A9VGI0 A9VHJ1 A9VJ15 A9VJQ7 A9VKG1
A9VKI1 A9VM05 A9VPW5 A9VQ76 A9VRA8 A9VS22 A9VS91 A9VS94 A9VS98 A9VSA7 A9VSA8
A9VSC2 A9VST4 A9VTD3 A9VTH6 A9VTH7 A9VTX6 A9VU75 A9VUA6 A9VUA7 A9VYW7 A9VYW8
A9VZA2 A9VZA4 A9W2K2 A9W350 A9W565 A9W590 A9W5W6 A9W739 A9W7L6 A9WAQ6 A9WEE0
A9WFC1 A9WG67 A9WG76 A9WGD1 A9WGS8 A9WGS9 A9WHY7 A9WJI9 A9WJM3 A9WK61 A9WMF7

A9WN51 A9WNC4 A9WPV3 A9WRE1 A9WSJ0 A9WT38 A9WU27 A9WW03 A9WW52 A9WWT8 A9WYS0
A9WZ33 A9X1C8 A9X4U2 A9XVA1 A9XVQ4 A9Y006 A9YWR6 A9YWS7 A9Z1Z3 A9ZM27 A9ZPI1
A9ZSY2 A9ZSY3 B0B7J4 B0B7J6 B0B7J7 B0B7U1 B0B7U3 B0B7X8 B0B8F4 B0B8Y3 B0B970
B0B999 B0B9A0 B0B9M3 B0B9M4 B0B9M5 B0B9V8 B0BAL2 B0BAX8 B0BAX9 B0BBQ9 B0BBR1
B0BBR2 B0BC06 B0BC08 B0BC43 B0BLS0 B0BLW3 B0BM39 B0BMG8 B0BMW8 B0BMY1 B0BN44
B0BN70 B0BN86 B0BNF0 B0BNG2 B0BNK7 B0BNL4 B0BP47 B0BP81 B0BPA2 B0BPQ8 B0BPR4
B0BPX8 B0BQN7 B0BQT0 B0BQV7 B0BR23 B0BRC0 B0BRH4 B0BRI0 B0BRR8 B0BRX6 B0BRX7
B0BRX8 B0BS59 B0BS70 B0BS74 B0BT71 B0BTB8 B0BTH4 B0BTX5 B0BU10 B0BU11 B0BV41
B0BVB1 B0BVP1 B0BVU2 B0BVZ1 B0BVZ4 B0BWT1 B0BX58 B0BXD2 B0BYD8 B0BYF0 B0BYP3
B0BYQ1 B0BYW9 B0BZ70 B0BZB1 B0BZI7 B0BZK7 B0BZK9 B0BZL0 B0BZW2 B0C0A5 B0C0E7
B0C0E8 B0C0S7 B0C1T8 B0C1V6 B0C3E3 B0C3P0 B0C474 B0C4B4 B0C5Z6 B0C6C4 B0C6G0
B0C6T1 B0C6T2 B0C6T5 B0C6Z7 B0C7S6 B0C9E9 B0CB14 B0CBA0 B0CCQ8 B0CDA5 B0CDZ1
B0CEA6 B0CEA7 B0CF38 B0CFM0 B0CFX9 B0CG09 B0CGI5 B0CHH0 B0CHM3 B0CHW7 B0CIQ7
B0CIT9 B0CJU8 B0CK70 B0CK72 B0CK73 B0CK83 B0CKF4 B0CKF6 B0CKT7 B0CKW5 B0CLB4
B0CLD6 B0CLD9 B0CLJ1 B0CLK7 B0CLX4 B0CM95 B0CMA4 B0CQL3 B0CQN7 B0CQN9 B0CXH5
B0D0N9 B0D0P0 B0D1L7 B0D4E6 B0D4J7 B0D4L0 B0D7T0 B0D8R3 B0DC53 B0DI84 B0DK33
B0DK57 B0E2U2 B0E843 B0EKR0 B0F2B4 B0F9L7 B0F9W3 B0FP48 B0FWC6 B0FWC7 B0FWC9
B0FWD1 B0FWD2 B0FWD3 B0FWD5 B0FWD6 B0FWD7 B0FWD8 B0FWK4 B0FYY4 B0G0Z9 B0G134
B0G143 B0G159 B0G162 B0G168 B0G180 B0JFF7 B0JFT3 B0JGA8 B0JGF6 B0JHI7 B0JHK5
B0JHQ5 B0JI44 B0JI68 B0JID4 B0JIS8 B0JJ94 B0JJA1 B0JKN9 B0JLL9 B0JLU8 B0JLU9
B0JLV0 B0JLV1 B0JLZ5 B0JM63 B0JM93 B0JM94 B0JNP7 B0JNP9 B0JPG4 B0JPN3 B0JR89
B0JS85 B0JSV4 B0JSW8 B0JT88 B0JTA8 B0JTK0 B0JUK8 B0JUV4 B0JWU6 B0JWU7 B0JWU8
B0JWU9 B0JWY5 B0JX58 B0JX68 B0JXB7 B0JXB8 B0JYB0 B0JYX6 B0JYZ2 B0JZD0 B0JZE1
B0JZG0 B0K004 B0K020 B0K025 B0K0H2 B0K165 B0K1B6 B0K2K2 B0K3E3 B0K3H3 B0K3X0
B0K4Z5 B0K5A3 B0K657 B0K897 B0K8K4 B0K8N4 B0K909 B0K9H8 B0K9M9 B0KB34 B0KCS0
B0KFS9 B0KFZ0 B0KG16 B0KHQ5 B0KHT9 B0KJ83 B0KLJ7 B0KLP3 B0KLW9 B0KM38 B0KM50
B0KME8 B0KMR7 B0KMX8 B0KMY4 B0KMY7 B0KN20 B0KN61 B0KNU2 B0KP20 B0KPE4 B0KPX4
B0KQ27 B0KQB7 B0KR91 B0KRB2 B0KRB3 B0KRB4 B0KRC1 B0KRX4 B0KT69 B0KTK9 B0KTL9
B0KUA6 B0KUP8 B0KUY1 B0KW95 B0KWE9 B0L3A2 B0LJC9 B0LPN4 B0R2U4 B0R367 B0R382
B0R3S1 B0R3W4 B0R461 B0R470 B0R474 B0R5G3 B0R5N7 B0R5T0 B0R610 B0R611 B0R632
B0R633 B0R6A6 B0R6A7 B0R6B1 B0R6I4 B0R737 B0R753 B0R8E4 B0R9L8 B0R9L9 B0R9M0
B0R9M1 B0R9X2 B0RCI3 B0RE27 B0RED8 B0RED9 B0REI2 B0RF30 B0RIJ0 B0RLE5 B0RLE6
B0RLE7 B0RLE8 B0RLZ0 B0RMM6 B0RMR4 B0RMT0 B0RNI5 B0RP10 B0RPI5 B0RQ88 B0RQA5
B0RRA5 B0RRA8 B0RS04 B0RU04 B0RUB3 B0RUI8 B0RUW5 B0RVA7 B0RVC5 B0RWC6 B0RXJ1
B0RYW0 B0RYY1 B0S1D0 B0S1F0 B0S1N1 B0S1S0 B0S4Q1 B0S5A7 B0S5D5 B0S5N4 B0S5Y3
B0S6R1 B0S6T2 B0S728 B0S977 B0S986 B0S9E8 B0SDA1 B0SE58 B0SFE6 B0SFU0 B0SFU2
B0SFU5 B0SLC4 B0SMP1 B0SNZ5 B0SP88 B0SP90 B0SR19 B0SRR1 B0SRS0 B0SS37 B0SVC7
B0SYJ5 B0SYJ7 B0SZ35 B0T010 B0T0S6 B0T0X6 B0T1Q6 B0T1Q7 B0T274 B0T2G5 B0T658
B0T669 B0T834 B0T872 B0T8A6 B0TA81 B0TBL5 B0TC89 B0TGB7 B0TH77 B0TH87 B0TI54
B0TI68 B0TII8 B0TIN9 B0TIP2 B0TJ18 B0TJ19 B0TJ20 B0TJB0 B0TJN4 B0TK05 B0TKB3
B0TKX6 B0TLA3 B0TNC7 B0TNS4 B0TQF8 B0TQF9 B0TQG0 B0TQH1 B0TQN4 B0TR00 B0TRE7
B0TRI8 B0TSP8 B0TUP5 B0TW03 B0TW73 B0TWK0 B0TWQ1 B0TWQ4 B0TWS3 B0TYI2 B0TYY2
B0TZ73 B0U042 B0U1E0 B0U1X2 B0U1Z0 B0U2A2 B0U2R2 B0U2V1 B0U3E5 B0U3J1 B0U3Q8
B0U458 B0U4Z9 B0U5A2 B0U5X0 B0U659 B0U6I1 B0U6S1 B0UDF9 B0UFH5 B0UFS2 B0UHI1

B0UHJ2 B0UI86 B0ULL4 B0ULY1 B0ULY3 B0ULY4 B0UMU6 B0UQ37 B0UR12 B0URU3 B0URV1
B0URX9 B0US64 B0US80 B0USG1 B0USN1 B0UT28 B0UT97 B0UTI5 B0UTW8 B0UUC9 B0UUJ2
B0UUL1 B0UUS0 B0UUY3 B0UV09 B0UVD4 B0UWF8 B0UWG9 B0UWH0 B0UWH1 B0UWZ1 B0UXR0
B0UYF2 B0UYT5 B0V1P1 B0V2N1 B0V4R4 B0V4S5 B0V4Z4 B0V7A3 B0V7G8 B0V7U8 B0V892
B0V8P2 B0V982 B0V9M4 B0VA17 B0VBM6 B0VBP7 B0VBP8 B0VCY6 B0VDQ0 B0VEC3 B0VEM6
B0VL58 B0VL66 B0VLD4 B0VM11 B0VM56 B0VMM0 B0VN34 B0VNJ8 B0VNJ9 B0VNK0 B0VPG4
B0VPY4 B0VR07 B0VRV0 B0VS31 B0VSQ2 B0VT08 B0VU47 B0VU56 B0VX73 B0W429 B0WC46
B0WH58 B0WN94 B0X4H5 B0XMZ5 B0XPG0 B0XPV4 B0XQ16 B0XQS8 B0XQZ0 B0XSI3 B0XUW3
B0XWT3 B0XXU1 B0XYF0 B0XYK8 B0Y240 B0Y384 B0Y3Z4 B0Y429 B0Y4J4 B0Y4N5 B0Y4Q5
B0Y4Q8 B0Y566 B0Y5H9 B0Y5Z6 B0Y606 B0Y6I4 B0Y6V2 B0Y7W2 B0YA61 B0YA89 B0YBR5
B0YCP1 B0YD09 B0YD91 B0YJ81 B0YPL7 B0YPM3 B0YPM4 B0YPM6 B0YPM8 B0YPN1 B0YPP2
B0YPP4 B0YPP7 B0YPP8 B0YPQ4 B0Z4K6 B0Z4L2 B0Z4M1 B0Z4M2 B0Z4M4 B0Z4M9 B0Z4N0
B0Z4N1 B0Z4N3 B0Z4N8 B0Z4N9 B0Z4P1 B0Z4P5 B0Z4P6 B0Z4S2 B0Z4S4 B0Z4S5 B0Z4S7
B0Z4S8 B0Z4U0 B0Z4U6 B0Z4V5 B0Z4V6 B0Z4V8 B0Z4W3 B0Z4W4 B0Z4W5 B0Z4W7 B0Z4X2
B0Z4X3 B0Z4X5 B0Z4X9 B0Z4Y0 B0Z4Y6 B0Z4Y8 B0Z506 B0Z508 B0Z509 B0Z511 B0Z512
B0Z524 B0Z530 B0Z539 B0Z540 B0Z542 B0Z547 B0Z548 B0Z549 B0Z551 B0Z556 B0Z557
B0Z559 B0Z563 B0Z564 B0Z570 B0Z572 B0Z590 B0Z592 B0Z593 B0Z595 B0Z596 B0Z5A8
B0Z5B4 B0Z5C3 B0Z5C4 B0Z5C6 B0Z5D1 B0Z5D2 B0Z5D3 B0Z5D5 B0Z5E0 B0Z5E1 B0Z5E3
B0Z5E7 B0Z5E8 B0Z5F4 B0Z5F6 B0Z5H4 B0Z5H6 B0Z5H7 B0Z5H9 B0Z5I0 B1A0U4 B1A0U5
B1A0U6 B1A0U7 B1A915 B1A919 B1A921 B1A922 B1A923 B1A928 B1A930 B1A931 B1A940
B1A947 B1A948 B1A950 B1A954 B1A955 B1A961 B1A981 B1A983 B1A984 B1A986 B1A987
B1AI60 B1AI70 B1AI94 B1AIC4 B1AIC5 B1AIQ4 B1AJ44 B1AL88 B1AQL3 B1AS29 B1AT66
B1ATG9 B1AUH1 B1AWJ5 B1AWN4 B1AWN6 B1AXP6 B1AXV0 B1AZA5 B1B1W5 B1B212 B1B3P3
B1B5P4 B1GT62 B1GVX2 B1GVX3 B1GYJ5 B1GZ50 B1GZE4 B1GZI2 B1GZK7 B1GZX3 B1H0B9
B1H0M3 B1H0M4 B1H134 B1H1G2 B1H1H3 B1H1P9 B1H234 B1H299 B1H2N3 B1H3B1 B1H3C9
B1H3G4 B1H3H9 B1HM50 B1HM52 B1HPV0 B1HPV1 B1HR37 B1HRA8 B1HST2 B1HUT5 B1HVI2
B1HVP3 B1HYA2 B1HYT6 B1I2Y8 B1I3H5 B1I461 B1I4C7 B1I4G8 B1I6G5 B1I6I5 B1I6K8
B1I6L8 B1I7K2 B1I892 B1I956 B1I9P6 B1IA22 B1IAY5 B1IB20 B1IC73 B1ICL2 B1ICT3
B1ICT4 B1ICT5 B1IDK9 B1IDY6 B1IE29 B1IE30 B1IGK6 B1II34 B1INB4 B1INP5 B1IP42
B1IPA0 B1IPB4 B1IPE3 B1IPL6 B1IQ09 B1IQ91 B1IQC2 B1IQC4 B1IQC7 B1IQG4 B1IQI5
B1IQN7 B1IQN8 B1IQN9 B1IQQ5 B1IQR5 B1IQZ0 B1IQZ1 B1IQZ8 B1IR08 B1IR91 B1IRC9
B1IRD6 B1IRE8 B1IRJ9 B1IRQ7 B1IRQ9 B1IRT1 B1IRT2 B1IRU5 B1IRU6 B1IRV8 B1IS43
B1IS55 B1IS59 B1ISB7 B1ISF1 B1ISR9 B1IST8 B1ITK0 B1ITK1 B1ITK7 B1ITP4 B1ITP5
B1IU08 B1IU09 B1IU16 B1IUA1 B1IUA6 B1IUI4 B1IUL2 B1IUL5 B1IUM1 B1IUT1 B1IV37
B1IV49 B1IV54 B1IVG1 B1IVG4 B1IVG9 B1IVH1 B1IVJ2 B1IW33 B1IW70 B1IWA6 B1IWE5
B1IWI7 B1IWS5 B1IWZ1 B1IX00 B1IX01 B1IX02 B1IX18 B1IX28 B1IX33 B1IX58 B1IX59
B1IX71 B1IX73 B1IXP7 B1IXQ4 B1IXR0 B1IXR3 B1IXR6 B1IXS8 B1IXS9 B1IXT0 B1IXT3
B1IY31 B1IY32 B1IY33 B1IY75 B1IYC8 B1IYI2 B1IYL5 B1IYQ9 B1IYZ7 B1IYZ8 B1IYZ9
B1IZQ8 B1IZU0 B1J021 B1J0A5 B1J0D8 B1J0Q9 B1J0R3 B1J0R7 B1J0R8 B1J1P0 B1J2E0
B1J2G5 B1J2K7 B1J2P4 B1J2R4 B1J2S6 B1J2X0 B1J3K9 B1J454 B1J467 B1J4W3 B1J6M8
B1J6N4 B1J6S8 B1J8R9 B1J9E8 B1J9Y3 B1JAB5 B1JAS8 B1JC07 B1JC48 B1JD87 B1JDU9
B1JDY4 B1JF19 B1JFI1 B1JFN5 B1JFU5 B1JFU6 B1JFU7 B1JFV4 B1JFW5 B1JFY9 B1JFZ6
B1JGF3 B1JGK5 B1JGL3 B1JGL9 B1JGM2 B1JGM5 B1JGZ7 B1JH81 B1JHF7 B1JHT2 B1JHV1
B1JHY8 B1JIC9 B1JII7 B1JIR6 B1JIU4 B1JJ25 B1JJ29 B1JJ32 B1JJ33 B1JJ34 B1JJ53

B1JJF9 B1JJM9 B1JJS9 B1JK21 B1JK84 B1JKF9 B1JKI1 B1JKI2 B1JKI3 B1JKN2 B1JKN8
B1JKS6 B1JKS9 B1JKT9 B1JKZ7 B1JL02 B1JL41 B1JL77 B1JLH7 B1JLJ7 B1JLJ8 B1JLM7
B1JLQ1 B1JLQ2 B1JM19 B1JM21 B1JMQ4 B1JNC3 B1JNE6 B1JNK6 B1JNR4 B1JP58 B1JPQ2
B1JPZ3 B1JPZ8 B1JQC1 B1JQC2 B1JQC9 B1JQG8 B1JQU5 B1JR04 B1JR26 B1JR35 B1JR36
B1JR96 B1JR97 B1JRE9 B1JRK3 B1JRM8 B1JS04 B1JSD2 B1JSD3 B1JSH0 B1JSH3 B1JSV3
B1JTT7 B1JTX9 B1JUB0 B1JV75 B1JVD0 B1JVN1 B1JVN4 B1JVP1 B1JVS7 B1JVZ5 B1JWU6
B1JX66 B1JXC4 B1JXG6 B1JZ29 B1JZ41 B1JZ77 B1K0J3 B1K0Y4 B1K3P2 B1KAE3 B1KD43
B1KG50 B1KH27 B1KHE3 B1KHE6 B1KIB5 B1KIT6 B1KJT6 B1KJV7 B1KJW0 B1KK11 B1KKU2
B1KKY0 B1KLY8 B1KM33 B1KM58 B1KNF6 B1KQ38 B1KQ39 B1KQ40 B1KQ65 B1KQ72 B1KR03
B1KR04 B1KR05 B1KSS3 B1KSS4 B1KVR4 B1KWJ2 B1KX15 B1L008 B1L108 B1L171 B1L1L6
B1L557 B1L5A4 B1L6Z7 B1L9R8 B1LAG5 B1LBC3 B1LBC4 B1LC41 B1LD43 B1LD47 B1LD52
B1LD53 B1LD87 B1LE19 B1LEG3 B1LEI0 B1LEM1 B1LEQ4 B1LEQ6 B1LEQ9 B1LET6 B1LET7
B1LEU5 B1LEV4 B1LEY8 B1LF36 B1LF51 B1LF84 B1LF85 B1LFA0 B1LFA1 B1LFB6 B1LFL8
B1LFV7 B1LFX3 B1LFY1 B1LG24 B1LG86 B1LGA6 B1LGH4 B1LGI9 B1LGK6 B1LGK7 B1LGP7
B1LGV8 B1LGX9 B1LH37 B1LH38 B1LH51 B1LHF1 B1LHG5 B1LHX5 B1LHY1 B1LI84 B1LIU5
B1LIV8 B1LIW5 B1LJ76 B1LJI2 B1LJW4 B1LKI4 B1LKL6 B1LKN8 B1LKU3 B1LL09 B1LL32
B1LL36 B1LL63 B1LL64 B1LL65 B1LL74 B1LLE0 B1LLE1 B1LLE2 B1LLK8 B1LLL1 B1LLL2
B1LLL3 B1LLM9 B1LLN2 B1LLN5 B1LLP1 B1LLP9 B1LLW7 B1LM23 B1LMI7 B1LMI9 B1LMK6
B1LMK7 B1LMS0 B1LMU8 B1LMZ9 B1LNC2 B1LNH1 B1LNL8 B1LNM1 B1LNW4 B1LNW5 B1LNW6
B1LPJ6 B1LPK1 B1LPK5 B1LPN2 B1LPV6 B1LQ68 B1LQH4 B1LQH5 B1LQN7 B1LR25 B1LR33
B1LR34 B1LSB3 B1LTW1 B1LU06 B1LWM0 B1LWM1 B1LXZ8 B1LZP1 B1M0R9 B1M282 B1M332
B1M4Y4 B1M595 B1M7D4 B1M7D7 B1MAF7 B1MAX9 B1MBV1 B1MC66 B1MCK4 B1MDL1 B1MEW8
B1MLM4 B1MLV8 B1MLV9 B1MP34 B1MP39 B1MPH4 B1MT31 B1MTH4 B1MTL0 B1MW89 B1MW96
B1MWU5 B1MXJ0 B1MXV8 B1MY16 B1MZ69 B1MZU9 B1NF18 B1NF19 B1NF20 B1NTN6 B1NTP4
B1NTR4 B1NWD0 B1NWD4 B1NWD6 B1NWD7 B1NWD8 B1NWE3 B1NWE5 B1NWE6 B1NWF5 B1NWG2
B1NWG3 B1NWG5 B1NWG9 B1NWH0 B1NWH6 B1NWJ6 B1NWJ8 B1NWJ9 B1NWK1 B1NWK2 B1PHQ8
B1PS78 B1PS79 B1Q005 B1Q006 B1Q236 B1Q257 B1V944 B1V9H9 B1VB61 B1VC86 B1VE23
B1VFY3 B1VHC7 B1VKB7 B1VKB8 B1VKC1 B1VKE3 B1VKE4 B1VKE8 B1VKF4 B1VKF9 B1VKG9
B1VKH0 B1VKH7 B1VKH8 B1VKH9 B1VKI2 B1VKI4 B1VKI6 B1VKI7 B1VKJ3 B1VSR0 B1W0A7
B1W0I7 B1W0K0 B1W0P3 B1W108 B1W1H9 B1W317 B1W3K2 B1W4P7 B1W4V1 B1W4V4 B1W506
B1W509 B1WB39 B1WBM3 B1WBT0 B1WBW4 B1WC86 B1WNA1 B1WNP9 B1WP68 B1WPK4 B1WPZ7
B1WQ89 B1WQP0 B1WR29 B1WRN9 B1WS17 B1WS50 B1WSD1 B1WU77 B1WUD5 B1WUE0 B1WUH7
B1WUH9 B1WUI0 B1WUL2 B1WUS5 B1WVQ2 B1WVR6 B1WVR7 B1WVR8 B1WVS0 B1WWA0 B1WWK9
B1WWL0 B1WXB2 B1WXB4 B1WXD2 B1WY29 B1WYX8 B1WZG2 B1WZK3 B1WZQ6 B1X0W5 B1X0Z7
B1X3F0 B1X3H4 B1X3V8 B1X3X2 B1X3Y1 B1X3Y2 B1X3Y3 B1X3Y4 B1X491 B1X495 B1X496
B1X4Q7 B1X4R1 B1X4T6 B1X5D6 B1X5F5 B1X5G6 B1X5P3 B1X5V5 B1X6C1 B1X6E2 B1X6K0
B1X6M7 B1X6M8 B1X6M9 B1X6T0 B1X712 B1X769 B1X7H1 B1X7H2 B1X7H3 B1X7L6 B1X7N0
B1X7U9 B1X7Y8 B1X838 B1X893 B1X8W7 B1X8X0 B1X8X1 B1X8X2 B1X8Z1 B1X8Z4 B1X900
B1X906 B1X9B5 B1X9G1 B1X9G6 B1X9H8 B1X9R1 B1X9R3 B1X9T4 B1X9T9 B1X9U9 B1X9W4
B1X9W5 B1X9W6 B1X9X4 B1XA73 B1XA78 B1XA83 B1XA84 B1XAE4 B1XAG5 B1XAJ9 B1XAT9
B1XAZ1 B1XB54 B1XB74 B1XB76 B1XB81 B1XB84 B1XBF2 B1XBG5 B1XBK3 B1XBK4 B1XC29
B1XC35 B1XC39 B1XC48 B1XC64 B1XCF0 B1XCH1 B1XCS4 B1XCV3 B1XD25 B1XD46 B1XDN4
B1XDP1 B1XDP2 B1XDQ6 B1XDQ7 B1XEA2 B1XEA3 B1XEB5 B1XEB6 B1XF47 B1XF57 B1XF64
B1XF65 B1XF92 B1XFF6 B1XFL6 B1XFU2 B1XFU4 B1XFX3 B1XG18 B1XG45 B1XG62 B1XG64

B1XGP9 B1XGT1 B1XH87 B1XH88 B1XH93 B1XH97 B1XHI3 B1XHJ7 B1XHL3 B1XHL4 B1XHP2
B1XHZ0 B1XHZ1 B1XHZ2 B1XHZ3 B1XIE1 B1XIE2 B1XIN0 B1XIP3 B1XJ18 B1XJC4 B1XJF2
B1XJL9 B1XJW1 B1XKB3 B1XLJ0 B1XLL7 B1XLQ0 B1XM24 B1XMC3 B1XMN4 B1XN58 B1XNK6
B1XNS2 B1XQC0 B1XQL3 B1XRK3 B1XRK5 B1XRV6 B1XS75 B1XSD0 B1XSN0 B1XSN7 B1XSV7
B1XT06 B1XT08 B1XTA2 B1XUJ8 B1XUL1 B1XUR7 B1XVJ0 B1XVV8 B1XW26 B1XWB1 B1XWE8
B1XWR5 B1XWS3 B1XX94 B1XYR0 B1XZY0 B1Y0B2 B1Y2I5 B1Y3T2 B1Y6Q4 B1Y818 B1Y827
B1Y854 B1Y8T2 B1YB28 B1YE76 B1YEC6 B1YEG1 B1YH82 B1YIT6 B1YIW5 B1YIW6 B1YIX1
B1YK60 B1YK80 B1YKA9 B1YKE7 B1YKK2 B1YL66 B1YMC0 B1YMR8 B1YMU9 B1YN86 B1YNL7
B1YPX4 B1YQJ7 B1YQL0 B1YR31 B1YRW6 B1YRY2 B1YS18 B1YSS1 B1YSX5 B1YTH7 B1YTP7
B1YTQ0 B1YTQ7 B1YTU3 B1YUI2 B1YUV9 B1YVA5 B1YVE9 B1Z1J6 B1Z4W3 B1Z8E7 B1Z8W9
B1Z9X9 B1Z9Y2 B1ZC68 B1ZE84 B1ZGD8 B1ZJN2 B1ZJN3 B1ZKN8 B1ZLL2 B1ZLT9 B1ZMG6
B1ZMT0 B1ZQ93 B1ZQR7 B1ZRS0 B1ZRS3 B1ZU32 B1ZUK1 B1ZUR4 B1ZWP0 B2A203 B2A2J1
B2A3G6 B2A3G7 B2A3Q4 B2A6J4 B2A951 B2AA26 B2AG73 B2AGL2 B2AGR8 B2ALT5 B2AMJ3
B2ANF9 B2AR67 B2ATB0 B2AU25 B2AVN3 B2AXJ5 B2B0S1 B2B1M4 B2B2N5 B2B3L4 B2B3T2
B2B585 B2B762 B2B9D9 B2B9E1 B2BNE5 B2C4J0 B2CPI4 B2CPI5 B2CPI6 B2CPI7 B2FHB6
B2FHZ2 B2FK89 B2FM90 B2FMB6 B2FN30 B2FN96 B2FNG8 B2FP54 B2FPA5 B2FPF9 B2FQ10
B2FSD7 B2FTT6 B2G5L7 B2G5U0 B2G5Z3 B2G685 B2G687 B2G6K3 B2G735 B2G7H6 B2GA61
B2GAT9 B2GAU0 B2GAU1 B2GB76 B2GDY8 B2GEU1 B2GI45 B2GIP7 B2GJQ1 B2GL41 B2GLY6
B2GUP8 B2GUY0 B2GUY2 B2GV22 B2GV36 B2GV46 B2GV54 B2GV87 B2HEA6 B2HEM2 B2HEY1
B2HFI5 B2HFV1 B2HFY7 B2HGD8 B2HGR9 B2HGX7 B2HI58 B2HM76 B2HPD3 B2HQK5 B2HQK6
B2HQK7 B2HQK8 B2HR11 B2HRQ7 B2HST3 B2HST4 B2HSU6 B2HSU8 B2HTU4 B2HU41 B2HU50
B2HVP6 B2HXB0 B2HXY4 B2HYF5 B2HZB0 B2HZL6 B2I0K2 B2I0Q1 B2I0Z6 B2I0Z7 B2I0Z8
B2I2E9 B2I2S5 B2I2V0 B2I3A5 B2I3B1 B2I583 B2I5D0 B2I611 B2I658 B2I675 B2I6F1
B2I6V1 B2I791 B2I7H3 B2I7I7 B2I864 B2I8J9 B2I938 B2I9B5 B2I9Z0 B2IA62 B2IAR7
B2ID03 B2IDV5 B2IFJ6 B2IGF7 B2IGJ6 B2IGK8 B2IGK9 B2IGL1 B2IGT8 B2IH44 B2IHV3
B2IHV6 B2II67 B2IIP5 B2IIP7 B2IJG4 B2IJX5 B2IK19 B2IM77 B2IM88 B2INX0 B2IQX4
B2IQX5 B2IQX6 B2ISB6 B2ISQ5 B2ITV2 B2ITV8 B2IU55 B2IUM6 B2IVV1 B2IWT4 B2IYZ8
B2IZT6 B2J053 B2J054 B2J055 B2J056 B2J0I9 B2J0R7 B2J1B5 B2J204 B2J266 B2J2W1
B2J3K1 B2J3K2 B2J490 B2J4T9 B2J4U1 B2J4U7 B2J565 B2J590 B2J591 B2J5G4 B2J5S9
B2J5T0 B2J6S7 B2J6T0 B2J6T1 B2J6T3 B2J7W4 B2J814 B2J8Q3 B2J9K4 B2JA12 B2JA13
B2JC54 B2JCE3 B2JCV1 B2JDL5 B2JDL8 B2JDM1 B2JDM8 B2JDU2 B2JDY8 B2JE25 B2JES9
B2JGG3 B2JGJ7 B2JGM9 B2JH09 B2JHG3 B2JHX4 B2JJK3 B2JJR8 B2JJU6 B2JLR2 B2JVU2
B2JYL3 B2JZ26 B2JZ67 B2JZ74 B2JZ75 B2JZA2 B2JZJ8 B2K059 B2K0F1 B2K0F2 B2K0R7
B2K0Y6 B2K137 B2K1U4 B2K1W7 B2K1Z0 B2K2I0 B2K2I2 B2K306 B2K336 B2K337 B2K3F9
B2K3G0 B2K3I4 B2K3M2 B2K3M7 B2K3Q0 B2K3U5 B2K3V5 B2K3V8 B2K3Y6 B2K436 B2K437
B2K438 B2K460 B2K4E3 B2K4J6 B2K4K2 B2K4Y4 B2K549 B2K576 B2K5F2 B2K5I9 B2K5K9
B2K5L0 B2K5L1 B2K5L4 B2K5S2 B2K5W4 B2K647 B2K660 B2K6A2 B2K6K2 B2K6M5 B2K787
B2K7E7 B2K7I6 B2K809 B2K812 B2K815 B2K821 B2K829 B2K841 B2K842 B2K843 B2K8K4
B2K911 B2K918 B2K942 B2K987 B2K990 B2K9M1 B2K9M2 B2K9Q1 B2K9W1 B2KA13 B2KA77
B2KA78 B2KBZ0 B2KBZ1 B2KD05 B2KE93 B2KED0 B2KEW6 B2KEW8 B2KG20 B2KI42 B2KI79
B2KKL4 B2KWH4 B2KWH5 B2KWH6 B2KWI0 B2KX91 B2LMH7 B2LMH8 B2LMI4 B2LMI5 B2LMI6
B2LMI8 B2LMI9 B2LMJ8 B2LMK5 B2LMK8 B2LML2 B2LML3 B2LML9 B2LMM1 B2LMP5 B2LMP6
B2LMP8 B2LMP9 B2LMQ1 B2LT48 B2LT61 B2LT62 B2LT64 B2LT65 B2LU20 B2LW77 B2LYG4
B2MVP8 B2MVX9 B2MZD0 B2RFN2 B2RI39 B2RI48 B2RI95 B2RIE8 B2RJ03 B2RJU3 B2RKS0

B2RLS8 B2RML6 B2RN74 B2RPY5 B2RQA1 B2RQR8 B2RTY4 B2RU40 B2RU80 B2RUZ4 B2RVW2
B2RVY9 B2RWJ3 B2RX12 B2RXF0 B2RXS4 B2RXV4 B2RXZ1 B2RYH9 B2RYW8 B2RZ37 B2RZ87
B2RZC9 B2RZD2 B2RZW9 B2S014 B2S0E5 B2S0H3 B2S2U2 B2S532 B2S553 B2S5A0 B2S5B5
B2S5P5 B2S6K8 B2S6Q7 B2S6Y9 B2S7N7 B2S888 B2S8E3 B2S9G8 B2S9M8 B2S9N0 B2S9N1
B2SA93 B2SAB9 B2SAT5 B2SB55 B2SC28 B2SD40 B2SDD0 B2SDR9 B2SE35 B2SE60 B2SEB4
B2SEL7 B2SEV6 B2SEX7 B2SF41 B2SF87 B2SG34 B2SHN5 B2SHQ4 B2SIH3 B2SNZ1 B2SPS7
B2SQB4 B2SQM8 B2SS70 B2ST34 B2STC5 B2SUA7 B2SUW0 B2SVK9 B2SVL2 B2SWL6 B2SX38
B2SXA0 B2SY66 B2SYS3 B2SYX9 B2SZ54 B2T152 B2T1K8 B2T2B4 B2T2E7 B2T2F4 B2T2F7
B2T349 B2T3W8 B2T419 B2T6I8 B2T7K4 B2T7U1 B2T9B0 B2TDV1 B2TJZ5 B2TJZ6 B2TMJ1
B2TMJ2 B2TMJ3 B2TPF2 B2TS25 B2TTF0 B2TTI6 B2TTJ6 B2TTJ7 B2TTK5 B2TTM5 B2TTZ9
B2TU32 B2TUN7 B2TUN8 B2TUN9 B2TUR5 B2TUS3 B2TUT7 B2TV39 B2TV73 B2TVI6 B2TVN0
B2TVP9 B2TVQ7 B2TVV4 B2TW40 B2TW41 B2TW57 B2TW60 B2TW63 B2TW69 B2TW76 B2TWL3
B2TWQ9 B2TWY6 B2TWY8 B2TWZ5 B2TWZ6 B2TX76 B2TXA0 B2TXP8 B2TXU1 B2TY29 B2TY30
B2TYA6 B2TYA7 B2TYA8 B2TYQ5 B2TYQ6 B2TYR3 B2TZB2 B2TZB6 B2TZI3 B2TZP2 B2TZQ8
B2TZZ4 B2TZZ9 B2U0A0 B2U0L5 B2U0Y9 B2U167 B2U1A7 B2U1G1 B2U1G3 B2U1Q2 B2U1Q8
B2U1Q9 B2U1U5 B2U1W0 B2U246 B2U255 B2U292 B2U2C6 B2U2C9 B2U2D1 B2U2G8 B2U2R0
B2U300 B2U320 B2U360 B2U3F5 B2U3H0 B2U3N2 B2U3Q2 B2U422 B2U460 B2U461 B2U467
B2U471 B2U4J7 B2U793 B2U7E0 B2U7Q1 B2U7Q4 B2U7R1 B2U820 B2U9C4 B2UAJ1 B2UAK3
B2UAT6 B2UB97 B2UBM4 B2UC26 B2UCY0 B2UDT8 B2UE66 B2UEE1 B2UGV3 B2UH25 B2UI31
B2UL32 B2ULY3 B2UMY1 B2UN92 B2UNX8 B2US19 B2USE9 B2UT00 B2UTZ3 B2UU75 B2UUA1
B2UUP4 B2UUX5 B2UUY7 B2UV31 B2UV34 B2UW72 B2UXE0 B2UZJ5 B2UZJ6 B2V2P2 B2V2P3
B2V2P4 B2V4V5 B2V5I7 B2V6F7 B2V6N8 B2V758 B2V983 B2V995 B2V9G8 B2V9H0 B2V9L9
B2VBI5 B2VBI6 B2VBI7 B2VBJ0 B2VBK2 B2VC50 B2VCA8 B2VCA9 B2VCB0 B2VCE6 B2VCK3
B2VCL2 B2VD87 B2VDB2 B2VDB4 B2VDY7 B2VE18 B2VE39 B2VE88 B2VEQ1 B2VEQ4 B2VEQ5
B2VEQ6 B2VET5 B2VFQ4 B2VFS7 B2VFU8 B2VFZ8 B2VG47 B2VG51 B2VGJ1 B2VGJ3 B2VGW0
B2VGW1 B2VH02 B2VH05 B2VH47 B2VHE5 B2VHL9 B2VHR9 B2VIF5 B2VIN4 B2VIN7 B2VIV3
B2VIV9 B2VJ02 B2VJ57 B2VJK2 B2VJK3 B2VJK4 B2VK30 B2VK47 B2VK97 B2VKE4 B2VKI2
B2VKR0 B2VKV4 B2VKV9 B2VS19 B2VSU4 B2VV00 B2VVB9 B2W0S3 B2W1P8 B2W244 B2W340
B2WAQ3 B2WBP3 B2WBQ6 B2WC36 B2WC78 B2WCU2 B2WDD8 B2WDP9 B2WEL2 B2WFD4 B2WMJ3
B2X1U6 B2X1U7 B2X1U8 B2X1V8 B2X1W4 B2X1W8 B2X1X3 B2X1X7 B2X1X9 B2X205 B2XT21
B2XT86 B2XT88 B2XT99 B2XTA0 B2XTC6 B2XTD8 B2XTF7 B2XTG4 B2XTH7 B2XTP2 B2XTP4
B2XTQ5 B2XTQ6 B2XTT2 B2XTX0 B2XWI8 B2XWJ0 B2XWJ3 B2XWJ4 B2XWK0 B2XWK1 B2XWK9
B2XWM0 B2XWM4 B2XWM5 B2XWN3 B2XWN5 B2XWN6 B2XWN7 B2XWP2 B2XWP9 B2XWQ1 B2Y1U5
B2Y1U6 B2Y1U7 B2Y1V3 B2Y1V4 B2Y1V9 B2Y1W0 B2Y1W1 B2Y1W3 B2Y1W4 B2Y1W5 B2Y1W9
B2Y1X0 B2Y1X1 B2Y1X2 B2Y1X3 B2Y1X7 B2Y1X8 B2Y1X9 B2Y1Y5 B2Y1Y8 B2Y1Y9 B2Y1Z0
B2Y206 B2ZHY2 B2ZI34 B2ZXD5 B3A0L9 B3A0M0 B3A0M1 B3A0M2 B3CLF8 B3CLG2 B3CLG3
B3CMI8 B3CMU6 B3CMZ0 B3CN58 B3CPW3 B3CQT8 B3CQT9 B3CRU7 B3CS08 B3CTF6 B3CUJ6
B3CUK2 B3CVD3 B3DFP2 B3DFW5 B3DH96 B3DHH5 B3DHU2 B3DHW5 B3DI94 B3DJK0 B3DK56
B3DLH6 B3DM66 B3DMA0 B3DN87 B3DQ87 B3DQM8 B3DRX3 B3DS08 B3DTV4 B3DTV5 B3DU18
B3DV46 B3DVW4 B3DWJ8 B3DWK0 B3DX74 B3DY14 B3DZT0 B3DZT3 B3E0Z6 B3E171 B3E340
B3E3S0 B3E3Y3 B3E3Y5 B3E4X5 B3E5K6 B3E8I3 B3E9V6 B3E9V9 B3E9W9 B3E9X3 B3E9X4
B3EA05 B3EA78 B3EAP8 B3EAQ1 B3EC54 B3ECE6 B3ECJ4 B3ED81 B3EDE6 B3EEG6 B3EEM7
B3EH31 B3EI29 B3EI93 B3EIJ6 B3EIL1 B3EIM7 B3EJP7 B3EKA1 B3EKJ5 B3EL80 B3ELA5
B3ELU5 B3EMS7 B3EPQ3 B3EPU8 B3EQ23 B3EQ96 B3EQ98 B3EQC1 B3EQF7 B3ER28 B3ER47

B3ETU2 B3EU56 B3EU96 B3EUQ9 B3EWY8 B3EWY9 B3EWZ1 B3EWZ3 B3EWZ9 B3FN88 B3FWR7
B3FWS2 B3FWT8 B3G515 B3GNI9 B3GXM1 B3GXV8 B3GXW4 B3GYC1 B3GYE7 B3GYF4 B3GYK8
B3GZD0 B3GZD1 B3GZK5 B3H011 B3H0G7 B3H0M8 B3H1E0 B3H1F9 B3H1S0 B3H276 B3H2D6
B3H2I9 B3H2P7 B3H2P8 B3H2P9 B3H2W4 B3H317 B3H4K7 B3H6K1 B3H707 B3H754 B3L2V1
B3LAJ9 B3LFA3 B3LGY4 B3LH09 B3LHC6 B3LHE1 B3LHF1 B3LHL3 B3LHR1 B3LHR7 B3LI04
B3LI56 B3LI73 B3LIC1 B3LIN5 B3LJ06 B3LJA1 B3LJJ8 B3LKQ3 B3LKX3 B3LLM2 B3LLQ4
B3LM17 B3LM36 B3LM44 B3LMP9 B3LN00 B3LN75 B3LNF5 B3LNR8 B3LNV9 B3LP27 B3LQ47
B3LQQ2 B3LR89 B3LRA0 B3LRH5 B3LRK1 B3LRL2 B3LRN4 B3LS11 B3LS51 B3LS52 B3LSR5
B3LTC4 B3LTC5 B3LTI7 B3LTW4 B3LTZ3 B3LU06 B3LUQ6 B3LUU7 B3LVB0 B3LVG7 B3LZ39
B3M1H7 B3M3X7 B3M8G0 B3M9A8 B3M9W1 B3MFK1 B3MFV7 B3MG58 B3MH43 B3MK83 B3MKS0
B3MR30 B3MRI9 B3MRS1 B3MTS2 B3MTS8 B3N5J3 B3N666 B3N7H1 B3N8M1 B3NBB6 B3NDM7
B3NGS7 B3NIN0 B3NS99 B3NSE1 B3NV41 B3NY03 B3NZM5 B3P048 B3P2E5 B3P3J9 B3P5J1
B3P6T8 B3P7K6 B3P9N0 B3PB31 B3PB33 B3PCL7 B3PCM1 B3PCM3 B3PE13 B3PFQ6 B3PH50
B3PHW0 B3PI58 B3PIC5 B3PIT1 B3PIT2 B3PIU1 B3PJ06 B3PJS2 B3PLV3 B3PLV4 B3PNH3
B3PNR8 B3PP14 B3PPV4 B3PQS4 B3PR08 B3PR74 B3PRF6 B3PRF8 B3PRF9 B3PS39 B3PSB2
B3PSB4 B3PTW3 B3PU61 B3PUW1 B3PVH0 B3PW09 B3PW35 B3PXU0 B3PY50 B3PYJ1 B3PZB2
B3Q040 B3Q0T2 B3Q5X8 B3Q6I3 B3Q6L4 B3Q6S6 B3Q6T5 B3Q7Z4 B3Q8C0 B3Q8S9 B3Q9X6
B3QBD2 B3QBQ3 B3QED3 B3QF26 B3QF34 B3QF35 B3QF37 B3QFN4 B3QI59 B3QKF4 B3QLQ4
B3QLV1 B3QLV3 B3QLW7 B3QLY4 B3QM18 B3QN57 B3QNH3 B3QNS4 B3QP51 B3QP54 B3QP58
B3QP89 B3QP93 B3QQ42 B3QQ95 B3QUX5 B3QW96 B3QWT4 B3QXL8 B3QXM3 B3QY38 B3QY48
B3QYV6 B3QZ04 B3QZE8 B3QZF0 B3QZS3 B3R057 B3R0R7 B3R0V4 B3R2M9 B3R3X4 B3R3X7
B3R3Y0 B3R523 B3R646 B3R6E9 B3R6W2 B3R789 B3R7L9 B3R883 B3R8Z5 B3RFJ6 B3RHM5
B3RML8 B3SGL0 B3SHH9 B3SRQ8 B3SRT2 B3SRV6 B3SRY0 B3TLD6 B3TN41 B3TN45 B3TN46
B3TN47 B3TN56 B3TN65 B3TN69 B3TN75 B3TN77 B3TN96 B3TN98 B3TNA1 B3TNA2 B3TNB7
B3TNB8 B3TNB9 B3TP03 B3VA58 B3VA59 B3VQ24 B3VSC2 B3W6P3 B3W6Y1 B3W705 B3WA65
B3WAS8 B3WCK5 B3WDL3 B3WDL4 B3WE66 B3WEB1 B3Y064 B3Y613 B3Y614 B3Y615 B3Y618
B4DJY2 B4DS77 B4DYI2 B4E5L2 B4E5L5 B4E5M2 B4E5Q9 B4E646 B4E6J5 B4E7D5 B4E7W0
B4E8G3 B4E8Z8 B4E9G3 B4E9G9 B4EA84 B4EBL0 B4EBU8 B4EC31 B4EEY5 B4EF95 B4ETJ0
B4ETL6 B4ETL9 B4ETM0 B4ETM1 B4ETN1 B4EUL1 B4EUT7 B4EUT8 B4EV00 B4EVU5 B4EVZ5
B4EW52 B4EWD2 B4EWI7 B4EWK1 B4EWN4 B4EWY4 B4EWY5 B4EX78 B4EXQ1 B4EXU4 B4EY22
B4EY74 B4EY98 B4EYA1 B4EYC2 B4EYY4 B4EZ47 B4EZ50 B4EZC4 B4EZD0 B4EZD8 B4F0E1
B4F0E2 B4F0E3 B4F114 B4F181 B4F1L3 B4F2G7 B4F2T1 B4F6I3 B4F6I5 B4F6J6 B4F753
B4F795 B4F7C5 B4F8I5 B4FAP1 B4FBQ7 B4FF80 B4FG96 B4FHU1 B4FIJ0 B4FNS3 B4FUS3
B4FVP3 B4G316 B4GAP7 B4GBH0 B4GD14 B4GE47 B4GF83 B4GHS8 B4GJC1 B4GKZ8 B4GMC3
B4GMC9 B4GPI0 B4GRI8 B4GVM9 B4GXS1 B4GZN1 B4H1F5 B4H3U8 B4HEB1 B4HEM8 B4HIJ8
B4HJV5 B4HNS0 B4HNS1 B4HNW4 B4HS00 B4HSS0 B4HVU2 B4HY03 B4HZ81 B4I1V5 B4I414
B4I9J6 B4IAB8 B4IKJ4 B4IL64 B4IXJ2 B4J043 B4J2W3 B4J780 B4J877 B4J913 B4JBN5
B4JEF2 B4JG34 B4JG39 B4JHV0 B4JLX2 B4JMC1 B4JTT7 B4JYJ2 B4JYU5 B4K4G5 B4K5X8
B4K835 B4KB36 B4KB41 B4KD90 B4KFW6 B4KJW1 B4KMZ1 B4KPU0 B4KQ08 B4KR05 B4L0H1
B4L184 B4L535 B4L7U0 B4L8V5 B4LC58 B4LCX4 B4LIH0 B4LNA8 B4LPX5 B4LQT7 B4LRN7
B4LY58 B4LYB8 B4LYC5 B4LZB5 B4M709 B4M7S0 B4MBU8 B4MDV2 B4MEG2 B4MGF8 B4MJN5
B4MLE8 B4MR28 B4MU83 B4MUQ2 B4MXW6 B4MYA4 B4N072 B4N1W9 B4N5D3 B4N937 B4NDL8
B4NE96 B4NFN4 B4NIY1 B4NK88 B4NKL0 B4NZY8 B4P0E1 B4P3A0 B4P417 B4P5Q9 B4P624
B4PD01 B4PD96 B4PET6 B4PF15 B4PQ50 B4PQC2 B4PST0 B4PUZ3 B4PVB0 B4PVC6 B4PZQ4

B4Q0P3 B4Q4M7 B4Q599 B4QBN2 B4QBN3 B4QC63 B4QH87 B4QJ33 B4QLP9 B4QMF4 B4QPR1
B4QVX4 B4QWH9 B4QWW7 B4QZI8 B4QZU6 B4R136 B4R313 B4R4T6 B4R8Q2 B4R910 B4RCI1
B4RCK7 B4RCL8 B4RCM1 B4RDI3 B4RDI4 B4RDV5 B4RF13 B4RFS3 B4RHR0 B4RIX6 B4RJ59
B4RJF6 B4RJJ2 B4RK00 B4RKA0 B4RLD8 B4RLE2 B4RLF2 B4RMD9 B4RNG2 B4RNM0 B4RNN7
B4RPH4 B4RQC6 B4RQC9 B4RQE0 B4RQT5 B4RS87 B4RU97 B4RUX1 B4RVP1 B4RVT2 B4RWY2
B4RY28 B4RYA6 B4RZU1 B4S170 B4S3H1 B4S3Q4 B4S5J8 B4S675 B4S6E4 B4S6E6 B4S6R2
B4S6X1 B4S751 B4S798 B4S7A0 B4S7I8 B4S891 B4S8V4 B4S9H4 B4SAH4 B4SBZ3 B4SCH3
B4SCV5 B4SD15 B4SDS7 B4SE06 B4SGK3 B4SGX7 B4SH38 B4SH40 B4SH93 B4SHE7 B4SHH0
B4SHI1 B4SI88 B4SJS3 B4SK34 B4SKS3 B4SNQ6 B4SNR8 B4SPG0 B4SPV6 B4SPY3 B4SQB7
B4SQS6 B4SR87 B4SRU3 B4STT2 B4SU47 B4SUC3 B4SUC4 B4SUC9 B4SUJ3 B4SUJ8 B4SUN3
B4SUR2 B4SUV7 B4SUY5 B4SV06 B4SV74 B4SV75 B4SV80 B4SV83 B4SVH5 B4SVM1 B4SWC6
B4SWG1 B4SWZ0 B4SXW0 B4SXW1 B4SXW2 B4SY13 B4SY75 B4SYB1 B4SYB3 B4SYD5 B4SYD6
B4SYD7 B4SYE8 B4SYI7 B4SYX0 B4SYX3 B4SYX4 B4SYX5 B4SYY7 B4SYZ0 B4SYZ3 B4SYZ9
B4SZ08 B4SZ39 B4SZ75 B4SZB0 B4SZB1 B4SZB2 B4SZP4 B4SZS7 B4SZU6 B4SZW9 B4SZZ5
B4T048 B4T0D1 B4T0K5 B4T0L1 B4T0Q0 B4T132 B4T1S0 B4T1S6 B4T1S9 B4T1W9 B4T2P9
B4T2Q0 B4T2Y0 B4T2Y5 B4T2Z8 B4T3W8 B4T458 B4T4E6 B4T4F0 B4T4G5 B4T4N5 B4T4Z4
B4T502 B4T503 B4T562 B4T591 B4T593 B4T596 B4T5B8 B4T5B9 B4T5E1 B4T5G1 B4T5G7
B4T5R6 B4T660 B4T675 B4T677 B4T695 B4T6B2 B4T6H3 B4T6J9 B4T6L2 B4T6P7 B4T6S8
B4T739 B4T749 B4T774 B4T775 B4T776 B4T8F0 B4T8W9 B4T8X4 B4T9U1 B4T9U2 B4T9U3
B4TA59 B4TAI2 B4TAK3 B4TAV2 B4TAV4 B4TAX6 B4TAX7 B4TAX8 B4TAY9 B4TB26 B4TBA5
B4TBA6 B4TBA7 B4TBG5 B4TBG8 B4TBG9 B4TBH0 B4TBI2 B4TBI5 B4TBJ4 B4TBK3 B4TBR5
B4TBV9 B4TBY5 B4TCD8 B4TCF7 B4TCJ9 B4TCW8 B4TD28 B4TD53 B4TD59 B4TD94 B4TDK9
B4TDL4 B4TDL7 B4TEB2 B4TES0 B4TES5 B4TET8 B4TF88 B4TF89 B4TFR0 B4TFW8 B4TGH8
B4TGQ7 B4TGR4 B4TGR5 B4TGX6 B4TGY0 B4TGZ4 B4TH65 B4THD1 B4THD3 B4THD6 B4THE9
B4THR3 B4THR4 B4THT6 B4THV6 B4TI02 B4TI40 B4TI56 B4TI58 B4TIE6 B4TIH3 B4TIS9
B4TIV3 B4TIX1 B4TJ42 B4TJ84 B4TJG4 B4TJL6 B4TJL7 B4TJM2 B4TJQ1 B4TJR2 B4TJT8
B4TJT9 B4TJU0 B4TJV2 B4TJY1 B4TK31 B4TK52 B4TKD1 B4TKD6 B4TKG4 B4TKG5 B4TKH1
B4TKH4 B4TKN2 B4TKP8 B4TKV0 B4TMX5 B4TN10 B4TN12 B4TN35 B4TN36 B4TN37 B4TN48
B4TNI6 B4TNI7 B4TNI8 B4TNN8 B4TNU6 B4TNY1 B4TPA0 B4TPI1 B4TPI4 B4TPI5 B4TPI6
B4TPJ8 B4TPK1 B4TPL0 B4TPN7 B4TPR1 B4TPS5 B4TQ21 B4TQ22 B4TQ23 B4TQ70 B4TQD9
B4TQE4 B4TQF9 B4TQP3 B4TQP9 B4TQQ2 B4TR50 B4TR56 B4TR96 B4TRK2 B4TRM3 B4TRS9
B4TSD5 B4TSD6 B4TSR0 B4TSR5 B4TTD0 B4TTW2 B4TU18 B4TU22 B4TU36 B4TU95 B4TUF7
B4TUM0 B4TUM8 B4TUM9 B4TUY3 B4TV14 B4TV16 B4TV19 B4TV35 B4TVE8 B4TVE9 B4TVH1
B4TVJ2 B4TVN6 B4TVS2 B4TVT9 B4TVU1 B4TVV9 B4TVX7 B4TWA8 B4TWH8 B4TWI9 B4TWN7
B4TWR7 B4TWT1 B4TWZ4 B4TX47 B4TX48 B4TX53 B4TX72 B4TX73 B4TX74 B4TX86 B4TXB4
B4TXF9 B4TXH5 B4TXQ7 B4TXX1 B4TXX6 B4TY03 B4TY04 B4TY09 B4TY12 B4TY33 B4TY77
B4TYD7 B4TZ13 B4TZP6 B4TZQ1 B4U1Z7 B4U2B4 B4U2D6 B4U2D7 B4U3D2 B4U3X6 B4U4K1
B4U4R2 B4U4T2 B4U4T7 B4U4X0 B4U7U4 B4U812 B4U8E3 B4U8E4 B4U8I1 B4U8I7 B4U8J0
B4U8V3 B4U8V5 B4U9Q9 B4U9R1 B4UCE4 B4UD86 B4UER8 B4UES6 B4UFX7 B4UIA6 B4UIV7
B4UIW0 B4UJU5 B4UJU7 B4UKG1 B4ULG0 B4UMI5 B4UN04 B4URD6 B4URD7 B4URD9 B4USZ0
B4WP19 B4XF06 B4YNF5 B4YQU1 B5BA35 B5BA93 B5BAN4 B5BAY4 B5BB21 B5BB26 B5BB70
B5BB71 B5BB80 B5BB84 B5BBC3 B5BBD5 B5BBE0 B5BBQ8 B5BBW7 B5BCA3 B5BCA4 B5BCG4
B5BCL0 B5BCL8 B5BCM7 B5BCP2 B5BCP3 B5BCP4 B5BCP7 B5BE54 B5BE74 B5BEY5 B5BF67
B5BF68 B5BF69 B5BFG7 B5BFH5 B5BFH6 B5BFN1 B5BFW5 B5BG02 B5BG16 B5BG18 B5BG54

B5BG59 B5BGE1 B5BGF7 B5BGN2 B5BGP4 B5BGR8 B5BGR9 B5BGS0 B5BGT2 B5BGV7 B5BH03
B5BH18 B5BHB2 B5BHB6 B5BHC3 B5BHH5 B5BHN6 B5BHQ7 B5BI47 B5BI52 B5BIA8 B5BIB2
B5BIB3 B5BIG3 B5BIJ0 B5BIL8 B5BIM0 B5BIP0 B5BIP1 B5BIP2 B5BIQ1 B5BIU2 B5BIY1
B5BJ45 B5BJ69 B5BJH0 B5BJI1 B5BJI4 B5BJU9 B5BJV4 B5BJV7 B5BJZ5 B5BK54 B5BK72
B5BK90 B5BKB0 B5BKB3 B5BKB5 B5BKE4 B5BKG4 B5BKG5 B5BKZ9 B5BL23 B5BL58 B5BL83
B5BLB9 B5BLL1 B5BP45 B5BP47 B5BP49 B5BSU1 B5BSU2 B5BSX1 B5D5N9 B5DE73 B5DEH2
B5DEK8 B5DF41 B5DF51 B5DFG1 B5DFH9 B5DFM7 B5DKS8 B5E1E8 B5E1R8 B5E1T5 B5E1Z4
B5E3I9 B5E3M4 B5E520 B5E5S0 B5E675 B5E676 B5E677 B5E6Z5 B5E921 B5E960 B5E964
B5E973 B5E9H5 B5E9Q0 B5EBP8 B5EDZ1 B5EFF3 B5EFG3 B5EFG8 B5EFG9 B5EFJ1 B5EGX9
B5EH78 B5EH80 B5ELC6 B5EN61 B5EN71 B5ENN2 B5EPU0 B5ER46 B5ER47 B5EWY8 B5EXV7
B5EXV8 B5EXV9 B5EY82 B5EYN1 B5EYX5 B5EYX7 B5EZ00 B5EZ01 B5EZ02 B5EZ13 B5EZ53
B5EZ69 B5EZE2 B5EZE3 B5EZE4 B5EZH7 B5EZI0 B5EZI1 B5EZI2 B5EZJ7 B5EZK0 B5EZK3
B5EZK9 B5EZL8 B5EZV0 B5EZZ9 B5F0D7 B5F0F6 B5F0K5 B5F0L1 B5F0N1 B5F0P5 B5F0Y9
B5F150 B5F199 B5F1P5 B5F1Q0 B5F1Q3 B5F2E9 B5F2L8 B5F2L9 B5F3P9 B5F3Q2 B5F3Q8
B5F3Q9 B5F412 B5F4E1 B5F4E6 B5F4L0 B5F4L5 B5F4L6 B5F4U6 B5F4V3 B5F4V4 B5F500
B5F504 B5F518 B5F588 B5F5B8 B5F5J1 B5F613 B5F641 B5F683 B5F6A1 B5F6A3 B5F6E1
B5F6G3 B5F6G4 B5F6I8 B5F6J1 B5F6J3 B5F6M2 B5F6N2 B5F6N4 B5F6Q0 B5F6X7 B5F725
B5F736 B5F753 B5F767 B5F7F5 B5F7J8 B5F7M4 B5F7M5 B5F7M6 B5F7N8 B5F7R7 B5F7W1
B5F873 B5F8G9 B5F8I5 B5F8P0 B5F8R6 B5F8T8 B5F941 B5F954 B5F959 B5F9I1 B5FA52
B5FAF8 B5FB38 B5FB78 B5FB81 B5FBI3 B5FBI4 B5FBS7 B5FBS8 B5FC41 B5FCB4 B5FCI6
B5FCI8 B5FCN0 B5FCN1 B5FCZ5 B5FCZ6 B5FCZ7 B5FDB4 B5FDK5 B5FDN8 B5FEZ3 B5FF80
B5FF81 B5FFD1 B5FFH5 B5FGC4 B5FGU8 B5FHE1 B5FHG8 B5FHN1 B5FHP9 B5FHR3 B5FHS1
B5FHS2 B5FHU0 B5FHU2 B5FHW0 B5FHX7 B5FI29 B5FI69 B5FIE5 B5FIE8 B5FIF0 B5FIH8
B5FII7 B5FIQ6 B5FIR7 B5FIU2 B5FIU3 B5FIU4 B5FIV6 B5FJ02 B5FJ24 B5FJA1 B5FJH1
B5FJI6 B5FJN1 B5FJP6 B5FKE3 B5FKN5 B5FKZ8 B5FL11 B5FL16 B5FLY5 B5FLZ0 B5FMC7
B5FMM5 B5FMM7 B5FMU6 B5FMU7 B5FMU8 B5FMZ7 B5FN13 B5FN15 B5FN37 B5FN38 B5FN39
B5FN50 B5FN87 B5FND9 B5FNE0 B5FNE1 B5FNT8 B5FNU1 B5FNW8 B5FP16 B5FPD7 B5FPE1
B5FPE2 B5FPF7 B5FPG0 B5FPG3 B5FPG9 B5FPH7 B5FPP7 B5FPR1 B5FQ40 B5FQA1 B5FQC0
B5FQG8 B5FQH4 B5FQQ0 B5FQQ5 B5FQQ8 B5FR64 B5FRF0 B5FRL0 B5FRL1 B5FTA1 B5FTA5
B5FTC0 B5FTJ0 B5FTJ3 B5FTJ9 B5FTK0 B5FTM8 B5FTN3 B5FTS6 B5FU52 B5FU57 B5FU58
B5FUB0 B5FUB8 B5FUB9 B5FUK9 B5FUN9 B5FV19 B5FV58 B5FVB3 B5FVG3 B5FXC9 B5FXE5
B5G2S6 B5LML1 B5LMM0 B5LMM1 B5LMM3 B5LMM8 B5LMM9 B5LMN0 B5LMP3 B5LMP4 B5LMP9
B5LMQ1 B5LMS3 B5LMS4 B5LMS9 B5MEV3 B5QU11 B5QU35 B5QUA4 B5QUC2 B5QUE3 B5QUE4
B5QUQ4 B5QUQ6 B5QV00 B5QV03 B5QV05 B5QV34 B5QVD6 B5QVD7 B5QVD8 B5QVE7 B5QVI5
B5QVN1 B5QVW1 B5QW19 B5QW75 B5QWE8 B5QWE9 B5QWF0 B5QWM0 B5QWT4 B5QWU1 B5QWU2
B5QWW1 B5QWY6 B5QWZ4 B5QWZ7 B5QXJ3 B5QXX4 B5QXZ9 B5QY11 B5QY16 B5QYC8 B5QYI9
B5QYJ4 B5QYP8 B5QYX4 B5QYX9 B5QZ27 B5QZ43 B5QZ59 B5QZ97 B5QZQ4 B5QZS1 B5R015
B5R016 B5R0C0 B5R0C1 B5R0C2 B5R0F4 B5R0J9 B5R0L1 B5R151 B5R1A7 B5R1A8 B5R1A9
B5R1C1 B5R1E7 B5R1N7 B5R1R3 B5R1S4 B5R271 B5R274 B5R275 B5R276 B5R2A8 B5R2C3
B5R2H8 B5R2M1 B5R2S7 B5R2T1 B5R2T8 B5R2W4 B5R2W9 B5R2Z7 B5R300 B5R309 B5R317
B5R383 B5R3G7 B5R3J0 B5R3N1 B5R3N5 B5R3N6 B5R3T7 B5R3U1 B5R3V5 B5R3V6 B5R3Y2
B5R3Z1 B5R415 B5R4A8 B5R4D6 B5R4I5 B5R4J1 B5R583 B5R669 B5R670 B5R671 B5R6L2
B5R6L6 B5R6L7 B5R7J1 B5R7P0 B5R7S5 B5R7T0 B5R7T3 B5R7X8 B5R857 B5R8I5 B5R8V8
B5R8W2 B5R8Z5 B5R8Z9 B5R937 B5R9A1 B5R9A2 B5R9T1 B5R9Z8 B5RAE4 B5RAG1 B5RAK0

B5RAK3 B5RAK5 B5RAN4 B5RAW6 B5RB26 B5RBD4 B5RBF1 B5RBK7 B5RBL2 B5RBW8 B5RBW9
B5RC02 B5RC20 B5RCC3 B5RCC6 B5RCC7 B5RCC8 B5RCE1 B5RCE4 B5RCF3 B5RCG1 B5RCN6
B5RCQ0 B5RCQ1 B5RCU8 B5RCV4 B5RCZ3 B5RDQ4 B5RDX8 B5RDY5 B5RDY6 B5RE27 B5REB2
B5REE9 B5REG5 B5REJ9 B5RES4 B5RET6 B5REW2 B5REW3 B5REW4 B5REX6 B5RFA7 B5RFB0
B5RFC1 B5RFE6 B5RFM6 B5RFR0 B5RFU8 B5RFV7 B5RFV8 B5RFV9 B5RFY3 B5RG43 B5RG82
B5RGA7 B5RGR2 B5RGT6 B5RH45 B5RH61 B5RHE1 B5RHG2 B5RHV8 B5RL84 B5RLC9 B5RLZ9
B5RM28 B5RR78 B5RRC2 B5RRP5 B5RRS4 B5RU30 B5RUL7 B5T267 B5TVM2 B5TYT3 B5UAQ8
B5VDL3 B5VDW2 B5VE39 B5VF36 B5VHD1 B5VHH5 B5VHP4 B5VI70 B5VIB8 B5VIN9 B5VIR6
B5VIU1 B5VJA0 B5VK80 B5VKG3 B5VKJ2 B5VLV9 B5VMF4 B5VMG3 B5VMP1 B5VMT6 B5VPC6
B5VPS9 B5VQN1 B5VQU2 B5VRJ9 B5VRU8 B5VS52 B5VSZ6 B5VT63 B5WWZ8 B5WWZ9 B5X0E4
B5X0N6 B5X1G3 B5X337 B5X3B2 B5X3I6 B5X3W7 B5X4H8 B5X570 B5X8S2 B5XB24 B5XJR7
B5XJV3 B5XJX2 B5XKG1 B5XKM5 B5XKP6 B5XKP7 B5XL22 B5XM58 B5XML0 B5XN31 B5XN50
B5XN76 B5XNC0 B5XNE5 B5XNL4 B5XNU5 B5XNW0 B5XNW3 B5XNW6 B5XP67 B5XPB7 B5XPB8
B5XPB9 B5XPI1 B5XQ41 B5XQ45 B5XQ52 B5XQ53 B5XQ77 B5XQ83 B5XQA9 B5XQW2 B5XR73
B5XR94 B5XRN5 B5XRW2 B5XSD1 B5XSP7 B5XSP8 B5XT15 B5XT16 B5XT62 B5XTD8 B5XTK8
B5XTL1 B5XTL2 B5XTL3 B5XTR5 B5XTX0 B5XU32 B5XU34 B5XU71 B5XUC1 B5XUP2 B5XUP3
B5XUQ1 B5XUV2 B5XV33 B5XVT3 B5XVU3 B5XVZ7 B5XWC8 B5XWL3 B5XWP3 B5XWP6 B5XWP8
B5XWQ1 B5XXJ2 B5XXK2 B5XXK5 B5XXT7 B5XXZ2 B5XY00 B5XYH9 B5XYN9 B5XZ41 B5XZ47
B5XZE5 B5XZE8 B5XZE9 B5XZF0 B5XZI5 B5XZK9 B5XZL8 B5XZL9 B5XZM0 B5XZP2 B5XZP5
B5XZS9 B5Y1F0 B5Y1J4 B5Y1L4 B5Y1V0 B5Y1Z8 B5Y234 B5Y280 B5Y298 B5Y348 B5Y349
B5YBQ1 B5YBQ2 B5YD74 B5YDL5 B5YEL6 B5YFT8 B5YGC8 B5YI20 B5YID8 B5YIG8 B5YIG9
B5YJT2 B5YKI9 B5YKJ2 B5YKM8 B5YL23 B5YL84 B5YPT7 B5YPX2 B5YPZ9 B5YQN8 B5YQN9
B5YQP0 B5YQS6 B5YQW1 B5YQW3 B5YQW8 B5YQX2 B5YR48 B5YR85 B5YR97 B5YR99 B5YS06
B5YSD8 B5YSJ8 B5YSS1 B5YSU1 B5YSV1 B5YSW6 B5YSW7 B5YT10 B5YT34 B5YTS2 B5YTX6
B5YUD3 B5YUD4 B5YUD5 B5YVH5 B5YVR9 B5YVT5 B5YW73 B5YWI0 B5YWY8 B5YX45 B5YXA9
B5YXB4 B5YXE0 B5YXE1 B5YXE2 B5YXL0 B5YXL5 B5YXQ0 B5YXQ1 B5YXQ2 B5YXR6 B5YXR9
B5YXS2 B5YXT5 B5YY05 B5YY42 B5YY84 B5YYB9 B5YYD4 B5YYF4 B5YYG0 B5YZ23 B5YZ47
B5YZ52 B5YZ56 B5YZ83 B5YZC3 B5YZU3 B5YZU5 B5YZV9 B5YZW0 B5Z009 B5Z0D5 B5Z0F6
B5Z0P2 B5Z0R2 B5Z0Y5 B5Z171 B5Z177 B5Z181 B5Z1C8 B5Z1Y4 B5Z2G2 B5Z2G3 B5Z364
B5Z3B0 B5Z428 B5Z435 B5Z436 B5Z461 B5Z464 B5Z466 B5Z497 B5Z4E5 B5Z4F3 B5Z4F4
B5Z4P2 B5Z4P6 B5Z4Q8 B5Z7J3 B5Z7V8 B5Z8D4 B5Z8H9 B5Z8K9 B5Z8M0 B5Z8R2 B5Z8R5
B5Z9I3 B5Z9W0 B5ZAQ4 B5ZAW9 B5ZBQ1 B5ZMF7 B5ZMX7 B5ZPK8 B5ZQP2 B5ZRH1 B5ZS16
B5ZS18 B5ZS19 B5ZSC5 B5ZSV2 B5ZSV4 B5ZTH0 B5ZU97 B5ZV34 B5ZVF6 B5ZW93 B5ZWJ7
B5ZXG4 B5ZXY9 B5ZY45 B5ZYG2 B5ZYM1 B5ZYM4 B6A8C7 B6A8R8 B6CZ46 B6CZ56 B6EGH2
B6EGH3 B6EGH4 B6EGH7 B6EGL4 B6EGQ7 B6EGU9 B6EH91 B6EHA6 B6EHA7 B6EHQ0 B6EHQ3
B6EHU1 B6EHU2 B6EHU3 B6EI58 B6EIC7 B6EIG5 B6EJ35 B6EJA1 B6EJQ7 B6EKC3 B6EKL7
B6ELH6 B6ELH8 B6EM11 B6EM14 B6EMS0 B6EMS1 B6EMX8 B6ENU2 B6EP40 B6EQE1 B6ETT4
B6GX67 B6H059 B6H065 B6H066 B6H067 B6H1I3 B6H1W0 B6H457 B6H465 B6H711 B6H7A4
B6HC89 B6HDM7 B6HFS8 B6HFX9 B6HIC2 B6HJA3 B6HJU0 B6HJU3 B6HL48 B6HN76 B6HN82
B6HNK5 B6HQ08 B6HR25 B6HR44 B6HS10 B6HTR9 B6HUQ4 B6HV31 B6HV35 B6HV36 B6HV37
B6HYQ4 B6HYQ5 B6HYQ6 B6HYS0 B6HYS1 B6HYS2 B6HYZ1 B6HZ22 B6HZ28 B6HZ64 B6HZD1
B6HZF1 B6I0P3 B6I117 B6I134 B6I195 B6I1T1 B6I1U2 B6I1V8 B6I1V9 B6I204 B6I260
B6I261 B6I2P2 B6I2P6 B6I2Q8 B6I2Q9 B6I2S3 B6I2Y7 B6I356 B6I394 B6I3R8 B6I3T8
B6I3U3 B6I3X3 B6I3X4 B6I3X5 B6I3Y4 B6I411 B6I428 B6I430 B6I491 B6I4D1 B6I4H7

B6I4M3 B6I4Q0 B6I4Q7 B6I4R1 B6I4Y9 B6I4Z0 B6I540 B6I588 B6I5P4 B6I5Q0 B6I5Q5
B6I5T5 B6I6D8 B6I6M3 B6I6U0 B6I6U2 B6I6W0 B6I6W1 B6I7F4 B6I7J7 B6I7K0 B6I7K1
B6I7K2 B6I7L7 B6I7M6 B6I7M9 B6I7N2 B6I829 B6I8G8 B6I8J1 B6I8R6 B6I8X0 B6I9C4
B6I9D0 B6I9E2 B6I9P7 B6I9W2 B6I9W8 B6I9W9 B6IA37 B6IA59 B6IAT1 B6IB16 B6IB33
B6IB34 B6IB62 B6IB68 B6IB70 B6IBA3 B6IBL2 B6IBP7 B6IBP8 B6IBQ3 B6IBQ7 B6ID01
B6IJ52 B6INE1 B6IQS5 B6IQX6 B6IQY7 B6IQY9 B6IRC8 B6IRG5 B6ISW8 B6ISX1 B6ISX4
B6ITJ7 B6IUZ7 B6IVR1 B6IWH9 B6IWJ6 B6IX46 B6IX47 B6IYW0 B6IZ52 B6IZ55 B6IZN9
B6J085 B6J0F8 B6J0H4 B6J2B4 B6J2D4 B6J2D6 B6J2Q9 B6J375 B6J454 B6J4N9 B6J517
B6J5K9 B6J612 B6J6A2 B6J6A5 B6J6V9 B6J6X5 B6J723 B6J736 B6J7P8 B6J8U7 B6J957
B6J959 B6JA60 B6JAN9 B6JB57 B6JBL9 B6JCF5 B6JCL3 B6JDB7 B6JDC7 B6JDC8 B6JDD0
B6JFP6 B6JGA5 B6JKP6 B6JL77 B6JM59 B6JMH5 B6JMX6 B6JN25 B6JN51 B6JN65 B6JNA7
B6JNB0 B6JP50 B6JPH7 B6JPT9 B6JWP7 B6JY29 B6JYT1 B6JZY9 B6K0N7 B6K2Z6 B6K327
B6K463 B6K6N3 B6K7U7 B6K9M7 B6KAM0 B6KEU8 B6KLP1 B6KV60 B6Q234 B6Q5G0 B6Q656
B6Q6B6 B6QEE0 B6QG32 B6QHK6 B6QHL8 B6QHL9 B6QIB3 B6QIM3 B6QMS8 B6QNP4 B6QQZ9
B6QVB2 B6QVW4 B6RAL1 B6SCF4 B6SEH8 B6SEH9 B6SGC5 B6SJQ0 B6SR79 B6SSW8 B6SZA7
B6SZU6 B6T959 B6T990 B6TAX2 B6TGJ8 B6TM88 B6TUB4 B6TUH4 B6TUW9 B6TVL4 B6TWJ1
B6TYV8 B6TZ45 B6U045 B6U300 B6U361 B6U769 B6U8R7 B6UBY6 B6VH76 B6VH77 B6VH79
B6VMU9 B6VQA5 B6YQ48 B6YQF9 B6YR08 B6YR09 B6YR61 B6YRY9 B6YS29 B6YSX5 B6YUC9
B6YV94 B6YVW0 B6YW71 B6YW76 B6ZJZ9 B6ZK76 B6ZK77 B7F138 B7F9G7 B7FA90 B7FXW1
B7GCG7 B7GF17 B7GFB0 B7GGI5 B7GGT5 B7GI41 B7GIK7 B7GJC8 B7GLN3 B7GLU2 B7GLW3
B7GME1 B7GME4 B7GME7 B7GMF8 B7GMF9 B7GPC7 B7GQ27 B7GR11 B7GRM7 B7GTW5 B7GTY8
B7GV24 B7GVI6 B7GWU1 B7GXE2 B7GXW9 B7GY42 B7GZB2 B7H0A3 B7H0K1 B7H0P1 B7H0W5
B7H133 B7H298 B7H299 B7H2A0 B7H305 B7H3P4 B7H502 B7H698 B7H6M9 B7H6P9 B7H6T1
B7H6T3 B7H705 B7H732 B7H7Y7 B7H9T4 B7HCE7 B7HCF8 B7HDF8 B7HDF9 B7HDG0 B7HEG3
B7HEV2 B7HFA9 B7HFI9 B7HFJ2 B7HFK6 B7HFK7 B7HFM2 B7HGA1 B7HGA2 B7HGR1 B7HKL5
B7HKP5 B7HLM7 B7HM34 B7HMC9 B7HMD0 B7HQ38 B7HRN5 B7HSG9 B7HSI6 B7HSK8 B7HU73
B7HWG0 B7HWG1 B7HWG2 B7HWY8 B7HXM3 B7HY28 B7HY51 B7HY54 B7HY69 B7HY70 B7HY84
B7HZC3 B7HZC4 B7HZS6 B7I1V8 B7I1V9 B7I1W0 B7I295 B7I4B4 B7I4U3 B7I4W9 B7I4Z5
B7I612 B7I748 B7I751 B7I7L7 B7I868 B7I8R1 B7IAW6 B7IB69 B7IBC0 B7IE18 B7IFP2
B7IFS1 B7IHD7 B7II08 B7II09 B7II10 B7IIB7 B7IIF0 B7IJZ3 B7IK49 B7IKT7 B7IL96
B7IPN4 B7IPS6 B7IQU6 B7IQU9 B7IQW2 B7IQW3 B7IQX7 B7IRQ1 B7IRX1 B7IRX2 B7ISM2
B7IUQ3 B7IUS3 B7IV78 B7IVI2 B7IVI3 B7IWR6 B7J0N5 B7J1I6 B7J1N1 B7J1T5 B7J208
B7J235 B7J3V5 B7J5J3 B7J5W3 B7J7T2 B7J7T5 B7J9Q0 B7JB88 B7JB89 B7JBL0 B7JDV6
B7JG51 B7JGI9 B7JGL8 B7JGM1 B7JGN4 B7JGN5 B7JGQ0 B7JIB5 B7JIF5 B7JIF6 B7JJY0
B7JK01 B7JK33 B7JK34 B7JMH1 B7JNH3 B7JQN4 B7JRB7 B7JRB8 B7JRB9 B7JRW8 B7JS25
B7JS48 B7JSD7 B7JUB6 B7JUD9 B7JUX4 B7JV08 B7JVF4 B7JVQ3 B7JVR2 B7JW07 B7JWT1
B7JX63 B7JX92 B7JY03 B7JYE8 B7JYG7 B7K085 B7K0U5 B7K1C1 B7K1G0 B7K1G1 B7K1G4
B7K1J7 B7K1L1 B7K2L3 B7K2V6 B7K336 B7K3S3 B7K3U2 B7K5I4 B7K5I5 B7K5Y2 B7K623
B7K624 B7K625 B7K626 B7K6Q5 B7K767 B7K968 B7K989 B7KA41 B7KAA7 B7KC47 B7KCV9
B7KDB2 B7KEZ9 B7KFG8 B7KFH4 B7KFK1 B7KFR0 B7KG11 B7KGD8 B7KH59 B7KH60 B7KH61
B7KH62 B7KHH8 B7KHH9 B7KHU9 B7KHV5 B7KHX2 B7KIS7 B7KIT4 B7KJP1 B7KK61 B7KKR7
B7KKR8 B7KL15 B7KPQ5 B7KTL8 B7KU72 B7KV80 B7KVJ7 B7KWA0 B7KX02 B7L1M9 B7L1N2
B7L2I5 B7L4E9 B7L4G3 B7L4H0 B7L4M7 B7L4N9 B7L4V0 B7L587 B7L5A9 B7L5D4 B7L5E4
B7L5F1 B7L5F2 B7L5I0 B7L5I3 B7L5I5 B7L5L6 B7L5X4 B7L612 B7L6I4 B7L6R4 B7L6U8

B7L6V0 B7L6V5 B7L7N1 B7L824 B7L849 B7L855 B7L883 B7L888 B7L897 B7L8H4 B7L8K7
B7L8Y9 B7L917 B7L952 B7L998 B7L9D9 B7L9F9 B7L9G9 B7L9U8 B7L9U9 B7L9V0 B7L9Z4
B7LA07 B7LA12 B7LA15 B7LAA3 B7LAA4 B7LAA5 B7LAM0 B7LAR9 B7LAS2 B7LAS3 B7LAS4
B7LAT8 B7LAU1 B7LAU4 B7LAX3 B7LAY3 B7LAY8 B7LB16 B7LBE9 B7LC12 B7LC13 B7LCE1
B7LCE3 B7LCF7 B7LCF8 B7LCK8 B7LD40 B7LD90 B7LDA8 B7LE10 B7LEG6 B7LF05 B7LF12
B7LF13 B7LFF7 B7LFG4 B7LFV7 B7LFZ9 B7LGE9 B7LGI2 B7LGL7 B7LGN8 B7LGU6 B7LGV1
B7LGZ1 B7LGZ5 B7LH53 B7LHJ4 B7LHK0 B7LHK1 B7LHS1 B7LHS8 B7LHU8 B7LHU9 B7LHV0
B7LHZ2 B7LJT8 B7LK48 B7LK52 B7LK81 B7LK82 B7LK83 B7LK92 B7LKC2 B7LKG9 B7LKR6
B7LKR7 B7LKR8 B7LL03 B7LL09 B7LL12 B7LL71 B7LL85 B7LL87 B7LLI4 B7LLI6 B7LLK9
B7LLT5 B7LLT6 B7LM37 B7LM50 B7LM53 B7LM72 B7LM73 B7LM74 B7LM77 B7LMN9 B7LMW8
B7LMY4 B7LN13 B7LN63 B7LNI3 B7LNJ1 B7LNJ2 B7LNW7 B7LPL6 B7LPM0 B7LPM5 B7LPM7
B7LPX2 B7LQ78 B7LQA4 B7LQB7 B7LQD5 B7LQD7 B7LQI2 B7LQN8 B7LQP0 B7LQP3 B7LRA1
B7LRC4 B7LRJ2 B7LRL5 B7LRL6 B7LRL7 B7LRM9 B7LRQ7 B7LRV3 B7LRX6 B7LS23 B7LS24
B7LS29 B7LS57 B7LSC6 B7LSJ3 B7LSJ8 B7LSP0 B7LSX5 B7LT01 B7LT82 B7LTZ9 B7LU62
B7LV39 B7LV40 B7LV41 B7LV89 B7LVA1 B7LVD0 B7LVD6 B7LVG3 B7LVN3 B7LVP2 B7LVR8
B7LVT8 B7LW74 B7LWB6 B7LWF0 B7LWK7 B7LWN1 B7LXA0 B7LXA5 B7LXG0 B7LXS2 B7LY04
B7LY10 B7LY11 B7LY84 B7LY91 B7LYI8 B7LYL2 B7LZD1 B7LZF6 B7LZI9 B7LZK7 B7LZK9
B7LZX4 B7LZY4 B7LZZ1 B7LZZ2 B7M023 B7M0C3 B7M0D7 B7M0E4 B7M0I4 B7M0I7 B7M0I9
B7M0M0 B7M0S6 B7M0T6 B7M0V2 B7M0V3 B7M0V4 B7M0Z6 B7M130 B7M196 B7M1C0 B7M1K3
B7M1Q2 B7M1R7 B7M1Y0 B7M294 B7M296 B7M2A1 B7M2A5 B7M2I4 B7M2Q4 B7M3B0 B7M3C8
B7M3E0 B7M458 B7M459 B7M460 B7M4H4 B7M4S3 B7M4X3 B7M4Y6 B7M547 B7M558 B7M562
B7M593 B7M594 B7M5A3 B7M5E1 B7M5F2 B7M5L2 B7M5L3 B7M5L4 B7M5T6 B7M5T9 B7M5U0
B7M5U1 B7M5V5 B7M5V8 B7M5W1 B7M5X4 B7M640 B7M685 B7M6Q9 B7M6R1 B7M6S2 B7M6S3
B7M6V9 B7M6W1 B7M6W6 B7M6W9 B7M7C2 B7M7H3 B7M7M0 B7M7U2 B7M7U8 B7M7V3 B7M7Y0
B7M826 B7M8R6 B7M8R7 B7M921 B7M928 B7M940 B7M9T5 B7M9V2 B7M9V3 B7M9Y1 B7M9Y4
B7M9Y6 B7MA17 B7MA67 B7MAB7 B7MAD4 B7MAE7 B7MAG3 B7MAH0 B7MAL0 B7MAS2 B7MAZ5
B7MB46 B7MBD8 B7MBF8 B7MBJ8 B7MBM6 B7MBM7 B7MBN2 B7MBN6 B7MBX7 B7MBY6 B7MC01
B7MC02 B7MC03 B7MCM5 B7MCQ6 B7MCW6 B7MCY1 B7MDB8 B7MDQ0 B7ME87 B7ME88 B7ME89
B7MEI9 B7MEN1 B7MF03 B7MF40 B7MF53 B7MFA9 B7MFW1 B7MFW2 B7MFW3 B7MG21 B7MG24
B7MG25 B7MG26 B7MG43 B7MG46 B7MG59 B7MGA5 B7MGC7 B7MGD1 B7MGF7 B7MGF8 B7MGG7
B7MGV4 B7MH49 B7MH51 B7MH58 B7MHC2 B7MHL0 B7MHR6 B7MHR7 B7MHW5 B7MI01 B7MI17
B7MI40 B7MI42 B7MI47 B7MI50 B7MII7 B7MIJ4 B7MIK6 B7MJ22 B7MJ27 B7MJ30 B7MJT5
B7MK83 B7MK88 B7MKM2 B7MKV8 B7MKV9 B7ML01 B7ML07 B7ML08 B7MLH4 B7MLI1 B7MLI2
B7MLZ8 B7MM18 B7MMZ1 B7MNB1 B7MNN2 B7MNP7 B7MNQ4 B7MNU6 B7MP17 B7MP37 B7MPJ9
B7MPK0 B7MPK1 B7MQV0 B7MR15 B7MRH2 B7MRM9 B7MRR2 B7MS16 B7MSH3 B7MSH4 B7MSH6
B7MTH8 B7MTI5 B7MTJ6 B7MTW4 B7MTW9 B7MU36 B7MU42 B7MU43 B7MUG2 B7MUI4 B7MUT7
B7MV12 B7MV28 B7MV39 B7MV43 B7MV45 B7MV46 B7MVA5 B7MVB0 B7MVJ0 B7MVK3 B7MVS1
B7MVV0 B7MVV5 B7MVV9 B7MW36 B7MWP9 B7MWY8 B7MWY9 B7MWZ0 B7MXF3 B7MXH4 B7MXM7
B7MXQ1 B7MXT5 B7MXT9 B7MXU0 B7MXU1 B7MXV5 B7MXV8 B7MXW1 B7MY49 B7MY66 B7MYB2
B7MYF1 B7MZ49 B7MZC7 B7MZD4 B7MZZ0 B7N061 B7N0B7 B7N0I9 B7N0K9 B7N0M6 B7N0T1
B7N112 B7N114 B7N156 B7N1D2 B7N1E7 B7N1S8 B7N1W8 B7N210 B7N241 B7N2D1 B7N2E3
B7N2F5 B7N2H6 B7N2I6 B7N2M7 B7N2N9 B7N2P4 B7N2P9 B7N2Q6 B7N2Q9 B7N364 B7N380
B7N3Z0 B7N3Z7 B7N460 B7N467 B7N468 B7N4C9 B7N4F0 B7N4V3 B7N549 B7N5F8 B7N5L9
B7N5M2 B7N5M3 B7N5M4 B7N5P1 B7N5P4 B7N5Q7 B7N5Z0 B7N5Z2 B7N604 B7N654 B7N6A4

B7N6X8 B7N760 B7N767 B7N768 B7N7Q1 B7N7R5 B7N7S2 B7N7W0 B7N824 B7N844 B7N9J8
B7N9N0 B7N9T9 B7N9U0 B7N9U1 B7NAJ3 B7NAP9 B7NAS2 B7NAS9 B7NAU0 B7NB32 B7NB42
B7NB49 B7NB81 B7NB84 B7NB86 B7NBB6 B7NBD4 B7NBG6 B7NBG8 B7NBH3 B7NBH7 B7NBW3
B7NC26 B7NCB1 B7NCB2 B7NCB3 B7NCH0 B7NCZ3 B7ND33 B7ND46 B7ND48 B7NDA4 B7NDJ2
B7NDL8 B7NDL9 B7NDM0 B7NDR2 B7NDV9 B7NDX4 B7NE28 B7NEC5 B7NEH6 B7NF02 B7NF23
B7NF27 B7NF53 B7NF54 B7NF63 B7NF99 B7NFD8 B7NFI5 B7NFK5 B7NFL2 B7NFL5 B7NFX5
B7NFY0 B7NFY4 B7NG10 B7NG90 B7NG91 B7NH28 B7NHD1 B7NHE4 B7NHF1 B7NHJ3 B7NHM5
B7NHP3 B7NIB8 B7NID9 B7NIN3 B7NJ05 B7NJF4 B7NJQ3 B7NJS0 B7NJS2 B7NJY3 B7NKS5
B7NKT5 B7NL69 B7NL82 B7NL87 B7NLF8 B7NLF9 B7NLG0 B7NLL0 B7NLV2 B7NLY4 B7NM77
B7NMA6 B7NMD3 B7NMI8 B7NMP9 B7NMQ0 B7NMQ1 B7NN09 B7NNB6 B7NNG2 B7NNT0 B7NNT3
B7NNT6 B7NNT7 B7NNT8 B7NNV5 B7NNV8 B7NNW1 B7NNX4 B7NPI5 B7NPS5 B7NPS9 B7NPU9
B7NQA9 B7NQB0 B7NQB1 B7NQL0 B7NQY5 B7NR08 B7NR12 B7NR39 B7NR40 B7NR50 B7NR82
B7NRB0 B7NRG6 B7NRZ5 B7NS00 B7NS04 B7NS90 B7NSA0 B7NSA1 B7NSM7 B7NSY7 B7NT65
B7NT92 B7NTE8 B7NTL2 B7NTL3 B7NTX0 B7NU01 B7NU03 B7NU06 B7NU89 B7NU92 B7NU97
B7NU99 B7NUP0 B7NUP7 B7NUQ7 B7NUV2 B7NUY5 B7NV31 B7NVC9 B7NVL8 B7NVM4 B7NVM5
B7NVX3 B7NVY1 B7NVY2 B7QIN3 B7SB91 B7SB92 B7T1R8 B7T1R9 B7T1V0 B7T1W7 B7TWW5
B7U540 B7UFE4 B7UFF7 B7UFL0 B7UFR6 B7UFR9 B7UFS0 B7UFS1 B7UFT4 B7UFT7 B7UFU0
B7UFV3 B7UG98 B7UGA0 B7UGB3 B7UGB4 B7UGM2 B7UGW2 B7UHG7 B7UHP5 B7UHQ3 B7UHQ4
B7UI72 B7UI86 B7UI93 B7UID7 B7UIK2 B7UIM2 B7UIQ5 B7UIV3 B7UIW5 B7UIW7 B7UJ20
B7UJU8 B7UJV7 B7UJX2 B7UJX3 B7UJX4 B7UK14 B7UK61 B7UK75 B7UKC8 B7UKN4 B7UKR6
B7UKX5 B7UKX6 B7UKX7 B7UL42 B7UL81 B7UM99 B7UMD5 B7UMF2 B7UMH2 B7UMH7 B7UMK2
B7UMK3 B7UML2 B7UMT9 B7UMY7 B7UNB2 B7UNG5 B7UNK3 B7UNM8 B7UNN0 B7UNN5 B7UNN8
B7UP62 B7UP67 B7UP79 B7UPJ2 B7UPJ7 B7UPK1 B7UPN5 B7UPX3 B7UPX4 B7UQ70 B7UQ75
B7UQD9 B7UQE5 B7UQE6 B7UQY8 B7UQZ2 B7UR04 B7URD0 B7URE7 B7URQ4 B7URS1 B7URT1
B7URT9 B7URU0 B7URW8 B7URX1 B7URX3 B7US04 B7US50 B7USG9 B7USJ6 B7USJ7 B7USK2
B7USK6 B7USY2 B7UT21 B7UTB3 B7UTB4 B7UTB5 B7UUZ6 B7UV86 B7UVJ4 B7UVW8 B7UVW9
B7UVX0 B7UW72 B7UWJ4 B7UXY2 B7UY17 B7UY90 B7UZG1 B7UZJ3 B7UZT8 B7UZU0 B7UZU1
B7V0B7 B7V0F9 B7V1A3 B7V2Q4 B7V3F8 B7V3G1 B7V3H0 B7V4G7 B7V5P9 B7V5T6 B7V6N7
B7V6V8 B7V719 B7V796 B7V797 B7V7A5 B7V7I2 B7V8H0 B7V9D2 B7VA62 B7VAB2 B7VAE6
B7VAQ6 B7VAQ9 B7VB57 B7VBM8 B7VBM9 B7VBN0 B7VBN3 B7VGH7 B7VH11 B7VHB5 B7VHC3
B7VHC4 B7VHE7 B7VHR5 B7VIK5 B7VIV7 B7VIW0 B7VJ00 B7VJ77 B7VK67 B7VKD5 B7VKD6
B7VL36 B7VLT3 B7VLT4 B7VLT5 B7VLT9 B7VLV8 B7VLW5 B7VMA0 B7VMM4 B7VPS1 B7VQF8
B7WFQ1 B7XIC4 B7YZT2 B7YZU2 B7Z031 B7Z0K8 B7Z0W9 B7Z8K6 B7ZAQ6 B7ZCC9 B7ZSK1
B7ZWI3 B7ZWR6 B7ZWR7 B8A4F0 B8A7Q8 B8A7Z5 B8A833 B8A927 B8A9X1 B8AAM2 B8AAV3
B8AC36 B8ACH9 B8AD72 B8AEC9 B8AF63 B8AFI3 B8AFI8 B8AFK5 B8AIW3 B8AIZ4 B8AJ09
B8AJL3 B8AJT9 B8ALI0 B8ANR3 B8APK3 B8ART0 B8ARW3 B8AT51 B8AV52 B8AY75 B8AYH1
B8AYM8 B8AYU8 B8B183 B8B2D0 B8B350 B8B4W4 B8B6G5 B8B6I2 B8B7Q4 B8BB68 B8BD96
B8BDK4 B8BDK8 B8BHF1 B8BIM2 B8BJ22 B8BKP4 B8BMY7 B8BPI2 B8CH68 B8CI02 B8CI03
B8CI04 B8CJ38 B8CJF5 B8CKC7 B8CLL1 B8CM53 B8CM54 B8CM55 B8CM58 B8CNG1 B8CNK8
B8CP83 B8CPD7 B8CQX8 B8CSC0 B8CTT5 B8CVV0 B8CVV1 B8CWG2 B8CWL9 B8D065 B8D0I7
B8D6S2 B8D6T3 B8D6X4 B8D718 B8D721 B8D723 B8D753 B8D765 B8D768 B8D7C2 B8D7H0
B8D7H2 B8D7H3 B8D7L3 B8D7W9 B8D7Z7 B8D860 B8D8G8 B8D8H9 B8D8M0 B8D8R4 B8D8R7
B8D8R9 B8D8U9 B8D8W1 B8D8W4 B8D917 B8D966 B8D968 B8D969 B8D9B1 B8D9L7 B8D9P5
B8D9V8 B8DAL8 B8DAW0 B8DAW1 B8DAW2 B8DBK8 B8DBM9 B8DC84 B8DDQ9 B8DDY1 B8DE85

B8DEH2 B8DF46 B8DF86 B8DG21 B8DG47 B8DGC6 B8DGI9 B8DGM3 B8DH65 B8DH71 B8DH72

B8DHE7 B8DHI0 B8DJU6 B8DNP8 B8DP11 B8DQF5 B8DR36 B8DR67 B8DRQ3 B8DSU2 B8DUF9

B8DWS7 B8DYT4 B8DZW4 B8E160 B8E2N2 B8E2N3 B8E328 B8E4B5 B8E548 B8E551 B8E552

B8E553 B8E613 B8E695 B8E6B2 B8E6B3 B8E6B4 B8E719 B8E7D2 B8E7U7 B8EAH3 B8EB95

B8EBH7 B8EBR4 B8EBV1 B8EBV4 B8EC46 B8ECI1 B8EDV5 B8EDV6 B8EDW5 B8EE41 B8EEF1

B8EET6 B8EFC6 B8EIM7 B8EJ45 B8EJK7 B8EJZ7 B8EK66 B8ELH2 B8ENH3 B8EPG5 B8EPK3

B8ETA0 B8ETD6 B8ETL9 B8F313 B8F381 B8F3B3 B8F3C3 B8F3F2 B8F486 B8F4W0 B8F543

B8F5U9 B8F6K1 B8F6K4 B8F6M1 B8F6M2 B8F6X1 B8F768 B8F769 B8F792 B8F7B0 B8F7B3

B8F7B7 B8F7T0 B8F8J1 B8F8N6 B8FAZ7 B8FBS1 B8FG65 B8FPI3 B8FRJ7 B8FRK0 B8FT17

B8FT59 B8FTI4 B8FX50 B8FYJ9 B8G014 B8G4Q6 B8G5X7 B8G5Y6 B8G5Z3 B8G6C2 B8G6H1

B8G6L3 B8G6Z6 B8GEB7 B8GEE2 B8GER0 B8GJG9 B8GLM2 B8GMB4 B8GMB9 B8GMM6 B8GMM8

B8GNZ5 B8GNZ8 B8GPF3 B8GPT8 B8GQ76 B8GRD1 B8GT83 B8GTM8 B8GTV4 B8GUR7 B8GVF8

B8GVG0 B8GWS7 B8GWU1 B8GX30 B8GX35 B8GX56 B8GX57 B8GX61 B8GX72 B8GXA0 B8GXB6

B8GXM1 B8GYX9 B8GZK4 B8GZV6 B8H051 B8H082 B8H092 B8H097 B8H0A0 B8H0N0 B8H0S5

B8H1E8 B8H1L5 B8H444 B8H551 B8H6A4 B8H6C3 B8H7N5 B8H8K7 B8HAZ5 B8HDM6 B8HGW7

B8HLB2 B8HMH3 B8HNR1 B8HPJ7 B8HQP5 B8HR15 B8HRF0 B8HRF1 B8HSK6 B8HTR1 B8HUA4

B8HUQ7 B8HUV5 B8HVD7 B8HVD8 B8HVD9 B8HVK3 B8HWC6 B8HWC7 B8HWF5 B8HWU5 B8HXG1

B8HY04 B8HYC0 B8I004 B8I0P7 B8I1G8 B8I378 B8I4B9 B8I574 B8I813 B8I9I5 B8I9I8

B8I9Q3 B8I9T7 B8IB41 B8IDA3 B8IHT8 B8IMM9 B8IMW7 B8ISP2 B8IUV6 B8IUV9 B8IUY7

B8IZT8 B8IZV2 B8J0P3 B8J1C7 B8J1D0 B8J1E9 B8J1I9 B8J229 B8J4L9 B8J670 B8J6V0

B8J708 B8J8E5 B8J8T6 B8J992 B8JAZ0 B8JBQ8 B8JBR1 B8JCU2 B8JDK5 B8JF01 B8JJV5

B8JK39 B8JKP6 B8JLV7 B8K1V7 B8K1W2 B8LQF9 B8LUE3 B8LWT1 B8LYF6 B8M5K5 B8M719

B8M853 B8M9J6 B8M9J8 B8MBN3 B8MD74 B8MG72 B8MJJ2 B8MJK3 B8MK20 B8MKT9 B8MTH6

B8MTT8 B8MY73 B8MYS7 B8MYS8 B8N076 B8N0E9 B8N0F1 B8N0Q7 B8N2C8 B8N6H2 B8N8R3

B8N9M0 B8NA66 B8NDR8 B8NDS8 B8NEM4 B8NFL5 B8NHD8 B8NHD9 B8NI03 B8NI21 B8NI27

B8NIM7 B8NJG7 B8NJG8 B8NJH0 B8NJL4 B8NM08 B8NM62 B8NM67 B8NM74 B8NM75 B8NNK9

B8NSP6 B8NTP7 B8NU01 B8NU02 B8NU03 B8NUK6 B8NWW3 B8NWW7 B8NX76 B8NYW8 B8P366

B8PL20 B8Q0B2 B8QHP1 B8QHP3 B8QHP5 B8R1K0 B8R1V5 B8RJM0 B8UU59 B8UU78 B8V7Q1

B8VIW9 B8XCH5 B8XH50 B8XX90 B8ZHC9 B8ZJU7 B8ZKX1 B8ZL54 B8ZLB4 B8ZLB5 B8ZLV4

B8ZLW2 B8ZNG7 B8ZNI6 B8ZNQ1 B8ZQP7 B8ZQQ2 B8ZR37 B8ZR39 B8ZR76 B8ZRC2 B8ZRD4

B8ZRJ4 B8ZRL0 B8ZS82 B8ZSY8 B8ZTP9 B8ZU13 B8ZUU1 B8ZYW3 B8ZZ34 B9A1Q4 B9DFI7

B9DFK5 B9DFU2 B9DFX7 B9DGY1 B9DHD7 B9DHG0 B9DIS4 B9DJL7 B9DJQ6 B9DK59 B9DLR8

B9DME9 B9DMF0 B9DML4 B9DMS4 B9DMY6 B9DP57 B9DP73 B9DP78 B9DPR2 B9DPV8 B9DPV9

B9DRJ4 B9DRR4 B9DTI6 B9DTP5 B9DTR8 B9DV75 B9E0Q4 B9E0W0 B9E4L8 B9E675 B9E8F1

B9E9M1 B9EA92 B9EB26 B9EB27 B9EB52 B9EBV2 B9EBV6 B9EJ80 B9EJ86 B9EJG8 B9EJI9

B9EJV3 B9EKR1 B9EKX1 B9EN89 B9EPI1 B9ES20 B9EXZ6 B9F6Z0 B9FCV3 B9FDT1 B9FGV7

B9FJ61 B9FKP6 B9FMS2 B9FMX4 B9G125 B9G2A8 B9G2E6 B9G300 B9G4M9 B9G5Y5 B9G934

B9GBJ9 B9GDE5 B9GFG6 B9GGL4 B9GHX8 B9GIE4 B9GXD6 B9H2V1 B9HBX2 B9HD38 B9HI87

B9HJR7 B9HMF8 B9HMP5 B9HMP6 B9HQ42 B9HTK2 B9HTL5 B9HVC2 B9I0G0 B9I0U9 B9I3X5

B9I534 B9IFI5 B9IH36 B9IIR4 B9IIR5 B9IK21 B9IM09 B9IQY6 B9IR99 B9IRS5 B9IRS8

B9IRU2 B9IRV7 B9ISM3 B9ISZ3 B9ISZ4 B9ITG0 B9IUJ0 B9IUK2 B9IVX0 B9IVZ0 B9IW24

B9IW25 B9IZP6 B9J095 B9J153 B9J176 B9J1Z7 B9J3N9 B9J4R6 B9J529 B9J8B4 B9J924

B9JA25 B9JA28 B9JAH9 B9JAP1 B9JBL3 B9JD11 B9JD51 B9JEG8 B9JEV2 B9JFT7 B9JG16

B9JH54 B9JKQ7 B9JKQ9 B9JSF9 B9JU45 B9JUJ0 B9JVB6 B9JVF3 B9JVI2 B9JW08 B9JWI7

B9JWY6 B9JXE1 B9JY57 B9JZ26 B9JZH6 B9JZJ6 B9JZK8 B9K2I1 B9K2I3 B9K2V4 B9K6P4
B9KCT5 B9KCX1 B9KD84 B9KDD4 B9KDV9 B9KEQ5 B9KEZ5 B9KF57 B9KG48 B9KGP6 B9KGU3
B9KH29 B9KH48 B9KH94 B9KHM9 B9KI88 B9KIT6 B9KLG9 B9KLZ5 B9KNF0 B9KNJ3 B9KNZ6
B9KPZ0 B9KQX4 B9KS10 B9KSX9 B9KW13 B9KXV3 B9KYE6 B9L173 B9L175 B9L178 B9L3S8
B9L7I2 B9L7V6 B9L8U9 B9LAV0 B9LBM5 B9LGS9 B9LK05 B9LKJ6 B9LRL5 B9LSA9 B9LSC9
B9LV21 B9LYY3 B9LZL2 B9LZL3 B9LZM7 B9LZU3 B9M0D1 B9M176 B9M1A8 B9M3Y9 B9M7R9
B9M837 B9M9E4 B9M9U5 B9MAB3 B9MB97 B9MB98 B9MBT5 B9MDW9 B9MDX6 B9ME24 B9MF14
B9MFI6 B9MFR5 B9MFV5 B9MH59 B9MIW0 B9MKR9 B9MPK5 B9MQ98 B9MS17 B9MS73 B9N2D0
B9N3F4 B9N5U6 B9NBE5 B9RA90 B9RH17 B9RK42 B9RQG7 B9RT03 B9RT04 B9RW00 B9RZ92
B9S2H4 B9S8Z3 B9SA89 B9SCX0 B9SLR1 B9SR15 B9SV63 B9SV84 B9SX12 B9SX13 B9SXY8
B9T3K6 B9T4E6 B9TRX0 B9U3F2 B9UIY3 B9VR26 B9VXD7 B9W5G6 B9W824 B9W8T5 B9W8Z2
B9W9N8 B9W9Z5 B9WA88 B9WAT8 B9WB34 B9WBE5 B9WCP3 B9WCV6 B9WD58 B9WE91 B9WFC9
B9WHT6 B9WJF2 B9WJJ7 B9WK46 B9WKT4 B9WL59 B9WLF1 B9WZX1 B9WZX4 B9WZX6 B9X187
B9X287 C0H3P8 C0H3Q1 C0H3Q4 C0H3Q7 C0H3T3 C0H3T7 C0H3T8 C0H3U0 C0H3U2 C0H3V0
C0H3V1 C0H3V4 C0H3V7 C0H3W0 C0H3X3 C0H3X6 C0H3X7 C0H3Y2 C0H3Y3 C0H3Y6 C0H3Y8
C0H409 C0H415 C0H417 C0H420 C0H423 C0H431 C0H433 C0H435 C0H438 C0H439 C0H451
C0H453 C0H463 C0H464 C0H465 C0H5F4 C0H9B6 C0HBB2 C0HJH3 C0HJH4 C0HJJ0 C0HJX2
C0HJX3 C0HK57 C0HK58 C0HK59 C0HK61 C0HK65 C0HK66 C0HK68 C0HK79 C0HK80 C0HKD1
C0HKD2 C0HKD3 C0HKD7 C0HKG7 C0HKG8 C0HKG9 C0HL12 C0HL13 C0HLB3 C0HLB4 C0HLB5
C0HLB6 C0HLM6 C0HLN0 C0HM28 C0J7L8 C0LGD6 C0LGD7 C0LGD8 C0LGD9 C0LGE0 C0LGE4
C0LGF4 C0LGF5 C0LGG3 C0LGG4 C0LGG6 C0LGG7 C0LGG8 C0LGG9 C0LGH2 C0LGH3 C0LGH8
C0LGI2 C0LGI5 C0LGJ1 C0LGJ9 C0LGK4 C0LGK9 C0LGL4 C0LGL9 C0LGN2 C0LGP2 C0LGP4
C0LGP9 C0LGQ4 C0LGQ5 C0LGQ7 C0LGQ9 C0LGR3 C0LGR6 C0LGR9 C0LGS2 C0LGS3 C0LGT1
C0LGT5 C0LGT6 C0LGU0 C0LGU1 C0LGU5 C0LGU7 C0LGV0 C0LGV1 C0LGW2 C0LGW6 C0LGX1
C0LGX3 C0M6K2 C0M715 C0M7A5 C0M7J7 C0M7Z3 C0M7Z7 C0M819 C0M824 C0M8J5 C0M958
C0MBF3 C0MCV2 C0MCZ9 C0ME59 C0MG10 C0MG44 C0MG49 C0MGG4 C0MGW1 C0MH74 C0MH92
C0NFC2 C0NIJ2 C0NJ57 C0NKP8 C0NKT8 C0NLX2 C0NU79 C0NUJ9 C0NUL6 C0NUQ8 C0NY51
C0NZU3 C0PEY7 C0PW59 C0PWC8 C0PWD2 C0PXA8 C0PXJ0 C0PXJ7 C0PXJ8 C0PXM8 C0PXT4
C0PY41 C0PYD4 C0PYM9 C0PYS4 C0PYW1 C0PYX8 C0PYZ8 C0PZ16 C0PZ64 C0PZB3 C0PZC8
C0PZM1 C0PZN3 C0PZQ9 C0PZR0 C0PZR1 C0PZS4 C0PZV3 C0Q030 C0Q045 C0Q048 C0Q051
C0Q065 C0Q066 C0Q067 C0Q070 C0Q0D3 C0Q0E8 C0Q0V6 C0Q135 C0Q145 C0Q1F2 C0Q1F3
C0Q1Q4 C0Q1Q9 C0Q212 C0Q247 C0Q2L3 C0Q2L5 C0Q2N7 C0Q2N8 C0Q2P7 C0Q2Z1 C0Q2Z5
C0Q301 C0Q328 C0Q332 C0Q393 C0Q398 C0Q399 C0Q3E3 C0Q3L6 C0Q3S8 C0Q3V4 C0Q413
C0Q416 C0Q4D1 C0Q4D6 C0Q4D9 C0Q4I6 C0Q4L6 C0Q4N0 C0Q4U6 C0Q4Y4 C0Q4Y5 C0Q506
C0Q511 C0Q552 C0Q5I3 C0Q5R6 C0Q5V7 C0Q6B2 C0Q6K0 C0Q711 C0Q7L8 C0Q842 C0Q855
C0Q859 C0Q8P0 C0QB81 C0QC85 C0QC99 C0QD59 C0QDP8 C0QEI1 C0QGT5 C0QJ09 C0QPE1
C0QR89 C0QR92 C0QU08 C0QYW6 C0R1X0 C0R2X3 C0R393 C0R3Z7 C0R4W3 C0R5T8 C0R5U2
C0R5U3 C0RE23 C0RE73 C0REG3 C0RF64 C0RFI0 C0RGK4 C0RH35 C0RH93 C0RHH6 C0RHH8
C0RHV4 C0RIC2 C0RIE6 C0RIE9 C0RIJ9 C0RIL5 C0RJ08 C0RLC2 C0RMD3 C0RMI1 C0RYV1
C0RYW2 C0RZV6 C0S345 C0S3S0 C0S6S4 C0S713 C0S7H1 C0S7J1 C0S7Q7 C0S9J4 C0SEQ8
C0SGJ2 C0SGL5 C0SJS2 C0SJS3 C0SJS4 C0SP78 C0SP79 C0SP89 C0SP90 C0SP95 C0SPA7
C0SPB2 C0SPB3 C0Z763 C0Z766 C0Z769 C0Z7G8 C0Z7H6 C0Z921 C0ZBY5 C0ZEP4 C0ZG49
C0ZGA2 C0ZIL1 C0ZL89 C0ZPH6 C0ZPK5 C0ZX54 C0ZYJ0 C0ZZU1 C1A1E9 C1A1Y4 C1A8A7
C1A8L5 C1A8P2 C1A8P3 C1A8Y1 C1A8Y4 C1ABH4 C1ABQ7 C1AFB2 C1AGR9 C1AGS2 C1AJ08

C1AKP3 C1ALX4 C1AM21 C1AM23 C1AML8 C1AMU9 C1AMV1 C1AN55 C1AN96 C1ANC7 C1ANJ0
C1ANN6 C1ANR3 C1AQ13 C1AQ52 C1AQ72 C1AQC3 C1AQK4 C1ARZ5 C1AU58 C1AUJ4 C1AZF7
C1AZG0 C1AZL6 C1B1K2 C1B7Z5 C1BEG3 C1BGG0 C1BI29 C1BKZ7 C1C524 C1C5J2 C1C5K9
C1C6F1 C1C6I5 C1C7R3 C1C811 C1C8A4 C1C8A5 C1C9E8 C1CBB5 C1CCM1 C1CCN0 C1CDG4
C1CDJ9 C1CEN8 C1CF00 C1CF98 C1CF99 C1CGA6 C1CIK1 C1CIV9 C1CIX7 C1CJQ4 C1CJU2
C1CL18 C1CLC0 C1CLL1 C1CLL2 C1CMJ2 C1CPL3 C1CPW6 C1CPZ1 C1CQU5 C1CR23 C1CS41
C1CS76 C1CSD3 C1CSD4 C1CTB2 C1CTN0 C1CVN6 C1CW54 C1CYN7 C1D0H8 C1D0I1 C1D186
C1D1H5 C1D4X7 C1D515 C1D5J3 C1D5L8 C1D5M0 C1D6F9 C1D6H8 C1D7X0 C1D896 C1D8R6
C1D8S5 C1D9H2 C1DAB4 C1DAK9 C1DBZ0 C1DCB3 C1DCB6 C1DCY1 C1DEB0 C1DEF3 C1DER3
C1DG34 C1DHS4 C1DHS7 C1DIY2 C1DK17 C1DK47 C1DKY9 C1DL20 C1DL23 C1DLV7 C1DMW2
C1DN28 C1DN39 C1DNF7 C1DP97 C1DQ96 C1DR03 C1DR06 C1DRP0 C1DRV2 C1DT39 C1EI34
C1EL62 C1ELN0 C1ENB1 C1ENC2 C1EPQ7 C1EPS7 C1EPV9 C1EPW0 C1ER33 C1ER34 C1ETB0
C1EUT9 C1EUZ7 C1EV23 C1EWH3 C1EWN1 C1EYJ9 C1EYK0 C1EYK1 C1EZE3 C1EZN6 C1F0L6
C1F0L9 C1F0N3 C1F0Q1 C1F0T4 C1F223 C1F454 C1F461 C1F686 C1F8X6 C1F9D0 C1F9D3
C1FA47 C1FL83 C1FPI3 C1FT66 C1FUP2 C1FYD2 C1FZI7 C1FZL3 C1G0X0 C1G2I2 C1G4S9
C1G520 C1G565 C1G784 C1G794 C1GC25 C1GG16 C1GJ63 C1GKM1 C1GP85 C1GT79 C1GTA5
C1GTI3 C1GU90 C1GVA2 C1GWM2 C1GXJ4 C1GYK6 C1GZK1 C1H0T6 C1H492 C1ITJ8 C1JJY3
C1KV55 C1KVL5 C1KVP1 C1KW75 C1KWE3 C1KWZ0 C1KX10 C1KX11 C1KX17 C1KXH9 C1KXM0
C1KYD5 C1KYP9 C1KYS0 C1KYV2 C1KZN4 C1KZN5 C1KZN6 C1L148 C1L1A5 C1L1N4 C1L254
C1L2J2 C1L2L3 C1L2Y0 C1N652 C1P5Z8 C1P5Z9 C1P602 C1P603 C1P609 C1P610 C1P611
C1P616 C1PGW0 C1PGW1 C3JY55 C3JY87 C3JY90 C3JYN4 C3JYP5 C3K0P6 C3K0Y6 C3K1C4
C3K1F1 C3K1F2 C3K1G2 C3K1X0 C3K222 C3K2C1 C3K2E3 C3K337 C3K3M0 C3K3Q9 C3K4C4
C3K4Y3 C3K4Z0 C3K6U0 C3K733 C3K816 C3K8T9 C3K8U2 C3KAC9 C3KAD3 C3KBV7 C3KCS7
C3KCS9 C3KDR5 C3KU54 C3KXR7 C3L101 C3L2A5 C3L712 C3L732 C3L8X4 C3L953 C3L9V6
C3L9Y0 C3LBH7 C3LBW9 C3LE57 C3LED1 C3LEN7 C3LF98 C3LF99 C3LFA0 C3LFG7 C3LFH0
C3LFI4 C3LFJ8 C3LGP7 C3LGP8 C3LH78 C3LH93 C3LI11 C3LI12 C3LJM3 C3LLJ2 C3LLS9
C3LLT4 C3LMV3 C3LN23 C3LNK5 C3LP04 C3LP77 C3LPQ2 C3LPS1 C3LPS7 C3LPT6 C3LPZ6
C3LQ63 C3LQ64 C3LQH1 C3LQU9 C3LS13 C3LS16 C3LS87 C3LS88 C3LSD1 C3LSG9 C3LSJ4
C3LSJ5 C3LSS9 C3LST8 C3LTL7 C3LTL8 C3LTR0 C3LTR1 C3LTR2 C3LU14 C3LUP2 C3LVE3
C3LVE6 C3LVM7 C3LWU0 C3M8M7 C3MA28 C3MA67 C3MA95 C3MAH1 C3MAH2 C3MAR8 C3MBT9
C3MCT9 C3MDQ7 C3MEK8 C3MEN2 C3MEY7 C3MF13 C3MF49 C3MFT5 C3MHB5 C3MHJ3 C3MHJ5
C3MJX7 C3MQ04 C3MQD0 C3MRS8 C3MVC3 C3MW09 C3MWB6 C3N0R7 C3N5M2 C3N652 C3N903
C3NBN0 C3NE86 C3NEK7 C3NH36 C3NHG3 C3NJZ2 C3NMG3 C3NMG6 C3NUE6 C3P0C9 C3P0L9
C3P0M1 C3P1B4 C3P1E2 C3P1E5 C3P1F9 C3P1N2 C3P286 C3P2J6 C3P2J7 C3P314 C3P3K4
C3P4P7 C3P663 C3P683 C3P6V2 C3P6V3 C3P8W4 C3PAI4 C3PBB5 C3PBD7 C3PC02 C3PD00
C3PE99 C3PFG5 C3PHA2 C3PHF9 C3PLM6 C3PLS6 C3PM50 C3PM51 C3PM81 C3PM83 C3PMV5
C3PN58 C3PNB2 C3PND1 C3PP25 C3PP35 C3S7F0 C3S7F1 C3U0S3 C3VEQ1 C3VMW3 C3YZ59
C3ZQF9 C3ZWH9 C4A0P0 C4IYS8 C4IZX0 C4JAF2 C4JDF8 C4JHR1 C4JHS3 C4JHY5 C4JI06
C4JLG3 C4JP24 C4JQ45 C4JQN4 C4JQN7 C4JT27 C4JTD3 C4JV00 C4JVW2 C4JYM4 C4K0P2
C4K0P3 C4K166 C4K168 C4K1G9 C4K1K2 C4K1S5 C4K222 C4K2C9 C4K2Q8 C4K3H9 C4K3W3
C4K4K3 C4K4T7 C4K6N1 C4KFQ2 C4KH98 C4KHL5 C4KJU4 C4KYQ8 C4L1J3 C4L2S7 C4L412
C4L477 C4L5D6 C4L5I5 C4L7X6 C4L804 C4L962 C4L975 C4L981 C4L9W1 C4LA22 C4LA24
C4LAI3 C4LAQ5 C4LB40 C4LB43 C4LB58 C4LBR0 C4LC87 C4LD03 C4LD11 C4LD18 C4LD54
C4LD55 C4LDD1 C4LDL6 C4LDL7 C4LDM1 C4LDW6 C4LDZ4 C4LE79 C4LEP4 C4LEP7 C4LES1

C4LI47 C4LIK6 C4LIK7 C4LTM1 C4M6U3 C4PLJ4 C4PLJ5 C4PQQ9 C4PQR0 C4QM85 C4QV79
C4QVD6 C4QVU3 C4QW04 C4QWJ4 C4QWY8 C4QX80 C4QXE9 C4QXN2 C4QXP7 C4QYK3 C4QYM9
C4QZ06 C4QZV7 C4R0M5 C4R127 C4R192 C4R3U0 C4R432 C4R4B3 C4R613 C4R628 C4R7Q4
C4R7S7 C4R7X8 C4R7X9 C4R7Z3 C4R7Z8 C4R892 C4R8D7 C4VBN2 C4XGW8 C4XH52 C4XK72
C4XKM8 C4XMN1 C4XMN4 C4XNJ6 C4XPM0 C4XQ07 C4XT96 C4XU20 C4XVQ6 C4XVW1 C4XW03
C4XXV8 C4XZH2 C4Y0Q6 C4Y1G1 C4Y206 C4Y2Z9 C4Y3C1 C4Y3D4 C4Y4X4 C4Y631 C4Y6F4
C4Y8B4 C4Y9H0 C4Y9R6 C4YBX9 C4YCC3 C4YDC4 C4YE34 C4YFU9 C4YG73 C4YI11 C4YIG3
C4YJ00 C4YJI1 C4YKP5 C4YL78 C4YLH0 C4YNJ0 C4YPM0 C4YQQ4 C4YRP9 C4YS59 C4YS65
C4YTG0 C4YTL7 C4YTQ0 C4ZPV3 C4ZPW6 C4ZPX3 C4ZQ24 C4ZQP2 C4ZQQ5 C4ZQS4 C4ZQV8
C4ZQX4 C4ZQX6 C4ZR30 C4ZRI2 C4ZRP8 C4ZRR9 C4ZRY8 C4ZRZ3 C4ZS05 C4ZSG3 C4ZSG4
C4ZSG5 C4ZSK9 C4ZSM1 C4ZSU7 C4ZSW2 C4ZSX8 C4ZSX9 C4ZSY0 C4ZT42 C4ZT46 C4ZT58
C4ZTM7 C4ZTN2 C4ZTU1 C4ZTU7 C4ZTU8 C4ZU39 C4ZU96 C4ZUB9 C4ZUC2 C4ZUC5 C4ZUE5
C4ZUK6 C4ZUM0 C4ZV37 C4ZV70 C4ZV90 C4ZVI8 C4ZVS6 C4ZVS8 C4ZVU3 C4ZVU4 C4ZVX5
C4ZW41 C4ZW75 C4ZWB3 C4ZWH2 C4ZWH3 C4ZWH4 C4ZWU5 C4ZWZ4 C4ZX47 C4ZX91 C4ZXQ4
C4ZY19 C4ZY54 C4ZY62 C4ZY63 C4ZY90 C4ZY93 C4ZY95 C4ZYC7 C4ZYH3 C4ZYW2 C4ZYY3
C4ZYY8 C4ZZ15 C4ZZ16 C4ZZ25 C4ZZ61 C4ZZE5 C4ZZH6 C4ZZH7 C4ZZI2 C4ZZI6 C4ZZQ2
C4ZZY0 C4ZZY8 C4ZZY9 C5A011 C5A054 C5A075 C5A077 C5A082 C5A085 C5A0W8 C5A130
C5A133 C5A160 C5A1E5 C5A1E6 C5A1J7 C5A2X0 C5A3H3 C5A3H4 C5A478 C5A4H9 C5A5K3
C5A7G3 C5B6Z4 C5B704 C5B7L9 C5B7R5 C5B828 C5B8H7 C5B8I0 C5B8J9 C5B9F3 C5B9W2
C5BB13 C5BB35 C5BC09 C5BC70 C5BC80 C5BCA6 C5BDA3 C5BDB0 C5BDE5 C5BDE9 C5BDL4
C5BDQ5 C5BDQ8 C5BDX6 C5BDY7 C5BE26 C5BED9 C5BF34 C5BF35 C5BF61 C5BFX0 C5BG86
C5BGE0 C5BGJ2 C5BGP5 C5BH44 C5BH58 C5BHG2 C5BHN3 C5BHX7 C5BII5 C5BJA8 C5BKK1
C5BKL7 C5BKU5 C5BMA2 C5BMS3 C5BNZ4 C5BP37 C5BQX3 C5BRP6 C5BW62 C5C0R4 C5C0R7
C5CA33 C5CA73 C5CB68 C5CBT8 C5CBV4 C5CDR2 C5CE71 C5CES8 C5CGT6 C5CIV2 C5CML1
C5CNF1 C5CNT0 C5CPD2 C5CPD4 C5CQK5 C5CQL2 C5CT10 C5CTD2 C5CTM8 C5CWY2 C5D3K0
C5D3Y4 C5D5W1 C5D657 C5D695 C5D6R3 C5D794 C5D862 C5D8M1 C5D978 C5D981 C5D984
C5D995 C5D9M1 C5D9N0 C5DA22 C5DAS3 C5DBT1 C5DCI0 C5DDH1 C5DDK8 C5DDY0 C5DDZ2
C5DE38 C5DE77 C5DEL5 C5DEP0 C5DG70 C5DGB6 C5DGX3 C5DH21 C5DH39 C5DHW0 C5DI25
C5DIL2 C5DJ31 C5DL56 C5DLL1 C5DLM9 C5DLZ7 C5DMK0 C5DNL7 C5DPA1 C5DQ08 C5DQ18
C5DQ25 C5DQ63 C5DQU6 C5DQY5 C5DRF7 C5DRK2 C5DRL9 C5DRQ1 C5DSA7 C5DSG3 C5DT30
C5DT72 C5DTA7 C5DTC6 C5DTJ3 C5DU19 C5DVU3 C5DWC4 C5DYQ1 C5DZR8 C5DZU3 C5DZY8
C5E006 C5E057 C5E0G6 C5E0J8 C5E109 C5E1G5 C5E1U4 C5E2E7 C5E2G4 C5E2Q0 C5E325
C5E368 C5E381 C5E398 C5E3S0 C5E3S7 C5E3W1 C5E4F2 C5E4T8 C5FDH0 C5FEL7 C5FGB1
C5FH98 C5FJT2 C5FMP1 C5FQ45 C5FRB0 C5FRT9 C5FSQ7 C5FTB4 C5FTV7 C5FUK3 C5FYX2
C5FYZ3 C5FZ62 C5G0E3 C5G729 C5G8H4 C5G8L5 C5GB89 C5GC75 C5GDJ2 C5GFG7 C5GIQ8
C5GL32 C5GLP2 C5GMK3 C5GN10 C5GNU1 C5GVC6 C5GVF3 C5H431 C5H884 C5H886 C5HGF3
C5IAW9 C5IGN8 C5J6A7 C5JC30 C5JC96 C5JCV0 C5JGE5 C5JIS0 C5JIT3 C5JJG5 C5JN54
C5JPM9 C5JVU4 C5JW43 C5JYV0 C5K0G3 C5K3E1 C5M1Z6 C5M3V6 C5M4D4 C5M4J6 C5M501
C5M545 C5M580 C5M5F1 C5M5S1 C5M696 C5M6Z4 C5M8B9 C5MA32 C5MAV2 C5MB30 C5MBE2
C5MC60 C5MCD3 C5MFP8 C5MK33 C5MSH2 C5NSL2 C5P0Z4 C5P311 C5P334 C5P436 C5P447
C5P635 C5P8K5 C5P998 C5PA33 C5PCN6 C5PEI5 C5W022 C5W4W1 C5W716 C5WNF5 C5WUP3
C5X4A5 C5XEK4 C5XIF2 C5XKI6 C5XTX2 C5XW97 C5XY39 C5Y376 C5Y494 C5Y7C6 C5Y7C7
C5Y7C8 C5Y9U6 C5YAP3 C5YDQ9 C5YHP6 C5YJG8 C5YLC9 C5YP66 C5YRU8 C5YVA2 C5Z782
C5Z7E3 C6A2C9 C6A335 C6ADP9 C6AMB3 C6AQL9 C6AR43 C6AUT0 C6BJN9 C6BRS7 C6BTV9

C6BUE1 C6BV90 C6BWF6 C6BXF6 C6BYG9 C6BYL0 C6C0J6 C6C4L7 C6C4L9 C6C9J7 C6C9P6
C6CAI2 C6CC69 C6CE57 C6CEF6 C6CJW3 C6CQJ7 C6CQK7 C6CU08 C6D5D2 C6D930 C6D9J9
C6D9P6 C6DA34 C6DA37 C6DA40 C6DA53 C6DAE8 C6DAJ1 C6DAW3 C6DAW6 C6DB46 C6DBC8
C6DBD0 C6DBD8 C6DBS3 C6DBU8 C6DBV4 C6DCB6 C6DDG3 C6DDL8 C6DDM0 C6DDT0 C6DE42
C6DE43 C6DEP2 C6DEU6 C6DEZ3 C6DF08 C6DFD4 C6DFG2 C6DFN7 C6DFR9 C6DFY1 C6DG20
C6DG45 C6DG97 C6DGG6 C6DGQ8 C6DGY3 C6DGY9 C6DGZ0 C6DH09 C6DH13 C6DH14 C6DH17
C6DHE3 C6DHK1 C6DI75 C6DIK6 C6DIM1 C6DIM2 C6DIM3 C6DIR2 C6DJ80 C6DJB7 C6DJF7
C6DJG6 C6DJG7 C6DJS3 C6DJS4 C6DJU8 C6DJX5 C6DK68 C6DK98 C6DKC9 C6DKE3 C6DKF1
C6DKV2 C6DS31 C6DTH3 C6DYP4 C6DYS1 C6DZK3 C6DZL0 C6E019 C6E0U8 C6E2F4 C6E8M5
C6E8P0 C6E8P1 C6E8Q5 C6E8U8 C6H1N5 C6H203 C6H220 C6H4B5 C6H7W0 C6HAY7 C6HB29
C6HFQ7 C6HRC7 C6K2K4 C6KEF6 C6KFA3 C6KI89 C6KTB7 C6SVQ5 C6SXZ3 C6SYW3 C6SZ04
C6SZP8 C6T1G0 C6T1K6 C6T1Z6 C6T2E7 C6T2J5 C6T4A0 C6T4E0 C6TBD0 C6TCJ2 C6TFL9
C6TG62 C6TH93 C6UJA3 C6ULT8 C6UX49 C6UYL8 C6V4R9 C6V5H7 C6V5K2 C6VKW6 C6VQL6
C6WX51 C6XAJ7 C6XAK0 C6XJX0 C6XMG3 C6Y4A3 C6Y4A4 C6Y4A7 C6Y4B3 C6Y4B6 C6Y4C7
C6Y4C8 C7C0H5 C7C1L2 C7EXK4 C7G3A0 C7GJ54 C7GJM5 C7GJY7 C7GK20 C7GKH8 C7GKW5
C7GLB5 C7GLH7 C7GLT4 C7GLU4 C7GMW8 C7GNE2 C7GNE3 C7GPP6 C7GQ65 C7GQ88 C7GQ96
C7GQI9 C7GR49 C7GR78 C7GRJ3 C7GRS7 C7GRY1 C7GSI5 C7GSI6 C7GT24 C7GT60 C7GTF0
C7GU92 C7GUR3 C7GUS4 C7GUZ7 C7GW31 C7GW39 C7GWV6 C7GWZ2 C7GX93 C7GXB2 C7J0P3
C7J0R5 C7J3A2 C7J4U3 C7J6G6 C7JGJ8 C7M0M0 C7MC16 C7N1I1 C7N914 C7NB69 C7NC37
C7P879 C7RCQ5 C7SFP6 C7T2J9 C7TC36 C7YGZ0 C7YIH6 C7YJ02 C7YQ31 C7YRS6 C7YYG9
C7Z274 C7Z504 C7Z7C3 C7ZHK5 C7ZIE1 C8AW46 C8AW47 C8UFK8 C8V0D4 C8V329 C8V3Y8
C8V4D5 C8V7P3 C8VBV0 C8VCL4 C8VCL5 C8VCP8 C8VE79 C8VI81 C8VJR0 C8VJR5 C8VJW1
C8VK15 C8VN86 C8VN91 C8VQ97 C8VQ98 C8VRV0 C8VTR5 C8W731 C8WEG0 C8WLM0 C8WQG6
C8WQG7 C8XPA8 C8XPB2 C8YUV0 C8Z3X9 C8Z6L4 C8Z6M6 C8Z6Z8 C8Z742 C8Z7B2 C8Z830
C8Z852 C8Z891 C8Z8G3 C8Z8H2 C8Z8J0 C8Z951 C8Z970 C8Z9U8 C8ZAC7 C8ZAS4 C8ZAV4
C8ZAW6 C8ZAZ9 C8ZBF1 C8ZCI0 C8ZCQ9 C8ZCR2 C8ZCS2 C8ZCS3 C8ZDW3 C8ZEH4 C8ZEK9
C8ZEY1 C8ZF05 C8ZF15 C8ZG52 C8ZGH0 C8ZHR0 C8ZI10 C8ZIT1 C8ZJM1 C9D7C2 C9E894
C9EIC7 C9JDP6 C9JG80 C9JH25 C9JI98 C9JQI7 C9JQL5 C9JVW0 C9K202 C9K7C6 C9S688
C9S8G0 C9SA05 C9SF29 C9SI22 C9SJ15 C9SM62 C9SXB4 C9Y3M8 C9YID6 D0B707 D0C9N4
D0CCT2 D0E0C2 D0G895 D0J948 D0JKF6 D0JNB0 D0JUX5 D0K261 D0K8E2 D0KEK2 D0KMH4
D0KWI6 D0KWS8 D0KXY3 D0KZ79 D0LB45 D0LWB8 D0MGK7 D0MGU8 D0MIN3 D0MIN4 D0MVS1
D0MZL5 D0NB60 D0NB65 D0PRN2 D0PRN3 D0PRN4 D0Q0Y7 D0QX25 D0U690 D0VFU8 D0VWR1
D0VWR3 D0VWR4 D0VWR5 D0VWR7 D0VWR8 D0VWU4 D0VWV4 D0Z7W2 D0ZCE7 D0ZE09 D0ZEW1
D0ZF88 D0ZGL2 D0ZHC6 D0ZK39 D0ZKX9 D0ZLQ7 D0ZQC5 D0ZSY8 D0ZTB2 D0ZV89 D0ZWT5
D0ZWU0 D0ZXP9 D0ZXQ1 D0ZXQ3 D0ZY51 D1A4G7 D1AFI6 D1AMK8 D1AMK9 D1AXT4 D1B5U2
D1BLD0 D1BTU8 D1C1U7 D1C2C6 D1C4U5 D1C8C0 D1CDJ5 D1CDJ6 D1CDT8 D1J722 D1LYT2
D1MEI6 D1MEI7 D1MEI8 D1MI46 D1MX85 D1MX86 D1MX87 D1MX88 D1MX89 D1Z4K7 D1Z5G1
D1Z9B4 D1Z9Q3 D1ZEM2 D1ZIW5 D1ZRK4 D1ZV85 D2A0H5 D2BGI6 D2BJS8 D2BKM6 D2BRP0
D2BUT3 D2BX50 D2C1Z6 D2CGS0 D2E9W6 D2E9W7 D2E9W8 D2H617 D2H6Z0 D2H788 D2HBV9
D2HFT7 D2HKB0 D2HSA6 D2HZB0 D2I2F3 D2I2M6 D2IYS2 D2K6F1 D2K762 D2KQI6 D2KUZ7
D2KX21 D2NQQ7 D2NT99 D2NWE2 D2NZ73 D2PAJ7 D2QZ34 D2RLC7 D2RLC8 D2RNY0 D2TBH1
D2TMS1 D2TN56 D2TN57 D2TNL4 D2TR04 D2TV88 D2WKD8 D2YW26 D2YZQ3 D2ZZC1 D3E045
D3E0P6 D3ERJ1 D3EZK2 D3F124 D3FA80 D3FFN2 D3FRP0 D3FSJ3 D3GE74 D3GS80 D3GSC3
D3HAJ5 D3INW7 D3K0R6 D3KZG3 D3NRE7 D3QW22 D3QY10 D3RNN6 D3RPC1 D3RVH4 D3RVH9

D3SD88 D3SD93 D3UEH0 D3UF10 D3UMA1 D3UTF3 D3UTF4 D3UW23 D3V3D1 D3V7P3 D3V7P4
D3W0D1 D3YTS9 D3YUK8 D3YVE8 D3YX43 D3YYI7 D3YZF7 D3Z1Q2 D3Z1U7 D3Z291 D3Z338
D3Z5L6 D3Z7H4 D3Z7Q2 D3Z813 D3Z9R8 D3ZAF6 D3ZAL8 D3ZAT9 D3ZAW2 D3ZB51 D3ZB94
D3ZBM4 D3ZCM3 D3ZE55 D3ZE85 D3ZEH5 D3ZF92 D3ZFB6 D3ZFW5 D3ZG27 D3ZGZ6 D3ZH22
D3ZHH1 D3ZHR2 D3ZI76 D3ZJ47 D3ZK93 D3ZLH5 D3ZLY0 D3ZNF5 D3ZNQ3 D3ZPG5 D3ZPX4
D3ZQJ0 D3ZQX2 D3ZR35 D3ZRC4 D3ZS74 D3ZTD8 D3ZTV3 D3ZTX0 D3ZUK3 D3ZVU9 D3ZWJ9
D3ZWT9 D3ZWZ9 D3ZXD8 D3ZYP5 D3ZZP4 D4A1J9 D4A2H2 D4A2Q0 D4A3T6 D4A3U0 D4A4Q2
D4A612 D4A6D8 D4A6L0 D4A723 D4A734 D4A7K7 D4A7P2 D4A9I3 D4A9R4 D4AA47 D4ABB8
D4ABL6 D4ABX8 D4AC13 D4ACN8 D4ACP2 D4ACX8 D4AD02 D4AD37 D4AD53 D4ADY9 D4AEP3
D4ALL0 D4AMV1 D4ANR0 D4AQT0 D4AQV3 D4ARB8 D4AT37 D4AU27 D4AVD4 D4AXV8 D4AYG1
D4AYS6 D4AYW0 D4AZD1 D4B086 D4B1G0 D4B4J2 D4B4K9 D4B5L8 D4B5N6 D4D4B1 D4D5I1
D4D802 D4D8C1 D4DA58 D4DCG0 D4DCV9 D4DDK2 D4DGR3 D4DHX2 D4G8R2 D4GCB2 D4GM14
D4GP35 D4GP36 D4GP37 D4GPW1 D4GPW2 D4GQ17 D4GSY8 D4GTK4 D4GTK5 D4GTV9 D4GU68
D4GU74 D4GUA0 D4GUZ4 D4GVK4 D4GWC8 D4GYE5 D4GYG3 D4GYG8 D4GYH0 D4GYH2 D4GYH3
D4GYH4 D4GZC9 D4GZD0 D4H031 D4HA34 D4I1C3 D4I6V4 D4PHA7 D5A7J3 D5A8E6 D5A972
D5ACW4 D5AGC8 D5ANZ2 D5ANZ3 D5ANZ4 D5AQD5 D5AQY7 D5AQY8 D5ARG8 D5ARG9 D5ARP7
D5ARY9 D5AT98 D5AUZ7 D5AUZ9 D5AV16 D5B5U5 D5BW25 D5CDA0 D5CHK9 D5CHY0 D5D8E3
D5DQW6 D5E7Y7 D5G4B1 D5GI81 D5GM60 D5GNC3 D5H7Z5 D5HA94 D5J9U8 D5JAJ1 D5JBW8
D5JBW9 D5JBX0 D5JBX1 D5K211 D5K8A9 D5KX81 D5LX53 D5RAW5 D5T166 D5WSC8 D5X4H4
D6CRB4 D6MZJ6 D6PVB4 D6PVB5 D6R8X8 D6VPM8 D6VTK4 D6W196 D6WJ77 D6WMX4 D6XTM9
D6XVS2 D6YXE8 D7A5Q8 D7AEL2 D7CEX8 D7DR00 D7DUI0 D7EBK7 D7GIS1 D7KBH3 D7KCH2
D7KFC7 D7KXP8 D7L342 D7L5G6 D7LAP2 D7LBN4 D7LD59 D7LD60 D7LGW9 D7LIK3 D7LIN7
D7LIR2 D7LTJ4 D7LZ50 D7M2M8 D7M7B3 D7M9V1 D7MAF5 D7MAF6 D7MAF7 D7MFJ8 D7MFW5
D7MGK0 D7MM00 D7MMW4 D7MUY4 D7PDD4 D7PHY8 D7PHZ6 D7PI13 D7RF80 D7SFH9 D7UPN3
D7UQ39 D8GR67 D8GR68 D8GR69 D8GR70 D8IIP4 D8ILJ0 D8JLE3 D8KFN3 D8KIE9 D8MEB8
D8MHM9 D8MMF7 D8MQN9 D8MXN7 D8PSG0 D8Q7Q5 D8QAM0 D8QGA7 D8QNI1 D8QNV6 D8QQW9
D8R354 D8R814 D8RQM9 D8S067 D8S069 D8S072 D8S6A8 D8SJ65 D8ST12 D8ST13 D8T2C0
D8T829 D8TC76 D8TFA8 D8TFB0 D8V075 D8X2C5 D9HP19 D9HP20 D9HP23 D9HP25 D9HP26
D9HP27 D9IA43 D9IA45 D9IL24 D9N164 D9PTN5 D9PXT1 D9QVP6 D9RQS7 D9RZP4 D9S0S1
D9SNZ5 D9U2B8 D9UBX4 D9UBX6 D9UC01 E0CX11 E0CYC6 E0CYR6 E0RGM4 E0SAK8 E0SCY2
E0SDG9 E0SIS2 E0SKT0 E0T2N0 E0T8U0 E0TW64 E0TW65 E0TW66 E0TW67 E0W1I1 E0W3E3
E1ACP7 E1ACQ2 E1ACQ3 E1ACQ6 E1ACR0 E1B2Z9 E1B9E5 E1BAR0 E1BBQ2 E1BD52 E1BE10
E1BHC3 E1BJD3 E1BLT8 E1BN97 E1BNE9 E1BPQ3 E1BWM5 E1BY51 E1BYA3 E1C2W7 E1C7U0
E1C8P7 E1JI63 E1JIB2 E1RHR1 E1RHR2 E1SFM6 E1SFR4 E1V6C5 E1V6K4 E1V825 E1V9L1
E1VBK2 E1VBK3 E1VBT7 E1W822 E1W8M5 E1W9W8 E1WAU4 E1WAU5 E1WF94 E1WFA0 E1WGF1
E1WIS2 E1WTB4 E2BJ30 E2IHA6 E2IUB0 E2JF22 E2R0A5 E2R4X3 E2R5I0 E2RA18 E2RD63
E2RDM9 E2RFJ3 E2RG47 E2RK30 E2RKN8 E2RP87 E2RQ08 E2RYF6 E3EYX6 E3F0U0 E3G1S2
E3G4M1 E3GC98 E3L8A5 E3LYX0 E3PJ86 E3PJ87 E3PJ89 E3PSD4 E3Q4R4 E3Q717 E3QDT3
E3QER5 E3QKD2 E3QNZ1 E3QWM6 E3QXY4 E3RE00 E3RFJ1 E3RIP0 E3RR70 E3S9K3 E3UBL5
E3UBL6 E3W9C4 E4MWF4 E4TJ18 E4TJ19 E4TN31 E4UPP6 E4UV76 E4UYL6 E4V094 E4V4X0
E4VP07 E4ZQC4 E4ZUB5 E4ZVE5 E4ZVX1 E5A7D6 E5A7D8 E5A8D2 E5AD87 E5AE35 E5AE43
E5DHB5 E5EZW6 E5EZW7 E5EZW9 E5EZX0 E5KCJ8 E5KK10 E5QVT2 E5QYX6 E5R540 E5RHQ5
E5RIL1 E5RP65 E6MXW0 E6MZM0 E6R427 E6R6G5 E6RCE9 E6RET3 E6UUM7 E6W4K8 E6W4K9
E6WE99 E6WMH4 E6Y2A0 E6ZZ11 E7A253 E7B4A5 E7DNB6 E7EAU8 E7EM37 E7ERA6 E7EXX2

E7EXZ6 E7EY42 E7EYQ9 E7EZG2 E7EZI4 E7F021 E7F3F0 E7F4N7 E7F4V6 E7F4Z4 E7F587

E7F594 E7F6D7 E7F6F7 E7F6V0 E7F7V7 E7FB98 E7FBS9 E7FBY6 E7FDE0 E7FDV5 E7FE13

E7FE40 E7FEC4 E7FFT2 E7FH11 E7FKV8 E7KBW4 E7KCH6 E7KDY6 E7KH03 E7KMS3 E7KNG0

E7KPQ3 E7KQ01 E7KTL7 E7LTJ6 E7LUD2 E7LVX4 E7LZK9 E7NHF8 E7NHN2 E7NIY5 E7NM81

E7NMC3 E7Q311 E7Q3U9 E7Q587 E7Q8M6 E7QE10 E7QET1 E7QG89 E7QJS8 E7QK11 E7R7C4

E7R7R2 E7RAI2 E8MGH9 E8N5I8 E8N5I9 E8SFP1 E8UEF9 E8WSG3 E8XVS9 E8XWL7 E8XY75

E8ZB61 E9AFX2 E9CS37 E9CUF4 E9CV02 E9CXH2 E9CZZ9 E9E1Z2 E9E6S9 E9E796 E9ED72

E9EM69 E9ESM3 E9ETL5 E9EYB5 E9F5E8 E9F8M0 E9F8V9 E9FCP5 E9FR69 E9KBR8 E9KID2

E9KID3 E9KMQ3 E9NA96 E9PQ53 E9PQX1 E9PSU6 E9PT23 E9PTA2 E9PTT0 E9PU17 E9PUL5

E9PV86 E9PVD3 E9PX95 E9PXX9 E9PZ19 E9PZ36 E9PZQ0 E9Q0B3 E9Q0N2 E9Q137 E9Q236

E9Q2Z6 E9Q343 E9Q355 E9Q3C1 E9Q3M5 E9Q401 E9Q4J9 E9Q612 E9Q649 E9Q6C8 E9Q6I0

E9Q7P9 E9Q7X6 E9Q7X7 E9Q876 E9Q8Q8 E9Q942 E9Q9F6 E9Q9K5 E9Q9R3 E9Q9W4 E9QAF0

E9QBI7 E9QCD3 E9QDC5 E9QJ73 E9QJQ6 E9QT42 E9QY26 E9R598 E9R5G2 E9R876 E9R9Y3

E9RBG1 E9RGS3 E9RGS4 F0P5S5 F0PBT2 F0T4D1 F0U7H7 F0UDD2 F0XJH4 F0XS04 F0ZBA6

F1CJ89 F1DBA7 F1DBA8 F1LMY4 F1LN46 F1LW30 F1LX07 F1LZW6 F1M2Z5 F1M391 F1M3J4

F1M8G0 F1MDB2 F1MJW3 F1MK05 F1MLX5 F1MMS9 F1MT22 F1MV99 F1MWM0 F1MYR9 F1N476

F1N4E5 F1N4M2 F1N5S9 F1NCD6 F1NHE9 F1NJ67 F1NNL1 F1NUK7 F1NVK6 F1NWE3 F1NXU8

F1NY98 F1NZP5 F1P4W9 F1PAA9 F1PCT7 F1PJP5 F1PZV2 F1Q4S1 F1Q7H8 F1Q8C3 F1Q8R9

F1Q930 F1QAM1 F1QF89 F1QFU0 F1QG30 F1QGD2 F1QHM7 F1QIK8 F1QLP1 F1QPC3 F1QR43

F1QUP1 F1QVU0 F1QX91 F1QXD3 F1QYC4 F1QZE9 F1R013 F1R0H0 F1R332 F1R4U0 F1RAX4

F1RBC8 F1RE08 F1RE57 F1RFX9 F1RMN2 F1RRW5 F1RT67 F1S584 F1S5L4 F1SAM7 F1SR90

F1SVE0 F1SVH9 F1SVK0 F1SVL2 F1SY49 F1SY52 F1SY62 F1SY66 F1SY68 F1SY70 F1SY71

F1SY73 F1SY74 F1SY75 F1SY77 F1SY82 F1SY83 F1SY91 F1SY95 F1SY96 F1SY99 F1SYA2

F1SYB2 F1SYB6 F1SYC2 F1SYD1 F1SYE5 F1SYF1 F1SYG6 F1SYH0 F1SYH1 F1SYH7 F1SYI7

F1SYI9 F2D276 F2E2E4 F2E5T1 F2EL82 F2EUZ7 F2EYE0 F2F702 F2I5T1 F2JVW9 F2M971

F2MHZ2 F2MNH8 F2PLH2 F2PRR1 F2Q5G0 F2QA93 F2QF13 F2QQ67 F2QZ45 F2QZ65 F2QZ66

F2RI42 F2RP52 F2RPA4 F2RSQ6 F2SG60 F2SH39 F2SHL1 F2SQT8 F2T0J9 F2T1C4 F2Z333

F2Z9C1 F2ZAF9 F3Y9Q5 F3YDF1 F4FSX0 F4HPH1 F4HPH2 F4HQ05 F4HQ17 F4HQ20 F4HQ22

F4HQ23 F4HSU3 F4HT41 F4HTB2 F4HTM3 F4HTV4 F4HVJ3 F4HVM3 F4HVS0 F4HVS6 F4HVW5

F4HVX7 F4HVY0 F4HW02 F4HW17 F4HW79 F4HWB6 F4HWL3 F4HXQ7 F4HXW9 F4HXY7 F4HYR3

F4HYR4 F4HZC3 F4HZH9 F4HZQ7 F4HZX7 F4HZZ4 F4HZZ5 F4I037 F4I0Z6 F4I111 F4I1G5

F4I214 F4I248 F4I2N7 F4I2Y3 F4I316 F4I340 F4I3V3 F4I3V6 F4I4K7 F4I4Q3 F4I5D5

F4I5Q2 F4I8H8 F4I8I0 F4I8K2 F4I8Q7 F4I8W6 F4I8X6 F4I8X8 F4I9A2 F4I9E1 F4I9S3

F4IAL1 F4IB81 F4IBD7 F4ICF4 F4IDU4 F4IDY5 F4IED2 F4IH21 F4IHS9 F4IHU9 F4IJ08

F4IJ77 F4IK45 F4IKF6 F4ILF8 F4ILY9 F4IME1 F4IME2 F4IMH3 F4IN58 F4INW9 F4IQG2

F4IQJ6 F4IQV7 F4IQX1 F4IR05 F4ISL7 F4ISV9 F4ITL6 F4ITM1 F4IU14 F4IUE7 F4IUU1

F4IUW3 F4IWA8 F4IXT6 F4IYM4 F4IYZ0 F4IZ80 F4IZC4 F4IZW8 F4J0A8 F4J0W4 F4J158

F4J1Q9 F4J220 F4J2C8 F4J3N2 F4J4C8 F4J4Y0 F4J7T6 F4J8G2 F4J9A3 F4J9A8 F4JAU3

F4JBM4 F4JCY2 F4JE35 F4JEQ1 F4JFU8 F4JGB6 F4JH46 F4JIE8 F4JIN3 F4JIP9 F4JK59

F4JKB9 F4JKK0 F4JKQ7 F4JLZ6 F4JMI5 F4JMI7 F4JN00 F4JN35 F4JNU8 F4JP36 F4JPE9

F4JPW1 F4JRE0 F4JS25 F4JSE7 F4JSH1 F4JTB3 F4JTN2 F4JTU7 F4JU70 F4JUU5 F4JW83

F4JY11 F4JYC8 F4JZ24 F4JZG9 F4JZN6 F4K1B4 F4K2U8 F4K4R6 F4K4T3 F4K5K4 F4K5L5

F4K6B8 F4K6F1 F4K956 F4KA50 F4KA51 F4KBM7 F4KCL7 F4KD68 F4KD71 F4KDA6 F4KF14

F4KF16 F4KFS7 F4KG58 F4KGN5 F4KHA2 F4KHA8 F4KHD8 F4KIB2 F4KID5 F4KIL8 F4P6T0

F5BHA2 F5H4A9 F5H8Q3 F5H8R0 F5H9I4 F5H9N9 F5H9T4 F5H9Z4 F5HA27 F5HAD1 F5HAE6
F5HAK6 F5HAK9 F5HAM0 F5HAR3 F5HAW0 F5HB41 F5HB52 F5HB53 F5HB81 F5HB98 F5HC14
F5HC33 F5HCK7 F5HD27 F5HD92 F5HDA4 F5HDD0 F5HDD3 F5HDE7 F5HDK1 F5HE05 F5HE44
F5HE69 F5HE74 F5HET1 F5HEU0 F5HF35 F5HF62 F5HFB4 F5HFG3 F5HFH0 F5HFJ7 F5HFQ0
F5HG68 F5HG95 F5HGH8 F5HGJ3 F5HGQ8 F5HGU6 F5HHH2 F5HHL7 F5HHQ0 F5HHS3 F5HHT6
F5HHZ3 F5HI25 F5HI32 F5HID8 F5HN69 F5WT83 F5ZBR5 F5ZP94 F6CD01 F6CFW7 F6D9Q8
F6H9N6 F6PTN1 F6Q1T7 F6QV99 F6RG56 F6RWY9 F6S3G9 F6UF99 F6USH3 F6V1J6 F6VAN0
F6Y5S8 F6ZQ54 F7B645 F7B909 F7BWT7 F7C0L1 F7C1E2 F7D3V9 F7E235 F7EQ49 F7J184
F7J186 F7J187 F7J188 F7J220 F7V995 F7V996 F7VJQ1 F7VJQ2 F7VJQ3 F7XXN6 F7XXN9
F7ZDD2 F8CLT8 F8DNI1 F8GAB4 F8GE40 F8HZP7 F8JJ22 F8LHW1 F8LR08 F8RT80 F8S1H3
F8S1H8 F8S1I0 F8VQ03 F8VQN3 F8W0I5 F8W3R9 F8W4H9 F9RBV9 F9UMX3 F9UQC5 F9USS6
F9USS7 F9UST0 F9UST3 F9USY3 F9UTW9 F9UUB3 F9UUD2 F9WWD1 F9X9V3 F9X9V4 F9ZZP5
F9ZZQ0 G0HV85 G0HV86 G0KYA8 G0KYA9 G0KYB2 G0LET5 G0LET6 G0LFX8 G0LXV8 G0R6T0
G0R6T1 G0REX9 G0S1H2 G0S235 G0S6T0 G0S7R3 G0SB44 G0Y287 G1FC92 G1FE62 G1QDE8
G1S9B8 G1SZD9 G1T7E7 G1TR84 G1TZA0 G1UB11 G1UB37 G1UBD1 G1UJF5 G1XU01 G1XU03
G2JZ15 G2JZ41 G2JZ43 G2QNH0 G2TRJ9 G2TRK0 G2TRL0 G2TRM3 G2TRN2 G2TRN3 G2TRN8
G2TRP0 G2TRP1 G2TRP4 G2TRP5 G2TRP7 G2TRP9 G2TRQ1 G2TRQ3 G2TRR0 G2TRR4 G2TRS3
G2TRT0 G2TRT3 G2TRT6 G2TRT7 G2TRT8 G2TRU0 G2TRU7 G3FEX6 G3GBK0 G3I6Z6 G3JUI6
G3M4F8 G3MTW7 G3MZR2 G3UVW3 G3V0H7 G3V7P1 G3V8V5 G3V909 G3V928 G3V9H8 G3V9Q9
G3X8R9 G3X939 G3X9R7 G3XA59 G3XCN6 G3XD12 G3XD24 G3XD29 G3XD30 G3XD43 G3XD46
G3XD51 G3XD94 G3XDA3 G3XDA7 G3XMC2 G3XMC9 G3XSI4 G3Y417 G3Y418 G3Y421 G3Y4N5
G4MS28 G4MVZ7 G4MWA8 G4MWA9 G4MWB3 G4MWY1 G4N2A8 G4N2B5 G4N2X2 G4N2Y3 G4N4W3
G4SDH4 G4SW86 G4V4G3 G4WJD4 G4WMX4 G4XV71 G5AY81 G5CTF8 G5CTF9 G5CTG0 G5CTG1
G5CTG2 G5CTG3 G5CTG4 G5CTG5 G5CTG6 G5CTG7 G5CTG8 G5DBJ0 G5E829 G5E8K6 G5E8Q8
G5EBF1 G5EBH1 G5EBI4 G5EBK1 G5EBL2 G5EBM5 G5EBN9 G5EBQ8 G5EBR3 G5EC63 G5EC84
G5ECB2 G5ECD6 G5ECD9 G5ECI1 G5ECJ0 G5ECL4 G5ECQ2 G5ECT0 G5ECX0 G5ED05 G5ED44
G5ED45 G5ED65 G5EDB8 G5EDE5 G5EDK5 G5EDN0 G5EDR5 G5EDW2 G5EE06 G5EE72 G5EE96
G5EEB1 G5EEE1 G5EEE5 G5EEE9 G5EEK9 G5EEP3 G5EEQ9 G5EEV9 G5EF33 G5EF48 G5EF60
G5EF96 G5EFC3 G5EFD4 G5EFD5 G5EFD9 G5EFE7 G5EFI7 G5EFJ9 G5EFP5 G5EFQ0 G5EG11
G5EG88 G5EGA5 G5EGH6 G5EGI7 G5EGJ5 G5EGJ9 G5EGK5 G5EGL9 G5EGM1 G5EGN2 G5EGP4
G5EGQ6 G5EGT9 G7CBF5 G7CBF6 G7IBJ4 G7IHF9 G7JG80 G7JIK2 G7JND3 G7KDA1 G7KGQ4
G7L0H3 G7L1W7 G7L218 G7LAA8 G7LG31 G8HTB6 G8QM60 G8QM62 G8QMC1 G8SD34 G8XYX6
G8YYA5 G8YYA6 G9LZD7 H0VDZ8 H0YL14 H1A981 H1A988 H1AE12 H1AFJ5 H1AFJ6 H1AFJ7
H1VM35 H2AM12 H2BXL6 H2CYR5 H2DF87 H2DF88 H2DH16 H2DH17 H2DH18 H2DH20 H2DH21
H2DH22 H2E7T5 H2E7T6 H2E7T7 H2E7T8 H2E7T9 H2E7U0 H2KWF1 H2KYE0 H2KYS3 H2KYS8
H2KZ60 H2KZB2 H2KZM6 H2KZU7 H2L002 H2L006 H2L0G5 H2L0H3 H2L0Q3 H2LNR5 H2LRU7
H2N4I1 H2Q5A1 H2QAR5 H2QAR6 H2VFV1 H3BR10 H3BS89 H3BV60 H3JU05 H6LC28 H6LC29
H6LC30 H6LC31 H6TB12 H6WS93 H6WS94 H7C241 H7C350 H7C6B6 H8EUF2 H8EVJ1 H8EVS9
H8EW46 H8EXN2 H8EZ77 H8F2P5 H8WEC1 H9A1V3 H9BZ66 H9GBX2 H9JAQ7 H9JBU5 H9JD76
H9JIQ1 I0HUL5 I0HUL6 I0HUL8 I1BRD6 I1FQB6 I1GQE7 I1H7R8 I1HL09 I1HTF7 I1IUJ6
I1MSF2 I1R9B3 I1RAK8 I1RAL7 I1RB03 I1RBR4 I1RD82 I1RE80 I1RF50 I1RF56 I1RF79
I1RFM2 I1RFN8 I1RJD6 I1RJR2 I1RL06 I1RL13 I1RL15 I1RL16 I1RMG9 I1RMK9 I1RNL0
I1RNT1 I1RPI4 I1RQ76 I1RR90 I1RT23 I1RV19 I1RV24 I1RXX1 I1RZZ3 I1S163 I1S1Q3
I1S2J9 I1S2M5 I1S9X9 I1TEM0 I1TEM1 I1WFC0 I1WSZ3 I1WUC2 I1WVY3 I1Z695 I2BAK7

I2C645 I2CYZ4 I3L273 I3LMB3 I3PFJ5 I3PLR0 I3PLR1 I3QBP4 I3R0S7 I3R635 I3R9N1
I3V6B1 I3V6B7 I6M4H4 I6V1W0 I6VSD2 I6X961 I6YA32 I6YA50 I6YB21 I6YC58 I6YET7
I6YF08 I7C6E8 I7CT85 I7LRH3 J3KML8 J3QMI4 J3QNX5 J3WAX0 J4UJ10 J5J924 J5JKC1
J7FIJ6 J7FIP9 J7FJH0 J7FJH1 J7FK10 J7H5K9 J7HBH4 J7QCA7 J7QLC0 J7RET5 J9RX10
J9VDQ4 J9VF17 J9VF33 J9VGT6 J9VJ99 J9VLH0 J9VLN4 J9VME1 J9VMX7 J9VNT4 J9VPA2
J9VQ06 J9VQQ3 J9VWT0 J9VWU3 J9VX37 J9VXM5 J9W453 K0DZ95 K0E3U9 K0E4D9 K0E684
K0E690 K2QVI4 K2RLM2 K2RU64 K3UII4 K3VFR5 K3VYH8 K4BVL1 K4CEE8 K4CF70 K4CI52
K4REQ6 K7EJ46 K7FQW8 K7FSQ4 K7GET2 K7K424 K7LC65 K7NA32 K7NAJ3 K7NBR2 K7NCS2
K7ND00 K7PEY4 K7R4D4 K7VYZ9 K7ZP88 K9JA28 K9N5Q8 K9N5R3 K9N638 K9N7A1 K9N7C7
K9N7D2 K9UJK2 L0E163 L0E2V1 L0E2Z4 L0E4F8 L0E4G6 L0GAZ8 L0GBQ6 L0GCI6 L0GCV8
L0MXJ1 L0MXJ3 L0MYS5 L0MYS8 L0N063 L0N065 L0R6Q1 L0T550 L0T911 L7HYA8 L7N697
L7T720 L7T8H2 L7V0S7 L7VG99 L7WGA7 L7WME6 L7WME9 L7WRR4 L7WRY5 L7WRZ2 L7WU90
L7X0L7 L7X3H5 L7X3S1 L7XCU0 L7XDS4 L7YAI7 L8EAY0 L8EBJ6 L8F4Z2 L8F618 L8FM50
M0L7V9 M0QZC1 M0R3Q7 M1C5M7 M1RNJ8 M1V1V5 M1VWN5 M1W266 M1W855 M1WCQ0 M1WEN7
M1WG91 M2R8W9 M2UCE5 M2UJ60 M2V933 M2WJF5 M2XHZ0 M2YI75 M2YIZ5 M2YJD1 M2YMU2
M3A333 M3APK4 M3XFH7 M3XWH1 M4VQY9 M5A7P9 M9M5N8 M9MRD1 M9MSG8 M9NDE3 M9PE65
N1NVB7 N1P212 N1PA11 N4WE43 N6Z5E2 O00124 O00144 O00155 O00168 O00180 O00198
O00206 O00219 O00220 O00222 O00237 O00238 O00241 O00254 O00258 O00264 O00270
O00299 O00322 O00337 O00341 O00391 O00398 O00400 O00421 O00445 O00453 O00461
O00468 O00476 O00478 O00481 O00483 O00501 O00519 O00526 O00533 O00548 O00555
O00559 O00574 O00587 O00590 O00591 O00592 O00623 O00624 O00631 O00767 O00844
O00933 O01346 O01393 O01509 O01538 O01578 O01608 O01609 O01634 O01635 O01668
O01670 O01705 O01735 O01811 O01840 O01916 O01925 O01971 O02051 O02100 O02164
O02194 O02213 O02228 O02241 O02275 O02300 O02351 O02464 O02465 O02466 O02495
O02662 O02663 O02664 O02666 O02667 O02670 O02671 O02695 O02703 O02713 O02718
O02721 O02733 O02740 O02757 O02765 O02766 O02769 O02773 O02777 O02781 O02813
O02822 O02824 O02835 O02836 O02839 O02840 O02858 O03060 O03061 O03076 O03166
O03167 O03168 O03169 O03170 O03171 O03172 O03173 O03174 O03175 O03176 O03197
O03198 O03199 O03200 O03201 O03202 O03203 O03204 O03205 O03206 O03207 O03252
O03359 O03363 O03376 O03473 O03474 O03478 O03503 O03515 O03520 O03521 O03522
O03523 O03524 O03538 O03539 O03543 O03545 O03546 O03548 O03553 O03554 O03570
O03692 O03695 O03697 O03698 O03699 O03700 O03701 O03702 O03703 O03704 O03707
O03710 O03713 O03726 O03727 O03733 O03763 O03772 O03773 O03778 O03780 O03792
O03807 O03810 O03811 O03812 O03847 O03848 O03850 O03851 O03889 O03890 O03891
O03892 O03893 O03895 O03978 O04006 O04023 O04034 O04036 O04086 O04089 O04133
O04161 O04164 O04195 O04200 O04249 O04253 O04289 O04331 O04348 O04353 O04354
O04378 O04472 O04478 O04508 O04514 O04523 O04532 O04533 O04534 O04536 O04567
O04584 O04616 O04619 O04621 O04660 O04714 O04722 O04925 O04928 O04940 O05000
O05011 O05012 O05083 O05087 O05097 O05098 O05131 O05215 O05216 O05219 O05224
O05227 O05228 O05229 O05241 O05245 O05248 O05249 O05250 O05254 O05255 O05256
O05257 O05259 O05260 O05318 O05329 O05330 O05331 O05332 O05333 O05390 O05395
O05399 O05401 O05402 O05403 O05406 O05407 O05442 O05458 O05461 O05467 O05493
O05500 O05507 O05522 O05523 O05524 O05525 O05543 O05560 O05731 O05883 O05938
O05962 O05963 O05976 O06005 O06079 O06081 O06136 O06291 O06342 O06349 O06432

O06433 O06434 O06473 O06475 O06481 O06493 O06640 O06653 O06659 O06725 O06730
O06740 O06748 O06750 O06751 O06752 O06753 O06754 O06758 O06836 O06844 O06873
O06900 O06912 O06923 O06967 O06979 O06981 O06984 O06990 O06991 O06992 O06999
O07002 O07003 O07011 O07017 O07080 O07084 O07086 O07091 O07107 O07134 O07295
O07344 O07350 O07366 O07380 O07395 O07443 O07515 O07516 O07518 O07520 O07523
O07533 O07536 O07543 O07549 O07550 O07553 O07556 O07557 O07559 O07560 O07563
O07568 O07569 O07576 O07577 O07580 O07581 O07585 O07589 O07590 O07591 O07594
O07596 O07599 O07601 O07605 O07620 O07625 O07626 O07628 O07630 O07632 O07639
O07650 O07668 O07777 O07801 O07802 O07837 O07838 O07881 O07886 O07887 O07888
O07899 O07923 O07940 O07942 O08310 O08387 O08455 O08462 O08520 O08522 O08530
O08532 O08547 O08556 O08562 O08564 O08565 O08579 O08581 O08589 O08590 O08595
O08597 O08617 O08625 O08644 O08650 O08675 O08680 O08703 O08705 O08707 O08715
O08721 O08722 O08725 O08726 O08734 O08742 O08747 O08770 O08775 O08786 O08790
O08812 O08832 O08835 O08843 O08852 O08858 O08878 O08888 O08889 O08890 O08892
O08893 O08912 O08914 O08962 O08966 O08984 O09005 O09008 O09009 O09010 O09014
O09027 O09028 O09030 O09047 O09101 O09111 O09117 O09126 O09127 O09160 O09198
O09495 O09800 O10236 O10620 O10683 O10684 O10685 O10686 O10687 O11283 O11457
O11982 O12000 O12163 O12164 O12674 O12947 O12948 O12971 O12972 O12976 O12977
O13001 O13018 O13076 O13082 O13085 O13092 O13097 O13113 O13134 O13146 O13147
O13227 O13262 O13263 O13306 O13317 O13332 O13353 O13356 O13359 O13375 O13394
O13395 O13444 O13498 O13511 O13512 O13514 O13518 O13520 O13522 O13523 O13529
O13530 O13533 O13537 O13540 O13543 O13544 O13545 O13550 O13554 O13560 O13561
O13562 O13566 O13567 O13568 O13570 O13572 O13579 O13583 O13584 O13597 O13619
O13621 O13633 O13636 O13644 O13656 O13657 O13660 O13663 O13666 O13679 O13681
O13689 O13695 O13709 O13712 O13714 O13717 O13718 O13726 O13729 O13737 O13742
O13746 O13752 O13760 O13761 O13770 O13779 O13780 O13785 O13787 O13789 O13793
O13805 O13809 O13813 O13814 O13818 O13825 O13844 O13846 O13861 O13879 O13880
O13883 O13898 O13901 O13904 O13909 O13912 O13918 O13920 O13931 O13932 O13933
O13934 O13938 O13941 O13942 O13946 O13950 O13954 O13960 O13961 O13965 O13967
O13968 O13980 O13981 O13987 O13989 O13994 O13995 O13999 O14001 O14020 O14022
O14031 O14035 O14038 O14042 O14046 O14050 O14065 O14067 O14072 O14073 O14084
O14090 O14091 O14103 O14107 O14119 O14123 O14136 O14142 O14166 O14169 O14173
O14175 O14193 O14194 O14197 O14199 O14200 O14201 O14202 O14217 O14223 O14225
O14234 O14237 O14238 O14239 O14244 O14252 O14255 O14260 O14264 O14267 O14281
O14286 O14287 O14290 O14300 O14320 O14322 O14324 O14329 O14333 O14345 O14346
O14355 O14357 O14364 O14493 O14494 O14495 O14511 O14514 O14520 O14521 O14522
O14523 O14524 O14525 O14569 O14581 O14609 O14626 O14638 O14649 O14653 O14662
O14668 O14669 O14672 O14678 O14681 O14683 O14684 O14718 O14735 O14763 O14764
O14786 O14788 O14804 O14817 O14828 O14836 O14842 O14843 O14863 O14880 O14894
O14917 O14925 O14931 O14944 O14949 O14957 O14967 O14975 O14983 O15031 O15079
O15118 O15120 O15121 O15126 O15127 O15146 O15155 O15162 O15165 O15173 O15197
O15218 O15229 O15239 O15243 O15244 O15245 O15247 O15258 O15260 O15269 O15270
O15303 O15321 O15342 O15354 O15374 O15375 O15389 O15393 O15394 O15399 O15400
O15403 O15427 O15431 O15432 O15438 O15439 O15440 O15455 O15466 O15482 O15503

O15529 O15533 O15547 O15551 O15552 O15554 O15943 O15973 O15974 O16000 O16005
O16017 O16018 O16019 O16020 O16110 O16242 O16262 O16374 O16406 O16444 O16452
O16544 O16715 O16796 O16881 O16887 O16974 O16975 O17040 O17185 O17204 O17240
O17287 O17386 O17405 O17444 O17528 O17554 O17645 O17699 O17798 O17800 O17816
O17818 O17819 O17820 O17821 O17822 O17823 O17842 O17844 O17846 O17847 O17956
O18016 O18023 O18037 O18175 O18276 O18280 O18303 O18304 O18312 O18315 O18405
O18412 O18423 O18424 O18425 O18481 O18485 O18486 O18497 O18498 O18689 O18690
O18735 O18738 O18756 O18757 O18765 O18766 O18784 O18793 O18796 O18821 O18824
O18839 O18851 O18866 O18867 O18868 O18875 O18882 O18894 O18906 O18910 O18911
O18912 O18913 O18914 O18917 O18926 O18935 O18951 O18956 O18965 O18968 O18982
O18983 O19005 O19012 O19014 O19024 O19025 O19032 O19037 O19054 O19058 O19071
O19091 O19103 O19105 O19131 O19133 O19179 O19182 O19477 O19882 O19893 O19895
O19901 O19902 O19913 O19916 O19922 O19925 O19926 O19927 O19928 O19930 O20030
O20272 O20433 O20434 O20435 O20539 O20540 O20544 O20546 O20547 O20549 O20603
O20604 O20611 O20612 O20613 O20644 O20645 O20672 O20964 O20965 O20968 O20998
O21001 O21003 O21004 O21039 O21042 O21047 O21048 O21049 O21069 O21070 O21077
O21078 O21079 O21159 O21163 O21166 O21167 O21175 O21205 O21217 O21220 O21223
O21224 O21243 O21266 O21273 O21298 O21325 O21326 O21327 O21329 O21330 O21331
O21332 O21333 O21334 O21335 O21336 O21337 O21366 O21397 O21398 O21399 O21400
O21401 O21402 O21403 O21405 O21406 O21407 O21408 O21409 O21411 O21414 O21416
O21424 O21425 O21513 O21514 O21515 O21517 O21519 O21520 O21521 O21535 O21540
O21543 O21546 O21549 O21554 O21558 O21560 O21563 O21566 O21567 O21569 O21572
O21575 O21578 O21581 O21584 O21587 O21590 O21592 O21595 O21598 O21601 O21604
O21607 O21610 O21613 O21616 O21619 O21764 O21783 O21787 O21789 O21795 O21798
O21804 O21807 O21816 O21942 O22007 O22008 O22048 O22049 O22141 O22151 O22187
O22203 O22208 O22218 O22252 O22255 O22261 O22267 O22285 O22342 O22397 O22476
O22552 O22588 O22622 O22704 O22719 O22752 O22755 O22757 O22775 O22780 O22784
O22802 O22807 O22808 O22815 O22824 O22833 O22834 O22881 O22912 O22920 O22925
O22938 O22943 O22960 O22993 O23035 O23039 O23040 O23051 O23053 O23066 O23081
O23086 O23087 O23144 O23164 O23342 O23373 O23386 O23413 O23429 O23461 O23482
O23492 O23596 O23680 O23693 O23913 O24011 O24060 O24088 O24303 O24312 O24367
O24381 O24509 O24585 O24594 O24723 O24817 O24849 O24866 O24867 O24958 O24972
O24982 O24989 O25004 O25088 O25170 O25235 O25291 O25300 O25369 O25396 O25425
O25455 O25526 O25551 O25578 O25582 O25609 O25700 O25701 O25722 O25754 O25755
O25788 O25792 O25797 O25823 O25832 O25847 O25857 O25860 O25879 O25888 O25889
O25897 O25898 O25899 O25901 O25917 O25986 O25989 O26018 O26024 O26039 O26073
O26074 O26075 O26076 O26134 O26206 O26234 O26240 O26317 O26478 O26484 O26521
O26618 O26669 O26757 O26759 O26788 O26830 O26878 O26920 O26929 O26936 O26937
O26973 O26990 O27041 O27101 O27106 O27134 O27170 O27184 O27225 O27226 O27227
O27228 O27229 O27230 O27231 O27279 O27318 O27329 O27435 O27460 O27479 O27564
O27713 O27726 O27741 O27840 O27961 O27993 O28044 O28047 O28052 O28069 O28078
O28092 O28100 O28116 O28118 O28122 O28124 O28147 O28148 O28149 O28150 O28156
O28158 O28166 O28169 O28178 O28221 O28231 O28238 O28243 O28251 O28254 O28269
O28280 O28288 O28330 O28377 O28435 O28452 O28474 O28476 O28482 O28498 O28517

O28519 O28525 O28528 O28534 O28619 O28640 O28652 O28655 O28656 O28674 O28675
O28688 O28690 O28693 O28694 O28695 O28699 O28703 O28710 O28712 O28720 O28726
O28739 O28748 O28773 O28785 O28790 O28797 O28810 O28812 O28815 O28817 O28838
O28846 O28851 O28852 O28933 O28934 O28942 O28968 O28985 O29000 O29002 O29019
O29043 O29047 O29105 O29106 O29169 O29246 O29249 O29285 O29292 O29316 O29338
O29371 O29391 O29397 O29398 O29416 O29422 O29423 O29470 O29487 O29495 O29496
O29497 O29499 O29506 O29522 O29529 O29530 O29546 O29561 O29565 O29573 O29600
O29670 O29709 O29710 O29714 O29727 O29740 O29747 O29749 O29750 O29762 O29777
O29793 O29843 O29848 O29853 O29867 O29878 O29883 O29918 O29939 O29972 O29973
O29976 O29981 O30003 O30004 O30049 O30065 O30074 O30079 O30083 O30085 O30088
O30089 O30091 O30092 O30114 O30131 O30135 O30138 O30139 O30141 O30143 O30151
O30195 O30198 O30248 O30270 O30278 O30318 O30387 O30417 O30499 O30513 O30646
O30795 O30990 O30993 O31000 O31100 O31101 O31126 O31127 O31128 O31185 O31214
O31215 O31216 O31250 O31285 O31289 O31421 O31425 O31428 O31431 O31433 O31442
O31444 O31445 O31446 O31447 O31448 O31453 O31460 O31462 O31464 O31467 O31469
O31479 O31485 O31487 O31488 O31491 O31493 O31507 O31514 O31515 O31516 O31519
O31520 O31522 O31533 O31539 O31540 O31543 O31553 O31554 O31555 O31559 O31561
O31563 O31568 O31569 O31577 O31580 O31582 O31583 O31593 O31600 O31603 O31615
O31623 O31634 O31638 O31645 O31651 O31652 O31655 O31657 O31658 O31660 O31670
O31671 O31674 O31686 O31688 O31695 O31696 O31707 O31708 O31709 O31710 O31712
O31715 O31729 O31752 O31754 O31762 O31771 O31773 O31774 O31780 O31785 O31789
O31793 O31804 O31808 O31809 O31811 O31813 O31814 O31815 O31816 O31820 O31821
O31823 O31830 O31833 O31845 O31855 O31859 O31861 O31862 O31867 O31872 O31873
O31884 O31888 O31919 O31923 O31929 O31930 O31933 O31934 O31937 O31941 O31974
O31975 O31984 O31989 O32015 O32022 O32031 O32032 O32045 O32047 O32050 O32051
O32052 O32060 O32064 O32071 O32074 O32077 O32081 O32082 O32084 O32086 O32087
O32095 O32100 O32101 O32104 O32105 O32107 O32109 O32111 O32115 O32131 O32134
O32139 O32140 O32154 O32155 O32168 O32182 O32193 O32198 O32199 O32202 O32203
O32204 O32205 O32206 O32209 O32212 O32216 O32217 O32219 O32220 O32225 O32226
O32227 O32233 O32239 O32241 O32244 O32246 O32247 O32256 O32257 O32261 O32262
O32269 O32270 O32273 O32274 O32278 O32282 O32327 O32328 O32332 O32333 O32521
O32522 O32591 O32617 O32619 O32629 O32859 O32864 O32865 O32866 O32867 O32868
O32870 O32871 O32882 O33006 O33021 O33057 O33071 O33192 O33369 O33407 O33507
O33517 O33518 O33533 O33551 O33568 O33641 O33654 O33683 O33787 O33789 O33814
O33821 O33854 O33953 O33980 O34206 O34238 O34245 O34307 O34311 O34315 O34319
O34322 O34329 O34341 O34343 O34349 O34356 O34358 O34367 O34368 O34375 O34378
O34382 O34383 O34416 O34424 O34427 O34428 O34430 O34431 O34434 O34436 O34437
O34440 O34442 O34447 O34451 O34456 O34460 O34470 O34472 O34474 O34486 O34500
O34502 O34505 O34506 O34507 O34509 O34518 O34521 O34524 O34525 O34535 O34545
O34546 O34551 O34560 O34572 O34573 O34575 O34578 O34586 O34597 O34606 O34610
O34616 O34618 O34624 O34636 O34638 O34648 O34649 O34653 O34654 O34659 O34663
O34671 O34674 O34684 O34686 O34691 O34694 O34706 O34708 O34713 O34718 O34719
O34724 O34726 O34733 O34734 O34738 O34739 O34741 O34742 O34745 O34752 O34753
O34755 O34757 O34758 O34765 O34774 O34784 O34792 O34798 O34811 O34830 O34832

| | | | | | | | | | | |
|---|---|---|---|---|---|---|---|---|---|---|
| O34840 | O34849 | O34853 | O34864 | O34876 | O34878 | O34881 | O34883 | O34894 | O34897 | O34898 |
| O34905 | O34908 | O34922 | O34927 | O34929 | O34931 | O34933 | O34952 | O34953 | O34961 | O34971 |
| O34978 | O34987 | O34989 | O34991 | O34997 | O35004 | O35005 | O35007 | O35018 | O35024 | O35027 |
| O35029 | O35036 | O35043 | O35044 | O35048 | O35052 | O35054 | O35074 | O35082 | O35083 | O35089 |
| O35092 | O35093 | O35111 | O35112 | O35114 | O35119 | O35136 | O35149 | O35152 | O35153 | O35158 |
| O35161 | O35165 | O35166 | O35173 | O35174 | O35181 | O35188 | O35210 | O35214 | O35219 | O35227 |
| O35235 | O35240 | O35245 | O35276 | O35296 | O35304 | O35305 | O35308 | O35316 | O35372 | O35375 |
| O35379 | O35393 | O35394 | O35405 | O35409 | O35425 | O35426 | O35433 | O35435 | O35439 | O35440 |
| O35445 | O35449 | O35451 | O35453 | O35454 | O35457 | O35458 | O35464 | O35465 | O35469 | O35476 |
| O35478 | O35488 | O35505 | O35516 | O35526 | O35532 | O35544 | O35547 | O35548 | O35566 | O35569 |
| O35587 | O35595 | O35598 | O35599 | O35600 | O35604 | O35607 | O35609 | O35623 | O35633 | O35659 |
| O35664 | O35674 | O35681 | O35682 | O35683 | O35696 | O35704 | O35714 | O35734 | O35764 | O35776 |
| O35777 | O35778 | O35786 | O35789 | O35790 | O35795 | O35799 | O35811 | O35874 | O35881 | O35899 |
| O35902 | O35910 | O35912 | O35913 | O35921 | O35930 | O35932 | O35949 | O35956 | O35988 | O36020 |
| O36022 | O36028 | O36359 | O36362 | O36364 | O36372 | O36377 | O36388 | O36400 | O36401 | O36403 |
| O36408 | O36410 | O36416 | O36417 | O36423 | O36424 | O36632 | O36633 | O36634 | O39519 | O39830 |
| O39842 | O39920 | O39927 | O39928 | O39929 | O40984 | O41517 | O41803 | O41933 | O42043 | O42127 |
| O42153 | O42154 | O42179 | O42204 | O42224 | O42266 | O42268 | O42281 | O42282 | O42294 | O42300 |
| O42301 | O42307 | O42327 | O42328 | O42329 | O42330 | O42334 | O42335 | O42384 | O42385 | O42398 |
| O42411 | O42412 | O42414 | O42422 | O42427 | O42431 | O42449 | O42451 | O42452 | O42466 | O42490 |
| O42574 | O42579 | O42580 | O42581 | O42582 | O42583 | O42596 | O42602 | O42603 | O42604 | O42633 |
| O42646 | O42656 | O42690 | O42701 | O42707 | O42826 | O42841 | O42843 | O42852 | O42885 | O42901 |
| O42907 | O42916 | O42922 | O42930 | O42933 | O42940 | O42944 | O42967 | O42968 | O42972 | O42976 |
| O42997 | O42999 | O43000 | O43002 | O43009 | O43019 | O43022 | O43023 | O43033 | O43043 | O43053 |
| O43062 | O43073 | O43080 | O43081 | O43085 | O43092 | O43108 | O43155 | O43157 | O43169 | O43173 |
| O43184 | O43193 | O43194 | O43246 | O43286 | O43291 | O43292 | O43300 | O43306 | O43315 | O43424 |
| O43451 | O43462 | O43464 | O43490 | O43493 | O43497 | O43505 | O43506 | O43508 | O43511 | O43520 |
| O43525 | O43526 | O43529 | O43556 | O43557 | O43561 | O43567 | O43570 | O43581 | O43603 | O43613 |
| O43614 | O43657 | O43674 | O43676 | O43677 | O43688 | O43699 | O43731 | O43736 | O43749 | O43752 |
| O43759 | O43760 | O43761 | O43772 | O43808 | O43819 | O43825 | O43826 | O43861 | O43868 | O43869 |
| O43889 | O43908 | O43909 | O43914 | O43916 | O43934 | O44017 | O44186 | O44252 | O44386 | O44390 |
| O44443 | O44477 | O44616 | O44827 | O44857 | O44953 | O45102 | O45145 | O45150 | O45166 | O45293 |
| O45300 | O45306 | O45363 | O45435 | O45503 | O45657 | O45731 | O45767 | O45870 | O45876 | O45923 |
| O45947 | O46077 | O46084 | O46101 | O46372 | O46373 | O46383 | O46420 | O46491 | O46502 | O46512 |
| O46516 | O46522 | O46547 | O46561 | O46577 | O46578 | O46579 | O46580 | O46581 | O46582 | O46584 |
| O46585 | O46586 | O46587 | O46588 | O46589 | O46590 | O46598 | O46600 | O46631 | O46634 | O46635 |
| O46639 | O46651 | O46669 | O46674 | O46680 | O46685 | O47040 | O47042 | O47043 | O47044 | O47414 |
| O47415 | O47420 | O47421 | O47423 | O47426 | O47427 | O47428 | O47430 | O47431 | O47434 | O47475 |
| O47476 | O47477 | O47478 | O47479 | O47488 | O47491 | O47492 | O47493 | O47494 | O47495 | O47496 |
| O47497 | O47498 | O47499 | O47554 | O47555 | O47556 | O47557 | O47558 | O47560 | O47561 | O47583 |
| O47584 | O47667 | O47668 | O47669 | O47670 | O47671 | O47672 | O47673 | O47674 | O47675 | O47676 |
| O47677 | O47678 | O47679 | O47680 | O47681 | O47685 | O47686 | O47687 | O47688 | O47689 | O47690 |
| O47691 | O47692 | O47693 | O47694 | O47695 | O47696 | O47697 | O47698 | O47699 | O47700 | O47701 |
| O47702 | O47705 | O47706 | O47708 | O47709 | O47710 | O47714 | O47716 | O47718 | O47721 | O47723 |

O47815 O47868 O47870 O47871 O47872 O47874 O47881 O47885 O47890 O47892 O47893
O47922 O47923 O47924 O47926 O47930 O47976 O47993 O48001 O48009 O48010 O48012
O48014 O48017 O48023 O48025 O48027 O48039 O48041 O48043 O48047 O48049 O48052
O48053 O48055 O48057 O48060 O48062 O48063 O48065 O48066 O48067 O48073 O48076
O48079 O48080 O48085 O48087 O48089 O48090 O48092 O48093 O48094 O48096 O48098
O48100 O48101 O48102 O48106 O48109 O48111 O48114 O48267 O48276 O48277 O48285
O48309 O48316 O48321 O48334 O48336 O48350 O48358 O48372 O48374 O48410 O48470
O48472 O48528 O48529 O48538 O48639 O48651 O48666 O48670 O48671 O48682 O48684
O48763 O48780 O48786 O48788 O48842 O48843 O48845 O48849 O48850 O48851 O48916
O48922 O48929 O48946 O48947 O48950 O48956 O48958 O48962 O49078 O49187 O49230
O49313 O49318 O49323 O49325 O49328 O49329 O49340 O49342 O49347 O49377 O49394
O49396 O49423 O49445 O49447 O49460 O49545 O49564 O49567 O49621 O49636 O49639
O49660 O49668 O49691 O49696 O49722 O49725 O49726 O49730 O49814 O49815 O49873
O49879 O49929 O49931 O50440 O50500 O50501 O50651 O51039 O51042 O51051 O51055
O51058 O51063 O51067 O51068 O51073 O51083 O51090 O51100 O51118 O51144 O51145
O51195 O51221 O51222 O51253 O51265 O51266 O51272 O51273 O51280 O51337 O51356
O51398 O51425 O51451 O51457 O51489 O51595 O51596 O51597 O51693 O51750 O51798
O51808 O51809 O51810 O51829 O51876 O51877 O51878 O51880 O51924 O52042 O52043
O52044 O52069 O52196 O52213 O52214 O52216 O52248 O52324 O52327 O52328 O52351
O52366 O52401 O52474 O52475 O52619 O52717 O52718 O52728 O52733 O52788 O52903
O53092 O53114 O53150 O53176 O53368 O53380 O53493 O53505 O53508 O53581 O53582
O53645 O53695 O53857 O53896 O53900 O53945 O54029 O54031 O54069 O54070 O54088
O54089 O54101 O54189 O54190 O54239 O54243 O54290 O54339 O54340 O54569 O54689
O54693 O54694 O54695 O54697 O54698 O54699 O54701 O54702 O54707 O54709 O54715
O54728 O54732 O54734 O54766 O54767 O54794 O54798 O54799 O54800 O54803 O54814
O54827 O54851 O54852 O54853 O54861 O54862 O54885 O54890 O54897 O54898 O54901
O54902 O54904 O54905 O54907 O54909 O54912 O54942 O54947 O54951 O54957 O54965
O54980 O54982 O54990 O54991 O55003 O55005 O55017 O55022 O55026 O55034 O55040
O55075 O55100 O55101 O55111 O55134 O55143 O55145 O55159 O55174 O55192 O55193
O55197 O55227 O55237 O55240 O55242 O55246 O55247 O55647 O55653 O55654 O55655
O55703 O55704 O55705 O55708 O55714 O55715 O55728 O55733 O55744 O55746 O55747
O55762 O56140 O56677 O56850 O56861 O57166 O57199 O57201 O57205 O57208 O57211
O57222 O57254 O57295 O57314 O57321 O57328 O57329 O57379 O57405 O57409 O57422
O57424 O57425 O57426 O57427 O57428 O57474 O57483 O57579 O57590 O57593 O57604
O57605 O57713 O57721 O57753 O57781 O57893 O57898 O57953 O58081 O58111 O58114
O58215 O58221 O58348 O58374 O58405 O58499 O58728 O58759 O58760 O58967 O58968
O58981 O58997 O59010 O59045 O59171 O59179 O59180 O59276 O59442 O59638 O59640
O59649 O59659 O59666 O59674 O59680 O59682 O59698 O59699 O59700 O59708 O59712
O59714 O59715 O59718 O59725 O59726 O59729 O59735 O59738 O59744 O59745 O59747
O59758 O59764 O59768 O59774 O59776 O59781 O59782 O59785 O59795 O59802 O59807
O59813 O59814 O59815 O59824 O59831 O59833 O59866 O59868 O59932 O59933 O59942
O60029 O60061 O60067 O60069 O60095 O60103 O60109 O60111 O60113 O60119 O60120
O60123 O60131 O60146 O60148 O60159 O60162 O60168 O60170 O60235 O60238 O60241
O60242 O60243 O60245 O60266 O60279 O60309 O60312 O60313 O60320 O60330 O60337

O60353 O60359 O60391 O60403 O60404 O60412 O60423 O60427 O60431 O60443 O60449
O60462 O60469 O60476 O60478 O60486 O60487 O60488 O60499 O60500 O60503 O60507
O60512 O60513 O60602 O60603 O60635 O60636 O60637 O60656 O60667 O60669 O60704
O60706 O60721 O60725 O60733 O60741 O60755 O60774 O60779 O60830 O60831 O60840
O60858 O60883 O60894 O60895 O60896 O60906 O60909 O60928 O60931 O60939 O60950
O61213 O61303 O61307 O61365 O61369 O61370 O61394 O61397 O61460 O61703 O61715
O61787 O61788 O61856 O61892 O61898 O62026 O62126 O62136 O62169 O62179 O62215
O62266 O62267 O62269 O62367 O62368 O62420 O62488 O62489 O62646 O62653 O62667
O62709 O62714 O62729 O62735 O62743 O62747 O62772 O62786 O62787 O62791 O62792
O62793 O62794 O62795 O62796 O62798 O62802 O62809 O62816 O62826 O62837 O62848
O62849 O62852 O62939 O63066 O63075 O63534 O63535 O63536 O63537 O63623 O63696
O63699 O63767 O63775 O63794 O63796 O63797 O63798 O63848 O63849 O63850 O63851
O63902 O63908 O63912 O63913 O63915 O64032 O64038 O64204 O64362 O64390 O64425
O64471 O64474 O64477 O64483 O64497 O64503 O64515 O64556 O64614 O64635 O64636
O64637 O64638 O64639 O64668 O64697 O64698 O64718 O64725 O64738 O64749 O64758
O64759 O64761 O64762 O64763 O64769 O64770 O64771 O64774 O64776 O64777 O64778
O64780 O64781 O64782 O64783 O64784 O64791 O64792 O64793 O64806 O64823 O64825
O64834 O64837 O64847 O64884 O64886 O64899 O64900 O64964 O64966 O64967 O64989
O65023 O65085 O65101 O65238 O65272 O65359 O65402 O65403 O65404 O65405 O65413
O65431 O65438 O65440 O65468 O65472 O65476 O65479 O65482 O65483 O65530 O65677
O65688 O65708 O65709 O65717 O65718 O65726 O65727 O65782 O65784 O65785 O65786
O65787 O65788 O65790 O65797 O65837 O65924 O65934 O65989 O66043 O66076 O66103
O66137 O66139 O66141 O66163 O66414 O66465 O66466 O66477 O66478 O66491 O66504
O66505 O66510 O66515 O66525 O66544 O66545 O66561 O66564 O66566 O66624 O66627
O66643 O66685 O66688 O66698 O66757 O66760 O66767 O66771 O66807 O66818 O66819
O66830 O66845 O66874 O66905 O66948 O66959 O66960 O66966 O66994 O67000 O67008
O67017 O67062 O67077 O67102 O67122 O67127 O67171 O67241 O67248 O67265 O67270
O67292 O67295 O67303 O67304 O67305 O67319 O67336 O67339 O67342 O67364 O67365
O67379 O67388 O67391 O67421 O67427 O67428 O67433 O67446 O67472 O67513 O67521
O67526 O67536 O67545 O67601 O67622 O67637 O67658 O67662 O67688 O67692 O67694
O67696 O67701 O67712 O67734 O67738 O67748 O67750 O67773 O67774 O67776 O67790
O67798 O67813 O67827 O67838 O67860 O67882 O67901 O67908 O67910 O67939 O68038
O68104 O68253 O68284 O68286 O68433 O68460 O68611 O68616 O68826 O68827 O68852
O68874 O68900 O69053 O69062 O69064 O69076 O69103 O69279 O69443 O69555 O69582
O69583 O69601 O69659 O69690 O69704 O69726 O69729 O70127 O70129 O70131 O70141
O70156 O70174 O70212 O70215 O70228 O70247 O70252 O70255 O70257 O70258 O70281
O70309 O70320 O70324 O70337 O70339 O70342 O70344 O70352 O70355 O70367 O70394
O70397 O70401 O70404 O70410 O70418 O70421 O70423 O70430 O70431 O70432 O70439
O70451 O70458 O70461 O70472 O70480 O70491 O70496 O70503 O70507 O70526 O70528
O70531 O70534 O70535 O70536 O70540 O70570 O70572 O70577 O70578 O70579 O70594
O70595 O70596 O70597 O70601 O70610 O70617 O70622 O70632 O70891 O70901 O70902
O71037 O71191 O71196 O71197 O71305 O72157 O72737 O72896 O72899 O73564 O73606
O73612 O73683 O73698 O73700 O73705 O73706 O73707 O73791 O73792 O73798 O73810
O73874 O73875 O73878 O73895 O73925 O74088 O74164 O74165 O74180 O74189 O74208

O74212 O74248 O74312 O74323 O74327 O74333 O74368 O74369 O74371 O74375 O74377

O74380 O74383 O74384 O74395 O74420 O74430 O74431 O74433 O74437 O74439 O74441

O74444 O74446 O74466 O74471 O74474 O74475 O74481 O74486 O74487 O74495 O74497

O74505 O74507 O74510 O74512 O74520 O74525 O74537 O74543 O74546 O74551 O74557

O74559 O74564 O74631 O74676 O74702 O74713 O74728 O74745 O74750 O74756 O74757

O74787 O74820 O74827 O74829 O74837 O74838 O74839 O74843 O74847 O74849 O74850

O74852 O74869 O74870 O74872 O74878 O74882 O74888 O74899 O74901 O74902 O74920

O74921 O74923 O74926 O74943 O74947 O74949 O74963 O74969 O74988 O74992 O75019

O75022 O75023 O75027 O75051 O75054 O75056 O75063 O75069 O75072 O75074 O75077

O75078 O75084 O75096 O75110 O75121 O75129 O75144 O75185 O75192 O75197 O75204

O75264 O75298 O75309 O75310 O75311 O75324 O75325 O75352 O75354 O75355 O75379

O75387 O75388 O75396 O75425 O75438 O75445 O75452 O75460 O75473 O75477 O75503

O75508 O75509 O75578 O75581 O75631 O75712 O75746 O75751 O75752 O75762 O75783

O75787 O75795 O75829 O75841 O75844 O75845 O75871 O75880 O75881 O75882 O75899

O75900 O75907 O75908 O75911 O75915 O75923 O75949 O75954 O75976 O76000 O76001

O76002 O76024 O76036 O76062 O76082 O76090 O76095 O76099 O76100 O76329 O76411

O76554 O76565 O76689 O76735 O76767 O76819 O76997 O77049 O77081 O77286 O77408

O77438 O77475 O77485 O77486 O77487 O77504 O77510 O77564 O77590 O77592 O77616

O77621 O77633 O77649 O77680 O77696 O77697 O77700 O77713 O77714 O77715 O77721

O77722 O77723 O77726 O77735 O77736 O77737 O77740 O77741 O77750 O77751 O77759

O77760 O77761 O77764 O77769 O77770 O77780 O77783 O77792 O77808 O77830 O77836

O78415 O78416 O78426 O78427 O78433 O78437 O78444 O78446 O78448 O78455 O78456

O78457 O78460 O78462 O78463 O78464 O78465 O78466 O78467 O78468 O78469 O78470

O78471 O78477 O78478 O78479 O78480 O78493 O78494 O78498 O78499 O78500 O78503

O78504 O78505 O78507 O78508 O78511 O78512 O78513 O78514 O78516 O78676 O78679

O78680 O78681 O78682 O78683 O78684 O78686 O78687 O78688 O78689 O78690 O78693

O78694 O78695 O78696 O78697 O78698 O78699 O78700 O78701 O78702 O78703 O78704

O78705 O78706 O78707 O78708 O78709 O78710 O78711 O78712 O78713 O78714 O78715

O78742 O78747 O78748 O78749 O78750 O78751 O78752 O78753 O78754 O78755 O78756

O78757 O78761 O78773 O78774 O78775 O78776 O78777 O78778 O78779 O78780 O78781

O78782 O78783 O78784 O78785 O78786 O78787 O78788 O78789 O78790 O78858 O78927

O78928 O78929 O78930 O78931 O78932 O78933 O78934 O78935 O78936 O78937 O79102

O79122 O79195 O79196 O79197 O79198 O79199 O79200 O79201 O79202 O79203 O79204

O79205 O79206 O79207 O79208 O79209 O79210 O79211 O79212 O79213 O79215 O79216

O79217 O79218 O79219 O79220 O79221 O79222 O79223 O79224 O79225 O79226 O79227

O79228 O79229 O79230 O79274 O79311 O79327 O79360 O79386 O79396 O79403 O79404

O79405 O79406 O79407 O79408 O79409 O79410 O79411 O79412 O79413 O79417 O79421

O79422 O79426 O79427 O79428 O79429 O79431 O79432 O79433 O79434 O79435 O79436

O79437 O79438 O79440 O79445 O79446 O79447 O79448 O79450 O79451 O79452 O79453

O79454 O79455 O79460 O79461 O79462 O79463 O79464 O79465 O79468 O79512 O79520

O79524 O79546 O79547 O79548 O79549 O79550 O79551 O79552 O79553 O79554 O79555

O79556 O79557 O79558 O79648 O79652 O79657 O79659 O79660 O79670 O79671 O79672

O79673 O79674 O79675 O79676 O79677 O79678 O79679 O79680 O79785 O79874 O79875

O79876 O79880 O79881 O79882 O79925 O79927 O79936 O79967 O79969 O79980 O79993

O79996 O79997 O80003 O80005 O80019 O80288 O80292 O80295 O80296 O80297 O80298
O80299 O80300 O80436 O80437 O80452 O80458 O80493 O80505 O80565 O80580 O80592
O80594 O80605 O80612 O80623 O80632 O80638 O80668 O80685 O80695 O80725 O80739
O80809 O80813 O80823 O80832 O80844 O80845 O80854 O80860 O80872 O80885 O80891
O80898 O80899 O80905 O80915 O80919 O80927 O80939 O80940 O80946 O80952 O80961
O80963 O80977 O80983 O81000 O81006 O81007 O81016 O81045 O81053 O81062 O81069
O81072 O81077 O81078 O81108 O81117 O81122 O81123 O81208 O81209 O81210 O81211
O81214 O81283 O81291 O81292 O81312 O81345 O81346 O81514 O81775 O81776 O81833
O81845 O81850 O81866 O81898 O81905 O81906 O81916 O81928 O81931 O81970 O81973
O82067 O82139 O82150 O82226 O82232 O82244 O82245 O82251 O82279 O82316 O82318
O82352 O82353 O82390 O82427 O82436 O82485 O82509 O82567 O82568 O82587 O82598
O82643 O82709 O82752 O82756 O82766 O82775 O82807 O82811 O82855 O82859 O82860
O83005 O83045 O83074 O83079 O83094 O83109 O83131 O83151 O83152 O83163 O83174
O83183 O83184 O83203 O83211 O83251 O83254 O83256 O83263 O83277 O83279 O83284
O83297 O83306 O83323 O83324 O83344 O83346 O83355 O83358 O83365 O83383 O83396
O83414 O83425 O83426 O83432 O83444 O83478 O83486 O83493 O83494 O83497 O83514
O83529 O83534 O83544 O83547 O83564 O83575 O83582 O83586 O83602 O83609 O83617
O83645 O83681 O83685 O83705 O83706 O83709 O83710 O83734 O83744 O83746 O83762
O83766 O83767 O83824 O83848 O83898 O83941 O83943 O83964 O83968 O83978 O83981
O83995 O84000 O84008 O84009 O84039 O84068 O84072 O84073 O84075 O84179 O84253
O84254 O84280 O84281 O84282 O84283 O84307 O84377 O84379 O84413 O84417 O84418
O84419 O84422 O84449 O84457 O84467 O84480 O84502 O84539 O84548 O84584 O84698
O84744 O84745 O84746 O84762 O84818 O84826 O84860 O84875 O84876 O84877 O84879
O84880 O84882 O84969 O85061 O85068 O85133 O85187 O85228 O85273 O85296 O85297
O85467 O86038 O86088 O86164 O86199 O86200 O86220 O86221 O86225 O86226 O86229
O86230 O86231 O86232 O86233 O86234 O86576 O86810 O86988 O87005 O87077 O87196
O87296 O87394 O87406 O87557 O87567 O87576 O87579 O87626 O87656 O87786 O87787
O87866 O87868 O87953 O88022 O88037 O88039 O88093 O88174 O88177 O88178 O88186
O88199 O88204 O88269 O88272 O88277 O88278 O88298 O88307 O88319 O88324 O88335
O88338 O88343 O88384 O88393 O88397 O88407 O88410 O88416 O88419 O88420 O88422
O88427 O88444 O88446 O88454 O88455 O88457 O88472 O88488 O88495 O88496 O88516
O88536 O88537 O88551 O88552 O88561 O88563 O88572 O88575 O88576 O88587 O88602
O88626 O88627 O88628 O88630 O88634 O88662 O88671 O88676 O88680 O88689 O88693
O88703 O88704 O88705 O88713 O88721 O88728 O88736 O88741 O88758 O88759 O88775
O88777 O88779 O88786 O88788 O88792 O88799 O88813 O88819 O88822 O88823 O88824
O88829 O88833 O88839 O88853 O88854 O88855 O88856 O88867 O88870 O88871 O88876
O88884 O88908 O88909 O88917 O88923 O88932 O88941 O88943 O88944 O88956 O88962
O88967 O88983 O89001 O89013 O89016 O89026 O89035 O89039 O89047 O89051 O89103
O89104 O89109 O89116 O89292 O89335 O89342 O89343 O89746 O90299 O90304 O90305
O90369 O90371 O91085 O91086 O91262 O91464 O91534 O91744 O91745 O91936 O92284
O92367 O92374 O92500 O92529 O92530 O92531 O92532 O92921 O92972 O93122 O93248
O93274 O93279 O93295 O93319 O93336 O93361 O93383 O93403 O93430 O93441 O93459
O93505 O93533 O93603 O93740 O93741 O93742 O93743 O93788 O93853 O93875 O93927
O93956 O93980 O94030 O94084 O94085 O94142 O94227 O94254 O94262 O94270 O94280

O94283 O94287 O94296 O94298 O94299 O94300 O94302 O94323 O94327 O94335 O94339
O94341 O94342 O94343 O94344 O94348 O94349 O94353 O94361 O94370 O94372 O94380
O94385 O94391 O94401 O94402 O94409 O94414 O94433 O94439 O94440 O94448 O94457
O94458 O94464 O94469 O94472 O94473 O94479 O94490 O94491 O94502 O94511 O94520
O94523 O94528 O94531 O94537 O94539 O94543 O94546 O94550 O94565 O94569 O94572
O94573 O94584 O94587 O94591 O94592 O94593 O94594 O94596 O94597 O94599 O94607
O94618 O94622 O94624 O94633 O94639 O94654 O94661 O94673 O94680 O94689 O94694
O94695 O94702 O94705 O94719 O94722 O94727 O94738 O94744 O94759 O94766 O94777
O94778 O94823 O94826 O94856 O94876 O94886 O94898 O94901 O94905 O94910 O94911
O94923 O94933 O94956 O94966 O94985 O94991 O95006 O95007 O95013 O95047 O95069
O95070 O95136 O95139 O95140 O95150 O95159 O95167 O95168 O95169 O95180 O95183
O95185 O95196 O95197 O95202 O95206 O95210 O95214 O95221 O95222 O95237 O95249
O95255 O95256 O95258 O95259 O95264 O95279 O95292 O95297 O95298 O95342 O95371
O95377 O95395 O95406 O95415 O95427 O95436 O95452 O95461 O95470 O95471 O95473
O95476 O95477 O95484 O95490 O95500 O95502 O95528 O95562 O95563 O95573 O95622
O95665 O95672 O95674 O95727 O95754 O95772 O95800 O95803 O95807 O95832 O95833
O95838 O95847 O95857 O95858 O95859 O95864 O95866 O95870 O95907 O95918 O95944
O95976 O95977 O95992 O96002 O96005 O96008 O96011 O96024 O96099 O96107 O96797
O96798 O96799 O96801 O96807 O96810 O96811 O96812 O96813 O97148 O97239 O97364
O97394 O97422 O97470 O97504 O97512 O97531 O97562 O97571 O97583 O97596 O97605
O97626 O97649 O97661 O97663 O97664 O97665 O97666 O97676 O97681 O97703 O97704
O97711 O97741 O97742 O97772 O97797 O97799 O97817 O97827 O97831 O97878 O97879
O97880 O97881 O97882 O97883 O97951 O97961 O97962 O97967 O97969 O97975 O98693
O98733 O98736 O99041 O99240 O99252 O99253 O99255 O99256 O99257 O99258 O99338
O99342 O99343 O99345 O99468 O99598 O99602 O99655 O99795 O99797 O99798 O99799
O99800 O99817 O99818 O99819 O99820 O99821 O99822 O99823 O99824 O99825 O99826
O99827 O99828 O99869 O99978 P00125 P00130 P00155 P00156 P00157 P00158 P00159
P00160 P00161 P00163 P00164 P00165 P00166 P00167 P00168 P00169 P00170 P00171
P00172 P00173 P00174 P00347 P00388 P00389 P00395 P00396 P00397 P00398 P00399
P00400 P00401 P00402 P00403 P00405 P00406 P00407 P00408 P00409 P00410 P00411
P00412 P00413 P00414 P00415 P00416 P00417 P00418 P00419 P00420 P00421 P00422
P00423 P00424 P00425 P00430 P00533 P00545 P00550 P00726 P00803 P00804 P00840
P00842 P00845 P00846 P00847 P00848 P00849 P00850 P00851 P00852 P00853 P00854
P00856 P00858 P01130 P01131 P01132 P01133 P01134 P01135 P01136 P01375 P01501
P01502 P01504 P01514 P01546 P01589 P01590 P01730 P01731 P01732 P01832 P01833
P01848 P01849 P01850 P01851 P01852 P01853 P01865 P01867 P01869 P01871 P01872
P01880 P01882 P01889 P01893 P01894 P01895 P01896 P01897 P01898 P01899 P01900
P01901 P01902 P01903 P01904 P01906 P01909 P01910 P01911 P01915 P01920 P01921
P02683 P02699 P02700 P02706 P02707 P02708 P02709 P02710 P02711 P02712 P02713
P02714 P02715 P02716 P02717 P02718 P02719 P02722 P02723 P02724 P02725 P02726
P02727 P02730 P02748 P02786 P02916 P02918 P02919 P02920 P02921 P02929 P02930
P02931 P02932 P02935 P02936 P02941 P02942 P02943 P02945 P02947 P02948 P02950
P02956 P02957 P02973 P02974 P02975 P02978 P02980 P02981 P02982 P02983 P03077
P03079 P03084 P03085 P03086 P03093 P03094 P03095 P03096 P03097 P03138 P03139

P03140 P03141 P03142 P03143 P03144 P03145 P03172 P03173 P03178 P03182 P03185
P03188 P03192 P03196 P03200 P03205 P03208 P03215 P03218 P03224 P03225 P03230
P03231 P03242 P03250 P03314 P03315 P03316 P03319 P03320 P03321 P03372 P03374
P03375 P03377 P03378 P03379 P03380 P03381 P03383 P03384 P03385 P03386 P03387
P03388 P03389 P03390 P03391 P03393 P03394 P03396 P03397 P03398 P03399 P03415
P03420 P03423 P03425 P03435 P03436 P03437 P03438 P03439 P03440 P03441 P03442
P03443 P03444 P03445 P03446 P03447 P03448 P03449 P03450 P03451 P03452 P03453
P03454 P03455 P03456 P03457 P03458 P03459 P03460 P03461 P03462 P03463 P03464
P03465 P03468 P03469 P03470 P03471 P03472 P03473 P03474 P03475 P03476 P03477
P03478 P03479 P03480 P03481 P03482 P03484 P03491 P03492 P03493 P03517 P03518
P03522 P03524 P03535 P03540 P03600 P03609 P03618 P03619 P03620 P03621 P03622
P03623 P03624 P03625 P03626 P03639 P03655 P03656 P03657 P03658 P03661 P03663
P03664 P03665 P03666 P03667 P03668 P03674 P03676 P03678 P03701 P03705 P03726
P03759 P03762 P03777 P03791 P03802 P03803 P03819 P03825 P03873 P03877 P03886
P03887 P03888 P03889 P03890 P03891 P03892 P03893 P03894 P03895 P03896 P03897
P03898 P03899 P03900 P03901 P03902 P03903 P03904 P03905 P03906 P03907 P03908
P03909 P03910 P03911 P03912 P03913 P03914 P03915 P03916 P03917 P03918 P03919
P03920 P03921 P03922 P03923 P03924 P03925 P03926 P03927 P03928 P03929 P03930
P03931 P03933 P03945 P03959 P03960 P03961 P03985 P03986 P03987 P03991 P03999
P04000 P04001 P04011 P04027 P04035 P04038 P04039 P04068 P04074 P04116 P04123
P04124 P04135 P04136 P04140 P04157 P04159 P04160 P04165 P04166 P04175 P04191
P04195 P04201 P04218 P04221 P04223 P04224 P04227 P04228 P04230 P04231 P04233
P04234 P04235 P04274 P04288 P04371 P04372 P04373 P04412 P04439 P04440 P04441
P04479 P04480 P04488 P04494 P04502 P04505 P04512 P04513 P04540 P04575 P04577
P04578 P04579 P04580 P04581 P04582 P04583 P04624 P04626 P04629 P04633 P04658
P04659 P04660 P04661 P04662 P04663 P04664 P04683 P04684 P04709 P04710 P04737
P04739 P04755 P04756 P04757 P04758 P04759 P04760 P04761 P04774 P04775 P04778
P04779 P04780 P04781 P04782 P04783 P04784 P04817 P04821 P04839 P04840 P04843
P04844 P04845 P04849 P04850 P04853 P04855 P04856 P04868 P04869 P04872 P04875
P04882 P04883 P04884 P04919 P04920 P04921 P04924 P04926 P04950 P04953 P04966
P04967 P04996 P04997 P04998 P04999 P05023 P05024 P05025 P05026 P05027 P05028
P05029 P05030 P05040 P05067 P05106 P05107 P05135 P05136 P05139 P05141 P05146
P05151 P05170 P05187 P05300 P05306 P05310 P05311 P05316 P05333 P05334 P05337
P05351 P05356 P05357 P05359 P05362 P05363 P05374 P05375 P05376 P05377 P05417
P05418 P05425 P05429 P05430 P05431 P05465 P05489 P05490 P05491 P05496 P05499
P05500 P05501 P05502 P05503 P05504 P05505 P05506 P05507 P05508 P05509 P05510
P05513 P05528 P05532 P05538 P05540 P05541 P05555 P05556 P05622 P05641 P05642
P05643 P05674 P05695 P05700 P05701 P05704 P05710 P05718 P05769 P05778 P05779
P05780 P05803 P05819 P05825 P05877 P05878 P05879 P05880 P05881 P05882 P05883
P05884 P05885 P05886 P05919 P05921 P05923 P05924 P05925 P05947 P05948 P05949
P05979 P05981 P05992 P06002 P06003 P06004 P06005 P06006 P06007 P06008 P06009
P06010 P06024 P06028 P06029 P06030 P06126 P06127 P06129 P06133 P06136 P06140
P06156 P06167 P06185 P06197 P06199 P06213 P06230 P06231 P06246 P06247 P06248
P06249 P06250 P06254 P06255 P06257 P06258 P06259 P06260 P06261 P06262 P06263

P06264 P06265 P06266 P06282 P06286 P06289 P06290 P06291 P06332 P06333 P06334
P06335 P06339 P06340 P06341 P06342 P06343 P06344 P06345 P06346 P06402 P06404
P06405 P06406 P06407 P06408 P06411 P06412 P06413 P06414 P06415 P06432 P06436
P06437 P06441 P06445 P06449 P06451 P06452 P06453 P06473 P06474 P06475 P06476
P06477 P06480 P06481 P06482 P06483 P06484 P06487 P06494 P06497 P06498 P06511
P06512 P06527 P06528 P06568 P06579 P06583 P06585 P06591 P06597 P06609 P06624
P06631 P06669 P06671 P06682 P06683 P06685 P06686 P06687 P06690 P06716 P06729
P06734 P06739 P06751 P06752 P06755 P06756 P06762 P06763 P06764 P06775 P06783
P06795 P06800 P06802 P06804 P06808 P06810 P06817 P06818 P06819 P06820 P06821
P06828 P06830 P06849 P06851 P06853 P06907 P06927 P06935 P06970 P06971 P06972
P06996 P07001 P07017 P07018 P07038 P07050 P07056 P07063 P07099 P07106 P07110
P07117 P07141 P07142 P07143 P07144 P07147 P07153 P07167 P07168 P07174 P07202
P07204 P07207 P07213 P07228 P07233 P07238 P07240 P07255 P07286 P07293 P07306
P07307 P07308 P07314 P07333 P07340 P07357 P07359 P07367 P07368 P07369 P07370
P07371 P07373 P07399 P07470 P07471 P07503 P07506 P07510 P07513 P07522 P07539
P07550 P07552 P07561 P07562 P07564 P07566 P07575 P07577 P07581 P07611 P07612
P07613 P07618 P07645 P07646 P07654 P07657 P07661 P07687 P07698 P07700 P07704
P07705 P07706 P07707 P07708 P07709 P07710 P07720 P07722 P07725 P07727 P07747
P07753 P07766 P07768 P07775 P07825 P07826 P07845 P07861 P07868 P07869 P07893
P07921 P07923 P07925 P07926 P07930 P07944 P07946 P07949 P07966 P07967 P07968
P07970 P07971 P07973 P07974 P07975 P07976 P07977 P08004 P08005 P08006 P08011
P08033 P08034 P08037 P08049 P08050 P08064 P08067 P08069 P08083 P08099 P08100
P08101 P08104 P08138 P08149 P08169 P08172 P08173 P08183 P08194 P08195 P08212
P08214 P08219 P08220 P08221 P08222 P08247 P08251 P08255 P08290 P08301 P08303
P08305 P08306 P08326 P08327 P08354 P08355 P08356 P08359 P08360 P08362 P08363
P08369 P08371 P08372 P08382 P08383 P08386 P08395 P08400 P08401 P08430 P08434
P08443 P08444 P08445 P08446 P08447 P08456 P08470 P08473 P08478 P08482 P08483
P08485 P08491 P08492 P08508 P08510 P08514 P08516 P08525 P08541 P08542 P08549
P08550 P08555 P08560 P08563 P08564 P08574 P08575 P08581 P08583 P08586 P08588
P08592 P08617 P08637 P08641 P08648 P08656 P08665 P08666 P08667 P08668 P08669
P08681 P08684 P08691 P08701 P08702 P08714 P08715 P08716 P08722 P08739 P08740
P08742 P08743 P08744 P08745 P08746 P08768 P08774 P08780 P08803 P08804 P08805
P08806 P08807 P08808 P08810 P08834 P08842 P08887 P08888 P08908 P08909 P08910
P08911 P08912 P08913 P08920 P08921 P08922 P08923 P08941 P08953 P08962 P08963
P08980 P08982 P08983 P08986 P08995 P08F94 P09003 P09004 P09011 P09045 P09047
P09048 P09055 P09127 P09130 P09131 P09138 P09144 P09146 P09166 P09167 P09169
P09172 P09175 P09182 P09190 P09191 P09192 P09193 P09208 P09218 P09221 P09231
P09241 P09247 P09256 P09257 P09258 P09259 P09260 P09261 P09264 P09266 P09273
P09280 P09290 P09298 P09312 P09323 P09343 P09344 P09345 P09348 P09354 P09362
P09391 P09438 P09447 P09458 P09460 P09470 P09478 P09479 P09480 P09481 P09482
P09483 P09484 P09500 P09501 P09506 P09507 P09545 P09549 P09564 P09572 P09573
P09574 P09581 P09592 P09601 P09603 P09610 P09612 P09613 P09619 P09620 P09626
P09627 P09628 P09660 P09669 P09688 P09689 P09690 P09691 P09692 P09693 P09703
P09705 P09712 P09713 P09716 P09717 P09718 P09719 P09720 P09721 P09723 P09724

P09727 P09728 P09729 P09730 P09731 P09732 P09752 P09755 P09756 P09758 P09759
P09765 P09766 P09767 P09776 P09779 P09781 P09783 P09790 P09793 P09803 P09817
P09829 P09835 P09836 P09843 P09848 P09849 P09866 P09875 P09888 P09912 P09925
P09927 P09933 P09958 P09962 P09969 P09973 P09974 P09983 P09986 P09990 P09991
P0A007 P0A008 P0A056 P0A057 P0A058 P0A059 P0A060 P0A063 P0A064 P0A065 P0A066
P0A067 P0A068 P0A069 P0A070 P0A0I1 P0A0I2 P0A0I3 P0A0I4 P0A0J4 P0A0J5 P0A0J6
P0A0J7 P0A0J8 P0A0J9 P0A0M1 P0A0M2 P0A0R8 P0A0R9 P0A0S0 P0A0S3 P0A0S4 P0A0U8
P0A0U9 P0A0V2 P0A0V3 P0A0V8 P0A0V9 P0A0Z3 P0A0Z4 P0A0Z5 P0A0Z6 P0A103 P0A104
P0A140 P0A141 P0A188 P0A189 P0A190 P0A197 P0A198 P0A1F4 P0A1F5 P0A1H1 P0A1H2
P0A1H7 P0A1H8 P0A1I3 P0A1I4 P0A1I5 P0A1I6 P0A1L1 P0A1L2 P0A1L3 P0A1L4 P0A1L5
P0A1L6 P0A1L7 P0A1L8 P0A1L9 P0A1M0 P0A1M1 P0A1M2 P0A1M3 P0A1M4 P0A1M5 P0A1M6
P0A1M7 P0A1M8 P0A1M9 P0A1U2 P0A1U3 P0A1W2 P0A1W3 P0A1W8 P0A1W9 P0A1X8 P0A1X9
P0A219 P0A220 P0A221 P0A222 P0A231 P0A232 P0A233 P0A234 P0A235 P0A236 P0A237
P0A238 P0A261 P0A262 P0A263 P0A264 P0A279 P0A280 P0A2D3 P0A2D4 P0A2F8 P0A2F9
P0A2G0 P0A2G3 P0A2G4 P0A2G5 P0A2G6 P0A2H3 P0A2H4 P0A2H5 P0A2H6 P0A2I7 P0A2I8
P0A2I9 P0A2J0 P0A2J1 P0A2J2 P0A2J3 P0A2J4 P0A2J5 P0A2J6 P0A2J8 P0A2L5 P0A2L6
P0A2N4 P0A2N5 P0A2N6 P0A2N7 P0A2R8 P0A2R9 P0A2V0 P0A2V1 P0A2Y8 P0A2Y9 P0A2Z0
P0A2Z1 P0A2Z2 P0A2Z3 P0A302 P0A303 P0A304 P0A305 P0A306 P0A307 P0A308 P0A309
P0A312 P0A313 P0A333 P0A334 P0A384 P0A385 P0A3G5 P0A3G6 P0A3M5 P0A3M6 P0A3Q7
P0A3Q8 P0A3R7 P0A3R8 P0A3S3 P0A3S4 P0A3S5 P0A3S6 P0A3U4 P0A3U5 P0A3V8 P0A3V9
P0A3Y1 P0A3Y2 P0A403 P0A404 P0A405 P0A406 P0A407 P0A408 P0A409 P0A425 P0A426
P0A427 P0A428 P0A429 P0A430 P0A444 P0A445 P0A446 P0A447 P0A4G8 P0A4G9 P0A4H0
P0A4H1 P0A4I5 P0A4I6 P0A4I7 P0A4I8 P0A4K4 P0A4K5 P0A4K6 P0A4K7 P0A4K8 P0A4M7
P0A4M8 P0A4M9 P0A4N0 P0A4N1 P0A4N2 P0A4N3 P0A4N4 P0A4N5 P0A4N6 P0A4N7 P0A4N8
P0A4N9 P0A4P0 P0A4P9 P0A4Q0 P0A4Q1 P0A4Q2 P0A4Q7 P0A4V9 P0A4W1 P0A4W5 P0A4Y1
P0A4Y3 P0A501 P0A503 P0A505 P0A561 P0A565 P0A5C2 P0A5D4 P0A5D6 P0A5E4 P0A5E8
P0A5F4 P0A5K1 P0A5K3 P0A5K9 P0A5M1 P0A5P5 P0A5S5 P0A5Z1 P0A5Z3 P0A5Z9 P0A601
P0A607 P0A615 P0A625 P0A627 P0A629 P0A631 P0A685 P0A695 P0A6A0 P0A6A1 P0A6A2
P0A6H8 P0A6H9 P0A6J3 P0A6J4 P0A6M2 P0A6M3 P0A6S5 P0A6S6 P0A6W3 P0A6W4 P0A710
P0A711 P0A712 P0A713 P0A714 P0A742 P0A743 P0A766 P0A767 P0A768 P0A769 P0A770
P0A771 P0A7C8 P0A7C9 P0A7D0 P0A830 P0A831 P0A843 P0A844 P0A845 P0A846 P0A8D9
P0A8E0 P0A8H3 P0A8H4 P0A8H5 P0A8K8 P0A8K9 P0A8L0 P0A8Q0 P0A8Q1 P0A8Q2 P0A8Q3
P0A8Q4 P0A8Q5 P0A8R7 P0A8R8 P0A8S5 P0A8S6 P0A8S7 P0A8S8 P0A910 P0A911 P0A915
P0A916 P0A917 P0A918 P0A919 P0A920 P0A921 P0A922 P0A923 P0A924 P0A925 P0A926
P0A927 P0A928 P0A929 P0A930 P0A931 P0A932 P0A933 P0A934 P0A940 P0A941 P0A942
P0A943 P0AA22 P0AA23 P0AA24 P0AA47 P0AA48 P0AA49 P0AA50 P0AA51 P0AA52 P0AA53
P0AA54 P0AA55 P0AA56 P0AA60 P0AA61 P0AA62 P0AA63 P0AA64 P0AA65 P0AA66 P0AA67
P0AA68 P0AA69 P0AA70 P0AA71 P0AA72 P0AA73 P0AA74 P0AA75 P0AA76 P0AA77 P0AA78
P0AA79 P0AA80 P0AA81 P0AA82 P0AA83 P0AA86 P0AA87 P0AA88 P0AA93 P0AA94 P0AAA1
P0AAA2 P0AAA5 P0AAA6 P0AAA7 P0AAA8 P0AAC4 P0AAC5 P0AAC6 P0AAC7 P0AAD2 P0AAD3
P0AAD4 P0AAD5 P0AAD6 P0AAD7 P0AAD8 P0AAD9 P0AAE0 P0AAE1 P0AAE2 P0AAE3 P0AAE4
P0AAE5 P0AAE6 P0AAE7 P0AAE8 P0AAE9 P0AAF0 P0AAF1 P0AAF2 P0AAG5 P0AAG6 P0AAG7
P0AAI3 P0AAI4 P0AAJ3 P0AAJ4 P0AAJ5 P0AAJ6 P0AAJ7 P0AAM1 P0AAM2 P0AAP1 P0AAP2
P0AAP7 P0AAP8 P0AAP9 P0AAQ0 P0AAQ1 P0AAR5 P0AAR6 P0AAR7 P0AAS0 P0AAS1 P0AAS2

P0AAS3 P0AAS4 P0AAT4 P0AAT5 P0AAU5 P0AAU6 P0AAW5 P0AAW6 P0AAW7 P0AAW8 P0AAW9
P0AAX0 P0AAX1 P0AAX2 P0AAZ0 P0AAZ1 P0AAZ2 P0AAZ3 P0AB01 P0AB02 P0AB12 P0AB13
P0AB35 P0AB36 P0AB37 P0AB58 P0AB59 P0AB60 P0AB67 P0AB68 P0AB69 P0AB70 P0AB93
P0AB94 P0AB95 P0AB98 P0AB99 P0ABA0 P0ABA1 P0ABA2 P0ABA3 P0ABB8 P0ABB9 P0ABC0
P0ABC1 P0ABC2 P0ABC3 P0ABC4 P0ABC5 P0ABC6 P0ABC7 P0ABC8 P0ABC9 P0ABD0 P0ABD1
P0ABD2 P0ABE5 P0ABE6 P0ABF8 P0ABF9 P0ABG0 P0ABG1 P0ABG2 P0ABG3 P0ABG4 P0ABG5
P0ABG6 P0ABG7 P0ABG8 P0ABG9 P0ABH4 P0ABH5 P0ABH6 P0ABI4 P0ABI5 P0ABI6 P0ABI7
P0ABI8 P0ABI9 P0ABJ0 P0ABJ1 P0ABJ2 P0ABJ3 P0ABJ4 P0ABJ5 P0ABJ6 P0ABJ7 P0ABJ8
P0ABJ9 P0ABK0 P0ABK1 P0ABK2 P0ABK3 P0ABK4 P0ABL5 P0ABL6 P0ABL7 P0ABL8 P0ABL9
P0ABM0 P0ABM1 P0ABM2 P0ABM3 P0ABM4 P0ABM5 P0ABM6 P0ABM7 P0ABM8 P0ABM9 P0ABN0
P0ABN1 P0ABN2 P0ABN3 P0ABN4 P0ABN5 P0ABN6 P0ABN7 P0ABN8 P0ABN9 P0ABP0 P0ABP1
P0ABP2 P0ABP3 P0ABP4 P0ABP5 P0ABP6 P0ABP7 P0ABT8 P0ABT9 P0ABU7 P0ABU8 P0ABU9
P0ABV0 P0ABV1 P0ABV2 P0ABV3 P0ABV4 P0ABV5 P0ABV6 P0ABV7 P0ABV8 P0ABV9 P0ABX8
P0ABX9 P0ABY0 P0ABY1 P0AC05 P0AC06 P0AC07 P0AC08 P0AC09 P0AC10 P0AC23 P0AC24
P0AC25 P0AC26 P0AC27 P0AC30 P0AC31 P0AC32 P0AC44 P0AC45 P0AC46 P0AC75 P0AC76
P0AC77 P0AC78 P0AC79 P0AC80 P0AC94 P0AC95 P0AC96 P0AC97 P0AC98 P0AC99 P0ACA0
P0ACB7 P0ACB8 P0ACB9 P0ACC0 P0ACG4 P0ACG5 P0ACG6 P0ACG7 P0ACV0 P0ACV1 P0ACV2
P0ACV3 P0ACV4 P0ACV5 P0ACX0 P0ACX1 P0ACX2 P0ACY6 P0ACY7 P0ACY8 P0AD03 P0AD04
P0AD14 P0AD15 P0AD16 P0AD17 P0AD18 P0AD19 P0AD20 P0AD27 P0AD28 P0AD29 P0AD30
P0AD31 P0AD32 P0AD42 P0AD43 P0AD47 P0AD48 P0AD65 P0AD66 P0AD67 P0AD68 P0AD69
P0AD99 P0ADA0 P0ADC3 P0ADC4 P0ADC5 P0ADC6 P0ADC7 P0ADD2 P0ADD3 P0ADD4 P0ADD5
P0ADD6 P0ADE4 P0ADE5 P0ADI9 P0ADJ0 P0ADJ1 P0ADJ2 P0ADJ5 P0ADJ6 P0ADJ7 P0ADJ8
P0ADJ9 P0ADK4 P0ADK5 P0ADL1 P0ADL2 P0ADL6 P0ADL7 P0ADL8 P0ADL9 P0ADM0 P0ADM1
P0ADM2 P0ADM3 P0ADM6 P0ADM7 P0ADP5 P0ADP6 P0ADQ7 P0ADQ8 P0ADQ9 P0ADR0 P0ADR1
P0ADR2 P0ADR3 P0ADR4 P0ADR5 P0ADT8 P0ADT9 P0ADU0 P0ADU1 P0ADV9 P0ADW0 P0ADW1
P0ADW2 P0ADW3 P0ADW4 P0ADW5 P0ADY1 P0ADY2 P0ADZ7 P0ADZ8 P0ADZ9 P0AE00 P0AE14
P0AE15 P0AE16 P0AE17 P0AE24 P0AE25 P0AE26 P0AE27 P0AE30 P0AE31 P0AE32 P0AE33
P0AE34 P0AE35 P0AE36 P0AE42 P0AE43 P0AE44 P0AE45 P0AE46 P0AE47 P0AE74 P0AE75
P0AE82 P0AE83 P0AE84 P0AEA5 P0AEA6 P0AEA7 P0AEB0 P0AEB1 P0AEB5 P0AEB6 P0AEC0
P0AEC1 P0AEC2 P0AEC3 P0AEC4 P0AEC5 P0AEC6 P0AEC7 P0AEC8 P0AEC9 P0AEE3 P0AEE4
P0AEF8 P0AEF9 P0AEG0 P0AEG1 P0AEG2 P0AEG3 P0AEH1 P0AEH2 P0AEH5 P0AEH6 P0AEH7
P0AEH8 P0AEH9 P0AEI0 P0AEJ0 P0AEJ1 P0AEJ4 P0AEJ5 P0AEK7 P0AEK8 P0AEK9 P0AEL0
P0AEL1 P0AEL2 P0AEN4 P0AEN5 P0AEN6 P0AEN7 P0AEP1 P0AEP2 P0AEQ6 P0AEQ7 P0AEQ8
P0AEQ9 P0AER0 P0AER1 P0AER2 P0AER3 P0AER4 P0AER5 P0AER6 P0AER7 P0AER8 P0AER9
P0AET5 P0AET6 P0AET7 P0AEU3 P0AEU4 P0AEU5 P0AEU6 P0AEW1 P0AEW2 P0AEW3 P0AEX3
P0AEX4 P0AEX7 P0AEX8 P0AEY1 P0AEY2 P0AEY8 P0AEY9 P0AEZ0 P0AF01 P0AF02 P0AF06
P0AF07 P0AF16 P0AF17 P0AF32 P0AF33 P0AF40 P0AF41 P0AF42 P0AF43 P0AF44 P0AF52
P0AF53 P0AF54 P0AF55 P0AF73 P0AF74 P0AF75 P0AF80 P0AF81 P0AF98 P0AF99 P0AFA0
P0AFA1 P0AFA2 P0AFA3 P0AFA4 P0AFA5 P0AFA6 P0AFA7 P0AFA8 P0AFA9 P0AFB0 P0AFC3
P0AFC4 P0AFC5 P0AFC6 P0AFD4 P0AFD5 P0AFE0 P0AFE1 P0AFE2 P0AFE3 P0AFE4 P0AFE5
P0AFE6 P0AFE7 P0AFE8 P0AFE9 P0AFF0 P0AFF1 P0AFF2 P0AFF3 P0AFF4 P0AFF5 P0AFH2
P0AFH3 P0AFH4 P0AFH5 P0AFH6 P0AFH7 P0AFI9 P0AFJ0 P0AFJ7 P0AFJ8 P0AFJ9 P0AFK2
P0AFK3 P0AFK4 P0AFK5 P0AFK6 P0AFK7 P0AFK8 P0AFL1 P0AFL2 P0AFL9 P0AFM0 P0AFM1
P0AFM9 P0AFN0 P0AFN1 P0AFN2 P0AFN3 P0AFN4 P0AFN5 P0AFN6 P0AFN7 P0AFN8 P0AFN9

P0AFP0 P0AFP1 P0AFP9 P0AFQ0 P0AFQ1 P0AFQ2 P0AFQ3 P0AFQ4 P0AFR2 P0AFR3 P0AFR7
P0AFR8 P0AFR9 P0AFS0 P0AFS1 P0AFS2 P0AFS5 P0AFS6 P0AFS7 P0AFS8 P0AFS9 P0AFT0
P0AFT1 P0AFT2 P0AFT3 P0AFT4 P0AFU0 P0AFU1 P0AFU2 P0AFU3 P0AFV0 P0AFV1 P0AFV2
P0AFV3 P0AFX7 P0AFX8 P0AFY2 P0AFY3 P0AFY4 P0AFY5 P0AFY6 P0AFY7 P0AFZ7 P0AFZ8
P0AFZ9 P0AG00 P0AG01 P0AG02 P0AG14 P0AG15 P0AG34 P0AG35 P0AG36 P0AG37 P0AG38
P0AG39 P0AG90 P0AG91 P0AG92 P0AG93 P0AG94 P0AG95 P0AG96 P0AG97 P0AG98 P0AG99
P0AGA0 P0AGA1 P0AGA2 P0AGA3 P0AGA4 P0AGA5 P0AGC0 P0AGC1 P0AGC2 P0AGC7 P0AGC8
P0AGC9 P0AGD0 P0AGE4 P0AGE5 P0AGF4 P0AGF5 P0AGH1 P0AGH2 P0AGH3 P0AGH4 P0AGH5
P0AGH6 P0AGH7 P0AGH8 P0AGH9 P0AGI0 P0AGI1 P0AGI2 P0AGI3 P0AGI4 P0AGI5 P0AGI6
P0AGI7 P0AGK1 P0AGK2 P0AGK3 P0AGM0 P0AGM1 P0AGM2 P0AGM3 P0AGM4 P0AGM7 P0AGM8
P0AGM9 P0AGN0 P0AGN1 P0AGN2 P0C029 P0C034 P0C035 P0C041 P0C048 P0C049 P0C052
P0C053 P0C070 P0C071 P0C081 P0C082 P0C083 P0C084 P0C085 P0C086 P0C087 P0C0F6
P0C0F7 P0C0G9 P0C0H0 P0C0H1 P0C0I3 P0C0K1 P0C0K2 P0C0K5 P0C0K6 P0C0K7 P0C0L6
P0C0L7 P0C0L8 P0C0N4 P0C0N5 P0C0S1 P0C0S2 P0C0S3 P0C0T8 P0C0V9 P0C0W0 P0C0W8
P0C0X4 P0C0X5 P0C0X9 P0C0Y0 P0C0Y1 P0C0Y2 P0C0Y7 P0C0Y8 P0C0Y9 P0C0Z0 P0C102
P0C105 P0C112 P0C117 P0C120 P0C147 P0C148 P0C150 P0C152 P0C155 P0C156 P0C157
P0C158 P0C179 P0C184 P0C187 P0C188 P0C190 P0C191 P0C192 P0C197 P0C1D2 P0C1D4
P0C1F6 P0C1F7 P0C1F8 P0C1G1 P0C1G7 P0C1H0 P0C1H1 P0C1H2 P0C1J4 P0C1J5 P0C1N9
P0C1P0 P0C1P1 P0C1P7 P0C1Q3 P0C1R8 P0C1S9 P0C1T0 P0C1U4 P0C1V1 P0C1V4 P0C1X9
P0C1Y6 P0C1Y7 P0C1Y8 P0C204 P0C212 P0C215 P0C217 P0C219 P0C261 P0C289 P0C2C9
P0C2E2 P0C2E9 P0C2G8 P0C2H1 P0C2H2 P0C2H3 P0C2H4 P0C2J9 P0C2L2 P0C2L3 P0C2M3
P0C2M9 P0C2N9 P0C2Q1 P0C2Q2 P0C2Q3 P0C2Q4 P0C2Q5 P0C2Q6 P0C2Q7 P0C2Q8 P0C2Q9
P0C2R0 P0C2S0 P0C2U2 P0C2U3 P0C2U8 P0C2V3 P0C2W0 P0C2W5 P0C2X0 P0C2Y5 P0C2Y6
P0C2Y7 P0C2Y8 P0C2Y9 P0C2Z0 P0C2Z9 P0C300 P0C301 P0C302 P0C303 P0C315 P0C316
P0C317 P0C318 P0C319 P0C320 P0C321 P0C322 P0C323 P0C324 P0C325 P0C326 P0C327
P0C328 P0C329 P0C330 P0C331 P0C332 P0C333 P0C334 P0C344 P0C345 P0C353 P0C354
P0C355 P0C356 P0C357 P0C358 P0C362 P0C363 P0C364 P0C365 P0C366 P0C367 P0C368
P0C369 P0C370 P0C371 P0C372 P0C373 P0C374 P0C375 P0C376 P0C377 P0C387 P0C388
P0C389 P0C390 P0C391 P0C392 P0C393 P0C394 P0C395 P0C396 P0C399 P0C400 P0C401
P0C405 P0C406 P0C407 P0C408 P0C409 P0C410 P0C411 P0C412 P0C413 P0C414 P0C415
P0C416 P0C417 P0C418 P0C419 P0C420 P0C421 P0C422 P0C423 P0C427 P0C428 P0C429
P0C430 P0C432 P0C433 P0C434 P0C435 P0C436 P0C437 P0C513 P0C514 P0C515 P0C518
P0C519 P0C523 P0C524 P0C526 P0C527 P0C528 P0C529 P0C531 P0C532 P0C546 P0C548
P0C550 P0C566 P0C572 P0C575 P0C582 P0C584 P0C588 P0C5A8 P0C5A9 P0C5B8 P0C5E0
P0C5E2 P0C5E4 P0C5H6 P0C5I1 P0C5J4 P0C5L2 P0C5M2 P0C5M3 P0C5M7 P0C5N0 P0C5N8
P0C5N9 P0C5P3 P0C5P5 P0C5Q3 P0C5Q4 P0C5R1 P0C5R7 P0C5R8 P0C5R9 P0C5S6 P0C5T0
P0C5T1 P0C5T2 P0C5T3 P0C5T4 P0C5T5 P0C5T6 P0C5X8 P0C5X9 P0C604 P0C617 P0C623
P0C626 P0C628 P0C629 P0C645 P0C646 P0C648 P0C649 P0C655 P0C672 P0C673 P0C684
P0C6A1 P0C6B1 P0C6B2 P0C6B3 P0C6B7 P0C6C9 P0C6D5 P0C6D9 P0C6E0 P0C6F3 P0C6F4
P0C6F5 P0C6F6 P0C6F7 P0C6F8 P0C6K4 P0C6K7 P0C6Q6 P0C6R0 P0C6R1 P0C6S8 P0C6T2
P0C6T3 P0C6T4 P0C6T5 P0C6T6 P0C6T7 P0C6T8 P0C6T9 P0C6U0 P0C6U1 P0C6U2 P0C6U3
P0C6U4 P0C6U5 P0C6U6 P0C6U7 P0C6U8 P0C6U9 P0C6V0 P0C6V1 P0C6V2 P0C6V3 P0C6V4
P0C6V5 P0C6V6 P0C6V7 P0C6V8 P0C6V9 P0C6W0 P0C6W1 P0C6W2 P0C6W3 P0C6W4 P0C6W5
P0C6W6 P0C6W7 P0C6W8 P0C6W9 P0C6X0 P0C6X1 P0C6X2 P0C6X3 P0C6X4 P0C6X5 P0C6X6

P0C6X7 P0C6X8 P0C6X9 P0C6Y0 P0C6Y1 P0C6Y2 P0C6Y3 P0C6Y4 P0C6Y5 P0C6Y6 P0C6Z1
P0C6Z3 P0C6Z4 P0C6Z5 P0C701 P0C729 P0C736 P0C737 P0C738 P0C739 P0C741 P0C742
P0C748 P0C751 P0C762 P0C763 P0C764 P0C778 P0C781 P0C782 P0C786 P0C787 P0C788
P0C7A3 P0C7A6 P0C7B2 P0C7B3 P0C7B7 P0C7H7 P0C7I9 P0C7J6 P0C7L9 P0C7M0 P0C7M1
P0C7M8 P0C7M9 P0C7N1 P0C7N4 P0C7N5 P0C7N8 P0C7P4 P0C7Q5 P0C7Q6 P0C7T2 P0C7T3
P0C7T8 P0C7U0 P0C7U3 P0C7U4 P0C7U5 P0C7U9 P0C7V7 P0C7W5 P0C851 P0C874 P0C889
P0C897 P0C8E5 P0C8E6 P0C8K7 P0C8K9 P0C8L0 P0C8M6 P0C8M7 P0C8N0 P0C8N2 P0C8N3
P0C8N4 P0C8N5 P0C8N6 P0C8N7 P0C8N8 P0C8Q9 P0C8R9 P0C8W2 P0C8Z2 P0C931 P0C941
P0C947 P0C948 P0C9A2 P0C9A3 P0C9A4 P0C9E0 P0C9E1 P0C9E2 P0C9G1 P0C9G2 P0C9G3
P0C9G4 P0C9G9 P0C9I9 P0C9J0 P0C9J1 P0C9J2 P0C9J3 P0C9J4 P0C9J5 P0C9J6 P0C9J7
P0C9J8 P0C9J9 P0C9K0 P0C9K1 P0C9K4 P0C9K5 P0C9K6 P0C9K7 P0C9P3 P0C9P4 P0C9P5
P0C9V6 P0C9V7 P0C9V8 P0C9V9 P0C9W0 P0C9W1 P0C9W2 P0C9W3 P0C9W4 P0C9W5 P0C9W6
P0C9W7 P0C9W8 P0C9W9 P0C9X0 P0C9Y1 P0C9Y2 P0C9Y3 P0C9Y4 P0C9Y8 P0C9Y9 P0C9Z0
P0C9Z8 P0C9Z9 P0CA00 P0CA13 P0CA14 P0CA15 P0CA19 P0CA20 P0CA21 P0CA26 P0CA27
P0CA28 P0CA29 P0CA52 P0CA53 P0CA54 P0CA63 P0CA64 P0CA65 P0CA66 P0CA67 P0CA68
P0CA69 P0CA70 P0CA71 P0CA72 P0CA94 P0CA95 P0CA96 P0CAB0 P0CAB1 P0CAB2 P0CAB7
P0CAB8 P0CAB9 P0CAC0 P0CAE3 P0CAE4 P0CAE5 P0CAE6 P0CAH7 P0CAH8 P0CAH9 P0CAJ1
P0CAJ2 P0CAJ3 P0CAJ4 P0CAL0 P0CAL4 P0CAL5 P0CAL6 P0CAL7 P0CAL8 P0CAL9 P0CAM0
P0CAM1 P0CAM2 P0CAM3 P0CAN6 P0CAP6 P0CAP7 P0CAT2 P0CAU8 P0CB17 P0CB28 P0CB29
P0CB30 P0CB31 P0CB39 P0CB40 P0CB59 P0CB62 P0CB63 P0CB64 P0CB69 P0CB70 P0CB71
P0CB72 P0CB73 P0CB74 P0CB85 P0CB86 P0CB93 P0CB94 P0CC10 P0CC20 P0CC21 P0CC22
P0CC23 P0CC24 P0CC25 P0CC26 P0CC27 P0CC28 P0CC29 P0CC30 P0CC31 P0CC32 P0CC33
P0CC34 P0CC35 P0CC36 P0CC37 P0CC38 P0CC39 P0CC40 P0CC41 P0CC42 P0CC43 P0CC44
P0CC45 P0CC46 P0CC47 P0CC48 P0CC49 P0CC50 P0CC51 P0CC52 P0CC53 P0CC54 P0CC55
P0CC56 P0CC57 P0CC58 P0CC59 P0CC60 P0CC61 P0CC62 P0CC63 P0CC64 P0CC65 P0CC66
P0CC67 P0CC68 P0CC69 P0CC70 P0CC71 P0CC72 P0CC73 P0CC74 P0CC75 P0CC76 P0CC77
P0CC78 P0CC79 P0CC80 P0CC81 P0CC82 P0CC83 P0CC84 P0CC85 P0CC86 P0CC87 P0CC88
P0CC89 P0CC90 P0CC91 P0CC92 P0CC93 P0CC94 P0CC95 P0CC96 P0CC97 P0CC98 P0CC99
P0CD00 P0CD01 P0CD02 P0CD03 P0CD04 P0CD05 P0CD06 P0CD07 P0CD08 P0CD09 P0CD10
P0CD11 P0CD12 P0CD13 P0CD14 P0CD15 P0CD16 P0CD17 P0CD18 P0CD19 P0CD20 P0CD21
P0CD22 P0CD23 P0CD24 P0CD25 P0CD26 P0CD27 P0CD28 P0CD29 P0CD30 P0CD31 P0CD32
P0CD33 P0CD34 P0CD35 P0CD36 P0CD37 P0CD38 P0CD39 P0CD40 P0CD41 P0CD42 P0CD43
P0CD44 P0CD45 P0CD46 P0CD47 P0CD48 P0CD49 P0CD50 P0CD51 P0CD52 P0CD53 P0CD54
P0CD55 P0CD56 P0CD57 P0CD58 P0CD59 P0CD61 P0CD64 P0CD89 P0CD90 P0CD91 P0CD92
P0CD94 P0CD97 P0CD98 P0CD99 P0CE00 P0CE11 P0CE14 P0CE44 P0CE45 P0CE46 P0CE68
P0CE70 P0CF19 P0CF20 P0CF38 P0CF51 P0CF98 P0CG08 P0CG16 P0CG27 P0CG41 P0CG44
P0CH01 P0CH02 P0CH03 P0CH61 P0CH79 P0CI03 P0CI27 P0CI36 P0CI37 P0CI38 P0CI39
P0CI40 P0CI70 P0CI71 P0CI89 P0CI90 P0CJ36 P0CJ37 P0CJ38 P0CJ39 P0CJ48 P0CJ82
P0CK16 P0CK21 P0CK22 P0CK23 P0CK27 P0CK28 P0CK29 P0CK30 P0CK96 P0CK97 P0CK99
P0CL04 P0CL13 P0CL14 P0CL27 P0CL28 P0CL29 P0CL30 P0CL31 P0CL32 P0CL33 P0CL34
P0CL35 P0CL36 P0CL37 P0CL38 P0CL39 P0CL40 P0CL41 P0CL42 P0CL48 P0CL49 P0CL92
P0CL93 P0CM10 P0CM11 P0CM32 P0CM33 P0CM40 P0CM41 P0CM42 P0CM43 P0CM50 P0CM51
P0CM56 P0CM57 P0CM68 P0CM69 P0CM72 P0CM73 P0CM84 P0CM85 P0CM92 P0CM93 P0CN16
P0CN17 P0CN66 P0CN67 P0CN70 P0CN71 P0CN72 P0CN73 P0CN92 P0CN93 P0CO60 P0CO61

P0CO66 P0CO67 P0CO70 P0CO71 P0CO78 P0CO79 P0CO80 P0CO81 P0CP14 P0CP15 P0CP32
P0CP33 P0CP36 P0CP37 P0CP48 P0CP49 P0CP52 P0CP53 P0CP62 P0CP63 P0CP64 P0CP65
P0CQ04 P0CQ05 P0CQ38 P0CQ39 P0CQ46 P0CQ47 P0CR34 P0CR35 P0CR36 P0CR37 P0CR46
P0CR47 P0CR88 P0CR89 P0CR90 P0CR91 P0CS02 P0CS03 P0CS04 P0CS05 P0CS18 P0CS19
P0CS24 P0CS25 P0CS58 P0CS59 P0CS66 P0CS67 P0CS68 P0CS69 P0CS86 P0CS87 P0CT18
P0CT44 P0CT94 P0CT95 P0CT96 P0CT97 P0CT98 P0CT99 P0CU06 P0CU07 P0CU10 P0CU11
P0CU14 P0CU15 P0CU16 P0CU19 P0CU20 P0CU22 P0CU26 P0CU30 P0CU31 P0CU66 P0CU70
P0CU77 P0CU83 P0CU86 P0CU87 P0CU88 P0CU91 P0CV71 P0CW39 P0CW70 P0CW77 P0CW78
P0CW80 P0CW81 P0CW82 P0CW83 P0CW87 P0CW97 P0CW98 P0CX12 P0CX13 P0CX92 P0CX93
P0CX94 P0CX95 P0CX96 P0CX97 P0CX98 P0CX99 P0CY00 P0CY01 P0CY41 P0CY42 P0CY43
P0CY44 P0CY45 P0CY46 P0CY47 P0CY48 P0CY49 P0CY56 P0CZ13 P0CZ21 P0CZ90 P0CZ91
P0CZ92 P0CZ93 P0DA10 P0DA11 P0DA98 P0DA99 P0DB16 P0DB17 P0DB42 P0DB43 P0DB50
P0DB51 P0DB56 P0DB57 P0DB60 P0DB61 P0DC20 P0DC21 P0DC24 P0DC25 P0DC86 P0DC87
P0DC88 P0DC89 P0DD10 P0DD11 P0DD30 P0DD31 P0DD32 P0DD33 P0DF20 P0DF21 P0DF78
P0DF79 P0DF96 P0DF97 P0DG86 P0DG87 P0DH04 P0DH05 P0DH30 P0DH31 P0DH58 P0DH61
P0DH63 P0DH64 P0DH65 P0DH67 P0DH69 P0DH79 P0DH80 P0DH81 P0DH82 P0DH83 P0DH84
P0DH86 P0DH87 P0DH92 P0DH93 P0DH94 P0DI20 P0DI21 P0DI22 P0DI23 P0DI24 P0DI25
P0DI26 P0DI27 P0DI28 P0DI29 P0DI30 P0DI31 P0DI32 P0DI33 P0DI34 P0DI35 P0DI36
P0DI37 P0DI38 P0DI39 P0DI40 P0DI41 P0DI42 P0DI43 P0DI44 P0DI45 P0DI46 P0DI47
P0DI48 P0DI49 P0DI50 P0DI51 P0DI52 P0DI53 P0DI54 P0DI55 P0DI56 P0DI57 P0DI58
P0DI59 P0DI60 P0DI61 P0DI62 P0DI63 P0DI64 P0DI65 P0DI66 P0DI67 P0DI68 P0DI69
P0DI70 P0DI71 P0DI72 P0DI73 P0DI74 P0DI75 P0DI78 P0DI79 P0DI80 P0DI93 P0DI94
P0DI95 P0DI96 P0DI97 P0DJ07 P0DJ90 P0DJ91 P0DJ92 P0DJ93 P0DJ97 P0DJ98 P0DJE0
P0DJE3 P0DJE4 P0DJE5 P0DJE6 P0DJE7 P0DJF3 P0DJI3 P0DJI4 P0DJI5 P0DJI7 P0DJO0
P0DJO1 P0DJO2 P0DJO3 P0DJO4 P0DJO6 P0DJP2 P0DJQ9 P0DJX9 P0DJY0 P0DJZ4 P0DJZ5
P0DJZ6 P0DJZ8 P0DJZ9 P0DKB5 P0DKB6 P0DKC0 P0DKC1 P0DKC2 P0DKC5 P0DKC6 P0DKC7
P0DKG2 P0DKG9 P0DKH0 P0DKH9 P0DKI0 P0DKI1 P0DKI2 P0DKI7 P0DKJ3 P0DKJ4 P0DKJ5
P0DKL2 P0DKM0 P0DKQ8 P0DKS0 P0DKS1 P0DKV0 P0DKX4 P0DKX5 P0DKX6 P0DKX7 P0DKX9
P0DKY0 P0DL10 P0DL12 P0DL55 P0DL56 P0DM37 P0DM40 P0DM44 P0DM64 P0DM79 P0DM80
P0DMA5 P0DMC5 P0DMC6 P0DME1 P0DMH7 P0DMH8 P0DMI3 P0DMI4 P0DMI5 P0DMJ8 P0DMK6
P0DMK9 P0DMQ5 P0DMQ8 P0DMS8 P0DMS9 P0DMT0 P0DMU2 P0DMV3 P0DMX2 P0DMX3 P0DMX4
P0DN25 P0DN34 P0DN35 P0DN66 P0DN67 P0DN77 P0DN78 P0DN80 P0DN81 P0DN82 P0DN83
P0DN84 P0DN89 P0DN90 P0DN91 P0DO05 P0DO06 P0DO13 P0DO14 P0DO17 P0DO18 P0DO19
P0DO22 P0DO27 P0DO28 P0DO32 P0DO34 P0DO35 P0DO36 P0DO37 P0DO38 P0DO39 P0DO40
P0DO41 P0DO42 P0DO44 P0DO45 P0DO53 P0DO54 P0DO99 P0DOA0 P0DOB2 P0DOC9 P0DOE4
P0DOE5 P0DOF5 P0DOF6 P0DOF8 P0DOF9 P0DOG8 P0DOG9 P0DOH0 P0DOH1 P0DOH2 P0DOH3
P0DOH4 P0DOH5 P0DOH6 P0DOH7 P0DOH8 P0DOH9 P0DOJ7 P0DOJ8 P0DOJ9 P0DOK0 P0DOK1
P0DOK8 P0DOL5 P0DOL6 P0DOM3 P0DOM4 P0DOM9 P0DON0 P0DON1 P0DON2 P0DON7 P0DON8
P0DOP9 P0DOQ0 P0DOR3 P0DOR4 P0DOR5 P0DOR6 P0DOS7 P0DOS8 P0DOT7 P0DOT8 P0DOU1
P0DOU2 P0DOV3 P0DOW2 P0DOW3 P0DP42 P0DP43 P0DP69 P0DP70 P0DP72 P0DP83 P0DP88
P0DP99 P0DPA2 P0DPA4 P0DPB1 P0DPB2 P0DPC4 P0DPD0 P0DPD1 P0DPD6 P0DPD8 P0DPD9
P0DPE2 P0DPE3 P0DPI0 P0DPI1 P0DPN0 P0DPN1 P0DPN4 P0DPO1 P0DPO4 P0DPO5 P0DPO8
P0DPP0 P0DPP7 P0DPQ5 P0DPR2 P0DPR4 P0DPR5 P0DPR6 P0DPR7 P0DPR9 P0DPS5 P0DPS6
P0DQD5 P0DQD7 P0DQH0 P0DQS0 P0DQU2 P0DSE1 P0DSE2 P0DSN6 P0DSP1 P0DSQ3 P0DSQ4

P0DSR7 P0DSR8 P0DSS1 P0DSS2 P0DST5 P0DST6 P0DSW1 P0DSW2 P0DSX7 P0DSX8 P0DSY1
P0DSY2 P0DSY5 P0DSY6 P0DSY7 P0DSY8 P0DSY9 P0DSZ0 P0DSZ3 P0DSZ4 P0DTB3 P0DTB4
P0DTB5 P0DTB6 P0DTC1 P0DTC2 P0DTC3 P0DTC4 P0DTC5 P0DTC7 P0DTD1 P0DTD3 P0DTD8
P0DTE0 P0DTE4 P0DTE5 P0DTF9 P0DTH6 P0DTH9 P0DTI0 P0DTI2 P0DTI4 P0DTJ0 P0DTJ1
P0DTK2 P0DTL5 P0DTU3 P0DTU4 P0DTW8 P0DUB1 P0DUD7 P0DUH5 P0DUK3 P0DUL3 P0DUM2
P0DUM3 P0DUR6 P0DUT8 P0DUT9 P0DUW6 P0DUW8 P0DUW9 P0DV10 P0DV25 P0DV27 P0DV39
P0DV43 P0DV44 P0DV46 P0DV48 P0DV50 P0DV52 P0DV84 P0DV85 P0DW01 P0DW02 P0DW09
P0DW10 P0DW45 P0DW86 P0DW88 P0DW90 P0DW93 P10033 P10040 P10041 P10047 P10049
P10063 P10064 P10089 P10090 P10170 P10171 P10174 P10175 P10176 P10204 P10211
P10218 P10224 P10227 P10228 P10229 P10240 P10247 P10250 P10259 P10260 P10261
P10269 P10287 P10288 P10300 P10305 P10307 P10321 P10329 P10330 P10332 P10347
P10348 P10379 P10384 P10393 P10404 P10415 P10417 P10438 P10439 P10448 P10466
P10499 P10502 P10510 P10513 P10522 P10566 P10586 P10603 P10608 P10611 P10613
P10614 P10615 P10620 P10641 P10681 P10707 P10708 P10716 P10717 P10718 P10721
P10730 P10731 P10747 P10757 P10761 P10799 P10803 P10804 P10817 P10818 P10820
P10844 P10852 P10858 P10861 P10864 P10865 P10866 P10867 P10870 P10888 P10897
P10898 P10900 P10903 P10905 P10906 P10912 P10920 P10921 P10955 P10966 P10980
P11004 P11005 P11022 P11024 P11049 P11072 P11093 P11095 P11117 P11127 P11128
P11132 P11133 P11134 P11135 P11162 P11166 P11167 P11168 P11169 P11170 P11188
P11209 P11215 P11222 P11223 P11224 P11225 P11229 P11230 P11235 P11236 P11261
P11267 P11268 P11279 P11286 P11293 P11296 P11297 P11306 P11311 P11313 P11316
P11317 P11318 P11323 P11339 P11344 P11350 P11362 P11432 P11438 P11456 P11461
P11483 P11485 P11491 P11505 P11506 P11507 P11511 P11528 P11545 P11551 P11557
P11584 P11599 P11607 P11608 P11609 P11610 P11613 P11614 P11615 P11616 P11617
P11622 P11623 P11627 P11628 P11629 P11631 P11636 P11646 P11647 P11654 P11655
P11658 P11661 P11662 P11667 P11669 P11688 P11695 P11696 P11710 P11716 P11717
P11718 P11764 P11835 P11836 P11847 P11848 P11849 P11869 P11881 P11895 P11898
P11902 P11911 P11912 P11933 P11942 P11947 P11948 P11950 P11951 P11991 P11992
P11993 P12012 P12023 P12050 P12060 P12062 P12074 P12080 P12117 P12119 P12120
P12122 P12123 P12124 P12154 P12160 P12163 P12164 P12168 P12170 P12177 P12179
P12180 P12182 P12184 P12185 P12186 P12187 P12188 P12189 P12191 P12192 P12193
P12194 P12205 P12207 P12211 P12212 P12213 P12214 P12216 P12221 P12222 P12228
P12229 P12234 P12235 P12236 P12238 P12239 P12240 P12241 P12242 P12287 P12312
P12313 P12314 P12318 P12319 P12327 P12328 P12329 P12330 P12331 P12332 P12333
P12336 P12342 P12357 P12360 P12361 P12363 P12371 P12377 P12384 P12389 P12390
P12391 P12392 P12403 P12404 P12407 P12409 P12410 P12428 P12430 P12439 P12440
P12441 P12442 P12443 P12446 P12449 P12463 P12469 P12470 P12471 P12487 P12488
P12489 P12492 P12515 P12516 P12517 P12518 P12522 P12526 P12553 P12554 P12555
P12556 P12557 P12558 P12559 P12561 P12562 P12563 P12564 P12565 P12566 P12567
P12568 P12569 P12570 P12571 P12572 P12574 P12575 P12581 P12582 P12583 P12584
P12585 P12586 P12587 P12588 P12589 P12590 P12605 P12606 P12607 P12608 P12640
P12641 P12649 P12650 P12651 P12655 P12657 P12667 P12673 P12681 P12683 P12684
P12685 P12691 P12696 P12697 P12700 P12701 P12702 P12719 P12722 P12771 P12772
P12773 P12774 P12775 P12776 P12777 P12778 P12786 P12820 P12821 P12822 P12823

P12824 P12830 P12857 P12866 P12890 P12906 P12908 P12909 P12910 P12911 P12924
P12934 P12946 P12983 P12984 P12989 P13011 P13025 P13036 P13073 P13090 P13101
P13102 P13103 P13112 P13128 P13134 P13155 P13156 P13160 P13164 P13165 P13180
P13181 P13182 P13183 P13184 P13198 P13201 P13224 P13253 P13271 P13272 P13285
P13290 P13291 P13293 P13296 P13347 P13348 P13352 P13355 P13368 P13369 P13379
P13386 P13387 P13388 P13415 P13422 P13423 P13436 P13464 P13467 P13473 P13504
P13508 P13509 P13510 P13511 P13512 P13514 P13516 P13518 P13536 P13547 P13554
P13555 P13563 P13568 P13569 P13583 P13585 P13586 P13587 P13588 P13590 P13591
P13595 P13596 P13597 P13598 P13599 P13607 P13612 P13625 P13626 P13627 P13633
P13637 P13638 P13679 P13688 P13701 P13709 P13721 P13726 P13738 P13747 P13748
P13749 P13750 P13751 P13752 P13753 P13762 P13765 P13773 P13794 P13801 P13806
P13808 P13838 P13841 P13843 P13847 P13850 P13851 P13865 P13866 P13869 P13881
P13882 P13890 P13897 P13901 P13908 P13910 P13924 P13945 P13970 P14009 P14020
P14058 P14060 P14075 P14089 P14092 P14093 P14094 P14142 P14151 P14154 P14176
P14180 P14185 P14186 P14198 P14209 P14220 P14221 P14222 P14224 P14225 P14231
P14236 P14237 P14246 P14271 P14273 P14274 P14275 P14276 P14277 P14278 P14279
P14283 P14285 P14319 P14334 P14335 P14336 P14337 P14338 P14339 P14340 P14351
P14359 P14377 P14378 P14403 P14406 P14410 P14413 P14414 P14415 P14416 P14426
P14427 P14428 P14429 P14430 P14432 P14434 P14435 P14436 P14437 P14438 P14439
P14483 P14494 P14495 P14496 P14544 P14545 P14546 P14548 P14551 P14553 P14562
P14569 P14570 P14571 P14573 P14574 P14575 P14578 P14579 P14584 P14585 P14599
P14600 P14616 P14617 P14622 P14623 P14627 P14650 P14660 P14672 P14677 P14679
P14698 P14719 P14740 P14745 P14753 P14763 P14769 P14770 P14772 P14773 P14778
P14784 P14787 P14790 P14791 P14801 P14813 P14835 P14842 P14852 P14853 P14862
P14867 P14891 P14893 P14895 P14896 P14897 P14901 P14906 P14925 P14992 P14997
P14998 P15011 P15012 P15013 P15014 P15015 P15029 P15030 P15068 P15073 P15078
P15083 P15097 P15098 P15101 P15116 P15127 P15133 P15134 P15136 P15144 P15145
P15151 P15191 P15192 P15193 P15194 P15195 P15199 P15208 P15209 P15260 P15261
P15278 P15286 P15291 P15306 P15321 P15325 P15347 P15362 P15365 P15367 P15377
P15378 P15379 P15380 P15381 P15382 P15383 P15384 P15385 P15387 P15388 P15389
P15390 P15391 P15396 P15400 P15404 P15405 P15409 P15413 P15414 P15415 P15416
P15418 P15420 P15421 P15422 P15423 P15425 P15431 P15464 P15481 P15484 P15492
P15499 P15509 P15514 P15529 P15530 P15535 P15541 P15544 P15545 P15546 P15547
P15548 P15549 P15550 P15551 P15552 P15553 P15554 P15575 P15576 P15577 P15578
P15579 P15580 P15581 P15582 P15583 P15584 P15585 P15589 P15643 P15644 P15647
P15658 P15684 P15685 P15686 P15688 P15701 P15702 P15710 P15715 P15725 P15727
P15729 P15745 P15746 P15747 P15748 P15749 P15750 P15751 P15752 P15753 P15754
P15773 P15775 P15776 P15777 P15779 P15795 P15798 P15811 P15812 P15813 P15819
P15823 P15831 P15853 P15877 P15878 P15892 P15893 P15896 P15897 P15898 P15899
P15900 P15901 P15902 P15903 P15907 P15910 P15911 P15914 P15920 P15921 P15928
P15939 P15941 P15943 P15952 P15953 P15954 P15956 P15957 P15958 P15959 P15963
P15978 P15979 P15980 P15981 P15982 P15983 P15993 P15994 P15995 P15996 P15997
P16000 P16001 P16003 P16004 P16005 P16013 P16033 P16036 P16056 P16060 P16065
P16066 P16067 P16070 P16071 P16082 P16090 P16092 P16109 P16141 P16144 P16145

P16150 P16151 P16153 P16155 P16170 P16177 P16191 P16192 P16195 P16196 P16199
P16200 P16201 P16202 P16203 P16204 P16205 P16206 P16207 P16208 P16209 P16210
P16211 P16212 P16215 P16221 P16232 P16234 P16235 P16237 P16256 P16257 P16260
P16261 P16262 P16264 P16265 P16283 P16284 P16288 P16297 P16305 P16310 P16313
P16336 P16357 P16358 P16359 P16360 P16362 P16363 P16364 P16366 P16367 P16382
P16388 P16389 P16390 P16391 P16393 P16395 P16406 P16410 P16422 P16429 P16430
P16435 P16442 P16445 P16449 P16454 P16465 P16471 P16473 P16477 P16482 P16483
P16492 P16493 P16496 P16517 P16532 P16534 P16535 P16536 P16537 P16547 P16549
P16552 P16561 P16567 P16573 P16575 P16581 P16582 P16599 P16603 P16610 P16615
P16620 P16621 P16645 P16646 P16651 P16652 P16655 P16661 P16662 P16670 P16671
P16672 P16673 P16674 P16701 P16710 P16718 P16719 P16720 P16721 P16733 P16739
P16740 P16741 P16742 P16743 P16744 P16745 P16751 P16754 P16757 P16758 P16778
P16791 P16795 P16808 P16809 P16815 P16843 P16844 P16849 P16853 P16863 P16864
P16869 P16871 P16872 P16873 P16875 P16876 P16877 P16882 P16893 P16899 P16926
P16960 P16967 P16970 P16994 P16995 P16996 P16997 P16998 P16999 P17000 P17001
P17002 P17003 P17004 P17005 P17046 P17047 P17057 P17064 P17101 P17124 P17152
P17154 P17155 P17159 P17176 P17181 P17194 P17195 P17200 P17226 P17227 P17228
P17229 P17230 P17254 P17261 P17266 P17281 P17286 P17301 P17302 P17315 P17326
P17327 P17332 P17334 P17342 P17344 P17345 P17397 P17398 P17399 P17400 P17404
P17413 P17416 P17417 P17438 P17451 P17452 P17471 P17472 P17501 P17503 P17504
P17517 P17519 P17520 P17527 P17528 P17549 P17551 P17583 P17590 P17592 P17605
P17635 P17636 P17643 P17644 P17645 P17646 P17658 P17659 P17665 P17693 P17717
P17747 P17755 P17763 P17781 P17787 P17790 P17792 P17793 P17796 P17798 P17800
P17801 P17809 P17810 P17811 P17813 P17822 P17823 P17824 P17825 P17826 P17827
P17836 P17837 P17838 P17852 P17861 P17880 P17898 P17927 P17942 P17948 P17970
P17971 P17972 P18006 P18011 P18033 P18034 P18040 P18050 P18052 P18084 P18089
P18090 P18094 P18125 P18130 P18141 P18156 P18163 P18168 P18194 P18195 P18200
P18211 P18238 P18239 P18246 P18257 P18263 P18275 P18290 P18293 P18337 P18345
P18347 P18348 P18356 P18357 P18392 P18396 P18397 P18398 P18399 P18405 P18409
P18410 P18411 P18413 P18414 P18433 P18434 P18438 P18439 P18450 P18460 P18461
P18466 P18467 P18468 P18469 P18470 P18475 P18489 P18490 P18502 P18505 P18506
P18507 P18508 P18519 P18521 P18522 P18535 P18536 P18537 P18538 P18540 P18551
P18560 P18563 P18564 P18572 P18577 P18581 P18587 P18594 P18596 P18597 P18598
P18599 P18614 P18627 P18630 P18631 P18696 P18762 P18767 P18774 P18777 P18780
P18790 P18795 P18799 P18806 P18812 P18814 P18817 P18825 P18827 P18828 P18841
P18845 P18850 P18860 P18861 P18871 P18875 P18876 P18877 P18878 P18879 P18880
P18881 P18892 P18901 P18907 P18910 P18916 P18924 P18929 P18930 P18931 P18932
P18933 P18934 P18935 P18936 P18937 P18938 P18939 P18940 P18941 P18942 P18943
P18944 P18945 P18946 P18962 P19019 P19020 P19021 P19022 P19024 P19030 P19040
P19041 P19042 P19043 P19044 P19045 P19046 P19047 P19048 P19049 P19052 P19053
P19054 P19056 P19057 P19070 P19072 P19075 P19101 P19106 P19110 P19145 P19150
P19152 P19153 P19154 P19155 P19156 P19173 P19224 P19235 P19238 P19240 P19255
P19256 P19258 P19320 P19327 P19328 P19331 P19334 P19357 P19370 P19377 P19381
P19390 P19391 P19397 P19407 P19423 P19430 P19431 P19437 P19438 P19439 P19440

P19444 P19445 P19449 P19456 P19462 P19467 P19485 P19488 P19490 P19491 P19492
P19493 P19503 P19516 P19518 P19526 P19528 P19534 P19535 P19542 P19549 P19550
P19551 P19554 P19557 P19568 P19579 P19580 P19581 P19586 P19621 P19634 P19638
P19642 P19643 P19657 P19694 P19695 P19696 P19697 P19698 P19699 P19700 P19701
P19702 P19711 P19712 P19716 P19720 P19725 P19739 P19741 P19742 P19744 P19754
P19756 P19758 P19762 P19783 P19807 P19809 P19811 P19814 P19819 P19845 P19903
P19934 P19941 P19965 P19969 P20020 P20021 P20023 P20036 P20037 P20040 P20048
P20063 P20093 P20103 P20107 P20113 P20114 P20120 P20121 P20138 P20143 P20148
P20149 P20150 P20166 P20169 P20175 P20176 P20223 P20225 P20236 P20237 P20239
P20241 P20242 P20262 P20272 P20273 P20274 P20288 P20291 P20292 P20303 P20309
P20310 P20333 P20334 P20350 P20352 P20374 P20375 P20386 P20387 P20388 P20395
P20401 P20411 P20420 P20431 P20454 P20471 P20488 P20489 P20490 P20491 P20494
P20496 P20497 P20500 P20508 P20539 P20540 P20594 P20598 P20600 P20601 P20603
P20610 P20611 P20645 P20647 P20648 P20649 P20657 P20672 P20679 P20680 P20681
P20682 P20684 P20686 P20687 P20688 P20693 P20701 P20702 P20715 P20716 P20720
P20725 P20735 P20755 P20756 P20781 P20782 P20786 P20788 P20789 P20792 P20796
P20806 P20826 P20843 P20864 P20865 P20866 P20871 P20872 P20882 P20888 P20895
P20896 P20898 P20905 P20916 P20917 P20923 P20924 P20928 P20934 P20937 P20938
P20944 P20953 P20954 P20963 P20966 P20978 P20987 P20990 P20991 P20993 P20996
P21014 P21030 P21049 P21050 P21057 P21061 P21063 P21065 P21083 P21093 P21109
P21115 P21129 P21138 P21145 P21147 P21178 P21183 P21188 P21217 P21245 P21303
P21334 P21345 P21365 P21395 P21396 P21397 P21398 P21400 P21401 P21405 P21409
P21412 P21415 P21420 P21430 P21436 P21439 P21440 P21441 P21443 P21445 P21447
P21448 P21449 P21450 P21451 P21452 P21453 P21458 P21462 P21463 P21482 P21503
P21521 P21526 P21530 P21534 P21535 P21536 P21537 P21548 P21554 P21555 P21556
P21559 P21560 P21564 P21579 P21581 P21583 P21592 P21602 P21608 P21616 P21627
P21628 P21634 P21641 P21647 P21653 P21707 P21709 P21714 P21715 P21716 P21717
P21718 P21719 P21720 P21721 P21722 P21723 P21727 P21728 P21729 P21730 P21731
P21754 P21757 P21758 P21761 P21783 P21796 P21802 P21803 P21804 P21817 P21822
P21823 P21825 P21849 P21854 P21855 P21860 P21865 P21903 P21904 P21905 P21906
P21917 P21918 P21926 P21932 P21958 P21960 P21964 P21994 P21995 P22001 P22002
P22036 P22056 P22067 P22071 P22072 P22083 P22086 P22089 P22092 P22094 P22109
P22110 P22111 P22112 P22113 P22140 P22152 P22167 P22170 P22171 P22179 P22180
P22182 P22185 P22189 P22214 P22215 P22223 P22235 P22236 P22261 P22263 P22269
P22270 P22272 P22273 P22289 P22290 P22292 P22300 P22309 P22310 P22316 P22328
P22329 P22330 P22331 P22332 P22338 P22340 P22380 P22407 P22408 P22409 P22411
P22413 P22426 P22427 P22428 P22429 P22430 P22432 P22440 P22455 P22456 P22459
P22460 P22462 P22475 P22476 P22481 P22483 P22484 P22485 P22519 P22520 P22525
P22545 P22554 P22575 P22576 P22586 P22593 P22596 P22597 P22598 P22607 P22609
P22610 P22621 P22622 P22638 P22646 P22648 P22650 P22651 P22652 P22653 P22655
P22656 P22666 P22671 P22680 P22686 P22700 P22708 P22709 P22710 P22723 P22729
P22732 P22733 P22734 P22739 P22747 P22756 P22760 P22770 P22771 P22794 P22804
P22815 P22819 P22821 P22825 P22831 P22833 P22888 P22897 P22905 P22909 P22930
P22933 P22934 P22940 P22961 P23000 P23024 P23036 P23041 P23052 P23054 P23064

P23067 P23068 P23073 P23075 P23079 P23080 P23088 P23114 P23126 P23134 P23135
P23136 P23150 P23169 P23173 P23174 P23188 P23200 P23220 P23229 P23230 P23231
P23242 P23265 P23266 P23267 P23268 P23269 P23270 P23271 P23272 P23273 P23274
P23275 P23276 P23289 P23290 P23294 P23299 P23316 P23317 P23318 P23319 P23336
P23338 P23347 P23348 P23355 P23372 P23373 P23377 P23380 P23383 P23385 P23387
P23414 P23415 P23416 P23421 P23422 P23423 P23438 P23447 P23452 P23462 P23467
P23468 P23469 P23470 P23471 P23482 P23491 P23500 P23501 P23502 P23510 P23516
P23525 P23531 P23545 P23562 P23563 P23564 P23574 P23576 P23577 P23585 P23586
P23596 P23597 P23598 P23621 P23622 P23631 P23633 P23634 P23641 P23642 P23644
P23645 P23648 P23649 P23650 P23654 P23662 P23685 P23702 P23705 P23711 P23723
P23728 P23732 P23735 P23739 P23749 P23763 P23791 P23797 P23811 P23818 P23819
P23820 P23833 P23837 P23842 P23849 P23876 P23877 P23886 P23889 P23890 P23894
P23895 P23897 P23900 P23910 P23913 P23930 P23936 P23942 P23944 P23945 P23956
P23957 P23958 P23968 P23975 P23977 P23978 P23979 P23980 P23988 P23993 P24003
P24006 P24008 P24009 P24010 P24011 P24012 P24013 P24016 P24017 P24025 P24038
P24045 P24046 P24053 P24055 P24062 P24063 P24065 P24071 P24077 P24083 P24084
P24105 P24126 P24128 P24129 P24138 P24139 P24140 P24144 P24145 P24161 P24177
P24181 P24205 P24207 P24241 P24244 P24262 P24282 P24305 P24306 P24310 P24311
P24326 P24328 P24338 P24348 P24358 P24361 P24380 P24389 P24390 P24391 P24392
P24394 P24400 P24412 P24413 P24414 P24422 P24430 P24458 P24464 P24465 P24469
P24485 P24499 P24503 P24505 P24506 P24507 P24520 P24524 P24530 P24545 P24557
P24584 P24585 P24603 P24608 P24612 P24614 P24616 P24625 P24628 P24638 P24643
P24668 P24687 P24688 P24689 P24690 P24721 P24725 P24726 P24754 P24755 P24761
P24763 P24765 P24786 P24794 P24797 P24798 P24811 P24815 P24849 P24872 P24873
P24874 P24875 P24876 P24877 P24878 P24879 P24880 P24881 P24882 P24883 P24884
P24885 P24886 P24887 P24888 P24889 P24890 P24891 P24892 P24893 P24894 P24895
P24896 P24904 P24905 P24935 P24942 P24943 P24944 P24945 P24946 P24947 P24948
P24949 P24950 P24951 P24952 P24953 P24954 P24955 P24956 P24957 P24958 P24959
P24960 P24962 P24964 P24965 P24966 P24967 P24968 P24969 P24970 P24971 P24972
P24974 P24975 P24976 P24978 P24979 P24980 P24981 P24982 P24983 P24984 P24985
P24986 P24987 P24988 P24989 P24992 P24993 P24994 P24995 P24996 P24997 P24998
P24999 P25000 P25001 P25002 P25003 P25004 P25005 P25014 P25021 P25023 P25024
P25025 P25045 P25049 P25050 P25057 P25060 P25061 P25062 P25076 P25083 P25088
P25089 P25090 P25092 P25094 P25095 P25099 P25100 P25101 P25102 P25103 P25104
P25105 P25106 P25107 P25108 P25109 P25110 P25115 P25116 P25117 P25118 P25119
P25122 P25123 P25129 P25130 P25131 P25149 P25162 P25169 P25178 P25180 P25181
P25184 P25185 P25189 P25190 P25191 P25192 P25193 P25194 P25218 P25221 P25222
P25235 P25253 P25254 P25276 P25286 P25297 P25304 P25305 P25312 P25334 P25338
P25340 P25346 P25351 P25353 P25354 P25358 P25360 P25365 P25371 P25376 P25385
P25396 P25412 P25445 P25446 P25465 P25466 P25467 P25489 P25504 P25505 P25506
P25507 P25515 P25527 P25560 P25565 P25568 P25573 P25574 P25580 P25584 P25585
P25587 P25594 P25596 P25610 P25613 P25618 P25619 P25621 P25625 P25628 P25639
P25641 P25656 P25706 P25707 P25710 P25714 P25722 P25737 P25743 P25744 P25747
P25761 P25794 P25798 P25818 P25874 P25876 P25878 P25902 P25917 P25918 P25926

P25929 P25930 P25931 P25936 P25937 P25942 P25945 P25954 P25955 P25956 P25957
P25960 P25961 P25962 P25965 P25966 P26006 P26007 P26008 P26009 P26010 P26011
P26012 P26013 P26016 P26021 P26025 P26028 P26031 P26032 P26048 P26049 P26051
P26094 P26095 P26096 P26097 P26098 P26099 P26100 P26101 P26102 P26103 P26129
P26134 P26135 P26136 P26137 P26138 P26139 P26140 P26141 P26142 P26143 P26149
P26150 P26151 P26152 P26153 P26170 P26171 P26176 P26178 P26185 P26201 P26207
P26217 P26218 P26219 P26235 P26246 P26255 P26260 P26261 P26266 P26273 P26274
P26279 P26280 P26286 P26287 P26288 P26289 P26290 P26291 P26292 P26293 P26303
P26310 P26313 P26338 P26342 P26361 P26362 P26363 P26376 P26381 P26382 P26400
P26405 P26406 P26417 P26429 P26430 P26431 P26432 P26433 P26434 P26435 P26439
P26453 P26455 P26456 P26457 P26458 P26459 P26466 P26467 P26468 P26471 P26489
P26502 P26522 P26523 P26524 P26562 P26572 P26587 P26591 P26618 P26619 P26628
P26629 P26660 P26661 P26662 P26663 P26664 P26671 P26677 P26678 P26681 P26682
P26684 P26710 P26714 P26715 P26717 P26718 P26725 P26756 P26757 P26760 P26761
P26762 P26769 P26770 P26789 P26790 P26803 P26804 P26824 P26835 P26842 P26844
P26845 P26846 P26847 P26848 P26849 P26850 P26851 P26852 P26853 P26855 P26856
P26857 P26858 P26896 P26897 P26898 P26903 P26904 P26936 P26937 P26949 P26951
P26952 P26954 P26955 P27021 P27022 P27023 P27024 P27025 P27026 P27037 P27038
P27039 P27040 P27041 P27046 P27062 P27080 P27081 P27112 P27113 P27115 P27125
P27135 P27148 P27168 P27171 P27173 P27174 P27175 P27178 P27181 P27182 P27183
P27196 P27200 P27201 P27212 P27218 P27219 P27243 P27277 P27284 P27285 P27286
P27303 P27312 P27315 P27331 P27332 P27333 P27338 P27340 P27358 P27359 P27360
P27364 P27365 P27367 P27370 P27372 P27380 P27382 P27386 P27387 P27388 P27389
P27390 P27392 P27395 P27399 P27415 P27416 P27426 P27427 P27434 P27449 P27455
P27465 P27471 P27487 P27489 P27490 P27491 P27492 P27493 P27494 P27495 P27496
P27497 P27512 P27514 P27516 P27517 P27518 P27520 P27521 P27522 P27523 P27524
P27525 P27544 P27545 P27548 P27572 P27573 P27589 P27591 P27593 P27597 P27599
P27611 P27614 P27615 P27645 P27648 P27650 P27655 P27662 P27668 P27669 P27670
P27674 P27681 P27688 P27689 P27690 P27691 P27701 P27712 P27716 P27732 P27757
P27799 P27808 P27809 P27810 P27811 P27812 P27814 P27824 P27825 P27835 P27837
P27843 P27844 P27869 P27884 P27896 P27904 P27906 P27907 P27908 P27909 P27912
P27913 P27914 P27915 P27922 P27930 P27931 P27945 P27953 P27954 P27955 P27956
P27957 P27958 P27959 P27960 P27961 P27977 P28008 P28056 P28057 P28058 P28059
P28067 P28068 P28078 P28081 P28082 P28083 P28085 P28086 P28087 P28088 P28164
P28190 P28192 P28221 P28222 P28223 P28228 P28229 P28230 P28231 P28232 P28233
P28234 P28235 P28236 P28238 P28246 P28251 P28253 P28254 P28265 P28282 P28285
P28286 P28288 P28294 P28303 P28306 P28328 P28334 P28335 P28336 P28471 P28472
P28473 P28476 P28494 P28496 P28517 P28527 P28529 P28530 P28539 P28540 P28541
P28542 P28564 P28565 P28566 P28568 P28569 P28570 P28571 P28572 P28573 P28584
P28611 P28612 P28620 P28626 P28628 P28642 P28646 P28647 P28648 P28649 P28660
P28662 P28678 P28679 P28680 P28681 P28682 P28683 P28684 P28687 P28693 P28721
P28728 P28729 P28730 P28731 P28737 P28754 P28774 P28785 P28791 P28795 P28824
P28825 P28826 P28827 P28828 P28843 P28845 P28847 P28864 P28873 P28876 P28877
P28882 P28883 P28884 P28885 P28886 P28898 P28906 P28907 P28908 P28930 P28948

P28954 P28959 P28967 P28968 P28971 P28977 P28980 P28981 P28986 P28987 P28991
P28992 P28994 P28995 P29000 P29001 P29016 P29017 P29018 P29033 P29040 P29041
P29057 P29058 P29070 P29075 P29076 P29086 P29089 P29101 P29109 P29110 P29111
P29113 P29119 P29131 P29149 P29150 P29154 P29163 P29165 P29254 P29255 P29270
P29274 P29275 P29276 P29285 P29317 P29318 P29319 P29320 P29322 P29323 P29328
P29329 P29362 P29370 P29371 P29376 P29388 P29402 P29403 P29404 P29414 P29415
P29431 P29463 P29465 P29470 P29481 P29485 P29491 P29518 P29525 P29526 P29527
P29528 P29529 P29530 P29531 P29533 P29534 P29553 P29563 P29610 P29630 P29631
P29632 P29633 P29634 P29635 P29636 P29637 P29638 P29639 P29640 P29641 P29642
P29643 P29644 P29645 P29646 P29647 P29648 P29649 P29650 P29651 P29652 P29653
P29654 P29655 P29656 P29657 P29658 P29659 P29660 P29661 P29662 P29663 P29664
P29665 P29666 P29667 P29668 P29669 P29670 P29671 P29672 P29704 P29705 P29739
P29754 P29755 P29757 P29791 P29801 P29812 P29816 P29823 P29824 P29826 P29837
P29838 P29846 P29856 P29859 P29860 P29865 P29867 P29868 P29869 P29870 P29871
P29872 P29873 P29874 P29875 P29876 P29877 P29878 P29879 P29880 P29881 P29895
P29896 P29897 P29905 P29907 P29919 P29920 P29922 P29923 P29924 P29925 P29926
P29936 P29938 P29939 P29940 P29942 P29945 P29960 P29961 P29963 P29965 P29972
P29973 P29974 P29975 P29984 P29988 P29990 P29991 P29993 P29994 P29995 P30000
P30019 P30025 P30026 P30029 P30030 P30031 P30082 P30083 P30098 P30130 P30143
P30144 P30145 P30149 P30160 P30191 P30193 P30194 P30203 P30204 P30215 P30246
P30247 P30248 P30273 P30296 P30301 P30302 P30323 P30329 P30336 P30345 P30353
P30354 P30355 P30356 P30357 P30358 P30361 P30372 P30375 P30376 P30377 P30378
P30379 P30380 P30381 P30382 P30383 P30385 P30386 P30387 P30388 P30391 P30393
P30394 P30395 P30396 P30398 P30408 P30409 P30411 P30413 P30430 P30432 P30433
P30434 P30511 P30515 P30516 P30517 P30518 P30519 P30530 P30531 P30532 P30535
P30536 P30540 P30542 P30543 P30544 P30545 P30546 P30547 P30548 P30549 P30550
P30551 P30552 P30553 P30554 P30555 P30556 P30557 P30558 P30559 P30560 P30572
P30573 P30576 P30577 P30578 P30579 P30580 P30581 P30582 P30583 P30584 P30585
P30586 P30587 P30588 P30589 P30590 P30591 P30592 P30593 P30594 P30595 P30596
P30597 P30598 P30599 P30600 P30601 P30602 P30603 P30604 P30605 P30606 P30607
P30608 P30609 P30612 P30628 P30638 P30639 P30650 P30673 P30680 P30686 P30687
P30688 P30689 P30690 P30691 P30692 P30704 P30705 P30714 P30715 P30716 P30728
P30729 P30730 P30731 P30777 P30796 P30803 P30804 P30816 P30823 P30825 P30833
P30834 P30836 P30839 P30844 P30845 P30847 P30855 P30872 P30873 P30874 P30875
P30878 P30926 P30931 P30932 P30935 P30936 P30937 P30938 P30939 P30940 P30944
P30951 P30953 P30954 P30955 P30957 P30959 P30961 P30962 P30964 P30966 P30968
P30969 P30974 P30975 P30980 P30987 P30988 P30989 P30992 P30993 P30994 P30995
P30996 P31024 P31041 P31042 P31043 P31048 P31053 P31064 P31069 P31077 P31084
P31088 P31091 P31092 P31094 P31095 P31109 P31119 P31122 P31125 P31126 P31135
P31140 P31141 P31157 P31167 P31172 P31213 P31214 P31224 P31242 P31243 P31302
P31303 P31309 P31348 P31349 P31355 P31356 P31377 P31379 P31381 P31382 P31387
P31388 P31389 P31390 P31391 P31392 P31397 P31403 P31413 P31414 P31421 P31422
P31423 P31424 P31431 P31435 P31436 P31437 P31440 P31442 P31446 P31448 P31452
P31462 P31466 P31468 P31474 P31479 P31480 P31481 P31489 P31499 P31510 P31512

P31513 P31527 P31547 P31549 P31553 P31555 P31557 P31559 P31560 P31563 P31564
P31565 P31567 P31585 P31586 P31587 P31588 P31589 P31590 P31591 P31596 P31597
P31601 P31602 P31603 P31606 P31613 P31614 P31615 P31619 P31621 P31626 P31627
P31636 P31637 P31639 P31640 P31641 P31643 P31644 P31645 P31646 P31647 P31648
P31649 P31650 P31651 P31652 P31661 P31662 P31675 P31679 P31680 P31691 P31692
P31694 P31695 P31696 P31698 P31699 P31700 P31701 P31704 P31705 P31706 P31707
P31708 P31709 P31710 P31711 P31712 P31733 P31734 P31735 P31736 P31737 P31738
P31739 P31740 P31743 P31744 P31755 P31760 P31776 P31780 P31783 P31785 P31789
P31791 P31793 P31794 P31796 P31801 P31809 P31819 P31826 P31830 P31831 P31833
P31836 P31839 P31840 P31841 P31842 P31844 P31845 P31853 P31868 P31869 P31872
P31873 P31875 P31898 P31955 P31964 P31971 P31994 P31995 P31996 P32004 P32007
P32011 P32014 P32015 P32022 P32036 P32037 P32071 P32082 P32089 P32095 P32113
P32129 P32135 P32136 P32137 P32154 P32166 P32198 P32207 P32211 P32212 P32214
P32215 P32225 P32229 P32236 P32237 P32238 P32239 P32240 P32241 P32244 P32245
P32246 P32247 P32248 P32249 P32250 P32251 P32266 P32291 P32297 P32299 P32300
P32301 P32302 P32303 P32304 P32305 P32306 P32307 P32308 P32309 P32310 P32311
P32312 P32313 P32319 P32331 P32332 P32334 P32339 P32341 P32343 P32352 P32353
P32360 P32361 P32363 P32368 P32369 P32378 P32386 P32394 P32399 P32417 P32418
P32421 P32462 P32465 P32466 P32467 P32476 P32482 P32486 P32487 P32500 P32506
P32507 P32510 P32513 P32514 P32515 P32522 P32536 P32541 P32550 P32554 P32563
P32564 P32568 P32584 P32592 P32595 P32596 P32612 P32616 P32621 P32629 P32633
P32646 P32650 P32660 P32672 P32696 P32701 P32703 P32705 P32709 P32710 P32714
P32715 P32716 P32720 P32722 P32739 P32745 P32746 P32747 P32748 P32758 P32784
P32791 P32798 P32799 P32802 P32803 P32804 P32808 P32819 P32837 P32839 P32842
P32843 P32847 P32850 P32851 P32854 P32856 P32857 P32858 P32867 P32870 P32876
P32884 P32886 P32897 P32900 P32901 P32903 P32906 P32907 P32915 P32926 P32927
P32940 P32942 P32958 P32970 P32971 P32972 P32974 P32977 P33011 P33015 P33021
P33024 P33026 P33027 P33032 P33033 P33055 P33059 P33085 P33108 P33113 P33116
P33121 P33124 P33129 P33135 P33145 P33146 P33147 P33148 P33151 P33152 P33209
P33212 P33218 P33226 P33231 P33236 P33250 P33251 P33256 P33258 P33274 P33281
P33296 P33300 P33302 P33303 P33310 P33311 P33326 P33328 P33335 P33336 P33338
P33341 P33359 P33361 P33366 P33369 P33379 P33388 P33389 P33390 P33391 P33392
P33396 P33397 P33408 P33410 P33413 P33421 P33448 P33449 P33450 P33455 P33464
P33465 P33468 P33470 P33478 P33481 P33495 P33496 P33498 P33500 P33501 P33502
P33503 P33504 P33505 P33506 P33507 P33508 P33509 P33510 P33511 P33512 P33513
P33515 P33517 P33518 P33527 P33532 P33533 P33534 P33535 P33543 P33545 P33550
P33553 P33554 P33555 P33560 P33566 P33591 P33607 P33612 P33613 P33614 P33615
P33617 P33620 P33639 P33650 P33659 P33669 P33671 P33681 P33691 P33692 P33698
P33699 P33704 P33705 P33706 P33725 P33729 P33730 P33733 P33742 P33743 P33754
P33766 P33767 P33775 P33790 P33807 P33835 P33841 P33851 P33852 P33865 P33879
P33894 P33896 P33897 P33908 P33915 P33924 P33927 P33934 P33941 P33946 P33947
P33948 P33951 P33969 P33970 P33971 P33972 P34001 P34010 P34026 P34027 P34054
P34080 P34082 P34158 P34163 P34186 P34187 P34188 P34189 P34190 P34191 P34192
P34193 P34194 P34195 P34196 P34197 P34198 P34199 P34200 P34201 P34202 P34210

P34224 P34230 P34237 P34240 P34248 P34251 P34261 P34271 P34272 P34273 P34276
P34285 P34290 P34298 P34299 P34311 P34315 P34317 P34319 P34322 P34352 P34356
P34358 P34364 P34374 P34380 P34389 P34410 P34413 P34422 P34434 P34444 P34446
P34454 P34465 P34479 P34488 P34490 P34519 P34525 P34530 P34548 P34551 P34557
P34571 P34576 P34577 P34579 P34586 P34595 P34608 P34616 P34639 P34641 P34644
P34652 P34655 P34660 P34669 P34672 P34678 P34679 P34693 P34698 P34702 P34709
P34711 P34712 P34713 P34740 P34741 P34810 P34834 P34836 P34838 P34840 P34842
P34843 P34844 P34845 P34846 P34847 P34848 P34849 P34850 P34851 P34852 P34853
P34854 P34855 P34856 P34857 P34858 P34859 P34860 P34861 P34862 P34863 P34864
P34865 P34866 P34867 P34868 P34869 P34870 P34871 P34872 P34873 P34874 P34875
P34876 P34886 P34891 P34892 P34900 P34901 P34902 P34903 P34904 P34907 P34910
P34925 P34927 P34941 P34956 P34957 P34958 P34959 P34968 P34969 P34970 P34971
P34972 P34973 P34974 P34975 P34976 P34978 P34979 P34980 P34981 P34982 P34983
P34984 P34985 P34986 P34987 P34989 P34992 P34993 P34994 P34995 P34996 P34997
P34998 P34999 P35000 P35008 P35011 P35012 P35013 P35054 P35070 P35071 P35075
P35077 P35088 P35089 P35090 P35091 P35092 P35093 P35094 P35095 P35096 P35097
P35098 P35099 P35101 P35102 P35103 P35104 P35105 P35106 P35107 P35108 P35109
P35113 P35114 P35118 P35157 P35158 P35160 P35161 P35162 P35164 P35171 P35179
P35180 P35206 P35212 P35253 P35254 P35272 P35315 P35316 P35317 P35329 P35330
P35331 P35342 P35343 P35344 P35345 P35346 P35347 P35348 P35349 P35350 P35351
P35352 P35353 P35356 P35357 P35358 P35359 P35360 P35361 P35362 P35363 P35364
P35365 P35366 P35367 P35368 P35370 P35371 P35372 P35373 P35374 P35375 P35376
P35377 P35378 P35379 P35382 P35383 P35384 P35400 P35402 P35403 P35404 P35405
P35406 P35407 P35408 P35409 P35410 P35411 P35412 P35413 P35414 P35419 P35436
P35438 P35439 P35449 P35462 P35463 P35464 P35498 P35499 P35500 P35503 P35504
P35522 P35523 P35524 P35525 P35526 P35528 P35535 P35536 P35537 P35538 P35546
P35560 P35561 P35563 P35564 P35565 P35575 P35576 P35589 P35590 P35595 P35597
P35598 P35610 P35613 P35620 P35645 P35646 P35647 P35652 P35654 P35655 P35656
P35670 P35720 P35721 P35723 P35724 P35735 P35737 P35739 P35740 P35741 P35742
P35743 P35762 P35763 P35764 P35765 P35766 P35771 P35801 P35802 P35803 P35818
P35822 P35832 P35839 P35841 P35848 P35855 P35860 P35865 P35867 P35874 P35894
P35895 P35896 P35897 P35898 P35899 P35906 P35916 P35917 P35918 P35930 P35936
P35938 P35949 P35950 P35951 P35952 P35953 P35954 P35961 P35966 P35968 P35969
P35971 P35973 P35992 P36021 P36027 P36029 P36032 P36033 P36034 P36035 P36039
P36044 P36046 P36051 P36057 P36060 P36062 P36066 P36081 P36092 P36096 P36098
P36101 P36107 P36112 P36125 P36137 P36142 P36147 P36148 P36163 P36164 P36169
P36172 P36173 P36209 P36253 P36269 P36291 P36299 P36300 P36301 P36312 P36318
P36319 P36320 P36323 P36329 P36330 P36331 P36332 P36334 P36335 P36336 P36337
P36342 P36343 P36344 P36346 P36348 P36365 P36366 P36367 P36370 P36371 P36372
P36380 P36381 P36382 P36383 P36423 P36438 P36442 P36491 P36492 P36494 P36495
P36497 P36498 P36510 P36511 P36512 P36513 P36514 P36537 P36544 P36549 P36554
P36555 P36557 P36561 P36581 P36587 P36596 P36606 P36619 P36640 P36641 P36642
P36643 P36646 P36647 P36655 P36672 P36678 P36690 P36691 P36696 P36705 P36707
P36836 P36837 P36886 P36888 P36890 P36894 P36895 P36896 P36897 P36898 P36920

P36937 P36939 P36940 P36941 P36948 P36956 P36963 P36964 P36965 P37002 P37019
P37020 P37023 P37034 P37039 P37040 P37059 P37067 P37068 P37069 P37070 P37071
P37072 P37082 P37083 P37088 P37089 P37090 P37091 P37103 P37114 P37115 P37116
P37117 P37118 P37131 P37147 P37148 P37169 P37172 P37173 P37176 P37180 P37201
P37212 P37217 P37255 P37256 P37264 P37265 P37267 P37268 P37277 P37278 P37279
P37287 P37288 P37289 P37295 P37296 P37298 P37299 P37301 P37305 P37327 P37340
P37367 P37385 P37386 P37409 P37432 P37433 P37439 P37443 P37446 P37447 P37451
P37458 P37460 P37461 P37475 P37476 P37482 P37483 P37484 P37489 P37498 P37508
P37511 P37514 P37555 P37559 P37575 P37592 P37593 P37594 P37597 P37602 P37608
P37617 P37619 P37621 P37624 P37630 P37637 P37642 P37643 P37645 P37649 P37652
P37653 P37659 P37660 P37661 P37662 P37665 P37669 P37674 P37675 P37683 P37703
P37711 P37716 P37726 P37729 P37730 P37731 P37737 P37738 P37739 P37746 P37748
P37758 P37772 P37781 P37784 P37785 P37792 P37813 P37814 P37815 P37816 P37827
P37828 P37841 P37868 P37873 P37894 P37908 P37915 P37916 P37917 P37918 P37919
P37924 P37943 P37948 P37953 P37972 P37973 P37989 P37990 P37998 P38006 P38008
P38044 P38047 P38050 P38054 P38055 P38069 P38072 P38079 P38083 P38084 P38085
P38087 P38090 P38097 P38101 P38102 P38107 P38114 P38124 P38125 P38127 P38130
P38131 P38134 P38142 P38150 P38152 P38156 P38166 P38168 P38173 P38176 P38179
P38185 P38186 P38189 P38190 P38192 P38196 P38206 P38211 P38212 P38220 P38221
P38224 P38225 P38227 P38228 P38233 P38240 P38242 P38244 P38247 P38250 P38252
P38253 P38258 P38264 P38267 P38268 P38269 P38279 P38286 P38287 P38292 P38297
P38298 P38301 P38307 P38310 P38311 P38312 P38317 P38318 P38320 P38325 P38327
P38329 P38335 P38341 P38342 P38343 P38353 P38355 P38356 P38357 P38358 P38359
P38360 P38361 P38362 P38368 P38369 P38374 P38375 P38376 P38377 P38379 P38381
P38382 P38383 P38385 P38386 P38387 P38388 P38389 P38392 P38396 P38397 P38399
P38428 P38435 P38438 P38441 P38444 P38445 P38453 P38454 P38459 P38461 P38462
P38477 P38484 P38551 P38570 P38572 P38573 P38574 P38575 P38591 P38592 P38593
P38594 P38595 P38596 P38597 P38598 P38599 P38600 P38601 P38602 P38603 P38626
P38631 P38649 P38670 P38680 P38695 P38702 P38703 P38717 P38718 P38721 P38723
P38724 P38726 P38727 P38728 P38731 P38734 P38735 P38736 P38739 P38740 P38745
P38750 P38752 P38756 P38757 P38767 P38770 P38775 P38776 P38778 P38799 P38800
P38813 P38819 P38825 P38834 P38837 P38842 P38843 P38848 P38849 P38851 P38857
P38865 P38868 P38869 P38875 P38878 P38880 P38881 P38921 P38925 P38928 P38929
P38943 P38954 P38958 P38962 P38967 P38969 P38971 P38988 P38992 P38993 P38995
P39003 P39004 P39006 P39007 P39010 P39012 P39038 P39063 P39064 P39086 P39087
P39088 P39098 P39103 P39106 P39107 P39109 P39128 P39129 P39137 P39139 P39141
P39150 P39154 P39180 P39196 P39209 P39210 P39214 P39215 P39216 P39217 P39263
P39270 P39276 P39277 P39282 P39284 P39285 P39301 P39308 P39310 P39314 P39321
P39328 P39338 P39344 P39349 P39352 P39355 P39357 P39365 P39367 P39379 P39381
P39382 P39385 P39386 P39396 P39398 P39401 P39414 P39438 P39453 P39465 P39479
P39480 P39481 P39504 P39515 P39524 P39532 P39535 P39540 P39541 P39542 P39543
P39547 P39548 P39551 P39552 P39557 P39559 P39563 P39569 P39570 P39578 P39580
P39582 P39584 P39589 P39590 P39591 P39595 P39599 P39600 P39602 P39603 P39604
P39608 P39612 P39613 P39618 P39619 P39627 P39636 P39637 P39642 P39649 P39656

P39668 P39685 P39690 P39694 P39695 P39704 P39709 P39718 P39719 P39722 P39727
P39734 P39755 P39764 P39766 P39767 P39775 P39793 P39794 P39798 P39803 P39816
P39817 P39823 P39830 P39832 P39835 P39838 P39843 P39850 P39853 P39854 P39855
P39886 P39898 P39920 P39924 P39925 P39926 P39928 P39932 P39952 P39953 P39980
P39981 P39982 P39986 P39991 P39996 P39997 P40004 P40006 P40009 P40015 P40030
P40035 P40039 P40046 P40056 P40071 P40073 P40074 P40075 P40085 P40086 P40088
P40097 P40098 P40100 P40107 P40113 P40126 P40137 P40144 P40145 P40146 P40163
P40169 P40180 P40189 P40190 P40197 P40198 P40200 P40207 P40219 P40223 P40237
P40238 P40239 P40240 P40241 P40259 P40260 P40293 P40305 P40309 P40310 P40312
P40318 P40319 P40321 P40330 P40341 P40345 P40350 P40351 P40358 P40365 P40367
P40401 P40410 P40411 P40416 P40419 P40420 P40438 P40440 P40441 P40445 P40451
P40452 P40458 P40464 P40474 P40475 P40476 P40478 P40497 P40499 P40500 P40501
P40502 P40504 P40508 P40514 P40515 P40519 P40524 P40526 P40527 P40528 P40532
P40533 P40534 P40538 P40540 P40544 P40548 P40549 P40550 P40556 P40557 P40570
P40576 P40578 P40587 P40605 P40606 P40614 P40682 P40684 P40700 P40701 P40702
P40709 P40719 P40727 P40729 P40730 P40736 P40739 P40748 P40749 P40758 P40760
P40779 P40785 P40786 P40800 P40812 P40857 P40862 P40866 P40867 P40868 P40879
P40885 P40886 P40890 P40897 P40900 P40901 P40913 P40931 P40934 P40941 P40948
P40967 P40970 P40975 P40979 P40980 P40986 P40988 P40989 P41006 P41013 P41014
P41015 P41025 P41026 P41027 P41028 P41029 P41032 P41036 P41047 P41083 P41085
P41086 P41132 P41143 P41144 P41145 P41146 P41147 P41149 P41155 P41172 P41173
P41180 P41181 P41216 P41217 P41231 P41232 P41233 P41234 P41237 P41248 P41251
P41265 P41266 P41272 P41273 P41274 P41280 P41281 P41282 P41283 P41285 P41286
P41287 P41288 P41289 P41290 P41291 P41292 P41293 P41294 P41295 P41296 P41297
P41298 P41299 P41300 P41301 P41303 P41304 P41305 P41306 P41307 P41308 P41309
P41310 P41311 P41312 P41313 P41314 P41315 P41322 P41355 P41356 P41360 P41406
P41413 P41438 P41440 P41441 P41442 P41443 P41468 P41484 P41507 P41543 P41544
P41586 P41587 P41588 P41590 P41591 P41592 P41593 P41594 P41595 P41596 P41597
P41598 P41599 P41600 P41601 P41603 P41604 P41608 P41611 P41613 P41614 P41615
P41616 P41617 P41618 P41619 P41620 P41624 P41625 P41626 P41627 P41628 P41639
P41640 P41642 P41643 P41644 P41647 P41650 P41688 P41690 P41698 P41701 P41705
P41717 P41718 P41722 P41731 P41732 P41733 P41740 P41771 P41774 P41775 P41783
P41786 P41796 P41800 P41806 P41809 P41815 P41820 P41821 P41823 P41834 P41849
P41850 P41851 P41852 P41909 P41930 P41941 P41943 P41948 P41950 P41951 P41955
P41956 P41968 P41983 P41984 P41985 P41986 P41987 P41990 P42010 P42011 P42015
P42032 P42049 P42050 P42051 P42054 P42055 P42056 P42057 P42062 P42063 P42067
P42069 P42070 P42071 P42072 P42081 P42082 P42086 P42087 P42098 P42099 P42102
P42107 P42108 P42109 P42112 P42116 P42117 P42159 P42167 P42177 P42194 P42196
P42197 P42200 P42201 P42205 P42237 P42243 P42245 P42247 P42248 P42252 P42257
P42258 P42259 P42260 P42261 P42262 P42263 P42264 P42287 P42288 P42289 P42290
P42291 P42292 P42306 P42308 P42312 P42314 P42333 P42334 P42361 P42362 P42394
P42396 P42397 P42399 P42401 P42417 P42422 P42424 P42432 P42459 P42501 P42512
P42513 P42514 P42519 P42523 P42531 P42532 P42533 P42536 P42548 P42557 P42590
P42598 P42601 P42603 P42615 P42619 P42628 P42640 P42658 P42659 P42667 P42668

P42670 P42701 P42702 P42703 P42707 P42726 P42767 P42781 P42782 P42787 P42792
P42833 P42836 P42838 P42839 P42840 P42857 P42864 P42866 P42867 P42872 P42881
P42891 P42892 P42893 P42905 P42910 P42911 P42915 P42925 P42934 P42946 P42953
P42956 P42959 P42964 P42970 P42971 P42978 P43002 P43003 P43004 P43005 P43006
P43007 P43008 P43009 P43010 P43023 P43024 P43040 P43042 P43045 P43054 P43059
P43071 P43083 P43088 P43109 P43111 P43114 P43115 P43116 P43117 P43118 P43119
P43121 P43140 P43141 P43142 P43143 P43144 P43146 P43150 P43189 P43190 P43191
P43192 P43193 P43194 P43195 P43196 P43197 P43198 P43199 P43200 P43201 P43202
P43203 P43204 P43205 P43206 P43207 P43208 P43218 P43219 P43220 P43221 P43240
P43245 P43252 P43253 P43256 P43257 P43258 P43259 P43260 P43261 P43266 P43286
P43287 P43298 P43303 P43307 P43308 P43322 P43369 P43370 P43372 P43373 P43374
P43375 P43376 P43377 P43406 P43407 P43416 P43427 P43428 P43439 P43440 P43454
P43457 P43466 P43470 P43481 P43488 P43489 P43524 P43531 P43537 P43542 P43548
P43549 P43552 P43553 P43555 P43557 P43560 P43561 P43562 P43564 P43566 P43571
P43575 P43578 P43581 P43585 P43590 P43595 P43608 P43613 P43614 P43617 P43620
P43624 P43626 P43627 P43628 P43629 P43630 P43631 P43632 P43636 P43650 P43657
P43662 P43666 P43671 P43676 P43679 P43681 P43682 P43719 P43720 P43721 P43756
P43768 P43769 P43787 P43804 P43805 P43810 P43838 P43839 P43840 P43842 P43931
P43932 P43939 P43940 P43947 P43952 P43957 P43958 P43960 P43969 P43975 P43984
P43989 P43991 P43993 P43999 P44005 P44010 P44016 P44018 P44019 P44020 P44023
P44026 P44028 P44035 P44038 P44043 P44051 P44054 P44056 P44060 P44063 P44067
P44069 P44070 P44079 P44080 P44081 P44087 P44089 P44092 P44095 P44097 P44110
P44112 P44113 P44114 P44115 P44127 P44129 P44130 P44132 P44133 P44135 P44136
P44139 P44145 P44157 P44163 P44170 P44185 P44186 P44201 P44202 P44203 P44219
P44220 P44222 P44236 P44250 P44252 P44253 P44263 P44265 P44266 P44270 P44272
P44273 P44274 P44275 P44277 P44278 P44284 P44285 P44287 P44288 P44289 P44291
P44292 P44300 P44301 P44302 P44407 P44408 P44424 P44450 P44451 P44454 P44468
P44469 P44472 P44476 P44477 P44483 P44484 P44520 P44523 P44528 P44530 P44535
P44543 P44545 P44546 P44555 P44560 P44567 P44570 P44577 P44578 P44579 P44581
P44590 P44591 P44592 P44596 P44600 P44601 P44603 P44604 P44607 P44608 P44610
P44614 P44615 P44620 P44621 P44623 P44626 P44629 P44640 P44646 P44653 P44655
P44660 P44661 P44678 P44691 P44693 P44699 P44706 P44707 P44713 P44714 P44717
P44720 P44727 P44736 P44739 P44741 P44742 P44747 P44768 P44772 P44776 P44783
P44789 P44791 P44795 P44799 P44806 P44809 P44826 P44836 P44840 P44843 P44849
P44855 P44872 P44885 P44890 P44891 P44892 P44903 P44908 P44917 P44919 P44927
P44928 P44930 P44933 P44935 P44936 P44937 P44943 P44944 P44947 P44958 P44963
P44969 P44970 P44973 P44974 P44975 P44985 P44993 P44994 P44998 P45002 P45014
P45019 P45020 P45021 P45023 P45029 P45030 P45033 P45034 P45035 P45036 P45037
P45038 P45039 P45045 P45053 P45054 P45058 P45059 P45062 P45064 P45067 P45075
P45079 P45081 P45082 P45089 P45090 P45096 P45108 P45114 P45117 P45122 P45123
P45130 P45145 P45146 P45169 P45170 P45174 P45190 P45191 P45221 P45239 P45240
P45243 P45246 P45268 P45272 P45280 P45286 P45287 P45290 P45315 P45320 P45322
P45332 P45333 P45335 P45336 P45341 P45356 P45357 P45380 P45384 P45385 P45386
P45387 P45394 P45399 P45400 P45401 P45402 P45403 P45404 P45405 P45406 P45407

P45408 P45417 P45419 P45420 P45428 P45433 P45434 P45442 P45459 P45468 P45490
P45495 P45503 P45518 P45522 P45537 P45539 P45546 P45562 P45566 P45570 P45598
P45604 P45608 P45609 P45610 P45650 P45675 P45700 P45701 P45706 P45708 P45710
P45711 P45751 P45755 P45756 P45757 P45758 P45760 P45761 P45762 P45763 P45767
P45768 P45772 P45773 P45774 P45775 P45776 P45777 P45778 P45779 P45780 P45781
P45782 P45783 P45784 P45789 P45790 P45791 P45793 P45794 P45800 P45804 P45817
P45820 P45827 P45828 P45829 P45830 P45843 P45844 P45860 P45861 P45862 P45865
P45866 P45869 P45874 P45879 P45880 P45895 P45901 P45919 P45946 P45963 P45971
P45996 P45997 P45998 P46000 P46002 P46005 P46009 P46014 P46022 P46023 P46024
P46025 P46026 P46027 P46029 P46030 P46032 P46059 P46069 P46078 P46089 P46090
P46091 P46092 P46093 P46094 P46095 P46096 P46097 P46098 P46101 P46104 P46105
P46116 P46125 P46133 P46136 P46139 P46141 P46187 P46194 P46197 P46231 P46242
P46249 P46269 P46270 P46274 P46304 P46305 P46306 P46310 P46312 P46313 P46317
P46322 P46324 P46332 P46333 P46335 P46339 P46340 P46344 P46348 P46349 P46359
P46383 P46408 P46411 P46455 P46469 P46478 P46481 P46482 P46490 P46491 P46492
P46499 P46506 P46530 P46531 P46555 P46556 P46564 P46565 P46566 P46567 P46568
P46569 P46570 P46571 P46572 P46576 P46591 P46592 P46612 P46613 P46616 P46617
P46620 P46621 P46626 P46627 P46628 P46630 P46636 P46641 P46642 P46659 P46663
P46691 P46694 P46695 P46720 P46721 P46785 P46826 P46827 P46832 P46838 P46839
P46840 P46889 P46895 P46896 P46904 P46907 P46909 P46912 P46913 P46921 P46925
P46928 P46950 P46956 P46961 P46964 P46965 P46967 P46971 P46972 P46975 P46977
P46978 P46985 P46986 P46989 P46991 P46993 P46996 P46998 P47003 P47010 P47013
P47014 P47020 P47022 P47026 P47028 P47034 P47040 P47045 P47046 P47055 P47056
P47062 P47069 P47075 P47080 P47081 P47082 P47088 P47090 P47091 P47094 P47106
P47111 P47114 P47124 P47131 P47144 P47153 P47154 P47155 P47159 P47161 P47185
P47187 P47190 P47191 P47192 P47193 P47194 P47211 P47234 P47260 P47261 P47274
P47279 P47289 P47290 P47301 P47307 P47308 P47310 P47315 P47317 P47321 P47322
P47323 P47324 P47332 P47344 P47351 P47355 P47360 P47366 P47367 P47369 P47376
P47377 P47379 P47381 P47390 P47393 P47416 P47427 P47434 P47435 P47467 P47468
P47475 P47483 P47484 P47485 P47489 P47497 P47498 P47509 P47521 P47522 P47523
P47528 P47533 P47544 P47548 P47558 P47561 P47562 P47564 P47573 P47629 P47637
P47643 P47644 P47645 P47651 P47653 P47655 P47662 P47679 P47681 P47685 P47690
P47694 P47695 P47702 P47735 P47740 P47741 P47742 P47743 P47745 P47746 P47747
P47748 P47749 P47750 P47751 P47752 P47758 P47759 P47774 P47775 P47787 P47789
P47790 P47798 P47799 P47800 P47801 P47802 P47803 P47804 P47818 P47820 P47824
P47842 P47843 P47861 P47862 P47863 P47864 P47865 P47866 P47869 P47870 P47871
P47872 P47881 P47883 P47884 P47887 P47888 P47890 P47893 P47898 P47899 P47900
P47901 P47918 P47936 P47937 P47946 P47951 P47983 P47984 P47985 P47987 P48016
P48017 P48018 P48020 P48021 P48022 P48023 P48027 P48029 P48030 P48039 P48040
P48042 P48043 P48044 P48048 P48050 P48051 P48055 P48056 P48057 P48058 P48060
P48065 P48066 P48067 P48079 P48084 P48085 P48086 P48094 P48102 P48103 P48104
P48105 P48106 P48107 P48108 P48109 P48112 P48113 P48115 P48116 P48117 P48121
P48122 P48123 P48145 P48146 P48165 P48167 P48168 P48169 P48170 P48171 P48172
P48173 P48174 P48175 P48176 P48177 P48178 P48180 P48181 P48182 P48183 P48184

P48185 P48186 P48187 P48188 P48189 P48192 P48201 P48230 P48231 P48236 P48241
P48244 P48245 P48256 P48257 P48258 P48265 P48266 P48268 P48269 P48272 P48278
P48280 P48302 P48303 P48305 P48306 P48356 P48357 P48366 P48410 P48421 P48422
P48439 P48440 P48442 P48503 P48505 P48509 P48517 P48518 P48519 P48520 P48521
P48522 P48537 P48542 P48543 P48544 P48545 P48546 P48547 P48548 P48549 P48550
P48551 P48558 P48565 P48583 P48613 P48622 P48623 P48624 P48625 P48626 P48630
P48631 P48632 P48651 P48652 P48653 P48654 P48655 P48656 P48657 P48658 P48659
P48660 P48661 P48662 P48663 P48664 P48665 P48746 P48747 P48748 P48751 P48761
P48762 P48763 P48764 P48765 P48766 P48767 P48768 P48770 P48771 P48772 P48777
P48813 P48829 P48830 P48831 P48832 P48833 P48834 P48866 P48867 P48868 P48869
P48870 P48871 P48872 P48873 P48874 P48875 P48876 P48877 P48878 P48879 P48880
P48881 P48882 P48884 P48885 P48886 P48887 P48888 P48889 P48890 P48891 P48892
P48893 P48894 P48895 P48896 P48897 P48898 P48899 P48900 P48901 P48902 P48903
P48904 P48905 P48906 P48907 P48909 P48910 P48911 P48912 P48913 P48914 P48915
P48916 P48917 P48918 P48919 P48920 P48921 P48922 P48923 P48924 P48925 P48926
P48927 P48928 P48929 P48930 P48931 P48934 P48935 P48960 P48962 P48974 P48990
P48994 P48995 P49015 P49018 P49019 P49020 P49022 P49049 P49059 P49069 P49070
P49096 P49097 P49098 P49099 P49100 P49109 P49111 P49129 P49130 P49131 P49132
P49133 P49134 P49144 P49145 P49146 P49159 P49160 P49161 P49173 P49175 P49179
P49190 P49191 P49217 P49218 P49219 P49220 P49238 P49253 P49256 P49257 P49259
P49260 P49279 P49280 P49281 P49282 P49283 P49285 P49286 P49287 P49288 P49300
P49301 P49326 P49328 P49333 P49334 P49341 P49346 P49374 P49380 P49382 P49390
P49414 P49415 P49416 P49430 P49446 P49447 P49460 P49461 P49470 P49471 P49472
P49473 P49474 P49475 P49476 P49478 P49479 P49480 P49483 P49484 P49485 P49486
P49487 P49488 P49489 P49509 P49511 P49512 P49513 P49514 P49515 P49516 P49523
P49524 P49526 P49527 P49528 P49529 P49531 P49538 P49539 P49542 P49543 P49573
P49578 P49579 P49580 P49581 P49582 P49599 P49600 P49603 P49641 P49650 P49651
P49652 P49653 P49654 P49655 P49656 P49658 P49660 P49681 P49682 P49683 P49684
P49685 P49727 P49728 P49729 P49755 P49768 P49769 P49771 P49772 P49779 P49780
P49781 P49782 P49783 P49784 P49785 P49788 P49810 P49825 P49853 P49856 P49857
P49858 P49892 P49894 P49895 P49896 P49897 P49898 P49899 P49912 P49922 P49936
P49937 P49939 P49940 P49950 P49961 P49976 P49977 P49978 P50012 P50013 P50017
P50052 P50054 P50076 P50077 P50078 P50080 P50083 P50085 P50087 P50088 P50093
P50108 P50110 P50123 P50126 P50127 P50128 P50129 P50130 P50131 P50132 P50133
P50155 P50156 P50170 P50172 P50232 P50253 P50267 P50268 P50269 P50270 P50271
P50276 P50279 P50281 P50283 P50284 P50285 P50333 P50334 P50363 P50364 P50365
P50366 P50367 P50368 P50369 P50370 P50391 P50402 P50406 P50407 P50416 P50427
P50443 P50466 P50482 P50487 P50488 P50489 P50490 P50491 P50492 P50493 P50494
P50501 P50505 P50515 P50523 P50537 P50555 P50571 P50572 P50573 P50591 P50592
P50598 P50599 P50600 P50605 P50636 P50637 P50638 P50644 P50645 P50653 P50654
P50655 P50656 P50657 P50658 P50659 P50660 P50661 P50662 P50663 P50664 P50665
P50666 P50667 P50668 P50669 P50671 P50672 P50673 P50675 P50676 P50677 P50678
P50679 P50680 P50681 P50682 P50683 P50684 P50685 P50686 P50687 P50688 P50689
P50690 P50691 P50692 P50693 P50726 P50730 P50731 P50732 P50738 P50741 P50742

P50744 P50847 P50851 P50852 P50860 P50876 P50895 P50939 P50940 P50943 P50944
P50945 P50973 P50974 P50975 P50976 P50992 P50993 P50996 P50997 P51000 P51021
P51027 P51046 P51047 P51048 P51049 P51050 P51051 P51052 P51055 P51057 P51095
P51096 P51097 P51098 P51099 P51100 P51130 P51131 P51132 P51133 P51134 P51135
P51143 P51144 P51164 P51165 P51167 P51168 P51169 P51170 P51171 P51172 P51180
P51184 P51193 P51194 P51197 P51206 P51212 P51220 P51221 P51222 P51232 P51236
P51239 P51244 P51245 P51246 P51247 P51264 P51265 P51274 P51275 P51276 P51284
P51285 P51297 P51316 P51317 P51318 P51321 P51322 P51323 P51324 P51325 P51327
P51340 P51341 P51356 P51357 P51360 P51363 P51364 P51369 P51370 P51385 P51387
P51388 P51389 P51390 P51391 P51392 P51393 P51395 P51435 P51436 P51470 P51471
P51472 P51473 P51474 P51475 P51476 P51488 P51489 P51490 P51491 P51511 P51512
P51515 P51519 P51520 P51533 P51542 P51559 P51563 P51564 P51571 P51572 P51574
P51575 P51576 P51577 P51578 P51579 P51582 P51639 P51648 P51651 P51658 P51674
P51675 P51676 P51677 P51678 P51679 P51680 P51681 P51682 P51683 P51684 P51685
P51686 P51693 P51700 P51712 P51727 P51728 P51729 P51742 P51743 P51749 P51750
P51751 P51753 P51759 P51761 P51762 P51763 P51764 P51765 P51766 P51769 P51770
P51773 P51787 P51788 P51789 P51790 P51791 P51792 P51793 P51794 P51795 P51796
P51797 P51798 P51799 P51800 P51801 P51802 P51803 P51804 P51805 P51809 P51810
P51811 P51828 P51829 P51830 P51839 P51840 P51841 P51842 P51866 P51867 P51869
P51881 P51882 P51898 P51899 P51905 P51906 P51907 P51912 P51914 P51915 P51916
P51926 P51930 P51931 P51932 P51937 P51938 P51940 P51941 P51973 P51975 P51993
P51995 P52002 P52005 P52047 P52059 P52067 P52071 P52094 P52101 P52112 P52138
P52143 P52146 P52159 P52166 P52177 P52178 P52185 P52186 P52187 P52188 P52189
P52190 P52191 P52192 P52194 P52202 P52204 P52205 P52219 P52220 P52221 P52222
P52223 P52225 P52226 P52237 P52280 P52352 P52353 P52366 P52370 P52371 P52372
P52373 P52380 P52381 P52382 P52383 P52429 P52443 P52444 P52445 P52449 P52465
P52466 P52500 P52542 P52543 P52547 P52569 P52583 P52586 P52587 P52591 P52592
P52599 P52600 P52625 P52636 P52638 P52649 P52687 P52702 P52703 P52764 P52765
P52768 P52770 P52785 P52795 P52796 P52799 P52800 P52848 P52849 P52850 P52853
P52867 P52869 P52870 P52871 P52872 P52875 P52876 P52879 P52880 P52881 P52883
P52887 P52923 P52924 P53006 P53007 P53008 P53012 P53039 P53045 P53047 P53048
P53049 P53053 P53059 P53062 P53069 P53073 P53074 P53083 P53087 P53089 P53093
P53099 P53106 P53108 P53109 P53113 P53116 P53117 P53120 P53121 P53134 P53142
P53146 P53151 P53153 P53154 P53157 P53159 P53160 P53161 P53171 P53173 P53174
P53176 P53181 P53182 P53190 P53192 P53198 P53203 P53206 P53209 P53214 P53217
P53220 P53223 P53224 P53226 P53229 P53239 P53241 P53245 P53247 P53253 P53257
P53259 P53262 P53266 P53268 P53269 P53273 P53279 P53282 P53283 P53285 P53288
P53293 P53306 P53308 P53310 P53311 P53320 P53322 P53324 P53325 P53337 P53339
P53342 P53343 P53344 P53380 P53387 P53388 P53389 P53390 P53391 P53392 P53393
P53394 P53403 P53425 P53426 P53427 P53431 P53432 P53439 P53452 P53453 P53454
P53507 P53512 P53513 P53514 P53517 P53525 P53541 P53561 P53563 P53584 P53601
P53624 P53629 P53631 P53633 P53660 P53690 P53694 P53697 P53706 P53708 P53710
P53711 P53712 P53713 P53714 P53717 P53721 P53723 P53726 P53730 P53735 P53740
P53744 P53745 P53746 P53747 P53748 P53751 P53752 P53756 P53767 P53785 P53786

P53788 P53790 P53791 P53792 P53793 P53794 P53798 P53799 P53800 P53801 P53815
P53816 P53817 P53822 P53825 P53832 P53835 P53838 P53842 P53845 P53856 P53862
P53868 P53878 P53880 P53882 P53884 P53895 P53896 P53898 P53903 P53906 P53908
P53918 P53925 P53926 P53928 P53932 P53938 P53943 P53947 P53950 P53954 P53956
P53961 P53964 P53966 P53969 P53975 P53979 P53983 P53985 P53986 P53987 P53988
P53993 P54002 P54003 P54069 P54070 P54072 P54074 P54078 P54079 P54085 P54086
P54104 P54126 P54127 P54128 P54129 P54131 P54133 P54134 P54141 P54142 P54144
P54145 P54146 P54147 P54148 P54158 P54163 P54172 P54175 P54176 P54179 P54180
P54181 P54209 P54210 P54211 P54219 P54244 P54245 P54246 P54247 P54248 P54249
P54250 P54251 P54265 P54267 P54281 P54282 P54289 P54290 P54302 P54323 P54360
P54371 P54373 P54392 P54393 P54396 P54417 P54425 P54428 P54437 P54438 P54442
P54444 P54445 P54447 P54449 P54463 P54465 P54466 P54467 P54469 P54478 P54481
P54483 P54485 P54486 P54487 P54488 P54493 P54496 P54500 P54502 P54505 P54506
P54510 P54513 P54514 P54515 P54516 P54529 P54536 P54538 P54544 P54558 P54559
P54567 P54571 P54576 P54580 P54582 P54585 P54587 P54589 P54593 P54595 P54596
P54603 P54642 P54660 P54678 P54679 P54683 P54700 P54702 P54707 P54708 P54709
P54710 P54715 P54718 P54719 P54720 P54723 P54736 P54738 P54743 P54744 P54745
P54751 P54753 P54754 P54755 P54756 P54757 P54758 P54759 P54760 P54761 P54762
P54763 P54764 P54773 P54781 P54813 P54815 P54820 P54825 P54829 P54830 P54833
P54834 P54837 P54848 P54849 P54850 P54851 P54852 P54854 P54855 P54856 P54862
P54867 P54880 P54881 P54883 P54900 P54908 P54909 P54913 P54914 P54917 P54932
P54942 P54946 P54953 P54957 P54960 P54982 P55003 P55011 P55012 P55013 P55014
P55015 P55016 P55017 P55018 P55019 P55024 P55025 P55026 P55027 P55028 P55033
P55061 P55062 P55064 P55069 P55073 P55082 P55085 P55086 P55087 P55088 P55096
P55098 P55117 P55118 P55122 P55123 P55125 P55128 P55129 P55130 P55131 P55140
P55144 P55146 P55154 P55160 P55161 P55162 P55167 P55182 P55183 P55189 P55190
P55202 P55203 P55204 P55205 P55237 P55244 P55245 P55270 P55277 P55280 P55281
P55283 P55284 P55285 P55286 P55287 P55288 P55289 P55291 P55292 P55340 P55344
P55368 P55375 P55377 P55384 P55396 P55397 P55402 P55403 P55406 P55416 P55421
P55431 P55432 P55439 P55442 P55448 P55449 P55450 P55452 P55469 P55471 P55475
P55478 P55492 P55493 P55505 P55506 P55522 P55527 P55554 P55561 P55569 P55571
P55572 P55574 P55580 P55581 P55583 P55585 P55602 P55603 P55647 P55652 P55660
P55661 P55671 P55679 P55682 P55698 P55699 P55705 P55712 P55714 P55720 P55721
P55722 P55723 P55725 P55726 P55727 P55734 P55738 P55772 P55777 P55778 P55779
P55780 P55781 P55782 P55783 P55803 P55808 P55815 P55849 P55850 P55851 P55891
P55892 P55899 P55908 P55910 P55916 P55917 P55919 P55920 P55926 P55934 P55981
P55989 P56079 P56085 P56086 P56087 P56096 P56098 P56100 P56117 P56134 P56135
P56136 P56180 P56190 P56199 P56295 P56296 P56297 P56305 P56306 P56307 P56308
P56309 P56310 P56312 P56313 P56314 P56315 P56316 P56318 P56319 P56321 P56322
P56323 P56324 P56325 P56326 P56327 P56328 P56338 P56339 P56340 P56341 P56342
P56343 P56348 P56349 P56369 P56370 P56373 P56378 P56379 P56383 P56384 P56392
P56393 P56395 P56400 P56401 P56402 P56403 P56404 P56405 P56411 P56412 P56416
P56427 P56433 P56434 P56436 P56439 P56440 P56442 P56443 P56444 P56445 P56446
P56447 P56448 P56449 P56450 P56451 P56475 P56476 P56479 P56481 P56482 P56483

P56484 P56485 P56486 P56488 P56489 P56490 P56491 P56492 P56493 P56494 P56495
P56496 P56497 P56498 P56499 P56500 P56501 P56502 P56508 P56509 P56510 P56513
P56514 P56515 P56516 P56528 P56557 P56560 P56563 P56564 P56579 P56580 P56583
P56589 P56603 P56621 P56627 P56629 P56630 P56631 P56632 P56677 P56695 P56696
P56697 P56698 P56699 P56718 P56719 P56720 P56722 P56726 P56729 P56731 P56745
P56746 P56747 P56748 P56749 P56750 P56751 P56752 P56758 P56759 P56760 P56761
P56766 P56767 P56768 P56769 P56770 P56771 P56773 P56774 P56775 P56776 P56777
P56778 P56779 P56780 P56781 P56782 P56783 P56785 P56788 P56790 P56817 P56818
P56819 P56833 P56834 P56856 P56857 P56874 P56880 P56882 P56895 P56911 P56937
P56938 P56939 P56940 P56941 P56949 P56962 P56971 P56974 P56975 P56976 P56982
P56983 P56984 P57003 P57004 P57012 P57019 P57021 P57022 P57034 P57039 P57041
P57042 P57046 P57047 P57048 P57051 P57054 P57057 P57061 P57062 P57063 P57083
P57087 P57088 P57097 P57103 P57105 P57118 P57119 P57120 P57131 P57152 P57170
P57175 P57184 P57185 P57186 P57189 P57213 P57216 P57218 P57223 P57234 P57239
P57248 P57252 P57258 P57260 P57261 P57262 P57263 P57264 P57296 P57312 P57314
P57317 P57318 P57331 P57334 P57335 P57347 P57355 P57361 P57363 P57364 P57368
P57370 P57382 P57392 P57402 P57406 P57408 P57414 P57415 P57437 P57440 P57460
P57462 P57496 P57514 P57524 P57527 P57538 P57540 P57541 P57542 P57543 P57544
P57550 P57551 P57552 P57571 P57601 P57630 P57631 P57635 P57647 P57648 P57663
P57679 P57680 P57683 P57684 P57685 P57686 P57687 P57688 P57695 P57698 P57699
P57700 P57701 P57709 P57716 P57727 P57738 P57739 P57757 P57758 P57764 P57773
P57787 P57788 P57789 P57791 P57792 P57800 P57816 P57839 P57846 P57864 P57929
P57950 P57959 P57970 P57999 P58007 P58008 P58010 P58021 P58027 P58028 P58029
P58030 P58034 P58045 P58046 P58047 P58048 P58049 P58050 P58051 P58061 P58118
P58119 P58120 P58126 P58155 P58158 P58162 P58163 P58164 P58165 P58170 P58173
P58180 P58181 P58182 P58191 P58195 P58196 P58198 P58199 P58200 P58214 P58216
P58219 P58229 P58239 P58244 P58246 P58247 P58266 P58273 P58274 P58281 P58295
P58307 P58308 P58309 P58310 P58311 P58312 P58325 P58335 P58341 P58342 P58344
P58345 P58351 P58352 P58353 P58354 P58355 P58356 P58359 P58361 P58363 P58365
P58366 P58367 P58368 P58369 P58377 P58383 P58385 P58386 P58387 P58390 P58391
P58392 P58400 P58401 P58402 P58406 P58411 P58412 P58414 P58418 P58419 P58420
P58421 P58428 P58492 P58511 P58529 P58530 P58531 P58549 P58550 P58560 P58564
P58565 P58566 P58568 P58569 P58576 P58577 P58583 P58593 P58594 P58595 P58597
P58603 P58612 P58650 P58651 P58655 P58658 P58659 P58662 P58681 P58682 P58684
P58689 P58690 P58691 P58701 P58702 P58710 P58720 P58721 P58722 P58725 P58727
P58734 P58735 P58740 P58741 P58743 P58749 P58762 P58764 P58765 P58767 P58769
P58819 P58821 P58825 P58826 P58872 P58873 P58905 P58931 P58932 P58933 P58934
P58950 P58951 P58952 P58953 P58954 P58955 P58956 P58957 P58958 P58959 P58960
P58961 P58962 P58985 P58997 P59017 P59025 P59034 P59035 P59041 P59053 P59087
P59111 P59112 P59158 P59194 P59195 P59198 P59219 P59221 P59222 P59228 P59229
P59246 P59247 P59248 P59249 P59250 P59251 P59252 P59253 P59255 P59256 P59257
P59261 P59262 P59266 P59267 P59268 P59269 P59270 P59277 P59333 P59334 P59335
P59340 P59341 P59342 P59343 P59344 P59345 P59346 P59347 P59363 P59364 P59365
P59366 P59382 P59383 P59436 P59440 P59481 P59494 P59523 P59528 P59529 P59530

P59532 P59533 P59534 P59535 P59536 P59537 P59538 P59539 P59540 P59541 P59542
P59543 P59544 P59551 P59559 P59585 P59594 P59596 P59632 P59635 P59637 P59638
P59639 P59645 P59646 P59647 P59648 P59649 P59652 P59653 P59654 P59662 P59668
P59670 P59676 P59684 P59693 P59694 P59695 P59699 P59702 P59703 P59705 P59706
P59709 P59710 P59711 P59737 P59741 P59762 P59763 P59770 P59771 P59773 P59777
P59783 P59789 P59798 P59809 P59810 P59811 P59822 P59823 P59824 P59832 P59836
P59841 P59844 P59845 P59852 P59862 P59889 P59901 P59902 P59903 P59904 P59905
P59907 P59908 P59909 P59917 P59922 P59926 P59947 P59949 P59950 P59960 P59963
P59972 P59977 P59983 P59985 P59988 P59994 P59995 P60015 P60018 P60019 P60020
P60021 P60025 P60026 P60029 P60033 P60034 P60035 P60036 P60037 P60058 P60059
P60060 P60061 P60062 P60063 P60064 P60065 P60066 P60067 P60112 P60113 P60114
P60115 P60116 P60117 P60118 P60124 P60125 P60126 P60127 P60128 P60129 P60130
P60131 P60132 P60133 P60134 P60135 P60136 P60137 P60138 P60139 P60140 P60141
P60142 P60143 P60144 P60145 P60146 P60147 P60148 P60149 P60150 P60151 P60152
P60159 P60160 P60161 P60162 P60181 P60182 P60183 P60201 P60202 P60203 P60247
P60248 P60249 P60250 P60251 P60300 P60363 P60459 P60460 P60461 P60462 P60463
P60464 P60465 P60466 P60467 P60468 P60497 P60498 P60503 P60505 P60506 P60507
P60508 P60509 P60528 P60570 P60571 P60572 P60573 P60574 P60588 P60602 P60603
P60606 P60612 P60613 P60614 P60620 P60621 P60632 P60633 P60634 P60635 P60636
P60637 P60638 P60639 P60640 P60641 P60642 P60643 P60644 P60645 P60646 P60647
P60648 P60649 P60650 P60671 P60672 P60674 P60675 P60676 P60677 P60678 P60679
P60680 P60681 P60682 P60683 P60684 P60685 P60686 P60687 P60688 P60689 P60690
P60691 P60692 P60693 P60694 P60695 P60696 P60697 P60698 P60699 P60752 P60753
P60777 P60778 P60779 P60782 P60783 P60784 P60815 P60823 P60844 P60845 P60852
P60869 P60870 P60871 P60872 P60873 P60874 P60882 P60883 P60893 P60894 P60895
P60925 P60926 P60927 P60928 P60929 P60932 P60933 P60934 P60935 P60936 P60937
P60938 P60939 P60940 P60941 P60942 P60943 P60944 P60945 P60946 P60947 P60949
P60950 P60951 P60955 P60956 P60957 P60958 P60959 P60960 P60961 P60962 P60963
P60964 P60965 P60966 P60967 P60968 P60969 P60970 P60971 P60972 P60973 P60974
P61008 P61009 P61012 P61013 P61014 P61015 P61016 P61032 P61033 P61034 P61035
P61036 P61037 P61038 P61039 P61040 P61041 P61042 P61045 P61046 P61047 P61048
P61049 P61051 P61052 P61053 P61072 P61073 P61130 P61131 P61132 P61165 P61166
P61168 P61169 P61170 P61171 P61172 P61228 P61252 P61264 P61265 P61266 P61267
P61268 P61315 P61382 P61383 P61384 P61385 P61386 P61387 P61388 P61389 P61390
P61391 P61392 P61393 P61394 P61395 P61402 P61403 P61415 P61483 P61484 P61507
P61543 P61544 P61545 P61546 P61547 P61550 P61552 P61553 P61554 P61555 P61556
P61557 P61558 P61561 P61562 P61563 P61564 P61565 P61566 P61567 P61570 P61619
P61620 P61621 P61622 P61625 P61636 P61637 P61638 P61639 P61642 P61643 P61644
P61645 P61646 P61647 P61648 P61649 P61752 P61755 P61756 P61757 P61776 P61777
P61784 P61785 P61793 P61794 P61795 P61796 P61803 P61804 P61805 P61806 P61807
P61808 P61813 P61814 P61815 P61828 P61829 P61837 P61838 P61839 P61840 P61900
P61901 P61902 P61903 P61914 P61915 P61943 P62021 P62079 P62080 P62095 P62096
P62097 P62098 P62099 P62100 P62101 P62102 P62103 P62104 P62105 P62106 P62107
P62108 P62109 P62111 P62112 P62113 P62114 P62115 P62116 P62341 P62342 P62447

P62461 P62462 P62463 P62464 P62465 P62466 P62467 P62480 P62481 P62513 P62514
P62517 P62518 P62519 P62530 P62531 P62586 P62587 P62595 P62596 P62648 P62651
P62670 P62671 P62719 P62720 P62721 P62723 P62724 P62725 P62812 P62813 P62816
P62817 P62950 P62951 P62952 P62953 P62954 P62955 P62956 P62957 P62967 P63024
P63025 P63026 P63027 P63030 P63031 P63044 P63045 P63079 P63080 P63081 P63082
P63115 P63116 P63137 P63138 P63141 P63142 P63160 P63161 P63183 P63184 P63232
P63235 P63236 P63250 P63251 P63252 P63253 P63304 P63305 P63306 P63307 P63308
P63332 P63333 P63340 P63341 P63342 P63343 P63344 P63350 P63359 P63360 P63392
P63394 P63398 P63400 P63520 P63528 P63619 P63620 P63655 P63657 P63682 P63683
P63684 P63686 P63688 P63690 P63692 P63694 P63696 P63701 P63702 P63714 P63725
P63726 P63752 P63754 P63755 P63756 P63757 P63759 P63761 P63763 P63798 P63799
P63800 P63801 P63802 P63853 P63855 P63857 P63858 P63859 P63860 P63861 P63863
P63865 P63866 P63867 P63886 P63888 P63912 P63993 P63994 P64002 P64003 P64099
P64100 P64101 P64102 P64160 P64161 P64162 P64163 P64164 P64165 P64166 P64167
P64169 P64176 P64177 P64255 P64256 P64257 P64258 P64260 P64293 P64423 P64424
P64425 P64429 P64430 P64431 P64432 P64433 P64434 P64439 P64440 P64441 P64445
P64446 P64447 P64453 P64454 P64471 P64472 P64473 P64481 P64482 P64485 P64486
P64487 P64499 P64500 P64501 P64502 P64512 P64513 P64514 P64515 P64516 P64536
P64537 P64538 P64539 P64545 P64546 P64547 P64550 P64551 P64552 P64553 P64564
P64565 P64566 P64581 P64582 P64583 P64584 P64585 P64586 P64587 P64590 P64591
P64592 P64593 P64604 P64605 P64606 P64607 P64608 P64609 P64619 P64620 P64621
P64622 P64623 P64627 P64628 P64629 P64630 P64634 P64635 P64646 P64647 P64648
P64651 P64652 P64661 P64662 P64682 P64684 P64690 P64696 P64700 P64712 P64716
P64730 P64736 P64738 P64744 P64752 P64756 P64758 P64772 P64776 P64782 P64784
P64802 P64812 P64816 P64830 P64838 P64848 P64850 P64856 P64858 P64884 P64890
P64892 P64894 P64896 P64904 P64932 P64934 P64940 P64948 P64950 P64952 P64954
P64958 P64962 P64970 P64972 P64998 P65008 P65016 P65020 P65022 P65024 P65026
P65030 P65072 P65082 P65195 P65196 P65197 P65198 P65207 P65208 P65210 P65212
P65213 P65214 P65253 P65254 P65263 P65264 P65265 P65266 P65267 P65268 P65317
P65371 P65373 P65375 P65377 P65379 P65381 P65410 P65411 P65524 P65525 P65527
P65540 P65541 P65542 P65543 P65544 P65545 P65562 P65564 P65566 P65594 P65622
P65623 P65624 P65625 P65627 P65628 P65629 P65630 P65631 P65638 P65651 P65713
P65727 P65731 P65733 P65811 P65812 P65813 P65814 P65816 P65817 P65818 P65822
P66772 P66792 P66884 P66887 P66888 P66890 P66891 P66892 P66896 P66965 P66967
P67053 P67054 P67116 P67118 P67127 P67128 P67129 P67131 P67143 P67144 P67145
P67147 P67148 P67149 P67150 P67153 P67154 P67155 P67156 P67158 P67161 P67162
P67163 P67164 P67165 P67166 P67167 P67168 P67170 P67244 P67245 P67277 P67278
P67279 P67280 P67281 P67282 P67283 P67284 P67288 P67289 P67290 P67291 P67292
P67293 P67294 P67295 P67297 P67377 P67378 P67379 P67384 P67385 P67386 P67387
P67388 P67389 P67390 P67391 P67392 P67394 P67439 P67444 P67445 P67446 P67470
P67471 P67559 P67560 P67601 P67602 P67638 P67639 P67644 P67645 P67646 P67686
P67687 P67729 P67730 P67780 P67781 P67782 P67783 P67784 P67793 P67794 P67795
P67796 P67797 P67798 P67799 P67810 P67811 P67812 P67866 P67867 P67907 P67908
P67909 P67923 P68002 P68003 P68088 P68089 P68090 P68091 P68092 P68093 P68094

P68095 P68159 P68160 P68161 P68162 P68178 P68179 P68183 P68184 P68185 P68186
P68269 P68270 P68294 P68295 P68296 P68297 P68298 P68299 P68300 P68307 P68308
P68309 P68310 P68323 P68324 P68325 P68326 P68327 P68328 P68329 P68330 P68331
P68332 P68333 P68334 P68343 P68347 P68348 P68407 P68408 P68409 P68434 P68452
P68453 P68458 P68459 P68513 P68514 P68526 P68527 P68528 P68529 P68530 P68531
P68532 P68533 P68534 P68537 P68538 P68539 P68540 P68548 P68549 P68553 P68554
P68557 P68558 P68571 P68572 P68579 P68580 P68592 P68593 P68604 P68605 P68606
P68616 P68617 P68618 P68619 P68622 P68623 P68632 P68633 P68667 P68668 P68670
P68672 P68674 P68699 P68700 P68701 P68702 P68703 P68704 P68705 P68706 P68711
P68712 P68755 P68756 P68757 P68758 P68759 P68760 P68761 P68762 P68782 P68783
P68803 P68804 P68805 P68806 P68827 P68832 P68833 P68853 P68854 P68855 P68856
P68857 P68858 P68859 P68860 P68914 P68920 P68921 P68978 P68979 P68980 P68981
P69034 P69051 P69052 P69054 P69055 P69168 P69169 P69187 P69188 P69194 P69195
P69210 P69211 P69212 P69213 P69214 P69215 P69216 P69217 P69218 P69219 P69220
P69221 P69232 P69233 P69234 P69235 P69236 P69237 P69238 P69239 P69302 P69303
P69304 P69305 P69306 P69307 P69332 P69333 P69334 P69335 P69336 P69337 P69338
P69339 P69340 P69345 P69350 P69351 P69353 P69354 P69355 P69356 P69357 P69358
P69359 P69360 P69367 P69368 P69371 P69372 P69373 P69374 P69375 P69378 P69379
P69380 P69381 P69382 P69383 P69384 P69385 P69386 P69387 P69388 P69390 P69393
P69394 P69395 P69396 P69397 P69398 P69399 P69400 P69401 P69402 P69420 P69421
P69422 P69423 P69424 P69425 P69426 P69427 P69428 P69429 P69430 P69431 P69432
P69433 P69434 P69435 P69447 P69448 P69449 P69453 P69454 P69455 P69456 P69457
P69458 P69460 P69461 P69462 P69463 P69464 P69465 P69478 P69490 P69491 P69492
P69493 P69518 P69519 P69520 P69521 P69523 P69524 P69525 P69526 P69528 P69529
P69530 P69531 P69532 P69533 P69534 P69535 P69536 P69537 P69538 P69539 P69540
P69541 P69550 P69551 P69552 P69553 P69554 P69555 P69556 P69557 P69558 P69559
P69560 P69561 P69562 P69563 P69564 P69565 P69599 P69600 P69601 P69602 P69603
P69604 P69605 P69606 P69607 P69610 P69611 P69612 P69613 P69616 P69617 P69669
P69670 P69671 P69672 P69673 P69674 P69675 P69676 P69677 P69680 P69681 P69685
P69686 P69690 P69691 P69692 P69693 P69694 P69695 P69696 P69699 P69700 P69703
P69704 P69705 P69739 P69740 P69744 P69786 P69787 P69788 P69789 P69790 P69801
P69802 P69803 P69804 P69805 P69806 P69807 P69826 P69827 P69831 P69832 P69833
P69849 P69856 P69857 P69858 P69927 P69937 P69938 P69955 P69956 P69972 P69973
P69980 P69981 P69982 P69983 P69984 P69985 P69986 P69987 P69998 P69999 P70031
P70057 P70083 P70097 P70105 P70106 P70110 P70115 P70126 P70170 P70172 P70174
P70180 P70187 P70193 P70194 P70205 P70206 P70207 P70208 P70211 P70224 P70225
P70227 P70232 P70245 P70259 P70263 P70277 P70280 P70289 P70302 P70310 P70312
P70387 P70403 P70406 P70407 P70408 P70412 P70414 P70419 P70423 P70424 P70428
P70452 P70500 P70502 P70505 P70526 P70536 P70539 P70545 P70549 P70551 P70553
P70555 P70579 P70580 P70585 P70596 P70597 P70604 P70605 P70606 P70612 P70617
P70627 P70628 P70658 P70669 P70673 P70689 P70691 P70704 P70705 P70758 P70775
P70786 P70841 P70845 P70864 P70952 P70954 P70963 P70966 P70972 P70995 P70997
P71005 P71008 P71010 P71012 P71013 P71038 P71040 P71041 P71042 P71044 P71045
P71050 P71052 P71056 P71060 P71062 P71067 P71070 P71082 P71083 P71238 P71241

P71338 P71342 P71345 P71355 P71356 P71360 P71364 P71367 P71369 P71375 P71377
P71380 P71395 P71399 P71408 P71411 P71485 P71486 P71707 P72109 P72179 P72202
P72203 P72204 P72205 P72267 P72281 P72287 P72292 P72295 P72296 P72302 P72331
P72336 P72358 P72367 P72482 P72575 P72604 P72637 P72640 P72655 P72660 P72701
P72705 P72712 P72714 P72717 P72732 P72752 P72771 P72793 P72804 P72823 P72831
P72836 P72846 P72855 P72926 P72929 P72935 P72941 P72942 P72958 P72973 P72978
P72986 P72991 P73009 P73070 P73087 P73157 P73179 P73182 P73241 P73354 P73387
P73437 P73451 P73517 P73528 P73540 P73548 P73553 P73555 P73633 P73676 P73689
P73714 P73726 P73771 P73795 P73801 P73809 P73863 P73866 P73867 P73869 P73888
P73893 P73894 P73912 P73920 P73933 P73962 P74055 P74068 P74072 P74078 P74149
P74155 P74165 P74167 P74168 P74180 P74311 P74343 P74369 P74393 P74409 P74436
P74475 P74505 P74511 P74547 P74564 P74625 P74635 P74653 P74714 P74787 P74805
P74810 P74852 P74856 P74884 P74890 P74891 P74930 P74931 P74932 P74985 P75024
P75028 P75039 P75040 P75041 P75047 P75048 P75057 P75058 P75071 P75083 P75094
P75095 P75109 P75112 P75120 P75121 P75125 P75130 P75134 P75146 P75149 P75160
P75166 P75174 P75185 P75191 P75201 P75204 P75207 P75208 P75213 P75251 P75262
P75263 P75291 P75295 P75307 P75314 P75318 P75323 P75328 P75329 P75330 P75343
P75346 P75357 P75369 P75377 P75379 P75383 P75384 P75385 P75387 P75397 P75405
P75408 P75421 P75422 P75428 P75439 P75440 P75441 P75443 P75445 P75449 P75450
P75459 P75462 P75463 P75468 P75472 P75484 P75501 P75503 P75506 P75507 P75513
P75514 P75515 P75520 P75528 P75535 P75547 P75553 P75554 P75555 P75556 P75569
P75582 P75585 P75586 P75588 P75596 P75597 P75603 P75606 P75607 P75608 P75613
P75681 P75683 P75685 P75693 P75712 P75733 P75742 P75747 P75750 P75757 P75763
P75769 P75770 P75780 P75783 P75785 P75788 P75798 P75799 P75800 P75801 P75806
P75810 P75822 P75826 P75830 P75831 P75835 P75839 P75851 P75857 P75870 P75885
P75892 P75901 P75905 P75908 P75910 P75916 P75920 P75925 P75955 P75958 P75961
P75962 P75968 P75971 P75979 P75995 P76007 P76011 P76037 P76045 P76090 P76091
P76093 P76103 P76111 P76115 P76136 P76146 P76147 P76169 P76173 P76175 P76180
P76182 P76185 P76186 P76197 P76198 P76208 P76219 P76221 P76224 P76226 P76228
P76230 P76236 P76237 P76239 P76242 P76249 P76261 P76264 P76269 P76272 P76278
P76298 P76299 P76330 P76334 P76335 P76339 P76343 P76345 P76350 P76352 P76361
P76372 P76387 P76389 P76398 P76399 P76417 P76425 P76445 P76446 P76460 P76470
P76473 P76474 P76482 P76507 P76521 P76538 P76552 P76576 P76577 P76628 P76630
P76655 P76657 P76773 P77031 P77091 P77156 P77172 P77173 P77179 P77180 P77196
P77211 P77219 P77221 P77228 P77265 P77272 P77285 P77293 P77304 P77306 P77307
P77308 P77315 P77328 P77335 P77338 P77354 P77377 P77389 P77400 P77409 P77416
P77437 P77463 P77468 P77473 P77485 P77494 P77504 P77506 P77510 P77529 P77538
P77549 P77579 P77589 P77610 P77619 P77672 P77682 P77716 P77726 P77733 P77747
P77757 P77799 P77858 P77868 P77871 P77921 P77964 P78036 P78283 P78310 P78324
P78325 P78334 P78348 P78357 P78363 P78369 P78380 P78381 P78382 P78383 P78410
P78423 P78504 P78508 P78536 P78552 P78562 P78575 P78588 P78589 P78591 P78595
P78611 P78741 P78746 P78763 P78768 P78810 P78817 P78831 P78898 P78966 P78970
P78979 P78990 P79003 P79082 P79088 P79098 P79099 P79100 P79107 P79108 P79110
P79113 P79138 P79143 P79144 P79148 P79164 P79165 P79166 P79168 P79169 P79171

P79175 P79176 P79177 P79178 P79184 P79185 P79188 P79189 P79190 P79191 P79194
P79197 P79200 P79211 P79213 P79217 P79218 P79222 P79229 P79234 P79235 P79236
P79237 P79240 P79241 P79242 P79243 P79250 P79266 P79282 P79291 P79292 P79307
P79336 P79337 P79350 P79365 P79368 P79374 P79381 P79382 P79391 P79393 P79394
P79399 P79400 P79483 P79701 P79747 P79748 P79749 P79750 P79755 P79756 P79762
P79763 P79784 P79785 P79795 P79798 P79801 P79802 P79803 P79807 P79808 P79809
P79812 P79826 P79848 P79863 P79883 P79898 P79901 P79902 P79903 P79911 P79914
P79928 P79945 P79948 P79949 P79960 P79995 P80065 P80086 P80100 P80101 P80102
P80103 P80104 P80105 P80106 P80107 P80143 P80150 P80172 P80183 P80184 P80185
P80186 P80201 P80202 P80203 P80204 P80228 P80259 P80260 P80285 P80332 P80333
P80368 P80369 P80370 P80431 P80432 P80433 P80439 P80440 P80441 P80442 P80443
P80444 P80451 P80452 P80453 P80454 P80470 P80478 P80479 P80481 P80482 P80497
P80498 P80500 P80560 P80586 P80587 P80588 P80589 P80590 P80591 P80595 P80603
P80651 P80652 P80653 P80654 P80655 P80656 P80667 P80672 P80696 P80746 P80747
P80883 P80896 P80897 P80943 P80967 P80971 P80975 P80977 P80979 P81004 P81103
P81134 P81155 P81230 P81231 P81232 P81233 P81234 P81265 P81301 P81304 P81305
P81306 P81310 P81311 P81313 P81314 P81316 P81317 P81318 P81320 P81322 P81323
P81324 P81327 P81328 P81329 P81331 P81376 P81378 P81379 P81380 P81381 P81408
P81425 P81450 P81451 P81549 P81605 P81662 P81721 P81780 P81785 P81794 P81861
P81862 P81863 P81864 P81865 P81866 P81891 P81909 P81910 P81911 P81912 P81913
P81914 P81915 P81916 P81917 P81918 P81919 P81921 P81922 P81923 P81924 P81928
P82008 P82009 P82013 P82045 P82046 P82047 P82048 P82049 P82149 P82179 P82251
P82252 P82279 P82295 P82347 P82348 P82349 P82350 P82352 P82451 P82539 P82542
P82543 P82805 P82872 P82873 P82874 P82889 P82945 P82982 P82983 P82984 P82985
P82986 P83009 P83079 P83088 P83093 P83094 P83097 P83118 P83119 P83120 P83149
P83151 P83239 P83240 P83247 P83260 P83292 P83293 P83294 P83295 P83296 P83297
P83312 P83313 P83314 P83362 P83502 P83528 P83555 P83556 P83626 P83650 P83681
P83740 P83755 P83756 P83781 P83791 P83792 P83793 P83794 P83795 P83796 P83797
P83798 P83854 P83858 P83861 P83936 P83970 P83978 P83981 P84039 P84157 P84180
P84181 P84201 P84202 P84285 P84287 P84288 P84289 P84290 P84294 P84295 P84300
P84301 P84302 P84303 P84304 P84305 P84309 P84344 P84345 P84392 P84393 P84400
P84406 P84449 P84628 P84817 P84889 P84903 P84988 P84990 P84991 P85007 P85118
P85226 P85296 P85410 P85521 P85828 P85928 P85967 P86044 P86045 P86046 P86050
P86091 P86148 P86176 P86218 P86223 P86229 P86243 P86275 P86789 P86889 P86905
P86911 P86935 P86936 P87014 P87023 P87024 P87041 P87055 P87056 P87059 P87073
P87110 P87113 P87119 P87122 P87124 P87130 P87131 P87136 P87139 P87146 P87148
P87149 P87155 P87170 P87178 P87179 P87200 P87207 P87234 P87236 P87264 P87266
P87267 P87270 P87272 P87273 P87274 P87275 P87280 P87281 P87284 P87285 P87287
P87293 P87320 P87365 P87366 P87367 P87368 P87369 P87410 P87416 P87419 P87421
P87506 P87666 P87671 P87691 P87730 P87745 P88825 P88838 P89035 P89053 P89059
P89061 P89063 P89433 P89443 P89445 P89457 P89464 P89465 P89475 P89477 P89946
P90471 P90489 P90495 P90522 P90680 P90747 P90754 P90755 P90771 P90837 P90859
P90920 P90970 P90992 P91133 P91193 P91200 P91209 P91210 P91211 P91275 P91390
P91406 P91409 P91454 P91535 P91536 P91550 P91645 P91657 P91660 P91679 P91682

P91685 P91730 P91766 P91799 P91927 P91928 P91994 P92001 P92002 P92015 P92029
P92127 P92230 P92248 P92251 P92276 P92277 P92417 P92432 P92472 P92475 P92476
P92477 P92478 P92479 P92480 P92481 P92482 P92483 P92484 P92485 P92486 P92487
P92492 P92493 P92494 P92503 P92507 P92509 P92511 P92514 P92527 P92528 P92533
P92536 P92541 P92547 P92550 P92553 P92556 P92558 P92559 P92564 P92565 P92583
P92584 P92592 P92599 P92600 P92604 P92605 P92613 P92623 P92649 P92651 P92657
P92658 P92659 P92660 P92661 P92662 P92663 P92664 P92665 P92666 P92667 P92668
P92669 P92670 P92671 P92681 P92690 P92691 P92692 P92693 P92694 P92695 P92696
P92697 P92698 P92699 P92700 P92701 P92717 P92759 P92792 P92793 P92794 P92811
P92817 P92845 P92846 P92847 P92848 P92849 P92850 P92852 P92853 P92857 P92870
P92916 P92919 P92934 P92935 P92939 P92941 P92942 P92943 P92946 P92949 P92960
P92961 P92962 P92994 P93004 P93026 P93030 P93039 P93042 P93050 P93051 P93149
P93194 P93278 P93280 P93283 P93284 P93285 P93286 P93296 P93298 P93303 P93312
P93313 P93314 P93316 P93319 P93332 P93401 P93472 P93484 P93604 P93654 P93664
P93690 P93703 P93735 P93756 P93761 P93766 P93823 P93825 P93829 P93834 P93846
P94024 P94131 P94139 P94177 P94178 P94185 P94199 P94250 P94287 P94304 P94311
P94312 P94336 P94346 P94353 P94357 P94362 P94363 P94364 P94365 P94366 P94367
P94369 P94371 P94372 P94373 P94375 P94376 P94381 P94383 P94392 P94394 P94395
P94408 P94412 P94414 P94418 P94419 P94422 P94431 P94434 P94438 P94441 P94442
P94456 P94469 P94476 P94485 P94488 P94493 P94497 P94499 P94507 P94513 P94515
P94516 P94519 P94520 P94529 P94530 P94532 P94543 P94546 P94571 P94572 P94574
P94575 P94577 P94585 P94606 P94608 P94633 P94665 P94700 P94774 P94851 P94853
P94854 P95160 P95200 P95357 P95374 P95375 P95503 P95504 P95505 P95522 P95549
P95557 P95577 P95654 P95655 P95656 P95673 P95674 P95695 P95730 P95745 P95783
P95784 P95785 P95822 P95827 P96000 P96064 P96065 P96068 P96069 P96118 P96119
P96169 P96335 P96368 P96419 P96454 P96499 P96517 P96554 P96581 P96585 P96587
P96589 P96593 P96594 P96601 P96603 P96604 P96606 P96607 P96610 P96615 P96616
P96617 P96619 P96634 P96640 P96641 P96644 P96646 P96650 P96656 P96661 P96664
P96665 P96668 P96672 P96675 P96678 P96680 P96685 P96687 P96689 P96694 P96696
P96697 P96702 P96704 P96706 P96709 P96710 P96712 P96715 P96787 P96790 P96792
P96796 P96949 P96989 P97029 P97041 P97046 P97054 P97055 P97207 P97215 P97224
P97253 P97259 P97260 P97266 P97267 P97281 P97287 P97288 P97291 P97292 P97295
P97300 P97321 P97325 P97326 P97333 P97353 P97363 P97370 P97378 P97402 P97414
P97435 P97438 P97441 P97445 P97449 P97464 P97467 P97468 P97484 P97490 P97501
P97520 P97521 P97523 P97524 P97525 P97544 P97546 P97557 P97564 P97570 P97583
P97603 P97607 P97610 P97612 P97629 P97639 P97675 P97677 P97678 P97685 P97686
P97687 P97689 P97698 P97700 P97707 P97708 P97710 P97714 P97717 P97718 P97739
P97742 P97751 P97772 P97776 P97792 P97793 P97794 P97797 P97798 P97799 P97808
P97819 P97829 P97846 P97849 P97858 P97872 P97881 P97887 P97926 P97943 P97952
P97998 P98000 P98001 P98002 P98003 P98004 P98005 P98008 P98009 P98012 P98019
P98020 P98021 P98023 P98024 P98025 P98027 P98030 P98031 P98032 P98033 P98034
P98035 P98036 P98037 P98038 P98039 P98042 P98043 P98044 P98046 P98047 P98048
P98049 P98052 P98053 P98054 P98055 P98056 P98057 P98059 P98072 P98073 P98074
P98105 P98106 P98107 P98109 P98110 P98131 P98135 P98136 P98153 P98154 P98155

P98156 P98157 P98158 P98161 P98162 P98163 P98164 P98165 P98166 P98172 P98183
P98187 P98188 P98191 P98194 P98195 P98196 P98197 P98198 P98199 P98200 P98204
P98205 P98999 P9WEP1 P9WEP4 P9WEP5 P9WEQ1 P9WEQ3 P9WEQ4 P9WEQ6 P9WEQ7 P9WEQ9
P9WER2 P9WER4 P9WES3 P9WEU1 P9WEU3 P9WEU4 P9WEV3 P9WEV5 P9WEV9 P9WEW7 P9WEX1
P9WEX2 P9WEX4 P9WEX7 P9WEX8 P9WEY0 P9WEY1 P9WEZ1 P9WEZ6 P9WFF8 P9WFF9 P9WFG0
P9WFG1 P9WFG2 P9WFG3 P9WFG4 P9WFG5 P9WFJ4 P9WFJ5 P9WFJ6 P9WFJ7 P9WFL2 P9WFL3
P9WFL4 P9WFL5 P9WFM2 P9WFM3 P9WFM4 P9WFM5 P9WFN6 P9WFN7 P9WFN8 P9WFN9 P9WFP0
P9WFP1 P9WFP2 P9WFP3 P9WFR4 P9WFR5 P9WFR6 P9WFR7 P9WFS2 P9WFS3 P9WFS4 P9WFS5
P9WFU6 P9WFU7 P9WFZ6 P9WFZ7 P9WFZ8 P9WFZ9 P9WG00 P9WG01 P9WG02 P9WG03 P9WG04
P9WG05 P9WG06 P9WG07 P9WG08 P9WG09 P9WG10 P9WG11 P9WG12 P9WG13 P9WG14 P9WG15
P9WG16 P9WG17 P9WG18 P9WG19 P9WG20 P9WG21 P9WG22 P9WG23 P9WG26 P9WG27 P9WG50
P9WG51 P9WG62 P9WG63 P9WG84 P9WG85 P9WG86 P9WG87 P9WG88 P9WG89 P9WG90 P9WG91
P9WG92 P9WG93 P9WG96 P9WG97 P9WG98 P9WG99 P9WGA0 P9WGA1 P9WGA6 P9WGA7 P9WGE4
P9WGE5 P9WGF0 P9WGF1 P9WGF4 P9WGF5 P9WGF6 P9WGF7 P9WGJ1 P9WGJ3 P9WGK4 P9WGK5
P9WGK6 P9WGK7 P9WGK8 P9WGK9 P9WGL0 P9WGL1 P9WGL2 P9WGL3 P9WGN2 P9WGN3 P9WGN4
P9WGN5 P9WGN6 P9WGN7 P9WGN8 P9WGN9 P9WGP0 P9WGP1 P9WGU4 P9WGU5 P9WGX4 P9WGX5
P9WH22 P9WH23 P9WHR0 P9WHR1 P9WHR8 P9WHR9 P9WHS2 P9WHS3 P9WHS4 P9WHS5 P9WHS6
P9WHS7 P9WHW4 P9WHW5 P9WHX0 P9WHX1 P9WHZ0 P9WHZ1 P9WI20 P9WI21 P9WI34 P9WI35
P9WI42 P9WI43 P9WI48 P9WI49 P9WI52 P9WI53 P9WI62 P9WI63 P9WI66 P9WI67 P9WI68
P9WI69 P9WI70 P9WI71 P9WI74 P9WI75 P9WI76 P9WI77 P9WI78 P9WI79 P9WI80 P9WI81
P9WI82 P9WI83 P9WIA4 P9WIA5 P9WIA6 P9WIA7 P9WIF2 P9WIF3 P9WIG8 P9WIG9 P9WIM6
P9WIM7 P9WIP2 P9WIP3 P9WIQ6 P9WIQ7 P9WIR8 P9WIR9 P9WIT4 P9WIT5 P9WIU4 P9WIU5
P9WIW0 P9WIW1 P9WIW2 P9WIW3 P9WIW4 P9WIW5 P9WIW6 P9WIW7 P9WIW8 P9WIW9 P9WIX0
P9WIX1 P9WIX2 P9WIX3 P9WIZ4 P9WIZ5 P9WJ64 P9WJ65 P9WJ66 P9WJ67 P9WJ70 P9WJ71
P9WJ82 P9WJ83 P9WJ98 P9WJ99 P9WJB0 P9WJB1 P9WJB2 P9WJB3 P9WJB4 P9WJB5 P9WJD2
P9WJD3 P9WJE2 P9WJE3 P9WJE4 P9WJE5 P9WJE6 P9WJE7 P9WJE8 P9WJE9 P9WJF2 P9WJF3
P9WJG4 P9WJG5 P9WJG6 P9WJG7 P9WJI2 P9WJI3 P9WJK2 P9WJK3 P9WJN4 P9WJN5 P9WJQ0
P9WJQ1 P9WJS6 P9WJS7 P9WJS8 P9WJS9 P9WJT0 P9WJT1 P9WJT2 P9WJT3 P9WJT4 P9WJT5
P9WJT6 P9WJT7 P9WJT8 P9WJT9 P9WJU0 P9WJU1 P9WJU2 P9WJU3 P9WJU4 P9WJU5 P9WJU6
P9WJU7 P9WJU8 P9WJU9 P9WJV0 P9WJV1 P9WJV2 P9WJV3 P9WJV4 P9WJV5 P9WJV6 P9WJV7
P9WJV8 P9WJV9 P9WJW6 P9WJW7 P9WJW8 P9WJW9 P9WJX0 P9WJX1 P9WJX2 P9WJX3 P9WJX4
P9WJX5 P9WJX6 P9WJX7 P9WJX8 P9WJX9 P9WJY0 P9WJY1 P9WJY2 P9WJY3 P9WJY4 P9WJY5
P9WJY6 P9WJY7 P9WK30 P9WK31 P9WK32 P9WK33 P9WK42 P9WK43 P9WK50 P9WK51 P9WK92
P9WK93 P9WK98 P9WK99 P9WKA0 P9WKA1 P9WKD0 P9WKD1 P9WKF0 P9WKF1 P9WKF2 P9WKF3
P9WKL6 P9WKL7 P9WKM8 P9WKM9 P9WKN2 P9WKN3 P9WKP6 P9WKP7 P9WKQ0 P9WKQ1 P9WKQ4
P9WKQ5 P9WKQ8 P9WKQ9 P9WKR0 P9WKR1 P9WKR4 P9WKR5 P9WKR8 P9WKR9 P9WKS6 P9WKS7
P9WKU2 P9WKU3 P9WKV6 P9WKV7 P9WKW0 P9WKW1 P9WKW4 P9WKW5 P9WKX8 P9WKX9 P9WKZ4
P9WKZ5 P9WKZ8 P9WKZ9 P9WL38 P9WL39 P9WL68 P9WL69 P9WL72 P9WL73 P9WL74 P9WL75
P9WL78 P9WL79 P9WL82 P9WL83 P9WL84 P9WL85 P9WL88 P9WL89 P9WLA0 P9WLA1 P9WLC6
P9WLC7 P9WLE0 P9WLE1 P9WLF2 P9WLF3 P9WLF4 P9WLF5 P9WLG2 P9WLG3 P9WLG6 P9WLG7
P9WLH0 P9WLH1 P9WLI0 P9WLI1 P9WLI2 P9WLI3 P9WLI4 P9WLI5 P9WLI6 P9WLI7 P9WLI8
P9WLI9 P9WLJ4 P9WLJ5 P9WLK2 P9WLK3 P9WLK4 P9WLK5 P9WLL0 P9WLL1 P9WLL2 P9WLL3
P9WLL4 P9WLL5 P9WLQ6 P9WLQ7 P9WLQ8 P9WLQ9 P9WLR6 P9WLR7 P9WLS4 P9WLS5 P9WLS8
P9WLS9 P9WLT4 P9WLT5 P9WLV8 P9WLV9 P9WLW0 P9WLW1 P9WLX0 P9WLX1 P9WLX8 P9WLX9

P9WLY0 P9WLY1 P9WLZ8 P9WLZ9 P9WM00 P9WM01 P9WM06 P9WM07 P9WM18 P9WM19 P9WM20
P9WM21 P9WM22 P9WM23 P9WM28 P9WM29 P9WM30 P9WM31 P9WM34 P9WM35 P9WM60 P9WM61
P9WM62 P9WM63 P9WM68 P9WM69 P9WM70 P9WM71 P9WM74 P9WM75 P9WM86 P9WM87 P9WM88
P9WM89 P9WMA2 P9WMA3 P9WMA6 P9WMA7 P9WMC8 P9WMC9 P9WMN6 P9WMN7 P9WMS8 P9WMS9
P9WMW4 P9WMW5 P9WMW6 P9WMW7 P9WMX0 P9WMX1 P9WMX4 P9WMX5 P9WMZ6 P9WMZ7 P9WMZ8
P9WMZ9 P9WN00 P9WN01 P9WN02 P9WN03 P9WN04 P9WN05 P9WN96 P9WN97 P9WN98 P9WN99
P9WNA0 P9WNA1 P9WNA2 P9WNA3 P9WNA4 P9WNA5 P9WNA6 P9WNA7 P9WNA8 P9WNA9 P9WNB2
P9WNB3 P9WNB4 P9WNB5 P9WNB6 P9WNB7 P9WNK6 P9WNK7 P9WNL4 P9WNL5 P9WNL6 P9WNL7
P9WNL8 P9WNL9 P9WNP8 P9WNP9 P9WNQ0 P9WNQ1 P9WNQ2 P9WNQ3 P9WNQ4 P9WNQ5 P9WNQ6
P9WNQ7 P9WNQ8 P9WNQ9 P9WNR0 P9WNR1 P9WNR2 P9WNR3 P9WNR4 P9WNR5 P9WNR6 P9WNR7
P9WNZ8 P9WNZ9 P9WP06 P9WP07 P9WP08 P9WP09 P9WP34 P9WP35 P9WP36 P9WP37 P9WP44
P9WP45 P9WP46 P9WP47 P9WP56 P9WP57 P9WP60 P9WP61 P9WP62 P9WP63 P9WP66 P9WP67
P9WP68 P9WP69 P9WP70 P9WP71 P9WP90 P9WP91 P9WP92 P9WP93 P9WPD6 P9WPD7 P9WPD8
P9WPD9 P9WPF6 P9WPF7 P9WPG0 P9WPG1 P9WPG2 P9WPG3 P9WPG4 P9WPG5 P9WPG6 P9WPG7
P9WPG8 P9WPG9 P9WPI6 P9WPI7 P9WPL6 P9WPL7 P9WPN6 P9WPN7 P9WPR6 P9WPR7 P9WPR8
P9WPR9 P9WPS0 P9WPS1 P9WPS2 P9WPS3 P9WPS4 P9WPS5 P9WPS6 P9WPS7 P9WPS8 P9WPS9
P9WPT0 P9WPT1 P9WPT2 P9WPT3 P9WPT4 P9WPT5 P9WPT6 P9WPT7 P9WPT8 P9WPT9 P9WPU0
P9WPU1 P9WPU2 P9WPU3 P9WPV2 P9WPV3 P9WPV4 P9WPV5 P9WPV6 P9WPV7 P9WQ28 P9WQ29
P9WQ30 P9WQ31 P9WQ32 P9WQ33 P9WQ34 P9WQ35 P9WQ44 P9WQ45 P9WQ46 P9WQ47 P9WQ52
P9WQ53 P9WQ64 P9WQ65 P9WQI6 P9WQI7 P9WQI8 P9WQI9 P9WQJ0 P9WQJ1 P9WQJ2 P9WQJ3
P9WQJ6 P9WQJ7 P9WQJ8 P9WQJ9 P9WQM2 P9WQM3 P9WQM4 P9WQM5 P9WQM6 P9WQM7 P9WQM8
P9WQM9 P9WQN2 P9WQN3 Q00045 Q00046 Q00058 Q00087 Q000K2 Q00102 Q00104 Q00133
Q00134 Q00135 Q00136 Q00137 Q00138 Q00141 Q00184 Q00185 Q00193 Q00194 Q00195
Q00224 Q00227 Q00228 Q00229 Q00230 Q00231 Q00232 Q00238 Q00242 Q00243 Q00244
Q00262 Q00269 Q00275 Q00276 Q00310 Q00319 Q00320 Q00325 Q00327 Q00330 Q00336
Q00342 Q00357 Q00383 Q00449 Q00471 Q00474 Q00478 Q00495 Q00496 Q00500 Q00502
Q00505 Q00506 Q00513 Q00514 Q00515 Q00516 Q00517 Q00518 Q00521 Q00522 Q00527
Q00528 Q00529 Q00530 Q00538 Q00540 Q00541 Q00542 Q00543 Q00552 Q00553 Q00554
Q00555 Q00560 Q00564 Q00568 Q00570 Q00576 Q00583 Q00589 Q00590 Q00595 Q00607
Q00608 Q00609 Q00619 Q00650 Q00651 Q00657 Q00660 Q00690 Q00702 Q00707 Q00709
Q00710 Q00716 Q00718 Q00748 Q00750 Q00751 Q00756 Q00757 Q00758 Q00765 Q00779
Q00788 Q00799 Q00804 Q00822 Q00823 Q00824 Q00825 Q00827 Q00842 Q00848 Q00860
Q008X3 Q008X4 Q008X5 Q008X6 Q00910 Q00912 Q00941 Q00951 Q00954 Q00959 Q00960
Q00961 Q00968 Q00973 Q00974 Q00975 Q00977 Q00986 Q00988 Q00991 Q00993 Q00GF0
Q00GQ9 Q00HT4 Q00M95 Q00PJ0 Q00PJ2 Q00PJ8 Q00U99 Q01016 Q01017 Q01035 Q01038
Q01045 Q01049 Q01083 Q01097 Q01098 Q01102 Q01107 Q01113 Q01114 Q01118 Q01148
Q01151 Q01218 Q01227 Q01231 Q01237 Q01245 Q01251 Q01255 Q01260 Q01266 Q01279
Q01285 Q01290 Q01299 Q01300 Q01319 Q01321 Q01337 Q01338 Q01341 Q01344 Q01345
Q01355 Q01356 Q01362 Q01366 Q01403 Q01404 Q01440 Q01441 Q01443 Q01447 Q01453
Q01455 Q01467 Q01481 Q01482 Q01497 Q01499 Q01501 Q01554 Q01555 Q01556 Q01559
Q01561 Q01562 Q01563 Q01564 Q01565 Q01580 Q01590 Q01602 Q01605 Q01606 Q01608
Q01625 Q01628 Q01629 Q01638 Q01650 Q01667 Q01668 Q01674 Q01685 Q01705 Q01717
Q01718 Q01725 Q01726 Q01727 Q01728 Q01740 Q01755 Q01776 Q01812 Q01814 Q01815
Q01818 Q01827 Q01854 Q01857 Q01887 Q01888 Q01895 Q01896 Q01926 Q01931 Q01956

Q01959 Q01961 Q01964 Q01965 Q01969 Q01973 Q01974 Q01977 Q01996 Q01H83 Q01H84
Q01IH3 Q01IU9 Q01JR9 Q01MI9 Q01MW8 Q01Q45 Q01QP2 Q01SK1 Q01UN2 Q01UN3 Q01UN6
Q01VL4 Q01WE6 Q01YC2 Q02004 Q02006 Q02012 Q02013 Q02060 Q02092 Q02094 Q020J1
Q020Q4 Q02115 Q02122 Q02127 Q02152 Q02161 Q02203 Q02211 Q02212 Q02217 Q02221
Q02223 Q02226 Q02242 Q02280 Q02282 Q02284 Q02294 Q02297 Q02317 Q02331 Q02334
Q02335 Q02342 Q02343 Q02353 Q02365 Q02367 Q02371 Q02372 Q02376 Q02377 Q02378
Q02380 Q02385 Q02391 Q02401 Q02409 Q02413 Q02432 Q02443 Q02465 Q02478 Q02482
Q02485 Q02487 Q02496 Q02500 Q02505 Q02527 Q02541 Q02563 Q02581 Q02585 Q02592
Q02643 Q02644 Q02653 Q02654 Q02655 Q02725 Q02729 Q02730 Q02731 Q02734 Q02738
Q02739 Q02742 Q02745 Q02755 Q02760 Q02761 Q02762 Q02763 Q02766 Q02768 Q02769
Q02774 Q02776 Q02783 Q02785 Q02789 Q02795 Q02820 Q02827 Q02837 Q02846 Q02847
Q02850 Q02851 Q02852 Q02854 Q02858 Q02883 Q02888 Q02889 Q02896 Q02920 Q02921
Q02929 Q02938 Q02955 Q02969 Q02978 Q02989 Q02B01 Q02BU5 Q02CT1 Q02CT4 Q02CT6
Q02CU2 Q02CX5 Q02CX7 Q02DE0 Q02DE8 Q02DE9 Q02DF0 Q02DM7 Q02DS7 Q02DW8 Q02E04
Q02EA7 Q02EU0 Q02EU8 Q02EU9 Q02EV2 Q02EZ6 Q02FY2 Q02G63 Q02GA3 Q02GB8 Q02H25
Q02H57 Q02I05 Q02I77 Q02IB2 Q02J03 Q02JC2 Q02JG2 Q02JW1 Q02K11 Q02KC0 Q02KW3
Q02L30 Q02LA5 Q02LE0 Q02LI9 Q02MI4 Q02NC9 Q02ND5 Q02ND8 Q02NZ3 Q02P72 Q02PF8
Q02PF9 Q02PG0 Q02PG2 Q02Q08 Q02QA7 Q02QX8 Q02QY1 Q02QY2 Q02QY3 Q02R24 Q02R27
Q02R28 Q02R29 Q02SF2 Q02SS7 Q02TI2 Q02TM7 Q02U74 Q02UV9 Q02UW5 Q02VT8 Q02VY0
Q02W23 Q02WP0 Q02XA1 Q02YQ2 Q02ZK6 Q02ZM1 Q02ZY6 Q03001 Q03024 Q03025 Q03027
Q03028 Q03031 Q03032 Q03035 Q03041 Q03064 Q03069 Q03073 Q03075 Q03079 Q03100
Q03101 Q03102 Q03105 Q03137 Q03142 Q03145 Q03146 Q03152 Q03155 Q03157 Q03163
Q03167 Q03185 Q03186 Q03187 Q03190 Q03193 Q03194 Q031B1 Q031B2 Q031Z8 Q03203
Q03212 Q03218 Q03227 Q03228 Q03236 Q03263 Q03268 Q03281 Q03289 Q03307 Q03322
Q03327 Q03343 Q03345 Q03348 Q03351 Q03364 Q03367 Q03370 Q03391 Q03395 Q03411
Q03412 Q03418 Q03419 Q03431 Q03438 Q03439 Q03440 Q03445 Q03446 Q03455 Q03463
Q03476 Q03477 Q03478 Q03480 Q03482 Q03499 Q034S0 Q03516 Q03518 Q03519 Q03521
Q03529 Q03530 Q03554 Q03566 Q03567 Q03574 Q03579 Q03583 Q03584 Q035B4 Q035P5
Q035X6 Q03600 Q03611 Q03613 Q03614 Q03640 Q03648 Q03650 Q03669 Q03671 Q03672
Q03673 Q03687 Q03691 Q03696 Q03697 Q03707 Q03713 Q03714 Q03717 Q03719 Q03720
Q03721 Q03723 Q03727 Q03730 Q03736 Q03755 Q03763 Q03770 Q03783 Q03788 Q03795
Q03798 Q037U3 Q03804 Q03816 Q03817 Q03824 Q03829 Q03845 Q03860 Q03880 Q03884
Q03885 Q03904 Q03909 Q03913 Q03944 Q03945 Q03965 Q039D7 Q039H8 Q039R9 Q03A22
Q03A23 Q03AP9 Q03AR6 Q03BE8 Q03D26 Q03D58 Q03D91 Q03DK0 Q03DY7 Q03E93 Q03EK8
Q03EK9 Q03EL0 Q03EQ6 Q03EY0 Q03EY3 Q03F77 Q03FU1 Q03GY4 Q03J12 Q03LB2 Q03LE6
Q03LM3 Q03LU4 Q03LX7 Q03M43 Q03MC3 Q03MC5 Q03MN6 Q03NQ6 Q03PY7 Q03QH3 Q03QR4
Q03QY3 Q03QY4 Q03R31 Q03S07 Q03S76 Q03SF6 Q03T13 Q03TH5 Q03US7 Q03V25 Q03VG1
Q03W33 Q03WN0 Q03XG1 Q03YG9 Q03YI6 Q03YJ0 Q03Z46 Q03ZL4 Q03ZT0 Q03ZT9 Q04013
Q04018 Q04050 Q04064 Q04069 Q04080 Q04082 Q04083 Q04100 Q04119 Q04121 Q04162
Q04164 Q04172 Q04174 Q04182 Q041C3 Q041L7 Q04203 Q04210 Q04216 Q04228 Q04261
Q042C3 Q042D3 Q042L1 Q042P7 Q04301 Q04302 Q04338 Q04350 Q04370 Q04396 Q04398
Q043I1 Q04400 Q04440 Q04441 Q04442 Q04443 Q04444 Q04452 Q04454 Q04458 Q04463
Q04464 Q04468 Q04472 Q04473 Q04487 Q044D6 Q04501 Q04507 Q04508 Q04516 Q04538
Q04547 Q04548 Q04561 Q04562 Q04565 Q04566 Q04567 Q04568 Q04569 Q04573 Q04582

Q04589 Q04592 Q04594 Q04602 Q04604 Q04609 Q04612 Q04613 Q04614 Q04629 Q04630
Q04645 Q04646 Q04651 Q04654 Q04656 Q04658 Q04662 Q04664 Q04671 Q04679 Q04680
Q04683 Q04684 Q04685 Q04686 Q04687 Q04697 Q046Q7 Q04703 Q04705 Q04721 Q04733
Q04734 Q04746 Q04767 Q04771 Q04785 Q04790 Q04799 Q04800 Q04804 Q04833 Q04835
Q04838 Q04844 Q04850 Q04854 Q04866 Q04874 Q04875 Q04876 Q04877 Q04878 Q04879
Q04880 Q04881 Q04882 Q04883 Q04884 Q04885 Q04895 Q048Q1 Q04900 Q04909 Q04910
Q04911 Q04912 Q04935 Q04941 Q04943 Q04947 Q04952 Q04954 Q04956 Q04958 Q04959
Q04969 Q04976 Q04991 Q04993 Q04995 Q04B74 Q04BA7 Q04BI8 Q04BJ7 Q04BL0 Q04CF3
Q04D15 Q04ES8 Q04ET1 Q04FK1 Q04FK3 Q04G24 Q04G25 Q04HT3 Q04HT4 Q04HT5 Q04IN8
Q04IS6 Q04J36 Q04J43 Q04JT8 Q04K41 Q04L63 Q04L94 Q04LE4 Q04M08 Q04M23 Q04MC4
Q04NG2 Q04PG4 Q04Q03 Q04QT4 Q04S14 Q04SA1 Q04TV3 Q04UZ1 Q04V64 Q04V66 Q04V92
Q04W30 Q04WP4 Q04XE2 Q04Y84 Q04YB0 Q04YB2 Q04YI7 Q04ZS2 Q04ZU1 Q05001 Q05005
Q05017 Q05030 Q05032 Q05033 Q05034 Q05037 Q05047 Q05052 Q05057 Q05059 Q05070
Q05078 Q05085 Q05098 Q050C7 Q05131 Q05143 Q05146 Q05152 Q05181 Q05199 Q05202
Q05204 Q05207 Q05217 Q05296 Q05312 Q05319 Q05320 Q05347 Q05359 Q05360 Q05364
Q05365 Q05366 Q05367 Q05376 Q05389 Q05394 Q053U5 Q05438 Q05490 Q05496 Q05497
Q05503 Q05511 Q05521 Q05567 Q05572 Q05577 Q05585 Q05586 Q05594 Q05595 Q05598
Q055W1 Q05600 Q05602 Q05605 Q05606 Q05611 Q05612 Q05613 Q05637 Q05648 Q05659
Q05688 Q05695 Q05740 Q05754 Q05787 Q05794 Q057F2 Q057W3 Q057W6 Q05807 Q05811
Q05867 Q05888 Q05889 Q05898 Q05899 Q058N4 Q058P6 Q05901 Q05902 Q05909 Q05910
Q05928 Q05934 Q05938 Q05939 Q05940 Q05941 Q05962 Q05973 Q05996 Q05998 Q059Y8
Q05A56 Q05AK6 Q05AM4 Q05AQ3 Q05AU4 Q05B21 Q05B45 Q05B54 Q05B67 Q05B71 Q05B73
Q05B81 Q05BQ1 Q05EQ4 Q05F77 Q05F85 Q05JG2 Q06001 Q06003 Q06011 Q06019 Q06034
Q06051 Q06055 Q06056 Q06057 Q06058 Q06059 Q06067 Q06089 Q06094 Q06104 Q06107
Q06114 Q06116 Q06119 Q06127 Q06136 Q06143 Q06144 Q06148 Q06154 Q06158 Q06159
Q06169 Q06170 Q06175 Q06186 Q06200 Q06220 Q06221 Q06222 Q06235 Q06236 Q06240
Q06247 Q06274 Q06303 Q06304 Q06305 Q06317 Q06325 Q06328 Q06335 Q06346 Q06379
Q06381 Q06393 Q06397 Q06400 Q06417 Q06418 Q06432 Q06438 Q06451 Q06466 Q06473
Q06474 Q06475 Q06480 Q06481 Q06485 Q06493 Q06495 Q06496 Q06497 Q06502 Q06508
Q06536 Q06537 Q06538 Q06541 Q06551 Q06563 Q06564 Q06567 Q06572 Q06593 Q06598
Q06599 Q06611 Q06636 Q06643 Q06644 Q06645 Q06646 Q06668 Q06676 Q06681 Q06686
Q06689 Q06693 Q06694 Q06799 Q06805 Q06806 Q06807 Q06810 Q06815 Q06833 Q06838
Q06885 Q06910 Q06916 Q06922 Q06987 Q06991 Q06AA5 Q06AV4 Q06BH3 Q06BI3 Q06BR2
Q06DU1 Q06DU9 Q06FK6 Q06FK7 Q06FK8 Q06FK9 Q06FL0 Q06FL1 Q06FL2 Q06FL3 Q06FL4
Q06FL5 Q06FL6 Q06FL7 Q06FN7 Q06FN8 Q06FN9 Q06FP0 Q06FP1 Q06FP2 Q06FP6 Q06FP9
Q06FQ1 Q06FQ2 Q06FQ4 Q06FQ5 Q06FV5 Q06FV9 Q06FW0 Q06FW1 Q06FW4 Q06FW5 Q06FW7
Q06FW8 Q06FX3 Q06FX4 Q06FX5 Q06FX7 Q06FX8 Q06FY1 Q06GK1 Q06GK2 Q06GK4 Q06GK7
Q06GK8 Q06GL0 Q06GM9 Q06GN0 Q06GN1 Q06GN2 Q06GN3 Q06GN4 Q06GP0 Q06GP1 Q06GP2
Q06GP3 Q06GP4 Q06GP5 Q06GP6 Q06GP7 Q06GP8 Q06GP9 Q06GQ0 Q06GQ5 Q06GR0 Q06GR1
Q06GR3 Q06GR4 Q06GR5 Q06GR6 Q06GR7 Q06GS2 Q06GS3 Q06GS4 Q06GS6 Q06GS7 Q06GT0
Q06GT8 Q06GU1 Q06GU3 Q06GU4 Q06GU6 Q06GU7 Q06GU9 Q06GW6 Q06GW7 Q06GW8 Q06GW9
Q06GX0 Q06GX1 Q06GX7 Q06GX8 Q06GX9 Q06GY0 Q06GY1 Q06GY2 Q06GY3 Q06GY4 Q06GY5
Q06GY6 Q06GY7 Q06GZ2 Q06GZ7 Q06GZ8 Q06H00 Q06H01 Q06H02 Q06H03 Q06H04 Q06H09
Q06H10 Q06H11 Q06H13 Q06H14 Q06H17 Q06HQ7 Q06J11 Q06J12 Q06J13 Q06J14 Q06J16

Q06J21 Q06J22 Q06J23 Q06J24 Q06J30 Q06J31 Q06J40 Q06J41 Q06J42 Q06J43 Q06J44
Q06J45 Q06J46 Q06J52 Q06J63 Q06J64 Q06J66 Q06J67 Q06J69 Q06J70 Q06J71 Q06R72
Q06R75 Q06R77 Q06R78 Q06R80 Q06R81 Q06R83 Q06RA1 Q06RA2 Q06RA3 Q06RA4 Q06RA5
Q06RA6 Q06RB2 Q06RB3 Q06RB4 Q06RB5 Q06RB6 Q06RB7 Q06RB8 Q06RB9 Q06RC0 Q06RC4
Q06RC9 Q06RD0 Q06RD1 Q06RD2 Q06RD4 Q06RD5 Q06RD6 Q06RD7 Q06RD8 Q06RE3 Q06RE4
Q06RE5 Q06RE7 Q06RE8 Q06RF1 Q06SC5 Q06SC6 Q06SC8 Q06SD2 Q06SD3 Q06SD4 Q06SD5
Q06SD7 Q06SE0 Q06SE1 Q06SE2 Q06SE6 Q06SF0 Q06SF1 Q06SF8 Q06SG1 Q06SG3 Q06SG8
Q06SH1 Q06SH2 Q06SH5 Q06SI0 Q06SI1 Q06SI7 Q06SI9 Q06SJ0 Q06SJ6 Q06SK2 Q06VD4
Q07001 Q07005 Q07008 Q07011 Q07060 Q07065 Q07066 Q07074 Q07075 Q07108 Q07113
Q07175 Q07185 Q07249 Q07252 Q07259 Q07263 Q07282 Q07287 Q072K5 Q07303 Q07307
Q07310 Q07326 Q07335 Q07349 Q07355 Q07376 Q07383 Q07404 Q07407 Q07421 Q07423
Q07429 Q07430 Q07434 Q07435 Q07441 Q07444 Q07451 Q07473 Q07494 Q07496 Q07497
Q07498 Q074N0 Q07509 Q07519 Q07520 Q07534 Q07537 Q07541 Q07549 Q07553 Q07560
Q07563 Q07564 Q07565 Q07570 Q07571 Q07572 Q07573 Q07574 Q07575 Q07576 Q07577
Q07578 Q07579 Q07580 Q07581 Q07582 Q07583 Q07584 Q07585 Q07591 Q07599 Q07609
Q07629 Q07631 Q07633 Q07646 Q07647 Q07651 Q07652 Q07660 Q07699 Q07730 Q07737
Q07757 Q07763 Q07764 Q07782 Q07788 Q07790 Q07798 Q07806 Q07811 Q07812 Q07813
Q07816 Q07817 Q07820 Q07824 Q07828 Q07829 Q07830 Q07835 Q07837 Q07865 Q07867
Q07868 Q07881 Q07891 Q07895 Q07904 Q07914 Q07920 Q07925 Q07926 Q07950 Q07954
Q07959 Q07969 Q07977 Q07984 Q07986 Q07994 Q07DV2 Q07DV5 Q07DV8 Q07DW5 Q07DW9
Q07DX5 Q07DX8 Q07DY1 Q07DY5 Q07DY8 Q07DZ1 Q07DZ6 Q07E01 Q07E08 Q07E16 Q07E24
Q07E37 Q07E42 Q07E45 Q07E48 Q07FI2 Q07FI4 Q07FI5 Q07FZ2 Q07FZ4 Q07H86 Q07H88
Q07H89 Q07HB0 Q07HB2 Q07HN8 Q07MP8 Q07N50 Q07N51 Q07NL6 Q07PQ2 Q07PT6 Q07Q28
Q07Q80 Q07QW3 Q07QW6 Q07QW9 Q07QX5 Q07QX6 Q07RP9 Q07T77 Q07T82 Q07UC1 Q07V52
Q07VS6 Q07VT8 Q07VU0 Q07VY9 Q07WI2 Q07WY2 Q07X68 Q07XN2 Q07YT7 Q07YY4 Q07ZL7
Q08001 Q08011 Q08013 Q08017 Q08022 Q08023 Q08027 Q08076 Q08084 Q08089 Q08098
Q080K8 Q080T2 Q08100 Q08101 Q08102 Q08103 Q08109 Q08120 Q08144 Q08156 Q08157
Q08174 Q08176 Q08179 Q08180 Q081P2 Q081P4 Q08201 Q08205 Q08207 Q08210 Q08215
Q08218 Q08219 Q08223 Q08232 Q08234 Q08238 Q08240 Q08268 Q08269 Q08272 Q08280
Q08299 Q08315 Q08322 Q08334 Q08336 Q08338 Q08339 Q08340 Q08345 Q08351 Q08357
Q08362 Q08363 Q08373 Q08382 Q08388 Q083B7 Q083H3 Q083H4 Q083Z6 Q08408 Q08409
Q08410 Q08416 Q08417 Q08428 Q08430 Q08435 Q08436 Q08448 Q08451 Q08460 Q08462
Q08463 Q08464 Q08469 Q08477 Q08481 Q08485 Q08498 Q084P1 Q08501 Q08520 Q08533
Q08537 Q08542 Q08548 Q08552 Q08554 Q08559 Q08560 Q08579 Q08584 Q08585 Q08586
Q08587 Q08604 Q08630 Q08633 Q08650 Q08651 Q08656 Q08676 Q08684 Q08693 Q086A9
Q086L0 Q086T5 Q08708 Q08717 Q08722 Q08733 Q08734 Q08736 Q08741 Q08743 Q08749
Q08750 Q08755 Q08760 Q08777 Q08786 Q08791 Q08793 Q08794 Q08811 Q08828 Q08832
Q08834 Q08844 Q08849 Q08850 Q08851 Q08853 Q08855 Q08857 Q088I0 Q088I1 Q088I2
Q088X1 Q08900 Q08902 Q08905 Q08908 Q08913 Q08915 Q08919 Q08922 Q08927 Q08929
Q08931 Q08955 Q08958 Q08959 Q08967 Q08970 Q08974 Q08980 Q08986 Q08991 Q089F2
Q089I2 Q089M7 Q08AI6 Q08AU7 Q08B22 Q08B29 Q08B84 Q08BA4 Q08BG1 Q08BI9 Q08BL3
Q08BY2 Q08C60 Q08C75 Q08C93 Q08CB3 Q08CD5 Q08CE6 Q08CG9 Q08CI8 Q08CS6 Q08CW8
Q08D11 Q08D19 Q08D64 Q08D68 Q08D83 Q08D99 Q08DA1 Q08DA4 Q08DB5 Q08DE1 Q08DE2
Q08DE5 Q08DE6 Q08DF2 Q08DG6 Q08DJ5 Q08DK0 Q08DK1 Q08DK4 Q08DK7 Q08DL4 Q08DM5

Q08DM6 Q08DP3 Q08DV9 Q08DW9 Q08DX7 Q08DZ5 Q08E08 Q08E15 Q08E24 Q08E25 Q08E36
Q08E40 Q08E50 Q08EA8 Q08ET2 Q08FX8 Q08GM0 Q08GU6 Q08GV9 Q08GY5 Q08GZ8 Q08H50
Q08H76 Q08HF4 Q08HG7 Q08HI0 Q08HJ3 Q08HK6 Q08HL9 Q08IG9 Q08IT1 Q08JR8 Q09013
Q09037 Q09039 Q09049 Q09056 Q09062 Q09068 Q09073 Q09081 Q09092 Q09104 Q09105
Q09106 Q09108 Q09118 Q09120 Q09143 Q09147 Q09154 Q09163 Q09174 Q09175 Q09188
Q09195 Q09199 Q09200 Q09201 Q09202 Q09203 Q09204 Q09205 Q09206 Q09208 Q09209
Q09210 Q09212 Q09213 Q09216 Q09218 Q09223 Q09225 Q09226 Q09231 Q09232 Q09251
Q09271 Q09272 Q09273 Q09274 Q09276 Q09285 Q09303 Q09304 Q09308 Q09320 Q09322
Q09323 Q09324 Q09325 Q09326 Q09327 Q09328 Q09333 Q09334 Q09341 Q09343 Q09344
Q09351 Q09363 Q09366 Q09368 Q09369 Q09371 Q09372 Q09388 Q09417 Q09422 Q09426
Q09427 Q09428 Q09429 Q09435 Q09453 Q09460 Q09461 Q09466 Q09470 Q09473 Q09488
Q09499 Q09500 Q09502 Q09554 Q09562 Q09572 Q09573 Q09606 Q09614 Q09621 Q09624
Q09630 Q09638 Q09653 Q09679 Q09680 Q09681 Q09684 Q09701 Q09710 Q09712 Q09714
Q09729 Q09730 Q09736 Q09743 Q09749 Q09752 Q09758 Q09764 Q09766 Q09782 Q09809
Q09812 Q09825 Q09827 Q09834 Q09835 Q09837 Q09850 Q09852 Q09875 Q09877 Q09887
Q09891 Q09895 Q09896 Q09906 Q09910 Q09917 Q09919 Q09920 Q09925 Q09951 Q09952
Q09954 Q09964 Q09965 Q09966 Q09967 Q09969 Q09993 Q09EG3 Q09FQ2 Q09FQ5 Q09FQ7
Q09FQ8 Q09FR0 Q09FR1 Q09FR3 Q09FT3 Q09FT5 Q09FU1 Q09FU2 Q09FU3 Q09FU6 Q09FU7
Q09FU8 Q09FU9 Q09FV0 Q09FV6 Q09FW1 Q09FW2 Q09FW5 Q09FW6 Q09FW7 Q09FW8 Q09FX3
Q09FX4 Q09FX5 Q09FX7 Q09FY1 Q09FY8 Q09FZ1 Q09FZ3 Q09FZ4 Q09FZ6 Q09FZ7 Q09FZ9
Q09G20 Q09G26 Q09G27 Q09G28 Q09G31 Q09G32 Q09G33 Q09G34 Q09G35 Q09G41 Q09G46
Q09G47 Q09G50 Q09G51 Q09G52 Q09G53 Q09G58 Q09G59 Q09G60 Q09G62 Q09G66 Q09J78
Q09J79 Q09JL7 Q09MB8 Q09MC1 Q09MC3 Q09MC4 Q09MC6 Q09MC7 Q09MC9 Q09ME6 Q09ME7
Q09ME8 Q09ME9 Q09MF0 Q09MF1 Q09MF8 Q09MF9 Q09MG0 Q09MG1 Q09MG2 Q09MG3 Q09MG4
Q09MG5 Q09MG6 Q09MG7 Q09MG8 Q09MH3 Q09MH8 Q09MH9 Q09MI1 Q09MI2 Q09MI3 Q09MI4
Q09MI5 Q09MJ0 Q09MJ1 Q09MJ2 Q09MJ4 Q09MJ5 Q09MJ8 Q09PK2 Q09QM4 Q09SZ7 Q09TM2
Q09TM4 Q09WW0 Q09WW3 Q09WW5 Q09WW6 Q09WW8 Q09WW9 Q09WX1 Q09WY6 Q09WY7 Q09WY8
Q09WY9 Q09WZ0 Q09WZ1 Q09WZ7 Q09WZ8 Q09WZ9 Q09X00 Q09X01 Q09X02 Q09X03 Q09X04
Q09X05 Q09X06 Q09X07 Q09X12 Q09X17 Q09X18 Q09X20 Q09X21 Q09X22 Q09X23 Q09X24
Q09X29 Q09X30 Q09X31 Q09X33 Q09X34 Q09X37 Q09YH0 Q09YH4 Q09YH7 Q09YI5 Q09YI9
Q09YJ4 Q09YJ7 Q09YK0 Q09YK5 Q09YK8 Q09YL1 Q09YN2 Q09YN5 Q0A2D6 Q0A2E4 Q0A2G2
Q0A2G4 Q0A2H5 Q0A2I4 Q0A2I6 Q0A2R1 Q0A3Q8 Q0A3Y1 Q0A415 Q0A436 Q0A447 Q0A448
Q0A456 Q0A458 Q0A4L5 Q0A4M2 Q0A4M4 Q0A4N8 Q0A4T1 Q0A4U4 Q0A5D5 Q0A5D6 Q0A5V9
Q0A6J9 Q0A6K1 Q0A782 Q0A785 Q0A788 Q0A7K5 Q0A805 Q0AAG2 Q0AAG7 Q0AAR8 Q0ABB2
Q0ABY1 Q0ADH9 Q0ADK7 Q0ADQ5 Q0AE56 Q0AET6 Q0AET7 Q0AFV3 Q0AFX4 Q0AGZ9 Q0AHJ0
Q0AHJ3 Q0AIG6 Q0AIM8 Q0AIQ0 Q0AJ92 Q0AJB4 Q0AJD8 Q0AJJ0 Q0AJY9 Q0AK30 Q0AK33
Q0AM65 Q0AMG9 Q0AMJ4 Q0AMW4 Q0AND7 Q0ANY0 Q0APH6 Q0APX7 Q0APY3 Q0AR54 Q0ASI6
Q0ATH6 Q0AUC1 Q0AUC8 Q0AUC9 Q0AWD5 Q0AXC2 Q0AXF8 Q0AXH9 Q0AXJ1 Q0AXT4 Q0AZ32
Q0AZJ4 Q0B3W2 Q0B8F7 Q0BAH4 Q0BAQ2 Q0BAT8 Q0BBI9 Q0BBM3 Q0BCG3 Q0BCK7 Q0BCQ8
Q0BD93 Q0BDD0 Q0BDD7 Q0BDE0 Q0BEC9 Q0BEG9 Q0BEY0 Q0BGG1 Q0BGK5 Q0BHC9 Q0BHM0
Q0BI45 Q0BIE9 Q0BIK4 Q0BIV8 Q0BIV9 Q0BJL9 Q0BK20 Q0BK58 Q0BK61 Q0BK80 Q0BKJ3
Q0BKM7 Q0BMC8 Q0BML9 Q0BNB6 Q0BNI0 Q0BNW5 Q0BNY1 Q0BP14 Q0BP56 Q0BPL2 Q0BPU7
Q0BQ20 Q0BQ81 Q0BQY4 Q0BQY5 Q0BQY6 Q0BQY7 Q0BRW3 Q0BSK9 Q0BSL2 Q0BSL5 Q0BSQ0
Q0BSV4 Q0BTT3 Q0BTZ8 Q0BU78 Q0BV22 Q0BVS2 Q0BW71 Q0BW85 Q0BWV3 Q0BXT9 Q0BZQ0

Q0BZT0 Q0C099 Q0C0V9 Q0C0X0 Q0C0X2 Q0C0X3 Q0C1D1 Q0C1D7 Q0C1N8 Q0C2L6 Q0C3D1
Q0C3G9 Q0C5R9 Q0C618 Q0C7E3 Q0C8A1 Q0C8A6 Q0C8A7 Q0C8L9 Q0C8M4 Q0C9L4 Q0C9L5
Q0CA27 Q0CCR9 Q0CEJ5 Q0CEN9 Q0CF72 Q0CFC0 Q0CGN5 Q0CH31 Q0CHZ8 Q0CI96 Q0CIV4
Q0CJ00 Q0CJ58 Q0CJ61 Q0CJ62 Q0CJC4 Q0CJD0 Q0CJH2 Q0CL24 Q0CL45 Q0CLP4 Q0CMM0
Q0CNC7 Q0CNZ5 Q0CQC5 Q0CQS4 Q0CRD8 Q0CRP6 Q0CRQ3 Q0CRW7 Q0CS59 Q0CS63 Q0CS68
Q0CS98 Q0CSG1 Q0CSZ7 Q0CT01 Q0CT23 Q0CT66 Q0CUP0 Q0CUT8 Q0CVD7 Q0CXB1 Q0CXF5
Q0CXU6 Q0CY11 Q0CY37 Q0CZG6 Q0CZG9 Q0CZY7 Q0D0W7 Q0D135 Q0D153 Q0D1P6 Q0D1P7
Q0D1P9 Q0D219 Q0D285 Q0D289 Q0D2D7 Q0D2E8 Q0D2G3 Q0D2K0 Q0D3C8 Q0D4G3 Q0D519
Q0D576 Q0D5P3 Q0D5R2 Q0D5R3 Q0D7E4 Q0D8I9 Q0D9S3 Q0DAK7 Q0DBF4 Q0DC45 Q0DET3
Q0DG05 Q0DHE3 Q0DHJ5 Q0DHM7 Q0DJC5 Q0DJY3 Q0DK16 Q0DKM0 Q0DKW8 Q0DLG0 Q0DS59
Q0DWA9 Q0DWC7 Q0DXZ1 Q0DY59 Q0E0I1 Q0E4J6 Q0E558 Q0E588 Q0E589 Q0E961 Q0EAB5
Q0EAB6 Q0EEE2 Q0G9G1 Q0G9G4 Q0G9G6 Q0G9G7 Q0G9G9 Q0G9H0 Q0G9H2 Q0G9I8 Q0G9I9
Q0G9J0 Q0G9J3 Q0G9J9 Q0G9K0 Q0G9K1 Q0G9K2 Q0G9K5 Q0G9K6 Q0G9K7 Q0G9K8 Q0G9K9
Q0G9L4 Q0G9L9 Q0G9M0 Q0G9M3 Q0G9M4 Q0G9M5 Q0G9M6 Q0G9N1 Q0G9N2 Q0G9N3 Q0G9N5
Q0G9N6 Q0G9N9 Q0G9Q4 Q0G9Q7 Q0G9Q9 Q0G9R0 Q0G9R2 Q0G9R3 Q0G9R5 Q0G9T1 Q0G9T2
Q0G9T3 Q0G9T4 Q0G9T5 Q0G9T6 Q0G9U2 Q0G9U3 Q0G9U4 Q0G9U5 Q0G9U6 Q0G9U7 Q0G9U8
Q0G9U9 Q0G9V0 Q0G9V1 Q0G9V2 Q0G9V7 Q0G9W2 Q0G9W3 Q0G9W5 Q0G9W6 Q0G9W7 Q0G9W8
Q0G9W9 Q0G9X4 Q0G9X5 Q0G9X6 Q0G9X8 Q0G9X9 Q0G9Y2 Q0GA42 Q0GBX6 Q0GBY1 Q0GBZ5
Q0GC71 Q0GE19 Q0GK35 Q0GQS6 Q0GXS4 Q0H8B9 Q0H8C3 Q0H8W8 Q0H8W9 Q0H8X0 Q0H8X1
Q0H8X2 Q0H8X3 Q0H8X4 Q0H8X5 Q0H8X6 Q0H8X8 Q0H8Y0 Q0H8Y1 Q0H8Y2 Q0H8Y3 Q0H8Y4
Q0H8Y5 Q0H8Y6 Q0H8Y7 Q0H8Y8 Q0H8Y9 Q0HD57 Q0HD59 Q0HD60 Q0HD64 Q0HD73 Q0HD74
Q0HD75 Q0HDH7 Q0HDK5 Q0HE80 Q0HE97 Q0HE98 Q0HE99 Q0HFK8 Q0HFZ0 Q0HG43 Q0HG46
Q0HG84 Q0HG86 Q0HGS5 Q0HHH4 Q0HHP5 Q0HIH5 Q0HIH7 Q0HII0 Q0HIS8 Q0HJ63 Q0HJX2
Q0HJX3 Q0HK32 Q0HK90 Q0HKG6 Q0HL71 Q0HLP8 Q0HLZ4 Q0HMJ1 Q0HMW6 Q0HN13 Q0HNH1
Q0HNQ9 Q0HPE6 Q0HPF5 Q0HPF6 Q0HPF7 Q0HPR0 Q0HPV6 Q0HQB7 Q0HQR8 Q0HQW4 Q0HR97
Q0HRU2 Q0HS83 Q0HSD6 Q0HSD9 Q0HSH7 Q0HSH9 Q0HT31 Q0HTS8 Q0HTZ7 Q0HUS0 Q0HV56
Q0HVF5 Q0HVF8 Q0HVG0 Q0HW68 Q0HW69 Q0HWD3 Q0HWJ2 Q0HWR8 Q0HXG9 Q0HY29 Q0HYC2
Q0HZP9 Q0HZQ0 Q0HZQ1 Q0HZR9 Q0I078 Q0I0W3 Q0I0Z1 Q0I185 Q0I1D6 Q0I1V9 Q0I206
Q0I207 Q0I208 Q0I2F6 Q0I2I2 Q0I2K9 Q0I2U0 Q0I309 Q0I337 Q0I372 Q0I3A2 Q0I455
Q0I487 Q0I4C5 Q0I4T6 Q0I4W6 Q0I5I2 Q0I5K7 Q0I5W7 Q0I5W8 Q0I5W9 Q0I5Y0 Q0I5Y1
Q0I604 Q0I6V9 Q0I6X0 Q0I6Z7 Q0I733 Q0I740 Q0I7J2 Q0I7P7 Q0I7Q7 Q0I7Q8 Q0I7R0
Q0I867 Q0I872 Q0I873 Q0I891 Q0I8A7 Q0I8V6 Q0I910 Q0I912 Q0IA94 Q0IAI7 Q0IAJ3
Q0IAK1 Q0IAM4 Q0IBJ2 Q0IBQ6 Q0IBR1 Q0ICP7 Q0ICP8 Q0ICR0 Q0ICR3 Q0ID47 Q0ID48
Q0IDC8 Q0IDC9 Q0IDD1 Q0IDF4 Q0IDJ4 Q0IDJ7 Q0IDJ9 Q0IDK0 Q0IFA9 Q0IH10 Q0IH22
Q0IH28 Q0IH73 Q0IHA5 Q0IHC4 Q0IHC5 Q0IHD6 Q0IHF1 Q0IHF9 Q0IHI1 Q0IHJ0 Q0IHM1
Q0IHQ3 Q0IHQ9 Q0IHU6 Q0IHY5 Q0II41 Q0II44 Q0II64 Q0II71 Q0II74 Q0II76 Q0II78
Q0IIE5 Q0IIE8 Q0IIF9 Q0IIH7 Q0III2 Q0IIK4 Q0IIL1 Q0IIL2 Q0IIL4 Q0IIS3 Q0IIV2
Q0IIY2 Q0IIY7 Q0IJ12 Q0IJ20 Q0IJ33 Q0ILJ3 Q0ILZ7 Q0IN16 Q0IQA5 Q0IUB5 Q0IWF3
Q0IZQ3 Q0J185 Q0J349 Q0J361 Q0J705 Q0J8R9 Q0J932 Q0JA29 Q0JAI9 Q0JAW2 Q0JBZ6
Q0JCR9 Q0JD42 Q0JDK9 Q0JEF7 Q0JEQ2 Q0JEU6 Q0JF01 Q0JFI2 Q0JJV0 Q0JJV1 Q0JJZ6
Q0JKV1 Q0JLC5 Q0JLS6 Q0JNB6 Q0JPT5 Q0JQS5 Q0JR55 Q0K020 Q0K2K6 Q0K437 Q0K5C1
Q0K5M3 Q0K698 Q0K699 Q0K6A4 Q0K6M1 Q0K739 Q0K799 Q0K7G0 Q0K7S4 Q0K843 Q0K9W2
Q0KB74 Q0KBH3 Q0KBR7 Q0KCE1 Q0KCS0 Q0KCS3 Q0KE35 Q0KER8 Q0KEX1 Q0KFB3 Q0KHU5
Q0KHV6 Q0KHY3 Q0KK59 Q0KL00 Q0MQ88 Q0MQ89 Q0MQ90 Q0MQ94 Q0MQ95 Q0MQ96 Q0MQ97

Q0MQ98 Q0MQ99 Q0MQB8 Q0MQB9 Q0MQC0 Q0MQC5 Q0MQC6 Q0MQD0 Q0MQD1 Q0MQD2 Q0MQD4
Q0MQD5 Q0MQD6 Q0MQD7 Q0MQD8 Q0MQE0 Q0MQE1 Q0MQE6 Q0MQE7 Q0MQF5 Q0MQF6 Q0MQF7
Q0MQF8 Q0MQF9 Q0MQJ3 Q0MQJ4 Q0MQJ5 Q0MS45 Q0MVP0 Q0P3J1 Q0P3J2 Q0P3J4 Q0P3J5
Q0P3J6 Q0P3J8 Q0P3J9 Q0P3K0 Q0P3K1 Q0P3K2 Q0P3K3 Q0P3K6 Q0P3K7 Q0P3K8 Q0P3M1
Q0P3N3 Q0P3N4 Q0P3N6 Q0P3N7 Q0P3N8 Q0P3N9 Q0P3P0 Q0P3P6 Q0P3P8 Q0P3P9 Q0P3Q0
Q0P3Q1 Q0P436 Q0P442 Q0P467 Q0P483 Q0P487 Q0P496 Q0P4A4 Q0P4G6 Q0P4G7 Q0P4J6
Q0P4J9 Q0P4L9 Q0P4P2 Q0P4Q4 Q0P4W3 Q0P4Y4 Q0P4Y8 Q0P4Z1 Q0P543 Q0P583 Q0P5A0
Q0P5B1 Q0P5B7 Q0P5C0 Q0P5C7 Q0P5D9 Q0P5E4 Q0P5F0 Q0P5H8 Q0P5J1 Q0P5L7 Q0P5M9
Q0P5V9 Q0P670 Q0P6D2 Q0P6H9 Q0P7X3 Q0P7X4 Q0P886 Q0P952 Q0P9C4 Q0P9C6 Q0P9C8
Q0P9D0 Q0P9D4 Q0P9F7 Q0P9W3 Q0P9Y2 Q0PA12 Q0PA25 Q0PAR0 Q0PBE9 Q0PMD2 Q0PV50
Q0PW25 Q0PW40 Q0Q2J4 Q0Q2J5 Q0Q2J6 Q0Q2J7 Q0Q2J8 Q0Q2J9 Q0Q2K0 Q0Q460 Q0Q463
Q0Q464 Q0Q465 Q0Q466 Q0Q470 Q0Q472 Q0Q473 Q0Q474 Q0Q475 Q0Q4E7 Q0Q4E8 Q0Q4E9
Q0Q4F2 Q0Q4Z0 Q0Q7H1 Q0Q7U7 Q0Q7U8 Q0Q7U9 Q0Q7V0 Q0QZ08 Q0R3M2 Q0R5Q9 Q0RD23
Q0RDA8 Q0RDB0 Q0RDW7 Q0RF16 Q0RFY2 Q0RG46 Q0RH20 Q0RHR9 Q0RHS2 Q0RJJ1 Q0RKT0
Q0RNP4 Q0RPD5 Q0RRV8 Q0RRW1 Q0RYC2 Q0S0I5 Q0S254 Q0S2P8 Q0S2P9 Q0S2S0 Q0S440
Q0S443 Q0S8L7 Q0SAA3 Q0SG15 Q0SGP5 Q0SHD0 Q0SHR8 Q0SHT6 Q0SHZ8 Q0SIC2 Q0SIG6
Q0SIZ9 Q0SN03 Q0SN37 Q0SN67 Q0SNF2 Q0SNK7 Q0SNQ3 Q0SQZ0 Q0SQZ1 Q0SRL7 Q0SRW4
Q0SSE8 Q0SSN5 Q0SSQ8 Q0SST2 Q0ST20 Q0STN1 Q0STX5 Q0SU39 Q0SVL9 Q0SWK0 Q0SWX2
Q0SX48 Q0SXC4 Q0SXC5 Q0SXE3 Q0SXP7 Q0SXQ2 Q0SXQ7 Q0SY25 Q0SYK8 Q0SYM8 Q0SYP9
Q0SYQ7 Q0SYR5 Q0SYT8 Q0SYT9 Q0SYU0 Q0SYU9 Q0SYY4 Q0SZ27 Q0SZ29 Q0SZ72 Q0SZA5
Q0SZC4 Q0SZH0 Q0SZM0 Q0SZM1 Q0SZP2 Q0SZV2 Q0T046 Q0T047 Q0T048 Q0T066 Q0T0F2
Q0T0H9 Q0T0K4 Q0T0K6 Q0T0L8 Q0T128 Q0T129 Q0T1H9 Q0T1S8 Q0T203 Q0T296 Q0T2A8
Q0T2B0 Q0T2J2 Q0T2J9 Q0T2K5 Q0T2K8 Q0T2M4 Q0T2M5 Q0T2M6 Q0T2M9 Q0T2S7 Q0T2Y2
Q0T2Z4 Q0T333 Q0T353 Q0T354 Q0T3C3 Q0T3F9 Q0T3L5 Q0T3Y9 Q0T4A9 Q0T4E5 Q0T4E7
Q0T4F0 Q0T4H7 Q0T4H8 Q0T4K0 Q0T4K1 Q0T4L7 Q0T4L8 Q0T4M5 Q0T4S1 Q0T513 Q0T515
Q0T519 Q0T522 Q0T579 Q0T5E1 Q0T5E2 Q0T5K9 Q0T5L5 Q0T5L6 Q0T5W7 Q0T5Z5 Q0T618
Q0T6D1 Q0T6N5 Q0T6U0 Q0T832 Q0T851 Q0T8B0 Q0T8E8 Q0T8F4 Q0T8G7 Q0T8H4 Q0T8J4
Q0T8M0 Q0T8T7 Q0T8U9 Q0T8V3 Q0T8X4 Q0T962 Q0T9N7 Q0T9N8 Q0T9Q5 Q0T9Y2 Q0TA22
Q0TA25 Q0TA30 Q0TA90 Q0TAE9 Q0TAF4 Q0TAF6 Q0TAH6 Q0TAL7 Q0TAL9 Q0TAT0 Q0TAW2
Q0TAX1 Q0TAX2 Q0TAX3 Q0TAY9 Q0TAZ2 Q0TAZ8 Q0TB02 Q0TB22 Q0TB40 Q0TBS5 Q0TBW9
Q0TC08 Q0TC09 Q0TC44 Q0TC96 Q0TCH1 Q0TCM3 Q0TCM4 Q0TCM5 Q0TCP2 Q0TCQ1 Q0TCY8
Q0TD17 Q0TD43 Q0TD47 Q0TD49 Q0TD62 Q0TD64 Q0TDA9 Q0TDZ6 Q0TDZ7 Q0TE04 Q0TEB0
Q0TER0 Q0TEW9 Q0TF04 Q0TF53 Q0TF54 Q0TF69 Q0TF71 Q0TFF1 Q0TFF8 Q0TFG4 Q0TFG7
Q0TFH0 Q0TFI3 Q0TFI4 Q0TFI5 Q0TFI8 Q0TFP4 Q0TFV0 Q0TFW4 Q0TFY6 Q0TG13 Q0TG14
Q0TG15 Q0TGE0 Q0TGM2 Q0TGS6 Q0TH03 Q0TH12 Q0TH13 Q0TH31 Q0TH40 Q0THB7 Q0THG4
Q0THH1 Q0THJ5 Q0THJ7 Q0THK0 Q0THM8 Q0THM9 Q0THP2 Q0THR0 Q0THR1 Q0THS6 Q0TI25
Q0TI43 Q0TIB5 Q0TIB6 Q0TIC2 Q0TII4 Q0TII9 Q0TIJ0 Q0TJ09 Q0TJ20 Q0TJ26 Q0TJ50
Q0TJD9 Q0TJF2 Q0TJH0 Q0TJK1 Q0TJL6 Q0TJL7 Q0TJY8 Q0TJY9 Q0TJZ0 Q0TK48 Q0TK80
Q0TKL4 Q0TLF6 Q0TLH6 Q0TLQ2 Q0TLU2 Q0TLU9 Q0TLW4 Q0TLW9 Q0TNB9 Q0TNC0 Q0TNZ8
Q0TP97 Q0TPT1 Q0TQ20 Q0TQG9 Q0TRA5 Q0TRK3 Q0TRL0 Q0TRQ9 Q0TRT2 Q0TRT3 Q0TRT4
Q0TTK8 Q0TTS0 Q0TU56 Q0TUQ9 Q0TUS0 Q0TV56 Q0TVB2 Q0TVF9 Q0TWF4 Q0TWV0 Q0TYM0
Q0U103 Q0U2R3 Q0U3N7 Q0U9W5 Q0UDX8 Q0UEY4 Q0UG89 Q0UI03 Q0UIR3 Q0UJ42 Q0UK49
Q0UK51 Q0ULB7 Q0UQV6 Q0URQ5 Q0USX0 Q0UTI3 Q0UUE1 Q0UUH1 Q0UV26 Q0UV43 Q0UVH1
Q0UVK7 Q0UYL2 Q0V0G4 Q0V1P1 Q0V302 Q0V3D6 Q0V4H8 Q0V4P1 Q0V4Z6 Q0V6M5 Q0V6Q0

Q0V6Q8 Q0V7M4 Q0V7N0 Q0V7R1 Q0V7T5 Q0V881 Q0V882 Q0V8E7 Q0V8G3 Q0V8L2 Q0V8N6
Q0V8S9 Q0V8T0 Q0V8T3 Q0V8T4 Q0V8T5 Q0V8T6 Q0V8T7 Q0V8T8 Q0V8T9 Q0V947 Q0V982
Q0V990 Q0V9A8 Q0V9E0 Q0V9G6 Q0V9J0 Q0V9U2 Q0V9V2 Q0V9V9 Q0V9Z3 Q0VA06 Q0VA36
Q0VA77 Q0VA82 Q0VAQ4 Q0VAX3 Q0VAX9 Q0VAY3 Q0VBD0 Q0VBF2 Q0VBF8 Q0VBN2 Q0VBP7
Q0VBU9 Q0VC00 Q0VC03 Q0VC16 Q0VC33 Q0VC54 Q0VC58 Q0VC81 Q0VC84 Q0VC89 Q0VCA3
Q0VCA9 Q0VCB1 Q0VCC1 Q0VCF5 Q0VCH4 Q0VCH6 Q0VCH8 Q0VCK9 Q0VCM2 Q0VCM6 Q0VCN0
Q0VCN6 Q0VCP2 Q0VCP9 Q0VCQ8 Q0VCR2 Q0VCR6 Q0VCS0 Q0VCS6 Q0VCT2 Q0VCT4 Q0VCU7
Q0VCY0 Q0VCY1 Q0VCY6 Q0VCZ0 Q0VD05 Q0VD07 Q0VD15 Q0VD38 Q0VD42 Q0VD51 Q0VD59
Q0VDE8 Q0VDI3 Q0VDU3 Q0VF94 Q0VFE3 Q0VFE7 Q0VFF9 Q0VFH9 Q0VFK3 Q0VFL4 Q0VFM0
Q0VFM5 Q0VFP2 Q0VFV6 Q0VG18 Q0VGK4 Q0VGT4 Q0VGV9 Q0VGW4 Q0VGW6 Q0VKU7 Q0VKW8
Q0VKX0 Q0VKZ3 Q0VLM2 Q0VLR6 Q0VM99 Q0VMA0 Q0VN94 Q0VP38 Q0VP41 Q0VPA7 Q0VPS7
Q0VQ74 Q0VQC0 Q0VQH0 Q0VQP5 Q0VQR4 Q0VRZ9 Q0VS03 Q0VS05 Q0VSY8 Q0VT01 Q0VTA7
Q0VTH3 Q0VTI9 Q0VTN6 Q0W1P5 Q0W4K2 Q0W5V8 Q0W6Q1 Q0WCB5 Q0WDX2 Q0WEG0 Q0WF87
Q0WFU0 Q0WJ82 Q0WJC2 Q0WLC7 Q0WMJ8 Q0WML0 Q0WMZ5 Q0WNG6 Q0WNW4 Q0WNY5 Q0WP01
Q0WP36 Q0WP96 Q0WPA5 Q0WPE9 Q0WPN7 Q0WPN8 Q0WPQ2 Q0WPS0 Q0WPS2 Q0WPW5 Q0WQ07
Q0WQ63 Q0WQD2 Q0WQG8 Q0WQJ3 Q0WQK2 Q0WQX7 Q0WR59 Q0WRB9 Q0WRJ2 Q0WRW8 Q0WS06
Q0WSX8 Q0WSZ6 Q0WT31 Q0WT40 Q0WT48 Q0WTF4 Q0WTI8 Q0WU02 Q0WUA3 Q0WUC5 Q0WUG6
Q0WUU6 Q0WUZ5 Q0WV13 Q0WV96 Q0WVE9 Q0WVM4 Q0WVN5 Q0WVZ1 Q0WW17 Q0WWW9 Q0X0A5
Q0Z7W6 Q0Z8I9 Q0ZAH6 Q0ZAH7 Q0ZAI3 Q0ZCA7 Q0ZHH6 Q0ZIW0 Q0ZIW3 Q0ZIW5 Q0ZIW6
Q0ZIW8 Q0ZIW9 Q0ZIX1 Q0ZIY8 Q0ZIY9 Q0ZIZ0 Q0ZIZ1 Q0ZIZ2 Q0ZIZ3 Q0ZJ00 Q0ZJ01
Q0ZJ03 Q0ZJ04 Q0ZJ06 Q0ZJ07 Q0ZJ08 Q0ZJ09 Q0ZJ10 Q0ZJ15 Q0ZJ20 Q0ZJ21 Q0ZJ23
Q0ZJ24 Q0ZJ25 Q0ZJ26 Q0ZJ27 Q0ZJ32 Q0ZJ33 Q0ZJ34 Q0ZJ36 Q0ZJ37 Q0ZJ40 Q0ZM14
Q0ZME4 Q0ZME5 Q0ZME7 Q0ZME8 Q0ZS23 Q0ZS24 Q0ZUL9 Q0ZUP0 Q0ZUP1 Q10000 Q10010
Q10015 Q10017 Q10022 Q10023 Q10025 Q10028 Q10029 Q10032 Q10037 Q10042 Q10044
Q10045 Q10046 Q10049 Q10058 Q10063 Q10065 Q10070 Q10071 Q10072 Q10073 Q10074
Q10084 Q10085 Q10087 Q10097 Q10109 Q10128 Q10130 Q10142 Q10151 Q10153 Q10176
Q10177 Q10185 Q10196 Q10211 Q10227 Q10244 Q10245 Q10248 Q10250 Q10252 Q10254
Q10255 Q10259 Q10260 Q10263 Q10268 Q10269 Q10279 Q10283 Q10286 Q10287 Q10309
Q10320 Q10323 Q10351 Q10354 Q10358 Q10359 Q10361 Q10408 Q10413 Q10418 Q10419
Q10436 Q10442 Q10449 Q10468 Q10469 Q10470 Q10471 Q10472 Q10473 Q10478 Q10479
Q10482 Q10484 Q10487 Q10493 Q10495 Q10496 Q104Y0 Q104Y1 Q104Y3 Q10589 Q10651
Q10656 Q10659 Q10710 Q10737 Q10739 Q10741 Q10743 Q10751 Q10836 Q108U0 Q108U3
Q108U6 Q10901 Q10904 Q10907 Q10909 Q10914 Q10917 Q10934 Q10935 Q10936 Q10937
Q10939 Q10941 Q10949 Q10979 Q10980 Q10981 Q10982 Q10983 Q10984 Q10BH4 Q10CV4
Q10D38 Q10EJ2 Q10EK7 Q10HD0 Q10I20 Q10KT6 Q10LI8 Q10LJ2 Q10LN5 Q10M80 Q10MK2
Q10MN2 Q10MN3 Q10MQ0 Q10MR5 Q10MX1 Q10MX2 Q10MX4 Q10N05 Q10NT7 Q10P01 Q10PI5
Q10PP8 Q10PW9 Q10Q65 Q10Q78 Q10Q99 Q10QE9 Q10QH3 Q10R54 Q10R93 Q10RR9 Q10RX7
Q10S25 Q10S27 Q10SM7 Q10SX7 Q10SY9 Q10T43 Q10VH5 Q10VK7 Q10VT7 Q10VT8 Q10VU1
Q10W93 Q10WA3 Q10WK0 Q10XM6 Q10XU6 Q10YS7 Q10YS8 Q10YS9 Q10YT0 Q10YT3 Q10YZ5
Q10ZF7 Q10ZG8 Q10ZK6 Q10ZX6 Q11068 Q11069 Q11071 Q11073 Q11080 Q11082 Q11085
Q11087 Q11095 Q11098 Q110N2 Q110N3 Q110N4 Q11106 Q11117 Q11122 Q11124 Q11126
Q11127 Q11128 Q11130 Q11131 Q11177 Q11180 Q11186 Q111U1 Q11200 Q11201 Q11203
Q11204 Q11205 Q11206 Q112P6 Q112Z1 Q112Z2 Q112Z3 Q112Z4 Q113E9 Q114L6 Q114L7
Q114N4 Q115B5 Q115G5 Q116J3 Q116L4 Q116S5 Q116S6 Q117J2 Q118H6 Q118J9 Q118K5

Q118V7 Q119Z5 Q11A00 Q11CA6 Q11CS6 Q11CT7 Q11CW2 Q11DP3 Q11GS2 Q11HC5 Q11HC6
Q11HT1 Q11IG3 Q11J69 Q11JJ6 Q11JJ9 Q11JY0 Q11KD0 Q11KD2 Q11KH6 Q11KH7 Q11KH9
Q11PK1 Q11PU6 Q11RH2 Q11RU2 Q11S04 Q11SZ7 Q11TN7 Q11TX2 Q11UI4 Q11VB2 Q11VB9
Q11VC2 Q11VC5 Q11VI0 Q11W14 Q11WJ0 Q11YP3 Q12001 Q12006 Q12010 Q12013 Q12015
Q12016 Q12023 Q12025 Q12029 Q12042 Q12057 Q12063 Q12066 Q12067 Q12077 Q12078
Q12079 Q12096 Q12106 Q12111 Q12116 Q12117 Q12119 Q12121 Q12133 Q12135 Q12142
Q12144 Q12147 Q12150 Q12155 Q12160 Q12164 Q12169 Q12171 Q12174 Q12185 Q12187
Q12200 Q12205 Q12207 Q12209 Q12210 Q12215 Q12225 Q12232 Q12235 Q12239 Q12240
Q12241 Q12249 Q12251 Q12252 Q12253 Q12255 Q12256 Q12270 Q12274 Q12275 Q12284
Q12289 Q12296 Q122S7 Q12300 Q12301 Q12312 Q12324 Q12325 Q12328 Q12333 Q12336
Q12346 Q12350 Q12352 Q12359 Q12361 Q12372 Q12374 Q12375 Q12382 Q12394 Q12396
Q123Z8 Q12402 Q12403 Q12407 Q12412 Q12424 Q12431 Q12436 Q12437 Q12442 Q12443
Q12445 Q12450 Q12465 Q12466 Q12467 Q12473 Q12479 Q12482 Q12489 Q12492 Q12498
Q12516 Q12519 Q12520 Q12528 Q12530 Q12564 Q12573 Q12581 Q12585 Q125P4 Q12612
Q12618 Q12632 Q12635 Q12649 Q12664 Q12674 Q12675 Q12691 Q12697 Q12732 Q12743
Q12746 Q12767 Q12770 Q12772 Q12791 Q12797 Q127N8 Q127X9 Q12809 Q12836 Q12846
Q12864 Q12866 Q12879 Q12884 Q12887 Q12893 Q12907 Q12908 Q12912 Q12913 Q12918
Q12981 Q12983 Q12999 Q12AY6 Q12BS8 Q12C33 Q12E37 Q12EL8 Q12EQ7 Q12F57 Q12FC1
Q12FC2 Q12FJ6 Q12GA4 Q12GQ4 Q12GQ5 Q12HF9 Q12HM8 Q12HP5 Q12HP6 Q12HP7 Q12HU1
Q12I78 Q12IC7 Q12IK3 Q12KB8 Q12KC1 Q12KG9 Q12L71 Q12L81 Q12M46 Q12MB4 Q12N25
Q12N27 Q12N30 Q12NE6 Q12NQ8 Q12NQ9 Q12P61 Q12P74 Q12Q34 Q12Q70 Q12QK1 Q12QK2
Q12QK3 Q12S01 Q12S25 Q12S26 Q12S27 Q12S48 Q12SC9 Q12TZ6 Q12U86 Q12UB0 Q12UC1
Q12UQ2 Q12VF3 Q12VU4 Q12X42 Q13002 Q13003 Q13018 Q13021 Q13061 Q13113 Q13145
Q13183 Q13190 Q13224 Q13241 Q13255 Q13258 Q13261 Q13277 Q13286 Q13291 Q132Z6
Q13304 Q13308 Q13323 Q13324 Q13326 Q13332 Q13336 Q13349 Q13370 Q133W8 Q133Z9
Q13410 Q13423 Q13433 Q13443 Q13444 Q13445 Q13454 Q13467 Q13477 Q13478 Q13488
Q13491 Q13505 Q13507 Q13520 Q13530 Q13563 Q13571 Q13585 Q13586 Q13591 Q135C9
Q135I3 Q135S8 Q135T0 Q135Y3 Q13606 Q13607 Q13621 Q13634 Q13635 Q13639 Q13641
Q13651 Q13683 Q13698 Q136H1 Q136Q9 Q136R8 Q136W9 Q13705 Q13724 Q13733 Q13740
Q13797 Q137R8 Q13873 Q13936 Q13948 Q139K2 Q13BG6 Q13BG9 Q13BH5 Q13BH6 Q13CX1
Q13CX3 Q13CX4 Q13CY3 Q13CY5 Q13D27 Q13DD7 Q13ED9 Q13ES7 Q13PK0 Q13SH4 Q13SP8
Q13TR5 Q13TR6 Q13TY9 Q13U46 Q13UQ8 Q13V13 Q13V94 Q13W85 Q13WE0 Q13WP2 Q13WP3
Q13X42 Q13YZ8 Q13Z35 Q14003 Q14028 Q140Y6 Q14108 Q14114 Q14118 Q14126 Q14162
Q14165 Q141I7 Q14242 Q14246 Q142G3 Q142G6 Q142H3 Q142K7 Q142P6 Q14318 Q14330
Q14332 Q14392 Q143Y3 Q14410 Q14416 Q14432 Q14435 Q14439 Q14494 Q14500 Q14517
Q14524 Q14542 Q14571 Q14573 Q14574 Q145H3 Q145N7 Q14626 Q14627 Q14643 Q14654
Q14656 Q146A1 Q146D6 Q14703 Q14714 Q14721 Q14728 Q14739 Q14761 Q14773 Q14789
Q147U7 Q14802 Q14831 Q14832 Q14833 Q14849 Q148F2 Q148G2 Q148G8 Q148H0 Q148I3
Q148K5 Q148L1 Q148M6 Q148R9 Q148W0 Q14916 Q14940 Q14943 Q14952 Q14953 Q14954
Q14956 Q14957 Q14973 Q149C3 Q149F5 Q149L6 Q149L7 Q149M0 Q149R9 Q14AK4 Q14AT5
Q14B02 Q14B62 Q14B80 Q14BI2 Q14BN4 Q14BT6 Q14C59 Q14C87 Q14CH0 Q14CN2 Q14CX5
Q14CZ8 Q14D33 Q14DG7 Q14DK4 Q14EA7 Q14EA8 Q14EB0 Q14EB1 Q14FA0 Q14FA3 Q14FA5
Q14FA6 Q14FA8 Q14FA9 Q14FB0 Q14FC6 Q14FC7 Q14FC8 Q14FC9 Q14FD0 Q14FD1 Q14FD7
Q14FD8 Q14FD9 Q14FE0 Q14FE1 Q14FE2 Q14FE3 Q14FE4 Q14FE5 Q14FE6 Q14FE7 Q14FF2

Q14FF7 Q14FF8 Q14FG0 Q14FG1 Q14FG2 Q14FG3 Q14FG4 Q14FG9 Q14FH0 Q14FH1 Q14FH3
Q14FH4 Q14FH6 Q14FP4 Q14H08 Q14H98 Q14HT3 Q14HY6 Q14J06 Q14JF9 Q14JI2 Q14JK8
Q14JW6 Q14K10 Q14K28 Q14K31 Q14SY0 Q15005 Q15011 Q15012 Q15035 Q15041 Q15043
Q15049 Q15053 Q15070 Q15077 Q15109 Q15116 Q15125 Q15155 Q15223 Q15256 Q15262
Q15303 Q15363 Q15375 Q15388 Q15391 Q15392 Q15399 Q15413 Q15526 Q15546 Q15612
Q15615 Q15617 Q15619 Q15620 Q15622 Q15629 Q15722 Q15738 Q15743 Q15758 Q15760
Q15761 Q15762 Q15768 Q15800 Q15822 Q15825 Q15836 Q15842 Q15849 Q15858 Q15878
Q15884 Q15904 Q15B89 Q15FB1 Q15FB4 Q15FB6 Q15GM1 Q15GP7 Q15GR0 Q15GS3 Q15KI9
Q15MS7 Q15MT8 Q15MU0 Q15N01 Q15N42 Q15N95 Q15NA2 Q15NT4 Q15P32 Q15P45 Q15Q14
Q15Q16 Q15QT0 Q15R05 Q15RL0 Q15RL1 Q15RL2 Q15RL5 Q15S31 Q15S32 Q15SF2 Q15UY7
Q15V35 Q15WB8 Q15WS4 Q15X16 Q15X19 Q15YQ1 Q15YQ2 Q15YQ3 Q15YQ5 Q15YY4 Q15Z93
Q15ZS2 Q15ZT9 Q16099 Q16280 Q16281 Q16288 Q162V9 Q16322 Q16348 Q16394 Q163C2
Q163R6 Q163T8 Q163W5 Q16445 Q16478 Q16515 Q16538 Q16549 Q16558 Q16563 Q16570
Q16572 Q16581 Q16585 Q16586 Q165T0 Q16602 Q16611 Q16617 Q16620 Q16623 Q16625
Q16647 Q16651 Q16653 Q16655 Q16671 Q16706 Q16720 Q16739 Q16790 Q16799 Q167V9
Q167W1 Q16819 Q16820 Q16821 Q16827 Q16832 Q16842 Q16849 Q16850 Q16853 Q16873
Q16880 Q16891 Q16928 Q16932 Q16943 Q16950 Q16951 Q16983 Q16AA1 Q16AB5 Q16AM5
Q16AM6 Q16AM7 Q16AM8 Q16BG3 Q16DZ2 Q16EI9 Q16N38 Q16QL3 Q16TM5 Q16WU7 Q16YE7
Q17046 Q17053 Q17094 Q17232 Q17239 Q17292 Q17296 Q17298 Q172W0 Q17307 Q17320
Q17328 Q17330 Q17374 Q17388 Q17405 Q17428 Q17433 Q17438 Q17528 Q17529 Q17586
Q17598 Q17638 Q17667 Q17678 Q17758 Q17760 Q17766 Q17833 Q17851 Q178U6 Q17938
Q17990 Q17A75 Q17A88 Q17CE7 Q17ED3 Q17JQ7 Q17NV8 Q17Q87 Q17QC0 Q17QD6 Q17QD8
Q17QE0 Q17QF8 Q17QI5 Q17QI7 Q17QJ2 Q17QJ5 Q17QL5 Q17QL9 Q17QM4 Q17QN4 Q17QN8
Q17QN9 Q17QQ5 Q17QR6 Q17QW0 Q17QW2 Q17QZ3 Q17QZ8 Q17R16 Q17R55 Q17RQ9 Q17SB3
Q17ST2 Q17VL4 Q17VM5 Q17VS8 Q17WM4 Q17WP6 Q17WX4 Q17X31 Q17X55 Q17XN0 Q17Y14
Q17Y82 Q17YP7 Q17YZ5 Q17Z59 Q17Z62 Q17Z65 Q17ZJ0 Q18007 Q18071 Q18078 Q18079
Q18081 Q18090 Q180E3 Q180T5 Q180X0 Q180X1 Q18120 Q18163 Q18179 Q18253 Q18297
Q182T8 Q182W4 Q182Y8 Q182Z4 Q18319 Q18321 Q18331 Q18411 Q18416 Q18428 Q18449
Q18473 Q18484 Q184H7 Q18515 Q18534 Q18593 Q18600 Q18673 Q18695 Q186E8 Q18775
Q18779 Q18788 Q18807 Q18812 Q18864 Q18881 Q188Z4 Q18904 Q18936 Q18BJ3 Q18BJ5
Q18DH5 Q18DH8 Q18EC3 Q18EC4 Q18FD0 Q18FQ7 Q18FQ8 Q18J00 Q18J48 Q18JU9 Q18KB3
Q18KL5 Q18KS3 Q18LE5 Q18LE9 Q18LF1 Q18P97 Q18PF5 Q18PF6 Q18PI6 Q19038 Q19053
Q19061 Q19076 Q19084 Q19127 Q19187 Q19189 Q19192 Q19200 Q19202 Q19238 Q19258
Q19267 Q19293 Q19319 Q19337 Q19351 Q19417 Q19425 Q19426 Q19443 Q19459 Q19468
Q19473 Q19474 Q19505 Q19507 Q19508 Q19509 Q19529 Q19549 Q19550 Q19552 Q19617
Q196T4 Q196T6 Q196U8 Q196W4 Q196X5 Q196Y7 Q196Z4 Q19707 Q19714 Q19746 Q19766
Q19768 Q197B3 Q197B7 Q197C3 Q197C7 Q197D6 Q197D8 Q197E7 Q197F4 Q197F6 Q19821
Q19897 Q19907 Q19951 Q19958 Q19967 Q19975 Q19978 Q19981 Q19992 Q19AE6 Q19KA1
Q19KC1 Q19KH7 Q19KI0 Q19LH4 Q19PY9 Q19T08 Q19V55 Q19V56 Q19V58 Q19V60 Q19V61
Q19V67 Q19V68 Q19V69 Q19V70 Q19V73 Q19V77 Q19V78 Q19V80 Q19V81 Q19V90 Q19V91
Q19V93 Q19V96 Q19V97 Q19VA2 Q19VA3 Q19VA4 Q19VC2 Q19VC4 Q19VC5 Q19VC6 Q19VC7
Q19VE6 Q1A243 Q1A244 Q1A261 Q1A262 Q1A3B0 Q1ACB3 Q1ACE4 Q1ACE6 Q1ACE7 Q1ACE9
Q1ACF3 Q1ACH0 Q1ACH1 Q1ACH2 Q1ACH3 Q1ACH4 Q1ACH5 Q1ACI1 Q1ACI4 Q1ACI5 Q1ACI7
Q1ACI8 Q1ACI9 Q1ACJ0 Q1ACJ1 Q1ACJ2 Q1ACJ5 Q1ACJ6 Q1ACJ7 Q1ACK4 Q1ACL3 Q1ACL4

Q1ACL6 Q1ACL7 Q1ACL9 Q1ACM1 Q1ACM2 Q1ACM5 Q1ACM9 Q1ACN0 Q1ACN1 Q1ACN8 Q1ACP0
Q1ACP1 Q1AE95 Q1AGV6 Q1AGV7 Q1AGX7 Q1AL67 Q1AU71 Q1AUL2 Q1AV13 Q1AVH3 Q1AVH5
Q1AVI7 Q1AVI9 Q1AVJ2 Q1AVX0 Q1AW41 Q1AWM0 Q1AWQ0 Q1AYN1 Q1AYN2 Q1B3Q7 Q1B3X9
Q1B453 Q1B4T8 Q1B548 Q1B549 Q1B550 Q1B6S1 Q1B6W8 Q1B769 Q1B7P4 Q1B913 Q1B952
Q1B971 Q1B9B1 Q1B9W9 Q1BBQ0 Q1BBQ3 Q1BC57 Q1BDZ2 Q1BES7 Q1BG50 Q1BGW2 Q1BH72
Q1BHB3 Q1BM23 Q1BPZ6 Q1BRA6 Q1BS36 Q1BS37 Q1BSF7 Q1BSJ6 Q1BT95 Q1BTD0 Q1BU57
Q1BUA0 Q1BUF3 Q1BUV6 Q1BUX7 Q1BV13 Q1BV20 Q1BV23 Q1BVU9 Q1BXA8 Q1BXE3 Q1BY52
Q1BYA7 Q1BYC6 Q1BYF5 Q1BYZ6 Q1BZB0 Q1BZG6 Q1C089 Q1C090 Q1C091 Q1C0I3 Q1C0J2
Q1C0J7 Q1C0M8 Q1C0T8 Q1C0W4 Q1C0X4 Q1C0Y6 Q1C136 Q1C137 Q1C158 Q1C190 Q1C1B3
Q1C1G0 Q1C1L1 Q1C1L2 Q1C1L3 Q1C1N6 Q1C211 Q1C246 Q1C296 Q1C2L2 Q1C2Q4 Q1C2S9
Q1C2X5 Q1C306 Q1C367 Q1C369 Q1C3K8 Q1C3X2 Q1C479 Q1C4D3 Q1C4D5 Q1C4J8 Q1C585
Q1C586 Q1C5I7 Q1C5L8 Q1C5L9 Q1C5Q9 Q1C5R1 Q1C5V2 Q1C5W7 Q1C5X8 Q1C5Y6 Q1C651
Q1C6A2 Q1C6A9 Q1C6B5 Q1C6B8 Q1C6C1 Q1C6Z1 Q1C741 Q1C744 Q1C745 Q1C746 Q1C769
Q1C7B0 Q1C7K1 Q1C7K7 Q1C7P8 Q1C7Q1 Q1C7R0 Q1C7V3 Q1C7V4 Q1C803 Q1C804 Q1C833
Q1C8L0 Q1C8M3 Q1C8X8 Q1C8X9 Q1C9U5 Q1CA68 Q1CA99 Q1CAD7 Q1CAM6 Q1CAS0 Q1CAS7
Q1CAS8 Q1CAY4 Q1CBF6 Q1CBF8 Q1CBH3 Q1CBH4 Q1CBP1 Q1CBU1 Q1CC20 Q1CC26 Q1CC44
Q1CCG9 Q1CCH0 Q1CCH1 Q1CCK6 Q1CCQ2 Q1CCS7 Q1CCX2 Q1CD32 Q1CD76 Q1CDG1 Q1CDI5
Q1CDU3 Q1CDW5 Q1CDW6 Q1CDW7 Q1CE18 Q1CE35 Q1CE95 Q1CEB8 Q1CEC8 Q1CEE0 Q1CEY6
Q1CF51 Q1CFA8 Q1CFB4 Q1CFF5 Q1CG99 Q1CGD7 Q1CGH0 Q1CGT7 Q1CH33 Q1CH34 Q1CHD3
Q1CHE5 Q1CHJ3 Q1CHP4 Q1CHQ1 Q1CHQ7 Q1CHR0 Q1CHR3 Q1CIC7 Q1CIH6 Q1CIH9 Q1CII0
Q1CII1 Q1CIK5 Q1CIP4 Q1CIY7 Q1CIZ3 Q1CJ34 Q1CJ37 Q1CJ46 Q1CJ89 Q1CJ90 Q1CJF4
Q1CJF5 Q1CJI3 Q1CJV0 Q1CJV8 Q1CJW8 Q1CK20 Q1CK22 Q1CK61 Q1CK62 Q1CK92 Q1CKH1
Q1CKH2 Q1CL75 Q1CLD8 Q1CLD9 Q1CLE0 Q1CLR7 Q1CLU6 Q1CMD9 Q1CME1 Q1CMJ5 Q1CMN0
Q1CMU0 Q1CMV0 Q1CMV5 Q1CMZ5 Q1CN16 Q1CN17 Q1CN43 Q1CNB0 Q1CNB2 Q1CNC7 Q1CNC8
Q1CNJ0 Q1CNP0 Q1CNR7 Q1CNS3 Q1CNU1 Q1CR54 Q1CRF2 Q1CRZ1 Q1CRZ4 Q1CS38 Q1CS52
Q1CS80 Q1CSD1 Q1CSR6 Q1CT41 Q1CTB0 Q1CU36 Q1CUB8 Q1CUI2 Q1CUX2 Q1CV80 Q1CY86
Q1CZI3 Q1D0A0 Q1D0S7 Q1D1K4 Q1D445 Q1D491 Q1D5J5 Q1D8S2 Q1DB00 Q1DCA9 Q1DDD0
Q1DDD3 Q1DF98 Q1DFA0 Q1DFF8 Q1DFX4 Q1DH70 Q1DHX9 Q1DJR8 Q1DK03 Q1DKX4 Q1DL22
Q1DLC7 Q1DME3 Q1DNC5 Q1DNE5 Q1DPX9 Q1DRJ3 Q1DTI4 Q1DWN4 Q1DWP9 Q1DXN1 Q1DXR6
Q1DZE0 Q1DZS0 Q1E1E0 Q1E2D2 Q1E432 Q1E4N0 Q1E4T3 Q1E5A9 Q1E6Q3 Q1E7G6 Q1E7P0
Q1E8Z0 Q1E9R5 Q1E9X9 Q1EBV7 Q1EC69 Q1ECT8 Q1ECW7 Q1EG05 Q1EG09 Q1EHB4 Q1EPG6
Q1EPG7 Q1ERH8 Q1ERP8 Q1G1A2 Q1G3K7 Q1G3L1 Q1G8W8 Q1GAT4 Q1GAT7 Q1GAW9 Q1GB06
Q1GB07 Q1GB50 Q1GB63 Q1GBZ5 Q1GCR3 Q1GD14 Q1GDE1 Q1GDE2 Q1GDE3 Q1GDE9 Q1GE49
Q1GE51 Q1GF14 Q1GFN7 Q1GGZ8 Q1GHK9 Q1GIN0 Q1GJX6 Q1GJZ3 Q1GMR9 Q1GNB5 Q1GQK7
Q1GR76 Q1GRB2 Q1GRX6 Q1GSZ2 Q1GSZ7 Q1GT53 Q1GTA5 Q1GTK6 Q1GTK9 Q1GTL2 Q1GU13
Q1GU76 Q1GU79 Q1GUG3 Q1GVA0 Q1GVK1 Q1GWK6 Q1GXB3 Q1GXL6 Q1GXM6 Q1GY68 Q1GY95
Q1GYH8 Q1GYU3 Q1GYZ8 Q1GZI0 Q1GZL6 Q1H012 Q1H042 Q1H058 Q1H1K6 Q1H1S8 Q1H1Z8
Q1H2C8 Q1H3M0 Q1H432 Q1H4H7 Q1H5C9 Q1H5D2 Q1H5H1 Q1H8Y8 Q1HAQ0 Q1HCL0 Q1HDT2
Q1HDT3 Q1HG43 Q1HG44 Q1HK43 Q1HK79 Q1HK80 Q1HK97 Q1HKA1 Q1HKB1 Q1HQF8 Q1HR20
Q1HRS5 Q1HRV8 Q1HUU1 Q1HUV6 Q1HUW0 Q1HUX2 Q1HUX8 Q1HUY4 Q1HUZ6 Q1HUZ9 Q1HV02
Q1HV05 Q1HV11 Q1HV14 Q1HV23 Q1HV29 Q1HV35 Q1HV38 Q1HV41 Q1HV53 Q1HV56 Q1HV59
Q1HVB3 Q1HVD2 Q1HVE9 Q1HVG2 Q1HVH3 Q1HVJ2 Q1I2H4 Q1I2I1 Q1I2I2 Q1I2I3 Q1I2L0
Q1I2R3 Q1I3C6 Q1I3R6 Q1I3S8 Q1I4K7 Q1I4R8 Q1I5B5 Q1I6D2 Q1I6X0 Q1I788 Q1I7I9
Q1I7N9 Q1I7P1 Q1I7Z1 Q1I7Z4 Q1I800 Q1I836 Q1I8U1 Q1I966 Q1I9H0 Q1IB51 Q1IB53

Q1IBG0 Q1ICK1 Q1ID18 Q1IDJ5 Q1IDN5 Q1IEP0 Q1IEP6 Q1IF21 Q1IG30 Q1IG71 Q1IGD6
Q1IGS0 Q1IGX0 Q1IGZ4 Q1IIG4 Q1IJ81 Q1IKG7 Q1IKZ6 Q1IN69 Q1INF0 Q1IP20 Q1IPI1
Q1IPR1 Q1IPW7 Q1IS40 Q1IS42 Q1IS45 Q1IS49 Q1IS52 Q1IS55 Q1IS58 Q1IUD3 Q1IUD5
Q1IVE8 Q1IVE9 Q1IXF1 Q1IY26 Q1IZI4 Q1IZW3 Q1IZW6 Q1J1C3 Q1J1H5 Q1J219 Q1J232
Q1J2Y8 Q1J3F4 Q1J5G1 Q1J5I5 Q1J5X4 Q1J722 Q1J798 Q1J7G3 Q1J7G4 Q1J7I7 Q1J7Q5
Q1J8B6 Q1J8D6 Q1J8E1 Q1J8H6 Q1JAG8 Q1JAJ3 Q1JC74 Q1JCF3 Q1JCL7 Q1JCL8 Q1JCP0
Q1JCV9 Q1JDD8 Q1JDG4 Q1JDJ9 Q1JFL1 Q1JFN6 Q1JG12 Q1JG56 Q1JHA3 Q1JHH8 Q1JHN9
Q1JHP0 Q1JHQ8 Q1JHX9 Q1JIG2 Q1JII1 Q1JII6 Q1JIM0 Q1JKM0 Q1JKP5 Q1JL33 Q1JM58
Q1JMD2 Q1JMJ3 Q1JMJ4 Q1JML7 Q1JMT5 Q1JNB1 Q1JND2 Q1JND7 Q1JNH1 Q1JP63 Q1JPA3
Q1JPD2 Q1JPD8 Q1JPG0 Q1JPM5 Q1JPS6 Q1JPY5 Q1JQ66 Q1JQA4 Q1JQB3 Q1JQC1 Q1JQC2
Q1JQC5 Q1JQE1 Q1JQE6 Q1JRP2 Q1JUP6 Q1JUZ1 Q1JUZ2 Q1JV40 Q1K528 Q1K722 Q1K7M0
Q1K9B6 Q1K9Q1 Q1KKR9 Q1KKT4 Q1KKV8 Q1KL86 Q1KLR6 Q1KPV0 Q1KTB0 Q1KVQ9 Q1KVR5
Q1KVR7 Q1KVS1 Q1KVS5 Q1KVS7 Q1KVS8 Q1KVT2 Q1KVU1 Q1KVU2 Q1KVU8 Q1KVV2 Q1KVV3
Q1KVV4 Q1KVV5 Q1KVW0 Q1KVW3 Q1KVW6 Q1KVW7 Q1KVW8 Q1KVW9 Q1KVX1 Q1KVX2 Q1KVX3
Q1KVY0 Q1KVY1 Q1KVY2 Q1KVY3 Q1KXG6 Q1KXH4 Q1KXK8 Q1KXQ2 Q1KXQ3 Q1KXQ5 Q1KXQ6
Q1KXQ8 Q1KXR1 Q1KXS8 Q1KXS9 Q1KXT0 Q1KXT1 Q1KXT2 Q1KXT3 Q1KXT9 Q1KXU0 Q1KXU1
Q1KXU2 Q1KXU3 Q1KXU4 Q1KXU5 Q1KXU6 Q1KXU7 Q1KXU8 Q1KXU9 Q1KXV4 Q1KXV9 Q1KXW0
Q1KXW2 Q1KXW3 Q1KXW4 Q1KXW6 Q1KXW7 Q1KXW8 Q1KXX3 Q1KXX4 Q1KXX5 Q1KXX6 Q1KXX9
Q1KZ54 Q1KZ56 Q1KZG0 Q1L864 Q1L867 Q1L8D2 Q1L8F4 Q1L8L6 Q1L8X9 Q1L911 Q1L990
Q1LFB1 Q1LG26 Q1LH91 Q1LHK6 Q1LIB7 Q1LIB8 Q1LIC3 Q1LIM3 Q1LIV1 Q1LJ64 Q1LJK1
Q1LJU8 Q1LLA9 Q1LLY6 Q1LM81 Q1LNJ0 Q1LP09 Q1LPI4 Q1LPV3 Q1LPV6 Q1LQD3 Q1LR23
Q1LRS0 Q1LRW2 Q1LS97 Q1LSL8 Q1LSU1 Q1LT89 Q1LT95 Q1LT98 Q1LTA1 Q1LTF8 Q1LTJ4
Q1LTM2 Q1LTU8 Q1LTU9 Q1LTV0 Q1LU19 Q1LU52 Q1LUD1 Q1LUK1 Q1LUQ4 Q1LV46 Q1LVN1
Q1LVS8 Q1LVW0 Q1LVZ2 Q1LW89 Q1LWC2 Q1LWG4 Q1LX59 Q1LX78 Q1LXV8 Q1LY46 Q1LY80
Q1LYG4 Q1LZ71 Q1LZ86 Q1LZA0 Q1LZA4 Q1LZB3 Q1LZB5 Q1LZD0 Q1LZD1 Q1LZE6 Q1LZF7
Q1LZF8 Q1LZI1 Q1LZI2 Q1M0V6 Q1M1A0 Q1M2P4 Q1M651 Q1M799 Q1M7A2 Q1M8M5 Q1MA32
Q1MA49 Q1MAB5 Q1MAC8 Q1MAL3 Q1MBJ9 Q1MDR5 Q1ME30 Q1MEX2 Q1MFK3 Q1MFY9 Q1MG67
Q1MGK1 Q1MGQ9 Q1MHM4 Q1MHM5 Q1MIH8 Q1MIK4 Q1MIK7 Q1MIM3 Q1MIP9 Q1MJ35 Q1MJ94
Q1MJ95 Q1MJ96 Q1MJ97 Q1MKI3 Q1MKI5 Q1MKS9 Q1MKT0 Q1MKT2 Q1MKT5 Q1MLH5 Q1MLQ8
Q1MM58 Q1MMI4 Q1MPC2 Q1MPF3 Q1MPG5 Q1MPG6 Q1MQ71 Q1MQW6 Q1MRX2 Q1MTD4 Q1MTQ5
Q1MTR5 Q1MTR7 Q1MTR8 Q1MWD9 Q1MWF2 Q1MWG5 Q1MWH4 Q1MWH8 Q1MWJ1 Q1MWK0 Q1MWK4
Q1MX30 Q1PCQ8 Q1PD50 Q1PDC7 Q1PDI1 Q1PDW5 Q1PDX9 Q1PE15 Q1PE39 Q1PE48 Q1PEH6
Q1PEM5 Q1PEN0 Q1PER9 Q1PET6 Q1PEX8 Q1PFB8 Q1PFD9 Q1PFG9 Q1PFJ4 Q1PG50 Q1PG51
Q1PG54 Q1PG55 Q1PG56 Q1PG57 Q1PG58 Q1PS23 Q1PUD6 Q1PUD9 Q1Q7Z7 Q1Q7Z8 Q1Q851
Q1Q893 Q1Q895 Q1Q8C2 Q1Q8V3 Q1Q8W7 Q1Q993 Q1Q996 Q1QAM6 Q1QB31 Q1QB33 Q1QBW0
Q1QD88 Q1QD91 Q1QD97 Q1QDA8 Q1QDU6 Q1QE84 Q1QEI0 Q1QES0 Q1QES7 Q1QEW2 Q1QF31
Q1QH37 Q1QH68 Q1QHV7 Q1QHV9 Q1QHY7 Q1QIP9 Q1QIW5 Q1QJ48 Q1QJT0 Q1QKJ7 Q1QL21
Q1QL44 Q1QL92 Q1QL95 Q1QLU4 Q1QME4 Q1QME5 Q1QN78 Q1QND7 Q1QNU6 Q1QQI5 Q1QR53
Q1QRH8 Q1QRI0 Q1QRI1 Q1QRL0 Q1QRN0 Q1QRR2 Q1QSC4 Q1QSC6 Q1QSG2 Q1QST2 Q1QST8
Q1QSU1 Q1QSU4 Q1QTI8 Q1QTW5 Q1QU58 Q1QU84 Q1QUC2 Q1QUD8 Q1QVG4 Q1QWB1 Q1QX69
Q1QX82 Q1QX85 Q1QXV6 Q1QYT5 Q1QYX9 Q1QYZ3 Q1QZ83 Q1QZ92 Q1QZP1 Q1R003 Q1R004
Q1R175 Q1R268 Q1R2E2 Q1R2E6 Q1R2G9 Q1R3A5 Q1R3A6 Q1R3C4 Q1R3H8 Q1R3J9 Q1R3P6
Q1R3Q0 Q1R3Q5 Q1R3U1 Q1R412 Q1R432 Q1R473 Q1R475 Q1R4E2 Q1R4I5 Q1R4J4 Q1R4J5
Q1R4J6 Q1R4L6 Q1R4L9 Q1R4M4 Q1R4M9 Q1R4Q2 Q1R4R9 Q1R575 Q1R5B5 Q1R5C2 Q1R5H6

Q1R5H7 Q1R5L1 Q1R5R8 Q1R5T2 Q1R642 Q1R697 Q1R698 Q1R699 Q1R6C6 Q1R6L7 Q1R6P1
Q1R6R8 Q1R6S2 Q1R6S4 Q1R6U2 Q1R7H5 Q1R7H6 Q1R7I5 Q1R7U3 Q1R8F4 Q1R8L7 Q1R8Q5
Q1R8U3 Q1R8V8 Q1R8W0 Q1R8X4 Q1R8X7 Q1R9C0 Q1R9C9 Q1R9D5 Q1R9D8 Q1R9E1 Q1R9F6
Q1R9F7 Q1R9F8 Q1R9G1 Q1R9M2 Q1R9U6 Q1R9W6 Q1R9Z4 Q1R9Z5 Q1R9Z6 Q1RAH0 Q1RAI4
Q1RAM8 Q1RAV8 Q1RAW2 Q1RAW9 Q1RAX0 Q1RAZ2 Q1RB03 Q1RB04 Q1RB84 Q1RBD2 Q1RBG4
Q1RBG6 Q1RBG9 Q1RBJ8 Q1RBJ9 Q1RBL7 Q1RBN8 Q1RBN9 Q1RBS2 Q1RC23 Q1RC43 Q1RCH9
Q1RCI0 Q1RCI7 Q1RCQ5 Q1RCR0 Q1RCR1 Q1RD92 Q1RDA5 Q1RDB2 Q1RDK0 Q1RDU4 Q1RDV7
Q1RE44 Q1RE78 Q1RE93 Q1RE94 Q1REC0 Q1REL9 Q1REM0 Q1RET6 Q1REX1 Q1RFB2 Q1RG12
Q1RG33 Q1RGA8 Q1RGF1 Q1RGF8 Q1RGH5 Q1RGI2 Q1RGL5 Q1RGM3 Q1RGP0 Q1RGS8 Q1RGT7
Q1RGU3 Q1RGU8 Q1RGZ0 Q1RGZ2 Q1RGZ3 Q1RH18 Q1RH19 Q1RH20 Q1RH21 Q1RH22 Q1RH27
Q1RH99 Q1RHA1 Q1RHB5 Q1RHB6 Q1RHC0 Q1RHC4 Q1RHH9 Q1RHK8 Q1RHP1 Q1RHU3 Q1RHX3
Q1RHX7 Q1RI01 Q1RI42 Q1RI44 Q1RI47 Q1RI77 Q1RI79 Q1RI93 Q1RIA4 Q1RIA5 Q1RIF2
Q1RIG3 Q1RII7 Q1RIL0 Q1RIL2 Q1RIN2 Q1RIN3 Q1RIN5 Q1RJ56 Q1RJ91 Q1RJB3 Q1RJI4
Q1RJI5 Q1RJM4 Q1RJQ9 Q1RJX8 Q1RK79 Q1RK92 Q1RKC5 Q1RKD1 Q1RKE5 Q1RKE6 Q1RKE7
Q1RKE8 Q1RKF1 Q1RKF6 Q1RKL1 Q1RKL2 Q1RKL9 Q1RKM2 Q1RKM3 Q1RLK6 Q1RLQ5 Q1RLT2
Q1RLU8 Q1RLY6 Q1RMB5 Q1RMH3 Q1RMH4 Q1RMI1 Q1RMK9 Q1RMP9 Q1RMQ3 Q1RMS4 Q1RMT2
Q1RMT9 Q1RMU4 Q1RMU8 Q1RMV9 Q1RMW2 Q1RPP5 Q1RPR6 Q1T7C2 Q1W0Y2 Q1W3A5 Q1W675
Q1WDN2 Q1WER1 Q1WIM1 Q1WIM2 Q1WIM3 Q1WLP9 Q1WM27 Q1WS51 Q1WS52 Q1WSN8 Q1WSX7
Q1WT14 Q1WT98 Q1WTL2 Q1WTS7 Q1WTX7 Q1WUD0 Q1WUI4 Q1WV87 Q1WVR1 Q1X6J6 Q1X6K1
Q1X880 Q1X881 Q1XA12 Q1XA32 Q1XA76 Q1XD92 Q1XD94 Q1XD95 Q1XD96 Q1XD97 Q1XD98
Q1XD99 Q1XDA0 Q1XDA2 Q1XDB7 Q1XDB8 Q1XDC3 Q1XDC4 Q1XDC7 Q1XDD0 Q1XDD1 Q1XDE6
Q1XDE7 Q1XDF9 Q1XDG1 Q1XDG2 Q1XDG3 Q1XDG4 Q1XDG5 Q1XDG9 Q1XDH0 Q1XDH1 Q1XDJ1
Q1XDK3 Q1XDK4 Q1XDL0 Q1XDL1 Q1XDL2 Q1XDM2 Q1XDM3 Q1XDP0 Q1XDP1 Q1XDP2 Q1XDP3
Q1XDP8 Q1XDQ1 Q1XDQ5 Q1XDR5 Q1XDR6 Q1XDR7 Q1XDS5 Q1XDT1 Q1XDU0 Q1XDU3 Q1XDU4
Q1XF11 Q1XHX8 Q1XI86 Q1XII1 Q1XII7 Q1XII9 Q1XIJ0 Q1XIJ4 Q1XIK7 Q1XIK9 Q1XIL4
Q1XIM0 Q1XIM1 Q1XIM5 Q1XIM8 Q1XIN0 Q1XIN1 Q1XIN6 Q1XIP1 Q1XIP3 Q1XIP4 Q1XIP7
Q1XIP8 Q1XIQ2 Q1XIQ3 Q1XIQ4 Q1ZXA1 Q1ZXA4 Q1ZXA7 Q1ZXA9 Q1ZXB0 Q1ZXC4 Q1ZXE4
Q1ZXF3 Q1ZXF5 Q1ZXG3 Q1ZXG6 Q1ZXH4 Q1ZXI7 Q1ZXI8 Q1ZXI9 Q1ZXJ0 Q1ZXL2 Q1ZXL7
Q1ZXN4 Q1ZXQ5 Q1ZXQ7 Q1ZY22 Q1ZYR0 Q1ZYR2 Q1ZZH0 Q1ZZH1 Q20024 Q20026 Q20027
Q20035 Q20086 Q20106 Q20123 Q20144 Q20157 Q20249 Q20263 Q202B1 Q20300 Q20303
Q20332 Q20405 Q20410 Q20411 Q20432 Q20435 Q20516 Q20574 Q20591 Q20593 Q20605
Q20618 Q20657 Q20687 Q20696 Q20735 Q20745 Q20748 Q20762 Q20779 Q20787 Q20797
Q20798 Q20799 Q20800 Q20806 Q20826 Q20924 Q20929 Q20961 Q20963 Q20BM9 Q20CR3
Q20ET2 Q20ET5 Q20ET6 Q20EU1 Q20EU3 Q20EU6 Q20EU7 Q20EU8 Q20EV3 Q20EV6 Q20EV7
Q20EV8 Q20EW2 Q20EW3 Q20EW4 Q20EW5 Q20EW6 Q20EW7 Q20EX4 Q20EX5 Q20EX6 Q20EY1
Q20EY3 Q20EY4 Q20EZ8 Q20EZ9 Q20F15 Q20F18 Q20F20 Q20F21 Q20F22 Q20FQ1 Q20FQ2
Q20FQ7 Q20FR3 Q20FR8 Q20FS0 Q20JQ7 Q20K40 Q20K43 Q20NP0 Q20NV8 Q20NW0 Q20P03
Q20PM2 Q20WZ8 Q20X00 Q20X01 Q20X23 Q20X25 Q20XR6 Q20Z38 Q20Z39 Q20Z45 Q20Z48
Q21005 Q21038 Q21041 Q21054 Q21121 Q21123 Q21153 Q21154 Q21190 Q211W6 Q21219
Q21281 Q21286 Q21339 Q21353 Q21389 Q213M9 Q21415 Q21433 Q21434 Q21455 Q214E7
Q214E8 Q214F0 Q214K6 Q214X1 Q214X2 Q21534 Q21540 Q21565 Q215H3 Q215M7 Q215Y7
Q21629 Q21642 Q216E1 Q216N0 Q21706 Q21751 Q21752 Q21767 Q21795 Q217L2 Q217M1
Q21802 Q21808 Q21815 Q21828 Q21874 Q21898 Q21973 Q21974 Q21997 Q21C73 Q21C99
Q21DG0 Q21DK2 Q21DK4 Q21DY7 Q21EC5 Q21EN9 Q21FI6 Q21FL0 Q21FP7 Q21FP8 Q21II7

Q21J49 Q21JJ9 Q21JR8 Q21JV7 Q21K23 Q21LC1 Q21MB6 Q21MH0 Q21MH2 Q21N99 Q21NS8
Q21P45 Q21PS1 Q21S73 Q21ST3 Q21SW6 Q21U51 Q21U85 Q21U86 Q21UZ2 Q21V77 Q21VA4
Q21VE7 Q21VT9 Q21WM5 Q21WN9 Q21XA2 Q21XE6 Q21XT4 Q21Y62 Q21YB7 Q21YC0 Q21YC7
Q21YI0 Q21ZA0 Q21ZA2 Q220N9 Q22141 Q22165 Q22169 Q22180 Q22181 Q22252 Q22267
Q22271 Q22295 Q22329 Q22361 Q22397 Q223D2 Q223K4 Q22472 Q22495 Q22505 Q22523
Q22549 Q22566 Q22638 Q22647 Q22682 Q22700 Q22701 Q22702 Q22747 Q22851 Q22931
Q22949 Q22957 Q23022 Q23027 Q23064 Q23125 Q23157 Q23221 Q23310 Q23316 Q23361
Q23369 Q23387 Q23424 Q23435 Q23444 Q23469 Q23497 Q23529 Q23534 Q23544 Q23593
Q23681 Q23682 Q23808 Q23868 Q24025 Q24046 Q24048 Q24093 Q24139 Q24157 Q24188
Q24238 Q24247 Q24270 Q24278 Q24292 Q24298 Q24319 Q24322 Q24332 Q24342 Q24352
Q24418 Q24451 Q24488 Q24498 Q24509 Q24547 Q24563 Q24738 Q24739 Q24760 Q24808
Q24810 Q24JN5 Q24JP1 Q24JP5 Q24JQ0 Q24JY6 Q24JY7 Q24MN7 Q24NH4 Q24PU6 Q24SP9
Q24TE3 Q24TI7 Q24UB5 Q24UB9 Q24UC2 Q24V86 Q24VA3 Q24VN7 Q24VQ5 Q24VQ6 Q24W62
Q24WB0 Q24WQ6 Q24X23 Q24XD0 Q24Y14 Q24Y16 Q24Y92 Q25011 Q25074 Q25147 Q25157
Q25158 Q25188 Q25190 Q25197 Q25263 Q25321 Q25322 Q25338 Q253G9 Q253X2 Q253X4
Q25410 Q25414 Q25452 Q25479 Q25489 Q25540 Q255H9 Q255I1 Q255W8 Q255Y6 Q25605
Q25AG2 Q25AG3 Q25BG1 Q25BG3 Q25BG7 Q25BH0 Q25BH1 Q25BH2 Q25BH5 Q25BH8 Q25BI2
Q25BI5 Q25CE8 Q25CF7 Q26250 Q26261 Q26365 Q26418 Q26419 Q26420 Q26421 Q26473
Q26495 Q26499 Q26614 Q26627 Q26710 Q26721 Q26896 Q26972 Q27002 Q27003 Q27115
Q27151 Q27218 Q27236 Q27238 Q27256 Q27257 Q27295 Q27324 Q27333 Q27367 Q27433
Q27533 Q27559 Q27591 Q27597 Q27675 Q27874 Q27946 Q27960 Q27963 Q27977 Q27979
Q27981 Q27987 Q27994 Q27GI3 Q27HK4 Q27ID8 Q27RZ6 Q27S35 Q27S45 Q27S50 Q27S51
Q27S64 Q27YE4 Q27YG9 Q28001 Q28003 Q28005 Q28030 Q28031 Q28036 Q28039 Q28041
Q28043 Q28044 Q28053 Q28056 Q28060 Q28067 Q28071 Q28072 Q28073 Q28074 Q28105
Q28110 Q28113 Q28125 Q28132 Q28139 Q28142 Q28143 Q28146 Q28172 Q28173 Q28181
Q28193 Q28203 Q28204 Q28235 Q28250 Q28256 Q28260 Q28265 Q28266 Q28269 Q28270
Q28279 Q28280 Q28284 Q28293 Q28296 Q28298 Q28309 Q28317 Q28362 Q28371 Q28422
Q28426 Q28427 Q28433 Q28446 Q28468 Q28474 Q28475 Q28478 Q28481 Q28483 Q28487
Q28505 Q28509 Q28519 Q28524 Q28527 Q28550 Q28553 Q28558 Q28560 Q28565 Q28575
Q28585 Q28589 Q28596 Q28597 Q28602 Q28610 Q28611 Q28612 Q28614 Q28615 Q28616
Q28620 Q28623 Q28634 Q28635 Q28642 Q28644 Q28645 Q28660 Q28677 Q28685 Q28686
Q28689 Q28691 Q28705 Q28709 Q28718 Q28722 Q28727 Q28728 Q28730 Q28735 Q28738
Q28740 Q28748 Q28756 Q28757 Q28768 Q28793 Q287T7 Q28806 Q28807 Q28808 Q28812
Q28813 Q28814 Q28820 Q28832 Q28838 Q28849 Q28851 Q28886 Q28889 Q28891 Q28892
Q288H1 Q288H2 Q288H4 Q288H5 Q288Z5 Q28905 Q28913 Q28914 Q28915 Q28927 Q28928
Q28929 Q28942 Q28983 Q28992 Q28997 Q28998 Q289M4 Q289M6 Q289M7 Q28BP2 Q28BQ6
Q28C41 Q28C60 Q28C88 Q28CE7 Q28CF8 Q28CH8 Q28CL4 Q28CV2 Q28CY9 Q28D16 Q28DA8
Q28DF1 Q28DG6 Q28DI5 Q28DS3 Q28DT3 Q28DX1 Q28E01 Q28E13 Q28ED6 Q28EH9 Q28EM2
Q28EM8 Q28ES4 Q28EW0 Q28EX7 Q28F21 Q28F36 Q28FA9 Q28FF3 Q28FG4 Q28FL7 Q28FY5
Q28GF4 Q28GF5 Q28GF8 Q28GL2 Q28GL3 Q28GR4 Q28H54 Q28H62 Q28HE5 Q28HF6 Q28HF8
Q28HR4 Q28HV5 Q28HW9 Q28I33 Q28I38 Q28I47 Q28I80 Q28IK8 Q28IL7 Q28IU1 Q28J34
Q28J44 Q28KD9 Q28L40 Q28LC5 Q28MJ1 Q28NN1 Q28NP3 Q28PL3 Q28PW0 Q28T49 Q28T60
Q28UC5 Q28UC6 Q28UC7 Q28UQ8 Q28US2 Q28VR9 Q28VU8 Q29000 Q29003 Q29005 Q29006
Q29010 Q29026 Q29030 Q29036 Q29037 Q29043 Q290J8 Q290M9 Q290N5 Q290X1 Q29101

Q29102 Q29121 Q29149 Q29154 Q29175 Q291H5 Q291H8 Q29238 Q29243 Q29244 Q29257
Q29259 Q292N4 Q292Q0 Q29380 Q29381 Q29397 Q29422 Q29423 Q29435 Q29441 Q29449
Q29450 Q29466 Q29512 Q29559 Q295A8 Q295B2 Q295N5 Q29608 Q29611 Q29612 Q29626
Q29627 Q296F7 Q296J9 Q297K8 Q297U2 Q298G6 Q298L5 Q298N4 Q298S5 Q29980 Q29983
Q29BL9 Q29BX8 Q29DG0 Q29EY2 Q29FE9 Q29G54 Q29GB8 Q29HG0 Q29I63 Q29IL2 Q29J90
Q29JH0 Q29JP8 Q29JX6 Q29M88 Q29Q28 Q29RH7 Q29RI0 Q29RK8 Q29RM1 Q29RM3 Q29RM6
Q29RN8 Q29RQ9 Q29RR6 Q29RT8 Q29RT9 Q29RU0 Q29RU6 Q29S03 Q29S14 Q29ZQ1 Q2A069
Q2A1A6 Q2A1F7 Q2A1G0 Q2A1H8 Q2A1U9 Q2A1Z8 Q2A3Z1 Q2A485 Q2A4Z5 Q2A564 Q2A5L1
Q2A5M8 Q2A5R2 Q2A5W5 Q2A7B8 Q2A7F4 Q2A865 Q2AAF6 Q2AB83 Q2ABP2 Q2ABP3 Q2AC31
Q2ACH7 Q2EEM0 Q2EES1 Q2EES6 Q2EET2 Q2EEX5 Q2EEX6 Q2EEX7 Q2EEY0 Q2EF88 Q2EG98
Q2EHL7 Q2EHL8 Q2EMT4 Q2EMW0 Q2EN82 Q2EY13 Q2EY14 Q2EY15 Q2F4V2 Q2F7I8 Q2F7J1
Q2F7Z7 Q2FDH6 Q2FDU2 Q2FDU4 Q2FDV0 Q2FDW5 Q2FDW8 Q2FE19 Q2FEB0 Q2FED4 Q2FED6
Q2FEE1 Q2FEF7 Q2FEG1 Q2FEW4 Q2FF18 Q2FF19 Q2FF20 Q2FF36 Q2FF48 Q2FF50 Q2FF88
Q2FFH9 Q2FFI4 Q2FFJ3 Q2FFK6 Q2FFM1 Q2FFM9 Q2FFP9 Q2FFV9 Q2FFZ9 Q2FG20 Q2FGF0
Q2FGW1 Q2FH03 Q2FH04 Q2FH24 Q2FH31 Q2FH37 Q2FH55 Q2FH56 Q2FH82 Q2FH87 Q2FHF2
Q2FHP2 Q2FHQ5 Q2FHU7 Q2FHW4 Q2FHW5 Q2FHX5 Q2FI17 Q2FI18 Q2FI19 Q2FI20 Q2FI55
Q2FI61 Q2FIA6 Q2FIC3 Q2FIC4 Q2FIC5 Q2FIC6 Q2FIC7 Q2FIC8 Q2FIC9 Q2FIN2 Q2FIP5
Q2FIS2 Q2FIT5 Q2FIV6 Q2FIX9 Q2FJ09 Q2FJ10 Q2FJ11 Q2FJ12 Q2FJ13 Q2FJ14 Q2FJ15
Q2FJ60 Q2FJS9 Q2FK07 Q2FK08 Q2FK10 Q2FK46 Q2FK59 Q2FK70 Q2FK73 Q2FKI2 Q2FKN7
Q2FLB3 Q2FLB7 Q2FLB8 Q2FLQ6 Q2FN44 Q2FNZ0 Q2FPH2 Q2FQC0 Q2FRN2 Q2FUR9 Q2FUT1
Q2FUW2 Q2FV54 Q2FV56 Q2FV58 Q2FV64 Q2FV87 Q2FV99 Q2FVD0 Q2FVE8 Q2FVE9 Q2FVH6
Q2FVN1 Q2FVQ8 Q2FVR0 Q2FVS6 Q2FVT1 Q2FW71 Q2FW84 Q2FWG4 Q2FWH7 Q2FWH9 Q2FWI1
Q2FWM7 Q2FWM8 Q2FWY4 Q2FWY7 Q2FX03 Q2FX99 Q2FXE5 Q2FXE6 Q2FXK8 Q2FXZ9 Q2FY80
Q2FYF1 Q2FYJ5 Q2FYJ6 Q2FYM4 Q2FYN3 Q2FYQ5 Q2FYQ6 Q2FYS6 Q2FZ08 Q2FZ79 Q2FZ91
Q2FZ93 Q2FZ95 Q2FZE3 Q2FZE5 Q2FZJ9 Q2FZK0 Q2FZK1 Q2FZK2 Q2FZP2 Q2FZP8 Q2FZT3
Q2FZV1 Q2FZV2 Q2FZV3 Q2FZW3 Q2G026 Q2G044 Q2G061 Q2G093 Q2G0B4 Q2G0D9 Q2G0J5
Q2G140 Q2G184 Q2G185 Q2G188 Q2G1E0 Q2G1F3 Q2G1G5 Q2G1G8 Q2G1N0 Q2G210 Q2G212
Q2G213 Q2G214 Q2G247 Q2G2B9 Q2G2C0 Q2G2E1 Q2G2F4 Q2G2F6 Q2G2F8 Q2G2G3 Q2G2H6
Q2G2H7 Q2G2H8 Q2G2H9 Q2G2I1 Q2G2M2 Q2G2M9 Q2G2R9 Q2G2U1 Q2G2U4 Q2G2U8 Q2G350
Q2G3F4 Q2G4F1 Q2G4Z2 Q2G506 Q2G5Z3 Q2G7B5 Q2G7D3 Q2G870 Q2G8I5 Q2G8J0 Q2G8R3
Q2G991 Q2G998 Q2G9D3 Q2G9D4 Q2G9W3 Q2G9Z8 Q2GBB8 Q2GBJ9 Q2GCS4 Q2GCU8 Q2GD32
Q2GDE4 Q2GDY1 Q2GE12 Q2GE13 Q2GEZ1 Q2GFB2 Q2GFJ2 Q2GG46 Q2GGK6 Q2GHG8 Q2GI73
Q2GIB5 Q2GIS5 Q2GIV1 Q2GJ17 Q2GK09 Q2GKI0 Q2GKQ9 Q2GLK1 Q2GLU7 Q2GMG9 Q2GMW8
Q2GN49 Q2GNR9 Q2GSI6 Q2GSS4 Q2GUT7 Q2GUU0 Q2GXG8 Q2GYB7 Q2H047 Q2H0D3 Q2H0U8
Q2H0X9 Q2H102 Q2H1P3 Q2H1V7 Q2H3I7 Q2H4Y0 Q2H608 Q2H6M8 Q2H740 Q2H7Q1 Q2H9Y1
Q2HA14 Q2HA54 Q2HA88 Q2HAB1 Q2HAR0 Q2HCW8 Q2HDJ0 Q2HDV5 Q2HEW1 Q2HF90 Q2HFE0
Q2HFL6 Q2HG02 Q2HIE9 Q2HIM5 Q2HIW2 Q2HJ10 Q2HJ17 Q2HJ22 Q2HJ40 Q2HJ55 Q2HJ59
Q2HJ61 Q2HJ63 Q2HJ66 Q2HJ69 Q2HJ95 Q2HJ96 Q2HJA4 Q2HJA8 Q2HJB9 Q2HJD1 Q2HJE5
Q2HJE9 Q2HJF8 Q2HJM9 Q2HQL6 Q2HRD4 Q2HRD5 Q2HWD6 Q2HWK7 Q2HXK9 Q2HXL0 Q2HXU8
Q2HZ96 Q2I0M4 Q2I3F2 Q2I3F4 Q2I3F5 Q2I3G3 Q2I3G4 Q2I3G5 Q2I3G6 Q2I3G7 Q2I3G8
Q2I3G9 Q2I3H0 Q2I3H1 Q2I3H2 Q2I3H3 Q2I3H4 Q2IAL6 Q2IBA1 Q2IBA6 Q2IBA8 Q2IBB3
Q2IBC0 Q2IBD0 Q2IBD8 Q2IBE0 Q2IBE4 Q2IBE8 Q2IBF2 Q2IBF6 Q2IBG7 Q2ICQ7 Q2ICR0
Q2IE76 Q2IF90 Q2IG26 Q2IH98 Q2IHA4 Q2IHP4 Q2IHP6 Q2IHR0 Q2IJN7 Q2IKX9 Q2IL11
Q2IL15 Q2IMA5 Q2IPD5 Q2IPE5 Q2IPI8 Q2IQM4 Q2IR91 Q2IR93 Q2IRA3 Q2IRA4 Q2IRA6

Q2IUG5 Q2IVP8 Q2IWG3 Q2IWG4 Q2IWP3 Q2IWV4 Q2IWX6 Q2IX43 Q2IX45 Q2IXD3 Q2IXI5
Q2IXX0 Q2IYH8 Q2IYL1 Q2J0E4 Q2J0E7 Q2J0F3 Q2J0F4 Q2J2D8 Q2J2T4 Q2J4I7 Q2J6B3
Q2J6G0 Q2J6M7 Q2J6M9 Q2J766 Q2J8A4 Q2J8M1 Q2J987 Q2J997 Q2J9S4 Q2J9U7 Q2JAC3
Q2JAK0 Q2JAK2 Q2JAL9 Q2JCG7 Q2JD53 Q2JFL0 Q2JFL3 Q2JHG9 Q2JHK3 Q2JHS8 Q2JI23
Q2JI37 Q2JIF4 Q2JIF6 Q2JIF7 Q2JIF8 Q2JJ02 Q2JJ29 Q2JJ68 Q2JJB0 Q2JJF6 Q2JJG7
Q2JJZ7 Q2JJZ9 Q2JK43 Q2JKT8 Q2JL31 Q2JL39 Q2JL40 Q2JLE7 Q2JLE8 Q2JLE9 Q2JLF0
Q2JLH4 Q2JLN5 Q2JLQ8 Q2JLQ9 Q2JMC1 Q2JN39 Q2JN82 Q2JN84 Q2JNL3 Q2JNP0 Q2JNP5
Q2JP67 Q2JPC2 Q2JPG8 Q2JPJ1 Q2JPK2 Q2JPR2 Q2JPX1 Q2JQ31 Q2JQ89 Q2JQ90 Q2JQK9
Q2JQR6 Q2JR85 Q2JR95 Q2JRH8 Q2JRJ5 Q2JRN5 Q2JRP9 Q2JRR7 Q2JRR9 Q2JRU9 Q2JRV0
Q2JS34 Q2JS35 Q2JS36 Q2JS37 Q2JS99 Q2JSN8 Q2JSV5 Q2JSV7 Q2JSV8 Q2JSV9 Q2JT51
Q2JT70 Q2JT76 Q2JTD6 Q2JTJ2 Q2JTM8 Q2JTM9 Q2JTN9 Q2JTP0 Q2JTT0 Q2JU34 Q2JUG5
Q2JUG6 Q2JUP0 Q2JUP1 Q2JUV4 Q2JUV5 Q2JV01 Q2JV29 Q2JVQ8 Q2JWB3 Q2JWK3 Q2JWU5
Q2JWW3 Q2JXH1 Q2JXK6 Q2JXZ7 Q2JXZ9 Q2JYF0 Q2K2I9 Q2K2T3 Q2K2U9 Q2K330 Q2K342
Q2K390 Q2K3T6 Q2K3V1 Q2K5E1 Q2K5Z8 Q2K6B8 Q2K711 Q2K7G5 Q2K7S4 Q2K819 Q2K969
Q2K970 Q2K9P7 Q2K9R9 Q2K9S2 Q2K9S5 Q2K9W9 Q2KA78 Q2KAC5 Q2KBM0 Q2KBM1 Q2KBP6
Q2KBP7 Q2KBV8 Q2KBV9 Q2KBW1 Q2KBW4 Q2KCN1 Q2KCQ1 Q2KDF4 Q2KEW0 Q2KHK3 Q2KHK6
Q2KHS5 Q2KHT4 Q2KHV2 Q2KHV4 Q2KHV9 Q2KHX3 Q2KHX4 Q2KHY2 Q2KHY8 Q2KHY9 Q2KI08
Q2KI11 Q2KI18 Q2KI29 Q2KI30 Q2KI76 Q2KI80 Q2KI92 Q2KI97 Q2KI99 Q2KIA4 Q2KIB3
Q2KIB5 Q2KIC8 Q2KID7 Q2KIG5 Q2KIG8 Q2KII7 Q2KIJ5 Q2KIK2 Q2KIK3 Q2KIN1 Q2KIN6
Q2KIP5 Q2KIP6 Q2KIP8 Q2KIQ4 Q2KIS9 Q2KIU0 Q2KIV1 Q2KIX3 Q2KIX5 Q2KIY1 Q2KIY2
Q2KJ11 Q2KJ37 Q2KJ53 Q2KJ84 Q2KJ92 Q2KJ98 Q2KJA5 Q2KJC3 Q2KJD2 Q2KJD9 Q2KJG4
Q2KJI2 Q2KJI7 Q2KJJ8 Q2KKX7 Q2KL21 Q2KN22 Q2KNB4 Q2KNB5 Q2KNB7 Q2KNE5 Q2KP58
Q2KTE1 Q2KTI5 Q2KTS6 Q2KTS7 Q2KTX0 Q2KU30 Q2KUU2 Q2KUY6 Q2KUZ4 Q2KUZ9 Q2KVC1
Q2KVS6 Q2KWC8 Q2KX31 Q2KXU0 Q2KYS6 Q2KYU4 Q2KZ03 Q2KZC5 Q2L004 Q2L113 Q2L4I7
Q2L897 Q2L8Y9 Q2L8Z0 Q2L8Z2 Q2L8Z3 Q2L8Z7 Q2L8Z8 Q2L900 Q2L901 Q2L902 Q2L903
Q2L905 Q2L907 Q2L912 Q2L913 Q2L914 Q2L918 Q2L919 Q2L920 Q2L921 Q2L922 Q2L923
Q2L924 Q2L925 Q2L930 Q2L931 Q2L932 Q2L933 Q2L934 Q2L935 Q2L939 Q2L951 Q2L952
Q2L955 Q2L957 Q2L958 Q2L960 Q2L962 Q2LA85 Q2LAM0 Q2LCP6 Q2LCP7 Q2LCP8 Q2LCP9
Q2LCQ6 Q2LCQ8 Q2LCR0 Q2LCR1 Q2LCR2 Q2LCR3 Q2LCR6 Q2LCV6 Q2LD37 Q2LFV4 Q2LK54
Q2LL16 Q2LQZ9 Q2LR51 Q2LRA0 Q2LRB9 Q2LSF9 Q2LTG5 Q2LUI9 Q2LUQ1 Q2LVL0 Q2LVM4
Q2LVX0 Q2LXE8 Q2LY33 Q2LZ37 Q2LZ62 Q2M146 Q2M1K6 Q2M2E3 Q2M2H8 Q2M2S2 Q2M2T3
Q2M2T6 Q2M2T8 Q2M385 Q2M3C6 Q2M3G0 Q2M3M2 Q2M3R5 Q2M3T9 Q2M591 Q2M5U1 Q2MH05
Q2MHH0 Q2MI42 Q2MI45 Q2MI47 Q2MI48 Q2MI51 Q2MI70 Q2MI71 Q2MI72 Q2MI73 Q2MI74
Q2MI75 Q2MI80 Q2MI81 Q2MI82 Q2MI83 Q2MI86 Q2MI87 Q2MI88 Q2MI89 Q2MI90 Q2MI95
Q2MIA0 Q2MIA1 Q2MIA3 Q2MIA4 Q2MIA5 Q2MIA6 Q2MIA7 Q2MIB2 Q2MIB3 Q2MIB4 Q2MIB6
Q2MIB7 Q2MIC0 Q2MIC9 Q2MID2 Q2MID4 Q2MID5 Q2MID8 Q2MIE0 Q2MIF7 Q2MIF8 Q2MIF9
Q2MIG0 Q2MIG1 Q2MIG2 Q2MIG7 Q2MIG8 Q2MIG9 Q2MIH0 Q2MIH1 Q2MIH3 Q2MIH4 Q2MIH5
Q2MIH6 Q2MIH7 Q2MII2 Q2MII7 Q2MII8 Q2MIJ0 Q2MIJ1 Q2MIJ2 Q2MIJ3 Q2MIJ4 Q2MIJ9
Q2MIK0 Q2MIK1 Q2MIK3 Q2MIK4 Q2MIK7 Q2MJ19 Q2MJ20 Q2MJ21 Q2MJQ7 Q2MKA5 Q2MV57
Q2N2B5 Q2N2B8 Q2N2J6 Q2N2J7 Q2N2K0 Q2N2K1 Q2N2K2 Q2N2K3 Q2N2K4 Q2N6M9 Q2N736
Q2N786 Q2N9P2 Q2N9P5 Q2NA73 Q2NA79 Q2NB90 Q2NBS1 Q2NCF2 Q2NCP2 Q2NCZ3 Q2ND29
Q2ND30 Q2ND79 Q2NDA1 Q2NEV8 Q2NGB4 Q2NGM1 Q2NHA4 Q2NHB7 Q2NHI7 Q2NI29 Q2NIA2
Q2NIN5 Q2NJ28 Q2NJF6 Q2NKB3 Q2NKH9 Q2NKR7 Q2NKS0 Q2NKS2 Q2NKU9 Q2NKV5 Q2NKV8
Q2NKY9 Q2NL01 Q2NL17 Q2NL23 Q2NL33 Q2NNR5 Q2NQ90 Q2NQ91 Q2NQ92 Q2NQ98 Q2NQF9

Q2NQH3 Q2NQQ0 Q2NR07 Q2NRH2 Q2NRL5 Q2NRV6 Q2NRV9 Q2NRW0 Q2NRW1 Q2NS78 Q2NS80
Q2NSC1 Q2NSJ5 Q2NSJ9 Q2NSK5 Q2NSK8 Q2NSL1 Q2NSY1 Q2NSZ1 Q2NSZ5 Q2NSZ8 Q2NSZ9
Q2NT00 Q2NT67 Q2NT68 Q2NT72 Q2NTA7 Q2NTB0 Q2NTC6 Q2NTC7 Q2NTD3 Q2NTK5 Q2NTS8
Q2NUA5 Q2NV87 Q2NVC0 Q2NVC1 Q2NVM5 Q2NVV4 Q2NVW1 Q2NVY5 Q2NVY9 Q2NW07 Q2NW73
Q2NW99 Q2NWA9 Q2NWE3 Q2NWE5 Q2NWN7 Q2NWT7 Q2NWT8 Q2NWT9 Q2NX52 Q2NXJ3 Q2NXS0
Q2NXS1 Q2NXY9 Q2NY86 Q2NYG1 Q2NYU0 Q2NZA0 Q2NZB6 Q2NZH8 Q2NZV5 Q2P0I7 Q2P0W2
Q2P0W5 Q2P1G3 Q2P1K9 Q2P2A1 Q2P2S3 Q2P352 Q2P3E7 Q2P3V2 Q2P4J0 Q2P4R3 Q2P4S8
Q2P576 Q2P5A9 Q2P6Q8 Q2P728 Q2P7Q8 Q2P8F5 Q2P8Q0 Q2P8R3 Q2PBL8 Q2PBL9 Q2PBM1
Q2PBM2 Q2PBY6 Q2PCF1 Q2PE60 Q2PG08 Q2PG42 Q2PG45 Q2PG55 Q2PI09 Q2PIK5 Q2PIM1
Q2PIM5 Q2PIY2 Q2PMM9 Q2PMN1 Q2PMN2 Q2PMN4 Q2PMN5 Q2PMN7 Q2PMP0 Q2PMQ4 Q2PMQ5
Q2PMQ6 Q2PMQ7 Q2PMQ8 Q2PMQ9 Q2PMR4 Q2PMR5 Q2PMR6 Q2PMR7 Q2PMR8 Q2PMR9 Q2PMS0
Q2PMS6 Q2PMS7 Q2PMS9 Q2PMT1 Q2PMT6 Q2PMT7 Q2PMT8 Q2PMT9 Q2PMU0 Q2PMU2 Q2PMU3
Q2PMU8 Q2PP52 Q2PS20 Q2PZH4 Q2PZI1 Q2PZL6 Q2Q1V7 Q2Q421 Q2Q426 Q2Q5T5 Q2QD36
Q2QD38 Q2QD39 Q2QD41 Q2QD43 Q2QD63 Q2QD69 Q2QD70 Q2QD75 Q2QD76 Q2QD77 Q2QD78
Q2QD89 Q2QD93 Q2QDA8 Q2QDF6 Q2QEI3 Q2QI47 Q2QKR2 Q2QL34 Q2QL74 Q2QL83 Q2QL86
Q2QL89 Q2QLA3 Q2QLA6 Q2QLA9 Q2QLB4 Q2QLB7 Q2QLC0 Q2QLC5 Q2QLD7 Q2QLE0 Q2QLE5
Q2QLE8 Q2QLF1 Q2QLF9 Q2QLG2 Q2QLG5 Q2QLH0 Q2QLH6 Q2QM69 Q2QMH2 Q2QMX9 Q2QN30
Q2QNE3 Q2QNK7 Q2QNS6 Q2QP53 Q2QQ55 Q2QR07 Q2QRN7 Q2QUC5 Q2QUP1 Q2QV81 Q2QWW7
Q2QWX8 Q2QXM3 Q2QXP0 Q2QXY1 Q2QY07 Q2QY12 Q2QYH7 Q2QYU2 Q2QYW3 Q2QZ14 Q2R041
Q2R0D1 Q2R0D2 Q2R0D4 Q2R224 Q2R237 Q2R2B1 Q2R2D5 Q2R2T4 Q2R2W8 Q2R2Z4 Q2R3K3
Q2R3P9 Q2R435 Q2R4J1 Q2R712 Q2RAP0 Q2RAP4 Q2RAR6 Q2RAS0 Q2RBL8 Q2RCH3 Q2RCH5
Q2RFA3 Q2RFA5 Q2RFW3 Q2RFX3 Q2RFX4 Q2RFX5 Q2RIS7 Q2RIV5 Q2RIZ7 Q2RJ53 Q2RJ58
Q2RJD6 Q2RJR6 Q2RJT7 Q2RJU3 Q2RJU7 Q2RK54 Q2RK82 Q2RK86 Q2RKK7 Q2RKY8 Q2RL34
Q2RL35 Q2RLE0 Q2RM80 Q2RNC2 Q2RPA4 Q2RPA5 Q2RPA6 Q2RPA7 Q2RPB4 Q2RPF8 Q2RQ23
Q2RR23 Q2RSB3 Q2RSB4 Q2RTE2 Q2RTH2 Q2RTH3 Q2RU27 Q2RU33 Q2RUF7 Q2RV87 Q2RV89
Q2RVU1 Q2RVU8 Q2RWQ0 Q2RWU7 Q2RXQ8 Q2RXV4 Q2RY62 Q2RY71 Q2RYF4 Q2RYG3 Q2RYH3
Q2RYV5 Q2RZS8 Q2S012 Q2S0F1 Q2S2H7 Q2S2K9 Q2S2L3 Q2S434 Q2S526 Q2S531 Q2S5P8
Q2S6C2 Q2S6M5 Q2S6N5 Q2S6N7 Q2S8G1 Q2S8V8 Q2S9Y9 Q2S9Z1 Q2S9Z5 Q2SE32 Q2SE53
Q2SF13 Q2SGY2 Q2SIN5 Q2SIP7 Q2SJ43 Q2SJB9 Q2SJQ5 Q2SK65 Q2SKU4 Q2SKU7 Q2SL19
Q2SMN9 Q2SN14 Q2SN17 Q2SN18 Q2SNG4 Q2SNV5 Q2SQ17 Q2SQU8 Q2SRU2 Q2SSZ9 Q2ST38
Q2ST39 Q2STE5 Q2STM0 Q2SUB3 Q2SUB4 Q2SUU5 Q2SUY6 Q2SWD1 Q2SWP0 Q2SWU6 Q2SY47
Q2SYA5 Q2SYD4 Q2SYE1 Q2SZ16 Q2SZD0 Q2SZI4 Q2SZI6 Q2SZM5 Q2SZM8 Q2SZN5 Q2SZS2
Q2SZW0 Q2T0B4 Q2T0H4 Q2T0L4 Q2T0V9 Q2T1F6 Q2T1J2 Q2T2A9 Q2T4B3 Q2T711 Q2T713
Q2T7Q0 Q2T9K0 Q2T9P5 Q2T9Q1 Q2T9R2 Q2T9S6 Q2T9W7 Q2TA01 Q2TA03 Q2TA21 Q2TA24
Q2TA35 Q2TA44 Q2TA57 Q2TA63 Q2TA68 Q2TAA5 Q2TAD5 Q2TAV2 Q2TB54 Q2TBF2 Q2TBG9
Q2TBH5 Q2TBK5 Q2TBM7 Q2TBM9 Q2TBP5 Q2TBQ4 Q2TBQ7 Q2TBS1 Q2TBU2 Q2TBU3 Q2TBX5
Q2TGI5 Q2TGI8 Q2TGJ1 Q2TGJ4 Q2TGK3 Q2TGY1 Q2TGY2 Q2TGY3 Q2TGY4 Q2TGY5 Q2TGY6
Q2THP9 Q2THQ0 Q2THW0 Q2THW7 Q2THW8 Q2THW9 Q2THX0 Q2THX1 Q2TL32 Q2TLY1 Q2TLY2
Q2TLZ1 Q2TLZ2 Q2TLZ3 Q2TLZ4 Q2TLZ5 Q2TQ12 Q2TQ13 Q2TQ14 Q2TQ15 Q2TQ17 Q2TQ18
Q2TQ19 Q2TQ20 Q2TV84 Q2TVN0 Q2TVN1 Q2TVY7 Q2TWP5 Q2TXA2 Q2TXA7 Q2TXB8 Q2TXF2
Q2TXG1 Q2TZ20 Q2U0J6 Q2U0J8 Q2U0K0 Q2U0K1 Q2U0M6 Q2U0S9 Q2U2Y9 Q2U492 Q2U4X9
Q2U5H8 Q2U5V7 Q2U6A4 Q2U7N9 Q2U8Y3 Q2U910 Q2U9A7 Q2U9J2 Q2UAP8 Q2UAZ7 Q2UAZ8
Q2UB01 Q2UB20 Q2UBI2 Q2UBW3 Q2UCW8 Q2UD41 Q2UD74 Q2UDE5 Q2UDH2 Q2UE23 Q2UEH7
Q2UEK5 Q2UEK7 Q2UEK9 Q2UET3 Q2UF96 Q2UFN3 Q2UG70 Q2UGQ3 Q2UH15 Q2UHA7 Q2UJF5

| | | | | | | | | | | |
|---|---|---|---|---|---|---|---|---|---|---|
| Q2UJS0 | Q2UJS7 | Q2UJY4 | Q2UKB8 | Q2UKE6 | Q2ULC6 | Q2ULH4 | Q2UM43 | Q2UMJ2 | Q2UMV7 | Q2UNJ0 |
| Q2UNK6 | Q2UPA8 | Q2UPB0 | Q2UPB8 | Q2UPC0 | Q2UPC1 | Q2UPC4 | Q2UPI1 | Q2UPW4 | Q2UQH4 | Q2URN2 |
| Q2USD9 | Q2USI0 | Q2USJ2 | Q2UTL1 | Q2UTP0 | Q2UTX5 | Q2UU23 | Q2UU67 | Q2UUI0 | Q2UUJ9 | Q2UUK2 |
| Q2UUT6 | Q2UVD8 | Q2UVE0 | Q2UVJ5 | Q2V091 | Q2V0A4 | Q2V2K5 | Q2V2P2 | Q2V2P3 | Q2V2P5 | Q2V2P6 |
| Q2V2P9 | Q2V3H2 | Q2V4B9 | Q2V4F9 | Q2V897 | Q2V898 | Q2V905 | Q2VC90 | Q2VC95 | Q2VEC3 | Q2VEC6 |
| Q2VEC8 | Q2VED0 | Q2VED1 | Q2VED3 | Q2VEE9 | Q2VEF0 | Q2VEF1 | Q2VEF2 | Q2VEF3 | Q2VEF4 | Q2VEF8 |
| Q2VEF9 | Q2VEG0 | Q2VEG1 | Q2VEG2 | Q2VEG3 | Q2VEG4 | Q2VEG5 | Q2VEG6 | Q2VEG7 | Q2VEH9 | Q2VEI0 |
| Q2VEI1 | Q2VEI2 | Q2VEI3 | Q2VEI8 | Q2VEI9 | Q2VEJ2 | Q2VEJ3 | Q2VEJ6 | Q2VJ42 | Q2VJ60 | Q2VL90 |
| Q2VLG4 | Q2VLG6 | Q2VLH6 | Q2VND0 | Q2VND2 | Q2VNF0 | Q2VNF2 | Q2VPW6 | Q2VPW7 | Q2VWP7 | Q2VWP9 |
| Q2VYF4 | Q2VZH2 | Q2W025 | Q2W027 | Q2W028 | Q2W031 | Q2W0H6 | Q2W158 | Q2W1R5 | Q2W252 | Q2W3J1 |
| Q2W3J7 | Q2W403 | Q2W4H6 | Q2W4T8 | Q2W548 | Q2W552 | Q2W612 | Q2W724 | Q2W8J4 | Q2W8K3 | Q2W8K4 |
| Q2W8K5 | Q2W8K7 | Q2W8L4 | Q2W8L6 | Q2W8M7 | Q2W8P3 | Q2W8P5 | Q2W8P6 | Q2W8P9 | Q2W8Q1 | Q2W8Q2 |
| Q2W8Q3 | Q2W8Q4 | Q2W8Q5 | Q2W8Q8 | Q2W8Q9 | Q2W8R0 | Q2W8R4 | Q2W8R5 | Q2W8R7 | Q2W8R8 | Q2W8R9 |
| Q2W8S0 | Q2W905 | Q2W906 | Q2W9D1 | Q2W9S6 | Q2W9Z8 | Q2WEA5 | Q2WED0 | Q2WF71 | Q2WFL8 | Q2WGC5 |
| Q2WGC7 | Q2WGD3 | Q2WGI0 | Q2WGJ8 | Q2WGJ9 | Q2WGK2 | Q2XVR3 | Q2XVR4 | Q2XVR5 | Q2XVR6 | Q2XVR7 |
| Q2XWK0 | Q2XWL3 | Q2Y060 | Q2Y067 | Q2Y0B9 | Q2Y0W8 | Q2Y2L9 | Q2Y2M0 | Q2Y2M3 | Q2Y2P0 | Q2Y5A8 |
| Q2Y5S3 | Q2Y635 | Q2Y642 | Q2Y6P3 | Q2Y767 | Q2Y7B5 | Q2Y8G6 | Q2Y8G8 | Q2Y8Q6 | Q2Y8T1 | Q2Y9Q8 |
| Q2Y9R4 | Q2YA20 | Q2YA85 | Q2YA93 | Q2YA96 | Q2YA99 | Q2YAA5 | Q2YAV4 | Q2YAV5 | Q2YBM4 | Q2YBW1 |
| Q2YC62 | Q2YC65 | Q2YCA7 | Q2YCL2 | Q2YCM2 | Q2YDD3 | Q2YDD4 | Q2YDD6 | Q2YDD9 | Q2YDE3 | Q2YDG0 |
| Q2YDG1 | Q2YDG7 | Q2YDH0 | Q2YDJ2 | Q2YDJ5 | Q2YDL5 | Q2YDM0 | Q2YDM3 | Q2YDM8 | Q2YDN1 | Q2YDR3 |
| Q2YDU8 | Q2YEF9 | Q2YEG0 | Q2YEG2 | Q2YFS1 | Q2YFS2 | Q2YFS3 | Q2YG58 | Q2YHF0 | Q2YHF2 | Q2YHT5 |
| Q2YHT7 | Q2YIG0 | Q2YIJ8 | Q2YIK1 | Q2YIT6 | Q2YIT7 | Q2YJ76 | Q2YJ78 | Q2YJ81 | Q2YJB2 | Q2YJB4 |
| Q2YJK0 | Q2YJK1 | Q2YJP6 | Q2YK65 | Q2YKD6 | Q2YKM0 | Q2YKR6 | Q2YKR7 | Q2YKY9 | Q2YL00 | Q2YLH3 |
| Q2YLU9 | Q2YLW7 | Q2YLY3 | Q2YM68 | Q2YM92 | Q2YMB3 | Q2YMC4 | Q2YMC5 | Q2YMU7 | Q2YMY7 | Q2YMY8 |
| Q2YNB0 | Q2YNF3 | Q2YNF6 | Q2YNK8 | Q2YNM0 | Q2YNM1 | Q2YNY8 | Q2YPJ2 | Q2YQ73 | Q2YQM9 | Q2YQS9 |
| Q2YR68 | Q2YRG7 | Q2YRJ8 | Q2YRP9 | Q2YS64 | Q2YSF9 | Q2YSG7 | Q2YSL2 | Q2YSM6 | Q2YSP8 | Q2YSS1 |
| Q2YST2 | Q2YSV1 | Q2YSV2 | Q2YSV3 | Q2YSV4 | Q2YSV5 | Q2YSV6 | Q2YT06 | Q2YTH2 | Q2YTJ7 | Q2YTK4 |
| Q2YTK5 | Q2YTZ1 | Q2YU13 | Q2YU20 | Q2YU38 | Q2YU56 | Q2YU61 | Q2YU74 | Q2YUD1 | Q2YUH6 | Q2YUH7 |
| Q2YUH8 | Q2YUI9 | Q2YUJ5 | Q2YUJ6 | Q2YUJ7 | Q2YUQ2 | Q2YUZ1 | Q2YV12 | Q2YV55 | Q2YV65 | Q2YV66 |
| Q2YV68 | Q2YVH4 | Q2YWA3 | Q2YWC1 | Q2YWE4 | Q2YWE6 | Q2YWT4 | Q2YWT5 | Q2YWT6 | Q2YWT7 | Q2YWT8 |
| Q2YWT9 | Q2YWU0 | Q2YWW2 | Q2YWW7 | Q2YX06 | Q2YX14 | Q2YX15 | Q2YX16 | Q2YX17 | Q2YX60 | Q2YX69 |
| Q2YX83 | Q2YX91 | Q2YXE0 | Q2YXH2 | Q2YXN1 | Q2YXS4 | Q2YXW4 | Q2YXW9 | Q2YXY7 | Q2YXY8 | Q2YXZ9 |
| Q2YY04 | Q2YY11 | Q2YY17 | Q2YY45 | Q2YY76 | Q2YYH7 | Q2YYX7 | Q2YYX9 | Q2YYZ8 | Q2YZ23 | Q2YZ25 |
| Q2YZ45 | Q2YZ90 | Q2YZA7 | Q30154 | Q30201 | Q304A0 | Q30631 | Q306W5 | Q306W7 | Q307W7 | Q30C86 |
| Q30DW6 | Q30NP8 | Q30NQ0 | Q30NQ1 | Q30NW9 | Q30PI6 | Q30PI9 | Q30PJ2 | Q30PL2 | Q30PM3 | Q30Q39 |
| Q30QW4 | Q30SA7 | Q30ST6 | Q30TB5 | Q30TH1 | Q30UH8 | Q30UT6 | Q30X24 | Q30XM9 | Q30XU9 | Q30XV0 |
| Q30YE7 | Q30YQ5 | Q30ZC2 | Q30ZF6 | Q30ZJ7 | Q310A3 | Q310M2 | Q31125 | Q312S1 | Q313Q6 | Q313V6 |
| Q313W7 | Q31651 | Q31652 | Q31684 | Q31696 | Q31720 | Q31721 | Q31792 | Q317C3 | Q317T2 | Q31849 |
| Q318A9 | Q318B0 | Q318L9 | Q318M0 | Q318T2 | Q318T6 | Q318T7 | Q318T8 | Q318T9 | Q31909 | Q31910 |
| Q31952 | Q319G0 | Q319Y1 | Q319Y5 | Q31A60 | Q31A92 | Q31AA8 | Q31AG7 | Q31B42 | Q31BB2 | Q31BD7 |
| Q31BW4 | Q31BZ4 | Q31C66 | Q31C72 | Q31C73 | Q31C74 | Q31C87 | Q31CA0 | Q31CK2 | Q31CK3 | Q31CL1 |
| Q31CL4 | Q31CN2 | Q31CN3 | Q31CN4 | Q31CN5 | Q31CN8 | Q31CR0 | Q31CS9 | Q31CT0 | Q31CT1 | Q31D21 |
| Q31D31 | Q31DC0 | Q31DL5 | Q31DL6 | Q31DM4 | Q31E60 | Q31E61 | Q31EL5 | Q31EM0 | Q31F55 | Q31FG2 |
| Q31G96 | Q31HC4 | Q31HE7 | Q31HF3 | Q31HL0 | Q31I56 | Q31I60 | Q31I62 | Q31ID1 | Q31JY9 | Q31KG3 |

Q31KZ4 Q31LI2 Q31LJ0 Q31LJ1 Q31LM3 Q31LP8 Q31LR3 Q31LS2 Q31LS7 Q31MA6 Q31MI4
Q31NA0 Q31NJ6 Q31NU0 Q31NV7 Q31NV8 Q31NV9 Q31P91 Q31PJ8 Q31QD8 Q31QE1 Q31QI3
Q31QX6 Q31R12 Q31R74 Q31RF3 Q31RF4 Q31RF5 Q31RF6 Q31RG5 Q31RR1 Q31RR2 Q31RR3
Q31RY5 Q31SX3 Q31T09 Q31T72 Q31T86 Q31T87 Q31TC4 Q31TR5 Q31TS7 Q31TU8 Q31U72
Q31U75 Q31U82 Q31UA7 Q31UF1 Q31UH1 Q31UJ0 Q31UM6 Q31UN6 Q31UN7 Q31US4 Q31UU4
Q31UV9 Q31UW4 Q31V97 Q31VD1 Q31VK9 Q31VS4 Q31VU0 Q31VY6 Q31W84 Q31WA8 Q31WA9
Q31WQ2 Q31WR5 Q31WU4 Q31WX1 Q31WX6 Q31WX8 Q31XB0 Q31XF2 Q31XF9 Q31XQ6 Q31XX4
Q31Y19 Q31Y50 Q31Y66 Q31Y67 Q31Y80 Q31Y82 Q31YG5 Q31YH3 Q31YH9 Q31YI2 Q31YI5
Q31YJ9 Q31YK0 Q31YK3 Q31YT6 Q31Z20 Q31ZA3 Q31ZC0 Q31ZM6 Q31ZM7 Q31ZN1 Q31ZU1
Q31ZU8 Q31ZU9 Q32007 Q32091 Q320A9 Q320D7 Q320K3 Q320K4 Q320T9 Q320V5 Q320V6
Q320Y4 Q320Y7 Q320Y9 Q32126 Q32131 Q321C2 Q321G8 Q321L5 Q321S5 Q321S6 Q321T8
Q321Y6 Q321Z1 Q321Z7 Q32238 Q322B3 Q322L9 Q322R9 Q322V1 Q32384 Q32389 Q323D8
Q323D9 Q323E0 Q323M3 Q323Q4 Q323U7 Q323W2 Q323W3 Q32440 Q324K9 Q324L0 Q324R8
Q324V2 Q32516 Q32539 Q32551 Q325H3 Q325W3 Q325Y4 Q32617 Q32618 Q32643 Q32644
Q326E8 Q326I6 Q326J6 Q326K4 Q32722 Q327N3 Q327T6 Q327U7 Q32880 Q328D2 Q328H2
Q328H3 Q328K4 Q328Y4 Q328Z8 Q32902 Q32904 Q32905 Q329B2 Q329C8 Q329F4 Q329S5
Q329S6 Q329S7 Q329T6 Q329X2 Q329Z2 Q32A13 Q32A56 Q32A72 Q32A79 Q32AE6 Q32AN6
Q32AT4 Q32AT5 Q32AW3 Q32AW5 Q32B02 Q32B97 Q32B98 Q32B99 Q32BC5 Q32BK1 Q32BN2
Q32BQ4 Q32BQ6 Q32BU5 Q32C68 Q32C86 Q32C93 Q32CI2 Q32D46 Q32D97 Q32DC5 Q32DE0
Q32DE1 Q32DF6 Q32DP4 Q32DQ1 Q32DQ7 Q32DR0 Q32DR3 Q32DT0 Q32DT1 Q32DT4 Q32DZ9
Q32E16 Q32E34 Q32E75 Q32E98 Q32ED1 Q32EI1 Q32EK8 Q32EM1 Q32ES6 Q32F28 Q32F32
Q32F38 Q32F39 Q32FA7 Q32FE1 Q32FE3 Q32FE6 Q32FI8 Q32FT6 Q32FU7 Q32FU8 Q32FX5
Q32G29 Q32G46 Q32G65 Q32G66 Q32GI7 Q32GL1 Q32GT9 Q32GU0 Q32GV1 Q32H24 Q32H30
Q32H31 Q32HD9 Q32HJ0 Q32HK6 Q32HZ7 Q32IB7 Q32IB8 Q32IC5 Q32IN4 Q32IV5 Q32IX8
Q32JT2 Q32JV3 Q32JZ9 Q32K05 Q32K09 Q32K57 Q32K70 Q32KA0 Q32KH8 Q32KL9 Q32KN2
Q32KP1 Q32KQ5 Q32KR6 Q32KU6 Q32L10 Q32L53 Q32L57 Q32L59 Q32L60 Q32L65 Q32L77
Q32L83 Q32L85 Q32L86 Q32L94 Q32L97 Q32LC4 Q32LF0 Q32LG5 Q32LJ5 Q32LK2 Q32LM8
Q32LN6 Q32LP4 Q32LQ0 Q32LQ6 Q32LS6 Q32LT7 Q32LU1 Q32M21 Q32M26 Q32M45 Q32MD9
Q32N32 Q32NG5 Q32NI8 Q32NJ7 Q32NM7 Q32NX4 Q32NY4 Q32NZ6 Q32P84 Q32PD8 Q32PD9
Q32PE3 Q32PF0 Q32PG7 Q32PH2 Q32PI9 Q32PK2 Q32PR0 Q32Q07 Q32RF7 Q32RF9 Q32RG3
Q32RG4 Q32RG6 Q32RG7 Q32RG8 Q32RG9 Q32RH5 Q32RH6 Q32RH7 Q32RH9 Q32RI2 Q32RJ3
Q32RJ5 Q32RJ6 Q32RJ7 Q32RJ8 Q32RK0 Q32RK2 Q32RK5 Q32RK9 Q32RL0 Q32RL2 Q32RL7
Q32RL8 Q32RL9 Q32RM1 Q32RM3 Q32RM5 Q32RM6 Q32RP1 Q32RP2 Q32RP3 Q32RP5 Q32RP6
Q32RQ3 Q32RQ4 Q32RQ5 Q32RQ6 Q32RS3 Q32RS4 Q32RS6 Q32RS7 Q32RS9 Q32RT0 Q32RT1
Q32RT2 Q32RT3 Q32RT4 Q32RT5 Q32RT6 Q32RU1 Q32RU2 Q32RU3 Q32RU4 Q32RU5 Q32RU6
Q32RW1 Q32RW2 Q32RW4 Q32RW6 Q32RW7 Q32RW8 Q32RW9 Q32RX1 Q32RX2 Q32RX4 Q32RX5
Q32RX6 Q32RX7 Q32RX8 Q32RY0 Q32RZ0 Q32RZ9 Q32S02 Q32S04 Q32S05 Q32S06 Q32S08
Q32US1 Q32UT4 Q32ZD4 Q32ZD5 Q32ZD7 Q32ZE0 Q32ZE1 Q32ZE2 Q32ZL2 Q32ZZ4 Q32ZZ5
Q32ZZ9 Q33066 Q330A0 Q330A3 Q330A4 Q330A5 Q330A7 Q330A8 Q330B1 Q330B2 Q330B3
Q330B4 Q330C3 Q330C4 Q330D6 Q330D8 Q330E4 Q330E5 Q330E6 Q330E9 Q330F2 Q330G8
Q330G9 Q330H0 Q330H1 Q330H2 Q330H3 Q33113 Q33282 Q332S2 Q332S4 Q332S5 Q332S8
Q332U6 Q332U7 Q332U8 Q332U9 Q332V0 Q332V1 Q332V8 Q332V9 Q332W0 Q332W1 Q332W2
Q332W4 Q332W5 Q332W6 Q332W7 Q332W8 Q332X3 Q332X8 Q332X9 Q332Y1 Q332Y2 Q33300
Q33302 Q33332 Q33375 Q33377 Q333S9 Q33401 Q33439 Q33452 Q33487 Q33500 Q33537

Q33568 Q33607 Q33629 Q33635 Q33636 Q33645 Q33688 Q33697 Q33706 Q33723 Q33752
Q33753 Q33754 Q33756 Q33772 Q33782 Q33784 Q33798 Q337C0 Q33800 Q33817 Q33818
Q33819 Q33820 Q33821 Q33822 Q33823 Q33824 Q33842 Q33845 Q33865 Q33884 Q33887
Q33941 Q33950 Q33954 Q33976 Q33987 Q33989 Q339M6 Q339N5 Q33AH8 Q33BW4 Q33BW7
Q33BW9 Q33BX0 Q33BX2 Q33BX3 Q33BX5 Q33C01 Q33C02 Q33C03 Q33C04 Q33C05 Q33C06
Q33C12 Q33C13 Q33C14 Q33C18 Q33C20 Q33C21 Q33C22 Q33C23 Q33C29 Q33C36 Q33C37
Q33C39 Q33C41 Q33C42 Q33C43 Q33C44 Q33C50 Q33C51 Q33C52 Q33C55 Q33C56 Q33C59
Q33DX9 Q33DY0 Q34028 Q34045 Q34048 Q34049 Q34050 Q34052 Q34070 Q34076 Q34101
Q34106 Q34107 Q34108 Q34161 Q34169 Q34172 Q34179 Q34214 Q34235 Q34253 Q34254
Q34260 Q34261 Q34279 Q34289 Q34300 Q34301 Q34302 Q34306 Q34313 Q34321 Q34340
Q34341 Q34345 Q34376 Q34382 Q34388 Q34391 Q34399 Q34409 Q34459 Q34462 Q34471
Q34473 Q34501 Q34522 Q34534 Q34571 Q34572 Q34573 Q34607 Q34653 Q34677 Q34683
Q34717 Q34724 Q34732 Q34760 Q34799 Q34800 Q34801 Q34814 Q34876 Q34878 Q34879
Q34889 Q34890 Q34891 Q34893 Q34900 Q34902 Q34916 Q34941 Q34942 Q34943 Q34944
Q34945 Q34946 Q34947 Q34948 Q34949 Q34950 Q34951 Q34952 Q34957 Q34972 Q34973
Q35000 Q35019 Q35020 Q35038 Q35054 Q35065 Q35066 Q35075 Q35076 Q35078 Q35099
Q35100 Q35101 Q35111 Q35113 Q35117 Q35118 Q35127 Q35128 Q35130 Q35131 Q35134
Q35135 Q35136 Q35137 Q35138 Q35140 Q35141 Q35142 Q35157 Q35160 Q35172 Q35196
Q35242 Q35262 Q35273 Q35350 Q35377 Q35380 Q35409 Q35416 Q35425 Q35438 Q35457
Q35459 Q35467 Q35468 Q35485 Q35505 Q35506 Q35516 Q35518 Q35522 Q35532 Q35533
Q35534 Q35535 Q35536 Q35537 Q35538 Q35539 Q35540 Q35541 Q35542 Q35543 Q35544
Q35553 Q35558 Q35584 Q35587 Q35588 Q35614 Q35618 Q35621 Q35630 Q35639 Q35647
Q35648 Q35653 Q35673 Q35675 Q35677 Q35695 Q35727 Q35805 Q35810 Q35813 Q35819
Q35826 Q35832 Q35861 Q35873 Q35877 Q35886 Q35895 Q35914 Q35915 Q35916 Q35920
Q35924 Q35925 Q35929 Q35930 Q35942 Q35966 Q35968 Q35994 Q36012 Q36035 Q36055
Q36058 Q36082 Q36089 Q36090 Q36096 Q36097 Q36098 Q36099 Q36118 Q36128 Q36150
Q36151 Q36192 Q36194 Q36201 Q36209 Q36222 Q36227 Q36229 Q36257 Q36258 Q36262
Q36266 Q36284 Q36291 Q36294 Q36309 Q36311 Q36319 Q36324 Q36346 Q36347 Q36362
Q36368 Q36421 Q36422 Q36423 Q36424 Q36425 Q36426 Q36427 Q36428 Q36445 Q36451
Q36452 Q36453 Q36454 Q36455 Q36456 Q36457 Q36458 Q36459 Q36460 Q36461 Q36507
Q36518 Q36548 Q36549 Q36551 Q36572 Q36590 Q36632 Q36644 Q36654 Q36659 Q36664
Q36675 Q36718 Q36724 Q36775 Q36814 Q36832 Q36834 Q36835 Q36836 Q36837 Q36838
Q36852 Q36860 Q36903 Q36918 Q36922 Q36952 Q36964 Q36967 Q36976 Q37006 Q37024
Q37050 Q37064 Q37078 Q37080 Q37108 Q37131 Q37143 Q37145 Q37165 Q37304 Q37311
Q37312 Q37313 Q37314 Q37315 Q37355 Q37369 Q37370 Q37371 Q37372 Q37374 Q37375
Q37376 Q37377 Q37378 Q37379 Q37381 Q37382 Q37385 Q37395 Q37399 Q37403 Q37411
Q37416 Q37419 Q37430 Q37440 Q37472 Q37476 Q37545 Q37546 Q37547 Q37548 Q37550
Q37556 Q37595 Q37596 Q37598 Q37600 Q37601 Q37603 Q37604 Q37605 Q37617 Q37620
Q37625 Q37626 Q37627 Q37643 Q37649 Q37676 Q37677 Q37679 Q37680 Q37684 Q37685
Q37691 Q37695 Q37704 Q37705 Q37706 Q37707 Q37708 Q37709 Q37710 Q37711 Q37712
Q37713 Q37714 Q37717 Q37718 Q37741 Q37751 Q37809 Q37875 Q37895 Q37958 Q37972
Q37975 Q37977 Q38008 Q38065 Q38066 Q38134 Q38623 Q38646 Q38674 Q38798 Q38820
Q38833 Q38846 Q38849 Q38856 Q38860 Q38885 Q38890 Q38898 Q388F1 Q38906 Q38932
Q38944 Q38954 Q38956 Q38967 Q38998 Q38BX0 Q38E53 Q38E54 Q38E55 Q38E56 Q38HS3

Q38IC7 Q38IC8 Q38J84 Q38J85 Q38PR1 Q38PR2 Q38PR3 Q38PR4 Q38PR5 Q38PR6 Q38PR7
Q38PR8 Q38PR9 Q38PS0 Q38PS1 Q38PS2 Q38PU2 Q38PU3 Q38PU4 Q38PU5 Q38PU6 Q38PU7
Q38PU8 Q38Q84 Q38Q87 Q38RU1 Q38RU4 Q38RU7 Q38RV3 Q38RV6 Q38RW5 Q38RW8 Q38RX1
Q38RY0 Q38RY6 Q38RY9 Q38RZ2 Q38RZ7 Q38S00 Q38S02 Q38S05 Q38S08 Q38S11 Q38S14
Q38S17 Q38S20 Q38S23 Q38S26 Q38S29 Q38S35 Q38S38 Q38S41 Q38S47 Q38S50 Q38S53
Q38S56 Q38S59 Q38S62 Q38S65 Q38SQ5 Q38SQ7 Q38SQ8 Q38UX8 Q38V39 Q38V59 Q38VP0
Q38W56 Q38WG4 Q38WK0 Q38WK1 Q38WU8 Q38X02 Q38X31 Q38XN0 Q38YB0 Q38YD8 Q38ZE4
Q38ZE5 Q38ZZ5 Q39002 Q39017 Q39041 Q39065 Q39079 Q39080 Q39085 Q39086 Q39102
Q39128 Q39134 Q39165 Q39191 Q39194 Q39196 Q39202 Q39203 Q39208 Q39219 Q39221
Q39228 Q39231 Q39232 Q39233 Q39253 Q39254 Q39287 Q393T3 Q39444 Q39524 Q39525
Q39593 Q39608 Q39609 Q39654 Q39709 Q39817 Q39994 Q399M3 Q39BQ2 Q39BU7 Q39CK7
Q39CQ5 Q39DI4 Q39DM7 Q39DT1 Q39E73 Q39EE5 Q39EF2 Q39EF5 Q39FB0 Q39FG1 Q39FS2
Q39HK8 Q39HP5 Q39II6 Q39IQ1 Q39IS2 Q39IU8 Q39JF0 Q39JR9 Q39JX3 Q39K79 Q39K80
Q39KY0 Q39N99 Q39PQ8 Q39Q52 Q39QA2 Q39QA3 Q39QA7 Q39QB4 Q39QB7 Q39QP4 Q39QX7
Q39R89 Q39R93 Q39SN1 Q39SW4 Q39SW6 Q39T32 Q39WF6 Q39WP5 Q39YE5 Q39YM0 Q39YM2
Q39YS9 Q39Z26 Q39ZA5 Q39ZB0 Q39ZB6 Q39ZC5 Q39ZN5 Q39ZN6 Q39ZT7 Q3A077 Q3A1R2
Q3A1R4 Q3A1T4 Q3A218 Q3A2G3 Q3A2H0 Q3A2U7 Q3A4S1 Q3A579 Q3A5X3 Q3A602 Q3A603
Q3A6U7 Q3A6V2 Q3A7W6 Q3A7Y9 Q3A815 Q3A818 Q3A821 Q3A826 Q3A8D5 Q3A8L3 Q3A8L4
Q3A931 Q3A941 Q3A942 Q3AAB1 Q3AAB2 Q3AAE3 Q3AAU7 Q3AC77 Q3AC81 Q3AC87 Q3ACP3
Q3ACX1 Q3ACY1 Q3ADS0 Q3AE01 Q3AE25 Q3AFC6 Q3AG75 Q3AGY8 Q3AGZ9 Q3AH25 Q3AH68
Q3AHI9 Q3AHK0 Q3AHK1 Q3AHK3 Q3AHU3 Q3AI50 Q3AJN5 Q3AJZ9 Q3AK05 Q3AK79 Q3AKT2
Q3AKY6 Q3ALM2 Q3ALX5 Q3ALY0 Q3ALY1 Q3ALZ8 Q3AM89 Q3AMC2 Q3AMC3 Q3AMD6 Q3AMS4
Q3AMS5 Q3AMZ3 Q3AMZ4 Q3AMZ6 Q3AN53 Q3AN56 Q3AN57 Q3AN58 Q3AN59 Q3ANG3 Q3ANG4
Q3ANR9 Q3ANV6 Q3ANW4 Q3ANW6 Q3ANZ7 Q3AP02 Q3APS6 Q3APU9 Q3AQC9 Q3AQW9 Q3ARQ6
Q3AS27 Q3ASV8 Q3ASW4 Q3ASW8 Q3ATC0 Q3ATR5 Q3AU90 Q3AUE0 Q3AUQ6 Q3AUQ9 Q3AUT5
Q3AUT6 Q3AUX3 Q3AUY3 Q3AV04 Q3AV15 Q3AVL6 Q3AW25 Q3AWF0 Q3AWF1 Q3AWJ7 Q3AWL5
Q3AWL6 Q3AWL7 Q3AWM2 Q3AWT0 Q3AXK1 Q3AXS1 Q3AXS7 Q3AY07 Q3AYB8 Q3AYG7 Q3AZ35
Q3AZ40 Q3AZD7 Q3AZM3 Q3AZM4 Q3AZM5 Q3AZM6 Q3AZN7 Q3AZZ7 Q3B038 Q3B063 Q3B075
Q3B0C3 Q3B0C6 Q3B0C7 Q3B0C8 Q3B0C9 Q3B0L7 Q3B0L8 Q3B110 Q3B126 Q3B141 Q3B143
Q3B181 Q3B199 Q3B1C1 Q3B2G6 Q3B399 Q3B3W2 Q3B400 Q3B402 Q3B410 Q3B4G1 Q3B4V9
Q3B4W3 Q3B4Y9 Q3B5J7 Q3B5L1 Q3B5P9 Q3B6R3 Q3B724 Q3B7D2 Q3B7D3 Q3B7N4 Q3B7S5
Q3B7T3 Q3B7T6 Q3B7U9 Q3B8G4 Q3B8G7 Q3B8H0 Q3B8H3 Q3B8P0 Q3B8Q3 Q3B8R1 Q3BAH8
Q3BAK6 Q3BAK7 Q3BAK8 Q3BAK9 Q3BAL0 Q3BAL2 Q3BAM0 Q3BAM1 Q3BAM2 Q3BAM3 Q3BAM4
Q3BAM5 Q3BAM6 Q3BAM8 Q3BAM9 Q3BAN1 Q3BAN2 Q3BAP0 Q3BAP1 Q3BAP3 Q3BAP4 Q3BAP5
Q3BAP6 Q3BAP7 Q3BAQ4 Q3BAQ5 Q3BAQ6 Q3BAQ8 Q3BAQ9 Q3BAR4 Q3BBX7 Q3BBX8 Q3BCQ4
Q3BCQ5 Q3BCQ7 Q3BCU0 Q3BCU4 Q3BDI7 Q3BKD5 Q3BLZ7 Q3BMG1 Q3BMG2 Q3BMJ1 Q3BND5
Q3BP11 Q3BPI3 Q3BQ08 Q3BQF3 Q3BQN4 Q3BQP8 Q3BRN6 Q3BRN9 Q3BRP2 Q3BSD6 Q3BSK6
Q3BTC8 Q3BTX1 Q3BUS9 Q3BUW9 Q3BUZ5 Q3BV17 Q3BVS6 Q3BW41 Q3BWV6 Q3BX14 Q3BX88
Q3BXF4 Q3BXV7 Q3BYI6 Q3BYM7 Q3BYT6 Q3BZ34 Q3BZ37 Q3BZ95 Q3C1F9 Q3C1G0 Q3C1G3
Q3C1G4 Q3C1G5 Q3C1H1 Q3C1H2 Q3C1H3 Q3C1H8 Q3C1H9 Q3C1I1 Q3C1I3 Q3C1I4 Q3C1I6
Q3C1I8 Q3C1I9 Q3C1J0 Q3C1J1 Q3C1J7 Q3C1K1 Q3C1K2 Q3C1K3 Q3C1K9 Q3C1L0 Q3C1L1
Q3C1L3 Q3C1M4 Q3C1M5 Q3C1M6 Q3C1N9 Q3C1P6 Q3C1P9 Q3C1Q1 Q3C1Q2 Q3C1Q4 Q3C1Q5
Q3C1V0 Q3C2E2 Q3E6R5 Q3E6T0 Q3E735 Q3E739 Q3E743 Q3E750 Q3E751 Q3E760 Q3E765
Q3E776 Q3E790 Q3E796 Q3E798 Q3E7A3 Q3E7B0 Q3E7B2 Q3E7B6 Q3E7B9 Q3E7D0 Q3E7Q9

Q3E7Z4 Q3E7Z8 Q3E810 Q3E814 Q3E815 Q3E820 Q3E824 Q3E828 Q3E834 Q3E842 Q3E884
Q3E8L0 Q3E8W4 Q3E8X3 Q3E8Z8 Q3E912 Q3E954 Q3E956 Q3E965 Q3E991 Q3E9A0 Q3E9A4
Q3E9B5 Q3E9B8 Q3E9X6 Q3E9Z9 Q3EA54 Q3EAQ5 Q3EAR7 Q3EAV6 Q3EB59 Q3EBC2 Q3EBF7
Q3EBR4 Q3EBY6 Q3EC11 Q3EC77 Q3ECH2 Q3ECI7 Q3ECP7 Q3ECR7 Q3ECT8 Q3ED15 Q3EDD7
Q3EDG3 Q3EDG5 Q3EDL4 Q3I5I9 Q3I5J0 Q3I5J2 Q3I5J3 Q3I5J4 Q3I5J5 Q3IDA4 Q3IDN2
Q3IDQ7 Q3IE16 Q3IE23 Q3IEA1 Q3IG01 Q3IG03 Q3IGX5 Q3IH20 Q3IHM5 Q3IHN8 Q3IHN9
Q3IHV1 Q3IHW5 Q3IHX2 Q3IJ43 Q3IJQ0 Q3IJV4 Q3IK44 Q3IK45 Q3IK46 Q3IK55 Q3IKD8
Q3IKL0 Q3ILE8 Q3INH7 Q3INR7 Q3IRC9 Q3IS61 Q3ITE8 Q3ITN6 Q3IUS6 Q3IUS7 Q3IUT9
Q3IW22 Q3IXC2 Q3IXC6 Q3IXW9 Q3IYD8 Q3IYQ0 Q3IYX7 Q3IYY6 Q3IZ12 Q3IZ13 Q3IZ14
Q3IZ15 Q3IZ47 Q3IZY4 Q3J015 Q3J0F7 Q3J144 Q3J145 Q3J170 Q3J1A3 Q3J1A4 Q3J1A5
Q3J1A6 Q3J1Q3 Q3J278 Q3J3D5 Q3J3E7 Q3J3F1 Q3J4L7 Q3J4M8 Q3J5F7 Q3J5F9 Q3J5J7
Q3J5P3 Q3J661 Q3J6C1 Q3J6L8 Q3J6M6 Q3J6M7 Q3J6P7 Q3J6R6 Q3J6Z7 Q3J786 Q3J7B5
Q3J7R8 Q3J7Z4 Q3J835 Q3J841 Q3J8K3 Q3J8V3 Q3JBS8 Q3JC15 Q3JC21 Q3JC24 Q3JC27
Q3JCH4 Q3JCJ0 Q3JCT0 Q3JDT9 Q3JDU1 Q3JE43 Q3JEE4 Q3JF14 Q3JGG7 Q3JIV3 Q3JL20
Q3JL23 Q3JMH0 Q3JN07 Q3JN08 Q3JND5 Q3JNX1 Q3JPB5 Q3JPL1 Q3JPP0 Q3JR90 Q3JRR7
Q3JTI8 Q3JTV2 Q3JU99 Q3JUA2 Q3JUA9 Q3JUE8 Q3JUI6 Q3JV10 Q3JV70 Q3JVC1 Q3JVZ2
Q3JWF7 Q3JWJ9 Q3JXD9 Q3JXI3 Q3JXV4 Q3JZQ1 Q3JZQ2 Q3JZQ9 Q3JZT4 Q3K059 Q3K0E6
Q3K0S9 Q3K0U5 Q3K107 Q3K1J9 Q3K1K0 Q3K1W8 Q3K2B6 Q3K374 Q3K391 Q3K3N4 Q3K428
Q3K435 Q3K436 Q3K437 Q3K4G3 Q3K4T4 Q3K543 Q3K584 Q3K5S2 Q3K6I0 Q3K6L6 Q3K741
Q3K767 Q3K773 Q3K8D1 Q3K8R2 Q3K8Y4 Q3K933 Q3K9J0 Q3K9V7 Q3KA54 Q3KA57 Q3KA63
Q3KA83 Q3KC07 Q3KCB7 Q3KCB8 Q3KCB9 Q3KCC2 Q3KCC9 Q3KD12 Q3KD25 Q3KE48 Q3KF57
Q3KFB3 Q3KFR3 Q3KFY2 Q3KG77 Q3KGX9 Q3KH21 Q3KHX7 Q3KJ31 Q3KJ66 Q3KJC7 Q3KJD0
Q3KJD1 Q3KJD8 Q3KK91 Q3KKB8 Q3KKG8 Q3KKG9 Q3KKT5 Q3KKV3 Q3KLQ8 Q3KLU5 Q3KLY0
Q3KLY4 Q3KM81 Q3KM82 Q3KM84 Q3KMF4 Q3KNA1 Q3KNM2 Q3KNS1 Q3KNT9 Q3KNW5 Q3KP22
Q3KPI0 Q3KPP4 Q3KPU7 Q3KPV4 Q3KQ18 Q3KQE5 Q3KQG9 Q3KQJ0 Q3KQZ1 Q3KR37 Q3KR51
Q3KR56 Q3KR82 Q3KR86 Q3KR92 Q3KRC4 Q3KRC6 Q3KRE0 Q3KSQ3 Q3KSQ7 Q3KSR7 Q3KST4
Q3KSU2 Q3KTM0 Q3KTM2 Q3L2T5 Q3L6L8 Q3L6L9 Q3L6P5 Q3L6Q6 Q3L6R7 Q3L6S8 Q3L6T9
Q3L6V0 Q3L6W1 Q3L6W3 Q3L6W7 Q3L6W9 Q3L6X2 Q3L6X7 Q3L6Y4 Q3L6Y5 Q3L6Y6 Q3L6Y8
Q3L6Z3 Q3L6Z6 Q3L701 Q3L708 Q3L7M0 Q3L890 Q3L8P3 Q3LFN0 Q3LR84 Q3LRP1 Q3LRP3
Q3LRV9 Q3LS21 Q3LSL6 Q3LTW2 Q3LUH2 Q3LX99 Q3LZS3 Q3LZW9 Q3LZX0 Q3LZX1 Q3LZX6
Q3LZX7 Q3LZX9 Q3M278 Q3M4K0 Q3M4N7 Q3M4N8 Q3M4V0 Q3M5L6 Q3M5W2 Q3M6F7 Q3M6T7
Q3M6U8 Q3M6Y1 Q3M8S5 Q3M918 Q3M999 Q3M9C7 Q3M9K9 Q3M9V5 Q3M9V6 Q3M9V7 Q3M9V8
Q3MA04 Q3MA59 Q3MA96 Q3MAB1 Q3MAR0 Q3MB01 Q3MB02 Q3MB63 Q3MB78 Q3MBD4 Q3MBP7
Q3MC09 Q3MC10 Q3MC11 Q3MC12 Q3MCB9 Q3MCF9 Q3MCT0 Q3MD07 Q3MD92 Q3MDP5 Q3MDS1
Q3MDY0 Q3MFH9 Q3MFT6 Q3MFU4 Q3MFU9 Q3MH22 Q3MHG1 Q3MHI3 Q3MHK4 Q3MHM8 Q3MHN2
Q3MHP8 Q3MHP9 Q3MHQ7 Q3MHU5 Q3MHV9 Q3MHW6 Q3MHW7 Q3MHW9 Q3MHZ1 Q3MHZ5 Q3MHZ7
Q3MID2 Q3MIP1 Q3MIR4 Q3MIW9 Q3MIX3 Q3MUY2 Q3S2T8 Q3S2U3 Q3S2U5 Q3S8M4 Q3SAG9
Q3SC83 Q3SF38 Q3SF60 Q3SF61 Q3SF62 Q3SF89 Q3SFE7 Q3SFG5 Q3SFQ0 Q3SFZ6 Q3SFZ8
Q3SG48 Q3SGB5 Q3SGQ5 Q3SGQ6 Q3SH71 Q3SI71 Q3SI72 Q3SJP4 Q3SJP7 Q3SJQ0 Q3SK07
Q3SLJ4 Q3SLW5 Q3SMH6 Q3SNN8 Q3SP57 Q3SPN4 Q3SQ28 Q3SQZ1 Q3SR23 Q3SRA3 Q3SRF1
Q3SRQ1 Q3SRQ2 Q3SSM1 Q3SSR2 Q3ST91 Q3STD7 Q3STT1 Q3SUL3 Q3SVC8 Q3SW35 Q3SW37
Q3SW38 Q3SW48 Q3SW84 Q3SWA8 Q3SWD2 Q3SWT0 Q3SWT5 Q3SWX0 Q3SWX5 Q3SWY0 Q3SWY4
Q3SWZ3 Q3SX00 Q3SX17 Q3SX46 Q3SXG2 Q3SXP7 Q3SXY7 Q3SY17 Q3SY77 Q3SYC2 Q3SYS8
Q3SYT0 Q3SYT6 Q3SYT8 Q3SYU3 Q3SYV1 Q3SYV3 Q3SYV5 Q3SYX1 Q3SYY2 Q3SYY9 Q3SYZ3

Q3SZ26 Q3SZ27 Q3SZ36 Q3SZ48 Q3SZ72 Q3SZ87 Q3SZ89 Q3SZ97 Q3SZB3 Q3SZE3 Q3SZI5
Q3SZI6 Q3SZK0 Q3SZL3 Q3SZL5 Q3SZL9 Q3SZM3 Q3SZN3 Q3SZQ2 Q3SZR6 Q3SZR9 Q3SZT1
Q3SZT4 Q3SZU5 Q3SZU9 Q3SZV6 Q3SZV8 Q3SZW3 Q3SZX0 Q3SZX8 Q3SZZ2 Q3SZZ5 Q3T013
Q3T021 Q3T033 Q3T073 Q3T075 Q3T0A0 Q3T0A9 Q3T0C6 Q3T0C9 Q3T0E3 Q3T0E5 Q3T0E9
Q3T0I5 Q3T0K8 Q3T0L5 Q3T0M2 Q3T0N3 Q3T0P7 Q3T0Q2 Q3T0R4 Q3T0S0 Q3T0S3 Q3T0T0
Q3T0W0 Q3T0Y8 Q3T0Y9 Q3T0Z4 Q3T100 Q3T104 Q3T110 Q3T113 Q3T124 Q3T126 Q3T130
Q3T133 Q3T134 Q3T144 Q3T181 Q3T196 Q3T1H8 Q3T1I5 Q3T1J2 Q3T1L5 Q3T4C0 Q3T4C2
Q3T4C8 Q3T4E5 Q3T4F1 Q3T4F3 Q3T4L9 Q3T4M7 Q3T904 Q3T906 Q3T9M1 Q3T9X0 Q3TA38
Q3TAE8 Q3TAS6 Q3TB48 Q3TB92 Q3TBN1 Q3TBT3 Q3TCT4 Q3TD49 Q3TDN0 Q3TDQ1 Q3TEW6
Q3TFD2 Q3TH73 Q3TIT8 Q3TIW9 Q3TMP8 Q3TMX7 Q3TNL8 Q3TNW5 Q3TP92 Q3TPR7 Q3TQB2
Q3TQR0 Q3TR08 Q3TR54 Q3TRM4 Q3TS39 Q3TT99 Q3TTE0 Q3TTY0 Q3TUA9 Q3TUD9 Q3TUL7
Q3TVA9 Q3TWI9 Q3TWL2 Q3TWN3 Q3TXX3 Q3TXX4 Q3TYD6 Q3TYE7 Q3TYP4 Q3TYS2 Q3TYU2
Q3TYX2 Q3TZ87 Q3TZM9 Q3TZW0 Q3TZX3 Q3TZZ7 Q3U095 Q3U0P5 Q3U0X8 Q3U0Y2 Q3U128
Q3U145 Q3U1F9 Q3U1J0 Q3U1N2 Q3U213 Q3U284 Q3U2C5 Q3U2S8 Q3U3D7 Q3U3F9 Q3U3R4
Q3U435 Q3U481 Q3U497 Q3U4G3 Q3U4N7 Q3U507 Q3U6B2 Q3U7R1 Q3U7U4 Q3U827 Q3U9G9
Q3U9N9 Q3UBX0 Q3UDF0 Q3UDR8 Q3UDW8 Q3UEZ8 Q3UF25 Q3UF64 Q3UFB7 Q3UFD7 Q3UFJ6
Q3UG50 Q3UG61 Q3UGP8 Q3UGX3 Q3UGY8 Q3UH53 Q3UH93 Q3UH99 Q3UHD1 Q3UHH2 Q3UHH8
Q3UHK1 Q3UHK3 Q3UHK6 Q3UHN9 Q3UIU2 Q3UJ81 Q3UJD6 Q3UJF0 Q3UJU9 Q3ULF4 Q3UM83
Q3UME2 Q3UMF9 Q3UMR5 Q3UMW8 Q3UMZ3 Q3UN02 Q3UN04 Q3UN16 Q3UN70 Q3UNB8 Q3UNN8
Q3UP23 Q3UP75 Q3UPI1 Q3UPL6 Q3UPR0 Q3UPR7 Q3UQ41 Q3UQS2 Q3UR32 Q3UR85 Q3URD2
Q3URD3 Q3URE8 Q3URE9 Q3URJ8 Q3URS9 Q3USF0 Q3USQ7 Q3USY0 Q3UTD9 Q3UU41 Q3UU67
Q3UU94 Q3UUA0 Q3UUQ7 Q3UUY6 Q3UV16 Q3UV31 Q3UV71 Q3UV74 Q3UVD5 Q3UVK0 Q3UVU3
Q3UVX5 Q3UVY1 Q3UVY5 Q3UW12 Q3UWA6 Q3UY51 Q3UZP0 Q3UZV7 Q3V009 Q3V050 Q3V0B4
Q3V0I2 Q3V0J1 Q3V0Q7 Q3V0T4 Q3V0X1 Q3V132 Q3V172 Q3V1F8 Q3V1H3 Q3V1N9 Q3V209
Q3V2G4 Q3V3F6 Q3V3G7 Q3V3K7 Q3V3R4 Q3V3Z3 Q3V4P9 Q3V4Q5 Q3V4S7 Q3V4U0 Q3V4U3
Q3V4U5 Q3V4V0 Q3V4V4 Q3V4V6 Q3V4W4 Q3V4W5 Q3V4X6 Q3V4X9 Q3V4Y1 Q3V4Y2 Q3V4Y4
Q3V4Y5 Q3V4Y7 Q3V503 Q3V508 Q3V515 Q3V516 Q3V517 Q3V518 Q3V521 Q3V522 Q3V523
Q3V524 Q3V529 Q3V534 Q3V535 Q3V538 Q3V539 Q3V540 Q3V541 Q3V546 Q3V547 Q3V548
Q3V550 Q3V554 Q3V5L5 Q3V7G2 Q3V7K4 Q3V7M0 Q3V7N8 Q3V7N9 Q3V7R3 Q3V815 Q3Y4E2
Q3Y6B8 Q3YAJ5 Q3YAW7 Q3YBM2 Q3YL57 Q3YL68 Q3YL95 Q3YL97 Q3YLA8 Q3YLC2 Q3YPZ4
Q3YQU4 Q3YQV2 Q3YR13 Q3YRB4 Q3YRZ4 Q3YS40 Q3YSH2 Q3YT97 Q3YU17 Q3YU30 Q3YUI6
Q3YUI7 Q3YUK4 Q3YUQ3 Q3YUR7 Q3YUU4 Q3YUU9 Q3YUV5 Q3YV10 Q3YV60 Q3YV63 Q3YV70
Q3YV90 Q3YVD0 Q3YVE9 Q3YVG9 Q3YVL0 Q3YVL2 Q3YVP0 Q3YVP1 Q3YVP2 Q3YVR0 Q3YW37
Q3YW59 Q3YWA4 Q3YWA8 Q3YWC7 Q3YWE6 Q3YWJ8 Q3YWK3 Q3YWQ7 Q3YWS1 Q3YWW9 Q3YX05
Q3YX06 Q3YX07 Q3YX33 Q3YXB3 Q3YXE2 Q3YXI0 Q3YXI3 Q3YXI5 Q3YXK1 Q3YXN2 Q3YY21
Q3YY22 Q3YY32 Q3YYB4 Q3YYT7 Q3YZ36 Q3YZ80 Q3YZC9 Q3YZD0 Q3YZE0 Q3YZE3 Q3YZR6
Q3YZS2 Q3YZS8 Q3YZT1 Q3YZT4 Q3YZU8 Q3YZU9 Q3YZV2 Q3Z010 Q3Z064 Q3Z077 Q3Z0A4
Q3Z0C7 Q3Z0C8 Q3Z0C9 Q3Z0K3 Q3Z0L7 Q3Z0N3 Q3Z0W3 Q3Z0Y2 Q3Z0Y3 Q3Z166 Q3Z192
Q3Z1P4 Q3Z1R4 Q3Z1R5 Q3Z1V2 Q3Z1V3 Q3Z1Y2 Q3Z1Y5 Q3Z1Y7 Q3Z218 Q3Z259 Q3Z269
Q3Z2C8 Q3Z2C9 Q3Z2D8 Q3Z2G1 Q3Z2G2 Q3Z2G6 Q3Z2G8 Q3Z2T6 Q3Z2V7 Q3Z2W2 Q3Z2W3
Q3Z351 Q3Z376 Q3Z398 Q3Z3K7 Q3Z3M2 Q3Z3Q4 Q3Z3T7 Q3Z3V1 Q3Z3V2 Q3Z4A5 Q3Z4A6
Q3Z4A7 Q3Z4G8 Q3Z4K2 Q3Z4X5 Q3Z5I1 Q3Z5K2 Q3Z5S2 Q3Z5W2 Q3Z5W8 Q3Z5Y3 Q3Z5Z4
Q3Z643 Q3Z646 Q3Z652 Q3Z6F1 Q3Z6K1 Q3Z6P4 Q3Z6R4 Q3Z7F0 Q3Z7W0 Q3Z7W1 Q3Z7Z7
Q3Z8Z6 Q3Z8Z7 Q3ZAI6 Q3ZAQ7 Q3ZAV1 Q3ZBE6 Q3ZBE9 Q3ZBF8 Q3ZBG6 Q3ZBG9 Q3ZBH3

Q3ZBI5 Q3ZBI7 Q3ZBI9 Q3ZBJ3 Q3ZBJ8 Q3ZBK9 Q3ZBN8 Q3ZBP2 Q3ZBR1 Q3ZBT5 Q3ZBU2
Q3ZBV0 Q3ZBX1 Q3ZBY0 Q3ZBY7 Q3ZC23 Q3ZC24 Q3ZC25 Q3ZC26 Q3ZC33 Q3ZC48 Q3ZC50
Q3ZC71 Q3ZC75 Q3ZC80 Q3ZCB8 Q3ZCD0 Q3ZCD2 Q3ZCD7 Q3ZCD8 Q3ZCG2 Q3ZCG8 Q3ZCI1
Q3ZCI8 Q3ZCQ0 Q3ZCQ3 Q3ZCQ8 Q3ZDR4 Q3ZEC1 Q3ZEC3 Q3ZEC9 Q3ZED1 Q3ZEG3 Q3ZIZ3
Q3ZIZ4 Q3ZIZ6 Q3ZIZ8 Q3ZJ04 Q3ZJ07 Q3ZJ10 Q3ZJ11 Q3ZJ14 Q3ZJ15 Q3ZJ17 Q3ZJ18
Q3ZJ19 Q3ZJ20 Q3ZJ21 Q3ZJ22 Q3ZJ26 Q3ZJ27 Q3ZJ28 Q3ZJ43 Q3ZJ46 Q3ZJ60 Q3ZJ61
Q3ZJ62 Q3ZJ64 Q3ZJ65 Q3ZJ66 Q3ZJ70 Q3ZJ76 Q3ZK22 Q3ZK64 Q3ZKN0 Q3ZMH1 Q3ZN06
Q3ZN07 Q3ZTK5 Q3ZU82 Q3ZW47 Q3ZW79 Q3ZW82 Q3ZWB4 Q3ZX56 Q3ZXR7 Q3ZXW0 Q3ZXW1
Q3ZY72 Q3ZYR8 Q3ZYT1 Q3ZYV4 Q3ZZ39 Q3ZZU1 Q3ZZU2 Q401C0 Q401N2 Q401Y8 Q40235
Q40294 Q40296 Q40297 Q40298 Q40299 Q402F3 Q402F4 Q402U7 Q40300 Q40301 Q40578
Q40585 Q40607 Q40608 Q40609 Q40746 Q40902 Q41009 Q41050 Q41093 Q41094 Q41131
Q41144 Q41224 Q41266 Q41342 Q41364 Q41385 Q41387 Q41437 Q41438 Q41629 Q41630
Q41706 Q41745 Q41760 Q41773 Q41870 Q41951 Q41963 Q41975 Q42093 Q42191 Q42342
Q42371 Q42374 Q42395 Q42400 Q42431 Q42496 Q42525 Q42556 Q42564 Q42569 Q42574
Q42593 Q42600 Q42602 Q42626 Q42627 Q42670 Q42716 Q42797 Q42841 Q42868 Q42870
Q42883 Q42969 Q42980 Q43033 Q43054 Q43067 Q43128 Q43135 Q43147 Q43240 Q43246
Q43250 Q43255 Q43257 Q43284 Q43348 Q43362 Q43402 Q43434 Q43469 Q43531 Q43715
Q43804 Q43857 Q43923 Q43924 Q43953 Q43961 Q43967 Q43975 Q44007 Q44091 Q44123
Q44287 Q44344 Q44346 Q44351 Q44360 Q44456 Q44470 Q44550 Q44576 Q44585 Q44586
Q44587 Q44600 Q44601 Q44619 Q44620 Q44664 Q44665 Q44756 Q44760 Q44763 Q44775
Q44776 Q44902 Q44906 Q44907 Q44909 Q44912 Q44930 Q45064 Q45067 Q45068 Q45078
Q45079 Q45104 Q45105 Q45110 Q45130 Q45132 Q45247 Q45298 Q45321 Q45324 Q45325
Q45326 Q45330 Q45331 Q45335 Q45340 Q45390 Q45391 Q45400 Q45407 Q45409 Q45411
Q45412 Q45461 Q45479 Q45514 Q45539 Q45544 Q45577 Q45588 Q45589 Q45592 Q45594
Q45605 Q45612 Q45614 Q45632 Q45658 Q45659 Q45689 Q45827 Q45832 Q45885 Q45894
Q45974 Q45975 Q45979 Q45980 Q45FN5 Q45R42 Q45U18 Q45UF1 Q45UF2 Q45W86 Q45W87
Q46065 Q46072 Q460G9 Q460M5 Q46136 Q46170 Q46203 Q46222 Q46378 Q463X4 Q46407
Q46409 Q46476 Q46493 Q464W5 Q46525 Q465Q6 Q465T0 Q465W3 Q46628 Q46629 Q46631
Q46633 Q46639 Q46646 Q466B1 Q46717 Q46755 Q46817 Q46821 Q46824 Q46831 Q46832
Q46833 Q46834 Q46836 Q46839 Q46841 Q46867 Q46892 Q46909 Q46916 Q46927 Q46943
Q46970 Q46AL8 Q46BD9 Q46BP2 Q46D59 Q46DN2 Q46EU5 Q46F66 Q46F67 Q46FW1 Q46GV5
Q46GW1 Q46GW2 Q46GX1 Q46GY3 Q46H42 Q46H51 Q46H53 Q46H79 Q46H80 Q46H81 Q46HA2
Q46HC1 Q46HC2 Q46HL3 Q46HM4 Q46HW5 Q46I69 Q46IH7 Q46IH8 Q46IV5 Q46IV6 Q46J48
Q46J52 Q46J53 Q46J54 Q46J55 Q46JD9 Q46JP5 Q46JS1 Q46JS5 Q46JV2 Q46JW4 Q46JW5
Q46JW8 Q46JW9 Q46JX0 Q46JZ3 Q46KE1 Q46L46 Q46L84 Q46LC1 Q46LG5 Q46LS0 Q46LY0
Q46LZ1 Q46M01 Q46M04 Q46RW7 Q46TZ4 Q46UZ7 Q46VL6 Q46VX6 Q46WM3 Q46WM4 Q46WM9
Q46WZ1 Q46XB8 Q46XG9 Q46XM6 Q46Y89 Q46ZT1 Q46ZY7 Q47085 Q47086 Q47108 Q47125
Q47142 Q47153 Q47162 Q47208 Q47258 Q472H9 Q47377 Q473G3 Q473T3 Q473T6 Q473U3
Q47421 Q47422 Q47455 Q47457 Q47490 Q474P6 Q47500 Q47502 Q47503 Q47539 Q475B0
Q475H6 Q47689 Q47692 Q476H8 Q476M4 Q47706 Q47745 Q47746 Q477C7 Q477Q4 Q477Z5
Q47825 Q47856 Q47866 Q47877 Q478K9 Q478U2 Q478U7 Q47908 Q47952 Q47A46 Q47AA1
Q47AF6 Q47AM7 Q47AR9 Q47BA6 Q47BQ7 Q47BZ8 Q47CR9 Q47DI2 Q47EA7 Q47G19 Q47GU1
Q47H38 Q47H39 Q47H40 Q47HG3 Q47HG6 Q47HG9 Q47I87 Q47IP1 Q47IT9 Q47J11 Q47JC1
Q47JR8 Q47KF9 Q47KG0 Q47LF1 Q47LF4 Q47LF7 Q47M78 Q47MI6 Q47ND5 Q47NU8 Q47QX2

Q47RS4 Q47SI8 Q47VQ6 Q47W31 Q47W34 Q47YD3 Q47YH9 Q47ZC7 Q480N3 Q480Q1 Q48153
Q48216 Q48217 Q48218 Q48219 Q48220 Q48221 Q48245 Q48247 Q48248 Q48253 Q48258
Q48262 Q48269 Q482I0 Q482U3 Q482U6 Q48314 Q48315 Q48334 Q483B6 Q48414 Q48445
Q48450 Q48452 Q48456 Q48457 Q48460 Q48465 Q48473 Q48475 Q48478 Q485F0 Q48624
Q48630 Q48724 Q48729 Q487D7 Q487F0 Q487I6 Q487R3 Q48815 Q489A5 Q48AB7 Q48AL9
Q48AW4 Q48AW6 Q48BF2 Q48BF9 Q48BG0 Q48BG1 Q48BQ5 Q48CC8 Q48CL6 Q48DU1 Q48E39
Q48EF5 Q48FA9 Q48FG6 Q48FK1 Q48FR0 Q48FX8 Q48FZ3 Q48GL4 Q48H45 Q48H48 Q48H54
Q48H73 Q48HY7 Q48HY8 Q48HY9 Q48HZ2 Q48JI5 Q48K34 Q48KB2 Q48KR3 Q48LG7 Q48M92
Q48M97 Q48N46 Q48N63 Q48NU7 Q48P40 Q48P74 Q48PJ6 Q48PJ9 Q48PK0 Q48PK6 Q48PL9
Q48PW5 Q48QE1 Q48RZ3 Q48S17 Q48SQ3 Q48U03 Q48U75 Q48UD7 Q48UD8 Q48UF9 Q48UM6
Q48V51 Q48V70 Q48V75 Q48VA9 Q49123 Q49125 Q49126 Q49135 Q49176 Q491V8 Q491Z9
Q492G8 Q492H6 Q492I2 Q492I5 Q492I8 Q492K8 Q492M6 Q492P7 Q492S9 Q49310 Q49378
Q49379 Q49397 Q49399 Q493G3 Q493Q2 Q493Q4 Q493S2 Q493X5 Q493X7 Q49401 Q49403
Q49409 Q49417 Q49430 Q49431 Q49432 Q49433 Q49460 Q494A7 Q494C7 Q494C8 Q494C9
Q494P0 Q494Q2 Q494T4 Q494W8 Q495A1 Q495M3 Q495N2 Q495T6 Q495W5 Q49606 Q49619
Q49626 Q49642 Q49647 Q496F6 Q496J9 Q496Z2 Q496Z4 Q49760 Q497B2 Q497B3 Q497J1
Q497K7 Q497L8 Q497L9 Q49894 Q498C7 Q498C8 Q498D5 Q498D6 Q498S8 Q498T9 Q498U0
Q498U3 Q498W5 Q499N4 Q499P3 Q499P8 Q499S9 Q499U1 Q499U8 Q49A17 Q49B93 Q49BZ4
Q49CA8 Q49CB2 Q49HH9 Q49HI0 Q49KI5 Q49KU0 Q49KU3 Q49KU5 Q49KU6 Q49KU8 Q49KU9
Q49KV1 Q49KW7 Q49KW8 Q49KW9 Q49KX0 Q49KX1 Q49KX2 Q49KX9 Q49KY0 Q49KY1 Q49KY2
Q49KY3 Q49KY4 Q49KY5 Q49KY6 Q49KY7 Q49KY8 Q49KZ3 Q49KZ8 Q49KZ9 Q49L01 Q49L02
Q49L03 Q49L04 Q49L05 Q49L10 Q49L11 Q49L12 Q49L14 Q49L15 Q49L18 Q49LR9 Q49LS0
Q49LS1 Q49LS3 Q49LS4 Q49LS5 Q49LS6 Q49LS7 Q49LS8 Q49LS9 Q49LX5 Q49LX6 Q49P94
Q49SH1 Q49SP8 Q49SQ1 Q49SQ2 Q49SQ3 Q49VC5 Q49VD3 Q49VG9 Q49VH0 Q49VH1 Q49VH2
Q49VH3 Q49VH4 Q49VH5 Q49VK4 Q49VN0 Q49VR4 Q49VU5 Q49VV9 Q49W06 Q49W85 Q49W86
Q49W87 Q49W88 Q49W89 Q49W90 Q49W91 Q49WA9 Q49WE5 Q49WF1 Q49WI1 Q49WI2 Q49WI3
Q49WI4 Q49WM9 Q49WP0 Q49WP1 Q49WW4 Q49X46 Q49X74 Q49XD7 Q49XE2 Q49XE7 Q49XK9
Q49XM6 Q49XN1 Q49XT4 Q49Y16 Q49YH0 Q49YL2 Q49YL3 Q49YN9 Q49YR9 Q49YT0 Q49YT4
Q49YU6 Q49YV5 Q49YW0 Q49YZ1 Q49Z38 Q49Z54 Q49Z55 Q49Z56 Q49ZA3 Q49ZB2 Q49ZN7
Q49ZR0 Q49ZT3 Q49ZT7 Q49ZT9 Q4A009 Q4A012 Q4A070 Q4A0C4 Q4A0G1 Q4A159 Q4A3R3
Q4A5F0 Q4A5Z9 Q4A600 Q4A729 Q4A8W3 Q4A8W4 Q4AAW1 Q4AAW2 Q4ACW4 Q4D3E8 Q4DHA1
Q4E4I4 Q4EW11 Q4F7G0 Q4F883 Q4FBI3 Q4FFM9 Q4FFN0 Q4FFN5 Q4FFP1 Q4FFP2 Q4FFP4
Q4FGE9 Q4FGF0 Q4FGF2 Q4FGF5 Q4FGF6 Q4FGF8 Q4FGG1 Q4FGG2 Q4FGG7 Q4FGG9 Q4FJU9
Q4FLQ2 Q4FLV1 Q4FLX6 Q4FM05 Q4FM78 Q4FM84 Q4FM91 Q4FNF1 Q4FNI5 Q4FNY3 Q4FPB0
Q4FPD2 Q4FPE7 Q4FPE8 Q4FPE9 Q4FPK2 Q4FPV2 Q4FPV3 Q4FQ10 Q4FQ31 Q4FQ33 Q4FQ51
Q4FQ60 Q4FQ61 Q4FQM2 Q4FQN5 Q4FR05 Q4FR08 Q4FS42 Q4FT89 Q4FU52 Q4FU55 Q4FU58
Q4FU64 Q4FU75 Q4FUU9 Q4FV70 Q4FVF4 Q4FVP3 Q4FVQ0 Q4FVS9 Q4FZ58 Q4FZ76 Q4FZC9
Q4FZH1 Q4FZQ0 Q4FZU9 Q4FZV2 Q4FZV7 Q4FZW7 Q4FZX7 Q4G019 Q4G068 Q4G072 Q4G0I0
Q4G0N0 Q4G0N8 Q4G0T1 Q4G148 Q4G1C9 Q4G1G8 Q4G2J3 Q4G2J4 Q4G2J5 Q4G2J6 Q4G2S9
Q4G2T3 Q4G351 Q4G371 Q4G373 Q4G378 Q4G380 Q4G381 Q4G382 Q4G383 Q4G385 Q4G395
Q4G399 Q4G3A0 Q4G3A1 Q4G3A9 Q4G3B0 Q4G3B2 Q4G3B3 Q4G3B5 Q4G3B8 Q4G3C0 Q4G3C2
Q4G3C3 Q4G3C4 Q4G3C6 Q4G3C7 Q4G3D2 Q4G3D7 Q4G3D8 Q4G3E7 Q4G3F2 Q4G3F5 Q4G3F6
Q4G3F7 Q4G3F8 Q4G3R0 Q4G4A7 Q4GZT3 Q4GZT4 Q4H3K6 Q4H425 Q4HVS2 Q4HVU7 Q4HX89
Q4HXT5 Q4HXT6 Q4HY20 Q4HZ95 Q4HZE0 Q4I073 Q4I099 Q4I0K3 Q4I1J3 Q4I298 Q4I2M7

Q4I2W2 Q4I4A4 Q4I5G1 Q4I5R9 Q4I785 Q4I7T5 Q4I8B6 Q4I8E9 Q4I8P5 Q4IA62 Q4IB63
Q4IBD0 Q4IBU4 Q4ICM9 Q4IJ79 Q4IJT0 Q4IK03 Q4ILH3 Q4IM48 Q4IMZ7 Q4IPX8 Q4IQ08
Q4J710 Q4J711 Q4J8E4 Q4J8K2 Q4J8L5 Q4J9J9 Q4J9K4 Q4J9L2 Q4JAC0 Q4JAE2 Q4JAQ5
Q4JB68 Q4JB69 Q4JB70 Q4JBJ9 Q4JBK2 Q4JIM4 Q4JLT5 Q4JM44 Q4JQH5 Q4JQH6 Q4JQH7
Q4JQH8 Q4JQH9 Q4JQI0 Q4JQI1 Q4JQI2 Q4JQI3 Q4JQI4 Q4JQI5 Q4JQI6 Q4JQI7 Q4JQV7
Q4JQW0 Q4JQW3 Q4JQX0 Q4JQX4 Q4JR05 Q4JSG6 Q4JSG7 Q4JTT0 Q4JUF6 Q4JUJ6 Q4JVK2
Q4JVN6 Q4JVQ1 Q4JW98 Q4JXD8 Q4JYC3 Q4K395 Q4K3A3 Q4K3A4 Q4K3A5 Q4K3D9 Q4K3L9
Q4K3X7 Q4K479 Q4K4C2 Q4K4W5 Q4K5Q6 Q4K5U5 Q4K6J0 Q4K6L6 Q4K6M1 Q4K878 Q4K884
Q4K8C3 Q4K8K4 Q4K8U3 Q4K909 Q4K9A4 Q4K9I6 Q4K9S7 Q4K9T0 Q4K9V6 Q4KBY6 Q4KC13
Q4KC78 Q4KC79 Q4KC80 Q4KC83 Q4KDD4 Q4KES7 Q4KFH2 Q4KG23 Q4KGB1 Q4KH68 Q4KHA4
Q4KI58 Q4KJB2 Q4KJF1 Q4KJL4 Q4KJL7 Q4KJL8 Q4KJM5 Q4KKI8 Q4KKL2 Q4KL18 Q4KL25
Q4KL91 Q4KLH9 Q4KLL3 Q4KLL4 Q4KLR8 Q4KLT0 Q4KLV1 Q4KLY3 Q4KLZ1 Q4KM64 Q4KM74
Q4KM77 Q4KM93 Q4KM98 Q4KMG0 Q4KMG9 Q4KMI4 Q4KMQ2 Q4KMZ8 Q4KRV1 Q4KUS1 Q4KWL2
Q4L208 Q4L2X2 Q4L2X3 Q4L3G8 Q4L3Q4 Q4L3Q9 Q4L443 Q4L444 Q4L445 Q4L446 Q4L447
Q4L448 Q4L449 Q4L482 Q4L4A6 Q4L4D7 Q4L4H1 Q4L4I8 Q4L4K9 Q4L4W1 Q4L4W2 Q4L4W3
Q4L4W4 Q4L4W5 Q4L4W6 Q4L4W7 Q4L4Y6 Q4L523 Q4L530 Q4L562 Q4L563 Q4L564 Q4L565
Q4L5B9 Q4L5C9 Q4L5D0 Q4L5N3 Q4L5P8 Q4L5W3 Q4L5Y9 Q4L652 Q4L656 Q4L661 Q4L667
Q4L6B4 Q4L6C5 Q4L6D0 Q4L6I5 Q4L6S3 Q4L783 Q4L7C3 Q4L7C4 Q4L7F0 Q4L7I0 Q4L7J6
Q4L7K3 Q4L7L6 Q4L7M5 Q4L7S0 Q4L7X2 Q4L7Y8 Q4L7Y9 Q4L7Z0 Q4L843 Q4L869 Q4L8H5
Q4L8J4 Q4L8J9 Q4L8L4 Q4L8L8 Q4L8M0 Q4L8N9 Q4L8Q5 Q4L8V3 Q4L945 Q4L970 Q4L977
Q4L979 Q4L9N9 Q4L9Y1 Q4LAB1 Q4LAI1 Q4LAI2 Q4LAI3 Q4LAJ8 Q4LBB6 Q4LBB9 Q4LCA6
Q4LDF9 Q4LDG0 Q4LDR2 Q4LE88 Q4N280 Q4N4N8 Q4N4T9 Q4P0H0 Q4P0S6 Q4P2W6 Q4P333
Q4P3I9 Q4P4U7 Q4P622 Q4P683 Q4P6L3 Q4P782 Q4P7J4 Q4P7Y8 Q4P8B7 Q4P8D2 Q4P8U8
Q4P937 Q4P958 Q4P983 Q4P996 Q4P9K6 Q4P9K9 Q4P9Q7 Q4P9R2 Q4PB75 Q4PDL5 Q4PE27
Q4PEP6 Q4PEQ0 Q4PEW9 Q4PF83 Q4PFA7 Q4PFQ1 Q4PGW7 Q4PH16 Q4PHZ2 Q4PIF8 Q4PIK6
Q4PIP8 Q4PJW3 Q4PJX1 Q4PKH3 Q4PNI0 Q4PNJ2 Q4PPC4 Q4PSF4 Q4PT07 Q4PT23 Q4PT37
Q4PZA2 Q4Q5P8 Q4QIU7 Q4QJQ3 Q4QJQ4 Q4QJQ5 Q4QJQ8 Q4QK50 Q4QKQ5 Q4QL94 Q4QLG1
Q4QLI8 Q4QLI9 Q4QLJ5 Q4QLL5 Q4QLP6 Q4QLQ6 Q4QLR0 Q4QLR4 Q4QM05 Q4QM23 Q4QM68
Q4QM74 Q4QMJ5 Q4QMJ9 Q4QMP3 Q4QMV0 Q4QMW0 Q4QN39 Q4QN58 Q4QN59 Q4QN60 Q4QNB7
Q4QNB8 Q4QNR3 Q4QNS2 Q4QNS5 Q4QNS6 Q4QNW1 Q4QP03 Q4QP19 Q4QP20 Q4QP21 Q4QP23
Q4QP40 Q4QP52 Q4QP67 Q4QPI4 Q4QPK9 Q4QQM4 Q4QQM5 Q4QQS1 Q4QQU5 Q4QQV3 Q4QQV7
Q4QQW1 Q4QR83 Q4QRH7 Q4QSC6 Q4QTJ1 Q4QTJ2 Q4QXJ7 Q4QXK3 Q4QXU0 Q4QXU2 Q4QXU3
Q4QXU4 Q4QXU5 Q4QXU6 Q4QXX9 Q4QYS6 Q4QYS8 Q4QYT1 Q4QYT2 Q4QYT4 Q4R1H1 Q4R1I4
Q4R1R8 Q4R1S6 Q4R2Y9 Q4R309 Q4R335 Q4R3C7 Q4R3I0 Q4R3L1 Q4R3U7 Q4R3Y4 Q4R445
Q4R495 Q4R4A8 Q4R4D5 Q4R4E0 Q4R4E1 Q4R4I5 Q4R4I9 Q4R4M1 Q4R4P4 Q4R4Q4 Q4R4R4
Q4R4S7 Q4R4T0 Q4R4V5 Q4R4X3 Q4R4Z3 Q4R504 Q4R510 Q4R512 Q4R516 Q4R518 Q4R532
Q4R534 Q4R540 Q4R542 Q4R566 Q4R581 Q4R583 Q4R589 Q4R5B1 Q4R5B4 Q4R5C3 Q4R5E3
Q4R5F4 Q4R5G7 Q4R5M4 Q4R5N9 Q4R5Q3 Q4R5T7 Q4R5V2 Q4R613 Q4R648 Q4R690 Q4R691
Q4R6B2 Q4R6E8 Q4R6F2 Q4R6H1 Q4R6K2 Q4R6L9 Q4R6N0 Q4R6P1 Q4R6W2 Q4R6Y5 Q4R717
Q4R749 Q4R763 Q4R766 Q4R794 Q4R7B7 Q4R7E2 Q4R7G8 Q4R7R1 Q4R7S2 Q4R7U0 Q4R7W6
Q4R7X9 Q4R7Z5 Q4R854 Q4R863 Q4R866 Q4R877 Q4R8A8 Q4R8C8 Q4R8M0 Q4R8M1 Q4R8P0
Q4R8S6 Q4R8T0 Q4R8V4 Q4R8W8 Q4R8X1 Q4R8Y9 Q4R910 Q4R942 Q4R9E0 Q4RSM6 Q4RY26
Q4TU93 Q4TVS0 Q4TWH8 Q4TWH9 Q4TZY1 Q4U116 Q4U2R8 Q4U3U4 Q4U3U6 Q4U4D2 Q4U9I8
Q4UDS9 Q4UEQ6 Q4UJ67 Q4UJ69 Q4UJM4 Q4UJQ7 Q4UJY1 Q4UK08 Q4UK23 Q4UK26 Q4UK27

Q4UK28 Q4UK29 Q4UK32 Q4UK37 Q4UK88 Q4UKA5 Q4UKA7 Q4UKC1 Q4UKC3 Q4UKC4 Q4UKC8
Q4UKK0 Q4UKP2 Q4UKP6 Q4UKQ2 Q4UKR6 Q4UKW1 Q4UKW3 Q4UKW4 Q4UL68 Q4UL85 Q4UL88
Q4ULJ8 Q4ULR7 Q4ULU0 Q4ULU3 Q4ULU5 Q4ULV9 Q4ULW4 Q4ULY0 Q4ULZ6 Q4UM04 Q4UM05
Q4UM22 Q4UM75 Q4UMD4 Q4UMI7 Q4UMJ9 Q4UMQ9 Q4UMU2 Q4UMZ3 Q4UN32 Q4UN46 Q4UN68
Q4UN74 Q4UN76 Q4UN77 Q4UN85 Q4UNA3 Q4UNB4 Q4UNB5 Q4UNB6 Q4UNB7 Q4UNB8 Q4UNC7
Q4UND5 Q4UNE0 Q4UNE6 Q4UNE7 Q4UNH7 Q4UNH9 Q4UNI0 Q4UNI5 Q4UNK8 Q4UP01 Q4UP02
Q4UP19 Q4UPT7 Q4UQF0 Q4UQV0 Q4UQW8 Q4UR34 Q4URA9 Q4URG5 Q4US41 Q4USI7 Q4UTA8
Q4UTC7 Q4UTK2 Q4UTP3 Q4UV65 Q4UVG6 Q4UVN7 Q4UWA8 Q4UWB1 Q4UXB0 Q4UXC5 Q4UXL9
Q4UXZ0 Q4UYH8 Q4UYZ8 Q4UZN6 Q4V052 Q4V056 Q4V093 Q4V398 Q4V3B8 Q4V3C7 Q4V3C9
Q4V5R4 Q4V622 Q4V786 Q4V7A5 Q4V7D3 Q4V7F2 Q4V7N7 Q4V7Q2 Q4V7Q8 Q4V7R2 Q4V7T3
Q4V7T7 Q4V7T9 Q4V7U1 Q4V885 Q4V887 Q4V888 Q4V892 Q4V897 Q4V899 Q4V8A1 Q4V8A9
Q4V8B1 Q4V8D1 Q4V8E0 Q4V8E5 Q4V8F3 Q4V8F8 Q4V8I1 Q4V8I7 Q4V8J4 Q4V8K1 Q4V8K4
Q4V8S3 Q4V8U5 Q4V8X0 Q4V8Y6 Q4V921 Q4V922 Q4V9F0 Q4V9L6 Q4V9P0 Q4V9Y1 Q4V9Z5
Q4VA61 Q4VA82 Q4VAE3 Q4VAH7 Q4VBD2 Q4VBG5 Q4VBI2 Q4VBI7 Q4VBT2 Q4VC39 Q4VCM1
Q4VGP8 Q4VHE7 Q4VKI6 Q4VKI7 Q4VKV3 Q4VKV4 Q4VNC0 Q4VNC1 Q4VUI0 Q4VUT8 Q4VUU0
Q4VUU2 Q4VUU4 Q4VUV4 Q4VUV8 Q4VUV9 Q4VUY0 Q4VUY6 Q4VUY8 Q4VUY9 Q4VUZ1 Q4VV71
Q4VWH4 Q4VWH9 Q4VWQ3 Q4VXA5 Q4VXF1 Q4VY51 Q4VZH0 Q4VZH5 Q4VZH6 Q4VZH7 Q4VZI2
Q4VZI6 Q4VZI7 Q4VZI9 Q4VZJ0 Q4VZJ9 Q4VZK4 Q4VZL0 Q4VZL8 Q4VZN4 Q4VZN6 Q4VZN8
Q4VZN9 Q4VZP0 Q4VZP5 Q4VZP7 Q4VZP9 Q4VZQ0 Q4W1W1 Q4W5P6 Q4W815 Q4W8A2 Q4W8A3
Q4W8E7 Q4W9C7 Q4W9I3 Q4W9R7 Q4WA15 Q4WA92 Q4WAG4 Q4WAH2 Q4WAW5 Q4WAW8 Q4WAX0
Q4WAZ6 Q4WB51 Q4WBI8 Q4WBL2 Q4WBM1 Q4WC37 Q4WC50 Q4WCL2 Q4WCV3 Q4WD46 Q4WD56
Q4WDD4 Q4WDE0 Q4WDL3 Q4WDM5 Q4WDV4 Q4WDZ0 Q4WED5 Q4WEE0 Q4WF31 Q4WF45 Q4WFQ4
Q4WGE2 Q4WGL2 Q4WGM4 Q4WGS5 Q4WH97 Q4WHE1 Q4WHZ1 Q4WI16 Q4WI88 Q4WI89 Q4WIQ0
Q4WIX5 Q4WJ38 Q4WJ59 Q4WJH4 Q4WJJ9 Q4WJM7 Q4WJS4 Q4WJU9 Q4WJW8 Q4WK25 Q4WK77
Q4WKA1 Q4WKA5 Q4WKN3 Q4WKX3 Q4WLC8 Q4WLD0 Q4WLD1 Q4WLD2 Q4WLI9 Q4WLN7 Q4WLP9
Q4WLT9 Q4WLW9 Q4WM67 Q4WMJ0 Q4WMP0 Q4WN24 Q4WN54 Q4WNB5 Q4WNN2 Q4WNT5 Q4WP49
Q4WP59 Q4WP81 Q4WPG0 Q4WPP2 Q4WPP6 Q4WQ08 Q4WQ14 Q4WQ21 Q4WQ82 Q4WQC5 Q4WQG9
Q4WQJ2 Q4WQJ5 Q4WQL0 Q4WQY6 Q4WR17 Q4WR59 Q4WR75 Q4WRQ4 Q4WRW0 Q4WS70 Q4WSI1
Q4WSY2 Q4WT65 Q4WTK9 Q4WTQ6 Q4WTT7 Q4WTT9 Q4WTW3 Q4WU12 Q4WUG9 Q4WUK1 Q4WUK5
Q4WUS1 Q4WV30 Q4WVG2 Q4WVT3 Q4WVV6 Q4WW43 Q4WWN2 Q4WWW3 Q4WX13 Q4WX68 Q4WXT2
Q4WYA5 Q4WYF4 Q4WYG7 Q4WYN3 Q4WYN4 Q4WZ65 Q4WZG1 Q4WZL8 Q4WZS1 Q4WZY1 Q4X006
Q4X022 Q4X0B5 Q4X0I8 Q4X0S3 Q4X117 Q4X162 Q4X180 Q4X1I8 Q4X251 Q4XZY3 Q4YCF6
Q4ZG55 Q4ZGE1 Q4ZHA6 Q4ZIN3 Q4ZJI4 Q4ZJY7 Q4ZJZ0 Q4ZL11 Q4ZL18 Q4ZL19 Q4ZL20
Q4ZL66 Q4ZLB4 Q4ZLV4 Q4ZM52 Q4ZMF5 Q4ZNG6 Q4ZNI5 Q4ZNN2 Q4ZNY7 Q4ZPY9 Q4ZQ33
Q4ZQ68 Q4ZQC4 Q4ZR01 Q4ZRI4 Q4ZRI7 Q4ZRJ3 Q4ZRL4 Q4ZS30 Q4ZSY8 Q4ZSY9 Q4ZSZ0
Q4ZSZ3 Q4ZT65 Q4ZV10 Q4ZVF7 Q4ZVP9 Q4ZWM7 Q4ZXC3 Q4ZXC8 Q4ZY49 Q4ZY63 Q4ZZ16
Q4ZZ50 Q4ZZG5 Q4ZZG8 Q4ZZG9 Q4ZZH4 Q4ZZI6 Q50097 Q50098 Q500P0 Q500V2 Q500W7
Q500Z2 Q500Z4 Q50103 Q50173 Q50186 Q501D6 Q501D8 Q501F8 Q501G6 Q501I9 Q501J2
Q50205 Q50230 Q50231 Q50232 Q50292 Q502B3 Q502E0 Q502E5 Q502F0 Q502G2 Q502I1
Q502J0 Q502L1 Q502L2 Q50325 Q50326 Q50327 Q50333 Q50334 Q50341 Q50361 Q50393
Q50394 Q50395 Q503G8 Q503I2 Q503M4 Q503P5 Q503U3 Q503V1 Q503V3 Q503W7 Q50414
Q50416 Q50418 Q504C1 Q504G0 Q504N2 Q504P2 Q504Y0 Q50568 Q505J3 Q505J6 Q505J9
Q50681 Q50773 Q50774 Q50799 Q50830 Q50862 Q508J0 Q508J1 Q508J2 Q508J3 Q508J4
Q508J5 Q508J7 Q508J9 Q508K1 Q508K2 Q508K4 Q508K5 Q508K6 Q508K7 Q508K8 Q508K9

Q508L0 Q508L1 Q508L2 Q508L3 Q508L4 Q508L6 Q508L8 Q508M1 Q508M2 Q508M3 Q508M4
Q508M6 Q508M8 Q508M9 Q508N0 Q508N1 Q508N2 Q508N3 Q508N6 Q508P2 Q50919 Q50DL2
Q50DL8 Q50DM5 Q50DM6 Q50DM7 Q50DM8 Q50EK0 Q50EK1 Q50EK3 Q50EK4 Q50EK5 Q50EK6
Q50JE5 Q50L44 Q50LE0 Q50LG9 Q50LH3 Q50LH4 Q51005 Q51330 Q51368 Q51389 Q51392
Q51397 Q51417 Q51463 Q51467 Q51468 Q51483 Q51485 Q51487 Q51575 Q51662 Q51699
Q51752 Q51769 Q51876 Q51881 Q51887 Q51910 Q51955 Q51IZ9 Q51MB1 Q51VY4 Q51WZ9
Q52106 Q52280 Q52356 Q52402 Q52454 Q52463 Q52488 Q52527 Q52581 Q525S7 Q52657
Q52664 Q52665 Q52689 Q52715 Q52718 Q52778 Q52813 Q52814 Q52819 Q52893 Q52910
Q52923 Q52938 Q52952 Q52965 Q52968 Q52969 Q52978 Q52980 Q52981 Q52982 Q52983
Q52E66 Q52KD1 Q52KD7 Q52KK3 Q52KL1 Q52KR2 Q52LC2 Q52MA5 Q52N47 Q52NJ0 Q52PA5
Q52PG9 Q52QI2 Q52RG7 Q52S86 Q52T38 Q52ZA1 Q53016 Q53020 Q53047 Q53151 Q53174
Q53178 Q53185 Q53191 Q53192 Q53229 Q53461 Q53496 Q534B1 Q534D9 Q534E0 Q53584
Q53588 Q53590 Q53630 Q53683 Q53705 Q53765 Q53866 Q53868 Q53902 Q53922 Q53955
Q53956 Q53957 Q539B4 Q539C2 Q539C6 Q539Z1 Q53AJ7 Q53AK6 Q53AK8 Q53AL7 Q53AM3
Q53AQ4 Q53AV2 Q53CF6 Q53CF9 Q53CG2 Q53EL9 Q53EP0 Q53EU6 Q53F39 Q53FP2 Q53FV1
Q53GD3 Q53GQ0 Q53HI1 Q53JG7 Q53P54 Q53P98 Q53PN2 Q53R12 Q53RH0 Q53RT3 Q53RY4
Q53S58 Q53TN4 Q53WK1 Q53WP9 Q53YF3 Q54001 Q54114 Q54151 Q54155 Q54337 Q54462
Q54471 Q54519 Q54697 Q54710 Q54711 Q54752 Q54753 Q54794 Q54875 Q54AX6 Q54B29
Q54B49 Q54B51 Q54B67 Q54BF2 Q54BF6 Q54BI3 Q54BM3 Q54BN0 Q54BN8 Q54BP9 Q54BR1
Q54BT3 Q54BT5 Q54BU4 Q54C03 Q54C27 Q54C37 Q54C55 Q54C67 Q54C78 Q54C80 Q54CE6
Q54CG0 Q54CH1 Q54CH8 Q54CJ4 Q54CK6 Q54CL3 Q54CL4 Q54CM6 Q54CS3 Q54CX4 Q54D07
Q54D10 Q54D36 Q54D38 Q54DA4 Q54DD7 Q54DE8 Q54DE9 Q54DM9 Q54DP2 Q54DP3 Q54DP7
Q54DQ8 Q54DR1 Q54DR6 Q54DS3 Q54DT1 Q54DT2 Q54DU1 Q54DV5 Q54DW2 Q54DY5 Q54DY9
Q54E04 Q54E62 Q54EB4 Q54EG5 Q54EG9 Q54EJ6 Q54EK2 Q54EM5 Q54ER8 Q54ES5 Q54ET0
Q54ET2 Q54EU7 Q54EV4 Q54EW2 Q54EX5 Q54F08 Q54F25 Q54F34 Q54F44 Q54F47 Q54F54
Q54F97 Q54FB6 Q54FC8 Q54FE6 Q54FE7 Q54FH7 Q54FM9 Q54FN7 Q54FP9 Q54FR4 Q54FU9
Q54FX7 Q54G02 Q54G85 Q54GA9 Q54GB3 Q54GC1 Q54GD8 Q54GD9 Q54GE8 Q54GG1 Q54GL3
Q54GN6 Q54GU0 Q54GV3 Q54GV8 Q54GY9 Q54H37 Q54H45 Q54H46 Q54H80 Q54H86 Q54HC6
Q54HG9 Q54HH9 Q54HI2 Q54HM0 Q54HM2 Q54HM6 Q54HY8 Q54I48 Q54I54 Q54I86 Q54I92
Q54IA4 Q54IC9 Q54IE8 Q54IJ2 Q54IK2 Q54IM2 Q54IN7 Q54IV7 Q54IX6 Q54J19 Q54J24
Q54J33 Q54J38 Q54J42 Q54J71 Q54J77 Q54J78 Q54JA3 Q54JA4 Q54JA5 Q54JB1 Q54JB3
Q54JE1 Q54JH4 Q54JN2 Q54JQ0 Q54JR2 Q54JT7 Q54JV6 Q54JW5 Q54JY7 Q54K09 Q54K11
Q54K24 Q54K35 Q54K38 Q54K40 Q54K74 Q54K87 Q54K99 Q54KB9 Q54KD1 Q54KD4 Q54KF7
Q54KH2 Q54KH5 Q54KI2 Q54KL9 Q54KV0 Q54KZ0 Q54L52 Q54L53 Q54L75 Q54L85 Q54L86
Q54L87 Q54L98 Q54LA2 Q54LA8 Q54LB5 Q54LC9 Q54LE6 Q54LG8 Q54LH8 Q54LJ7 Q54LP7
Q54LQ4 Q54LQ9 Q54LR6 Q54LT0 Q54LT5 Q54LT7 Q54LY1 Q54LY6 Q54LY7 Q54M40 Q54M59
Q54M75 Q54M86 Q54MA0 Q54MB9 Q54MC0 Q54MC6 Q54ME0 Q54ME7 Q54MI3 Q54MJ9 Q54MK0
Q54MV1 Q54MW0 Q54MY4 Q54MZ4 Q54N33 Q54N40 Q54N73 Q54N80 Q54N94 Q54NA3 Q54NC1
Q54NC9 Q54ND0 Q54ND5 Q54NH7 Q54NL1 Q54NL4 Q54NM7 Q54NM9 Q54NN1 Q54NQ9 Q54NS7
Q54NT4 Q54NT8 Q54NV2 Q54NW7 Q54NY3 Q54NZ2 Q54P00 Q54P08 Q54P13 Q54P44 Q54P45
Q54P59 Q54P68 Q54P69 Q54PA5 Q54PB1 Q54PF8 Q54PG8 Q54PI4 Q54PK8 Q54PN1 Q54PP1
Q54PP8 Q54PR7 Q54PT3 Q54PT9 Q54PW9 Q54PY7 Q54Q31 Q54Q42 Q54Q53 Q54Q69 Q54Q92
Q54QA8 Q54QD0 Q54QD2 Q54QE8 Q54QG6 Q54QG7 Q54QH3 Q54QI8 Q54QJ7 Q54QM0 Q54QM5
Q54QN1 Q54QN2 Q54QN4 Q54QP7 Q54QR5 Q54QR8 Q54QU8 Q54QV5 Q54QV8 Q54QY3 Q54QY9

Q54R31 Q54R34 Q54R36 Q54R37 Q54R45 Q54R52 Q54R82 Q54RB9 Q54RE1 Q54RJ1 Q54RJ4
Q54RJ6 Q54RN1 Q54RQ1 Q54RQ3 Q54RU1 Q54RV9 Q54RZ2 Q54RZ7 Q54S10 Q54S12 Q54S31
Q54S52 Q54SD5 Q54SG3 Q54SH2 Q54SH7 Q54SH8 Q54SJ4 Q54SK0 Q54SK5 Q54SN0 Q54ST6
Q54SV9 Q54SW3 Q54T01 Q54T02 Q54T06 Q54T09 Q54T60 Q54T74 Q54T82 Q54TA1 Q54TA2
Q54TB2 Q54TC9 Q54TL0 Q54TM2 Q54TS4 Q54TU8 Q54TV1 Q54TV2 Q54TW0 Q54TW5 Q54U35
Q54U44 Q54U65 Q54U71 Q54U75 Q54U86 Q54U87 Q54U89 Q54UA5 Q54UB0 Q54UD3 Q54UD4
Q54UF5 Q54UF8 Q54UI0 Q54UJ6 Q54UK4 Q54UP0 Q54UQ1 Q54UW8 Q54V07 Q54V53 Q54V54
Q54V76 Q54V79 Q54V86 Q54V96 Q54VC1 Q54VF9 Q54VG6 Q54VH7 Q54VI6 Q54VI8 Q54VJ0
Q54VP1 Q54VS2 Q54VS4 Q54VS7 Q54VW1 Q54VX4 Q54VY0 Q54W11 Q54W19 Q54W20 Q54W24
Q54W90 Q54WB9 Q54WH4 Q54WM0 Q54WN6 Q54WR6 Q54WS5 Q54WT7 Q54WT8 Q54WZ5 Q54X25
Q54X86 Q54XG9 Q54XK1 Q54XK2 Q54XM0 Q54XM6 Q54XS3 Q54XX1 Q54XX9 Q54XY6 Q54Y83
Q54YC4 Q54YH4 Q54YK1 Q54YM7 Q54YN3 Q54YR4 Q54ZG7 Q54ZM4 Q54ZW0 Q54ZX5 Q55026
Q55027 Q550A6 Q550C1 Q550R7 Q550S9 Q550T0 Q550T1 Q550W1 Q550W6 Q55106 Q55115
Q55143 Q55147 Q55153 Q55179 Q55190 Q551M4 Q551X6 Q55274 Q55282 Q552L9 Q552M6
Q552P3 Q552S0 Q55329 Q55330 Q55339 Q55354 Q55364 Q553K4 Q553P5 Q553P6 Q553Y0
Q553Y1 Q553Y7 Q55403 Q55415 Q55429 Q55438 Q55461 Q55472 Q55473 Q55487 Q554E2
Q554E7 Q554S4 Q55500 Q55518 Q555E8 Q555I8 Q555N7 Q555U5 Q555X4 Q555Z5 Q55610
Q55611 Q55684 Q556C6 Q556J4 Q556J9 Q556K9 Q556M4 Q556M5 Q556N7 Q556S2 Q556W1
Q556W2 Q556Z9 Q55700 Q55705 Q55714 Q55717 Q55720 Q55721 Q55736 Q55774 Q557E9
Q557H5 Q55827 Q55834 Q558K8 Q558Y6 Q558Z2 Q55909 Q55932 Q55939 Q55979 Q55986
Q559A0 Q559K0 Q559M5 Q559W3 Q55A00 Q55A09 Q55AJ4 Q55AP1 Q55AP3 Q55AP8 Q55AQ3
Q55AW9 Q55B06 Q55B97 Q55B99 Q55BC0 Q55BF4 Q55BH9 Q55BP8 Q55BU8 Q55BU9 Q55C17
Q55C66 Q55C70 Q55CD6 Q55CN6 Q55CN8 Q55CT1 Q55CV4 Q55CV5 Q55CV7 Q55CW2 Q55CW7
Q55CY2 Q55CY9 Q55CZ1 Q55D52 Q55D54 Q55D55 Q55D90 Q55DA0 Q55DA7 Q55DM5 Q55DQ0
Q55DQ2 Q55DR1 Q55DT7 Q55DU7 Q55DV2 Q55DW4 Q55DY8 Q55E32 Q55E35 Q55E44 Q55E45
Q55E69 Q55E85 Q55E98 Q55EA1 Q55EF8 Q55EH8 Q55EK2 Q55EX3 Q55F11 Q55F68 Q55F69
Q55FD5 Q55FD8 Q55FL1 Q55FL5 Q55FP0 Q55FQ9 Q55FR8 Q55FS3 Q55FV8 Q55FX9 Q55FY6
Q55G23 Q55G31 Q55G32 Q55G45 Q55GB1 Q55GE2 Q55GH9 Q55GI8 Q55GJ2 Q55GJ3 Q55GK8
Q55GS3 Q55GT2 Q55GT5 Q55GU3 Q55GU4 Q55GW4 Q56019 Q56023 Q56036 Q56072 Q56110
Q56111 Q56113 Q56119 Q56128 Q56134 Q561R8 Q561T9 Q561X0 Q56217 Q56225 Q56226
Q56227 Q56228 Q56229 Q562B5 Q56331 Q56338 Q56347 Q56348 Q56352 Q56408 Q56460
Q56461 Q56463 Q564G3 Q564G7 Q564N6 Q564N8 Q564P1 Q564P2 Q56577 Q56578 Q56580
Q56582 Q56584 Q56587 Q56589 Q56652 Q566D0 Q566E6 Q566G2 Q566J8 Q566M8 Q566N0
Q566R2 Q566S6 Q566Y9 Q56714 Q56724 Q56725 Q56740 Q56763 Q56770 Q567V2 Q567X1
Q567X9 Q56828 Q56850 Q56886 Q56887 Q568B8 Q568I2 Q568J8 Q568N3 Q568T5 Q568Y3
Q568Y5 Q568Z4 Q56902 Q56954 Q56955 Q56956 Q56957 Q56983 Q56989 Q56992 Q569A2
Q569A3 Q569A6 Q569C0 Q569T7 Q56A06 Q56A07 Q56A20 Q56A55 Q56EB3 Q56H28 Q56H79
Q56NL1 Q56P05 Q56P07 Q56P08 Q56P09 Q56P14 Q56P15 Q56P16 Q56P17 Q56P28 Q56RZ1
Q56RZ2 Q56RZ5 Q56RZ6 Q56UD9 Q56UJ5 Q56VS4 Q56WD3 Q56X46 Q56X52 Q56X72 Q56X75
Q56X95 Q56XE8 Q56XP4 Q56XQ0 Q56XQ6 Q56XR7 Q56Y01 Q56ZN6 Q56ZQ3 Q56ZZ7 Q57007
Q57017 Q57020 Q57038 Q57048 Q57071 Q57095 Q57097 Q570B4 Q570Q8 Q570S7 Q570X5
Q57101 Q57124 Q57130 Q57147 Q57180 Q57208 Q57251 Q57256 Q57286 Q57288 Q57306
Q57320 Q57321 Q57335 Q57341 Q57347 Q57350 Q57362 Q57380 Q57389 Q573B7 Q573C0
Q573C2 Q573C7 Q573D3 Q573E0 Q573E1 Q573F0 Q573F1 Q57408 Q57424 Q57425 Q57442

Q57484 Q57486 Q57491 Q57493 Q57500 Q57506 Q57523 Q57538 Q57552 Q57556 Q57561
Q57570 Q57571 Q57574 Q57575 Q57593 Q57595 Q57601 Q57603 Q57604 Q57613 Q57627
Q57629 Q57634 Q57643 Q57644 Q57674 Q57675 Q57683 Q57685 Q576B4 Q576B5 Q576B8
Q576D7 Q576D9 Q576S3 Q57708 Q57718 Q57719 Q57723 Q57727 Q57736 Q57737 Q57740
Q57753 Q57769 Q57772 Q57789 Q577J7 Q57806 Q57817 Q57837 Q57848 Q57856 Q57858
Q57861 Q57862 Q57863 Q57873 Q57879 Q57882 Q57883 Q57891 Q57898 Q578A0 Q578E7
Q578E8 Q578M9 Q578U0 Q578U1 Q57904 Q57915 Q57928 Q57938 Q57939 Q57940 Q57941
Q57942 Q57943 Q57944 Q57945 Q57946 Q57947 Q57948 Q57965 Q57966 Q57974 Q57985
Q57986 Q57994 Q57996 Q579K4 Q57A43 Q57AW3 Q57B74 Q57BC3 Q57BM1 Q57BY9 Q57C75
Q57CC4 Q57CD6 Q57CD7 Q57D97 Q57DM5 Q57DP0 Q57DU1 Q57DU4 Q57DW5 Q57ED2 Q57EX8
Q57EY0 Q57F40 Q57FM7 Q57G46 Q57G59 Q57G63 Q57GJ9 Q57GN6 Q57GN7 Q57GQ5 Q57GV4
Q57GZ3 Q57GZ6 Q57H01 Q57H87 Q57HD7 Q57HE1 Q57HE2 Q57HF4 Q57HG6 Q57HI9 Q57HN4
Q57HN6 Q57HQ5 Q57HS3 Q57HW3 Q57HX3 Q57HX4 Q57HX5 Q57HZ5 Q57HZ7 Q57I28 Q57I71
Q57IK8 Q57IN7 Q57IS1 Q57IS2 Q57IV1 Q57IW8 Q57J00 Q57J15 Q57J60 Q57J90 Q57JA2
Q57JA3 Q57JA4 Q57JD0 Q57JE2 Q57JL7 Q57JN4 Q57JQ2 Q57JQ4 Q57JR8 Q57JS2 Q57JW1
Q57K48 Q57KA7 Q57KA8 Q57KB5 Q57KJ3 Q57L56 Q57LI5 Q57LM7 Q57LN2 Q57LT0 Q57LU5
Q57LU9 Q57LY2 Q57M19 Q57M28 Q57M34 Q57M37 Q57M40 Q57M52 Q57M53 Q57M54 Q57M56
Q57MA1 Q57MH8 Q57MM4 Q57MM5 Q57MW3 Q57MX5 Q57MY0 Q57N14 Q57N57 Q57NG5 Q57NG9
Q57NH7 Q57NH8 Q57NK5 Q57NK6 Q57NL1 Q57NS0 Q57NS4 Q57NS5 Q57NX8 Q57P04 Q57P83
Q57PB5 Q57PB6 Q57PD4 Q57PF4 Q57PF5 Q57PH8 Q57PI1 Q57PI3 Q57PL1 Q57PU6 Q57Q12
Q57Q14 Q57Q25 Q57QC4 Q57QJ1 Q57QK4 Q57QK9 Q57R14 Q57R33 Q57R58 Q57R91 Q57RA9
Q57RB0 Q57RM6 Q57RM9 Q57RN0 Q57RN1 Q57RU6 Q57RY1 Q57SC4 Q57SD7 Q57SD8 Q57T31
Q57T52 Q57TD3 Q57TH4 Q57TI7 Q57TL4 Q57TM1 Q57VW6 Q58004 Q58005 Q58007 Q58023
Q58026 Q58028 Q58047 Q58061 Q58076 Q58086 Q58091 Q58107 Q58111 Q58115 Q58117
Q58119 Q58121 Q58128 Q58159 Q58160 Q58169 Q58172 Q58186 Q58189 Q58190 Q58191
Q58200 Q58202 Q58203 Q58204 Q58205 Q58207 Q58213 Q58236 Q58237 Q58245 Q58257
Q58258 Q58259 Q58260 Q58261 Q58262 Q58263 Q58281 Q58286 Q58287 Q58290 Q58308
Q58311 Q58312 Q58313 Q58314 Q58333 Q58343 Q58348 Q58353 Q58355 Q58363 Q58381
Q58382 Q58386 Q58389 Q58402 Q58404 Q58419 Q58420 Q58421 Q58423 Q58430 Q58437
Q58438 Q58439 Q58442 Q58467 Q58472 Q58474 Q58478 Q58479 Q58480 Q58489 Q58490
Q58491 Q58492 Q58507 Q58513 Q58537 Q58538 Q58541 Q58543 Q58547 Q58561 Q58578
Q58580 Q58589 Q58596 Q58607 Q58609 Q58616 Q58618 Q58620 Q58623 Q58650 Q58651
Q58665 Q58666 Q58671 Q58676 Q58679 Q58681 Q58682 Q58685 Q58686 Q58689 Q58702
Q58703 Q58704 Q58705 Q58706 Q58710 Q58713 Q58714 Q58715 Q58739 Q58749 Q58752
Q58753 Q58756 Q58757 Q58760 Q58763 Q58768 Q58770 Q58781 Q58788 Q58789 Q58790
Q58791 Q58795 Q58798 Q587I9 Q58812 Q58828 Q58833 Q58837 Q58838 Q58852 Q58863
Q58864 Q58865 Q58866 Q58872 Q58878 Q58880 Q58886 Q58887 Q58890 Q588Y6 Q58901
Q58902 Q58911 Q58916 Q58918 Q58920 Q58922 Q58930 Q58948 Q58951 Q58955 Q58956
Q58957 Q58961 Q58964 Q58965 Q58966 Q58971 Q58972 Q58975 Q58983 Q58984 Q58985
Q58990 Q58995 Q589A5 Q589B1 Q589B2 Q589G5 Q58A42 Q58CP9 Q58CR4 Q58CT4 Q58CT8
Q58CU4 Q58CU5 Q58CV5 Q58CW4 Q58CW5 Q58CY8 Q58D72 Q58D78 Q58D85 Q58D91 Q58DA4
Q58DA6 Q58DA8 Q58DD0 Q58DD2 Q58DD4 Q58DF6 Q58DF9 Q58DH2 Q58DH7 Q58DI5 Q58DK1
Q58DM3 Q58DN3 Q58DR2 Q58DR5 Q58DR6 Q58DS3 Q58DS4 Q58DT3 Q58DX5 Q58DZ9 Q58E03
Q58E26 Q58E76 Q58EA0 Q58EC8 Q58EG2 Q58EG3 Q58EJ9 Q58EK4 Q58EL2 Q58EX2 Q58F56

Q58F60 Q58F69 Q58F73 Q58F82 Q58F86 Q58G33 Q58G35 Q58HT5 Q58HT7 Q58P69 Q58P71

Q58Y75 Q59012 Q59016 Q59038 Q59071 Q59076 Q59086 Q59167 Q591M5 Q591M6 Q591P2

Q591V2 Q591Y0 Q591Y7 Q591Y8 Q59207 Q59297 Q592E4 Q59327 Q59385 Q59465 Q59467

Q594P3 Q59548 Q59550 Q59584 Q59589 Q595Z2 Q59640 Q59646 Q59647 Q59659 Q59660

Q597C5 Q597C7 Q597E2 Q597E3 Q597E4 Q597E6 Q597E7 Q597E8 Q597E9 Q597F0 Q597F1

Q597F2 Q59835 Q598S9 Q598T9 Q598U3 Q598V2 Q598Z5 Q59912 Q59916 Q59947 Q59986

Q59998 Q599A0 Q599A1 Q599A4 Q59E83 Q59I64 Q59J82 Q59KC4 Q59KJ7 Q59LF2 Q59LL4

Q59LP6 Q59LV5 Q59LX9 Q59M70 Q59MA6 Q59MC8 Q59MJ1 Q59MJ2 Q59MZ9 Q59N74 Q59NP1

Q59NR8 Q59NY7 Q59P03 Q59Q30 Q59Q36 Q59Q43 Q59Q79 Q59QC6 Q59QL0 Q59R09 Q59R28

Q59RA2 Q59RF7 Q59RG0 Q59RQ2 Q59S72 Q59S85 Q59SI2 Q59SI5 Q59SU1 Q59UQ7 Q59V93

Q59VG6 Q59VL7 Q59VN0 Q59VP0 Q59VW7 Q59W44 Q59W55 Q59WB3 Q59WF4 Q59WG7 Q59WU0

Q59WU8 Q59WV0 Q59X23 Q59XM0 Q59XW9 Q59YF0 Q59YF4 Q59YK4 Q59YS7 Q59YT1 Q59YV2

Q59ZI3 Q59ZX6 Q5A044 Q5A061 Q5A0L7 Q5A0L9 Q5A1L6 Q5A2J7 Q5A302 Q5A368 Q5A3M6

Q5A3Z6 Q5A446 Q5A477 Q5A4E3 Q5A4J4 Q5A4M8 Q5A4N0 Q5A5N6 Q5A5S7 Q5A663 Q5A687

Q5A688 Q5A6K2 Q5A6N8 Q5A6P6 Q5A762 Q5A846 Q5A861 Q5A869 Q5A872 Q5A888 Q5A970

Q5AA40 Q5AB93 Q5ABC5 Q5ABP8 Q5ABR2 Q5ABT8 Q5ABU7 Q5ABU8 Q5ACH7 Q5ACL7 Q5ACU3

Q5ACW6 Q5AD51 Q5AD72 Q5ADM9 Q5ADN8 Q5ADN9 Q5ADQ9 Q5AEC6 Q5AEK8 Q5AEM8 Q5AFF7

Q5AFH3 Q5AFI4 Q5AFP8 Q5AFT2 Q5AG77 Q5AGA0 Q5AGA9 Q5AGV7 Q5AH11 Q5AH12 Q5AHC2

Q5AHD6 Q5AI21 Q5AJD2 Q5AJX2 Q5AK24 Q5AKW3 Q5AKZ2 Q5AL27 Q5ALN1 Q5ALW2 Q5AM72

Q5AMH3 Q5AMQ4 Q5AMR5 Q5ANA3 Q5ANC9 Q5AND1 Q5ANE3 Q5ANH2 Q5AP90 Q5APA2 Q5APC0

Q5APD4 Q5APQ8 Q5AQ36 Q5AQA2 Q5AQY2 Q5AR21 Q5AR23 Q5AR27 Q5AR29 Q5AR32 Q5AR50

Q5AS50 Q5AS82 Q5ASG4 Q5ATG7 Q5ATG9 Q5ATH2 Q5ATK1 Q5AUX8 Q5AUY2 Q5AUZ9 Q5AV01

Q5AV03 Q5AVI2 Q5AVT9 Q5AVW1 Q5AVZ7 Q5AW93 Q5AWM9 Q5AXD1 Q5AXJ9 Q5AXV1 Q5AY94

Q5AYC8 Q5AYH8 Q5AZB4 Q5AZQ5 Q5B0L1 Q5B0M8 Q5B0R9 Q5B0V6 Q5B1G6 Q5B1Q2 Q5B1X5

Q5B2K4 Q5B3W7 Q5B430 Q5B433 Q5B4H1 Q5B4H2 Q5B4P0 Q5B538 Q5B5L3 Q5B5L5 Q5B5W2

Q5B6I7 Q5B6J9 Q5B6U6 Q5B717 Q5B7A1 Q5B7I9 Q5B7W0 Q5B8A2 Q5B8K4 Q5B905 Q5B934

Q5BAE9 Q5BAW7 Q5BAX7 Q5BAY0 Q5BB01 Q5BB40 Q5BB76 Q5BBC6 Q5BBM5 Q5BC23 Q5BCB9

Q5BCK8 Q5BD15 Q5BD67 Q5BDG8 Q5BDJ6 Q5BE65 Q5BEE5 Q5BEG1 Q5BEJ7 Q5BEJ9 Q5BF53

Q5BFT5 Q5BG98 Q5BGC2 Q5BGS2 Q5BI42 Q5BIM5 Q5BIM9 Q5BIN4 Q5BIN6 Q5BIR3 Q5BIY5

Q5BIZ0 Q5BJ68 Q5BJC1 Q5BJC2 Q5BJD5 Q5BJF2 Q5BJH2 Q5BJH7 Q5BJI9 Q5BJK8 Q5BJM8

Q5BJN9 Q5BJP5 Q5BJQ0 Q5BJS2 Q5BJS4 Q5BJT4 Q5BJU5 Q5BJW3 Q5BJW9 Q5BK09 Q5BK13

Q5BK17 Q5BK26 Q5BK38 Q5BK39 Q5BK49 Q5BK54 Q5BK62 Q5BK65 Q5BK75 Q5BK83 Q5BK84

Q5BK85 Q5BKD0 Q5BKJ7 Q5BKR2 Q5BKT4 Q5BKW8 Q5BKX0 Q5BKX6 Q5BL21 Q5BL29 Q5BL33

Q5BL38 Q5BL44 Q5BL50 Q5BL63 Q5BL72 Q5BL81 Q5BLB7 Q5BLC7 Q5BLE2 Q5BLG4 Q5BMR2

Q5BPJ0 Q5BPL5 Q5BPZ5 Q5BQ04 Q5BQE6 Q5BQG8 Q5BQG9 Q5BU39 Q5BU73 Q5BU77 Q5BU90

Q5BVD1 Q5C9H5 Q5CAZ6 Q5CB03 Q5CB04 Q5CC92 Q5CC93 Q5CCK1 Q5CCK3 Q5CD18 Q5CL93

Q5CZ37 Q5CZ51 Q5CZL8 Q5CZN0 Q5CZQ0 Q5CZV0 Q5D0K2 Q5D1Z7 Q5DC12 Q5DID0 Q5DID3

Q5DJV7 Q5DNB7 Q5DRA2 Q5DRA3 Q5DRA4 Q5DRA5 Q5DRA6 Q5DRA7 Q5DRA8 Q5DRA9 Q5DRB0

Q5DRB1 Q5DRB2 Q5DRB3 Q5DRB4 Q5DRB5 Q5DRB6 Q5DRB7 Q5DRB8 Q5DRB9 Q5DRC0 Q5DRC1

Q5DRC2 Q5DRC3 Q5DRC4 Q5DRC6 Q5DRC7 Q5DRC8 Q5DRC9 Q5DRD0 Q5DRD1 Q5DRD2 Q5DRD3

Q5DRD4 Q5DRD5 Q5DRD6 Q5DRD7 Q5DRD8 Q5DRD9 Q5DRE0 Q5DRE1 Q5DRE2 Q5DRE3 Q5DRE4

Q5DRE5 Q5DRE6 Q5DRE7 Q5DRE8 Q5DRE9 Q5DRF0 Q5DRF1 Q5DRF2 Q5DRF3 Q5DRF4 Q5DRF5

Q5DRK1 Q5DT36 Q5DT37 Q5DT39 Q5DTI8 Q5DTK1 Q5DTL9 Q5DTZ0 Q5DTZ6 Q5DU28 Q5DU41

Q5DUB1 Q5DUB2 Q5DUB3 Q5DUY0 Q5DWG5 Q5DWV1 Q5DWV2 Q5DX17 Q5DX21 Q5DZC3 Q5E0F2

Q5E1N1 Q5E1N2 Q5E1N3 Q5E207 Q5E2B5 Q5E2K3 Q5E2K6 Q5E2P3 Q5E2P7 Q5E2Q2 Q5E327
Q5E3L0 Q5E3T1 Q5E4B6 Q5E4B7 Q5E4I5 Q5E4Y6 Q5E5M9 Q5E5P9 Q5E5T6 Q5E626 Q5E684
Q5E6B6 Q5E6C0 Q5E6C1 Q5E6C7 Q5E6G2 Q5E6G4 Q5E6L3 Q5E6X2 Q5E6X3 Q5E733 Q5E7P2
Q5E861 Q5E8Q1 Q5E8V2 Q5E8V3 Q5E930 Q5E943 Q5E960 Q5E965 Q5E971 Q5E972 Q5E975
Q5E978 Q5E980 Q5E9B7 Q5E9C2 Q5E9E4 Q5E9E7 Q5E9E8 Q5E9F2 Q5E9H5 Q5E9H7 Q5E9H8
Q5E9I3 Q5E9J2 Q5E9J5 Q5E9K8 Q5E9L0 Q5E9M1 Q5E9M8 Q5E9M9 Q5E9P3 Q5E9P6 Q5E9Q3
Q5E9R1 Q5E9R2 Q5E9R6 Q5E9S9 Q5E9T5 Q5E9U1 Q5E9U3 Q5E9V4 Q5E9W3 Q5E9W5 Q5E9X4
Q5E9Y2 Q5E9Z5 Q5E9Z9 Q5EA01 Q5EA03 Q5EA06 Q5EA10 Q5EA26 Q5EA41 Q5EA42 Q5EA43
Q5EA45 Q5EA46 Q5EA48 Q5EA65 Q5EA68 Q5EA70 Q5EA85 Q5EA86 Q5EA87 Q5EA90 Q5EA91
Q5EA97 Q5EAA5 Q5EAB0 Q5EAB6 Q5EAC0 Q5EAE0 Q5EAE9 Q5EAJ6 Q5EAK8 Q5EAL3 Q5EAL6
Q5EAT6 Q5EAU0 Q5EAU3 Q5EAU9 Q5EAX9 Q5EAY8 Q5EB14 Q5EB52 Q5EB62 Q5EB63 Q5EB66
Q5EB73 Q5EB76 Q5EB81 Q5EBA8 Q5EBF8 Q5EBI0 Q5ECR9 Q5EG65 Q5EG71 Q5EGZ1 Q5ENN5
Q5ENP6 Q5EXK5 Q5EZ72 Q5F226 Q5F267 Q5F285 Q5F289 Q5F297 Q5F2L1 Q5F2L2 Q5F2N2
Q5F2N3 Q5F2N4 Q5F334 Q5F349 Q5F362 Q5F364 Q5F380 Q5F381 Q5F383 Q5F384 Q5F3A1
Q5F3A4 Q5F3C1 Q5F3F5 Q5F3G7 Q5F3I6 Q5F3L3 Q5F3L7 Q5F3N0 Q5F3W2 Q5F3X0 Q5F407
Q5F409 Q5F410 Q5F433 Q5F448 Q5F468 Q5F499 Q5F4B8 Q5F4C0 Q5F4E4 Q5F4E8 Q5F4F1
Q5F4F4 Q5F4F5 Q5F4G0 Q5F4G2 Q5F4G3 Q5F4G7 Q5F4G9 Q5F4W6 Q5F4X8 Q5F4Z4 Q5F5G9
Q5F600 Q5F622 Q5F625 Q5F629 Q5F688 Q5F6A1 Q5F6F6 Q5F6K4 Q5F6K8 Q5F6L4 Q5F6M1
Q5F6V6 Q5F6X5 Q5F6X6 Q5F6X7 Q5F791 Q5F8C7 Q5F8D0 Q5F8D6 Q5F8Z3 Q5F9J4 Q5F9S8
Q5FA49 Q5FA50 Q5FAC7 Q5FAF3 Q5FB95 Q5FBC3 Q5FBC4 Q5FC15 Q5FC64 Q5FC78 Q5FC79
Q5FF11 Q5FG54 Q5FGC3 Q5FGG9 Q5FGK0 Q5FGK8 Q5FGW5 Q5FHG0 Q5FI75 Q5FJL6 Q5FK09
Q5FKD1 Q5FKD2 Q5FKJ5 Q5FKJ8 Q5FKV4 Q5FL71 Q5FL81 Q5FLF5 Q5FLW0 Q5FMI9 Q5FMK7
Q5FMS7 Q5FNB7 Q5FNE6 Q5FNY7 Q5FPU4 Q5FQF4 Q5FQL0 Q5FRH0 Q5FRJ4 Q5FRU4 Q5FRW5
Q5FRW6 Q5FRW8 Q5FU11 Q5FUJ8 Q5FUS6 Q5FV41 Q5FVF4 Q5FVG1 Q5FVH2 Q5FVI4 Q5FVJ3
Q5FVJ4 Q5FVJ6 Q5FVL3 Q5FVL6 Q5FVM1 Q5FVM3 Q5FVM5 Q5FVM7 Q5FVN0 Q5FVN1 Q5FVN2
Q5FVP8 Q5FVQ0 Q5FVQ4 Q5FVQ5 Q5FVQ7 Q5FVR0 Q5FVR1 Q5FVR3 Q5FW56 Q5FWC3 Q5FWE3
Q5FWH7 Q5FWI2 Q5FWI3 Q5FWK6 Q5FWL4 Q5FWL7 Q5FWM4 Q5FWM8 Q5FWR8 Q5FWS4 Q5FWU3
Q5FWV6 Q5FY69 Q5FYA8 Q5G235 Q5G2C6 Q5G5D5 Q5G7X0 Q5G8Z3 Q5GA86 Q5GCA5 Q5GF25
Q5GFD5 Q5GH22 Q5GH56 Q5GH57 Q5GH59 Q5GH60 Q5GH61 Q5GH62 Q5GH64 Q5GH66 Q5GH67
Q5GH68 Q5GH70 Q5GH72 Q5GH73 Q5GH76 Q5GH77 Q5GIT4 Q5GJ04 Q5GJ77 Q5GKZ7 Q5GRV8
Q5GRZ3 Q5GS15 Q5GS17 Q5GSH6 Q5GSH7 Q5GSI0 Q5GSL3 Q5GSR1 Q5GSX9 Q5GTK3 Q5GTT3
Q5GU80 Q5GV02 Q5GV87 Q5GW39 Q5GW40 Q5GWB4 Q5GWQ2 Q5GXD6 Q5GXT9 Q5GXU2 Q5GYG0
Q5GYK5 Q5GZ91 Q5GZS7 Q5H077 Q5H0H0 Q5H0W6 Q5H1N2 Q5H1V9 Q5H1X6 Q5H2A5 Q5H2E0
Q5H3V0 Q5H488 Q5H4Y8 Q5H5R3 Q5H616 Q5H629 Q5H879 Q5H8A4 Q5H8A5 Q5H8A6 Q5H943
Q5H9E4 Q5H9R4 Q5H9S7 Q5HA45 Q5HA73 Q5HAA6 Q5HAH7 Q5HB99 Q5HBM4 Q5HBS1 Q5HCG2
Q5HCL2 Q5HCN0 Q5HCN1 Q5HCN3 Q5HCP4 Q5HCY3 Q5HCY5 Q5HCY7 Q5HCZ3 Q5HD01 Q5HD10
Q5HD11 Q5HD13 Q5HD69 Q5HDB2 Q5HDG7 Q5HDJ3 Q5HDJ5 Q5HDJ9 Q5HDL7 Q5HDM2 Q5HDV2
Q5HDX8 Q5HE15 Q5HE37 Q5HE48 Q5HE91 Q5HE92 Q5HE93 Q5HEA9 Q5HEB2 Q5HEC3 Q5HEC4
Q5HEC5 Q5HEG5 Q5HEG6 Q5HEK4 Q5HEL0 Q5HEM0 Q5HEN5 Q5HEN9 Q5HEQ0 Q5HEQ8 Q5HES9
Q5HEZ2 Q5HEZ3 Q5HF02 Q5HF35 Q5HF56 Q5HFI7 Q5HFT1 Q5HFU2 Q5HFY7 Q5HFY8 Q5HG01
Q5HG05 Q5HG12 Q5HG19 Q5HG38 Q5HG39 Q5HG57 Q5HG59 Q5HG66 Q5HG71 Q5HG76 Q5HGA3
Q5HGE5 Q5HGE8 Q5HGF5 Q5HGG9 Q5HGH0 Q5HGN6 Q5HGP9 Q5HGV0 Q5HGW8 Q5HGW9 Q5HGX9
Q5HH23 Q5HH24 Q5HH25 Q5HH26 Q5HH63 Q5HH70 Q5HHB1 Q5HHB5 Q5HHB9 Q5HHC0 Q5HHD3
Q5HHD4 Q5HHD5 Q5HHD6 Q5HHD7 Q5HHD8 Q5HHD9 Q5HHN9 Q5HHQ7 Q5HHS4 Q5HHV4 Q5HHW5

Q5HHX4 Q5HHY5 Q5HI08 Q5HI39 Q5HI40 Q5HI41 Q5HI42 Q5HI43 Q5HI44 Q5HI45 Q5HI93
Q5HIA2 Q5HIE0 Q5HIV4 Q5HIW0 Q5HJ86 Q5HJ87 Q5HJ89 Q5HJ90 Q5HJA6 Q5HJB3 Q5HJB4
Q5HJB6 Q5HJF6 Q5HJG9 Q5HJI0 Q5HJI3 Q5HJX6 Q5HK19 Q5HK63 Q5HK64 Q5HK65 Q5HKB0
Q5HKE1 Q5HKP7 Q5HKP9 Q5HKQ0 Q5HL56 Q5HL70 Q5HL71 Q5HL73 Q5HL85 Q5HLG0 Q5HLG1
Q5HLG3 Q5HLG6 Q5HLK7 Q5HLN1 Q5HLN3 Q5HLN7 Q5HLR3 Q5HLT2 Q5HLZ2 Q5HM19 Q5HM40
Q5HM62 Q5HMB3 Q5HMB4 Q5HMB5 Q5HMD1 Q5HMD3 Q5HMY5 Q5HN24 Q5HN32 Q5HN45 Q5HN49
Q5HN60 Q5HN90 Q5HND0 Q5HND1 Q5HNH0 Q5HNI8 Q5HNX4 Q5HP65 Q5HPA2 Q5HPB7 Q5HPC4
Q5HPD5 Q5HPI1 Q5HPI7 Q5HPJ2 Q5HPJ7 Q5HPM5 Q5HPQ5 Q5HPQ8 Q5HPR5 Q5HPT0 Q5HPZ5
Q5HQ10 Q5HQ51 Q5HQ52 Q5HQ64 Q5HQA9 Q5HQB0 Q5HQB1 Q5HQB2 Q5HQE2 Q5HQE8 Q5HQJ2
Q5HQL0 Q5HQL1 Q5HQL2 Q5HQL3 Q5HQL4 Q5HQL5 Q5HQL6 Q5HQU8 Q5HQW7 Q5HQY3 Q5HR16
Q5HR29 Q5HR54 Q5HR80 Q5HRA6 Q5HRA7 Q5HRA8 Q5HRA9 Q5HRB0 Q5HRB1 Q5HRB2 Q5HRG3
Q5HRH0 Q5HRI3 Q5HRL7 Q5HRT2 Q5HSD8 Q5HSD9 Q5HSL9 Q5HSM5 Q5HTQ5 Q5HTR0 Q5HTT8
Q5HTX9 Q5HU07 Q5HUM3 Q5HUT1 Q5HV55 Q5HVG3 Q5HVG7 Q5HVJ1 Q5HVP6 Q5HW33 Q5HW60
Q5HWA6 Q5HWB0 Q5HWW4 Q5HWZ9 Q5HXN6 Q5HYA8 Q5HYJ1 Q5HYL7 Q5HZA9 Q5HZB0 Q5HZD4
Q5HZE0 Q5HZE2 Q5HZE5 Q5HZE8 Q5HZH7 Q5HZI9 Q5HZJ5 Q5HZK2 Q5HZL5 Q5HZN7 Q5HZP7
Q5HZP8 Q5HZQ9 Q5HZS2 Q5HZT8 Q5HZT9 Q5HZW3 Q5HZW5 Q5I012 Q5I020 Q5I027 Q5I030
Q5I047 Q5I0B2 Q5I0C4 Q5I0E7 Q5I0E9 Q5I0H4 Q5I0I2 Q5I0I8 Q5I130 Q5I191 Q5I195
Q5I2B1 Q5I2M3 Q5I2M4 Q5I2M5 Q5I2M7 Q5I2M8 Q5I2N3 Q5I396 Q5I3B1 Q5I3B2 Q5I4B8
Q5I4F8 Q5I4F9 Q5I7T1 Q5IBK3 Q5IBK4 Q5IFJ9 Q5IFP1 Q5IGR6 Q5IGR7 Q5IGR8 Q5IHA3
Q5IHA8 Q5IHA9 Q5IHB0 Q5IHC2 Q5IHC3 Q5IHD7 Q5IHD9 Q5IHI1 Q5IJ48 Q5IJ49 Q5IRJ6
Q5IS35 Q5IS37 Q5IS39 Q5IS45 Q5IS46 Q5IS50 Q5IS51 Q5IS52 Q5IS53 Q5IS59 Q5IS62
Q5IS64 Q5IS65 Q5IS66 Q5IS72 Q5IS73 Q5IS75 Q5IS76 Q5IS77 Q5IS80 Q5IS82 Q5IS84
Q5IS98 Q5ISE2 Q5IXF8 Q5IZC8 Q5J1T0 Q5J1T3 Q5J1T4 Q5J1T5 Q5J316 Q5J3E5 Q5J3F6
Q5J3G9 Q5J3K5 Q5J3K9 Q5J3L4 Q5J3L6 Q5J3L7 Q5J3M3 Q5J3M4 Q5J3M9 Q5J3N1 Q5J583
Q5J6M8 Q5J7P3 Q5J8M3 Q5J8X5 Q5JCK5 Q5JCK6 Q5JCS9 Q5JCT0 Q5JDA9 Q5JDK7 Q5JDN5
Q5JDS2 Q5JEB3 Q5JEX0 Q5JEZ8 Q5JF03 Q5JH32 Q5JHX4 Q5JI24 Q5JJI4 Q5JJY5 Q5JK32
Q5JK39 Q5JKN1 Q5JLM1 Q5JLY8 Q5JM04 Q5JM57 Q5JMH0 Q5JN63 Q5JPE7 Q5JQD7 Q5JQS5
Q5JRA6 Q5JRM2 Q5JRS4 Q5JRV8 Q5JTH9 Q5JTV8 Q5JUK3 Q5JW98 Q5JX69 Q5JX71 Q5JXA9
Q5JXX7 Q5JZQ7 Q5JZQ8 Q5JZQ9 Q5JZY3 Q5K027 Q5K0D9 Q5K2K4 Q5K2N1 Q5K3R9 Q5K3S0
Q5K3S1 Q5K3S2 Q5K3S7 Q5K3S8 Q5K3S9 Q5K3T2 Q5K3T3 Q5K3T5 Q5K3T8 Q5K3U3 Q5K3U4
Q5K3U6 Q5K3V1 Q5K3V3 Q5K3V4 Q5K4H9 Q5K4L6 Q5K6N0 Q5K8R6 Q5K8S8 Q5KA65 Q5KH67
Q5KQH7 Q5KQN0 Q5KR61 Q5KS41 Q5KSK8 Q5KSU9 Q5KTN1 Q5KTN2 Q5KTN3 Q5KTQ9 Q5KU26
Q5KUH7 Q5KUI7 Q5KUI8 Q5KUI9 Q5KUJ5 Q5KUJ9 Q5KUK3 Q5KUK6 Q5KUV3 Q5KVC0 Q5KW53
Q5KWB4 Q5KWE8 Q5KWE9 Q5KXL9 Q5KXP1 Q5KXR9 Q5KZ35 Q5KZL2 Q5L081 Q5L0F4 Q5L0J5
Q5L0V0 Q5L0X5 Q5L0X8 Q5L114 Q5L115 Q5L1S5 Q5L248 Q5L2A6 Q5L2R6 Q5L3H7 Q5L3P4
Q5L520 Q5L5E6 Q5L5E8 Q5L6C0 Q5L6C2 Q5L6Q8 Q5L6T1 Q5L7B9 Q5L9Y0 Q5LC41 Q5LC87
Q5LD84 Q5LD85 Q5LE03 Q5LG62 Q5LG69 Q5LG75 Q5LGV7 Q5LH48 Q5LH51 Q5LHU5 Q5LHU7
Q5LIJ4 Q5LJ30 Q5LL36 Q5LLI6 Q5LLZ3 Q5LMR7 Q5LNH0 Q5LNH2 Q5LNI0 Q5LNX8 Q5LNX9
Q5LPS7 Q5LPT3 Q5LQ14 Q5LRS1 Q5LS91 Q5LU74 Q5LUA8 Q5LUA9 Q5LUZ4 Q5LW11 Q5LX62
Q5LY94 Q5M0E9 Q5M0J6 Q5M0R8 Q5M0X5 Q5M108 Q5M171 Q5M1F2 Q5M1F4 Q5M1R0 Q5M245
Q5M2U9 Q5M4V4 Q5M4Z6 Q5M532 Q5M5A4 Q5M5G2 Q5M5J5 Q5M5Q8 Q5M5Y6 Q5M5Z0 Q5M614
Q5M6A1 Q5M757 Q5M7A3 Q5M7C7 Q5M7E0 Q5M7E9 Q5M7F4 Q5M7K0 Q5M7K3 Q5M7N9 Q5M7R3
Q5M7S0 Q5M7T4 Q5M7U7 Q5M7V1 Q5M7W4 Q5M7W6 Q5M7X9 Q5M7Z0 Q5M807 Q5M826 Q5M828
Q5M836 Q5M843 Q5M844 Q5M848 Q5M854 Q5M869 Q5M871 Q5M880 Q5M883 Q5M8E3 Q5M8F7

Q5M8F9 Q5M8H5 Q5M8I8 Q5M8L8 Q5M8M9 Q5M8S2 Q5M8T2 Q5M8U1 Q5M8Y1 Q5M900 Q5M962
Q5M9A7 Q5M9B7 Q5M9I1 Q5MB13 Q5MD61 Q5MD62 Q5MD89 Q5MJS3 Q5MK06 Q5MK23 Q5MK24
Q5MNI1 Q5MNU5 Q5MNV6 Q5MNY4 Q5MPP0 Q5MQC7 Q5MQC8 Q5MQD0 Q5MQD1 Q5MR23 Q5MWV9
Q5MY90 Q5MY95 Q5MZ55 Q5MZ82 Q5MZD9 Q5MZM0 Q5MZZ8 Q5N047 Q5N052 Q5N060 Q5N075
Q5N0A1 Q5N0D5 Q5N0D6 Q5N0E4 Q5N0X7 Q5N159 Q5N160 Q5N1F0 Q5N1X0 Q5N2C7 Q5N2J1
Q5N2J2 Q5N2J3 Q5N326 Q5N337 Q5N366 Q5N386 Q5N3C1 Q5N3Q6 Q5N3Q7 Q5N3U6 Q5N3U9
Q5N4D2 Q5N4M6 Q5N4X8 Q5N553 Q5N554 Q5N555 Q5N556 Q5N559 Q5N5A8 Q5N5A9 Q5N5B0
Q5N5C7 Q5N5L8 Q5N5R2 Q5N5T8 Q5N5W1 Q5N675 Q5N794 Q5N7A7 Q5N7W3 Q5N800 Q5N808
Q5N855 Q5N892 Q5N8J1 Q5N8Q3 Q5N941 Q5N9A1 Q5N9J9 Q5N9N2 Q5NA18 Q5NAY7 Q5NAZ9
Q5NC32 Q5NCE8 Q5NCH9 Q5NCM1 Q5NCP0 Q5NCS9 Q5ND28 Q5ND56 Q5NDE3 Q5NDE4 Q5NDE5
Q5NDE6 Q5NDE7 Q5NDE8 Q5NDE9 Q5NDF0 Q5NDF1 Q5NDF2 Q5NDM9 Q5NDN0 Q5NE91 Q5NFK6
Q5NFU6 Q5NGD1 Q5NGI4 Q5NGK7 Q5NGW7 Q5NGY0 Q5NHK4 Q5NI07 Q5NI30 Q5NI56 Q5NIG3
Q5NIK7 Q5NIM5 Q5NIM8 Q5NKN2 Q5NKN4 Q5NKT8 Q5NKU6 Q5NKV1 Q5NKV2 Q5NKV4 Q5NKV6
Q5NKV9 Q5NLV5 Q5NN30 Q5NN66 Q5NN67 Q5NN77 Q5NN91 Q5NNP2 Q5NPR9 Q5NRB1 Q5NRN4
Q5NU32 Q5NUL3 Q5NUR5 Q5NUR7 Q5NV96 Q5NVA3 Q5NVB3 Q5NVB9 Q5NVC1 Q5NVC3 Q5NVF6
Q5NVI6 Q5NVJ6 Q5NVM7 Q5NVN3 Q5NVQ1 Q5NVQ2 Q5NVS6 Q5NVS8 Q5NXM9 Q5NXX6 Q5NYM6
Q5NYX9 Q5NZ68 Q5P1E0 Q5P1E3 Q5P260 Q5P2E3 Q5P2N0 Q5P2R7 Q5P2S7 Q5P2Z1 Q5P3E3
Q5P3K6 Q5P4E6 Q5P4E7 Q5P4P4 Q5P4W4 Q5P539 Q5P5M0 Q5P6D5 Q5P6E9 Q5P6J8 Q5P6K4
Q5P6Z3 Q5P6Z5 Q5P799 Q5P7A0 Q5P9D8 Q5P9I4 Q5P9K3 Q5P9R8 Q5P9U0 Q5P9Y8 Q5PAQ2
Q5PB35 Q5PBP7 Q5PC53 Q5PC58 Q5PC79 Q5PC81 Q5PCA7 Q5PCA8 Q5PCJ7 Q5PCJ8 Q5PCQ2
Q5PCR8 Q5PCR9 Q5PCS9 Q5PD19 Q5PD20 Q5PD50 Q5PD65 Q5PDC3 Q5PDE4 Q5PDL4 Q5PDL8
Q5PDS9 Q5PDU1 Q5PDU6 Q5PDW7 Q5PDW8 Q5PDW9 Q5PE66 Q5PEG0 Q5PEN5 Q5PEN7 Q5PEN8
Q5PEQ4 Q5PFP5 Q5PFQ5 Q5PFQ6 Q5PGH0 Q5PGH4 Q5PGK9 Q5PGN8 Q5PGP5 Q5PGP6 Q5PGW7
Q5PGX5 Q5PGY0 Q5PH16 Q5PH87 Q5PHE0 Q5PHE1 Q5PHF4 Q5PHH0 Q5PHJ7 Q5PHN9 Q5PHP3
Q5PHR6 Q5PHY0 Q5PI18 Q5PI76 Q5PI77 Q5PIC1 Q5PIC6 Q5PIC8 Q5PIL3 Q5PIN5 Q5PIQ0
Q5PIR4 Q5PIR7 Q5PIT4 Q5PJ05 Q5PJ35 Q5PJ48 Q5PJE1 Q5PJE5 Q5PJE6 Q5PJI9 Q5PJK9
Q5PJL0 Q5PJP1 Q5PJR1 Q5PJT2 Q5PJT7 Q5PJT8 Q5PJT9 Q5PJV6 Q5PJX7 Q5PK28 Q5PK67
Q5PKD5 Q5PKI0 Q5PKI8 Q5PKK8 Q5PKM1 Q5PKP2 Q5PKQ9 Q5PKS7 Q5PKU2 Q5PKV6 Q5PKW6
Q5PKW7 Q5PKW8 Q5PKZ7 Q5PL02 Q5PL17 Q5PL21 Q5PL70 Q5PL71 Q5PLF0 Q5PLI4 Q5PLR6
Q5PLW2 Q5PLZ1 Q5PLZ8 Q5PM72 Q5PM94 Q5PME0 Q5PMJ0 Q5PMS0 Q5PMU6 Q5PMU9 Q5PMV3
Q5PN10 Q5PN46 Q5PN55 Q5PN59 Q5PN65 Q5PN71 Q5PN94 Q5PN95 Q5PNA5 Q5PNA8 Q5PNA9
Q5PNB0 Q5PNB4 Q5PND3 Q5PND8 Q5PNE6 Q5PNF0 Q5PNI3 Q5PNK5 Q5PNP1 Q5PNZ1 Q5PP23
Q5PP32 Q5PPI4 Q5PPI7 Q5PPK1 Q5PPL3 Q5PPM7 Q5PPM8 Q5PPN7 Q5PPS7 Q5PPX5 Q5PQ27
Q5PQ35 Q5PQ63 Q5PQA4 Q5PQL3 Q5PQL5 Q5PQL7 Q5PQM0 Q5PQM9 Q5PQN2 Q5PQP2 Q5PQQ4
Q5PQQ5 Q5PQS5 Q5PQV5 Q5PQX0 Q5PQX1 Q5PQZ3 Q5PQZ7 Q5PR00 Q5PR34 Q5PR61 Q5PRC0
Q5PRC1 Q5PSV5 Q5PT50 Q5PT53 Q5PT54 Q5PT55 Q5PT56 Q5PXD3 Q5PXE2 Q5PXE3 Q5Q995
Q5QA78 Q5QA79 Q5QA80 Q5QA82 Q5QD04 Q5QD05 Q5QD06 Q5QD07 Q5QD08 Q5QD09 Q5QD10
Q5QD11 Q5QD12 Q5QD13 Q5QD14 Q5QD15 Q5QD16 Q5QD17 Q5QD21 Q5QD23 Q5QD24 Q5QD25
Q5QD28 Q5QD29 Q5QD74 Q5QD76 Q5QF96 Q5QFB9 Q5QFW4 Q5QFW5 Q5QFW6 Q5QGT7 Q5QGU6
Q5QGZ2 Q5QGZ9 Q5QJU3 Q5QM25 Q5QM84 Q5QN13 Q5QNI1 Q5QNI2 Q5QNP2 Q5QNS5 Q5QQ37
Q5QQ49 Q5QQ50 Q5QQ51 Q5QQ52 Q5QQ53 Q5QQ54 Q5QQ55 Q5QQ56 Q5QQ57 Q5QQX7 Q5QR91
Q5QRZ3 Q5QS02 Q5QS06 Q5QS15 Q5QS19 Q5QS28 Q5QS32 Q5QS41 Q5QS45 Q5QS58 Q5QS67
Q5QS71 Q5QS84 Q5QS97 Q5QSA4 Q5QSK2 Q5QT56 Q5QU36 Q5QUB1 Q5QUC2 Q5QUP3 Q5QUY5
Q5QV33 Q5QVU3 Q5QVY3 Q5QWN6 Q5QWR6 Q5QX91 Q5QY47 Q5QYQ7 Q5QYQ8 Q5QZ20 Q5QZI0

Q5QZI1 Q5QZI2 Q5QZQ1 Q5R065 Q5R0A9 Q5R0E0 Q5R0M2 Q5R0M4 Q5R0Y6 Q5R1P0 Q5R1P3
Q5R1Q1 Q5R231 Q5R2I8 Q5R2J4 Q5R3F8 Q5R3K3 Q5R419 Q5R422 Q5R440 Q5R443 Q5R447
Q5R454 Q5R460 Q5R461 Q5R482 Q5R491 Q5R4B0 Q5R4B2 Q5R4C3 Q5R4D0 Q5R4D7 Q5R4D8
Q5R4G0 Q5R4H0 Q5R4I4 Q5R4J5 Q5R4L2 Q5R4L9 Q5R4M0 Q5R4P1 Q5R4Q3 Q5R4Q6 Q5R4S2
Q5R4U5 Q5R4X4 Q5R4X8 Q5R4Y7 Q5R4Z3 Q5R501 Q5R504 Q5R533 Q5R540 Q5R542 Q5R563
Q5R570 Q5R574 Q5R589 Q5R597 Q5R598 Q5R5A1 Q5R5A8 Q5R5B0 Q5R5B8 Q5R5C5 Q5R5E8
Q5R5G4 Q5R5H4 Q5R5H7 Q5R5J8 Q5R5K5 Q5R5L5 Q5R5L9 Q5R5M0 Q5R5M4 Q5R5M5 Q5R5T8
Q5R5U4 Q5R5V2 Q5R5V8 Q5R5X2 Q5R5X4 Q5R5Y3 Q5R5Z8 Q5R601 Q5R602 Q5R603 Q5R608
Q5R611 Q5R617 Q5R621 Q5R634 Q5R660 Q5R662 Q5R668 Q5R676 Q5R687 Q5R692 Q5R6B1
Q5R6B2 Q5R6B8 Q5R6C0 Q5R6E6 Q5R6E9 Q5R6F5 Q5R6H1 Q5R6H3 Q5R6I6 Q5R6J1 Q5R6J3
Q5R6J7 Q5R6K7 Q5R6L8 Q5R6N3 Q5R6Q2 Q5R6R3 Q5R6R6 Q5R6S0 Q5R6T0 Q5R6T5 Q5R6U1
Q5R6U3 Q5R6W5 Q5R6X6 Q5R6Z1 Q5R6Z4 Q5R703 Q5R705 Q5R751 Q5R757 Q5R764 Q5R770
Q5R780 Q5R795 Q5R7A2 Q5R7A3 Q5R7B1 Q5R7B4 Q5R7C1 Q5R7C5 Q5R7D0 Q5R7E2 Q5R7E3
Q5R7E4 Q5R7E6 Q5R7F5 Q5R7H4 Q5R7J0 Q5R7J7 Q5R7K2 Q5R7K6 Q5R7M3 Q5R7Q1 Q5R7Q3
Q5R7R1 Q5R7R7 Q5R7U6 Q5R7V3 Q5R7V4 Q5R7W2 Q5R7W8 Q5R7Y0 Q5R7Z3 Q5R7Z6 Q5R800
Q5R803 Q5R804 Q5R809 Q5R819 Q5R826 Q5R831 Q5R834 Q5R838 Q5R839 Q5R866 Q5R868
Q5R875 Q5R876 Q5R880 Q5R885 Q5R891 Q5R8B1 Q5R8B5 Q5R8C7 Q5R8C8 Q5R8D5 Q5R8D6
Q5R8E0 Q5R8E5 Q5R8F1 Q5R8F6 Q5R8G3 Q5R8G5 Q5R8H1 Q5R8H3 Q5R8H8 Q5R8I8 Q5R8M2
Q5R8M3 Q5R8N4 Q5R8N9 Q5R8Q2 Q5R8S8 Q5R8V1 Q5R8X3 Q5R8X5 Q5R8Y5 Q5R8Y6 Q5R921
Q5R946 Q5R949 Q5R957 Q5R960 Q5R987 Q5R991 Q5R996 Q5R9A1 Q5R9A6 Q5R9A7 Q5R9B0
Q5R9C2 Q5R9C4 Q5R9C7 Q5R9E4 Q5R9F5 Q5R9I0 Q5R9I4 Q5R9J3 Q5R9K1 Q5R9K2 Q5R9K4
Q5R9L5 Q5R9M4 Q5R9M9 Q5R9Q3 Q5R9Q8 Q5R9Q9 Q5R9R3 Q5R9S1 Q5R9S6 Q5R9S8 Q5R9T5
Q5R9U1 Q5R9U7 Q5R9W1 Q5R9W3 Q5R9X1 Q5RA25 Q5RA31 Q5RA41 Q5RA57 Q5RA70 Q5RA76
Q5RA79 Q5RAB7 Q5RAC0 Q5RAC1 Q5RAC4 Q5RAC8 Q5RAE3 Q5RAF8 Q5RAG2 Q5RAG4 Q5RAG7
Q5RAH0 Q5RAH7 Q5RAI2 Q5RAI9 Q5RAJ4 Q5RAJ8 Q5RAK3 Q5RAL1 Q5RAL2 Q5RAL3 Q5RAL7
Q5RAL8 Q5RAM3 Q5RAN0 Q5RAP2 Q5RAP3 Q5RAP8 Q5RAP9 Q5RAQ1 Q5RAS5 Q5RAS6 Q5RAS8
Q5RAT9 Q5RAU6 Q5RAV0 Q5RAX1 Q5RAY6 Q5RAY9 Q5RAZ3 Q5RB09 Q5RB22 Q5RB29 Q5RB31
Q5RB84 Q5RB85 Q5RBB4 Q5RBC3 Q5RBC8 Q5RBD7 Q5RBD8 Q5RBG7 Q5RBJ7 Q5RBK3 Q5RBK4
Q5RBL0 Q5RBL6 Q5RBL9 Q5RBM1 Q5RBM3 Q5RBN0 Q5RBN1 Q5RBP9 Q5RBQ2 Q5RBQ8 Q5RBQ9
Q5RBS4 Q5RBT7 Q5RBT8 Q5RBU0 Q5RBW6 Q5RBX2 Q5RBY5 Q5RBY7 Q5RBZ6 Q5RBZ8 Q5RC10
Q5RC27 Q5RC29 Q5RC30 Q5RC33 Q5RC35 Q5RC38 Q5RC45 Q5RC74 Q5RC98 Q5RCA9 Q5RCB6
Q5RCB9 Q5RCC0 Q5RCC5 Q5RCC7 Q5RCD5 Q5RCD7 Q5RCD8 Q5RCE2 Q5RCF0 Q5RCG1 Q5RCH6
Q5RCI4 Q5RCJ4 Q5RCJ9 Q5RCK6 Q5RCL0 Q5RCM7 Q5RCM9 Q5RCN4 Q5RCN6 Q5RCN7 Q5RCN9
Q5RCP4 Q5RCQ5 Q5RCS0 Q5RCS3 Q5RCV1 Q5RCV8 Q5RCY3 Q5RCY6 Q5RCY9 Q5RCZ2 Q5RCZ8
Q5RD07 Q5RD09 Q5RD16 Q5RD28 Q5RD30 Q5RD34 Q5RD44 Q5RD64 Q5RD67 Q5RD81 Q5RD93
Q5RDA5 Q5RDB4 Q5RDB5 Q5RDB8 Q5RDC9 Q5RDD0 Q5RDE9 Q5RDG5 Q5RDH6 Q5RDJ4 Q5RDJ5
Q5RDJ7 Q5RDM3 Q5RDP8 Q5RDQ8 Q5RDR3 Q5RDR6 Q5RDS0 Q5RDV3 Q5RDV6 Q5RDV7 Q5RDY2
Q5RDY4 Q5RDY9 Q5RDZ8 Q5RE11 Q5RE31 Q5RE32 Q5RE33 Q5RE35 Q5RE43 Q5RE51 Q5RE57
Q5RE60 Q5RE69 Q5RE72 Q5RE85 Q5RE87 Q5RE98 Q5RE99 Q5REA1 Q5REA7 Q5REB4 Q5RED0
Q5REE3 Q5REF6 Q5REH2 Q5REH6 Q5REI5 Q5REI9 Q5REK0 Q5REK8 Q5REL0 Q5REM8 Q5REP2
Q5REP3 Q5REP8 Q5REQ5 Q5RER2 Q5RES1 Q5RET5 Q5RET6 Q5RET7 Q5REV9 Q5REW1 Q5REX0
Q5REZ0 Q5REZ1 Q5REZ6 Q5RF01 Q5RF07 Q5RF08 Q5RF19 Q5RF34 Q5RF46 Q5RF50 Q5RF53
Q5RF57 Q5RF58 Q5RF60 Q5RF73 Q5RF74 Q5RF75 Q5RF94 Q5RF96 Q5RFB6 Q5RFB7 Q5RFC1
Q5RFD2 Q5RFD5 Q5RFE0 Q5RFF0 Q5RFI0 Q5RFI9 Q5RFJ3 Q5RFJ6 Q5RFK9 Q5RFL2 Q5RFL5

Q5RFL7 Q5RFN1 Q5RFN8 Q5RFP3 Q5RFQ9 Q5RFR2 Q5RFR4 Q5RFR6 Q5RFS5 Q5RFT1 Q5RFV4
Q5RFW0 Q5RGJ8 Q5RGQ8 Q5RGS3 Q5RGU1 Q5RH02 Q5RH51 Q5RH71 Q5RHB5 Q5RHZ2 Q5RI15
Q5RII3 Q5RIU9 Q5RIV7 Q5RJF7 Q5RJH2 Q5RJH3 Q5RJI2 Q5RJK0 Q5RJM1 Q5RJQ0 Q5RJQ8
Q5RJR8 Q5RJS6 Q5RJS8 Q5RJX2 Q5RJY4 Q5RK23 Q5RK27 Q5RKH7 Q5RKI7 Q5RKI8 Q5RKJ2
Q5RKL5 Q5RKL7 Q5RKR3 Q5RKS2 Q5RL79 Q5RLM2 Q5S1V3 Q5S248 Q5S589 Q5S7T7 Q5S7W5
Q5SCW1 Q5SCW2 Q5SCW5 Q5SCW8 Q5SCX0 Q5SCX1 Q5SCX4 Q5SCX5 Q5SCX7 Q5SCY2 Q5SCY3
Q5SCY5 Q5SCY7 Q5SCZ0 Q5SCZ2 Q5SCZ6 Q5SCZ8 Q5SCZ9 Q5SD04 Q5SD06 Q5SD09 Q5SD11
Q5SD22 Q5SD23 Q5SD24 Q5SD29 Q5SD31 Q5SD33 Q5SD34 Q5SD35 Q5SD38 Q5SD39 Q5SD40
Q5SD41 Q5SD42 Q5SD43 Q5SDA5 Q5SGD2 Q5SHB3 Q5SHQ8 Q5SI82 Q5SIG4 Q5SJ11 Q5SJ79
Q5SJ80 Q5SJG0 Q5SK58 Q5SKE6 Q5SKV3 Q5SLA9 Q5SLF8 Q5SLI3 Q5SLX1 Q5SMG8 Q5SML4
Q5SMQ9 Q5SNQ7 Q5SNT2 Q5SP46 Q5SPB1 Q5SPC5 Q5SPF7 Q5SPX3 Q5SQ64 Q5SR56 Q5SRD1
Q5SRI9 Q5SRN2 Q5SS91 Q5SSE9 Q5SSG8 Q5SSN7 Q5SSZ7 Q5STR5 Q5SUC9 Q5SV17 Q5SVR0
Q5SVS4 Q5SWH9 Q5SWK7 Q5SWT3 Q5SWX8 Q5SWY8 Q5SWZ9 Q5SX19 Q5SY80 Q5SYH2 Q5SZA1
Q5SZI1 Q5SZK8 Q5SZV5 Q5T0T0 Q5T197 Q5T1A1 Q5T1Q4 Q5T1S8 Q5T292 Q5T2D2 Q5T2E6
Q5T3F8 Q5T3U5 Q5T442 Q5T4D3 Q5T4F4 Q5T4S7 Q5T4T1 Q5T601 Q5T6L9 Q5T6X4 Q5T6X5
Q5T700 Q5T7M9 Q5T7P6 Q5T7P8 Q5T7R7 Q5T848 Q5T8D3 Q5T9L3 Q5T9Z0 Q5TAH2 Q5TAT6
Q5TEA6 Q5TF21 Q5TF39 Q5TFQ8 Q5TGI0 Q5TGU0 Q5TGY1 Q5TGZ0 Q5TH69 Q5TJ59 Q5TJE4
Q5TJE8 Q5TJF6 Q5TKG3 Q5TM22 Q5TM23 Q5TM67 Q5TYP8 Q5TYV0 Q5TZ07 Q5TZ20 Q5TZ24
Q5TZ51 Q5TZA3 Q5TZE2 Q5TZJ5 Q5U1V9 Q5U1W4 Q5U1X7 Q5U1Y0 Q5U220 Q5U239 Q5U258
Q5U2P1 Q5U2P2 Q5U2P6 Q5U2Q4 Q5U2S1 Q5U2T1 Q5U2V5 Q5U2V8 Q5U2V9 Q5U2W7 Q5U2X6
Q5U2X7 Q5U305 Q5U308 Q5U310 Q5U395 Q5U3A7 Q5U3C3 Q5U3G6 Q5U3H3 Q5U3I0 Q5U3T0
Q5U3T3 Q5U3U7 Q5U3V7 Q5U3W1 Q5U3Y5 Q5U3Y7 Q5U3Y9 Q5U405 Q5U419 Q5U430 Q5U431
Q5U462 Q5U4D8 Q5U4E0 Q5U4F4 Q5U4K5 Q5U4P2 Q5U4Q2 Q5U4S8 Q5U4T9 Q5U4U5 Q5U4V1
Q5U4X2 Q5U4X7 Q5U4X8 Q5U520 Q5U534 Q5U583 Q5U5A3 Q5U680 Q5U820 Q5U9D7 Q5U9D9
Q5U9X3 Q5UAW9 Q5UB51 Q5UBV8 Q5UCB8 Q5UCC4 Q5UE96 Q5UEM7 Q5UEM8 Q5UKY4 Q5ULB0
Q5UNU7 Q5UNU8 Q5UNV5 Q5UNW0 Q5UNW1 Q5UNZ1 Q5UP28 Q5UP55 Q5UP70 Q5UP90 Q5UPC1
Q5UPE0 Q5UPE7 Q5UPJ5 Q5UPL2 Q5UPL7 Q5UPR2 Q5UPT3 Q5UPV8 Q5UPW0 Q5UPW5 Q5UPX2
Q5UPY8 Q5UPZ1 Q5UPZ4 Q5UPZ5 Q5UQ28 Q5UQ30 Q5UQ33 Q5UQ44 Q5UQ56 Q5UQ82 Q5UQ86
Q5UQ88 Q5UQ90 Q5UQA0 Q5UQB7 Q5UQD3 Q5UQE2 Q5UQE4 Q5UQF9 Q5UQG5 Q5UQG7 Q5UQG8
Q5UQI2 Q5UQI4 Q5UQI9 Q5UQJ5 Q5UQJ8 Q5UQL5 Q5UQN2 Q5UQN5 Q5UQN8 Q5UQP0 Q5UQP2
Q5UQP7 Q5UQR8 Q5UQS3 Q5UQT2 Q5UQT8 Q5UQV2 Q5UQV6 Q5UQW3 Q5UQW4 Q5UQW6 Q5UQX9
Q5UQY3 Q5UQZ8 Q5UR00 Q5UR15 Q5UR31 Q5UR46 Q5UR61 Q5UR63 Q5UR77 Q5UR78 Q5UR80
Q5UR99 Q5UVI3 Q5UVI4 Q5UVI5 Q5UVI6 Q5UVI8 Q5UVJ0 Q5UXM8 Q5UXM9 Q5UXT1 Q5UXY6
Q5UY54 Q5UZH5 Q5UZJ2 Q5V069 Q5V070 Q5V0J1 Q5V0R5 Q5V1N0 Q5V456 Q5V4H7 Q5V4H8
Q5V5S6 Q5V5V3 Q5V5V4 Q5V7F4 Q5V9L7 Q5VAN0 Q5VI41 Q5VJ36 Q5VJ37 Q5VJ39 Q5VJ41
Q5VJ42 Q5VJ44 Q5VJ45 Q5VJ46 Q5VJ49 Q5VJ50 Q5VJ52 Q5VJ55 Q5VJ57 Q5VJ58 Q5VJ59
Q5VJ60 Q5VJ61 Q5VJ65 Q5VJ67 Q5VJ70 Q5VJL3 Q5VJS9 Q5VKJ7 Q5VKN9 Q5VKP3 Q5VNW5
Q5VP70 Q5VQK9 Q5VQL3 Q5VQY3 Q5VR89 Q5VRB2 Q5VRD7 Q5VRI5 Q5VRJ8 Q5VRM7 Q5VSB5
Q5VSB6 Q5VSG8 Q5VT66 Q5VT99 Q5VTJ3 Q5VTT2 Q5VTY9 Q5VU36 Q5VU65 Q5VU97 Q5VUB5
Q5VUD6 Q5VUY2 Q5VV42 Q5VV43 Q5VV63 Q5VVB8 Q5VVP1 Q5VW36 Q5VW38 Q5VWC8 Q5VWK5
Q5VX71 Q5VXT5 Q5VXU1 Q5VY43 Q5VYJ5 Q5VYP0 Q5VZ72 Q5VZI3 Q5VZR4 Q5VZY2 Q5W0B7
Q5W0N0 Q5W0Z9 Q5W264 Q5W270 Q5W274 Q5W676 Q5W6B9 Q5W6F1 Q5W6H5 Q5W6M3 Q5W6Y3
Q5W7C1 Q5W7F1 Q5W7P8 Q5W8I7 Q5W8I8 Q5W8W0 Q5W9T5 Q5WA50 Q5WB61 Q5WB72 Q5WB73
Q5WB74 Q5WCX5 Q5WDH2 Q5WDX4 Q5WDX6 Q5WDX9 Q5WEF9 Q5WFD1 Q5WFG7 Q5WFI5 Q5WFT5

Q5WFX6 Q5WG37 Q5WGU3 Q5WGV5 Q5WGY8 Q5WH44 Q5WH46 Q5WHE8 Q5WHV2 Q5WIE4 Q5WJP2
Q5WJQ1 Q5WJQ2 Q5WLH2 Q5WN23 Q5WNW9 Q5WNX1 Q5WNX2 Q5WPU5 Q5WQY5 Q5WRN1 Q5WSE9
Q5WSG4 Q5WSQ5 Q5WSU4 Q5WT27 Q5WT30 Q5WT33 Q5WT71 Q5WTE1 Q5WTI4 Q5WTR2 Q5WTT0
Q5WU67 Q5WU69 Q5WU90 Q5WUW3 Q5WVN2 Q5WWR8 Q5WX43 Q5WXA2 Q5WY48 Q5WZ32 Q5WZC7
Q5WZY7 Q5X0M2 Q5X0N9 Q5X0X3 Q5X118 Q5X1B4 Q5X1B7 Q5X1F8 Q5X1N2 Q5X1S3 Q5X1Y7
Q5X204 Q5X2E5 Q5X2E7 Q5X2Q6 Q5X2T2 Q5X3H2 Q5X498 Q5X5D5 Q5X5R4 Q5X5Y0 Q5X6P3
Q5X7N1 Q5X7X6 Q5X8K6 Q5X9A4 Q5X9A9 Q5X9U6 Q5XAL6 Q5XAP0 Q5XB99 Q5XBB2 Q5XCC9
Q5XCK1 Q5XCS3 Q5XCY4 Q5XCY5 Q5XD04 Q5XD68 Q5XD70 Q5XDI2 Q5XDQ0 Q5XDQ1 Q5XDQ5
Q5XDS0 Q5XDS5 Q5XDV8 Q5XDY9 Q5XE40 Q5XEP9 Q5XEZ0 Q5XEZ1 Q5XEZ5 Q5XF03 Q5XF04
Q5XF09 Q5XF36 Q5XF85 Q5XF89 Q5XF90 Q5XFV8 Q5XG04 Q5XG41 Q5XG64 Q5XG73 Q5XG99
Q5XGD7 Q5XGE0 Q5XGF7 Q5XGI1 Q5XGK0 Q5XGR0 Q5XGR4 Q5XGS4 Q5XGU1 Q5XGZ9 Q5XH03
Q5XH39 Q5XH57 Q5XH84 Q5XH90 Q5XH94 Q5XH95 Q5XHA0 Q5XHA2 Q5XHB4 Q5XHF9 Q5XHG1
Q5XHG6 Q5XHH7 Q5XHI3 Q5XHM7 Q5XI00 Q5XI05 Q5XI31 Q5XI41 Q5XI43 Q5XI52 Q5XI60
Q5XI64 Q5XI83 Q5XI99 Q5XIA1 Q5XIA8 Q5XID0 Q5XID7 Q5XIE5 Q5XIE8 Q5XIF1 Q5XIF5
Q5XIF9 Q5XIH2 Q5XII8 Q5XIJ4 Q5XIK2 Q5XIL0 Q5XIN6 Q5XIN7 Q5XIP9 Q5XIS8 Q5XIT3
Q5XIU9 Q5XIX0 Q5XIY2 Q5XJP0 Q5XJR6 Q5XJS0 Q5XJV6 Q5XJX0 Q5XJY4 Q5XK03 Q5XK91
Q5XK92 Q5XK94 Q5XKA2 Q5XKA6 Q5XKN4 Q5XKP0 Q5XM32 Q5XNR9 Q5XPH7 Q5XPT3 Q5XTS1
Q5XV40 Q5XV67 Q5XVI1 Q5XXA6 Q5XXP2 Q5XXP3 Q5Y388 Q5Y4N8 Q5Y4Q0 Q5Y4Q1 Q5Y4Q2
Q5Y4Q3 Q5Y4Q4 Q5Y4Q6 Q5Y4Q8 Q5Y4Q9 Q5Y4R0 Q5Y4Y6 Q5Y4Y8 Q5Y4Y9 Q5Y4Z0 Q5Y4Z2
Q5Y4Z5 Q5Y4Z8 Q5Y500 Q5Y5T1 Q5Y5T2 Q5Y5T3 Q5Y5T5 Q5Y750 Q5Y944 Q5YA25 Q5YB40
Q5YB65 Q5YBK0 Q5YCC5 Q5YCC7 Q5YCW8 Q5YF91 Q5YJ58 Q5YJ78 Q5YJ93 Q5YJC1 Q5YKK4
Q5YKK9 Q5YLM1 Q5YPB8 Q5YQF7 Q5YR14 Q5YR85 Q5YR86 Q5YRP8 Q5YTG0 Q5YTS1 Q5YU15
Q5YUY0 Q5YVL4 Q5YWD4 Q5YWD6 Q5YYN2 Q5YYY2 Q5YZ19 Q5YZ36 Q5YZ37 Q5YZH9 Q5Z0K2
Q5Z0Y5 Q5Z2Z5 Q5Z3P6 Q5Z413 Q5Z5F2 Q5Z5R4 Q5Z5R7 Q5Z5S6 Q5Z653 Q5Z6B1 Q5Z6E5
Q5Z6F0 Q5Z6K9 Q5Z880 Q5Z8N6 Q5Z8P0 Q5Z8T8 Q5Z922 Q5Z974 Q5Z9E2 Q5Z9I2 Q5Z9N5
Q5Z9P8 Q5Z9S8 Q5ZAJ0 Q5ZBY9 Q5ZC87 Q5ZCC5 Q5ZD08 Q5ZEG0 Q5ZEJ0 Q5ZHM5 Q5ZHQ5
Q5ZHQ6 Q5ZHU0 Q5ZHX6 Q5ZHX7 Q5ZI05 Q5ZI20 Q5ZI25 Q5ZI82 Q5ZIE4 Q5ZIF1 Q5ZIF5
Q5ZIG3 Q5ZIH3 Q5ZII3 Q5ZII6 Q5ZIL6 Q5ZIN0 Q5ZIR9 Q5ZIS0 Q5ZIS9 Q5ZIT9 Q5ZIU0
Q5ZIU9 Q5ZJ06 Q5ZJ41 Q5ZJ75 Q5ZJC7 Q5ZJD7 Q5ZJD8 Q5ZJD9 Q5ZJE4 Q5ZJH2 Q5ZJH7
Q5ZJH8 Q5ZJI0 Q5ZJL8 Q5ZJN8 Q5ZJP5 Q5ZJQ8 Q5ZJR3 Q5ZJR8 Q5ZJU0 Q5ZJW4 Q5ZJX0
Q5ZJY9 Q5ZJZ4 Q5ZK32 Q5ZK33 Q5ZK43 Q5ZK55 Q5ZK82 Q5ZKB7 Q5ZKD1 Q5ZKD5 Q5ZKH8
Q5ZKI6 Q5ZKJ0 Q5ZKJ5 Q5ZKK0 Q5ZKL6 Q5ZKN2 Q5ZKN3 Q5ZKP7 Q5ZKQ5 Q5ZKS8 Q5ZKU4
Q5ZKX9 Q5ZKY0 Q5ZKZ4 Q5ZKZ9 Q5ZL00 Q5ZL05 Q5ZL19 Q5ZL36 Q5ZL67 Q5ZL74 Q5ZL84
Q5ZLA9 Q5ZLD4 Q5ZLF4 Q5ZLG8 Q5ZLH4 Q5ZLJ4 Q5ZLK4 Q5ZLL0 Q5ZLL6 Q5ZLL8 Q5ZLR1
Q5ZLR5 Q5ZLR7 Q5ZLW2 Q5ZM31 Q5ZM57 Q5ZM60 Q5ZM65 Q5ZM72 Q5ZM73 Q5ZM83 Q5ZMD4
Q5ZMJ7 Q5ZML6 Q5ZML7 Q5ZMP3 Q5ZMT9 Q5ZN98 Q5ZNA1 Q5ZNA2 Q5ZPR3 Q5ZR81 Q5ZR97
Q5ZRL2 Q5ZRU5 Q5ZRU8 Q5ZS62 Q5ZSA2 Q5ZSA4 Q5ZSH7 Q5ZSJ6 Q5ZSV1 Q5ZSY2 Q5ZT09
Q5ZT21 Q5ZTN5 Q5ZU17 Q5ZUH9 Q5ZV56 Q5ZVL0 Q5ZVZ5 Q5ZW98 Q5ZWN4 Q5ZWR1 Q5ZX72
Q5ZXT8 Q5ZY60 Q5ZYC6 Q5ZYC7 Q5ZYG0 Q5ZZ31 Q60010 Q60019 Q60048 Q60153 Q60175
Q601Z9 Q60236 Q60263 Q60293 Q60294 Q602A0 Q602K4 Q602M6 Q602S2 Q60304 Q60309
Q60318 Q60324 Q60333 Q60336 Q60362 Q60366 Q60371 Q603S3 Q603T3 Q603U6 Q603U8
Q60409 Q60411 Q60414 Q60416 Q60417 Q60429 Q60441 Q60457 Q60472 Q60474 Q60475
Q60476 Q60483 Q60484 Q60489 Q60490 Q60492 Q60495 Q604V4 Q604V6 Q604Z9 Q60513
Q60519 Q60522 Q60528 Q60534 Q60538 Q60553 Q60555 Q60565 Q605A0 Q605R1 Q605R3

Q60603 Q60612 Q60613 Q60614 Q60625 Q60629 Q60651 Q60652 Q60653 Q60654 Q60660
Q60664 Q60673 Q60677 Q60682 Q60696 Q606P3 Q60714 Q60718 Q60720 Q60738 Q60748
Q60750 Q60751 Q60754 Q60755 Q60767 Q60771 Q60774 Q607G3 Q60805 Q60813 Q60819
Q60825 Q60837 Q60846 Q60850 Q60856 Q60857 Q60860 Q60870 Q60878 Q60879 Q60880
Q60881 Q60882 Q60883 Q60884 Q60885 Q60886 Q60887 Q60888 Q60889 Q60890 Q60891
Q60892 Q60893 Q60894 Q60895 Q608K2 Q608N5 Q608X7 Q608Y3 Q608Y6 Q608Y9 Q608Z9
Q60928 Q60930 Q60931 Q60932 Q60934 Q60936 Q60943 Q60961 Q60991 Q60997 Q60A92
Q60AA3 Q60AK1 Q60B18 Q60BP6 Q60CP3 Q60CR8 Q60CS0 Q60D27 Q60DN5 Q60E70 Q60EA5
Q60EC2 Q60EN8 Q60EW9 Q60F97 Q60GF7 Q60HC0 Q60HC1 Q60HC5 Q60HD1 Q60HD2 Q60HE1
Q60HE7 Q60HE8 Q60HF7 Q60HG0 Q60HH0 Q60HH8 Q60M68 Q60MW2 Q60NC0 Q60QP4 Q60S81
Q60SZ2 Q60T32 Q60T34 Q60T36 Q60V90 Q60WL8 Q60WT2 Q60WU2 Q60ZN5 Q60ZR7 Q61003
Q61006 Q61009 Q61029 Q61038 Q61041 Q61056 Q61070 Q61072 Q61086 Q61088 Q61089
Q61090 Q61091 Q61093 Q61098 Q610N4 Q61102 Q61115 Q61121 Q61124 Q61125 Q61137
Q61139 Q61143 Q61144 Q61146 Q61153 Q61165 Q61168 Q61180 Q61184 Q61190 Q611C8
Q61212 Q61224 Q61263 Q61271 Q61285 Q61288 Q61290 Q61327 Q61334 Q61335 Q61391
Q61409 Q61418 Q61420 Q61423 Q61450 Q61451 Q61469 Q61470 Q61476 Q61483 Q61490
Q61495 Q61500 Q61521 Q61526 Q61527 Q61542 Q61543 Q61549 Q61559 Q61586 Q61595
Q615A2 Q61603 Q61606 Q61609 Q61614 Q61616 Q61618 Q61625 Q61626 Q61627 Q61647
Q61672 Q61694 Q616J1 Q61712 Q61727 Q61730 Q61735 Q61738 Q61739 Q61743 Q61762
Q61767 Q61772 Q61790 Q617M0 Q617Y0 Q61809 Q61817 Q61824 Q61826 Q61830 Q61835
Q61847 Q61851 Q61885 Q618G2 Q61907 Q61923 Q61941 Q61982 Q61983 Q61985 Q619N4
Q61C05 Q61CA3 Q61CQ8 Q61G93 Q61H86 Q61IS6 Q61JS7 Q61LC0 Q61Z83 Q61ZW5 Q62005
Q62007 Q62011 Q62028 Q62035 Q62052 Q62053 Q62092 Q620A6 Q62132 Q62147 Q62151
Q62165 Q62170 Q62178 Q62179 Q62186 Q62190 Q62192 Q62205 Q62217 Q62230 Q62273
Q62277 Q62283 Q622X0 Q62302 Q62312 Q62313 Q62314 Q62351 Q62371 Q62398 Q623S8
Q62413 Q62425 Q62442 Q62452 Q62463 Q62469 Q62470 Q62556 Q62611 Q62633 Q62634
Q62638 Q62640 Q62645 Q62656 Q62666 Q62668 Q62674 Q62687 Q626H5 Q626N3 Q62720
Q62730 Q62733 Q62745 Q62746 Q62747 Q62758 Q62760 Q62765 Q62773 Q62786 Q62789
Q62795 Q62799 Q627G6 Q62805 Q62838 Q62849 Q62850 Q62865 Q62866 Q62875 Q62876
Q62878 Q62888 Q62889 Q62893 Q62896 Q62897 Q628Z2 Q62902 Q62904 Q62927 Q62928
Q62929 Q62930 Q62931 Q62941 Q62953 Q62956 Q62959 Q62962 Q62968 Q62969 Q62976
Q62994 Q629K1 Q62B05 Q62B07 Q62EM4 Q62F24 Q62F61 Q62FR9 Q62GF0 Q62GF1 Q62GS4
Q62H69 Q62HL3 Q62HT6 Q62HU2 Q62HW8 Q62I47 Q62IG3 Q62IJ6 Q62IJ8 Q62IN5 Q62IP2
Q62IP5 Q62JH4 Q62JM5 Q62JX8 Q62L95 Q62LG5 Q62LW6 Q62M01 Q62M79 Q62MP2 Q62MT2
Q63008 Q63016 Q63085 Q63089 Q63099 Q630G3 Q630G4 Q630S4 Q630T8 Q630T9 Q630U8
Q630V2 Q630V5 Q630Y6 Q63113 Q63118 Q63120 Q63132 Q63135 Q63149 Q63151 Q63155
Q63175 Q63180 Q63190 Q63191 Q63199 Q631J1 Q631P3 Q631P4 Q63202 Q63203 Q63226
Q63257 Q63258 Q63259 Q63269 Q63273 Q63315 Q63344 Q63345 Q63371 Q63372 Q63373
Q63374 Q63376 Q63377 Q63378 Q63384 Q633B8 Q633E6 Q633K2 Q63400 Q63418 Q63424
Q63447 Q63472 Q63474 Q63475 Q63488 Q63493 Q63495 Q634T9 Q63511 Q63524 Q63563
Q63564 Q63584 Q635Y0 Q635Y1 Q63604 Q63621 Q63632 Q63633 Q63634 Q63635 Q63645
Q63652 Q63661 Q63664 Q63666 Q63690 Q636B3 Q636D3 Q636P8 Q63704 Q63707 Q63722
Q63734 Q637E7 Q637F8 Q637M1 Q63881 Q63886 Q638L6 Q63931 Q63959 Q63961 Q63965
Q63994 Q63AK0 Q63AN4 Q63AU4 Q63BB0 Q63BU9 Q63DQ8 Q63DZ3 Q63DZ4 Q63EL0 Q63FE8

Q63FQ9 Q63FR0 Q63FR1 Q63FW2 Q63GA7 Q63GU6 Q63HM2 Q63I54 Q63K32 Q63K34 Q63MM6
Q63PH6 Q63Q97 Q63Q98 Q63QJ0 Q63QJ4 Q63QK0 Q63QP9 Q63R13 Q63RM9 Q63RN4 Q63RQ7
Q63RY4 Q63S57 Q63T62 Q63US0 Q63V24 Q63V58 Q63VA1 Q63VM3 Q63VM6 Q63VN3 Q63VS1
Q63VS3 Q63VX7 Q63WB7 Q63WI4 Q63WM5 Q63X97 Q63XS7 Q63XW5 Q63YR4 Q63YW1 Q63ZE4
Q63ZG0 Q63ZI0 Q63ZL3 Q63ZR0 Q63ZR6 Q63ZR7 Q63ZU3 Q63ZZ0 Q64017 Q64018 Q64077
Q64093 Q640B3 Q640B5 Q640K3 Q640L2 Q640L3 Q640M6 Q640P4 Q640R3 Q640S1 Q640U0
Q640U3 Q640X6 Q64121 Q64133 Q64151 Q64159 Q64198 Q641C8 Q641E3 Q641M3 Q641S4
Q641Y0 Q641Y1 Q64230 Q64232 Q64237 Q64244 Q64255 Q64264 Q64273 Q64281 Q64294
Q642A2 Q642A3 Q642A7 Q642B3 Q642F4 Q64302 Q64310 Q64314 Q64319 Q64322 Q64323
Q64326 Q64329 Q64335 Q64343 Q64347 Q64349 Q64359 Q64373 Q64385 Q64392 Q643R3
Q64420 Q64421 Q64430 Q64435 Q64436 Q64445 Q64446 Q64448 Q64449 Q64455 Q64487
Q64505 Q64518 Q64519 Q64535 Q64541 Q64542 Q64548 Q64550 Q64566 Q64568 Q64578
Q645J3 Q645S2 Q645S5 Q645T0 Q645T2 Q645T3 Q645T4 Q645T6 Q645T7 Q645T9 Q645U0
Q645U1 Q645U2 Q645U4 Q645U5 Q645U6 Q645U7 Q645U8 Q645U9 Q645V0 Q645V1 Q645V2
Q645V3 Q645V4 Q645V6 Q645V7 Q645V8 Q645V9 Q645Y3 Q645Y4 Q645Y5 Q645Y7 Q645Y8
Q645Y9 Q645Z0 Q645Z1 Q645Z2 Q645Z3 Q645Z4 Q645Z5 Q645Z6 Q645Z7 Q645Z9 Q64604
Q64605 Q64612 Q64632 Q64633 Q64634 Q64637 Q64638 Q64654 Q64663 Q64676 Q64685
Q64686 Q64687 Q64689 Q64690 Q64692 Q64695 Q64697 Q646A0 Q646A1 Q646A2 Q646A3
Q646A4 Q646A5 Q646A7 Q646A9 Q646B0 Q646B1 Q646B2 Q646B3 Q646B4 Q646B5 Q646B6
Q646B7 Q646B8 Q646B9 Q646C0 Q646C1 Q646C2 Q646C3 Q646C4 Q646C5 Q646C6 Q646C7
Q646C8 Q646D0 Q646D1 Q646D2 Q646D3 Q646D4 Q646D5 Q646D6 Q646D7 Q646D8 Q646D9
Q646E0 Q646E1 Q646E2 Q646E3 Q646E4 Q646E5 Q646E6 Q646E7 Q646E8 Q646E9 Q646F0
Q646F1 Q646F2 Q646F3 Q646F4 Q646F5 Q646F6 Q646F7 Q646F8 Q646F9 Q646G0 Q646G1
Q646G2 Q646G3 Q646G4 Q646G5 Q646G6 Q646G7 Q646G9 Q646H0 Q64704 Q64716 Q64729
Q64735 Q64764 Q647G3 Q647G4 Q647G5 Q64964 Q64965 Q64968 Q64GI8 Q64GJ1 Q64H34
Q64J97 Q64JA0 Q64JA4 Q64L89 Q64MG9 Q64PN7 Q64Q86 Q64SU9 Q64T26 Q64T76 Q64TC3
Q64UA6 Q64UA7 Q64V41 Q64VB3 Q64X36 Q64X43 Q64X50 Q64XQ8 Q64Y06 Q64Y09 Q64Y15
Q64YU7 Q64YU9 Q64ZL8 Q650L9 Q650U0 Q65107 Q65134 Q65139 Q65145 Q65150 Q65158
Q65164 Q65166 Q65167 Q65172 Q65176 Q65178 Q65188 Q65194 Q65197 Q65198 Q651D5
Q651J5 Q651V6 Q651X6 Q651X7 Q65200 Q65214 Q65223 Q65233 Q65240 Q65242 Q65244
Q65272 Q652J4 Q652N5 Q652U9 Q65387 Q653B6 Q653F4 Q653P0 Q653S9 Q653V6 Q65475
Q654D9 Q655S1 Q655Y0 Q65675 Q65676 Q65706 Q657S5 Q657W3 Q658G7 Q658H5 Q658I5
Q658N2 Q658P3 Q65984 Q65999 Q659K9 Q659L0 Q659L1 Q659X0 Q65AC2 Q65CI5 Q65DJ4
Q65DM3 Q65DW1 Q65DW9 Q65DX0 Q65E27 Q65E87 Q65EF1 Q65FP0 Q65G47 Q65G86 Q65H64
Q65HX8 Q65I04 Q65IG9 Q65J17 Q65J36 Q65JB1 Q65JB2 Q65JF1 Q65JJ2 Q65JY3 Q65K13
Q65K14 Q65KL0 Q65KV0 Q65KV1 Q65LE3 Q65LI4 Q65LU9 Q65LX3 Q65LX4 Q65M31 Q65M79
Q65MC6 Q65MT6 Q65P50 Q65Q01 Q65Q03 Q65Q27 Q65Q77 Q65QC7 Q65QF7 Q65QT7 Q65RA1
Q65RI0 Q65RK2 Q65RN5 Q65RY3 Q65RZ8 Q65S74 Q65S86 Q65SL4 Q65SY9 Q65T32 Q65TG9
Q65TH4 Q65U21 Q65U34 Q65UC1 Q65UL6 Q65UZ8 Q65VA2 Q65VA3 Q65VC2 Q65VH3 Q65VH4
Q65VJ7 Q65VT3 Q65VU8 Q65VU9 Q65W61 Q65WL3 Q65WM1 Q65X71 Q65XS5 Q65YU5 Q65YU7
Q65Z14 Q65ZA5 Q65ZG0 Q66000 Q66001 Q66165 Q66181 Q661B6 Q661F0 Q661H9 Q661Q7
Q661W1 Q66209 Q66262 Q66285 Q662A7 Q663Q2 Q663Q3 Q663Q4 Q663W8 Q66405 Q664D7
Q664F8 Q664J1 Q664N1 Q664Q5 Q664V0 Q664X0 Q664X6 Q665E9 Q665H0 Q665H1 Q665H2
Q665P2 Q665U6 Q665U8 Q66608 Q66613 Q66615 Q66625 Q66630 Q66631 Q66655 Q66660

Q66662 Q66669 Q66673 Q66676 Q666Q4 Q666Z8 Q66798 Q66799 Q667F1 Q667F2 Q667F8
Q667J7 Q667Q9 Q667R0 Q667S4 Q667S5 Q667Z8 Q66810 Q66814 Q668C4 Q668C5 Q668C6
Q668G1 Q668G3 Q668K2 Q668L6 Q668M5 Q668N2 Q668U2 Q668Z2 Q669A0 Q669A6 Q669A9
Q669B2 Q669I8 Q669V7 Q669Z9 Q66A06 Q66A07 Q66A27 Q66A67 Q66A75 Q66AG7 Q66AK9
Q66AL1 Q66AL9 Q66AQ9 Q66AR0 Q66AS8 Q66AS9 Q66AV8 Q66BN4 Q66BP6 Q66BY5 Q66BY6
Q66C77 Q66CI3 Q66CJ8 Q66CL2 Q66CP5 Q66D85 Q66DF7 Q66DU5 Q66E01 Q66E03 Q66EC4
Q66EK8 Q66EQ5 Q66ER4 Q66ES3 Q66ES8 Q66EW4 Q66EZ0 Q66EZ1 Q66FC9 Q66FD1 Q66FE1
Q66FF0 Q66FH0 Q66FN0 Q66FS6 Q66FS8 Q66FU5 Q66FU6 Q66FY7 Q66G06 Q66G57 Q66GA9
Q66GF5 Q66GH5 Q66GI1 Q66GI9 Q66GN2 Q66GQ5 Q66GS2 Q66GV0 Q66H15 Q66H21 Q66H23
Q66H28 Q66H29 Q66H44 Q66H50 Q66H52 Q66H69 Q66H78 Q66H88 Q66H95 Q66H96 Q66HA2
Q66HE2 Q66HF0 Q66HF2 Q66HG0 Q66HG5 Q66HG9 Q66HH1 Q66HH2 Q66HH4 Q66HR0 Q66HV9
Q66HW4 Q66HX0 Q66I08 Q66I12 Q66I51 Q66I79 Q66IE4 Q66IF1 Q66IJ4 Q66IM1 Q66IQ1
Q66IR0 Q66IS8 Q66IV1 Q66IV3 Q66IZ4 Q66J01 Q66J05 Q66J17 Q66J27 Q66J44 Q66J52
Q66J54 Q66JC0 Q66JF2 Q66JT1 Q66JT7 Q66K14 Q66K66 Q66KA5 Q66KF2 Q66KG0 Q66KH2
Q66KI8 Q66KL0 Q66KU2 Q66KX2 Q66KY5 Q66KZ9 Q66L44 Q66L49 Q66LN0 Q66PF2 Q66PG1
Q66PG2 Q66PG3 Q66PG4 Q66RM2 Q66T47 Q66T62 Q67018 Q67143 Q67146 Q67160 Q67162
Q67166 Q67168 Q67170 Q67172 Q67176 Q67179 Q67186 Q67201 Q67203 Q67205 Q67206
Q67207 Q67210 Q67211 Q67212 Q67214 Q67282 Q672I5 Q672J9 Q672K1 Q672V4 Q67333
Q67344 Q67369 Q67370 Q67371 Q67372 Q67373 Q67374 Q67375 Q67376 Q67377 Q67378
Q67379 Q67380 Q67381 Q67387 Q673U1 Q674R7 Q67593 Q675B7 Q675B8 Q675B9 Q675C0
Q67726 Q67825 Q67867 Q67875 Q678S1 Q678S2 Q678S3 Q678S5 Q678S8 Q678S9 Q67926
Q679A0 Q679A1 Q679A2 Q679A3 Q679A5 Q679A7 Q679A8 Q679B0 Q679B1 Q679B2 Q679B3
Q679B6 Q679B8 Q679B9 Q679C1 Q67BJ4 Q67BT3 Q67EQ0 Q67EQ1 Q67ER8 Q67ER9 Q67ES0
Q67ES1 Q67ES2 Q67ES3 Q67ES5 Q67ES6 Q67ES7 Q67ES9 Q67ET0 Q67ET1 Q67ET2 Q67ET3
Q67ET4 Q67ET5 Q67ET7 Q67GH9 Q67H84 Q67H85 Q67H89 Q67H92 Q67H93 Q67H97 Q67HA0
Q67HA1 Q67HB2 Q67HB3 Q67HC0 Q67HC1 Q67HD2 Q67HD3 Q67HD6 Q67HD7 Q67HH2 Q67HH3
Q67HJ6 Q67HJ7 Q67HM5 Q67HN6 Q67HN7 Q67HR8 Q67HR9 Q67HS2 Q67HS3 Q67HT0 Q67HT1
Q67HT8 Q67HT9 Q67HV0 Q67HV1 Q67HV8 Q67HV9 Q67HX0 Q67HX1 Q67HX4 Q67HX5 Q67I10
Q67I33 Q67I34 Q67I70 Q67I71 Q67IA0 Q67IA1 Q67IA3 Q67IA6 Q67IA9 Q67IB2 Q67IB5
Q67IC4 Q67ID0 Q67ID9 Q67IE2 Q67IG8 Q67IK8 Q67IL7 Q67JH0 Q67KP0 Q67KP3 Q67KP6
Q67L13 Q67L43 Q67LC0 Q67LG6 Q67MD6 Q67ML5 Q67MN1 Q67NH9 Q67NS8 Q67P10 Q67P12
Q67P15 Q67PL7 Q67Q16 Q67Q50 Q67S16 Q67S17 Q67S38 Q67SE9 Q67T82 Q67TC1 Q67TC3
Q67UC7 Q67UL3 Q67UQ7 Q67UX7 Q67VS5 Q67VS7 Q67VZ8 Q67W83 Q67WJ2 Q67WJ8 Q67WQ7
Q67X31 Q67X40 Q67X45 Q67XC4 Q67XQ0 Q67XW5 Q67YF8 Q67YI6 Q67YS6 Q67YT8 Q67ZE1
Q67ZF6 Q67ZM7 Q681I0 Q681K7 Q682H0 Q68396 Q684M3 Q685J3 Q68749 Q68772 Q68798
Q687X5 Q68801 Q688J2 Q688R1 Q688S6 Q689D1 Q68CI2 Q68CJ9 Q68CP4 Q68CQ1 Q68CQ7
Q68CR1 Q68CR7 Q68D42 Q68D85 Q68DH5 Q68DV7 Q68ED2 Q68EF4 Q68EK2 Q68EL3 Q68EQ9
Q68ES4 Q68EU0 Q68EU6 Q68EU8 Q68EV0 Q68EV1 Q68EV7 Q68EV8 Q68EW1 Q68EY2 Q68F00
Q68F18 Q68F33 Q68F35 Q68F37 Q68F62 Q68F72 Q68F79 Q68FA7 Q68FB2 Q68FB8 Q68FD9
Q68FE2 Q68FE7 Q68FF9 Q68FM6 Q68FN7 Q68FQ2 Q68FR5 Q68FS7 Q68FT6 Q68FU0 Q68FV0
Q68FV1 Q68FV2 Q68FV3 Q68FV4 Q68FV6 Q68FW3 Q68FW4 Q68G12 Q68G75 Q68J47 Q68KI4
Q68RJ7 Q68RU8 Q68RV1 Q68RV3 Q68RV4 Q68RV6 Q68RV7 Q68RV9 Q68RX6 Q68RX7 Q68RX8
Q68RX9 Q68RY0 Q68RY1 Q68RY6 Q68RY7 Q68RY8 Q68RY9 Q68RZ0 Q68RZ1 Q68RZ2 Q68RZ3
Q68RZ4 Q68RZ5 Q68RZ6 Q68S01 Q68S06 Q68S07 Q68S09 Q68S10 Q68S11 Q68S18 Q68S19

Q68S20 Q68S22 Q68S23 Q68S26 Q68SN8 Q68US5 Q68UT4 Q68VK5 Q68VS6 Q68VU2 Q68VV3
Q68VV6 Q68VV7 Q68VV8 Q68VV9 Q68VW2 Q68VW8 Q68W01 Q68W11 Q68W42 Q68W49 Q68W71
Q68W98 Q68WC5 Q68WD6 Q68WF0 Q68WF2 Q68WF3 Q68WJ7 Q68WQ3 Q68WQ5 Q68WV5 Q68WX1
Q68WY5 Q68WZ7 Q68X09 Q68X15 Q68X16 Q68X99 Q68XA0 Q68XB6 Q68XB9 Q68XN4 Q68XN8
Q68XP0 Q68XP1 Q68XP8 Q68XQ0 Q68XQ1 Q68XR9 Q68XS2 Q68XS4 Q68XS5 Q68XS7 Q68XU9
Q68XV0 Q68XV2 Q68XV7 Q68XW4 Q68XX2 Q68XX3 Q68XX4 Q68XX5 Q68XX6 Q68XY2 Q68Y21
Q68Y56 Q68Y62 Q69024 Q69091 Q69150 Q692W3 Q69384 Q69422 Q69467 Q69489 Q69505
Q69550 Q69552 Q69553 Q69555 Q69556 Q69557 Q69558 Q69559 Q69566 Q69569 Q695P6
Q695T7 Q695T8 Q695T9 Q695U0 Q69603 Q69606 Q697L1 Q697L2 Q697L3 Q697L4 Q697L5
Q697L6 Q698P7 Q69AY8 Q69B66 Q69B76 Q69B78 Q69B80 Q69B84 Q69B86 Q69BJ6 Q69BJ7
Q69BJ8 Q69BJ9 Q69BK0 Q69BK1 Q69BK2 Q69BK3 Q69BK4 Q69BK5 Q69BK6 Q69D47 Q69D48
Q69DJ1 Q69F95 Q69FB3 Q69FB6 Q69FH1 Q69GM3 Q69GY7 Q69HU0 Q69JN6 Q69JW3 Q69L19
Q69L87 Q69L93 Q69LA1 Q69MT2 Q69P51 Q69P80 Q69PA8 Q69RI8 Q69S39 Q69SA9 Q69SP5
Q69SV0 Q69T22 Q69T29 Q69T94 Q69TN4 Q69U49 Q69V23 Q69VR7 Q69X58 Q69XK5 Q69XM6
Q69XV0 Q69YG0 Q69YU5 Q69YW2 Q69YZ2 Q69Z08 Q69Z14 Q69ZF7 Q69ZK9 Q69ZN6 Q69ZN7
Q69ZQ1 Q69ZS6 Q69ZU6 Q69ZZ6 Q6A044 Q6A051 Q6A058 Q6A152 Q6A155 Q6A198 Q6A1G2
Q6A1G3 Q6A329 Q6A4L0 Q6A4L1 Q6A553 Q6A5N5 Q6A5S0 Q6A666 Q6A6G5 Q6A6G8 Q6A6H1
Q6A7G0 Q6A8C3 Q6A8N5 Q6A8U1 Q6A900 Q6A9N9 Q6A9P0 Q6A9Q5 Q6AB77 Q6ABJ2 Q6AE61
Q6AE72 Q6AF34 Q6AF65 Q6AFM6 Q6AFW1 Q6AG62 Q6AHI4 Q6AHI5 Q6AI14 Q6AJ39 Q6AJ51
Q6AJ81 Q6AJW3 Q6AK46 Q6AKL2 Q6ALU7 Q6AMM5 Q6ANM8 Q6ANN6 Q6AP26 Q6APL9 Q6APV5
Q6APZ3 Q6AQ14 Q6AQ28 Q6AQ29 Q6AQH5 Q6AQV6 Q6ARA9 Q6AS08 Q6AT26 Q6AT32 Q6ATV4
Q6ATY4 Q6AU53 Q6AU90 Q6AVD0 Q6AWJ9 Q6AWW5 Q6AWX0 Q6AX23 Q6AX31 Q6AX42 Q6AX44
Q6AX53 Q6AX57 Q6AX73 Q6AX80 Q6AXF6 Q6AXL3 Q6AXM1 Q6AXM5 Q6AXN3 Q6AXN4 Q6AXP6
Q6AXR5 Q6AXR8 Q6AXS0 Q6AXS2 Q6AXS4 Q6AXT9 Q6AXU5 Q6AXV4 Q6AXV7 Q6AXW8 Q6AXY3
Q6AY01 Q6AY05 Q6AY06 Q6AY19 Q6AY20 Q6AY25 Q6AY39 Q6AY41 Q6AY47 Q6AY76 Q6AY78
Q6AY82 Q6AY85 Q6AY92 Q6AY94 Q6AYA5 Q6AYA6 Q6AYD4 Q6AYF4 Q6AYF6 Q6AYF7 Q6AYG0
Q6AYH6 Q6AYL2 Q6AYM7 Q6AYM9 Q6AYN2 Q6AYP2 Q6AYP5 Q6AYQ4 Q6AYR5 Q6AYR9 Q6AYS5
Q6AYT7 Q6AYT8 Q6AYY8 Q6AZ51 Q6AZ61 Q6AZ83 Q6AZB0 Q6AZD1 Q6AZD4 Q6AZG1 Q6AZH0
Q6AZI2 Q6AZM3 Q6AZN8 Q6AZR3 Q6AZV0 Q6AZW1 Q6AZY7 Q6B0R2 Q6B0V8 Q6B0W0 Q6B0X2
Q6B0Y1 Q6B0Y5 Q6B0Y6 Q6B0Z2 Q6B2I9 Q6B3P0 Q6B457 Q6B4J2 Q6B6R8 Q6B8K0 Q6B8K4
Q6B8K5 Q6B8K6 Q6B8K7 Q6B8K8 Q6B8L0 Q6B8M4 Q6B8P4 Q6B8P5 Q6B8P8 Q6B8Q2 Q6B8Q5
Q6B8R0 Q6B8R1 Q6B8R2 Q6B8S9 Q6B8T0 Q6B8U0 Q6B8U6 Q6B8U7 Q6B8U9 Q6B8V0 Q6B8V1
Q6B8Y5 Q6B8Y6 Q6B8Y7 Q6B903 Q6B904 Q6B917 Q6B918 Q6B923 Q6B926 Q6B929 Q6B935
Q6B947 Q6B948 Q6B959 Q6BBL6 Q6BD04 Q6BEA0 Q6BF25 Q6BF87 Q6BH02 Q6BH37 Q6BH42
Q6BH65 Q6BHI9 Q6BHK4 Q6BHT4 Q6BHZ8 Q6BI17 Q6BI99 Q6BIG0 Q6BIT1 Q6BIV4 Q6BJ94
Q6BJ96 Q6BK00 Q6BK59 Q6BKG0 Q6BKQ1 Q6BKU8 Q6BKW6 Q6BKZ1 Q6BKZ8 Q6BL60 Q6BLG8
Q6BLM0 Q6BLY8 Q6BM09 Q6BM18 Q6BMD0 Q6BMD6 Q6BMV2 Q6BMV3 Q6BMY0 Q6BMZ7 Q6BN08
Q6BN11 Q6BNJ2 Q6BNJ4 Q6BNL9 Q6BP15 Q6BP23 Q6BP80 Q6BP97 Q6BPD6 Q6BPF8 Q6BPR6
Q6BPV1 Q6BPW0 Q6BQ54 Q6BQJ1 Q6BQK1 Q6BQK9 Q6BRE5 Q6BRG1 Q6BRZ9 Q6BS98 Q6BSA9
Q6BSB9 Q6BSD5 Q6BSK8 Q6BSU1 Q6BSY9 Q6BT35 Q6BTC2 Q6BTN1 Q6BTS4 Q6BTT3 Q6BTX9
Q6BUS8 Q6BUS9 Q6BUX2 Q6BUZ2 Q6BVA4 Q6BVB2 Q6BVC2 Q6BVE3 Q6BVH1 Q6BVK1 Q6BVM4
Q6BVN0 Q6BVY9 Q6BW42 Q6BW46 Q6BW58 Q6BW93 Q6BWE3 Q6BWE7 Q6BWH8 Q6BWK4 Q6BWY8
Q6BX67 Q6BXC8 Q6BXD7 Q6BXM0 Q6BXM9 Q6BXQ6 Q6BY05 Q6BY28 Q6BYG4 Q6BYM1 Q6BYU3
Q6BYY8 Q6BZ39 Q6BZ57 Q6BZQ6 Q6BZU7 Q6BZW3 Q6BZY4 Q6C060 Q6C0L2 Q6C0R5 Q6C0U0

Q6C0Z3 Q6C105 Q6C107 Q6C188 Q6C216 Q6C2F5 Q6C2J1 Q6C2N7 Q6C2P2 Q6C2Z2 Q6C3B0
Q6C3K2 Q6C3V7 Q6C427 Q6C457 Q6C4Q0 Q6C4R5 Q6C4W5 Q6C4X5 Q6C503 Q6C520 Q6C555
Q6C5K1 Q6C5S0 Q6C6I3 Q6C6L1 Q6C6N0 Q6C6S3 Q6C6U9 Q6C741 Q6C7D1 Q6C7F7 Q6C7Q0
Q6C7Q6 Q6C7V5 Q6C7W0 Q6C7Y5 Q6C880 Q6C890 Q6C8F0 Q6C8H7 Q6C8S0 Q6C8U1 Q6C8W4
Q6C938 Q6C994 Q6C9G8 Q6C9T3 Q6CA06 Q6CA86 Q6CAB8 Q6CAC5 Q6CAG4 Q6CAG9 Q6CAP9
Q6CAQ9 Q6CAW5 Q6CAW6 Q6CB37 Q6CBQ8 Q6CBS9 Q6CC06 Q6CCF6 Q6CD56 Q6CDD0 Q6CDE6
Q6CDK3 Q6CDV6 Q6CDV7 Q6CE07 Q6CE86 Q6CEA4 Q6CEA5 Q6CEE7 Q6CEU0 Q6CF02 Q6CF18
Q6CF60 Q6CFD4 Q6CFJ0 Q6CFP3 Q6CFU5 Q6CFW6 Q6CG20 Q6CGU8 Q6CHP1 Q6CHT7 Q6CIB3
Q6CID0 Q6CII3 Q6CIJ3 Q6CIN9 Q6CIY2 Q6CIY7 Q6CJ97 Q6CJA0 Q6CJC5 Q6CJG3 Q6CJH4
Q6CJI9 Q6CJM3 Q6CJR2 Q6CJU9 Q6CJX2 Q6CJX5 Q6CK18 Q6CK73 Q6CKC6 Q6CKK4 Q6CKU5
Q6CKX6 Q6CKY2 Q6CL16 Q6CLD6 Q6CLF2 Q6CLI2 Q6CLJ5 Q6CLL5 Q6CLM9 Q6CLN0 Q6CLN2
Q6CLN9 Q6CLP4 Q6CLU6 Q6CLZ3 Q6CLZ9 Q6CM45 Q6CMA4 Q6CMF1 Q6CMG6 Q6CMH6 Q6CMK7
Q6CMM1 Q6CMQ1 Q6CMR5 Q6CMW6 Q6CN26 Q6CN27 Q6CN76 Q6CNN5 Q6CNR4 Q6CNW2 Q6CNY8
Q6CP77 Q6CP93 Q6CPU8 Q6CPW5 Q6CPX6 Q6CPY8 Q6CPY9 Q6CQB5 Q6CQP9 Q6CQR3 Q6CQW1
Q6CR04 Q6CR06 Q6CRE7 Q6CRJ6 Q6CRM6 Q6CRV3 Q6CRX0 Q6CRY8 Q6CS27 Q6CS34 Q6CS73
Q6CSA1 Q6CSB8 Q6CSN0 Q6CSN2 Q6CT08 Q6CT28 Q6CT96 Q6CTB6 Q6CTE6 Q6CTK9 Q6CTT3
Q6CTW6 Q6CTX8 Q6CTY0 Q6CU02 Q6CU37 Q6CU99 Q6CUA2 Q6CUA4 Q6CUB5 Q6CUC1 Q6CUG3
Q6CUV7 Q6CUZ1 Q6CUZ6 Q6CV11 Q6CVE0 Q6CVS4 Q6CVU2 Q6CVX2 Q6CW19 Q6CW36 Q6CW96
Q6CWC2 Q6CWD4 Q6CWK9 Q6CWQ0 Q6CWT4 Q6CWT7 Q6CWW7 Q6CX96 Q6CXA2 Q6CXJ8 Q6CXK7
Q6CXQ8 Q6CXR8 Q6CXX9 Q6CXZ7 Q6CY37 Q6CY85 Q6CYC1 Q6CYD0 Q6CYG2 Q6CYI9 Q6CYJ0
Q6CYJ1 Q6CYR0 Q6CYV4 Q6CYZ1 Q6CZ32 Q6CZ33 Q6CZ44 Q6CZ45 Q6CZB5 Q6CZF1 Q6CZH4
Q6CZL3 Q6CZM7 Q6CZU5 Q6CZZ8 Q6D029 Q6D090 Q6D0B9 Q6D0D4 Q6D0E4 Q6D0I0 Q6D107
Q6D108 Q6D157 Q6D159 Q6D1B2 Q6D275 Q6D2A1 Q6D2A9 Q6D2B0 Q6D2B1 Q6D2C3 Q6D2F0
Q6D2F3 Q6D2F4 Q6D2F5 Q6D2Q7 Q6D2R7 Q6D2S3 Q6D2S6 Q6D2S9 Q6D364 Q6D3B1 Q6D3B2
Q6D3B6 Q6D3R1 Q6D437 Q6D469 Q6D4G3 Q6D4K1 Q6D4K2 Q6D4M8 Q6D4M9 Q6D4S2 Q6D4S7
Q6D4T5 Q6D4W0 Q6D4W1 Q6D4W4 Q6D5P8 Q6D5Q4 Q6D5W6 Q6D602 Q6D656 Q6D6A7 Q6D7E5
Q6D7I3 Q6D7I5 Q6D7N0 Q6D7N6 Q6D7R0 Q6D836 Q6D8D5 Q6D8I7 Q6D8S6 Q6D8S7 Q6D8T5
Q6D913 Q6D9G4 Q6D9I8 Q6D9J6 Q6D9Z3 Q6DA13 Q6DAH4 Q6DAH5 Q6DAH6 Q6DAJ0 Q6DAQ3
Q6DAQ5 Q6DB52 Q6DB92 Q6DBG8 Q6DBH0 Q6DBM8 Q6DBN4 Q6DBP3 Q6DBW9 Q6DBX0 Q6DBY9
Q6DC58 Q6DC66 Q6DC75 Q6DC77 Q6DCC7 Q6DCD5 Q6DCE3 Q6DCE8 Q6DCG4 Q6DCG9 Q6DCK1
Q6DCL6 Q6DCP1 Q6DCP8 Q6DCQ3 Q6DCT2 Q6DCT3 Q6DCU6 Q6DCW7 Q6DCX5 Q6DD32 Q6DD44
Q6DD53 Q6DD88 Q6DDA1 Q6DDB3 Q6DDG2 Q6DDK2 Q6DDK3 Q6DDK5 Q6DDL7 Q6DDR3 Q6DDW2
Q6DDW6 Q6DE06 Q6DE15 Q6DE21 Q6DE23 Q6DE37 Q6DE40 Q6DE75 Q6DE92 Q6DED0 Q6DED9
Q6DEJ6 Q6DEL1 Q6DEL2 Q6DEX3 Q6DF08 Q6DF19 Q6DF34 Q6DF55 Q6DF80 Q6DF87 Q6DFC0
Q6DFE7 Q6DFI2 Q6DFJ3 Q6DFJ8 Q6DFK0 Q6DFK2 Q6DFM9 Q6DFQ5 Q6DFQ7 Q6DFR1 Q6DFR5
Q6DFS0 Q6DFT4 Q6DFT6 Q6DFV6 Q6DFW5 Q6DFX2 Q6DG19 Q6DG32 Q6DG36 Q6DG38 Q6DGA7
Q6DGF5 Q6DGF8 Q6DGG9 Q6DGL7 Q6DGM2 Q6DGP4 Q6DGU5 Q6DGV7 Q6DGW9 Q6DGY3 Q6DH46
Q6DH66 Q6DH69 Q6DH87 Q6DH88 Q6DH94 Q6DHB5 Q6DHC3 Q6DHI1 Q6DHJ3 Q6DHK8 Q6DHN0
Q6DHN3 Q6DHP1 Q6DHQ1 Q6DHR8 Q6DHS9 Q6DHU1 Q6DI48 Q6DIB5 Q6DID7 Q6DIE4 Q6DIS1
Q6DIT7 Q6DIV6 Q6DIY8 Q6DIZ8 Q6DJ08 Q6DJ35 Q6DJ83 Q6DJ87 Q6DJC8 Q6DJM3 Q6DJR8
Q6DKC0 Q6DKI7 Q6DKM1 Q6DLD8 Q6DLD9 Q6DLH1 Q6DLH8 Q6DN07 Q6DN12 Q6DN14 Q6DN61
Q6DN72 Q6DNA2 Q6DNG6 Q6DPH9 Q6DPI6 Q6DPJ8 Q6DPJ9 Q6DPK0 Q6DPK1 Q6DPK2 Q6DPQ1
Q6DPQ7 Q6DPT5 Q6DPT7 Q6DPT9 Q6DPU1 Q6DPU3 Q6DPZ9 Q6DQ15 Q6DQ18 Q6DQ19 Q6DQ20
Q6DQ21 Q6DQ22 Q6DQX6 Q6DR03 Q6DR04 Q6DR10 Q6DR98 Q6DRM0 Q6DST1 Q6DUL2 Q6DV88

Q6DVA0 Q6DWJ6 Q6DYE5 Q6DYE8 Q6E0V4 Q6E0W7 Q6E1M8 Q6E213 Q6E240 Q6E263 Q6E279
Q6EBV9 Q6ECI6 Q6ED58 Q6EGH7 Q6EGH8 Q6EGH9 Q6EGI0 Q6EGI2 Q6EGI3 Q6EGI4 Q6EGI5
Q6EGJ0 Q6EGJ1 Q6EIG7 Q6ELU6 Q6ELU7 Q6ELU8 Q6ELU9 Q6ELV2 Q6ELV4 Q6ELV5 Q6ELV7
Q6ELV8 Q6ELV9 Q6ELW1 Q6ELW2 Q6ELW3 Q6ELW6 Q6EMK4 Q6EMS3 Q6EMS7 Q6EMS9 Q6EMT0
Q6EMT1 Q6ENA2 Q6ENA4 Q6ENA5 Q6ENA7 Q6ENA8 Q6ENB0 Q6ENE3 Q6ENE4 Q6ENE5 Q6ENE6
Q6ENE7 Q6ENE8 Q6ENF3 Q6ENF4 Q6ENF5 Q6ENF6 Q6ENF7 Q6ENF8 Q6ENF9 Q6ENG0 Q6ENG1
Q6ENG3 Q6ENG4 Q6ENG9 Q6ENH4 Q6ENH5 Q6ENH8 Q6ENH9 Q6ENI0 Q6ENI5 Q6ENI6 Q6ENI9
Q6ENJ0 Q6ENJ1 Q6ENJ3 Q6ENJ4 Q6ENJ7 Q6ENP2 Q6ENP4 Q6ENP5 Q6ENP7 Q6ENP8 Q6ENQ0
Q6ENT3 Q6ENT4 Q6ENT5 Q6ENT6 Q6ENT7 Q6ENT8 Q6ENU4 Q6ENU5 Q6ENU6 Q6ENU7 Q6ENU8
Q6ENU9 Q6ENV0 Q6ENV1 Q6ENV2 Q6ENV3 Q6ENV4 Q6ENV8 Q6ENW3 Q6ENW4 Q6ENW7 Q6ENW8
Q6ENW9 Q6ENX4 Q6ENX5 Q6ENX9 Q6ENY0 Q6ENY1 Q6ENY3 Q6ENY5 Q6EP58 Q6EPN8 Q6EPQ3
Q6EQ60 Q6ERE5 Q6ESG1 Q6ETK9 Q6ETL8 Q6ETN2 Q6ETQ7 Q6EU94 Q6EUN0 Q6EV76 Q6EV77
Q6EVZ2 Q6EVZ5 Q6EVZ7 Q6EVZ8 Q6EW00 Q6EW01 Q6EW03 Q6EW21 Q6EW22 Q6EW23 Q6EW24
Q6EW25 Q6EW26 Q6EW33 Q6EW34 Q6EW35 Q6EW36 Q6EW37 Q6EW38 Q6EW39 Q6EW40 Q6EW41
Q6EW42 Q6EW43 Q6EW48 Q6EW49 Q6EW51 Q6EW52 Q6EW53 Q6EW54 Q6EW55 Q6EW60 Q6EW61
Q6EW62 Q6EW64 Q6EW65 Q6EW68 Q6EW69 Q6EWG9 Q6EYB9 Q6EYC0 Q6EYC3 Q6EYC4 Q6EYD5
Q6EYD6 Q6EYE3 Q6EYE4 Q6EYF1 Q6EYF2 Q6EYG3 Q6EYG4 Q6EYH4 Q6EYH5 Q6EYH8 Q6EYH9
Q6EYJ0 Q6EYJ1 Q6EYJ4 Q6EYJ5 Q6EYJ8 Q6EYJ9 Q6EYK6 Q6EYK7 Q6EYL4 Q6EYL5 Q6EYL8
Q6EYL9 Q6EYM6 Q6EYM7 Q6EYN0 Q6EYN1 Q6EYP1 Q6EYP2 Q6EYP8 Q6EYR0 Q6EYS0 Q6EYS3
Q6EYS6 Q6EYT6 Q6EYT9 Q6EYU0 Q6EYU3 Q6EYU4 Q6EYV1 Q6EYV8 Q6EYV9 Q6EYW2 Q6EYW3
Q6F055 Q6F056 Q6F0E5 Q6F0N9 Q6F1C9 Q6F1Q9 Q6F206 Q6F207 Q6F2Z1 Q6F3F9 Q6F4F5
Q6F5E0 Q6F5E3 Q6F6K2 Q6F6S7 Q6F6X5 Q6F754 Q6F793 Q6F7D6 Q6F7F9 Q6F7T9 Q6F7V8
Q6F813 Q6F8Q1 Q6F9N8 Q6F9R1 Q6F9V4 Q6F9X0 Q6FA90 Q6FAW0 Q6FB65 Q6FBS3 Q6FCI0
Q6FDF5 Q6FDF6 Q6FDP8 Q6FE62 Q6FE65 Q6FE71 Q6FE93 Q6FEB4 Q6FER0 Q6FER4 Q6FEY7
Q6FF10 Q6FFC1 Q6FFK4 Q6FFK5 Q6FFK6 Q6FFS6 Q6FG03 Q6FIK2 Q6FIK3 Q6FIK6 Q6FIN8
Q6FIQ1 Q6FIR8 Q6FIW0 Q6FIY2 Q6FJ31 Q6FJ70 Q6FJ81 Q6FJD1 Q6FJG2 Q6FJH4 Q6FJJ0
Q6FJJ9 Q6FJT9 Q6FK15 Q6FK23 Q6FK33 Q6FKA1 Q6FKH5 Q6FKI5 Q6FKJ1 Q6FKL3 Q6FKM3
Q6FL09 Q6FL34 Q6FLC5 Q6FLC9 Q6FLH2 Q6FLL2 Q6FLM2 Q6FLT3 Q6FLY9 Q6FLZ2 Q6FM91
Q6FMA9 Q6FMB2 Q6FMP5 Q6FMQ0 Q6FMU3 Q6FMZ2 Q6FN06 Q6FN38 Q6FNE2 Q6FNJ6 Q6FNP6
Q6FNQ2 Q6FNV5 Q6FNX1 Q6FP25 Q6FPB2 Q6FPB6 Q6FPC0 Q6FPE8 Q6FPF1 Q6FPI5 Q6FPI6
Q6FPJ3 Q6FPU1 Q6FQ03 Q6FQ14 Q6FQ15 Q6FQ42 Q6FQ45 Q6FQE0 Q6FQK7 Q6FQN0 Q6FQN3
Q6FQT7 Q6FR11 Q6FR88 Q6FRE5 Q6FRM0 Q6FRR4 Q6FRT5 Q6FRT6 Q6FRU5 Q6FRU9 Q6FRX4
Q6FRY3 Q6FRZ3 Q6FS06 Q6FS52 Q6FS62 Q6FS98 Q6FSD1 Q6FSF8 Q6FSG0 Q6FSQ7 Q6FSS4
Q6FSU7 Q6FSW5 Q6FSZ7 Q6FT25 Q6FT37 Q6FT92 Q6FT93 Q6FTB1 Q6FTC4 Q6FTE5 Q6FTI0
Q6FTM9 Q6FTN2 Q6FTY5 Q6FU40 Q6FU42 Q6FUG4 Q6FUX5 Q6FUY5 Q6FV22 Q6FV75 Q6FV98
Q6FVB6 Q6FVE6 Q6FVS6 Q6FVT7 Q6FW26 Q6FW43 Q6FW70 Q6FWD1 Q6FWD4 Q6FWE8 Q6FWN4
Q6FWS5 Q6FWT0 Q6FWV0 Q6FX62 Q6FX92 Q6FX96 Q6FXC6 Q6FXH5 Q6FXJ3 Q6FXQ5 Q6FXS2
Q6FXV6 Q6FYG1 Q6FYL0 Q6FYV9 Q6FYW1 Q6FYW3 Q6FYW5 Q6FYW8 Q6FZF2 Q6FZS3 Q6FZY1
Q6FZY4 Q6G011 Q6G039 Q6G0D0 Q6G0F4 Q6G0H0 Q6G0H1 Q6G0H3 Q6G0H9 Q6G121 Q6G147
Q6G160 Q6G1A8 Q6G1V5 Q6G2A8 Q6G2B2 Q6G2B4 Q6G2L7 Q6G2Q2 Q6G2Z5 Q6G395 Q6G398
Q6G3F5 Q6G3L5 Q6G422 Q6G494 Q6G4C6 Q6G4F8 Q6G4V3 Q6G5A2 Q6G5K1 Q6G5K9 Q6G5L0
Q6G5L2 Q6G5S9 Q6G5Z0 Q6G605 Q6G607 Q6G608 Q6G610 Q6G6A7 Q6G6A9 Q6G6B1 Q6G6B7
Q6G6C4 Q6G6D3 Q6G6D4 Q6G6D6 Q6G6I7 Q6G6M9 Q6G6T3 Q6G6V8 Q6G6W0 Q6G6W4 Q6G6Y2
Q6G6Y6 Q6G765 Q6G791 Q6G7C4 Q6G7E6 Q6G7F6 Q6G7K1 Q6G7K2 Q6G7K3 Q6G7M0 Q6G7M2

Q6G7N2 Q6G7N3 Q6G7N4 Q6G7S1 Q6G7S2 Q6G816 Q6G821 Q6G831 Q6G845 Q6G849 Q6G860
Q6G868 Q6G889 Q6G8E7 Q6G8E8 Q6G8F6 Q6G8I8 Q6G8L0 Q6G8Z4 Q6G973 Q6G983 Q6G9C6
Q6G9E1 Q6G9E7 Q6G9F4 Q6G9G0 Q6G9H8 Q6G9H9 Q6G9J7 Q6G9J9 Q6G9K6 Q6G9L1 Q6G9L5
Q6G9S7 Q6G9T0 Q6G9T7 Q6G9V1 Q6G9V2 Q6GA17 Q6GA30 Q6GA81 Q6GA98 Q6GA99 Q6GAA9
Q6GAF2 Q6GAF3 Q6GAF4 Q6GAF5 Q6GAJ1 Q6GAR1 Q6GAV2 Q6GAV7 Q6GAW1 Q6GAW2 Q6GAX4
Q6GAX5 Q6GAX6 Q6GAX7 Q6GAX8 Q6GAX9 Q6GAY0 Q6GB52 Q6GB69 Q6GB84 Q6GBB1 Q6GBC5
Q6GBD5 Q6GBE7 Q6GBH0 Q6GBK1 Q6GBK2 Q6GBK3 Q6GBK4 Q6GBK5 Q6GBK6 Q6GBK7 Q6GBQ4
Q6GBR4 Q6GBV2 Q6GCD1 Q6GCD7 Q6GCI5 Q6GCI6 Q6GCI8 Q6GCI9 Q6GCK2 Q6GCK9 Q6GCL0
Q6GCL2 Q6GCQ2 Q6GCR5 Q6GCT4 Q6GCT7 Q6GD71 Q6GDB9 Q6GDD5 Q6GDD7 Q6GDD8 Q6GDE0
Q6GDN2 Q6GDN4 Q6GDN6 Q6GDP1 Q6GDP8 Q6GDQ7 Q6GDQ8 Q6GDR0 Q6GDV3 Q6GDZ1 Q6GE44
Q6GE72 Q6GE74 Q6GE78 Q6GE96 Q6GEA0 Q6GEH7 Q6GEK3 Q6GEN9 Q6GER1 Q6GES1 Q6GEW6
Q6GEW7 Q6GEW8 Q6GEY5 Q6GEY7 Q6GEZ6 Q6GEZ7 Q6GEZ8 Q6GF36 Q6GF37 Q6GFE0 Q6GFE5
Q6GFF5 Q6GFG8 Q6GFH2 Q6GFI3 Q6GFJ1 Q6GFL0 Q6GFR9 Q6GFS0 Q6GFT0 Q6GFW9 Q6GFZ0
Q6GGC7 Q6GGK7 Q6GGT1 Q6GGX2 Q6GGX3 Q6GGY8 Q6GGZ4 Q6GH01 Q6GH07 Q6GH25 Q6GH26
Q6GH43 Q6GH45 Q6GH52 Q6GH57 Q6GH63 Q6GH88 Q6GHE9 Q6GHF2 Q6GHF9 Q6GHH3 Q6GHH4
Q6GHN9 Q6GHQ1 Q6GHQ3 Q6GHV2 Q6GHW9 Q6GHX0 Q6GHY0 Q6GI23 Q6GI24 Q6GI25 Q6GI26
Q6GI62 Q6GI68 Q6GIB3 Q6GIB9 Q6GIC3 Q6GIC4 Q6GID6 Q6GID7 Q6GID8 Q6GID9 Q6GIE0
Q6GIE1 Q6GIE2 Q6GIL2 Q6GIN0 Q6GIP5 Q6GIS3 Q6GIT7 Q6GIU7 Q6GIV9 Q6GIX1 Q6GIX4
Q6GJ10 Q6GJ42 Q6GJ43 Q6GJ44 Q6GJ45 Q6GJ46 Q6GJ47 Q6GJ86 Q6GJ96 Q6GJD3 Q6GJX7
Q6GJY2 Q6GK24 Q6GK25 Q6GK27 Q6GK28 Q6GK41 Q6GK49 Q6GK50 Q6GK52 Q6GK92 Q6GKB4
Q6GKB7 Q6GKN3 Q6GKN4 Q6GKS6 Q6GL33 Q6GL42 Q6GLA2 Q6GLC5 Q6GLE1 Q6GLJ0 Q6GLL2
Q6GLN5 Q6GLN7 Q6GLU0 Q6GLW8 Q6GLX2 Q6GLX4 Q6GLZ9 Q6GM04 Q6GM05 Q6GM19 Q6GM22
Q6GM29 Q6GM42 Q6GM44 Q6GM59 Q6GMB1 Q6GMF8 Q6GMG6 Q6GMI9 Q6GMK0 Q6GMK6 Q6GMR7
Q6GMZ9 Q6GN01 Q6GN29 Q6GN30 Q6GN42 Q6GN58 Q6GNB5 Q6GND7 Q6GNG3 Q6GNL1 Q6GNM0
Q6GNR9 Q6GNS1 Q6GNT2 Q6GNT7 Q6GNV7 Q6GNX5 Q6GNY6 Q6GNY8 Q6GP51 Q6GP81 Q6GPA5
Q6GPA8 Q6GPH6 Q6GPL4 Q6GPN9 Q6GPP7 Q6GPQ3 Q6GPW4 Q6GPY1 Q6GPZ5 Q6GQ22 Q6GQ26
Q6GQ39 Q6GQ49 Q6GQ62 Q6GQ70 Q6GQ81 Q6GQE1 Q6GQF5 Q6GQI7 Q6GQI8 Q6GQK9 Q6GQS1
Q6GQT5 Q6GQT6 Q6GQT9 Q6GQU6 Q6GQW0 Q6GR21 Q6GR43 Q6GR51 Q6GTX8 Q6GUG4 Q6GV12
Q6GV17 Q6GV23 Q6GV27 Q6GV28 Q6GX83 Q6GZQ6 Q6GZS3 Q6GZT3 Q6GZU2 Q6GZV5 Q6GZW4
Q6GZW5 Q6GZX1 Q6GZX3 Q6H1V1 Q6H2Y3 Q6H3X3 Q6H3Z3 Q6H3Z6 Q6H4L9 Q6H4M2 Q6H4N0
Q6H4R6 Q6H501 Q6H5U3 Q6H641 Q6H658 Q6H6R9 Q6H765 Q6H7D2 Q6H7J6 Q6H7M3 Q6H7U3
Q6H8M7 Q6H941 Q6H952 Q6H956 Q6HAJ6 Q6HAJ7 Q6HAV9 Q6HAX4 Q6HAX5 Q6HAY5 Q6HAY9
Q6HAZ2 Q6HBC7 Q6HBH8 Q6HBI2 Q6HBI3 Q6HCK7 Q6HCM8 Q6HCT7 Q6HDR9 Q6HEL8 Q6HEL9
Q6HEQ1 Q6HES1 Q6HF37 Q6HFC1 Q6HFD1 Q6HFJ3 Q6HG49 Q6HHY7 Q6HI21 Q6HI77 Q6HIP2
Q6HJB9 Q6HL81 Q6HLG8 Q6HLG9 Q6HM31 Q6HMW0 Q6HN77 Q6HN78 Q6HN79 Q6HND2 Q6HNQ4
Q6HNS1 Q6HP15 Q6HPB7 Q6HQX5 Q6I628 Q6I681 Q6I7B9 Q6IA17 Q6IAN0 Q6IC98 Q6ICH7
Q6ICI0 Q6ICL7 Q6ICY4 Q6ID99 Q6IDL4 Q6IE08 Q6IE14 Q6IE26 Q6IE63 Q6IE75 Q6IED8
Q6IEE6 Q6IEE7 Q6IEE8 Q6IEU7 Q6IEV9 Q6IEY1 Q6IEZ7 Q6IF00 Q6IF36 Q6IF42 Q6IF63
Q6IF82 Q6IF99 Q6IFG1 Q6IFH4 Q6IFN5 Q6IFT6 Q6IFY7 Q6IM80 Q6IM82 Q6IM84 Q6IM86
Q6IM87 Q6IM89 Q6IM91 Q6IM92 Q6IM93 Q6IM94 Q6IM95 Q6IMP4 Q6INB3 Q6INC8 Q6INE8
Q6INN0 Q6INP0 Q6INQ6 Q6INU6 Q6INU7 Q6INX1 Q6INX3 Q6IP19 Q6IP59 Q6IQ69 Q6IQ85
Q6IQA2 Q6IQC7 Q6IQH6 Q6IQX7 Q6IR37 Q6IR55 Q6IR74 Q6IR76 Q6IRC8 Q6IRI9 Q6IRP4
Q6IS24 Q6IS41 Q6ISU1 Q6ITQ4 Q6ITT3 Q6IUF7 Q6IUU3 Q6IUY1 Q6IV13 Q6IVV8 Q6IWH7
Q6IWY1 Q6IYF8 Q6IYF9 Q6IYG6 Q6IYH0 Q6IZB5 Q6J163 Q6J1J8 Q6J1Y9 Q6J2K5 Q6J329

Q6J3P1 Q6J4G7 Q6J4K2 Q6J540 Q6J541 Q6J8D5 Q6J8E4 Q6J8E7 Q6J8F6 Q6J8I9 Q6J9G0
Q6J9G1 Q6JAM9 Q6JAN0 Q6JD68 Q6JDN5 Q6JDP8 Q6JDQ6 Q6JDS2 Q6JDS4 Q6JDS5 Q6JDS6
Q6JDS8 Q6JDT0 Q6JDT3 Q6JDT4 Q6JDT8 Q6JDU0 Q6JDU8 Q6JN46 Q6JN47 Q6JQN2 Q6JTA8
Q6JTJ0 Q6JWR2 Q6JWV8 Q6JWX5 Q6K0P5 Q6K1C4 Q6K1J1 Q6K215 Q6K3D8 Q6K3R5 Q6K3T2
Q6K478 Q6K4T4 Q6K4V2 Q6K548 Q6K602 Q6K739 Q6K7S7 Q6K8E7 Q6K8S7 Q6K965 Q6K991
Q6K9F6 Q6K9G1 Q6K9G3 Q6KB54 Q6KB55 Q6KB58 Q6KB59 Q6KCM7 Q6KEQ9 Q6KF39 Q6KF53
Q6KFX9 Q6KHA4 Q6KHJ9 Q6KI76 Q6KI77 Q6KIH2 Q6KZG0 Q6L013 Q6L0N2 Q6L147 Q6L174
Q6L2Q7 Q6L371 Q6L372 Q6L373 Q6L374 Q6L375 Q6L376 Q6L381 Q6L382 Q6L383 Q6L384
Q6L385 Q6L386 Q6L387 Q6L388 Q6L389 Q6L390 Q6L394 Q6L398 Q6L399 Q6L3A2 Q6L3A3
Q6L3A8 Q6L3A9 Q6L3B0 Q6L3B1 Q6L3B2 Q6L3B3 Q6L3B5 Q6L3D5 Q6L3D7 Q6L3D8 Q6L3E0
Q6L3E1 Q6L3E3 Q6L4D2 Q6L538 Q6L568 Q6L5F5 Q6L5F6 Q6L5I5 Q6L602 Q6L603 Q6L708
Q6L786 Q6L8F7 Q6L8F9 Q6L8G0 Q6L8G1 Q6L8S8 Q6L9W6 Q6LA55 Q6LEJ4 Q6LG82 Q6LIN8
Q6LK12 Q6LL02 Q6LLG2 Q6LLG4 Q6LLR8 Q6LLU3 Q6LM12 Q6LM13 Q6LMF3 Q6LMF5 Q6LNA1
Q6LNF7 Q6LNY7 Q6LNY8 Q6LP61 Q6LP88 Q6LPF8 Q6LPH2 Q6LPK6 Q6LPY5 Q6LQ01 Q6LQ49
Q6LQ76 Q6LQ84 Q6LTL4 Q6LTU6 Q6LTY5 Q6LTY6 Q6LUJ8 Q6LUL9 Q6LUM2 Q6LV09 Q6LV37
Q6LVR2 Q6LVS3 Q6LVW7 Q6LVW8 Q6LVY3 Q6LW55 Q6LWL1 Q6LWM4 Q6LWN2 Q6LWY8 Q6LWZ0
Q6LWZ1 Q6LWZ2 Q6LWZ3 Q6LWZ4 Q6LX42 Q6LX70 Q6LXA4 Q6LXB7 Q6LXR6 Q6LYN9 Q6LYT4
Q6LZR9 Q6LZU5 Q6LZY8 Q6M2F0 Q6M8W7 Q6MAD8 Q6MAK1 Q6MAK2 Q6MAK3 Q6MBS3 Q6MBY2
Q6MC92 Q6MCB9 Q6MDI5 Q6MDQ3 Q6MDQ6 Q6MDQ9 Q6ME04 Q6MG14 Q6MG22 Q6MG55 Q6MG56
Q6MG59 Q6MG61 Q6MG71 Q6MG82 Q6MG97 Q6MGM3 Q6MGN3 Q6MGN5 Q6MGN8 Q6MIF7 Q6MJV1
Q6ML26 Q6ML54 Q6MLJ0 Q6MLL0 Q6MLS7 Q6MN25 Q6MN26 Q6MNQ3 Q6MPX9 Q6MR11 Q6MRC6
Q6MRR3 Q6MS89 Q6MS90 Q6MTI6 Q6MUC0 Q6MWE5 Q6MWG9 Q6MYT0 Q6MYX6 Q6MZM0 Q6N022
Q6N075 Q6N1M9 Q6N1Y7 Q6N1Y8 Q6N1Z4 Q6N1Z7 Q6N356 Q6N408 Q6N435 Q6N5F0 Q6N5F2
Q6N5G6 Q6N5H0 Q6N5H2 Q6N5N1 Q6N5N4 Q6N5U3 Q6N5X1 Q6N660 Q6N6D0 Q6N715 Q6N893
Q6N8I3 Q6N8Y1 Q6N9G6 Q6NB79 Q6NBI2 Q6NBI4 Q6NBI5 Q6NBJ5 Q6NCE7 Q6NE49 Q6NE56
Q6NE58 Q6NE62 Q6NE63 Q6NE74 Q6NE75 Q6NE76 Q6NE78 Q6NFM3 Q6NG92 Q6NG98 Q6NG99
Q6NGA0 Q6NGA1 Q6NGA2 Q6NGA3 Q6NGC5 Q6NH44 Q6NH98 Q6NHP1 Q6NHT3 Q6NHY4 Q6NIE7
Q6NIN5 Q6NJ43 Q6NKQ9 Q6NKR1 Q6NKS4 Q6NKU9 Q6NKZ8 Q6NKZ9 Q6NLA5 Q6NLC1 Q6NM25
Q6NM73 Q6NMB6 Q6NMB7 Q6NMD6 Q6NMK1 Q6NML0 Q6NMM1 Q6NMM8 Q6NMN6 Q6NN03 Q6NN55
Q6NP60 Q6NPD8 Q6NPF8 Q6NPR7 Q6NPT7 Q6NQ51 Q6NQ87 Q6NQA8 Q6NQB7 Q6NQG7 Q6NQI8
Q6NQK9 Q6NQL6 Q6NQN5 Q6NRB7 Q6NRB9 Q6NRI1 Q6NRI4 Q6NRL6 Q6NRM1 Q6NRQ1 Q6NRQ2
Q6NRS2 Q6NRS6 Q6NRV4 Q6NRX0 Q6NS09 Q6NS19 Q6NS26 Q6NS65 Q6NS82 Q6NSJ0 Q6NSJ5
Q6NSN2 Q6NSQ9 Q6NSR3 Q6NSU3 Q6NT16 Q6NT55 Q6NTF9 Q6NTJ7 Q6NTL1 Q6NTT6 Q6NTV1
Q6NTW5 Q6NTZ6 Q6NU09 Q6NU91 Q6NUB3 Q6NUI2 Q6NUJ2 Q6NUK1 Q6NUK4 Q6NUQ4 Q6NUS6
Q6NUS8 Q6NUT2 Q6NUT3 Q6NUT7 Q6NUZ2 Q6NV04 Q6NV12 Q6NV38 Q6NV75 Q6NVC5 Q6NVD0
Q6NVG1 Q6NVP8 Q6NVQ1 Q6NVR9 Q6NVS2 Q6NVV3 Q6NWD4 Q6NWF1 Q6NWG0 Q6NWG1 Q6NWH5
Q6NWW9 Q6NX28 Q6NXD8 Q6NXH2 Q6NXK7 Q6NXK8 Q6NXM3 Q6NXN4 Q6NXT4 Q6NXT6 Q6NYD1
Q6NYE4 Q6NYE7 Q6NYF1 Q6NYG4 Q6NYI0 Q6NYK8 Q6NYN7 Q6NYP0 Q6NYS8 Q6NYV8 Q6NYY9
Q6NYZ6 Q6NZ07 Q6NZ21 Q6NZ63 Q6NZH5 Q6NZI6 Q6NZL0 Q6NZQ8 Q6NZZ4 Q6P011 Q6P036
Q6P073 Q6P0C6 Q6P0C7 Q6P0D1 Q6P0E8 Q6P0F0 Q6P0S3 Q6P135 Q6P179 Q6P1A2 Q6P1B3
Q6P1C6 Q6P1D5 Q6P1H1 Q6P1H6 Q6P1I6 Q6P1J0 Q6P1J6 Q6P1K1 Q6P1M0 Q6P1Q0 Q6P1S2
Q6P1U2 Q6P1V1 Q6P257 Q6P2H8 Q6P2L7 Q6P2T0 Q6P2X9 Q6P301 Q6P312 Q6P316 Q6P317
Q6P342 Q6P355 Q6P360 Q6P3A4 Q6P3L6 Q6P3N5 Q6P3N9 Q6P3P5 Q6P418 Q6P499 Q6P4A7
Q6P4E1 Q6P4F1 Q6P4H8 Q6P4M0 Q6P4N1 Q6P4P2 Q6P4Q7 Q6P4S8 Q6P4U0 Q6P4Y0 Q6P501

Q6P531 Q6P5B0 Q6P5F6 Q6P5F7 Q6P5I8 Q6P5P5 Q6P5S7 Q6P5W5 Q6P5X7 Q6P606 Q6P689
Q6P6G2 Q6P6G5 Q6P6J9 Q6P6M5 Q6P6Q9 Q6P6R0 Q6P6S2 Q6P6S5 Q6P6T5 Q6P6V1 Q6P6V6
Q6P719 Q6P726 Q6P742 Q6P752 Q6P767 Q6P768 Q6P7A1 Q6P7B2 Q6P7B4 Q6P7B7 Q6P7B9
Q6P7C4 Q6P7C7 Q6P7D4 Q6P7G9 Q6P7K5 Q6P7L4 Q6P7N7 Q6P7R8 Q6P806 Q6P825 Q6P8F8
Q6P8G8 Q6P8H8 Q6P8I6 Q6P995 Q6P9A2 Q6P9B9 Q6P9F7 Q6P9G4 Q6P9J9 Q6P9K9 Q6P9S7
Q6P9X9 Q6P9Z6 Q6PA06 Q6PA35 Q6PA90 Q6PAB8 Q6PAQ9 Q6PAX2 Q6PAZ3 Q6PB03 Q6PB15
Q6PB26 Q6PB70 Q6PB90 Q6PB93 Q6PBD1 Q6PBE5 Q6PBF7 Q6PBH5 Q6PBI8 Q6PBK8 Q6PBS6
Q6PBT7 Q6PBX9 Q6PC24 Q6PC64 Q6PC78 Q6PCB7 Q6PCB8 Q6PCF8 Q6PCF9 Q6PCG6 Q6PCP5
Q6PCP7 Q6PCS6 Q6PCT8 Q6PD19 Q6PD26 Q6PD82 Q6PDC8 Q6PDD0 Q6PDE7 Q6PDE8 Q6PDF3
Q6PDJ1 Q6PDU4 Q6PE13 Q6PE84 Q6PEC3 Q6PEH1 Q6PEH9 Q6PEM8 Q6PEX7 Q6PEY0 Q6PEY1
Q6PF45 Q6PFC5 Q6PFE7 Q6PFJ7 Q6PFM1 Q6PFM4 Q6PFS5 Q6PFT6 Q6PFX6 Q6PG34 Q6PGC7
Q6PGD0 Q6PGE7 Q6PGK7 Q6PGS5 Q6PGW3 Q6PGX3 Q6PGY3 Q6PH58 Q6PH61 Q6PHB0 Q6PHL4
Q6PHN7 Q6PHS9 Q6PHU5 Q6PHW0 Q6PI25 Q6PI62 Q6PI73 Q6PI78 Q6PIC6 Q6PIE5 Q6PIS1
Q6PIU1 Q6PIU2 Q6PIV7 Q6PIX5 Q6PIX9 Q6PIZ9 Q6PJF5 Q6PJG9 Q6PJW8 Q6PK18 Q6PKC3
Q6PL45 Q6PLK3 Q6PML9 Q6PNM1 Q6PP77 Q6PQZ1 Q6PQZ2 Q6PQZ3 Q6PRD1 Q6PST4 Q6PT86
Q6PT90 Q6PWL6 Q6PXP3 Q6PYT3 Q6PZD2 Q6Q0C1 Q6Q0N0 Q6Q1P3 Q6Q1R9 Q6Q1S0 Q6Q1S1
Q6Q1S2 Q6Q2J0 Q6Q304 Q6Q3F5 Q6Q473 Q6Q477 Q6Q487 Q6Q4G3 Q6Q595 Q6Q5X2 Q6Q629
Q6Q760 Q6Q7K0 Q6Q876 Q6Q879 Q6Q881 Q6Q884 Q6Q889 Q6Q8B3 Q6Q972 Q6QA32 Q6QA33
Q6QA74 Q6QAJ8 Q6QBN9 Q6QBQ4 Q6QD59 Q6QDP7 Q6QHC5 Q6QI25 Q6QI68 Q6QIX3 Q6QIY3
Q6QJE2 Q6QLL4 Q6QLQ4 Q6QMG1 Q6QMT2 Q6QN11 Q6QN12 Q6QN13 Q6QNF3 Q6QNI4 Q6QNK2
Q6QNU9 Q6QQT1 Q6QRN8 Q6QRN9 Q6QT02 Q6QU67 Q6QUN5 Q6R0J2 Q6R2J8 Q6R2K1 Q6R2K2
Q6R2K3 Q6R3K4 Q6R3K6 Q6R3K8 Q6R3K9 Q6R3L0 Q6R4A8 Q6R4Q5 Q6R567 Q6R5A3 Q6R5J1
Q6R5J2 Q6R5N8 Q6R5P0 Q6R653 Q6R6I6 Q6R6I7 Q6R6L8 Q6R6U4 Q6R755 Q6R7B9 Q6R7C6
Q6R7E1 Q6R7E5 Q6R7E9 Q6R7F2 Q6R7F6 Q6R7G0 Q6R7G6 Q6R7G8 Q6R7H1 Q6R7I3 Q6R7I8
Q6R7J8 Q6R7K1 Q6R8F2 Q6R8G0 Q6R8G2 Q6R8G3 Q6R8G4 Q6R8G5 Q6R8G6 Q6R8G7 Q6R8G8
Q6R8J2 Q6RET6 Q6RH31 Q6RH51 Q6RH59 Q6RHR6 Q6RI86 Q6RI88 Q6RJQ3 Q6RJR6 Q6RKD8
Q6RS95 Q6RS96 Q6RSH5 Q6RSH6 Q6RUT7 Q6RVG2 Q6RW13 Q6RWA9 Q6RX08 Q6RYS9 Q6RYW5
Q6RZ07 Q6RZG6 Q6RZI9 Q6RZJ7 Q6S5G3 Q6S5G4 Q6S5H8 Q6S6Q5 Q6S6R5 Q6S6T9 Q6S6V5
Q6S6V7 Q6S6W0 Q6S9C0 Q6S9C4 Q6SJE0 Q6SJP2 Q6SJQ0 Q6SJQ5 Q6SJQ7 Q6SLE7 Q6SQI7
Q6SVX3 Q6SVZ0 Q6SVZ5 Q6SW00 Q6SW10 Q6SW14 Q6SW31 Q6SW43 Q6SW67 Q6SW81 Q6SW94
Q6SW98 Q6SWB7 Q6SWB9 Q6SWC0 Q6SWC3 Q6SWC4 Q6SWC7 Q6SWC8 Q6SWC9 Q6SWD0 Q6SWD1
Q6SWN0 Q6SZ61 Q6SZ87 Q6SZS5 Q6SZS6 Q6T3U3 Q6T3U4 Q6T423 Q6T4P5 Q6T5K2 Q6T5K3
Q6T707 Q6T752 Q6TA49 Q6TAC4 Q6TAC8 Q6TAW2 Q6TCG2 Q6TCG5 Q6TCG8 Q6TCH4 Q6TCH7
Q6TEK3 Q6TEK4 Q6TEK5 Q6TGJ4 Q6TL19 Q6TLE4 Q6TLE6 Q6TLI7 Q6TLI9 Q6TLJ0 Q6TMJ1
Q6TMJ3 Q6TMJ9 Q6TNT2 Q6TUD4 Q6TV02 Q6TXB9 Q6TXF5 Q6TYA7 Q6TYA8 Q6TYA9 Q6TYI6
Q6U1P6 Q6U1Z4 Q6U6J0 Q6U736 Q6U7R4 Q6U7Y2 Q6U841 Q6UC88 Q6UD73 Q6UDF0 Q6UDF4
Q6UDF5 Q6UDF6 Q6UDG4 Q6UDH4 Q6UDI5 Q6UDI7 Q6UDI8 Q6UDJ1 Q6UDJ7 Q6UDK4 Q6UDL0
Q6UDL6 Q6UDL9 Q6UDM3 Q6UE05 Q6UE39 Q6UEF4 Q6UEH3 Q6UEH4 Q6UG62 Q6UGB2 Q6UJB9
Q6UJY2 Q6UK00 Q6UKI2 Q6UN27 Q6UNA4 Q6UPR8 Q6UQ04 Q6UQ17 Q6UR05 Q6URK6 Q6US81
Q6UUW5 Q6UUW6 Q6UVK1 Q6UVM3 Q6UVM4 Q6UVW9 Q6UVY6 Q6UW02 Q6UW56 Q6UW60 Q6UW68
Q6UW78 Q6UW88 Q6UWB1 Q6UWD8 Q6UWF3 Q6UWF5 Q6UWH4 Q6UWH6 Q6UWI2 Q6UWI4 Q6UWJ1
Q6UWJ8 Q6UWL2 Q6UWL6 Q6UWM7 Q6UWM9 Q6UWP7 Q6UWU4 Q6UWV2 Q6UWV6 Q6UWV7 Q6UWW9
Q6UX01 Q6UX15 Q6UX27 Q6UX34 Q6UX40 Q6UX41 Q6UX65 Q6UX68 Q6UX71 Q6UX72 Q6UX98
Q6UXB4 Q6UXC1 Q6UXD1 Q6UXD5 Q6UXD7 Q6UXE8 Q6UXF1 Q6UXG2 Q6UXG3 Q6UXG8 Q6UXK2

Q6UXK5 Q6UXL0 Q6UXM1 Q6UXN7 Q6UXN8 Q6UXP3 Q6UXU4 Q6UXU6 Q6UXV0 Q6UXV1 Q6UXV4
Q6UXY8 Q6UXZ0 Q6UXZ3 Q6UXZ4 Q6UY09 Q6UY11 Q6UY18 Q6UY66 Q6UY87 Q6V0I7 Q6V0L0
Q6V1P9 Q6V3W9 Q6V4S5 Q6V7K3 Q6V8Y0 Q6V8Y1 Q6V915 Q6V9D6 Q6V9D9 Q6V9E0 Q6VB83
Q6VBQ5 Q6VE48 Q6VMN6 Q6VNS1 Q6VPV2 Q6VV64 Q6VVA6 Q6VVW5 Q6VZT9 Q6VZZ7 Q6W049
Q6W3B0 Q6W3E1 Q6W3E5 Q6W3F4 Q6W5G4 Q6W5P4 Q6W630 Q6W6K7 Q6W6L7 Q6W6Q8 Q6W6Q9
Q6W6R2 Q6W6R3 Q6W8W4 Q6W8W5 Q6WAY2 Q6WB94 Q6WB95 Q6WB98 Q6WEB5 Q6WG24 Q6WG30
Q6WGK6 Q6WKY9 Q6WKZ0 Q6WKZ1 Q6WL85 Q6WN19 Q6WN98 Q6WNG7 Q6WNQ8 Q6WNQ9 Q6WNR0
Q6WP49 Q6WP51 Q6WQJ1 Q6WQW2 Q6WRG2 Q6WRG3 Q6WRG4 Q6WRG7 Q6WRH2 Q6WRH4 Q6WRH5
Q6WRS2 Q6WRU0 Q6WRY0 Q6WV16 Q6WV17 Q6WV19 Q6WV20 Q6WWW3 Q6WXV7 Q6WYY1 Q6WZB0
Q6X0I2 Q6X1D1 Q6X1D5 Q6X1Y6 Q6X2M0 Q6X2U1 Q6X2U3 Q6X5T8 Q6X5U0 Q6X5U1 Q6X5V0
Q6X632 Q6X862 Q6X893 Q6X936 Q6X9S4 Q6X9T7 Q6X9T8 Q6XAT2 Q6XBV1 Q6XBV6 Q6XBW1
Q6XBW4 Q6XBW5 Q6XBW7 Q6XBW8 Q6XCF2 Q6XFR6 Q6XJV4 Q6XJV6 Q6XK22 Q6XKD3 Q6XL41
Q6XL69 Q6XPS3 Q6XPU4 Q6XQ14 Q6XQ84 Q6XQ97 Q6XQ98 Q6XQ99 Q6XQG8 Q6XQG9 Q6XQH0
Q6XR72 Q6XT33 Q6XT43 Q6XTM7 Q6XTN2 Q6XTV0 Q6XU12 Q6XUA7 Q6XUD6 Q6XUE4 Q6XV23
Q6XV24 Q6XV26 Q6XV27 Q6XV28 Q6XV43 Q6XV52 Q6XXX8 Q6XXX9 Q6XXY0 Q6XYQ8 Q6XZH2
Q6XZH3 Q6XZN8 Q6XZN9 Q6XZP4 Q6Y1E2 Q6Y1H2 Q6Y1R5 Q6Y288 Q6Y290 Q6Y2X3 Q6Y306
Q6Y8A2 Q6Y8J2 Q6Y8J6 Q6Y8J7 Q6YAT4 Q6YBV0 Q6YC49 Q6YDK5 Q6YDK7 Q6YDL2 Q6YDL6
Q6YDL7 Q6YI21 Q6YI46 Q6YI48 Q6YK44 Q6YLN1 Q6YLT3 Q6YLT4 Q6YLT7 Q6YLV5 Q6YLX9
Q6YMS3 Q6YMS4 Q6YNB6 Q6YNI2 Q6YQV1 Q6YRJ5 Q6YRM6 Q6YSA9 Q6YT98 Q6YTF1 Q6YTF5
Q6YUU5 Q6YV88 Q6YVM4 Q6YW53 Q6YW62 Q6YWK8 Q6YWQ4 Q6YWS8 Q6YXK0 Q6YXK1 Q6YXK2
Q6YXL3 Q6YXL6 Q6YXL7 Q6YXL8 Q6YXL9 Q6YXM0 Q6YXM1 Q6YXM2 Q6YXM8 Q6YXN1 Q6YXN2
Q6YXN3 Q6YXN4 Q6YXN8 Q6YXN9 Q6YXP5 Q6YXP8 Q6YXQ0 Q6YXQ1 Q6YXQ3 Q6YXQ6 Q6YXR0
Q6YXR1 Q6YXR4 Q6YXR5 Q6YXR8 Q6YXR9 Q6YXZ1 Q6YZF3 Q6Z0E2 Q6Z0Q9 Q6Z1G3 Q6Z1Y7
Q6Z2T3 Q6Z2T9 Q6Z2U5 Q6Z3Y6 Q6Z401 Q6Z4P2 Q6Z4U4 Q6Z5I7 Q6Z5M3 Q6Z6D6 Q6Z7I3
Q6Z7K5 Q6Z8D0 Q6Z8F5 Q6ZBF6 Q6ZCX1 Q6ZCZ2 Q6ZDE3 Q6ZF85 Q6ZF86 Q6ZF89 Q6ZFH6
Q6ZFR0 Q6ZFZ4 Q6ZG25 Q6ZG85 Q6ZGL9 Q6ZGM7 Q6ZH45 Q6ZH92 Q6ZI50 Q6ZIB5 Q6ZIB9
Q6ZJ91 Q6ZJJ1 Q6ZKB2 Q6ZKF0 Q6ZL17 Q6ZLD3 Q6ZLK0 Q6ZM28 Q6ZM51 Q6ZMB0 Q6ZMB5
Q6ZMC9 Q6ZMD2 Q6ZMG9 Q6ZMH5 Q6ZMI3 Q6ZMJ2 Q6ZMQ8 Q6ZMR5 Q6ZMZ0 Q6ZMZ3 Q6ZN44
Q6ZN68 Q6ZNA5 Q6ZNB6 Q6ZNB7 Q6ZNC8 Q6ZNI0 Q6ZNR0 Q6ZP29 Q6ZP80 Q6ZPB5 Q6ZPD8
Q6ZPD9 Q6ZPJ0 Q6ZPR4 Q6ZPR5 Q6ZQ11 Q6ZQ29 Q6ZQ89 Q6ZQA6 Q6ZQE4 Q6ZQI3 Q6ZQM8
Q6ZQN7 Q6ZQQ2 Q6ZRH7 Q6ZRP7 Q6ZRR5 Q6ZS10 Q6ZS62 Q6ZS82 Q6ZSA7 Q6ZSJ9 Q6ZSM3
Q6ZSS7 Q6ZSY5 Q6ZT12 Q6ZT21 Q6ZT89 Q6ZTQ4 Q6ZU64 Q6ZU69 Q6ZUB0 Q6ZUB1 Q6ZUK4
Q6ZUT9 Q6ZUX7 Q6ZV29 Q6ZVE7 Q6ZVK1 Q6ZVL6 Q6ZVX9 Q6ZW05 Q6ZWK4 Q6ZWK6 Q6ZWL3
Q6ZWQ0 Q6ZWQ7 Q6ZWR6 Q6ZWS4 Q6ZWT7 Q6ZWX0 Q6ZXA0 Q6ZXC8 Q6ZXC9 Q6ZXD2 Q6ZXV5
Q701P2 Q701R0 Q701R1 Q701R2 Q701R3 Q701R4 Q703F0 Q704E8 Q704V6 Q704Y3 Q70625
Q70626 Q708S3 Q708S4 Q708S5 Q708S6 Q708S7 Q708S8 Q709R6 Q70AG8 Q70AK3 Q70AN6
Q70BM6 Q70CQ3 Q70CT4 Q70D51 Q70EX6 Q70FG9 Q70HW3 Q70JA7 Q70JP0 Q70KP1 Q70LB2
Q70LB3 Q70LB8 Q70LC2 Q70LC9 Q70LD2 Q70RD5 Q70RM6 Q70RN9 Q70RQ2 Q70RR5 Q70RS8
Q70RT7 Q70RU1 Q70RV0 Q70RV4 Q70RW3 Q70RW7 Q70RY0 Q70S06 Q70S19 Q70S32 Q70SY1
Q70UQ0 Q70UT9 Q70VB1 Q70VZ7 Q70VZ8 Q70XG0 Q70XG4 Q70XH7 Q70XI6 Q70XJ0 Q70XU8
Q70XU9 Q70XV2 Q70XV6 Q70XV9 Q70XW1 Q70XW2 Q70XW4 Q70XW6 Q70XX7 Q70XX8 Q70XX9
Q70XY1 Q70XY6 Q70XY7 Q70XY8 Q70XY9 Q70XZ0 Q70XZ1 Q70XZ2 Q70XZ3 Q70XZ8 Q70Y03
Q70Y04 Q70Y06 Q70Y07 Q70Y08 Q70Y09 Q70Y10 Q70Y12 Q70Y13 Q70Y14 Q70Y17 Q70Y22
Q70Y23 Q70Y24 Q70Y26 Q70Y27 Q70Y28 Q70Y29 Q70Y30 Q70Z44 Q710D3 Q710E6 Q710S3

Q710U6 Q711L6 Q712G6 Q712P7 Q717C1 Q717C2 Q718F4 Q719N3 Q71B07 Q71E53 Q71E87
Q71E89 Q71E95 Q71E97 Q71E99 Q71EA6 Q71EB0 Q71ED1 Q71FH1 Q71FH2 Q71FI6 Q71FI8
Q71FI9 Q71FJ0 Q71FJ2 Q71H61 Q71KN3 Q71KN4 Q71KN5 Q71KP0 Q71KP3 Q71KP4 Q71KP6
Q71KP7 Q71KQ5 Q71KQ6 Q71KQ7 Q71KQ8 Q71KT5 Q71L51 Q71L52 Q71L58 Q71L59 Q71L70
Q71L71 Q71L78 Q71L79 Q71L82 Q71L83 Q71L86 Q71L87 Q71L94 Q71L95 Q71LA6 Q71LA7
Q71LB5 Q71LB6 Q71LB9 Q71LC0 Q71M36 Q71M42 Q71MB6 Q71MR3 Q71MR6 Q71MR7 Q71MY2
Q71MY6 Q71NS7 Q71NV7 Q71P68 Q71PB7 Q71RC9 Q71RG4 Q71RH2 Q71RJ2 Q71RS6 Q71RU1
Q71RU5 Q71S46 Q71SG7 Q71SP5 Q71SS4 Q71SV0 Q71U07 Q71VQ8 Q71W89 Q71W90 Q71W91
Q71WP3 Q71WP5 Q71WS5 Q71WU5 Q71XA1 Q71XE1 Q71XL9 Q71XT1 Q71XT2 Q71XV0 Q71XV6
Q71XV7 Q71XX6 Q71YH0 Q71YH2 Q71YP2 Q71Z73 Q71Z99 Q71ZP6 Q71ZS1 Q71ZU1 Q720B5
Q720D7 Q720N7 Q720T3 Q721K3 Q721Y1 Q722D6 Q723S2 Q724L1 Q725B6 Q725T6 Q726E5
Q727E5 Q728D2 Q728U5 Q729H8 Q72A64 Q72AR4 Q72B27 Q72BN6 Q72CB6 Q72CX3 Q72D53
Q72DI6 Q72DL1 Q72DL2 Q72DL5 Q72E00 Q72E86 Q72EF4 Q72EN4 Q72FU0 Q72G45 Q72GD5
Q72GD8 Q72GL2 Q72H22 Q72HM1 Q72IU3 Q72JC0 Q72JC2 Q72JC3 Q72JC4 Q72JT2 Q72LP2
Q72M34 Q72MD8 Q72NT0 Q72NT2 Q72PB6 Q72PQ2 Q72PS8 Q72Q29 Q72QX8 Q72R82 Q72SI0
Q72SY3 Q72TF8 Q72TG0 Q72TM5 Q72TM6 Q72TM7 Q72V37 Q72V65 Q72VU2 Q72VY8 Q72X07
Q72XC7 Q72XE3 Q72XE4 Q72XF3 Q72XF7 Q72XG0 Q72XU4 Q72XV7 Q72Y02 Q72Z61 Q72Z83
Q72ZE2 Q730U2 Q732C1 Q732C2 Q732F5 Q732H6 Q732U7 Q73372 Q733E2 Q733F5 Q733N4
Q736X4 Q737I1 Q737Z5 Q738X1 Q73B22 Q73BB8 Q73C11 Q73CW2 Q73DA2 Q73DA4 Q73DE0
Q73DW7 Q73E41 Q73G61 Q73GB4 Q73GI2 Q73H08 Q73HV8 Q73HW2 Q73HW3 Q73I75 Q73IK1
Q73IK8 Q73JG0 Q73JM1 Q73K39 Q73R54 Q73UD3 Q73UD4 Q73US5 Q73V05 Q73V08 Q73V87
Q73VC3 Q73WV6 Q73X55 Q73X56 Q73Y38 Q73YK6 Q73YL5 Q73YL6 Q73YM3 Q73YQ5 Q73YX3
Q740J9 Q740Y5 Q74126 Q741B3 Q741T7 Q742C0 Q744R9 Q746Q2 Q746S3 Q746T2 Q746T6
Q747C1 Q747K6 Q747Y0 Q748D0 Q748D3 Q748D5 Q748D9 Q748J7 Q749C5 Q749E9 Q74AA5
Q74AA8 Q74AB0 Q74AK4 Q74AM0 Q74AP9 Q74E27 Q74EE6 Q74G98 Q74GA1 Q74GA8 Q74GB2
Q74GB3 Q74GM3 Q74GY4 Q74HV9 Q74I63 Q74IR1 Q74JC2 Q74JJ0 Q74JY6 Q74K19 Q74K20
Q74KB1 Q74L41 Q74LN2 Q74LN3 Q74NQ3 Q74Q23 Q74Q30 Q74RF8 Q74RF9 Q74RU7 Q74RZ6
Q74SD5 Q74TF8 Q74U12 Q74Z23 Q74Z34 Q74Z81 Q74ZD2 Q74ZD5 Q74ZP5 Q74ZS2 Q74ZU2
Q74ZX0 Q74ZZ2 Q75007 Q75008 Q750C9 Q750M8 Q750Q7 Q750R7 Q750S6 Q750Y9 Q750Z6
Q751C1 Q751J2 Q751K5 Q751M6 Q751N2 Q751P7 Q751Q6 Q751T2 Q751U2 Q751V6 Q751W1
Q751Z0 Q752F7 Q752G2 Q752H7 Q752I1 Q752M5 Q752Q2 Q752X9 Q753C1 Q753C3 Q753C5
Q753H5 Q753I3 Q753J1 Q753L2 Q753M9 Q753T9 Q753V6 Q754B9 Q754F0 Q754F4 Q754G0
Q754G4 Q754I2 Q754J7 Q754N9 Q754Q4 Q754Q7 Q754R6 Q754Z2 Q755A9 Q755C1 Q755G4
Q755H7 Q755H8 Q755J8 Q755Q0 Q755S6 Q755Y0 Q755Y8 Q756D4 Q756F2 Q756G1 Q756I2
Q756X9 Q756Z0 Q757A5 Q757F0 Q757K7 Q757X5 Q757Z8 Q758B5 Q758B6 Q758B8 Q758C3
Q758C9 Q758G4 Q758T7 Q758W8 Q758X6 Q759B8 Q759D2 Q759E9 Q759H4 Q759J5 Q759K4
Q759K9 Q759P7 Q759R3 Q759T8 Q759W4 Q759Y0 Q759Y2 Q75A48 Q75A56 Q75A60 Q75A69
Q75A73 Q75A82 Q75AH6 Q75AI5 Q75AL4 Q75AQ7 Q75AQ8 Q75AW7 Q75B12 Q75BA5 Q75BD5
Q75BG6 Q75BG9 Q75BY0 Q75C24 Q75C38 Q75C62 Q75C82 Q75CB4 Q75CC0 Q75CC8 Q75CE1
Q75CF5 Q75CQ8 Q75CX4 Q75CY3 Q75D07 Q75D19 Q75D23 Q75D26 Q75D30 Q75D33 Q75E41
Q75E80 Q75EB9 Q75EG5 Q75EI3 Q75EN3 Q75EP3 Q75EU0 Q75EY7 Q75F43 Q75F69 Q75F70
Q75F81 Q75FA5 Q75FA7 Q75FP1 Q75FR2 Q75G34 Q75G38 Q75G39 Q75G40 Q75G84 Q75GA5
Q75GK3 Q75HB1 Q75HP9 Q75IC7 Q75IQ4 Q75IW1 Q75J39 Q75JB5 Q75JF3 Q75JN3 Q75JP4
Q75JP9 Q75JQ3 Q75JT4 Q75KA9 Q75L84 Q75LD5 Q75N23 Q75N73 Q75NA5 Q75NE6 Q75PL7

Q75Q39 Q75Q40 Q75Q41 Q75QW1 Q75R30 Q75R59 Q75R60 Q75R61 Q75R62 Q75R63 Q75T13
Q75UR0 Q75V66 Q75VQ0 Q75VQ4 Q75VR0 Q75VR1 Q75VT8 Q75W84 Q75XW3 Q75Z89 Q75ZH0
Q75ZY9 Q761V0 Q761X5 Q762B4 Q762D5 Q764M8 Q764M9 Q764N2 Q765A7 Q765H6 Q765N9
Q765P1 Q76638 Q766D5 Q767I8 Q767L9 Q769E5 Q769E8 Q769F9 Q769I5 Q76B49 Q76CA2
Q76CU2 Q76EC5 Q76EI6 Q76EJ3 Q76HL1 Q76HP2 Q76HP3 Q76I25 Q76IC3 Q76IC6 Q76II0
Q76JU8 Q76JU9 Q76JV1 Q76K24 Q76K27 Q76KB1 Q76KB2 Q76KF0 Q76KJ8 Q76KP1 Q76L40
Q76LB1 Q76LL8 Q76LN2 Q76LN6 Q76LT9 Q76LW2 Q76M72 Q76M99 Q76MJ5 Q76MS7 Q76MX5
Q76NM1 Q76NQ1 Q76NT6 Q76P23 Q76PC3 Q76R36 Q76R62 Q76RD1 Q76RD2 Q76TT7 Q76V04
Q76V06 Q76V11 Q76V12 Q76VD9 Q76WJ1 Q76ZQ3 Q76ZQ4 Q77376 Q77377 Q775J3 Q779L9
Q77CE4 Q77GW2 Q77IM6 Q77JN0 Q77MP7 Q77MR0 Q77MR4 Q77MS0 Q77MS3 Q77MS4 Q77NN4
Q77NN6 Q77NP2 Q77NU1 Q77PA8 Q77Q91 Q77SK0 Q77TI4 Q77TL4 Q77W49 Q77XR9 Q77ZJ9
Q77ZK8 Q77ZL3 Q77ZL5 Q783Y1 Q786F2 Q786F3 Q788J2 Q788J5 Q78DX7 Q78EA7 Q78HU7
Q78IK2 Q78IK4 Q78IQ7 Q78IS1 Q78KK3 Q78PG9 Q78RX3 Q78S06 Q78T54 Q78XF5 Q78YY6
Q791F6 Q791T5 Q791V5 Q792S6 Q793U7 Q794C0 Q794F9 Q794W0 Q795K2 Q795M8 Q795Q5
Q79669 Q79670 Q796K8 Q796Q1 Q797A7 Q797B3 Q797E3 Q797S1 Q799A3 Q79AV1 Q79BT1
Q79EF2 Q79IF2 Q79K00 Q79VC4 Q79VD7 Q79VE0 Q79VE8 Q79VE9 Q79VG9 Q79ZV2 Q79ZV3
Q79ZY2 Q7A040 Q7A047 Q7A086 Q7A092 Q7A0A4 Q7A0C2 Q7A0C3 Q7A0C4 Q7A0H2 Q7A0H9
Q7A0I6 Q7A0J1 Q7A0M4 Q7A0W5 Q7A0W9 Q7A0X4 Q7A0Y0 Q7A0Y4 Q7A121 Q7A160 Q7A166
Q7A181 Q7A182 Q7A183 Q7A184 Q7A1B1 Q7A1B5 Q7A1F4 Q7A1G8 Q7A1I3 Q7A1J2 Q7A1M9
Q7A1N0 Q7A1N1 Q7A1P4 Q7A1V2 Q7A1Y1 Q7A215 Q7A2K4 Q7A2M6 Q7A2Q0 Q7A2R3 Q7A2R7
Q7A2S8 Q7A2V2 Q7A305 Q7A341 Q7A348 Q7A350 Q7A351 Q7A363 Q7A3D6 Q7A3D8 Q7A3E0
Q7A3E6 Q7A3F2 Q7A3G4 Q7A3L2 Q7A3L7 Q7A3Q5 Q7A3U8 Q7A3X0 Q7A3X2 Q7A3X6 Q7A3Z2
Q7A468 Q7A4A6 Q7A4B3 Q7A4E5 Q7A4E6 Q7A4E7 Q7A4G4 Q7A4P8 Q7A4Q7 Q7A4R6 Q7A4S6
Q7A4T3 Q7A4V2 Q7A531 Q7A5C5 Q7A5H7 Q7A5I6 Q7A5M0 Q7A5M9 Q7A5N3 Q7A5N9 Q7A5P4
Q7A5Q6 Q7A5Q7 Q7A5R9 Q7A5Y4 Q7A651 Q7A664 Q7A665 Q7A698 Q7A699 Q7A6A0 Q7A6A1
Q7A6C9 Q7A6D3 Q7A6G7 Q7A6H2 Q7A6Q1 Q7A6U1 Q7A6V4 Q7A6Z3 Q7A722 Q7A723 Q7A724
Q7A725 Q7A726 Q7A727 Q7A763 Q7A771 Q7A7M5 Q7A7N0 Q7A7S0 Q7A7S2 Q7A7S3 Q7A7X8
Q7A804 Q7A807 Q7A8E0 Q7A937 Q7AA80 Q7ABM8 Q7ACM1 Q7ADY8 Q7AFA1 Q7AHS5 Q7AKA4
Q7AKE5 Q7ANN4 Q7ANN5 Q7AP31 Q7APG1 Q7B6T4 Q7BCK4 Q7BHI8 Q7BHL7 Q7BS42 Q7BSW5
Q7C1M5 Q7CB38 Q7CCJ4 Q7CEG0 Q7CEG1 Q7CEG3 Q7CFM3 Q7CFM4 Q7CFX8 Q7CG48 Q7CG49
Q7CG59 Q7CG77 Q7CGA7 Q7CGB1 Q7CGD8 Q7CGD9 Q7CGZ2 Q7CH99 Q7CHI2 Q7CIC6 Q7CJ84
Q7CJ87 Q7CJG3 Q7CJL1 Q7CJM7 Q7CKP5 Q7CKV5 Q7CKV6 Q7CLR2 Q7CLW2 Q7CND5 Q7CNS4
Q7CNU3 Q7CP67 Q7CP73 Q7CP98 Q7CPB4 Q7CPC0 Q7CPE3 Q7CPE4 Q7CPG0 Q7CPI7 Q7CPI8
Q7CPM7 Q7CPM8 Q7CQ24 Q7CQ37 Q7CQ49 Q7CQ53 Q7CQ90 Q7CQB9 Q7CQD5 Q7CQK0 Q7CQK1
Q7CQP8 Q7CQV4 Q7CQY0 Q7CQY4 Q7CR81 Q7CR84 Q7CR86 Q7CRU3 Q7CZN9 Q7D0U9 Q7D0Y8
Q7D745 Q7DB77 Q7DD66 Q7DDB6 Q7DDJ2 Q7DDQ4 Q7DDS0 Q7DGD4 Q7DJ60 Q7DLB7 Q7DM58
Q7DMA9 Q7DNC3 Q7EYH7 Q7EZD7 Q7EZR4 Q7EZW6 Q7F0Q2 Q7F239 Q7F4F8 Q7F613 Q7F8Q9
Q7FA29 Q7FAZ0 Q7FAZ2 Q7FAZ3 Q7FB56 Q7FK82 Q7FMW4 Q7FNR8 Q7FNS2 Q7FNS3 Q7FNS4
Q7FNS7 Q7FNS8 Q7FNT0 Q7FNT1 Q7FNT3 Q7FZR1 Q7G4P2 Q7G602 Q7G768 Q7G7C7 Q7GB25
Q7GCL7 Q7GEN7 Q7GEV4 Q7GF83 Q7GGQ9 Q7GI23 Q7GQD2 Q7GQE8 Q7GQG6 Q7GUC9 Q7GUD2
Q7GZ99 Q7GZA1 Q7GZA2 Q7GZA5 Q7GZA7 Q7H823 Q7H824 Q7H825 Q7H826 Q7H827 Q7H828
Q7H833 Q7H834 Q7H8A5 Q7H8E4 Q7H8E5 Q7H8E6 Q7H8E7 Q7H8I7 Q7H8I8 Q7H8J7 Q7H8J8
Q7H8J9 Q7H8K0 Q7H8K6 Q7H8K7 Q7H8K8 Q7H8K9 Q7H8L0 Q7H8L1 Q7H8L2 Q7H8L3 Q7H8L5
Q7H8L6 Q7HEI2 Q7HFF9 Q7HFG0 Q7HIT4 Q7HIT5 Q7HIT6 Q7HIT7 Q7HIT8 Q7HIT9 Q7HIU0

Q7HIU1 Q7HIU5 Q7HIU6 Q7HIU7 Q7HIU8 Q7HIU9 Q7HIV0 Q7HIV1 Q7HIV8 Q7HIV9 Q7HIW2
Q7HIW3 Q7HIW4 Q7HIW5 Q7HIW6 Q7HJJ2 Q7HKW1 Q7HKX5 Q7HKX6 Q7HKX7 Q7HKX8 Q7HKX9
Q7HKY0 Q7HKY1 Q7HKY2 Q7HLD2 Q7HML5 Q7HML6 Q7HP56 Q7I5M1 Q7IC90 Q7IKU8 Q7ISH5
Q7IT44 Q7IV45 Q7IV55 Q7IVV2 Q7IVV3 Q7IW39 Q7IW40 Q7IW41 Q7IW42 Q7IW43 Q7IY93
Q7IY99 Q7IYN8 Q7IYZ1 Q7IZ01 Q7IZ08 Q7IZ11 Q7IZ14 Q7IZ15 Q7IZ16 Q7IZ21 Q7J185
Q7J186 Q7J187 Q7J188 Q7J189 Q7J190 Q7J191 Q7J192 Q7J193 Q7J194 Q7J195 Q7J196
Q7J197 Q7J198 Q7J199 Q7J1A0 Q7J1A1 Q7J1A2 Q7J1A3 Q7J1A4 Q7J1A5 Q7J1A6 Q7J1A7
Q7J1A8 Q7J1A9 Q7J1B1 Q7J1B2 Q7J1B3 Q7J1B4 Q7J1B5 Q7J1B6 Q7J1B7 Q7J1B8 Q7J1B9
Q7J1C0 Q7J1C1 Q7J1C2 Q7J1C3 Q7J1C4 Q7J1C5 Q7J370 Q7J3B5 Q7J3C3 Q7J6G2 Q7J6G4
Q7J6I1 Q7JE02 Q7JGL3 Q7JGX4 Q7JII7 Q7JII8 Q7JK24 Q7JK25 Q7JK26 Q7JKY3 Q7JQ32
Q7JQF1 Q7JR49 Q7JRE4 Q7JUN3 Q7JVE7 Q7JW12 Q7JX57 Q7JYV2 Q7K0P4 Q7K1V5 Q7K237
Q7K4B6 Q7K4W1 Q7K4Y6 Q7K566 Q7K755 Q7KIN0 Q7KQM4 Q7KRW1 Q7KSC4 Q7KT91 Q7KVA1
Q7KVW5 Q7KVY7 Q7KWM5 Q7KWN2 Q7KWS2 Q7KWU5 Q7KYR7 Q7KZN9 Q7L0J3 Q7L0L9 Q7L0X0
Q7L1I2 Q7L1S5 Q7L1W4 Q7L211 Q7L311 Q7L4E1 Q7L4S7 Q7L5A8 Q7L5L3 Q7L5N7 Q7L8C5
Q7L985 Q7LBE3 Q7LFX5 Q7LGA3 Q7LGC8 Q7LKX0 Q7LL00 Q7LXU6 Q7LYX6 Q7M157 Q7M1T6
Q7M290 Q7M370 Q7M3I4 Q7M6G6 Q7M707 Q7M709 Q7M710 Q7M711 Q7M712 Q7M713 Q7M715
Q7M717 Q7M718 Q7M720 Q7M721 Q7M722 Q7M723 Q7M724 Q7M725 Q7M729 Q7M730 Q7M732
Q7M734 Q7M750 Q7M758 Q7M7A9 Q7M7S6 Q7M7X0 Q7M8U0 Q7M8Z9 Q7M915 Q7M922 Q7M9C0
Q7M9M1 Q7MA22 Q7MA37 Q7MA40 Q7MA43 Q7MAD5 Q7MAR3 Q7MAR9 Q7MAV0 Q7MBS2 Q7MD16
Q7MDC8 Q7MDF0 Q7MFC1 Q7MFC2 Q7MFH2 Q7MGA3 Q7MGH4 Q7MGH6 Q7MGM2 Q7MGV6 Q7MGX4
Q7MGX5 Q7MHQ2 Q7MHR9 Q7MIC8 Q7MIC9 Q7MID0 Q7MID1 Q7MID2 Q7MIR4 Q7MIV9 Q7MJ07
Q7MJM9 Q7MJN7 Q7MJN8 Q7MJT4 Q7MK21 Q7MKP8 Q7MLE7 Q7MLF2 Q7MLF6 Q7MLI9 Q7MLJ3
Q7MLU5 Q7MM60 Q7MM81 Q7MM84 Q7MM85 Q7MM86 Q7MMF0 Q7MMT6 Q7MN04 Q7MNM7 Q7MNN7
Q7MNV2 Q7MNV4 Q7MNZ4 Q7MNZ7 Q7MP82 Q7MQ07 Q7MQ29 Q7MQ30 Q7MQ31 Q7MQ42 Q7MQ87
Q7MQD2 Q7MQK5 Q7MQL4 Q7MRQ2 Q7MRX1 Q7MSR8 Q7MTF7 Q7MTR7 Q7MUD1 Q7MUH6 Q7MUS4
Q7MUZ1 Q7MV19 Q7MVL1 Q7MW41 Q7MWM6 Q7MWN8 Q7MX21 Q7MYE3 Q7MYM5 Q7MYP0 Q7MZ83
Q7MZ84 Q7MZ85 Q7MZB6 Q7MZX2 Q7MZY0 Q7MZY2 Q7N087 Q7N0A2 Q7N0B7 Q7N0B9 Q7N0K3
Q7N144 Q7N183 Q7N1G0 Q7N283 Q7N2D7 Q7N2D8 Q7N2I2 Q7N2I7 Q7N2J3 Q7N2J6 Q7N2J9
Q7N2S6 Q7N365 Q7N3E1 Q7N3E2 Q7N3J0 Q7N3J2 Q7N3L4 Q7N3L5 Q7N3N4 Q7N3Q3 Q7N3Q6
Q7N3Q9 Q7N3R0 Q7N3R1 Q7N3V2 Q7N3Y1 Q7N3Y5 Q7N3Z3 Q7N3Z4 Q7N471 Q7N475 Q7N4F7
Q7N4G1 Q7N536 Q7N537 Q7N568 Q7N5C1 Q7N5C8 Q7N6C6 Q7N6F9 Q7N6K1 Q7N6W5 Q7N6W6
Q7N6W7 Q7N6Y5 Q7N750 Q7N768 Q7N7A8 Q7N7A9 Q7N7F0 Q7N7F1 Q7N8K8 Q7N8N9 Q7N8U5
Q7N8V3 Q7N8X1 Q7N8X6 Q7N983 Q7N984 Q7N987 Q7N9D9 Q7N9W4 Q7NA72 Q7NA89 Q7NA90
Q7NAE3 Q7NAR5 Q7NB09 Q7NB93 Q7NBQ3 Q7NCE0 Q7NCE1 Q7NCH7 Q7NCH8 Q7NCR8 Q7NCR9
Q7NCS0 Q7NCS1 Q7ND43 Q7NDN8 Q7NDQ8 Q7NEY9 Q7NF27 Q7NF92 Q7NFT5 Q7NFT6 Q7NG92
Q7NGP8 Q7NH05 Q7NH81 Q7NH97 Q7NHY3 Q7NIE7 Q7NIH2 Q7NIL7 Q7NJ10 Q7NJ11 Q7NJB3
Q7NJB4 Q7NJF8 Q7NJV1 Q7NJX5 Q7NK94 Q7NLN8 Q7NMA7 Q7NMA8 Q7NMA9 Q7NMB0 Q7NML7
Q7NMQ4 Q7NMV2 Q7NN39 Q7NN41 Q7NNA0 Q7NNA7 Q7NNP3 Q7NP39 Q7NPH6 Q7NPI3 Q7NPT8
Q7NPZ6 Q7NQI3 Q7NQZ3 Q7NRM5 Q7NRS3 Q7NRU2 Q7NS41 Q7NS58 Q7NSE1 Q7NSG6 Q7NT68
Q7NTX4 Q7NU39 Q7NUG7 Q7NUI5 Q7NUJ3 Q7NWP1 Q7NWQ2 Q7NXM7 Q7NXN3 Q7NXQ1 Q7NYB4
Q7NYE7 Q7NYJ8 Q7NZD1 Q7NZH1 Q7NZH4 Q7NZU6 Q7NZV9 Q7P006 Q7P0A1 Q7P0E4 Q7P0G0
Q7P0J4 Q7PC67 Q7PC69 Q7PC70 Q7PC71 Q7PC73 Q7PC76 Q7PC80 Q7PC81 Q7PC82 Q7PC83
Q7PC84 Q7PC85 Q7PC86 Q7PC87 Q7PC88 Q7PH91 Q7PIR5 Q7PMG3 Q7PPA5 Q7PPU9 Q7PQV7
Q7PRJ0 Q7PWV1 Q7PXN1 Q7PYF4 Q7Q297 Q7Q3N5 Q7Q5D4 Q7Q5R7 Q7Q6A7 Q7Q6R1 Q7Q732

Q7QC49 Q7QCC7 Q7QDY3 Q7QHP6 Q7QIQ6 Q7REK3 Q7RRM4 Q7RTM1 Q7RTP0 Q7RTR8 Q7RTS5
Q7RTS6 Q7RTT9 Q7RTX0 Q7RTX1 Q7RTX7 Q7RTX9 Q7RTY0 Q7RTY1 Q7RTY8 Q7RVX9 Q7RWM9
Q7RX59 Q7RX75 Q7RXL1 Q7RXP5 Q7RY44 Q7RY98 Q7RYE5 Q7RYI0 Q7RYM7 Q7RYY1 Q7RZA2
Q7RZK9 Q7RZR2 Q7S0A1 Q7S130 Q7S158 Q7S2H8 Q7S3M5 Q7S3S2 Q7S455 Q7S4D7 Q7S4V5
Q7S4Z3 Q7S4Z6 Q7S5E7 Q7S5M7 Q7S5W9 Q7S693 Q7S7C5 Q7S7U5 Q7S7Z6 Q7S8J1 Q7S8M1
Q7S8S5 Q7S9I1 Q7SA35 Q7SAJ6 Q7SAM0 Q7SAP1 Q7SB51 Q7SB92 Q7SBA0 Q7SBB6 Q7SBC5
Q7SBE0 Q7SC45 Q7SCL1 Q7SCY6 Q7SCY7 Q7SDN3 Q7SDU5 Q7SEE1 Q7SFD8 Q7SFL7 Q7SFN5
Q7SFQ9 Q7SFY2 Q7SG38 Q7SGB6 Q7SGL6 Q7SGT7 Q7SGY6 Q7SH60 Q7SHI2 Q7SHQ1 Q7SHU8
Q7SI11 Q7SXB3 Q7SXB7 Q7SXE4 Q7SXF1 Q7SXP0 Q7SXS2 Q7SXW0 Q7SXW4 Q7SXZ1 Q7SY06
Q7SY09 Q7SY10 Q7SY29 Q7SY31 Q7SY73 Q7SYC7 Q7SYC9 Q7SYH5 Q7SYI5 Q7SYR6 Q7SZ56
Q7SZC5 Q7SZN2 Q7SZP9 Q7SZQ0 Q7SZQ4 Q7SZQ7 Q7SZU9 Q7T047 Q7T070 Q7T0R0 Q7T0U6
Q7T0V6 Q7T0Z7 Q7T163 Q7T199 Q7T1N4 Q7T207 Q7T277 Q7T278 Q7T292 Q7T2B0 Q7T2D1
Q7T2D4 Q7T2H2 Q7T2L7 Q7T2P6 Q7T2Z5 Q7T310 Q7T312 Q7T316 Q7T326 Q7T352 Q7T381
Q7T384 Q7T392 Q7T3B2 Q7T3Q2 Q7T3Q7 Q7T3R4 Q7T3S3 Q7T3S5 Q7T3T4 Q7T6D2 Q7T6S7
Q7T6S9 Q7T6T0 Q7T6T1 Q7T6T3 Q7T6X2 Q7T6Y2 Q7T9D9 Q7TD19 Q7TDX0 Q7TFA1 Q7TM99
Q7TMA4 Q7TMB0 Q7TMB7 Q7TMD7 Q7TME0 Q7TMJ8 Q7TML3 Q7TMM8 Q7TMP6 Q7TMS5 Q7TMV1
Q7TN37 Q7TN38 Q7TN39 Q7TN40 Q7TN41 Q7TN44 Q7TN45 Q7TN49 Q7TN50 Q7TN51 Q7TN58
Q7TN60 Q7TN73 Q7TN84 Q7TN88 Q7TNC8 Q7TNI2 Q7TNJ0 Q7TNJ2 Q7TNJ4 Q7TNK0 Q7TNK7
Q7TNN2 Q7TNN9 Q7TNS2 Q7TNS7 Q7TNT2 Q7TNU7 Q7TNV1 Q7TP48 Q7TP65 Q7TPB0 Q7TPB4
Q7TPD3 Q7TPF1 Q7TPJ0 Q7TPN3 Q7TQ16 Q7TQ65 Q7TQ69 Q7TQ79 Q7TQ95 Q7TQA1 Q7TQA3
Q7TQA4 Q7TQA5 Q7TQA6 Q7TQA7 Q7TQA8 Q7TQA9 Q7TQB0 Q7TQB8 Q7TQB9 Q7TQE6 Q7TQH7
Q7TQI0 Q7TQJ1 Q7TQM3 Q7TQM4 Q7TQN7 Q7TQN8 Q7TQN9 Q7TQP0 Q7TQP2 Q7TQP3 Q7TQP4
Q7TQQ0 Q7TR96 Q7TRF3 Q7TS48 Q7TS55 Q7TSA0 Q7TSA3 Q7TSF0 Q7TSF1 Q7TSF2 Q7TSH7
Q7TSH8 Q7TSK2 Q7TSK3 Q7TSN2 Q7TSN4 Q7TSN5 Q7TSN6 Q7TSN7 Q7TST0 Q7TST5 Q7TSU7
Q7TSW6 Q7TSX5 Q7TSY2 Q7TT15 Q7TT36 Q7TT41 Q7TT47 Q7TTH6 Q7TTI2 Q7TTJ7 Q7TUF3
Q7TUF5 Q7TUF8 Q7TUM3 Q7TUP7 Q7TUQ1 Q7TVK7 Q7TVL0 Q7TVN3 Q7TVN4 Q7TVN5 Q7TX22
Q7TX91 Q7TY95 Q7TYX4 Q7TYY6 Q7TZ67 Q7TZN3 Q7U021 Q7U042 Q7U095 Q7U0X3 Q7U1Z7
Q7U2X8 Q7U351 Q7U3B6 Q7U402 Q7U413 Q7U482 Q7U4E4 Q7U4E5 Q7U4T0 Q7U4T3 Q7U4U6
Q7U4U7 Q7U4Z3 Q7U522 Q7U538 Q7U550 Q7U564 Q7U565 Q7U566 Q7U571 Q7U5H9 Q7U5S4
Q7U620 Q7U669 Q7U6N5 Q7U6P8 Q7U6W9 Q7U6X4 Q7U772 Q7U7A6 Q7U8A1 Q7U8E3 Q7U8N7
Q7U8W7 Q7U8W9 Q7U8X0 Q7U8Y8 Q7U9C4 Q7U9I7 Q7U9I8 Q7U9J0 Q7U9L1 Q7U9P6 Q7U9P9
Q7U9Q0 Q7U9Q1 Q7U9Q2 Q7UA04 Q7UA05 Q7UAH8 Q7UAJ5 Q7UBE9 Q7UBJ3 Q7UBK4 Q7UBP8
Q7UC61 Q7UC63 Q7UC94 Q7UCD5 Q7UCG7 Q7UCH6 Q7UCL3 Q7UDF9 Q7UDG0 Q7UDT6 Q7UF32
Q7UFB9 Q7UFZ2 Q7UGJ8 Q7UH06 Q7UH36 Q7UHW8 Q7UJF4 Q7ULB5 Q7ULI3 Q7UM18 Q7UMF7
Q7UNE3 Q7UNQ8 Q7UQD2 Q7UQJ2 Q7URM7 Q7UTY3 Q7UUZ7 Q7UWD1 Q7UWS1 Q7UZF8 Q7UZM6
Q7UZM7 Q7UZX6 Q7V028 Q7V032 Q7V033 Q7V034 Q7V035 Q7V0D9 Q7V0G7 Q7V0R8 Q7V0U6
Q7V129 Q7V1C3 Q7V1U0 Q7V1W2 Q7V2B1 Q7V2E0 Q7V2K8 Q7V2L4 Q7V2L5 Q7V2L6 Q7V2M7
Q7V2N8 Q7V2X6 Q7V2Y5 Q7V2Z7 Q7V2Z8 Q7V2Z9 Q7V300 Q7V323 Q7V339 Q7V340 Q7V341
Q7V365 Q7V3C1 Q7V3C8 Q7V3L3 Q7V3S7 Q7V4D7 Q7V4E4 Q7V4P9 Q7V4Q0 Q7V4Q1 Q7V4Q2
Q7V4T1 Q7V4T2 Q7V4U9 Q7V4V1 Q7V4V2 Q7V510 Q7V511 Q7V5A6 Q7V5B9 Q7V5H0 Q7V5R7
Q7V5S2 Q7V5S3 Q7V5S4 Q7V5S5 Q7V5Y3 Q7V610 Q7V626 Q7V638 Q7V652 Q7V653 Q7V654
Q7V658 Q7V6I0 Q7V6I1 Q7V6M3 Q7V6U4 Q7V7J0 Q7V7J6 Q7V7R5 Q7V806 Q7V872 Q7V897
Q7V8K1 Q7V8Z1 Q7V937 Q7V938 Q7V9F5 Q7V9N5 Q7V9N6 Q7VA54 Q7VA58 Q7VA59 Q7VA60
Q7VA61 Q7VAG9 Q7VAP9 Q7VAT8 Q7VB00 Q7VB45 Q7VB49 Q7VBC2 Q7VBC8 Q7VBF3 Q7VC24

Q7VC50 Q7VC57 Q7VCA3 Q7VCF7 Q7VCH2 Q7VCH3 Q7VCK5 Q7VD03 Q7VD17 Q7VDC0 Q7VDC1
Q7VDC2 Q7VDD5 Q7VDE7 Q7VDK8 Q7VDK9 Q7VDL9 Q7VDM3 Q7VDN7 Q7VDN8 Q7VDN9 Q7VDP0
Q7VDP3 Q7VDR4 Q7VDR6 Q7VDT3 Q7VDT4 Q7VDT5 Q7VE30 Q7VE41 Q7VED7 Q7VEN1 Q7VEN6
Q7VEQ2 Q7VEQ4 Q7VEV7 Q7VEW5 Q7VEZ5 Q7VFJ7 Q7VFN0 Q7VFN1 Q7VFP7 Q7VFS9 Q7VFT2
Q7VFT5 Q7VG27 Q7VG35 Q7VGZ9 Q7VH06 Q7VHP1 Q7VIA0 Q7VIL1 Q7VIL7 Q7VIU4 Q7VJ25
Q7VJY0 Q7VKA0 Q7VKU4 Q7VKX4 Q7VKY2 Q7VKY3 Q7VKY8 Q7VKZ1 Q7VL38 Q7VL52 Q7VLT4
Q7VLY3 Q7VM56 Q7VMB5 Q7VMB8 Q7VMF9 Q7VMJ8 Q7VMX6 Q7VMZ4 Q7VN08 Q7VN39 Q7VN59
Q7VN62 Q7VN63 Q7VN64 Q7VNT1 Q7VNT3 Q7VNT6 Q7VNU1 Q7VNU4 Q7VNU5 Q7VNU6 Q7VNU7
Q7VNU8 Q7VNX2 Q7VP55 Q7VP57 Q7VPB5 Q7VPK5 Q7VPM2 Q7VPN2 Q7VPP4 Q7VPP5 Q7VPP6
Q7VQ46 Q7VQI6 Q7VQJ0 Q7VQK9 Q7VQQ8 Q7VQU0 Q7VQW0 Q7VQW1 Q7VQW2 Q7VQX5 Q7VQZ2
Q7VR12 Q7VR44 Q7VRA5 Q7VRF2 Q7VRH6 Q7VRM7 Q7VRV2 Q7VRV8 Q7VRW1 Q7VRW4 Q7VSE8
Q7VSX4 Q7VSX6 Q7VSX8 Q7VSY1 Q7VSY2 Q7VT22 Q7VTK0 Q7VU48 Q7VU50 Q7VUN5 Q7VUP7
Q7VUQ0 Q7VUQ6 Q7VV78 Q7VVZ9 Q7VW80 Q7VWD8 Q7VWH2 Q7VXA0 Q7VXL6 Q7VXN3 Q7VYU0
Q7VZP5 Q7VZP8 Q7W039 Q7W079 Q7W1Y0 Q7W2K1 Q7W2T8 Q7W2U0 Q7W2U2 Q7W2U4 Q7W2U5
Q7W2W9 Q7W2X4 Q7W2X5 Q7W316 Q7W3A4 Q7W3A6 Q7W3V2 Q7W496 Q7W4B1 Q7W4D4 Q7W536
Q7W5B4 Q7W5B7 Q7W5H7 Q7W5P0 Q7W5V1 Q7W669 Q7W740 Q7W7H9 Q7W7R8 Q7W866 Q7W8J2
Q7W917 Q7W9D1 Q7W9N7 Q7WA16 Q7WAK9 Q7WCL8 Q7WCU6 Q7WCU9 Q7WD07 Q7WD13 Q7WD76
Q7WDT6 Q7WDT8 Q7WDU0 Q7WDU2 Q7WDU3 Q7WDW9 Q7WDX5 Q7WE16 Q7WEM3 Q7WEM5 Q7WF82
Q7WFQ4 Q7WFR9 Q7WFU2 Q7WGK2 Q7WH20 Q7WHD3 Q7WI35 Q7WIP4 Q7WJ43 Q7WJR0 Q7WKG2
Q7WKW8 Q7WL55 Q7WLL4 Q7WM55 Q7WMN7 Q7WQV8 Q7WTB2 Q7WTR3 Q7WVD3 Q7WWR6 Q7WXB8
Q7WXK8 Q7WY60 Q7WY64 Q7WY71 Q7WY74 Q7WY78 Q7WYU3 Q7X660 Q7X6L2 Q7X6P3 Q7X7C4
Q7X7K8 Q7X7S8 Q7X7V2 Q7X7X4 Q7X843 Q7X8B5 Q7X8C5 Q7X911 Q7X9A6 Q7XA06 Q7XA61
Q7XA64 Q7XA72 Q7XA86 Q7XA87 Q7XAB8 Q7XB13 Q7XB14 Q7XBS0 Q7XCJ7 Q7XDI3 Q7XDZ7
Q7XEK4 Q7XEM4 Q7XFE1 Q7XGU4 Q7XHV0 Q7XHW5 Q7XI45 Q7XIF5 Q7XIT1 Q7XIV8 Q7XJ98
Q7XJL2 Q7XJR2 Q7XK12 Q7XKC8 Q7XKF4 Q7XKI5 Q7XLC6 Q7XLD4 Q7XLG3 Q7XLR1 Q7XLY8
Q7XMK3 Q7XN54 Q7XP59 Q7XPF7 Q7XPF8 Q7XPL3 Q7XPU9 Q7XPY2 Q7XPZ4 Q7XQ12 Q7XQ88
Q7XQQ1 Q7XR51 Q7XRH8 Q7XRV1 Q7XRV2 Q7XSQ9 Q7XSR9 Q7XSZ4 Q7XT08 Q7XTB2 Q7XTF8
Q7XTL7 Q7XTQ5 Q7XU31 Q7XU38 Q7XU74 Q7XUA6 Q7XUH5 Q7XUJ2 Q7XUK4 Q7XUN6 Q7XUT9
Q7XUU0 Q7XUV2 Q7XUV7 Q7XUW4 Q7XV05 Q7XVB3 Q7XVN7 Q7XX84 Q7XXN8 Q7XYM4 Q7XZF5
Q7XZP5 Q7Y030 Q7Y1E6 Q7Y1V5 Q7Y201 Q7Y208 Q7Y228 Q7Y2B9 Q7Y2C0 Q7Y2C1 Q7Y4H2
Q7Y5X9 Q7Y778 Q7Y779 Q7Y7B2 Q7Y7G0 Q7Y8D2 Q7Y8D5 Q7Y8E7 Q7Y8F6 Q7Y8G0 Q7Y8I2
Q7Y8I6 Q7Y8J5 Q7Y8J9 Q7Y8K8 Q7Y8L0 Q7Y8L4 Q7Y8L5 Q7Y8L6 Q7Y8L9 Q7Y8M0 Q7YA49
Q7YBC1 Q7YBY4 Q7YBY6 Q7YC63 Q7YC71 Q7YC73 Q7YC74 Q7YC75 Q7YC76 Q7YCA5 Q7YD12
Q7YD13 Q7YD16 Q7YD72 Q7YD73 Q7YD75 Q7YEL1 Q7YEV2 Q7YFV7 Q7YGD4 Q7YJS6 Q7YJS9
Q7YJT1 Q7YJT2 Q7YJT3 Q7YJT4 Q7YJT6 Q7YJU7 Q7YJU8 Q7YJU9 Q7YJV0 Q7YJV1 Q7YJV6
Q7YJV7 Q7YJV8 Q7YJV9 Q7YJW0 Q7YJW1 Q7YJW2 Q7YJW3 Q7YJW7 Q7YJX2 Q7YJX3 Q7YJX5
Q7YJX6 Q7YJX7 Q7YJY2 Q7YJY3 Q7YJY5 Q7YJY8 Q7YMA2 Q7YN52 Q7YN54 Q7YNG3 Q7YNH0
Q7YNH4 Q7YNI0 Q7YNI1 Q7YNT5 Q7YNT6 Q7YNV4 Q7YNW2 Q7YQE1 Q7YQE5 Q7YQI4 Q7YQK3
Q7YQK4 Q7YQK9 Q7YQL0 Q7YQL9 Q7YR42 Q7YR43 Q7YR73 Q7YRB2 Q7YRB7 Q7YRC0 Q7YRF4
Q7YRH6 Q7YRJ8 Q7YRJ9 Q7YRK0 Q7YRK1 Q7YRK2 Q7YRK3 Q7YRK5 Q7YRK7 Q7YRL5 Q7YRN2
Q7YRP1 Q7YRP2 Q7YRP3 Q7YRU6 Q7YRU7 Q7YRZ9 Q7YS44 Q7YS54 Q7YS88 Q7YS92 Q7YSJ4
Q7YTA6 Q7YTM7 Q7YTM8 Q7YU24 Q7YW31 Q7YWV6 Q7YWX7 Q7YXD4 Q7YZW5 Q7Z020 Q7Z0Q2
Q7Z0V9 Q7Z118 Q7Z139 Q7Z1B8 Q7Z1V1 Q7Z1Z1 Q7Z2C4 Q7Z2D5 Q7Z2H8 Q7Z2K6 Q7Z2Q7
Q7Z2W7 Q7Z388 Q7Z3B0 Q7Z3C6 Q7Z3D4 Q7Z3F1 Q7Z3J2 Q7Z3Q1 Q7Z3S7 Q7Z3T1 Q7Z402

Q7Z403 Q7Z404 Q7Z407 Q7Z408 Q7Z410 Q7Z412 Q7Z418 Q7Z419 Q7Z429 Q7Z434 Q7Z442
Q7Z443 Q7Z449 Q7Z4F1 Q7Z4J2 Q7Z4L0 Q7Z4N2 Q7Z4T8 Q7Z5B4 Q7Z5H4 Q7Z5H5 Q7Z5J8
Q7Z5M5 Q7Z5N4 Q7Z5S9 Q7Z601 Q7Z602 Q7Z692 Q7Z695 Q7Z6A9 Q7Z6J6 Q7Z6L0 Q7Z6M3
Q7Z6W1 Q7Z769 Q7Z7B1 Q7Z7D3 Q7Z7H5 Q7Z7J7 Q7Z7M0 Q7Z7M1 Q7Z7M8 Q7Z7M9 Q7Z7N9
Q7Z859 Q7Z877 Q7Z888 Q7Z8R5 Q7Z8U2 Q7Z9I0 Q7Z9I5 Q7ZAK5 Q7ZTL7 Q7ZTM6 Q7ZTV5
Q7ZTY7 Q7ZU13 Q7ZU80 Q7ZUA6 Q7ZUB3 Q7ZUC7 Q7ZUU1 Q7ZV14 Q7ZV15 Q7ZV50 Q7ZV60
Q7ZVH1 Q7ZVN4 Q7ZVP8 Q7ZVW1 Q7ZVX5 Q7ZW02 Q7ZW11 Q7ZW24 Q7ZW46 Q7ZW77 Q7ZWF4
Q7ZWG6 Q7ZWG9 Q7ZWJ3 Q7ZWK8 Q7ZWN0 Q7ZWN9 Q7ZWS5 Q7ZWU7 Q7ZWW7 Q7ZX53 Q7ZX75
Q7ZXF5 Q7ZXH1 Q7ZXJ7 Q7ZXQ3 Q7ZXQ8 Q7ZXS5 Q7ZXS7 Q7ZXS8 Q7ZXX1 Q7ZY07 Q7ZY36
Q7ZY85 Q7ZY86 Q7ZYA0 Q7ZYD5 Q7ZYQ3 Q7ZZ25 Q7ZZL8 Q800H9 Q800K9 Q800L1 Q801D8
Q801G2 Q801M1 Q802F2 Q802T2 Q802V6 Q803C7 Q803C9 Q803F5 Q803G5 Q803I2 Q803I8
Q803X0 Q803Z2 Q804E1 Q804Q2 Q804X9 Q805E5 Q805R1 Q806Y7 Q809U7 Q809V0 Q809V2
Q809V4 Q80A22 Q80A26 Q80A28 Q80A30 Q80DN6 Q80DU8 Q80HT7 Q80HV6 Q80HX1 Q80IU3
Q80IU6 Q80KJ7 Q80KJ8 Q80RC7 Q80S27 Q80SS6 Q80SS9 Q80SU6 Q80SV1 Q80SX5 Q80SZ5
Q80T02 Q80T14 Q80T22 Q80T32 Q80T41 Q80T62 Q80T79 Q80T91 Q80TA1 Q80TE3 Q80TF3
Q80TG9 Q80TI0 Q80TL1 Q80TN4 Q80TN5 Q80TR1 Q80TS3 Q80TS8 Q80U23 Q80U56 Q80U63
Q80UA9 Q80UC6 Q80UC8 Q80UF9 Q80UG1 Q80UG2 Q80UJ1 Q80UK8 Q80UL9 Q80UM7 Q80UP8
Q80UU9 Q80UW0 Q80UX8 Q80V03 Q80V26 Q80V53 Q80VA0 Q80VJ8 Q80VM9 Q80VP8 Q80VP9
Q80W04 Q80W35 Q80W54 Q80W57 Q80W66 Q80W68 Q80W89 Q80W94 Q80W99 Q80WD0 Q80WE5
Q80WF4 Q80WG5 Q80WJ6 Q80WJ7 Q80WK2 Q80WK7 Q80WL1 Q80WM9 Q80WN7 Q80WN8 Q80WN9
Q80WQ6 Q80WR1 Q80WT0 Q80WT4 Q80WV3 Q80WY6 Q80X45 Q80X59 Q80X71 Q80X72 Q80X80
Q80X89 Q80XA0 Q80XD9 Q80XF7 Q80XG9 Q80XH2 Q80XH4 Q80XL7 Q80XM3 Q80XM9 Q80XQ8
Q80XR2 Q80XU8 Q80XZ4 Q80YA8 Q80YD3 Q80YE4 Q80YF6 Q80YN4 Q80Z10 Q80Z39 Q80Z70
Q80Z96 Q80ZA4 Q80ZA5 Q80ZA7 Q80ZC8 Q80ZD3 Q80ZD5 Q80ZD7 Q80ZD8 Q80ZD9 Q80ZE3
Q80ZE4 Q80ZE5 Q80ZF7 Q80ZF8 Q80ZI2 Q80ZM8 Q80ZU4 Q80ZU9 Q810B7 Q810B8 Q810B9
Q810C0 Q810C1 Q810F0 Q810F1 Q810F5 Q810K3 Q810K9 Q810L4 Q810M5 Q810S1 Q810S2
Q810U2 Q810U3 Q810U4 Q810W6 Q810Y4 Q81162 Q811A2 Q811B1 Q811P0 Q811Q0 Q811Q2
Q811Q4 Q811U4 Q81258 Q812E9 Q812F8 Q812G0 Q812J5 Q812P5 Q812Y8 Q812Z6 Q813A6
Q813Y0 Q81487 Q81495 Q814F4 Q814J0 Q814J2 Q814J3 Q814U5 Q814V7 Q814V8 Q814W6
Q814X0 Q814X3 Q815A2 Q815I7 Q815R5 Q815R6 Q815V9 Q816V0 Q816Z0 Q81754 Q817A9
Q81870 Q818K7 Q819N0 Q819N1 Q819P9 Q819Q1 Q819Q5 Q81A03 Q81A17 Q81A38 Q81AB0
Q81B38 Q81BL7 Q81CK9 Q81CP1 Q81CW7 Q81DD1 Q81DG8 Q81DZ9 Q81EJ6 Q81FU5 Q81G27
Q81G28 Q81GK4 Q81GP1 Q81HM5 Q81HP9 Q81HQ0 Q81HQ1 Q81HT3 Q81HV4 Q81I00 Q81I77
Q81IX5 Q81JH1 Q81JL2 Q81JL4 Q81JL5 Q81JX6 Q81JZ0 Q81JZ1 Q81K00 Q81K04 Q81K07
Q81K10 Q81K33 Q81KM7 Q81KR6 Q81KT8 Q81KZ4 Q81L64 Q81LY8 Q81MT7 Q81MT8 Q81MZ9
Q81N43 Q81PQ8 Q81PZ9 Q81QD6 Q81QF8 Q81QL7 Q81R17 Q81SZ9 Q81T84 Q81T85 Q81TR2
Q81TV2 Q81UK6 Q81UU9 Q81UW5 Q81UW7 Q81V01 Q81V21 Q81V75 Q81VC9 Q81VJ7 Q81WC4
Q81WC8 Q81WD2 Q81WE6 Q81WH6 Q81WP2 Q81WQ4 Q81WT3 Q81X30 Q81X52 Q81XB1 Q81XB2
Q81XB6 Q81XC0 Q81XC1 Q81XH4 Q81Y17 Q81Y27 Q81Y92 Q81YU3 Q82028 Q82029 Q82030
Q82033 Q82034 Q82035 Q82055 Q820A4 Q820B4 Q820B8 Q820C8 Q820C9 Q821Q3 Q821S0
Q822F2 Q822F4 Q822U2 Q823E7 Q823P2 Q823P3 Q823P4 Q823P5 Q824E2 Q824Q5 Q82509
Q82559 Q82634 Q82683 Q82706 Q827A4 Q827U4 Q82857 Q828H7 Q828X4 Q829U3 Q82A79
Q82AC7 Q82AD9 Q82AK5 Q82AK8 Q82AK9 Q82AL0 Q82DT1 Q82DX1 Q82DX4 Q82E60 Q82EB2
Q82EJ8 Q82EL6 Q82EU4 Q82IL9 Q82IS8 Q82IY3 Q82J80 Q82K93 Q82LR8 Q82NI1 Q82NI3

Q82NI4 Q82PI3 Q82PI5 Q82SA8 Q82SE7 Q82TF3 Q82TP2 Q82TU3 Q82TV0 Q82TV3 Q82TV6
Q82U09 Q82U73 Q82UR0 Q82VB9 Q82VG8 Q82VK1 Q82VQ6 Q82VS4 Q82VS6 Q82W83 Q82WC5
Q82WF6 Q82WM9 Q82WN0 Q82X64 Q82XN3 Q82XQ2 Q82XQ4 Q82XW5 Q82Y34 Q82Y49 Q82YV1
Q82Z75 Q82Z99 Q82ZB4 Q82ZN5 Q82ZT8 Q83017 Q83034 Q83046 Q830V1 Q830Z9 Q831A1
Q831R1 Q83207 Q832R4 Q83356 Q83470 Q834C0 Q834D2 Q834D8 Q834K7 Q834N5 Q835X3
Q836A6 Q836V6 Q837B5 Q837C1 Q837G9 Q83883 Q83887 Q838R5 Q83953 Q83AF9 Q83AG1
Q83AR1 Q83AR2 Q83B45 Q83BB2 Q83BD3 Q83BG0 Q83BQ5 Q83BR2 Q83BR5 Q83BR8 Q83BT9
Q83C89 Q83CV6 Q83D36 Q83D55 Q83D84 Q83EC8 Q83EK8 Q83F15 Q83F26 Q83F34 Q83F93
Q83FV7 Q83G22 Q83G86 Q83G87 Q83GF8 Q83GH4 Q83GN4 Q83H47 Q83HJ8 Q83HQ2 Q83HQ7
Q83HY4 Q83HY5 Q83I41 Q83IG0 Q83IH3 Q83IH7 Q83II9 Q83IP5 Q83IP7 Q83IQ8 Q83IS4
Q83IU1 Q83IX6 Q83IZ9 Q83J01 Q83JE2 Q83JJ9 Q83JV7 Q83JY0 Q83K22 Q83K56 Q83KB1
Q83KB5 Q83KB6 Q83KF5 Q83KI4 Q83KI8 Q83KM2 Q83KM9 Q83KS5 Q83KY5 Q83KY7 Q83L10
Q83L69 Q83L78 Q83LB2 Q83LC9 Q83LI8 Q83LK5 Q83LP0 Q83LR7 Q83LX0 Q83MH1 Q83MI4
Q83MJ2 Q83MN6 Q83MP1 Q83MS8 Q83N53 Q83N58 Q83NC0 Q83NJ2 Q83NK0 Q83P38 Q83P81
Q83P86 Q83P87 Q83P94 Q83PD6 Q83PD8 Q83PJ2 Q83PJ8 Q83PL6 Q83PS4 Q83PU6 Q83PU7
Q83PV6 Q83PW3 Q83PY0 Q83Q03 Q83Q28 Q83Q34 Q83Q35 Q83QB9 Q83QD0 Q83QK7 Q83QN3
Q83QN8 Q83QN9 Q83QP2 Q83QS9 Q83QT2 Q83QV3 Q83QZ0 Q83R13 Q83R47 Q83RD1 Q83RD8
Q83RM8 Q83RQ1 Q83RQ5 Q83RR1 Q83RR7 Q83RW9 Q83RZ5 Q83S00 Q83S20 Q83S25 Q83S26
Q83S83 Q83S84 Q83SA2 Q83SB6 Q83SF7 Q83SQ3 Q83SR3 Q83ST5 Q83VR1 Q83W27 Q83XX3
Q83Z42 Q840P7 Q84168 Q84568 Q84706 Q84710 Q848I3 Q848J1 Q849R0 Q849R1 Q84GK0
Q84I33 Q84JA6 Q84JB8 Q84JH9 Q84JN1 Q84JQ4 Q84JW1 Q84K00 Q84K11 Q84K47 Q84K71
Q84KI1 Q84KJ6 Q84KJ7 Q84L08 Q84L09 Q84M24 Q84M43 Q84M47 Q84M97 Q84MA5 Q84MA8
Q84MA9 Q84MB5 Q84MS3 Q84MS4 Q84N34 Q84NP7 Q84NQ7 Q84P95 Q84P97 Q84PB7 Q84Q83
Q84QC0 Q84QU8 Q84R16 Q84R21 Q84R49 Q84RL6 Q84RL7 Q84S07 Q84S11 Q84S18 Q84T21
Q84T91 Q84TH5 Q84TI7 Q84TJ0 Q84TU6 Q84TV3 Q84TW8 Q84UB8 Q84UB9 Q84UC0 Q84UC7
Q84UP7 Q84UT5 Q84V22 Q84VK7 Q84VQ4 Q84VT2 Q84W04 Q84W06 Q84W40 Q84W41 Q84W54
Q84WB7 Q84WE9 Q84WF5 Q84WF6 Q84WG0 Q84WG1 Q84WG3 Q84WN3 Q84WP5 Q84WU0 Q84WU8
Q84WV9 Q84WW5 Q84X12 Q84XI3 Q84XU2 Q84YJ9 Q84Z01 Q84ZM7 Q84ZN6 Q85041 Q850M0
Q851F9 Q851G4 Q851L8 Q851M9 Q851X7 Q85213 Q85230 Q85281 Q852F6 Q852K0 Q852M4
Q85320 Q85444 Q85646 Q859V1 Q859W0 Q85A24 Q85A51 Q85A78 Q85A91 Q85AC9 Q85AE6
Q85AG5 Q85AH1 Q85AI0 Q85AI7 Q85AJ6 Q85AJ9 Q85AL8 Q85AP4 Q85AP7 Q85AQ3 Q85AR2
Q85AU0 Q85AV3 Q85AV8 Q85AY5 Q85B27 Q85B66 Q85BB6 Q85BG0 Q85BG2 Q85BG5 Q85BH5
Q85BI5 Q85BK0 Q85BP9 Q85BV1 Q85C06 Q85C23 Q85C42 Q85C43 Q85CD4 Q85CI4 Q85CN0
Q85CN1 Q85CT7 Q85DA0 Q85DA4 Q85DA7 Q85DD9 Q85DE4 Q85DF2 Q85DF5 Q85DF8 Q85FG7
Q85FH0 Q85FH2 Q85FH3 Q85FH5 Q85FH6 Q85FH9 Q85FJ2 Q85FJ3 Q85FJ4 Q85FJ5 Q85FJ6
Q85FJ7 Q85FK2 Q85FK3 Q85FK4 Q85FK5 Q85FK6 Q85FK7 Q85FK8 Q85FK9 Q85FL0 Q85FL1
Q85FL2 Q85FL5 Q85FM0 Q85FM2 Q85FM3 Q85FM4 Q85FM5 Q85FM6 Q85FN1 Q85FN2 Q85FN3
Q85FN5 Q85FN6 Q85FN7 Q85FP1 Q85FP4 Q85FP7 Q85FP8 Q85FP9 Q85FQ1 Q85FQ2 Q85FQ3
Q85FQ4 Q85FQ5 Q85FR0 Q85FR1 Q85FR2 Q85FS8 Q85FU6 Q85FX0 Q85FX7 Q85FX8 Q85FY1
Q85FY2 Q85FY3 Q85FY6 Q85FY7 Q85FZ2 Q85FZ3 Q85FZ4 Q85G15 Q85G16 Q85G27 Q85G30
Q85G59 Q85G61 Q85G86 Q85IN3 Q85IN4 Q85IN5 Q85IN6 Q85IN7 Q85IN8 Q85IN9 Q85PK7
Q85PM6 Q85PN0 Q85PN4 Q85PN6 Q85PN7 Q85PN8 Q85PP0 Q85PP1 Q85PP8 Q85PQ0 Q85PQ1
Q85PQ2 Q85PQ3 Q85PQ7 Q85PQ8 Q85PR2 Q85PS1 Q85Q58 Q85Q96 Q85Q98 Q85Q99 Q85QA3
Q85QA4 Q85RV3 Q85T01 Q85TA0 Q85UK4 Q85V00 Q85WS3 Q85WS7 Q85WU2 Q85WW5 Q85WW6

Q85WX0 Q85WX3 Q85X03 Q85X07 Q85X08 Q85X09 Q85X12 Q85X26 Q85X29 Q85X30 Q85X31
Q85X32 Q85X34 Q85X47 Q85X66 Q85X70 Q85X72 Q86198 Q861W5 Q86283 Q862A8 Q862A9
Q862Z7 Q862Z8 Q862Z9 Q863C4 Q863F8 Q863F9 Q863G0 Q863G1 Q863G2 Q863G3 Q863G4
Q863G5 Q863G6 Q863G7 Q863G8 Q863G9 Q863H2 Q863H3 Q863H8 Q863I4 Q863T6 Q863Y7
Q863Y8 Q863Y9 Q863Z5 Q864F4 Q864F5 Q864F6 Q864F7 Q864F8 Q864F9 Q864G0 Q864G1
Q864G2 Q864G3 Q864G4 Q864G6 Q864G7 Q864G9 Q864H0 Q864H1 Q864H2 Q864H3 Q864H4
Q864H5 Q864H6 Q864H7 Q864H8 Q864H9 Q864I0 Q864I1 Q864I2 Q864I3 Q864I4 Q864I5
Q864I6 Q864I7 Q864I8 Q864I9 Q864J0 Q864J1 Q864J2 Q864J3 Q864J4 Q864J5 Q864J6
Q864J7 Q864J8 Q864J9 Q864K0 Q864K1 Q864K2 Q864K3 Q864K4 Q864K5 Q864K6 Q864K7
Q864K8 Q864K9 Q864L0 Q864L3 Q864R5 Q864R9 Q864U6 Q864U8 Q864V4 Q864V5 Q864Z3
Q86516 Q86539 Q865E5 Q865E8 Q865E9 Q865K8 Q865K9 Q865S0 Q86607 Q86695 Q866A2
Q866C5 Q866C7 Q866C9 Q866D2 Q866D6 Q866D9 Q866E1 Q866E4 Q866E6 Q866E7 Q866E8
Q866F0 Q866F1 Q866G7 Q866S3 Q866T7 Q866X0 Q866Y9 Q866Z4 Q866Z5 Q866Z6 Q867B2
Q867C0 Q867X6 Q868T3 Q86917 Q86925 Q869R1 Q869V1 Q869W7 Q86A22 Q86A83 Q86AE3
Q86AF0 Q86AQ7 Q86AV5 Q86AZ6 Q86B61 Q86B91 Q86BE9 Q86BJ3 Q86CZ2 Q86DA7 Q86FQ0
Q86GL4 Q86GV3 Q86H13 Q86H31 Q86H65 Q86HH3 Q86HN8 Q86HQ2 Q86HQ3 Q86HT1 Q86I81
Q86I95 Q86ID5 Q86IG3 Q86IJ0 Q86IV0 Q86IW5 Q86IX2 Q86IZ2 Q86J02 Q86J18 Q86JD4
Q86JE4 Q86JE5 Q86JF2 Q86JG6 Q86JM5 Q86JM7 Q86K80 Q86KL7 Q86KP5 Q86KR1 Q86L28
Q86LG1 Q86ME2 Q86PM2 Q86PM4 Q86SF2 Q86SJ2 Q86SJ6 Q86SK9 Q86SM5 Q86SM8 Q86SP6
Q86SQ3 Q86SQ4 Q86SQ6 Q86SR1 Q86SS6 Q86SU0 Q86T03 Q86T13 Q86T20 Q86T26 Q86T96
Q86TG1 Q86TL2 Q86TM6 Q86TY3 Q86U02 Q86UB9 Q86UD3 Q86UD5 Q86UE4 Q86UE6 Q86UF1
Q86UF2 Q86UG4 Q86UK0 Q86UK5 Q86UL3 Q86UP0 Q86UP2 Q86UP6 Q86UP9 Q86UQ4 Q86UQ5
Q86UW1 Q86UW2 Q86V24 Q86V35 Q86V40 Q86V85 Q86VB7 Q86VD7 Q86VD9 Q86VE9 Q86VF5
Q86VH4 Q86VH5 Q86VI4 Q86VL8 Q86VR2 Q86VR7 Q86VU5 Q86VW1 Q86VY9 Q86VZ1 Q86VZ4
Q86VZ5 Q86W10 Q86W33 Q86W47 Q86W74 Q86WA9 Q86WB7 Q86WC4 Q86WI0 Q86WI1 Q86WK6
Q86WK7 Q86WK9 Q86WR0 Q86WS5 Q86WV6 Q86X19 Q86X29 Q86X52 Q86XE3 Q86XJ0 Q86XK7
Q86XL3 Q86XM0 Q86XQ3 Q86XR5 Q86XS8 Q86XT9 Q86XX4 Q86Y07 Q86Y22 Q86Y34 Q86Y38
Q86Y39 Q86Y82 Q86YA3 Q86YC3 Q86YD3 Q86YD5 Q86YJ5 Q86YL7 Q86YN1 Q86YT5 Q86YT9
Q86YW5 Q86Z14 Q86ZU7 Q87008 Q87041 Q87088 Q870I1 Q870L2 Q870L3 Q870V7 Q870W0
Q871R7 Q871U9 Q871V2 Q873L8 Q873N0 Q873N1 Q873N2 Q874C0 Q874C1 Q874I4 Q874I5
Q874K0 Q874L4 Q874L8 Q874X5 Q875A7 Q875C2 Q875M2 Q875S9 Q876A6 Q876K6 Q876L2
Q876L3 Q876L4 Q876L5 Q876N4 Q876Q0 Q879S3 Q879Z5 Q87A36 Q87A87 Q87AD2 Q87AD3
Q87AD4 Q87AF7 Q87AV3 Q87AW7 Q87B79 Q87B80 Q87BH3 Q87BH5 Q87BL6 Q87BT3 Q87BZ9
Q87C89 Q87C90 Q87CC5 Q87CI7 Q87CN1 Q87CN8 Q87D01 Q87DK8 Q87DL2 Q87DT0 Q87DY5
Q87E55 Q87E86 Q87EF0 Q87EF5 Q87EI0 Q87EJ9 Q87EN2 Q87EP5 Q87EP8 Q87F82 Q87FA1
Q87FD5 Q87FP8 Q87FR1 Q87FV4 Q87G36 Q87GB7 Q87GB8 Q87GU5 Q87GZ9 Q87IH5 Q87IR2
Q87J97 Q87JM4 Q87KA3 Q87KA4 Q87KE9 Q87KM9 Q87KN9 Q87KV8 Q87KW0 Q87KY1 Q87KY2
Q87LQ1 Q87LR6 Q87LZ1 Q87M69 Q87MA7 Q87MA8 Q87MA9 Q87MB0 Q87ML2 Q87MQ5 Q87MW9
Q87MX0 Q87MX1 Q87MX4 Q87MZ5 Q87N03 Q87N04 Q87NA5 Q87NG3 Q87NJ8 Q87NZ5 Q87PP5
Q87Q39 Q87Q45 Q87Q50 Q87QB8 Q87QJ4 Q87QN1 Q87QP4 Q87R16 Q87R62 Q87RJ5 Q87RJ6
Q87RP8 Q87S89 Q87SA1 Q87SG7 Q87SL0 Q87SL3 Q87ST2 Q87TE5 Q87TH0 Q87TH1 Q87TH2
Q87TI7 Q87TJ1 Q87TN7 Q87TR5 Q87TS1 Q87TS8 Q87TS9 Q87TT0 Q87TY7 Q87U39 Q87UL3
Q87UW8 Q87UY1 Q87UY7 Q87UZ0 Q87VB6 Q87VF3 Q87VX7 Q87WB2 Q87WD2 Q87WG7 Q87WL3
Q87WY2 Q87Y19 Q87YP0 Q87YY5 Q87Z07 Q87ZQ0 Q87ZQ3 Q87ZQ9 Q87ZS5 Q87ZS8 Q880L3

Q881Q1 Q88261 Q882Z2 Q883V4 Q883V6 Q88416 Q88417 Q88418 Q88421 Q88422 Q88424
Q88425 Q88426 Q88427 Q884D4 Q885M2 Q885Q3 Q885W4 Q885Z9 Q886F1 Q887G7 Q887H2
Q887P9 Q887Q6 Q887V6 Q88893 Q888L8 Q888N1 Q88938 Q88940 Q889E3 Q88A56 Q88AF9
Q88BA2 Q88BW8 Q88BW9 Q88BX0 Q88C19 Q88C65 Q88CB9 Q88CN8 Q88CS3 Q88CW4 Q88D04
Q88D09 Q88D92 Q88DN4 Q88DZ3 Q88E10 Q88E23 Q88EX9 Q88F17 Q88F24 Q88F88 Q88FD7
Q88FD8 Q88FG8 Q88FH1 Q88FH7 Q88FQ6 Q88FS8 Q88FT1 Q88FW9 Q88FX8 Q88FY6 Q88GQ7
Q88IY7 Q88IY8 Q88JK6 Q88K49 Q88K63 Q88KP1 Q88LQ8 Q88M93 Q88N31 Q88N32 Q88N45
Q88N79 Q88NC5 Q88ND2 Q88NI1 Q88NK7 Q88NL9 Q88NS2 Q88PN3 Q88Q16 Q88Q91 Q88QS0
Q88QU5 Q88R14 Q88RJ4 Q88RL9 Q88RX1 Q88RZ3 Q88S59 Q88SK7 Q88SV0 Q88TT1 Q88UB3
Q88UI3 Q88UT8 Q88UT9 Q88UX2 Q88UZ5 Q88V72 Q88V79 Q88VI8 Q88VY3 Q88W28 Q88W75
Q88WR8 Q88X65 Q88X97 Q88XQ3 Q88YK2 Q88Z31 Q88Z42 Q88Z48 Q88ZT7 Q88ZT8 Q88ZZ1
Q89182 Q89186 Q891X1 Q89277 Q893D9 Q893H9 Q89424 Q89443 Q89448 Q894B9 Q894C1
Q894D4 Q89501 Q89504 Q895I4 Q895I8 Q89607 Q89609 Q89669 Q89687 Q89705 Q89730
Q89738 Q89784 Q897L1 Q897L3 Q897L5 Q89829 Q89843 Q89846 Q89853 Q89881 Q89894
Q898B8 Q898D6 Q898L6 Q898Q9 Q89904 Q89905 Q899D2 Q899S4 Q899T2 Q89A13 Q89A23
Q89A24 Q89A36 Q89A39 Q89A40 Q89A60 Q89A85 Q89A96 Q89A97 Q89A98 Q89AA3 Q89AA4
Q89AA5 Q89AA6 Q89AA7 Q89AA9 Q89AB5 Q89AB7 Q89AC7 Q89AF2 Q89AG6 Q89AI1 Q89AI6
Q89AI9 Q89AJ1 Q89AL0 Q89AL2 Q89AL3 Q89AL5 Q89AL9 Q89AM6 Q89AN5 Q89AN6 Q89AP9
Q89AQ0 Q89AQ3 Q89AR2 Q89AT1 Q89AT3 Q89AT4 Q89AT5 Q89AT6 Q89AT7 Q89AT8 Q89AU0
Q89AU6 Q89AV0 Q89AV4 Q89AV7 Q89AW1 Q89AW5 Q89AW6 Q89AW7 Q89AX0 Q89AX2 Q89AY7
Q89AZ0 Q89AZ1 Q89AZ2 Q89B00 Q89B14 Q89B34 Q89B43 Q89B44 Q89B45 Q89B48 Q89BQ0
Q89BU5 Q89DJ5 Q89DV4 Q89E33 Q89EA0 Q89EG9 Q89FC2 Q89FC4 Q89FU4 Q89FV1 Q89FY8
Q89G85 Q89I98 Q89IC2 Q89K17 Q89K46 Q89K67 Q89K83 Q89KD9 Q89KJ5 Q89KZ7 Q89KZ8
Q89L53 Q89LF9 Q89M95 Q89NN5 Q89NN6 Q89NW0 Q89NX6 Q89PK4 Q89PK5 Q89PK9 Q89Q75
Q89QD9 Q89RX3 Q89RX4 Q89UT8 Q89V40 Q89V68 Q89V70 Q89V71 Q89VU8 Q89W72 Q89WA1
Q89WH1 Q89WN1 Q89WR2 Q89YX1 Q89Z03 Q89Z16 Q89Z22 Q89ZG0 Q89ZV7 Q8A0F4 Q8A0F7
Q8A1G1 Q8A255 Q8A294 Q8A2U3 Q8A337 Q8A519 Q8A521 Q8A5Z4 Q8A6G5 Q8A6M3 Q8A7J8
Q8A7J9 Q8A7K1 Q8A7K2 Q8A8D8 Q8A8H2 Q8A9S9 Q8A9U9 Q8A9V0 Q8AA31 Q8AA76 Q8AAG5
Q8AIH5 Q8AIH6 Q8AV57 Q8AV58 Q8AV61 Q8AVC3 Q8AVF4 Q8AVI9 Q8AVJ9 Q8AVM3 Q8AVM5
Q8AVS8 Q8AVY0 Q8AVY8 Q8AW42 Q8AWB5 Q8AWB6 Q8AWC7 Q8AWW4 Q8AWW5 Q8AXB3 Q8AXC6
Q8AXC7 Q8AXS6 Q8AXU4 Q8AXY6 Q8AY29 Q8AY31 Q8AY32 Q8AY33 Q8AY34 Q8AY35 Q8AY36
Q8AYD0 Q8AYM7 Q8AYM8 Q8AYN0 Q8AYS8 Q8AYW1 Q8AYY5 Q8B0H1 Q8B0H6 Q8B0I1 Q8B115
Q8B119 Q8B120 Q8B121 Q8B8X3 Q8B912 Q8BAC4 Q8BB24 Q8BB25 Q8BB26 Q8BB27 Q8BDV6
Q8BFQ2 Q8BFQ3 Q8BFQ6 Q8BFT9 Q8BFU1 Q8BFU7 Q8BFU8 Q8BFW6 Q8BFW9 Q8BFY0 Q8BFZ1
Q8BFZ2 Q8BFZ9 Q8BG07 Q8BG09 Q8BG16 Q8BG19 Q8BG21 Q8BG22 Q8BG28 Q8BG39 Q8BG47
Q8BG50 Q8BG51 Q8BG54 Q8BG55 Q8BG58 Q8BG75 Q8BG84 Q8BGA2 Q8BGA3 Q8BGA9 Q8BGC3
Q8BGD4 Q8BGD5 Q8BGD6 Q8BGE4 Q8BGE9 Q8BGF8 Q8BGF9 Q8BGG0 Q8BGG6 Q8BGH2 Q8BGH4
Q8BGI1 Q8BGI3 Q8BGI5 Q8BGJ0 Q8BGK5 Q8BGK6 Q8BGK9 Q8BGM5 Q8BGN5 Q8BGN6 Q8BGN8
Q8BGN9 Q8BGP5 Q8BGP6 Q8BGQ4 Q8BGR2 Q8BGS7 Q8BGT0 Q8BGT1 Q8BGT9 Q8BGV3 Q8BGV8
Q8BGW2 Q8BGX2 Q8BGX3 Q8BGY5 Q8BGY6 Q8BGY7 Q8BGY9 Q8BGZ8 Q8BH01 Q8BH02 Q8BH06
Q8BH07 Q8BH10 Q8BH18 Q8BH24 Q8BH26 Q8BH27 Q8BH31 Q8BH32 Q8BH47 Q8BH51 Q8BH52
Q8BH59 Q8BH66 Q8BH73 Q8BH79 Q8BH98 Q8BHA1 Q8BHA9 Q8BHB3 Q8BHB9 Q8BHC0 Q8BHC7
Q8BHC9 Q8BHD4 Q8BHE4 Q8BHG3 Q8BHH1 Q8BHH8 Q8BHH9 Q8BHI7 Q8BHJ6 Q8BHJ7 Q8BHK1
Q8BHK2 Q8BHK3 Q8BHK6 Q8BHL4 Q8BHM9 Q8BHN0 Q8BHS6 Q8BHS8 Q8BHT6 Q8BHW5 Q8BHY2

Q8BHY3 Q8BHY8 Q8BI08 Q8BI36 Q8BI41 Q8BI84 Q8BIF0 Q8BIG7 Q8BIK6 Q8BIV7 Q8BIX9
Q8BIZ0 Q8BJ03 Q8BJ48 Q8BJ51 Q8BJ52 Q8BJ56 Q8BJ83 Q8BJA2 Q8BJE2 Q8BJI1 Q8BJM5
Q8BJN4 Q8BJQ9 Q8BJS4 Q8BJU2 Q8BJZ3 Q8BK03 Q8BK08 Q8BKD6 Q8BKE6 Q8BKG3 Q8BKG4
Q8BKN6 Q8BKT6 Q8BKU8 Q8BL00 Q8BL03 Q8BL07 Q8BLA8 Q8BLD9 Q8BLE7 Q8BLF1 Q8BLG2
Q8BLH5 Q8BLI4 Q8BLN6 Q8BLQ7 Q8BLQ9 Q8BLU0 Q8BLV3 Q8BLX4 Q8BLY3 Q8BM55 Q8BM81
Q8BM86 Q8BM92 Q8BM96 Q8BMC0 Q8BMD6 Q8BMD8 Q8BMF5 Q8BMF8 Q8BMG8 Q8BMK4 Q8BML2
Q8BMN3 Q8BMP4 Q8BMT4 Q8BMT9 Q8BMW7 Q8BN21 Q8BN57 Q8BN82 Q8BNA6 Q8BND5 Q8BNI4
Q8BNJ6 Q8BNQ3 Q8BNX1 Q8BNX7 Q8BP07 Q8BP31 Q8BP97 Q8BPE4 Q8BPM6 Q8BPS4 Q8BPT6
Q8BPX9 Q8BQ86 Q8BQC3 Q8BQM7 Q8BQQ1 Q8BQS5 Q8BQT2 Q8BQU6 Q8BQU7 Q8BQX5 Q8BQZ8
Q8BR26 Q8BR70 Q8BR76 Q8BR86 Q8BRG8 Q8BRH0 Q8BRJ3 Q8BRK9 Q8BRU4 Q8BRU6 Q8BRX9
Q8BS35 Q8BS95 Q8BSD4 Q8BSE0 Q8BSF4 Q8BSL4 Q8BSM7 Q8BSU2 Q8BSY0 Q8BT18 Q8BT35
Q8BT42 Q8BTC1 Q8BTE5 Q8BTG6 Q8BTJ4 Q8BTP0 Q8BTP3 Q8BTV1 Q8BTY2 Q8BTZ5 Q8BU14
Q8BU33 Q8BU91 Q8BUB6 Q8BUD0 Q8BUE1 Q8BUE4 Q8BUH7 Q8BUJ9 Q8BUM6 Q8BUN9 Q8BUV8
Q8BUX5 Q8BUY5 Q8BVA2 Q8BVF7 Q8BVG5 Q8BVG8 Q8BVI5 Q8BVM2 Q8BVN3 Q8BVN7 Q8BW41
Q8BW66 Q8BW75 Q8BW93 Q8BWB6 Q8BWC0 Q8BWF2 Q8BWG9 Q8BWH0 Q8BWJ4 Q8BWM0 Q8BWP8
Q8BWQ1 Q8BWQ6 Q8BWY2 Q8BWY7 Q8BWY9 Q8BX10 Q8BX35 Q8BX43 Q8BX51 Q8BX79 Q8BX90
Q8BXA0 Q8BXA1 Q8BXA5 Q8BXA6 Q8BXB6 Q8BXJ9 Q8BXK9 Q8BXN9 Q8BXQ2 Q8BXQ3 Q8BXR1
Q8BXR5 Q8BXS7 Q8BXT9 Q8BXV2 Q8BXZ0 Q8BXZ1 Q8BY79 Q8BY89 Q8BYC4 Q8BYF6 Q8BYG9
Q8BYI6 Q8BYI8 Q8BYM5 Q8BYR8 Q8BYU6 Q8BZ00 Q8BZ09 Q8BZ10 Q8BZ39 Q8BZ81 Q8BZA7
Q8BZB3 Q8BZF2 Q8BZG5 Q8BZH0 Q8BZK4 Q8BZL4 Q8BZN2 Q8BZP8 Q8BZR0 Q8BZT5 Q8C010
Q8C013 Q8C025 Q8C031 Q8C0G2 Q8C0K5 Q8C0L0 Q8C0N2 Q8C0T0 Q8C0T7 Q8C0X2 Q8C0X7
Q8C0Z1 Q8C102 Q8C110 Q8C129 Q8C131 Q8C138 Q8C145 Q8C160 Q8C170 Q8C172 Q8C1E7
Q8C1F4 Q8C1Q6 Q8C1T8 Q8C206 Q8C261 Q8C2L6 Q8C2R7 Q8C2S7 Q8C341 Q8C351 Q8C353
Q8C398 Q8C3B8 Q8C3K5 Q8C3K6 Q8C3Q5 Q8C3S2 Q8C3X2 Q8C3X8 Q8C407 Q8C419 Q8C428
Q8C432 Q8C4A0 Q8C4G9 Q8C4N4 Q8C4Q9 Q8C4U2 Q8C4X7 Q8C503 Q8C561 Q8C567 Q8C581
Q8C5C9 Q8C5H1 Q8C5U4 Q8C5W0 Q8C6N3 Q8C6U2 Q8C6V3 Q8C6Z1 Q8C708 Q8C784 Q8C7E7
Q8C7N7 Q8C7U7 Q8C7X2 Q8C811 Q8C817 Q8C838 Q8C850 Q8C8C1 Q8C8N2 Q8C8S3 Q8C8T7
Q8C985 Q8C996 Q8C9E8 Q8CA03 Q8CA71 Q8CAA5 Q8CAC8 Q8CAE3 Q8CAE9 Q8CAQ8 Q8CB12
Q8CB19 Q8CB44 Q8CB65 Q8CB67 Q8CB93 Q8CBB2 Q8CBC4 Q8CBC6 Q8CBF3 Q8CBG9 Q8CBH5
Q8CBH7 Q8CBR1 Q8CBX0 Q8CC12 Q8CC84 Q8CCM6 Q8CCS2 Q8CD26 Q8CD54 Q8CD78 Q8CDN1
Q8CE47 Q8CE72 Q8CE93 Q8CE94 Q8CEF8 Q8CEF9 Q8CEG0 Q8CEK7 Q8CES1 Q8CEZ1 Q8CF82
Q8CF93 Q8CF94 Q8CFA6 Q8CFC4 Q8CFD0 Q8CFD9 Q8CFE6 Q8CFG5 Q8CFG6 Q8CFJ7 Q8CFM6
Q8CFS6 Q8CFW1 Q8CFY5 Q8CFZ5 Q8CG09 Q8CG64 Q8CGA3 Q8CGF5 Q8CGK5 Q8CGL2 Q8CGM1
Q8CGQ8 Q8CGU6 Q8CH34 Q8CH36 Q8CH60 Q8CH87 Q8CHI9 Q8CHJ0 Q8CHJ1 Q8CHJ2 Q8CHK3
Q8CHL0 Q8CHM1 Q8CHM9 Q8CHN6 Q8CHQ6 Q8CHQ9 Q8CHS2 Q8CHT3 Q8CHX6 Q8CI52 Q8CI59
Q8CI66 Q8CI85 Q8CIA5 Q8CIA9 Q8CIB6 Q8CIF6 Q8CII8 Q8CII9 Q8CIM5 Q8CIP3 Q8CIP5
Q8CIQ6 Q8CIR4 Q8CIV2 Q8CIW6 Q8CIY2 Q8CIZ5 Q8CIZ9 Q8CJ11 Q8CJ12 Q8CJ26 Q8CJ44
Q8CJ52 Q8CJ58 Q8CJ61 Q8CJ91 Q8CJC7 Q8CJH3 Q8CJI3 Q8CM25 Q8CMK4 Q8CMK8 Q8CMM5
Q8CMU8 Q8CN02 Q8CN16 Q8CN53 Q8CN54 Q8CN76 Q8CN95 Q8CNB2 Q8CNF2 Q8CNF3 Q8CNF7
Q8CNJ1 Q8CNJ2 Q8CNJ3 Q8CNK3 Q8CNP2 Q8CNQ3 Q8CNV5 Q8CNW5 Q8CP59 Q8CP76 Q8CP86
Q8CPC0 Q8CPC4 Q8CPD8 Q8CPF7 Q8CPK0 Q8CPK7 Q8CPM1 Q8CPM2 Q8CPM6 Q8CPP6 Q8CPP7
Q8CPP8 Q8CPP9 Q8CPR4 Q8CPU8 Q8CPU9 Q8CPV0 Q8CPV1 Q8CPV2 Q8CPV3 Q8CPV4 Q8CPY2
Q8CQ10 Q8CQ45 Q8CQ46 Q8CQ47 Q8CQ48 Q8CQ49 Q8CQ50 Q8CQ86 Q8CQF4 Q8CQF7 Q8CQT3
Q8CR38 Q8CR39 Q8CR79 Q8CRA8 Q8CRA9 Q8CRD1 Q8CRK7 Q8CRT4 Q8CRU8 Q8CRV2 Q8CRX9

Q8CS28 Q8CS29 Q8CSD1 Q8CSK8 Q8CSL7 Q8CSP3 Q8CSS7 Q8CST9 Q8CT52 Q8CT77 Q8CTE6
Q8CTF5 Q8CTI3 Q8CTJ6 Q8CTL4 Q8CTM8 Q8CTQ5 Q8CU03 Q8CU36 Q8CU87 Q8CUL5 Q8CUT6
Q8CV14 Q8CV51 Q8CVG4 Q8CVH5 Q8CVH8 Q8CVI4 Q8CVJ0 Q8CVJ9 Q8CVU7 Q8CVW1 Q8CW90
Q8CWA4 Q8CWC7 Q8CWZ1 Q8CX05 Q8CX16 Q8CX97 Q8CXF3 Q8CXI8 Q8CXK0 Q8CYJ8 Q8D074
Q8D124 Q8D1E4 Q8D1U8 Q8D281 Q8D2A8 Q8D2I8 Q8D2I9 Q8D2N5 Q8D2N9 Q8D2Q7 Q8D2R1
Q8D2T1 Q8D2U8 Q8D2Z3 Q8D2Z5 Q8D339 Q8D342 Q8D350 Q8D396 Q8D3D5 Q8D3I8 Q8D3J7
Q8D3J8 Q8D3J9 Q8D3U7 Q8D3U8 Q8D3Z8 Q8D4Q1 Q8D4Z3 Q8D5Z6 Q8D666 Q8D6H2 Q8D6J0
Q8D825 Q8D885 Q8D887 Q8D890 Q8D8M2 Q8D8Y2 Q8D8Y7 Q8D922 Q8D927 Q8D9N8 Q8DA60
Q8DAD7 Q8DAG9 Q8DAH7 Q8DAV2 Q8DB17 Q8DB58 Q8DBJ1 Q8DBJ2 Q8DBJ3 Q8DBJ4 Q8DBJ5
Q8DBX0 Q8DC44 Q8DC61 Q8DCX3 Q8DCX4 Q8DCZ0 Q8DD41 Q8DD49 Q8DD90 Q8DDH2 Q8DDH3
Q8DDH4 Q8DDI2 Q8DDL3 Q8DDQ1 Q8DDQ2 Q8DDQ3 Q8DDS2 Q8DED6 Q8DEG8 Q8DEK7 Q8DER8
Q8DES8 Q8DFE4 Q8DFK4 Q8DG81 Q8DGA3 Q8DGB4 Q8DGM9 Q8DH03 Q8DHA7 Q8DHB0 Q8DHJ2
Q8DHQ2 Q8DHX4 Q8DIC3 Q8DIF8 Q8DIH3 Q8DIN8 Q8DIN9 Q8DIP0 Q8DIQ0 Q8DIQ1 Q8DIV4
Q8DJ02 Q8DJ15 Q8DJ41 Q8DJ42 Q8DJ43 Q8DJ44 Q8DJ91 Q8DJI1 Q8DJM9 Q8DJZ6 Q8DK20
Q8DK95 Q8DKI2 Q8DKM3 Q8DKN2 Q8DKW4 Q8DKY0 Q8DKZ3 Q8DL16 Q8DL32 Q8DL96 Q8DLP5
Q8DLP6 Q8DLP7 Q8DLP8 Q8DMR6 Q8DMX7 Q8DMY2 Q8DMY3 Q8DN93 Q8DNB6 Q8DNE1 Q8DNS0
Q8DNY1 Q8DP63 Q8DPA3 Q8DPG6 Q8DPG7 Q8DPH5 Q8DPQ6 Q8DQ64 Q8DQE5 Q8DQM0 Q8DQN5
Q8DQY6 Q8DR34 Q8DR55 Q8DR69 Q8DR70 Q8DRB1 Q8DRN6 Q8DRY7 Q8DRZ4 Q8DSP8 Q8DSQ4
Q8DSX7 Q8DTQ3 Q8DUQ0 Q8DUY3 Q8DUY4 Q8DV98 Q8DVB6 Q8DVK7 Q8DVM4 Q8DVX3 Q8DWQ7
Q8DXQ8 Q8DY58 Q8DY59 Q8DY66 Q8DY84 Q8DY91 Q8DYV7 Q8DZH6 Q8DZL1 Q8E076 Q8E077
Q8E0J4 Q8E0U6 Q8E1R5 Q8E218 Q8E260 Q8E2K6 Q8E3C7 Q8E3S4 Q8E3S5 Q8E3T2 Q8E3U9
Q8E3V6 Q8E418 Q8E480 Q8E4G5 Q8E541 Q8E575 Q8E5N2 Q8E5V2 Q8E5V3 Q8E669 Q8E6G4
Q8E6W4 Q8E779 Q8E7H4 Q8E7M0 Q8E8B4 Q8E8B5 Q8E8B6 Q8E8F7 Q8E8P7 Q8E985 Q8E9K1
Q8E9P1 Q8E9P5 Q8E9P9 Q8E9R3 Q8E9R4 Q8E9R5 Q8EA87 Q8EAB0 Q8EAM8 Q8EAX8 Q8EAZ0
Q8EB02 Q8EB99 Q8EBI5 Q8EBR1 Q8ECL5 Q8ED56 Q8ED69 Q8ED79 Q8EDF0 Q8EDL5 Q8EE76
Q8EE77 Q8EE78 Q8EE81 Q8EER0 Q8EER3 Q8EES3 Q8EF50 Q8EF78 Q8EG03 Q8EH93 Q8EH95
Q8EHC1 Q8EHD3 Q8EHP1 Q8EHV6 Q8EI17 Q8EI29 Q8EI35 Q8EI38 Q8EIL8 Q8EIX5 Q8EJE5
Q8EJJ5 Q8EJM6 Q8EK44 Q8EKT6 Q8EKU1 Q8EL25 Q8EM16 Q8EM65 Q8EM77 Q8EM78 Q8EM79
Q8EMX2 Q8ENF3 Q8ENQ1 Q8ENS1 Q8EPB2 Q8EPX2 Q8EQA1 Q8EQL9 Q8EQQ2 Q8EQR7 Q8EQS7
Q8EQU7 Q8ER40 Q8ER51 Q8ER78 Q8ERW3 Q8ERY3 Q8ESA3 Q8ETN0 Q8EU91 Q8EUA6 Q8EVM0
Q8EWA4 Q8EWS0 Q8EWZ2 Q8EWZ3 Q8EX10 Q8EXN4 Q8EXQ8 Q8EYK8 Q8EYY4 Q8EZP1 Q8EZS4
Q8F1M1 Q8F1M2 Q8F1W7 Q8F1X4 Q8F2J0 Q8F332 Q8F397 Q8F4J3 Q8F4W2 Q8F641 Q8F6H0
Q8F724 Q8F7Q5 Q8F7Q7 Q8F7R0 Q8F8F1 Q8F9F8 Q8FA58 Q8FA73 Q8FA95 Q8FAJ3 Q8FAL7
Q8FAL8 Q8FAX0 Q8FAX1 Q8FAX2 Q8FAZ0 Q8FB38 Q8FBC8 Q8FBD1 Q8FBD6 Q8FBI6 Q8FBI7
Q8FBI8 Q8FBP8 Q8FBS8 Q8FBU5 Q8FBV0 Q8FBW6 Q8FBX9 Q8FCI8 Q8FCK7 Q8FCN5 Q8FCQ0
Q8FCQ1 Q8FCS5 Q8FCT7 Q8FCY0 Q8FD11 Q8FD49 Q8FD50 Q8FD51 Q8FD59 Q8FD68 Q8FDC3
Q8FDE0 Q8FDE1 Q8FDG7 Q8FDI3 Q8FDL6 Q8FDU9 Q8FDW4 Q8FDZ8 Q8FE55 Q8FEA6 Q8FEA7
Q8FEJ4 Q8FF11 Q8FF56 Q8FF81 Q8FFA9 Q8FFB9 Q8FFC0 Q8FFD3 Q8FFD5 Q8FFJ2 Q8FFK0
Q8FFK3 Q8FFL7 Q8FFL8 Q8FFL9 Q8FFM2 Q8FFU1 Q8FFV6 Q8FFX9 Q8FG02 Q8FG03 Q8FG04
Q8FGA4 Q8FGI6 Q8FGJ7 Q8FGT7 Q8FGU0 Q8FGV4 Q8FGW2 Q8FH26 Q8FH68 Q8FH91 Q8FH92
Q8FH94 Q8FHB4 Q8FHB5 Q8FHC1 Q8FHC6 Q8FHE5 Q8FHE6 Q8FHF9 Q8FHG6 Q8FHW3 Q8FI24
Q8FIB8 Q8FIR9 Q8FIS3 Q8FJ36 Q8FJB1 Q8FJC1 Q8FJC7 Q8FJK8 Q8FJK9 Q8FJV3 Q8FJV4
Q8FJV5 Q8FJY4 Q8FJZ8 Q8FK20 Q8FK36 Q8FK37 Q8FL15 Q8FL49 Q8FL94 Q8FLA1 Q8FLB6
Q8FLC1 Q8FLS3 Q8FMF7 Q8FMR6 Q8FMT1 Q8FNP2 Q8FNQ1 Q8FNQ7 Q8FNQ8 Q8FNQ9 Q8FNR0

Q8FNR1 Q8FNR2 Q8FNT7 Q8FP03 Q8FPC1 Q8FQ24 Q8FQD9 Q8FRF9 Q8FS11 Q8FT77 Q8FTC9

Q8FTE9 Q8FTH1 Q8FTK0 Q8FUI7 Q8FUN0 Q8FUN2 Q8FUS3 Q8FUW9 Q8FUX0 Q8FV29 Q8FV59

Q8FVL9 Q8FVM0 Q8FVS6 Q8FVU0 Q8FW08 Q8FW09 Q8FWN8 Q8FWN9 Q8FWV9 Q8FXM5 Q8FXT9

Q8FY85 Q8FYI0 Q8FYQ1 Q8FYT4 Q8FYV0 Q8FZ43 Q8FZF5 Q8FZT8 Q8FZV0 Q8FZV1 Q8G022

Q8G0E0 Q8G0E1 Q8G0Q5 Q8G0T8 Q8G0Y6 Q8G156 Q8G1A7 Q8G1B0 Q8G1D1 Q8G1E6 Q8G1U5

Q8G261 Q8G2C7 Q8G2D8 Q8G2D9 Q8G2E1 Q8G2I1 Q8G2K1 Q8G2M6 Q8G308 Q8G3C1 Q8G4H3

Q8G4Q7 Q8G5C0 Q8G5C2 Q8G5C3 Q8G5I3 Q8G5R2 Q8G692 Q8G6C4 Q8G6J6 Q8G6P5 Q8G6T7

Q8G6U1 Q8G6U3 Q8G782 Q8G7A8 Q8G7A9 Q8G7Q3 Q8G7W3 Q8G838 Q8G845 Q8G846 Q8GAI5

Q8GAI6 Q8GFA2 Q8GH68 Q8GHM3 Q8GJ26 Q8GJ30 Q8GLC5 Q8GLH3 Q8GMS2 Q8GNS3 Q8GP19

Q8GQ96 Q8GRI8 Q8GRT8 Q8GRU6 Q8GSD9 Q8GSG4 Q8GSJ6 Q8GSQ1 Q8GT66 Q8GT74 Q8GT75

Q8GTZ9 Q8GU82 Q8GU83 Q8GU84 Q8GU85 Q8GU86 Q8GU87 Q8GU88 Q8GU89 Q8GU92 Q8GUH2

Q8GUH7 Q8GUJ1 Q8GUJ2 Q8GUK1 Q8GUK6 Q8GUK7 Q8GUM0 Q8GUM3 Q8GUM4 Q8GUM5 Q8GUN6

Q8GUQ5 Q8GUU2 Q8GVE3 Q8GW16 Q8GW19 Q8GW20 Q8GW38 Q8GW61 Q8GW96 Q8GWA7 Q8GWB7

Q8GWC3 Q8GWD2 Q8GWD5 Q8GWG0 Q8GWG6 Q8GWH5 Q8GWJ7 Q8GWP3 Q8GWR0 Q8GWT1 Q8GWT5

Q8GWW4 Q8GWX2 Q8GX23 Q8GX37 Q8GX78 Q8GX86 Q8GX92 Q8GX93 Q8GXA4 Q8GXB4 Q8GXE6

Q8GXG6 Q8GXJ4 Q8GXK5 Q8GXM8 Q8GXN2 Q8GXQ3 Q8GXR2 Q8GXT7 Q8GXU8 Q8GXV7 Q8GY03

Q8GY97 Q8GYA4 Q8GYB4 Q8GYC7 Q8GYE0 Q8GYF4 Q8GYF5 Q8GYG1 Q8GYH5 Q8GYH6 Q8GYH8

Q8GYK7 Q8GYP8 Q8GYS1 Q8GYS4 Q8GYT9 Q8GYW9 Q8GZ52 Q8GZ79 Q8GZ81 Q8GZ99 Q8GZA4

Q8GZC2 Q8GZC3 Q8GZD4 Q8GZR2 Q8H038 Q8H074 Q8H0B2 Q8H0B6 Q8H0T6 Q8H0Y1 Q8H112

Q8H116 Q8H118 Q8H157 Q8H184 Q8H199 Q8H1D8 Q8H1E6 Q8H1F2 Q8H1G3 Q8H1R3 Q8H1X1

Q8H1Z0 Q8H329 Q8H384 Q8H3P9 Q8H4H5 Q8H569 Q8H5N9 Q8H5T6 Q8H5X5 Q8H6G7 Q8H6G8

Q8H6G9 Q8H6H0 Q8H6H2 Q8H6H4 Q8H7L6 Q8H7N9 Q8H8C7 Q8H8K7 Q8H8U0 Q8H8V7 Q8H930

Q8HB52 Q8HBG1 Q8HBG2 Q8HC87 Q8HC89 Q8HCJ4 Q8HD57 Q8HD60 Q8HEB9 Q8HEC0 Q8HEC1

Q8HEC2 Q8HEC3 Q8HEC4 Q8HEC5 Q8HEM1 Q8HF67 Q8HFZ6 Q8HG02 Q8HG55 Q8HHD2 Q8HHD3

Q8HL81 Q8HL93 Q8HLB9 Q8HLD2 Q8HLR2 Q8HMZ7 Q8HQ92 Q8HQB1 Q8HQE6 Q8HQE7 Q8HQE8

Q8HQE9 Q8HQF0 Q8HQF1 Q8HQF2 Q8HRZ5 Q8HRZ6 Q8HS08 Q8HS09 Q8HS12 Q8HS15 Q8HS16

Q8HS19 Q8HS25 Q8HS28 Q8HS29 Q8HS36 Q8HS39 Q8HS40 Q8HTL5 Q8HU50 Q8HWB0 Q8HXE3

Q8HXG5 Q8HXG6 Q8HXI3 Q8HXJ7 Q8HXQ5 Q8HXW0 Q8HXW2 Q8HXW5 Q8HXW7 Q8HXX3 Q8HXY2

Q8HXZ7 Q8HXZ8 Q8HY00 Q8HY01 Q8HY02 Q8HY03 Q8HY04 Q8HY06 Q8HY10 Q8HY11 Q8HY12

Q8HY24 Q8HY39 Q8HY46 Q8HY56 Q8HY59 Q8HY69 Q8HYB2 Q8HYB7 Q8HYC0 Q8HYC3 Q8HYJ3

Q8HYJ4 Q8HYJ5 Q8HYJ6 Q8HYJ7 Q8HYJ9 Q8HYN8 Q8HYS5 Q8HYW2 Q8HYX8 Q8HYZ0 Q8HYZ1

Q8HZ64 Q8HZ69 Q8HZ70 Q8HZD9 Q8HZJ2 Q8HZJ6 Q8HZJ8 Q8HZK2 Q8HZK3 Q8HZN9 Q8HZP1

Q8HZP2 Q8HZP3 Q8HZR2 Q8HZR3 Q8HZR8 Q8HZT6 Q8HZT9 Q8HZU0 Q8HZU1 Q8HZV3 Q8I0G4

Q8I0L6 Q8I136 Q8I1F0 Q8I2A6 Q8I3H7 Q8I3Z1 Q8I480 Q8I4B0 Q8I4N3 Q8I4N4 Q8I5I1

Q8I5V4 Q8I5Z5 Q8I6V3 Q8I6Z5 Q8I6Z7 Q8I7D9 Q8I7F8 Q8I7I5 Q8I7J4 Q8I7T3 Q8I7Z2

Q8I9S1 Q8I9S2 Q8IA41 Q8IA42 Q8IA43 Q8IA44 Q8IA98 Q8IBP1 Q8IBZ9 Q8IDA0 Q8IDG7

Q8IDX6 Q8IE14 Q8IFN0 Q8IGJ0 Q8IHY1 Q8IHZ5 Q8IIG1 Q8IIG4 Q8IKD3 Q8ILG2 Q8ILR9

Q8ILT5 Q8IM15 Q8IM16 Q8IM77 Q8IMN5 Q8IMN6 Q8IMQ6 Q8IMX7 Q8IMZ5 Q8IMZ9 Q8IN23

Q8IN35 Q8IN58 Q8IN78 Q8INM9 Q8INQ7 Q8INZ1 Q8INZ2 Q8IPU5 Q8IQ56 Q8IQ70 Q8IQ72

Q8IQC1 Q8IRI6 Q8IRL8 Q8IRZ5 Q8IS44 Q8IS64 Q8ISS3 Q8ITC7 Q8ITC9 Q8IU57 Q8IU68

Q8IU80 Q8IU89 Q8IU99 Q8IUA7 Q8IUC8 Q8IUH4 Q8IUH5 Q8IUH8 Q8IUK5 Q8IUN9 Q8IUR5

Q8IUS5 Q8IUW5 Q8IUX1 Q8IUY3 Q8IV01 Q8IV08 Q8IV31 Q8IV45 Q8IV77 Q8IVB4 Q8IVJ1

Q8IVJ8 Q8IVM8 Q8IVP5 Q8IVQ6 Q8IVU1 Q8IVV8 Q8IVW8 Q8IVY1 Q8IW00 Q8IW52 Q8IW70

Q8IWA4 Q8IWA5 Q8IWB1 Q8IWB4 Q8IWB9 Q8IWD5 Q8IWK6 Q8IWR1 Q8IWT1 Q8IWT6 Q8IWU2

Q8IWU4 Q8IWV1 Q8IWX5 Q8IWY9 Q8IX05 Q8IX19 Q8IX94 Q8IXA5 Q8IXB3 Q8IXE1 Q8IXF9
Q8IXH8 Q8IXI1 Q8IXI2 Q8IXK2 Q8IXL6 Q8IXM6 Q8IXU6 Q8IXX5 Q8IY17 Q8IY26 Q8IY34
Q8IY49 Q8IY50 Q8IY95 Q8IYJ0 Q8IYL9 Q8IYP9 Q8IYR6 Q8IYS0 Q8IYS2 Q8IYS5 Q8IYV9
Q8IZ08 Q8IZ52 Q8IZ57 Q8IZ96 Q8IZA0 Q8IZD6 Q8IZF0 Q8IZF2 Q8IZF3 Q8IZF4 Q8IZF5
Q8IZF6 Q8IZF7 Q8IZJ1 Q8IZK6 Q8IZM9 Q8IZN3 Q8IZP7 Q8IZP9 Q8IZR5 Q8IZS7 Q8IZS8
Q8IZT8 Q8IZU8 Q8IZU9 Q8IZV2 Q8IZV5 Q8IZY2 Q8J025 Q8J0F3 Q8J0F4 Q8J0F6 Q8J0I8
Q8J0L4 Q8J127 Q8J1F5 Q8J1M7 Q8J1Z2 Q8J207 Q8J2P4 Q8J2Q1 Q8J2Q2 Q8J2Q9 Q8JFB7
Q8JFG3 Q8JFN7 Q8JFR5 Q8JG38 Q8JG69 Q8JG70 Q8JGM4 Q8JGT4 Q8JH92 Q8JHE9 Q8JHF0
Q8JHF2 Q8JI14 Q8JIL9 Q8JIY1 Q8JJX0 Q8JL88 Q8JMY6 Q8JMZ6 Q8JN07 Q8JNB2 Q8JPR1
Q8JSD9 Q8JSP5 Q8JSP8 Q8JSP9 Q8JSZ3 Q8JTH0 Q8JTR0 Q8JU56 Q8JUX5 Q8JV21 Q8JXB9
Q8JZL1 Q8JZL2 Q8JZM4 Q8JZM8 Q8JZN3 Q8JZN7 Q8JZQ0 Q8JZQ2 Q8JZR0 Q8JZR4 Q8JZR6
Q8JZU2 Q8JZZ0 Q8JZZ7 Q8K021 Q8K064 Q8K071 Q8K072 Q8K078 Q8K087 Q8K093 Q8K097
Q8K099 Q8K0B2 Q8K0B3 Q8K0C4 Q8K0D7 Q8K0E3 Q8K0E7 Q8K0H1 Q8K0H7 Q8K0J2 Q8K0L2
Q8K0S5 Q8K0T0 Q8K0U3 Q8K0Z9 Q8K129 Q8K135 Q8K158 Q8K174 Q8K177 Q8K182 Q8K190
Q8K1A5 Q8K1B9 Q8K1C7 Q8K1E0 Q8K1E3 Q8K1F9 Q8K1G0 Q8K1G2 Q8K1N1 Q8K1P8 Q8K1P9
Q8K1Q3 Q8K1S2 Q8K1S3 Q8K1S4 Q8K1S7 Q8K1T0 Q8K1T1 Q8K1X4 Q8K1Z6 Q8K201 Q8K209
Q8K211 Q8K249 Q8K273 Q8K299 Q8K2A8 Q8K2C8 Q8K2C9 Q8K2I3 Q8K2J7 Q8K2P7 Q8K2T4
Q8K2W3 Q8K2X1 Q8K2Y3 Q8K304 Q8K348 Q8K353 Q8K358 Q8K377 Q8K385 Q8K3A6 Q8K3C0
Q8K3D6 Q8K3F6 Q8K3J9 Q8K3K7 Q8K3L6 Q8K3P1 Q8K3P6 Q8K3T4 Q8K3V3 Q8K3V5 Q8K3V7
Q8K3Y7 Q8K3Z9 Q8K404 Q8K410 Q8K415 Q8K418 Q8K424 Q8K440 Q8K441 Q8K442 Q8K445
Q8K448 Q8K449 Q8K451 Q8K458 Q8K467 Q8K480 Q8K4B4 Q8K4B7 Q8K4C0 Q8K4C2 Q8K4C8
Q8K4D3 Q8K4D6 Q8K4F0 Q8K4G2 Q8K4G9 Q8K4P7 Q8K4Q8 Q8K4Q9 Q8K4R8 Q8K4S3 Q8K4T1
Q8K4V2 Q8K4X7 Q8K4Y7 Q8K4Z2 Q8K4Z4 Q8K4Z6 Q8K558 Q8K592 Q8K593 Q8K595 Q8K596
Q8K5A9 Q8K5B1 Q8K5E0 Q8K902 Q8K903 Q8K911 Q8K914 Q8K915 Q8K942 Q8K969 Q8K984
Q8K985 Q8K987 Q8K993 Q8K994 Q8K995 Q8K996 Q8K997 Q8K999 Q8K9B1 Q8K9B4 Q8K9D5
Q8K9G8 Q8K9G9 Q8K9I8 Q8K9J0 Q8K9L3 Q8K9L4 Q8K9M0 Q8K9M1 Q8K9M4 Q8K9M7 Q8K9M9
Q8K9N7 Q8K9N8 Q8K9P8 Q8K9P9 Q8K9Q4 Q8K9R0 Q8K9S0 Q8K9S1 Q8K9S5 Q8K9T3 Q8K9X4
Q8K9X5 Q8K9X6 Q8K9X7 Q8K9X8 Q8K9X9 Q8K9Y1 Q8K9Y7 Q8K9Z3 Q8KA02 Q8KA08 Q8KA12
Q8KA17 Q8KA18 Q8KA19 Q8KA21 Q8KA32 Q8KA35 Q8KA36 Q8KA37 Q8KA45 Q8KA52 Q8KA64
Q8KAQ4 Q8KAT2 Q8KBH7 Q8KBK0 Q8KBX2 Q8KC14 Q8KCC4 Q8KCT5 Q8KD14 Q8KDG0 Q8KDL8
Q8KDT8 Q8KDU6 Q8KDV4 Q8KE45 Q8KEB6 Q8KEB9 Q8KEC3 Q8KEF5 Q8KFE9 Q8KFJ7 Q8KG75
Q8KGD1 Q8KGE7 Q8KGE9 Q8KGG2 Q8KHU8 Q8KKH0 Q8KLH5 Q8KPP0 Q8KPP1 Q8KPP2 Q8KPP3
Q8KPP6 Q8KQC3 Q8KR68 Q8KRV2 Q8KRV3 Q8KS19 Q8KSV0 Q8KTA9 Q8KTB5 Q8KTC0 Q8KU73
Q8KWS7 Q8KWT2 Q8KX34 Q8KX40 Q8KX53 Q8KX56 Q8KX60 Q8KY01 Q8KY50 Q8KYH8 Q8L171
Q8L1D5 Q8L2J7 Q8L3A1 Q8L3K8 Q8L481 Q8L4B0 Q8L4H4 Q8L4K5 Q8L4R0 Q8L4S2 Q8L4X4
Q8L4Y2 Q8L540 Q8L586 Q8L5Y5 Q8L615 Q8L616 Q8L636 Q8L638 Q8L6Z8 Q8L706 Q8L707
Q8L709 Q8L710 Q8L718 Q8L725 Q8L731 Q8L778 Q8L783 Q8L7A0 Q8L7C8 Q8L7D0 Q8L7D5
Q8L7E3 Q8L7E5 Q8L7E9 Q8L7F9 Q8L7G3 Q8L7H8 Q8L7M0 Q8L7M1 Q8L7N4 Q8L7Q7 Q8L7R3
Q8L7R5 Q8L7R8 Q8L7V3 Q8L7W1 Q8L7W5 Q8L7Z0 Q8L7Z9 Q8L856 Q8L883 Q8L884 Q8L899
Q8L8U9 Q8L8W0 Q8L8Z1 Q8L924 Q8L928 Q8L970 Q8L986 Q8L9B5 Q8L9C4 Q8L9G7 Q8L9J7
Q8L9S0 Q8L9T5 Q8L9W3 Q8LAA6 Q8LAI1 Q8LAL0 Q8LAN3 Q8LB08 Q8LB17 Q8LBA0 Q8LBA6
Q8LBB5 Q8LBF7 Q8LBI9 Q8LBL1 Q8LBP4 Q8LBX7 Q8LC45 Q8LC69 Q8LCA1 Q8LCL8 Q8LCQ4
Q8LD38 Q8LD98 Q8LDB8 Q8LDD3 Q8LDG7 Q8LDL0 Q8LDS3 Q8LE26 Q8LE59 Q8LE84 Q8LEB6
Q8LED0 Q8LED3 Q8LEE7 Q8LEJ9 Q8LEK2 Q8LEQ4 Q8LES0 Q8LET2 Q8LF05 Q8LF65 Q8LFD1

Q8LFH5 Q8LFN2 Q8LFP1 Q8LFP7 Q8LFP9 Q8LFY8 Q8LG30 Q8LG50 Q8LG53 Q8LG88 Q8LGA5
Q8LGE9 Q8LGN0 Q8LGN1 Q8LGU1 Q8LIB3 Q8LIF2 Q8LIN5 Q8LIY0 Q8LKI3 Q8LKV5 Q8LKZ1
Q8LM92 Q8LPB4 Q8LPF4 Q8LPF8 Q8LPG1 Q8LPK0 Q8LPK2 Q8LPK5 Q8LPL2 Q8LPL8 Q8LPQ5
Q8LPQ6 Q8LPQ8 Q8LPR9 Q8LPS2 Q8LPS5 Q8LPT1 Q8LPT3 Q8LQ68 Q8LQX2 Q8LR09 Q8LTZ5
Q8LV29 Q8LW54 Q8LW83 Q8LWM2 Q8LWM4 Q8LWN0 Q8LWN4 Q8LWP6 Q8LWS1 Q8LWT4 Q8LWU4
Q8LWU8 Q8LWW8 Q8LX23 Q8LX26 Q8LX27 Q8LX28 Q8LX30 Q8LX31 Q8LX40 Q8LX57 Q8LX87
Q8LZ88 Q8LZ94 Q8LZ97 Q8M0A6 Q8M0E7 Q8M0K7 Q8M0K9 Q8M154 Q8M354 Q8M3J6 Q8M4C6
Q8M4C7 Q8M4D6 Q8M4E2 Q8M4E6 Q8M5M3 Q8M6Z3 Q8M700 Q8M703 Q8M706 Q8M708 Q8M867
Q8M873 Q8M880 Q8M881 Q8M888 Q8M889 Q8M891 Q8M892 Q8M893 Q8M894 Q8M896 Q8M897
Q8M899 Q8M9T2 Q8M9T6 Q8M9T8 Q8M9T9 Q8M9U1 Q8M9U5 Q8M9V9 Q8M9W0 Q8M9W3 Q8M9W4
Q8M9W5 Q8M9W6 Q8M9W8 Q8M9W9 Q8M9X0 Q8M9X2 Q8M9X3 Q8M9X4 Q8M9X5 Q8M9X8 Q8M9Y3
Q8M9Y4 Q8M9Y5 Q8M9Z0 Q8M9Z1 Q8M9Z2 Q8M9Z3 Q8M9Z4 Q8M9Z5 Q8M9Z7 Q8M9Z8 Q8M9Z9
Q8MA00 Q8MA06 Q8MA07 Q8MA08 Q8MA13 Q8MA15 Q8MA16 Q8MAV7 Q8MAW2 Q8MAW5 Q8MFA2
Q8MFA3 Q8MH63 Q8MHW5 Q8MHY9 Q8MI01 Q8MI05 Q8MI28 Q8MIA3 Q8MIB3 Q8MIB6 Q8MII5
Q8MII8 Q8MIQ9 Q8MIS5 Q8MIT8 Q8MIT9 Q8MIU0 Q8MJ02 Q8MJ08 Q8MJ88 Q8MJ89 Q8MJD7
Q8MJG0 Q8MJH1 Q8MJI6 Q8MJJ2 Q8MJN0 Q8MJS1 Q8MJU6 Q8MJV2 Q8MJV3 Q8MJW8 Q8MJZ2
Q8MJZ7 Q8MK44 Q8MK48 Q8MKG8 Q8MKK4 Q8MLV1 Q8MMJ7 Q8MNZ1 Q8MPP0 Q8MPP3 Q8MRC9
Q8MRQ4 Q8MS59 Q8MSF5 Q8MSU3 Q8MT80 Q8MTI2 Q8MUJ1 Q8MV48 Q8MVS5 Q8MXJ9 Q8MXL9
Q8MXN7 Q8MXQ3 Q8MXQ7 Q8MXY0 Q8MY85 Q8MY86 Q8MYY6 Q8MZC4 Q8MZR5 Q8MZR6 Q8N0U2
Q8N0U8 Q8N0V5 Q8N0W4 Q8N0W7 Q8N0Y3 Q8N0Y5 Q8N0Z9 Q8N109 Q8N111 Q8N112 Q8N114
Q8N118 Q8N126 Q8N127 Q8N130 Q8N131 Q8N138 Q8N139 Q8N144 Q8N146 Q8N148 Q8N149
Q8N162 Q8N1C3 Q8N1L4 Q8N1M1 Q8N1N0 Q8N1N2 Q8N1S5 Q8N1Y9 Q8N201 Q8N205 Q8N271
Q8N292 Q8N2A8 Q8N2C7 Q8N2F6 Q8N2H4 Q8N2K0 Q8N2K1 Q8N2M4 Q8N2Q7 Q8N2U0 Q8N2U9
Q8N326 Q8N349 Q8N350 Q8N357 Q8N370 Q8N386 Q8N387 Q8N394 Q8N3F9 Q8N3G9 Q8N3J6
Q8N3S3 Q8N3T1 Q8N3T6 Q8N3Y3 Q8N3Y7 Q8N413 Q8N423 Q8N428 Q8N434 Q8N441 Q8N468
Q8N4A0 Q8N4C9 Q8N4F4 Q8N4F7 Q8N4H5 Q8N4K4 Q8N4L1 Q8N4L2 Q8N4L4 Q8N4M1 Q8N4S7
Q8N4S9 Q8N4V1 Q8N4V2 Q8N4W6 Q8N511 Q8N539 Q8N5B7 Q8N5C1 Q8N5D6 Q8N5G0 Q8N5G2
Q8N5K1 Q8N5M9 Q8N5S1 Q8N5U1 Q8N5Y8 Q8N608 Q8N609 Q8N614 Q8N628 Q8N661 Q8N682
Q8N695 Q8N697 Q8N6C5 Q8N6D2 Q8N6F1 Q8N6G5 Q8N6I4 Q8N6K0 Q8N6L0 Q8N6L1 Q8N6L7
Q8N6M3 Q8N6P7 Q8N6Q1 Q8N6R1 Q8N6S5 Q8N6U8 Q8N6Y1 Q8N743 Q8N755 Q8N766 Q8N7C0
Q8N7C4 Q8N7C7 Q8N7P1 Q8N7P3 Q8N7S6 Q8N7X8 Q8N808 Q8N816 Q8N8D7 Q8N8F6 Q8N8F7
Q8N8J7 Q8N8N0 Q8N8Q1 Q8N8Q8 Q8N8Q9 Q8N8R3 Q8N8V8 Q8N8Z6 Q8N912 Q8N966 Q8N967
Q8N9A8 Q8N9F0 Q8N9F7 Q8N9I0 Q8N9I5 Q8N9M5 Q8N9R8 Q8N9W7 Q8N9X5 Q8NA29 Q8NA58
Q8NAC3 Q8NAN2 Q8NAT1 Q8NAU1 Q8NB49 Q8NB59 Q8NBD8 Q8NBF6 Q8NBI2 Q8NBI5 Q8NBI6
Q8NBJ4 Q8NBJ9 Q8NBL3 Q8NBM4 Q8NBN3 Q8NBP5 Q8NBQ7 Q8NBR0 Q8NBS3 Q8NBT3 Q8NBU5
Q8NBV4 Q8NBV8 Q8NBW4 Q8NBZ7 Q8NC01 Q8NC24 Q8NC42 Q8NC44 Q8NC54 Q8NC56 Q8NC67
Q8NCC5 Q8NCG5 Q8NCG7 Q8NCH0 Q8NCK7 Q8NCL4 Q8NCL8 Q8NCL9 Q8NCM2 Q8NCQ3 Q8NCR0
Q8NCR9 Q8NCS4 Q8NCS7 Q8NCU8 Q8NCW0 Q8NCW6 Q8ND61 Q8ND94 Q8NDB6 Q8NDH2 Q8NDV1
Q8NDV2 Q8NDX2 Q8NDY8 Q8NDZ6 Q8NE00 Q8NE01 Q8NE79 Q8NE86 Q8NEA5 Q8NEB5 Q8NEC5
Q8NEN9 Q8NEQ5 Q8NER1 Q8NER5 Q8NES3 Q8NET5 Q8NET6 Q8NET8 Q8NEW0 Q8NEW7 Q8NF37
Q8NF91 Q8NFB2 Q8NFF2 Q8NFJ5 Q8NFJ6 Q8NFK1 Q8NFL0 Q8NFM4 Q8NFM7 Q8NFN8 Q8NFQ8
Q8NFR3 Q8NFR9 Q8NFT2 Q8NFT8 Q8NFU0 Q8NFU1 Q8NFY4 Q8NFZ3 Q8NFZ4 Q8NFZ6 Q8NFZ8
Q8NG04 Q8NG11 Q8NG75 Q8NG76 Q8NG77 Q8NG78 Q8NG80 Q8NG81 Q8NG83 Q8NG84 Q8NG85
Q8NG92 Q8NG94 Q8NG95 Q8NG97 Q8NG98 Q8NG99 Q8NGA0 Q8NGA1 Q8NGA2 Q8NGA4 Q8NGA5

Q8NGA6 Q8NGA8 Q8NGB2 Q8NGB4 Q8NGB6 Q8NGB8 Q8NGB9 Q8NGC0 Q8NGC1 Q8NGC2 Q8NGC3
Q8NGC4 Q8NGC5 Q8NGC6 Q8NGC7 Q8NGC8 Q8NGC9 Q8NGD0 Q8NGD1 Q8NGD2 Q8NGD3 Q8NGD4
Q8NGD5 Q8NGE0 Q8NGE1 Q8NGE2 Q8NGE3 Q8NGE5 Q8NGE7 Q8NGE8 Q8NGE9 Q8NGF0 Q8NGF1
Q8NGF3 Q8NGF4 Q8NGF6 Q8NGF7 Q8NGF8 Q8NGF9 Q8NGG0 Q8NGG1 Q8NGG2 Q8NGG3 Q8NGG4
Q8NGG5 Q8NGG6 Q8NGG7 Q8NGG8 Q8NGH3 Q8NGH5 Q8NGH6 Q8NGH7 Q8NGH8 Q8NGH9 Q8NGI0
Q8NGI1 Q8NGI2 Q8NGI3 Q8NGI4 Q8NGI6 Q8NGI7 Q8NGI8 Q8NGI9 Q8NGJ0 Q8NGJ1 Q8NGJ2
Q8NGJ3 Q8NGJ4 Q8NGJ5 Q8NGJ6 Q8NGJ7 Q8NGJ8 Q8NGJ9 Q8NGK0 Q8NGK1 Q8NGK2 Q8NGK3
Q8NGK4 Q8NGK5 Q8NGK6 Q8NGK9 Q8NGL0 Q8NGL1 Q8NGL2 Q8NGL3 Q8NGL4 Q8NGL6 Q8NGL7
Q8NGL9 Q8NGM1 Q8NGM8 Q8NGM9 Q8NGN0 Q8NGN1 Q8NGN2 Q8NGN3 Q8NGN4 Q8NGN5 Q8NGN6
Q8NGN7 Q8NGN8 Q8NGP0 Q8NGP2 Q8NGP3 Q8NGP4 Q8NGP6 Q8NGP8 Q8NGP9 Q8NGQ1 Q8NGQ2
Q8NGQ3 Q8NGQ4 Q8NGQ5 Q8NGQ6 Q8NGR1 Q8NGR2 Q8NGR3 Q8NGR4 Q8NGR5 Q8NGR6 Q8NGR8
Q8NGR9 Q8NGS0 Q8NGS1 Q8NGS2 Q8NGS3 Q8NGS4 Q8NGS5 Q8NGS6 Q8NGS7 Q8NGS8 Q8NGS9
Q8NGT0 Q8NGT1 Q8NGT2 Q8NGT5 Q8NGT7 Q8NGT9 Q8NGU1 Q8NGU2 Q8NGU4 Q8NGU9 Q8NGV0
Q8NGV5 Q8NGV6 Q8NGV7 Q8NGW1 Q8NGW6 Q8NGX0 Q8NGX1 Q8NGX2 Q8NGX3 Q8NGX5 Q8NGX6
Q8NGX8 Q8NGX9 Q8NGY0 Q8NGY1 Q8NGY2 Q8NGY3 Q8NGY5 Q8NGY6 Q8NGY7 Q8NGY9 Q8NGZ0
Q8NGZ2 Q8NGZ3 Q8NGZ4 Q8NGZ5 Q8NGZ6 Q8NGZ9 Q8NH00 Q8NH01 Q8NH02 Q8NH03 Q8NH04
Q8NH05 Q8NH06 Q8NH07 Q8NH08 Q8NH09 Q8NH10 Q8NH16 Q8NH18 Q8NH19 Q8NH21 Q8NH37
Q8NH40 Q8NH41 Q8NH42 Q8NH43 Q8NH48 Q8NH49 Q8NH50 Q8NH51 Q8NH53 Q8NH54 Q8NH55
Q8NH56 Q8NH57 Q8NH59 Q8NH60 Q8NH61 Q8NH63 Q8NH64 Q8NH67 Q8NH69 Q8NH70 Q8NH72
Q8NH73 Q8NH74 Q8NH76 Q8NH79 Q8NH80 Q8NH81 Q8NH83 Q8NH85 Q8NH87 Q8NH89 Q8NH90
Q8NH92 Q8NH93 Q8NH94 Q8NH95 Q8NHA4 Q8NHA6 Q8NHA8 Q8NHB1 Q8NHB7 Q8NHB8 Q8NHC4
Q8NHC5 Q8NHC6 Q8NHC7 Q8NHC8 Q8NHE4 Q8NHH9 Q8NHJ6 Q8NHK3 Q8NHL6 Q8NHP6 Q8NHS1
Q8NHS3 Q8NHU3 Q8NHV5 Q8NHX9 Q8NHY0 Q8NI17 Q8NI28 Q8NI60 Q8NIP8 Q8NIQ1 Q8NJ57
Q8NJ59 Q8NKE2 Q8NKG8 Q8NKG9 Q8NKS6 Q8NKW5 Q8NKX1 Q8NL19 Q8NL44 Q8NLB7 Q8NLK2
Q8NLN8 Q8NLQ9 Q8NLR0 Q8NLR5 Q8NMM8 Q8NMM9 Q8NN75 Q8NNI8 Q8NNJ7 Q8NNK2 Q8NNK3
Q8NNK5 Q8NNM0 Q8NNN2 Q8NNU2 Q8NP09 Q8NP53 Q8NPB7 Q8NPZ2 Q8NQ66 Q8NQ73 Q8NQC3
Q8NQC8 Q8NQE4 Q8NQG6 Q8NQK0 Q8NRD0 Q8NRZ6 Q8NS49 Q8NS93 Q8NSD6 Q8NSG3 Q8NSP8
Q8NSS8 Q8NTW4 Q8NU99 Q8NUH7 Q8NUI7 Q8NUQ2 Q8NUQ4 Q8NUQ9 Q8NUR5 Q8NUS2 Q8NUV7
Q8NUZ4 Q8NV31 Q8NV46 Q8NV48 Q8NVE1 Q8NVI1 Q8NVI2 Q8NVI3 Q8NVK5 Q8NVK6 Q8NVS5
Q8NVT9 Q8NVV4 Q8NW03 Q8NW04 Q8NW09 Q8NW47 Q8NWB3 Q8NWF3 Q8NWM5 Q8NWQ5 Q8NWQ6
Q8NWR2 Q8NWT4 Q8NWU7 Q8NWY8 Q8NWZ4 Q8NX36 Q8NX64 Q8NX68 Q8NXB8 Q8NXC4 Q8NXF6
Q8NXL8 Q8NXQ4 Q8NXR5 Q8NXS9 Q8NXT0 Q8NXT1 Q8NXT2 Q8NXW9 Q8NYA5 Q8NYB0 Q8NYF3
Q8NYF4 Q8NYF5 Q8NYG6 Q8NYH2 Q8NYJ8 Q8NYK8 Q8NYM1 Q8NYZ3 Q8NZB3 Q8NZY2 Q8P000
Q8P0C8 Q8P171 Q8P1G4 Q8P1K8 Q8P1M3 Q8P276 Q8P2D4 Q8P2D8 Q8P2K0 Q8P2K3 Q8P2P8
Q8P2T7 Q8P338 Q8P3H8 Q8P3H9 Q8P3J6 Q8P487 Q8P4E6 Q8P532 Q8P5J5 Q8P5M6 Q8P626
Q8P6E6 Q8P6T5 Q8P6V1 Q8P7T9 Q8P8F0 Q8P8L7 Q8P8R8 Q8P8W4 Q8P8Z6 Q8P993 Q8P9Z2
Q8PA33 Q8PA99 Q8PAB7 Q8PB24 Q8PBG5 Q8PC32 Q8PC75 Q8PCE8 Q8PCJ7 Q8PCK2 Q8PCK6
Q8PCM0 Q8PCM1 Q8PCZ9 Q8PDD7 Q8PDE8 Q8PDW1 Q8PDZ9 Q8PEH7 Q8PEX2 Q8PEX3 Q8PEZ6
Q8PFU4 Q8PG07 Q8PGG3 Q8PGZ1 Q8PH20 Q8PHE9 Q8PHV1 Q8PI34 Q8PI51 Q8PJ47 Q8PJX8
Q8PK49 Q8PKQ5 Q8PKS5 Q8PKX1 Q8PKZ5 Q8PL00 Q8PLR9 Q8PLW2 Q8PLY9 Q8PM10 Q8PMS3
Q8PN18 Q8PNQ6 Q8PNV2 Q8PP47 Q8PPB0 Q8PPC8 Q8PPC9 Q8PPR7 Q8PQA7 Q8PQD5 Q8PQI6
Q8PQT0 Q8PQW6 Q8PSE5 Q8PU58 Q8PU59 Q8PU61 Q8PV96 Q8PVB4 Q8PVI7 Q8PVW4 Q8PX50
Q8PXI2 Q8PY55 Q8PYN2 Q8PYN3 Q8PYZ7 Q8PYZ8 Q8PZ51 Q8Q0J5 Q8QFN3 Q8QFP1 Q8QFP9
Q8QFV0 Q8QFX6 Q8QGH3 Q8QHJ9 Q8QHL3 Q8QL14 Q8QL22 Q8QL30 Q8QL32 Q8QL45 Q8QL47

Q8QL52 Q8QMS5 Q8QPK2 Q8QPL1 Q8QV59 Q8QVV0 Q8QZ72 Q8QZR0 Q8QZS5 Q8QZT4 Q8QZV2
Q8QZW3 Q8QZW7 Q8QZY6 Q8R000 Q8R007 Q8R043 Q8R070 Q8R079 Q8R087 Q8R090 Q8R092
Q8R0C3 Q8R0G7 Q8R0I0 Q8R0I4 Q8R0J4 Q8R0M8 Q8R0N9 Q8R0S9 Q8R0T6 Q8R0W6 Q8R0X7
Q8R0Y8 Q8R0Z5 Q8R100 Q8R115 Q8R138 Q8R139 Q8R143 Q8R173 Q8R182 Q8R183 Q8R189
Q8R191 Q8R1C0 Q8R1C3 Q8R1E7 Q8R1I1 Q8R1J1 Q8R1K1 Q8R1L4 Q8R1P5 Q8R1R5 Q8R1S9
Q8R1T4 Q8R1V4 Q8R1W2 Q8R1Z9 Q8R207 Q8R218 Q8R235 Q8R239 Q8R2E6 Q8R2F2 Q8R2H2
Q8R2H3 Q8R2N1 Q8R2Q6 Q8R2Q8 Q8R2R1 Q8R2S8 Q8R2Y0 Q8R2Y2 Q8R2Y3 Q8R2Z3 Q8R310
Q8R314 Q8R366 Q8R373 Q8R3G9 Q8R3H7 Q8R3I2 Q8R3I9 Q8R3L0 Q8R3L5 Q8R3N2 Q8R3Q0
Q8R3R5 Q8R3S2 Q8R3U1 Q8R400 Q8R404 Q8R411 Q8R412 Q8R415 Q8R416 Q8R420 Q8R428
Q8R429 Q8R454 Q8R455 Q8R456 Q8R464 Q8R493 Q8R4C1 Q8R4C4 Q8R4C5 Q8R4C6 Q8R4D1
Q8R4D5 Q8R4E1 Q8R4F0 Q8R4G1 Q8R4G6 Q8R4G9 Q8R4H9 Q8R4I7 Q8R4P4 Q8R4P5 Q8R4P9
Q8R4S8 Q8R4T9 Q8R4U0 Q8R4V1 Q8R4X1 Q8R4Y0 Q8R4Y4 Q8R4Z9 Q8R500 Q8R502 Q8R507
Q8R508 Q8R523 Q8R526 Q8R534 Q8R553 Q8R5I0 Q8R5J9 Q8R5M3 Q8R5M8 Q8R5S4 Q8R5T5
Q8R664 Q8R666 Q8R6A3 Q8R6K6 Q8R6L9 Q8R7S2 Q8R8I5 Q8R8I6 Q8R8I7 Q8R936 Q8R980
Q8R9G3 Q8R9J2 Q8R9R0 Q8RA57 Q8RB29 Q8RC19 Q8RCN8 Q8RCX1 Q8RDQ0 Q8RFE0 Q8RFY9
Q8RFZ3 Q8RGD7 Q8RGD8 Q8RH46 Q8RHA4 Q8RHJ2 Q8RIA6 Q8RII2 Q8RKH1 Q8RKH4 Q8RKT5
Q8RPN9 Q8RQL4 Q8RQL5 Q8RQM4 Q8RQS4 Q8RR17 Q8RSW0 Q8RSW1 Q8RSW2 Q8RSW3 Q8RU61
Q8RUN1 Q8RUS5 Q8RV04 Q8RVC7 Q8RVL1 Q8RVL2 Q8RWA5 Q8RWA7 Q8RWB7 Q8RWE9 Q8RWF4
Q8RWF5 Q8RWG3 Q8RWH8 Q8RWI9 Q8RWL5 Q8RWM6 Q8RWM7 Q8RWN2 Q8RWQ5 Q8RWS1 Q8RWS6
Q8RWS9 Q8RWU6 Q8RWW0 Q8RWW1 Q8RWW7 Q8RWZ1 Q8RWZ5 Q8RWZ6 Q8RWZ7 Q8RX29 Q8RX55
Q8RX63 Q8RX67 Q8RX77 Q8RX80 Q8RX83 Q8RX88 Q8RXE1 Q8RXE8 Q8RXF8 Q8RXK1 Q8RXL7
Q8RXL8 Q8RXN0 Q8RXN3 Q8RXP6 Q8RXQ1 Q8RXQ2 Q8RXR2 Q8RXS1 Q8RXW0 Q8RXX9 Q8RXZ9
Q8RY00 Q8RY17 Q8RY46 Q8RY60 Q8RY65 Q8RY67 Q8RY74 Q8RY81 Q8RY83 Q8RYL0 Q8RYL1
Q8RZQ8 Q8RZV7 Q8S0F0 Q8S1N1 Q8S1X7 Q8S1X8 Q8S1X9 Q8S230 Q8S233 Q8S237 Q8S2A7
Q8S339 Q8S396 Q8S397 Q8S3C0 Q8S3C1 Q8S3J3 Q8S3R1 Q8S3R2 Q8S3W4 Q8S403 Q8S4F6
Q8S626 Q8S628 Q8S7W0 Q8S8A0 Q8S8K8 Q8S8K9 Q8S8M0 Q8S8N4 Q8S8N9 Q8S8P3 Q8S8Q6
Q8S8S1 Q8S8S2 Q8S8S3 Q8S8T4 Q8S8T8 Q8S8U2 Q8S8U5 Q8S8U6 Q8S8U7 Q8S8U8 Q8S8V0
Q8S8W0 Q8S8W1 Q8S8W3 Q8S8W4 Q8S8W5 Q8S8W6 Q8S8X0 Q8S8X4 Q8S8X5 Q8S8X7 Q8S8X8
Q8S8Y2 Q8S8Y4 Q8S9J0 Q8S9K7 Q8S9L6 Q8SAA5 Q8SE00 Q8SE72 Q8SE77 Q8SE89 Q8SEH7
Q8SEJ1 Q8SEL4 Q8SEM9 Q8SEP4 Q8SET9 Q8SEW8 Q8SFK5 Q8SFK7 Q8SG72 Q8SGA0 Q8SGQ9
Q8SHP7 Q8SHP8 Q8SJ54 Q8SJB6 Q8SJG7 Q8SJH9 Q8SJJ1 Q8SJK5 Q8SJK6 Q8SJK7 Q8SJK8
Q8SJK9 Q8SJL1 Q8SJL2 Q8SJL3 Q8SJX4 Q8SJX5 Q8SJZ1 Q8SJZ2 Q8SJZ3 Q8SJZ4 Q8SL87
Q8SL89 Q8SL91 Q8SL93 Q8SL95 Q8SPE8 Q8SPE9 Q8SPF0 Q8SPI1 Q8SPI4 Q8SPI5 Q8SPJ9
Q8SPK0 Q8SPK1 Q8SPN1 Q8SPN2 Q8SPP9 Q8SPQ7 Q8SPR2 Q8SPS2 Q8SPU6 Q8SPU7 Q8SPV8
Q8SPV9 Q8SPW1 Q8SPW2 Q8SPZ1 Q8SQ19 Q8SQ20 Q8SQ34 Q8SQ54 Q8SQ72 Q8SQ78 Q8SQ79
Q8SQ80 Q8SQ81 Q8SQA4 Q8SQB8 Q8SQC1 Q8SQH5 Q8SQI5 Q8SQU5 Q8SRA2 Q8SRK2 Q8SRW9
Q8SS86 Q8SSI7 Q8ST66 Q8ST87 Q8STK5 Q8SUE5 Q8SUF9 Q8SUG0 Q8SUG7 Q8SUL4 Q8SUS0
Q8SV25 Q8SVN2 Q8SVN8 Q8SVP6 Q8SVT0 Q8SW90 Q8SWK6 Q8SWR3 Q8SWV6 Q8SX37 Q8SXQ7
Q8SXY6 Q8SY72 Q8SYV9 Q8SZ63 Q8T062 Q8T0B1 Q8T0Z3 Q8T197 Q8T2Q0 Q8T2Q6 Q8T2T5
Q8T3C8 Q8T5G8 Q8T5S1 Q8T5Z7 Q8T673 Q8T674 Q8T675 Q8T683 Q8T685 Q8T686 Q8T689
Q8T690 Q8T6H3 Q8T6H8 Q8T6J0 Q8T6J1 Q8T6J2 Q8T6J5 Q8T8L8 Q8T8P3 Q8T9W1 Q8T9W2
Q8T9W4 Q8TAA9 Q8TAB3 Q8TAC9 Q8TAD4 Q8TAE7 Q8TAF8 Q8TAQ9 Q8TAV4 Q8TAX9 Q8TAZ6
Q8TB36 Q8TB61 Q8TB68 Q8TB96 Q8TBA6 Q8TBB6 Q8TBE1 Q8TBE3 Q8TBE7 Q8TBF5 Q8TBG9
Q8TBJ4 Q8TBM7 Q8TBM8 Q8TBP5 Q8TBP6 Q8TBQ9 Q8TBR7 Q8TC12 Q8TC26 Q8TC27 Q8TC36

Q8TC41 Q8TCB6 Q8TCC7 Q8TCG1 Q8TCG5 Q8TCJ2 Q8TCP9 Q8TCQ1 Q8TCT6 Q8TCT7 Q8TCT8
Q8TCT9 Q8TCU3 Q8TCU5 Q8TCW7 Q8TCW9 Q8TCY5 Q8TCZ2 Q8TD07 Q8TD20 Q8TD22 Q8TD43
Q8TD46 Q8TD84 Q8TDB4 Q8TDB8 Q8TDD5 Q8TDF5 Q8TDI7 Q8TDI8 Q8TDN1 Q8TDN2 Q8TDN7
Q8TDQ0 Q8TDQ1 Q8TDS4 Q8TDS5 Q8TDS7 Q8TDT2 Q8TDU5 Q8TDU6 Q8TDU9 Q8TDV0 Q8TDV2
Q8TDV5 Q8TDW0 Q8TDW4 Q8TDW7 Q8TDX6 Q8TDX9 Q8TDY8 Q8TE23 Q8TE54 Q8TE99 Q8TEB7
Q8TEB9 Q8TED1 Q8TED4 Q8TEF2 Q8TEM1 Q8TEQ8 Q8TEY5 Q8TEZ7 Q8TF08 Q8TF62 Q8TF66
Q8TF71 Q8TF80 Q8TFD3 Q8TFG0 Q8TFG1 Q8TFG3 Q8TFG8 Q8TFH2 Q8TFH3 Q8TFH6 Q8TGB2
Q8TGG5 Q8TGG8 Q8TGH6 Q8TGJ3 Q8TGK5 Q8TGM9 Q8TGN7 Q8TGN9 Q8TGP9 Q8TGQ0 Q8TGQ7
Q8TGR3 Q8TGR7 Q8TGS0 Q8TGS1 Q8TGS7 Q8TGT2 Q8TGT4 Q8TGT9 Q8TGU2 Q8TGU7 Q8THH5
Q8TI84 Q8TIQ3 Q8TIQ4 Q8TJ52 Q8TJ54 Q8TJA8 Q8TJA9 Q8TJN1 Q8TKT8 Q8TL61 Q8TMG0
Q8TP15 Q8TPA6 Q8TQP1 Q8TS44 Q8TS66 Q8TSG7 Q8TSV8 Q8TSX9 Q8TSY0 Q8TSY1 Q8TSY2
Q8TSY3 Q8TTZ9 Q8TU00 Q8TU01 Q8TU02 Q8TU03 Q8TU04 Q8TU05 Q8TUT2 Q8TVA7 Q8TW25
Q8TW26 Q8TWG2 Q8TWM4 Q8TWM5 Q8TWS9 Q8TX54 Q8TY29 Q8TYT4 Q8TYX0 Q8TZH7 Q8TZK2
Q8TZM7 Q8U019 Q8U0T6 Q8U1H3 Q8U1S0 Q8U1Z6 Q8U230 Q8U271 Q8U3P6 Q8U3Q8 Q8U400
Q8U403 Q8U4B4 Q8U4B5 Q8U4D2 Q8U4K4 Q8U526 Q8U527 Q8U530 Q8U7I2 Q8U9D8 Q8U9D9
Q8U9E0 Q8UB30 Q8UBM5 Q8UBX8 Q8UC06 Q8UDB4 Q8UDF5 Q8UDM5 Q8UEM1 Q8UEP9 Q8UEQ0
Q8UFC8 Q8UFL7 Q8UFP1 Q8UFU1 Q8UFX0 Q8UG67 Q8UGH9 Q8UGR1 Q8UHB2 Q8UHH1 Q8UHY0
Q8UI24 Q8UIB3 Q8UID7 Q8UJ37 Q8UJL0 Q8UJW4 Q8UKE4 Q8UN02 Q8UUG8 Q8UUY8 Q8UVJ6
Q8UVJ7 Q8UVR8 Q8UVX8 Q8UWF0 Q8UWJ4 Q8UZB4 Q8UZK5 Q8V0N6 Q8V3T9 Q8V3U0 Q8V433
Q8V436 Q8V437 Q8V4U9 Q8V4Y0 Q8V5E0 Q8V790 Q8V9S1 Q8V9T2 Q8V9T3 Q8V9T7 Q8V9U2
Q8VA10 Q8VBS7 Q8VBT0 Q8VBT3 Q8VBV9 Q8VBW1 Q8VBW9 Q8VBX4 Q8VBZ3 Q8VC04 Q8VC26
Q8VC48 Q8VC49 Q8VC65 Q8VC69 Q8VC74 Q8VC82 Q8VC90 Q8VCA0 Q8VCA5 Q8VCA6 Q8VCB1
Q8VCD3 Q8VCD6 Q8VCF0 Q8VCF1 Q8VCF5 Q8VCH2 Q8VCH6 Q8VCH7 Q8VCH9 Q8VCJ6 Q8VCK6
Q8VCL2 Q8VCL5 Q8VCM5 Q8VCM8 Q8VCN6 Q8VCP9 Q8VCQ6 Q8VCR3 Q8VCS3 Q8VCV9 Q8VCW4
Q8VCX2 Q8VCX5 Q8VCY8 Q8VCZ2 Q8VD00 Q8VD26 Q8VD31 Q8VD53 Q8VD57 Q8VD58 Q8VD98
Q8VDA1 Q8VDB2 Q8VDB9 Q8VDI9 Q8VDN2 Q8VDP6 Q8VDR5 Q8VDS8 Q8VDT1 Q8VDX6 Q8VDZ4
Q8VE49 Q8VE91 Q8VE96 Q8VE98 Q8VEA7 Q8VEC3 Q8VEC4 Q8VEF1 Q8VEG4 Q8VEH0 Q8VEK0
Q8VEK2 Q8VEL0 Q8VEM1 Q8VEM8 Q8VES2 Q8VEW2 Q8VEW5 Q8VEW6 Q8VEX5 Q8VEX6 Q8VEY3
Q8VEZ0 Q8VF12 Q8VF13 Q8VF65 Q8VF66 Q8VF76 Q8VFB9 Q8VFC9 Q8VFD0 Q8VFD1 Q8VFD2
Q8VFD3 Q8VFK1 Q8VFK2 Q8VFK7 Q8VFL5 Q8VFL9 Q8VFM9 Q8VFR8 Q8VFT4 Q8VFV4 Q8VFX2
Q8VG02 Q8VG03 Q8VG04 Q8VG05 Q8VG06 Q8VG07 Q8VG08 Q8VG09 Q8VG13 Q8VG42 Q8VG43
Q8VG44 Q8VGD6 Q8VGI1 Q8VGI4 Q8VGI5 Q8VGI6 Q8VGK5 Q8VGQ7 Q8VGR8 Q8VGR9 Q8VGS1
Q8VGS3 Q8VH49 Q8VHB5 Q8VHD6 Q8VHD7 Q8VHE0 Q8VHF2 Q8VHG0 Q8VHH7 Q8VHI8 Q8VHJ4
Q8VHK5 Q8VHK6 Q8VHK8 Q8VHL0 Q8VHN2 Q8VHN7 Q8VHP6 Q8VHR0 Q8VHS2 Q8VHV8 Q8VHW1
Q8VHW2 Q8VHW3 Q8VHW4 Q8VHW5 Q8VHW7 Q8VHW8 Q8VHW9 Q8VHX0 Q8VHY0 Q8VHZ8 Q8VI16
Q8VI21 Q8VI23 Q8VI38 Q8VI47 Q8VI51 Q8VI56 Q8VI59 Q8VI64 Q8VI88 Q8VIB3 Q8VIC6
Q8VIC7 Q8VIC9 Q8VIH3 Q8VIH7 Q8VIH9 Q8VII6 Q8VIK2 Q8VIK5 Q8VIM0 Q8VIM4 Q8VM97
Q8VNJ8 Q8VNU8 Q8VNW3 Q8VNZ2 Q8VPC3 Q8VPZ0 Q8VQ25 Q8VQ71 Q8VQK4 Q8VQK5 Q8VRH0
Q8VRZ1 Q8VRZ2 Q8VRZ3 Q8VSL2 Q8VWI1 Q8VWJ1 Q8VWZ7 Q8VWZ8 Q8VXB1 Q8VXB5 Q8VXX9
Q8VXY7 Q8VXZ5 Q8VY13 Q8VY16 Q8VY22 Q8VY39 Q8VY40 Q8VY49 Q8VY63 Q8VY69 Q8VY77
Q8VY92 Q8VY97 Q8VY98 Q8VYA3 Q8VYC8 Q8VYD4 Q8VYD8 Q8VYE2 Q8VYE4 Q8VYF4 Q8VYF9
Q8VYG0 Q8VYH2 Q8VYH6 Q8VYI1 Q8VYJ1 Q8VYL8 Q8VYM2 Q8VYP0 Q8VYP5 Q8VYP9 Q8VYR0
Q8VYR3 Q8VYR4 Q8VYR7 Q8VYR9 Q8VYS5 Q8VYS8 Q8VYT3 Q8VYY4 Q8VYY5 Q8VYZ7 Q8VZ10
Q8VZ13 Q8VZ14 Q8VZ48 Q8VZ52 Q8VZ80 Q8VZ87 Q8VZ95 Q8VZ96 Q8VZB2 Q8VZC0 Q8VZE2

Q8VZF6 Q8VZG2 Q8VZG8 Q8VZI2 Q8VZI8 Q8VZJ0 Q8VZJ7 Q8VZK9 Q8VZL4 Q8VZM5 Q8VZM7
Q8VZN4 Q8VZN9 Q8VZP6 Q8VZQ3 Q8VZQ5 Q8VZR6 Q8VZR7 Q8VZS0 Q8VZT3 Q8VZU2 Q8VZU9
Q8VZV7 Q8VZW1 Q8VZZ4 Q8W036 Q8W037 Q8W0H5 Q8W0K2 Q8W110 Q8W115 Q8W245 Q8W246
Q8W3F9 Q8W404 Q8W405 Q8W469 Q8W472 Q8W486 Q8W488 Q8W496 Q8W4A7 Q8W4B2 Q8W4D4
Q8W4E6 Q8W4E7 Q8W4G3 Q8W4G6 Q8W4I4 Q8W4K3 Q8W4P8 Q8W4Q5 Q8W4R9 Q8W4S4 Q8W4S5
Q8W4Y5 Q8W4Z5 Q8W571 Q8W585 Q8W7G4 Q8W822 Q8W855 Q8W8H5 Q8W8Q1 Q8W8W1 Q8W8Y1
Q8W926 Q8W9B6 Q8W9E4 Q8W9F5 Q8W9F9 Q8W9G2 Q8W9G4 Q8W9G7 Q8W9G8 Q8W9J6 Q8W9L2
Q8W9M5 Q8W9M6 Q8W9M7 Q8W9M8 Q8W9M9 Q8W9N0 Q8W9N1 Q8W9N2 Q8W9N3 Q8W9N4 Q8W9N5
Q8W9N6 Q8W9N7 Q8WA39 Q8WA47 Q8WAB4 Q8WB03 Q8WB05 Q8WBP9 Q8WBV2 Q8WCJ7 Q8WCK9
Q8WCL1 Q8WDK6 Q8WDU1 Q8WDV0 Q8WDV2 Q8WDV6 Q8WE97 Q8WEK2 Q8WEK7 Q8WEW3 Q8WEW4
Q8WEW8 Q8WFX3 Q8WGF6 Q8WGF7 Q8WGF8 Q8WGG5 Q8WGG8 Q8WGL0 Q8WHC6 Q8WHX1 Q8WHX4
Q8WHX6 Q8WHX7 Q8WHX8 Q8WHZ2 Q8WHZ3 Q8WHZ4 Q8WHZ5 Q8WHZ6 Q8WI01 Q8WI02 Q8WI03
Q8WI04 Q8WI05 Q8WI06 Q8WI07 Q8WI08 Q8WI09 Q8WI10 Q8WI13 Q8WI19 Q8WI20 Q8WI21
Q8WI22 Q8WI23 Q8WI28 Q8WI29 Q8WI31 Q8WI33 Q8WI34 Q8WI36 Q8WMG3 Q8WMG7 Q8WML4
Q8WMN1 Q8WMQ3 Q8WMR7 Q8WMS0 Q8WMU5 Q8WMV1 Q8WMV3 Q8WMW0 Q8WMW2 Q8WMX5 Q8WN57
Q8WN91 Q8WN92 Q8WN93 Q8WN95 Q8WN96 Q8WN97 Q8WNQ5 Q8WNQ6 Q8WNQ8 Q8WNQ9 Q8WNR0
Q8WNR1 Q8WNY2 Q8WP17 Q8WPA2 Q8WQA4 Q8WQG1 Q8WR62 Q8WSR4 Q8WTE6 Q8WTE7 Q8WTR4
Q8WTT0 Q8WTV0 Q8WTX9 Q8WU17 Q8WU67 Q8WUD6 Q8WUG5 Q8WUH6 Q8WUM9 Q8WUR0 Q8WUS8
Q8WUT4 Q8WUT9 Q8WUU8 Q8WUX1 Q8WUY8 Q8WV15 Q8WV19 Q8WV48 Q8WV83 Q8WVE6 Q8WVE7
Q8WVI0 Q8WVN6 Q8WVP7 Q8WVQ1 Q8WVV5 Q8WVX3 Q8WVX9 Q8WVZ1 Q8WVZ7 Q8WW34 Q8WW43
Q8WW52 Q8WW62 Q8WWA1 Q8WWB7 Q8WWF3 Q8WWF5 Q8WWG1 Q8WWG9 Q8WWI5 Q8WWP7 Q8WWQ8
Q8WWT9 Q8WWU5 Q8WWV6 Q8WWX8 Q8WWZ4 Q8WWZ7 Q8WXA8 Q8WXD0 Q8WXF7 Q8WXG9 Q8WXH0
Q8WXH2 Q8WXI7 Q8WXI8 Q8WXS4 Q8WXS5 Q8WY07 Q8WY21 Q8WY22 Q8WY98 Q8WYK1 Q8WZ04
Q8WZ55 Q8WZ59 Q8WZ71 Q8WZ84 Q8WZ92 Q8WZ94 Q8WZA1 Q8WZA6 Q8WZJ9 Q8WZK2 Q8X092
Q8X0H8 Q8X0Z1 Q8X170 Q8X1N9 Q8X1Z7 Q8X226 Q8X2V8 Q8X3J5 Q8X3T0 Q8X3T6 Q8X3T7
Q8X3U5 Q8X400 Q8X479 Q8X490 Q8X492 Q8X4J8 Q8X4L7 Q8X4N7 Q8X4P0 Q8X4Q7 Q8X4S4
Q8X4V6 Q8X504 Q8X524 Q8X534 Q8X564 Q8X5H9 Q8X5K7 Q8X5K8 Q8X5L7 Q8X5L8 Q8X5M2
Q8X5R2 Q8X5R8 Q8X5R9 Q8X5T7 Q8X5W7 Q8X5X8 Q8X5Z7 Q8X614 Q8X625 Q8X638 Q8X651
Q8X653 Q8X659 Q8X691 Q8X6J8 Q8X6K6 Q8X6P4 Q8X6V3 Q8X6V6 Q8X6V7 Q8X6Z6 Q8X714
Q8X739 Q8X794 Q8X7A5 Q8X7A6 Q8X7I3 Q8X7J4 Q8X7L9 Q8X7U9 Q8X7W7 Q8X800 Q8X826
Q8X845 Q8X878 Q8X8F2 Q8X8K9 Q8X8U9 Q8X968 Q8X9A9 Q8X9D3 Q8X9E0 Q8X9E5 Q8X9E6
Q8X9F8 Q8X9F9 Q8X9G0 Q8X9G8 Q8X9I1 Q8X9I3 Q8X9I6 Q8X9L0 Q8XA19 Q8XA20 Q8XA30
Q8XA47 Q8XA48 Q8XA63 Q8XA81 Q8XAF5 Q8XAJ0 Q8XAQ5 Q8XAR5 Q8XAS7 Q8XAS8 Q8XAW9
Q8XAX0 Q8XAY8 Q8XAZ3 Q8XAZ4 Q8XB24 Q8XB33 Q8XB59 Q8XB73 Q8XB84 Q8XB92 Q8XB94
Q8XBJ0 Q8XBJ1 Q8XBK2 Q8XBK4 Q8XBP8 Q8XBW9 Q8XBX0 Q8XBY1 Q8XBY3 Q8XBY4 Q8XC17
Q8XC28 Q8XC49 Q8XCB7 Q8XCC6 Q8XCN6 Q8XCN8 Q8XCP1 Q8XCP5 Q8XD10 Q8XD24 Q8XDD1
Q8XDJ5 Q8XDQ5 Q8XDS6 Q8XDT4 Q8XDT5 Q8XDY7 Q8XDZ1 Q8XDZ5 Q8XE41 Q8XEB7 Q8XED0
Q8XED4 Q8XEK3 Q8XEK5 Q8XEM4 Q8XEN3 Q8XEV9 Q8XEW6 Q8XF02 Q8XF83 Q8XF88 Q8XFA3
Q8XFA5 Q8XFD0 Q8XFE5 Q8XFG0 Q8XFK6 Q8XFT0 Q8XFY2 Q8XG16 Q8XG35 Q8XG71 Q8XGC2
Q8XGD7 Q8XGS2 Q8XGT8 Q8XGZ7 Q8XH28 Q8XHH1 Q8XIC9 Q8XID0 Q8XJ01 Q8XJA1 Q8XJR2
Q8XJS8 Q8XJT1 Q8XK19 Q8XK42 Q8XL56 Q8XLK2 Q8XLK5 Q8XLL2 Q8XM19 Q8XMG8 Q8XN03
Q8XNY9 Q8XP36 Q8XPC4 Q8XPF2 Q8XPH4 Q8XQC5 Q8XQC8 Q8XQF2 Q8XR16 Q8XR66 Q8XRH0
Q8XSF6 Q8XTT4 Q8XTY1 Q8XU10 Q8XU11 Q8XU12 Q8XU72 Q8XUB6 Q8XV41 Q8XV51 Q8XV89
Q8XV90 Q8XVA1 Q8XVC2 Q8XVI4 Q8XVQ4 Q8XVY9 Q8XWC8 Q8XWL5 Q8XWR3 Q8XWS4 Q8XWX9

Q8XXB6 Q8XXQ1 Q8XXQ8 Q8XXR1 Q8XYL2 Q8XYY5 Q8XZ47 Q8XZI4 Q8XZM2 Q8XZR2 Q8Y0B4
Q8Y0D3 Q8Y1I9 Q8Y210 Q8Y275 Q8Y2G4 Q8Y2K5 Q8Y3A6 Q8Y3H6 Q8Y3I2 Q8Y3Z6 Q8Y3Z7
Q8Y3Z8 Q8Y4B6 Q8Y4B8 Q8Y4E3 Q8Y4G2 Q8Y513 Q8Y552 Q8Y588 Q8Y5B8 Q8Y5E4 Q8Y5I0
Q8Y5I1 Q8Y5J6 Q8Y5K2 Q8Y5K3 Q8Y5M0 Q8Y5M3 Q8Y5W0 Q8Y652 Q8Y654 Q8Y656 Q8Y6B5
Q8Y6T8 Q8Y6W3 Q8Y773 Q8Y7A3 Q8Y7A9 Q8Y7G3 Q8Y7I6 Q8Y7J3 Q8Y7S9 Q8Y7X1 Q8Y8E1
Q8Y8H5 Q8Y8Q5 Q8Y8Q9 Q8Y8W3 Q8Y9K8 Q8Y9U6 Q8Y9V5 Q8YAF3 Q8YAF7 Q8YAR6 Q8YB48
Q8YBN8 Q8YBN9 Q8YC64 Q8YCA8 Q8YCB0 Q8YCQ8 Q8YCQ9 Q8YD73 Q8YDA3 Q8YDG7 Q8YDG8
Q8YDK9 Q8YDM4 Q8YDR8 Q8YDS0 Q8YDX1 Q8YDZ0 Q8YDZ2 Q8YEA6 Q8YEG4 Q8YES8 Q8YFB7
Q8YFD7 Q8YFH6 Q8YFH7 Q8YFI5 Q8YFQ7 Q8YG29 Q8YG56 Q8YGH4 Q8YGI7 Q8YGK7 Q8YGL0
Q8YGQ8 Q8YH20 Q8YH52 Q8YHH1 Q8YHH2 Q8YHT9 Q8YI11 Q8YI12 Q8YI23 Q8YIF6 Q8YIS0
Q8YIT2 Q8YJ03 Q8YJ15 Q8YJ50 Q8YJB8 Q8YJF3 Q8YLI4 Q8YLK8 Q8YLP3 Q8YM13 Q8YM86
Q8YMK0 Q8YMQ0 Q8YMW5 Q8YMZ8 Q8YN71 Q8YNB0 Q8YNI8 Q8YP83 Q8YPA9 Q8YPE8 Q8YPE9
Q8YPF1 Q8YQ35 Q8YQ44 Q8YQ64 Q8YQ78 Q8YQ90 Q8YQ91 Q8YQH9 Q8YQI0 Q8YQI1 Q8YQI2
Q8YQS7 Q8YRM9 Q8YRV2 Q8YSD5 Q8YSD7 Q8YSU5 Q8YUS1 Q8YVQ2 Q8YWE0 Q8YXD6 Q8YXJ9
Q8YY27 Q8YYA3 Q8YYB0 Q8YYB4 Q8YYG5 Q8YYG7 Q8YYI9 Q8YYK1 Q8YYK2 Q8YZG8 Q8YZH6
Q8YZS8 Q8YZV7 Q8Z075 Q8Z076 Q8Z0F9 Q8Z0T9 Q8Z0W0 Q8Z0W2 Q8Z172 Q8Z1A8 Q8Z1R2
Q8Z1T9 Q8Z1U2 Q8Z1U3 Q8Z1Y7 Q8Z1Z8 Q8Z229 Q8Z246 Q8Z247 Q8Z257 Q8Z287 Q8Z290
Q8Z291 Q8Z2E3 Q8Z2J2 Q8Z2L7 Q8Z2N7 Q8Z2N9 Q8Z2R2 Q8Z2T0 Q8Z2T8 Q8Z2V6 Q8Z2W4
Q8Z2W6 Q8Z2X6 Q8Z2X7 Q8Z2Y1 Q8Z314 Q8Z332 Q8Z386 Q8Z396 Q8Z3B1 Q8Z3B3 Q8Z3B4
Q8Z3C0 Q8Z3D4 Q8Z3D8 Q8Z3E4 Q8Z3E5 Q8Z3G0 Q8Z3L4 Q8Z3L5 Q8Z3P2 Q8Z3Q8 Q8Z3W2
Q8Z406 Q8Z407 Q8Z4J7 Q8Z4M8 Q8Z4P3 Q8Z4S7 Q8Z4W3 Q8Z4X5 Q8Z4X8 Q8Z4Y8 Q8Z4Z9
Q8Z521 Q8Z527 Q8Z530 Q8Z537 Q8Z538 Q8Z541 Q8Z579 Q8Z5F5 Q8Z5F6 Q8Z5F7 Q8Z5G4
Q8Z5G6 Q8Z5M7 Q8Z5N3 Q8Z5N8 Q8Z5R0 Q8Z670 Q8Z671 Q8Z684 Q8Z6E5 Q8Z6F3 Q8Z6I5
Q8Z6N7 Q8Z6P7 Q8Z6Q5 Q8Z6Q7 Q8Z6Q8 Q8Z6Y0 Q8Z6Z2 Q8Z706 Q8Z727 Q8Z784 Q8Z7A9
Q8Z7E2 Q8Z7E3 Q8Z7H3 Q8Z7L0 Q8Z7L7 Q8Z7L8 Q8Z7L9 Q8Z7S0 Q8Z810 Q8Z814 Q8Z824
Q8Z851 Q8Z862 Q8Z8B6 Q8Z8E4 Q8Z8E5 Q8Z8E6 Q8Z8G7 Q8Z8H3 Q8Z8L4 Q8Z8S4 Q8Z8V5
Q8Z8W9 Q8Z8X0 Q8Z991 Q8Z9A3 Q8Z9A4 Q8Z9B3 Q8Z9H1 Q8Z9K0 Q8Z9L1 Q8Z9M5 Q8Z9M6
Q8Z9N1 Q8Z9N5 Q8Z9U3 Q8Z9V7 Q8ZA28 Q8ZA34 Q8ZA50 Q8ZAA1 Q8ZAE1 Q8ZAF1 Q8ZAG5
Q8ZAM1 Q8ZAM2 Q8ZAM3 Q8ZAS3 Q8ZAS9 Q8ZAU8 Q8ZAU9 Q8ZAV0 Q8ZAX0 Q8ZB72 Q8ZBM0
Q8ZBP5 Q8ZBZ1 Q8ZBZ2 Q8ZBZ3 Q8ZBZ5 Q8ZCA7 Q8ZCD6 Q8ZCH3 Q8ZCK2 Q8ZCV9 Q8ZCW0
Q8ZD52 Q8ZD54 Q8ZD96 Q8ZD97 Q8ZD98 Q8ZDB1 Q8ZDB2 Q8ZDH1 Q8ZDH2 Q8ZDJ8 Q8ZDL3
Q8ZDX4 Q8ZDX7 Q8ZDX9 Q8ZDY0 Q8ZDZ8 Q8ZE38 Q8ZE46 Q8ZEB3 Q8ZEC8 Q8ZED2 Q8ZED3
Q8ZED4 Q8ZEH4 Q8ZEH5 Q8ZEI2 Q8ZEM1 Q8ZEW3 Q8ZF51 Q8ZFF7 Q8ZFF8 Q8ZFG9 Q8ZFJ7
Q8ZG77 Q8ZGA9 Q8ZGC3 Q8ZGC7 Q8ZGH1 Q8ZGJ0 Q8ZGS9 Q8ZGY6 Q8ZH39 Q8ZH58 Q8ZH59
Q8ZHH6 Q8ZHP0 Q8ZHV0 Q8ZI65 Q8ZI67 Q8ZIF2 Q8ZIL9 Q8ZIQ1 Q8ZIX7 Q8ZIX8 Q8ZIY9
Q8ZIZ8 Q8ZJ73 Q8ZJ83 Q8ZJC4 Q8ZJE6 Q8ZJL7 Q8ZJR1 Q8ZJT0 Q8ZJV0 Q8ZJX4 Q8ZJX6
Q8ZK55 Q8ZK90 Q8ZKC3 Q8ZKF8 Q8ZKL0 Q8ZKP8 Q8ZKQ2 Q8ZKQ3 Q8ZKR4 Q8ZKT5 Q8ZKW1
Q8ZKY1 Q8ZKY4 Q8ZL04 Q8ZL63 Q8ZL64 Q8ZLD6 Q8ZLE4 Q8ZLF2 Q8ZLF3 Q8ZLH5 Q8ZLK7
Q8ZLL3 Q8ZLP1 Q8ZLP5 Q8ZLR7 Q8ZLW1 Q8ZLX1 Q8ZLX2 Q8ZLZ9 Q8ZM13 Q8ZM68 Q8ZM94
Q8ZMA4 Q8ZMA5 Q8ZML6 Q8ZMX5 Q8ZN39 Q8ZN53 Q8ZN80 Q8ZNA5 Q8ZNB8 Q8ZND7 Q8ZNE4
Q8ZNF0 Q8ZNK7 Q8ZNQ0 Q8ZNQ1 Q8ZNQ2 Q8ZNQ9 Q8ZNT5 Q8ZP00 Q8ZP01 Q8ZP14 Q8ZP49
Q8ZP50 Q8ZP64 Q8ZPI0 Q8ZPJ2 Q8ZPK1 Q8ZPK3 Q8ZPK5 Q8ZPM3 Q8ZPM4 Q8ZPM6 Q8ZPN4
Q8ZPP2 Q8ZPP6 Q8ZPS8 Q8ZPW6 Q8ZPW7 Q8ZQ19 Q8ZQ25 Q8ZQ26 Q8ZQ27 Q8ZQD2 Q8ZQD5
Q8ZQE4 Q8ZQK4 Q8ZQM2 Q8ZQT3 Q8ZQW1 Q8ZQW2 Q8ZQW3 Q8ZQZ4 Q8ZR35 Q8ZR95 Q8ZRC4

Q8ZRJ9 Q8ZRN0 Q8ZRP0 Q8ZRP1 Q8ZRP8 Q8ZRU5 Q8ZRV1 Q8ZRW2 Q8ZRX1 Q8ZRY4 Q8ZRY9
Q8ZRZ3 Q8ZSB0 Q8ZSL3 Q8ZST3 Q8ZT30 Q8ZV98 Q8ZWI8 Q8ZXD4 Q8ZYR4 Q8ZYX0 Q8ZZ89
Q90021 Q90037 Q90214 Q90215 Q90218 Q90240 Q90245 Q90252 Q90275 Q902F8 Q902F9
Q90304 Q90305 Q90308 Q90309 Q90314 Q90328 Q90330 Q90334 Q90343 Q90344 Q90352
Q90372 Q90373 Q90374 Q90375 Q90413 Q90456 Q90460 Q90478 Q90479 Q90515 Q90584
Q90592 Q90610 Q90615 Q90617 Q90631 Q90632 Q90669 Q90670 Q90674 Q90693 Q90762
Q90763 Q90805 Q90812 Q90845 Q90854 Q90972 Q90980 Q90997 Q90999 Q90VY2 Q90WI4
Q90WT7 Q90WY4 Q90WY5 Q90WY6 Q90X07 Q90X46 Q90X48 Q90X64 Q90XY5 Q90Y50 Q90Y54
Q90Y57 Q90Y90 Q90YH8 Q90YK3 Q90YL4 Q90YM4 Q90YM5 Q90Z00 Q90Z04 Q90Z05 Q90ZB9
Q90ZC7 Q90ZE4 Q90ZK6 Q90ZM2 Q90ZN1 Q91009 Q91044 Q91048 Q91049 Q91081 Q91085
Q91094 Q910E2 Q910M3 Q91147 Q91175 Q91178 Q911P0 Q91280 Q91285 Q91286 Q91287
Q91288 Q913A6 Q913D4 Q913V3 Q91498 Q91499 Q914F8 Q914H5 Q914H7 Q914H9 Q914I0
Q914I1 Q914I3 Q914I9 Q914M0 Q91502 Q91513 Q91514 Q91559 Q91571 Q91577 Q91590
Q91664 Q91687 Q91694 Q91735 Q91736 Q91743 Q91755 Q91756 Q91827 Q91828 Q91838
Q91845 Q91892 Q918W0 Q918W1 Q91909 Q91987 Q91A23 Q91A26 Q91AV0 Q91AV1 Q91B85
Q91C28 Q91C35 Q91DD8 Q91DM1 Q91DS0 Q91E93 Q91F60 Q91F63 Q91F68 Q91FG3 Q91FI7
Q91FJ9 Q91FL7 Q91FM4 Q91FM9 Q91FR3 Q91FT8 Q91FU5 Q91FU7 Q91FV8 Q91FW7 Q91G04
Q91G40 Q91G50 Q91G63 Q91G84 Q91MA2 Q91MA7 Q91MG0 Q91MG4 Q91PP5 Q91QS9 Q91T40
Q91UL0 Q91V01 Q91V04 Q91V08 Q91V14 Q91V24 Q91V37 Q91V45 Q91V61 Q91V79 Q91V87
Q91V95 Q91V98 Q91VA1 Q91VC4 Q91VC9 Q91VE0 Q91VH1 Q91VK4 Q91VM4 Q91VN0 Q91VP7
Q91VS7 Q91VT8 Q91VU1 Q91VX9 Q91W10 Q91W20 Q91W29 Q91W34 Q91W52 Q91W98 Q91WA9
Q91WB2 Q91WC3 Q91WC7 Q91WC9 Q91WD0 Q91WD2 Q91WD8 Q91WE6 Q91WF3 Q91WG1 Q91WM6
Q91WN2 Q91WN3 Q91WN4 Q91WR6 Q91WS0 Q91WU2 Q91WU4 Q91WU6 Q91WV7 Q91WW2 Q91WW3
Q91WW4 Q91WW5 Q91WW7 Q91X05 Q91X21 Q91X49 Q91X56 Q91X60 Q91X78 Q91X85 Q91X86
Q91X88 Q91XA2 Q91XB7 Q91XC9 Q91XD3 Q91XD7 Q91XD8 Q91XE7 Q91XE8 Q91XE9 Q91XF4
Q91XL3 Q91XN4 Q91XP5 Q91XQ5 Q91XQ6 Q91XT9 Q91XU7 Q91XU8 Q91XV6 Q91XV7 Q91XY4
Q91XZ2 Q91XZ4 Q91Y02 Q91Y11 Q91Y13 Q91Y20 Q91Y55 Q91Y57 Q91Y63 Q91Y74 Q91Y77
Q91Y94 Q91YD1 Q91YD4 Q91YH5 Q91YL7 Q91YN1 Q91YQ5 Q91YT2 Q91YT8 Q91YV9 Q91YX5
Q91YY2 Q91YY5 Q91Z22 Q91Z92 Q91ZB4 Q91ZB5 Q91ZB7 Q91ZB8 Q91ZB9 Q91ZC0 Q91ZC1
Q91ZC4 Q91ZC5 Q91ZC6 Q91ZC7 Q91ZD4 Q91ZE5 Q91ZF1 Q91ZH7 Q91ZI0 Q91ZI8 Q91ZM8
Q91ZN5 Q91ZP6 Q91ZQ0 Q91ZR5 Q91ZR7 Q91ZT1 Q91ZU6 Q91ZU9 Q91ZV0 Q91ZV2 Q91ZV3
Q91ZV4 Q91ZV7 Q91ZV8 Q91ZW7 Q91ZW8 Q91ZX1 Q91ZX7 Q91ZY1 Q91ZY2 Q91ZZ5 Q92030
Q92036 Q92038 Q92052 Q92075 Q920A1 Q920A7 Q920B6 Q920C4 Q920E0 Q920E1 Q920E3
Q920G3 Q920G8 Q920H4 Q920H8 Q920L1 Q920L5 Q920L6 Q920L7 Q920N7 Q920P9 Q920R3
Q920R6 Q920V1 Q92105 Q92123 Q92126 Q92154 Q92179 Q92180 Q92182 Q92183 Q92184
Q92185 Q92186 Q92187 Q92197 Q921A2 Q921C1 Q921I0 Q921L3 Q921L8 Q921N7 Q921Q3
Q921R4 Q921R7 Q921R8 Q921T2 Q921V5 Q921V7 Q921W8 Q921Y4 Q92206 Q92253 Q92254
Q92262 Q92266 Q92275 Q922G0 Q922J6 Q922J9 Q922P8 Q922Q5 Q922Q6 Q922Q8 Q922T2
Q922Y2 Q922Z1 Q92316 Q92337 Q92339 Q92341 Q92356 Q92367 Q923B6 Q923D3 Q923G5
Q923I7 Q923J1 Q923J4 Q923K1 Q923L3 Q923S2 Q923U9 Q923W9 Q923X1 Q923X5 Q923X6
Q923X8 Q923X9 Q923Y0 Q923Y1 Q923Y2 Q923Y3 Q923Y4 Q923Y5 Q923Y6 Q923Y7 Q923Y8
Q923Y9 Q923Z0 Q92444 Q92459 Q92478 Q92482 Q924A5 Q924C3 Q924C9 Q924H0 Q924I3
Q924L1 Q924N4 Q924P2 Q924S1 Q924S4 Q924T4 Q924T8 Q924T9 Q924U0 Q924U1 Q924V1
Q924V4 Q924X1 Q924X2 Q924X6 Q924Y8 Q924Z4 Q924Z5 Q92504 Q92508 Q92521 Q92523

Q92535 Q92536 Q92537 Q92542 Q92544 Q92545 Q92581 Q925B3 Q925B4 Q925B5 Q925C0
Q925D4 Q925D8 Q925E8 Q925F2 Q925F3 Q925G2 Q925H0 Q925I1 Q925I4 Q925N0 Q925N1
Q925N2 Q925N4 Q925N6 Q925N7 Q925P2 Q925Q3 Q925R7 Q925S8 Q925U2 Q925U4 Q92604
Q92611 Q92617 Q92629 Q92633 Q92637 Q92643 Q92667 Q92673 Q92685 Q92692 Q926K7
Q926Q5 Q92729 Q92736 Q92781 Q927F9 Q927G0 Q927G1 Q927P0 Q927U4 Q927V8 Q927W0
Q927Z0 Q92806 Q92813 Q92819 Q92823 Q92824 Q92838 Q92839 Q92847 Q92854 Q92859
Q92887 Q92896 Q928B1 Q92903 Q92911 Q92932 Q92935 Q92952 Q92953 Q92956 Q92959
Q92968 Q92982 Q929B1 Q929E9 Q929M4 Q929T6 Q929T7 Q929V3 Q929V9 Q929W0 Q929X7
Q929Y0 Q92AG0 Q92AG2 Q92AG4 Q92AN2 Q92BB4 Q92BE7 Q92BT1 Q92BW9 Q92BX6 Q92C36
Q92C68 Q92CL6 Q92CQ9 Q92D58 Q92DI5 Q92DP0 Q92EE1 Q92EL4 Q92EP8 Q92ES8 Q92EU0
Q92EU1 Q92F91 Q92F94 Q92FD1 Q92FR2 Q92FX1 Q92FX2 Q92G50 Q92G78 Q92G93 Q92G96
Q92G97 Q92G98 Q92G99 Q92GA4 Q92GA9 Q92GG3 Q92GI5 Q92GL0 Q92GP9 Q92GV3 Q92GY6
Q92H24 Q92H61 Q92H75 Q92H77 Q92H78 Q92HH5 Q92HP9 Q92HQ3 Q92HV4 Q92HW0 Q92I31
Q92I62 Q92I65 Q92I67 Q92I81 Q92I85 Q92I98 Q92IC5 Q92ID4 Q92ID5 Q92IF1 Q92IL2
Q92IR1 Q92IR2 Q92IR4 Q92IS7 Q92IT3 Q92IT6 Q92J29 Q92J66 Q92J92 Q92J96 Q92J98
Q92J99 Q92JB1 Q92JB3 Q92JC1 Q92JC2 Q92JC3 Q92JC4 Q92JC5 Q92JD6 Q92JD8 Q92JE9
Q92JF7 Q92JF8 Q92JG1 Q92JG2 Q92JH1 Q92JH8 Q92JI6 Q92JJ2 Q92JJ3 Q92JJ5 Q92JJ9
Q92JL6 Q92JM2 Q92JP0 Q92JP1 Q92JP3 Q92JP8 Q92KX3 Q92L23 Q92L49 Q92L50 Q92L89
Q92N77 Q92NM3 Q92NU9 Q92P50 Q92P97 Q92PF5 Q92PZ0 Q92Q24 Q92Q25 Q92Q49 Q92QE1
Q92QE8 Q92QL7 Q92QN9 Q92QP2 Q92QP5 Q92QR8 Q92R43 Q92R68 Q92RG6 Q92RG8 Q92RM5
Q92RM6 Q92RM8 Q92RN0 Q92RV0 Q92S72 Q92SF5 Q92SJ3 Q92SP2 Q92UW5 Q92WD7 Q92WD8
Q92X78 Q92XI9 Q92XJ0 Q92XJ1 Q92Y37 Q92Y93 Q92YC7 Q92ZW9 Q93033 Q93038 Q93050
Q93063 Q93084 Q93086 Q93105 Q93109 Q93126 Q93127 Q93149 Q931E4 Q931G6 Q931R6
Q931S1 Q931T9 Q93212 Q93235 Q93249 Q932F5 Q932J9 Q932K6 Q93373 Q93380 Q933E0
Q933I3 Q934G3 Q93561 Q93655 Q93704 Q93834 Q93841 Q93890 Q93971 Q93A44 Q93AK1
Q93AR8 Q93AR9 Q93AS0 Q93AS1 Q93CB7 Q93D93 Q93D98 Q93DY1 Q93DY2 Q93DY4 Q93DY8
Q93DZ0 Q93DZ1 Q93F90 Q93FD1 Q93FG6 Q93FH0 Q93FH2 Q93FH3 Q93FH6 Q93HS4 Q93HU4
Q93IM6 Q93IN1 Q93IN2 Q93JL8 Q93KD5 Q93LK8 Q93LK9 Q93MV4 Q93MV5 Q93N69 Q93N70
Q93NB7 Q93PD4 Q93PD9 Q93PU3 Q93PU4 Q93Q23 Q93QX2 Q93QY7 Q93RR6 Q93S26 Q93SE0
Q93SM8 Q93SW6 Q93SW8 Q93SW9 Q93SX0 Q93SX1 Q93TE2 Q93V04 Q93V66 Q93V85 Q93VH1
Q93VJ2 Q93VM8 Q93VV0 Q93VV5 Q93W54 Q93W57 Q93W95 Q93WD2 Q93WG3 Q93XI5 Q93XM7
Q93XX4 Q93XY5 Q93Y06 Q93Y07 Q93Y08 Q93Y37 Q93Y38 Q93Y91 Q93YP7 Q93YP9 Q93YR2
Q93YS2 Q93YS4 Q93YT1 Q93YT3 Q93YV7 Q93YW7 Q93YX4 Q93Z13 Q93Z16 Q93Z20 Q93Z29
Q93Z32 Q93Z38 Q93Z75 Q93Z79 Q93Z80 Q93Z81 Q93Z92 Q93ZA0 Q93ZB2 Q93ZD7 Q93ZF5
Q93ZH9 Q93ZQ5 Q93ZR6 Q93ZR8 Q93ZS4 Q93ZX7 Q93ZY3 Q940E8 Q940G0 Q940H3 Q940J9
Q940M4 Q940N9 Q940P5 Q940Q3 Q940Q4 Q940Q8 Q940S0 Q940U6 Q940V4 Q940Y5 Q940Y9
Q94175 Q94177 Q941A0 Q941L0 Q941R4 Q941V3 Q94225 Q94263 Q942F3 Q942X4 Q942X8
Q94316 Q943K1 Q94421 Q94494 Q944A9 Q944G5 Q944H2 Q944H5 Q944J0 Q944K2 Q944M5
Q944N8 Q944P0 Q944W2 Q94515 Q94526 Q945F0 Q945L4 Q945M8 Q945S5 Q946Y7 Q94741
Q947B7 Q94887 Q948L0 Q948R8 Q948T9 Q948U0 Q948Y1 Q94900 Q94920 Q949G3 Q949N0
Q949P1 Q949Q1 Q949Q5 Q949R9 Q949U1 Q949U4 Q949X0 Q949Y5 Q94A02 Q94A03 Q94A05
Q94A15 Q94A18 Q94A20 Q94A32 Q94A76 Q94A87 Q94A99 Q94AA1 Q94AA9 Q94AF9 Q94AG2
Q94AG6 Q94AH1 Q94AH3 Q94AL1 Q94AL8 Q94AN2 Q94AP3 Q94AS5 Q94AS9 Q94AU2 Q94AV5
Q94AX5 Q94AZ2 Q94B38 Q94B65 Q94B77 Q94BP3 Q94BX4 Q94BY4 Q94BY6 Q94BZ1 Q94C25

Q94C26 Q94C49 Q94C70 Q94C77 Q94CA0 Q94CC0 Q94CG2 Q94CI6 Q94CI7 Q94CJ5 Q94CK0
Q94CK3 Q94CS9 Q94DB8 Q94DD4 Q94DG6 Q94EC3 Q94EC4 Q94EE4 Q94EG9 Q94EH2 Q94EI9
Q94EJ6 Q94F09 Q94F27 Q94F62 Q94FB9 Q94FM7 Q94G16 Q94G17 Q94GB1 Q94GM9 Q94HV7
Q94I55 Q94IA6 Q94IB8 Q94IB9 Q94II3 Q94IP1 Q94IW5 Q94JG1 Q94JQ6 Q94JS0 Q94JS4
Q94JU2 Q94K00 Q94K18 Q94K66 Q94K78 Q94K98 Q94KB1 Q94KB2 Q94KB4 Q94KB7 Q94KB9
Q94KE0 Q94KE1 Q94KI8 Q94KK5 Q94KK6 Q94KK7 Q94LW6 Q94N24 Q94ND7 Q94NF7 Q94NM3
Q94NW9 Q94NY9 Q94P21 Q94P36 Q94PA3 Q94PA4 Q94PB4 Q94Q33 Q94Q38 Q94Q45 Q94Q84
Q94QD3 Q94QP2 Q94QQ7 Q94RJ2 Q94S37 Q94S78 Q94SA4 Q94SD0 Q94SJ5 Q94SK8 Q94SZ7
Q94T21 Q94T42 Q94T68 Q94TE4 Q94TI3 Q94UY0 Q94UZ9 Q94V00 Q94V01 Q94V02 Q94V92
Q94VC3 Q94VE5 Q94VF8 Q94VI4 Q94VJ5 Q94VJ9 Q94VK0 Q94VK4 Q94VK5 Q94WP7 Q94WQ8
Q94WR7 Q94WW5 Q94WW7 Q94WW9 Q94WX2 Q94WX4 Q94WX5 Q94WY6 Q94WY7 Q94XZ1 Q94XZ5
Q94Y46 Q94Y47 Q94Y50 Q94Y51 Q94Y56 Q94Y62 Q94Y64 Q94Y66 Q94Y71 Q94Y72 Q94Y87
Q94YC5 Q94YD6 Q94YE5 Q94YL4 Q94YL5 Q94YL9 Q94YT7 Q94Z64 Q94ZD1 Q94ZI2 Q94ZJ2
Q94ZK8 Q94ZL2 Q95050 Q95078 Q950B9 Q950C2 Q950C6 Q950C7 Q950S3 Q950U6 Q950U8
Q95102 Q95118 Q95119 Q95123 Q95125 Q95126 Q95132 Q95135 Q95136 Q95137 Q95141
Q95152 Q95154 Q95155 Q95156 Q95157 Q95158 Q95165 Q95167 Q95170 Q95179 Q95194
Q95195 Q95198 Q951T4 Q951T6 Q95204 Q95207 Q95209 Q95220 Q95235 Q95237 Q95241
Q95242 Q95245 Q95247 Q95250 Q95252 Q95254 Q952R8 Q95334 Q95339 Q953G9 Q953H3
Q953I3 Q953I4 Q953I6 Q953J5 Q953J9 Q953K8 Q953L2 Q95460 Q955G5 Q955G6 Q955T4
Q95695 Q956S6 Q956Y7 Q956Y9 Q956Z1 Q956Z2 Q956Z3 Q956Z5 Q956Z6 Q956Z7 Q956Z8
Q956Z9 Q95704 Q95705 Q95706 Q95707 Q95708 Q95709 Q95710 Q95711 Q95718 Q95719
Q95723 Q95726 Q95727 Q95728 Q95729 Q95730 Q95731 Q95734 Q95735 Q95736 Q95737
Q95738 Q95739 Q95741 Q95744 Q95745 Q95750 Q95776 Q957A1 Q957A2 Q957A4 Q957A7
Q957A8 Q957A9 Q957B0 Q957B1 Q957B2 Q957B3 Q957B4 Q957B5 Q957B6 Q957B7 Q957B8
Q957B9 Q957C1 Q957C3 Q957C6 Q957C7 Q957C8 Q957D1 Q957T2 Q95840 Q95881 Q95885
Q95891 Q95897 Q95910 Q95911 Q95912 Q95913 Q95914 Q95915 Q95916 Q95917 Q95918
Q95919 Q95920 Q95921 Q95928 Q959V6 Q95A26 Q95AG0 Q95H43 Q95H44 Q95H46 Q95H59
Q95IT1 Q95IT3 Q95J56 Q95J75 Q95J79 Q95J96 Q95JB9 Q95JC7 Q95JD3 Q95JD4 Q95JF2
Q95JG1 Q95JH0 Q95JH2 Q95JI2 Q95JJ0 Q95JJ6 Q95JK0 Q95JK4 Q95JN5 Q95JP5 Q95JR3
Q95JR4 Q95JR7 Q95JW2 Q95JX4 Q95K04 Q95K73 Q95K74 Q95KA5 Q95KB4 Q95KC3 Q95KC8
Q95KD5 Q95KE2 Q95KG7 Q95KH4 Q95KI3 Q95KI5 Q95KM5 Q95KM6 Q95KU1 Q95KV7 Q95KW8
Q95KZ0 Q95L01 Q95L04 Q95L11 Q95L14 Q95L55 Q95L74 Q95L97 Q95LA0 Q95LA1 Q95LA2
Q95LA9 Q95LB4 Q95LC6 Q95LF0 Q95LF2 Q95LF3 Q95LF4 Q95LF5 Q95LF7 Q95LF9 Q95LG1
Q95LG5 Q95LH0 Q95LH1 Q95LI5 Q95LI7 Q95LI8 Q95LL7 Q95LN3 Q95LU3 Q95LV7 Q95M19
Q95M53 Q95M54 Q95M74 Q95M75 Q95M77 Q95MD2 Q95MF9 Q95MG6 Q95MH6 Q95MI1 Q95MI4
Q95MI5 Q95MI7 Q95ML5 Q95MM9 Q95MN6 Q95MQ5 Q95N00 Q95N02 Q95N03 Q95N18 Q95NC0
Q95NC1 Q95NC2 Q95NC3 Q95NC4 Q95NC5 Q95NC6 Q95NC7 Q95NC8 Q95NC9 Q95ND0 Q95ND1
Q95ND2 Q95ND4 Q95NE8 Q95NQ0 Q95NT6 Q95PI2 Q95PJ8 Q95PZ2 Q95Q39 Q95QD1 Q95QW4
Q95QZ5 Q95R48 Q95RU0 Q95SP2 Q95SS8 Q95ST2 Q95T12 Q95T64 Q95TU8 Q95US5 Q95V11
Q95V25 Q95WY5 Q95X35 Q95XG8 Q95XZ6 Q95Y52 Q95Y82 Q95YI3 Q95YI5 Q95YM9 Q95ZE3
Q95ZJ1 Q95ZV7 Q95ZW1 Q95ZY0 Q95ZY1 Q95ZY2 Q95ZY4 Q96000 Q96007 Q96061 Q96062
Q96063 Q96064 Q96065 Q96066 Q96067 Q96068 Q96069 Q96070 Q96071 Q960A0 Q960W6
Q96114 Q96126 Q96127 Q96133 Q96182 Q96183 Q96186 Q96189 Q96190 Q961C9 Q961J5
Q96243 Q96247 Q96282 Q96285 Q96290 Q962X7 Q96303 Q96326 Q96328 Q96423 Q96473

Q96495 Q96514 Q96543 Q965M2 Q965Q4 Q965Q8 Q965T1 Q96662 Q966A0 Q966W3 Q967D7
Q967F4 Q968A5 Q968Y9 Q969E2 Q969F0 Q969F8 Q969I6 Q969K7 Q969L2 Q969M1 Q969M2
Q969M3 Q969N2 Q969N4 Q969P0 Q969S0 Q969S6 Q969V1 Q969V3 Q969V5 Q969W0 Q969W1
Q969W9 Q969X1 Q969X2 Q969X5 Q969Z4 Q96A11 Q96A25 Q96A26 Q96A28 Q96A29 Q96A33
Q96A46 Q96A54 Q96A57 Q96A59 Q96AA3 Q96AD5 Q96AG3 Q96AG4 Q96AJ9 Q96AM1 Q96AN5
Q96AP7 Q96AQ2 Q96AQ8 Q96AW1 Q96B21 Q96B33 Q96B42 Q96B77 Q96B96 Q96BA8 Q96BD0
Q96BF3 Q96BI1 Q96BI3 Q96BM0 Q96BY9 Q96BZ4 Q96BZ9 Q96C03 Q96CC6 Q96CE8 Q96CH1
Q96CP6 Q96CP7 Q96CQ1 Q96CU9 Q96D05 Q96D31 Q96D42 Q96D53 Q96D59 Q96D96 Q96DA6
Q96DB9 Q96DC7 Q96DD7 Q96DL1 Q96DS6 Q96DU3 Q96DW6 Q96DX8 Q96DY5 Q96DZ7 Q96DZ9
Q96E16 Q96E22 Q96E52 Q96E93 Q96EC8 Q96EP9 Q96ER9 Q96ES6 Q96ET8 Q96EU7 Q96EX1
Q96EX2 Q96F05 Q96F15 Q96F25 Q96F46 Q96F81 Q96FB5 Q96FE5 Q96FE7 Q96FL8 Q96FL9
Q96FM1 Q96FT7 Q96FV3 Q96FX8 Q96FZ5 Q96G23 Q96G30 Q96G79 Q96G91 Q96G97 Q96GC9
Q96GE9 Q96GF1 Q96GL9 Q96GM1 Q96GP6 Q96GQ5 Q96GR4 Q96GX1 Q96GZ6 Q96H15 Q96H72
Q96H78 Q96H96 Q96HA1 Q96HA4 Q96HA9 Q96HD1 Q96HE8 Q96HG1 Q96HH4 Q96HH6 Q96HJ5
Q96HP8 Q96HR9 Q96HS1 Q96HV5 Q96I36 Q96I45 Q96IK0 Q96IQ7 Q96IV6 Q96IW7 Q96IX5
Q96IZ2 Q96J42 Q96J65 Q96J66 Q96J84 Q96J86 Q96JA1 Q96JA4 Q96JF0 Q96JJ6 Q96JJ7
Q96JN2 Q96JP9 Q96JQ0 Q96JQ2 Q96JQ5 Q96JT2 Q96JW4 Q96JX3 Q96K12 Q96K19 Q96K37
Q96K49 Q96K78 Q96KA5 Q96KC8 Q96KF7 Q96KG7 Q96KJ4 Q96KJ9 Q96KK3 Q96KK4 Q96KN9
Q96KR6 Q96KT7 Q96KV6 Q96L08 Q96L42 Q96L58 Q96LA5 Q96LA6 Q96LA9 Q96LB0 Q96LB1
Q96LB2 Q96LC7 Q96LD1 Q96LL3 Q96LL9 Q96LR9 Q96LT4 Q96LU7 Q96LW7 Q96LZ7 Q96M19
Q96MC6 Q96MH6 Q96MM7 Q96MS0 Q96MT1 Q96MU8 Q96MV1 Q96MV8 Q96MX0 Q96N19 Q96N35
Q96N66 Q96N68 Q96N87 Q96NA8 Q96NB2 Q96ND0 Q96NI6 Q96NL1 Q96NR3 Q96NT5 Q96NU0
Q96NY7 Q96NY8 Q96P31 Q96P56 Q96P65 Q96P66 Q96P67 Q96P68 Q96P69 Q96P88 Q96PB1
Q96PB8 Q96PC5 Q96PD2 Q96PD6 Q96PD7 Q96PE1 Q96PE5 Q96PG1 Q96PG2 Q96PH1 Q96PJ5
Q96PL5 Q96PQ0 Q96PQ1 Q96PR1 Q96PS6 Q96PS8 Q96PX8 Q96PZ7 Q96Q04 Q96Q45 Q96Q80
Q96Q91 Q96QA5 Q96QD8 Q96QE2 Q96QE4 Q96QI5 Q96QK8 Q96QS1 Q96QT4 Q96QU1 Q96QZ0
Q96R08 Q96R09 Q96R27 Q96R28 Q96R30 Q96R45 Q96R47 Q96R48 Q96R54 Q96R67 Q96R69
Q96R72 Q96R84 Q96RA2 Q96RB7 Q96RC9 Q96RD0 Q96RD1 Q96RD2 Q96RD3 Q96RD6 Q96RD7
Q96RD9 Q96RI0 Q96RI8 Q96RI9 Q96RJ0 Q96RJ3 Q96RL6 Q96RN1 Q96RP7 Q96RP8 Q96RQ1
Q96RT6 Q96RV3 Q96S06 Q96S37 Q96S52 Q96S66 Q96S97 Q96SA4 Q96SE0 Q96SJ8 Q96SK2
Q96SL1 Q96SN7 Q96T52 Q96T53 Q96T54 Q96T55 Q96T83 Q96TA0 Q96TA2 Q96TC7 Q96TW8
Q96TW9 Q96U08 Q96UB6 Q96UR9 Q96UY1 Q96UY2 Q96VG2 Q96VG6 Q96VH1 Q96VH3 Q96VH5
Q96VT7 Q96WS1 Q96WT3 Q96WV4 Q96WW4 Q96WW6 Q96Y22 Q96Y25 Q96ZM1 Q970T4 Q971A3
Q971I8 Q973R2 Q975E2 Q975W7 Q977V3 Q979K5 Q979X0 Q979Z2 Q97BF5 Q97BF6 Q97BF7
Q97BG1 Q97BY3 Q97CB5 Q97CS1 Q97CW0 Q97D63 Q97G69 Q97GR5 Q97H86 Q97H88 Q97H98
Q97I14 Q97I38 Q97I41 Q97I57 Q97IQ3 Q97IQ4 Q97JA2 Q97KF6 Q97KG9 Q97LH9 Q97LQ3
Q97M68 Q97MU9 Q97MW3 Q97N28 Q97NI6 Q97NP5 Q97P79 Q97PA9 Q97Q19 Q97QC4 Q97QC5
Q97QD6 Q97QN5 Q97QP7 Q97RK0 Q97RU7 Q97SA4 Q97SH4 Q97SJ6 Q97SR2 Q97T80 Q97TA9
Q97TN5 Q97TZ6 Q97TZ9 Q97UY9 Q97UZ0 Q97UZ1 Q97X94 Q97X95 Q980C6 Q98146 Q982X5
Q98300 Q98329 Q983A3 Q983H5 Q984U2 Q985G2 Q985W8 Q985X0 Q986D1 Q986D2 Q986D4
Q986R8 Q98803 Q98855 Q98864 Q98880 Q98894 Q98895 Q98902 Q98907 Q98910 Q98913
Q98914 Q98931 Q98936 Q98949 Q98980 Q98982 Q98998 Q98AV2 Q98B79 Q98BJ6 Q98C38
Q98CU0 Q98D15 Q98D88 Q98DG5 Q98DH7 Q98DM7 Q98E66 Q98EB3 Q98EB7 Q98EH3 Q98ET0
Q98FG8 Q98FL3 Q98FL4 Q98FM7 Q98GN8 Q98GR1 Q98GX5 Q98GX6 Q98GX7 Q98HW8 Q98I99

Q98IV9 Q98KB0 Q98KL7 Q98KL8 Q98KP0 Q98KR3 Q98KR6 Q98LC5 Q98LF1 Q98M18 Q98M84
Q98MC1 Q98MN3 Q98N26 Q98NJ1 Q98PE4 Q98PL0 Q98QR6 Q98QU0 Q98QU1 Q98R58 Q98R64
Q98SH2 Q98SN8 Q98SN9 Q98ST7 Q98UF7 Q98VG9 Q99027 Q99062 Q99063 Q99075 Q99087
Q99088 Q99102 Q99126 Q99127 Q99155 Q99161 Q99163 Q99165 Q99173 Q99190 Q99191
Q99234 Q99244 Q99246 Q99250 Q99252 Q99271 Q99279 Q99280 Q99287 Q99297 Q99325
Q99332 Q99340 Q99376 Q99380 Q99382 Q99385 Q99393 Q99396 Q99401 Q99437 Q99442
Q99463 Q99466 Q99467 Q99500 Q99518 Q99523 Q99527 Q99571 Q99572 Q99595 Q99624
Q99643 Q99650 Q99665 Q99677 Q99678 Q99679 Q99680 Q99705 Q99706 Q99712 Q99720
Q99726 Q99735 Q99758 Q99788 Q99795 Q997C2 Q99805 Q99808 Q99835 Q99884 Q998L9
Q998M2 Q99928 Q99941 Q99942 Q99943 Q99946 Q99965 Q99999 Q99CW3 Q99D35 Q99HR4
Q99HS3 Q99IB8 Q99IE6 Q99J19 Q99J21 Q99J27 Q99J47 Q99J56 Q99J59 Q99J85 Q99J86
Q99J93 Q99J94 Q99JA4 Q99JB3 Q99JB4 Q99JB7 Q99JD3 Q99JG2 Q99JH7 Q99JH8 Q99JP7
Q99JR1 Q99JS0 Q99JT6 Q99JW5 Q99JY6 Q99JY8 Q99K10 Q99K24 Q99KC7 Q99KF1 Q99KG5
Q99KI3 Q99KK1 Q99KU0 Q99KU6 Q99KW9 Q99L43 Q99LE2 Q99LE3 Q99LG1 Q99LH2 Q99LI2
Q99LJ5 Q99LJ8 Q99LL3 Q99LP8 Q99LQ7 Q99LR1 Q99LV7 Q99LX8 Q99LY2 Q99M02 Q99M08
Q99M66 Q99M80 Q99M96 Q99MB1 Q99MB3 Q99ME6 Q99MF4 Q99MI6 Q99ML4 Q99MQ3 Q99MQ5
Q99MR3 Q99MR9 Q99MS3 Q99MT6 Q99MT7 Q99MT8 Q99MU1 Q99MZ4 Q99N01 Q99N02 Q99N03
Q99N05 Q99N07 Q99N08 Q99N09 Q99N10 Q99N16 Q99N28 Q99N32 Q99N43 Q99N46 Q99N99
Q99NB2 Q99NB5 Q99NH7 Q99NH8 Q99P47 Q99P50 Q99P55 Q99P65 Q99P66 Q99P72 Q99P81
Q99P82 Q99P91 Q99PA3 Q99PA5 Q99PD7 Q99PE3 Q99PE4 Q99PE5 Q99PE6 Q99PE7 Q99PE8
Q99PF4 Q99PG0 Q99PG6 Q99PH1 Q99PJ1 Q99PL5 Q99PL7 Q99PL8 Q99PN0 Q99PR0 Q99PW6
Q99Q02 Q99QV6 Q99QX3 Q99R71 Q99R72 Q99R74 Q99R80 Q99R86 Q99R97 Q99RE7 Q99RF2
Q99RJ0 Q99RP1 Q99RR5 Q99RR7 Q99RS0 Q99RT8 Q99S39 Q99S77 Q99S97 Q99S98 Q99SA3
Q99SE9 Q99SF0 Q99SF1 Q99SH9 Q99SX1 Q99SY5 Q99SZ7 Q99T05 Q99T13 Q99T32 Q99TC5
Q99TS3 Q99TZ9 Q99U09 Q99U52 Q99U54 Q99U67 Q99U80 Q99U85 Q99UA0 Q99UA1 Q99UB9
Q99UL1 Q99UX0 Q99UY7 Q99UZ7 Q99V36 Q99V37 Q99V38 Q99V39 Q99V70 Q99V76 Q99VB6
Q99VC1 Q99VK3 Q99VM8 Q99VQ4 Q99VR8 Q99VW1 Q99VY6 Q99VY7 Q99VY8 Q99VY9 Q99VZ0
Q99VZ1 Q99VZ2 Q99W28 Q99W36 Q99WN6 Q99WP2 Q99WT9 Q99WU0 Q99WU2 Q99WU3 Q99WZ9
Q99X12 Q99X29 Q99X32 Q99XY3 Q99YK2 Q99Z39 Q99ZV5 Q9A0D2 Q9A0J1 Q9A0J2 Q9A0K9
Q9A155 Q9A1B9 Q9A1C3 Q9A1D5 Q9A1E0 Q9A262 Q9A288 Q9A2A3 Q9A2G0 Q9A2H6 Q9A301
Q9A4R9 Q9A4T3 Q9A5E1 Q9A5K1 Q9A6R5 Q9A6T0 Q9A6T1 Q9A6V2 Q9A6V7 Q9A6Y1 Q9A6Y6
Q9A710 Q9A7X6 Q9A7X7 Q9A7X8 Q9A8H9 Q9A8J0 Q9AA40 Q9AAA6 Q9AAC9 Q9AAH2 Q9AAV1
Q9AAZ2 Q9AB09 Q9AB66 Q9ABA6 Q9ABR0 Q9ABT9 Q9ABX9 Q9AC16 Q9ADP8 Q9AE87 Q9AEL8
Q9AEU0 Q9AFF7 Q9AFI0 Q9AGA7 Q9AGD5 Q9AGF2 Q9AIS6 Q9AIV3 Q9AKD8 Q9AKI9 Q9AKP2
Q9AL93 Q9ALA2 Q9ANR2 Q9APE0 Q9AQU5 Q9AR14 Q9AR74 Q9ARD5 Q9ARX2 Q9ASI1 Q9ASQ2
Q9ASQ6 Q9ASQ7 Q9ASR3 Q9ASS4 Q9ASS7 Q9AST3 Q9AST6 Q9ASU1 Q9ASW1 Q9ASY8 Q9AT72
Q9AT74 Q9AT75 Q9ATL2 Q9ATL3 Q9ATL4 Q9ATL5 Q9ATL6 Q9ATL7 Q9ATL8 Q9ATL9 Q9ATM0
Q9ATM1 Q9ATM2 Q9ATM3 Q9ATM4 Q9ATM5 Q9ATM6 Q9ATM7 Q9ATM8 Q9ATN0 Q9ATN1 Q9ATN2
Q9ATN4 Q9AUE0 Q9AUK4 Q9AV23 Q9AV39 Q9AV71 Q9AVE6 Q9AXH9 Q9AXI0 Q9AXI1 Q9AY27
Q9B0S2 Q9B0Y1 Q9B106 Q9B161 Q9B167 Q9B1C5 Q9B1D2 Q9B1N0 Q9B1P9 Q9B1X4 Q9B1Z4
Q9B203 Q9B205 Q9B229 Q9B2C4 Q9B2F7 Q9B2G1 Q9B365 Q9B506 Q9B5Q1 Q9B5Q3 Q9B5Q4
Q9B5Q5 Q9B5Q6 Q9B5Q7 Q9B5Q8 Q9B5Q9 Q9B5R0 Q9B5R1 Q9B5R2 Q9B5R3 Q9B5R7 Q9B5R8
Q9B5S0 Q9B5S1 Q9B6C7 Q9B6C8 Q9B6D0 Q9B6D3 Q9B6D4 Q9B6D5 Q9B6D6 Q9B6D8 Q9B6E7
Q9B6E8 Q9B6E9 Q9B6F0 Q9B6H6 Q9B7V7 Q9B7W1 Q9B7W3 Q9B840 Q9B8C8 Q9B8C9 Q9B8D0

Q9B8D1 Q9B8D2 Q9B8D3 Q9B8D4 Q9B8D5 Q9B8D6 Q9B8D7 Q9B8D8 Q9B8T5 Q9B8T9 Q9B985
Q9B987 Q9B996 Q9B9E4 Q9B9E8 Q9B9F5 Q9B9F6 Q9B9F9 Q9B9G0 Q9B9Z1 Q9BA02 Q9BAC7
Q9BAE0 Q9BBC1 Q9BBN6 Q9BBN9 Q9BBP1 Q9BBP2 Q9BBP3 Q9BBP4 Q9BBP6 Q9BBQ5 Q9BBQ6
Q9BBQ7 Q9BBQ8 Q9BBR3 Q9BBR4 Q9BBR5 Q9BBR6 Q9BBR7 Q9BBR8 Q9BBR9 Q9BBS0 Q9BBS2
Q9BBS4 Q9BBS5 Q9BBT0 Q9BBT1 Q9BBT2 Q9BBT8 Q9BBU3 Q9BCU3 Q9BCZ4 Q9BD50 Q9BDE0
Q9BDK6 Q9BDM3 Q9BDM7 Q9BDN0 Q9BDN1 Q9BDN3 Q9BDN4 Q9BDP2 Q9BDP4 Q9BDQ5 Q9BDR0
Q9BDR7 Q9BDS6 Q9BDS7 Q9BDS8 Q9BE53 Q9BE63 Q9BE65 Q9BE71 Q9BE72 Q9BE74 Q9BEA0
Q9BEA1 Q9BEA8 Q9BEG2 Q9BEG5 Q9BEG8 Q9BG77 Q9BG98 Q9BGI7 Q9BGL2 Q9BGL8 Q9BGN4
Q9BGN6 Q9BGP6 Q9BGS7 Q9BGX9 Q9BGY6 Q9BH10 Q9BH11 Q9BH13 Q9BH41 Q9BKJ9 Q9BKS0
Q9BLG3 Q9BLG4 Q9BLM6 Q9BLY8 Q9BMK9 Q9BMN8 Q9BPN8 Q9BPQ5 Q9BPV8 Q9BPX6 Q9BQ31
Q9BQ49 Q9BQ51 Q9BQA9 Q9BQB6 Q9BQD7 Q9BQE4 Q9BQG1 Q9BQI7 Q9BQJ4 Q9BQQ7 Q9BQS2
Q9BQS7 Q9BQT8 Q9BQT9 Q9BR10 Q9BR26 Q9BR39 Q9BRB3 Q9BRI3 Q9BRK0 Q9BRK3 Q9BRL7
Q9BRN9 Q9BRQ5 Q9BRQ8 Q9BRR3 Q9BRV3 Q9BRY0 Q9BS91 Q9BSA4 Q9BSA9 Q9BSE2 Q9BSE4
Q9BSF4 Q9BSJ5 Q9BSJ8 Q9BSK0 Q9BSK2 Q9BSN7 Q9BSR8 Q9BT22 Q9BT67 Q9BT76 Q9BT88
Q9BTD3 Q9BTN0 Q9BTV4 Q9BTX1 Q9BTX3 Q9BU23 Q9BU79 Q9BUB7 Q9BUF7 Q9BUJ0 Q9BUM1
Q9BUN8 Q9BUR5 Q9BUV8 Q9BV10 Q9BV23 Q9BV35 Q9BV40 Q9BV81 Q9BV87 Q9BVA6 Q9BVC6
Q9BVG9 Q9BVH7 Q9BVI4 Q9BVK2 Q9BVK6 Q9BVK8 Q9BVT8 Q9BVV7 Q9BVV8 Q9BVW6 Q9BVX2
Q9BW60 Q9BW72 Q9BWH2 Q9BWL3 Q9BWM7 Q9BWQ6 Q9BWQ8 Q9BWV1 Q9BWV2 Q9BX59 Q9BX67
Q9BX73 Q9BX74 Q9BX79 Q9BX84 Q9BX95 Q9BX97 Q9BXA5 Q9BXB1 Q9BXC0 Q9BXC1 Q9BXE9
Q9BXI2 Q9BXJ7 Q9BXJ8 Q9BXK5 Q9BXM7 Q9BXN2 Q9BXP2 Q9BXR5 Q9BXS0 Q9BXS4 Q9BXS9
Q9BXT2 Q9BXU9 Q9BY07 Q9BY08 Q9BY10 Q9BY15 Q9BY19 Q9BY21 Q9BY50 Q9BY64 Q9BY67
Q9BY71 Q9BY78 Q9BY79 Q9BYC5 Q9BYE2 Q9BYE9 Q9BYF1 Q9BYG0 Q9BYG8 Q9BYH1 Q9BYT1
Q9BYT9 Q9BYW1 Q9BZ11 Q9BZ76 Q9BZ97 Q9BZA7 Q9BZA8 Q9BZC7 Q9BZD2 Q9BZD6 Q9BZD7
Q9BZF1 Q9BZG2 Q9BZJ4 Q9BZJ6 Q9BZJ7 Q9BZJ8 Q9BZL3 Q9BZV2 Q9BZV3 Q9BZW2 Q9BZW4
Q9BZW5 Q9BZW8 Q9BZZ2 Q9C091 Q9C0A0 Q9C0B5 Q9C0B7 Q9C0C4 Q9C0D9 Q9C0E8 Q9C0H2
Q9C0I4 Q9C0J1 Q9C0K1 Q9C0L9 Q9C0Q7 Q9C0U9 Q9C0V0 Q9C0V1 Q9C0V2 Q9C0V8 Q9C0W3
Q9C0W5 Q9C0W9 Q9C0X1 Q9C0X4 Q9C0X5 Q9C0X7 Q9C0Y5 Q9C0Z0 Q9C0Z1 Q9C100 Q9C101
Q9C112 Q9C113 Q9C121 Q9C1B3 Q9C1B8 Q9C1E8 Q9C1K8 Q9C1W3 Q9C1W4 Q9C1W7 Q9C1W8
Q9C206 Q9C2D4 Q9C2M6 Q9C447 Q9C449 Q9C450 Q9C4B4 Q9C4Z5 Q9C508 Q9C509 Q9C512
Q9C521 Q9C533 Q9C540 Q9C565 Q9C595 Q9C5D6 Q9C5G5 Q9C5H6 Q9C5L3 Q9C5M0 Q9C5N2
Q9C5S2 Q9C5S8 Q9C5S9 Q9C5U0 Q9C5U1 Q9C5U2 Q9C5V5 Q9C5W7 Q9C5Y2 Q9C5Z6 Q9C5Z7
Q9C637 Q9C639 Q9C654 Q9C660 Q9C664 Q9C699 Q9C6A6 Q9C6A8 Q9C6B2 Q9C6B8 Q9C6D3
Q9C6K9 Q9C6L5 Q9C6L8 Q9C6M1 Q9C6M2 Q9C6S4 Q9C6S5 Q9C6S7 Q9C6T0 Q9C6T7 Q9C6W5
Q9C6X2 Q9C720 Q9C733 Q9C757 Q9C788 Q9C7A2 Q9C7C1 Q9C7D7 Q9C7E9 Q9C7F2 Q9C7F8
Q9C7F9 Q9C7H4 Q9C7I1 Q9C7S0 Q9C7S5 Q9C7T7 Q9C7U1 Q9C809 Q9C821 Q9C823 Q9C835
Q9C884 Q9C889 Q9C8C1 Q9C8E7 Q9C8G5 Q9C8G9 Q9C8H0 Q9C8H1 Q9C8H3 Q9C8I6 Q9C8J8
Q9C8K2 Q9C8M1 Q9C8M9 Q9C8W3 Q9C8X2 Q9C900 Q9C919 Q9C992 Q9C994 Q9C999 Q9C9B9
Q9C9H6 Q9C9H7 Q9C9J0 Q9C9K1 Q9C9K4 Q9C9K5 Q9C9L5 Q9C9M8 Q9C9M9 Q9C9N5 Q9C9Q5
Q9C9Q6 Q9C9Q8 Q9C9R4 Q9C9U1 Q9C9Y8 Q9C9Z1 Q9C9Z2 Q9C9Z3 Q9C9Z5 Q9CA25 Q9CA34
Q9CA60 Q9CA61 Q9CA71 Q9CA75 Q9CA79 Q9CA93 Q9CAC4 Q9CAC5 Q9CAD6 Q9CAH5 Q9CAK4
Q9CAL2 Q9CAL3 Q9CAL8 Q9CAN1 Q9CAN8 Q9CAR9 Q9CAT6 Q9CAX1 Q9CAX3 Q9CAY3 Q9CB43
Q9CBA4 Q9CBJ9 Q9CBL9 Q9CBP1 Q9CBQ5 Q9CBU1 Q9CBU2 Q9CBU4 Q9CC08 Q9CC23 Q9CC42
Q9CC52 Q9CCF0 Q9CCF1 Q9CCF6 Q9CCJ1 Q9CCL1 Q9CCM6 Q9CCN4 Q9CCP7 Q9CD58 Q9CDA3
Q9CDA6 Q9CDA7 Q9CDA8 Q9CDA9 Q9CDE7 Q9CDF7 Q9CDG9 Q9CDM7 Q9CDR6 Q9CDT3 Q9CDW0

Q9CE02 Q9CE03 Q9CEE2 Q9CEU8 Q9CEY5 Q9CF25 Q9CF93 Q9CFU9 Q9CG19 Q9CG27 Q9CGU5
Q9CGW4 Q9CH70 Q9CHL6 Q9CHL8 Q9CHU4 Q9CHU5 Q9CHU9 Q9CHZ9 Q9CII5 Q9CIN2 Q9CJ32
Q9CJ66 Q9CJ67 Q9CJ72 Q9CJB0 Q9CJB2 Q9CJB5 Q9CJB7 Q9CJB8 Q9CJL2 Q9CJW3 Q9CJZ2
Q9CK96 Q9CKC7 Q9CKC8 Q9CKC9 Q9CKD3 Q9CKD4 Q9CKD8 Q9CKI7 Q9CKJ9 Q9CKK5 Q9CKN5
Q9CKU1 Q9CKW4 Q9CKW5 Q9CKW6 Q9CKX3 Q9CKY2 Q9CKY9 Q9CL11 Q9CL24 Q9CL26 Q9CL57
Q9CLA6 Q9CLA7 Q9CLA8 Q9CLA9 Q9CLB0 Q9CLC4 Q9CLF4 Q9CLJ2 Q9CLN1 Q9CLQ2 Q9CLT6
Q9CLV1 Q9CLV6 Q9CLX8 Q9CLY8 Q9CM14 Q9CM47 Q9CM87 Q9CMC3 Q9CMD9 Q9CME9 Q9CMG7
Q9CMJ4 Q9CMP4 Q9CMS3 Q9CMV2 Q9CMZ9 Q9CN32 Q9CN97 Q9CNC5 Q9CNG8 Q9CNJ5 Q9CNJ6
Q9CNL3 Q9CNM5 Q9CNN9 Q9CNP0 Q9CNP3 Q9CNP4 Q9CNP5 Q9CNV0 Q9CP13 Q9CP40 Q9CP52
Q9CP58 Q9CP59 Q9CPA9 Q9CPG2 Q9CPJ1 Q9CPK9 Q9CPL9 Q9CPM6 Q9CPM8 Q9CPQ1 Q9CPQ3
Q9CPU4 Q9CPU9 Q9CPV0 Q9CPV7 Q9CPV9 Q9CPW0 Q9CPW5 Q9CPX8 Q9CPZ3 Q9CPZ6 Q9CQ00
Q9CQ35 Q9CQ44 Q9CQ54 Q9CQ56 Q9CQ61 Q9CQ69 Q9CQ74 Q9CQ85 Q9CQ88 Q9CQ91 Q9CQ92
Q9CQB5 Q9CQC3 Q9CQC4 Q9CQC7 Q9CQD6 Q9CQD8 Q9CQE2 Q9CQE7 Q9CQG0 Q9CQG6 Q9CQG9
Q9CQH0 Q9CQH3 Q9CQJ1 Q9CQM1 Q9CQM2 Q9CQN6 Q9CQQ0 Q9CQR7 Q9CQS4 Q9CQS8 Q9CQT9
Q9CQU3 Q9CQV4 Q9CQV7 Q9CQW0 Q9CQW9 Q9CQX2 Q9CQX5 Q9CQY5 Q9CQY7 Q9CQY8 Q9CQY9
Q9CQZ0 Q9CQZ6 Q9CR20 Q9CR22 Q9CR23 Q9CR33 Q9CR48 Q9CR58 Q9CR60 Q9CR62 Q9CR63
Q9CR64 Q9CR67 Q9CR68 Q9CR75 Q9CR76 Q9CR84 Q9CR89 Q9CR96 Q9CRA0 Q9CRA4 Q9CRB8
Q9CRC0 Q9CRG1 Q9CRY7 Q9CRZ8 Q9CS84 Q9CSV6 Q9CTG6 Q9CTN8 Q9CTY5 Q9CUS9 Q9CV60
Q9CW42 Q9CW73 Q9CWG1 Q9CWP6 Q9CWR7 Q9CWU2 Q9CX13 Q9CX19 Q9CX30 Q9CX83 Q9CX92
Q9CX98 Q9CXB2 Q9CXD6 Q9CXE7 Q9CXI3 Q9CXJ4 Q9CXK4 Q9CXL1 Q9CXL7 Q9CXM0 Q9CXR4
Q9CXT7 Q9CXV1 Q9CXY1 Q9CXY9 Q9CY24 Q9CY27 Q9CY50 Q9CYB0 Q9CYC3 Q9CYN2 Q9CYN9
Q9CYV5 Q9CZ16 Q9CZ52 Q9CZ69 Q9CZB0 Q9CZB9 Q9CZH7 Q9CZJ9 Q9CZL2 Q9CZM9 Q9CZN4
Q9CZP5 Q9CZR2 Q9CZR3 Q9CZR4 Q9CZT5 Q9CZW4 Q9CZW5 Q9CZX7 Q9CZX9 Q9D023 Q9D024
Q9D035 Q9D081 Q9D099 Q9D0F3 Q9D0I4 Q9D0K1 Q9D0L6 Q9D0L7 Q9D0M3 Q9D0N8 Q9D0P0
Q9D0U9 Q9D0V7 Q9D0Z3 Q9D103 Q9D123 Q9D132 Q9D136 Q9D159 Q9D162 Q9D164 Q9D173
Q9D176 Q9D1D1 Q9D1D4 Q9D1E5 Q9D1E8 Q9D1G3 Q9D1I2 Q9D1M7 Q9D1N4 Q9D1Q4 Q9D1R1
Q9D1T0 Q9D1X9 Q9D1Y9 Q9D1Z3 Q9D232 Q9D241 Q9D270 Q9D2A5 Q9D2C7 Q9D2E9 Q9D2F0
Q9D2F7 Q9D2J4 Q9D2L6 Q9D2N8 Q9D2R4 Q9D2R6 Q9D2V8 Q9D2W0 Q9D2Y9 Q9D2Z1 Q9D304
Q9D306 Q9D311 Q9D321 Q9D328 Q9D342 Q9D379 Q9D387 Q9D3A9 Q9D3B1 Q9D3G2 Q9D3H0
Q9D3L0 Q9D3P8 Q9D3R5 Q9D3X9 Q9D4B1 Q9D4D7 Q9D4F2 Q9D4J3 Q9D4M9 Q9D4R2 Q9D4V6
Q9D4Y8 Q9D517 Q9D563 Q9D5K1 Q9D5T0 Q9D5U0 Q9D611 Q9D659 Q9D665 Q9D666 Q9D676
Q9D687 Q9D6B9 Q9D6F4 Q9D6G5 Q9D6G9 Q9D6H5 Q9D6I7 Q9D6J1 Q9D6J5 Q9D6K5 Q9D6K8
Q9D6K9 Q9D6M3 Q9D6U8 Q9D6W7 Q9D6X5 Q9D701 Q9D702 Q9D709 Q9D710 Q9D771 Q9D780
Q9D7B7 Q9D7D1 Q9D7D7 Q9D7I0 Q9D7J4 Q9D7L8 Q9D7N9 Q9D7P7 Q9D7R2 Q9D7S1 Q9D7U6
Q9D7W4 Q9D806 Q9D808 Q9D844 Q9D850 Q9D856 Q9D871 Q9D880 Q9D882 Q9D8B1 Q9D8B4
Q9D8B6 Q9D8B7 Q9D8C2 Q9D8D0 Q9D8F3 Q9D8H7 Q9D8I2 Q9D8K3 Q9D8K8 Q9D8M3 Q9D8N1
Q9D8N3 Q9D8T2 Q9D8T4 Q9D8U2 Q9D8U6 Q9D8V0 Q9D8V7 Q9D8X0 Q9D8Y1 Q9D938 Q9D956
Q9D958 Q9D992 Q9D9D5 Q9D9E0 Q9D9H2 Q9D9I4 Q9D9J7 Q9D9M0 Q9D9R3 Q9D9S2 Q9D9V2
Q9D9X8 Q9DA04 Q9DA16 Q9DA21 Q9DA32 Q9DA37 Q9DA39 Q9DA47 Q9DA48 Q9DA57 Q9DA75
Q9DA77 Q9DAC0 Q9DAH2 Q9DAL0 Q9DAM5 Q9DAM7 Q9DAR1 Q9DAS1 Q9DAV9 Q9DAX2 Q9DAZ5
Q9DB10 Q9DB25 Q9DB41 Q9DB43 Q9DB70 Q9DB73 Q9DBD2 Q9DBE8 Q9DBG6 Q9DBH5 Q9DBI0
Q9DBM0 Q9DBN1 Q9DBP0 Q9DBS1 Q9DBT4 Q9DBU0 Q9DBV4 Q9DBW0 Q9DBX3 Q9DBY1 Q9DBY4
Q9DBZ1 Q9DC11 Q9DC16 Q9DC26 Q9DC29 Q9DC37 Q9DC58 Q9DC60 Q9DCA7 Q9DCC3 Q9DCC8
Q9DCF1 Q9DCF9 Q9DCG2 Q9DCI3 Q9DCK3 Q9DCN7 Q9DCP2 Q9DCQ3 Q9DCS1 Q9DCU0 Q9DCU2

Q9DCV3 Q9DCW2 Q9DCX7 Q9DCX8 Q9DCZ4 Q9DCZ9 Q9DD78 Q9DDD0 Q9DDD1 Q9DDK4 Q9DDN6
Q9DE49 Q9DEB5 Q9DER5 Q9DEX7 Q9DF69 Q9DFS4 Q9DG23 Q9DG25 Q9DGB6 Q9DGD1 Q9DGG4
Q9DGG6 Q9DGY2 Q9DHD5 Q9DHD6 Q9DHJ5 Q9DHV7 Q9DIC6 Q9DK03 Q9DLD4 Q9DR81 Q9DUD9
Q9DWR1 Q9E006 Q9E2H5 Q9E6L5 Q9E6L6 Q9E6M3 Q9E6M9 Q9E6N5 Q9E6P6 Q9E6Q6 Q9E6R4
Q9E6S4 Q9EA42 Q9ENL3 Q9EP51 Q9EP64 Q9EP66 Q9EP69 Q9EP72 Q9EP73 Q9EP78 Q9EP79
Q9EP86 Q9EP93 Q9EP96 Q9EPB7 Q9EPB8 Q9EPE9 Q9EPF2 Q9EPH0 Q9EPI0 Q9EPI9 Q9EPK8
Q9EPL0 Q9EPL2 Q9EPQ0 Q9EPQ1 Q9EPR4 Q9EPR5 Q9EPS3 Q9EPT5 Q9EPT8 Q9EPU5 Q9EPW9
Q9EPX4 Q9EPZ7 Q9EPZ8 Q9EQ09 Q9EQ16 Q9EQ25 Q9EQ31 Q9EQ44 Q9EQ45 Q9EQ46 Q9EQ47
Q9EQ48 Q9EQ51 Q9EQ52 Q9EQ60 Q9EQ76 Q9EQC0 Q9EQC1 Q9EQC4 Q9EQD0 Q9EQD2 Q9EQF2
Q9EQG5 Q9EQG6 Q9EQG7 Q9EQH2 Q9EQH7 Q9EQJ0 Q9EQK7 Q9EQL5 Q9EQN9 Q9EQQ2 Q9EQQ3
Q9EQQ4 Q9EQR3 Q9EQR5 Q9EQS5 Q9EQS9 Q9EQT6 Q9EQU3 Q9EQW8 Q9EQY0 Q9EQY6 Q9ER00
Q9ER04 Q9ER18 Q9ER26 Q9ER31 Q9ER47 Q9ER60 Q9ER62 Q9ER64 Q9ER65 Q9ER80 Q9ERB5
Q9ERC5 Q9ERC8 Q9ERH8 Q9ERK7 Q9ERK9 Q9ERM2 Q9ERM3 Q9ERN0 Q9ERQ6 Q9ERS0 Q9ERS1
Q9ERS2 Q9ERS6 Q9ERS7 Q9ERY9 Q9ERZ3 Q9ERZ4 Q9ERZ8 Q9ES07 Q9ES08 Q9ES14 Q9ES21
Q9ES38 Q9ES40 Q9ES43 Q9ES44 Q9ES45 Q9ES57 Q9ES58 Q9ES81 Q9ES82 Q9ES83 Q9ES87
Q9ES88 Q9ES89 Q9ES90 Q9ES97 Q9ESC1 Q9ESC2 Q9ESD1 Q9ESD6 Q9ESD7 Q9ESE1 Q9ESE2
Q9ESF1 Q9ESF4 Q9ESG3 Q9ESG4 Q9ESG6 Q9ESG8 Q9ESK8 Q9ESM6 Q9ESM7 Q9ESN3 Q9ESP4
Q9ESQ4 Q9ESQ5 Q9ESR9 Q9ESS6 Q9EST1 Q9ESY6 Q9ET09 Q9ET20 Q9ET30 Q9ET32 Q9ET35
Q9ET37 Q9ET38 Q9ET39 Q9ET43 Q9ET61 Q9ET64 Q9ET67 Q9ET77 Q9ET78 Q9ET80 Q9ETX2
Q9EV27 Q9EV29 Q9EV30 Q9EV31 Q9EV32 Q9EV33 Q9EV34 Q9EV66 Q9EV71 Q9EVN2 Q9EVN3
Q9EVN4 Q9EXD0 Q9EXD1 Q9EXD2 Q9EXD7 Q9EXN5 Q9EXN6 Q9EYX5 Q9EYY3 Q9EZE7 Q9F1K9
Q9F1N3 Q9F1R6 Q9F2V2 Q9F2V6 Q9F487 Q9F488 Q9F495 Q9F4E2 Q9F4G9 Q9F5A5 Q9F5N7
Q9F5Y1 Q9F721 Q9F722 Q9F7B1 Q9F7P4 Q9F7Y9 Q9F8D7 Q9F9B1 Q9FA52 Q9FB58 Q9FBK8
Q9FBW3 Q9FC36 Q9FC37 Q9FC39 Q9FCI4 Q9FDM1 Q9FDW8 Q9FDZ5 Q9FE29 Q9FE38 Q9FE59
Q9FE70 Q9FE98 Q9FEC1 Q9FED2 Q9FEG2 Q9FEL6 Q9FEL7 Q9FEL8 Q9FEP7 Q9FF17 Q9FF18
Q9FF29 Q9FF31 Q9FF46 Q9FF53 Q9FF75 Q9FF78 Q9FF99 Q9FFB7 Q9FFB8 Q9FFD0 Q9FFD9
Q9FFE8 Q9FFH5 Q9FFK1 Q9FFK3 Q9FFK8 Q9FFL1 Q9FFN2 Q9FFR9 Q9FFS0 Q9FFT2 Q9FFU6
Q9FFW5 Q9FFX0 Q9FFZ4 Q9FFZ7 Q9FG00 Q9FG14 Q9FG21 Q9FG24 Q9FG33 Q9FG35 Q9FG39
Q9FG65 Q9FG70 Q9FG72 Q9FG87 Q9FG89 Q9FG93 Q9FGE9 Q9FGG3 Q9FGH6 Q9FGJ6 Q9FGK0
Q9FGK4 Q9FGL0 Q9FGL2 Q9FGL5 Q9FGM0 Q9FGN1 Q9FGP9 Q9FGQ2 Q9FGS5 Q9FGS8 Q9FGT9
Q9FGU8 Q9FGW0 Q9FH02 Q9FH16 Q9FH18 Q9FH21 Q9FH27 Q9FH28 Q9FH36 Q9FH66 Q9FH67
Q9FH75 Q9FH76 Q9FH86 Q9FH90 Q9FH92 Q9FHB6 Q9FHD4 Q9FHD5 Q9FHD8 Q9FHF1 Q9FHG4
Q9FHG8 Q9FHH5 Q9FHK7 Q9FHM0 Q9FHT4 Q9FHX2 Q9FHX3 Q9FI00 Q9FI03 Q9FI10 Q9FI17
Q9FI39 Q9FI43 Q9FI46 Q9FI73 Q9FI79 Q9FIB0 Q9FID5 Q9FID6 Q9FID8 Q9FID9 Q9FIF0
Q9FIF1 Q9FIG9 Q9FIH5 Q9FII5 Q9FIJ0 Q9FIJ6 Q9FIM2 Q9FIQ5 Q9FIR0 Q9FIS2 Q9FIU5
Q9FIZ3 Q9FIZ9 Q9FJ06 Q9FJ57 Q9FJ62 Q9FJ72 Q9FJ73 Q9FJ74 Q9FJ75 Q9FJ76 Q9FJ77
Q9FJ78 Q9FJ81 Q9FJ87 Q9FJB4 Q9FJD1 Q9FJD2 Q9FJD6 Q9FJH7 Q9FJH8 Q9FJI4 Q9FJV7
Q9FJW3 Q9FJX3 Q9FJX6 Q9FK04 Q9FK05 Q9FK10 Q9FK13 Q9FK28 Q9FK29 Q9FK30 Q9FK43
Q9FK63 Q9FK66 Q9FK89 Q9FK96 Q9FKC1 Q9FKE9 Q9FKF2 Q9FKF4 Q9FKH6 Q9FKM2 Q9FKP1
Q9FKP2 Q9FKQ1 Q9FKS5 Q9FKS8 Q9FKT8 Q9FKV1 Q9FKX5 Q9FKY3 Q9FKY4 Q9FKY5 Q9FL07
Q9FL08 Q9FL25 Q9FL28 Q9FL41 Q9FL42 Q9FL44 Q9FL51 Q9FL59 Q9FL63 Q9FLA5 Q9FLB6
Q9FLB7 Q9FLC6 Q9FLC8 Q9FLD5 Q9FLD7 Q9FLG2 Q9FLG8 Q9FLJ8 Q9FLK2 Q9FLL4 Q9FLM3
Q9FLN5 Q9FLS8 Q9FLT4 Q9FLT5 Q9FLT8 Q9FLT9 Q9FLV4 Q9FLV9 Q9FLW0 Q9FLW2 Q9FLX5
Q9FM10 Q9FM20 Q9FM67 Q9FM86 Q9FMA5 Q9FMD5 Q9FMD7 Q9FME8 Q9FMF7 Q9FMF8 Q9FMR5

Q9FMU6 Q9FMV3 Q9FMV7 Q9FMW3 Q9FMW9 Q9FMX3 Q9FMY1 Q9FMY4 Q9FN04 Q9FN15 Q9FN18
Q9FN37 Q9FN48 Q9FN50 Q9FN51 Q9FN75 Q9FN77 Q9FN92 Q9FN93 Q9FN94 Q9FNA5 Q9FNA9
Q9FNB2 Q9FNB5 Q9FNC1 Q9FNC9 Q9FNE0 Q9FNE1 Q9FNE8 Q9FNH6 Q9FNI7 Q9FNL3 Q9FNL7
Q9FNL8 Q9FNU2 Q9FP13 Q9FPD5 Q9FPG2 Q9FPN0 Q9FPP4 Q9FPP7 Q9FPP8 Q9FPS0 Q9FR82
Q9FRK5 Q9FRL3 Q9FRR1 Q9FRS6 Q9FT51 Q9FT88 Q9FTT3 Q9FTU1 Q9FTZ2 Q9FUJ9 Q9FUT3
Q9FUY7 Q9FV68 Q9FVN0 Q9FVQ4 Q9FVQ5 Q9FVR3 Q9FVS9 Q9FVU0 Q9FVV9 Q9FVZ7 Q9FVZ9
Q9FWA4 Q9FWR2 Q9FWV6 Q9FWX6 Q9FWX7 Q9FWX8 Q9FWY9 Q9FX01 Q9FX21 Q9FX97 Q9FX99
Q9FXA7 Q9FXF2 Q9FXG3 Q9FXH5 Q9FXH6 Q9FXT6 Q9FXW4 Q9FY06 Q9FY14 Q9FY46 Q9FY51
Q9FY68 Q9FY75 Q9FY84 Q9FY94 Q9FYB9 Q9FYC0 Q9FYC1 Q9FYC7 Q9FYE5 Q9FYG3 Q9FYH7
Q9FYK0 Q9FYL2 Q9FYL3 Q9FYL6 Q9FYQ8 Q9FZ22 Q9FZ33 Q9FZ37 Q9FZ45 Q9FZ49 Q9FZ59
Q9FZ63 Q9FZ81 Q9FZ95 Q9FZ96 Q9FZ97 Q9FZB1 Q9FZB8 Q9FZF1 Q9FZI8 Q9FZJ1 Q9G049
Q9G0M4 Q9G0S9 Q9G0U1 Q9G0W6 Q9G156 Q9G185 Q9G1B9 Q9G1C4 Q9G1F7 Q9G1N5 Q9G1R4
Q9G1Z0 Q9G210 Q9G215 Q9G240 Q9G250 Q9G273 Q9G2D7 Q9G2I2 Q9G2R8 Q9G2S5 Q9G2U4
Q9G2U7 Q9G2W6 Q9G2W7 Q9G2W8 Q9G2W9 Q9G2X0 Q9G2X1 Q9G2X2 Q9G385 Q9G396 Q9G3M2
Q9G3M7 Q9G3M9 Q9G3N1 Q9G3N2 Q9G3N4 Q9G3R1 Q9G3R5 Q9G3S4 Q9G3S5 Q9G3S8 Q9G3T7
Q9G4Q1 Q9G5S9 Q9G6G2 Q9G6G3 Q9G6J1 Q9G6M3 Q9G6M4 Q9G6M7 Q9G7T9 Q9G7U1 Q9G946
Q9G964 Q9G9J4 Q9G9J6 Q9G9J7 Q9GA18 Q9GAM6 Q9GAM7 Q9GAM9 Q9GAN2 Q9GAN4 Q9GAP0
Q9GAR5 Q9GAR6 Q9GAW0 Q9GAW3 Q9GAW6 Q9GBG9 Q9GBY1 Q9GBY4 Q9GBY5 Q9GBY6 Q9GBY8
Q9GBZ0 Q9GBZ1 Q9GBZ2 Q9GBZ4 Q9GBZ5 Q9GBZ6 Q9GBZ7 Q9GCE7 Q9GDV1 Q9GDV2 Q9GE29
Q9GE34 Q9GF77 Q9GF82 Q9GF83 Q9GF86 Q9GF87 Q9GF90 Q9GF93 Q9GF98 Q9GFA1 Q9GFA4
Q9GFA7 Q9GFC2 Q9GFC3 Q9GIP4 Q9GJR5 Q9GJS9 Q9GJT6 Q9GJU1 Q9GJX5 Q9GJY3 Q9GK74
Q9GK76 Q9GKE2 Q9GKE7 Q9GKE8 Q9GKN7 Q9GKQ8 Q9GKS6 Q9GKT2 Q9GKU6 Q9GKU7 Q9GKY1
Q9GKY7 Q9GKY8 Q9GKZ0 Q9GKZ4 Q9GL01 Q9GL41 Q9GL42 Q9GL43 Q9GL50 Q9GL59 Q9GL65
Q9GL76 Q9GL77 Q9GLC3 Q9GLD3 Q9GLD9 Q9GLE4 Q9GLF3 Q9GLF5 Q9GLJ8 Q9GLJ9 Q9GLL1
Q9GLN7 Q9GLN9 Q9GLP0 Q9GLX0 Q9GLX8 Q9GM01 Q9GMB0 Q9GMC9 Q9GME8 Q9GMF1 Q9GMS6
Q9GMU6 Q9GMZ1 Q9GN97 Q9GNK5 Q9GNL3 Q9GNN8 Q9GPH7 Q9GPH8 Q9GPP8 Q9GPR3 Q9GQ00
Q9GQQ0 Q9GSB0 Q9GSD3 Q9GSD7 Q9GSD8 Q9GT50 Q9GTN7 Q9GUM2 Q9GV72 Q9GV77 Q9GYF0
Q9GYK2 Q9GYL2 Q9GYQ4 Q9GZC8 Q9GZE9 Q9GZH4 Q9GZK3 Q9GZK4 Q9GZK6 Q9GZK7 Q9GZM5
Q9GZM6 Q9GZN0 Q9GZN6 Q9GZP1 Q9GZP7 Q9GZP9 Q9GZQ4 Q9GZQ6 Q9GZR5 Q9GZS9 Q9GZT6
Q9GZU1 Q9GZU3 Q9GZV3 Q9GZW8 Q9GZX3 Q9GZY4 Q9GZY6 Q9GZY8 Q9GZZ6 Q9H013 Q9H015
Q9H061 Q9H0A3 Q9H0C2 Q9H0C3 Q9H0H0 Q9H0Q3 Q9H0R3 Q9H0U3 Q9H0V1 Q9H0V9 Q9H0X4
Q9H0X9 Q9H156 Q9H158 Q9H159 Q9H172 Q9H195 Q9H1B5 Q9H1C0 Q9H1C3 Q9H1C4 Q9H1C7
Q9H1D0 Q9H1E5 Q9H1K4 Q9H1N7 Q9H1U4 Q9H1U9 Q9H1V8 Q9H1X3 Q9H1Y3 Q9H1Z9 Q9H205
Q9H207 Q9H208 Q9H209 Q9H210 Q9H221 Q9H222 Q9H228 Q9H237 Q9H244 Q9H251 Q9H252
Q9H255 Q9H295 Q9H2A7 Q9H2A9 Q9H2B2 Q9H2B4 Q9H2C2 Q9H2C5 Q9H2C8 Q9H2D1 Q9H2E6
Q9H2F3 Q9H2H9 Q9H2J7 Q9H2L4 Q9H2S1 Q9H2S6 Q9H2U9 Q9H2V7 Q9H2W1 Q9H2X3 Q9H2X8
Q9H2X9 Q9H2Y9 Q9H300 Q9H310 Q9H313 Q9H330 Q9H339 Q9H340 Q9H341 Q9H342 Q9H343
Q9H344 Q9H346 Q9H354 Q9H3H5 Q9H3K2 Q9H3M0 Q9H3N1 Q9H3N8 Q9H3Q3 Q9H3R1 Q9H3R2
Q9H3S1 Q9H3S3 Q9H3S5 Q9H3T2 Q9H3T3 Q9H3U5 Q9H3V2 Q9H3W5 Q9H400 Q9H427 Q9H461
Q9H490 Q9H4D0 Q9H4F1 Q9H4I9 Q9H553 Q9H598 Q9H5I5 Q9H5J4 Q9H5K3 Q9H5V8 Q9H5Y7
Q9H665 Q9H6A9 Q9H6B4 Q9H6B9 Q9H6D3 Q9H6D8 Q9H6F2 Q9H6H4 Q9H6L2 Q9H6L5 Q9H6R6
Q9H6U8 Q9H6X2 Q9H6X4 Q9H6Y7 Q9H720 Q9H741 Q9H756 Q9H799 Q9H7F0 Q9H7F4 Q9H7M9
Q9H7T0 Q9H7V2 Q9H7X2 Q9H7Z7 Q9H813 Q9H819 Q9H841 Q9H8J5 Q9H8M5 Q9H8M9 Q9H8P0
Q9H8X9 Q9H902 Q9H920 Q9H936 Q9H9B4 Q9H9K5 Q9H9P2 Q9H9S3 Q9H9S5 Q9H9V4 Q9HA72

Q9HA82 Q9HAB3 Q9HAR2 Q9HAS3 Q9HAT1 Q9HAV5 Q9HAW7 Q9HAW8 Q9HAW9 Q9HB03 Q9HB14
Q9HB15 Q9HB29 Q9HB89 Q9HBA0 Q9HBB8 Q9HBE5 Q9HBG4 Q9HBG7 Q9HBI6 Q9HBJ8 Q9HBL6
Q9HBL7 Q9HBM0 Q9HBR0 Q9HBT6 Q9HBU9 Q9HBV1 Q9HBV2 Q9HBW0 Q9HBW1 Q9HBW9 Q9HBX8
Q9HBX9 Q9HBY0 Q9HC07 Q9HC10 Q9HC21 Q9HC24 Q9HC47 Q9HC56 Q9HC58 Q9HC73 Q9HC97
Q9HCC8 Q9HCE9 Q9HCF6 Q9HCJ1 Q9HCJ2 Q9HCK4 Q9HCL0 Q9HCL2 Q9HCM2 Q9HCM3 Q9HCN3
Q9HCN6 Q9HCP6 Q9HCQ5 Q9HCS2 Q9HCU0 Q9HCU4 Q9HCU5 Q9HCX4 Q9HD20 Q9HD23 Q9HD36
Q9HD43 Q9HD45 Q9HD87 Q9HDB5 Q9HDC5 Q9HDC9 Q9HDD0 Q9HDG2 Q9HDU0 Q9HDU1 Q9HDU8
Q9HDV1 Q9HDV2 Q9HDV8 Q9HDW5 Q9HDW7 Q9HDW9 Q9HDX1 Q9HDX4 Q9HDX5 Q9HDY3 Q9HDZ2
Q9HDZ8 Q9HE10 Q9HE12 Q9HE13 Q9HEK4 Q9HEM6 Q9HFC7 Q9HFE1 Q9HFF0 Q9HFV1 Q9HGM2
Q9HGM3 Q9HGM4 Q9HGM6 Q9HGN3 Q9HGN7 Q9HGP8 Q9HGQ3 Q9HGZ6 Q9HHN1 Q9HJ89 Q9HJ91
Q9HJV2 Q9HL05 Q9HLW7 Q9HLZ7 Q9HM61 Q9HMD1 Q9HND8 Q9HNE5 Q9HNE6 Q9HNG1 Q9HNW1
Q9HNW2 Q9HP81 Q9HP84 Q9HPB1 Q9HPD9 Q9HPH5 Q9HPL2 Q9HPU7 Q9HQ19 Q9HQ63 Q9HR72
Q9HRJ8 Q9HRP0 Q9HS72 Q9HSC3 Q9HSQ2 Q9HT06 Q9HT13 Q9HT14 Q9HT15 Q9HT16 Q9HT88
Q9HTH0 Q9HTI1 Q9HTK0 Q9HTR0 Q9HU16 Q9HU17 Q9HU20 Q9HUA6 Q9HUB1 Q9HUB4 Q9HUB5
Q9HUB8 Q9HUE8 Q9HUG6 Q9HUG8 Q9HUH5 Q9HUI3 Q9HUW5 Q9HUW6 Q9HUX3 Q9HUX5 Q9HUX6
Q9HV31 Q9HV52 Q9HVA5 Q9HVD1 Q9HVH7 Q9HVL0 Q9HVM5 Q9HVZ6 Q9HVZ8 Q9HW19 Q9HW50
Q9HW91 Q9HW93 Q9HWC3 Q9HWF5 Q9HWR7 Q9HWZ3 Q9HXE9 Q9HXI1 Q9HXI2 Q9HXY3 Q9HXZ6
Q9HY23 Q9HY59 Q9HY60 Q9HY61 Q9HY64 Q9HYB5 Q9HYB6 Q9HYB7 Q9HYC0 Q9HYK9 Q9HYT3
Q9HYW6 Q9HZ39 Q9HZ47 Q9HZA6 Q9HZK7 Q9HZK8 Q9HZK9 Q9HZL0 Q9HZL1 Q9HZT7 Q9HZX3
Q9I013 Q9I055 Q9I061 Q9I068 Q9I069 Q9I0I2 Q9I0I4 Q9I0I6 Q9I0I9 Q9I0J0 Q9I0J1
Q9I0J2 Q9I0J3 Q9I0J5 Q9I0K1 Q9I0M3 Q9I0R2 Q9I104 Q9I190 Q9I1K1 Q9I273 Q9I2B6
Q9I2E5 Q9I2L4 Q9I2S5 Q9I310 Q9I3F6 Q9I3I5 Q9I3M9 Q9I3N0 Q9I3N1 Q9I3N2 Q9I3N3
Q9I3N4 Q9I3N5 Q9I3N6 Q9I3Y3 Q9I423 Q9I424 Q9I425 Q9I426 Q9I427 Q9I463 Q9I469
Q9I4F5 Q9I4F8 Q9I507 Q9I540 Q9I595 Q9I5G7 Q9I5V6 Q9I5W9 Q9I633 Q9I6B7 Q9I6F2
Q9I6Q3 Q9I6V2 Q9I703 Q9I713 Q9I719 Q9I739 Q9I755 Q9I7K5 Q9I816 Q9I829 Q9I830
Q9I859 Q9I8C7 Q9I8D0 Q9I8D8 Q9I8I2 Q9I8K8 Q9I8N6 Q9I8U0 Q9I8W5 Q9I8X3 Q9I918
Q9I919 Q9I9B9 Q9I9C3 Q9I9M5 Q9I9R3 Q9IA02 Q9IA03 Q9IA05 Q9IA06 Q9IA79 Q9IAL7
Q9IAL8 Q9IAT6 Q9IAX2 Q9IB61 Q9IB86 Q9IBD8 Q9IBI3 Q9IDV2 Q9IDV3 Q9IFX2 Q9IFX3
Q9IGQ0 Q9IGQ6 Q9IGY9 Q9IH62 Q9IH63 Q9IKB5 Q9IKB7 Q9IKC7 Q9IKC8 Q9IKD1 Q9IKD2
Q9ILI5 Q9ILI6 Q9IPJ6 Q9IQ49 Q9IV52 Q9IV53 Q9IZK2 Q9J3M8 Q9J3N8 Q9J529 Q9J541
Q9J548 Q9J551 Q9J552 Q9J554 Q9J555 Q9J560 Q9J586 Q9J590 Q9J5D2 Q9J5H4 Q9J5I0
Q9J6K8 Q9JEB4 Q9JF44 Q9JFA1 Q9JFA9 Q9JFC1 Q9JFH4 Q9JGF2 Q9JGK8 Q9JGT4 Q9JH66
Q9JH67 Q9JH69 Q9JH70 Q9JHB2 Q9JHE3 Q9JHE4 Q9JHE5 Q9JHE8 Q9JHF3 Q9JHG1 Q9JHG3
Q9JHH2 Q9JHI3 Q9JHI4 Q9JHI8 Q9JHI9 Q9JHJ1 Q9JHJ3 Q9JHJ5 Q9JHJ8 Q9JHJ9 Q9JHL0
Q9JHL3 Q9JHS6 Q9JHW1 Q9JHW5 Q9JHX3 Q9JHY1 Q9JHY2 Q9JHZ2 Q9JHZ9 Q9JI12 Q9JI18
Q9JI35 Q9JI39 Q9JI51 Q9JI59 Q9JI60 Q9JI61 Q9JI66 Q9JI67 Q9JI71 Q9JI76 Q9JI93
Q9JI97 Q9JI99 Q9JIF3 Q9JIG1 Q9JIG4 Q9JIG8 Q9JII9 Q9JIK1 Q9JIL6 Q9JIM1 Q9JIM5
Q9JIN6 Q9JIP0 Q9JIP3 Q9JIP4 Q9JIP6 Q9JIP7 Q9JIQ8 Q9JIS3 Q9JIS4 Q9JIS5 Q9JIS7
Q9JIS8 Q9JIT0 Q9JIY5 Q9JIY6 Q9JIY7 Q9JIY8 Q9JIZ0 Q9JIZ9 Q9JJ00 Q9JJ04 Q9JJ05
Q9JJ06 Q9JJ09 Q9JJ16 Q9JJ22 Q9JJ46 Q9JJ59 Q9JJ61 Q9JJ72 Q9JJ73 Q9JJ74 Q9JJB1
Q9JJB9 Q9JJC8 Q9JJC9 Q9JJE4 Q9JJE7 Q9JJF2 Q9JJF9 Q9JJG6 Q9JJG7 Q9JJH2 Q9JJH3
Q9JJH7 Q9JJI5 Q9JJI6 Q9JJJ3 Q9JJJ7 Q9JJK1 Q9JJK3 Q9JJK4 Q9JJK5 Q9JJL3 Q9JJL9
Q9JJM7 Q9JJP0 Q9JJQ0 Q9JJR8 Q9JJS7 Q9JJV3 Q9JJV4 Q9JJV5 Q9JJV7 Q9JJV8 Q9JJV9
Q9JJW0 Q9JJW1 Q9JJW3 Q9JJX6 Q9JJZ1 Q9JJZ4 Q9JJZ8 Q9JJZ9 Q9JK00 Q9JK11 Q9JK39

Q9JK41 Q9JK45 Q9JK79 Q9JK82 Q9JK95 Q9JK96 Q9JK97 Q9JKA3 Q9JKA5 Q9JKA7 Q9JKA8
Q9JKA9 Q9JKB0 Q9JKC6 Q9JKD3 Q9JKD6 Q9JKE1 Q9JKE2 Q9JKE3 Q9JKE7 Q9JKE8 Q9JKE9
Q9JKF0 Q9JKF4 Q9JKF6 Q9JKJ9 Q9JKK1 Q9JKL1 Q9JKM5 Q9JKN1 Q9JKQ2 Q9JKR5 Q9JKT2
Q9JKT3 Q9JKT4 Q9JKT5 Q9JKT7 Q9JKT8 Q9JKT9 Q9JKU0 Q9JKU1 Q9JKV5 Q9JKV7 Q9JKW0
Q9JKW1 Q9JKX3 Q9JKX8 Q9JKZ2 Q9JL06 Q9JL18 Q9JL21 Q9JL27 Q9JL55 Q9JL56 Q9JL58
Q9JL59 Q9JL89 Q9JL99 Q9JLB5 Q9JLB9 Q9JLC4 Q9JLF1 Q9JLF7 Q9JLG4 Q9JLI3 Q9JLJ1
Q9JLJ4 Q9JLJ5 Q9JLL0 Q9JLL3 Q9JLM2 Q9JLN5 Q9JLN6 Q9JLR1 Q9JLR9 Q9JLS0 Q9JLS3
Q9JLS8 Q9JLU8 Q9JLZ8 Q9JM10 Q9JM15 Q9JM51 Q9JM61 Q9JM62 Q9JM63 Q9JM72 Q9JM80
Q9JM95 Q9JMA9 Q9JMD7 Q9JME2 Q9JME9 Q9JMF7 Q9JMG2 Q9JMG3 Q9JMG4 Q9JMH2 Q9JMH4
Q9JMI0 Q9JMI9 Q9JMK0 Q9JMS3 Q9JMS4 Q9JMS5 Q9JMS7 Q9JMT4 Q9JMT6 Q9JRX2 Q9JSY6
Q9JSZ3 Q9JTA9 Q9JTK4 Q9JTL9 Q9JU31 Q9JUK4 Q9JUK9 Q9JUL1 Q9JUL4 Q9JV19 Q9JV27
Q9JV93 Q9JVA8 Q9JVK1 Q9JVP9 Q9JVQ0 Q9JVQ1 Q9JVQ2 Q9JVQ3 Q9JVQ5 Q9JVR5 Q9JVT9
Q9JW48 Q9JW59 Q9JWA2 Q9JWA8 Q9JX23 Q9JX32 Q9JX59 Q9JX88 Q9JX92 Q9JXB8 Q9JXK7
Q9JXM0 Q9JXM5 Q9JXR3 Q9JXS4 Q9JYC6 Q9JYD0 Q9JYM0 Q9JZ36 Q9JZF9 Q9JZG4 Q9JZG6
Q9JZG9 Q9JZN9 Q9K006 Q9K015 Q9K083 Q9K0A2 Q9K0J6 Q9K0M4 Q9K0M5 Q9K0M6 Q9K0M7
Q9K0M8 Q9K0N0 Q9K0N7 Q9K0R0 Q9K0U7 Q9K0U9 Q9K0X9 Q9K0Y6 Q9K0Z3 Q9K165 Q9K1B0
Q9K1B4 Q9K1E1 Q9K1G9 Q9K1H0 Q9K1H6 Q9K1M2 Q9K266 Q9K275 Q9K2S2 Q9K2Y1 Q9K3A9
Q9K407 Q9K451 Q9K489 Q9K490 Q9K4D7 Q9K537 Q9K5Z9 Q9K618 Q9K620 Q9K678 Q9K6F9
Q9K6H0 Q9K6H1 Q9K6Q8 Q9K6Y5 Q9K794 Q9K802 Q9K809 Q9K846 Q9K8L9 Q9K8M0 Q9K8M3
Q9K8Z4 Q9K920 Q9K923 Q9K997 Q9K9A5 Q9K9E6 Q9K9G6 Q9K9K2 Q9K9M8 Q9K9M9 Q9K9S6
Q9K9V2 Q9KA70 Q9KA95 Q9KAB2 Q9KAD9 Q9KB63 Q9KBI6 Q9KCD3 Q9KCE6 Q9KCH9 Q9KCI3
Q9KCJ4 Q9KCP8 Q9KD29 Q9KD62 Q9KDI4 Q9KDL8 Q9KDP2 Q9KDR3 Q9KE91 Q9KEA2 Q9KEE9
Q9KEF0 Q9KEJ2 Q9KF65 Q9KF90 Q9KFL5 Q9KG49 Q9KG78 Q9KGA4 Q9KGC7 Q9KGE8 Q9KHL7
Q9KHT6 Q9KHT8 Q9KI10 Q9KIN5 Q9KIV0 Q9KIV1 Q9KIV2 Q9KJ74 Q9KJ75 Q9KJC1 Q9KJC2
Q9KJN3 Q9KJY8 Q9KJZ3 Q9KK91 Q9KKA3 Q9KKQ7 Q9KKU0 Q9KL06 Q9KL07 Q9KLK7 Q9KLS6
Q9KLX9 Q9KM59 Q9KM62 Q9KM66 Q9KM72 Q9KNC9 Q9KNE4 Q9KNE5 Q9KNG8 Q9KNG9 Q9KNH0
Q9KNH1 Q9KNJ6 Q9KNM9 Q9KNN1 Q9KNS2 Q9KNS3 Q9KNU5 Q9KPG0 Q9KPG4 Q9KPG9 Q9KPI5
Q9KPS2 Q9KPV9 Q9KPZ7 Q9KQE7 Q9KQE9 Q9KQS0 Q9KQS1 Q9KQU6 Q9KQU7 Q9KQW9 Q9KR19
Q9KR65 Q9KR66 Q9KRE2 Q9KRU4 Q9KS42 Q9KSA1 Q9KSG1 Q9KSG9 Q9KSL2 Q9KSL8 Q9KSM3
Q9KSW1 Q9KSY9 Q9KT06 Q9KT86 Q9KT89 Q9KT90 Q9KT91 Q9KTD2 Q9KTD3 Q9KTE4 Q9KTJ6
Q9KTU4 Q9KTU5 Q9KU26 Q9KU46 Q9KUC0 Q9KUH2 Q9KUJ3 Q9KUJ4 Q9KUJ7 Q9KUR8 Q9KV08
Q9KV09 Q9KV36 Q9KVD8 Q9KVD9 Q9KVF1 Q9KVI9 Q9KVK7 Q9KVP7 Q9KVQ4 Q9KVR0 Q9KVS9
Q9KVY4 Q9KWK6 Q9KWK8 Q9KWL2 Q9KWM8 Q9KWV3 Q9KWV4 Q9KYS0 Q9L091 Q9L3P8 Q9L3P9
Q9L3Q0 Q9L4N4 Q9L523 Q9L598 Q9L6B3 Q9L6L2 Q9L6N6 Q9L6N7 Q9L6P1 Q9L6R7 Q9L772
Q9L7M1 Q9L7Q2 Q9L7R3 Q9L7R4 Q9L7R5 Q9L8J3 Q9L8M1 Q9L8M2 Q9L8M5 Q9LA01 Q9LA06
Q9LA29 Q9LA53 Q9LA56 Q9LA60 Q9LCB5 Q9LCJ0 Q9LD18 Q9LD37 Q9LD40 Q9LD45 Q9LD83
Q9LDC0 Q9LDF2 Q9LDG2 Q9LDH0 Q9LDH3 Q9LDM5 Q9LDN1 Q9LDQ3 Q9LDR2 Q9LDR4 Q9LDS6
Q9LDT0 Q9LDU0 Q9LDU6 Q9LDX3 Q9LE20 Q9LE59 Q9LEA3 Q9LEM9 Q9LEN0 Q9LEQ3 Q9LET3
Q9LEV7 Q9LEX1 Q9LF09 Q9LF50 Q9LF61 Q9LF64 Q9LF79 Q9LF80 Q9LF97 Q9LFB2 Q9LFB8
Q9LFC6 Q9LFE3 Q9LFG1 Q9LFG8 Q9LFH0 Q9LFH9 Q9LFL0 Q9LFN3 Q9LFN5 Q9LFN8 Q9LFP1
Q9LFP3 Q9LFP6 Q9LFQ0 Q9LFR0 Q9LFR1 Q9LFS2 Q9LFS4 Q9LFT0 Q9LFT1 Q9LFV3 Q9LFX9
Q9LFZ8 Q9LGB4 Q9LH39 Q9LH74 Q9LH79 Q9LH82 Q9LH83 Q9LH84 Q9LHA1 Q9LHD1 Q9LHD2
Q9LHD9 Q9LHE5 Q9LHJ3 Q9LHK3 Q9LHK4 Q9LHL6 Q9LHN4 Q9LHN7 Q9LHP4 Q9LHQ6 Q9LHS7
Q9LHZ7 Q9LI17 Q9LI61 Q9LI65 Q9LI83 Q9LIC2 Q9LIC6 Q9LIC7 Q9LIE4 Q9LIG2 Q9LIH7

Q9LIK6 Q9LIK7 Q9LIL4 Q9LIN9 Q9LIP3 Q9LIP4 Q9LIP5 Q9LIP6 Q9LIR9 Q9LIS3 Q9LJF3
Q9LJI2 Q9LJJ7 Q9LJK1 Q9LJK2 Q9LJK3 Q9LJK4 Q9LJL6 Q9LJM4 Q9LJN2 Q9LJP4 Q9LJQ5
Q9LJS0 Q9LJS2 Q9LJU2 Q9LJU6 Q9LJV9 Q9LJW0 Q9LJW7 Q9LJX0 Q9LJX5 Q9LJY0 Q9LJY5
Q9LJY7 Q9LK03 Q9LK09 Q9LK16 Q9LK25 Q9LK29 Q9LK31 Q9LK35 Q9LK43 Q9LK50 Q9LK62
Q9LK64 Q9LK65 Q9LK90 Q9LK94 Q9LKA9 Q9LKG8 Q9LKH3 Q9LKU2 Q9LKW9 Q9LLC6 Q9LLY4
Q9LM24 Q9LM25 Q9LM60 Q9LM69 Q9LMB9 Q9LMC8 Q9LMI3 Q9LMI4 Q9LMJ1 Q9LMM0 Q9LMM1
Q9LMM2 Q9LMN6 Q9LMN7 Q9LMN8 Q9LMP1 Q9LMP7 Q9LMT9 Q9LMU1 Q9LMX7 Q9LMZ9 Q9LN20
Q9LN27 Q9LN49 Q9LN50 Q9LN59 Q9LN62 Q9LN68 Q9LN71 Q9LND8 Q9LND9 Q9LNH5 Q9LNH6
Q9LNJ0 Q9LNJ4 Q9LNK7 Q9LNL1 Q9LNQ1 Q9LNQ4 Q9LNV3 Q9LNV9 Q9LP10 Q9LP11 Q9LP24
Q9LP74 Q9LP77 Q9LPC1 Q9LPF1 Q9LPF6 Q9LPG3 Q9LPQ8 Q9LPR6 Q9LPS1 Q9LPT1 Q9LPU2
Q9LPU9 Q9LPV4 Q9LPV5 Q9LPZ3 Q9LPZ9 Q9LQ32 Q9LQ73 Q9LQC9 Q9LQL2 Q9LQM2 Q9LQU2
Q9LQU4 Q9LQY3 Q9LQZ0 Q9LR57 Q9LR68 Q9LR87 Q9LRB5 Q9LRB7 Q9LRI7 Q9LRP1 Q9LRP2
Q9LRP3 Q9LRS2 Q9LRS5 Q9LRT1 Q9LRW9 Q9LS19 Q9LS22 Q9LS23 Q9LS42 Q9LS43 Q9LS44
Q9LS45 Q9LS46 Q9LS79 Q9LS80 Q9LS95 Q9LS99 Q9LSB1 Q9LSC4 Q9LSE4 Q9LSE7 Q9LSE8
Q9LSF0 Q9LSF6 Q9LSF8 Q9LSG3 Q9LSI9 Q9LSJ2 Q9LSJ5 Q9LSJ6 Q9LSJ8 Q9LSK9 Q9LSL5
Q9LSP7 Q9LSR8 Q9LSR9 Q9LSS0 Q9LSS4 Q9LSV3 Q9LSW9 Q9LSX7 Q9LSZ9 Q9LT02 Q9LT15
Q9LT68 Q9LT71 Q9LT84 Q9LT96 Q9LTD8 Q9LTD9 Q9LTG0 Q9LTG5 Q9LTH9 Q9LTI3 Q9LTL0
Q9LTL2 Q9LTL8 Q9LTM0 Q9LTM1 Q9LTM2 Q9LTM3 Q9LTM4 Q9LTM6 Q9LTM7 Q9LTP6 Q9LTR2
Q9LTW0 Q9LTX4 Q9LTZ9 Q9LU01 Q9LU40 Q9LU41 Q9LU76 Q9LU77 Q9LUC5 Q9LUC6 Q9LUC8
Q9LUC9 Q9LUD7 Q9LUE3 Q9LUE4 Q9LUF2 Q9LUH2 Q9LUH3 Q9LUL1 Q9LUL4 Q9LUL6 Q9LUL8
Q9LUN2 Q9LUN4 Q9LUR4 Q9LUS2 Q9LUZ6 Q9LUZ9 Q9LV10 Q9LV11 Q9LV16 Q9LV20 Q9LV48
Q9LV72 Q9LV81 Q9LV85 Q9LVD2 Q9LVD6 Q9LVD9 Q9LVE0 Q9LVE4 Q9LVF1 Q9LVF4 Q9LVH5
Q9LVI6 Q9LVK9 Q9LVM0 Q9LVM1 Q9LVN2 Q9LVP0 Q9LVP9 Q9LVR3 Q9LVR4 Q9LVR5 Q9LVU1
Q9LVV3 Q9LVY7 Q9LVZ3 Q9LVZ6 Q9LVZ7 Q9LW27 Q9LW56 Q9LW77 Q9LW83 Q9LW86 Q9LWR0
Q9LWR2 Q9LX29 Q9LX66 Q9LX93 Q9LXA5 Q9LXD4 Q9LXF3 Q9LXF8 Q9LXH0 Q9LXJ2 Q9LXL9
Q9LXM3 Q9LXR8 Q9LXS1 Q9LXT9 Q9LXV3 Q9LXX4 Q9LXX8 Q9LY03 Q9LY08 Q9LY09 Q9LY28
Q9LY32 Q9LY41 Q9LY46 Q9LY50 Q9LY62 Q9LY74 Q9LY77 Q9LY99 Q9LYD5 Q9LYG1 Q9LYG9
Q9LYK2 Q9LYM2 Q9LYN0 Q9LYN3 Q9LYN6 Q9LYN8 Q9LYP1 Q9LYQ4 Q9LYR2 Q9LYR6 Q9LYS2
Q9LYS6 Q9LYS9 Q9LYT3 Q9LYU7 Q9LYX1 Q9LYY2 Q9LZ20 Q9LZ31 Q9LZ39 Q9LZ72 Q9LZ76
Q9LZA4 Q9LZB8 Q9LZD0 Q9LZH8 Q9LZI2 Q9LZJ3 Q9LZJ5 Q9LZJ6 Q9LZM4 Q9LZM5 Q9LZM7
Q9LZN2 Q9LZQ0 Q9LZQ1 Q9LZR3 Q9LZR8 Q9LZT4 Q9LZU4 Q9LZU9 Q9LZV7 Q9LZV8 Q9M001
Q9M007 Q9M008 Q9M012 Q9M020 Q9M021 Q9M024 Q9M028 Q9M038 Q9M066 Q9M092 Q9M095
Q9M0A4 Q9M0B6 Q9M0B7 Q9M0B8 Q9M0C3 Q9M0G0 Q9M0G5 Q9M0G7 Q9M0G9 Q9M0L3 Q9M0M2
Q9M0R4 Q9M0R5 Q9M0R6 Q9M0R7 Q9M0X5 Q9M0Y1 Q9M0Y2 Q9M0Y3 Q9M0Z3 Q9M0Z9 Q9M115
Q9M126 Q9M129 Q9M130 Q9M131 Q9M145 Q9M146 Q9M158 Q9M171 Q9M172 Q9M173 Q9M174
Q9M175 Q9M1B5 Q9M1C7 Q9M1E1 Q9M1E2 Q9M1E7 Q9M1G3 Q9M1G4 Q9M1I2 Q9M1I6 Q9M1J0
Q9M1J1 Q9M1K3 Q9M1K5 Q9M1L7 Q9M1P7 Q9M1Q9 Q9M1S8 Q9M1V9 Q9M1Z9 Q9M271 Q9M2A0
Q9M2C0 Q9M2C3 Q9M2D2 Q9M2G7 Q9M2J9 Q9M2L4 Q9M2P2 Q9M2R0 Q9M2S4 Q9M2S6 Q9M2U0
Q9M2U2 Q9M2V5 Q9M2V6 Q9M2V7 Q9M2W3 Q9M2W6 Q9M2Y3 Q9M2Y6 Q9M2Z1 Q9M2Z4 Q9M2Z8
Q9M304 Q9M306 Q9M313 Q9M325 Q9M331 Q9M342 Q9M345 Q9M353 Q9M363 Q9M376 Q9M390
Q9M392 Q9M393 Q9M3B0 Q9M3B1 Q9M3B2 Q9M3B3 Q9M3B9 Q9M3D6 Q9M3D7 Q9M3D8 Q9M3E5
Q9M3H5 Q9M3I6 Q9M3I8 Q9M3I9 Q9M3J0 Q9M3J1 Q9M3J4 Q9M3J5 Q9M3L0 Q9M3L2 Q9M3L4
Q9M3L5 Q9M3L9 Q9M3M6 Q9M3T9 Q9M3W5 Q9M4G3 Q9M5A9 Q9M5P2 Q9M5P3 Q9M5Q1 Q9M622
Q9M643 Q9M647 Q9M670 Q9M6A3 Q9M6A7 Q9M6N7 Q9M7B7 Q9M7B8 Q9M7I9 Q9M7J9 Q9M7K4

Q9M7M1 Q9M812 Q9M815 Q9M817 Q9M841 Q9M883 Q9M895 Q9M896 Q9M897 Q9M8J2 Q9M8L9
Q9M8S6 Q9M8T0 Q9M8W5 Q9M8W7 Q9M903 Q9M9A5 Q9M9A8 Q9M9C5 Q9M9E0 Q9M9E1 Q9M9H7
Q9M9L3 Q9M9M4 Q9M9N9 Q9M9R9 Q9M9S4 Q9M9S6 Q9M9U0 Q9M9V7 Q9M9X0 Q9M9Y5 Q9M9Z2
Q9MA44 Q9MA60 Q9MA83 Q9MA87 Q9MA88 Q9MA90 Q9MA96 Q9MAG3 Q9MAH4 Q9MAM3 Q9MAP8
Q9MAS5 Q9MAX3 Q9MBD8 Q9MCI8 Q9MDB7 Q9MDF2 Q9MDJ1 Q9MDZ9 Q9ME31 Q9ME79 Q9MED6
Q9MEI2 Q9MEI6 Q9MES8 Q9MET2 Q9MEV6 Q9MFN3 Q9MFN9 Q9MFP7 Q9MG84 Q9MGD7 Q9MGL5
Q9MGM1 Q9MGM4 Q9MI31 Q9MI32 Q9MI35 Q9MI36 Q9MI37 Q9MI38 Q9MI39 Q9MI64 Q9MI97
Q9MIX8 Q9MIX9 Q9MIY0 Q9MIY1 Q9MIY2 Q9MIY3 Q9MIY4 Q9MIY5 Q9MIY6 Q9MIY7 Q9MIY8
Q9MIY9 Q9MIZ0 Q9MIZ2 Q9MJ99 Q9MJA1 Q9MJA2 Q9MJA8 Q9MJB2 Q9MJC0 Q9MJV1 Q9MLD6
Q9MLD7 Q9MLI3 Q9MLI6 Q9MLI7 Q9MLI8 Q9MLI9 Q9MLJ0 Q9MLJ1 Q9MLJ2 Q9MLJ3 Q9MLJ4
Q9MLJ5 Q9MLJ6 Q9MLJ7 Q9MLJ8 Q9MLJ9 Q9MLK0 Q9MLK1 Q9MLK2 Q9MLK3 Q9MLK4 Q9MLK5
Q9MLK6 Q9MLK7 Q9MLK8 Q9MLK9 Q9MLL0 Q9MLL3 Q9MLL4 Q9MLL5 Q9MLL6 Q9MLQ5 Q9MMZ2
Q9MP77 Q9MQJ9 Q9MQK2 Q9MQX1 Q9MQX8 Q9MQY0 Q9MQY1 Q9MQY3 Q9MQY5 Q9MQY6 Q9MR49
Q9MS53 Q9MS54 Q9MS55 Q9MS57 Q9MS58 Q9MS59 Q9MS60 Q9MS61 Q9MS62 Q9MS63 Q9MS64
Q9MS65 Q9MS66 Q9MS67 Q9MS68 Q9MS69 Q9MS70 Q9MS71 Q9MS72 Q9MS73 Q9MS75 Q9MS76
Q9MS77 Q9MS78 Q9MS79 Q9MSC0 Q9MSC1 Q9MSC2 Q9MSQ0 Q9MSR1 Q9MSR2 Q9MTH5 Q9MTH7
Q9MTH9 Q9MTI0 Q9MTI2 Q9MTI4 Q9MTJ4 Q9MTJ5 Q9MTJ7 Q9MTK3 Q9MTK4 Q9MTK5 Q9MTK6
Q9MTK7 Q9MTK9 Q9MTL1 Q9MTL2 Q9MTL6 Q9MTL8 Q9MTM0 Q9MTM8 Q9MTN1 Q9MTN2 Q9MTN5
Q9MTN7 Q9MTN8 Q9MTP5 Q9MTQ3 Q9MTQ4 Q9MUC0 Q9MUJ0 Q9MUJ1 Q9MUJ2 Q9MUJ3 Q9MUJ4
Q9MUJ5 Q9MUJ6 Q9MUJ7 Q9MUJ8 Q9MUJ9 Q9MUK0 Q9MUK1 Q9MUK2 Q9MUK3 Q9MUK4 Q9MUK5
Q9MUK8 Q9MUL1 Q9MUL3 Q9MUL4 Q9MUL9 Q9MUM0 Q9MUM3 Q9MUM4 Q9MUM8 Q9MUN3 Q9MUN4
Q9MUN6 Q9MUN7 Q9MUN8 Q9MUP4 Q9MUP9 Q9MUQ0 Q9MUQ1 Q9MUQ2 Q9MUQ3 Q9MUQ4 Q9MUQ6
Q9MUQ7 Q9MUQ9 Q9MUR7 Q9MUR8 Q9MUS0 Q9MUS1 Q9MUS2 Q9MUS3 Q9MUS4 Q9MUS9 Q9MUT0
Q9MUT1 Q9MUV2 Q9MUV3 Q9MUV4 Q9MUV5 Q9MUV6 Q9MUV7 Q9MUV9 Q9MUW0 Q9MUW1 Q9MUW2
Q9MVK2 Q9MVL6 Q9MYL0 Q9MYM7 Q9MYS6 Q9MYU4 Q9MYV9 Q9MYW0 Q9MYW3 Q9MYW9 Q9MYX0
Q9MYX7 Q9MYY3 Q9MYY9 Q9MYZ3 Q9MYZ9 Q9MZ01 Q9MZ03 Q9MZ08 Q9MZ13 Q9MZ15 Q9MZ16
Q9MZ19 Q9MZ30 Q9MZ34 Q9MZ37 Q9MZ41 Q9MZD1 Q9MZF4 Q9MZI6 Q9MZJ3 Q9MZJ7 Q9MZK6
Q9MZK9 Q9MZS1 Q9MZS9 Q9MZT1 Q9MZT2 Q9MZV8 Q9MZZ8 Q9N008 Q9N092 Q9N0A4 Q9N0B3
Q9N0B5 Q9N0D0 Q9N0J9 Q9N0K1 Q9N0V3 Q9N0W2 Q9N0W7 Q9N0Y0 Q9N0Z0 Q9N0Z4 Q9N0Z6
Q9N119 Q9N126 Q9N1E4 Q9N1E5 Q9N1E6 Q9N1F0 Q9N1F2 Q9N1Q4 Q9N1R2 Q9N1R5 Q9N1R6
Q9N234 Q9N281 Q9N285 Q9N291 Q9N293 Q9N294 Q9N295 Q9N296 Q9N297 Q9N298 Q9N2A2
Q9N2A3 Q9N2A4 Q9N2A7 Q9N2B0 Q9N2B1 Q9N2B2 Q9N2B6 Q9N2B7 Q9N2D9 Q9N2G5 Q9N2I5
Q9N2I7 Q9N2I9 Q9N2J1 Q9N2J4 Q9N2J8 Q9N2K0 Q9N3T5 Q9N3X8 Q9N491 Q9N4K0 Q9N4M4
Q9N4U2 Q9N4Y9 Q9N587 Q9N598 Q9N5N8 Q9N5Y2 Q9N623 Q9N6P4 Q9NA13 Q9NA25 Q9NA64
Q9NA73 Q9NA75 Q9NAQ9 Q9NBW1 Q9NDH7 Q9NDM2 Q9NDV2 Q9NEQ0 Q9NFA0 Q9NGJ7 Q9NGP5
Q9NHW7 Q9NIV0 Q9NIV1 Q9NIW4 Q9NJ15 Q9NJP2 Q9NL29 Q9NLA1 Q9NNX6 Q9NNZ3 Q9NP58
Q9NP59 Q9NP60 Q9NP78 Q9NP80 Q9NP84 Q9NP91 Q9NP94 Q9NP99 Q9NPA0 Q9NPA1 Q9NPB0
Q9NPB9 Q9NPC1 Q9NPC2 Q9NPC4 Q9NPD5 Q9NPE6 Q9NPF0 Q9NPF2 Q9NPG1 Q9NPG4 Q9NPG8
Q9NPH3 Q9NPH5 Q9NPI0 Q9NPI9 Q9NPL8 Q9NPR2 Q9NPR9 Q9NPU4 Q9NPY3 Q9NPZ5 Q9NQ11
Q9NQ25 Q9NQ34 Q9NQ40 Q9NQ60 Q9NQ84 Q9NQ90 Q9NQA5 Q9NQC3 Q9NQG1 Q9NQG6 Q9NQN1
Q9NQQ7 Q9NQR9 Q9NQS3 Q9NQS5 Q9NQW8 Q9NQX5 Q9NQX7 Q9NQZ7 Q9NR16 Q9NR34 Q9NR61
Q9NR71 Q9NR77 Q9NR82 Q9NR96 Q9NR97 Q9NRA2 Q9NRB3 Q9NRC1 Q9NRD8 Q9NRD9 Q9NRJ7
Q9NRK6 Q9NRM0 Q9NRM6 Q9NRP0 Q9NRQ2 Q9NRQ5 Q9NRR2 Q9NRS4 Q9NRU3 Q9NRX5 Q9NRX6
Q9NRY6 Q9NRY7 Q9NRZ5 Q9NRZ7 Q9NS00 Q9NS40 Q9NS62 Q9NS64 Q9NS66 Q9NS67 Q9NS68

Q9NS69 Q9NS75 Q9NS82 Q9NS84 Q9NS93 Q9NSA0 Q9NSA2 Q9NSC7 Q9NSD5 Q9NSD7 Q9NSI5
Q9NSK7 Q9NST1 Q9NT68 Q9NT99 Q9NTG1 Q9NTI2 Q9NTJ5 Q9NTN3 Q9NTN9 Q9NTQ9 Q9NU53
Q9NUB4 Q9NUD9 Q9NUE0 Q9NUH8 Q9NUM3 Q9NUM4 Q9NUN5 Q9NUN7 Q9NUQ2 Q9NUR3 Q9NUT2
Q9NUV7 Q9NV12 Q9NV29 Q9NV58 Q9NV64 Q9NV92 Q9NV96 Q9NVA4 Q9NVC3 Q9NVH0 Q9NVI7
Q9NVM1 Q9NVV0 Q9NVV5 Q9NW15 Q9NW97 Q9NWC5 Q9NWD8 Q9NWF4 Q9NWH2 Q9NWQ8 Q9NWR8
Q9NWS6 Q9NWW5 Q9NWW9 Q9NX00 Q9NX14 Q9NX47 Q9NX52 Q9NX61 Q9NX62 Q9NX76 Q9NX77
Q9NX78 Q9NX94 Q9NX95 Q9NXB9 Q9NXE4 Q9NXF8 Q9NXG6 Q9NXH8 Q9NXI6 Q9NXJ0 Q9NXK6
Q9NXL6 Q9NXS2 Q9NXW2 Q9NY15 Q9NY25 Q9NY26 Q9NY28 Q9NY35 Q9NY37 Q9NY46 Q9NY47
Q9NY64 Q9NY72 Q9NY91 Q9NY97 Q9NYB5 Q9NYG2 Q9NYG8 Q9NYJ7 Q9NYK1 Q9NYL4 Q9NYL5
Q9NYM4 Q9NYM9 Q9NYP7 Q9NYQ6 Q9NYQ7 Q9NYQ8 Q9NYR8 Q9NYV7 Q9NYV8 Q9NYV9 Q9NYW0
Q9NYW1 Q9NYW2 Q9NYW3 Q9NYW4 Q9NYW5 Q9NYW6 Q9NYW7 Q9NYX4 Q9NYZ1 Q9NYZ2 Q9NYZ4
Q9NZ01 Q9NZ08 Q9NZ42 Q9NZ43 Q9NZ45 Q9NZ53 Q9NZ94 Q9NZA1 Q9NZC2 Q9NZC3 Q9NZD1
Q9NZG7 Q9NZH0 Q9NZJ5 Q9NZJ7 Q9NZM1 Q9NZM6 Q9NZN1 Q9NZP0 Q9NZP2 Q9NZP5 Q9NZQ7
Q9NZQ8 Q9NZR2 Q9NZS2 Q9NZS9 Q9NZU0 Q9NZU1 Q9NZV1 Q9NZV8 Q9P003 Q9P035 Q9P055
Q9P0B6 Q9P0I2 Q9P0J0 Q9P0K1 Q9P0K9 Q9P0L0 Q9P0L9 Q9P0N5 Q9P0N8 Q9P0S2 Q9P0S3
Q9P0S9 Q9P0T7 Q9P0U1 Q9P0V8 Q9P0X4 Q9P109 Q9P126 Q9P1P4 Q9P1P5 Q9P1Q5 Q9P1W3
Q9P1W8 Q9P1Z3 Q9P241 Q9P244 Q9P246 Q9P273 Q9P283 Q9P290 Q9P291 Q9P296 Q9P298
Q9P2B2 Q9P2C4 Q9P2D8 Q9P2E5 Q9P2E7 Q9P2E8 Q9P2E9 Q9P2J2 Q9P2K9 Q9P2P1 Q9P2S2
Q9P2U7 Q9P2U8 Q9P2V4 Q9P2W7 Q9P2W9 Q9P2X0 Q9P353 Q9P375 Q9P377 Q9P380 Q9P381
Q9P3B2 Q9P3E5 Q9P3E6 Q9P3K8 Q9P3S1 Q9P3T7 Q9P3T9 Q9P3U6 Q9P3U7 Q9P3V0 Q9P3V3
Q9P3V5 Q9P3V7 Q9P414 Q9P429 Q9P4U6 Q9P4W1 Q9P4X1 Q9P544 Q9P5L0 Q9P5N0 Q9P5N2
Q9P5N4 Q9P5X6 Q9P6I3 Q9P6J0 Q9P6J2 Q9P6J5 Q9P6J7 Q9P6J9 Q9P6K1 Q9P6K6 Q9P6L4
Q9P6L7 Q9P6M5 Q9P6M7 Q9P6N3 Q9P6N5 Q9P6N6 Q9P6N9 Q9P6P1 Q9P6P8 Q9P6Q3 Q9P6R3
Q9P6R5 Q9P6S2 Q9P6S6 Q9P768 Q9P771 Q9P781 Q9P787 Q9P7B3 Q9P7C1 Q9P7C2 Q9P7C3
Q9P7D0 Q9P7D8 Q9P7E0 Q9P7E9 Q9P7F3 Q9P7F9 Q9P7G3 Q9P7G4 Q9P7G5 Q9P7G8 Q9P7H7
Q9P7I1 Q9P7I9 Q9P7K7 Q9P7L7 Q9P7M0 Q9P7P0 Q9P7Q0 Q9P7Q9 Q9P7R8 Q9P7S8 Q9P7T9
Q9P7U3 Q9P7V2 Q9P7V8 Q9P7X0 Q9P7X9 Q9P7Y3 Q9P824 Q9P838 Q9P854 Q9P864 Q9P8E3
Q9P904 Q9P959 Q9P9U1 Q9PA34 Q9PA93 Q9PAG1 Q9PAN4 Q9PAN6 Q9PAS8 Q9PB12 Q9PB87
Q9PBK1 Q9PBK2 Q9PC78 Q9PCE0 Q9PCE8 Q9PCR3 Q9PCY1 Q9PDC5 Q9PDC9 Q9PDL9 Q9PDT7
Q9PE46 Q9PE81 Q9PEE7 Q9PEF2 Q9PEI1 Q9PEL3 Q9PEQ2 Q9PF83 Q9PFB2 Q9PFB3 Q9PFB4
Q9PFU3 Q9PFU4 Q9PGD4 Q9PGG2 Q9PGI5 Q9PGI8 Q9PH76 Q9PHA5 Q9PHR3 Q9PHS3 Q9PHT7
Q9PHT8 Q9PHZ4 Q9PI72 Q9PI98 Q9PIE4 Q9PIN2 Q9PIQ8 Q9PIS4 Q9PIW1 Q9PJB9 Q9PJG0
Q9PJJ8 Q9PJK8 Q9PJN1 Q9PJP6 Q9PJT2 Q9PJU1 Q9PJV1 Q9PJX8 Q9PJY1 Q9PJY2 Q9PJY3
Q9PJY8 Q9PK20 Q9PK22 Q9PK88 Q9PKB3 Q9PKB4 Q9PKB5 Q9PKB6 Q9PKE2 Q9PKE3 Q9PKI5
Q9PKL7 Q9PKW7 Q9PKW9 Q9PKX0 Q9PKX5 Q9PL02 Q9PL34 Q9PL35 Q9PL41 Q9PL44 Q9PL45
Q9PL46 Q9PL47 Q9PL67 Q9PLA1 Q9PLB0 Q9PLG6 Q9PLI2 Q9PLI3 Q9PLI7 Q9PLL3 Q9PLN5
Q9PMA3 Q9PMA7 Q9PMQ9 Q9PN86 Q9PNB9 Q9PNJ9 Q9PNM6 Q9PNU1 Q9PNX7 Q9PPI1 Q9PPN7
Q9PQ85 Q9PQM6 Q9PQW0 Q9PQW5 Q9PQX8 Q9PQY5 Q9PQY6 Q9PR08 Q9PR09 Q9PR13 Q9PR17
Q9PR60 Q9PR70 Q9PR81 Q9PR84 Q9PR89 Q9PR97 Q9PR99 Q9PRA1 Q9PRA3 Q9PRA4 Q9PRB5
Q9PRD7 Q9PRD8 Q9PSZ3 Q9PT62 Q9PT84 Q9PTE6 Q9PTS8 Q9PTW1 Q9PTW3 Q9PU16 Q9PU17
Q9PUB8 Q9PUF6 Q9PUI7 Q9PUK8 Q9PUN1 Q9PUQ8 Q9PUU6 Q9PUU8 Q9PVZ4 Q9PW88 Q9PWH2
Q9PWR4 Q9PWW3 Q9PYC7 Q9PYC8 Q9PYD0 Q9PYD1 Q9PYD2 Q9Q0L9 Q9Q0U6 Q9Q0U7 Q9Q2W5
Q9Q6P4 Q9Q714 Q9Q8T2 Q9Q8W2 Q9Q9G3 Q9QAB7 Q9QAQ8 Q9QAQ9 Q9QAR5 Q9QAR6 Q9QAR9
Q9QAX1 Q9QBF0 Q9QBY2 Q9QBZ0 Q9QBZ4 Q9QBZ8 Q9QIZ6 Q9QJ14 Q9QJ17 Q9QJ35 Q9QJ36

Q9QJ41 Q9QJ42 Q9QJ43 Q9QJ44 Q9QJ45 Q9QJ46 Q9QJ48 Q9QJ51 Q9QJ54 Q9QJ59 Q9QJT6
Q9QMI0 Q9QNA6 Q9QR69 Q9QSP1 Q9QSQ7 Q9QSQ8 Q9QUI6 Q9QUJ0 Q9QUJ7 Q9QUK3 Q9QUK6
Q9QUL7 Q9QUM0 Q9QUM4 Q9QUN7 Q9QUP4 Q9QUQ5 Q9QUT0 Q9QW30 Q9QWE9 Q9QWJ9 Q9QWS8
Q9QX01 Q9QX05 Q9QX29 Q9QX75 Q9QX82 Q9QXA6 Q9QXF7 Q9QXH4 Q9QXI3 Q9QXM0 Q9QXP0
Q9QXS4 Q9QXS7 Q9QXS8 Q9QXT3 Q9QXT4 Q9QXU2 Q9QXW9 Q9QXX0 Q9QXX4 Q9QXY7 Q9QXY9
Q9QXZ6 Q9QXZ9 Q9QY00 Q9QY30 Q9QY33 Q9QY40 Q9QY42 Q9QY44 Q9QY73 Q9QY76 Q9QY80
Q9QY81 Q9QY96 Q9QYA2 Q9QYB1 Q9QYC1 Q9QYC5 Q9QYC6 Q9QYC7 Q9QYE2 Q9QYE5 Q9QYE6
Q9QYE7 Q9QYF1 Q9QYH9 Q9QYI4 Q9QYJ1 Q9QYJ4 Q9QYK4 Q9QYK5 Q9QYM0 Q9QYM9 Q9QYN8
Q9QYP0 Q9QYP1 Q9QYP2 Q9QYS2 Q9QYT7 Q9QYU3 Q9QYV0 Q9QYV1 Q9QYW1 Q9QYW5 Q9QZ03
Q9QZ09 Q9QZ15 Q9QZ26 Q9QZ39 Q9QZ49 Q9QZ65 Q9QZ70 Q9QZ71 Q9QZA6 Q9QZC1 Q9QZC2
Q9QZC8 Q9QZD8 Q9QZE9 Q9QZG1 Q9QZH0 Q9QZH8 Q9QZI8 Q9QZI9 Q9QZL2 Q9QZM4 Q9QZM6
Q9QZN9 Q9QZS6 Q9QZS7 Q9QZT0 Q9QZU4 Q9QZW0 Q9QZX8 Q9QZY5 Q9QZY6 Q9QZY7 Q9QZY8
Q9QZY9 Q9QZZ0 Q9QZZ1 Q9QZZ2 Q9R007 Q9R024 Q9R031 Q9R032 Q9R044 Q9R049 Q9R053
Q9R066 Q9R069 Q9R0A1 Q9R0C5 Q9R0C9 Q9R0E0 Q9R0H2 Q9R0K7 Q9R0L9 Q9R0M0 Q9R0M1
Q9R0M4 Q9R0M8 Q9R0N3 Q9R0N4 Q9R0N5 Q9R0N6 Q9R0N7 Q9R0N8 Q9R0N9 Q9R0P6 Q9R0Q2
Q9R0Q3 Q9R0Q8 Q9R0Q9 Q9R0S2 Q9R0S5 Q9R0T4 Q9R0W2 Q9R0W5 Q9R0W9 Q9R0Y1 Q9R0Y8
Q9R100 Q9R110 Q9R141 Q9R154 Q9R155 Q9R157 Q9R158 Q9R159 Q9R160 Q9R172 Q9R186
Q9R187 Q9R193 Q9R1C6 Q9R1C8 Q9R1F8 Q9R1I1 Q9R1J0 Q9R1K2 Q9R1K6 Q9R1M7 Q9R1N2
Q9R1N3 Q9R1N9 Q9R1Q6 Q9R1Q7 Q9R1Q9 Q9R1S3 Q9R1S4 Q9R1S7 Q9R1T7 Q9R1T9 Q9R1U7
Q9R1U9 Q9R1V4 Q9R1V6 Q9R1V7 Q9R1W3 Q9R1W5 Q9R1X5 Q9R1Z0 Q9R216 Q9R233 Q9R244
Q9R261 Q9R279 Q9R283 Q9R2B6 Q9R2C1 Q9R3F2 Q9R6T0 Q9R6T9 Q9R6U0 Q9R6W9 Q9R6X1
Q9R6X2 Q9R6Y2 Q9R9I1 Q9R9S0 Q9R9U8 Q9R9Y4 Q9RA45 Q9RA46 Q9RB37 Q9RB65 Q9RBJ2
Q9RC07 Q9RC08 Q9RC53 Q9RCA5 Q9RCG7 Q9RCG8 Q9RDV0 Q9REB3 Q9REM8 Q9RF98 Q9RFA0
Q9RFC8 Q9RFV6 Q9RFV7 Q9RFV8 Q9RFV9 Q9RFW0 Q9RGG2 Q9RGG3 Q9RGS4 Q9RGY5 Q9RGY9
Q9RGZ0 Q9RGZ1 Q9RGZ2 Q9RGZ3 Q9RGZ4 Q9RGZ5 Q9RH13 Q9RJ68 Q9RKM3 Q9RKN3 Q9RM98
Q9RMT1 Q9RMT3 Q9RMW9 Q9RMX8 Q9RMZ1 Q9RN02 Q9RN05 Q9RN06 Q9RN08 Q9RN14 Q9RN18
Q9RN19 Q9RN20 Q9RN21 Q9RN27 Q9RND2 Q9RNL1 Q9RNL5 Q9RNV1 Q9RPF2 Q9RPF3 Q9RPF4
Q9RPP2 Q9RPX5 Q9RPX7 Q9RPX9 Q9RPY2 Q9RPY3 Q9RQ29 Q9RQ30 Q9RQB4 Q9RQP6 Q9RQP8
Q9RQP9 Q9RQQ1 Q9RQQ9 Q9RR45 Q9RR78 Q9RRF5 Q9RRG7 Q9RRM6 Q9RRU7 Q9RRW7 Q9RS57
Q9RSH5 Q9RSV1 Q9RTD0 Q9RTP8 Q9RU85 Q9RU94 Q9RU97 Q9RUA0 Q9RUR0 Q9RVX8 Q9RW62
Q9RW63 Q9RWB5 Q9RWH3 Q9RX61 Q9RXG6 Q9RXX0 Q9RY44 Q9RYR9 Q9RZN6 Q9RZN7 Q9RZP0
Q9S1E4 Q9S1E6 Q9S1H4 Q9S1X8 Q9S2C8 Q9S2H4 Q9S2U8 Q9S2X7 Q9S341 Q9S342 Q9S3J9
Q9S3K0 Q9S3N1 Q9S3R8 Q9S3R9 Q9S3W3 Q9S3W5 Q9S3W9 Q9S5X0 Q9S6Z8 Q9S735 Q9S760
Q9S784 Q9S7D9 Q9S7G2 Q9S7I6 Q9S7J7 Q9S7J8 Q9S7M0 Q9S7M9 Q9S7N7 Q9S7V4 Q9S7W1
Q9S7Z8 Q9S814 Q9S836 Q9S837 Q9S845 Q9S972 Q9S9M1 Q9S9M2 Q9S9M3 Q9S9M5 Q9S9N4
Q9S9N5 Q9S9N8 Q9S9U0 Q9S9U1 Q9S9U3 Q9SA00 Q9SA05 Q9SA23 Q9SA25 Q9SA35 Q9SA36
Q9SA37 Q9SA38 Q9SA63 Q9SA71 Q9SA72 Q9SA77 Q9SAA4 Q9SAA9 Q9SAB0 Q9SAB6 Q9SAC0
Q9SAE3 Q9SAE4 Q9SAF5 Q9SAF8 Q9SAH0 Q9SAH3 Q9SAH5 Q9SAH8 Q9SAI4 Q9SAJ3 Q9SAJ7
Q9SAK7 Q9SAK8 Q9SAY1 Q9SB41 Q9SB48 Q9SB51 Q9SB52 Q9SB58 Q9SB67 Q9SB75 Q9SBA7
Q9SBI0 Q9SBM5 Q9SBM8 Q9SBN3 Q9SBQ9 Q9SCG2 Q9SCK3 Q9SCK6 Q9SCN0 Q9SCN2 Q9SCS6
Q9SCS7 Q9SCS8 Q9SCS9 Q9SCT4 Q9SCW7 Q9SCX5 Q9SCZ4 Q9SD00 Q9SD39 Q9SD42 Q9SD53
Q9SD62 Q9SD67 Q9SD80 Q9SD85 Q9SD88 Q9SDI4 Q9SDM1 Q9SDQ4 Q9SDW9 Q9SE51 Q9SE97
Q9SEK2 Q9SEK3 Q9SEL5 Q9SEL6 Q9SEZ9 Q9SF09 Q9SF29 Q9SF33 Q9SF57 Q9SFB0 Q9SFC4
Q9SFE9 Q9SFG0 Q9SFU6 Q9SFX3 Q9SG77 Q9SG79 Q9SG86 Q9SG91 Q9SG96 Q9SGD2 Q9SGG3

Q9SGI7 Q9SGP2 Q9SGY1 Q9SGY7 Q9SH30 Q9SH31 Q9SH59 Q9SH71 Q9SH76 Q9SH89 Q9SH93
Q9SHC8 Q9SHE8 Q9SHG5 Q9SHH0 Q9SHI2 Q9SHI3 Q9SHI4 Q9SHI7 Q9SHJ3 Q9SHJ5 Q9SHJ6
Q9SHL8 Q9SHM1 Q9SHR7 Q9SHU7 Q9SHV1 Q9SHV2 Q9SHY2 Q9SHZ3 Q9SI03 Q9SI06 Q9SI09
Q9SI56 Q9SI62 Q9SIA1 Q9SIA3 Q9SIA4 Q9SIA5 Q9SID2 Q9SIH4 Q9SIJ9 Q9SIK7 Q9SIL6
Q9SIN2 Q9SIQ9 Q9SIT1 Q9SIT5 Q9SIT6 Q9SIX1 Q9SIY5 Q9SIY7 Q9SIZ3 Q9SIZ4 Q9SIZ8
Q9SJ02 Q9SJ22 Q9SJ52 Q9SJ54 Q9SJ74 Q9SJA1 Q9SJA2 Q9SJA4 Q9SJB3 Q9SJE8 Q9SJE9
Q9SJG2 Q9SJH6 Q9SJJ7 Q9SJK6 Q9SJL6 Q9SJM0 Q9SJP2 Q9SJP4 Q9SJP9 Q9SJQ1 Q9SJQ5
Q9SJT0 Q9SJT7 Q9SJV5 Q9SJY5 Q9SK50 Q9SK92 Q9SK96 Q9SK99 Q9SKA9 Q9SKB2 Q9SKD3
Q9SKD6 Q9SKD7 Q9SKG5 Q9SKJ5 Q9SKJ7 Q9SKK2 Q9SKK5 Q9SKK8 Q9SKN3 Q9SKR2 Q9SKT0
Q9SKT3 Q9SKT6 Q9SKX0 Q9SKX9 Q9SKY1 Q9SKZ5 Q9SL03 Q9SL29 Q9SL56 Q9SL78 Q9SL95
Q9SLA2 Q9SLC3 Q9SLC4 Q9SLD2 Q9SLD9 Q9SLF3 Q9SLG3 Q9SLI6 Q9SLK2 Q9SLK6 Q9SLP1
Q9SLV0 Q9SM02 Q9SM23 Q9SM57 Q9SMB4 Q9SMC4 Q9SME8 Q9SME9 Q9SMM5 Q9SMN4 Q9SMP2
Q9SMP5 Q9SMS0 Q9SMU4 Q9SMX3 Q9SMY6 Q9SN26 Q9SN27 Q9SN28 Q9SN36 Q9SN38 Q9SNA3
Q9SNE5 Q9SNV9 Q9SP35 Q9SP61 Q9SP62 Q9SPI9 Q9SPK6 Q9SPM4 Q9SPM5 Q9SQ76 Q9SQG2
Q9SQH6 Q9SQH9 Q9SQL2 Q9SQN2 Q9SQR5 Q9SQU2 Q9SQZ0 Q9SR05 Q9SR44 Q9SR64 Q9SR87
Q9SR92 Q9SR93 Q9SRD3 Q9SRE4 Q9SRH5 Q9SRI2 Q9SRK7 Q9SRL2 Q9SRL3 Q9SRL7 Q9SRM0
Q9SRM9 Q9SRN0 Q9SRN1 Q9SRQ1 Q9SRQ8 Q9SRR2 Q9SRS3 Q9SRT3 Q9SRU2 Q9SRV7 Q9SRW9
Q9SRY4 Q9SS37 Q9SS39 Q9SS40 Q9SS43 Q9SS86 Q9SS98 Q9SSD1 Q9SSE8 Q9SSG5 Q9SSL9
Q9SSN0 Q9SSR0 Q9SSY6 Q9STE8 Q9STG2 Q9STI6 Q9STJ8 Q9STK7 Q9STK8 Q9STK9 Q9STL0
Q9STL1 Q9STL2 Q9STP2 Q9STP8 Q9STT5 Q9STT6 Q9STT7 Q9STT8 Q9STW3 Q9STW8 Q9STX9
Q9STY0 Q9STY7 Q9STY8 Q9SU50 Q9SU56 Q9SU58 Q9SU64 Q9SUA4 Q9SUC5 Q9SUD4 Q9SUD5
Q9SUE0 Q9SUF1 Q9SUI0 Q9SUI4 Q9SUI5 Q9SUI6 Q9SUI7 Q9SUJ1 Q9SUM3 Q9SUN3 Q9SUP0
Q9SUQ3 Q9SUQ7 Q9SUR3 Q9SUT8 Q9SUT9 Q9SUU1 Q9SUV1 Q9SUV2 Q9SUW1 Q9SUW4 Q9SUY9
Q9SV07 Q9SV13 Q9SV31 Q9SV71 Q9SV84 Q9SVC2 Q9SVC4 Q9SVE6 Q9SVE7 Q9SVF3 Q9SVG0
Q9SVL6 Q9SVM3 Q9SVN2 Q9SVT8 Q9SVU4 Q9SVV6 Q9SW07 Q9SW08 Q9SW17 Q9SW25 Q9SW31
Q9SW97 Q9SWH4 Q9SWH5 Q9SWQ9 Q9SWR5 Q9SWS1 Q9SWW6 Q9SX20 Q9SX24 Q9SX26 Q9SX28
Q9SX31 Q9SX33 Q9SX48 Q9SX55 Q9SX83 Q9SX90 Q9SX93 Q9SX98 Q9SXA5 Q9SXB0 Q9SXB3
Q9SXB4 Q9SXB5 Q9SXB6 Q9SXB8 Q9SXC4 Q9SXC9 Q9SXE1 Q9SXF8 Q9SXJ8 Q9SXL4 Q9SXS2
Q9SXS3 Q9SXX7 Q9SY14 Q9SY29 Q9SY55 Q9SY57 Q9SY89 Q9SY95 Q9SY97 Q9SYA0 Q9SYB0
Q9SYB5 Q9SYC8 Q9SYD6 Q9SYG9 Q9SYH3 Q9SYI2 Q9SYI3 Q9SYJ2 Q9SYM0 Q9SYM1 Q9SYM9
Q9SYQ1 Q9SYQ8 Q9SYS3 Q9SYS7 Q9SYW8 Q9SYX1 Q9SYX7 Q9SYY4 Q9SYZ0 Q9SYZ7 Q9SYZ8
Q9SZ11 Q9SZ13 Q9SZ42 Q9SZ46 Q9SZ74 Q9SZ96 Q9SZC9 Q9SZD5 Q9SZE2 Q9SZG0 Q9SZG1
Q9SZL4 Q9SZL9 Q9SZR1 Q9SZR9 Q9SZT0 Q9SZT4 Q9SZU1 Q9SZW4 Q9SZX8 Q9SZY4 Q9SZY7
Q9SZZ8 Q9T020 Q9T042 Q9T050 Q9T058 Q9T071 Q9T093 Q9T095 Q9T0E0 Q9T0F7 Q9T0G0
Q9T0H9 Q9T0J1 Q9T0K0 Q9T0K2 Q9T0L2 Q9T1D6 Q9T1D7 Q9T1X2 Q9T2L6 Q9T351 Q9T3H9
Q9T3Q2 Q9T3R3 Q9T3S9 Q9T3Y0 Q9T3Y2 Q9T3Y3 Q9T443 Q9T469 Q9T475 Q9T4A7 Q9T4B2
Q9T4N0 Q9T4P2 Q9T4R0 Q9T4R3 Q9T4U3 Q9T5N4 Q9T6H8 Q9T6I4 Q9T6J0 Q9T6R2 Q9T6R3
Q9T6R5 Q9T6R8 Q9T6R9 Q9T7L5 Q9T7L6 Q9T7L7 Q9T7L8 Q9T7L9 Q9T7M0 Q9T7M3 Q9T7N2
Q9T7N5 Q9T7P0 Q9T7P1 Q9T7Q0 Q9T7Q5 Q9T7Q6 Q9T7Q8 Q9T7Q9 Q9T7S2 Q9T7T7 Q9T7T8
Q9T7T9 Q9T9A5 Q9T9A6 Q9T9A7 Q9T9A8 Q9T9B2 Q9T9B6 Q9T9B7 Q9T9B9 Q9T9C0 Q9T9C1
Q9T9C8 Q9T9D5 Q9T9I2 Q9T9I7 Q9T9J2 Q9T9J5 Q9T9V5 Q9T9V6 Q9T9V8 Q9T9V9 Q9T9W0
Q9T9W1 Q9T9W3 Q9T9W4 Q9T9W5 Q9T9W6 Q9T9W7 Q9T9W8 Q9T9W9 Q9T9X0 Q9T9X1 Q9T9X3
Q9T9X5 Q9T9X6 Q9T9X8 Q9T9X9 Q9T9Y3 Q9T9Y5 Q9T9Y6 Q9T9Y7 Q9T9Y9 Q9T9Z0 Q9TA00
Q9TA07 Q9TA18 Q9TA19 Q9TA21 Q9TA24 Q9TA25 Q9TA26 Q9TA27 Q9TA28 Q9TA29 Q9TAB1

Q9TAD3 Q9TBI5 Q9TBI6 Q9TBI7 Q9TBI8 Q9TBJ1 Q9TBJ9 Q9TC94 Q9TDJ6 Q9TDJ9 Q9TDK1
Q9TDK2 Q9TDK4 Q9TDK6 Q9TDK7 Q9TDK9 Q9TDL0 Q9TDL1 Q9TDL2 Q9TDL3 Q9TDL4 Q9TDL5
Q9TDL6 Q9TDL7 Q9TDL8 Q9TDL9 Q9TDM0 Q9TDM1 Q9TDM3 Q9TDM5 Q9TDM6 Q9TDM7 Q9TDN0
Q9TDN1 Q9TDN2 Q9TDQ5 Q9TDQ7 Q9TDR1 Q9TDT5 Q9TEB4 Q9TEB5 Q9TEC1 Q9TEG7 Q9TEU7
Q9TEX8 Q9TEY4 Q9TEY5 Q9TEY6 Q9TEY7 Q9TEY8 Q9TEZ5 Q9TF10 Q9TF12 Q9TF14 Q9TF17
Q9TF20 Q9TF27 Q9TF30 Q9TF31 Q9TF32 Q9TF34 Q9TF37 Q9TF45 Q9TF46 Q9TF47 Q9TF49
Q9TF50 Q9TF51 Q9TF52 Q9TF55 Q9TF57 Q9TF61 Q9TF73 Q9TF74 Q9TF80 Q9TF86 Q9TF89
Q9TF91 Q9TF93 Q9TF95 Q9TFA0 Q9TFX4 Q9TFX5 Q9TFX6 Q9TG16 Q9TGD8 Q9TGE6 Q9TGH6
Q9TGH8 Q9TH43 Q9TH45 Q9TH46 Q9TH47 Q9TH52 Q9TH56 Q9THD6 Q9THZ1 Q9THZ2 Q9THZ3
Q9TJ82 Q9TJ83 Q9TJR3 Q9TJR4 Q9TKU7 Q9TKU8 Q9TKV2 Q9TKV3 Q9TKV5 Q9TKV7 Q9TKV8
Q9TKW1 Q9TKW2 Q9TKW3 Q9TKW4 Q9TKW5 Q9TKW6 Q9TKW7 Q9TKX2 Q9TKX9 Q9TKY1 Q9TKY2
Q9TKY3 Q9TKY4 Q9TKY6 Q9TKY7 Q9TKY8 Q9TKY9 Q9TKZ1 Q9TKZ2 Q9TKZ3 Q9TKZ8 Q9TKZ9
Q9TL00 Q9TL01 Q9TL07 Q9TL12 Q9TL13 Q9TL14 Q9TL15 Q9TL31 Q9TL32 Q9TL35 Q9TL36
Q9TL37 Q9TL56 Q9TLA3 Q9TLC2 Q9TLQ5 Q9TLQ6 Q9TLQ9 Q9TLR0 Q9TLR5 Q9TLR6 Q9TLS4
Q9TLS5 Q9TLW5 Q9TLW6 Q9TLX0 Q9TLX5 Q9TLZ7 Q9TLZ8 Q9TM16 Q9TM17 Q9TM18 Q9TM19
Q9TM20 Q9TM21 Q9TM22 Q9TM23 Q9TM24 Q9TM28 Q9TM29 Q9TM30 Q9TM31 Q9TM46 Q9TM47
Q9TM69 Q9TM72 Q9TNF7 Q9TNF8 Q9TQX4 Q9TQZ3 Q9TR28 Q9TR29 Q9TR30 Q9TRZ8 Q9TS33
Q9TSL6 Q9TSN4 Q9TSP2 Q9TSP5 Q9TSQ8 Q9TST4 Q9TST5 Q9TST6 Q9TSY7 Q9TSZ3 Q9TSZ6
Q9TT13 Q9TT15 Q9TT16 Q9TT17 Q9TT23 Q9TT37 Q9TT94 Q9TT96 Q9TTC0 Q9TTC5 Q9TTC7
Q9TTF3 Q9TTI8 Q9TTJ7 Q9TTK6 Q9TTK7 Q9TTM9 Q9TTN0 Q9TTQ9 Q9TTT5 Q9TTT7 Q9TTT8
Q9TTU3 Q9TTY3 Q9TTY5 Q9TU05 Q9TU17 Q9TU31 Q9TU32 Q9TU34 Q9TU45 Q9TU69 Q9TU72
Q9TU77 Q9TU84 Q9TU86 Q9TU88 Q9TU89 Q9TU90 Q9TU92 Q9TU93 Q9TU94 Q9TU95 Q9TU97
Q9TU99 Q9TUA0 Q9TUA1 Q9TUA2 Q9TUA4 Q9TUA6 Q9TUA7 Q9TUA8 Q9TUA9 Q9TUD4 Q9TUD6
Q9TUE1 Q9TUF8 Q9TUF9 Q9TUH9 Q9TUI4 Q9TUI8 Q9TUK4 Q9TUN9 Q9TUP7 Q9TUQ2 Q9TV16
Q9TV42 Q9TV43 Q9TV45 Q9TV47 Q9TV48 Q9TV49 Q9TV52 Q9TV66 Q9TVC8 Q9TVY6 Q9TX43
Q9TYV2 Q9TYW1 Q9TYY7 Q9U0M8 Q9U1G3 Q9U1H8 Q9U1M2 Q9U1S7 Q9U280 Q9U298 Q9U2B7
Q9U2C4 Q9U2G5 Q9U320 Q9U357 Q9U358 Q9U3A0 Q9U3D4 Q9U3H8 Q9U3I4 Q9U3K5 Q9U3N4
Q9U3P2 Q9U3U2 Q9U4F3 Q9U4G1 Q9U4H5 Q9U4L6 Q9U505 Q9U516 Q9U539 Q9U5V3 Q9U5Y0
Q9U641 Q9U6A0 Q9U6B8 Q9U6L4 Q9U6X1 Q9U720 Q9U7D5 Q9U8P7 Q9U8Q1 Q9U969 Q9UA35
Q9UBD6 Q9UBG0 Q9UBH6 Q9UBI4 Q9UBJ2 Q9UBK5 Q9UBL9 Q9UBM1 Q9UBM7 Q9UBM8 Q9UBN1
Q9UBN4 Q9UBN6 Q9UBQ6 Q9UBR5 Q9UBS5 Q9UBS9 Q9UBU6 Q9UBV2 Q9UBV7 Q9UBX3 Q9UBX8
Q9UBY0 Q9UBY5 Q9UBY8 Q9UDW1 Q9UDX5 Q9UEF7 Q9UEU0 Q9UEW3 Q9UF02 Q9UF33 Q9UG56
Q9UGF5 Q9UGF6 Q9UGF7 Q9UGH3 Q9UGI6 Q9UGM1 Q9UGN4 Q9UGP8 Q9UGQ2 Q9UGQ3 Q9UGT4
Q9UH62 Q9UH99 Q9UHC3 Q9UHC6 Q9UHC9 Q9UHE5 Q9UHE8 Q9UHF3 Q9UHF4 Q9UHI5 Q9UHI7
Q9UHJ9 Q9UHM6 Q9UHN6 Q9UHP7 Q9UHQ4 Q9UHQ9 Q9UHW9 Q9UHX3 Q9UI14 Q9UI33 Q9UI40
Q9UIB8 Q9UIG8 Q9UIJ5 Q9UIK5 Q9UIQ6 Q9UIR0 Q9UIW2 Q9UIX4 Q9UJ14 Q9UJ37 Q9UJ42
Q9UJ71 Q9UJ90 Q9UJ96 Q9UJ99 Q9UJA2 Q9UJA9 Q9UJG1 Q9UJQ1 Q9UJS0 Q9UK00 Q9UK17
Q9UK23 Q9UK28 Q9UKB5 Q9UKF2 Q9UKF5 Q9UKG4 Q9UKH3 Q9UKJ0 Q9UKJ1 Q9UKJ8 Q9UKL2
Q9UKL4 Q9UKM7 Q9UKN1 Q9UKP6 Q9UKQ2 Q9UKR5 Q9UKR8 Q9UKU0 Q9UKU6 Q9UKV5 Q9UKX5
Q9UKY4 Q9UKZ4 Q9UL01 Q9UL19 Q9UL51 Q9UL52 Q9UL54 Q9UL62 Q9ULB1 Q9ULB4 Q9ULB5
Q9ULC0 Q9ULC5 Q9ULC8 Q9ULD8 Q9ULF5 Q9ULG6 Q9ULH0 Q9ULH4 Q9ULI3 Q9ULK0 Q9ULK5
Q9ULK6 Q9ULL4 Q9ULQ1 Q9ULS5 Q9ULS6 Q9ULT6 Q9ULV1 Q9ULW2 Q9ULX5 Q9ULX7 Q9ULY5
Q9UM00 Q9UM01 Q9UM21 Q9UM44 Q9UM47 Q9UM73 Q9UMD9 Q9UMF0 Q9UMR7 Q9UMS5 Q9UMX3
Q9UMX9 Q9UMZ3 Q9UN42 Q9UN66 Q9UN67 Q9UN70 Q9UN71 Q9UN72 Q9UN73 Q9UN74 Q9UN75

Q9UN76 Q9UN88 Q9UNA3 Q9UNE0 Q9UNG2 Q9UNK0 Q9UNL2 Q9UNN8 Q9UNP4 Q9UNQ0 Q9UNU6
Q9UNW8 Q9UNX9 Q9UP38 Q9UP52 Q9UP95 Q9UPC5 Q9UPI3 Q9UPQ8 Q9UPR5 Q9UPU3 Q9UPX0
Q9UPX6 Q9UPY5 Q9UPZ6 Q9UQ05 Q9UQ53 Q9UQ90 Q9UQC9 Q9UQD0 Q9UQF0 Q9UQQ1 Q9UQV4
Q9UR17 Q9UR35 Q9URX1 Q9URY6 Q9URY8 Q9URZ3 Q9URZ4 Q9URZ8 Q9US00 Q9US04 Q9US09
Q9US10 Q9US11 Q9US20 Q9US27 Q9US37 Q9US40 Q9US44 Q9US48 Q9US57 Q9USD4 Q9USG8
Q9USH0 Q9USH7 Q9USJ2 Q9USJ3 Q9USJ4 Q9USK3 Q9USM6 Q9USM7 Q9USN0 Q9USN1 Q9USN4
Q9USP2 Q9USP5 Q9USQ0 Q9USS9 Q9UST2 Q9USV0 Q9USV6 Q9USV7 Q9USW2 Q9USW3 Q9USW4
Q9USW6 Q9USX0 Q9USY7 Q9USZ8 Q9UT00 Q9UT11 Q9UT18 Q9UT20 Q9UT29 Q9UT30 Q9UT35
Q9UT37 Q9UT41 Q9UT43 Q9UT62 Q9UT67 Q9UT91 Q9UT92 Q9UTA0 Q9UTA3 Q9UTB1 Q9UTB5
Q9UTD3 Q9UTD6 Q9UTD8 Q9UTE0 Q9UTF7 Q9UTF8 Q9UTI9 Q9UTK1 Q9UTK7 Q9UTK8 Q9UTL4
Q9UTM2 Q9UTN1 Q9UTN7 Q9UTP5 Q9UTQ9 Q9UTR0 Q9UTR6 Q9UU82 Q9UU86 Q9UU91 Q9UUA4
Q9UUA6 Q9UUC9 Q9UUD1 Q9UUD4 Q9UUF4 Q9UUH0 Q9UUH1 Q9UUH4 Q9UUH8 Q9UUI8 Q9UUJ0
Q9UUJ8 Q9UUM6 Q9UUN5 Q9UUQ6 Q9UV71 Q9UVF6 Q9UVG5 Q9UVH3 Q9UW00 Q9UW81 Q9UW82
Q9UW95 Q9UWW3 Q9UWY6 Q9UX46 Q9UX84 Q9UXG3 Q9UXQ8 Q9UXU2 Q9UY86 Q9UYC6 Q9UYS2
Q9UYV6 Q9UZ49 Q9UZK3 Q9V0X2 Q9V0Y1 Q9V1V8 Q9V274 Q9V2C1 Q9V2M8 Q9V2N1 Q9V2P5
Q9V2S1 Q9V2S5 Q9V3A4 Q9V3C5 Q9V3L1 Q9V3N2 Q9V3Q2 Q9V3R8 Q9V3W6 Q9V3X4 Q9V3X5
Q9V422 Q9V427 Q9V447 Q9V477 Q9V492 Q9V498 Q9V4A7 Q9V4K2 Q9V4N3 Q9V4Q0 Q9V568
Q9V576 Q9V589 Q9V5E1 Q9V5N8 Q9V5Z7 Q9V668 Q9V6A9 Q9V6H2 Q9V6K3 Q9V730 Q9V784
Q9V7A4 Q9V7H4 Q9V7S5 Q9V7W1 Q9V7Y2 Q9V817 Q9V818 Q9V8Y7 Q9V969 Q9V9A8 Q9V9I2
Q9V9I4 Q9V9N1 Q9V9P3 Q9V9Y4 Q9VA73 Q9VAC5 Q9VAF0 Q9VAF5 Q9VAJ3 Q9VAL0 Q9VAM6
Q9VAP3 Q9VAS7 Q9VAW0 Q9VAY3 Q9VAZ3 Q9VB11 Q9VB26 Q9VB30 Q9VB46 Q9VBN7 Q9VBW3
Q9VC27 Q9VC35 Q9VC57 Q9VCA2 Q9VCA9 Q9VCG4 Q9VCI5 Q9VCR7 Q9VCS8 Q9VCS9 Q9VCY8
Q9VCZ3 Q9VD26 Q9VD73 Q9VD74 Q9VD76 Q9VDD3 Q9VDD7 Q9VDH6 Q9VDM1 Q9VDQ0 Q9VE46
Q9VE50 Q9VED0 Q9VEG9 Q9VEI3 Q9VES1 Q9VEU0 Q9VEU1 Q9VEV4 Q9VF81 Q9VFJ3 Q9VFN2
Q9VFS6 Q9VFX3 Q9VGA2 Q9VGG5 Q9VGG8 Q9VGH7 Q9VGN7 Q9VGW1 Q9VH58 Q9VH78 Q9VHB6
Q9VHE6 Q9VHG4 Q9VHN5 Q9VHQ2 Q9VHQ6 Q9VHQ7 Q9VHR8 Q9VHS2 Q9VHS4 Q9VHS7 Q9VHT4
Q9VHX7 Q9VI20 Q9VIK2 Q9VIP7 Q9VJ83 Q9VJB6 Q9VJF2 Q9VJJ7 Q9VK50 Q9VKA4 Q9VKA5
Q9VKD7 Q9VKJ7 Q9VKX7 Q9VL32 Q9VL84 Q9VLA1 Q9VLC1 Q9VLE5 Q9VLM5 Q9VLP9 Q9VLQ1
Q9VLR9 Q9VM08 Q9VM09 Q9VM51 Q9VM64 Q9VMA7 Q9VMD5 Q9VMH0 Q9VMK1 Q9VML9 Q9VMU8
Q9VMW8 Q9VN12 Q9VN13 Q9VN97 Q9VNA0 Q9VNA4 Q9VNB3 Q9VNB5 Q9VNJ5 Q9VNK9 Q9VNM1
Q9VP04 Q9VP76 Q9VPB1 Q9VPD3 Q9VPE4 Q9VPF8 Q9VPI2 Q9VPN2 Q9VPT1 Q9VPT5 Q9VPT7
Q9VQ37 Q9VQ57 Q9VQE9 Q9VQG2 Q9VQG4 Q9VQH2 Q9VQP9 Q9VR07 Q9VR50 Q9VR82 Q9VRC9
Q9VRG7 Q9VRJ2 Q9VRJ8 Q9VRL2 Q9VRN2 Q9VRX6 Q9VS29 Q9VS60 Q9VS77 Q9VSB9 Q9VSE7
Q9VSH2 Q9VSH3 Q9VSL7 Q9VSU7 Q9VT04 Q9VT08 Q9VT20 Q9VT90 Q9VT92 Q9VTG7 Q9VTJ8
Q9VTK2 Q9VTN0 Q9VTX8 Q9VU27 Q9VU52 Q9VUK5 Q9VUL9 Q9VUN0 Q9VUT6 Q9VUT8 Q9VUW9
Q9VV87 Q9VVA8 Q9VVD9 Q9VVF3 Q9VVH3 Q9VVH9 Q9VVL1 Q9VVT2 Q9VVT9 Q9VVV2 Q9VVW3
Q9VVX3 Q9VVX5 Q9VW12 Q9VW60 Q9VW71 Q9VW87 Q9VWE0 Q9VWE9 Q9VWG0 Q9VWH8 Q9VWK6
Q9VWL5 Q9VWR8 Q9VWS4 Q9VWV9 Q9VX31 Q9VX39 Q9VXD1 Q9VXD6 Q9VXD9 Q9VXL0 Q9VXT2
Q9VXV9 Q9VY86 Q9VYB7 Q9VYF2 Q9VYN8 Q9VYV5 Q9VYY2 Q9VYZ1 Q9VYZ2 Q9VZA4 Q9VZE7
Q9VZJ6 Q9VZJ9 Q9VZL7 Q9VZM5 Q9VZS6 Q9VZU2 Q9VZW5 Q9VZW8 Q9W056 Q9W0A0 Q9W0C1
Q9W0H3 Q9W0I6 Q9W0M2 Q9W0R5 Q9W0R6 Q9W0S3 Q9W0T5 Q9W0V7 Q9W0Y6 Q9W0Y8 Q9W145
Q9W1C5 Q9W1K0 Q9W1M8 Q9W1N5 Q9W1N6 Q9W1P7 Q9W1P8 Q9W1U5 Q9W266 Q9W2B1 Q9W2B2
Q9W2E1 Q9W2H1 Q9W2U9 Q9W345 Q9W367 Q9W3F7 Q9W3I5 Q9W3T5 Q9W3V8 Q9W429 Q9W436
Q9W462 Q9W495 Q9W497 Q9W4C5 Q9W4V8 Q9W4Y2 Q9W534 Q9W594 Q9W5A5 Q9W5D4 Q9W5G6

Q9W5Y0 Q9W6A5 Q9W6A6 Q9W6A7 Q9W6A8 Q9W6A9 Q9W6T7 Q9W6V5 Q9W6V6 Q9W701 Q9W720
Q9W725 Q9W735 Q9W742 Q9W754 Q9W7C5 Q9W7R3 Q9W7R4 Q9W7Y7 Q9WAA1 Q9WAI7 Q9WAI8
Q9WAL8 Q9WC60 Q9WC69 Q9WCD1 Q9WCD8 Q9WCD9 Q9WCE1 Q9WCE3 Q9WCE8 Q9WDA6 Q9WFX3
Q9WIJ7 Q9WJB2 Q9WKC4 Q9WMX2 Q9WNW0 Q9WRL5 Q9WT38 Q9WT46 Q9WT55 Q9WTJ5 Q9WTK3
Q9WTL3 Q9WTL4 Q9WTM3 Q9WTN3 Q9WTN5 Q9WTN6 Q9WTP5 Q9WTQ2 Q9WTQ8 Q9WTR0 Q9WTR1
Q9WTR6 Q9WTR7 Q9WTS2 Q9WTS4 Q9WTS5 Q9WTS6 Q9WTU3 Q9WTV0 Q9WTV8 Q9WTV9 Q9WTW2
Q9WTW3 Q9WTW5 Q9WTW7 Q9WTW8 Q9WTX4 Q9WTY0 Q9WTY4 Q9WTZ2 Q9WTZ3 Q9WU02 Q9WU03
Q9WU08 Q9WU25 Q9WU38 Q9WU39 Q9WU40 Q9WU45 Q9WU47 Q9WU60 Q9WU61 Q9WU65 Q9WU72
Q9WU74 Q9WU81 Q9WUB6 Q9WUB7 Q9WUB9 Q9WUD2 Q9WUE3 Q9WUF1 Q9WUF4 Q9WUH1 Q9WUH7
Q9WUK7 Q9WUL5 Q9WUP0 Q9WUP1 Q9WUP4 Q9WUQ2 Q9WUS4 Q9WUT7 Q9WUV2 Q9WUX5 Q9WV08
Q9WV18 Q9WV26 Q9WV27 Q9WV38 Q9WV55 Q9WV57 Q9WV78 Q9WV82 Q9WV87 Q9WV91 Q9WVC5
Q9WVC8 Q9WVD0 Q9WVD4 Q9WVD5 Q9WVF9 Q9WVJ0 Q9WVJ4 Q9WVK0 Q9WVK5 Q9WVK8 Q9WVL3
Q9WVL6 Q9WVM4 Q9WVR6 Q9WVS0 Q9WVT0 Q9WVT6 Q9WWM4 Q9WWR1 Q9WWR2 Q9WWR3 Q9WWR4
Q9WWR5 Q9WWW6 Q9WWZ8 Q9WX61 Q9WX62 Q9WX70 Q9WX75 Q9WXM8 Q9WXU1 Q9WXX9 Q9WY05
Q9WY77 Q9WYC4 Q9WYR0 Q9WYU9 Q9WZ31 Q9WZ43 Q9WZ49 Q9WZQ6 Q9WZS2 Q9WZW3 Q9WZZ2
Q9WZZ5 Q9X0M7 Q9X0N0 Q9X0R9 Q9X109 Q9X170 Q9X1B7 Q9X1E2 Q9X1E3 Q9X1F9 Q9X1G6
Q9X1H2 Q9X1I9 Q9X1N5 Q9X1U9 Q9X1V0 Q9X2H2 Q9X2I1 Q9X2N9 Q9X2S4 Q9X2W0 Q9X3P3
Q9X442 Q9X447 Q9X480 Q9X4A5 Q9X4G9 Q9X4M3 Q9X4Q6 Q9X4Q7 Q9X4Q8 Q9X5A6 Q9X5H9
Q9X5V3 Q9X5X3 Q9X6B2 Q9X722 Q9X756 Q9X772 Q9X7E7 Q9X7I4 Q9X7I5 Q9X7K6 Q9X7P0
Q9X806 Q9X807 Q9X809 Q9X812 Q9X813 Q9X814 Q9X889 Q9X8S2 Q9X8Z8 Q9X8Z9 Q9X900
Q9X913 Q9X929 Q9X972 Q9XA10 Q9XA14 Q9XA15 Q9XA86 Q9XAC2 Q9XAR1 Q9XAR4 Q9XAR5
Q9XB52 Q9XBF4 Q9XBM7 Q9XCY0 Q9XD50 Q9XD78 Q9XD84 Q9XDB2 Q9XDP0 Q9XE11 Q9XEA1
Q9XEC6 Q9XEC7 Q9XEC8 Q9XEE9 Q9XEL8 Q9XES1 Q9XF43 Q9XF58 Q9XF59 Q9XF63 Q9XF87
Q9XF88 Q9XF89 Q9XF91 Q9XF92 Q9XFJ8 Q9XFM6 Q9XFR0 Q9XFW4 Q9XGM8 Q9XGT9 Q9XGY6
Q9XH46 Q9XH57 Q9XH58 Q9XH75 Q9XHC6 Q9XHE6 Q9XHE7 Q9XHE8 Q9XHL5 Q9XHP2 Q9XI23
Q9XI46 Q9XI60 Q9XI62 Q9XI72 Q9XI74 Q9XI77 Q9XI78 Q9XI80 Q9XI81 Q9XI96 Q9XIA4
Q9XIA7 Q9XIC5 Q9XIC7 Q9XID2 Q9XID3 Q9XIE2 Q9XIE6 Q9XIG2 Q9XIH6 Q9XIH7 Q9XII1
Q9XIM7 Q9XIN1 Q9XIN7 Q9XIP7 Q9XIQ7 Q9XJJ6 Q9XJR0 Q9XJR2 Q9XJR4 Q9XJR5 Q9XJR6
Q9XJR8 Q9XJS7 Q9XJY3 Q9XK21 Q9XK52 Q9XK70 Q9XK79 Q9XKC0 Q9XKK0 Q9XKK2 Q9XKK5
Q9XKZ5 Q9XL36 Q9XL45 Q9XL48 Q9XL62 Q9XL66 Q9XLE0 Q9XLE1 Q9XLE2 Q9XLW8 Q9XLW9
Q9XLX0 Q9XM10 Q9XMC1 Q9XMC2 Q9XMI9 Q9XMJ3 Q9XMN9 Q9XN27 Q9XN35 Q9XNM0 Q9XNM1
Q9XNM5 Q9XNN0 Q9XNN1 Q9XNN3 Q9XNN4 Q9XNT9 Q9XNU0 Q9XNU2 Q9XNU4 Q9XNU5 Q9XNU6
Q9XNU9 Q9XNV3 Q9XNV4 Q9XNV7 Q9XNW3 Q9XNW4 Q9XNW5 Q9XNW7 Q9XNW8 Q9XNW9 Q9XNX0
Q9XNX1 Q9XNX3 Q9XNX4 Q9XNX6 Q9XNX7 Q9XNX8 Q9XP34 Q9XP68 Q9XP73 Q9XP76 Q9XP77
Q9XP78 Q9XP81 Q9XP82 Q9XP83 Q9XP84 Q9XP86 Q9XP87 Q9XP88 Q9XP89 Q9XPI2 Q9XPS4
Q9XPS6 Q9XPT0 Q9XQ96 Q9XQA8 Q9XQR2 Q9XQR3 Q9XQR4 Q9XQU4 Q9XQU7 Q9XQV1 Q9XQV2
Q9XQV3 Q9XS59 Q9XS72 Q9XS78 Q9XSA7 Q9XSC2 Q9XSD4 Q9XSD7 Q9XSI1 Q9XSJ9 Q9XSK2
Q9XSL6 Q9XSM3 Q9XSM7 Q9XSP1 Q9XST2 Q9XSV3 Q9XSZ1 Q9XSZ4 Q9XSZ6 Q9XT10 Q9XT45
Q9XT47 Q9XT49 Q9XT50 Q9XT55 Q9XT56 Q9XT57 Q9XT58 Q9XT74 Q9XT76 Q9XT77 Q9XT82
Q9XT90 Q9XT96 Q9XT97 Q9XT98 Q9XTA0 Q9XTA8 Q9XTB5 Q9XTG1 Q9XTQ6 Q9XTQ7 Q9XTW2
Q9XTY1 Q9XU42 Q9XU63 Q9XU84 Q9XU98 Q9XUC4 Q9XUU9 Q9XUY5 Q9XV66 Q9XVE5 Q9XVM9
Q9XVQ9 Q9XVV5 Q9XVW1 Q9XWD6 Q9XWE9 Q9XWF0 Q9XWL9 Q9XWU2 Q9XWU8 Q9XWV0 Q9XWW1
Q9XX00 Q9XX85 Q9XXD1 Q9XXN2 Q9XXN3 Q9XXQ9 Q9XXR3 Q9XXX1 Q9XY35 Q9XYH7 Q9XYN0
Q9XYN1 Q9XYS3 Q9XYS8 Q9XZ08 Q9XZ14 Q9XZC0 Q9Y006 Q9Y008 Q9Y068 Q9Y0A1 Q9Y0E4

Q9Y115 Q9Y117 Q9Y123 Q9Y140 Q9Y169 Q9Y194 Q9Y1U9 Q9Y1W0 Q9Y210 Q9Y219 Q9Y225
Q9Y226 Q9Y227 Q9Y228 Q9Y231 Q9Y241 Q9Y256 Q9Y257 Q9Y267 Q9Y271 Q9Y274 Q9Y275
Q9Y276 Q9Y277 Q9Y278 Q9Y279 Q9Y282 Q9Y284 Q9Y286 Q9Y287 Q9Y289 Q9Y2A7 Q9Y2A9
Q9Y2B1 Q9Y2B2 Q9Y2C2 Q9Y2C3 Q9Y2C5 Q9Y2C9 Q9Y2D2 Q9Y2E8 Q9Y2G1 Q9Y2G3 Q9Y2G8
Q9Y2H6 Q9Y2P4 Q9Y2P5 Q9Y2Q0 Q9Y2R0 Q9Y2T5 Q9Y2T6 Q9Y2U2 Q9Y2U8 Q9Y2W3 Q9Y2Y6
Q9Y320 Q9Y328 Q9Y336 Q9Y342 Q9Y345 Q9Y385 Q9Y397 Q9Y3A6 Q9Y3B3 Q9Y3D6 Q9Y3E0
Q9Y3N9 Q9Y3P4 Q9Y3P8 Q9Y3Q0 Q9Y3Q3 Q9Y3Q4 Q9Y3Q7 Q9Y426 Q9Y442 Q9Y487 Q9Y493
Q9Y4A9 Q9Y4C0 Q9Y4C5 Q9Y4D2 Q9Y4D7 Q9Y4D8 Q9Y4W6 Q9Y512 Q9Y519 Q9Y548 Q9Y561
Q9Y584 Q9Y585 Q9Y5E1 Q9Y5E2 Q9Y5E3 Q9Y5E4 Q9Y5E5 Q9Y5E6 Q9Y5E7 Q9Y5E8 Q9Y5E9
Q9Y5F0 Q9Y5F1 Q9Y5F2 Q9Y5F3 Q9Y5F6 Q9Y5F7 Q9Y5F8 Q9Y5F9 Q9Y5G0 Q9Y5G1 Q9Y5G2
Q9Y5G3 Q9Y5G4 Q9Y5G5 Q9Y5G6 Q9Y5G7 Q9Y5G8 Q9Y5G9 Q9Y5H0 Q9Y5H1 Q9Y5H2 Q9Y5H3
Q9Y5H4 Q9Y5H5 Q9Y5H6 Q9Y5H7 Q9Y5H8 Q9Y5H9 Q9Y5I0 Q9Y5I1 Q9Y5I2 Q9Y5I3 Q9Y5I4
Q9Y5I7 Q9Y5L2 Q9Y5L3 Q9Y5M8 Q9Y5N1 Q9Y5P0 Q9Y5P1 Q9Y5Q0 Q9Y5Q5 Q9Y5R2 Q9Y5S1
Q9Y5S8 Q9Y5T4 Q9Y5U4 Q9Y5U5 Q9Y5U8 Q9Y5U9 Q9Y5W7 Q9Y5X5 Q9Y5Y0 Q9Y5Y3 Q9Y5Y4
Q9Y5Y5 Q9Y5Y6 Q9Y5Y7 Q9Y5Y9 Q9Y5Z0 Q9Y5Z6 Q9Y5Z9 Q9Y619 Q9Y624 Q9Y639 Q9Y644
Q9Y653 Q9Y661 Q9Y662 Q9Y663 Q9Y666 Q9Y672 Q9Y673 Q9Y691 Q9Y693 Q9Y694 Q9Y696
Q9Y698 Q9Y6A1 Q9Y6A2 Q9Y6A9 Q9Y6C5 Q9Y6C9 Q9Y6D0 Q9Y6F6 Q9Y6G1 Q9Y6H6 Q9Y6H8
Q9Y6I8 Q9Y6I9 Q9Y6J6 Q9Y6K0 Q9Y6L6 Q9Y6M5 Q9Y6M7 Q9Y6N1 Q9Y6N7 Q9Y6N8 Q9Y6Q6
Q9Y6R1 Q9Y6U7 Q9Y6W8 Q9Y6X1 Q9Y6X5 Q9Y706 Q9Y711 Q9Y753 Q9Y761 Q9Y7C8 Q9Y7D2
Q9Y7I4 Q9Y7K7 Q9Y7K8 Q9Y7L0 Q9Y7L7 Q9Y7M7 Q9Y7N5 Q9Y7P2 Q9Y7P7 Q9Y7P8 Q9Y7Q9
Q9Y7R5 Q9Y7R6 Q9Y7S4 Q9Y7S5 Q9Y7T0 Q9Y7T6 Q9Y7T9 Q9Y7U1 Q9Y7V0 Q9Y7X4 Q9Y7Y0
Q9Y7Y4 Q9Y7Y9 Q9Y7Z5 Q9Y800 Q9Y806 Q9Y810 Q9Y819 Q9Y874 Q9Y876 Q9Y8G1 Q9Y8G2
Q9Y8H5 Q9Y8K0 Q9Y8K1 Q9Y8K2 Q9Y8K3 Q9Y8K4 Q9Y8K5 Q9Y8K6 Q9YAA0 Q9YAR5 Q9YBS3
Q9YD67 Q9YDD0 Q9YDF8 Q9YDX4 Q9YDX6 Q9YDX7 Q9YEA0 Q9YEC1 Q9YF05 Q9YGC3 Q9YGL9
Q9YGY9 Q9YGZ0 Q9YGZ1 Q9YGZ2 Q9YGZ3 Q9YGZ4 Q9YGZ5 Q9YGZ6 Q9YGZ7 Q9YGZ8 Q9YGZ9
Q9YH00 Q9YH01 Q9YH02 Q9YH03 Q9YH04 Q9YH05 Q9YH26 Q9YH90 Q9YH91 Q9YH92 Q9YHB3
Q9YHX4 Q9YI66 Q9YJN7 Q9YJQ8 Q9YJU5 Q9YMJ7 Q9YN02 Q9YPD6 Q9YRV3 Q9YS17 Q9YTU2
Q9YWQ3 Q9Z0C8 Q9Z0D9 Q9Z0E8 Q9Z0F0 Q9Z0F3 Q9Z0F5 Q9Z0F8 Q9Z0G9 Q9Z0H5 Q9Z0K5
Q9Z0L0 Q9Z0L1 Q9Z0L5 Q9Z0M3 Q9Z0M4 Q9Z0M6 Q9Z0N7 Q9Z0R7 Q9Z0R8 Q9Z0R9 Q9Z0S3
Q9Z0S4 Q9Z0S5 Q9Z0S6 Q9Z0S7 Q9Z0S8 Q9Z0S9 Q9Z0T6 Q9Z0T9 Q9Z0U0 Q9Z0U4 Q9Z0U9
Q9Z0V1 Q9Z0V2 Q9Z0V7 Q9Z0V8 Q9Z0W0 Q9Z0W1 Q9Z0W7 Q9Z0X2 Q9Z0X4 Q9Z0Y8 Q9Z0Z4
Q9Z0Z5 Q9Z0Z6 Q9Z109 Q9Z122 Q9Z123 Q9Z125 Q9Z127 Q9Z137 Q9Z138 Q9Z139 Q9Z142
Q9Z143 Q9Z158 Q9Z173 Q9Z186 Q9Z1E9 Q9Z1F2 Q9Z1F6 Q9Z1G4 Q9Z1K8 Q9Z1K9 Q9Z1L1
Q9Z1L2 Q9Z1L5 Q9Z1M0 Q9Z1M7 Q9Z1N3 Q9Z1P1 Q9Z1P3 Q9Z1P4 Q9Z1P5 Q9Z1Q2 Q9Z1Q5
Q9Z1S9 Q9Z1V0 Q9Z1W5 Q9Z1W6 Q9Z1W8 Q9Z1X1 Q9Z1X2 Q9Z1Y3 Q9Z1Y9 Q9Z1Z1 Q9Z209
Q9Z210 Q9Z211 Q9Z218 Q9Z222 Q9Z239 Q9Z258 Q9Z260 Q9Z261 Q9Z262 Q9Z269 Q9Z270
Q9Z282 Q9Z2A7 Q9Z2A8 Q9Z2A9 Q9Z2B1 Q9Z2B2 Q9Z2B3 Q9Z2B5 Q9Z2C6 Q9Z2D3 Q9Z2D5
Q9Z2E3 Q9Z2E9 Q9Z2F7 Q9Z2G6 Q9Z2H4 Q9Z2H6 Q9Z2I0 Q9Z2I3 Q9Z2I4 Q9Z2I6 Q9Z2I7
Q9Z2J0 Q9Z2J5 Q9Z2J6 Q9Z2L0 Q9Z2P3 Q9Z2P8 Q9Z2Q7 Q9Z2T1 Q9Z2T2 Q9Z2V2 Q9Z2W8
Q9Z2W9 Q9Z2Y2 Q9Z2Y9 Q9Z2Z6 Q9Z2Z8 Q9Z306 Q9Z307 Q9Z319 Q9Z324 Q9Z325 Q9Z329
Q9Z351 Q9Z393 Q9Z398 Q9Z3A1 Q9Z3G1 Q9Z3Q3 Q9Z3Q5 Q9Z3R6 Q9Z3R7 Q9Z3U1 Q9Z429
Q9Z469 Q9Z4H7 Q9Z4S7 Q9Z521 Q9Z5D6 Q9Z5G4 Q9Z5G7 Q9Z645 Q9Z647 Q9Z670 Q9Z6C1
Q9Z6M6 Q9Z6M8 Q9Z6P9 Q9Z6U5 Q9Z706 Q9Z722 Q9Z723 Q9Z726 Q9Z752 Q9Z7H5 Q9Z7M4
Q9Z7N9 Q9Z7Q1 Q9Z7S5 Q9Z7U0 Q9Z7W1 Q9Z7X6 Q9Z7Y6 Q9Z7Z6 Q9Z809 Q9Z812 Q9Z813

Q9Z817 Q9Z880 Q9Z882 Q9Z883 Q9Z895 Q9Z896 Q9Z898 Q9Z899 Q9Z8A0 Q9Z8A1 Q9Z8A3
Q9Z8B3 Q9Z8B4 Q9Z8B5 Q9Z8B6 Q9Z8J2 Q9Z8J6 Q9Z8J7 Q9Z8V7 Q9Z949 Q9Z990 Q9Z9G5
Q9Z9P4 Q9ZA21 Q9ZAA6 Q9ZAA8 Q9ZAF3 Q9ZB71 Q9ZB72 Q9ZB74 Q9ZB80 Q9ZB81 Q9ZBM2
Q9ZC45 Q9ZC85 Q9ZC92 Q9ZC95 Q9ZC99 Q9ZCA5 Q9ZCA6 Q9ZCA7 Q9ZCC5 Q9ZCD4 Q9ZCE9
Q9ZCF7 Q9ZCG0 Q9ZCG1 Q9ZCG2 Q9ZCG3 Q9ZCG6 Q9ZCG7 Q9ZCI0 Q9ZCJ1 Q9ZCJ8 Q9ZCK7
Q9ZCM8 Q9ZCN4 Q9ZCN5 Q9ZCQ1 Q9ZCS5 Q9ZCT3 Q9ZCU7 Q9ZCV6 Q9ZCW0 Q9ZCW3 Q9ZCW4
Q9ZCW6 Q9ZCW8 Q9ZCW9 Q9ZCY3 Q9ZCY5 Q9ZD13 Q9ZD18 Q9ZD41 Q9ZD47 Q9ZD57 Q9ZD67
Q9ZD69 Q9ZD71 Q9ZD72 Q9ZDA5 Q9ZDA8 Q9ZDA9 Q9ZDB2 Q9ZDC4 Q9ZDC6 Q9ZDD7 Q9ZDD9
Q9ZDE9 Q9ZDF2 Q9ZDF9 Q9ZDG1 Q9ZDG3 Q9ZDH0 Q9ZDH1 Q9ZDI2 Q9ZDK4 Q9ZDL0 Q9ZDM3
Q9ZDN8 Q9ZDN9 Q9ZDQ5 Q9ZDQ7 Q9ZDR8 Q9ZDS0 Q9ZDS2 Q9ZDS3 Q9ZDS5 Q9ZDU3 Q9ZDX1
Q9ZDX2 Q9ZDX3 Q9ZDZ4 Q9ZE02 Q9ZE15 Q9ZE32 Q9ZE34 Q9ZE40 Q9ZE41 Q9ZE42 Q9ZE43
Q9ZE44 Q9ZE49 Q9ZE51 Q9ZE57 Q9ZE58 Q9ZE68 Q9ZE70 Q9ZE71 Q9ZE80 Q9ZE96 Q9ZE97
Q9ZE99 Q9ZEA2 Q9ZEC0 Q9ZEC1 Q9ZEC2 Q9ZEC4 Q9ZEC9 Q9ZEF9 Q9ZEL6 Q9ZEM5 Q9ZEP3
Q9ZEZ0 Q9ZEZ4 Q9ZF37 Q9ZFB4 Q9ZFB5 Q9ZFB6 Q9ZFB7 Q9ZFF8 Q9ZFU5 Q9ZG88 Q9ZGG0
Q9ZH29 Q9ZH79 Q9ZHC7 Q9ZHC9 Q9ZHD2 Q9ZHD4 Q9ZHG0 Q9ZHR4 Q9ZHV9 Q9ZI86 Q9ZIJ1
Q9ZJ31 Q9ZJ48 Q9ZJ65 Q9ZJ66 Q9ZJ67 Q9ZJ68 Q9ZJB1 Q9ZJC8 Q9ZJD1 Q9ZJD5 Q9ZJG8
Q9ZJH1 Q9ZJP2 Q9ZJP4 Q9ZJP5 Q9ZJP6 Q9ZJQ3 Q9ZJQ4 Q9ZJS9 Q9ZJV7 Q9ZJW0 Q9ZJX2
Q9ZJZ2 Q9ZK01 Q9ZK12 Q9ZK31 Q9ZK41 Q9ZK47 Q9ZK77 Q9ZK83 Q9ZK84 Q9ZKG7 Q9ZKP6
Q9ZKS4 Q9ZKT1 Q9ZKW2 Q9ZKW5 Q9ZKW7 Q9ZKX9 Q9ZL15 Q9ZL29 Q9ZL53 Q9ZL73 Q9ZL83
Q9ZLA0 Q9ZLC0 Q9ZLE0 Q9ZLF3 Q9ZLQ5 Q9ZLR4 Q9ZLY1 Q9ZM40 Q9ZM58 Q9ZM59 Q9ZM66
Q9ZM68 Q9ZM69 Q9ZM74 Q9ZM87 Q9ZMB8 Q9ZMH8 Q9ZML7 Q9ZMN9 Q9ZMP2 Q9ZMP3 Q9ZMQ0
Q9ZMZ3 Q9ZN40 Q9ZNB0 Q9ZNE7 Q9ZNG6 Q9ZNQ7 Q9ZNQ8 Q9ZNR0 Q9ZNS0 Q9ZNS6 Q9ZNT1
Q9ZNT7 Q9ZNV4 Q9ZNW2 Q9ZP16 Q9ZPE9 Q9ZPH4 Q9ZPH9 Q9ZPJ8 Q9ZPR4 Q9ZPR6 Q9ZPR7
Q9ZPS7 Q9ZPS9 Q9ZPV7 Q9ZQ88 Q9ZQ89 Q9ZQ94 Q9ZQB9 Q9ZQC6 Q9ZQI2 Q9ZQN8 Q9ZQP6
Q9ZQQ7 Q9ZQR3 Q9ZQR4 Q9ZQR5 Q9ZQX7 Q9ZQZ8 Q9ZQZ9 Q9ZR03 Q9ZR08 Q9ZR72 Q9ZR79
Q9ZRA3 Q9ZRF9 Q9ZRP7 Q9ZS51 Q9ZS88 Q9ZSD4 Q9ZSJ0 Q9ZSJ2 Q9ZSR7 Q9ZST8 Q9ZT05
Q9ZT07 Q9ZT29 Q9ZT49 Q9ZT63 Q9ZT81 Q9ZT82 Q9ZTN2 Q9ZTP3 Q9ZTW3 Q9ZTX0 Q9ZTZ7
Q9ZU07 Q9ZU35 Q9ZU43 Q9ZU46 Q9ZU49 Q9ZU50 Q9ZU82 Q9ZU94 Q9ZU96 Q9ZUA0 Q9ZUA5
Q9ZUE0 Q9ZUH3 Q9ZUH8 Q9ZUI0 Q9ZUI8 Q9ZUK3 Q9ZUK7 Q9ZUM2 Q9ZUN3 Q9ZUN5 Q9ZUS1
Q9ZUT0 Q9ZUT3 Q9ZUT8 Q9ZUU9 Q9ZUV3 Q9ZUV9 Q9ZUX1 Q9ZUZ0 Q9ZV02 Q9ZV07 Q9ZV24
Q9ZV28 Q9ZV29 Q9ZV51 Q9ZV53 Q9ZV57 Q9ZV71 Q9ZV87 Q9ZV89 Q9ZVD2 Q9ZVE7 Q9ZVF7
Q9ZVH5 Q9ZVH7 Q9ZVI7 Q9ZVI9 Q9ZVK6 Q9ZVL6 Q9ZVR7 Q9ZVS8 Q9ZVT9 Q9ZVX8 Q9ZW09
Q9ZW11 Q9ZW22 Q9ZW75 Q9ZW86 Q9ZW95 Q9ZWB9 Q9ZWC8 Q9ZWD1 Q9ZWG1 Q9ZWI7 Q9ZWL6
Q9ZWQ7 Q9ZWT2 Q9ZWT3 Q9ZXB7 Q9ZXD8 Q9ZXX5 Q9ZXX6 Q9ZXX7 Q9ZXX8 Q9ZXX9 Q9ZXY0
Q9ZXY2 Q9ZXY3 Q9ZXY4 Q9ZY14 Q9ZY15 Q9ZY46 Q9ZY49 Q9ZYM7 Q9ZZ08 Q9ZZ38 Q9ZZ40
Q9ZZ42 Q9ZZ43 Q9ZZ44 Q9ZZ45 Q9ZZ46 Q9ZZ47 Q9ZZ48 Q9ZZ49 Q9ZZ50 Q9ZZ51 Q9ZZ52
Q9ZZ53 Q9ZZ54 Q9ZZ56 Q9ZZ57 Q9ZZ58 Q9ZZ59 Q9ZZ60 Q9ZZ61 Q9ZZ62 Q9ZZ63 Q9ZZ64
Q9ZZ65 Q9ZZD2 Q9ZZD4 Q9ZZD8 Q9ZZE3 Q9ZZE4 Q9ZZF6 Q9ZZM2 Q9ZZM3 Q9ZZM4 Q9ZZM5
Q9ZZM6 Q9ZZR4 Q9ZZT5 Q9ZZT6 Q9ZZT7 Q9ZZT8 Q9ZZU1 Q9ZZW7 Q9ZZX1 Q9ZZY0 Q9ZZY1
Q9ZZY2 Q9ZZY3 Q9ZZY4 Q9ZZY5 Q9ZZY6 Q9ZZY7 Q9ZZY8 Q9ZZY9 Q9ZZZ0 Q9ZZZ1 R4V2Q5
R8NBR0 S0ARX1 S0AU91 S0DPM1 S0DPY2 S0DQM8 S0DS64 S0DW25 S0DZN4 S0E2X6 S0E2Y2
S0E3D0 S0E9M0 S0ECK8 S0EE84 S0EEY7 S0EGU4 S0ELQ3 S3D778 S3DQN8 S4R2P9 S4UX02
S4W283 S4W288 S4X0Q8 S5VBU1 S5ZH89 S7UQV8 S7VVK4 S7WCF5 S7WII9 S7ZK48 S7ZK63
S8ASK3 S8B3I0 S8B3I8 S8EZA7 S8GJB7 T0DVE4 T1NXB5 T2KM18 T2KMF4 T2KMI3 T2KPJ3

T2KPJ7 T2MBC4 U3H0P2 U3JA75 U3KPV4 U5HKE8 U5JCC6 U5NDT8 U5NE19 U9W802 V5IP95
V5IRD7 V5JFY4 V5NAL9 V5XZS6 V6F235 V6F244 V6F2B6 V6F2C2 V6F2Z8 V6F4W4 V6F510
V6F5F3 V6QX21 V6RPN0 W0FKI0 W0RYD3 W0SKS3 W0TIW1 W3X9K4 W5P3P0 W5PFI3 W5PLZ6
W5PSH7 W5XKT8 W6KHH6 W6Q3Z9 W6Q403 W6Q4S2 W6Q4S4 W6Q9B3 W6QB15 W6QIT8 W6QP06
W6QRN8 W6R4H8 W6VBF4 W7JPD9 W7LKY5 W7MC48 W7MLD1 W7MLD3 W7MM11 W7MSH6 W7MTI3
W7N2B4 W7NDQ0 W8DXL4 W8JDE2 W8JIS5 W8JIT4 W8JMU7 W8JMV1 W8JWW3 W8W138 X1WBB5
X1WEM4 X2ESR4 X2EVQ5 X2EVV9 X2EW55 X2JE85 X2JI34 X5M8U1

#### 2.3 The UniProt_Sol dataset (371069 sequences)

A0A011QK89 A0A017SE81 A0A017SE85 A0A017SEF3 A0A017SEX7 A0A017SEY2 A0A017SP50
A0A017SPL2 A0A017SQ41 A0A023GPI8 A0A023GPJ0 A0A023GS28 A0A023GS29 A0A023I4F1
A0A023IWD9 A0A023IWE0 A0A023IWE1 A0A023IWE2 A0A023IWE3 A0A023IWG1 A0A023IWG2
A0A023IWG3 A0A023IWG4 A0A023IWI4 A0A023IWI5 A0A023IWI6 A0A023IWI8 A0A023IWK3
A0A023IWK4 A0A023IWK5 A0A023IWK6 A0A023IWK7 A0A023IWM4 A0A023IWM5 A0A023IWM6
A0A023IWM7 A0A023IWM8 A0A023PXA5 A0A023PXB0 A0A023PXD3 A0A023PXE5 A0A023PXE8
A0A023PXF2 A0A023PXF5 A0A023PXH5 A0A023PXI5 A0A023PXI8 A0A023PXM2 A0A023PXN3
A0A023PXP4 A0A023PXP7 A0A023PYD3 A0A023PYF7 A0A023PYG5 A0A023PYH5 A0A023PYH8
A0A023PYI2 A0A023PYJ4 A0A023PZA4 A0A023PZA9 A0A023PZB3 A0A023PZC3 A0A023PZC7
A0A023PZD5 A0A023PZE4 A0A023PZE8 A0A023PZE9 A0A023PZF2 A0A023PZF5 A0A023PZG4
A0A023PZH9 A0A023PZI1 A0A023PZI9 A0A023PZJ9 A0A023PZK4 A0A023PZL7 A0A023UA23
A0A023UBA8 A0A023UBX6 A0A023UBY3 A0A023UCA6 A0A023UCC1 A0A023W421 A0A024B2V6
A0A024B3D0 A0A024B3G5 A0A024B404 A0A024B4E4 A0A024B7I0 A0A024F8Y4 A0A024F910
A0A024F9J0 A0A024SMV2 A0A031WDE4 A0A059TC02 A0A059WLZ7 A0A059WYP6 A0A059ZV61
A0A060KY90 A0A060X6Z0 A0A061AYY2 A0A061B0Q2 A0A061DFA8 A0A061DFM8 A0A061DM17
A0A061FKL9 A0A061FKM4 A0A061FLA2 A0A061FMF5 A0A061FTC2 A0A063C1W0 A0A067SEC9
A0A067SLB9 A0A067XG71 A0A067XGX8 A0A067XH53 A0A067XMK8 A0A067XML2 A0A067XMN9
A0A067XMT3 A0A067XMT7 A0A067XMV2 A0A067XMV3 A0A067XNH7 A0A067XNI2 A0A067XNI6
A0A067XNJ0 A0A067XR63 A0A067YB04 A0A067YBQ3 A0A068A9T3 A0A068AA98 A0A068ABB7
A0A068ACU9 A0A068BGA5 A0A068BIF1 A0A068FIK2 A0A068J840 A0A069AMK2 A0A072TLV8
A0A072VIM5 A0A073CEA3 A0A075BSX9 A0A075DVI9 A0A075F9Z3 A0A075FA51 A0A075FAK4
A0A075FBG7 A0A075QQ08 A0A075TXZ8 A0A077ES70 A0A077ESS0 A0A077EW86 A0A077EWA5
A0A077EYL8 A0A077LPS9 A0A078BQN7 A0A084AFG6 A0A084AFG7 A0A084AFG8 A0A084AFG9
A0A084API2 A0A084API3 A0A084API4 A0A084API5 A0A084API6 A0A084B9Y9 A0A084B9Z0
A0A084B9Z1 A0A084B9Z3 A0A084B9Z6 A0A084B9Z7 A0A084B9Z8 A0A084R1H6 A0A084R1I1
A0A084R1I4 A0A084R1J2 A0A084R1J7 A0A084R1K1 A0A084R1K2 A0A084R1K6 A0A084R1K7
A0A084R1N2 A0A084R1N8 A0A087WNH6 A0A087WUL8 A0A087WV53 A0A087WWA1 A0A087WXM9
A0A089FQI6 A0A089FS99 A0A089FSA4 A0A089GT80 A0A089GT84 A0A089QKZ7 A0A095AMW7
A0A096LP49 A0A096LPI5 A0A096MJN4 A0A096MJY4 A0A096MJZ0 A0A096P8D3 A0A096SRM5
A0A096VHX1 A0A096VHX6 A0A096VHY7 A0A096VHZ1 A0A096VHZ6 A0A096ZEC9 A0A096ZED0
A0A097I1R9 A0A097I2B5 A0A097I2D0 A0A097SRV6 A0A097SRX3 A0A097SRX8 A0A097ZPD4
A0A097ZPD5 A0A097ZPD9 A0A097ZPE0 A0A097ZPE2 A0A097ZPE8 A0A097ZPG2 A0A098CZ12
A0A098D1N7 A0A098D1P1 A0A098D6U0 A0A098D8A0 A0A098DDI1 A0A098DPY0 A0A098E171
A0A0A0RM07 A0A0A1EA36 A0A0A1EQ07 A0A0A1GKA2 A0A0A1H7N4 A0A0A1HA03 A0A0A2IBN3
A0A0A2IDH4 A0A0A2IJP3 A0A0A2J5V5 A0A0A2JB87 A0A0A2JP58 A0A0A2JW88 A0A0A2JW91

A0A0A2JW93 A0A0A2JWD0 A0A0A2JWD5 A0A0A2JY25 A0A0A2JY30 A0A0A2JYK3 A0A0A2KMZ4
A0A0A2KT65 A0A0A6YY25 A0A0A6YYL3 A0A0A7AAW9 A0A0A7EPL0 A0A0A7GEY4 A0A0A7HB61
A0A0A7HF73 A0A0A7HF79 A0A0A7HFE1 A0A0A7HFE6 A0A0A7HGA1 A0A0A7HIF0 A0A0A7HIX1
A0A0A7HIX6 A0A0A7HJC7 A0A0A7HMS2 A0A0B0QJN8 A0A0B0QJR1 A0A0B0SG80 A0A0B2XG51
A0A0B4EB91 A0A0B4ESU9 A0A0B4FAW4 A0A0B4FLB2 A0A0B4FP77 A0A0B4FTU6 A0A0B4FWC3
A0A0B4G3Q7 A0A0B4G504 A0A0B4G9F9 A0A0B4GDU5 A0A0B4GN88 A0A0B4GT47 A0A0B4GU97
A0A0B4H2J9 A0A0B4H4F0 A0A0B4H8J1 A0A0B4HM53 A0A0B4HVU2 A0A0B4I1J8 A0A0B4I1L1
A0A0B4IF96 A0A0B4J1W7 A0A0B4J2D5 A0A0B4K859 A0A0B4KGY6 A0A0B5AC19 A0A0B5KU17
A0A0B5L585 A0A0B5L781 A0A0B5L7R4 A0A0B5LB55 A0A0B6CGH9 A0A0B6VIJ5 A0A0B6VQ48
A0A0B7P221 A0A0C1E5J2 A0A0C1E5J8 A0A0C1EFK1 A0A0C2S258 A0A0C2SRU0 A0A0C2SRU5
A0A0C2W6G2 A0A0C2WKN7 A0A0C3FBR2 A0A0C5DM37 A0A0C5URS1 A0A0C5XKJ0 A0A0C5XL88
A0A0C6DRT7 A0A0C6DRW4 A0A0C6DWS6 A0A0C6E0I7 A0A0C9VD04 A0A0D1C1H8 A0A0D1C8Z4
A0A0D1CVS5 A0A0D1CY03 A0A0D1DRJ3 A0A0D1DWZ5 A0A0D1DXB8 A0A0D1E015 A0A0D1E6L2
A0A0D2L718 A0A0D2UG83 A0A0D2YFZ7 A0A0D2YFZ9 A0A0D2YG00 A0A0D2YG01 A0A0D2YG02
A0A0D2YG03 A0A0D2YG05 A0A0D2YG06 A0A0D2YG09 A0A0D2YG10 A0A0D3MJQ5 A0A0D4BS77
A0A0D4BSN8 A0A0D4BSP3 A0A0D4D912 A0A0D5ZDC8 A0A0D8BWP6 A0A0D9MRV9 A0A0E0RXA7
A0A0E0RXA9 A0A0E2IV13 A0A0E3D8L2 A0A0E3D8L9 A0A0E3D8M4 A0A0E3D8M5 A0A0E3D8M9
A0A0E3D8N0 A0A0E3D8N4 A0A0E3D8N6 A0A0E3D8P4 A0A0E3D8P9 A0A0E3D8Q1 A0A0E3JT70
A0A0E3KBH3 A0A0E3NEE1 A0A0E3T552 A0A0E3U2K2 A0A0E3URH8 A0A0E3VJW8 A0A0E4AVP3
A0A0E4AY46 A0A0E4AYE7 A0A0E4AZF8 A0A0E4AZP0 A0A0E4FKF7 A0A0F0I5G4 A0A0F0IP79
A0A0F5HNC1 A0A0F6B244 A0A0F6B4W4 A0A0F6B8A2 A0A0F6MY85 A0A0F7CUE9 A0A0F7GFI5
A0A0F7GFS4 A0A0F7GG11 A0A0F7GGT4 A0A0F7GH49 A0A0F7RJ52 A0A0F7TN06 A0A0F7TN16
A0A0F7TN65 A0A0F7TN69 A0A0F7TQP2 A0A0F7TRU0 A0A0F7TSZ1 A0A0F7TXA5 A0A0F7TXA8
A0A0F7TZD6 A0A0F7U103 A0A0F7U1Z1 A0A0F7U1Z8 A0A0F8VPX6 A0A0F9XIF1 A0A0F9XJT1
A0A0F9ZXE3 A0A0G0AID6 A0A0G0MF47 A0A0G2JEB6 A0A0G2JMD5 A0A0G2JTZ2 A0A0G2JXN2
A0A0G2K0D3 A0A0G2KIZ8 A0A0G2Q9D6 A0A0G2Q9F8 A0A0G2UGT2 A0A0G3VTN5 A0A0G4DBR5
A0A0H2UPF7 A0A0H2UR93 A0A0H2UR96 A0A0H2URA3 A0A0H2URB1 A0A0H2URH2 A0A0H2URH7
A0A0H2URJ6 A0A0H2V660 A0A0H2V760 A0A0H2VA12 A0A0H2VA68 A0A0H2VBH0 A0A0H2VFI8
A0A0H2VKG8 A0A0H2WWB5 A0A0H2WWV6 A0A0H2WZY4 A0A0H2XCS3 A0A0H2XEA6 A0A0H2XFI6
A0A0H2XHV5 A0A0H2XI17 A0A0H2ZDG9 A0A0H2ZDT7 A0A0H2ZF83 A0A0H2ZF87 A0A0H2ZGB9
A0A0H2ZGJ4 A0A0H2ZGR9 A0A0H2ZHV3 A0A0H2ZIC3 A0A0H2ZJC1 A0A0H2ZKA1 A0A0H2ZMB4
A0A0H2ZNL3 A0A0H2ZPS7 A0A0H2ZQL5 A0A0H3AFM6 A0A0H3AL67 A0A0H3CC29 A0A0H3CC30
A0A0H3CCP8 A0A0H3CEP9 A0A0H3G0N3 A0A0H3GEZ8 A0A0H3H456 A0A0H3JGH6 A0A0H3JNB0
A0A0H3JPC6 A0A0H3JQ59 A0A0H3JR16 A0A0H3JRU9 A0A0H3JT43 A0A0H3JT80 A0A0H3JU78
A0A0H3JUG6 A0A0H3JVA1 A0A0H3JX98 A0A0H3JXA8 A0A0H3K9R3 A0A0H3KB22 A0A0H3KB98
A0A0H3KP73 A0A0H3KZS3 A0A0H3L116 A0A0H3L1B8 A0A0H3L952 A0A0H3LQK8 A0A0H3LX82
A0A0H3M0L1 A0A0H3M776 A0A0H3MAB5 A0A0H3N8G7 A0A0H3NBY9 A0A0H3NGF1 A0A0H3NGK0
A0A0H3NM73 A0A0H3PEK7 A0A0H3PIP6 A0A0H4ADX3 A0A0H4LJX8 A0A0H4TXY1 A0A0H5AXT3
A0A0H5BB10 A0A0H5BB17 A0A0H5BMX5 A0A0H5BN57 A0A0H5BN61 A0A0H5SJ89 A0A0I9QGZ2
A0A0I9QGZ7 A0A0I9RJ56 A0A0J1BDH3 A0A0J1I1I3 A0A0J5WTU0 A0A0J6G7P5 A0A0J9U3L6
A0A0J9UBD6 A0A0J9WZF0 A0A0J9X1Q5 A0A0J9YWL9 A0A0J9YX94 A0A0J9YXQ4 A0A0J9YY54
A0A0K0JFP3 A0A0K0MCJ4 A0A0K0PVL0 A0A0K0PVL3 A0A0K0PVM5 A0A0K0QSV4 A0A0K0XCY8
A0A0K1SC59 A0A0K1TPY7 A0A0K1TQ05 A0A0K2G9F6 A0A0K2H5X8 A0A0K2H9D8 A0A0K3AXH1
A0A0K9RL25 A0A0L0P4F8 A0A0L0P6P7 A0A0M3KKZ0 A0A0M3Q1Q3 A0A0M3U1B0 A0A0M4FMK2

A0A0M4KDW1 A0A0M4KE44 A0A0M4L8I7 A0A0M4LAF6 A0A0M4M0T9 A0A0M5KK13 A0A0M5L832
A0A0M7LBC1 A0A0M9ADP1 A0A0M9AFS9 A0A0M9ER62 A0A0M9ER64 A0A0N0BLP2 A0A0N0BLS5
A0A0N0DCA3 A0A0N6WJC7 A0A0N7D483 A0A0N7D4P6 A0A0N7D745 A0A0N7KIY3 A0A0N8YGA2
A0A0N9HMN6 A0A0N9HN32 A0A0N9HNS9 A0A0N9HP11 A0A0N9HQ36 A0A0N9HTA1 A0A0P0E482
A0A0P0UZP7 A0A0P0V4R0 A0A0P0V5U9 A0A0P0VI36 A0A0P0VUY4 A0A0P0WFC8 A0A0P0WGX7
A0A0P0WIY3 A0A0P0WQ90 A0A0P0X0C0 A0A0P0X9Z7 A0A0P0XB70 A0A0P0XCU3 A0A0P0ZBS7
A0A0P0ZD79 A0A0P0ZEM1 A0A0P1DJE3 A0A0P9JFY5 A0A0R0IHP4 A0A0R3K2G2 A0A0R4I987
A0A0R4IBL7 A0A0R4IEW8 A0A0R4IFG5 A0A0R4IKJ1 A0A0R4IVA4 A0A0R4IWG9 A0A0R4IWI1
A0A0R4IXF6 A0A0R4IYX6 A0A0R4IZ84 A0A0R6Y3I5 A0A0R8YWJ7 A0A0S0FTT9 A0A0S0MVI5
A0A0S0MX59 A0A0S0N8M3 A0A0S0N9S9 A0A0S1RUN4 A0A0S1RVB0 A0A0S1X9S7 A0A0S2DN66
A0A0S2E7V8 A0A0S2E7W3 A0A0S2E7W7 A0A0S2E7X0 A0A0S2E7Z1 A0A0S2PM82 A0A0S2PM92
A0A0S2PMA8 A0A0S2SZW1 A0A0S2SZX3 A0A0S2T050 A0A0S2UQQ5 A0A0S2UWA5 A0A0S2UWC9
A0A0S2UWT1 A0A0S3QTC6 A0A0S3QTD0 A0A0S4IJL0 A0A0S6XAW4 A0A0S6XAX9 A0A0S6XGG4
A0A0S6XGG5 A0A0S6XGJ8 A0A0S6XH49 A0A0S6XHF8 A0A0S6XHH0 A0A0S6XHQ0 A0A0T7AQA7
A0A0U1LQE6 A0A0U1LSP0 A0A0U1RGY0 A0A0U1RQE8 A0A0U1RQF7 A0A0U1RQI7 A0A0U1RR11
A0A0U1RR37 A0A0U1RRE5 A0A0U1RRI6 A0A0U1RRK4 A0A0U1RRL7 A0A0U1WZ18 A0A0U2D9C5
A0A0U2JT80 A0A0U2WFX7 A0A0U3AL34 A0A0U3BRC5 A0A0U3C228 A0A0U3LQ20 A0A0U4CDK4
A0A0U4ZPL6 A0A0U4ZX08 A0A0U5AF03 A0A0U5CJU2 A0A0U5CNN8 A0A0U5CNP2 A0A0U5G0B1
A0A0U5GFQ6 A0A0U5GFR4 A0A0U5GFS8 A0A0U5GGX2 A0A0U5GHC1 A0A0U5GHD4 A0A0U5GHG9
A0A0U5GHU6 A0A0U5GIM7 A0A0U5GJ41 A0A0U5GJY1 A0A0U5GK88 A0A0U5GMR5 A0A0U5GNT1
A0A0U5GQ05 A0A0U5HAP0 A0A0U5HAQ4 A0A0U8TRG9 A0A0U8UFF4 A0A0Y0M151 A0A110A2W7
A0A120HUS7 A0A120HYZ1 A0A120KVR5 A0A120KVR8 A0A120KVW2 A0A125R003 A0A140C435
A0A140F3K6 A0A140F3K7 A0A140JWS2 A0A140JWS4 A0A140JWS5 A0A140JWS6 A0A140JWS7
A0A140JWS8 A0A140JWT5 A0A140JWT7 A0A140JWU0 A0A140KFG9 A0A140LI88 A0A140N890
A0A140NAP5 A0A140NCB4 A0A140ND86 A0A140NDS5 A0A142BX70 A0A142BX74 A0A142C799
A0A142C7A0 A0A142C7A1 A0A142C7A2 A0A142C7A3 A0A142C7A4 A0A142C7A5 A0A142PTM2
A0A144A134 A0A144KPJ6 A0A144Y7G4 A0A145P7T2 A0A150XSC5 A0A151ZEA6 A0A159BP93
A0A161CAI1 A0A161CEU9 A0A161CEV4 A0A161CKG1 A0A162J3X8 A0A162J448 A0A162J452
A0A163UT06 A0A165U5V5 A0A166B1A6 A0A166U5H3 A0A166YZR3 A0A166YZT6 A0A166YZW0
A0A166YZY4 A0A166YZY9 A0A167LUQ4 A0A167V661 A0A167V873 A0A168WVR6 A0A169T193
A0A172J1R7 A0A172J1S0 A0A172J1V3 A0A172J2D0 A0A172J2G3 A0A173GP47 A0A178U9Y5
A0A178VBJ0 A0A178VEK7 A0A179G9G5 A0A179GC70 A0A179H0J7 A0A179H0U5 A0A179H164
A0A179H2I8 A0A179H2P2 A0A179H324 A0A179HJB8 A0A179HKZ8 A0A179HLJ8 A0A179HLW2
A0A179HM40 A0A179HM59 A0A179HME7 A0A183 A0A193AU77 A0A193AUF6 A0A193PS46
A0A193PS74 A0A1A9TAK5 A0A1B0GTD5 A0A1B0GTH6 A0A1B0GTI1 A0A1B0GTK5 A0A1B0GTR3
A0A1B0GTR4 A0A1B0GTS1 A0A1B0GTU1 A0A1B0GTZ2 A0A1B0GU33 A0A1B0GU71 A0A1B0GUA6
A0A1B0GUA9 A0A1B0GUC4 A0A1B0GUI7 A0A1B0GUJ8 A0A1B0GUQ0 A0A1B0GUS0 A0A1B0GUS4
A0A1B0GUT2 A0A1B0GUU1 A0A1B0GUV1 A0A1B0GUV7 A0A1B0GUX0 A0A1B0GV03 A0A1B0GV22
A0A1B0GVG6 A0A1B0GVH6 A0A1B0GVH7 A0A1B0GVK7 A0A1B0GVM5 A0A1B0GVM6 A0A1B0GVQ3
A0A1B0GVR7 A0A1B0GVS7 A0A1B0GVZ2 A0A1B0GVZ6 A0A1B0GWH4 A0A1B0GWK0 A0A1B1FHP3
A0A1B1WAJ0 A0A1B1ZGB5 A0A1B2CTA7 A0A1B2CTA9 A0A1B2CTB2 A0A1B2CTB3 A0A1B2CTB6
A0A1B2CTB7 A0A1B2CTB8 A0A1B2CTB9 A0A1B2CTC1 A0A1B2CTC5 A0A1B2CTC6 A0A1B2LS07
A0A1B4XBG9 A0A1B4XBH2 A0A1B4XBH3 A0A1B4XBH4 A0A1B4XBH5 A0A1B4XBH9 A0A1B4XBK0
A0A1B4XBK1 A0A1B4XBK2 A0A1B5L6A0 A0A1B5L8S2 A0A1B7YCC9 A0A1B8YAB1 A0A1C3YKE0

A0A1C5G2V9 A0A1C7AAN2 A0A1C7D1B7 A0A1C9CX56 A0A1C9CX66 A0A1C9II22 A0A1C9J6A7
A0A1C9U5X5 A0A1C9U5X7 A0A1D1V3Z0 A0A1D3PCI9 A0A1D3PCJ5 A0A1D3PCK2 A0A1D3PCM3
A0A1D3PDD4 A0A1D5NS60 A0A1D5P556 A0A1D5PPP7 A0A1D5PRR9 A0A1D5XGW0 A0A1D6EFT8
A0A1D6F7U7 A0A1D6GDY8 A0A1D6HQ92 A0A1D6K6U5 A0A1D6KUI6 A0A1D6LAB7 A0A1D6LAG9
A0A1D6LTV0 A0A1D8EJF9 A0A1D8PL26 A0A1D8PNZ7 A0A1D8PPK1 A0A1D9BZF0 A0A1E1FFL0
A0A1E1FFL1 A0A1E1FFN5 A0A1E1FFN7 A0A1E1FFN8 A0A1E3B0S1 A0A1E3B0T2 A0A1E3B0T7
A0A1E3P8S6 A0A1E3P8S8 A0A1E4S2N7 A0A1E4S2P1 A0A1E5RIW9 A0A1E5RUL9 A0A1E7MYN1
A0A1F3 A0A1F5UTC9 A0A1F6CTG8 A0A1G2PK14 A0A1G6LGU2 A0A1H0NKS3 A0A1H7TQR5
A0A1I5E523 A0A1I8M2I8 A0A1I8MUL8 A0A1I9LM04 A0A1I9LQ12 A0A1J0HSK7 A0A1J0HSL4
A0A1J0HSL5 A0A1J0HSL6 A0A1J0HSL8 A0A1J0HSP5 A0A1J0HSQ1 A0A1J0HSR1 A0A1J0KKC6
A0A1J1DL12 A0A1J1EM40 A0A1J6KGJ9 A0A1L1SUL6 A0A1L2D3T3 A0A1L4BKP6 A0A1L4BKQ4
A0A1L4BKS3 A0A1L5BTC1 A0A1L5BU05 A0A1L5BUG8 A0A1L5BUX8 A0A1L5YKS7 A0A1L5YRA2
A0A1L6Z3A0 A0A1L6Z3A5 A0A1L7NQ96 A0A1L7U8F2 A0A1L7VZE7 A0A1L8EV45 A0A1L8EXB5
A0A1L8EYB2 A0A1L8FFY5 A0A1L8G016 A0A1L8GR68 A0A1L8GSA2 A0A1L8GUX5 A0A1L8GXY4
A0A1L8GXY6 A0A1L8H579 A0A1L8H8C0 A0A1L8HCK2 A0A1L8HFX9 A0A1L8HU22 A0A1L8HV70
A0A1L8HX76 A0A1L9NGU5 A0A1L9U7P9 A0A1L9UCG2 A0A1L9UKS1 A0A1L9UR19 A0A1L9URE0
A0A1L9WLD2 A0A1L9WLD5 A0A1L9WLD6 A0A1L9WLD7 A0A1L9WLD9 A0A1L9WLE6 A0A1L9WLF1
A0A1L9WLF2 A0A1L9WLF6 A0A1L9WLG2 A0A1L9WLH9 A0A1L9WLI9 A0A1L9WLK0 A0A1L9WLK2
A0A1L9WLL5 A0A1L9WN23 A0A1L9WN37 A0A1L9WN49 A0A1L9WN60 A0A1L9WQM9 A0A1L9WQN2
A0A1L9WQN9 A0A1L9WQQ1 A0A1L9WQW4 A0A1L9WR63 A0A1L9WUI2 A0A1L9WUI4 A0A1L9WUM2
A0A1L9WUW3 A0A1L9WUX8 A0A1M3T4K3 A0A1M7D0R2 A0A1P8ARG1 A0A1P8AS03 A0A1P8AW69
A0A1P8B760 A0A1P8BH59 A0A1P8NTI3 A0A1P8YT88 A0A1Q9P359 A0A1Q9PA05 A0A1R3RGJ2
A0A1R3RGJ9 A0A1R3RGK0 A0A1R3RGK1 A0A1R3RGK4 A0A1S1YUU1 A0A1S3C4H6 A0A1S3THR8
A0A1S3WYR7 A0A1S3X835 A0A1S3YCW2 A0A1S3YEG8 A0A1S4A695 A0A1S4AUX8 A0A1S4BJT3
A0A1S4BZI5 A0A1S4CB73 A0A1S4CL59 A0A1S4D475 A0A1S4DF18 A0A1S4DFD3 A0A1S4NYE4
A0A1S5RW73 A0A1U8F5V2 A0A1U8GR65 A0A1U8IPT1 A0A1U8QGE6 A0A1U8QGV2 A0A1U8QH20
A0A1U8QHE3 A0A1U8QIH0 A0A1U8QJR1 A0A1U8QK63 A0A1U8QL22 A0A1U8QLG8 A0A1U8QLK0
A0A1U8QN18 A0A1U8QNG8 A0A1U8QP15 A0A1U8QTA2 A0A1U8QVN4 A0A1U8QW91 A0A1U8QXK4
A0A1U8QZJ7 A0A1U9X9N3 A0A1U9YHZ1 A0A1U9YHZ2 A0A1U9YHZ6 A0A1U9YI02 A0A1U9YI04
A0A1U9YI06 A0A1U9YI11 A0A1U9YI21 A0A1U9YI27 A0A1V0E492 A0A1V0E4A6 A0A1V0HUX5
A0A1V0QS34 A0A1V0QSA8 A0A1V0QSC6 A0A1V0QSE8 A0A1V1FVQ6 A0A1V6NVX3 A0A1V6NWD3
A0A1V6NXN7 A0A1V6NYI6 A0A1V6NYM5 A0A1V6NYZ6 A0A1V6P9U2 A0A1V6PAA5 A0A1V6PAF7
A0A1V6PAN1 A0A1V6PAP3 A0A1V6PAU0 A0A1W2P884 A0A1W2PP81 A0A1W2PPE2 A0A1W2PPF3
A0A1W2PPH5 A0A1W2PPK0 A0A1W2PPL8 A0A1W2PPM1 A0A1W2PPW3 A0A1W2PQ09 A0A1W2PQ72
A0A1W2PQ73 A0A1W2PQL4 A0A1W2PR19 A0A1W2PR48 A0A1W2PR64 A0A1W2PR82 A0A1W2PRP0
A0A1W2PRV1 A0A1W5SKE9 A0A1W5SKT4 A0A1W5SMT6 A0A1W5SR39 A0A1W5T1T1 A0A1W5T1T3
A0A1W5T1U1 A0A1W5T1Y4 A0A1W5T1Y5 A0A1W5T1Y7 A0A1W5T312 A0A1W5T323 A0A1W5T363
A0A1W5T3K7 A0A1W5T4X6 A0A1W6BT46 A0A1W6BT53 A0A1W6GW06 A0A1W6GW18 A0A1W6GW32
A0A1W6I162 A0A1W6I186 A0A1W6I187 A0A1W6QDI6 A0A1W6QDI7 A0A1W6QDJ1 A0A1W7M1U5
A0A1X9IRP7 A0A1X9IRQ7 A0A1X9IRT6 A0A1X9ISH5 A0A1X9ISN9 A0A1X9ISP7 A0A1X9PY88
A0A1X9QHJ0 A0A1X9WEP1 A0A1Y0BRF1 A0A1Y0BRF3 A0A1Y0BRF4 A0A1Y0BRF8 A0A1Y0BRG0
A0A1Y1C7Q5 A0A1Y3DYH2 A0A1Z3GBK8 A0A1Z3GC64 A0A1Z3GCD1 A0A217EJJ0 A0A217ER65
A0A221J5P3 A0A221J5W8 A0A221J5X1 A0A221J5X3 A0A221J5X6 A0A224AKZ9 A0A224AM54
A0A244CMP0 A0A248AFK6 A0A248QE08 A0A251PW43 A0A256XLS3 A0A284RE13 A0A284RNH4

A0A286LEZ6 A0A286LEZ7 A0A286LEZ8 A0A286LEZ9 A0A286R227 A0A286R7K5 A0A286Y9D1
A0A286YDK6 A0A286YEV6 A0A286YEX9 A0A286YEY9 A0A286YF01 A0A286YF46 A0A286YF60
A0A286YF77 A0A286YFB4 A0A286YFG1 A0A291LSD6 A0A291SJC7 A0A292G9J6 A0A2C9EVE6
A0A2D1VNJ8 A0A2D1VNM2 A0A2G3D6U1 A0A2G5I8P4 A0A2G5I8W0 A0A2G5IC53 A0A2G5IC96
A0A2G5ICA0 A0A2G5ICB4 A0A2G5ICC7 A0A2G5ICG8 A0A2G5ICZ9 A0A2H1VE33 A0A2H3CNT9
A0A2H3CSB7 A0A2H3CTK0 A0A2H3D8Y2 A0A2H3D905 A0A2H3E4G0 A0A2H5AIW0 A0A2H5AIX5
A0A2H5AIY0 A0A2H5AIY2 A0A2H5AIY4 A0A2H5AIY6 A0A2H5AIZ1 A0A2H5AIZ6 A0A2H5BHG5
A0A2H5BHG9 A0A2H5BHJ5 A0A2H5BHJ6 A0A2H5Q1B8 A0A2I1BSV7 A0A2I1BSV9 A0A2I1BSW6
A0A2I1BSX0 A0A2I1BSZ6 A0A2I1BT15 A0A2I1BT56 A0A2I1C3T0 A0A2I1C3T1 A0A2I1C3T5
A0A2I1C3T9 A0A2I1C3U0 A0A2I1C3U1 A0A2I1C3U2 A0A2I1C3V0 A0A2I1C3V7 A0A2I1C3W6
A0A2I1C3W7 A0A2I1C3W8 A0A2I1C3X5 A0A2I1C3X9 A0A2I1C3Y3 A0A2I1C3Z1 A0A2I2F262
A0A2I2F272 A0A2I6B3P0 A0A2I6PIZ1 A0A2I6PIZ6 A0A2I6PIZ7 A0A2I6PIZ8 A0A2I6PJ02
A0A2I6PJ05 A0A2I7G3B0 A0A2I7G3B2 A0A2I7G3B3 A0A2I9 A0A2J6L8Y7 A0A2K3CNL6
A0A2K3D5Z7 A0A2K3DDJ2 A0A2K3DZC4 A0A2K4Z9G8 A0A2K5X3B6 A0A2K8K5C5 A0A2K9M484
A0A2K9RFZ2 A0A2K9RFZ7 A0A2K9RFZ8 A0A2K9RFZ9 A0A2K9RG07 A0A2L0P0J8 A0A2L0P0K1
A0A2L0P0K3 A0A2L0P0L5 A0A2L0V130 A0A2L0V156 A0A2L0V161 A0A2L0V166 A0A2L0VXR0
A0A2L0VXR5 A0A2P1BT06 A0A2P1GIW2 A0A2P1GIW3 A0A2P1GIW4 A0A2P1GIW7 A0A2P1GIY1
A0A2P1GIY2 A0A2P1GIY9 A0A2R4LMF9 A0A2R4QKX7 A0A2R6P624 A0A2R6PI27 A0A2R6PLK2
A0A2R6PN00 A0A2R6Q324 A0A2R6Q326 A0A2R6Q8R5 A0A2R6QE26 A0A2R6QXF8 A0A2R6S148
A0A2R8Q1W5 A0A2R8QFQ6 A0A2R8RWN9 A0A2R8VHR8 A0A2R8Y2Y2 A0A2R8Y4L2 A0A2R8Y619
A0A2R8Y7D0 A0A2S1WBY6 A0A2S1WC15 A0A2T5Y4G4 A0A2U1Q018 A0A2U7XUH7 A0A2U8NET4
A0A2U8NEV6 A0A2U8U2K7 A0A2U8U2K8 A0A2U8U2L2 A0A2U8U2L3 A0A2U8U2L4 A0A2U8U2L5
A0A2U8U2L8 A0A2U9GGW3 A0A2U9GHG9 A0A2V5GSC6 A0A2V5GWU4 A0A2V5GX43 A0A2V5HCN7
A0A2V5HP22 A0A2W1NDJ7 A0A2Z4HPY1 A0A2Z4HPY5 A0A2Z4HPZ6 A0A2Z4HPZ8 A0A2Z4LIS9
A0A2Z5CV93 A0A2Z5CVA1 A0A2Z5CVK5 A0A2Z5CVQ4 A0A2Z5TIQ0 A0A2Z5TIR0 A0A2Z5TJB0
A0A2Z5TM64 A0A2Z5TWF0 A0A2Z5U1Z1 A0A2Z5VKM8 A0A2Z5WA18 A0A2Z5XAK4 A0A2Z5XAK6
A0A2Z5XAK7 A0A2Z5XAL7 A0A2Z5XAS1 A0A2Z5XAU0 A0A2Z6AQX6 A0A2Z6AQX7 A0A2Z6FZ31
A0A2Z6FZX4 A0A314 A0A314KSQ4 A0A314LG79 A0A316 A0A317 A0A317VEE1 A0A318Z3U0
A0A319A6V2 A0A319BQC1 A0A319DV72 A0A324 A0A325 A0A326 A0A327 A0A333
A0A336U965 A0A337 A0A338P6K9 A0A343W969 A0A343W970 A0A345BJN0 A0A345BJN1
A0A345BJN2 A0A345BJN3 A0A345BJN4 A0A345BJN5 A0A345BJN6 A0A345BJN7 A0A345BJN9
A0A345BJP0 A0A345BJP2 A0A345BJP6 A0A345BJP7 A0A345MJY6 A0A348AUV5 A0A348AUV6
A0A348AUV7 A0A348AUV9 A0A348AUW0 A0A348AUW1 A0A348AXX3 A0A348AXX4 A0A348AXX5
A0A348AXX6 A0A348AXX8 A0A348AXY0 A0A348AXY2 A0A348B779 A0A348B780 A0A348B781
A0A348B782 A0A348B784 A0A348B785 A0A348B788 A0A348B790 A0A348B793 A0A348B794
A0A348DU52 A0A348FUE1 A0A356 A0A357 A0A358 A0A360 A0A364LXP7 A0A367 A0A368
A0A369 A0A370 A0A371 A0A372 A0A373 A0A374 A0A375 A0A376 A0A377 A0A378 A0A379
A0A381 A0A381HAP5 A0A381HBN1 A0A383 A0A384E126 A0A384JAD8 A0A384JIP2
A0A384JK99 A0A384JQC9 A0A384JQF5 A0A384JRP0 A0A384KG77 A0A384LP51 A0A384XC21
A0A384XHA3 A0A384XLH1 A0A384XR80 A0A385AJM2 A0A385AJM4 A0A385AJM5 A0A385XJ53
A0A385XJE6 A0A385XJK5 A0A385XJL2 A0A385XJL4 A0A386CAB9 A0A386JV86 A0A386JVA3
A0A386KZ50 A0A386KZ95 A0A386L1Q6 A0A391NTR7 A0A392 A0A395I3F8 A0A396GMX6
A0A396IUP1 A0A398 A0A3B1EFP8 A0A3B1EFP9 A0A3B1EFQ1 A0A3B1EFQ2 A0A3B3IRV3
A0A3B3IT33 A0A3B3IT52 A0A3B3IU63 A0A3B6KF13 A0A3B6MPP5 A0A3B6UES5 A0A3G1DJE9

A0A3G1DJF0 A0A3G1DJF3 A0A3G1DJF4 A0A3G1DJF8 A0A3G1DJG8 A0A3G1DJG9 A0A3G1DJH7
A0A3G1DJI9 A0A3G1DJJ7 A0A3G1DJJ8 A0A3G1DJK5 A0A3G1QTS7 A0A3G1QTT3 A0A3G1QTU5
A0A3G5BB24 A0A3G9GQ29 A0A3G9GR23 A0A3G9H185 A0A3G9H990 A0A3G9H9H8 A0A3G9H9I5
A0A3G9HAL8 A0A3G9HCC8 A0A3G9HHK2 A0A3G9HN61 A0A3G9HPE6 A0A3G9HRC2 A0A3G9HRC8
A0A3G9JYJ6 A0A3G9K3K9 A0A3L6E0R4 A0A3L6G2C1 A0A3L6G998 A0A3Q0KR05 A0A3Q1LFK7
A0A3Q1N1R0 A0A3Q2TTB3 A0A3Q2UEI8 A0A3Q7ELQ2 A0A3Q7FGP1 A0A3Q7GFB5 A0A3Q7H216
A0A3Q7HES4 A0A3Q7HH64 A0A3Q7HRZ6 A0A3Q7I5Y9 A0A3Q7I7R4 A0A3Q7I9E2 A0A3Q7JC00
A0A3Q8GL18 A0A3Q8GLE8 A0A3Q8GYY4 A0A3Q9U4A9 A0A3Q9U4G3 A0A3Q9U4Z5 A0A3S7WQS5
A0A3T0E4B9 A0A3T0QHS4 A0A3T0QHT6 A0A411KUP5 A0A411KUP8 A0A411KUP9 A0A411KUQ4
A0A411KUQ7 A0A411KUQ8 A0A411KUQ9 A0A411KUU5 A0A411KZU9 A0A411KZZ8 A0A411L027
A0A411L030 A0A411PQL6 A0A411PQM3 A0A411PQN2 A0A411PQN4 A0A411PQN6 A0A411PQN7
A0A411PQN8 A0A411PQN9 A0A411PQP1 A0A411PQP6 A0A411PQP9 A0A443HJY8 A0A443HJZ3
A0A443HK05 A0A443HK11 A0A443HK66 A0A443HK81 A0A445AGS0 A0A447CCJ8 A0A455LLW5
A0A455LXK6 A0A455R5H4 A0A455R5K2 A0A455R5P9 A0A455R614 A0A455R7E6 A0A455R7M0
A0A455ZAR2 A0A480NP79 A0A481NV25 A0A481WNL2 A0A481WNM5 A0A481WNM7 A0A481WNM8
A0A481WNP4 A0A481WNP6 A0A481WP25 A0A481WP37 A0A481WPJ6 A0A481WQ01 A0A481WQ18
A0A481WQB6 A0A481WQL4 A0A481WR96 A0A482GDX1 A0A482N8M8 A0A482N9T9 A0A482N9V7
A0A482NAR8 A0A482PU20 A0A486WWJ9 A0A494C086 A0A494C0N9 A0A494C0Y3 A0A494C0Z2
A0A494C191 A0A494C1R9 A0A494TJG7 A0A498KE69 A0A498KFL4 A0A498KIQ3 A0A498KJ69
A0A4D6IA24 A0A4D6IA61 A0A4D6Q411 A0A4D6Q440 A0A4D6Q4S0 A0A4D6Q4T7 A0A4D6Q5I0
A0A4D6Q9B0 A0A4D6QCQ2 A0A4P8DJD9 A0A4P8DJF6 A0A4P8DJG9 A0A4P8DJH6 A0A4P8DJU1
A0A4P8DJU2 A0A4P8DJU7 A0A4P8DJV0 A0A4P8DJV2 A0A4P8DJW5 A0A4P8DJW8 A0A4P8DJY5
A0A4P8DJZ3 A0A4P8DK24 A0A4P8DY91 A0A4P8GEE8 A0A4P8GF19 A0A4P8GFV8 A0A4P8GG75
A0A4P8GG91 A0A4P8GHJ1 A0A4P8W733 A0A4P8W7L7 A0A4P8W7Y2 A0A4P8W851 A0A4P8WAD3
A0A4P8WAE5 A0A4S8MAF3 A0A4V1DXC3 A0A4V1DXC4 A0A4V2JTK3 A0A4W3 A0A4X1T4U3
A0A4Y5UJ70 A0A4Y6HUD7 A0A4Y6HUI7 A0A4Y6UQ63 A0A509ADV9 A0A509AF89 A0A509AFG4
A0A509AIR7 A0A509AJL3 A0A509AJP9 A0A509ALD0 A0A516F180 A0A516F183 A0A516KNH9
A0A5B8YU33 A0A5B8YU39 A0A5B8YU67 A0A5B8YU68 A0A5B8YU69 A0A5B8YU81 A0A5B8YUW5
A0A5B8YUX3 A0A5B8YUX5 A0A5B8YUY5 A0A5B8YVD7 A0A5B8YVD8 A0A5B8YW48 A0A5B8YWJ2
A0A5B8YWJ9 A0A5B8YXH5 A0A5B8YXI2 A0A5C1RD96 A0A5C1RDA3 A0A5C1RDA6 A0A5C1RDG7
A0A5C1REZ4 A0A5C1RF03 A0A5C1RFE1 A0A5C1RFS2 A0A5C1RHX3 A0A5E4M3Q4 A0A5F8AH41
A0A5F8MPE6 A0A5F9ZHS7 A0A5K4F1D0 A0A5K7RLP0 A0A5M3Z5R2 A0A5N1I561 A0A5P8DHZ6
A0A5P8DI06 A0A5P8DI07 A0A5P8DI15 A0A5Q0QMX0 A0A5Q0QMX1 A0A5Q0QMX6 A0A5Q0QN66
A0A5Q0QNH9 A0A5Q0QP64 A0A5Q0QRJ3 A0A5Q0QRK8 A0A5Q0QU70 A0A5S9I252 A0A5S9MMK5
A0A646QV53 A0A663DJA2 A0A6C0WW36 A0A6C0WW38 A0A6C0WX00 A0A6C7EEG6 A0A6G9KH54
A0A6G9KH61 A0A6G9KIF9 A0A6G9KJC3 A0A6G9KJL7 A0A6J4B487 A0A6J4B4M6 A0A6J4B4P7
A0A6J4B5J2 A0A6J4B898 A0A6M3VWZ7 A0A6M3VXD3 A0A6M3VZT9 A0A6M7H989 A0A6P6W6H5
A0A6S5ZY48 A0A6S6AAU0 A0A6S6QJ62 A0A6S6QR11 A0A7D0AGU9 A0A7G5KLV0 A0A7G5KLV1
A0A7G5KLV2 A0A7G5KLV3 A0A7L8UVC6 A0A7L8UVC9 A0A7L8UWS6 A0A7L8UXK7 A0A7L9EYL1
A0A7L9EZ90 A0A7L9EZZ4 A0A7L9F0X4 A0A7M4BDQ2 A0A7N9VSG0 A0A7T1FRB0 A0A7U3TBV3
A0A7U3TBV6 A0A7U3TCA2 A0A7U9P668 A0A848M4Z0 A0A858E4Y6 A0A858E4Y8 A0A858E6N7
A0A858E7G0 A0A858E7J4 A0A858E899 A0A858EAD5 A0A868BQY3 A0A8C2MDK8 A0A8D9PCQ4
A0A8D9PH56 A0A8D9UGQ7 A0A8D9UGU6 A0A8F4PN06 A0A8F4SKJ7 A0A8F5SIS3 A0A8F5XX49
A0A8G1LR08 A0A8K1AWG4 A0A8K1AY78 A0A8M2 A0AAL4 A0AAR7 A0ACX8 A0AEI7 A0AEJ1

A0AEM0 A0AEM1 A0AEM3 A0AEN0 A0AEN2 A0AEY1 A0AEY3 A0AEZ3 A0AEZ5 A0AEZ6 A0AF06
A0AF11 A0AF20 A0AF21 A0AF28 A0AF31 A0AF36 A0AF38 A0AF43 A0AF47 A0AF48 A0AF49
A0AF50 A0AF63 A0AF64 A0AF79 A0AFA6 A0AFC3 A0AFC5 A0AFI3 A0AFT6 A0AFT8 A0AFU3
A0AFU8 A0AFY0 A0AG13 A0AG14 A0AG15 A0AG16 A0AG17 A0AG18 A0AG20 A0AG21 A0AGV7
A0AGY4 A0AH04 A0AH12 A0AH18 A0AH37 A0AH42 A0AH75 A0AH92 A0AHA7 A0AHD1 A0AHD2
A0AHE8 A0AHF9 A0AHG0 A0AHG3 A0AHH1 A0AHH9 A0AHI4 A0AHK4 A0AHK7 A0AHS0 A0AHT7
A0AHV1 A0AHW2 A0AHX3 A0AHX5 A0AHX8 A0AHX9 A0AI64 A0AI65 A0AI70 A0AI71 A0AI76
A0AI79 A0AI80 A0AI81 A0AI82 A0AI83 A0AI95 A0AIA4 A0AIB4 A0AIB5 A0AIB8 A0AIC0
A0AIC1 A0AIC2 A0AIC6 A0AIC8 A0AIC9 A0AID1 A0AID2 A0AID6 A0AIE9 A0AIF0 A0AIF1
A0AIF6 A0AIG0 A0AIG1 A0AIG2 A0AIG3 A0AIG6 A0AIG8 A0AIH1 A0AII3 A0AII5 A0AII6
A0AIJ9 A0AIK0 A0AIK5 A0AIK6 A0AIM3 A0AIN8 A0AIQ0 A0AIQ1 A0AIQ7 A0AIQ8 A0AIQ9
A0AIR0 A0AIR1 A0AIR6 A0AIS0 A0AIS2 A0AIS3 A0AIS4 A0AIS5 A0AIS6 A0AIS9 A0AIT7
A0AIU3 A0AIU5 A0AIU6 A0AIV0 A0AIV2 A0AIV3 A0AIW1 A0AIW8 A0AIX1 A0AIX3 A0AIX9
A0AIY0 A0AIY1 A0AIY2 A0AIY6 A0AIY7 A0AIY9 A0AIZ1 A0AIZ4 A0AIZ8 A0AJ02 A0AJ05
A0AJ06 A0AJ07 A0AJ08 A0AJ11 A0AJ20 A0AJ26 A0AJ38 A0AJ40 A0AJ41 A0AJ45 A0AJ48
A0AJ57 A0AJ60 A0AJ67 A0AJ79 A0AJ80 A0AJ81 A0AJ82 A0AJ83 A0AJB0 A0AJB1 A0AJB3
A0AJB5 A0AJB9 A0AJC9 A0AJD5 A0AJD9 A0AJE1 A0AJE7 A0AJG3 A0AJJ5 A0AJK7 A0AJK8
A0AJK9 A0AJL1 A0AJL9 A0AJM1 A0AJM3 A0AJM4 A0AJM6 A0AJN4 A0AJP1 A0AJP5 A0AJP7
A0AJP8 A0AJQ2 A0AJR0 A0AJS1 A0AJS8 A0AJT1 A0AJT7 A0AJT8 A0AJT9 A0AJU0 A0AJU2
A0AJU3 A0AJU5 A0AJW1 A0AJW2 A0AJW4 A0AJW5 A0AJX1 A0AJY0 A0AJY2 A0AJY3 A0AJY7
A0AJZ0 A0AJZ3 A0AJZ4 A0AJZ6 A0AK01 A0AK05 A0AK06 A0AK07 A0AK10 A0AK11 A0AK12
A0AK33 A0AK35 A0AK37 A0AK39 A0AK40 A0AK41 A0AK43 A0AK45 A0AK49 A0AK51 A0AK62
A0AK63 A0AK66 A0AK73 A0AK80 A0AK82 A0AK88 A0AK91 A0AK92 A0AK94 A0AK95 A0AKA1
A0AKC4 A0AKC5 A0AKD0 A0AKE1 A0AKE2 A0AKE7 A0AKE9 A0AKF8 A0AKG0 A0AKH5 A0AKH6
A0AKH9 A0AKI2 A0AKJ7 A0AKJ8 A0AKK8 A0AKK9 A0AKL8 A0AKM3 A0AKU3 A0AKV8 A0AKW5
A0AKX3 A0AKX6 A0AL18 A0AL19 A0AL52 A0AL74 A0AL76 A0ALA8 A0ALD2 A0ALE0 A0ALE2
A0ALF5 A0ALF6 A0ALF8 A0ALG5 A0ALG7 A0ALM3 A0ALM4 A0ALM8 A0ALN0 A0ALN3 A0ALP6
A0ALP7 A0ALT2 A0ALT3 A0ALT4 A0ALU1 A0ALU2 A0ALU3 A0ALU4 A0ALU5 A0ALU6 A0ALU7
A0ALU9 A0ALV0 A0ALV1 A0ALV2 A0ALV3 A0ALV4 A0ALV5 A0ALV6 A0ALV7 A0ALV8 A0ALV9
A0ALW0 A0ALW1 A0ALW2 A0ALW3 A0ALW4 A0ALW5 A0ALW6 A0ALW7 A0ALW8 A0ALW9 A0ALY8
A0ALY9 A0ALZ0 A0ALZ1 A0AM28 A0AM39 A0AM40 A0AM81 A0AMA3 A0AMC3 A0AMC4 A0AMC5
A0AMD1 A0AMD2 A0AMG4 A0AMG5 A0AMG7 A0AMG8 A0AMG9 A0AQW4 A0AUJ5 A0AUP1 A0AUQ6
A0AUS0 A0AUT1 A0AUZ9 A0AV96 A0AVF1 A0AVK6 A0AVT1 A0AWY9 A0AXI7 A0AXK2 A0AXR0
A0AXY5 A0AY13 A0AYH4 A0AYX8 A0AYY8 A0AYZ0 A0AYZ1 A0AYZ2 A0AZ43 A0AZ60 A0AZ67
A0AZ71 A0AZQ0 A0B0B8 A0B0M1 A0B0V3 A0B137 A0B1L8 A0B2F5 A0B2F6 A0B2F7 A0B2Q6
A0B2Y5 A0B2Z3 A0B338 A0B3M8 A0B3Q5 A0B3W7 A0B459 A0B466 A0B471 A0B532 A0B533
A0B555 A0B560 A0B562 A0B563 A0B571 A0B577 A0B5A7 A0B5B1 A0B5B7 A0B5C3 A0B5C6
A0B5C7 A0B5D0 A0B5E0 A0B5E6 A0B5E9 A0B5H8 A0B5I5 A0B5I7 A0B5J1 A0B5K3 A0B5K5
A0B5K9 A0B5L3 A0B5N0 A0B5N2 A0B5P0 A0B5P4 A0B5P5 A0B5Q5 A0B5R0 A0B5R2 A0B5S0
A0B5T8 A0B5U4 A0B601 A0B603 A0B624 A0B632 A0B635 A0B645 A0B688 A0B6A6 A0B6C1
A0B6C9 A0B6D7 A0B6E1 A0B6E5 A0B6E6 A0B6E7 A0B6E8 A0B6F6 A0B6F9 A0B6I9 A0B6K6
A0B6K7 A0B6L2 A0B6S6 A0B6U3 A0B6X3 A0B6X7 A0B6X8 A0B6Y9 A0B6Z6 A0B738 A0B742
A0B747 A0B748 A0B796 A0B7A6 A0B7B7 A0B7C2 A0B7D5 A0B7D6 A0B7D7 A0B7E1 A0B7F2
A0B7F9 A0B7H6 A0B7J8 A0B7K2 A0B7N4 A0B7P3 A0B7Q2 A0B7U9 A0B7V0 A0B7V2 A0B7V3

A0B7V4 A0B7W4 A0B7X3 A0B7Y8 A0B7Z0 A0B870 A0B879 A0B880 A0B885 A0B886 A0B8A1
A0B8B0 A0B8B3 A0B8D4 A0B8F1 A0B8J2 A0B8J6 A0B8J7 A0B8K5 A0B8K9 A0B8L1 A0B8L2
A0B8M6 A0B8N3 A0B8P7 A0B8Q1 A0B8Q6 A0B8Q8 A0B8S7 A0B8S8 A0B8S9 A0B8V5 A0B8V8
A0B8W0 A0B8Z1 A0B913 A0B922 A0B923 A0B966 A0B976 A0B977 A0B993 A0B999 A0B9A2
A0B9D8 A0B9G1 A0B9G7 A0B9L1 A0B9L2 A0B9L3 A0B9M7 A0B9M9 A0B9N3 A0B9N7 A0B9P3
A0B9P8 A0B9Q9 A0B9S8 A0B9U1 A0B9U5 A0B9U9 A0B9V0 A0B9V1 A0B9V4 A0B9V5 A0B9V9
A0B9W0 A0B9W3 A0B9W4 A0B9W5 A0B9W6 A0B9W7 A0B9W9 A0B9X0 A0B9X4 A0BD73 A0BD92
A0BV78 A0C1E4 A0C574 A0CC46 A0CCD2 A0CEY2 A0CH87 A0CIS3 A0CNL9 A0CRL1 A0CXT3
A0CY51 A0CYG2 A0DB19 A0DJ90 A0E3W6 A0FGR0 A0FIN4 A0FKE6 A0FKG7 A0FLQ6 A0JC51
A0JC76 A0JC77 A0JJU0 A0JJU1 A0JJZ6 A0JLY1 A0JM08 A0JM23 A0JM24 A0JM49 A0JM51
A0JM56 A0JM64 A0JM80 A0JM83 A0JM98 A0JMA0 A0JMA6 A0JMA8 A0JMA9 A0JMD0 A0JME2
A0JME6 A0JMF1 A0JMF6 A0JMF8 A0JMG1 A0JMH2 A0JMI9 A0JMK8 A0JMK9 A0JML8 A0JMM9
A0JMP0 A0JMQ0 A0JMR6 A0JMT0 A0JMU5 A0JMU8 A0JMV4 A0JMW2 A0JMY4 A0JMZ1 A0JMZ3
A0JMZ4 A0JN27 A0JN40 A0JN41 A0JN51 A0JN52 A0JN53 A0JN71 A0JN74 A0JN76 A0JN86
A0JN92 A0JN95 A0JNA3 A0JNB1 A0JNC2 A0JND2 A0JND4 A0JNE3 A0JNF3 A0JNG6 A0JNG7
A0JNH1 A0JNH6 A0JNH9 A0JNI1 A0JNI4 A0JNI5 A0JNI8 A0JNJ4 A0JNJ5 A0JNL1 A0JNN3
A0JNN6 A0JNQ6 A0JNT9 A0JP26 A0JP43 A0JP70 A0JP75 A0JP82 A0JP85 A0JP94 A0JPB4
A0JPD7 A0JPE9 A0JPF5 A0JPF9 A0JPG1 A0JPH7 A0JPI9 A0JPK3 A0JPL0 A0JPM9 A0JPN1
A0JPN6 A0JPP1 A0JPP5 A0JPP7 A0JPP8 A0JPQ1 A0JPQ7 A0JQT5 A0JQW8 A0JR24 A0JR71
A0JR91 A0JRF5 A0JRH2 A0JRH3 A0JRH4 A0JRH5 A0JRH6 A0JRH8 A0JSA1 A0JSD3 A0JSM6
A0JSP6 A0JSU9 A0JSV8 A0JT07 A0JT43 A0JT53 A0JTA3 A0JTE5 A0JTZ0 A0JU87 A0JU88
A0JU94 A0JUH2 A0JUI0 A0JUI2 A0JUI7 A0JUP1 A0JUP2 A0JUP3 A0JUP4 A0JUS3 A0JUS8
A0JUT1 A0JUT7 A0JUU0 A0JUU1 A0JUU8 A0JUV5 A0JUV7 A0JUV8 A0JUY4 A0JUY6 A0JUY7
A0JUZ5 A0JUZ7 A0JV02 A0JV03 A0JV20 A0JV22 A0JV26 A0JV33 A0JV85 A0JV86 A0JV95
A0JVA0 A0JVB0 A0JVB5 A0JVC9 A0JVG9 A0JVJ5 A0JVK2 A0JVK3 A0JVK4 A0JVK5 A0JVK7
A0JVL1 A0JWN0 A0JWP2 A0JWP3 A0JWP5 A0JWP8 A0JWQ6 A0JWT7 A0JWV0 A0JWX8 A0JWY3
A0JWY6 A0JWY7 A0JWY8 A0JX18 A0JX36 A0JX41 A0JX46 A0JX52 A0JX57 A0JX72 A0JX76
A0JX79 A0JX80 A0JX82 A0JX84 A0JX88 A0JX91 A0JXA1 A0JXB1 A0JXB2 A0JXB3 A0JXB4
A0JXB5 A0JXC7 A0JXJ4 A0JXJ8 A0JXJ9 A0JXK0 A0JXL2 A0JXL5 A0JXN9 A0JXQ4 A0JXT3
A0JXT5 A0JXT8 A0JXT9 A0JXU0 A0JXW1 A0JXW3 A0JXX3 A0JXX7 A0JXX8 A0JXY5 A0JXZ6
A0JXZ9 A0JY49 A0JY58 A0JY78 A0JYC1 A0JYC5 A0JYI6 A0JYM2 A0JYN1 A0JYN2 A0JYP1
A0JYU5 A0JYZ5 A0JYZ6 A0JZ00 A0JZ01 A0JZ09 A0JZ10 A0JZ25 A0JZ26 A0JZ27 A0JZ48
A0JZ49 A0JZ50 A0JZ51 A0JZ52 A0JZ53 A0JZ67 A0JZ68 A0JZ70 A0JZ71 A0JZ72 A0JZ73
A0JZ74 A0JZ77 A0JZ78 A0JZ79 A0JZ80 A0JZ81 A0JZ82 A0JZ84 A0JZ85 A0JZ88 A0JZ90
A0JZ91 A0JZ92 A0JZ93 A0JZ99 A0JZA2 A0JZA3 A0JZB9 A0JZC2 A0JZC7 A0JZP6 A0JZZ4
A0K010 A0K0C0 A0K0Y1 A0K1I7 A0K1L3 A0K1R7 A0K1U2 A0K1X4 A0K1X5 A0K1X6 A0K258
A0K2D5 A0K2H2 A0K2H3 A0K2H5 A0K2M2 A0K2M6 A0K2M7 A0K2M8 A0K2U3 A0K2U6 A0K2V4
A0K2X0 A0K2X1 A0K321 A0K322 A0K3G7 A0K3L0 A0K3L3 A0K3L4 A0K3L5 A0K3L6 A0K3L7
A0K3L8 A0K3M0 A0K3M1 A0K3M4 A0K3M5 A0K3M6 A0K3M7 A0K3M8 A0K3M9 A0K3N0 A0K3N1
A0K3N2 A0K3N3 A0K3N4 A0K3N5 A0K3N6 A0K3N7 A0K3N8 A0K3N9 A0K3P0 A0K3P1 A0K3P2
A0K3P3 A0K3P4 A0K3P6 A0K3P7 A0K3P8 A0K3P9 A0K3Q0 A0K3Q1 A0K3Q2 A0K3Q6 A0K3R3
A0K3S0 A0K3V1 A0K3V4 A0K3V7 A0K3V8 A0K3V9 A0K3W0 A0K437 A0K438 A0K458 A0K459
A0K477 A0K478 A0K484 A0K487 A0K488 A0K4A1 A0K4A8 A0K4A9 A0K4B0 A0K4B1 A0K4B3
A0K4B4 A0K4C9 A0K4H7 A0K4I0 A0K4I3 A0K4K6 A0K4K7 A0K4K8 A0K4L0 A0K4L4 A0K4L5

A0K4L6 A0K4L7 A0K4M4 A0K4P9 A0K4Q2 A0K4S0 A0K4S1 A0K4S2 A0K4S6 A0K4S8 A0K4S9
A0K4T1 A0K4T7 A0K4U0 A0K4Y3 A0K4Z6 A0K536 A0K540 A0K541 A0K543 A0K573 A0K574
A0K575 A0K576 A0K577 A0K578 A0K579 A0K594 A0K5D1 A0K5D2 A0K5D7 A0K5F5 A0K5F9
A0K5G1 A0K5H3 A0K5H6 A0K5J0 A0K5L4 A0K5L8 A0K5M1 A0K5M2 A0K5M4 A0K5M9 A0K5P0
A0K5P5 A0K5P6 A0K5P7 A0K5P8 A0K5Q4 A0K5Q6 A0K5S6 A0K5T4 A0K5U4 A0K5W6 A0K5W8
A0K608 A0K609 A0K610 A0K648 A0K6C1 A0K6C4 A0K6D2 A0K6N4 A0K6Q3 A0K6T5 A0K6V2
A0K6V3 A0K6X3 A0K6X5 A0K6X6 A0K724 A0K730 A0K731 A0K733 A0K7D5 A0K7F1 A0K7Q6
A0K7S6 A0K7T1 A0K7T2 A0K7T5 A0K7T8 A0K7T9 A0K7U1 A0K7U6 A0K7U9 A0K7V0 A0K7V7
A0K7Z5 A0K7Z6 A0K7Z8 A0K817 A0K832 A0K833 A0K846 A0K847 A0K848 A0K866 A0K867
A0K8B3 A0K8C3 A0K8C7 A0K8C9 A0K8D7 A0K8E0 A0K8E1 A0K8E2 A0K8E3 A0K8E8 A0K8F4
A0K8F6 A0K8I0 A0K8K2 A0K8L2 A0K8N3 A0K8P2 A0K8P6 A0K8R4 A0K8T1 A0K8U2 A0K8U6
A0K8Z7 A0K927 A0K928 A0K933 A0K937 A0K965 A0K968 A0K973 A0K988 A0K989 A0K993
A0K9A9 A0K9B9 A0K9C0 A0K9K2 A0K9K6 A0K9L0 A0K9L5 A0K9L6 A0K9N9 A0K9S4 A0K9S5
A0K9S6 A0K9S9 A0K9T5 A0K9T8 A0K9U2 A0K9W9 A0K9X1 A0K9X4 A0K9X6 A0K9X7 A0K9X8
A0K9Z1 A0KA00 A0KA02 A0KA61 A0KA72 A0KAA1 A0KAB6 A0KAB8 A0KAD1 A0KAE8 A0KAF5
A0KAG0 A0KAJ8 A0KAK7 A0KAL7 A0KAL8 A0KAM1 A0KAM4 A0KAM6 A0KAM7 A0KAN0 A0KAS4
A0KAS7 A0KAV9 A0KAX6 A0KAZ0 A0KAZ7 A0KB03 A0KB04 A0KB05 A0KB51 A0KBA2 A0KBF2
A0KBG2 A0KBG3 A0KBG7 A0KBH0 A0KBH8 A0KBH9 A0KBI0 A0KBJ0 A0KBJ7 A0KBN1 A0KBN6
A0KCF0 A0KCQ9 A0KCS0 A0KCV5 A0KDG4 A0KDV0 A0KDV2 A0KE38 A0KE56 A0KEC3 A0KEG6
A0KEG8 A0KEG9 A0KEH8 A0KEJ1 A0KEJ2 A0KEJ9 A0KEK0 A0KEK5 A0KEL3 A0KEM1 A0KEM2
A0KEM6 A0KEM8 A0KEM9 A0KEN0 A0KEN2 A0KEN4 A0KEP8 A0KEQ6 A0KER2 A0KEW9 A0KEX3
A0KEX6 A0KEX7 A0KEX8 A0KEY0 A0KEY2 A0KEZ2 A0KF07 A0KF09 A0KF11 A0KF21 A0KF22
A0KF23 A0KF24 A0KF25 A0KF26 A0KF27 A0KF28 A0KF30 A0KF31 A0KF32 A0KF33 A0KF34
A0KF35 A0KF36 A0KF37 A0KF38 A0KF39 A0KF40 A0KF42 A0KF43 A0KF44 A0KF45 A0KF46
A0KF84 A0KF85 A0KFC0 A0KFE0 A0KFH9 A0KFI1 A0KFJ6 A0KFN8 A0KFQ0 A0KFR1 A0KFT8
A0KFU8 A0KFV0 A0KFV1 A0KFV3 A0KFV4 A0KG11 A0KG16 A0KG23 A0KG24 A0KG25 A0KG26
A0KG29 A0KG32 A0KG36 A0KG39 A0KG42 A0KG67 A0KG68 A0KG69 A0KGD2 A0KGD3 A0KGG9
A0KGH2 A0KGH5 A0KGH7 A0KGH8 A0KGH9 A0KGI3 A0KGI4 A0KGJ2 A0KGL0 A0KGL1 A0KGL5
A0KGM6 A0KGM7 A0KGM9 A0KGR9 A0KGS0 A0KGS1 A0KGS7 A0KGS8 A0KGS9 A0KGT6 A0KGT8
A0KGZ0 A0KH11 A0KHA2 A0KHB9 A0KHC4 A0KHD5 A0KHE0 A0KHE3 A0KHF9 A0KHG0 A0KHG3
A0KHG4 A0KHG5 A0KHG6 A0KHG9 A0KHH7 A0KHL9 A0KHR8 A0KHU1 A0KHV1 A0KHV5 A0KHY2
A0KHZ9 A0KI46 A0KI53 A0KI76 A0KI80 A0KIB5 A0KIC3 A0KIC6 A0KIC7 A0KIC9 A0KIF5
A0KIF6 A0KIF9 A0KIG3 A0KIH3 A0KIK2 A0KIK4 A0KIK9 A0KIL8 A0KIN6 A0KIN9 A0KIP7
A0KIP8 A0KIQ4 A0KIT3 A0KIV0 A0KIY0 A0KIZ1 A0KJ03 A0KJ04 A0KJ19 A0KJ20 A0KJ32
A0KJ35 A0KJ36 A0KJ40 A0KJ41 A0KJ45 A0KJ48 A0KJ91 A0KJ92 A0KJ94 A0KJA0 A0KJD4
A0KJD6 A0KJD8 A0KJD9 A0KJE0 A0KJE8 A0KJG4 A0KJH4 A0KJI0 A0KJJ0 A0KJJ1 A0KJJ9
A0KJK9 A0KJL8 A0KJM0 A0KJP1 A0KJP3 A0KJS0 A0KJT2 A0KJU0 A0KJU2 A0KK26 A0KK38
A0KK42 A0KK55 A0KK57 A0KK58 A0KK62 A0KKB0 A0KKB4 A0KKB7 A0KKB9 A0KKD3 A0KKD5
A0KKE6 A0KKE7 A0KKF0 A0KKF3 A0KKF6 A0KKF8 A0KKF9 A0KKH0 A0KKH9 A0KKI5 A0KKK7
A0KKK8 A0KKL4 A0KKM7 A0KKN1 A0KKP3 A0KKP5 A0KKP6 A0KKP7 A0KKP9 A0KKS7 A0KKS9
A0KKW8 A0KL51 A0KL52 A0KL53 A0KL54 A0KL76 A0KLC8 A0KLD7 A0KLD9 A0KLN7 A0KLP0
A0KLP9 A0KLT5 A0KLY0 A0KM23 A0KM24 A0KM45 A0KM75 A0KMC8 A0KMD0 A0KMD1 A0KMH0
A0KMI5 A0KMI6 A0KMI7 A0KMM4 A0KMP9 A0KMV4 A0KMY0 A0KMZ1 A0KMZ8 A0KMZ9 A0KN01
A0KN04 A0KN18 A0KN28 A0KN29 A0KN65 A0KN67 A0KN81 A0KN83 A0KN87 A0KN91 A0KN93

A0KNA1 A0KNA3 A0KND9 A0KNE0 A0KNE2 A0KNE3 A0KNE5 A0KNE8 A0KNF1 A0KNF9 A0KNG6
A0KNH2 A0KNH4 A0KNH6 A0KNI2 A0KNI3 A0KNJ1 A0KNK6 A0KNK7 A0KNL0 A0KNQ8 A0KNT0
A0KNT4 A0KNU5 A0KNV3 A0KNV5 A0KNY4 A0KNY6 A0KNY9 A0KP02 A0KP07 A0KP08 A0KP33
A0KP35 A0KP36 A0KP37 A0KPA2 A0KPA3 A0KPA5 A0KPB6 A0KPC2 A0KPC4 A0KPE2 A0KPE3
A0KPE4 A0KPF3 A0KPF9 A0KPG1 A0KPG3 A0KPG5 A0KPI3 A0KPL4 A0KPN1 A0KPN5 A0KPP9
A0KPQ1 A0KPW2 A0KPX1 A0KPY0 A0KPY4 A0KPZ2 A0KPZ3 A0KQ08 A0KQ23 A0KQ25 A0KQ95
A0KQ96 A0KQ97 A0KQ98 A0KQA4 A0KQA5 A0KQA6 A0KQA7 A0KQA8 A0KQA9 A0KQF1 A0KQG0
A0KQG1 A0KQG4 A0KQH4 A0KQH8 A0KQI7 A0KQI8 A0KQJ4 A0KQK0 A0KQR3 A0KQS4 A0KQV3
A0KQV6 A0KQY8 A0KQY9 A0KQZ6 A0KQZ9 A0KR00 A0KR23 A0KR27 A0KR28 A0KR31 A0KR34
A0KR35 A0KR37 A0KR42 A0KR43 A0KR47 A0KR51 A0KR60 A0KR64 A0KR66 A0KR67 A0KR69
A0KR77 A0KR79 A0KRB5 A0KRC4 A0KRC6 A0KRF0 A0KRF2 A0KRF9 A0KRH2 A0KRH3 A0KRK0
A0KRK1 A0KRK4 A0KRK5 A0KRK7 A0KRK9 A0KRL0 A0KRL3 A0KRL4 A0KRL5 A0KRL6 A0KRL7
A0KRL8 A0KRL9 A0KRM0 A0KRM3 A0KRM4 A0KRM5 A0KRM7 A0KRM8 A0KRM9 A0KRN0 A0KRN1
A0KRN2 A0KRN3 A0KRN4 A0KRN5 A0KRN6 A0KRN7 A0KRN8 A0KRN9 A0KRP0 A0KRP1 A0KRP2
A0KRP3 A0KRP5 A0KRP6 A0KRP7 A0KRP8 A0KRP9 A0KRQ0 A0KRT7 A0KRU9 A0KRY8 A0KRZ0
A0KS32 A0KS60 A0KS62 A0KS65 A0KS67 A0KS72 A0KS74 A0KS84 A0KS88 A0KSA9 A0KSI2
A0KSL6 A0KSN0 A0KSP4 A0KSQ0 A0KSR0 A0KSR5 A0KSR6 A0KSR7 A0KST2 A0KSW7 A0KSX4
A0KSY8 A0KT03 A0KT11 A0KT12 A0KT20 A0KT22 A0KT23 A0KT33 A0KT36 A0KT69 A0KT85
A0KT87 A0KT88 A0KTG8 A0KTH4 A0KTH5 A0KTI2 A0KTI5 A0KTL9 A0KTP8 A0KTR5 A0KTS5
A0KTS6 A0KTT2 A0KTU3 A0KTU4 A0KTV4 A0KTV5 A0KTV6 A0KTW2 A0KTW4 A0KTW7 A0KTX2
A0KTX4 A0KTX7 A0KTY8 A0KTZ1 A0KTZ2 A0KTZ4 A0KTZ6 A0KTZ7 A0KTZ8 A0KTZ9 A0KU01
A0KU03 A0KU07 A0KU08 A0KU09 A0KU20 A0KU60 A0KU61 A0KU66 A0KU76 A0KU81 A0KU85
A0KU87 A0KU88 A0KU91 A0KU92 A0KU94 A0KU96 A0KUE6 A0KUE8 A0KUF0 A0KUF7 A0KUG6
A0KUJ2 A0KUJ5 A0KUJ6 A0KUJ9 A0KUK0 A0KUK2 A0KV13 A0KV33 A0KV49 A0KV50 A0KV82
A0KV84 A0KV89 A0KV91 A0KV94 A0KVD1 A0KVD3 A0KVG7 A0KVI4 A0KVL6 A0KVM4 A0KVN6
A0KVN8 A0KVP0 A0KVP1 A0KVS0 A0KVW5 A0KVX6 A0KVZ2 A0KVZ9 A0KW11 A0KW16 A0KW18
A0KW19 A0KW20 A0KW24 A0KW26 A0KW27 A0KW30 A0KW34 A0KWB0 A0KWB1 A0KWB6 A0KWC7
A0KWF4 A0KWH4 A0KWK2 A0KWL2 A0KWL3 A0KWL5 A0KWL7 A0KWL8 A0KWL9 A0KWN3 A0KWN5
A0KWN7 A0KWN8 A0KWP0 A0KWV4 A0KWV6 A0KWV7 A0KWW5 A0KWX7 A0KX28 A0KX39 A0KX43
A0KX52 A0KX58 A0KX70 A0KX75 A0KXH1 A0KXI2 A0KXI4 A0KXI6 A0KXI7 A0KXJ0 A0KXK2
A0KXK3 A0KXL6 A0KXL7 A0KXS6 A0KXS9 A0KXU1 A0KXU2 A0KXU4 A0KXU5 A0KXY6 A0KY01
A0KY03 A0KY04 A0KY07 A0KY73 A0KY79 A0KY83 A0KY98 A0KYA2 A0KYA3 A0KYC0 A0KYD1
A0KYE0 A0KYF8 A0KYL8 A0KYL9 A0KYM0 A0KYM1 A0KYM2 A0KYQ5 A0KYQ6 A0KYQ7 A0KYY0
A0KYY8 A0KZ02 A0KZ09 A0KZ17 A0KZ20 A0KZ21 A0KZ22 A0KZ23 A0KZ26 A0KZ89 A0KZ94
A0KZ99 A0KZA4 A0KZA9 A0KZB0 A0KZB1 A0KZD8 A0KZF9 A0KZH8 A0KZL7 A0KZM0 A0KZM1
A0KZM2 A0KZM3 A0KZN2 A0KZP5 A0KZP9 A0KZQ0 A0KZQ7 A0KZR5 A0KZS2 A0KZS6 A0KZT1
A0KZT8 A0KZT9 A0KZZ1 A0KZZ5 A0KZZ8 A0L002 A0L003 A0L052 A0L064 A0L065 A0L066
A0L072 A0L073 A0L074 A0L0B8 A0L0D3 A0L0F1 A0L0G4 A0L0I8 A0L0K6 A0L0K7 A0L0K9
A0L0M2 A0L0Q7 A0L0R2 A0L0R3 A0L0V1 A0L0W9 A0L0Y0 A0L103 A0L104 A0L105 A0L113
A0L114 A0L170 A0L171 A0L1C5 A0L1C8 A0L1D3 A0L1H0 A0L1H3 A0L1J8 A0L1J9 A0L1M4
A0L1P1 A0L1Q0 A0L1Q1 A0L1Q5 A0L1Q7 A0L1Q8 A0L1R0 A0L1R7 A0L1R8 A0L1R9 A0L1S1
A0L1S2 A0L1S4 A0L1S6 A0L1W0 A0L1Y5 A0L1Z0 A0L230 A0L231 A0L242 A0L243 A0L250
A0L251 A0L253 A0L254 A0L257 A0L2B4 A0L2F1 A0L2I5 A0L2M8 A0L2N2 A0L2N9 A0L2Q3
A0L2Q8 A0L2R5 A0L3I7 A0L3I9 A0L3J9 A0L3K1 A0L3K3 A0L3K8 A0L3L1 A0L3L7 A0L3L9

A0L3M0 A0L3M5 A0L3M6 A0L3N2 A0L3R7 A0L3S9 A0L3T9 A0L3X0 A0L3X7 A0L3Z9 A0L403
A0L407 A0L408 A0L480 A0L481 A0L482 A0L486 A0L4B0 A0L4B2 A0L4C9 A0L4J3 A0L4J6
A0L4J8 A0L4J9 A0L4K0 A0L4K5 A0L4K6 A0L4N8 A0L4P8 A0L4R4 A0L4R8 A0L4V5 A0L4V9
A0L4W0 A0L4W1 A0L4X9 A0L4Y6 A0L4Y7 A0L4Y8 A0L4Y9 A0L4Z0 A0L4Z2 A0L4Z4 A0L509
A0L510 A0L516 A0L576 A0L579 A0L5E8 A0L5G0 A0L5G1 A0L5I2 A0L5J6 A0L5K4 A0L5L8
A0L5M8 A0L5M9 A0L5N0 A0L5N9 A0L5T9 A0L5U1 A0L5U5 A0L5V8 A0L5W2 A0L5W3 A0L5W5
A0L5W6 A0L5W7 A0L5W8 A0L5W9 A0L5X0 A0L5X2 A0L5X3 A0L5X4 A0L5X5 A0L5X6 A0L5X7
A0L5X8 A0L5X9 A0L5Y0 A0L5Y1 A0L5Y2 A0L5Y3 A0L5Y4 A0L5Y5 A0L5Y6 A0L5Y7 A0L5Y8
A0L5Z0 A0L5Z1 A0L5Z2 A0L5Z4 A0L5Z5 A0L5Z6 A0L5Z7 A0L5Z8 A0L5Z9 A0L626 A0L633
A0L636 A0L667 A0L669 A0L673 A0L678 A0L6E2 A0L6E9 A0L6F0 A0L6F1 A0L6F2 A0L6F4
A0L6F5 A0L6F7 A0L6H3 A0L6H5 A0L6J8 A0L6W5 A0L6X0 A0L6Y9 A0L7K3 A0L7K6 A0L7K9
A0L7L5 A0L7P3 A0L7R3 A0L7R4 A0L7S8 A0L7T9 A0L7X0 A0L7X6 A0L837 A0L859 A0L886
A0L887 A0L8B0 A0L8B5 A0L8B9 A0L8F5 A0L8G8 A0L8H9 A0L8I1 A0L8I2 A0L8J0 A0L8J4
A0L8J5 A0L8Q3 A0L8Q5 A0L8Q6 A0L8Q7 A0L8Q8 A0L8S1 A0L8S5 A0L8T5 A0L8U7 A0L8V0
A0L900 A0L901 A0L909 A0L9S5 A0LA23 A0LA32 A0LA38 A0LA39 A0LA84 A0LAG5 A0LAG7
A0LAG9 A0LAN9 A0LBT6 A0LBW3 A0LBZ1 A0LC14 A0LC15 A0LC19 A0LC90 A0LCB4 A0LCF2
A0LCF3 A0LCF4 A0LCF5 A0LCI7 A0LCQ7 A0LCU4 A0LCX5 A0LCX6 A0LCY8 A0LCZ1 A0LCZ3
A0LCZ4 A0LCZ5 A0LD64 A0LD65 A0LD67 A0LD71 A0LDB0 A0LDB5 A0LDB8 A0LDC6 A0LDF3
A0LDN3 A0LDQ0 A0LDR1 A0LDT3 A0LDT4 A0LDT5 A0LDT6 A0LDT7 A0LDU6 A0LDX4 A0LDY1
A0LE04 A0LE14 A0LE15 A0LE17 A0LE19 A0LE22 A0LE31 A0LE33 A0LE34 A0LE46 A0LE47
A0LE48 A0LE52 A0LE53 A0LE92 A0LE95 A0LEA2 A0LEA5 A0LEA6 A0LEA7 A0LEB1 A0LEB2
A0LEB5 A0LEB7 A0LEB8 A0LEE9 A0LEF1 A0LEF2 A0LEF5 A0LEH2 A0LEJ7 A0LEL7 A0LF21
A0LF54 A0LF65 A0LF66 A0LF67 A0LFB5 A0LFB7 A0LFC4 A0LFC5 A0LFC6 A0LFF8 A0LFI1
A0LFP6 A0LFQ9 A0LFR2 A0LG38 A0LG61 A0LG64 A0LG76 A0LG91 A0LGB7 A0LGJ7 A0LGM1
A0LGY6 A0LGY7 A0LGY9 A0LGZ0 A0LH11 A0LH26 A0LH27 A0LH38 A0LH56 A0LH88 A0LHE3
A0LHG0 A0LHK5 A0LHL6 A0LHL8 A0LHM0 A0LHM2 A0LHM3 A0LHM4 A0LHQ3 A0LHQ7 A0LHR1
A0LHS2 A0LI05 A0LI08 A0LI16 A0LI20 A0LI21 A0LI55 A0LI59 A0LI63 A0LI66 A0LI67
A0LIB0 A0LIH6 A0LIH7 A0LII0 A0LII1 A0LII2 A0LII3 A0LII4 A0LII5 A0LII6 A0LII7
A0LII9 A0LIJ0 A0LIJ1 A0LIJ2 A0LIJ3 A0LIJ4 A0LIJ5 A0LIJ6 A0LIJ7 A0LIJ8 A0LIJ9
A0LIK0 A0LIK1 A0LIK2 A0LIK3 A0LIK4 A0LIK5 A0LIK7 A0LIK8 A0LIL1 A0LIL3 A0LIL4
A0LIL5 A0LIL6 A0LIL7 A0LIL8 A0LIM0 A0LIM5 A0LIN6 A0LIN8 A0LIP3 A0LIR9 A0LIS1
A0LIS2 A0LIV2 A0LIV3 A0LIV4 A0LIW0 A0LIY8 A0LIZ1 A0LIZ7 A0LIZ8 A0LJ00 A0LJ16
A0LJ24 A0LJ41 A0LJ43 A0LJ55 A0LJ58 A0LJ59 A0LJ63 A0LJ64 A0LJ65 A0LJ66 A0LJ92
A0LJA4 A0LJA8 A0LJK2 A0LJK3 A0LJR4 A0LJR5 A0LJZ1 A0LJZ7 A0LK12 A0LK14 A0LK15
A0LK18 A0LK20 A0LK21 A0LK22 A0LK48 A0LK95 A0LK96 A0LKS4 A0LKV6 A0LL24 A0LL48
A0LL53 A0LL58 A0LLA3 A0LLA4 A0LLA6 A0LLA7 A0LLH0 A0LLH1 A0LLH5 A0LLH7 A0LLI8
A0LLL8 A0LLR2 A0LLR3 A0LLU6 A0LLU7 A0LLU8 A0LLW3 A0LM16 A0LM17 A0LM45 A0LM89
A0LMC0 A0LMC1 A0LMD8 A0LMH0 A0LMI3 A0LMM1 A0LMM5 A0LN56 A0LN58 A0LN59 A0LN73
A0LNE5 A0LNI2 A0LNQ8 A0LNR3 A0LNR4 A0LNW5 A0LNY1 A0LNY2 A0LNY9 A0LNZ0 A0LP01
A0LP22 A0LP26 A0LPA9 A0LPE9 A0LPF7 A0LPF8 A0LPF9 A0LPG2 A0LPG9 A0LPH1 A0LQ64
A0LQG8 A0LQM0 A0LQM1 A0LQP0 A0LQR3 A0LQR5 A0LQR9 A0LQX9 A0LQY9 A0LQZ7 A0LQZ8
A0LR17 A0LR22 A0LR32 A0LR93 A0LRC5 A0LRC6 A0LRC9 A0LRE3 A0LRE5 A0LRF1 A0LRF2
A0LRH8 A0LRK0 A0LRK1 A0LRK9 A0LRL0 A0LRL1 A0LRL2 A0LRL3 A0LRL4 A0LRL5 A0LRL6
A0LRL7 A0LRL8 A0LRL9 A0LRM0 A0LRM2 A0LRM3 A0LRM4 A0LRM5 A0LRM6 A0LRM7 A0LRN0

A0LRN1 A0LRN2 A0LRN3 A0LRN4 A0LRN5 A0LRN6 A0LRN7 A0LRN8 A0LRP1 A0LRP3 A0LRP4
A0LRP5 A0LRP6 A0LRP7 A0LRP8 A0LRQ7 A0LRS4 A0LRS7 A0LRU4 A0LRU9 A0LS98 A0LSK3
A0LSK4 A0LSQ2 A0LSQ7 A0LSQ9 A0LSR0 A0LSR8 A0LSS7 A0LSU9 A0LSV2 A0LSW9 A0LSX0
A0LSX1 A0LSX5 A0LSY4 A0LSZ6 A0LT03 A0LT79 A0LT86 A0LTD0 A0LTD9 A0LTE0 A0LTH9
A0LTK3 A0LTL4 A0LTL5 A0LTM4 A0LTN7 A0LTP8 A0LTS3 A0LTS5 A0LTT2 A0LTT7 A0LTX2
A0LTX3 A0LTX5 A0LTY8 A0LU34 A0LU46 A0LU48 A0LU50 A0LU51 A0LU53 A0LUA5 A0LUB6
A0LUB8 A0LUB9 A0LUC1 A0LUC3 A0LUC5 A0LUC9 A0LUD0 A0LUD3 A0LUE0 A0LUE1 A0LUF4
A0LUG3 A0LUG6 A0LUG7 A0LUH0 A0LUH1 A0LUJ2 A0LUJ6 A0LUK1 A0LUK5 A0LUK6 A0LUK7
A0LUK8 A0LUK9 A0LUP4 A0LUY9 A0LUZ5 A0LUZ6 A0LV00 A0LV04 A0LV09 A0LV11 A0LV15
A0LV20 A0LV21 A0LV25 A0LV27 A0LV30 A0LV31 A0LV34 A0LV48 A0LV49 A0LV51 A0LV52
A0LV53 A0LV54 A0LV62 A0LV66 A0LV69 A0LV70 A0LV71 A0LV73 A0LV84 A0LV85 A0LV87
A0LV90 A0LV94 A0LVA2 A0LVA3 A0LVG0 A0LVJ8 A0LVP0 A0LVP4 A0LVX1 A0LVY3 A0LW20
A0LW30 A0LW37 A0LW38 A0LWH7 A0LWI8 A0LWR6 A0LWS1 A0LWS4 A0LWS7 A0LWT8 A0LWU2
A0LWU3 A0LWU5 A0LWW8 A0LWX2 A0LWX3 A0LXG7 A0LXJ3 A0LXJ4 A0LXJ9 A0LXL4 A0LXL9
A0LXM6 A0LXM7 A0LXM8 A0LXM9 A0LXP9 A0LXQ1 A0LXR1 A0LXS4 A0LXS6 A0LXV8 A0LXW0
A0LXX3 A0LXY3 A0LXZ7 A0LY29 A0LY55 A0LY71 A0LY86 A0LY87 A0LY94 A0LYC0 A0LYC2
A0LYD2 A0LYK2 A0LYL5 A0LYL6 A0LYZ0 A0LYZ1 A0LZ02 A0LZ03 A0LZ23 A0LZ30 A0LZ43
A0LZ51 A0LZ81 A0LZD4 A0LZE0 A0LZF7 A0LZH1 A0LZH4 A0LZH6 A0LZH8 A0LZI4 A0LZM9
A0M0A2 A0M0C9 A0M0D0 A0M0D7 A0M0D8 A0M0D9 A0M0E2 A0M0J2 A0M1H4 A0M1V1 A0M1V5
A0M1X3 A0M283 A0M284 A0M287 A0M298 A0M2C1 A0M2D8 A0M2G0 A0M2G8 A0M2N6 A0M345
A0M353 A0M360 A0M367 A0M376 A0M380 A0M3I8 A0M3J2 A0M3J8 A0M3K2 A0M3K8 A0M3L3
A0M3L4 A0M3L5 A0M3M1 A0M3N2 A0M3P0 A0M3P3 A0M3P4 A0M3T2 A0M3Y8 A0M3Y9 A0M3Z0
A0M3Z1 A0M3Z2 A0M3Z3 A0M3Z6 A0M4S0 A0M4T1 A0M4T5 A0M4U0 A0M4U5 A0M4U6 A0M4W4
A0M4X2 A0M4Y1 A0M4Z1 A0M4Z2 A0M4Z3 A0M500 A0M526 A0M529 A0M534 A0M535 A0M537
A0M554 A0M572 A0M573 A0M574 A0M575 A0M576 A0M577 A0M579 A0M580 A0M581 A0M583
A0M584 A0M585 A0M586 A0M587 A0M588 A0M590 A0M591 A0M592 A0M593 A0M594 A0M595
A0M597 A0M598 A0M599 A0M5A0 A0M5A1 A0M5A2 A0M5C6 A0M5D4 A0M5H5 A0M5H7 A0M5H8
A0M5I5 A0M5L6 A0M5N3 A0M5P1 A0M5T0 A0M5T1 A0M5U1 A0M5U7 A0M5W6 A0M5Z7 A0M5Z9
A0M665 A0M672 A0M684 A0M6C1 A0M6C4 A0M6J7 A0M6J8 A0M6K0 A0M6M4 A0M6N1 A0M6N7
A0M6W9 A0M6X6 A0M745 A0M764 A0M786 A0M787 A0M7A5 A0M7A9 A0M7D0 A0M7D3 A0M7D9
A0M8R8 A0M8S4 A0M8S9 A0M8T3 A0M8T5 A0M8U6 A0M8V0 A0MD36 A0MDQ1 A0MEB5 A0MFS5
A0MH68 A0MQH0 A0MS83 A0MTA1 A0MWD1 A0MZA0 A0MZE1 A0NEF7 A0NGY0 A0NLY7 A0NXQ7
A0NXQ8 A0NXQ9 A0P8Z4 A0P8Z5 A0P9L2 A0PA85 A0PDV5 A0PFK7 A0PGB2 A0PJ24 A0PJ26
A0PJE2 A0PJS5 A0PJW8 A0PJX0 A0PJX2 A0PJY2 A0PJZ0 A0PKB2 A0PKB4 A0PKB5 A0PKF8
A0PKG2 A0PKH7 A0PKH8 A0PKQ3 A0PKS1 A0PKZ7 A0PL16 A0PL52 A0PL81 A0PLB8 A0PLC0
A0PLC7 A0PLD8 A0PLI0 A0PLM7 A0PLQ1 A0PLS0 A0PLU7 A0PLU9 A0PLV5 A0PLW6 A0PLZ1
A0PM01 A0PM07 A0PM08 A0PM24 A0PM25 A0PM27 A0PM39 A0PM40 A0PM41 A0PM42 A0PM48
A0PM49 A0PM50 A0PM51 A0PM62 A0PM63 A0PM64 A0PM65 A0PM66 A0PM67 A0PM68 A0PM69
A0PM70 A0PM71 A0PM72 A0PM75 A0PM76 A0PM78 A0PM79 A0PM80 A0PM81 A0PM82 A0PM83
A0PM84 A0PM87 A0PM88 A0PM89 A0PM98 A0PMB2 A0PMB3 A0PMB4 A0PMB5 A0PMB6 A0PMB7
A0PMB8 A0PMD1 A0PMD2 A0PMD3 A0PME2 A0PME7 A0PME8 A0PME9 A0PMI0 A0PMQ6 A0PMV4
A0PMW8 A0PMX1 A0PMY7 A0PN13 A0PN72 A0PNG6 A0PNH7 A0PNH8 A0PNJ1 A0PNK2 A0PNK6
A0PNP3 A0PNQ1 A0PNR5 A0PNU6 A0PP02 A0PP03 A0PP05 A0PP15 A0PP16 A0PP18 A0PP20
A0PP21 A0PP28 A0PP30 A0PP48 A0PP59 A0PP60 A0PP75 A0PP77 A0PP78 A0PP79 A0PP80

A0PPA8 A0PPD1 A0PPD2 A0PPD3 A0PPE4 A0PPE8 A0PPG6 A0PPG7 A0PPH0 A0PPH1 A0PPH6
A0PPI7 A0PPI8 A0PPJ3 A0PPJ8 A0PPK9 A0PPL6 A0PPL7 A0PPM2 A0PPM4 A0PPN3 A0PPN4
A0PPN8 A0PPP5 A0PPW5 A0PPW9 A0PPX3 A0PPX4 A0PPY3 A0PPY6 A0PPY8 A0PPZ6 A0PPZ7
A0PQ00 A0PQ01 A0PQ03 A0PQ07 A0PQ11 A0PQ17 A0PQ49 A0PQ63 A0PQ65 A0PQ68 A0PQ70
A0PQ75 A0PQ79 A0PQ80 A0PQ87 A0PQ89 A0PQ95 A0PQB8 A0PQC1 A0PQC4 A0PQC5 A0PQD9
A0PQE4 A0PQE7 A0PQF8 A0PQG7 A0PQH6 A0PQT1 A0PQT2 A0PQT3 A0PQT9 A0PQV0 A0PQV5
A0PQZ1 A0PQZ6 A0PR18 A0PR19 A0PRD3 A0PRF5 A0PRG8 A0PRJ3 A0PRT4 A0PRV0 A0PRX9
A0PRY2 A0PS01 A0PSA3 A0PSA4 A0PSD4 A0PSG0 A0PSG2 A0PSG3 A0PSG4 A0PSI5 A0PST9
A0PSX2 A0PSY4 A0PSY6 A0PSZ4 A0PT34 A0PT40 A0PT52 A0PT73 A0PT74 A0PT80 A0PT81
A0PT87 A0PT88 A0PT89 A0PTC0 A0PTG1 A0PTH3 A0PTI1 A0PTI6 A0PTJ6 A0PTJ7 A0PTM6
A0PTP2 A0PTP8 A0PTP9 A0PTS1 A0PTT2 A0PTT6 A0PTU2 A0PTY0 A0PTY6 A0PU04 A0PU14
A0PU15 A0PU16 A0PU17 A0PU19 A0PU27 A0PU31 A0PU35 A0PU57 A0PU74 A0PUG0 A0PUH6
A0PUI6 A0PUJ0 A0PUL1 A0PUL2 A0PUL4 A0PUL8 A0PUT6 A0PUX2 A0PUX3 A0PV23 A0PV26
A0PV27 A0PV29 A0PV48 A0PV49 A0PV50 A0PV57 A0PVE6 A0PVF2 A0PVM7 A0PVN0 A0PVN7
A0PVP6 A0PVU7 A0PVV1 A0PVW4 A0PVW7 A0PVY3 A0PVY7 A0PVZ1 A0PVZ3 A0PW28 A0PW29
A0PW66 A0PW69 A0PW79 A0PWB5 A0PWC5 A0PWD1 A0PWG2 A0PWG3 A0PWG4 A0PWH2 A0PWI9
A0PWL9 A0PWM0 A0PWM1 A0PX13 A0PX17 A0PX66 A0PX70 A0PX71 A0PX72 A0PX73 A0PX77
A0PX78 A0PX79 A0PX87 A0PX89 A0PX93 A0PXA4 A0PXA5 A0PXA8 A0PXD7 A0PXE8 A0PXK5
A0PXK7 A0PXL3 A0PXN0 A0PXN1 A0PXN3 A0PXN4 A0PXN6 A0PXP2 A0PXP3 A0PXP5 A0PXP6
A0PXP7 A0PXP9 A0PXQ3 A0PXQ4 A0PXQ5 A0PXR8 A0PXS4 A0PXS5 A0PXT1 A0PXT4 A0PXT5
A0PXT7 A0PXT8 A0PXT9 A0PXU1 A0PXU2 A0PXU3 A0PXU5 A0PXU6 A0PXU7 A0PXU8 A0PXU9
A0PXV0 A0PXV1 A0PXV2 A0PXV3 A0PXV4 A0PXV5 A0PXV6 A0PXV7 A0PXV8 A0PXV9 A0PXW0
A0PXW1 A0PXW2 A0PXW3 A0PXW4 A0PXW5 A0PXW7 A0PXX0 A0PXX1 A0PXX2 A0PXX3 A0PXX4
A0PXX5 A0PXX6 A0PXY1 A0PXY2 A0PXZ6 A0PY42 A0PY64 A0PY66 A0PY67 A0PY73 A0PY95
A0PYA5 A0PYB2 A0PYB3 A0PYB4 A0PYB5 A0PYB7 A0PYC8 A0PYE7 A0PYH3 A0PYH7 A0PYK3
A0PYP1 A0PYP2 A0PYP3 A0PYP8 A0PYU7 A0PYU9 A0PYW1 A0PYX0 A0PYX5 A0PYX6 A0PYZ1
A0PZ93 A0PZD5 A0PZF8 A0PZG3 A0PZG4 A0PZG9 A0PZH1 A0PZH3 A0PZK9 A0PZL5 A0PZM2
A0PZM3 A0PZM7 A0PZN1 A0PZN3 A0PZN4 A0PZN8 A0PZP4 A0PZS6 A0PZU0 A0PZV4 A0PZW4
A0PZW6 A0PZW9 A0PZX0 A0PZX4 A0PZY2 A0PZZ5 A0PZZ6 A0Q055 A0Q067 A0Q074 A0Q087
A0Q099 A0Q0A0 A0Q0A1 A0Q0A2 A0Q0A4 A0Q0A7 A0Q0E8 A0Q0E9 A0Q0K2 A0Q0M5 A0Q0M6
A0Q0M7 A0Q0M8 A0Q0M9 A0Q0P4 A0Q0P6 A0Q0Q1 A0Q0Q2 A0Q0Q4 A0Q0Q6 A0Q0Q7 A0Q0R0
A0Q0R4 A0Q0R7 A0Q0R8 A0Q0R9 A0Q0S0 A0Q0X6 A0Q0X7 A0Q0X9 A0Q0Y0 A0Q0Y1 A0Q0Y3
A0Q0Z3 A0Q0Z5 A0Q0Z7 A0Q101 A0Q106 A0Q109 A0Q112 A0Q115 A0Q119 A0Q121 A0Q125
A0Q128 A0Q140 A0Q145 A0Q149 A0Q150 A0Q151 A0Q152 A0Q1A0 A0Q1B0 A0Q1D1 A0Q1F0
A0Q1F1 A0Q1F7 A0Q1J0 A0Q1J2 A0Q1J4 A0Q1N5 A0Q1P9 A0Q1Q3 A0Q1Q8 A0Q1R2 A0Q1R3
A0Q1R4 A0Q1S2 A0Q1T2 A0Q1T4 A0Q1T6 A0Q1T8 A0Q1U7 A0Q1U9 A0Q1Z1 A0Q1Z2 A0Q217
A0Q222 A0Q299 A0Q2A0 A0Q2A8 A0Q2B1 A0Q2B3 A0Q2E1 A0Q2I3 A0Q2J7 A0Q2K7 A0Q2L0
A0Q2L1 A0Q2L2 A0Q2L7 A0Q2N3 A0Q2N5 A0Q2P2 A0Q2P9 A0Q2Q0 A0Q2Q3 A0Q2Q9 A0Q2R0
A0Q2S8 A0Q2T1 A0Q2T2 A0Q2U4 A0Q2U7 A0Q2W0 A0Q2Y4 A0Q306 A0Q312 A0Q316 A0Q323
A0Q330 A0Q345 A0Q355 A0Q380 A0Q388 A0Q3B0 A0Q3B8 A0Q3H1 A0Q3H6 A0Q3H7 A0Q3I1
A0Q3I7 A0Q3K0 A0Q3L2 A0Q3M5 A0Q3Q9 A0Q3R3 A0Q3R4 A0Q3T2 A0Q3U3 A0Q3U6 A0Q3U7
A0Q3W9 A0Q402 A0Q406 A0Q414 A0Q415 A0Q422 A0Q425 A0Q444 A0Q452 A0Q454 A0Q467
A0Q468 A0Q481 A0Q492 A0Q4A6 A0Q4C3 A0Q4G6 A0Q4H1 A0Q4H2 A0Q4H3 A0Q4H4 A0Q4H7
A0Q4H9 A0Q4I0 A0Q4I1 A0Q4I2 A0Q4I3 A0Q4I4 A0Q4I5 A0Q4I6 A0Q4I7 A0Q4I8 A0Q4I9

A0Q4J0 A0Q4J1 A0Q4J2 A0Q4J3 A0Q4J4 A0Q4J5 A0Q4J6 A0Q4J7 A0Q4J8 A0Q4J9 A0Q4K0
A0Q4K1 A0Q4K2 A0Q4K4 A0Q4K5 A0Q4K6 A0Q4K7 A0Q4K8 A0Q4K9 A0Q4L0 A0Q4L3 A0Q4L4
A0Q4M1 A0Q4Q1 A0Q4S0 A0Q4S3 A0Q4S4 A0Q4T9 A0Q4U9 A0Q4X5 A0Q4Z0 A0Q505 A0Q512
A0Q526 A0Q529 A0Q534 A0Q549 A0Q556 A0Q585 A0Q587 A0Q588 A0Q593 A0Q595 A0Q596
A0Q599 A0Q5C2 A0Q5C7 A0Q5D7 A0Q5E0 A0Q5E2 A0Q5H4 A0Q5I1 A0Q5I2 A0Q5I5 A0Q5I8
A0Q5I9 A0Q5J0 A0Q5J3 A0Q5J5 A0Q5K3 A0Q5K6 A0Q5M5 A0Q5M6 A0Q5P0 A0Q5P2 A0Q5Q2
A0Q5Q3 A0Q5Q7 A0Q5Q9 A0Q5S7 A0Q5W0 A0Q5X2 A0Q5Y0 A0Q5Y3 A0Q5Y4 A0Q5Y5 A0Q5Y6
A0Q604 A0Q614 A0Q621 A0Q626 A0Q628 A0Q635 A0Q638 A0Q656 A0Q669 A0Q689 A0Q690
A0Q693 A0Q696 A0Q6A6 A0Q6B4 A0Q6B9 A0Q6C0 A0Q6C2 A0Q6D2 A0Q6E3 A0Q6F0 A0Q6H0
A0Q6H2 A0Q6H3 A0Q6H4 A0Q6H6 A0Q6K3 A0Q6K9 A0Q6L9 A0Q6M6 A0Q6N0 A0Q6P7 A0Q6P9
A0Q6Q0 A0Q6Q2 A0Q6R5 A0Q6R7 A0Q6S3 A0Q6S4 A0Q6S9 A0Q6T0 A0Q6T5 A0Q6T7 A0Q6U8
A0Q6W0 A0Q6X1 A0Q6Y1 A0Q6Z1 A0Q706 A0Q707 A0Q713 A0Q735 A0Q736 A0Q753 A0Q756
A0Q759 A0Q760 A0Q762 A0Q767 A0Q768 A0Q7A0 A0Q7A9 A0Q7B4 A0Q7B6 A0Q7C5 A0Q7F0
A0Q7F1 A0Q7F4 A0Q7F9 A0Q7G4 A0Q7I7 A0Q7J1 A0Q7J5 A0Q7L0 A0Q7L2 A0Q7L3 A0Q7L4
A0Q7M7 A0Q7N3 A0Q7Q2 A0Q7Q5 A0Q7Q7 A0Q7R6 A0Q7W5 A0Q7W6 A0Q7Y5 A0Q7Y8 A0Q804
A0Q809 A0Q810 A0Q814 A0Q838 A0Q839 A0Q841 A0Q842 A0Q844 A0Q845 A0Q858 A0Q859
A0Q860 A0Q861 A0Q862 A0Q864 A0Q866 A0Q867 A0Q868 A0Q869 A0Q870 A0Q871 A0Q874
A0Q880 A0Q892 A0Q893 A0Q8A2 A0Q8C4 A0Q8C5 A0Q8C6 A0Q8F1 A0Q8F2 A0Q8F3 A0Q8F5
A0Q8F6 A0Q8I1 A0Q8I2 A0Q8K2 A0Q8L5 A0Q8M6 A0Q8N2 A0Q8R1 A0Q8R8 A0Q8W4 A0Q8W7
A0Q8Y7 A0Q8Y8 A0Q981 A0Q994 A0Q999 A0Q9F3 A0Q9K2 A0Q9T5 A0QA85 A0QA93 A0QA94
A0QA95 A0QAA9 A0QAB4 A0QAB5 A0QAB6 A0QAC2 A0QAG8 A0QAG9 A0QAK7 A0QAT1 A0QAU0
A0QAU8 A0QAW3 A0QB57 A0QBE6 A0QBI3 A0QBI8 A0QBM5 A0QBN4 A0QBP0 A0QBQ8 A0QBT9
A0QBW0 A0QBW1 A0QBW3 A0QC23 A0QC35 A0QC69 A0QC96 A0QCA6 A0QCB7 A0QCH0 A0QCI6
A0QCW6 A0QCW7 A0QCZ8 A0QD02 A0QD04 A0QDA8 A0QDF1 A0QDF5 A0QDF8 A0QDG6 A0QDG8
A0QDG9 A0QDH0 A0QDH1 A0QDH4 A0QDI7 A0QDK7 A0QDL1 A0QDN6 A0QEA3 A0QEA7 A0QEC2
A0QEI3 A0QEJ0 A0QEV5 A0QEY6 A0QEY7 A0QEZ6 A0QEZ9 A0QF43 A0QF44 A0QF47 A0QF50
A0QF53 A0QF59 A0QF76 A0QFA4 A0QFA5 A0QFB2 A0QFB5 A0QFB6 A0QFB7 A0QFS9 A0QG83
A0QGA4 A0QGE9 A0QGF1 A0QGF2 A0QGI9 A0QGK3 A0QGM8 A0QGN2 A0QGN7 A0QH59 A0QH60
A0QH68 A0QHA7 A0QHA8 A0QHA9 A0QHB0 A0QHB2 A0QHB4 A0QHB6 A0QHC1 A0QHC2 A0QHG8
A0QHH0 A0QHH5 A0QHH6 A0QHH8 A0QHI0 A0QHI1 A0QHI6 A0QHJ1 A0QHJ8 A0QHJ9 A0QHT7
A0QHY0 A0QHY3 A0QHY4 A0QI01 A0QI03 A0QI08 A0QI09 A0QI16 A0QI26 A0QI28 A0QI32
A0QI37 A0QI38 A0QI54 A0QI55 A0QI58 A0QI60 A0QI61 A0QI75 A0QI91 A0QIA9 A0QIB7
A0QIB8 A0QIB9 A0QIC5 A0QIC6 A0QIC8 A0QID4 A0QIL2 A0QIL6 A0QIM8 A0QIQ0 A0QIQ1
A0QIQ8 A0QIQ9 A0QIR5 A0QIR6 A0QIT4 A0QIU1 A0QIW5 A0QIW6 A0QIX1 A0QIY1 A0QIY2
A0QIY5 A0QIY9 A0QJ14 A0QJ18 A0QJ19 A0QJ27 A0QJ36 A0QJ41 A0QJ43 A0QJ46 A0QJ48
A0QJ63 A0QJ66 A0QJ93 A0QJ99 A0QJA9 A0QJB2 A0QJB3 A0QJB6 A0QJB7 A0QJC2 A0QJC6
A0QJD1 A0QJD4 A0QJD5 A0QJE3 A0QJE5 A0QJI1 A0QJJ4 A0QJL0 A0QJS1 A0QKC7 A0QKC8
A0QKD9 A0QKJ4 A0QKL5 A0QKM1 A0QKN5 A0QKR2 A0QKR3 A0QKR4 A0QKR9 A0QKT1 A0QKT2
A0QKT3 A0QKU4 A0QKU5 A0QKU6 A0QKU7 A0QKU8 A0QKU9 A0QKV0 A0QKY2 A0QKY8 A0QKY9
A0QKZ0 A0QKZ3 A0QKZ4 A0QKZ6 A0QKZ7 A0QKZ8 A0QKZ9 A0QL00 A0QL01 A0QL02 A0QL10
A0QL11 A0QL12 A0QL13 A0QL14 A0QL15 A0QL16 A0QL17 A0QL18 A0QL19 A0QL30 A0QL35
A0QL36 A0QL37 A0QL38 A0QL48 A0QL49 A0QL54 A0QL66 A0QL67 A0QL73 A0QL81 A0QL99
A0QLA0 A0QLB4 A0QLB6 A0QLC9 A0QLG2 A0QLI0 A0QLI1 A0QLK3 A0QLK5 A0QLK9 A0QLL2
A0QLR0 A0QLS0 A0QLT3 A0QLT6 A0QLU8 A0QLV0 A0QLV3 A0QLW6 A0QLX9 A0QLZ6 A0QM15

A0QM44 A0QM57 A0QM60 A0QMB9 A0QMC8 A0QME9 A0QMH2 A0QMX8 A0QMX9 A0QN14 A0QND0
A0QND4 A0QND5 A0QND6 A0QND8 A0QND9 A0QNE0 A0QNF3 A0QNH4 A0QNH5 A0QNH6 A0QNI5
A0QNI7 A0QNJ3 A0QNM1 A0QNM2 A0QNX1 A0QP32 A0QP90 A0QPN1 A0QPN2 A0QQ37 A0QQ45
A0QQ53 A0QQ72 A0QQ98 A0QQH7 A0QQJ4 A0QQK1 A0QQK3 A0QQK7 A0QQL0 A0QQP8 A0QQQ2
A0QQV6 A0QQZ3 A0QQZ8 A0QR17 A0QR33 A0QR54 A0QRD3 A0QRF3 A0QRG0 A0QRG5 A0QRH1
A0QRM0 A0QRN3 A0QRN6 A0QRX8 A0QRX9 A0QRY0 A0QRY1 A0QRY5 A0QRY6 A0QS28 A0QS29
A0QS30 A0QS39 A0QS45 A0QS46 A0QS62 A0QS66 A0QS96 A0QS97 A0QS98 A0QSB6 A0QSB7
A0QSB8 A0QSB9 A0QSC0 A0QSD0 A0QSD1 A0QSD2 A0QSD3 A0QSD4 A0QSD5 A0QSD6 A0QSD7
A0QSD8 A0QSD9 A0QSE0 A0QSF9 A0QSG0 A0QSG1 A0QSG2 A0QSG3 A0QSG4 A0QSG6 A0QSG7
A0QSH3 A0QSH4 A0QSH5 A0QSH6 A0QSH8 A0QSK5 A0QSK6 A0QSL3 A0QSL4 A0QSL5 A0QSL6
A0QSL7 A0QSL8 A0QSL9 A0QSP8 A0QSP9 A0QSQ1 A0QSR6 A0QSS1 A0QSS3 A0QSS4 A0QST8
A0QSU4 A0QSY0 A0QSY5 A0QSZ0 A0QT53 A0QTE5 A0QTF8 A0QTG1 A0QTG2 A0QTG3 A0QTL6
A0QTP2 A0QTP3 A0QTU7 A0QTU8 A0QTU9 A0QTV0 A0QTV4 A0QU58 A0QU63 A0QU88 A0QUA1
A0QUB2 A0QUP1 A0QUV5 A0QUW3 A0QUW7 A0QUW9 A0QUX1 A0QUX7 A0QUX8 A0QUY7 A0QUZ2
A0QUZ4 A0QUZ7 A0QV01 A0QV09 A0QV10 A0QV16 A0QV20 A0QV21 A0QV24 A0QV37 A0QV39
A0QV42 A0QV44 A0QV47 A0QVA9 A0QVB4 A0QVB8 A0QVB9 A0QVD9 A0QVE0 A0QVE4 A0QVH7
A0QVI7 A0QVL2 A0QVM0 A0QVM3 A0QVM8 A0QVQ3 A0QVQ5 A0QVQ8 A0QVR3 A0QVR9 A0QVT0
A0QVT2 A0QVU1 A0QVX5 A0QVX6 A0QVX9 A0QVY0 A0QVY8 A0QW08 A0QW19 A0QW23 A0QW25
A0QWG2 A0QWG8 A0QWH0 A0QWH1 A0QWH4 A0QWH5 A0QWH6 A0QWJ5 A0QWN3 A0QWQ4 A0QWQ5
A0QWQ9 A0QWR4 A0QWR5 A0QWS1 A0QWT1 A0QWT2 A0QWT3 A0QWU2 A0QWV6 A0QWV7 A0QWV9
A0QWW2 A0QWW3 A0QWX4 A0QWX9 A0QX20 A0QX29 A0QX65 A0QX66 A0QX70 A0QX82 A0QX83
A0QX86 A0QX87 A0QX88 A0QX93 A0QX95 A0QXD8 A0QXE4 A0QXE5 A0QXJ6 A0QXX7 A0QY28
A0QYC2 A0QYE0 A0QYE2 A0QYE8 A0QYG2 A0QYG3 A0QYP1 A0QYQ7 A0QYS5 A0QYS6 A0QYS8
A0QYS9 A0QYT0 A0QYU6 A0QYU7 A0QYY6 A0QYZ8 A0QZ11 A0QZ42 A0QZ46 A0QZ47 A0QZ49
A0QZ54 A0QZA1 A0QZE3 A0QZU1 A0QZU2 A0QZX2 A0QZY0 A0QZZ6 A0R006 A0R008 A0R015
A0R018 A0R024 A0R025 A0R048 A0R066 A0R074 A0R075 A0R079 A0R080 A0R083 A0R0B0
A0R0N3 A0R0N4 A0R0S5 A0R0T0 A0R0T9 A0R0W2 A0R0W9 A0R102 A0R109 A0R112 A0R115
A0R148 A0R149 A0R150 A0R151 A0R152 A0R191 A0R196 A0R199 A0R1B5 A0R1C2 A0R1T3
A0R1T4 A0R1V9 A0R1W8 A0R1X0 A0R1Y1 A0R1Y4 A0R1Y7 A0R1Z0 A0R213 A0R215 A0R219
A0R220 A0R2B1 A0R2D4 A0R2D5 A0R2E1 A0R2E2 A0R2E6 A0R2G4 A0R2G5 A0R2I7 A0R2K7
A0R2N3 A0R2V7 A0R2V9 A0R2W9 A0R2X1 A0R2Y2 A0R316 A0R3C4 A0R3D3 A0R3D7 A0R3F9
A0R3H2 A0R3I9 A0R3M4 A0R3N9 A0R3Q8 A0R3R7 A0R3S7 A0R3T8 A0R420 A0R443 A0R480
A0R4C9 A0R4D0 A0R4D2 A0R4I0 A0R4M9 A0R4Q0 A0R4S6 A0R4S7 A0R4Z6 A0R518 A0R549
A0R550 A0R551 A0R557 A0R558 A0R561 A0R563 A0R564 A0R566 A0R567 A0R574 A0R579
A0R580 A0R5C5 A0R5D9 A0R5E1 A0R5K4 A0R5M9 A0R5N0 A0R5N1 A0R5R0 A0R5R7 A0R5T2
A0R5T3 A0R5X8 A0R5Z0 A0R609 A0R610 A0R635 A0R638 A0R643 A0R647 A0R652 A0R664
A0R666 A0R692 A0R6D0 A0R6D9 A0R6E0 A0R6E3 A0R6J8 A0R6Q0 A0R756 A0R757 A0R758
A0R7F6 A0R7F7 A0R7F9 A0R7G6 A0R7H5 A0R7H8 A0R7J5 A0R7K0 A0R7K1 A0R883 A0R890
A0R891 A0R898 A0R899 A0R8A3 A0R8B4 A0R8B7 A0R8D8 A0R8D9 A0R8E9 A0R8F7 A0R8F8
A0R8F9 A0R8G1 A0R8G8 A0R8H0 A0R8H2 A0R8H3 A0R8H5 A0R8H6 A0R8H8 A0R8H9 A0R8I0
A0R8I1 A0R8I2 A0R8I3 A0R8I4 A0R8I5 A0R8I6 A0R8I7 A0R8I9 A0R8J0 A0R8J1 A0R8J2
A0R8J3 A0R8J4 A0R8J7 A0R8J8 A0R8J9 A0R8K1 A0R8K3 A0R8K4 A0R8K5 A0R8K6 A0R8K7
A0R8L2 A0R8L3 A0R8M4 A0R8S0 A0R8U6 A0R8V9 A0R8W1 A0R8W4 A0R8W7 A0R8Z5 A0R8Z6
A0R8Z8 A0R900 A0R906 A0R909 A0R920 A0R921 A0R922 A0R946 A0R951 A0R969 A0R979

A0R980 A0R981 A0R9A0 A0R9B0 A0R9F4 A0R9G2 A0R9G6 A0R9J0 A0R9V2 A0RA08 A0RAB4
A0RAB9 A0RAL4 A0RAN0 A0RAP3 A0RAQ6 A0RAS3 A0RAX6 A0RAX7 A0RB09 A0RB27 A0RB32
A0RB53 A0RB61 A0RB62 A0RB63 A0RB64 A0RB65 A0RBC7 A0RBC8 A0RBC9 A0RBE8 A0RBE9
A0RBG1 A0RBJ0 A0RBK5 A0RBL2 A0RBL4 A0RBL6 A0RBL9 A0RBM0 A0RBM1 A0RBM2 A0RBM3
A0RBM4 A0RBN2 A0RBN3 A0RBS0 A0RBV2 A0RBV4 A0RBV9 A0RBW5 A0RBX0 A0RBX1 A0RBX4
A0RBY6 A0RBY7 A0RBZ2 A0RBZ3 A0RBZ4 A0RC02 A0RC08 A0RC09 A0RC17 A0RCF8 A0RCH7
A0RCL3 A0RCM7 A0RCP2 A0RCV1 A0RCX4 A0RCZ8 A0RD27 A0RD31 A0RD64 A0RD69 A0RD70
A0RD97 A0RDG3 A0RDL9 A0RDW1 A0RE53 A0RE96 A0REB4 A0REB5 A0REB6 A0REB7 A0REC6
A0REX0 A0REX1 A0REX2 A0RFD1 A0RFD4 A0RFE8 A0RFP3 A0RGR8 A0RH02 A0RH37 A0RH38
A0RH39 A0RH40 A0RH87 A0RH88 A0RHB6 A0RHE0 A0RHE1 A0RHE4 A0RHF1 A0RHF2 A0RHH8
A0RHH9 A0RHI1 A0RHI2 A0RHI4 A0RHI8 A0RHJ2 A0RHJ7 A0RHJ9 A0RHK0 A0RHK1 A0RHK3
A0RHK5 A0RHK9 A0RHL0 A0RHL3 A0RHL4 A0RHL5 A0RHL7 A0RHN1 A0RHN7 A0RHN9 A0RHP3
A0RHP5 A0RHQ6 A0RHQ7 A0RHQ8 A0RHR0 A0RHR1 A0RHR3 A0RHS0 A0RHT3 A0RHT9 A0RHU0
A0RHU2 A0RHU5 A0RHU9 A0RHV8 A0RHW7 A0RHX6 A0RHX9 A0RHY8 A0RHZ2 A0RHZ3 A0RI12
A0RI38 A0RI39 A0RI42 A0RI43 A0RI44 A0RI45 A0RI59 A0RI62 A0RI78 A0RI91 A0RIB3
A0RIB4 A0RIB6 A0RIB9 A0RIC0 A0RID0 A0RID9 A0RIH1 A0RIH2 A0RIH3 A0RIH4 A0RII7
A0RIK9 A0RIL0 A0RIL1 A0RIQ2 A0RIQ8 A0RIR1 A0RIR7 A0RIS3 A0RIT1 A0RIT3 A0RIT5
A0RIU9 A0RIV1 A0RIY7 A0RIY8 A0RIZ3 A0RIZ8 A0RIZ9 A0RJ00 A0RJ05 A0RJ11 A0RJ14
A0RJ16 A0RJ24 A0RJ26 A0RJ27 A0RJ35 A0RJ47 A0RJ49 A0RJ51 A0RJ56 A0RJ78 A0RJ81
A0RJ83 A0RJ86 A0RJ87 A0RJ90 A0RJC3 A0RJE3 A0RJF2 A0RJH2 A0RJH3 A0RJJ0 A0RJJ6
A0RJK8 A0RJM3 A0RJM4 A0RJN9 A0RJP7 A0RJS6 A0RJT1 A0RJT7 A0RJV9 A0RJX2 A0RJZ5
A0RJZ7 A0RJZ8 A0RK73 A0RK76 A0RK86 A0RK94 A0RKB5 A0RKB9 A0RKC7 A0RKF7 A0RKM3
A0RKR7 A0RKS4 A0RKS6 A0RKU0 A0RKU2 A0RKU8 A0RL17 A0RLA2 A0RLA3 A0RLB6 A0RLC3
A0RLC4 A0RLG5 A0RLH0 A0RLI1 A0RLI8 A0RLN5 A0RLN8 A0RLP6 A0RLP8 A0RLQ1 A0RLQ9
A0RLR0 A0RLR1 A0RLR2 A0RLX8 A0RM10 A0RM12 A0RM13 A0RM14 A0RM15 A0RM16 A0RM17
A0RM18 A0RM21 A0RM22 A0RM24 A0RM25 A0RM26 A0RM27 A0RM28 A0RM32 A0RM33 A0RM34
A0RM35 A0RM36 A0RM37 A0RM41 A0RM43 A0RM81 A0RM82 A0RM89 A0RM90 A0RM91 A0RM99
A0RME8 A0RMG7 A0RMH3 A0RMI1 A0RMI6 A0RMI8 A0RMJ2 A0RMK4 A0RMK6 A0RMM1 A0RMM6
A0RMN5 A0RMN9 A0RMP5 A0RMQ6 A0RMQ7 A0RMR3 A0RMR7 A0RMU4 A0RMV6 A0RMY9 A0RN28
A0RN29 A0RN42 A0RN82 A0RN84 A0RN89 A0RNF8 A0RNG2 A0RNH4 A0RNJ6 A0RNK3 A0RNK4
A0RNM0 A0RNN3 A0RNS2 A0RNS3 A0RNT0 A0RNT1 A0RNT3 A0RNU2 A0RNU3 A0RNU4 A0RNV4
A0RNX3 A0RNY9 A0RP16 A0RP26 A0RP30 A0RP36 A0RP49 A0RP54 A0RP84 A0RP96 A0RPB3
A0RPC8 A0RPD1 A0RPD2 A0RPD3 A0RPE8 A0RPI2 A0RPK6 A0RPL1 A0RPL5 A0RPL8 A0RPN3
A0RPN4 A0RPR0 A0RPR5 A0RPR6 A0RPR7 A0RPR9 A0RPS8 A0RPU6 A0RPV3 A0RPV9 A0RPW7
A0RPW8 A0RPX5 A0RPY4 A0RPY7 A0RPZ6 A0RQ02 A0RQ06 A0RQ08 A0RQ16 A0RQ21 A0RQ26
A0RQ44 A0RQ49 A0RQ51 A0RQ64 A0RQ71 A0RQ72 A0RQ83 A0RQ89 A0RQ93 A0RQC0 A0RQC4
A0RQC5 A0RQD0 A0RQD7 A0RQI0 A0RQI1 A0RQI2 A0RQI4 A0RQI5 A0RQI6 A0RQI7 A0RQI8
A0RQI9 A0RQJ2 A0RQJ3 A0RQJ5 A0RQJ6 A0RQJ7 A0RQK2 A0RQK5 A0RQK7 A0RQL1 A0RQL2
A0RQM7 A0RQM8 A0RQM9 A0RQN5 A0RQR9 A0RQT4 A0RQT6 A0RQU0 A0RQU7 A0RQU8 A0RQV2
A0RQV9 A0RQX6 A0RQY3 A0RQZ2 A0RQZ4 A0RQZ6 A0RR35 A0RR37 A0RR38 A0RR39 A0RR46
A0RR59 A0RR62 A0RR66 A0RR72 A0RR73 A0RR75 A0RR76 A0RRF8 A0RRG4 A0RRI0 A0RRI8
A0RRK2 A0RRK3 A0RRK4 A0RRM6 A0RRM7 A0RRN7 A0RRR2 A0RRR3 A0RRT6 A0RRT8 A0RRU0
A0RRU6 A0RTP4 A0RTS8 A0RTT1 A0RTU3 A0RTV6 A0RU27 A0RU64 A0RU67 A0RU76 A0RU82
A0RU86 A0RU95 A0RUA5 A0RUB4 A0RUE6 A0RUE7 A0RUE9 A0RUF1 A0RUG0 A0RUI8 A0RUM4

A0RUM5 A0RUP0 A0RUP3 A0RUR1 A0RUR2 A0RUR7 A0RUR8 A0RUZ0 A0RV25 A0RV28 A0RV30
A0RVQ7 A0RVQ8 A0RVX7 A0RVX8 A0RVX9 A0RVY3 A0RVY4 A0RVY6 A0RVY8 A0RW02 A0RW30
A0RW81 A0RW96 A0RWD1 A0RWD6 A0RWI9 A0RWW0 A0RWW1 A0RWW9 A0RX07 A0RX10 A0RX34
A0RXB3 A0RXE1 A0RXF8 A0RXH7 A0RXJ5 A0RXJ6 A0RXQ5 A0RXT6 A0RXU1 A0RXV1 A0RY01
A0RY02 A0RY06 A0RY14 A0RY20 A0RY85 A0RYC3 A0RYD1 A0RYF4 A0RYF8 A0RYM0 A0RYN2
A0RYP2 A0RYQ9 A0RYU6 A0RYW7 A0RYW9 A0RZ01 A0RZ09 A0RZ37 A0RZ60 A0RZ75 A0RZ77
A0RZ79 A0RZA3 A0SJA5 A0SPJ3 A0SPJ4 A0SPJ6 A0SPJ8 A0SPJ9 A0SQM0 A0ST41 A0ST43
A0ST44 A0ST45 A0ST46 A0SVK0 A0SWL0 A0SXL6 A0T0B3 A0T0C0 A0T0C1 A0T0C2 A0T0C7
A0T0D4 A0T0D5 A0T0D6 A0T0D7 A0T0D8 A0T0D9 A0T0E0 A0T0F5 A0T0F6 A0T0G1 A0T0G2
A0T0G3 A0T0G4 A0T0G6 A0T0H2 A0T0H4 A0T0H6 A0T0H7 A0T0H8 A0T0H9 A0T0I0 A0T0I1
A0T0I2 A0T0I4 A0T0I5 A0T0I6 A0T0I7 A0T0I8 A0T0I9 A0T0J0 A0T0J1 A0T0J2 A0T0J3
A0T0J4 A0T0J5 A0T0J7 A0T0J8 A0T0J9 A0T0K0 A0T0K2 A0T0K3 A0T0K4 A0T0K5 A0T0K6
A0T0K7 A0T0L0 A0T0L1 A0T0N1 A0T0N6 A0T0Q1 A0T0Q2 A0T0Q5 A0T0Q6 A0T0Q7 A0T0Q8
A0T0Q9 A0T0R0 A0T0R1 A0T0R2 A0T0R3 A0T0R4 A0T0S2 A0T0V4 A0T0V5 A0T0V6 A0T0V7
A0T0V9 A0T0W6 A0T0W7 A0T0X0 A0T0X1 A0T0X2 A0T0X3 A0T0X4 A0T0X5 A0T0X6 A0T0X8
A0T0X9 A0T0Y0 A0T0Y2 A0T0Y3 A0T0Y4 A0T0Y5 A0T0Y6 A0T0Y7 A0T0Y8 A0T0Y9 A0T0Z1
A0T0Z2 A0T0Z3 A0T0Z4 A0T0Z6 A0T0Z7 A0T0Z8 A0T0Z9 A0T100 A0T101 A0T103 A0T104
A0T2M3 A0ZWU1 A0ZZ16 A0ZZ17 A0ZZ24 A0ZZ25 A0ZZ26 A0ZZ27 A0ZZ33 A0ZZ37 A0ZZ43
A0ZZ56 A0ZZ57 A0ZZ58 A0ZZ59 A0ZZ60 A0ZZ67 A0ZZ68 A0ZZ69 A0ZZ70 A0ZZ71 A0ZZ72
A0ZZ73 A0ZZ74 A0ZZ75 A0ZZ76 A0ZZ77 A0ZZ78 A0ZZ80 A0ZZ83 A0ZZ92 A0ZZA1 A0ZZG5
A0ZZH5 A0ZZH6 A0ZZM2 A0ZZN6 A0ZZR0 A0ZZS4 A0ZZS7 A0ZZS8 A0ZZT4 A0ZZV1 A0ZZV4
A0ZZV5 A0ZZV7 A0ZZX2 A0ZZY0 A0ZZY8 A0ZZY9 A0ZZZ0 A0ZZZ1 A0ZZZ5 A0ZZZ6 A1A007
A1A008 A1A032 A1A033 A1A040 A1A041 A1A048 A1A057 A1A058 A1A068 A1A069 A1A070
A1A072 A1A073 A1A074 A1A075 A1A076 A1A078 A1A079 A1A080 A1A081 A1A082 A1A083
A1A084 A1A085 A1A086 A1A087 A1A088 A1A091 A1A092 A1A093 A1A094 A1A095 A1A096
A1A0A2 A1A0A3 A1A0A9 A1A0B7 A1A0E0 A1A0H0 A1A0I8 A1A0I9 A1A0K4 A1A0L0 A1A0L5
A1A0M5 A1A0R8 A1A0S2 A1A0S3 A1A0S8 A1A0S9 A1A0T0 A1A0T1 A1A0T4 A1A0T7 A1A0T8
A1A0T9 A1A0U2 A1A0U4 A1A0U7 A1A0Z4 A1A141 A1A171 A1A190 A1A195 A1A198 A1A199
A1A1A2 A1A1B4 A1A1B9 A1A1E3 A1A1F4 A1A1F7 A1A1G3 A1A1H9 A1A1I0 A1A1I1 A1A1I2
A1A1I4 A1A1I5 A1A1I6 A1A1J2 A1A1J3 A1A1K1 A1A1K2 A1A1K3 A1A1K5 A1A1M6 A1A1N2
A1A1N3 A1A1N4 A1A1N5 A1A1Q5 A1A1R2 A1A1U9 A1A1V0 A1A1W2 A1A1W7 A1A1W8 A1A1X1
A1A1X6 A1A200 A1A216 A1A236 A1A279 A1A280 A1A2C2 A1A2C3 A1A2E8 A1A2F1 A1A2F7
A1A2H2 A1A2H5 A1A2H6 A1A2I3 A1A2J2 A1A2K3 A1A2L6 A1A2Z1 A1A2Z2 A1A316 A1A317
A1A321 A1A399 A1A3A4 A1A3A5 A1A3A8 A1A3B6 A1A3C3 A1A3D2 A1A3F7 A1A3H8 A1A3H9
A1A3N3 A1A3P4 A1A3P5 A1A3S8 A1A3T0 A1A3W3 A1A3W4 A1A3X4 A1A3X8 A1A4G5 A1A4J7
A1A4K3 A1A4K8 A1A4L5 A1A4L8 A1A4M4 A1A4P5 A1A4Q2 A1A4Q3 A1A4Q4 A1A4Q8 A1A4Q9
A1A4R1 A1A4R8 A1A4R9 A1A4V9 A1A519 A1A546 A1A580 A1A5B6 A1A5D9 A1A5F2 A1A5H6
A1A5I1 A1A5K2 A1A5P5 A1A5Q0 A1A5Q4 A1A5Q5 A1A5Q7 A1A5R1 A1A5R7 A1A5R8 A1A5R9
A1A5S1 A1A5V9 A1A5X2 A1A5Y5 A1A5Z3 A1A600 A1A619 A1A653 A1A6H3 A1A6K6 A1A708
A1A756 A1A759 A1A766 A1A774 A1A777 A1A778 A1A780 A1A785 A1A787 A1A788 A1A789
A1A792 A1A798 A1A799 A1A7A0 A1A7A6 A1A7A9 A1A7B0 A1A7B8 A1A7B9 A1A7C0 A1A7C2
A1A7C7 A1A7D6 A1A7E5 A1A7E7 A1A7G7 A1A7H7 A1A7H9 A1A7I0 A1A7J2 A1A7K0 A1A7K2
A1A7K5 A1A7K6 A1A7K9 A1A7L0 A1A7L1 A1A7L3 A1A7L5 A1A7M7 A1A7N8 A1A7Q3 A1A7U2
A1A7V0 A1A7V1 A1A7V3 A1A7V4 A1A7X7 A1A831 A1A834 A1A835 A1A836 A1A854 A1A859

A1A860 A1A863 A1A875 A1A876 A1A883 A1A890 A1A892 A1A8A5 A1A8A6 A1A8A7 A1A8B3
A1A8B5 A1A8D5 A1A8D7 A1A8D8 A1A8D9 A1A8E1 A1A8E2 A1A8G9 A1A8H0 A1A8H1 A1A8H2
A1A8H7 A1A8J2 A1A8J4 A1A8L2 A1A8P2 A1A8P3 A1A8Q3 A1A8Q5 A1A8Q6 A1A8R1 A1A8R4
A1A8R7 A1A8R9 A1A8S7 A1A8S9 A1A8T7 A1A8U7 A1A8W9 A1A8X0 A1A8Z3 A1A8Z8 A1A900
A1A909 A1A917 A1A918 A1A921 A1A925 A1A944 A1A947 A1A974 A1A991 A1A998 A1A9A5
A1A9B1 A1A9B8 A1A9C8 A1A9D6 A1A9I4 A1A9I5 A1A9I7 A1A9J9 A1A9K1 A1A9L2 A1A9L8
A1A9M5 A1A9M7 A1A9N5 A1A9N6 A1A9N7 A1A9Q6 A1A9Q7 A1A9Q9 A1A9R2 A1A9R3 A1A9R4
A1A9R5 A1A9R6 A1A9R7 A1A9T2 A1A9V1 A1A9V4 A1A9V7 A1A9W2 A1A9Z2 A1AA00 A1AA01
A1AA13 A1AA21 A1AA27 A1AA28 A1AA51 A1AA91 A1AA96 A1AA98 A1AAA0 A1AAA3 A1AAD5
A1AAD7 A1AAD8 A1AAF7 A1AAH4 A1AAN0 A1AAN1 A1AAQ4 A1AAV5 A1AB46 A1AB75 A1AB99
A1ABA1 A1ABF1 A1ABG8 A1ABI2 A1ABI3 A1ABI5 A1ABJ5 A1ABJ9 A1ABL7 A1ABL8 A1ABM9
A1ABN0 A1ABP0 A1ABP2 A1ABQ0 A1ABQ2 A1ABR2 A1ABR4 A1ABS1 A1ABS3 A1ABS5 A1ABS6
A1ABS8 A1ABS9 A1ABW3 A1ABY1 A1ABZ7 A1AC00 A1AC10 A1AC21 A1AC22 A1AC24 A1AC27
A1AC31 A1AC32 A1AC35 A1AC37 A1AC51 A1AC53 A1AC65 A1AC86 A1ACA3 A1ACB2 A1ACG4
A1ACI8 A1ACN1 A1ACN3 A1ACN6 A1ACN7 A1ACS1 A1ACW1 A1ACW2 A1ACX0 A1ACX8 A1AD04
A1AD13 A1AD14 A1AD18 A1AD20 A1AD29 A1AD43 A1AD63 A1AD97 A1AD99 A1ADA3 A1ADB3
A1ADB5 A1ADB6 A1ADC0 A1ADE0 A1ADG7 A1ADG9 A1ADH3 A1ADH8 A1ADJ6 A1ADP3 A1ADQ1
A1ADR4 A1ADS2 A1ADS8 A1ADT5 A1ADU3 A1ADU8 A1ADV0 A1ADV4 A1ADX2 A1ADX7 A1ADY9
A1ADZ1 A1AE43 A1AE45 A1AE52 A1AE53 A1AE55 A1AE56 A1AE61 A1AE64 A1AE65 A1AE68
A1AE69 A1AE82 A1AE95 A1AE97 A1AEA5 A1AEA9 A1AEB0 A1AED6 A1AED9 A1AEE5 A1AEE9
A1AEK4 A1AEL9 A1AEN7 A1AEN8 A1AEQ0 A1AEQ1 A1AES5 A1AET7 A1AET8 A1AET9 A1AEU0
A1AEU1 A1AEU4 A1AEU5 A1AEU6 A1AEU8 A1AEU9 A1AEV0 A1AEW8 A1AEX2 A1AEY1 A1AEY5
A1AEY9 A1AEZ1 A1AEZ3 A1AF39 A1AF43 A1AF64 A1AF79 A1AF88 A1AF90 A1AF92 A1AF93
A1AFA2 A1AFB0 A1AFB1 A1AFC5 A1AFD0 A1AFD4 A1AFD5 A1AFD9 A1AFE6 A1AFE8 A1AFV9
A1AFX0 A1AFX4 A1AFX8 A1AFY6 A1AFY9 A1AFZ3 A1AG02 A1AG09 A1AG22 A1AG35 A1AG39
A1AG46 A1AG53 A1AG54 A1AG59 A1AG60 A1AG69 A1AG70 A1AG71 A1AG72 A1AG73 A1AG75
A1AG76 A1AG79 A1AG82 A1AG86 A1AG91 A1AGA6 A1AGB6 A1AGB7 A1AGB9 A1AGC0 A1AGC3
A1AGC4 A1AGC9 A1AGD0 A1AGF9 A1AGG1 A1AGH3 A1AGH4 A1AGH6 A1AGH8 A1AGH9 A1AGI0
A1AGI5 A1AGI6 A1AGI7 A1AGI8 A1AGI9 A1AGJ1 A1AGJ2 A1AGJ3 A1AGJ4 A1AGJ5 A1AGJ6
A1AGJ7 A1AGJ8 A1AGJ9 A1AGK0 A1AGK1 A1AGK2 A1AGK3 A1AGK4 A1AGK5 A1AGK6 A1AGK7
A1AGK8 A1AGM6 A1AGM7 A1AGM8 A1AGM9 A1AGN0 A1AGN1 A1AGN2 A1AGQ2 A1AGR6 A1AGS9
A1AGT6 A1AGT8 A1AGU2 A1AGU3 A1AGU8 A1AGW2 A1AGW4 A1AGW5 A1AH17 A1AH34 A1AH61
A1AHA0 A1AHA1 A1AHB7 A1AHD0 A1AHE6 A1AHE9 A1AHF3 A1AHF5 A1AHG8 A1AHG9 A1AHH1
A1AHH2 A1AHH4 A1AHH8 A1AHI0 A1AHK5 A1AHM2 A1AHM5 A1AHM7 A1AHN5 A1AHN8 A1AHP0
A1AHS0 A1AHS1 A1AHS4 A1AHS5 A1AHS6 A1AHS8 A1AHU3 A1AHU6 A1AHU9 A1AHV0 A1AHX3
A1AHX9 A1AHZ7 A1AI22 A1AI34 A1AI47 A1AI48 A1AI64 A1AI70 A1AI75 A1AI78 A1AI79
A1AI80 A1AI81 A1AI82 A1AI83 A1AI95 A1AIA6 A1AIA7 A1AIC1 A1AID4 A1AID7 A1AID8
A1AIE2 A1AIE7 A1AIF2 A1AIF3 A1AIF7 A1AIF9 A1AIG0 A1AIG4 A1AIG5 A1AIG6 A1AIG7
A1AIG8 A1AIG9 A1AIH7 A1AII3 A1AII6 A1AIJ2 A1AIK3 A1AIL6 A1AJ51 A1AJ55 A1AJ76
A1AJ77 A1AJ78 A1AJ82 A1AJ83 A1AJ96 A1AJA2 A1AJA3 A1AJA5 A1AJA6 A1AJA7 A1AJC0
A1AJD7 A1AJD9 A1AJE9 A1AJF0 A1AJF5 A1AJG3 A1AJI8 A1AJJ1 A1AJL8 A1AJN5 A1AJP2
A1AJU2 A1AJU8 A1AJV0 A1AJV3 A1AJW2 A1AJW4 A1AJX2 A1AJY3 A1AK01 A1AKP9 A1AKQ8
A1AKR3 A1AKS6 A1AKW2 A1AKW3 A1AL40 A1AL91 A1ALA7 A1ALC1 A1ALC4 A1ALC9 A1ALD8
A1ALE8 A1ALG7 A1ALM4 A1ALM7 A1ALN0 A1ALN7 A1ALN8 A1ALR6 A1ALS6 A1ALT0 A1ALT1

A1ALT2 A1ALT3 A1ALT4 A1ALT5 A1ALT6 A1ALT7 A1ALU0 A1ALU1 A1ALU2 A1ALU3 A1ALU4
A1ALU5 A1ALU6 A1ALU7 A1ALU8 A1ALV0 A1ALV1 A1ALV2 A1ALV3 A1ALV4 A1ALV5 A1ALV6
A1ALV7 A1ALV8 A1ALV9 A1ALW0 A1ALW2 A1ALW4 A1ALW5 A1ALW6 A1ALW7 A1ALW8 A1ALX3
A1ALZ9 A1AM04 A1AM05 A1AM18 A1AM19 A1AM21 A1AME1 A1AMK6 A1AML8 A1AMM1 A1AMM2
A1AMM5 A1AMM6 A1AMN5 A1AMT5 A1AMT7 A1AMW9 A1AMZ7 A1AMZ8 A1AN01 A1AN03 A1AN65
A1AN77 A1AN84 A1AN85 A1AN86 A1AN88 A1ANI8 A1ANJ2 A1ANJ5 A1ANJ6 A1ANJ7 A1ANL2
A1ANL4 A1ANP7 A1ANP9 A1ANS1 A1ANS7 A1ANT3 A1ANT6 A1ANV0 A1ANV1 A1APE2 A1APF9
A1APK9 A1APM0 A1APN7 A1APN8 A1APP8 A1APP9 A1APQ8 A1APR6 A1APR9 A1APS3 A1APT2
A1APU0 A1APU5 A1APV4 A1APW6 A1APW8 A1APZ3 A1AQ29 A1AQJ8 A1AQL5 A1AQL6 A1AQN2
A1AQN3 A1AQN4 A1AQN5 A1AQN8 A1AQS8 A1AQT1 A1AR93 A1AR94 A1ARE1 A1ARE2 A1ARE3
A1ARE5 A1ARF8 A1ARJ6 A1ARJ7 A1ARN3 A1ARN6 A1ARP1 A1ARX8 A1AS07 A1AS12 A1AS13
A1AS39 A1AS43 A1AS76 A1AS90 A1ASB3 A1ASD1 A1ASD4 A1ASE9 A1ASF4 A1ASF5 A1ASF6
A1ASJ7 A1ASL7 A1ASL8 A1ASM0 A1AST1 A1AST2 A1ASW6 A1AT10 A1AT17 A1AT30 A1AT66
A1AT71 A1AT72 A1AT80 A1AT81 A1AT85 A1AT86 A1AT89 A1AT90 A1ATG7 A1ATH8 A1ATH9
A1ATI1 A1ATI5 A1ATI6 A1ATI7 A1ATI8 A1ATJ1 A1ATJ6 A1ATK5 A1ATL0 A1ATL1 A1ATL2
A1ATL3 A1ATL4 A1ATL8 A1ATL9 A1ATU1 A1ATU4 A1ATU5 A1ATU7 A1ATU9 A1AU17 A1AU18
A1AU47 A1AU48 A1AU58 A1AU59 A1AU60 A1AU63 A1AU69 A1AU70 A1AUE4 A1AUE7 A1AUE8
A1AUE9 A1AUF4 A1AUF5 A1AUQ6 A1AUS1 A1AUS2 A1AUT0 A1AUV5 A1AUV7 A1AUV8 A1AUZ4
A1AV03 A1AV12 A1AV13 A1AV24 A1AV29 A1AV33 A1AV35 A1AV41 A1AV42 A1AV43 A1AV47
A1AV52 A1AV54 A1AV71 A1AV74 A1AV91 A1AV97 A1AV99 A1AVA0 A1AVA8 A1AVB1 A1AVB2
A1AVC2 A1AVC7 A1AVD5 A1AVE9 A1AVF0 A1AVF1 A1AVF9 A1AVH0 A1AVH5 A1AVI3 A1AVI4
A1AVI7 A1AVJ5 A1AVJ7 A1AVJ8 A1AVJ9 A1AVK0 A1AVK1 A1AVK2 A1AVK3 A1AVK4 A1AVK5
A1AVK6 A1AVK7 A1AVK8 A1AVK9 A1AVL0 A1AVL1 A1AVL2 A1AVL3 A1AVL4 A1AVL5 A1AVL6
A1AVL7 A1AVL8 A1AVM1 A1AVM2 A1AVM3 A1AVM4 A1AVM5 A1AVM6 A1AVM7 A1AVN1 A1AVN5
A1AVN6 A1AVP9 A1AVQ1 A1AVQ5 A1AVQ7 A1AVT4 A1AVT6 A1AVT7 A1AVU5 A1AVU9 A1AVV3
A1AVV5 A1AVW2 A1AVW3 A1AVW6 A1AVX1 A1AVX5 A1AW02 A1AW09 A1AW10 A1AW12 A1AW13
A1AW14 A1AW17 A1AW21 A1AW22 A1AW32 A1AW35 A1AW36 A1AW38 A1AW41 A1AW44 A1AW46
A1AW54 A1AW56 A1AW62 A1AW65 A1AW67 A1AW70 A1AW71 A1AW72 A1AW83 A1AW86 A1AW87
A1AW99 A1AWA1 A1AWA3 A1AWB1 A1AWC3 A1AWC8 A1AWD0 A1AWD4 A1AWE2 A1AWE7 A1AWF3
A1AWH6 A1AWH8 A1AWI6 A1AWI7 A1AWI8 A1AWJ5 A1AWK6 A1AWK7 A1AWK9 A1AWL4 A1AWL7
A1AWM1 A1AWN8 A1AWP4 A1AWP5 A1AWQ8 A1AWR5 A1AWS0 A1AWS1 A1AWS2 A1AWS4 A1AWT1
A1AWT3 A1AWT6 A1AWU9 A1AWV0 A1AWV1 A1AWV4 A1AWV6 A1AWW7 A1AWX5 A1AWZ5 A1AX01
A1AX02 A1AX14 A1AX17 A1AX20 A1AX25 A1AX27 A1AX29 A1AX35 A1AX38 A1AX51 A1AX52
A1AX74 A1AX75 A1AX76 A1AX77 A1AX78 A1AX79 A1AX82 A1AXA0 A1AXA6 A1AXB0 A1AXD1
A1AXD2 A1AXE0 A1AXG6 A1AXH0 A1AXH4 A1AXH5 A1AXH9 A1AXI6 A1AXJ2 A1AXM7 A1AXM9
A1AXN4 A1AXN8 A1AXN9 A1AXP2 A1AXP3 A1AXP4 A1AXP6 A1AXR3 A1AXS1 A1AXS3 A1AXS4
A1AXS9 A1AXV2 A1AXV8 A1AXV9 A1AXW4 A1AXW5 A1AXW7 A1AXW9 A1AXX0 A1AXX3 A1AXX6
A1AXX7 A1AXX8 A1AXY8 A1AXZ2 A1AY12 A1AY28 A1AYS1 A1AYU1 A1AYV3 A1AZ43 A1AZ46
A1AZ56 A1AZB7 A1AZC8 A1AZF2 A1AZF6 A1AZF8 A1AZG3 A1AZH7 A1AZJ2 A1AZJ3 A1AZK1
A1AZK7 A1AZL6 A1AZM1 A1AZQ0 A1AZR0 A1AZW1 A1AZW3 A1AZW4 A1AZX8 A1AZX9 A1B002
A1B011 A1B012 A1B013 A1B014 A1B015 A1B017 A1B021 A1B022 A1B023 A1B026 A1B027
A1B028 A1B029 A1B030 A1B031 A1B032 A1B033 A1B034 A1B036 A1B037 A1B038 A1B039
A1B040 A1B041 A1B042 A1B043 A1B044 A1B045 A1B047 A1B049 A1B050 A1B051 A1B052
A1B053 A1B072 A1B095 A1B0C6 A1B0E2 A1B0E7 A1B0F1 A1B0F4 A1B0F6 A1B0F7 A1B0F8

A1B0G1 A1B0G4 A1B0G5 A1B0I7 A1B0J4 A1B0M2 A1B195 A1B196 A1B198 A1B1B6 A1B1B7
A1B1B8 A1B1B9 A1B1C2 A1B1C3 A1B1H5 A1B1H7 A1B1I1 A1B1J4 A1B1L0 A1B1M8 A1B1T3
A1B1V8 A1B1Z0 A1B2B8 A1B2I7 A1B2K9 A1B2L0 A1B2L1 A1B2N1 A1B2P8 A1B2Q2 A1B2U5
A1B2U9 A1B2W3 A1B2Z3 A1B316 A1B323 A1B362 A1B366 A1B373 A1B374 A1B384 A1B387
A1B388 A1B389 A1B3A6 A1B3B3 A1B3B5 A1B3C5 A1B3D5 A1B3F6 A1B3F9 A1B3I3 A1B3J6
A1B3K8 A1B3M0 A1B3N7 A1B3P3 A1B3V0 A1B3Y5 A1B416 A1B422 A1B431 A1B467 A1B474
A1B475 A1B477 A1B4A2 A1B4B5 A1B4B8 A1B4C1 A1B4C2 A1B4E9 A1B4F0 A1B4F1 A1B4L2
A1B4L5 A1B4S0 A1B4S2 A1B4S6 A1B4Z0 A1B4Z2 A1B4Z8 A1B4Z9 A1B502 A1B507 A1B510
A1B513 A1B515 A1B548 A1B584 A1B587 A1B5A7 A1B5K8 A1B5M0 A1B5M1 A1B5P9 A1B5Q0
A1B5Q5 A1B5S4 A1B5T0 A1B5T1 A1B5T9 A1B5U1 A1B5V3 A1B5X6 A1B5X7 A1B5Y2 A1B5Z0
A1B5Z2 A1B656 A1B673 A1B684 A1B6A9 A1B6P4 A1B840 A1B842 A1B843 A1B852 A1B853
A1B870 A1B877 A1B890 A1B8B6 A1B8C4 A1B8D4 A1B8D6 A1B8E0 A1B8E8 A1B8E9 A1B8K9
A1B8L1 A1B8L3 A1B8V3 A1B8V4 A1B8V6 A1B8V7 A1B8Z0 A1B8Z1 A1B8Z2 A1B8Z3 A1B8Z4
A1B937 A1B943 A1B945 A1B962 A1B964 A1B965 A1B966 A1B969 A1B9C7 A1B9C8 A1B9D5
A1B9E9 A1B9F0 A1B9F2 A1B9F5 A1B9H0 A1B9H4 A1B9H5 A1B9J4 A1B9Q2 A1B9Q5 A1B9Z3
A1BA12 A1BA17 A1BA23 A1BA52 A1BA96 A1BAE6 A1BAJ6 A1BAL1 A1BAL4 A1BAL9 A1BAM3
A1BAN1 A1BAN2 A1BAN4 A1BAT3 A1BB44 A1BB78 A1BBM5 A1BBS1 A1BCG1 A1BCI0 A1BCI1
A1BCQ5 A1BCU6 A1BCW2 A1BCW4 A1BD19 A1BD20 A1BD21 A1BD22 A1BD23 A1BD24 A1BD27
A1BD33 A1BD85 A1BDC1 A1BDE6 A1BDE9 A1BDF1 A1BDF2 A1BDF5 A1BDG1 A1BDG7 A1BDH1
A1BDH2 A1BDI1 A1BDJ5 A1BDJ6 A1BDK4 A1BDK8 A1BDQ5 A1BDQ9 A1BDR8 A1BDS2 A1BDT2
A1BDW1 A1BDW8 A1BDW9 A1BDY0 A1BDY1 A1BDY2 A1BDZ0 A1BDZ5 A1BDZ8 A1BE02 A1BE03
A1BE04 A1BE10 A1BE11 A1BE13 A1BE20 A1BE30 A1BE51 A1BE55 A1BE59 A1BE85 A1BE95
A1BEA5 A1BEB4 A1BED6 A1BED8 A1BEE6 A1BEF7 A1BEH0 A1BEI1 A1BEM6 A1BEN1 A1BEN4
A1BEN9 A1BEP5 A1BEP9 A1BER1 A1BES5 A1BES8 A1BET8 A1BEV6 A1BEZ1 A1BEZ2 A1BEZ3
A1BF00 A1BF67 A1BF69 A1BF81 A1BF85 A1BF89 A1BFE5 A1BFF0 A1BFF9 A1BFI0 A1BFM9
A1BFN1 A1BFN2 A1BFP7 A1BFY3 A1BG26 A1BG28 A1BG29 A1BG31 A1BGB1 A1BGB4 A1BGL6
A1BGL8 A1BGL9 A1BGM7 A1BGN2 A1BGN4 A1BGN6 A1BGP1 A1BGS9 A1BGY2 A1BGZ4 A1BH02
A1BH06 A1BH19 A1BH65 A1BH76 A1BHA0 A1BHA1 A1BHA5 A1BHD5 A1BHD6 A1BHE5 A1BHJ1
A1BHJ3 A1BHJ9 A1BHK0 A1BHL1 A1BHL3 A1BHL6 A1BHM3 A1BHN0 A1BHN5 A1BHN8 A1BHN9
A1BHP4 A1BHP6 A1BHS2 A1BHS4 A1BHS5 A1BHS6 A1BHZ3 A1BHZ4 A1BHZ5 A1BHZ6 A1BHZ7
A1BHZ8 A1BI02 A1BI09 A1BI15 A1BI19 A1BI20 A1BI58 A1BI59 A1BI65 A1BI66 A1BI67
A1BI74 A1BI80 A1BI83 A1BI85 A1BIP2 A1BIP3 A1BIQ7 A1BIW8 A1BIX2 A1BIX9 A1BIY8
A1BIZ7 A1BIZ8 A1BIZ9 A1BJ01 A1BJ02 A1BJ05 A1BJ06 A1BJ07 A1BJ08 A1BJ09 A1BJ10
A1BJ11 A1BJ12 A1BJ15 A1BJ16 A1BJ17 A1BJ18 A1BJ19 A1BJ20 A1BJ21 A1BJ22 A1BJ23
A1BJ24 A1BJ27 A1BJ28 A1BJ29 A1BJ30 A1BJ31 A1BJ32 A1BJ33 A1BJ34 A1BJ35 A1BJ36
A1BJ37 A1BJ38 A1BJ39 A1BJ56 A1BJ57 A1BJ58 A1BJ68 A1BJ99 A1BJA7 A1BJB3 A1BJB4
A1BJB5 A1BJB6 A1BJC0 A1BJC1 A1BJC3 A1BJF2 A1BJF6 A1BJF9 A1BJG0 A1BJG7 A1BJG8
A1BJI2 A1BJI4 A1BJR8 A1BJT7 A1BJT9 A1BJV5 A1BJV9 A1BJW0 A1BJX7 A1BJY0 A1BJY3
A1BJY6 A1BJY7 A1BJZ1 A1BJZ9 A1BK00 A1C3U1 A1C3V7 A1C408 A1C4N6 A1C4R7 A1C4X8
A1C4Z4 A1C595 A1C5V3 A1C5W6 A1C602 A1C603 A1C646 A1C688 A1C690 A1C6B4 A1C6C4
A1C6F8 A1C6J2 A1C6L9 A1C6R8 A1C712 A1C777 A1C7E4 A1C7F7 A1C7P9 A1C7R6 A1C7T4
A1C7Y5 A1C807 A1C874 A1C8C3 A1C8E1 A1C8E3 A1C8R3 A1C8Y9 A1C928 A1C987 A1C9M5
A1C9M6 A1C9R2 A1C9W6 A1CA18 A1CAF0 A1CAP4 A1CAQ1 A1CAU2 A1CAU3 A1CAZ7 A1CB55
A1CB93 A1CBC9 A1CBG9 A1CBI6 A1CBI9 A1CCP2 A1CD56 A1CD85 A1CDD4 A1CDT9 A1CE19

A1CE66 A1CE73 A1CE78 A1CE98 A1CED0 A1CEK7 A1CEY5 A1CEY6 A1CF18 A1CF47 A1CFB5
A1CFJ6 A1CFL9 A1CFY8 A1CGN9 A1CGW7 A1CH02 A1CH27 A1CH36 A1CH57 A1CH75 A1CH78
A1CHC0 A1CHC3 A1CHD1 A1CHE9 A1CHL0 A1CHL3 A1CHT0 A1CIC2 A1CIF1 A1CIJ6 A1CIK4
A1CIQ5 A1CJ09 A1CJ10 A1CJ18 A1CJ34 A1CJ75 A1CJ92 A1CJC1 A1CJC4 A1CJJ9 A1CJM9
A1CJT5 A1CJY4 A1CK40 A1CKE0 A1CKI4 A1CKJ0 A1CKL4 A1CKN7 A1CKP5 A1CL59 A1CL63
A1CL70 A1CL82 A1CL91 A1CL96 A1CLD2 A1CLT5 A1CLY6 A1CLY7 A1CLY8 A1CLZ0 A1CLZ1
A1CLZ2 A1CLZ5 A1CM73 A1CM94 A1CMA3 A1CMB9 A1CMQ7 A1CMX6 A1CNH7 A1CNK1 A1CNR3
A1CNV1 A1CNV8 A1CNW6 A1CNY3 A1CP31 A1CP80 A1CP85 A1CPW7 A1CQ63 A1CQA9 A1CQB4
A1CQH7 A1CQH8 A1CQI3 A1CQI9 A1CQL6 A1CQN6 A1CQP0 A1CQR5 A1CQS5 A1CQZ0 A1CQZ4
A1CR32 A1CR70 A1CRE5 A1CRF7 A1CRI1 A1CRM1 A1CS00 A1CS92 A1CSI0 A1CSR1 A1CT57
A1CT75 A1CTE6 A1CTF4 A1CTL8 A1CTZ2 A1CTZ6 A1CU75 A1CUD6 A1CUK5 A1CV59 A1CVQ6
A1CVY3 A1CW14 A1CW59 A1CW67 A1CWA9 A1CWD8 A1CWY1 A1CX72 A1CX75 A1CXD6 A1CXF4
A1CXG4 A1CXG7 A1CXK7 A1CXL0 A1CXQ2 A1CXR0 A1CXT5 A1CXW3 A1CY38 A1CYJ1 A1CYM1
A1CYR9 A1CYY6 A1CZ18 A1CZ92 A1CZE5 A1CZJ5 A1CZL7 A1CZP6 A1CZX3 A1D071 A1D1B7
A1D1E3 A1D1M2 A1D1R1 A1D1R8 A1D1S6 A1D1U0 A1D1Y5 A1D240 A1D244 A1D2R0 A1D327
A1D373 A1D379 A1D3E1 A1D3E2 A1D3F5 A1D3I2 A1D3K3 A1D3M8 A1D3N8 A1D3V4 A1D3V8
A1D405 A1D434 A1D4A7 A1D4E3 A1D4K4 A1D4V5 A1D558 A1D5T7 A1D611 A1D619 A1D624
A1D688 A1D695 A1D699 A1D6S8 A1D6T6 A1D6V7 A1D6X9 A1D6Y7 A1D731 A1D7C4 A1D7I5
A1D7N3 A1D7X4 A1D7X8 A1D855 A1D858 A1D887 A1D8A4 A1D8E4 A1D8G1 A1D8G8 A1D8G9
A1D8I8 A1D8I9 A1D8J1 A1D8J2 A1D9C9 A1D9E6 A1D9I5 A1D9P1 A1D9Z6 A1D9Z7 A1DA59
A1DA61 A1DA62 A1DA64 A1DA66 A1DA84 A1DAL6 A1DAY1 A1DB13 A1DB29 A1DB76 A1DC33
A1DC62 A1DCB2 A1DCU5 A1DDV0 A1DDV3 A1DDX0 A1DE13 A1DE49 A1DE84 A1DED8 A1DEK2
A1DEK3 A1DES4 A1DF20 A1DF27 A1DFN6 A1DFW7 A1DG00 A1DG01 A1DG38 A1DG51 A1DG66
A1DGF2 A1DGS2 A1DGW4 A1DGW6 A1DGY9 A1DGZ7 A1DH31 A1DH62 A1DH89 A1DHD8 A1DHP1
A1DHV3 A1DHW6 A1DI25 A1DIC0 A1DIL2 A1DIN9 A1DJ20 A1DL11 A1DL85 A1DLP3 A1DLU6
A1DLV0 A1DLW4 A1DM27 A1DM75 A1DM82 A1DM87 A1DMA6 A1DMG9 A1DMI8 A1DMJ3 A1DMT9
A1DN09 A1DN10 A1DN11 A1DN30 A1DN59 A1DNF9 A1DNG2 A1DNV8 A1DP19 A1DP85 A1DPK7
A1DQP9 A1DZZ3 A1E260 A1E261 A1E2V0 A1E952 A1E9H0 A1E9I1 A1E9I2 A1E9I3 A1E9I4
A1E9I9 A1E9J3 A1E9L1 A1E9L2 A1E9L3 A1E9M3 A1E9M4 A1E9M5 A1E9M6 A1E9M7 A1E9M8
A1E9M9 A1E9N0 A1E9N5 A1E9N6 A1E9N8 A1E9Q2 A1E9Q3 A1E9Q6 A1E9R4 A1E9R5 A1E9R7
A1E9S2 A1E9S6 A1E9T2 A1E9T3 A1E9U5 A1E9U6 A1E9U7 A1E9U9 A1E9V6 A1E9V7 A1E9V8
A1E9V9 A1E9W0 A1E9W1 A1E9W2 A1E9W3 A1E9W4 A1E9W6 A1E9W7 A1E9W9 A1E9X0 A1E9X2
A1E9Y6 A1E9Y7 A1E9Z0 A1E9Z8 A1E9Z9 A1EA00 A1EA01 A1EA06 A1EA10 A1EA16 A1EA17
A1EA29 A1EA30 A1EA31 A1EA33 A1EA40 A1EA41 A1EA42 A1EA43 A1EA44 A1EA45 A1EA46
A1EA47 A1EA48 A1EA50 A1EA51 A1EA53 A1EA54 A1EA57 A1EC95 A1EGX6 A1IGA8 A1IGU3
A1IGU4 A1IGU5 A1IHE6 A1IHE7 A1IIX2 A1IIX3 A1IIX4 A1IIX5 A1IP83 A1IPH6 A1IPQ8
A1IQ34 A1IQ68 A1IQ95 A1IQG0 A1IQR7 A1IQS5 A1IR09 A1IRQ3 A1IRY3 A1IS67 A1ISB3
A1ISU7 A1ITK1 A1JHR0 A1JHR2 A1JHR3 A1JHR4 A1JHR5 A1JHR7 A1JHR8 A1JHR9 A1JHT4
A1JHT5 A1JHU4 A1JHV7 A1JHV9 A1JHW7 A1JHW8 A1JHW9 A1JHX0 A1JHX2 A1JHX6 A1JHX8
A1JHY1 A1JHY4 A1JHZ3 A1JHZ7 A1JI05 A1JI06 A1JI08 A1JI09 A1JI13 A1JI14 A1JI20
A1JI23 A1JI24 A1JI37 A1JI40 A1JI42 A1JI47 A1JI53 A1JI57 A1JI66 A1JI86 A1JI88
A1JI90 A1JIB4 A1JIE5 A1JIF2 A1JIG5 A1JIH1 A1JIH3 A1JIH6 A1JIH7 A1JIH8 A1JIH9
A1JII0 A1JII1 A1JII4 A1JII9 A1JIJ0 A1JIJ1 A1JIJ2 A1JIJ7 A1JIK3 A1JIM9 A1JIP2
A1JIP3 A1JIP6 A1JIQ4 A1JIQ7 A1JIQ8 A1JIR3 A1JIR4 A1JIR5 A1JIS0 A1JIS1 A1JIS8

A1JIS9 A1JIT0 A1JIT1 A1JIU0 A1JIU6 A1JIU9 A1JIV0 A1JIV3 A1JIV4 A1JIV6 A1JIW2
A1JIW5 A1JIW7 A1JIW9 A1JIX0 A1JIX1 A1JIX2 A1JIX3 A1JIY2 A1JIY7 A1JIZ7 A1JJ31
A1JJ50 A1JJ51 A1JJ57 A1JJ89 A1JJ92 A1JJ98 A1JJ99 A1JJA0 A1JJB6 A1JJB7 A1JJB8
A1JJC7 A1JJC9 A1JJD0 A1JJD5 A1JJD6 A1JJD9 A1JJE1 A1JJE4 A1JJE5 A1JJF2 A1JJF3
A1JJF4 A1JJF5 A1JJG0 A1JJG7 A1JJH3 A1JJH5 A1JJH7 A1JJI4 A1JJI5 A1JJJ4 A1JJK4
A1JJK6 A1JJK8 A1JJM4 A1JJM7 A1JJM8 A1JJN6 A1JJN7 A1JJN8 A1JJP3 A1JJQ1 A1JJQ3
A1JJQ6 A1JJQ7 A1JJR0 A1JJR3 A1JJS2 A1JJS3 A1JJS4 A1JJS8 A1JJS9 A1JJT0 A1JJT5
A1JJT6 A1JJT7 A1JJT8 A1JJU1 A1JJW5 A1JK01 A1JK02 A1JK08 A1JK09 A1JK11 A1JK14
A1JK22 A1JK24 A1JK25 A1JK26 A1JK38 A1JK48 A1JKA7 A1JKA9 A1JKB1 A1JKB8 A1JKD7
A1JKD8 A1JKD9 A1JKE0 A1JKE1 A1JKE2 A1JKE3 A1JKF8 A1JKI0 A1JKI6 A1JKI8 A1JKI9
A1JKJ4 A1JKK3 A1JKK5 A1JKL6 A1JKP3 A1JKQ0 A1JKQ1 A1JKQ4 A1JKQ5 A1JKQ7 A1JKR7
A1JKR9 A1JKS2 A1JKS3 A1JKS6 A1JKT0 A1JKT2 A1JKT8 A1JKU0 A1JKU9 A1JKZ7 A1JKZ8
A1JL00 A1JL07 A1JL09 A1JL12 A1JL17 A1JL31 A1JL36 A1JL37 A1JL38 A1JL55 A1JL75
A1JLA0 A1JLA4 A1JLB3 A1JLD7 A1JLD8 A1JLF1 A1JLK6 A1JLK7 A1JLK9 A1JLM5 A1JLM8
A1JLM9 A1JLN0 A1JLS2 A1JLS4 A1JLV6 A1JLX8 A1JM26 A1JM34 A1JM38 A1JM74 A1JMA3
A1JMA4 A1JMB6 A1JMC0 A1JMC5 A1JMD0 A1JMD2 A1JME4 A1JME5 A1JMF5 A1JMG0 A1JMI5
A1JMJ0 A1JMJ9 A1JMK2 A1JMK4 A1JMK7 A1JMK8 A1JML2 A1JML4 A1JML7 A1JML8 A1JMM2
A1JMM4 A1JMN8 A1JMP2 A1JMQ6 A1JMT1 A1JMT8 A1JMU2 A1JMV3 A1JMW1 A1JMW4 A1JMW5
A1JMW8 A1JMX0 A1JMX1 A1JMX4 A1JMX8 A1JMY1 A1JN27 A1JN32 A1JN45 A1JN60 A1JN84
A1JN89 A1JNA4 A1JNB1 A1JNB3 A1JNB5 A1JNB8 A1JNC2 A1JND8 A1JNG6 A1JNH0 A1JNH5
A1JNH6 A1JNL2 A1JNM3 A1JNN1 A1JNN2 A1JNN4 A1JNP9 A1JNQ7 A1JNR2 A1JNR4 A1JNR7
A1JNS2 A1JNS3 A1JNS5 A1JNT0 A1JNT4 A1JNV0 A1JNV1 A1JNV2 A1JNV3 A1JNV9 A1JNW9
A1JNX2 A1JNX6 A1JNX7 A1JNX8 A1JNX9 A1JNY1 A1JNY3 A1JNZ8 A1JP02 A1JP08 A1JP09
A1JP10 A1JP11 A1JP13 A1JP36 A1JP52 A1JP53 A1JP58 A1JP66 A1JP78 A1JP79 A1JP80
A1JP81 A1JP82 A1JP85 A1JP86 A1JP87 A1JP94 A1JP96 A1JPA5 A1JPB6 A1JPC1 A1JPC9
A1JPE1 A1JPE3 A1JPI0 A1JPK8 A1JPL4 A1JPL5 A1JPM8 A1JPN3 A1JPN4 A1JPN6 A1JPP0
A1JPP4 A1JPP6 A1JPR0 A1JPR1 A1JPR7 A1JPR8 A1JPR9 A1JPS4 A1JPT0 A1JPT5 A1JPT6
A1JPU6 A1JPV4 A1JPV6 A1JPW3 A1JPW6 A1JPX6 A1JPY2 A1JQ25 A1JQ46 A1JQ74 A1JQA3
A1JQB6 A1JQE8 A1JQG8 A1JQH0 A1JQJ2 A1JQJ6 A1JQN9 A1JQU7 A1JQV2 A1JQV6 A1JQW2
A1JQW9 A1JQX1 A1JR10 A1JR24 A1JR25 A1JR34 A1JR39 A1JR73 A1JR74 A1JR92 A1JR94
A1JRA1 A1JRA2 A1JRB1 A1JRB5 A1JRB6 A1JRC1 A1JRD2 A1JRD5 A1JRD6 A1JRH2 A1JRJ5
A1JRJ9 A1JRK0 A1JRK2 A1JRK5 A1JRL1 A1JRL5 A1JRL7 A1JRM2 A1JRM4 A1JRM5 A1JRM9
A1JRP2 A1JRR0 A1JRT1 A1JRU0 A1JRU4 A1JRU5 A1JRV9 A1JRW1 A1JRX0 A1JRX5 A1JRX9
A1JRY4 A1JRY6 A1JRZ1 A1JRZ2 A1JRZ3 A1JS00 A1JS01 A1JS02 A1JS03 A1JS04 A1JS06
A1JS08 A1JS10 A1JS11 A1JS13 A1JS15 A1JS16 A1JS18 A1JS20 A1JS21 A1JS23 A1JS24
A1JS25 A1JS26 A1JS27 A1JS29 A1JS31 A1JS32 A1JS33 A1JS34 A1JS35 A1JS43 A1JS44
A1JS45 A1JS46 A1JS47 A1JS48 A1JS52 A1JS56 A1JS57 A1JS58 A1JS59 A1JS60 A1JS64
A1JS67 A1JS69 A1JS71 A1JS94 A1JSB7 A1JSC3 A1JSD4 A1JSE7 A1JSF2 A1JSF6 A1JSF8
A1JSG1 A1JSH7 A1JSI5 A1JSI8 A1JSJ0 A1JSK6 A1JSK7 A1JSN4 A1JSN5 A1JSN7 A1JSQ2
A1JSU2 A1JSW6 A1JSW8 A1JT10 A1JT25 A1JT26 A1JT31 A1JT32 A1JT33 A1JT46 A1JT50
A1JT51 A1JT78 A1JT80 A1JT83 A1JT84 A1JT87 A1JTB7 A1JTC4 A1JTD7 A1JTD9 A1JTF2
A1JTI8 A1JTL6 A1JTP8 A1JTQ2 A1JTT9 A1JTU7 A1JTV7 A1JTV9 A1JTW2 A1JTW3 A1JTX3
A1JTX4 A1JTX6 A1JTY5 A1JU09 A1JU33 A1JU34 A1JU40 A1JU66 A1JU75 A1JU77 A1JU91
A1JU92 A1JUB1 A1K1B4 A1K1F6 A1K1J9 A1K1K9 A1K1M2 A1K1N5 A1K1Q4 A1K1Q5 A1K1T3

A1K1T5 A1K1T6 A1K1U6 A1K1U7 A1K279 A1K280 A1K2E2 A1K2G1 A1K2H5 A1K2I5 A1K2I6
A1K2N7 A1K2P0 A1K2P7 A1K2Q5 A1K2R3 A1K2R9 A1K2Y3 A1K2Y4 A1K2Y6 A1K301 A1K306
A1K313 A1K314 A1K320 A1K339 A1K344 A1K3B9 A1K3D0 A1K3D3 A1K3E2 A1K3E5 A1K3G5
A1K3G6 A1K3G8 A1K3H4 A1K3J0 A1K3J2 A1K3J8 A1K3M9 A1K3Q2 A1K3T2 A1K3T3 A1K3T5
A1K3T7 A1K3T8 A1K3U7 A1K3U8 A1K3W2 A1K3W9 A1K3X0 A1K3X8 A1K3Y5 A1K3Y6 A1K3Z0
A1K3Z3 A1K3Z4 A1K3Z9 A1K400 A1K421 A1K430 A1K436 A1K442 A1K443 A1K451 A1K454
A1K471 A1K487 A1K498 A1K4A1 A1K4A2 A1K4B0 A1K4C4 A1K4C5 A1K4C6 A1K4D5 A1K4E1
A1K4E3 A1K4E4 A1K4E5 A1K4E7 A1K4E9 A1K4F0 A1K4F8 A1K4G2 A1K4J5 A1K4J7 A1K4J8
A1K4J9 A1K4K1 A1K4K7 A1K4P9 A1K4Q9 A1K4R0 A1K4R2 A1K4R4 A1K4R7 A1K4Y1 A1K4Z8
A1K524 A1K581 A1K589 A1K598 A1K5A1 A1K5A6 A1K5I3 A1K5I5 A1K5J1 A1K5J4 A1K5J9
A1K5N0 A1K5N3 A1K5Q9 A1K5T6 A1K5T8 A1K5U3 A1K5Y3 A1K606 A1K607 A1K616 A1K639
A1K644 A1K645 A1K6F9 A1K6L8 A1K6L9 A1K6P9 A1K6Q1 A1K6Q7 A1K6R8 A1K6R9 A1K6S0
A1K6S1 A1K6Y4 A1K6Y5 A1K6Y7 A1K713 A1K717 A1K719 A1K724 A1K725 A1K764 A1K782
A1K783 A1K784 A1K7B5 A1K7B6 A1K7B7 A1K7B8 A1K7B9 A1K7C1 A1K7F8 A1K7I0 A1K7J5
A1K7J6 A1K7J8 A1K7J9 A1K7K2 A1K7M0 A1K7T8 A1K7V8 A1K828 A1K833 A1K893 A1K8P2
A1K8P4 A1K8Q1 A1K8R4 A1K8U5 A1K922 A1K962 A1K969 A1K970 A1K971 A1K975 A1K976
A1K983 A1K991 A1K9B2 A1K9B3 A1K9B9 A1K9C3 A1K9C8 A1K9D3 A1K9D7 A1K9D9 A1K9F0
A1K9I4 A1K9J6 A1K9K9 A1K9L0 A1K9L2 A1K9L5 A1K9L8 A1K9S9 A1KA92 A1KA97 A1KAA5
A1KAA6 A1KAB7 A1KAC8 A1KAC9 A1KAD0 A1KAD1 A1KAF7 A1KAG0 A1KAG3 A1KAG5 A1KAH5
A1KAH8 A1KAI4 A1KAI5 A1KAJ9 A1KAK4 A1KAT2 A1KAT3 A1KAU3 A1KAV2 A1KAV3 A1KAV4
A1KAV5 A1KAV7 A1KAY0 A1KB00 A1KB01 A1KB02 A1KB03 A1KB04 A1KB05 A1KB06 A1KB08
A1KB09 A1KB10 A1KB11 A1KB12 A1KB13 A1KB14 A1KB15 A1KB16 A1KB17 A1KB18 A1KB19
A1KB20 A1KB21 A1KB22 A1KB23 A1KB24 A1KB25 A1KB26 A1KB27 A1KB28 A1KB29 A1KB30
A1KB31 A1KB32 A1KB33 A1KB34 A1KB35 A1KB36 A1KB37 A1KB38 A1KB59 A1KB60 A1KB69
A1KB70 A1KB72 A1KB74 A1KBB1 A1KBB2 A1KBB3 A1KBB4 A1KBB5 A1KBB7 A1KBB8 A1KBB9
A1KBC7 A1KBD3 A1KBH0 A1KBH4 A1KBL9 A1KBQ3 A1KBQ4 A1KBR9 A1KBU9 A1KC59 A1KCI5
A1KCL2 A1KCL7 A1KCM3 A1KCM8 A1KCM9 A1KCN1 A1KCN9 A1KCP8 A1KCQ1 A1KEG3 A1KEK6
A1KEK9 A1KEM1 A1KEM3 A1KEM4 A1KET4 A1KEU8 A1KEV7 A1KEW5 A1KEW6 A1KF28 A1KF31
A1KF44 A1KF54 A1KF76 A1KFA4 A1KFB7 A1KFE2 A1KFE4 A1KFH3 A1KFH8 A1KFK3 A1KFN6
A1KFN9 A1KFP5 A1KFQ1 A1KFQ5 A1KFU9 A1KFV2 A1KFV6 A1KFW0 A1KFW3 A1KFY5 A1KFY6
A1KG00 A1KG30 A1KG32 A1KG35 A1KG37 A1KG43 A1KG98 A1KGB4 A1KGC0 A1KGC1 A1KGD1
A1KGE7 A1KGE8 A1KGF0 A1KGG2 A1KGG3 A1KGG4 A1KGG5 A1KGI1 A1KGI2 A1KGI3 A1KGI4
A1KGI5 A1KGI6 A1KGI7 A1KGI8 A1KGI9 A1KGJ0 A1KGJ5 A1KGJ6 A1KGJ7 A1KGJ8 A1KGJ9
A1KGK0 A1KGK1 A1KGK2 A1KGK3 A1KGK4 A1KGK6 A1KGK7 A1KGL2 A1KGL4 A1KGR3 A1KGT3
A1KGT6 A1KGU6 A1KGV2 A1KGW3 A1KGY5 A1KH25 A1KH80 A1KH85 A1KH91 A1KHA2 A1KHB5
A1KHD9 A1KHE8 A1KHE9 A1KHF2 A1KHL8 A1KHN2 A1KHN9 A1KHP7 A1KHP8 A1KHS4 A1KHV0
A1KHW1 A1KI01 A1KI08 A1KI28 A1KI36 A1KI72 A1KI80 A1KI86 A1KI87 A1KIB6 A1KIC1
A1KIC8 A1KID0 A1KIG2 A1KIG8 A1KIG9 A1KIH4 A1KIH9 A1KII1 A1KIJ5 A1KIK4 A1KIK9
A1KIL0 A1KIL2 A1KIM6 A1KIM7 A1KIN7 A1KIS5 A1KJ00 A1KJ01 A1KJ05 A1KJ10 A1KJ16
A1KJ17 A1KJ19 A1KJ21 A1KJ22 A1KJ27 A1KJ29 A1KJ59 A1KJ60 A1KJ66 A1KJ69 A1KJ71
A1KJ73 A1KJ74 A1KJ75 A1KJ76 A1KJA4 A1KJB5 A1KJC9 A1KJD0 A1KJE6 A1KJE8 A1KJG2
A1KJN8 A1KJP4 A1KJP9 A1KJR0 A1KJR1 A1KJR2 A1KJR3 A1KJR4 A1KJW1 A1KJW2 A1KJW6
A1KJX3 A1KKA1 A1KKA2 A1KKA3 A1KKE3 A1KKF2 A1KKF3 A1KKF8 A1KKG4 A1KKG5 A1KKH3
A1KKJ0 A1KKJ5 A1KKJ8 A1KKK8 A1KKK9 A1KKN4 A1KKP9 A1KKQ5 A1KKQ9 A1KKR0 A1KKR8

A1KKZ1 A1KL43 A1KL52 A1KL57 A1KL64 A1KL82 A1KL99 A1KLA4 A1KLB3 A1KLC0 A1KLC4
A1KLD5 A1KLD6 A1KLD7 A1KLE1 A1KLE9 A1KLF3 A1KLF8 A1KLK6 A1KLN0 A1KLN1 A1KLN4
A1KLN6 A1KLN7 A1KLQ2 A1KLQ3 A1KLS7 A1KLS8 A1KLT2 A1KLU0 A1KLU1 A1KLU2 A1KLV3
A1KLV4 A1KLV6 A1KLV7 A1KLW4 A1KM16 A1KM20 A1KM35 A1KM51 A1KM55 A1KM56 A1KM64
A1KM65 A1KM71 A1KM74 A1KM94 A1KM95 A1KMA6 A1KMB5 A1KMC6 A1KMC8 A1KMD6 A1KMI1
A1KMI2 A1KMI5 A1KMI8 A1KMJ5 A1KML3 A1KML5 A1KMM4 A1KMM6 A1KMM7 A1KMN3 A1KMN8
A1KMP2 A1KMP6 A1KMP8 A1KMQ0 A1KMQ1 A1KMQ3 A1KMR7 A1KMR9 A1KMV1 A1KMV2 A1KMV6
A1KMV9 A1KMW6 A1KMX0 A1KMX5 A1KMX7 A1KMX8 A1KMY1 A1KMY2 A1KMY6 A1KMY9 A1KMZ6
A1KN04 A1KN06 A1KN07 A1KN09 A1KN20 A1KN49 A1KN60 A1KN98 A1KNJ7 A1KNQ0 A1KNR3
A1KNR4 A1KNT4 A1KNZ5 A1KP00 A1KP32 A1KP49 A1KP65 A1KP87 A1KP90 A1KPA8 A1KPA9
A1KPB0 A1KPC8 A1KPC9 A1KPD0 A1KPE2 A1KPE3 A1KPE4 A1KPE5 A1KPE6 A1KPE7 A1KPE8
A1KPH5 A1KPL9 A1KPM0 A1KPR5 A1KPR6 A1KPR7 A1KPR8 A1KPS2 A1KPT4 A1KPT5 A1KPT6
A1KPT7 A1KPU4 A1KQ47 A1KQ48 A1KQ98 A1KQA5 A1KQC1 A1KQG9 A1KQH3 A1KQI8 A1KQY3
A1KQY9 A1KR57 A1KR91 A1KRE1 A1KRE2 A1KRE5 A1KRE6 A1KRF2 A1KRF9 A1KRG2 A1KRG3
A1KRG4 A1KRG5 A1KRG6 A1KRG7 A1KRG8 A1KRG9 A1KRH0 A1KRH2 A1KRH3 A1KRH4 A1KRH5
A1KRH6 A1KRH7 A1KRH8 A1KRH9 A1KRI0 A1KRI1 A1KRI2 A1KRI3 A1KRI4 A1KRI5 A1KRI6
A1KRI7 A1KRI8 A1KRI9 A1KRJ0 A1KRJ1 A1KRJ2 A1KRJ4 A1KRJ5 A1KRJ6 A1KRJ7 A1KRJ8
A1KRJ9 A1KRK0 A1KRK1 A1KRK3 A1KRK7 A1KRL9 A1KRM2 A1KRM4 A1KRM5 A1KRN3 A1KRN6
A1KRR2 A1KRU1 A1KRU2 A1KRU4 A1KRU9 A1KRV4 A1KRY5 A1KRZ4 A1KS00 A1KS01 A1KS02
A1KS07 A1KS25 A1KS31 A1KS32 A1KS33 A1KS34 A1KS35 A1KS36 A1KS57 A1KS60 A1KS62
A1KS64 A1KS84 A1KS92 A1KS96 A1KS98 A1KSA7 A1KSA9 A1KSD4 A1KSD9 A1KSE7 A1KSG3
A1KSG6 A1KSH0 A1KSH8 A1KSI4 A1KSI5 A1KSJ8 A1KSJ9 A1KSK0 A1KSK1 A1KSL1 A1KSL7
A1KSM7 A1KSN4 A1KSN5 A1KSN6 A1KSN7 A1KSP4 A1KSS7 A1KST1 A1KST5 A1KSU5 A1KSU6
A1KSU7 A1KSV1 A1KSW0 A1KSW2 A1KSY0 A1KSY1 A1KSY3 A1KSY4 A1KSY5 A1KSZ1 A1KSZ5
A1KSZ9 A1KT00 A1KT06 A1KT11 A1KT19 A1KT34 A1KT40 A1KT53 A1KT54 A1KT57 A1KT61
A1KT64 A1KT70 A1KT71 A1KT73 A1KT74 A1KT78 A1KT90 A1KT92 A1KT93 A1KT95 A1KTB1
A1KTB2 A1KTB5 A1KTB7 A1KTD5 A1KTJ9 A1KTK4 A1KTK6 A1KTL0 A1KTL1 A1KTL3 A1KTM6
A1KTN4 A1KTP9 A1KTU6 A1KTV4 A1KTV5 A1KTV6 A1KTW0 A1KU01 A1KU04 A1KU05 A1KU06
A1KU10 A1KU16 A1KU21 A1KU22 A1KU26 A1KU46 A1KU50 A1KU52 A1KU57 A1KU60 A1KU62
A1KU78 A1KU80 A1KU82 A1KU93 A1KUB3 A1KUC8 A1KUD0 A1KUD5 A1KUD6 A1KUD8 A1KUD9
A1KUE0 A1KUE7 A1KUE8 A1KUE9 A1KUF0 A1KUF5 A1KUG1 A1KUG2 A1KUG4 A1KUH8 A1KUI0
A1KUI2 A1KUI5 A1KUJ4 A1KUJ5 A1KUK1 A1KUK4 A1KUL0 A1KUM5 A1KUN1 A1KUN6 A1KUP6
A1KUP8 A1KUQ0 A1KUQ1 A1KUQ3 A1KUS7 A1KUT0 A1KUT9 A1KUU4 A1KUU8 A1KUU9 A1KUV0
A1KUW3 A1KUW7 A1KUX8 A1KUY0 A1KUZ7 A1KUZ8 A1KV03 A1KV06 A1KV07 A1KV20 A1KV25
A1KV29 A1KV38 A1KV41 A1KV48 A1KV49 A1KV51 A1KV56 A1KV59 A1KV60 A1KV69 A1KV71
A1KV85 A1KV88 A1KV89 A1KV92 A1KV93 A1KV95 A1KV99 A1KVB3 A1KVF6 A1KVF9 A1KVG2
A1KVG4 A1KVG8 A1KVH8 A1KVH9 A1KVL1 A1KVL2 A1KVL5 A1KVM4 A1KVM5 A1KVN9 A1KVQ5
A1KVS3 A1KVS6 A1KVT3 A1KVT5 A1KVT7 A1KVU9 A1KVV4 A1KVV8 A1KVV9 A1KVW1 A1KVW2
A1KVW5 A1KW08 A1KW09 A1KW26 A1KW52 A1KW58 A1KW66 A1KW76 A1KW97 A1KW99 A1KWA0
A1KWA2 A1KWB6 A1KWC0 A1KWD5 A1KWD6 A1KWE0 A1KWE2 A1KWF0 A1KWF4 A1KWG2 A1KWG5
A1KWG6 A1KWH2 A1KWH6 A1KWH7 A1KWH8 A1KWJ8 A1KWK6 A1KWL6 A1KWM6 A1KWM9 A1KWN6
A1KWN7 A1KXM5 A1KXP4 A1KYZ2 A1L020 A1L0Y2 A1L0Z0 A1L0Z1 A1L0Z4 A1L0Z6 A1L108
A1L112 A1L162 A1L168 A1L188 A1L190 A1L1C7 A1L1F0 A1L1F4 A1L1F6 A1L1G1 A1L1G9
A1L1H3 A1L1J6 A1L1K3 A1L1L1 A1L1L7 A1L1M4 A1L1N5 A1L1P7 A1L1R0 A1L1R5 A1L1R6

A1L1S5 A1L209 A1L251 A1L253 A1L258 A1L2E4 A1L2F3 A1L2H3 A1L2H9 A1L2L5 A1L2P5
A1L2P7 A1L2Q8 A1L2T6 A1L2U9 A1L2W7 A1L2Y1 A1L317 A1L3C1 A1L3F4 A1L3F5 A1L3H4
A1L3I5 A1L3K1 A1L3L1 A1L3N6 A1L3P1 A1L3T7 A1L3X4 A1L429 A1L443 A1L4K1 A1L4L8
A1L4Q6 A1L4T4 A1L4V7 A1L4W5 A1L4Y1 A1L4Y2 A1L595 A1QNY1 A1QQ42 A1QYG5 A1QYG8
A1QYH6 A1QYH7 A1QYH8 A1QYK3 A1QYK6 A1QYL0 A1QYL4 A1QYN6 A1QYR2 A1QYR4 A1QYS0
A1QYS1 A1QYS2 A1QYS8 A1QYT1 A1QYX2 A1QYX4 A1QYX8 A1QYX9 A1QYY3 A1QYY4 A1QYY5
A1QYZ0 A1QYZ4 A1QZ05 A1QZ08 A1QZ26 A1QZ46 A1QZ92 A1QZA2 A1QZD0 A1QZD1 A1QZD3
A1QZE2 A1QZF9 A1QZH4 A1QZH5 A1QZH6 A1QZH7 A1QZH8 A1QZH9 A1QZI0 A1QZI4 A1QZK3
A1QZK9 A1QZM2 A1QZM6 A1QZM7 A1QZN3 A1QZP2 A1QZP7 A1QZR2 A1QZR3 A1QZR4 A1QZR8
A1QZR9 A1QZS0 A1QZS1 A1QZS2 A1QZS3 A1QZS4 A1QZS5 A1QZS6 A1QZS7 A1QZS8 A1QZS9
A1QZT0 A1QZT5 A1QZT6 A1QZT9 A1QZU9 A1QZY4 A1R019 A1R025 A1R030 A1R032 A1R037
A1R039 A1R043 A1R050 A1R074 A1R083 A1R0A7 A1R0B9 A1R0C2 A1R0C7 A1R0C8 A1R0E4
A1R0H8 A1R0J1 A1R0K3 A1R0K4 A1R0K5 A1R0K8 A1R0K9 A1R0L0 A1R0M2 A1R0M4 A1R0M9
A1R0N0 A1R0N4 A1R0P6 A1R0Q9 A1R0R8 A1R0S5 A1R120 A1R163 A1R1C3 A1R1C4 A1R1C5
A1R1C6 A1R1C7 A1R1C9 A1R1K8 A1R1M6 A1R228 A1R2A8 A1R2B5 A1R2K7 A1R2V3 A1R2Y2
A1R317 A1R383 A1R3H9 A1R3M0 A1R3R7 A1R3U5 A1R451 A1R4F7 A1R4F8 A1R4G4 A1R4P6
A1R4R0 A1R4R1 A1R4R2 A1R4S6 A1R4W0 A1R4W1 A1R4W2 A1R4Z8 A1R501 A1R508 A1R513
A1R516 A1R517 A1R520 A1R524 A1R525 A1R548 A1R550 A1R551 A1R558 A1R559 A1R562
A1R567 A1R568 A1R585 A1R587 A1R591 A1R5C7 A1R5E9 A1R5F0 A1R5F9 A1R5G4 A1R5H3
A1R5H8 A1R5J6 A1R5N7 A1R5R2 A1R5R8 A1R5R9 A1R5S0 A1R5S1 A1R5S3 A1R5S7 A1R665
A1R674 A1R6F3 A1R6G5 A1R6G6 A1R6G8 A1R6H1 A1R6L8 A1R6P9 A1R6Q4 A1R6Q7 A1R6Q8
A1R6Q9 A1R6T5 A1R6V4 A1R6V9 A1R6X0 A1R6X5 A1R6X6 A1R6Z3 A1R6Z7 A1R700 A1R701
A1R703 A1R705 A1R709 A1R712 A1R724 A1R726 A1R727 A1R728 A1R735 A1R783 A1R787
A1R788 A1R7A5 A1R7D2 A1R7F9 A1R7G2 A1R7G3 A1R7G4 A1R7I5 A1R7I7 A1R7J5 A1R7J9
A1R7K0 A1R7K6 A1R7M9 A1R7X2 A1R807 A1R863 A1R898 A1R8A6 A1R8M1 A1R8M2 A1R8M6
A1R8M8 A1R8N0 A1R8N8 A1R8N9 A1R8P8 A1R8Q0 A1R8Q1 A1R8Q2 A1R8R4 A1R8R5 A1R8R6
A1R8R7 A1R8R8 A1R8R9 A1R8S8 A1R8S9 A1R8T1 A1R8T2 A1R8T3 A1R8T4 A1R8T5 A1R8T8
A1R8T9 A1R8U0 A1R8U1 A1R8U2 A1R8U3 A1R8U5 A1R8U6 A1R8U9 A1R8V0 A1R8V1 A1R8V2
A1R8V3 A1R8V4 A1R8V8 A1R8W1 A1R8W2 A1R8X8 A1R8Y2 A1R8Y7 A1R990 A1R996 A1R9J0
A1RAI8 A1RAU1 A1RAW1 A1RAY5 A1RAZ4 A1RB22 A1RB23 A1RB24 A1RBC4 A1RC67 A1RC68
A1RC70 A1RCA8 A1RCB2 A1RDH3 A1RDW2 A1RDW4 A1RDW5 A1RDW7 A1RDX6 A1RDX7 A1RDX9
A1RDY5 A1RDY7 A1RDY8 A1RDZ0 A1RDZ6 A1RE02 A1RE07 A1RE18 A1RE25 A1RE29 A1RE77
A1RE99 A1REA3 A1REA4 A1REA5 A1REA6 A1REA7 A1REA8 A1REA9 A1REB0 A1REB3 A1REB4
A1REB5 A1REB7 A1REB8 A1REB9 A1REC0 A1REC1 A1REC2 A1REC3 A1REC4 A1REC5 A1REC6
A1REC7 A1REC8 A1REC9 A1RED0 A1RED1 A1RED2 A1RED3 A1RED5 A1RED6 A1RED7 A1RED8
A1RED9 A1REE0 A1REI6 A1REK0 A1REM5 A1REM9 A1REN3 A1RET8 A1REU0 A1REU1 A1REU2
A1REU4 A1REU5 A1REU6 A1REY0 A1REY2 A1REY3 A1REY5 A1REY7 A1REY8 A1REZ7 A1RFA1
A1RFA3 A1RFB3 A1RFB7 A1RFE3 A1RFJ9 A1RFM0 A1RFN3 A1RFP5 A1RFQ1 A1RFR0 A1RFR6
A1RFR7 A1RFX7 A1RFX8 A1RFY6 A1RFY7 A1RFY8 A1RG35 A1RG66 A1RG67 A1RG72 A1RGD1
A1RGD7 A1RGE6 A1RGF1 A1RGF7 A1RGG0 A1RGJ2 A1RGK8 A1RGN1 A1RGN2 A1RGP1 A1RGQ6
A1RGR7 A1RGR8 A1RGT2 A1RGT3 A1RGT4 A1RGU0 A1RGU2 A1RGU5 A1RGV2 A1RGV4 A1RGW7
A1RGX0 A1RGX1 A1RGX3 A1RGX5 A1RGX6 A1RGX7 A1RGX8 A1RGY0 A1RH82 A1RH87 A1RH88
A1RH89 A1RHD0 A1RHD1 A1RHD5 A1RHD6 A1RHE4 A1RHF0 A1RHF2 A1RHF6 A1RHF8 A1RHF9
A1RHG2 A1RHG3 A1RHG5 A1RHG7 A1RHK8 A1RHL0 A1RHL2 A1RHL9 A1RHM8 A1RHQ0 A1RHQ3

A1RHQ4 A1RHQ7 A1RHR0 A1RI33 A1RI52 A1RI66 A1RI67 A1RI98 A1RIA0 A1RIA5 A1RIA7
A1RIB0 A1RIE3 A1RIE5 A1RIH0 A1RII7 A1RIK1 A1RIK5 A1RIL5 A1RIP7 A1RIP9 A1RIQ0
A1RIQ1 A1RIQ3 A1RIQ6 A1RIR2 A1RIS2 A1RIT3 A1RIU9 A1RIV6 A1RIW8 A1RIX2 A1RIX4
A1RIX6 A1RIY0 A1RIY2 A1RIY3 A1RIZ0 A1RJ16 A1RJ17 A1RJ52 A1RJ55 A1RJ56 A1RJ58
A1RJ60 A1RJ71 A1RJ78 A1RJD1 A1RJD3 A1RJD6 A1RJF8 A1RJF9 A1RJG1 A1RJI3 A1RJI6
A1RJJ4 A1RJP0 A1RJQ2 A1RJQ3 A1RJQ4 A1RJQ6 A1RJQ8 A1RJQ9 A1RJR5 A1RJU6 A1RJV7
A1RJW6 A1RJX1 A1RJY8 A1RK12 A1RK14 A1RK52 A1RK54 A1RK75 A1RK78 A1RK88 A1RK89
A1RK91 A1RKF2 A1RKK6 A1RKK8 A1RKL0 A1RKL4 A1RKM2 A1RKQ1 A1RKQ4 A1RKQ5 A1RKS3
A1RL10 A1RL19 A1RL35 A1RL88 A1RL89 A1RL90 A1RL91 A1RL92 A1RLI0 A1RLI8 A1RLK6
A1RLL3 A1RLM4 A1RLM5 A1RLM6 A1RLM7 A1RLN0 A1RLT1 A1RLT6 A1RLU3 A1RLU8 A1RLV3
A1RLV4 A1RLV5 A1RLX5 A1RLX8 A1RLZ8 A1RM77 A1RMB2 A1RMB5 A1RMB6 A1RMB7 A1RMB8
A1RMC7 A1RME0 A1RME4 A1RME5 A1RMF2 A1RMG0 A1RMG3 A1RMG7 A1RMH9 A1RMM6 A1RMM9
A1RMN2 A1RMN6 A1RMN7 A1RMT4 A1RMU8 A1RMU9 A1RMV0 A1RMV6 A1RMV7 A1RMV8 A1RMZ5
A1RN12 A1RN54 A1RN68 A1RN69 A1RN71 A1RNE1 A1RNE2 A1RNE4 A1RNF8 A1RNI3 A1RNI6
A1RNI7 A1RNI8 A1RNI9 A1RNJ7 A1RNJ8 A1RNL3 A1RNM8 A1RNN6 A1RNN7 A1RNQ0 A1RNR2
A1RNU4 A1RNX8 A1RNY7 A1RP36 A1RP57 A1RP78 A1RP91 A1RP94 A1RP95 A1RPC1 A1RPC2
A1RPD0 A1RPD2 A1RPD5 A1RPD7 A1RPG2 A1RPI1 A1RPK5 A1RPL0 A1RPP5 A1RPP6 A1RPQ6
A1RPQ7 A1RPR5 A1RPR6 A1RPR8 A1RPS2 A1RPV7 A1RPZ7 A1RQ03 A1RQ05 A1RQ25 A1RQ57
A1RQ59 A1RQ67 A1RQ93 A1RQ95 A1RQC0 A1RQC1 A1RQE8 A1RQF1 A1RQF2 A1RQH3 A1RQK2
A1RQK4 A1RQK6 A1RQP5 A1RQP7 A1RQQ2 A1RQQ5 A1RQR5 A1RR50 A1RR73 A1RR74 A1RR78
A1RRA0 A1RRA5 A1RRE2 A1RRG7 A1RRI0 A1RRJ3 A1RRJ4 A1RRJ5 A1RRJ7 A1RRJ9 A1RRK0
A1RRK1 A1RRN0 A1RRN3 A1RRN7 A1RRN8 A1RRQ3 A1RRQ5 A1RRQ6 A1RRR1 A1RRS5 A1RRT7
A1RRU0 A1RRV1 A1RRV2 A1RRV7 A1RRV9 A1RRX4 A1RRX6 A1RRY6 A1RRZ4 A1RRZ8 A1RS05
A1RS06 A1RS16 A1RS21 A1RS24 A1RS27 A1RS32 A1RS33 A1RS34 A1RS58 A1RS86 A1RSA1
A1RSA2 A1RSA3 A1RSC6 A1RSC7 A1RSD0 A1RSD4 A1RSD7 A1RSD9 A1RSE1 A1RSE3 A1RSE4
A1RSE6 A1RSH9 A1RSI5 A1RSJ7 A1RSJ8 A1RSK3 A1RSL5 A1RSM4 A1RSN2 A1RSP7 A1RSQ1
A1RSQ5 A1RSR6 A1RSS5 A1RSS8 A1RST0 A1RSV1 A1RSY7 A1RT03 A1RT04 A1RT11 A1RT13
A1RT34 A1RT54 A1RT57 A1RT97 A1RTB8 A1RTD5 A1RTF7 A1RTG5 A1RTI0 A1RTI3 A1RTI4
A1RTI5 A1RTI7 A1RTJ3 A1RTJ4 A1RTJ7 A1RTK4 A1RTL6 A1RTL7 A1RTU8 A1RTU9 A1RTX9
A1RU12 A1RU15 A1RU25 A1RU26 A1RU36 A1RU37 A1RU38 A1RU43 A1RU67 A1RU92 A1RUA9
A1RUC0 A1RUG4 A1RUV2 A1RUV7 A1RUW1 A1RUX0 A1RUX2 A1RUZ2 A1RV13 A1RV26 A1RV28
A1RV37 A1RV38 A1RV54 A1RV74 A1RV79 A1RV82 A1RV83 A1RV98 A1RVA1 A1RVD1 A1RVD2
A1RVE1 A1RVG3 A1RVH0 A1RVH3 A1RVH4 A1RVI0 A1RVJ1 A1RVN1 A1RVN4 A1RVP3 A1RVQ8
A1RVT0 A1RVT1 A1RVT3 A1RVT4 A1RVX2 A1RW13 A1RW47 A1RWC3 A1RWC8 A1RWQ1 A1RWQ3
A1RWQ5 A1RWQ7 A1RWR4 A1RWR5 A1RWR6 A1RWR7 A1RWS0 A1RWS4 A1RWT7 A1RWT8 A1RWT9
A1RWU6 A1RWU7 A1RWV4 A1RWW1 A1RWW7 A1RWX0 A1RWX6 A1RWY2 A1RWY6 A1RX33 A1RX40
A1RX59 A1RX71 A1RX88 A1RX99 A1RXA6 A1RXA9 A1RXD1 A1RXE1 A1RXF0 A1RXF3 A1RXG2
A1RXG4 A1RXG6 A1RXG7 A1RXH2 A1RXH6 A1RXH7 A1RXH8 A1RXJ1 A1RXJ4 A1RXM2 A1RXN2
A1RXN7 A1RXQ3 A1RXQ7 A1RXS9 A1RXT9 A1RXU2 A1RXU8 A1RXV1 A1RXW9 A1RXY3 A1RY04
A1RY08 A1RY17 A1RY30 A1RY31 A1RY32 A1RY70 A1RY72 A1RY76 A1RYD7 A1RYD8 A1RYE4
A1RYM0 A1RYM3 A1RYS1 A1RYX4 A1RYX5 A1RZ22 A1RZC5 A1RZC6 A1RZN3 A1RZT8 A1RZT9
A1S0B0 A1S0R2 A1S0T1 A1S1G4 A1S1G7 A1S1G8 A1S1G9 A1S1H1 A1S1I1 A1S1I2 A1S1I9
A1S1J8 A1S1K2 A1S1K5 A1S1K6 A1S1K8 A1S1L6 A1S1L8 A1S1N0 A1S1N6 A1S1P1 A1S1P8
A1S1Q3 A1S1S3 A1S1S7 A1S1X0 A1S1Z4 A1S1Z8 A1S203 A1S204 A1S207 A1S208 A1S209

A1S210 A1S211 A1S212 A1S213 A1S214 A1S217 A1S218 A1S219 A1S220 A1S221 A1S222
A1S223 A1S224 A1S225 A1S226 A1S227 A1S228 A1S229 A1S230 A1S231 A1S232 A1S233
A1S234 A1S235 A1S236 A1S237 A1S239 A1S240 A1S241 A1S242 A1S243 A1S244 A1S257
A1S262 A1S264 A1S265 A1S270 A1S2C5 A1S2C7 A1S2C9 A1S2D0 A1S2D2 A1S2D3 A1S2D4
A1S2E0 A1S2E2 A1S2E3 A1S2E6 A1S2F0 A1S2F1 A1S2G0 A1S2G9 A1S2J2 A1S2J9 A1S2P4
A1S2P5 A1S2P7 A1S2Q1 A1S2T9 A1S2U5 A1S2Z7 A1S321 A1S351 A1S382 A1S397 A1S3C4
A1S3I3 A1S3L3 A1S3L6 A1S3M6 A1S3M7 A1S3N0 A1S3N2 A1S3S7 A1S3S8 A1S3U0 A1S3U4
A1S3U9 A1S3V6 A1S3W3 A1S3W8 A1S3Y1 A1S3Z0 A1S3Z1 A1S3Z2 A1S3Z3 A1S3Z6 A1S413
A1S414 A1S418 A1S422 A1S425 A1S454 A1S457 A1S458 A1S460 A1S462 A1S463 A1S464
A1S465 A1S467 A1S474 A1S475 A1S476 A1S477 A1S487 A1S4B5 A1S4B6 A1S4C0 A1S4C1
A1S4D3 A1S4D6 A1S4D9 A1S4E0 A1S4E2 A1S4E3 A1S4E6 A1S4E7 A1S4E9 A1S4F1 A1S4F9
A1S4N7 A1S4P0 A1S4P1 A1S4P2 A1S4P3 A1S4P6 A1S4Q4 A1S4R2 A1S4X2 A1S4X3 A1S4X4
A1S4X5 A1S4X6 A1S528 A1S544 A1S547 A1S548 A1S550 A1S552 A1S556 A1S560 A1S562
A1S563 A1S564 A1S566 A1S572 A1S579 A1S590 A1S5C5 A1S5I0 A1S5I9 A1S5J3 A1S5P6
A1S5R7 A1S5V3 A1S5V8 A1S5Y5 A1S601 A1S605 A1S615 A1S647 A1S684 A1S699 A1S6A5
A1S6A8 A1S6C9 A1S6D1 A1S6D3 A1S6D4 A1S6D6 A1S6G4 A1S6G6 A1S6G7 A1S6H5 A1S6K0
A1S6M5 A1S6N8 A1S6N9 A1S6P0 A1S6P2 A1S6P4 A1S6P5 A1S6Q5 A1S6Q9 A1S6T0 A1S6T3
A1S6V2 A1S6X6 A1S6X8 A1S6X9 A1S6Y1 A1S6Z2 A1S6Z5 A1S6Z6 A1S700 A1S702 A1S710
A1S712 A1S716 A1S723 A1S733 A1S788 A1S792 A1S795 A1S796 A1S798 A1S7A2 A1S7A3
A1S7A4 A1S7A6 A1S7B1 A1S7C2 A1S7C8 A1S7E5 A1S7E6 A1S7I0 A1S7I2 A1S7I3 A1S7J9
A1S7K2 A1S7K4 A1S7K8 A1S7P2 A1S7P3 A1S7Q6 A1S7S0 A1S7T2 A1S7U2 A1S856 A1S858
A1S859 A1S862 A1S863 A1S866 A1S886 A1S8B8 A1S8C4 A1S8C9 A1S8D4 A1S8D5 A1S8E8
A1S8H7 A1S8I4 A1S8I6 A1S8I8 A1S8K1 A1S8K6 A1S8K7 A1S8L1 A1S8L2 A1S8M3 A1S8N5
A1S8P4 A1S8Q3 A1S8R2 A1S8R3 A1S8R5 A1S8R8 A1S8S1 A1S8S3 A1S8S8 A1S8T1 A1S8T3
A1S8T9 A1S8U0 A1S8U1 A1S8W7 A1S965 A1S966 A1S967 A1S982 A1S987 A1S9A4 A1S9A6
A1S9F3 A1S9G5 A1S9H3 A1S9H4 A1S9H5 A1S9I4 A1S9L6 A1S9M3 A1S9N4 A1S9N7 A1S9W1
A1SA21 A1SA22 A1SA23 A1SA28 A1SA53 A1SA57 A1SA58 A1SA60 A1SA65 A1SA66 A1SA67
A1SA71 A1SA87 A1SAC0 A1SAC1 A1SAC7 A1SAD4 A1SAE2 A1SAF6 A1SAJ8 A1SAL1 A1SAM0
A1SAM2 A1SAP3 A1SAP5 A1SAR6 A1SAS5 A1SAS8 A1SAW5 A1SB01 A1SB02 A1SB13 A1SB14
A1SB44 A1SB48 A1SB70 A1SBA6 A1SBE8 A1SBK3 A1SBL0 A1SBR7 A1SBR9 A1SBV0 A1SBV1
A1SCL9 A1SCM2 A1SCV3 A1SD28 A1SD34 A1SDD5 A1SDH7 A1SDH8 A1SDP0 A1SDW7 A1SDW8
A1SDX1 A1SE01 A1SE06 A1SE17 A1SE26 A1SE71 A1SED2 A1SEF9 A1SEI6 A1SEI7 A1SEI9
A1SEK1 A1SEK2 A1SEM0 A1SEM4 A1SEQ0 A1SER2 A1SET2 A1SF13 A1SF27 A1SF28 A1SF29
A1SF33 A1SFK7 A1SFT8 A1SG03 A1SG51 A1SGA2 A1SGG5 A1SGT4 A1SH77 A1SH78 A1SHH8
A1SHH9 A1SHP5 A1SHP7 A1SHV9 A1SHW4 A1SHX8 A1SHY8 A1SI05 A1SI37 A1SIB6 A1SIM3
A1SIM4 A1SJ19 A1SJ31 A1SJ39 A1SJ61 A1SJ90 A1SJA2 A1SJA3 A1SJA5 A1SJA6 A1SJA7
A1SJB1 A1SJB8 A1SJC7 A1SJC8 A1SJD2 A1SJD7 A1SJD8 A1SJE8 A1SJF3 A1SJF7 A1SJF9
A1SJG2 A1SJG9 A1SJH0 A1SJH9 A1SJI0 A1SJI9 A1SJJ3 A1SJJ4 A1SJK4 A1SJP6 A1SJP8
A1SJZ6 A1SK07 A1SK13 A1SK14 A1SK16 A1SKI3 A1SKL2 A1SKM6 A1SKY7 A1SL44 A1SL49
A1SL56 A1SL57 A1SL60 A1SL74 A1SL79 A1SL88 A1SL89 A1SLE8 A1SLG6 A1SLJ4 A1SLJ5
A1SLK6 A1SLK7 A1SLL0 A1SLL4 A1SLQ4 A1SLQ7 A1SLQ8 A1SLQ9 A1SLR0 A1SLR5 A1SLS0
A1SLS2 A1SLS4 A1SLU7 A1SLV0 A1SLV4 A1SLW4 A1SLW5 A1SLX8 A1SM03 A1SM80 A1SM84
A1SMA0 A1SMA5 A1SMB0 A1SMB4 A1SMB5 A1SMB6 A1SMD1 A1SME0 A1SME5 A1SMP3 A1SMP7
A1SMP8 A1SMQ5 A1SMU5 A1SMW1 A1SMW2 A1SMX2 A1SMY9 A1SN17 A1SNC7 A1SNF6 A1SNG0

A1SNG8 A1SNI7 A1SNI8 A1SNI9 A1SNJ0 A1SNJ1 A1SNJ2 A1SNJ3 A1SNJ7 A1SNJ9 A1SNK0
A1SNK1 A1SNK2 A1SNK3 A1SNK4 A1SNK5 A1SNK7 A1SNK8 A1SNL1 A1SNL2 A1SNL3 A1SNL4
A1SNL5 A1SNL8 A1SNL9 A1SNN2 A1SNN5 A1SNN6 A1SNN7 A1SNN8 A1SNU4 A1SNW4 A1SNY3
A1SNY6 A1SP05 A1SP12 A1SPL0 A1SPL4 A1SPN2 A1SPT8 A1SPV1 A1SPV6 A1SPW6 A1SPX5
A1SPZ7 A1SQ09 A1SQ24 A1SQH3 A1SQN1 A1SQS1 A1SQS2 A1SQS4 A1SQV5 A1SQV9 A1SQW0
A1SQW3 A1SQW4 A1SQW6 A1SQX0 A1SQZ9 A1SR01 A1SR02 A1SR12 A1SR13 A1SR14 A1SR15
A1SR17 A1SR30 A1SR31 A1SR38 A1SR48 A1SR71 A1SR75 A1SR92 A1SRB6 A1SRB7 A1SRC9
A1SRG3 A1SRG8 A1SRH8 A1SRI1 A1SRI3 A1SRN6 A1SRP6 A1SRP8 A1SRP9 A1SRR4 A1SRR5
A1SRS4 A1SRS5 A1SRT6 A1SRU1 A1SRU7 A1SRV2 A1SRX1 A1SRX3 A1SRX7 A1SRZ1 A1SS81
A1SS83 A1SS86 A1SS88 A1SS92 A1SS96 A1SSA8 A1SSB4 A1SSB5 A1SSB6 A1SSB8 A1SSC2
A1SSC4 A1SSC6 A1SSG7 A1SSG8 A1SSG9 A1SSI8 A1SSJ0 A1SSL6 A1SSM8 A1SSN0 A1SSQ6
A1SSR3 A1SST4 A1SST5 A1SST6 A1SSU3 A1SSU8 A1SSV2 A1SSV3 A1SSV5 A1SSV6 A1SSY1
A1SSY3 A1SSZ1 A1SSZ3 A1ST25 A1ST27 A1ST34 A1ST36 A1ST38 A1ST41 A1ST43 A1ST45
A1ST46 A1ST47 A1ST52 A1ST53 A1ST72 A1ST89 A1ST92 A1ST94 A1STA5 A1STC9 A1STD6
A1STD7 A1STD9 A1STE3 A1STE4 A1STE5 A1STE7 A1STF0 A1STH1 A1STI2 A1STI3 A1STI9
A1STJ9 A1STP2 A1STQ3 A1STQ6 A1STS0 A1STS1 A1STT0 A1STT2 A1STT9 A1STV0 A1STV3
A1STV6 A1STV9 A1STX3 A1STZ0 A1SU08 A1SU10 A1SU11 A1SU20 A1SU36 A1SU39 A1SU43
A1SU47 A1SU57 A1SU60 A1SU80 A1SUA7 A1SUA8 A1SUE3 A1SUE5 A1SUI4 A1SUI7 A1SUJ5
A1SUQ7 A1SUU0 A1SUU1 A1SUU5 A1SUU6 A1SUW6 A1SUW7 A1SUW8 A1SUX2 A1SUX5 A1SUY2
A1SUY9 A1SV84 A1SV86 A1SV88 A1SV89 A1SV95 A1SVA5 A1SVA7 A1SVA9 A1SVC5 A1SVD0
A1SVD5 A1SVE4 A1SVE6 A1SVE7 A1SVE8 A1SVJ9 A1SVN9 A1SVW5 A1SVX2 A1SVX4 A1SW15
A1SW31 A1SW33 A1SW72 A1SW90 A1SW94 A1SW96 A1SWB8 A1SWF0 A1SWM1 A1SWP5 A1SWQ3
A1SWQ8 A1SWR4 A1SWS4 A1SWS5 A1SWU0 A1SWU2 A1SWU5 A1SWV0 A1SWV3 A1SWV9 A1SWW0
A1SWW4 A1SWW6 A1SWX6 A1SWY4 A1SWY7 A1SWZ6 A1SWZ8 A1SX02 A1SX04 A1SX09 A1SX20
A1SX22 A1SX24 A1SX61 A1SX95 A1SXA4 A1SXF9 A1SXK4 A1SXW4 A1SXW5 A1SXW6 A1SXW7
A1SXW8 A1SY50 A1SY66 A1SY73 A1SY74 A1SY89 A1SY91 A1SYB5 A1SYB8 A1SYD3 A1SYG9
A1SYH7 A1SYJ9 A1SYK3 A1SYK4 A1SYK5 A1SYL6 A1SYM0 A1SYN3 A1SYT6 A1SYT9 A1SYU9
A1SYV7 A1SYW0 A1SYW1 A1SYW2 A1SYW3 A1SYX8 A1SYX9 A1SYY0 A1SYY1 A1SYY2 A1SYY3
A1SYY4 A1SYZ6 A1SYZ9 A1SZ16 A1SZ17 A1SZ19 A1SZ24 A1SZ30 A1SZA2 A1SZA3 A1SZE1
A1SZG4 A1SZG8 A1SZH4 A1SZH7 A1SZI0 A1SZI3 A1SZI8 A1SZJ2 A1SZJ4 A1SZJ9 A1SZK6
A1SZL0 A1SZL1 A1SZL2 A1SZP0 A1SZP3 A1SZP6 A1SZR1 A1SZR6 A1SZR7 A1SZR8 A1SZR9
A1SZU8 A1SZY2 A1SZY5 A1SZY6 A1SZY7 A1SZY8 A1SZZ9 A1T001 A1T004 A1T007 A1T026
A1T038 A1T040 A1T041 A1T047 A1T054 A1T056 A1T058 A1T059 A1T064 A1T065 A1T066
A1T067 A1T068 A1T086 A1T097 A1T099 A1T0B7 A1T0B8 A1T0C3 A1T0C4 A1T0C5 A1T0C6
A1T0C7 A1T0C8 A1T0C9 A1T0D0 A1T0D1 A1T0D2 A1T0D4 A1T0D5 A1T0D6 A1T0D7 A1T0D8
A1T0D9 A1T0E0 A1T0E1 A1T0E2 A1T0E3 A1T0H3 A1T0M6 A1T0M7 A1T0N0 A1T0P0 A1T0P2
A1T0R0 A1T0U7 A1T0X4 A1T0X6 A1T0Y5 A1T0Y6 A1T0Z9 A1T100 A1T102 A1T105 A1T165
A1T1A5 A1T1J8 A1T1K1 A1T1P0 A1T225 A1T235 A1T273 A1T297 A1T2M8 A1T2N5 A1T2N6
A1T2S3 A1T2W5 A1T2Y9 A1T2Z5 A1T320 A1T324 A1T3B5 A1T3D2 A1T3D3 A1T3F2 A1T3J8
A1T3R3 A1T3S1 A1T3T5 A1T4G4 A1T4H0 A1T4H1 A1T4I7 A1T4I8 A1T4J2 A1T4J3 A1T4L3
A1T4L4 A1T4L5 A1T4L6 A1T4N8 A1T4N9 A1T4P1 A1T4P2 A1T4P3 A1T4P4 A1T4P5 A1T4P6
A1T4P7 A1T4R6 A1T4R7 A1T4R9 A1T4S0 A1T4S1 A1T4S4 A1T4S8 A1T4S9 A1T4T0 A1T4T2
A1T515 A1T516 A1T517 A1T518 A1T519 A1T520 A1T521 A1T551 A1T554 A1T566 A1T571
A1T576 A1T577 A1T5D0 A1T5G2 A1T5H1 A1T5Q3 A1T5Z3 A1T639 A1T6B0 A1T6D1 A1T6D6

A1T6P2 A1T6Q8 A1T6X1 A1T6X7 A1T6X9 A1T6Y0 A1T6Y9 A1T6Z4 A1T6Z8 A1T700 A1T701
A1T705 A1T708 A1T717 A1T726 A1T732 A1T737 A1T739 A1T741 A1T752 A1T754 A1T757
A1T759 A1T766 A1T773 A1T774 A1T784 A1T785 A1T787 A1T7G4 A1T7H2 A1T7H5 A1T7H8
A1T7H9 A1T7L3 A1T7L4 A1T7N8 A1T7P4 A1T7Q0 A1T7T3 A1T7T4 A1T7U2 A1T7U7 A1T7U8
A1T7V6 A1T7Y0 A1T7Z0 A1T7Z4 A1T868 A1T876 A1T879 A1T882 A1T883 A1T884 A1T894
A1T898 A1T8E8 A1T8F3 A1T8F5 A1T8F9 A1T8G0 A1T8G7 A1T8H1 A1T8H2 A1T8H5 A1T8H7
A1T8J4 A1T8K1 A1T8K2 A1T8K4 A1T8K5 A1T8K7 A1T8L1 A1T8L2 A1T8R1 A1T8R2 A1T8U6
A1T8U7 A1T8V0 A1T8W2 A1T8W3 A1T8W5 A1T8W7 A1T8W8 A1T8X3 A1T903 A1T9D9 A1T9F4
A1T9L9 A1T9N2 A1T9N3 A1T9N4 A1T9N5 A1T9N6 A1T9P9 A1T9T8 A1TA09 A1TA72 A1TA81
A1TAA5 A1TAA6 A1TAB0 A1TAB5 A1TAC3 A1TAC4 A1TAP0 A1TAP3 A1TAP4 A1TAQ3 A1TAR6
A1TAS3 A1TAW2 A1TAW7 A1TAX0 A1TAX6 A1TAX7 A1TB00 A1TB25 A1TBP6 A1TBP9 A1TBR2
A1TBR8 A1TBY9 A1TC01 A1TC03 A1TC11 A1TC20 A1TC21 A1TC22 A1TC25 A1TC40 A1TC97
A1TCB3 A1TCB8 A1TCX1 A1TCX2 A1TD14 A1TD35 A1TD66 A1TD67 A1TD73 A1TD77 A1TDB7
A1TDB8 A1TDE9 A1TDH1 A1TDK2 A1TDM2 A1TDM8 A1TDR0 A1TDS4 A1TE06 A1TE33 A1TE43
A1TE53 A1TEG1 A1TEH2 A1TEH3 A1TEL8 A1TEP3 A1TEQ0 A1TEV2 A1TF46 A1TF55 A1TFB7
A1TFC6 A1TFE0 A1TFF9 A1TFL3 A1TFQ5 A1TFQ6 A1TFU3 A1TFU9 A1TG04 A1TG08 A1TG09
A1TG10 A1TG33 A1TG34 A1TG35 A1TG42 A1TGD7 A1TGF9 A1TGG0 A1TGG1 A1TGJ3 A1TGJ4
A1TGS6 A1TGX4 A1TGX7 A1TGZ7 A1TH24 A1TH50 A1THY4 A1THY5 A1TI06 A1TI09 A1TI25
A1TI29 A1TI30 A1TI78 A1TI79 A1TI97 A1TIF0 A1TIG3 A1TIS2 A1TIW2 A1TJ02 A1TJ03
A1TJ04 A1TJ05 A1TJ06 A1TJ07 A1TJ08 A1TJ10 A1TJ11 A1TJ12 A1TJ13 A1TJ14 A1TJ15
A1TJ16 A1TJ31 A1TJ32 A1TJ83 A1TJA3 A1TJQ1 A1TJQ2 A1TJR8 A1TJS7 A1TJS8 A1TJS9
A1TJT0 A1TJT1 A1TJT2 A1TJT3 A1TJT4 A1TJT5 A1TJT6 A1TJT8 A1TJT9 A1TJU0 A1TJU1
A1TJU2 A1TJU3 A1TJU6 A1TKA9 A1TKC1 A1TKC3 A1TKD2 A1TKD3 A1TKD9 A1TKE3 A1TKF1
A1TKI5 A1TKK8 A1TKL7 A1TKM0 A1TKQ3 A1TKQ5 A1TKQ6 A1TKT9 A1TKV4 A1TKW2 A1TKY2
A1TKY3 A1TKZ0 A1TKZ1 A1TKZ3 A1TL06 A1TL07 A1TL08 A1TL40 A1TL44 A1TL98 A1TLA3
A1TLB1 A1TLB7 A1TLC1 A1TLC4 A1TLC6 A1TLD5 A1TLF2 A1TLG8 A1TLH0 A1TLH5 A1TLH6
A1TLH8 A1TLH9 A1TLI0 A1TLI1 A1TLI3 A1TLI4 A1TLL1 A1TLL2 A1TLL4 A1TLP7 A1TLT0
A1TLY4 A1TM23 A1TM24 A1TM31 A1TM32 A1TM36 A1TM37 A1TM53 A1TM61 A1TM62 A1TM63
A1TMA7 A1TMF3 A1TMP4 A1TMP7 A1TN69 A1TN70 A1TN71 A1TN72 A1TN83 A1TN89 A1TNA0
A1TNA2 A1TNA3 A1TNB1 A1TND0 A1TND1 A1TND2 A1TND3 A1TND7 A1TND9 A1TNF7 A1TNJ5
A1TNR1 A1TNU7 A1TNY6 A1TP41 A1TP96 A1TP97 A1TPC1 A1TPK8 A1TPW5 A1TPY4 A1TPZ6
A1TQ19 A1TQ21 A1TQ53 A1TQ96 A1TQ98 A1TQC1 A1TQF0 A1TQF3 A1TQI0 A1TQI7 A1TQY3
A1TR21 A1TR35 A1TR36 A1TR37 A1TR39 A1TRD3 A1TRE0 A1TRE4 A1TRG1 A1TRG5 A1TRH1
A1TRH2 A1TRH4 A1TRL2 A1TRL4 A1TRM3 A1TRM6 A1TRM7 A1TRN6 A1TRP8 A1TRT1 A1TRT4
A1TRT6 A1TRV8 A1TS23 A1TS27 A1TS63 A1TSA0 A1TSA3 A1TSE0 A1TSJ4 A1TSK2 A1TSK3
A1TSK5 A1TSN1 A1TSP7 A1TSP8 A1TSQ7 A1TSX9 A1TSY0 A1TSZ3 A1TSZ5 A1TSZ6 A1TSZ8
A1TSZ9 A1TT41 A1TT47 A1TT56 A1TT64 A1TT66 A1TT72 A1TT74 A1TT75 A1TT99 A1TTC3
A1TTC4 A1TTD0 A1TTD1 A1TTD2 A1TTD3 A1TTD4 A1TTD5 A1TTM6 A1TTN2 A1TTU1 A1TTV0
A1TTV9 A1TTW0 A1TTW2 A1TTW5 A1TUF2 A1TUF5 A1TUF6 A1TUG3 A1TUP2 A1TUS2 A1TUT0
A1TUT1 A1TUV3 A1TUW0 A1TUY3 A1TUY9 A1TUZ2 A1TUZ4 A1TUZ5 A1TV10 A1TVC1 A1TVM0
A1TVN0 A1TVP2 A1TVS9 A1TVT0 A1TVT1 A1TVT2 A1TVT3 A1TVT4 A1TVU4 A1TVU7 A1TVV5
A1TW81 A1TW87 A1TW90 A1TW93 A1TWB8 A1TWF5 A1TWI0 A1TWI4 A1TWI9 A1TWJ0 A1TWJ3
A1TWM6 A1TWN0 A1TWN4 A1TWN6 A1TWU4 A1TWZ3 A1TXB6 A1TXM4 A1TXT7 A1TXW9 A1TY02
A1TY08 A1TY14 A1TY28 A1TY30 A1TYB4 A1TYD9 A1TYE0 A1TYE5 A1TYE6 A1TYF4 A1TYG5

A1TYG6 A1TYH0 A1TYH3 A1TYI2 A1TYI6 A1TYI7 A1TYI8 A1TYI9 A1TYJ0 A1TYJ1 A1TYJ2
A1TYJ3 A1TYJ4 A1TYJ5 A1TYJ6 A1TYJ7 A1TYJ8 A1TYJ9 A1TYK0 A1TYK1 A1TYK2 A1TYK3
A1TYK4 A1TYK5 A1TYK6 A1TYK7 A1TYK8 A1TYK9 A1TYL0 A1TYL1 A1TYL2 A1TYL3 A1TYL4
A1TYL5 A1TYL6 A1TYL8 A1TYL9 A1TYM0 A1TYM1 A1TYM2 A1TYM3 A1TYQ0 A1TYR4 A1TYR6
A1TYT7 A1TYT9 A1TYU2 A1TYU5 A1TYU6 A1TYU8 A1TYV1 A1TYW3 A1TYW8 A1TYW9 A1TYX3
A1TYY2 A1TYY3 A1TYY4 A1TYY6 A1TYY9 A1TYZ2 A1TZ11 A1TZ34 A1TZ35 A1TZ36 A1TZ46
A1TZ51 A1TZ52 A1TZ53 A1TZ54 A1TZ66 A1TZ68 A1TZ73 A1TZ92 A1TZ94 A1TZA3 A1TZF1
A1TZF4 A1TZG2 A1TZH7 A1TZH9 A1TZN6 A1TZN7 A1TZP6 A1TZP7 A1TZQ0 A1TZQ4 A1TZS9
A1TZT9 A1TZU2 A1TZV5 A1U0K1 A1U0K2 A1U0R2 A1U0U9 A1U0V1 A1U0W2 A1U0X2 A1U0X4
A1U0X9 A1U0Y0 A1U0Y3 A1U123 A1U124 A1U126 A1U127 A1U1A4 A1U1A6 A1U1B1 A1U1B9
A1U1C0 A1U1C1 A1U1C2 A1U1E2 A1U1F0 A1U1F4 A1U1G9 A1U1H0 A1U1H1 A1U1H5 A1U1I8
A1U1J4 A1U1K6 A1U1P2 A1U1Q2 A1U1Q4 A1U1Q5 A1U1Q6 A1U1Q7 A1U1T3 A1U1U7 A1U1V3
A1U1Y1 A1U1Y5 A1U204 A1U241 A1U260 A1U292 A1U293 A1U2B7 A1U2B9 A1U2C0 A1U2C1
A1U2C3 A1U2D4 A1U2E3 A1U2E8 A1U2K1 A1U2L4 A1U2M9 A1U2P6 A1U2S9 A1U2T9 A1U2U2
A1U2U4 A1U2U9 A1U2V3 A1U2Y6 A1U2Y8 A1U2Y9 A1U324 A1U326 A1U339 A1U342 A1U344
A1U360 A1U367 A1U368 A1U371 A1U373 A1U374 A1U380 A1U388 A1U390 A1U392 A1U393
A1U3B4 A1U3C0 A1U3C2 A1U3C3 A1U3D8 A1U3F6 A1U3F7 A1U3G0 A1U3G6 A1U3G7 A1U3H0
A1U3H7 A1U3I4 A1U3J9 A1U3K1 A1U3K5 A1U3L1 A1U3L2 A1U3P0 A1U3Q2 A1U3Q3 A1U3Q4
A1U3R0 A1U3R1 A1U3U4 A1U3X8 A1U3Y0 A1U458 A1U459 A1U474 A1U485 A1U486 A1U487
A1U488 A1U490 A1U494 A1U496 A1U4A0 A1U4A9 A1U4B1 A1U4B2 A1U4B8 A1U4B9 A1U4C0
A1U4C1 A1U4C5 A1U4C6 A1U4D2 A1U4D3 A1U4H0 A1U4L8 A1U4P7 A1U4R4 A1U4Z6 A1U4Z7
A1U4Z8 A1U4Z9 A1U500 A1U501 A1U543 A1U547 A1U560 A1U578 A1U5D9 A1U5G5 A1U5G9
A1U5H5 A1U5H6 A1U5H8 A1U5I2 A1U5W7 A1U5W8 A1U5Z6 A1U5Z7 A1U5Z8 A1U5Z9 A1U600
A1U602 A1U604 A1U605 A1U613 A1U614 A1U622 A1U670 A1U671 A1U689 A1U695 A1U698
A1U6F3 A1U6F8 A1U6F9 A1U6H0 A1U6K2 A1U6L0 A1U6L4 A1U6L5 A1U6L6 A1U6M5 A1U6P4
A1U6R9 A1U6U9 A1U749 A1U754 A1U761 A1U763 A1U7G1 A1U7I4 A1U7I5 A1U7J3 A1U7J6
A1U7J7 A1U8R4 A1U8R8 A1U8R9 A1U8S0 A1U8S3 A1U908 A1U909 A1U992 A1U995 A1U9D7
A1U9T8 A1U9V4 A1U9Z8 A1UA31 A1UA84 A1UAA8 A1UAB4 A1UAD9 A1UAE3 A1UAF0 A1UAM4
A1UAM8 A1UAP7 A1UAR1 A1UAX3 A1UAY5 A1UBE0 A1UBE6 A1UBE7 A1UBJ0 A1UBJ3 A1UBJ4
A1UBK8 A1UBK9 A1UBL0 A1UBL1 A1UBN5 A1UBN6 A1UBN7 A1UBN8 A1UBN9 A1UBP0 A1UBP1
A1UBP2 A1UBP3 A1UBP4 A1UBP8 A1UBP9 A1UBQ0 A1UBQ1 A1UBQ2 A1UBQ3 A1UBQ6 A1UBR5
A1UBR6 A1UBR7 A1UBR9 A1UBY0 A1UBY1 A1UBY2 A1UBY3 A1UBY4 A1UBY5 A1UBY6 A1UC03
A1UC06 A1UC23 A1UC26 A1UC75 A1UC78 A1UCA4 A1UCG3 A1UCJ5 A1UCK8 A1UCN4 A1UDD2
A1UDD7 A1UDY9 A1UE06 A1UE63 A1UE68 A1UE81 A1UE95 A1UE98 A1UEA6 A1UEA8 A1UEA9
A1UEB3 A1UEB6 A1UEC0 A1UEC5 A1UEC6 A1UED2 A1UED7 A1UED9 A1UEF2 A1UEF4 A1UEF7
A1UEF9 A1UEH2 A1UEH7 A1UEI0 A1UEI1 A1UEJ0 A1UEJ1 A1UEJ3 A1UEN7 A1UEQ5 A1UER2
A1UER5 A1UER8 A1UER9 A1UET1 A1UET5 A1UET6 A1UEU3 A1UEU8 A1UEV1 A1UEY3 A1UEY4
A1UEZ3 A1UEZ8 A1UEZ9 A1UF06 A1UF27 A1UF44 A1UF48 A1UF79 A1UF87 A1UF88 A1UFA2
A1UFA3 A1UFA4 A1UFB6 A1UFE5 A1UFI3 A1UFI4 A1UFI9 A1UFJ5 A1UFJ6 A1UFK0 A1UFK4
A1UFL0 A1UFM0 A1UFM8 A1UFN2 A1UFN3 A1UFN5 A1UFP0 A1UFP1 A1UFU2 A1UFV1 A1UGG4
A1UGI7 A1UGT3 A1UGT5 A1UGT6 A1UGT7 A1UGU7 A1UGX5 A1UH74 A1UH85 A1UH86 A1UH95
A1UH96 A1UHA0 A1UHA9 A1UHB0 A1UHJ6 A1UHK1 A1UHK2 A1UHK4 A1UHK6 A1UHK7 A1UHL4
A1UHL9 A1UHM2 A1UHM3 A1UHS6 A1UHS7 A1UHT0 A1UHU3 A1UHU4 A1UHV1 A1UI48 A1UI53
A1UI62 A1UI63 A1UI88 A1UIA8 A1UIB3 A1UIB4 A1UID0 A1UIP2 A1UIP9 A1UIQ4 A1UIQ9

A1UIW4 A1UIW8 A1UIY9 A1UIZ7 A1UJ07 A1UJ08 A1UJ09 A1UJ35 A1UJ38 A1UJ99 A1UJP3
A1UJP4 A1UJV2 A1UJW6 A1UJW9 A1UJZ5 A1UJZ6 A1UK02 A1UK06 A1UK27 A1UK81 A1UKA3
A1UKE3 A1UKF3 A1UKL5 A1UKL7 A1UKP2 A1UKU2 A1UL23 A1UL32 A1UL71 A1UL88 A1UL96
A1ULE3 A1ULM8 A1ULN4 A1ULP7 A1ULU4 A1ULV3 A1ULY1 A1UM30 A1UM80 A1UM81 A1UME8
A1UMF4 A1UMF5 A1UMH8 A1UMH9 A1UMI0 A1UMX3 A1UMX4 A1UN51 A1UN98 A1UNA3 A1UNC5
A1UNG1 A1UNJ0 A1UNS0 A1UP83 A1UP84 A1UPA5 A1UQA0 A1UQA1 A1UQU2 A1UQU4 A1UQU6
A1UQU7 A1UQU8 A1UQV5 A1UQV8 A1UQV9 A1UQX3 A1UQY9 A1UQZ7 A1UR04 A1UR09 A1UR12
A1UR20 A1UR21 A1UR22 A1UR36 A1UR39 A1UR41 A1UR54 A1UR57 A1UR66 A1UR75 A1UR84
A1UR85 A1UR86 A1UR93 A1UR96 A1UR98 A1URA6 A1URB3 A1URD8 A1URE4 A1URK5 A1URL6
A1URM1 A1URM5 A1URM6 A1URM7 A1URN3 A1URN4 A1URN6 A1URN9 A1URR0 A1URS1 A1URS5
A1URT5 A1URU0 A1URV3 A1URV4 A1URW0 A1URW3 A1URW5 A1URW6 A1URX2 A1URX4 A1US08
A1US11 A1US27 A1US28 A1US30 A1US32 A1US36 A1US38 A1US39 A1US40 A1US43 A1US53
A1US55 A1US56 A1US65 A1US76 A1US81 A1US90 A1US92 A1USA0 A1USA2 A1USA7 A1USA8
A1USB2 A1USB3 A1USB4 A1USC1 A1USC4 A1USC5 A1USC6 A1USC7 A1USC8 A1USC9 A1USD9
A1USF5 A1USF7 A1USG2 A1USH6 A1USI0 A1USI1 A1USI4 A1USJ0 A1USJ2 A1USJ5 A1USK9
A1USL0 A1USL2 A1USL4 A1USL7 A1USL8 A1USM0 A1USM1 A1USM4 A1USM5 A1USQ6 A1USQ7
A1USQ8 A1USQ9 A1USR0 A1USR1 A1USR2 A1USR3 A1USR5 A1USR6 A1USR7 A1USR8 A1USR9
A1USS3 A1USS4 A1USU3 A1USY3 A1UT10 A1UT11 A1UT12 A1UT20 A1UT21 A1UT22 A1UT24
A1UT31 A1UT47 A1UT52 A1UT80 A1UT83 A1UT84 A1UT97 A1UTA5 A1UTB5 A1UTC2 A1UTC3
A1UTC4 A1UTD3 A1UTE7 A1UTE8 A1UTG0 A1UTH4 A1UTH5 A1UTJ1 A1UTK7 A1UTL4 A1UTL8
A1UTM1 A1UTM3 A1UTM4 A1UTQ0 A1UTQ4 A1UTQ7 A1UTX7 A1UTX8 A1UTY3 A1UU01 A1UU20
A1UU35 A1UU36 A1UU39 A1UU41 A1UU46 A1UU47 A1UU50 A1UU51 A1UU52 A1UU53 A1UU54
A1UU56 A1UU59 A1UU72 A1UU77 A1UU92 A1UU95 A1UU99 A1UUA2 A1UUA8 A1UUA9 A1UUB0
A1UUB4 A1UUB5 A1UUB6 A1UUB8 A1UUC1 A1UUC3 A1UUC9 A1UUD2 A1UUD5 A1UUE1 A1UUF5
A1UUF7 A1UUS6 A1UVC0 A1UVN3 A1UVS4 A1UW43 A1UW99 A1UWA0 A1UWC8 A1UWC9 A1UWN5
A1UXP8 A1UY21 A1UYM9 A1UYQ2 A1UYQ5 A1UZ10 A1UZ32 A1UZ33 A1UZ38 A1UZ42 A1UZE5
A1UZH3 A1UZL8 A1UZM8 A1UZM9 A1UZN2 A1UZN5 A1UZN7 A1UZN8 A1UZP1 A1UZX6 A1UZX9
A1V016 A1V017 A1V034 A1V051 A1V052 A1V053 A1V060 A1V064 A1V065 A1V068 A1V076
A1V0A3 A1V0A6 A1V0B1 A1V0G7 A1V0H0 A1V0I2 A1V0M1 A1V0N6 A1V0N7 A1V0P0 A1V0P1
A1V0P2 A1V0P3 A1V0Q3 A1V0R6 A1V0R7 A1V0S0 A1V0S6 A1V0S7 A1V0T9 A1V0U1 A1V0U4
A1V0U6 A1V0U8 A1V0V0 A1V0V7 A1V0V9 A1V115 A1V117 A1V120 A1V193 A1V194 A1V195
A1V1A2 A1V1B6 A1V1B7 A1V1C0 A1V1G3 A1V1G4 A1V1G5 A1V1G6 A1V1G7 A1V1G8 A1V1H0
A1V1K4 A1V1K5 A1V1L0 A1V1N6 A1V1P0 A1V1P2 A1V1Q6 A1V1S2 A1V1U3 A1V1Z9 A1V203
A1V204 A1V206 A1V209 A1V292 A1V2A0 A1V2C2 A1V2D0 A1V2F7 A1V2G0 A1V2G2 A1V2G4
A1V2G6 A1V2K2 A1V2L1 A1V2L2 A1V2P7 A1V2S9 A1V319 A1V322 A1V3F2 A1V3G1 A1V3K9
A1V3N3 A1V3N4 A1V3N6 A1V3R0 A1V3R1 A1V3R3 A1V4A4 A1V4B2 A1V4H3 A1V4I9 A1V4J0
A1V4J5 A1V4K0 A1V4K1 A1V4K3 A1V4K4 A1V4L2 A1V4L3 A1V4L5 A1V4L6 A1V4M5 A1V4Q8
A1V4Q9 A1V4R1 A1V4T2 A1V4U8 A1V4V8 A1V4X0 A1V4X1 A1V4X2 A1V4Z9 A1V500 A1V517
A1V534 A1V545 A1V552 A1V554 A1V563 A1V566 A1V567 A1V568 A1V569 A1V572 A1V575
A1V582 A1V584 A1V591 A1V5A8 A1V5B3 A1V5G8 A1V5I2 A1V5K4 A1V5L6 A1V5M2 A1V5M3
A1V5P2 A1V5R9 A1V5U0 A1V5U2 A1V5V1 A1V5V5 A1V5W0 A1V5W8 A1V5W9 A1V5Z0 A1V5Z5
A1V674 A1V675 A1V6A8 A1V6B0 A1V6D3 A1V6I4 A1V6I6 A1V6J1 A1V6J2 A1V6J3 A1V6J4
A1V6K0 A1V6L4 A1V6M0 A1V6M2 A1V6R3 A1V6R5 A1V6U0 A1V6U2 A1V6U5 A1V6V5 A1V6W8
A1V713 A1V727 A1V745 A1V753 A1V760 A1V775 A1V776 A1V777 A1V788 A1V7D3 A1V7D7

A1V7D8 A1V7D9 A1V7J8 A1V7K1 A1V7K6 A1V7L3 A1V7L5 A1V7N8 A1V7S0 A1V7S8 A1V7X3
A1V7Z5 A1V817 A1V818 A1V819 A1V827 A1V839 A1V876 A1V877 A1V878 A1V879 A1V880
A1V881 A1V882 A1V884 A1V885 A1V886 A1V887 A1V888 A1V889 A1V890 A1V891 A1V892
A1V893 A1V894 A1V895 A1V896 A1V897 A1V898 A1V899 A1V8A0 A1V8A1 A1V8A2 A1V8A3
A1V8A4 A1V8A5 A1V8A7 A1V8A8 A1V8B1 A1V8B2 A1V8B3 A1V8B5 A1V8B6 A1V8C9 A1V8D7
A1V8G8 A1V8H1 A1V8H4 A1V8H5 A1V8H6 A1V8H7 A1V8N7 A1V8S5 A1V8U2 A1V8U3 A1V8X0
A1V8X2 A1V8Z2 A1V905 A1V915 A1V9B1 A1V9B4 A1V9B8 A1V9E1 A1V9F1 A1V9I9 A1V9L9
A1V9N9 A1V9Q2 A1V9Q3 A1V9Q4 A1V9S2 A1V9U1 A1V9V1 A1V9X6 A1V9Z2 A1VA10 A1VA24
A1VA26 A1VA64 A1VAA0 A1VAE6 A1VAE7 A1VAI9 A1VAJ5 A1VAJ6 A1VAJ7 A1VAJ9 A1VAK0
A1VAK3 A1VAK4 A1VAK8 A1VAL0 A1VAL8 A1VAM1 A1VAN0 A1VAN1 A1VB03 A1VB97 A1VBA3
A1VBA6 A1VBB4 A1VBB5 A1VBB6 A1VBB7 A1VBB8 A1VBC3 A1VBD2 A1VBD3 A1VBD9 A1VBE0
A1VBE6 A1VBE9 A1VBJ1 A1VBJ2 A1VBJ4 A1VBV4 A1VBW6 A1VBY0 A1VC40 A1VC41 A1VC42
A1VC43 A1VC44 A1VC53 A1VC54 A1VC77 A1VC86 A1VC99 A1VCE4 A1VCE6 A1VCP5 A1VCP9
A1VCQ0 A1VCR9 A1VCU3 A1VCV9 A1VCW7 A1VCW8 A1VCW9 A1VCX0 A1VCX6 A1VCX9 A1VCY2
A1VCZ9 A1VD15 A1VD23 A1VD24 A1VD34 A1VD37 A1VD65 A1VD68 A1VD83 A1VDB2 A1VDC3
A1VDC5 A1VDD3 A1VDD9 A1VDK3 A1VDL2 A1VDM6 A1VDM7 A1VDN0 A1VDN2 A1VDN4 A1VDQ5
A1VDQ7 A1VDQ8 A1VDR0 A1VDR1 A1VDU9 A1VDV0 A1VDW3 A1VDW4 A1VDW9 A1VDX6 A1VDX8
A1VE01 A1VE02 A1VE41 A1VE69 A1VE71 A1VE74 A1VE84 A1VE85 A1VE86 A1VE87 A1VE89
A1VE90 A1VE91 A1VE92 A1VE93 A1VE94 A1VE97 A1VE98 A1VE99 A1VEA0 A1VEA1 A1VEA2
A1VEA3 A1VEA4 A1VEA5 A1VEA6 A1VEA7 A1VEA8 A1VEA9 A1VEB0 A1VEB1 A1VEB2 A1VEB3
A1VEB4 A1VEB5 A1VEB6 A1VEB7 A1VEB9 A1VEC0 A1VEC1 A1VED4 A1VEG0 A1VEG3 A1VEG5
A1VEI5 A1VEJ7 A1VEJ9 A1VEK5 A1VEL0 A1VEL1 A1VEL2 A1VEL6 A1VEP0 A1VEP9 A1VEQ0
A1VEQ1 A1VEQ5 A1VEQ6 A1VER5 A1VES0 A1VEV0 A1VEV3 A1VEV5 A1VEW4 A1VEW5 A1VEW8
A1VF04 A1VF25 A1VF28 A1VF29 A1VF30 A1VF31 A1VF55 A1VF56 A1VF57 A1VF58 A1VF73
A1VF78 A1VF79 A1VF83 A1VF85 A1VF90 A1VFA8 A1VFA9 A1VFB1 A1VFB2 A1VFB6 A1VFE2
A1VFE4 A1VFE6 A1VFE7 A1VFE8 A1VFF3 A1VFG1 A1VFG4 A1VFG6 A1VFG8 A1VFG9 A1VFH0
A1VFP3 A1VFP6 A1VFS4 A1VFZ7 A1VG02 A1VG69 A1VG70 A1VG80 A1VG83 A1VG87 A1VG88
A1VGB0 A1VGB6 A1VGE5 A1VGE6 A1VGU5 A1VGV4 A1VGV6 A1VGY9 A1VGZ0 A1VH35 A1VHE4
A1VHE5 A1VHH2 A1VHK1 A1VHP4 A1VI55 A1VI67 A1VIB1 A1VIB2 A1VIH8 A1VIL5 A1VIL9
A1VIP5 A1VIP6 A1VIP8 A1VIP9 A1VIQ0 A1VIQ1 A1VIQ2 A1VIQ3 A1VIQ4 A1VIQ5 A1VIQ6
A1VIQ7 A1VIQ8 A1VIQ9 A1VIT9 A1VIU0 A1VIW8 A1VJ15 A1VJ23 A1VJ25 A1VJ26 A1VJ27
A1VJ28 A1VJ29 A1VJ30 A1VJ31 A1VJ32 A1VJ33 A1VJ34 A1VJ36 A1VJ37 A1VJ38 A1VJ39
A1VJ40 A1VJ41 A1VJ58 A1VJ84 A1VJD7 A1VJD8 A1VJF1 A1VJG0 A1VJJ3 A1VJJ4 A1VJJ5
A1VJK0 A1VJK1 A1VJK5 A1VJK9 A1VJX7 A1VJZ5 A1VJZ6 A1VJZ8 A1VJZ9 A1VK01 A1VK35
A1VK36 A1VK38 A1VK39 A1VK41 A1VK42 A1VK43 A1VK86 A1VK87 A1VK89 A1VK90 A1VK91
A1VK92 A1VKB6 A1VKB7 A1VKF1 A1VKF6 A1VKF9 A1VKN7 A1VKN9 A1VKP2 A1VKP5 A1VKP7
A1VKP8 A1VKR6 A1VKR9 A1VKS2 A1VKS3 A1VKS5 A1VKS9 A1VKW0 A1VKW2 A1VKW3 A1VKW4
A1VKW6 A1VKX0 A1VKZ3 A1VL56 A1VL69 A1VL71 A1VL72 A1VL83 A1VL96 A1VLE3 A1VLH3
A1VLZ6 A1VM24 A1VM52 A1VM55 A1VM56 A1VM97 A1VM98 A1VMA0 A1VMA8 A1VMB2 A1VMD5
A1VMD7 A1VMG1 A1VMG2 A1VMG3 A1VMH6 A1VMK3 A1VMN2 A1VMR1 A1VMV7 A1VMW5 A1VN04
A1VN06 A1VN09 A1VN11 A1VN15 A1VN17 A1VN40 A1VN41 A1VN42 A1VN43 A1VN54 A1VN58
A1VN60 A1VN92 A1VN98 A1VNB7 A1VNF0 A1VNF1 A1VNF8 A1VNG5 A1VNK2 A1VNK4 A1VNP5
A1VNQ9 A1VNR3 A1VNU1 A1VNU2 A1VNU4 A1VP76 A1VP90 A1VP99 A1VPP2 A1VPX1 A1VPX2
A1VPX4 A1VPX9 A1VPZ7 A1VQ02 A1VQ03 A1VQ12 A1VQ18 A1VQD4 A1VQF1 A1VQF4 A1VQI3

A1VQI6 A1VQK2 A1VQW9 A1VQZ6 A1VR16 A1VR25 A1VR71 A1VR74 A1VR75 A1VR77 A1VR98
A1VRB1 A1VRB4 A1VRC8 A1VRC9 A1VRD9 A1VRF5 A1VRF7 A1VRH7 A1VRH9 A1VRL0 A1VRN0
A1VRN9 A1VRP1 A1VRQ1 A1VRR0 A1VRR1 A1VRR5 A1VRR7 A1VRR8 A1VRT0 A1VRU9 A1VRW9
A1VS34 A1VS36 A1VS84 A1VS85 A1VS93 A1VSK4 A1VSK8 A1VSL4 A1VSM6 A1VSP2 A1VSS8
A1VST4 A1VST5 A1VSU4 A1VSY7 A1VSZ8 A1VT30 A1VT89 A1VTA5 A1VTF7 A1VTF8 A1VTF9
A1VTG0 A1VTG1 A1VTG2 A1VTG7 A1VTI2 A1VTL9 A1VTR9 A1VTV4 A1VTW0 A1VTY5 A1VU11
A1VU36 A1VU45 A1VUC0 A1VUC3 A1VUJ4 A1VUJ6 A1VUN7 A1VUS4 A1VUS8 A1VUV0 A1VUV1
A1VUV2 A1VW83 A1VWK8 A1VX79 A1VX88 A1VX91 A1VX95 A1VXA6 A1VXD7 A1VXF1 A1VXG9
A1VXH7 A1VXH8 A1VXH9 A1VXI1 A1VXK4 A1VXK9 A1VXL7 A1VXL9 A1VXM1 A1VXN2 A1VXN9
A1VXP5 A1VXP9 A1VXQ2 A1VXS0 A1VXS1 A1VXS2 A1VXS5 A1VXT5 A1VXT6 A1VXU6 A1VXU8
A1VXV5 A1VXV6 A1VXW3 A1VXW6 A1VXW7 A1VXX6 A1VY04 A1VY10 A1VY14 A1VY17 A1VY27
A1VY30 A1VY31 A1VY36 A1VY40 A1VY43 A1VY44 A1VY45 A1VY49 A1VY51 A1VY69 A1VY70
A1VY80 A1VY90 A1VY95 A1VYA1 A1VYA2 A1VYA3 A1VYA6 A1VYA9 A1VYB8 A1VYC1 A1VYC2
A1VYE9 A1VYF1 A1VYG6 A1VYG9 A1VYH4 A1VYI6 A1VYI7 A1VYJ0 A1VYJ1 A1VYJ2 A1VYJ3
A1VYJ4 A1VYJ5 A1VYJ6 A1VYJ7 A1VYJ8 A1VYK3 A1VYL5 A1VYL8 A1VYL9 A1VYM4 A1VYN0
A1VYP2 A1VYQ1 A1VYQ2 A1VYQ4 A1VYR0 A1VYR7 A1VYT7 A1VYU1 A1VYU6 A1VYV0 A1VYV7
A1VYZ2 A1VYZ5 A1VYZ7 A1VYZ9 A1VZ00 A1VZ01 A1VZ10 A1VZ21 A1VZ22 A1VZ23 A1VZ24
A1VZ41 A1VZ44 A1VZ48 A1VZ59 A1VZ65 A1VZ67 A1VZ68 A1VZ69 A1VZB3 A1VZB4 A1VZB5
A1VZE6 A1VZE7 A1VZF0 A1VZF4 A1VZF8 A1VZH1 A1VZH5 A1VZI4 A1VZI8 A1VZJ6 A1VZJ8
A1VZK1 A1VZL9 A1VZM8 A1VZM9 A1VZN1 A1VZN9 A1VZR0 A1VZR4 A1VZR5 A1VZU4 A1VZU6
A1VZU7 A1VZV1 A1VZV2 A1VZW5 A1VZY1 A1VZY2 A1VZY5 A1VZZ6 A1VZZ8 A1W023 A1W035
A1W038 A1W043 A1W048 A1W057 A1W059 A1W062 A1W068 A1W078 A1W085 A1W089 A1W091
A1W092 A1W094 A1W096 A1W0A4 A1W0D6 A1W0F6 A1W0F9 A1W0G5 A1W0G6 A1W0H2 A1W0I1
A1W0I5 A1W0K3 A1W0K4 A1W0M1 A1W0M2 A1W0M7 A1W0N0 A1W0N3 A1W0N8 A1W0P5 A1W0P8
A1W0Q3 A1W0Q8 A1W0Q9 A1W0R3 A1W0R9 A1W0S0 A1W0S2 A1W0U6 A1W0U7 A1W0U8 A1W0U9
A1W0W0 A1W0W2 A1W0W6 A1W0X7 A1W0X9 A1W0Z2 A1W0Z5 A1W116 A1W118 A1W162 A1W163
A1W165 A1W187 A1W1A5 A1W1D0 A1W1D6 A1W1E0 A1W1J2 A1W1J3 A1W1J4 A1W1J5 A1W1J6
A1W1J7 A1W1J8 A1W1J9 A1W1K1 A1W1K3 A1W1K9 A1W1L3 A1W1L4 A1W1N6 A1W1N8 A1W1S5
A1W1T3 A1W1U3 A1W1U4 A1W1U5 A1W1U6 A1W1U7 A1W1U8 A1W1U9 A1W1V0 A1W1V1 A1W1V2
A1W1V3 A1W1V4 A1W1V5 A1W1V6 A1W1V7 A1W1V8 A1W1V9 A1W1W1 A1W1W6 A1W1W9 A1W1X0
A1W1X2 A1W1X5 A1W1X7 A1W1Y1 A1W207 A1W208 A1W247 A1W2F6 A1W2K2 A1W2K3 A1W2Q2
A1W2Q3 A1W2Q4 A1W2Q5 A1W2Q6 A1W2Q7 A1W2Q8 A1W2R0 A1W2R1 A1W2R2 A1W2R3 A1W2R4
A1W2R5 A1W2R6 A1W2S7 A1W2Y5 A1W2Y8 A1W2Z7 A1W2Z8 A1W309 A1W318 A1W319 A1W320
A1W321 A1W322 A1W323 A1W324 A1W325 A1W326 A1W327 A1W329 A1W330 A1W331 A1W332
A1W333 A1W334 A1W341 A1W3A1 A1W3C0 A1W3D0 A1W3D7 A1W3F2 A1W3F3 A1W3F9 A1W3L0
A1W3P0 A1W3P7 A1W3P8 A1W3Q0 A1W3R6 A1W3W5 A1W3W8 A1W3W9 A1W3X4 A1W3Z7 A1W428
A1W431 A1W432 A1W434 A1W444 A1W445 A1W446 A1W4A8 A1W4A9 A1W4B0 A1W4B1 A1W4B2
A1W4B3 A1W4B8 A1W4B9 A1W4E1 A1W4G0 A1W4G1 A1W4G3 A1W4G6 A1W4G9 A1W4H1 A1W4H2
A1W4I7 A1W4L3 A1W4L7 A1W4L8 A1W4R3 A1W4S3 A1W4U7 A1W4U9 A1W538 A1W551 A1W573
A1W574 A1W578 A1W581 A1W582 A1W587 A1W588 A1W5A4 A1W5B5 A1W5B6 A1W5B7 A1W631
A1W6C8 A1W6E1 A1W6H4 A1W6H5 A1W6H6 A1W6I6 A1W6N2 A1W6Q8 A1W6R1 A1W6T4 A1W6T7
A1W6V2 A1W6V9 A1W6W1 A1W748 A1W753 A1W757 A1W776 A1W7D0 A1W7D2 A1W7D8 A1W7E4
A1W7J6 A1W7K9 A1W7L8 A1W7M0 A1W7N4 A1W7S2 A1W7T4 A1W838 A1W893 A1W895 A1W8C6
A1W8D4 A1W8G7 A1W8H0 A1W8I2 A1W8I3 A1W8I4 A1W8I6 A1W8J9 A1W8S8 A1W8Y5 A1W8Z3

A1W8Z4 A1W8Z6 A1W904 A1W915 A1W916 A1W917 A1W918 A1W9B0 A1W9F0 A1W9F8 A1W9F9
A1W9G0 A1W9G6 A1W9H4 A1W9H6 A1W9I3 A1W9I4 A1W9K6 A1W9K7 A1W9P4 A1W9S4 A1W9U1
A1W9U6 A1W9V0 A1WA22 A1WA60 A1WAG1 A1WAJ7 A1WAM6 A1WAR1 A1WAR2 A1WAR4 A1WAR5
A1WAR6 A1WAR7 A1WAS7 A1WAT0 A1WAT5 A1WAT8 A1WAW3 A1WAY0 A1WAZ1 A1WB09 A1WB23
A1WB43 A1WB49 A1WB51 A1WB52 A1WB79 A1WB80 A1WB87 A1WBD4 A1WBD9 A1WBH1 A1WBI2
A1WBI7 A1WBI8 A1WBJ0 A1WBJ3 A1WBK4 A1WBM9 A1WBQ7 A1WBR0 A1WBR8 A1WBW2 A1WBZ2
A1WBZ4 A1WBZ7 A1WC04 A1WC05 A1WC11 A1WC14 A1WC15 A1WC16 A1WC30 A1WC36 A1WC38
A1WC42 A1WC79 A1WCB1 A1WCJ3 A1WCK2 A1WCM8 A1WCM9 A1WCN0 A1WCN1 A1WCN2 A1WCN3
A1WCN6 A1WCP4 A1WCP8 A1WCQ6 A1WD26 A1WD32 A1WD35 A1WD47 A1WD86 A1WDB0 A1WDB4
A1WDB9 A1WDC8 A1WDU6 A1WDW0 A1WDW4 A1WDW5 A1WDX2 A1WDX7 A1WE11 A1WE12 A1WE19
A1WE20 A1WE26 A1WEE4 A1WEE7 A1WEL1 A1WEN1 A1WF48 A1WF49 A1WF62 A1WF86 A1WFG0
A1WFG9 A1WFH2 A1WFH8 A1WFK6 A1WFL2 A1WFS9 A1WFT0 A1WG50 A1WGH1 A1WGH3 A1WGI4
A1WGJ9 A1WGK0 A1WGK1 A1WGK3 A1WGK4 A1WGM5 A1WGR0 A1WGR9 A1WGS0 A1WGS2 A1WGT2
A1WGT3 A1WGU0 A1WGU4 A1WH11 A1WH14 A1WH73 A1WH74 A1WH84 A1WH85 A1WH89 A1WHC0
A1WHC1 A1WHC2 A1WHC3 A1WHC4 A1WHC5 A1WHC6 A1WHC8 A1WHC9 A1WHD0 A1WHD1 A1WHD2
A1WHD3 A1WHD4 A1WHI6 A1WHJ5 A1WHP7 A1WHU2 A1WHU3 A1WHU4 A1WHU5 A1WHV6 A1WHW6
A1WHX2 A1WI17 A1WI20 A1WI55 A1WI56 A1WIC3 A1WID5 A1WIF4 A1WIG8 A1WIH0 A1WIJ3
A1WIL9 A1WIM2 A1WIM3 A1WIM4 A1WIM6 A1WIQ3 A1WIQ4 A1WIS4 A1WIS6 A1WJ05 A1WJ68
A1WJJ2 A1WJL0 A1WJP5 A1WJR4 A1WJV9 A1WJY8 A1WK51 A1WK52 A1WK53 A1WK54 A1WK55
A1WK56 A1WK91 A1WK92 A1WK93 A1WK94 A1WK95 A1WK97 A1WK98 A1WK99 A1WKA0 A1WKA1
A1WKA2 A1WKA3 A1WKA4 A1WKA5 A1WKA6 A1WKB4 A1WKD3 A1WKG6 A1WKI1 A1WKI2 A1WL02
A1WL04 A1WL05 A1WL18 A1WL60 A1WL74 A1WL75 A1WL78 A1WL89 A1WLD5 A1WLG5 A1WLI1
A1WLI2 A1WLI3 A1WLI5 A1WLL2 A1WLL3 A1WLP8 A1WLP9 A1WM26 A1WM72 A1WMA0 A1WME3
A1WMG0 A1WMG5 A1WMG7 A1WMG8 A1WMG9 A1WMN5 A1WMN6 A1WMU5 A1WMV5 A1WMV6 A1WMW1
A1WMX7 A1WN06 A1WN86 A1WNH8 A1WNY2 A1WNY5 A1WP07 A1WP10 A1WPC7 A1WPQ3 A1WPR4
A1WPR5 A1WPR6 A1WPR7 A1WPT9 A1WPU0 A1WQ64 A1WQ66 A1WQ68 A1WQ79 A1WQF4 A1WQL7
A1WQM5 A1WQN5 A1WQN6 A1WQP4 A1WQP5 A1WQW2 A1WQY3 A1WR02 A1WR07 A1WR16 A1WR17
A1WR21 A1WR23 A1WR24 A1WR98 A1WRA7 A1WRC4 A1WRC7 A1WRJ3 A1WRK0 A1WRK1 A1WRK2
A1WRK3 A1WRL2 A1WRL3 A1WS08 A1WS32 A1WS74 A1WS75 A1WS77 A1WS95 A1WSA8 A1WSC8
A1WSF0 A1WSF1 A1WSF3 A1WSH6 A1WSK3 A1WSL6 A1WSR5 A1WSU0 A1WSU4 A1WSY4 A1WSY5
A1WSY9 A1WSZ8 A1WT15 A1WT61 A1WT89 A1WT90 A1WT98 A1WTG1 A1WTK1 A1WTK7 A1WTL2
A1WTL3 A1WTM6 A1WU53 A1WU55 A1WU56 A1WU59 A1WU91 A1WUF2 A1WUF3 A1WUF4 A1WUG8
A1WUH0 A1WUH1 A1WUI3 A1WUI6 A1WUI9 A1WUJ7 A1WUL5 A1WUM6 A1WUM7 A1WUM8 A1WUN2
A1WUN4 A1WUS2 A1WUS4 A1WUS6 A1WUS7 A1WUT0 A1WUT6 A1WUT7 A1WUU3 A1WUU4 A1WUU5
A1WUV1 A1WUW3 A1WUX1 A1WUX3 A1WUY3 A1WUY7 A1WUZ5 A1WUZ6 A1WV00 A1WV94 A1WV96
A1WV97 A1WV98 A1WV99 A1WVA0 A1WVA1 A1WVA3 A1WVA4 A1WVA5 A1WVA6 A1WVA7 A1WVA8
A1WVA9 A1WVB0 A1WVB1 A1WVB2 A1WVB3 A1WVB4 A1WVB5 A1WVB6 A1WVB7 A1WVB8 A1WVB9
A1WVC0 A1WVC1 A1WVC2 A1WVC4 A1WVC5 A1WVC6 A1WVC7 A1WVC8 A1WVC9 A1WVD0 A1WVD1
A1WVD2 A1WVD3 A1WVD6 A1WVD8 A1WVE3 A1WVE4 A1WVE9 A1WVF5 A1WVF7 A1WVG0 A1WVG1
A1WVG5 A1WVG6 A1WVI0 A1WVK5 A1WVL3 A1WVM4 A1WVM6 A1WVM7 A1WVN0 A1WVN4 A1WVN6
A1WVN7 A1WVQ1 A1WVQ2 A1WVQ4 A1WVQ7 A1WVQ8 A1WVR6 A1WVR7 A1WVR8 A1WVS9 A1WVT7
A1WVT8 A1WVT9 A1WVV5 A1WVW0 A1WVW8 A1WVX2 A1WW04 A1WW05 A1WW06 A1WW08 A1WW09
A1WW12 A1WW18 A1WW41 A1WW56 A1WWA3 A1WWA4 A1WWA5 A1WWB3 A1WWB6 A1WWC6 A1WWE0
A1WWE1 A1WWE2 A1WWE4 A1WWF0 A1WWG6 A1WWH7 A1WWH9 A1WWI7 A1WWP8 A1WWU7 A1WWV2

A1WWV3 A1WWV4 A1WWV6 A1WWV9 A1WWY5 A1WWY6 A1WWY9 A1WWZ0 A1WWZ4 A1WX09 A1WX18
A1WX19 A1WX20 A1WX21 A1WX26 A1WX29 A1WX30 A1WX31 A1WX32 A1WX34 A1WX38 A1WX40
A1WXD7 A1WXD9 A1WXE1 A1WXE4 A1WXH5 A1WXI6 A1WXI8 A1WXI9 A1WXK3 A1WXK5 A1WXK6
A1WXK7 A1WXK9 A1WXL3 A1WXM0 A1WXM7 A1WXS8 A1WXU7 A1WXU8 A1WXU9 A1WXV0 A1WXV1
A1WXV3 A1WXW9 A1WXX0 A1WXX3 A1WXX6 A1WXY4 A1WXY6 A1WXZ0 A1WXZ3 A1WXZ4 A1WY05
A1WY07 A1WY14 A1WY16 A1WY18 A1WY38 A1WY45 A1WY47 A1WY48 A1WY49 A1WY50 A1WY69
A1WY80 A1WY83 A1WY97 A1WYA9 A1WYB5 A1WYD1 A1WYD5 A1WYI1 A1WYI6 A1WYI8 A1WYK5
A1WYL3 A1WYL4 A1WYP9 A1WYS1 A1WYS6 A1WYS8 A1WYS9 A1WYT0 A1WYT2 A1WYU0 A1WYU2
A1WYV1 A1WYV2 A1WYV5 A1WYV6 A1WYW5 A1WYW6 A1WYY2 A1WYZ1 A1WYZ3 A1WYZ4 A1WYZ7
A1WZ13 A1WZ50 A1WZ62 A1WZ63 A1WZ64 A1WZ65 A1WZ92 A1WZ95 A1WZA2 A1WZA9 A1WZB0
A1WZB3 A1WZE3 A1WZE7 A1WZE9 A1WZF2 A1WZG0 A1WZG2 A1WZG4 A1WZG5 A1WZH0 A1WZH3
A1WZH6 A1WZH8 A1WZH9 A1WZI2 A1WZI8 A1WZJ1 A1WZJ2 A1WZK4 A1WZP9 A1WZR2 A1WZR7
A1X151 A1X154 A1X157 A1XBB7 A1XDC0 A1XFT4 A1XFT6 A1XFU6 A1XFY0 A1XFZ0 A1XFZ3
A1XFZ5 A1XFZ7 A1XFZ9 A1XGL7 A1XGL9 A1XGM7 A1XGM8 A1XGM9 A1XGR3 A1XGS0 A1XGS5
A1XGS8 A1XGT0 A1XIQ0 A1XIQ1 A1XPK1 A1XQR9 A1XQU1 A1XQU7 A1XQV4 A1XSY8 A1XWY7
A1Y2B7 A1YCA5 A1YEP8 A1YEQ3 A1YER0 A1YER2 A1YER7 A1YEV8 A1YEV9 A1YEW3 A1YEW9
A1YEX3 A1YF12 A1YF15 A1YF16 A1YF22 A1YF56 A1YFA5 A1YFA7 A1YFC1 A1YFD8 A1YFI3
A1YFT7 A1YFU7 A1YFW2 A1YFW6 A1YFX5 A1YFX9 A1YFY3 A1YG01 A1YG22 A1YG25 A1YG26
A1YG31 A1YG48 A1YG60 A1YG61 A1YG88 A1YG91 A1YG92 A1YG93 A1YG99 A1YGA2 A1YGA4
A1YGI6 A1YGJ4 A1YGK1 A1YGK6 A1YGK7 A1YGK9 A1YIZ1 A1YKT1 A1YKW8 A1YKY7 A1YPR0
A1YPR2 A1YPR3 A1YTM8 A1YTU9 A1YVX4 A1Z1Q3 A1Z6J5 A1Z6K7 A1Z6M6 A1Z7A8 A1Z7K9
A1Z897 A1Z8D0 A1Z8J0 A1Z8X3 A1Z9A2 A1Z9A8 A1Z9E2 A1Z9G2 A1Z9J4 A1Z9L3 A1Z9R4
A1ZA92 A1ZAJ2 A1ZAK1 A1ZAU8 A1ZAX1 A1ZB42 A1ZB86 A1ZB91 A1ZBR5 A1ZBT5 A1ZBY1
A2A1A0 A2A288 A2A2Z9 A2A368 A2A3K4 A2A3L6 A2A3V1 A2A591 A2A5E6 A2A5K6 A2A5N8
A2A5X4 A2A5X5 A2A690 A2A6A1 A2A6H3 A2A6Q5 A2A6T1 A2A791 A2A7S8 A2A7Y5 A2A935
A2A9A2 A2A9I7 A2A9R3 A2A9T0 A2AA28 A2AAX3 A2ABU4 A2ABV5 A2ACG1 A2ACJ2 A2ACP1
A2ADZ8 A2AEV7 A2AEY4 A2AFE9 A2AFR3 A2AG06 A2AG50 A2AG58 A2AGB2 A2AGH6 A2AGX3
A2AHC3 A2AHG0 A2AHJ4 A2AI05 A2AIG8 A2AIP0 A2AIV2 A2AIV8 A2AJ77 A2AJB1 A2AJI0
A2AJK6 A2AJL3 A2AKB9 A2AKE7 A2AKQ8 A2AL36 A2ALV5 A2AM05 A2AMZ4 A2ANX9 A2APB8
A2APC3 A2APF3 A2AQ19 A2AQ25 A2ARM1 A2ARS0 A2ARZ3 A2AS55 A2AS89 A2ASS6 A2AT37
A2ATU0 A2AU37 A2AU72 A2AUU0 A2AUY4 A2AV36 A2AV37 A2AVJ0 A2AVQ5 A2AWA9 A2AWL7
A2AWP0 A2AWP8 A2AWT3 A2AXG5 A2AXI2 A2BD05 A2BD66 A2BD83 A2BD89 A2BD94 A2BDA5
A2BDB7 A2BDR7 A2BDX3 A2BE28 A2BEA6 A2BF66 A2BFC9 A2BFL2 A2BGA0 A2BGD5 A2BGM5
A2BGU8 A2BGU9 A2BHB7 A2BHJ4 A2BID7 A2BIG9 A2BIJ3 A2BIL7 A2BIL8 A2BIR6 A2BIT1
A2BIU4 A2BIW2 A2BIX0 A2BIX4 A2BIZ8 A2BJ23 A2BJ24 A2BJ25 A2BJ90 A2BJB7 A2BJD9
A2BJL3 A2BJL6 A2BJP9 A2BJU0 A2BJU2 A2BJX6 A2BJY6 A2BJY7 A2BJY9 A2BJZ0 A2BJZ2
A2BJZ7 A2BJZ8 A2BK08 A2BK57 A2BK76 A2BK77 A2BK78 A2BK83 A2BK84 A2BK91 A2BK92
A2BK95 A2BK96 A2BK98 A2BKC0 A2BKH8 A2BKJ6 A2BKJ8 A2BKX3 A2BL26 A2BL27 A2BL34
A2BL58 A2BL68 A2BL74 A2BL78 A2BL82 A2BL86 A2BL93 A2BLB9 A2BLC0 A2BLC6 A2BLD4
A2BLE8 A2BLF3 A2BLL5 A2BLL6 A2BLM0 A2BLQ8 A2BLU2 A2BLV6 A2BLW2 A2BLW3 A2BLW4
A2BLY5 A2BM05 A2BM26 A2BM70 A2BM73 A2BMB2 A2BMB3 A2BMC4 A2BMC5 A2BMC7 A2BMC9
A2BMD0 A2BMD1 A2BMD4 A2BMD5 A2BMD8 A2BMD9 A2BME0 A2BME1 A2BME5 A2BME7 A2BMH3
A2BMI0 A2BMI7 A2BMW0 A2BMY4 A2BN08 A2BN21 A2BN24 A2BN27 A2BN31 A2BN32 A2BN38
A2BN41 A2BN42 A2BN51 A2BN55 A2BN56 A2BN57 A2BN58 A2BN59 A2BN63 A2BN64 A2BN65

A2BN82 A2BN93 A2BN94 A2BN97 A2BNA6 A2BNB6 A2BNC9 A2BND8 A2BNE2 A2BNE6 A2BNF2
A2BNG3 A2BNH2 A2BNH3 A2BNH4 A2BNH6 A2BNK7 A2BNP5 A2BNQ7 A2BNS4 A2BNV9 A2BNW6
A2BNW8 A2BNX7 A2BNY4 A2BNZ0 A2BNZ6 A2BNZ7 A2BNZ8 A2BNZ9 A2BP15 A2BP19 A2BP22
A2BP31 A2BP33 A2BP36 A2BP39 A2BP40 A2BP49 A2BP55 A2BP56 A2BP57 A2BP61 A2BP63
A2BP65 A2BP70 A2BP81 A2BPB2 A2BPB4 A2BPC1 A2BPD1 A2BPL5 A2BPL9 A2BPM3 A2BPM9
A2BPN3 A2BPN7 A2BPP7 A2BPR2 A2BPS1 A2BPT4 A2BPT7 A2BPV2 A2BPV5 A2BPW3 A2BPW6
A2BPX4 A2BPX7 A2BPY1 A2BPY8 A2BPZ0 A2BQ02 A2BQ03 A2BQ06 A2BQ24 A2BQ25 A2BQ26
A2BQ31 A2BQ40 A2BQ41 A2BQ46 A2BQ64 A2BQ65 A2BQ71 A2BQ78 A2BQ88 A2BQ90 A2BQ94
A2BQA4 A2BQE5 A2BQG2 A2BQN3 A2BQN9 A2BQP0 A2BQQ2 A2BQQ3 A2BQQ9 A2BQS5 A2BQS8
A2BQT3 A2BQW2 A2BQW3 A2BQW4 A2BQW5 A2BQW6 A2BQW7 A2BQW8 A2BQY1 A2BQY7 A2BQZ0
A2BQZ1 A2BQZ8 A2BR00 A2BR01 A2BR27 A2BR32 A2BR42 A2BR44 A2BR45 A2BR50 A2BR65
A2BR66 A2BR89 A2BR98 A2BRA1 A2BRB4 A2BRE1 A2BRG6 A2BRN2 A2BRP4 A2BRP7 A2BRS2
A2BRS3 A2BRS6 A2BRT6 A2BRU5 A2BRV3 A2BS39 A2BS59 A2BS60 A2BS76 A2BS99 A2BSA0
A2BSA4 A2BSA6 A2BSA7 A2BSJ2 A2BSJ5 A2BSK6 A2BSL2 A2BSL8 A2BSM1 A2BSM6 A2BSM7
A2BSN1 A2BSN6 A2BSQ0 A2BSQ4 A2BSR8 A2BSS0 A2BSS2 A2BSS6 A2BT05 A2BT10 A2BT11
A2BT35 A2BT49 A2BT57 A2BT59 A2BT60 A2BT61 A2BT63 A2BT67 A2BT70 A2BT75 A2BT77
A2BT82 A2BT83 A2BT84 A2BT85 A2BT86 A2BTA9 A2BTB0 A2BTB1 A2BTB2 A2BTB3 A2BTB4
A2BTB5 A2BTB6 A2BTB7 A2BTB8 A2BTC0 A2BTC1 A2BTC2 A2BTC4 A2BTC5 A2BTC6 A2BTC7
A2BTC8 A2BTC9 A2BTD0 A2BTD1 A2BTD2 A2BTD3 A2BTD4 A2BTD5 A2BTD6 A2BTD7 A2BTE1
A2BTF4 A2BTG1 A2BTH0 A2BTH1 A2BTH3 A2BTH4 A2BTI7 A2BTJ3 A2BTJ7 A2BTJ8 A2BTL7
A2BTL8 A2BTM4 A2BTM6 A2BTN4 A2BTN6 A2BTN8 A2BTP2 A2BTP3 A2BTP5 A2BTQ0 A2BTQ4
A2BTQ6 A2BTR7 A2BTR9 A2BTS0 A2BTS1 A2BTS3 A2BTT7 A2BTT9 A2BTV0 A2BTV4 A2BTV8
A2BTV9 A2BTX3 A2BTY3 A2BTY4 A2BTZ9 A2BU00 A2BU01 A2BU03 A2BU25 A2BU62 A2BU74
A2BU86 A2BUA6 A2BUE1 A2BUE8 A2BUF0 A2BUF9 A2BUG6 A2BUH2 A2BUH8 A2BUH9 A2BUI0
A2BUI1 A2BUJ6 A2BUK0 A2BUK4 A2BUL3 A2BUL5 A2BUL8 A2BUM1 A2BUM2 A2BUN1 A2BUN7
A2BUN8 A2BUN9 A2BUP3 A2BUP5 A2BUP7 A2BUQ2 A2BUQ6 A2BUR2 A2BUU2 A2BUU4 A2BUV0
A2BUV8 A2BV47 A2BV51 A2BV55 A2BV61 A2BV65 A2BV69 A2BV78 A2BV90 A2BVB2 A2BVB5
A2BVD2 A2BVD5 A2BVE2 A2BVE5 A2BVF3 A2BVF7 A2BVG1 A2BVG8 A2BVH0 A2BVI1 A2BVI3
A2BVI4 A2BVI7 A2BVK5 A2BVK6 A2BVK7 A2BVL2 A2BVM2 A2BVM7 A2BVP6 A2BVP7 A2BVQ3
A2BVQ8 A2BVR0 A2BVS0 A2BVS2 A2BVS6 A2BVT6 A2BVY6 A2BW03 A2BW33 A2BW38 A2BW57
A2BW63 A2BW64 A2BW76 A2BW77 A2BW83 A2BWF0 A2BWG3 A2BWG6 A2BWH6 A2BWJ6 A2BWJ7
A2BWL2 A2BWL7 A2BWL8 A2BWM0 A2BWN0 A2BWN6 A2BWR1 A2BWR2 A2BWR4 A2BWS2 A2BWS5
A2BWT1 A2BWW2 A2BWY5 A2BX41 A2BX52 A2BX55 A2BX80 A2BX81 A2BX84 A2BX94 A2BXA3
A2BXB1 A2BXC2 A2BXK1 A2BXK2 A2BXM3 A2BXM8 A2BXN1 A2BXQ1 A2BXQ2 A2BXQ6 A2BXQ9
A2BXX8 A2BXZ2 A2BXZ8 A2BY04 A2BY07 A2BY12 A2BY13 A2BY17 A2BY22 A2BY35 A2BY50
A2BY51 A2BY52 A2BY54 A2BY56 A2BY60 A2BY75 A2BYF5 A2BYG1 A2BYG2 A2BYI6 A2BYK1
A2BYK7 A2BYK9 A2BYL0 A2BYL1 A2BYL3 A2BYL7 A2BYM0 A2BYM5 A2BYM6 A2BYM8 A2BYN3
A2BYN4 A2BYN5 A2BYN6 A2BYN7 A2BYQ7 A2BYQ8 A2BYQ9 A2BYR0 A2BYR1 A2BYR2 A2BYR3
A2BYR4 A2BYR5 A2BYR6 A2BYR7 A2BYR9 A2BYS0 A2BYS1 A2BYS2 A2BYS3 A2BYS4 A2BYS5
A2BYS6 A2BYS7 A2BYS8 A2BYS9 A2BYT0 A2BYT1 A2BYT2 A2BYT3 A2BYT4 A2BYT5 A2BYT6
A2BYU0 A2BYV8 A2BYW5 A2BYX5 A2BYX6 A2BYY9 A2BYZ5 A2BZ00 A2BZ01 A2BZ03 A2BZ20
A2BZ21 A2BZ29 A2BZ39 A2BZ41 A2BZ43 A2BZ47 A2BZ48 A2BZ50 A2BZ55 A2BZ59 A2BZ61
A2BZ72 A2BZ74 A2BZ75 A2BZ76 A2BZ78 A2BZ91 A2BZ92 A2BZ93 A2BZ94 A2BZA6 A2BZB5
A2BZB9 A2BZC2 A2BZC3 A2BZC5 A2BZC9 A2BZD8 A2BZD9 A2BZG2 A2BZG4 A2BZG6 A2BZH4

A2BZN6 A2BZQ6 A2BZT9 A2BZV2 A2BZV6 A2BZZ2 A2C000 A2C002 A2C016 A2C018 A2C025
A2C030 A2C031 A2C032 A2C033 A2C034 A2C050 A2C055 A2C058 A2C066 A2C068 A2C071
A2C073 A2C082 A2C088 A2C089 A2C090 A2C094 A2C096 A2C098 A2C0A9 A2C0B3 A2C0B5
A2C0E4 A2C0F4 A2C0F7 A2C0G6 A2C0H4 A2C0J2 A2C0J7 A2C0K1 A2C0K7 A2C0L1 A2C0L5
A2C0L8 A2C0M6 A2C0N7 A2C0Q8 A2C0S7 A2C0T1 A2C0T9 A2C0U2 A2C0V4 A2C0V9 A2C0W6
A2C0W8 A2C0X8 A2C0Y0 A2C0Y1 A2C0Y4 A2C0Z9 A2C100 A2C101 A2C106 A2C116 A2C117
A2C122 A2C129 A2C141 A2C142 A2C148 A2C153 A2C155 A2C165 A2C167 A2C172 A2C175
A2C182 A2C1C0 A2C1E0 A2C1I7 A2C1I8 A2C1K3 A2C1K4 A2C1M9 A2C1N4 A2C1X5 A2C1Y1
A2C1Y4 A2C1Y5 A2C1Z3 A2C1Z5 A2C1Z6 A2C220 A2C226 A2C228 A2C236 A2C239 A2C241
A2C242 A2C243 A2C247 A2C249 A2C260 A2C261 A2C283 A2C296 A2C2A5 A2C2B2 A2C2S8
A2C3G4 A2C3H1 A2C3I3 A2C3I4 A2C3L3 A2C3L7 A2C3N0 A2C3T6 A2C3T8 A2C3V6 A2C3V9
A2C3W6 A2C3W9 A2C404 A2C405 A2C409 A2C411 A2C412 A2C436 A2C451 A2C457 A2C462
A2C466 A2C471 A2C472 A2C476 A2C482 A2C4A1 A2C4B2 A2C4B7 A2C4B8 A2C4C0 A2C4C2
A2C4H7 A2C4I0 A2C4I2 A2C4I3 A2C4M0 A2C4M8 A2C4N0 A2C4N1 A2C4N2 A2C4N4 A2C4N6
A2C4N8 A2C4P1 A2C4R6 A2C4R9 A2C4S0 A2C4S1 A2C4S2 A2C4T1 A2C4T7 A2C4T9 A2C4U4
A2C4U5 A2C4U6 A2C4U7 A2C4U8 A2C4X1 A2C4X2 A2C4X4 A2C4X6 A2C4X7 A2C4X8 A2C4X9
A2C4Y0 A2C4Y2 A2C4Y3 A2C4Y4 A2C4Y5 A2C4Y6 A2C4Y7 A2C4Y8 A2C4Y9 A2C4Z0 A2C4Z1
A2C4Z2 A2C4Z3 A2C4Z4 A2C4Z5 A2C4Z6 A2C4Z7 A2C4Z9 A2C503 A2C524 A2C535 A2C536
A2C538 A2C539 A2C552 A2C559 A2C563 A2C564 A2C585 A2C586 A2C5A7 A2C5A9 A2C5B3
A2C5B5 A2C5C0 A2C5C2 A2C5C6 A2C5C7 A2C5C9 A2C5D4 A2C5D5 A2C5D9 A2C5E1 A2C5E6
A2C5F4 A2C5F7 A2C5F9 A2C5G1 A2C5I5 A2C5I6 A2C5J9 A2C5K9 A2C5L7 A2C5M0 A2C5M1
A2C5M2 A2C5M3 A2C5M7 A2C5P1 A2C5P2 A2C5Q6 A2C5Q7 A2C5R5 A2C5R7 A2C5R9 A2C5S6
A2C5T5 A2C5T7 A2C600 A2C601 A2C620 A2C629 A2C637 A2C638 A2C642 A2C650 A2C661
A2C698 A2C6N6 A2C6Q5 A2C6Q8 A2C6S4 A2C6S6 A2C6S8 A2C6S9 A2C6T1 A2C6T3 A2C6U2
A2C6W2 A2C6Z6 A2C700 A2C717 A2C725 A2C730 A2C732 A2C733 A2C735 A2C740 A2C764
A2C794 A2C795 A2C798 A2C7A3 A2C7A4 A2C7E2 A2C7G2 A2C7H0 A2C7I7 A2C7K4 A2C7L8
A2C7M5 A2C7N3 A2C7N5 A2C7Q3 A2C7Q7 A2C7T4 A2C7T5 A2C7T6 A2C7U9 A2C7W3 A2C7X1
A2C7X7 A2C7X9 A2C808 A2C835 A2C859 A2C873 A2C897 A2C8B2 A2C8D5 A2C8D6 A2C8E1
A2C8K0 A2C9H8 A2C9I6 A2C9M6 A2C9P3 A2C9R4 A2C9R5 A2C9T2 A2C9U0 A2C9U9 A2C9V7
A2C9W0 A2C9X1 A2C9Z1 A2C9Z7 A2CA16 A2CA19 A2CA37 A2CA48 A2CA51 A2CA52 A2CA92
A2CA98 A2CA99 A2CAB7 A2CAB8 A2CAC6 A2CAH8 A2CAR8 A2CAR9 A2CAS7 A2CAU0 A2CAU4
A2CAW2 A2CAW5 A2CAX1 A2CAX3 A2CB17 A2CB21 A2CB37 A2CB44 A2CB59 A2CB74 A2CB75
A2CB80 A2CB87 A2CBC7 A2CBH2 A2CBJ1 A2CBJ6 A2CBL8 A2CBM2 A2CBN5 A2CBQ3 A2CBR5
A2CBU0 A2CBU3 A2CBW8 A2CBW9 A2CBX2 A2CBX4 A2CC22 A2CC27 A2CC28 A2CC29 A2CC30
A2CC31 A2CC32 A2CC33 A2CC34 A2CC35 A2CC36 A2CC37 A2CC38 A2CC40 A2CC41 A2CC43
A2CC44 A2CC47 A2CC48 A2CC49 A2CC50 A2CC51 A2CC52 A2CC53 A2CC54 A2CC55 A2CC56
A2CC84 A2CC85 A2CC86 A2CC87 A2CC89 A2CC94 A2CC97 A2CCA2 A2CCB6 A2CCF2 A2CCG0
A2CCG3 A2CCI1 A2CCJ0 A2CCJ3 A2CCJ9 A2CCK3 A2CCK6 A2CCL1 A2CCL7 A2CCN3 A2CCR1
A2CCX2 A2CCY5 A2CD15 A2CD19 A2CD93 A2CDA0 A2CDB7 A2CDC4 A2CDE8 A2CDE9 A2CDG7
A2CDJ8 A2CDK0 A2CDK1 A2CDK7 A2CDR0 A2CDR1 A2CDR6 A2CDR8 A2CDS6 A2CDU0 A2CDU4
A2CDX0 A2CDX3 A2CDX5 A2CDX8 A2CDY7 A2CDZ5 A2CDZ8 A2CDZ9 A2CE00 A2CE01 A2CE27
A2CE29 A2CE36 A2CE44 A2CEH0 A2CEI4 A2CEI6 A2CEM9 A2CG63 A2CHX2 A2CI44 A2CI45
A2CI46 A2CI50 A2CI56 A2CI66 A2CI69 A2CI70 A2CIR7 A2D4P8 A2D4R4 A2D4U8 A2D4Z7
A2D5H2 A2D5I1 A2D5K9 A2D5N1 A2D5V0 A2D5Y4 A2D635 A2D649 A2D671 A2E5K9 A2E9H3

A2EF58 A2EYC4 A2I3Y6 A2I3Z2 A2I469 A2I8Z7 A2IB53 A2IBH5 A2IBY7 A2ICC6 A2ICN5
A2ICR5 A2P2R3 A2PYH4 A2PZF9 A2PZN8 A2Q0R8 A2Q127 A2Q1V6 A2Q7T1 A2Q848 A2Q879
A2Q8B5 A2Q8I1 A2Q8R2 A2Q908 A2Q9A9 A2Q9E4 A2Q9L0 A2Q9M4 A2Q9T6 A2QA23 A2QA83
A2QAB5 A2QAC0 A2QAF9 A2QAP8 A2QAQ3 A2QAU6 A2QAW7 A2QAX7 A2QAY5 A2QB52 A2QBE8
A2QBF0 A2QBF1 A2QBH6 A2QC41 A2QC74 A2QCC7 A2QCJ9 A2QCR5 A2QCU8 A2QCW6 A2QD49
A2QDB9 A2QDI9 A2QE29 A2QE32 A2QE38 A2QEN5 A2QEP2 A2QEV8 A2QF55 A2QF59 A2QFG8
A2QFH1 A2QFK0 A2QFL3 A2QGA1 A2QGF0 A2QGR5 A2QGS5 A2QGZ1 A2QHE5 A2QHG9 A2QHV0
A2QHX0 A2QI22 A2QI25 A2QI73 A2QI77 A2QID5 A2QIK9 A2QIL2 A2QJI5 A2QJX5 A2QJZ4
A2QK64 A2QK65 A2QK66 A2QK67 A2QKF6 A2QMS4 A2QMS5 A2QMY6 A2QN07 A2QNQ8 A2QNW5
A2QP30 A2QPG3 A2QPM6 A2QPN9 A2QPT2 A2QPW4 A2QPZ4 A2QQ09 A2QQ10 A2QQA2 A2QQA8
A2QRR5 A2QRY2 A2QS00 A2QTE3 A2QTE5 A2QTE9 A2QTF1 A2QU58 A2QU71 A2QU77 A2QUI2
A2QUJ2 A2QUS7 A2QVS9 A2QVV2 A2QW83 A2QWA6 A2QWI7 A2QWW0 A2QX22 A2QX23 A2QX24
A2QX25 A2QX26 A2QX45 A2QXL3 A2QY22 A2QY39 A2QY49 A2QY54 A2QYG2 A2QYX4 A2QZ71
A2R0B5 A2R156 A2R1F6 A2R1P9 A2R2E3 A2R345 A2R346 A2R3A7 A2R3H4 A2R472 A2R475
A2R4I0 A2R4I6 A2R4J9 A2R4K5 A2R4R1 A2R5W7 A2R692 A2R6G7 A2R6G8 A2R6H0 A2R6H1
A2R6M7 A2R7B4 A2R7P5 A2R7S7 A2R7T0 A2R7Z2 A2R8S7 A2R919 A2R9B9 A2R9H9 A2R9P6
A2R9S1 A2RA31 A2RA55 A2RA56 A2RA63 A2RAF3 A2RAW3 A2RB17 A2RB71 A2RB75 A2RBA1
A2RBB7 A2RBC7 A2RBX3 A2RBX7 A2RBY9 A2RBZ5 A2RC05 A2RC13 A2RC14 A2RC15 A2RC16
A2RC17 A2RC18 A2RC19 A2RC20 A2RC21 A2RC22 A2RC23 A2RC24 A2RC25 A2RC26 A2RC27
A2RC28 A2RC29 A2RC30 A2RC31 A2RC32 A2RC33 A2RC35 A2RC36 A2RC37 A2RC38 A2RC39
A2RC40 A2RC41 A2RC49 A2RC51 A2RC52 A2RC62 A2RC73 A2RCA2 A2RCB2 A2RCC9 A2RCD9
A2RCF4 A2RCF8 A2RCG1 A2RCG2 A2RCH2 A2RCI0 A2RCI1 A2RCI2 A2RCI9 A2RCK6 A2RCK9
A2RCL1 A2RCL2 A2RCL6 A2RCM1 A2RCM6 A2RCM9 A2RCN0 A2RCN4 A2RCN9 A2RCP2 A2RCQ1
A2RCQ5 A2RCQ8 A2RCR4 A2RCR6 A2RCS2 A2RCS4 A2RCT3 A2RCV3 A2RCV6 A2RCV7 A2RCV8
A2RCW4 A2RCW6 A2RCY1 A2RCZ6 A2RCZ7 A2RD01 A2RD02 A2RD12 A2RD23 A2RD24 A2RD25
A2RD27 A2RD32 A2RD38 A2RD40 A2RD51 A2RD54 A2RD55 A2RD56 A2RD68 A2RD70 A2RD86
A2RD87 A2RDH0 A2RDJ1 A2RDJ5 A2RDJ7 A2RDK1 A2RDK9 A2RDL5 A2RDM9 A2RDN0 A2RDN1
A2RDV3 A2RDV4 A2RDV9 A2RDW0 A2RDW3 A2RDW6 A2RDX5 A2RDX6 A2RDY3 A2RDZ9 A2RE06
A2RE13 A2RE14 A2RE19 A2RE30 A2RE43 A2RE45 A2RE65 A2RE90 A2RE93 A2REA5 A2REA6
A2REC4 A2RED2 A2RED8 A2REE2 A2REE5 A2REF2 A2REF7 A2REG0 A2REG9 A2REH5 A2REH9
A2REI4 A2REI5 A2REJ8 A2REK0 A2REL5 A2REL9 A2REM4 A2REM5 A2REM7 A2REN0 A2REP8
A2REX9 A2REY2 A2REY5 A2REZ0 A2RF00 A2RF04 A2RF05 A2RF09 A2RF12 A2RF13 A2RF16
A2RF17 A2RF20 A2RF34 A2RF47 A2RF48 A2RF49 A2RF55 A2RF60 A2RF62 A2RF64 A2RF70
A2RF72 A2RF77 A2RF78 A2RF79 A2RF82 A2RF83 A2RF84 A2RF85 A2RF89 A2RFA6 A2RFA8
A2RFA9 A2RFD0 A2RFE9 A2RFF4 A2RFM2 A2RFM4 A2RFM5 A2RFN2 A2RFQ3 A2RFQ4 A2RFQ7
A2RFR7 A2RFS6 A2RFW8 A2RFY5 A2RFZ5 A2RFZ7 A2RG00 A2RG22 A2RG27 A2RG28 A2RG30
A2RG31 A2RG32 A2RG33 A2RG55 A2RG67 A2RG70 A2RG71 A2RG72 A2RG73 A2RG75 A2RG81
A2RG91 A2RG92 A2RG93 A2RG97 A2RGA4 A2RGA7 A2RGB0 A2RGB3 A2RGB9 A2RGC8 A2RGC9
A2RGE8 A2RGF2 A2RGF8 A2RGH2 A2RGH7 A2RGI0 A2RGI3 A2RGI4 A2RGJ5 A2RGK2 A2RGQ2
A2RGR1 A2RGR2 A2RGR8 A2RGS2 A2RGS5 A2RGS6 A2RGS9 A2RGT0 A2RGT7 A2RGU4 A2RGU5
A2RGU6 A2RGU8 A2RGU9 A2RGV1 A2RGV3 A2RGX2 A2RGX5 A2RGY0 A2RGY1 A2RGY2 A2RGY3
A2RGZ8 A2RH01 A2RH03 A2RH05 A2RH21 A2RH29 A2RH74 A2RH76 A2RH77 A2RH85 A2RH97
A2RHD8 A2RHG9 A2RHH0 A2RHI1 A2RHI6 A2RHJ9 A2RHL0 A2RHL3 A2RHL6 A2RHM1 A2RHM2
A2RHM3 A2RHN5 A2RHP2 A2RHP4 A2RHP5 A2RHP6 A2RHR3 A2RHS0 A2RHS3 A2RHW0 A2RHW1

A2RHW2 A2RHW9 A2RHX1 A2RHX9 A2RI05 A2RI06 A2RI10 A2RI17 A2RI22 A2RI67 A2RI71
A2RI72 A2RI79 A2RI85 A2RI94 A2RIA0 A2RIA6 A2RIB1 A2RIB7 A2RIC4 A2RIC7 A2RIF3
A2RIG0 A2RIH1 A2RIH4 A2RIM1 A2RIM2 A2RIM4 A2RIM8 A2RIP3 A2RIR4 A2RIS0 A2RIS2
A2RIT6 A2RIU4 A2RIV7 A2RIW0 A2RIW1 A2RIW3 A2RIX0 A2RIY6 A2RJ76 A2RJ85 A2RJB4
A2RJC6 A2RJC8 A2RJD1 A2RJD5 A2RJK2 A2RJK5 A2RJL6 A2RJM8 A2RJP7 A2RJQ3 A2RJQ6
A2RJR5 A2RJR7 A2RJS1 A2RJS3 A2RJS5 A2RJS9 A2RJT1 A2RJT9 A2RJW3 A2RJY0 A2RK17
A2RK21 A2RK24 A2RK25 A2RK41 A2RK45 A2RK93 A2RKA2 A2RKH0 A2RKH4 A2RKH5 A2RKH6
A2RKI8 A2RKI9 A2RKJ5 A2RKJ7 A2RKK4 A2RKP9 A2RKQ0 A2RKQ6 A2RKQ9 A2RKR1 A2RKR3
A2RKR4 A2RKR7 A2RKR8 A2RKS1 A2RKS3 A2RKS4 A2RKS5 A2RKU2 A2RKV4 A2RKV6 A2RKW7
A2RKX0 A2RL29 A2RL40 A2RL43 A2RL47 A2RL82 A2RL87 A2RL88 A2RL98 A2RLA2 A2RLA4
A2RLC0 A2RLC3 A2RLC4 A2RLD8 A2RLF5 A2RLF8 A2RLG3 A2RLG5 A2RLI8 A2RLK1 A2RLK2
A2RLS5 A2RLT4 A2RLU3 A2RLU4 A2RLU5 A2RLW6 A2RLX1 A2RLY6 A2RLZ8 A2RM00 A2RM21
A2RM36 A2RM37 A2RM41 A2RM49 A2RM54 A2RM55 A2RM59 A2RM73 A2RMB7 A2RMC2 A2RME2
A2RME6 A2RMG0 A2RMH2 A2RMH3 A2RMH5 A2RMH6 A2RML8 A2RML9 A2RMM5 A2RMN0 A2RMP4
A2RMQ1 A2RMR1 A2RMR2 A2RMS1 A2RMS5 A2RMT1 A2RMV9 A2RN27 A2RN34 A2RN40 A2RN48
A2RN61 A2RN75 A2RN80 A2RN83 A2RN94 A2RN96 A2RNA8 A2RNF3 A2RNF4 A2RNF7 A2RNG1
A2RNH9 A2RNI8 A2RNI9 A2RNJ0 A2RNK1 A2RNK5 A2RNM6 A2RNM7 A2RNM8 A2RNM9 A2RNN0
A2RNN1 A2RNN2 A2RNN4 A2RNN5 A2RNN6 A2RNN7 A2RNN8 A2RNN9 A2RNP2 A2RNP3 A2RNP4
A2RNP5 A2RNP6 A2RNP7 A2RNP8 A2RNP9 A2RNQ0 A2RNQ1 A2RNQ2 A2RNQ4 A2RNQ5 A2RNQ6
A2RNR2 A2RNS2 A2RNT2 A2RNT8 A2RNU9 A2RNV0 A2RNW2 A2RNW8 A2RNY5 A2RNZ2 A2RNZ4
A2RP06 A2RP07 A2RP08 A2RP10 A2RP20 A2RP28 A2RP37 A2RP61 A2RP65 A2RP72 A2RP73
A2RP74 A2RPW9 A2RQ29 A2RRD8 A2RRH5 A2RRS8 A2RRT3 A2RRU1 A2RRU3 A2RRV3 A2RRV9
A2RRW4 A2RRX6 A2RRY8 A2RSQ1 A2RSX4 A2RSX7 A2RSY6 A2RT62 A2RTL5 A2RTY3 A2RU37
A2RU49 A2RU54 A2RUB1 A2RUB6 A2RUC4 A2RUH7 A2RUQ5 A2RUR9 A2RUV5 A2RV01 A2RV06
A2RV18 A2RV29 A2RVA7 A2RVK7 A2RVQ5 A2RVR7 A2RVS4 A2RVS6 A2RVV7 A2RWD6 A2RWD7
A2RWH4 A2RWR9 A2RXI5 A2RXU4 A2RYI2 A2RYI3 A2RYL1 A2RYL2 A2RYW5 A2RZW5 A2RZY1
A2S0L3 A2S0N4 A2S0N7 A2S105 A2S127 A2S129 A2S134 A2S138 A2S233 A2S242 A2S244
A2S245 A2S278 A2S286 A2S287 A2S290 A2S291 A2S299 A2S2A0 A2S2A2 A2S2A3 A2S2A6
A2S2A7 A2S2B2 A2S2B3 A2S2C9 A2S2I6 A2S2L0 A2S2L1 A2S2L3 A2S2N4 A2S2N6 A2S370
A2S373 A2S381 A2S3C6 A2S3D4 A2S3Q4 A2S3Q7 A2S3Z6 A2S430 A2S463 A2S464 A2S472
A2S4A9 A2S4B1 A2S4B3 A2S4B5 A2S4B8 A2S4E7 A2S4F3 A2S4I0 A2S4M9 A2S4N1 A2S4N7
A2S4N8 A2S4N9 A2S4P0 A2S4P8 A2S4R1 A2S4R7 A2S4R9 A2S4V8 A2S4W0 A2S4Y8 A2S4Y9
A2S4Z0 A2S4Z4 A2S504 A2S514 A2S516 A2S519 A2S552 A2S554 A2S561 A2S563 A2S565
A2S567 A2S570 A2S581 A2S582 A2S583 A2S587 A2S590 A2S594 A2S595 A2S597 A2S5A6
A2S5D2 A2S5D5 A2S5J4 A2S5J7 A2S5L0 A2S5P8 A2S5R3 A2S5R4 A2S5R7 A2S5R8 A2S5R9
A2S5S0 A2S5S9 A2S5U3 A2S5U4 A2S5U7 A2S5V3 A2S5V4 A2S5X1 A2S5Y8 A2S5Y9 A2S625
A2S628 A2S6B1 A2S6B4 A2S6B7 A2S6C0 A2S6C3 A2S6C4 A2S6D4 A2S6F4 A2S6F6 A2S6J1
A2S6K9 A2S6L0 A2S6N7 A2S6N9 A2S6T3 A2S6U9 A2S6W2 A2S703 A2S718 A2S747 A2S750
A2S753 A2S754 A2S755 A2S756 A2S7F1 A2S7F9 A2S7G2 A2S7G3 A2S7G4 A2S7G5 A2S7G6
A2S7G7 A2S7H1 A2S7H2 A2S7H3 A2S7H5 A2S7H6 A2S7H7 A2S7H8 A2S7H9 A2S7I0 A2S7I1
A2S7I2 A2S7I3 A2S7I4 A2S7I5 A2S7I6 A2S7I7 A2S7I8 A2S7I9 A2S7J0 A2S7J1 A2S7J2
A2S7J3 A2S7J4 A2S7J5 A2S7J7 A2S7J8 A2S7J9 A2S7K0 A2S7K1 A2S7K2 A2S7K3 A2S7P3
A2S7Q4 A2S7R1 A2S7R2 A2S7R3 A2S7T4 A2S7V6 A2S800 A2S808 A2S836 A2S860 A2S862
A2S869 A2S874 A2S877 A2S8D2 A2S8D3 A2S8D4 A2S8D8 A2S8H7 A2S8I8 A2S8I9 A2S8J0

A2S8K4 A2S8L1 A2S8L9 A2S8N7 A2S8Q1 A2S8U3 A2S924 A2S925 A2S926 A2S933 A2S947
A2S948 A2S951 A2S993 A2S994 A2S995 A2S996 A2S997 A2S998 A2S999 A2S9D3 A2S9D4
A2S9D9 A2S9G4 A2S9G8 A2S9H0 A2S9I2 A2S9I5 A2S9K2 A2S9M1 A2S9S5 A2S9S9 A2S9T0
A2S9T2 A2S9T5 A2S9Y0 A2SA02 A2SA04 A2SA47 A2SA96 A2SAC2 A2SAC7 A2SAE9 A2SAF0
A2SAF7 A2SAG2 A2SAG6 A2SAH4 A2SAH5 A2SAN1 A2SAQ8 A2SAS9 A2SAT3 A2SAU5 A2SAW7
A2SAX8 A2SB31 A2SB35 A2SB51 A2SB58 A2SB60 A2SB66 A2SB69 A2SB72 A2SB73 A2SB74
A2SB75 A2SB78 A2SB87 A2SB89 A2SB97 A2SB99 A2SBA7 A2SBC3 A2SBD6 A2SBD7 A2SBG2
A2SBG3 A2SBG4 A2SBH6 A2SBI6 A2SBI7 A2SBK1 A2SBM4 A2SBU8 A2SBV7 A2SBW1 A2SC07
A2SC26 A2SC77 A2SC91 A2SC97 A2SCE2 A2SCE3 A2SCF3 A2SCF7 A2SCG5 A2SCI8 A2SCK0
A2SCV0 A2SCV1 A2SCX6 A2SCX7 A2SCY6 A2SCY7 A2SD05 A2SD22 A2SD34 A2SD35 A2SD36
A2SD37 A2SD39 A2SD40 A2SD46 A2SD47 A2SD53 A2SD55 A2SD77 A2SDC3 A2SDF7 A2SDI6
A2SDI7 A2SDI8 A2SDJ0 A2SDJ1 A2SDJ2 A2SDN1 A2SDN6 A2SDN9 A2SDR5 A2SDV6 A2SE03
A2SE05 A2SE06 A2SE08 A2SE09 A2SE10 A2SE11 A2SEP2 A2SES6 A2SES7 A2SES8 A2SES9
A2SET3 A2SET8 A2SET9 A2SEX7 A2SEX8 A2SEY3 A2SF60 A2SF75 A2SF76 A2SF90 A2SF93
A2SFB4 A2SFB5 A2SFB6 A2SFG0 A2SFG3 A2SFG5 A2SFJ5 A2SFL0 A2SFL8 A2SFM2 A2SFP8
A2SFS2 A2SFU1 A2SFU2 A2SFV1 A2SFV4 A2SFW9 A2SFZ3 A2SFZ9 A2SG01 A2SG02 A2SG83
A2SG87 A2SG89 A2SG95 A2SGN7 A2SGQ6 A2SGS4 A2SGT2 A2SGT8 A2SH03 A2SH04 A2SH05
A2SH07 A2SH39 A2SH40 A2SH42 A2SH61 A2SH83 A2SH85 A2SH95 A2SH96 A2SH97 A2SHA4
A2SHB1 A2SHB8 A2SHB9 A2SHF3 A2SHH0 A2SHK9 A2SHL0 A2SHL8 A2SHM0 A2SHP8 A2SHQ1
A2SHS3 A2SHS4 A2SHS5 A2SHS6 A2SHT0 A2SHT9 A2SI25 A2SI33 A2SI34 A2SI71 A2SI90
A2SIL9 A2SIM2 A2SIQ2 A2SIQ4 A2SIR1 A2SIR4 A2SIR5 A2SIT7 A2SIU8 A2SIU9 A2SIX7
A2SIY4 A2SJ45 A2SJ46 A2SJA7 A2SJA8 A2SJB0 A2SJB9 A2SJD1 A2SJJ0 A2SJJ7 A2SJK3
A2SJQ7 A2SJR5 A2SJS7 A2SJZ5 A2SK03 A2SK12 A2SK91 A2SK99 A2SKA3 A2SKF8 A2SKF9
A2SKK9 A2SKL7 A2SKM0 A2SKM5 A2SKM7 A2SKQ7 A2SKR5 A2SKU1 A2SKU2 A2SKU4 A2SKV0
A2SKV4 A2SKV9 A2SKW6 A2SKX8 A2SL00 A2SL35 A2SL36 A2SL47 A2SL48 A2SL50 A2SL57
A2SL82 A2SL95 A2SLA3 A2SLA5 A2SLD0 A2SLD1 A2SLD2 A2SLD3 A2SLD4 A2SLD5 A2SLD6
A2SLD8 A2SLD9 A2SLE0 A2SLE1 A2SLE2 A2SLE3 A2SLE4 A2SLE6 A2SLE8 A2SLE9 A2SLF0
A2SLF1 A2SLF2 A2SLF3 A2SLF4 A2SLF6 A2SLF7 A2SLF8 A2SLF9 A2SLG0 A2SLG1 A2SLG2
A2SLG4 A2SLG5 A2SLG6 A2SLG7 A2SLG8 A2SLG9 A2SLH5 A2SLJ8 A2SLS6 A2SLT0 A2SLT3
A2SLU8 A2SLW2 A2SLX7 A2SLY9 A2SM29 A2SM89 A2SM97 A2SMA4 A2SMC9 A2SMD0 A2SMF0
A2SMF1 A2SMH6 A2SMI8 A2SMJ2 A2SMZ3 A2SPC3 A2SPE0 A2SPG1 A2SPI9 A2SPJ9 A2SPK0
A2SPK1 A2SPK3 A2SPK6 A2SPK7 A2SPK8 A2SPK9 A2SPL1 A2SPL2 A2SPL3 A2SPL5 A2SPL7
A2SPM1 A2SPM2 A2SPM3 A2SPM8 A2SPS3 A2SPT0 A2SPT1 A2SPV6 A2SPY9 A2SPZ6 A2SQ45
A2SQ53 A2SQ57 A2SQ60 A2SQ85 A2SQ86 A2SQ87 A2SQ91 A2SQB8 A2SQC6 A2SQD1 A2SQE9
A2SQF6 A2SQG5 A2SQG6 A2SQG9 A2SQH8 A2SQI5 A2SQJ8 A2SQM1 A2SQM4 A2SQM6 A2SQR0
A2SQR6 A2SQS1 A2SQT3 A2SQU1 A2SQU3 A2SQU7 A2SQU8 A2SQV5 A2SQV7 A2SQW1 A2SQW3
A2SQZ4 A2SR20 A2SR38 A2SR44 A2SR54 A2SR62 A2SR70 A2SR71 A2SR73 A2SR93 A2SRC6
A2SRE2 A2SRG2 A2SRI7 A2SRK7 A2SRK8 A2SRK9 A2SRZ6 A2SS66 A2SS72 A2SS78 A2SS82
A2SS93 A2SSJ1 A2SSJ7 A2SSL7 A2SSM7 A2SSN1 A2SSN5 A2SSS7 A2SST9 A2SSV0 A2SSV2
A2SSV3 A2SSV8 A2SSV9 A2SSW0 A2SSW2 A2SSW4 A2SSW6 A2SSW7 A2ST12 A2ST15 A2ST18
A2ST39 A2ST45 A2ST53 A2ST55 A2ST56 A2ST69 A2ST72 A2ST81 A2ST82 A2ST90 A2ST94
A2ST96 A2STA4 A2STA7 A2STB6 A2STB8 A2STE9 A2STF0 A2STI6 A2STI7 A2STI8 A2STI9
A2STJ1 A2STJ3 A2STJ6 A2STK8 A2STL6 A2STM4 A2STM7 A2STM8 A2STQ3 A2STQ8 A2STS4
A2STS6 A2STT9 A2STU0 A2STU3 A2STX0 A2STX1 A2STX4 A2STY4 A2STZ4 A2STZ6 A2SU05

A2SU08 A2SU11 A2SU30 A2SU54 A2SU66 A2SU89 A2SU98 A2SUH3 A2SW84 A2SXS5 A2SZS3
A2T308 A2T310 A2T312 A2T321 A2T322 A2T323 A2T324 A2T331 A2T335 A2T354 A2T355
A2T356 A2T357 A2T358 A2T365 A2T366 A2T367 A2T368 A2T369 A2T370 A2T371 A2T372
A2T373 A2T374 A2T375 A2T378 A2T381 A2T390 A2T391 A2T392 A2T3M0 A2T3M2 A2T3M4
A2T3N5 A2T3P0 A2T3Q9 A2T3R0 A2T3R5 A2T3S0 A2T3S5 A2T3T1 A2T6E3 A2T6F8 A2T6H5
A2T6K9 A2T6V8 A2T6W2 A2T6X6 A2T6Z0 A2T711 A2T712 A2T713 A2T715 A2T736 A2T737
A2T748 A2T759 A2T762 A2T763 A2T764 A2T771 A2T777 A2T779 A2T7D1 A2T7D2 A2T7D7
A2T7D9 A2T7E6 A2T7F2 A2T7F3 A2T7F4 A2T7G6 A2T7G9 A2T7H5 A2T7H7 A2T7J2 A2T7L5
A2T7L7 A2T7L8 A2T7M0 A2T7P4 A2T7S4 A2T7T2 A2T806 A2T812 A2T833 A2T928 A2TBT9
A2TBU0 A2TBU1 A2TBU2 A2TBU4 A2TC44 A2TC45 A2TED3 A2TIL1 A2V735 A2VCW9 A2VCX1
A2VCZ5 A2VD00 A2VD13 A2VD33 A2VD82 A2VD83 A2VD92 A2VD93 A2VD95 A2VDB3 A2VDC2
A2VDD2 A2VDM0 A2VDN6 A2VDP0 A2VDP1 A2VDP4 A2VDQ5 A2VDQ7 A2VDR2 A2VDR9 A2VDS0
A2VDS1 A2VDT4 A2VDW6 A2VDW9 A2VDX7 A2VDY4 A2VDY6 A2VDZ3 A2VE01 A2VE14 A2VE40
A2VE44 A2VE45 A2VE52 A2VE56 A2VE78 A2VE99 A2VEA3 A2VEA7 A2WJP3 A2WKK5 A2WKP3
A2WKP5 A2WKS3 A2WKS5 A2WKS8 A2WKT1 A2WKT4 A2WL19 A2WLS0 A2WM14 A2WMH2 A2WN93
A2WNB0 A2WNH1 A2WNM0 A2WPN7 A2WPU3 A2WQ39 A2WQE4 A2WQG7 A2WSI7 A2WSZ6 A2WTQ5
A2WV68 A2WW87 A2WWU2 A2WWU7 A2WX30 A2WXB2 A2WXR5 A2WXT0 A2WXU2 A2WXV8 A2WY46
A2WYE9 A2WYS7 A2WYS8 A2WZI4 A2X052 A2X0M1 A2X0Q0 A2X0Q3 A2X0R4 A2X1A1 A2X254
A2X2G3 A2X2K3 A2X4M8 A2X5H5 A2X635 A2X674 A2X6S3 A2X6X1 A2X7F7 A2X7Q3 A2X7U1
A2X899 A2X8Q3 A2X980 A2X9L8 A2X9W8 A2XAM0 A2XAM1 A2XAZ3 A2XB18 A2XBT1 A2XCH8
A2XCT8 A2XD08 A2XD15 A2XD35 A2XDD6 A2XDK5 A2XDY1 A2XE76 A2XEV1 A2XEX2 A2XF66
A2XFC1 A2XFC7 A2XFC8 A2XFF4 A2XFI3 A2XFU4 A2XFU5 A2XG55 A2XG77 A2XGP6 A2XGQ1
A2XHJ3 A2XHJ5 A2XHU0 A2XI04 A2XIA8 A2XIK9 A2XK00 A2XK56 A2XK57 A2XKG2 A2XKR7
A2XKU9 A2XLF2 A2XLF4 A2XLG5 A2XLL2 A2XLV9 A2XM23 A2XMJ1 A2XMK6 A2XMN1 A2XMN2
A2XMV1 A2XN40 A2XNF0 A2XNK3 A2XNL6 A2XNR6 A2XNS1 A2XQV4 A2XQX0 A2XRZ0 A2XRZ6
A2XSX6 A2XT03 A2XTW9 A2XU53 A2XU85 A2XUN8 A2XUW1 A2XV58 A2XVF7 A2XW22 A2XW69
A2XWA9 A2XWL6 A2XX39 A2XY47 A2XYW4 A2XYY8 A2XZI2 A2XZL0 A2XZN0 A2XZR3 A2Y007
A2Y040 A2Y053 A2Y0H2 A2Y0J7 A2Y0Q2 A2Y0T4 A2Y1D7 A2Y205 A2Y3Q5 A2Y4B6 A2Y4R8
A2Y533 A2Y5G8 A2Y5L9 A2Y5T7 A2Y609 A2Y720 A2Y7R3 A2Y7R5 A2Y8B9 A2Y8S6 A2Y8X2
A2Y931 A2Y953 A2Y9M4 A2Y9P0 A2Y9X7 A2YAA5 A2YAG8 A2YAW8 A2YB34 A2YCH5 A2YEB4
A2YEQ6 A2YEZ6 A2YG67 A2YGL9 A2YGP6 A2YH41 A2YH60 A2YII8 A2YIW7 A2YL07 A2YM35
A2YMC5 A2YMC6 A2YMR0 A2YMU2 A2YMV6 A2YN17 A2YN56 A2YNH4 A2YNI2 A2YNP0 A2YNY4
A2YP56 A2YPT7 A2YQ58 A2YQ76 A2YQD9 A2YQK9 A2YQL4 A2YQT7 A2YQU8 A2YR10 A2YST1
A2YUL5 A2YUQ6 A2YV85 A2YVE5 A2YVR7 A2YW03 A2YWA6 A2YWC0 A2YWI3 A2YXJ7 A2YXQ1
A2YXQ7 A2YXS5 A2YXU2 A2Z0C0 A2Z0Q0 A2Z182 A2Z1U1 A2Z1W9 A2Z259 A2Z2J6 A2Z388
A2Z389 A2Z3C4 A2Z3E5 A2Z3I9 A2Z4C4 A2Z4C5 A2Z5S8 A2Z6W1 A2Z730 A2Z734 A2Z7B3
A2Z7C4 A2Z8F9 A2Z8S0 A2Z928 A2Z9B8 A2Z9Q7 A2ZAC2 A2ZAX5 A2ZB00 A2ZBV1 A2ZCP0
A2ZCQ7 A2ZDX4 A2ZDX6 A2ZDX8 A2ZDX9 A2ZET6 A2ZEX7 A2ZF53 A2ZH20 A2ZH47 A2ZHL0
A2ZIW7 A2ZK26 A2ZK29 A2ZLC1 A2ZLS4 A2ZLU6 A2ZM73 A2ZMH2 A2ZMR9 A2ZP58 A2ZRG4
A2ZRY8 A2ZVY5 A3A463 A3A8W2 A3ABE1 A3ACF3 A3AGM4 A3AHG5 A3AJ77 A3ANB5 A3AVH5
A3AWH5 A3AZ88 A3AZ89 A3AZW5 A3B0Y1 A3B6V0 A3B9A0 A3BDI8 A3BFT0 A3BH85 A3BH91
A3BKM2 A3BLC3 A3BLS0 A3BMN9 A3BN26 A3BNA1 A3BT52 A3BUD2 A3BYC1 A3C039 A3C4N5
A3C4S4 A3C6D7 A3C6G9 A3CEM4 A3CJX1 A3CJX7 A3CJY8 A3CJZ1 A3CJZ4 A3CK22 A3CK30
A3CK60 A3CK62 A3CK63 A3CK64 A3CK65 A3CK66 A3CK67 A3CK68 A3CK69 A3CK70 A3CK71

A3CK72 A3CK73 A3CK74 A3CK75 A3CK76 A3CK77 A3CK79 A3CK80 A3CK81 A3CK82 A3CK83
A3CK85 A3CK86 A3CK87 A3CK88 A3CK89 A3CK90 A3CK91 A3CKD3 A3CKD4 A3CKF4 A3CKG4
A3CKI0 A3CKI1 A3CKK5 A3CKK8 A3CKP1 A3CKQ4 A3CKR5 A3CKS0 A3CKS8 A3CKT0 A3CKU4
A3CKV4 A3CL11 A3CL24 A3CL27 A3CL31 A3CL32 A3CL34 A3CL43 A3CL56 A3CL74 A3CL77
A3CL78 A3CL81 A3CLF4 A3CLF7 A3CLF8 A3CLF9 A3CLH1 A3CLH4 A3CLI1 A3CLJ1 A3CLL0
A3CLL3 A3CLL8 A3CLL9 A3CLM2 A3CLN5 A3CLP1 A3CLP4 A3CLQ4 A3CLQ7 A3CLQ8 A3CLS0
A3CLS2 A3CLS3 A3CLS9 A3CLU1 A3CLW6 A3CLX2 A3CLX4 A3CLY3 A3CLY4 A3CLY6 A3CM01
A3CM02 A3CM04 A3CM30 A3CM33 A3CM36 A3CM38 A3CM39 A3CM46 A3CM69 A3CMF2 A3CMF5
A3CMJ2 A3CMJ3 A3CMP2 A3CMP3 A3CMP6 A3CMQ4 A3CMQ6 A3CMR4 A3CMR5 A3CMR7 A3CMS0
A3CMS1 A3CMT0 A3CMT1 A3CMU3 A3CMU4 A3CMU9 A3CMV1 A3CMV2 A3CMW1 A3CMW2 A3CMY9
A3CN05 A3CN08 A3CN22 A3CN32 A3CN38 A3CN43 A3CN49 A3CN67 A3CN70 A3CN85 A3CN88
A3CN89 A3CNA8 A3CNA9 A3CNB0 A3CNB2 A3CNB9 A3CNE1 A3CNE6 A3CNE7 A3CNF4 A3CNH0
A3CNI5 A3CNI7 A3CNI9 A3CNJ9 A3CNM0 A3CNN8 A3CNR2 A3CNR3 A3CNR9 A3CNS2 A3CNS8
A3CNS9 A3CNT0 A3CNT1 A3CNT2 A3CNT3 A3CNT5 A3CNT6 A3CNT7 A3CNT9 A3CNU0 A3CNV3
A3CNV4 A3CNV6 A3CNV7 A3CNV8 A3CNV9 A3CNX3 A3CNY4 A3CNY7 A3CNY8 A3CNZ0 A3CP09
A3CP10 A3CP17 A3CP33 A3CP36 A3CP46 A3CP56 A3CP85 A3CP96 A3CPA0 A3CPA2 A3CPA3
A3CPA5 A3CPA6 A3CPH0 A3CPH5 A3CPH6 A3CPJ2 A3CPJ7 A3CPJ9 A3CPK7 A3CPK8 A3CPK9
A3CPL6 A3CPM5 A3CPQ6 A3CPQ7 A3CPQ8 A3CPR2 A3CPS3 A3CPS7 A3CPT0 A3CPT4 A3CPV3
A3CPV8 A3CPV9 A3CPX4 A3CPX6 A3CPY3 A3CPY4 A3CPY5 A3CPY9 A3CPZ9 A3CQ04 A3CQ05
A3CQ06 A3CQ17 A3CQ18 A3CQ22 A3CQ23 A3CQ45 A3CQ78 A3CQ86 A3CQ97 A3CQB0 A3CQB4
A3CQB5 A3CQB8 A3CQC2 A3CQC3 A3CQC4 A3CQD3 A3CQF1 A3CQG6 A3CQH4 A3CQI2 A3CQI5
A3CQM2 A3CQM3 A3CQM4 A3CQN5 A3CQP8 A3CQQ3 A3CQQ4 A3CQQ6 A3CQT4 A3CQT5 A3CQU5
A3CQV0 A3CQW2 A3CQW3 A3CQZ6 A3CQZ7 A3CQZ8 A3CR02 A3CR11 A3CR14 A3CR17 A3CR40
A3CR42 A3CR43 A3CRA2 A3CRA9 A3CRB1 A3CRB2 A3CRC5 A3CRD3 A3CRD6 A3CRE5 A3CRI3
A3CRJ3 A3CRK8 A3CRP6 A3CRT6 A3CRT9 A3CS13 A3CS45 A3CS46 A3CS56 A3CSG3 A3CSI6
A3CSJ1 A3CSJ5 A3CSJ6 A3CSK8 A3CSL0 A3CSL3 A3CSP4 A3CSP5 A3CSQ2 A3CSQ6 A3CSU7
A3CSX5 A3CSY4 A3CSZ5 A3CSZ7 A3CSZ8 A3CT00 A3CT01 A3CT02 A3CT03 A3CT06 A3CT07
A3CT08 A3CT10 A3CT12 A3CT16 A3CT17 A3CT18 A3CT21 A3CT23 A3CT71 A3CT76 A3CT77
A3CT78 A3CTA1 A3CTF1 A3CTG2 A3CTG3 A3CTG9 A3CTJ9 A3CTK0 A3CTP4 A3CTR4 A3CTR8
A3CTR9 A3CTT1 A3CTT5 A3CTU1 A3CTY6 A3CU20 A3CU49 A3CU51 A3CU63 A3CU65 A3CU68
A3CU73 A3CU82 A3CU83 A3CU84 A3CUA0 A3CUB9 A3CUF2 A3CUG3 A3CUG8 A3CUS9 A3CUX9
A3CV15 A3CV22 A3CV25 A3CV28 A3CV31 A3CV35 A3CVB4 A3CVF0 A3CVI3 A3CVI6 A3CVI7
A3CVI8 A3CVJ2 A3CVJ3 A3CVP2 A3CVP8 A3CVQ1 A3CVQ3 A3CVR8 A3CVT5 A3CVT7 A3CVV0
A3CVW1 A3CVW4 A3CVZ3 A3CW49 A3CW55 A3CW56 A3CW58 A3CW66 A3CW67 A3CW75 A3CWF5
A3CWF9 A3CWH6 A3CWH7 A3CWH8 A3CWI3 A3CWI6 A3CWJ3 A3CWL0 A3CWL3 A3CWL4 A3CWL5
A3CWM4 A3CWP1 A3CWS5 A3CWS6 A3CWS8 A3CWU4 A3CWU5 A3CWV2 A3CWW3 A3CWX7 A3CWY7
A3CWZ2 A3CWZ4 A3CWZ5 A3CWZ6 A3CWZ8 A3CX01 A3CX23 A3CX25 A3CX32 A3CX39 A3CX68
A3CX76 A3CXE7 A3CXH4 A3CXI2 A3CXJ3 A3CXJ5 A3CXJ6 A3CXM8 A3CXN0 A3CXN3 A3CXN8
A3CXP8 A3CXR7 A3CXS0 A3CXS2 A3CXS4 A3CXU2 A3CXW9 A3CXX4 A3CY14 A3CY20 A3CY32
A3CYB6 A3CYG8 A3CYG9 A3CYH5 A3CYH7 A3CYI6 A3CYI7 A3CYJ2 A3CYJ5 A3CYK4 A3CYK8
A3CYL0 A3CYL1 A3CYL3 A3CYL9 A3CYM5 A3CYN0 A3CYP1 A3CYP7 A3CYQ1 A3CYW3 A3CYZ9
A3CZ93 A3CZA5 A3CZD8 A3CZE0 A3CZH4 A3CZI9 A3CZJ1 A3CZJ3 A3CZJ4 A3CZJ6 A3CZJ7
A3CZJ8 A3CZK5 A3CZK7 A3CZK8 A3CZL0 A3CZL2 A3CZL3 A3CZM2 A3CZN2 A3CZT7 A3CZX1
A3CZY6 A3D001 A3D007 A3D017 A3D022 A3D023 A3D024 A3D039 A3D074 A3D075 A3D083

A3D084 A3D085 A3D0C6 A3D0E1 A3D0F8 A3D0G0 A3D0G1 A3D0H5 A3D0P6 A3D0T8 A3D0U1
A3D0V3 A3D0Y0 A3D0Y5 A3D0Y7 A3D0Z9 A3D130 A3D147 A3D178 A3D179 A3D180 A3D186
A3D187 A3D188 A3D1A0 A3D1F4 A3D1F5 A3D1F9 A3D1G2 A3D1G5 A3D1Q3 A3D1Q4 A3D1R1
A3D1R5 A3D1R9 A3D1S4 A3D1T1 A3D1T8 A3D1T9 A3D1U3 A3D1V6 A3D1W5 A3D1W6 A3D1W7
A3D1W8 A3D1X1 A3D262 A3D286 A3D289 A3D2B0 A3D2B1 A3D2B2 A3D2B7 A3D2C2 A3D2C6
A3D2D1 A3D2K2 A3D2K5 A3D2K6 A3D2K7 A3D2K8 A3D2L1 A3D2L9 A3D2M7 A3D2P0 A3D2P8
A3D302 A3D303 A3D304 A3D305 A3D306 A3D367 A3D383 A3D392 A3D3B1 A3D3D1 A3D3D2
A3D3D5 A3D3E8 A3D3F2 A3D3F9 A3D3G2 A3D3G4 A3D3G5 A3D3Q9 A3D3R1 A3D3R2 A3D3R4
A3D3R8 A3D3S2 A3D3U4 A3D3V7 A3D437 A3D440 A3D465 A3D474 A3D475 A3D477 A3D479
A3D480 A3D481 A3D499 A3D4A1 A3D4A6 A3D4A7 A3D4A9 A3D4I2 A3D4I4 A3D4I5 A3D4J4
A3D4P4 A3D547 A3D558 A3D569 A3D571 A3D573 A3D574 A3D577 A3D595 A3D5A5 A3D5B0
A3D5B1 A3D5D5 A3D5E2 A3D5E3 A3D5E5 A3D5E9 A3D5F1 A3D5F3 A3D5F8 A3D5G9 A3D5H7
A3D5J3 A3D5K4 A3D5L4 A3D5L9 A3D5M3 A3D5M6 A3D5M8 A3D5N2 A3D5N8 A3D5P1 A3D5Q0
A3D5Q2 A3D5Q5 A3D5Y2 A3D600 A3D628 A3D630 A3D631 A3D666 A3D669 A3D671 A3D676
A3D678 A3D6B1 A3D6B2 A3D6C6 A3D6H7 A3D6V2 A3D6V4 A3D6V5 A3D6V8 A3D6V9 A3D6W2
A3D6Z7 A3D705 A3D707 A3D709 A3D738 A3D781 A3D783 A3D785 A3D786 A3D789 A3D790
A3D792 A3D796 A3D798 A3D7B5 A3D7C4 A3D7C5 A3D7C9 A3D7D0 A3D7I1 A3D7J1 A3D7J2
A3D7J3 A3D7J4 A3D7J9 A3D7K1 A3D7K3 A3D7K4 A3D7K5 A3D7K6 A3D7K8 A3D7L0 A3D7L1
A3D7L4 A3D7M5 A3D7M8 A3D7N0 A3D7N6 A3D7N9 A3D7P1 A3D7P7 A3D7P8 A3D7P9 A3D7R0
A3D7R1 A3D7S3 A3D7T3 A3D7T4 A3D7V9 A3D7X8 A3D819 A3D822 A3D829 A3D837 A3D8A3
A3D8A9 A3D8E4 A3D8P6 A3D8P9 A3D8Q2 A3D8Q3 A3D8Q4 A3D8Q5 A3D8R3 A3D8R4 A3D8R8
A3D8T3 A3D8W0 A3D8W9 A3D8X0 A3D944 A3D958 A3D9A0 A3D9A3 A3D9C5 A3D9C6 A3D9F2
A3D9I1 A3D9I5 A3D9J5 A3D9J7 A3D9Q0 A3D9Q3 A3D9Q4 A3D9T2 A3D9W1 A3D9W6 A3DA02
A3DA03 A3DA14 A3DA15 A3DA22 A3DA23 A3DA25 A3DA26 A3DA29 A3DA46 A3DA47 A3DA48
A3DA49 A3DA50 A3DA51 A3DA53 A3DA54 A3DA55 A3DA56 A3DA57 A3DA58 A3DA59 A3DA60
A3DA61 A3DA62 A3DA63 A3DA64 A3DA65 A3DA66 A3DA67 A3DA68 A3DA69 A3DA71 A3DA72
A3DA73 A3DA74 A3DA75 A3DA79 A3DA80 A3DA84 A3DA85 A3DAB1 A3DAB2 A3DAC1 A3DAC9
A3DAF4 A3DAF6 A3DAG6 A3DAM8 A3DAN0 A3DAS4 A3DAS5 A3DAS7 A3DAT0 A3DAT1 A3DB95
A3DB98 A3DBA0 A3DBD3 A3DBD5 A3DBD7 A3DBE8 A3DBI2 A3DBJ9 A3DBK4 A3DBK6 A3DBL4
A3DBL9 A3DBM3 A3DBM4 A3DBM5 A3DBM8 A3DBQ1 A3DBQ2 A3DBR3 A3DBR5 A3DBS2 A3DBS4
A3DBS5 A3DBU0 A3DBU1 A3DBU2 A3DBU3 A3DBU4 A3DBX6 A3DBX9 A3DC22 A3DCC8 A3DCD4
A3DCF7 A3DCG9 A3DCH6 A3DCH7 A3DCK9 A3DCM0 A3DCX4 A3DCX5 A3DCX9 A3DD04 A3DD27
A3DD54 A3DD79 A3DD83 A3DD94 A3DDA0 A3DDC1 A3DDC2 A3DDC8 A3DDC9 A3DDD8 A3DDG4
A3DDG7 A3DDH1 A3DDH3 A3DDH5 A3DDI1 A3DDI3 A3DDI9 A3DDJ2 A3DDJ7 A3DDM1 A3DDM2
A3DDP0 A3DDQ3 A3DDS6 A3DDS7 A3DDS8 A3DDV5 A3DDV6 A3DDV7 A3DDX7 A3DDZ7 A3DE07
A3DE08 A3DE09 A3DE13 A3DE14 A3DE17 A3DE34 A3DE35 A3DE43 A3DE44 A3DE48 A3DE52
A3DE56 A3DE57 A3DE58 A3DE59 A3DE67 A3DE77 A3DE79 A3DE86 A3DE88 A3DE94 A3DEA5
A3DEB1 A3DEB6 A3DEB9 A3DEC2 A3DEE6 A3DEL6 A3DEN2 A3DEQ2 A3DER9 A3DES1 A3DES6
A3DES7 A3DET1 A3DEU0 A3DEU4 A3DEU9 A3DEV1 A3DEX9 A3DEY2 A3DF00 A3DF04 A3DF15
A3DF19 A3DF23 A3DF24 A3DF25 A3DF27 A3DF35 A3DF46 A3DF48 A3DF51 A3DF80 A3DF94
A3DFH7 A3DGC4 A3DGF4 A3DGF7 A3DGF8 A3DGF9 A3DGG0 A3DGI7 A3DGJ4 A3DGK5 A3DGK6
A3DGR3 A3DGS4 A3DGS7 A3DH19 A3DH20 A3DH43 A3DH53 A3DH61 A3DHB7 A3DHB8 A3DHF8
A3DHH6 A3DHI4 A3DHL2 A3DHL3 A3DHM4 A3DHN0 A3DHU9 A3DHY6 A3DHY7 A3DHY8 A3DHZ3
A3DHZ4 A3DHZ7 A3DI04 A3DI22 A3DI26 A3DIE1 A3DIE8 A3DIF0 A3DIF3 A3DIH0 A3DIJ8

A3DIK9 A3DIL4 A3DIL9 A3DIP6 A3DIP8 A3DIY3 A3DIY7 A3DIZ0 A3DIZ1 A3DIZ4 A3DIZ5
A3DIZ7 A3DIZ8 A3DIZ9 A3DJ00 A3DJ07 A3DJ09 A3DJ10 A3DJ11 A3DJ13 A3DJ18 A3DJ77
A3DJE3 A3DJE9 A3DJF0 A3DJF3 A3DJF6 A3DJF7 A3DJG6 A3DJH1 A3DJH2 A3DJH3 A3DJH4
A3DJH5 A3DJH6 A3DJH7 A3DJH8 A3DJH9 A3DJI0 A3DJI1 A3DJI2 A3DJI3 A3DJI5 A3DJI6
A3DJI8 A3DJI9 A3DJJ0 A3DJJ1 A3DJJ3 A3DJJ6 A3DJJ7 A3DJJ8 A3DJJ9 A3DJK0 A3DJK1
A3DJK2 A3DJK8 A3DJL6 A3DJP1 A3DJX6 A3DK03 A3DK09 A3DK16 A3DK17 A3DK21 A3DK82
A3DKB6 A3DKL1 A3DKS1 A3DL27 A3DL40 A3DL59 A3DLD9 A3DLI4 A3DLP6 A3DLR9 A3DLT5
A3DLU8 A3DLV7 A3DLW5 A3DM32 A3DM47 A3DM48 A3DM49 A3DM71 A3DM74 A3DM76 A3DM85
A3DM89 A3DM90 A3DMD0 A3DMF7 A3DMF9 A3DMG2 A3DMG3 A3DMH6 A3DML6 A3DMM2 A3DMP2
A3DMP8 A3DMQ1 A3DMQ3 A3DMQ4 A3DMQ5 A3DMR0 A3DMR1 A3DMR5 A3DMR6 A3DMR8 A3DMR9
A3DMS0 A3DMS1 A3DMS6 A3DMS9 A3DMT2 A3DMT5 A3DMU2 A3DMV6 A3DMX7 A3DMX9 A3DMY1
A3DMZ6 A3DN06 A3DN07 A3DN21 A3DN24 A3DN27 A3DN33 A3DN67 A3DNA4 A3DNA6 A3DNA7
A3DNB1 A3DNB2 A3DNB3 A3DNB4 A3DNB5 A3DNB7 A3DNB8 A3DNB9 A3DNC2 A3DNC3 A3DNC6
A3DNC7 A3DNC8 A3DNC9 A3DND2 A3DND3 A3DND4 A3DNG5 A3DNG9 A3DNH3 A3DNH4 A3DNH5
A3DNH6 A3DNI1 A3DNI2 A3DNI3 A3DNI8 A3DNJ6 A3DNJ9 A3DNK0 A3DNK3 A3DNN7 A3DNT5
A3DNV8 A3DNV9 A3DNW6 A3DNX9 A3DNY5 A3DNY7 A3DP40 A3DP41 A3DP49 A3DP85 A3DP93
A3DPB2 A3DSK5 A3DSK6 A3DSK8 A3DSK9 A3DSL0 A3E3H0 A3E4D8 A3E4F9 A3E7Z6 A3EX96
A3EX97 A3EXD2 A3EXD3 A3EXH2 A3F4S0 A3F5F0 A3F5L9 A3FEM2 A3FEW8 A3FJ64 A3FPJ2
A3FPN7 A3GEV2 A3GF47 A3GF61 A3GFF4 A3GFI4 A3GFJ4 A3GFM6 A3GFV3 A3GFV6 A3GG03
A3GG12 A3GG51 A3GGB3 A3GGB4 A3GGE8 A3GGE9 A3GGG8 A3GGJ5 A3GGT2 A3GGU2 A3GGU3
A3GGV1 A3GH78 A3GHA4 A3GHN6 A3GHP2 A3GHR8 A3GHT9 A3GHW9 A3GIA4 A3KFF6 A3KFM7
A3KFX0 A3KG59 A3KGA4 A3KGB4 A3KGF9 A3KGZ2 A3KI17 A3KI18 A3KMH1 A3KMI0 A3KMN5
A3KMP2 A3KMR8 A3KMT2 A3KMU5 A3KMV1 A3KMV2 A3KMV5 A3KMW7 A3KMX0 A3KMX7 A3KMX8
A3KN12 A3KN22 A3KN24 A3KN27 A3KN32 A3KN36 A3KNA5 A3KND5 A3KNI7 A3KNJ8 A3KNK7
A3KNL6 A3KNM4 A3KNX5 A3KNX6 A3KP40 A3KP59 A3KP74 A3KP77 A3KP85 A3KPE8 A3KPF2
A3KPF8 A3KPK0 A3KPN8 A3KPW7 A3KQ55 A3KQ58 A3KQA5 A3KQH2 A3LN13 A3LN21 A3LN53
A3LNC4 A3LNI7 A3LNJ3 A3LNL1 A3LNL5 A3LNM0 A3LNR4 A3LNR6 A3LNW4 A3LP13 A3LP19
A3LP72 A3LP87 A3LP95 A3LPA1 A3LPG0 A3LPQ8 A3LPR4 A3LPV8 A3LPW2 A3LPX1 A3LQ01
A3LQ10 A3LQ44 A3LQ55 A3LQ86 A3LQ93 A3LQC5 A3LQD7 A3LQF9 A3LQG3 A3LQI7 A3LQL9
A3LQV7 A3LQW7 A3LQX9 A3LQY1 A3LQZ8 A3LR41 A3LRF0 A3LRH4 A3LRM2 A3LRM6 A3LRV8
A3LRW2 A3LRX6 A3LRZ0 A3LS22 A3LS24 A3LS77 A3LSJ2 A3LSN3 A3LST5 A3LSY0 A3LSY7
A3LSZ2 A3LT90 A3LTE2 A3LTS5 A3LTV8 A3LU10 A3LU29 A3LU54 A3LU56 A3LUB9 A3LUK5
A3LUM8 A3LUT0 A3LUW5 A3LUY1 A3LV29 A3LV40 A3LV51 A3LVD5 A3LVK6 A3LVM1 A3LVM9
A3LVX0 A3LVY7 A3LW29 A3LW52 A3LWC0 A3LWH3 A3LWH8 A3LWI0 A3LWJ7 A3LWK3 A3LWR2
A3LWX3 A3LWY2 A3LX02 A3LX18 A3LX67 A3LX72 A3LX99 A3LXD5 A3LXE6 A3LXE7 A3LXF0
A3LXG0 A3LXL0 A3LXM4 A3LXS6 A3LXV1 A3LXX5 A3LY29 A3LY88 A3LY92 A3LYE4 A3LYF7
A3LYI0 A3LYL7 A3LYQ7 A3LYR2 A3LYR9 A3LYS9 A3LYY9 A3LZ54 A3LZD1 A3LZG5 A3LZM4
A3LZT3 A3LZU7 A3LZU8 A3LZW1 A3LZY1 A3LZZ0 A3LZZ1 A3M022 A3M056 A3M0B1 A3M0C9
A3M0E6 A3M0F4 A3M0J3 A3M0M0 A3M0Q4 A3M0Q6 A3M0R4 A3M0U3 A3M0W5 A3M0Z0 A3M154
A3M1A1 A3M1A8 A3M1C6 A3M1D2 A3M1D8 A3M1E0 A3M1F6 A3M1F9 A3M1G0 A3M1G1 A3M1G2
A3M1G3 A3M1G4 A3M1J3 A3M1K0 A3M1K2 A3M1K4 A3M1M6 A3M1M7 A3M1P7 A3M1Q6 A3M1S3
A3M1S5 A3M1S8 A3M1S9 A3M1T2 A3M1T4 A3M1V8 A3M1V9 A3M1W6 A3M1X9 A3M1Y2 A3M1Y4
A3M1Y6 A3M1Y8 A3M1Z8 A3M207 A3M208 A3M211 A3M212 A3M215 A3M237 A3M254 A3M294
A3M295 A3M299 A3M2A4 A3M2B0 A3M2I8 A3M2K0 A3M2N7 A3M2N9 A3M2R1 A3M2T0 A3M2T7

A3M2U3 A3M2W0 A3M2W9 A3M2X2 A3M2X4 A3M2X7 A3M2Y4 A3M304 A3M305 A3M306 A3M311
A3M317 A3M319 A3M324 A3M326 A3M361 A3M365 A3M394 A3M3B1 A3M3B7 A3M3C8 A3M3D6
A3M3E9 A3M3F0 A3M3F1 A3M3F3 A3M3F5 A3M3G1 A3M3G7 A3M3I7 A3M3K3 A3M3K6 A3M3N7
A3M3S7 A3M3W2 A3M3X6 A3M419 A3M451 A3M452 A3M4A9 A3M4Q3 A3M4Q8 A3M4Q9 A3M4T0
A3M4U4 A3M4U7 A3M4V2 A3M4W0 A3M4W5 A3M4X6 A3M4Y3 A3M500 A3M504 A3M508 A3M550
A3M561 A3M562 A3M5B7 A3M5C1 A3M5C7 A3M5X8 A3M5Y3 A3M5Z6 A3M5Z8 A3M645 A3M654
A3M657 A3M658 A3M659 A3M663 A3M665 A3M684 A3M692 A3M6H8 A3M6J6 A3M6N7 A3M6P6
A3M6Q4 A3M6Q5 A3M6Q6 A3M6R5 A3M6R7 A3M6X4 A3M6Z4 A3M6Z6 A3M736 A3M741 A3M750
A3M767 A3M775 A3M784 A3M785 A3M795 A3M7A4 A3M7E3 A3M7E7 A3M7F7 A3M7G3 A3M7I8
A3M7M1 A3M7M2 A3M7P2 A3M7Q1 A3M7R2 A3M7W0 A3M7W1 A3M7W9 A3M7X8 A3M7Y8 A3M7Z0
A3M804 A3M810 A3M828 A3M836 A3M837 A3M840 A3M846 A3M851 A3M887 A3M899 A3M8A0
A3M8F1 A3M8H2 A3M8I1 A3M8J9 A3M8N2 A3M8Q6 A3M8S1 A3M8S3 A3M8S9 A3M8T4 A3M8W8
A3M8W9 A3M8Y8 A3M8Z3 A3M907 A3M928 A3M934 A3M938 A3M958 A3M959 A3M960 A3M961
A3M962 A3M963 A3M965 A3M966 A3M967 A3M968 A3M969 A3M970 A3M971 A3M972 A3M973
A3M974 A3M975 A3M976 A3M977 A3M978 A3M980 A3M981 A3M982 A3M983 A3M984 A3M990
A3M9B0 A3M9B2 A3M9C2 A3M9D1 A3M9D2 A3M9G4 A3M9G5 A3M9G7 A3M9H2 A3M9H4 A3M9I8
A3M9K5 A3M9N4 A3M9P1 A3M9W4 A3M9W5 A3M9X9 A3M9Y0 A3M9Y5 A3MA02 A3MA25 A3MA28
A3MA34 A3MA35 A3MA48 A3MA70 A3MA87 A3MA88 A3MA94 A3MAA0 A3MAA1 A3MAC8 A3MAE3
A3MAQ6 A3MBA7 A3MBD0 A3MBD3 A3MBR2 A3MBT5 A3MBT7 A3MBU2 A3MBU6 A3MCV7 A3MD35
A3MDY1 A3ME84 A3MEC5 A3MEC6 A3MF00 A3MFQ3 A3MFQ4 A3MFT3 A3MFT4 A3MG40 A3MGM0
A3MH48 A3MH60 A3MHE1 A3MHE2 A3MHE3 A3MHF0 A3MHG5 A3MHG6 A3MHG9 A3MHJ7 A3MHJ9
A3MHN9 A3MHP1 A3MHP7 A3MHR1 A3MHR7 A3MHR8 A3MHR9 A3MHS0 A3MHS6 A3MHS8 A3MHX8
A3MHY5 A3MI07 A3MI14 A3MI42 A3MI45 A3MI47 A3MI49 A3MI51 A3MI87 A3MI96 A3MI97
A3MID2 A3MIG6 A3MIQ3 A3MIQ6 A3MJ36 A3MJ45 A3MJ92 A3MJ99 A3MJA2 A3MJT8 A3MJU0
A3MJU1 A3MJU2 A3MJW2 A3MJW4 A3MJW5 A3MJW6 A3MJY8 A3MK46 A3MK63 A3MK64 A3MK69
A3MK70 A3MK74 A3MK75 A3MK77 A3MK78 A3MK86 A3MK87 A3MK89 A3MK90 A3MK98 A3MKD3
A3MKD4 A3MKD6 A3MKF7 A3MKI3 A3MKJ7 A3MKJ8 A3MKJ9 A3MKM3 A3MKM4 A3MKP1 A3MKQ7
A3MKR8 A3MKS6 A3MKS8 A3MKT7 A3MKU0 A3MKU1 A3MKU2 A3MKU3 A3MKU6 A3MKU9 A3MKV6
A3MKV8 A3MKW5 A3MKY3 A3MKY8 A3ML43 A3ML57 A3ML79 A3ML91 A3ML96 A3MLB7 A3MLE3
A3MLK5 A3MLK7 A3MLL6 A3MLM0 A3MLM5 A3MLN2 A3MLN3 A3MLQ5 A3MLR1 A3MLT7 A3MLY7
A3MM33 A3MM35 A3MM58 A3MMA4 A3MMA7 A3MMA9 A3MMB0 A3MMB4 A3MMH2 A3MMJ2 A3MML1
A3MML4 A3MMM6 A3MMM8 A3MMN2 A3MMQ8 A3MMR3 A3MMR4 A3MMU8 A3MMV0 A3MMV1 A3MMV2
A3MMV3 A3MMV4 A3MMV5 A3MMZ4 A3MMZ6 A3MN00 A3MN10 A3MN46 A3MN48 A3MN51 A3MN86
A3MN88 A3MN95 A3MN97 A3MN99 A3MNA1 A3MNA4 A3MNA5 A3MNB7 A3MND8 A3MNG1 A3MNG2
A3MNG3 A3MNH1 A3MNI3 A3MNQ3 A3MNR5 A3MNR6 A3MNR7 A3MNT1 A3MNT8 A3MNU6 A3MNW4
A3MNX8 A3MP59 A3MP60 A3MP61 A3MP68 A3MP72 A3MP73 A3MP74 A3MP76 A3MP84 A3MPB4
A3MPB8 A3MPH9 A3MPI2 A3MPU2 A3MPU3 A3MPU4 A3MPU5 A3MPU8 A3MPV1 A3MPY1 A3MQ23
A3MQ26 A3MQB0 A3MQB3 A3MQB4 A3MQB6 A3MQC2 A3MQC3 A3MQF7 A3MQF9 A3MQI7 A3MQI8
A3MQK6 A3MQP3 A3MQP5 A3MQV3 A3MQW7 A3MR08 A3MR17 A3MR18 A3MR36 A3MR54 A3MR55
A3MR61 A3MR64 A3MR65 A3MR78 A3MR88 A3MR89 A3MR90 A3MR91 A3MR94 A3MR95 A3MRB1
A3MRF1 A3MRJ1 A3MRJ9 A3MRM9 A3MRQ4 A3MRQ6 A3MRR3 A3MRR8 A3MRS1 A3MRS9 A3MRT8
A3MRU1 A3MRU2 A3MRU4 A3MRU5 A3MRU6 A3MRU9 A3MRV0 A3MRV3 A3MRV4 A3MRV5 A3MRV6
A3MRV7 A3MRV8 A3MRV9 A3MRW0 A3MRW1 A3MRW2 A3MRW3 A3MRW4 A3MRW5 A3MRW6 A3MRW7
A3MRW8 A3MRW9 A3MRX0 A3MRX1 A3MRX2 A3MRX3 A3MRX5 A3MRX6 A3MRX7 A3MRX8 A3MRX9

A3MRY0 A3MS17 A3MS22 A3MS23 A3MS27 A3MS28 A3MS37 A3MS41 A3MS42 A3MS44 A3MS50
A3MS51 A3MS54 A3MS62 A3MS76 A3MS77 A3MS85 A3MSA1 A3MSE0 A3MSE7 A3MSF2 A3MSF5
A3MSF6 A3MSG0 A3MSH9 A3MSJ3 A3MSJ6 A3MSM9 A3MSN0 A3MSN3 A3MSP1 A3MSS2 A3MSX5
A3MSX6 A3MT22 A3MT36 A3MT40 A3MT41 A3MT44 A3MTA0 A3MTA8 A3MTA9 A3MTB5 A3MTI8
A3MTK6 A3MTL3 A3MTL6 A3MTM9 A3MTQ3 A3MTQ5 A3MTT6 A3MTU1 A3MTU4 A3MTU5 A3MTU7
A3MTW6 A3MU00 A3MU09 A3MU14 A3MU81 A3MU87 A3MU88 A3MU96 A3MUA4 A3MUC6 A3MUD0
A3MUE4 A3MUI7 A3MUI8 A3MUK4 A3MUK8 A3MUN9 A3MUP1 A3MUQ5 A3MUS9 A3MUT0 A3MUU4
A3MUU9 A3MUV2 A3MUX9 A3MUY9 A3MUZ0 A3MUZ1 A3MV08 A3MV29 A3MV32 A3MV36 A3MV40
A3MV50 A3MV61 A3MV63 A3MV65 A3MV67 A3MV68 A3MV69 A3MV81 A3MV95 A3MVB7 A3MVC9
A3MVD1 A3MVE0 A3MVG6 A3MVS2 A3MVZ5 A3MW04 A3MW09 A3MW83 A3MWA6 A3MWC5 A3MWD6
A3MWE7 A3MWF6 A3MWI4 A3MWJ1 A3MWJ4 A3MWJ5 A3MWK3 A3MWK6 A3MWM4 A3MWN4 A3MWN5
A3MWN6 A3MWN7 A3MWQ7 A3MWR8 A3MWS2 A3MWU9 A3MWV7 A3MWW7 A3MWX9 A3MWY2 A3MWZ7
A3MWZ8 A3MX05 A3MX07 A3MX10 A3MX13 A3MX14 A3MX26 A3MX27 A3MX34 A3MX52 A3MX54
A3MX63 A3MX72 A3MX75 A3MX77 A3MX78 A3MX83 A3MX99 A3MXD4 A3MXE4 A3MXE6 A3MXN2
A3MXP1 A3MXP3 A3MXQ6 A3MXQ7 A3MXT2 A3MXT8 A3MXW0 A3MXX9 A3MXZ2 A3MXZ4 A3MXZ5
A3MXZ9 A3MY01 A3MY04 A3MY12 A3MY15 A3MY16 A3MY45 A3MY52 A3MY75 A3MY81 A3MY82
A3MY85 A3MY91 A3MY92 A3MYA2 A3MYA5 A3MYB2 A3MYB6 A3MYC7 A3MYD2 A3MYD4 A3MYE6
A3MYE7 A3MYF1 A3MYG3 A3MYG4 A3MYG9 A3MYI2 A3MYI4 A3MYI9 A3MYK1 A3MYK2 A3MYK8
A3MYL0 A3MYL1 A3MYL5 A3MYM1 A3MYM2 A3MYN0 A3MYN3 A3MYQ4 A3MYR5 A3MYR6 A3MYS3
A3MYS9 A3MYT3 A3MYU1 A3MYU6 A3MYU7 A3MYV3 A3MYV4 A3MYW5 A3MYX4 A3MYY8 A3MYZ9
A3MZ05 A3MZ06 A3MZ07 A3MZ32 A3MZ36 A3MZ45 A3MZ52 A3MZ57 A3MZ64 A3MZ67 A3MZ69
A3MZ70 A3MZ76 A3MZ79 A3MZ80 A3MZ85 A3MZA2 A3MZA3 A3MZB6 A3MZB7 A3MZB9 A3MZC0
A3MZC1 A3MZC2 A3MZC4 A3MZD3 A3MZE1 A3MZE7 A3MZH0 A3MZI3 A3MZI4 A3MZI7 A3MZI8
A3MZK5 A3MZQ9 A3MZR1 A3MZT2 A3MZT3 A3MZT5 A3MZT6 A3MZT8 A3MZT9 A3MZU3 A3MZU5
A3MZU7 A3MZV0 A3MZV8 A3MZW0 A3MZW6 A3MZW7 A3MZW8 A3MZX4 A3MZX6 A3MZY9 A3N003
A3N005 A3N006 A3N007 A3N011 A3N017 A3N018 A3N031 A3N039 A3N042 A3N048 A3N049
A3N054 A3N055 A3N058 A3N059 A3N062 A3N063 A3N073 A3N075 A3N076 A3N087 A3N094
A3N0A0 A3N0A3 A3N0B0 A3N0B6 A3N0B9 A3N0C2 A3N0C7 A3N0D8 A3N0F8 A3N0G0 A3N0G2
A3N0G3 A3N0G6 A3N0G9 A3N0H4 A3N0H9 A3N0J2 A3N0J5 A3N0K4 A3N0L8 A3N0M3 A3N0M7
A3N0M8 A3N0N1 A3N0N2 A3N0N5 A3N0Q6 A3N0Q7 A3N0Q9 A3N0R2 A3N0R3 A3N0S3 A3N0S5
A3N0T5 A3N0X3 A3N0X6 A3N0Y3 A3N0Z0 A3N0Z5 A3N103 A3N113 A3N120 A3N121 A3N122
A3N123 A3N148 A3N181 A3N186 A3N1A1 A3N1A3 A3N1A7 A3N1B2 A3N1B5 A3N1B7 A3N1B8
A3N1C3 A3N1C4 A3N1C8 A3N1D7 A3N1D8 A3N1E1 A3N1E3 A3N1E4 A3N1E6 A3N1F3 A3N1F5
A3N1F7 A3N1F9 A3N1G7 A3N1H1 A3N1H2 A3N1H3 A3N1H8 A3N1I1 A3N1M8 A3N1N1 A3N1Q1
A3N1S4 A3N1S5 A3N1T0 A3N1T6 A3N1U5 A3N1U6 A3N1U8 A3N1V2 A3N1W3 A3N1W5 A3N1X3
A3N1Z1 A3N1Z6 A3N211 A3N228 A3N231 A3N234 A3N246 A3N247 A3N248 A3N249 A3N262
A3N298 A3N2A1 A3N2A2 A3N2B0 A3N2C1 A3N2D3 A3N2D8 A3N2D9 A3N2E6 A3N2F0 A3N2F5
A3N2F7 A3N2G1 A3N2G8 A3N2H9 A3N2I3 A3N2I4 A3N2I5 A3N2I9 A3N2J4 A3N2K3 A3N2K6
A3N2L0 A3N2N5 A3N2P1 A3N2P2 A3N2P8 A3N2Q0 A3N2Q3 A3N2Q9 A3N2R0 A3N2R1 A3N2R4
A3N2R5 A3N2R6 A3N2S7 A3N2T3 A3N2T4 A3N2T5 A3N2T7 A3N2V2 A3N2V3 A3N2W7 A3N2Y4
A3N308 A3N316 A3N317 A3N318 A3N319 A3N325 A3N326 A3N333 A3N334 A3N335 A3N338
A3N348 A3N350 A3N352 A3N353 A3N357 A3N358 A3N359 A3N361 A3N362 A3N363 A3N364
A3N365 A3N366 A3N367 A3N368 A3N369 A3N370 A3N371 A3N372 A3N373 A3N374 A3N375
A3N376 A3N378 A3N379 A3N380 A3N381 A3N382 A3N384 A3N385 A3N386 A3N387 A3N398

A3N3A1 A3N3B3 A3N3B7 A3N3B8 A3N3C2 A3N3C6 A3N3D3 A3N3D7 A3N3D8 A3N3E9 A3N3F7
A3N3F8 A3N3G7 A3N3G9 A3N3J1 A3N3J4 A3N3J5 A3N3J9 A3N3K0 A3N3K2 A3N3L1 A3N3L9
A3N3M1 A3N3M9 A3N3N0 A3N3P1 A3N3P3 A3N3P8 A3N3P9 A3N3Q5 A3N3Q6 A3N3Q8 A3N3Q9
A3N3R0 A3N3R7 A3N3S8 A3N3T7 A3N3T8 A3N3U3 A3N3U7 A3N3V7 A3N3W1 A3N3W4 A3N3Y5
A3N3Y7 A3N473 A3N474 A3N475 A3N480 A3N4D4 A3N4E3 A3N4F3 A3N4F4 A3N4F5 A3N4G4
A3N4G7 A3N4H2 A3N4H9 A3N4I1 A3N4P2 A3N4T7 A3N4V9 A3N520 A3N521 A3N522 A3N530
A3N542 A3N559 A3N575 A3N576 A3N5B0 A3N5B3 A3N5J6 A3N5J9 A3N5K0 A3N5K2 A3N5K5
A3N5K8 A3N5K9 A3N5L9 A3N5U2 A3N5U8 A3N5V6 A3N5X6 A3N5Z1 A3N693 A3N6H2 A3N6H3
A3N6H4 A3N6I1 A3N6J5 A3N6J6 A3N6J9 A3N6K4 A3N6K6 A3N6N9 A3N6P4 A3N6Q4 A3N6Q7
A3N6Q8 A3N6Q9 A3N6U4 A3N6U9 A3N6X1 A3N6X2 A3N6Y1 A3N6Y6 A3N6Z8 A3N700 A3N759
A3N786 A3N794 A3N7B6 A3N7C4 A3N7F4 A3N7F7 A3N7F9 A3N7G1 A3N7G3 A3N7K4 A3N7K7
A3N7L2 A3N7L3 A3N7P8 A3N7W7 A3N853 A3N856 A3N8B3 A3N8K7 A3N8L4 A3N8Q9 A3N8T1
A3N8T3 A3N8T4 A3N8V6 A3N8V8 A3N8V9 A3N8W0 A3N919 A3N926 A3N927 A3NA26 A3NA42
A3NA43 A3NA48 A3NA49 A3NA53 A3NA54 A3NA56 A3NA57 A3NA64 A3NA65 A3NA68 A3NA69
A3NA78 A3NAB3 A3NAB4 A3NAB6 A3NAE5 A3NAH2 A3NAI4 A3NAI5 A3NAI6 A3NAL8 A3NAL9
A3NAR3 A3NAS3 A3NAT0 A3NAT3 A3NAU2 A3NAU5 A3NAU6 A3NAU7 A3NAU8 A3NAV0 A3NAV3
A3NAW0 A3NAW2 A3NAY8 A3NAZ3 A3NB50 A3NB64 A3NB96 A3NBA0 A3NBA1 A3NBC0 A3NBE8
A3NBG6 A3NBH0 A3NBM6 A3NBM7 A3NBR9 A3NBS1 A3NBU4 A3NBZ6 A3NBZ8 A3NC04 A3NC05
A3NC06 A3NC07 A3NC12 A3NC24 A3NC29 A3NC31 A3NC70 A3NC72 A3NC92 A3NC95 A3NCA9
A3NCB1 A3NCC8 A3NCF3 A3NCF9 A3NCG0 A3NCL3 A3NCL5 A3NCL6 A3NCL7 A3NCL8 A3NCL9
A3NCM0 A3NCP7 A3NCQ0 A3NCQ2 A3NCQ3 A3NCQ8 A3NCY6 A3ND55 A3ND57 A3ND64 A3ND66
A3ND68 A3ND70 A3ND73 A3ND74 A3NDB2 A3NDB6 A3NDE4 A3NDF2 A3NDF4 A3NDF5 A3NDF6
A3NDG6 A3NDG7 A3NDG8 A3NDK5 A3NDK8 A3NDL7 A3NDQ7 A3NDS2 A3NDS3 A3NDS6 A3NDS7
A3NDS9 A3NDT0 A3NDU9 A3NDW2 A3NDW3 A3NDW6 A3NDX2 A3NDX3 A3NDZ2 A3NDZ3 A3NE23
A3NE24 A3NE91 A3NE92 A3NE93 A3NE94 A3NE97 A3NEA0 A3NED2 A3NED9 A3NEF2 A3NEF3
A3NEF4 A3NEF5 A3NEF6 A3NEF7 A3NEF8 A3NEG0 A3NEG1 A3NEG2 A3NEG3 A3NEG4 A3NEG5
A3NEG6 A3NEG7 A3NEG8 A3NEG9 A3NEH0 A3NEH1 A3NEH2 A3NEH3 A3NEH4 A3NEH5 A3NEH6
A3NEH7 A3NEH8 A3NEH9 A3NEI0 A3NEI1 A3NEI3 A3NEI4 A3NEI6 A3NEI7 A3NEI8 A3NEJ0
A3NEJ1 A3NEP2 A3NER1 A3NEZ8 A3NF00 A3NF35 A3NF53 A3NF54 A3NF91 A3NGW6 A3NH07
A3NHB6 A3NIZ0 A3NJS6 A3NK14 A3NKA8 A3NKK4 A3NKP7 A3NKP8 A3NKT2 A3NM66 A3NM70
A3NM75 A3NM77 A3NM97 A3NMF5 A3NMF6 A3NMF8 A3NMH8 A3NML2 A3NML3 A3NN77 A3NNQ0
A3NNW7 A3NPL5 A3NPL7 A3NPW7 A3NPW8 A3NPW9 A3NPX5 A3NQ23 A3NQ33 A3NQ44 A3NQ45
A3NQ46 A3NQ54 A3NQ57 A3NQ62 A3NQ63 A3NQ71 A3NQ73 A3NQ97 A3NQD4 A3NQI1 A3NQK2
A3NQS1 A3NQS2 A3NQS3 A3NQT1 A3NQU2 A3NQV8 A3NQX5 A3NQX6 A3NR09 A3NR12 A3NR15
A3NR92 A3NR95 A3NR98 A3NRA1 A3NRA4 A3NRA5 A3NRB5 A3NRI7 A3NRJ4 A3NRK2 A3NRM0
A3NRN5 A3NRX7 A3NS55 A3NS56 A3NS57 A3NS64 A3NS79 A3NS80 A3NS83 A3NS87 A3NS89
A3NSC3 A3NSC8 A3NSD8 A3NSE2 A3NSE3 A3NSE4 A3NSI1 A3NSI6 A3NSK7 A3NSK8 A3NSL6
A3NSM2 A3NSN6 A3NSU7 A3NSX5 A3NSY2 A3NT04 A3NT12 A3NT43 A3NT46 A3NT48 A3NT50
A3NT52 A3NT91 A3NT94 A3NTA0 A3NTA2 A3NTD7 A3NTK3 A3NTW3 A3NTW6 A3NU16 A3NUB0
A3NUB7 A3NUG3 A3NUI4 A3NUI6 A3NUI7 A3NUI8 A3NUK8 A3NUL0 A3NUL1 A3NUL2 A3NUR6
A3NUS2 A3NUS3 A3NVU2 A3NVV9 A3NVW0 A3NVW5 A3NVW6 A3NVX0 A3NVX1 A3NVX3 A3NVX4
A3NVY1 A3NVY2 A3NVY5 A3NVY6 A3NVZ5 A3NW31 A3NW32 A3NW34 A3NW63 A3NW79 A3NW90
A3NWA5 A3NWA6 A3NWA7 A3NWD9 A3NWE0 A3NWJ3 A3NWK5 A3NWL4 A3NWL6 A3NWM5 A3NWM8
A3NWM9 A3NWN0 A3NWN1 A3NWN4 A3NWN7 A3NWP3 A3NWP5 A3NWR9 A3NWS4 A3NWX8 A3NWZ4

A3NX26 A3NX27 A3NX33 A3NX34 A3NX53 A3NX81 A3NX99 A3NXA3 A3NXG1 A3NXG2 A3NXK5
A3NXK9 A3NXN1 A3NXT2 A3NXT4 A3NXU0 A3NXU1 A3NXU2 A3NXU3 A3NXU9 A3NXW3 A3NXW8
A3NXX0 A3NY08 A3NY10 A3NY34 A3NY37 A3NY50 A3NY52 A3NY56 A3NY82 A3NY88 A3NY89
A3NYC5 A3NYC7 A3NYC8 A3NYC9 A3NYD0 A3NYD1 A3NYD2 A3NYF9 A3NYG2 A3NYG4 A3NYG5
A3NYH0 A3NYN0 A3NYQ0 A3NYW4 A3NYW6 A3NYX3 A3NYX5 A3NYX7 A3NYX9 A3NYY2 A3NYY3
A3NZ22 A3NZ26 A3NZ55 A3NZ64 A3NZ65 A3NZ66 A3NZ67 A3NZ74 A3NZ78 A3NZ79 A3NZ80
A3NZB6 A3NZB9 A3NZD0 A3NZI6 A3NZI7 A3NZJ0 A3NZJ1 A3NZJ2 A3NZK0 A3NZL3 A3NZL4
A3NZL7 A3NZM3 A3NZM4 A3NZP5 A3NZP6 A3NZS3 A3NZS4 A3P025 A3P026 A3P027 A3P028
A3P031 A3P034 A3P066 A3P073 A3P082 A3P086 A3P087 A3P088 A3P089 A3P090 A3P091
A3P092 A3P094 A3P095 A3P096 A3P097 A3P098 A3P099 A3P0A0 A3P0A1 A3P0A2 A3P0A3
A3P0A4 A3P0A5 A3P0A6 A3P0A7 A3P0A8 A3P0A9 A3P0B0 A3P0B1 A3P0B2 A3P0B3 A3P0B4
A3P0B5 A3P0B7 A3P0B8 A3P0C4 A3P0C5 A3P0C6 A3P0C8 A3P0C9 A3P0J9 A3P0L8 A3P0N1
A3P0U5 A3P0U7 A3P0Y4 A3P103 A3P104 A3P139 A3P2G7 A3P2K7 A3P2X2 A3P4L6 A3P5E2
A3P5N3 A3P5X2 A3P669 A3P6A9 A3P6B0 A3P6E0 A3P6S9 A3P7M8 A3P7N2 A3P7N7 A3P7N9
A3P7Q9 A3P7W1 A3P7W3 A3P7W4 A3P7W6 A3P7Z0 A3P825 A3P826 A3P8N2 A3P975 A3PA50
A3PA59 A3PA63 A3PA67 A3PA73 A3PA84 A3PA95 A3PA96 A3PA97 A3PAG5 A3PAH7 A3PAI9
A3PAJ7 A3PAN2 A3PAN9 A3PAP1 A3PAQ0 A3PAQ7 A3PAR3 A3PAR9 A3PAS0 A3PAS1 A3PAS2
A3PAT7 A3PAU1 A3PAU4 A3PAV3 A3PAV5 A3PAV8 A3PAW2 A3PAX1 A3PAX7 A3PAX8 A3PAX9
A3PAY3 A3PAY5 A3PAY7 A3PAZ2 A3PB03 A3PB34 A3PB36 A3PB44 A3PB53 A3PBA5 A3PBA9
A3PBB3 A3PBB9 A3PBC3 A3PBC7 A3PBD7 A3PBF2 A3PBG1 A3PBH4 A3PBH7 A3PBJ3 A3PBJ6
A3PBK4 A3PBK8 A3PBL6 A3PBM0 A3PBM4 A3PBN1 A3PBN3 A3PBP3 A3PBP6 A3PBP9 A3PBR7
A3PBR8 A3PBR9 A3PBS4 A3PBT3 A3PBT4 A3PBT9 A3PBV7 A3PBV8 A3PBW4 A3PBW9 A3PBX1
A3PBY1 A3PBY3 A3PBY7 A3PBZ7 A3PC66 A3PC83 A3PCF5 A3PCG1 A3PCG2 A3PCH5 A3PCH6
A3PCI2 A3PCJ7 A3PCK0 A3PCK5 A3PCN5 A3PCN6 A3PCN7 A3PCN8 A3PCN9 A3PCP0 A3PCP1
A3PCQ4 A3PCR0 A3PCR3 A3PCR4 A3PCS1 A3PCS3 A3PCS4 A3PCV0 A3PCV5 A3PCW5 A3PCW7
A3PCW8 A3PCX3 A3PCY8 A3PCY9 A3PD13 A3PD22 A3PD39 A3PD66 A3PD91 A3PDF8 A3PDH0
A3PDH3 A3PDJ8 A3PDJ9 A3PDK2 A3PDM1 A3PDX8 A3PDZ9 A3PE00 A3PE10 A3PE32 A3PE33
A3PE37 A3PE39 A3PE40 A3PE89 A3PEA4 A3PEB8 A3PEC4 A3PED0 A3PED3 A3PED9 A3PEE0
A3PEE4 A3PEE9 A3PEG2 A3PEG6 A3PEH8 A3PEH9 A3PEI1 A3PEI3 A3PEI7 A3PER9 A3PES4
A3PES5 A3PEU9 A3PEW3 A3PEX0 A3PEX2 A3PEX3 A3PEX4 A3PEX6 A3PEY0 A3PEY3 A3PEY9
A3PEZ1 A3PEZ6 A3PEZ7 A3PEZ8 A3PEZ9 A3PF00 A3PF19 A3PF20 A3PF21 A3PF22 A3PF23
A3PF24 A3PF25 A3PF26 A3PF27 A3PF28 A3PF30 A3PF31 A3PF32 A3PF34 A3PF35 A3PF36
A3PF37 A3PF38 A3PF39 A3PF40 A3PF41 A3PF42 A3PF43 A3PF44 A3PF45 A3PF46 A3PF47
A3PF51 A3PF64 A3PF71 A3PF79 A3PF80 A3PF82 A3PF83 A3PF96 A3PFA2 A3PFA5 A3PFA6
A3PFA8 A3PFB5 A3PFC4 A3PFC5 A3PFD1 A3PFD3 A3PFD7 A3PFE1 A3PFE3 A3PFE5 A3PFE9
A3PFF0 A3PFF2 A3PFF7 A3PFG1 A3PFG3 A3PFH4 A3PFH6 A3PFH7 A3PFH8 A3PFI0 A3PFJ4
A3PFJ6 A3PFL1 A3PFL2 A3PFL4 A3PFL5 A3PFN0 A3PFQ4 A3PFR1 A3PFS0 A3PFX4 A3PFZ5
A3PG14 A3PG34 A3PG35 A3PGE9 A3PGF2 A3PGF5 A3PGF8 A3PGG1 A3PGG3 A3PGG7 A3PGI1
A3PGI6 A3PGI9 A3PGJ0 A3PGJ2 A3PGJ3 A3PGL0 A3PGL2 A3PGL3 A3PGL4 A3PGL5 A3PGL6
A3PGL7 A3PGL8 A3PGM0 A3PGM1 A3PGM2 A3PGM3 A3PGM4 A3PGM5 A3PGM6 A3PGM7 A3PGM8
A3PGM9 A3PGN0 A3PGN3 A3PGN4 A3PGN5 A3PGN6 A3PGQ2 A3PGQ3 A3PGQ9 A3PGR0 A3PGR1
A3PGR7 A3PGS3 A3PGS6 A3PGW1 A3PGY5 A3PGZ4 A3PH01 A3PH07 A3PH12 A3PH14 A3PH28
A3PH34 A3PH74 A3PH80 A3PHH4 A3PHJ3 A3PHJ6 A3PHK2 A3PHK6 A3PHK7 A3PHK8 A3PHL6
A3PHM3 A3PHR6 A3PHR7 A3PHS4 A3PHS9 A3PHT1 A3PHT2 A3PI00 A3PI15 A3PI61 A3PI62

A3PI63 A3PI66 A3PI75 A3PI87 A3PI89 A3PI90 A3PIA3 A3PIA4 A3PIB4 A3PIC9 A3PID0
A3PIG0 A3PIG5 A3PII0 A3PIN5 A3PIP7 A3PIQ5 A3PIR7 A3PIU6 A3PIU9 A3PIY7 A3PIZ8
A3PIZ9 A3PJ63 A3PJ66 A3PJ71 A3PJ74 A3PJ78 A3PJ80 A3PJ90 A3PJB1 A3PJB4 A3PJD6
A3PJF0 A3PJF5 A3PJF6 A3PJF7 A3PJG0 A3PJH4 A3PJI8 A3PJK8 A3PJL9 A3PJM6 A3PJM7
A3PJP5 A3PJQ3 A3PJQ4 A3PJS2 A3PJT9 A3PJX6 A3PJY2 A3PJZ3 A3PJZ6 A3PK22 A3PK23
A3PK32 A3PK36 A3PK37 A3PK50 A3PK61 A3PK79 A3PK84 A3PK94 A3PK95 A3PKG3 A3PKL5
A3PKL6 A3PKL7 A3PKL8 A3PKR6 A3PKS0 A3PKS1 A3PL07 A3PL16 A3PL17 A3PL19 A3PL34
A3PL35 A3PL38 A3PL41 A3PL42 A3PL43 A3PL91 A3PL98 A3PLA0 A3PLC6 A3PLD1 A3PLD4
A3PLG8 A3PLI0 A3PLJ1 A3PLJ2 A3PLM1 A3PLP3 A3PLR7 A3PLS9 A3PLT9 A3PLU2 A3PLU4
A3PLU5 A3PLV0 A3PLW1 A3PLX6 A3PM14 A3PM19 A3PM22 A3PM38 A3PM44 A3PM50 A3PM52
A3PM71 A3PMA3 A3PMA9 A3PMB7 A3PMC8 A3PMF8 A3PMH0 A3PMI4 A3PMI6 A3PMJ1 A3PMK8
A3PML8 A3PML9 A3PMM1 A3PMM5 A3PMN4 A3PMP8 A3PMQ7 A3PMQ9 A3PMR2 A3PMR3 A3PMS1
A3PMS2 A3PMS9 A3PMX6 A3PMX9 A3PN13 A3PN14 A3PN16 A3PN31 A3PN45 A3PN48 A3PN90
A3PN91 A3PN94 A3PN95 A3PNA6 A3PNA9 A3PNB5 A3PNE9 A3PNF3 A3PNF4 A3PNF6 A3PNF9
A3PNG0 A3PNH5 A3PNH8 A3PNK9 A3PNL2 A3PNM0 A3PNM1 A3PNM3 A3PNM4 A3PNQ9 A3PNR5
A3PNR8 A3PNS5 A3PNS6 A3PNS7 A3PNT1 A3PNT4 A3PNX3 A3PNX7 A3PNY0 A3PP56 A3PPJ8
A3PPV2 A3PPV4 A3PPV7 A3PPW0 A3PPW5 A3PQ70 A3PQD8 A3PQI7 A3PQI8 A3PQJ0 A3PQJ2
A3PRB5 A3PRF6 A3PRP9 A3PRU3 A3PRW1 A3PRW6 A3PRX7 A3PSD7 A3PSE0 A3PSL4 A3PSL5
A3PSS2 A3PSV1 A3PSZ4 A3PTE4 A3PTG0 A3PTJ4 A3PTK2 A3PTN4 A3PTT9 A3PTW3 A3PTW9
A3PTZ4 A3PTZ8 A3PU05 A3PU80 A3PU84 A3PUA3 A3PUB7 A3PUH9 A3PUJ1 A3PV27 A3PV33
A3PV34 A3PV75 A3PV78 A3PV79 A3PV93 A3PV94 A3PV95 A3PV96 A3PVC0 A3PVC1 A3PVC2
A3PVC3 A3PVC4 A3PVC5 A3PVC6 A3PVC7 A3PVC8 A3PVC9 A3PVD5 A3PVD6 A3PVD7 A3PVD8
A3PVD9 A3PVE0 A3PVE3 A3PVF1 A3PVF2 A3PVF3 A3PVF5 A3PVL3 A3PVL4 A3PVL5 A3PVL6
A3PVL7 A3PVL8 A3PVL9 A3PVN4 A3PVN7 A3PVQ4 A3PVQ7 A3PVV6 A3PVY4 A3PW53 A3PW85
A3PW97 A3PWC1 A3PWV0 A3PWV5 A3PXE2 A3PXF8 A3PXL7 A3PXM2 A3PXN5 A3PXP9 A3PXQ2
A3PXR0 A3PXR2 A3PXR3 A3PXR7 A3PXS0 A3PXS4 A3PXS9 A3PXT0 A3PXT6 A3PXU1 A3PXU3
A3PXV6 A3PXV8 A3PXW1 A3PXW3 A3PXX6 A3PXY1 A3PXY4 A3PXY5 A3PXZ4 A3PXZ5 A3PXZ7
A3PY44 A3PY62 A3PY72 A3PY75 A3PY76 A3PY88 A3PY92 A3PY93 A3PYA0 A3PYA8 A3PYB1
A3PYE3 A3PYE4 A3PYF3 A3PYF8 A3PYF9 A3PYG8 A3PYI9 A3PYK6 A3PYL0 A3PYU0 A3PYU8
A3PYU9 A3PYW3 A3PYW4 A3PYW5 A3PYX7 A3PZ05 A3PZ44 A3PZ45 A3PZ50 A3PZ56 A3PZ57
A3PZ61 A3PZ65 A3PZ71 A3PZ81 A3PZ89 A3PZ93 A3PZ94 A3PZ96 A3PZA2 A3PZA3 A3PZF2
A3PZG1 A3Q016 A3Q040 A3Q0D3 A3Q0D4 A3Q0D5 A3Q0D6 A3Q0D7 A3Q0E7 A3Q0H8 A3Q0R1
A3Q0S2 A3Q0T2 A3Q0T3 A3Q0T7 A3Q0U6 A3Q0U7 A3Q119 A3Q124 A3Q125 A3Q127 A3Q129
A3Q130 A3Q137 A3Q142 A3Q145 A3Q146 A3Q192 A3Q193 A3Q196 A3Q1A9 A3Q1B0 A3Q1B7
A3Q1L2 A3Q1L7 A3Q1M6 A3Q1M7 A3Q1Q2 A3Q1S2 A3Q1S7 A3Q1S8 A3Q1U4 A3Q255 A3Q262
A3Q267 A3Q272 A3Q2B8 A3Q2C2 A3Q2D5 A3Q2E3 A3Q2F3 A3Q2F4 A3Q2F5 A3Q2I1 A3Q2I4
A3Q2P5 A3Q338 A3Q339 A3Q379 A3Q393 A3Q396 A3Q3A2 A3Q3C2 A3Q3C3 A3Q3C9 A3Q3D3
A3Q3I2 A3Q3N5 A3Q3Q7 A3Q4B8 A3Q4B9 A3Q4F8 A3Q4G7 A3Q4N0 A3Q4N2 A3Q4Q8 A3Q5H1
A3Q5I0 A3Q5M0 A3Q5N7 A3Q5P5 A3Q5U2 A3Q628 A3Q635 A3Q648 A3Q695 A3Q6A4 A3Q6D1
A3Q6H9 A3Q6M9 A3Q6N0 A3Q6V2 A3Q6V9 A3Q6W0 A3Q6Y3 A3Q6Y4 A3Q6Y5 A3Q7A7 A3Q7A8
A3Q7J9 A3Q7P6 A3Q7Q1 A3Q7S1 A3Q7V7 A3Q7Y6 A3Q868 A3Q8M9 A3Q8N0 A3Q8Q1 A3Q8S0
A3Q8S4 A3Q8S5 A3Q8S6 A3Q8S8 A3Q8T4 A3Q8T5 A3Q8U0 A3Q8U5 A3Q8W5 A3Q913 A3Q930
A3Q958 A3Q962 A3Q963 A3Q967 A3Q968 A3Q971 A3Q972 A3Q973 A3Q974 A3Q975 A3Q976
A3Q977 A3Q978 A3Q980 A3Q981 A3Q982 A3Q983 A3Q984 A3Q985 A3Q986 A3Q987 A3Q988

A3Q989 A3Q990 A3Q991 A3Q992 A3Q993 A3Q994 A3Q995 A3Q996 A3Q997 A3Q998 A3Q999
A3Q9A0 A3Q9A1 A3Q9A3 A3Q9A4 A3Q9A5 A3Q9A6 A3Q9A7 A3Q9A8 A3Q9C2 A3Q9C4 A3Q9C6
A3Q9C7 A3Q9C9 A3Q9D0 A3Q9D7 A3Q9D8 A3Q9E9 A3Q9F0 A3Q9I6 A3Q9J1 A3Q9L6 A3Q9P7
A3Q9Q3 A3Q9Q5 A3Q9R3 A3Q9U8 A3Q9U9 A3Q9V2 A3Q9V5 A3QA17 A3QA31 A3QA45 A3QA46
A3QAD3 A3QAD8 A3QAD9 A3QAE0 A3QAN3 A3QAP0 A3QAT8 A3QAV2 A3QAX5 A3QAX7 A3QAX8
A3QAZ2 A3QAZ7 A3QB25 A3QB36 A3QB39 A3QB49 A3QB62 A3QB90 A3QB91 A3QB93 A3QB94
A3QB95 A3QBA1 A3QBA2 A3QBA3 A3QBG2 A3QBM2 A3QBM3 A3QBN1 A3QBN5 A3QBP0 A3QBP3
A3QBQ0 A3QBR1 A3QBR4 A3QBS7 A3QBT6 A3QBT7 A3QBT8 A3QBT9 A3QBU2 A3QBW1 A3QBW2
A3QBW6 A3QBW9 A3QBX2 A3QBZ5 A3QBZ9 A3QC07 A3QC17 A3QC57 A3QC58 A3QC62 A3QC63
A3QC76 A3QC79 A3QC80 A3QC82 A3QC83 A3QC86 A3QC87 A3QC89 A3QC90 A3QCE0 A3QCF9
A3QCG2 A3QCG3 A3QCG6 A3QCG8 A3QCH0 A3QCT9 A3QCU1 A3QCV5 A3QCW8 A3QD32 A3QD33
A3QD52 A3QD67 A3QD79 A3QD84 A3QD86 A3QD88 A3QD92 A3QD95 A3QD98 A3QDA2 A3QDB8
A3QDC9 A3QDD6 A3QDD8 A3QDE1 A3QDE6 A3QDF9 A3QDH6 A3QDN8 A3QDU9 A3QDY0 A3QDZ3
A3QDZ8 A3QE30 A3QE68 A3QE80 A3QE83 A3QEC0 A3QEC1 A3QEC3 A3QEG3 A3QEI8 A3QEJ7
A3QEJ8 A3QEK0 A3QEN0 A3QEP3 A3QEP4 A3QEP5 A3QEP7 A3QEP9 A3QEQ0 A3QER0 A3QER4
A3QET0 A3QET2 A3QEV1 A3QF01 A3QF02 A3QF04 A3QF15 A3QF19 A3QF20 A3QF21 A3QF32
A3QF43 A3QF48 A3QF50 A3QF51 A3QF52 A3QF54 A3QF58 A3QF71 A3QF73 A3QFC4 A3QFC7
A3QFC8 A3QFD1 A3QFD2 A3QFD5 A3QFF5 A3QFF6 A3QFI1 A3QFJ5 A3QFL1 A3QFM9 A3QFN4
A3QFR4 A3QFX5 A3QFX6 A3QFX7 A3QFX8 A3QFX9 A3QG19 A3QG64 A3QG89 A3QG97 A3QGA0
A3QGA1 A3QGA2 A3QGA3 A3QGA6 A3QGJ4 A3QGL9 A3QGM4 A3QGM9 A3QGN4 A3QGN9 A3QGP0
A3QGP1 A3QGR1 A3QGR7 A3QGT0 A3QGT1 A3QGT2 A3QGT3 A3QGT8 A3QGU0 A3QGU2 A3QGU3
A3QGU4 A3QGU5 A3QGU7 A3QGU9 A3QGV0 A3QGV3 A3QGV8 A3QGW1 A3QGW2 A3QGY9 A3QH04
A3QH11 A3QH30 A3QH31 A3QH33 A3QH38 A3QH41 A3QH43 A3QH47 A3QH52 A3QH58 A3QH59
A3QH60 A3QH80 A3QHI0 A3QHI1 A3QHI2 A3QHP6 A3QHQ1 A3QHQ2 A3QHZ8 A3QI29 A3QI45
A3QI49 A3QI50 A3QI52 A3QI59 A3QI61 A3QI66 A3QI82 A3QID1 A3QIE1 A3QII0 A3QIJ0
A3QIM0 A3QIM9 A3QIN0 A3QIN4 A3QIN7 A3QIN8 A3QIP0 A3QIP7 A3QIP8 A3QIP9 A3QIQ1
A3QIQ2 A3QIQ4 A3QIQ6 A3QIR6 A3QIU7 A3QIW0 A3QJ17 A3QJ40 A3QJ41 A3QJ65 A3QJ82
A3QJA4 A3QJA8 A3QJB0 A3QJB2 A3QJC8 A3QJD3 A3QJD9 A3QJE8 A3QJF1 A3QJF5 A3QJF9
A3QJG1 A3QJH7 A3QJI9 A3QJM2 A3QJM4 A3QJS0 A3QJS1 A3QJT0 A3QJT4 A3QJZ6 A3QJZ7
A3QMC8 A3QMD7 A3QVV1 A3R052 A3RCV9 A3RGC1 A3RK74 A3RK75 A3RLD7 A3RLD8 A3RLE1
A3RLE2 A3RLT6 A3RM19 A3RM23 A3RXB7 A3SK19 A3SQG3 A3SR25 A3U3H1 A3UZK3 A4AQI7
A4D0Y5 A4D126 A4D161 A4D1E1 A4D1E9 A4D1N5 A4D1U4 A4D1Z8 A4D250 A4D263 A4D2B8
A4D7Q3 A4D7S9 A4D7T3 A4D998 A4D9J5 A4D9U3 A4DA05 A4DA84 A4F1Q8 A4F2N8 A4F4L4
A4F5G6 A4F5N5 A4F6D0 A4F6D1 A4F6L9 A4F6T0 A4F6T1 A4F6T2 A4F6W3 A4F6W7 A4F710
A4F727 A4F749 A4F7B2 A4F7F5 A4F7F6 A4F7F9 A4F7G0 A4F7G2 A4F7P2 A4F7P5 A4F7R8
A4F7R9 A4F7S0 A4F7S3 A4F7S4 A4F7W6 A4F7X3 A4F866 A4F870 A4F891 A4F8D5 A4F8G5
A4F8R2 A4F8R5 A4F910 A4F913 A4F929 A4F9F6 A4F9J8 A4F9L1 A4F9L2 A4F9L3 A4F9M1
A4F9M5 A4F9N6 A4F9T1 A4F9T9 A4F9Z0 A4FA00 A4FA49 A4FA70 A4FAB4 A4FAC5 A4FAH9
A4FAI5 A4FAJ1 A4FAJ5 A4FAK5 A4FAQ5 A4FAS3 A4FBA2 A4FBA3 A4FBA6 A4FBA7 A4FBA8
A4FBB2 A4FBB7 A4FBC3 A4FBD3 A4FBE6 A4FBE7 A4FBE8 A4FBE9 A4FBF7 A4FBF8 A4FBG1
A4FBI2 A4FBJ4 A4FBK5 A4FBL9 A4FBM0 A4FBM2 A4FBM5 A4FBM6 A4FBN0 A4FBP0 A4FBV6
A4FBW7 A4FBW8 A4FBX6 A4FBY1 A4FBY2 A4FBY4 A4FCP0 A4FCP1 A4FDR2 A4FDS0 A4FDY3
A4FF33 A4FFJ9 A4FFQ4 A4FFQ6 A4FFX3 A4FG06 A4FG18 A4FG19 A4FH01 A4FHB5 A4FI92
A4FID1 A4FIM1 A4FIQ1 A4FIR1 A4FIS6 A4FK61 A4FK76 A4FKA9 A4FKC1 A4FKC5 A4FKC8

A4FKE3 A4FKE4 A4FL80 A4FLF6 A4FLG1 A4FLL0 A4FLL5 A4FLL9 A4FLN6 A4FLU1 A4FLV9
A4FLW9 A4FLX5 A4FLX6 A4FLZ0 A4FM22 A4FM23 A4FM32 A4FM34 A4FM36 A4FM88 A4FMC5
A4FMD4 A4FMD5 A4FMD6 A4FME3 A4FME5 A4FME7 A4FME8 A4FME9 A4FMF1 A4FMH0 A4FMJ7
A4FMJ9 A4FMP1 A4FMP2 A4FMP6 A4FMQ2 A4FMQ5 A4FMS5 A4FMT5 A4FMU3 A4FN40 A4FN47
A4FND4 A4FNE8 A4FNF1 A4FNI0 A4FNQ1 A4FNR2 A4FP42 A4FP60 A4FPA5 A4FPG9 A4FPH0
A4FPH1 A4FPJ1 A4FPJ2 A4FPJ3 A4FPJ4 A4FPJ5 A4FPJ6 A4FPK6 A4FPK7 A4FPK8 A4FPL0
A4FPL1 A4FPL2 A4FPL3 A4FPL4 A4FPL5 A4FPL6 A4FPL7 A4FPL8 A4FPL9 A4FPM0 A4FPM1
A4FPM2 A4FPM3 A4FPM4 A4FPM5 A4FPM6 A4FPM7 A4FPM8 A4FPM9 A4FPN0 A4FPP2 A4FPP3
A4FPQ6 A4FPQ7 A4FPQ9 A4FPR0 A4FPR5 A4FPR6 A4FPS1 A4FPV1 A4FPX3 A4FPY4 A4FQ08
A4FQ15 A4FQ42 A4FQD4 A4FQE4 A4FQI3 A4FQK6 A4FQV7 A4FR29 A4FR30 A4FR32 A4FR54
A4FR78 A4FU01 A4FU69 A4FUA8 A4FUB0 A4FUB7 A4FUD1 A4FUD3 A4FUE7 A4FUF0 A4FUG7
A4FUH0 A4FUI1 A4FUI2 A4FUM1 A4FUX8 A4FUY7 A4FUZ0 A4FUZ6 A4FV08 A4FV14 A4FV29
A4FV42 A4FV57 A4FV58 A4FV61 A4FV72 A4FV84 A4FV97 A4FV98 A4FVI0 A4FVJ7 A4FVP2
A4FVR1 A4FVT7 A4FVT8 A4FVV0 A4FVV8 A4FVX8 A4FVX9 A4FVY0 A4FVY1 A4FVY2 A4FW11
A4FW14 A4FW27 A4FW32 A4FW36 A4FW46 A4FW52 A4FW54 A4FW56 A4FW79 A4FW85 A4FW91
A4FW92 A4FW93 A4FW97 A4FW98 A4FW99 A4FWA1 A4FWA2 A4FWA4 A4FWA5 A4FWA6 A4FWA8
A4FWA9 A4FWB0 A4FWB1 A4FWB4 A4FWB5 A4FWB6 A4FWB7 A4FWB9 A4FWC6 A4FWD2 A4FWE5
A4FWE6 A4FWE8 A4FWE9 A4FWF0 A4FWF1 A4FWF2 A4FWF4 A4FWF5 A4FWF9 A4FWG7 A4FWK1
A4FWK8 A4FWK9 A4FWL3 A4FWL4 A4FWM4 A4FWM6 A4FWP5 A4FWP8 A4FWQ2 A4FWR5 A4FWR6
A4FWS3 A4FWS4 A4FWT8 A4FWU0 A4FWU2 A4FWV5 A4FWV6 A4FWW1 A4FWW2 A4FWW6 A4FWW9
A4FWX0 A4FWX2 A4FWX4 A4FWY5 A4FWY7 A4FWY9 A4FWZ9 A4FX24 A4FX32 A4FX33 A4FX39
A4FX65 A4FX97 A4FX98 A4FXC3 A4FXC7 A4FXE1 A4FXE4 A4FXE7 A4FXF3 A4FXG8 A4FXH2
A4FXH5 A4FXH7 A4FXR1 A4FXR3 A4FXR4 A4FXT1 A4FXT9 A4FXV0 A4FXV4 A4FXV6 A4FXV7
A4FXW1 A4FXW8 A4FXX6 A4FXX9 A4FXY0 A4FXY2 A4FXY5 A4FY19 A4FY32 A4FY35 A4FY69
A4FY92 A4FY94 A4FYA3 A4FYA4 A4FYA5 A4FYB1 A4FYB3 A4FYB9 A4FYC1 A4FYC7 A4FYC9
A4FYD1 A4FYD7 A4FYE4 A4FYI4 A4FYI6 A4FYI7 A4FYK0 A4FYK1 A4FYK2 A4FYK8 A4FYL0
A4FYL2 A4FYL8 A4FYL9 A4FYM2 A4FYM5 A4FYN1 A4FYP1 A4FYQ2 A4FYQ3 A4FYQ8 A4FYS5
A4FYS8 A4FYX5 A4FZ57 A4FZ58 A4FZ59 A4FZ71 A4FZ74 A4FZ85 A4FZ86 A4FZ87 A4FZ97
A4FZ98 A4FZA5 A4FZB0 A4FZB7 A4FZG7 A4FZL0 A4FZL1 A4FZL2 A4FZL6 A4FZL8 A4FZM0
A4FZN7 A4FZN9 A4FZP0 A4FZP2 A4FZP7 A4FZQ1 A4FZQ3 A4FZQ7 A4FZS6 A4FZS7 A4FZS8
A4FZT6 A4FZT8 A4FZU2 A4FZX1 A4FZY1 A4FZY3 A4FZY5 A4G002 A4G005 A4G010 A4G029
A4G031 A4G037 A4G043 A4G060 A4G064 A4G068 A4G074 A4G075 A4G086 A4G087 A4G096
A4G0A1 A4G0A4 A4G0B2 A4G0B9 A4G0C7 A4G0C8 A4G0C9 A4G0D0 A4G0D4 A4G0E2 A4G0E3
A4G0F2 A4G0F7 A4G0F9 A4G0G9 A4G0H7 A4G0I4 A4G0J1 A4G0J7 A4G0K8 A4G0L3 A4G0L4
A4G0N3 A4G0P1 A4G0R3 A4G0R4 A4G0R6 A4G0S4 A4G0T1 A4G0V5 A4G0V7 A4G0Y3 A4G0Y9
A4G0Z0 A4G0Z3 A4G0Z4 A4G154 A4G168 A4G1E2 A4G1F1 A4G1G4 A4G1G8 A4G1K0 A4G1L0
A4G1L3 A4G1L4 A4G1L5 A4G1M4 A4G1N3 A4G1P4 A4G1P5 A4G1Q7 A4G1T2 A4G1T3 A4G1T4
A4G1T7 A4G1U1 A4G1U6 A4G1U7 A4G1U8 A4G1U9 A4G1V0 A4G1V2 A4G1V8 A4G1W7 A4G1X0
A4G1Z2 A4G1Z3 A4G213 A4G215 A4G224 A4G230 A4G255 A4G256 A4G262 A4G265 A4G267
A4G288 A4G289 A4G2A7 A4G2B9 A4G2M2 A4G2M4 A4G2N0 A4G2N3 A4G2P0 A4G2S9 A4G2T5
A4G2T6 A4G2T7 A4G2T8 A4G2V0 A4G310 A4G341 A4G352 A4G3E3 A4G3E6 A4G3G5 A4G3H0
A4G3H1 A4G3H2 A4G3L8 A4G3R1 A4G3R6 A4G3S0 A4G3S6 A4G3S7 A4G3U8 A4G3W7 A4G3Y9
A4G3Z0 A4G413 A4G416 A4G419 A4G422 A4G431 A4G433 A4G4B0 A4G4B3 A4G4F3 A4G4F5
A4G4G0 A4G4G1 A4G4G3 A4G4H8 A4G4J2 A4G4J3 A4G4J8 A4G4J9 A4G4K6 A4G4K7 A4G4L1

A4G4L2 A4G4L7 A4G4M2 A4G4P0 A4G4Q7 A4G4R0 A4G4R4 A4G4S1 A4G4S2 A4G4S3 A4G4S4
A4G4S7 A4G4T5 A4G4T7 A4G4U7 A4G4X3 A4G4Y1 A4G5N9 A4G5P1 A4G5V4 A4G5V5 A4G5X0
A4G5X2 A4G5Z9 A4G615 A4G616 A4G617 A4G619 A4G646 A4G647 A4G648 A4G6D4 A4G6R2
A4G6T4 A4G6Y1 A4G709 A4G712 A4G717 A4G724 A4G738 A4G778 A4G782 A4G7G3 A4G7G5
A4G7M3 A4G7M4 A4G7P5 A4G7S6 A4G7S7 A4G7S9 A4G7T5 A4G7T6 A4G7T9 A4G7U3 A4G7V9
A4G7W0 A4G7W1 A4G7X8 A4G7Y0 A4G7Y7 A4G807 A4G836 A4G837 A4G858 A4G861 A4G865
A4G883 A4G895 A4G8B2 A4G8B3 A4G8D1 A4G8D2 A4G8D3 A4G8D5 A4G8E2 A4G8E6 A4G8E9
A4G8L5 A4G8L8 A4G8N0 A4G8N3 A4G8P1 A4G8P4 A4G8R0 A4G8R1 A4G8R3 A4G8R4 A4G8R5
A4G8R6 A4G8T6 A4G8T7 A4G8U6 A4G8U7 A4G8X1 A4G912 A4G916 A4G919 A4G921 A4G927
A4G964 A4G969 A4G975 A4G977 A4G980 A4G989 A4G995 A4G9B1 A4G9B3 A4G9C5 A4G9C6
A4G9I4 A4G9I5 A4G9I6 A4G9I7 A4G9I9 A4G9J2 A4G9J3 A4G9P9 A4G9Q7 A4G9R2 A4G9R3
A4G9R4 A4G9R5 A4G9R6 A4G9R7 A4G9R9 A4G9S0 A4G9S1 A4G9S2 A4G9S3 A4G9S4 A4G9S5
A4G9S6 A4G9S7 A4G9S8 A4G9T0 A4G9T1 A4G9T2 A4G9T3 A4G9T4 A4G9T5 A4G9T6 A4G9T7
A4G9T8 A4G9U0 A4G9U1 A4G9U2 A4G9U3 A4G9U4 A4G9U5 A4G9U6 A4G9U7 A4G9U8 A4G9U9
A4G9W3 A4GAF0 A4GAG3 A4GAI1 A4GAI2 A4GAN2 A4GAN6 A4GCR5 A4GCR6 A4GCR7 A4GCR8
A4GD50 A4GD54 A4GD55 A4GD56 A4GD58 A4GDQ6 A4GDQ8 A4GDR1 A4GDR3 A4GDR4 A4GDR6
A4GDR7 A4GDR8 A4GDR9 A4GDS0 A4GDS6 A4GDS7 A4GDS9 A4GDT0 A4GDT1 A4GDT3 A4GDT4
A4GDT5 A4GDT9 A4GDU0 A4GDU2 A4GDU3 A4GDU5 A4GDU6 A4GE38 A4GE39 A4GE42 A4GE44
A4GE45 A4GE47 A4GE48 A4GE49 A4GE50 A4GE53 A4GE54 A4GE55 A4GE69 A4GE70 A4GFB7
A4GFB9 A4GFC0 A4GFC2 A4GFC3 A4GFC4 A4GG84 A4GG85 A4GG86 A4GG89 A4GG95 A4GG99
A4GGA5 A4GGA6 A4GGA7 A4GGA8 A4GGC7 A4GGC8 A4GGC9 A4GGD0 A4GGD7 A4GGD8 A4GGD9
A4GGE0 A4GGE1 A4GGE4 A4GGF3 A4GGF5 A4GGF8 A4GGF9 A4GVE9 A4GWX9 A4GXA9 A4GYN8
A4GYP0 A4GYP7 A4GYP8 A4GYP9 A4GYQ0 A4GYQ8 A4GYR2 A4GYR8 A4GYT2 A4GYT3 A4GYT4
A4GYT5 A4GYT6 A4GYU3 A4GYU4 A4GYU5 A4GYU6 A4GYU7 A4GYU8 A4GYU9 A4GYV0 A4GYV1
A4GYV2 A4GYV4 A4GYV5 A4GYV8 A4GYX3 A4GYZ0 A4GZ95 A4GZ97 A4GZV0 A4H3Z2 A4H4W7
A4H5X5 A4H6F7 A4H7A3 A4H7G5 A4H7T9 A4H804 A4H823 A4H824 A4H8F7 A4H9H8 A4H9Z3
A4HE59 A4HFE4 A4HGN7 A4HH79 A4HM77 A4HQ10 A4HQ28 A4HQI9 A4HRK0 A4HS78 A4HT41
A4HU70 A4HUV2 A4HVP7 A4HVU6 A4HW65 A4HWD2 A4HWE9 A4HWF0 A4HWT0 A4HXU4 A4HY57
A4HYT9 A4I1F7 A4I1H7 A4I2L4 A4I3R0 A4I4C5 A4IAU0 A4IAU1 A4ICE5 A4IDS4 A4IF62
A4IF63 A4IF69 A4IF78 A4IF93 A4IF97 A4IF98 A4IFA3 A4IFA8 A4IFB1 A4IFB6 A4IFC4
A4IFD0 A4IFD2 A4IFD8 A4IFE3 A4IFF3 A4IFG2 A4IFH4 A4IFH5 A4IFI1 A4IFJ0 A4IFJ6
A4IFM7 A4IFR3 A4IFR8 A4IFU7 A4IFU8 A4IG62 A4IGH2 A4IGH4 A4IGK4 A4IGL8 A4IGN3
A4IGQ8 A4IGU3 A4IGU9 A4IGY3 A4IGY9 A4IGZ4 A4IH24 A4IH68 A4IH95 A4IHD2 A4IHD9
A4IHF8 A4IHG1 A4IHJ3 A4IHR0 A4IHR1 A4IHR5 A4IHS0 A4IHS2 A4IHT0 A4IHT9 A4IHU6
A4IHU7 A4IHW6 A4IHY1 A4II09 A4II15 A4II20 A4II29 A4II40 A4II60 A4II71 A4II73
A4II96 A4IIB1 A4III8 A4IIJ8 A4IIM2 A4IIN5 A4IIW0 A4IIW5 A4IIX9 A4IIY2 A4IIZ9
A4IJ15 A4IJ21 A4IJ27 A4IJ84 A4IJ87 A4IJ94 A4IJ95 A4IJ96 A4IJA0 A4IJA1 A4IJA6
A4IJB8 A4IJC2 A4IJC5 A4IJC8 A4IJC9 A4IJE6 A4IJG0 A4IJG4 A4IJG5 A4IJG6 A4IJG8
A4IJH3 A4IJH6 A4IJH7 A4IJH8 A4IJH9 A4IJI1 A4IJI2 A4IJI3 A4IJI4 A4IJI5 A4IJI6
A4IJI7 A4IJI8 A4IJI9 A4IJJ0 A4IJJ1 A4IJJ2 A4IJJ3 A4IJJ4 A4IJJ5 A4IJJ6 A4IJJ7
A4IJJ8 A4IJJ9 A4IJK0 A4IJK1 A4IJK2 A4IJK4 A4IJK5 A4IJK6 A4IJK7 A4IJK9 A4IJL1
A4IJL2 A4IJL3 A4IJL4 A4IJL5 A4IJL9 A4IJM0 A4IJM1 A4IJN4 A4IJP9 A4IJS8 A4IJU4
A4IJU6 A4IJU7 A4IJV2 A4IJV3 A4IJX2 A4IJX3 A4IJX5 A4IJY6 A4IJZ1 A4IJZ2 A4IJZ3
A4IK74 A4IK89 A4IK90 A4IK94 A4IK95 A4IK96 A4IKB1 A4IKB5 A4IKB7 A4IKB8 A4IKI6

A4IKI9 A4IKK1 A4IKL1 A4IKL5 A4IKQ7 A4IKR6 A4IKS9 A4IKT9 A4IKU7 A4IKU8 A4IKW6
A4IKW7 A4IL06 A4IL46 A4IL48 A4IL64 A4IL84 A4IL88 A4ILB9 A4ILI2 A4ILL1 A4ILL2
A4ILL5 A4ILL6 A4ILL7 A4ILL8 A4ILN7 A4ILP0 A4ILT5 A4ILT6 A4ILU2 A4ILW6 A4ILY5
A4ILY8 A4ILZ2 A4ILZ4 A4ILZ9 A4IM00 A4IM04 A4IM21 A4IM25 A4IM28 A4IM30 A4IM31
A4IM34 A4IM35 A4IM36 A4IM41 A4IM47 A4IM49 A4IM52 A4IM55 A4IM67 A4IM72 A4IM75
A4IM80 A4IM83 A4IM88 A4IM89 A4IM90 A4IMB9 A4IMC3 A4IMC4 A4IMC5 A4IMC6 A4IMD1
A4IMD3 A4IMD7 A4IMD9 A4IME0 A4IME2 A4IME3 A4IMH7 A4IMI0 A4IMI1 A4IMI5 A4IMK2
A4IMM1 A4IMM3 A4IMP1 A4IMP4 A4IMR6 A4IMR9 A4IMT1 A4IMT6 A4IMW8 A4IN19 A4IN26
A4IN28 A4IN31 A4IN50 A4IN56 A4IN63 A4IN74 A4IN75 A4IN84 A4IN85 A4IN93 A4INM5
A4INR2 A4INW9 A4INY1 A4INZ4 A4IP00 A4IP64 A4IP67 A4IP74 A4IP75 A4IPA1 A4IPA2
A4IPB2 A4IPB3 A4IPB4 A4IPB5 A4IPB6 A4IPF0 A4IPF5 A4IPI2 A4IPN3 A4IPN4 A4IPN9
A4IPP2 A4IPX2 A4IPX4 A4IQ49 A4IQ54 A4IQ59 A4IQ60 A4IQ61 A4IQ63 A4IQ66 A4IQ67
A4IQ78 A4IQ80 A4IQ81 A4IQ82 A4IQ83 A4IQ84 A4IQ85 A4IQ88 A4IQ89 A4IQA2 A4IQA8
A4IQG3 A4IQG7 A4IQI5 A4IQK7 A4IQR5 A4IQR7 A4IQR9 A4IQS0 A4IQS1 A4IQT4 A4IQU0
A4IQV4 A4IQV5 A4IQY1 A4IQZ4 A4IR02 A4IR05 A4IR12 A4IR14 A4IR19 A4IR26 A4IR29
A4IR30 A4IR31 A4IR32 A4IR39 A4IR49 A4IR66 A4IR67 A4IR71 A4IR72 A4IR77 A4IR78
A4IR79 A4IR80 A4IR87 A4IR95 A4IR99 A4IRA1 A4IRA9 A4IRB0 A4IRB2 A4IRB3 A4IRB5
A4IRC0 A4IRC6 A4IRC7 A4IRD1 A4IRD5 A4IRD9 A4IRG2 A4IRG5 A4IRG7 A4IRG9 A4IRH2
A4IRH3 A4IRH6 A4IRH8 A4IRH9 A4IRI5 A4IRL1 A4IRM4 A4IRN1 A4IRN2 A4IRP9 A4IRR6
A4IRS2 A4IRS3 A4IRT9 A4IRU5 A4IRW1 A4IRW4 A4IRW9 A4IRY3 A4IRZ1 A4IRZ2 A4IS14
A4IS20 A4IS22 A4IS33 A4IS40 A4ISB5 A4ISB8 A4ISB9 A4ISE0 A4ISE6 A4ISE8 A4ISE9
A4ISG5 A4ISJ7 A4ISN8 A4ISP5 A4ISP6 A4ISQ2 A4ISQ4 A4ISQ6 A4ISR2 A4ISR3 A4ISR4
A4ISR5 A4ISR7 A4ISR8 A4ISS3 A4ISU3 A4ISU9 A4ISV0 A4IT42 A4IT43 A4IT49 A4IT50
A4IT51 A4IT57 A4IT59 A4IT60 A4IT64 A4ITJ9 A4ITK8 A4ITL0 A4ITL1 A4ITL5 A4ITM9
A4ITP1 A4ITP5 A4ITQ2 A4ITQ8 A4ITU1 A4ITU9 A4ITV1 A4ITV8 A4ITW8 A4ITW9 A4ITX0
A4ITX1 A4ITX4 A4ITX5 A4IVR6 A4IVS4 A4IVS5 A4IVS8 A4IVW3 A4IVY3 A4IVY4 A4IW07
A4IW08 A4IW10 A4IW11 A4IW12 A4IW38 A4IW39 A4IW74 A4IW75 A4IW85 A4IW92 A4IW95
A4IW96 A4IW97 A4IW98 A4IW99 A4IWA0 A4IWA2 A4IWA4 A4IWA5 A4IWA6 A4IWA7 A4IWA8
A4IWB3 A4IWC0 A4IWC1 A4IWC4 A4IWC5 A4IWC7 A4IWF2 A4IWH3 A4IWH6 A4IWH7 A4IWI7
A4IWJ1 A4IWK9 A4IWL0 A4IWQ8 A4IWR6 A4IWT9 A4IWV0 A4IWW4 A4IWW5 A4IWW6 A4IWW8
A4IWY3 A4IWY7 A4IX14 A4IX19 A4IX24 A4IX27 A4IX28 A4IX29 A4IX62 A4IX63 A4IX68
A4IX70 A4IX71 A4IXB4 A4IXD7 A4IXE6 A4IXE8 A4IXG2 A4IXH0 A4IXH9 A4IXI6 A4IXK7
A4IXL1 A4IXL2 A4IXN4 A4IXN6 A4IXN8 A4IXP2 A4IXP5 A4IXR0 A4IXS1 A4IXT7 A4IXT8
A4IXU1 A4IXU4 A4IXV3 A4IXW0 A4IXW5 A4IXW6 A4IXW8 A4IXX8 A4IXY7 A4IXZ5 A4IY03
A4IY05 A4IY06 A4IY07 A4IY09 A4IY35 A4IY57 A4IY63 A4IY68 A4IY77 A4IY85 A4IY87
A4IY88 A4IY90 A4IYA1 A4IYA4 A4IYB0 A4IYB1 A4IYB6 A4IYB7 A4IYC1 A4IYC3 A4IYE6
A4IYF8 A4IYG6 A4IYH5 A4IYI9 A4IYJ0 A4IYJ6 A4IYL4 A4IYL5 A4IYN2 A4IYN5 A4IYP1
A4IYP4 A4IYQ8 A4IYR4 A4IYS7 A4IYU1 A4IYW2 A4IYZ0 A4IZ01 A4IZ18 A4IZ25 A4IZ40
A4IZ44 A4IZ45 A4IZ52 A4IZ54 A4IZ68 A4IZ69 A4IZ90 A4IZ92 A4IZA2 A4IZA5 A4IZA6
A4IZA7 A4IZB0 A4IZB4 A4IZB5 A4IZC2 A4IZE9 A4IZF1 A4IZF4 A4IZG9 A4IZJ6 A4IZJ9
A4IZK0 A4IZK2 A4IZK5 A4IZK6 A4IZK8 A4IZP2 A4IZP3 A4IZQ6 A4IZQ8 A4IZQ9 A4IZR0
A4IZR1 A4IZR2 A4IZR3 A4IZR5 A4IZR6 A4IZR7 A4IZR8 A4IZR9 A4IZS0 A4IZS1 A4IZS2
A4IZS3 A4IZS4 A4IZS5 A4IZS6 A4IZS7 A4IZS8 A4IZS9 A4IZT0 A4IZT1 A4IZT2 A4IZT3
A4IZT4 A4IZT5 A4IZT6 A4IZT7 A4IZT8 A4IZU0 A4IZU3 A4IZU4 A4IZU5 A4IZU6 A4IZU9

A4IZY4 A4IZZ4 A4J011 A4J014 A4J019 A4J020 A4J037 A4J053 A4J072 A4J079 A4J081
A4J0B0 A4J0C0 A4J0C1 A4J0C7 A4J0E5 A4J0F0 A4J0F3 A4J0G0 A4J0G4 A4J0J0 A4J0J1
A4J0K6 A4J0M0 A4J0P1 A4J0Q5 A4J0Q6 A4J0S4 A4J0S6 A4J0X5 A4J0Y3 A4J0Y4 A4J0Y7
A4J0Y9 A4J0Z5 A4J0Z6 A4J0Z9 A4J100 A4J102 A4J103 A4J104 A4J106 A4J107 A4J108
A4J110 A4J111 A4J112 A4J113 A4J114 A4J115 A4J116 A4J117 A4J118 A4J119 A4J121
A4J122 A4J123 A4J124 A4J125 A4J126 A4J127 A4J128 A4J129 A4J130 A4J134 A4J135
A4J136 A4J137 A4J138 A4J139 A4J140 A4J146 A4J147 A4J148 A4J151 A4J152 A4J153
A4J167 A4J169 A4J173 A4J174 A4J179 A4J181 A4J190 A4J1M3 A4J233 A4J246 A4J247
A4J272 A4J2A2 A4J2A3 A4J2A9 A4J2B2 A4J2B3 A4J2C8 A4J2E9 A4J2F3 A4J2F4 A4J2F6
A4J2F8 A4J2F9 A4J2G8 A4J2H0 A4J2J1 A4J2J2 A4J2K1 A4J2K4 A4J2L0 A4J2L1 A4J2L3
A4J3A0 A4J3A1 A4J3A3 A4J3D2 A4J3D4 A4J3F1 A4J3F5 A4J3F7 A4J3G3 A4J3G5 A4J3L6
A4J3N2 A4J3N3 A4J3P1 A4J3P5 A4J3Q0 A4J3Q5 A4J412 A4J451 A4J474 A4J4E2 A4J4E3
A4J4I2 A4J4I4 A4J4X3 A4J4Y7 A4J4Y9 A4J4Z0 A4J4Z4 A4J510 A4J533 A4J535 A4J536
A4J537 A4J540 A4J541 A4J547 A4J555 A4J556 A4J563 A4J568 A4J573 A4J579 A4J582
A4J592 A4J5B2 A4J5C3 A4J5C7 A4J5D0 A4J5D2 A4J5G4 A4J5G8 A4J5N8 A4J5Q0 A4J5Q1
A4J5Q3 A4J5Q6 A4J5Q8 A4J5U0 A4J5U4 A4J5U5 A4J5V5 A4J5W6 A4J5W7 A4J5X1 A4J5X2
A4J5X6 A4J5Y2 A4J5Y5 A4J5Z0 A4J5Z1 A4J5Z2 A4J5Z3 A4J5Z5 A4J5Z6 A4J5Z8 A4J649
A4J661 A4J664 A4J665 A4J666 A4J669 A4J675 A4J678 A4J683 A4J698 A4J6A3 A4J6A4
A4J6A8 A4J6G9 A4J6H0 A4J6H7 A4J6H8 A4J6I7 A4J6S4 A4J6S5 A4J6Z8 A4J703 A4J707
A4J708 A4J710 A4J716 A4J726 A4J742 A4J774 A4J778 A4J7B3 A4J7C9 A4J7D0 A4J7D2
A4J7D9 A4J7E7 A4J7F1 A4J7F3 A4J7F6 A4J7G2 A4J7H2 A4J7I2 A4J7I8 A4J7I9 A4J7J1
A4J7J9 A4J7K6 A4J7K7 A4J7K8 A4J7L5 A4J7L6 A4J7L9 A4J7M0 A4J7Y6 A4J7Y7 A4J806
A4J829 A4J872 A4J898 A4J8C2 A4J8E6 A4J8G6 A4J8H4 A4J8J1 A4J8T4 A4J8U4 A4J903
A4J945 A4J9B1 A4J9C1 A4J9C4 A4J9D2 A4J9P0 A4J9P7 A4J9Q0 A4J9Q3 A4J9Q5 A4J9R9
A4J9S0 A4J9S1 A4J9S4 A4J9S5 A4J9S6 A4J9Z5 A4J9Z8 A4JA07 A4JA22 A4JA23 A4JA40
A4JA69 A4JA70 A4JA71 A4JAD2 A4JAG1 A4JAM5 A4JAM8 A4JAM9 A4JAN1 A4JAN2 A4JAN3
A4JAN5 A4JAN6 A4JAN9 A4JAP0 A4JAP1 A4JAP2 A4JAP3 A4JAP4 A4JAP5 A4JAP6 A4JAP7
A4JAP8 A4JAP9 A4JAQ0 A4JAQ1 A4JAQ2 A4JAQ3 A4JAQ4 A4JAQ5 A4JAQ6 A4JAQ7 A4JAQ8
A4JAQ9 A4JAR1 A4JAR2 A4JAR3 A4JAR4 A4JAR5 A4JAR6 A4JAR7 A4JAS1 A4JAS8 A4JAT5
A4JAW2 A4JAW5 A4JAW8 A4JAW9 A4JAX0 A4JAX1 A4JB50 A4JB51 A4JB67 A4JB68 A4JB85
A4JB86 A4JB92 A4JB95 A4JB96 A4JBA8 A4JBB6 A4JBB7 A4JBB8 A4JBB9 A4JBC1 A4JBC2
A4JBD7 A4JBH8 A4JBI1 A4JBI4 A4JBK4 A4JBK5 A4JBK6 A4JBK7 A4JBK9 A4JBL2 A4JBL3
A4JBL4 A4JBL5 A4JBM2 A4JBP3 A4JBP6 A4JBR4 A4JBR5 A4JBR6 A4JBR9 A4JBS1 A4JBS2
A4JBS4 A4JBT0 A4JBT2 A4JBU8 A4JBW6 A4JC03 A4JC07 A4JC08 A4JC10 A4JC39 A4JC40
A4JC41 A4JC42 A4JC43 A4JC44 A4JC45 A4JC58 A4JC81 A4JC82 A4JC86 A4JCA3 A4JCA7
A4JCA9 A4JCC2 A4JCC5 A4JCE2 A4JCG7 A4JCH1 A4JCH4 A4JCH5 A4JCH7 A4JCI2 A4JCJ3
A4JCJ8 A4JCJ9 A4JCK0 A4JCK6 A4JCM8 A4JCN6 A4JCP6 A4JCR8 A4JCS0 A4JCW0 A4JCW1
A4JCZ0 A4JDA6 A4JDA9 A4JDB7 A4JDM9 A4JDP6 A4JDS6 A4JDU4 A4JDU6 A4JDU7 A4JDU8
A4JDW9 A4JDX1 A4JDX2 A4JE17 A4JE23 A4JE24 A4JE26 A4JEA9 A4JEB6 A4JEL4 A4JEN0
A4JEN5 A4JEN6 A4JEN9 A4JEP2 A4JEP3 A4JEQ0 A4JEQ1 A4JEQ3 A4JEQ4 A4JER1 A4JEU8
A4JEU9 A4JEV1 A4JEW8 A4JEZ0 A4JEZ1 A4JF04 A4JF05 A4JF06 A4JF19 A4JF20 A4JF45
A4JF55 A4JF59 A4JF61 A4JF69 A4JF72 A4JF73 A4JF74 A4JF75 A4JF78 A4JF81 A4JF86
A4JF88 A4JFB2 A4JFD0 A4JFW7 A4JFY7 A4JFZ6 A4JG01 A4JG19 A4JG35 A4JG46 A4JGA4
A4JGD4 A4JGD5 A4JGE0 A4JGE3 A4JGH2 A4JGH4 A4JGH5 A4JGI0 A4JGJ9 A4JGK4 A4JGM1

A4JGV4 A4JGV5 A4JGV8 A4JGW2 A4JGW6 A4JGX1 A4JGX2 A4JGX5 A4JGZ2 A4JH21 A4JH22
A4JH23 A4JH32 A4JH35 A4JH39 A4JH78 A4JH80 A4JH83 A4JH86 A4JH87 A4JHA1 A4JHD7
A4JHJ1 A4JHJ8 A4JHM8 A4JHN3 A4JHP7 A4JHP8 A4JHR9 A4JHS6 A4JHT1 A4JHX7 A4JHY7
A4JHY8 A4JHZ1 A4JHZ4 A4JHZ6 A4JI00 A4JI42 A4JI45 A4JI72 A4JI88 A4JIA2 A4JIA9
A4JIB5 A4JIB6 A4JIB7 A4JIF6 A4JIK4 A4JIL2 A4JIN6 A4JIQ4 A4JIQ6 A4JIR2 A4JIR3
A4JIR7 A4JIS1 A4JIS9 A4JIT0 A4JIT1 A4JIU1 A4JJ22 A4JJ44 A4JJ49 A4JJE1 A4JJG4
A4JJG5 A4JJG6 A4JJS1 A4JK94 A4JKA9 A4JKE6 A4JKF7 A4JKG7 A4JKV8 A4JL57 A4JLD9
A4JLS0 A4JLT2 A4JLT5 A4JLT6 A4JM71 A4JMB6 A4JMC2 A4JN03 A4JNX5 A4JP25 A4JPD1
A4JPG9 A4JPX5 A4JPX7 A4JPX8 A4JPX9 A4JPY0 A4JPY1 A4JQH4 A4JR39 A4JRK7 A4JRL3
A4JRR6 A4JS50 A4JS72 A4JSU5 A4JT91 A4JW24 A4JW25 A4K2M3 A4K2P5 A4K2Q5 A4K2S1
A4K2U9 A4K2Y1 A4K9Z8 A4KA31 A4KA32 A4KA33 A4KA34 A4KA36 A4KA37 A4KA38 A4KA39
A4KA40 A4KA41 A4KA43 A4KA44 A4KA45 A4KA49 A4KA50 A4KA51 A4KA52 A4KA53 A4KA54
A4KA55 A4KA56 A4KA57 A4KA58 A4KA59 A4KA60 A4KA61 A4KAG7 A4KAG8 A4KX57 A4KX71
A4KXB1 A4KXB2 A4KXB4 A4KXK0 A4KXL9 A4KZ49 A4L691 A4L9I8 A4L9I9 A4LBC0 A4PB26
A4PBL4 A4PBP6 A4PBP7 A4PBP8 A4PBP9 A4PBQ0 A4PU48 A4Q8F7 A4Q8H2 A4Q998 A4Q9E4
A4Q9E5 A4Q9E8 A4Q9F0 A4Q9F1 A4Q9F3 A4Q9F4 A4Q9F6 A4Q9R9 A4Q9S2 A4Q9S3 A4QA10
A4QA21 A4QA22 A4QA24 A4QA25 A4QA26 A4QA27 A4QAA5 A4QAA6 A4QAA8 A4QAD0 A4QAF9
A4QAK7 A4QAL4 A4QAN2 A4QAN3 A4QAP0 A4QB37 A4QB40 A4QB41 A4QB58 A4QB59 A4QB64
A4QB65 A4QB76 A4QB78 A4QB97 A4QBE1 A4QBE5 A4QBF0 A4QBF1 A4QBF9 A4QBG0 A4QBG2
A4QBG3 A4QBG7 A4QBG8 A4QBG9 A4QBH0 A4QBH7 A4QBH9 A4QBI0 A4QBI2 A4QBI3 A4QBI5
A4QBI6 A4QBI7 A4QBI8 A4QBI9 A4QBJ1 A4QBJ4 A4QBJ5 A4QBJ6 A4QBL0 A4QBL1 A4QBL3
A4QBL5 A4QBL6 A4QBP4 A4QBQ3 A4QBQ4 A4QBQ5 A4QBQ6 A4QBQ7 A4QBQ8 A4QBS3 A4QBS4
A4QBS5 A4QBT0 A4QBT6 A4QBT9 A4QBU0 A4QBV0 A4QBY8 A4QC29 A4QC50 A4QC87 A4QC99
A4QCC4 A4QCD3 A4QCD6 A4QCG9 A4QCI7 A4QCJ4 A4QCK4 A4QCK8 A4QCK9 A4QCL0 A4QCL1
A4QCL4 A4QCL5 A4QCN5 A4QCP0 A4QCP1 A4QCR9 A4QCW5 A4QCW6 A4QCX4 A4QCY8 A4QCY9
A4QD64 A4QD83 A4QDA4 A4QDE1 A4QDE5 A4QDF8 A4QDG1 A4QDG2 A4QDH7 A4QDK1 A4QDK5
A4QDK7 A4QDK8 A4QDL6 A4QDM0 A4QDM9 A4QDN4 A4QDP9 A4QDQ6 A4QDR7 A4QDS8 A4QDS9
A4QDT4 A4QDU2 A4QDY1 A4QDY2 A4QDY3 A4QDY8 A4QDZ0 A4QDZ3 A4QDZ4 A4QDZ5 A4QE10
A4QE51 A4QE80 A4QE83 A4QE90 A4QE91 A4QEA0 A4QEC1 A4QEG0 A4QEG1 A4QEG3 A4QEG5
A4QEG6 A4QEG8 A4QEG9 A4QEH4 A4QEI3 A4QEI7 A4QEI8 A4QEJ4 A4QEJ6 A4QEJ8 A4QEK7
A4QEL0 A4QEM8 A4QEN3 A4QEN4 A4QEN6 A4QEQ9 A4QER2 A4QES6 A4QET6 A4QEV1 A4QEV2
A4QEV7 A4QEW3 A4QEY0 A4QEY2 A4QEY3 A4QEY6 A4QEZ1 A4QEZ2 A4QEZ5 A4QF01 A4QF02
A4QF06 A4QF08 A4QF22 A4QF26 A4QF29 A4QF30 A4QF31 A4QF32 A4QF37 A4QF39 A4QFA4
A4QFA8 A4QFB9 A4QFC1 A4QFD3 A4QFD9 A4QFF8 A4QFF9 A4QFG5 A4QFG6 A4QFK8 A4QFM2
A4QFN1 A4QFN2 A4QFR5 A4QFS2 A4QFS3 A4QFW4 A4QFY0 A4QFZ5 A4QFZ9 A4QG13 A4QG64
A4QG75 A4QG78 A4QG84 A4QG85 A4QG90 A4QGA0 A4QGA7 A4QGE8 A4QGM2 A4QGM5 A4QGQ5
A4QGQ6 A4QGR5 A4QGT2 A4QGT4 A4QGT5 A4QGU3 A4QGX4 A4QGX7 A4QGY6 A4QGZ4 A4QH04
A4QH19 A4QH41 A4QH44 A4QH48 A4QH59 A4QH60 A4QH66 A4QH94 A4QHB7 A4QHD8 A4QHF4
A4QHG1 A4QHG2 A4QHG9 A4QHI9 A4QHJ0 A4QHN8 A4QHQ4 A4QHQ6 A4QHT1 A4QHW5 A4QHY3
A4QI23 A4QI35 A4QI41 A4QI59 A4QI74 A4QI76 A4QI77 A4QIE3 A4QIE5 A4QIE6 A4QJ97
A4QJA4 A4QJA5 A4QJA6 A4QJA7 A4QJB3 A4QJB7 A4QJC3 A4QJD6 A4QJD7 A4QJD8 A4QJD9
A4QJE0 A4QJE7 A4QJE8 A4QJE9 A4QJF0 A4QJF1 A4QJF2 A4QJF3 A4QJF4 A4QJF5 A4QJF6
A4QJF8 A4QJG0 A4QJG4 A4QJH3 A4QJI8 A4QJI9 A4QJJ0 A4QJJ1 A4QJJ7 A4QJK1 A4QJK7
A4QJM0 A4QJM1 A4QJM2 A4QJM3 A4QJM4 A4QJN1 A4QJN2 A4QJN3 A4QJN4 A4QJN5 A4QJN6

A4QJN7 A4QJN8 A4QJN9 A4QJP0 A4QJP2 A4QJP4 A4QJP8 A4QJQ7 A4QJS2 A4QJS3 A4QJS4
A4QJS5 A4QJT1 A4QJT5 A4QJU1 A4QJV2 A4QJV3 A4QJV4 A4QJV5 A4QJV6 A4QJW3 A4QJW4
A4QJW5 A4QJW6 A4QJW7 A4QJW8 A4QJW9 A4QJX0 A4QJX1 A4QJX2 A4QJX4 A4QJX6 A4QJX7
A4QJY0 A4QJY9 A4QK00 A4QK07 A4QK08 A4QK09 A4QK10 A4QK16 A4QK20 A4QK26 A4QK39
A4QK40 A4QK41 A4QK42 A4QK43 A4QK50 A4QK51 A4QK52 A4QK53 A4QK54 A4QK55 A4QK56
A4QK57 A4QK58 A4QK59 A4QK61 A4QK62 A4QK64 A4QK68 A4QK77 A4QK86 A4QK94 A4QK95
A4QK96 A4QK97 A4QKA3 A4QKA7 A4QKB3 A4QKC6 A4QKC7 A4QKC8 A4QKC9 A4QKD0 A4QKD7
A4QKD8 A4QKD9 A4QKE0 A4QKE1 A4QKE2 A4QKE3 A4QKE4 A4QKE5 A4QKE6 A4QKE8 A4QKE9
A4QKF1 A4QKF5 A4QKG4 A4QKH3 A4QKI1 A4QKI2 A4QKI3 A4QKI4 A4QKJ0 A4QKJ4 A4QKK0
A4QKL3 A4QKL4 A4QKL5 A4QKL6 A4QKL7 A4QKM4 A4QKM5 A4QKM6 A4QKM7 A4QKM8 A4QKM9
A4QKN0 A4QKN1 A4QKN2 A4QKN3 A4QKN5 A4QKN6 A4QKN8 A4QKP3 A4QKQ2 A4QKR2 A4QKS0
A4QKS1 A4QKS2 A4QKS3 A4QKS9 A4QKT3 A4QKT9 A4QKV2 A4QKV3 A4QKV4 A4QKV5 A4QKV6
A4QKW3 A4QKW4 A4QKW5 A4QKW6 A4QKW7 A4QKW8 A4QKW9 A4QKX0 A4QKX1 A4QKX2 A4QKX4
A4QKX5 A4QKX7 A4QKY2 A4QKZ1 A4QKZ7 A4QL01 A4QL08 A4QL09 A4QL10 A4QL11 A4QL17
A4QL21 A4QL27 A4QL40 A4QL41 A4QL42 A4QL43 A4QL44 A4QL51 A4QL52 A4QL53 A4QL54
A4QL55 A4QL56 A4QL57 A4QL58 A4QL59 A4QL60 A4QL62 A4QL63 A4QL65 A4QL69 A4QL78
A4QL87 A4QL95 A4QL96 A4QL97 A4QL98 A4QLA4 A4QLA8 A4QLB4 A4QLC7 A4QLC8 A4QLC9
A4QLD0 A4QLD1 A4QLD8 A4QLD9 A4QLE0 A4QLE1 A4QLE2 A4QLE3 A4QLE4 A4QLE5 A4QLE6
A4QLE7 A4QLE9 A4QLF0 A4QLF2 A4QLF7 A4QLG6 A4QLH6 A4QLI4 A4QLI5 A4QLI6 A4QLJ2
A4QLJ6 A4QLK2 A4QLL5 A4QLL6 A4QLL7 A4QLL8 A4QLL9 A4QLM6 A4QLM7 A4QLM8 A4QLM9
A4QLN0 A4QLN1 A4QLN2 A4QLN3 A4QLN4 A4QLN5 A4QLN7 A4QLN8 A4QLP0 A4QLP5 A4QLQ4
A4QLR4 A4QLS2 A4QLS3 A4QLS4 A4QLS5 A4QLT1 A4QLT5 A4QLU1 A4QLV4 A4QLV5 A4QLV6
A4QLV7 A4QLV8 A4QLW5 A4QLW6 A4QLW7 A4QLW8 A4QLW9 A4QLX0 A4QLX1 A4QLX2 A4QLX3
A4QLX4 A4QLX6 A4QLX7 A4QLX9 A4QLY3 A4QLZ2 A4QMS7 A4QN01 A4QN59 A4QNB6 A4QNC6
A4QND5 A4QNE6 A4QNG3 A4QNG5 A4QNI8 A4QNL8 A4QNM5 A4QNP7 A4QNR3 A4QNR8 A4QNW7
A4QNX6 A4QNZ7 A4QP16 A4QP31 A4QP75 A4QP78 A4QPB2 A4QQ05 A4QQE0 A4QQN1 A4QR60
A4QRM8 A4QS18 A4QSS5 A4QSX4 A4QT78 A4QTA3 A4QTC6 A4QTQ9 A4QTR1 A4QTY2 A4QU31
A4QUC1 A4QUF0 A4QVH4 A4QVI3 A4QVP2 A4QW40 A4QX49 A4QXV8 A4QXX4 A4QYF9 A4QYG1
A4QYM6 A4QYU7 A4QZL9 A4R0E5 A4R0E6 A4R0J9 A4R0Q1 A4R0R0 A4R0T9 A4R188 A4R1G4
A4R1J6 A4R1T1 A4R227 A4R2Q6 A4R2R1 A4R2S1 A4R3M4 A4R4J0 A4R4R3 A4R5B8 A4R5F6
A4R606 A4R6X5 A4R713 A4R715 A4R796 A4R7U3 A4R8B5 A4R8D7 A4R8F9 A4R8G3 A4R962
A4R9U5 A4RA36 A4RAE9 A4RBS3 A4RBW8 A4RCW0 A4RCX7 A4RD09 A4RD36 A4RD79 A4RD93
A4RDD7 A4RDI6 A4RE46 A4REU9 A4REV8 A4RF35 A4RF51 A4RFH6 A4RGD1 A4RGE6 A4RGU2
A4RGU7 A4RHF1 A4RHN3 A4RHR8 A4RHU9 A4RI88 A4RJA0 A4RJH4 A4RK04 A4RK48 A4RK52
A4RK68 A4RK80 A4RK96 A4RKC3 A4RKF7 A4RLF4 A4RLI4 A4RM69 A4RM80 A4RMA9 A4RMB1
A4RMV8 A4RN08 A4RN46 A4RPM5 A4RQ11 A4RQ29 A4RS24 A4RSQ6 A4RUK2 A4RW69 A4RYD2
A4RZ86 A4RZA6 A4S051 A4S068 A4S1G4 A4S1K1 A4S3S3 A4S450 A4S6Z4 A4S779 A4SAD2
A4SC23 A4SC27 A4SC34 A4SC35 A4SCF1 A4SCF3 A4SCI7 A4SCJ5 A4SCJ7 A4SCJ8 A4SCK0
A4SCK1 A4SCK2 A4SCK3 A4SCK4 A4SCK8 A4SCL6 A4SCN7 A4SCP4 A4SCQ4 A4SCQ5 A4SCQ6
A4SCQ7 A4SCQ8 A4SCQ9 A4SCR0 A4SCR1 A4SCR3 A4SCR4 A4SCR6 A4SCR9 A4SCS0 A4SCS1
A4SCS2 A4SCS3 A4SCS4 A4SCS5 A4SCS6 A4SCS7 A4SCT1 A4SCT2 A4SCT3 A4SCT4 A4SCT5
A4SCT6 A4SCT7 A4SCT8 A4SCU1 A4SCU2 A4SCU5 A4SCU6 A4SCU9 A4SCW6 A4SCX0 A4SD12
A4SD47 A4SD50 A4SD53 A4SD60 A4SD65 A4SD66 A4SD71 A4SD72 A4SD82 A4SD83 A4SDA4
A4SDB4 A4SDB8 A4SDB9 A4SDC0 A4SDC1 A4SDC2 A4SDE2 A4SDE6 A4SDF7 A4SDG1 A4SDH2

A4SDK2 A4SDL1 A4SDL6 A4SDM0 A4SDM4 A4SDP8 A4SDP9 A4SDQ0 A4SDQ2 A4SDR0 A4SDU0
A4SE02 A4SE07 A4SE10 A4SE17 A4SE28 A4SE31 A4SE60 A4SE63 A4SE69 A4SE77 A4SE82
A4SE83 A4SE87 A4SEC8 A4SEG3 A4SEI5 A4SEI7 A4SEI8 A4SEN7 A4SEN9 A4SEP8 A4SEQ5
A4SEQ7 A4SER5 A4SER8 A4SEX9 A4SEY8 A4SEY9 A4SF00 A4SF04 A4SF07 A4SF37 A4SF66
A4SF76 A4SF77 A4SF78 A4SFB1 A4SFC2 A4SFC5 A4SFD1 A4SFD5 A4SFG0 A4SFG2 A4SFG4
A4SFG7 A4SFH7 A4SFH8 A4SFI9 A4SFJ7 A4SFJ8 A4SFK6 A4SFM6 A4SFM8 A4SFN0 A4SFP1
A4SFP2 A4SFP8 A4SFR5 A4SFR6 A4SFR7 A4SFS0 A4SFS4 A4SFT1 A4SFT2 A4SFT3 A4SFT4
A4SFT9 A4SFU1 A4SFU6 A4SFW2 A4SFY3 A4SFY4 A4SFZ0 A4SFZ6 A4SFZ8 A4SFZ9 A4SG05
A4SG06 A4SG07 A4SG11 A4SG14 A4SG19 A4SG25 A4SG26 A4SG27 A4SG33 A4SG34 A4SG41
A4SG66 A4SG77 A4SG83 A4SG85 A4SGA5 A4SGB1 A4SGC0 A4SGD2 A4SGE1 A4SGE2 A4SGE6
A4SGF2 A4SGF8 A4SGG0 A4SGG1 A4SGG2 A4SGG4 A4SGH4 A4SGJ0 A4SGJ6 A4SGK2 A4SGK5
A4SGK7 A4SGK8 A4SGL0 A4SGM5 A4SGM6 A4SGN0 A4SGN3 A4SGN4 A4SGN6 A4SGR9 A4SGT2
A4SGU2 A4SGU4 A4SGV9 A4SGW2 A4SGX9 A4SGY3 A4SGY4 A4SH01 A4SH04 A4SH10 A4SH11
A4SH15 A4SH23 A4SH46 A4SH73 A4SHB7 A4SHC0 A4SHC9 A4SHE9 A4SHF6 A4SHG2 A4SHK8
A4SHL4 A4SHM1 A4SHM2 A4SHP4 A4SHP7 A4SHP8 A4SHQ9 A4SHR6 A4SHU2 A4SHU5 A4SHU6
A4SHU7 A4SHU8 A4SHU9 A4SHV0 A4SHV6 A4SHV7 A4SHV8 A4SHW1 A4SHW3 A4SHW5 A4SHY0
A4SHZ4 A4SHZ5 A4SI48 A4SI57 A4SI66 A4SIB9 A4SIE1 A4SIE3 A4SIF7 A4SIG1 A4SIG6
A4SIJ4 A4SIL2 A4SIM0 A4SIM2 A4SIM3 A4SIM5 A4SIM6 A4SIU5 A4SIV0 A4SIV7 A4SIV8
A4SIV9 A4SIW0 A4SIW4 A4SIW7 A4SIX2 A4SIX5 A4SIX8 A4SIZ7 A4SIZ8 A4SIZ9 A4SJ23
A4SJ25 A4SJ26 A4SJ51 A4SJ60 A4SJ61 A4SJ66 A4SJ79 A4SJ82 A4SJ84 A4SJB3 A4SJC2
A4SJD0 A4SJD4 A4SJF6 A4SJK0 A4SJK1 A4SJL7 A4SJM5 A4SJM6 A4SJN2 A4SJN4 A4SJN7
A4SJP2 A4SJP9 A4SJQ1 A4SJQ7 A4SJR0 A4SJR3 A4SJR5 A4SJR6 A4SJR7 A4SJR8 A4SJR9
A4SJV5 A4SJV7 A4SJW3 A4SJW5 A4SJW8 A4SJX2 A4SJX4 A4SJY6 A4SJY8 A4SK24 A4SK25
A4SK35 A4SK61 A4SK63 A4SK64 A4SK71 A4SK74 A4SK82 A4SKA6 A4SKE9 A4SKG7 A4SKH7
A4SKI2 A4SKJ6 A4SKK7 A4SKL1 A4SKR0 A4SKS0 A4SKT0 A4SKT1 A4SKT3 A4SKY4 A4SL28
A4SL29 A4SL69 A4SLF3 A4SLF8 A4SLS4 A4SLS6 A4SLT2 A4SLV6 A4SLX9 A4SLY0 A4SLY1
A4SM13 A4SM23 A4SM33 A4SM41 A4SM42 A4SM43 A4SM83 A4SM99 A4SMA4 A4SMA6 A4SMA7
A4SMA8 A4SMC0 A4SMC4 A4SMD7 A4SME4 A4SMI9 A4SMK3 A4SML2 A4SML3 A4SML6 A4SML8
A4SMM1 A4SMM2 A4SMM4 A4SMN1 A4SMN3 A4SMP5 A4SMP7 A4SMQ0 A4SMQ4 A4SMV5 A4SMV9
A4SMW0 A4SMW2 A4SMX2 A4SMX4 A4SMX7 A4SMZ0 A4SN40 A4SNA7 A4SNB0 A4SNB5 A4SND2
A4SNE3 A4SNF0 A4SNF1 A4SNH3 A4SNI2 A4SNJ1 A4SNK6 A4SNL5 A4SNL6 A4SNL8 A4SNM0
A4SNT8 A4SNU1 A4SNV2 A4SNY1 A4SNZ8 A4SP01 A4SP04 A4SP05 A4SP09 A4SP10 A4SP14
A4SP26 A4SP27 A4SP34 A4SP35 A4SP53 A4SP64 A4SP90 A4SPA7 A4SPD6 A4SPD7 A4SPE5
A4SPE8 A4SPH9 A4SPI4 A4SPI6 A4SPM2 A4SPN4 A4SPN5 A4SPN7 A4SPN8 A4SPR4 A4SPR6
A4SPR7 A4SPS7 A4SPV7 A4SPX2 A4SQ09 A4SQ24 A4SQ25 A4SQ26 A4SQ27 A4SQ71 A4SQG8
A4SQH6 A4SQH9 A4SQI0 A4SQI1 A4SQI2 A4SQI5 A4SQI6 A4SQJ9 A4SQK3 A4SQK6 A4SQM2
A4SQM8 A4SQN8 A4SQW5 A4SR08 A4SR17 A4SR18 A4SR19 A4SR25 A4SR26 A4SR27 A4SR47
A4SR62 A4SR64 A4SR65 A4SR75 A4SR80 A4SR81 A4SR98 A4SRB0 A4SRB1 A4SRB5 A4SRB6
A4SRB7 A4SRC2 A4SRC5 A4SRF1 A4SRF2 A4SRG6 A4SRG8 A4SRG9 A4SRH0 A4SRR7 A4SRS3
A4SRS5 A4SRU2 A4SRU3 A4SRU4 A4SRW7 A4SRW9 A4SRY4 A4SS04 A4SS26 A4SS38 A4SS45
A4SS61 A4SS99 A4SSA1 A4SSA7 A4SSI2 A4SSJ2 A4SSN4 A4SSN5 A4SSY0 A4SSY1 A4SSY2
A4SSY3 A4SSY4 A4SSY5 A4SSY7 A4SSY8 A4SSY9 A4SSZ0 A4SSZ1 A4SSZ2 A4SSZ3 A4SSZ4
A4SSZ5 A4SSZ6 A4SSZ7 A4SSZ8 A4SSZ9 A4ST00 A4ST01 A4ST02 A4ST03 A4ST04 A4ST05
A4ST06 A4ST17 A4ST19 A4ST21 A4ST23 A4ST45 A4ST47 A4ST49 A4ST50 A4ST51 A4ST54

A4ST57 A4ST58 A4STA4 A4STB6 A4STC9 A4STD1 A4STD3 A4STD4 A4STD5 A4STD7 A4STE1
A4STE2 A4STF0 A4STF8 A4STG2 A4STG3 A4STH3 A4STH4 A4STJ7 A4STJ8 A4STK0 A4STQ3
A4STQ4 A4STS4 A4STS7 A4STS8 A4SUS3 A4SUS4 A4SUU7 A4SUV0 A4SUV1 A4SUV3 A4SUV4
A4SUV5 A4SUV6 A4SUV7 A4SUV8 A4SUW0 A4SUW1 A4SUW2 A4SUW3 A4SUW4 A4SUW5 A4SUW6
A4SUW7 A4SUW8 A4SUX0 A4SUX1 A4SUX2 A4SUX3 A4SUX4 A4SUX5 A4SUX6 A4SUX7 A4SUX8
A4SUX9 A4SUY0 A4SUY2 A4SUY3 A4SUY4 A4SUY5 A4SUY6 A4SUY7 A4SUZ2 A4SV13 A4SV17
A4SV19 A4SV20 A4SV43 A4SV44 A4SV52 A4SV53 A4SV63 A4SV66 A4SV75 A4SV76 A4SVA1
A4SVA2 A4SVA3 A4SVA6 A4SVA7 A4SVB8 A4SVD9 A4SVE2 A4SVE6 A4SVF3 A4SVH2 A4SVH9
A4SVI6 A4SVI7 A4SVI8 A4SVJ6 A4SVJ7 A4SVT7 A4SVT8 A4SVU3 A4SVV2 A4SVW4 A4SVW9
A4SVX6 A4SVX8 A4SVX9 A4SW55 A4SW73 A4SW74 A4SW75 A4SW76 A4SWE1 A4SWG7 A4SWH2
A4SWJ8 A4SWK6 A4SWK9 A4SWL6 A4SWL7 A4SWM3 A4SWN4 A4SWP6 A4SWQ5 A4SWQ6 A4SWT5
A4SWU6 A4SWV7 A4SWW0 A4SWW8 A4SWX0 A4SWX4 A4SWX7 A4SX33 A4SX35 A4SX36 A4SX51
A4SX62 A4SX83 A4SXC8 A4SXD4 A4SXD5 A4SXD7 A4SXE7 A4SXF2 A4SXH9 A4SXJ6 A4SXL3
A4SXL6 A4SXM1 A4SXM6 A4SXQ5 A4SXQ7 A4SXQ8 A4SXR0 A4SXR3 A4SXS9 A4SY41 A4SY42
A4SY43 A4SY44 A4SY47 A4SY48 A4SY77 A4SY78 A4SY80 A4SYB3 A4SYB4 A4SYD1 A4SYD2
A4SYD7 A4SYE2 A4SYE3 A4SYE9 A4SYF9 A4SYS2 A4SYS6 A4SYT7 A4SYU8 A4SYV0 A4SYV1
A4SYV5 A4SYV8 A4SYV9 A4SYX2 A4SYY9 A4SZ24 A4SZK0 A4SZK1 A4SZK2 A4SZL8 A4SZM1
A4SZN3 A4SZN4 A4SZP2 A4SZQ2 A4SZQ5 A4SZQ9 A4SZR8 A4SZR9 A4SZT7 A4SZU5 A4SZV4
A4SZV5 A4SZX3 A4SZY0 A4T007 A4T008 A4T011 A4T025 A4T026 A4T027 A4T031 A4T032
A4T034 A4T063 A4T067 A4T069 A4T070 A4T072 A4T096 A4T0B0 A4T0E0 A4T0E9 A4T0F0
A4T0F4 A4T0G0 A4T0G5 A4T0M5 A4T0N1 A4T0N5 A4T0R6 A4T0W0 A4T0W5 A4T0X9 A4T112
A4T142 A4T163 A4T166 A4T174 A4T183 A4T197 A4T1E2 A4T1L7 A4T1M1 A4T1M2 A4T1N5
A4T1P3 A4T1P4 A4T1P7 A4T1R2 A4T1R3 A4T1R4 A4T1T9 A4T1U0 A4T1U2 A4T1U3 A4T2B7
A4T2C0 A4T2C4 A4T2G0 A4T2G4 A4T2H9 A4T2I0 A4T2J0 A4T2J3 A4T2J4 A4T2J5 A4T2N4
A4T2N8 A4T2P3 A4T2R5 A4T2T4 A4T2U3 A4T318 A4T324 A4T340 A4T341 A4T358 A4T3S1
A4T3S4 A4T3V1 A4T3Z0 A4T413 A4T432 A4T4N9 A4T4P0 A4T4R2 A4T4R8 A4T4S9 A4T4T3
A4T4T9 A4T4U1 A4T4Y2 A4T523 A4T5K5 A4T5K9 A4T5L0 A4T5M9 A4T5N4 A4T5N5 A4T5P2
A4T5Y1 A4T5Z8 A4T5Z9 A4T600 A4T623 A4T6G3 A4T6N5 A4T6N7 A4T6P3 A4T6Q0 A4T6T4
A4T6V7 A4T6W8 A4T6Y5 A4T792 A4T7D6 A4T7D7 A4T7H0 A4T7H3 A4T7J3 A4T7K2 A4T7Q6
A4T7U3 A4T7U4 A4T8B6 A4T8B7 A4T8F7 A4T8H8 A4T8L2 A4T8L3 A4T8M4 A4T8Q2 A4T8S3
A4T8W9 A4T8X4 A4T907 A4T986 A4T9A9 A4T9D5 A4T9I1 A4T9L2 A4T9L3 A4T9L9 A4T9M4
A4T9M6 A4T9N1 A4T9N2 A4T9N5 A4T9S3 A4T9U9 A4T9V0 A4T9V2 A4T9V7 A4T9W3 A4T9W4
A4T9W6 A4T9Y6 A4TA35 A4TA86 A4TA90 A4TAC4 A4TAD3 A4TAK5 A4TB47 A4TB55 A4TB64
A4TB65 A4TB82 A4TB83 A4TB85 A4TBE2 A4TBE7 A4TBF5 A4TBF6 A4TBH5 A4TBJ9 A4TBK7
A4TBQ5 A4TBQ6 A4TBR2 A4TBU4 A4TBW4 A4TBX3 A4TBX4 A4TBX8 A4TBY0 A4TBY6 A4TBZ0
A4TC02 A4TC09 A4TC12 A4TC13 A4TC15 A4TC18 A4TC20 A4TC25 A4TC66 A4TC67 A4TC75
A4TC77 A4TCE3 A4TCF3 A4TCF7 A4TCF8 A4TCG8 A4TCH6 A4TCI6 A4TCI9 A4TCJ3 A4TCJ4
A4TCL7 A4TCM9 A4TCN3 A4TCN4 A4TCN7 A4TCR4 A4TCS1 A4TCS5 A4TD09 A4TD17 A4TDE5
A4TDF4 A4TDS2 A4TDU4 A4TDZ4 A4TDZ8 A4TE06 A4TE07 A4TE12 A4TE16 A4TE22 A4TE23
A4TE25 A4TE28 A4TE29 A4TE31 A4TE41 A4TE51 A4TE57 A4TE63 A4TE69 A4TE75 A4TE78
A4TE84 A4TEA7 A4TEB1 A4TEB6 A4TEB7 A4TEB8 A4TEB9 A4TEC0 A4TEC2 A4TEC3 A4TEC4
A4TEC6 A4TEC7 A4TEC9 A4TED3 A4TED4 A4TED9 A4TEE0 A4TEI1 A4TEI2 A4TEI3 A4TEJ2
A4TEJ4 A4TEL0 A4TEL3 A4TEN0 A4TEN6 A4TEN7 A4TET2 A4TEW1 A4TEW9 A4TF31 A4TFB8
A4TFF4 A4TGL5 A4TGM6 A4TGM7 A4TGN6 A4TGN7 A4TGP0 A4TGP7 A4TGQ2 A4TGQ3 A4TGR3

A4TGR4 A4TGR7 A4TGS1 A4TGS8 A4TGS9 A4TGU1 A4TGU2 A4TGU8 A4TGW5 A4TGX8 A4TGY1
A4TGY2 A4TGY3 A4TGY4 A4TGY5 A4TGY6 A4TGY7 A4TGZ1 A4TGZ2 A4TGZ3 A4TGZ5 A4TGZ6
A4TGZ7 A4TGZ8 A4TGZ9 A4TH00 A4TH01 A4TH02 A4TH03 A4TH04 A4TH05 A4TH06 A4TH07
A4TH08 A4TH09 A4TH10 A4TH12 A4TH13 A4TH14 A4TH15 A4TH16 A4TH21 A4TH22 A4TH23
A4TH25 A4TH27 A4TH28 A4TH30 A4TH34 A4TH39 A4TH89 A4THB7 A4THC4 A4THF5 A4THG5
A4THG6 A4THG8 A4THI5 A4THJ1 A4THJ2 A4THK2 A4THK3 A4THL9 A4THM0 A4THM8 A4THT0
A4THT1 A4THT5 A4THT9 A4THU2 A4TI59 A4TI61 A4TI69 A4TI76 A4TI77 A4TI83 A4TI90
A4TI95 A4TIA1 A4TIA5 A4TIA6 A4TIA7 A4TIB4 A4TIC5 A4TIL1 A4TIL3 A4TIL4 A4TIL5
A4TIL7 A4TIN0 A4TIN1 A4TIN3 A4TIP2 A4TIP3 A4TIQ4 A4TIR6 A4TIR8 A4TIS1 A4TIV5
A4TIW7 A4TJ19 A4TJ37 A4TJ45 A4TJ65 A4TJ67 A4TJ68 A4TJA0 A4TJB4 A4TJB9 A4TJC4
A4TJH7 A4TJI0 A4TJI1 A4TJI3 A4TJK0 A4TJK2 A4TJK3 A4TJK5 A4TJK8 A4TJK9 A4TJL0
A4TJL3 A4TJM2 A4TJN7 A4TJN8 A4TJP0 A4TJT7 A4TJU3 A4TJU4 A4TJU5 A4TJU6 A4TK49
A4TK60 A4TKC4 A4TKC7 A4TKC8 A4TKD0 A4TKG8 A4TKI3 A4TKJ3 A4TKK2 A4TKK4 A4TKK7
A4TKK8 A4TKM6 A4TKM7 A4TKM9 A4TKN7 A4TKN8 A4TKV4 A4TKV5 A4TKV8 A4TKW3 A4TKX6
A4TKX8 A4TKY7 A4TKZ1 A4TKZ2 A4TKZ3 A4TKZ6 A4TL15 A4TL28 A4TL30 A4TL31 A4TL32
A4TL34 A4TL54 A4TL56 A4TL66 A4TL70 A4TL77 A4TL87 A4TL88 A4TL89 A4TL91 A4TL92
A4TL94 A4TL95 A4TL96 A4TLA2 A4TLA4 A4TLA5 A4TLC5 A4TLC8 A4TLC9 A4TLE6 A4TLG7
A4TLL3 A4TLL5 A4TLN3 A4TLP0 A4TLQ4 A4TLR5 A4TLS5 A4TLT2 A4TLT6 A4TLT9 A4TLZ2
A4TLZ3 A4TM00 A4TM05 A4TM07 A4TM08 A4TM15 A4TM39 A4TM42 A4TM44 A4TM66 A4TM73
A4TM78 A4TMF1 A4TMF9 A4TMG4 A4TMJ3 A4TMJ5 A4TMK1 A4TMK2 A4TMK3 A4TMK5 A4TMM3
A4TMM9 A4TMN2 A4TMP0 A4TMP2 A4TMP3 A4TMS8 A4TMT0 A4TMT3 A4TMT6 A4TMT7 A4TMU0
A4TMU1 A4TMU7 A4TMV1 A4TMV2 A4TMV5 A4TMV6 A4TMW4 A4TMY1 A4TMZ0 A4TMZ6 A4TN03
A4TN09 A4TN15 A4TN17 A4TN18 A4TN19 A4TN29 A4TN37 A4TN38 A4TN41 A4TN50 A4TN73
A4TN86 A4TNA4 A4TNC5 A4TNC6 A4TNE3 A4TNE5 A4TNE6 A4TNI8 A4TNP0 A4TNP1 A4TNP2
A4TNP7 A4TNQ1 A4TNQ5 A4TNQ6 A4TNQ8 A4TNR8 A4TNS2 A4TNS5 A4TNT8 A4TNU6 A4TNV0
A4TNV2 A4TNX8 A4TNY0 A4TNY6 A4TNY8 A4TNZ7 A4TP00 A4TP02 A4TP07 A4TP08 A4TP09
A4TP60 A4TP63 A4TPA3 A4TPA4 A4TPA5 A4TPA7 A4TPA8 A4TPB0 A4TPB9 A4TPC0 A4TPD3
A4TPD5 A4TPE2 A4TPE3 A4TPE4 A4TPF5 A4TPF8 A4TPG2 A4TPG6 A4TPG7 A4TPG9 A4TPH6
A4TPJ2 A4TPJ3 A4TPJ4 A4TPK0 A4TPK1 A4TPK2 A4TPK3 A4TPK8 A4TPM1 A4TPM8 A4TPM9
A4TPN0 A4TPN1 A4TPN2 A4TPS3 A4TPS4 A4TPS6 A4TPU1 A4TPU3 A4TPU4 A4TPV3 A4TPV4
A4TPV5 A4TPW0 A4TPW6 A4TPW8 A4TPX2 A4TPX3 A4TPY0 A4TPY5 A4TPY6 A4TPZ1 A4TPZ2
A4TPZ7 A4TPZ8 A4TPZ9 A4TQ00 A4TQ01 A4TQ04 A4TQ50 A4TQ51 A4TQ52 A4TQ53 A4TQ62
A4TQ63 A4TQ64 A4TQ65 A4TQ81 A4TQ82 A4TQ91 A4TQ92 A4TQA2 A4TQA4 A4TQA5 A4TQA7
A4TQB3 A4TQD8 A4TQD9 A4TQE0 A4TQE9 A4TQF0 A4TQF3 A4TQF5 A4TQF8 A4TQF9 A4TQG9
A4TQH5 A4TQH6 A4TQJ0 A4TQJ1 A4TQJ9 A4TQK1 A4TQL2 A4TQL9 A4TQM0 A4TQP6 A4TQQ1
A4TQS2 A4TQT0 A4TQU4 A4TQU9 A4TQV0 A4TQV1 A4TQV2 A4TQY1 A4TR20 A4TR26 A4TR39
A4TR46 A4TRA3 A4TRA5 A4TRA8 A4TRC8 A4TRC9 A4TRD9 A4TRE8 A4TRF4 A4TRG1 A4TRG4
A4TRH3 A4TRH4 A4TRH7 A4TRI0 A4TRI1 A4TRI2 A4TRI3 A4TRI5 A4TRI7 A4TRJ0 A4TRJ6
A4TRJ8 A4TRJ9 A4TRK3 A4TRK4 A4TRK7 A4TRL2 A4TRL8 A4TRM3 A4TRM4 A4TRM5 A4TRN4
A4TRN9 A4TRP0 A4TRP1 A4TRP7 A4TRQ1 A4TRQ7 A4TRR0 A4TRR1 A4TRR5 A4TRS7 A4TRS8
A4TRT0 A4TRT1 A4TRT2 A4TRT4 A4TRU9 A4TS06 A4TS14 A4TS20 A4TS22 A4TS25 A4TS28
A4TS29 A4TS30 A4TS31 A4TS32 A4TS33 A4TS36 A4TS44 A4TS63 A4TS76 A4TS79 A4TS83
A4TS84 A4TS86 A4TS94 A4TSA1 A4TSA7 A4TSB9 A4TSC2 A4TSC6 A4TSC8 A4TSD3 A4TSD4
A4TSD5 A4TSD6 A4TSD7 A4TSE0 A4TSE1 A4TSE2 A4TSE5 A4TSE7 A4TSF7 A4TSG5 A4TSG6

A4TSH1 A4TSH7 A4TSH9 A4TSI0 A4TSI1 A4TSI4 A4TSI5 A4TSL0 A4TSL3 A4TSL4 A4UBV5
A4UGR9 A4UHC0 A4UHP8 A4UHQ2 A4UHQ3 A4UHQ7 A4UHT7 A4ULF8 A4URH3 A4UTP7 A4UTQ0
A4UTQ2 A4VCM0 A4VDN2 A4VFG0 A4VFG6 A4VFH7 A4VFH8 A4VFI1 A4VFI3 A4VFI4 A4VFL7
A4VFR3 A4VFW8 A4VFW9 A4VFY3 A4VFY4 A4VFZ6 A4VG81 A4VG93 A4VGC9 A4VGD5 A4VGE1
A4VGE2 A4VGE5 A4VGE9 A4VGK4 A4VGK9 A4VGL4 A4VGL9 A4VGR5 A4VGR8 A4VGS1 A4VGS4
A4VGS6 A4VGS8 A4VGT8 A4VGU2 A4VGW3 A4VGW5 A4VGW8 A4VGX6 A4VGZ1 A4VH02 A4VH14
A4VHG8 A4VHG9 A4VHH5 A4VHH7 A4VHH8 A4VHI1 A4VHK0 A4VHK1 A4VHK9 A4VHL6 A4VHL9
A4VHM0 A4VHM1 A4VHM2 A4VHM3 A4VHM4 A4VHM5 A4VHM6 A4VHM7 A4VHM8 A4VHM9 A4VHN0
A4VHN1 A4VHN2 A4VHN3 A4VHN4 A4VHN5 A4VHN6 A4VHN7 A4VHN8 A4VHN9 A4VHP0 A4VHP1
A4VHP2 A4VHP3 A4VHP4 A4VHP5 A4VHP7 A4VHP8 A4VHP9 A4VHQ1 A4VHQ2 A4VHQ3 A4VHQ4
A4VHQ5 A4VHQ6 A4VHT4 A4VHT5 A4VHV6 A4VI15 A4VI28 A4VI36 A4VI53 A4VI54 A4VI55
A4VI57 A4VI60 A4VI89 A4VI91 A4VIA8 A4VIC4 A4VIE0 A4VIE6 A4VIF7 A4VIF8 A4VIG6
A4VIH0 A4VIH9 A4VII0 A4VIL0 A4VIM1 A4VIR4 A4VIR7 A4VIT1 A4VIT5 A4VIT6 A4VIT7
A4VIT8 A4VIU5 A4VIU7 A4VIX7 A4VIX8 A4VIY1 A4VIY2 A4VJ05 A4VJ19 A4VJ35 A4VJ40
A4VJ70 A4VJ99 A4VJA8 A4VJB3 A4VJB4 A4VJB5 A4VJG1 A4VJG3 A4VJG5 A4VJH6 A4VJN9
A4VJR0 A4VJR9 A4VJS1 A4VJS2 A4VJS3 A4VJS7 A4VJT5 A4VJT9 A4VJV0 A4VJV1 A4VJV2
A4VJV3 A4VK13 A4VK43 A4VKA0 A4VKC2 A4VKC4 A4VKE7 A4VKE8 A4VKF4 A4VKH1 A4VKJ1
A4VKJ5 A4VKK1 A4VKK4 A4VKM0 A4VKN0 A4VKQ1 A4VKZ1 A4VL58 A4VL60 A4VL72 A4VL73
A4VL74 A4VL77 A4VL90 A4VL92 A4VLM9 A4VLN9 A4VLQ5 A4VLR0 A4VLU6 A4VLU7 A4VLV3
A4VLV4 A4VLV5 A4VLW2 A4VLX7 A4VLZ2 A4VLZ8 A4VM16 A4VM19 A4VM20 A4VM22 A4VM39
A4VM82 A4VM84 A4VMC4 A4VMF7 A4VMG3 A4VMP0 A4VMR6 A4VMS4 A4VMT4 A4VMU6 A4VMX4
A4VMY9 A4VN38 A4VN44 A4VN63 A4VN86 A4VN94 A4VNA3 A4VNA4 A4VNA5 A4VNB1 A4VNB4
A4VNE5 A4VNI2 A4VNJ1 A4VNP3 A4VNR4 A4VNT7 A4VNV0 A4VNV6 A4VNW7 A4VNX0 A4VNX4
A4VNX5 A4VNX8 A4VNX9 A4VNY2 A4VNZ7 A4VP05 A4VP32 A4VP33 A4VP82 A4VP83 A4VP90
A4VPB2 A4VPB6 A4VPB9 A4VPC0 A4VPC3 A4VPC5 A4VPI6 A4VPK2 A4VPK9 A4VPM6 A4VPM7
A4VPM8 A4VPM9 A4VPN6 A4VPN7 A4VPN8 A4VPN9 A4VPP0 A4VPP2 A4VPP4 A4VPP5 A4VPQ3
A4VPQ5 A4VPQ6 A4VPR2 A4VQE0 A4VQH4 A4VQH5 A4VQH6 A4VQH7 A4VQH8 A4VQH9 A4VQI5
A4VQM5 A4VQM7 A4VQM9 A4VQN0 A4VQN6 A4VQP0 A4VQP1 A4VQP2 A4VQS8 A4VQU8 A4VQU9
A4VQV0 A4VQV5 A4VQW0 A4VQX9 A4VQY0 A4VQY3 A4VQY5 A4VQZ0 A4VQZ6 A4VQZ7 A4VR04
A4VR07 A4VR15 A4VR38 A4VR47 A4VR87 A4VR88 A4VR98 A4VRA3 A4VRE8 A4VRF3 A4VRF7
A4VRH8 A4VRI1 A4VRJ2 A4VRK4 A4VRK5 A4VRM9 A4VRN4 A4VRT4 A4VRU9 A4VRV9 A4VRW0
A4VRW2 A4VRX0 A4VRY9 A4VRZ0 A4VS72 A4VS73 A4VS81 A4VS84 A4VS85 A4VS92 A4VSA6
A4VSB5 A4VSD3 A4VSE3 A4VSE5 A4VSE9 A4VSF0 A4VSF1 A4VSF4 A4VSF5 A4VSF6 A4VSF7
A4VSF8 A4VSF9 A4VSG0 A4VSG1 A4VSG2 A4VSG3 A4VSG4 A4VSG5 A4VSG6 A4VSG7 A4VSG8
A4VSG9 A4VSH0 A4VSH1 A4VSH4 A4VSH5 A4VSH6 A4VSH7 A4VSH8 A4VSK2 A4VSK3 A4VSL6
A4VSN1 A4VSN2 A4VSN3 A4VSN8 A4VSP7 A4VSS3 A4VSW4 A4VT05 A4VT06 A4VT07 A4VT25
A4VT28 A4VT29 A4VT40 A4VT43 A4VT57 A4VT63 A4VT69 A4VT80 A4VT82 A4VT83 A4VT84
A4VT88 A4VTD0 A4VTD4 A4VTE0 A4VTE3 A4VTE4 A4VTE5 A4VTF1 A4VTF3 A4VTH2 A4VTK3
A4VTL1 A4VTM4 A4VTP6 A4VTQ4 A4VTQ7 A4VTQ8 A4VTS0 A4VTT4 A4VTT5 A4VTX6 A4VTZ1
A4VTZ7 A4VTZ9 A4VU03 A4VU11 A4VU28 A4VU40 A4VU42 A4VU67 A4VU96 A4VUB1 A4VUC8
A4VUD1 A4VUD2 A4VUF4 A4VUF9 A4VUG6 A4VUG7 A4VUH1 A4VUI0 A4VUI1 A4VUJ2 A4VUJ6
A4VUJ7 A4VUK5 A4VUK6 A4VUL1 A4VUM9 A4VUQ8 A4VUS8 A4VV10 A4VV23 A4VV30 A4VV36
A4VV69 A4VVA3 A4VVB2 A4VVB5 A4VVB6 A4VVF5 A4VVH1 A4VVI4 A4VVK7 A4VVR3 A4VVS8
A4VVS9 A4VVT7 A4VVT9 A4VVU0 A4VVU5 A4VVU6 A4VVU8 A4VVY4 A4VVZ0 A4VVZ3 A4VVZ9

A4VW03 A4VW04 A4VW08 A4VW09 A4VW30 A4VW43 A4VW51 A4VW67 A4VW76 A4VW79 A4VWC2
A4VWC5 A4VWC6 A4VWD1 A4VWE5 A4VWG9 A4VWM6 A4VWM9 A4VWT8 A4VWU8 A4VWY2 A4VWZ2
A4VX00 A4VX07 A4VX28 A4VX29 A4VX33 A4VX43 A4VX47 A4VX53 A4VX57 A4VX71 A4VX91
A4VX92 A4VXA1 A4VXD9 A4VXF9 A4VXH2 A4VXH3 A4VXH7 A4VXH8 A4VXL3 A4VXL9 A4VXQ4
A4VXR1 A4VXS1 A4VXS3 A4VXT1 A4VXT8 A4VXU6 A4VXV6 A4VXV7 A4VXW3 A4VXZ3 A4VXZ4
A4VY04 A4VY21 A4VY24 A4VY58 A4VY70 A4VYA3 A4VYB8 A4VYD3 A4VYE0 A4VYE7 A4VYF9
A4VYG8 A4VYI0 A4VYJ7 A4VYK5 A4VYM0 A4VYN2 A4VYN4 A4VYN7 A4VYN8 A4VYN9 A4VYP1
A4VYP4 A4VYP5 A4VYP6 A4VYP7 A4VYP8 A4VYP9 A4VYQ0 A4VYQ1 A4VYQ2 A4VYQ3 A4VYQ4
A4VYQ5 A4VYQ6 A4VYQ7 A4VYQ8 A4VYQ9 A4VYR0 A4VYR4 A4VYR5 A4VYR6 A4VYR7 A4VYR8
A4VYR9 A4VYU1 A4VYU3 A4VYU4 A4VYX4 A4VYX5 A4VYX6 A4VYY1 A4VYZ0 A4VZ14 A4VZ21
A4VZ51 A4VZ90 A4VZ91 A4VZ92 A4VZB1 A4VZB4 A4VZB5 A4VZC6 A4VZC9 A4VZE3 A4VZE7
A4VZF0 A4VZG2 A4VZG4 A4VZG5 A4VZG6 A4VZI7 A4VZK6 A4VZL1 A4VZL7 A4VZM0 A4VZM1
A4VZM2 A4VZM9 A4VZN1 A4VZQ1 A4VZT8 A4VZU7 A4VZV9 A4VZY0 A4VZZ0 A4VZZ3 A4VZZ4
A4W027 A4W028 A4W069 A4W070 A4W083 A4W088 A4W090 A4W094 A4W0A3 A4W0C1 A4W0D2
A4W0D4 A4W0G0 A4W0I7 A4W0K2 A4W0K4 A4W0M2 A4W0M6 A4W0M7 A4W0P5 A4W0Q0 A4W0Q9
A4W0R0 A4W0R4 A4W0S2 A4W0S3 A4W0T4 A4W0T9 A4W0U7 A4W0U8 A4W0V3 A4W0W9 A4W104
A4W125 A4W1B6 A4W1C8 A4W1D5 A4W1D9 A4W1F7 A4W1H6 A4W1K3 A4W1K6 A4W1L5 A4W1L8
A4W1L9 A4W1Q9 A4W1S9 A4W1U2 A4W1W6 A4W221 A4W235 A4W236 A4W244 A4W246 A4W247
A4W251 A4W252 A4W254 A4W292 A4W299 A4W2A2 A4W2A7 A4W2B1 A4W2B2 A4W2B6 A4W2B7
A4W2D1 A4W2F0 A4W2F8 A4W2H3 A4W2I1 A4W2M2 A4W2M7 A4W2N6 A4W2Q0 A4W2X8 A4W2X9
A4W2Z7 A4W341 A4W351 A4W369 A4W384 A4W395 A4W3A3 A4W3A5 A4W3A8 A4W3D2 A4W3D3
A4W3D7 A4W3E8 A4W3F1 A4W3F7 A4W3G1 A4W3H4 A4W3J4 A4W3J5 A4W3N5 A4W3Q0 A4W3Q2
A4W3R6 A4W3R7 A4W3S1 A4W3S2 A4W3V8 A4W3W3 A4W3W8 A4W401 A4W410 A4W418 A4W420
A4W428 A4W435 A4W440 A4W443 A4W452 A4W453 A4W460 A4W483 A4W487 A4W488 A4W496
A4W4B4 A4W4B5 A4W4F3 A4W4G5 A4W4J7 A4W4J9 A4W4L2 A4W4M5 A4W4N0 A4W4N7 A4W4P9
A4W4R3 A4W4R9 A4W4S4 A4W500 A4W502 A4W506 A4W507 A4W508 A4W509 A4W511 A4W512
A4W513 A4W514 A4W515 A4W527 A4W529 A4W533 A4W536 A4W566 A4W572 A4W573 A4W599
A4W5A0 A4W5A3 A4W5A4 A4W5A5 A4W5A6 A4W5A7 A4W5A8 A4W5B3 A4W5B6 A4W5B8 A4W5B9
A4W5C0 A4W5C1 A4W5C7 A4W5D0 A4W5E0 A4W5F2 A4W5N7 A4W5N8 A4W5P1 A4W5Q2 A4W5Q6
A4W5R1 A4W5R2 A4W5R3 A4W5R8 A4W5R9 A4W5S9 A4W5T0 A4W5T1 A4W5T2 A4W5T7 A4W5V5
A4W5X6 A4W5X9 A4W604 A4W605 A4W687 A4W691 A4W698 A4W699 A4W6A0 A4W6A1 A4W6A2
A4W6B2 A4W6B3 A4W6C5 A4W6D5 A4W6D6 A4W6D9 A4W6E4 A4W6E7 A4W6F5 A4W6F6 A4W6F7
A4W6F8 A4W6G3 A4W6G6 A4W6G7 A4W6H6 A4W6H7 A4W6H9 A4W6I4 A4W6I5 A4W6I8 A4W6J4
A4W6K3 A4W6K4 A4W6K6 A4W6M4 A4W6M5 A4W6M6 A4W6N6 A4W6N7 A4W6N8 A4W6P3 A4W6Q1
A4W6Q3 A4W6Q6 A4W6Q7 A4W6R0 A4W6R1 A4W6R4 A4W6R5 A4W6R6 A4W6R7 A4W6S8 A4W6T3
A4W6T5 A4W6T9 A4W6U9 A4W6V2 A4W6V4 A4W6W5 A4W6W8 A4W6X1 A4W6X2 A4W6X4 A4W6X5
A4W757 A4W762 A4W763 A4W766 A4W767 A4W778 A4W779 A4W785 A4W787 A4W788 A4W791
A4W793 A4W797 A4W7A7 A4W7A9 A4W7B5 A4W7B7 A4W7D2 A4W7D6 A4W7D9 A4W7F3 A4W7F5
A4W7F6 A4W7F7 A4W7F8 A4W7F9 A4W7I4 A4W7J0 A4W7T6 A4W7Z0 A4W7Z1 A4W7Z2 A4W7Z3
A4W7Z4 A4W814 A4W816 A4W817 A4W822 A4W827 A4W829 A4W836 A4W838 A4W844 A4W846
A4W853 A4W870 A4W871 A4W879 A4W892 A4W897 A4W899 A4W8A9 A4W8B2 A4W8B4 A4W8B5
A4W8B7 A4W8B8 A4W8C2 A4W8C6 A4W8E5 A4W8F0 A4W8H0 A4W8L7 A4W8M4 A4W8N3 A4W8P1
A4W8P9 A4W8Q8 A4W8Q9 A4W8R7 A4W8S7 A4W8S8 A4W8S9 A4W8T1 A4W8T6 A4W8U3 A4W8U9
A4W8V0 A4W8V4 A4W8W0 A4W8W8 A4W8X0 A4W8X5 A4W8X9 A4W8Y0 A4W916 A4W920 A4W921

A4W922 A4W923 A4W924 A4W925 A4W949 A4W960 A4W968 A4W975 A4W980 A4W9A3 A4W9B2
A4W9C2 A4W9E5 A4W9G9 A4W9I2 A4W9J6 A4W9J7 A4W9J8 A4W9K3 A4W9L0 A4W9M3 A4W9M5
A4W9M6 A4W9M7 A4W9M9 A4W9N5 A4W9P3 A4W9R3 A4W9R4 A4W9U3 A4W9V7 A4W9V9 A4W9W0
A4W9X4 A4W9Y2 A4W9Y8 A4W9Z4 A4WA78 A4WAF3 A4WAF9 A4WAG4 A4WAJ3 A4WAM9 A4WAR9
A4WAT7 A4WAX5 A4WAX8 A4WAY2 A4WAY6 A4WB02 A4WB03 A4WB85 A4WB99 A4WBC6 A4WBC7
A4WBC9 A4WBD3 A4WBF2 A4WBF7 A4WBF9 A4WBG0 A4WBG2 A4WBG9 A4WBH1 A4WBJ0 A4WBJ4
A4WBL8 A4WBL9 A4WBM0 A4WBM1 A4WBM3 A4WBM7 A4WBM8 A4WBM9 A4WBN1 A4WBQ9 A4WBR1
A4WBW7 A4WBY2 A4WC00 A4WC14 A4WC16 A4WC68 A4WC70 A4WC73 A4WC74 A4WCB5 A4WCB6
A4WCE6 A4WCF6 A4WCI1 A4WCI8 A4WCJ5 A4WCK1 A4WCK2 A4WCL6 A4WCL7 A4WCL8 A4WCM3
A4WCN5 A4WCP6 A4WCP8 A4WCP9 A4WCQ3 A4WCS0 A4WCS3 A4WCV2 A4WCV4 A4WCV7 A4WCW2
A4WCW4 A4WD05 A4WD15 A4WD23 A4WD41 A4WD46 A4WD51 A4WD56 A4WD70 A4WD72 A4WD82
A4WD84 A4WD89 A4WD92 A4WD93 A4WD95 A4WD96 A4WDA4 A4WDA7 A4WDA8 A4WDA9 A4WDB1
A4WDB2 A4WDC0 A4WDD1 A4WDD6 A4WDD8 A4WDE6 A4WDE8 A4WDE9 A4WDH1 A4WDH2 A4WDH3
A4WDH4 A4WDH8 A4WDH9 A4WDI4 A4WDK7 A4WDN8 A4WDQ2 A4WDQ5 A4WDQ6 A4WDQ7 A4WDQ8
A4WDR6 A4WDR7 A4WDU5 A4WDU8 A4WDU9 A4WDV0 A4WDV1 A4WDV2 A4WDV5 A4WDV7 A4WDV9
A4WDW0 A4WDW1 A4WDW4 A4WDW8 A4WDX1 A4WDY6 A4WDZ5 A4WE03 A4WE06 A4WE07 A4WE30
A4WE41 A4WE42 A4WE49 A4WE55 A4WE56 A4WE57 A4WE66 A4WE71 A4WE72 A4WE75 A4WE78
A4WE80 A4WE85 A4WE91 A4WE99 A4WEA6 A4WEI2 A4WEI4 A4WEI8 A4WEJ2 A4WEJ3 A4WEJ4
A4WEJ5 A4WEJ6 A4WEJ7 A4WEJ8 A4WEJ9 A4WEK0 A4WEK3 A4WEQ6 A4WEQ9 A4WER0 A4WER5
A4WER6 A4WER8 A4WEU0 A4WEU6 A4WEV0 A4WEV6 A4WEW3 A4WEW4 A4WEW9 A4WEX0 A4WEX9
A4WEY0 A4WEY1 A4WEY2 A4WEY3 A4WEY5 A4WEY6 A4WEY7 A4WEZ0 A4WEZ6 A4WEZ8 A4WEZ9
A4WF02 A4WF18 A4WF25 A4WF38 A4WF39 A4WF42 A4WF48 A4WF49 A4WF69 A4WF74 A4WF77
A4WF90 A4WF91 A4WF93 A4WF95 A4WF96 A4WF97 A4WFA2 A4WFA3 A4WFA4 A4WFA5 A4WFA7
A4WFA9 A4WFB0 A4WFB1 A4WFB2 A4WFB3 A4WFB4 A4WFB5 A4WFB6 A4WFB7 A4WFB8 A4WFB9
A4WFC0 A4WFC1 A4WFC2 A4WFC3 A4WFC4 A4WFC5 A4WFC7 A4WFC8 A4WFC9 A4WFD4 A4WFD5
A4WFD6 A4WFD7 A4WFD8 A4WFE1 A4WFF7 A4WFH1 A4WFH7 A4WFH8 A4WFI9 A4WFJ1 A4WFJ8
A4WFK2 A4WFK6 A4WFK9 A4WFL2 A4WFL3 A4WFL4 A4WFL5 A4WFQ2 A4WFT5 A4WFX3 A4WFX9
A4WFY5 A4WFZ2 A4WG08 A4WG10 A4WG12 A4WG30 A4WG31 A4WG34 A4WG37 A4WG46 A4WG49
A4WG51 A4WG52 A4WG54 A4WG55 A4WG57 A4WG61 A4WG62 A4WG66 A4WG67 A4WG69 A4WG72
A4WG77 A4WG81 A4WG90 A4WG91 A4WG92 A4WG93 A4WG94 A4WGA0 A4WGB3 A4WGB6 A4WGC4
A4WGC5 A4WGD9 A4WGE2 A4WGE3 A4WGE6 A4WGE7 A4WGH1 A4WGH5 A4WGV2 A4WGV3 A4WGW1
A4WGX8 A4WGY5 A4WGZ8 A4WH01 A4WH02 A4WH03 A4WH05 A4WH11 A4WH12 A4WH15 A4WH24
A4WH36 A4WH37 A4WH44 A4WH96 A4WHB4 A4WHB8 A4WHH5 A4WHH6 A4WHI1 A4WHK6 A4WHN0
A4WHP6 A4WHP7 A4WHQ7 A4WHQ8 A4WHQ9 A4WHY5 A4WHY9 A4WI40 A4WIJ1 A4WIJ6 A4WIK0
A4WIK2 A4WIK5 A4WIN2 A4WIW2 A4WIW5 A4WIW6 A4WIY8 A4WIY9 A4WIZ3 A4WJ12 A4WJ22
A4WJ29 A4WJ34 A4WJ90 A4WJE1 A4WJG1 A4WJI0 A4WJI4 A4WJK0 A4WJK1 A4WJK5 A4WJL8
A4WJM7 A4WJP4 A4WJR7 A4WJU1 A4WK37 A4WK38 A4WKG3 A4WKI4 A4WKJ7 A4WKK8 A4WKL0
A4WKL2 A4WKM4 A4WKM8 A4WKQ6 A4WKQ9 A4WKR0 A4WKS9 A4WKT1 A4WKX3 A4WKY7 A4WL16
A4WL27 A4WL31 A4WL72 A4WL88 A4WL93 A4WL94 A4WL95 A4WL96 A4WL99 A4WLE0 A4WLE9
A4WLG6 A4WLG8 A4WLH7 A4WLH8 A4WLJ5 A4WLJ8 A4WLK1 A4WLK4 A4WLK6 A4WLL3 A4WLL4
A4WLM9 A4WLN2 A4WLN5 A4WLP0 A4WLP1 A4WLP2 A4WLQ5 A4WLR8 A4WLS1 A4WLS5 A4WLS6
A4WLV3 A4WLV9 A4WLY0 A4WLY6 A4WM07 A4WM20 A4WM27 A4WM28 A4WM67 A4WM72 A4WMA2
A4WMA6 A4WMB5 A4WMB7 A4WMD4 A4WMD6 A4WMD9 A4WMH5 A4WMI3 A4WMI6 A4WMK4 A4WMP8
A4WMQ7 A4WMR8 A4WMV7 A4WMX1 A4WMZ0 A4WN05 A4WN07 A4WN19 A4WN24 A4WN33 A4WN38

A4WN42 A4WN46 A4WN67 A4WN87 A4WNA3 A4WNA4 A4WNA8 A4WNB3 A4WNC1 A4WNC4 A4WNC5
A4WNE9 A4WNG4 A4WNI9 A4WNJ4 A4WNJ6 A4WNL5 A4WNL8 A4WNM6 A4WNM7 A4WNM9 A4WNS0
A4WNS1 A4WNU8 A4WNV1 A4WNV2 A4WNV5 A4WNV6 A4WNX7 A4WNX8 A4WNY1 A4WNY2 A4WP19
A4WP35 A4WPB6 A4WPB7 A4WPE1 A4WPE3 A4WPE4 A4WPE5 A4WPF7 A4WPH4 A4WPH6 A4WPI0
A4WPJ3 A4WPR4 A4WPT1 A4WPU4 A4WPW1 A4WPY6 A4WPZ2 A4WQ11 A4WQ13 A4WQ19 A4WQ25
A4WQ33 A4WQ36 A4WQ41 A4WQ67 A4WQ76 A4WQ81 A4WQ91 A4WQ93 A4WQA7 A4WQB2 A4WQC4
A4WQC5 A4WQD2 A4WQD6 A4WQD8 A4WQD9 A4WQT3 A4WQT6 A4WQU1 A4WQW2 A4WQW4 A4WQX2
A4WQY3 A4WQY6 A4WQY7 A4WQZ1 A4WR15 A4WR71 A4WR72 A4WR73 A4WR75 A4WR78 A4WR79
A4WR94 A4WR96 A4WR97 A4WRA6 A4WRC8 A4WRD4 A4WRE3 A4WRF5 A4WRI4 A4WRK0 A4WRK3
A4WRK9 A4WRL4 A4WRM1 A4WRM2 A4WRP6 A4WRQ0 A4WRQ2 A4WRT4 A4WRT7 A4WRW1 A4WRW7
A4WRW8 A4WRX0 A4WRZ6 A4WS82 A4WS83 A4WS84 A4WS85 A4WSC5 A4WSH8 A4WSH9 A4WSI3
A4WSL4 A4WSL5 A4WSM3 A4WSP0 A4WSP1 A4WSQ8 A4WTF5 A4WTH0 A4WTM7 A4WTP4 A4WTQ6
A4WTS8 A4WTT2 A4WTT7 A4WTU0 A4WU04 A4WU05 A4WU16 A4WU98 A4WUH5 A4WUH8 A4WUH9
A4WUI0 A4WUI6 A4WUJ2 A4WUK1 A4WUK2 A4WUL5 A4WUL6 A4WUN2 A4WUN9 A4WUP0 A4WUQ2
A4WUQ3 A4WUQ5 A4WUS5 A4WUS7 A4WUS8 A4WUS9 A4WUX2 A4WUY6 A4WV24 A4WV36 A4WV47
A4WV48 A4WV81 A4WVB1 A4WVB9 A4WVE5 A4WVE8 A4WVF5 A4WVG1 A4WVG2 A4WVH0 A4WVI3
A4WVI4 A4WVI5 A4WVI6 A4WVI9 A4WVJ0 A4WVJ1 A4WVJ2 A4WVJ3 A4WVJ4 A4WVJ5 A4WVJ6
A4WVJ7 A4WVJ8 A4WVJ9 A4WVK0 A4WVK1 A4WVK2 A4WVK3 A4WVK4 A4WVK5 A4WVK6 A4WVK7
A4WVK9 A4WVL0 A4WVL1 A4WVL2 A4WVL3 A4WVL5 A4WVL6 A4WVL7 A4WVL8 A4WVL9 A4WVM0
A4WVN6 A4WVP0 A4WVP2 A4WVP5 A4WVQ1 A4WVQ4 A4WVR6 A4WVR7 A4WVT8 A4WVX3 A4WVX8
A4WVY1 A4WVY8 A4WVY9 A4WVZ0 A4WVZ4 A4WVZ7 A4WW31 A4WW35 A4WW38 A4WW77 A4WW80
A4WW88 A4WW89 A4WW91 A4WWC0 A4WWC9 A4WWI7 A4WWJ3 A4WWL8 A4WWN6 A4WWP0 A4WWP1
A4WWP4 A4WWP6 A4WWP7 A4WWQ0 A4WWS0 A4WWS1 A4WWV3 A4WWV6 A4WX36 A4WX55 A4WX62
A4WX66 A4WX67 A4WX68 A4WX84 A4WX92 A4WXA5 A4WXK6 A4WXS6 A4WZ45 A4WZB3 A4WZQ4
A4WZV1 A4X0U0 A4X0V0 A4X0Z1 A4X148 A4X1K4 A4X1M6 A4X1P5 A4X1R6 A4X1U1 A4X1X0
A4X307 A4X310 A4X378 A4X388 A4X398 A4X3E5 A4X3K6 A4X3K9 A4X3P0 A4X3Q0 A4X3X5
A4X3X8 A4X484 A4X487 A4X489 A4X490 A4X491 A4X4A5 A4X4C8 A4X4C9 A4X4F2 A4X4F8
A4X4H5 A4X4H7 A4X4H8 A4X4H9 A4X4I2 A4X4J0 A4X4J2 A4X4J3 A4X4J5 A4X4J7 A4X4L9
A4X4N4 A4X4N7 A4X4N9 A4X4P7 A4X4P8 A4X4R3 A4X4S2 A4X4U1 A4X4V8 A4X4W1 A4X4W8
A4X4Y7 A4X548 A4X585 A4X5V3 A4X5V4 A4X5V5 A4X5V6 A4X5X2 A4X5X3 A4X5X7 A4X5Y3
A4X5Z0 A4X607 A4X608 A4X610 A4X611 A4X612 A4X615 A4X621 A4X624 A4X626 A4X631
A4X639 A4X642 A4X645 A4X646 A4X650 A4X652 A4X692 A4X6K7 A4X6P4 A4X6W9 A4X741
A4X744 A4X745 A4X747 A4X776 A4X7S0 A4X7W7 A4X7Y6 A4X8U4 A4X8U5 A4X8U8 A4X915
A4X981 A4X982 A4X9H1 A4X9N0 A4X9P7 A4X9Q0 A4X9Q4 A4X9R5 A4X9S3 A4X9S4 A4X9S8
A4X9Z2 A4XA20 A4XA62 A4XAD4 A4XAE7 A4XAF6 A4XAG1 A4XAG2 A4XAG3 A4XAH9 A4XAK7
A4XAK8 A4XAL2 A4XAM8 A4XAX7 A4XB58 A4XB85 A4XB95 A4XBC7 A4XBE5 A4XBH2 A4XBI0
A4XBI5 A4XBI6 A4XBI7 A4XBL2 A4XBL7 A4XBM0 A4XBM1 A4XBM2 A4XBM5 A4XBM7 A4XBM8
A4XBM9 A4XBN1 A4XBN2 A4XBN3 A4XBN4 A4XBN5 A4XBN6 A4XBN7 A4XBN8 A4XBN9 A4XBP0
A4XBP1 A4XBP2 A4XBP3 A4XBP6 A4XBP7 A4XBP8 A4XBP9 A4XBQ0 A4XBQ1 A4XBQ5 A4XBQ9
A4XBR0 A4XBR5 A4XC73 A4XCN1 A4XCQ3 A4XCQ4 A4XCT1 A4XD40 A4XDG5 A4XDG6 A4XDG8
A4XDK0 A4XDK4 A4XDK5 A4XDR3 A4XDR5 A4XDR7 A4XDT6 A4XDT7 A4XES9 A4XF23 A4XF35
A4XF36 A4XFR8 A4XFZ6 A4XG24 A4XG37 A4XG66 A4XGB6 A4XGB8 A4XGE5 A4XGE7 A4XH45
A4XHB0 A4XHE0 A4XHE2 A4XHH4 A4XHK5 A4XHL5 A4XHM5 A4XHN7 A4XHR9 A4XHT8 A4XHU4
A4XHV1 A4XHV2 A4XHV9 A4XHW0 A4XHW1 A4XHX9 A4XHY7 A4XHY9 A4XHZ5 A4XHZ6 A4XI02

A4XI30 A4XI31 A4XI35 A4XI36 A4XI37 A4XI43 A4XI64 A4XI71 A4XI74 A4XI79 A4XI81
A4XIA5 A4XIB6 A4XID4 A4XIH3 A4XIL7 A4XIM9 A4XIN0 A4XIN2 A4XIQ3 A4XIQ4 A4XIQ6
A4XIS3 A4XIT8 A4XIU2 A4XIW8 A4XIW9 A4XIX0 A4XJ08 A4XJ09 A4XJ18 A4XJ25 A4XJ31
A4XJ48 A4XJ49 A4XJ54 A4XJ59 A4XJ60 A4XJ89 A4XJH1 A4XJI0 A4XJI3 A4XJK7 A4XJL4
A4XJM8 A4XJN1 A4XJN7 A4XJN8 A4XJN9 A4XJS1 A4XJS2 A4XJS3 A4XJS5 A4XJS7 A4XJS8
A4XJU0 A4XJU4 A4XJV0 A4XJV1 A4XJV6 A4XK04 A4XK06 A4XK09 A4XK25 A4XK60 A4XK62
A4XK85 A4XKA4 A4XKA5 A4XKA6 A4XKB7 A4XKC0 A4XKC1 A4XKC3 A4XKC6 A4XKC7 A4XKE2
A4XKE6 A4XKG2 A4XKG3 A4XKG4 A4XKG5 A4XKG9 A4XKH4 A4XKI3 A4XKI9 A4XKJ4 A4XKJ8
A4XKL5 A4XKM4 A4XKM9 A4XKN9 A4XKP4 A4XKP6 A4XKP7 A4XKQ3 A4XKQ8 A4XKR2 A4XKS3
A4XKS9 A4XKT1 A4XKT2 A4XKT3 A4XKT6 A4XKT8 A4XKU9 A4XKV7 A4XKX8 A4XKY5 A4XKZ2
A4XKZ4 A4XL21 A4XL23 A4XL26 A4XL36 A4XL40 A4XL44 A4XL45 A4XL47 A4XL48 A4XL57
A4XL61 A4XL64 A4XL65 A4XL67 A4XL69 A4XL70 A4XL73 A4XL78 A4XL81 A4XL87 A4XLC1
A4XLD7 A4XLD9 A4XLE0 A4XLE5 A4XLE8 A4XLF0 A4XLF1 A4XLF2 A4XLF4 A4XLI3 A4XLJ3
A4XLK1 A4XLK2 A4XLK5 A4XLL8 A4XLM2 A4XLM3 A4XLM7 A4XLN1 A4XLN3 A4XLP5 A4XLQ1
A4XLQ2 A4XLQ3 A4XLQ4 A4XLQ5 A4XLQ6 A4XLQ7 A4XLR0 A4XLR2 A4XLR3 A4XLR4 A4XLR5
A4XLR6 A4XLR7 A4XLR9 A4XLS0 A4XLS1 A4XLS2 A4XLS3 A4XLS4 A4XLS5 A4XLS6 A4XLS7
A4XLS8 A4XLS9 A4XLT0 A4XLT1 A4XLW0 A4XLZ4 A4XLZ9 A4XM00 A4XM01 A4XM02 A4XM18
A4XMA9 A4XML6 A4XMY1 A4XMY4 A4XMZ2 A4XMZ3 A4XMZ4 A4XN49 A4XN50 A4XN51 A4XN56
A4XN62 A4XN72 A4XN73 A4XNB2 A4XNB3 A4XNB6 A4XNB8 A4XNB9 A4XNC8 A4XNJ7 A4XNL3
A4XNV3 A4XNV9 A4XNX4 A4XNX7 A4XNY5 A4XP02 A4XP32 A4XP44 A4XP61 A4XP63 A4XP70
A4XP74 A4XP75 A4XPA4 A4XPF3 A4XPF7 A4XPF8 A4XPG4 A4XPI5 A4XPI6 A4XPI7 A4XPL2
A4XPL8 A4XPM3 A4XPM7 A4XPN6 A4XPN7 A4XPN9 A4XPP1 A4XPQ2 A4XPQ6 A4XPR2 A4XPS6
A4XPU9 A4XPW1 A4XPX8 A4XPY0 A4XPY6 A4XPY7 A4XPZ2 A4XPZ3 A4XPZ7 A4XPZ8 A4XQ00
A4XQ16 A4XQ17 A4XQ18 A4XQ41 A4XQ45 A4XQ47 A4XQ48 A4XQ60 A4XQ64 A4XQ67 A4XQA3
A4XQE4 A4XQF5 A4XQI3 A4XQK5 A4XQK6 A4XQK7 A4XQL6 A4XQL9 A4XQN4 A4XQN5 A4XQP2
A4XQQ3 A4XQQ4 A4XQR2 A4XQR5 A4XQR6 A4XQR9 A4XQS5 A4XQS6 A4XQU1 A4XQU8 A4XQV2
A4XQV5 A4XQV8 A4XR11 A4XR34 A4XR36 A4XR54 A4XR55 A4XR57 A4XR60 A4XR61 A4XR64
A4XR78 A4XRA9 A4XRJ0 A4XRL4 A4XRN0 A4XRN2 A4XRN4 A4XRQ7 A4XRR1 A4XRR7 A4XRR9
A4XRS1 A4XRT0 A4XRT1 A4XS41 A4XS43 A4XS54 A4XS57 A4XS59 A4XS60 A4XS85 A4XSC4
A4XSC7 A4XSD0 A4XSD1 A4XSQ1 A4XSR3 A4XSS3 A4XST1 A4XSW1 A4XSY1 A4XSZ1 A4XT11
A4XT24 A4XT41 A4XT53 A4XT55 A4XT72 A4XT74 A4XT90 A4XTE5 A4XTE7 A4XTF3 A4XTH9
A4XTK0 A4XTK4 A4XTL9 A4XTM7 A4XTN0 A4XTR0 A4XTR1 A4XTR2 A4XTR4 A4XTS1 A4XTS3
A4XTS4 A4XTS5 A4XTS7 A4XTZ3 A4XTZ4 A4XTZ5 A4XTZ6 A4XU09 A4XU11 A4XU29 A4XU32
A4XU74 A4XU99 A4XUA5 A4XUP3 A4XUW4 A4XUX3 A4XUX4 A4XUY0 A4XUY1 A4XUY2 A4XUY4
A4XUY9 A4XV44 A4XV81 A4XV90 A4XVA3 A4XVB1 A4XVB5 A4XVG1 A4XVG2 A4XVK6 A4XVL7
A4XVN4 A4XVN5 A4XVW1 A4XVW2 A4XVX2 A4XVY5 A4XW05 A4XWE0 A4XWE2 A4XWE4 A4XWE6
A4XWE7 A4XWI3 A4XWK1 A4XWN5 A4XWP8 A4XWQ3 A4XWR1 A4XWR2 A4XWR3 A4XWR4 A4XWR9
A4XWS3 A4XWS7 A4XWT5 A4XWT8 A4XWT9 A4XWU0 A4XWU1 A4XWU3 A4XWV4 A4XXB7 A4XXF3
A4XXN1 A4XXN9 A4XXP5 A4XXP6 A4XXQ6 A4XXT4 A4XXT6 A4XXT7 A4XXZ2 A4XY05 A4XY17
A4XY24 A4XY28 A4XY31 A4XY32 A4XY35 A4XY36 A4XY39 A4XY40 A4XY43 A4XY53 A4XY67
A4XY81 A4XYB5 A4XYB8 A4XYC4 A4XYC5 A4XYC6 A4XYD6 A4XYD7 A4XYD8 A4XYD9 A4XYE0
A4XYE2 A4XYE4 A4XYE5 A4XYE8 A4XYF4 A4XYF5 A4XYF6 A4XYF7 A4XYG1 A4XYG4 A4XYH6
A4XYJ7 A4XYL1 A4XYL2 A4XYL3 A4XYL4 A4XYL9 A4XYM0 A4XYN8 A4XYV5 A4XYV6 A4XYW1
A4XYW3 A4XYW8 A4XYX3 A4XYX4 A4XYX5 A4XYY1 A4XYY3 A4XYY4 A4XZ25 A4XZ35 A4XZ39

A4XZ62 A4XZ64 A4XZ65 A4XZ66 A4XZ67 A4XZ68 A4XZ69 A4XZ71 A4XZ72 A4XZ73 A4XZ74
A4XZ75 A4XZ76 A4XZ77 A4XZ78 A4XZ79 A4XZ80 A4XZ81 A4XZ82 A4XZ83 A4XZ84 A4XZ85
A4XZ86 A4XZ87 A4XZ88 A4XZ89 A4XZ90 A4XZ91 A4XZ92 A4XZ93 A4XZ94 A4XZ95 A4XZ96
A4XZ97 A4XZ98 A4XZ99 A4XZA0 A4XZA1 A4XZA6 A4XZB1 A4XZB4 A4XZC5 A4XZC6 A4XZJ3
A4XZJ5 A4XZJ6 A4XZJ7 A4XZJ9 A4XZK2 A4XZK6 A4XZK7 A4XZQ4 A4XZR8 A4XZR9 A4XZX0
A4XZY4 A4XZY6 A4XZZ7 A4XZZ8 A4Y022 A4Y028 A4Y036 A4Y044 A4Y049 A4Y074 A4Y077
A4Y083 A4Y084 A4Y087 A4Y091 A4Y0F3 A4Y0K5 A4Y0K7 A4Y0K9 A4Y0L1 A4Y0L7 A4Y0M0
A4Y119 A4Y197 A4Y198 A4Y199 A4Y1A2 A4Y1A3 A4Y1A4 A4Y1A6 A4Y1A9 A4Y1B0 A4Y1B4
A4Y1B7 A4Y1C6 A4Y1D0 A4Y1D2 A4Y1D3 A4Y1D5 A4Y1E3 A4Y1E5 A4Y1H2 A4Y1I0 A4Y1I2
A4Y1K3 A4Y1N1 A4Y1N7 A4Y1V6 A4Y259 A4Y2C8 A4Y2D9 A4Y2G4 A4Y2G8 A4Y2H2 A4Y2K5
A4Y2K7 A4Y2K8 A4Y2K9 A4Y2L1 A4Y2L2 A4Y2L3 A4Y2M0 A4Y2M2 A4Y2M3 A4Y2M5 A4Y2M7
A4Y2M8 A4Y2N7 A4Y2Q5 A4Y2S7 A4Y2S8 A4Y2U9 A4Y2Z8 A4Y307 A4Y344 A4Y373 A4Y379
A4Y384 A4Y397 A4Y398 A4Y3A9 A4Y3C4 A4Y3E1 A4Y3E2 A4Y3F0 A4Y3F1 A4Y3F2 A4Y3F3
A4Y3F6 A4Y3I1 A4Y3J5 A4Y3J7 A4Y3J8 A4Y3R7 A4Y3R9 A4Y3S0 A4Y3T4 A4Y3Y0 A4Y425
A4Y426 A4Y427 A4Y433 A4Y434 A4Y435 A4Y447 A4Y493 A4Y494 A4Y498 A4Y4A1 A4Y4A4
A4Y4F2 A4Y4F3 A4Y4G4 A4Y4G8 A4Y4H1 A4Y4H9 A4Y4I6 A4Y4I7 A4Y4J1 A4Y4K4 A4Y4L3
A4Y4L4 A4Y4L5 A4Y4L6 A4Y4L9 A4Y4Q4 A4Y4S4 A4Y4U3 A4Y4U6 A4Y4W8 A4Y4W9 A4Y4X0
A4Y4X5 A4Y4Y0 A4Y4Y4 A4Y4Y9 A4Y540 A4Y543 A4Y544 A4Y545 A4Y546 A4Y557 A4Y564
A4Y581 A4Y589 A4Y5H9 A4Y5I0 A4Y5I1 A4Y5I2 A4Y5I3 A4Y5N5 A4Y5Q1 A4Y5R0 A4Y5S0
A4Y5T8 A4Y5T9 A4Y5U2 A4Y5X4 A4Y5Y2 A4Y5Y6 A4Y5Y9 A4Y5Z1 A4Y644 A4Y6A5 A4Y6A7
A4Y6A8 A4Y6B8 A4Y6C1 A4Y6E2 A4Y6E4 A4Y6G8 A4Y6H0 A4Y6J4 A4Y6L5 A4Y6M4 A4Y6N5
A4Y6R2 A4Y6R6 A4Y6S2 A4Y6S3 A4Y6S5 A4Y6S7 A4Y6S8 A4Y6S9 A4Y6T9 A4Y6Z5 A4Y703
A4Y704 A4Y706 A4Y729 A4Y731 A4Y732 A4Y757 A4Y759 A4Y7B1 A4Y7B8 A4Y7C9 A4Y7D1
A4Y7D3 A4Y7D4 A4Y7D7 A4Y7H1 A4Y7H2 A4Y7J8 A4Y7K5 A4Y7K6 A4Y7K8 A4Y7L2 A4Y7L3
A4Y7L4 A4Y7L6 A4Y7M1 A4Y7N2 A4Y7N9 A4Y7Q5 A4Y7R6 A4Y7S6 A4Y7T2 A4Y7T5 A4Y7T7
A4Y7T8 A4Y7T9 A4Y7U1 A4Y7X3 A4Y7Y3 A4Y7Y7 A4Y802 A4Y819 A4Y843 A4Y845 A4Y879
A4Y882 A4Y884 A4Y889 A4Y891 A4Y8C2 A4Y8C3 A4Y8D7 A4Y8F6 A4Y8T3 A4Y8T6 A4Y8T9
A4Y8U0 A4Y8U3 A4Y8W7 A4Y8X6 A4Y8Y3 A4Y8Y5 A4Y8Y7 A4Y929 A4Y931 A4Y933 A4Y934
A4Y937 A4Y938 A4Y940 A4Y944 A4Y946 A4Y952 A4Y960 A4Y961 A4Y965 A4Y966 A4Y9A6
A4Y9A7 A4Y9A8 A4Y9B3 A4Y9B5 A4Y9B7 A4Y9B8 A4Y9B9 A4Y9C0 A4Y9C2 A4Y9C4 A4Y9C5
A4Y9C8 A4Y9D9 A4Y9E1 A4Y9E3 A4Y9E9 A4Y9F2 A4Y9F4 A4Y9G0 A4Y9G1 A4Y9G2 A4Y9L3
A4Y9L4 A4Y9M3 A4Y9P3 A4Y9Q2 A4Y9Q3 A4Y9S5 A4Y9U2 A4Y9X0 A4Y9X3 A4Y9X9 A4Y9Y4
A4Y9Z3 A4Y9Z9 A4YA56 A4YA61 A4YA62 A4YA93 A4YAD8 A4YAD9 A4YAE0 A4YAE8 A4YAE9
A4YAK9 A4YAL0 A4YAL6 A4YAM5 A4YAN1 A4YAP3 A4YAQ6 A4YAS7 A4YAY4 A4YB05 A4YB09
A4YB19 A4YB21 A4YB86 A4YBB9 A4YBC0 A4YBE6 A4YBE7 A4YBF5 A4YBF7 A4YBG0 A4YBG2
A4YBI7 A4YBK6 A4YBM9 A4YBN4 A4YBR8 A4YBR9 A4YBT0 A4YBT1 A4YBT8 A4YBT9 A4YBU1
A4YBU5 A4YBV7 A4YBV8 A4YBV9 A4YBW0 A4YBW1 A4YBW2 A4YBW4 A4YBW5 A4YBW6 A4YBW7
A4YBW8 A4YBW9 A4YBX0 A4YBX1 A4YBX2 A4YBX3 A4YBX4 A4YBX5 A4YBX6 A4YBX7 A4YBX8
A4YBX9 A4YBY0 A4YBY2 A4YBY3 A4YBY4 A4YBY5 A4YBY7 A4YBY8 A4YBY9 A4YBZ0 A4YBZ1
A4YBZ2 A4YBZ3 A4YBZ4 A4YBZ8 A4YC18 A4YCA5 A4YCB1 A4YCC5 A4YCD0 A4YCD7 A4YCI8
A4YCI9 A4YCM1 A4YCM4 A4YCM5 A4YCN4 A4YCP2 A4YCQ1 A4YCQ2 A4YCQ3 A4YCQ4 A4YCQ5
A4YCQ7 A4YCR0 A4YCR6 A4YCR7 A4YCS3 A4YCT0 A4YCT6 A4YCU5 A4YCU9 A4YCV9 A4YCW6
A4YCW9 A4YCX0 A4YCX1 A4YCX2 A4YCX6 A4YCX7 A4YCX8 A4YCY1 A4YCY5 A4YCY6 A4YCY7
A4YD25 A4YD26 A4YD40 A4YD47 A4YD78 A4YD82 A4YD84 A4YD89 A4YD91 A4YD92 A4YE96

A4YEA2 A4YEA7 A4YEC1 A4YEC3 A4YED8 A4YEH8 A4YEI1 A4YEM3 A4YEN2 A4YEN4 A4YEP4
A4YEQ8 A4YER6 A4YES9 A4YEU7 A4YF02 A4YF03 A4YF54 A4YF67 A4YGA7 A4YGN0 A4YGN2
A4YGQ6 A4YGR1 A4YH38 A4YH39 A4YH62 A4YH79 A4YH82 A4YH83 A4YH87 A4YH88 A4YH91
A4YH98 A4YH99 A4YHA3 A4YHA5 A4YHA6 A4YHB7 A4YHC9 A4YHD8 A4YHE2 A4YHE5 A4YHH3
A4YHH8 A4YHH9 A4YHI8 A4YHJ0 A4YHJ3 A4YHK0 A4YHK6 A4YHK7 A4YHK9 A4YHL5 A4YHL6
A4YHP0 A4YHP1 A4YHV0 A4YHV3 A4YHV8 A4YHW1 A4YI01 A4YI09 A4YI11 A4YI12 A4YI15
A4YI28 A4YI31 A4YI33 A4YI34 A4YI36 A4YI40 A4YI51 A4YI52 A4YI68 A4YI75 A4YI77
A4YIC2 A4YIC4 A4YIE0 A4YIG5 A4YIK8 A4YIL0 A4YIL1 A4YIL2 A4YIL7 A4YIL8 A4YIL9
A4YIM3 A4YIM6 A4YIM7 A4YIM8 A4YIN3 A4YIN4 A4YIP0 A4YIP3 A4YIQ9 A4YIU6 A4YIV7
A4YIV8 A4YIW0 A4YIX4 A4YIX5 A4YIY0 A4YIY3 A4YIY8 A4YIZ5 A4YIZ6 A4YJ03 A4YJ04
A4YJ14 A4YJ15 A4YJ98 A4YJD7 A4YJE6 A4YJE9 A4YJF0 A4YJF2 A4YJF3 A4YJG9 A4YJH0
A4YJH3 A4YJI5 A4YJI6 A4YJI7 A4YJI9 A4YJJ2 A4YJJ3 A4YJM8 A4YJM9 A4YJN0 A4YJP8
A4YJP9 A4YJQ1 A4YJR1 A4YJR2 A4YJR3 A4YJR8 A4YJS8 A4YJT3 A4YJT4 A4YJT5 A4YJT8
A4YJU2 A4YJU7 A4YJV0 A4YJV1 A4YJV4 A4YJV5 A4YJV7 A4YJX9 A4YJY2 A4YJY8 A4YK09
A4YK33 A4YK44 A4YK54 A4YK73 A4YK97 A4YK99 A4YKA0 A4YKA5 A4YKC5 A4YKC6 A4YKE4
A4YKE8 A4YKF0 A4YKF1 A4YKF3 A4YKF6 A4YKP3 A4YKQ0 A4YKQ8 A4YKR8 A4YKS8 A4YKT6
A4YKZ2 A4YL90 A4YL97 A4YLB3 A4YLB4 A4YLB9 A4YLC0 A4YLC1 A4YLC3 A4YLC4 A4YLC7
A4YLD2 A4YLK4 A4YLX6 A4YM24 A4YM26 A4YM31 A4YM32 A4YM47 A4YMC4 A4YMC5 A4YMC6
A4YMC8 A4YMG9 A4YN80 A4YND6 A4YNH1 A4YNK6 A4YNK7 A4YNK8 A4YNP1 A4YNP2 A4YNP4
A4YNQ1 A4YNQ7 A4YNR1 A4YNR8 A4YNU5 A4YPC0 A4YPM4 A4YPP8 A4YPQ3 A4YPV1 A4YQ21
A4YQ28 A4YQ36 A4YQ37 A4YQD3 A4YQS3 A4YQV7 A4YR32 A4YRD9 A4YRI5 A4YRL1 A4YRR9
A4YRT2 A4YRU4 A4YS25 A4YSB5 A4YSH5 A4YSH6 A4YSH9 A4YSI0 A4YSI1 A4YSI2 A4YSI7
A4YSJ0 A4YSJ1 A4YSJ2 A4YSJ3 A4YSJ5 A4YSJ6 A4YSJ7 A4YSJ8 A4YSJ9 A4YSK1 A4YSK2
A4YSK3 A4YSK4 A4YSK6 A4YSK7 A4YSK8 A4YSK9 A4YSL0 A4YSL1 A4YSL4 A4YSL5 A4YSL6
A4YSL7 A4YSR9 A4YSS3 A4YSU1 A4YT21 A4YT30 A4YT74 A4YT76 A4YT77 A4YT90 A4YTA2
A4YTA9 A4YTC3 A4YTC4 A4YTQ2 A4YTY9 A4YUE1 A4YUI3 A4YUJ5 A4YUL6 A4YUN8 A4YUP3
A4YUQ7 A4YUR5 A4YV19 A4YV23 A4YV54 A4YV66 A4YV68 A4YV70 A4YV78 A4YV79 A4YV90
A4YVC1 A4YVC9 A4YVD3 A4YVD5 A4YVD6 A4YVF1 A4YVG0 A4YVG3 A4YVG4 A4YVG5 A4YVG6
A4YVI0 A4YVM3 A4YVM6 A4YVP2 A4YVP3 A4YW83 A4YW92 A4YW96 A4YW97 A4YWA5 A4YWC5
A4YWD1 A4YWD3 A4YWD7 A4YWF1 A4YWF3 A4YWF9 A4YWG4 A4YWH5 A4YWV8 A4YXJ9 A4YXK4
A4YXM5 A4YXN2 A4YXQ6 A4YXQ9 A4YXZ8 A4YYA7 A4YYB6 A4YYB9 A4YYF1 A4YYT0 A4YZ37
A4YZ76 A4YZA6 A4YZB3 A4YZC4 A4YZJ3 A4YZK1 A4YZK2 A4YZL1 A4YZQ4 A4YZQ6 A4YZY5
A4Z005 A4Z012 A4Z027 A4Z049 A4Z057 A4Z082 A4Z0B8 A4Z0D8 A4Z0I5 A4Z0Q5 A4Z0T3
A4Z0U1 A4Z0W1 A4Z0W6 A4Z112 A4Z1C6 A4Z1S1 A4Z1S4 A4Z1Z9 A4Z216 A4Z217 A4Z264
A4Z2C2 A4Z2J6 A4Z3G9 A4Z3J9 A4Z6H0 A4Z6H1 A4Z943 A4Z944 A4Z945 A4ZCW2 A4ZCW3
A4ZCW4 A4ZCW5 A4ZCW6 A4ZCW8 A4ZDG8 A4ZGQ8 A4ZHR8 A4ZHT6 A4ZU87 A4ZU88 A4ZU89
A4ZU90 A4ZU91 A4ZU93 A4ZU94 A4ZU96 A4ZU97 A4ZU98 A4ZU99 A4ZUA0 A4ZUA1 A4ZUA2
A4ZUA3 A4ZUA4 A4ZUA5 A4ZUA6 A4ZUA8 A4ZUA9 A4ZUB0 A4ZUB2 A4ZUB8 A4ZUC1 A4ZUC2
A4ZUC5 A4ZUD2 A4ZUD3 A4ZUD4 A4ZUD6 A4ZUD7 A4ZUD8 A4ZUD9 A4ZUE0 A4ZUE1 A4ZUE2
A4ZZ93 A5A3E0 A5A607 A5A611 A5A612 A5A613 A5A614 A5A619 A5A625 A5A6H4 A5A6H9
A5A6I1 A5A6I5 A5A6I8 A5A6J5 A5A6J9 A5A6K1 A5A6K2 A5A6K3 A5A6K4 A5A6K8 A5A6L0
A5A6L7 A5A6M0 A5A6M4 A5A6M5 A5A6M8 A5A6N0 A5A6N2 A5A6N4 A5A6P1 A5A6P3 A5A6P5
A5A779 A5AA68 A5AAA4 A5AAE5 A5AAL8 A5AAW6 A5AB13 A5ABG0 A5ABG3 A5ABG4 A5ABG5
A5ABG7 A5ABG8 A5ABM5 A5ABU3 A5ABX6 A5AEC8 A5AEM3 A5ANT9 A5B4D2 A5B4K1 A5BJL8

A5BUU4 A5C9M2 A5CAL1 A5CB81 A5CC41 A5CC52 A5CC57 A5CC62 A5CC63 A5CC73 A5CC78
A5CC87 A5CC91 A5CC92 A5CC93 A5CC95 A5CC96 A5CCA0 A5CCA2 A5CCC1 A5CCC2 A5CCC4
A5CCC8 A5CCE0 A5CCI7 A5CCI8 A5CCI9 A5CCJ0 A5CCJ1 A5CCJ4 A5CCJ5 A5CCJ6 A5CCJ8
A5CCJ9 A5CCK1 A5CCK2 A5CCK3 A5CCK5 A5CCK6 A5CCK7 A5CCK8 A5CCK9 A5CCL1 A5CCL4
A5CCM4 A5CCM6 A5CCN0 A5CCQ4 A5CCS3 A5CCS8 A5CCT4 A5CCV1 A5CCV6 A5CCY5 A5CCY6
A5CCY7 A5CCZ1 A5CCZ2 A5CCZ3 A5CCZ4 A5CCZ6 A5CCZ7 A5CD00 A5CD02 A5CD03 A5CD09
A5CD12 A5CD16 A5CD21 A5CD22 A5CD80 A5CD85 A5CD86 A5CDH4 A5CDI3 A5CDI4 A5CDL9
A5CDM6 A5CDP1 A5CDP2 A5CDP5 A5CDS4 A5CDS8 A5CDS9 A5CDT2 A5CDU2 A5CDU5 A5CDU7
A5CE04 A5CE05 A5CE49 A5CE64 A5CEC8 A5CEC9 A5CEJ4 A5CEJ6 A5CEL3 A5CEN4 A5CEN6
A5CEP5 A5CEP8 A5CER5 A5CES8 A5CET3 A5CET5 A5CEY0 A5CF17 A5CF21 A5CF22 A5CF23
A5CF34 A5CF57 A5CF58 A5CF62 A5CF64 A5CF65 A5CF69 A5CF98 A5CFB7 A5CFB8 A5CFJ6
A5CFM7 A5CFN0 A5CFN1 A5CFN3 A5CFP2 A5CKB9 A5CLT3 A5CLT6 A5CLV7 A5CLW1 A5CM68
A5CM86 A5CNA2 A5CNC0 A5CNC6 A5CNH4 A5CNI7 A5CPA0 A5CPB7 A5CPC1 A5CPD5 A5CPD8
A5CPE3 A5CPG8 A5CPK0 A5CPY3 A5CPY6 A5CPZ4 A5CPZ8 A5CQ46 A5CQ48 A5CQ99 A5CQA0
A5CQK2 A5CQM6 A5CQN3 A5CQP6 A5CQP8 A5CQQ7 A5CQQ8 A5CQR0 A5CQR1 A5CQR4 A5CQR8
A5CQS1 A5CQS5 A5CQS6 A5CQS7 A5CQU4 A5CQU5 A5CQV0 A5CQY7 A5CR04 A5CR08 A5CR30
A5CR31 A5CR32 A5CR33 A5CR34 A5CR36 A5CR89 A5CR93 A5CRA6 A5CRB1 A5CRB9 A5CRI6
A5CRN7 A5CRQ9 A5CRT6 A5CRT9 A5CRU0 A5CRU3 A5CRV6 A5CRW1 A5CRW2 A5CRW4 A5CRW8
A5CRX0 A5CRX2 A5CRX5 A5CRX9 A5CRY5 A5CRY6 A5CRZ1 A5CRZ4 A5CRZ5 A5CRZ7 A5CS06
A5CS07 A5CS08 A5CS09 A5CS10 A5CS23 A5CS25 A5CS39 A5CS46 A5CS50 A5CS59 A5CS60
A5CSB9 A5CSI8 A5CSI9 A5CSJ0 A5CSJ2 A5CSJ6 A5CSJ9 A5CSK2 A5CSK8 A5CSL3 A5CSL4
A5CSL5 A5CSL7 A5CSN2 A5CSN9 A5CSP0 A5CSP7 A5CSW0 A5CSZ3 A5CSZ4 A5CSZ7 A5CT01
A5CT42 A5CT74 A5CTL3 A5CTX3 A5CTY3 A5CTY4 A5CTZ0 A5CTZ4 A5CTZ8 A5CU17 A5CU18
A5CU44 A5CU64 A5CU73 A5CU74 A5CU75 A5CU79 A5CU80 A5CU81 A5CU83 A5CU84 A5CU93
A5CU96 A5CU99 A5CUA0 A5CUA3 A5CUA4 A5CUA6 A5CUA7 A5CUA8 A5CUA9 A5CUB0 A5CUB1
A5CUB3 A5CUB4 A5CUB6 A5CUB7 A5CUB8 A5CUB9 A5CUC6 A5CUC7 A5CUT3 A5CUT4 A5CUU0
A5CUU9 A5CUV9 A5CV36 A5CV47 A5CV83 A5CV84 A5CV85 A5CVA8 A5CVA9 A5CVC1 A5CVC5
A5CVC6 A5CVE7 A5CVE8 A5CVF0 A5CVF3 A5CVF7 A5CVG7 A5CVH0 A5CVH1 A5CVH4 A5CVI0
A5CVJ3 A5CVK1 A5CVK4 A5CVK5 A5CVK6 A5CVL9 A5CVM3 A5CVN0 A5CVN3 A5CVN4 A5CVN5
A5CVQ5 A5CVR1 A5CVR5 A5CVS6 A5CVT9 A5CVU0 A5CVV1 A5CVV4 A5CVV8 A5CVV9 A5CVW8
A5CVW9 A5CVZ0 A5CVZ6 A5CVZ8 A5CW02 A5CW05 A5CW24 A5CW25 A5CW26 A5CW27 A5CW28
A5CW29 A5CW32 A5CW34 A5CW46 A5CW49 A5CW51 A5CW53 A5CW55 A5CW57 A5CW65 A5CW77
A5CW80 A5CW82 A5CW91 A5CW96 A5CW97 A5CWD1 A5CWD9 A5CWE0 A5CWE1 A5CWE3 A5CWE4
A5CWE8 A5CWF6 A5CWF7 A5CWF9 A5CWH5 A5CWI0 A5CWI3 A5CWI5 A5CWI9 A5CWJ0 A5CWK3
A5CWL4 A5CWL9 A5CWM2 A5CWM6 A5CWN5 A5CWP5 A5CWP6 A5CWP9 A5CWQ3 A5CWQ5 A5CWR4
A5CWR5 A5CWR6 A5CWS3 A5CWU3 A5CWU5 A5CWU9 A5CWV5 A5CWW0 A5CWW8 A5CWX9 A5CWY3
A5CWZ1 A5CWZ3 A5CX05 A5CX08 A5CX12 A5CX15 A5CX17 A5CX20 A5CX21 A5CX22 A5CX27
A5CX29 A5CX31 A5CX33 A5CX35 A5CX42 A5CX55 A5CX56 A5CX57 A5CX59 A5CX61 A5CX62
A5CX64 A5CX65 A5CX66 A5CX68 A5CX75 A5CX85 A5CX90 A5CX97 A5CXA1 A5CXB2 A5CXB8
A5CXC6 A5CXC7 A5CXD1 A5CXD6 A5CXD7 A5CXE4 A5CXE9 A5CXF1 A5CXH6 A5CXH8 A5CXI2
A5CXI5 A5CXI6 A5CXI7 A5CXI8 A5CXI9 A5CXJ0 A5CXJ1 A5CXJ4 A5CXJ7 A5CXJ8 A5CXK2
A5CXK3 A5CXK4 A5CXK5 A5CXK6 A5CXK7 A5CXK8 A5CXK9 A5CXL0 A5CXL1 A5CXL2 A5CXL3
A5CXL4 A5CXL5 A5CXL6 A5CXL7 A5CXL8 A5CXL9 A5CXM0 A5CXM1 A5CXM2 A5CXM3 A5CXM6
A5CXM9 A5CXN5 A5CXN7 A5CXP4 A5CXP9 A5CXQ6 A5CXR1 A5CXR3 A5CXS1 A5CXS6 A5CXT4

A5CXU9 A5CXV2 A5CXW2 A5CXW7 A5CXX4 A5CXX6 A5CXX7 A5CXY8 A5CY09 A5CY11 A5CY29
A5CY44 A5CY45 A5CY46 A5CY49 A5CY50 A5CY52 A5CY74 A5CY78 A5CY81 A5CYA0 A5CYB7
A5CYC3 A5CYC5 A5CYD0 A5CYJ7 A5CYK7 A5CYM8 A5CYN8 A5CYP3 A5CYQ6 A5CYR8 A5CYZ0
A5CZ03 A5CZ07 A5CZ28 A5CZ68 A5CZ71 A5CZ72 A5CZ73 A5CZ74 A5CZ77 A5CZ78 A5CZ96
A5D0T6 A5D0U1 A5D0U4 A5D0U5 A5D0W6 A5D113 A5D114 A5D143 A5D144 A5D147 A5D165
A5D172 A5D180 A5D186 A5D193 A5D197 A5D1A4 A5D1A5 A5D1B6 A5D1B9 A5D1C0 A5D1C3
A5D1C5 A5D1C7 A5D1G1 A5D1G6 A5D1G8 A5D1H6 A5D1I2 A5D1I3 A5D1J3 A5D1J4 A5D1J5
A5D1P3 A5D1P4 A5D1R6 A5D1R9 A5D1S5 A5D1T9 A5D1U6 A5D1V4 A5D243 A5D2A7 A5D2B7
A5D2C2 A5D2D8 A5D2E1 A5D2F9 A5D2G6 A5D2K1 A5D2K2 A5D2K5 A5D2K7 A5D2K8 A5D2M7
A5D2Q5 A5D2R3 A5D2R7 A5D2S0 A5D2S1 A5D2S3 A5D2S5 A5D2S6 A5D2T5 A5D2T8 A5D2U3
A5D2U6 A5D2U8 A5D2W5 A5D2W6 A5D2W7 A5D2W8 A5D2Z3 A5D2Z6 A5D2Z8 A5D300 A5D319
A5D323 A5D337 A5D339 A5D369 A5D372 A5D373 A5D374 A5D3C3 A5D3C4 A5D3E0 A5D3E2
A5D3E3 A5D3F0 A5D3F4 A5D3F8 A5D3F9 A5D3G1 A5D3G5 A5D3G6 A5D3H8 A5D3L4 A5D3L5
A5D3M2 A5D3M8 A5D3N0 A5D3N4 A5D3N6 A5D3W2 A5D3W7 A5D3W9 A5D3X2 A5D3X8 A5D3Y3
A5D407 A5D408 A5D410 A5D416 A5D426 A5D434 A5D446 A5D447 A5D451 A5D4I1 A5D4K1
A5D4K4 A5D4K5 A5D4R0 A5D4W0 A5D4W7 A5D4Y2 A5D4Y6 A5D504 A5D506 A5D508 A5D510
A5D522 A5D523 A5D5E3 A5D5E4 A5D5E5 A5D5E6 A5D5E8 A5D5F0 A5D5F6 A5D5F7 A5D5G1
A5D5G2 A5D5G3 A5D5G4 A5D5G5 A5D5G6 A5D5G8 A5D5G9 A5D5H0 A5D5H1 A5D5H3 A5D5H4
A5D5H7 A5D5H8 A5D5H9 A5D5I2 A5D5I3 A5D5I5 A5D5I6 A5D5I7 A5D5I8 A5D5I9 A5D5J0
A5D5J1 A5D5J2 A5D5J3 A5D5J4 A5D5J5 A5D5K0 A5D5K1 A5D5K4 A5D5K7 A5D5L4 A5D5L5
A5D5L8 A5D5M0 A5D5P9 A5D5Q4 A5D5R7 A5D5T5 A5D5T6 A5D5U8 A5D632 A5D654 A5D673
A5D690 A5D6B2 A5D6B3 A5D6D2 A5D6D5 A5D6E6 A5D789 A5D796 A5D7B1 A5D7C1 A5D7C2
A5D7D1 A5D7F6 A5D7F8 A5D7I0 A5D7J1 A5D7J3 A5D7M3 A5D7N9 A5D7P0 A5D7Q3 A5D7R8
A5D7S3 A5D7T0 A5D7U0 A5D7V9 A5D8M0 A5D8M6 A5D8N2 A5D8N7 A5D8Q0 A5D8Q8 A5D8R3
A5D8S0 A5D8S5 A5D8S8 A5D8V7 A5D8W1 A5D962 A5D989 A5D9C6 A5D9W7 A5DA73 A5DA88
A5DAC8 A5DAD0 A5DAF0 A5DAH9 A5DAK5 A5DAL3 A5DAQ7 A5DAR2 A5DAX7 A5DAY6 A5DB27
A5DB51 A5DB52 A5DB88 A5DB98 A5DBF4 A5DBG1 A5DBG4 A5DBG5 A5DBJ8 A5DBS1 A5DC23
A5DC72 A5DC85 A5DCB6 A5DCF5 A5DCP8 A5DCV3 A5DDB7 A5DDF4 A5DDH8 A5DDJ4 A5DDN0
A5DDU4 A5DE24 A5DE65 A5DE68 A5DEE9 A5DEF8 A5DEJ2 A5DEK6 A5DEV6 A5DEY7 A5DEZ5
A5DF03 A5DF43 A5DFC2 A5DFC5 A5DFH6 A5DFI4 A5DFT1 A5DFX7 A5DG41 A5DG57 A5DG59
A5DGF3 A5DGG9 A5DGI1 A5DGL8 A5DGM4 A5DGV3 A5DGY0 A5DGZ7 A5DH34 A5DH98 A5DHD2
A5DHL9 A5DHT2 A5DHV9 A5DHW0 A5DI11 A5DI69 A5DI74 A5DID3 A5DID7 A5DIP0 A5DIP4
A5DIR2 A5DIU9 A5DIV5 A5DIX5 A5DJ67 A5DJ85 A5DJJ1 A5DJJ2 A5DJK0 A5DJW8 A5DJX5
A5DK38 A5DK47 A5DK49 A5DKB1 A5DKB9 A5DKC0 A5DKC4 A5DKF1 A5DKH0 A5DKN0 A5DKQ3
A5DKW3 A5DL80 A5DL92 A5DLC6 A5DLD1 A5DLE0 A5DLF4 A5DLG8 A5DLJ8 A5DLL7 A5DLM5
A5DLN4 A5DLR3 A5DLU8 A5DLW3 A5DM21 A5DM91 A5DMB6 A5DMB8 A5DME6 A5DMM1 A5DN53
A5DN58 A5DN78 A5DNJ4 A5DNK9 A5DNL9 A5DNM7 A5DNT0 A5DNW6 A5DNX8 A5DNX9 A5DNZ1
A5DPE3 A5DPQ4 A5DPQ5 A5DPU0 A5DQ50 A5DQ88 A5DQA6 A5DQA8 A5DQF1 A5DQI5 A5DQJ8
A5DQN2 A5DQP9 A5DQS0 A5DR03 A5DR04 A5DR43 A5DR68 A5DR89 A5DRH5 A5DRJ2 A5DRM4
A5DRQ6 A5DRX1 A5DS77 A5DSB5 A5DSD3 A5DSI2 A5DSQ8 A5DSR2 A5DSS4 A5DSS9 A5DST9
A5DSU5 A5DSZ3 A5DT31 A5DT64 A5DTA8 A5DTF4 A5DTJ3 A5DTK7 A5DTP3 A5DTS1 A5DTV4
A5DTX8 A5DU46 A5DU73 A5DUA3 A5DUA7 A5DUB2 A5DUN2 A5DUP0 A5DUW8 A5DUX4 A5DVF8
A5DVG6 A5DVI0 A5DVL3 A5DVM3 A5DVY3 A5DW14 A5DWE2 A5DWF0 A5DWF1 A5DWF4 A5DWG1
A5DWJ1 A5DWJ3 A5DWN5 A5DWP3 A5DWY0 A5DX41 A5DXA0 A5DXC3 A5DXQ6 A5DXS8 A5DXV0

A5DXV6 A5DY33 A5DY34 A5DYN3 A5DYR5 A5DYS6 A5DYV8 A5DZ02 A5DZE6 A5DZI5 A5DZP0
A5DZP6 A5DZS4 A5DZT7 A5DZY1 A5E032 A5E058 A5E092 A5E094 A5E0B1 A5E0T7 A5E0U1
A5E0U9 A5E0W5 A5E0Z9 A5E121 A5E145 A5E181 A5E1A0 A5E1F6 A5E1N2 A5E1T3 A5E1W0
A5E1W4 A5E1Z4 A5E203 A5E287 A5E2H7 A5E2I8 A5E2R6 A5E2Y4 A5E2Z7 A5E2Z9 A5E3E3
A5E3J7 A5E3K3 A5E3Q2 A5E3Q3 A5E3R9 A5E3W5 A5E406 A5E450 A5E4A8 A5E4P1 A5E4V9
A5E4Y7 A5E572 A5E598 A5E5F7 A5E5I9 A5E5U8 A5E5Y8 A5E625 A5E654 A5E6K6 A5E6M3
A5E6W6 A5E713 A5E726 A5E748 A5E7B9 A5E7I7 A5E7M8 A5E7T4 A5E7U6 A5E812 A5E855
A5E857 A5E863 A5E866 A5E867 A5E869 A5E870 A5E887 A5E888 A5E891 A5E8A4 A5E8A5
A5E8A7 A5E8B1 A5E8E3 A5E8E5 A5E8E8 A5E8F1 A5E8F2 A5E8F5 A5E8G0 A5E8G4 A5E8G7
A5E8G8 A5E8G9 A5E8H4 A5E8I4 A5E8I8 A5E8I9 A5E8J0 A5E8J8 A5E8K0 A5E8K1 A5E8L7
A5E8L8 A5E8L9 A5E8P0 A5E8P3 A5E8P9 A5E8S0 A5E8T8 A5E8U9 A5E8V7 A5E8X6 A5E8Z8
A5E907 A5E909 A5E910 A5E915 A5E935 A5E936 A5E954 A5E958 A5E960 A5E961 A5E964
A5E965 A5E968 A5E969 A5E987 A5E993 A5EA34 A5EAG2 A5EAG4 A5EAL1 A5EAL2 A5EAM9
A5EB33 A5EB50 A5EB56 A5EBK6 A5EBP8 A5ECG6 A5ECH2 A5ECI7 A5ECJ5 A5ECL8 A5ECT2
A5ECX6 A5ED09 A5ED38 A5EDA7 A5EDB2 A5EDB3 A5EDY0 A5EE90 A5EEC1 A5EEC6 A5EED4
A5EEN4 A5EEP1 A5EEQ0 A5EEQ1 A5EES6 A5EES7 A5EES8 A5EET6 A5EF40 A5EF62 A5EFG5
A5EFG7 A5EFR6 A5EG60 A5EG87 A5EGD7 A5EGE2 A5EGF4 A5EGF9 A5EGH4 A5EHE7 A5EHJ0
A5EHP2 A5EI59 A5EI95 A5EI97 A5EI98 A5EIA8 A5EIB4 A5EIC0 A5EID1 A5EIE5 A5EIE6
A5EIM5 A5EIT5 A5EIY9 A5EIZ5 A5EJ03 A5EJ24 A5EJ46 A5EJ58 A5EJC6 A5EJD2 A5EJR4
A5EJR7 A5EJU4 A5EJV5 A5EJV7 A5EJV9 A5EJW7 A5EJW8 A5EJY2 A5EJY3 A5EK14 A5EK18
A5EK22 A5EK24 A5EK25 A5EK39 A5EK49 A5EK52 A5EK53 A5EK55 A5EK68 A5EKA7 A5EKA8
A5EKB0 A5EKC6 A5EKC7 A5EKG4 A5EKH8 A5EKI2 A5EKI3 A5EKJ1 A5EKL4 A5EKM2 A5EKM6
A5EKP2 A5EKP4 A5EKP6 A5EKQ4 A5EKQ8 A5EKS0 A5EKW3 A5EL70 A5EL79 A5ELK2 A5ELK3
A5ELK4 A5ELK5 A5ELK8 A5ELK9 A5ELL0 A5ELL1 A5ELL2 A5ELL3 A5ELL4 A5ELL5 A5ELL6
A5ELL7 A5ELL8 A5ELL9 A5ELM0 A5ELM1 A5ELM2 A5ELM3 A5ELM4 A5ELM6 A5ELM7 A5ELM8
A5ELM9 A5ELN0 A5ELN1 A5ELN2 A5ELN6 A5ELN7 A5ELN8 A5ELN9 A5ELP1 A5ELP2 A5ELY1
A5EM76 A5EMF5 A5EMG8 A5EMI1 A5EML9 A5EMM2 A5EMW5 A5EN78 A5EN81 A5ENA7 A5ENA8
A5ENR3 A5EP16 A5EP79 A5EPB5 A5EPC1 A5EPD3 A5EPJ2 A5EPK0 A5EPK1 A5EPL2 A5EPQ3
A5EPQ5 A5EPZ7 A5EQ55 A5EQ63 A5EQ67 A5EQ73 A5EQ80 A5EQ82 A5EQ83 A5EQB6 A5EQB7
A5EQB8 A5EQF2 A5EQH6 A5ER96 A5ERF9 A5ERG8 A5ERJ9 A5ERK1 A5ERK2 A5ERK3 A5ERN6
A5ERQ9 A5ERU7 A5ERU8 A5ERV2 A5ERV4 A5ES06 A5ESE1 A5ESP9 A5ESQ2 A5ESQ3 A5ESQ5
A5ESQ6 A5ESQ7 A5ESR4 A5ESR5 A5EST0 A5EST5 A5ETA3 A5ETC5 A5ETG1 A5ETH1 A5ETI5
A5ETJ3 A5ETJ9 A5EU55 A5EUT5 A5EUT6 A5EV06 A5EV13 A5EV26 A5EV27 A5EV28 A5EV29
A5EV41 A5EV42 A5EV43 A5EV63 A5EV65 A5EV74 A5EV80 A5EV96 A5EVA6 A5EVB4 A5EVB6
A5EVB8 A5EVE1 A5EVF4 A5EVG3 A5EVG5 A5EVH6 A5EVK4 A5EVK6 A5EVM0 A5EVM2 A5EVM4
A5EVN1 A5EVN8 A5EVP4 A5EVP6 A5EVP7 A5EVQ3 A5EVQ4 A5EVQ5 A5EVQ9 A5EVR0 A5EVR7
A5EVS5 A5EVU0 A5EVW7 A5EVX0 A5EVX5 A5EVY7 A5EVZ5 A5EW22 A5EW27 A5EW39 A5EW54
A5EW66 A5EW70 A5EW83 A5EW88 A5EW90 A5EW94 A5EWA3 A5EWD6 A5EWE5 A5EWF3 A5EWF5
A5EWF6 A5EWG6 A5EWH1 A5EWL5 A5EWL8 A5EWL9 A5EWQ0 A5EWQ2 A5EWR5 A5EWS0 A5EWS4
A5EWT0 A5EWT6 A5EWU9 A5EWV0 A5EWV4 A5EWV8 A5EWV9 A5EWY6 A5EWY8 A5EWY9 A5EWZ1
A5EWZ5 A5EX17 A5EX18 A5EX24 A5EX25 A5EX29 A5EX42 A5EX48 A5EX54 A5EX57 A5EX62
A5EX67 A5EX68 A5EX69 A5EX70 A5EX71 A5EX72 A5EX73 A5EX74 A5EX77 A5EX78 A5EX79
A5EX81 A5EX82 A5EX83 A5EX84 A5EX85 A5EX86 A5EX87 A5EX88 A5EX89 A5EX90 A5EX91
A5EX92 A5EX93 A5EX94 A5EX95 A5EX96 A5EX97 A5EX99 A5EXA0 A5EXA1 A5EXA2 A5EXA3

A5EXA4 A5EXA5 A5EXA7 A5EXB0 A5EXB2 A5EXB6 A5EXD5 A5EXE3 A5EXE7 A5EXG1 A5EXG2
A5EXH6 A5EXH7 A5EXH9 A5EXK3 A5EXK6 A5EXK9 A5EXM0 A5EXM4 A5EXM6 A5EXM7 A5EXM9
A5EXN0 A5EXN1 A5EXP1 A5EXP9 A5EXQ4 A5EXR1 A5EXR7 A5EXR9 A5EXS0 A5EXS3 A5EXT0
A5EXT4 A5EXU0 A5EXU1 A5EXU4 A5EXW8 A5EXX0 A5EXX1 A5EXX2 A5EXX6 A5EXY9 A5EXZ1
A5EXZ2 A5EXZ5 A5EY11 A5EY13 A5EY21 A5EY22 A5EY31 A5EY32 A5EY33 A5EY39 A5EY43
A5EY47 A5EY50 A5EY60 A5EY62 A5EY64 A5EY68 A5EY80 A5EY84 A5EY93 A5EYD6 A5EYE5
A5EYG1 A5EYG2 A5EYG3 A5EYI0 A5EYJ8 A5EYL9 A5EYT7 A5EYY8 A5EZ07 A5EZ15 A5EZ16
A5EZ18 A5EZN1 A5EZR2 A5EZS6 A5EZS7 A5EZT0 A5EZT5 A5EZV8 A5EZZ8 A5F028 A5F049
A5F051 A5F083 A5F089 A5F0G1 A5F0H9 A5F0I2 A5F0J0 A5F0M6 A5F0N6 A5F0U5 A5F0W1
A5F0Z4 A5F125 A5F155 A5F159 A5F176 A5F1A0 A5F1C1 A5F1F1 A5F1F9 A5F1N4 A5F1P8
A5F1U0 A5F1U8 A5F1V4 A5F1V7 A5F1W4 A5F1W7 A5F1X2 A5F1X6 A5F1X7 A5F1X8 A5F1Y1
A5F1Y2 A5F1Z0 A5F207 A5F208 A5F210 A5F263 A5F277 A5F278 A5F289 A5F294 A5F2A2
A5F2F2 A5F2H1 A5F2H3 A5F2L8 A5F2N2 A5F2N5 A5F2P8 A5F2Q5 A5F2Q7 A5F2T1 A5F2T4
A5F2T9 A5F2U0 A5F2W3 A5F2W6 A5F2X0 A5F2X4 A5F2X6 A5F2Y0 A5F2Y1 A5F2Y2 A5F2Y7
A5F331 A5F333 A5F340 A5F342 A5F343 A5F345 A5F359 A5F361 A5F362 A5F368 A5F369
A5F375 A5F376 A5F377 A5F379 A5F384 A5F3A2 A5F3A7 A5F3C2 A5F3C5 A5F3D0 A5F3E6
A5F3E9 A5F3F1 A5F3F4 A5F3F7 A5F3F8 A5F3G1 A5F3G4 A5F3G7 A5F3G8 A5F3H1 A5F3H2
A5F3I4 A5F3I5 A5F3I6 A5F3I8 A5F3J3 A5F3J8 A5F3K0 A5F3K3 A5F3K5 A5F3L3 A5F3L5
A5F3L6 A5F3L7 A5F3M2 A5F3M7 A5F3M9 A5F3N0 A5F3N4 A5F3P1 A5F3P2 A5F3P3 A5F3P4
A5F3P5 A5F3P6 A5F3P9 A5F3R3 A5F3R4 A5F3S1 A5F3S3 A5F3S6 A5F3U8 A5F3Z4 A5F402
A5F404 A5F405 A5F407 A5F408 A5F449 A5F468 A5F469 A5F482 A5F483 A5F485 A5F487
A5F493 A5F497 A5F4A1 A5F4B4 A5F4C3 A5F4C4 A5F4C5 A5F4C9 A5F4D2 A5F4E2 A5F4E5
A5F4G9 A5F4H7 A5F4I0 A5F4I4 A5F4I6 A5F4I9 A5F4K5 A5F4M4 A5F4P4 A5F4Q2 A5F4Q4
A5F4Q7 A5F4R9 A5F4S8 A5F4V7 A5F4W0 A5F4W4 A5F4X3 A5F4X4 A5F4X8 A5F4Y1 A5F4Y6
A5F4Z3 A5F4Z4 A5F4Z5 A5F4Z7 A5F514 A5F520 A5F523 A5F529 A5F534 A5F542 A5F543
A5F544 A5F545 A5F546 A5F547 A5F548 A5F549 A5F550 A5F553 A5F554 A5F555 A5F556
A5F557 A5F558 A5F559 A5F561 A5F562 A5F563 A5F564 A5F565 A5F566 A5F571 A5F572
A5F573 A5F574 A5F578 A5F579 A5F587 A5F599 A5F5A9 A5F5B4 A5F5C5 A5F5C9 A5F5D0
A5F5D8 A5F5E2 A5F5E3 A5F5F3 A5F5F7 A5F5G0 A5F5H6 A5F5H8 A5F5I2 A5F5I4 A5F5I8
A5F5J2 A5F5K1 A5F5M8 A5F5N1 A5F5N9 A5F5R2 A5F5R5 A5F5R6 A5F5R7 A5F5R9 A5F5S6
A5F5T1 A5F5U0 A5F5U6 A5F5X3 A5F5Y1 A5F5Z2 A5F5Z6 A5F601 A5F602 A5F604 A5F605
A5F606 A5F607 A5F615 A5F618 A5F619 A5F623 A5F626 A5F633 A5F635 A5F637 A5F642
A5F643 A5F660 A5F684 A5F686 A5F696 A5F697 A5F699 A5F6B5 A5F6C3 A5F6D7 A5F6E9
A5F6F4 A5F6G4 A5F6G6 A5F6M1 A5F6M2 A5F6P1 A5F6P9 A5F6R2 A5F6T1 A5F6T5 A5F6U8
A5F6U9 A5F6V4 A5F6X0 A5F6X1 A5F6Y1 A5F6Y4 A5F6Z1 A5F706 A5F727 A5F745 A5F760
A5F761 A5F762 A5F763 A5F769 A5F7B9 A5F7C0 A5F7F1 A5F7F2 A5F7F3 A5F7G0 A5F7G5
A5F7G9 A5F7H0 A5F7H8 A5F7I0 A5F7P6 A5F7R5 A5F7V7 A5F7X7 A5F818 A5F827 A5F836
A5F888 A5F890 A5F8E4 A5F8G4 A5F8G8 A5F8G9 A5F8N0 A5F8N1 A5F8N2 A5F8P0 A5F8P1
A5F8P7 A5F8U2 A5F8Y1 A5F8Y6 A5F8Z0 A5F8Z3 A5F8Z6 A5F904 A5F913 A5F924 A5F926
A5F927 A5F928 A5F929 A5F930 A5F937 A5F942 A5F945 A5F947 A5F964 A5F974 A5F975
A5F980 A5F986 A5F992 A5F997 A5F998 A5F9A7 A5F9A8 A5F9A9 A5F9B5 A5F9B6 A5F9B7
A5F9C0 A5F9C1 A5F9C4 A5F9D7 A5F9D8 A5F9D9 A5F9E6 A5F9E8 A5F9F1 A5F9F9 A5F9G1
A5F9G2 A5F9G8 A5F9H3 A5F9H4 A5F9H5 A5F9H9 A5F9I5 A5F9I8 A5F9R5 A5F9Y3 A5F9Z0
A5F9Z1 A5F9Z4 A5F9Z6 A5FA30 A5FA31 A5FA35 A5FA67 A5FA70 A5FA76 A5FAD0 A5FAD1

A5FAD2 A5FAD3 A5FAE2 A5FAQ5 A5FAX4 A5FB38 A5FB49 A5FB76 A5FB79 A5FC03 A5FC30
A5FC71 A5FD11 A5FDF9 A5FE24 A5FE44 A5FE53 A5FE75 A5FE76 A5FE77 A5FE82 A5FE87
A5FE96 A5FEA5 A5FEB8 A5FEC6 A5FEF4 A5FEP6 A5FF23 A5FFL9 A5FFP4 A5FFT1 A5FFT9
A5FFX6 A5FFX7 A5FFY0 A5FFY3 A5FG48 A5FG55 A5FG67 A5FG89 A5FG91 A5FG96 A5FGE0
A5FGL1 A5FGN1 A5FGN3 A5FGS7 A5FGU0 A5FGV5 A5FGX1 A5FGX2 A5FH28 A5FH89 A5FHA4
A5FHA9 A5FHB4 A5FHG4 A5FHI8 A5FHM5 A5FHM7 A5FHP3 A5FHR3 A5FHV5 A5FHV7 A5FHW9
A5FHX5 A5FHX8 A5FIF5 A5FIJ0 A5FIJ1 A5FIJ2 A5FIJ3 A5FIJ4 A5FIJ9 A5FIK2 A5FIL0
A5FIL6 A5FIL7 A5FIM3 A5FIP9 A5FIR0 A5FIU2 A5FIV1 A5FIX6 A5FIY4 A5FIZ1 A5FJ06
A5FJ26 A5FJ42 A5FJ63 A5FJ93 A5FJ95 A5FJE5 A5FJF7 A5FJF9 A5FJG9 A5FJI4 A5FJI6
A5FJJ1 A5FJJ8 A5FJK3 A5FJP5 A5FJP6 A5FJR8 A5FJT9 A5FJU1 A5FJV8 A5FJZ1 A5FJZ2
A5FK02 A5FK26 A5FK68 A5FKC6 A5FKD7 A5FKD8 A5FKD9 A5FKJ2 A5FKL6 A5FKV4 A5FKW3
A5FL10 A5FL22 A5FL73 A5FL86 A5FL87 A5FLE3 A5FLE7 A5FLM7 A5FLP4 A5FLT1 A5FLW1
A5FLZ0 A5FLZ4 A5FM03 A5FM31 A5FM90 A5FMF6 A5FMG2 A5FMI1 A5FMJ5 A5FML2 A5FMM4
A5FMM5 A5FMN0 A5FMP6 A5FMT0 A5FMV7 A5FMW5 A5FMX0 A5FMY0 A5FMY1 A5FMY2 A5FMY4
A5FMY5 A5FMY8 A5FMY9 A5FMZ0 A5FMZ2 A5FMZ3 A5FMZ4 A5FMZ5 A5FMZ6 A5FMZ7 A5FMZ9
A5FN00 A5FN02 A5FN03 A5FN06 A5FN07 A5FN08 A5FN14 A5FN16 A5FN17 A5FN18 A5FN24
A5FND6 A5FNE3 A5FNF9 A5FNG4 A5FNH2 A5FNH3 A5FNH5 A5FNN7 A5FNY2 A5FP01 A5FP03
A5FP04 A5FP31 A5FP49 A5FP50 A5FP73 A5FP84 A5FP88 A5FP90 A5FPB2 A5FPB5 A5FPC2
A5FPD9 A5FPF7 A5FPK5 A5FPL3 A5FPM3 A5FPN6 A5FPQ5 A5FPR2 A5FPR4 A5FPS8 A5FPT3
A5FPU4 A5FPV2 A5FPW5 A5FPX2 A5FPY7 A5FQ06 A5FQ07 A5FQ13 A5FQ25 A5FQ39 A5FQ48
A5FQ49 A5FQ51 A5FQ55 A5FQ60 A5FQ65 A5FQ73 A5FQ74 A5FQ84 A5FQB7 A5FQC0 A5FQC5
A5FQD3 A5FQE0 A5FQE1 A5FQE3 A5FQI9 A5FQJ4 A5FQL1 A5FQM9 A5FQQ4 A5FQQ5 A5FQQ9
A5FQR0 A5FQR1 A5FQR2 A5FQR4 A5FQR5 A5FQR6 A5FQS4 A5FQT1 A5FQT2 A5FQT3 A5FQT5
A5FQW8 A5FR28 A5FR29 A5FR32 A5FR35 A5FR44 A5FR63 A5FR64 A5FR68 A5FR69 A5FR71
A5FR82 A5FR84 A5FR85 A5FR86 A5FR87 A5FR94 A5FRA7 A5FRB0 A5FRB1 A5FRB3 A5FRB5
A5FRB6 A5FRB9 A5FRC5 A5FRE3 A5FRE4 A5FRE5 A5FRF1 A5FRG9 A5FRK5 A5FRK6 A5FRK7
A5FRL3 A5FRL4 A5FRL6 A5FRM7 A5FRN4 A5FRN5 A5FRN7 A5FRP5 A5FRP7 A5FRP8 A5FRT0
A5FRU8 A5FRV2 A5FRV6 A5FRV8 A5FRV9 A5FRW2 A5FRW3 A5FRW5 A5FRW7 A5FRW8 A5FRW9
A5FRX0 A5FRX2 A5FRX3 A5FRX4 A5FRX5 A5FRX6 A5FRX7 A5FRX9 A5FRY0 A5FRY1 A5FRY2
A5FRY3 A5FRY4 A5FRY5 A5FRY6 A5FRY7 A5FRY8 A5FRY9 A5FRZ0 A5FRZ1 A5FRZ2 A5FRZ3
A5FRZ4 A5FRZ5 A5FRZ6 A5FRZ7 A5FS00 A5FS02 A5FS06 A5FS18 A5FS25 A5FS26 A5FS31
A5FS34 A5FS52 A5FS53 A5FS65 A5FS74 A5FS79 A5FS82 A5FS84 A5FS85 A5FS95 A5FSB3
A5FSB5 A5FSB6 A5FSB7 A5FSC0 A5FSC8 A5FSN4 A5FSN9 A5FSQ2 A5FSR7 A5FSS1 A5FSU1
A5FSU3 A5FUG4 A5FUG5 A5FUG6 A5FUH4 A5FUH7 A5FUH8 A5FUI6 A5FUJ7 A5FUK2 A5FUL1
A5FUL2 A5FUL3 A5FUM6 A5FUS8 A5FUV6 A5FUV7 A5FV18 A5FV21 A5FV22 A5FV27 A5FV28
A5FV29 A5FV42 A5FV67 A5FVE7 A5FVG2 A5FVG6 A5FVI6 A5FVJ4 A5FVK3 A5FVK4 A5FVN0
A5FVN2 A5FVN5 A5FVN9 A5FVT7 A5FVX2 A5FVZ7 A5FW68 A5FW78 A5FWA3 A5FWG7 A5FWH1
A5FWI3 A5FWI5 A5FWJ5 A5FWK8 A5FWQ4 A5FWQ8 A5FWR1 A5FWR6 A5FWV7 A5FWV9 A5FWW0
A5FWY8 A5FWZ7 A5FX03 A5FX05 A5FX23 A5FX26 A5FX43 A5FX97 A5FX98 A5FX99 A5FXB8
A5FXC2 A5FXD0 A5FXD8 A5FXD9 A5FXQ8 A5FXT9 A5FXW0 A5FXW5 A5FXW7 A5FY40 A5FY49
A5FY57 A5FY58 A5FY68 A5FY70 A5FY82 A5FYD2 A5FYD8 A5FYE1 A5FYE4 A5FYE5 A5FYE6
A5FYH8 A5FYN3 A5FYN4 A5FYN6 A5FYN7 A5FYN8 A5FYQ7 A5FYS4 A5FYS5 A5FYZ9 A5FZ18
A5FZ19 A5FZ25 A5FZ26 A5FZ47 A5FZ48 A5FZ67 A5FZ68 A5FZ78 A5FZ90 A5FZ94 A5FZ95
A5FZ96 A5FZ98 A5FZB8 A5FZB9 A5FZC1 A5FZC4 A5FZD5 A5FZE9 A5FZF9 A5FZG9 A5FZN7

A5FZN8 A5FZN9 A5FZS6 A5FZS7 A5FZU0 A5FZU1 A5FZU2 A5FZU3 A5FZU6 A5FZU7 A5FZU8
A5FZU9 A5FZV0 A5FZV1 A5FZV2 A5FZV4 A5FZV5 A5FZV6 A5FZV7 A5FZV8 A5FZV9 A5FZW0
A5FZW1 A5FZW2 A5FZW4 A5FZW5 A5FZW6 A5FZW7 A5FZW8 A5FZW9 A5FZX0 A5FZX1 A5FZX2
A5FZX3 A5FZX4 A5FZX5 A5FZX8 A5G009 A5G016 A5G017 A5G020 A5G029 A5G068 A5G0D9
A5G0E0 A5G0F5 A5G0G0 A5G0G6 A5G0G7 A5G0G9 A5G0H0 A5G0L2 A5G0M4 A5G0S4 A5G0S8
A5G0V3 A5G0Y2 A5G126 A5G127 A5G146 A5G167 A5G168 A5G169 A5G1D5 A5G1D6 A5G1F7
A5G1G2 A5G1G3 A5G1J3 A5G1L0 A5G1L2 A5G1L3 A5G1L8 A5G1U2 A5G1W5 A5G1X4 A5G1Y4
A5G1Y9 A5G201 A5G209 A5G211 A5G222 A5G270 A5G276 A5G293 A5G2D2 A5G2E2 A5G2H9
A5G2L3 A5G2L4 A5G2M7 A5G2N0 A5G2N1 A5G2T9 A5G320 A5G360 A5G381 A5G382 A5G3A0
A5G3A1 A5G3A2 A5G3A5 A5G3G6 A5G3H0 A5G3H3 A5G3J6 A5G3J8 A5G3J9 A5G3K3 A5G3L2
A5G3L5 A5G3P2 A5G3Q7 A5G3S1 A5G3W1 A5G3W5 A5G3X1 A5G439 A5G447 A5G460 A5G4G0
A5G4G8 A5G4S5 A5G4S7 A5G4S8 A5G4T3 A5G4T4 A5G4T5 A5G535 A5G536 A5G5A2 A5G5C1
A5G5F6 A5G5G9 A5G5H3 A5G5R2 A5G5S2 A5G5T3 A5G670 A5G674 A5G677 A5G692 A5G696
A5G6A1 A5G6A4 A5G6F4 A5G6I5 A5G6I6 A5G6I9 A5G6M1 A5G6S0 A5G733 A5G742 A5G743
A5G7C7 A5G7P4 A5G7Q7 A5G7Q8 A5G7Q9 A5G7R0 A5G7R2 A5G7R3 A5G7R5 A5G7R6 A5G7R8
A5G7S1 A5G7S3 A5G7U4 A5G7V3 A5G7V6 A5G7W2 A5G7W5 A5G7W7 A5G7W8 A5G7Y7 A5G7Y9
A5G7Z1 A5G7Z2 A5G7Z3 A5G7Z5 A5G7Z6 A5G7Z8 A5G863 A5G879 A5G8G3 A5G8J7 A5G8J8
A5G8J9 A5G8K2 A5G8K8 A5G8K9 A5G8P0 A5G8S1 A5G8S2 A5G8S7 A5G8S9 A5G8T0 A5G8T2
A5G8T5 A5G8T7 A5G8U0 A5G8X1 A5G8X2 A5G903 A5G905 A5G906 A5G913 A5G937 A5G938
A5G939 A5G949 A5G962 A5G964 A5G9C0 A5G9C7 A5G9G1 A5G9G5 A5G9G7 A5G9I1 A5G9I2
A5G9J7 A5G9J8 A5G9J9 A5G9K0 A5G9K4 A5G9T7 A5G9V1 A5G9V2 A5G9V3 A5G9V6 A5G9V7
A5G9Y2 A5G9Y4 A5G9Y5 A5GA37 A5GA66 A5GA97 A5GA98 A5GAC4 A5GAC6 A5GAM5 A5GAS3
A5GAU3 A5GAU5 A5GAU7 A5GAV3 A5GAV4 A5GAV5 A5GAV6 A5GAV7 A5GAV8 A5GAW0 A5GAW1
A5GAW3 A5GAW4 A5GAW5 A5GAW6 A5GAW7 A5GAW8 A5GAW9 A5GAX0 A5GAX1 A5GAX2 A5GAX4
A5GAX7 A5GAX8 A5GAX9 A5GAY0 A5GAY1 A5GAY2 A5GAY3 A5GAY4 A5GAY7 A5GAY8 A5GB00
A5GB52 A5GBB7 A5GBF9 A5GBI7 A5GBL0 A5GBR0 A5GBW0 A5GBX4 A5GBX9 A5GBY8 A5GC19
A5GC79 A5GC94 A5GCS6 A5GCW2 A5GCW3 A5GCZ8 A5GD27 A5GD28 A5GD29 A5GD47 A5GD54
A5GD74 A5GD89 A5GD90 A5GD93 A5GD98 A5GDA1 A5GDA4 A5GDA7 A5GDB2 A5GDB4 A5GDC7
A5GDC8 A5GDD6 A5GDG7 A5GDW7 A5GDX1 A5GDX3 A5GE08 A5GE10 A5GE36 A5GE43 A5GEC1
A5GEC2 A5GEF7 A5GEQ5 A5GER7 A5GES2 A5GES5 A5GEV5 A5GEV6 A5GF03 A5GF05 A5GF08
A5GF41 A5GF42 A5GF66 A5GF75 A5GF83 A5GF86 A5GF87 A5GF89 A5GF90 A5GF91 A5GF99
A5GFA0 A5GFA1 A5GFC7 A5GFN6 A5GFQ0 A5GFT6 A5GFW7 A5GFX0 A5GFY4 A5GFY8 A5GFZ6
A5GHL4 A5GHM4 A5GHN3 A5GHN7 A5GHN8 A5GHN9 A5GHP3 A5GHR6 A5GHR9 A5GHS0 A5GHS5
A5GHT2 A5GHT4 A5GHU1 A5GHU5 A5GHU9 A5GI06 A5GI07 A5GI30 A5GI38 A5GI45 A5GI46
A5GI51 A5GI62 A5GIA4 A5GID0 A5GIF4 A5GIF7 A5GIF9 A5GIG4 A5GIG6 A5GIG7 A5GII9
A5GIJ2 A5GIJ9 A5GIK5 A5GIL9 A5GIN1 A5GIN4 A5GIN9 A5GIP0 A5GIP1 A5GIP2 A5GIP3
A5GIR6 A5GIR7 A5GIR9 A5GIS1 A5GIS2 A5GIS3 A5GIS4 A5GIS5 A5GIS6 A5GIS8 A5GIS9
A5GIT0 A5GIT1 A5GIT2 A5GIT3 A5GIT4 A5GIT5 A5GIT6 A5GIT7 A5GIT8 A5GIT9 A5GIU0
A5GIU1 A5GIU2 A5GIU3 A5GIU4 A5GIU5 A5GIV0 A5GIW2 A5GIW5 A5GIY3 A5GIY5 A5GIZ3
A5GJ24 A5GJ39 A5GJ48 A5GJ60 A5GJ71 A5GJ79 A5GJ83 A5GJ87 A5GJ90 A5GJ95 A5GJA2
A5GJA5 A5GJC5 A5GJD1 A5GJD3 A5GJE9 A5GJF0 A5GJF5 A5GJI1 A5GJI2 A5GJI3 A5GJK5
A5GJK6 A5GJL3 A5GJR6 A5GJR7 A5GJS5 A5GJT4 A5GJT7 A5GJV3 A5GKI2 A5GKI7 A5GKJ3
A5GKJ5 A5GKM1 A5GKM7 A5GKN0 A5GKN1 A5GKP6 A5GKR0 A5GKR3 A5GKS6 A5GKX4 A5GL34
A5GL48 A5GL51 A5GL71 A5GL79 A5GLA2 A5GLA3 A5GLA9 A5GLB0 A5GLD9 A5GLE0 A5GLH6

A5GLH7 A5GLI2 A5GLJ3 A5GLK1 A5GLP6 A5GLQ5 A5GLV1 A5GLY6 A5GLZ1 A5GM25 A5GM31
A5GM47 A5GM62 A5GM79 A5GMB7 A5GMB9 A5GME5 A5GMH4 A5GMH7 A5GMI5 A5GMJ1 A5GMJ9
A5GMK4 A5GML6 A5GML7 A5GMM6 A5GMN1 A5GMR6 A5GMT2 A5GMU7 A5GMV4 A5GMW9 A5GMX4
A5GN09 A5GN10 A5GN16 A5GN18 A5GN19 A5GN53 A5GN62 A5GN71 A5GN76 A5GN78 A5GN79
A5GN82 A5GN83 A5GN87 A5GNA0 A5GNA9 A5GNB0 A5GNG2 A5GNG9 A5GNH1 A5GNH3 A5GNH5
A5GNH7 A5GNI7 A5GNJ0 A5GNK6 A5GNP9 A5GNU1 A5GNW7 A5GNX9 A5GP01 A5GP05 A5GP57
A5GP60 A5GP74 A5GP76 A5GP97 A5GPA1 A5GPB2 A5GPD2 A5GPD3 A5GPD4 A5GPD5 A5GPE6
A5GPE9 A5GPF9 A5GPH4 A5GPH9 A5GPI1 A5GPI9 A5GPL8 A5GPL9 A5GPM2 A5GPN0 A5GPS8
A5GPT0 A5GPT7 A5GPV1 A5GPW9 A5GPX0 A5GPX1 A5GPX5 A5GPX7 A5GQ03 A5GQ04 A5GQ09
A5GQ11 A5GQ13 A5GQ22 A5GQ24 A5GQ34 A5GQ35 A5GQ58 A5GQ67 A5GQ68 A5GQ73 A5GQ79
A5GQA2 A5GQJ8 A5GQK2 A5GQM5 A5GQN5 A5GQP4 A5GQQ5 A5GQT3 A5GQU6 A5GQU9 A5GR03
A5GR07 A5GR10 A5GR30 A5GR42 A5GR50 A5GR53 A5GR60 A5GR64 A5GRD7 A5GRD9 A5GRE1
A5GRE3 A5GRI8 A5GRJ3 A5GRJ7 A5GRJ8 A5GRL0 A5GRL5 A5GRN3 A5GRN4 A5GRX2 A5GRZ0
A5GRZ3 A5GS00 A5GS37 A5GS43 A5GS45 A5GS63 A5GS82 A5GS98 A5GSA3 A5GSC0 A5GSC3
A5GSD0 A5GSE0 A5GSG3 A5GSG4 A5GSH0 A5GSI7 A5GSL0 A5GSL5 A5GSL8 A5GSQ4 A5GT34
A5GT38 A5GT44 A5GT49 A5GT82 A5GT90 A5GTB8 A5GTB9 A5GTE2 A5GTE8 A5GTE9 A5GTG7
A5GTG8 A5GTK8 A5GTK9 A5GTL6 A5GTL7 A5GTM0 A5GTM5 A5GTM6 A5GTR3 A5GTS7 A5GTT4
A5GTZ1 A5GU17 A5GU20 A5GU32 A5GU85 A5GU97 A5GUA8 A5GUA9 A5GUB0 A5GUJ2 A5GUL0
A5GUP3 A5GUP6 A5GUQ3 A5GUS1 A5GUT4 A5GUY6 A5GV03 A5GV19 A5GV21 A5GV22 A5GV27
A5GV37 A5GV53 A5GVE0 A5GVE8 A5GVF0 A5GVF1 A5GVF3 A5GVF8 A5GVG1 A5GVG4 A5GVQ6
A5GVR3 A5GVR5 A5GVW0 A5GVW2 A5GVW3 A5GVW4 A5GVW5 A5GVW6 A5GVW7 A5GVW8 A5GVW9
A5GVX0 A5GVX1 A5GVX2 A5GVX3 A5GVX4 A5GVX5 A5GVX6 A5GVX7 A5GVX9 A5GVY0 A5GVY1
A5GVY2 A5GVY3 A5GVY4 A5GVY6 A5GVY8 A5GVY9 A5GW06 A5GW11 A5GW12 A5GW13 A5GW14
A5GW15 A5GW20 A5GW23 A5GW34 A5GW48 A5GW52 A5GW63 A5GW73 A5GW82 A5GW85 A5GW90
A5GWA0 A5GWA2 A5GWB6 A5GWD4 A5GWG3 A5GWG7 A5GWH2 A5GWH4 A5GWH5 A5GWJ9 A5GWK0
A5GWK1 A5GWK2 A5GWM2 A5GWM4 A5GWN4 A5GWN5 A5GWP0 A5GWP3 A5GWQ3 A5GWT1 A5GWT2
A5GWT6 A5GWT8 A5GWV3 A5GWV5 A5GWV6 A5GWV7 A5GX14 A5GX16 A5GX22 A5H0J2 A5H0J3
A5H0J4 A5H2P4 A5H2Q8 A5HBD7 A5HBK6 A5HBL2 A5HC98 A5HEH4 A5HEI1 A5HNV6 A5HWA8
A5HXR1 A5HXS7 A5HXS8 A5HXT1 A5HXW3 A5HXX7 A5HY17 A5HY31 A5HY32 A5HY34 A5HY35
A5HY41 A5HY62 A5HY79 A5HYC0 A5HYC6 A5HYC7 A5HYL0 A5HYU6 A5HYX0 A5HYY0 A5HYY2
A5HYZ3 A5HYZ4 A5HYZ6 A5HZ44 A5HZ60 A5HZE1 A5HZP0 A5HZY2 A5I0A5 A5I0B2 A5I0C8
A5I0I2 A5I0M6 A5I0R5 A5I0T6 A5I0Y0 A5I112 A5I123 A5I133 A5I174 A5I1A3 A5I1A6
A5I1C9 A5I1H8 A5I1K5 A5I1T3 A5I1T6 A5I1U5 A5I1Z6 A5I231 A5I232 A5I237 A5I242
A5I244 A5I245 A5I246 A5I247 A5I248 A5I2A9 A5I2R9 A5I2S0 A5I2S2 A5I2S3 A5I2T0
A5I2T7 A5I2X9 A5I302 A5I308 A5I309 A5I310 A5I321 A5I322 A5I323 A5I348 A5I365
A5I389 A5I3B0 A5I3D3 A5I3F5 A5I3X9 A5I421 A5I426 A5I427 A5I4I1 A5I4I7 A5I4I8
A5I4J2 A5I4J3 A5I4J7 A5I4K2 A5I4K9 A5I4L0 A5I4L2 A5I4L3 A5I4L4 A5I4L8 A5I4L9
A5I4M2 A5I4M3 A5I4M4 A5I4N7 A5I4R8 A5I4S1 A5I4S6 A5I4T1 A5I4T4 A5I4U1 A5I4U3
A5I4V0 A5I4W1 A5I4X1 A5I4Z2 A5I4Z3 A5I4Z4 A5I4Z5 A5I4Z9 A5I524 A5I526 A5I540
A5I5A3 A5I5A4 A5I5H1 A5I5H2 A5I5I5 A5I5M0 A5I5P4 A5I5R9 A5I5U3 A5I5U9 A5I5V9
A5I5W3 A5I5Y9 A5I5Z5 A5I603 A5I629 A5I634 A5I638 A5I640 A5I648 A5I656 A5I664
A5I666 A5I667 A5I669 A5I683 A5I684 A5I689 A5I6D5 A5I6D6 A5I6D9 A5I6E1 A5I6E9
A5I6F0 A5I6F1 A5I6F2 A5I6F4 A5I6L1 A5I6L5 A5I6L6 A5I6L8 A5I6M3 A5I6N5 A5I6W0
A5I6W1 A5I6W2 A5I6W5 A5I6W8 A5I6X0 A5I6X1 A5I6Z2 A5I6Z3 A5I6Z4 A5I6Z7 A5I705

A5I720 A5I723 A5I724 A5I731 A5I738 A5I748 A5I768 A5I788 A5I798 A5I7A0 A5I7A2
A5I7A3 A5I7A4 A5I7A5 A5I7D0 A5I7E5 A5I7H0 A5I7H6 A5I7H7 A5I7H8 A5I7H9 A5I7I0
A5I7I1 A5I7I2 A5I7I5 A5I7I7 A5I7I8 A5I7I9 A5I7J0 A5I7J1 A5I7J2 A5I7J3 A5I7J4
A5I7J5 A5I7J6 A5I7J7 A5I7J8 A5I7J9 A5I7K0 A5I7K2 A5I7K3 A5I7K4 A5I7K5 A5I7K6
A5I7K7 A5I7K8 A5I7K9 A5I7L0 A5I7L1 A5I7L3 A5I7L4 A5I7L5 A5I7L7 A5I7L8 A5I7M1
A5I7M8 A5I7N1 A5I7P2 A5I7P4 A5I7P5 A5I7P8 A5I7P9 A5I7R5 A5I7S1 A5I7S3 A5I7T7
A5I7U4 A5I7Y4 A5I7Y9 A5I7Z5 A5I7Z9 A5I801 A5I814 A5I815 A5I816 A5I820 A5I821
A5I9F1 A5I9G0 A5I9H3 A5I9P3 A5I9Q7 A5I9R2 A5I9T3 A5I9T5 A5I9T6 A5I9T7 A5I9U4
A5I9V1 A5I9V7 A5IA69 A5IA87 A5IAB4 A5IAG1 A5IAI0 A5IAK4 A5IAK9 A5IAL1 A5IAL2
A5IAL3 A5IAL5 A5IAL7 A5IAM8 A5IAR4 A5IAR5 A5IAS3 A5IAS4 A5IAS5 A5IAS6 A5IAS7
A5IAT1 A5IAU0 A5IAU5 A5IAV2 A5IAW2 A5IAW3 A5IAW4 A5IAY3 A5IB64 A5IB79 A5IB92
A5IBA2 A5IBB3 A5IBD5 A5IBD8 A5IBD9 A5IBE0 A5IBE1 A5IBE7 A5IBF0 A5IBF5 A5IBF6
A5IBF7 A5IBF8 A5IBG0 A5IBH7 A5IBI8 A5IBJ1 A5IBK8 A5IBL1 A5IBL5 A5IBL8 A5IBM2
A5IBM5 A5IBN0 A5IBN1 A5IBN3 A5IBP1 A5IBP5 A5IBQ3 A5IBR0 A5IBR1 A5IBR7 A5IBT2
A5IBU7 A5IBW2 A5IBW4 A5IBW5 A5IBW8 A5IC06 A5IC23 A5IC27 A5IC29 A5IC36 A5IC42
A5IC43 A5IC64 A5IC73 A5IC87 A5IC89 A5IC90 A5ICA4 A5ICB9 A5ICK3 A5ICK4 A5ICK7
A5ICK8 A5ICK9 A5ICL0 A5ICM0 A5ICM2 A5ICN0 A5ICP2 A5ICP3 A5ICS3 A5ICS5 A5ICT2
A5ICV6 A5ICV7 A5ICV8 A5ICW2 A5ICZ5 A5ID02 A5ID16 A5ID17 A5ID18 A5ID21 A5ID38
A5ID69 A5ID78 A5ID85 A5IDI1 A5IDI6 A5IDJ2 A5IDJ5 A5IDK5 A5IDK7 A5IDK8 A5IDK9
A5IDL1 A5IDL4 A5IDM1 A5IDN6 A5IE48 A5IE87 A5IEA1 A5IEA6 A5IEB3 A5IEC9 A5IED6
A5IED7 A5IEF4 A5IEF9 A5IEG0 A5IEW9 A5IEX1 A5IEY2 A5IF70 A5IFN1 A5IFV1 A5IFW5
A5IFW8 A5IFX5 A5IFX9 A5IFY0 A5IFZ9 A5IG00 A5IG02 A5IG23 A5IG27 A5IG28 A5IG41
A5IG48 A5IG81 A5IG82 A5IG90 A5IG97 A5IGB3 A5IGC8 A5IGG3 A5IGG9 A5IGI0 A5IGI2
A5IGJ7 A5IGK7 A5IGL1 A5IGL5 A5IGM1 A5IGM2 A5IGM4 A5IGQ8 A5IGR7 A5IGR8 A5IGU1
A5IGU3 A5IGX6 A5IGX8 A5IH02 A5IH08 A5IH19 A5IH21 A5IH84 A5IH96 A5IHA2 A5IHA3
A5IHA9 A5IHB4 A5IHB6 A5IHB7 A5IHD0 A5IHD3 A5IHG9 A5IHI0 A5IHJ5 A5IHJ6 A5IHJ8
A5IHM9 A5IHP0 A5IHP1 A5IHP2 A5IHP3 A5IHP4 A5IHP6 A5IHP7 A5IHP8 A5IHP9 A5IHQ0
A5IHQ2 A5IHQ3 A5IHQ4 A5IHQ5 A5IHQ6 A5IHQ7 A5IHQ8 A5IHQ9 A5IHR0 A5IHR1 A5IHR2
A5IHR3 A5IHR4 A5IHR5 A5IHR6 A5IHR7 A5IHR8 A5IHR9 A5IHS0 A5IHS1 A5IHS2 A5IHS3
A5IHS4 A5IHS5 A5IHS9 A5IHT0 A5IHU3 A5IHU4 A5IHU5 A5IHU6 A5IHU7 A5IHU9 A5IHW5
A5IHW7 A5IHX0 A5IHY1 A5IHY4 A5IHZ5 A5IHZ7 A5II06 A5II09 A5II10 A5II41 A5II45
A5II55 A5II59 A5II62 A5II63 A5II90 A5IIB8 A5IIC0 A5IIJ5 A5IIK3 A5IIK5 A5IIK6
A5IIK7 A5IIL8 A5IIM2 A5IIN8 A5IIP2 A5IIP4 A5IIQ2 A5IIR1 A5IIS0 A5IIS7 A5IIS8
A5IIT2 A5IIT3 A5IIT4 A5IIT9 A5IIW2 A5IIX7 A5IIY7 A5IIZ4 A5IIZ7 A5IJ00 A5IJ09
A5IJ21 A5IJ44 A5IJ45 A5IJ51 A5IJ55 A5IJ64 A5IJ65 A5IJ66 A5IJ79 A5IJ83 A5IJ91
A5IJ92 A5IJ98 A5IJA0 A5IJD2 A5IJD3 A5IJD4 A5IJG9 A5IJH1 A5IJJ8 A5IJJ9 A5IJK0
A5IJK9 A5IJL8 A5IJM2 A5IJM4 A5IJM5 A5IJN0 A5IJP6 A5IJP8 A5IJP9 A5IJQ2 A5IJQ3
A5IJQ4 A5IJQ9 A5IJR6 A5IJR7 A5IJU8 A5IJU9 A5IJV5 A5IJW2 A5IJW3 A5IJW4 A5IJW5
A5IJW6 A5IJW7 A5IJX0 A5IJX2 A5IK18 A5IK42 A5IK44 A5IK54 A5IK57 A5IK69 A5IK72
A5IK73 A5IK80 A5IK83 A5IK84 A5IK86 A5IKB4 A5IKB6 A5IKE5 A5IKE7 A5IKF4 A5IKF6
A5IKF8 A5IKG2 A5IKG7 A5IKG8 A5IKG9 A5IKH0 A5IKI1 A5IKI7 A5IKJ0 A5IKK4 A5IKK5
A5IKK7 A5IKK8 A5IKL0 A5IKL2 A5IKM1 A5IKN6 A5IKQ2 A5IKQ3 A5IKQ4 A5IKQ6 A5IKR2
A5IKS1 A5IKS6 A5IKT2 A5IKT4 A5IKW2 A5IKX0 A5IKX2 A5IKX3 A5IKY3 A5IKY9 A5IL01
A5IL06 A5IL29 A5IL32 A5IL67 A5IL75 A5IL77 A5IL80 A5IL83 A5IL84 A5ILC5 A5ILC9

A5ILD1 A5ILE6 A5ILE7 A5ILE8 A5ILF1 A5ILF5 A5ILF6 A5ILF8 A5ILG0 A5ILH0 A5ILH3
A5ILI6 A5ILJ3 A5ILJ5 A5ILJ7 A5ILJ8 A5ILJ9 A5ILK0 A5ILK1 A5ILK2 A5ILK6 A5ILL0
A5ILL1 A5ILL3 A5ILL5 A5ILP6 A5ILP7 A5ILQ1 A5ILR5 A5ILS0 A5ILS1 A5ILS4 A5ILS5
A5ILT8 A5ILV5 A5ILV6 A5ILX8 A5ILX9 A5ILZ2 A5ILZ3 A5IM04 A5IM09 A5IM12 A5IM14
A5IM15 A5IM17 A5IM19 A5IM22 A5IM24 A5IM26 A5IM27 A5IM62 A5IM63 A5IM64 A5IM70
A5IM74 A5IM78 A5IM79 A5IM80 A5IM81 A5IM82 A5IM83 A5IM84 A5IM85 A5IM86 A5IM87
A5IM88 A5IM89 A5IM90 A5IM91 A5IM92 A5IM93 A5IM94 A5IM96 A5IM97 A5IM98 A5IM99
A5IMA0 A5IMA1 A5IMA2 A5IMA4 A5IMA6 A5IMA7 A5IMA8 A5IMA9 A5IMB0 A5IMB1 A5IMB2
A5IMB8 A5IMC0 A5IMC1 A5IMC2 A5IMC7 A5IMC9 A5IMD1 A5IMD2 A5IMD9 A5IME1 A5IME2
A5IME8 A5IMH5 A5IMH8 A5IMI0 A5IMI1 A5IMI9 A5IMJ0 A5IMJ4 A5IML2 A5IMM2 A5IMM8
A5IMM9 A5IMS4 A5IMS8 A5IMT3 A5IMW3 A5IMZ6 A5IN10 A5IN20 A5IN74 A5IN77 A5IN78
A5IN82 A5IN85 A5IN97 A5IN99 A5IND6 A5INE0 A5INE2 A5INE3 A5INE5 A5INE6 A5INL0
A5INL8 A5INN0 A5INP2 A5INP5 A5INP9 A5INQ0 A5INQ6 A5INR5 A5INY7 A5IP09 A5IP51
A5IP80 A5IPA9 A5IPC8 A5IPD8 A5IPI2 A5IPI5 A5IPT7 A5IPU6 A5IPU8 A5IPW7 A5IPX0
A5IQ31 A5IQ32 A5IQ34 A5IQ38 A5IQ45 A5IQ47 A5IQ50 A5IQ55 A5IQ65 A5IQ72 A5IQ73
A5IQ81 A5IQ83 A5IQ88 A5IQ91 A5IQ92 A5IQ93 A5IQ94 A5IQ96 A5IQ98 A5IQ99 A5IQA0
A5IQA1 A5IQA2 A5IQA5 A5IQC0 A5IQC3 A5IQD5 A5IQE1 A5IQE7 A5IQG1 A5IQM3 A5IQN5
A5IQR6 A5IQS7 A5IQT1 A5IQT4 A5IQU2 A5IQU7 A5IQW4 A5IQW9 A5IQX1 A5IQX2 A5IQX7
A5IQX8 A5IQY5 A5IR07 A5IR11 A5IR97 A5IRA1 A5IRB0 A5IRB7 A5IRD8 A5IRD9 A5IRE4
A5IRE5 A5IRE8 A5IRH5 A5IRI1 A5IRI3 A5IRJ1 A5IRJ6 A5IRS9 A5IRV0 A5IRV3 A5IRV6
A5IRV8 A5IRW0 A5IRX7 A5IRY4 A5IRZ3 A5IRZ9 A5IS08 A5IS11 A5IS12 A5IS14 A5IS24
A5IS26 A5IS30 A5IS32 A5IS36 A5IS63 A5IS64 A5IS65 A5IS69 A5IS75 A5IS79 A5IS83
A5IS85 A5IS88 A5IS95 A5ISA1 A5ISA3 A5ISA9 A5ISB8 A5ISC3 A5ISC4 A5ISC5 A5ISC6
A5ISC9 A5ISD0 A5ISD5 A5ISD6 A5ISD7 A5ISD8 A5ISD9 A5ISE0 A5ISE1 A5ISE3 A5ISE7
A5ISE9 A5ISF3 A5ISF4 A5ISF5 A5ISF7 A5ISF8 A5ISG9 A5ISH6 A5ISH9 A5ISI2 A5ISI5
A5ISL2 A5ISL7 A5ISL8 A5ISL9 A5ISM8 A5ISM9 A5ISP0 A5ISQ1 A5ISQ2 A5ISQ4 A5ISQ5
A5ISS7 A5ISS8 A5ISS9 A5ISU5 A5ISV2 A5ISV4 A5ISV8 A5ISX6 A5ISX7 A5ISX9 A5ISY4
A5ISZ2 A5ISZ4 A5ISZ5 A5ISZ6 A5ISZ7 A5ISZ9 A5IT03 A5IT05 A5IT22 A5IT23 A5IT32
A5IT49 A5IT51 A5IT52 A5IT53 A5IT57 A5IT63 A5IT64 A5IT65 A5IT66 A5IT79 A5IT85
A5IT91 A5IT93 A5IT98 A5ITA3 A5ITA6 A5ITA7 A5ITA8 A5ITA9 A5ITB0 A5ITB3 A5ITC1
A5ITC3 A5ITC6 A5ITD0 A5ITD5 A5ITD6 A5ITE0 A5ITE1 A5ITE2 A5ITE3 A5ITE6 A5ITF4
A5ITF5 A5ITF7 A5ITF9 A5ITG3 A5ITG4 A5ITG5 A5ITG6 A5ITG7 A5ITG8 A5ITG9 A5ITH1
A5ITI8 A5ITJ2 A5ITJ5 A5ITJ7 A5ITJ8 A5ITJ9 A5ITK0 A5ITK3 A5ITK4 A5ITK7 A5ITL0
A5ITM2 A5ITN0 A5ITN4 A5ITN8 A5ITP2 A5ITQ1 A5ITQ4 A5ITR2 A5ITR5 A5ITS0 A5ITT0
A5ITT7 A5ITU6 A5ITV6 A5ITV7 A5ITW2 A5ITZ9 A5IU10 A5IU30 A5IU33 A5IU39 A5IU61
A5IU66 A5IU67 A5IU68 A5IU71 A5IU80 A5IU87 A5IUH6 A5IUH7 A5IUJ2 A5IUJ5 A5IUJ9
A5IUK2 A5IUK3 A5IUK5 A5IUK6 A5IUK8 A5IUL1 A5IUM9 A5IUN7 A5IUN8 A5IUQ7 A5IUQ8
A5IUQ9 A5IUR3 A5IUR5 A5IUS2 A5IUS3 A5IUU9 A5IUV1 A5IUX8 A5IUY1 A5IUY2 A5IUY3
A5IUY4 A5IV01 A5IV02 A5IV03 A5IV07 A5IV09 A5IV10 A5IV11 A5IV12 A5IV15 A5IV16
A5IV17 A5IV18 A5IV19 A5IV20 A5IV21 A5IV22 A5IV23 A5IV24 A5IV25 A5IV26 A5IV27
A5IV29 A5IV30 A5IV31 A5IV32 A5IV33 A5IV34 A5IV35 A5IV50 A5IV51 A5IV56 A5IV62
A5IV69 A5IV70 A5IV71 A5IV72 A5IV73 A5IV74 A5IV75 A5IVB2 A5IVB5 A5IVB6 A5IVC5
A5IVC7 A5IVF3 A5IVH3 A5IVJ6 A5IVK3 A5IVK5 A5IVU3 A5IVY0 A5IVY4 A5IW22 A5IW23
A5IW24 A5IW27 A5IW31 A5IW37 A5IW58 A5IW62 A5IWA0 A5IWA1 A5IWA3 A5IWA6 A5IWB6

A5IWB7 A5IWD5 A5IWD6 A5IWD7 A5IWD8 A5IXG3 A5IXI1 A5IXJ1 A5IXJ4 A5IXK9 A5IXM0
A5IXN6 A5IXR8 A5IXS0 A5IXT1 A5IXT5 A5IXT6 A5IXW3 A5IXW4 A5IXY4 A5IXZ0 A5IY02
A5IY09 A5IY12 A5IY18 A5IY26 A5IY48 A5IY49 A5IY65 A5IY67 A5IY71 A5IY97 A5IYA9
A5IYB2 A5IYF2 A5IYF5 A5IYG2 A5IYH4 A5IYK4 A5IYK5 A5IYK6 A5IYP4 A5IYR5 A5IYR6
A5IYT8 A5IYU8 A5IYV3 A5IYV8 A5IYW1 A5IYW8 A5IYW9 A5IYX2 A5IYX3 A5IYX4 A5IYX5
A5IYX6 A5IYX7 A5IYX8 A5IYX9 A5IYY0 A5IYY1 A5IYY2 A5IYY3 A5IYY6 A5IYZ5 A5IYZ6
A5IZ19 A5IZ33 A5IZ34 A5IZ35 A5IZ52 A5IZ59 A5IZ66 A5IZ67 A5IZ68 A5IZ80 A5IZA0
A5IZB9 A5IZC6 A5IZD7 A5IZF7 A5IZG5 A5IZG6 A5IZH0 A5IZH1 A5IZH6 A5IZI0 A5IZI8
A5J1W9 A5J277 A5JSS4 A5JST6 A5JSW9 A5JTM5 A5JTM6 A5JV83 A5JYS0 A5JYW9 A5JZ19
A5JZQ2 A5K2F3 A5K3F9 A5K464 A5K5K4 A5K6L0 A5K6W4 A5K7J8 A5K9M4 A5K9S0 A5KAL1
A5KB67 A5KE01 A5KE24 A5KUH4 A5LFW4 A5LFW8 A5LFY3 A5LHX3 A5MZ33 A5MZ34 A5MZ48
A5MZ75 A5MZ76 A5MZ82 A5MZJ0 A5MZP4 A5MZX7 A5MZY0 A5MZZ9 A5N0Q0 A5N0Q2 A5N0Q4
A5N0V1 A5N0Z3 A5N106 A5N127 A5N1C9 A5N1D3 A5N1Y8 A5N1Y9 A5N1Z2 A5N1Z4 A5N204
A5N205 A5N206 A5N217 A5N219 A5N245 A5N254 A5N256 A5N257 A5N258 A5N2K5 A5N2K7
A5N2K8 A5N2K9 A5N2L5 A5N2L8 A5N2N2 A5N2N6 A5N2N7 A5N2N8 A5N2R5 A5N2U5 A5N2V9
A5N2W5 A5N2X7 A5N2Y5 A5N2Y9 A5N2Z2 A5N359 A5N360 A5N361 A5N376 A5N399 A5N3J1
A5N3J7 A5N3K1 A5N3K4 A5N3L3 A5N3P1 A5N3R3 A5N3V4 A5N3V7 A5N3V8 A5N434 A5N438
A5N440 A5N449 A5N450 A5N451 A5N455 A5N456 A5N457 A5N460 A5N469 A5N4D3 A5N4F7
A5N4H0 A5N4H7 A5N4I2 A5N4I4 A5N4J0 A5N4K7 A5N4L0 A5N4L1 A5N4L2 A5N4L3 A5N4L9
A5N4M5 A5N4M6 A5N4N1 A5N4N2 A5N4N5 A5N4N6 A5N4N8 A5N4N9 A5N4P0 A5N4P2 A5N4P3
A5N4P4 A5N4P6 A5N4P7 A5N4P8 A5N4P9 A5N4Q0 A5N4Q1 A5N4Q2 A5N4Q3 A5N4Q4 A5N4Q5
A5N4Q6 A5N4Q7 A5N4Q8 A5N4Q9 A5N4R0 A5N4R1 A5N4R2 A5N4R3 A5N4R5 A5N4R6 A5N4R8
A5N4S0 A5N4S1 A5N4S2 A5N4S3 A5N4S5 A5N4S6 A5N4T0 A5N4T1 A5N4T2 A5N4Y9 A5N505
A5N597 A5N5B3 A5N5B5 A5N5B9 A5N5C6 A5N5D6 A5N5D7 A5N5D9 A5N5W0 A5N5W4 A5N5Y0
A5N5Y1 A5N628 A5N643 A5N646 A5N670 A5N680 A5N6A8 A5N6A9 A5N6D9 A5N6F5 A5N6H8
A5N6I0 A5N6I4 A5N6I5 A5N6J0 A5N6J2 A5N6J9 A5N6K1 A5N6K3 A5N6L4 A5N6M2 A5N6M3
A5N6M4 A5N6M8 A5N6N3 A5N6N7 A5N6Q7 A5N6S3 A5N6T4 A5N6U2 A5N6U3 A5N6V8 A5N6V9
A5N6W8 A5N6Y3 A5N749 A5N7A5 A5N7G2 A5N7H7 A5N7I8 A5N7I9 A5N7J0 A5N7J2 A5N7J4
A5N7J5 A5N7K3 A5N7N7 A5N7N8 A5N7N9 A5N7P0 A5N7P1 A5N7P5 A5N7Q4 A5N7Q5 A5N7Q7
A5N7Q8 A5N7R0 A5N7T5 A5N7T6 A5N7T7 A5N7T8 A5N7V5 A5N7W7 A5N7X5 A5N7X7 A5N7Y7
A5N7Z1 A5N7Z7 A5N800 A5N802 A5N807 A5N813 A5N815 A5N816 A5N817 A5N820 A5N821
A5N827 A5N828 A5N829 A5N831 A5N834 A5N838 A5N842 A5N843 A5N845 A5N847 A5N848
A5N854 A5N867 A5N8A0 A5N8F9 A5N8G4 A5N8G6 A5N8H9 A5N8I5 A5N8I7 A5N8I8 A5N8M0
A5N8N4 A5N8P8 A5N8V4 A5N934 A5N937 A5N960 A5PF62 A5PHD6 A5PJA1 A5PJC4 A5PJD0
A5PJD8 A5PJF6 A5PJI7 A5PJJ5 A5PJK7 A5PJM5 A5PJM7 A5PJN1 A5PJN8 A5PJP6 A5PJR4
A5PJU6 A5PJU8 A5PJU9 A5PJV3 A5PJV8 A5PJX4 A5PK19 A5PK21 A5PK23 A5PK29 A5PK30
A5PK39 A5PK42 A5PK51 A5PK61 A5PK63 A5PK65 A5PK71 A5PK74 A5PKB7 A5PKC7 A5PKE4
A5PKF5 A5PKG6 A5PKG7 A5PKG8 A5PKK7 A5PKK9 A5PKL1 A5PKL6 A5PKN5 A5PKP9 A5PKR8
A5PLC8 A5PLI4 A5PLK2 A5PLL1 A5PLN7 A5PLN9 A5PMF7 A5PMU4 A5PN52 A5TY69 A5TY71
A5TY72 A5TY73 A5TY79 A5TY80 A5TYA9 A5TYB2 A5TYC4 A5TYC6 A5TYC7 A5TYI7 A5TYK2
A5TYL2 A5TYM0 A5TYM1 A5TYT3 A5TYT6 A5TYU9 A5TYV9 A5TYY1 A5TZ08 A5TZ21 A5TZ46
A5TZ48 A5TZ77 A5TZ78 A5TZ84 A5TZA9 A5TZE0 A5TZE3 A5TZE9 A5TZF5 A5TZF9 A5TZK3
A5TZK6 A5TZL0 A5TZL4 A5TZL7 A5TZN9 A5TZP0 A5TZQ4 A5TZT3 A5TZT5 A5TZT8 A5TZU0
A5TZU6 A5U003 A5U019 A5U025 A5U026 A5U036 A5U052 A5U053 A5U056 A5U068 A5U069

A5U070 A5U071 A5U086 A5U087 A5U088 A5U089 A5U090 A5U091 A5U092 A5U093 A5U094
A5U095 A5U0A0 A5U0A1 A5U0A2 A5U0A3 A5U0A4 A5U0A5 A5U0A6 A5U0A7 A5U0A8 A5U0A9
A5U0B1 A5U0B2 A5U0B7 A5U0B9 A5U0G8 A5U0I9 A5U0J2 A5U0K2 A5U0K8 A5U0L8 A5U0P1
A5U0T2 A5U0Y6 A5U0Z1 A5U0Z7 A5U121 A5U144 A5U153 A5U154 A5U157 A5U1C6 A5U1D9
A5U1E6 A5U1F4 A5U1F5 A5U1I0 A5U1K8 A5U1L9 A5U1R1 A5U1R8 A5U1T8 A5U1U6 A5U1Y3
A5U1Z1 A5U1Z7 A5U1Z8 A5U226 A5U231 A5U238 A5U240 A5U275 A5U281 A5U282 A5U287
A5U292 A5U294 A5U2A8 A5U2B7 A5U2C2 A5U2C3 A5U2C5 A5U2D9 A5U2E0 A5U2F1 A5U2I9
A5U2S6 A5U2S7 A5U2U5 A5U2V0 A5U2V6 A5U2V7 A5U2V9 A5U2W1 A5U2W2 A5U2W7 A5U2W9
A5U2Z9 A5U300 A5U306 A5U309 A5U311 A5U313 A5U314 A5U315 A5U316 A5U346 A5U357
A5U371 A5U372 A5U388 A5U390 A5U3A5 A5U3J3 A5U3J9 A5U3K4 A5U3L5 A5U3L6 A5U3L7
A5U3L8 A5U3L9 A5U3R5 A5U3R6 A5U3S0 A5U3S7 A5U406 A5U480 A5U481 A5U482 A5U4C3
A5U4D5 A5U4D6 A5U4E1 A5U4E7 A5U4E8 A5U4F6 A5U4H4 A5U4H9 A5U4I2 A5U4J2 A5U4J3
A5U4M0 A5U4N0 A5U4N5 A5U4P1 A5U4P5 A5U4P6 A5U4Y2 A5U533 A5U555 A5U560 A5U567
A5U586 A5U5A1 A5U5A7 A5U5B6 A5U5C2 A5U5C5 A5U5D5 A5U5D6 A5U5D7 A5U5E1 A5U5E9
A5U5F3 A5U5F8 A5U5K8 A5U5N2 A5U5N3 A5U5N6 A5U5N8 A5U5N9 A5U5Q4 A5U5Q5 A5U5S9
A5U5T0 A5U5T4 A5U5U2 A5U5U3 A5U5U4 A5U5V4 A5U5V5 A5U5V7 A5U5V8 A5U5W5 A5U630
A5U634 A5U649 A5U654 A5U665 A5U669 A5U670 A5U678 A5U679 A5U683 A5U685 A5U688
A5U6A6 A5U6A7 A5U6B7 A5U6C6 A5U6D6 A5U6D8 A5U6E6 A5U6J0 A5U6J1 A5U6J4 A5U6K4
A5U6M2 A5U6M4 A5U6N5 A5U6N7 A5U6N8 A5U6P4 A5U6P9 A5U6Q3 A5U6Q7 A5U6Q9 A5U6R1
A5U6R2 A5U6R4 A5U6S3 A5U6S8 A5U6T0 A5U6W5 A5U6W6 A5U6X0 A5U6X4 A5U6Y1 A5U6Y7
A5U6Z2 A5U6Z4 A5U6Z5 A5U6Z8 A5U6Z9 A5U703 A5U706 A5U713 A5U721 A5U723 A5U724
A5U726 A5U737 A5U766 A5U776 A5U7B5 A5U7L9 A5U7Q1 A5U7S2 A5U7T5 A5U7T6 A5U7V6
A5U7Y3 A5U7Y8 A5U815 A5U830 A5U846 A5U871 A5U874 A5U892 A5U893 A5U894 A5U8B7
A5U8B8 A5U8B9 A5U8D2 A5U8D3 A5U8D4 A5U8D5 A5U8D6 A5U8D7 A5U8D8 A5U8G5 A5U8K8
A5U8K9 A5U8Q4 A5U8Q5 A5U8Q6 A5U8Q7 A5U8R1 A5U8S4 A5U8S5 A5U8S6 A5U8S7 A5U8T4
A5U908 A5U920 A5U941 A5U942 A5U994 A5U9A1 A5U9B6 A5U9G3 A5U9G7 A5U9I2 A5U9P7
A5U9Q2 A5U9Q7 A5U9Q8 A5U9Q9 A5U9R0 A5U9R1 A5U9R7 A5U9S9 A5U9U1 A5U9U3 A5U9U6
A5U9U8 A5U9U9 A5U9V0 A5U9V1 A5U9V2 A5U9V3 A5U9V4 A5U9V5 A5U9V6 A5U9V7 A5U9V8
A5U9W0 A5U9W4 A5U9X0 A5U9X5 A5U9X6 A5U9X7 A5U9X8 A5U9X9 A5U9Y3 A5U9Z0 A5U9Z5
A5U9Z6 A5UA03 A5UA15 A5UA16 A5UA18 A5UA19 A5UA21 A5UA25 A5UA30 A5UA33 A5UA46
A5UA47 A5UA56 A5UA57 A5UA66 A5UA71 A5UA73 A5UA77 A5UA83 A5UA84 A5UA85 A5UA93
A5UAA8 A5UAB3 A5UAB7 A5UAB9 A5UAC2 A5UAD6 A5UAE5 A5UAF5 A5UAF9 A5UAG1 A5UAG5
A5UAG7 A5UAH5 A5UAH6 A5UAH7 A5UAH8 A5UAI4 A5UAI5 A5UAJ6 A5UAK7 A5UAL2 A5UAL5
A5UAL6 A5UAL7 A5UAN5 A5UAP2 A5UAP3 A5UAQ6 A5UAR3 A5UAS4 A5UAS6 A5UAS8 A5UAT0
A5UAU5 A5UAU8 A5UAU9 A5UAV1 A5UAV2 A5UAV3 A5UAV4 A5UAV8 A5UAX1 A5UAZ2 A5UB07
A5UB10 A5UB16 A5UB17 A5UB28 A5UB32 A5UB42 A5UB44 A5UB54 A5UB70 A5UB71 A5UB83
A5UB85 A5UBA5 A5UBB0 A5UBB1 A5UBB2 A5UBB4 A5UBB9 A5UBC0 A5UBC1 A5UBC4 A5UBC5
A5UBC8 A5UBD4 A5UBD9 A5UBE2 A5UBE8 A5UBF2 A5UBF3 A5UBF4 A5UBF7 A5UBF9 A5UBG1
A5UBG2 A5UBH1 A5UBH4 A5UBK4 A5UBK5 A5UBM3 A5UBN1 A5UBN2 A5UBN4 A5UBN7 A5UBP1
A5UBP3 A5UBP8 A5UBQ1 A5UBQ2 A5UBS8 A5UBT4 A5UBT6 A5UBU1 A5UBU2 A5UBU4 A5UBV1
A5UBV7 A5UBV9 A5UBW0 A5UBW4 A5UBW8 A5UBX3 A5UBX5 A5UBX9 A5UBY0 A5UBZ6 A5UBZ9
A5UC01 A5UC39 A5UC40 A5UC41 A5UC48 A5UC49 A5UC51 A5UC54 A5UC55 A5UC58 A5UC59
A5UC76 A5UC84 A5UC89 A5UC92 A5UCA8 A5UCB0 A5UCB3 A5UCB7 A5UCB8 A5UCB9 A5UCC3
A5UCC6 A5UCC9 A5UCE4 A5UCG6 A5UCH1 A5UCH8 A5UCJ5 A5UCK0 A5UCK4 A5UCM3 A5UCM6

A5UCM7 A5UCM9 A5UCN0 A5UCP1 A5UCP4 A5UCQ0 A5UCQ1 A5UCQ2 A5UCQ6 A5UCR4 A5UCR5
A5UCR9 A5UCS8 A5UCT6 A5UCU0 A5UCU6 A5UCV6 A5UCW1 A5UCW6 A5UCW7 A5UCX0 A5UCX3
A5UCX6 A5UCX7 A5UCY0 A5UCY6 A5UCY8 A5UCZ3 A5UD04 A5UD24 A5UD28 A5UD40 A5UD45
A5UD62 A5UD65 A5UD69 A5UD71 A5UD74 A5UD82 A5UD83 A5UD85 A5UD87 A5UD89 A5UD90
A5UD91 A5UD93 A5UDA0 A5UDA4 A5UDA6 A5UDA9 A5UDB0 A5UDB2 A5UDB3 A5UDB4 A5UDB6
A5UDB7 A5UDB8 A5UDC3 A5UDC6 A5UDE0 A5UDE3 A5UDF4 A5UDG1 A5UDG2 A5UDH0 A5UDH3
A5UDH4 A5UDI1 A5UDI2 A5UDI4 A5UDJ4 A5UDJ5 A5UDJ7 A5UDJ8 A5UDL7 A5UDM5 A5UDM6
A5UDN2 A5UDN3 A5UDN8 A5UDP4 A5UDQ5 A5UDQ7 A5UDR0 A5UDR1 A5UDR3 A5UDR4 A5UDR5
A5UDR7 A5UDS1 A5UDS2 A5UDS3 A5UDS4 A5UDS5 A5UDS6 A5UDS8 A5UDS9 A5UDT0 A5UDT1
A5UDT2 A5UDT3 A5UDT4 A5UDT5 A5UDT6 A5UDT7 A5UDT8 A5UDT9 A5UDU0 A5UDU1 A5UDU2
A5UDU3 A5UDU4 A5UDU5 A5UDU6 A5UDU7 A5UDW2 A5UDW5 A5UDW7 A5UDW8 A5UDX5 A5UDY2
A5UDY7 A5UDZ4 A5UDZ6 A5UE01 A5UE04 A5UE11 A5UE12 A5UE18 A5UE20 A5UE24 A5UE25
A5UE34 A5UE38 A5UE39 A5UE41 A5UE44 A5UE59 A5UE60 A5UE61 A5UE63 A5UE77 A5UE78
A5UE83 A5UE95 A5UE96 A5UEB3 A5UEB4 A5UEC1 A5UEC2 A5UED2 A5UEH1 A5UEH4 A5UEH8
A5UEI5 A5UEI7 A5UEI8 A5UEJ1 A5UEJ2 A5UEJ5 A5UEK0 A5UEK1 A5UEK6 A5UEK8 A5UEN6
A5UEN9 A5UEU3 A5UEU5 A5UEU6 A5UEV0 A5UEV3 A5UEV4 A5UEV6 A5UEV9 A5UEW0 A5UEW3
A5UEW4 A5UEW7 A5UEY4 A5UEY9 A5UEZ1 A5UF06 A5UF10 A5UF14 A5UF15 A5UF16 A5UF19
A5UF26 A5UF28 A5UF34 A5UF36 A5UF42 A5UF43 A5UF66 A5UF67 A5UF68 A5UF70 A5UF74
A5UF76 A5UF83 A5UF86 A5UF87 A5UF93 A5UFB3 A5UFB4 A5UFD9 A5UFE2 A5UFF1 A5UFF2
A5UFF5 A5UFF7 A5UFF8 A5UFG0 A5UFG3 A5UFG5 A5UFH0 A5UFH3 A5UFH8 A5UFI3 A5UFI6
A5UFI8 A5UFJ0 A5UFJ1 A5UFJ6 A5UFJ8 A5UFJ9 A5UFK0 A5UFK3 A5UFL7 A5UFM4 A5UFN4
A5UFP3 A5UFQ1 A5UFQ7 A5UFQ9 A5UFR9 A5UFS2 A5UFT8 A5UFT9 A5UFU3 A5UFU9 A5UFW3
A5UFW8 A5UFX6 A5UFZ9 A5UG02 A5UG03 A5UG04 A5UG05 A5UG06 A5UG09 A5UG10 A5UG14
A5UG27 A5UG34 A5UG47 A5UG48 A5UG62 A5UG90 A5UG91 A5UG92 A5UG95 A5UGA0 A5UGC3
A5UGC4 A5UGC9 A5UGD0 A5UGD1 A5UGD3 A5UGD5 A5UGD7 A5UGE0 A5UGE3 A5UGF0 A5UGF8
A5UGH1 A5UGH2 A5UGH6 A5UGI1 A5UGR0 A5UGR1 A5UGR9 A5UGS0 A5UGS5 A5UGS9 A5UGT0
A5UGT6 A5UGU3 A5UGU4 A5UGV2 A5UGV3 A5UGW7 A5UGX0 A5UGX5 A5UGY0 A5UGY2 A5UGY3
A5UGY5 A5UGZ7 A5UH05 A5UH08 A5UH15 A5UH19 A5UH20 A5UH21 A5UH22 A5UH23 A5UH30
A5UH36 A5UH41 A5UH43 A5UH44 A5UH45 A5UH46 A5UH47 A5UH48 A5UH49 A5UH50 A5UH52
A5UH53 A5UH57 A5UH58 A5UH70 A5UH79 A5UH91 A5UH92 A5UH99 A5UHA0 A5UHA3 A5UHB4
A5UHB8 A5UHB9 A5UHC0 A5UHC1 A5UHD1 A5UHD2 A5UHE3 A5UHE8 A5UHE9 A5UHF7 A5UHF9
A5UHG0 A5UHG1 A5UHG7 A5UHG8 A5UHH1 A5UHH7 A5UHI7 A5UHJ5 A5UHJ6 A5UHK3 A5UHL4
A5UHL6 A5UHM6 A5UHN1 A5UHN3 A5UHP2 A5UHP8 A5UHQ4 A5UHR1 A5UHR2 A5UHR4 A5UHR8
A5UHS9 A5UHT0 A5UHT1 A5UHT2 A5UHT3 A5UHT4 A5UHT5 A5UHT6 A5UHT7 A5UHT8 A5UHT9
A5UHU1 A5UHU2 A5UHU3 A5UHU4 A5UHU5 A5UHU6 A5UHU7 A5UHU8 A5UHU9 A5UHV0 A5UHV2
A5UHV4 A5UHV5 A5UHV6 A5UHV9 A5UHW0 A5UHW1 A5UHW2 A5UHW3 A5UHW5 A5UHW6 A5UHW8
A5UHX0 A5UHY0 A5UHY5 A5UHY8 A5UHY9 A5UHZ6 A5UI04 A5UI22 A5UI23 A5UI24 A5UI25
A5UI33 A5UI35 A5UI36 A5UI44 A5UI45 A5UI47 A5UI55 A5UI56 A5UI62 A5UI65 A5UI68
A5UI87 A5UI92 A5UI93 A5UI95 A5UI96 A5UI99 A5UIA3 A5UIB2 A5UIB7 A5UIB9 A5UIC0
A5UIC3 A5UIC6 A5UID2 A5UID3 A5UID5 A5UID9 A5UIF3 A5UIG1 A5UIG4 A5UII7 A5UIJ1
A5UIK2 A5UIK7 A5UIM6 A5UIN7 A5UIN9 A5UIP2 A5UIP9 A5UIQ2 A5UIQ3 A5UIQ7 A5UIR0
A5UIR3 A5UIR4 A5UIS4 A5UIT6 A5UIU0 A5UIU8 A5UIV9 A5UIW0 A5UIW8 A5UIX2 A5UIX3
A5UIX4 A5UIX8 A5UIY2 A5UIY9 A5UIZ0 A5UIZ2 A5UIZ5 A5UJ18 A5UJ23 A5UJ35 A5UJ40
A5UJ44 A5UJ82 A5UJ98 A5UJA2 A5UJB5 A5UJB8 A5UJD5 A5UJF2 A5UJF3 A5UJL1 A5UJM1

A5UJM2 A5UJM8 A5UJM9 A5UJN0 A5UJN1 A5UJN3 A5UJQ8 A5UJR5 A5UJS0 A5UJS2 A5UJS3
A5UJV0 A5UJW2 A5UJW4 A5UJW5 A5UK11 A5UK12 A5UK20 A5UK38 A5UK48 A5UK52 A5UK79
A5UK92 A5UKA1 A5UKA4 A5UKD0 A5UKG5 A5UKI3 A5UKI5 A5UKI8 A5UKJ0 A5UKJ7 A5UKS9
A5UKT8 A5UKU4 A5UKU6 A5UKU8 A5UKU9 A5UKV0 A5UKV8 A5UKW7 A5UKX2 A5UKY0 A5UKY3
A5UKY4 A5UKY8 A5UL29 A5UL31 A5UL33 A5UL35 A5UL38 A5UL47 A5UL52 A5UL61 A5UL62
A5UL66 A5UL67 A5UL68 A5UL69 A5UL72 A5UL73 A5UL75 A5UL76 A5UL77 A5UL78 A5UL80
A5UL82 A5UL83 A5UL84 A5UL85 A5UL86 A5UL87 A5UL88 A5UL89 A5ULA6 A5ULF7 A5ULF8
A5ULG0 A5ULG2 A5ULG4 A5ULI2 A5ULI6 A5ULI7 A5ULI9 A5ULK1 A5ULK4 A5ULL0 A5ULL8
A5ULM4 A5ULM5 A5ULM6 A5ULM8 A5ULN4 A5ULP3 A5ULP4 A5ULP6 A5ULQ9 A5ULR3 A5ULS3
A5ULT9 A5ULU4 A5ULV0 A5ULV3 A5ULV6 A5ULW4 A5UM08 A5UM10 A5UM12 A5UM14 A5UM16
A5UM67 A5UM78 A5UM80 A5UM85 A5UMA9 A5UMB0 A5UMB4 A5UMC1 A5UMF3 A5UMF4 A5UMF6
A5UMF9 A5UMH1 A5UMH5 A5UMI1 A5UMI3 A5UMJ1 A5UMJ2 A5UMJ5 A5UMJ9 A5UMK3 A5UMK4
A5UMK8 A5UML0 A5UML4 A5UMN0 A5UMN4 A5UMN6 A5UMP0 A5UMS3 A5UMW0 A5UMW4 A5UMW7
A5UMW9 A5UMZ1 A5UN04 A5UN52 A5UN53 A5UN54 A5UN55 A5UN64 A5UN65 A5UN79 A5UN82
A5UN95 A5UNA1 A5UNB3 A5UNH4 A5UNI4 A5UNI8 A5UNK4 A5UNK6 A5UNQ5 A5UNQ7 A5UNR1
A5UP00 A5UP91 A5UPB5 A5UPG5 A5UPG6 A5UPS2 A5UPS3 A5UPS5 A5UPV2 A5UPW5 A5UPY6
A5UQ40 A5UQB7 A5UQF4 A5UQK6 A5UQL0 A5UQQ1 A5UQQ2 A5UQR2 A5UQS8 A5UQV0 A5UQV9
A5UQW9 A5URA4 A5URA7 A5URA8 A5URC6 A5URD0 A5URD5 A5URF5 A5URM3 A5URV4 A5URV5
A5URX3 A5URZ1 A5URZ5 A5US07 A5US14 A5US40 A5US42 A5US48 A5US49 A5US58 A5US83
A5USB5 A5USC4 A5USE3 A5USE7 A5USF9 A5USG0 A5USG2 A5USG3 A5USG4 A5USG5 A5USG6
A5USG7 A5USH0 A5USH1 A5USH2 A5USH3 A5USH4 A5USH5 A5USH6 A5USH7 A5USH8 A5USH9
A5USI0 A5USI1 A5USI2 A5USI3 A5USI4 A5USI5 A5USI6 A5USI7 A5USI9 A5USJ0 A5USJ1
A5USJ2 A5USJ3 A5USJ4 A5USK5 A5USQ4 A5USR6 A5USR7 A5UST8 A5USU0 A5USV2 A5USV3
A5UT22 A5UT23 A5UT95 A5UTB6 A5UTG1 A5UTH3 A5UTH4 A5UTJ6 A5UTY2 A5UU10 A5UU33
A5UU35 A5UU37 A5UU40 A5UU55 A5UU62 A5UUG7 A5UUJ4 A5UUJ5 A5UUJ6 A5UUL2 A5UUL3
A5UUN1 A5UUN7 A5UUP4 A5UUP5 A5UUR3 A5UUU9 A5UUV0 A5UUV2 A5UUW9 A5UUZ9 A5UV64
A5UVB2 A5UVB3 A5UVE3 A5UVE4 A5UVV6 A5UVX7 A5UVZ9 A5UW19 A5UWC5 A5UWE5 A5UWE6
A5UWI3 A5UWI8 A5UWR6 A5UWR7 A5UWX7 A5UWZ9 A5UX01 A5UX78 A5UX79 A5UXC5 A5UXF9
A5UXG2 A5UXT6 A5UXX5 A5UXX8 A5UY13 A5UY19 A5UY26 A5UY50 A5UY52 A5UY81 A5UY83
A5UY90 A5UY93 A5UY94 A5UYA1 A5UYI1 A5UYI4 A5UYI6 A5UYJ2 A5UYQ5 A5UYR6 A5UYU9
A5UYW4 A5UYW7 A5UYX1 A5UZ80 A5UZ98 A5UZF9 A5UZG3 A5UZK7 A5UZQ2 A5UZQ3 A5UZS3
A5UZS8 A5UZS9 A5UZT4 A5UZT9 A5UZV5 A5UZW2 A5UZW4 A5UZX0 A5V022 A5V070 A5V080
A5V094 A5V0B3 A5V0J4 A5V0J9 A5V0P4 A5V0S2 A5V1D8 A5V1E4 A5V1E5 A5V1H2 A5V1U3
A5V1W3 A5V230 A5V243 A5V280 A5V282 A5V287 A5V2D8 A5V2E0 A5V2E1 A5V2F8 A5V2S4
A5V2U6 A5V2U9 A5V3A1 A5V3E1 A5V3G2 A5V3G3 A5V3G4 A5V3G5 A5V3H3 A5V3I6 A5V3J3
A5V3L0 A5V3L8 A5V3L9 A5V3M9 A5V3R9 A5V3S4 A5V3S9 A5V3T8 A5V3U4 A5V3U6 A5V3U9
A5V3V0 A5V3V1 A5V3V3 A5V3V4 A5V3W0 A5V3W3 A5V3Y9 A5V3Z0 A5V3Z4 A5V4F9 A5V4L0
A5V4M8 A5V4U1 A5V592 A5V5D1 A5V5D2 A5V5E0 A5V5P9 A5V5Q2 A5V5Q4 A5V5U9 A5V5X4
A5V5X5 A5V5X6 A5V5Y3 A5V5Y4 A5V5Y5 A5V5Y6 A5V5Y7 A5V5Y8 A5V5Y9 A5V5Z0 A5V5Z1
A5V5Z2 A5V5Z3 A5V5Z4 A5V5Z5 A5V5Z6 A5V5Z7 A5V5Z8 A5V5Z9 A5V601 A5V602 A5V604
A5V605 A5V606 A5V607 A5V625 A5V626 A5V649 A5V652 A5V656 A5V667 A5V6A9 A5V6F9
A5V6G8 A5V6T7 A5V6T8 A5V6Z0 A5V7I1 A5V807 A5V856 A5V881 A5V883 A5V884 A5V885
A5V8R9 A5V8T0 A5V8U0 A5V8U1 A5V8Y0 A5V8Y2 A5V8Y6 A5V922 A5V924 A5V925 A5V932
A5V939 A5V9C3 A5V9D8 A5V9E5 A5V9F5 A5V9F6 A5V9F7 A5V9J6 A5V9P7 A5V9P8 A5V9Q2

A5V9Q3 A5V9R3 A5V9T6 A5V9V7 A5V9V9 A5V9W0 A5V9W3 A5V9W7 A5VA00 A5VA16 A5VA19
A5VA26 A5VA34 A5VA37 A5VA47 A5VA56 A5VA69 A5VA70 A5VA82 A5VA83 A5VA84 A5VA89
A5VAA0 A5VAA8 A5VAC6 A5VAE6 A5VAG4 A5VAJ8 A5VAK5 A5VAL5 A5VAN5 A5VAZ8 A5VB15
A5VB78 A5VB82 A5VB94 A5VBA0 A5VBQ3 A5VBQ5 A5VBQ6 A5VBZ7 A5VBZ8 A5VC06 A5VC07
A5VC58 A5VC84 A5VC87 A5VCB6 A5VCE4 A5VCF0 A5VCW6 A5VCY8 A5VCY9 A5VCZ4 A5VCZ5
A5VCZ9 A5VD08 A5VD53 A5VD70 A5VD71 A5VD74 A5VDC2 A5VDC3 A5VDC5 A5VDD4 A5VDL9
A5VDM0 A5VDM1 A5VDM3 A5VDM4 A5VDM8 A5VDT3 A5VDT7 A5VE44 A5VEJ8 A5VEM2 A5VEW5
A5VF37 A5VF46 A5VFA2 A5VFA5 A5VFF3 A5VFI9 A5VFJ8 A5VFK5 A5VFM2 A5VFM5 A5VFP1
A5VFP2 A5VFX8 A5VG00 A5VG06 A5VG07 A5VGU4 A5VGU5 A5VHF3 A5VHF6 A5VHF9 A5VHG1
A5VHG3 A5VHH9 A5VHK1 A5VHK2 A5VHM0 A5VHN6 A5VHP5 A5VHQ2 A5VHR0 A5VHR2 A5VHS1
A5VHS3 A5VHS9 A5VHT7 A5VHT8 A5VHU0 A5VHU2 A5VHU6 A5VI09 A5VI11 A5VI25 A5VI29
A5VI51 A5VI57 A5VI81 A5VI94 A5VI99 A5VIA4 A5VIA9 A5VIB0 A5VIC4 A5VIC6 A5VID8
A5VIE0 A5VIE7 A5VIE8 A5VIE9 A5VIG4 A5VII1 A5VII3 A5VII4 A5VIJ0 A5VIJ2 A5VIJ8
A5VIK0 A5VIK4 A5VIL5 A5VIM0 A5VIN9 A5VIP9 A5VIQ0 A5VIS1 A5VIT7 A5VIU4 A5VIU7
A5VIW6 A5VIW7 A5VIW9 A5VIX0 A5VIX1 A5VIX2 A5VIX8 A5VIX9 A5VIY0 A5VIY5 A5VIZ6
A5VJ12 A5VJ17 A5VJ27 A5VJ28 A5VJ32 A5VJ37 A5VJ48 A5VJ56 A5VJ57 A5VJ58 A5VJ69
A5VJ71 A5VJ82 A5VJ88 A5VJ89 A5VJ92 A5VJ93 A5VJ94 A5VJ95 A5VJ98 A5VJ99 A5VJA0
A5VJC3 A5VJC5 A5VJC6 A5VJC7 A5VJC8 A5VJD4 A5VJD5 A5VJD6 A5VJE0 A5VJE1 A5VJE2
A5VJE5 A5VJE6 A5VJE7 A5VJE8 A5VJF8 A5VJG2 A5VJG4 A5VJG8 A5VJH1 A5VJH2 A5VJH6
A5VJH9 A5VJI1 A5VJK2 A5VJK4 A5VJK7 A5VJK9 A5VJL5 A5VJL7 A5VJX0 A5VK00 A5VK09
A5VK10 A5VK11 A5VK28 A5VK99 A5VKA1 A5VKB2 A5VKE8 A5VKG0 A5VKN2 A5VKN4 A5VKN5
A5VKN7 A5VKP2 A5VKP8 A5VKQ1 A5VKQ8 A5VKR5 A5VKR6 A5VKR7 A5VKR8 A5VKS3 A5VKS5
A5VKT1 A5VKT4 A5VKT6 A5VKU3 A5VKU8 A5VKV1 A5VKW2 A5VKW4 A5VKX2 A5VKX3 A5VKX7
A5VKY3 A5VKZ3 A5VL01 A5VL03 A5VL28 A5VL29 A5VL77 A5VLA3 A5VLG2 A5VLG3 A5VLG6
A5VLG9 A5VLH3 A5VLH4 A5VLH8 A5VLH9 A5VLI0 A5VLI1 A5VLI2 A5VLI3 A5VLI6 A5VLI7
A5VLI8 A5VLI9 A5VLJ0 A5VLJ1 A5VLJ2 A5VLJ3 A5VLJ4 A5VLJ5 A5VLJ7 A5VLJ8 A5VLJ9
A5VLK0 A5VLK1 A5VLK2 A5VLK3 A5VLK4 A5VLK5 A5VLK6 A5VLK8 A5VLK9 A5VLL0 A5VLL3
A5VLL4 A5VLS3 A5VLU9 A5VLY3 A5VLY5 A5VM22 A5VM65 A5VM67 A5VM68 A5VM70 A5VM87
A5VMA5 A5VMB3 A5VME2 A5VML0 A5VMP8 A5VMP9 A5VMU5 A5VMV1 A5VMV2 A5VMX5 A5VMY2
A5VMY3 A5VN19 A5VN22 A5VN26 A5VN43 A5VN81 A5VN83 A5VN89 A5VNA6 A5VNA8 A5VNC5
A5VNE4 A5VNK0 A5VNK4 A5VNL0 A5VNL5 A5VNL6 A5VNM7 A5VNP2 A5VNS0 A5VNS7 A5VNV9
A5VNW6 A5VNZ1 A5VP02 A5VP09 A5VP24 A5VP26 A5VP46 A5VP47 A5VP50 A5VP58 A5VP72
A5VPA1 A5VPA5 A5VPA6 A5VPI5 A5VPI6 A5VPI8 A5VPI9 A5VPJ0 A5VPL3 A5VPL7 A5VPM3
A5VPM8 A5VPP4 A5VPT5 A5VPT9 A5VPU7 A5VPU8 A5VPV2 A5VPX1 A5VPX4 A5VPX5 A5VQ02
A5VQ09 A5VQ10 A5VQ15 A5VQ20 A5VQ53 A5VQ54 A5VQ63 A5VQ76 A5VQ80 A5VQ81 A5VQC8
A5VQC9 A5VQD7 A5VQF9 A5VQI8 A5VQL0 A5VQL9 A5VQM2 A5VQN3 A5VQN6 A5VQP4 A5VQP8
A5VQQ9 A5VQR4 A5VQR5 A5VQR6 A5VQR9 A5VQT0 A5VQT2 A5VQT3 A5VQV8 A5VQW7 A5VQX4
A5VQY1 A5VQY2 A5VQY3 A5VQY4 A5VQY5 A5VQY7 A5VQY8 A5VQY9 A5VQZ0 A5VQZ1 A5VQZ2
A5VQZ3 A5VQZ4 A5VQZ5 A5VQZ6 A5VQZ7 A5VQZ8 A5VQZ9 A5VR00 A5VR02 A5VR03 A5VR04
A5VR05 A5VR06 A5VR07 A5VR08 A5VR09 A5VR10 A5VR11 A5VR14 A5VR15 A5VR16 A5VR18
A5VR19 A5VR48 A5VR62 A5VR64 A5VRB1 A5VRB2 A5VRC9 A5VRD3 A5VRD9 A5VRE7 A5VRF8
A5VRG5 A5VRH5 A5VRH6 A5VRI5 A5VRJ7 A5VRM1 A5VRR6 A5VRR9 A5VRW1 A5VS09 A5VS40
A5VS47 A5VS49 A5VS58 A5VS59 A5VS60 A5VS65 A5VS71 A5VS72 A5VS98 A5VSA3 A5VSC1
A5VSD5 A5VSE7 A5VSE9 A5VSF3 A5VSF8 A5VSG4 A5VSH6 A5VSI0 A5VSI3 A5VSI4 A5VSI5

A5VSI8 A5VSK3 A5VSL5 A5VSL6 A5VSN3 A5VSN4 A5VSP1 A5VSP2 A5VSQ0 A5VSQ3 A5VSQ4
A5VSQ7 A5VSR1 A5VSU2 A5VSU9 A5VSV7 A5VSY2 A5VSZ3 A5VT18 A5VT19 A5VT20 A5VT24
A5VT25 A5VT27 A5VT30 A5VT37 A5VT38 A5VT39 A5VT42 A5VT43 A5VT44 A5VT45 A5VT52
A5VT61 A5VT72 A5VT73 A5VTA1 A5VTA9 A5VTB2 A5VTB3 A5VTB4 A5VTB5 A5VTB6 A5VTD0
A5VTJ7 A5VTK6 A5VTN1 A5VTQ7 A5VTT4 A5VTT5 A5VTT6 A5VTU1 A5VTU2 A5VTW1 A5VU41
A5VU45 A5VU71 A5VUC1 A5VUF9 A5VUG1 A5VUQ0 A5VUS6 A5VUS7 A5VUS8 A5VUS9 A5VUT1
A5VUU1 A5VUZ8 A5VV25 A5VV55 A5VV73 A5VV94 A5VVE8 A5VVG4 A5VVH9 A5VVI9 A5VVJ0
A5VVJ5 A5VVJ6 A5VVJ8 A5VVL2 A5VVM8 A5VVS7 A5VVT1 A5VVT4 A5VVT7 A5VVT9 A5VVU0
A5VVV1 A5VVV5 A5VVW9 A5VVX1 A5VVZ5 A5VWB8 A5VWC0 A5VWJ1 A5VWJ2 A5VWJ8 A5VWK1
A5VWK4 A5VWL2 A5VWL3 A5VWQ6 A5VWX3 A5VX17 A5VX27 A5VX31 A5VX32 A5VX45 A5VX68
A5VX72 A5VX75 A5VX76 A5VXF1 A5VXF2 A5VXG4 A5VXG6 A5VXG7 A5VXI3 A5VXI4 A5VXI8
A5VXJ0 A5VXJ2 A5VXJ3 A5VXJ4 A5VXL4 A5VXL5 A5VXM5 A5VXN1 A5VXN3 A5VXN6 A5VXN7
A5VXN9 A5VXP0 A5VXP1 A5VXP2 A5VXP3 A5VXP6 A5VXP7 A5VXP8 A5VXP9 A5VXQ0 A5VXQ1
A5VXQ2 A5VXQ3 A5VXQ4 A5VXQ5 A5VXQ6 A5VXQ7 A5VXQ8 A5VXQ9 A5VXR0 A5VXR1 A5VXR2
A5VXR3 A5VXR4 A5VXR5 A5VXR6 A5VXR8 A5VXR9 A5VXS0 A5VXS1 A5VXS2 A5VXS3 A5VXT7
A5VXV6 A5VXW0 A5VXW1 A5VXW5 A5VXW9 A5VXX1 A5VXY9 A5VY49 A5VY52 A5VY55 A5VY79
A5VY92 A5VYC4 A5VYC5 A5VYC6 A5VYC7 A5VYF8 A5VYF9 A5VYG1 A5VYG4 A5VYG5 A5VYG8
A5VYH6 A5VYH8 A5VYJ3 A5VYQ1 A5VYR4 A5VYS4 A5VYS7 A5VYT1 A5VYT2 A5VYT5 A5VYT6
A5VYT9 A5VYU4 A5VZ21 A5VZ22 A5VZ23 A5VZ32 A5VZ39 A5VZ54 A5VZ55 A5VZ57 A5VZ63
A5VZ65 A5VZ69 A5VZB6 A5VZC3 A5VZG7 A5VZH7 A5VZH8 A5VZI0 A5VZI8 A5VZJ1 A5VZK6
A5VZL6 A5VZN0 A5VZS9 A5VZT3 A5VZT5 A5VZT8 A5VZU3 A5VZU5 A5VZU6 A5VZU7 A5VZV6
A5VZW1 A5VZX1 A5VZY6 A5VZZ2 A5VZZ9 A5W002 A5W024 A5W059 A5W0C1 A5W0D5 A5W0D6
A5W0D8 A5W0D9 A5W0J4 A5W0S6 A5W0U0 A5W0U6 A5W0U7 A5W0U8 A5W0W6 A5W114 A5W167
A5W169 A5W1B2 A5W1E8 A5W1F2 A5W1G2 A5W1G7 A5W1G9 A5W1H0 A5W1H1 A5W1H6 A5W1H7
A5W241 A5W2C0 A5W2U1 A5W348 A5W384 A5W454 A5W458 A5W488 A5W4B2 A5W4B3 A5W4B5
A5W4B6 A5W4B8 A5W4B9 A5W4E5 A5W4E9 A5W4F0 A5W4F1 A5W4F2 A5W4F9 A5W4G5 A5W4Y8
A5W516 A5W5B3 A5W5D5 A5W5D7 A5W5D8 A5W5D9 A5W5E0 A5W5E1 A5W5E7 A5W5F5 A5W5G8
A5W5N3 A5W634 A5W635 A5W636 A5W638 A5W695 A5W696 A5W6E6 A5W6F5 A5W6I8 A5W6I9
A5W6J0 A5W6K7 A5W6M0 A5W6M3 A5W6U4 A5W6Y0 A5W6Z7 A5W709 A5W717 A5W727 A5W756
A5W759 A5W770 A5W794 A5W7B0 A5W7F5 A5W7G2 A5W7G3 A5W7H0 A5W7H7 A5W7I1 A5W7K1
A5W7K3 A5W7Q4 A5W7Q6 A5W7R7 A5W7U0 A5W7U2 A5W7V1 A5W7V2 A5W816 A5W820 A5W821
A5W822 A5W823 A5W827 A5W831 A5W836 A5W844 A5W847 A5W848 A5W849 A5W850 A5W852
A5W863 A5W873 A5W880 A5W881 A5W890 A5W8C0 A5W8C1 A5W8C2 A5W8C3 A5W8E4 A5W8E5
A5W8E9 A5W8F0 A5W8F2 A5W8H2 A5W8J7 A5W8M6 A5W8M7 A5W8N5 A5W8P9 A5W8Q8 A5W8Q9
A5W8R2 A5W8R3 A5W8S0 A5W8S1 A5W8T2 A5W8U6 A5W8Z1 A5W920 A5W926 A5W952 A5W965
A5W968 A5W974 A5W975 A5W976 A5W983 A5W984 A5W985 A5W986 A5W987 A5W989 A5W991
A5W992 A5W995 A5W9A1 A5W9A2 A5W9A3 A5W9A4 A5W9A9 A5W9G9 A5W9H0 A5W9H4 A5W9H9
A5W9I5 A5W9I6 A5W9I7 A5W9J3 A5W9J6 A5W9K3 A5W9K7 A5W9L9 A5W9N5 A5W9N6 A5W9Q6
A5W9Q7 A5W9R5 A5W9R7 A5W9R8 A5W9S5 A5W9S6 A5W9T0 A5W9T1 A5W9T8 A5W9T9 A5W9V8
A5W9W6 A5W9Z6 A5WA00 A5WA01 A5WA09 A5WA29 A5WA32 A5WA35 A5WA45 A5WA48 A5WA49
A5WA61 A5WA64 A5WA67 A5WA74 A5WA86 A5WA89 A5WA96 A5WA97 A5WAA1 A5WAA7 A5WAB1
A5WAB2 A5WAC2 A5WAD0 A5WAD6 A5WAD7 A5WAF7 A5WAF9 A5WAG2 A5WAK1 A5WAL1 A5WAL5
A5WAR6 A5WAU2 A5WAU7 A5WAY3 A5WAY8 A5WB00 A5WB02 A5WB04 A5WB06 A5WB07 A5WB10
A5WB18 A5WB73 A5WBB3 A5WBB4 A5WBB6 A5WBB9 A5WBC0 A5WBC6 A5WBF8 A5WBG1 A5WBH9

A5WBP5 A5WBQ0 A5WBQ1 A5WBR2 A5WBR3 A5WBS1 A5WBS3 A5WBS5 A5WBS6 A5WBT1 A5WBT2
A5WBU9 A5WBW8 A5WBW9 A5WBX0 A5WBX3 A5WBY3 A5WC33 A5WC53 A5WC54 A5WC55 A5WC58
A5WC69 A5WC71 A5WC75 A5WC94 A5WCA0 A5WCB3 A5WCC4 A5WCC7 A5WCG4 A5WCI2 A5WCI9
A5WCJ0 A5WCJ1 A5WCJ2 A5WCJ3 A5WCJ4 A5WCJ5 A5WCJ6 A5WCJ7 A5WCJ8 A5WCJ9 A5WCK0
A5WCK1 A5WCK2 A5WCK3 A5WCK4 A5WCK5 A5WCK6 A5WCK7 A5WCK8 A5WCK9 A5WCL1 A5WCL2
A5WCL3 A5WCL4 A5WCL5 A5WCL9 A5WCN7 A5WCQ0 A5WCQ5 A5WCR1 A5WCR5 A5WCS1 A5WCT9
A5WCU0 A5WCU8 A5WCX0 A5WCX2 A5WCX3 A5WCZ0 A5WCZ2 A5WD02 A5WD03 A5WD25 A5WD51
A5WD52 A5WD53 A5WD56 A5WDG2 A5WDG7 A5WDI5 A5WDJ2 A5WDJ3 A5WDK9 A5WDL8 A5WDN1
A5WDP0 A5WDP1 A5WDP3 A5WDR1 A5WDU0 A5WDV0 A5WDV1 A5WDV6 A5WDV8 A5WDY2 A5WDY8
A5WE06 A5WE20 A5WE50 A5WE72 A5WEG6 A5WEH0 A5WEW5 A5WEW6 A5WEW9 A5WFA2 A5WFC8
A5WFD0 A5WFD7 A5WFF6 A5WFG0 A5WFG1 A5WFM3 A5WFN9 A5WFR0 A5WFU8 A5WFU9 A5WFV1
A5WFW1 A5WFW3 A5WFX2 A5WFX4 A5WFY9 A5WFZ0 A5WFZ1 A5WFZ6 A5WG08 A5WG19 A5WG49
A5WG58 A5WG70 A5WG72 A5WG77 A5WG78 A5WG99 A5WGA8 A5WGB3 A5WGB5 A5WGC0 A5WGD0
A5WGE7 A5WGF0 A5WGF1 A5WGF2 A5WGF9 A5WGG4 A5WGH5 A5WGH7 A5WGH9 A5WGK5 A5WGK6
A5WGK9 A5WGL0 A5WGL1 A5WGL2 A5WGM2 A5WGM4 A5WGQ0 A5WGQ4 A5WGQ5 A5WGS3 A5WGS5
A5WGS6 A5WGT1 A5WGT4 A5WGT5 A5WGY0 A5WGY8 A5WH04 A5WH34 A5WH35 A5WH36 A5WH37
A5WH38 A5WH39 A5WH42 A5WH45 A5WH56 A5WH78 A5WH82 A5WH93 A5WHA3 A5WHA5 A5WHA7
A5WHC3 A5WHC5 A5WHC8 A5WHC9 A5WHD0 A5WHD5 A5WHI6 A5WHJ0 A5WHK7 A5WHT5 A5WHW6
A5WHW7 A5WHX7 A5WHY0 A5WI04 A5WI20 A5WI35 A5WI36 A5WI39 A5WII2 A5WJJ8 A5WKI8
A5WLC4 A5WLJ9 A5WMQ5 A5WMR0 A5WP74 A5WP83 A5WP84 A5WP89 A5WPA3 A5WRN6 A5WUL3
A5WUN7 A5WUT8 A5WUX7 A5WVX0 A5WVX1 A5WW08 A5WW21 A5WWC6 A5X7A0 A5Y5L5 A5YBJ3
A5YBJ4 A5YBJ5 A5YBJ6 A5YBJ7 A5YBJ9 A5YC49 A5YM69 A5YM72 A5YVF1 A6AMN2 A6B4U8
A6BLW4 A6BLY7 A6BM06 A6BM07 A6BM42 A6BM44 A6BM45 A6BM46 A6BM47 A6BM48 A6BM50
A6BM51 A6BM54 A6BM55 A6BM56 A6BM57 A6BM60 A6GVJ9 A6GVK0 A6GVN8 A6GVP1 A6GVQ2
A6GVQ6 A6GVR5 A6GVV2 A6GVV3 A6GVW9 A6GVY8 A6GVZ1 A6GW04 A6GW32 A6GW45 A6GW65
A6GW77 A6GW80 A6GW82 A6GW89 A6GW90 A6GWA9 A6GWB0 A6GWB6 A6GWD9 A6GWH3 A6GWI4
A6GWI5 A6GWI6 A6GWN3 A6GWP9 A6GWQ2 A6GWR9 A6GWS0 A6GWS3 A6GWS4 A6GWT7 A6GWU1
A6GWU2 A6GWU3 A6GWV6 A6GWV8 A6GWZ0 A6GWZ4 A6GX80 A6GXB2 A6GXC2 A6GXD5 A6GXD6
A6GXG2 A6GXJ8 A6GXP5 A6GXW2 A6GXW3 A6GXW4 A6GXZ1 A6GY19 A6GY20 A6GY22 A6GY30
A6GY75 A6GY76 A6GY79 A6GY96 A6GYG9 A6GYI0 A6GYJ4 A6GYR1 A6GYT9 A6GYU0 A6GYU1
A6GYU2 A6GYU3 A6GYU4 A6GYU7 A6GYV0 A6GYW0 A6GYW1 A6GYW4 A6GYW8 A6GYX5 A6GZ60
A6GZ72 A6GZ73 A6GZ74 A6GZ75 A6GZ76 A6GZ77 A6GZ78 A6GZ80 A6GZ81 A6GZ82 A6GZ84
A6GZ85 A6GZ86 A6GZ87 A6GZ88 A6GZ89 A6GZ91 A6GZ92 A6GZ93 A6GZ94 A6GZ95 A6GZ96
A6GZ98 A6GZ99 A6GZA0 A6GZA2 A6GZA3 A6GZB7 A6GZB9 A6GZF2 A6GZF3 A6GZF6 A6GZG1
A6GZG2 A6GZH5 A6GZI2 A6GZI5 A6GZI6 A6GZI7 A6GZJ6 A6GZL9 A6GZM5 A6GZP8 A6GZS1
A6H064 A6H070 A6H098 A6H0A6 A6H0C4 A6H0E5 A6H0E6 A6H0E7 A6H0G2 A6H0G9 A6H0H4
A6H0I8 A6H0L2 A6H0M8 A6H0N9 A6H0P1 A6H0P2 A6H0P3 A6H0Q2 A6H0R5 A6H0R6 A6H0S6
A6H0T6 A6H0U1 A6H0U4 A6H0V1 A6H0V8 A6H0W3 A6H0Y3 A6H103 A6H119 A6H125 A6H141
A6H143 A6H148 A6H162 A6H184 A6H185 A6H194 A6H1A2 A6H1A3 A6H1A5 A6H1B8 A6H1D1
A6H1G0 A6H1G2 A6H1G3 A6H1H2 A6H1N2 A6H1P4 A6H1P7 A6H1P8 A6H213 A6H233 A6H244
A6H247 A6H261 A6H294 A6H2B4 A6H2H7 A6H4Q1 A6H5E5 A6H5E7 A6H5F6 A6H5F7 A6H5F8
A6H5F9 A6H5H0 A6H5H5 A6H5I5 A6H5K3 A6H5K4 A6H5K5 A6H5K6 A6H5L3 A6H5L4 A6H5L5
A6H5L6 A6H5L7 A6H5L8 A6H5L9 A6H5M0 A6H5M1 A6H5M2 A6H5M3 A6H5M5 A6H5M8 A6H5P9
A6H5Q1 A6H5Q2 A6H5Y3 A6H603 A6H611 A6H619 A6H630 A6H639 A6H684 A6H687 A6H690

A6H694 A6H6E9 A6H6Q4 A6H6W9 A6H6X4 A6H6Z7 A6H712 A6H739 A6H750 A6H754 A6H759
A6H768 A6H791 A6H795 A6H7B4 A6H7D1 A6H7E1 A6H7E2 A6H7F2 A6H7F7 A6H7H7 A6H7I3
A6H7I7 A6H7I8 A6H7J1 A6H7J2 A6H8I2 A6H8J1 A6H8T2 A6H8T7 A6H8Y1 A6H8Z2 A6H907
A6H909 A6KWC4 A6KWD5 A6KWD6 A6KWD7 A6KWL1 A6KWL7 A6KWQ7 A6KWR2 A6KWR4 A6KX92
A6KXA0 A6KXA1 A6KXA6 A6KXB3 A6KXB4 A6KXF7 A6KXG4 A6KXG8 A6KXG9 A6KXH3 A6KXK1
A6KXL2 A6KXL8 A6KXM3 A6KXS0 A6KXS5 A6KXS6 A6KXU3 A6KXU8 A6KXY5 A6KXY8 A6KY05
A6KYG9 A6KYH0 A6KYH1 A6KYH2 A6KYH3 A6KYH6 A6KYH7 A6KYH8 A6KYH9 A6KYI0 A6KYI1
A6KYI2 A6KYI3 A6KYI4 A6KYI5 A6KYI8 A6KYI9 A6KYJ0 A6KYJ1 A6KYJ2 A6KYJ4 A6KYJ5
A6KYJ6 A6KYJ7 A6KYJ8 A6KYJ9 A6KYK2 A6KYK3 A6KYK4 A6KYK6 A6KYK7 A6KYK9 A6KYL2
A6KYN9 A6KYP1 A6KYR3 A6KYS7 A6KZ28 A6KZ38 A6KZ76 A6KZA8 A6KZH6 A6KZJ2 A6KZJ9
A6KZK0 A6KZL1 A6KZL8 A6KZM1 A6KZN7 A6KZN9 A6KZP0 A6KZP3 A6KZR2 A6KZV1 A6KZW5
A6KZZ5 A6L011 A6L012 A6L013 A6L028 A6L030 A6L046 A6L048 A6L050 A6L064 A6L065
A6L070 A6L075 A6L078 A6L079 A6L083 A6L089 A6L091 A6L0A5 A6L0E8 A6L0U5 A6L0V0
A6L0V1 A6L0V2 A6L0V6 A6L0W0 A6L0W6 A6L0W8 A6L0X5 A6L0Y7 A6L0Z9 A6L100 A6L175
A6L1E7 A6L1H7 A6L1H8 A6L1I4 A6L1J0 A6L1L8 A6L1M6 A6L1N4 A6L1P5 A6L1P6 A6L1Q8
A6L1U8 A6L1V8 A6L1X1 A6L1Z3 A6L239 A6L2G2 A6L2I3 A6L2J2 A6L2N4 A6L2R5 A6L2T0
A6L2V6 A6L2V8 A6L2W9 A6L2X7 A6L313 A6L330 A6L391 A6L3C0 A6L3C5 A6L3D0 A6L3D5
A6L3E7 A6L3F8 A6L3G7 A6L3H3 A6L3I9 A6L3K7 A6L3K9 A6L3Y2 A6L3Y9 A6L404 A6L4C3
A6L4C4 A6L4C6 A6L4J3 A6L4J6 A6L4N0 A6L4P0 A6L4P5 A6L4U4 A6L4U5 A6L4Y5 A6L518
A6L532 A6L585 A6L5A6 A6L5E2 A6L5E4 A6L5E5 A6L5E7 A6L5G7 A6L5G8 A6L5G9 A6L5H1
A6L5I2 A6L5J5 A6L5J9 A6L5K0 A6L5K3 A6L5X6 A6L5X9 A6L645 A6L646 A6L683 A6L686
A6L6D1 A6L6D2 A6L6E8 A6L6X0 A6L6X1 A6L6X5 A6L6Z1 A6L6Z2 A6L792 A6L7E4 A6L7E5
A6L7J5 A6L7J6 A6L7J7 A6L7J9 A6L7K0 A6L7M5 A6L7M8 A6L7N0 A6L7N1 A6L7P5 A6L7P7
A6L7Q8 A6L7T5 A6L849 A6L852 A6L8C4 A6L8C6 A6L8D1 A6L8E5 A6L8G0 A6L8H3 A6L8H5
A6L8I6 A6L8I7 A6L8J2 A6L8P3 A6L8Q7 A6L8S9 A6L8T0 A6L8T2 A6L8T3 A6L8U2 A6L8U3
A6L8V3 A6L8V4 A6L903 A6L927 A6L982 A6L991 A6L9A0 A6L9B7 A6L9E1 A6L9J8 A6L9J9
A6L9K4 A6L9K8 A6L9R7 A6L9R8 A6L9S2 A6L9S3 A6L9S4 A6L9S5 A6L9U9 A6L9V0 A6L9X1
A6LA51 A6LA55 A6LA57 A6LA68 A6LA76 A6LA92 A6LA96 A6LA97 A6LAJ6 A6LAM0 A6LAM2
A6LAP3 A6LAR6 A6LB11 A6LB29 A6LB35 A6LB99 A6LBC3 A6LBD0 A6LBD8 A6LBG7 A6LBQ5
A6LBU3 A6LBU6 A6LBX4 A6LBX7 A6LBY4 A6LC32 A6LC43 A6LC86 A6LCB6 A6LCP4 A6LCQ4
A6LCS5 A6LCY1 A6LCZ1 A6LD04 A6LD45 A6LD46 A6LD68 A6LD69 A6LD84 A6LDA7 A6LDB6
A6LDG8 A6LDI6 A6LDI7 A6LDN1 A6LDP0 A6LDP9 A6LDQ1 A6LDR9 A6LDS0 A6LDS8 A6LDT9
A6LE09 A6LE74 A6LE80 A6LE81 A6LE82 A6LE84 A6LE85 A6LE88 A6LE91 A6LE92 A6LEA1
A6LEF2 A6LEG4 A6LEG5 A6LEG6 A6LEG7 A6LEG8 A6LEG9 A6LEH2 A6LEH3 A6LEH4 A6LEH5
A6LEH6 A6LEH7 A6LEH8 A6LEH9 A6LEI0 A6LEI1 A6LEI3 A6LEI4 A6LEI5 A6LEI6 A6LEI7
A6LEI8 A6LEJ0 A6LEJ1 A6LEJ2 A6LEJ3 A6LEJ4 A6LEJ5 A6LEJ8 A6LEL4 A6LEL6 A6LEL7
A6LEM6 A6LEP5 A6LEU2 A6LEU7 A6LEV0 A6LEZ7 A6LEZ8 A6LF20 A6LF29 A6LF39 A6LF43
A6LF90 A6LFA6 A6LFB1 A6LFB5 A6LFB9 A6LFD0 A6LFE8 A6LFF0 A6LFI7 A6LFK9 A6LFL9
A6LFP0 A6LFQ4 A6LFQ8 A6LFT6 A6LFV1 A6LG11 A6LG55 A6LG59 A6LG60 A6LG62 A6LG92
A6LG93 A6LGA2 A6LGA4 A6LGF1 A6LGR3 A6LGR5 A6LHA9 A6LHB0 A6LHB5 A6LHB6 A6LHM6
A6LHM7 A6LHM8 A6LHN0 A6LHR9 A6LHS1 A6LHT2 A6LHT8 A6LHV2 A6LHW2 A6LHY5 A6LI30
A6LI52 A6LI67 A6LI95 A6LIA1 A6LIB2 A6LIB3 A6LIC2 A6LIF1 A6LIF5 A6LIF7 A6LIF9
A6LIG0 A6LIG7 A6LIS9 A6LIT0 A6LIW6 A6LIX7 A6LIX8 A6LIX9 A6LIY2 A6LJ30 A6LJ31
A6LJ38 A6LJ39 A6LJ40 A6LJ41 A6LJ42 A6LJ47 A6LJ53 A6LJ63 A6LJ74 A6LJ76 A6LJD8

A6LJG1 A6LJG3 A6LJI7 A6LJK3 A6LJK7 A6LJK8 A6LJK9 A6LJL4 A6LJL7 A6LJL9 A6LJM6
A6LJM7 A6LJM9 A6LJP2 A6LJQ5 A6LJS3 A6LJS5 A6LJT4 A6LJT6 A6LJT9 A6LJZ0 A6LJZ1
A6LJZ9 A6LK77 A6LK89 A6LK98 A6LKA2 A6LKB7 A6LKB8 A6LKB9 A6LKC0 A6LKD2 A6LKE7
A6LKE9 A6LKF8 A6LKI7 A6LKJ3 A6LKP0 A6LKP9 A6LKQ1 A6LKQ2 A6LKQ5 A6LKQ6 A6LKU9
A6LKW9 A6LKX4 A6LKY1 A6LL06 A6LL16 A6LL24 A6LL30 A6LL44 A6LL47 A6LL84 A6LL86
A6LL96 A6LLA5 A6LLA7 A6LLH1 A6LLI5 A6LLI6 A6LLI7 A6LLJ7 A6LLK6 A6LLK8 A6LLK9
A6LLL0 A6LLL1 A6LLL2 A6LLL3 A6LLL4 A6LLL6 A6LLL7 A6LLL8 A6LLL9 A6LLM0 A6LLM1
A6LLM2 A6LLM3 A6LLM4 A6LLM5 A6LLM6 A6LLM7 A6LLM8 A6LLM9 A6LLN0 A6LLN1 A6LLN2
A6LLN4 A6LLN6 A6LLN7 A6LLN8 A6LLN9 A6LLP0 A6LLP1 A6LLP2 A6LLP5 A6LLR1 A6LLR7
A6LLT7 A6LLU6 A6LLU8 A6LLV6 A6LM26 A6LM32 A6LM33 A6LM59 A6LM62 A6LM69 A6LM79
A6LMG3 A6LMG9 A6LMH6 A6LMH9 A6LMK5 A6LML3 A6LMN3 A6LMN4 A6LMP4 A6LMR3 A6LMR9
A6LMS2 A6LMS3 A6LMS4 A6LMU0 A6LMU4 A6LMU8 A6LMV5 A6LMV9 A6LMW2 A6LMW5 A6LMW8
A6LN42 A6LN47 A6LN55 A6LN73 A6LN82 A6LN85 A6LN93 A6LNA3 A6LNA6 A6LNF3 A6LNF5
A6LNG2 A6LNG3 A6LNG5 A6LNG7 A6LNH1 A6LNM3 A6LNR5 A6LNR9 A6LNU3 A6LNU4 A6LNU8
A6LNY4 A6LNY6 A6LNY7 A6LNY9 A6LP26 A6LP38 A6LP41 A6LP48 A6LP58 A6LP59 A6LP60
A6LP67 A6LP69 A6LP75 A6LP88 A6LP90 A6LP91 A6LPA6 A6LPB1 A6LPB4 A6LPC3 A6LPC4
A6LPD5 A6LPD6 A6LPD9 A6LPF0 A6LPI2 A6LPI6 A6LPI8 A6LPJ0 A6LPJ5 A6LPL1 A6LPL2
A6LPL4 A6LPL6 A6LPL7 A6LPM3 A6LPN9 A6LPP0 A6LPP1 A6LPP6 A6LPP7 A6LPQ0 A6LPQ1
A6LPQ3 A6LPQ4 A6LPQ5 A6LPQ6 A6LPQ7 A6LPQ8 A6LPR0 A6LPR1 A6LPR2 A6LPR3 A6LPR4
A6LPR5 A6LPR6 A6LPR7 A6LPR8 A6LPR9 A6LPS0 A6LPS1 A6LPS2 A6LPS3 A6LPS4 A6LPS5
A6LPS6 A6LPS7 A6LPS9 A6LPT0 A6LPT2 A6LPT5 A6LPT6 A6LPT7 A6LPT8 A6LPT9 A6LPU0
A6LPU1 A6LPU5 A6LPU6 A6LPU7 A6LPX0 A6LPX4 A6LPX5 A6LPX8 A6LQ55 A6LQ78 A6LQ86
A6LQ87 A6LQ90 A6LQ99 A6LQB0 A6LQD5 A6LQF2 A6LQF6 A6LQF9 A6LQG2 A6LQG6 A6LQK4
A6LQL0 A6LQP7 A6LQP8 A6LQQ3 A6LQQ6 A6LQR6 A6LQR8 A6LQR9 A6LQS1 A6LQV9 A6LQX4
A6LQY8 A6LR05 A6LR06 A6LR07 A6LR41 A6LRL7 A6LRM5 A6LRN2 A6LRN3 A6LRN4 A6LRN5
A6LRN7 A6LRN8 A6LRP7 A6LS00 A6LS54 A6LS60 A6LS61 A6LS73 A6LSA9 A6LSB3 A6LSF2
A6LSF6 A6LSF7 A6LSF8 A6LSF9 A6LSH5 A6LSI6 A6LSJ2 A6LSJ4 A6LSJ8 A6LSK1 A6LSK4
A6LSK7 A6LSL2 A6LSL7 A6LSM2 A6LSM8 A6LSN0 A6LSN1 A6LSN2 A6LSN4 A6LSN5 A6LSN9
A6LSP0 A6LSP3 A6LSP7 A6LSQ0 A6LSQ4 A6LSQ5 A6LSQ9 A6LSR0 A6LSR6 A6LSS9 A6LSV5
A6LSX2 A6LSX9 A6LSY3 A6LSY5 A6LSY8 A6LSZ4 A6LSZ5 A6LT19 A6LT21 A6LT22 A6LT23
A6LT26 A6LT28 A6LT31 A6LTA6 A6LTC5 A6LTK6 A6LTL5 A6LTM6 A6LTM7 A6LTM8 A6LTN5
A6LTN7 A6LTP1 A6LTR4 A6LTR5 A6LTR7 A6LTR9 A6LTS0 A6LTT3 A6LTU0 A6LU43 A6LU44
A6LU45 A6LU46 A6LU48 A6LU50 A6LU51 A6LU79 A6LU81 A6LU82 A6LU93 A6LU95 A6LU96
A6LU97 A6LUA1 A6LUD2 A6LUD4 A6LUE1 A6LUF3 A6LUK9 A6LUQ4 A6LUQ7 A6LVB6 A6LVU2
A6LVY2 A6LW30 A6LW55 A6LW95 A6LWD4 A6LWG6 A6LWG7 A6LWI1 A6LWI9 A6LWJ0 A6LWJ1
A6LWJ5 A6LWN4 A6LWN6 A6LX30 A6LX31 A6LXM3 A6LXU8 A6LY33 A6LZF4 A6M020 A6M0F0
A6M0L7 A6M0X2 A6M152 A6M163 A6M1C5 A6M1C6 A6M1N9 A6M1P1 A6M1P8 A6M1T6 A6M1W6
A6M1W7 A6M1Y8 A6M1Z4 A6M1Z5 A6M1Z6 A6M1Z8 A6M220 A6M225 A6M229 A6M241 A6M249
A6M250 A6M251 A6M254 A6M255 A6M268 A6M273 A6M293 A6M2W4 A6M2Y1 A6M2Y3 A6M2Y4
A6M2Y5 A6M2Z0 A6M317 A6M320 A6M327 A6M331 A6M334 A6M378 A6M3G5 A6M3I3 A6M3K0
A6M3K7 A6M3L0 A6M3L2 A6M3M3 A6M3M4 A6M3M5 A6M3M9 A6M3N0 A6M931 A6MM15 A6MM17
A6MM25 A6MM26 A6MM27 A6MM28 A6MM34 A6MM38 A6MM44 A6MM57 A6MM58 A6MM59 A6MM61
A6MM68 A6MM69 A6MM70 A6MM71 A6MM72 A6MM73 A6MM74 A6MM75 A6MM76 A6MM77 A6MM78
A6MM80 A6MM82 A6MM94 A6MMA1 A6MMA3 A6MMB1 A6MMB2 A6MMB3 A6MMB4 A6MMC0 A6MMC4

A6MMD0 A6MME3 A6MME4 A6MME5 A6MME7 A6MMF4 A6MMF5 A6MMF6 A6MMF7 A6MMF8 A6MMF9
A6MMG0 A6MMG1 A6MMG2 A6MMG3 A6MMG4 A6MMG6 A6MMG8 A6MMH0 A6MMI6 A6MMI7 A6MMI9
A6MMJ6 A6MMJ7 A6MMJ8 A6MMJ9 A6MMK5 A6MMK9 A6MML5 A6MMM8 A6MMM9 A6MMN0 A6MMN2
A6MMN9 A6MMP0 A6MMP1 A6MMP2 A6MMP3 A6MMP4 A6MMP5 A6MMP6 A6MMP8 A6MMP9 A6MMQ1
A6MMQ2 A6MMQ4 A6MMQ6 A6MMS3 A6MMS5 A6MMT3 A6MMT4 A6MMT5 A6MMT6 A6MMU2 A6MMU6
A6MMV2 A6MMW5 A6MMW6 A6MMW7 A6MMW9 A6MMX6 A6MMX7 A6MMX8 A6MMX9 A6MMY0 A6MMY1
A6MMY2 A6MMY3 A6MMY4 A6MMY5 A6MMY6 A6MMY8 A6MMZ0 A6MN02 A6MVP4 A6MVP8 A6MVT7
A6MVT9 A6MVX0 A6MVX1 A6MVX2 A6MVX3 A6MVX4 A6MVX6 A6MVX7 A6MVX8 A6MVY4 A6MVZ5
A6MW01 A6MW04 A6MW05 A6MW06 A6MW07 A6MW08 A6MW11 A6MW12 A6MW13 A6MW14 A6MW17
A6MW19 A6MW20 A6MW21 A6MW22 A6MW27 A6MW28 A6MZM2 A6N0M9 A6N6J5 A6N7Y9 A6N9N3
A6NA00 A6NAF9 A6NC05 A6NC42 A6NC57 A6NC62 A6NCF5 A6NCF6 A6NCI8 A6NCJ1 A6NCK2
A6NCL1 A6NCL7 A6NCM1 A6NCN2 A6NCN8 A6NCS4 A6NCS6 A6NCW7 A6ND91 A6NDE4 A6NDE8
A6NDI0 A6NDL7 A6NDN8 A6NDR6 A6NDU8 A6NDX5 A6NDY0 A6NDY2 A6NDZ8 A6NE01 A6NE52
A6NE82 A6NEC2 A6NEE1 A6NEF3 A6NEH8 A6NEK1 A6NEL2 A6NEL3 A6NEM1 A6NEN9 A6NEQ0
A6NEQ2 A6NER3 A6NES4 A6NEV1 A6NEY3 A6NEY8 A6NF36 A6NF83 A6NFD8 A6NFE3 A6NFF2
A6NFK2 A6NFN3 A6NFN9 A6NFQ7 A6NFT4 A6NFU8 A6NFY7 A6NGB9 A6NGD5 A6NGE4 A6NGG3
A6NGH7 A6NGH8 A6NGJ6 A6NGK3 A6NGN4 A6NGQ2 A6NGR9 A6NGS2 A6NGU7 A6NGY1 A6NGY3
A6NH13 A6NHG4 A6NHJ4 A6NHP3 A6NHQ2 A6NHQ4 A6NHR8 A6NHR9 A6NHS1 A6NHT5 A6NHZ5
A6NI03 A6NI15 A6NI47 A6NI56 A6NI72 A6NI86 A6NI87 A6NIE6 A6NIJ5 A6NIK2 A6NIN4
A6NIR3 A6NIU2 A6NIY4 A6NJ08 A6NJ46 A6NJ64 A6NJ78 A6NJ88 A6NJB7 A6NJG2 A6NJG6
A6NJI1 A6NJI9 A6NJJ6 A6NJL1 A6NJQ4 A6NJR5 A6NJT0 A6NJV1 A6NJZ7 A6NK02 A6NK06
A6NK44 A6NK53 A6NK58 A6NK75 A6NKC0 A6NKC9 A6NKD2 A6NKH3 A6NKN8 A6NKP2 A6NKT7
A6NKU9 A6NKX1 A6NL46 A6NL82 A6NLC5 A6NLC8 A6NLI5 A6NLP5 A6NLU0 A6NLW8 A6NLX3
A6NM28 A6NM43 A6NM66 A6NMA1 A6NMB9 A6NMK7 A6NMK8 A6NMT0 A6NMX2 A6NN06 A6NN14
A6NN90 A6NNA2 A6NNA5 A6NNF4 A6NNH2 A6NNJ1 A6NNL0 A6NNM3 A6NNM8 A6NNP5 A6NNT2
A6NNV3 A6NNW6 A6NNX1 A6NP11 A6NP61 A6P7L8 A6PVC2 A6PVI3 A6PVS8 A6PVY3 A6PWD2
A6PWY4 A6Q0K5 A6Q0W4 A6Q102 A6Q115 A6Q117 A6Q128 A6Q160 A6Q162 A6Q163 A6Q164
A6Q167 A6Q168 A6Q169 A6Q182 A6Q186 A6Q193 A6Q194 A6Q1C0 A6Q1C1 A6Q1C9 A6Q1H0
A6Q1H1 A6Q1H7 A6Q1H9 A6Q1I0 A6Q1I1 A6Q1I2 A6Q1I3 A6Q1I4 A6Q1I5 A6Q1I6 A6Q1I7
A6Q1I8 A6Q1J1 A6Q1J2 A6Q1J3 A6Q1J4 A6Q1J5 A6Q1J6 A6Q1J9 A6Q1K0 A6Q1K1 A6Q1K2
A6Q1K3 A6Q1K4 A6Q1K5 A6Q1L1 A6Q1L5 A6Q1L9 A6Q1M0 A6Q1M1 A6Q1M2 A6Q1M3 A6Q1M4
A6Q1M5 A6Q1M6 A6Q1M7 A6Q1P1 A6Q1Q8 A6Q1S4 A6Q1S6 A6Q1U2 A6Q1W0 A6Q1W3 A6Q1W6
A6Q1W8 A6Q1X7 A6Q1X8 A6Q1Y1 A6Q1Y9 A6Q1Z5 A6Q1Z6 A6Q1Z8 A6Q200 A6Q202 A6Q225
A6Q226 A6Q228 A6Q232 A6Q234 A6Q235 A6Q239 A6Q291 A6Q2B4 A6Q2C1 A6Q2G5 A6Q2K0
A6Q2K1 A6Q2R2 A6Q2R6 A6Q2R7 A6Q2U7 A6Q2V4 A6Q2V7 A6Q2W2 A6Q2W6 A6Q2X6 A6Q2Y9
A6Q311 A6Q312 A6Q322 A6Q330 A6Q339 A6Q342 A6Q349 A6Q354 A6Q367 A6Q371 A6Q377
A6Q388 A6Q391 A6Q3B4 A6Q3C7 A6Q3D1 A6Q3D2 A6Q3D6 A6Q3E8 A6Q3K8 A6Q3N0 A6Q3P0
A6Q3P9 A6Q3Q0 A6Q3Q1 A6Q3R4 A6Q3S1 A6Q3S4 A6Q3S5 A6Q3T0 A6Q3U3 A6Q3V1 A6Q3Y3
A6Q3Y5 A6Q3Y6 A6Q3Z7 A6Q3Z8 A6Q401 A6Q418 A6Q421 A6Q422 A6Q424 A6Q429 A6Q433
A6Q441 A6Q446 A6Q461 A6Q464 A6Q466 A6Q469 A6Q478 A6Q485 A6Q486 A6Q496 A6Q4C9
A6Q4D2 A6Q4D3 A6Q4D4 A6Q4D7 A6Q4I0 A6Q4I1 A6Q4L2 A6Q4L7 A6Q4N2 A6Q4N8 A6Q4Q9
A6Q4R0 A6Q4R8 A6Q4S8 A6Q4V5 A6Q4V8 A6Q4V9 A6Q4Z5 A6Q4Z6 A6Q504 A6Q512 A6Q522
A6Q524 A6Q526 A6Q531 A6Q538 A6Q539 A6Q541 A6Q543 A6Q548 A6Q549 A6Q550 A6Q552
A6Q559 A6Q573 A6Q574 A6Q576 A6Q581 A6Q582 A6Q584 A6Q585 A6Q596 A6Q599 A6Q5A7

A6Q5D0 A6Q5D5 A6Q5F6 A6Q5F7 A6Q5H2 A6Q5H7 A6Q5I8 A6Q5J0 A6Q5J3 A6Q5K0 A6Q5K5
A6Q5L0 A6Q5L6 A6Q638 A6Q639 A6Q652 A6Q674 A6Q675 A6Q687 A6Q688 A6Q692 A6Q693
A6Q6B4 A6Q6B5 A6Q6B6 A6Q6C1 A6Q6D3 A6Q6D8 A6Q6E1 A6Q6F9 A6Q6H4 A6Q6H5 A6Q6H8
A6Q6H9 A6Q6I0 A6Q6I1 A6Q6I2 A6Q6I3 A6Q6I4 A6Q6I5 A6Q6I6 A6Q6J0 A6Q6J3 A6Q6J8
A6Q6K6 A6Q6K7 A6Q6L9 A6Q6M5 A6Q6M8 A6Q6N9 A6Q6P0 A6Q6Q7 A6Q6R2 A6Q6S1 A6Q6S7
A6Q6T2 A6Q6T6 A6Q715 A6Q716 A6Q718 A6Q719 A6Q747 A6Q760 A6Q769 A6Q771 A6Q772
A6Q773 A6Q776 A6Q782 A6Q784 A6Q7C5 A6Q7C7 A6Q7C9 A6Q7E9 A6Q7F6 A6Q7G2 A6Q7G6
A6Q7H8 A6Q7I8 A6Q7J0 A6Q7M3 A6Q7P5 A6Q7P7 A6Q7P9 A6Q7Q0 A6Q7Q7 A6Q7S7 A6Q7V3
A6Q7Y2 A6Q7Y4 A6Q7Y7 A6Q7Y8 A6Q809 A6Q815 A6Q826 A6Q8F8 A6Q8G9 A6Q8H2 A6Q8K5
A6Q906 A6Q908 A6Q936 A6Q941 A6Q992 A6Q9C9 A6Q9D4 A6Q9I6 A6Q9M2 A6Q9N6 A6Q9P9
A6Q9Q9 A6Q9R0 A6Q9U8 A6Q9U9 A6Q9V0 A6Q9W7 A6Q9Z8 A6Q9Z9 A6QAB4 A6QAE8 A6QAE9
A6QAK6 A6QAL0 A6QAL4 A6QAM1 A6QAM9 A6QAQ4 A6QAR3 A6QAR9 A6QAX2 A6QAX3 A6QAX6
A6QAX8 A6QB12 A6QB13 A6QB34 A6QB35 A6QB48 A6QB68 A6QB69 A6QB70 A6QB71 A6QB72
A6QB84 A6QBB6 A6QBB7 A6QBC0 A6QBG0 A6QBG1 A6QBG2 A6QBG7 A6QBH4 A6QBI2 A6QBI5
A6QBK6 A6QBK7 A6QBL4 A6QBN6 A6QBN7 A6QBP2 A6QBP7 A6QBP8 A6QBQ3 A6QBQ5 A6QBQ7
A6QBT0 A6QBY2 A6QBY4 A6QBY6 A6QBY8 A6QBZ2 A6QBZ9 A6QC00 A6QC07 A6QC22 A6QC28
A6QC54 A6QC65 A6QC86 A6QC89 A6QCC6 A6QCD0 A6QCD3 A6QCD6 A6QCE8 A6QCI0 A6QCI9
A6QCJ0 A6QCJ1 A6QCJ3 A6QCK9 A6QCP7 A6QCQ0 A6QCQ1 A6QCQ2 A6QCQ3 A6QCQ4 A6QCQ5
A6QCQ6 A6QCQ8 A6QCQ9 A6QCR0 A6QCR1 A6QCR2 A6QCR3 A6QCR4 A6QCR5 A6QCR9 A6QCS0
A6QCS1 A6QCS2 A6QCS3 A6QCS4 A6QCS5 A6QCU4 A6QCU6 A6QCX3 A6QD00 A6QD02 A6QD40
A6QD41 A6QD43 A6QD47 A6QD48 A6QD54 A6QD63 A6QD96 A6QDA0 A6QDC3 A6QDF1 A6QDG6
A6QDL6 A6QDN3 A6QDP3 A6QDU7 A6QDV0 A6QE38 A6QE47 A6QE49 A6QE68 A6QE71 A6QED3
A6QED4 A6QEE0 A6QEE6 A6QEE8 A6QEF1 A6QEF5 A6QEG4 A6QEH1 A6QEH2 A6QEI0 A6QEI2
A6QEJ0 A6QEJ1 A6QEJ2 A6QEJ4 A6QEJ6 A6QEJ7 A6QEJ8 A6QEJ9 A6QEK0 A6QEK3 A6QEK4
A6QEL9 A6QEM2 A6QEN4 A6QEP0 A6QEP6 A6QER9 A6QES8 A6QEX8 A6QEZ0 A6QF21 A6QF32
A6QF37 A6QF39 A6QF47 A6QF52 A6QF68 A6QF73 A6QF75 A6QF76 A6QF82 A6QF83 A6QF90
A6QFB2 A6QFB6 A6QFD2 A6QFD6 A6QFE3 A6QFF0 A6QFH1 A6QFH2 A6QFH3 A6QFH7 A6QFH8
A6QFI1 A6QFK8 A6QFL5 A6QFL7 A6QFM5 A6QFN0 A6QFQ3 A6QFS2 A6QFS5 A6QFS7 A6QFS9
A6QFT1 A6QFV3 A6QFW2 A6QFW7 A6QFX5 A6QFX8 A6QFY1 A6QG37 A6QG39 A6QG41 A6QG46
A6QG48 A6QG53 A6QG68 A6QG77 A6QG78 A6QG79 A6QG83 A6QG89 A6QG93 A6QG99 A6QGA1
A6QGA4 A6QGA5 A6QGB6 A6QGB8 A6QGC4 A6QGD3 A6QGD8 A6QGD9 A6QGE0 A6QGE1 A6QGE4
A6QGE5 A6QGF1 A6QGF2 A6QGF3 A6QGF4 A6QGF5 A6QGF6 A6QGF7 A6QGF8 A6QGF9 A6QGG3
A6QGG4 A6QGG5 A6QGG8 A6QGG9 A6QGH0 A6QGH2 A6QGH3 A6QGI4 A6QGJ1 A6QGJ4 A6QGJ5
A6QGJ8 A6QGK1 A6QGN2 A6QGN7 A6QGN8 A6QGN9 A6QGP7 A6QGP8 A6QGR0 A6QGS1 A6QGS2
A6QGS4 A6QGS5 A6QGU6 A6QGU7 A6QGU8 A6QGV5 A6QGW6 A6QGX4 A6QGX8 A6QGZ6 A6QGZ7
A6QH05 A6QH08 A6QH13 A6QH15 A6QH16 A6QH17 A6QH18 A6QH20 A6QH24 A6QH26 A6QH42
A6QH51 A6QH66 A6QH68 A6QH69 A6QH73 A6QH79 A6QH81 A6QH82 A6QH94 A6QHA0 A6QHA6
A6QHA8 A6QHA9 A6QHB3 A6QHB8 A6QHC1 A6QHC2 A6QHC3 A6QHC4 A6QHC5 A6QHC8 A6QHD6
A6QHD8 A6QHE1 A6QHE6 A6QHF1 A6QHF2 A6QHF6 A6QHF7 A6QHF8 A6QHF9 A6QHG3 A6QHH2
A6QHH3 A6QHH5 A6QHH7 A6QHI1 A6QHI2 A6QHI3 A6QHI4 A6QHI5 A6QHI6 A6QHI7 A6QHI9
A6QHJ5 A6QHJ8 A6QHK1 A6QHK4 A6QHK6 A6QHK7 A6QHK8 A6QHK9 A6QHL2 A6QHL3 A6QHL4
A6QHL6 A6QHL9 A6QHN3 A6QHP1 A6QHP5 A6QHP9 A6QHQ3 A6QHR2 A6QHR5 A6QHS3 A6QHS6
A6QHT1 A6QHU1 A6QHU9 A6QHV6 A6QHV8 A6QHX0 A6QHX1 A6QHX6 A6QI15 A6QI27 A6QI46
A6QI49 A6QI55 A6QIC3 A6QIC7 A6QIC8 A6QIC9 A6QID2 A6QIE0 A6QIM7 A6QIM8 A6QIP3

A6QIP6 A6QIQ0 A6QIQ2 A6QIQ3 A6QIQ5 A6QIQ6 A6QIQ8 A6QIR1 A6QIR4 A6QIR5 A6QIS7
A6QIT5 A6QIT6 A6QIV6 A6QIV7 A6QIV8 A6QIW2 A6QIW4 A6QIX1 A6QIX2 A6QJ00 A6QJ02
A6QJ33 A6QJ36 A6QJ37 A6QJ38 A6QJ39 A6QJ59 A6QJ60 A6QJ61 A6QJ65 A6QJ66 A6QJ67
A6QJ68 A6QJ69 A6QJ70 A6QJ73 A6QJ74 A6QJ75 A6QJ76 A6QJ77 A6QJ78 A6QJ79 A6QJ80
A6QJ81 A6QJ82 A6QJ83 A6QJ85 A6QJ86 A6QJ87 A6QJ88 A6QJ89 A6QJ90 A6QJ91 A6QJ92
A6QJ93 A6QJA8 A6QJA9 A6QJB4 A6QJC0 A6QJC8 A6QJC9 A6QJD0 A6QJD1 A6QJD2 A6QJD3
A6QJD4 A6QJH1 A6QJH4 A6QJH5 A6QJI5 A6QJI7 A6QJJ8 A6QJL4 A6QJN2 A6QJQ5 A6QJR2
A6QJR4 A6QK04 A6QK44 A6QK48 A6QK84 A6QK85 A6QK86 A6QK89 A6QK93 A6QK99 A6QKC0
A6QKC4 A6QKF4 A6QKG1 A6QKG2 A6QKG4 A6QKG7 A6QKG8 A6QKH7 A6QKH8 A6QKK0 A6QKK1
A6QKK2 A6QKK4 A6QL29 A6QL50 A6QL70 A6QL72 A6QL79 A6QLA4 A6QLA6 A6QLC7 A6QLD5
A6QLE1 A6QLH5 A6QLH6 A6QLI5 A6QLI8 A6QLJ8 A6QLK5 A6QLP2 A6QLR3 A6QLS2 A6QLU1
A6QLU5 A6QLU8 A6QLW2 A6QLW9 A6QLY4 A6QLZ5 A6QM00 A6QNM2 A6QNM3 A6QNM8 A6QNP3
A6QNQ6 A6QNR0 A6QNT4 A6QNX3 A6QNX5 A6QNZ0 A6QNZ4 A6QP05 A6QP24 A6QP35 A6QP66
A6QP81 A6QP92 A6QPA3 A6QPC0 A6QPC8 A6QPE1 A6QPE7 A6QPH1 A6QPH8 A6QPH9 A6QPL4
A6QPM3 A6QPM6 A6QPR6 A6QPR9 A6QPT6 A6QPY8 A6QQ14 A6QQ21 A6QQ66 A6QQ68 A6QQ70
A6QQ71 A6QQ74 A6QQ94 A6QQA5 A6QQC0 A6QQD2 A6QQF5 A6QQL3 A6QQL5 A6QQM4 A6QQM8
A6QQQ9 A6QQS3 A6QQV9 A6QQW0 A6QQW8 A6QQX5 A6QQY2 A6QQY4 A6QR00 A6QR06 A6QR16
A6QR20 A6QR31 A6QR40 A6QR44 A6QR56 A6QRA1 A6QRQ7 A6QSQ0 A6QUM7 A6QV55 A6QX61
A6QXB4 A6QXC1 A6QXC6 A6QXU0 A6QYC6 A6QYF6 A6QYH8 A6QYX4 A6R035 A6R040 A6R104
A6R213 A6R2L6 A6R363 A6R364 A6R371 A6R3K5 A6R3L3 A6R3R5 A6R4H9 A6R603 A6R687
A6R7B8 A6R7F0 A6R7H2 A6R7I3 A6R7S1 A6R853 A6R8I2 A6R8Q8 A6R8V2 A6R918 A6R989
A6R9C3 A6R9K6 A6R9U4 A6RAS3 A6RBB0 A6RBC0 A6RBE9 A6RBX7 A6RC73 A6RCJ9 A6RCT2
A6RDW5 A6RDX3 A6REV3 A6RF09 A6RF22 A6RG85 A6RGC8 A6RGE3 A6RGF3 A6RHI1 A6RHT9
A6RHW0 A6RI12 A6RIE1 A6RIS1 A6RJ45 A6RJA2 A6RJP7 A6RK67 A6RKG5 A6RL85 A6RLH0
A6RMI1 A6RMY2 A6RMY5 A6RMZ2 A6RNR4 A6RPB6 A6RPU0 A6RPU8 A6RQZ9 A6RR15 A6RRD4
A6RSH5 A6RSP5 A6RT32 A6RT94 A6RTU0 A6RUH2 A6RUL1 A6RVT8 A6RVU0 A6RW56 A6RW62
A6RW79 A6RWR8 A6RY31 A6RYJ8 A6RZS5 A6S043 A6S0T8 A6S146 A6S1A3 A6S374 A6S4N4
A6S5C9 A6S5X1 A6S694 A6S6H9 A6S6V7 A6S8E7 A6S936 A6S9T6 A6SCT6 A6SCX6 A6SDA8
A6SDE9 A6SDR8 A6SDT7 A6SDU6 A6SED8 A6SEH9 A6SFG0 A6SFQ6 A6SFV4 A6SFW7 A6SG85
A6SGN8 A6SIZ0 A6SJW6 A6SK19 A6SK81 A6SL16 A6SLT9 A6SM77 A6SMQ7 A6SNX1 A6SP81
A6SRT0 A6SRX6 A6SSW1 A6SSW5 A6SSW6 A6SSW9 A6STW2 A6SU56 A6SU64 A6SU65 A6SU66
A6SU83 A6SU87 A6SU94 A6SUC4 A6SUD4 A6SUD6 A6SUD7 A6SUD8 A6SUE7 A6SUF6 A6SUF7
A6SUH5 A6SUH6 A6SUL1 A6SUN7 A6SUN8 A6SUN9 A6SUP2 A6SUP6 A6SUQ1 A6SUQ2 A6SUQ3
A6SUQ4 A6SUQ5 A6SUQ8 A6SUR1 A6SUR4 A6SUS2 A6SUS5 A6SUU3 A6SUU4 A6SUW6 A6SUX7
A6SUY3 A6SV12 A6SV13 A6SV18 A6SV19 A6SV22 A6SV24 A6SV54 A6SV55 A6SV83 A6SVC2
A6SVD2 A6SVD7 A6SVE7 A6SVH6 A6SVI5 A6SVI6 A6SVI7 A6SVJ9 A6SVL3 A6SVP5 A6SVR5
A6SW68 A6SW71 A6SW83 A6SW89 A6SW90 A6SWY0 A6SWZ4 A6SWZ8 A6SX05 A6SX06 A6SX19
A6SX31 A6SX50 A6SX59 A6SX66 A6SX69 A6SX71 A6SX77 A6SXA4 A6SXE9 A6SXF3 A6SXF4
A6SXG2 A6SXI2 A6SXJ4 A6SXJ7 A6SXM0 A6SXP7 A6SXR9 A6SXS8 A6SXU1 A6SXY6 A6SY01
A6SY02 A6SY03 A6SY30 A6SY33 A6SY34 A6SY47 A6SY68 A6SY73 A6SY74 A6SY75 A6SY97
A6SZ04 A6SZ06 A6SZ07 A6SZ08 A6SZ09 A6SZ10 A6SZ52 A6SZH9 A6SZI7 A6SZM5 A6SZN5
A6SZN7 A6SZP5 A6SZP8 A6SZP9 A6SZQ0 A6SZQ1 A6SZQ8 A6SZR2 A6SZR5 A6SZT5 A6SZV1
A6SZV5 A6SZW1 A6SZW5 A6SZW6 A6SZX3 A6SZX4 A6SZX9 A6SZY0 A6SZZ1 A6T006 A6T009
A6T015 A6T017 A6T078 A6T081 A6T0F1 A6T0H1 A6T0H2 A6T0T5 A6T0T6 A6T0V4 A6T0Y7

A6T0Y8 A6T0Z0 A6T104 A6T105 A6T108 A6T112 A6T144 A6T146 A6T169 A6T171 A6T178
A6T1A0 A6T1E4 A6T1E5 A6T1G0 A6T1G3 A6T1G7 A6T1X9 A6T1Z0 A6T203 A6T204 A6T221
A6T225 A6T226 A6T227 A6T228 A6T235 A6T239 A6T242 A6T287 A6T290 A6T2A2 A6T2A5
A6T2B3 A6T2B6 A6T2D1 A6T2D2 A6T2D4 A6T2D5 A6T2D6 A6T2D7 A6T2E3 A6T2F6 A6T2F7
A6T2G3 A6T2G6 A6T2G7 A6T2R3 A6T2R7 A6T2S0 A6T2S2 A6T2S8 A6T2X9 A6T2Y3 A6T2Y9
A6T2Z1 A6T306 A6T312 A6T328 A6T332 A6T343 A6T344 A6T345 A6T374 A6T375 A6T376
A6T377 A6T379 A6T382 A6T383 A6T3G3 A6T3H0 A6T3H7 A6T3H8 A6T3H9 A6T3I0 A6T3I1
A6T3I2 A6T3I3 A6T3I5 A6T3I6 A6T3I7 A6T3I8 A6T3I9 A6T3J0 A6T3J1 A6T3J2 A6T3J3
A6T3J4 A6T3J6 A6T3J7 A6T3J8 A6T3J9 A6T3K0 A6T3K1 A6T3K2 A6T3K3 A6T3K4 A6T3K6
A6T3K7 A6T3K8 A6T3K9 A6T3L2 A6T3L3 A6T3L4 A6T3L5 A6T3L6 A6T3L7 A6T3N1 A6T450
A6T461 A6T463 A6T481 A6T482 A6T4D6 A6T4E0 A6T4E1 A6T4E3 A6T4E6 A6T4F4 A6T4F5
A6T4G3 A6T4G4 A6T4H4 A6T4H8 A6T4I5 A6T4I7 A6T4J7 A6T4K0 A6T4K1 A6T4L2 A6T4L6
A6T4L7 A6T4L9 A6T4M4 A6T4M5 A6T4N4 A6T4P3 A6T4P4 A6T4P6 A6T4R4 A6T4R5 A6T4R6
A6T4S9 A6T4T0 A6T4T3 A6T4T5 A6T4V8 A6T4W0 A6T4W3 A6T4W4 A6T4W7 A6T4W8 A6T4X1
A6T4X2 A6T4X3 A6T4Y5 A6T4Z7 A6T510 A6T512 A6T518 A6T521 A6T533 A6T541 A6T561
A6T562 A6T571 A6T5A7 A6T5B1 A6T5B6 A6T5B7 A6T5D7 A6T5D8 A6T5E6 A6T5E8 A6T5E9
A6T5F3 A6T5F5 A6T5F6 A6T5G0 A6T5H9 A6T5I1 A6T5I7 A6T5I9 A6T5L7 A6T5N2 A6T5N4
A6T5N5 A6T5N6 A6T5N7 A6T5R2 A6T5R4 A6T5Z2 A6T613 A6T644 A6T655 A6T656 A6T663
A6T672 A6T673 A6T689 A6T691 A6T692 A6T697 A6T6A0 A6T6A3 A6T6B3 A6T6B5 A6T6C1
A6T6C3 A6T6E6 A6T6E8 A6T6F6 A6T6I3 A6T6J6 A6T6K7 A6T6K8 A6T6L0 A6T6L1 A6T6L5
A6T6L6 A6T6L9 A6T6M5 A6T6M7 A6T6Q1 A6T6Q6 A6T6T1 A6T6X9 A6T6Y1 A6T6Y2 A6T6Z0
A6T700 A6T701 A6T702 A6T704 A6T709 A6T714 A6T716 A6T726 A6T742 A6T750 A6T752
A6T765 A6T766 A6T769 A6T797 A6T798 A6T799 A6T7A0 A6T7A1 A6T7A2 A6T7B9 A6T7D6
A6T7E0 A6T7E9 A6T7F8 A6T7G7 A6T7K3 A6T7Q4 A6T7R0 A6T7T5 A6T7U7 A6T7W1 A6T7W5
A6T7W6 A6T7Y5 A6T7Z3 A6T7Z4 A6T886 A6T8M8 A6T8N3 A6T8R9 A6T8X0 A6T8Z6 A6T900
A6T923 A6T962 A6T9C8 A6T9H0 A6T9H1 A6T9H2 A6T9K7 A6T9Q5 A6T9S2 A6T9W8 A6T9Y5
A6T9Y6 A6T9Y8 A6TA01 A6TA06 A6TAC4 A6TAC5 A6TAC6 A6TAC7 A6TAC8 A6TAC9 A6TAE3
A6TAE4 A6TAG8 A6TAH1 A6TAI1 A6TAI3 A6TAI4 A6TAJ8 A6TAN9 A6TAP0 A6TAP2 A6TAP7
A6TAW4 A6TAX0 A6TAX2 A6TAX3 A6TAX5 A6TAY2 A6TAY4 A6TB02 A6TB06 A6TB30 A6TB31
A6TB32 A6TB33 A6TB35 A6TB39 A6TB40 A6TB41 A6TB43 A6TB47 A6TB69 A6TB83 A6TBB2
A6TBC2 A6TBC4 A6TBC5 A6TBC7 A6TBC8 A6TBG5 A6TBG6 A6TBJ8 A6TBK1 A6TBK7 A6TBN1
A6TBN8 A6TBP4 A6TBQ0 A6TBR5 A6TBS0 A6TBS1 A6TBT7 A6TBU6 A6TBU8 A6TBV3 A6TBV5
A6TBV7 A6TBV8 A6TBW1 A6TBX7 A6TC04 A6TC10 A6TC15 A6TC17 A6TC33 A6TC43 A6TC47
A6TC68 A6TC90 A6TC94 A6TC99 A6TCA1 A6TCA8 A6TCB0 A6TCB3 A6TCC2 A6TCC4 A6TCD0
A6TCD4 A6TCD6 A6TCD7 A6TCE4 A6TCE7 A6TCE8 A6TCE9 A6TCF1 A6TCF2 A6TCF3 A6TCG5
A6TCH9 A6TCI1 A6TCI9 A6TCJ1 A6TCJ2 A6TCL4 A6TCL5 A6TCL6 A6TCL7 A6TCM1 A6TCM2
A6TCM7 A6TCU0 A6TCV4 A6TCV8 A6TCV9 A6TCW0 A6TCW1 A6TCX5 A6TCX6 A6TCZ3 A6TD24
A6TD35 A6TD37 A6TD38 A6TD39 A6TD42 A6TD44 A6TD45 A6TD47 A6TD48 A6TD49 A6TD54
A6TD57 A6TD67 A6TD79 A6TD83 A6TD85 A6TD87 A6TDB1 A6TDC4 A6TDD9 A6TDE9 A6TDG3
A6TDG6 A6TDG7 A6TDP3 A6TDQ4 A6TDR5 A6TDR6 A6TDR7 A6TDS1 A6TDS5 A6TDS6 A6TDT2
A6TDU9 A6TDV1 A6TDV3 A6TDV7 A6TDV8 A6TDW3 A6TDW8 A6TDX0 A6TE25 A6TE29 A6TE31
A6TE35 A6TE39 A6TE40 A6TE41 A6TE42 A6TE43 A6TE44 A6TE45 A6TE46 A6TE47 A6TE50
A6TEB2 A6TEB3 A6TEB4 A6TEB9 A6TEC0 A6TEC2 A6TEE9 A6TEF4 A6TEF8 A6TEG7 A6TEG8
A6TEH3 A6TEH4 A6TEI3 A6TEI4 A6TEI5 A6TEI6 A6TEI7 A6TEI9 A6TEJ0 A6TEJ5 A6TEJ8

A6TEK2 A6TEK4 A6TEK5 A6TEK8 A6TEM4 A6TEN5 A6TEN8 A6TEN9 A6TEQ3 A6TEQ4 A6TES6
A6TES8 A6TET6 A6TEU0 A6TEU1 A6TEU2 A6TEU7 A6TEU8 A6TEU9 A6TEV0 A6TEV1 A6TEV3
A6TEV4 A6TEV5 A6TEV6 A6TEV7 A6TEV8 A6TEW0 A6TEW1 A6TEW2 A6TEW3 A6TEW4 A6TEW5
A6TEW6 A6TEW7 A6TEW8 A6TEW9 A6TEX1 A6TEX2 A6TEX3 A6TEX7 A6TEX9 A6TEY0 A6TEY1
A6TEY2 A6TEY3 A6TEY6 A6TEZ3 A6TF07 A6TF13 A6TF33 A6TF35 A6TF37 A6TF41 A6TF44
A6TF48 A6TF49 A6TF50 A6TF51 A6TF89 A6TFA8 A6TFB5 A6TFD9 A6TFH4 A6TFH5 A6TFJ1
A6TFJ4 A6TFK4 A6TFL2 A6TFL4 A6TFM5 A6TFM6 A6TFM7 A6TFM8 A6TFM9 A6TFN1 A6TFN2
A6TFN4 A6TFN5 A6TFP6 A6TFP8 A6TFR2 A6TFW2 A6TFW3 A6TFY9 A6TFZ4 A6TG02 A6TG05
A6TG09 A6TG44 A6TG45 A6TG48 A6TG52 A6TG68 A6TG77 A6TG82 A6TG86 A6TGA0 A6TGA6
A6TGA7 A6TGA8 A6TGA9 A6TGB0 A6TGB8 A6TGB9 A6TGC1 A6TGC2 A6TGC6 A6TGC7 A6TGD7
A6TGE3 A6TGE4 A6TGE6 A6TGE9 A6TGF8 A6TGG1 A6TGG8 A6TGG9 A6TGI8 A6TGJ0 A6TGK9
A6TGL3 A6TGM5 A6TGN1 A6TGN5 A6TGN6 A6TGN8 A6TGN9 A6TGP0 A6TGP1 A6TGP7 A6TGQ0
A6TGQ1 A6TGQ3 A6TGQ4 A6TGQ5 A6TGR3 A6TGR5 A6TGR8 A6TGT4 A6TGU0 A6TGU8 A6TH13
A6TH52 A6TH53 A6TH65 A6TH74 A6TH78 A6TH79 A6TH85 A6TH86 A6TH87 A6TH93 A6TH94
A6THB1 A6THB3 A6THB4 A6THB9 A6THE3 A6THE7 A6THH3 A6THH4 A6THI2 A6THJ2 A6THJ4
A6THU5 A6THX4 A6THY6 A6THZ0 A6THZ7 A6TI00 A6TI07 A6TI08 A6TI09 A6TI29 A6TJ76
A6TJ79 A6TJ92 A6TJ99 A6TJA7 A6TJA9 A6TJD3 A6TJD5 A6TJD8 A6TJE5 A6TJK9 A6TJM2
A6TJM4 A6TJM7 A6TJX7 A6TK31 A6TK36 A6TK43 A6TK51 A6TKH4 A6TKJ0 A6TKJ3 A6TKN0
A6TKP1 A6TKR6 A6TKS4 A6TKS7 A6TKS9 A6TKT0 A6TKT3 A6TKT5 A6TKT6 A6TL04 A6TL06
A6TL07 A6TL09 A6TL10 A6TL31 A6TL60 A6TL87 A6TL88 A6TLJ0 A6TLJ1 A6TLM6 A6TLR3
A6TLS2 A6TLS3 A6TLS5 A6TLS7 A6TLU4 A6TLX6 A6TM00 A6TM05 A6TM20 A6TM60 A6TM61
A6TM62 A6TM67 A6TM69 A6TM73 A6TM74 A6TM76 A6TM77 A6TMH9 A6TMJ6 A6TMN5 A6TMN6
A6TMY6 A6TN07 A6TN94 A6TNI2 A6TNN7 A6TNP3 A6TNP4 A6TNP8 A6TNW2 A6TNW5 A6TNX0
A6TNX2 A6TNZ5 A6TPR4 A6TPX1 A6TQ57 A6TQ96 A6TQC4 A6TQH7 A6TQH8 A6TQI5 A6TQI6
A6TQJ2 A6TQJ6 A6TQK3 A6TQL2 A6TQM3 A6TQM4 A6TQM5 A6TQM7 A6TQN8 A6TQP0 A6TQQ1
A6TQZ4 A6TQZ5 A6TR01 A6TR08 A6TR25 A6TR38 A6TR39 A6TR45 A6TR59 A6TR67 A6TR76
A6TR77 A6TR78 A6TR79 A6TR80 A6TRF9 A6TRG1 A6TRH0 A6TRI3 A6TRJ4 A6TRK1 A6TRK2
A6TRK6 A6TRK7 A6TRL1 A6TRM0 A6TRM1 A6TRM2 A6TRM3 A6TRS2 A6TRS3 A6TRS5 A6TRS6
A6TRS7 A6TRS9 A6TRU4 A6TRU6 A6TRU7 A6TRV0 A6TRV6 A6TRW0 A6TRW3 A6TRW7 A6TRW8
A6TRX1 A6TRX3 A6TRX5 A6TRX7 A6TRY5 A6TS03 A6TS39 A6TS50 A6TS63 A6TS66 A6TS69
A6TS70 A6TSC8 A6TSF6 A6TSF8 A6TSH1 A6TSI5 A6TSJ2 A6TSJ4 A6TSJ6 A6TSK3 A6TSL4
A6TSL8 A6TSM0 A6TSM2 A6TSM8 A6TSW9 A6TSX0 A6TT15 A6TT16 A6TT61 A6TTD6 A6TTJ1
A6TTJ3 A6TTJ8 A6TTL2 A6TTY3 A6TU24 A6TU32 A6TU46 A6TU70 A6TU88 A6TU96 A6TU99
A6TUA0 A6TUB2 A6TUE9 A6TUM7 A6TVD3 A6TVF1 A6TVF5 A6TVF6 A6TVF9 A6TVI5 A6TVI7
A6TVN2 A6TVN9 A6TVP5 A6TVQ2 A6TVR2 A6TVS0 A6TVS9 A6TVU0 A6TVU5 A6TVU7 A6TVW5
A6TVX6 A6TW06 A6TWC9 A6TWD0 A6TWD9 A6TWE6 A6TWE7 A6TWF2 A6TWF3 A6TWF4 A6TWF5
A6TWF6 A6TWF7 A6TWF8 A6TWG3 A6TWG4 A6TWG5 A6TWG6 A6TWG7 A6TWG8 A6TWG9 A6TWH0
A6TWH1 A6TWH2 A6TWH3 A6TWH4 A6TWH5 A6TWH6 A6TWH7 A6TWH8 A6TWH9 A6TWI0 A6TWI1
A6TWI2 A6TWI3 A6TWI4 A6TWI5 A6TWI6 A6TWI7 A6TWI9 A6TWJ0 A6TWJ1 A6TWJ2 A6TWJ3
A6TWJ4 A6TWJ7 A6TWJ8 A6TWK5 A6TWK7 A6TWK9 A6TWL0 A6TWN2 A6TWV7 A6TWW1 A6TWW9
A6TWX8 A6TX17 A6TX97 A6TXB3 A6TXB7 A6TXC7 A6TXC8 A6TXE3 A6TXE4 A6TXE5 A6TXE9
A6TXF0 A6TXF1 A6TXF4 A6TXF8 A6TXF9 A6TXG5 A6TXH4 A6TXQ1 A6TXS4 A6TXW6 A6TXZ6
A6TY25 A6TY44 A6TY54 A6TY99 A6TYA2 A6TYK8 A6TYL8 A6TYM0 A6TYN9 A6TYP2 A6TYV3
A6TYV4 A6TYV6 A6TYW0 A6TYW7 A6TYW9 A6TYX2 A6TYX7 A6TYY7 A6TYZ5 A6TYZ6 A6TZ04

A6TZ06 A6TZ11 A6TZ14 A6TZ15 A6TZ16 A6TZ17 A6TZ19 A6TZ21 A6TZ22 A6TZ23 A6TZ24
A6TZ25 A6TZ28 A6TZ43 A6TZ46 A6TZ58 A6TZ65 A6TZ71 A6TZ85 A6TZE7 A6TZF9 A6TZJ0
A6TZK2 A6TZK7 A6TZK9 A6TZL7 A6TZM2 A6TZN3 A6TZN9 A6TZP4 A6TZP6 A6TZP7 A6TZQ2
A6TZQ3 A6TZR0 A6TZT3 A6TZT7 A6U025 A6U030 A6U039 A6U046 A6U067 A6U068 A6U073
A6U074 A6U077 A6U0A4 A6U0B0 A6U0B2 A6U0C0 A6U0C5 A6U0L1 A6U0N1 A6U0N4 A6U0N7
A6U0N9 A6U0P1 A6U0Q8 A6U0R5 A6U0S4 A6U0T0 A6U0T9 A6U0U2 A6U0U3 A6U0U5 A6U0V5
A6U0V7 A6U0W1 A6U0W3 A6U0W7 A6U0Z7 A6U0Z8 A6U0Z9 A6U103 A6U109 A6U113 A6U117
A6U119 A6U122 A6U129 A6U135 A6U137 A6U143 A6U152 A6U157 A6U158 A6U159 A6U160
A6U163 A6U164 A6U169 A6U170 A6U171 A6U172 A6U173 A6U174 A6U176 A6U178 A6U182
A6U184 A6U188 A6U189 A6U190 A6U192 A6U193 A6U1A4 A6U1B1 A6U1B4 A6U1B5 A6U1B8
A6U1C1 A6U1F0 A6U1F5 A6U1F6 A6U1F7 A6U1G6 A6U1G7 A6U1H8 A6U1J1 A6U1J2 A6U1J4
A6U1J5 A6U1L6 A6U1L7 A6U1L8 A6U1N4 A6U1P1 A6U1P3 A6U1P7 A6U1R5 A6U1R6 A6U1R8
A6U1S3 A6U1T1 A6U1T3 A6U1T4 A6U1T5 A6U1T7 A6U1T9 A6U1U3 A6U1U6 A6U1W3 A6U1W4
A6U1X3 A6U1Z2 A6U1Z4 A6U1Z5 A6U1Z6 A6U200 A6U206 A6U207 A6U208 A6U209 A6U223
A6U229 A6U235 A6U237 A6U242 A6U247 A6U250 A6U251 A6U252 A6U253 A6U254 A6U258
A6U266 A6U268 A6U271 A6U274 A6U279 A6U280 A6U284 A6U285 A6U286 A6U287 A6U290
A6U298 A6U299 A6U2A1 A6U2A3 A6U2A7 A6U2A8 A6U2A9 A6U2B0 A6U2B1 A6U2B2 A6U2B3
A6U2B5 A6U2D1 A6U2D5 A6U2D8 A6U2E0 A6U2E1 A6U2E2 A6U2E3 A6U2E6 A6U2E7 A6U2F0
A6U2F3 A6U2G5 A6U2H3 A6U2H8 A6U2I2 A6U2I6 A6U2J5 A6U2J8 A6U2K6 A6U2K9 A6U2L4
A6U2M4 A6U2N1 A6U2P0 A6U2Q1 A6U2Q2 A6U2Q7 A6U2T7 A6U2U8 A6U2W8 A6U2X1 A6U2X7
A6U2Z9 A6U304 A6U305 A6U306 A6U309 A6U318 A6U325 A6U3B5 A6U3B6 A6U3D1 A6U3D4
A6U3D8 A6U3E1 A6U3E2 A6U3E4 A6U3E5 A6U3E7 A6U3F0 A6U3G9 A6U3H7 A6U3H8 A6U3J7
A6U3J8 A6U3J9 A6U3K3 A6U3K5 A6U3L2 A6U3L3 A6U3N9 A6U3P1 A6U3R9 A6U3S2 A6U3S3
A6U3S4 A6U3S5 A6U3U2 A6U3U3 A6U3U4 A6U3U8 A6U3V0 A6U3V1 A6U3V2 A6U3V3 A6U3V6
A6U3V7 A6U3V8 A6U3V9 A6U3W0 A6U3W1 A6U3W2 A6U3W3 A6U3W4 A6U3W5 A6U3W6 A6U3W7
A6U3W8 A6U3X0 A6U3X1 A6U3X2 A6U3X3 A6U3X4 A6U3X5 A6U3X6 A6U3Z2 A6U3Z3 A6U3Z8
A6U404 A6U412 A6U413 A6U414 A6U415 A6U416 A6U417 A6U418 A6U455 A6U458 A6U459
A6U471 A6U473 A6U499 A6U4C1 A6U4E6 A6U4F3 A6U4F5 A6U4P6 A6U4T5 A6U4T9 A6U4X7
A6U4X8 A6U4X9 A6U4Y2 A6U4Y6 A6U4Z2 A6U515 A6U519 A6U558 A6U559 A6U561 A6U564
A6U565 A6U574 A6U575 A6U593 A6U594 A6U595 A6U596 A6U597 A6U5D1 A6U5E0 A6U5E1
A6U5E2 A6U5F1 A6U5G0 A6U5G1 A6U5H0 A6U5H2 A6U5H5 A6U5H7 A6U5I5 A6U5I6 A6U5J1
A6U5J9 A6U5K5 A6U5K8 A6U5L1 A6U5N6 A6U5N7 A6U5N9 A6U5R0 A6U5X9 A6U5Y5 A6U5Z0
A6U5Z2 A6U608 A6U619 A6U636 A6U659 A6U669 A6U6F0 A6U6I1 A6U6I2 A6U6I5 A6U6K5
A6U6M8 A6U6N3 A6U6P3 A6U6S1 A6U6T5 A6U6U1 A6U6V4 A6U6V5 A6U6V8 A6U6Y8 A6U6Y9
A6U707 A6U753 A6U7A0 A6U7A2 A6U7A8 A6U7B9 A6U7C5 A6U7D9 A6U7G4 A6U7G6 A6U7G7
A6U7I6 A6U7J2 A6U7K1 A6U7L8 A6U7M8 A6U7N0 A6U7N6 A6U7P3 A6U7P8 A6U7Q6 A6U7T4
A6U7T6 A6U7T7 A6U7U8 A6U7Z3 A6U802 A6U811 A6U816 A6U817 A6U818 A6U819 A6U820
A6U821 A6U825 A6U828 A6U839 A6U842 A6U847 A6U848 A6U850 A6U851 A6U852 A6U854
A6U855 A6U856 A6U858 A6U859 A6U860 A6U861 A6U862 A6U863 A6U864 A6U865 A6U866
A6U867 A6U868 A6U869 A6U870 A6U871 A6U872 A6U873 A6U874 A6U875 A6U876 A6U877
A6U878 A6U880 A6U881 A6U882 A6U883 A6U884 A6U899 A6U8A1 A6U8C1 A6U8C2 A6U8C7
A6U8E2 A6U8F5 A6U8F8 A6U8G6 A6U8K2 A6U8K3 A6U8K4 A6U8K5 A6U8M8 A6U8M9 A6U8N8
A6U8N9 A6U8P9 A6U8Q2 A6U8Q3 A6U8Q4 A6U8X4 A6U8Y7 A6U901 A6U9C4 A6U9C5 A6U9C6
A6U9H2 A6U9H4 A6U9I7 A6U9L2 A6U9L9 A6U9M6 A6U9M8 A6U9N0 A6U9N4 A6U9N5 A6U9R1

A6U9T4 A6U9U3 A6U9W4 A6UAC0 A6UAW6 A6UB14 A6UB24 A6UB62 A6UB63 A6UB73 A6UB80
A6UB83 A6UB84 A6UB93 A6UB95 A6UB96 A6UB98 A6UBM6 A6UBR5 A6UBS2 A6UBU5 A6UBV0
A6UC29 A6UC30 A6UC35 A6UC38 A6UC40 A6UC41 A6UCB5 A6UCC1 A6UCF2 A6UCF6 A6UCK6
A6UCL4 A6UCQ0 A6UCR6 A6UCS2 A6UCT7 A6UCT8 A6UCT9 A6UCU1 A6UCV3 A6UCW1 A6UCX4
A6UCY0 A6UD23 A6UD82 A6UD89 A6UD99 A6UDA0 A6UDC0 A6UDD3 A6UDH2 A6UDH9 A6UDM7
A6UDP2 A6UDP3 A6UDR4 A6UDV2 A6UDV3 A6UDV6 A6UDV7 A6UDV8 A6UDW1 A6UDW8 A6UDX5
A6UDZ3 A6UE05 A6UE09 A6UE14 A6UE17 A6UE43 A6UE44 A6UE47 A6UE65 A6UE79 A6UED3
A6UED8 A6UEE2 A6UEE7 A6UEE8 A6UEE9 A6UEF2 A6UEF3 A6UEF5 A6UEF8 A6UEI0 A6UEI1
A6UEI2 A6UEJ1 A6UEJ7 A6UEK1 A6UEK2 A6UEK3 A6UEK4 A6UEK7 A6UEK8 A6UEL0 A6UEL4
A6UET3 A6UET4 A6UET5 A6UET6 A6UEW0 A6UEW2 A6UEW3 A6UEY0 A6UEY1 A6UF02 A6UF23
A6UF26 A6UF29 A6UF30 A6UF31 A6UF33 A6UF34 A6UF45 A6UF66 A6UF72 A6UF73 A6UF74
A6UF86 A6UFA3 A6UFC6 A6UFD1 A6UFD9 A6UFF7 A6UFF8 A6UH06 A6UH73 A6UKC3 A6ULQ4
A6UM48 A6UN45 A6UN58 A6UN66 A6UN70 A6UN73 A6UN84 A6UN85 A6UN87 A6UNL1 A6UNM1
A6UNQ2 A6UNQ4 A6UNQ5 A6UNQ9 A6UNR2 A6UNS4 A6UNT0 A6UNT3 A6UNU0 A6UNU1 A6UNY0
A6UP00 A6UP03 A6UP14 A6UP16 A6UP19 A6UP25 A6UP37 A6UP41 A6UP44 A6UP47 A6UP61
A6UP65 A6UP68 A6UP85 A6UP86 A6UPB5 A6UPD8 A6UPD9 A6UPE5 A6UPE6 A6UPF4 A6UPH7
A6UPI8 A6UPJ0 A6UPJ2 A6UPK3 A6UPK5 A6UPK8 A6UPL1 A6UPL5 A6UPL6 A6UPM1 A6UPN5
A6UPN7 A6UPN9 A6UPQ3 A6UPQ4 A6UPR3 A6UPR4 A6UPS6 A6UPT0 A6UPV2 A6UPV4 A6UPW5
A6UPW6 A6UPX0 A6UPX1 A6UPX8 A6UPZ7 A6UQ14 A6UQ18 A6UQ31 A6UQ39 A6UQ46 A6UQ48
A6UQ49 A6UQ52 A6UQ54 A6UQ55 A6UQ56 A6UQ57 A6UQ59 A6UQ60 A6UQ61 A6UQ63 A6UQ64
A6UQ65 A6UQ66 A6UQ70 A6UQ71 A6UQ72 A6UQ76 A6UQ77 A6UQ78 A6UQ84 A6UQ90 A6UQB4
A6UQB6 A6UQC1 A6UQD0 A6UQD4 A6UQD9 A6UQF3 A6UQF6 A6UQJ0 A6UQJ1 A6UQJ3 A6UQJ4
A6UQL3 A6UQL8 A6UQL9 A6UQM0 A6UQM3 A6UQM4 A6UQM7 A6UQM8 A6UQN4 A6UQQ6 A6UQQ8
A6UQS6 A6UQT3 A6UQT7 A6UQT9 A6UQU0 A6UQW0 A6UQY8 A6UQY9 A6UQZ4 A6UR05 A6UR12
A6UR14 A6UR43 A6UR44 A6UR49 A6UR53 A6UR54 A6UR55 A6UR56 A6UR69 A6UR75 A6UR76
A6UR78 A6UR83 A6UR90 A6UR91 A6UR99 A6URA0 A6URA6 A6URB0 A6URB4 A6URC0 A6URC7
A6URD2 A6URD8 A6URE0 A6URG2 A6URG5 A6URH2 A6URH7 A6URI4 A6URJ1 A6URK3 A6URL2
A6URL5 A6URN3 A6URN7 A6URN9 A6URP5 A6URP9 A6URQ1 A6URR6 A6URS1 A6URS3 A6URS6
A6URT1 A6URT3 A6URT7 A6URV4 A6URV6 A6URV8 A6URW1 A6URW2 A6URW3 A6URY5 A6URZ7
A6US04 A6US09 A6US15 A6US17 A6US18 A6US27 A6US28 A6US29 A6US36 A6US39 A6US51
A6US52 A6US53 A6US67 A6US71 A6US72 A6US81 A6USA0 A6USA3 A6USA4 A6USB0 A6USB4
A6USC0 A6USC1 A6USC2 A6USC8 A6USD4 A6USD6 A6USE4 A6USE6 A6USF2 A6USF9 A6USH1
A6USH4 A6USH5 A6USJ0 A6USJ3 A6USJ4 A6USJ5 A6USK4 A6USK6 A6USL5 A6USM7 A6USP4
A6USY3 A6USY4 A6USZ5 A6USZ9 A6UT01 A6UT04 A6UT10 A6UT12 A6UT20 A6UT21 A6UT22
A6UT27 A6UT49 A6UT50 A6UT51 A6UT74 A6UT94 A6UT95 A6UT96 A6UT97 A6UTA2 A6UTA7
A6UTA8 A6UTB1 A6UTB9 A6UTD8 A6UTD9 A6UTE4 A6UTE5 A6UTE9 A6UTF2 A6UTF3 A6UTF6
A6UTF8 A6UTF9 A6UTG8 A6UTH4 A6UTK3 A6UTK4 A6UTL4 A6UTL8 A6UTM2 A6UTP0 A6UTP5
A6UTP9 A6UTQ0 A6UTQ8 A6UTT2 A6UTV8 A6UTV9 A6UTW9 A6UTZ1 A6UU47 A6UU49 A6UU55
A6UU57 A6UU64 A6UU75 A6UU96 A6UU98 A6UUG9 A6UUH1 A6UUK5 A6UUK8 A6UUL6 A6UUL7
A6UUL9 A6UUM1 A6UUN0 A6UUN1 A6UUN3 A6UUN6 A6UUN9 A6UUQ4 A6UUS0 A6UUS4 A6UUS5
A6UUS7 A6UUT0 A6UUT1 A6UUT2 A6UUU2 A6UUU3 A6UUV1 A6UUW0 A6UUW5 A6UUY1 A6UUY8
A6UUZ4 A6UUZ8 A6UUZ9 A6UV02 A6UV05 A6UV11 A6UV17 A6UV20 A6UV22 A6UV33 A6UV34
A6UV42 A6UV43 A6UV44 A6UV45 A6UV46 A6UV49 A6UV53 A6UV59 A6UV64 A6UV65 A6UV66
A6UV67 A6UV68 A6UV93 A6UVA1 A6UVA8 A6UVC4 A6UVC9 A6UVD8 A6UVE3 A6UVF4 A6UVG0

A6UVG7 A6UVG8 A6UVH0 A6UVH2 A6UVH4 A6UVI4 A6UVJ0 A6UVK2 A6UVK7 A6UVN4 A6UVQ1
A6UVQ4 A6UVQ6 A6UVR4 A6UVR5 A6UVT6 A6UVU3 A6UVV2 A6UVW6 A6UW00 A6UW04 A6UW05
A6UW07 A6UW15 A6UW21 A6UW24 A6UW25 A6UW32 A6UW33 A6UW35 A6UW53 A6UW55 A6UW60
A6UW69 A6UW73 A6UW87 A6UW89 A6UW94 A6UWA5 A6UWB3 A6UWC6 A6UWF5 A6UWF6 A6UWI0
A6UWK8 A6UWL8 A6UWM0 A6UWM1 A6UWN1 A6UWP2 A6UWP3 A6UWR5 A6UWR8 A6UWS1 A6UWT0
A6UWT1 A6UWT5 A6UWU0 A6UWU2 A6UWU3 A6UWU4 A6UWU6 A6UWU8 A6UWU9 A6UWV0 A6UWV1
A6UWV3 A6UWV4 A6UWV7 A6UWV8 A6UWV9 A6UWW0 A6UWW4 A6UWW5 A6UWW6 A6UWX3 A6UWX4
A6UWZ3 A6UX13 A6UX30 A6UX37 A6UX46 A6UX50 A6UX54 A6UX55 A6UX64 A6UX70 A6UX80
A6UX84 A6UX86 A6UX87 A6UX98 A6UXB5 A6UXT7 A6UY04 A6UY05 A6UY93 A6UYD2 A6UYD9
A6UYE6 A6UYF6 A6UYG0 A6UYJ0 A6UYJ1 A6UYJ9 A6UYL2 A6UYL3 A6UYL4 A6UYL5 A6UYU5
A6UYW0 A6UYW1 A6UYW3 A6UYW4 A6UZ09 A6UZ15 A6UZ82 A6UZ83 A6UZ87 A6UZ90 A6UZ93
A6UZ94 A6UZ95 A6UZF1 A6UZF2 A6UZF5 A6UZG3 A6UZG6 A6UZG7 A6UZH2 A6UZH4 A6UZH7
A6UZH8 A6UZH9 A6UZI0 A6UZI1 A6UZI2 A6UZI3 A6UZI4 A6UZI7 A6UZI8 A6UZI9 A6UZJ0
A6UZJ1 A6UZJ2 A6UZJ3 A6UZJ4 A6UZJ5 A6UZJ6 A6UZJ7 A6UZJ8 A6UZJ9 A6UZK0 A6UZK1
A6UZK2 A6UZK3 A6UZK4 A6UZK5 A6UZK6 A6UZK7 A6UZK9 A6UZL0 A6UZL1 A6UZL2 A6UZL3
A6UZL4 A6UZV3 A6V049 A6V055 A6V058 A6V060 A6V082 A6V0A3 A6V0A4 A6V0A6 A6V0B2
A6V0B3 A6V0B4 A6V0C2 A6V0C7 A6V0C9 A6V0D2 A6V0D3 A6V0J0 A6V0K2 A6V0Q7 A6V0S3
A6V0T2 A6V0T8 A6V0U8 A6V0V1 A6V0V2 A6V0V6 A6V0V7 A6V0W0 A6V0W1 A6V0W4 A6V0X2
A6V123 A6V124 A6V125 A6V126 A6V187 A6V1A0 A6V1A1 A6V1A8 A6V1C2 A6V1D0 A6V1D2
A6V1D3 A6V1D4 A6V1D5 A6V1D8 A6V1E6 A6V1F1 A6V1F5 A6V1G1 A6V1G2 A6V1G3 A6V1G4
A6V1G8 A6V1H2 A6V1H3 A6V1I9 A6V1J0 A6V1K5 A6V1Q6 A6V1R8 A6V1R9 A6V1S1 A6V1U3
A6V1U5 A6V1V5 A6V1W3 A6V1X0 A6V1X3 A6V255 A6V262 A6V271 A6V291 A6V2F6 A6V2H4
A6V2H6 A6V2H8 A6V2N1 A6V2Q0 A6V2Q4 A6V2Q6 A6V2Q8 A6V2R2 A6V2R4 A6V2T9 A6V2V3
A6V2V4 A6V2W1 A6V2Y8 A6V328 A6V333 A6V365 A6V3A6 A6V3B8 A6V3C8 A6V3D6 A6V3I3
A6V3L5 A6V3N9 A6V3Q8 A6V3R3 A6V3T3 A6V3U3 A6V485 A6V487 A6V488 A6V489 A6V491
A6V4G0 A6V4G5 A6V4G7 A6V4G8 A6V4G9 A6V4H5 A6V4H6 A6V4J2 A6V4P3 A6V5Z4 A6V616
A6V628 A6V669 A6V681 A6V6B1 A6V6F5 A6V6I1 A6V6I3 A6V6I4 A6V6X0 A6V6X8 A6V6Z8
A6V705 A6V707 A6V718 A6V720 A6V726 A6V727 A6V753 A6V7A6 A6V7A7 A6V7A8 A6V7B0
A6V7B8 A6V7E9 A6V7F4 A6V7J7 A6V7K5 A6V7S3 A6V7T5 A6V7T6 A6V7V8 A6V7W8 A6V7X3
A6V7Y9 A6V7Z0 A6V8H8 A6V8H9 A6V8P9 A6V8R1 A6V8R5 A6V8R6 A6V8T2 A6V906 A6V910
A6V936 A6V990 A6V9E9 A6V9K1 A6V9L0 A6V9R6 A6V9Z6 A6VA05 A6VA06 A6VA07 A6VA08
A6VA12 A6VA16 A6VA18 A6VA50 A6VA58 A6VA71 A6VA73 A6VA75 A6VAA4 A6VAC0 A6VAE3
A6VAK1 A6VAK2 A6VAK3 A6VAK4 A6VAK6 A6VAM7 A6VAQ5 A6VAU7 A6VB57 A6VB58 A6VB67
A6VB84 A6VB93 A6VB97 A6VB98 A6VBA5 A6VBA6 A6VBB6 A6VBC2 A6VBC3 A6VBD9 A6VBI8
A6VBJ7 A6VBJ8 A6VBJ9 A6VBN0 A6VBR6 A6VBU7 A6VBV0 A6VBV2 A6VBV3 A6VBV4 A6VBV5
A6VC08 A6VC20 A6VC42 A6VC51 A6VC58 A6VC61 A6VC62 A6VC65 A6VC67 A6VC68 A6VCE7
A6VCH5 A6VCH6 A6VCH9 A6VCI6 A6VCI7 A6VCI8 A6VCJ6 A6VCJ8 A6VCJ9 A6VCK0 A6VCK1
A6VCK3 A6VCK5 A6VCK6 A6VCK9 A6VCL6 A6VCL7 A6VCL8 A6VCL9 A6VCM3 A6VCM5 A6VCN1
A6VCR3 A6VCV6 A6VCW0 A6VCX1 A6VCX2 A6VCX3 A6VCX4 A6VCY0 A6VD00 A6VD01 A6VD02
A6VD32 A6VD45 A6VD47 A6VD48 A6VD51 A6VD52 A6VD57 A6VD58 A6VD59 A6VD64 A6VD86
A6VDB5 A6VDE4 A6VDF5 A6VDF9 A6VDG0 A6VDH0 A6VDH2 A6VDH4 A6VDH5 A6VDI6 A6VDI9
A6VDJ0 A6VDK2 A6VDL0 A6VDL7 A6VDM7 A6VDN5 A6VDP6 A6VDP9 A6VDQ9 A6VDR0 A6VDR3
A6VDR9 A6VDW7 A6VDW9 A6VDX3 A6VDX9 A6VDY0 A6VDZ9 A6VE23 A6VE37 A6VE40 A6VE49
A6VE52 A6VE54 A6VEA3 A6VEC3 A6VEC6 A6VEC8 A6VED0 A6VED8 A6VEE1 A6VEE4 A6VEI3

A6VEI4 A6VEI5 A6VEW7 A6VEZ4 A6VF17 A6VF42 A6VF43 A6VF44 A6VF46 A6VF47 A6VFD6
A6VFG8 A6VFH0 A6VFH1 A6VFI6 A6VFI9 A6VFJ7 A6VFK8 A6VFL2 A6VFL4 A6VFL5 A6VFL9
A6VFM6 A6VFM7 A6VFM8 A6VFN4 A6VFN7 A6VFN8 A6VFP0 A6VFP3 A6VFQ9 A6VFS2 A6VFS5
A6VFT8 A6VFU0 A6VFU3 A6VFU7 A6VFW2 A6VFX9 A6VFY2 A6VFY5 A6VFZ9 A6VG03 A6VG07
A6VG23 A6VG24 A6VG49 A6VG76 A6VG82 A6VG83 A6VG90 A6VGC7 A6VGC9 A6VGD1 A6VGE3
A6VGE5 A6VGE7 A6VGE8 A6VGF1 A6VGF5 A6VGF6 A6VGG1 A6VGG2 A6VGH5 A6VGH7 A6VGH9
A6VGJ3 A6VGJ4 A6VGK4 A6VGK5 A6VGL9 A6VGM3 A6VGM6 A6VGP5 A6VGP7 A6VGQ7 A6VGQ8
A6VGR2 A6VGR3 A6VGS0 A6VGT9 A6VGU6 A6VGV0 A6VGV1 A6VGV3 A6VGV4 A6VGV5 A6VGV6
A6VGV7 A6VGV9 A6VGW0 A6VGX3 A6VGY0 A6VGY7 A6VGY9 A6VGZ0 A6VGZ1 A6VGZ2 A6VGZ5
A6VGZ6 A6VGZ7 A6VGZ8 A6VH00 A6VH01 A6VH02 A6VH04 A6VH05 A6VH06 A6VH07 A6VH11
A6VH12 A6VH13 A6VH17 A6VH18 A6VH19 A6VH25 A6VH31 A6VH54 A6VH56 A6VH58 A6VH63
A6VH72 A6VH76 A6VH81 A6VH94 A6VH97 A6VHC3 A6VHD2 A6VHD3 A6VHD4 A6VHD5 A6VHD6
A6VHG3 A6VHG6 A6VHG7 A6VHH0 A6VHH1 A6VHH7 A6VHK3 A6VHK5 A6VHQ4 A6VHR1 A6VHR9
A6VHS1 A6VHS2 A6VHU3 A6VHV2 A6VHX1 A6VHX2 A6VHX7 A6VHY8 A6VHZ4 A6VI01 A6VI08
A6VI14 A6VI22 A6VI24 A6VI28 A6VI37 A6VI38 A6VI43 A6VI47 A6VI48 A6VI49 A6VI50
A6VI60 A6VI67 A6VI74 A6VI77 A6VI82 A6VI91 A6VI92 A6VIA5 A6VIA6 A6VIB2 A6VIB6
A6VIC0 A6VID5 A6VIE1 A6VIE7 A6VIE9 A6VIG8 A6VIH3 A6VIH6 A6VIK1 A6VIL1 A6VIL3
A6VIL5 A6VIN4 A6VIN8 A6VIP0 A6VIP7 A6VIP9 A6VIQ0 A6VIQ1 A6VIS0 A6VIS4 A6VIS6
A6VIT0 A6VIT5 A6VIT7 A6VIT8 A6VIU0 A6VIV7 A6VIV9 A6VIW1 A6VIW4 A6VIW5 A6VIW6
A6VJ05 A6VJ20 A6VJ27 A6VJ32 A6VJ39 A6VJ40 A6VJ50 A6VJ51 A6VJ52 A6VJ61 A6VJ64
A6VJ76 A6VJ77 A6VJ78 A6VJJ9 A6VJK4 A6VJM2 A6VJM9 A6VJN0 A6VJP1 A6VJQ1 A6VJQ7
A6VJR0 A6VJR3 A6VJR4 A6VJS2 A6VJS4 A6VJT0 A6VJT1 A6VJT2 A6VJU5 A6VJU6 A6VJU8
A6VJW0 A6VJW6 A6VJX1 A6VJX3 A6VJX5 A6VJY2 A6VJY4 A6VJZ0 A6VJZ5 A6VK02 A6VK03
A6VK04 A6VK13 A6VK15 A6VK38 A6VK86 A6VK88 A6VK96 A6VK98 A6VK99 A6VKA0 A6VKA1
A6VKA4 A6VKC3 A6VKC4 A6VKC5 A6VKC6 A6VKC7 A6VKC8 A6VKC9 A6VKD5 A6VKD6 A6VKH3
A6VKH7 A6VKH8 A6VKJ6 A6VKK5 A6VKM8 A6VKQ3 A6VKQ4 A6VKR5 A6VKS6 A6VKS7 A6VKT5
A6VKU4 A6VKU5 A6VKV4 A6VKV5 A6VKX6 A6VKX9 A6VKY5 A6VKY6 A6VKY9 A6VKZ5 A6VKZ6
A6VL11 A6VL12 A6VL27 A6VL65 A6VL66 A6VL68 A6VL69 A6VL72 A6VL74 A6VL76 A6VL77
A6VL80 A6VL81 A6VL83 A6VL87 A6VLB1 A6VLB7 A6VLC4 A6VLD2 A6VLD6 A6VLD8 A6VLE0
A6VLE1 A6VLE6 A6VLF1 A6VLF2 A6VLF5 A6VLF9 A6VLG0 A6VLI7 A6VLI8 A6VLI9 A6VLJ1
A6VLJ2 A6VLJ3 A6VLJ4 A6VLJ5 A6VLJ6 A6VLJ7 A6VLJ8 A6VLJ9 A6VLK0 A6VLK1 A6VLK2
A6VLK3 A6VLK4 A6VLK5 A6VLK6 A6VLK7 A6VLK9 A6VLL1 A6VLL2 A6VLL3 A6VLL4 A6VLM2
A6VLM8 A6VLN1 A6VLN4 A6VLN8 A6VLP4 A6VLP5 A6VLP6 A6VLP7 A6VLP8 A6VLQ3 A6VLR0
A6VLR3 A6VLR7 A6VLR9 A6VLS1 A6VLT3 A6VLU0 A6VLU1 A6VLV0 A6VLV3 A6VLV4 A6VLV5
A6VLV8 A6VLW0 A6VLX6 A6VLY4 A6VM01 A6VM02 A6VM11 A6VM22 A6VM24 A6VM31 A6VM32
A6VM35 A6VM38 A6VM39 A6VM51 A6VM63 A6VM72 A6VM78 A6VM79 A6VM96 A6VMA0 A6VMA3
A6VMA4 A6VMA6 A6VMA9 A6VMB8 A6VMD3 A6VMD4 A6VMD5 A6VMD6 A6VMD8 A6VME0 A6VME1
A6VME2 A6VME7 A6VME8 A6VMF1 A6VMF2 A6VMF8 A6VMG5 A6VMI0 A6VMI2 A6VMK7 A6VMK8
A6VML0 A6VML2 A6VMN7 A6VMP1 A6VMP5 A6VMP9 A6VMQ1 A6VMQ9 A6VMR0 A6VMR3 A6VMR4
A6VMR9 A6VMS2 A6VMT0 A6VMT8 A6VMT9 A6VMU1 A6VMU2 A6VMU5 A6VMV1 A6VMV2 A6VMV4
A6VMX3 A6VMY4 A6VMY7 A6VMZ1 A6VN03 A6VN04 A6VN06 A6VN08 A6VN09 A6VN10 A6VN13
A6VN14 A6VN18 A6VN19 A6VN24 A6VN35 A6VN44 A6VN49 A6VN75 A6VN85 A6VN92 A6VN94
A6VNB0 A6VNB1 A6VNB3 A6VNC3 A6VND5 A6VND6 A6VND8 A6VNE0 A6VNE1 A6VNE2 A6VNE4
A6VNE5 A6VNF1 A6VNI7 A6VNK4 A6VNL0 A6VNL1 A6VNL8 A6VNS2 A6VNS4 A6VNT3 A6VNV7

A6VNZ2 A6VP06 A6VP11 A6VP15 A6VP17 A6VP18 A6VP23 A6VP30 A6VP37 A6VP50 A6VP52
A6VP66 A6VP67 A6VP68 A6VP74 A6VP80 A6VP83 A6VP84 A6VP88 A6VPA9 A6VPB0 A6VPC2
A6VPC4 A6VPD9 A6VPE0 A6VPE4 A6VPE8 A6VPE9 A6VPF4 A6VPF5 A6VPG3 A6VPG7 A6VPG9
A6VPH1 A6VPH6 A6VPH7 A6VPK5 A6VPK8 A6VPL0 A6VPL3 A6VPM2 A6VPR2 A6VPS2 A6VPY1
A6VPY2 A6VPZ1 A6VPZ6 A6VQ01 A6VQ16 A6VQ17 A6VQ29 A6VQ53 A6VQ56 A6VQ66 A6VQ73
A6VQ74 A6VQ76 A6VQ79 A6VQ84 A6VQA3 A6VQA4 A6VQA5 A6VQA7 A6VQA9 A6VQB0 A6VQB6
A6VQC3 A6VQD4 A6VQD5 A6VQG4 A6VQH1 A6VQH4 A6VQH8 A6VQJ4 A6VQK2 A6VQL0 A6VQL1
A6VQL8 A6VQN1 A6VQN2 A6VQP1 A6VQP2 A6VQP6 A6VQQ3 A6VQR8 A6VQS4 A6VQS6 A6VQS7
A6VQT1 A6VQT2 A6VQU2 A6VQV4 A6VQW2 A6VQW4 A6VQW6 A6VQX5 A6VQX6 A6VQX9 A6VQY0
A6VQY1 A6VR01 A6VR02 A6VR10 A6VR16 A6VR54 A6VR57 A6VR60 A6VR64 A6VR65 A6VR67
A6VRA1 A6VRG3 A6VSA9 A6VSD2 A6VSP6 A6VSP9 A6VSX4 A6VSX5 A6VSX6 A6VSX9 A6VSY4
A6VSY5 A6VSY6 A6VT70 A6VT87 A6VT94 A6VTA1 A6VTA6 A6VTA8 A6VTB4 A6VTC3 A6VTQ2
A6VTV5 A6VTV6 A6VTV7 A6VTV8 A6VTV9 A6VTW0 A6VTW1 A6VU13 A6VU24 A6VU27 A6VU29
A6VU30 A6VU31 A6VU32 A6VU33 A6VU46 A6VU47 A6VU51 A6VU53 A6VU64 A6VUD3 A6VUE5
A6VUE7 A6VUG0 A6VUP1 A6VUP6 A6VUQ2 A6VUQ5 A6VUQ6 A6VUQ9 A6VUR3 A6VUR6 A6VUS2
A6VUS5 A6VUT6 A6VUV4 A6VUX8 A6VV02 A6VV03 A6VV06 A6VV07 A6VV10 A6VV11 A6VVD9
A6VVI2 A6VVJ7 A6VVN4 A6VVU9 A6VVZ9 A6VW08 A6VW16 A6VW19 A6VW20 A6VW21 A6VW36
A6VWH1 A6VWL7 A6VWM5 A6VWM9 A6VWN0 A6VWS0 A6VWW9 A6VWX6 A6VWX9 A6VWY0 A6VWY4
A6VX14 A6VX18 A6VX34 A6VX35 A6VX48 A6VX53 A6VX63 A6VX72 A6VX88 A6VX96 A6VX99
A6VXA6 A6VXC5 A6VXD0 A6VXD2 A6VXD3 A6VXF8 A6VXH3 A6VXH5 A6VXI7 A6VXK1 A6VXL0
A6VXM3 A6VXM6 A6VXM7 A6VXT1 A6VXU5 A6VXW3 A6VXW9 A6VXX9 A6VXY2 A6VXY8 A6VXZ2
A6VXZ3 A6VY00 A6VYA4 A6VYC0 A6VYC3 A6VYC5 A6VYH5 A6VYH7 A6VYH8 A6VYH9 A6VYI1
A6VYJ6 A6VYJ7 A6VYK6 A6VYK7 A6VYL0 A6VYL1 A6VYL6 A6VYL7 A6VYN2 A6VZ12 A6VZ14
A6VZ30 A6VZ75 A6VZ82 A6VZ84 A6VZ85 A6VZ89 A6VZ92 A6VZA1 A6VZB3 A6VZE1 A6VZE3
A6VZE7 A6VZF8 A6VZL5 A6VZN8 A6W097 A6W0A0 A6W0C9 A6W0E1 A6W0Q2 A6W0S5 A6W0S9
A6W0X8 A6W0Y0 A6W0Y1 A6W0Y6 A6W0Y8 A6W0Y9 A6W0Z9 A6W133 A6W1C0 A6W1C2 A6W1C3
A6W1C6 A6W1C7 A6W1C8 A6W1C9 A6W1F1 A6W1F2 A6W1K0 A6W1Q1 A6W1Q6 A6W1Q9 A6W1R4
A6W1S0 A6W1T1 A6W1T9 A6W1U0 A6W1U1 A6W1U2 A6W1V6 A6W1W3 A6W1W5 A6W1X1 A6W1X2
A6W216 A6W284 A6W2C1 A6W2D1 A6W2D2 A6W2E0 A6W2I3 A6W2I4 A6W2I5 A6W2K8 A6W2L6
A6W2L8 A6W2M9 A6W2P7 A6W2T3 A6W2T4 A6W2U2 A6W2U5 A6W2U6 A6W2V1 A6W2V3 A6W2V5
A6W2V6 A6W2V7 A6W2W4 A6W346 A6W349 A6W351 A6W352 A6W353 A6W354 A6W366 A6W368
A6W369 A6W370 A6W371 A6W373 A6W374 A6W376 A6W377 A6W378 A6W379 A6W380 A6W381
A6W382 A6W383 A6W384 A6W385 A6W387 A6W388 A6W389 A6W390 A6W391 A6W392 A6W393
A6W394 A6W396 A6W397 A6W398 A6W399 A6W3A0 A6W3A1 A6W3A2 A6W3A3 A6W3A7 A6W3C7
A6W3C9 A6W3D3 A6W3D6 A6W3E8 A6W3H7 A6W3K1 A6W3T8 A6W3T9 A6W3V0 A6W3V2 A6W3V3
A6W3V4 A6W3V7 A6W4E9 A6W4H8 A6W4I5 A6W4N5 A6W4N6 A6W4N7 A6W504 A6W572 A6W5D4
A6W5D5 A6W5H6 A6W5K2 A6W5L7 A6W5M6 A6W5P6 A6W5R3 A6W5S6 A6W5S7 A6W5S8 A6W5S9
A6W5T0 A6W5T1 A6W5T2 A6W5T3 A6W5T4 A6W5T5 A6W5T7 A6W5T8 A6W5U1 A6W5U2 A6W5U3
A6W5U4 A6W5U5 A6W5U7 A6W5U8 A6W5U9 A6W5V0 A6W5V1 A6W5V2 A6W5V3 A6W5V4 A6W5V5
A6W5V6 A6W5W1 A6W5W2 A6W5W3 A6W5W4 A6W5W5 A6W5W6 A6W5X0 A6W5X1 A6W5X2 A6W5Y5
A6W5Y6 A6W614 A6W6A1 A6W6A2 A6W6C7 A6W6D1 A6W6D7 A6W6D9 A6W6E9 A6W6G0 A6W6P9
A6W6U1 A6W6U4 A6W6U5 A6W6V3 A6W733 A6W7E9 A6W7F6 A6W7K7 A6W7L4 A6W7L6 A6W7L7
A6W7L8 A6W7N6 A6W7Q5 A6W7Q7 A6W7Q8 A6W7R2 A6W7S4 A6W7S6 A6W7U7 A6W7U9 A6W7V0
A6W7V1 A6W7V4 A6W7V6 A6W7W4 A6W7W5 A6W7W7 A6W7W9 A6W7Y5 A6W7Y9 A6W7Z2 A6W7Z4

A6W814 A6W815 A6W828 A6W833 A6W841 A6W848 A6W849 A6W857 A6W933 A6W961 A6W963
A6W970 A6W971 A6W972 A6WC54 A6WC55 A6WC57 A6WC59 A6WC91 A6WC96 A6WCA2 A6WCA9
A6WCB6 A6WCC6 A6WCD0 A6WCD3 A6WCF6 A6WCF7 A6WCG5 A6WCH4 A6WCI2 A6WCI3 A6WCS8
A6WCT0 A6WCT4 A6WCU2 A6WCU3 A6WCV0 A6WCW8 A6WCX3 A6WCY2 A6WCY3 A6WD63 A6WD69
A6WDF2 A6WDG1 A6WDI1 A6WDJ6 A6WDL7 A6WDM4 A6WDM6 A6WDM9 A6WDN0 A6WDN1 A6WDT9
A6WDU2 A6WE21 A6WE36 A6WE60 A6WEK3 A6WEK6 A6WF54 A6WF55 A6WFC5 A6WFJ3 A6WFU4
A6WFV7 A6WG40 A6WG60 A6WG61 A6WGA4 A6WGF9 A6WGM8 A6WGN2 A6WGN3 A6WH85 A6WH87
A6WH92 A6WH93 A6WH97 A6WHA1 A6WHB0 A6WHB4 A6WHB6 A6WHB7 A6WHB9 A6WHC7 A6WHC9
A6WHG9 A6WHH8 A6WHI0 A6WHI5 A6WHK9 A6WHL6 A6WHM7 A6WHM8 A6WHQ4 A6WHQ5 A6WHQ8
A6WHQ9 A6WHR3 A6WHR4 A6WHR7 A6WHR8 A6WHR9 A6WHS0 A6WHS1 A6WHS2 A6WHS3 A6WHS4
A6WHS7 A6WHS8 A6WHS9 A6WHT1 A6WHT2 A6WHT3 A6WHT4 A6WHT5 A6WHT6 A6WHT7 A6WHT8
A6WHT9 A6WHU0 A6WHU1 A6WHU2 A6WHU3 A6WHU4 A6WHU5 A6WHU6 A6WHU7 A6WHU9 A6WHV0
A6WHV1 A6WHV2 A6WHV3 A6WHV4 A6WHZ7 A6WI10 A6WI48 A6WI50 A6WI83 A6WI99 A6WIA1
A6WIA3 A6WIA4 A6WIA6 A6WIA7 A6WIA8 A6WIB5 A6WIB7 A6WIB8 A6WIC0 A6WIC2 A6WIC3
A6WID2 A6WIE9 A6WIH5 A6WIH6 A6WIK3 A6WIK6 A6WIP6 A6WIQ6 A6WJ16 A6WJ17 A6WJ51
A6WJ66 A6WJ70 A6WJ71 A6WJ79 A6WJ80 A6WJ81 A6WJ82 A6WJ85 A6WJ89 A6WJH4 A6WJK9
A6WJL5 A6WJS7 A6WJT5 A6WJU3 A6WJU6 A6WJZ1 A6WK08 A6WK49 A6WK50 A6WK51 A6WK57
A6WK58 A6WK59 A6WK72 A6WKC3 A6WKC4 A6WKC8 A6WKD1 A6WKD4 A6WKK3 A6WKK4 A6WKL1
A6WKL6 A6WKM0 A6WKM4 A6WKN2 A6WKN9 A6WKP0 A6WKP4 A6WKQ7 A6WKR6 A6WKR7 A6WKR8
A6WKR9 A6WKS2 A6WKV9 A6WKY2 A6WKY5 A6WL02 A6WL03 A6WL04 A6WL09 A6WL14 A6WL18
A6WL23 A6WLA3 A6WLA6 A6WLA7 A6WLA8 A6WLA9 A6WLC0 A6WLC8 A6WLE1 A6WLE9 A6WLP8
A6WLP9 A6WLQ0 A6WLQ1 A6WLQ2 A6WLV9 A6WLX6 A6WLY5 A6WM16 A6WM34 A6WM35 A6WM39
A6WM52 A6WM55 A6WM62 A6WM65 A6WM67 A6WM68 A6WMJ7 A6WMJ9 A6WMK0 A6WMK2 A6WMK4
A6WMK6 A6WMK8 A6WMN1 A6WMP4 A6WMW5 A6WMW8 A6WMZ6 A6WN68 A6WN87 A6WN92 A6WNA1
A6WNB3 A6WNG1 A6WNG9 A6WNH0 A6WNH1 A6WNH2 A6WNM7 A6WNM9 A6WNN0 A6WNN5 A6WNN7
A6WNQ2 A6WNQ3 A6WNQ4 A6WNQ6 A6WNQ8 A6WNQ9 A6WNR8 A6WNV3 A6WNW5 A6WNX6 A6WNX9
A6WNY1 A6WNY2 A6WNY5 A6WP02 A6WP16 A6WP21 A6WP22 A6WP46 A6WP53 A6WP54 A6WP56
A6WP60 A6WP62 A6WP64 A6WP69 A6WP80 A6WP88 A6WPA4 A6WPB5 A6WPC5 A6WPD0 A6WPH7
A6WPI0 A6WPI2 A6WPI6 A6WPJ2 A6WPJ5 A6WPK6 A6WPK9 A6WPL2 A6WPS5 A6WPT9 A6WPW9
A6WPX1 A6WPX2 A6WQ04 A6WQ07 A6WQ09 A6WQ14 A6WQ16 A6WQ52 A6WQ53 A6WQ67 A6WQB8
A6WQD3 A6WQP0 A6WQP2 A6WQP3 A6WQP6 A6WQP7 A6WQQ0 A6WQT5 A6WQU2 A6WQU4 A6WQU6
A6WQX2 A6WR14 A6WR16 A6WR18 A6WR19 A6WR22 A6WR23 A6WR25 A6WR29 A6WR31 A6WR37
A6WR46 A6WR47 A6WR51 A6WR52 A6WRA3 A6WRB5 A6WRB6 A6WRB7 A6WRB8 A6WRG3 A6WRG5
A6WRG6 A6WRG7 A6WRG8 A6WRH0 A6WRH2 A6WRH3 A6WRH7 A6WRI8 A6WRJ1 A6WRJ3 A6WRJ9
A6WRK2 A6WRK4 A6WRL0 A6WRL1 A6WRL2 A6WRS7 A6WRS8 A6WRU0 A6WRU8 A6WRU9 A6WRW5
A6WRY2 A6WS13 A6WS27 A6WS29 A6WS34 A6WS64 A6WS76 A6WS79 A6WS92 A6WSD7 A6WSF0
A6WSF1 A6WSF3 A6WSH0 A6WSI3 A6WSL1 A6WSL2 A6WSL3 A6WSM1 A6WSM2 A6WSQ6 A6WST7
A6WSV2 A6WSV3 A6WSV4 A6WSV9 A6WSW9 A6WSX5 A6WSY9 A6WT04 A6WT39 A6WTA5 A6WTC8
A6WTC9 A6WTD3 A6WTE5 A6WTE7 A6WTF5 A6WTF8 A6WTG0 A6WTI9 A6WTL2 A6WTL7 A6WTP9
A6WTQ0 A6WTR1 A6WTR2 A6WTR9 A6WTS0 A6WTS2 A6WTS3 A6WTS6 A6WTY4 A6WU19 A6WU68
A6WUE5 A6WUE9 A6WUF6 A6WUG6 A6WUH1 A6WUH7 A6WUK0 A6WUK1 A6WUK3 A6WUK6 A6WUK7
A6WUS9 A6WUU9 A6WUV0 A6WUV8 A6WV13 A6WV17 A6WV21 A6WV39 A6WV52 A6WV84 A6WV86
A6WV92 A6WVA7 A6WVA9 A6WVB4 A6WVC7 A6WVK9 A6WVQ0 A6WVQ2 A6WVQ8 A6WVR3 A6WVR4
A6WVR5 A6WVU6 A6WVX4 A6WW03 A6WW16 A6WW65 A6WW84 A6WW98 A6WWA8 A6WWB6 A6WWC4

A6WWE3 A6WWE5 A6WWE6 A6WWG6 A6WWG7 A6WWG8 A6WWI2 A6WWR5 A6WWW1 A6WWW2 A6WWW3
A6WWW4 A6WWW5 A6WWW8 A6WWW9 A6WWX4 A6WWX6 A6WX07 A6WX16 A6WX18 A6WX19 A6WX21
A6WX28 A6WX30 A6WX40 A6WX47 A6WX50 A6WX51 A6WX52 A6WX53 A6WX56 A6WX57 A6WX59
A6WX66 A6WX71 A6WX72 A6WX76 A6WX77 A6WX78 A6WXC7 A6WXD5 A6WXE0 A6WXE4 A6WXE7
A6WXE8 A6WXF0 A6WXG0 A6WXG3 A6WXG4 A6WXJ1 A6WXJ2 A6WXL6 A6WXL9 A6WXM7 A6WXQ0
A6WXQ3 A6WXR2 A6WXR6 A6WXR7 A6WXS0 A6WXS1 A6WXS2 A6WXS5 A6WXT0 A6WXU5 A6WXV2
A6WXV3 A6WXV8 A6WXW1 A6WXW5 A6WXX7 A6WXY9 A6WXZ4 A6WY25 A6WY31 A6WY63 A6WY64
A6WY74 A6WY75 A6WY76 A6WY85 A6WY87 A6WY94 A6WYD2 A6WYH5 A6WYH7 A6WYH9 A6WYI0
A6WYJ2 A6WYJ6 A6WYK8 A6WYK9 A6WYL3 A6WYL7 A6WYM1 A6WYM5 A6WYM6 A6WYQ4 A6WYS5
A6WYU6 A6WYU8 A6WYU9 A6WYV7 A6WYX0 A6WZ88 A6WZC6 A6WZC8 A6WZD8 A6WZE3 A6WZH2
A6WZJ5 A6WZN1 A6WZP8 A6WZQ7 A6WZQ8 A6WZQ9 A6WZR6 A6WZS8 A6WZV4 A6WZW3 A6WZX2
A6WZX5 A6X034 A6X049 A6X051 A6X071 A6X073 A6X0A2 A6X0A5 A6X0A6 A6X0A7 A6X0A8
A6X0A9 A6X0B0 A6X0B3 A6X0B4 A6X0B5 A6X0B7 A6X0B8 A6X0B9 A6X0C0 A6X0C1 A6X0C2
A6X0C4 A6X0C5 A6X0C7 A6X0C8 A6X0C9 A6X0D0 A6X0D1 A6X0D2 A6X0D3 A6X0D4 A6X0D5
A6X0D6 A6X0D7 A6X0D9 A6X0E0 A6X0E1 A6X0E2 A6X0E3 A6X0E8 A6X0E9 A6X0F8 A6X0G9
A6X0J1 A6X0J2 A6X0J3 A6X0J4 A6X0K5 A6X0K9 A6X0L0 A6X0L1 A6X0L6 A6X0M7 A6X0N1
A6X0R0 A6X0U8 A6X0X0 A6X0X1 A6X0Y9 A6X0Z2 A6X117 A6X120 A6X187 A6X191 A6X192
A6X1E3 A6X1F1 A6X1G1 A6X1G2 A6X1H0 A6X1J2 A6X1J9 A6X1K4 A6X1K6 A6X1L5 A6X1P9
A6X1Q4 A6X1V4 A6X1V5 A6X1V9 A6X1X7 A6X1Y4 A6X1Y8 A6X1Y9 A6X1Z7 A6X200 A6X235
A6X253 A6X259 A6X265 A6X270 A6X294 A6X296 A6X299 A6X2G6 A6X2G7 A6X2G8 A6X2S6
A6X2S8 A6X307 A6X357 A6X374 A6X376 A6X391 A6X3A0 A6X3C2 A6X3C3 A6X3D0 A6X3F3
A6X3G3 A6X3M9 A6X3N5 A6X438 A6X449 A6X4G3 A6X4W8 A6X529 A6X554 A6X562 A6X564
A6X565 A6X566 A6X568 A6X578 A6X5D7 A6X5L4 A6X5N3 A6X610 A6X6N5 A6X6Q6 A6X6V1
A6X6W2 A6X6Y1 A6X792 A6X798 A6X8E9 A6X8Z9 A6X942 A6X970 A6X972 A6X974 A6X978
A6X980 A6X992 A6XH05 A6XH06 A6XIG6 A6XIG7 A6XMY9 A6XNC6 A6Y9S5 A6YB01 A6YG63
A6YG72 A6YG96 A6YGA0 A6YGA1 A6YGA2 A6YGA3 A6YGA4 A6YGA5 A6YGB1 A6YGB4 A6YGC4
A6YGC5 A6YGC7 A6YGC8 A6YGC9 A6YGD0 A6YGD2 A6YGD3 A6YGD8 A6YGD9 A6YGE0 A6YPJ8
A6YRN9 A6YSL1 A6ZJ55 A6ZKI3 A6ZKL2 A6ZKM2 A6ZKR6 A6ZKW8 A6ZKX7 A6ZL17 A6ZL57
A6ZL85 A6ZL94 A6ZLA6 A6ZLD4 A6ZLF4 A6ZLG6 A6ZLH6 A6ZLH8 A6ZM02 A6ZM17 A6ZM60
A6ZM67 A6ZM92 A6ZMA9 A6ZMC1 A6ZMC3 A6ZME0 A6ZME2 A6ZMG4 A6ZMG6 A6ZMK0 A6ZMK5
A6ZMM3 A6ZMQ6 A6ZMZ8 A6ZN26 A6ZN46 A6ZN74 A6ZND9 A6ZNN4 A6ZNQ5 A6ZNR2 A6ZNU5
A6ZNW6 A6ZP42 A6ZP43 A6ZP47 A6ZP58 A6ZP88 A6ZPA6 A6ZPA9 A6ZPB3 A6ZPC2 A6ZPE1
A6ZPE5 A6ZPJ1 A6ZPQ2 A6ZPT1 A6ZPU3 A6ZPV9 A6ZPY2 A6ZPY9 A6ZPZ1 A6ZQ26 A6ZQ37
A6ZQ47 A6ZQ59 A6ZQF3 A6ZQH0 A6ZQJ1 A6ZQJ2 A6ZQJ9 A6ZQK9 A6ZQM3 A6ZQM4 A6ZQR2
A6ZQU2 A6ZQU8 A6ZQX9 A6ZQY2 A6ZR01 A6ZR25 A6ZR27 A6ZR36 A6ZR60 A6ZR64 A6ZRG7
A6ZRI9 A6ZRK3 A6ZRK4 A6ZRL6 A6ZRM0 A6ZRM7 A6ZRN4 A6ZRQ0 A6ZRQ4 A6ZRR9 A6ZRW0
A6ZRW1 A6ZRW2 A6ZRW4 A6ZRW5 A6ZRW7 A6ZRX0 A6ZRY1 A6ZRZ0 A6ZRZ4 A6ZS02 A6ZS33
A6ZS79 A6ZS83 A6ZSB3 A6ZSM1 A6ZSX1 A6ZSZ4 A6ZT11 A6ZT19 A6ZT44 A6ZT54 A6ZT67
A6ZT71 A6ZT77 A6ZT79 A6ZTA3 A6ZTA4 A6ZTA5 A6ZTB4 A6ZTR3 A6ZTT5 A6ZTU2 A6ZTX8
A6ZU08 A6ZU15 A6ZU34 A6ZU71 A6ZUA0 A6ZUA1 A6ZUB2 A6ZUC0 A6ZUG2 A6ZUM5 A6ZUS8
A6ZUT6 A6ZV04 A6ZV63 A6ZV70 A6ZV85 A6ZVB1 A6ZVE9 A6ZVF4 A6ZVI4 A6ZVI6 A6ZVL3
A6ZVR0 A6ZVS0 A6ZVS1 A6ZVT0 A6ZW01 A6ZW87 A6ZW95 A6ZWC8 A6ZWD3 A6ZWK1 A6ZWN8
A6ZWX0 A6ZWZ7 A6ZWZ9 A6ZX45 A6ZX59 A6ZX64 A6ZXB7 A6ZXD2 A6ZXD3 A6ZXG9 A6ZXP4
A6ZXT5 A6ZXU0 A6ZXY5 A6ZY30 A6ZY43 A6ZY89 A6ZYC3 A6ZYI0 A6ZYM0 A6ZYQ3 A6ZYV1

A6ZYV6 A6ZYX7 A6ZYX9 A6ZZ25 A6ZZE3 A6ZZF6 A6ZZJ1 A6ZZK4 A6ZZL7 A6ZZL8 A6ZZM4
A6ZZQ5 A6ZZR2 A6ZZR4 A6ZZS7 A6ZZW1 A6ZZY2 A6ZZY8 A7A018 A7A024 A7A031 A7A061
A7A075 A7A084 A7A0M4 A7A0P8 A7A0V3 A7A0X4 A7A0X8 A7A0Z5 A7A134 A7A136 A7A1A3
A7A1D7 A7A1G0 A7A1I2 A7A1L2 A7A1S5 A7A1T1 A7A1V6 A7A1V9 A7A1W1 A7A237 A7A239
A7A241 A7A261 A7A270 A7A283 A7AMD1 A7ANL7 A7AU38 A7AWN0 A7AX58 A7B1V0 A7BHQ9
A7BJC4 A7BJC5 A7DTG3 A7DY56 A7DZP8 A7E2S9 A7E2U8 A7E2Y6 A7E2Z2 A7E305 A7E320
A7E321 A7E346 A7E369 A7E371 A7E379 A7E3D8 A7E3Q8 A7E3S5 A7E3Z4 A7E436 A7E449
A7E471 A7E4H3 A7E4K0 A7E4S9 A7E4T8 A7E564 A7E5J5 A7E5P6 A7E664 A7E7B3 A7E7C4
A7E8H8 A7EAA1 A7EAG1 A7EAY2 A7EB95 A7EC69 A7ECP3 A7EDG9 A7EF03 A7EF41 A7EFH4
A7EFS3 A7EGB5 A7EGG4 A7EGL7 A7EGU0 A7EGZ3 A7EH92 A7EHE5 A7EHM8 A7EI75 A7EI93
A7EIX7 A7EJ50 A7EJY3 A7EKL8 A7EKM8 A7EM78 A7EM88 A7EML8 A7EMM3 A7ENE0 A7ENP9
A7ENU3 A7EPT0 A7EPT5 A7EQA8 A7ER98 A7ERG1 A7ESB8 A7ESL7 A7ESL8 A7ESR0 A7ETU4
A7EUB3 A7EUE6 A7EV21 A7EVF4 A7EVV6 A7EVV9 A7EWH6 A7EWL8 A7EWN6 A7EWR0 A7EWX9
A7EXU7 A7EXZ2 A7EYK3 A7EYW0 A7EYZ4 A7EZ71 A7EZ86 A7EZJ5 A7EZP8 A7F075 A7F080
A7F190 A7F1E9 A7F1L5 A7F2D1 A7F2R3 A7F2R6 A7F2S3 A7F3L0 A7F3V4 A7F4L5 A7F582
A7F5M4 A7F7H0 A7F8K7 A7F8R0 A7F8U1 A7F8V8 A7F996 A7F9K1 A7FA72 A7FCM9 A7FCN0
A7FCN1 A7FCN3 A7FCP0 A7FCP5 A7FCP6 A7FCQ4 A7FCR3 A7FCR5 A7FCS9 A7FCT0 A7FCT1
A7FCT2 A7FCT4 A7FCT5 A7FCT6 A7FCT7 A7FCT8 A7FCU3 A7FCU5 A7FCU8 A7FCV1 A7FCW1
A7FCW8 A7FCX5 A7FCY2 A7FCY4 A7FCY5 A7FCZ0 A7FCZ2 A7FCZ7 A7FCZ8 A7FD00 A7FD01
A7FD10 A7FD13 A7FD15 A7FD20 A7FD26 A7FD32 A7FD46 A7FD47 A7FD67 A7FD72 A7FD74
A7FDA1 A7FDD3 A7FDE0 A7FDF3 A7FDF8 A7FDG2 A7FDG8 A7FDL9 A7FDQ0 A7FDQ3 A7FDQ8
A7FDU0 A7FDV0 A7FDV1 A7FDV3 A7FDW9 A7FDX4 A7FDX5 A7FDY8 A7FDY9 A7FE06 A7FE15
A7FE70 A7FE71 A7FE75 A7FE79 A7FE82 A7FEX4 A7FEX6 A7FEY3 A7FEY8 A7FEZ0 A7FEZ1
A7FEZ6 A7FEZ7 A7FF05 A7FF10 A7FF15 A7FF19 A7FF20 A7FF21 A7FF28 A7FF38 A7FF39
A7FFD6 A7FFD7 A7FFE0 A7FFE9 A7FFF7 A7FFF8 A7FFG0 A7FFG5 A7FFG6 A7FFG7 A7FFG9
A7FFH0 A7FFH1 A7FFH2 A7FFH3 A7FFI3 A7FFI7 A7FFJ0 A7FFJ4 A7FFK3 A7FFK5 A7FFK7
A7FFL7 A7FFN0 A7FFN1 A7FFN2 A7FFN3 A7FFN4 A7FFN6 A7FFP8 A7FFS0 A7FFS3 A7FFS4
A7FFS5 A7FFT0 A7FFT9 A7FFU1 A7FFU6 A7FFW1 A7FFX0 A7FFX3 A7FFX4 A7FFX8 A7FFY5
A7FFY6 A7FFY9 A7FFZ0 A7FFZ3 A7FFZ6 A7FFZ9 A7FG37 A7FG38 A7FG40 A7FG49 A7FG51
A7FG57 A7FG78 A7FG81 A7FG82 A7FG95 A7FG96 A7FGC2 A7FGC6 A7FGD3 A7FGK4 A7FGK6
A7FGL1 A7FGL9 A7FGM1 A7FGP4 A7FGP6 A7FGQ0 A7FGT2 A7FGT3 A7FGT5 A7FGU1 A7FGU7
A7FGU8 A7FH09 A7FH13 A7FH16 A7FH23 A7FH32 A7FH41 A7FH42 A7FH52 A7FH54 A7FH61
A7FH78 A7FH79 A7FH83 A7FHC0 A7FHE3 A7FHG1 A7FHG2 A7FHG3 A7FHG4 A7FHG5 A7FHG7
A7FHI0 A7FHI1 A7FHI3 A7FHJ2 A7FHJ3 A7FHK2 A7FHK6 A7FHL4 A7FHL7 A7FHL9 A7FHM0
A7FHP9 A7FHR1 A7FHW9 A7FHX5 A7FHY4 A7FI01 A7FI10 A7FI31 A7FI33 A7FI34 A7FI51
A7FI67 A7FI81 A7FI87 A7FI92 A7FIA1 A7FIA4 A7FIA5 A7FIA7 A7FIC5 A7FIC6 A7FIC7
A7FIC8 A7FID0 A7FID3 A7FID4 A7FID5 A7FID8 A7FIE3 A7FIE8 A7FIG1 A7FIG2 A7FIG4
A7FIG9 A7FIK8 A7FIL2 A7FIL3 A7FIL6 A7FIZ3 A7FJ02 A7FJ17 A7FJ89 A7FJ93 A7FJ95
A7FJD3 A7FJE9 A7FJF8 A7FJG9 A7FJH1 A7FJH4 A7FJH5 A7FJJ3 A7FJJ4 A7FJJ6 A7FJK3
A7FJK4 A7FJL1 A7FJR7 A7FJR8 A7FJR9 A7FJS2 A7FJS7 A7FJT1 A7FJT9 A7FJU4 A7FJU5
A7FJV2 A7FJV9 A7FJW6 A7FJW8 A7FJW9 A7FJX0 A7FJY0 A7FJZ0 A7FJZ1 A7FJZ4 A7FK04
A7FK27 A7FK39 A7FK62 A7FK85 A7FK86 A7FKA7 A7FKA8 A7FKA9 A7FKF4 A7FKL4 A7FKL5
A7FKL6 A7FKM0 A7FKM4 A7FKM8 A7FKM9 A7FKN1 A7FKP2 A7FKP6 A7FKQ0 A7FKR4 A7FKS2
A7FKS6 A7FKS8 A7FKU1 A7FKU3 A7FKU9 A7FKV1 A7FKW0 A7FKW3 A7FKY2 A7FKY7 A7FKY8

A7FKY9 A7FL44 A7FL46 A7FL85 A7FL86 A7FL89 A7FL90 A7FL92 A7FLA1 A7FLA2 A7FLB5
A7FLB7 A7FLC3 A7FLC4 A7FLC5 A7FLD6 A7FLE0 A7FLE2 A7FLE4 A7FLE7 A7FLE8 A7FLF0
A7FLF5 A7FLF6 A7FLH3 A7FLH4 A7FLH5 A7FLI1 A7FLI2 A7FLI3 A7FLI4 A7FLI9 A7FLJ8
A7FLK2 A7FLL0 A7FLL1 A7FLL2 A7FLL3 A7FLL4 A7FLQ4 A7FLQ5 A7FLQ6 A7FLQ7 A7FLR6
A7FLR7 A7FLR8 A7FLR9 A7FLX1 A7FLX4 A7FLX5 A7FLX6 A7FLX7 A7FLX8 A7FLY2 A7FLY3
A7FLY4 A7FLY8 A7FLY9 A7FLZ0 A7FLZ6 A7FM03 A7FM04 A7FM07 A7FM09 A7FM16 A7FM22
A7FM23 A7FM24 A7FM33 A7FM34 A7FM37 A7FM52 A7FM54 A7FM55 A7FM64 A7FM65 A7FM74
A7FM75 A7FM84 A7FM86 A7FM87 A7FM96 A7FMC0 A7FMC1 A7FMC2 A7FMC3 A7FMD2 A7FMD4
A7FMD7 A7FMD9 A7FME2 A7FME3 A7FME9 A7FMF0 A7FMF2 A7FMF8 A7FMF9 A7FMH2 A7FMH3
A7FMH4 A7FMI1 A7FMI4 A7FMJ5 A7FMK1 A7FMK2 A7FML9 A7FMM5 A7FMP6 A7FMQ4 A7FMR8
A7FMR9 A7FMS0 A7FMS1 A7FMS2 A7FMS4 A7FMS6 A7FMS9 A7FMT5 A7FMT7 A7FMT8 A7FMU2
A7FMU3 A7FMU7 A7FMV2 A7FMV8 A7FMW2 A7FMW3 A7FMW4 A7FMW5 A7FMX3 A7FMX4 A7FMX9
A7FMY0 A7FMY1 A7FMY7 A7FMY8 A7FMZ1 A7FMZ8 A7FN01 A7FN02 A7FN06 A7FN79 A7FN80
A7FN82 A7FN83 A7FN85 A7FNA0 A7FNG1 A7FNG6 A7FNH1 A7FNH2 A7FNH4 A7FNH5 A7FNH7
A7FNH9 A7FNI2 A7FNI3 A7FNI4 A7FNI5 A7FNI6 A7FNI7 A7FNJ0 A7FNJ3 A7FNJ7 A7FNJ8
A7FNK0 A7FNK2 A7FNK3 A7FNK4 A7FNK9 A7FNL0 A7FNL1 A7FNL2 A7FNL3 A7FNL5 A7FNL6
A7FNL7 A7FNL8 A7FNL9 A7FNM0 A7FNM1 A7FNM2 A7FNM3 A7FNM4 A7FNM6 A7FNM7 A7FNM8
A7FNM9 A7FNN0 A7FNN1 A7FNN2 A7FNN4 A7FNN5 A7FNN6 A7FNN8 A7FNN9 A7FNP0 A7FNP1
A7FNP2 A7FNP3 A7FNP4 A7FNP7 A7FNR1 A7FNS9 A7FNT5 A7FNT6 A7FNU7 A7FNU8 A7FNV6
A7FNV8 A7FNW0 A7FNW4 A7FNW5 A7FNX2 A7FNX3 A7FNX4 A7FNX5 A7FP06 A7FP52 A7FP68
A7FP80 A7FP81 A7FP85 A7FP92 A7FP95 A7FP96 A7FPA7 A7FPA8 A7FPB5 A7FPB7 A7FPB8
A7FPB9 A7FPC2 A7FPE8 A7FPF0 A7FPF3 A7FPG0 A7FPG2 A7FPH4 A7FPH5 A7FPH8 A7FPI7
A7FPJ7 A7FPK3 A7FPL5 A7FPL6 A7FPL8 A7FPL9 A7FPM0 A7FPP5 A7FPR6 A7FPT0 A7FPT1
A7FPZ5 A7FPZ6 A7FPZ7 A7FPZ8 A7FPZ9 A7FQ31 A7FQ39 A7FQ40 A7FQ41 A7FQ53 A7FQ66
A7FQ67 A7FQ70 A7FQ95 A7FQA9 A7FQF0 A7FQF8 A7FQF9 A7FQG2 A7FQG8 A7FQI8 A7FQL2
A7FQM1 A7FQN7 A7FQN8 A7FQN9 A7FQP3 A7FQP4 A7FQU9 A7FQY1 A7FR34 A7FR58 A7FR59
A7FR60 A7FR61 A7FRB3 A7FRC9 A7FRK8 A7FRV3 A7FRV4 A7FS46 A7FSG1 A7FSG8 A7FSI4
A7FSN7 A7FSS9 A7FSW7 A7FSY9 A7FT29 A7FT56 A7FT69 A7FT79 A7FTB7 A7FTE0 A7FTE3
A7FTE6 A7FTH1 A7FTM1 A7FTP7 A7FTX8 A7FTY1 A7FU32 A7FU67 A7FU68 A7FU73 A7FU76
A7FU78 A7FU80 A7FU81 A7FU82 A7FU83 A7FU84 A7FUE5 A7FUK7 A7FUK8 A7FUK9 A7FUL0
A7FUL1 A7FUL8 A7FUM5 A7FUR1 A7FUT4 A7FUT5 A7FUT9 A7FUU0 A7FUU1 A7FUV2 A7FUV3
A7FUV4 A7FUX9 A7FUZ6 A7FV20 A7FV42 A7FV66 A7FVM3 A7FVR9 A7FVS5 A7FVS6 A7FVY1
A7FVY7 A7FVY8 A7FVZ2 A7FVZ3 A7FVZ7 A7FW01 A7FW04 A7FW05 A7FW08 A7FW09 A7FW10
A7FW12 A7FW24 A7FW56 A7FW59 A7FW64 A7FW69 A7FW72 A7FW79 A7FW81 A7FW89 A7FW99
A7FWA9 A7FWJ2 A7FWJ3 A7FWJ4 A7FWJ5 A7FWJ9 A7FWM4 A7FWM6 A7FWP0 A7FWU5 A7FWU6
A7FWY7 A7FWY8 A7FX01 A7FX30 A7FX38 A7FX54 A7FX73 A7FX96 A7FXC4 A7FXD3 A7FXD5
A7FXD7 A7FXG3 A7FXG9 A7FXH7 A7FXK4 A7FXK9 A7FXL3 A7FXL5 A7FXM3 A7FXT6 A7FXU4
A7FXU6 A7FXU7 A7FXU9 A7FXV6 A7FXW3 A7FXW4 A7FXW9 A7FY03 A7FY04 A7FY07 A7FY09
A7FY17 A7FY18 A7FY19 A7FY20 A7FY22 A7FY72 A7FY80 A7FY84 A7FY85 A7FY87 A7FY92
A7FYA5 A7FYI1 A7FYI2 A7FYI3 A7FYI6 A7FYI9 A7FYJ1 A7FYJ2 A7FYL2 A7FYL3 A7FYL4
A7FYL7 A7FYM5 A7FYP0 A7FYP3 A7FYP4 A7FYQ1 A7FYQ8 A7FYR9 A7FYU0 A7FYV9 A7FYW7
A7FYW9 A7FYX1 A7FYX2 A7FYX3 A7FYX4 A7FYY1 A7FZ00 A7FZ14 A7FZ37 A7FZ38 A7FZ43
A7FZ44 A7FZ45 A7FZ46 A7FZ47 A7FZ48 A7FZ49 A7FZ52 A7FZ53 A7FZ54 A7FZ55 A7FZ56
A7FZ57 A7FZ58 A7FZ59 A7FZ60 A7FZ61 A7FZ62 A7FZ63 A7FZ65 A7FZ66 A7FZ67 A7FZ68

A7FZ69 A7FZ70 A7FZ71 A7FZ72 A7FZ73 A7FZ74 A7FZ76 A7FZ77 A7FZ78 A7FZ80 A7FZ81
A7FZ84 A7FZ91 A7FZ94 A7FZA3 A7FZA5 A7FZA6 A7FZA9 A7FZB0 A7FZC4 A7FZC7 A7FZD1
A7FZF4 A7FZF9 A7FZG5 A7FZG9 A7FZH1 A7G9B0 A7G9B3 A7G9C5 A7G9D4 A7G9D5 A7G9D8
A7G9H1 A7G9I5 A7G9M4 A7G9N8 A7G9N9 A7G9P2 A7G9P8 A7G9S1 A7G9T9 A7G9V5 A7G9W4
A7G9Y0 A7G9Y1 A7G9Y2 A7G9Y6 A7G9Y7 A7GA53 A7GA73 A7GA85 A7GAG3 A7GAI4 A7GAI5
A7GAI6 A7GAI7 A7GAJ8 A7GAK0 A7GAL0 A7GAL1 A7GAL4 A7GAQ4 A7GAS1 A7GB01 A7GB83
A7GB84 A7GBI7 A7GBV6 A7GBW3 A7GBX9 A7GC28 A7GC78 A7GCB9 A7GCD9 A7GCI1 A7GCK7
A7GCL0 A7GCM3 A7GCN5 A7GCS6 A7GCV1 A7GCV4 A7GCV8 A7GCY3 A7GD48 A7GD74 A7GDE2
A7GDF4 A7GDK3 A7GDP1 A7GDP2 A7GDP8 A7GDQ0 A7GDQ1 A7GDQ3 A7GDQ5 A7GDQ6 A7GDQ7
A7GDW6 A7GE42 A7GE43 A7GE44 A7GE45 A7GE46 A7GE53 A7GE60 A7GE89 A7GEA6 A7GEJ0
A7GEJ1 A7GEJ6 A7GEJ7 A7GEJ8 A7GEK9 A7GEL0 A7GEL1 A7GEN6 A7GEQ5 A7GES9 A7GEU9
A7GEX3 A7GFE5 A7GFJ3 A7GFJ8 A7GFJ9 A7GFZ4 A7GG00 A7GG01 A7GG03 A7GG05 A7GG06
A7GG10 A7GG14 A7GG20 A7GG21 A7GG22 A7GG23 A7GG24 A7GG28 A7GG29 A7GG32 A7GG33
A7GG34 A7GG36 A7GG48 A7GG76 A7GG79 A7GG84 A7GG89 A7GG92 A7GG99 A7GGA1 A7GGA7
A7GGB7 A7GGC7 A7GGE1 A7GGE8 A7GGE9 A7GGF0 A7GGF1 A7GGI0 A7GGI2 A7GGJ6 A7GGQ8
A7GGQ9 A7GGV5 A7GGV6 A7GGW9 A7GH02 A7GH10 A7GH18 A7GH26 A7GH51 A7GH77 A7GH84
A7GH96 A7GH98 A7GHA0 A7GHC7 A7GHD2 A7GHE0 A7GHG5 A7GHH0 A7GHH4 A7GHH6 A7GHI4
A7GHJ2 A7GHK0 A7GHK2 A7GHK3 A7GHK5 A7GHL2 A7GHL9 A7GHM0 A7GHM5 A7GHS2 A7GHS3
A7GHS6 A7GHS8 A7GHT6 A7GHT7 A7GHT8 A7GHT9 A7GHU1 A7GHZ0 A7GHZ8 A7GI02 A7GI03
A7GI05 A7GI10 A7GI22 A7GIH1 A7GIH2 A7GIH3 A7GIH6 A7GIH9 A7GII1 A7GII2 A7GIK1
A7GIK2 A7GIK3 A7GIK6 A7GIL4 A7GIN0 A7GIN3 A7GIN4 A7GIP1 A7GIP8 A7GIQ9 A7GIT0
A7GIV2 A7GIW9 A7GIX1 A7GIX3 A7GIX4 A7GIX5 A7GIX6 A7GIY3 A7GJ02 A7GJ15 A7GJ38
A7GJ44 A7GJ45 A7GJ46 A7GJ47 A7GJ48 A7GJ49 A7GJ50 A7GJ53 A7GJ55 A7GJ56 A7GJ57
A7GJ58 A7GJ59 A7GJ60 A7GJ61 A7GJ62 A7GJ63 A7GJ64 A7GJ65 A7GJ66 A7GJ67 A7GJ68
A7GJ70 A7GJ71 A7GJ72 A7GJ73 A7GJ74 A7GJ75 A7GJ76 A7GJ77 A7GJ78 A7GJ79 A7GJ81
A7GJ82 A7GJ83 A7GJ85 A7GJ86 A7GJ89 A7GJ96 A7GJ99 A7GJB0 A7GJB3 A7GJB4 A7GJB7
A7GJB8 A7GJD4 A7GJE0 A7GJE2 A7GJE7 A7GJF6 A7GJG3 A7GJK7 A7GJL2 A7GJL8 A7GJM2
A7GJM4 A7GJN7 A7GJN8 A7GJN9 A7GJP3 A7GJP4 A7GJR9 A7GJS2 A7GJS8 A7GJS9 A7GJT0
A7GJT6 A7GJT7 A7GJU2 A7GJU5 A7GJV3 A7GJV7 A7GJW0 A7GJW3 A7GJX8 A7GJX9 A7GJY9
A7GJZ2 A7GJZ7 A7GJZ8 A7GJZ9 A7GK01 A7GK08 A7GK09 A7GK10 A7GK11 A7GK13 A7GK15
A7GK16 A7GK17 A7GK18 A7GK19 A7GK20 A7GK21 A7GK22 A7GK23 A7GK24 A7GK25 A7GK26
A7GK27 A7GK28 A7GK29 A7GK30 A7GK31 A7GK32 A7GK33 A7GK34 A7GK36 A7GK37 A7GK38
A7GK39 A7GK41 A7GK43 A7GK44 A7GK45 A7GK46 A7GK47 A7GK48 A7GK53 A7GK54 A7GK66
A7GKE0 A7GKF0 A7GKF6 A7GKF9 A7GKG0 A7GKG1 A7GKH5 A7GKH7 A7GKH8 A7GKI0 A7GKI2
A7GKJ1 A7GKJ4 A7GKK2 A7GKK3 A7GKK4 A7GKN4 A7GKP8 A7GKR3 A7GKR4 A7GKT9 A7GKX3
A7GKY4 A7GKY7 A7GKZ6 A7GL00 A7GL12 A7GL73 A7GLA2 A7GLA6 A7GLF8 A7GLK9 A7GLN2
A7GLP0 A7GLX6 A7GLY7 A7GLZ1 A7GM02 A7GM36 A7GM37 A7GM53 A7GM70 A7GM89 A7GM94
A7GMB9 A7GMD4 A7GMJ2 A7GMJ3 A7GMJ4 A7GML9 A7GMM0 A7GMN2 A7GMN5 A7GMR0 A7GMT6
A7GMU0 A7GMU1 A7GMU3 A7GMU4 A7GMU5 A7GMU7 A7GMU8 A7GMU9 A7GMV0 A7GMV1 A7GMV2
A7GMW0 A7GMW1 A7GN01 A7GN34 A7GN36 A7GN41 A7GN48 A7GN50 A7GN53 A7GN54 A7GN55
A7GN56 A7GN58 A7GN70 A7GN71 A7GN74 A7GN76 A7GN77 A7GN78 A7GN87 A7GN94 A7GN95
A7GNB6 A7GNL0 A7GNN8 A7GNQ7 A7GNR9 A7GNS3 A7GNS8 A7GNV5 A7GNW5 A7GNY4 A7GNY8
A7GP55 A7GP90 A7GPD9 A7GPH3 A7GPN5 A7GPY2 A7GPY3 A7GQ01 A7GQB6 A7GQB8 A7GQI9
A7GQQ6 A7GQU9 A7GQV6 A7GQV9 A7GQZ8 A7GQZ9 A7GR00 A7GR01 A7GR30 A7GR70 A7GR71

A7GR91 A7GR98 A7GR99 A7GRA2 A7GRA9 A7GRB0 A7GRD7 A7GRD8 A7GRE0 A7GRE1 A7GRE3
A7GRE7 A7GRE9 A7GRF4 A7GRF6 A7GRF7 A7GRF8 A7GRF9 A7GRG0 A7GRG2 A7GRG6 A7GRG7
A7GRH0 A7GRH2 A7GRH4 A7GRH9 A7GRI8 A7GRJ1 A7GRJ4 A7GRJ6 A7GRK0 A7GRK2 A7GRK7
A7GRK8 A7GRK9 A7GRL0 A7GRL1 A7GRL3 A7GRL4 A7GRL6 A7GRM0 A7GRM4 A7GRN8 A7GRP4
A7GRP5 A7GRP9 A7GRV4 A7GRV8 A7GRW1 A7GRX7 A7GRY8 A7GRY9 A7GRZ1 A7GS04 A7GS08
A7GS09 A7GS30 A7GS56 A7GS57 A7GS60 A7GS61 A7GS62 A7GS63 A7GS78 A7GS81 A7GS82
A7GS97 A7GSD5 A7GSD8 A7GSD9 A7GSE1 A7GSF2 A7GSF4 A7GSG2 A7GSI3 A7GSJ5 A7GSJ7
A7GSJ8 A7GSJ9 A7GSK0 A7GSL5 A7GSL7 A7GSM1 A7GSN6 A7GSN7 A7GSN8 A7GSP5 A7GSW3
A7GSX5 A7GSY1 A7GSY4 A7GSZ0 A7GSZ1 A7GSZ3 A7GSZ7 A7GT03 A7GT06 A7GT07 A7GT08
A7GT09 A7GT10 A7GT16 A7GT17 A7GT26 A7GT28 A7GT35 A7GT52 A7GT53 A7GT58 A7GT59
A7GT65 A7GT66 A7GT67 A7GT74 A7GT84 A7GT85 A7GT88 A7GT90 A7GT99 A7GTA0 A7GTA2
A7GTA3 A7GTA8 A7GTC2 A7GTC4 A7GTC9 A7GTD6 A7GTE2 A7GTE5 A7GTE7 A7GTE8 A7GTF1
A7GTF2 A7GTG6 A7GTK1 A7GTK8 A7GTL0 A7GTL2 A7GTM0 A7GTM1 A7GTM7 A7GTM8 A7GTN7
A7GTP4 A7GTQ5 A7GTR4 A7GTR5 A7GTR9 A7GTS8 A7GTT6 A7GTV6 A7GTW1 A7GTW9 A7GTZ9
A7GU01 A7GU19 A7GU36 A7GU40 A7GU41 A7GU88 A7GU91 A7GU98 A7GUA0 A7GUA1 A7GUA6
A7GUC8 A7GUD5 A7GUE1 A7GUG3 A7GUH9 A7GUR1 A7GUR8 A7GUR9 A7GUS0 A7GUT5 A7GUT7
A7GUU5 A7GUZ5 A7GV65 A7GV66 A7GV79 A7GV80 A7GV88 A7GVB2 A7GVD1 A7GVD5 A7GVE5
A7GVF0 A7GVM2 A7GVN0 A7GVN2 A7GVN5 A7GVN7 A7GVP4 A7GVP5 A7GVP6 A7GVP7 A7GVQ0
A7GVQ1 A7GVR3 A7GVT2 A7GVW3 A7GVW5 A7GVW6 A7GVX1 A7GVX8 A7GVX9 A7GVZ2 A7GVZ3
A7GVZ7 A7GW10 A7GW11 A7GW12 A7GW36 A7GW42 A7GW44 A7GW77 A7GW82 A7GW99 A7GWA5
A7GWB1 A7GWC2 A7GWM5 A7GWR7 A7GWR8 A7GWS5 A7GWT1 A7GWT2 A7GWX9 A7GWY9 A7GWZ2
A7GWZ6 A7GWZ7 A7GWZ8 A7GX26 A7GX27 A7GX33 A7GX37 A7GX39 A7GX43 A7GX48 A7GX79
A7GX97 A7GXA8 A7GXB4 A7GXB5 A7GXE4 A7GXE5 A7GXF6 A7GXF7 A7GXG2 A7GXH1 A7GXH6
A7GXI1 A7GXJ2 A7GXK2 A7GXK4 A7GXP7 A7GXQ2 A7GXQ4 A7GXQ8 A7GXS4 A7GXS7 A7GXT2
A7GXT8 A7GXU2 A7GXU4 A7GXW0 A7GXW7 A7GY40 A7GYB0 A7GYB4 A7GYC2 A7GYC8 A7GYD5
A7GYE1 A7GYG9 A7GYH0 A7GYI5 A7GYJ8 A7GYK7 A7GYL9 A7GYN4 A7GYN5 A7GYN6 A7GYQ7
A7GYS1 A7GYU4 A7GYU6 A7GYU7 A7GYX6 A7GYY6 A7GYY9 A7GYZ1 A7GZ43 A7GZ44 A7GZ54
A7GZ69 A7GZ81 A7GZ93 A7GZ96 A7GZA4 A7GZB0 A7GZB1 A7GZB2 A7GZB9 A7GZE1 A7GZE9
A7GZF6 A7GZI1 A7GZJ4 A7GZJ5 A7GZJ6 A7GZK0 A7GZK1 A7GZK2 A7GZK3 A7GZK6 A7GZL7
A7GZL8 A7GZM1 A7GZM8 A7GZP8 A7GZP9 A7GZU8 A7GZX4 A7GZZ0 A7GZZ1 A7GZZ3 A7H000
A7H026 A7H029 A7H030 A7H051 A7H084 A7H088 A7H097 A7H0A1 A7H0A8 A7H0F9 A7H0G1
A7H0H0 A7H0I4 A7H0K1 A7H0K4 A7H0L8 A7H0Y1 A7H0Y2 A7H0Y9 A7H0Z0 A7H0Z1 A7H0Z2
A7H0Z3 A7H0Z4 A7H0Z5 A7H0Z6 A7H0Z7 A7H0Z9 A7H100 A7H101 A7H102 A7H103 A7H104
A7H105 A7H106 A7H107 A7H108 A7H109 A7H110 A7H121 A7H147 A7H149 A7H185 A7H186
A7H187 A7H193 A7H194 A7H1A3 A7H1A6 A7H1A8 A7H1B7 A7H1D7 A7H1F1 A7H1G3 A7H1G8
A7H1G9 A7H1H0 A7H1H2 A7H1K5 A7H1L3 A7H1L5 A7H1L7 A7H1M6 A7H1N3 A7H1N9 A7H1P3
A7H1R0 A7H1R1 A7H1R4 A7H1S0 A7H1S1 A7H1S3 A7H1S5 A7H1T2 A7H1T3 A7H1U1 A7H1U2
A7H1U7 A7H1W2 A7H1W5 A7H1X6 A7H220 A7H239 A7H243 A7H245 A7H291 A7H295 A7H2A2
A7H2A6 A7H2A7 A7H2B2 A7H2B7 A7H2C0 A7H2C7 A7H2D2 A7H2D5 A7H2D8 A7H2E1 A7H2F8
A7H2F9 A7H2H6 A7H2I0 A7H2I7 A7H2J2 A7H2J3 A7H2J9 A7H2K2 A7H2L7 A7H2P1 A7H2P8
A7H2Q0 A7H2Q2 A7H2Q3 A7H2Q5 A7H2Q9 A7H2R6 A7H2S9 A7H2T2 A7H2T4 A7H2U1 A7H2V3
A7H2V9 A7H2Z6 A7H303 A7H331 A7H333 A7H342 A7H345 A7H346 A7H361 A7H367 A7H368
A7H372 A7H373 A7H375 A7H3B9 A7H3C0 A7H3C4 A7H3E3 A7H3F0 A7H3F2 A7H3F3 A7H3L9
A7H3M3 A7H3N1 A7H3N3 A7H3N6 A7H3P9 A7H3U1 A7H3U4 A7H3U9 A7H422 A7H426 A7H439

A7H443 A7H447 A7H450 A7H451 A7H475 A7H476 A7H477 A7H484 A7H485 A7H491 A7H492
A7H4B1 A7H4B2 A7H4B3 A7H4B5 A7H4C1 A7H4D2 A7H4D5 A7H4D8 A7H4E9 A7H4F0 A7H4F1
A7H4F2 A7H4G2 A7H4H1 A7H4H2 A7H4H3 A7H4H5 A7H4H6 A7H4H9 A7H4J1 A7H4J7 A7H4J9
A7H4K0 A7H4K9 A7H4M0 A7H4M4 A7H4N2 A7H4N3 A7H4N6 A7H4P0 A7H4P5 A7H4P6 A7H4P7
A7H4Q6 A7H4Q7 A7H4Q8 A7H4Q9 A7H4R2 A7H4R3 A7H4S5 A7H4T0 A7H4T2 A7H4U3 A7H4U5
A7H4X6 A7H4X7 A7H4Y0 A7H4Y8 A7H4Z1 A7H4Z4 A7H4Z5 A7H4Z6 A7H502 A7H506 A7H514
A7H522 A7H523 A7H537 A7H539 A7H543 A7H544 A7H552 A7H556 A7H563 A7H564 A7H574
A7H575 A7H576 A7H578 A7H585 A7H590 A7H591 A7H5C4 A7H5C5 A7H5G3 A7H5G4 A7H5G6
A7H5K0 A7H5L7 A7H5P5 A7H5Q5 A7H5U1 A7H5U2 A7H5U3 A7H5U4 A7H5U5 A7H5U6 A7H5U7
A7H5U9 A7H5V3 A7H5V4 A7H5V8 A7H5W2 A7H5W3 A7H5Y1 A7H5Y2 A7H623 A7H632 A7H639
A7H640 A7H641 A7H642 A7H643 A7H644 A7H645 A7H646 A7H647 A7H648 A7H649 A7H650
A7H651 A7H652 A7H653 A7H654 A7H655 A7H657 A7H662 A7H664 A7H665 A7H667 A7H668
A7H674 A7H677 A7H6E5 A7H6G8 A7H6G9 A7H6H5 A7H6J2 A7H6J3 A7H6J5 A7H6J6 A7H6J8
A7H6J9 A7H6L1 A7H6M4 A7H6N1 A7H6Q3 A7H6S6 A7H6S7 A7H6S8 A7H6U9 A7H6X6 A7H715
A7H716 A7H722 A7H724 A7H725 A7H750 A7H771 A7H773 A7H774 A7H783 A7H793 A7H796
A7H7H6 A7H7J7 A7H7Z8 A7H804 A7H8A2 A7H8A3 A7H8C1 A7H8C3 A7H8C5 A7H8F4 A7H8F6
A7H8K2 A7H8W2 A7H8W4 A7H905 A7H906 A7H910 A7H9A5 A7H9D8 A7H9E0 A7H9E8 A7H9F0
A7H9F3 A7H9F5 A7H9F7 A7H9F8 A7H9F9 A7H9S6 A7H9Y1 A7HA08 A7HA27 A7HA35 A7HA60
A7HAA4 A7HAB8 A7HAD3 A7HAE1 A7HAH8 A7HAL6 A7HB50 A7HB82 A7HBF2 A7HBF8 A7HBI2
A7HBI3 A7HBI7 A7HBI8 A7HBI9 A7HBJ3 A7HBL4 A7HBL5 A7HBL7 A7HBL8 A7HBL9 A7HBM0
A7HBM1 A7HBM2 A7HBM3 A7HBM4 A7HBM5 A7HBM7 A7HBM8 A7HBM9 A7HBN0 A7HBN1 A7HBN2
A7HBN3 A7HBN4 A7HBN5 A7HBN6 A7HBN7 A7HBN9 A7HBP1 A7HBP2 A7HBP3 A7HBP4 A7HBP5
A7HBP6 A7HBP8 A7HBQ0 A7HBQ1 A7HBQ2 A7HBQ4 A7HBU9 A7HC22 A7HC32 A7HC45 A7HC64
A7HCA3 A7HCC3 A7HCD6 A7HCD8 A7HCE9 A7HCH6 A7HCH7 A7HCH8 A7HCH9 A7HCI0 A7HCI1
A7HCI4 A7HCN7 A7HCP4 A7HCQ8 A7HCR6 A7HCR8 A7HCU0 A7HCV7 A7HD67 A7HD93 A7HDA1
A7HDA2 A7HDA9 A7HDB0 A7HDB7 A7HDC0 A7HDC5 A7HDC9 A7HDD6 A7HDE0 A7HDS7 A7HDU3
A7HDU6 A7HDW6 A7HDX2 A7HDY2 A7HDY3 A7HDY7 A7HDY8 A7HEI5 A7HFV9 A7HFW0 A7HFW2
A7HFW3 A7HG34 A7HG96 A7HG97 A7HGP4 A7HGP8 A7HGS8 A7HGU3 A7HGU4 A7HGV2 A7HGZ0
A7HGZ2 A7HGZ6 A7HH57 A7HH59 A7HH68 A7HH69 A7HH70 A7HH87 A7HHA2 A7HHM9 A7HHN1
A7HHN3 A7HHV3 A7HI45 A7HI47 A7HI48 A7HIA1 A7HID3 A7HIF0 A7HIF8 A7HIF9 A7HIG6
A7HIG8 A7HIH1 A7HIJ4 A7HIL1 A7HIL2 A7HIX3 A7HIY6 A7HJ26 A7HJ56 A7HJ58 A7HJ69
A7HJ78 A7HJ80 A7HJA5 A7HJA9 A7HJB0 A7HJD6 A7HJF3 A7HJG6 A7HJI0 A7HJI3 A7HJQ1
A7HJQ9 A7HJR3 A7HJT8 A7HJX5 A7HJZ8 A7HJZ9 A7HK07 A7HK24 A7HK34 A7HK39 A7HK58
A7HK70 A7HK86 A7HKC0 A7HKM4 A7HKM5 A7HKP2 A7HKQ7 A7HKR8 A7HKS0 A7HKS1 A7HKS3
A7HKT4 A7HKU5 A7HKU7 A7HKV0 A7HKV1 A7HKV2 A7HKV8 A7HKW8 A7HL08 A7HL11 A7HL14
A7HL16 A7HL30 A7HL47 A7HL71 A7HL98 A7HLA6 A7HLB0 A7HLB6 A7HLB7 A7HLC0 A7HLE2
A7HLE9 A7HLG3 A7HLH4 A7HLL7 A7HLM0 A7HLP1 A7HLU0 A7HLU6 A7HLV7 A7HLW9 A7HLZ4
A7HLZ9 A7HM20 A7HM23 A7HM24 A7HM25 A7HM26 A7HM27 A7HM28 A7HM29 A7HM31 A7HM33
A7HM34 A7HM35 A7HM36 A7HM37 A7HM38 A7HM39 A7HM40 A7HM41 A7HM42 A7HM43 A7HM44
A7HM45 A7HM46 A7HM47 A7HM48 A7HM49 A7HM51 A7HM52 A7HM53 A7HM54 A7HM55 A7HM56
A7HM57 A7HM68 A7HM70 A7HM73 A7HM80 A7HM89 A7HMA1 A7HME3 A7HME4 A7HME6 A7HMF2
A7HMG4 A7HMG8 A7HMJ2 A7HMJ7 A7HMJ8 A7HMK2 A7HMM1 A7HMP1 A7HMQ2 A7HMY3 A7HMY6
A7HN01 A7HN13 A7HN21 A7HN55 A7HN59 A7HN62 A7HN65 A7HN68 A7HN69 A7HN71 A7HN78
A7HN79 A7HN82 A7HNA3 A7HNA4 A7HND1 A7HNE6 A7HNE8 A7HNK6 A7HNL1 A7HNL5 A7HNN0

A7HNQ1 A7HNR3 A7HNS0 A7HNS2 A7HNT0 A7HNV0 A7HNV7 A7HNX2 A7HNX8 A7HNY1 A7HNY2
A7HNY3 A7HNY4 A7HNY7 A7HNZ1 A7HP03 A7HP26 A7HP27 A7HP29 A7HP87 A7HP99 A7HPD3
A7HPD4 A7HPD7 A7HPE1 A7HPJ7 A7HPK8 A7HPK9 A7HPP8 A7HPQ2 A7HPQ3 A7HPQ6 A7HPQ7
A7HPT7 A7HPU1 A7HQ17 A7HQ48 A7HQF9 A7HQG6 A7HQP6 A7HQQ0 A7HQQ1 A7HQR0 A7HQX4
A7HQX5 A7HQX9 A7HQY0 A7HQZ3 A7HR11 A7HR75 A7HR80 A7HRI8 A7HS32 A7HS42 A7HSA5
A7HSG6 A7HSG7 A7HSH0 A7HSH1 A7HSH4 A7HSH6 A7HSH7 A7HSI7 A7HSJ1 A7HSK9 A7HSL1
A7HSL2 A7HSL7 A7HSQ6 A7HSR8 A7HSV9 A7HSW1 A7HSW4 A7HSW7 A7HSX5 A7HT06 A7HT07
A7HT09 A7HT10 A7HT11 A7HT37 A7HT39 A7HT46 A7HT62 A7HT64 A7HT65 A7HT67 A7HT69
A7HT70 A7HT94 A7HTA7 A7HTE5 A7HTI8 A7HTU2 A7HTX8 A7HTY3 A7HU16 A7HU86 A7HU87
A7HU88 A7HUC7 A7HUQ6 A7HUR5 A7HUU8 A7HUX6 A7HUY3 A7HUZ5 A7HUZ6 A7HUZ7 A7HUZ8
A7HV12 A7HV25 A7HV30 A7HV58 A7HV69 A7HVA3 A7HVC7 A7HVD7 A7HVD8 A7HVT8 A7HVT9
A7HVU8 A7HVU9 A7HVV0 A7HVV8 A7HVX4 A7HVY9 A7HVZ2 A7HW48 A7HWA4 A7HWA5 A7HWA8
A7HWB1 A7HWB4 A7HWJ0 A7HWJ3 A7HWM6 A7HWP7 A7HWQ0 A7HWQ1 A7HWQ2 A7HWQ3 A7HWQ4
A7HWQ5 A7HWQ6 A7HWQ7 A7HWQ8 A7HWR0 A7HWR1 A7HWR2 A7HWR4 A7HWR5 A7HWR7 A7HWR8
A7HWR9 A7HWS0 A7HWS1 A7HWS2 A7HWS3 A7HWS4 A7HWS5 A7HWS6 A7HWS7 A7HWS8 A7HWS9
A7HWT0 A7HWT2 A7HWT3 A7HWT4 A7HWT5 A7HWT6 A7HWU9 A7HWX2 A7HWZ1 A7HX10 A7HX17
A7HX36 A7HX37 A7HX41 A7HX43 A7HX57 A7HX83 A7HXB2 A7HXI4 A7HXJ5 A7HXJ6 A7HXK4
A7HXK5 A7HXK6 A7HXL5 A7HXM4 A7HXM7 A7HXU8 A7HXV6 A7HXW0 A7HXX1 A7HXX4 A7HXX9
A7HXZ5 A7HXZ6 A7HXZ9 A7HY03 A7HY16 A7HY17 A7HY18 A7HY19 A7HY28 A7HY53 A7HY57
A7HYA5 A7HYC4 A7HYE5 A7HYF2 A7HYF4 A7HYI2 A7HYI3 A7HYI4 A7HYS8 A7HYV8 A7HYW1
A7HZ35 A7HZ36 A7HZ38 A7HZ39 A7HZ43 A7HZ45 A7HZ51 A7HZ64 A7HZ77 A7HZ82 A7HZ84
A7HZ88 A7HZ90 A7HZ93 A7HZ94 A7HZ96 A7HZ97 A7HZC0 A7HZC4 A7HZE0 A7HZE7 A7HZE8
A7HZG0 A7HZG1 A7HZG2 A7HZH0 A7HZI3 A7HZK5 A7HZK7 A7HZK8 A7HZK9 A7HZL0 A7HZL1
A7HZL2 A7HZL3 A7HZL5 A7HZL6 A7HZL7 A7HZL8 A7HZL9 A7HZM0 A7HZM1 A7HZM2 A7HZM3
A7HZN3 A7HZN6 A7HZP2 A7HZP3 A7HZP6 A7HZW6 A7HZW7 A7HZX0 A7HZX1 A7HZX2 A7HZX3
A7HZX4 A7HZX5 A7HZX6 A7HZX7 A7HZY2 A7HZY9 A7I002 A7I005 A7I006 A7I008 A7I010
A7I057 A7I058 A7I064 A7I0A8 A7I0C2 A7I0D4 A7I0I5 A7I0M0 A7I0N0 A7I0N1 A7I0N5
A7I0N8 A7I0P0 A7I0P1 A7I0P7 A7I0P8 A7I0P9 A7I0Q3 A7I0T3 A7I0V4 A7I0V6 A7I0V7
A7I0V8 A7I0W4 A7I0W5 A7I0X0 A7I0X6 A7I0Y0 A7I107 A7I108 A7I131 A7I140 A7I145
A7I157 A7I162 A7I166 A7I168 A7I198 A7I199 A7I1A0 A7I1B8 A7I1C5 A7I1D0 A7I1F3
A7I1G1 A7I1G2 A7I1I7 A7I1J3 A7I1L2 A7I1L3 A7I1M0 A7I1P3 A7I1R7 A7I1R8 A7I1S0
A7I1S7 A7I1S8 A7I1U0 A7I1U3 A7I1U4 A7I1U5 A7I1V2 A7I1V7 A7I1W4 A7I1W6 A7I1Y4
A7I206 A7I226 A7I232 A7I236 A7I241 A7I243 A7I252 A7I254 A7I261 A7I279 A7I280
A7I281 A7I291 A7I2C6 A7I2D4 A7I2F2 A7I2G3 A7I2H6 A7I2I6 A7I2L9 A7I2M4 A7I2S8
A7I2T7 A7I2U3 A7I2V3 A7I2V7 A7I2V8 A7I2W3 A7I2W4 A7I2W9 A7I2X0 A7I2X3 A7I2Y2
A7I2Z8 A7I2Z9 A7I316 A7I328 A7I332 A7I338 A7I377 A7I3C8 A7I3D2 A7I3D4 A7I3J2
A7I3Q0 A7I3Q1 A7I3S0 A7I3S8 A7I3S9 A7I3T6 A7I3T7 A7I3T8 A7I3U0 A7I3U1 A7I3U2
A7I3U3 A7I3U6 A7I3U7 A7I3U8 A7I3V0 A7I3V2 A7I3W7 A7I3W8 A7I3X0 A7I3X1 A7I3X8
A7I3Y5 A7I3Z5 A7I414 A7I416 A7I417 A7I439 A7I444 A7I456 A7I458 A7I463 A7I464
A7I472 A7I4D1 A7I4D4 A7I4F7 A7I4H6 A7I4I0 A7I4M1 A7I4Q7 A7I4S0 A7I4T3 A7I4T5
A7I4T7 A7I4U9 A7I4V4 A7I4W9 A7I4X4 A7I4X5 A7I4X6 A7I4Y2 A7I4Y4 A7I5I7 A7I5I8
A7I5J0 A7I5K0 A7I5N0 A7I5N7 A7I5N9 A7I5P0 A7I5P2 A7I5P3 A7I5P4 A7I5P5 A7I5P8
A7I5P9 A7I5Q0 A7I5Q2 A7I5Q4 A7I5Q8 A7I5Q9 A7I5R0 A7I5R5 A7I615 A7I616 A7I637
A7I644 A7I655 A7I656 A7I661 A7I6B7 A7I6C5 A7I6C9 A7I6H3 A7I6H7 A7I6J1 A7I6J7

A7I6K3 A7I6L5 A7I6N9 A7I6P0 A7I6S5 A7I6S6 A7I6W6 A7I728 A7I757 A7I771 A7I773
A7I774 A7I781 A7I798 A7I7A4 A7I7L0 A7I7N1 A7I7P2 A7I7P5 A7I7S4 A7I7T1 A7I7T4
A7I7T6 A7I7Y6 A7I807 A7I809 A7I841 A7I848 A7I849 A7I854 A7I890 A7I8A7 A7I8B8
A7I8L5 A7I8L8 A7I8M6 A7I8N1 A7I8P9 A7I8Q1 A7I8R6 A7I8W5 A7I8Y0 A7I912 A7I937
A7I940 A7I9C7 A7I9D0 A7I9D1 A7I9E0 A7I9E9 A7I9H8 A7I9I3 A7I9I6 A7I9I7 A7I9J0
A7I9J3 A7I9K5 A7I9K6 A7I9M0 A7I9N4 A7I9P7 A7I9Q6 A7I9Q9 A7I9R3 A7I9R7 A7I9Z8
A7IA08 A7IA09 A7IA21 A7IA25 A7IA28 A7IA59 A7IA69 A7IA72 A7IA92 A7IA93 A7IAA9
A7IAB9 A7IAE1 A7IAF1 A7IAF4 A7IAF5 A7IAG1 A7IAG4 A7IAG8 A7IAH1 A7IAH4 A7IAH6
A7IAH7 A7IAI8 A7IAJ1 A7IAK4 A7IAK5 A7IAL9 A7IAM2 A7IAP7 A7IAP8 A7IAR0 A7IAR4
A7IAS7 A7IAU3 A7IAZ7 A7IB57 A7IB61 A7IB67 A7IBD6 A7IBL9 A7IC03 A7IC04 A7IC07
A7IC11 A7IC40 A7IC65 A7IC67 A7IC75 A7ICA9 A7ICB3 A7ICB9 A7ICI5 A7IDJ5 A7IDK1
A7IDU0 A7IDU5 A7IDU6 A7IDY1 A7IE31 A7IEB7 A7IEC3 A7IEJ1 A7IEJ8 A7IEN8 A7IEQ6
A7IES2 A7IF36 A7IF97 A7IFB7 A7IFB8 A7IFD4 A7IFE5 A7IFF8 A7IFW1 A7IFX6 A7IFX7
A7IFX8 A7IFY0 A7IFY1 A7IFY2 A7IFY4 A7IFY5 A7IFY7 A7IFY8 A7IFY9 A7IFZ0 A7IFZ1
A7IFZ2 A7IG40 A7IG64 A7IGA7 A7IGB2 A7IGE1 A7IGF4 A7IGF5 A7IGG9 A7IGL7 A7IGM0
A7IGM5 A7IGN4 A7IGN8 A7IGW4 A7IH06 A7IH12 A7IH13 A7IH44 A7IH93 A7IH94 A7IHD3
A7IHE7 A7IHE9 A7IHF1 A7IHH0 A7IHN7 A7IHN9 A7IHP0 A7IHP3 A7IHP6 A7IHQ4 A7IHS8
A7IHU0 A7IHV6 A7II77 A7IIA9 A7IIB1 A7IID0 A7IID4 A7IIE0 A7IIE4 A7IIG5 A7IIJ3
A7IIK5 A7IIK6 A7IIM2 A7IIX3 A7IJ56 A7IJ80 A7IJ81 A7IJF6 A7IJG7 A7IJU9 A7IJV2
A7IJV4 A7IJW3 A7IJY7 A7IJZ8 A7IK05 A7IK06 A7IK08 A7IK14 A7IK18 A7IK27 A7IKI9
A7IKJ3 A7IKP6 A7IKP7 A7IKP8 A7IKP9 A7IKQ0 A7IKQ1 A7IKQ7 A7IL64 A7ILC5 A7ILC6
A7ILC7 A7ILW0 A7ILW8 A7IM13 A7IM18 A7IM41 A7IM59 A7IM67 A7IM78 A7IMA0 A7IMA6
A7IMA7 A7IMA9 A7IMR0 A7IMS4 A7IMW3 A7IMW7 A7IMY3 A7IMY7 A7IN73 A7IN97 A7INA4
A7INA7 A7INB9 A7INE1 A7INH1 A7INH2 A7INJ6 A7INK0 A7INK2 A7INK3 A7INK7 A7INL3
A7INN1 A7INP1 A7INR3 A7INR5 A7INR6 A7INT8 A7INU8 A7INU9 A7INV0 A7IPJ0 A7IPJ5
A7IPK6 A7IPP5 A7IPP6 A7IPP7 A7IPP8 A7IPP9 A7IPQ1 A7IPQ2 A7IPQ3 A7IPQ4 A7IPQ5
A7IPQ6 A7IPQ7 A7IPQ8 A7IPQ9 A7IPX7 A7IQE5 A7IQE6 A7IQF2 A7IQH5 A7IQW5 A7ISP5
A7ISP6 A7IXI8 A7IZZ1 A7IZZ2 A7J390 A7J391 A7J392 A7J398 A7J399 A7J3A0 A7J3A2
A7J3A3 A7J3A4 A7J3A6 A7K6Y8 A7K6Y9 A7KAI6 A7KAJ8 A7KAL8 A7KAM4 A7KAN7 A7KBS4
A7KE01 A7KGA5 A7KII6 A7KWF8 A7L2Z6 A7L6A0 A7L6A2 A7L9Z4 A7LBL2 A7LFZ6 A7LKB2
A7LN87 A7M7B9 A7M7C7 A7M8Z3 A7M8Z9 A7M903 A7M906 A7M918 A7M919 A7M920 A7M922
A7M929 A7M930 A7M931 A7M932 A7M933 A7M934 A7M935 A7M936 A7M937 A7M939 A7M943
A7M948 A7M955 A7M956 A7M957 A7M958 A7M964 A7M968 A7M984 A7M985 A7M986 A7M988
A7M996 A7M997 A7M998 A7M999 A7M9A0 A7M9A1 A7M9A2 A7M9A3 A7M9A4 A7M9B1 A7MAS3
A7MAS5 A7MAZ5 A7MB12 A7MB16 A7MB34 A7MB35 A7MB40 A7MB47 A7MB54 A7MB62 A7MB76
A7MB78 A7MB89 A7MBC0 A7MBD1 A7MBE4 A7MBF6 A7MBG0 A7MBH5 A7MBJ1 A7MBJ2 A7MBJ5
A7MBP4 A7MBU6 A7MCB7 A7MD48 A7ME27 A7ME43 A7ME44 A7ME45 A7ME47 A7ME52 A7ME53
A7ME54 A7ME70 A7ME76 A7ME80 A7MEA0 A7MEA1 A7MEA2 A7MEA3 A7MEB0 A7MEB1 A7MEB6
A7MEB8 A7MEC0 A7MEE9 A7MEF1 A7MEG3 A7MEG4 A7MEG5 A7MEJ8 A7MEL5 A7MEN4 A7MEQ6
A7MEQ8 A7MER7 A7MES7 A7MES8 A7MET6 A7MEU5 A7MEV9 A7MEW3 A7MEW8 A7MEX5 A7MEY6
A7MEZ1 A7MF19 A7MF48 A7MF54 A7MF58 A7MF59 A7MFA6 A7MFA8 A7MFC6 A7MFD3 A7MFG0
A7MFG4 A7MFG5 A7MFH0 A7MFH4 A7MFI7 A7MFI9 A7MFJ6 A7MFJ8 A7MFN6 A7MFQ6 A7MFR1
A7MFT1 A7MFV2 A7MFW4 A7MFW6 A7MFX7 A7MFX8 A7MFY0 A7MG20 A7MG30 A7MG41 A7MGA3
A7MGA7 A7MGA8 A7MGA9 A7MGB0 A7MGC5 A7MGD1 A7MGD2 A7MGD4 A7MGE0 A7MGF3 A7MGF4

A7MGG4 A7MGP2 A7MGP3 A7MGP6 A7MGP7 A7MGP8 A7MGQ7 A7MGQ9 A7MGR3 A7MGR5 A7MGS0
A7MGS1 A7MGS2 A7MGS4 A7MGS5 A7MGT0 A7MGT1 A7MGT2 A7MGU1 A7MGU7 A7MGV1 A7MGV2
A7MGV3 A7MGV7 A7MGW6 A7MGW9 A7MGX5 A7MGX6 A7MGX8 A7MGY0 A7MGY5 A7MGZ1 A7MH01
A7MH06 A7MH19 A7MH28 A7MH30 A7MH33 A7MH59 A7MH63 A7MH65 A7MH70 A7MHB0 A7MHE1
A7MHE2 A7MHG8 A7MHG9 A7MHJ8 A7MHL3 A7MHT9 A7MHU4 A7MHU5 A7MHW7 A7MHX2 A7MHX6
A7MHX8 A7MI07 A7MI15 A7MI22 A7MI25 A7MI34 A7MI37 A7MI45 A7MI49 A7MI52 A7MI96
A7MIA7 A7MIB1 A7MIB2 A7MIC1 A7MIC5 A7MIC7 A7MID0 A7MID1 A7MID6 A7MIE1 A7MIF0
A7MIG0 A7MIG6 A7MIG7 A7MIH6 A7MIJ0 A7MIJ1 A7MIJ5 A7MIJ7 A7MIK3 A7MIK5 A7MIM0
A7MIM3 A7MIM5 A7MIM8 A7MIN6 A7MIN8 A7MIP1 A7MIP9 A7MIR9 A7MIS8 A7MIU0 A7MIX5
A7MIX7 A7MIZ2 A7MJ02 A7MJ03 A7MJ08 A7MJ10 A7MJ27 A7MJ32 A7MJ38 A7MJ39 A7MJ41
A7MJ43 A7MJ57 A7MJ58 A7MJ59 A7MJ60 A7MJ61 A7MJ63 A7MJ64 A7MJ65 A7MJ67 A7MJ68
A7MJ69 A7MJ70 A7MJ78 A7MJ80 A7MJ81 A7MJ89 A7MJ90 A7MJA9 A7MJB9 A7MJD3 A7MJD4
A7MJF1 A7MJF4 A7MJG4 A7MJJ7 A7MJM0 A7MJM9 A7MJN1 A7MJP4 A7MJQ1 A7MJQ2 A7MJQ4
A7MJQ6 A7MJR0 A7MJR8 A7MJR9 A7MJS4 A7MJU0 A7MJU1 A7MJU3 A7MJV7 A7MJW4 A7MJW5
A7MJX1 A7MJZ3 A7MJZ6 A7MK00 A7MK09 A7MK11 A7MK12 A7MK40 A7MK41 A7MK78 A7MKB5
A7MKC0 A7MKC1 A7MKD3 A7MKE3 A7MKE5 A7MKE8 A7MKF0 A7MKG2 A7MKH5 A7MKI5 A7MKI9
A7MKJ0 A7MKJ3 A7MKJ5 A7MKJ6 A7MKK2 A7MKK9 A7MKL4 A7MKM8 A7MKR9 A7MKS3 A7MKV5
A7MKW4 A7MKX3 A7MKX5 A7MKY0 A7MKY7 A7ML00 A7ML04 A7ML10 A7ML12 A7ML17 A7ML45
A7ML56 A7ML57 A7ML61 A7ML62 A7ML65 A7ML67 A7ML69 A7ML79 A7ML80 A7ML81 A7ML84
A7ML85 A7ML88 A7ML95 A7MLI7 A7MLJ4 A7MLL2 A7MLL8 A7MLL9 A7MLM8 A7MLM9 A7MLN0
A7MLV0 A7MLV5 A7MLV7 A7MLW9 A7MLX1 A7MLY0 A7MLY2 A7MM35 A7MM48 A7MM56 A7MM70
A7MM76 A7MM77 A7MM78 A7MM80 A7MM88 A7MM89 A7MM94 A7MMA2 A7MMB9 A7MMC0 A7MMC3
A7MMD8 A7MMF2 A7MMF7 A7MMG1 A7MMG7 A7MMK0 A7MMK1 A7MMK2 A7MMM5 A7MMT5 A7MMV1
A7MMV4 A7MMW2 A7MMW3 A7MMW6 A7MMZ8 A7MN01 A7MN03 A7MN07 A7MN58 A7MN74 A7MN76
A7MNA3 A7MNB3 A7MNE5 A7MNG1 A7MNI5 A7MNK0 A7MNK1 A7MNK3 A7MNM8 A7MNP0 A7MNP9
A7MNR3 A7MNR4 A7MNS2 A7MNU8 A7MNV4 A7MNV7 A7MNV8 A7MNW9 A7MNY7 A7MNY9 A7MNZ1
A7MNZ2 A7MNZ6 A7MP11 A7MP52 A7MP70 A7MP89 A7MP93 A7MP95 A7MPA0 A7MPA9 A7MPC2
A7MPD4 A7MPD7 A7MPE1 A7MPE7 A7MPE8 A7MPE9 A7MPF3 A7MPF5 A7MPF7 A7MPF8 A7MPG2
A7MPG4 A7MPG5 A7MPG6 A7MPG7 A7MPG8 A7MPH1 A7MPH2 A7MPH4 A7MPH5 A7MPH6 A7MPH8
A7MPI3 A7MPI4 A7MPI5 A7MPI6 A7MPI7 A7MPJ5 A7MPJ7 A7MPN9 A7MPR2 A7MPS3 A7MQ53
A7MQ61 A7MQ83 A7MQ91 A7MQ93 A7MQ95 A7MQ96 A7MQ97 A7MQ98 A7MQA9 A7MQB0 A7MQB3
A7MQB5 A7MQD6 A7MQD9 A7MQE0 A7MQE1 A7MQE3 A7MQG0 A7MQG1 A7MQH1 A7MQI2 A7MQI3
A7MQJ1 A7MQK2 A7MQK4 A7MQL7 A7MQM1 A7MQN1 A7MQN9 A7MQP3 A7MQP6 A7MQP7 A7MQP8
A7MQQ0 A7MQQ2 A7MQQ8 A7MQQ9 A7MQS0 A7MQS3 A7MQS5 A7MQS7 A7MQT2 A7MQT6 A7MQV7
A7MQW6 A7MQX5 A7MQY9 A7MQZ8 A7MR02 A7MR03 A7MR08 A7MR28 A7MR31 A7MR65 A7MR71
A7MR73 A7MR82 A7MR83 A7MR85 A7MR91 A7MRX1 A7MRY0 A7MRY3 A7MRY9 A7MRZ7 A7MS04
A7MS57 A7MS71 A7MS76 A7MS81 A7MS83 A7MS84 A7MS85 A7MSA8 A7MSB2 A7MSC7 A7MSD7
A7MSD8 A7MSE0 A7MSE1 A7MSE5 A7MSM1 A7MSN3 A7MSN5 A7MSP5 A7MSP7 A7MSP8 A7MSP9
A7MST1 A7MST3 A7MSW1 A7MSW8 A7MSX5 A7MSX6 A7MSX8 A7MSY2 A7MSY3 A7MSY9 A7MSZ6
A7MSZ7 A7MT09 A7MT10 A7MT26 A7MT45 A7MT47 A7MT54 A7MT82 A7MT83 A7MT89 A7MT92
A7MTC2 A7MTC9 A7MTD6 A7MTL8 A7MTM7 A7MTN6 A7MTN8 A7MTP0 A7MTP2 A7MTQ1 A7MTQ2
A7MTQ7 A7MTQ9 A7MTS4 A7MTS5 A7MTS6 A7MTS9 A7MTT6 A7MTT7 A7MTT8 A7MTT9 A7MTV7
A7MTX1 A7MU00 A7MU26 A7MU31 A7MU37 A7MU38 A7MU39 A7MU42 A7MU46 A7MU47 A7MU48
A7MU49 A7MU79 A7MUE1 A7MUE4 A7MUE5 A7MUE7 A7MUF3 A7MUJ1 A7MUJ3 A7MUL2 A7MUN4

A7MUN7 A7MUP0 A7MUP2 A7MUT5 A7MUU6 A7MUU9 A7MUV0 A7MUV2 A7MUW4 A7MUW7 A7MUW8
A7MV01 A7MV11 A7MV27 A7MV82 A7MV84 A7MV87 A7MV92 A7MVD3 A7MVE1 A7MVE7 A7MVG2
A7MVH4 A7MVH5 A7MVH6 A7MVH9 A7MVI6 A7MVJ3 A7MVJ5 A7MVK1 A7MVK3 A7MVN0 A7MVW6
A7MVW7 A7MVY7 A7MW43 A7MWB9 A7MWC3 A7MWC5 A7MWC6 A7MWD3 A7MWD9 A7MWE0 A7MWE4
A7MWE9 A7MWG4 A7MWH2 A7MWH3 A7MWH4 A7MWH5 A7MWH6 A7MWH7 A7MWH9 A7MWI1 A7MWI2
A7MWI3 A7MWI4 A7MWI5 A7MWI6 A7MWI7 A7MWI8 A7MWJ1 A7MWJ3 A7MWJ4 A7MWK7 A7MWM0
A7MWM9 A7MWN0 A7MWN3 A7MWN9 A7MWP7 A7MWS7 A7MWT6 A7MWU0 A7MWU1 A7MWW0 A7MWW1
A7MWW3 A7MWX4 A7MX07 A7MX10 A7MX14 A7MX17 A7MX30 A7MX33 A7MX36 A7MX54 A7MX55
A7MX59 A7MX66 A7MX70 A7MX74 A7MX76 A7MX77 A7MX86 A7MX95 A7MX96 A7MXA2 A7MXB2
A7MXB9 A7MXC0 A7MXC2 A7MXD3 A7MXE3 A7MXE4 A7MXE5 A7MXF0 A7MXF1 A7MXF2 A7MXG0
A7MXI3 A7MXJ9 A7MXK6 A7MXK7 A7MXL2 A7MXM2 A7MXP2 A7MXR0 A7MXR2 A7MXR3 A7MXT2
A7MXT4 A7MXU9 A7MXX7 A7MXY6 A7MXZ4 A7MXZ7 A7MXZ8 A7MY01 A7MY07 A7MY11 A7MY21
A7MY25 A7MY37 A7MY61 A7MY68 A7MY77 A7MY79 A7MY81 A7MY83 A7MY86 A7MY91 A7MY96
A7MY97 A7MYB6 A7MYB8 A7MYC1 A7MYC5 A7MYC6 A7MYG2 A7MYT3 A7MYT5 A7MYT6 A7MYU9
A7MYV0 A7MYV1 A7MYV2 A7MYW9 A7MYX2 A7MYX3 A7MYX4 A7MZ01 A7MZ47 A7MZ48 A7MZ63
A7MZ64 A7MZ92 A7MZA1 A7MZA8 A7MZB2 A7MZB3 A7MZE2 A7MZE5 A7MZG9 A7MZH4 A7MZI2
A7MZI3 A7MZI4 A7MZI5 A7MZM6 A7MZN5 A7MZR6 A7MZS0 A7MZT3 A7MZU8 A7MZY1 A7N009
A7N011 A7N012 A7N025 A7N060 A7N075 A7N080 A7N0F0 A7N0G1 A7N0G2 A7N0G6 A7N0H6
A7N0H7 A7N0H8 A7N0I3 A7N0I4 A7N0I5 A7N0I6 A7N0I7 A7N0I8 A7N0I9 A7N0J0 A7N0J7
A7N0K6 A7N0M1 A7N0M8 A7N0R3 A7N0T3 A7N0V8 A7N0W0 A7N0W3 A7N0X5 A7N0X6 A7N0X8
A7N118 A7N121 A7N122 A7N127 A7N128 A7N129 A7N137 A7N141 A7N156 A7N158 A7N178
A7N1D8 A7N1D9 A7N1E2 A7N1E4 A7N1E5 A7N1E9 A7N1F1 A7N1G7 A7N1G9 A7N1H7 A7N1I0
A7N1I8 A7N1I9 A7N1J1 A7N1J2 A7N1J4 A7N1J9 A7N1K3 A7N1L2 A7N1L5 A7N1L7 A7N1L8
A7N1L9 A7N1R1 A7N1U8 A7N1W0 A7N1W2 A7N1X1 A7N1X3 A7N1X4 A7N1X5 A7N1X6 A7N1X7
A7N1X9 A7N1Y0 A7N1Z7 A7N204 A7N212 A7N2P9 A7N2Q0 A7N2S3 A7N2T8 A7N333 A7N3D6
A7N3R7 A7N3S9 A7N3Y9 A7N3Z0 A7N3Z2 A7N4I5 A7N4V5 A7N512 A7N5C3 A7N5C4 A7N5E9
A7N5N6 A7N5Q6 A7N5W3 A7N609 A7N654 A7N655 A7N681 A7N6L2 A7N721 A7N787 A7N7P0
A7N7S2 A7N7S3 A7N7T1 A7N805 A7N819 A7N8I1 A7N8K5 A7N926 A7N938 A7N939 A7N949
A7N982 A7N992 A7N9A3 A7N9A5 A7N9C6 A7N9D1 A7N9D2 A7N9H6 A7N9I8 A7N9L2 A7N9L5
A7N9M0 A7N9M1 A7N9Q8 A7N9R4 A7N9R5 A7N9R6 A7N9R7 A7N9S0 A7N9S2 A7N9S3 A7N9S4
A7N9S5 A7N9S6 A7N9S7 A7N9S8 A7N9S9 A7N9T0 A7N9T1 A7N9T2 A7N9T3 A7N9T4 A7N9T5
A7N9T6 A7N9T7 A7N9T8 A7N9T9 A7N9U0 A7N9U1 A7N9U2 A7N9U3 A7N9U4 A7N9U6 A7N9U7
A7N9U8 A7N9U9 A7N9V0 A7N9V7 A7N9Y1 A7N9Y5 A7N9Y6 A7NA22 A7NA32 A7NA52 A7NA74
A7NA91 A7NAA1 A7NAB1 A7NAC9 A7NAD2 A7NAE2 A7NAH6 A7NAH8 A7NAI2 A7NAI4 A7NAI5
A7NAI8 A7NAM0 A7NAM3 A7NAM4 A7NAN4 A7NAP0 A7NAR0 A7NAR1 A7NAX2 A7NAY2 A7NAY4
A7NB07 A7NB16 A7NB31 A7NB32 A7NB34 A7NB66 A7NB76 A7NB78 A7NB92 A7NBA0 A7NBA3
A7NBC6 A7NBC7 A7NBG3 A7NBG9 A7NBH0 A7NBJ3 A7NBK5 A7NBL6 A7NBM9 A7NBQ8 A7NBR0
A7NBR5 A7NBR6 A7NBS1 A7NBS2 A7NBT0 A7NBV0 A7NBV2 A7NBV3 A7NBV5 A7NBW2 A7NBX3
A7NBY0 A7NBY8 A7NC07 A7NC13 A7NC51 A7NC55 A7NC56 A7NC58 A7NC76 A7NC90 A7NCA0
A7NCA3 A7NCA4 A7NCA9 A7NCD9 A7NCF1 A7NCH9 A7NCI3 A7NCL5 A7NCM0 A7NCM5 A7NCM8
A7NCM9 A7NCN7 A7NCP9 A7NCQ3 A7NCQ7 A7NCQ8 A7NCT0 A7NCU4 A7NCW7 A7NCX0 A7NCX8
A7NCY0 A7NCY2 A7ND07 A7ND08 A7ND27 A7ND36 A7ND52 A7NDA9 A7NDB3 A7NDB4 A7NDB6
A7NDC1 A7NDC2 A7NDF7 A7NDG1 A7NDG2 A7NDI1 A7NDI3 A7NDJ7 A7NDJ8 A7NDM9 A7NDN9
A7NDP2 A7NDP3 A7NDP4 A7NDP7 A7NDQ2 A7NDQ3 A7NDR1 A7NDV1 A7NDV3 A7NDV6 A7NDW6

A7NDX4 A7NE12 A7NE28 A7NE29 A7NE53 A7NE88 A7NE89 A7NE94 A7NE96 A7NE97 A7NEA4
A7NEB0 A7NEB1 A7NEB2 A7NEB3 A7NEB4 A7NEB6 A7NEC0 A7NEC1 A7NEC2 A7NEC3 A7NEC4
A7NEC7 A7NEF8 A7NEF9 A7NEG0 A7NEI6 A7NEI7 A7NEI8 A7NEJ0 A7NEJ1 A7NEM0 A7NEM1
A7NEQ3 A7NES0 A7NET9 A7NEW1 A7NF00 A7NF26 A7NF27 A7NF38 A7NF69 A7NFA0 A7NFA1
A7NFB8 A7NFC5 A7NFC6 A7NFF1 A7NFF4 A7NFG3 A7NFJ2 A7NFR2 A7NFR6 A7NFV2 A7NG29
A7NG41 A7NG77 A7NG78 A7NG79 A7NGC5 A7NGG1 A7NGH6 A7NGQ9 A7NH01 A7NH10 A7NH25
A7NH85 A7NH92 A7NHA9 A7NHN1 A7NHP6 A7NHQ4 A7NI01 A7NI03 A7NI09 A7NI71 A7NI79
A7NI92 A7NI97 A7NIA2 A7NIA3 A7NIB1 A7NID0 A7NID1 A7NIF2 A7NIG2 A7NIK8 A7NIS7
A7NIS8 A7NJ21 A7NJ53 A7NJ65 A7NJ67 A7NJA5 A7NJB8 A7NJE1 A7NJF1 A7NJF9 A7NJG0
A7NJG1 A7NJH1 A7NJH2 A7NJH8 A7NJM0 A7NJT6 A7NJT8 A7NK01 A7NK07 A7NK64 A7NK74
A7NK79 A7NKB4 A7NKL7 A7NKM0 A7NKR9 A7NKS0 A7NKS5 A7NKU4 A7NKU6 A7NKU8 A7NKV1
A7NKX0 A7NKY6 A7NL64 A7NLC3 A7NLL2 A7NLR3 A7NLU2 A7NLU3 A7NLW5 A7NLW6 A7NM80
A7NM84 A7NM86 A7NMS9 A7NMZ8 A7NN19 A7NN26 A7NN50 A7NN87 A7NN90 A7NNA3 A7NNA4
A7NND5 A7NNH1 A7NNW3 A7NNY4 A7NP08 A7NP12 A7NP88 A7NPB0 A7NPB5 A7NPB9 A7NPC0
A7NPG2 A7NPM9 A7NPT5 A7NPY6 A7NPZ0 A7NPZ1 A7NQ20 A7NQA5 A7NQB4 A7NQB7 A7NQB8
A7NQC5 A7NQH9 A7NQM2 A7NQM8 A7NQR0 A7NQU3 A7NQW2 A7NQW5 A7NQY6 A7NQZ5 A7NR17
A7NR21 A7NR33 A7NR34 A7NR36 A7NR37 A7NR38 A7NR39 A7NR40 A7NR41 A7NR44 A7NR45
A7NR47 A7NR48 A7NR49 A7NR50 A7NR51 A7NR52 A7NR53 A7NR54 A7NR55 A7NR56 A7NR58
A7NR59 A7NR60 A7NR61 A7NR63 A7NR64 A7NR65 A7NR66 A7NR67 A7NR68 A7NR80 A7NRA6
A7NRF7 A7NRF8 A7NRP8 A7NRU6 A7NRW4 A7NRZ0 A7NRZ6 A7NRZ8 A7NS01 A7NS25 A7NS27
A7NS28 A7NS62 A7NS65 A7NS83 A7NSB7 A7NSC5 A7NVX9 A7P514 A7PQS0 A7PRJ6 A7Q0V4
A7Q5X9 A7QJG1 A7RF00 A7RFA1 A7RFT2 A7RG82 A7RH72 A7RHB3 A7RHG8 A7RHL5 A7RIT9
A7RK30 A7RKF6 A7RKS5 A7RL75 A7RLE5 A7RM20 A7RNG8 A7RP64 A7RPT4 A7RR04 A7RRG3
A7RRJ0 A7RRZ8 A7RS11 A7RSH7 A7RT29 A7RTB3 A7RU46 A7RUD5 A7RV13 A7RVK7 A7RWC9
A7RWD2 A7RWP6 A7RX26 A7RX34 A7RZW4 A7S1A4 A7S1H9 A7S338 A7S3I2 A7S3J7 A7S6A1
A7S6A5 A7S6M8 A7S710 A7S7F2 A7S8T5 A7SA47 A7SBF0 A7SBN6 A7SCH8 A7SD85 A7SDA8
A7SDW5 A7SE05 A7SE07 A7SEP9 A7SG48 A7SG73 A7SGU3 A7SGU6 A7SGZ5 A7SJ66 A7SK48
A7SKE9 A7SKJ3 A7SL20 A7SLC8 A7SLW1 A7SLX5 A7SM54 A7SMR0 A7SMR1 A7SMW7 A7SN31
A7SNN5 A7SNN9 A7SP74 A7SPE8 A7SPW6 A7SPX9 A7STV9 A7SUU7 A7SWD3 A7SWH1 A7SXZ6
A7T0T1 A7T0W1 A7T167 A7TDI2 A7TDL7 A7TDM3 A7TDZ7 A7TDZ8 A7TE16 A7TE19 A7TE38
A7TE77 A7TE86 A7TEF4 A7TEG8 A7TEH5 A7TEM0 A7TEN6 A7TER7 A7TES6 A7TET7 A7TEU3
A7TEW6 A7TEY0 A7TF50 A7TFN8 A7TFW4 A7TFZ9 A7TG30 A7TGA7 A7TGD2 A7TGD9 A7TGI3
A7TGI5 A7TGL9 A7TGQ2 A7TGR0 A7TGR2 A7TGU7 A7TGW7 A7TH19 A7TH34 A7TH62 A7TH64
A7TH93 A7THE0 A7THE5 A7THG7 A7THH1 A7THU5 A7THU9 A7THV0 A7THV5 A7THW2 A7THY5
A7TI13 A7TI41 A7TIC4 A7TID0 A7TIF5 A7TIJ7 A7TIN6 A7TJ36 A7TJ59 A7TJ71 A7TJ85
A7TJI3 A7TJK8 A7TJL1 A7TJM9 A7TJP2 A7TJS7 A7TJT3 A7TJT7 A7TJV5 A7TJY9 A7TK55
A7TK63 A7TK72 A7TK94 A7TKC6 A7TKF2 A7TKN1 A7TKR8 A7TKU9 A7TKY1 A7TL17 A7TL18
A7TL22 A7TL64 A7TL79 A7TL97 A7TLA0 A7TLF1 A7TLF7 A7TLL0 A7TLU2 A7TLZ8 A7TM88
A7TM98 A7TMF9 A7TMH4 A7TMJ6 A7TMK6 A7TML4 A7TMW6 A7TMZ9 A7TN25 A7TN35 A7TN63
A7TNP9 A7TNQ2 A7TNS4 A7TNT1 A7TP22 A7TP28 A7TPB3 A7TPC9 A7TPF8 A7TPS5 A7TPS8
A7TPV3 A7TPW4 A7TQ03 A7TQ21 A7TQ35 A7TQ73 A7TQ77 A7TQB2 A7TQJ9 A7TQK0 A7TQL5
A7TQN2 A7TQP0 A7TQR2 A7TQS6 A7TQT8 A7TQZ1 A7TR10 A7TR75 A7TR79 A7TR87 A7TR90
A7TRH1 A7TRI3 A7TRR5 A7TRU5 A7TRX9 A7TRZ8 A7TS37 A7TS67 A7TSA5 A7TSA8 A7TSF4
A7TSJ7 A7TSL7 A7TSM3 A7TSQ8 A7TSV4 A7TSV6 A7TT02 A7TT36 A7TT46 A7TT88 A7TTE2

A7TTE6 A7TTL0 A7TTQ1 A7TTT5 A7TTT7 A7TTV0 A7TX81 A7TZT2 A7U6E7 A7U6E9 A7U6F0
A7U6F1 A7U6F2 A7U6F3 A7UFC2 A7UKY7 A7UL74 A7UMC0 A7UMW1 A7UNK4 A7UVN1 A7UW97
A7UWD5 A7UWE6 A7UWH7 A7UX13 A7UXD4 A7UXE4 A7UXI1 A7VLV1 A7VLV2 A7WLI0 A7WNA9
A7WNB2 A7WQL9 A7WWM4 A7WWN1 A7WWP0 A7WWQ0 A7WWR5 A7WX68 A7WXA5 A7WXE5 A7WXH2
A7WXH5 A7WXK6 A7WXP7 A7WXS1 A7WXT5 A7WXZ2 A7WXZ6 A7WY46 A7WY58 A7WY61 A7WY89
A7WY93 A7WYL7 A7WYM0 A7WYM2 A7WYN2 A7WYP0 A7WYP4 A7WYQ5 A7WYS1 A7WYS8 A7WYT1
A7WYT2 A7WYT9 A7WYU4 A7WYU7 A7WYW0 A7WYW1 A7WYW2 A7WYW4 A7WYW7 A7WYX0 A7WYX1
A7WYX3 A7WYX4 A7WYX6 A7WYY0 A7WYY2 A7WZ02 A7WZ06 A7WZ22 A7WZ30 A7WZ38 A7WZ56
A7WZF1 A7WZJ4 A7WZL0 A7WZL5 A7WZL9 A7WZM9 A7WZN6 A7WZQ8 A7WZR5 A7WZR7 A7WZR9
A7WZS6 A7WZS8 A7WZT9 A7X013 A7X019 A7X0C7 A7X0D6 A7X0E5 A7X0H4 A7X0H5 A7X0H6
A7X0I1 A7X0I2 A7X0I7 A7X0M2 A7X0M6 A7X0M9 A7X0N2 A7X0P2 A7X0P6 A7X0S7 A7X0V3
A7X0V7 A7X0W1 A7X0W4 A7X0Z9 A7X113 A7X122 A7X136 A7X140 A7X141 A7X145 A7X160
A7X163 A7X169 A7X172 A7X179 A7X1B5 A7X1B6 A7X1B8 A7X1C4 A7X1D2 A7X1D8 A7X1E5
A7X1E9 A7X1F4 A7X1F5 A7X1G4 A7X1H4 A7X1H8 A7X1I6 A7X1J9 A7X1K6 A7X1K9 A7X1L0
A7X1L2 A7X1L7 A7X1L8 A7X1M6 A7X1M7 A7X1M9 A7X1N1 A7X1N3 A7X1N5 A7X1N6 A7X1N7
A7X1P3 A7X1P4 A7X1P6 A7X1Q1 A7X1Q2 A7X1Q5 A7X1Q7 A7X1Q8 A7X1S3 A7X1T3 A7X1T6
A7X1T8 A7X1U3 A7X1U8 A7X1Y2 A7X1Y9 A7X1Z0 A7X1Z1 A7X202 A7X203 A7X215 A7X232
A7X234 A7X235 A7X238 A7X240 A7X271 A7X273 A7X274 A7X283 A7X295 A7X2A5 A7X2A9
A7X2B3 A7X2D9 A7X2E0 A7X2E3 A7X2E9 A7X2F4 A7X2G1 A7X2G5 A7X2G6 A7X2G8 A7X2H3
A7X2H6 A7X2I1 A7X2I6 A7X2K7 A7X2M2 A7X2P6 A7X2P9 A7X2Q2 A7X2Q3 A7X2Q9 A7X2R9
A7X2S0 A7X2S3 A7X2S4 A7X2U5 A7X2V2 A7X2W0 A7X2W2 A7X2W3 A7X2W8 A7X2X5 A7X2X9
A7X2Y0 A7X2Y1 A7X2Y2 A7X2Y4 A7X2Y8 A7X2Z9 A7X302 A7X306 A7X313 A7X319 A7X320
A7X325 A7X327 A7X328 A7X333 A7X344 A7X345 A7X347 A7X350 A7X354 A7X356 A7X357
A7X358 A7X359 A7X361 A7X362 A7X364 A7X383 A7X388 A7X391 A7X393 A7X395 A7X396
A7X397 A7X3A1 A7X3A2 A7X3A6 A7X3B0 A7X3D5 A7X3E0 A7X3E5 A7X3F0 A7X3G2 A7X3G5
A7X3H5 A7X3H9 A7X3I5 A7X3J9 A7X3K7 A7X3L7 A7X3N2 A7X3N3 A7X3P0 A7X3T8 A7X3V3
A7X3X9 A7X3Y3 A7X3Z1 A7X400 A7X420 A7X426 A7X427 A7X428 A7X432 A7X442 A7X451
A7X4J1 A7X4J3 A7X4L6 A7X4L9 A7X4M5 A7X4M9 A7X4N1 A7X4N3 A7X4N4 A7X4N7 A7X4P1
A7X4P5 A7X4P6 A7X4R6 A7X4S8 A7X4S9 A7X4V6 A7X4V7 A7X4V8 A7X4W4 A7X4W8 A7X4X7
A7X4Y0 A7X522 A7X524 A7X569 A7X574 A7X575 A7X578 A7X579 A7X5B3 A7X5B4 A7X5B5
A7X5C3 A7X5C4 A7X5C5 A7X5C7 A7X5C8 A7X5D0 A7X5D4 A7X5D6 A7X5D8 A7X5D9 A7X5E1
A7X5E2 A7X5E4 A7X5E5 A7X5E7 A7X5E9 A7X5F0 A7X5F2 A7X5F4 A7X5F5 A7X5F6 A7X5F7
A7X5G0 A7X5G1 A7X5G2 A7X5G3 A7X5G6 A7X5J1 A7X5J2 A7X5J9 A7X5K8 A7X5L9 A7X5M0
A7X5M3 A7X5M4 A7X5M5 A7X5M7 A7X5T3 A7X5T8 A7X5U0 A7X5V7 A7X5W0 A7X5Z9 A7X624
A7X656 A7X666 A7X669 A7X6K5 A7X6R7 A7X6S3 A7X6X6 A7X6X7 A7X6Y1 A7X6Y6 A7X6Z3
A7X715 A7X719 A7X763 A7X764 A7X766 A7X770 A7X771 A7X781 A7X782 A7X7A5 A7X7A7
A7X7A8 A7X7A9 A7X7B0 A7XCC9 A7XCD9 A7XCE1 A7XCE4 A7XCE8 A7XGN8 A7XN92 A7XRY0
A7XRY3 A7XRY6 A7XRY8 A7XRZ1 A7XUK7 A7XXB6 A7XXB7 A7XXB9 A7XXC1 A7XXC2 A7XYH5
A7XYH9 A7XYI6 A7XYJ6 A7XYQ1 A7Y397 A7Y3B3 A7Y3B4 A7Y3B6 A7Y3B9 A7Y3D0 A7Y3D8
A7Y3G7 A7Y3G8 A7Y3G9 A7Y3I1 A7Y3I2 A7Y3I4 A7Y3I5 A7Y3I6 A7Y3I8 A7Y3I9 A7Y3J0
A7Y3J1 A7Y3J3 A7Y3J6 A7Y3J9 A7Y3K4 A7Y521 A7Y7W2 A7Y7W3 A7Y7X5 A7YDW0 A7YE96
A7YH32 A7YSY2 A7YVH9 A7YVI8 A7YW45 A7YW98 A7YWC8 A7YWD2 A7YWE4 A7YWH3 A7YWI9
A7YWK3 A7YWL5 A7YWM2 A7YWP4 A7YWS7 A7YWU3 A7YY28 A7YY35 A7YY45 A7YY53 A7YY54
A7YY62 A7YY73 A7YY97 A7Z017 A7Z019 A7Z026 A7Z052 A7Z061 A7Z0C3 A7Z0C6 A7Z0D3

A7Z0D4 A7Z0D5 A7Z0E4 A7Z0F1 A7Z0F6 A7Z0F8 A7Z0G9 A7Z0H2 A7Z0H5 A7Z0H6 A7Z0J3
A7Z0J4 A7Z0K7 A7Z0L2 A7Z0L3 A7Z0L4 A7Z0L6 A7Z0M1 A7Z0M4 A7Z0M5 A7Z0M6 A7Z0M7
A7Z0M9 A7Z0N0 A7Z0N2 A7Z0N3 A7Z0N4 A7Z0N5 A7Z0N7 A7Z0N8 A7Z0N9 A7Z0P0 A7Z0P1
A7Z0P2 A7Z0P3 A7Z0P4 A7Z0P5 A7Z0P6 A7Z0P7 A7Z0P8 A7Z0P9 A7Z0Q0 A7Z0Q1 A7Z0Q2
A7Z0Q3 A7Z0Q4 A7Z0Q5 A7Z0Q6 A7Z0Q7 A7Z0Q9 A7Z0R1 A7Z0R2 A7Z0R3 A7Z0R4 A7Z0R5
A7Z0R6 A7Z0S0 A7Z0S1 A7Z0S2 A7Z0U6 A7Z0V3 A7Z0Z2 A7Z0Z7 A7Z119 A7Z152 A7Z159
A7Z171 A7Z1E8 A7Z1F1 A7Z1F3 A7Z1L3 A7Z1M9 A7Z1N4 A7Z1T2 A7Z200 A7Z201 A7Z206
A7Z207 A7Z235 A7Z248 A7Z249 A7Z251 A7Z261 A7Z266 A7Z267 A7Z268 A7Z282 A7Z2E2
A7Z2G6 A7Z2G8 A7Z2J3 A7Z2L7 A7Z2M2 A7Z2N5 A7Z2N8 A7Z2P5 A7Z2Q3 A7Z2Q4 A7Z2R4
A7Z2U3 A7Z2Z1 A7Z305 A7Z311 A7Z323 A7Z354 A7Z367 A7Z368 A7Z391 A7Z3A4 A7Z3A7
A7Z3A9 A7Z3C9 A7Z3E8 A7Z3F2 A7Z3F8 A7Z3H9 A7Z3M5 A7Z3T1 A7Z3U1 A7Z3X0 A7Z3X1
A7Z3X4 A7Z3X5 A7Z3X6 A7Z3X7 A7Z3Y6 A7Z3Y9 A7Z432 A7Z433 A7Z434 A7Z439 A7Z463
A7Z465 A7Z487 A7Z488 A7Z4A7 A7Z4C3 A7Z4C5 A7Z4C9 A7Z4D1 A7Z4D3 A7Z4D5 A7Z4D6
A7Z4D7 A7Z4E1 A7Z4E3 A7Z4E6 A7Z4F9 A7Z4G7 A7Z4G9 A7Z4H3 A7Z4H4 A7Z4H7 A7Z4H9
A7Z4I0 A7Z4I7 A7Z4I9 A7Z4J3 A7Z4J5 A7Z4J8 A7Z4K2 A7Z4L3 A7Z4L9 A7Z4M2 A7Z4M3
A7Z4M4 A7Z4M6 A7Z4M9 A7Z4N3 A7Z4N5 A7Z4N6 A7Z4N7 A7Z4R7 A7Z4S0 A7Z4S1 A7Z4S2
A7Z4S3 A7Z4S6 A7Z4S8 A7Z4T0 A7Z4T4 A7Z4T6 A7Z4T7 A7Z4T9 A7Z4U0 A7Z4U8 A7Z4W4
A7Z4W5 A7Z4X0 A7Z4X1 A7Z4X2 A7Z4X5 A7Z4X6 A7Z4X7 A7Z4X8 A7Z4X9 A7Z4Y0 A7Z500
A7Z501 A7Z504 A7Z522 A7Z553 A7Z555 A7Z565 A7Z566 A7Z570 A7Z571 A7Z5B2 A7Z5B3
A7Z5B4 A7Z5B6 A7Z5E4 A7Z5J9 A7Z5M8 A7Z5N6 A7Z5S1 A7Z5S2 A7Z5T3 A7Z5T7 A7Z5U1
A7Z5W2 A7Z5W9 A7Z5X0 A7Z5Y2 A7Z5Y7 A7Z5Z2 A7Z5Z3 A7Z5Z4 A7Z5Z6 A7Z5Z9 A7Z600
A7Z610 A7Z612 A7Z614 A7Z615 A7Z616 A7Z617 A7Z618 A7Z619 A7Z622 A7Z623 A7Z627
A7Z629 A7Z630 A7Z636 A7Z638 A7Z640 A7Z670 A7Z671 A7Z674 A7Z675 A7Z692 A7Z6C4
A7Z6E7 A7Z6E9 A7Z6F0 A7Z6F3 A7Z6J3 A7Z6J5 A7Z6J7 A7Z6J9 A7Z6K0 A7Z6L3 A7Z6L9
A7Z6M0 A7Z6M2 A7Z6M3 A7Z6M4 A7Z6Q6 A7Z6S2 A7Z6S8 A7Z6T1 A7Z6T7 A7Z6T8 A7Z6U1
A7Z6U2 A7Z6U6 A7Z6V5 A7Z6V9 A7Z6W0 A7Z6W1 A7Z6W2 A7Z6W3 A7Z6W8 A7Z6W9 A7Z6X8
A7Z721 A7Z722 A7Z726 A7Z727 A7Z732 A7Z733 A7Z734 A7Z735 A7Z745 A7Z750 A7Z751
A7Z754 A7Z756 A7Z766 A7Z767 A7Z768 A7Z769 A7Z774 A7Z780 A7Z781 A7Z783 A7Z785
A7Z789 A7Z793 A7Z794 A7Z7A2 A7Z7A5 A7Z7A7 A7Z7A9 A7Z7B2 A7Z7B3 A7Z7B5 A7Z7B6
A7Z7B8 A7Z7B9 A7Z7C6 A7Z7C8 A7Z7D7 A7Z7E7 A7Z7F1 A7Z7F5 A7Z7H3 A7Z7H4 A7Z7I1
A7Z7I6 A7Z7I7 A7Z7J8 A7Z7K6 A7Z7L6 A7Z7L9 A7Z7M0 A7Z7M2 A7Z7N3 A7Z7P1 A7Z7R1
A7Z7S3 A7Z7S6 A7Z7T3 A7Z7V7 A7Z7Y9 A7Z7Z0 A7Z803 A7Z809 A7Z812 A7Z848 A7Z8B5
A7Z8C1 A7Z8C3 A7Z8D3 A7Z8D5 A7Z8E0 A7Z8E8 A7Z8F5 A7Z8J3 A7Z8R5 A7Z8S7 A7Z8U4
A7Z8Y2 A7Z8Y3 A7Z900 A7Z901 A7Z902 A7Z939 A7Z951 A7Z953 A7Z962 A7Z963 A7Z964
A7Z965 A7Z967 A7Z968 A7Z971 A7Z973 A7Z975 A7Z985 A7Z9A2 A7Z9A6 A7Z9A7 A7Z9G4
A7Z9I5 A7Z9N4 A7Z9N5 A7Z9P0 A7Z9Q8 A7Z9Q9 A7Z9R0 A7Z9S5 A7Z9S6 A7Z9T0 A7Z9T4
A7Z9T5 A7Z9T8 A7Z9V1 A7Z9X5 A7Z9Y5 A7Z9Y9 A7ZA24 A7ZA58 A7ZA59 A7ZA74 A7ZA84
A7ZAB2 A7ZAE3 A7ZAE4 A7ZAE5 A7ZAE6 A7ZAF1 A7ZAH2 A7ZAH3 A7ZAH5 A7ZAH7 A7ZAH8
A7ZAH9 A7ZAI0 A7ZAI1 A7ZAI5 A7ZAK3 A7ZAL8 A7ZAP4 A7ZAQ7 A7ZAR2 A7ZAU7 A7ZAU9
A7ZAV8 A7ZAV9 A7ZAW0 A7ZAW1 A7ZAW4 A7ZAW5 A7ZAW7 A7ZAY0 A7ZAY1 A7ZAZ4 A7ZAZ7
A7ZAZ8 A7ZB08 A7ZB09 A7ZB10 A7ZB34 A7ZB36 A7ZB70 A7ZB75 A7ZB82 A7ZB87 A7ZB92
A7ZBD9 A7ZBK0 A7ZBP3 A7ZBQ0 A7ZBR7 A7ZBS1 A7ZBS6 A7ZBS7 A7ZC03 A7ZC05 A7ZC11
A7ZC60 A7ZC67 A7ZC69 A7ZC71 A7ZC85 A7ZC90 A7ZCA7 A7ZCA8 A7ZCB2 A7ZCB6 A7ZCD0
A7ZCD9 A7ZCE9 A7ZCF3 A7ZCG7 A7ZCH3 A7ZCH5 A7ZCL1 A7ZCL5 A7ZCL8 A7ZCL9 A7ZCN0

A7ZCN3 A7ZCN4 A7ZCN5 A7ZCN6 A7ZCP1 A7ZCQ2 A7ZCQ6 A7ZCT9 A7ZCV1 A7ZCV2 A7ZCY6
A7ZCZ1 A7ZCZ7 A7ZD05 A7ZD34 A7ZD35 A7ZD37 A7ZD85 A7ZD89 A7ZD95 A7ZDA4 A7ZDF2
A7ZDF3 A7ZDF4 A7ZDH6 A7ZDJ9 A7ZDK0 A7ZDK1 A7ZDN6 A7ZDP4 A7ZDQ0 A7ZDQ1 A7ZDQ5
A7ZDV3 A7ZDX6 A7ZDX8 A7ZDY6 A7ZE07 A7ZE19 A7ZE20 A7ZE21 A7ZE25 A7ZE26 A7ZE39
A7ZE50 A7ZE53 A7ZE60 A7ZE62 A7ZEA2 A7ZEB5 A7ZEB6 A7ZEC2 A7ZEC8 A7ZED5 A7ZED8
A7ZEF1 A7ZEF6 A7ZEF8 A7ZEG5 A7ZEH3 A7ZEH4 A7ZEJ1 A7ZEJ3 A7ZEJ7 A7ZEK1 A7ZEL6
A7ZEM1 A7ZEQ3 A7ZES2 A7ZES6 A7ZES8 A7ZEX0 A7ZEX8 A7ZEY3 A7ZEY4 A7ZEZ4 A7ZEZ5
A7ZF21 A7ZF27 A7ZF28 A7ZF95 A7ZFA0 A7ZFA4 A7ZFB7 A7ZFB9 A7ZFC1 A7ZFJ4 A7ZFP0
A7ZFR4 A7ZFS7 A7ZFX5 A7ZFX6 A7ZFY6 A7ZFY7 A7ZFY8 A7ZFY9 A7ZFZ0 A7ZFZ1 A7ZFZ4
A7ZFZ5 A7ZFZ6 A7ZFZ7 A7ZFZ8 A7ZFZ9 A7ZG00 A7ZG01 A7ZG02 A7ZG03 A7ZG04 A7ZG05
A7ZG06 A7ZG07 A7ZG08 A7ZG09 A7ZG10 A7ZG11 A7ZG16 A7ZG18 A7ZGB0 A7ZGB2 A7ZGB4
A7ZGD3 A7ZGE5 A7ZH92 A7ZH96 A7ZHA4 A7ZHA5 A7ZHB2 A7ZHB5 A7ZHB8 A7ZHB9 A7ZHC0
A7ZHC6 A7ZHC8 A7ZHC9 A7ZHD0 A7ZHD4 A7ZHD5 A7ZHE2 A7ZHE3 A7ZHE4 A7ZHE5 A7ZHF0
A7ZHF3 A7ZHF4 A7ZHG3 A7ZHG4 A7ZHG6 A7ZHH2 A7ZHH3 A7ZHI2 A7ZHJ2 A7ZHJ3 A7ZHJ5
A7ZHK8 A7ZHK9 A7ZHL0 A7ZHM0 A7ZHM3 A7ZHM4 A7ZHN7 A7ZHP6 A7ZHP8 A7ZHQ1 A7ZHQ2
A7ZHQ5 A7ZHQ6 A7ZHQ7 A7ZHQ9 A7ZHR0 A7ZHR1 A7ZHR2 A7ZHS3 A7ZHT0 A7ZHT6 A7ZHU5
A7ZHU8 A7ZHV1 A7ZHY1 A7ZHZ2 A7ZHZ8 A7ZHZ9 A7ZI01 A7ZI02 A7ZI28 A7ZI30 A7ZI50
A7ZI51 A7ZI52 A7ZI88 A7ZI91 A7ZI94 A7ZI95 A7ZI96 A7ZI97 A7ZI98 A7ZI99 A7ZIA3
A7ZIA4 A7ZIA5 A7ZID1 A7ZID6 A7ZID7 A7ZIE0 A7ZIE2 A7ZIF7 A7ZIF8 A7ZIG6 A7ZIG8
A7ZIG9 A7ZIH3 A7ZIH5 A7ZIH7 A7ZII0 A7ZIJ4 A7ZIJ5 A7ZIJ6 A7ZIK2 A7ZIK4 A7ZIM8
A7ZIN1 A7ZIN2 A7ZIN3 A7ZIN4 A7ZIN5 A7ZIN6 A7ZIR0 A7ZIR2 A7ZIR8 A7ZIT5 A7ZIT8
A7ZIW5 A7ZIZ9 A7ZJ00 A7ZJ03 A7ZJ15 A7ZJ18 A7ZJ19 A7ZJ25 A7ZJ28 A7ZJ31 A7ZJ42
A7ZJ51 A7ZJ54 A7ZJ60 A7ZJ63 A7ZJ94 A7ZJ95 A7ZJA8 A7ZJC5 A7ZJD0 A7ZJD2 A7ZJE2
A7ZJG7 A7ZJI4 A7ZJI5 A7ZJI8 A7ZJJ0 A7ZJJ2 A7ZJL8 A7ZJM2 A7ZJM7 A7ZJQ2 A7ZJS0
A7ZJS7 A7ZJT5 A7ZJU2 A7ZJU9 A7ZJV1 A7ZJV2 A7ZJW2 A7ZJZ6 A7ZJZ7 A7ZJZ9 A7ZK01
A7ZK06 A7ZK12 A7ZK14 A7ZK21 A7ZK22 A7ZK27 A7ZK52 A7ZK59 A7ZK62 A7ZK67 A7ZK71
A7ZKA2 A7ZKA4 A7ZKA5 A7ZKA9 A7ZKB2 A7ZKB3 A7ZKB4 A7ZKB5 A7ZKB6 A7ZKB7 A7ZKD6
A7ZKF9 A7ZKG4 A7ZKG7 A7ZKH2 A7ZKK1 A7ZKL0 A7ZKL1 A7ZKL2 A7ZKM3 A7ZKR7 A7ZKS3
A7ZKS4 A7ZKT2 A7ZKU1 A7ZKU5 A7ZKU7 A7ZKV0 A7ZKV2 A7ZKW0 A7ZKX7 A7ZKY2 A7ZKY4
A7ZKY5 A7ZL06 A7ZL19 A7ZL78 A7ZL79 A7ZL97 A7ZLA2 A7ZLA9 A7ZLH4 A7ZLK8 A7ZLN7
A7ZLQ4 A7ZLS1 A7ZLW9 A7ZLX0 A7ZLX4 A7ZLX5 A7ZLX7 A7ZM66 A7ZM70 A7ZM83 A7ZM99
A7ZMA0 A7ZMA2 A7ZMA4 A7ZMB5 A7ZMC4 A7ZME4 A7ZME5 A7ZMF7 A7ZMF8 A7ZMH0 A7ZMH3
A7ZMI0 A7ZMI2 A7ZMI4 A7ZMI6 A7ZMJ8 A7ZMK0 A7ZMK7 A7ZMK9 A7ZML2 A7ZML3 A7ZML4
A7ZML5 A7ZML6 A7ZMN3 A7ZMP8 A7ZMQ4 A7ZMT4 A7ZMT6 A7ZMV4 A7ZMV8 A7ZMY4 A7ZMY5
A7ZMY7 A7ZMZ0 A7ZMZ5 A7ZMZ6 A7ZN00 A7ZN02 A7ZN19 A7ZN22 A7ZN37 A7ZN60 A7ZN75
A7ZN77 A7ZN88 A7ZNE5 A7ZNF5 A7ZNH9 A7ZNJ1 A7ZNJ3 A7ZNJ6 A7ZNJ7 A7ZNN9 A7ZNR4
A7ZNR5 A7ZNS3 A7ZNS4 A7ZNT3 A7ZNW5 A7ZNX6 A7ZNX7 A7ZNY3 A7ZNY5 A7ZNZ5 A7ZP09
A7ZP10 A7ZP12 A7ZP32 A7ZP50 A7ZP69 A7ZP80 A7ZP82 A7ZP83 A7ZP87 A7ZP88 A7ZPA5
A7ZPA8 A7ZPD3 A7ZPD6 A7ZPE0 A7ZPE5 A7ZPE7 A7ZPH2 A7ZPI2 A7ZPJ8 A7ZPK7 A7ZPL2
A7ZPL9 A7ZPM7 A7ZPM8 A7ZPN6 A7ZPP0 A7ZPR0 A7ZPR5 A7ZPS1 A7ZPS4 A7ZPT9 A7ZPU1
A7ZPU2 A7ZPV1 A7ZPV4 A7ZPV7 A7ZPV8 A7ZPW0 A7ZPW1 A7ZPW6 A7ZPW9 A7ZPX0 A7ZPX4
A7ZPX5 A7ZPY2 A7ZPY3 A7ZPY4 A7ZPY5 A7ZPZ4 A7ZQ09 A7ZQ11 A7ZQ20 A7ZQ24 A7ZQ25
A7ZQ46 A7ZQ47 A7ZQ48 A7ZQ49 A7ZQ54 A7ZQ55 A7ZQ58 A7ZQ60 A7ZQ90 A7ZQA5 A7ZQC1
A7ZQC4 A7ZQC5 A7ZQC7 A7ZQC8 A7ZQD9 A7ZQE0 A7ZQH3 A7ZQI7 A7ZQI8 A7ZQI9 A7ZQJ0

A7ZQJ1 A7ZQJ4 A7ZQJ5 A7ZQJ6 A7ZQK5 A7ZQK6 A7ZQK7 A7ZQM3 A7ZQN7 A7ZQP2 A7ZQP6
A7ZQP8 A7ZQQ1 A7ZQT3 A7ZQT6 A7ZQT7 A7ZQV1 A7ZQY1 A7ZQZ8 A7ZR07 A7ZR09 A7ZR12
A7ZR13 A7ZR14 A7ZR19 A7ZR23 A7ZR24 A7ZR34 A7ZR35 A7ZR59 A7ZR64 A7ZR67 A7ZR71
A7ZR72 A7ZR85 A7ZR88 A7ZRR9 A7ZRS4 A7ZRT0 A7ZRT3 A7ZRT7 A7ZRU6 A7ZRU7 A7ZRV0
A7ZRV6 A7ZRW7 A7ZRX6 A7ZS00 A7ZS18 A7ZS26 A7ZS33 A7ZS44 A7ZS45 A7ZS49 A7ZS50
A7ZS61 A7ZS62 A7ZS63 A7ZS64 A7ZS65 A7ZS68 A7ZS69 A7ZS72 A7ZS75 A7ZS79 A7ZS81
A7ZS83 A7ZS86 A7ZSA4 A7ZSB5 A7ZSB6 A7ZSB8 A7ZSC4 A7ZSC5 A7ZSD0 A7ZSD1 A7ZSF3
A7ZSF6 A7ZSH0 A7ZSH1 A7ZSH3 A7ZSH6 A7ZSH7 A7ZSI3 A7ZSI4 A7ZSI5 A7ZSI6 A7ZSI8
A7ZSJ0 A7ZSJ1 A7ZSJ2 A7ZSJ3 A7ZSJ4 A7ZSJ5 A7ZSJ6 A7ZSJ7 A7ZSJ8 A7ZSJ9 A7ZSK0
A7ZSK1 A7ZSK2 A7ZSK3 A7ZSK4 A7ZSK5 A7ZSK6 A7ZSK8 A7ZSK9 A7ZSL0 A7ZSL4 A7ZSL5
A7ZSL6 A7ZSL7 A7ZSL8 A7ZSL9 A7ZSM0 A7ZSM3 A7ZSN0 A7ZSP4 A7ZSR4 A7ZSR5 A7ZST1
A7ZST9 A7ZSU1 A7ZSU3 A7ZSU8 A7ZSU9 A7ZSV5 A7ZSW2 A7ZSW3 A7ZSW4 A7ZSW5 A7ZT06
A7ZT19 A7ZT33 A7ZT60 A7ZTA5 A7ZTA6 A7ZTB2 A7ZTC2 A7ZTD9 A7ZTE9 A7ZTG3 A7ZTG6
A7ZTH0 A7ZTH2 A7ZTI5 A7ZTI6 A7ZTI7 A7ZTI8 A7ZTI9 A7ZTJ1 A7ZTJ2 A7ZTJ3 A7ZTJ4
A7ZTJ9 A7ZTK1 A7ZTM0 A7ZTP0 A7ZTP1 A7ZTP5 A7ZTP7 A7ZTQ6 A7ZTQ8 A7ZTQ9 A7ZTR0
A7ZTR2 A7ZTR3 A7ZTR6 A7ZTV2 A7ZTV5 A7ZTV6 A7ZTV7 A7ZTV9 A7ZTW7 A7ZTX3 A7ZTX6
A7ZTY0 A7ZTY2 A7ZU09 A7ZU14 A7ZU16 A7ZU36 A7ZU40 A7ZU52 A7ZU66 A7ZU67 A7ZU92
A7ZUA1 A7ZUB2 A7ZUB3 A7ZUB4 A7ZUB5 A7ZUB7 A7ZUB8 A7ZUC9 A7ZUD3 A7ZUE0 A7ZUE3
A7ZUE4 A7ZUE6 A7ZUE7 A7ZUF1 A7ZUF7 A7ZUG1 A7ZUH4 A7ZUH5 A7ZUH7 A7ZUH8 A7ZUI2
A7ZUI5 A7ZUI7 A7ZUJ0 A7ZUJ2 A7ZUJ6 A7ZUJ7 A7ZUJ8 A7ZUK0 A7ZUK1 A7ZUK2 A7ZUK5
A7ZUK8 A7ZUK9 A7ZUL0 A7ZUL1 A7ZUL3 A7ZUL4 A7ZUM3 A7ZUM6 A7ZUN0 A7ZUN5 A7ZUP3
A7ZUR1 A7ZV06 A7ZV11 A7ZV12 A7ZV18 A7ZV28 A7ZV32 A7ZV33 A7ZV39 A7ZV40 A7ZV41
A7ZV47 A7ZV48 A7ZV62 A7ZV63 A7ZV66 A7ZV68 A7ZV69 A7ZV71 A7ZV72 A7ZV73 A7ZV74
A7ZV87 A7ZV98 A7ZVB0 A7ZVB3 A7ZVC6 A7ZVC7 A7ZVD3 A7ZVE0 A7ZVL9 A7ZVM4 A7ZVN0
A7ZVQ2 A7ZVR0 A7ZVR5 A7ZVS4 A7ZVS5 A7ZVS6 A7ZVS7 A7ZVS8 A7ZVT5 A7ZVT7 A7ZVU9
A7ZVV7 A7ZVV8 A7ZVW0 A7ZVW2 A7ZVX1 A7ZVX4 A7ZVX7 A7ZVX8 A7ZVX9 A7ZVY5 A7ZVY7
A7ZVY8 A7ZVY9 A7ZVZ3 A7ZVZ4 A7ZW01 A7ZW02 A7ZW03 A7ZW04 A7ZW09 A7ZW12 A7ZW13
A7ZW22 A7ZW23 A7ZW25 A7ZW33 A7ZW34 A7ZW43 A7ZW52 A7ZW53 A7ZW55 A7ZW68 A7ZW69
A7ZW70 A7ZW80 A7ZW82 A7ZW83 A7ZW93 A7ZWA2 A7ZWA4 A7ZWA7 A7ZWA8 A7ZWB1 A7ZWB2
A7ZWB3 A7ZWB5 A7ZWB6 A7ZWB7 A7ZWB8 A7ZWC9 A7ZWD6 A7ZWE2 A7ZWF4 A7ZWF6 A7ZWI6
A7ZWJ6 A7ZWK2 A7ZWK3 A7ZWK5 A7ZWK6 A7ZWT2 A7ZWT3 A7ZWT4 A7ZWV4 A7ZWV5 A7ZWV6
A7ZWY8 A7ZWZ1 A7ZWZ4 A7ZWZ5 A7ZWZ6 A7ZWZ7 A7ZWZ8 A7ZWZ9 A7ZX03 A7ZX04 A7ZX05
A7ZX32 A7ZX37 A7ZX38 A7ZX42 A7ZX56 A7ZX57 A7ZX65 A7ZX67 A7ZX68 A7ZX72 A7ZX74
A7ZX76 A7ZX79 A7ZX94 A7ZX95 A7ZX96 A7ZXA2 A7ZXA4 A7ZXC7 A7ZXC9 A7ZXD0 A7ZXD1
A7ZXD2 A7ZXD3 A7ZXD4 A7ZXF7 A7ZXF9 A7ZXG5 A7ZXI1 A7ZXI3 A7ZXK4 A7ZXP0 A7ZXP1
A7ZXP4 A7ZXQ4 A7ZXQ7 A7ZXQ8 A7ZXR2 A7ZXR5 A7ZXR8 A7ZXS0 A7ZXS8 A7ZXT0 A7ZXT7
A7ZXU0 A7ZXX4 A7ZXX5 A7ZXY8 A7ZY06 A7ZY11 A7ZY13 A7ZY23 A7ZY31 A7ZY32 A7ZY35
A7ZY37 A7ZY39 A7ZY61 A7ZY64 A7ZY70 A7ZY97 A7ZYF5 A7ZYG2 A7ZYH0 A7ZYH7 A7ZYI5
A7ZYI7 A7ZYI8 A7ZYJ7 A7ZYL0 A7ZYL1 A7ZYL3 A7ZYL5 A7ZYL9 A7ZYM4 A7ZYM6 A7ZYN3
A7ZYN4 A7ZYN8 A7ZYQ0 A7ZYQ8 A7ZYR0 A7ZYR5 A7ZYR9 A7ZYU9 A7ZYV1 A7ZYV2 A7ZYV7
A7ZYW2 A7ZYW3 A7ZYW4 A7ZYW5 A7ZYW6 A7ZYW7 A7ZYY7 A7ZZ13 A7ZZ17 A7ZZ20 A7ZZ25
A7ZZ46 A7ZZ55 A7ZZ64 A7ZZ65 A7ZZ66 A7ZZ76 A7ZZ82 A7ZZ88 A7ZZ89 A7ZZ97 A7ZZA4
A7ZZA8 A7ZZB3 A7ZZB5 A7ZZC3 A7ZZE0 A7ZZE4 A7ZZE6 A7ZZE7 A7ZZH0 A7ZZI4 A7ZZJ6
A7ZZJ7 A7ZZL5 A7ZZM0 A7ZZM6 A7ZZT7 A7ZZX1 A8A002 A8A018 A8A036 A8A064 A8A065

A8A070 A8A071 A8A072 A8A074 A8A0E8 A8A0F2 A8A0G5 A8A0I0 A8A0I1 A8A0I3 A8A0I5
A8A0J6 A8A0K5 A8A0M5 A8A0M6 A8A0N8 A8A0N9 A8A0P5 A8A0P8 A8A0Q4 A8A0Q6 A8A0Q9
A8A0R0 A8A0R2 A8A0S2 A8A0S5 A8A0T1 A8A0T3 A8A0T6 A8A0T7 A8A0T8 A8A0T9 A8A0U0
A8A0V7 A8A0X0 A8A0X7 A8A110 A8A129 A8A133 A8A160 A8A161 A8A163 A8A166 A8A171
A8A172 A8A176 A8A178 A8A195 A8A1A0 A8A1B5 A8A1C2 A8A1D9 A8A1F4 A8A1G6 A8A1J6
A8A1K5 A8A1N0 A8A1P3 A8A1P5 A8A1P8 A8A1P9 A8A1T7 A8A1W0 A8A1W1 A8A1W9 A8A1X0
A8A1X8 A8A204 A8A214 A8A215 A8A222 A8A225 A8A234 A8A248 A8A249 A8A251 A8A273
A8A296 A8A2B1 A8A2B3 A8A2B6 A8A2B8 A8A2C9 A8A2D1 A8A2D2 A8A2D6 A8A2G0 A8A2I7
A8A2I9 A8A2J3 A8A2J9 A8A2K1 A8A2M0 A8A2M8 A8A2P2 A8A2Q2 A8A2Q7 A8A2R4 A8A2S4
A8A2S5 A8A2T4 A8A2T8 A8A2V7 A8A2W3 A8A2W8 A8A2X1 A8A2Y8 A8A2Z0 A8A2Z1 A8A313
A8A315 A8A317 A8A320 A8A321 A8A323 A8A324 A8A329 A8A332 A8A333 A8A336 A8A337
A8A344 A8A345 A8A346 A8A347 A8A348 A8A359 A8A375 A8A377 A8A386 A8A391 A8A392
A8A3B3 A8A3B4 A8A3B5 A8A3B6 A8A3C0 A8A3C1 A8A3C4 A8A3C6 A8A3E0 A8A3F6 A8A3H0
A8A3H3 A8A3H4 A8A3H5 A8A3H6 A8A3I7 A8A3I8 A8A3L2 A8A3M2 A8A3M3 A8A3M4 A8A3M5
A8A3M6 A8A3M9 A8A3N0 A8A3N1 A8A3P3 A8A3P4 A8A3P5 A8A3R5 A8A3S9 A8A3T3 A8A3T7
A8A3T9 A8A3U1 A8A3W0 A8A3W3 A8A3W5 A8A3X7 A8A415 A8A430 A8A432 A8A439 A8A441
A8A444 A8A445 A8A446 A8A450 A8A454 A8A456 A8A466 A8A467 A8A477 A8A481 A8A484
A8A488 A8A489 A8A4A3 A8A4J3 A8A4J8 A8A4K4 A8A4K6 A8A4L0 A8A4M1 A8A4M2 A8A4M5
A8A4N0 A8A4P1 A8A4Q0 A8A4S3 A8A4T9 A8A4U6 A8A4V3 A8A4W5 A8A4X0 A8A4X1 A8A4Y0
A8A4Y1 A8A4Y2 A8A4Y3 A8A4Y4 A8A4Y6 A8A4Y7 A8A4Z0 A8A4Z3 A8A4Z8 A8A500 A8A501
A8A504 A8A520 A8A532 A8A533 A8A535 A8A539 A8A540 A8A545 A8A546 A8A570 A8A573
A8A586 A8A587 A8A589 A8A591 A8A592 A8A593 A8A599 A8A5A0 A8A5A1 A8A5A2 A8A5A4
A8A5A6 A8A5A7 A8A5A8 A8A5A9 A8A5B0 A8A5B1 A8A5B2 A8A5B3 A8A5B4 A8A5B5 A8A5B6
A8A5B7 A8A5B8 A8A5B9 A8A5C0 A8A5C1 A8A5C2 A8A5C4 A8A5C5 A8A5C6 A8A5E6 A8A5E7
A8A5E8 A8A5E9 A8A5F0 A8A5F1 A8A5F2 A8A5F5 A8A5G2 A8A5H5 A8A5J6 A8A5J7 A8A5L1
A8A5L8 A8A5M0 A8A5M2 A8A5M7 A8A5M8 A8A5N3 A8A5N9 A8A5P0 A8A5P1 A8A5P2 A8A5S8
A8A5T9 A8A5V4 A8A5X9 A8A616 A8A617 A8A623 A8A634 A8A651 A8A660 A8A674 A8A677
A8A681 A8A683 A8A696 A8A697 A8A698 A8A699 A8A6A0 A8A6A2 A8A6A3 A8A6A4 A8A6A5
A8A6A9 A8A6B1 A8A6C8 A8A6E6 A8A6E7 A8A6F0 A8A6F2 A8A6G1 A8A6G3 A8A6G4 A8A6G5
A8A6G8 A8A6H0 A8A6H3 A8A6K3 A8A6K4 A8A6K7 A8A6K8 A8A6L1 A8A6M0 A8A6M7 A8A6N0
A8A6N3 A8A6N4 A8A6R1 A8A6R7 A8A6R9 A8A6T6 A8A6U0 A8A6V2 A8A6W6 A8A6Y8 A8A6Z4
A8A704 A8A705 A8A706 A8A707 A8A709 A8A710 A8A720 A8A724 A8A731 A8A734 A8A735
A8A737 A8A738 A8A742 A8A746 A8A750 A8A764 A8A765 A8A767 A8A768 A8A770 A8A773
A8A775 A8A778 A8A779 A8A782 A8A783 A8A784 A8A785 A8A786 A8A787 A8A790 A8A793
A8A794 A8A795 A8A796 A8A797 A8A798 A8A7A6 A8A7A9 A8A7B3 A8A7B7 A8A7C4 A8A7D9
A8A7N3 A8A7N8 A8A7N9 A8A7P4 A8A7Q4 A8A7Q8 A8A7Q9 A8A7R4 A8A7R5 A8A7R6 A8A7S2
A8A7S3 A8A7T7 A8A7T8 A8A7U2 A8A7U3 A8A7U4 A8A7U6 A8A7U7 A8A7U8 A8A7U9 A8A7W5
A8A7X5 A8A7Y7 A8A7Y9 A8A7Z8 A8A802 A8A803 A8A810 A8A816 A8A847 A8A853 A8A858
A8A876 A8A891 A8A899 A8A8A3 A8A8B0 A8A8B1 A8A8B2 A8A8B3 A8A8B4 A8A8C2 A8A8C4
A8A8D3 A8A8G1 A8A8H1 A8A8H5 A8A8H6 A8A8J3 A8A8L4 A8A8L8 A8A8L9 A8A8M4 A8A8N6
A8A8T2 A8A8T7 A8A8V6 A8A8W1 A8A8W3 A8A8W4 A8A8X0 A8A8X8 A8A8Z7 A8A906 A8A912
A8A929 A8A935 A8A948 A8A952 A8A953 A8A954 A8A962 A8A968 A8A990 A8A9B8 A8A9E3
A8A9E4 A8A9H3 A8A9M3 A8A9P0 A8A9U0 A8A9U3 A8A9W1 A8A9W2 A8A9X7 A8A9Z2 A8AA18
A8AA19 A8AA20 A8AA22 A8AA46 A8AA47 A8AA51 A8AA61 A8AA65 A8AAB3 A8AAB4 A8AAB6

A8AAC1 A8AAF5 A8AAF8 A8AAG3 A8AAI4 A8AAI6 A8AAJ1 A8AAJ4 A8AAJ7 A8AAK3 A8AAK8
A8AAN4 A8AAQ5 A8AAR8 A8AAU0 A8AAU3 A8AAV7 A8AB06 A8AB20 A8AB26 A8AB39 A8AB48
A8AB57 A8AB58 A8AB61 A8AB67 A8AB68 A8AB69 A8AB83 A8ABB4 A8ABE2 A8ABI5 A8ABJ8
A8ABJ9 A8ABK3 A8ABM5 A8ABM7 A8ABN0 A8ABQ7 A8ABX6 A8ABX8 A8ABY5 A8ABZ6 A8ABZ9
A8AC00 A8AC01 A8AC04 A8AC05 A8AC24 A8AC69 A8AC70 A8AC96 A8AC98 A8ACA7 A8ACB1
A8ACD2 A8ACD3 A8ACD4 A8ACD7 A8ACE0 A8ACG6 A8ACI7 A8ACJ9 A8ACK0 A8ACL2 A8ACL6
A8ACL8 A8ACM1 A8ACP6 A8ACP7 A8ACQ1 A8ACQ4 A8ACS4 A8ACT2 A8ACT3 A8ACV5 A8ACX7
A8ACY2 A8ACY8 A8ACZ6 A8AD07 A8AD08 A8AD10 A8AD38 A8AD50 A8AD51 A8AD55 A8AD56
A8AD60 A8AD67 A8AD69 A8AD71 A8AD75 A8AD78 A8AD80 A8AD93 A8AD95 A8ADB1 A8ADB3
A8ADB6 A8ADC6 A8ADF0 A8ADG3 A8ADI2 A8ADJ2 A8ADK6 A8ADP7 A8ADP9 A8ADQ4 A8ADQ9
A8ADR1 A8ADU4 A8ADU7 A8ADW5 A8ADW9 A8ADX0 A8ADX2 A8ADY5 A8AE07 A8AE28 A8AE29
A8AE31 A8AE51 A8AE66 A8AE79 A8AE87 A8AEB2 A8AEC1 A8AEF5 A8AEJ9 A8AEK0 A8AEK3
A8AEL4 A8AEL8 A8AEM8 A8AEP5 A8AEQ8 A8AF20 A8AF43 A8AF74 A8AFB5 A8AFD4 A8AFG5
A8AFG8 A8AFH0 A8AFH1 A8AFH5 A8AFH7 A8AFH8 A8AFI0 A8AFI1 A8AFL6 A8AFM0 A8AFP5
A8AFP7 A8AFQ7 A8AFR0 A8AFR2 A8AFS0 A8AFY6 A8AFZ9 A8AG01 A8AG02 A8AG49 A8AG60
A8AG61 A8AG83 A8AG96 A8AGE0 A8AGG0 A8AGJ9 A8AGN6 A8AGR0 A8AGR2 A8AGY5 A8AGY9
A8AH18 A8AH19 A8AH33 A8AH79 A8AH80 A8AH93 A8AHA5 A8AHA8 A8AHC2 A8AHC4 A8AHD1
A8AHD3 A8AHD6 A8AHD7 A8AHD8 A8AHD9 A8AHE0 A8AHK2 A8AHU0 A8AHU6 A8AHW2 A8AHX4
A8AI08 A8AI14 A8AI19 A8AI24 A8AI35 A8AI48 A8AI59 A8AI78 A8AI79 A8AIA9 A8AIB0
A8AIB1 A8AIB5 A8AIC1 A8AIC3 A8AID1 A8AID7 A8AIE5 A8AIF5 A8AIF7 A8AIG4 A8AIH1
A8AIH4 A8AIH5 A8AIH6 A8AII9 A8AIJ8 A8AIJ9 A8AIK9 A8AIP7 A8AIQ7 A8AIR5 A8AIT5
A8AIW9 A8AIX5 A8AJ02 A8AJ07 A8AJ11 A8AJ12 A8AJ15 A8AJ16 A8AJ18 A8AJ27 A8AJ37
A8AJ40 A8AJA5 A8AJA6 A8AJA9 A8AJD6 A8AJE0 A8AJE9 A8AJF1 A8AJF8 A8AJG0 A8AJG3
A8AJG7 A8AJH3 A8AJH4 A8AJH6 A8AJI8 A8AJJ1 A8AJJ2 A8AJL0 A8AJN0 A8AJN1 A8AJS4
A8AJW9 A8AJX0 A8AJX1 A8AJX2 A8AJX4 A8AJY9 A8AJZ0 A8AK07 A8AK09 A8AK15 A8AK17
A8AK28 A8AK31 A8AK32 A8AK34 A8AK38 A8AK39 A8AK48 A8AK49 A8AK73 A8AK77 A8AK82
A8AKB8 A8AKE2 A8AKK2 A8AKP4 A8AKP5 A8AKP8 A8AKP9 A8AKQ2 A8AKR0 A8AKR2 A8AKS0
A8AKS5 A8AKS6 A8AKS7 A8AKS9 A8AKT0 A8AKT3 A8AKT8 A8AKT9 A8AKU0 A8AKU1 A8AKU2
A8AKU4 A8AKV0 A8AKV5 A8AKW0 A8AKW2 A8AKW3 A8AKW6 A8AKX8 A8AKY5 A8AKZ1 A8AKZ4
A8AKZ7 A8AL00 A8AL13 A8AL27 A8AL30 A8AL31 A8AL34 A8AL48 A8AL53 A8AL94 A8AL98
A8ALA5 A8ALB8 A8ALC0 A8ALC1 A8ALC3 A8ALC4 A8ALC5 A8ALC8 A8ALC9 A8ALD2 A8ALD5
A8ALE4 A8ALE6 A8ALE9 A8ALF1 A8ALG8 A8ALG9 A8ALH0 A8ALJ2 A8ALJ5 A8ALJ6 A8ALK5
A8ALL4 A8ALL5 A8ALM5 A8ALM7 A8ALM8 A8ALN9 A8ALP0 A8ALP3 A8ALP8 A8ALP9 A8ALQ0
A8ALQ2 A8ALR4 A8ALR6 A8ALR7 A8ALR8 A8ALS4 A8ALT3 A8ALT6 A8ALU3 A8ALV0 A8ALV4
A8ALW1 A8ALW2 A8ALX2 A8ALX7 A8ALY7 A8ALZ1 A8AM04 A8AMA5 A8AMC8 A8AMC9 A8AMG1
A8AMG4 A8AMI9 A8AMJ4 A8AMJ6 A8AMJ7 A8AMJ8 A8AMM3 A8AMM9 A8AMN0 A8AMN1 A8AMN6
A8AMP1 A8AMQ1 A8AMQ6 A8AMR3 A8AN29 A8AN89 A8ANA6 A8ANB2 A8ANB7 A8ANC0 A8AND9
A8ANE1 A8ANE2 A8ANF0 A8ANF2 A8ANI0 A8ANI1 A8ANI2 A8ANI3 A8ANI4 A8ANL5 A8ANN2
A8ANP6 A8ANQ0 A8ANQ1 A8ANQ3 A8ANR7 A8ANR8 A8ANU8 A8ANV7 A8ANV8 A8ANV9 A8ANW0
A8ANW1 A8ANW4 A8ANW5 A8ANW6 A8ANW8 A8ANX0 A8ANX1 A8ANY8 A8ANZ1 A8AP02 A8AP08
A8AP13 A8AP17 A8AP21 A8AP42 A8AP50 A8AP75 A8AP95 A8AP96 A8APA8 A8APB1 A8APB3
A8APB4 A8APC5 A8APE1 A8APF8 A8APG1 A8APG7 A8APG8 A8APH3 A8API1 A8APS2 A8APS7
A8APT1 A8APT7 A8APU7 A8APU8 A8APU9 A8APV0 A8APV1 A8APV2 A8APV3 A8APV4 A8APV6
A8APW0 A8APX8 A8APY0 A8AQ02 A8AQ07 A8AQ14 A8AQ21 A8AQ28 A8AQ33 A8AQ34 A8AQ39

A8AQ40 A8AQ53 A8AQ54 A8AQ55 A8AQ56 A8AQ58 A8AQ61 A8AQ63 A8AQ65 A8AQ68 A8AQ74
A8AQ77 A8AQ80 A8AQ96 A8AQB2 A8AQB6 A8AQC0 A8AQC1 A8AQC8 A8AQC9 A8AQF2 A8AQF7
A8AQG0 A8AQH6 A8AQH7 A8AQI1 A8AQI3 A8AQJ0 A8AQJ1 A8AQJ5 A8AQJ6 A8AQJ8 A8AQJ9
A8AQK0 A8AQK2 A8AQK3 A8AQK4 A8AQK5 A8AQK6 A8AQK7 A8AQK8 A8AQK9 A8AQL0 A8AQL2
A8AQL3 A8AQL8 A8AQM0 A8AQM1 A8AQM8 A8AQM9 A8AQN2 A8AQN3 A8AQN6 A8AQQ4 A8AQS4
A8AQU3 A8AQU4 A8AQV7 A8AQW4 A8AQW5 A8AQW7 A8AQX2 A8AQX5 A8AQY0 A8AQY2 A8AQY3
A8AR31 A8AR62 A8AR83 A8ARE4 A8ARE6 A8ARG3 A8ARJ6 A8ARK6 A8ARK8 A8ARM1 A8ARM2
A8ARM4 A8ARM5 A8ARM7 A8ARM8 A8ARN1 A8ARN4 A8ARN6 A8AU71 A8AU84 A8AU87 A8AU89
A8AU93 A8AU96 A8AU99 A8AUA2 A8AUH0 A8AUI0 A8AUL3 A8AUL5 A8AUL9 A8AUM0 A8AUN5
A8AUN6 A8AUN7 A8AUN9 A8AUP4 A8AUQ4 A8AUR5 A8AUR6 A8AUR9 A8AUS9 A8AUT2 A8AUU3
A8AUV1 A8AUV7 A8AUW5 A8AUW9 A8AV19 A8AV30 A8AV42 A8AV53 A8AV68 A8AV96 A8AVA6
A8AVA7 A8AVA8 A8AVB3 A8AVB7 A8AVB8 A8AVB9 A8AVC4 A8AVD7 A8AVE1 A8AVE2 A8AVE3
A8AVF8 A8AVG0 A8AVI2 A8AVJ7 A8AVL4 A8AVL8 A8AVM1 A8AVM5 A8AVN4 A8AVP7 A8AVP8
A8AVQ2 A8AVQ3 A8AVS4 A8AVS5 A8AVS6 A8AVS9 A8AVT9 A8AVU3 A8AVU4 A8AVU5 A8AVV3
A8AVW8 A8AVW9 A8AVX5 A8AVZ4 A8AVZ7 A8AW02 A8AW03 A8AW04 A8AW06 A8AW14 A8AW20
A8AW24 A8AW32 A8AW35 A8AW36 A8AW46 A8AW52 A8AW61 A8AW75 A8AW92 A8AW99 A8AWA0
A8AWA1 A8AWB7 A8AWC2 A8AWD2 A8AWE1 A8AWE3 A8AWF1 A8AWF2 A8AWF4 A8AWF7 A8AWF9
A8AWG9 A8AWM5 A8AWM9 A8AWP5 A8AWP6 A8AWT2 A8AWT5 A8AWT7 A8AWW6 A8AWX1 A8AWX4
A8AWY0 A8AWY1 A8AX10 A8AX12 A8AX13 A8AX17 A8AX28 A8AX31 A8AX56 A8AX59 A8AX76
A8AX77 A8AX83 A8AX86 A8AX88 A8AX95 A8AX96 A8AXA5 A8AXB8 A8AXB9 A8AXC2 A8AXC8
A8AXD5 A8AXD6 A8AXG8 A8AXG9 A8AXH0 A8AXH8 A8AXI2 A8AXJ8 A8AXK6 A8AXK9 A8AXM4
A8AXM7 A8AXM8 A8AXN4 A8AXN8 A8AXN9 A8AXP0 A8AXP8 A8AXQ6 A8AXS8 A8AXT3 A8AXT6
A8AXT7 A8AXU4 A8AXW1 A8AXY8 A8AXY9 A8AXZ1 A8AXZ4 A8AXZ5 A8AXZ6 A8AY03 A8AY10
A8AY22 A8AY23 A8AY24 A8AY26 A8AY27 A8AY29 A8AY30 A8AY31 A8AY33 A8AY34 A8AY61
A8AY66 A8AY67 A8AY70 A8AY71 A8AY74 A8AY75 A8AYF6 A8AYF8 A8AYH0 A8AYH2 A8AYH3
A8AYI5 A8AYI7 A8AYI8 A8AYK4 A8AYK5 A8AYL1 A8AYN4 A8AYP9 A8AYQ0 A8AYT7 A8AYT8
A8AYT9 A8AYU3 A8AYV0 A8AYY6 A8AYZ9 A8AZ08 A8AZ43 A8AZ63 A8AZ70 A8AZ72 A8AZ77
A8AZ80 A8AZA7 A8AZB9 A8AZD5 A8AZD7 A8AZD8 A8AZE1 A8AZE2 A8AZH2 A8AZI2 A8AZI3
A8AZI5 A8AZJ8 A8AZJ9 A8AZK0 A8AZK1 A8AZK2 A8AZK3 A8AZK6 A8AZK7 A8AZK8 A8AZK9
A8AZL0 A8AZL1 A8AZL2 A8AZL3 A8AZL4 A8AZL5 A8AZL6 A8AZL7 A8AZL8 A8AZL9 A8AZM0
A8AZM1 A8AZM2 A8AZM3 A8AZM4 A8AZM5 A8AZM6 A8AZM9 A8AZN1 A8AZP0 A8AZP1 A8AZS8
A8AZS9 A8AZT0 A8AZT4 A8AZT8 A8AZU1 A8AZU4 A8AZV0 A8AZV3 A8AZV4 A8AZY6 A8B000
A8B006 A8B011 A8B2Z8 A8B4Q0 A8B672 A8B6A2 A8BCK6 A8BE86 A8BFN4 A8BKZ6 A8BM50
A8BQB4 A8BQJ3 A8BQZ5 A8BS79 A8C750 A8C752 A8C754 A8C7R3 A8C7R6 A8C7S0 A8C7S1
A8C8X2 A8C927 A8C984 A8CAG3 A8CEP3 A8CVF3 A8CVX7 A8D7K2 A8D888 A8D8P8 A8D8X1
A8DEZ8 A8DRH7 A8DT10 A8DYH2 A8DYP0 A8DYP7 A8DYP9 A8DYY5 A8DZE6 A8DZJ1 A8E0R9
A8E1G1 A8E264 A8E283 A8E4L3 A8E4M4 A8E4N3 A8E4S7 A8E4V2 A8E534 A8E5T6 A8E5W8
A8E5Y3 A8E604 A8E653 A8E657 A8E7D2 A8E7G4 A8E7I5 A8EQT0 A8EQW4 A8EQW5 A8EQW8
A8EQW9 A8EQY4 A8EQY5 A8EQZ0 A8EQZ2 A8EQZ3 A8EQZ4 A8ER05 A8ER06 A8ER37 A8ER43
A8ER62 A8ER63 A8ER74 A8ER78 A8ERB7 A8ERB8 A8ERD0 A8ERD1 A8ERD8 A8ERE4 A8ERE6
A8ERE9 A8ERF6 A8ERG2 A8ERI4 A8ERI7 A8ERJ2 A8ERJ4 A8ERP6 A8ERS0 A8ERS1 A8ERT8
A8ERT9 A8ERV3 A8ERX9 A8ERY0 A8ERY3 A8ERZ9 A8ES07 A8ES24 A8ES35 A8ES36 A8ES39
A8ES62 A8ES83 A8ES84 A8ESC9 A8ESF7 A8ESG0 A8ESH8 A8ESH9 A8ESI0 A8ESI8 A8ESJ1
A8ESJ5 A8ESP0 A8ESR5 A8EST5 A8ESU2 A8ESU4 A8ESU5 A8ESU6 A8ESU7 A8ESU8 A8ESU9

A8ESV0 A8ESV1 A8ESV2 A8ESV3 A8ESV4 A8ESV6 A8ESV7 A8ESV8 A8ESW0 A8ESW4 A8ESW7
A8ESX7 A8ESX8 A8ESZ6 A8ESZ8 A8ESZ9 A8ET01 A8ET36 A8ET60 A8ET67 A8ETA7 A8ETF2
A8ETF6 A8ETF7 A8ETG3 A8ETG7 A8ETJ2 A8ETJ8 A8ETK2 A8ETK3 A8ETK4 A8ETK5 A8ETK6
A8ETK7 A8ETK8 A8ETQ3 A8ETY6 A8EU19 A8EU50 A8EU55 A8EU69 A8EU90 A8EUB2 A8EUC7
A8EUD5 A8EUE8 A8EUF3 A8EUZ4 A8EV04 A8EV17 A8EV23 A8EV24 A8EV83 A8EV84 A8EV85
A8EV88 A8EV89 A8EV95 A8EV99 A8EVA1 A8EVH5 A8EVH6 A8EVN0 A8EVP8 A8EVP9 A8EVQ3
A8EVR0 A8EVR4 A8EVR5 A8EVV4 A8EVW0 A8EVW3 A8EVW4 A8EVZ3 A8EVZ4 A8EVZ5 A8EVZ6
A8EVZ7 A8EVZ8 A8EW01 A8EW02 A8EW05 A8EW70 A8EW72 A8EW78 A8EW86 A8EW87 A8EW88
A8EWB9 A8EWD5 A8EWD7 A8EWD9 A8EWE5 A8EWF5 A8EWG2 A8EWG5 A8EWH6 A8EWI3 A8EWI6
A8EWJ4 A8EWM0 A8EWM2 A8EWM4 A8EWM9 A8EWN0 A8EWN8 A8EWR5 A8EWR6 A8EWT6 A8EWT7
A8EWU8 A8EWV0 A8EWW1 A8EWX7 A8EWY1 A8EX03 A8EX06 A8EX12 A8EX66 A8EX70 A8EX71
A8EX95 A8EX98 A8EXA5 A8EXA7 A8EXA8 A8EXA9 A8EXC2 A8EXC3 A8EXD1 A8EXD2 A8EXD6
A8EXD7 A8EXD9 A8EXE9 A8EXF0 A8EXF1 A8EXF8 A8EXG2 A8EXG4 A8EXH8 A8EXH9 A8EXI3
A8EXJ2 A8EXJ9 A8EXK0 A8EXK1 A8EXK4 A8EXK5 A8EXK6 A8EXK7 A8EXK8 A8EXK9 A8EXL1
A8EXL4 A8EXL6 A8EXM0 A8EXM1 A8EXM2 A8EXM3 A8EXM4 A8EXN1 A8EXP6 A8EXS4 A8EXS5
A8EXS6 A8EXT7 A8EXV2 A8EXV6 A8EXW3 A8EXW5 A8EXY1 A8EXY2 A8EXY4 A8EXY5 A8EXY7
A8EXZ6 A8EXZ7 A8EXZ8 A8EY32 A8EY33 A8EY35 A8EY36 A8EY39 A8EY40 A8EY49 A8EY51
A8EY59 A8EY60 A8EY61 A8EY62 A8EY64 A8EY65 A8EY77 A8EYA6 A8EYA7 A8EYB9 A8EYC4
A8EYC5 A8EYE9 A8EYF2 A8EYF4 A8EYF7 A8EYH9 A8EYJ9 A8EYL5 A8EYL6 A8EYM2 A8EYM5
A8EYM7 A8EYR1 A8EYU0 A8EYU1 A8EYU2 A8EYU8 A8EYV4 A8EYV5 A8EYX8 A8EYY2 A8EYY5
A8EYY8 A8EYZ3 A8EYZ4 A8EZ02 A8EZ06 A8EZ08 A8EZ30 A8EZ44 A8EZ47 A8EZ51 A8EZ58
A8EZ69 A8EZ91 A8EZ93 A8EZ98 A8EZA9 A8EZB2 A8EZB8 A8EZD2 A8EZD3 A8EZE6 A8EZE8
A8EZG1 A8EZJ0 A8EZJ1 A8EZJ2 A8EZJ3 A8EZJ5 A8EZJ7 A8EZJ8 A8EZJ9 A8EZK0 A8EZK1
A8EZK2 A8EZK3 A8EZK4 A8EZK5 A8EZK6 A8EZK7 A8EZK8 A8EZK9 A8EZL0 A8EZL1 A8EZL2
A8EZL3 A8EZL4 A8EZL5 A8EZL6 A8EZL7 A8EZL8 A8EZN3 A8EZN6 A8EZN9 A8EZP4 A8EZP5
A8EZQ1 A8EZQ5 A8EZR2 A8EZR4 A8EZT5 A8EZT7 A8EZU0 A8EZU1 A8EZU2 A8EZU3 A8EZV1
A8EZV2 A8EZV9 A8EZW1 A8EZX4 A8EZY3 A8F011 A8F030 A8F031 A8F033 A8F059 A8F060
A8F062 A8F063 A8F071 A8F072 A8F078 A8F079 A8F083 A8F088 A8F095 A8F0A0 A8F0A2
A8F0A3 A8F0A4 A8F0B2 A8F0B3 A8F0B8 A8F0D4 A8F0D7 A8F0E3 A8F0E6 A8F0E7 A8F0E8
A8F0G3 A8F0G4 A8F0H7 A8F0H9 A8F0I8 A8F0I9 A8F0J0 A8F0K6 A8F0L7 A8F0L8 A8F0M2
A8F0M4 A8F0N3 A8F0N9 A8F0P0 A8F0P3 A8F0P5 A8F0P7 A8F0P8 A8F0Q0 A8F0R2 A8F0R3
A8F0S0 A8F0U0 A8F0V7 A8F0V8 A8F0V9 A8F0W5 A8F0X7 A8F0Z2 A8F0Z8 A8F108 A8F109
A8F110 A8F164 A8F172 A8F187 A8F197 A8F1A5 A8F1B5 A8F1B9 A8F1C1 A8F1D8 A8F1F2
A8F1F7 A8F1H0 A8F1I9 A8F1J5 A8F1K3 A8F1K4 A8F1K7 A8F1L4 A8F1N1 A8F1N2 A8F1N7
A8F1P3 A8F1P4 A8F1S1 A8F1X5 A8F1Y8 A8F204 A8F223 A8F225 A8F228 A8F247 A8F259
A8F284 A8F291 A8F2B0 A8F2B5 A8F2C2 A8F2C3 A8F2C4 A8F2C5 A8F2C8 A8F2C9 A8F2D0
A8F2D2 A8F2D3 A8F2D4 A8F2D6 A8F2D7 A8F2D9 A8F2E0 A8F2E1 A8F2E2 A8F2E3 A8F2E4
A8F2E7 A8F2E8 A8F2E9 A8F2G9 A8F2H0 A8F2H5 A8F2I3 A8F2I5 A8F2K5 A8F2K6 A8F2L0
A8F2M1 A8F2M4 A8F2M5 A8F2N7 A8F2P7 A8F2Q0 A8F2S3 A8F2U6 A8F2V3 A8F2V6 A8F2Y0
A8F2Y2 A8F2Y3 A8F2Z1 A8F2Z7 A8F2Z8 A8F300 A8F303 A8F323 A8F331 A8F346 A8F397
A8F3A3 A8F3F9 A8F3G2 A8F3G4 A8F3M5 A8F3R6 A8F3S3 A8F3S6 A8F3T2 A8F3W0 A8F3W8
A8F3X9 A8F401 A8F402 A8F446 A8F448 A8F450 A8F453 A8F465 A8F478 A8F479 A8F498
A8F4D7 A8F4E1 A8F4E2 A8F4E8 A8F4F3 A8F4F6 A8F4F7 A8F4F9 A8F4G0 A8F4G1 A8F4K1
A8F4K3 A8F4K6 A8F4M2 A8F4P7 A8F4Q6 A8F4Q7 A8F4Q8 A8F4Q9 A8F4R0 A8F4R1 A8F4R2

A8F4R3 A8F4R4 A8F4R5 A8F4R6 A8F4R7 A8F4R8 A8F4R9 A8F4S0 A8F4S1 A8F4S2 A8F4S3
A8F4S4 A8F4S5 A8F4S6 A8F4S7 A8F4S8 A8F4S9 A8F4T0 A8F4T2 A8F4T4 A8F4T5 A8F4T6
A8F4T7 A8F4T8 A8F4T9 A8F4W4 A8F4W9 A8F4X2 A8F4Y8 A8F4Z0 A8F4Z8 A8F4Z9 A8F501
A8F502 A8F516 A8F522 A8F523 A8F524 A8F525 A8F526 A8F531 A8F541 A8F560 A8F593
A8F595 A8F596 A8F597 A8F5A0 A8F5A7 A8F5A8 A8F5B3 A8F5B4 A8F5B8 A8F5C4 A8F5E8
A8F5G0 A8F5H5 A8F5H6 A8F5H8 A8F5K4 A8F5M9 A8F5R0 A8F5X7 A8F5X9 A8F5Y1 A8F5Z3
A8F670 A8F679 A8F681 A8F6A5 A8F6I7 A8F6M1 A8F6N8 A8F6Q8 A8F6X2 A8F701 A8F716
A8F717 A8F720 A8F724 A8F725 A8F732 A8F753 A8F754 A8F755 A8F758 A8F791 A8F7A0
A8F7B1 A8F7B3 A8F7B4 A8F7D2 A8F7D4 A8F7D5 A8F7D6 A8F7F8 A8F7L9 A8F7P0 A8F7P3
A8F7P9 A8F7Q7 A8F7Q8 A8F7R7 A8F7R9 A8F7S0 A8F7S2 A8F7S8 A8F7S9 A8F7T6 A8F7V1
A8F7W0 A8F7Z9 A8F801 A8F811 A8F821 A8F840 A8F851 A8F863 A8F866 A8F8B7 A8F8C2
A8F8C8 A8F8F7 A8F8G5 A8F8I1 A8F8I3 A8F8I4 A8F8L8 A8F8M6 A8F8M9 A8F8N4 A8F8P5
A8F8P6 A8F8P9 A8F8Q3 A8F8Q8 A8F8T5 A8F8W3 A8F8W9 A8F8Y4 A8F8Y7 A8F8Z5 A8F900
A8F902 A8F909 A8F913 A8F916 A8F919 A8F920 A8F938 A8F953 A8F958 A8F959 A8F960
A8F961 A8F962 A8F967 A8F971 A8F972 A8F973 A8F976 A8F977 A8F978 A8F979 A8F980
A8F981 A8F982 A8F984 A8F985 A8F986 A8F987 A8F988 A8F989 A8F990 A8F991 A8F992
A8F994 A8F995 A8F996 A8F997 A8F998 A8F999 A8F9A1 A8F9A2 A8F9A3 A8F9A4 A8F9A6
A8F9A8 A8F9A9 A8F9B0 A8F9B1 A8F9B2 A8F9B3 A8F9B7 A8F9B8 A8F9B9 A8F9E0 A8F9E6
A8F9I7 A8F9Q8 A8F9S0 A8F9S1 A8F9T1 A8F9Z9 A8FA07 A8FA09 A8FA13 A8FA55 A8FA73
A8FAD5 A8FAD6 A8FAD7 A8FAE6 A8FAF5 A8FAF6 A8FAG2 A8FAG3 A8FAH5 A8FAK7 A8FAM6
A8FAM7 A8FAM9 A8FAQ1 A8FAR9 A8FAS0 A8FAS1 A8FAY5 A8FB03 A8FB12 A8FB45 A8FB51
A8FB62 A8FB65 A8FB84 A8FB92 A8FBJ5 A8FBK3 A8FBL1 A8FBM5 A8FBM6 A8FBR0 A8FBR1
A8FBU5 A8FBW1 A8FBY4 A8FC05 A8FC15 A8FC34 A8FCD2 A8FCG5 A8FCG6 A8FCG8 A8FCG9
A8FCH0 A8FCH1 A8FCH9 A8FCI2 A8FCN1 A8FCN2 A8FCN3 A8FCN9 A8FCR4 A8FCR6 A8FCS8
A8FCS9 A8FCU1 A8FCV9 A8FCW1 A8FCW7 A8FCW9 A8FCX1 A8FCX3 A8FCX9 A8FCY2 A8FD04
A8FD12 A8FD14 A8FD15 A8FD18 A8FD19 A8FD20 A8FD21 A8FD22 A8FD24 A8FD25 A8FD32
A8FD34 A8FD38 A8FD40 A8FD43 A8FD47 A8FD63 A8FD66 A8FD67 A8FD68 A8FD69 A8FD70
A8FD73 A8FD77 A8FD78 A8FD79 A8FD80 A8FD81 A8FDB4 A8FDB6 A8FDB9 A8FDC1 A8FDC7
A8FDD1 A8FDD3 A8FDD4 A8FDD6 A8FDD7 A8FDG2 A8FDG3 A8FDG9 A8FDH0 A8FDH3 A8FDH4
A8FDI9 A8FDJ0 A8FDN4 A8FDN5 A8FDN6 A8FDP8 A8FDQ0 A8FDQ8 A8FDQ9 A8FDR3 A8FDR4
A8FDV4 A8FDY3 A8FE33 A8FE66 A8FE93 A8FEA3 A8FEA4 A8FEB9 A8FEC5 A8FEC9 A8FED0
A8FEE7 A8FEF4 A8FEF5 A8FEG6 A8FEH5 A8FEH6 A8FEH7 A8FEH9 A8FEI2 A8FEI3 A8FEJ2
A8FEJ4 A8FEJ6 A8FEJ7 A8FEJ8 A8FEK0 A8FEK1 A8FEK4 A8FEK5 A8FEK9 A8FEL2 A8FEL8
A8FEM3 A8FEM5 A8FEQ7 A8FEQ8 A8FER2 A8FET0 A8FET4 A8FEW4 A8FF09 A8FF11 A8FF13
A8FF14 A8FF15 A8FF16 A8FF29 A8FF31 A8FF36 A8FF40 A8FF41 A8FF72 A8FF87 A8FFA1
A8FFA7 A8FFB1 A8FFB2 A8FFB6 A8FFC6 A8FFD0 A8FFD1 A8FFD2 A8FFD3 A8FFD4 A8FFD9
A8FFE0 A8FFF0 A8FFL1 A8FFL2 A8FFL6 A8FFL8 A8FFM6 A8FFM7 A8FFM8 A8FFM9 A8FFP0
A8FFP8 A8FFQ0 A8FFQ9 A8FFR0 A8FFR2 A8FFR3 A8FFR8 A8FFS2 A8FFS7 A8FFS8 A8FFT0
A8FFT2 A8FFT6 A8FFU0 A8FFU1 A8FFU9 A8FFV2 A8FFV4 A8FFV6 A8FFV9 A8FFW0 A8FFW2
A8FFW3 A8FFW5 A8FFW6 A8FFY6 A8FFY8 A8FG01 A8FG15 A8FG17 A8FG25 A8FG26 A8FG47
A8FG55 A8FG66 A8FG69 A8FG70 A8FG94 A8FGA1 A8FGA6 A8FGB7 A8FGC0 A8FGG0 A8FGH7
A8FGH8 A8FGK6 A8FGK9 A8FGR8 A8FGT0 A8FGU0 A8FH04 A8FH11 A8FH15 A8FH17 A8FH24
A8FH34 A8FH75 A8FHC4 A8FHG7 A8FHJ1 A8FHJ2 A8FHJ3 A8FHN9 A8FHQ0 A8FHQ2 A8FHQ9
A8FHR0 A8FHR1 A8FHR2 A8FHR4 A8FHR5 A8FHS1 A8FHS4 A8FHS6 A8FHW9 A8FHX3 A8FHX4

A8FI32 A8FI51 A8FI96 A8FI98 A8FIA1 A8FIC0 A8FIC1 A8FID1 A8FID5 A8FID6 A8FIE1
A8FIE5 A8FIE6 A8FIG1 A8FIG8 A8FII0 A8FIM7 A8FIR6 A8FIR7 A8FIY2 A8FJ14 A8FJA3
A8FJC0 A8FJD2 A8FJD5 A8FJE5 A8FJE7 A8FJF7 A8FJF8 A8FJF9 A8FJG0 A8FJG3 A8FJG4
A8FJG5 A8FJH3 A8FJH6 A8FJH9 A8FJJ0 A8FJL3 A8FJM2 A8FJP1 A8FJP9 A8FJQ0 A8FJQ1
A8FJQ2 A8FJQ3 A8FJT1 A8FJT9 A8FJU1 A8FJU2 A8FJU3 A8FJV7 A8FJW4 A8FJX0 A8FJX4
A8FJZ3 A8FJZ4 A8FJZ5 A8FJZ8 A8FK07 A8FK08 A8FK14 A8FK16 A8FK23 A8FK24 A8FK31
A8FK33 A8FK34 A8FK41 A8FK69 A8FK75 A8FK82 A8FK85 A8FK86 A8FK87 A8FK97 A8FKA0
A8FKA1 A8FKA6 A8FKB0 A8FKB3 A8FKB4 A8FKB5 A8FKB9 A8FKC1 A8FKD6 A8FKD7 A8FKD8
A8FKE8 A8FKF8 A8FKG2 A8FKG8 A8FKG9 A8FKH0 A8FKH3 A8FKH6 A8FKI5 A8FKI7 A8FKI8
A8FKI9 A8FKL9 A8FKM1 A8FKN5 A8FKN8 A8FKP3 A8FKQ5 A8FKQ6 A8FKQ9 A8FKR0 A8FKR1
A8FKR2 A8FKR3 A8FKR4 A8FKR5 A8FKR6 A8FKR7 A8FKS2 A8FKS9 A8FKT2 A8FKT3 A8FKT8
A8FKU4 A8FKV6 A8FKW5 A8FKW6 A8FKW8 A8FKX4 A8FL01 A8FL05 A8FL10 A8FL14 A8FL53
A8FL56 A8FL58 A8FL60 A8FL61 A8FL62 A8FL71 A8FL80 A8FL81 A8FL82 A8FL83 A8FLB6
A8FLB9 A8FLC2 A8FLD3 A8FLD9 A8FLE1 A8FLE2 A8FLE3 A8FLH1 A8FLH2 A8FLH8 A8FLH9
A8FLI0 A8FLL1 A8FLL2 A8FLL5 A8FLL9 A8FLM3 A8FLN6 A8FLP0 A8FLP9 A8FLQ4 A8FLR2
A8FLR4 A8FLR7 A8FLT4 A8FLU3 A8FLU4 A8FLU6 A8FLV4 A8FLX6 A8FLY0 A8FLY1 A8FM07
A8FM09 A8FM10 A8FM14 A8FM15 A8FM27 A8FM42 A8FM43 A8FM46 A8FM57 A8FM59 A8FM84
A8FM94 A8FMA1 A8FMA8 A8FMB2 A8FMC3 A8FMC5 A8FMC8 A8FMD4 A8FME4 A8FMF1 A8FMF5
A8FMF7 A8FMF8 A8FMG0 A8FMG2 A8FMH0 A8FMK7 A8FMK8 A8FMM8 A8FMN1 A8FMN7 A8FMN8
A8FMP4 A8FMQ3 A8FMQ7 A8FMS5 A8FMS6 A8FMU3 A8FMU4 A8FMU8 A8FMV1 A8FMV3 A8FMV9
A8FMW6 A8FMW9 A8FMX4 A8FMX9 A8FMY0 A8FMY4 A8FMZ1 A8FMZ3 A8FMZ7 A8FN22 A8FN24
A8FN28 A8FN39 A8FN42 A8FN54 A8FN57 A8FN77 A8FN79 A8FNB6 A8FNC1 A8FNC2 A8FNC4
A8FNE5 A8FNG4 A8FNH9 A8FNJ1 A8FNJ7 A8FNK1 A8FNQ2 A8FNQ3 A8FNQ4 A8FNQ5 A8FNQ6
A8FNQ7 A8FNQ8 A8FNQ9 A8FNR1 A8FNR3 A8FNR4 A8FNR8 A8FNS2 A8FNS3 A8FNU6 A8FNU7
A8FNU8 A8FNY4 A8FNZ2 A8FP04 A8FP05 A8FP06 A8FP07 A8FP08 A8FP09 A8FP10 A8FP11
A8FP12 A8FP13 A8FP14 A8FP15 A8FP16 A8FP17 A8FP18 A8FP19 A8FP20 A8FP22 A8FP27
A8FP32 A8FP33 A8FP35 A8FP38 A8FP40 A8FP41 A8FP45 A8FP46 A8FP48 A8FP54 A8FP60
A8FP64 A8FP74 A8FP78 A8FP81 A8FP82 A8FPD5 A8FPE0 A8FPF5 A8FPG1 A8FPK4 A8FPL9
A8FPR6 A8FPZ5 A8FPZ7 A8FQ25 A8FQ68 A8FQ72 A8FQ73 A8FQ75 A8FQ76 A8FQ77 A8FQ83
A8FQ85 A8FQ86 A8FQ89 A8FQ91 A8FQ92 A8FQA1 A8FQB0 A8FQD0 A8FQD4 A8FQL9 A8FQM0
A8FQM2 A8FQM5 A8FQV1 A8FQV9 A8FQW4 A8FQY0 A8FQY1 A8FR61 A8FR77 A8FR86 A8FR95
A8FR96 A8FR97 A8FR98 A8FRA1 A8FRD3 A8FRD4 A8FRD5 A8FRE9 A8FRF1 A8FRH9 A8FRI2
A8FRL1 A8FRL4 A8FRM9 A8FRN6 A8FRQ4 A8FRT9 A8FRU0 A8FRU2 A8FRU3 A8FRU4 A8FRV0
A8FRV1 A8FRV2 A8FS02 A8FS63 A8FS64 A8FS72 A8FS76 A8FS90 A8FSA3 A8FSC0 A8FSC3
A8FSD6 A8FSE7 A8FSE8 A8FSE9 A8FSF0 A8FSF3 A8FSH2 A8FSH3 A8FSH6 A8FSH9 A8FSI2
A8FSK6 A8FSK8 A8FSL7 A8FSM5 A8FSQ9 A8FSR0 A8FSR4 A8FSR5 A8FSS4 A8FSS7 A8FST0
A8FST1 A8FST3 A8FST4 A8FST7 A8FST8 A8FSV6 A8FSV7 A8FSY8 A8FT45 A8FT67 A8FT70
A8FT71 A8FT74 A8FTB0 A8FTH6 A8FTH7 A8FTH8 A8FTH9 A8FTI0 A8FTP1 A8FTS6 A8FTT1
A8FTT7 A8FTU0 A8FU26 A8FU90 A8FU94 A8FU97 A8FUD3 A8FUD4 A8FUF1 A8FUF8 A8FUG9
A8FUH3 A8FUH5 A8FUH7 A8FUK8 A8FUS8 A8FUU9 A8FUV3 A8FUW3 A8FUW4 A8FUW6 A8FUW8
A8FUW9 A8FUX0 A8FUY4 A8FV56 A8FV58 A8FV59 A8FV67 A8FVF7 A8FVN0 A8FVN3 A8FVN5
A8FVN6 A8FVN8 A8FVR6 A8FVS5 A8FVT2 A8FVU8 A8FW04 A8FW35 A8FW40 A8FW59 A8FWA0
A8FWA2 A8FWA3 A8FWA5 A8FWC1 A8FWC5 A8FWC9 A8FWD0 A8FWD1 A8FWI0 A8FWI2 A8FWJ0
A8FWJ3 A8FWJ5 A8FWK2 A8FWL2 A8FWM8 A8FWN7 A8FWP4 A8FX10 A8FX24 A8FX46 A8FX58

A8FX70 A8FX76 A8FX83 A8FX84 A8FX85 A8FX87 A8FX91 A8FX96 A8FXA1 A8FXA5 A8FXC6
A8FXE0 A8FXH3 A8FXJ5 A8FXM2 A8FY04 A8FY06 A8FY28 A8FY36 A8FY39 A8FY41 A8FY42
A8FY45 A8FYI7 A8FYJ2 A8FYK0 A8FYK5 A8FYL0 A8FYL1 A8FYQ7 A8FYQ8 A8FYQ9 A8FYR3
A8FYR5 A8FYR7 A8FYR9 A8FYS0 A8FYS2 A8FYS4 A8FYS5 A8FYS8 A8FYT6 A8FYT9 A8FYU0
A8FYU6 A8FYU7 A8FYV3 A8FYW7 A8FYX4 A8FYZ0 A8FYZ1 A8FYZ3 A8FYZ9 A8FZ02 A8FZ04
A8FZ10 A8FZ13 A8FZ15 A8FZ21 A8FZ22 A8FZ23 A8FZ49 A8FZC6 A8FZE2 A8FZK4 A8FZK5
A8FZK6 A8FZR5 A8FZY7 A8FZZ3 A8G079 A8G084 A8G085 A8G094 A8G097 A8G0F9 A8G0I7
A8G0J1 A8G0R9 A8G0S5 A8G0S7 A8G0U0 A8G0U8 A8G0V0 A8G0W0 A8G0W4 A8G0Z3 A8G183
A8G184 A8G195 A8G196 A8G1A3 A8G1A4 A8G1A6 A8G1A7 A8G1A9 A8G1B1 A8G1C4 A8G1C6
A8G1C7 A8G1C8 A8G1D0 A8G1D2 A8G1D3 A8G1D4 A8G1D5 A8G1D6 A8G1D7 A8G1D8 A8G1D9
A8G1E0 A8G1E1 A8G1E3 A8G1E4 A8G1E5 A8G1E6 A8G1E7 A8G1E8 A8G1E9 A8G1F0 A8G1F2
A8G1F3 A8G1F4 A8G1F5 A8G1F6 A8G1F7 A8G1F8 A8G1F9 A8G1G2 A8G1G4 A8G1G7 A8G1H1
A8G1J0 A8G1J1 A8G1K1 A8G1M9 A8G1Q8 A8G1S5 A8G1S6 A8G1S8 A8G1T3 A8G1U7 A8G1U9
A8G1X5 A8G1X6 A8G1Z0 A8G1Z9 A8G203 A8G207 A8G213 A8G222 A8G223 A8G242 A8G243
A8G244 A8G2A6 A8G2B8 A8G2D5 A8G2H0 A8G2H7 A8G2H9 A8G2I8 A8G2J5 A8G2K1 A8G2K8
A8G2K9 A8G2L0 A8G2L1 A8G2M6 A8G2N0 A8G2N2 A8G2P1 A8G2P3 A8G2P6 A8G2P9 A8G2Q0
A8G2Q9 A8G2R6 A8G2R7 A8G2R8 A8G2S2 A8G2S6 A8G2T1 A8G2U2 A8G2X2 A8G2X4 A8G2Y1
A8G2Y9 A8G399 A8G3A3 A8G3A7 A8G3B3 A8G3B7 A8G3C1 A8G3D1 A8G3E4 A8G3F3 A8G3G6
A8G3G9 A8G3I5 A8G3I8 A8G3J6 A8G3J9 A8G3K7 A8G3L1 A8G3L5 A8G3M2 A8G3M4 A8G3N6
A8G3N7 A8G3P0 A8G3Q8 A8G3Q9 A8G3R0 A8G3R5 A8G3S5 A8G3T0 A8G3U9 A8G3V0 A8G3V6
A8G3W1 A8G3W3 A8G3X3 A8G3X5 A8G3X9 A8G434 A8G451 A8G4C5 A8G4D1 A8G4D2 A8G4E5
A8G4E6 A8G4F2 A8G4G7 A8G4H0 A8G4K3 A8G4K4 A8G4K5 A8G4K6 A8G4K7 A8G4K8 A8G4K9
A8G4M3 A8G4M8 A8G4N1 A8G4N2 A8G4N9 A8G4P1 A8G4P2 A8G4R9 A8G4S4 A8G4T4 A8G4T6
A8G4T7 A8G4U2 A8G4V7 A8G4V8 A8G4Y1 A8G4Z1 A8G4Z4 A8G507 A8G532 A8G5C3 A8G5D5
A8G5D8 A8G5G3 A8G5G4 A8G5G7 A8G5H7 A8G5I6 A8G5S9 A8G5U9 A8G5V0 A8G5V8 A8G5W1
A8G5Y1 A8G5Y2 A8G5Y6 A8G5Y8 A8G5Y9 A8G657 A8G678 A8G687 A8G690 A8G695 A8G6A1
A8G6A7 A8G6B6 A8G6B7 A8G6C1 A8G6C6 A8G6D9 A8G6E3 A8G6F5 A8G6F6 A8G6F8 A8G6G0
A8G6G4 A8G6T6 A8G6T7 A8G6W1 A8G6X5 A8G6Y2 A8G6Y4 A8G6Y5 A8G6Y6 A8G6Y8 A8G6Z2
A8G6Z5 A8G702 A8G708 A8G709 A8G710 A8G711 A8G714 A8G733 A8G734 A8G735 A8G736
A8G737 A8G738 A8G739 A8G740 A8G741 A8G742 A8G744 A8G745 A8G746 A8G748 A8G749
A8G750 A8G751 A8G752 A8G753 A8G754 A8G756 A8G757 A8G758 A8G759 A8G760 A8G761
A8G762 A8G766 A8G779 A8G786 A8G794 A8G795 A8G797 A8G798 A8G7B1 A8G7B7 A8G7C1
A8G7C2 A8G7C4 A8G7D1 A8G7E1 A8G7E2 A8G7E8 A8G7F8 A8G7G0 A8G7G2 A8G7G6 A8G7G7
A8G7G9 A8G7H4 A8G7H8 A8G7I0 A8G7J0 A8G7J2 A8G7J3 A8G7J4 A8G7J6 A8G7L0 A8G7L2
A8G7P7 A8G7Q0 A8G7Q1 A8G7Q2 A8G7Q4 A8G7R4 A8G7R8 A8G7R9 A8G7T9 A8G7U0 A8G7U3
A8G7V7 A8G7W0 A8G7W8 A8G827 A8G828 A8G848 A8G850 A8G852 A8G875 A8G8B2 A8G8B8
A8G8C4 A8G8D3 A8G8D9 A8G8E0 A8G8E3 A8G8E4 A8G8E5 A8G8E6 A8G8E7 A8G8E8 A8G8F0
A8G8F3 A8G8F7 A8G8F8 A8G8G3 A8G8G7 A8G8S6 A8G8S7 A8G8T0 A8G8T8 A8G8U1 A8G8U2
A8G8U7 A8G8U8 A8G8U9 A8G8V4 A8G8V5 A8G8W4 A8G8W5 A8G8W6 A8G8W7 A8G8X6 A8G8Y2
A8G8Y6 A8G8Y7 A8G8Z1 A8G8Z2 A8G8Z4 A8G900 A8G903 A8G905 A8G907 A8G908 A8G909
A8G910 A8G911 A8G925 A8G928 A8G960 A8G961 A8G978 A8G9G5 A8G9G8 A8G9H6 A8G9H7
A8G9H8 A8G9J2 A8G9J3 A8G9J4 A8G9K0 A8G9K2 A8G9K8 A8G9K9 A8G9L2 A8G9L4 A8G9L7
A8G9M8 A8G9N8 A8G9N9 A8G9P0 A8G9P1 A8G9P6 A8G9Q8 A8G9Q9 A8G9R1 A8G9R8 A8G9R9
A8G9S8 A8G9T9 A8G9U1 A8G9U7 A8G9U9 A8G9V3 A8G9V4 A8G9W0 A8G9X6 A8G9X7 A8G9X8

A8G9Y3 A8G9Y4 A8G9Z2 A8G9Z3 A8G9Z4 A8G9Z5 A8G9Z6 A8G9Z9 A8GA07 A8GA08 A8GA09
A8GA13 A8GA18 A8GA19 A8GA20 A8GA21 A8GA75 A8GA77 A8GAA8 A8GAA9 A8GAB0 A8GAB1
A8GAB2 A8GAB6 A8GAB9 A8GAC8 A8GAD1 A8GAD2 A8GAD3 A8GAD4 A8GAI2 A8GAJ0 A8GAJ1
A8GAM4 A8GAM5 A8GAN5 A8GAN7 A8GAN8 A8GAP2 A8GAP4 A8GAP5 A8GAP8 A8GAQ8 A8GAR0
A8GAR6 A8GAR8 A8GAT0 A8GAT8 A8GAT9 A8GAU8 A8GAV0 A8GAV1 A8GAV2 A8GAV3 A8GAV7
A8GAY0 A8GAY2 A8GB09 A8GB10 A8GB11 A8GB16 A8GB18 A8GB21 A8GB31 A8GB34 A8GB41
A8GB43 A8GB71 A8GB83 A8GB96 A8GBA2 A8GBA5 A8GBC1 A8GBC5 A8GBC6 A8GBD1 A8GBD6
A8GBT4 A8GBU5 A8GBX7 A8GBX8 A8GBX9 A8GBY8 A8GBZ6 A8GBZ7 A8GC24 A8GC31 A8GC32
A8GC37 A8GC40 A8GC41 A8GC74 A8GC75 A8GC78 A8GC80 A8GCA7 A8GCB4 A8GCC2 A8GCD7
A8GCD9 A8GCE0 A8GCF0 A8GCG3 A8GCH0 A8GCH1 A8GCH2 A8GCH4 A8GCI2 A8GCI7 A8GCI9
A8GCJ5 A8GCJ6 A8GCK1 A8GCK7 A8GCL2 A8GCL9 A8GCM4 A8GCM7 A8GCM8 A8GCM9 A8GCS6
A8GCT2 A8GCT3 A8GCT4 A8GCT5 A8GCT6 A8GCW2 A8GCZ9 A8GD09 A8GD15 A8GD24 A8GD38
A8GD47 A8GD97 A8GDA0 A8GDA2 A8GDA3 A8GDA9 A8GDC2 A8GDD5 A8GDJ0 A8GDM6 A8GDM9
A8GDQ5 A8GDQ7 A8GDQ8 A8GDQ9 A8GDR1 A8GDS3 A8GDU4 A8GDV6 A8GDW2 A8GDX5 A8GDX7
A8GDX9 A8GDY0 A8GE04 A8GE05 A8GE12 A8GES6 A8GEX8 A8GF18 A8GF39 A8GF49 A8GF80
A8GF82 A8GF83 A8GFA1 A8GFC3 A8GFD0 A8GFD8 A8GFE3 A8GFF7 A8GFG2 A8GFG6 A8GFG7
A8GFG9 A8GFH0 A8GFI7 A8GFI8 A8GFI9 A8GFJ0 A8GFJ2 A8GFJ7 A8GFJ8 A8GFJ9 A8GFK2
A8GFK5 A8GFL3 A8GFL7 A8GFQ1 A8GFQ2 A8GFQ3 A8GFZ8 A8GG07 A8GG50 A8GG79 A8GG82
A8GG83 A8GG86 A8GG94 A8GG95 A8GGH0 A8GGN0 A8GGN3 A8GGQ0 A8GGR6 A8GGT5 A8GGU5
A8GGU6 A8GGW2 A8GGW3 A8GGX8 A8GGZ0 A8GGZ1 A8GH24 A8GH44 A8GH77 A8GH82 A8GH84
A8GHC5 A8GHF2 A8GHH2 A8GHJ1 A8GHK9 A8GHL5 A8GHT4 A8GHV1 A8GHV3 A8GHW1 A8GHW4
A8GHW5 A8GHW8 A8GHW9 A8GHX6 A8GHX9 A8GHY0 A8GHY1 A8GHY3 A8GHY4 A8GHZ4 A8GHZ8
A8GI17 A8GI23 A8GI25 A8GI34 A8GI38 A8GI39 A8GI40 A8GI41 A8GI44 A8GI46 A8GI78
A8GIB4 A8GIB8 A8GIC1 A8GID2 A8GIE2 A8GIE3 A8GIE4 A8GIE5 A8GIE6 A8GIE8 A8GIE9
A8GIF0 A8GIF8 A8GIF9 A8GIG8 A8GIH7 A8GII0 A8GII1 A8GIL6 A8GIP4 A8GIQ4 A8GIR9
A8GIS0 A8GIS1 A8GIS5 A8GIT0 A8GIV0 A8GIW3 A8GIX7 A8GIX9 A8GIY9 A8GIZ1 A8GIZ7
A8GIZ8 A8GJ06 A8GJ07 A8GJ12 A8GJ28 A8GJ32 A8GJ33 A8GJ38 A8GJ45 A8GJ47 A8GJ49
A8GJM1 A8GJS8 A8GJU0 A8GJU3 A8GJU7 A8GJV1 A8GJV2 A8GJV5 A8GJW6 A8GJX5 A8GJZ1
A8GJZ2 A8GK02 A8GK03 A8GK08 A8GK24 A8GK41 A8GK75 A8GK78 A8GKB8 A8GKD1 A8GKF5
A8GKF8 A8GKG0 A8GKG1 A8GKG3 A8GKG5 A8GKG6 A8GKG7 A8GKH2 A8GKH3 A8GKH4 A8GKH5
A8GKH6 A8GKH8 A8GKH9 A8GKI0 A8GKI1 A8GKI2 A8GKI3 A8GKI4 A8GKI5 A8GKI6 A8GKI7
A8GKI8 A8GKI9 A8GKJ0 A8GKJ1 A8GKJ2 A8GKJ4 A8GKJ6 A8GKJ7 A8GKJ8 A8GKK1 A8GKK3
A8GKK4 A8GKK5 A8GKK6 A8GKK7 A8GKN0 A8GKQ1 A8GKQ7 A8GKQ8 A8GKR9 A8GKT3 A8GKT5
A8GKT7 A8GKU0 A8GKU1 A8GKU7 A8GKU8 A8GKU9 A8GKV0 A8GKV9 A8GKW1 A8GL03 A8GL43
A8GL54 A8GL60 A8GL72 A8GL74 A8GL77 A8GL81 A8GL82 A8GL83 A8GL86 A8GL90 A8GL91
A8GL95 A8GL96 A8GL98 A8GL99 A8GLA8 A8GLB0 A8GLB9 A8GLC2 A8GLC6 A8GLC8 A8GLE1
A8GLE2 A8GLE3 A8GLE4 A8GLE5 A8GLE7 A8GLE8 A8GLE9 A8GLF0 A8GLH0 A8GLH9 A8GLI7
A8GLI8 A8GLJ4 A8GLK3 A8GLK6 A8GLK7 A8GLL0 A8GLL1 A8GLR3 A8GLU3 A8GLV2 A8GLX7
A8GLX8 A8GLX9 A8GLZ8 A8GLZ9 A8GM00 A8GM08 A8GM09 A8GM16 A8GM17 A8GM19 A8GM31
A8GM32 A8GM33 A8GM49 A8GM54 A8GM58 A8GM60 A8GM71 A8GM73 A8GM74 A8GM75 A8GM79
A8GM81 A8GM90 A8GM98 A8GM99 A8GMA0 A8GMA3 A8GMA4 A8GMA5 A8GMA6 A8GMA7 A8GMA8
A8GMB3 A8GMB6 A8GMC0 A8GMC1 A8GMC3 A8GMC4 A8GMD2 A8GMD3 A8GMF8 A8GMF9 A8GMH0
A8GMI8 A8GMI9 A8GMJ0 A8GMJ1 A8GMJ8 A8GMK3 A8GML2 A8GML4 A8GMM9 A8GMN0 A8GMN4
A8GMN6 A8GMP6 A8GMP7 A8GMP8 A8GMX2 A8GMY3 A8GMY5 A8GMZ9 A8GN00 A8GN05 A8GN11

A8GN18 A8GN21 A8GN35 A8GN44 A8GN45 A8GN71 A8GN82 A8GN89 A8GN90 A8GN94 A8GN97
A8GNB1 A8GND7 A8GND9 A8GNE3 A8GNF1 A8GNF2 A8GNF5 A8GNF9 A8GNG2 A8GNH0 A8GNK0
A8GNK2 A8GNL2 A8GNL8 A8GNL9 A8GNM0 A8GNQ4 A8GNR5 A8GNU0 A8GNV8 A8GNW1 A8GNW3
A8GNW7 A8GNY2 A8GNY3 A8GNY6 A8GNZ4 A8GNZ6 A8GP28 A8GP29 A8GP33 A8GP45 A8GP46
A8GP75 A8GP84 A8GP85 A8GP88 A8GP89 A8GP90 A8GP91 A8GP98 A8GPA1 A8GPB1 A8GPB2
A8GPB6 A8GPB7 A8GPB9 A8GPC0 A8GPC4 A8GPC6 A8GPC7 A8GPC8 A8GPC9 A8GPD1 A8GPD2
A8GPD3 A8GPD4 A8GPD5 A8GPD6 A8GPD7 A8GPD8 A8GPD9 A8GPE0 A8GPE1 A8GPE2 A8GPE3
A8GPE4 A8GPE5 A8GPE6 A8GPE7 A8GPE8 A8GPE9 A8GPF0 A8GPF1 A8GPF2 A8GPG7 A8GPH1
A8GPH5 A8GPI0 A8GPJ0 A8GPJ3 A8GPK1 A8GPK3 A8GPM9 A8GPN2 A8GPN6 A8GPP8 A8GPP9
A8GPR4 A8GPR5 A8GPR8 A8GPR9 A8GPT6 A8GPW2 A8GPW6 A8GPX3 A8GQ03 A8GQ12 A8GQ13
A8GQ15 A8GQ50 A8GQ51 A8GQ54 A8GQ55 A8GQ64 A8GQ65 A8GQ73 A8GQ80 A8GQ89 A8GQ98
A8GQA5 A8GQA7 A8GQA8 A8GQA9 A8GQB7 A8GQC1 A8GQC2 A8GQG2 A8GQH5 A8GQJ1 A8GQJ4
A8GQJ5 A8GQJ6 A8GQJ7 A8GQL6 A8GQL7 A8GQL8 A8GQM7 A8GQM8 A8GQN3 A8GQN4 A8GQN6
A8GQP6 A8GQP7 A8GQP8 A8GQR1 A8GQR9 A8GQS1 A8GQT2 A8GQT3 A8GQT4 A8GQU0 A8GQU9
A8GQV5 A8GQV6 A8GQV7 A8GQW0 A8GQW1 A8GQW2 A8GQW4 A8GQW5 A8GQW7 A8GQX3 A8GQX6
A8GQX7 A8GQX8 A8GQY0 A8GQY1 A8GQY9 A8GQZ0 A8GR21 A8GR22 A8GR33 A8GR49 A8GR50
A8GR51 A8GR53 A8GR69 A8GR81 A8GR83 A8GRA0 A8GRA1 A8GRA7 A8GRA8 A8GRB9 A8GRC0
A8GRC1 A8GRI6 A8GRJ5 A8GRL7 A8GRL8 A8GRM5 A8GRM9 A8GRN5 A8GRN9 A8GRQ0 A8GRQ6
A8GRQ8 A8GRR0 A8GRT7 A8GRV2 A8GRV6 A8GRW0 A8GRW1 A8GRW4 A8GRW7 A8GRY1 A8GS10
A8GS12 A8GS16 A8GS25 A8GS26 A8GS29 A8GS39 A8GS42 A8GS48 A8GS71 A8GS72 A8GS87
A8GS96 A8GS97 A8GS98 A8GSC7 A8GSG4 A8GSH4 A8GSH5 A8GSH6 A8GSI6 A8GSK8 A8GSK9
A8GSP0 A8GSP4 A8GSP6 A8GSQ1 A8GSS6 A8GSS7 A8GST1 A8GSU9 A8GSV0 A8GSY1 A8GSZ4
A8GSZ5 A8GSZ7 A8GSZ8 A8GSZ9 A8GT00 A8GT12 A8GT27 A8GT30 A8GT31 A8GT33 A8GT43
A8GT44 A8GT45 A8GT46 A8GT47 A8GT50 A8GT51 A8GT52 A8GT53 A8GT54 A8GT55 A8GT56
A8GT57 A8GT58 A8GT59 A8GT60 A8GT61 A8GT62 A8GT63 A8GT64 A8GT65 A8GT66 A8GT67
A8GT68 A8GT69 A8GT70 A8GT71 A8GT85 A8GT93 A8GT99 A8GTA0 A8GTB0 A8GTB4 A8GTC4
A8GTC6 A8GTF2 A8GTF4 A8GTI4 A8GTI5 A8GTI8 A8GTI9 A8GTK6 A8GTK7 A8GTM1 A8GTP3
A8GTQ7 A8GTT8 A8GTV5 A8GTV6 A8GTV8 A8GTZ4 A8GTZ5 A8GTZ8 A8GTZ9 A8GU09 A8GU16
A8GU17 A8GU26 A8GU44 A8GU54 A8GU56 A8GU57 A8GU58 A8GU63 A8GU66 A8GU77 A8GU80
A8GU86 A8GU95 A8GUA5 A8GUB6 A8GUB7 A8GUE4 A8GUF0 A8GUF3 A8GUG0 A8GUH1 A8GUH2
A8GUH3 A8GUH4 A8GUI1 A8GUI2 A8GUI6 A8GUI7 A8GUI9 A8GUK0 A8GUK1 A8GUK2 A8GUK4
A8GUK6 A8GUM4 A8GUN3 A8GUN4 A8GUQ0 A8GUQ3 A8GUQ4 A8GUR0 A8GUR1 A8GUR2 A8GUT7
A8GUU1 A8GUU4 A8GUV0 A8GUW0 A8GUZ9 A8GV08 A8GV15 A8GV17 A8GV20 A8GV21 A8GV22
A8GV23 A8GV24 A8GV25 A8GV42 A8GV45 A8GV46 A8GV47 A8GV48 A8GV49 A8GV60 A8GV61
A8GV67 A8GV86 A8GVB2 A8GVB3 A8GVB4 A8GVB5 A8GVB6 A8GVB7 A8GVB8 A8GVB9 A8GVC0
A8GVC1 A8GVC2 A8GVC3 A8GVC4 A8GVC6 A8GVC7 A8GVC8 A8GVC9 A8GVD0 A8GVD2 A8GVD3
A8GVD5 A8GVD6 A8GVD7 A8GVD8 A8GVD9 A8GVG9 A8GVJ9 A8GVK6 A8GVL0 A8GVL1 A8GVL2
A8GVL3 A8GVL5 A8GVL6 A8GVN1 A8GVN7 A8GVR7 A8GVR9 A8GVS7 A8GVS9 A8GVT0 A8GVU7
A8GVU9 A8GVV0 A8GVV1 A8GVV5 A8GVV6 A8GVY2 A8GVY3 A8GW07 A8GW08 A8GW18 A8GW24
A8GW25 A8GW29 A8GW31 A8GW63 A8GW64 A8GW74 A8GW75 A8GW83 A8GW85 A8GW94 A8GW95
A8GW96 A8GWB2 A8GWC2 A8GWC4 A8GWH2 A8GWH4 A8GWI0 A8GWJ3 A8GWJ4 A8GWP0 A8GWP2
A8GWR2 A8GWS0 A8GWS1 A8GWU6 A8GWV0 A8GWV1 A8GWV2 A8GWV8 A8GWX0 A8GX08 A8GX11
A8GX62 A8GX84 A8GX86 A8GXC3 A8GXC5 A8GXC8 A8GXE5 A8GXI1 A8GXI7 A8GXN6 A8GXN7
A8GXQ1 A8GXR0 A8GXR2 A8GXR7 A8GXS3 A8GXS7 A8GXT8 A8GXV3 A8GXY8 A8GXY9 A8GY07

A8GY12 A8GY13 A8GY14 A8GY17 A8GY24 A8GY58 A8GY79 A8GY80 A8GY81 A8GY89 A8GY99
A8GYB3 A8GYB4 A8GYB5 A8GYC8 A8GYD7 A8GYD8 A8GYE2 A8GYE3 A8GYE5 A8GYE8 A8GYF4
A8GYG1 A8GYH2 A8GYH6 A8GYH9 A8GYI0 A8GYI2 A8GYP3 A8GYP8 A8GYS3 A8GYT2 A8GYV2
A8GYV6 A8GYV7 A8GYW1 A8GYW2 A8GYW5 A8GYW6 A8GYW7 A8GYW8 A8GYW9 A8GYX0 A8GYX1
A8GYX2 A8GYX5 A8GYX6 A8GYX7 A8GYX8 A8GYX9 A8GYY0 A8GYY1 A8GYY3 A8GYY4 A8GYY5
A8GYY6 A8GYY7 A8GYY8 A8GYY9 A8GYZ0 A8GYZ1 A8GYZ2 A8GYZ4 A8GYZ5 A8GYZ7 A8GYZ8
A8GYZ9 A8GZ01 A8GZ14 A8GZ16 A8GZ18 A8GZ19 A8GZ21 A8GZ22 A8GZ29 A8GZ30 A8GZ41
A8GZ42 A8GZ99 A8GZD9 A8GZG4 A8GZI0 A8GZI2 A8GZJ0 A8GZJ9 A8GZL8 A8GZM3 A8GZV5
A8H029 A8H038 A8H086 A8H094 A8H0A3 A8H0A6 A8H0D3 A8H0D4 A8H0D9 A8H0H1 A8H0J7
A8H0M7 A8H0M9 A8H0N0 A8H0P3 A8H0P9 A8H0Q6 A8H0R6 A8H0T9 A8H0U0 A8H0U2 A8H0U3
A8H0U4 A8H0V1 A8H0V2 A8H151 A8H152 A8H161 A8H164 A8H175 A8H183 A8H191 A8H1B1
A8H1B4 A8H1C7 A8H1D9 A8H1E0 A8H1E1 A8H1E2 A8H1E5 A8H1G3 A8H1G4 A8H1G8 A8H1H0
A8H1H3 A8H1H6 A8H1J7 A8H1J9 A8H1Q0 A8H1Q1 A8H1Q5 A8H1Q6 A8H1S1 A8H1S4 A8H1S7
A8H1S8 A8H1T0 A8H1T1 A8H1T4 A8H1T5 A8H1T6 A8H1T7 A8H1U8 A8H222 A8H242 A8H245
A8H246 A8H249 A8H252 A8H254 A8H2H4 A8H2I7 A8H2J4 A8H2K7 A8H2K8 A8H2M6 A8H2M9
A8H2N0 A8H2N6 A8H2P5 A8H2P7 A8H2R3 A8H2R4 A8H2S2 A8H2X4 A8H2Z3 A8H305 A8H356
A8H376 A8H392 A8H3E4 A8H3F0 A8H3F1 A8H3F8 A8H3H6 A8H3I7 A8H3V2 A8H3V4 A8H3W2
A8H3W9 A8H3Y1 A8H3Y9 A8H3Z4 A8H415 A8H424 A8H467 A8H479 A8H482 A8H499 A8H4A2
A8H4A4 A8H4A5 A8H4A7 A8H4D4 A8H4E9 A8H4F0 A8H4Q7 A8H4R2 A8H4T0 A8H4U3 A8H4U4
A8H4U6 A8H532 A8H545 A8H546 A8H547 A8H549 A8H551 A8H552 A8H562 A8H566 A8H589
A8H5A5 A8H5C4 A8H5C6 A8H5C9 A8H5D9 A8H5E4 A8H5E5 A8H5E7 A8H5F1 A8H5G0 A8H5G3
A8H5G5 A8H5H1 A8H5I1 A8H5K7 A8H5L2 A8H5L4 A8H5L6 A8H5M1 A8H5N3 A8H5P1 A8H5Q9
A8H5R0 A8H5U9 A8H613 A8H614 A8H615 A8H616 A8H617 A8H6B7 A8H6J5 A8H6K2 A8H6L0
A8H6L3 A8H6L6 A8H6L9 A8H6S3 A8H6T8 A8H6V1 A8H6V6 A8H6W1 A8H6W6 A8H6X1 A8H6X2
A8H726 A8H727 A8H728 A8H733 A8H735 A8H737 A8H738 A8H739 A8H740 A8H742 A8H744
A8H745 A8H748 A8H756 A8H759 A8H760 A8H764 A8H765 A8H771 A8H786 A8H793 A8H7A5
A8H7A6 A8H7A8 A8H7B6 A8H7B8 A8H7C2 A8H7C5 A8H7C7 A8H7D3 A8H7D4 A8H7D5 A8H7G5
A8H7J4 A8H7S9 A8H7T0 A8H7T1 A8H7X2 A8H819 A8H8E0 A8H8G6 A8H8G7 A8H8G8 A8H8K6
A8H8K9 A8H8L0 A8H8L1 A8H8L2 A8H8L8 A8H8M0 A8H8M9 A8H8Q9 A8H8R3 A8H8U2 A8H8W3
A8H8W4 A8H8X8 A8H8Z4 A8H950 A8H953 A8H955 A8H956 A8H966 A8H983 A8H992 A8H993
A8H995 A8H998 A8H999 A8H9A1 A8H9A8 A8H9A9 A8H9B0 A8H9B2 A8H9B3 A8H9B7 A8H9F1
A8H9H9 A8H9P4 A8H9T3 A8H9X0 A8HA20 A8HA27 A8HA30 A8HA69 A8HA91 A8HA92 A8HA94
A8HAA3 A8HAD8 A8HAE0 A8HAF6 A8HAH3 A8HAH4 A8HAH9 A8HAI3 A8HME3 A8HMZ4 A8HN58
A8HNV0 A8HQ27 A8HQ40 A8HQ54 A8HQ60 A8HQP5 A8HR94 A8HRJ3 A8HRP6 A8HRQ8 A8HRS7
A8HRT0 A8HRT6 A8HRT9 A8HS37 A8HS48 A8HS51 A8HS67 A8HS70 A8HS90 A8HT01 A8HT29
A8HTM2 A8HTN2 A8HTW6 A8HTX5 A8HTX8 A8HTY1 A8HTY4 A8HTY8 A8HTZ1 A8HU60 A8HUA1
A8HUG2 A8HUR7 A8HV31 A8HV62 A8HV93 A8HVC4 A8HVE8 A8HVF1 A8HVI9 A8HVL5 A8HVS0
A8HVS4 A8HWL0 A8HWM0 A8HWM3 A8HX69 A8HXU1 A8HYF0 A8HYF2 A8HYF4 A8HYF7 A8HYK3
A8HYP5 A8HYS9 A8HYT3 A8HYT7 A8HYT9 A8HYU5 A8HYU9 A8HZ68 A8HZ71 A8HZ95 A8HZA5
A8HZA8 A8HZS2 A8I060 A8I066 A8I0D5 A8I0F9 A8I0G1 A8I0I5 A8I0J5 A8I0R2 A8I1Q0
A8I1V7 A8I1W4 A8I264 A8I266 A8I267 A8I275 A8I2D3 A8I2M2 A8I2V9 A8I3B5 A8I438
A8I460 A8I464 A8I467 A8I468 A8I4E9 A8I4G3 A8I4J4 A8I4L8 A8I4M5 A8I4R6 A8I4S6
A8I4T0 A8I4X6 A8I541 A8I5B0 A8I5C0 A8I5F1 A8I5K8 A8I5N7 A8I5P5 A8I5Q2 A8I5R5
A8I5S8 A8I655 A8I679 A8I6A9 A8I6D2 A8I6E6 A8I711 A8I7Z7 A8I7Z9 A8I819 A8I831

A8I836 A8I839 A8I889 A8I8B0 A8I8V7 A8I9E8 A8I9L7 A8I9M0 A8I9W7 A8I9X4 A8I9X6
A8IA58 A8IAC9 A8IAD2 A8IAD8 A8IAN3 A8IAN5 A8IAN8 A8IAP0 A8IAP5 A8IAP6 A8IAP8
A8IAP9 A8IAQ0 A8IAQ2 A8IAQ3 A8IAQ5 A8IAQ6 A8IAQ7 A8IAR0 A8IAR2 A8IAR6 A8IAR8
A8IAS6 A8IAS9 A8IAT3 A8IAT5 A8IAT8 A8IAX3 A8IB22 A8IB25 A8IBI0 A8IBL6 A8IBS1
A8IC06 A8IC17 A8IC24 A8IC48 A8ICF8 A8ICS9 A8ICV0 A8ICW0 A8ICW8 A8ID68 A8ID74
A8ID95 A8IEF3 A8IEI6 A8IES7 A8IEU9 A8IEY6 A8IF44 A8IFI2 A8IFX1 A8IFY6 A8IG00
A8IG05 A8IG13 A8IG16 A8IG20 A8IG22 A8IGA3 A8IGA5 A8IGE6 A8IGK2 A8IGX4 A8IGY0
A8IGZ9 A8IH47 A8IHN8 A8IHT2 A8II77 A8IIZ0 A8IJI4 A8IJU8 A8IKD2 A8IKN6 A8IKU6
A8IKV0 A8IKV6 A8IL81 A8ILC6 A8ILE7 A8ILK1 A8ILM7 A8ILN3 A8ILP8 A8ILV4 A8ILY5
A8IM50 A8IM57 A8IM75 A8IM90 A8IM96 A8IM99 A8IMA7 A8IMG3 A8IMH3 A8IMK0 A8IMT6
A8IMV5 A8IMX7 A8IN83 A8INH8 A8IPQ8 A8IPT0 A8IPT1 A8IPV3 A8IPW5 A8IQ54 A8IQ73
A8IQD2 A8IQE0 A8IQL3 A8IQM5 A8IQT2 A8IR43 A8IRJ7 A8IRK7 A8ITB0 A8ITV9 A8IU92
A8IUG5 A8IVJ1 A8IVX2 A8IW34 A8IYS6 A8IZG4 A8J0N6 A8J1V4 A8J2Z9 A8J353 A8J3A0
A8J3F6 A8J637 A8J666 A8J6X1 A8J6X7 A8J8F6 A8J9T5 A8JA42 A8JAF2 A8JAM0 A8JAN3
A8JB22 A8JBB2 A8JC00 A8JF70 A8JF71 A8JGF7 A8JGV8 A8JHD7 A8JID5 A8JJB2 A8JNV4
A8JPF9 A8JQX3 A8JR14 A8JUV0 A8JYX6 A8K010 A8K0R7 A8K0S8 A8K554 A8K830 A8K8V0
A8K979 A8KB34 A8KB59 A8KBE0 A8KBF3 A8KBL7 A8KBY2 A8KRL3 A8KYR0 A8KYR1 A8KYR3
A8KYR5 A8KYS5 A8KYU3 A8KYV9 A8KYW3 A8KZA9 A8KZB0 A8KZE2 A8KZE6 A8KZE8 A8KZE9
A8KZF0 A8L0K9 A8L175 A8L177 A8L188 A8L1A5 A8L1J2 A8L1V4 A8L1V6 A8L1V7 A8L1V8
A8L1W8 A8L1X0 A8L1X3 A8L2B6 A8L2C4 A8L2N0 A8L2N1 A8L3U5 A8L3V0 A8L3V1 A8L3X7
A8L536 A8L538 A8L548 A8L551 A8L554 A8L567 A8L568 A8L585 A8L588 A8L591 A8L594
A8L5A4 A8L5A5 A8L6C4 A8L6C6 A8L6C8 A8L6D6 A8L6D8 A8L6E2 A8L6F1 A8L6F4 A8L6G0
A8L6G1 A8L6J8 A8L6K0 A8L6K6 A8L764 A8L766 A8L7B0 A8L7B6 A8L7B7 A8L893 A8L8M0
A8L8S7 A8L8T1 A8L8T2 A8L8W2 A8L8W3 A8LAA1 A8LAA2 A8LAW3 A8LAX2 A8LAY7 A8LAZ9
A8LB01 A8LB04 A8LB07 A8LB08 A8LB14 A8LB15 A8LBC6 A8LC40 A8LC41 A8LC42 A8LC43
A8LC44 A8LC45 A8LC46 A8LC49 A8LC50 A8LC51 A8LC52 A8LC53 A8LC54 A8LC56 A8LC57
A8LC58 A8LC59 A8LC60 A8LC61 A8LC63 A8LC64 A8LC65 A8LC66 A8LC67 A8LC68 A8LC73
A8LC74 A8LC81 A8LCA4 A8LCD0 A8LCI3 A8LCM4 A8LCM7 A8LCV4 A8LCW4 A8LD98 A8LDG7
A8LDJ8 A8LDK0 A8LDL3 A8LDR3 A8LDU1 A8LE02 A8LE05 A8LE09 A8LE17 A8LE19 A8LE21
A8LE22 A8LE32 A8LE33 A8LE44 A8LE46 A8LE67 A8LEP8 A8LEP9 A8LF85 A8LFB7 A8LFH1
A8LGQ5 A8LGR0 A8LGR8 A8LH49 A8LH53 A8LH54 A8LH57 A8LH67 A8LHA4 A8LHP8 A8LHP9
A8LHV2 A8LHW1 A8LHX5 A8LHX7 A8LHY2 A8LI17 A8LI32 A8LI68 A8LI72 A8LI73 A8LI79
A8LI95 A8LIH2 A8LIH3 A8LIJ7 A8LIJ9 A8LIN0 A8LIN4 A8LIN6 A8LIP0 A8LIR1 A8LIR3
A8LIR8 A8LIV9 A8LIX8 A8LJ00 A8LJ19 A8LJ23 A8LJ24 A8LJ26 A8LJ36 A8LJ68 A8LJA7
A8LJA8 A8LJB4 A8LJI8 A8LJK6 A8LJP9 A8LJQ0 A8LJS9 A8LJT5 A8LJU6 A8LJY0 A8LK04
A8LK09 A8LK10 A8LK12 A8LK18 A8LK35 A8LK37 A8LK49 A8LK52 A8LK58 A8LK91 A8LK92
A8LK96 A8LK98 A8LKA2 A8LKA5 A8LKC5 A8LKE4 A8LKE7 A8LKI9 A8LKJ1 A8LKJ4 A8LKJ5
A8LKJ8 A8LKM4 A8LKN5 A8LKP0 A8LKQ3 A8LKQ5 A8LKS9 A8LKV5 A8LKW5 A8LKX1 A8LKY0
A8LL20 A8LL24 A8LL25 A8LL26 A8LL46 A8LL57 A8LL75 A8LLC0 A8LLC1 A8LLC7 A8LLD2
A8LLD9 A8LLE2 A8LLE8 A8LLF2 A8LLG2 A8LLG9 A8LLH0 A8LLH4 A8LLI5 A8LLJ9 A8LLK0
A8LLK4 A8LLR2 A8LLT2 A8LLT5 A8LLU7 A8LLX6 A8LLZ7 A8LM40 A8LM41 A8LM43 A8LM44
A8LM45 A8LM48 A8LM49 A8LM50 A8LM57 A8LM58 A8LM59 A8LM60 A8LM61 A8LM65 A8LM66
A8LM67 A8LM68 A8LM69 A8LM70 A8LM72 A8LM73 A8LM74 A8LM75 A8LM76 A8LM80 A8LM82
A8LM96 A8LMA2 A8LMA5 A8LMB2 A8LMB3 A8LMB6 A8LMC0 A8LMD0 A8LME1 A8LME6 A8LMF7

A8LMJ6 A8LMJ8 A8LMK2 A8LMT3 A8LMZ2 A8LN36 A8LN54 A8LNB8 A8LNF0 A8LNH2 A8LNK7
A8LNK8 A8LNL0 A8LNY7 A8LNZ7 A8LP53 A8LP83 A8LPA8 A8LPA9 A8LPB5 A8LPB8 A8LPC2
A8LPC3 A8LPC4 A8LPD1 A8LPD4 A8LPE0 A8LPN3 A8LPV7 A8LPW1 A8LQ20 A8LQ27 A8LQ28
A8LQ30 A8LQ43 A8LQ53 A8LQ56 A8LQ63 A8LQA8 A8LQI0 A8LQK5 A8LQK6 A8LQM0 A8LQM4
A8LQQ5 A8LQQ6 A8LQQ7 A8LQQ9 A8LQU6 A8LQX9 A8LQZ6 A8LR16 A8LR70 A8LRQ1 A8LRR5
A8LRR7 A8LRS0 A8LRS3 A8LRS4 A8LRS5 A8LRS7 A8LRT3 A8LRT6 A8LRW0 A8LRY5 A8LRY8
A8LS12 A8LS15 A8LS17 A8LS39 A8LS40 A8LS45 A8LS52 A8LS60 A8LS61 A8LS63 A8LS71
A8LS92 A8LSB5 A8LSE3 A8LSE7 A8LSF0 A8LSF1 A8LSF5 A8LSF9 A8LSK4 A8LSL0 A8LSV9
A8LTG4 A8LUR1 A8LVF4 A8LVH1 A8LVI6 A8LW18 A8LW20 A8LW22 A8LW31 A8LWH4 A8LWI0
A8LWM4 A8LWZ4 A8LWZ5 A8LWZ6 A8LX37 A8LX55 A8LX58 A8LX62 A8LX76 A8LX80 A8LX88
A8LX89 A8LX93 A8LXG0 A8LXG1 A8LXG3 A8LXK0 A8LXK4 A8LXW7 A8LXW8 A8LXW9 A8LXX3
A8LXY1 A8LXY8 A8LY07 A8LY09 A8LY10 A8LY11 A8LY14 A8LY20 A8LY23 A8LY25 A8LY30
A8LY38 A8LY39 A8LY44 A8LY45 A8LY49 A8LY51 A8LYC4 A8LYE7 A8LYF4 A8LYF8 A8LYN0
A8LYQ4 A8LYQ9 A8LYU3 A8LZ53 A8LZD6 A8LZF8 A8LZG1 A8LZI9 A8M064 A8M065 A8M0C0
A8M0L5 A8M0W3 A8M0X6 A8M0Y6 A8M0Y9 A8M0Z0 A8M0Z1 A8M1D3 A8M1E8 A8M1K7 A8M1L0
A8M1N0 A8M1N1 A8M218 A8M223 A8M264 A8M2A0 A8M2A3 A8M2A4 A8M2A6 A8M2K8 A8M2V8
A8M2W8 A8M3D4 A8M3H2 A8M3K2 A8M3K6 A8M3R1 A8M3R4 A8M449 A8M469 A8M497 A8M4B3
A8M4B8 A8M4B9 A8M4C0 A8M4E3 A8M4E6 A8M4I1 A8M4I4 A8M4S6 A8M4X3 A8M4Z9 A8M502
A8M503 A8M504 A8M505 A8M508 A8M510 A8M511 A8M512 A8M514 A8M515 A8M516 A8M517
A8M518 A8M519 A8M520 A8M521 A8M522 A8M523 A8M524 A8M525 A8M526 A8M529 A8M530
A8M531 A8M532 A8M533 A8M534 A8M538 A8M542 A8M543 A8M548 A8M5E1 A8M5E3 A8M5E5
A8M5E6 A8M5E7 A8M5F5 A8M5F9 A8M5I3 A8M5I4 A8M5I9 A8M5J6 A8M661 A8M677 A8M679
A8M680 A8M681 A8M684 A8M6A9 A8M6B1 A8M6B2 A8M6B4 A8M6B6 A8M6K7 A8M6L0 A8M6L1
A8M6S5 A8M6W7 A8M6W8 A8M723 A8M743 A8M746 A8M748 A8M757 A8M758 A8M773 A8M783
A8M7S7 A8M7V5 A8M7W6 A8M7W9 A8M7X6 A8M8E8 A8M8F2 A8M8F3 A8M8I3 A8M8Q8 A8M8Q9
A8M8R2 A8M8R5 A8M8T7 A8M8Y9 A8M8Z5 A8M909 A8M919 A8M9D7 A8M9G2 A8M9I3 A8M9J5
A8M9L3 A8M9N4 A8M9N5 A8M9S4 A8M9U2 A8M9Z7 A8M9Z9 A8MA42 A8MA52 A8MA54 A8MA77
A8MA81 A8MA86 A8MA91 A8MAC1 A8MAC7 A8MAG0 A8MAH5 A8MAH6 A8MAJ1 A8MAJ5 A8MAL5
A8MAV7 A8MAX6 A8MAY1 A8MB19 A8MB20 A8MB21 A8MB38 A8MB46 A8MB58 A8MB69 A8MB70
A8MB73 A8MB75 A8MB76 A8MB78 A8MB89 A8MB97 A8MBA2 A8MBB6 A8MBI2 A8MBK0 A8MBL4
A8MBL5 A8MBP6 A8MBQ3 A8MBU4 A8MBV3 A8MBV9 A8MBW0 A8MBW8 A8MBZ5 A8MC04 A8MC09
A8MC33 A8MC54 A8MC60 A8MC64 A8MC69 A8MC80 A8MCA3 A8MCC8 A8MCC9 A8MCD3 A8MCJ8
A8MCK2 A8MCN3 A8MCT2 A8MCU7 A8MCV2 A8MCX6 A8MCY4 A8MD44 A8MD73 A8MDD0 A8MDE4
A8MDI3 A8MDL8 A8MDN0 A8MDP6 A8MDS5 A8MDU3 A8MDU6 A8MDY8 A8MDY9 A8ME30 A8ME45
A8ME61 A8ME99 A8MEA0 A8MEA3 A8MEB3 A8MED4 A8MED6 A8MEE1 A8MEG4 A8MEG5 A8MEG7
A8MEH2 A8MEI9 A8MET3 A8MEV2 A8MEV9 A8MEX7 A8MF35 A8MF40 A8MF41 A8MF64 A8MF65
A8MF68 A8MFA2 A8MFA8 A8MFA9 A8MFB1 A8MFB3 A8MFB4 A8MFD1 A8MFD2 A8MFD4 A8MFD5
A8MFE2 A8MFE4 A8MFG6 A8MFH2 A8MFI5 A8MFI7 A8MFJ0 A8MFJ1 A8MFJ3 A8MFK4 A8MFK5
A8MFQ5 A8MFQ6 A8MFY0 A8MFY7 A8MG11 A8MG12 A8MG25 A8MG50 A8MG53 A8MG57 A8MG66
A8MG71 A8MG73 A8MG75 A8MGE1 A8MGE4 A8MGH9 A8MGI2 A8MGI3 A8MGJ3 A8MGJ5 A8MGK0
A8MGL7 A8MGM7 A8MGN0 A8MGN2 A8MGS9 A8MGY8 A8MH14 A8MH15 A8MH16 A8MH18 A8MH27
A8MH28 A8MH31 A8MH34 A8MH47 A8MH56 A8MH65 A8MH75 A8MH77 A8MH79 A8MH81 A8MH85
A8MH89 A8MH95 A8MHA0 A8MHA3 A8MHC1 A8MHC3 A8MHC4 A8MHC5 A8MHC7 A8MHC8 A8MHG9
A8MHH0 A8MHH1 A8MHH2 A8MHH5 A8MHI0 A8MHI3 A8MHI4 A8MHI5 A8MHJ4 A8MHJ6 A8MHK8

A8MHL1 A8MHL7 A8MHM4 A8MHM5 A8MHN6 A8MHN7 A8MHS1 A8MHU4 A8MI45 A8MI49 A8MI61
A8MI62 A8MI73 A8MI78 A8MI82 A8MI83 A8MIA8 A8MIB3 A8MIF4 A8MIM4 A8MIN1 A8MIS4
A8MIS7 A8MIS8 A8MIS9 A8MIU1 A8MIU4 A8MIX7 A8MJ06 A8MJ41 A8MJ51 A8MJ53 A8MJ87
A8MJ91 A8MJA9 A8MJD2 A8MJJ7 A8MJJ8 A8MJP4 A8MJT0 A8MJX1 A8MJX9 A8MJY6 A8MJY9
A8MJZ5 A8MK43 A8MK46 A8MK48 A8MK56 A8MK92 A8MK93 A8MKA7 A8MKA8 A8MKD9 A8MKH4
A8MKJ7 A8MKQ1 A8MKQ2 A8MKR7 A8MKR8 A8MKR9 A8MKS3 A8MKS4 A8MKZ5 A8ML24 A8ML31
A8ML45 A8ML82 A8MLB1 A8MLB2 A8MLB3 A8MLB5 A8MLB7 A8MLC4 A8MLC5 A8MLC8 A8MLC9
A8MLD0 A8MLD1 A8MLD2 A8MLD3 A8MLD5 A8MLD6 A8MLD7 A8MLD9 A8MLE0 A8MLE1 A8MLE2
A8MLE3 A8MLE4 A8MLE5 A8MLE6 A8MLE7 A8MLE8 A8MLE9 A8MLF0 A8MLF1 A8MLF2 A8MLF3
A8MLF4 A8MLF5 A8MLF6 A8MLF7 A8MLF8 A8MLF9 A8MLG1 A8MLG4 A8MLG5 A8MLG6 A8MLG7
A8MLG8 A8MLG9 A8MLH0 A8MLH5 A8MLH6 A8MLI7 A8MLI9 A8MLJ8 A8MLM7 A8MLM9 A8MLU1
A8MLU4 A8MLW5 A8MLW8 A8MLW9 A8MLX3 A8MLX7 A8MLY0 A8MM16 A8MM35 A8MPH9 A8MPP1
A8MPS7 A8MPX8 A8MQ03 A8MQ11 A8MQ14 A8MQA3 A8MQB3 A8MQG7 A8MR21 A8MR89 A8MRP2
A8MRX0 A8MS20 A8MS41 A8MS68 A8MS85 A8MSI8 A8MT19 A8MT33 A8MT65 A8MT66 A8MT69
A8MT70 A8MTA8 A8MTJ6 A8MTL0 A8MTL3 A8MTQ0 A8MTY0 A8MTY7 A8MTZ7 A8MU10 A8MU46
A8MU93 A8MUA0 A8MUI8 A8MUL3 A8MUM7 A8MUP2 A8MUU1 A8MUU9 A8MUV8 A8MUX0 A8MUZ8
A8MV24 A8MV57 A8MV65 A8MV72 A8MVA2 A8MVJ9 A8MVM7 A8MVU1 A8MVX0 A8MW92 A8MW95
A8MW99 A8MWA4 A8MWA6 A8MWD9 A8MWE9 A8MWP4 A8MX19 A8MX34 A8MX76 A8MX80 A8MXD5
A8MXJ8 A8MXQ7 A8MXT2 A8MXY4 A8MXZ1 A8MXZ3 A8MY62 A8MYA2 A8MYJ7 A8MYP8 A8MYX2
A8MYZ0 A8MYZ5 A8MYZ6 A8MZ25 A8MZ59 A8MZA4 A8MZF0 A8MZG2 A8N142 A8N1T1 A8N1X3
A8N2M6 A8N2Y6 A8N2Z9 A8N339 A8N374 A8N4R7 A8N4R8 A8N5J5 A8N767 A8N7C6 A8N7N4
A8N8S3 A8N8Z0 A8N936 A8NA43 A8NAN2 A8NCK5 A8NDT2 A8NE23 A8NEG8 A8NEP3 A8NF97
A8NF99 A8NFF0 A8NHT8 A8NIX5 A8NJ91 A8NMB4 A8NQC2 A8NQV9 A8NS27 A8NS61 A8NS89
A8NSD1 A8NU13 A8NU66 A8NV38 A8NVB7 A8NWK4 A8NWP2 A8NWR2 A8NX92 A8NY27 A8NYG2
A8NYM5 A8NZ80 A8NZM5 A8NZY7 A8P008 A8P1W0 A8P353 A8P5H7 A8P7F7 A8P7J8 A8P7Y3
A8PAA1 A8PB32 A8PCG3 A8PD13 A8PDE3 A8PF69 A8PGQ3 A8PHP4 A8PJ38 A8PJJ2 A8PJX4
A8PKH2 A8PR17 A8PRT2 A8PS95 A8PT44 A8PTG4 A8PU71 A8PUI7 A8PV03 A8PVM6 A8PW87
A8PWB6 A8PWF6 A8PWG8 A8PWH1 A8PWQ8 A8PXR7 A8PXY6 A8PZI1 A8PZS4 A8Q0E3 A8Q1F0
A8Q2H5 A8Q2R5 A8Q513 A8Q6P6 A8Q8J2 A8Q8M5 A8Q9M0 A8QA64 A8QB65 A8QBB1 A8QBF3
A8QBZ2 A8QC60 A8QCB2 A8QCE4 A8QCH0 A8QCY3 A8QD31 A8QD49 A8QDN3 A8QE42 A8QE76
A8QFF6 A8QFQ3 A8QHQ0 A8QMS7 A8QW39 A8QW51 A8QW52 A8QW53 A8R4Q8 A8RG82 A8RQK7
A8SE44 A8SE74 A8SE78 A8SE82 A8SE86 A8SEA1 A8SEA5 A8SEC4 A8SEC5 A8SEC6 A8SED5
A8SED6 A8SED7 A8SED8 A8SED9 A8SEE0 A8SEE1 A8SEE2 A8SEE3 A8SEE4 A8SEE5 A8SEE7
A8SEG3 A8SMG2 A8STG1 A8T6P4 A8TSS9 A8W3A6 A8W3B3 A8W3B4 A8W3B5 A8W3B6 A8W3C2
A8W3C6 A8W3D6 A8W3E2 A8W3E3 A8W3E4 A8W3F3 A8W3F4 A8W3F5 A8W3F6 A8W3F7 A8W3F8
A8W3F9 A8W3G0 A8W3G1 A8W3G2 A8W3G3 A8W3G4 A8W3G8 A8W3H9 A8W3I5 A8W3I9 A8W3J2
A8W3K4 A8W3K5 A8W3K6 A8W3L5 A8W3L6 A8W3L7 A8W3L8 A8W3L9 A8W3M0 A8W3M1 A8W3M2
A8W3M5 A8W3M7 A8W7M4 A8W7M6 A8W7M7 A8W7M9 A8W7N0 A8W7N1 A8W7N2 A8W7N3 A8W7N4
A8W7N5 A8W7N6 A8W7N7 A8W7N8 A8W7N9 A8W7P0 A8W7P1 A8W7P2 A8W7P3 A8W995 A8WAT2
A8WE67 A8WFF7 A8WFT6 A8WFU8 A8WG43 A8WGF4 A8WGP3 A8WH18 A8WHT1 A8WJ41 A8WJ66
A8WJR8 A8WK63 A8WL06 A8WL28 A8WLG6 A8WLV5 A8WM57 A8WMD5 A8WMM4 A8WP66 A8WPC5
A8WPG6 A8WQQ5 A8WQV9 A8WQY8 A8WR63 A8WRE3 A8WRG3 A8WS58 A8WS92 A8WSQ9 A8WSU9
A8WSV7 A8WT19 A8WTA7 A8WTE8 A8WTH5 A8WTI8 A8WTJ7 A8WTM7 A8WTV9 A8WVJ9 A8WVR2
A8WVX8 A8WVY9 A8WWC0 A8WWH5 A8WWQ7 A8WWU0 A8WX15 A8WX27 A8WX89 A8WXF6 A8WXM1

A8WXX7 A8WY26 A8WZU5 A8X0C4 A8X0L9 A8X0P0 A8X0Z1 A8X2R1 A8X2U9 A8X2W2 A8X354
A8X371 A8X379 A8X3V8 A8X409 A8X419 A8X485 A8X493 A8X4H1 A8X5H6 A8X633 A8X679
A8X6I9 A8X7H5 A8X7X9 A8X7Z3 A8X811 A8X858 A8X8C6 A8X8D0 A8X8I6 A8X9U4 A8X9V4
A8XA58 A8XA80 A8XAA9 A8XAC4 A8XAC6 A8XAF4 A8XBR9 A8XDI8 A8XDR5 A8XDT0 A8XE00
A8XEA2 A8XEG9 A8XEN7 A8XEQ8 A8XFF4 A8XFH3 A8XFZ3 A8XG63 A8XGH1 A8XGS6 A8XGY6
A8XGZ9 A8XHQ1 A8XHX1 A8XI07 A8XI75 A8XJ98 A8XJD0 A8XJN7 A8XKG6 A8XKM2 A8XKT0
A8XKV0 A8XL02 A8XL25 A8XLL3 A8XLS0 A8XLU5 A8XMF2 A8XND8 A8XNK6 A8XP91 A8XPH5
A8XPL7 A8XRF5 A8XSQ8 A8XSS1 A8XSW2 A8XSX8 A8XT16 A8XT81 A8XU68 A8XV40 A8XV95
A8XW44 A8XW52 A8XWB7 A8XX92 A8XXB0 A8XXC0 A8XXC7 A8XY47 A8XY95 A8XYW9 A8XYX2
A8XYX3 A8XYZ2 A8XZB0 A8XZJ0 A8XZJ9 A8XZP2 A8XZU0 A8Y0L5 A8Y183 A8Y197 A8Y1E9
A8Y1R8 A8Y3Q1 A8Y3R9 A8Y3X9 A8Y461 A8Y5T1 A8Y5U1 A8Y985 A8Y986 A8Y9A7 A8Y9A8
A8Y9A9 A8Y9B8 A8Y9B9 A8Y9C0 A8Y9C1 A8Y9C2 A8Y9C3 A8Y9C4 A8Y9C5 A8Y9C6 A8Y9C7
A8Y9C8 A8Y9D1 A8Y9D2 A8Y9F1 A8Y9G0 A8Y9G1 A8Y9G2 A8Y9G3 A8Y9G8 A8Y9H8 A8Y9I2
A8YD90 A8YER2 A8YIE1 A8YMJ4 A8YTB0 A8YTB1 A8YTD2 A8YTD3 A8YTD9 A8YTE3 A8YTF1
A8YTF2 A8YTF6 A8YTF7 A8YTF9 A8YTH7 A8YTH8 A8YTH9 A8YTI0 A8YTI1 A8YTI2 A8YTJ0
A8YTJ1 A8YTJ2 A8YTJ4 A8YTK4 A8YTK7 A8YTN8 A8YTQ6 A8YTQ7 A8YTQ9 A8YTR0 A8YTR6
A8YTT2 A8YTT9 A8YTZ3 A8YTZ6 A8YTZ7 A8YU29 A8YU37 A8YU42 A8YU90 A8YUA3 A8YUC7
A8YUD6 A8YUD8 A8YUD9 A8YUE3 A8YUE4 A8YUE7 A8YUF1 A8YUF8 A8YUH0 A8YUI5 A8YUJ1
A8YUJ4 A8YUL2 A8YUL5 A8YUN5 A8YUN6 A8YUN8 A8YUP3 A8YUP7 A8YUQ2 A8YUQ8 A8YUR0
A8YUR4 A8YUR9 A8YUS2 A8YUS3 A8YUS4 A8YUS5 A8YUT1 A8YUT2 A8YUT3 A8YUT4 A8YUW0
A8YUY2 A8YUZ6 A8YUZ8 A8YUZ9 A8YV13 A8YV14 A8YV16 A8YV19 A8YV22 A8YV35 A8YV39
A8YV42 A8YV45 A8YV48 A8YV51 A8YVD9 A8YVH1 A8YVJ2 A8YVJ3 A8YVJ7 A8YVK0 A8YVK1
A8YVM4 A8YVM6 A8YVM8 A8YVN1 A8YVN2 A8YVP7 A8YVQ3 A8YVQ6 A8YVQ7 A8YVR1 A8YVR3
A8YVR6 A8YVR7 A8YVR8 A8YVR9 A8YVT1 A8YVT2 A8YVT3 A8YVT4 A8YVV4 A8YVV8 A8YVW4
A8YVW8 A8YVW9 A8YVX0 A8YVX6 A8YVX7 A8YVY9 A8YVZ3 A8YVZ5 A8YVZ6 A8YVZ9 A8YW41
A8YW44 A8YW47 A8YW49 A8YW53 A8YW74 A8YW78 A8YW83 A8YW85 A8YWA5 A8YWB0 A8YWB7
A8YWC6 A8YWD3 A8YWE1 A8YWE4 A8YWE6 A8YWG1 A8YWH0 A8YWL2 A8YWL3 A8YWM6 A8YWP6
A8YWQ7 A8YWU4 A8YWU7 A8YWU9 A8YX24 A8YX30 A8YX55 A8YXD7 A8YXF1 A8YXF8 A8YXG4
A8YXH9 A8YXI3 A8YXI6 A8YXJ3 A8YXJ8 A8YXJ9 A8YXK1 A8YXK2 A8YXK3 A8YXK4 A8YXK5
A8YXK6 A8YXK7 A8YXK8 A8YXK9 A8YXL0 A8YXL1 A8YXL2 A8YXL3 A8YXL4 A8YXL5 A8YXL6
A8YXL7 A8YXL9 A8YXM0 A8YXM2 A8YXM3 A8YXM4 A8YXM6 A8YXM7 A8YXM8 A8YXM9 A8YXN0
A8YXN1 A8YXZ5 A8YY02 A8YY15 A8YY36 A8YY39 A8YY45 A8YY79 A8YY80 A8YY81 A8YY83
A8YY85 A8YY92 A8YY93 A8YYC5 A8YYC6 A8YYF6 A8YYF9 A8YYG0 A8YYG1 A8YYG4 A8YYH5
A8YYR9 A8YYS0 A8YYS1 A8YYS2 A8YYS3 A8YYS4 A8YYS7 A8YYT1 A8YYT2 A8YYT8 A8YYU7
A8YZ41 A8YZ65 A8YZA3 A8YZD9 A8YZE2 A8YZI1 A8YZJ1 A8YZJ3 A8YZL0 A8YZL5 A8YZL6
A8YZM5 A8YZM7 A8YZN2 A8YZN5 A8YZN6 A8YZN7 A8YZN8 A8YZP0 A8YZP3 A8YZP4 A8YZP5
A8YZP8 A8YZR4 A8YZR7 A8YZS8 A8YZT4 A8YZU0 A8YZV2 A8YZY3 A8YZZ5 A8Z000 A8Z002
A8Z012 A8Z018 A8Z036 A8Z042 A8Z044 A8Z045 A8Z066 A8Z067 A8Z072 A8Z073 A8Z076
A8Z0A3 A8Z0A8 A8Z0B0 A8Z0B9 A8Z0C4 A8Z0D7 A8Z0H0 A8Z0K3 A8Z0M2 A8Z0N3 A8Z0Q8
A8Z0R1 A8Z0V0 A8Z0V1 A8Z0X3 A8Z0X4 A8Z0Y8 A8Z0Z1 A8Z117 A8Z190 A8Z1A1 A8Z1A2
A8Z1A9 A8Z1D4 A8Z1D8 A8Z1H7 A8Z1I3 A8Z1J1 A8Z1J8 A8Z1K5 A8Z1K8 A8Z1L1 A8Z1L3
A8Z1M9 A8Z1N7 A8Z1P7 A8Z1Q5 A8Z1Q8 A8Z1Q9 A8Z1R5 A8Z1R9 A8Z1S5 A8Z1S8 A8Z1T2
A8Z1V6 A8Z1W3 A8Z1W6 A8Z1W7 A8Z1X0 A8Z1X3 A8Z213 A8Z216 A8Z219 A8Z228 A8Z242
A8Z244 A8Z245 A8Z2D7 A8Z2F7 A8Z2F8 A8Z2G0 A8Z2G2 A8Z2G7 A8Z2G8 A8Z2G9 A8Z2H0

A8Z2H1 A8Z2H2 A8Z2H3 A8Z2H5 A8Z2I4 A8Z2I8 A8Z2J1 A8Z2J3 A8Z2J4 A8Z2J5 A8Z2J6
A8Z2J8 A8Z2K2 A8Z2L4 A8Z2M2 A8Z2M7 A8Z2N1 A8Z2N6 A8Z2P6 A8Z2Q0 A8Z2R3 A8Z2R4
A8Z2R5 A8Z2R8 A8Z2S7 A8Z2T4 A8Z305 A8Z324 A8Z325 A8Z326 A8Z330 A8Z332 A8Z333
A8Z334 A8Z335 A8Z338 A8Z339 A8Z340 A8Z341 A8Z342 A8Z343 A8Z344 A8Z345 A8Z346
A8Z347 A8Z348 A8Z349 A8Z366 A8Z367 A8Z372 A8Z378 A8Z385 A8Z386 A8Z387 A8Z388
A8Z389 A8Z390 A8Z391 A8Z3B1 A8Z3F5 A8Z3F9 A8Z3J8 A8Z3J9 A8Z3K5 A8Z3L8 A8Z3L9
A8Z3M0 A8Z3M4 A8Z3M7 A8Z3N1 A8Z3N5 A8Z3N7 A8Z3P0 A8Z3P7 A8Z3Q2 A8Z3Q4 A8Z3R9
A8Z3S4 A8Z3S5 A8Z3T1 A8Z3T2 A8Z3T3 A8Z3T4 A8Z3T5 A8Z3T6 A8Z3T8 A8Z3U3 A8Z3U5
A8Z3U9 A8Z3V0 A8Z3V1 A8Z3V3 A8Z3V4 A8Z3X3 A8Z3X4 A8Z3X5 A8Z3Z0 A8Z400 A8Z402
A8Z406 A8Z425 A8Z426 A8Z429 A8Z434 A8Z437 A8Z442 A8Z444 A8Z445 A8Z446 A8Z448
A8Z450 A8Z454 A8Z457 A8Z462 A8Z463 A8Z464 A8Z468 A8Z474 A8Z476 A8Z477 A8Z491
A8Z497 A8Z4A3 A8Z4A4 A8Z4A5 A8Z4A9 A8Z4B4 A8Z4B7 A8Z4B8 A8Z4B9 A8Z4C0 A8Z4C1
A8Z4C5 A8Z4D3 A8Z4D5 A8Z4D8 A8Z4E3 A8Z4E8 A8Z4E9 A8Z4F3 A8Z4F4 A8Z4F5 A8Z4F6
A8Z4F9 A8Z4G7 A8Z4H5 A8Z4I6 A8Z4J3 A8Z4K0 A8Z4K2 A8Z4L2 A8Z4L3 A8Z4L9 A8Z4T2
A8Z4T3 A8Z4V2 A8Z4V6 A8Z4V9 A8Z4W0 A8Z4W2 A8Z4W3 A8Z4W5 A8Z4W8 A8Z4Y6 A8Z519
A8Z522 A8Z523 A8Z534 A8Z536 A8Z563 A8Z581 A8Z5A4 A8Z5D0 A8Z5H3 A8Z5H4 A8Z5H6
A8Z5H9 A8Z5I0 A8Z5J0 A8Z5J1 A8Z5Q4 A8Z5Q5 A8Z5R9 A8Z5S7 A8Z5S8 A8Z5T2 A8Z5T3
A8Z5T7 A8Z5T8 A8Z5V2 A8Z5V5 A8Z5V7 A8Z5V8 A8Z5V9 A8Z5W0 A8Z5W4 A8Z5X0 A8Z5X3
A8Z5Y2 A8Z5Z0 A8Z5Z2 A8Z608 A8Z609 A8Z618 A8Z619 A8Z621 A8Z624 A8Z629 A8Z631
A8Z640 A8Z642 A8Z651 A8Z653 A8Z655 A8Z660 A8Z661 A8Z664 A8Z665 A8Z666 A8Z668
A8Z671 A8Z672 A8Z673 A8Z674 A8Z675 A8Z676 A8Z677 A8Z678 A8Z680 A8Z681 A8Z683
A8Z687 A8Z688 A8Z689 A8Z690 A8Z694 A8Z6A5 A8Z6D0 A8Z6D5 A8Z6D9 A8Z6F5 A8Z6F6
A8Z6F7 A8Z6G0 A8Z6G1 A8Z6I5 A8Z6I6 A8Z6J5 A8Z6K1 A8Z6L3 A8Z6M6 A8Z6P5 A8Z6P6
A8ZNW7 A8ZQ66 A8ZRQ0 A8ZRQ2 A8ZRQ3 A8ZRV0 A8ZRW3 A8ZRW7 A8ZRX7 A8ZRX9 A8ZRY0
A8ZRY1 A8ZRY3 A8ZRZ2 A8ZRZ3 A8ZS81 A8ZSB6 A8ZSE7 A8ZSH8 A8ZSP8 A8ZSP9 A8ZT17
A8ZT18 A8ZT68 A8ZT78 A8ZTF0 A8ZTL5 A8ZTL7 A8ZTL8 A8ZTM1 A8ZTM3 A8ZTM6 A8ZTP9
A8ZTQ4 A8ZTQ7 A8ZTR0 A8ZTS7 A8ZTU6 A8ZTU7 A8ZTV1 A8ZTV3 A8ZU05 A8ZU12 A8ZU38
A8ZU47 A8ZU48 A8ZUC6 A8ZUC7 A8ZUH7 A8ZUH8 A8ZUJ6 A8ZUJ9 A8ZUK5 A8ZUK6 A8ZUN0
A8ZUR6 A8ZUR7 A8ZUS4 A8ZUS7 A8ZUS9 A8ZUU5 A8ZUU6 A8ZV12 A8ZV18 A8ZV23 A8ZV25
A8ZV50 A8ZV51 A8ZV52 A8ZV53 A8ZV54 A8ZV57 A8ZV58 A8ZV59 A8ZV60 A8ZV61 A8ZV62
A8ZV63 A8ZV64 A8ZV65 A8ZV66 A8ZV67 A8ZV68 A8ZV69 A8ZV70 A8ZV72 A8ZV73 A8ZV75
A8ZV76 A8ZV79 A8ZV80 A8ZV81 A8ZV82 A8ZV83 A8ZV98 A8ZVC5 A8ZVD2 A8ZVH2 A8ZVH5
A8ZVI6 A8ZVP3 A8ZVP4 A8ZVQ3 A8ZW25 A8ZW56 A8ZW65 A8ZWA3 A8ZWE7 A8ZWG2 A8ZWG5
A8ZWM4 A8ZWP0 A8ZWQ4 A8ZWU1 A8ZWU7 A8ZX50 A8ZXB6 A8ZXB7 A8ZXB8 A8ZXG6 A8ZXG7
A8ZXG8 A8ZXH5 A8ZXL1 A8ZXL3 A8ZXR7 A8ZXR8 A8ZXS1 A8ZXV5 A8ZXW1 A8ZXW2 A8ZXW5
A8ZXX1 A8ZXX2 A8ZXZ4 A8ZY67 A8ZYA5 A8ZYC8 A8ZYF3 A8ZZ59 A8ZZ60 A8ZZ63 A8ZZ65
A8ZZ67 A8ZZ70 A8ZZ71 A8ZZ72 A8ZZG8 A8ZZH7 A8ZZJ0 A8ZZU9 A8ZZW7 A8ZZX0 A8ZZX1
A8ZZX4 A9A043 A9A059 A9A078 A9A079 A9A099 A9A0A5 A9A0B2 A9A0B5 A9A0E0 A9A0F0
A9A0H0 A9A0L3 A9A0L5 A9A0L9 A9A0S9 A9A0U3 A9A110 A9A115 A9A135 A9A141 A9A148
A9A1A0 A9A1A1 A9A1D3 A9A1F0 A9A1F7 A9A1K6 A9A1K7 A9A1T2 A9A1Y1 A9A215 A9A217
A9A231 A9A233 A9A235 A9A298 A9A2B1 A9A2E4 A9A2G6 A9A2Q5 A9A2Q6 A9A2T9 A9A2U7
A9A2X4 A9A2Y3 A9A2Y8 A9A2Z3 A9A305 A9A323 A9A331 A9A389 A9A3A9 A9A3J7 A9A3K1
A9A3L9 A9A3M9 A9A3N4 A9A3N5 A9A3T2 A9A3X2 A9A3Y9 A9A413 A9A423 A9A428 A9A432
A9A436 A9A450 A9A458 A9A492 A9A498 A9A4A1 A9A4B0 A9A4N5 A9A4S0 A9A4V0 A9A4V3

A9A541 A9A555 A9A557 A9A565 A9A582 A9A592 A9A597 A9A5C8 A9A5C9 A9A5G9 A9A5I0
A9A5I2 A9A5I4 A9A5I5 A9A5I6 A9A5I7 A9A5J0 A9A5J1 A9A5K6 A9A5K8 A9A5S1 A9A5W3
A9A5W9 A9A5X0 A9A5X1 A9A5Y7 A9A623 A9A624 A9A655 A9A656 A9A661 A9A697 A9A698
A9A699 A9A6A5 A9A6B5 A9A6B6 A9A6B9 A9A6C2 A9A6C3 A9A6C8 A9A6D8 A9A6I9 A9A6J0
A9A6J1 A9A6K3 A9A6K6 A9A6L5 A9A6L6 A9A6L7 A9A6M7 A9A6M9 A9A6N2 A9A6N4 A9A6N6
A9A6P4 A9A6P5 A9A6Q2 A9A6Q7 A9A6R4 A9A756 A9A787 A9A788 A9A794 A9A799 A9A7A5
A9A7A8 A9A7B4 A9A7B6 A9A7B8 A9A7D0 A9A7E4 A9A7E6 A9A7E7 A9A7M3 A9A7Q1 A9A7S5
A9A7T0 A9A7T1 A9A802 A9A807 A9A811 A9A813 A9A817 A9A828 A9A836 A9A837 A9A838
A9A840 A9A846 A9A848 A9A852 A9A871 A9A873 A9A875 A9A885 A9A8A9 A9A8B2 A9A8D5
A9A8D7 A9A8E3 A9A8E9 A9A8H9 A9A8I0 A9A8I8 A9A8I9 A9A8J8 A9A8K3 A9A8K6 A9A8L2
A9A8L9 A9A8M9 A9A8N0 A9A8N1 A9A8N2 A9A8N5 A9A8P0 A9A8P1 A9A8Q0 A9A8Q7 A9A8R5
A9A8S1 A9A8S8 A9A8T5 A9A8U1 A9A8V2 A9A8V7 A9A8V8 A9A8X7 A9A8Y5 A9A906 A9A907
A9A909 A9A916 A9A923 A9A945 A9A947 A9A972 A9A978 A9A979 A9A983 A9A986 A9A9B4
A9A9B5 A9A9B6 A9A9B7 A9A9B8 A9A9F2 A9A9F5 A9A9H3 A9A9J1 A9A9J3 A9A9J5 A9A9L8
A9A9M4 A9A9N0 A9A9N1 A9A9N2 A9A9N6 A9A9N7 A9A9N8 A9A9P2 A9A9P4 A9A9P5 A9A9P7
A9A9P8 A9A9P9 A9A9Q1 A9A9Q2 A9A9Q4 A9A9Q7 A9A9Q9 A9A9R0 A9A9R2 A9A9R9 A9A9S6
A9A9T9 A9A9U0 A9A9U2 A9A9U3 A9A9U4 A9A9U5 A9A9U8 A9A9U9 A9A9V3 A9A9W1 A9A9Y7
A9A9Z3 A9AA03 A9AA06 A9AA25 A9AA28 A9AA32 A9AA46 A9AA47 A9AA58 A9AA59 A9AA73
A9AA75 A9AA77 A9AA90 A9AA91 A9AA96 A9AA97 A9AAA1 A9AAA4 A9AAA5 A9AAA7 A9AAB0
A9AAC4 A9AAG2 A9AAG9 A9AAH0 A9AAH5 A9AAK2 A9AAN8 A9AAP2 A9AAP6 A9AAR1 A9AAR4
A9AAR7 A9AAS2 A9AAT7 A9AAU1 A9AAU4 A9AAU6 A9AAW1 A9AAX4 A9AAZ2 A9AAZ7 A9AAZ8
A9AB01 A9AB07 A9AB08 A9AB09 A9AB16 A9AB20 A9AB21 A9AB23 A9AB27 A9AB37 A9AB46
A9AB64 A9AB65 A9AB67 A9AB97 A9ABB5 A9ABD4 A9ABD5 A9ABD7 A9ABF6 A9ABF9 A9ABG9
A9ABP3 A9ABQ1 A9ABS7 A9ABZ7 A9AC16 A9AC32 A9AC42 A9AC50 A9AC51 A9AC62 A9AC69
A9ACI3 A9ACK7 A9ACL5 A9ACL8 A9ACU3 A9ACU4 A9ACU5 A9ACX9 A9AD18 A9AD19 A9ADC9
A9ADD1 A9ADF3 A9ADI1 A9ADI2 A9ADI3 A9ADI4 A9ADI5 A9ADI6 A9ADI8 A9ADI9 A9ADJ2
A9ADJ3 A9ADJ4 A9ADJ5 A9ADJ6 A9ADJ7 A9ADJ8 A9ADJ9 A9ADK0 A9ADK1 A9ADK2 A9ADK3
A9ADK4 A9ADK5 A9ADK6 A9ADK7 A9ADK8 A9ADK9 A9ADL0 A9ADL1 A9ADL2 A9ADL5 A9ADL6
A9ADL7 A9ADL8 A9ADL9 A9ADM0 A9ADM4 A9ADN1 A9ADP0 A9ADS8 A9ADS9 A9ADT0 A9ADT1
A9ADT7 A9ADU8 A9ADV3 A9ADV5 A9ADV6 A9ADV9 A9ADW3 A9AE44 A9AE45 A9AE46 A9AE52
A9AE58 A9AEF0 A9AEG7 A9AEK0 A9AEN5 A9AEQ4 A9AEQ7 A9AER8 A9AES0 A9AES3 A9AEU0
A9AEW9 A9AEY6 A9AEY7 A9AEY9 A9AEZ2 A9AEZ5 A9AEZ6 A9AF06 A9AF24 A9AF25 A9AF70
A9AF71 A9AF72 A9AF74 A9AFB4 A9AFC0 A9AFC7 A9AFE4 A9AFF8 A9AFH2 A9AFI4 A9AFI6
A9AFI7 A9AFM6 A9AFP3 A9AFT0 A9AFT3 A9AFU0 A9AG13 A9AG80 A9AGA7 A9AGA9 A9AGB5
A9AGB8 A9AGC0 A9AGC4 A9AGC5 A9AGE5 A9AGH1 A9AGH3 A9AGI1 A9AGJ4 A9AGJ5 A9AGJ8
A9AGK1 A9AGK3 A9AGM9 A9AGN3 A9AGN6 A9AGP7 A9AGQ1 A9AGR3 A9AGS9 A9AGU6 A9AGU7
A9AGV1 A9AGW0 A9AGY0 A9AGY1 A9AGZ3 A9AGZ7 A9AH00 A9AH01 A9AH05 A9AH21 A9AH63
A9AH70 A9AH71 A9AH72 A9AH73 A9AH76 A9AH79 A9AH80 A9AHA9 A9AHB2 A9AHD4 A9AHD5
A9AHG9 A9AHJ2 A9AHJ3 A9AHJ8 A9AHK2 A9AHN6 A9AHQ3 A9AHQ6 A9AHS8 A9AHT0 A9AHY0
A9AI41 A9AI56 A9AI57 A9AI59 A9AI60 A9AI61 A9AI62 A9AI82 A9AI94 A9AI95 A9AI97
A9AIJ8 A9AIK9 A9AIL5 A9AIL7 A9AIM0 A9AIN0 A9AIP4 A9AIT5 A9AIT6 A9AJ14 A9AJ16
A9AJ18 A9AJ24 A9AJ25 A9AJ43 A9AJ67 A9AJ68 A9AJC5 A9AJC7 A9AJD5 A9AJF1 A9AJF2
A9AJJ5 A9AJJ8 A9AJK0 A9AJK1 A9AJK3 A9AJN1 A9AJN4 A9AJN9 A9AJP0 A9AJQ9 A9AJR1
A9AJR8 A9AJS6 A9AJS7 A9AJW1 A9AJW3 A9AJW4 A9AJX3 A9AKB0 A9AKG8 A9AKH1 A9AL79

A9ALD1 A9ALD3 A9AMA5 A9AMB3 A9AMD5 A9AMZ9 A9AN00 A9AN01 A9AN40 A9ANA0 A9ANA1
A9ANA4 A9ANP0 A9AQ22 A9AQJ3 A9AQW9 A9AR15 A9AR24 A9AR64 A9ARG7 A9AST0 A9AT43
A9AUB8 A9AUH0 A9AUN9 A9AUP1 A9AUT1 A9AUX0 A9AV71 A9AVM2 A9AW17 A9AW39 A9AW42
A9AWD5 A9AWD6 A9AWD7 A9AWF4 A9AWL1 A9AWY7 A9AXA6 A9AXC6 A9AXD6 A9AXD9 A9AXI5
A9AXJ1 A9AXK0 A9AXP4 A9AXS2 A9AXY4 A9AY15 A9AY55 A9AY83 A9AY92 A9AYA7 A9AYB7
A9AYE8 A9AYL1 A9AYM5 A9AYX1 A9AZ03 A9AZ11 A9AZ32 A9AZ72 A9AZD9 A9AZE0 A9AZE6
A9AZE7 A9AZE8 A9AZE9 A9AZL0 A9AZM5 A9AZP9 A9AZS3 A9AZY8 A9AZZ2 A9B055 A9B076
A9B090 A9B0A2 A9B0I2 A9B0N2 A9B0N4 A9B0X2 A9B119 A9B129 A9B148 A9B168 A9B182
A9B1D4 A9B1E0 A9B1H0 A9B1N9 A9B1Q2 A9B2B7 A9B2N4 A9B2Q5 A9B2Q6 A9B2U4 A9B2Z8
A9B2Z9 A9B302 A9B312 A9B327 A9B356 A9B392 A9B3C5 A9B3N9 A9B3P8 A9B3R2 A9B3S3
A9B3W0 A9B406 A9B412 A9B414 A9B415 A9B416 A9B417 A9B419 A9B421 A9B422 A9B423
A9B424 A9B426 A9B427 A9B429 A9B430 A9B434 A9B435 A9B436 A9B437 A9B440 A9B457
A9B458 A9B461 A9B464 A9B496 A9B4H8 A9B4J6 A9B4R2 A9B4R4 A9B4S4 A9B4U5 A9B518
A9B519 A9B527 A9B543 A9B546 A9B550 A9B562 A9B567 A9B5C4 A9B5E6 A9B5E9 A9B5G7
A9B5G8 A9B5H2 A9B5I2 A9B5I3 A9B5I5 A9B5N1 A9B5N7 A9B5T5 A9B5V4 A9B5Y6 A9B660
A9B689 A9B6A4 A9B6D5 A9B6H3 A9B6H5 A9B6J1 A9B6J3 A9B6M7 A9B6Q3 A9B6S7 A9B6X1
A9B6X5 A9B732 A9B745 A9B746 A9B775 A9B7I9 A9B805 A9B890 A9B891 A9B8B2 A9B8B9
A9B8G1 A9B8G2 A9B9E5 A9B9E7 A9B9H1 A9B9I2 A9B9L0 A9B9L4 A9B9L9 A9B9W1 A9B9W7
A9B9X0 A9B9X1 A9B9X8 A9B9Y0 A9B9Z7 A9BA44 A9BA66 A9BAC1 A9BAC7 A9BAD0 A9BAE1
A9BAE3 A9BAE4 A9BAE9 A9BAF1 A9BAG3 A9BAH6 A9BAI8 A9BAJ3 A9BAK7 A9BAN4 A9BAN9
A9BAP2 A9BAQ7 A9BAR3 A9BAS8 A9BAS9 A9BAW0 A9BAX2 A9BAY7 A9BAY8 A9BAZ9 A9BB12
A9BB39 A9BB43 A9BB49 A9BB56 A9BBH5 A9BBH9 A9BBI9 A9BBJ0 A9BBJ6 A9BBJ7 A9BBN4
A9BBQ1 A9BBQ7 A9BBR3 A9BBS2 A9BBT2 A9BBV1 A9BBV5 A9BBW2 A9BBW6 A9BBW8 A9BC43
A9BCB8 A9BCC5 A9BCH2 A9BCH4 A9BCH5 A9BCH6 A9BCH8 A9BCI0 A9BCI2 A9BCI5 A9BCJ1
A9BCJ9 A9BCK0 A9BCK1 A9BCK2 A9BCK3 A9BCL8 A9BCL9 A9BCM0 A9BCM2 A9BCM3 A9BCM5
A9BCM6 A9BCM7 A9BCM8 A9BCN2 A9BCN3 A9BCN4 A9BCN5 A9BCN7 A9BCN9 A9BCP0 A9BCP1
A9BCP2 A9BCP3 A9BCP4 A9BCP6 A9BCP7 A9BCQ0 A9BCR1 A9BCS9 A9BCT2 A9BCT3 A9BCV9
A9BCW6 A9BCX0 A9BCY2 A9BCY7 A9BD24 A9BD32 A9BD34 A9BD43 A9BD47 A9BD54 A9BD59
A9BD62 A9BD69 A9BD80 A9BD81 A9BD87 A9BD93 A9BD97 A9BD99 A9BDA1 A9BDA5 A9BDA6
A9BDB8 A9BDC5 A9BDD8 A9BDD9 A9BDF0 A9BDF8 A9BDG4 A9BDG5 A9BDG6 A9BDG7 A9BDI4
A9BDJ2 A9BDK0 A9BDK3 A9BDK6 A9BDM7 A9BDM9 A9BDN4 A9BDN6 A9BDN8 A9BDP3 A9BDP7
A9BDR2 A9BDR3 A9BDS9 A9BDW6 A9BDW9 A9BDX7 A9BDY5 A9BE04 A9BE09 A9BE13 A9BE20
A9BE25 A9BE29 A9BE38 A9BE47 A9BE76 A9BE95 A9BEA2 A9BEA5 A9BEB6 A9BEC8 A9BED0
A9BEE2 A9BEE4 A9BEE9 A9BEG6 A9BEG7 A9BEG8 A9BEH3 A9BEI3 A9BEI4 A9BEI9 A9BEK6
A9BEN2 A9BER7 A9BER9 A9BEU2 A9BEU9 A9BEV2 A9BEX0 A9BEX2 A9BEX3 A9BEZ8 A9BF00
A9BF03 A9BF05 A9BF21 A9BF22 A9BF32 A9BF33 A9BF34 A9BF35 A9BF36 A9BF37 A9BF40
A9BF44 A9BF51 A9BFB6 A9BFB9 A9BFC0 A9BFC2 A9BFH2 A9BFQ1 A9BFS6 A9BFT1 A9BFT2
A9BFU0 A9BFU2 A9BFW1 A9BFY9 A9BFZ1 A9BFZ2 A9BFZ3 A9BFZ4 A9BFZ5 A9BFZ7 A9BFZ9
A9BG00 A9BG02 A9BG04 A9BG05 A9BG28 A9BG54 A9BG68 A9BGA8 A9BGL0 A9BGL2 A9BGN7
A9BGP6 A9BGS3 A9BGS7 A9BGT8 A9BGT9 A9BGU1 A9BGV2 A9BGV6 A9BGV7 A9BGW2 A9BGW7
A9BGZ9 A9BH07 A9BH33 A9BH64 A9BH65 A9BH72 A9BH73 A9BH95 A9BH96 A9BH97 A9BH98
A9BHA0 A9BHA1 A9BHA2 A9BHA4 A9BHA5 A9BHA7 A9BHA8 A9BHA9 A9BHB0 A9BHB5 A9BHB6
A9BHC0 A9BHC2 A9BHC5 A9BHF0 A9BHG1 A9BHH8 A9BHH9 A9BHI9 A9BHJ0 A9BHK3 A9BHK4
A9BHK7 A9BHK9 A9BHL8 A9BHL9 A9BHP7 A9BHQ1 A9BHQ2 A9BHQ4 A9BHQ5 A9BHQ6 A9BHR2

A9BHR6 A9BHU3 A9BHV0 A9BHV4 A9BHV5 A9BHW9 A9BHX2 A9BHX3 A9BHX8 A9BHZ7 A9BI29
A9BI45 A9BI46 A9BI93 A9BI99 A9BIA3 A9BIC0 A9BID3 A9BIE9 A9BIH4 A9BIJ5 A9BIJ7
A9BIK8 A9BIM4 A9BIS3 A9BIS4 A9BIS5 A9BIU9 A9BIV8 A9BIV9 A9BIX4 A9BIX9 A9BJ07
A9BJ15 A9BJ18 A9BJ54 A9BJ81 A9BJB2 A9BJB9 A9BJC3 A9BJC4 A9BJD6 A9BJE0 A9BJF6
A9BJH4 A9BJH6 A9BJK2 A9BJN0 A9BJQ0 A9BJV3 A9BJX5 A9BJY3 A9BJZ4 A9BJZ9 A9BK05
A9BK06 A9BM04 A9BM51 A9BM59 A9BM60 A9BM65 A9BME3 A9BME9 A9BMK7 A9BML0 A9BML8
A9BMM9 A9BMN1 A9BMN2 A9BMP0 A9BMR1 A9BMU3 A9BMU4 A9BMU8 A9BMU9 A9BMV6 A9BMV7
A9BMZ9 A9BN39 A9BN87 A9BNB6 A9BNB8 A9BNB9 A9BNG0 A9BNG1 A9BNG3 A9BNG4 A9BNG5
A9BNG6 A9BNH1 A9BNH5 A9BNH8 A9BNI6 A9BNJ3 A9BNJ5 A9BNK8 A9BNL7 A9BNL9 A9BNR6
A9BNS8 A9BNS9 A9BNU9 A9BNV0 A9BNV7 A9BP03 A9BP04 A9BP06 A9BP16 A9BP18 A9BP19
A9BP37 A9BP60 A9BP65 A9BP66 A9BP67 A9BP68 A9BP69 A9BP70 A9BPN0 A9BPR3 A9BPR4
A9BPR7 A9BPR8 A9BPR9 A9BPS1 A9BPS2 A9BPS3 A9BPS5 A9BPS6 A9BPS7 A9BPT7 A9BPT8
A9BPZ7 A9BQ97 A9BQA5 A9BQB0 A9BQJ1 A9BQN8 A9BQU2 A9BQY2 A9BR02 A9BR94 A9BR95
A9BR96 A9BR97 A9BR98 A9BR99 A9BRD7 A9BRF0 A9BRF4 A9BRT1 A9BRV2 A9BRW1 A9BRW2
A9BRW3 A9BRW4 A9BRW5 A9BRW6 A9BRW7 A9BRW8 A9BRW9 A9BRX0 A9BRX2 A9BRX3 A9BRX4
A9BRX5 A9BRX6 A9BS05 A9BS06 A9BS40 A9BS60 A9BS67 A9BSM4 A9BTA2 A9BTA7 A9BTN3
A9BU40 A9BUB4 A9BUC1 A9BUC2 A9BUC4 A9BUC7 A9BUC8 A9BUE9 A9BUG5 A9BUG7 A9BUG8
A9BUJ5 A9BUJ7 A9BUJ8 A9BUK7 A9BUK8 A9BUL6 A9BUQ1 A9BUZ3 A9BV01 A9BV02 A9BV10
A9BV13 A9BVD2 A9BVE1 A9BVK0 A9BVK1 A9BVN7 A9BVZ2 A9BVZ6 A9BVZ7 A9BW85 A9BWD4
A9BWE1 A9BWI1 A9BWI3 A9BWI7 A9BWT7 A9BWU9 A9BX71 A9BX80 A9BXA1 A9BXA2 A9BXD6
A9BXF7 A9BXL2 A9BXL3 A9BXM7 A9BXV6 A9BXZ7 A9BXZ8 A9BY30 A9BY74 A9BY83 A9BY86
A9BYK1 A9BYK5 A9BZ45 A9BZG3 A9BZW2 A9BZW4 A9BZY6 A9BZY8 A9C011 A9C020 A9C021
A9C0A8 A9C0B9 A9C177 A9C178 A9C179 A9C184 A9C1G5 A9C1G8 A9C1M4 A9C1U7 A9C1U9
A9C1Z0 A9C1Z3 A9C2L1 A9C2R2 A9C3B1 A9C3C8 A9C3D0 A9C3D1 A9C3D2 A9C3D4 A9C3F6
A9C3G0 A9C3G2 A9C3H6 A9C3W3 A9CB27 A9CB34 A9CB42 A9CB60 A9CB82 A9CB83 A9CB84
A9CB85 A9CB86 A9CB88 A9CB89 A9CB90 A9CB91 A9CB92 A9CB93 A9CB94 A9CB95 A9CB96
A9CB97 A9CB98 A9CB99 A9CBA1 A9CEQ7 A9CEQ8 A9CES3 A9CES4 A9CES5 A9CES6 A9CF16
A9CFA8 A9CFX5 A9CG70 A9CH01 A9CH28 A9CH39 A9CHB2 A9CHM9 A9CHR3 A9CHW6 A9CIB3
A9CIG1 A9CJ88 A9CJC9 A9CK16 A9CK30 A9CK39 A9CKA8 A9CKB4 A9CKC4 A9CKM5 A9CKQ7
A9CKY2 A9CPT4 A9CRA7 A9CRB8 A9CRJ7 A9CS50 A9CS74 A9CSU5 A9CSZ1 A9CTE0 A9CTP8
A9D2P7 A9EN94 A9ENT2 A9EPV3 A9ER52 A9ERU2 A9ESS5 A9ESS9 A9ESU7 A9ET18 A9ETA6
A9ETB7 A9ETC1 A9ETD1 A9ETE3 A9ETE8 A9ETF4 A9ETZ9 A9EUF9 A9EVN4 A9EXF3 A9EXH2
A9EXM5 A9EXT3 A9EYT5 A9EYY7 A9EZV5 A9F0C1 A9F1Y8 A9F3N2 A9F3N7 A9F3P4 A9F3W3
A9F8E5 A9F8L0 A9F8Z2 A9F8Z6 A9F907 A9FBU8 A9FCX7 A9FCY0 A9FCY5 A9FD62 A9FD66
A9FD89 A9FDL0 A9FDP5 A9FDP9 A9FFI1 A9FFJ6 A9FG63 A9FG66 A9FGA8 A9FGB5 A9FGB8
A9FGC9 A9FGD2 A9FGF3 A9FGF8 A9FGG6 A9FGH7 A9FGI0 A9FGI9 A9FGJ3 A9FGJ7 A9FGK2
A9FHA5 A9FI02 A9FI06 A9FI25 A9FI59 A9FJ48 A9FJF8 A9FK50 A9FL08 A9FM88 A9FML0
A9FNE0 A9FQE0 A9FQE4 A9FQM4 A9FR00 A9FRC1 A9FRF5 A9FRP1 A9FST8 A9FU01 A9FYZ4
A9FZ87 A9FZA0 A9FZW7 A9G0D6 A9G109 A9G209 A9G5S0 A9G6C7 A9G6E5 A9G6T8 A9G6U6
A9G7W0 A9G7Z8 A9G862 A9G882 A9G896 A9G8M9 A9G9K6 A9G9L1 A9G9R0 A9GB11 A9GB86
A9GBF1 A9GBI6 A9GBQ2 A9GBR2 A9GCQ1 A9GD65 A9GD69 A9GD73 A9GDD3 A9GEX7 A9GF63
A9GFF3 A9GFF6 A9GFL5 A9GFW6 A9GGK7 A9GHU4 A9GHW2 A9GI16 A9GIM9 A9GIN6 A9GIS9
A9GIT1 A9GK58 A9GKM6 A9GL99 A9GM36 A9GN39 A9GN91 A9GP79 A9GP84 A9GP90 A9GPH2
A9GQU6 A9GR94 A9GRA1 A9GRA4 A9GRA5 A9GRA8 A9GRB1 A9GRB4 A9GS95 A9GT78 A9GU24

A9GUD0 A9GUD3 A9GVA4 A9GVS4 A9GVS8 A9GW22 A9GW78 A9GWT4 A9GWV1 A9GWW0 A9GXJ8
A9GXK1 A9GXK6 A9GZI5 A9GZJ4 A9GZR7 A9GZS1 A9GZW9 A9GZY1 A9H0B5 A9H0E5 A9H0E8
A9H0F1 A9H0G8 A9H0G9 A9H0J3 A9H0L6 A9H0U2 A9H167 A9H170 A9H1M3 A9H1M6 A9H1N0
A9H253 A9H259 A9H271 A9H298 A9H2B4 A9H311 A9H3A6 A9H3E2 A9H3I6 A9H3I9 A9H3J3
A9H3J7 A9H3K5 A9H3L2 A9H3L5 A9H3L8 A9H3M1 A9H3M4 A9H3N1 A9H3N4 A9H3N9 A9H3P2
A9H3P5 A9H3P8 A9H3Q2 A9H3Q5 A9H3Q8 A9H3R2 A9H3R5 A9H3R6 A9H3R7 A9H3R8 A9H3R9
A9H3S0 A9H3S4 A9H3S7 A9H3T0 A9H3T4 A9H481 A9H4F6 A9H5B7 A9H6E1 A9H863 A9H867
A9H8D2 A9H8E4 A9H8G1 A9H8G4 A9H8G7 A9H9B7 A9H9C9 A9H9I9 A9H9N2 A9HAC0 A9HAV5
A9HAV7 A9HB20 A9HBI8 A9HBJ2 A9HC01 A9HC14 A9HC19 A9HDB1 A9HDQ7 A9HDX7 A9HE10
A9HE13 A9HE16 A9HE41 A9HE84 A9HE98 A9HE99 A9HEA1 A9HEA3 A9HEC4 A9HEC6 A9HEL1
A9HEW1 A9HF10 A9HF18 A9HF21 A9HF26 A9HF29 A9HF35 A9HFM1 A9HFN0 A9HFP5 A9HFY5
A9HGR3 A9HHS1 A9HHY9 A9HI30 A9HI31 A9HI56 A9HIM2 A9HIQ4 A9HIR6 A9HIW2 A9HIW5
A9HJ12 A9HJ48 A9HJ49 A9HJ50 A9HJ57 A9HJ69 A9HJ81 A9HJ86 A9HJF3 A9HJH4 A9HJV3
A9HK37 A9HK92 A9HKQ4 A9HKR2 A9HKR8 A9HKS8 A9HKW6 A9HKY5 A9HL98 A9HLD9 A9HLG2
A9HLQ1 A9HLS1 A9HLY4 A9HMI3 A9HPH6 A9HRF2 A9HRF6 A9HRH6 A9HRJ1 A9HRJ3 A9HRQ0
A9HRQ4 A9HRQ9 A9HRV3 A9HRV7 A9HS05 A9HS49 A9HS54 A9HS58 A9HS61 A9HS64 A9HS66
A9HS91 A9HVA1 A9HVE9 A9HVN4 A9HVX4 A9HW07 A9HW20 A9HWB8 A9HWC3 A9HWC8 A9HWD9
A9HWU4 A9HWU6 A9HWW6 A9HWX2 A9HX42 A9HY11 A9HY12 A9HY25 A9HY51 A9HYE0 A9HYU0
A9HYV0 A9HYZ1 A9HZ10 A9HZ13 A9HZF7 A9HZV6 A9HZV9 A9HZZ2 A9I000 A9I023 A9I042
A9I097 A9I0D6 A9I0E6 A9I0G7 A9I0H0 A9I0H5 A9I0H9 A9I0K4 A9I0M2 A9I0N5 A9I0P4
A9I0U7 A9I174 A9I1G8 A9I1I9 A9I1R3 A9I1S7 A9I1T0 A9I1T6 A9I1U7 A9I1W8 A9I219
A9I237 A9I240 A9I242 A9I246 A9I280 A9I292 A9I295 A9I3K3 A9I3W2 A9I3Y9 A9I3Z9
A9I4I5 A9I4K2 A9I4S5 A9I4V6 A9I4V9 A9I518 A9I5F2 A9I602 A9I682 A9I685 A9I6E7
A9I6G5 A9I6L9 A9I6W1 A9I6W5 A9I6Y3 A9I714 A9I734 A9I7K9 A9I7L6 A9I7R5 A9I7T3
A9I7X7 A9I847 A9I8B8 A9I8C3 A9I8F2 A9I8F5 A9I8M3 A9I8M6 A9IE54 A9IER8 A9IFD9
A9IFF5 A9IFF7 A9IFF9 A9IFG1 A9IFG8 A9IFJ0 A9IFJ5 A9IFL0 A9IFP1 A9IFQ1 A9IGC0
A9IGC3 A9IGC5 A9IGG4 A9IGI5 A9IGJ3 A9IGJ8 A9IGJ9 A9IGW1 A9IGY5 A9IH21 A9IH51
A9IH53 A9IH67 A9IH79 A9IHE2 A9IHH4 A9IHJ7 A9IHP8 A9IHR4 A9IHR6 A9IHR7 A9IHR9
A9IHS0 A9IHS2 A9IHS7 A9IHS8 A9IHT0 A9IHT2 A9IHT3 A9IHT5 A9IHT7 A9IHT8 A9IHU0
A9IHU2 A9II44 A9II47 A9II65 A9II69 A9IIC5 A9IIE0 A9IIG9 A9IIJ2 A9IIK3 A9IIP5
A9IIS3 A9IIY6 A9IIY8 A9IIY9 A9IIZ1 A9IIZ3 A9IIZ4 A9IIZ6 A9IIZ8 A9IIZ9 A9IJ01
A9IJ03 A9IJ08 A9IJ10 A9IJ22 A9IJ25 A9IJ27 A9IJ29 A9IJ30 A9IJ32 A9IJ46 A9IJ48
A9IJ97 A9IJC1 A9IJC3 A9IJI3 A9IJI7 A9IJJ1 A9IJJ7 A9IJN0 A9IJP4 A9IJV4 A9IJZ8
A9IK31 A9IK34 A9IK36 A9IK55 A9IK57 A9IK66 A9IK70 A9IK79 A9IK82 A9IK87 A9IKD2
A9IKD8 A9IKE3 A9IKF3 A9IKK0 A9IKM2 A9IKU8 A9IKX8 A9IKY5 A9IL25 A9IL44 A9IL50
A9IL83 A9ILA7 A9ILC8 A9ILD7 A9ILE7 A9ILE9 A9ILF0 A9ILH7 A9ILI4 A9ILJ1 A9ILJ3
A9ILJ7 A9ILK1 A9ILK4 A9ILL7 A9ILM1 A9ILN1 A9ILQ3 A9ILQ7 A9ILR1 A9ILS3 A9ILT1
A9ILY6 A9IM00 A9IM39 A9IM41 A9IM54 A9IM57 A9IMA1 A9IMA6 A9IMB2 A9IMC0 A9IMC9
A9IMI9 A9IML0 A9IMM3 A9IMR3 A9IMR9 A9IMS2 A9IMS8 A9IMT2 A9IMT5 A9IMU4 A9IMW1
A9IMW7 A9IMX8 A9IMY0 A9IN26 A9IN82 A9INL0 A9INU2 A9INV3 A9INV9 A9IP04 A9IP06
A9IP11 A9IP97 A9IPU1 A9IPU4 A9IQ03 A9IQ32 A9IQ39 A9IQ52 A9IQ57 A9IQ90 A9IQD2
A9IQD9 A9IQF4 A9IQH4 A9IQK6 A9IQK8 A9IQL1 A9IQN4 A9IQP0 A9IQP2 A9IQQ8 A9IQQ9
A9IQR2 A9IQR5 A9IQR9 A9IQY2 A9IQZ6 A9IQZ9 A9IR19 A9IR50 A9IR54 A9IR57 A9IR78
A9IR81 A9IR82 A9IR85 A9IRC2 A9IRC3 A9IRH0 A9IRH6 A9IRH7 A9IRJ9 A9IRK3 A9IRK6

A9IRL0 A9IRL2 A9IRL8 A9IRN0 A9IRN6 A9IRN9 A9IRP2 A9IRQ9 A9IRT7 A9IRU1 A9IRU5
A9IRW8 A9IS02 A9IS21 A9IS38 A9IS43 A9IS79 A9IS87 A9ISA4 A9ISA8 A9ISB8 A9ISC1
A9ISC3 A9ISD9 A9ISE8 A9ISF1 A9ISF5 A9ISF8 A9ISG1 A9ISG4 A9ISJ8 A9ISK1 A9ISK9
A9ISQ6 A9ISS8 A9ISZ5 A9IT47 A9IT50 A9IT77 A9IT92 A9IT96 A9ITA2 A9ITA7 A9ITB0
A9ITB6 A9ITM2 A9ITM7 A9ITN0 A9ITN7 A9ITX2 A9ITX6 A9ITY3 A9IU26 A9IU27 A9IU29
A9IU46 A9IU51 A9IU58 A9IU63 A9IU91 A9IU96 A9IU99 A9IUA2 A9IUA4 A9IUL1 A9IUY4
A9IVB0 A9IVB2 A9IVB9 A9IVC0 A9IVC4 A9IVC5 A9IVD2 A9IVF6 A9IVH1 A9IVH5 A9IVI2
A9IVK2 A9IVK4 A9IVT5 A9IVW5 A9IVX0 A9IVX4 A9IVX6 A9IVY8 A9IVY9 A9IVZ4 A9IVZ5
A9IVZ7 A9IVZ8 A9IVZ9 A9IW02 A9IW03 A9IW04 A9IW06 A9IW07 A9IW08 A9IW11 A9IW12
A9IW13 A9IW16 A9IW17 A9IW18 A9IW20 A9IW23 A9IW24 A9IW25 A9IW26 A9IW27 A9IW28
A9IW31 A9IW33 A9IW34 A9IW67 A9IW77 A9IW82 A9IW95 A9IWA2 A9IWA3 A9IWA4 A9IWB6
A9IWF5 A9IWF8 A9IWH0 A9IWV1 A9IXA1 A9IXC8 A9IXD6 A9IXE1 A9IXE5 A9IXE7 A9IXQ8
A9IXS0 A9IXV7 A9IY10 A9IY12 A9IYI0 A9IYI2 A9IYI8 A9IYK5 A9IYK7 A9IYK9 A9IYN9
A9IYQ7 A9IYR6 A9IYV7 A9IYW1 A9IYY3 A9IYY8 A9IYZ3 A9IZA0 A9IZA2 A9IZB5 A9IZD0
A9IZE1 A9IZF6 A9IZG7 A9IZS4 A9IZV3 A9IZV5 A9IZW1 A9IZW8 A9IZX8 A9IZX9 A9IZY1
A9JPE1 A9JQL9 A9JR23 A9JR78 A9JRC7 A9JRD8 A9JRE2 A9JRI0 A9JRL3 A9JRM0 A9JS51
A9JSB3 A9JSQ8 A9JSR5 A9JTP3 A9JTS5 A9JTS9 A9JTY4 A9JX05 A9JX06 A9JX07 A9JX08
A9KB94 A9KB96 A9KBA0 A9KBB0 A9KBB1 A9KBC1 A9KBD3 A9KBI0 A9KBI1 A9KBI4 A9KBJ2
A9KBJ3 A9KBK2 A9KBL9 A9KBM1 A9KBM2 A9KBM3 A9KBN4 A9KBN5 A9KBN6 A9KBQ4 A9KBR3
A9KBR4 A9KBR6 A9KBS7 A9KBS8 A9KBS9 A9KBT3 A9KBT4 A9KBT7 A9KBT8 A9KBW2 A9KBY0
A9KBY1 A9KBY6 A9KBY7 A9KBZ3 A9KC01 A9KC14 A9KC15 A9KC18 A9KC19 A9KC20 A9KC25
A9KC67 A9KC74 A9KC82 A9KC83 A9KCC7 A9KCD5 A9KCK3 A9KCL1 A9KCP2 A9KCP4 A9KCQ4
A9KCQ7 A9KCR0 A9KCS0 A9KCT0 A9KCT5 A9KCT6 A9KCU2 A9KCX4 A9KCZ5 A9KD05 A9KD06
A9KD07 A9KD08 A9KD09 A9KD10 A9KD12 A9KD13 A9KD15 A9KD16 A9KD17 A9KD19 A9KD20
A9KD21 A9KD22 A9KD23 A9KD24 A9KD25 A9KD26 A9KD27 A9KD28 A9KD29 A9KD30 A9KD31
A9KD32 A9KD33 A9KD34 A9KD35 A9KD36 A9KD37 A9KD39 A9KD40 A9KD41 A9KD42 A9KD43
A9KD63 A9KD67 A9KD75 A9KD87 A9KD88 A9KDA0 A9KDC4 A9KDC5 A9KDD3 A9KDE5 A9KDE7
A9KDE8 A9KDE9 A9KDF8 A9KDG3 A9KDH0 A9KDH9 A9KDJ2 A9KDN5 A9KDP4 A9KDQ8 A9KDS6
A9KDS7 A9KDT8 A9KE14 A9KE31 A9KE32 A9KE65 A9KE66 A9KE87 A9KE95 A9KEA3 A9KEC8
A9KED8 A9KEE7 A9KEE8 A9KEE9 A9KEF0 A9KEG1 A9KEG7 A9KEG8 A9KEI8 A9KEJ4 A9KEJ7
A9KEJ8 A9KEJ9 A9KEL6 A9KEP6 A9KER1 A9KER2 A9KER6 A9KET2 A9KET3 A9KET4 A9KET6
A9KEU8 A9KEV0 A9KEY5 A9KEY9 A9KEZ9 A9KF12 A9KF13 A9KF39 A9KF41 A9KF42 A9KF43
A9KF44 A9KF61 A9KF70 A9KF84 A9KFA0 A9KFA2 A9KFA7 A9KFF3 A9KFG7 A9KFK2 A9KFK5
A9KFK6 A9KFK7 A9KFL5 A9KFL6 A9KFL8 A9KFM0 A9KFP2 A9KFP4 A9KFQ0 A9KFQ4 A9KFS0
A9KFT9 A9KFU3 A9KFU6 A9KFV0 A9KFV6 A9KFW4 A9KFW5 A9KFZ5 A9KG10 A9KG24 A9KG28
A9KG71 A9KG87 A9KG88 A9KG91 A9KG94 A9KGA1 A9KGA3 A9KGB5 A9KGB7 A9KGB8 A9KGB9
A9KGD2 A9KGD3 A9KGD5 A9KGE3 A9KGE6 A9KGL7 A9KGN4 A9KGN7 A9KGQ0 A9KGQ5 A9KGQ8
A9KGR6 A9KGR9 A9KGS5 A9KGS6 A9KGS7 A9KGU5 A9KGX9 A9KGY3 A9KGY4 A9KGY8 A9KGZ8
A9KH04 A9KH12 A9KH14 A9KH17 A9KH21 A9KH27 A9KH30 A9KH31 A9KH99 A9KHA0 A9KHD4
A9KHD9 A9KHE4 A9KHG8 A9KHG9 A9KHK4 A9KHL4 A9KHL6 A9KHN1 A9KHN3 A9KHU4 A9KHX3
A9KHY1 A9KHY6 A9KI94 A9KIA0 A9KIA5 A9KIA6 A9KIB0 A9KIE9 A9KIP3 A9KIR8 A9KIT8
A9KIW5 A9KJ19 A9KJ25 A9KJ26 A9KJ27 A9KJ36 A9KJF4 A9KJF5 A9KJG7 A9KJG8 A9KJG9
A9KJH0 A9KJH1 A9KJH3 A9KJH5 A9KJH6 A9KJH8 A9KJH9 A9KJI0 A9KJI2 A9KJI3 A9KJI4
A9KJI5 A9KJI6 A9KJI7 A9KJI8 A9KJI9 A9KJJ0 A9KJJ1 A9KJJ2 A9KJJ3 A9KJJ4 A9KJJ5

A9KJL3 A9KJL7 A9KJM0 A9KJM3 A9KJR5 A9KJR6 A9KJR7 A9KJR8 A9KJR9 A9KJS0 A9KJS1
A9KK15 A9KK69 A9KK74 A9KK75 A9KKB3 A9KKC8 A9KKE9 A9KKR4 A9KKR5 A9KKS6 A9KKS7
A9KKT0 A9KKT8 A9KKT9 A9KKU0 A9KKU1 A9KKU2 A9KKU7 A9KKZ0 A9KKZ8 A9KL08 A9KL10
A9KL35 A9KL42 A9KL43 A9KL44 A9KL72 A9KL97 A9KLG6 A9KLJ9 A9KLK7 A9KLL5 A9KLL7
A9KLM0 A9KLM2 A9KLM3 A9KLM5 A9KLS2 A9KLS3 A9KLU2 A9KLV5 A9KLX7 A9KLX8 A9KLX9
A9KLY3 A9KLY4 A9KLZ5 A9KM25 A9KM34 A9KM56 A9KM86 A9KM87 A9KM95 A9KM99 A9KMA3
A9KMB4 A9KMB5 A9KMB6 A9KMB8 A9KMC1 A9KMC2 A9KMD6 A9KMD8 A9KMF3 A9KMF5 A9KMF6
A9KMF8 A9KMJ6 A9KMP5 A9KMS8 A9KMT3 A9KMU9 A9KMV0 A9KMV9 A9KMW6 A9KMW7 A9KMX2
A9KMX5 A9KMZ5 A9KN01 A9KN28 A9KN62 A9KNC4 A9KND1 A9KNG9 A9KNI2 A9KNI6 A9KNI7
A9KNJ3 A9KNJ5 A9KNK6 A9KNL0 A9KNN3 A9KNS9 A9KNU8 A9KNV3 A9KNV9 A9KNW1 A9KNW3
A9KNW4 A9KNW8 A9KNX3 A9KNX4 A9KNX6 A9KNY7 A9KP29 A9KP30 A9KP49 A9KP81 A9KP86
A9KPF1 A9KPF4 A9KPF8 A9KPG3 A9KPG4 A9KPH7 A9KPI9 A9KPK9 A9KPL0 A9KPP1 A9KPP4
A9KPQ3 A9KPW1 A9KPX7 A9KQ51 A9KQ65 A9KQ75 A9KQ87 A9KQ90 A9KQ99 A9KQC2 A9KQC8
A9KQC9 A9KQI8 A9KQK0 A9KQM9 A9KQX0 A9KR30 A9KR32 A9KR74 A9KRX7 A9KRZ1 A9KRZ2
A9KRZ3 A9KRZ4 A9KS30 A9KS68 A9KSC2 A9KSC7 A9KSE6 A9KSH6 A9KSJ1 A9KSJ2 A9KSM5
A9KSQ1 A9KSQ2 A9KSS0 A9KSS1 A9KSS3 A9KSS4 A9KSS5 A9KSU3 A9KSU6 A9KSW8 A9KSW9
A9KSX1 A9KT32 A9KT71 A9KT72 A9KT78 A9KT79 A9KTA4 A9KTB9 A9KTE0 A9KTE6 A9KTE7
A9KTG8 A9KTJ1 A9KTJ4 A9KTL1 A9KTL2 A9KTL5 A9KTM0 A9KTM5 A9KTM9 A9KTN4 A9KTR5
A9KTR7 A9KTR8 A9KTU7 A9KTV0 A9KTV2 A9KTV7 A9KTV9 A9KTY6 A9KTY7 A9KTY9 A9KTZ0
A9KTZ3 A9KU65 A9KU72 A9KU74 A9KU83 A9KU84 A9KU88 A9KU92 A9KUA1 A9KUA5 A9KUA7
A9KUB0 A9KUB8 A9KUC0 A9KUK2 A9KUK5 A9KUK6 A9KUK8 A9KUL1 A9KUL9 A9KUN0 A9KUN1
A9KUU5 A9KUY3 A9KV34 A9KV69 A9KV78 A9KV80 A9KV86 A9KVB0 A9KVB7 A9KVC7 A9KVC8
A9KVD6 A9KVE9 A9KVF7 A9KVR8 A9KVT4 A9KW43 A9KW47 A9KW78 A9KW79 A9KW82 A9KW83
A9KW85 A9KW87 A9KW88 A9KW91 A9KW92 A9KW93 A9KW94 A9KW96 A9KW97 A9KW98 A9KWA0
A9KWA1 A9KWA2 A9KWA3 A9KWA5 A9KWA6 A9KWA7 A9KWA8 A9KWA9 A9KWB0 A9KWB1 A9KWB2
A9KWB3 A9KWB4 A9KWB5 A9KWB6 A9KWB7 A9KWB8 A9KWB9 A9KWC0 A9KWC1 A9KWC3 A9KWC4
A9KWC5 A9KWC7 A9KWH4 A9KWH5 A9KWH6 A9KWH7 A9KWH8 A9KWW6 A9KWW8 A9KWW9 A9KWX0
A9KWY0 A9KWY5 A9KWZ3 A9KX16 A9KX17 A9KX19 A9KX22 A9KX23 A9KX59 A9KX72 A9KXA7
A9KXA9 A9KXC4 A9KXE1 A9KXF0 A9KXI7 A9KXK7 A9KXK8 A9KXL1 A9KXN7 A9KXP6 A9KXQ3
A9KXQ5 A9KXQ7 A9KXT3 A9KXX5 A9KXZ7 A9KXZ9 A9KY01 A9KY02 A9KY04 A9KY05 A9KY06
A9KY13 A9KY15 A9KY16 A9KY18 A9KY20 A9KY21 A9KY30 A9KY38 A9KY39 A9KY56 A9KY60
A9KY67 A9KY70 A9KY72 A9KY73 A9KYF6 A9KYF8 A9KYG0 A9KYG1 A9KYG4 A9KYG5 A9KYG7
A9KYH1 A9KYH3 A9KYI1 A9KYJ0 A9KYJ1 A9KYJ5 A9KYJ6 A9KYL8 A9KYP8 A9KYP9 A9KYS9
A9KYT2 A9KYY3 A9KYY5 A9KYY6 A9KYY8 A9KYZ2 A9KYZ4 A9KZ17 A9KZ30 A9KZ67 A9KZ78
A9KZ79 A9KZ80 A9KZF7 A9KZH0 A9KZW6 A9KZW8 A9KZW9 A9KZX0 A9KZX1 A9KZX3 A9KZX5
A9KZX6 A9KZX9 A9KZZ0 A9KZZ3 A9KZZ5 A9L001 A9L004 A9L006 A9L012 A9L013 A9L014
A9L0C0 A9L0E9 A9L0F2 A9L0H9 A9L0P5 A9L0P6 A9L0Q8 A9L0R7 A9L0R8 A9L0T3 A9L0V0
A9L0Z2 A9L107 A9L111 A9L112 A9L120 A9L121 A9L122 A9L123 A9L126 A9L129 A9L185
A9L1A8 A9L1B3 A9L1C2 A9L1D4 A9L1G4 A9L1H8 A9L1I0 A9L1I5 A9L1L1 A9L1M3 A9L1M6
A9L1P1 A9L1V0 A9L269 A9L2C1 A9L2D4 A9L2D5 A9L2D7 A9L2F4 A9L2H4 A9L2I0 A9L2P0
A9L2P8 A9L2Q6 A9L2Q7 A9L2Q9 A9L2X4 A9L2X6 A9L2X7 A9L2Y2 A9L2Y4 A9L330 A9L331
A9L332 A9L340 A9L341 A9L375 A9L389 A9L392 A9L3D8 A9L3F5 A9L3H7 A9L3H8 A9L3H9
A9L3I1 A9L3I3 A9L3I4 A9L3J3 A9L3M8 A9L3N9 A9L3Q1 A9L3Q4 A9L3Q6 A9L3Q7 A9L3R0
A9L3U4 A9L3V9 A9L3W0 A9L3W1 A9L3X6 A9L3Y2 A9L3Z6 A9L428 A9L429 A9L430 A9L436

A9L437 A9L438 A9L4A0 A9L4B4 A9L4B9 A9L4C0 A9L4E6 A9L4F3 A9L4F4 A9L4F6 A9L4G0
A9L4G2 A9L4G4 A9L4G9 A9L4I0 A9L4I9 A9L4M4 A9L4T5 A9L4T6 A9L4U0 A9L4U3 A9L4U7
A9L514 A9L516 A9L532 A9L543 A9L555 A9L560 A9L564 A9L567 A9L569 A9L573 A9L579
A9L582 A9L591 A9L594 A9L5A8 A9L5D1 A9L5D5 A9L5E5 A9L5E7 A9L5F5 A9L5F8 A9L5G0
A9L5I3 A9L5J0 A9L5J5 A9L5J9 A9L5K3 A9L5L1 A9L5L8 A9L5L9 A9L5M3 A9L5N6 A9L5P5
A9L5P6 A9L5P7 A9L5P8 A9L5Q1 A9L5S5 A9L5Z1 A9L605 A9L621 A9L650 A9L682 A9L683
A9L694 A9L695 A9L913 A9L937 A9L948 A9L985 A9L986 A9L987 A9L988 A9L994 A9L9A4
A9L9B7 A9L9B8 A9L9B9 A9L9C9 A9L9D0 A9L9D2 A9L9D3 A9L9D5 A9L9D6 A9L9D7 A9L9D9
A9L9E1 A9L9E3 A9LLI7 A9LLI8 A9LMC0 A9LNK9 A9LS46 A9LYA9 A9LYD5 A9LYD8 A9LYE0
A9LYE2 A9LYE4 A9LYE6 A9LYG6 A9LYH4 A9LYH6 A9LYH7 A9LYX4 A9LYX9 A9LYY9 A9LYZ5
A9LZ69 A9LZ72 A9LZ73 A9LZ77 A9LZ86 A9LZ95 A9LZ96 A9LZA2 A9LZB7 A9LZC1 A9LZC3
A9LZD2 A9LZD5 A9LZF5 A9LZF7 A9LZF9 A9LZH0 A9LZH6 A9LZI3 A9LZI5 A9LZN6 A9LZN8
A9LZN9 A9LZP0 A9LZQ0 A9LZQ1 A9LZQ2 A9LZQ3 A9LZQ8 A9LZR4 A9LZR5 A9LZS9 A9LZV9
A9M003 A9M005 A9M007 A9M020 A9M021 A9M029 A9M033 A9M041 A9M061 A9M068 A9M072
A9M081 A9M087 A9M088 A9M0A4 A9M0B8 A9M0C4 A9M0D0 A9M0D2 A9M0D3 A9M0D4 A9M0G1
A9M0G3 A9M0K2 A9M0M3 A9M0N5 A9M0Q0 A9M0Q5 A9M0Q6 A9M0S4 A9M0T5 A9M0T9 A9M0U5
A9M0U6 A9M0U7 A9M0W1 A9M0W6 A9M0Y3 A9M0Y5 A9M105 A9M125 A9M127 A9M138 A9M148
A9M159 A9M162 A9M177 A9M185 A9M186 A9M198 A9M1A2 A9M1A6 A9M1D3 A9M1D5 A9M1F6
A9M1G2 A9M1G3 A9M1G4 A9M1G6 A9M1H2 A9M1J1 A9M1J5 A9M1J7 A9M1J9 A9M1M1 A9M1N1
A9M1N3 A9M1P7 A9M1Q1 A9M1Q2 A9M1Q5 A9M1Q6 A9M1Q8 A9M1R2 A9M1U4 A9M1U5 A9M1V2
A9M1V8 A9M1W7 A9M1W9 A9M1Y9 A9M1Z4 A9M251 A9M254 A9M257 A9M260 A9M276 A9M296
A9M2A3 A9M2B7 A9M2B8 A9M2D0 A9M2D1 A9M2D2 A9M2D3 A9M2F1 A9M2H2 A9M2H5 A9M2I4
A9M2J9 A9M2L4 A9M2N2 A9M2N9 A9M2P1 A9M2P6 A9M2Q5 A9M2Q6 A9M2R4 A9M2U7 A9M2V6
A9M2W6 A9M2W7 A9M2W8 A9M2Y6 A9M2Z1 A9M2Z6 A9M2Z9 A9M300 A9M342 A9M352 A9M354
A9M374 A9M375 A9M378 A9M383 A9M388 A9M392 A9M393 A9M3A0 A9M3B7 A9M3C0 A9M3F3
A9M3H4 A9M3H7 A9M3I9 A9M3J0 A9M3K6 A9M3L2 A9M3M7 A9M3M8 A9M3N1 A9M3N5 A9M3N9
A9M3P3 A9M3P6 A9M3P7 A9M3Q2 A9M3Q3 A9M3Q7 A9M3R4 A9M3R5 A9M3R8 A9M3S1 A9M3T2
A9M3T6 A9M3T9 A9M3U1 A9M3U2 A9M3U3 A9M3U4 A9M3U5 A9M3U7 A9M3U8 A9M3U9 A9M3V0
A9M3V1 A9M3V2 A9M3V3 A9M3V4 A9M3V5 A9M3V6 A9M3V7 A9M3V8 A9M3V9 A9M3W0 A9M3W1
A9M3W2 A9M3W3 A9M3W5 A9M3W6 A9M3W8 A9M3X0 A9M3X1 A9M3X2 A9M3X3 A9M3X5 A9M3X7
A9M3X8 A9M3Z7 A9M3Z9 A9M402 A9M421 A9M423 A9M439 A9M441 A9M456 A9M463 A9M464
A9M469 A9M478 A9M479 A9M485 A9M489 A9M490 A9M4B1 A9M4C4 A9M4C8 A9M4D3 A9M4D6
A9M4E0 A9M4E1 A9M4E3 A9M4F8 A9M4G6 A9M4I2 A9M4K6 A9M4L8 A9M560 A9M582 A9M586
A9M587 A9M5A4 A9M5A7 A9M5C1 A9M5C2 A9M5C4 A9M5D7 A9M5E7 A9M5F3 A9M5F4 A9M5F5
A9M5G0 A9M5H1 A9M5H3 A9M5H4 A9M5I2 A9M5J8 A9M5L1 A9M5L7 A9M5L8 A9M5M4 A9M5M5
A9M5M6 A9M5M7 A9M5M9 A9M5N1 A9M5N2 A9M5N3 A9M5N4 A9M5N5 A9M5N6 A9M5N7 A9M5N8
A9M5N9 A9M5P0 A9M5P1 A9M5P2 A9M5P3 A9M5P4 A9M5P6 A9M5P7 A9M5P8 A9M5P9 A9M5Q0
A9M5Q1 A9M5Q2 A9M5Q3 A9M5Q4 A9M5Q5 A9M5Q8 A9M5R0 A9M5R1 A9M5R2 A9M5R3 A9M5R4
A9M5U3 A9M5U5 A9M5W0 A9M5X3 A9M616 A9M617 A9M618 A9M637 A9M642 A9M648 A9M657
A9M670 A9M676 A9M679 A9M688 A9M689 A9M698 A9M6C1 A9M6E4 A9M6F1 A9M6H1 A9M6K0
A9M6K1 A9M6K5 A9M6L4 A9M6N0 A9M6N1 A9M6N7 A9M6N8 A9M6S7 A9M6S9 A9M6T4 A9M6V2
A9M700 A9M752 A9M757 A9M792 A9M798 A9M7B6 A9M7B7 A9M7B8 A9M7C3 A9M7D7 A9M7F3
A9M7H5 A9M7I3 A9M7I9 A9M7J1 A9M7K0 A9M7K1 A9M7K2 A9M7K7 A9M7L3 A9M7L4 A9M7P1
A9M7P7 A9M7T6 A9M7T8 A9M7U1 A9M7V2 A9M7V7 A9M7V8 A9M7X1 A9M7Z1 A9M817 A9M820

A9M832 A9M843 A9M845 A9M849 A9M854 A9M862 A9M874 A9M880 A9M881 A9M882 A9M886
A9M887 A9M8B2 A9M8B3 A9M8C0 A9M8F4 A9M8G2 A9M8H7 A9M8J1 A9M8K5 A9M8K8 A9M8M2
A9M8M3 A9M8P2 A9M8P3 A9M8Q0 A9M8Q1 A9M8Q9 A9M8R2 A9M8R3 A9M8R8 A9M8S2 A9M8V2
A9M8W0 A9M8X5 A9M8X7 A9M8X8 A9M8Z9 A9M903 A9M916 A9M929 A9M959 A9M966 A9M968
A9M9A0 A9M9B3 A9M9B4 A9M9E3 A9M9E4 A9M9E5 A9M9E9 A9M9F0 A9M9H3 A9M9H7 A9M9H8
A9M9H9 A9M9Q4 A9M9Q7 A9M9R4 A9M9R5 A9M9R8 A9M9R9 A9M9S0 A9M9S1 A9M9S3 A9M9T1
A9M9U0 A9M9U2 A9M9U3 A9M9V1 A9M9V2 A9M9V4 A9M9V9 A9M9W1 A9M9Y3 A9M9Y5 A9M9Z1
A9M9Z4 A9M9Z5 A9M9Z6 A9M9Z7 A9M9Z8 A9MA22 A9MA26 A9MA28 A9MA35 A9MA50 A9MA55
A9MA67 A9MA84 A9MAB2 A9MAB4 A9MAD3 A9MAD7 A9MAE5 A9MAE6 A9MAF0 A9MAF7 A9MAG5
A9MAG7 A9MAG8 A9MAJ9 A9MAK7 A9MAK8 A9MAL4 A9MAL9 A9MAP0 A9MAP9 A9MAQ0 A9MAQ8
A9MAQ9 A9MAR7 A9MAS2 A9MAS9 A9MAU6 A9MAX5 A9MB09 A9MB31 A9MB89 A9MBD0 A9MBD4
A9MBD6 A9MBK6 A9MBN4 A9MBN5 A9MBN6 A9MBN7 A9MBN9 A9MBP9 A9MBW8 A9MBZ4 A9MC11
A9MC50 A9MC67 A9MC92 A9MC99 A9MCA1 A9MCF4 A9MCG4 A9MCH2 A9MCJ0 A9MCK3 A9MCK4
A9MCL0 A9MCL2 A9MCM7 A9MCP6 A9MCU7 A9MCU8 A9MCV1 A9MCV4 A9MCV6 A9MCV8 A9MCV9
A9MCX4 A9MCX8 A9MCZ2 A9MCZ3 A9MCZ4 A9MCZ5 A9MD56 A9MDJ3 A9MDK1 A9MDR2 A9MDU4
A9MDU5 A9MDU6 A9MDV1 A9MDV2 A9MDX3 A9ME65 A9ME73 A9ME88 A9MEB0 A9MEH2 A9MEH5
A9MEJ1 A9MEP2 A9MEP3 A9MEQ9 A9MET9 A9MEV5 A9MEW2 A9MEW3 A9MEY0 A9MEY3 A9MEZ2
A9MF07 A9MF10 A9MF16 A9MF17 A9MF18 A9MF23 A9MF24 A9MF25 A9MF28 A9MF29 A9MF30
A9MF31 A9MF32 A9MF43 A9MFB2 A9MFB7 A9MFC0 A9MFC1 A9MFE4 A9MFF4 A9MFF8 A9MFF9
A9MFG0 A9MFG1 A9MFH4 A9MFI1 A9MFK9 A9MFL0 A9MFL1 A9MFM3 A9MFN5 A9MFP0 A9MFP1
A9MFP2 A9MFQ6 A9MFR5 A9MFR9 A9MFS0 A9MFX5 A9MFX6 A9MFY9 A9MFZ1 A9MFZ2 A9MFZ5
A9MG06 A9MG80 A9MG86 A9MG95 A9MGA5 A9MGB6 A9MGD1 A9MGI9 A9MGS5 A9MGT6 A9MGT8
A9MGV9 A9MGW0 A9MGW2 A9MGZ5 A9MH00 A9MH03 A9MH08 A9MH26 A9MH45 A9MH53 A9MH54
A9MH59 A9MH91 A9MHA7 A9MHB2 A9MHC0 A9MHC3 A9MHD1 A9MHD2 A9MHD5 A9MHD6 A9MHD7
A9MHE0 A9MHE9 A9MHF1 A9MHF3 A9MHF4 A9MHF5 A9MHF7 A9MHG0 A9MHG7 A9MHH3 A9MHI3
A9MHJ3 A9MHJ4 A9MHJ6 A9MHJ7 A9MHJ8 A9MHK1 A9MHL2 A9MHL3 A9MHL5 A9MHL6 A9MHM3
A9MHM5 A9MHP1 A9MHP3 A9MHQ2 A9MHQ4 A9MHQ8 A9MHR4 A9MHS1 A9MHS2 A9MHS3 A9MHS6
A9MHT1 A9MHT3 A9MHU0 A9MHU8 A9MHV9 A9MHW1 A9MHW6 A9MHX0 A9MHX2 A9MHX5 A9MHX6
A9MHZ1 A9MHZ9 A9MI00 A9MI06 A9MI11 A9MI13 A9MI14 A9MI17 A9MI19 A9MI34 A9MI36
A9MI38 A9MI40 A9MI50 A9MI63 A9MI64 A9MI67 A9MI70 A9MI98 A9MIA3 A9MIC6 A9MID0
A9MIE1 A9MIG0 A9MIG3 A9MIH3 A9MIK8 A9MIM3 A9MIP6 A9MIS0 A9MIS5 A9MIU5 A9MIV9
A9MIX1 A9MIY3 A9MIY8 A9MJ08 A9MJ10 A9MJ29 A9MJ30 A9MJ39 A9MJ42 A9MJ47 A9MJ55
A9MJ57 A9MJ94 A9MJB6 A9MJB7 A9MJB9 A9MJC8 A9MJE1 A9MJE5 A9MJE6 A9MJE9 A9MJF0
A9MJF2 A9MJF3 A9MJH1 A9MJI2 A9MJL9 A9MJM8 A9MJN1 A9MJN4 A9MJQ1 A9MJQ5 A9MJQ9
A9MJR0 A9MJT6 A9MJT8 A9MJT9 A9MJU2 A9MJV6 A9MJY3 A9MK24 A9MK26 A9MK27 A9MK40
A9MK49 A9MK63 A9MKA7 A9MKA9 A9MKB7 A9MKB9 A9MKC6 A9MKC8 A9MKD1 A9MKD5 A9MKE1
A9MKE4 A9MKF6 A9MKG0 A9MKH2 A9MKM3 A9MKM6 A9MKN2 A9MKN3 A9MKN4 A9MKN5 A9MKN7
A9MKN8 A9MKN9 A9MKP0 A9MKQ6 A9MKQ8 A9MKR5 A9MKS5 A9MKV0 A9MKV9 A9MKX9 A9ML11
A9ML12 A9ML15 A9ML30 A9ML47 A9MLA4 A9MLA7 A9MLB3 A9MLE0 A9MLF6 A9MLF8 A9MLK7
A9MLL7 A9MLN2 A9MLN4 A9MLP6 A9MLR2 A9MLS5 A9MLS8 A9MLV4 A9MLY4 A9MLY7 A9MLY8
A9MLY9 A9MM02 A9MM14 A9MM16 A9MM22 A9MM24 A9MM39 A9MM40 A9MM42 A9MM45 A9MM57
A9MM58 A9MM66 A9MMA0 A9MMA2 A9MMA3 A9MMA6 A9MMA9 A9MMB3 A9MMB5 A9MMB8 A9MMC5
A9MMD9 A9MME5 A9MMJ6 A9MML1 A9MMQ2 A9MMQ4 A9MMR9 A9MMS3 A9MMS6 A9MN11 A9MN31
A9MN34 A9MN35 A9MN36 A9MN38 A9MN39 A9MN45 A9MN46 A9MN48 A9MN51 A9MN52 A9MN53

A9MN54 A9MN55 A9MN56 A9MN57 A9MN58 A9MN59 A9MN60 A9MN61 A9MN62 A9MN63 A9MN64
A9MN66 A9MN67 A9MN68 A9MN70 A9MN71 A9MN78 A9MN80 A9MN84 A9MN98 A9MNA2 A9MNC4
A9MNC7 A9MNC8 A9MND2 A9MND4 A9MND5 A9MND7 A9MNE8 A9MNH4 A9MNH8 A9MNJ9 A9MNR4
A9MNR5 A9MNR7 A9MNR8 A9MNS1 A9MNX4 A9MNX5 A9MNY0 A9MNY1 A9MNY5 A9MNY6 A9MNY8
A9MNY9 A9MP02 A9MP18 A9MP21 A9MP23 A9MP28 A9MP31 A9MP33 A9MP34 A9MP36 A9MP37
A9MP38 A9MP39 A9MP47 A9MP50 A9MP52 A9MP60 A9MP98 A9MPA0 A9MPA1 A9MPD4 A9MPF1
A9MPF3 A9MPG5 A9MPH8 A9MPI9 A9MPJ3 A9MPJ5 A9MPJ6 A9MPK1 A9MPK2 A9MPK5 A9MPK7
A9MPL5 A9MPM1 A9MPM2 A9MPM3 A9MPN3 A9MPN4 A9MPN7 A9MPN8 A9MPP3 A9MPP7 A9MPQ0
A9MPQ4 A9MPR6 A9MPU2 A9MPU4 A9MPV1 A9MPV4 A9MPV5 A9MPV9 A9MPW3 A9MPW6 A9MPX4
A9MPY6 A9MPY7 A9MQ20 A9MQ82 A9MQA3 A9MQA5 A9MQA6 A9MQC1 A9MQD1 A9MQD8 A9MQE0
A9MQE1 A9MQF0 A9MQF1 A9MQF6 A9MQG1 A9MQG2 A9MQG3 A9MQG4 A9MQH5 A9MQH7 A9MQL9
A9MQM1 A9MQN1 A9MQR4 A9MQR6 A9MQS2 A9MQS7 A9MQT5 A9MQU9 A9MQY3 A9MR05 A9MR28
A9MR41 A9MR43 A9MR48 A9MR76 A9MR77 A9MR84 A9MR86 A9MR94 A9MR95 A9MR96 A9MRA3
A9MRA4 A9MRA5 A9MRA6 A9MRB6 A9MRB9 A9MRC8 A9MRD1 A9MRF0 A9MRF1 A9MRG1 A9MRG2
A9MRH0 A9MRH1 A9MRH2 A9MRH4 A9MRH6 A9MRI4 A9MRI5 A9MRK3 A9MRQ5 A9MRV0 A9MRX5
A9MS78 A9MS82 A9MS91 A9MSA2 A9MSA4 A9MSA6 A9MSB5 A9MSC2 A9MSK6 A9MSM9 A9MSQ9
A9MST4 A9MSU1 A9MSX9 A9MSY0 A9MSY1 A9MSY2 A9MSY3 A9MSY4 A9MSY5 A9MSY6 A9MSY7
A9MSY8 A9MSY9 A9MSZ0 A9MSZ1 A9MSZ2 A9MSZ3 A9MSZ4 A9MSZ5 A9MT00 A9MT05 A9MT06
A9MT07 A9MT08 A9MT09 A9MT10 A9MT13 A9MT26 A9MT45 A9MT55 A9MT72 A9MT88 A9MTA4
A9MTF1 A9MTH5 A9MTH7 A9MTI2 A9MTI7 A9MTI8 A9MTJ1 A9MTJ2 A9MTJ4 A9MTJ5 A9MTJ8
A9MTL0 A9MTL3 A9MTQ2 A9MTR2 A9MTS5 A9MTT1 A9MTT7 A9MTU1 A9MTV1 A9MTV2 A9MTV3
A9MTV4 A9MU21 A9MU40 A9MU62 A9MU69 A9MU86 A9MU90 A9MUA9 A9MUB2 A9MUB3 A9MUB4
A9MUB9 A9MUC2 A9MUG5 A9MUG8 A9MUH8 A9MUI2 A9MUJ0 A9MUK3 A9MUK6 A9MUL2 A9MUU1
A9MUV0 A9MUW2 A9MUX3 A9MUX4 A9MUY8 A9MV57 A9MV61 A9MV74 A9MVB6 A9MVD3 A9MVI1
A9MVI4 A9MVK3 A9MVK6 A9MVL0 A9MVL2 A9MVM9 A9MVN0 A9MVN1 A9MVN5 A9MVN6 A9MVN7
A9MVP4 A9MVP7 A9MVU1 A9MVU3 A9MVV1 A9MVV3 A9MVV4 A9MVV6 A9MVW7 A9MVZ4 A9MW00
A9MW03 A9MW04 A9MW26 A9MW31 A9MW45 A9MW53 A9MW55 A9MW81 A9MW82 A9MW88 A9MW89
A9MW90 A9MW92 A9MWA7 A9MWA8 A9MWC0 A9MWC2 A9MWI4 A9MWJ6 A9MWJ7 A9MWL1 A9MWL9
A9MWP8 A9MWR3 A9MWR4 A9MWU4 A9MWU8 A9MWX5 A9MWX9 A9MWZ5 A9MWZ8 A9MWZ9 A9MX06
A9MX07 A9MX09 A9MX13 A9MX16 A9MX29 A9MX31 A9MX46 A9MX47 A9MX50 A9MX51 A9MX56
A9MX74 A9MX77 A9MX79 A9MX84 A9MXB7 A9MXB8 A9MXC1 A9MXC2 A9MXC3 A9MXC5 A9MXD5
A9MXE2 A9MXE7 A9MXF2 A9MXF3 A9MXH1 A9MXH3 A9MXI2 A9MXI3 A9MXN4 A9MXQ2 A9MY02
A9MY03 A9MY08 A9MY13 A9MY16 A9MY64 A9MY70 A9MY90 A9MY97 A9MYB1 A9MYC7 A9MYC8
A9MYG8 A9MYH1 A9MYI6 A9MYJ4 A9MYJ5 A9MYJ6 A9MYJ9 A9MYK3 A9MYM2 A9MYM3 A9MYM4
A9MYN1 A9MYN8 A9MYN9 A9MYQ4 A9MYU8 A9MYZ4 A9MZ19 A9MZ21 A9MZ47 A9MZ50 A9MZ51
A9MZ63 A9MZA2 A9MZB0 A9MZC2 A9MZC4 A9MZC5 A9MZC6 A9MZC8 A9MZC9 A9MZE7 A9MZF8
A9MZF9 A9MZG5 A9MZG6 A9MZG8 A9MZH4 A9MZH7 A9MZI1 A9MZI2 A9MZI7 A9MZJ3 A9MZK0
A9MZL0 A9MZL1 A9MZM0 A9MZN0 A9MZN3 A9MZQ5 A9MZR1 A9MZR2 A9MZS9 A9MZT0 A9MZU5
A9N011 A9N034 A9N035 A9N037 A9N078 A9N099 A9N0B5 A9N0C0 A9N0C1 A9N0C4 A9N0C5
A9N0D9 A9N0E0 A9N0F0 A9N0F4 A9N0G6 A9N0G7 A9N0G9 A9N0H0 A9N0H2 A9N0H5 A9N0I2
A9N0I9 A9N0J1 A9N0J3 A9N0J4 A9N0J5 A9N0K2 A9N0K5 A9N0K6 A9N0K7 A9N0K8 A9N0K9
A9N0L0 A9N0L8 A9N0M3 A9N0M7 A9N0P9 A9N0Q2 A9N0Q5 A9N0Q9 A9N0R2 A9N0R7 A9N0S0
A9N0T3 A9N0U0 A9N0U3 A9N0U7 A9N0W3 A9N0W4 A9N0W9 A9N0Y2 A9N141 A9N142 A9N158
A9N159 A9N171 A9N1F8 A9N1H6 A9N1I0 A9N1J6 A9N1L2 A9N1T3 A9N1U1 A9N1W0 A9N1X3

A9N1X5 A9N1X7 A9N1X8 A9N1X9 A9N1Y3 A9N1Z8 A9N201 A9N202 A9N205 A9N216 A9N218
A9N229 A9N230 A9N236 A9N238 A9N240 A9N241 A9N243 A9N261 A9N269 A9N274 A9N275
A9N276 A9N277 A9N278 A9N292 A9N2A1 A9N2D0 A9N2D1 A9N2D2 A9N2D3 A9N2D7 A9N2D8
A9N2D9 A9N2E4 A9N2E5 A9N2E6 A9N2F5 A9N2H7 A9N2I4 A9N2J1 A9N2J4 A9N2J6 A9N2L7
A9N2M1 A9N2M4 A9N2Y1 A9N2Y3 A9N2Z5 A9N2Z9 A9N308 A9N329 A9N336 A9N362 A9N372
A9N3I7 A9N3J4 A9N3L5 A9N3L6 A9N3M5 A9N3M7 A9N3N1 A9N3N2 A9N3N3 A9N3P6 A9N3P8
A9N3Q5 A9N3Q6 A9N3R6 A9N3Z7 A9N413 A9N420 A9N432 A9N457 A9N460 A9N465 A9N474
A9N476 A9N4B8 A9N4K8 A9N4M6 A9N4N1 A9N4P2 A9N4P3 A9N4P8 A9N4Q8 A9N4T4 A9N4U5
A9N4U7 A9N4Y4 A9N4Y5 A9N4Y6 A9N4Z1 A9N4Z2 A9N507 A9N508 A9N512 A9N513 A9N514
A9N518 A9N520 A9N521 A9N529 A9N540 A9N548 A9N563 A9N564 A9N578 A9N598 A9N5A2
A9N5A3 A9N5A5 A9N5B7 A9N5C2 A9N5C4 A9N5J5 A9N5J6 A9N5J7 A9N5K6 A9N5P3 A9N5P9
A9N5Q2 A9N5Q7 A9N5T2 A9N5W3 A9N5X2 A9N5X8 A9N5Y2 A9N5Y7 A9N5Y8 A9N5Z1 A9N5Z8
A9N601 A9N623 A9N658 A9N659 A9N680 A9N6A3 A9N6F2 A9N6F3 A9N6F6 A9N6H3 A9N6I5
A9N6K0 A9N6L3 A9N6R4 A9N6S2 A9N6S3 A9N6T0 A9N6W0 A9N6W1 A9N6W4 A9N6X0 A9N6X2
A9N6Y0 A9N6Y6 A9N6Z0 A9N6Z8 A9N702 A9N712 A9N716 A9N717 A9N728 A9N729 A9N730
A9N731 A9N732 A9N734 A9N735 A9N740 A9N744 A9N750 A9N752 A9N753 A9N756 A9N773
A9N7B3 A9N7C4 A9N7C9 A9N7E1 A9N7E3 A9N7E6 A9N7F3 A9N7F4 A9N7F5 A9N7I2 A9N7J2
A9N7M0 A9N7M1 A9N7S0 A9N7S1 A9N7T7 A9N7T9 A9N7U5 A9N7V2 A9N7V4 A9N7V6 A9N7V7
A9N7X6 A9N7Y5 A9N7Y7 A9N7Z5 A9N804 A9N812 A9N824 A9N830 A9N831 A9N833 A9N839
A9N840 A9N855 A9N889 A9N8A7 A9N8A9 A9N8B1 A9N8B2 A9N8B3 A9N8B8 A9N8C1 A9N8H1
A9N8H2 A9N8H5 A9N8H8 A9N8I3 A9N8I5 A9N8J9 A9N8K1 A9N8K2 A9N8K3 A9N8L3 A9N8L5
A9N8L6 A9N8M1 A9N8M5 A9N8Q7 A9N8Q9 A9N8R0 A9N8R8 A9N8T6 A9N8T7 A9N8T8 A9N8V4
A9N8V5 A9N8V6 A9N8V8 A9N8X5 A9N8Z4 A9N8Z5 A9N900 A9N902 A9N909 A9N910 A9N917
A9N924 A9N925 A9N936 A9N940 A9N942 A9N957 A9N970 A9N980 A9N999 A9N9A0 A9N9A1
A9N9A2 A9N9A3 A9N9C7 A9N9E2 A9N9F4 A9N9F8 A9N9F9 A9N9G2 A9N9G8 A9N9H2 A9N9H5
A9N9I1 A9N9I7 A9N9J6 A9N9K1 A9N9K2 A9N9M4 A9N9M9 A9N9S2 A9N9U7 A9N9V5 A9N9Z2
A9NA21 A9NA23 A9NA24 A9NA25 A9NA40 A9NA45 A9NA46 A9NA54 A9NA60 A9NA67 A9NA71
A9NA76 A9NA77 A9NA78 A9NA82 A9NA83 A9NA93 A9NA98 A9NAA4 A9NAA5 A9NAB0 A9NAB1
A9NAD8 A9NAF8 A9NAF9 A9NAH7 A9NAJ3 A9NAJ6 A9NAK7 A9NAL0 A9NAL1 A9NAL2 A9NAL3
A9NAL4 A9NAL6 A9NAL9 A9NAM0 A9NAM1 A9NAM3 A9NAM4 A9NAM5 A9NAM6 A9NAM7 A9NAM8
A9NAM9 A9NAN0 A9NAN1 A9NAQ7 A9NAR1 A9NAS1 A9NAT6 A9NAT7 A9NAW7 A9NAW9 A9NAX3
A9NAX8 A9NAX9 A9NAY0 A9NAY1 A9NAY2 A9NAY4 A9NAY5 A9NAY6 A9NAY8 A9NAY9 A9NAZ1
A9NAZ2 A9NAZ3 A9NAZ4 A9NAZ5 A9NAZ6 A9NB09 A9NB23 A9NB26 A9NB28 A9NB33 A9NB36
A9NB44 A9NB47 A9NB52 A9NB65 A9NB80 A9NB81 A9NB98 A9NB99 A9NBA2 A9NBA3 A9NBA7
A9NBA8 A9NBA9 A9NBB7 A9NBF7 A9NBH0 A9NBI0 A9NBL3 A9NBL4 A9NBL5 A9NBL7 A9NBM5
A9NBP1 A9NBP7 A9NBQ8 A9NBQ9 A9NBR0 A9NBR1 A9NBS5 A9NBT9 A9NBU1 A9NBU2 A9NBU3
A9NBU5 A9NBV6 A9NBY1 A9NC09 A9NC17 A9NC18 A9NC23 A9NC33 A9NC46 A9NC49 A9NC56
A9NC58 A9NC75 A9NCD4 A9NCE2 A9NCE9 A9NCF0 A9NCJ8 A9NCK3 A9NCK4 A9NCM4 A9NCN3
A9NCN7 A9NCS0 A9NCS8 A9NCX7 A9NCY5 A9ND11 A9ND24 A9ND48 A9ND50 A9ND52 A9ND53
A9ND61 A9ND62 A9ND63 A9ND66 A9NDA0 A9NDB3 A9NDE8 A9NDF9 A9NDG0 A9NDK6 A9NDK7
A9NDN1 A9NDN6 A9NDQ6 A9NDR1 A9NDR7 A9NDS1 A9NDT3 A9NDV1 A9NDV6 A9NDV9 A9NDW2
A9NDW8 A9NDX7 A9NDX8 A9NE10 A9NE14 A9NE34 A9NE58 A9NE63 A9NE64 A9NE65 A9NE68
A9NE72 A9NE80 A9NE82 A9NE95 A9NE96 A9NEA6 A9NEA9 A9NEC4 A9NEC5 A9NED3 A9NED4
A9NED6 A9NED7 A9NED8 A9NED9 A9NEE0 A9NEE1 A9NEE2 A9NEE3 A9NEE4 A9NEE5 A9NEE6

A9NEE7 A9NEE8 A9NEE9 A9NEF1 A9NEF2 A9NEF4 A9NEF6 A9NEF7 A9NEF8 A9NEF9 A9NEG1
A9NEH3 A9NEI3 A9NEI5 A9NEI9 A9NEJ0 A9NEJ5 A9NEL3 A9NEL4 A9NEL5 A9NEL7 A9NEL8
A9NEN1 A9NEN2 A9NEN3 A9NEN4 A9NEN9 A9NEP1 A9NEP7 A9NEQ2 A9NEQ8 A9NET8 A9NEU5
A9NEU7 A9NEV1 A9NEX8 A9NF20 A9NF23 A9NF30 A9NF46 A9NF54 A9NF57 A9NF61 A9NF62
A9NF91 A9NF94 A9NF97 A9NFF6 A9NFG5 A9NFH1 A9NFI4 A9NFK6 A9NFM5 A9NFN6 A9NFN7
A9NFP2 A9NFP3 A9NFP5 A9NFP7 A9NG16 A9NG27 A9NG35 A9NG38 A9NGC2 A9NGD7 A9NGE2
A9NGE3 A9NGE6 A9NGF8 A9NGG3 A9NGH5 A9NGH6 A9NGI4 A9NGI6 A9NGK2 A9NGK9 A9NGL1
A9NGL4 A9NGL6 A9NGM3 A9NGM5 A9NGN4 A9NGN9 A9NGP8 A9NGT0 A9NGT2 A9NHA2 A9NHC3
A9NHC5 A9NHC6 A9NHD0 A9NHF4 A9NHF7 A9NHF8 A9NHG0 A9NHG7 A9NHG9 A9NHH8 A9NHL6
A9NHW0 A9NHW6 A9NHX0 A9NK39 A9NKD9 A9NNH7 A9NUH8 A9NYY0 A9P2J1 A9P2P4 A9P822
A9PBH9 A9PCL4 A9PDZ7 A9PEK8 A9PHC5 A9PHE9 A9Q1D5 A9Q1J7 A9Q1K7 A9Q1K8 A9Q1K9
A9Q1L2 A9Q751 A9QA56 A9QC39 A9QC40 A9QC56 A9QCA5 A9QGV6 A9QM73 A9QT41 A9QXE0
A9QYG3 A9QYG4 A9QYG5 A9QYG7 A9QYH5 A9QYI0 A9QYI1 A9QYL0 A9QYL3 A9QYL4 A9QYL5
A9QYM6 A9QYN1 A9QYN2 A9QYN3 A9QYN9 A9QYP2 A9QYP8 A9QYQ1 A9QYQ2 A9QYQ6 A9QYR6
A9QYR8 A9QYS1 A9QYS2 A9QYS3 A9QYS5 A9QYT2 A9QYU7 A9QYV1 A9QYV2 A9QYV4 A9QYX2
A9QYX3 A9QYX5 A9QYX6 A9QYX8 A9QYY1 A9QYY2 A9QYY3 A9QYY6 A9QYZ5 A9QZ08 A9QZ09
A9QZ11 A9QZ55 A9QZ59 A9QZ60 A9QZ61 A9QZ62 A9QZ63 A9QZ95 A9QZ96 A9QZA1 A9QZB3
A9QZC8 A9QZC9 A9QZG0 A9QZG6 A9QZH0 A9QZJ5 A9QZK6 A9QZK7 A9QZM4 A9QZN1 A9QZP3
A9QZP5 A9QZQ2 A9QZQ4 A9QZR6 A9QZR9 A9QZS1 A9QZS3 A9QZW5 A9QZW6 A9QZX0 A9QZY9
A9QZZ0 A9QZZ1 A9QZZ2 A9R003 A9R005 A9R008 A9R010 A9R014 A9R015 A9R021 A9R023
A9R025 A9R032 A9R033 A9R046 A9R047 A9R055 A9R058 A9R072 A9R087 A9R0A2 A9R0A4
A9R0A5 A9R0A6 A9R0A8 A9R0C4 A9R0E4 A9R0G0 A9R0H5 A9R0H6 A9R0H7 A9R0H8 A9R0H9
A9R0I6 A9R0J7 A9R0J8 A9R0L5 A9R0M2 A9R0Q1 A9R0Q3 A9R0Q4 A9R0Q7 A9R0Q8 A9R0S6
A9R0S7 A9R0U2 A9R0U3 A9R0U4 A9R0U5 A9R0V7 A9R0V9 A9R111 A9R115 A9R116 A9R117
A9R118 A9R119 A9R122 A9R132 A9R133 A9R142 A9R144 A9R145 A9R154 A9R1B1 A9R1B9
A9R1C3 A9R1D2 A9R1D9 A9R1E0 A9R1E3 A9R1E6 A9R1G2 A9R1G3 A9R1G4 A9R1H4 A9R1H5
A9R1H8 A9R1J4 A9R1J6 A9R1J7 A9R1N3 A9R1P0 A9R1Q7 A9R1Q8 A9R1R9 A9R1S0 A9R1S6
A9R1U0 A9R1U2 A9R1U3 A9R1V4 A9R1Y4 A9R1Z3 A9R253 A9R256 A9R284 A9R2E0 A9R2E4
A9R2F2 A9R2H1 A9R2H7 A9R2J1 A9R2J3 A9R2J6 A9R2J8 A9R2J9 A9R2K3 A9R2K5 A9R2K8
A9R2M4 A9R2M5 A9R2M7 A9R2N0 A9R2N5 A9R2N8 A9R2N9 A9R2P6 A9R2Q4 A9R2R6 A9R2R7
A9R2W1 A9R2W2 A9R2W3 A9R2X0 A9R2X1 A9R2X2 A9R2X3 A9R2Y6 A9R2Y9 A9R2Z4 A9R2Z5
A9R2Z6 A9R2Z7 A9R2Z8 A9R307 A9R308 A9R314 A9R351 A9R352 A9R361 A9R371 A9R375
A9R382 A9R392 A9R393 A9R395 A9R396 A9R397 A9R398 A9R399 A9R3A9 A9R3B3 A9R3B5
A9R3C8 A9R3C9 A9R3D1 A9R3D3 A9R3D8 A9R3I1 A9R3I2 A9R3I4 A9R3P0 A9R3Q0 A9R3V0
A9R3V1 A9R3V2 A9R3W4 A9R3Y2 A9R3Y5 A9R3Y6 A9R3Y7 A9R3Z3 A9R402 A9R404 A9R410
A9R431 A9R462 A9R463 A9R464 A9R465 A9R466 A9R467 A9R470 A9R484 A9R4A9 A9R4B0
A9R4C1 A9R4C2 A9R4D0 A9R4D2 A9R4D5 A9R4E4 A9R4E5 A9R4E9 A9R4F6 A9R4F9 A9R4G0
A9R4H1 A9R4H2 A9R4I4 A9R4I7 A9R4J2 A9R4J7 A9R4K2 A9R4K6 A9R4K7 A9R4K8 A9R4L4
A9R4M4 A9R4P9 A9R4S1 A9R4U4 A9R4V6 A9R4X7 A9R4Y5 A9R4Y8 A9R514 A9R515 A9R536
A9R541 A9R568 A9R574 A9R579 A9R583 A9R584 A9R587 A9R588 A9R591 A9R594 A9R599
A9R5A1 A9R5A3 A9R5A4 A9R5A6 A9R5A9 A9R5C3 A9R5D1 A9R5F5 A9R5L9 A9R5M0 A9R5Q1
A9R5R3 A9R5R5 A9R5R6 A9R5S1 A9R5U7 A9R5U9 A9R5V2 A9R5W6 A9R5W9 A9R5X0 A9R5X2
A9R622 A9R633 A9R655 A9R664 A9R666 A9R668 A9R670 A9R671 A9R672 A9R674 A9R675
A9R676 A9R677 A9R678 A9R683 A9R685 A9R690 A9R6A3 A9R6B0 A9R6B7 A9R6C5 A9R6C7

A9R6C8 A9R6H5 A9R6H6 A9R6I2 A9R6I7 A9R6I9 A9R6J0 A9R6J7 A9R6M4 A9R6M7 A9R6M9
A9R6N5 A9R6N8 A9R6R1 A9R6X8 A9R6Y0 A9R6Z0 A9R6Z3 A9R6Z5 A9R701 A9R703 A9R755
A9R7D2 A9R7D5 A9R7D9 A9R7E3 A9R7E4 A9R7G5 A9R7I1 A9R7I2 A9R7I3 A9R7I5 A9R7J2
A9R7J9 A9R7L5 A9R7R8 A9R7S4 A9R7V0 A9R7V8 A9R7W3 A9R7W5 A9R7Y6 A9R7Z2 A9R7Z4
A9R7Z8 A9R801 A9R802 A9R805 A9R806 A9R811 A9R815 A9R816 A9R819 A9R8C1 A9R8D2
A9R8D3 A9R8D5 A9R8E1 A9R8E7 A9R8F3 A9R8G1 A9R8H6 A9R8K0 A9R8K3 A9R8K5 A9R8K9
A9R8Q5 A9R8T1 A9R8T9 A9R8V1 A9R8Z5 A9R8Z6 A9R8Z7 A9R8Z9 A9R900 A9R901 A9R902
A9R903 A9R904 A9R905 A9R906 A9R907 A9R908 A9R909 A9R910 A9R911 A9R913 A9R915
A9R916 A9R917 A9R919 A9R925 A9R926 A9R927 A9R931 A9R933 A9R934 A9R941 A9R995
A9R9A9 A9R9C4 A9R9D3 A9RA82 A9RAG8 A9RBS1 A9RFQ5 A9RHW6 A9RHX1 A9RWC9 A9RY03
A9S0B8 A9S5R3 A9S6X4 A9S9Q8 A9SCJ6 A9SCV9 A9SDL4 A9SDW6 A9SLE5 A9SMC8 A9SME1
A9SNJ1 A9SR33 A9SS00 A9SUX2 A9SVH7 A9T142 A9T6G5 A9TF79 A9TIK2 A9TND9 A9TWD9
A9U328 A9UHW6 A9UL13 A9UL14 A9UL70 A9UL91 A9ULH0 A9ULR1 A9ULY7 A9ULZ2 A9UMG5
A9UMM1 A9UMP7 A9UMQ3 A9UMU8 A9UMV8 A9UNU6 A9UP22 A9UPA2 A9UQ62 A9UQN0 A9UQS1
A9UR29 A9URZ4 A9US33 A9USH8 A9UTP3 A9UVV1 A9UXG6 A9UYK7 A9UZ24 A9UZS7 A9V196
A9V2C1 A9V2Y9 A9V3C0 A9V3P1 A9V549 A9V767 A9V790 A9V7A1 A9VB27 A9VCM7 A9VCQ2
A9VCY6 A9VDW7 A9VE54 A9VEU6 A9VF06 A9VFA6 A9VFD5 A9VFD6 A9VFD9 A9VFE0 A9VFE1
A9VFE2 A9VFF7 A9VFG0 A9VFG1 A9VFH8 A9VFJ2 A9VFL4 A9VFV1 A9VFW9 A9VG50 A9VG51
A9VG53 A9VG56 A9VG70 A9VG72 A9VG79 A9VGB0 A9VGC9 A9VGD1 A9VGD3 A9VGD4 A9VGE7
A9VGI1 A9VGI8 A9VGU0 A9VGW3 A9VGW4 A9VGY0 A9VGY2 A9VGY7 A9VH10 A9VH11 A9VH12
A9VH19 A9VH49 A9VH61 A9VHG0 A9VHP7 A9VHP8 A9VHP9 A9VHR4 A9VHR7 A9VHS3 A9VHS5
A9VHT0 A9VHT6 A9VHU0 A9VHU1 A9VHU3 A9VHU8 A9VHU9 A9VHV9 A9VHW1 A9VHZ5 A9VHZ6
A9VI01 A9VI07 A9VI08 A9VI09 A9VI37 A9VI58 A9VI69 A9VI76 A9VI77 A9VI78 A9VI98
A9VI99 A9VID8 A9VIG8 A9VIM2 A9VIM8 A9VIN2 A9VIN4 A9VIP3 A9VIP4 A9VIP6 A9VIP7
A9VIQ3 A9VIR5 A9VIR7 A9VIR9 A9VIS4 A9VIS8 A9VIS9 A9VIT5 A9VIT8 A9VIU0 A9VIU3
A9VIU7 A9VIV4 A9VJ01 A9VJ02 A9VJ21 A9VJ45 A9VJ67 A9VJG1 A9VJI0 A9VJK2 A9VJL2
A9VJL6 A9VJM4 A9VJM6 A9VJN6 A9VJN7 A9VJP3 A9VJP4 A9VJQ4 A9VJR1 A9VJS4 A9VJS5
A9VJU8 A9VJV9 A9VJW0 A9VJW1 A9VJW2 A9VJW3 A9VJX9 A9VK31 A9VK32 A9VK81 A9VKA7
A9VKB0 A9VKE0 A9VKE1 A9VKE9 A9VKG5 A9VKJ7 A9VKK3 A9VKK9 A9VKN0 A9VKN1 A9VKN2
A9VKQ2 A9VKQ3 A9VKR8 A9VKS1 A9VKV6 A9VKX8 A9VL86 A9VLA2 A9VLC2 A9VLC6 A9VLC7
A9VLG7 A9VLG9 A9VLH1 A9VLH2 A9VLH4 A9VLH5 A9VLH6 A9VLH7 A9VLH9 A9VLI9 A9VLJ0
A9VLN1 A9VM74 A9VM77 A9VM89 A9VM90 A9VM93 A9VM99 A9VMA0 A9VMA5 A9VMA7 A9VMB3
A9VMC0 A9VMC2 A9VMC6 A9VMD0 A9VME3 A9VME4 A9VME7 A9VME9 A9VMF0 A9VMF1 A9VMG0
A9VMG7 A9VMG8 A9VMH5 A9VML1 A9VMS6 A9VMV8 A9VMW5 A9VMY8 A9VMZ3 A9VN01 A9VN02
A9VN37 A9VN38 A9VN43 A9VN46 A9VN54 A9VN58 A9VN61 A9VN79 A9VN80 A9VN90 A9VN93
A9VN98 A9VN99 A9VNA0 A9VNA2 A9VNA7 A9VNB0 A9VNB1 A9VNB2 A9VNB3 A9VNN2 A9VNW3
A9VNX6 A9VNZ4 A9VP41 A9VP69 A9VP70 A9VP72 A9VP73 A9VP74 A9VP75 A9VP76 A9VP77
A9VP78 A9VP79 A9VP80 A9VP81 A9VP82 A9VP84 A9VP85 A9VP86 A9VP87 A9VP88 A9VP89
A9VP90 A9VP91 A9VP93 A9VP95 A9VP96 A9VP98 A9VPA1 A9VPA2 A9VPA3 A9VPA5 A9VPB0
A9VPB1 A9VPC3 A9VPH9 A9VPK1 A9VPN7 A9VPT6 A9VPT7 A9VPT8 A9VPT9 A9VQ41 A9VQ50
A9VQ51 A9VQ67 A9VQ69 A9VQ75 A9VQ77 A9VQG4 A9VQG7 A9VQG8 A9VQG9 A9VQI6 A9VQJ9
A9VQK2 A9VQT3 A9VQW6 A9VQW7 A9VR02 A9VR03 A9VR36 A9VR68 A9VRE8 A9VRF0 A9VRF1
A9VRF3 A9VRF5 A9VRG3 A9VRG6 A9VRH6 A9VRH7 A9VRH8 A9VRK3 A9VRK8 A9VRN3 A9VRN4
A9VRQ8 A9VRT5 A9VRT9 A9VS12 A9VS13 A9VS16 A9VS23 A9VS24 A9VSB3 A9VSB4 A9VSC7

A9VSC8 A9VSD6 A9VSE5 A9VSG3 A9VSH6 A9VSI0 A9VSJ0 A9VSL6 A9VSP6 A9VSR3 A9VSR6
A9VSS3 A9VSS7 A9VT44 A9VT45 A9VT48 A9VT50 A9VT54 A9VT57 A9VT62 A9VT64 A9VT65
A9VT66 A9VT67 A9VT68 A9VT70 A9VT74 A9VT75 A9VT78 A9VT79 A9VT80 A9VT82 A9VT87
A9VT96 A9VTA2 A9VTA4 A9VTA8 A9VTB0 A9VTC2 A9VTC3 A9VTC4 A9VTC5 A9VTC6 A9VTC8
A9VTC9 A9VTD1 A9VTD5 A9VTE3 A9VTJ5 A9VTK5 A9VTK8 A9VTL0 A9VTL7 A9VTL8 A9VTL9
A9VTM0 A9VTM3 A9VTM4 A9VTN6 A9VU29 A9VU61 A9VU74 A9VU80 A9VU81 A9VU83 A9VU85
A9VU87 A9VU91 A9VU94 A9VUB0 A9VUC1 A9VUC2 A9VUC4 A9VUD8 A9VUE2 A9VUE3 A9VW37
A9VWA4 A9VWE9 A9VWV4 A9VWV7 A9VWV8 A9VWZ8 A9VX03 A9VX42 A9VXC3 A9VXD6 A9VXT4
A9VY48 A9VY49 A9VY56 A9VY77 A9VY95 A9VYM4 A9VYM7 A9VYM8 A9VYP7 A9VYV8 A9VYW5
A9VYW6 A9VYY7 A9VYY9 A9VYZ2 A9VYZ9 A9VZF3 A9VZG4 A9VZN6 A9VZP1 A9VZR7 A9VZS2
A9VZT0 A9W0A8 A9W0F2 A9W0F3 A9W0G1 A9W0G2 A9W0N8 A9W103 A9W106 A9W185 A9W1A4
A9W1B1 A9W1B3 A9W1C8 A9W1H7 A9W1K0 A9W1L5 A9W1Z5 A9W222 A9W265 A9W296 A9W297
A9W2W3 A9W317 A9W327 A9W370 A9W371 A9W376 A9W377 A9W381 A9W382 A9W383 A9W386
A9W3H7 A9W3H8 A9W3H9 A9W3I0 A9W3I1 A9W3L4 A9W3L5 A9W3R0 A9W3R2 A9W427 A9W480
A9W4A4 A9W4C5 A9W4E5 A9W4G3 A9W4G4 A9W4H6 A9W4L9 A9W4P6 A9W4P7 A9W4P8 A9W4Q0
A9W4Q1 A9W4Q2 A9W4Q3 A9W4Q4 A9W4Q5 A9W4Q6 A9W4Q8 A9W4R0 A9W4R4 A9W4R7 A9W4R8
A9W4R9 A9W4S1 A9W4S2 A9W4S3 A9W4S4 A9W4S5 A9W4S7 A9W4S9 A9W4T0 A9W4X1 A9W4X2
A9W4X6 A9W528 A9W537 A9W592 A9W5A7 A9W5B4 A9W5F4 A9W5F5 A9W5F6 A9W5L3 A9W5L4
A9W5P5 A9W5T6 A9W5U0 A9W5V0 A9W5Y4 A9W605 A9W606 A9W613 A9W640 A9W681 A9W685
A9W6A5 A9W6C6 A9W6G8 A9W6I4 A9W6K8 A9W6Q2 A9W6Q3 A9W6R7 A9W6R8 A9W6T5 A9W6U7
A9W6X0 A9W6X1 A9W6X7 A9W6X8 A9W714 A9W731 A9W737 A9W747 A9W748 A9W7L3 A9W7T2
A9W8D2 A9W8G1 A9W8G3 A9W8G4 A9W8M0 A9W8M1 A9W8M8 A9W8N6 A9W8N7 A9W8N8 A9W8N9
A9W8P8 A9W8P9 A9W8R7 A9W8T7 A9W8W7 A9W900 A9W910 A9W963 A9W966 A9W982 A9W9E2
A9W9G7 A9W9J2 A9W9L9 A9W9Q2 A9W9Q7 A9W9U2 A9W9U3 A9WA87 A9WA88 A9WA95 A9WA97
A9WAF9 A9WAG2 A9WAH5 A9WAI7 A9WAJ7 A9WAL0 A9WAR0 A9WAX3 A9WB43 A9WB45 A9WB97
A9WBA4 A9WBG7 A9WBM7 A9WBM9 A9WBS1 A9WBS4 A9WC26 A9WC30 A9WC34 A9WC35 A9WC36
A9WC39 A9WC40 A9WC41 A9WC63 A9WCA1 A9WCC8 A9WCD8 A9WCD9 A9WCH7 A9WCR2 A9WCR4
A9WCU2 A9WCV7 A9WD25 A9WD26 A9WD80 A9WD89 A9WDA2 A9WDA3 A9WDC2 A9WDD4 A9WDJ3
A9WDJ6 A9WDL8 A9WDL9 A9WDM8 A9WDU6 A9WDU7 A9WDZ1 A9WE23 A9WE34 A9WE39 A9WE59
A9WE79 A9WEA8 A9WEA9 A9WEH7 A9WEJ6 A9WEJ7 A9WEK3 A9WEK5 A9WEK8 A9WEM1 A9WEP1
A9WEX3 A9WF68 A9WFA3 A9WFA4 A9WFH2 A9WFI5 A9WFJ2 A9WFJ8 A9WFP3 A9WFP6 A9WFP7
A9WFP8 A9WFP9 A9WFR1 A9WFR6 A9WFR7 A9WFT9 A9WFU0 A9WFY6 A9WFZ7 A9WFZ9 A9WG10
A9WG69 A9WG70 A9WG80 A9WG82 A9WGB5 A9WGE3 A9WGI4 A9WGI6 A9WGJ4 A9WGJ5 A9WGJ7
A9WGP6 A9WGP7 A9WGQ3 A9WGQ8 A9WH11 A9WH12 A9WH61 A9WH62 A9WH65 A9WH66 A9WH69
A9WH70 A9WH71 A9WH72 A9WH73 A9WH74 A9WH76 A9WH77 A9WH78 A9WH79 A9WH80 A9WH81
A9WH82 A9WH83 A9WH84 A9WH85 A9WH88 A9WH89 A9WH90 A9WH92 A9WH94 A9WH95 A9WH96
A9WH99 A9WHB1 A9WHB7 A9WHF8 A9WHG7 A9WHH4 A9WHH9 A9WHR4 A9WHT8 A9WHV4 A9WI34
A9WI57 A9WI58 A9WI69 A9WIA0 A9WIH1 A9WIH5 A9WIH6 A9WII8 A9WIJ8 A9WIN9 A9WIQ7
A9WIS2 A9WIS6 A9WIS7 A9WIW3 A9WJ10 A9WJ21 A9WJ28 A9WJ45 A9WJ51 A9WJ69 A9WJ74
A9WJ75 A9WJ95 A9WJJ9 A9WJN7 A9WJR8 A9WJV5 A9WK04 A9WK05 A9WK38 A9WK62 A9WK98
A9WKE3 A9WKE4 A9WKF8 A9WKG4 A9WKI1 A9WKL7 A9WKM1 A9WKM8 A9WKU9 A9WKW2 A9WL07
A9WL08 A9WL12 A9WL37 A9WL73 A9WLN5 A9WLY4 A9WM40 A9WMA0 A9WMF8 A9WMF9 A9WMG0
A9WML8 A9WMS3 A9WMV4 A9WMW3 A9WMY0 A9WMZ3 A9WN14 A9WN15 A9WNA7 A9WNB3 A9WNI5
A9WNI9 A9WNU3 A9WP05 A9WP08 A9WP26 A9WP46 A9WP47 A9WP48 A9WP53 A9WPH1 A9WPI0

A9WPN1 A9WPN2 A9WPT6 A9WPU0 A9WPV8 A9WPV9 A9WPX3 A9WPX4 A9WQ47 A9WQ48 A9WQ57
A9WQ58 A9WQ84 A9WQ86 A9WQ88 A9WQ89 A9WQ90 A9WQ91 A9WQA3 A9WQB0 A9WQB1 A9WQH7
A9WQR3 A9WQR6 A9WQS5 A9WQT7 A9WR14 A9WR32 A9WR59 A9WR62 A9WR65 A9WR74 A9WRA9
A9WRD2 A9WRD7 A9WRE6 A9WRE7 A9WRT1 A9WRT6 A9WS39 A9WS75 A9WS85 A9WS93 A9WSA8
A9WSB2 A9WSC4 A9WSE5 A9WSE6 A9WSE7 A9WSE8 A9WSH9 A9WSI0 A9WSI1 A9WSI4 A9WSI9
A9WSN5 A9WSR1 A9WSR2 A9WSR3 A9WSR4 A9WSR5 A9WST7 A9WSU0 A9WSU3 A9WSU4 A9WSU5
A9WSU6 A9WSU7 A9WSV2 A9WSV3 A9WSV4 A9WSV5 A9WSV6 A9WSV9 A9WSW5 A9WSW6 A9WSW7
A9WSW8 A9WSX9 A9WSY0 A9WT08 A9WT19 A9WT40 A9WT63 A9WT66 A9WT68 A9WT69 A9WT85
A9WTE8 A9WTR8 A9WTS8 A9WTS9 A9WTZ3 A9WU36 A9WUU2 A9WUU3 A9WUW1 A9WUW4 A9WV65
A9WV79 A9WVL2 A9WVL3 A9WVP1 A9WVP8 A9WVQ6 A9WVS4 A9WVT5 A9WVT6 A9WVU2 A9WVU3
A9WVU5 A9WVV8 A9WVX6 A9WW32 A9WW33 A9WW36 A9WW39 A9WW41 A9WW43 A9WW44 A9WW58
A9WW62 A9WW74 A9WW75 A9WW76 A9WW77 A9WWC9 A9WWF2 A9WWG0 A9WWG7 A9WWG9 A9WWH9
A9WWI0 A9WWI1 A9WWI6 A9WWJ2 A9WWJ3 A9WWM1 A9WWM7 A9WWP9 A9WWQ2 A9WWR5 A9WWS9
A9WWT1 A9WWT6 A9WWU1 A9WWU9 A9WWW1 A9WWW7 A9WWW8 A9WWW9 A9WWX2 A9WWX3 A9WX00
A9WXE2 A9WXF1 A9WXL2 A9WXP3 A9WXP4 A9WXP5 A9WXQ0 A9WXQ1 A9WXS2 A9WY13 A9WY20
A9WY35 A9WY54 A9WYC6 A9WYG6 A9WYH0 A9WYH2 A9WYN5 A9WYR3 A9WYR4 A9WYR5 A9WYR6
A9WYR8 A9WYS8 A9WYY2 A9WZ04 A9WZ23 A9WZ60 A9WZ75 A9WZ97 A9WZA1 A9WZA3 A9WZF8
A9X185 A9X192 A9X1A5 A9X1A9 A9X1C6 A9X1D0 A9X4T1 A9X6P9 A9X7L0 A9XLE1 A9XLE2
A9XLG1 A9Y0K8 A9Y0K9 A9YDN6 A9YF60 A9YTQ3 A9YUB5 A9YWH3 A9ZLL8 A9ZND1 A9ZPC9
A9ZPH9 A9ZPJ6 A9ZPJ7 A9ZSY0 A9ZSZ2 B0B0A5 B0B0T7 B0B7F6 B0B7I8 B0B7J8 B0B7K3
B0B7K8 B0B7L3 B0B7L8 B0B7N0 B0B7N1 B0B7N2 B0B7N3 B0B7N4 B0B7N5 B0B7N8 B0B7P4
B0B7P5 B0B7P6 B0B7P7 B0B7P9 B0B7Q0 B0B7Q3 B0B7R0 B0B7R1 B0B7R5 B0B7R6 B0B7R8
B0B7S2 B0B7S3 B0B7T5 B0B7T7 B0B7T8 B0B7U5 B0B7U7 B0B7V8 B0B7W0 B0B7W3 B0B7W4
B0B7W5 B0B7W6 B0B7X6 B0B7Y8 B0B7Y9 B0B7Z0 B0B7Z4 B0B800 B0B802 B0B803 B0B804
B0B806 B0B808 B0B809 B0B810 B0B818 B0B825 B0B830 B0B832 B0B835 B0B836 B0B843
B0B858 B0B872 B0B874 B0B875 B0B876 B0B879 B0B880 B0B881 B0B882 B0B883 B0B885
B0B887 B0B889 B0B890 B0B891 B0B892 B0B893 B0B894 B0B895 B0B896 B0B897 B0B898
B0B899 B0B8A0 B0B8A1 B0B8A2 B0B8A4 B0B8B6 B0B8B7 B0B8D2 B0B8E9 B0B8F5 B0B8I0
B0B8I2 B0B8J2 B0B8K0 B0B8K1 B0B8K2 B0B8K9 B0B8M5 B0B8N8 B0B8Q3 B0B8Q5 B0B8R9
B0B8S4 B0B8S9 B0B8T1 B0B8T3 B0B8U8 B0B8V5 B0B8V8 B0B8W9 B0B8X3 B0B8X5 B0B8Y4
B0B8Z2 B0B8Z8 B0B907 B0B910 B0B911 B0B912 B0B913 B0B918 B0B923 B0B925 B0B926
B0B927 B0B928 B0B929 B0B930 B0B931 B0B944 B0B950 B0B960 B0B961 B0B964 B0B969
B0B971 B0B972 B0B991 B0B998 B0B9A9 B0B9B0 B0B9B1 B0B9B5 B0B9C6 B0B9C9 B0B9D0
B0B9D1 B0B9D3 B0B9D4 B0B9D5 B0B9D6 B0B9D9 B0B9E6 B0B9E8 B0B9F3 B0B9G5 B0B9H0
B0B9H9 B0B9I2 B0B9I6 B0B9J4 B0B9K2 B0B9K3 B0B9K4 B0B9L8 B0B9L9 B0B9N3 B0B9N4
B0B9N6 B0B9Q4 B0B9Q8 B0B9S0 B0B9T3 B0B9T4 B0B9T8 B0B9U3 B0B9U7 B0B9V9 B0B9W0
B0B9W1 B0B9W3 B0B9W8 B0B9Y3 B0B9Z5 B0B9Z6 B0BA11 B0BA28 B0BA34 B0BA59 B0BA61
B0BA71 B0BA79 B0BA80 B0BA81 B0BA88 B0BAA4 B0BAB7 B0BAD2 B0BAD4 B0BAE8 B0BAF3
B0BAF8 B0BAG0 B0BAG2 B0BAH7 B0BAI4 B0BAI7 B0BAK2 B0BAK4 B0BAL3 B0BAM1 B0BAM7
B0BAN6 B0BAN9 B0BAP0 B0BAP1 B0BAP2 B0BAP7 B0BAQ4 B0BAQ5 B0BAQ6 B0BAQ7 B0BAQ8
B0BAQ9 B0BAR0 B0BAS3 B0BAS9 B0BAT9 B0BAU0 B0BAU3 B0BAU8 B0BAV0 B0BAV1 B0BAX0
B0BAX7 B0BAY8 B0BAY9 B0BAZ0 B0BAZ4 B0BB05 B0BB08 B0BB09 B0BB12 B0BB13 B0BB14
B0BB15 B0BB18 B0BB25 B0BB27 B0BB32 B0BB44 B0BB49 B0BB58 B0BB61 B0BB65 B0BB73
B0BB81 B0BB82 B0BB83 B0BB97 B0BB98 B0BBB2 B0BBB3 B0BBB5 B0BBD3 B0BBD7 B0BBF0

B0BBG3 B0BBG4 B0BBG8 B0BBH3 B0BBH7 B0BBI9 B0BBJ0 B0BBJ1 B0BBJ3 B0BBJ8 B0BBM1
B0BBQ3 B0BBR3 B0BBR8 B0BBS3 B0BBS8 B0BBT2 B0BBT3 B0BBU5 B0BBU6 B0BBU7 B0BBU8
B0BBU9 B0BBV0 B0BBV3 B0BBV9 B0BBW0 B0BBW1 B0BBW2 B0BBW4 B0BBW8 B0BBX5 B0BBX6
B0BBY0 B0BBY1 B0BBY3 B0BBY7 B0BBY8 B0BC00 B0BC02 B0BC03 B0BC10 B0BC12 B0BC23
B0BC25 B0BC28 B0BC29 B0BC30 B0BC31 B0BC41 B0BC53 B0BC54 B0BC55 B0BC59 B0BC65
B0BC67 B0BC69 B0BC71 B0BC73 B0BC74 B0BC75 B0BC83 B0BC90 B0BC95 B0BC97 B0BCA0
B0BCA1 B0BCA8 B0BCC3 B0BCD7 B0BCD9 B0BCE0 B0BCE1 B0BCE5 B0BCE6 B0BCE7 B0BCE8
B0BCF0 B0BCF2 B0BCF3 B0BCF4 B0BCF5 B0BCF6 B0BCF7 B0BCF8 B0BCF9 B0BCG0 B0BCG1
B0BCG2 B0BCG3 B0BCG4 B0BCG5 B0BCG6 B0BCG7 B0BCM0 B0BCM1 B0BCM2 B0BCM3 B0BCM4
B0BCM5 B0BK70 B0BLT0 B0BLU1 B0BM16 B0BM20 B0BM24 B0BM28 B0BM40 B0BM95 B0BMJ2
B0BML1 B0BML7 B0BMX9 B0BMZ3 B0BMZ6 B0BN13 B0BN18 B0BN49 B0BN56 B0BN72 B0BN85
B0BN93 B0BN94 B0BN95 B0BNA7 B0BNA9 B0BNC9 B0BND4 B0BNE2 B0BNE4 B0BNE5 B0BNK8
B0BNK9 B0BNN3 B0BNN9 B0BNR3 B0BNR5 B0BNR6 B0BNS1 B0BNS7 B0BNS8 B0BNV4 B0BNW4
B0BNW7 B0BNX2 B0BNX3 B0BNY0 B0BNY1 B0BNZ0 B0BNZ1 B0BNZ2 B0BP03 B0BP10 B0BP16
B0BP19 B0BP27 B0BP33 B0BP37 B0BP40 B0BP45 B0BP57 B0BP79 B0BP80 B0BP82 B0BP83
B0BP87 B0BP90 B0BP95 B0BP99 B0BPB3 B0BPB6 B0BPC7 B0BPE2 B0BPE7 B0BPF3 B0BPF6
B0BPF7 B0BPG0 B0BPI1 B0BPI2 B0BPI4 B0BPI7 B0BPI8 B0BPJ8 B0BPK8 B0BPQ7 B0BPR0
B0BPR8 B0BPS5 B0BPS7 B0BPT5 B0BPU4 B0BPV1 B0BPV2 B0BPV3 B0BPV4 B0BPX9 B0BQ12
B0BQ17 B0BQ30 B0BQ32 B0BQ37 B0BQ46 B0BQ49 B0BQ51 B0BQ52 B0BQ57 B0BQ60 B0BQ64
B0BQ73 B0BQ77 B0BQ79 B0BQ80 B0BQ90 B0BQ92 B0BQ96 B0BQA6 B0BQB0 B0BQB1 B0BQB2
B0BQB3 B0BQB9 B0BQC2 B0BQG0 B0BQG3 B0BQI3 B0BQK6 B0BQK7 B0BQL3 B0BQM0 B0BQM9
B0BQN0 B0BQN2 B0BQN6 B0BQP9 B0BQQ2 B0BQR0 B0BQT2 B0BQT7 B0BQV2 B0BQW9 B0BQX5
B0BQZ1 B0BQZ3 B0BQZ4 B0BQZ5 B0BR13 B0BR42 B0BR43 B0BR51 B0BR65 B0BR82 B0BR83
B0BR90 B0BR94 B0BRA0 B0BRA2 B0BRA6 B0BRB3 B0BRC5 B0BRC9 B0BRD0 B0BRD1 B0BRD5
B0BRE1 B0BRF0 B0BRF3 B0BRG1 B0BRG8 B0BRG9 B0BRH8 B0BRH9 B0BRI9 B0BRJ2 B0BRJ9
B0BRL3 B0BRL8 B0BRM0 B0BRN3 B0BRR3 B0BRR9 B0BRS0 B0BRS6 B0BRS8 B0BRT1 B0BRT7
B0BRT8 B0BRT9 B0BRU2 B0BRU3 B0BRU4 B0BRV5 B0BRW2 B0BRW3 B0BRW5 B0BRY0 B0BRY1
B0BRY5 B0BS03 B0BS15 B0BS17 B0BS22 B0BS34 B0BS35 B0BS41 B0BS43 B0BS44 B0BS54
B0BS55 B0BS63 B0BS66 B0BS95 B0BSB3 B0BSD7 B0BSE7 B0BSE8 B0BSF5 B0BSF6 B0BSG4
B0BSG5 B0BSG6 B0BSG9 B0BSH8 B0BSI0 B0BSI2 B0BSJ7 B0BSK4 B0BSL0 B0BSL4 B0BSM2
B0BSM7 B0BSM8 B0BSN8 B0BSN9 B0BSQ0 B0BSS1 B0BSS6 B0BST0 B0BST1 B0BST2 B0BST4
B0BST5 B0BST6 B0BST8 B0BST9 B0BSU0 B0BSU1 B0BSU2 B0BSU3 B0BSU4 B0BSU5 B0BSU6
B0BSU7 B0BSU9 B0BSV0 B0BSV2 B0BSV4 B0BSV5 B0BSV7 B0BSV8 B0BSV9 B0BSW0 B0BSW1
B0BSW9 B0BSX2 B0BSX5 B0BSY9 B0BSZ3 B0BSZ4 B0BSZ8 B0BT03 B0BT12 B0BT16 B0BT17
B0BT25 B0BT63 B0BT67 B0BT84 B0BT96 B0BTA1 B0BTA2 B0BTA9 B0BTB3 B0BTB4 B0BTB9
B0BTC2 B0BTD3 B0BTE1 B0BTF1 B0BTF3 B0BTH6 B0BTI0 B0BTI6 B0BTI7 B0BTI9 B0BTJ8
B0BTK6 B0BTK8 B0BTL7 B0BTL8 B0BTN8 B0BTN9 B0BTQ3 B0BTQ4 B0BTQ6 B0BTQ8 B0BTQ9
B0BTR0 B0BTR9 B0BTT2 B0BTV5 B0BTW8 B0BTX7 B0BTX9 B0BTY3 B0BTY5 B0BTY6 B0BTZ2
B0BTZ3 B0BTZ5 B0BTZ6 B0BTZ7 B0BU06 B0BU16 B0BU26 B0BU33 B0BU37 B0BU47 B0BU51
B0BU54 B0BU72 B0BU73 B0BU91 B0BUA6 B0BUA8 B0BUC4 B0BUC5 B0BUC8 B0BUD0 B0BUD1
B0BUD5 B0BUD7 B0BUD9 B0BUE2 B0BUF0 B0BUF3 B0BUF9 B0BUG0 B0BUG1 B0BUG7 B0BUI6
B0BUI7 B0BUJ0 B0BUJ1 B0BUJ2 B0BUK2 B0BUL7 B0BUM0 B0BUM3 B0BUN5 B0BUN6 B0BUN7
B0BUN8 B0BUP1 B0BUP2 B0BUP3 B0BUP4 B0BUP5 B0BUP6 B0BUP7 B0BUP8 B0BUP9 B0BUQ0
B0BUQ1 B0BUQ2 B0BUQ3 B0BUQ4 B0BUQ5 B0BUQ6 B0BUQ7 B0BUQ8 B0BUQ9 B0BUR0 B0BUR1

B0BUR2 B0BUT3 B0BUT9 B0BUU0 B0BUU8 B0BUV1 B0BUW3 B0BUW5 B0BUZ1 B0BUZ2 B0BV24
B0BV26 B0BV27 B0BV43 B0BV44 B0BV57 B0BV81 B0BV96 B0BVC7 B0BVE3 B0BVE4 B0BVE6
B0BVI5 B0BVI6 B0BVJ0 B0BVJ1 B0BVK1 B0BVK9 B0BVL0 B0BVL9 B0BVN5 B0BVP3 B0BVP5
B0BVP6 B0BVP7 B0BVQ3 B0BVQ6 B0BVQ7 B0BVU8 B0BVW3 B0BVX7 B0BVY1 B0BVY2 B0BVY4
B0BW02 B0BW03 B0BW04 B0BW16 B0BW17 B0BW24 B0BW26 B0BW37 B0BW38 B0BW39 B0BW55
B0BW65 B0BW78 B0BW79 B0BW80 B0BW84 B0BW86 B0BW95 B0BWA1 B0BWA2 B0BWA5 B0BWA6
B0BWA7 B0BWA9 B0BWB0 B0BWB2 B0BWC2 B0BWC3 B0BWC4 B0BWC7 B0BWD7 B0BWD8 B0BWH0
B0BWH1 B0BWJ7 B0BWJ8 B0BWJ9 B0BWK0 B0BWL1 B0BWL5 B0BWM7 B0BWQ0 B0BWQ1 B0BWQ7
B0BWQ9 B0BWS0 B0BWS1 B0BWS2 B0BWY8 B0BWZ7 B0BWZ9 B0BX21 B0BX22 B0BX32 B0BX35
B0BX42 B0BX57 B0BX63 B0BX65 B0BX67 B0BX92 B0BXA8 B0BXB3 B0BXB8 B0BXB9 B0BXC2
B0BXE1 B0BXH3 B0BXI2 B0BXJ1 B0BXJ2 B0BXJ5 B0BXK5 B0BXK8 B0BXL4 B0BXN7 B0BXN8
B0BXQ1 B0BXR0 B0BXR1 B0BXR2 B0BXT9 B0BXU8 B0BXX6 B0BXY6 B0BXZ0 B0BY00 B0BY23
B0BY57 B0BY61 B0BY63 B0BY69 B0BYA0 B0BYA1 B0BYA6 B0BYC5 B0BYC6 B0BYF7 B0BYK8
B0BYM7 B0BYP7 B0BYP8 B0BYR8 B0BYW4 B0BZ26 B0BZ38 B0BZ87 B0BZ94 B0BZ98 B0BZF8
B0BZF9 B0BZH9 B0BZI8 B0BZP2 B0BZP6 B0BZQ6 B0BZS8 B0BZS9 B0BZY6 B0C074 B0C075
B0C079 B0C090 B0C0B6 B0C0B9 B0C0E2 B0C118 B0C132 B0C142 B0C146 B0C1B5 B0C1D3
B0C1D4 B0C1D5 B0C1D6 B0C1D7 B0C1D8 B0C1D9 B0C1E0 B0C1E1 B0C1E2 B0C1E3 B0C1E4
B0C1E5 B0C1E6 B0C1E7 B0C1E8 B0C1E9 B0C1F0 B0C1F3 B0C1M5 B0C1P2 B0C1R1 B0C1T2
B0C1U8 B0C1Y1 B0C1Z3 B0C205 B0C218 B0C230 B0C250 B0C253 B0C269 B0C2C8 B0C2D1
B0C2D5 B0C2J3 B0C2J4 B0C2K7 B0C2T9 B0C2W2 B0C2W4 B0C2Z9 B0C308 B0C312 B0C321
B0C384 B0C385 B0C390 B0C3C2 B0C3C3 B0C3J3 B0C3N6 B0C424 B0C425 B0C426 B0C427
B0C429 B0C431 B0C432 B0C4F3 B0C4G1 B0C4I0 B0C4N8 B0C4Q9 B0C512 B0C5D4 B0C5Q2
B0C6E1 B0C6F8 B0C6I2 B0C6R3 B0C6S1 B0C6V3 B0C6V8 B0C6Z1 B0C790 B0C791 B0C793
B0C7G7 B0C7R9 B0C7S1 B0C7S8 B0C7T1 B0C7T2 B0C811 B0C872 B0C882 B0C899 B0C8B9
B0C8D6 B0C8E3 B0C8E5 B0C8H2 B0C8J2 B0C8J3 B0C8Q6 B0C904 B0C915 B0C921 B0C929
B0C977 B0C9F4 B0C9F5 B0C9F6 B0C9G3 B0C9N7 B0C9S0 B0C9Y4 B0CA72 B0CA77 B0CAA5
B0CAB9 B0CAC6 B0CAC9 B0CAD0 B0CAD1 B0CAD2 B0CAD9 B0CAE7 B0CAI0 B0CAI3 B0CAL3
B0CAM1 B0CAM2 B0CAM3 B0CAV7 B0CAX3 B0CAZ0 B0CB11 B0CB57 B0CB70 B0CB83 B0CBB0
B0CBC7 B0CBC9 B0CBD5 B0CBE9 B0CBF7 B0CBS9 B0CBU2 B0CBX6 B0CC57 B0CC69 B0CC88
B0CC89 B0CCC1 B0CCC9 B0CCD0 B0CCD2 B0CCD3 B0CCD5 B0CCD6 B0CCF6 B0CCG9 B0CCH1
B0CCJ3 B0CCQ2 B0CCR3 B0CCT9 B0CD11 B0CD21 B0CD47 B0CD57 B0CD68 B0CD77 B0CD95
B0CDA0 B0CDA2 B0CDE2 B0CDF8 B0CDH5 B0CDK8 B0CDM2 B0CDQ3 B0CDX9 B0CDZ6 B0CE09
B0CE23 B0CE31 B0CE35 B0CE36 B0CE39 B0CED7 B0CED9 B0CEI9 B0CEJ2 B0CEJ4 B0CEK4
B0CEL5 B0CEN4 B0CEN8 B0CEQ4 B0CEU2 B0CEV0 B0CEX1 B0CEZ1 B0CEZ5 B0CF01 B0CF30
B0CF78 B0CFL0 B0CFM3 B0CFQ6 B0CFX4 B0CG28 B0CG35 B0CG96 B0CGE4 B0CGG7 B0CGJ9
B0CGM0 B0CGM4 B0CGM5 B0CGP2 B0CGP5 B0CGR0 B0CGR1 B0CGR3 B0CGS4 B0CGT4 B0CGT8
B0CGT9 B0CGU0 B0CGU5 B0CGV6 B0CGV8 B0CGV9 B0CGW7 B0CGY1 B0CGZ3 B0CGZ9 B0CH00
B0CH06 B0CH07 B0CH08 B0CH09 B0CH11 B0CH13 B0CH14 B0CH15 B0CH16 B0CH17 B0CH18
B0CH19 B0CH20 B0CH21 B0CH22 B0CH23 B0CH24 B0CH25 B0CH26 B0CH27 B0CH28 B0CH29
B0CH30 B0CH31 B0CH32 B0CH33 B0CH34 B0CH35 B0CH36 B0CH37 B0CH40 B0CH41 B0CH42
B0CH43 B0CH44 B0CH45 B0CH77 B0CHA6 B0CHF1 B0CHG8 B0CHH2 B0CHH8 B0CHI7 B0CHJ9
B0CHK5 B0CHK8 B0CHL8 B0CHL9 B0CHM8 B0CHP1 B0CHR3 B0CHS0 B0CHU1 B0CHX4 B0CHX5
B0CHX9 B0CI45 B0CI46 B0CI52 B0CI53 B0CI68 B0CIC3 B0CIC6 B0CID9 B0CIE0 B0CIF7
B0CIF8 B0CII7 B0CII9 B0CIJ4 B0CIL1 B0CIM5 B0CIQ3 B0CIQ5 B0CIR1 B0CIR8 B0CIR9

B0CIS7 B0CIT0 B0CIT1 B0CIT6 B0CIU0 B0CIX5 B0CIY2 B0CIY4 B0CJ30 B0CJ31 B0CJ32
B0CJ37 B0CJ51 B0CJ70 B0CJB7 B0CJD0 B0CJD1 B0CJG0 B0CJG1 B0CJG2 B0CJG6 B0CJG9
B0CJH2 B0CJH9 B0CJI0 B0CJI2 B0CJI5 B0CJI6 B0CJI7 B0CJI8 B0CJJ7 B0CJK6 B0CJK8
B0CJK9 B0CJL7 B0CJL8 B0CJM7 B0CJM9 B0CJN2 B0CJP3 B0CJP9 B0CJQ0 B0CJR3 B0CJT2
B0CJW0 B0CJW1 B0CJW8 B0CJX0 B0CJX5 B0CJX7 B0CK00 B0CK02 B0CK08 B0CK11 B0CK12
B0CK13 B0CK14 B0CK15 B0CK30 B0CK66 B0CK74 B0CK89 B0CKA2 B0CKB2 B0CKC0 B0CKD6
B0CKD8 B0CKD9 B0CKG1 B0CKG2 B0CKH7 B0CKJ0 B0CKM9 B0CKN3 B0CKN4 B0CKN5 B0CKX3
B0CKX7 B0CKX9 B0CKY7 B0CL02 B0CL06 B0CL18 B0CL36 B0CL77 B0CL81 B0CL90 B0CL91
B0CL95 B0CLA3 B0CLB5 B0CLB7 B0CLB8 B0CLE9 B0CLF6 B0CLF8 B0CLG4 B0CLG9 B0CLJ2
B0CLK1 B0CLK2 B0CLL0 B0CLL1 B0CLM0 B0CLM5 B0CLN2 B0CLQ0 B0CLS9 B0CLT0 B0CM26
B0CM32 B0CM36 B0CM52 B0CN28 B0CN31 B0CN39 B0CPQ8 B0CQH8 B0CQL7 B0CR45 B0CRJ3
B0CRK4 B0CRL4 B0CRU6 B0CS49 B0CXE1 B0CXQ2 B0CY45 B0CZ32 B0D0T8 B0D0U4 B0D174
B0D1Q8 B0D316 B0D360 B0D442 B0D4E7 B0D4J6 B0D6H2 B0DAD3 B0DK66 B0DN41 B0DOB4
B0DOB5 B0DQB9 B0DSK4 B0DSN9 B0DUI8 B0DWM8 B0DWN3 B0DX25 B0DZL3 B0E412 B0EAV2
B0EIW5 B0EMY3 B0EN90 B0EXJ8 B0F481 B0FGA9 B0FIH3 B0FIH7 B0FII2 B0FIL5 B0FIN6
B0FRF9 B0FXI0 B0FXI7 B0FXQ5 B0FYK7 B0FZN7 B0FZN8 B0FZN9 B0FZP0 B0FZP1 B0FZP2
B0FZP3 B0G0Y5 B0G0Z3 B0G100 B0G101 B0G102 B0G103 B0G104 B0G107 B0G124 B0G125
B0G132 B0G138 B0G170 B0G172 B0G186 B0G188 B0G195 B0I1G7 B0JFL6 B0JFP4 B0JFQ0
B0JFQ9 B0JFW8 B0JFX8 B0JFY0 B0JFY2 B0JGA0 B0JGA6 B0JGE2 B0JGG3 B0JGK2 B0JGK4
B0JGK8 B0JGL3 B0JGQ4 B0JGW5 B0JGZ3 B0JH56 B0JH61 B0JH94 B0JHB1 B0JHC1 B0JHH4
B0JHT4 B0JHV0 B0JHV3 B0JHX2 B0JHY6 B0JHY7 B0JHY8 B0JHY9 B0JHZ0 B0JHZ1 B0JHZ2
B0JHZ3 B0JHZ4 B0JHZ5 B0JHZ6 B0JHZ7 B0JHZ8 B0JHZ9 B0JI00 B0JI02 B0JI03 B0JI21
B0JI39 B0JI84 B0JIA6 B0JID6 B0JII8 B0JIQ1 B0JIR2 B0JIY7 B0JJ52 B0JJ58 B0JJC4
B0JJJ4 B0JJJ5 B0JJJ7 B0JJJ8 B0JJL5 B0JJP7 B0JJS5 B0JK88 B0JK91 B0JKA1 B0JKN4
B0JKN5 B0JKQ1 B0JKS7 B0JKT7 B0JL12 B0JL16 B0JL23 B0JL55 B0JL88 B0JL91 B0JL93
B0JL96 B0JLH2 B0JLI8 B0JLJ8 B0JLM0 B0JLN5 B0JLP1 B0JLR5 B0JLU4 B0JM14 B0JM53
B0JMT8 B0JN42 B0JN62 B0JN63 B0JN99 B0JNA9 B0JNB0 B0JNC2 B0JNJ4 B0JNQ1 B0JNQ4
B0JNQ5 B0JNU3 B0JNW2 B0JNW9 B0JPD9 B0JPG5 B0JPJ2 B0JPM4 B0JPM6 B0JPS9 B0JPT8
B0JPW6 B0JPX8 B0JPY2 B0JQ94 B0JQ95 B0JQG3 B0JQG4 B0JQG5 B0JQJ1 B0JQL4 B0JQV0
B0JQZ0 B0JR14 B0JR45 B0JR63 B0JR86 B0JRF1 B0JRM9 B0JRN7 B0JRP2 B0JRP8 B0JRV0
B0JRV5 B0JS32 B0JS46 B0JS91 B0JSC7 B0JSD7 B0JSD9 B0JSE0 B0JSE2 B0JSE3 B0JSJ8
B0JSJ9 B0JSK4 B0JSN9 B0JSP4 B0JSQ9 B0JSW2 B0JSX3 B0JT33 B0JT86 B0JTA6 B0JTA9
B0JTD7 B0JTF4 B0JTF5 B0JTF9 B0JTG1 B0JTL2 B0JTL3 B0JTM2 B0JTP1 B0JTP2 B0JTT9
B0JTU1 B0JTW2 B0JTZ2 B0JU07 B0JU67 B0JU70 B0JU82 B0JUF7 B0JUG8 B0JUH1 B0JUI1
B0JUK5 B0JUK9 B0JUM0 B0JUP8 B0JV09 B0JV88 B0JVA7 B0JVB1 B0JVE7 B0JVG7 B0JVL8
B0JVM6 B0JVN3 B0JVN4 B0JVQ9 B0JVR4 B0JVV0 B0JVY2 B0JVZ6 B0JW10 B0JW21 B0JW23
B0JW81 B0JWQ2 B0JWQ4 B0JWQ8 B0JWW5 B0JWW6 B0JWX2 B0JX00 B0JX03 B0JX07 B0JX47
B0JXE8 B0JXF5 B0JXF6 B0JXI2 B0JXU3 B0JY33 B0JY34 B0JY35 B0JY37 B0JY39 B0JY40
B0JY53 B0JY70 B0JY77 B0JY78 B0JY97 B0JYC2 B0JYC3 B0JYC5 B0JYE6 B0JYF4 B0JYS7
B0JYW5 B0JZ65 B0JZ85 B0JZ89 B0JZV4 B0K012 B0K035 B0K036 B0K0D9 B0K0F7 B0K0I7
B0K0J1 B0K0J7 B0K0L3 B0K0L7 B0K0L8 B0K0M0 B0K0M1 B0K0N0 B0K0N1 B0K0N4 B0K0N5
B0K0P8 B0K0Q1 B0K0Q3 B0K0S9 B0K0T1 B0K0T2 B0K0T3 B0K0T5 B0K0U6 B0K0U8 B0K0U9
B0K0V4 B0K0V5 B0K0W8 B0K0X1 B0K0Y0 B0K0Y2 B0K0Y9 B0K0Z0 B0K0Z5 B0K100 B0K101
B0K119 B0K150 B0K158 B0K172 B0K177 B0K189 B0K197 B0K199 B0K1A1 B0K1A3 B0K1A4

B0K1A5 B0K1A7 B0K1B9 B0K1C1 B0K1C7 B0K1D0 B0K1D1 B0K1D6 B0K1E0 B0K1E5 B0K1F3
B0K1F4 B0K1F7 B0K1G2 B0K1L3 B0K1M3 B0K1N2 B0K1P6 B0K1P8 B0K1P9 B0K1T5 B0K1T6
B0K1T7 B0K1U6 B0K1U8 B0K1V0 B0K1X1 B0K1X4 B0K1Y2 B0K1Y9 B0K212 B0K213 B0K226
B0K241 B0K242 B0K250 B0K272 B0K280 B0K292 B0K295 B0K297 B0K2C2 B0K2C3 B0K2E4
B0K2E5 B0K2E7 B0K2E8 B0K2F0 B0K2J6 B0K2K5 B0K2T8 B0K2T9 B0K2U0 B0K2U1 B0K2U2
B0K2V6 B0K2W0 B0K2W1 B0K2W6 B0K346 B0K363 B0K364 B0K365 B0K3E4 B0K3F9 B0K3H2
B0K3H8 B0K3H9 B0K3I5 B0K3J8 B0K3L3 B0K3N1 B0K3P5 B0K3P6 B0K3Q1 B0K3Q4 B0K3Q6
B0K3Q8 B0K3R0 B0K3S1 B0K3S2 B0K3S3 B0K3S4 B0K3T4 B0K3T5 B0K3V0 B0K3V4 B0K3V6
B0K3W4 B0K3X5 B0K3X8 B0K3Y0 B0K3Y2 B0K403 B0K413 B0K414 B0K416 B0K418 B0K421
B0K427 B0K429 B0K431 B0K437 B0K460 B0K466 B0K474 B0K485 B0K4D2 B0K4D4 B0K4D7
B0K4D8 B0K4D9 B0K4E2 B0K4I3 B0K4I5 B0K4M4 B0K4N6 B0K4N7 B0K530 B0K532 B0K533
B0K534 B0K544 B0K568 B0K569 B0K584 B0K590 B0K5A5 B0K5B1 B0K5B9 B0K5C0 B0K5C1
B0K5C5 B0K5C8 B0K5F2 B0K5F4 B0K5F6 B0K5F7 B0K5G4 B0K5G5 B0K5G6 B0K5G7 B0K5G8
B0K5G9 B0K5H2 B0K5I0 B0K5L4 B0K5L7 B0K5L9 B0K5N2 B0K5N3 B0K5N4 B0K5N8 B0K5N9
B0K5P0 B0K5P2 B0K5P3 B0K5P4 B0K5P5 B0K5P6 B0K5P7 B0K5P8 B0K5Q0 B0K5Q1 B0K5Q2
B0K5Q3 B0K5Q4 B0K5Q5 B0K5Q6 B0K5Q7 B0K5Q8 B0K5Q9 B0K5R0 B0K5R1 B0K5R3 B0K5R7
B0K5R8 B0K5R9 B0K5S0 B0K5S1 B0K5S2 B0K5S6 B0K5S7 B0K5S8 B0K5X4 B0K5Y2 B0K5Z9
B0K622 B0K623 B0K625 B0K626 B0K628 B0K629 B0K631 B0K643 B0K664 B0K680 B0K682
B0K683 B0K6D2 B0K6G7 B0K6J6 B0K6K7 B0K6L2 B0K6M6 B0K6X4 B0K6Y4 B0K6Z9 B0K706
B0K709 B0K710 B0K714 B0K718 B0K732 B0K733 B0K735 B0K736 B0K742 B0K754 B0K775
B0K791 B0K793 B0K794 B0K7G1 B0K7G6 B0K7G9 B0K7H0 B0K7K7 B0K7S0 B0K7T0 B0K7U2
B0K7U7 B0K7V7 B0K800 B0K801 B0K880 B0K889 B0K8A4 B0K8B2 B0K8B4 B0K8B5 B0K8D0
B0K8D7 B0K8F8 B0K8G2 B0K8G4 B0K8H7 B0K8H8 B0K8H9 B0K8I3 B0K8I4 B0K8J2 B0K8J8
B0K8J9 B0K8K5 B0K8L8 B0K8N3 B0K8Q5 B0K8R2 B0K8R7 B0K8R8 B0K8S2 B0K8T3 B0K8T4
B0K8T5 B0K8T6 B0K8T7 B0K8Z7 B0K924 B0K928 B0K934 B0K951 B0K955 B0K956 B0K958
B0K959 B0K969 B0K971 B0K977 B0K978 B0K991 B0K994 B0K996 B0K9C2 B0K9C4 B0K9C5
B0K9C6 B0K9C8 B0K9D9 B0K9E1 B0K9E2 B0K9E7 B0K9E8 B0K9G3 B0K9H1 B0K9I5 B0K9J0
B0K9K2 B0K9L0 B0K9L2 B0K9L4 B0K9L6 B0K9L7 B0K9L8 B0K9M0 B0K9N3 B0K9N5 B0K9P1
B0K9P4 B0K9P5 B0K9Q0 B0K9Q4 B0K9Q9 B0K9R3 B0K9R5 B0K9R6 B0K9V2 B0K9V3 B0K9V4
B0K9W3 B0K9W5 B0K9W7 B0K9Y6 B0K9Y8 B0K9Z1 B0K9Z9 B0KA06 B0KA11 B0KA14 B0KA16
B0KA31 B0KA32 B0KA34 B0KA35 B0KA38 B0KA44 B0KA48 B0KA52 B0KA53 B0KA56 B0KA63
B0KA79 B0KA81 B0KA83 B0KAA6 B0KAB6 B0KAB8 B0KAC0 B0KAC2 B0KAC5 B0KAD1 B0KAD3
B0KAD5 B0KAE1 B0KAG0 B0KAG3 B0KAH3 B0KAH5 B0KAI2 B0KAI3 B0KAI8 B0KAL7 B0KAL9
B0KAQ8 B0KAS0 B0KAS1 B0KB08 B0KB09 B0KBA1 B0KBA3 B0KBA4 B0KBA5 B0KBA7 B0KBD3
B0KBF4 B0KBG1 B0KBH0 B0KBI1 B0KBM4 B0KBM5 B0KBN3 B0KBN4 B0KBN5 B0KBN6 B0KBP8
B0KBQ1 B0KBQ3 B0KBQ5 B0KBQ8 B0KBR3 B0KBR4 B0KBS9 B0KBV9 B0KBW1 B0KBW4 B0KBW5
B0KBW6 B0KBW9 B0KC14 B0KC20 B0KC36 B0KC42 B0KC64 B0KC73 B0KC90 B0KC91 B0KC92
B0KCA9 B0KCB7 B0KCB9 B0KCE3 B0KCE4 B0KCE5 B0KCE9 B0KCF2 B0KCH5 B0KCH7 B0KCH9
B0KCI5 B0KCI8 B0KCI9 B0KCJ0 B0KCJ1 B0KCJ2 B0KCJ3 B0KCJ5 B0KCJ6 B0KCJ7 B0KCJ9
B0KCK0 B0KCK1 B0KCK2 B0KCK3 B0KCK4 B0KCK5 B0KCK7 B0KCK8 B0KCK9 B0KCL0 B0KCL1
B0KCL2 B0KCL3 B0KCL4 B0KCL5 B0KCL6 B0KCL7 B0KCL8 B0KCM0 B0KCM4 B0KCM5 B0KCM6
B0KCM7 B0KCM8 B0KCM9 B0KCN3 B0KCN4 B0KCN5 B0KCS6 B0KCW3 B0KCX1 B0KD39 B0KD48
B0KD66 B0KD87 B0KD95 B0KD96 B0KDA4 B0KDB7 B0KDB8 B0KDE7 B0KEU7 B0KEU8 B0KEU9
B0KEV1 B0KF22 B0KF23 B0KF29 B0KF32 B0KF35 B0KF42 B0KF52 B0KF60 B0KF73 B0KF81

B0KF82 B0KF94 B0KFA9 B0KFQ2 B0KFQ3 B0KFR1 B0KFS4 B0KFS5 B0KFT4 B0KFT5 B0KFT8
B0KFT9 B0KFU7 B0KFU8 B0KFW2 B0KFW3 B0KG04 B0KG76 B0KG79 B0KG92 B0KGD7 B0KGD8
B0KGD9 B0KGK3 B0KGN7 B0KGP8 B0KGQ1 B0KGU6 B0KH03 B0KH05 B0KH56 B0KH90 B0KH99
B0KHA7 B0KHQ8 B0KHR5 B0KHU2 B0KHV5 B0KHV8 B0KHW5 B0KHW6 B0KHX3 B0KHX5 B0KHX6
B0KHX7 B0KHX8 B0KHY0 B0KHY2 B0KHY3 B0KHY6 B0KI00 B0KI04 B0KI05 B0KI18 B0KI37
B0KI41 B0KI44 B0KI45 B0KIG1 B0KIS3 B0KIS4 B0KIS5 B0KIS6 B0KIT1 B0KJ25 B0KJ53
B0KJ54 B0KJ68 B0KJ69 B0KJ81 B0KJ82 B0KJ86 B0KJ88 B0KJ90 B0KJ91 B0KJ92 B0KJB2
B0KJG3 B0KJG5 B0KJG6 B0KJG7 B0KJV8 B0KJV9 B0KJW3 B0KJW8 B0KJX5 B0KJX6 B0KJY2
B0KJY4 B0KJY5 B0KJZ2 B0KJZ6 B0KK08 B0KK25 B0KK26 B0KK35 B0KK45 B0KK51 B0KK53
B0KK56 B0KK57 B0KK58 B0KK59 B0KK60 B0KK61 B0KK62 B0KK63 B0KK66 B0KK67 B0KK68
B0KK69 B0KK70 B0KK71 B0KK72 B0KK74 B0KK75 B0KK76 B0KK77 B0KK78 B0KK79 B0KK80
B0KK81 B0KK82 B0KK83 B0KK84 B0KK85 B0KK86 B0KK88 B0KK89 B0KK90 B0KK92 B0KKB9
B0KKI2 B0KKK0 B0KKR2 B0KKR4 B0KKR5 B0KKR6 B0KKR7 B0KKR8 B0KKS4 B0KKX3 B0KKX4
B0KKY0 B0KKY2 B0KKY3 B0KKZ0 B0KKZ1 B0KKZ5 B0KKZ6 B0KKZ7 B0KL04 B0KL24 B0KL32
B0KL66 B0KL70 B0KL71 B0KL75 B0KL79 B0KL81 B0KL94 B0KLE3 B0KLM9 B0KLP4 B0KLW8
B0KLX3 B0KLX4 B0KLX5 B0KLX7 B0KLY7 B0KLZ1 B0KLZ2 B0KM00 B0KM20 B0KM22 B0KM23
B0KM26 B0KM36 B0KM39 B0KM40 B0KM52 B0KM55 B0KM58 B0KM65 B0KM81 B0KM84 B0KM87
B0KMD1 B0KMF4 B0KMF5 B0KMF6 B0KMF7 B0KMQ6 B0KMR5 B0KMS1 B0KN00 B0KN02 B0KN08
B0KN11 B0KN18 B0KN19 B0KN22 B0KN28 B0KN41 B0KN43 B0KN51 B0KN57 B0KN58 B0KN77
B0KN79 B0KN82 B0KND4 B0KND5 B0KND7 B0KNE3 B0KNE4 B0KNE7 B0KNF5 B0KNF7 B0KNH3
B0KNK9 B0KNW8 B0KP24 B0KP28 B0KP70 B0KP86 B0KPA6 B0KPF4 B0KPG6 B0KPH6 B0KPH9
B0KPI3 B0KPI4 B0KPI7 B0KPI8 B0KPJ1 B0KPJ6 B0KPL8 B0KPU9 B0KPW6 B0KPW7 B0KPW8
B0KPX3 B0KPY7 B0KQ38 B0KQ41 B0KQ43 B0KQ69 B0KQ74 B0KQ85 B0KQ87 B0KQ89 B0KQ91
B0KQ92 B0KQ94 B0KQA1 B0KQD8 B0KQF9 B0KQG0 B0KQG1 B0KQH0 B0KQH8 B0KQJ3 B0KQJ4
B0KQJ6 B0KQK2 B0KQK4 B0KQK8 B0KQS4 B0KQS7 B0KQW4 B0KQZ5 B0KR43 B0KR44 B0KR46
B0KR47 B0KR83 B0KR89 B0KRB8 B0KRB9 B0KRC0 B0KRD9 B0KRE6 B0KRH6 B0KRI1 B0KRI2
B0KRI3 B0KRI4 B0KRL5 B0KRW3 B0KS44 B0KS63 B0KS64 B0KS71 B0KS83 B0KS97 B0KS99
B0KSA0 B0KSA2 B0KSA5 B0KSB3 B0KSB7 B0KSC1 B0KSC5 B0KSC6 B0KSC7 B0KSC8 B0KSD2
B0KSS1 B0KSS4 B0KST1 B0KSY0 B0KT20 B0KT21 B0KT30 B0KT32 B0KT52 B0KT65 B0KT67
B0KTG8 B0KTH8 B0KTI3 B0KTJ2 B0KTJ3 B0KTJ4 B0KTJ6 B0KTK1 B0KTK4 B0KTK6 B0KTL0
B0KTR8 B0KTU0 B0KTU2 B0KTV7 B0KTW1 B0KTW7 B0KTX4 B0KTX5 B0KTY0 B0KTY3 B0KU95
B0KUA9 B0KUC0 B0KUD5 B0KUD8 B0KUE7 B0KUE9 B0KUG0 B0KUJ6 B0KUL0 B0KUN5 B0KUP7
B0KUQ0 B0KUT6 B0KUU0 B0KUX1 B0KUZ3 B0KUZ4 B0KUZ6 B0KUZ7 B0KUZ8 B0KUZ9 B0KV00
B0KV19 B0KV20 B0KV24 B0KV25 B0KV27 B0KV50 B0KV77 B0KW86 B0KW94 B0KWC3 B0KWC9
B0KWQ2 B0KWR6 B0KWU8 B0KZ40 B0LDU5 B0LDU6 B0LKP1 B0LL23 B0LXM0 B0M0P5 B0M1H3
B0M2T2 B0M3E8 B0QZF7 B0R061 B0R0B1 B0R0I6 B0R0T1 B0R0W9 B0R160 B0R1D5 B0R2L5
B0R2M2 B0R2M5 B0R2Q7 B0R2R5 B0R2S6 B0R2T0 B0R2T2 B0R2T7 B0R2U1 B0R2U6 B0R2V5
B0R2X8 B0R2Y0 B0R2Y1 B0R2Y4 B0R2Y5 B0R2Y7 B0R328 B0R332 B0R333 B0R336 B0R338
B0R349 B0R384 B0R386 B0R3A3 B0R3D6 B0R3D9 B0R3H3 B0R3H4 B0R3H6 B0R3L9 B0R3M0
B0R3M1 B0R3M8 B0R3R4 B0R3T7 B0R3Y0 B0R3Y1 B0R400 B0R403 B0R414 B0R434 B0R450
B0R457 B0R4B1 B0R4B2 B0R4B5 B0R4C0 B0R4C8 B0R4C9 B0R4D2 B0R4D3 B0R4J5 B0R4J8
B0R4P0 B0R4Q3 B0R4Q4 B0R4Q9 B0R4U1 B0R4U8 B0R4W1 B0R4W2 B0R4W3 B0R4W9 B0R4X4
B0R4Y0 B0R4Y2 B0R4Y3 B0R4Y6 B0R4Y7 B0R4Y9 B0R4Z6 B0R4Z9 B0R500 B0R501 B0R502
B0R506 B0R509 B0R511 B0R515 B0R518 B0R541 B0R542 B0R543 B0R544 B0R548 B0R558

B0R567 B0R569 B0R583 B0R591 B0R594 B0R596 B0R5A8 B0R5A9 B0R5F0 B0R5F5 B0R5I4
B0R5I7 B0R5J9 B0R5L0 B0R5L2 B0R5L3 B0R5M0 B0R5N8 B0R5P1 B0R5P8 B0R5R1 B0R5R9
B0R5S5 B0R5U5 B0R5W9 B0R5X2 B0R5Y6 B0R5Y7 B0R5Y8 B0R601 B0R621 B0R622 B0R636
B0R639 B0R642 B0R653 B0R656 B0R657 B0R659 B0R660 B0R664 B0R665 B0R666 B0R668
B0R669 B0R671 B0R675 B0R677 B0R678 B0R685 B0R686 B0R6B4 B0R6B9 B0R6C0 B0R6C7
B0R6D5 B0R6D7 B0R6F1 B0R6G1 B0R6G2 B0R6G6 B0R6H4 B0R6I6 B0R6K3 B0R6L8 B0R6L9
B0R6P3 B0R6P7 B0R6P8 B0R6S2 B0R6S4 B0R6T1 B0R6T4 B0R6U5 B0R6U7 B0R6V1 B0R6V2
B0R6V4 B0R6V9 B0R6W9 B0R6Y1 B0R6Y2 B0R6Y3 B0R6Y7 B0R6Z8 B0R700 B0R701 B0R713
B0R717 B0R731 B0R748 B0R7B4 B0R7C0 B0R7C4 B0R7D8 B0R7E0 B0R7E2 B0R7E9 B0R7F2
B0R7F5 B0R7F7 B0R7F9 B0R7G3 B0R7H0 B0R7H7 B0R7I0 B0R7I1 B0R7J1 B0R7K0 B0R7L9
B0R7P9 B0R7Q0 B0R7Q2 B0R7Q4 B0R7Q6 B0R7R3 B0R7U1 B0R7U7 B0R7V3 B0R7V5 B0R7W8
B0R7X6 B0R7X7 B0R7X8 B0R7X9 B0R7Y4 B0R806 B0R807 B0R815 B0R841 B0R858 B0R868
B0R870 B0R871 B0R877 B0R879 B0R884 B0R888 B0R889 B0R8A2 B0R8C2 B0R8C3 B0R8C8
B0R8D0 B0R8D1 B0R8D2 B0R8D3 B0R8D4 B0R8D5 B0R8D6 B0R8D7 B0R9H2 B0R9L6 B0R9M2
B0R9V8 B0R9W5 B0R9W6 B0R9W7 B0R9X5 B0R9X6 B0R9X9 B0RAE3 B0RAP7 B0RAQ5 B0RAS0
B0RAS4 B0RAV5 B0RAV9 B0RAW4 B0RAX1 B0RAY1 B0RB25 B0RB26 B0RB33 B0RB34 B0RB35
B0RB36 B0RB38 B0RB39 B0RB41 B0RB42 B0RB43 B0RB44 B0RB45 B0RB46 B0RB49 B0RB50
B0RB53 B0RB56 B0RB59 B0RB68 B0RB69 B0RB71 B0RB72 B0RB73 B0RB77 B0RB78 B0RBB1
B0RBI1 B0RBL3 B0RBM5 B0RC26 B0RC50 B0RC64 B0RCF1 B0RCQ8 B0RCR5 B0RCZ0 B0RCZ5
B0RD07 B0RD37 B0RDA1 B0RDA6 B0RDF9 B0RDG2 B0RDG3 B0RDG4 B0RDG5 B0RDK9 B0RDL0
B0RDL1 B0RDN5 B0RDN6 B0RDP8 B0RDQ4 B0RDR2 B0RDR3 B0RDY8 B0RDY9 B0RDZ1 B0RDZ5
B0RE23 B0RE24 B0REA1 B0REB6 B0REB7 B0REB8 B0REB9 B0REG5 B0REJ2 B0REJ5 B0REJ6
B0REK0 B0REK2 B0REK3 B0REL6 B0REL8 B0REM7 B0REM8 B0REN0 B0REN1 B0REN4 B0REN8
B0REP4 B0REP5 B0REP7 B0RES6 B0REU1 B0REU6 B0REU9 B0REV2 B0REV4 B0REV6 B0REV9
B0REW3 B0REX0 B0REX1 B0REY1 B0RF94 B0RG24 B0RGA6 B0RGA7 B0RGB2 B0RGG4 B0RGI6
B0RGV9 B0RGW6 B0RGW7 B0RGX5 B0RGZ1 B0RGZ2 B0RH22 B0RH69 B0RH73 B0RH97 B0RHB0
B0RHB5 B0RHC1 B0RHC2 B0RHD0 B0RHD2 B0RHD4 B0RHD5 B0RHD6 B0RHY9 B0RI48 B0RI49
B0RIB7 B0RIC6 B0RIE3 B0RIF1 B0RIG4 B0RIH2 B0RIJ4 B0RIJ8 B0RIK5 B0RIN3 B0RIN6
B0RIP4 B0RIP8 B0RIR7 B0RIR8 B0RLD0 B0RLF1 B0RLI8 B0RLK6 B0RLW8 B0RM75 B0RMB4
B0RMM4 B0RMM7 B0RMM8 B0RMN0 B0RMP9 B0RMQ9 B0RMR1 B0RMR2 B0RMW5 B0RMY6 B0RMZ8
B0RN02 B0RN05 B0RN06 B0RN07 B0RN15 B0RN27 B0RN47 B0RN51 B0RN52 B0RN85 B0RN90
B0RN93 B0RNI9 B0RNK1 B0RNK3 B0RNK7 B0RNP8 B0RNP9 B0RNU6 B0RNU9 B0RNV0 B0RNW7
B0RNZ3 B0RP28 B0RP32 B0RP37 B0RP51 B0RP52 B0RP74 B0RP81 B0RP90 B0RP96 B0RP99
B0RPA3 B0RPF1 B0RPG5 B0RPG6 B0RPG8 B0RPJ1 B0RPP2 B0RPQ9 B0RPU7 B0RPU9 B0RPX9
B0RPY2 B0RPY3 B0RPY4 B0RPY7 B0RPZ6 B0RQ25 B0RQ26 B0RQ27 B0RQ28 B0RQ55 B0RQ92
B0RQB0 B0RQC7 B0RQG9 B0RQH0 B0RQI0 B0RQI6 B0RQK3 B0RQM8 B0RQM9 B0RQN0 B0RQR6
B0RQR7 B0RQU0 B0RQZ0 B0RR06 B0RR12 B0RR22 B0RR28 B0RR56 B0RR62 B0RR76 B0RR81
B0RR82 B0RR84 B0RR86 B0RR88 B0RRB2 B0RRB4 B0RRB5 B0RRB6 B0RRB7 B0RRB8 B0RRD9
B0RRG9 B0RRH1 B0RRH2 B0RRH3 B0RRH5 B0RRL3 B0RRP8 B0RRQ3 B0RRR0 B0RRR1 B0RRT4
B0RRV9 B0RRW2 B0RRX4 B0RRZ3 B0RRZ8 B0RS12 B0RS13 B0RS24 B0RS27 B0RS35 B0RS54
B0RS55 B0RS59 B0RS70 B0RSA2 B0RSA3 B0RSA5 B0RSB3 B0RSB6 B0RSD4 B0RSF9 B0RSI3
B0RSK6 B0RSL3 B0RSL5 B0RSL6 B0RSL8 B0RSL9 B0RSM3 B0RSM4 B0RSM5 B0RSN5 B0RSP5
B0RSQ8 B0RSS2 B0RSS7 B0RT32 B0RT36 B0RT37 B0RT49 B0RT51 B0RT56 B0RTD8 B0RTE0
B0RTE9 B0RTF5 B0RTF7 B0RTF8 B0RTH4 B0RTH5 B0RTI2 B0RTS4 B0RTS9 B0RTT8 B0RTT9

B0RTU0 B0RTU1 B0RTU4 B0RTW1 B0RTY0 B0RTY1 B0RTY2 B0RTY3 B0RTY9 B0RTZ0 B0RTZ3
B0RTZ8 B0RTZ9 B0RU01 B0RU02 B0RU03 B0RU31 B0RU49 B0RU57 B0RU58 B0RU59 B0RU60
B0RU61 B0RU64 B0RU65 B0RU66 B0RU67 B0RU68 B0RU69 B0RU70 B0RU71 B0RU72 B0RU73
B0RU74 B0RU75 B0RU77 B0RU78 B0RU79 B0RU80 B0RU81 B0RU82 B0RU83 B0RU84 B0RU85
B0RU86 B0RU88 B0RU89 B0RU90 B0RU91 B0RU92 B0RU93 B0RU96 B0RU99 B0RUA1 B0RUA4
B0RUA5 B0RUE4 B0RUE7 B0RUG0 B0RUI2 B0RUI3 B0RUI4 B0RUK3 B0RUV8 B0RUW1 B0RUW3
B0RUX3 B0RUX9 B0RUY2 B0RUY7 B0RUY8 B0RUY9 B0RUZ7 B0RUZ9 B0RV00 B0RV07 B0RV66
B0RVA4 B0RVB2 B0RVB3 B0RVB9 B0RVD1 B0RVD2 B0RVE0 B0RVE1 B0RVE9 B0RVI0 B0RVI4
B0RVJ2 B0RVK2 B0RVK5 B0RVK6 B0RVN9 B0RVR1 B0RVU1 B0RVU2 B0RVV0 B0RW53 B0RW57
B0RW65 B0RW66 B0RW69 B0RW72 B0RW80 B0RWA1 B0RWE6 B0RWG2 B0RWR1 B0RWR5 B0RWT2
B0RWT6 B0RWU0 B0RWU6 B0RX17 B0RX25 B0RXD3 B0RXD6 B0RXD7 B0RXD8 B0RXD9 B0RXI2
B0RXI3 B0RXL2 B0RXM3 B0RXM6 B0RXN7 B0RXV1 B0RXV6 B0RXX4 B0RY26 B0RY28 B0RY31
B0RY32 B0RY33 B0RY34 B0RY43 B0RY47 B0RY61 B0RY63 B0RY89 B0RYD5 B0RYI8 B0RYK3
B0RYM2 B0RYP5 B0RYR2 B0RYR3 B0RYS0 B0RYX3 B0RYX4 B0RYY7 B0RYY8 B0RZ13 B0RZ30
B0RZ32 B0RZ50 B0RZJ0 B0RZJ9 B0RZL3 B0RZP8 B0RZQ3 B0RZR9 B0RZS0 B0RZS2 B0RZS3
B0RZS4 B0RZS5 B0RZS7 B0RZS8 B0RZT0 B0RZT3 B0RZT6 B0RZT8 B0RZU0 B0RZU1 B0RZU2
B0RZU3 B0RZU4 B0RZU5 B0RZU6 B0RZU7 B0RZU8 B0RZV0 B0RZW0 B0RZW3 B0RZY8 B0S032
B0S044 B0S046 B0S047 B0S048 B0S053 B0S054 B0S057 B0S061 B0S062 B0S064 B0S065
B0S093 B0S0D9 B0S0G1 B0S0I2 B0S0I3 B0S0I4 B0S0K3 B0S0K7 B0S0K8 B0S0L4 B0S0L6
B0S0M3 B0S0N1 B0S0R2 B0S0S7 B0S0W2 B0S0W3 B0S0Y8 B0S0Y9 B0S0Z8 B0S101 B0S104
B0S115 B0S134 B0S136 B0S139 B0S140 B0S143 B0S150 B0S155 B0S168 B0S169 B0S174
B0S175 B0S177 B0S184 B0S185 B0S187 B0S196 B0S198 B0S1A1 B0S1B1 B0S1C0 B0S1C1
B0S1C3 B0S1D8 B0S1D9 B0S1E0 B0S1E4 B0S1E5 B0S1E8 B0S1F2 B0S1F7 B0S1F8 B0S1G4
B0S1G9 B0S1H0 B0S1I1 B0S1I7 B0S1J2 B0S1J4 B0S1J5 B0S1L6 B0S1L9 B0S1M1 B0S1M9
B0S1N0 B0S1N2 B0S1N3 B0S1P2 B0S1Q1 B0S1R1 B0S1R4 B0S1R5 B0S1S1 B0S1S2 B0S216
B0S218 B0S230 B0S255 B0S259 B0S264 B0S273 B0S296 B0S2A9 B0S2B2 B0S2B6 B0S2B9
B0S2E1 B0S2E5 B0S2F7 B0S2F8 B0S2G9 B0S2I6 B0S2N4 B0S2N7 B0S2Q1 B0S2Q7 B0S341
B0S3A6 B0S3F3 B0S3F6 B0S3F7 B0S3H8 B0S3R4 B0S3U9 B0S3V0 B0S3V1 B0S3V2 B0S3V6
B0S3Y4 B0S3Z1 B0S3Z4 B0S415 B0S4Q5 B0S6S9 B0S733 B0S794 B0S8I0 B0S8M8 B0S8U6
B0S8V2 B0S8V3 B0S8X7 B0S8X8 B0S8Z4 B0S902 B0S904 B0S907 B0S909 B0S912 B0S919
B0S959 B0S989 B0S9A0 B0S9A2 B0S9A3 B0S9A5 B0S9E2 B0S9F1 B0S9F2 B0S9H9 B0S9I4
B0S9J3 B0S9J5 B0S9J6 B0S9S9 B0S9U5 B0SA19 B0SA20 B0SA21 B0SA23 B0SA24 B0SA26
B0SA29 B0SA31 B0SA32 B0SA33 B0SA34 B0SA35 B0SA36 B0SA37 B0SA40 B0SA41 B0SA42
B0SA43 B0SA75 B0SA95 B0SAB8 B0SAC5 B0SAE4 B0SAF1 B0SAF2 B0SAF3 B0SAF4 B0SAF5
B0SAF6 B0SAF8 B0SAF9 B0SAG0 B0SAG1 B0SAG3 B0SAG4 B0SAG5 B0SAP0 B0SAR6 B0SAT2
B0SB16 B0SB21 B0SB22 B0SB28 B0SB29 B0SB31 B0SB32 B0SB35 B0SB37 B0SB38 B0SB40
B0SB76 B0SB77 B0SB85 B0SB88 B0SBG1 B0SBH6 B0SBH8 B0SBH9 B0SBK8 B0SBL3 B0SBW4
B0SBX0 B0SC03 B0SC16 B0SC36 B0SCA8 B0SCB9 B0SCC0 B0SCD1 B0SCE2 B0SCH2 B0SCH3
B0SCH4 B0SCM4 B0SCQ5 B0SCT0 B0SCV6 B0SCX2 B0SDC5 B0SDE2 B0SDL3 B0SDL8 B0SDM7
B0SDM8 B0SDM9 B0SDN1 B0SDN4 B0SDN6 B0SDS4 B0SDS6 B0SDY7 B0SDZ6 B0SE38 B0SE82
B0SE83 B0SEC2 B0SEF8 B0SEH0 B0SEH6 B0SEH8 B0SEI3 B0SF41 B0SF60 B0SF64 B0SF75
B0SFD6 B0SFE8 B0SFG5 B0SFJ3 B0SFJ4 B0SFU6 B0SG67 B0SG84 B0SGA8 B0SGD3 B0SGD4
B0SGD8 B0SGE9 B0SGN8 B0SGN9 B0SGR0 B0SGR1 B0SGV6 B0SGW0 B0SGW3 B0SGZ5 B0SH16
B0SH18 B0SH19 B0SH21 B0SH22 B0SH39 B0SH49 B0SH84 B0SHA8 B0SHG0 B0SHH1 B0SHJ0

B0SHK4 B0SHL8 B0SHT0 B0SHT1 B0SHT2 B0SHT3 B0SHV8 B0SI62 B0SI93 B0SIB4 B0SIF3
B0SII1 B0SII2 B0SJ24 B0SJ25 B0SJ27 B0SJ28 B0SJ52 B0SJ57 B0SJA9 B0SJC4 B0SJF2
B0SJL7 B0SJM4 B0SJP0 B0SJQ8 B0SJR1 B0SJR2 B0SJT4 B0SK31 B0SK33 B0SK44 B0SK77
B0SKC1 B0SKG9 B0SKQ2 B0SKS9 B0SKU1 B0SKU2 B0SKV3 B0SKW4 B0SKZ5 B0SKZ6 B0SKZ7
B0SL24 B0SL33 B0SL58 B0SLE9 B0SLQ3 B0SLS0 B0SLU7 B0SLU9 B0SM42 B0SM47 B0SM57
B0SM58 B0SM60 B0SM63 B0SM65 B0SMB9 B0SMD0 B0SMF0 B0SMI6 B0SMJ9 B0SMK5 B0SMK7
B0SMR8 B0SMR9 B0SMW9 B0SN74 B0SNJ9 B0SNK3 B0SNL6 B0SNY3 B0SNZ7 B0SP16 B0SP94
B0SPA9 B0SPB4 B0SPC6 B0SPF0 B0SPF1 B0SPQ8 B0SPT4 B0SPT5 B0SPT9 B0SPV0 B0SQ03
B0SQ07 B0SQ10 B0SQA1 B0SQA2 B0SQF0 B0SQH2 B0SQH4 B0SQH5 B0SQH7 B0SQH8 B0SQL1
B0SQM2 B0SQM8 B0SQQ4 B0SQR4 B0SQV0 B0SQX5 B0SQZ8 B0SR12 B0SR27 B0SR57 B0SRF0
B0SRF1 B0SRF2 B0SRH8 B0SRN4 B0SRP0 B0SRP1 B0SRS3 B0SRT4 B0SRT6 B0SRT7 B0SRT9
B0SRX5 B0SRX6 B0SRZ1 B0SRZ9 B0SS01 B0SS31 B0SS40 B0SS41 B0SS68 B0SS77 B0SSF1
B0SSF2 B0SSF3 B0SSF5 B0SSF7 B0SSG0 B0SSG2 B0SSG3 B0SSG4 B0SSG5 B0SSG6 B0SSG7
B0SSH0 B0SSH1 B0SSH2 B0SSH3 B0SSH4 B0SSH5 B0SSH6 B0SSH7 B0SSH8 B0SSH9 B0SSI1
B0SSI2 B0SSI3 B0SSI4 B0SSI6 B0SSI7 B0SSI8 B0SST8 B0SSU3 B0SSU4 B0SSV0 B0SSV1
B0SSV3 B0SSV4 B0SSV7 B0SSV9 B0SSW0 B0SSW2 B0ST01 B0ST02 B0ST13 B0ST16 B0STE5
B0STF1 B0STJ3 B0STK1 B0STS6 B0STV8 B0SU49 B0SU50 B0SUK3 B0SUL0 B0SUL4 B0SUL9
B0SUN3 B0SUN7 B0SUN8 B0SUN9 B0SUP2 B0SUP3 B0SUP7 B0SUP8 B0SUQ4 B0SUQ5 B0SUQ6
B0SUQ7 B0SV28 B0SVF8 B0SVG5 B0SVG8 B0SW01 B0SW60 B0SW61 B0SW62 B0SW63 B0SW81
B0SWD8 B0SWK9 B0SWL0 B0SWL1 B0SWN7 B0SWQ6 B0SWX9 B0SX45 B0SX50 B0SXA5 B0SXA8
B0SXB0 B0SXB7 B0SXC3 B0SXF2 B0SXF9 B0SXP0 B0SXR2 B0SXR3 B0SXZ7 B0SXZ8 B0SXZ9
B0SY03 B0SY17 B0SY33 B0SY51 B0SYU4 B0SYU6 B0SYU7 B0SYY1 B0SYZ3 B0SYZ4 B0SZ02
B0SZ18 B0SZ20 B0SZ21 B0SZ28 B0SZ33 B0SZ62 B0SZ84 B0SZC3 B0SZF7 B0SZN2 B0SZN4
B0SZQ2 B0SZS4 B0SZS9 B0SZT0 B0SZU3 B0T014 B0T019 B0T024 B0T025 B0T064 B0T065
B0T0B1 B0T0C5 B0T0G9 B0T0I3 B0T0R1 B0T0S3 B0T0T0 B0T0V2 B0T0Z7 B0T101 B0T130
B0T134 B0T135 B0T138 B0T142 B0T152 B0T155 B0T157 B0T162 B0T164 B0T167 B0T172
B0T173 B0T174 B0T175 B0T179 B0T190 B0T191 B0T192 B0T193 B0T1G7 B0T1I4 B0T1I5
B0T1K8 B0T1M3 B0T1N5 B0T1Q1 B0T1Q2 B0T1Q3 B0T1R2 B0T1S6 B0T1S7 B0T1S9 B0T1T2
B0T1T3 B0T1V6 B0T1Y4 B0T202 B0T207 B0T229 B0T270 B0T271 B0T273 B0T287 B0T2A6
B0T2B5 B0T2C1 B0T2C2 B0T2C3 B0T2C4 B0T2C5 B0T2C6 B0T2C8 B0T2C9 B0T2D1 B0T2D2
B0T2D3 B0T2D4 B0T2D5 B0T2D6 B0T2D7 B0T2D8 B0T2D9 B0T2E0 B0T2E1 B0T2E3 B0T2E4
B0T2E5 B0T2E6 B0T2E7 B0T2L1 B0T2Q1 B0T2R0 B0T2T7 B0T2Y6 B0T310 B0T315 B0T316
B0T320 B0T360 B0T367 B0T368 B0T369 B0T377 B0T387 B0T395 B0T3B7 B0T3B9 B0T3D1
B0T3F7 B0T3H0 B0T3H5 B0T3H6 B0T3I0 B0T3I3 B0T3J0 B0T3J3 B0T3X7 B0T3Y0 B0T3Z9
B0T410 B0T4C1 B0T4L2 B0T4U9 B0T4Y1 B0T542 B0T549 B0T562 B0T563 B0T583 B0T5D0
B0T660 B0T663 B0T692 B0T694 B0T6C1 B0T6E0 B0T6E1 B0T6E2 B0T6E7 B0T6F4 B0T725
B0T734 B0T791 B0T797 B0T798 B0T799 B0T7A1 B0T7A8 B0T7B0 B0T7D0 B0T7D1 B0T7D5
B0T7V9 B0T816 B0T824 B0T826 B0T827 B0T833 B0T839 B0T866 B0T874 B0T890 B0T899
B0T8D1 B0T8F3 B0T8H5 B0T8I8 B0T8J6 B0T8W6 B0TA38 B0TA39 B0TA52 B0TA56 B0TAB5
B0TAB6 B0TAC7 B0TAD4 B0TAD7 B0TAD9 B0TAE8 B0TAF3 B0TAF7 B0TAH2 B0TAK6 B0TAK8
B0TAL0 B0TAN9 B0TAQ4 B0TAQ9 B0TAU6 B0TAY4 B0TAZ6 B0TB07 B0TB10 B0TB20 B0TB90
B0TB98 B0TBA3 B0TBI8 B0TBI9 B0TBJ3 B0TBM6 B0TBM7 B0TBM8 B0TBN6 B0TBP1 B0TBP5
B0TBP6 B0TBU3 B0TBV8 B0TBW1 B0TBW5 B0TBW7 B0TBY0 B0TBY2 B0TC10 B0TC35 B0TC40
B0TC43 B0TC44 B0TC46 B0TC48 B0TC49 B0TC51 B0TC52 B0TC53 B0TC54 B0TC55 B0TC56

B0TC57 B0TC58 B0TC59 B0TC60 B0TC61 B0TC63 B0TC64 B0TC65 B0TC66 B0TC67 B0TC68
B0TC69 B0TC70 B0TC71 B0TC72 B0TC73 B0TC74 B0TC75 B0TC77 B0TC81 B0TC82 B0TC84
B0TC85 B0TC86 B0TC91 B0TC92 B0TC99 B0TCA0 B0TCA6 B0TCA8 B0TCJ8 B0TCL9 B0TCQ9
B0TCR1 B0TCR2 B0TCR3 B0TCS1 B0TD37 B0TD39 B0TDI0 B0TDI1 B0TDK7 B0TDK8 B0TDL0
B0TDM7 B0TDM8 B0TDN0 B0TDZ7 B0TDZ8 B0TE53 B0TE71 B0TE72 B0TEC5 B0TEH1 B0TEI6
B0TEJ2 B0TEJ5 B0TEJ8 B0TEK0 B0TEP6 B0TER7 B0TEV3 B0TEV4 B0TEV9 B0TF65 B0TF68
B0TF70 B0TF76 B0TF85 B0TF88 B0TF89 B0TFA7 B0TFC9 B0TFI7 B0TFI9 B0TFJ0 B0TFR3
B0TFV0 B0TFW3 B0TFY7 B0TFZ0 B0TFZ1 B0TG92 B0TGB1 B0TGB2 B0TGC2 B0TGD0 B0TGK8
B0TGL0 B0TGL6 B0TGN6 B0TGP1 B0TGQ3 B0TGQ8 B0TGR9 B0TGS2 B0TGS4 B0TGS8 B0TGS9
B0TGT1 B0TGU2 B0TGU9 B0TGV2 B0TGW4 B0TH41 B0TH58 B0TH64 B0TH66 B0TH67 B0TH69
B0TH93 B0THD7 B0THD8 B0THE0 B0THN7 B0THP0 B0THQ9 B0THR5 B0THR7 B0THS1 B0THS2
B0THT2 B0THU1 B0TI09 B0TI15 B0TI16 B0TI63 B0TI64 B0TI71 B0TI73 B0TI77 B0TI89
B0TI92 B0TIB8 B0TIF2 B0TIH8 B0TII6 B0TIM6 B0TIN6 B0TIN7 B0TIQ0 B0TIR0 B0TIR7
B0TIS5 B0TIU1 B0TIU4 B0TIV7 B0TIW4 B0TIW9 B0TIX7 B0TIY9 B0TIZ6 B0TJ03 B0TJ05
B0TJ16 B0TJ29 B0TJ37 B0TJ39 B0TJ54 B0TJ75 B0TJ76 B0TJ77 B0TJ78 B0TJ99 B0TJA0
B0TJA4 B0TJA6 B0TJA9 B0TJB2 B0TJD1 B0TJD3 B0TJN8 B0TJP8 B0TJR3 B0TJY5 B0TJY6
B0TJZ0 B0TJZ1 B0TK00 B0TK03 B0TK06 B0TK07 B0TK09 B0TK10 B0TK13 B0TK14 B0TK15
B0TK16 B0TK23 B0TK42 B0TK50 B0TK55 B0TKB2 B0TKD2 B0TKD7 B0TKZ4 B0TKZ7 B0TL07
B0TL40 B0TL64 B0TL65 B0TL76 B0TL77 B0TL84 B0TL85 B0TL87 B0TL88 B0TL90 B0TL92
B0TLA4 B0TLA6 B0TLA9 B0TLB0 B0TLB5 B0TLC0 B0TLC9 B0TLD3 B0TLD6 B0TLD7 B0TLD8
B0TLE6 B0TLE8 B0TLG2 B0TLI1 B0TLI4 B0TLI5 B0TLI8 B0TLJ2 B0TLJ4 B0TLU8 B0TLU9
B0TLV0 B0TLV1 B0TLV2 B0TLY7 B0TLY9 B0TLZ0 B0TLZ1 B0TLZ3 B0TLZ4 B0TLZ6 B0TLZ7
B0TLZ8 B0TLZ9 B0TM00 B0TM01 B0TM02 B0TM03 B0TM04 B0TM05 B0TM07 B0TM08 B0TM09
B0TM10 B0TM11 B0TM12 B0TM13 B0TM14 B0TM16 B0TM17 B0TM18 B0TM20 B0TM21 B0TM22
B0TM23 B0TM27 B0TM50 B0TM54 B0TM56 B0TM80 B0TMA7 B0TMM7 B0TMS3 B0TMV5 B0TMV6
B0TMW0 B0TMY0 B0TMY9 B0TMZ9 B0TN04 B0TN10 B0TN76 B0TN99 B0TNB2 B0TNB9 B0TND3
B0TND4 B0TNJ5 B0TNM4 B0TNM9 B0TNS0 B0TNX0 B0TNY1 B0TNY4 B0TNY5 B0TNY9 B0TNZ8
B0TP00 B0TP04 B0TP05 B0TP07 B0TP13 B0TP63 B0TP69 B0TP77 B0TP80 B0TP82 B0TP83
B0TP86 B0TPD4 B0TPE7 B0TPF3 B0TPG2 B0TPG7 B0TPH1 B0TPR8 B0TPU5 B0TPW7 B0TPW8
B0TPY4 B0TPZ1 B0TQ02 B0TQ07 B0TQ11 B0TQ31 B0TQ36 B0TQ37 B0TQ38 B0TQ89 B0TQ90
B0TQ91 B0TQ95 B0TQ97 B0TQ99 B0TQA0 B0TQA1 B0TQA2 B0TQA4 B0TQA6 B0TQA7 B0TQB0
B0TQB8 B0TQC1 B0TQC2 B0TQC4 B0TQC5 B0TQD1 B0TQE1 B0TQG4 B0TQG5 B0TQH0 B0TQH4
B0TQK4 B0TQK8 B0TQK9 B0TQL1 B0TQL2 B0TQL3 B0TQM0 B0TQM2 B0TQM3 B0TQM6 B0TQM8
B0TQM9 B0TQN8 B0TQP7 B0TQU3 B0TQV3 B0TQX1 B0TQX3 B0TQX6 B0TQY4 B0TQY6 B0TQY8
B0TQY9 B0TR07 B0TR14 B0TR27 B0TR28 B0TR30 B0TR35 B0TR38 B0TR40 B0TR44 B0TR47
B0TR49 B0TR55 B0TR56 B0TR57 B0TR87 B0TR91 B0TRH1 B0TRI1 B0TRI3 B0TRI4 B0TRI6
B0TRK1 B0TRL8 B0TRM1 B0TRN5 B0TRP7 B0TRQ2 B0TRS6 B0TS18 B0TS36 B0TS89 B0TSA0
B0TSA1 B0TSA3 B0TSA5 B0TSA6 B0TSA7 B0TSC0 B0TSG5 B0TSG6 B0TSG7 B0TSQ4 B0TSS5
B0TST3 B0TSU8 B0TT20 B0TT22 B0TT23 B0TT38 B0TT41 B0TT43 B0TT44 B0TT46 B0TT63
B0TT68 B0TT69 B0TT70 B0TT71 B0TT72 B0TT73 B0TT74 B0TT81 B0TTD2 B0TTD5 B0TTE4
B0TTF2 B0TTI1 B0TTI2 B0TTI7 B0TTL6 B0TTP2 B0TTP5 B0TTR5 B0TTS9 B0TTT0 B0TU38
B0TU64 B0TU65 B0TU66 B0TUB8 B0TUC0 B0TUC1 B0TUD3 B0TUD9 B0TUH7 B0TUH8 B0TUI0
B0TUI1 B0TUI2 B0TUN8 B0TUP4 B0TUS0 B0TUS6 B0TUS9 B0TUU1 B0TUU6 B0TUU7 B0TUU8
B0TUU9 B0TUV4 B0TUV6 B0TUW5 B0TUY1 B0TUZ2 B0TUZ6 B0TV29 B0TV51 B0TV52 B0TVE5

B0TVI4 B0TVL3 B0TVL4 B0TVM8 B0TVP3 B0TVU8 B0TVV1 B0TVV3 B0TVV4 B0TVV8 B0TVX5
B0TVZ0 B0TW04 B0TW08 B0TW17 B0TW32 B0TW34 B0TW44 B0TW67 B0TW70 B0TW77 B0TW84
B0TW90 B0TWF2 B0TWF7 B0TWG4 B0TWI3 B0TWJ4 B0TWJ9 B0TWK7 B0TWN0 B0TWR0 B0TWR1
B0TWR3 B0TWR4 B0TWR5 B0TWU0 B0TWU1 B0TWU2 B0TWY5 B0TWY6 B0TWZ7 B0TX03 B0TX06
B0TX07 B0TX08 B0TX09 B0TX10 B0TX11 B0TX13 B0TX15 B0TX16 B0TX17 B0TX18 B0TX19
B0TX27 B0TX30 B0TX36 B0TX37 B0TX39 B0TX40 B0TX64 B0TX65 B0TX95 B0TX99 B0TXA4
B0TXF5 B0TXH8 B0TXH9 B0TXQ5 B0TXQ6 B0TXR2 B0TXS4 B0TXS6 B0TXX4 B0TXY0 B0TXZ9
B0TY02 B0TY17 B0TY21 B0TY34 B0TY35 B0TY41 B0TY42 B0TY45 B0TY78 B0TY81 B0TY86
B0TY87 B0TY89 B0TY94 B0TY95 B0TYA1 B0TYB0 B0TYD1 B0TYD6 B0TYE8 B0TYF1 B0TYF2
B0TYF3 B0TYH3 B0TYH9 B0TYI1 B0TYI6 B0TYK1 B0TYK2 B0TYL2 B0TYL4 B0TYM8 B0TYM9
B0TYR5 B0TYR8 B0TYS6 B0TYX5 B0TYY3 B0TZ05 B0TZ08 B0TZ09 B0TZ10 B0TZ13 B0TZ16
B0TZ17 B0TZ23 B0TZ52 B0TZ54 B0TZ67 B0TZ72 B0TZ91 B0TZA4 B0TZA5 B0TZA7 B0TZB0
B0TZB1 B0TZB8 B0TZC0 B0TZD6 B0TZD8 B0TZD9 B0TZF1 B0TZF5 B0TZG2 B0TZH7 B0TZI2
B0TZI3 B0TZI5 B0TZJ5 B0TZJ6 B0TZJ8 B0TZP1 B0TZQ0 B0TZQ5 B0TZQ8 B0TZS5 B0TZT2
B0TZW0 B0TZY8 B0U003 B0U009 B0U040 B0U054 B0U064 B0U079 B0U080 B0U081 B0U083
B0U092 B0U0B1 B0U0B2 B0U0B3 B0U0B6 B0U0C4 B0U0D4 B0U0E8 B0U0E9 B0U0F0 B0U0F9
B0U0G0 B0U0G3 B0U0I3 B0U0L6 B0U0L7 B0U0M7 B0U0Q5 B0U0S3 B0U0S6 B0U0T6 B0U0T8
B0U0U4 B0U0W4 B0U0W6 B0U0W7 B0U0W8 B0U0X0 B0U0X1 B0U0X3 B0U0X4 B0U0X5 B0U0X6
B0U0X7 B0U0X8 B0U0X9 B0U0Y0 B0U0Y1 B0U0Y2 B0U0Y4 B0U0Y5 B0U0Y6 B0U0Y7 B0U0Y8
B0U0Y9 B0U0Z0 B0U0Z1 B0U0Z2 B0U0Z3 B0U0Z5 B0U0Z8 B0U100 B0U101 B0U106 B0U111
B0U150 B0U151 B0U169 B0U172 B0U178 B0U1D6 B0U1E7 B0U1E8 B0U1F2 B0U1G5 B0U1G7
B0U1H5 B0U1H6 B0U1H9 B0U1I2 B0U1J0 B0U1J8 B0U1K8 B0U1K9 B0U1L0 B0U1L1 B0U1L5
B0U1N3 B0U1N9 B0U1P0 B0U1P6 B0U1Q2 B0U1Q3 B0U1Q4 B0U1Q5 B0U1Q6 B0U1Q8 B0U1Q9
B0U1R1 B0U1R2 B0U1U0 B0U1W3 B0U1Z4 B0U207 B0U208 B0U219 B0U232 B0U235 B0U242
B0U262 B0U269 B0U270 B0U281 B0U288 B0U289 B0U291 B0U296 B0U2A0 B0U2A9 B0U2B7
B0U2D9 B0U2E1 B0U2E3 B0U2F2 B0U2I4 B0U2I5 B0U2S7 B0U2T5 B0U2T9 B0U2W8 B0U2Y2
B0U328 B0U354 B0U365 B0U376 B0U379 B0U399 B0U3A4 B0U3A5 B0U3A6 B0U3A8 B0U3A9
B0U3B1 B0U3B2 B0U3B4 B0U3B6 B0U3B8 B0U3C1 B0U3C2 B0U3C6 B0U3D8 B0U3E0 B0U3E1
B0U3F2 B0U3F5 B0U3G5 B0U3J7 B0U3J8 B0U3K0 B0U3K6 B0U3L0 B0U3M5 B0U3M6 B0U3N1
B0U3Q7 B0U3Q9 B0U3R3 B0U3R4 B0U3R8 B0U3S8 B0U3S9 B0U3V7 B0U3W1 B0U3W3 B0U3W7
B0U3Y1 B0U410 B0U417 B0U418 B0U431 B0U447 B0U453 B0U454 B0U456 B0U466 B0U468
B0U472 B0U489 B0U494 B0U495 B0U498 B0U4A7 B0U4B5 B0U4B6 B0U4D9 B0U4E6 B0U4E7
B0U4I5 B0U4J3 B0U4K2 B0U4K3 B0U4K7 B0U4K9 B0U4L2 B0U4M3 B0U4M4 B0U4M7 B0U4N0
B0U4P1 B0U4P3 B0U4U5 B0U4U6 B0U4V3 B0U4X0 B0U4X9 B0U4Z6 B0U504 B0U509 B0U565
B0U568 B0U572 B0U5A6 B0U5A7 B0U5A8 B0U5A9 B0U5B0 B0U5C4 B0U5C6 B0U5D5 B0U5D7
B0U5D9 B0U5E0 B0U5E1 B0U5E3 B0U5E6 B0U5F8 B0U5G3 B0U5G7 B0U5H0 B0U5H1 B0U5H2
B0U5I2 B0U5I9 B0U5K3 B0U5K4 B0U5K5 B0U5K6 B0U5K8 B0U5K9 B0U5L0 B0U5L1 B0U5L2
B0U5L3 B0U5L4 B0U5L5 B0U5L6 B0U5M0 B0U5M1 B0U5M2 B0U5M3 B0U5M4 B0U5N0 B0U5N1
B0U5N2 B0U5P7 B0U5P8 B0U5Q1 B0U5U4 B0U5V3 B0U5V4 B0U5X3 B0U5X4 B0U5X5 B0U5X6
B0U5X7 B0U5X8 B0U5X9 B0U5Y0 B0U5Y6 B0U5Y7 B0U5Z2 B0U5Z3 B0U5Z5 B0U620 B0U641
B0U656 B0U660 B0U6A9 B0U6B4 B0U6B6 B0U6C5 B0U6C7 B0U6C8 B0U6D0 B0U6I3 B0U6I9
B0U6J0 B0U6J4 B0U6K0 B0U6K4 B0U6K5 B0U6K6 B0U6K7 B0U6M8 B0U6M9 B0U6N8 B0U6Q7
B0U6Q9 B0U6R3 B0U6T2 B0U6U5 B0U6U9 B0U6V1 B0U6V9 B0U6W0 B0U6W4 B0U6W7 B0U6Z3
B0U6Z5 B0U6Z6 B0U6Z8 B0U741 B0U767 B0U772 B0U794 B0U7A0 B0U7B1 B0U7B2 B0U7F9

B0U7H1 B0U7Z3 B0U7Z5 B0U802 B0U811 B0U833 B0U881 B0U8L6 B0U8M0 B0U8N5 B0U8Y5
B0U8Y9 B0U9V5 B0UA56 B0UA74 B0UAJ5 B0UAJ8 B0UAK2 B0UAP4 B0UAV0 B0UAV4 B0UB90
B0UBD0 B0UBD4 B0UBI3 B0UBI6 B0UBI8 B0UBJ1 B0UC46 B0UC53 B0UC54 B0UCF0 B0UCG5
B0UCI8 B0UCJ9 B0UCL3 B0UCL4 B0UCS0 B0UCS1 B0UD11 B0UD12 B0UD14 B0UD17 B0UD19
B0UDF4 B0UDG6 B0UDI8 B0UDK6 B0UDK8 B0UE10 B0UE13 B0UEE1 B0UEK2 B0UET5 B0UEX4
B0UEX5 B0UF36 B0UFA1 B0UFC6 B0UFE2 B0UFH4 B0UFI0 B0UFU2 B0UGC4 B0UGM0 B0UGN1
B0UGP0 B0UGY1 B0UGY2 B0UH14 B0UH52 B0UHC4 B0UHC7 B0UHD1 B0UHH9 B0UHP4 B0UHR4
B0UHR5 B0UHR6 B0UHS5 B0UHU4 B0UHU5 B0UHU6 B0UHU7 B0UHV0 B0UHV1 B0UHV2 B0UHV3
B0UHV4 B0UHV6 B0UHV8 B0UHV9 B0UHW0 B0UHW2 B0UHW4 B0UHW5 B0UHW6 B0UHW7 B0UHW8
B0UHW9 B0UHX0 B0UHX2 B0UHX3 B0UHX4 B0UHX5 B0UHX6 B0UHX7 B0UHX8 B0UHY1 B0UHY2
B0UI66 B0UI81 B0UI96 B0UIA8 B0UIS8 B0UIW7 B0UIY3 B0UJ26 B0UJ71 B0UJ78 B0UJ99
B0UJB1 B0UJH5 B0UJI7 B0UJI8 B0UJI9 B0UKC4 B0UKC8 B0UKF2 B0UKH0 B0UKV2 B0UL29
B0UL30 B0UL32 B0UL34 B0UL49 B0UL53 B0UL61 B0UL68 B0UL80 B0ULF6 B0ULM2 B0ULP6
B0ULQ6 B0ULY8 B0ULZ9 B0UM25 B0UM37 B0UML4 B0UML5 B0UMS1 B0UMS2 B0UMS4 B0UMS9
B0UN04 B0UN75 B0UN77 B0UN89 B0UNU2 B0UNY0 B0UNY7 B0UP23 B0UP34 B0UP37 B0UP40
B0UP41 B0UP43 B0UP55 B0UP59 B0UPC1 B0UPC4 B0UPD3 B0UPF4 B0UPF9 B0UPH0 B0UPJ5
B0UPJ6 B0UPJ7 B0UPN1 B0UPZ4 B0UQ09 B0UQ15 B0UQ16 B0UQ51 B0UQE6 B0UQF5 B0UQR1
B0UQS5 B0UQT7 B0UQW5 B0UQW6 B0UR46 B0UR68 B0UR82 B0UR84 B0URH4 B0URI0 B0URI5
B0URI7 B0URJ9 B0URL9 B0URM0 B0URM7 B0URM8 B0URM9 B0URN5 B0URN6 B0URR1 B0URT3
B0URU2 B0URU5 B0URU6 B0URU7 B0URU8 B0URV5 B0URV9 B0URW9 B0URX4 B0URX8 B0URZ0
B0URZ3 B0URZ6 B0URZ7 B0US05 B0US06 B0US07 B0US22 B0US27 B0US28 B0US32 B0US43
B0US44 B0US45 B0US46 B0US54 B0US58 B0US59 B0US68 B0US69 B0USD9 B0USE8 B0USF1
B0USF3 B0USF4 B0USF6 B0USG8 B0USG9 B0USH2 B0USH5 B0USI0 B0USI5 B0USJ0 B0USJ3
B0USK5 B0USK6 B0USK7 B0USL1 B0USL5 B0USM2 B0USP0 B0USP1 B0USP2 B0USR6 B0USS1
B0USS3 B0USS4 B0UST3 B0USU0 B0USU1 B0USU2 B0UT10 B0UT19 B0UT47 B0UT51 B0UT65
B0UT66 B0UT69 B0UT70 B0UT76 B0UT78 B0UT80 B0UT85 B0UT89 B0UT98 B0UT99 B0UTA0
B0UTC6 B0UTE1 B0UTE2 B0UTE6 B0UTE7 B0UTF5 B0UTH6 B0UTI0 B0UTI4 B0UTI6 B0UTI7
B0UTJ5 B0UTS2 B0UTT1 B0UTU0 B0UTU4 B0UTU5 B0UTU8 B0UTX7 B0UTZ3 B0UTZ5 B0UTZ7
B0UU01 B0UU02 B0UU11 B0UU13 B0UU14 B0UU15 B0UU16 B0UU19 B0UU22 B0UU23 B0UU27
B0UU33 B0UU34 B0UU36 B0UU42 B0UU49 B0UU57 B0UU60 B0UU61 B0UU62 B0UU63 B0UU73
B0UU75 B0UU76 B0UU87 B0UU89 B0UUA1 B0UUA4 B0UUA9 B0UUD2 B0UUD3 B0UUD4 B0UUE6
B0UUF1 B0UUF3 B0UUG0 B0UUH4 B0UUI5 B0UUJ3 B0UUJ4 B0UUJ6 B0UUK6 B0UUM0 B0UUM5
B0UUN2 B0UUN5 B0UUN7 B0UUP7 B0UUR2 B0UUR6 B0UUR7 B0UUT2 B0UUT7 B0UUU5 B0UUU6
B0UUU9 B0UUV6 B0UUW1 B0UUW5 B0UUW7 B0UUW8 B0UUW9 B0UUX0 B0UUY0 B0UUY8 B0UUZ4
B0UUZ5 B0UUZ6 B0UUZ7 B0UUZ9 B0UV08 B0UV12 B0UV21 B0UV22 B0UV30 B0UV33 B0UV42
B0UV47 B0UV54 B0UV55 B0UV57 B0UV71 B0UV72 B0UV80 B0UV84 B0UV86 B0UVH6 B0UVH9
B0UVI0 B0UVI2 B0UVK3 B0UVK4 B0UVK6 B0UVL5 B0UVL9 B0UVM2 B0UVM6 B0UVM7 B0UVM8
B0UVM9 B0UVN8 B0UVN9 B0UVP3 B0UVP7 B0UVP8 B0UVQ3 B0UVX4 B0UVY2 B0UVY6 B0UVZ4
B0UVZ8 B0UW05 B0UW11 B0UW18 B0UW19 B0UW20 B0UW21 B0UW56 B0UW58 B0UW63 B0UW70
B0UW86 B0UW92 B0UW94 B0UWA6 B0UWA7 B0UWA8 B0UWA9 B0UWB7 B0UWC4 B0UWC5 B0UWC6
B0UWC7 B0UWD5 B0UWD6 B0UWE3 B0UWE5 B0UWE8 B0UWF1 B0UWF2 B0UWG2 B0UWH2 B0UWH3
B0UWH5 B0UWH8 B0UWJ9 B0UWK6 B0UWK7 B0UWK9 B0UWL1 B0UWL5 B0UWL8 B0UWM2 B0UWM5
B0UWP6 B0UWQ4 B0UWQ8 B0UWR0 B0UWR3 B0UWR4 B0UWS7 B0UWS8 B0UWT1 B0UWT3 B0UWT4
B0UWT7 B0UWU3 B0UWU6 B0UWV3 B0UWV7 B0UWV8 B0UWV9 B0UWW1 B0UWW2 B0UWY1 B0UWZ4

B0UWZ5 B0UWZ8 B0UX12 B0UX13 B0UX14 B0UX16 B0UX17 B0UX18 B0UX20 B0UX21 B0UX22
B0UX24 B0UX25 B0UX26 B0UX27 B0UX28 B0UX29 B0UX30 B0UX31 B0UX32 B0UX33 B0UX35
B0UX37 B0UX38 B0UX40 B0UX44 B0UXH9 B0UXL7 B0UXP9 B0UXU6 B0UYH6 B0UYI1 B0V207
B0V2S2 B0V3F8 B0V3H4 B0V494 B0V495 B0V496 B0V497 B0V498 B0V4K1 B0V4L0 B0V4L7
B0V4N1 B0V4N2 B0V4P7 B0V4R6 B0V4R8 B0V4S0 B0V4S6 B0V4T7 B0V4U0 B0V4U1 B0V4V4
B0V4V6 B0V4V8 B0V4Y6 B0V4Z1 B0V529 B0V550 B0V5E6 B0V5G6 B0V5G7 B0V5G8 B0V5H2
B0V5M5 B0V5N2 B0V5N7 B0V5P1 B0V5P2 B0V5P6 B0V5P7 B0V5Q4 B0V5Q6 B0V5Q9 B0V5R2
B0V5R3 B0V5S5 B0V5U3 B0V5U4 B0V5W8 B0V5X4 B0V625 B0V628 B0V634 B0V636 B0V669
B0V675 B0V681 B0V684 B0V6E0 B0V6E8 B0V6F0 B0V6F3 B0V6G3 B0V6G5 B0V6J2 B0V6N4
B0V6R4 B0V6R7 B0V6T5 B0V6U3 B0V6U4 B0V6U5 B0V6U6 B0V6U7 B0V6U8 B0V6V5 B0V6V6
B0V6V7 B0V6V8 B0V6V9 B0V6W0 B0V6W1 B0V6W2 B0V6W3 B0V6W4 B0V6W8 B0V6W9 B0V6X0
B0V6X1 B0V6X2 B0V6X3 B0V6X4 B0V6X5 B0V6X6 B0V6X7 B0V707 B0V709 B0V710 B0V732
B0V744 B0V782 B0V7B1 B0V7E0 B0V7F2 B0V7G5 B0V7G6 B0V7G7 B0V7H8 B0V7I0 B0V7Q2
B0V7R3 B0V7W2 B0V7X3 B0V7X4 B0V810 B0V811 B0V876 B0V8B1 B0V8B3 B0V8C7 B0V8E4
B0V8F1 B0V8G0 B0V8G1 B0V8G6 B0V8H2 B0V8I3 B0V8J1 B0V8J2 B0V8K4 B0V8L2 B0V8M2
B0V8M8 B0V8M9 B0V8N7 B0V8Q3 B0V8R7 B0V8S1 B0V8W3 B0V8X8 B0V8Y3 B0V8Y4 B0V8Y6
B0V8Y8 B0V8Z3 B0V8Z5 B0V944 B0V945 B0V946 B0V9C8 B0V9C9 B0V9F6 B0V9F7 B0V9I9
B0V9L0 B0V9L5 B0V9M3 B0V9N5 B0V9P2 B0V9P4 B0V9P6 B0V9P7 B0V9P8 B0V9Q3 B0V9Q9
B0V9T8 B0V9V8 B0V9X4 B0V9X5 B0V9X6 B0VA00 B0VA24 B0VA26 B0VA98 B0VAC1 B0VAE0
B0VAE1 B0VAE9 B0VAF3 B0VAF5 B0VAI1 B0VAI3 B0VAI5 B0VAL3 B0VAL5 B0VAU1 B0VAU7
B0VAY4 B0VAY5 B0VB44 B0VBB3 B0VBC6 B0VBD8 B0VBE4 B0VBN5 B0VBS1 B0VBS2 B0VBT2
B0VBU8 B0VBX6 B0VBY5 B0VBZ1 B0VC45 B0VC63 B0VC84 B0VC98 B0VCA5 B0VCA8 B0VCB7
B0VCN6 B0VCR5 B0VCR6 B0VCU0 B0VCV4 B0VCW7 B0VCX5 B0VCZ2 B0VCZ6 B0VD03 B0VD17
B0VD21 B0VD25 B0VD42 B0VD51 B0VD54 B0VD55 B0VDB2 B0VDC8 B0VDE4 B0VDE5 B0VDF1
B0VDG3 B0VDG4 B0VDG5 B0VDG6 B0VDH9 B0VDI0 B0VDI5 B0VDI6 B0VDJ1 B0VDK0 B0VDK5
B0VDM7 B0VDN4 B0VDQ2 B0VDR1 B0VDR5 B0VDR6 B0VDW7 B0VDX7 B0VDY4 B0VE41 B0VE54
B0VE65 B0VE72 B0VE80 B0VE81 B0VE83 B0VEA3 B0VEA4 B0VEC1 B0VEC9 B0VEE3 B0VEE9
B0VEF2 B0VEG5 B0VEJ4 B0VEJ5 B0VEN3 B0VH76 B0VHH0 B0VKA4 B0VKC6 B0VKC7 B0VKJ4
B0VKK2 B0VKK3 B0VKK4 B0VKL3 B0VKP5 B0VKP9 B0VKR1 B0VKR4 B0VKR7 B0VKR8 B0VKS2
B0VKS3 B0VKT1 B0VKU4 B0VKU6 B0VKV0 B0VKW9 B0VL04 B0VL27 B0VL32 B0VL49 B0VL51
B0VL72 B0VL79 B0VL80 B0VLA5 B0VLB1 B0VLB6 B0VLD2 B0VLE0 B0VLF5 B0VLH0 B0VLH2
B0VLL0 B0VLL2 B0VLR7 B0VLR8 B0VLU1 B0VLU2 B0VLU4 B0VLU6 B0VLV6 B0VLV9 B0VLX1
B0VLZ5 B0VLZ6 B0VLZ7 B0VLZ8 B0VLZ9 B0VM08 B0VM09 B0VM10 B0VM18 B0VM26 B0VM64
B0VM84 B0VMA3 B0VMC7 B0VMC8 B0VMD7 B0VMG3 B0VMH0 B0VMK0 B0VMK2 B0VML0 B0VMN8
B0VMP4 B0VMS2 B0VMT6 B0VMU3 B0VMU5 B0VMU8 B0VMV7 B0VMX3 B0VMY0 B0VMZ7 B0VN00
B0VN04 B0VNC8 B0VNF5 B0VNF6 B0VNJ1 B0VNL3 B0VNL8 B0VNU2 B0VNV8 B0VNV9 B0VNW2
B0VNX9 B0VNZ7 B0VNZ8 B0VP26 B0VP57 B0VPB7 B0VPC2 B0VPF6 B0VPF9 B0VPL6 B0VPM5
B0VPN2 B0VPT9 B0VPU0 B0VPV2 B0VPX5 B0VPZ8 B0VQ00 B0VQ11 B0VQ12 B0VQ33 B0VQ34
B0VQ51 B0VQ53 B0VQ56 B0VQ58 B0VQ59 B0VQ88 B0VQ89 B0VQA1 B0VQB5 B0VQB7 B0VQB8
B0VQD3 B0VQD4 B0VQD5 B0VQD6 B0VQD7 B0VQH8 B0VQI6 B0VQK3 B0VQK6 B0VQP7 B0VQP8
B0VQR8 B0VQR9 B0VQS0 B0VQS1 B0VQS2 B0VQS3 B0VQS4 B0VQS5 B0VQS6 B0VQS7 B0VQS8
B0VQS9 B0VQT0 B0VQT1 B0VQT2 B0VQT3 B0VQT4 B0VQT5 B0VQT6 B0VQT7 B0VQT9 B0VQU0
B0VQU1 B0VQU2 B0VQU3 B0VQU4 B0VQX0 B0VQX2 B0VQX5 B0VQZ2 B0VR13 B0VR22 B0VR30
B0VR37 B0VR41 B0VR58 B0VR80 B0VR86 B0VR94 B0VRA8 B0VRB1 B0VRC7 B0VRF8 B0VRJ4

B0VRK7 B0VRR2 B0VRV3 B0VRW9 B0VRY3 B0VS13 B0VS71 B0VSA3 B0VSA9 B0VSB5 B0VSB7
B0VSC1 B0VSC2 B0VSC3 B0VSD7 B0VSE0 B0VSH5 B0VSH6 B0VSK0 B0VSL0 B0VSL8 B0VSM4
B0VSN6 B0VSP4 B0VSP5 B0VSR5 B0VST2 B0VST3 B0VT10 B0VT23 B0VT30 B0VT55 B0VT70
B0VT80 B0VT89 B0VT90 B0VTA6 B0VTE3 B0VTE5 B0VTF0 B0VTF2 B0VTF8 B0VTG3 B0VTG4
B0VTG5 B0VTH7 B0VTL6 B0VTM5 B0VTQ1 B0VTQ2 B0VTW7 B0VTX2 B0VTX3 B0VTY5 B0VU02
B0VU08 B0VU22 B0VU43 B0VU92 B0VUC9 B0VUD6 B0VUE2 B0VUE9 B0VUQ6 B0VUQ9 B0VUS0
B0VUS1 B0VUU0 B0VUW1 B0VUX1 B0VV21 B0VV43 B0VV83 B0VV85 B0VV91 B0VV92 B0VV99
B0VX69 B0VXE6 B0VXK3 B0VXM6 B0VXM7 B0VZR4 B0VZS1 B0W010 B0W041 B0W0S3 B0W1V2
B0W2S0 B0W2W6 B0W377 B0W3H3 B0W3L6 B0W3R7 B0W3S2 B0W517 B0W562 B0W6M9 B0W6N3
B0W730 B0W8L4 B0W8M5 B0W939 B0W9B4 B0W9F4 B0WAE3 B0WAM5 B0WAN0 B0WAQ0 B0WAU6
B0WAU8 B0WC25 B0WC36 B0WCT9 B0WCZ5 B0WD26 B0WDA9 B0WG73 B0WGM1 B0WGN4 B0WI30
B0WII7 B0WK43 B0WMG3 B0WN96 B0WPU9 B0WQ75 B0WQG0 B0WQV1 B0WR18 B0WS18 B0WSB8
B0WSW8 B0WSX1 B0WTN3 B0WTU5 B0WU24 B0WU52 B0WV73 B0WVC4 B0WWP2 B0WYR6 B0WYS3
B0X0K1 B0X1V5 B0X2G0 B0X2V9 B0X4N1 B0X4N8 B0X560 B0X5E3 B0X6E8 B0X6P7 B0X6V0
B0X755 B0X911 B0X9H6 B0X9V1 B0XAF3 B0XDC4 B0XDJ0 B0XEA7 B0XFQ9 B0XFT7 B0XHW8
B0XK69 B0XKF3 B0XL39 B0XM00 B0XMC1 B0XMV6 B0XN37 B0XNF8 B0XNR0 B0XNT4 B0XP13
B0XPE4 B0XPE7 B0XPW9 B0XPZ4 B0XPZ9 B0XQ15 B0XQ41 B0XQ42 B0XQ54 B0XQ55 B0XQB9
B0XQT1 B0XR80 B0XRA0 B0XRG7 B0XRM8 B0XS28 B0XS72 B0XS74 B0XS99 B0XSH8 B0XSK6
B0XSW5 B0XTJ7 B0XTS1 B0XTT5 B0XU47 B0XU60 B0XUC7 B0XUL1 B0XUR7 B0XV83 B0XVP0
B0XVV1 B0XW47 B0XWG9 B0XXK5 B0XXL3 B0XXN5 B0XXN8 B0XXS1 B0XY69 B0XYC8 B0XYE0
B0XYQ4 B0XYY2 B0XZ33 B0XZU3 B0Y0F3 B0Y0P7 B0Y0Y6 B0Y1D1 B0Y1E7 B0Y2U0 B0Y356
B0Y3V7 B0Y3W4 B0Y462 B0Y4D5 B0Y4D6 B0Y4F9 B0Y4I9 B0Y4N9 B0Y567 B0Y568 B0Y581
B0Y5C3 B0Y5N4 B0Y5Q9 B0Y5R7 B0Y5V6 B0Y601 B0Y604 B0Y612 B0Y691 B0Y6H2 B0Y6R2
B0Y762 B0Y7H6 B0Y7I1 B0Y8S0 B0Y8W7 B0Y9Q4 B0Y9Z9 B0YA45 B0YAK0 B0YAX5 B0YC65
B0YCA6 B0YCF6 B0YCZ3 B0YD89 B0YDH7 B0YEH1 B0YLW6 B0YLW7 B0YLW8 B0YPL5 B0YPL9
B0YPM0 B0YPM1 B0YPN0 B0YPN8 B0YPP9 B0YPQ0 B0YPQ1 B0YPQ8 B0YPQ9 B0YPR0 B0YPR2
B0YPR3 B0YPR5 B0YPR6 B0YPR7 B0YPS2 B0Z4K7 B0Z4K9 B0Z4L9 B0Z4M6 B0Z4M7 B0Z4M8
B0Z4P7 B0Z4P8 B0Z4P9 B0Z4Q9 B0Z4R0 B0Z4R2 B0Z4R3 B0Z4R5 B0Z4R6 B0Z4R7 B0Z4R9
B0Z4S1 B0Z4T2 B0Z4U1 B0Z4U3 B0Z4V3 B0Z4W0 B0Z4W1 B0Z4W2 B0Z4Y1 B0Z4Y2 B0Z4Y3
B0Z4Z3 B0Z4Z4 B0Z4Z6 B0Z4Z7 B0Z4Z9 B0Z500 B0Z501 B0Z503 B0Z505 B0Z516 B0Z525
B0Z527 B0Z537 B0Z544 B0Z545 B0Z546 B0Z565 B0Z566 B0Z567 B0Z577 B0Z578 B0Z580
B0Z581 B0Z583 B0Z584 B0Z585 B0Z587 B0Z589 B0Z5A0 B0Z5A9 B0Z5B1 B0Z5C1 B0Z5C8
B0Z5C9 B0Z5D0 B0Z5E9 B0Z5F0 B0Z5F1 B0Z5G1 B0Z5G2 B0Z5G4 B0Z5G5 B0Z5G7 B0Z5G8
B0Z5G9 B0Z5H1 B0Z5H3 B0Z5I4 B0ZB55 B0ZB56 B0ZB57 B0ZSF2 B0ZSF3 B0ZSH1 B0ZT44
B0ZT45 B0ZT46 B0ZT47 B0ZTE1 B0ZTE2 B1A0U8 B1A612 B1A7S3 B1A916 B1A924 B1A925
B1A926 B1A933 B1A956 B1A957 B1A958 B1A968 B1A969 B1A971 B1A972 B1A974 B1A975
B1A976 B1A978 B1A980 B1A991 B1AAA7 B1ABR6 B1ABS0 B1AHY7 B1AHY9 B1AHZ6 B1AHZ7
B1AI04 B1AI06 B1AI12 B1AI24 B1AI25 B1AI43 B1AI47 B1AI63 B1AI69 B1AI74 B1AI75
B1AI95 B1AIA5 B1AIA6 B1AIA9 B1AIB0 B1AIG8 B1AIG9 B1AIH0 B1AIH5 B1AIH6 B1AIJ1
B1AIJ7 B1AIJ8 B1AIK0 B1AIK5 B1AIL7 B1AIL8 B1AIL9 B1AIM0 B1AIM1 B1AIM2 B1AIM3
B1AIM4 B1AIM5 B1AIM6 B1AIM7 B1AIM8 B1AIM9 B1AIN0 B1AIN1 B1AIN2 B1AIN3 B1AIN4
B1AIN6 B1AIN7 B1AIN8 B1AIP3 B1AIP4 B1AIP5 B1AIP7 B1AIQ5 B1AIQ8 B1AIR7 B1AIR8
B1AIS5 B1AIS8 B1AIT3 B1AIT6 B1AIU2 B1AIU8 B1AIV8 B1AIV9 B1AIX1 B1AIX2 B1AIX8
B1AIY1 B1AIY6 B1AIY7 B1AIZ3 B1AJ02 B1AJ04 B1AJ08 B1AJ09 B1AJ10 B1AJ20 B1AJ22

B1AJ24 B1AJ25 B1AJ36 B1AJ41 B1AJ46 B1AJ51 B1AJ58 B1AJ61 B1AJ69 B1AJ71 B1AJ72
B1AJ73 B1AJ74 B1AJ75 B1AJ79 B1AJ89 B1AJ90 B1AJ93 B1AJ99 B1AJA2 B1AJA4 B1AJA6
B1AJC8 B1AJD0 B1AJD3 B1AJD7 B1AJF3 B1AJF5 B1AJG3 B1AJG4 B1AJG5 B1AJG6 B1AJH4
B1AJH5 B1AJH7 B1AJI1 B1AJI2 B1AJI4 B1AJI8 B1AJJ1 B1AJJ2 B1AJJ4 B1AJK7 B1AJK8
B1AJL7 B1AJL8 B1AJM6 B1AJN5 B1AJP0 B1AJP1 B1AJP2 B1AJP4 B1AJP5 B1AJZ9 B1AK53
B1AK76 B1AL46 B1AMM8 B1ANH7 B1ANS9 B1ANY3 B1APH4 B1AQ75 B1AR13 B1ARD6 B1ARD8
B1ARI9 B1ARW8 B1AS42 B1ASB6 B1ATL7 B1AUF7 B1AUR6 B1AUS7 B1AWL2 B1AX39 B1AXH1
B1AYB6 B1B0V2 B1B1A0 B1B1U3 B1B1U4 B1B557 B1ELW6 B1G889 B1G894 B1GVX4 B1GVX5
B1GVX6 B1GVX7 B1GYK8 B1GYM8 B1GYN1 B1GYP6 B1GYQ3 B1GYQ9 B1GYR2 B1GYR7 B1GYS7
B1GYT2 B1GYT3 B1GYT4 B1GYU9 B1GYV6 B1GYV7 B1GYV8 B1GYW9 B1GYY3 B1GYY7 B1GYY8
B1GYY9 B1GYZ9 B1GZ10 B1GZ11 B1GZ18 B1GZ20 B1GZ23 B1GZ25 B1GZ34 B1GZ43 B1GZ47
B1GZ68 B1GZ70 B1GZ74 B1GZ76 B1GZ77 B1GZ79 B1GZ80 B1GZ83 B1GZ85 B1GZ86 B1GZ87
B1GZ88 B1GZ89 B1GZ90 B1GZ92 B1GZ93 B1GZ94 B1GZ96 B1GZ97 B1GZ99 B1GZA6 B1GZA7
B1GZA8 B1GZA9 B1GZB1 B1GZB9 B1GZC0 B1GZC1 B1GZD2 B1GZD3 B1GZG7 B1GZG8 B1GZH1
B1GZH8 B1GZI5 B1GZJ0 B1GZJ3 B1GZJ4 B1GZJ8 B1GZJ9 B1GZM9 B1GZN4 B1GZN9 B1GZP2
B1GZQ6 B1GZS8 B1GZT3 B1GZU1 B1GZU2 B1GZV6 B1GZW3 B1GZX0 B1GZX5 B1GZY6 B1GZY8
B1GZY9 B1GZZ1 B1H009 B1H013 B1H014 B1H028 B1H052 B1H053 B1H056 B1H060 B1H063
B1H069 B1H070 B1H072 B1H084 B1H0A5 B1H0A6 B1H0A7 B1H0B1 B1H0B2 B1H0B3 B1H0C6
B1H0C7 B1H0D2 B1H0E6 B1H0E7 B1H0E9 B1H0H0 B1H0H2 B1H0H8 B1H0H9 B1H0I0 B1H0I4
B1H0I6 B1H0I8 B1H0J3 B1H0L0 B1H0L3 B1H0L5 B1H0Q9 B1H0R2 B1H116 B1H158 B1H1E4
B1H1S4 B1H1W9 B1H1X1 B1H1X4 B1H1Z8 B1H222 B1H224 B1H228 B1H268 B1H278 B1H283
B1H285 B1H2P7 B1H2Q6 B1H349 B1H369 B1H3B4 B1H3D5 B1HM45 B1HM47 B1HMT3 B1HMU8
B1HMV0 B1HMV4 B1HMV6 B1HMV7 B1HMV9 B1HMW1 B1HMW2 B1HMW3 B1HMW4 B1HMW6 B1HMW8
B1HMW9 B1HMX0 B1HMX2 B1HMX3 B1HMX4 B1HMX5 B1HMX6 B1HMX8 B1HMX9 B1HMY0 B1HMZ0
B1HMZ1 B1HMZ3 B1HMZ5 B1HMZ6 B1HMZ8 B1HMZ9 B1HN47 B1HN64 B1HN89 B1HN90 B1HNL5
B1HNL6 B1HNM3 B1HNM5 B1HNM6 B1HNM7 B1HNS5 B1HNU2 B1HNX3 B1HNY1 B1HNY3 B1HPA1
B1HPF6 B1HPJ4 B1HPK4 B1HPK6 B1HPM0 B1HPM1 B1HPM2 B1HPM3 B1HPM8 B1HPQ8 B1HPR6
B1HPR7 B1HPT4 B1HPT5 B1HPU6 B1HPW8 B1HPX1 B1HPX3 B1HPX8 B1HPY0 B1HQB9 B1HQC1
B1HQC2 B1HQC3 B1HQE0 B1HQE4 B1HQE6 B1HQI0 B1HQI2 B1HQI4 B1HQJ3 B1HQJ5 B1HQK2
B1HQT3 B1HQV3 B1HQZ0 B1HQZ1 B1HQZ7 B1HQZ9 B1HR01 B1HR05 B1HR10 B1HR13 B1HR35
B1HR59 B1HR71 B1HR96 B1HR98 B1HR99 B1HRG8 B1HRH3 B1HRK0 B1HRT3 B1HRT5 B1HRX4
B1HRX6 B1HRX7 B1HRX9 B1HRY0 B1HS10 B1HS22 B1HS35 B1HS47 B1HS62 B1HS82 B1HSN5
B1HSN6 B1HSN7 B1HSS4 B1HSU6 B1HSV6 B1HSY5 B1HT11 B1HT15 B1HT93 B1HTA3 B1HTA6
B1HTD1 B1HTD2 B1HTD4 B1HTF0 B1HTF1 B1HTF6 B1HTG5 B1HTI4 B1HTI6 B1HTJ4 B1HTK2
B1HTK8 B1HTT1 B1HTV1 B1HTV3 B1HTV6 B1HTV8 B1HTW6 B1HTW8 B1HTW9 B1HTX0 B1HU19
B1HU20 B1HUD0 B1HUD1 B1HUH0 B1HUI4 B1HUJ1 B1HUK4 B1HUL3 B1HUT1 B1HUU8 B1HUY7
B1HV71 B1HV72 B1HV77 B1HV79 B1HV83 B1HVA1 B1HVA2 B1HVA4 B1HVB0 B1HVB2 B1HVB6
B1HVC0 B1HVD3 B1HVD6 B1HVD8 B1HVE5 B1HVE6 B1HVI3 B1HVK6 B1HVP6 B1HVP7 B1HVQ1
B1HVQ6 B1HVQ8 B1HVR0 B1HVS2 B1HVX5 B1HVX6 B1HW35 B1HW77 B1HW86 B1HW99 B1HWA2
B1HWC8 B1HWC9 B1HWD3 B1HWE9 B1HX23 B1HX32 B1HX34 B1HX36 B1HX43 B1HX90 B1HXA1
B1HXC1 B1HXC6 B1HXC8 B1HXY7 B1HXZ6 B1HY21 B1HY35 B1HY37 B1HYB1 B1HYR5 B1HYT4
B1HYT5 B1HZ05 B1HZ48 B1HZE2 B1HZE3 B1HZE5 B1HZE7 B1HZE8 B1HZH4 B1HZH6 B1HZM2
B1HZM7 B1HZY7 B1I012 B1I022 B1I024 B1I057 B1I067 B1I069 B1I0S9 B1I0T1 B1I0W9
B1I0X4 B1I0X5 B1I0X7 B1I0X9 B1I0Y8 B1I158 B1I161 B1I162 B1I165 B1I190 B1I193

B1I196 B1I197 B1I1B1 B1I1B2 B1I1B3 B1I1B4 B1I1B5 B1I1B6 B1I1C5 B1I1D1 B1I1E6
B1I1G4 B1I1H6 B1I1I3 B1I1I4 B1I1I5 B1I1I7 B1I1I8 B1I1I9 B1I1J0 B1I1J1 B1I1J2
B1I1J3 B1I1J4 B1I1J5 B1I1J6 B1I1J7 B1I1J8 B1I1J9 B1I1K0 B1I1K1 B1I1K2 B1I1K9
B1I1L2 B1I1L3 B1I1L4 B1I1L5 B1I1L7 B1I1L8 B1I1M3 B1I1M4 B1I1M6 B1I1M9 B1I1N9
B1I1P4 B1I1P6 B1I1S3 B1I1S7 B1I1S8 B1I1T4 B1I1Y1 B1I1Y5 B1I241 B1I250 B1I258
B1I260 B1I293 B1I295 B1I296 B1I2D4 B1I2D5 B1I2E4 B1I2I8 B1I2J1 B1I2K0 B1I2N0
B1I2N1 B1I2N2 B1I2P1 B1I2P4 B1I2P9 B1I310 B1I313 B1I337 B1I359 B1I363 B1I370
B1I373 B1I374 B1I383 B1I392 B1I393 B1I3B8 B1I3C0 B1I3D5 B1I3D6 B1I3I9 B1I3J0
B1I3J6 B1I3J9 B1I3K2 B1I3M7 B1I3P6 B1I3R5 B1I3T1 B1I3V7 B1I3W2 B1I3W6 B1I3Z0
B1I3Z6 B1I3Z8 B1I462 B1I477 B1I493 B1I494 B1I4A3 B1I4D9 B1I4F5 B1I4F9 B1I4G4
B1I4H2 B1I4I5 B1I4I8 B1I4I9 B1I4J1 B1I4K2 B1I4K8 B1I4L1 B1I4L8 B1I4M2 B1I4M6
B1I4M8 B1I4N1 B1I4P7 B1I4P8 B1I4Q9 B1I4R5 B1I4S0 B1I4S1 B1I4W9 B1I4X2 B1I4Z1
B1I501 B1I504 B1I509 B1I516 B1I544 B1I545 B1I551 B1I556 B1I557 B1I559 B1I5R2
B1I5R7 B1I5V8 B1I5Z0 B1I661 B1I672 B1I677 B1I693 B1I699 B1I6C4 B1I6D9 B1I6E6
B1I6H9 B1I6L2 B1I6L4 B1I6M4 B1I6P8 B1I6Q1 B1I6Q5 B1I6S1 B1I6S2 B1I6S7 B1I6S9
B1I6T1 B1I6X3 B1I6X7 B1I6Y3 B1I700 B1I703 B1I710 B1I720 B1I721 B1I726 B1I733
B1I734 B1I755 B1I766 B1I770 B1I7I6 B1I7J8 B1I7K0 B1I7K5 B1I7P5 B1I7S6 B1I7S7
B1I7S8 B1I7S9 B1I7T0 B1I7V1 B1I7V5 B1I7V8 B1I7Y6 B1I7Z3 B1I814 B1I815 B1I833
B1I835 B1I837 B1I843 B1I857 B1I863 B1I885 B1I886 B1I887 B1I891 B1I8A9 B1I8B0
B1I8B2 B1I8B3 B1I8C4 B1I8F2 B1I8F3 B1I8F7 B1I8I4 B1I8I5 B1I8I6 B1I8J7 B1I8J8
B1I8J9 B1I8K0 B1I8K1 B1I8K2 B1I8K3 B1I8K4 B1I8K5 B1I8K6 B1I8K7 B1I8K8 B1I8K9
B1I8L0 B1I8L1 B1I8L2 B1I8L3 B1I8L4 B1I8L6 B1I8L7 B1I8L9 B1I8M1 B1I8M2 B1I8M3
B1I8M5 B1I8M7 B1I8Q1 B1I8Q2 B1I8R2 B1I8R3 B1I8R4 B1I8S2 B1I8S4 B1I8T7 B1I8U6
B1I8Y2 B1I8Y6 B1I8Z1 B1I8Z7 B1I8Z8 B1I8Z9 B1I918 B1I922 B1I953 B1I960 B1I998
B1I9A0 B1I9B4 B1I9D7 B1I9D8 B1I9E5 B1I9E7 B1I9G2 B1I9G3 B1I9G6 B1I9H0 B1I9H1
B1I9H2 B1I9J3 B1I9J5 B1I9J8 B1I9K1 B1I9K5 B1I9M4 B1I9M6 B1I9M7 B1I9M8 B1I9M9
B1I9N0 B1I9N1 B1I9N6 B1I9S9 B1I9T7 B1I9U2 B1I9V0 B1I9V1 B1I9W0 B1I9W1 B1I9W8
B1I9Z6 B1I9Z8 B1IA16 B1IA31 B1IA36 B1IA50 B1IA51 B1IA52 B1IA75 B1IA76 B1IA80
B1IA81 B1IAA1 B1IAA7 B1IAA9 B1IAC7 B1IAC8 B1IAD9 B1IAG1 B1IAH3 B1IAH4 B1IAK1
B1IAK9 B1IAL2 B1IAM3 B1IAM6 B1IAN6 B1IAN9 B1IAP6 B1IAS4 B1IAS5 B1IAT8 B1IAU3
B1IAU9 B1IAV2 B1IAV3 B1IAZ7 B1IAZ8 B1IB05 B1IB07 B1IB08 B1IB13 B1IB28 B1IB49
B1IB64 B1IB66 B1IB77 B1IB89 B1IB99 B1IBA0 B1IBA2 B1IBA4 B1IBA6 B1IBA8 B1IBB0
B1IBB9 B1IBC0 B1IBC1 B1IBC7 B1IBD3 B1IBD6 B1IBG7 B1IBG8 B1IBH4 B1IBH5 B1IBH9
B1IBI3 B1IBL9 B1IBM3 B1IBM8 B1IBN0 B1IBN7 B1IBP0 B1IBP2 B1IBP4 B1IBQ3 B1IBQ8
B1IBR0 B1IBR5 B1IBR6 B1IBW1 B1IBW3 B1IBW9 B1IBZ5 B1IBZ6 B1IBZ7 B1IC08 B1IC09
B1IC10 B1IC17 B1IC26 B1IC35 B1IC43 B1IC44 B1IC62 B1IC65 B1IC67 B1IC68 B1ICA7
B1ICB3 B1ICF4 B1ICF5 B1ICG1 B1ICG3 B1ICH2 B1ICH3 B1ICH5 B1ICH6 B1ICH7 B1ICH8
B1ICI3 B1ICI9 B1ICK4 B1ICL3 B1ICL4 B1ICL5 B1ICL6 B1ICL9 B1ICM9 B1ICN7 B1ICP8
B1ICP9 B1ICQ4 B1ICQ5 B1ICQ6 B1ICR4 B1ICT9 B1ICU3 B1ICV4 B1ICV9 B1ICW1 B1ICW8
B1ICX7 B1ICY0 B1ICY3 B1ICY4 B1ICY6 B1ICY8 B1ICZ4 B1ICZ6 B1ICZ7 B1ID02 B1ID11
B1ID16 B1ID41 B1ID54 B1ID91 B1IDA0 B1IDB6 B1IDB7 B1IDC2 B1IDC3 B1IDU3 B1IDU6
B1IDV4 B1IDW7 B1IDW8 B1IDX1 B1IE03 B1IE17 B1IE18 B1IE21 B1IE45 B1IE69 B1IEF5
B1IEG6 B1IEG7 B1IEG9 B1IEK9 B1IEL4 B1IEL5 B1IEL6 B1IEL7 B1IEN0 B1IEV3 B1IEV4
B1IEY8 B1IF28 B1IFD1 B1IFD4 B1IFD5 B1IFE2 B1IFE9 B1IFG0 B1IFI2 B1IFK1 B1IFV9

B1IFW1 B1IFW3 B1IFW4 B1IFW5 B1IFW6 B1IFX3 B1IG08 B1IGB8 B1IGC4 B1IGC5 B1IGC7
B1IGC8 B1IGC9 B1IGD3 B1IGD5 B1IGD6 B1IGD8 B1IGD9 B1IGE0 B1IGE1 B1IGE2 B1IGE3
B1IGE4 B1IGE5 B1IGE6 B1IGE7 B1IGE8 B1IGF0 B1IGF1 B1IGF2 B1IGF3 B1IGF4 B1IGF5
B1IGF6 B1IGF7 B1IGF8 B1IGF9 B1IGG1 B1IGG2 B1IGG3 B1IGG5 B1IGG6 B1IGG9 B1IGH6
B1IGH9 B1IGJ0 B1IGJ2 B1IGJ3 B1IGV8 B1IGY0 B1IGY1 B1IGZ7 B1IH03 B1IH05 B1IH09
B1IH18 B1IH25 B1IHC4 B1IHD1 B1IHE5 B1IHH3 B1IHH8 B1IHH9 B1IHN7 B1IHP2 B1IHP8
B1IHQ2 B1IHQ4 B1IHR7 B1IHR8 B1IHR9 B1IHS3 B1IHS4 B1IHX4 B1II37 B1II43 B1II44
B1II48 B1II49 B1II53 B1II57 B1II63 B1II65 B1II66 B1II67 B1II71 B1II72 B1II75
B1II76 B1II79 B1II90 B1IIA6 B1IIE6 B1IIG7 B1IIK1 B1IIK4 B1IIK9 B1IIL4 B1IIL7
B1IIM4 B1IIM6 B1IIN2 B1IIP2 B1IIW9 B1IIY2 B1IJ68 B1IJ78 B1IJE7 B1IJF0 B1IJG4
B1IJG5 B1IJG6 B1IJG7 B1IJJ6 B1IJJ8 B1IJL2 B1IJR0 B1IK07 B1IK08 B1IK65 B1IK66
B1IK79 B1IKA6 B1IKG3 B1IKJ7 B1IKK5 B1IKM0 B1IKR0 B1IKX1 B1IKY4 B1IL14 B1IL45
B1IL57 B1IL59 B1IL61 B1IL94 B1IL95 B1ILA1 B1ILA3 B1ILA4 B1ILA6 B1ILA8 B1ILA9
B1ILB0 B1ILB1 B1ILB2 B1ILH4 B1ILH9 B1ILL2 B1ILL7 B1ILM1 B1ILM3 B1ILN1 B1ILR0
B1ILX5 B1ILY3 B1ILY5 B1ILY6 B1ILY8 B1ILZ5 B1IM02 B1IM03 B1IM08 B1IM65 B1IM66
B1IM67 B1IM68 B1IM69 B1IM76 B1IM83 B1IMC4 B1IMD7 B1IMD8 B1IME1 B1IME3 B1IMF1
B1IMF2 B1IMF3 B1IMF4 B1IMF6 B1IMK5 B1IML3 B1IMN1 B1IMN2 B1IMN7 B1IMN8 B1IMN9
B1IMQ0 B1IMQ1 B1IMQ2 B1IMT7 B1IMV0 B1IMV1 B1IMV3 B1IMV8 B1IMX1 B1IN35 B1IN59
B1IN79 B1INA3 B1IND6 B1IND7 B1IND8 B1INE1 B1INE4 B1INE6 B1INE7 B1INF6 B1INF7
B1INF8 B1ING1 B1ING9 B1IP32 B1IP33 B1IP34 B1IP35 B1IP41 B1IP46 B1IP47 B1IP51
B1IP53 B1IP55 B1IP62 B1IP75 B1IP76 B1IP96 B1IPB0 B1IPB7 B1IPC0 B1IPC1 B1IPE7
B1IPF3 B1IPI2 B1IPI3 B1IPI4 B1IPI5 B1IPI6 B1IPI8 B1IPJ0 B1IPJ7 B1IPK9 B1IPL0
B1IPL1 B1IPL2 B1IPL3 B1IPL5 B1IPP6 B1IPU4 B1IPU6 B1IPU9 B1IPV5 B1IPV6 B1IPV7
B1IPV8 B1IPV9 B1IPW0 B1IPX8 B1IPX9 B1IPY0 B1IPY2 B1IPY3 B1IPY4 B1IPY5 B1IPY6
B1IPY7 B1IPY8 B1IPY9 B1IPZ0 B1IPZ1 B1IPZ2 B1IPZ3 B1IPZ4 B1IPZ5 B1IPZ6 B1IPZ7
B1IPZ8 B1IQ00 B1IQ02 B1IQ03 B1IQ05 B1IQ11 B1IQ12 B1IQ13 B1IQ15 B1IQ17 B1IQ18
B1IQ31 B1IQ33 B1IQ50 B1IQ53 B1IQ63 B1IQ64 B1IQ76 B1IQ77 B1IQ96 B1IQB4 B1IQB6
B1IQB7 B1IQD2 B1IQE6 B1IQF1 B1IQF8 B1IQG9 B1IQH1 B1IQH2 B1IQH4 B1IQH5 B1IQH7
B1IQI0 B1IQI1 B1IQI4 B1IQI6 B1IQJ4 B1IQK6 B1IQK7 B1IQK9 B1IQL9 B1IQM0 B1IQM1
B1IQN5 B1IQP2 B1IQP3 B1IQP8 B1IQP9 B1IQQ3 B1IQQ4 B1IQQ6 B1IQQ7 B1IQR8 B1IQT4
B1IQT7 B1IQT8 B1IQU0 B1IQU4 B1IQU7 B1IQV0 B1IQV1 B1IQV3 B1IQV4 B1IQV5 B1IQV6
B1IQV7 B1IQW7 B1IQX2 B1IQX3 B1IQY0 B1IQY4 B1IR77 B1IR78 B1IR87 B1IR96 B1IR97
B1IRA4 B1IRA6 B1IRA7 B1IRB5 B1IRB6 B1IRB9 B1IRC4 B1IRC5 B1IRC6 B1IRC7 B1IRD4
B1IRD5 B1IRD7 B1IRD8 B1IRD9 B1IRE0 B1IRE1 B1IRE5 B1IRE6 B1IRE9 B1IRF1 B1IRF9
B1IRG0 B1IRG7 B1IRH0 B1IRH9 B1IRI6 B1IRJ2 B1IRK5 B1IRM5 B1IRN3 B1IRP4 B1IRP8
B1IRQ1 B1IRQ2 B1IRR0 B1IRR4 B1IRR7 B1IRT9 B1IRU1 B1IRU2 B1IRU3 B1IRU8 B1IRU9
B1IRX9 B1IRZ6 B1IS15 B1IS24 B1IS25 B1IS26 B1IS33 B1IS35 B1IS36 B1IS37 B1IS38
B1IS45 B1IS49 B1IS56 B1IS70 B1IS82 B1ISA3 B1ISB1 B1ISB5 B1ISC0 B1ISK7 B1ISN3
B1ISR7 B1ISV4 B1ISW0 B1ISW1 B1ISX1 B1ISX3 B1ISY2 B1ISZ1 B1IT03 B1IT04 B1IT05
B1IT06 B1IT08 B1IT09 B1IT10 B1IT14 B1IT15 B1IT28 B1IT29 B1IT34 B1IT35 B1IT36
B1IT42 B1IT47 B1IT54 B1IT58 B1IT65 B1IT67 B1IT76 B1IT77 B1IT87 B1IT89 B1IT97
B1IT98 B1IT99 B1ITA2 B1ITA4 B1ITB1 B1ITB3 B1ITC8 B1ITG7 B1ITG9 B1ITH3 B1ITH7
B1ITJ4 B1ITJ5 B1ITK8 B1ITM1 B1ITP1 B1ITQ1 B1ITQ5 B1ITQ6 B1ITR1 B1IU01 B1IU13
B1IU14 B1IU17 B1IU35 B1IU37 B1IU39 B1IU47 B1IU61 B1IU77 B1IU78 B1IU82 B1IU83

B1IU85 B1IU89 B1IUA3 B1IUB2 B1IUB5 B1IUB7 B1IUB8 B1IUC4 B1IUD3 B1IUD4 B1IUE0
B1IUE6 B1IUF7 B1IUF8 B1IUF9 B1IUG7 B1IUL3 B1IUM7 B1IUN1 B1IUN6 B1IUN9 B1IUP1
B1IUP9 B1IUQ0 B1IUQ1 B1IUQ2 B1IUQ3 B1IUQ4 B1IUQ7 B1IUQ9 B1IUR0 B1IUR1 B1IUR2
B1IUR3 B1IUR4 B1IUR7 B1IUS7 B1IUS8 B1IUS9 B1IUT2 B1IUT3 B1IUT4 B1IUT5 B1IUT6
B1IUU6 B1IUW8 B1IUW9 B1IUY0 B1IUY1 B1IUY2 B1IUY3 B1IUY6 B1IUZ9 B1IV14 B1IV35
B1IV40 B1IV43 B1IV47 B1IV67 B1IV85 B1IV86 B1IV87 B1IV88 B1IV89 B1IV90 B1IV93
B1IV97 B1IV98 B1IV99 B1IVA7 B1IVB3 B1IVB6 B1IVB8 B1IVB9 B1IVC1 B1IVC2 B1IVD5
B1IVD9 B1IVE3 B1IVE7 B1IVE8 B1IVF0 B1IVF1 B1IVF3 B1IVG0 B1IVG3 B1IVH3 B1IVH4
B1IVH5 B1IVH6 B1IVH7 B1IVI3 B1IVI7 B1IVJ1 B1IVJ9 B1IVL4 B1IVL6 B1IVL9 B1IVM0
B1IVM4 B1IVM5 B1IVM6 B1IVM7 B1IVP7 B1IVP8 B1IVQ2 B1IVR0 B1IVR2 B1IVS6 B1IVT5
B1IVT6 B1IVT8 B1IVW5 B1IVW6 B1IVW9 B1IVX4 B1IVX6 B1IVY4 B1IVZ6 B1IW01 B1IW07
B1IW09 B1IW14 B1IW19 B1IW21 B1IW23 B1IW24 B1IW48 B1IW60 B1IW72 B1IW76 B1IW93
B1IW95 B1IW97 B1IWC0 B1IWC1 B1IWC4 B1IWD0 B1IWD1 B1IWD3 B1IWD4 B1IWD5 B1IWD8
B1IWE3 B1IWE4 B1IWE6 B1IWE7 B1IWF0 B1IWF2 B1IWF4 B1IWG1 B1IWG2 B1IWG4 B1IWI1
B1IWI3 B1IWI8 B1IWJ3 B1IWM2 B1IWM6 B1IWN0 B1IWN8 B1IWP1 B1IWP9 B1IWQ6 B1IWR3
B1IWS0 B1IWX5 B1IWY2 B1IWZ0 B1IWZ2 B1IWZ3 B1IWZ4 B1IWZ7 B1IWZ8 B1IX27 B1IX30
B1IX32 B1IX36 B1IX44 B1IX45 B1IX53 B1IX60 B1IX65 B1IX74 B1IX88 B1IX96 B1IXB4
B1IXB6 B1IXC1 B1IXD2 B1IXF6 B1IXG1 B1IXG4 B1IXI5 B1IXI9 B1IXJ2 B1IXJ3 B1IXL9
B1IXM2 B1IXM4 B1IXP8 B1IXQ1 B1IXR8 B1IXS2 B1IXS3 B1IXS5 B1IXT6 B1IXT8 B1IXU0
B1IXU2 B1IXV6 B1IXX9 B1IXY1 B1IXY6 B1IY02 B1IY15 B1IY16 B1IY48 B1IY50 B1IY62
B1IY82 B1IY83 B1IY84 B1IY98 B1IYA8 B1IYB0 B1IYB5 B1IYB6 B1IYC6 B1IYE7 B1IYE9
B1IYF7 B1IYG6 B1IYG9 B1IYH2 B1IYH7 B1IYH8 B1IYI0 B1IYI9 B1IYJ2 B1IYJ3 B1IYM7
B1IYP2 B1IYP4 B1IYQ2 B1IYQ7 B1IYQ8 B1IYS4 B1IYV1 B1IYV3 B1IYV7 B1IYV8 B1IYV9
B1IYW0 B1IYW7 B1IYX7 B1IYY4 B1IYY5 B1IZ09 B1IZ49 B1IZ50 B1IZ53 B1IZ55 B1IZ72
B1IZ75 B1IZ89 B1IZ95 B1IZ97 B1IZB8 B1IZB9 B1IZC0 B1IZC1 B1IZC2 B1IZC3 B1IZC5
B1IZE7 B1IZF6 B1IZF7 B1IZF8 B1IZF9 B1IZH2 B1IZH4 B1IZH8 B1IZI1 B1IZJ2 B1IZK2
B1IZL7 B1IZM7 B1IZN3 B1IZN4 B1IZQ6 B1IZQ9 B1IZT7 B1IZU5 B1IZV5 B1IZV7 B1IZY5
B1IZZ1 B1J004 B1J010 B1J011 B1J012 B1J023 B1J026 B1J027 B1J029 B1J033 B1J034
B1J036 B1J043 B1J044 B1J055 B1J057 B1J060 B1J061 B1J084 B1J085 B1J086 B1J0B4
B1J0D5 B1J0E9 B1J0G0 B1J0J4 B1J0J6 B1J0L2 B1J0L4 B1J0L7 B1J0L8 B1J0M2 B1J0M5
B1J0M7 B1J0M8 B1J0Q3 B1J0Q7 B1J0S5 B1J0S7 B1J0S8 B1J0T5 B1J0T9 B1J0W4 B1J0W5
B1J0W6 B1J0X7 B1J0X8 B1J0Z1 B1J0Z2 B1J0Z4 B1J0Z5 B1J100 B1J133 B1J134 B1J136
B1J142 B1J143 B1J144 B1J150 B1J153 B1J166 B1J173 B1J176 B1J195 B1J1A3 B1J1B4
B1J1C4 B1J1D1 B1J1F5 B1J1K1 B1J1K3 B1J1M3 B1J1N6 B1J1N8 B1J1S3 B1J1S5 B1J1U3
B1J1V0 B1J1V2 B1J1V8 B1J1W0 B1J1W8 B1J1W9 B1J1X7 B1J1X8 B1J1Y1 B1J1Y2 B1J1Y6
B1J1Y7 B1J201 B1J247 B1J253 B1J254 B1J255 B1J256 B1J262 B1J265 B1J266 B1J268
B1J2A2 B1J2A9 B1J2B0 B1J2B2 B1J2B3 B1J2C0 B1J2C1 B1J2C2 B1J2C8 B1J2D7 B1J2E3
B1J2E9 B1J2G8 B1J2G9 B1J2H5 B1J2I3 B1J2I5 B1J2J5 B1J2J9 B1J2K5 B1J2K9 B1J2M0
B1J2M4 B1J2P0 B1J2P7 B1J2Q5 B1J2R2 B1J2S4 B1J2S5 B1J2S8 B1J2T7 B1J2T8 B1J2V4
B1J2W7 B1J2Y8 B1J2Z7 B1J2Z8 B1J302 B1J331 B1J347 B1J355 B1J356 B1J358 B1J390
B1J3F1 B1J3F6 B1J3G0 B1J3G4 B1J3G6 B1J3J6 B1J3K4 B1J3K5 B1J3L3 B1J3Q4 B1J3Y2
B1J3Y4 B1J425 B1J426 B1J432 B1J499 B1J4A2 B1J4B0 B1J4B1 B1J4E0 B1J4E2 B1J4E3
B1J4E4 B1J4E5 B1J4I2 B1J4I4 B1J4J3 B1J4L4 B1J4L6 B1J4L7 B1J4L9 B1J4M1 B1J4M3
B1J4N3 B1J4P3 B1J4S1 B1J4V1 B1J4W2 B1J4W5 B1J4Y7 B1J4Z5 B1J507 B1J522 B1J530

B1J531 B1J538 B1J554 B1J593 B1J596 B1J5B7 B1J5D5 B1J5D6 B1J5D7 B1J5F1 B1J5F7
B1J5G4 B1J5G5 B1J5H0 B1J5H2 B1J5Q0 B1J5T3 B1J5U4 B1J5W6 B1J5W7 B1J5X5 B1J606
B1J615 B1J623 B1J649 B1J675 B1J691 B1J692 B1J693 B1J6F5 B1J6H3 B1J6H6 B1J6T8
B1J6U4 B1J6U5 B1J6U6 B1J6U7 B1J6U8 B1J6V0 B1J6Y1 B1J6Y2 B1J6Y6 B1J6Y7 B1J7Q1
B1J805 B1J812 B1J813 B1J814 B1J815 B1J816 B1J818 B1J819 B1J865 B1J9C7 B1J9E4
B1J9G3 B1J9R7 B1JA05 B1JAB6 B1JAD0 B1JAD4 B1JAD5 B1JAD6 B1JAE3 B1JAG3 B1JAG8
B1JAH0 B1JAH1 B1JAI0 B1JAI6 B1JAI8 B1JAI9 B1JAJ0 B1JAJ1 B1JAJ3 B1JAJ4 B1JAJ5
B1JAJ6 B1JAJ7 B1JAJ8 B1JAJ9 B1JAK0 B1JAK1 B1JAV3 B1JB25 B1JB29 B1JB30 B1JB31
B1JB32 B1JB36 B1JB85 B1JBA9 B1JBB3 B1JBB5 B1JBC0 B1JBC2 B1JBC3 B1JBD9 B1JBE0
B1JBE5 B1JBE6 B1JBE9 B1JBG6 B1JBH6 B1JBJ1 B1JBM5 B1JBM6 B1JBN1 B1JBN3 B1JBP2
B1JBP6 B1JBQ4 B1JBQ7 B1JBQ9 B1JBR0 B1JBR2 B1JBS2 B1JBT2 B1JBX3 B1JBZ0 B1JBZ1
B1JBZ2 B1JC06 B1JC20 B1JC95 B1JCG9 B1JCH0 B1JCH2 B1JCH3 B1JCI6 B1JCS1 B1JCV3
B1JCW0 B1JD29 B1JD39 B1JD61 B1JD70 B1JD71 B1JD72 B1JD74 B1JD79 B1JD82 B1JD88
B1JDA1 B1JDA2 B1JDB4 B1JDB7 B1JDE2 B1JDE3 B1JDE4 B1JDE5 B1JDF0 B1JDH6 B1JDI2
B1JDM3 B1JDN1 B1JDP0 B1JDP1 B1JDP2 B1JDQ5 B1JDQ6 B1JDR0 B1JDR3 B1JDS3 B1JDU1
B1JDV4 B1JDV7 B1JDV8 B1JDV9 B1JDW0 B1JDW1 B1JDW2 B1JDW3 B1JDW4 B1JDW5 B1JDW6
B1JDW8 B1JDW9 B1JDX0 B1JDX1 B1JDX2 B1JDX3 B1JDX4 B1JDX5 B1JDX6 B1JDX7 B1JDX9
B1JDY1 B1JDY5 B1JDY6 B1JDZ6 B1JDZ7 B1JDZ8 B1JDZ9 B1JE08 B1JE09 B1JE10 B1JE12
B1JE13 B1JE18 B1JE24 B1JE34 B1JE35 B1JE54 B1JE55 B1JE82 B1JED1 B1JED2 B1JED5
B1JED9 B1JEN1 B1JEN3 B1JEP3 B1JEP6 B1JEP7 B1JEQ0 B1JEQ2 B1JEQ3 B1JER5 B1JET1
B1JET2 B1JET4 B1JET8 B1JEV9 B1JEZ1 B1JF56 B1JF57 B1JF58 B1JF59 B1JF64 B1JF80
B1JF83 B1JF86 B1JFB0 B1JFD6 B1JFJ3 B1JFM7 B1JFN3 B1JFP4 B1JFV1 B1JFV2 B1JFV3
B1JFV9 B1JFW2 B1JFZ0 B1JFZ4 B1JG01 B1JG57 B1JG65 B1JG69 B1JG85 B1JG88 B1JG95
B1JG97 B1JGA5 B1JGA8 B1JGB0 B1JGB5 B1JGB6 B1JGB7 B1JGD5 B1JGG7 B1JGH2 B1JGI0
B1JGK4 B1JGK6 B1JGL0 B1JGN4 B1JGZ5 B1JGZ6 B1JH06 B1JH07 B1JH11 B1JH20 B1JH25
B1JH26 B1JH40 B1JH60 B1JH88 B1JH89 B1JH91 B1JH95 B1JHA1 B1JHA2 B1JHG7 B1JHH1
B1JHH4 B1JHI1 B1JHI9 B1JHJ1 B1JHN0 B1JHN1 B1JHN4 B1JHN6 B1JHP7 B1JHP8 B1JHR2
B1JHR4 B1JHS0 B1JHS2 B1JHU7 B1JHX8 B1JHX9 B1JHY0 B1JHY9 B1JHZ0 B1JHZ3 B1JHZ5
B1JHZ7 B1JI04 B1JI05 B1JI38 B1JI47 B1JI48 B1JI57 B1JI59 B1JI67 B1JI86 B1JI88
B1JI92 B1JIC7 B1JID2 B1JID5 B1JID6 B1JID8 B1JIE1 B1JIE2 B1JIE4 B1JIE9 B1JIF0
B1JIG5 B1JIG6 B1JIG7 B1JIH2 B1JIH3 B1JIH4 B1JIH5 B1JIJ3 B1JIK2 B1JIK3 B1JIK4
B1JIK5 B1JIK6 B1JIQ3 B1JIQ4 B1JIT3 B1JIU7 B1JIV0 B1JIV1 B1JIV2 B1JIV3 B1JIV4
B1JIV5 B1JIW0 B1JIW1 B1JIW2 B1JIW4 B1JIW5 B1JIW6 B1JIW7 B1JIW8 B1JIW9 B1JIX0
B1JIX1 B1JIX2 B1JIX3 B1JIX4 B1JIX5 B1JIX6 B1JIX7 B1JIX9 B1JIY0 B1JIY2 B1JIY3
B1JJ01 B1JJ17 B1JJ19 B1JJ20 B1JJ21 B1JJ23 B1JJ36 B1JJ37 B1JJ39 B1JJ48 B1JJ49
B1JJ59 B1JJ73 B1JJ75 B1JJ88 B1JJ89 B1JJ90 B1JJ91 B1JJ99 B1JJA0 B1JJA2 B1JJF1
B1JJF4 B1JJF5 B1JJF6 B1JJF7 B1JJF8 B1JJG3 B1JJG7 B1JJG8 B1JJG9 B1JJH1 B1JJH6
B1JJH7 B1JJH8 B1JJI0 B1JJI2 B1JJI3 B1JJI8 B1JJJ4 B1JJJ5 B1JJJ6 B1JJJ7 B1JJJ9
B1JJK0 B1JJK2 B1JJK7 B1JJK8 B1JJL0 B1JJL6 B1JJL9 B1JJM7 B1JJR1 B1JJU2 B1JK04
B1JK10 B1JK16 B1JK17 B1JK20 B1JK22 B1JK35 B1JK36 B1JK37 B1JK45 B1JK46 B1JK49
B1JK64 B1JK67 B1JK68 B1JK79 B1JK89 B1JK90 B1JK98 B1JKA0 B1JKA1 B1JKC0 B1JKF0
B1JKF2 B1JKF8 B1JKI7 B1JKJ7 B1JKJ8 B1JKK0 B1JKL9 B1JKM7 B1JKP2 B1JKQ0 B1JKS0
B1JKS2 B1JKS3 B1JKV7 B1JKY2 B1JKY3 B1JKY4 B1JKY5 B1JKZ4 B1JKZ5 B1JKZ8 B1JL00
B1JL03 B1JL04 B1JL09 B1JL10 B1JL13 B1JL20 B1JL21 B1JL34 B1JL35 B1JL36 B1JL43

B1JL46 B1JL60 B1JL66 B1JL67 B1JL79 B1JL80 B1JL96 B1JLA4 B1JLE7 B1JLG2 B1JLG9
B1JLH4 B1JLI4 B1JLI7 B1JLI8 B1JLJ0 B1JLL0 B1JLL1 B1JLL2 B1JLL3 B1JLL5 B1JLL8
B1JLL9 B1JLM0 B1JLM2 B1JLN0 B1JLP8 B1JLQ4 B1JLQ5 B1JLS9 B1JLV4 B1JLW2 B1JLX6
B1JLX7 B1JLX9 B1JLY0 B1JLY2 B1JM17 B1JM18 B1JM22 B1JM26 B1JM30 B1JM85 B1JM88
B1JMC6 B1JMC9 B1JMD0 B1JMD1 B1JMD2 B1JMH4 B1JMH7 B1JMI3 B1JMJ6 B1JMJ7 B1JMK1
B1JMK2 B1JMK6 B1JML1 B1JML7 B1JMM1 B1JMM2 B1JMM4 B1JMN3 B1JMN9 B1JMP0 B1JMP1
B1JMP8 B1JMQ1 B1JMQ7 B1JMR1 B1JMR2 B1JMR6 B1JNA9 B1JNC4 B1JNC5 B1JNC9 B1JND0
B1JND2 B1JNE7 B1JNM8 B1JNN7 B1JNP5 B1JNP6 B1JNQ2 B1JNR1 B1JNR6 B1JNS1 B1JNS6
B1JNS7 B1JNS8 B1JNT4 B1JP35 B1JP38 B1JP39 B1JP41 B1JP62 B1JP69 B1JP75 B1JPB6
B1JPE1 B1JPE3 B1JPE6 B1JPH6 B1JPS4 B1JPT9 B1JPU8 B1JPV9 B1JPW1 B1JPW4 B1JPW5
B1JPY5 B1JPY6 B1JPY8 B1JPZ5 B1JPZ6 B1JQ11 B1JQ12 B1JQ26 B1JQ32 B1JQ39 B1JQ44
B1JQ46 B1JQ50 B1JQ59 B1JQ60 B1JQ63 B1JQ68 B1JQ70 B1JQ75 B1JQ76 B1JQ78 B1JQ83
B1JQ90 B1JQC6 B1JQC7 B1JQD9 B1JQE8 B1JQE9 B1JQF1 B1JQF6 B1JQF7 B1JQG0 B1JQG1
B1JQG3 B1JQG4 B1JQH4 B1JQI1 B1JQI5 B1JQJ4 B1JQP6 B1JQP7 B1JQQ0 B1JQQ5 B1JQQ8
B1JQR7 B1JQT1 B1JQT7 B1JQU4 B1JQV6 B1JQW2 B1JQW4 B1JQW9 B1JQX0 B1JQX1 B1JQX2
B1JQX3 B1JQX6 B1JQX7 B1JQX8 B1JQZ3 B1JQZ5 B1JR03 B1JR11 B1JR12 B1JR17 B1JR25
B1JR27 B1JR28 B1JR29 B1JR32 B1JR33 B1JR44 B1JR46 B1JR56 B1JR69 B1JR71 B1JR72
B1JR73 B1JR75 B1JR86 B1JRA7 B1JRB0 B1JRB1 B1JRB2 B1JRB8 B1JRC9 B1JRD6 B1JRD8
B1JRD9 B1JRE0 B1JRF0 B1JRF9 B1JRG2 B1JRH0 B1JRJ4 B1JRK6 B1JRM5 B1JRQ1 B1JRQ5
B1JRW1 B1JRX7 B1JRZ0 B1JRZ1 B1JRZ6 B1JS01 B1JS02 B1JS05 B1JS06 B1JS27 B1JS28
B1JS45 B1JS46 B1JS47 B1JS96 B1JSA4 B1JSA8 B1JSB0 B1JSF4 B1JSF5 B1JSF7 B1JSG7
B1JSH5 B1JSJ3 B1JSJ5 B1JSJ6 B1JSQ9 B1JSR0 B1JSR4 B1JSR9 B1JSS3 B1JSS4 B1JSS6
B1JST8 B1JSU1 B1JSU6 B1JSZ2 B1JSZ3 B1JSZ4 B1JT65 B1JT82 B1JT87 B1JT88 B1JT91
B1JT94 B1JT95 B1JTA2 B1JTA5 B1JTA6 B1JTB3 B1JTF0 B1JTF1 B1JTF3 B1JTI2 B1JTS0
B1JTT5 B1JTT6 B1JTU4 B1JTU9 B1JTV1 B1JTX0 B1JU10 B1JU11 B1JU12 B1JU13 B1JU14
B1JU15 B1JU17 B1JU18 B1JU21 B1JU22 B1JU23 B1JU24 B1JU25 B1JU26 B1JU27 B1JU28
B1JU29 B1JU30 B1JU31 B1JU32 B1JU33 B1JU34 B1JU35 B1JU36 B1JU37 B1JU38 B1JU39
B1JU40 B1JU41 B1JU44 B1JU45 B1JU46 B1JU47 B1JU48 B1JU49 B1JU53 B1JU60 B1JU67
B1JU98 B1JUA1 B1JUA4 B1JUA5 B1JUA6 B1JUA7 B1JUC0 B1JUD4 B1JUD6 B1JUE4 B1JUE7
B1JUE9 B1JUF0 B1JUF2 B1JUF5 B1JUG0 B1JUG2 B1JUI6 B1JUK8 B1JUS4 B1JUS5 B1JUU5
B1JUU6 B1JUW3 B1JUW4 B1JUW8 B1JUY8 B1JUZ7 B1JV00 B1JV19 B1JV36 B1JV46 B1JV50
B1JV76 B1JV79 B1JV80 B1JV92 B1JV99 B1JVA0 B1JVA1 B1JVA2 B1JVA4 B1JVA5 B1JVC0
B1JVG6 B1JVG9 B1JVH2 B1JVL5 B1JVP5 B1JVP6 B1JVQ1 B1JVQ4 B1JVU4 B1JVU5 B1JVU6
B1JVU7 B1JVU9 B1JVV3 B1JVV4 B1JVV5 B1JVV6 B1JVW3 B1JVY9 B1JVZ2 B1JW12 B1JW13
B1JW14 B1JW17 B1JW19 B1JW20 B1JW28 B1JW31 B1JW38 B1JW40 B1JW41 B1JW43 B1JW46
B1JW61 B1JW62 B1JW82 B1JWI0 B1JWJ3 B1JWN9 B1JWP0 B1JWP2 B1JWR0 B1JWR5 B1JWR9
B1JWS3 B1JWS8 B1JWS9 B1JWV2 B1JX26 B1JX27 B1JX29 B1JX30 B1JX31 B1JX75 B1JX76
B1JX81 B1JX98 B1JXA1 B1JXA3 B1JXB2 B1JXB3 B1JXB4 B1JXB7 B1JXC3 B1JXC6 B1JXD0
B1JXF6 B1JXF8 B1JXG1 B1JXG4 B1JXG5 B1JXH8 B1JXI7 B1JXI9 B1JXM2 B1JXM5 B1JXN8
B1JXR2 B1JXR6 B1JXR9 B1JXS0 B1JXS2 B1JXS7 B1JXT8 B1JXU3 B1JXU4 B1JXU5 B1JXU6
B1JXV4 B1JY03 B1JY14 B1JY43 B1JY57 B1JY71 B1JY80 B1JY88 B1JY98 B1JYC0 B1JYC2
B1JYI9 B1JYJ6 B1JYK1 B1JYM2 B1JYN1 B1JYP2 B1JYP8 B1JYQ0 B1JYQ1 B1JYQ4 B1JYT5
B1JYT6 B1JYT7 B1JYX0 B1JZ58 B1JZ61 B1JZ91 B1JZB2 B1JZC6 B1JZD3 B1JZD9 B1JZE0
B1JZE1 B1JZE9 B1JZF2 B1JZG0 B1JZU7 B1JZZ7 B1K026 B1K045 B1K077 B1K091 B1K093

B1K094 B1K0B1 B1K0F2 B1K0G1 B1K0G2 B1K0G5 B1K0G8 B1K0H6 B1K0H7 B1K0H8 B1K0I8
B1K0J5 B1K0L8 B1K0M0 B1K0M1 B1K0R9 B1K0S5 B1K0S6 B1K0S8 B1K0Y2 B1K0Y7 B1K0Y8
B1K153 B1K168 B1K1H7 B1K1H9 B1K1I7 B1K1K2 B1K259 B1K264 B1K269 B1K2U6 B1K2X7
B1K366 B1K370 B1K375 B1K377 B1K394 B1K3J6 B1K3S7 B1K3S8 B1K3S9 B1K3T1 B1K3U1
B1K3Y3 B1K4Q5 B1K568 B1K5H3 B1K5N3 B1K5S8 B1K5T6 B1K6C3 B1K6I9 B1K707 B1K708
B1K709 B1K759 B1K774 B1K7A6 B1K943 B1KAV4 B1KAV6 B1KB47 B1KBY1 B1KC83 B1KCV0
B1KCW3 B1KCW7 B1KCX0 B1KCX1 B1KCX3 B1KCX5 B1KCY4 B1KCY7 B1KCZ4 B1KD01 B1KD27
B1KD28 B1KD33 B1KD54 B1KD73 B1KDF2 B1KDK2 B1KDK4 B1KDM9 B1KDN0 B1KDP2 B1KDS3
B1KDU3 B1KDU4 B1KDU6 B1KDV1 B1KDV4 B1KDV6 B1KDW0 B1KDW3 B1KDW5 B1KDX1 B1KDX2
B1KDX3 B1KE86 B1KEG7 B1KEH1 B1KEH7 B1KEI0 B1KEK5 B1KEL4 B1KEP1 B1KEP6 B1KEP7
B1KF18 B1KF21 B1KF36 B1KF45 B1KF47 B1KF48 B1KF50 B1KFN0 B1KFR6 B1KFR7 B1KFS0
B1KFT2 B1KFU3 B1KFY6 B1KG49 B1KG58 B1KG59 B1KG60 B1KG61 B1KG87 B1KG88 B1KG89
B1KGG6 B1KGG7 B1KGG9 B1KGH0 B1KGH1 B1KGH8 B1KGH9 B1KGX9 B1KGY0 B1KHA5 B1KHA6
B1KHE1 B1KHE2 B1KHF1 B1KHF6 B1KHG5 B1KHG7 B1KHG9 B1KHH0 B1KHI0 B1KHI4 B1KHK0
B1KHR8 B1KHS7 B1KHT1 B1KHU8 B1KHU9 B1KHV0 B1KHY4 B1KHY7 B1KHY8 B1KHY9 B1KHZ0
B1KI25 B1KI32 B1KI41 B1KI42 B1KI57 B1KI68 B1KI69 B1KI70 B1KI71 B1KI74 B1KI99
B1KIC1 B1KIC5 B1KID0 B1KIE1 B1KIG8 B1KIG9 B1KIH9 B1KII1 B1KII7 B1KIK3 B1KIR6
B1KIR7 B1KIS3 B1KIS4 B1KIS8 B1KIT2 B1KIT5 B1KIW2 B1KIW4 B1KIX3 B1KIY1 B1KJ42
B1KJ47 B1KJ76 B1KJA5 B1KJC0 B1KJJ9 B1KJK0 B1KJK4 B1KJK5 B1KJK9 B1KJL0 B1KJM2
B1KJM4 B1KJM7 B1KJR3 B1KJT5 B1KJX4 B1KK02 B1KK45 B1KK48 B1KK50 B1KK51 B1KKG8
B1KKI9 B1KKJ2 B1KKJ3 B1KKN8 B1KKP1 B1KKR3 B1KKS1 B1KKX6 B1KKY5 B1KKY6 B1KKY8
B1KKZ1 B1KKZ2 B1KKZ4 B1KL03 B1KL04 B1KL05 B1KL07 B1KL08 B1KL12 B1KL24 B1KL36
B1KL38 B1KL44 B1KLA4 B1KLB1 B1KLB9 B1KLC1 B1KLM0 B1KLT6 B1KLT7 B1KLT8 B1KLT9
B1KLU0 B1KLV2 B1KLY5 B1KM34 B1KM36 B1KM44 B1KM47 B1KM97 B1KMH3 B1KMH4 B1KMJ3
B1KMK3 B1KML5 B1KMU6 B1KMW0 B1KMW1 B1KMW2 B1KMW4 B1KMW5 B1KMW7 B1KMW8 B1KMW9
B1KMX0 B1KMX1 B1KMX2 B1KMX3 B1KMX4 B1KMX5 B1KMX6 B1KMX8 B1KMX9 B1KMY0 B1KMY1
B1KMY2 B1KMY3 B1KMY4 B1KMY7 B1KMY8 B1KMY9 B1KMZ0 B1KMZ1 B1KMZ2 B1KMZ3 B1KMZ4
B1KMZ8 B1KN02 B1KN03 B1KN07 B1KN30 B1KN33 B1KN91 B1KNA0 B1KNA4 B1KNC9 B1KND3
B1KND6 B1KND9 B1KNG3 B1KNI3 B1KNI6 B1KNI7 B1KNJ0 B1KNJ9 B1KNK1 B1KNK2 B1KNK5
B1KNK9 B1KNL5 B1KNM8 B1KNR3 B1KNS7 B1KNS9 B1KNT7 B1KNU0 B1KNU2 B1KNU3 B1KNY2
B1KP09 B1KP42 B1KP55 B1KP56 B1KP58 B1KP64 B1KP71 B1KP72 B1KP81 B1KP82 B1KPD1
B1KPH0 B1KPJ7 B1KPS4 B1KPS5 B1KPS6 B1KPS7 B1KPT0 B1KPT1 B1KPT3 B1KPT7 B1KPU0
B1KPZ1 B1KQ11 B1KQ25 B1KQ44 B1KQ45 B1KQ64 B1KQ67 B1KQ68 B1KQ88 B1KQC6 B1KQD0
B1KQE6 B1KQE8 B1KQF6 B1KQL8 B1KQM2 B1KQN1 B1KQX1 B1KQX8 B1KQY3 B1KQY8 B1KQY9
B1KQZ0 B1KR07 B1KR25 B1KRB5 B1KRC5 B1KRD2 B1KRD4 B1KRD7 B1KRE3 B1KRE5 B1KRF9
B1KRG4 B1KRG9 B1KRI9 B1KRJ1 B1KRP6 B1KRP7 B1KRP8 B1KRQ3 B1KRQ5 B1KRQ7 B1KRQ8
B1KRQ9 B1KRR0 B1KRR2 B1KRR4 B1KRR8 B1KRS8 B1KRT1 B1KRT2 B1KRT6 B1KRT7 B1KRU0
B1KRX5 B1KRY8 B1KS25 B1KSA0 B1KSA1 B1KSA2 B1KSA3 B1KSA4 B1KSB1 B1KSB9 B1KSG4
B1KSI9 B1KSJ0 B1KSJ5 B1KSJ6 B1KSJ8 B1KSJ9 B1KSK0 B1KSK4 B1KSK6 B1KSK7 B1KSK9
B1KSL0 B1KSL1 B1KSL2 B1KSL3 B1KSL4 B1KSL5 B1KSL6 B1KSL7 B1KSL8 B1KSL9 B1KSM1
B1KSM2 B1KSM3 B1KSM4 B1KSM5 B1KSM6 B1KSM7 B1KSM8 B1KSM9 B1KSN0 B1KSN2 B1KSN3
B1KSN4 B1KSN6 B1KSP3 B1KSQ7 B1KSQ8 B1KSR1 B1KSR7 B1KST9 B1KSX3 B1KSY5 B1KT07
B1KT22 B1KT46 B1KT47 B1KT52 B1KT53 B1KT54 B1KT65 B1KT66 B1KT67 B1KT95 B1KT98
B1KTA5 B1KTA8 B1KTB9 B1KTC1 B1KTC2 B1KTC5 B1KTC6 B1KTE2 B1KTE8 B1KTF0 B1KTF5

B1KTG4 B1KTH1 B1KTJ5 B1KTJ6 B1KTK1 B1KTK2 B1KTS4 B1KU02 B1KU20 B1KU80 B1KU85
B1KU91 B1KU95 B1KU97 B1KUB0 B1KUB1 B1KUB2 B1KUB6 B1KUB7 B1KUD0 B1KUN4 B1KUQ8
B1KUY5 B1KUY6 B1KUY7 B1KUY8 B1KUZ8 B1KV00 B1KV12 B1KV13 B1KV16 B1KV69 B1KVJ4
B1KVV5 B1KW01 B1KW07 B1KW08 B1KWD5 B1KWD6 B1KWJ5 B1KWK1 B1KWK2 B1KWK6 B1KWK7
B1KWL1 B1KWL5 B1KWM1 B1KWM4 B1KWM5 B1KWM6 B1KWN0 B1KWN1 B1KWN4 B1KWN5 B1KWN8
B1KWP9 B1KX40 B1KX43 B1KX48 B1KX53 B1KX56 B1KX63 B1KX65 B1KX71 B1KX81 B1KX91
B1KXA7 B1KXB4 B1KXB5 B1KXB6 B1KXB7 B1KXQ3 B1KXQ5 B1KXR9 B1KXY0 B1KXY1 B1KY08
B1KY15 B1KY30 B1KY93 B1KYF3 B1KYF4 B1KYG7 B1KYK0 B1KYK8 B1KYP3 B1KYT1 B1KYV1
B1KYX4 B1KYX5 B1KYY3 B1KZ07 B1KZ37 B1KZ46 B1KZ55 B1KZ57 B1KZ59 B1KZC8 B1KZE1
B1KZF5 B1KZJ2 B1KZJ7 B1KZM6 B1KZN1 B1KZN5 B1KZN7 B1KZP5 B1KZQ3 B1KZR1 B1KZR3
B1KZR4 B1KZR6 B1KZS3 B1KZT0 B1KZT1 B1KZT7 B1KZY0 B1KZY3 B1L010 B1L096 B1L097
B1L0A0 B1L0A2 B1L0B0 B1L0B1 B1L0B2 B1L0B3 B1L0B5 B1L0G9 B1L0J7 B1L0S3 B1L0T0
B1L0T4 B1L0T5 B1L0T7 B1L0U2 B1L0V4 B1L164 B1L1D6 B1L1D7 B1L1D8 B1L1E1 B1L1E4
B1L1E6 B1L1E7 B1L1G8 B1L1G9 B1L1H0 B1L1H3 B1L1I1 B1L1J7 B1L1K0 B1L1K1 B1L1K6
B1L1K9 B1L1M0 B1L1N3 B1L1S1 B1L1S2 B1L1S5 B1L1Y6 B1L1Z3 B1L203 B1L224 B1L242
B1L266 B1L268 B1L270 B1L271 B1L273 B1L280 B1L384 B1L385 B1L3E4 B1L3K8 B1L3L5
B1L3V2 B1L3W1 B1L3X7 B1L400 B1L402 B1L448 B1L4Z3 B1L550 B1L5G0 B1L5G9 B1L5H2
B1L5K9 B1L5N1 B1L5N5 B1L5R5 B1L5U2 B1L5U5 B1L5W7 B1L5W9 B1L640 B1L657 B1L659
B1L6B4 B1L6E9 B1L6L2 B1L6L3 B1L6M8 B1L6M9 B1L6N8 B1L6R7 B1L6R9 B1L6S6 B1L6S7
B1L6T7 B1L6T9 B1L6U5 B1L6X8 B1L707 B1L708 B1L712 B1L719 B1L726 B1L735 B1L762
B1L765 B1L776 B1L777 B1L778 B1L781 B1L784 B1L785 B1L787 B1L793 B1L795 B1L796
B1L7A5 B1L7A7 B1L7A8 B1L7C0 B1L7D5 B1L7I3 B1L7I8 B1L7J8 B1L7J9 B1L7Q0 B1L7Q7
B1L7R6 B1L7S2 B1L7T1 B1L7U3 B1L7W5 B1L7W6 B1L7X2 B1L7X6 B1L7Y5 B1L7Y6 B1L7Y7
B1L800 B1L804 B1L812 B1L813 B1L821 B1L826 B1L829 B1L841 B1L843 B1L862 B1L867
B1L869 B1L870 B1L872 B1L873 B1L8B8 B1L8F1 B1L8F3 B1L8K5 B1L8K7 B1L8M5 B1L8P0
B1L8R5 B1L8R9 B1L8S0 B1L8S9 B1L8T8 B1L8U2 B1L8U4 B1L8U5 B1L8W7 B1L8W9 B1L8X0
B1L8X3 B1L8X4 B1L8X5 B1L8Y1 B1L8Y8 B1L8Y9 B1L920 B1L921 B1L927 B1L934 B1L935
B1L936 B1L937 B1L938 B1L939 B1L942 B1L944 B1L989 B1L993 B1L9B4 B1L9B6 B1L9C6
B1L9E1 B1L9E3 B1L9E4 B1L9E5 B1L9F1 B1L9F7 B1L9J1 B1L9J9 B1L9M7 B1L9M9 B1L9N6
B1L9N8 B1L9P0 B1L9P4 B1L9P8 B1L9P9 B1L9Q0 B1L9Q1 B1L9Q2 B1L9R3 B1L9R9 B1L9S2
B1L9T5 B1L9T6 B1L9T8 B1L9T9 B1L9U1 B1L9U3 B1L9W8 B1LA07 B1LA13 B1LA15 B1LA43
B1LA51 B1LA53 B1LA54 B1LA64 B1LA70 B1LA82 B1LA87 B1LAA9 B1LAB2 B1LAE7 B1LAF5
B1LAF7 B1LAG0 B1LAG3 B1LAG4 B1LAJ8 B1LAK2 B1LAK3 B1LAP0 B1LAP2 B1LAP4 B1LAP5
B1LAP9 B1LAQ3 B1LAQ5 B1LAQ6 B1LAQ7 B1LAQ8 B1LAR0 B1LAR2 B1LAU0 B1LAU3 B1LAU5
B1LAW3 B1LAW7 B1LAW8 B1LAX2 B1LAX5 B1LAX6 B1LAZ1 B1LAZ3 B1LAZ7 B1LB08 B1LB13
B1LB14 B1LB17 B1LB18 B1LB31 B1LB32 B1LB65 B1LB68 B1LB70 B1LB75 B1LB78 B1LB80
B1LB83 B1LB88 B1LBA0 B1LBA1 B1LBB4 B1LBD6 B1LBD7 B1LBH5 B1LBH9 B1LBI1 B1LBI2
B1LBI4 B1LBJ1 B1LBJ2 B1LBJ4 B1LBJ6 B1LBK1 B1LBK2 B1LBK3 B1LBK5 B1LBL1 B1LBL3
B1LBL4 B1LBL5 B1LBL6 B1LBL9 B1LBM1 B1LBM2 B1LBM3 B1LBM4 B1LBM5 B1LBM6 B1LBM7
B1LBM8 B1LBM9 B1LBN0 B1LBN1 B1LBN2 B1LBN3 B1LBN4 B1LBN5 B1LBN6 B1LBN7 B1LBN8
B1LBN9 B1LBP0 B1LBP1 B1LBP2 B1LBP3 B1LBP4 B1LBP5 B1LBP9 B1LBQ9 B1LBR0 B1LBR1
B1LBS6 B1LBS9 B1LBT1 B1LBT2 B1LBU1 B1LBU2 B1LBU6 B1LBX4 B1LBY0 B1LBY1 B1LC08
B1LC19 B1LC23 B1LC39 B1LC43 B1LC63 B1LC69 B1LC74 B1LCA5 B1LCE8 B1LCF0 B1LCF9
B1LCG8 B1LCH6 B1LCI0 B1LCI1 B1LCI2 B1LCI7 B1LCK9 B1LCN9 B1LCP2 B1LCQ5 B1LCS9

B1LCY9 B1LCZ5 B1LCZ7 B1LD00 B1LD01 B1LD05 B1LD08 B1LD10 B1LD11 B1LD37 B1LD41
B1LD60 B1LD62 B1LD92 B1LD99 B1LDA1 B1LDA3 B1LDA4 B1LDA5 B1LDB5 B1LDC0 B1LDD2
B1LDE6 B1LDE8 B1LDF1 B1LDF3 B1LDF7 B1LDF8 B1LDG2 B1LDH2 B1LDQ8 B1LDR4 B1LDT4
B1LDV3 B1LDY3 B1LDY4 B1LDY5 B1LDY6 B1LDY7 B1LDY9 B1LDZ1 B1LDZ8 B1LE00 B1LE12
B1LE14 B1LE15 B1LE16 B1LE18 B1LE24 B1LE27 B1LE43 B1LE44 B1LE56 B1LE57 B1LEG6
B1LEH4 B1LEH8 B1LEI6 B1LEI7 B1LEI8 B1LEI9 B1LEJ1 B1LEK0 B1LEK2 B1LEM6 B1LEP4
B1LEP6 B1LEP8 B1LEP9 B1LER3 B1LET0 B1LF30 B1LF38 B1LF40 B1LF43 B1LF48 B1LF56
B1LF57 B1LF61 B1LF65 B1LF94 B1LF96 B1LF97 B1LF98 B1LFA3 B1LFA4 B1LFD8 B1LFF6
B1LFH3 B1LFI2 B1LFJ0 B1LFL2 B1LFM7 B1LFN5 B1LFP2 B1LFP9 B1LFQ0 B1LFQ5 B1LFQ6
B1LFR6 B1LFR7 B1LFR9 B1LFS0 B1LFS2 B1LFS3 B1LFS6 B1LFS9 B1LFT3 B1LFT4 B1LFT7
B1LFU5 B1LFV4 B1LFV6 B1LFV9 B1LFW0 B1LFW1 B1LFW8 B1LFW9 B1LFX0 B1LFX1 B1LFX2
B1LFX5 B1LFX6 B1LFY5 B1LFY6 B1LFY7 B1LFY8 B1LFZ3 B1LFZ6 B1LFZ7 B1LG08 B1LG09
B1LG11 B1LG18 B1LG19 B1LG28 B1LG37 B1LG38 B1LG40 B1LG84 B1LGE7 B1LGE9 B1LGF1
B1LGF2 B1LGF5 B1LGH1 B1LGI7 B1LGI8 B1LGJ0 B1LGJ6 B1LGJ7 B1LGK2 B1LGK3 B1LGM4
B1LGM6 B1LGN8 B1LGN9 B1LGP1 B1LGP3 B1LGP4 B1LGP5 B1LGQ0 B1LGS0 B1LGS1 B1LGS2
B1LGT2 B1LGT5 B1LGT6 B1LGU8 B1LGV7 B1LGV9 B1LGW2 B1LGW3 B1LGW6 B1LGW7 B1LGW8
B1LGX0 B1LGX2 B1LGX4 B1LGY5 B1LGZ2 B1LGZ7 B1LH07 B1LH12 B1LH31 B1LH32 B1LH52
B1LH89 B1LH90 B1LH92 B1LH96 B1LHB0 B1LHB1 B1LHB3 B1LHB5 B1LHB6 B1LHB8 B1LHB9
B1LHC0 B1LHC1 B1LHC2 B1LHC3 B1LHC4 B1LHC5 B1LHC6 B1LHC7 B1LHC8 B1LHC9 B1LHD0
B1LHD1 B1LHD3 B1LHD4 B1LHD5 B1LHE0 B1LHE1 B1LHE2 B1LHE3 B1LHE4 B1LHE5 B1LHE8
B1LHF5 B1LHG9 B1LHI4 B1LHJ8 B1LHK1 B1LHL0 B1LHL2 B1LHL4 B1LHL8 B1LHM1 B1LHM3
B1LHM8 B1LHT5 B1LHT6 B1LHT8 B1LHT9 B1LHW0 B1LHW3 B1LHW4 B1LHY6 B1LHY8 B1LHZ1
B1LHZ3 B1LHZ6 B1LI03 B1LI09 B1LI10 B1LI25 B1LI36 B1LI37 B1LI38 B1LI47 B1LI56
B1LI79 B1LI80 B1LI85 B1LI90 B1LI91 B1LI92 B1LID3 B1LIE4 B1LIG5 B1LIJ7 B1LIJ8
B1LIJ9 B1LIM5 B1LIM9 B1LIN2 B1LIN3 B1LIN5 B1LIN6 B1LIN7 B1LIP0 B1LIP1 B1LIP2
B1LIR6 B1LIS1 B1LIS2 B1LIS5 B1LIS7 B1LIU3 B1LIU8 B1LIV1 B1LIV6 B1LIY0 B1LIZ3
B1LIZ4 B1LIZ5 B1LIZ6 B1LIZ7 B1LIZ8 B1LJ00 B1LJ04 B1LJ05 B1LJ06 B1LJ30 B1LJ31
B1LJ35 B1LJ40 B1LJ43 B1LJ63 B1LJC0 B1LJC1 B1LJC7 B1LJD7 B1LJF3 B1LJF4 B1LJG3
B1LJG5 B1LJG6 B1LJH0 B1LJH2 B1LJH3 B1LJH6 B1LJJ3 B1LJJ5 B1LJK1 B1LJK3 B1LJM6
B1LJM8 B1LJM9 B1LJN2 B1LJN3 B1LJN4 B1LJR4 B1LJS7 B1LJT2 B1LJT9 B1LJU1 B1LJU6
B1LJV1 B1LJV3 B1LJV6 B1LJV7 B1LJX0 B1LJX9 B1LK21 B1LK35 B1LK48 B1LK55 B1LK58
B1LK71 B1LK72 B1LK73 B1LK74 B1LK75 B1LK77 B1LK78 B1LK79 B1LK80 B1LK84 B1LK86
B1LKC2 B1LKC4 B1LKD0 B1LKE5 B1LKE8 B1LKH2 B1LKK5 B1LKK6 B1LKK9 B1LKL9 B1LKM1
B1LKM2 B1LKM9 B1LKP0 B1LKQ0 B1LKQ1 B1LKQ6 B1LKQ9 B1LKS0 B1LKT4 B1LKT5 B1LKT7
B1LKV7 B1LKZ0 B1LL11 B1LL12 B1LL17 B1LL25 B1LL27 B1LL30 B1LL31 B1LL33 B1LL35
B1LL67 B1LL68 B1LL71 B1LL72 B1LL73 B1LL75 B1LL85 B1LL88 B1LL91 B1LLA1 B1LLB0
B1LLB3 B1LLC0 B1LLC4 B1LLF0 B1LLF1 B1LLG1 B1LLI3 B1LLJ9 B1LLK1 B1LLK3 B1LLK5
B1LLL6 B1LLL8 B1LLL9 B1LLP5 B1LLP8 B1LLS3 B1LLS6 B1LLT0 B1LLT5 B1LLT7 B1LLU0
B1LLU6 B1LLU9 B1LLV2 B1LLV3 B1LLX7 B1LLY0 B1LLY2 B1LM00 B1LM21 B1LM33 B1LM42
B1LM46 B1LM48 B1LM58 B1LM66 B1LM67 B1LM70 B1LM72 B1LM74 B1LM96 B1LM99 B1LMA4
B1LMD0 B1LMH0 B1LMI6 B1LMJ5 B1LMK5 B1LML3 B1LMM1 B1LMM2 B1LMN0 B1LMN4 B1LMP7
B1LMP8 B1LMS1 B1LMS6 B1LMU1 B1LMU2 B1LMU3 B1LMU4 B1LMU6 B1LN04 B1LN12 B1LN20
B1LN27 B1LN54 B1LN84 B1LN92 B1LN93 B1LNB6 B1LNC1 B1LNC6 B1LNC8 B1LNE6 B1LNE8
B1LNF0 B1LNF9 B1LNG1 B1LNG6 B1LNG9 B1LNH0 B1LNH2 B1LNH3 B1LNH8 B1LNI1 B1LNI2

B1LNI4 B1LNI6 B1LNI7 B1LNJ5 B1LNJ7 B1LNJ8 B1LNK7 B1LNL9 B1LNM2 B1LNM9 B1LNN1
B1LNN4 B1LNN6 B1LNN7 B1LNP1 B1LNP3 B1LNQ3 B1LNR6 B1LNR7 B1LNS2 B1LNS5 B1LNT0
B1LNT5 B1LNT6 B1LNT7 B1LNT8 B1LNT9 B1LNU0 B1LNU6 B1LNU7 B1LNU8 B1LNU9 B1LNV0
B1LNX6 B1LP16 B1LP17 B1LP20 B1LP22 B1LP36 B1LP78 B1LP80 B1LP88 B1LP92 B1LP93
B1LPB3 B1LPB4 B1LPB5 B1LPB6 B1LPC1 B1LPC2 B1LPC5 B1LPC7 B1LPD2 B1LPE9 B1LPG3
B1LPG6 B1LPH0 B1LPH3 B1LPH7 B1LPI9 B1LPK4 B1LPR3 B1LPS5 B1LPV7 B1LPW8 B1LQ13
B1LQ15 B1LQ16 B1LQ17 B1LQ28 B1LQ29 B1LQ55 B1LQ63 B1LQ65 B1LQ66 B1LQ67 B1LQ71
B1LQ72 B1LQ73 B1LQ85 B1LQ86 B1LQB3 B1LQF8 B1LQG3 B1LQG4 B1LQG8 B1LQH8 B1LQI4
B1LQJ0 B1LQJ1 B1LQJ2 B1LQJ7 B1LQJ8 B1LQL1 B1LQL2 B1LQL6 B1LQL7 B1LQL8 B1LQM0
B1LQM1 B1LQP2 B1LQQ4 B1LQQ6 B1LQT7 B1LQU4 B1LQW0 B1LQW3 B1LQY7 B1LQZ6 B1LQZ8
B1LR00 B1LR27 B1LR29 B1LR41 B1LR71 B1LR73 B1LR79 B1LR94 B1LRA3 B1LRB5 B1LRB8
B1LRD1 B1LRD2 B1LRE5 B1LS97 B1LSA1 B1LSF9 B1LSG1 B1LSI6 B1LSM6 B1LSP9 B1LSQ8
B1LTD9 B1LTE0 B1LTE4 B1LTL1 B1LTQ4 B1LTQ7 B1LTQ8 B1LU98 B1LUA2 B1LUA9 B1LUB2
B1LUB9 B1LUE7 B1LUE9 B1LUJ9 B1LUL8 B1LUP6 B1LUP7 B1LUQ0 B1LUQ4 B1LUS0 B1LUU4
B1LUU9 B1LUW6 B1LV15 B1LV19 B1LV66 B1LV84 B1LV85 B1LVA0 B1LVA1 B1LVB8 B1LVC3
B1LVR4 B1LVU1 B1LVU2 B1LW27 B1LW28 B1LW29 B1LW44 B1LWE8 B1LWL6 B1LWN4 B1LWN6
B1LWP8 B1LWP9 B1LWQ1 B1LWQ2 B1LWQ5 B1LWQ7 B1LWQ8 B1LWQ9 B1LWR0 B1LWR1 B1LWR3
B1LWR5 B1LWR6 B1LWS1 B1LWS2 B1LWS3 B1LWS5 B1LWS6 B1LWS7 B1LWS8 B1LWS9 B1LWT0
B1LWT1 B1LWT3 B1LWT5 B1LWT6 B1LWY6 B1LWZ6 B1LX00 B1LXE5 B1LXF3 B1LXF6 B1LXK9
B1LXR9 B1LXV0 B1LXZ1 B1LXZ3 B1LY35 B1LY42 B1LY43 B1LY44 B1LY45 B1LY47 B1LY48
B1LY53 B1LYB4 B1LYI1 B1LYI3 B1LYI4 B1LYJ9 B1LYP9 B1LYT9 B1LZ51 B1LZ52 B1LZ78
B1LZ88 B1LZ89 B1LZ99 B1LZB2 B1LZG0 B1LZN1 B1LZP2 B1LZQ1 B1LZQ2 B1LZR8 B1LZS5
B1LZY1 B1M0D1 B1M0E0 B1M0I6 B1M0M1 B1M0N0 B1M0N9 B1M0Q0 B1M186 B1M187 B1M1C0
B1M1K0 B1M1U6 B1M1V6 B1M1W2 B1M1W4 B1M200 B1M201 B1M272 B1M2B1 B1M325 B1M335
B1M3B2 B1M3E0 B1M3G3 B1M3X3 B1M3X4 B1M445 B1M4N1 B1M4X4 B1M4Z8 B1M509 B1M537
B1M565 B1M575 B1M584 B1M597 B1M5H7 B1M5I5 B1M5I6 B1M5N6 B1M5Q2 B1M5Q9 B1M692
B1M695 B1M697 B1M699 B1M6A0 B1M6A3 B1M6A7 B1M6B8 B1M6B9 B1M6H4 B1M6N5 B1M6N6
B1M6N9 B1M6P0 B1M6P3 B1M6P4 B1M6P5 B1M6P9 B1M760 B1M766 B1M787 B1M7L0 B1M7M3
B1M860 B1M869 B1M9D6 B1M9D7 B1MAH2 B1MAI2 B1MAI3 B1MAJ3 B1MAS5 B1MAY2 B1MAY3
B1MAZ6 B1MB03 B1MB62 B1MB69 B1MB83 B1MB85 B1MB87 B1MBD0 B1MBV2 B1MBV4 B1MBX3
B1MBX5 B1MBZ3 B1MBZ5 B1MC29 B1MC85 B1MC86 B1MC89 B1MC91 B1MC92 B1MCA3 B1MCA4
B1MCB9 B1MCD9 B1MCE2 B1MCE5 B1MCE8 B1MCF1 B1MCF8 B1MCH3 B1MCI5 B1MCJ1 B1MCJ6
B1MCJ8 B1MCK6 B1MCU7 B1MCZ3 B1MD00 B1MD01 B1MD05 B1MD16 B1MD17 B1MD42 B1MD62
B1MD64 B1MD72 B1MD86 B1MD87 B1MD92 B1MD96 B1MDE3 B1MDF1 B1MDF2 B1MDH6 B1MDH9
B1MDI1 B1MDI4 B1MDI6 B1MDL2 B1MDL6 B1MDP0 B1MDP6 B1MDQ0 B1MDQ1 B1MDQ5 B1MDU8
B1MDU9 B1MDV2 B1MDW3 B1ME21 B1ME26 B1ME65 B1ME90 B1MEG8 B1MEI9 B1MEP9 B1MES1
B1MEU3 B1MEU8 B1MFC0 B1MFL8 B1MFN1 B1MFN2 B1MFN5 B1MFT1 B1MFZ1 B1MFZ9 B1MG21
B1MG63 B1MG81 B1MG98 B1MGA0 B1MGA1 B1MGA2 B1MGA3 B1MGB3 B1MGB7 B1MGC3 B1MGC4
B1MGC5 B1MGC7 B1MGD4 B1MGD5 B1MGD6 B1MGD7 B1MGE2 B1MGE3 B1MGE4 B1MGE5 B1MGE6
B1MGE7 B1MGE8 B1MGF1 B1MGH7 B1MGH8 B1MGH9 B1MGI0 B1MGU9 B1MGV8 B1MGY4 B1MGY6
B1MH39 B1MH40 B1MH61 B1MH69 B1MH70 B1MH79 B1MH85 B1MH86 B1MH92 B1MHA5 B1MHC3
B1MHC6 B1MHH1 B1MHI5 B1MHI6 B1MHQ0 B1MHW4 B1MHY1 B1MI06 B1MI19 B1MIM7 B1MIN9
B1MIP8 B1MIT2 B1MIV1 B1MIX5 B1MJJ0 B1MJK9 B1MJP3 B1MJT8 B1MJT9 B1MJX0 B1MK43
B1MK49 B1MKD8 B1MKI7 B1MKJ7 B1MKN0 B1ML08 B1ML09 B1ML10 B1ML11 B1ML30 B1ML31

B1MLG8 B1MLH7 B1MLL3 B1MLM0 B1MLU2 B1MLU6 B1MLV0 B1MLV1 B1MLW9 B1MLZ3 B1MLZ4
B1MM05 B1MM06 B1MMH9 B1MMI2 B1MML1 B1MMV3 B1MMV6 B1MMX9 B1MMY3 B1MMY4 B1MMY5
B1MN07 B1MN34 B1MN38 B1MN42 B1MN48 B1MN96 B1MN97 B1MP28 B1MP29 B1MP38 B1MP43
B1MPE6 B1MPG1 B1MT51 B1MTG4 B1MTI8 B1MTJ4 B1MVJ0 B1MVK9 B1MVM6 B1MVN7 B1MVQ9
B1MVR0 B1MVT4 B1MVT6 B1MVU1 B1MVU2 B1MVU3 B1MVW7 B1MVW8 B1MVY2 B1MVY3 B1MVY9
B1MVZ2 B1MVZ4 B1MVZ6 B1MVZ7 B1MVZ9 B1MW00 B1MW01 B1MW02 B1MW03 B1MW04 B1MW06
B1MW07 B1MW08 B1MW09 B1MW10 B1MW11 B1MW12 B1MW13 B1MW14 B1MW15 B1MW21 B1MW22
B1MW23 B1MW26 B1MW28 B1MW32 B1MW54 B1MW68 B1MW69 B1MW95 B1MWD1 B1MWL9 B1MWS3
B1MWS4 B1MWU1 B1MWV2 B1MWV4 B1MWX1 B1MWZ3 B1MWZ8 B1MX30 B1MX39 B1MX40 B1MX52
B1MX61 B1MX63 B1MX68 B1MX70 B1MXB1 B1MXB4 B1MXE0 B1MXE3 B1MXG8 B1MXG9 B1MXI0
B1MXL9 B1MXM6 B1MXP1 B1MXP8 B1MXP9 B1MXQ0 B1MXQ8 B1MXR1 B1MXS2 B1MXT7 B1MXT8
B1MXT9 B1MXU5 B1MXU9 B1MXV4 B1MXV5 B1MXV9 B1MXY4 B1MXZ5 B1MXZ9 B1MY00 B1MY04
B1MY11 B1MY28 B1MY30 B1MY42 B1MY45 B1MY57 B1MY62 B1MY98 B1MYA3 B1MYB3 B1MYB5
B1MYC1 B1MYC9 B1MYD0 B1MYD1 B1MYE1 B1MYE4 B1MYF2 B1MYF5 B1MYJ5 B1MZ17 B1MZ27
B1MZ49 B1MZ55 B1MZ60 B1MZ61 B1MZ62 B1MZ64 B1MZ65 B1MZ74 B1MZ77 B1MZ79 B1MZ82
B1MZE7 B1MZG6 B1MZG7 B1MZG8 B1MZH3 B1MZH4 B1MZH7 B1MZH8 B1MZH9 B1MZI4 B1MZJ0
B1MZK4 B1MZM2 B1MZM3 B1MZM6 B1MZM8 B1MZM9 B1MZN0 B1MZN8 B1MZP3 B1MZP4 B1MZP8
B1MZQ2 B1MZQ4 B1MZT4 B1MZT5 B1MZU5 B1MZV1 B1MZW8 B1N002 B1N009 B1N016 B1N017
B1N024 B1N025 B1N044 B1N057 B1N064 B1N065 B1N066 B1N079 B1N092 B1N094 B1N0C1
B1N1A2 B1N1A3 B1NKQ6 B1NKQ9 B1NKR0 B1NKR1 B1NKR3 B1NKR4 B1NKR5 B1NKR8 B1NKS7
B1NKS9 B1NKT0 B1NKT1 B1NKT7 B1NKT8 B1NKT9 B1NKU1 B1NKU2 B1NKU3 B1NWD1 B1NWD9
B1NWE0 B1NWE1 B1NWE8 B1NWF8 B1NWH1 B1NWH2 B1NWH3 B1NWI3 B1NWI4 B1NWI6 B1NWI7
B1NWI9 B1NWJ0 B1NWJ1 B1NWJ3 B1NWJ5 B1NWK6 B1P123 B1P1W2 B1PK17 B1PM81 B1PS76
B1PS77 B1PS80 B1PS81 B1PVZ9 B1Q2A8 B1Q2B6 B1Q3J6 B1V8K7 B1V8W6 B1V8X4 B1V8Z1
B1V8Z6 B1V8Z9 B1V910 B1V924 B1V932 B1V950 B1V955 B1V975 B1V9I4 B1V9I5 B1V9R1
B1VA46 B1VA58 B1VA59 B1VA75 B1VA77 B1VA79 B1VA80 B1VA93 B1VA97 B1VAA7 B1VAA9
B1VAB0 B1VAB8 B1VAC0 B1VAC2 B1VAC3 B1VAC4 B1VAC7 B1VAC9 B1VAD1 B1VAD2 B1VAD4
B1VAD5 B1VAD7 B1VAD8 B1VAD9 B1VAE1 B1VAE2 B1VAE4 B1VAE5 B1VAE6 B1VAE7 B1VAE8
B1VAE9 B1VAF0 B1VAF2 B1VAF3 B1VAF5 B1VAG0 B1VAH1 B1VAH8 B1VAH9 B1VAI9 B1VAJ8
B1VAJ9 B1VAK6 B1VAM2 B1VAM3 B1VAM4 B1VAM6 B1VAM7 B1VAM8 B1VAM9 B1VAN0 B1VAN5
B1VAN7 B1VAN8 B1VAN9 B1VAX7 B1VAY1 B1VAZ2 B1VB63 B1VB67 B1VB68 B1VB70 B1VB71
B1VB72 B1VB74 B1VB75 B1VB76 B1VB80 B1VB81 B1VB82 B1VDC0 B1VDC1 B1VDC2 B1VDC4
B1VDD6 B1VDD8 B1VDE1 B1VDE8 B1VDF7 B1VDJ3 B1VDJ5 B1VDJ7 B1VDK9 B1VDL1 B1VDL4
B1VDM5 B1VDM8 B1VDN5 B1VDN7 B1VDP0 B1VDP6 B1VDQ2 B1VDQ4 B1VDU4 B1VDV2 B1VDV5
B1VEE1 B1VEI4 B1VEJ5 B1VEK3 B1VEN4 B1VEQ2 B1VEQ8 B1VEQ9 B1VER8 B1VES1 B1VES2
B1VES9 B1VET0 B1VET1 B1VET5 B1VET6 B1VET7 B1VET9 B1VEU0 B1VEU1 B1VEU2 B1VEU3
B1VEU4 B1VEU5 B1VEW4 B1VEW5 B1VEW6 B1VEW9 B1VEX0 B1VEX2 B1VEX3 B1VEX4 B1VEX6
B1VEX9 B1VEY0 B1VEY1 B1VEY3 B1VEZ9 B1VF43 B1VF60 B1VF77 B1VFF9 B1VFG0 B1VFG1
B1VFG2 B1VFG3 B1VFG4 B1VFG5 B1VFI1 B1VFK0 B1VFM5 B1VFT1 B1VFW6 B1VFZ0 B1VFZ9
B1VG01 B1VG26 B1VG35 B1VG41 B1VG49 B1VG60 B1VG70 B1VG72 B1VG80 B1VG89 B1VG90
B1VG92 B1VGA4 B1VGB6 B1VGB7 B1VGC2 B1VGC3 B1VGK8 B1VGL9 B1VGM0 B1VGM1 B1VGM4
B1VGW8 B1VH27 B1VH30 B1VH34 B1VH40 B1VH41 B1VH82 B1VH88 B1VH93 B1VHC3 B1VHC6
B1VHD2 B1VHD3 B1VHG8 B1VHK2 B1VHM9 B1VHR9 B1VHT0 B1VHT2 B1VHT3 B1VHT5 B1VHT9
B1VHX4 B1VI61 B1VI76 B1VI88 B1VIF7 B1VII5 B1VII7 B1VIK2 B1VIQ0 B1VIS5 B1VIZ6

B1VIZ8 B1VJ37 B1VJ38 B1VKC8 B1VKC9 B1VKD2 B1VKD3 B1VKD5 B1VKD6 B1VKD7 B1VKE0
B1VKE1 B1VKE2 B1VKG7 B1VKH4 B1VKH5 B1VKH6 B1VKJ1 B1VKJ4 B1VKZ4 B1VLT7 B1VM97
B1VMC3 B1VMF2 B1VMF3 B1VMF9 B1VMX8 B1VN49 B1VN93 B1VN94 B1VNF2 B1VNN9 B1VNP0
B1VNY5 B1VP80 B1VP90 B1VP97 B1VPA3 B1VPC1 B1VPC2 B1VPC4 B1VPE8 B1VPE9 B1VPF0
B1VPF3 B1VQH0 B1VQW8 B1VR40 B1VRF5 B1VRF7 B1VRF8 B1VRG0 B1VRH5 B1VRH6 B1VS15
B1VS45 B1VS48 B1VS51 B1VS68 B1VS80 B1VS88 B1VS94 B1VSF5 B1VSG4 B1VSI6 B1VSL0
B1VSQ0 B1VSW8 B1VSW9 B1VT93 B1VTB7 B1VTD1 B1VTI5 B1VTI7 B1VTI8 B1VTN7 B1VTW2
B1VU78 B1VUC2 B1VUF9 B1VUR5 B1VUR8 B1VUV4 B1VUW6 B1VV08 B1VV57 B1VVV6 B1VWE7
B1VWF9 B1VWH6 B1VWJ8 B1VXA5 B1VXA8 B1VXB3 B1VXD6 B1VXD7 B1VXD8 B1VXE5 B1VXF6
B1VXM7 B1VXN6 B1VXR4 B1VXS2 B1VXS9 B1VXU2 B1VXU5 B1VXY2 B1VXZ4 B1VXZ6 B1VXZ7
B1VY27 B1VY37 B1VY42 B1VY47 B1VY58 B1VYN2 B1VYN3 B1VYN5 B1VYN8 B1VYP2 B1VYP5
B1VYT2 B1VYT6 B1VYT7 B1VYV2 B1VYV9 B1VYW1 B1VYW3 B1VYY1 B1VYY3 B1VYY6 B1VYZ5
B1VZ02 B1VZ03 B1VZ06 B1VZ57 B1VZ72 B1VZM3 B1VZM4 B1VZN5 B1VZR5 B1VZU6 B1VZV9
B1VZW1 B1VZW7 B1VZY6 B1VZY7 B1W019 B1W052 B1W089 B1W0B6 B1W0B7 B1W0D5 B1W0I2
B1W0M1 B1W0M4 B1W0M6 B1W0N3 B1W0N8 B1W0P0 B1W0X1 B1W0Y8 B1W0Y9 B1W0Z1 B1W0Z3
B1W0Z7 B1W1K9 B1W1P9 B1W1R5 B1W1W0 B1W1Y7 B1W203 B1W221 B1W2D7 B1W2D9 B1W2E1
B1W2G7 B1W2P8 B1W2Y7 B1W303 B1W306 B1W307 B1W310 B1W353 B1W354 B1W3A9 B1W3B1
B1W3B3 B1W3B4 B1W3F0 B1W3G0 B1W3G1 B1W3G2 B1W3G3 B1W3G4 B1W3H3 B1W3H7 B1W3I1
B1W3N4 B1W3N7 B1W3P6 B1W3T1 B1W3U3 B1W3X3 B1W3X4 B1W3X5 B1W3X8 B1W3X9 B1W3Y0
B1W3Y1 B1W3Y2 B1W3Y3 B1W3Y6 B1W3Y8 B1W3Y9 B1W3Z0 B1W3Z1 B1W3Z2 B1W3Z3 B1W3Z4
B1W3Z5 B1W3Z6 B1W3Z7 B1W3Z8 B1W3Z9 B1W400 B1W401 B1W402 B1W403 B1W404 B1W407
B1W408 B1W417 B1W418 B1W419 B1W441 B1W443 B1W444 B1W446 B1W447 B1W455 B1W456
B1W459 B1W463 B1W468 B1W470 B1W473 B1W4A5 B1W4A6 B1W4G3 B1W4R8 B1W4R9 B1W4U4
B1W525 B1W5A8 B1W5F4 B1W5G7 B1W5G8 B1W5H1 B1W5H2 B1W5J2 B1W5Q9 B1W5S5 B1WAV2
B1WAZ8 B1WB17 B1WB65 B1WBB4 B1WBP0 B1WBU4 B1WBU5 B1WBU8 B1WBV0 B1WC15 B1WC37
B1WC39 B1WC40 B1WC58 B1WC68 B1WN69 B1WN93 B1WNA0 B1WNP1 B1WNP3 B1WNP5 B1WNQ1
B1WNQ4 B1WP07 B1WP72 B1WP79 B1WPH9 B1WPL5 B1WPP6 B1WPY4 B1WPZ0 B1WQ07 B1WQ09
B1WQ23 B1WQ52 B1WQB7 B1WQD7 B1WQE4 B1WQF1 B1WQH2 B1WQL2 B1WQQ4 B1WQQ9 B1WQR2
B1WQR3 B1WQR4 B1WQR5 B1WQR6 B1WQR7 B1WQR8 B1WQS0 B1WQS1 B1WQS4 B1WQS5 B1WQS6
B1WQS7 B1WQS8 B1WQT3 B1WQT6 B1WQT8 B1WQT9 B1WQW1 B1WQY3 B1WQY4 B1WQY6 B1WQY7
B1WQZ8 B1WR00 B1WR01 B1WR40 B1WR77 B1WR90 B1WR94 B1WRJ4 B1WRJ7 B1WRN0 B1WRP7
B1WRQ9 B1WRV7 B1WRV9 B1WRW8 B1WS35 B1WS51 B1WSG7 B1WSJ0 B1WSK6 B1WSL8 B1WSM0
B1WSW1 B1WSY4 B1WT08 B1WT19 B1WT39 B1WT42 B1WT69 B1WTF7 B1WTI3 B1WTK3 B1WTN1
B1WTN3 B1WTW9 B1WTZ2 B1WU11 B1WU13 B1WU69 B1WU96 B1WUB0 B1WUD1 B1WUD2 B1WUD4
B1WUG7 B1WUL8 B1WUP2 B1WUT9 B1WUV7 B1WV53 B1WV74 B1WVA3 B1WVE8 B1WVR2 B1WVY1
B1WVZ0 B1WW40 B1WW41 B1WW59 B1WW94 B1WW99 B1WWG9 B1WWJ3 B1WWL8 B1WWM7 B1WX40
B1WX86 B1WXF7 B1WXH3 B1WXT3 B1WXY6 B1WY20 B1WY30 B1WY56 B1WY59 B1WY64 B1WYB2
B1WYG5 B1WYI6 B1WYK7 B1WYP3 B1WYQ5 B1WYT6 B1WYU1 B1WYU3 B1WYX6 B1WZ70 B1WZH9
B1WZK0 B1WZL6 B1WZM0 B1WZN0 B1WZP0 B1WZR3 B1WZT6 B1WZT9 B1WZV3 B1WZW5 B1X018
B1X023 B1X058 B1X067 B1X073 B1X074 B1X076 B1X082 B1X090 B1X0B0 B1X0D2 B1X0G3
B1X0L5 B1X0M2 B1X0N5 B1X0Q1 B1X0U7 B1X0X6 B1X0X7 B1X0Z4 B1X109 B1X3H9 B1X3I0
B1X3K4 B1X3M8 B1X3M9 B1X3S1 B1X416 B1X437 B1X4D6 B1X4K3 B1X4W8 B1X4X1 B1X4Y8
B1X4Z6 B1X4Z7 B1X4Z8 B1X501 B1X503 B1X504 B1X508 B1X536 B1X559 B1X580 B1X5S1
B1X631 B1X654 B1X656 B1X658 B1X671 B1X677 B1X691 B1X6A6 B1X6A8 B1X6B7 B1X6C4

B1X6D4 B1X6D5 B1X6D7 B1X6D9 B1X6E0 B1X6E1 B1X6E6 B1X6E8 B1X6E9 B1X6F1 B1X6F3
B1X6F4 B1X6F6 B1X6F7 B1X6F8 B1X6F9 B1X6G0 B1X6G1 B1X6G2 B1X6G3 B1X6G4 B1X6G5
B1X6G6 B1X6G7 B1X6G8 B1X6H0 B1X6H1 B1X6H2 B1X6J0 B1X6J1 B1X6J2 B1X6J3 B1X6J4
B1X6J7 B1X6K4 B1X6L1 B1X6L3 B1X6P4 B1X6P5 B1X6Q8 B1X6S4 B1X6S9 B1X6V6 B1X6V8
B1X6W1 B1X6W2 B1X701 B1X716 B1X736 B1X737 B1X748 B1X750 B1X757 B1X758 B1X760
B1X764 B1X765 B1X770 B1X774 B1X775 B1X776 B1X786 B1X797 B1X7A5 B1X7A6 B1X7A9
B1X7B1 B1X7B3 B1X7D4 B1X7D7 B1X7E2 B1X7I7 B1X7J5 B1X7K4 B1X7N2 B1X7P1 B1X7P2
B1X7S8 B1X7T9 B1X7V2 B1X7X4 B1X7X6 B1X7Z3 B1X800 B1X807 B1X814 B1X822 B1X825
B1X833 B1X847 B1X848 B1X850 B1X852 B1X857 B1X860 B1X869 B1X883 B1X884 B1X885
B1X8A6 B1X8C6 B1X8H5 B1X8I1 B1X8J1 B1X8L4 B1X8M1 B1X8M6 B1X8M8 B1X8N5 B1X8N9
B1X8Q2 B1X8R0 B1X8R2 B1X8R7 B1X8S0 B1X8S1 B1X8V3 B1X8V4 B1X8V5 B1X8V8 B1X8W0
B1X8W2 B1X8W4 B1X8X5 B1X8X7 B1X8X8 B1X903 B1X905 B1X929 B1X931 B1X935 B1X943
B1X950 B1X953 B1X968 B1X969 B1X970 B1X971 B1X972 B1X974 B1X975 B1X976 B1X980
B1X982 B1X998 B1X9B6 B1X9B7 B1X9C1 B1X9C5 B1X9C8 B1X9C9 B1X9D0 B1X9D1 B1X9D2
B1X9D3 B1X9E8 B1X9G8 B1X9H2 B1X9H5 B1X9I0 B1X9K2 B1X9K4 B1X9N9 B1X9P6 B1X9R0
B1X9R9 B1X9S9 B1X9T1 B1X9T2 B1X9T5 B1X9T7 B1X9U0 B1X9W8 B1X9W9 B1X9X2 B1X9X3
B1X9X5 B1X9Y2 B1X9Z0 B1X9Z3 B1X9Z4 B1X9Z9 B1XA08 B1XA16 B1XA17 B1XA18 B1XA29
B1XA37 B1XA43 B1XA44 B1XA47 B1XA53 B1XA58 B1XA61 B1XA63 B1XA66 B1XA68 B1XA76
B1XA82 B1XA89 B1XA98 B1XA99 B1XAA7 B1XAB1 B1XAD8 B1XAE3 B1XAE8 B1XAH2 B1XAH3
B1XAH6 B1XAH7 B1XAJ3 B1XAJ7 B1XAK9 B1XAM2 B1XAP5 B1XAP9 B1XAQ1 B1XAQ2 B1XAS5
B1XAT8 B1XAX1 B1XAX3 B1XAX4 B1XAY2 B1XAY4 B1XAY6 B1XAY9 B1XAZ0 B1XAZ2 B1XAZ3
B1XAZ8 B1XB01 B1XB02 B1XB03 B1XB05 B1XB06 B1XB14 B1XB16 B1XB17 B1XB26 B1XB39
B1XB41 B1XB55 B1XB59 B1XB63 B1XB68 B1XB69 B1XB70 B1XB71 B1XB72 B1XB82 B1XB85
B1XB92 B1XB94 B1XB95 B1XB97 B1XB98 B1XBA2 B1XBA4 B1XBA8 B1XBC1 B1XBC2 B1XBC4
B1XBC5 B1XBC6 B1XBC8 B1XBD1 B1XBE0 B1XBE8 B1XBF1 B1XBF4 B1XBF5 B1XBF6 B1XBG0
B1XBG2 B1XBG3 B1XBG4 B1XBG6 B1XBG7 B1XBI8 B1XBJ4 B1XBJ5 B1XBJ6 B1XBK9 B1XBL0
B1XBM8 B1XBN2 B1XBN7 B1XBQ2 B1XBQ6 B1XBQ7 B1XBS8 B1XBS9 B1XBT0 B1XBT1 B1XBT4
B1XBT5 B1XBT8 B1XBU0 B1XBW5 B1XBX3 B1XBX6 B1XBX8 B1XBY1 B1XBY5 B1XBY6 B1XBY7
B1XBY8 B1XBY9 B1XBZ0 B1XBZ3 B1XBZ7 B1XBZ8 B1XBZ9 B1XC00 B1XC01 B1XC09 B1XC12
B1XC15 B1XC20 B1XC24 B1XC38 B1XC50 B1XC51 B1XC52 B1XC53 B1XC58 B1XC59 B1XC68
B1XC77 B1XC78 B1XC80 B1XC94 B1XC95 B1XC96 B1XCA6 B1XCA8 B1XCA9 B1XCC1 B1XCH3
B1XCL1 B1XCM1 B1XCM4 B1XCM5 B1XCM6 B1XCN7 B1XCN8 B1XCR0 B1XCR9 B1XCS0 B1XCS1
B1XCS2 B1XCS3 B1XCS6 B1XCS7 B1XCS8 B1XD17 B1XD24 B1XD26 B1XD29 B1XD30 B1XD33
B1XD35 B1XD36 B1XD38 B1XD39 B1XD41 B1XD52 B1XD59 B1XD64 B1XD73 B1XD76 B1XD78
B1XD84 B1XDF5 B1XDG9 B1XDH5 B1XDJ0 B1XDK3 B1XDK7 B1XDL1 B1XDL3 B1XDL5 B1XDN6
B1XDN7 B1XDP6 B1XDP7 B1XDQ0 B1XDQ9 B1XDR5 B1XDS1 B1XDS2 B1XDS3 B1XDS8 B1XDS9
B1XDU2 B1XDU3 B1XDU7 B1XDU8 B1XDU9 B1XDV1 B1XDV2 B1XDV3 B1XDV9 B1XDW8 B1XDY2
B1XDY3 B1XDY5 B1XDY6 B1XE36 B1XE38 B1XE39 B1XE52 B1XE70 B1XE99 B1XEA0 B1XEA5
B1XEA6 B1XEA7 B1XED7 B1XEG2 B1XEH6 B1XEH8 B1XEI5 B1XEI7 B1XEJ0 B1XEJ1 B1XEJ5
B1XEJ9 B1XEK1 B1XEL0 B1XEL4 B1XEL6 B1XEM5 B1XEM6 B1XEN9 B1XET3 B1XET7 B1XET8
B1XEU8 B1XEX1 B1XEX6 B1XEX7 B1XEY0 B1XEY2 B1XEZ3 B1XEZ4 B1XF01 B1XF03 B1XF04
B1XF08 B1XF10 B1XF11 B1XF75 B1XF88 B1XF99 B1XFA4 B1XFA5 B1XFA8 B1XFA9 B1XFB3
B1XFC2 B1XFH3 B1XFI0 B1XFI4 B1XFJ1 B1XFJ2 B1XFJ3 B1XFJ4 B1XFJ6 B1XFK3 B1XFK4
B1XFK5 B1XFL4 B1XFM4 B1XFM6 B1XFN2 B1XFN4 B1XFQ6 B1XFQ8 B1XFQ9 B1XFR1 B1XFR2

B1XFR3 B1XFT8 B1XFU0 B1XFU9 B1XFV0 B1XFV2 B1XFV4 B1XFY7 B1XFY8 B1XG00 B1XG01
B1XG10 B1XG13 B1XG19 B1XG21 B1XG23 B1XG24 B1XG46 B1XG54 B1XG57 B1XG61 B1XG69
B1XG70 B1XG73 B1XG77 B1XG88 B1XG96 B1XGB0 B1XGC4 B1XGC7 B1XGI1 B1XGJ4 B1XGK1
B1XGK3 B1XGK5 B1XGK6 B1XGK7 B1XGK8 B1XGK9 B1XGN8 B1XGP4 B1XGS5 B1XGT9 B1XGU5
B1XGU9 B1XGW0 B1XGW1 B1XGW6 B1XGW7 B1XGX6 B1XGX7 B1XGX9 B1XGY0 B1XGY2 B1XGY3
B1XGY6 B1XGY9 B1XH79 B1XH80 B1XH99 B1XHA3 B1XHB7 B1XHC8 B1XHC9 B1XHD1 B1XHD4
B1XHD8 B1XHD9 B1XHE2 B1XHE4 B1XHF8 B1XHF9 B1XHG1 B1XHG2 B1XHI0 B1XHJ5 B1XHJ6
B1XHJ8 B1XHJ9 B1XHK3 B1XHK4 B1XHK9 B1XHL0 B1XHM8 B1XHN0 B1XHQ1 B1XHW1 B1XHW6
B1XHZ7 B1XI06 B1XI09 B1XI10 B1XI62 B1XI63 B1XI65 B1XI66 B1XI84 B1XI94 B1XI96
B1XIA1 B1XIC0 B1XIE7 B1XIF6 B1XIG4 B1XIH9 B1XIL4 B1XIL5 B1XIM6 B1XIQ9 B1XIR0
B1XIT5 B1XIV9 B1XIZ3 B1XJ08 B1XJ13 B1XJ17 B1XJ47 B1XJ88 B1XJ90 B1XJD1 B1XJD2
B1XJF0 B1XJG8 B1XJG9 B1XJH0 B1XJH1 B1XJH2 B1XJH5 B1XJH8 B1XJH9 B1XJI0 B1XJI1
B1XJI2 B1XJI3 B1XJI6 B1XJJ0 B1XJJ1 B1XJJ2 B1XJJ3 B1XJK0 B1XJM0 B1XJM1 B1XJP0
B1XJS6 B1XJS7 B1XJS8 B1XJS9 B1XJT1 B1XJT2 B1XJT3 B1XJT4 B1XJT5 B1XJT6 B1XJT9
B1XJU0 B1XJV9 B1XJY4 B1XJZ5 B1XK11 B1XK17 B1XK39 B1XK45 B1XK79 B1XK80 B1XK83
B1XKA0 B1XKC3 B1XKC5 B1XKE8 B1XKF6 B1XKH1 B1XKM7 B1XKN8 B1XKQ0 B1XKV1 B1XKY8
B1XKZ0 B1XL11 B1XL12 B1XL18 B1XL20 B1XL23 B1XL26 B1XL39 B1XL79 B1XLA4 B1XLD1
B1XLE7 B1XLF1 B1XLH8 B1XLJ1 B1XLK5 B1XLK6 B1XLM3 B1XLP1 B1XLP7 B1XLQ6 B1XLQ9
B1XLR4 B1XLR5 B1XLV0 B1XLW1 B1XM04 B1XM11 B1XM20 B1XM22 B1XM75 B1XM86 B1XMA0
B1XMC5 B1XMC7 B1XMM7 B1XMP0 B1XN05 B1XN07 B1XN08 B1XN09 B1XN32 B1XN45 B1XN65
B1XN66 B1XN71 B1XNB2 B1XND1 B1XNE1 B1XNE8 B1XNE9 B1XNG1 B1XNH7 B1XNI3 B1XNN1
B1XNQ0 B1XNQ1 B1XNV4 B1XNV8 B1XNY9 B1XP08 B1XP15 B1XP37 B1XP39 B1XP49 B1XP50
B1XP53 B1XP68 B1XP73 B1XP88 B1XP90 B1XP99 B1XPA6 B1XPB0 B1XPD9 B1XPE0 B1XPE2
B1XPG7 B1XPL6 B1XPL7 B1XPN9 B1XPQ6 B1XPQ9 B1XPT1 B1XPT2 B1XPT3 B1XPV1 B1XPV8
B1XPX4 B1XPX8 B1XPZ2 B1XPZ7 B1XPZ8 B1XPZ9 B1XQ35 B1XQ41 B1XQ51 B1XQ70 B1XQE1
B1XQE7 B1XQF6 B1XQH8 B1XQI2 B1XQI5 B1XQK7 B1XQL0 B1XQQ0 B1XQQ1 B1XRS6 B1XRS9
B1XRT3 B1XRU1 B1XRU2 B1XRV8 B1XRX1 B1XRX2 B1XRZ4 B1XS06 B1XS10 B1XS23 B1XS24
B1XS25 B1XS32 B1XS59 B1XS61 B1XS63 B1XS65 B1XS66 B1XS68 B1XS92 B1XSA5 B1XSC4
B1XSD8 B1XSE8 B1XSE9 B1XSF4 B1XSG8 B1XSH8 B1XSH9 B1XSI4 B1XSJ0 B1XSN1 B1XSN2
B1XSP0 B1XSP1 B1XSP2 B1XSP3 B1XSP4 B1XSP6 B1XSP7 B1XSP8 B1XSQ0 B1XSQ1 B1XSQ2
B1XSQ3 B1XSQ4 B1XSQ5 B1XSQ6 B1XSQ8 B1XSQ9 B1XSR0 B1XSR1 B1XSR2 B1XSR3 B1XSR4
B1XSR5 B1XSR6 B1XSR7 B1XSR9 B1XSS0 B1XSS3 B1XSS4 B1XSS5 B1XSS6 B1XSS7 B1XSS8
B1XST2 B1XSU7 B1XSV0 B1XSV2 B1XSV3 B1XSY0 B1XSZ1 B1XSZ2 B1XT01 B1XT10 B1XT11
B1XT33 B1XT34 B1XT35 B1XT37 B1XT38 B1XT48 B1XT51 B1XT71 B1XT72 B1XT79 B1XT83
B1XTA7 B1XTC3 B1XTC4 B1XTC5 B1XTC7 B1XTD4 B1XTK4 B1XTL6 B1XTM0 B1XTM5 B1XTM7
B1XTM8 B1XTR3 B1XTS4 B1XTT6 B1XTU0 B1XTU3 B1XTU4 B1XTU6 B1XTV7 B1XTW6 B1XU55
B1XU63 B1XU64 B1XU70 B1XU72 B1XU78 B1XU84 B1XU85 B1XU95 B1XU96 B1XUA6 B1XUH5
B1XUJ0 B1XUJ3 B1XUJ4 B1XUJ5 B1XUJ6 B1XUM3 B1XUP3 B1XUR6 B1XUS0 B1XUS8 B1XUT0
B1XUT6 B1XUX7 B1XUY8 B1XV11 B1XV12 B1XV14 B1XV46 B1XV49 B1XV53 B1XV55 B1XV63
B1XV66 B1XV76 B1XV88 B1XV89 B1XV91 B1XVE1 B1XVE9 B1XVF7 B1XVG9 B1XVH8 B1XVK0
B1XVN0 B1XVN1 B1XVN2 B1XVN3 B1XVP6 B1XVQ2 B1XVZ1 B1XVZ2 B1XW11 B1XW20 B1XW21
B1XW28 B1XW54 B1XW69 B1XW95 B1XW99 B1XWA3 B1XWC5 B1XWG6 B1XWG9 B1XWK4 B1XWM2
B1XWM8 B1XWQ5 B1XWR8 B1XWS5 B1XWZ1 B1XX30 B1XX46 B1XX60 B1XX71 B1XX72 B1XX77
B1XX97 B1XXG0 B1XXG3 B1XXH2 B1XXH3 B1XXH5 B1XXH9 B1XXJ2 B1XXJ4 B1XXJ5 B1XXK8

B1XXK9 B1XXL6 B1XXL7 B1XXP1 B1XXX9 B1XXY5 B1XXY8 B1XXY9 B1XY02 B1XY11 B1XY19
B1XY20 B1XY29 B1XY42 B1XY47 B1XY48 B1XY49 B1XY56 B1XY65 B1XY67 B1XY68 B1XYA8
B1XYB1 B1XYE2 B1XYE7 B1XYE8 B1XYF6 B1XYL1 B1XYL2 B1XYP5 B1XYQ4 B1XYR4 B1XYW4
B1XYW5 B1XYX1 B1XYX5 B1XYX6 B1XYZ3 B1XYZ5 B1XYZ6 B1XYZ7 B1XZ11 B1XZ34 B1XZP2
B1Y070 B1Y0F6 B1Y0G0 B1Y0H5 B1Y0H6 B1Y0H7 B1Y0H8 B1Y0I4 B1Y0I6 B1Y0I9 B1Y0J8
B1Y0M2 B1Y0P9 B1Y0T0 B1Y0T1 B1Y0U3 B1Y0V7 B1Y148 B1Y149 B1Y1E3 B1Y1E6 B1Y1E7
B1Y1G9 B1Y1J9 B1Y1L9 B1Y1M1 B1Y222 B1Y223 B1Y225 B1Y234 B1Y279 B1Y280 B1Y281
B1Y282 B1Y2E5 B1Y2E6 B1Y2G9 B1Y2H6 B1Y2L3 B1Y2X5 B1Y2Y7 B1Y2Y8 B1Y382 B1Y3C2
B1Y3E4 B1Y3E6 B1Y3G7 B1Y3K6 B1Y3K7 B1Y3N8 B1Y3P1 B1Y3P5 B1Y3Q0 B1Y3Q1 B1Y3R5
B1Y3V1 B1Y3V2 B1Y3V4 B1Y3W7 B1Y3Z2 B1Y4K6 B1Y4K7 B1Y4K8 B1Y4W6 B1Y500 B1Y502
B1Y547 B1Y580 B1Y5G6 B1Y5I3 B1Y5U2 B1Y5W3 B1Y629 B1Y634 B1Y647 B1Y650 B1Y654
B1Y6D4 B1Y6D6 B1Y6E5 B1Y6E7 B1Y6H2 B1Y6H3 B1Y6H4 B1Y6I5 B1Y6I7 B1Y6N9 B1Y6R3
B1Y6R5 B1Y6S3 B1Y6S6 B1Y708 B1Y785 B1Y786 B1Y787 B1Y7F0 B1Y7G7 B1Y7G8 B1Y7G9
B1Y7H1 B1Y7H2 B1Y7H4 B1Y7H5 B1Y7H6 B1Y7H7 B1Y7I2 B1Y7I3 B1Y7L9 B1Y7M8 B1Y7M9
B1Y7N0 B1Y7P2 B1Y7P3 B1Y7V8 B1Y816 B1Y822 B1Y823 B1Y839 B1Y847 B1Y852 B1Y856
B1Y8A3 B1Y8B3 B1Y8B4 B1Y8B5 B1Y8B8 B1Y8B9 B1Y8C0 B1Y8C1 B1Y8C2 B1Y8C3 B1Y8C4
B1Y8C6 B1Y8C7 B1Y8E1 B1Y8E2 B1Y8E7 B1Y8G2 B1Y8H8 B1Y8H9 B1Y8I0 B1Y8I1 B1Y8I2
B1Y8I3 B1Y8I4 B1Y8I5 B1Y8I6 B1Y8I7 B1Y8J4 B1Y8P7 B1Y8P8 B1Y8Q5 B1Y8T9 B1Y8V2
B1Y8V5 B1Y8Z1 B1Y901 B1Y919 B1Y945 B1Y949 B1Y953 B1Y963 B1Y973 B1Y976 B1Y978
B1Y992 B1Y9B8 B1Y9C5 B1Y9D1 B1Y9H0 B1Y9H2 B1Y9M7 B1Y9M9 B1Y9N2 B1Y9N5 B1Y9N6
B1Y9R0 B1Y9S4 B1Y9T4 B1Y9T9 B1Y9U2 B1Y9U5 B1Y9V2 B1Y9V3 B1Y9V4 B1Y9W1 B1Y9W7
B1Y9Y4 B1Y9Y6 B1YA13 B1YA16 B1YA29 B1YA32 B1YA33 B1YA34 B1YA36 B1YA42 B1YA45
B1YA52 B1YA63 B1YA64 B1YAA9 B1YAC8 B1YAD4 B1YAD6 B1YAF1 B1YAF2 B1YAF6 B1YAF9
B1YAJ2 B1YAJ4 B1YAJ6 B1YAJ8 B1YAJ9 B1YAL1 B1YAM9 B1YAN0 B1YAS9 B1YAT9 B1YAU2
B1YAW1 B1YAX4 B1YAZ7 B1YB29 B1YB34 B1YB49 B1YB53 B1YB54 B1YB61 B1YB67 B1YB76
B1YB90 B1YBD4 B1YBH2 B1YBK5 B1YBL3 B1YBM2 B1YBM5 B1YBN0 B1YBN2 B1YBN8 B1YC14
B1YC27 B1YC30 B1YC32 B1YC34 B1YC36 B1YC39 B1YC43 B1YC46 B1YC47 B1YC67 B1YC69
B1YCG8 B1YCL7 B1YCN7 B1YCP4 B1YCQ5 B1YCQ7 B1YCS2 B1YCT7 B1YCV9 B1YCW1 B1YCX0
B1YCY3 B1YD00 B1YD06 B1YD09 B1YD10 B1YD24 B1YD56 B1YD57 B1YD64 B1YD88 B1YD95
B1YD98 B1YDA5 B1YDC4 B1YDE0 B1YDH4 B1YDH6 B1YDJ0 B1YDJ1 B1YDK8 B1YDP0 B1YDP2
B1YDP4 B1YDX0 B1YE08 B1YE74 B1YED2 B1YEE3 B1YEG7 B1YEH3 B1YEH9 B1YEJ3 B1YEK4
B1YEL0 B1YEP6 B1YEP7 B1YEQ1 B1YES2 B1YF09 B1YFI1 B1YFL0 B1YFM6 B1YFM9 B1YFS8
B1YG36 B1YG43 B1YG64 B1YG85 B1YG93 B1YG95 B1YG99 B1YGA6 B1YGA7 B1YGA8 B1YGB0
B1YGB1 B1YGB2 B1YGB5 B1YGC2 B1YGC4 B1YGC9 B1YGD0 B1YGD3 B1YGP2 B1YGP4 B1YGP7
B1YGQ4 B1YGQ7 B1YGS7 B1YGS9 B1YGT4 B1YGT7 B1YGU3 B1YGU5 B1YGU6 B1YGU7 B1YGU8
B1YGU9 B1YGV0 B1YGV1 B1YGV2 B1YGV3 B1YGV4 B1YGV5 B1YGV6 B1YGV7 B1YGV8 B1YGV9
B1YGW0 B1YGW1 B1YGW2 B1YGW3 B1YGW4 B1YGW5 B1YGW6 B1YGW8 B1YGW9 B1YGX1 B1YGX4
B1YGX5 B1YGX6 B1YGX8 B1YGY0 B1YH73 B1YH83 B1YH84 B1YH85 B1YH95 B1YHD4 B1YHD6
B1YHH3 B1YHJ8 B1YHQ7 B1YHQ8 B1YHQ9 B1YHS1 B1YHT1 B1YHU0 B1YI19 B1YI20 B1YI22
B1YI33 B1YI57 B1YI58 B1YI66 B1YI68 B1YI70 B1YI73 B1YI75 B1YI76 B1YIA8 B1YIA9
B1YIB2 B1YIH8 B1YIL7 B1YIM3 B1YIM4 B1YIM5 B1YIM6 B1YIP3 B1YIQ5 B1YIR0 B1YIR7
B1YIR9 B1YIS7 B1YIT1 B1YIT8 B1YIU3 B1YIU4 B1YIU7 B1YIU9 B1YIV1 B1YIX9 B1YIY0
B1YIY1 B1YIY2 B1YIY3 B1YIY4 B1YIZ1 B1YJ02 B1YJ04 B1YJ07 B1YJ17 B1YJ30 B1YJ35
B1YJ36 B1YJ39 B1YJ40 B1YJ41 B1YJ65 B1YJ90 B1YJ94 B1YJC0 B1YJD6 B1YJE1 B1YJE5

B1YJE9 B1YJF7 B1YJF9 B1YJG1 B1YJJ2 B1YJK3 B1YJL9 B1YJP1 B1YJR0 B1YJR1 B1YJR2
B1YJR9 B1YJS1 B1YJS3 B1YJT1 B1YJU8 B1YJV1 B1YJV2 B1YJV8 B1YJW0 B1YJW1 B1YJW5
B1YJW6 B1YJW7 B1YJY5 B1YJY9 B1YJZ8 B1YK03 B1YK58 B1YK68 B1YK70 B1YK83 B1YK88
B1YK95 B1YK97 B1YKA0 B1YKC5 B1YKD2 B1YKE8 B1YKF9 B1YKH4 B1YKI7 B1YKL9 B1YKM7
B1YKM8 B1YKR3 B1YKR5 B1YKS4 B1YKS7 B1YKS8 B1YKS9 B1YKT0 B1YKT1 B1YKT7 B1YKV4
B1YKW2 B1YKZ4 B1YL09 B1YL11 B1YL27 B1YL34 B1YL70 B1YL74 B1YL75 B1YL84 B1YL88
B1YL94 B1YLA2 B1YLB5 B1YLE0 B1YLE1 B1YLE8 B1YLF0 B1YLF7 B1YLH0 B1YLK2 B1YLN6
B1YLN7 B1YLN8 B1YLP0 B1YLP7 B1YLQ1 B1YLQ2 B1YLQ5 B1YLQ7 B1YLR4 B1YLS1 B1YLS2
B1YLS4 B1YLU2 B1YLU8 B1YM01 B1YM02 B1YM12 B1YM95 B1YMA3 B1YMB8 B1YMB9 B1YMC6
B1YMD7 B1YMD8 B1YMH0 B1YMH1 B1YMW2 B1YMW5 B1YMX3 B1YMY4 B1YMY7 B1YN34 B1YN45
B1YN48 B1YN51 B1YN53 B1YN54 B1YN57 B1YNA3 B1YNA6 B1YNJ6 B1YNM7 B1YNP3 B1YNQ7
B1YNR4 B1YNS0 B1YNS1 B1YNS2 B1YNW1 B1YNZ5 B1YP13 B1YP15 B1YP16 B1YP34 B1YP36
B1YP37 B1YP61 B1YPB3 B1YPC1 B1YPI1 B1YPI7 B1YPI8 B1YPJ0 B1YPQ4 B1YPR2 B1YPR3
B1YPR7 B1YPU9 B1YPV7 B1YPV8 B1YPV9 B1YPW9 B1YPX6 B1YPZ0 B1YQ54 B1YQ56 B1YQ64
B1YQC8 B1YQE3 B1YQJ5 B1YQK0 B1YQK2 B1YQK3 B1YQQ1 B1YQQ2 B1YQQ3 B1YR17 B1YR34
B1YR39 B1YR40 B1YR43 B1YR44 B1YR46 B1YR47 B1YRB8 B1YRB9 B1YRC0 B1YRC1 B1YRC2
B1YRC3 B1YRC5 B1YRC6 B1YRC9 B1YRD0 B1YRD1 B1YRD2 B1YRD3 B1YRD4 B1YRD5 B1YRD6
B1YRD7 B1YRE4 B1YRE7 B1YRE8 B1YRF5 B1YRJ1 B1YRJ2 B1YRJ4 B1YRL3 B1YRN5 B1YRN6
B1YRN7 B1YRN8 B1YRN9 B1YRP0 B1YRP1 B1YRP2 B1YRP3 B1YRP4 B1YRP5 B1YRP6 B1YRP7
B1YRP8 B1YRQ1 B1YRQ2 B1YRQ3 B1YRQ4 B1YRQ5 B1YRQ6 B1YRR0 B1YRR7 B1YRS4 B1YRV4
B1YRV7 B1YRW0 B1YRW1 B1YRW2 B1YRW3 B1YRY9 B1YRZ4 B1YRZ6 B1YS10 B1YS44 B1YS58
B1YS60 B1YS68 B1YS71 B1YS73 B1YS74 B1YS76 B1YSD5 B1YSD6 B1YSF2 B1YSF5 B1YSF6
B1YSH2 B1YSH7 B1YSH9 B1YSK3 B1YSM8 B1YSP3 B1YSR5 B1YSR6 B1YSS2 B1YSS5 B1YSS6
B1YST8 B1YSU5 B1YSU6 B1YSU7 B1YSU8 B1YSV0 B1YSV1 B1YSW5 B1YT08 B1YT11 B1YT20
B1YT24 B1YT42 B1YT59 B1YT76 B1YT80 B1YTA7 B1YTB0 B1YTD2 B1YTD3 B1YTD4 B1YTD6
B1YTD9 B1YTE0 B1YTE1 B1YTE2 B1YTE9 B1YTH0 B1YTH4 B1YTJ3 B1YTJ4 B1YTJ5 B1YTK0
B1YTK1 B1YTN1 B1YTR1 B1YTR2 B1YTR7 B1YTS0 B1YTV3 B1YTV7 B1YTX4 B1YTZ1 B1YU27
B1YU28 B1YU30 B1YU46 B1YU48 B1YU49 B1YU52 B1YU55 B1YU72 B1YU93 B1YUF6 B1YUF7
B1YUF9 B1YUG0 B1YUG1 B1YUJ1 B1YUJ2 B1YUJ7 B1YUL5 B1YUL9 B1YUM1 B1YUN3 B1YUS8
B1YUT2 B1YUT9 B1YUU0 B1YUW5 B1YUY0 B1YUZ7 B1YV26 B1YV30 B1YV33 B1YV34 B1YV36
B1YV41 B1YV52 B1YV57 B1YV58 B1YV59 B1YV60 B1YV68 B1YV93 B1YV94 B1YV95 B1YV98
B1YVA4 B1YVA7 B1YVB1 B1YVD8 B1YVE0 B1YVE4 B1YVE7 B1YVE8 B1YVG1 B1YVH0 B1YVH2
B1YVJ0 B1YVJ8 B1YVK9 B1YVN1 B1YVN3 B1YVX4 B1YVY6 B1YW19 B1YW20 B1YW60 B1YWB2
B1YWB9 B1YWD5 B1YWF2 B1YWF9 B1YWG4 B1YWR8 B1YX25 B1YXE9 B1YXP0 B1YXQ0 B1YY34
B1YYR8 B1YZG1 B1YZP8 B1YZQ0 B1Z032 B1Z033 B1Z034 B1Z0E2 B1Z0R0 B1Z0R8 B1Z0T3
B1Z151 B1Z181 B1Z1G0 B1Z1G2 B1Z1G3 B1Z1G4 B1Z1N0 B1Z1N2 B1Z1N7 B1Z1P1 B1Z1Y2
B1Z2C1 B1Z3G9 B1Z405 B1Z5U9 B1Z700 B1Z757 B1Z759 B1Z761 B1Z767 B1Z768 B1Z770
B1Z771 B1Z772 B1Z774 B1Z775 B1Z776 B1Z777 B1Z778 B1Z780 B1Z782 B1Z783 B1Z7C0
B1Z7C1 B1Z7C5 B1Z7L0 B1Z7Y5 B1Z7Y8 B1Z888 B1Z896 B1Z8E9 B1Z8R3 B1Z8R4 B1Z8U3
B1Z914 B1Z922 B1Z924 B1Z939 B1Z942 B1Z962 B1Z970 B1Z980 B1Z987 B1Z9C6 B1Z9C7
B1Z9C8 B1Z9F8 B1Z9P1 B1Z9R0 B1Z9R9 B1Z9T1 B1Z9X0 B1Z9Y8 B1ZA19 B1ZA28 B1ZA86
B1ZAD1 B1ZAD6 B1ZAJ4 B1ZAK8 B1ZAP4 B1ZAP8 B1ZAR5 B1ZAU5 B1ZAU6 B1ZB15 B1ZB16
B1ZB17 B1ZB18 B1ZB19 B1ZB59 B1ZBP2 B1ZBR8 B1ZBT6 B1ZBW7 B1ZC86 B1ZC90 B1ZCB3
B1ZCB4 B1ZCM6 B1ZDL3 B1ZDN3 B1ZDQ7 B1ZDQ8 B1ZDR1 B1ZDT1 B1ZDV2 B1ZE36 B1ZEG3

B1ZEJ7 B1ZEL4 B1ZEN4 B1ZES7 B1ZEV4 B1ZFB1 B1ZFJ0 B1ZFQ5 B1ZFQ7 B1ZFQ8 B1ZFW9
B1ZFX0 B1ZFX6 B1ZFX7 B1ZG57 B1ZG93 B1ZG98 B1ZG99 B1ZGA0 B1ZGA6 B1ZGA7 B1ZGB4
B1ZGB5 B1ZGB8 B1ZGG1 B1ZGG2 B1ZGG8 B1ZGG9 B1ZGP5 B1ZGP6 B1ZGP9 B1ZGR1 B1ZGR2
B1ZGR8 B1ZGR9 B1ZGS0 B1ZGS1 B1ZGS7 B1ZGS8 B1ZGU8 B1ZGW7 B1ZGZ6 B1ZHG6 B1ZHG7
B1ZHJ4 B1ZHK6 B1ZHL0 B1ZHM8 B1ZHN6 B1ZHN7 B1ZHP3 B1ZI53 B1ZI64 B1ZIB2 B1ZIB4
B1ZIG8 B1ZIK6 B1ZIL2 B1ZIP6 B1ZJ09 B1ZJ12 B1ZJ42 B1ZJ47 B1ZJ73 B1ZJ78 B1ZJ88
B1ZJM3 B1ZJM9 B1ZJN1 B1ZJQ3 B1ZJQ5 B1ZJQ8 B1ZJR5 B1ZJU3 B1ZJW1 B1ZJX9 B1ZKA1
B1ZKB3 B1ZKE5 B1ZKG7 B1ZKI7 B1ZKR9 B1ZKT6 B1ZKX6 B1ZL15 B1ZL17 B1ZL59 B1ZL67
B1ZL68 B1ZL76 B1ZLB5 B1ZLB6 B1ZLC8 B1ZLG6 B1ZLG8 B1ZLJ9 B1ZLK0 B1ZLK1 B1ZLK3
B1ZLK4 B1ZLK5 B1ZLK6 B1ZLK7 B1ZLK8 B1ZLL0 B1ZLM0 B1ZLM1 B1ZLX0 B1ZLY7 B1ZMC1
B1ZMD5 B1ZMQ8 B1ZMR3 B1ZMW8 B1ZN03 B1ZN17 B1ZN99 B1ZNA6 B1ZNB4 B1ZNC2 B1ZNC4
B1ZNC6 B1ZND0 B1ZND3 B1ZND4 B1ZND5 B1ZND7 B1ZND8 B1ZND9 B1ZNE0 B1ZNE1 B1ZNE2
B1ZNE3 B1ZNE4 B1ZNE5 B1ZNE6 B1ZNE9 B1ZNF1 B1ZNF2 B1ZNL9 B1ZNY5 B1ZNZ6 B1ZNZ7
B1ZP06 B1ZP07 B1ZP08 B1ZP17 B1ZP50 B1ZP77 B1ZPB7 B1ZPC2 B1ZPC5 B1ZPF0 B1ZQ51
B1ZQM8 B1ZQN2 B1ZQN4 B1ZQS6 B1ZQW3 B1ZQZ5 B1ZR75 B1ZRZ1 B1ZS98 B1ZS99 B1ZSC4
B1ZSD3 B1ZSR1 B1ZSR5 B1ZSW3 B1ZSW8 B1ZSX8 B1ZT05 B1ZT49 B1ZT51 B1ZTJ1 B1ZTR5
B1ZU25 B1ZU74 B1ZUD7 B1ZUE3 B1ZUE6 B1ZUH7 B1ZUM3 B1ZUS0 B1ZUS4 B1ZUV7 B1ZUW6
B1ZUX5 B1ZUZ1 B1ZUZ4 B1ZVE0 B1ZVF7 B1ZVI7 B1ZVM5 B1ZVU0 B1ZVV8 B1ZW06 B1ZW12
B1ZW79 B1ZW80 B1ZW93 B1ZWC4 B1ZWF5 B1ZWG7 B1ZWH2 B1ZWL2 B1ZWP4 B1ZWP5 B1ZX44
B1ZX46 B1ZX57 B1ZXD4 B1ZXD5 B1ZXJ5 B1ZY08 B1ZYG2 B1ZYH0 B1ZYW1 B1ZZ32 B1ZZ37
B1ZZ40 B1ZZ94 B1ZZH6 B1ZZL7 B1ZZL8 B1ZZL9 B1ZZW8 B1ZZY2 B2A0B9 B2A0H0 B2A0K0
B2A0V2 B2A157 B2A158 B2A159 B2A170 B2A1A8 B2A1E8 B2A1G4 B2A1G7 B2A1H4 B2A1I5
B2A1I6 B2A1L9 B2A1M7 B2A1N7 B2A1Y5 B2A1Y6 B2A242 B2A250 B2A252 B2A259 B2A2C4
B2A2D7 B2A2F7 B2A2F9 B2A2G0 B2A2G3 B2A2G4 B2A2H0 B2A2I4 B2A2I8 B2A2J6 B2A2K1
B2A2K2 B2A2K6 B2A2L1 B2A2L7 B2A2M1 B2A2M3 B2A2N1 B2A2N7 B2A2N9 B2A2P0 B2A2P1
B2A2P2 B2A2P3 B2A2T4 B2A2U4 B2A2U7 B2A2U8 B2A2Y6 B2A2Y9 B2A2Z6 B2A2Z7 B2A2Z8
B2A302 B2A334 B2A335 B2A337 B2A380 B2A383 B2A390 B2A393 B2A397 B2A398 B2A3A2
B2A3A3 B2A3B2 B2A3C0 B2A3C4 B2A3D8 B2A3D9 B2A3H5 B2A3H6 B2A3J0 B2A3J2 B2A3J5
B2A3K5 B2A3L2 B2A3L9 B2A3N5 B2A3Q6 B2A3X6 B2A3X9 B2A3Y0 B2A449 B2A451 B2A454
B2A456 B2A469 B2A470 B2A474 B2A475 B2A499 B2A4B2 B2A4B3 B2A4C3 B2A4C6 B2A4C8
B2A4C9 B2A4D2 B2A4D4 B2A4D5 B2A4D6 B2A4D8 B2A4D9 B2A4E0 B2A4E1 B2A4E2 B2A4E3
B2A4E4 B2A4E5 B2A4E6 B2A4E7 B2A4E8 B2A4E9 B2A4F0 B2A4F1 B2A4F2 B2A4F3 B2A4F4
B2A4F6 B2A4F7 B2A4F8 B2A4G1 B2A4G2 B2A4J4 B2A4J5 B2A4M9 B2A4P7 B2A4P8 B2A4P9
B2A4Q0 B2A4Q1 B2A4Q5 B2A4Q6 B2A4Q7 B2A4R9 B2A4X5 B2A524 B2A526 B2A531 B2A532
B2A534 B2A549 B2A578 B2A5J5 B2A5J7 B2A5K1 B2A5K8 B2A5L1 B2A5L5 B2A5L6 B2A5L8
B2A5M4 B2A5N5 B2A5N9 B2A5P0 B2A5P2 B2A5S2 B2A5T8 B2A5V2 B2A5V3 B2A5V5 B2A5W1
B2A5W5 B2A5W6 B2A5W7 B2A5Z1 B2A610 B2A611 B2A618 B2A663 B2A685 B2A6B3 B2A6B5
B2A6B7 B2A6C2 B2A6E4 B2A6I7 B2A6V4 B2A6X0 B2A6Y0 B2A6Z3 B2A6Z7 B2A700 B2A713
B2A7A5 B2A7A8 B2A826 B2A827 B2A875 B2A8C9 B2A8E3 B2A8I5 B2A8L6 B2A8L8 B2A8M8
B2A8X1 B2AAJ5 B2ACC7 B2ACV0 B2AEZ5 B2AFA8 B2AFW1 B2AFZ7 B2AG17 B2AG22 B2AG25
B2AG26 B2AG37 B2AG42 B2AG45 B2AG52 B2AG76 B2AG85 B2AG98 B2AGA0 B2AGB6 B2AGB7
B2AGC8 B2AGE5 B2AGF3 B2AGF6 B2AGG2 B2AGH1 B2AGH6 B2AGP0 B2AGP7 B2AGQ0 B2AGT3
B2AGT6 B2AGT7 B2AGT9 B2AGU2 B2AGU6 B2AGU8 B2AGV9 B2AGY8 B2AH07 B2AH15 B2AH54
B2AH57 B2AH58 B2AH60 B2AH66 B2AH72 B2AH73 B2AH75 B2AH76 B2AH89 B2AH98 B2AHA4

B2AHB8 B2AHC1 B2AHC7 B2AHE3 B2AHM5 B2AI46 B2AIF3 B2AIJ5 B2AIJ6 B2AL40 B2AL82
B2AM55 B2AML6 B2AND4 B2AR36 B2AR91 B2ASU5 B2AT43 B2ATX1 B2AW39 B2AWF0 B2AWV6
B2AWZ9 B2AXG6 B2AY28 B2B0E2 B2B1Z3 B2B223 B2B2C5 B2B4I8 B2B4S6 B2B4X6 B2B5P2
B2B5Q3 B2B5V0 B2B674 B2B720 B2B738 B2B763 B2B766 B2B7W0 B2BMF7 B2BMF8 B2BMF9
B2BMQ6 B2BND9 B2BNE0 B2BNE1 B2BNE2 B2BNE3 B2BNE6 B2BNE7 B2BNE8 B2BNE9 B2BNF0
B2BRG2 B2BRG3 B2BRG4 B2BRG5 B2C4D0 B2C4J1 B2C4J3 B2C6F2 B2C6R6 B2CAZ2 B2CML6
B2CNY4 B2CQK0 B2CQK1 B2CW77 B2D6K9 B2D6M2 B2D6P4 B2DBE8 B2DBE9 B2DBF0 B2DBF1
B2DEU7 B2DEU8 B2DEV1 B2DFG5 B2FDA8 B2FHD7 B2FHH6 B2FHI7 B2FHW7 B2FI02 B2FI06
B2FI28 B2FI29 B2FIA5 B2FIA8 B2FIA9 B2FIC0 B2FIQ2 B2FIR3 B2FIR4 B2FIR7 B2FIS1
B2FIU9 B2FIV0 B2FIV8 B2FJ58 B2FJ68 B2FJ87 B2FJ92 B2FJE7 B2FJN5 B2FJP0 B2FJP9
B2FJS1 B2FJT4 B2FJU2 B2FJU3 B2FJU4 B2FJW0 B2FJW1 B2FJX2 B2FK05 B2FK06 B2FK26
B2FK84 B2FK85 B2FK90 B2FK94 B2FK95 B2FKA1 B2FKE2 B2FKE4 B2FKF0 B2FKG1 B2FKI7
B2FKJ1 B2FKJ7 B2FKJ8 B2FKK7 B2FKL1 B2FKL2 B2FKM0 B2FKN7 B2FKR1 B2FKV6 B2FL25
B2FL26 B2FL31 B2FL32 B2FL33 B2FL34 B2FL61 B2FL67 B2FL68 B2FL70 B2FL71 B2FL74
B2FL80 B2FL93 B2FL97 B2FL98 B2FLM4 B2FLM5 B2FLM6 B2FLQ1 B2FLR2 B2FLW4 B2FLX2
B2FM87 B2FM88 B2FMC1 B2FMD6 B2FMD7 B2FMF6 B2FMH4 B2FMH7 B2FMR5 B2FMS2 B2FMS4
B2FMX5 B2FMY4 B2FMY5 B2FMY6 B2FN00 B2FN41 B2FN57 B2FN73 B2FN75 B2FN76 B2FN79
B2FN86 B2FN87 B2FN89 B2FN90 B2FN92 B2FN98 B2FND0 B2FNE2 B2FNE3 B2FNF9 B2FNK2
B2FNK6 B2FNL3 B2FNL4 B2FNN0 B2FNN8 B2FNP3 B2FNP4 B2FNP6 B2FNQ5 B2FNQ6 B2FNR1
B2FNS1 B2FNY1 B2FNY7 B2FNZ1 B2FNZ5 B2FNZ6 B2FP01 B2FP10 B2FP14 B2FP36 B2FP40
B2FP48 B2FPA7 B2FPC7 B2FPG4 B2FPG6 B2FPH5 B2FPL6 B2FPL8 B2FPM0 B2FPM1 B2FPM3
B2FPM4 B2FPP2 B2FPP3 B2FPP4 B2FPR2 B2FPR4 B2FPR7 B2FPT0 B2FPW9 B2FPX2 B2FPX7
B2FQ14 B2FQ15 B2FQ21 B2FQ34 B2FQ35 B2FQ36 B2FQ37 B2FQ38 B2FQ39 B2FQ40 B2FQ41
B2FQ42 B2FQ44 B2FQ45 B2FQ46 B2FQ47 B2FQ48 B2FQ49 B2FQ50 B2FQ52 B2FQ53 B2FQ58
B2FQ70 B2FQ89 B2FQ90 B2FQA1 B2FQE7 B2FQI7 B2FQJ3 B2FQJ4 B2FQJ5 B2FQJ6 B2FQJ7
B2FQJ8 B2FQJ9 B2FQK1 B2FQK2 B2FQK5 B2FQK6 B2FQK7 B2FQK8 B2FQK9 B2FQL8 B2FQN4
B2FQR1 B2FQR2 B2FQR3 B2FQX2 B2FQX5 B2FR00 B2FR20 B2FR21 B2FR57 B2FR75 B2FR85
B2FRM5 B2FRN4 B2FRN6 B2FRN7 B2FRN8 B2FRP1 B2FRS3 B2FRV1 B2FRV3 B2FRW0 B2FRW1
B2FRX1 B2FRX8 B2FRZ7 B2FSC4 B2FSD0 B2FSD1 B2FSF3 B2FSI8 B2FSQ5 B2FSQ8 B2FT05
B2FT17 B2FT23 B2FT30 B2FT35 B2FT48 B2FT70 B2FT78 B2FT82 B2FTA6 B2FTB9 B2FTC1
B2FTD1 B2FTD2 B2FTD3 B2FTI7 B2FTJ0 B2FTY5 B2FU10 B2FU49 B2FU56 B2FU58 B2FU61
B2FU83 B2FUB3 B2FUB4 B2FUB5 B2FUB6 B2FUM0 B2FUP5 B2FUS8 B2FUT0 B2FUU6 B2FUV7
B2FUW1 B2G493 B2G4Y5 B2G4Y8 B2G4Z1 B2G4Z3 B2G4Z5 B2G510 B2G531 B2G532 B2G550
B2G552 B2G564 B2G575 B2G582 B2G590 B2G592 B2G5A1 B2G5A3 B2G5A9 B2G5B7 B2G5B8
B2G5C0 B2G5C2 B2G5I8 B2G5J0 B2G5K4 B2G5K8 B2G5M9 B2G5N5 B2G5Q7 B2G5S0 B2G5S5
B2G5S8 B2G5T1 B2G5T2 B2G5T6 B2G5T7 B2G5V1 B2G5V3 B2G5W6 B2G5W8 B2G5X5 B2G5X6
B2G5X7 B2G5Z2 B2G609 B2G611 B2G613 B2G616 B2G624 B2G626 B2G632 B2G634 B2G638
B2G649 B2G653 B2G669 B2G676 B2G679 B2G680 B2G6B4 B2G6C0 B2G6C3 B2G6E2 B2G6E3
B2G6E5 B2G6E6 B2G6E7 B2G6E8 B2G6F4 B2G6F5 B2G6F6 B2G6G1 B2G6H2 B2G6I6 B2G6J1
B2G6K0 B2G6K1 B2G6K4 B2G6K9 B2G6L2 B2G6L9 B2G6M7 B2G6M8 B2G6M9 B2G6N9 B2G6P1
B2G6Q2 B2G6Q8 B2G6Q9 B2G6R2 B2G6R3 B2G6R5 B2G6R8 B2G6R9 B2G6S0 B2G6U1 B2G6U2
B2G6U4 B2G6V0 B2G6V2 B2G6V6 B2G6V7 B2G6W1 B2G6W2 B2G6W3 B2G6W4 B2G6X4 B2G6X7
B2G6X9 B2G6Y3 B2G6Y6 B2G6Y7 B2G6Z1 B2G6Z4 B2G6Z6 B2G716 B2G718 B2G721 B2G729
B2G731 B2G7B1 B2G7E3 B2G7F2 B2G7F3 B2G7H0 B2G7Q2 B2G7Q4 B2G7U6 B2G7V8 B2G816

B2G818 B2G819 B2G821 B2G826 B2G832 B2G835 B2G842 B2G848 B2G849 B2G850 B2G851
B2G856 B2G858 B2G868 B2G870 B2G877 B2G882 B2G885 B2G896 B2G8A7 B2G8A8 B2G8B2
B2G8C8 B2G8D6 B2G8D8 B2G8G2 B2G8G3 B2G8K6 B2G8N1 B2G8T5 B2G8T6 B2G8T9 B2G8U2
B2G8U6 B2G8U7 B2G8V1 B2G8V2 B2G8V3 B2G8V4 B2G8V5 B2G8V7 B2G8V9 B2G8W0 B2G8W3
B2G8W4 B2G8W5 B2G8W6 B2G8W7 B2G8W8 B2G8X0 B2G8X1 B2G8X3 B2G8X4 B2G8X5 B2G8X6
B2G8X7 B2G8X8 B2G8X9 B2G8Y0 B2G8Y1 B2G8Y2 B2G8Y5 B2G944 B2G967 B2G994 B2G996
B2G9D0 B2G9H2 B2G9H4 B2G9H5 B2G9H7 B2G9J4 B2G9P6 B2G9W3 B2GA02 B2GA53 B2GA54
B2GA55 B2GA58 B2GA65 B2GA68 B2GA76 B2GA78 B2GA98 B2GAB3 B2GAB5 B2GAB9 B2GAC2
B2GAC3 B2GAC5 B2GAC6 B2GAD9 B2GAE1 B2GAF4 B2GAG0 B2GAH8 B2GAH9 B2GAI0 B2GAL0
B2GAL2 B2GAL4 B2GAL8 B2GAM5 B2GAM7 B2GAN2 B2GAN5 B2GAT1 B2GAT4 B2GAT5 B2GAV5
B2GAZ1 B2GAZ4 B2GB14 B2GB15 B2GB17 B2GB18 B2GB19 B2GB27 B2GB28 B2GB29 B2GB35
B2GB45 B2GB54 B2GB72 B2GB73 B2GB77 B2GB82 B2GB86 B2GB92 B2GBA3 B2GBB0 B2GBB7
B2GBB8 B2GBC2 B2GBC3 B2GBC5 B2GBC9 B2GBD0 B2GBD1 B2GBL8 B2GBL9 B2GBM5 B2GBN0
B2GBN3 B2GBN7 B2GBN8 B2GBP8 B2GBQ4 B2GBQ5 B2GBQ6 B2GBQ9 B2GBR2 B2GBR5 B2GBR7
B2GBR8 B2GBS2 B2GBV5 B2GBV7 B2GBW0 B2GBW2 B2GBW6 B2GBY6 B2GBY7 B2GBY9 B2GBZ2
B2GBZ5 B2GBZ7 B2GC09 B2GC10 B2GC11 B2GC12 B2GC19 B2GC33 B2GC35 B2GC39 B2GC51
B2GCB7 B2GCB8 B2GCD0 B2GCM0 B2GCN4 B2GCQ8 B2GCR0 B2GCR2 B2GCV0 B2GCV4 B2GCY7
B2GD06 B2GD08 B2GD23 B2GD25 B2GD26 B2GD28 B2GD42 B2GD48 B2GD51 B2GD55 B2GD58
B2GD65 B2GD67 B2GD68 B2GD71 B2GD73 B2GD80 B2GD87 B2GD90 B2GDA3 B2GDA5 B2GDB3
B2GDB4 B2GDB8 B2GDC4 B2GDC8 B2GDD5 B2GDD7 B2GDF8 B2GDG2 B2GDS5 B2GDS6 B2GDS9
B2GDT2 B2GDT7 B2GDT8 B2GDU2 B2GDU4 B2GDU5 B2GDU6 B2GDU8 B2GDV0 B2GDV1 B2GDV4
B2GDV5 B2GDV6 B2GDV7 B2GDV8 B2GDV9 B2GDW1 B2GDW2 B2GDW4 B2GDW5 B2GDW6 B2GDW7
B2GDW8 B2GDW9 B2GDX0 B2GDX1 B2GDX2 B2GDX3 B2GDX6 B2GE56 B2GE66 B2GE78 B2GE89
B2GEA0 B2GEF7 B2GEJ1 B2GEP6 B2GER9 B2GES0 B2GEU6 B2GEU7 B2GEU8 B2GEV1 B2GEV4
B2GEV6 B2GEV8 B2GEW8 B2GEY3 B2GEY4 B2GEZ1 B2GEZ3 B2GF13 B2GF15 B2GF22 B2GF64
B2GF71 B2GF73 B2GF74 B2GF76 B2GFD7 B2GFH1 B2GFJ0 B2GFJ7 B2GFK2 B2GFK7 B2GFL1
B2GFL8 B2GFM0 B2GFS8 B2GFU4 B2GFY1 B2GFY3 B2GFY4 B2GFY5 B2GG36 B2GG84 B2GG85
B2GG86 B2GGA2 B2GGA3 B2GGB4 B2GGB7 B2GGD7 B2GGD8 B2GGD9 B2GGG9 B2GGP0 B2GGU6
B2GGU7 B2GGU8 B2GGU9 B2GH80 B2GH81 B2GH83 B2GH92 B2GHC3 B2GHI3 B2GHI7 B2GHR6
B2GHT7 B2GHU5 B2GHV1 B2GI02 B2GI03 B2GI04 B2GI05 B2GI06 B2GI07 B2GI08 B2GI63
B2GI87 B2GI94 B2GI97 B2GIA1 B2GIA9 B2GIH3 B2GII3 B2GII4 B2GIJ3 B2GIJ4 B2GIK0
B2GIK9 B2GIL0 B2GIL1 B2GIL2 B2GIL3 B2GIL4 B2GIL9 B2GIM1 B2GIN9 B2GIP2 B2GIP6
B2GIZ5 B2GIZ6 B2GIZ7 B2GIZ8 B2GIZ9 B2GJ00 B2GJ03 B2GJ04 B2GJ05 B2GJ06 B2GJ07
B2GJ10 B2GJ13 B2GJ15 B2GJ17 B2GJ18 B2GJ19 B2GJ23 B2GJ26 B2GJ56 B2GJ87 B2GJ88
B2GJB5 B2GJD5 B2GJD6 B2GJD8 B2GJF5 B2GJF6 B2GJP2 B2GJQ6 B2GJQ7 B2GK12 B2GKC1
B2GKC8 B2GKD7 B2GKF3 B2GKF4 B2GKG9 B2GKH0 B2GKI3 B2GKI4 B2GKJ1 B2GKK6 B2GKN0
B2GKR5 B2GKR8 B2GKS1 B2GKS3 B2GKT3 B2GKT5 B2GKT6 B2GL12 B2GL19 B2GL27 B2GL48
B2GL64 B2GLJ3 B2GLQ7 B2GLR9 B2GLU6 B2GLW8 B2GLX4 B2GLZ4 B2GM30 B2GM31 B2GM49
B2GU58 B2GUB3 B2GUE2 B2GUN4 B2GUS6 B2GUV7 B2GUZ2 B2GV05 B2GV17 B2GV47 B2GV50
B2GV72 B2GV77 B2GV91 B2GVM7 B2HCT1 B2HCT2 B2HCT4 B2HCT5 B2HCT6 B2HCT7 B2HCT8
B2HCT9 B2HCU0 B2HCU3 B2HCU4 B2HCU5 B2HCV2 B2HCV3 B2HCV5 B2HCX0 B2HCX1 B2HCX2
B2HCX3 B2HCX5 B2HCZ0 B2HCZ1 B2HCZ2 B2HD09 B2HD59 B2HD96 B2HD97 B2HDA1 B2HDJ5
B2HDJ9 B2HDL4 B2HDP0 B2HDP8 B2HDR8 B2HDS6 B2HDU8 B2HDV9 B2HE69 B2HE73 B2HEB1
B2HEC3 B2HED1 B2HED8 B2HEF0 B2HEK5 B2HEN2 B2HEP6 B2HES1 B2HFA4 B2HFV4 B2HFV5

B2HFV6 B2HFW2 B2HFX5 B2HFY1 B2HGA4 B2HGH0 B2HGH1 B2HGJ2 B2HGL4 B2HGM7 B2HGN2
B2HGR0 B2HGR5 B2HGS5 B2HGW0 B2HGX6 B2HGY2 B2HGZ9 B2HH13 B2HH20 B2HH58 B2HHB4
B2HHE1 B2HHL3 B2HHL4 B2HHM7 B2HHY2 B2HI25 B2HI26 B2HI46 B2HI48 B2HI49 B2HI98
B2HIA2 B2HIB9 B2HIC0 B2HIE6 B2HIF0 B2HIF6 B2HIF7 B2HIH1 B2HIH4 B2HII1 B2HII2
B2HII5 B2HII6 B2HIJ2 B2HIJ9 B2HIK6 B2HJ17 B2HJ20 B2HJ21 B2HJ22 B2HJ23 B2HJ44
B2HJ53 B2HJJ6 B2HJL2 B2HJL4 B2HJL7 B2HJL9 B2HJM5 B2HJN3 B2HJP0 B2HJP3 B2HJP9
B2HJU4 B2HJU9 B2HK09 B2HK79 B2HKQ1 B2HKQ2 B2HKQ3 B2HKR9 B2HKS2 B2HKS3 B2HKV0
B2HKV3 B2HKW6 B2HKZ0 B2HL06 B2HL07 B2HL08 B2HL15 B2HL16 B2HLB2 B2HLB7 B2HLJ5
B2HLJ8 B2HLP4 B2HLR5 B2HLS4 B2HLS5 B2HLS6 B2HLY1 B2HM03 B2HM23 B2HM74 B2HM80
B2HMC3 B2HMC9 B2HME5 B2HME9 B2HMG0 B2HMG1 B2HMG2 B2HMG3 B2HMG5 B2HMM1 B2HMM5
B2HMS9 B2HMX9 B2HMZ9 B2HN48 B2HN50 B2HN61 B2HN62 B2HN63 B2HN75 B2HN79 B2HNC1
B2HNC2 B2HNC6 B2HNC7 B2HND2 B2HNE5 B2HNE6 B2HNF8 B2HNG2 B2HNG6 B2HNJ6 B2HNT4
B2HNT8 B2HNT9 B2HNW1 B2HNX5 B2HP18 B2HP20 B2HP45 B2HP50 B2HP60 B2HP67 B2HP68
B2HP72 B2HP73 B2HP75 B2HP85 B2HP86 B2HP98 B2HPD6 B2HPS1 B2HPW4 B2HPZ3 B2HPZ9
B2HQ02 B2HQ90 B2HQ91 B2HQ93 B2HQA3 B2HQA4 B2HQA8 B2HQA9 B2HQH3 B2HQI2 B2HQJ2
B2HQL3 B2HQL4 B2HQM5 B2HQN4 B2HQU3 B2HQU8 B2HQV2 B2HQV4 B2HQX7 B2HQX9 B2HR13
B2HR14 B2HR28 B2HR31 B2HR32 B2HR33 B2HR69 B2HR81 B2HRH5 B2HRJ2 B2HRL5 B2HRP4
B2HRP6 B2HRQ2 B2HRR5 B2HRU9 B2HS47 B2HS55 B2HS63 B2HSB3 B2HSF4 B2HSG5 B2HSH1
B2HSH2 B2HSJ3 B2HSJ4 B2HSJ8 B2HSL0 B2HSL1 B2HSL2 B2HSL3 B2HSN0 B2HSN1 B2HSN2
B2HSN3 B2HSN4 B2HSN5 B2HSN6 B2HSN7 B2HSN8 B2HSN9 B2HSP0 B2HSY2 B2HSZ1 B2HSZ2
B2HSZ3 B2HSZ4 B2HT44 B2HT62 B2HTB2 B2HTC4 B2HTC8 B2HTF7 B2HTG4 B2HTG5 B2HTG6
B2HTH0 B2HTH5 B2HTH6 B2HTI2 B2HTR5 B2HTT5 B2HTW5 B2HTX7 B2HU54 B2HU82 B2HUA3
B2HUA4 B2HUB2 B2HUI0 B2HUI9 B2HUJ5 B2HUL1 B2HUM1 B2HUM2 B2HUM5 B2HUM8 B2HUN4
B2HUN5 B2HUQ2 B2HUQ4 B2HUQ9 B2HUR7 B2HUR9 B2HUS7 B2HUU9 B2HUV1 B2HUY9 B2HUZ6
B2HV06 B2HV23 B2HV28 B2HV30 B2HV78 B2HV79 B2HV80 B2HVA8 B2HVC6 B2HVD2 B2HVE7
B2HVF8 B2HVG6 B2HVL6 B2HVM2 B2HVM9 B2HVN5 B2HVR7 B2HVR8 B2HVR9 B2HVS2 B2HVS4
B2HVT0 B2HVT7 B2HW19 B2HWE2 B2HWF6 B2HWK5 B2HWK7 B2HWK9 B2HWM8 B2HWM9 B2HWQ7
B2HWR7 B2HWT2 B2HWT3 B2HWU2 B2HWV2 B2HWW1 B2HWW3 B2HWX8 B2HWY0 B2HX34 B2HX50
B2HX82 B2HX89 B2HXA8 B2HXB6 B2HXC1 B2HXD3 B2HXG0 B2HXI0 B2HXI1 B2HXU5 B2HXU8
B2HXU9 B2HY00 B2HY23 B2HY33 B2HYF8 B2HYJ0 B2HYL7 B2HYN4 B2HYN7 B2HYP8 B2HYT2
B2HYT7 B2HYW1 B2HYX9 B2HYY2 B2HYY9 B2HYZ7 B2HZ03 B2HZ29 B2HZ52 B2HZ55 B2HZ57
B2HZ61 B2HZ83 B2HZ84 B2HZ85 B2HZ86 B2HZ87 B2HZ89 B2HZ90 B2HZ91 B2HZ95 B2HZ97
B2HZ98 B2HZ99 B2HZA0 B2HZA1 B2HZA2 B2HZA5 B2HZA7 B2HZB6 B2HZC3 B2HZE0 B2HZE4
B2HZE8 B2HZH0 B2HZI1 B2HZI2 B2HZI5 B2HZL7 B2HZL8 B2HZM0 B2HZM1 B2HZM2 B2HZM3
B2HZQ0 B2HZQ2 B2HZR3 B2HZS2 B2HZS3 B2HZV2 B2HZV3 B2HZV5 B2HZW3 B2HZW5 B2HZZ7
B2HZZ8 B2I023 B2I029 B2I056 B2I0A5 B2I0B1 B2I0B7 B2I0I2 B2I0I3 B2I0I4 B2I0I5
B2I0J2 B2I0K7 B2I0L0 B2I0N7 B2I0P3 B2I0Y7 B2I111 B2I1H7 B2I1H8 B2I1J3 B2I1J4
B2I1J9 B2I1L5 B2I1P1 B2I1P4 B2I1Q0 B2I1T3 B2I1U0 B2I1V6 B2I1W2 B2I1W8 B2I1W9
B2I1Y7 B2I1Y9 B2I1Z0 B2I1Z1 B2I1Z2 B2I2A2 B2I2A7 B2I2B2 B2I2C1 B2I2C4 B2I2E3
B2I2G4 B2I2G6 B2I2H6 B2I2H7 B2I2I7 B2I2L3 B2I2L6 B2I2M3 B2I2M5 B2I2M7 B2I2M8
B2I2P9 B2I2Q0 B2I2R9 B2I2S7 B2I2V4 B2I2V7 B2I316 B2I325 B2I330 B2I336 B2I352
B2I354 B2I359 B2I360 B2I367 B2I391 B2I392 B2I399 B2I3B3 B2I3B6 B2I3B8 B2I3C0
B2I3C2 B2I3D3 B2I3E1 B2I3E2 B2I3E5 B2I3E6 B2I3F0 B2I3G6 B2I3H4 B2I4L2 B2I4L7
B2I4M8 B2I4M9 B2I4P8 B2I4Q1 B2I4R3 B2I4S5 B2I4S7 B2I4S9 B2I4T1 B2I4U0 B2I4Y4

B2I4Y5 B2I596 B2I598 B2I5A7 B2I5B5 B2I5C0 B2I5K8 B2I5M3 B2I5R5 B2I5R6 B2I5S5
B2I5T5 B2I5T8 B2I5U9 B2I5W7 B2I5X2 B2I5X3 B2I5X4 B2I5X6 B2I5X7 B2I5X9 B2I5Y0
B2I5Y2 B2I5Y4 B2I5Y6 B2I5Y9 B2I5Z0 B2I5Z2 B2I604 B2I606 B2I607 B2I618 B2I621
B2I631 B2I637 B2I656 B2I663 B2I669 B2I672 B2I680 B2I691 B2I692 B2I696 B2I699
B2I6A6 B2I6A7 B2I6A8 B2I6A9 B2I6B4 B2I6C2 B2I6F5 B2I6F6 B2I6F7 B2I6F8 B2I6G4
B2I6G9 B2I6I5 B2I6I6 B2I6I8 B2I6K2 B2I6K3 B2I6K9 B2I6L3 B2I6M0 B2I6M1 B2I6M4
B2I6M7 B2I6N0 B2I6P5 B2I6Q0 B2I6Q1 B2I6R9 B2I6S4 B2I6S5 B2I6T4 B2I6V0 B2I6V2
B2I6V6 B2I6V7 B2I6W2 B2I6X0 B2I6X2 B2I6X3 B2I700 B2I705 B2I707 B2I720 B2I721
B2I723 B2I724 B2I726 B2I727 B2I729 B2I730 B2I757 B2I782 B2I7B1 B2I7C8 B2I7D3
B2I7D4 B2I7E8 B2I7G4 B2I7H1 B2I7H2 B2I7H7 B2I7J1 B2I7K9 B2I7L1 B2I7L2 B2I7M5
B2I7N4 B2I7N7 B2I7P4 B2I7Q9 B2I7R5 B2I7R6 B2I7S0 B2I7V0 B2I7V5 B2I7V6 B2I7V8
B2I7W6 B2I7X4 B2I7X5 B2I841 B2I844 B2I848 B2I858 B2I891 B2I897 B2I898 B2I8D4
B2I8F9 B2I8G8 B2I8G9 B2I8H0 B2I8H1 B2I8H2 B2I8H3 B2I8H4 B2I8H5 B2I8H6 B2I8H8
B2I8H9 B2I8I0 B2I8I1 B2I8I2 B2I8I3 B2I8I4 B2I8I5 B2I8I6 B2I8I7 B2I8J0 B2I8J1
B2I8J2 B2I8J3 B2I8J4 B2I8K2 B2I8K3 B2I8K4 B2I8L7 B2I8L8 B2I8M1 B2I8Q4 B2I8Q5
B2I8Q9 B2I8R0 B2I8R2 B2I8S3 B2I8S4 B2I8S7 B2I8S9 B2I8U1 B2I8U3 B2I8X7 B2I8X8
B2I8Y3 B2I8Y4 B2I916 B2I923 B2I925 B2I935 B2I939 B2I961 B2I965 B2I967 B2I985
B2I994 B2I9B2 B2I9C0 B2I9C6 B2I9E4 B2I9E6 B2I9F0 B2I9F9 B2I9G0 B2I9G1 B2I9H6
B2I9H7 B2I9I0 B2I9I6 B2I9J0 B2I9J1 B2I9J2 B2I9J3 B2I9L4 B2I9L5 B2I9M4 B2I9N9
B2I9P2 B2I9P4 B2I9P5 B2I9P6 B2I9P8 B2I9Q1 B2I9R4 B2I9S0 B2I9S4 B2I9S7 B2I9S8
B2I9S9 B2I9T6 B2I9U3 B2I9W5 B2I9W6 B2I9X1 B2I9X8 B2I9Y0 B2I9Y3 B2IA03 B2IA16
B2IA18 B2IA19 B2IA20 B2IA21 B2IA30 B2IA31 B2IA38 B2IA64 B2IA65 B2IA67 B2IA68
B2IA69 B2IA70 B2IA72 B2IA78 B2IA79 B2IA84 B2IA85 B2IA87 B2IAB2 B2IAE5 B2IAE7
B2IAE8 B2IAF0 B2IAI0 B2IAI3 B2IAK4 B2IAR9 B2IAW0 B2IAW3 B2IAW4 B2IAX9 B2IB12
B2IB44 B2IB58 B2IB74 B2IBC4 B2IBG2 B2IBG3 B2IBH3 B2IBH8 B2IBH9 B2IBR4 B2IBR5
B2IBR8 B2IBR9 B2IBT9 B2IBU5 B2IC04 B2IC30 B2IC83 B2ICH3 B2ICK3 B2ICL0 B2ICL3
B2ICL6 B2ICL8 B2ICM1 B2ICM2 B2ICM8 B2ICM9 B2ICQ8 B2ICR0 B2ICU3 B2ICU4 B2ICY4
B2ICY8 B2ICZ5 B2ID54 B2ID97 B2IDA4 B2IDB5 B2IDD9 B2IDE7 B2IDI9 B2IDK3 B2IDQ8
B2IDQ9 B2IDT5 B2IDU5 B2IDV3 B2IDV9 B2IDW3 B2IDX7 B2IDX8 B2IE09 B2IE16 B2IE42
B2IE44 B2IE46 B2IE47 B2IE54 B2IE63 B2IE81 B2IEA3 B2IEA4 B2IEA5 B2IEC1 B2IED0
B2IEE1 B2IEF8 B2IEM1 B2IEN5 B2IEP2 B2IEV6 B2IEW0 B2IEX5 B2IEZ6 B2IF46 B2IF48
B2IF93 B2IF94 B2IF96 B2IF97 B2IFA4 B2IFY8 B2IG63 B2IG64 B2IG85 B2IGB3 B2IGF1
B2IGF2 B2IGG1 B2IGG2 B2IGG3 B2IGH4 B2IGI6 B2IGK2 B2IGK3 B2IGL6 B2IGP2 B2IGP4
B2IGR2 B2IGT0 B2IGT1 B2IGU6 B2IGY0 B2IGZ5 B2IH12 B2IH14 B2IH50 B2IHC3 B2IHD1
B2IHD2 B2IHD4 B2IHG2 B2IHG4 B2IHJ7 B2IHL7 B2IHR4 B2IHR5 B2IHX3 B2IHY4 B2IHZ4
B2II34 B2II71 B2II88 B2IIA1 B2III9 B2IIJ1 B2IIJ6 B2IIJ7 B2IIK0 B2IIK5 B2IIK7
B2IJ38 B2IJF4 B2IJG0 B2IJG1 B2IJG9 B2IJH6 B2IJH9 B2IJJ3 B2IJK8 B2IJM5 B2IJQ3
B2IJQ4 B2IJR2 B2IJT2 B2IK51 B2IK52 B2IK53 B2IK54 B2IK56 B2IK57 B2IK58 B2IK59
B2IK61 B2IK62 B2IK63 B2IK65 B2IK66 B2IK67 B2IK68 B2IK69 B2IK71 B2IK72 B2IK73
B2IK75 B2IK76 B2IK77 B2IK78 B2IK79 B2IK80 B2IK81 B2IK83 B2IK87 B2IKA4 B2IKE2
B2IKJ9 B2IKL1 B2IKL7 B2IKL8 B2IKP4 B2IKQ2 B2IKT2 B2IKV4 B2IKV6 B2IKW1 B2IKW6
B2ILR2 B2ILR3 B2ILS5 B2ILU3 B2ILU5 B2ILU9 B2ILV2 B2ILX6 B2ILX7 B2ILX8 B2ILY0
B2ILY3 B2ILY8 B2ILZ1 B2ILZ2 B2ILZ5 B2ILZ6 B2IM11 B2IM28 B2IM29 B2IM38 B2IM39
B2IM47 B2IM49 B2IM67 B2IM97 B2IMA2 B2IMB5 B2IMD9 B2IME0 B2IME4 B2IMG7 B2IML6

B2IMN3 B2IMQ9 B2IMT4 B2IMU7 B2IMU8 B2IMY3 B2IMY5 B2IMY9 B2IMZ0 B2IMZ8 B2IN14
B2IN15 B2IN18 B2IN32 B2IN41 B2IN52 B2IN60 B2IN61 B2IN76 B2IN86 B2IN95 B2IN99
B2INA6 B2INC9 B2IND1 B2INE5 B2INF0 B2INF5 B2INF6 B2ING0 B2ING9 B2INH0 B2INH7
B2INK5 B2INK7 B2INM2 B2INN1 B2INN2 B2INP4 B2INV3 B2INW0 B2INZ9 B2IP11 B2IP51
B2IP53 B2IP67 B2IP79 B2IP88 B2IP90 B2IP98 B2IPA0 B2IPA2 B2IPB4 B2IPB5 B2IPC1
B2IPC9 B2IPG2 B2IPG3 B2IPH2 B2IPH6 B2IPI0 B2IPN4 B2IPN5 B2IPP4 B2IPQ3 B2IPS2
B2IPS4 B2IPT5 B2IPT6 B2IPT7 B2IPW6 B2IPW8 B2IPX3 B2IPX4 B2IPY4 B2IPY6 B2IPZ1
B2IQ02 B2IQ06 B2IQ16 B2IQ29 B2IQ55 B2IQ63 B2IQ67 B2IQ69 B2IQ70 B2IQ88 B2IQH7
B2IQJ0 B2IQJ2 B2IQJ3 B2IQJ4 B2IQJ5 B2IQK0 B2IQK6 B2IQM0 B2IQN8 B2IQR1 B2IQS1
B2IQT4 B2IQU0 B2IQU1 B2IQU2 B2IQY8 B2IQY9 B2IQZ3 B2IQZ8 B2IR26 B2IR37 B2IR53
B2IR58 B2IR60 B2IR66 B2IR74 B2IR75 B2IR78 B2IR81 B2IR82 B2IR84 B2IR86 B2IR87
B2IR95 B2IRA0 B2IRB1 B2IRB6 B2IRD4 B2IRG4 B2IRH1 B2IRK2 B2IRK6 B2IRL3 B2IRP8
B2IRR1 B2IRR4 B2IRR9 B2IRS6 B2IRS7 B2IRW4 B2IS10 B2IS11 B2IS15 B2IS27 B2IS28
B2IS29 B2IS39 B2IS40 B2IS41 B2IS42 B2IS43 B2IS44 B2IS45 B2IS46 B2IS47 B2IS48
B2IS49 B2IS50 B2IS51 B2IS52 B2IS53 B2IS54 B2IS55 B2IS58 B2IS59 B2IS63 B2IS65
B2IS66 B2IS68 B2IS85 B2IS88 B2IS93 B2ISD7 B2ISI0 B2ISI4 B2ISI9 B2ISJ7 B2ISJ8
B2ISJ9 B2ISL9 B2ISM4 B2ISQ2 B2ISQ9 B2ISS1 B2ISS3 B2ISS4 B2ISU6 B2ISV1 B2ISX3
B2ISX4 B2ISX5 B2ISX9 B2IT24 B2IT63 B2IT64 B2IT65 B2IT66 B2IT70 B2IT82 B2IT87
B2IT89 B2IT91 B2ITL1 B2ITL6 B2ITM5 B2ITM6 B2ITM7 B2ITN0 B2ITN2 B2ITN3 B2ITN4
B2ITN8 B2ITN9 B2ITP0 B2ITP1 B2ITP2 B2ITP3 B2ITP4 B2ITP5 B2ITP7 B2ITP8 B2ITP9
B2ITQ0 B2ITQ1 B2ITQ2 B2ITQ3 B2ITQ4 B2ITQ5 B2ITR9 B2ITU1 B2ITX3 B2IU18 B2IU46
B2IU50 B2IUA1 B2IUI4 B2IUK8 B2IUL7 B2IUM7 B2IUR7 B2IUT0 B2IUW7 B2IUY3 B2IV09
B2IV10 B2IVH7 B2IVH8 B2IVK6 B2IVM2 B2IVR7 B2IVU1 B2IVV0 B2IVV2 B2IVW4 B2IVW8
B2IVX7 B2IVZ4 B2IW30 B2IWE2 B2IWG6 B2IWH8 B2IWI8 B2IWJ2 B2IWJ8 B2IWK4 B2IWL4
B2IX22 B2IX27 B2IX28 B2IX33 B2IX35 B2IX54 B2IXH4 B2IY34 B2IY45 B2IY66 B2IY74
B2IY96 B2IYC1 B2IYD7 B2IYE4 B2IYI1 B2IYW1 B2IYX1 B2IYY1 B2IZ22 B2IZ57 B2IZ83
B2IZ89 B2IZE7 B2IZP5 B2IZP9 B2IZV5 B2J068 B2J0A5 B2J0A6 B2J0A9 B2J0C5 B2J0D0
B2J0D1 B2J0D3 B2J0D4 B2J0J6 B2J0Q5 B2J0Q6 B2J0V5 B2J116 B2J183 B2J191 B2J199
B2J1A0 B2J1A3 B2J1D2 B2J1G1 B2J1K8 B2J1L2 B2J1L6 B2J1L7 B2J1M1 B2J1M7 B2J1M8
B2J1S6 B2J1T0 B2J1W1 B2J1W7 B2J224 B2J2A2 B2J2A3 B2J2B2 B2J2D9 B2J2G1 B2J2U3
B2J2U6 B2J2V3 B2J2X2 B2J323 B2J384 B2J397 B2J3A7 B2J3C7 B2J3F1 B2J3G8 B2J3J3
B2J3M9 B2J3P0 B2J3X9 B2J427 B2J428 B2J434 B2J442 B2J448 B2J456 B2J471 B2J4A4
B2J4E5 B2J4H5 B2J528 B2J573 B2J588 B2J593 B2J5A8 B2J5A9 B2J5B0 B2J5B1 B2J5B2
B2J5E4 B2J5F1 B2J5F8 B2J5G0 B2J5J5 B2J5L9 B2J5M3 B2J5P1 B2J5P6 B2J5Q6 B2J5U5
B2J5Z9 B2J6C6 B2J6D0 B2J6D3 B2J6F7 B2J6I0 B2J6I3 B2J6K8 B2J6L1 B2J6L3 B2J6P8
B2J6T8 B2J6U2 B2J6U8 B2J6U9 B2J6X9 B2J707 B2J711 B2J717 B2J718 B2J719 B2J722
B2J764 B2J7E8 B2J7M9 B2J7T9 B2J7Y8 B2J876 B2J8B7 B2J8B8 B2J8F0 B2J8T2 B2J908
B2J913 B2J914 B2J918 B2J938 B2J955 B2J958 B2J997 B2J9G3 B2J9H6 B2J9P0 B2J9Q1
B2J9R3 B2J9R9 B2J9T1 B2JA15 B2JA20 B2JA48 B2JA49 B2JA71 B2JA74 B2JA77 B2JC42
B2JC45 B2JC95 B2JC98 B2JCG4 B2JCH8 B2JCI0 B2JCJ1 B2JCN4 B2JCN7 B2JCN8 B2JCP0
B2JCP3 B2JCP6 B2JCP7 B2JCQ7 B2JCU2 B2JCU8 B2JCV5 B2JCX3 B2JD61 B2JD86 B2JD88
B2JD95 B2JDC4 B2JDJ8 B2JDN2 B2JDN3 B2JDN6 B2JDN9 B2JDS4 B2JDT0 B2JDU3 B2JDU4
B2JDU7 B2JDV1 B2JDV3 B2JDY3 B2JDY5 B2JDY9 B2JDZ5 B2JDZ8 B2JDZ9 B2JE33 B2JE34
B2JE39 B2JE74 B2JED0 B2JED1 B2JED5 B2JEF5 B2JEG8 B2JEH0 B2JEI0 B2JEI3 B2JEJ7

B2JEN9 B2JEP3 B2JEP7 B2JEQ7 B2JEQ8 B2JER3 B2JET5 B2JEZ6 B2JF00 B2JF04 B2JF05
B2JF07 B2JF12 B2JF25 B2JF29 B2JF30 B2JF31 B2JF32 B2JF36 B2JF65 B2JF66 B2JF69
B2JF70 B2JFC8 B2JFD3 B2JFD4 B2JFE0 B2JFE1 B2JFE9 B2JFI7 B2JFL0 B2JFN9 B2JFP2
B2JFR4 B2JG18 B2JG20 B2JG21 B2JG54 B2JG59 B2JG60 B2JG67 B2JG80 B2JG93 B2JGD1
B2JGD2 B2JGD9 B2JGE1 B2JGE2 B2JGE4 B2JGE7 B2JGE8 B2JGE9 B2JGG8 B2JGI9 B2JGK2
B2JGK3 B2JGL4 B2JGL6 B2JGM3 B2JGP7 B2JGU1 B2JGU2 B2JGU4 B2JGU7 B2JGV0 B2JGV1
B2JGX3 B2JGX6 B2JGX9 B2JH19 B2JHD3 B2JHD4 B2JHD6 B2JHD7 B2JHD8 B2JHD9 B2JHE6
B2JHF8 B2JHF9 B2JHG2 B2JHG8 B2JHI7 B2JHI8 B2JHJ1 B2JHK7 B2JHK8 B2JHX9 B2JHY0
B2JHY3 B2JHY7 B2JI26 B2JI34 B2JI38 B2JI39 B2JI40 B2JI42 B2JI43 B2JI46 B2JI47
B2JI48 B2JI49 B2JI50 B2JI51 B2JI52 B2JI53 B2JI54 B2JI55 B2JI57 B2JI59 B2JI60
B2JI61 B2JI62 B2JIA0 B2JIA4 B2JIC0 B2JIC3 B2JIC5 B2JIC6 B2JID1 B2JID7 B2JID9
B2JIG2 B2JIG3 B2JIG4 B2JIG5 B2JIG6 B2JIG7 B2JIH0 B2JIH1 B2JIH3 B2JIH4 B2JIH5
B2JIH6 B2JIH7 B2JIH8 B2JIL8 B2JIM3 B2JIN1 B2JIP4 B2JIQ8 B2JIT0 B2JIU0 B2JIU1
B2JIU6 B2JIU7 B2JIV0 B2JIV3 B2JIV4 B2JIX2 B2JIY1 B2JIY6 B2JJ71 B2JJ72 B2JJ97
B2JJC9 B2JJF7 B2JJJ4 B2JJJ5 B2JJJ7 B2JJL1 B2JJN6 B2JJS1 B2JJT8 B2JJU4 B2JJV1
B2JJW1 B2JJW2 B2JJW3 B2JJW9 B2JJX2 B2JJX7 B2JJX8 B2JJY4 B2JJY6 B2JK28 B2JK72
B2JKA8 B2JKH4 B2JKH5 B2JKH6 B2JKI4 B2JKJ2 B2JKK7 B2JKL5 B2JKL7 B2JKL8 B2JKP8
B2JKT4 B2JKT6 B2JLI5 B2JLI8 B2JLM6 B2JLM7 B2JLS3 B2JMF6 B2JML0 B2JMM1 B2JN27
B2JND6 B2JNF0 B2JNZ4 B2JP23 B2JP66 B2JP67 B2JP68 B2JPU0 B2JQD2 B2JQE1 B2JQE2
B2JQE6 B2JQF0 B2JQI5 B2JQV7 B2JQV8 B2JQV9 B2JQW0 B2JQW2 B2JRI2 B2JRS8 B2JS24
B2JS87 B2JS88 B2JS89 B2JU32 B2JVN8 B2JX47 B2JX50 B2JYI8 B2JYK0 B2JYK5 B2JYK6
B2JYL4 B2JYM2 B2JYM4 B2JYM8 B2JYM9 B2JYN0 B2JYN3 B2JYN4 B2JYN5 B2JYN6 B2JYN7
B2JYP2 B2JYP4 B2JYQ0 B2JYQ7 B2JYS1 B2JYT3 B2JYT5 B2JYU0 B2JYU3 B2JYU4 B2JZ02
B2JZ09 B2JZ13 B2JZ20 B2JZ30 B2JZ31 B2JZ33 B2JZ34 B2JZ37 B2JZ38 B2JZ44 B2JZ46
B2JZ47 B2JZ56 B2JZ69 B2JZ70 B2JZA3 B2JZB1 B2JZB8 B2JZC3 B2JZC5 B2JZC6 B2JZD1
B2JZD3 B2JZD9 B2JZE3 B2JZE4 B2JZF3 B2JZF7 B2JZF9 B2JZG4 B2JZH0 B2JZH8 B2JZJ5
B2JZJ6 B2JZK3 B2JZK5 B2JZK6 B2JZM4 B2JZM5 B2JZM8 B2JZN0 B2JZP2 B2JZQ2 B2K045
B2K046 B2K065 B2K070 B2K072 B2K098 B2K0C8 B2K0H0 B2K0H2 B2K0H3 B2K0H6 B2K0N6
B2K0P7 B2K0Q3 B2K0Q4 B2K0Q5 B2K0R0 B2K0R4 B2K0R5 B2K0S0 B2K0S7 B2K0S8 B2K0T3
B2K0T4 B2K0V1 B2K0Y4 B2K0Z7 B2K103 B2K108 B2K109 B2K110 B2K111 B2K113 B2K114
B2K116 B2K121 B2K122 B2K124 B2K130 B2K198 B2K1U3 B2K1V8 B2K1W0 B2K1W1 B2K1W5
B2K1W6 B2K1X9 B2K1Y3 B2K1Y4 B2K1Y7 B2K1Z3 B2K1Z6 B2K202 B2K203 B2K204 B2K217
B2K295 B2K296 B2K297 B2K298 B2K2H0 B2K2H4 B2K2H9 B2K2I3 B2K2I4 B2K2K4 B2K2L3
B2K2L5 B2K2L6 B2K2M0 B2K2M6 B2K2N1 B2K2N4 B2K2N5 B2K2N9 B2K2P0 B2K2P2 B2K2P8
B2K2Q1 B2K2Q3 B2K2Q5 B2K2Q6 B2K2Q8 B2K2Q9 B2K2S3 B2K2T1 B2K2Y7 B2K2Z0 B2K303
B2K308 B2K312 B2K314 B2K315 B2K316 B2K319 B2K321 B2K322 B2K323 B2K324 B2K3B8
B2K3C6 B2K3E9 B2K3F6 B2K3F7 B2K3G3 B2K3H9 B2K3I2 B2K3J0 B2K3J1 B2K3K4 B2K3K5
B2K3L2 B2K3L4 B2K3L5 B2K3M0 B2K3M1 B2K3M4 B2K3M6 B2K3M9 B2K3N1 B2K3N7 B2K3N9
B2K3P0 B2K3P3 B2K3Q3 B2K3Q8 B2K3R4 B2K3S9 B2K3W1 B2K3W2 B2K3W4 B2K3Y3 B2K3Y4
B2K3Z6 B2K3Z7 B2K403 B2K419 B2K421 B2K422 B2K432 B2K461 B2K467 B2K469 B2K485
B2K486 B2K487 B2K488 B2K4B3 B2K4C6 B2K4C7 B2K4C9 B2K4D7 B2K4D8 B2K4E8 B2K4F9
B2K4G0 B2K4G3 B2K4H8 B2K4I1 B2K4I2 B2K4I4 B2K4J2 B2K4K7 B2K4L5 B2K4U5 B2K4Y6
B2K4Z5 B2K4Z9 B2K500 B2K502 B2K504 B2K505 B2K506 B2K511 B2K513 B2K514 B2K515
B2K516 B2K518 B2K519 B2K521 B2K522 B2K523 B2K524 B2K532 B2K533 B2K534 B2K547

B2K550 B2K553 B2K554 B2K560 B2K566 B2K567 B2K572 B2K577 B2K578 B2K579 B2K580
B2K581 B2K584 B2K5E0 B2K5H0 B2K5I4 B2K5J3 B2K5J4 B2K5K4 B2K5K6 B2K5K7 B2K5L8
B2K5L9 B2K5M0 B2K5M1 B2K5M2 B2K5M3 B2K5M4 B2K5M5 B2K5M6 B2K5M7 B2K5M9 B2K5N0
B2K5N1 B2K5N5 B2K5N6 B2K5N7 B2K5N8 B2K5N9 B2K5P0 B2K5P3 B2K5Q7 B2K5T4 B2K5T5
B2K5U7 B2K5U8 B2K5V5 B2K5V7 B2K5V9 B2K5W2 B2K5W3 B2K5X7 B2K5X9 B2K5Y0 B2K5Y8
B2K5Y9 B2K5Z0 B2K5Z1 B2K630 B2K631 B2K632 B2K633 B2K634 B2K659 B2K662 B2K664
B2K665 B2K666 B2K668 B2K682 B2K6A4 B2K6E0 B2K6E4 B2K6E6 B2K6F8 B2K6F9 B2K6G0
B2K6K6 B2K6P6 B2K6Q2 B2K6Q3 B2K6Q4 B2K6Q9 B2K6R0 B2K6R1 B2K6S5 B2K6S6 B2K6T1
B2K6T3 B2K6T4 B2K6T7 B2K6T9 B2K6U1 B2K6U4 B2K6V6 B2K6V8 B2K6W4 B2K6W6 B2K6Y0
B2K6Y1 B2K6Z1 B2K6Z3 B2K6Z6 B2K6Z7 B2K711 B2K719 B2K721 B2K730 B2K731 B2K751
B2K760 B2K767 B2K770 B2K776 B2K7D2 B2K7E5 B2K7E6 B2K7F7 B2K7F8 B2K7G3 B2K7H2
B2K7H7 B2K7H8 B2K7I5 B2K7I7 B2K7I8 B2K7I9 B2K7J2 B2K7J3 B2K7N2 B2K7N4 B2K7X9
B2K7Y0 B2K7Y6 B2K7Z0 B2K7Z2 B2K7Z3 B2K800 B2K824 B2K828 B2K830 B2K868 B2K872
B2K874 B2K875 B2K876 B2K881 B2K883 B2K886 B2K894 B2K896 B2K8A2 B2K8A4 B2K8D9
B2K8E3 B2K8F1 B2K8H7 B2K8I4 B2K8I9 B2K8Q8 B2K8R3 B2K8R6 B2K8S8 B2K8T0 B2K8T1
B2K8T5 B2K8U0 B2K8U4 B2K8U5 B2K8U6 B2K906 B2K913 B2K917 B2K945 B2K948 B2K962
B2K963 B2K985 B2K993 B2K9A4 B2K9F0 B2K9F1 B2K9F2 B2K9G8 B2K9G9 B2K9J1 B2K9J7
B2K9J9 B2K9K0 B2K9P0 B2K9P2 B2K9P6 B2K9P9 B2K9Q0 B2K9Q3 B2K9Q4 B2K9R0 B2K9R4
B2K9R5 B2K9S8 B2K9V8 B2K9X0 B2K9Z2 B2KA00 B2KA03 B2KA12 B2KA22 B2KA23 B2KA24
B2KA26 B2KA33 B2KA40 B2KA45 B2KA56 B2KA62 B2KA63 B2KA64 B2KA67 B2KA89 B2KAA0
B2KAA2 B2KAA3 B2KAA4 B2KAA6 B2KAB9 B2KAC9 B2KAD1 B2KAM4 B2KAQ0 B2KAR1 B2KAT6
B2KAU3 B2KAV3 B2KAW0 B2KAW2 B2KAX0 B2KAZ5 B2KB05 B2KB17 B2KB18 B2KB59 B2KB67
B2KB83 B2KB84 B2KB85 B2KBB9 B2KBC0 B2KBD0 B2KBD7 B2KBL5 B2KBN0 B2KBT7 B2KBU4
B2KBX0 B2KC30 B2KC44 B2KC76 B2KC93 B2KCB2 B2KCC0 B2KCD9 B2KCE9 B2KCF7 B2KCF8
B2KCI5 B2KCJ9 B2KCM6 B2KCN3 B2KCN5 B2KCP2 B2KCQ1 B2KCQ4 B2KCS5 B2KCS8 B2KCX0
B2KD56 B2KD65 B2KD81 B2KD82 B2KDH2 B2KDK9 B2KDP9 B2KDR9 B2KDU6 B2KDV7 B2KDX2
B2KDX3 B2KE25 B2KE31 B2KE32 B2KE60 B2KE70 B2KE80 B2KE94 B2KE95 B2KEA9 B2KEC4
B2KEF9 B2KEG0 B2KEG2 B2KEH0 B2KEJ2 B2KEJ4 B2KEJ6 B2KEJ7 B2KEK1 B2KEK3 B2KEK8
B2KEL0 B2KEL1 B2KEL4 B2KEL5 B2KEL6 B2KEL7 B2KEL8 B2KEM8 B2KEM9 B2KEN0 B2KEN2
B2KER5 B2KER6 B2KES3 B2KEU6 B2KEU8 B2KEV5 B2KEW4 B2KF05 B2KFW1 B2KGE5 B2KHZ4
B2KI57 B2KI88 B2KIQ4 B2KPR3 B2KSJ5 B2KSJ6 B2KWH7 B2KWH8 B2KWH9 B2KWI1 B2KWI4
B2LMH4 B2LMI1 B2LMI2 B2LMI3 B2LMJ1 B2LMK1 B2LML4 B2LML5 B2LML6 B2LMM6 B2LMM7
B2LMN0 B2LMN1 B2LMN3 B2LMN4 B2LMN7 B2LMN9 B2LMP1 B2LSD2 B2MUU6 B2MW49 B2MW51
B2NI93 B2NID7 B2RBV5 B2RC85 B2RD01 B2RFS9 B2RFT0 B2RGM5 B2RGP4 B2RGP8 B2RGQ9
B2RGR2 B2RGR5 B2RH08 B2RH13 B2RH32 B2RH34 B2RH41 B2RH62 B2RH69 B2RH81 B2RHB4
B2RHB7 B2RHC2 B2RHD7 B2RHE0 B2RHF2 B2RHF3 B2RHF5 B2RHI3 B2RHK7 B2RHL6 B2RHM7
B2RHM9 B2RHN4 B2RHN9 B2RHP0 B2RHR8 B2RHS7 B2RHV8 B2RI67 B2RI74 B2RI92 B2RID6
B2RIE4 B2RIE9 B2RIF2 B2RIF8 B2RIF9 B2RIG0 B2RIG3 B2RIG4 B2RIG5 B2RIG7 B2RIH0
B2RII6 B2RII9 B2RIJ1 B2RIJ4 B2RIJ5 B2RIL8 B2RIL9 B2RIR0 B2RIS4 B2RIT4 B2RIU1
B2RIU4 B2RIV3 B2RIV6 B2RIW6 B2RIY7 B2RIY8 B2RIZ0 B2RIZ3 B2RJ01 B2RJ07 B2RJ15
B2RJ75 B2RJ90 B2RJD6 B2RJD7 B2RJD8 B2RJE9 B2RJG1 B2RJH1 B2RJH4 B2RJI6 B2RJJ6
B2RJM1 B2RJN1 B2RJS2 B2RJT7 B2RJV4 B2RJW0 B2RJX5 B2RJY1 B2RK02 B2RK20 B2RK25
B2RK28 B2RK42 B2RK53 B2RKG6 B2RKJ1 B2RKL2 B2RKL4 B2RKS1 B2RKS4 B2RKS5 B2RKS6
B2RKT6 B2RKU2 B2RKW6 B2RKY9 B2RL24 B2RL29 B2RL41 B2RL44 B2RL45 B2RL46 B2RL48

B2RL49 B2RL52 B2RL54 B2RL61 B2RL62 B2RL63 B2RL67 B2RL77 B2RL89 B2RLA0 B2RLA8
B2RLC1 B2RLC2 B2RLG7 B2RLG9 B2RLH2 B2RLI5 B2RLI9 B2RLJ0 B2RLS5 B2RLV1 B2RLV5
B2RLV6 B2RLW4 B2RLW6 B2RLW7 B2RLW8 B2RLW9 B2RLX3 B2RLX4 B2RLX6 B2RLX7 B2RLX8
B2RLX9 B2RLY0 B2RLY1 B2RLY2 B2RLY3 B2RLY4 B2RLY5 B2RLY6 B2RLY7 B2RLY8 B2RLY9
B2RLZ1 B2RLZ2 B2RLZ3 B2RLZ4 B2RLZ5 B2RLZ6 B2RM14 B2RM15 B2RM62 B2RM63 B2RM78
B2RM92 B2RMA5 B2RMB4 B2RMB9 B2RMD0 B2RME4 B2RMG0 B2RMI0 B2RMJ1 B2RMK4 B2RPK0
B2RPU2 B2RQC6 B2RR83 B2RRE4 B2RRF6 B2RRL2 B2RS91 B2RU33 B2RUI1 B2RUR4 B2RV13
B2RV46 B2RVL6 B2RW38 B2RW88 B2RX14 B2RX88 B2RXB0 B2RXB2 B2RXC1 B2RXC5 B2RXH2
B2RXH4 B2RXH8 B2RXM4 B2RXR6 B2RY50 B2RY56 B2RY71 B2RYD2 B2RYD6 B2RYF1 B2RYG1
B2RYG8 B2RYJ4 B2RYJ8 B2RYL1 B2RYM0 B2RYM5 B2RYN2 B2RYR0 B2RYT9 B2RYU6 B2RYU8
B2RYW9 B2RYZ5 B2RZ17 B2RZ39 B2RZ50 B2RZ86 B2RZC4 B2RZD7 B2RZI6 B2RZI7 B2RZI8
B2RZJ4 B2RZJ7 B2RZL5 B2RZN8 B2RZP4 B2RZP5 B2RZP9 B2RZQ0 B2RZQ1 B2RZQ6 B2RZR0
B2RZS4 B2RZT0 B2RZU2 B2RZV0 B2RZW1 B2RZW7 B2S007 B2S017 B2S045 B2S046 B2S048
B2S057 B2S074 B2S075 B2S076 B2S089 B2S090 B2S091 B2S092 B2S093 B2S094 B2S095
B2S099 B2S0D7 B2S0D9 B2S0E3 B2S0E4 B2S0F0 B2S0F9 B2S0G4 B2S0H9 B2S0I0 B2S0I1
B2S0I5 B2S0I6 B2S0I7 B2S0I8 B2S0I9 B2S0J0 B2S0J1 B2S0J2 B2S0J3 B2S0J4 B2S0J5
B2S0J6 B2S0J7 B2S0J9 B2S0K2 B2S0K3 B2S0K6 B2S0L6 B2S0M0 B2S0M1 B2S0M7 B2S0Q2
B2S0T6 B2S0U2 B2S0U7 B2S0U9 B2S0V7 B2S0W1 B2S0W8 B2S100 B2S101 B2S125 B2S137
B2S140 B2S145 B2S146 B2S162 B2S197 B2S1B0 B2S1C2 B2S1C3 B2S1C4 B2S1C7 B2S1C8
B2S1C9 B2S1E1 B2S1E3 B2S1E6 B2S1E8 B2S1E9 B2S1F3 B2S1G5 B2S1H8 B2S1K1 B2S1K4
B2S1L1 B2S1L2 B2S1L3 B2S1N3 B2S1N8 B2S1P0 B2S1P1 B2S1P5 B2S1P9 B2S1S1 B2S1U6
B2S1U7 B2S1U9 B2S1V1 B2S1W3 B2S1W7 B2S1X8 B2S1Z2 B2S1Z3 B2S1Z9 B2S208 B2S209
B2S211 B2S216 B2S239 B2S264 B2S270 B2S2A7 B2S2B5 B2S2D1 B2S2D5 B2S2D6 B2S2D7
B2S2D8 B2S2D9 B2S2E0 B2S2E1 B2S2E2 B2S2E3 B2S2E4 B2S2E5 B2S2E6 B2S2E7 B2S2E8
B2S2E9 B2S2F0 B2S2F1 B2S2F2 B2S2F3 B2S2F4 B2S2F5 B2S2F7 B2S2F8 B2S2F9 B2S2G1
B2S2G4 B2S2I1 B2S2I4 B2S2I5 B2S2I6 B2S2I7 B2S2I9 B2S2J0 B2S2J1 B2S2J8 B2S2K3
B2S2L2 B2S2N1 B2S2Q3 B2S2Q4 B2S2S5 B2S2T4 B2S2T8 B2S2U7 B2S2U8 B2S2V0 B2S2V9
B2S2W8 B2S2W9 B2S2X0 B2S2Y0 B2S2Y1 B2S2Z5 B2S303 B2S328 B2S334 B2S337 B2S348
B2S360 B2S367 B2S369 B2S370 B2S371 B2S3A0 B2S3A2 B2S3B1 B2S3B8 B2S3C9 B2S3D0
B2S3D5 B2S3D9 B2S3E2 B2S3E8 B2S3F1 B2S3H7 B2S3I2 B2S3I6 B2S3J4 B2S3J5 B2S3J6
B2S3J9 B2S3K6 B2S3M4 B2S3M6 B2S3N0 B2S3N6 B2S3N9 B2S3P6 B2S3P7 B2S3Q9 B2S3R1
B2S3R2 B2S3R9 B2S3S0 B2S3S1 B2S3S8 B2S3T1 B2S3X1 B2S3Y0 B2S3Y1 B2S3Y2 B2S3Y4
B2S3Z3 B2S3Z5 B2S3Z6 B2S3Z7 B2S433 B2S446 B2S448 B2S457 B2S462 B2S468 B2S469
B2S475 B2S486 B2S487 B2S488 B2S4C2 B2S4C3 B2S4C4 B2S4C6 B2S4C7 B2S4E2 B2S4E6
B2S4F0 B2S4I3 B2S4I8 B2S4L0 B2S4M2 B2S4P7 B2S4P8 B2S4Q0 B2S4Q2 B2S4Q7 B2S4Q8
B2S4R0 B2S4R1 B2S4R2 B2S4S6 B2S4T7 B2S4U1 B2S4V2 B2S4W8 B2S501 B2S504 B2S513
B2S514 B2S518 B2S525 B2S533 B2S535 B2S536 B2S561 B2S567 B2S568 B2S573 B2S578
B2S5A1 B2S5B0 B2S5B8 B2S5B9 B2S5D1 B2S5D2 B2S5D7 B2S5E0 B2S5I0 B2S5I1 B2S5K6
B2S5M8 B2S5Q9 B2S5S6 B2S5T0 B2S5T1 B2S5U5 B2S5U8 B2S5W0 B2S5W1 B2S5W3 B2S5X5
B2S5Y5 B2S5Z0 B2S5Z1 B2S5Z2 B2S5Z7 B2S608 B2S610 B2S611 B2S618 B2S630 B2S641
B2S648 B2S654 B2S655 B2S656 B2S657 B2S658 B2S660 B2S661 B2S662 B2S663 B2S664
B2S665 B2S666 B2S667 B2S668 B2S669 B2S670 B2S671 B2S672 B2S673 B2S675 B2S676
B2S677 B2S678 B2S679 B2S680 B2S682 B2S683 B2S684 B2S686 B2S687 B2S688 B2S689
B2S690 B2S691 B2S6B8 B2S6C0 B2S6D5 B2S6E8 B2S6K5 B2S6L0 B2S6L5 B2S6P1 B2S6P4

B2S6Q2 B2S6Q3 B2S6R2 B2S6U9 B2S6W5 B2S6Z6 B2S701 B2S710 B2S746 B2S793 B2S7B5
B2S7D1 B2S7D9 B2S7E0 B2S7E1 B2S7E6 B2S7F2 B2S7H9 B2S7K3 B2S7K6 B2S7L7 B2S7M9
B2S7N1 B2S7N5 B2S7P0 B2S7P8 B2S7T0 B2S7T1 B2S7T7 B2S7X3 B2S7X6 B2S7X9 B2S7Z5
B2S7Z7 B2S801 B2S805 B2S806 B2S810 B2S811 B2S825 B2S828 B2S842 B2S843 B2S860
B2S861 B2S868 B2S869 B2S877 B2S880 B2S881 B2S885 B2S889 B2S8A9 B2S8D8 B2S8E6
B2S8G3 B2S8G4 B2S8H0 B2S8J9 B2S8M9 B2S8N8 B2S8R7 B2S8S9 B2S903 B2S906 B2S928
B2S946 B2S959 B2S965 B2S968 B2S971 B2S978 B2S979 B2S980 B2S983 B2S984 B2S985
B2S986 B2S994 B2S9A3 B2S9A5 B2S9A6 B2S9B5 B2S9B6 B2S9B8 B2S9C2 B2S9C4 B2S9E5
B2S9F3 B2S9F7 B2S9F9 B2S9G0 B2S9H8 B2S9H9 B2S9I5 B2S9M3 B2S9Q7 B2S9R7 B2S9T2
B2S9T4 B2S9T6 B2S9V1 B2S9V3 B2S9V4 B2S9W9 B2S9X0 B2S9Y5 B2SA00 B2SA37 B2SA42
B2SAC5 B2SAC9 B2SAD1 B2SAD9 B2SAE8 B2SAF6 B2SAG4 B2SAM1 B2SAP4 B2SAR1 B2SAU5
B2SAW2 B2SAY0 B2SB47 B2SB57 B2SB59 B2SB60 B2SB61 B2SB62 B2SB89 B2SBF0 B2SBF2
B2SBF6 B2SBN5 B2SBQ3 B2SBS0 B2SBS2 B2SBS5 B2SC12 B2SC15 B2SC17 B2SC20 B2SC33
B2SC34 B2SC37 B2SC50 B2SC51 B2SC52 B2SC53 B2SCB2 B2SCP3 B2SCQ3 B2SCV8 B2SCY7
B2SCY8 B2SCY9 B2SCZ4 B2SCZ5 B2SD15 B2SD64 B2SD81 B2SD94 B2SD96 B2SD97 B2SDA2
B2SDB5 B2SDD4 B2SDE2 B2SDE3 B2SDE7 B2SDE8 B2SDF5 B2SDF7 B2SDH2 B2SDI9 B2SDJ2
B2SDK3 B2SDK6 B2SDK7 B2SDK9 B2SDL2 B2SDL4 B2SDL5 B2SDM1 B2SDP9 B2SDQ1 B2SDQ4
B2SDR3 B2SDR8 B2SDS9 B2SDV9 B2SDW1 B2SDW2 B2SDW3 B2SDW4 B2SDW6 B2SDW7 B2SDW9
B2SDX0 B2SDX1 B2SDX2 B2SDX3 B2SDX4 B2SDX5 B2SDX6 B2SDX7 B2SDX8 B2SDX9 B2SDY0
B2SDY1 B2SDY2 B2SDY3 B2SDY4 B2SDY5 B2SDY6 B2SDY7 B2SDY8 B2SDY9 B2SDZ1 B2SDZ4
B2SDZ6 B2SE01 B2SE34 B2SE44 B2SE51 B2SE57 B2SE64 B2SE78 B2SEA8 B2SEE2 B2SEF6
B2SEJ5 B2SEJ7 B2SEK7 B2SEK8 B2SEM6 B2SEM7 B2SEQ2 B2SEQ6 B2SES8 B2SEW4 B2SEW5
B2SEW7 B2SEW8 B2SEW9 B2SEZ5 B2SEZ6 B2SEZ7 B2SF22 B2SF23 B2SF32 B2SF43 B2SF54
B2SF67 B2SF73 B2SF85 B2SF88 B2SF94 B2SFA2 B2SFC9 B2SFD2 B2SFD3 B2SFD4 B2SFD5
B2SFD6 B2SFD7 B2SFD9 B2SFE1 B2SFE2 B2SFE3 B2SFE4 B2SFE5 B2SFE8 B2SFF5 B2SFF6
B2SFF8 B2SFF9 B2SFG1 B2SFH9 B2SFJ3 B2SFK0 B2SFK1 B2SFK9 B2SFL4 B2SFL7 B2SFM1
B2SFM3 B2SFM4 B2SFM7 B2SFM9 B2SFN0 B2SFN3 B2SFQ6 B2SFR3 B2SFS8 B2SFS9 B2SFT1
B2SFU4 B2SFU7 B2SFV9 B2SFW5 B2SFW7 B2SFY4 B2SG03 B2SG11 B2SG13 B2SG17 B2SG18
B2SG23 B2SG24 B2SG30 B2SG31 B2SG47 B2SG50 B2SG83 B2SG84 B2SGA3 B2SGC2 B2SGC8
B2SGC9 B2SGE0 B2SGE3 B2SGE5 B2SGG3 B2SGG5 B2SGG7 B2SGG9 B2SGH0 B2SGH2 B2SGH6
B2SGI4 B2SGJ2 B2SGK2 B2SGK4 B2SGK5 B2SGL0 B2SGL8 B2SGM6 B2SGM9 B2SGN1 B2SGN2
B2SGQ7 B2SGS0 B2SGS1 B2SGS3 B2SGS9 B2SGT0 B2SGV8 B2SGV9 B2SGW2 B2SGW7 B2SH29
B2SH34 B2SH45 B2SH51 B2SH55 B2SH67 B2SH69 B2SH70 B2SH72 B2SH83 B2SH90 B2SH99
B2SHB2 B2SHB9 B2SHD4 B2SHF6 B2SHI9 B2SHJ1 B2SHP3 B2SHP8 B2SHQ8 B2SHR2 B2SHR3
B2SHS9 B2SHW2 B2SHW6 B2SI03 B2SI05 B2SI06 B2SI21 B2SI43 B2SIG7 B2SIG8 B2SII2
B2SIL3 B2SIN8 B2SIS8 B2SIT6 B2SIT8 B2SIV0 B2SIV1 B2SIV2 B2SIW1 B2SIZ4 B2SIZ5
B2SJ67 B2SJF5 B2SJG3 B2SJG4 B2SJI1 B2SJJ1 B2SJJ3 B2SJL4 B2SJL8 B2SJL9 B2SJM1
B2SJV1 B2SJV2 B2SJV3 B2SJY0 B2SJY7 B2SJZ7 B2SJZ8 B2SK35 B2SK49 B2SK75 B2SK80
B2SKA9 B2SKC6 B2SKI8 B2SKN4 B2SKN6 B2SKN9 B2SKP3 B2SKP5 B2SKP7 B2SKS0 B2SL02
B2SL03 B2SL06 B2SL09 B2SL10 B2SL46 B2SL50 B2SL54 B2SLA2 B2SLA3 B2SLC0 B2SLG1
B2SLG3 B2SLH4 B2SLH6 B2SLH7 B2SLL1 B2SLM6 B2SLQ8 B2SLT7 B2SM13 B2SM18 B2SM33
B2SM82 B2SM83 B2SM84 B2SMB3 B2SMD7 B2SMI2 B2SMI3 B2SMI4 B2SMP6 B2SMV3 B2SN02
B2SNG3 B2SNG5 B2SNG9 B2SNH3 B2SNH4 B2SNJ3 B2SNS8 B2SNV6 B2SNW4 B2SNW5 B2SNY1
B2SNY6 B2SNZ4 B2SPE7 B2SPF0 B2SPG9 B2SPN6 B2SPP5 B2SPT1 B2SPT2 B2SPT3 B2SPT4

B2SPV7 B2SQ45 B2SQ77 B2SQI9 B2SQJ4 B2SQN5 B2SQN6 B2SQP2 B2SQP7 B2SQP8 B2SQP9
B2SQQ0 B2SQQ1 B2SQQ2 B2SQQ3 B2SQQ4 B2SQQ9 B2SQR0 B2SQR1 B2SQR2 B2SQR4 B2SQR5
B2SQR7 B2SQR8 B2SQR9 B2SQS0 B2SQS1 B2SQS2 B2SQS3 B2SQS6 B2SQS7 B2SQS8 B2SQT1
B2SQT2 B2SQT3 B2SQT4 B2SQU3 B2SQU4 B2SQU5 B2SQV2 B2SQV8 B2SQY5 B2SQZ9 B2SR02
B2SR05 B2SR68 B2SRB2 B2SRE3 B2SRH1 B2SRI5 B2SRM7 B2SRM8 B2SRQ4 B2SRX5 B2SS19
B2SS21 B2SS22 B2SS56 B2SS66 B2SS67 B2SS96 B2SS98 B2SSC6 B2SSI8 B2SSM9 B2SSQ0
B2SSQ1 B2SSW1 B2SSW2 B2SSW3 B2ST87 B2STA8 B2STB9 B2STC0 B2STC1 B2STC6 B2STI7
B2STK0 B2STK2 B2STK3 B2STK4 B2STK7 B2STN5 B2STN6 B2STP6 B2STU3 B2SU21 B2SU25
B2SUA3 B2SUA8 B2SUA9 B2SUB2 B2SUB3 B2SUB8 B2SUC1 B2SUC8 B2SUC9 B2SUD0 B2SUD1
B2SUK8 B2SUL9 B2SUM0 B2SUM1 B2SUN1 B2SUV8 B2SUW2 B2SUW3 B2SUX4 B2SUX7 B2SV55
B2SV69 B2SV97 B2SVJ9 B2SVK0 B2SVK3 B2SVK5 B2SVM1 B2SVN3 B2SVN5 B2SVN7 B2SVN9
B2SVP0 B2SVP7 B2SVT3 B2SW86 B2SWC0 B2SWC5 B2SWE7 B2SWF4 B2SWG5 B2SWH2 B2SWK7
B2SWS5 B2SWS7 B2SWU4 B2SWV4 B2SWX0 B2SWY1 B2SWY4 B2SX15 B2SX18 B2SX29 B2SX35
B2SX42 B2SX63 B2SX74 B2SX83 B2SXB6 B2SXB7 B2SXB8 B2SXC3 B2SXC5 B2SXC6 B2SXC7
B2SXG0 B2SXG3 B2SXG4 B2SXG6 B2SXG9 B2SXH2 B2SXH3 B2SXI3 B2SXN2 B2SXN5 B2SXQ9
B2SXR9 B2SXS0 B2SXS5 B2SXS6 B2SXS9 B2SXT2 B2SXT3 B2SXV1 B2SXV7 B2SXY6 B2SXZ9
B2SY60 B2SY63 B2SY64 B2SY65 B2SY78 B2SY84 B2SY86 B2SYE3 B2SYE8 B2SYH7 B2SYI5
B2SYI7 B2SYJ7 B2SYJ9 B2SYK1 B2SYK2 B2SYK5 B2SYK8 B2SYK9 B2SYL0 B2SYN3 B2SYT3
B2SYU8 B2SYV1 B2SYV2 B2SYV3 B2SYV4 B2SYW1 B2SYX3 B2SYX4 B2SYX8 B2SYY3 B2SYY4
B2SZ04 B2SZ26 B2SZ27 B2SZ59 B2SZ60 B2SZ63 B2SZ71 B2SZ75 B2SZF8 B2SZF9 B2SZG0
B2SZG2 B2SZH3 B2SZN0 B2SZQ7 B2SZR6 B2SZR9 B2SZU5 B2SZU7 B2T067 B2T068 B2T084
B2T0H9 B2T0I0 B2T0I4 B2T0I5 B2T0I7 B2T0P6 B2T0P7 B2T146 B2T167 B2T171 B2T172
B2T181 B2T1A6 B2T1B6 B2T1B8 B2T1C7 B2T1D1 B2T1K6 B2T1L3 B2T1M3 B2T1M4 B2T1M5
B2T1N1 B2T1N4 B2T1N8 B2T1N9 B2T1P5 B2T1P7 B2T236 B2T256 B2T2A0 B2T2A3 B2T2A5
B2T2A6 B2T2D1 B2T2E3 B2T2H9 B2T2S1 B2T2S2 B2T2S5 B2T2Y2 B2T2Y6 B2T2Z5 B2T314
B2T319 B2T379 B2T381 B2T382 B2T3F4 B2T3H8 B2T3H9 B2T3I1 B2T3J1 B2T3K5 B2T3N3
B2T3N5 B2T3N6 B2T3R7 B2T3S4 B2T3S5 B2T3U5 B2T3X2 B2T3X3 B2T3Z9 B2T404 B2T411
B2T421 B2T436 B2T4H0 B2T4N1 B2T4U5 B2T4Y1 B2T5G8 B2T5H8 B2T5I0 B2T5I8 B2T5J1
B2T5J3 B2T5J4 B2T5K0 B2T5K6 B2T5K8 B2T5L6 B2T5L9 B2T5N8 B2T5R7 B2T5U4 B2T600
B2T604 B2T605 B2T606 B2T607 B2T612 B2T625 B2T629 B2T630 B2T634 B2T637 B2T641
B2T672 B2T699 B2T6B2 B2T6B5 B2T6C2 B2T6D0 B2T6D1 B2T6H6 B2T6H7 B2T6H8 B2T6I7
B2T6J1 B2T6J4 B2T6M2 B2T6M4 B2T6U6 B2T6V1 B2T6V9 B2T6X2 B2T711 B2T719 B2T724
B2T725 B2T726 B2T728 B2T729 B2T732 B2T733 B2T734 B2T735 B2T736 B2T737 B2T738
B2T739 B2T740 B2T741 B2T742 B2T743 B2T744 B2T745 B2T746 B2T747 B2T748 B2T749
B2T750 B2T751 B2T752 B2T755 B2T756 B2T758 B2T759 B2T760 B2T761 B2T762 B2T763
B2T792 B2T7I7 B2T7I8 B2T7J6 B2T7L1 B2T7M8 B2T7N6 B2T7N8 B2T7U4 B2T7X5 B2T8L2
B2T929 B2T9B1 B2T9I7 B2T9N0 B2T9N4 B2T9P8 B2TA88 B2TBH5 B2TBS1 B2TBS5 B2TBV3
B2TCB2 B2TCJ8 B2TCJ9 B2TCK0 B2TCR1 B2TD41 B2TDB3 B2TDV9 B2TEK6 B2TFB8 B2TFJ4
B2TFM4 B2TFM7 B2TFQ5 B2TG13 B2TGB8 B2THB4 B2THB7 B2THC5 B2THC7 B2THC8 B2THG5
B2THH3 B2THM0 B2THM7 B2THN0 B2THN1 B2THN2 B2THN3 B2THP9 B2THQ9 B2THR6 B2THR8
B2THS2 B2THS4 B2THS7 B2THS9 B2THT3 B2THU3 B2THZ1 B2TI01 B2TI04 B2TI06 B2TI10
B2TI22 B2TI25 B2TI29 B2TIA8 B2TIB6 B2TIB7 B2TIB8 B2TIC3 B2TIC9 B2TIE4 B2TIF4
B2TIF5 B2TIF6 B2TIG4 B2TIG5 B2TIG7 B2TIG8 B2TIG9 B2TIH0 B2TIH1 B2TIH2 B2TIH4
B2TIH5 B2TIH6 B2TIH7 B2TIH8 B2TIH9 B2TII1 B2TII2 B2TII3 B2TII4 B2TII5 B2TII6

B2TII7 B2TII8 B2TII9 B2TIJ0 B2TIJ1 B2TIJ3 B2TIJ4 B2TIJ6 B2TIK0 B2TIK1 B2TIK3
B2TIK4 B2TIK5 B2TIK9 B2TIL0 B2TIL1 B2TIN4 B2TIN7 B2TIQ9 B2TIR0 B2TIR2 B2TIR4
B2TIT5 B2TIU0 B2TIV9 B2TIX0 B2TIX3 B2TIX6 B2TIZ1 B2TIZ7 B2TIZ8 B2TIZ9 B2TJ00
B2TJ05 B2TJ12 B2TJ17 B2TJ22 B2TJ28 B2TJ30 B2TJ31 B2TJ32 B2TJ35 B2TJ39 B2TJ40
B2TJ41 B2TJ43 B2TJ47 B2TJ51 B2TJ55 B2TJ56 B2TJ60 B2TJ61 B2TJ67 B2TJ69 B2TJ78
B2TJ82 B2TJ86 B2TJ88 B2TJE9 B2TJH6 B2TJK1 B2TJM0 B2TJM9 B2TJY1 B2TJY5 B2TJY9
B2TK10 B2TK54 B2TK55 B2TK60 B2TK62 B2TK64 B2TK69 B2TK70 B2TK75 B2TK87 B2TK93
B2TKE1 B2TKE5 B2TKI3 B2TKP6 B2TL11 B2TL32 B2TL64 B2TL69 B2TL73 B2TL74 B2TL86
B2TLN8 B2TLQ3 B2TLR6 B2TLR7 B2TLY1 B2TLY9 B2TLZ5 B2TLZ7 B2TLZ8 B2TM00 B2TM01
B2TM06 B2TM09 B2TM49 B2TML4 B2TML8 B2TMY2 B2TMZ1 B2TMZ2 B2TMZ3 B2TMZ9 B2TN01
B2TN05 B2TN12 B2TN28 B2TN52 B2TN72 B2TN76 B2TNB1 B2TNB5 B2TND7 B2TNF7 B2TNG1
B2TNG3 B2TNM9 B2TP27 B2TP76 B2TPB6 B2TPB8 B2TPB9 B2TPC0 B2TPC6 B2TPC7 B2TPD4
B2TPD7 B2TPD9 B2TPE3 B2TPE7 B2TPG6 B2TPM3 B2TPV9 B2TPW8 B2TPX3 B2TPX4 B2TPY4
B2TQ00 B2TQ13 B2TQ14 B2TQ23 B2TQ24 B2TQ25 B2TQ47 B2TQ51 B2TQ57 B2TQ95 B2TQR6
B2TQR9 B2TQS1 B2TQS2 B2TQS8 B2TQW1 B2TQW6 B2TQZ9 B2TR25 B2TR28 B2TR53 B2TR54
B2TR90 B2TRC7 B2TRE4 B2TRG2 B2TRH8 B2TRI3 B2TRI4 B2TRM1 B2TRM3 B2TRM5 B2TRM7
B2TRM8 B2TRN5 B2TRN6 B2TRQ5 B2TRR9 B2TS13 B2TS30 B2TS31 B2TS45 B2TS55 B2TS59
B2TS60 B2TS64 B2TS69 B2TS78 B2TS86 B2TTA2 B2TTA4 B2TTB0 B2TTB3 B2TTH6 B2TTH7
B2TTI0 B2TTI9 B2TTK3 B2TTK9 B2TTL2 B2TTN5 B2TTP8 B2TTP9 B2TTQ0 B2TTS7 B2TTS8
B2TTT1 B2TTT7 B2TTT9 B2TTU8 B2TTU9 B2TTV0 B2TTV1 B2TTV2 B2TTV4 B2TTV5 B2TTV6
B2TTV7 B2TTX9 B2TU12 B2TU13 B2TU17 B2TU20 B2TU26 B2TU31 B2TU51 B2TU53 B2TU59
B2TU61 B2TU69 B2TU77 B2TU80 B2TU82 B2TU87 B2TU88 B2TU96 B2TU97 B2TUB2 B2TUD1
B2TUE4 B2TUF5 B2TUF7 B2TUG2 B2TUG9 B2TUH1 B2TUH3 B2TUH4 B2TUI5 B2TUJ4 B2TUJ7
B2TUK6 B2TUL3 B2TUM0 B2TUM8 B2TUN0 B2TUN1 B2TUN4 B2TUN5 B2TUR4 B2TUR7 B2TUS2
B2TUS4 B2TUS5 B2TUS6 B2TUS8 B2TUT5 B2TUT6 B2TUW2 B2TUW5 B2TUW7 B2TUY1 B2TUY6
B2TUY8 B2TV25 B2TV60 B2TV62 B2TV63 B2TV93 B2TVB6 B2TVC2 B2TVC6 B2TVE7 B2TVE9
B2TVF1 B2TVF4 B2TVF5 B2TVG3 B2TVH5 B2TVI4 B2TVJ6 B2TVL1 B2TVL2 B2TVN1 B2TVN6
B2TVP4 B2TVP7 B2TVQ8 B2TVR1 B2TVS0 B2TVT4 B2TVT9 B2TVV2 B2TVX7 B2TW20 B2TW47
B2TW49 B2TW50 B2TW54 B2TW72 B2TW75 B2TWA0 B2TWA2 B2TWA6 B2TWB1 B2TWB3 B2TWC2
B2TWC4 B2TWC5 B2TWC7 B2TWC8 B2TWD3 B2TWD5 B2TWE0 B2TWF1 B2TWF2 B2TWF4 B2TWF5
B2TWF7 B2TWG0 B2TWG4 B2TWG9 B2TWH0 B2TWH1 B2TWH2 B2TWH3 B2TWH4 B2TWH8 B2TWI0
B2TWI1 B2TWI2 B2TWI3 B2TWI4 B2TWI5 B2TWJ3 B2TWJ5 B2TWJ8 B2TWL8 B2TWQ1 B2TWQ2
B2TWR0 B2TWS7 B2TWX3 B2TWY5 B2TX02 B2TX08 B2TX16 B2TX17 B2TX24 B2TX41 B2TX50
B2TX62 B2TX77 B2TXF7 B2TXG4 B2TXI8 B2TXJ0 B2TXL2 B2TXL5 B2TXP7 B2TXQ2 B2TXQ4
B2TXS1 B2TXS3 B2TXS4 B2TXT2 B2TXT4 B2TXT6 B2TXT9 B2TXU0 B2TXU2 B2TXU3 B2TXU8
B2TXV1 B2TXV2 B2TXV5 B2TXV6 B2TXW4 B2TXX8 B2TXY0 B2TY12 B2TY17 B2TY18 B2TY23
B2TY33 B2TY39 B2TY45 B2TY46 B2TY47 B2TY52 B2TY53 B2TY64 B2TY65 B2TY68 B2TY69
B2TY70 B2TY73 B2TY74 B2TY75 B2TY76 B2TY84 B2TY92 B2TY93 B2TYB7 B2TYF5 B2TYF6
B2TYF9 B2TYG1 B2TYH1 B2TYH5 B2TYI8 B2TYK1 B2TYK2 B2TYM8 B2TYM9 B2TYN0 B2TYN1
B2TYN5 B2TYN7 B2TYP0 B2TYP2 B2TYP7 B2TYR0 B2TYR1 B2TYT0 B2TYV0 B2TYW4 B2TYY4
B2TYZ8 B2TYZ9 B2TZ10 B2TZ12 B2TZ80 B2TZ86 B2TZ97 B2TZ99 B2TZA6 B2TZA8 B2TZB5
B2TZC9 B2TZD0 B2TZD5 B2TZD9 B2TZF3 B2TZH1 B2TZH2 B2TZH9 B2TZI0 B2TZI1 B2TZI4
B2TZI5 B2TZI6 B2TZI7 B2TZI8 B2TZJ7 B2TZL3 B2TZN1 B2TZP5 B2TZQ2 B2TZQ6 B2TZR4
B2TZR5 B2TZR6 B2TZR7 B2TZR8 B2TZS6 B2TZS8 B2TZV2 B2TZV6 B2TZV8 B2TZV9 B2TZX7

B2TZZ3 B2U036 B2U047 B2U048 B2U049 B2U050 B2U053 B2U069 B2U084 B2U092 B2U0A8
B2U0F1 B2U0F2 B2U0G9 B2U0H4 B2U0I4 B2U0Q6 B2U0Q7 B2U0R5 B2U0R7 B2U0S0 B2U0S1
B2U0S2 B2U0S6 B2U0T0 B2U0T2 B2U0U3 B2U0V8 B2U0W2 B2U0W4 B2U0W8 B2U0W9 B2U0Z1
B2U177 B2U195 B2U1E4 B2U1F0 B2U1F7 B2U1G0 B2U1G7 B2U1G8 B2U1M1 B2U1R9 B2U1S3
B2U1T3 B2U1U1 B2U1U8 B2U1U9 B2U1V4 B2U1V5 B2U1V9 B2U1W1 B2U1W2 B2U1X2 B2U1Y8
B2U1Z1 B2U1Z2 B2U1Z4 B2U1Z8 B2U201 B2U204 B2U205 B2U207 B2U208 B2U210 B2U211
B2U217 B2U219 B2U221 B2U233 B2U242 B2U245 B2U248 B2U249 B2U250 B2U257 B2U258
B2U259 B2U260 B2U265 B2U268 B2U269 B2U278 B2U279 B2U281 B2U286 B2U287 B2U296
B2U2A6 B2U2A7 B2U2A9 B2U2C1 B2U2D7 B2U2D8 B2U2E0 B2U2E2 B2U2G3 B2U2I3 B2U2I4
B2U2J6 B2U2J7 B2U2K8 B2U2L2 B2U2L3 B2U2L8 B2U2L9 B2U2N5 B2U2N7 B2U2P9 B2U2Q0
B2U2Q2 B2U2Q4 B2U2Q5 B2U2Q6 B2U2R3 B2U2R5 B2U2R6 B2U2R9 B2U2S0 B2U2S2 B2U2S3
B2U2S4 B2U2S5 B2U2S6 B2U2S7 B2U2S8 B2U2S9 B2U2T0 B2U2T1 B2U2T2 B2U2T3 B2U2T4
B2U2T5 B2U2T7 B2U2T8 B2U2T9 B2U2U7 B2U2U8 B2U2U9 B2U2V0 B2U2V1 B2U2V2 B2U2V5
B2U2W8 B2U2W9 B2U2X0 B2U2X8 B2U2Y0 B2U2Y1 B2U2Z0 B2U2Z9 B2U301 B2U303 B2U304
B2U306 B2U308 B2U309 B2U311 B2U313 B2U315 B2U326 B2U333 B2U338 B2U350 B2U352
B2U355 B2U390 B2U392 B2U394 B2U395 B2U397 B2U3C0 B2U3C2 B2U3C5 B2U3C8 B2U3C9
B2U3D0 B2U3F9 B2U3H4 B2U3J5 B2U3J6 B2U3L1 B2U3M0 B2U3M2 B2U3M4 B2U3M8 B2U3N3
B2U3P9 B2U3R4 B2U3S3 B2U3T0 B2U3T1 B2U3T3 B2U3T4 B2U3Y5 B2U3Y9 B2U3Z0 B2U3Z3
B2U3Z6 B2U416 B2U447 B2U449 B2U473 B2U477 B2U494 B2U4B0 B2U4C2 B2U4G1 B2U4G2
B2U4G3 B2U4I7 B2U4K6 B2U4K7 B2U4L6 B2U4L8 B2U4L9 B2U4M3 B2U4M5 B2U4M7 B2U4N0
B2U4P1 B2U4P3 B2U4P9 B2U4Q1 B2U4S2 B2U4S4 B2U4S5 B2U4S7 B2U4S8 B2U4S9 B2U4X1
B2U4X2 B2U4X5 B2U4X7 B2U512 B2U513 B2U523 B2U532 B2U560 B2U565 B2U566 B2U5A3
B2U5B1 B2U5C8 B2U5D5 B2U5D7 B2U6Q7 B2U6V1 B2U6V5 B2U6Y7 B2U6Z0 B2U6Z1 B2U705
B2U764 B2U774 B2U795 B2U7A8 B2U7A9 B2U7B1 B2U7E2 B2U7E9 B2U7F0 B2U7G6 B2U7G7
B2U7H6 B2U7J7 B2U7J9 B2U7L1 B2U7M8 B2U7M9 B2U7N2 B2U7N4 B2U7R6 B2U7R7 B2U7S3
B2U7U6 B2U7V6 B2U7V7 B2U7W0 B2U7W2 B2U7X2 B2U7X3 B2U7Y8 B2U804 B2U814 B2U822
B2U823 B2U826 B2U846 B2U882 B2U886 B2U887 B2U894 B2U8B2 B2U8C4 B2U8D3 B2U8D8
B2U8L3 B2U8L7 B2U8S6 B2U8U5 B2U8U9 B2U8V1 B2U8V2 B2U8X6 B2U8Z0 B2U8Z2 B2U8Z4
B2U8Z7 B2U928 B2U930 B2U932 B2U942 B2U956 B2U966 B2U981 B2U984 B2U987 B2U9C0
B2U9D2 B2U9E4 B2U9P8 B2U9R2 B2U9R5 B2U9S6 B2U9T8 B2U9U3 B2U9U5 B2U9U6 B2U9U9
B2U9V3 B2U9V4 B2U9V9 B2U9W0 B2U9W5 B2U9W6 B2U9W7 B2U9Y5 B2UA10 B2UA12 B2UA33
B2UAA1 B2UAA3 B2UAA4 B2UAA5 B2UAG4 B2UAG7 B2UAK8 B2UAP0 B2UAP1 B2UAP2 B2UAP5
B2UAQ4 B2UAR0 B2UAR2 B2UAU3 B2UAX2 B2UAZ1 B2UAZ4 B2UB00 B2UB06 B2UB07 B2UB09
B2UB12 B2UB83 B2UB85 B2UBA0 B2UBA2 B2UBA3 B2UBA4 B2UBA6 B2UBB5 B2UBC4 B2UBD1
B2UBJ3 B2UBJ4 B2UBL0 B2UBN8 B2UBP2 B2UBP3 B2UBP7 B2UBQ0 B2UBZ3 B2UC74 B2UC75
B2UC85 B2UC88 B2UC98 B2UCA1 B2UCA2 B2UCA8 B2UCD6 B2UCE8 B2UCF1 B2UCS1 B2UCU9
B2UCV0 B2UCV2 B2UCV3 B2UCV4 B2UCV5 B2UCW2 B2UCW3 B2UCX5 B2UCX6 B2UCY5 B2UCY6
B2UD22 B2UD25 B2UD27 B2UD46 B2UDA1 B2UDK8 B2UDK9 B2UDL1 B2UDM7 B2UDV1 B2UDW0
B2UE01 B2UE02 B2UE35 B2UEE3 B2UEE4 B2UEE5 B2UEE6 B2UEE9 B2UEG8 B2UEG9 B2UEH7
B2UEI8 B2UEJ2 B2UEJ3 B2UEJ4 B2UEJ5 B2UEJ6 B2UEJ7 B2UEK0 B2UEK1 B2UEK2 B2UEK3
B2UEK4 B2UEK5 B2UEK6 B2UEK7 B2UEK8 B2UEK9 B2UEL0 B2UEL1 B2UEL2 B2UEL3 B2UEL4
B2UEL5 B2UEL6 B2UEL7 B2UEL8 B2UEL9 B2UEM0 B2UEM3 B2UEM4 B2UEN6 B2UEN7 B2UEN8
B2UER1 B2UER4 B2UER5 B2UER7 B2UES0 B2UES4 B2UES6 B2UF65 B2UF70 B2UF71 B2UF87
B2UFG8 B2UFJ8 B2UFK1 B2UFK2 B2UFK5 B2UFL1 B2UFL2 B2UFL4 B2UFL5 B2UFN5 B2UFP3

B2UFQ3 B2UFT6 B2UFU5 B2UFW4 B2UFW5 B2UFY4 B2UG56 B2UG65 B2UG80 B2UG82 B2UG86
B2UGE0 B2UGE2 B2UGE9 B2UGJ4 B2UGJ5 B2UGJ6 B2UGJ8 B2UGK8 B2UGM5 B2UGP7 B2UGQ0
B2UGT5 B2UGW0 B2UGY6 B2UGZ7 B2UGZ9 B2UHU5 B2UHV5 B2UIG7 B2UIZ4 B2UJ29 B2UJ42
B2UJ73 B2UJ81 B2UJF1 B2UJF2 B2UJY9 B2UK94 B2UKL9 B2UKN1 B2UKT4 B2UKT9 B2UKV6
B2UKV8 B2UKW8 B2UKY5 B2UKZ6 B2UL21 B2UL23 B2UL25 B2UL54 B2UL69 B2UL92 B2UL96
B2UL99 B2ULE2 B2ULJ1 B2ULM9 B2ULP2 B2ULP4 B2ULP6 B2ULR3 B2ULR4 B2ULZ9 B2UM17
B2UM23 B2UM93 B2UMF1 B2UMG0 B2UMG5 B2UMH5 B2UML6 B2UMP4 B2UMP5 B2UMS9 B2UMT2
B2UMT3 B2UMT4 B2UMT6 B2UMT7 B2UMT8 B2UMU0 B2UMU1 B2UMU3 B2UMU4 B2UMU5 B2UMV0
B2UMV5 B2UN87 B2UN91 B2UN97 B2UNA9 B2UNB0 B2UNB6 B2UNC5 B2UND0 B2UNE0 B2UNE2
B2UNE3 B2UNF2 B2UNF6 B2UNG3 B2UNG5 B2UNH4 B2UNJ3 B2UNJ5 B2UNS0 B2UNV3 B2UNW0
B2UP00 B2UP25 B2UP55 B2UP63 B2UPE6 B2UPE7 B2UPI9 B2UPK2 B2UPK4 B2UPK5 B2UPN5
B2UPP6 B2UPQ2 B2UPR9 B2UPW8 B2UQ11 B2UQ30 B2UQA2 B2UQD6 B2UQE3 B2UQE7 B2UQM2
B2UQM5 B2UQM8 B2UQQ2 B2UQR9 B2UQS1 B2UQS5 B2UQY1 B2UQY3 B2UQY6 B2UQY9 B2UQZ3
B2UR09 B2UR37 B2UR52 B2URB8 B2URC9 B2URE3 B2URE7 B2URG7 B2URH4 B2URH6 B2URI3
B2URJ0 B2URN8 B2URQ7 B2URQ8 B2URR4 B2URR5 B2URT8 B2URT9 B2URV1 B2URV3 B2URV4
B2URY3 B2URY7 B2URZ2 B2US00 B2US06 B2US13 B2US14 B2US24 B2US27 B2US28 B2US30
B2US41 B2US42 B2US45 B2US50 B2US69 B2US70 B2US71 B2US87 B2US91 B2USA0 B2USB2
B2USB4 B2USC5 B2USC7 B2USC8 B2USD4 B2USD7 B2USE8 B2USG0 B2USG1 B2USG3 B2USG5
B2USJ6 B2USL4 B2USM0 B2USM3 B2USM4 B2USN4 B2USN5 B2USN6 B2USP1 B2USP3 B2USR1
B2USR7 B2UST7 B2USV3 B2USW4 B2USW5 B2USX2 B2USX3 B2USX4 B2USY3 B2USY5 B2USZ4
B2USZ6 B2USZ8 B2UT06 B2UT08 B2UT29 B2UT33 B2UT36 B2UT37 B2UT38 B2UT54 B2UT94
B2UT98 B2UTC1 B2UTG2 B2UTG3 B2UTG4 B2UTH9 B2UTI4 B2UTI7 B2UTK0 B2UTK2 B2UTK9
B2UTL0 B2UTL2 B2UTM1 B2UTP7 B2UTQ6 B2UTQ8 B2UTR2 B2UTR6 B2UTS9 B2UTT1 B2UTX2
B2UTX3 B2UTX4 B2UTX8 B2UTZ5 B2UU34 B2UU37 B2UU54 B2UU73 B2UU74 B2UU76 B2UU78
B2UU89 B2UU96 B2UUA6 B2UUB6 B2UUE3 B2UUF1 B2UUF7 B2UUF9 B2UUG0 B2UUH2 B2UUI4
B2UUI8 B2UUJ8 B2UUK7 B2UUQ6 B2UUQ7 B2UUQ8 B2UUR0 B2UUR9 B2UUS4 B2UUU1 B2UUU2
B2UUV2 B2UUV6 B2UUV7 B2UUV8 B2UUV9 B2UUW1 B2UUW2 B2UUW3 B2UUW4 B2UUW7 B2UUW8
B2UUX6 B2UUZ0 B2UV01 B2UV03 B2UV04 B2UV09 B2UV11 B2UV20 B2UV21 B2UV41 B2UV44
B2UV55 B2UV57 B2UV58 B2UV59 B2UV60 B2UV64 B2UV65 B2UV66 B2UV67 B2UV68 B2UV69
B2UV70 B2UV71 B2UV72 B2UV73 B2UV74 B2UV75 B2UV76 B2UV77 B2UV78 B2UV79 B2UV80
B2UV81 B2UV83 B2UV86 B2UV99 B2UVA2 B2UVB0 B2UVE8 B2UVF4 B2UVF6 B2UVG2 B2UVG6
B2UVG9 B2UVI7 B2UVI9 B2UVJ2 B2UVJ3 B2UVJ7 B2UVL9 B2UVM0 B2UVM6 B2UVM8 B2UVP1
B2UVP2 B2UVP3 B2UVS4 B2UVU0 B2UVU9 B2UVV4 B2UVV5 B2UVX4 B2UVY4 B2UVZ8 B2UVZ9
B2UW00 B2UW05 B2UW06 B2UW09 B2UW10 B2UW12 B2UW13 B2UW31 B2UW36 B2UW37 B2UW64
B2UW65 B2UW67 B2UW70 B2UW71 B2UW73 B2UW74 B2UW87 B2UW91 B2UW93 B2UWE6 B2UWS9
B2UWW9 B2UX10 B2UX12 B2UX13 B2UX14 B2UX19 B2UX20 B2UX21 B2UX23 B2UX29 B2UX32
B2UX33 B2UX41 B2UX43 B2UX46 B2UX54 B2UX56 B2UX57 B2UX78 B2UX86 B2UX89 B2UX99
B2UXC4 B2UXC6 B2UXD0 B2UXD4 B2UXF7 B2UXL6 B2UXQ2 B2UXS0 B2UXS3 B2UXS5 B2UXS9
B2UXU5 B2UXU6 B2UXU9 B2UXV7 B2UY12 B2UY22 B2UY23 B2UY33 B2UY48 B2UY65 B2UY66
B2UY89 B2UY90 B2UY96 B2UY99 B2UYA0 B2UYA2 B2UYA3 B2UYA4 B2UYA5 B2UYA6 B2UYA7
B2UYA9 B2UYB0 B2UYB1 B2UYB2 B2UYB3 B2UYB4 B2UYB6 B2UYB7 B2UYB8 B2UYB9 B2UYC0
B2UYC1 B2UYC2 B2UYC3 B2UYC4 B2UYC5 B2UYC6 B2UYC8 B2UYC9 B2UYD1 B2UYD5 B2UYD6
B2UYD8 B2UYD9 B2UYE0 B2UYE4 B2UYE5 B2UYE6 B2UYH0 B2UYH3 B2UYH6 B2UYI0 B2UYI1
B2UYI2 B2UYK3 B2UYK7 B2UYK8 B2UYM8 B2UYT4 B2UYT6 B2UYT8 B2UYW3 B2UYW8 B2UYY8

B2UZ01 B2UZ02 B2UZ05 B2UZ17 B2UZ23 B2UZ25 B2UZ26 B2UZ27 B2UZ31 B2UZH3 B2UZI1
B2UZI5 B2UZI9 B2UZL0 B2UZT8 B2UZU4 B2UZX8 B2UZY1 B2UZY3 B2UZY4 B2UZZ0 B2V017
B2V046 B2V049 B2V089 B2V090 B2V095 B2V097 B2V099 B2V0A5 B2V0A7 B2V0A8 B2V0B0
B2V0F0 B2V0J5 B2V0J9 B2V0W0 B2V0W1 B2V110 B2V158 B2V163 B2V167 B2V168 B2V184
B2V1L8 B2V1M0 B2V1M2 B2V1Q9 B2V1S6 B2V1T2 B2V1U8 B2V1V3 B2V1V4 B2V1Y4 B2V1Z7
B2V1Z8 B2V267 B2V268 B2V276 B2V2G9 B2V2H7 B2V2I3 B2V2I5 B2V2I6 B2V2I8 B2V2I9
B2V2J3 B2V2J8 B2V2S0 B2V2S2 B2V2T4 B2V307 B2V327 B2V328 B2V337 B2V338 B2V339
B2V345 B2V347 B2V351 B2V358 B2V377 B2V398 B2V3B9 B2V3C3 B2V3P0 B2V3T7 B2V3V9
B2V3W2 B2V3W3 B2V3W4 B2V3W5 B2V3Y1 B2V3Z1 B2V430 B2V479 B2V486 B2V4A6 B2V4A8
B2V4B1 B2V4B2 B2V4B5 B2V4B8 B2V4C1 B2V4C6 B2V4D1 B2V4D6 B2V4E2 B2V4E4 B2V4E5
B2V4E6 B2V4E9 B2V4F3 B2V4F4 B2V4F5 B2V4F7 B2V4G1 B2V4G5 B2V4G9 B2V4H0 B2V4H4
B2V4H5 B2V4I1 B2V4I3 B2V4J4 B2V4J8 B2V4K0 B2V4K2 B2V4R0 B2V4R1 B2V4R3 B2V4R5
B2V4R6 B2V4R7 B2V4R8 B2V4S9 B2V4U3 B2V4W0 B2V4W1 B2V515 B2V525 B2V536 B2V583
B2V594 B2V599 B2V5A0 B2V5A4 B2V5A9 B2V5B7 B2V5C5 B2V5E8 B2V5F2 B2V5H4 B2V5I6
B2V5J1 B2V5J5 B2V5L9 B2V5M0 B2V5M9 B2V5N9 B2V5P9 B2V5T9 B2V5U0 B2V5U1 B2V5W6
B2V630 B2V652 B2V693 B2V6B0 B2V6B4 B2V6C3 B2V6E2 B2V6F1 B2V6F5 B2V6M2 B2V6N0
B2V6N2 B2V6T1 B2V6U1 B2V6V0 B2V6V1 B2V6V2 B2V6V4 B2V6V8 B2V6W5 B2V6W9 B2V6Y7
B2V6Y9 B2V709 B2V713 B2V726 B2V751 B2V752 B2V753 B2V771 B2V790 B2V795 B2V7A4
B2V7C0 B2V7E1 B2V7F0 B2V7F8 B2V7G0 B2V7G7 B2V7H6 B2V7H9 B2V7I4 B2V7I6 B2V7I9
B2V7J4 B2V7J5 B2V7J6 B2V7J9 B2V7K0 B2V7K2 B2V7K3 B2V7K5 B2V7K6 B2V7K7 B2V7K8
B2V7K9 B2V7L0 B2V7L1 B2V7L3 B2V7L4 B2V7L6 B2V7L7 B2V7L8 B2V7L9 B2V7M1 B2V7M3
B2V7M4 B2V7P6 B2V7Q4 B2V7S8 B2V7T1 B2V7T6 B2V7W7 B2V7X0 B2V7Y0 B2V7Y2 B2V7Y5
B2V842 B2V844 B2V847 B2V855 B2V890 B2V895 B2V8B3 B2V8C0 B2V8C9 B2V8D8 B2V8E1
B2V8F1 B2V8G9 B2V8I4 B2V8N8 B2V8N9 B2V8P5 B2V8P8 B2V8W5 B2V8X3 B2V8X4 B2V900
B2V901 B2V910 B2V930 B2V935 B2V937 B2V939 B2V955 B2V959 B2V963 B2V966 B2V968
B2V969 B2V9E3 B2V9E6 B2V9F3 B2V9H1 B2V9I5 B2V9M1 B2V9M8 B2V9P4 B2V9P8 B2V9P9
B2V9Q7 B2V9R3 B2V9T1 B2V9T6 B2V9U2 B2VA33 B2VA52 B2VA65 B2VA83 B2VAA3 B2VAA6
B2VAB3 B2VAC3 B2VAE4 B2VBD1 B2VBD5 B2VBJ9 B2VBK0 B2VBK1 B2VBL4 B2VBL6 B2VBM4
B2VBM9 B2VBN3 B2VBN5 B2VBP1 B2VBP4 B2VBR6 B2VBS6 B2VBT3 B2VBT8 B2VBT9 B2VBU3
B2VBV2 B2VBW5 B2VBW9 B2VBZ3 B2VC08 B2VC34 B2VC55 B2VC63 B2VC66 B2VC72 B2VC76
B2VC78 B2VC79 B2VC80 B2VC88 B2VC92 B2VC99 B2VCB2 B2VCE1 B2VCE3 B2VCE4 B2VCE5
B2VCE7 B2VCF8 B2VCH0 B2VCH1 B2VCH7 B2VCL6 B2VCL7 B2VCU8 B2VCV1 B2VCV6 B2VCV8
B2VCV9 B2VCW0 B2VCW1 B2VCW8 B2VCX3 B2VCX4 B2VCY1 B2VCZ2 B2VD05 B2VD17 B2VD18
B2VD19 B2VD21 B2VD27 B2VD29 B2VD52 B2VD53 B2VD79 B2VD83 B2VD93 B2VDA2 B2VDA7
B2VDB1 B2VDC9 B2VDD5 B2VDE5 B2VDF6 B2VDG5 B2VDG6 B2VDH0 B2VDJ2 B2VDJ8 B2VDJ9
B2VDK3 B2VDK9 B2VDM3 B2VDN8 B2VDQ9 B2VDU2 B2VDV1 B2VDW1 B2VDY5 B2VDZ9 B2VE05
B2VE09 B2VE11 B2VE12 B2VE16 B2VE25 B2VE26 B2VE28 B2VE30 B2VE40 B2VE50 B2VE56
B2VE61 B2VE71 B2VE89 B2VE90 B2VE93 B2VE96 B2VE97 B2VE98 B2VEB1 B2VEC0 B2VEC2
B2VEC5 B2VEC6 B2VEC7 B2VED1 B2VEH7 B2VEH8 B2VEI4 B2VEI9 B2VEK0 B2VEK5 B2VEK6
B2VEL2 B2VEL4 B2VEL5 B2VEL7 B2VEL8 B2VEM1 B2VEM5 B2VEN3 B2VEN4 B2VEN5 B2VEQ9
B2VER0 B2VES1 B2VES2 B2VES3 B2VEZ3 B2VEZ9 B2VF06 B2VF08 B2VF12 B2VF17 B2VF18
B2VF23 B2VF24 B2VF34 B2VF35 B2VF41 B2VF45 B2VF49 B2VF53 B2VF62 B2VF68 B2VF69
B2VF70 B2VF71 B2VF76 B2VF77 B2VF78 B2VF80 B2VFA0 B2VFA6 B2VFC7 B2VFE3 B2VFG0
B2VFG2 B2VFL2 B2VFL7 B2VFM6 B2VFR1 B2VFR2 B2VFV0 B2VFX1 B2VFX4 B2VFY9 B2VFZ2

B2VFZ5 B2VFZ6 B2VG00 B2VG01 B2VG02 B2VG05 B2VG06 B2VG22 B2VG25 B2VG26 B2VG41
B2VG45 B2VG55 B2VG58 B2VG69 B2VG72 B2VG76 B2VG77 B2VG81 B2VG82 B2VG89 B2VG92
B2VG93 B2VG94 B2VG95 B2VG96 B2VG97 B2VGA0 B2VGA2 B2VGA3 B2VGA5 B2VGA6 B2VGA8
B2VGA9 B2VGH5 B2VGJ0 B2VGJ4 B2VGJ7 B2VGJ9 B2VGK1 B2VGN3 B2VGN5 B2VGN7 B2VGP5
B2VGP6 B2VGP9 B2VGQ6 B2VGQ8 B2VGR2 B2VGR4 B2VGR6 B2VGR9 B2VGS0 B2VGS4 B2VGS9
B2VGT0 B2VGT1 B2VGT5 B2VGT8 B2VGU0 B2VGU9 B2VGV3 B2VGW6 B2VGW7 B2VGX3 B2VGX5
B2VH04 B2VH06 B2VH10 B2VH13 B2VH20 B2VH43 B2VH50 B2VH51 B2VH52 B2VH53 B2VH94
B2VHD0 B2VHD3 B2VHD6 B2VHE7 B2VHF1 B2VHI1 B2VHI6 B2VHJ5 B2VHJ7 B2VHM6 B2VHM7
B2VHN2 B2VHN8 B2VHQ1 B2VHQ4 B2VHS0 B2VHS1 B2VHS3 B2VHS6 B2VHS8 B2VHS9 B2VHT2
B2VHT7 B2VHU9 B2VHV7 B2VHX1 B2VHX2 B2VHY0 B2VHZ5 B2VI13 B2VI25 B2VI31 B2VI32
B2VI36 B2VI46 B2VI48 B2VI49 B2VI65 B2VI72 B2VI89 B2VI92 B2VIA0 B2VIE6 B2VIF2
B2VIF3 B2VIF6 B2VIG6 B2VIH1 B2VII2 B2VII6 B2VII8 B2VIL6 B2VIP6 B2VIQ7 B2VIQ9
B2VIR1 B2VIR2 B2VIS2 B2VIS3 B2VIT0 B2VIT2 B2VIU4 B2VIV0 B2VIX9 B2VIY1 B2VJ29
B2VJ36 B2VJ42 B2VJ44 B2VJ45 B2VJ51 B2VJ52 B2VJ55 B2VJ68 B2VJ83 B2VJ90 B2VJ92
B2VJ93 B2VJ94 B2VJ95 B2VJA4 B2VJB6 B2VJB9 B2VJC1 B2VJC2 B2VJD3 B2VJN9 B2VJP2
B2VJP4 B2VJR7 B2VJU2 B2VJU7 B2VJU8 B2VJU9 B2VJW0 B2VJW7 B2VJW8 B2VJX8 B2VK01
B2VK11 B2VK22 B2VK34 B2VK36 B2VK37 B2VK38 B2VK39 B2VK41 B2VK44 B2VK51 B2VK52
B2VK53 B2VK54 B2VK55 B2VK56 B2VK57 B2VK58 B2VK59 B2VK60 B2VK61 B2VK63 B2VK65
B2VK70 B2VK72 B2VK73 B2VK74 B2VK77 B2VK78 B2VK80 B2VK81 B2VK82 B2VK85 B2VK88
B2VK93 B2VK94 B2VK95 B2VKA5 B2VKB1 B2VKN8 B2VKS8 B2VKT1 B2VKT2 B2VKT4 B2VKV3
B2VKX1 B2VL10 B2VL12 B2VL34 B2VL36 B2VL42 B2VL47 B2VL52 B2VL59 B2VL73 B2VL75
B2VL81 B2VL84 B2VL85 B2VR76 B2VTT3 B2VUE4 B2VUU7 B2VVA8 B2VW14 B2VWG7 B2VXY5
B2VY36 B2VZH2 B2VZQ8 B2W0B1 B2W0D9 B2W0K4 B2W1N6 B2W1T4 B2W2N2 B2W2Y7 B2W6W8
B2W722 B2W8X8 B2W978 B2W9N9 B2WB15 B2WBA7 B2WBE8 B2WBM8 B2WBW4 B2WCB2 B2WGR9
B2WH84 B2WIC3 B2WIX9 B2WKR4 B2WKU1 B2WM34 B2WME0 B2WME4 B2WMQ2 B2WMX8 B2WSM8
B2WSN0 B2WSN1 B2X050 B2X1Y2 B2X1Y4 B2X1Y5 B2X1Y6 B2X1Y7 B2X1Z3 B2X1Z4 B2X201
B2XBQ5 B2XTD5 B2XTE3 B2XTE4 B2XTU1 B2XTU9 B2XTV0 B2XWJ8 B2XWK3 B2XWM6 B2XWM8
B2XWN0 B2XWN9 B2XWP0 B2XWQ6 B2XWR0 B2XWR1 B2XWR3 B2XWR4 B2Y1U8 B2Y1V2 B2Y1V6
B2Y1V7 B2Y1Y0 B2Y1Y2 B2Y1Z2 B2Y1Z3 B2Y1Z5 B2Y1Z6 B2Y1Z7 B2Y1Z9 B2Y203 B2Y208
B2Z449 B2ZCQ0 B2ZCQ1 B2ZCQ2 B2ZCQ3 B2ZDY2 B2ZFP3 B2ZGJ1 B2ZRS9 B2ZX90 B2ZYY5
B2ZYZ1 B2ZZS9 B3A0L6 B3A0N1 B3BM81 B3CKG1 B3CL66 B3CL69 B3CL70 B3CL71 B3CL80
B3CL83 B3CL86 B3CL87 B3CLA3 B3CLA4 B3CLA5 B3CLB0 B3CLB1 B3CLB3 B3CLC0 B3CLF0
B3CLF3 B3CLF5 B3CLG9 B3CLI3 B3CLJ0 B3CLJ5 B3CLK1 B3CLK2 B3CLK3 B3CLL6 B3CLN1
B3CLQ6 B3CLR5 B3CLR6 B3CLT1 B3CLT4 B3CLT6 B3CLT9 B3CLU1 B3CLU4 B3CLV0 B3CLX0
B3CLX1 B3CLX6 B3CLY6 B3CLZ0 B3CM01 B3CM06 B3CM19 B3CM20 B3CM21 B3CM25 B3CM26
B3CM33 B3CM44 B3CM46 B3CM50 B3CM56 B3CM59 B3CM64 B3CM65 B3CM66 B3CM71 B3CM94
B3CMB1 B3CMB5 B3CMB8 B3CMC0 B3CMC1 B3CME9 B3CMH0 B3CMH5 B3CMH6 B3CMJ4 B3CMJ7
B3CMJ8 B3CML3 B3CMP5 B3CMS0 B3CMS4 B3CMT8 B3CMV5 B3CMX3 B3CMZ6 B3CN01 B3CN07
B3CN15 B3CN25 B3CN27 B3CN29 B3CN30 B3CN31 B3CN32 B3CN33 B3CN34 B3CN36 B3CN37
B3CN38 B3CN39 B3CN40 B3CN42 B3CN44 B3CN46 B3CN47 B3CN48 B3CN50 B3CN54 B3CN64
B3CN77 B3CN84 B3CN89 B3CNB0 B3CNB5 B3CNB7 B3CNE0 B3CNF3 B3CNG9 B3CNH0 B3CNH2
B3CNH7 B3CNK5 B3CNN4 B3CNP2 B3CP03 B3CP11 B3CP16 B3CPG9 B3CPJ5 B3CPK8 B3CPS1
B3CPS8 B3CPU0 B3CPU1 B3CPU7 B3CPW2 B3CPX8 B3CPY0 B3CPY6 B3CPY7 B3CQ06 B3CQ25
B3CQ26 B3CQ33 B3CQC3 B3CQC6 B3CQC7 B3CQD1 B3CQD5 B3CQF1 B3CQN9 B3CQP3 B3CQP5

B3CQQ8 B3CQQ9 B3CQR0 B3CQX7 B3CQX9 B3CQY2 B3CR58 B3CR61 B3CR62 B3CR66 B3CR92
B3CRA8 B3CRA9 B3CRE6 B3CRT6 B3CRX8 B3CRX9 B3CRY6 B3CRY7 B3CRZ0 B3CRZ1 B3CRZ3
B3CRZ7 B3CRZ8 B3CRZ9 B3CSG3 B3CSI0 B3CSN9 B3CSP0 B3CSQ9 B3CSR8 B3CSS5 B3CSS7
B3CST3 B3CST4 B3CST6 B3CT06 B3CT08 B3CT09 B3CT10 B3CT11 B3CT12 B3CT14 B3CT15
B3CT16 B3CT18 B3CT19 B3CT21 B3CT24 B3CT27 B3CT28 B3CT29 B3CT30 B3CT31 B3CTD7
B3CTE5 B3CTE6 B3CTE7 B3CTF9 B3CTK8 B3CTQ6 B3CTT3 B3CTT4 B3CTT8 B3CTU1 B3CTU9
B3CTW0 B3CTW5 B3CTX1 B3CTX2 B3CTY9 B3CTZ1 B3CTZ4 B3CU37 B3CU42 B3CUJ5 B3CUK4
B3CUK6 B3CUK7 B3CUN5 B3CUN6 B3CUN9 B3CUP9 B3CV07 B3CV08 B3CV32 B3CV38 B3CV48
B3CV53 B3CV54 B3CV57 B3CV58 B3CV64 B3CVD8 B3CVD9 B3CVG2 B3CVK1 B3D6W2 B3DF93
B3DFA5 B3DFA8 B3DFB2 B3DFV0 B3DGJ2 B3DH20 B3DIV9 B3DIY3 B3DJM5 B3DJT0 B3DK16
B3DL37 B3DL53 B3DL65 B3DL84 B3DLA6 B3DLB3 B3DLD3 B3DLE8 B3DLF3 B3DM23 B3DM25
B3DM43 B3DM47 B3DNN5 B3DNP9 B3DNT3 B3DNU3 B3DP22 B3DP23 B3DP24 B3DP27 B3DP37
B3DP38 B3DP64 B3DPH4 B3DPH5 B3DPI2 B3DPI3 B3DPJ1 B3DPK2 B3DPK4 B3DPM0 B3DPR8
B3DPS3 B3DPS4 B3DPS5 B3DPS9 B3DPT0 B3DPW9 B3DPX0 B3DPX3 B3DPX4 B3DPY3 B3DPY6
B3DPZ9 B3DQ00 B3DQ12 B3DQ13 B3DQ14 B3DQ15 B3DQ16 B3DQ17 B3DQ29 B3DQ30 B3DQ31
B3DQ32 B3DQ51 B3DQ62 B3DQ65 B3DQ80 B3DQ84 B3DQA1 B3DQA4 B3DQB9 B3DQC0 B3DQC1
B3DQC3 B3DQC4 B3DQC5 B3DQC6 B3DQC7 B3DQC9 B3DQD0 B3DQD1 B3DQD2 B3DQD3 B3DQD7
B3DQD8 B3DQD9 B3DQE1 B3DQE2 B3DQF0 B3DQF1 B3DQF2 B3DQF7 B3DQG7 B3DQI6 B3DQM3
B3DQM9 B3DQN2 B3DQN7 B3DQQ9 B3DQT3 B3DQT4 B3DQV1 B3DQX6 B3DR08 B3DR12 B3DR33
B3DR50 B3DR51 B3DR79 B3DRA9 B3DRD0 B3DRM6 B3DRM7 B3DRM9 B3DRN4 B3DRN8 B3DRT0
B3DRT1 B3DRT4 B3DRT8 B3DRT9 B3DRV5 B3DRV7 B3DRV8 B3DRV9 B3DRW0 B3DRX0 B3DRX5
B3DRX7 B3DRX8 B3DRX9 B3DRY0 B3DRY2 B3DRY6 B3DS05 B3DS06 B3DS13 B3DS55 B3DS61
B3DS65 B3DSB0 B3DSB4 B3DSC2 B3DSC7 B3DSD9 B3DSE2 B3DSF0 B3DSF5 B3DSL6 B3DSQ0
B3DSS3 B3DSV5 B3DSX1 B3DSX9 B3DSY6 B3DSY7 B3DSY8 B3DSZ1 B3DSZ3 B3DT29 B3DT30
B3DT31 B3DT37 B3DT56 B3DT80 B3DT97 B3DTB5 B3DTC2 B3DTC8 B3DTE1 B3DTE2 B3DTE6
B3DTF8 B3DTJ5 B3DTK7 B3DTL4 B3DTL7 B3DTM9 B3DTN1 B3DTQ9 B3DTR2 B3DTR5 B3DTT9
B3DTU8 B3DU23 B3DU31 B3DU32 B3DU42 B3DU87 B3DUA1 B3DUB4 B3DUF9 B3DUG9 B3DUR6
B3DV19 B3DV36 B3DVB4 B3DVC8 B3DVG5 B3DVG8 B3DVG9 B3DVH5 B3DVH7 B3DVI3 B3DVI6
B3DVI7 B3DVR2 B3DVT5 B3DVV9 B3DVX0 B3DVY9 B3DVZ3 B3DVZ4 B3DVZ6 B3DW88 B3DWG1
B3DWG6 B3DWH2 B3DWQ3 B3DWR3 B3DWS5 B3DWT4 B3DWT5 B3DWU2 B3DWU5 B3DWZ0 B3DX01
B3DX06 B3DX49 B3DX68 B3DX88 B3DXF7 B3DXI7 B3DXK2 B3DXK4 B3DXL0 B3DXL1 B3DXN2
B3DXP7 B3DXQ2 B3DXV4 B3DY05 B3DYE4 B3DYH9 B3DYI1 B3DYJ8 B3DYQ1 B3DYQ6 B3DYW1
B3DYW2 B3DYW5 B3DYX1 B3DZ86 B3DZ87 B3DZB3 B3DZN7 B3DZP5 B3DZP6 B3DZP7 B3DZZ7
B3E044 B3E045 B3E058 B3E073 B3E099 B3E0A0 B3E0I5 B3E0I7 B3E0I8 B3E0I9 B3E0J0
B3E0J1 B3E0J5 B3E0J8 B3E0J9 B3E0K2 B3E0M0 B3E0Q7 B3E0U0 B3E0U1 B3E0U2 B3E147
B3E156 B3E159 B3E163 B3E164 B3E177 B3E1H7 B3E1K3 B3E1K5 B3E1K9 B3E1P0 B3E1T2
B3E1T7 B3E1T8 B3E1T9 B3E1X5 B3E1Z3 B3E1Z4 B3E1Z5 B3E1Z8 B3E252 B3E278 B3E280
B3E2A4 B3E2D1 B3E2H7 B3E2H8 B3E2K0 B3E2K5 B3E2L0 B3E2M3 B3E2S1 B3E325 B3E330
B3E331 B3E339 B3E394 B3E399 B3E3D2 B3E3H5 B3E3I1 B3E3I2 B3E3K3 B3E3K5 B3E3M5
B3E3M7 B3E3P0 B3E3Q8 B3E3R0 B3E3S3 B3E3X0 B3E3X9 B3E3Y0 B3E3Y1 B3E3Y4 B3E3Z0
B3E3Z1 B3E421 B3E424 B3E441 B3E458 B3E468 B3E479 B3E4K8 B3E4L0 B3E4R2 B3E4R5
B3E4T4 B3E4U4 B3E4X8 B3E4Y5 B3E590 B3E599 B3E5I0 B3E5K9 B3E5L0 B3E5M9 B3E5N0
B3E5S3 B3E5S5 B3E5V8 B3E5X4 B3E5X8 B3E5Z1 B3E5Z2 B3E5Z5 B3E5Z6 B3E600 B3E609
B3E610 B3E615 B3E616 B3E617 B3E618 B3E621 B3E622 B3E625 B3E628 B3E677 B3E683

B3E692 B3E693 B3E6F9 B3E6G0 B3E6H3 B3E6H4 B3E6H6 B3E6I4 B3E6L6 B3E6L7 B3E6P2
B3E6X0 B3E6X7 B3E6X8 B3E6Y9 B3E6Z6 B3E716 B3E717 B3E719 B3E754 B3E757 B3E758
B3E787 B3E789 B3E790 B3E7B0 B3E7B3 B3E7F6 B3E7F7 B3E7G1 B3E7K2 B3E7P9 B3E7Q7
B3E7S4 B3E7S5 B3E7S6 B3E7S7 B3E7S8 B3E7S9 B3E7T0 B3E7T1 B3E7T2 B3E7T4 B3E7T5
B3E7T6 B3E7T7 B3E7T8 B3E7T9 B3E7U0 B3E7U1 B3E7U2 B3E7U3 B3E7U4 B3E7U5 B3E7U6
B3E7U7 B3E7U8 B3E7U9 B3E7W9 B3E7X0 B3E845 B3E846 B3E847 B3E848 B3E849 B3E851
B3E853 B3E854 B3E856 B3E858 B3E8C4 B3E8F8 B3E8F9 B3E8G0 B3E8N9 B3E8X9 B3E8Z1
B3E933 B3E935 B3E936 B3E940 B3E966 B3E9B8 B3E9H6 B3E9N0 B3E9N5 B3E9Q7 B3E9Q8
B3E9R5 B3E9X0 B3E9X6 B3E9Y3 B3E9Z0 B3E9Z4 B3EA22 B3EA23 B3EA24 B3EA25 B3EA29
B3EA77 B3EAE0 B3EAE4 B3EAE7 B3EAE8 B3EAF1 B3EAF2 B3EAF3 B3EAK5 B3EAM2 B3EAP0
B3EAQ9 B3EAR8 B3EAS4 B3EB70 B3EB76 B3EB83 B3EBB7 B3EBH2 B3EBZ3 B3EBZ4 B3EBZ5
B3EBZ6 B3ECA6 B3ECB1 B3ECB9 B3ECG2 B3ECG5 B3ECH5 B3ECH9 B3ECI3 B3ECK5 B3ECN9
B3ECP1 B3ECP2 B3ECP3 B3ED30 B3ED33 B3ED41 B3ED46 B3ED49 B3ED51 B3EDD6 B3EDE5
B3EDF1 B3EDF4 B3EDG7 B3EDH8 B3EDK3 B3EDL2 B3EDN1 B3EDP4 B3EDX3 B3EDY2 B3EDY3
B3EE09 B3EE12 B3EE18 B3EE19 B3EE24 B3EE31 B3EE33 B3EE36 B3EE40 B3EE55 B3EE57
B3EE77 B3EEA0 B3EEB2 B3EEB5 B3EEB7 B3EEB8 B3EEB9 B3EEC0 B3EEC1 B3EEC8 B3EED6
B3EEE1 B3EEE4 B3EEE5 B3EEF0 B3EEF2 B3EEI0 B3EEJ7 B3EEL6 B3EEL7 B3EEM6 B3EEP4
B3EEQ8 B3EER0 B3EER1 B3EER3 B3EER6 B3EEV3 B3EEX3 B3EF00 B3EFA6 B3EFA9 B3EFB1
B3EFB2 B3EFB5 B3EFC1 B3EFC7 B3EFD1 B3EFD2 B3EFE2 B3EFF5 B3EFG3 B3EFG9 B3EFK8
B3EFL3 B3EFN5 B3EFQ8 B3EFS3 B3EFT2 B3EFT6 B3EFU4 B3EFU5 B3EFW0 B3EFW1 B3EFX1
B3EFX7 B3EFY1 B3EFY2 B3EFY3 B3EFY5 B3EFY7 B3EG28 B3EG35 B3EG36 B3EG59 B3EG61
B3EG68 B3EG74 B3EGF3 B3EGF4 B3EGF5 B3EGF7 B3EGG9 B3EGH6 B3EGI2 B3EGK5 B3EGL2
B3EGN4 B3EGN5 B3EGP4 B3EGP8 B3EGT4 B3EGU1 B3EGV0 B3EGV3 B3EGV4 B3EGV7 B3EGV8
B3EGW1 B3EGW2 B3EGW3 B3EGW4 B3EGW5 B3EGW6 B3EGW7 B3EGX2 B3EGX3 B3EGX4 B3EGX5
B3EGX6 B3EGX7 B3EGX8 B3EGX9 B3EGY0 B3EGY3 B3EGY4 B3EGY5 B3EGY6 B3EGY8 B3EGY9
B3EGZ0 B3EH05 B3EH06 B3EH07 B3EH20 B3EH23 B3EH27 B3EH28 B3EH29 B3EH35 B3EH36
B3EH38 B3EH68 B3EH88 B3EH92 B3EH93 B3EH94 B3EH95 B3EH96 B3EHE6 B3EHE8 B3EHF8
B3EHI9 B3EHJ0 B3EHP0 B3EHP3 B3EHR1 B3EHS7 B3EHV0 B3EHV1 B3EHV5 B3EI06 B3EI07
B3EI21 B3EI42 B3EIC2 B3EIC4 B3EIC5 B3EID0 B3EII6 B3EIJ1 B3EIJ2 B3EIK7 B3EIL0
B3EIL6 B3EIL7 B3EIM1 B3EIN0 B3EIN1 B3EIN2 B3EIP1 B3EIT1 B3EIU4 B3EIU9 B3EJ23
B3EJ24 B3EJ26 B3EJ62 B3EJ63 B3EJF5 B3EJF7 B3EJG6 B3EJG9 B3EJI1 B3EJJ5 B3EJP4
B3EJS3 B3EK13 B3EK39 B3EK52 B3EKC2 B3EKE9 B3EKF9 B3EKG2 B3EKG5 B3EKG6 B3EKG7
B3EKG8 B3EKG9 B3EKI0 B3EKJ7 B3EKJ8 B3EKM6 B3EKT9 B3EKV1 B3EKW5 B3EKW7 B3EL29
B3EL33 B3EL38 B3EL42 B3EL44 B3EL57 B3EL59 B3EL60 B3EL62 B3EL75 B3EL81 B3EL96
B3ELA4 B3ELC6 B3ELD9 B3ELE3 B3ELG8 B3ELI1 B3ELJ9 B3ELN3 B3ELN6 B3ELN8 B3ELN9
B3ELP2 B3ELP8 B3ELQ7 B3ELQ8 B3ELU3 B3ELV3 B3ELV8 B3EM22 B3EM42 B3EM49 B3EM52
B3EM61 B3EM81 B3EMG1 B3EMG7 B3EMI1 B3EMI3 B3EMI6 B3EMI7 B3EMK1 B3EML3 B3EMM7
B3EMN5 B3EMN6 B3EMN7 B3EMR0 B3EMR5 B3EMS2 B3EMS4 B3EMT1 B3EMT2 B3EMT4 B3EMU1
B3EMW0 B3EMY1 B3EMY2 B3EMY7 B3EMY8 B3EMZ5 B3EN04 B3EN08 B3EN11 B3EN95 B3EN99
B3ENA3 B3ENJ1 B3ENJ2 B3ENK5 B3ENP4 B3ENQ2 B3ENS3 B3ENV6 B3ENV7 B3ENV8 B3ENW0
B3ENZ9 B3EP05 B3EP06 B3EP07 B3EP08 B3EP11 B3EP19 B3EP23 B3EP24 B3EP28 B3EP33
B3EP34 B3EP35 B3EP36 B3EP37 B3EP38 B3EP42 B3EP43 B3EP44 B3EP45 B3EP46 B3EP47
B3EP48 B3EP49 B3EP50 B3EP51 B3EP52 B3EP54 B3EP55 B3EP56 B3EP57 B3EP58 B3EP59
B3EP60 B3EP61 B3EP62 B3EP63 B3EP64 B3EP65 B3EP66 B3EP74 B3EP77 B3EPA8 B3EPC5

B3EPC6 B3EPD4 B3EPG5 B3EPG6 B3EPH3 B3EPH5 B3EPH6 B3EPI9 B3EPK0 B3EPK1 B3EPL0
B3EPT4 B3EPT5 B3EPT7 B3EPW2 B3EPX3 B3EPX5 B3EPY3 B3EPY6 B3EPZ5 B3EQ14 B3EQ18
B3EQ26 B3EQ34 B3EQ72 B3EQ83 B3EQ84 B3EQB7 B3EQC0 B3EQC6 B3EQC7 B3EQD1 B3EQK4
B3EQL0 B3EQL4 B3EQM3 B3EQQ7 B3EQT0 B3EQT2 B3EQT3 B3ER35 B3ER55 B3ER62 B3ER64
B3ER68 B3ER80 B3ER84 B3ER86 B3ER92 B3ERE1 B3ERG3 B3ERG6 B3ERM8 B3ERN3 B3ERN8
B3ERN9 B3ERQ7 B3ERW0 B3ERW1 B3ERW9 B3ERX0 B3ERY6 B3ERZ2 B3ERZ8 B3ES11 B3ES13
B3ES20 B3ES21 B3ES47 B3ES89 B3ES99 B3ESB4 B3ESC6 B3ESD7 B3ESF6 B3ESS3 B3ESU1
B3ESU3 B3ET45 B3ETF1 B3ETH8 B3ETH9 B3ETM6 B3ETT4 B3ETT5 B3ETU4 B3ETV4 B3ETW5
B3ETY9 B3ETZ1 B3ETZ2 B3ETZ3 B3ETZ4 B3ETZ7 B3EU33 B3EU50 B3EU51 B3EU68 B3EU83
B3EU84 B3EU85 B3EUC9 B3EUD6 B3EUD7 B3EUD8 B3EUF2 B3EUF3 B3EUF4 B3EUF5 B3EUG6
B3EUG8 B3EUH5 B3EUJ5 B3EUJ7 B3EUJ9 B3EUK0 B3EUK4 B3EUK5 B3EUK8 B3EUK9 B3EUL0
B3EUL2 B3EUL3 B3EUL5 B3EUL6 B3EUL8 B3EUL9 B3EUM0 B3EUM2 B3EUM3 B3EUR5 B3EWC5
B3EWC6 B3EWC7 B3EWC8 B3EWC9 B3EWD0 B3EWD1 B3EWD2 B3EWD3 B3EWD4 B3EWD5 B3EWD6
B3EWD7 B3EWD8 B3EWD9 B3EWE0 B3EWE1 B3EWE2 B3EWE3 B3EWE4 B3EWE5 B3EWE7 B3EWE8
B3EWF0 B3EWF1 B3EWF7 B3EWG3 B3EWG5 B3EWG6 B3EWG7 B3EWG9 B3EWH5 B3EWH7 B3EWI0
B3EWI1 B3EWI5 B3EWI6 B3EWI7 B3EWI8 B3EWJ2 B3EWK7 B3EWK9 B3EWM8 B3EWN4 B3EWN6
B3EWN7 B3EWN8 B3EWN9 B3EWP0 B3EWP1 B3EWP5 B3EWP8 B3EWQ1 B3EWQ2 B3EWQ3 B3EWQ7
B3EWQ9 B3EWR5 B3EWR7 B3EWR8 B3EWR9 B3EWS0 B3EX35 B3EX63 B3EY95 B3EYM8 B3F209
B3F2X5 B3F2X6 B3F2X7 B3FHP0 B3FHT4 B3FIW8 B3FK34 B3FL73 B3FWT6 B3FWT9 B3FWU0
B3GNI6 B3GW58 B3GX29 B3GX42 B3GX43 B3GX46 B3GX47 B3GX65 B3GX82 B3GX87 B3GXB0
B3GXB1 B3GXB4 B3GXB6 B3GXB7 B3GXC1 B3GXC3 B3GXC5 B3GXC6 B3GXC9 B3GXD3 B3GXD4
B3GXE4 B3GXE6 B3GXE7 B3GXF8 B3GXG6 B3GXH2 B3GXH5 B3GXI4 B3GXJ0 B3GXJ4 B3GXJ7
B3GXL9 B3GXP2 B3GXP3 B3GXP5 B3GXP8 B3GXP9 B3GXR0 B3GXR2 B3GXS0 B3GXV7 B3GXW0
B3GXW8 B3GXX5 B3GXX7 B3GXY6 B3GXZ2 B3GXZ7 B3GXZ9 B3GY04 B3GY07 B3GY11 B3GY20
B3GY24 B3GY26 B3GY27 B3GY30 B3GY37 B3GY39 B3GY41 B3GY43 B3GY52 B3GY57 B3GY58
B3GY59 B3GY63 B3GY64 B3GY67 B3GY94 B3GY96 B3GYA4 B3GYC3 B3GYC8 B3GYE2 B3GYF9
B3GYG5 B3GYI3 B3GYI8 B3GYJ0 B3GYJ4 B3GYK1 B3GYL3 B3GYL7 B3GYL8 B3GYL9 B3GYM3
B3GYM9 B3GYN8 B3GYP1 B3GYP5 B3GYS0 B3GYT6 B3GYT7 B3GYU4 B3GYU5 B3GYU6 B3GYU7
B3GYV5 B3GYW2 B3GYW3 B3GYW4 B3GYW7 B3GZ01 B3GZ03 B3GZ05 B3GZ06 B3GZ10 B3GZ11
B3GZ12 B3GZ14 B3GZ15 B3GZ16 B3GZ18 B3GZ19 B3GZ20 B3GZ21 B3GZ22 B3GZ23 B3GZ24
B3GZ25 B3GZ26 B3GZ27 B3GZ29 B3GZ30 B3GZ34 B3GZ35 B3GZ37 B3GZ38 B3GZ39 B3GZ40
B3GZ41 B3GZ49 B3GZ52 B3GZ55 B3GZ58 B3GZ69 B3GZ73 B3GZ78 B3GZ82 B3GZ90 B3GZ94
B3GZ95 B3GZA0 B3GZB8 B3GZC5 B3GZC6 B3GZD6 B3GZE5 B3GZE6 B3GZF1 B3GZF5 B3GZG5
B3GZG9 B3GZH2 B3GZH3 B3GZJ0 B3GZJ3 B3GZJ9 B3GZK0 B3GZK9 B3GZL0 B3GZM0 B3GZM3
B3GZN0 B3GZN4 B3GZP5 B3GZQ0 B3GZQ2 B3GZR5 B3GZR9 B3GZT0 B3GZT1 B3GZT6 B3GZV2
B3GZV8 B3GZX1 B3GZX2 B3GZX8 B3GZY0 B3GZY1 B3GZZ1 B3GZZ2 B3H003 B3H035 B3H044
B3H050 B3H053 B3H061 B3H066 B3H067 B3H073 B3H074 B3H084 B3H0A8 B3H0B9 B3H0C5
B3H0C8 B3H0F9 B3H0G3 B3H0H9 B3H0J1 B3H0J4 B3H0J6 B3H0J7 B3H0M0 B3H0M3 B3H0M4
B3H0M9 B3H0P8 B3H0P9 B3H0R2 B3H0R3 B3H0R5 B3H0R7 B3H0R8 B3H0S9 B3H108 B3H120
B3H123 B3H129 B3H130 B3H131 B3H137 B3H139 B3H152 B3H161 B3H163 B3H164 B3H169
B3H176 B3H177 B3H198 B3H1A1 B3H1A7 B3H1B0 B3H1B7 B3H1D8 B3H1D9 B3H1E1 B3H1E2
B3H1E5 B3H1E8 B3H1F1 B3H1F6 B3H1G9 B3H1H2 B3H1I1 B3H1K2 B3H1K6 B3H1K7 B3H1L0
B3H1L1 B3H1L4 B3H1M3 B3H1N7 B3H1P4 B3H1P5 B3H1P6 B3H1P7 B3H1V8 B3H1W3 B3H1X8
B3H202 B3H222 B3H248 B3H252 B3H259 B3H268 B3H269 B3H271 B3H275 B3H282 B3H285

B3H286 B3H296 B3H2A8 B3H2E6 B3H2G5 B3H2G6 B3H2H2 B3H2H8 B3H2I5 B3H2J0 B3H2J1
B3H2J7 B3H2J9 B3H2K8 B3H2K9 B3H2L3 B3H2L4 B3H2L5 B3H2M6 B3H2N1 B3H2N2 B3H2N3
B3H2N4 B3H2N6 B3H2Q1 B3H2Q2 B3H2R4 B3H2T2 B3H2U3 B3H2W6 B3H2W9 B3H2X0 B3H2X7
B3H2X9 B3H2Y8 B3H2Z6 B3H2Z8 B3H307 B3H308 B3H319 B3H321 B3H330 B3H331 B3H337
B3H338 B3H340 B3H341 B3H342 B3H469 B3H4F1 B3H4P1 B3H4X8 B3H5A8 B3H5K9 B3H5R8
B3H615 B3H672 B3H6C3 B3H6Y2 B3H6Z8 B3H7G2 B3IUN8 B3KS81 B3L014 B3L145 B3L230
B3L2G0 B3L2R5 B3L3W7 B3L4K9 B3L781 B3L9G8 B3LC82 B3LF83 B3LFA4 B3LFU5 B3LFZ1
B3LG61 B3LGB9 B3LGE9 B3LGJ2 B3LGK5 B3LH81 B3LH96 B3LHE5 B3LHQ7 B3LI22 B3LI79
B3LIS9 B3LIY9 B3LJ84 B3LJA2 B3LJA9 B3LJG1 B3LJM6 B3LJN9 B3LJT5 B3LJV1 B3LJX0
B3LK55 B3LK80 B3LK82 B3LKH7 B3LKR9 B3LKV0 B3LL20 B3LLJ2 B3LLK7 B3LLP9 B3LLV5
B3LLV7 B3LLX4 B3LLX6 B3LLZ8 B3LM82 B3LMH0 B3LMS3 B3LMT5 B3LMW8 B3LMZ5 B3LN41
B3LN76 B3LNB2 B3LNL2 B3LNM2 B3LNU5 B3LNX6 B3LP36 B3LP57 B3LP64 B3LP78 B3LPN5
B3LPY7 B3LQ10 B3LQ11 B3LQ40 B3LQ66 B3LQ92 B3LQB9 B3LQG6 B3LQP5 B3LQY3 B3LQZ6
B3LR01 B3LR30 B3LR49 B3LRD9 B3LRE5 B3LRH6 B3LRI8 B3LRJ8 B3LRM5 B3LRX9 B3LS63
B3LS90 B3LSM6 B3LSR0 B3LSS3 B3LSS7 B3LSV8 B3LSV9 B3LT19 B3LT39 B3LT42 B3LT59
B3LT96 B3LT98 B3LTD9 B3LTJ8 B3LTK6 B3LTL7 B3LTR3 B3LTT8 B3LTX4 B3LU13 B3LU25
B3LUP1 B3LUQ4 B3LVF9 B3LW62 B3LWP6 B3LWU3 B3LXF2 B3LY22 B3LY71 B3LYC0 B3LYP1
B3LZN3 B3M011 B3M070 B3M098 B3M0F3 B3M0Y8 B3M123 B3M173 B3M1B7 B3M1E1 B3M1F2
B3M1G0 B3M268 B3M269 B3M343 B3M383 B3M3R5 B3M611 B3M6I4 B3M7M6 B3M7W0 B3M881
B3M996 B3M9R3 B3M9V8 B3MA91 B3MB79 B3MC02 B3MC74 B3MCF3 B3MCQ5 B3MCZ5 B3MDA3
B3MDR0 B3MDU3 B3ME25 B3MET8 B3MEY6 B3MEZ6 B3MF31 B3MF39 B3MG50 B3MGT3 B3MGU5
B3MGZ0 B3MHX6 B3MI37 B3MI72 B3MI77 B3MIF1 B3MIY7 B3MJ61 B3MJ69 B3MJV4 B3MJV8
B3MLA8 B3MLC5 B3MLI0 B3MLX7 B3MM49 B3MMA5 B3MN22 B3MN57 B3MP94 B3MPC0 B3MQ24
B3MQP7 B3MQV3 B3MRC6 B3MRT7 B3MRX2 B3MS75 B3MSG8 B3MSI4 B3MSP2 B3MT31 B3MTI8
B3MTI9 B3MU92 B3MUX9 B3MVL6 B3MW09 B3MYX2 B3MYY4 B3MZN7 B3MZY6 B3N018 B3N0Q8
B3N1G9 B3N3F7 B3N449 B3N4C7 B3N4F1 B3N534 B3N538 B3N5N3 B3N6D9 B3N6U7 B3N7D5
B3N7L9 B3N899 B3N8H0 B3N8L3 B3N8S9 B3N9E4 B3NA01 B3NA78 B3NAN8 B3NAN9 B3NB67
B3NCH1 B3NDN0 B3NDY8 B3NE99 B3NEF0 B3NFA2 B3NFQ7 B3NGA1 B3NH71 B3NHQ1 B3NIL9
B3NIP2 B3NJY0 B3NK72 B3NKH7 B3NLK7 B3NLX1 B3NM45 B3NMI5 B3NML0 B3NN00 B3NNJ9
B3NP10 B3NP26 B3NP61 B3NPW0 B3NPZ0 B3NQ53 B3NQ86 B3NQR5 B3NRC6 B3NRK6 B3NRP1
B3NSK1 B3NSW1 B3NU20 B3NU52 B3NUB6 B3NUC9 B3NVC6 B3NVZ7 B3NW64 B3NWW9 B3NXQ7
B3NY01 B3NY06 B3NY08 B3NY19 B3NYF7 B3NYS4 B3NYY7 B3NZ68 B3P0D7 B3P0Q4 B3P0R4
B3P1F9 B3P1R1 B3P211 B3P230 B3P239 B3P2A7 B3P2S2 B3P321 B3P3E8 B3P3W1 B3P4F8
B3P4I7 B3P4M4 B3P4N5 B3P538 B3P5D2 B3P5D3 B3P6R4 B3P6R5 B3P773 B3P851 B3P8G6
B3P8M3 B3P8V8 B3P935 B3P9J3 B3P9W1 B3PB55 B3PB58 B3PB59 B3PBB9 B3PBD6 B3PBH2
B3PBH5 B3PBK5 B3PBL9 B3PBM0 B3PBP6 B3PBP7 B3PBQ4 B3PBV0 B3PBY5 B3PBZ8 B3PC08
B3PC09 B3PC16 B3PC19 B3PC22 B3PC77 B3PCD4 B3PCJ2 B3PCK0 B3PCM8 B3PCM9 B3PCR4
B3PCR5 B3PCS0 B3PCS1 B3PCS2 B3PCV7 B3PCZ8 B3PD15 B3PD57 B3PDB1 B3PDB6 B3PDC1
B3PDC2 B3PDC3 B3PDH6 B3PDL7 B3PDM1 B3PDM5 B3PDM9 B3PDN1 B3PDU1 B3PDW7 B3PE25
B3PEB5 B3PEI8 B3PEM4 B3PER3 B3PER6 B3PEU8 B3PEY6 B3PF20 B3PF22 B3PF24 B3PF33
B3PF48 B3PFH5 B3PFK3 B3PFN2 B3PFN3 B3PFN8 B3PFP8 B3PFQ2 B3PFU1 B3PFU2 B3PG51
B3PG60 B3PG61 B3PG68 B3PG71 B3PG74 B3PGA1 B3PGA3 B3PGA4 B3PGC8 B3PGD6 B3PGE8
B3PGF2 B3PGF6 B3PGN4 B3PGN5 B3PGP3 B3PGQ8 B3PGV8 B3PGW2 B3PGY7 B3PGZ1 B3PGZ3
B3PH48 B3PHF4 B3PHG5 B3PHG9 B3PHH8 B3PHI3 B3PHK5 B3PHK7 B3PHR3 B3PI65 B3PI66

B3PI81 B3PI82 B3PI87 B3PI88 B3PI89 B3PI90 B3PI94 B3PI96 B3PI97 B3PI99 B3PIA0
B3PIB4 B3PID5 B3PIE4 B3PIG3 B3PIG8 B3PII1 B3PIK9 B3PIL6 B3PIL7 B3PIT7 B3PIT8
B3PIU4 B3PIU8 B3PIU9 B3PIV4 B3PIW6 B3PJ20 B3PJ64 B3PJ74 B3PJA8 B3PJA9 B3PJB0
B3PJM0 B3PJN6 B3PJN8 B3PJP4 B3PK10 B3PK14 B3PK17 B3PK21 B3PK26 B3PK27 B3PK29
B3PK30 B3PK31 B3PK33 B3PK36 B3PK37 B3PK38 B3PK39 B3PK40 B3PK41 B3PK42 B3PK44
B3PK46 B3PK47 B3PK48 B3PK49 B3PK50 B3PK51 B3PK52 B3PK53 B3PK56 B3PK58 B3PK59
B3PK60 B3PK61 B3PK62 B3PK63 B3PK83 B3PKA4 B3PKA5 B3PKB3 B3PKE3 B3PKF2 B3PKG0
B3PKG8 B3PKG9 B3PKM9 B3PKN1 B3PKN9 B3PKU5 B3PKV6 B3PL12 B3PL13 B3PL16 B3PL38
B3PL49 B3PL51 B3PL52 B3PL62 B3PL65 B3PL68 B3PL73 B3PLF9 B3PLM6 B3PLN2 B3PLP1
B3PLQ1 B3PLQ4 B3PLS0 B3PLS2 B3PLW7 B3PLX9 B3PLZ0 B3PLZ2 B3PM19 B3PM20 B3PM27
B3PM28 B3PM35 B3PM40 B3PM62 B3PM63 B3PM68 B3PM70 B3PM73 B3PM75 B3PM77 B3PM95
B3PM96 B3PMA0 B3PMA1 B3PMA2 B3PMA3 B3PMB4 B3PMB5 B3PMC1 B3PMC2 B3PMC6 B3PMD5
B3PME9 B3PMF0 B3PMF1 B3PMF7 B3PMF8 B3PMG6 B3PMH1 B3PMI7 B3PMJ0 B3PMJ8 B3PMK1
B3PMK3 B3PML9 B3PMM0 B3PMM2 B3PMM3 B3PMM4 B3PMM9 B3PMN2 B3PMN3 B3PMN4 B3PMN7
B3PMN9 B3PMP1 B3PMP2 B3PMP3 B3PMP4 B3PMP8 B3PMQ1 B3PMU1 B3PN01 B3PN08 B3PN19
B3PN30 B3PN46 B3PN57 B3PN58 B3PN60 B3PN94 B3PNC6 B3PNC7 B3PND6 B3PND9 B3PNG8
B3PNG9 B3PNH1 B3PNH7 B3PNI8 B3PNJ7 B3PNM4 B3PNT1 B3PNT4 B3PNT8 B3PNV4 B3PNV9
B3PP01 B3PP19 B3PP20 B3PP83 B3PPC1 B3PPE3 B3PPF1 B3PPK3 B3PPL9 B3PPP3 B3PPS1
B3PPT2 B3PPT3 B3PPU4 B3PPZ4 B3PQ08 B3PQ37 B3PQ90 B3PQ91 B3PQA4 B3PQB3 B3PQL4
B3PQS3 B3PQX7 B3PQY8 B3PR16 B3PR17 B3PR19 B3PR22 B3PR35 B3PR71 B3PR77 B3PRB0
B3PRB1 B3PRD9 B3PRG0 B3PRG5 B3PRH7 B3PRI2 B3PRW0 B3PRW7 B3PRY8 B3PRY9 B3PRZ3
B3PRZ4 B3PRZ5 B3PRZ6 B3PS03 B3PS18 B3PS36 B3PS41 B3PS46 B3PS51 B3PS53 B3PS68
B3PS72 B3PS94 B3PSA1 B3PSB9 B3PSC0 B3PSC4 B3PT89 B3PTD3 B3PTE0 B3PTE1 B3PTJ5
B3PTN1 B3PTP8 B3PTU0 B3PTU7 B3PTV8 B3PTW8 B3PU35 B3PUC6 B3PUI4 B3PUK3 B3PUN0
B3PUN2 B3PUN8 B3PUP2 B3PUS4 B3PUT4 B3PUT6 B3PUT7 B3PUU6 B3PUV3 B3PV10 B3PV29
B3PV52 B3PV59 B3PV77 B3PV93 B3PVB6 B3PVD4 B3PVE1 B3PVH2 B3PVJ6 B3PVT2 B3PVT4
B3PVT7 B3PVV1 B3PVV2 B3PVV3 B3PVW2 B3PVW3 B3PVW4 B3PVY5 B3PW17 B3PW29 B3PW37
B3PW55 B3PW56 B3PW57 B3PW58 B3PW59 B3PW60 B3PW62 B3PW63 B3PW72 B3PWH4 B3PWH7
B3PWH8 B3PWI0 B3PWI5 B3PWI6 B3PWI7 B3PWJ0 B3PWQ6 B3PWR8 B3PWS0 B3PWS1 B3PWS2
B3PWS3 B3PWS4 B3PWS5 B3PWS7 B3PWS8 B3PWS9 B3PWT0 B3PWT1 B3PWT2 B3PWT3 B3PWT4
B3PWT5 B3PWT6 B3PWT7 B3PWT8 B3PWT9 B3PWU0 B3PWU2 B3PWU3 B3PWU4 B3PWU5 B3PWU6
B3PWV1 B3PWZ9 B3PXA8 B3PXA9 B3PXB0 B3PXB3 B3PXB6 B3PXB8 B3PXD7 B3PXD9 B3PXE2
B3PXE3 B3PXE6 B3PXF0 B3PXG8 B3PXH2 B3PXH3 B3PXI8 B3PXK3 B3PXK6 B3PXK7 B3PXL5
B3PXN0 B3PXN1 B3PXT4 B3PXT5 B3PXU3 B3PXU4 B3PXV9 B3PXW5 B3PXX0 B3PXX1 B3PXX2
B3PXX5 B3PXX8 B3PXY4 B3PXY7 B3PY13 B3PYE0 B3PYE1 B3PYE2 B3PYF6 B3PYH7 B3PYJ5
B3PYP2 B3PYP3 B3PYP5 B3PYS0 B3PYS4 B3PYT5 B3PYU7 B3PYW4 B3PYX0 B3PYY8 B3PYZ1
B3PYZ4 B3PYZ5 B3PZ23 B3PZ58 B3PZ69 B3PZ92 B3PZ93 B3PZ95 B3PZA2 B3PZA3 B3PZA4
B3PZB4 B3PZB6 B3PZE1 B3PZF7 B3PZG1 B3PZQ8 B3PZR1 B3PZS1 B3PZT5 B3PZU8 B3Q000
B3Q001 B3Q031 B3Q039 B3Q042 B3Q0A1 B3Q0A6 B3Q0A9 B3Q0L2 B3Q0L3 B3Q0M2 B3Q0R5
B3Q0S3 B3Q0S4 B3Q0T5 B3Q0Y0 B3Q0Y6 B3Q1C3 B3Q1Q5 B3Q4F9 B3Q5X1 B3Q5X2 B3Q5X3
B3Q5Z7 B3Q5Z9 B3Q602 B3Q604 B3Q605 B3Q606 B3Q616 B3Q619 B3Q620 B3Q6A0 B3Q6H7
B3Q6H9 B3Q6L1 B3Q6M4 B3Q6M8 B3Q6M9 B3Q6P5 B3Q6P8 B3Q6Q0 B3Q6U1 B3Q6U2 B3Q6V6
B3Q6W6 B3Q6Y9 B3Q725 B3Q726 B3Q731 B3Q732 B3Q733 B3Q736 B3Q760 B3Q761 B3Q7C4
B3Q7C5 B3Q7D4 B3Q7E1 B3Q7H1 B3Q7H9 B3Q7J8 B3Q7K1 B3Q7K3 B3Q7K4 B3Q7L2 B3Q7P4

B3Q7P5 B3Q7P6 B3Q7Q5 B3Q7T8 B3Q7X3 B3Q7X4 B3Q7Y9 B3Q845 B3Q851 B3Q853 B3Q854
B3Q855 B3Q856 B3Q862 B3Q867 B3Q881 B3Q8A1 B3Q8A6 B3Q8A7 B3Q8A8 B3Q8B1 B3Q8B5
B3Q8B9 B3Q8C2 B3Q8J5 B3Q8X5 B3Q8Z5 B3Q950 B3Q951 B3Q953 B3Q968 B3Q969 B3Q970
B3Q972 B3Q973 B3Q976 B3Q978 B3Q986 B3Q999 B3Q9C1 B3Q9C2 B3Q9D3 B3Q9D7 B3Q9E4
B3Q9J8 B3Q9N8 B3Q9Q1 B3Q9Q2 B3Q9Q6 B3Q9R4 B3Q9R8 B3Q9S3 B3Q9S4 B3Q9T5 B3Q9T6
B3Q9T8 B3Q9V3 B3Q9V6 B3Q9W5 B3Q9W8 B3Q9X0 B3Q9X4 B3Q9Y7 B3QA01 B3QA32 B3QA35
B3QA57 B3QA91 B3QA92 B3QAA7 B3QAB0 B3QAB1 B3QAB3 B3QAB7 B3QAC6 B3QAG7 B3QAH7
B3QAR0 B3QAV6 B3QAW1 B3QB18 B3QB33 B3QB68 B3QB72 B3QB98 B3QBC6 B3QBD3 B3QBS6
B3QBS9 B3QBT0 B3QBT1 B3QBV5 B3QBV6 B3QBV7 B3QBW1 B3QBW2 B3QBW3 B3QBW4 B3QBW5
B3QBW6 B3QBW7 B3QBW8 B3QBW9 B3QBX0 B3QBX1 B3QBX2 B3QBX3 B3QBX4 B3QBX6 B3QBX7
B3QBX9 B3QBY0 B3QBY1 B3QBY3 B3QBY4 B3QBY5 B3QBZ9 B3QC00 B3QC01 B3QC02 B3QC04
B3QC05 B3QC92 B3QCB2 B3QCC6 B3QCE1 B3QCF3 B3QCF6 B3QCG6 B3QCG7 B3QCH1 B3QCH2
B3QCH4 B3QCH6 B3QCH9 B3QCI1 B3QCI4 B3QCQ7 B3QCR0 B3QCS4 B3QCV7 B3QCW2 B3QCX3
B3QCX5 B3QDE9 B3QE24 B3QE25 B3QEP7 B3QEW0 B3QEZ3 B3QF66 B3QFA1 B3QFB0 B3QFC6
B3QFD2 B3QFL8 B3QFM9 B3QFN0 B3QFN9 B3QFS7 B3QFS9 B3QFV9 B3QFY7 B3QFY8 B3QG72
B3QGC4 B3QGJ8 B3QGJ9 B3QGK1 B3QGK3 B3QGK4 B3QGV7 B3QH27 B3QH35 B3QH88 B3QHD0
B3QHD1 B3QHK8 B3QHR9 B3QHS1 B3QHS2 B3QHS4 B3QHV4 B3QIF6 B3QII5 B3QIK5 B3QIL0
B3QIL2 B3QIL5 B3QIM2 B3QIM6 B3QIN0 B3QIN8 B3QIV5 B3QIX7 B3QJ15 B3QJ47 B3QJF8
B3QJG5 B3QJG9 B3QJH1 B3QJH7 B3QJI8 B3QJK5 B3QJK9 B3QJM1 B3QJT3 B3QJZ1 B3QJZ9
B3QK65 B3QKG3 B3QKG4 B3QKH1 B3QKK5 B3QKL9 B3QKR4 B3QKS1 B3QKT6 B3QKU1 B3QKY6
B3QKY9 B3QL00 B3QL05 B3QL06 B3QL07 B3QL17 B3QL18 B3QL20 B3QL35 B3QL59 B3QL60
B3QL67 B3QL68 B3QL69 B3QLC5 B3QLE2 B3QLE4 B3QLF1 B3QLF4 B3QLF5 B3QLF6 B3QLF7
B3QLF8 B3QLI5 B3QLI9 B3QLJ0 B3QLJ9 B3QLK3 B3QLL5 B3QLQ8 B3QLR6 B3QLR7 B3QLU3
B3QLU4 B3QLU7 B3QLU8 B3QLW3 B3QLW6 B3QLX2 B3QLX3 B3QLX7 B3QLY7 B3QLZ4 B3QLZ5
B3QM08 B3QM09 B3QM10 B3QM19 B3QM22 B3QM27 B3QM31 B3QM32 B3QM33 B3QM41 B3QM42
B3QM44 B3QM48 B3QM51 B3QM53 B3QM61 B3QM89 B3QM90 B3QM94 B3QMA2 B3QMB2 B3QMC9
B3QMD7 B3QMF2 B3QMF5 B3QMF6 B3QMG8 B3QMJ7 B3QML1 B3QMM1 B3QMM2 B3QMM3 B3QMN0
B3QMN9 B3QMP2 B3QMP9 B3QMQ8 B3QMS4 B3QMS8 B3QMT1 B3QMU5 B3QMU7 B3QMU8 B3QN90
B3QN91 B3QN92 B3QN94 B3QNE9 B3QNL1 B3QNL4 B3QNM2 B3QNM5 B3QNN0 B3QNN5 B3QNS6
B3QNY9 B3QNZ0 B3QNZ9 B3QP02 B3QP08 B3QP11 B3QP81 B3QP85 B3QP90 B3QPB1 B3QPB3
B3QPB5 B3QPB6 B3QPI5 B3QPN4 B3QPN8 B3QPP2 B3QPR3 B3QPS3 B3QPT5 B3QPT6 B3QPT8
B3QPU5 B3QPW0 B3QPW8 B3QPW9 B3QPX0 B3QPX3 B3QPX7 B3QPY5 B3QPY8 B3QPY9 B3QPZ8
B3QQ00 B3QQ12 B3QQ33 B3QQ34 B3QQ73 B3QQ80 B3QQ83 B3QQ93 B3QQA5 B3QQA9 B3QQC6
B3QQD1 B3QQD9 B3QQE0 B3QQF5 B3QQG1 B3QQG2 B3QQG6 B3QQH1 B3QQH8 B3QQI1 B3QQI2
B3QQI4 B3QQI7 B3QQM1 B3QQR8 B3QQS1 B3QQS3 B3QQS4 B3QQS6 B3QQT5 B3QQT8 B3QQU5
B3QQU6 B3QQX9 B3QQY0 B3QQY5 B3QQY9 B3QQZ0 B3QQZ3 B3QQZ4 B3QR00 B3QR09 B3QR32
B3QR49 B3QR56 B3QR57 B3QR64 B3QR66 B3QR72 B3QR73 B3QR75 B3QR76 B3QR77 B3QR78
B3QR79 B3QR81 B3QR82 B3QR83 B3QR84 B3QR86 B3QR87 B3QR88 B3QR89 B3QR90 B3QR94
B3QR95 B3QR96 B3QR97 B3QR98 B3QR99 B3QRD2 B3QRD3 B3QRD6 B3QRD8 B3QRH7 B3QRL0
B3QRL5 B3QRL8 B3QRM0 B3QRM1 B3QRM2 B3QRM3 B3QRN9 B3QRP3 B3QRR1 B3QRS6 B3QRU2
B3QRV1 B3QS01 B3QS09 B3QS35 B3QS41 B3QS43 B3QS44 B3QS57 B3QS62 B3QS77 B3QS95
B3QSA6 B3QSB4 B3QSB6 B3QSD5 B3QSH0 B3QSH6 B3QSH7 B3QSH8 B3QSI0 B3QSI5 B3QSJ7
B3QSM8 B3QSM9 B3QSP0 B3QSP3 B3QSS3 B3QSU1 B3QSU2 B3QSZ9 B3QT42 B3QT85 B3QTC8
B3QTP2 B3QTP5 B3QTP6 B3QTQ3 B3QTR8 B3QTT2 B3QTV1 B3QTV3 B3QTV8 B3QTX2 B3QU22

B3QU24 B3QU88 B3QUD7 B3QUG2 B3QUL4 B3QUL7 B3QUN0 B3QUN2 B3QUN3 B3QUT2 B3QUT6
B3QUU7 B3QUU9 B3QUV1 B3QUY6 B3QUZ7 B3QV24 B3QV27 B3QV45 B3QV46 B3QV54 B3QV56
B3QV72 B3QV75 B3QV79 B3QV82 B3QVA0 B3QVA9 B3QVB0 B3QVE8 B3QVL0 B3QVR4 B3QVR7
B3QVU6 B3QW61 B3QW69 B3QW77 B3QW86 B3QWA4 B3QWC0 B3QWD9 B3QWF5 B3QWF8 B3QWH2
B3QWH6 B3QWI0 B3QWI1 B3QWJ0 B3QWJ5 B3QWK0 B3QWR1 B3QWS9 B3QWT5 B3QWU0 B3QWU7
B3QWX6 B3QX21 B3QX75 B3QXB5 B3QXE1 B3QXJ5 B3QXJ6 B3QXJ8 B3QXK8 B3QXK9 B3QXR3
B3QXY0 B3QXY8 B3QXZ3 B3QY08 B3QY13 B3QY14 B3QY19 B3QY20 B3QY21 B3QY22 B3QY23
B3QY24 B3QY25 B3QY26 B3QY27 B3QY28 B3QY29 B3QYB6 B3QYD2 B3QYD4 B3QYD5 B3QYD7
B3QYD8 B3QYD9 B3QYE0 B3QYE1 B3QYE2 B3QYE3 B3QYE6 B3QYE7 B3QYE8 B3QYE9 B3QYF0
B3QYF1 B3QYF4 B3QYG7 B3QYH5 B3QYH8 B3QYI4 B3QYJ3 B3QYJ4 B3QYL0 B3QYL2 B3QYL3
B3QYL5 B3QYM8 B3QYP1 B3QYQ7 B3QYR1 B3QYV9 B3QYX5 B3QYZ5 B3QZ05 B3QZ07 B3QZ38
B3QZ51 B3QZ92 B3QZ96 B3QZC7 B3QZC8 B3QZD6 B3QZE1 B3QZE2 B3QZE3 B3QZF2 B3QZF8
B3QZG1 B3QZG6 B3QZG8 B3QZG9 B3QZH0 B3QZH2 B3QZH3 B3QZH4 B3QZH5 B3QZK4 B3QZM3
B3QZN3 B3QZS0 B3QZS5 B3QZS8 B3QZT1 B3QZT2 B3QZT4 B3QZV0 B3QZV1 B3QZV8 B3QZW0
B3QZW6 B3QZW9 B3QZX0 B3QZZ1 B3QZZ2 B3QZZ5 B3QZZ6 B3QZZ7 B3QZZ8 B3R000 B3R001
B3R002 B3R005 B3R006 B3R007 B3R010 B3R014 B3R015 B3R016 B3R018 B3R022 B3R068
B3R075 B3R0A1 B3R0D7 B3R0E5 B3R0E7 B3R0E8 B3R0F3 B3R0F5 B3R0F6 B3R0F8 B3R0G4
B3R0G5 B3R0I1 B3R0I8 B3R0J0 B3R0J1 B3R0K3 B3R0L6 B3R0M3 B3R0N4 B3R0P0 B3R0P2
B3R0P5 B3R0Q2 B3R0X7 B3R0X9 B3R113 B3R117 B3R181 B3R1A0 B3R1E2 B3R1E3 B3R1E4
B3R1E6 B3R1I2 B3R1I3 B3R1I4 B3R1J2 B3R1J7 B3R1J8 B3R1K1 B3R1K6 B3R1L4 B3R1M6
B3R1M7 B3R1N6 B3R1N9 B3R1R9 B3R1S2 B3R1X2 B3R1Y0 B3R1Z3 B3R1Z6 B3R1Z9 B3R215
B3R259 B3R290 B3R297 B3R2B1 B3R2B4 B3R2B6 B3R2B7 B3R2C2 B3R2C8 B3R2D1 B3R2F9
B3R2H3 B3R2J3 B3R2K7 B3R2P4 B3R2P5 B3R2P7 B3R2Y2 B3R2Y3 B3R312 B3R368 B3R370
B3R376 B3R396 B3R397 B3R398 B3R399 B3R3A1 B3R3C0 B3R3D9 B3R3E4 B3R3G1 B3R3N0
B3R3N5 B3R3R6 B3R3T1 B3R3V4 B3R3W1 B3R3W3 B3R3W5 B3R3Z9 B3R401 B3R404 B3R447
B3R450 B3R474 B3R475 B3R478 B3R496 B3R4A7 B3R4D1 B3R4J1 B3R4J3 B3R4J4 B3R4J5
B3R4L6 B3R4T1 B3R4T2 B3R4U4 B3R4W0 B3R4W1 B3R4W2 B3R4X7 B3R508 B3R570 B3R5F6
B3R5H2 B3R5H4 B3R5J8 B3R5L0 B3R5L8 B3R5N3 B3R5Q1 B3R5Q8 B3R5S0 B3R5S1 B3R5T4
B3R5Y0 B3R605 B3R622 B3R647 B3R663 B3R678 B3R684 B3R6A3 B3R6B8 B3R6C0 B3R6D1
B3R6D3 B3R6D5 B3R6D6 B3R6D7 B3R6E8 B3R6F7 B3R6F8 B3R6G5 B3R6G6 B3R6G7 B3R6J3
B3R6J4 B3R6K3 B3R6L9 B3R6Q0 B3R6Q3 B3R6R0 B3R6R4 B3R6S4 B3R6U7 B3R6V7 B3R6V8
B3R6W7 B3R6W8 B3R703 B3R722 B3R793 B3R794 B3R795 B3R798 B3R7A1 B3R7A2 B3R7B8
B3R7C6 B3R7D7 B3R7E2 B3R7E3 B3R7E4 B3R7E5 B3R7E6 B3R7E9 B3R7F0 B3R7F1 B3R7F2
B3R7F3 B3R7F4 B3R7F5 B3R7F6 B3R7J7 B3R7K5 B3R7R2 B3R7R3 B3R7R4 B3R7R5 B3R7R6
B3R7R7 B3R7R8 B3R7R9 B3R7S0 B3R7S1 B3R7S2 B3R7S3 B3R7S9 B3R7T2 B3R7T3 B3R7T5
B3R7T6 B3R7T7 B3R7T8 B3R7T9 B3R7U0 B3R808 B3R886 B3R894 B3R895 B3R897 B3R898
B3R899 B3R8A0 B3R8A8 B3R8D4 B3R8D5 B3R8S4 B3R9F2 B3RAC1 B3RBH4 B3RBL3 B3RBL8
B3RD79 B3RDA4 B3RHD9 B3RHI0 B3RHK4 B3RHR9 B3RHU5 B3RHV0 B3RID0 B3RLE6 B3RNE8
B3RNR8 B3RNT0 B3RPX4 B3RPX6 B3RQN1 B3RTL9 B3RVF0 B3RVZ1 B3RYZ0 B3S0D3 B3S4I5
B3S6Y0 B3S866 B3SAE4 B3SBB5 B3SRQ2 B3SRQ6 B3SRR0 B3SRR2 B3SRR3 B3SRR4 B3SRR7
B3SRR8 B3SRS0 B3SRS1 B3SRS2 B3SRS5 B3SRS6 B3SRT4 B3SRU4 B3SRV0 B3SRV2 B3SRV3
B3SRV4 B3SRV7 B3SRV8 B3SRX0 B3SRX3 B3SRX4 B3SRX6 B3SRX8 B3SRY1 B3SRY2 B3STP6
B3STU3 B3TKC8 B3TMR8 B3TN43 B3TN49 B3TN59 B3TN70 B3TN72 B3TN82 B3TN86 B3TN87
B3TN89 B3TN90 B3TN95 B3TNB4 B3TPQ6 B3TPQ7 B3V4Q2 B3V4S7 B3VI55 B3VI56 B3VKQ2

B3VML1 B3VML2 B3VML3 B3VMP4 B3VTV7 B3W6N4 B3W6Q9 B3W6R4 B3W6R6 B3W6W6 B3W6W7
B3W6W8 B3W6W9 B3W782 B3W783 B3W789 B3W7C1 B3W7E5 B3W7E6 B3W7E7 B3W7F8 B3W7G1
B3W7G6 B3W7I3 B3W7I5 B3W7I7 B3W7M8 B3W8L5 B3W8N4 B3W8T2 B3W8V7 B3W8W8 B3W8Y1
B3W952 B3W953 B3W963 B3W9A1 B3W9A3 B3W9A8 B3W9F8 B3W9M6 B3W9T6 B3W9W2 B3W9W3
B3W9W4 B3W9W7 B3W9W8 B3W9X7 B3W9Y5 B3W9Y8 B3W9Z0 B3W9Z1 B3WA01 B3WA02 B3WA08
B3WA09 B3WA35 B3WA39 B3WA82 B3WAE3 B3WAH9 B3WAI0 B3WAI6 B3WAJ1 B3WAJ2 B3WAJ3
B3WAJ4 B3WAJ5 B3WAJ7 B3WAK0 B3WAK2 B3WAK3 B3WAK4 B3WAK5 B3WAK6 B3WAK7 B3WAK8
B3WAL0 B3WAL2 B3WAL3 B3WAL4 B3WAL5 B3WAL6 B3WAL7 B3WAL8 B3WAM2 B3WAM3 B3WAM4
B3WAM7 B3WAM8 B3WAN3 B3WAP7 B3WAR0 B3WAR8 B3WB42 B3WBI0 B3WBT3 B3WBV6 B3WBV7
B3WC55 B3WC56 B3WC69 B3WC70 B3WC71 B3WC74 B3WC77 B3WCB2 B3WCC9 B3WCD5 B3WCJ6
B3WCK9 B3WCM0 B3WCP4 B3WCQ5 B3WCQ8 B3WCR0 B3WCR2 B3WCW5 B3WCW6 B3WCX4 B3WCZ5
B3WD16 B3WD20 B3WD44 B3WD54 B3WD57 B3WD58 B3WDB0 B3WDC0 B3WDK7 B3WDL0 B3WDL1
B3WDP7 B3WDV5 B3WDV9 B3WDX7 B3WDY1 B3WDY6 B3WDZ1 B3WE10 B3WE13 B3WE28 B3WE33
B3WE34 B3WE38 B3WE40 B3WE42 B3WE47 B3WE52 B3WE54 B3WE58 B3WE64 B3WE80 B3WE84
B3WE94 B3WEA2 B3WEA7 B3WEA8 B3WEC9 B3WED0 B3WED1 B3WED2 B3WED3 B3WED4 B3WED6
B3WED7 B3WEF5 B3WEG3 B3WEI0 B3WEI1 B3WEI2 B3WEJ1 B3WEJ2 B3WEJ5 B3WEK6 B3WEL4
B3WEL7 B3WEL8 B3WEM2 B3WEM3 B3WEN3 B3WEN7 B3WEQ6 B3WEQ7 B3WEQ9 B3WER2 B3WER4
B3WER5 B3WER9 B3WES5 B3WES7 B3WES8 B3WET9 B3WEU0 B3WEU1 B3WEU4 B3WEW2 B3WEW9
B3WEX2 B3WEY3 B3WEY4 B3WEY5 B3WEZ5 B3WF00 B3WF19 B3WF30 B3WF43 B3WF44 B3WF49
B3WF53 B3WF58 B3WF63 B3WF72 B3WF81 B3WF93 B3WF98 B3WFA0 B3WFB1 B3WFB7 B3XPQ7
B3XZI5 B3Y679 B3Y681 B3Y682 B3Y683 B4DH59 B4DU55 B4DX44 B4DXR9 B4E2M5 B4E558
B4E566 B4E581 B4E582 B4E5A8 B4E5A9 B4E5B0 B4E5B1 B4E5B2 B4E5B3 B4E5B5 B4E5B6
B4E5B9 B4E5C0 B4E5C1 B4E5C2 B4E5C3 B4E5C4 B4E5C5 B4E5C6 B4E5C7 B4E5C8 B4E5C9
B4E5D0 B4E5D1 B4E5D2 B4E5D3 B4E5D4 B4E5D5 B4E5D6 B4E5D7 B4E5D8 B4E5D9 B4E5E2
B4E5E3 B4E5E4 B4E5E5 B4E5E6 B4E5E7 B4E5F3 B4E5F6 B4E5M6 B4E5M7 B4E5N2 B4E5N5
B4E5R3 B4E5R5 B4E5R6 B4E5R8 B4E5R9 B4E5S1 B4E5V2 B4E5W6 B4E5W7 B4E5W9 B4E5X0
B4E5X1 B4E5X2 B4E5X9 B4E5Y9 B4E5Z6 B4E603 B4E634 B4E637 B4E640 B4E641 B4E642
B4E643 B4E6D6 B4E6D7 B4E6F9 B4E6J0 B4E6J1 B4E6K0 B4E6K1 B4E6U6 B4E6U7 B4E7A8
B4E7D1 B4E7D2 B4E7H2 B4E7I6 B4E7I7 B4E7I8 B4E7I9 B4E7K9 B4E7L1 B4E7L2 B4E7V8
B4E7W5 B4E7X5 B4E7X6 B4E7X7 B4E7Y5 B4E7Y8 B4E7Z1 B4E7Z2 B4E800 B4E852 B4E8C4
B4E8D2 B4E8D6 B4E8E0 B4E8E5 B4E8E6 B4E8G9 B4E8K1 B4E8N5 B4E8N6 B4E8N8 B4E8S9
B4E8V9 B4E8Y7 B4E8Y8 B4E8Z7 B4E900 B4E904 B4E9B2 B4E9C7 B4E9F9 B4E9G1 B4E9G4
B4E9G6 B4E9G8 B4E9I1 B4E9I9 B4E9J1 B4E9L4 B4E9L5 B4E9L6 B4E9M9 B4E9P3 B4EA64
B4EA67 B4EAF5 B4EAH3 B4EAM5 B4EAN3 B4EAN4 B4EAN6 B4EAP1 B4EAQ8 B4EAR1 B4EAR4
B4EAR7 B4EAS0 B4EAS1 B4EAW4 B4EAW5 B4EAW8 B4EAW9 B4EAX1 B4EAX2 B4EAY2 B4EAY3
B4EAZ1 B4EB34 B4EB39 B4EB41 B4EB42 B4EB45 B4EB49 B4EB94 B4EBB9 B4EBC4 B4EBD1
B4EBD9 B4EBE8 B4EBG2 B4EBH0 B4EBH1 B4EBH3 B4EBJ2 B4EBK8 B4EBK9 B4EBL7 B4EBM2
B4EBM4 B4EBP5 B4EBP8 B4EBQ9 B4EBR1 B4EBR4 B4EBT3 B4EBT8 B4EBT9 B4EBY3 B4EBY6
B4EBZ2 B4EC22 B4EC23 B4EC68 B4EC82 B4ECC5 B4ECC6 B4ECC7 B4ECC9 B4ECL7 B4ECM5
B4ECM8 B4ECN0 B4ECN1 B4ECN6 B4ECP1 B4ECP3 B4ECR7 B4ECT0 B4ECT2 B4ECT3 B4ECX5
B4ECY8 B4ECZ8 B4ECZ9 B4ED00 B4ED40 B4ED73 B4ED80 B4ED83 B4EDA3 B4EDB1 B4EDB2
B4EDH3 B4EDH7 B4EDS3 B4EDS4 B4EDU1 B4EDW1 B4EDX1 B4EDX5 B4EDZ1 B4EDZ2 B4EDZ4
B4EDZ7 B4EDZ8 B4EE21 B4EE24 B4EE50 B4EEQ6 B4EES4 B4EES5 B4EES6 B4EES7 B4EET1
B4EET3 B4EET4 B4EET5 B4EET6 B4EEW1 B4EEW4 B4EEZ3 B4EF26 B4EF27 B4EF28 B4EFB0

B4EFI7 B4EFI8 B4EFK2 B4EFS7 B4EFZ7 B4EG04 B4EG09 B4EH54 B4EHE6 B4EHF8 B4EHJ0
B4EHJ1 B4EHJ2 B4EHL5 B4EIP2 B4EJ37 B4EJA6 B4EJC2 B4EJQ7 B4EK21 B4EK77 B4EK78
B4EKM1 B4EKS2 B4EKX6 B4ELA5 B4ELG6 B4EM48 B4EN17 B4EN27 B4EN29 B4EN31 B4ENA5
B4ENZ2 B4EPB7 B4EPK7 B4ESJ0 B4ESN7 B4ESR1 B4EST0 B4ESU3 B4ESU4 B4ESU7 B4ESV1
B4ESY2 B4ESY3 B4ESY6 B4ESY8 B4ESY9 B4ESZ8 B4ESZ9 B4ET04 B4ET09 B4ET17 B4ET24
B4ET25 B4ET26 B4ET28 B4ET33 B4ET62 B4ET91 B4ET92 B4ET93 B4ETA2 B4ETA5 B4ETB9
B4ETC5 B4ETI7 B4ETK6 B4ETK7 B4ETK8 B4ETK9 B4ETL0 B4ETL2 B4ETN5 B4ETN7 B4ETP0
B4ETP1 B4ETP3 B4ETP5 B4ETP6 B4ETP7 B4ETP8 B4ETX3 B4EU12 B4EU13 B4EU17 B4EU19
B4EU20 B4EU30 B4EU31 B4EU34 B4EU40 B4EU52 B4EU53 B4EU54 B4EU60 B4EU76 B4EU78
B4EU79 B4EUD1 B4EUD2 B4EUD3 B4EUD8 B4EUE1 B4EUE2 B4EUF0 B4EUF2 B4EUF6 B4EUG3
B4EUH7 B4EUH8 B4EUI8 B4EUQ7 B4EUR1 B4EUS0 B4EUS1 B4EUT1 B4EUV0 B4EUV1 B4EUV6
B4EUV8 B4EUV9 B4EUW4 B4EUW6 B4EUW7 B4EV01 B4EV08 B4EV14 B4EV26 B4EV28 B4EV71
B4EV85 B4EV93 B4EVC0 B4EVC4 B4EVC6 B4EVD2 B4EVE1 B4EVE7 B4EVF5 B4EVG2 B4EVM7
B4EVR1 B4EVR4 B4EVR6 B4EVR7 B4EVV5 B4EVV7 B4EVV8 B4EVW1 B4EVZ1 B4EW02 B4EW04
B4EW08 B4EW41 B4EW46 B4EW50 B4EW57 B4EW58 B4EWA1 B4EWA2 B4EWA3 B4EWA4 B4EWB0
B4EWC2 B4EWC9 B4EWE3 B4EWH5 B4EWH7 B4EWH8 B4EWL7 B4EWL8 B4EWM9 B4EWW4 B4EWY8
B4EWZ6 B4EX21 B4EX23 B4EX28 B4EX40 B4EX94 B4EX96 B4EXC9 B4EXE2 B4EXE3 B4EXL1
B4EXL9 B4EXM0 B4EXS5 B4EXU2 B4EXV4 B4EY23 B4EY24 B4EY25 B4EY26 B4EY51 B4EY52
B4EY59 B4EY61 B4EY70 B4EY76 B4EY85 B4EY88 B4EY89 B4EYB8 B4EYN9 B4EYQ7 B4EYR7
B4EYR8 B4EYS9 B4EYT2 B4EYT7 B4EYT8 B4EYU2 B4EYU5 B4EYU9 B4EYV0 B4EYV1 B4EYV2
B4EYV3 B4EYV7 B4EYV8 B4EYV9 B4EYW2 B4EYW6 B4EZ30 B4EZ39 B4EZ42 B4EZ57 B4EZ94
B4EZ95 B4EZA3 B4EZB4 B4EZD9 B4EZF9 B4EZG9 B4EZH2 B4EZM4 B4EZM6 B4EZM8 B4EZM9
B4EZQ3 B4EZQ7 B4EZR2 B4EZS7 B4EZS9 B4EZT2 B4EZT3 B4EZT6 B4EZT7 B4EZU4 B4EZU5
B4EZU6 B4EZU8 B4EZV5 B4EZY9 B4F016 B4F025 B4F026 B4F045 B4F047 B4F055 B4F057
B4F058 B4F059 B4F063 B4F065 B4F088 B4F0A6 B4F0B1 B4F0D4 B4F0D5 B4F0D8 B4F0M6
B4F0M7 B4F0N7 B4F0N8 B4F0N9 B4F0Q0 B4F0Q1 B4F0S0 B4F0S1 B4F0S5 B4F0U0 B4F0U3
B4F0U4 B4F0U5 B4F0U7 B4F0W3 B4F0W4 B4F0W5 B4F0W6 B4F0W9 B4F0X0 B4F0X6 B4F0X7
B4F0Y2 B4F0Z5 B4F0Z6 B4F110 B4F119 B4F120 B4F132 B4F134 B4F137 B4F139 B4F161
B4F168 B4F169 B4F171 B4F172 B4F176 B4F178 B4F183 B4F192 B4F194 B4F196 B4F197
B4F1A3 B4F1A4 B4F1A5 B4F1B5 B4F1C2 B4F1G5 B4F1G7 B4F1G9 B4F1H1 B4F1H4 B4F1H6
B4F1I3 B4F1I4 B4F1I5 B4F1I6 B4F1I7 B4F1I8 B4F1I9 B4F1J1 B4F1J2 B4F1J3 B4F1J4
B4F1J5 B4F1J6 B4F1J7 B4F1J8 B4F1J9 B4F1K0 B4F1K2 B4F1K3 B4F1K5 B4F1K7 B4F1K8
B4F1L0 B4F1L5 B4F1L6 B4F1M1 B4F1M6 B4F1Q1 B4F1Q2 B4F1Q4 B4F1U3 B4F1U6 B4F1V2
B4F1V3 B4F1X3 B4F1X7 B4F1Z6 B4F203 B4F220 B4F223 B4F224 B4F225 B4F226 B4F231
B4F233 B4F234 B4F249 B4F252 B4F256 B4F265 B4F266 B4F271 B4F272 B4F275 B4F276
B4F277 B4F278 B4F280 B4F298 B4F2A0 B4F2A1 B4F2A6 B4F2A7 B4F2A8 B4F2B2 B4F2B5
B4F2B7 B4F2B9 B4F2C0 B4F2C2 B4F2C3 B4F2C9 B4F2D0 B4F2D2 B4F2D3 B4F2D6 B4F2D7
B4F2E4 B4F2E7 B4F2F5 B4F2G9 B4F2H6 B4F2I2 B4F2I3 B4F2I4 B4F2M1 B4F2M2 B4F2M5
B4F2N1 B4F2T3 B4F2T5 B4F2T8 B4F2T9 B4F2U3 B4F2U6 B4F2V1 B4F2V2 B4F2V5 B4F2V6
B4F2X9 B4F6G1 B4F6K2 B4F6Q9 B4F6U4 B4F6V6 B4F6W9 B4F6X2 B4F721 B4F739 B4F769
B4F777 B4F7C4 B4F7E9 B4F7L9 B4F8Z1 B4F9A4 B4FAT0 B4FFK9 B4FGS2 B4FHF0 B4FHY0
B4FK49 B4FNX4 B4FR29 B4FT40 B4FXN6 B4FZ81 B4G072 B4G0F3 B4G2G5 B4G325 B4G3E2
B4G518 B4G5C9 B4G5J0 B4G652 B4G653 B4G7U3 B4G7Y6 B4G8N3 B4G8R5 B4G9L6 B4G9X6
B4GA28 B4GAJ1 B4GAY7 B4GBA9 B4GBN7 B4GBR1 B4GCP2 B4GD81 B4GDB3 B4GDK5 B4GDM5

B4GDM7 B4GDU3 B4GDX4 B4GE20 B4GEL3 B4GEU1 B4GEU5 B4GF49 B4GFE6 B4GFN6 B4GFS1
B4GG58 B4GGT6 B4GH42 B4GHY8 B4GIB1 B4GIF2 B4GIJ0 B4GIM3 B4GIU9 B4GJ61 B4GK79
B4GKQ3 B4GLK8 B4GMG4 B4GMI7 B4GN43 B4GN46 B4GNT0 B4GP48 B4GP93 B4GQJ7 B4GR63
B4GRJ2 B4GS98 B4GSC0 B4GSH1 B4GT01 B4GT53 B4GTK1 B4GU19 B4GUT1 B4GUY6 B4GUZ2
B4GW22 B4GWP5 B4GWY0 B4GX14 B4GXC8 B4GXH4 B4GY79 B4GYC7 B4GYU1 B4GZ07 B4GZ20
B4GZZ4 B4H0S8 B4H1X9 B4H2S0 B4H303 B4H4H5 B4H4I3 B4H4U3 B4H4X0 B4H538 B4H581
B4H732 B4H7P4 B4H957 B4H9N7 B4H9W2 B4HBD2 B4HBU3 B4HE12 B4HEN5 B4HEQ8 B4HFJ3
B4HFV9 B4HGE0 B4HGK2 B4HHG8 B4HI06 B4HIK1 B4HJA7 B4HJC0 B4HKF2 B4HKJ7 B4HL48
B4HLH4 B4HMH2 B4HMQ1 B4HMV9 B4HMY3 B4HN85 B4HND9 B4HPU1 B4HQ29 B4HQJ2 B4HQT6
B4HR14 B4HR78 B4HRL4 B4HRQ6 B4HSF8 B4HSI1 B4HSL3 B4HSL7 B4HT57 B4HTA1 B4HUE4
B4HVA2 B4HW93 B4HWD7 B4HWV2 B4HWV6 B4HXA6 B4HXL7 B4HYP0 B4HYQ2 B4HZE7 B4HZE8
B4I029 B4I0K4 B4I0W6 B4I100 B4I195 B4I1C2 B4I298 B4I2A8 B4I2S0 B4I3P3 B4I3R2
B4I3S1 B4I3X6 B4I4N0 B4I4Y1 B4I540 B4I5W3 B4I7J9 B4I7U3 B4I8G2 B4I9F6 B4I9N7
B4I9X1 B4IAA7 B4IAD1 B4IAQ8 B4IB36 B4IBA4 B4IBI9 B4IBY3 B4IC49 B4IDM2 B4IDV8
B4IF39 B4IFU5 B4IG10 B4IG61 B4IH30 B4II37 B4II46 B4II68 B4IIC7 B4IJG7 B4IJG8
B4IK21 B4IK33 B4IL62 B4IL68 B4IL76 B4IMF6 B4IMH3 B4IMI7 B4IMQ8 B4ISL0 B4ISV0
B4IUG3 B4IUH5 B4IWN1 B4IX08 B4IXD3 B4IYG8 B4J045 B4J184 B4J1U4 B4J1U5 B4J272
B4J340 B4J3A3 B4J3F1 B4J3F3 B4J497 B4J4E7 B4J4G8 B4J4Y2 B4J5B3 B4J5U3 B4J613
B4J672 B4J6D5 B4J6M4 B4J6N6 B4J6Q0 B4J6R7 B4J789 B4J8A0 B4J8H6 B4J9K1 B4J9U3
B4J9W6 B4JAS7 B4JAX0 B4JB43 B4JBC4 B4JBE6 B4JBI7 B4JCG4 B4JCS7 B4JCV8 B4JCX8
B4JDU5 B4JDZ5 B4JE52 B4JEY0 B4JF25 B4JGW7 B4JGX4 B4JHI7 B4JHJ7 B4JHK2 B4JHP4
B4JIY0 B4JJQ3 B4JKB7 B4JKK1 B4JLC3 B4JLL3 B4JLU7 B4JM29 B4JMQ2 B4JND4 B4JNS2
B4JPR2 B4JPT9 B4JQY3 B4JRX2 B4JSI3 B4JSL2 B4JSP3 B4JSW8 B4JT42 B4JTN0 B4JTS9
B4JTV9 B4JUE4 B4JUG8 B4JUM0 B4JUT1 B4JV25 B4JVG7 B4JVK9 B4JVN0 B4JW81 B4JW83
B4JWA1 B4JWF5 B4JWL5 B4JWR9 B4JWS7 B4JXG8 B4JXP7 B4JXU2 B4JXV2 B4JYN1 B4JYN3
B4JZG4 B4JZG8 B4JZQ9 B4K250 B4K4X6 B4K4Y6 B4K5R2 B4K5S6 B4K616 B4K617 B4K6T8
B4K7S5 B4K843 B4K892 B4K8A4 B4K8J8 B4K8W7 B4K934 B4K982 B4KA23 B4KA44 B4KBH3
B4KBI4 B4KBJ0 B4KBL1 B4KCD5 B4KCG1 B4KCL9 B4KDI2 B4KDK5 B4KE10 B4KEN8 B4KF66
B4KFU7 B4KG03 B4KGS4 B4KGX9 B4KH32 B4KHL5 B4KHX1 B4KI53 B4KID9 B4KIE3 B4KIW3
B4KJ11 B4KJS5 B4KKN1 B4KKN5 B4KLC0 B4KLL0 B4KLY1 B4KLY7 B4KMF8 B4KML2 B4KN00
B4KN44 B4KNA9 B4KNC5 B4KND9 B4KNM1 B4KNN9 B4KPG8 B4KPY6 B4KQQ4 B4KQU8 B4KRQ4
B4KS18 B4KSC0 B4KSL6 B4KSR4 B4KSY3 B4KT48 B4KT65 B4KTH6 B4KTK4 B4KTN5 B4KVI7
B4KX02 B4KXI8 B4KY56 B4KY72 B4KYT0 B4KYX8 B4KYY0 B4KZ45 B4KZN6 B4L0G9 B4L1K2
B4L1Z2 B4L1Z8 B4L2J8 B4L340 B4L529 B4L629 B4L6Q5 B4L6T9 B4L710 B4L760 B4L7F0
B4L7V4 B4L8C7 B4L8M2 B4LDA6 B4LDJ6 B4LE67 B4LEJ0 B4LF72 B4LFQ9 B4LFW2 B4LGB4
B4LIG6 B4LIK8 B4LJT7 B4LJT9 B4LKE5 B4LKS9 B4LL39 B4LM02 B4LM71 B4LM90 B4LMA2
B4LMK1 B4LMQ3 B4LMR9 B4LN42 B4LNA1 B4LNQ8 B4LNV5 B4LP08 B4LP65 B4LPB6 B4LPP8
B4LQ21 B4LQD0 B4LQD3 B4LQR5 B4LQW4 B4LQY8 B4LRB9 B4LRY2 B4LS78 B4LS82 B4LSE6
B4LTW0 B4LTY6 B4LUA5 B4LUF5 B4LVP8 B4LVR8 B4LVS8 B4LW09 B4LWK0 B4LWT7 B4LWZ8
B4LX81 B4LXW1 B4LY66 B4LYI3 B4LZ60 B4LZ88 B4LZW7 B4M140 B4M1W5 B4M205 B4M2R8
B4M357 B4M375 B4M3W0 B4M401 B4M416 B4M4L4 B4M4W4 B4M4X4 B4M5A7 B4M5L0 B4M5T5
B4M5T7 B4M686 B4M693 B4M6M6 B4M703 B4M7J1 B4M7P8 B4M7T6 B4M818 B4M891 B4M8J8
B4M8L8 B4M8Y0 B4M9A8 B4M9F7 B4MA02 B4MA12 B4MA85 B4MB32 B4MCL6 B4MDT2 B4MDZ0
B4MEW8 B4MFM2 B4MHR1 B4MIW0 B4MIX7 B4MKC7 B4MLV0 B4MM23 B4MNL1 B4MP81 B4MQL4

B4MQL8 B4MQT3 B4MQY1 B4MR33 B4MR46 B4MR59 B4MR74 B4MR84 B4MRE7 B4MRW2 B4MRZ8
B4MRZ9 B4MSH7 B4MTY2 B4MU54 B4MUM8 B4MV81 B4MVH6 B4MVR2 B4MX21 B4MX71 B4MXH8
B4MXR6 B4MXR8 B4MXW3 B4MY65 B4MY75 B4MY77 B4MYA1 B4MYI5 B4MYR2 B4MZ79 B4N002
B4N0L0 B4N0P7 B4N116 B4N1C3 B4N1G8 B4N1L0 B4N1V2 B4N208 B4N278 B4N3B0 B4N4D9
B4N4E1 B4N549 B4N665 B4N7R4 B4N7S9 B4N8G6 B4N8G7 B4N8Z4 B4N943 B4N984 B4N9D3
B4N9T4 B4NAB3 B4NAJ1 B4NAU8 B4NB54 B4NBB0 B4NBL5 B4NBL6 B4NC41 B4NDE5 B4NDG5
B4NE56 B4NE93 B4NEH6 B4NER4 B4NEU3 B4NEU8 B4NFJ7 B4NFS5 B4NFY1 B4NG41 B4NHI4
B4NI64 B4NIC3 B4NIM9 B4NJR8 B4NJW0 B4NKI9 B4NKT1 B4NLG2 B4NN33 B4NP05 B4NP88
B4NPT0 B4NSP6 B4NSS7 B4NTY9 B4NU50 B4NUB3 B4NW98 B4NWM2 B4NX24 B4NXB8 B4NXB9
B4NXF7 B4NXG9 B4NXN5 B4NY70 B4NYQ2 B4NYT3 B4NYU1 B4NYV0 B4P0U1 B4P116 B4P1N5
B4P238 B4P2P8 B4P2Y8 B4P3Q9 B4P3T9 B4P3W7 B4P4H2 B4P4K8 B4P528 B4P562 B4P5F7
B4P5U9 B4P6M6 B4P6P9 B4P6S9 B4P6V4 B4P6W7 B4P7H8 B4P7Q3 B4P8I0 B4P8V1 B4P925
B4P9A8 B4P9H8 B4PAP8 B4PAR6 B4PAZ6 B4PCA1 B4PD06 B4PDM5 B4PES4 B4PEU8 B4PEV9
B4PF83 B4PG99 B4PIP5 B4PIS2 B4PJ01 B4PKZ7 B4PL68 B4PLB3 B4PMC6 B4PNE2 B4PNN4
B4PNV2 B4PP90 B4PPG0 B4PPK2 B4PPK3 B4PPT4 B4PR18 B4PRE2 B4PRJ9 B4PRU6 B4PS24
B4PS83 B4PST7 B4PTS9 B4PU14 B4PU44 B4PU77 B4PUD0 B4PUD1 B4PUG5 B4PUM2 B4PV71
B4PVH6 B4PVI7 B4PXF5 B4PY37 B4PY93 B4PYA4 B4PYH5 B4PYR0 B4PYW0 B4PZ52 B4PZB4
B4Q034 B4Q068 B4Q0P1 B4Q0P5 B4Q0Y7 B4Q1B6 B4Q2F5 B4Q2J2 B4Q354 B4Q572 B4Q5Z1
B4Q785 B4Q7F5 B4Q865 B4Q8A7 B4Q8M1 B4Q9T2 B4Q9T6 B4Q9Z1 B4QB64 B4QB88 B4QBH8
B4QC10 B4QCR6 B4QD77 B4QE02 B4QEE3 B4QEJ9 B4QFD2 B4QFM2 B4QFP7 B4QFZ8 B4QGZ1
B4QHD6 B4QHG6 B4QHS6 B4QHU5 B4QI55 B4QID8 B4QIG6 B4QIJ9 B4QJ46 B4QK53 B4QKW3
B4QL99 B4QLQ2 B4QMP1 B4QPV0 B4QQE2 B4QR64 B4QRJ0 B4QS37 B4QSM8 B4QT07 B4QT24
B4QTL6 B4QU20 B4QUC0 B4QUC1 B4QUF6 B4QV17 B4QVI2 B4QVL3 B4QVR7 B4QVV3 B4QVW6
B4QW91 B4QX57 B4QYZ3 B4QZ45 B4QZV5 B4R089 B4R090 B4R0A5 B4R1Z3 B4R222 B4R338
B4R345 B4R3T1 B4R3W7 B4R4E9 B4R4H1 B4R694 B4R6G1 B4R7V3 B4R8K1 B4R8K2 B4R8K3
B4R8K4 B4R8K5 B4R8L1 B4R8L2 B4R8L3 B4R8L6 B4R8L7 B4R8L8 B4R8M0 B4R8M1 B4R8M2
B4R8M3 B4R8M4 B4R8M5 B4R8M6 B4R8M7 B4R8M9 B4R8N0 B4R8N2 B4R8N3 B4R8N4 B4R8N6
B4R8N8 B4R8P0 B4R8P1 B4R8P2 B4R8Q7 B4R8Q8 B4R8T3 B4R8T4 B4R8T7 B4R8T9 B4R8U3
B4R906 B4R947 B4R955 B4R966 B4R967 B4R968 B4R981 B4R9A9 B4R9C1 B4R9C4 B4R9C5
B4R9C6 B4R9L7 B4R9Q3 B4R9R7 B4R9Z1 B4R9Z2 B4RA25 B4RA70 B4RAD7 B4RAE8 B4RAL5
B4RB18 B4RB34 B4RB35 B4RB97 B4RBF7 B4RBG0 B4RBI9 B4RBJ4 B4RBL4 B4RBN1 B4RBP5
B4RBQ2 B4RBS4 B4RBV1 B4RBW5 B4RBX4 B4RBX5 B4RBX8 B4RBZ1 B4RBZ3 B4RBZ4 B4RBZ9
B4RC03 B4RC21 B4RC42 B4RC43 B4RC46 B4RC48 B4RC49 B4RC51 B4RC55 B4RC58 B4RC70
B4RC79 B4RC87 B4RCA4 B4RCC0 B4RCC2 B4RCC6 B4RCD1 B4RCD6 B4RCE2 B4RCH3 B4RCK3
B4RCL0 B4RCL1 B4RCN8 B4RCP4 B4RCQ8 B4RCR3 B4RCS1 B4RCS3 B4RCS6 B4RCT1 B4RCU1
B4RCU4 B4RCV1 B4RCW4 B4RCZ4 B4RD04 B4RD10 B4RD11 B4RD56 B4RD63 B4RD64 B4RD67
B4RDA2 B4RDH0 B4RDI0 B4RDI1 B4RDT4 B4RDT6 B4RDT8 B4RDU2 B4RDU4 B4RDV7 B4RDX2
B4RE11 B4RE90 B4RE95 B4RED8 B4REE0 B4REI0 B4RET2 B4REY7 B4RF16 B4RF17 B4RF90
B4RFE9 B4RFG1 B4RFG2 B4RFQ5 B4RFS2 B4RFS8 B4RFS9 B4RFX5 B4RFX6 B4RG33 B4RG67
B4RGF4 B4RGF5 B4RGP0 B4RGS2 B4RGV8 B4RGW0 B4RH30 B4RH54 B4RHE8 B4RHH0 B4RHN9
B4RHV8 B4RI01 B4RIW2 B4RIW3 B4RIW5 B4RIY4 B4RIY7 B4RJ05 B4RJ40 B4RJ66 B4RJ69
B4RJC7 B4RJC9 B4RJG2 B4RJG3 B4RJH7 B4RJI8 B4RJJ5 B4RJJ6 B4RJL0 B4RJM3 B4RJN1
B4RJN2 B4RJN3 B4RJN4 B4RJQ3 B4RJQ9 B4RJR3 B4RJR8 B4RJT2 B4RJT3 B4RJT4 B4RJT7
B4RJU3 B4RJV5 B4RJV8 B4RJY2 B4RJY6 B4RJZ4 B4RJZ8 B4RK02 B4RK03 B4RK11 B4RK29

B4RK50 B4RK59 B4RK78 B4RK79 B4RK82 B4RK90 B4RK97 B4RK98 B4RKA2 B4RKA3 B4RKC9
B4RKD1 B4RKD2 B4RKE5 B4RKF0 B4RKF2 B4RKH1 B4RKH3 B4RKR4 B4RKR9 B4RKY8 B4RL14
B4RL16 B4RL22 B4RL23 B4RL28 B4RL47 B4RL51 B4RL58 B4RL59 B4RL77 B4RL93 B4RLC9
B4RLD4 B4RLE6 B4RLE8 B4RLF4 B4RLI6 B4RLK3 B4RLK9 B4RLL0 B4RLL1 B4RLL6 B4RLV1
B4RLV5 B4RLV8 B4RLX4 B4RLX6 B4RLX8 B4RLX9 B4RM12 B4RM65 B4RM67 B4RM84 B4RM85
B4RM87 B4RM90 B4RMA9 B4RMB1 B4RMB8 B4RMC3 B4RMD5 B4RMG0 B4RMG2 B4RMG4 B4RMG6
B4RMH7 B4RMH8 B4RMI6 B4RMJ5 B4RMM2 B4RMZ1 B4RMZ3 B4RMZ8 B4RN20 B4RN40 B4RN45
B4RN46 B4RN50 B4RN52 B4RNE5 B4RNF8 B4RNG7 B4RNL5 B4RNN0 B4RNP1 B4RNP4 B4RNP9
B4RNT1 B4RNV9 B4RNW5 B4RNW6 B4RNW8 B4RNX2 B4RP08 B4RP13 B4RP57 B4RP71 B4RP93
B4RP98 B4RPA1 B4RPA5 B4RPF1 B4RPG0 B4RPG3 B4RPP9 B4RPQ3 B4RPR1 B4RPR2 B4RPW7
B4RPW8 B4RPX5 B4RPY2 B4RPY6 B4RPZ2 B4RQ17 B4RQ29 B4RQ33 B4RQ36 B4RQ47 B4RQ50
B4RQ61 B4RQ67 B4RQ94 B4RQC3 B4RQC7 B4RQD7 B4RQD8 B4RQI7 B4RQK5 B4RQL3 B4RQN5
B4RQP4 B4RQP8 B4RQP9 B4RQQ5 B4RQS8 B4RQU4 B4RQV8 B4RQV9 B4RQW1 B4RQW2 B4RQW3
B4RQX2 B4RQX4 B4RQX9 B4RQY1 B4RQY3 B4RQY4 B4RQY5 B4RQY6 B4RQY7 B4RR02 B4RR07
B4RR25 B4RR35 B4RR38 B4RR84 B4RR86 B4RRA1 B4RRA8 B4RRB9 B4RRF0 B4RRY0 B4RRY8
B4RS18 B4RS41 B4RS43 B4RS91 B4RS92 B4RS97 B4RSJ5 B4RSJ6 B4RSL5 B4RSL6 B4RSL8
B4RSS7 B4RST1 B4RSU5 B4RSU6 B4RSV6 B4RSW4 B4RSX7 B4RSX8 B4RSX9 B4RSY0 B4RSY2
B4RT38 B4RT39 B4RT40 B4RT41 B4RT42 B4RT43 B4RT45 B4RT46 B4RT47 B4RT49 B4RT50
B4RT51 B4RT52 B4RT54 B4RT72 B4RTK8 B4RTW4 B4RTX3 B4RU03 B4RU34 B4RU35 B4RU48
B4RU55 B4RU62 B4RU63 B4RU89 B4RUH2 B4RUJ5 B4RUL1 B4RUW0 B4RV70 B4RV73 B4RV80
B4RV85 B4RV88 B4RV89 B4RV94 B4RV95 B4RVA4 B4RVF4 B4RVH4 B4RVI3 B4RVI6 B4RVJ6
B4RVP7 B4RVU1 B4RVU4 B4RVU9 B4RVV4 B4RVY5 B4RVY8 B4RW95 B4RWC5 B4RWC7 B4RWE1
B4RWE6 B4RWG2 B4RWP8 B4RWR9 B4RWS2 B4RWY7 B4RX37 B4RX40 B4RX41 B4RXI5 B4RXK9
B4RXN1 B4RXT2 B4RXT6 B4RXT8 B4RXT9 B4RXU2 B4RXY7 B4RY33 B4RY34 B4RY96 B4RYA0
B4RYB0 B4RYE3 B4RYE6 B4RYG2 B4RYH7 B4RYN2 B4RYN8 B4RYN9 B4RYP8 B4RYQ8 B4RYR7
B4RYS5 B4RYS6 B4RZ25 B4RZ39 B4RZ48 B4RZB4 B4RZG5 B4RZG8 B4RZH3 B4RZH4 B4RZH5
B4RZM6 B4RZN1 B4RZN2 B4RZS0 B4RZU0 B4RZW3 B4RZX7 B4S010 B4S011 B4S012 B4S092
B4S094 B4S096 B4S098 B4S099 B4S0D0 B4S0F6 B4S0H4 B4S0H7 B4S0H8 B4S0I7 B4S0J8
B4S0J9 B4S0P6 B4S0P9 B4S0R8 B4S0S0 B4S0U1 B4S112 B4S113 B4S130 B4S133 B4S195
B4S1A6 B4S1J4 B4S1N2 B4S1N7 B4S1U9 B4S1X4 B4S200 B4S201 B4S253 B4S277 B4S278
B4S284 B4S291 B4S294 B4S298 B4S2B7 B4S2C3 B4S2C4 B4S2C6 B4S2C7 B4S2E4 B4S2E8
B4S2H6 B4S2K9 B4S2L3 B4S2L7 B4S2W0 B4S2W4 B4S2X2 B4S2X7 B4S385 B4S387 B4S3A5
B4S3B3 B4S3C6 B4S3C8 B4S3C9 B4S3D0 B4S3D1 B4S3D2 B4S3F3 B4S3F9 B4S3G9 B4S3H9
B4S3I3 B4S3N4 B4S3P2 B4S3Q6 B4S3Q7 B4S3W2 B4S3W8 B4S3X8 B4S404 B4S410 B4S417
B4S427 B4S429 B4S432 B4S433 B4S437 B4S462 B4S493 B4S495 B4S496 B4S498 B4S4A1
B4S4H1 B4S4H7 B4S4H8 B4S4I3 B4S4I4 B4S4I5 B4S4J4 B4S4K5 B4S4Q8 B4S4Q9 B4S4S0
B4S4S4 B4S4S6 B4S4S7 B4S4T0 B4S4T8 B4S4U7 B4S4U8 B4S4W1 B4S4X8 B4S4Y6 B4S4Y9
B4S4Z8 B4S517 B4S561 B4S564 B4S573 B4S584 B4S591 B4S592 B4S596 B4S5A0 B4S5A1
B4S5A2 B4S5A3 B4S5A4 B4S5A5 B4S5A9 B4S5B0 B4S5B1 B4S5B2 B4S5B3 B4S5B4 B4S5B5
B4S5B6 B4S5B7 B4S5B8 B4S5C1 B4S5C2 B4S5C3 B4S5F5 B4S5F6 B4S5G4 B4S5G5 B4S5G6
B4S5H3 B4S5H8 B4S5I1 B4S5J3 B4S5J4 B4S5J5 B4S5K2 B4S5K3 B4S5K5 B4S5L1 B4S5M3
B4S5M4 B4S5M5 B4S5M6 B4S5M7 B4S5M8 B4S5M9 B4S5N0 B4S5N1 B4S5N2 B4S5P1 B4S5P3
B4S5Q4 B4S5S1 B4S5W7 B4S5W8 B4S5Y8 B4S5Y9 B4S612 B4S616 B4S620 B4S665 B4S687
B4S690 B4S6D0 B4S6D8 B4S6D9 B4S6H1 B4S6H2 B4S6H3 B4S6H5 B4S6J0 B4S6J7 B4S6K6

B4S6L0 B4S6M3 B4S6M7 B4S6N3 B4S6P7 B4S6Q8 B4S6R7 B4S6R8 B4S6S2 B4S6X4 B4S715
B4S729 B4S787 B4S790 B4S7L7 B4S7L8 B4S7M0 B4S7P3 B4S7P9 B4S7R9 B4S7S0 B4S808
B4S810 B4S817 B4S820 B4S888 B4S893 B4S8A8 B4S8C4 B4S8K5 B4S8K9 B4S8L3 B4S8L6
B4S8U3 B4S8U7 B4S8V2 B4S8V9 B4S8W3 B4S8X6 B4S8Y3 B4S8Y8 B4S8Z6 B4S8Z9 B4S912
B4S921 B4S922 B4S930 B4S937 B4S950 B4S9A9 B4S9B0 B4S9B2 B4S9B8 B4S9B9 B4S9D0
B4S9D1 B4S9D2 B4S9D5 B4S9D9 B4S9E9 B4S9F0 B4S9F1 B4S9F8 B4S9G0 B4S9H5 B4S9J3
B4S9J4 B4S9K0 B4S9K2 B4S9T4 B4S9U5 B4S9U6 B4S9U8 B4S9V0 B4SA43 B4SA46 B4SA62
B4SA64 B4SA76 B4SA80 B4SAG3 B4SAG7 B4SAH1 B4SAH9 B4SAI2 B4SAJ4 B4SAK0 B4SAK8
B4SAM9 B4SAR5 B4SAR7 B4SAT4 B4SAT7 B4SAU0 B4SB22 B4SB42 B4SB45 B4SB89 B4SB90
B4SB91 B4SB93 B4SB96 B4SBB7 B4SBD5 B4SBF1 B4SBF6 B4SBF7 B4SBF9 B4SBG6 B4SBN2
B4SBR1 B4SBR3 B4SBU2 B4SBU3 B4SBU4 B4SBU5 B4SBU6 B4SBU7 B4SBU9 B4SBV1 B4SBV2
B4SBV3 B4SBV4 B4SBV7 B4SBV8 B4SBV9 B4SBW0 B4SBW1 B4SBW2 B4SBW3 B4SBW4 B4SBW6
B4SBX0 B4SBX1 B4SBX2 B4SBX3 B4SBX4 B4SC59 B4SC70 B4SC71 B4SC74 B4SC75 B4SC78
B4SC79 B4SC83 B4SC92 B4SCB6 B4SCC0 B4SCE2 B4SCE5 B4SCE7 B4SCE8 B4SCF0 B4SCF6
B4SCG2 B4SCH0 B4SCL0 B4SCQ5 B4SCQ7 B4SCR5 B4SCR7 B4SCS6 B4SCU0 B4SCU3 B4SCV2
B4SCV6 B4SCX5 B4SCX8 B4SCY8 B4SCZ7 B4SD14 B4SD35 B4SD42 B4SD43 B4SD44 B4SD86
B4SDA0 B4SDK2 B4SDK7 B4SDN7 B4SDP5 B4SDS5 B4SDT6 B4SDV8 B4SDW5 B4SDW6 B4SDX3
B4SDX4 B4SDX5 B4SDY9 B4SDZ2 B4SDZ4 B4SDZ7 B4SE01 B4SE02 B4SE03 B4SE11 B4SE13
B4SE23 B4SE30 B4SE31 B4SE58 B4SEC4 B4SEC6 B4SEC8 B4SEC9 B4SED0 B4SED4 B4SEH6
B4SEI0 B4SEI4 B4SEN0 B4SEN1 B4SEN7 B4SEQ2 B4SER5 B4SER6 B4SES7 B4SES8 B4SET5
B4SEU9 B4SEZ1 B4SEZ5 B4SEZ8 B4SF71 B4SF75 B4SFD3 B4SFD4 B4SFE7 B4SFH4 B4SFI2
B4SFI3 B4SFM7 B4SFQ3 B4SFQ8 B4SFQ9 B4SFU3 B4SFX7 B4SG07 B4SG10 B4SG12 B4SG13
B4SG15 B4SG33 B4SG51 B4SG54 B4SG55 B4SG68 B4SGC3 B4SGD1 B4SGD2 B4SGD6 B4SGL8
B4SGM1 B4SGN6 B4SGN7 B4SGP8 B4SGQ2 B4SGQ6 B4SGR0 B4SGR4 B4SGR7 B4SGR9 B4SGW0
B4SGW1 B4SGW9 B4SGZ7 B4SH30 B4SH34 B4SH35 B4SH48 B4SH62 B4SH92 B4SHB4 B4SHE6
B4SHF1 B4SHF2 B4SHF6 B4SHH3 B4SHI8 B4SHI9 B4SHM0 B4SHS5 B4SHT7 B4SHV0 B4SHV9
B4SHW0 B4SHW1 B4SHX1 B4SHX7 B4SI11 B4SI18 B4SI23 B4SI37 B4SI60 B4SI67 B4SIE4
B4SIF5 B4SIN4 B4SIN7 B4SIY9 B4SJ16 B4SJ34 B4SJ64 B4SJ70 B4SJ73 B4SJB0 B4SJB3
B4SJC0 B4SJC1 B4SJN0 B4SJP5 B4SJT3 B4SJT8 B4SJW7 B4SJW8 B4SJX6 B4SK03 B4SK42
B4SK51 B4SKG5 B4SKH6 B4SKH7 B4SKI0 B4SKL8 B4SKL9 B4SKN8 B4SKS8 B4SKS9 B4SKT5
B4SKV2 B4SKV3 B4SKV4 B4SKV5 B4SKV6 B4SKV8 B4SKV9 B4SKW0 B4SKW2 B4SKW3 B4SKW4
B4SKW5 B4SKW6 B4SKW7 B4SKW8 B4SKX0 B4SKX1 B4SKX2 B4SKX3 B4SKX4 B4SKX5 B4SKX6
B4SKX7 B4SKX8 B4SKY0 B4SKY1 B4SL67 B4SL82 B4SL92 B4SLB7 B4SLD1 B4SLD9 B4SLE0
B4SLE1 B4SLE4 B4SLE8 B4SLE9 B4SLF7 B4SLH2 B4SLH3 B4SLH4 B4SLH5 B4SLH6 B4SLI5
B4SLN0 B4SLN2 B4SLN7 B4SLQ3 B4SM23 B4SM81 B4SM82 B4SM83 B4SM96 B4SMG8 B4SMH0
B4SMH8 B4SMH9 B4SMR2 B4SMU1 B4SMU6 B4SMZ6 B4SN21 B4SN29 B4SN50 B4SNL7 B4SNM9
B4SNN0 B4SNP6 B4SNR2 B4SNS0 B4SNX3 B4SNY7 B4SP00 B4SP02 B4SP12 B4SP13 B4SP14
B4SP15 B4SP18 B4SP20 B4SP40 B4SP45 B4SP54 B4SP57 B4SP84 B4SP88 B4SP92 B4SP93
B4SP94 B4SPG1 B4SPG2 B4SPH1 B4SPH2 B4SPH3 B4SPH4 B4SPL6 B4SPM7 B4SPV3 B4SPV4
B4SQ19 B4SQ22 B4SQ23 B4SQ35 B4SQ79 B4SQ92 B4SQ93 B4SQB1 B4SQC8 B4SQG8 B4SQH0
B4SQH1 B4SQH2 B4SQH4 B4SQI3 B4SQK2 B4SQK7 B4SQQ3 B4SQR6 B4SQR7 B4SQR9 B4SQS0
B4SQS2 B4SQT8 B4SQU4 B4SQU8 B4SQV5 B4SQV6 B4SQW1 B4SQX0 B4SQX4 B4SQZ7 B4SR01
B4SR07 B4SR24 B4SR82 B4SR88 B4SR92 B4SR93 B4SRA0 B4SRA4 B4SRA5 B4SRB0 B4SRB1
B4SRB3 B4SRE2 B4SRE7 B4SRG1 B4SRK3 B4SRK6 B4SRK8 B4SRN0 B4SRN7 B4SRN9 B4SRP3

B4SRP4 B4SRX7 B4SRY3 B4SRY4 B4SRY6 B4SRY7 B4SRZ0 B4SRZ6 B4SS10 B4SS16 B4SS22
B4SS56 B4SS67 B4SS80 B4SSA3 B4SSC0 B4SSD9 B4SSE0 B4SSF8 B4SSQ5 B4SSQ6 B4SSQ7
B4SSS0 B4SSS1 B4SSV0 B4SSV1 B4SSV3 B4SSW2 B4SSW3 B4SSW8 B4SSX7 B4ST23 B4ST32
B4ST34 B4ST35 B4ST36 B4ST40 B4ST72 B4ST96 B4STE2 B4STE3 B4STF6 B4STN3 B4STN6
B4STN8 B4STN9 B4STP1 B4STP2 B4STR0 B4STR1 B4STR2 B4STS0 B4STS6 B4STS7 B4STV2
B4STZ0 B4STZ5 B4SU06 B4SU11 B4SU17 B4SU23 B4SU24 B4SU36 B4SU41 B4SU42 B4SU51
B4SU60 B4SU61 B4SU63 B4SU84 B4SU90 B4SU91 B4SUA6 B4SUA7 B4SUA8 B4SUB3 B4SUG2
B4SUG3 B4SUH1 B4SUJ5 B4SUK4 B4SUK6 B4SUK7 B4SUK9 B4SUL6 B4SUL8 B4SUN8 B4SUP1
B4SUQ6 B4SUQ8 B4SUQ9 B4SUR0 B4SUR5 B4SUR7 B4SUR8 B4SUS1 B4SUS2 B4SUS4 B4SUS5
B4SUS6 B4SUS7 B4SUS8 B4SUS9 B4SUT0 B4SUT1 B4SUT2 B4SUT3 B4SUT4 B4SUT5 B4SUT6
B4SUT7 B4SUT9 B4SUU0 B4SUU1 B4SUU6 B4SUU7 B4SUU8 B4SUU9 B4SUV0 B4SUV1 B4SUV4
B4SUW4 B4SUY4 B4SUY6 B4SUY9 B4SUZ3 B4SUZ5 B4SUZ6 B4SUZ8 B4SUZ9 B4SV01 B4SV12
B4SV18 B4SV20 B4SV24 B4SV34 B4SV37 B4SV39 B4SV85 B4SV89 B4SVD3 B4SVE4 B4SVE5
B4SVE8 B4SVF1 B4SVF5 B4SVF6 B4SVF7 B4SVG0 B4SVI1 B4SVI8 B4SVI9 B4SVK3 B4SVL0
B4SVL3 B4SVL5 B4SVL9 B4SVM2 B4SVN2 B4SVN3 B4SVN4 B4SVV2 B4SVW0 B4SVW1 B4SVW5
B4SVW6 B4SW22 B4SW27 B4SW28 B4SW31 B4SW34 B4SW54 B4SW60 B4SW74 B4SW90 B4SWA7
B4SWC5 B4SWD9 B4SWF1 B4SWK1 B4SWK2 B4SWK9 B4SWL6 B4SWN1 B4SWN2 B4SWP5 B4SWP7
B4SWQ7 B4SWQ9 B4SWR0 B4SWR4 B4SWR6 B4SWR7 B4SWS3 B4SWS4 B4SWS7 B4SWU0 B4SWU1
B4SWU2 B4SWU8 B4SWV0 B4SWW2 B4SWW3 B4SWX6 B4SWX8 B4SWX9 B4SWY1 B4SWY3 B4SWY4
B4SX00 B4SX27 B4SX40 B4SX42 B4SX45 B4SX46 B4SX92 B4SX95 B4SXB2 B4SXB9 B4SXC1
B4SXD6 B4SXD7 B4SXD8 B4SXD9 B4SXE0 B4SXE1 B4SXE2 B4SXE3 B4SXE4 B4SXE9 B4SXF1
B4SXL9 B4SXM1 B4SXM8 B4SXP2 B4SXP7 B4SXT5 B4SXV3 B4SXX8 B4SXY7 B4SY15 B4SY26
B4SY37 B4SY47 B4SY68 B4SY76 B4SY89 B4SY97 B4SYA6 B4SYA8 B4SYA9 B4SYB0 B4SYB2
B4SYB4 B4SYE1 B4SYE2 B4SYE5 B4SYE6 B4SYE7 B4SYE9 B4SYF7 B4SYJ0 B4SYJ4 B4SYJ9
B4SYK7 B4SYM0 B4SYM8 B4SYN0 B4SYN7 B4SYN9 B4SYR0 B4SYR2 B4SYR3 B4SYU8 B4SYW1
B4SYW3 B4SYW7 B4SYX8 B4SYY0 B4SYY1 B4SZ04 B4SZ17 B4SZ22 B4SZ25 B4SZ26 B4SZ48
B4SZ51 B4SZ67 B4SZ73 B4SZ86 B4SZ98 B4SZ99 B4SZB9 B4SZD3 B4SZE2 B4SZF9 B4SZH5
B4SZH7 B4SZI8 B4SZJ1 B4SZJ3 B4SZJ4 B4SZJ7 B4SZJ8 B4SZK1 B4SZK4 B4SZK6 B4SZP0
B4SZP2 B4SZQ7 B4SZQ8 B4SZS6 B4SZT6 B4SZU5 B4SZX0 B4SZX8 B4SZY1 B4SZY8 B4SZZ0
B4SZZ1 B4SZZ2 B4SZZ3 B4SZZ4 B4T049 B4T083 B4T089 B4T0B2 B4T0D6 B4T0E3 B4T0F2
B4T0F9 B4T0G6 B4T0I0 B4T0I5 B4T0L0 B4T0L4 B4T0M5 B4T0M7 B4T0N7 B4T0N9 B4T0P5
B4T0P8 B4T0P9 B4T0Q1 B4T0Q2 B4T0R5 B4T0R8 B4T0R9 B4T0S0 B4T0S2 B4T0S7 B4T0T2
B4T0T4 B4T0T6 B4T0T8 B4T0T9 B4T0U3 B4T0U7 B4T0V5 B4T0V8 B4T0W7 B4T0W8 B4T0X0
B4T0X1 B4T0X5 B4T0Y0 B4T0Y5 B4T0Y6 B4T0Y7 B4T0Y8 B4T0Y9 B4T0Z5 B4T0Z6 B4T0Z7
B4T0Z8 B4T0Z9 B4T100 B4T108 B4T110 B4T119 B4T128 B4T141 B4T142 B4T144 B4T146
B4T151 B4T156 B4T158 B4T172 B4T1C0 B4T1D1 B4T1E8 B4T1F6 B4T1M4 B4T1N0 B4T1R5
B4T1S8 B4T1Z1 B4T1Z9 B4T201 B4T206 B4T208 B4T209 B4T289 B4T290 B4T2B2 B4T2B3
B4T2B5 B4T2B9 B4T2C0 B4T2C3 B4T2C5 B4T2N5 B4T2P5 B4T2Q3 B4T2Q8 B4T2R5 B4T2R6
B4T2R7 B4T2S2 B4T2T9 B4T2U4 B4T2U5 B4T2V2 B4T2W7 B4T2X8 B4T2Y7 B4T2Z2 B4T2Z5
B4T300 B4T332 B4T340 B4T361 B4T381 B4T394 B4T397 B4T399 B4T3A0 B4T3B2 B4T3E2
B4T3E3 B4T3E7 B4T3F1 B4T3F2 B4T3F3 B4T3F4 B4T3G2 B4T3H1 B4T3H9 B4T3I1 B4T3K5
B4T3K6 B4T3M4 B4T3P6 B4T3P7 B4T3Q8 B4T3R6 B4T3X3 B4T3Y1 B4T3Z4 B4T3Z5 B4T3Z6
B4T3Z7 B4T436 B4T453 B4T454 B4T455 B4T456 B4T457 B4T460 B4T461 B4T462 B4T473
B4T474 B4T484 B4T4A3 B4T4E9 B4T4F9 B4T4G3 B4T4H1 B4T4H2 B4T4H3 B4T4I0 B4T4I7

B4T4I8 B4T4I9 B4T4K3 B4T4K7 B4T4L5 B4T4L7 B4T4M8 B4T4N0 B4T4N1 B4T4N2 B4T4N4
B4T4P0 B4T4P3 B4T4Q4 B4T4R6 B4T4R7 B4T4W1 B4T4W5 B4T4X1 B4T4X3 B4T4X5 B4T4Z7
B4T4Z8 B4T512 B4T534 B4T535 B4T543 B4T545 B4T548 B4T549 B4T550 B4T555 B4T567
B4T583 B4T585 B4T586 B4T5A0 B4T5A5 B4T5G4 B4T5H1 B4T5H2 B4T5I5 B4T5J3 B4T5J6
B4T5J9 B4T5K3 B4T5K4 B4T5K9 B4T5L9 B4T5P6 B4T5P8 B4T5R8 B4T5V6 B4T657 B4T665
B4T670 B4T674 B4T678 B4T679 B4T682 B4T688 B4T690 B4T6A7 B4T6B6 B4T6B9 B4T6C0
B4T6C6 B4T6C8 B4T6D6 B4T6D7 B4T6G9 B4T6H5 B4T6H8 B4T6J1 B4T6J4 B4T6J5 B4T6J6
B4T6J8 B4T6K2 B4T6L4 B4T6L5 B4T6L6 B4T6P5 B4T6U4 B4T6U6 B4T6V0 B4T6X1 B4T6X2
B4T6Y0 B4T6Y1 B4T6Y5 B4T6Z5 B4T6Z7 B4T6Z8 B4T700 B4T702 B4T705 B4T708 B4T714
B4T716 B4T717 B4T720 B4T736 B4T747 B4T750 B4T751 B4T754 B4T755 B4T769 B4T770
B4T7P3 B4T7Q1 B4T7Q2 B4T7Q5 B4T7Q6 B4T7Z2 B4T7Z3 B4T7Z6 B4T7Z9 B4T803 B4T804
B4T805 B4T808 B4T825 B4T842 B4T849 B4T867 B4T868 B4T869 B4T8C7 B4T8D8 B4T8M0
B4T8M6 B4T8M7 B4T8N0 B4T8N1 B4T8P4 B4T8P5 B4T8Q6 B4T8Q8 B4T8Q9 B4T8R3 B4T8R5
B4T8R6 B4T8S2 B4T8T9 B4T8V5 B4T8W8 B4T8X1 B4T8Y4 B4T911 B4T945 B4T946 B4T952
B4T959 B4T974 B4T977 B4T994 B4T9A1 B4T9A3 B4T9B9 B4T9C0 B4T9C1 B4T9C2 B4T9C3
B4T9C4 B4T9C5 B4T9C6 B4T9C7 B4T9D0 B4T9E2 B4T9E3 B4T9E4 B4T9F0 B4T9F2 B4T9G3
B4T9G4 B4T9H6 B4T9H8 B4T9H9 B4T9I1 B4T9I3 B4T9K6 B4T9K8 B4T9L5 B4T9N3 B4T9N5
B4T9N8 B4T9N9 B4T9T3 B4T9T4 B4T9W1 B4T9X0 B4T9Y5 B4T9Y6 B4T9Z1 B4T9Z3 B4TA53
B4TA60 B4TA73 B4TA81 B4TA89 B4TA94 B4TAD4 B4TAI5 B4TAI9 B4TAK5 B4TAL5 B4TAM6
B4TAN5 B4TAP8 B4TAQ0 B4TAQ1 B4TAU7 B4TAU9 B4TAV0 B4TAV1 B4TAV5 B4TAY2 B4TAY3
B4TAY6 B4TAY7 B4TAY8 B4TAZ0 B4TAZ8 B4TB04 B4TB09 B4TB12 B4TB13 B4TB34 B4TB37
B4TB44 B4TB51 B4TB65 B4TB72 B4TB74 B4TB82 B4TB84 B4TBB5 B4TBC8 B4TBD7 B4TBE3
B4TBF6 B4TBF8 B4TBG2 B4TBH3 B4TBH5 B4TBH6 B4TBI0 B4TBJ9 B4TBN7 B4TBN9 B4TBQ7
B4TBR3 B4TBS6 B4TBT8 B4TBT9 B4TBW0 B4TBW8 B4TBX1 B4TBX8 B4TBY0 B4TBY1 B4TBY2
B4TBY3 B4TBY4 B4TC11 B4TC26 B4TC28 B4TC39 B4TC41 B4TC42 B4TC44 B4TC45 B4TC48
B4TC49 B4TC52 B4TC55 B4TC57 B4TC80 B4TC86 B4TCA5 B4TCA6 B4TCD7 B4TCE7 B4TCF6
B4TCI1 B4TCI7 B4TCK0 B4TCL7 B4TCM2 B4TCM4 B4TCM6 B4TCM8 B4TCM9 B4TCN3 B4TCN8
B4TCP6 B4TCP9 B4TCQ2 B4TCQ3 B4TCQ5 B4TCQ6 B4TCR0 B4TCR5 B4TCS0 B4TCS1 B4TCS2
B4TCS3 B4TCS4 B4TCT2 B4TCT3 B4TCT4 B4TCT5 B4TCT6 B4TCT7 B4TCV0 B4TCX3 B4TCZ1
B4TCZ8 B4TD06 B4TD16 B4TD24 B4TD37 B4TD38 B4TD40 B4TD42 B4TD58 B4TD62 B4TD71
B4TD73 B4TD75 B4TD82 B4TD84 B4TD92 B4TD93 B4TD95 B4TD96 B4TDB2 B4TDB3 B4TDB4
B4TDB6 B4TDB7 B4TDC8 B4TDE5 B4TDF3 B4TDF7 B4TDG1 B4TDG7 B4TDK4 B4TDL6 B4TDQ4
B4TDQ9 B4TDR1 B4TDS5 B4TDY8 B4TDZ6 B4TDZ8 B4TE03 B4TE05 B4TE06 B4TE13 B4TE27
B4TE28 B4TE50 B4TE51 B4TE53 B4TE57 B4TE58 B4TE61 B4TE63 B4TEM9 B4TEN4 B4TEN5
B4TEP2 B4TEQ7 B4TES7 B4TET2 B4TET5 B4TEU0 B4TEZ3 B4TF06 B4TF10 B4TF12 B4TF13
B4TF25 B4TF74 B4TF79 B4TF80 B4TF84 B4TF92 B4TF97 B4TFA4 B4TFA5 B4TFA6 B4TFB1
B4TFB2 B4TFC5 B4TFC6 B4TFD0 B4TFD1 B4TFD5 B4TFD6 B4TFD7 B4TFD8 B4TFE3 B4TFF2
B4TFG0 B4TFG2 B4TFH5 B4TFI3 B4TFI4 B4TFI5 B4TFJ6 B4TFK5 B4TFR5 B4TFU6 B4TFW3
B4TFW5 B4TFW6 B4TFW7 B4TFX0 B4TFX1 B4TFX2 B4TFY3 B4TFY4 B4TFZ2 B4TG16 B4TG40
B4TG41 B4TG58 B4TG77 B4TGA3 B4TGC8 B4TGE0 B4TGE1 B4TGE2 B4TGE3 B4TGE4 B4TGE8
B4TGF6 B4TGF8 B4TGH1 B4TGH3 B4TGH4 B4TGH5 B4TGH7 B4TGI2 B4TGI5 B4TGJ6 B4TGJ7
B4TGK9 B4TGL0 B4TGM7 B4TGN3 B4TGN5 B4TGN7 B4TGQ9 B4TGR0 B4TGS4 B4TGV9 B4TGW0
B4TGW8 B4TGX0 B4TGX3 B4TGX4 B4TGX5 B4TGX9 B4TGY8 B4TGZ2 B4TGZ9 B4TH00 B4TH01
B4TH02 B4TH09 B4TH16 B4TH17 B4TH18 B4TH70 B4THC3 B4THC5 B4THC6 B4THE2 B4THE6

B4THF3 B4THF4 B4THG3 B4THH1 B4THH4 B4THH7 B4THI1 B4THI2 B4THI7 B4THJ7 B4THM6
B4THM8 B4THP5 B4THQ0 B4THV9 B4TI37 B4TI45 B4TI51 B4TI55 B4TI59 B4TI60 B4TI63
B4TIA4 B4TIA6 B4TIB4 B4TIB5 B4TIE3 B4TIE9 B4TIF1 B4TIG4 B4TIG8 B4TIG9 B4TIH0
B4TIH2 B4TII8 B4TIS7 B4TIU6 B4TIU8 B4TIW6 B4TIX5 B4TIX9 B4TIY0 B4TIY8 B4TIY9
B4TIZ3 B4TJ03 B4TJ05 B4TJ06 B4TJ08 B4TJ09 B4TJ12 B4TJ16 B4TJ21 B4TJ23 B4TJ24
B4TJ27 B4TJ45 B4TJ46 B4TJ47 B4TJ59 B4TJ60 B4TJ72 B4TJ78 B4TJ79 B4TJ88 B4TJ97
B4TJ98 B4TJA0 B4TJC7 B4TJD1 B4TJD2 B4TJI0 B4TJI2 B4TJK8 B4TJK9 B4TJP8 B4TJR0
B4TJR3 B4TJR8 B4TJR9 B4TJT3 B4TJT4 B4TJV7 B4TJV9 B4TJX2 B4TJX5 B4TJX7 B4TJX8
B4TJX9 B4TJY4 B4TJY6 B4TJY7 B4TJZ0 B4TJZ1 B4TJZ3 B4TJZ4 B4TK03 B4TK04 B4TK05
B4TK10 B4TK30 B4TK32 B4TK35 B4TK39 B4TK41 B4TK42 B4TK44 B4TK45 B4TK47 B4TK58
B4TK64 B4TK66 B4TK70 B4TK80 B4TK83 B4TK85 B4TKA4 B4TKA5 B4TKA8 B4TKB3 B4TKD3
B4TKE2 B4TKE4 B4TKE5 B4TKE7 B4TKF5 B4TKF7 B4TKH6 B4TKI0 B4TKK1 B4TKK2 B4TKK3
B4TKK4 B4TKK5 B4TKK6 B4TKK7 B4TKK8 B4TKK9 B4TKL0 B4TKL1 B4TKL2 B4TKL4 B4TKL5
B4TKL6 B4TKM1 B4TKM2 B4TKM3 B4TKM4 B4TKM5 B4TKM6 B4TKM9 B4TKN8 B4TKQ4 B4TKR2
B4TKR3 B4TKS6 B4TKT3 B4TKT6 B4TKT8 B4TKU2 B4TKU5 B4TKV1 B4TM95 B4TM97 B4TM98
B4TMA2 B4TMA4 B4TMA5 B4TMB1 B4TMB2 B4TMB5 B4TMC5 B4TMC6 B4TMC7 B4TMD3 B4TMD5
B4TME7 B4TME8 B4TMG1 B4TMG3 B4TMG4 B4TMG6 B4TMG7 B4TMG8 B4TMJ2 B4TMJ8 B4TMR4
B4TMR6 B4TMR9 B4TMS0 B4TMW9 B4TMX6 B4TMX7 B4TMY9 B4TN05 B4TN07 B4TN08 B4TN09
B4TN11 B4TN13 B4TN41 B4TN42 B4TN45 B4TN46 B4TN47 B4TN49 B4TN59 B4TN64 B4TNH8
B4TNH9 B4TNK3 B4TNL2 B4TNP0 B4TNQ0 B4TNR0 B4TNR5 B4TNS1 B4TNS6 B4TNT2 B4TNT3
B4TNV3 B4TNV5 B4TNV9 B4TNX3 B4TNX9 B4TNZ2 B4TP54 B4TP70 B4TPA3 B4TPA7 B4TPB4
B4TPC7 B4TPC8 B4TPC9 B4TPG0 B4TPH2 B4TPH4 B4TPH8 B4TPI9 B4TPJ1 B4TPJ2 B4TPL2
B4TPL3 B4TPN8 B4TPP6 B4TPP9 B4TPQ4 B4TPQ6 B4TPQ7 B4TPQ8 B4TPQ9 B4TPR0 B4TPS6
B4TPU2 B4TPU7 B4TPU9 B4TPV0 B4TPV2 B4TPV3 B4TPV7 B4TPW1 B4TPW8 B4TPX5 B4TPY1
B4TPY8 B4TPZ0 B4TPZ8 B4TQ00 B4TQ30 B4TQ44 B4TQ53 B4TQ66 B4TQA4 B4TQA6 B4TQB9
B4TQC0 B4TQD8 B4TQE9 B4TQF8 B4TQG7 B4TQG9 B4TQH2 B4TQH3 B4TQH5 B4TQH6 B4TQI0
B4TQI6 B4TQJ1 B4TQJ2 B4TQJ3 B4TQJ4 B4TQJ5 B4TQK0 B4TQK3 B4TQK4 B4TQK5 B4TQK6
B4TQK7 B4TQK8 B4TQL6 B4TQL9 B4TQM3 B4TQM9 B4TQN8 B4TQQ1 B4TQR7 B4TQR9 B4TQT0
B4TQT2 B4TQT3 B4TQT5 B4TQT6 B4TQT9 B4TQU0 B4TQU3 B4TQU6 B4TQU8 B4TQW9 B4TQX5
B4TQZ6 B4TR28 B4TR32 B4TR55 B4TR59 B4TR62 B4TR72 B4TR74 B4TR76 B4TR83 B4TR85
B4TR94 B4TR95 B4TR97 B4TR98 B4TRB1 B4TRM8 B4TRN5 B4TRP6 B4TRQ3 B4TRR1 B4TRR7
B4TRS5 B4TRT8 B4TRT9 B4TRU1 B4TRU3 B4TRU9 B4TRV4 B4TRV6 B4TRW6 B4TRX1 B4TRX2
B4TRX3 B4TRX5 B4TRX7 B4TRY8 B4TS04 B4TS12 B4TS28 B4TS29 B4TS53 B4TS54 B4TS56
B4TS61 B4TS62 B4TS65 B4TS67 B4TSC0 B4TSC5 B4TSC6 B4TSD0 B4TSD9 B4TSE3 B4TSF0
B4TSF1 B4TSF2 B4TSF7 B4TSF8 B4TSH1 B4TSH2 B4TSI2 B4TSI4 B4TSI9 B4TSJ1 B4TSJ2
B4TSL7 B4TSM2 B4TSM3 B4TSN0 B4TSP6 B4TSR7 B4TSS2 B4TSS5 B4TSY7 B4TT00 B4TT03
B4TT05 B4TT06 B4TT18 B4TT28 B4TT29 B4TT33 B4TT34 B4TT35 B4TT36 B4TT41 B4TT49
B4TT57 B4TT59 B4TT82 B4TT83 B4TTA0 B4TTB9 B4TTD2 B4TTH0 B4TTH8 B4TTH9 B4TTI0
B4TTJ1 B4TTK0 B4TTU0 B4TTV7 B4TTV8 B4TTV9 B4TTW0 B4TTW1 B4TTW4 B4TTW5 B4TTW6
B4TTX7 B4TTX8 B4TTY6 B4TU21 B4TU29 B4TU33 B4TU42 B4TU43 B4TU44 B4TU46 B4TU53
B4TU54 B4TU55 B4TUA0 B4TUA8 B4TUC2 B4TUC3 B4TUC4 B4TUC5 B4TUC9 B4TUD7 B4TUD9
B4TUF0 B4TUF2 B4TUF3 B4TUF4 B4TUF6 B4TUG2 B4TUG5 B4TUI7 B4TUJ1 B4TUJ7 B4TUJ9
B4TUK1 B4TUM2 B4TUM3 B4TUN8 B4TUQ7 B4TUQ8 B4TUR5 B4TUR7 B4TUS2 B4TUS3 B4TUT5
B4TUT6 B4TUY8 B4TV05 B4TV07 B4TV08 B4TV23 B4TV24 B4TV29 B4TV33 B4TV38 B4TV39

B4TV53 B4TV55 B4TV58 B4TV61 B4TV65 B4TV66 B4TV71 B4TV80 B4TVB1 B4TVB3 B4TVC7
B4TVD5 B4TVJ5 B4TVR9 B4TVS9 B4TVT4 B4TVT8 B4TVU2 B4TVU3 B4TVU6 B4TVV2 B4TVV4
B4TVX2 B4TVY2 B4TVY5 B4TVY7 B4TVZ5 B4TVZ6 B4TW15 B4TWA6 B4TWB0 B4TWB1 B4TWB9
B4TWC0 B4TWC4 B4TWD4 B4TWD6 B4TWD8 B4TWE0 B4TWE1 B4TWE3 B4TWE7 B4TWF3 B4TWF5
B4TWF6 B4TWF9 B4TWH5 B4TWI7 B4TWJ0 B4TWJ1 B4TWJ4 B4TWJ5 B4TWK9 B4TWL0 B4TWN4
B4TWP0 B4TWP3 B4TWQ6 B4TWR2 B4TWR3 B4TWR4 B4TWR6 B4TWT4 B4TWT5 B4TWT6 B4TWU3
B4TWU8 B4TWU9 B4TWV8 B4TWV9 B4TWW1 B4TX11 B4TX13 B4TX38 B4TX39 B4TX91 B4TX93
B4TXA5 B4TXA8 B4TXB0 B4TXB1 B4TXB2 B4TXB7 B4TXB9 B4TXC0 B4TXC3 B4TXC4 B4TXC6
B4TXC7 B4TXC8 B4TXC9 B4TXD0 B4TXD1 B4TXD2 B4TXD3 B4TXD4 B4TXD5 B4TXD6 B4TXD7
B4TXD8 B4TXD9 B4TXE1 B4TXE2 B4TXE3 B4TXE8 B4TXE9 B4TXF0 B4TXF1 B4TXF2 B4TXF3
B4TXF6 B4TXG5 B4TXG9 B4TXH0 B4TXH9 B4TXI7 B4TXI8 B4TXJ0 B4TXL1 B4TXL6 B4TXL7
B4TXN2 B4TXN3 B4TXP7 B4TXQ6 B4TXQ8 B4TXR1 B4TXR2 B4TXR6 B4TXR8 B4TXR9 B4TXS1
B4TXS2 B4TXU2 B4TXU3 B4TXU6 B4TXV1 B4TXX3 B4TXY1 B4TXY3 B4TXY4 B4TXY6 B4TXZ4
B4TXZ6 B4TY14 B4TY18 B4TY38 B4TY46 B4TY47 B4TY59 B4TY66 B4TY71 B4TY75 B4TY78
B4TY86 B4TY87 B4TY88 B4TYD2 B4TYE3 B4TYE9 B4TYF1 B4TYF5 B4TYG5 B4TYG8 B4TYH0
B4TYM6 B4TYQ6 B4TYR7 B4TYR8 B4TYS1 B4TYS4 B4TYS8 B4TYS9 B4TYT0 B4TYT3 B4TYV0
B4TYW7 B4TYX4 B4TYY1 B4TYZ1 B4TZ02 B4TZ49 B4TZ50 B4TZ55 B4TZ64 B4TZ86 B4TZ87
B4TZ90 B4TZ91 B4TZF4 B4TZG0 B4TZG1 B4TZG4 B4TZG5 B4TZH8 B4TZI0 B4TZJ7 B4TZN1
B4TZP5 B4TZP8 B4TZR1 B4TZT5 B4TZT8 B4TZV2 B4TZV9 B4TZW1 B4TZX6 B4TZX7 B4TZX8
B4TZX9 B4TZY0 B4TZY1 B4TZY2 B4TZY3 B4TZY4 B4TZZ8 B4U000 B4U041 B4U045 B4U047
B4U063 B4U080 B4U081 B4U0A1 B4U0D8 B4U0F0 B4U0H0 B4U0I1 B4U0I8 B4U0I9 B4U0J5
B4U0J7 B4U0K5 B4U0K6 B4U0K7 B4U0K8 B4U0N4 B4U0N8 B4U0P1 B4U0R8 B4U0S7 B4U0T5
B4U0U9 B4U0V7 B4U0V8 B4U0V9 B4U0W3 B4U0X4 B4U0X7 B4U0X9 B4U0Y6 B4U108 B4U113
B4U126 B4U143 B4U151 B4U153 B4U161 B4U178 B4U194 B4U196 B4U197 B4U198 B4U1A3
B4U1A7 B4U1C3 B4U1E2 B4U1E4 B4U1E8 B4U1F0 B4U1F8 B4U1F9 B4U1G6 B4U1J1 B4U1J4
B4U1S0 B4U1S2 B4U1S6 B4U1S8 B4U1T1 B4U1U3 B4U1W4 B4U1W7 B4U1W8 B4U1Y5 B4U1Z1
B4U1Z2 B4U206 B4U220 B4U228 B4U237 B4U238 B4U246 B4U247 B4U276 B4U278 B4U292
B4U295 B4U2A0 B4U2C7 B4U2E8 B4U2H0 B4U2H1 B4U2J8 B4U2J9 B4U2K0 B4U2K1 B4U2K6
B4U2K7 B4U2K8 B4U2M2 B4U2P9 B4U2S2 B4U2S5 B4U2T6 B4U2W6 B4U2X5 B4U2Y3 B4U2Z3
B4U309 B4U313 B4U317 B4U337 B4U360 B4U394 B4U395 B4U396 B4U3B0 B4U3B2 B4U3C0
B4U3D4 B4U3E1 B4U3E6 B4U3E7 B4U3F9 B4U3I0 B4U3I6 B4U3L7 B4U3M1 B4U3M4 B4U3P4
B4U3Q7 B4U3R2 B4U3U0 B4U3U1 B4U3X7 B4U3X9 B4U434 B4U460 B4U464 B4U465 B4U467
B4U475 B4U476 B4U4B1 B4U4F7 B4U4F8 B4U4H7 B4U4I5 B4U4I6 B4U4I9 B4U4K4 B4U4K6
B4U4K7 B4U4L6 B4U4M2 B4U4M6 B4U4M9 B4U4N8 B4U4Q5 B4U4Q6 B4U4Q7 B4U4S2 B4U4S8
B4U4T4 B4U4U3 B4U4U5 B4U4X8 B4U4Y3 B4U4Y6 B4U4Z9 B4U500 B4U501 B4U503 B4U504
B4U505 B4U506 B4U507 B4U508 B4U509 B4U510 B4U511 B4U512 B4U513 B4U514 B4U515
B4U516 B4U518 B4U519 B4U523 B4U524 B4U525 B4U527 B4U545 B4U561 B4U570 B4U573
B4U574 B4U576 B4U592 B4U5A5 B4U5E5 B4U5F8 B4U5G6 B4U5G8 B4U5H3 B4U5H9 B4U5L8
B4U5M3 B4U5Q9 B4U5Z8 B4U600 B4U605 B4U606 B4U609 B4U616 B4U626 B4U640 B4U6A9
B4U6B2 B4U6H0 B4U6H1 B4U6I9 B4U6J5 B4U6K0 B4U6K3 B4U6N3 B4U6N4 B4U6P0 B4U6R6
B4U6R7 B4U6R8 B4U6S1 B4U6T9 B4U6U1 B4U6Z8 B4U701 B4U714 B4U733 B4U736 B4U738
B4U739 B4U740 B4U741 B4U743 B4U744 B4U745 B4U746 B4U747 B4U748 B4U749 B4U750
B4U751 B4U752 B4U754 B4U755 B4U757 B4U759 B4U760 B4U761 B4U762 B4U763 B4U765
B4U768 B4U769 B4U772 B4U773 B4U7A6 B4U7D5 B4U7D7 B4U7E2 B4U7F1 B4U7F2 B4U7J9

B4U7L4 B4U7M0 B4U7M4 B4U7M7 B4U7P3 B4U7P8 B4U7S5 B4U7S7 B4U7T2 B4U7T8 B4U7U3
B4U7U5 B4U7Y7 B4U813 B4U831 B4U858 B4U872 B4U896 B4U8A0 B4U8B2 B4U8G5 B4U8H0
B4U8K0 B4U8K1 B4U8M6 B4U8P1 B4U8R9 B4U8S7 B4U8T1 B4U8T3 B4U8T5 B4U8T6 B4U8W2
B4U8W9 B4U8X0 B4U901 B4U930 B4U931 B4U950 B4U955 B4U969 B4U973 B4U9B9 B4U9C8
B4U9D9 B4U9I4 B4U9K4 B4U9K9 B4U9L1 B4U9N6 B4U9P0 B4U9S4 B4U9V3 B4U9X0 B4U9X4
B4U9Z9 B4UA20 B4UA87 B4UAM2 B4UAM8 B4UAN0 B4UAN1 B4UAN4 B4UAN5 B4UAN6 B4UAN7
B4UAP1 B4UAX9 B4UAY5 B4UAY6 B4UAY8 B4UB95 B4UB96 B4UB99 B4UBA0 B4UBA1 B4UBA2
B4UBA3 B4UBA4 B4UBA5 B4UBA6 B4UBA8 B4UBA9 B4UBB0 B4UBB1 B4UBB2 B4UBB3 B4UBB4
B4UBB5 B4UBB7 B4UBB8 B4UBC0 B4UBC2 B4UBC3 B4UBC5 B4UBC6 B4UBC7 B4UBC9 B4UBD2
B4UBD3 B4UBI2 B4UBJ2 B4UBV8 B4UC43 B4UCB1 B4UCB2 B4UCN6 B4UCP2 B4UCU5 B4UCU9
B4UCW3 B4UCY7 B4UD63 B4UD71 B4UD97 B4UDA8 B4UDH6 B4UDJ7 B4UDJ9 B4UDK2 B4UDM7
B4UDP2 B4UDP4 B4UDQ2 B4UDQ3 B4UDQ9 B4UDT1 B4UDT2 B4UDT3 B4UDT4 B4UDT5 B4UDT6
B4UDT9 B4UDY7 B4UDZ1 B4UDZ3 B4UDZ6 B4UE87 B4UE89 B4UEC9 B4UEJ2 B4UEJ9 B4UEK4
B4UEL1 B4UEL3 B4UEL6 B4UER1 B4UER3 B4UES2 B4UES3 B4UES4 B4UEZ9 B4UF77 B4UF79
B4UFJ8 B4UFU2 B4UFX9 B4UFY0 B4UFY1 B4UFZ1 B4UG87 B4UG88 B4UG90 B4UGB5 B4UGC3
B4UGH8 B4UGI0 B4UGJ6 B4UGK3 B4UGQ8 B4UGT5 B4UGU5 B4UGV4 B4UGW9 B4UGX0 B4UGX9
B4UGY2 B4UGZ2 B4UGZ3 B4UH04 B4UHA8 B4UHC8 B4UHD1 B4UHE7 B4UHF2 B4UHF5 B4UHF7
B4UHG0 B4UHG2 B4UHG4 B4UHG5 B4UHG6 B4UHX9 B4UHY1 B4UHY4 B4UI08 B4UI29 B4UI79
B4UID2 B4UIE3 B4UIG2 B4UIH9 B4UII2 B4UIK8 B4UIM1 B4UIM2 B4UIM6 B4UIM7 B4UIS1
B4UIT6 B4UIU2 B4UIU3 B4UIU4 B4UIU8 B4UIV0 B4UIV1 B4UIV2 B4UIX7 B4UIZ5 B4UJ23
B4UJ61 B4UJ96 B4UJT7 B4UJV1 B4UJX6 B4UJZ9 B4UK35 B4UKE6 B4UKF8 B4UKG4 B4UKM8
B4UKQ2 B4UKQ3 B4ULA3 B4ULB9 B4ULC0 B4ULC2 B4ULC3 B4ULC5 B4ULC6 B4ULD8 B4ULF0
B4ULF7 B4ULH7 B4ULJ1 B4ULJ2 B4ULJ3 B4ULR5 B4ULW4 B4UM59 B4UM78 B4UMB7 B4UMB8
B4UMC6 B4UMJ1 B4UMK8 B4UMQ6 B4UN34 B4UN38 B4UN83 B4UNA2 B4USW8 B4UT01 B4XMC6
B4XXY3 B4XY98 B4XY99 B4XYB8 B4YB07 B4YNE1 B4YNE2 B4YNE3 B4YNE4 B4YNE5 B4YNE6
B4YNE7 B4YNE8 B4YNE9 B4YNF0 B4YNF1 B4YNF2 B4YNF3 B4YNF4 B4YNF6 B4YNF7 B4YNF8
B4YNF9 B4YNG0 B4YNG1 B4YPV4 B4YPW6 B4YYA9 B4ZIX8 B5A434 B5A435 B5A7L9 B5APK2
B5B0J6 B5BA16 B5BA27 B5BA30 B5BA36 B5BA38 B5BA39 B5BA42 B5BA56 B5BA58 B5BA65
B5BA68 B5BA69 B5BA70 B5BA71 B5BA72 B5BA90 B5BAE8 B5BAF5 B5BAG6 B5BAG8 B5BAH5
B5BAI4 B5BAJ8 B5BAJ9 B5BAK0 B5BAK2 B5BAK9 B5BAL0 B5BAL1 B5BAM6 B5BAM7 B5BAM9
B5BAP0 B5BAQ2 B5BAR7 B5BAR8 B5BAS2 B5BAT0 B5BAT2 B5BAU0 B5BAV4 B5BAW5 B5BAW6
B5BAW8 B5BAW9 B5BAX0 B5BAY2 B5BAY3 B5BAY5 B5BAY6 B5BAY9 B5BAZ3 B5BAZ5 B5BB05
B5BB06 B5BB08 B5BB16 B5BB18 B5BB22 B5BB46 B5BB50 B5BB72 B5BB76 B5BB85 B5BBA3
B5BBA5 B5BBC1 B5BBC6 B5BBD3 B5BBF2 B5BBG6 B5BBH2 B5BBH3 B5BBH8 B5BBK0 B5BBK1
B5BBK4 B5BBK9 B5BBL1 B5BBL9 B5BBM4 B5BBN4 B5BBN6 B5BBP1 B5BBP5 B5BBP7 B5BBP9
B5BBR2 B5BBS0 B5BBT0 B5BBT7 B5BBU4 B5BBV5 B5BBW2 B5BBX8 B5BBZ6 B5BC01 B5BC22
B5BC24 B5BC27 B5BC30 B5BC31 B5BC34 B5BC36 B5BC37 B5BC39 B5BC50 B5BC52 B5BC59
B5BC72 B5BC82 B5BC95 B5BCC3 B5BCC5 B5BCD1 B5BCD3 B5BCD9 B5BCE6 B5BCE9 B5BCF3
B5BCF8 B5BCG1 B5BCH6 B5BCH8 B5BCL1 B5BCL4 B5BCN2 B5BCN6 B5BCN7 B5BCN9 B5BCQ0
B5BCQ4 B5BCQ6 B5BCS0 B5BCV2 B5BCW8 B5BCZ5 B5BCZ8 B5BD11 B5BD17 B5BD19 B5BD42
B5BD43 B5BD44 B5BD46 B5BD47 B5BD49 B5BD63 B5BD74 B5BD76 B5BD82 B5BD83 B5BD84
B5BDA0 B5BDA1 B5BDA7 B5BDA8 B5BDB0 B5BDB4 B5BDB5 B5BDB7 B5BDC8 B5BDC9 B5BDE2
B5BDE5 B5BDE6 B5BDF0 B5BDP7 B5BDP8 B5BDQ0 B5BDQ1 B5BDQ9 B5BDW5 B5BDW7 B5BE09
B5BE10 B5BE12 B5BE26 B5BE34 B5BE43 B5BE52 B5BE81 B5BE91 B5BE92 B5BE93 B5BE94

B5BE99 B5BEA1 B5BEA6 B5BEK3 B5BEM1 B5BEN1 B5BEN4 B5BEN6 B5BEN7 B5BEP9 B5BEY0
B5BEY1 B5BEY2 B5BEY3 B5BEY4 B5BEY7 B5BEY8 B5BEY9 B5BEZ7 B5BEZ8 B5BF03 B5BF23
B5BF27 B5BF31 B5BF33 B5BF35 B5BF36 B5BF73 B5BF74 B5BFB5 B5BFB6 B5BFB9 B5BFC1
B5BFD0 B5BFH0 B5BFH1 B5BFI4 B5BFK4 B5BFK5 B5BFK6 B5BFL3 B5BFL5 B5BFL9 B5BFM0
B5BFM8 B5BFM9 B5BFN4 B5BFN5 B5BFP3 B5BFP7 B5BFP9 B5BFQ3 B5BFQ4 B5BFQ9 B5BFR8
B5BFU8 B5BFV0 B5BFZ9 B5BG06 B5BG08 B5BG11 B5BG15 B5BG20 B5BG21 B5BG24 B5BG29
B5BG31 B5BG44 B5BG57 B5BG60 B5BG71 B5BGA1 B5BGB4 B5BGC1 B5BGF2 B5BGG5 B5BGG9
B5BGH7 B5BGH8 B5BGI2 B5BGJ1 B5BGJ2 B5BGJ4 B5BGJ5 B5BGJ7 B5BGK0 B5BGK3 B5BGK7
B5BGK9 B5BGL0 B5BGL3 B5BGM9 B5BGP2 B5BGP5 B5BGP6 B5BGP9 B5BGQ0 B5BGR3 B5BGR4
B5BGT7 B5BGT9 B5BGU8 B5BGV1 B5BGV3 B5BGV4 B5BGV5 B5BGW0 B5BGW2 B5BGW7 B5BGW8
B5BGX0 B5BGX1 B5BGX2 B5BGX3 B5BGX4 B5BGX5 B5BGX6 B5BGX7 B5BGX8 B5BGX9 B5BGY0
B5BGY1 B5BGY2 B5BGY3 B5BGY5 B5BGY6 B5BGY7 B5BGZ2 B5BGZ3 B5BGZ4 B5BGZ5 B5BGZ6
B5BGZ7 B5BH08 B5BH22 B5BH29 B5BH30 B5BH46 B5BH48 B5BH49 B5BH50 B5BH54 B5BH57
B5BH60 B5BH61 B5BH72 B5BHA6 B5BHB0 B5BHD1 B5BHD3 B5BHE0 B5BHE2 B5BHE3 B5BHE5
B5BHF6 B5BHF8 B5BHG5 B5BHG7 B5BHG9 B5BHH3 B5BHH6 B5BHH9 B5BHI0 B5BHI1 B5BHL9
B5BHM4 B5BHN9 B5BHU1 B5BHU2 B5BHV4 B5BHW8 B5BHX1 B5BHY4 B5BHZ1 B5BHZ3 B5BI08
B5BI09 B5BI10 B5BI11 B5BI12 B5BI14 B5BI15 B5BI16 B5BI17 B5BI22 B5BI24 B5BI50
B5BI72 B5BI74 B5BI75 B5BI93 B5BIA7 B5BIC0 B5BIC1 B5BID8 B5BIE3 B5BII6 B5BIJ1
B5BIJ2 B5BIK2 B5BIK9 B5BIL2 B5BIL4 B5BIL5 B5BIL6 B5BIL9 B5BIP4 B5BIP5 B5BIP8
B5BIQ0 B5BIQ2 B5BIR0 B5BIR7 B5BIS2 B5BIS8 B5BIS9 B5BIU8 B5BIV3 B5BIV6 B5BIX4
B5BIX9 B5BIZ1 B5BJ27 B5BJ71 B5BJF4 B5BJF7 B5BJG3 B5BJG5 B5BJG6 B5BJG7 B5BJG8
B5BJG9 B5BJI2 B5BJJ8 B5BJK3 B5BJK5 B5BJK6 B5BJK8 B5BJK9 B5BJL3 B5BJL6 B5BJM5
B5BJM7 B5BJN0 B5BJN1 B5BJN3 B5BJN4 B5BJN6 B5BJN9 B5BJP1 B5BJP4 B5BJP9 B5BJQ0
B5BJQ1 B5BJQ2 B5BJQ3 B5BJQ9 B5BJR2 B5BJR3 B5BJR4 B5BJR5 B5BJR6 B5BJR7 B5BJS5
B5BJS7 B5BJT4 B5BJU4 B5BJV6 B5BKA2 B5BKC0 B5BKC1 B5BKC3 B5BKC5 B5BKD9 B5BKE7
B5BKF3 B5BKF4 B5BKF8 B5BKG8 B5BKH9 B5BKI0 B5BKI1 B5BKI5 B5BKI6 B5BKJ9 B5BKK0
B5BKK4 B5BKK5 B5BKK6 B5BKK9 B5BKL0 B5BKL1 B5BKL2 B5BKL8 B5BKM8 B5BKN5 B5BKN6
B5BKQ8 B5BKQ9 B5BKS6 B5BKW3 B5BKZ8 B5BL07 B5BL11 B5BL14 B5BL25 B5BL26 B5BL27
B5BL33 B5BL41 B5BL43 B5BL44 B5BL53 B5BL54 B5BL55 B5BL56 B5BL57 B5BL59 B5BL63
B5BL64 B5BL65 B5BL68 B5BL70 B5BL82 B5BL87 B5BL88 B5BL91 B5BL93 B5BL94 B5BL96
B5BL97 B5BLB2 B5BLB3 B5BLB5 B5BLC3 B5BLC4 B5BLD4 B5BLD5 B5BLD7 B5BLF5 B5BLG3
B5BLG4 B5BLG7 B5BLG9 B5BLH1 B5BLH8 B5BLH9 B5BLK7 B5BLL0 B5BLL3 B5BLL5 B5BLW5
B5BP46 B5BP48 B5BT18 B5BUZ8 B5DDX6 B5DE03 B5DE69 B5DE70 B5DE93 B5DEB9 B5DEI4
B5DEL1 B5DEQ3 B5DF07 B5DF11 B5DF21 B5DF91 B5DFC8 B5DFI8 B5DFK3 B5DFW7 B5DG42
B5DG51 B5DG67 B5DGH9 B5DGI7 B5DGK1 B5DGL6 B5DGM3 B5DGM4 B5DGM7 B5DGQ7 B5DHB2
B5DHW4 B5DI68 B5DJJ2 B5DK05 B5DK31 B5DKJ8 B5DKM4 B5DLV1 B5DMC9 B5DPF1 B5DQK2
B5DR03 B5DRT7 B5DRY3 B5DS72 B5DUH6 B5DUR8 B5DX45 B5DXG8 B5DY99 B5DZ31 B5DZN7
B5E081 B5E0H4 B5E0K3 B5E0U2 B5E1I4 B5E1N0 B5E1P3 B5E1P5 B5E1P6 B5E1P7 B5E1P8
B5E1P9 B5E1Q0 B5E1Q6 B5E1U3 B5E1U9 B5E1V8 B5E1W4 B5E1Y7 B5E1Y8 B5E1Z3 B5E216
B5E217 B5E223 B5E224 B5E230 B5E231 B5E232 B5E261 B5E262 B5E266 B5E267 B5E298
B5E2A0 B5E2C6 B5E2E0 B5E2E1 B5E2F2 B5E2F3 B5E2G0 B5E2G2 B5E2H6 B5E2I5 B5E2L2
B5E2M5 B5E2M6 B5E2Q5 B5E2R3 B5E2R6 B5E2W2 B5E2W4 B5E2X9 B5E2Z8 B5E301 B5E302
B5E317 B5E320 B5E327 B5E349 B5E364 B5E369 B5E375 B5E378 B5E379 B5E385 B5E387
B5E3A3 B5E3A4 B5E3A7 B5E3C1 B5E3C8 B5E3D3 B5E3E2 B5E3E3 B5E3F7 B5E3F8 B5E3G0

B5E3G1 B5E3K1 B5E3K2 B5E3K8 B5E3L0 B5E3L1 B5E3L5 B5E3S7 B5E3S9 B5E3V0 B5E3W0
B5E3W1 B5E3Y2 B5E3Z6 B5E3Z8 B5E435 B5E437 B5E439 B5E446 B5E448 B5E455 B5E466
B5E469 B5E478 B5E479 B5E480 B5E486 B5E494 B5E4D5 B5E4D9 B5E4E3 B5E4J4 B5E4K3
B5E4K5 B5E4L2 B5E4L5 B5E4L9 B5E4M9 B5E4N4 B5E4N6 B5E4P4 B5E4P5 B5E4Q2 B5E4Q4
B5E4R3 B5E4T0 B5E4T1 B5E4T2 B5E4U4 B5E4U5 B5E4U6 B5E4V6 B5E4W4 B5E4X3 B5E4Y2
B5E4Y3 B5E508 B5E512 B5E514 B5E515 B5E522 B5E529 B5E535 B5E568 B5E572 B5E578
B5E5F0 B5E5F1 B5E5F6 B5E5F8 B5E5I5 B5E5J3 B5E5J6 B5E5J9 B5E5M9 B5E5N0 B5E5N2
B5E5N3 B5E5N4 B5E5N5 B5E5P6 B5E5R2 B5E5S1 B5E5S2 B5E5S3 B5E5S4 B5E5S7 B5E5U0
B5E5U9 B5E5W0 B5E5W1 B5E5W8 B5E5X4 B5E605 B5E609 B5E614 B5E641 B5E642 B5E643
B5E653 B5E681 B5E685 B5E699 B5E6A4 B5E6A6 B5E6B3 B5E6C5 B5E6C6 B5E6D0 B5E6E0
B5E6E1 B5E6E2 B5E6F4 B5E6F5 B5E6F6 B5E6F7 B5E6F8 B5E6F9 B5E6G0 B5E6G1 B5E6G2
B5E6G3 B5E6G4 B5E6G5 B5E6G6 B5E6G7 B5E6G8 B5E6G9 B5E6H0 B5E6H1 B5E6H3 B5E6H4
B5E6H6 B5E6H9 B5E6I1 B5E6I3 B5E6J9 B5E6K0 B5E6K3 B5E6K6 B5E6K9 B5E6L1 B5E6L2
B5E6L7 B5E6L9 B5E6M0 B5E6N8 B5E6P3 B5E6Q4 B5E6Q8 B5E6S0 B5E6T0 B5E6T3 B5E6T8
B5E6U3 B5E6U4 B5E6U5 B5E6W5 B5E6W9 B5E6Z2 B5E706 B5E709 B5E714 B5E721 B5E722
B5E745 B5E756 B5E760 B5E782 B5E785 B5E7B0 B5E7B1 B5E7B2 B5E7E0 B5E7E2 B5E7E5
B5E7E8 B5E7F2 B5E7F9 B5E7G3 B5E7M2 B5E7M3 B5E7M4 B5E7M5 B5E7M6 B5E7P6 B5E7P8
B5E7R1 B5E844 B5E845 B5E846 B5E853 B5E854 B5E858 B5E8F0 B5E8F6 B5E8I9 B5E8R2
B5E8U0 B5E956 B5E957 B5E958 B5E988 B5E9A4 B5E9C1 B5E9D0 B5E9M9 B5E9N4 B5E9N7
B5E9S9 B5E9T0 B5E9T1 B5E9W9 B5E9X4 B5E9X5 B5E9Y1 B5E9Y2 B5EA63 B5EAH0 B5EAH1
B5EAH2 B5EAH3 B5EAK6 B5EAS1 B5EAS6 B5EAS8 B5EAW7 B5EAX0 B5EB44 B5EB48 B5EB49
B5EB50 B5EB79 B5EBB8 B5EBC6 B5EBC7 B5EBC9 B5EBD2 B5EBF9 B5EBG1 B5EBG2 B5EBG7
B5EBG8 B5EBM3 B5EBP3 B5EBP9 B5EBQ2 B5EBQ4 B5EBU6 B5EBV0 B5EBX9 B5EBY1 B5EBY2
B5EBY3 B5EC38 B5EC42 B5EC43 B5EC56 B5EC81 B5EC90 B5EC98 B5ECF3 B5ECN1 B5ED01
B5ED03 B5ED05 B5ED10 B5ED13 B5ED16 B5ED18 B5ED19 B5ED20 B5ED21 B5ED52 B5ED57
B5ED60 B5ED61 B5ED77 B5ED83 B5ED84 B5ED95 B5EDQ6 B5EDQ7 B5EDR2 B5EDR4 B5EDR5
B5EDR7 B5EDS0 B5EDS6 B5EDX8 B5EDZ5 B5EE14 B5EE25 B5EE39 B5EE44 B5EE49 B5EE88
B5EEE6 B5EEE7 B5EEF0 B5EEI1 B5EEI3 B5EEI4 B5EEI5 B5EEK6 B5EEL9 B5EEM1 B5EEV8
B5EEZ1 B5EEZ9 B5EF22 B5EF40 B5EF41 B5EF56 B5EFG4 B5EFL1 B5EFM8 B5EFM9 B5EFN1
B5EFN6 B5EFN9 B5EFP1 B5EFP2 B5EFP3 B5EFP4 B5EFP5 B5EFP6 B5EFP7 B5EFP9 B5EFQ0
B5EFQ1 B5EFQ2 B5EFQ3 B5EFQ4 B5EFQ5 B5EFQ6 B5EFQ7 B5EFQ8 B5EFQ9 B5EFR0 B5EFR1
B5EFR2 B5EFR3 B5EFR4 B5EFR5 B5EFR6 B5EFR8 B5EFR9 B5EFS1 B5EFS3 B5EFS4 B5EFS6
B5EFU4 B5EFU7 B5EFV5 B5EFW2 B5EFX9 B5EFY8 B5EG09 B5EG40 B5EGD4 B5EGD5 B5EGN2
B5EGN6 B5EGQ4 B5EGX2 B5EGX3 B5EGX4 B5EGY0 B5EGY1 B5EH68 B5EHA6 B5EHA7 B5EHF2
B5EHF5 B5EHR3 B5EHR5 B5EHU2 B5EHV2 B5EHV5 B5EHV8 B5EHW0 B5EHW1 B5EHW6 B5EHW8
B5EHW9 B5EHX1 B5EHX2 B5EHY0 B5EI10 B5EI11 B5EI26 B5EI27 B5EI28 B5EI54 B5EI57
B5EI59 B5EI62 B5EI63 B5EI65 B5EIJ0 B5EIQ6 B5EIQ7 B5EIT8 B5EIT9 B5EIU8 B5EIV1
B5EIW4 B5EJ44 B5EJ75 B5EJ84 B5EJA0 B5EJC7 B5EJE0 B5EJE1 B5EJE8 B5EJE9 B5EJF3
B5EJF7 B5EJJ2 B5EJL3 B5EJL6 B5EJL8 B5EJT1 B5EK08 B5EK09 B5EK17 B5EK18 B5EK28
B5EKF7 B5EKH2 B5EKH7 B5EKJ0 B5EKW6 B5EKY1 B5EKZ2 B5EL11 B5EL42 B5EL74 B5EL82
B5EL84 B5EL87 B5ELA6 B5ELC2 B5ELD1 B5ELD2 B5ELF7 B5ELK7 B5ELM2 B5ELM5 B5ELR8
B5ELU2 B5ELU3 B5ELU4 B5ELU7 B5ELV1 B5ELV3 B5ELV4 B5ELV8 B5ELW8 B5ELW9 B5ELX0
B5ELX1 B5ELX2 B5ELX4 B5ELX5 B5ELX6 B5ELX8 B5ELX9 B5ELY0 B5ELY1 B5ELY2 B5ELY3
B5ELY4 B5ELY6 B5ELY7 B5ELY8 B5ELY9 B5ELZ0 B5ELZ2 B5ELZ3 B5ELZ4 B5ELZ5 B5ELZ6

B5ELZ7 B5ELZ8 B5EM95 B5EM96 B5EM98 B5EMA0 B5EMA3 B5EMB5 B5EMC5 B5EMC8 B5EMD1
B5EMD3 B5EMD4 B5EMD8 B5EMF9 B5EMG2 B5EMR4 B5EMS4 B5EMU4 B5EN11 B5EN19 B5EN20
B5EN39 B5EN53 B5EN54 B5EN55 B5EN57 B5EN98 B5ENA2 B5ENA3 B5ENA4 B5ENA6 B5ENA7
B5ENA8 B5ENG4 B5ENJ1 B5ENJ2 B5ENJ5 B5ENK9 B5ENM4 B5ENR5 B5ENR7 B5ENS6 B5ENT6
B5ENY5 B5ENY8 B5EP40 B5EP52 B5EP57 B5EP61 B5EP95 B5EP97 B5EPA0 B5EPA4 B5EPC7
B5EPD0 B5EPM5 B5EPM9 B5EPR5 B5EPW0 B5EPW2 B5EPX1 B5EQ20 B5EQ26 B5EQ29 B5EQ42
B5EQ47 B5EQ48 B5EQ61 B5EQ88 B5EQC6 B5EQE5 B5EQE6 B5EQE9 B5EQF1 B5EQF2 B5EQG3
B5EQH0 B5EQH6 B5EQH8 B5EQL5 B5EQL6 B5EQP9 B5EQQ4 B5EQW2 B5EQZ5 B5ER53 B5ER58
B5ER76 B5ERI2 B5ERI3 B5ERJ0 B5ERJ7 B5ERK4 B5ERM3 B5ERN6 B5ERN9 B5ERP0 B5ERR7
B5ES06 B5ES07 B5ES53 B5ESI0 B5ESS5 B5ET10 B5ET63 B5ET96 B5ETC9 B5ETI9 B5ETJ6
B5ETK6 B5ETN1 B5ETN4 B5ETU8 B5ETW5 B5ETZ2 B5ETZ4 B5ETZ6 B5EUF3 B5EUG3 B5EUH0
B5EUH1 B5EUP7 B5EUP9 B5EUQ0 B5EUS9 B5EUW2 B5EUW3 B5EV09 B5EV28 B5EVR2 B5EVS5
B5EVT2 B5EWJ3 B5EWK1 B5EWK2 B5EWK5 B5EWK6 B5EWR4 B5EWR9 B5EWS0 B5EWS3 B5EWS4
B5EWT8 B5EWT9 B5EWY2 B5EWY5 B5EWZ8 B5EX25 B5EX38 B5EX40 B5EX43 B5EX44 B5EX65
B5EX66 B5EX72 B5EX79 B5EX94 B5EX99 B5EXB6 B5EXC3 B5EXC5 B5EXD9 B5EXE0 B5EXE1
B5EXE2 B5EXE3 B5EXE4 B5EXE5 B5EXE6 B5EXE7 B5EXF2 B5EXF6 B5EXF8 B5EXF9 B5EXG3
B5EXG5 B5EXG6 B5EXH1 B5EXH2 B5EXH5 B5EXI7 B5EXI9 B5EXJ5 B5EXJ7 B5EXK9 B5EXL0
B5EXM5 B5EXM7 B5EXM8 B5EXN0 B5EXN2 B5EXN3 B5EXU9 B5EXV0 B5EXY6 B5EXZ5 B5EY10
B5EY75 B5EY83 B5EY97 B5EYB2 B5EYB4 B5EYB6 B5EYC3 B5EYD7 B5EYE2 B5EYH4 B5EYN3
B5EYP3 B5EYQ5 B5EYS6 B5EYS8 B5EYS9 B5EYW1 B5EYX2 B5EYX3 B5EYX4 B5EYX8 B5EZ06
B5EZ07 B5EZ10 B5EZ11 B5EZ14 B5EZ22 B5EZ28 B5EZ33 B5EZ39 B5EZ40 B5EZ64 B5EZ72
B5EZ76 B5EZ81 B5EZ89 B5EZA2 B5EZB0 B5EZB2 B5EZC1 B5EZC3 B5EZF1 B5EZF2 B5EZG1
B5EZG8 B5EZH0 B5EZH4 B5EZI5 B5EZI7 B5EZI8 B5EZJ2 B5EZL4 B5EZQ2 B5EZQ4 B5EZR6
B5EZR7 B5EZS3 B5EZU1 B5EZU8 B5EZW1 B5EZX3 B5EZX4 B5F000 B5F008 B5F011 B5F034
B5F050 B5F052 B5F063 B5F065 B5F066 B5F068 B5F069 B5F072 B5F073 B5F076 B5F079
B5F081 B5F0A4 B5F0B0 B5F0D6 B5F0E6 B5F0F5 B5F0I0 B5F0I6 B5F0L0 B5F0L3 B5F0M4
B5F0M6 B5F0M7 B5F0M8 B5F0M9 B5F0N0 B5F0P6 B5F0R2 B5F0R7 B5F0R9 B5F0S0 B5F0S2
B5F0S3 B5F0S7 B5F0T1 B5F0U2 B5F0U6 B5F0U8 B5F0U9 B5F0V3 B5F0V8 B5F0W3 B5F0W4
B5F0W5 B5F0W6 B5F0W7 B5F0X7 B5F0Z4 B5F101 B5F111 B5F118 B5F127 B5F138 B5F146
B5F159 B5F161 B5F163 B5F165 B5F170 B5F171 B5F173 B5F185 B5F187 B5F197 B5F198
B5F1A0 B5F1A1 B5F1B3 B5F1B6 B5F1B7 B5F1B8 B5F1C0 B5F1C1 B5F1D2 B5F1F1 B5F1F9
B5F1H3 B5F1H6 B5F1H7 B5F1H8 B5F1H9 B5F1I0 B5F1I1 B5F1I9 B5F1J1 B5F1J4 B5F1K4
B5F1P0 B5F1Q2 B5F1S4 B5F1S6 B5F1U0 B5F1U5 B5F1V3 B5F1V5 B5F1W0 B5F1W2 B5F1W3
B5F1Y9 B5F1Z4 B5F1Z5 B5F202 B5F227 B5F228 B5F286 B5F287 B5F289 B5F295 B5F298
B5F2A0 B5F2K4 B5F2K9 B5F2L0 B5F2L4 B5F2M2 B5F2S0 B5F2T4 B5F2U1 B5F333 B5F346
B5F349 B5F351 B5F352 B5F365 B5F378 B5F385 B5F386 B5F387 B5F392 B5F393 B5F3A7
B5F3A8 B5F3B2 B5F3B3 B5F3B4 B5F3B7 B5F3B8 B5F3B9 B5F3C0 B5F3C5 B5F3D4 B5F3E2
B5F3F3 B5F3F4 B5F3G6 B5F3I0 B5F3I3 B5F3I4 B5F3I5 B5F3I9 B5F3J2 B5F3J6 B5F3J7
B5F3P3 B5F3P7 B5F3R6 B5F3R8 B5F3S6 B5F3S8 B5F3S9 B5F3T1 B5F3Y8 B5F407 B5F409
B5F410 B5F411 B5F414 B5F415 B5F416 B5F429 B5F430 B5F445 B5F446 B5F465 B5F4C2
B5F4E4 B5F4G7 B5F4H2 B5F4H5 B5F4H6 B5F4M3 B5F4M4 B5F4P0 B5F4R2 B5F4R7 B5F4S3
B5F4S5 B5F4S7 B5F4U8 B5F4U9 B5F4W4 B5F503 B5F512 B5F516 B5F525 B5F526 B5F527
B5F534 B5F541 B5F542 B5F543 B5F565 B5F569 B5F571 B5F5C0 B5F5G3 B5F5G4 B5F5H2
B5F5H4 B5F5H7 B5F5H9 B5F5I0 B5F5I5 B5F5I9 B5F5J4 B5F5J5 B5F5K3 B5F5L1 B5F5L4

B5F5L7 B5F5M1 B5F5M2 B5F5M7 B5F5N7 B5F5W5 B5F610 B5F621 B5F623 B5F680 B5F690
B5F696 B5F6A0 B5F6A4 B5F6A5 B5F6A8 B5F6B4 B5F6B6 B5F6C2 B5F6C4 B5F6H7 B5F6I4
B5F6J8 B5F6J9 B5F6K1 B5F6L7 B5F6P5 B5F6P9 B5F6Q5 B5F6R0 B5F6R8 B5F6R9 B5F6S3
B5F6T5 B5F6T7 B5F6T8 B5F6U0 B5F6U1 B5F6U4 B5F6U7 B5F6V2 B5F6V4 B5F6V5 B5F6V8
B5F6X4 B5F6Y8 B5F6Y9 B5F721 B5F728 B5F731 B5F744 B5F748 B5F749 B5F750 B5F752
B5F756 B5F757 B5F769 B5F770 B5F771 B5F778 B5F783 B5F784 B5F7C3 B5F7C4 B5F7D6
B5F7D7 B5F7E7 B5F7F0 B5F7F6 B5F7F8 B5F7F9 B5F7G0 B5F7G2 B5F7H6 B5F7I4 B5F7I8
B5F7I9 B5F7J0 B5F7J6 B5F7J7 B5F7J9 B5F7K4 B5F7K5 B5F7L9 B5F7M0 B5F7P3 B5F7P5
B5F7R1 B5F7R3 B5F7R4 B5F7R5 B5F7S0 B5F7S2 B5F7S3 B5F7S6 B5F7S7 B5F7S9 B5F7T0
B5F7T1 B5F7T2 B5F7T3 B5F7T4 B5F7T5 B5F7T6 B5F7T7 B5F7T8 B5F7T9 B5F7U7 B5F7U9
B5F7V5 B5F7V6 B5F7W5 B5F7X5 B5F7X6 B5F7X8 B5F808 B5F814 B5F815 B5F831 B5F832
B5F833 B5F838 B5F845 B5F846 B5F860 B5F867 B5F8C6 B5F8D3 B5F8E4 B5F8E5 B5F8E6
B5F8E7 B5F8E8 B5F8E9 B5F8F1 B5F8F2 B5F8F3 B5F8F8 B5F8F9 B5F8G0 B5F8G1 B5F8G2
B5F8G3 B5F8G6 B5F8H5 B5F8J0 B5F8K0 B5F8K1 B5F8L6 B5F8M3 B5F8M6 B5F8M8 B5F8N2
B5F8N4 B5F8P1 B5F8Q1 B5F8Q2 B5F8Q3 B5F8R5 B5F8R7 B5F8S1 B5F8S5 B5F8S7 B5F8S8
B5F8T0 B5F8T1 B5F8T3 B5F8U4 B5F8V0 B5F8V2 B5F8V6 B5F8W7 B5F8X0 B5F8X2 B5F907
B5F939 B5F944 B5F947 B5F952 B5F961 B5F973 B5F981 B5F9F6 B5F9H0 B5F9M4 B5F9M8
B5F9P4 B5F9P6 B5F9P8 B5F9P9 B5F9Q1 B5F9Q2 B5F9Q6 B5F9R3 B5F9R5 B5F9R6 B5F9S6
B5F9S7 B5F9T2 B5F9T4 B5F9T5 B5F9T8 B5F9U5 B5F9U8 B5F9V3 B5F9W2 B5F9X0 B5F9X3
B5F9X5 B5F9X6 B5F9X8 B5F9X9 B5F9Y3 B5F9Y4 B5F9Z0 B5FA11 B5FA13 B5FA19 B5FA49
B5FA57 B5FA60 B5FA63 B5FA64 B5FA72 B5FA75 B5FA77 B5FA79 B5FA80 B5FA81 B5FA82
B5FA83 B5FA93 B5FA98 B5FA99 B5FAA0 B5FAC8 B5FAD0 B5FAD3 B5FAD9 B5FAE5 B5FAE6
B5FAE7 B5FAE8 B5FAF3 B5FAF5 B5FAF6 B5FAF7 B5FAG0 B5FAG4 B5FAG8 B5FAI0 B5FAI5
B5FAI8 B5FAJ2 B5FAJ7 B5FAJ9 B5FAK1 B5FAL1 B5FAL7 B5FAL8 B5FAM0 B5FAN7 B5FAP6
B5FAQ3 B5FAQ4 B5FAS8 B5FAT6 B5FAT7 B5FAU1 B5FAU4 B5FAW3 B5FAW4 B5FAW8 B5FAW9
B5FAX2 B5FAX3 B5FAX6 B5FAX9 B5FAY1 B5FAY4 B5FB05 B5FB06 B5FB26 B5FB34 B5FB37
B5FB43 B5FB46 B5FB54 B5FB55 B5FB67 B5FB77 B5FB82 B5FB83 B5FBA6 B5FBB1 B5FBB5
B5FBC3 B5FBC4 B5FBC5 B5FBC6 B5FBC7 B5FBC8 B5FBC9 B5FBD6 B5FBF0 B5FBF4 B5FBF8
B5FBF9 B5FBG6 B5FBG8 B5FBH1 B5FBH8 B5FBI0 B5FBJ5 B5FBJ6 B5FBK3 B5FBK7 B5FBL1
B5FBL3 B5FBL7 B5FBL8 B5FBM2 B5FBN1 B5FBN2 B5FBP4 B5FBP5 B5FBP7 B5FBP8 B5FBQ0
B5FBQ1 B5FBQ3 B5FBQ6 B5FBR4 B5FBR5 B5FBR6 B5FBT1 B5FBT9 B5FBU7 B5FBW5 B5FBW7
B5FBW9 B5FBX0 B5FBZ5 B5FBZ6 B5FBZ7 B5FBZ8 B5FBZ9 B5FC12 B5FC28 B5FC42 B5FC62
B5FC65 B5FC67 B5FC70 B5FC78 B5FC88 B5FC89 B5FC90 B5FC91 B5FC92 B5FCA3 B5FCA6
B5FCB8 B5FCD0 B5FCE6 B5FCE7 B5FCG1 B5FCJ8 B5FCK7 B5FCP1 B5FCP7 B5FCP8 B5FCP9
B5FCQ0 B5FCQ1 B5FCQ2 B5FCR3 B5FCV4 B5FCV8 B5FCW3 B5FCW6 B5FCW7 B5FCY6 B5FD98
B5FDA0 B5FDA3 B5FDA4 B5FDA8 B5FDB6 B5FDB7 B5FDJ8 B5FDS3 B5FDT8 B5FDV3 B5FDV4
B5FDX4 B5FDX6 B5FDX9 B5FE04 B5FE06 B5FE11 B5FE19 B5FE41 B5FEA5 B5FEB2 B5FEE3
B5FEF2 B5FEJ4 B5FEN2 B5FEP5 B5FEQ2 B5FEQ9 B5FEU9 B5FEV5 B5FEV9 B5FEW0 B5FEW3
B5FEY5 B5FEZ9 B5FF00 B5FF08 B5FF36 B5FF54 B5FF66 B5FF70 B5FF86 B5FF87 B5FF97
B5FFA6 B5FFA9 B5FFB3 B5FFC4 B5FFD2 B5FFD6 B5FFE2 B5FFE3 B5FFE9 B5FFF4 B5FFF6
B5FFF7 B5FFF8 B5FFG1 B5FFG3 B5FFJ0 B5FFK1 B5FFK8 B5FFM1 B5FFM4 B5FFM5 B5FFN7
B5FFP0 B5FFP4 B5FFP5 B5FFP7 B5FFR4 B5FFR9 B5FFS9 B5FFX7 B5FFX9 B5FFY0 B5FFY1
B5FFY9 B5FFZ7 B5FG04 B5FG05 B5FG08 B5FG09 B5FG10 B5FG11 B5FG12 B5FG13 B5FG14
B5FG15 B5FG16 B5FG17 B5FG19 B5FG20 B5FG21 B5FG22 B5FG23 B5FG24 B5FG25 B5FG26

B5FG27 B5FG34 B5FG43 B5FG51 B5FG53 B5FG54 B5FG57 B5FG59 B5FG60 B5FG61 B5FG65
B5FG68 B5FG69 B5FG80 B5FG83 B5FG96 B5FG97 B5FGA8 B5FGB1 B5FGD5 B5FGD7 B5FGD9
B5FGE0 B5FGE1 B5FGE2 B5FGE3 B5FGE9 B5FGF5 B5FGF7 B5FGF8 B5FGF9 B5FGG4 B5FGG5
B5FGH1 B5FGH2 B5FGH4 B5FGI2 B5FGI9 B5FGJ0 B5FGJ8 B5FGJ9 B5FGU2 B5FGU7 B5FGX0
B5FGX4 B5FGY4 B5FGY5 B5FH97 B5FH99 B5FHA6 B5FHA7 B5FHD7 B5FHE3 B5FHE5 B5FHF8
B5FHG3 B5FHG4 B5FHG5 B5FHG7 B5FHH0 B5FHJ6 B5FHM8 B5FHS9 B5FHT5 B5FHT9 B5FHU3
B5FHU4 B5FHU7 B5FHV3 B5FHV5 B5FHX2 B5FHY2 B5FHY5 B5FHZ3 B5FHZ4 B5FHZ8 B5FI10
B5FI12 B5FI13 B5FI15 B5FI16 B5FI19 B5FI23 B5FI32 B5FI33 B5FI34 B5FI41 B5FI45
B5FI46 B5FI58 B5FI63 B5FI64 B5FI73 B5FI83 B5FI84 B5FI86 B5FIA8 B5FIB3 B5FIB4
B5FID6 B5FIE1 B5FIF4 B5FIF5 B5FIF7 B5FIH3 B5FIM2 B5FIM3 B5FIN3 B5FIN4 B5FIN7
B5FIQ3 B5FIR5 B5FIR8 B5FIS2 B5FIS3 B5FIT7 B5FIT8 B5FIW1 B5FIW4 B5FIX4 B5FIX6
B5FIX7 B5FIX8 B5FIY3 B5FJ01 B5FJ03 B5FJ06 B5FJ11 B5FJ13 B5FJ14 B5FJ16 B5FJ17
B5FJ19 B5FJ30 B5FJ36 B5FJ38 B5FJ42 B5FJ53 B5FJ56 B5FJ58 B5FJ81 B5FJ82 B5FJ93
B5FJ96 B5FJA2 B5FJA4 B5FJA5 B5FJA6 B5FJA8 B5FJC0 B5FJC2 B5FJD0 B5FJD4 B5FJD5
B5FJD6 B5FJD7 B5FJD8 B5FJF9 B5FJG6 B5FJI0 B5FJI2 B5FJI3 B5FJI4 B5FJI9 B5FJJ1
B5FJJ2 B5FJJ5 B5FJJ6 B5FJJ8 B5FJJ9 B5FJK0 B5FJK1 B5FJK2 B5FJK3 B5FJK4 B5FJK5
B5FJK6 B5FJK7 B5FJK8 B5FJK9 B5FJL0 B5FJL1 B5FJL3 B5FJL4 B5FJL5 B5FJM0 B5FJM1
B5FJM2 B5FJM3 B5FJM4 B5FJM5 B5FJM8 B5FJN6 B5FJQ1 B5FJQ9 B5FJR0 B5FJS1 B5FJS8
B5FJT1 B5FJW2 B5FJX0 B5FJX1 B5FJX4 B5FJX5 B5FK32 B5FK77 B5FK86 B5FK97 B5FK98
B5FK99 B5FKA7 B5FKD2 B5FKD6 B5FKE4 B5FKF4 B5FKF5 B5FKF6 B5FKL3 B5FKM4 B5FKP2
B5FKP3 B5FKP6 B5FKP7 B5FKQ9 B5FKR0 B5FKS0 B5FKS2 B5FKS3 B5FKS7 B5FKS9 B5FKT0
B5FKT6 B5FKT7 B5FKU0 B5FKV2 B5FKV4 B5FKW0 B5FKW1 B5FKX3 B5FKX4 B5FKY7 B5FKY9
B5FKZ6 B5FL01 B5FL04 B5FL09 B5FL29 B5FL88 B5FLC9 B5FLD0 B5FLD6 B5FLE5 B5FLG0
B5FLG3 B5FLH9 B5FLI6 B5FLI8 B5FLJ4 B5FLJ7 B5FLJ9 B5FLK0 B5FLM4 B5FLM6 B5FLN3
B5FLP7 B5FLQ2 B5FLY4 B5FLY7 B5FM00 B5FM27 B5FM40 B5FM42 B5FM45 B5FM46 B5FM58
B5FM59 B5FM60 B5FM61 B5FM62 B5FM63 B5FM64 B5FM65 B5FM66 B5FM71 B5FM73 B5FMC1
B5FMC8 B5FME2 B5FMI4 B5FMM8 B5FMN2 B5FMN7 B5FMP4 B5FMU0 B5FMU1 B5FMW2 B5FMX1
B5FMZ9 B5FN00 B5FN08 B5FN10 B5FN11 B5FN12 B5FN16 B5FN43 B5FN44 B5FN47 B5FN48
B5FN49 B5FN51 B5FN59 B5FN65 B5FN70 B5FN73 B5FN74 B5FN93 B5FNA2 B5FNB0 B5FNB2
B5FNB9 B5FNC1 B5FNE9 B5FNF0 B5FNF9 B5FNJ7 B5FNK8 B5FNL7 B5FNN0 B5FNN2 B5FNN3
B5FNR7 B5FNS9 B5FNT1 B5FNT5 B5FNU2 B5FNU5 B5FNW1 B5FNW6 B5FNX9 B5FNZ2 B5FNZ3
B5FP17 B5FP25 B5FP28 B5FP36 B5FP38 B5FP39 B5FP41 B5FP52 B5FP54 B5FP55 B5FP57
B5FP58 B5FP61 B5FP62 B5FP65 B5FP68 B5FP70 B5FP91 B5FP96 B5FPB9 B5FPE5 B5FPE7
B5FPE8 B5FPF2 B5FPH3 B5FPL2 B5FPL4 B5FPM7 B5FPM8 B5FPP3 B5FPP4 B5FPP5 B5FPP6
B5FPR2 B5FPS9 B5FPT4 B5FPT6 B5FPT8 B5FPU0 B5FPU1 B5FPU5 B5FPU9 B5FPW2 B5FPX1
B5FPX2 B5FPX4 B5FPX5 B5FPX9 B5FPY5 B5FPY9 B5FPZ6 B5FQ06 B5FQ13 B5FQ21 B5FQ28
B5FQ36 B5FQ49 B5FQ51 B5FQ53 B5FQ55 B5FQ60 B5FQ65 B5FQ67 B5FQ81 B5FQA0 B5FQB0
B5FQB9 B5FQE4 B5FQF0 B5FQH3 B5FQH7 B5FQI8 B5FQJ0 B5FQJ2 B5FQJ5 B5FQJ6 B5FQJ7
B5FQJ8 B5FQJ9 B5FQK8 B5FQK9 B5FQL0 B5FQL1 B5FQL2 B5FQL3 B5FQM1 B5FQM3 B5FQM6
B5FQP5 B5FQQ7 B5FQZ1 B5FQZ9 B5FR01 B5FR06 B5FR08 B5FR09 B5FR34 B5FR39 B5FR40
B5FR47 B5FR52 B5FR54 B5FR62 B5FR63 B5FR66 B5FR67 B5FR78 B5FR81 B5FR82 B5FR83
B5FR85 B5FR87 B5FRB5 B5FRC3 B5FRD8 B5FRD9 B5FRJ6 B5FRK1 B5FRK2 B5FRK6 B5FRL4
B5FRL9 B5FRM6 B5FRM7 B5FRM8 B5FRN3 B5FRW4 B5FRX8 B5FRY5 B5FS06 B5FS07 B5FS09
B5FS14 B5FS17 B5FS93 B5FS94 B5FS97 B5FS98 B5FSA1 B5FSA2 B5FSA3 B5FSA4 B5FSB1

B5FSB9 B5FSC7 B5FSC9 B5FSF2 B5FSF3 B5FSH0 B5FSM1 B5FSM4 B5FSM5 B5FSM6 B5FSN0
B5FSN3 B5FSN7 B5FSN8 B5FSQ0 B5FSW2 B5FSX4 B5FSX7 B5FSX9 B5FSY0 B5FSZ2 B5FT52
B5FTA4 B5FTB3 B5FTB7 B5FTC5 B5FTC6 B5FTC7 B5FTC8 B5FTD0 B5FTD7 B5FTD8 B5FTD9
B5FTI4 B5FTI8 B5FTK8 B5FTL0 B5FTL7 B5FTL9 B5FTM0 B5FTM2 B5FTN1 B5FTQ5 B5FTS1
B5FTS3 B5FTS4 B5FTS5 B5FTS8 B5FTS9 B5FTT0 B5FTU1 B5FTU2 B5FTV0 B5FTX0 B5FTX5
B5FTY1 B5FTY3 B5FTY5 B5FU10 B5FU18 B5FU19 B5FU65 B5FU66 B5FU91 B5FU93 B5FUB2
B5FUB4 B5FUC8 B5FUF2 B5FUF3 B5FUG1 B5FUG3 B5FUG6 B5FUG7 B5FUG8 B5FUH3 B5FUH7
B5FUI2 B5FUI3 B5FUJ0 B5FUJ7 B5FUK0 B5FUP1 B5FUV5 B5FUV9 B5FUW0 B5FUW5 B5FUX4
B5FUZ9 B5FV01 B5FV55 B5FVI0 B5FW36 B5FWC0 B5FX50 B5FX56 B5FXA0 B5FXJ6 B5FXK1
B5FXN8 B5FXU9 B5FXZ4 B5FY35 B5FY93 B5FYC7 B5FZ19 B5FZ63 B5FZA8 B5FZS9 B5FZY7
B5G1C4 B5G1P1 B5G279 B5GCP6 B5GMG2 B5GRC8 B5GS26 B5GW45 B5H7H3 B5HDJ6 B5I9H4
B5IAJ6 B5IEE5 B5IEU6 B5KFL3 B5KFM4 B5KM66 B5KRH5 B5LML8 B5LMP5 B5LMQ6 B5LMQ9
B5LMR0 B5LMR2 B5LMR9 B5LVL2 B5M9L6 B5MCN3 B5MCY1 B5MD39 B5ME19 B5QBD4 B5QTD6
B5QTD7 B5QTE0 B5QTF3 B5QTF4 B5QTG3 B5QTG5 B5QTG6 B5QTH0 B5QTH2 B5QTH3 B5QTH9
B5QTI0 B5QTJ5 B5QTJ6 B5QTJ7 B5QTN6 B5QTS1 B5QTT8 B5QTU6 B5QTU8 B5QTV6 B5QTW0
B5QTW1 B5QTY1 B5QU31 B5QU45 B5QU47 B5QU58 B5QU72 B5QU74 B5QU75 B5QU77 B5QU78
B5QU79 B5QUA2 B5QUF5 B5QUG0 B5QUG1 B5QUG2 B5QUG3 B5QUG4 B5QUG9 B5QUH0 B5QUH3
B5QUH5 B5QUL6 B5QUN2 B5QUN9 B5QUP8 B5QUQ0 B5QUQ1 B5QUQ2 B5QUQ5 B5QUQ7 B5QUS9
B5QUT6 B5QUV0 B5QUV5 B5QUY5 B5QV10 B5QV11 B5QV13 B5QV29 B5QV72 B5QV75 B5QV77
B5QV78 B5QV90 B5QVE1 B5QVE4 B5QVE6 B5QVE8 B5QVG2 B5QVG7 B5QVH1 B5QVH2 B5QVJ3
B5QVK0 B5QVN4 B5QVN7 B5QVP2 B5QVP6 B5QVP9 B5QVR2 B5QVS8 B5QVS9 B5QVU2 B5QVU3
B5QVV3 B5QVV6 B5QVW2 B5QVW4 B5QVW5 B5QVW6 B5QVW8 B5QW14 B5QW15 B5QW16 B5QW17
B5QW18 B5QW21 B5QW22 B5QW23 B5QW34 B5QW35 B5QW41 B5QW68 B5QW73 B5QW86 B5QW98
B5QWB9 B5QWC1 B5QWC8 B5QWD0 B5QWF7 B5QWF8 B5QWG8 B5QWH4 B5QWH6 B5QWI3 B5QWI6
B5QWI7 B5QWI8 B5QWI9 B5QWJ0 B5QWL7 B5QWQ2 B5QWQ6 B5QWR0 B5QWR2 B5QWR4 B5QWT6
B5QWT7 B5QWV0 B5QWW2 B5QWW9 B5QWX3 B5QWX9 B5QWY1 B5QWY2 B5QWY3 B5QWY4 B5QWY5
B5QWZ5 B5QX14 B5QX27 B5QX43 B5QX45 B5QX56 B5QX58 B5QX59 B5QX61 B5QX62 B5QX65
B5QX66 B5QX69 B5QX73 B5QX75 B5QXA9 B5QXB6 B5QXC6 B5QXC7 B5QXC8 B5QXD6 B5QXE5
B5QXG6 B5QXG7 B5QXH5 B5QXH7 B5QXI0 B5QXI1 B5QXI2 B5QXI6 B5QXJ0 B5QXJ1 B5QXJ6
B5QXJ7 B5QXK5 B5QXL5 B5QXL7 B5QXL8 B5QXM0 B5QXM1 B5QXM5 B5QXM8 B5QXP2 B5QXQ1
B5QXQ2 B5QXQ4 B5QXQ5 B5QXQ7 B5QXR0 B5QXR2 B5QXR5 B5QXT1 B5QXT6 B5QXV6 B5QXZ7
B5QY02 B5QY05 B5QY09 B5QY29 B5QY66 B5QY68 B5QY72 B5QY73 B5QY78 B5QY87 B5QYA5
B5QYB0 B5QYB2 B5QYD4 B5QYD5 B5QYD6 B5QYD7 B5QYD8 B5QYE5 B5QYE8 B5QYE9 B5QYF0
B5QYF1 B5QYF2 B5QYF3 B5QYG1 B5QYG3 B5QYG6 B5QYI4 B5QYJ7 B5QYK4 B5QYL4 B5QYM1
B5QYM8 B5QYN6 B5QYP4 B5QYQ7 B5QYQ8 B5QYT3 B5QYX3 B5QYX6 B5QYY9 B5QZ24 B5QZ31
B5QZ33 B5QZ36 B5QZ40 B5QZ44 B5QZ45 B5QZ48 B5QZ53 B5QZ55 B5QZ58 B5QZB4 B5QZB6
B5QZC1 B5QZC6 B5QZC8 B5QZD8 B5QZF1 B5QZF9 B5QZG1 B5QZG6 B5QZG9 B5QZH0 B5QZJ9
B5QZL1 B5QZL3 B5QZL6 B5QZL7 B5QZR6 B5QZS6 B5QZS8 B5QZS9 B5QZT7 B5QZT8 B5QZU2
B5QZV4 B5QZV5 B5QZV7 B5QZV8 B5QZW0 B5QZW1 B5QZW4 B5QZZ9 B5R004 B5R005 B5R009
B5R019 B5R024 B5R043 B5R048 B5R049 B5R056 B5R0A0 B5R0B4 B5R0B5 B5R0D9 B5R0E7
B5R0H0 B5R0H4 B5R0H6 B5R0H7 B5R0I0 B5R0J6 B5R0K9 B5R0L2 B5R0L3 B5R0L7 B5R0L8
B5R0N2 B5R0N7 B5R0N8 B5R0N9 B5R0P3 B5R0P4 B5R0Q8 B5R0Q9 B5R0R3 B5R0R4 B5R0R8
B5R0R9 B5R0S0 B5R0S1 B5R0S8 B5R0T7 B5R0U4 B5R0U6 B5R0X8 B5R0Z0 B5R104 B5R110
B5R142 B5R144 B5R145 B5R146 B5R153 B5R163 B5R173 B5R181 B5R194 B5R196 B5R197

B5R1A3 B5R1C6 B5R1C8 B5R1D8 B5R1E1 B5R1E3 B5R1E4 B5R1E5 B5R1F1 B5R1F3 B5R1F8
B5R1F9 B5R1G1 B5R1G2 B5R1G3 B5R1I5 B5R1I6 B5R1K2 B5R1M7 B5R1N6 B5R1N9 B5R1P1
B5R1Q4 B5R1Q8 B5R1Q9 B5R1R0 B5R1R1 B5R1R2 B5R1S6 B5R1S7 B5R1S8 B5R1T4 B5R1T9
B5R1U0 B5R1V2 B5R1V5 B5R1V8 B5R249 B5R262 B5R264 B5R268 B5R279 B5R280 B5R281
B5R282 B5R283 B5R284 B5R285 B5R286 B5R287 B5R288 B5R290 B5R291 B5R292 B5R297
B5R298 B5R299 B5R2A0 B5R2A1 B5R2A2 B5R2A5 B5R2B3 B5R2C7 B5R2D5 B5R2D6 B5R2H7
B5R2I6 B5R2J0 B5R2J7 B5R2J8 B5R2J9 B5R2K1 B5R2K9 B5R2L5 B5R2L6 B5R2M5 B5R2N6
B5R2N7 B5R2N9 B5R2Q9 B5R2R4 B5R2R5 B5R2S1 B5R2S5 B5R2U5 B5R2U7 B5R2V4 B5R2V6
B5R2V7 B5R2V9 B5R2W7 B5R2Y7 B5R2Y9 B5R2Z0 B5R2Z4 B5R313 B5R316 B5R349 B5R360
B5R367 B5R369 B5R371 B5R375 B5R378 B5R384 B5R394 B5R395 B5R396 B5R3B5 B5R3B6
B5R3B7 B5R3D3 B5R3E4 B5R3E5 B5R3E6 B5R3F1 B5R3G6 B5R3G8 B5R3H1 B5R3H2 B5R3H6
B5R3H8 B5R3H9 B5R3I2 B5R3I3 B5R3I5 B5R3J2 B5R3J6 B5R3J8 B5R3J9 B5R3L7 B5R3P3
B5R3P4 B5R3Q1 B5R3R5 B5R3R7 B5R3T6 B5R3U6 B5R3V4 B5R3X9 B5R3Y5 B5R3Z4 B5R404
B5R422 B5R434 B5R447 B5R449 B5R470 B5R486 B5R490 B5R4D8 B5R4G0 B5R4G5 B5R4J0
B5R4J4 B5R4P2 B5R4P3 B5R4P8 B5R4Q5 B5R4S2 B5R4S3 B5R4S5 B5R4S6 B5R541 B5R558
B5R560 B5R569 B5R571 B5R578 B5R581 B5R582 B5R584 B5R585 B5R595 B5R598 B5R599
B5R5A0 B5R5A2 B5R5A3 B5R5B6 B5R5B9 B5R5D4 B5R5E1 B5R5E3 B5R5F8 B5R5F9 B5R5G0
B5R5G1 B5R5G2 B5R5G4 B5R5G5 B5R5G6 B5R5G7 B5R5H2 B5R5H5 B5R5H7 B5R5I2 B5R5I3
B5R5I6 B5R5J8 B5R5L1 B5R5L3 B5R5Q7 B5R5R5 B5R5R6 B5R5R8 B5R5R9 B5R5X9 B5R5Y2
B5R5Y3 B5R5Z3 B5R607 B5R609 B5R610 B5R612 B5R613 B5R637 B5R639 B5R646 B5R651
B5R678 B5R679 B5R689 B5R6A2 B5R6B4 B5R6C2 B5R6C4 B5R6C9 B5R6D2 B5R6D3 B5R6F7
B5R6G2 B5R6G3 B5R6H0 B5R6J8 B5R6M4 B5R6M5 B5R6P3 B5R6Q6 B5R6Q7 B5R6R6 B5R6R8
B5R6R9 B5R6S3 B5R6S5 B5R6S6 B5R6T2 B5R6T3 B5R6U8 B5R6V0 B5R6V6 B5R6V8 B5R6X0
B5R6X5 B5R705 B5R739 B5R741 B5R752 B5R754 B5R755 B5R757 B5R758 B5R761 B5R762
B5R765 B5R769 B5R771 B5R792 B5R797 B5R7E9 B5R7G4 B5R7H8 B5R7J6 B5R7J9 B5R7K3
B5R7K5 B5R7K7 B5R7L4 B5R7M7 B5R7M8 B5R7N6 B5R7P4 B5R7P7 B5R7Q1 B5R7Q4 B5R7Q9
B5R7S0 B5R7T2 B5R7Y1 B5R7Y4 B5R7Y9 B5R7Z3 B5R7Z6 B5R809 B5R815 B5R817 B5R824
B5R840 B5R861 B5R878 B5R885 B5R890 B5R896 B5R8C8 B5R8D0 B5R8D1 B5R8D2 B5R8D6
B5R8D9 B5R8E2 B5R8G5 B5R8H3 B5R8I1 B5R8J4 B5R8J5 B5R8J7 B5R8J9 B5R8K4 B5R8K9
B5R8L1 B5R8M1 B5R8V2 B5R8V6 B5R8X8 B5R8Y5 B5R8Y7 B5R8Y8 B5R8Z0 B5R917 B5R921
B5R923 B5R924 B5R985 B5R990 B5R991 B5R995 B5R9A5 B5R9B7 B5R9B8 B5R9B9 B5R9C3
B5R9C4 B5R9D9 B5R9E3 B5R9E4 B5R9E8 B5R9E9 B5R9F0 B5R9F1 B5R9F8 B5R9G7 B5R9H4
B5R9H6 B5R9J8 B5R9J9 B5R9L5 B5R9P0 B5R9T0 B5R9T9 B5R9U3 B5R9V0 B5R9V1 B5R9V2
B5R9V4 B5R9W1 B5R9W2 B5R9W3 B5R9Y0 B5RA25 B5RAB4 B5RAE1 B5RAG0 B5RAJ1 B5RAJ6
B5RAL0 B5RAL1 B5RAL3 B5RAM9 B5RAT3 B5RAT4 B5RAU7 B5RAU8 B5RAV8 B5RAW7 B5RAW9
B5RAX0 B5RAX1 B5RAX3 B5RAY5 B5RAY7 B5RAZ4 B5RAZ7 B5RAZ8 B5RAZ9 B5RB01 B5RB75
B5RB82 B5RB92 B5RB93 B5RB94 B5RBA2 B5RBB1 B5RBD7 B5RBE0 B5RBE4 B5RBF3 B5RBG7
B5RBK6 B5RBK9 B5RBP9 B5RBR1 B5RBR3 B5RBR6 B5RBR7 B5RBW2 B5RBW3 B5RBY7 B5RBZ5
B5RC22 B5RC32 B5RC42 B5RC50 B5RC63 B5RC65 B5RC66 B5RCA1 B5RCB4 B5RCB6 B5RCC0
B5RCD1 B5RCD3 B5RCD4 B5RCD8 B5RCF7 B5RCG0 B5RCJ3 B5RCJ5 B5RCK9 B5RCL1 B5RCN1
B5RCP1 B5RCP9 B5RCS3 B5RCS8 B5RCV3 B5RCV7 B5RCW8 B5RCX0 B5RCX9 B5RCY1 B5RCY8
B5RCZ1 B5RCZ2 B5RCZ4 B5RCZ5 B5RD05 B5RD08 B5RD09 B5RD10 B5RD12 B5RD13 B5RD39
B5RD47 B5RD49 B5RD57 B5RD61 B5RD62 B5RD82 B5RD83 B5RD84 B5RD85 B5RD90 B5RD91
B5RD94 B5RD96 B5RDD6 B5RDE8 B5RDF1 B5RDF3 B5RDF4 B5RDG6 B5RDN2 B5RDP9 B5RDQ0

B5RDQ1 B5RDQ2 B5RDQ3 B5RDQ6 B5RDQ7 B5RDQ8 B5RDR9 B5RDS0 B5RDS6 B5RDU6 B5RDV0
B5RDV4 B5RDV6 B5RDV8 B5RDY0 B5RDY1 B5RDZ4 B5RE00 B5RE01 B5RE09 B5RE11 B5RE14
B5RE15 B5RE16 B5RE20 B5RE24 B5RE25 B5RE30 B5RE31 B5RE39 B5RE50 B5RE52 B5RE56
B5RE57 B5RE62 B5RE71 B5RE89 B5RE94 B5RE96 B5REE6 B5REF3 B5REF8 B5REG2 B5REG6
B5REG7 B5REH0 B5REH5 B5REH7 B5REJ4 B5REK4 B5REK6 B5REK7 B5REL5 B5REL6 B5REM0
B5REN2 B5REN3 B5REN5 B5REN6 B5REN8 B5REP2 B5REP5 B5REP9 B5REQ1 B5REQ2 B5REQ5
B5RES1 B5RET4 B5RET7 B5RET8 B5REU3 B5REV7 B5REV8 B5REY1 B5REY3 B5REZ1 B5REZ3
B5REZ6 B5REZ7 B5REZ9 B5RF01 B5RF08 B5RF09 B5RF40 B5RF43 B5RF45 B5RF46 B5RF48
B5RF49 B5RF58 B5RF72 B5RF75 B5RF79 B5RF80 B5RF82 B5RF83 B5RF85 B5RF90 B5RFA9
B5RFC3 B5RFC4 B5RFC5 B5RFC6 B5RFC8 B5RFD4 B5RFD8 B5RFE5 B5RFH8 B5RFI6 B5RFI7
B5RFI8 B5RFI9 B5RFJ0 B5RFJ1 B5RFJ4 B5RFK1 B5RFK2 B5RFK3 B5RFK4 B5RFK5 B5RFK9
B5RFL5 B5RFM8 B5RFN3 B5RFQ2 B5RFS3 B5RFS4 B5RFS8 B5RFT3 B5RFU7 B5RFU9 B5RFV4
B5RFY2 B5RFY5 B5RFY6 B5RFY7 B5RFY9 B5RFZ8 B5RG05 B5RG23 B5RG73 B5RG75 B5RG80
B5RG81 B5RG84 B5RG86 B5RG98 B5RG99 B5RGA3 B5RGA4 B5RGA5 B5RGA6 B5RGA8 B5RGC0
B5RGC1 B5RGC2 B5RGD3 B5RGD4 B5RGE4 B5RGE6 B5RGE7 B5RGE9 B5RGF0 B5RGF1 B5RGF2
B5RGF3 B5RGG8 B5RGH0 B5RGH7 B5RGJ1 B5RGJ4 B5RGL6 B5RGM1 B5RGM2 B5RGS3 B5RGT3
B5RGU2 B5RGU8 B5RGZ3 B5RH01 B5RH04 B5RH05 B5RH06 B5RH07 B5RH08 B5RH09 B5RH14
B5RH15 B5RH16 B5RH18 B5RH19 B5RH20 B5RH21 B5RH22 B5RH23 B5RH24 B5RH25 B5RH26
B5RH27 B5RH29 B5RH30 B5RH31 B5RH33 B5RH34 B5RH38 B5RH39 B5RH41 B5RH47 B5RH48
B5RH49 B5RH55 B5RH56 B5RH65 B5RH76 B5RH77 B5RH79 B5RH99 B5RHA4 B5RHA5 B5RHB9
B5RHC0 B5RHC1 B5RHC6 B5RHE0 B5RHE2 B5RHE5 B5RHE6 B5RHE9 B5RHF1 B5RHF2 B5RHF4
B5RHF5 B5RHF7 B5RHS5 B5RI54 B5RKU7 B5RKV2 B5RKV4 B5RKV5 B5RKV9 B5RKW3 B5RL03
B5RL05 B5RL09 B5RL10 B5RL14 B5RL15 B5RL16 B5RL25 B5RL36 B5RL39 B5RL50 B5RL65
B5RL72 B5RL76 B5RL78 B5RL82 B5RLC2 B5RLD2 B5RLD9 B5RLH3 B5RLI6 B5RLI7 B5RLI9
B5RLJ5 B5RLJ6 B5RLJ7 B5RLK3 B5RLK6 B5RLM0 B5RLM3 B5RLN1 B5RLN2 B5RLQ9 B5RLR1
B5RLS3 B5RLS4 B5RLS6 B5RLT4 B5RLU6 B5RLU7 B5RLU8 B5RLU9 B5RLV0 B5RLV1 B5RLV2
B5RLV6 B5RLX5 B5RLY1 B5RLZ3 B5RLZ7 B5RLZ8 B5RM04 B5RM14 B5RM19 B5RM34 B5RM35
B5RM36 B5RM37 B5RM39 B5RM40 B5RM41 B5RM42 B5RM43 B5RM44 B5RM45 B5RM46 B5RM47
B5RM48 B5RM49 B5RM50 B5RM51 B5RM52 B5RM57 B5RM58 B5RM61 B5RM71 B5RM75 B5RMA6
B5RME0 B5RME6 B5RMF3 B5RMG0 B5RMG2 B5RMG4 B5RMH1 B5RMJ5 B5RMK4 B5RMM8 B5RMP0
B5RMP3 B5RMP6 B5RMP7 B5RMR3 B5RMU7 B5RMW0 B5RMX2 B5RMX3 B5RMX7 B5RMX8 B5RMX9
B5RMZ3 B5RMZ9 B5RN03 B5RN19 B5RN32 B5RN40 B5RPI0 B5RPI1 B5RPI2 B5RPI3 B5RPI5
B5RPI6 B5RPI7 B5RPI8 B5RPI9 B5RPJ0 B5RPJ1 B5RPJ2 B5RPJ3 B5RPJ4 B5RPJ5 B5RPJ6
B5RPJ7 B5RPJ8 B5RPK3 B5RPK4 B5RPK7 B5RPL7 B5RPM1 B5RPQ2 B5RPT6 B5RPU2 B5RPU9
B5RPV6 B5RPV8 B5RPW0 B5RPW7 B5RPZ1 B5RQ00 B5RQ24 B5RQ36 B5RQ39 B5RQ42 B5RQ43
B5RQ59 B5RQ92 B5RQA5 B5RQB7 B5RQB8 B5RQC2 B5RQC3 B5RQC4 B5RQD7 B5RQE3 B5RQE7
B5RQF8 B5RQH0 B5RQH8 B5RQJ5 B5RQJ8 B5RQK6 B5RQK7 B5RQM7 B5RQN2 B5RQN4 B5RQN5
B5RQN9 B5RQP3 B5RQS6 B5RQT9 B5RQU0 B5RQU2 B5RQU6 B5RQU7 B5RQU8 B5RQV4 B5RQZ6
B5RQZ8 B5RR02 B5RR03 B5RR07 B5RR08 B5RR09 B5RR18 B5RR29 B5RR32 B5RR43 B5RR63
B5RR67 B5RRB6 B5RRC5 B5RRD2 B5RRF2 B5RRF3 B5RRF5 B5RRG3 B5RRH9 B5RRI1 B5RRJ4
B5RRJ5 B5RRJ6 B5RRJ7 B5RRJ8 B5RRJ9 B5RRK0 B5RRK4 B5RRM3 B5RRM9 B5RRN9 B5RRP3
B5RRP4 B5RRQ0 B5RRR0 B5RRR5 B5RSM1 B5RSP9 B5RTE0 B5RTW5 B5RUN4 B5RUP5 B5RUV1
B5RV24 B5SNH4 B5UAT8 B5UBC1 B5VDQ0 B5VDR1 B5VDZ9 B5VE38 B5VEB9 B5VEH8 B5VEQ2
B5VFW2 B5VFX4 B5VG60 B5VGI4 B5VH97 B5VHI3 B5VHN4 B5VI57 B5VI68 B5VIP5 B5VJQ0

B5VK45 B5VK87 B5VK90 B5VKC6 B5VKC8 B5VKE8 B5VKI1 B5VL46 B5VLD2 B5VLI6 B5VLV2
B5VM59 B5VM60 B5VM65 B5VMH6 B5VML1 B5VMP2 B5VMR4 B5VN01 B5VN64 B5VN66 B5VNM2
B5VNY0 B5VP69 B5VP80 B5VPI9 B5VPJ0 B5VPK7 B5VQ50 B5VQF5 B5VQH0 B5VQJ1 B5VQJ9
B5VQM2 B5VQS7 B5VRU9 B5VRV8 B5VS81 B5VS96 B5VSG4 B5VST2 B5VT12 B5VT41 B5VTQ6
B5VTQ8 B5X0I6 B5X0W9 B5X165 B5X171 B5X1P9 B5X212 B5X277 B5X2D1 B5X2S3 B5X2Z0
B5X392 B5X3C4 B5X3I1 B5X3V2 B5X3X5 B5X3Z6 B5X441 B5X4Y9 B5X4Z4 B5X574 B5X582
B5X5B4 B5X5D0 B5X5L2 B5X5N3 B5X5U9 B5X601 B5X737 B5X7E4 B5X8A5 B5X8M4 B5X9P2
B5X9S3 B5XAM2 B5XB27 B5XBI1 B5XBP5 B5XC71 B5XCA1 B5XCB8 B5XCZ6 B5XD90 B5XDD3
B5XE27 B5XEX1 B5XF11 B5XFA7 B5XG19 B5XGE7 B5XGH3 B5XHH0 B5XHV2 B5XHV3 B5XHV7
B5XHV8 B5XHX4 B5XHZ0 B5XHZ8 B5XHZ9 B5XI00 B5XI03 B5XI21 B5XI30 B5XI32 B5XI38
B5XI40 B5XI46 B5XI49 B5XI61 B5XID8 B5XIE3 B5XIE6 B5XIF1 B5XIF6 B5XIG1 B5XIG6
B5XII4 B5XIJ1 B5XIK4 B5XIK9 B5XIL1 B5XIL5 B5XIM6 B5XIN3 B5XIP2 B5XIP6 B5XIW7
B5XIW8 B5XIX5 B5XIY2 B5XIY3 B5XIY5 B5XIY6 B5XIZ3 B5XJ00 B5XJ01 B5XJ02 B5XJ05
B5XJ07 B5XJ09 B5XJ12 B5XJ18 B5XJ20 B5XJ28 B5XJ35 B5XJ36 B5XJ37 B5XJ39 B5XJ40
B5XJ41 B5XJ43 B5XJ44 B5XJ45 B5XJ46 B5XJ47 B5XJ48 B5XJ49 B5XJ50 B5XJ51 B5XJ52
B5XJ54 B5XJ55 B5XJ57 B5XJ59 B5XJ60 B5XJ61 B5XJ63 B5XJ70 B5XJ72 B5XJ75 B5XJ77
B5XJ82 B5XJ83 B5XJ84 B5XJ99 B5XJA2 B5XJA6 B5XJC1 B5XJC8 B5XJD9 B5XJE9 B5XJH8
B5XJI9 B5XJL3 B5XJM0 B5XJN3 B5XJN7 B5XJP0 B5XJP1 B5XJR0 B5XJR1 B5XJR8 B5XJU1
B5XJU2 B5XJV1 B5XJV7 B5XJW0 B5XJW3 B5XJX3 B5XJX7 B5XJX8 B5XJX9 B5XJY9 B5XJZ7
B5XJZ8 B5XJZ9 B5XK00 B5XK03 B5XK14 B5XK61 B5XK62 B5XK64 B5XK66 B5XK67 B5XK72
B5XK94 B5XK96 B5XKA6 B5XKB7 B5XKG9 B5XKI1 B5XKI2 B5XKL0 B5XKL2 B5XKL7 B5XKM2
B5XKP3 B5XKQ6 B5XKR3 B5XKR4 B5XKR6 B5XKT3 B5XKT7 B5XKT8 B5XKT9 B5XKU0 B5XKU3
B5XKU4 B5XKU5 B5XKV0 B5XKV2 B5XKV8 B5XKW0 B5XKW2 B5XKW7 B5XKX3 B5XKX5 B5XL03
B5XL04 B5XL07 B5XL08 B5XL15 B5XL20 B5XL29 B5XL33 B5XL36 B5XL39 B5XLD8 B5XLE2
B5XLE8 B5XLF2 B5XLG6 B5XLG8 B5XLI1 B5XLI3 B5XLI8 B5XLJ2 B5XLJ8 B5XLK7 B5XLM1
B5XLM6 B5XLN3 B5XLN9 B5XLR5 B5XLR6 B5XLS8 B5XLV6 B5XLX5 B5XLX7 B5XLZ0 B5XM06
B5XM07 B5XM14 B5XM21 B5XM45 B5XM46 B5XM60 B5XM63 B5XM64 B5XM69 B5XM82 B5XM83
B5XM84 B5XMA3 B5XMB1 B5XMB5 B5XMB7 B5XMC1 B5XMC3 B5XMC5 B5XME6 B5XMF0 B5XMG6
B5XMG7 B5XMI3 B5XMI5 B5XMJ7 B5XMJ8 B5XMJ9 B5XML3 B5XML5 B5XML6 B5XMM1 B5XMM6
B5XMM8 B5XMM9 B5XMN0 B5XMV6 B5XMW1 B5XMX0 B5XMY6 B5XMY7 B5XN47 B5XN72 B5XN79
B5XN82 B5XN83 B5XN84 B5XN85 B5XN86 B5XN93 B5XN94 B5XN95 B5XN97 B5XN98 B5XN99
B5XNA0 B5XNA1 B5XNA2 B5XNA4 B5XNA5 B5XNA6 B5XNA7 B5XNA8 B5XNA9 B5XNB1 B5XNB2
B5XNB4 B5XNB5 B5XNB7 B5XNC2 B5XNC3 B5XNC4 B5XNC6 B5XND7 B5XND9 B5XNE4 B5XNF8
B5XNF9 B5XNG1 B5XNH0 B5XNH2 B5XNI6 B5XNJ6 B5XNJ7 B5XNJ9 B5XNK0 B5XNK1 B5XNK4
B5XNL1 B5XNL2 B5XNL5 B5XNL6 B5XNM2 B5XNM4 B5XNQ0 B5XNQ4 B5XNQ6 B5XNR7 B5XNR9
B5XNU6 B5XNU9 B5XNW7 B5XNX0 B5XNX1 B5XNX3 B5XNX5 B5XNY1 B5XNZ3 B5XP20 B5XP25
B5XP41 B5XP51 B5XP57 B5XP65 B5XP89 B5XP95 B5XP99 B5XPC8 B5XPE2 B5XPE3 B5XPE6
B5XPE8 B5XPH2 B5XPH8 B5XPI0 B5XPU4 B5XPV7 B5XPW3 B5XPW9 B5XPZ2 B5XPZ4 B5XPZ5
B5XPZ6 B5XQ00 B5XQ02 B5XQ03 B5XQ04 B5XQ05 B5XQ21 B5XQ35 B5XQ39 B5XQ60 B5XQ62
B5XQ69 B5XQ71 B5XQ72 B5XQ74 B5XQ81 B5XQ92 B5XQA8 B5XQC6 B5XQC8 B5XQC9 B5XQD0
B5XQD2 B5XQE2 B5XQE5 B5XQH2 B5XQH3 B5XQI9 B5XQJ0 B5XQJ3 B5XQJ4 B5XQT8 B5XQU5
B5XQU9 B5XQY2 B5XRA3 B5XRA9 B5XRE3 B5XRN3 B5XS03 B5XS05 B5XS09 B5XS13 B5XS71
B5XS77 B5XSN3 B5XSP2 B5XSQ5 B5XSQ7 B5XSS1 B5XSS2 B5XSS5 B5XST6 B5XSV2 B5XSV5
B5XSV6 B5XSV8 B5XSW2 B5XSW5 B5XSX1 B5XSX2 B5XSX4 B5XSX5 B5XSX7 B5XSX9 B5XSY9

B5XSZ4 B5XSZ5 B5XT02 B5XT03 B5XT12 B5XT26 B5XT28 B5XT51 B5XT53 B5XT60 B5XT61
B5XTD4 B5XTE8 B5XTF0 B5XTF9 B5XTG0 B5XTG2 B5XTG3 B5XTG5 B5XTG6 B5XTG7 B5XTG8
B5XTG9 B5XTI2 B5XTI4 B5XTJ1 B5XTJ9 B5XTL8 B5XTQ8 B5XTQ9 B5XTR0 B5XTR4 B5XTR7
B5XTS2 B5XTS4 B5XTS6 B5XTU7 B5XTW7 B5XTX7 B5XTY7 B5XU18 B5XU21 B5XU22 B5XU24
B5XU25 B5XU26 B5XU27 B5XU28 B5XU29 B5XU35 B5XU40 B5XU43 B5XU47 B5XU89 B5XU96
B5XUA1 B5XUA2 B5XUA6 B5XUA8 B5XUB1 B5XUB2 B5XUB7 B5XUB8 B5XUC3 B5XUC4 B5XUC9
B5XUD4 B5XUD5 B5XUE4 B5XUE6 B5XUF3 B5XUF5 B5XUP7 B5XUP9 B5XUR6 B5XUU2 B5XUV5
B5XUY0 B5XUY2 B5XUY5 B5XUZ0 B5XV05 B5XV15 B5XV18 B5XV25 B5XV26 B5XV29 B5XV31
B5XV34 B5XV35 B5XV36 B5XV37 B5XV38 B5XV49 B5XV79 B5XVA0 B5XVB6 B5XVB7 B5XVB9
B5XVC3 B5XVD7 B5XVJ3 B5XVJ5 B5XVJ8 B5XVJ9 B5XVK3 B5XVK4 B5XVK5 B5XVK6 B5XVM9
B5XVN4 B5XVR1 B5XVT4 B5XVT8 B5XVU8 B5XVW4 B5XVW6 B5XVZ8 B5XW18 B5XW43 B5XW48
B5XW50 B5XW51 B5XWL8 B5XWN4 B5XWN7 B5XWP0 B5XWQ7 B5XWV7 B5XWY0 B5XX25 B5XX58
B5XX59 B5XX60 B5XX61 B5XX62 B5XXG3 B5XXG4 B5XXH2 B5XXI1 B5XXJ0 B5XXJ5 B5XXK0
B5XXK9 B5XXL2 B5XXN0 B5XXN1 B5XXN2 B5XXN3 B5XXN4 B5XXN5 B5XXP0 B5XXW1 B5XXZ3
B5XY03 B5XY38 B5XY39 B5XY41 B5XY47 B5XY49 B5XY56 B5XY73 B5XY75 B5XY80 B5XY84
B5XY86 B5XY87 B5XY88 B5XY99 B5XYA8 B5XYB2 B5XYB9 B5XYC7 B5XYD0 B5XYD2 B5XYE0
B5XYE1 B5XYE2 B5XYE4 B5XYE8 B5XYF5 B5XYF6 B5XYF7 B5XYF8 B5XYF9 B5XYG3 B5XYG9
B5XYH5 B5XYI1 B5XYJ1 B5XYK9 B5XYL1 B5XYN4 B5XYQ3 B5XYT2 B5XYT7 B5XYW4 B5XYW6
B5XYX2 B5XYY9 B5XYZ0 B5XYZ8 B5XZ01 B5XZ09 B5XZ11 B5XZ14 B5XZ16 B5XZ17 B5XZ19
B5XZ20 B5XZ23 B5XZ31 B5XZ32 B5XZ36 B5XZ37 B5XZ39 B5XZ40 B5XZ49 B5XZ51 B5XZ52
B5XZ54 B5XZ60 B5XZ74 B5XZ75 B5XZ79 B5XZ80 B5XZ82 B5XZ83 B5XZA0 B5XZB2 B5XZD1
B5XZD9 B5XZE0 B5XZE1 B5XZG7 B5XZG9 B5XZH5 B5XZH9 B5XZI4 B5XZJ3 B5XZK8 B5XZL2
B5XZL6 B5XZP4 B5XZP6 B5XZP7 B5XZQ0 B5XZQ2 B5XZR2 B5XZR5 B5XZR8 B5XZS3 B5XZS4
B5XZS6 B5XZU3 B5XZV4 B5XZW2 B5XZW3 B5Y006 B5Y007 B5Y008 B5Y035 B5Y0K4 B5Y0K6
B5Y0N2 B5Y0N3 B5Y0N5 B5Y0N6 B5Y0N8 B5Y0Q3 B5Y0Q4 B5Y0T3 B5Y0T5 B5Y0U1 B5Y0U3
B5Y0W4 B5Y0W8 B5Y0W9 B5Y0X1 B5Y0X5 B5Y0X6 B5Y0X8 B5Y0Y6 B5Y0Y7 B5Y109 B5Y110
B5Y115 B5Y119 B5Y159 B5Y168 B5Y169 B5Y1B5 B5Y1D2 B5Y1D9 B5Y1F1 B5Y1F4 B5Y1G2
B5Y1G4 B5Y1G7 B5Y1H7 B5Y1I8 B5Y1J9 B5Y1K1 B5Y1K2 B5Y1K5 B5Y1K6 B5Y1K9 B5Y1L0
B5Y1L3 B5Y1L5 B5Y1P0 B5Y1P2 B5Y1P5 B5Y1P6 B5Y1R1 B5Y1R2 B5Y1R3 B5Y1T3 B5Y1T6
B5Y1T7 B5Y1U6 B5Y1V5 B5Y1V6 B5Y1W2 B5Y1W4 B5Y1W5 B5Y1W9 B5Y1Y0 B5Y1Y1 B5Y1Y4
B5Y1Z4 B5Y1Z5 B5Y1Z6 B5Y210 B5Y214 B5Y231 B5Y232 B5Y235 B5Y237 B5Y241 B5Y242
B5Y250 B5Y254 B5Y256 B5Y264 B5Y265 B5Y266 B5Y273 B5Y274 B5Y275 B5Y276 B5Y277
B5Y282 B5Y286 B5Y299 B5Y2D2 B5Y2S3 B5Y2S5 B5Y2T8 B5Y2U6 B5Y2W8 B5Y2X3 B5Y2Y3
B5Y301 B5Y305 B5Y306 B5Y307 B5Y324 B5Y325 B5Y332 B5Y333 B5Y334 B5Y341 B5Y345
B5Y354 B5Y368 B5Y369 B5Y5X6 B5Y646 B5Y659 B5Y6A0 B5Y6A9 B5Y6G9 B5Y6I0 B5Y6I1
B5Y6R9 B5Y6U9 B5Y6V5 B5Y714 B5Y795 B5Y7I0 B5Y7N9 B5Y7Q7 B5Y7R0 B5Y7R3 B5Y7S6
B5Y7T0 B5Y7T1 B5Y7V3 B5Y7V5 B5Y7V9 B5Y7W0 B5Y804 B5Y805 B5Y806 B5Y807 B5Y822
B5Y862 B5Y863 B5Y876 B5Y878 B5Y893 B5Y894 B5Y8A3 B5Y8C1 B5Y8C8 B5Y8D8 B5Y8F0
B5Y8F3 B5Y8F5 B5Y8G1 B5Y8G6 B5Y8H0 B5Y8H6 B5Y8I0 B5Y8I3 B5Y8I4 B5Y8I9 B5Y8J0
B5Y8J2 B5Y8J7 B5Y8K5 B5Y8K7 B5Y8L1 B5Y8M4 B5Y8N2 B5Y8Q4 B5Y8Q8 B5Y8R7 B5Y8R8
B5Y8S2 B5Y8S5 B5Y8Y0 B5Y8Y9 B5Y926 B5Y930 B5Y931 B5Y932 B5Y934 B5Y955 B5Y959
B5Y962 B5Y963 B5Y969 B5Y973 B5Y974 B5Y975 B5Y978 B5Y980 B5Y981 B5Y982 B5Y985
B5Y988 B5Y991 B5Y9B9 B5Y9D1 B5Y9D4 B5Y9H0 B5Y9P7 B5Y9T8 B5Y9V1 B5Y9X7 B5Y9Z4
B5YA17 B5YA73 B5YA97 B5YAD0 B5YAD3 B5YAK0 B5YAL8 B5YAL9 B5YAR3 B5YAR4 B5YB40

B5YB52 B5YBN8 B5YBN9 B5YCC8 B5YCT4 B5YD05 B5YD09 B5YD73 B5YD80 B5YD88 B5YD92
B5YDA8 B5YDB0 B5YDB2 B5YDB7 B5YDB8 B5YDD8 B5YDE1 B5YDN9 B5YDP7 B5YDR3 B5YDR9
B5YDS4 B5YDT0 B5YDT1 B5YDT2 B5YDT8 B5YDT9 B5YDU6 B5YDU7 B5YDU9 B5YDV3 B5YDV4
B5YDV5 B5YDV6 B5YDV7 B5YDV8 B5YDV9 B5YDW6 B5YDW7 B5YDW8 B5YDW9 B5YDX1 B5YDX5
B5YDX6 B5YDY1 B5YDY6 B5YDZ9 B5YE05 B5YE06 B5YE16 B5YE18 B5YE40 B5YE42 B5YE43
B5YE53 B5YE70 B5YE84 B5YE87 B5YE89 B5YE90 B5YE93 B5YE95 B5YE97 B5YEA6 B5YEA9
B5YEB0 B5YEB2 B5YEC2 B5YEC9 B5YEF3 B5YEF4 B5YEG8 B5YEG9 B5YEH1 B5YEH6 B5YEL2
B5YEM1 B5YEM2 B5YEP5 B5YEQ0 B5YEQ1 B5YEQ2 B5YEQ3 B5YEQ8 B5YER1 B5YER5 B5YER8
B5YES4 B5YES5 B5YET6 B5YEU3 B5YEU4 B5YEW7 B5YEW9 B5YEX7 B5YEX8 B5YEX9 B5YEY0
B5YEY8 B5YF00 B5YF11 B5YF42 B5YF45 B5YF46 B5YF48 B5YF50 B5YF65 B5YF73 B5YF77
B5YF84 B5YF85 B5YF86 B5YFA9 B5YFB8 B5YFC6 B5YFD1 B5YFD4 B5YFD5 B5YFD8 B5YFE5
B5YFF7 B5YFG2 B5YFH2 B5YFK5 B5YFL3 B5YFL4 B5YFN3 B5YFT1 B5YFU3 B5YFU5 B5YFV7
B5YFX9 B5YFY8 B5YFZ0 B5YFZ8 B5YFZ9 B5YG00 B5YG20 B5YG21 B5YG23 B5YG25 B5YG29
B5YG32 B5YG33 B5YG34 B5YG37 B5YG38 B5YG39 B5YG40 B5YG41 B5YG43 B5YG44 B5YG45
B5YG46 B5YG51 B5YG52 B5YG54 B5YG55 B5YG57 B5YG78 B5YG91 B5YGA0 B5YGC5 B5YGD9
B5YGE0 B5YGE4 B5YGR4 B5YGU2 B5YGX0 B5YH59 B5YH61 B5YH68 B5YH69 B5YH80 B5YHE5
B5YHH6 B5YHI3 B5YHI4 B5YHK1 B5YHN1 B5YHN2 B5YHP1 B5YHP2 B5YHQ1 B5YHQ2 B5YHS2
B5YHS7 B5YHS9 B5YHT8 B5YHT9 B5YHV2 B5YHX4 B5YHX5 B5YI09 B5YI18 B5YI28 B5YI37
B5YI38 B5YI39 B5YIA9 B5YIB4 B5YIB7 B5YID4 B5YIF8 B5YIF9 B5YIG1 B5YIH1 B5YIK6
B5YIL6 B5YIL7 B5YIM0 B5YIM4 B5YIM9 B5YIN3 B5YIQ4 B5YIQ5 B5YIQ7 B5YIQ8 B5YIR0
B5YIU7 B5YIU9 B5YIZ6 B5YJ09 B5YJ12 B5YJ13 B5YJ32 B5YJ35 B5YJ37 B5YJ55 B5YJ65
B5YJ66 B5YJ67 B5YJ68 B5YJ73 B5YJ75 B5YJ89 B5YJ91 B5YJA6 B5YJD3 B5YJE1 B5YJG7
B5YJG9 B5YJH1 B5YJH3 B5YJJ2 B5YJL3 B5YJN3 B5YJN4 B5YJP5 B5YJS1 B5YJS2 B5YJS4
B5YJT0 B5YJT7 B5YJT8 B5YJV0 B5YJV1 B5YJW3 B5YJW4 B5YJX5 B5YJY3 B5YJZ3 B5YK66
B5YK79 B5YK82 B5YK85 B5YK86 B5YK93 B5YK94 B5YK96 B5YK97 B5YKD1 B5YKD5 B5YKE0
B5YKE1 B5YKE5 B5YKE9 B5YKF0 B5YKG0 B5YKK4 B5YKK9 B5YKL4 B5YKN8 B5YKP5 B5YKS2
B5YKU0 B5YKW0 B5YKW2 B5YKX4 B5YKY0 B5YL10 B5YL53 B5YL59 B5YL60 B5YL83 B5YLB7
B5YLD0 B5YLE5 B5YM03 B5YPM0 B5YPM2 B5YPP1 B5YPP4 B5YPS4 B5YPU3 B5YPW2 B5YPW5
B5YPX9 B5YPY0 B5YPZ1 B5YPZ4 B5YQ00 B5YQ02 B5YQ04 B5YQ05 B5YQ07 B5YQ18 B5YQ20
B5YQ27 B5YQ29 B5YQ31 B5YQ32 B5YQ33 B5YQ34 B5YQ35 B5YQ71 B5YQ77 B5YQ81 B5YQ91
B5YQ93 B5YQ95 B5YQ96 B5YQ97 B5YQA3 B5YQA8 B5YQA9 B5YQB9 B5YQD4 B5YQD6 B5YQD9
B5YQE6 B5YQE7 B5YQF1 B5YQG1 B5YQH9 B5YQI1 B5YQI2 B5YQI8 B5YQJ1 B5YQJ4 B5YQK2
B5YQL0 B5YQL3 B5YQM0 B5YQM4 B5YQQ2 B5YQR7 B5YQV2 B5YQV4 B5YQX5 B5YQX9 B5YR05
B5YR06 B5YR08 B5YR11 B5YR15 B5YR16 B5YR19 B5YR22 B5YR82 B5YR87 B5YR89 B5YR91
B5YR96 B5YRA4 B5YRA5 B5YRA8 B5YRB3 B5YRC4 B5YRF2 B5YRF5 B5YRG5 B5YRL4 B5YRL5
B5YRL8 B5YRM0 B5YRM2 B5YRN8 B5YRT7 B5YRU5 B5YRW4 B5YRY4 B5YS00 B5YS14 B5YS22
B5YS30 B5YS38 B5YS39 B5YS44 B5YS45 B5YS54 B5YS55 B5YS57 B5YS58 B5YS60 B5YS62
B5YS65 B5YS68 B5YS72 B5YS74 B5YS75 B5YS78 B5YS96 B5YS99 B5YSA4 B5YSC6 B5YSE3
B5YSF1 B5YSG2 B5YSG9 B5YSH7 B5YSI2 B5YSI4 B5YSJ3 B5YSQ7 B5YSS0 B5YST8 B5YSU9
B5YSV0 B5YSV2 B5YSV6 B5YSV7 B5YSW2 B5YSW3 B5YSY4 B5YSY6 B5YT01 B5YT04 B5YT06
B5YT07 B5YT08 B5YT13 B5YT15 B5YT16 B5YT20 B5YT28 B5YT41 B5YT42 B5YT44 B5YT46
B5YT51 B5YT56 B5YT58 B5YT65 B5YT69 B5YT79 B5YT86 B5YT88 B5YT93 B5YT97 B5YT98
B5YTE1 B5YTM2 B5YTM3 B5YTM5 B5YTM6 B5YTM7 B5YTM8 B5YTM9 B5YTN0 B5YTN1 B5YTN2
B5YTN3 B5YTN4 B5YTN5 B5YTN6 B5YTN7 B5YTN8 B5YTP0 B5YTP1 B5YTP2 B5YTP7 B5YTP8

B5YTP9 B5YTQ0 B5YTQ1 B5YTQ2 B5YTQ5 B5YTR1 B5YTS6 B5YTT8 B5YTV3 B5YTW1 B5YTW3
B5YTW5 B5YTW9 B5YTX2 B5YU41 B5YU42 B5YU43 B5YU47 B5YU49 B5YU50 B5YU51 B5YU52
B5YU53 B5YU75 B5YU77 B5YU80 B5YU81 B5YUC5 B5YUH7 B5YUI5 B5YUI6 B5YUI7 B5YUM9
B5YUQ6 B5YUS8 B5YUX1 B5YUX2 B5YUX3 B5YUX4 B5YUX5 B5YUX6 B5YV42 B5YV43 B5YV51
B5YV64 B5YVG5 B5YVL1 B5YVL2 B5YVL8 B5YVQ4 B5YVS5 B5YVS9 B5YVT2 B5YVT7 B5YVV7
B5YVW6 B5YVX5 B5YVX6 B5YVX7 B5YW75 B5YW91 B5YW99 B5YWB1 B5YWB7 B5YWC0 B5YWD2
B5YWD3 B5YWD4 B5YWD5 B5YWD6 B5YWD8 B5YWD9 B5YWE0 B5YWE1 B5YWE6 B5YWE9 B5YWJ2
B5YWU3 B5YWV0 B5YWV3 B5YWW4 B5YWX8 B5YWX9 B5YWY1 B5YX02 B5YX17 B5YX72 B5YX92
B5YX93 B5YXA2 B5YXA4 B5YXA7 B5YXA8 B5YXB0 B5YXB2 B5YXB5 B5YXE4 B5YXE5 B5YXJ7
B5YXJ9 B5YXK3 B5YXM2 B5YXM6 B5YXM8 B5YXM9 B5YXQ5 B5YXQ7 B5YXQ8 B5YXT1 B5YXT4
B5YXV8 B5YXW1 B5YXX0 B5YXX2 B5YY02 B5YY03 B5YY04 B5YY06 B5YY14 B5YY20 B5YY23
B5YY27 B5YY28 B5YY53 B5YY56 B5YY58 B5YY78 B5YY82 B5YY94 B5YY97 B5YYA0 B5YYA8
B5YYB6 B5YYB8 B5YYC1 B5YYC2 B5YYC3 B5YYC9 B5YYD1 B5YYD2 B5YYD3 B5YYD6 B5YYF3
B5YYW7 B5YYX4 B5YZ00 B5YZ24 B5YZ28 B5YZ31 B5YZ38 B5YZ39 B5YZ40 B5YZ41 B5YZ43
B5YZ53 B5YZ57 B5YZ64 B5YZ66 B5YZ69 B5YZ71 B5YZ72 B5YZ77 B5YZ87 B5YZ88 B5YZ89
B5YZ90 B5YZ98 B5YZ99 B5YZA7 B5YZA8 B5YZB0 B5YZB7 B5YZB8 B5YZC7 B5YZD6 B5YZD7
B5YZD9 B5YZF4 B5YZF5 B5YZF6 B5YZG7 B5YZH0 B5YZH1 B5YZI3 B5YZP0 B5YZP1 B5YZQ8
B5YZR7 B5YZR8 B5YZU2 B5YZV2 B5YZV8 B5YZW5 B5YZX2 B5YZX3 B5YZY1 B5YZY5 B5Z008
B5Z013 B5Z015 B5Z033 B5Z035 B5Z036 B5Z040 B5Z058 B5Z059 B5Z061 B5Z062 B5Z064
B5Z069 B5Z079 B5Z080 B5Z081 B5Z082 B5Z083 B5Z089 B5Z090 B5Z091 B5Z092 B5Z093
B5Z094 B5Z0A3 B5Z0A5 B5Z0A8 B5Z0B3 B5Z0C4 B5Z0D4 B5Z0D6 B5Z0D9 B5Z0E0 B5Z0E3
B5Z0E4 B5Z0E5 B5Z0E7 B5Z0E9 B5Z0F1 B5Z0G2 B5Z0G9 B5Z0H4 B5Z0I3 B5Z0I6 B5Z0I8
B5Z0M1 B5Z0R4 B5Z0W0 B5Z0X6 B5Z0X8 B5Z0Y0 B5Z0Y3 B5Z0Y4 B5Z0Y6 B5Z0Y7 B5Z0Z6
B5Z0Z9 B5Z100 B5Z104 B5Z105 B5Z112 B5Z114 B5Z115 B5Z123 B5Z139 B5Z141 B5Z149
B5Z153 B5Z154 B5Z180 B5Z1I3 B5Z1I4 B5Z1I6 B5Z1I7 B5Z1N3 B5Z1R0 B5Z1R1 B5Z1R2
B5Z1S5 B5Z1T9 B5Z1X6 B5Z1X8 B5Z225 B5Z226 B5Z227 B5Z228 B5Z231 B5Z232 B5Z235
B5Z237 B5Z271 B5Z288 B5Z2A3 B5Z2A6 B5Z2A9 B5Z2B0 B5Z2E7 B5Z2F1 B5Z2F2 B5Z2F6
B5Z2G6 B5Z2H0 B5Z2H6 B5Z2H7 B5Z2H8 B5Z2I3 B5Z2I4 B5Z2J7 B5Z2J8 B5Z2K2 B5Z2K3
B5Z2K4 B5Z2K6 B5Z2K7 B5Z2K8 B5Z2P3 B5Z2P7 B5Z2Q2 B5Z2Q3 B5Z2Q5 B5Z2Q6 B5Z2Q7
B5Z2T4 B5Z2T9 B5Z2U0 B5Z2U3 B5Z2U5 B5Z2W1 B5Z2W2 B5Z372 B5Z397 B5Z3A5 B5Z3A6
B5Z3A7 B5Z3A8 B5Z3A9 B5Z3B2 B5Z3B3 B5Z3B4 B5Z3C5 B5Z3C6 B5Z3E4 B5Z3F9 B5Z3G5
B5Z3H5 B5Z3I7 B5Z3I9 B5Z3K2 B5Z3K3 B5Z3L7 B5Z3R0 B5Z3R6 B5Z3R7 B5Z3R9 B5Z3S0
B5Z3S5 B5Z3S7 B5Z3S9 B5Z3T2 B5Z3U4 B5Z3U5 B5Z3V1 B5Z3V4 B5Z3X9 B5Z3Y1 B5Z3Y2
B5Z3Y6 B5Z3Y7 B5Z457 B5Z471 B5Z472 B5Z474 B5Z476 B5Z492 B5Z4B2 B5Z4B3 B5Z4C2
B5Z4C4 B5Z4C6 B5Z4E7 B5Z4E9 B5Z4G1 B5Z4N0 B5Z4P5 B5Z4Q2 B5Z4Q6 B5Z4R3 B5Z4R4
B5Z4R5 B5Z4R6 B5Z4S5 B5Z4S7 B5Z670 B5Z671 B5Z672 B5Z674 B5Z677 B5Z679 B5Z680
B5Z683 B5Z684 B5Z6A3 B5Z6D1 B5Z6D2 B5Z6D7 B5Z6D8 B5Z6E3 B5Z6E6 B5Z6E7 B5Z6E8
B5Z6E9 B5Z6F4 B5Z6F6 B5Z6H4 B5Z6I0 B5Z6J9 B5Z6L6 B5Z6L9 B5Z6N2 B5Z6N7 B5Z6N8
B5Z6N9 B5Z6S2 B5Z6S4 B5Z6T3 B5Z6T7 B5Z6U1 B5Z6U3 B5Z6V2 B5Z6X6 B5Z6Y6 B5Z6Y8
B5Z6Y9 B5Z6Z7 B5Z711 B5Z716 B5Z721 B5Z735 B5Z736 B5Z737 B5Z772 B5Z796 B5Z7D2
B5Z7F5 B5Z7F6 B5Z7F7 B5Z7G0 B5Z7I6 B5Z7I8 B5Z7J5 B5Z7J7 B5Z7J9 B5Z7K9 B5Z7L6
B5Z7L8 B5Z7M6 B5Z7N7 B5Z7P2 B5Z7P5 B5Z7S9 B5Z7T0 B5Z7T2 B5Z7T4 B5Z7U5 B5Z7V2
B5Z7W3 B5Z7X4 B5Z809 B5Z842 B5Z848 B5Z850 B5Z863 B5Z875 B5Z879 B5Z887 B5Z889
B5Z8E3 B5Z8E7 B5Z8F0 B5Z8F5 B5Z8G1 B5Z8H2 B5Z8H3 B5Z8I6 B5Z8J0 B5Z8J2 B5Z8J6

B5Z8J7 B5Z8J8 B5Z8J9 B5Z8K3 B5Z8L0 B5Z8M3 B5Z8N3 B5Z8N5 B5Z8P0 B5Z8P2 B5Z8P5
B5Z8Q1 B5Z8Q2 B5Z8S2 B5Z8S5 B5Z8U1 B5Z8U3 B5Z8U4 B5Z8U5 B5Z8U9 B5Z8V1 B5Z8V2
B5Z8V3 B5Z8V4 B5Z8V5 B5Z8V7 B5Z8V8 B5Z8W0 B5Z8W1 B5Z8W2 B5Z8W3 B5Z8W4 B5Z8W5
B5Z8W6 B5Z8W8 B5Z8X1 B5Z8Y2 B5Z8Y5 B5Z8Z4 B5Z928 B5Z935 B5Z937 B5Z943 B5Z947
B5Z950 B5Z961 B5Z963 B5Z993 B5Z994 B5Z999 B5Z9A1 B5Z9B5 B5Z9B6 B5Z9E3 B5Z9F8
B5Z9G5 B5Z9I6 B5Z9I8 B5Z9J7 B5Z9L2 B5Z9L3 B5Z9L4 B5Z9P0 B5Z9P1 B5Z9Q1 B5Z9Q3
B5Z9Q4 B5Z9S8 B5Z9T2 B5Z9T6 B5Z9U4 B5Z9V0 B5Z9V6 B5Z9V7 B5Z9W5 B5Z9W8 B5Z9W9
B5Z9Y1 B5Z9Y2 B5Z9Y5 B5ZA04 B5ZA05 B5ZA06 B5ZA21 B5ZA25 B5ZA35 B5ZA47 B5ZA49
B5ZA60 B5ZA62 B5ZA63 B5ZA69 B5ZA72 B5ZA76 B5ZA86 B5ZA97 B5ZAA0 B5ZAA2 B5ZAB7
B5ZAD2 B5ZAF2 B5ZAF5 B5ZAF6 B5ZAG0 B5ZAH6 B5ZAI3 B5ZAI4 B5ZAJ9 B5ZAL6 B5ZAP2
B5ZAQ7 B5ZAR3 B5ZAS3 B5ZAS4 B5ZAT6 B5ZAU8 B5ZAU9 B5ZAV3 B5ZAY6 B5ZAY8 B5ZAZ3
B5ZAZ4 B5ZB10 B5ZB16 B5ZB18 B5ZB20 B5ZB25 B5ZB37 B5ZB38 B5ZB39 B5ZB40 B5ZB41
B5ZB42 B5ZB43 B5ZB44 B5ZB46 B5ZB47 B5ZB48 B5ZB49 B5ZB50 B5ZB51 B5ZB52 B5ZB53
B5ZB54 B5ZB56 B5ZB57 B5ZB58 B5ZB63 B5ZB64 B5ZB65 B5ZB67 B5ZB76 B5ZB79 B5ZB96
B5ZB99 B5ZBA5 B5ZBA7 B5ZBB2 B5ZBB4 B5ZBB8 B5ZBC9 B5ZBD0 B5ZBE2 B5ZBE3 B5ZBE9
B5ZBF2 B5ZBF7 B5ZBG4 B5ZBH5 B5ZBH7 B5ZBI2 B5ZBI3 B5ZBM7 B5ZBM9 B5ZBN1 B5ZBN2
B5ZBP8 B5ZBQ8 B5ZBR5 B5ZBR8 B5ZBS5 B5ZBS7 B5ZBS8 B5ZBS9 B5ZBT0 B5ZBT1 B5ZBT4
B5ZBT5 B5ZBT6 B5ZBT9 B5ZBV3 B5ZBV6 B5ZBV9 B5ZBZ7 B5ZC00 B5ZC02 B5ZC20 B5ZC22
B5ZC31 B5ZC32 B5ZC33 B5ZC34 B5ZC42 B5ZC44 B5ZC48 B5ZC49 B5ZC51 B5ZC55 B5ZC58
B5ZC59 B5ZC61 B5ZC77 B5ZC78 B5ZC79 B5ZC90 B5ZC91 B5ZCA0 B5ZCA9 B5ZCB4 B5ZCB5
B5ZCB6 B5ZCB8 B5ZCB9 B5ZMF1 B5ZMF2 B5ZMG0 B5ZMG1 B5ZMH5 B5ZMI0 B5ZMI7 B5ZMI8
B5ZMI9 B5ZMJ0 B5ZMJ6 B5ZMK3 B5ZMN5 B5ZMN6 B5ZMN7 B5ZMP0 B5ZMP3 B5ZMP5 B5ZMP6
B5ZMS8 B5ZMW8 B5ZMW9 B5ZMX0 B5ZMX9 B5ZMY1 B5ZN05 B5ZN20 B5ZN21 B5ZN23 B5ZN24
B5ZN33 B5ZN41 B5ZN44 B5ZN83 B5ZN84 B5ZN86 B5ZNB1 B5ZNB6 B5ZNC1 B5ZND2 B5ZNK8
B5ZNL4 B5ZNS1 B5ZNS6 B5ZNS9 B5ZNW9 B5ZP73 B5ZP76 B5ZP79 B5ZP80 B5ZPA0 B5ZPA7
B5ZPD5 B5ZPG3 B5ZPM0 B5ZPM3 B5ZPM7 B5ZPR5 B5ZPR8 B5ZPY6 B5ZPY7 B5ZPZ7 B5ZQ46
B5ZQ47 B5ZQC1 B5ZQD7 B5ZQG0 B5ZQK2 B5ZQK7 B5ZQP7 B5ZQP8 B5ZQT5 B5ZQU1 B5ZQU6
B5ZQV6 B5ZQW4 B5ZQW5 B5ZQY2 B5ZR38 B5ZR94 B5ZRD0 B5ZRD1 B5ZRD6 B5ZRD7 B5ZRF0
B5ZRF1 B5ZRG2 B5ZRL6 B5ZRM8 B5ZRN9 B5ZRR4 B5ZRR7 B5ZRX6 B5ZS00 B5ZS20 B5ZS25
B5ZS33 B5ZS37 B5ZS68 B5ZS73 B5ZS84 B5ZSC4 B5ZSI4 B5ZSI5 B5ZSR9 B5ZSS0 B5ZST4
B5ZSU2 B5ZSV9 B5ZSW0 B5ZSW5 B5ZT17 B5ZTC5 B5ZTD4 B5ZTF1 B5ZTF5 B5ZTH4 B5ZTH8
B5ZTH9 B5ZTI5 B5ZTJ9 B5ZU68 B5ZU74 B5ZU94 B5ZUA0 B5ZUD8 B5ZUD9 B5ZUE3 B5ZUE4
B5ZUE5 B5ZUE6 B5ZUF2 B5ZUF6 B5ZUG2 B5ZUG3 B5ZUL2 B5ZUP2 B5ZV14 B5ZV31 B5ZV36
B5ZV40 B5ZV45 B5ZV51 B5ZV56 B5ZV69 B5ZV70 B5ZV71 B5ZV80 B5ZV86 B5ZV89 B5ZV90
B5ZV92 B5ZV97 B5ZV98 B5ZV99 B5ZVA2 B5ZVD0 B5ZVQ5 B5ZVU9 B5ZW24 B5ZW25 B5ZW28
B5ZW32 B5ZW36 B5ZW65 B5ZW67 B5ZW73 B5ZW78 B5ZWB3 B5ZWC6 B5ZWC8 B5ZWC9 B5ZWD8
B5ZWH3 B5ZWH9 B5ZWJ2 B5ZWK2 B5ZWK3 B5ZWP8 B5ZWQ1 B5ZWQ2 B5ZWT2 B5ZWT5 B5ZWU4
B5ZWY7 B5ZWZ6 B5ZX12 B5ZX21 B5ZX28 B5ZXA0 B5ZXE7 B5ZXG2 B5ZXI2 B5ZXI3 B5ZXI4
B5ZXR7 B5ZXR8 B5ZXR9 B5ZXS0 B5ZXT9 B5ZXV2 B5ZXV9 B5ZXZ1 B5ZXZ5 B5ZXZ6 B5ZY09
B5ZY55 B5ZY62 B5ZY73 B5ZY95 B5ZYA2 B5ZYC6 B5ZYE9 B5ZYG6 B5ZYH2 B5ZYI7 B5ZYK8
B5ZYK9 B5ZYL1 B5ZYN2 B5ZYQ2 B5ZYS3 B5ZYS4 B5ZYS5 B5ZYS6 B5ZYS7 B5ZYT0 B5ZYT1
B5ZYT2 B5ZYT4 B5ZYT5 B5ZYT6 B5ZYT7 B5ZYT8 B5ZYT9 B5ZYU1 B5ZYU2 B5ZYU3 B5ZYU4
B5ZYU5 B5ZYU6 B5ZYU7 B5ZYU8 B5ZYU9 B5ZYV0 B5ZYV1 B5ZYV2 B5ZYV3 B5ZYV4 B5ZYX3
B5ZZ10 B5ZZ34 B5ZZ55 B5ZZ56 B5ZZ57 B5ZZ58 B5ZZ62 B5ZZA8 B5ZZC8 B6A3G5 B6AFP1

B6CDL6 B6CHA3 B6CJX2 B6CZ40 B6CZ45 B6CZ54 B6CZ61 B6D1N4 B6D5I7 B6D5P1 B6D5P3
B6D5P6 B6D9A7 B6D9A8 B6DDU1 B6DMK2 B6E107 B6E2X2 B6ECN9 B6EGB3 B6EGC0 B6EGC1
B6EGC2 B6EGC9 B6EGG2 B6EGG3 B6EGG7 B6EGH0 B6EGH8 B6EGI3 B6EGI9 B6EGJ0 B6EGJ1
B6EGJ2 B6EGJ3 B6EGJ4 B6EGN8 B6EGP6 B6EGP7 B6EGQ6 B6EGR4 B6EGT2 B6EGT3 B6EGT6
B6EGV1 B6EGV5 B6EGX4 B6EGX8 B6EGX9 B6EGY3 B6EGY4 B6EGY7 B6EGZ1 B6EGZ4 B6EGZ6
B6EGZ9 B6EH18 B6EH43 B6EH68 B6EH86 B6EH87 B6EHH0 B6EHH2 B6EHH4 B6EHH5 B6EHJ9
B6EHK0 B6EHK1 B6EHK2 B6EHK3 B6EHL7 B6EHU8 B6EHV6 B6EHW3 B6EHX0 B6EHZ5 B6EHZ6
B6EHZ8 B6EI32 B6EI57 B6EI79 B6EI86 B6EI95 B6EI99 B6EIA0 B6EIA7 B6EIA9 B6EIB2
B6EIC1 B6EIC3 B6EIF1 B6EIF3 B6EII1 B6EIJ8 B6EIJ9 B6EIL4 B6EIM4 B6EIM5 B6EIM8
B6EIN2 B6EIN6 B6EIN8 B6EIU9 B6EIV0 B6EIV6 B6EIV8 B6EIV9 B6EIW3 B6EIW6 B6EIX3
B6EIX7 B6EIX8 B6EIX9 B6EIY1 B6EIY4 B6EIY5 B6EIY7 B6EIZ6 B6EJ04 B6EJ41 B6EJ43
B6EJ44 B6EJ48 B6EJ59 B6EJ87 B6EJ89 B6EJ92 B6EJ93 B6EJA2 B6EJA3 B6EJQ3 B6EJQ6
B6EJS8 B6EJT2 B6EJU2 B6EJU3 B6EJV2 B6EJV4 B6EJW6 B6EJZ8 B6EK11 B6EK51 B6EK54
B6EK56 B6EK57 B6EK65 B6EK66 B6EK72 B6EKA0 B6EKA3 B6EKA7 B6EKA9 B6EKH2 B6EKK3
B6EKL2 B6EKL4 B6EKL5 B6EKL6 B6EKL9 B6EKM3 B6EKM7 B6EKN9 B6EKP3 B6EKP6 B6EKQ4
B6EKR0 B6EKU7 B6EKX9 B6EKY3 B6EKY5 B6EKY7 B6EKZ7 B6EL03 B6EL04 B6EL06 B6EL23
B6EL39 B6EL41 B6EL42 B6EL43 B6EL47 B6EL48 B6ELB8 B6ELC0 B6ELC5 B6ELD2 B6ELD3
B6ELD6 B6ELE1 B6ELE2 B6ELG5 B6ELG6 B6ELH4 B6ELH7 B6ELI3 B6ELI6 B6ELJ4 B6ELJ5
B6ELJ7 B6ELJ8 B6ELK0 B6ELK6 B6ELK8 B6ELM5 B6ELM6 B6ELP2 B6ELP3 B6ELS5 B6ELU0
B6ELZ7 B6EM10 B6EM15 B6EM43 B6EM44 B6EM99 B6EMB5 B6EMC2 B6EMJ7 B6EMK8 B6EML5
B6EML9 B6EMM4 B6EMN4 B6EMN5 B6EMN8 B6EMP3 B6EMP4 B6EMP6 B6EMP9 B6EMQ7 B6EMQ8
B6EMQ9 B6EMS4 B6EMS5 B6EMS7 B6EMT2 B6EMT9 B6EMU1 B6EMU2 B6EMV2 B6EMV3 B6EMV8
B6EMW1 B6EMW2 B6EMW5 B6EMX2 B6EMX5 B6EMY3 B6EMY6 B6EN03 B6EN06 B6EN08 B6EN31
B6EN32 B6EN33 B6EN68 B6EN78 B6EN80 B6ENA0 B6ENA3 B6ENA5 B6ENA8 B6ENB7 B6ENC7
B6ENC8 B6END5 B6END8 B6ENE0 B6ENE2 B6ENE3 B6ENE4 B6ENE5 B6ENE6 B6ENF7 B6ENG3
B6ENG4 B6ENG5 B6ENH6 B6ENH8 B6ENI1 B6ENI2 B6ENK8 B6ENN5 B6ENR3 B6ENR4 B6ENR5
B6ENR6 B6ENR7 B6ENT1 B6ENT4 B6ENU6 B6ENV8 B6ENX1 B6EP21 B6EP24 B6EP29 B6EP33
B6EP41 B6EP42 B6EP46 B6EP48 B6EP83 B6EP85 B6EPA4 B6EPI8 B6EPI9 B6EPJ3 B6EPK1
B6EPL9 B6EPM6 B6EPN7 B6EPP1 B6EPP2 B6EPP3 B6EPQ0 B6EPQ8 B6EPR5 B6EPR6 B6EPS4
B6EPS5 B6EPS6 B6EPS7 B6EPS8 B6EPS9 B6EPT0 B6EPT1 B6EPT2 B6EPT3 B6EPT4 B6EPT5
B6EPT6 B6EPT7 B6EPT8 B6EPT9 B6EPU0 B6EPU1 B6EPU2 B6EPU3 B6EPU4 B6EPU7 B6EPU8
B6EPU9 B6EPV0 B6EPV3 B6EPV7 B6EQ13 B6EQ59 B6EQC3 B6EQJ5 B6EQL7 B6ER09 B6ER23
B6ER81 B6ER91 B6ERB4 B6ERC3 B6ERE2 B6ERM4 B6ERM6 B6ERM8 B6ERS6 B6ERX5 B6ERX7
B6ERZ8 B6ES34 B6ES35 B6ES52 B6ES82 B6ESA4 B6ESC4 B6ESC6 B6ESC7 B6ESF2 B6EU02
B6EUA9 B6F0W8 B6F107 B6F1W5 B6F209 B6GVZ2 B6GWE7 B6GZA1 B6GZD3 B6H060 B6H061
B6H062 B6H063 B6H068 B6H069 B6H2G9 B6H2I7 B6H2M0 B6H3T4 B6H460 B6H464 B6H4C6
B6H4I3 B6H5L5 B6H5R4 B6H6F3 B6H6L7 B6H768 B6H7A3 B6H7F3 B6H9U8 B6H9W0 B6HAA7
B6HAN0 B6HBG5 B6HCY4 B6HDT2 B6HE56 B6HEM2 B6HFE0 B6HFQ1 B6HFX8 B6HGB5 B6HHJ8
B6HI95 B6HIM5 B6HJU1 B6HJU2 B6HJU4 B6HJU6 B6HK90 B6HLP3 B6HLP6 B6HLP7 B6HLP8
B6HLP9 B6HN75 B6HN77 B6HNK3 B6HNK4 B6HNY8 B6HP17 B6HP56 B6HQC9 B6HQD4 B6HQN4
B6HRA4 B6HRL5 B6HSQ3 B6HTQ4 B6HU70 B6HUC4 B6HUK8 B6HUQ5 B6HV32 B6HV33 B6HV34
B6HV38 B6HV39 B6HVR6 B6HYM8 B6HYN0 B6HYN6 B6HYN9 B6HYT6 B6HYU5 B6HYV5 B6HYX7
B6HYX8 B6HYX9 B6HYY8 B6HYY9 B6HYZ0 B6HYZ2 B6HYZ3 B6HYZ8 B6HZ00 B6HZ03 B6HZ11
B6HZ18 B6HZ21 B6HZ24 B6HZ30 B6HZ31 B6HZ32 B6HZ33 B6HZ39 B6HZ42 B6HZ43 B6HZ51

B6HZ52 B6HZ58 B6HZ59 B6HZ68 B6HZ78 B6HZ79 B6HZ81 B6HZ94 B6HZ96 B6HZA6 B6HZA8
B6HZA9 B6HZC1 B6HZD0 B6HZD2 B6HZD5 B6HZD6 B6HZD9 B6HZE0 B6HZE1 B6HZE3 B6HZE4
B6HZE6 B6HZF7 B6HZG4 B6HZI2 B6HZI7 B6HZI8 B6HZJ1 B6HZJ3 B6HZK5 B6HZK6 B6HZL4
B6HZL6 B6HZL7 B6HZM1 B6HZM5 B6HZM6 B6HZM9 B6HZP3 B6HZP5 B6HZQ1 B6HZQ3 B6HZR3
B6HZS5 B6HZS7 B6HZW0 B6HZW6 B6HZX3 B6HZX4 B6HZX5 B6HZX7 B6HZX8 B6I019 B6I020
B6I022 B6I023 B6I030 B6I074 B6I075 B6I082 B6I084 B6I093 B6I0B9 B6I0C1 B6I0C3
B6I0C4 B6I0C6 B6I0C7 B6I0F4 B6I0F6 B6I0G2 B6I0H1 B6I0H4 B6I0N0 B6I0S2 B6I0S3
B6I0S6 B6I0S7 B6I0U4 B6I0W0 B6I0Y4 B6I0Z9 B6I101 B6I112 B6I137 B6I139 B6I140
B6I146 B6I150 B6I153 B6I163 B6I184 B6I185 B6I186 B6I1A8 B6I1E8 B6I1E9 B6I1F2
B6I1J9 B6I1K5 B6I1L2 B6I1M3 B6I1M4 B6I1M9 B6I1N0 B6I1N9 B6I1P0 B6I1P2 B6I1P3
B6I1P5 B6I1P6 B6I1P9 B6I1Q2 B6I1Q6 B6I1Q8 B6I1Q9 B6I1R2 B6I1S8 B6I1U0 B6I1U1
B6I1U3 B6I1U4 B6I1U8 B6I1U9 B6I1V4 B6I1V5 B6I1X9 B6I1Y1 B6I1Z5 B6I1Z8 B6I200
B6I201 B6I202 B6I208 B6I210 B6I211 B6I215 B6I216 B6I218 B6I219 B6I220 B6I221
B6I222 B6I223 B6I224 B6I225 B6I226 B6I227 B6I228 B6I229 B6I230 B6I231 B6I233
B6I234 B6I235 B6I240 B6I241 B6I242 B6I243 B6I244 B6I245 B6I248 B6I254 B6I264
B6I268 B6I274 B6I275 B6I276 B6I281 B6I282 B6I296 B6I297 B6I2A1 B6I2A2 B6I2A3
B6I2A6 B6I2A7 B6I2A8 B6I2A9 B6I2C2 B6I2D2 B6I2E6 B6I2F4 B6I2F8 B6I2F9 B6I2H3
B6I2K2 B6I2L3 B6I2N1 B6I2P5 B6I2Q2 B6I2Q6 B6I2R3 B6I2S7 B6I2U7 B6I2W3 B6I2X3
B6I2X6 B6I2X8 B6I2Y2 B6I2Y8 B6I2Z5 B6I2Z6 B6I2Z7 B6I335 B6I346 B6I359 B6I385
B6I3C9 B6I3D0 B6I3D6 B6I3E8 B6I3G5 B6I3H6 B6I3I8 B6I3J6 B6I3J9 B6I3L1 B6I3L2
B6I3L3 B6I3L4 B6I3L7 B6I3L8 B6I3L9 B6I3M0 B6I3M8 B6I3P6 B6I3R9 B6I3S0 B6I3S5
B6I3T3 B6I3T5 B6I3T6 B6I3T7 B6I3T9 B6I3U1 B6I3U4 B6I3X7 B6I3X8 B6I3Y1 B6I3Y2
B6I3Y5 B6I408 B6I413 B6I419 B6I423 B6I427 B6I436 B6I437 B6I441 B6I445 B6I464
B6I485 B6I4A0 B6I4A8 B6I4B1 B6I4B5 B6I4B6 B6I4E0 B6I4E2 B6I4F2 B6I4H1 B6I4H5
B6I4I7 B6I4K3 B6I4M4 B6I4N1 B6I4N6 B6I4P3 B6I4P4 B6I4P5 B6I4P6 B6I4P8 B6I4Q8
B6I4R2 B6I4S1 B6I4S2 B6I4S4 B6I4S5 B6I4S9 B6I4T6 B6I4U6 B6I4U8 B6I4X6 B6I4X8
B6I4Y8 B6I4Z5 B6I503 B6I504 B6I512 B6I516 B6I539 B6I546 B6I548 B6I568 B6I570
B6I571 B6I579 B6I581 B6I583 B6I586 B6I587 B6I589 B6I590 B6I595 B6I598 B6I599
B6I5A0 B6I5A2 B6I5A3 B6I5B2 B6I5B4 B6I5B5 B6I5C4 B6I5D9 B6I5E1 B6I5E9 B6I5F4
B6I5F5 B6I5H2 B6I5H3 B6I5H5 B6I5H6 B6I5I0 B6I5I3 B6I5I5 B6I5J3 B6I5J4 B6I5J5
B6I5J6 B6I5J7 B6I5K1 B6I5K4 B6I5K5 B6I5K6 B6I5K7 B6I5K8 B6I5K9 B6I5L8 B6I5M1
B6I5M4 B6I5M9 B6I5N7 B6I5Q4 B6I609 B6I614 B6I615 B6I628 B6I629 B6I630 B6I631
B6I635 B6I636 B6I639 B6I641 B6I665 B6I679 B6I682 B6I684 B6I685 B6I697 B6I6B9
B6I6D3 B6I6D4 B6I6D5 B6I6D6 B6I6D7 B6I6E0 B6I6E1 B6I6E2 B6I6F6 B6I6F7 B6I6H6
B6I6J2 B6I6J6 B6I6K0 B6I6K2 B6I6K4 B6I6M8 B6I6M9 B6I6N0 B6I6N1 B6I6N3 B6I6P1
B6I6P2 B6I6P3 B6I6R7 B6I6S5 B6I6T9 B6I6U7 B6I6V5 B6I6V6 B6I6W8 B6I707 B6I722
B6I731 B6I733 B6I736 B6I737 B6I738 B6I742 B6I746 B6I748 B6I761 B6I775 B6I780
B6I782 B6I786 B6I787 B6I789 B6I799 B6I7H5 B6I7I7 B6I7J4 B6I7K5 B6I7K7 B6I7K8
B6I7L2 B6I7L3 B6I7L6 B6I7Q4 B6I7Q9 B6I7R1 B6I7S1 B6I7S9 B6I7T0 B6I7T4 B6I7T6
B6I7T8 B6I7W9 B6I7Y4 B6I7Y5 B6I7Z9 B6I815 B6I828 B6I833 B6I846 B6I848 B6I851
B6I896 B6I8D6 B6I8F5 B6I8H3 B6I8H9 B6I8I6 B6I8J3 B6I8K3 B6I8L0 B6I8L3 B6I8M3
B6I8Q5 B6I8Q7 B6I8Q8 B6I8R1 B6I8U4 B6I8V2 B6I8V6 B6I8W4 B6I8X9 B6I8Y0 B6I8Y2
B6I8Y4 B6I8Y9 B6I8Z4 B6I8Z6 B6I904 B6I909 B6I930 B6I933 B6I938 B6I941 B6I942
B6I974 B6I975 B6I976 B6I980 B6I983 B6I984 B6I985 B6I986 B6I987 B6I988 B6I9B0

B6I9D2 B6I9D6 B6I9D9 B6I9E4 B6I9G6 B6I9H5 B6I9I5 B6I9I6 B6I9J7 B6I9K6 B6I9L6
B6I9N0 B6I9N3 B6I9N5 B6I9N9 B6I9Q0 B6I9R9 B6I9S3 B6I9S5 B6I9S6 B6I9U8 B6I9W1
B6I9X4 B6I9X5 B6I9Z2 B6I9Z7 B6IA01 B6IA03 B6IA61 B6IAI9 B6IAL3 B6IAS2 B6IAS4
B6IB44 B6IB57 B6IB76 B6IB77 B6IB79 B6IB98 B6IBB8 B6IBB9 B6IBD2 B6IBD3 B6IBD4
B6IBD5 B6IBD7 B6IBF1 B6IBF3 B6IBG0 B6IBG2 B6IBG4 B6IBG5 B6IBG6 B6IBG7 B6IBG8
B6IBJ9 B6IBK5 B6IBN8 B6IBP0 B6IBQ9 B6IBR3 B6IBT9 B6IBU0 B6IBU2 B6IBU5 B6IDH3
B6IDH8 B6IFG5 B6IFN4 B6IMI3 B6IMQ2 B6IMQ3 B6IMR7 B6IMS9 B6IMT0 B6IMY9 B6IN07
B6IN13 B6IN23 B6IN24 B6IN25 B6IN31 B6IN38 B6IN46 B6IN60 B6IN79 B6IN81 B6INF7
B6INF8 B6INH0 B6INJ3 B6INM0 B6INM1 B6INN5 B6INP2 B6INQ0 B6IP33 B6IP39 B6IP72
B6IP82 B6IP92 B6IP93 B6IP95 B6IPE0 B6IPE2 B6IPF8 B6IPG6 B6IPH3 B6IPH8 B6IPI0
B6IPI1 B6IPJ0 B6IPJ7 B6IPJ8 B6IPK0 B6IPN3 B6IPQ9 B6IPU4 B6IQ15 B6IQ27 B6IQ36
B6IQI0 B6IQZ3 B6IQZ9 B6IR00 B6IR01 B6IRB5 B6IRC6 B6IRF1 B6IRG1 B6IRH0 B6IRJ4
B6IRP2 B6IRP3 B6IRP4 B6IRP5 B6IRP6 B6IRP7 B6IRQ0 B6IRQ1 B6IRQ2 B6IRQ5 B6IRQ6
B6IRQ7 B6IRQ9 B6IRR0 B6IRR1 B6IRR2 B6IRR3 B6IRR4 B6IRR5 B6IRR6 B6IRR7 B6IRR9
B6IRS0 B6IRS1 B6IRS2 B6IRS5 B6IRS7 B6IRS8 B6IRT0 B6IRT1 B6IRV8 B6IS04 B6IS93
B6ISD0 B6ISG3 B6ISG4 B6ISG8 B6ISM1 B6ISM3 B6ISP7 B6IST4 B6ISU5 B6ISU8 B6ISV0
B6ISV1 B6ISW6 B6ISY6 B6ISY8 B6IT90 B6ITA0 B6ITG2 B6ITH3 B6ITI4 B6ITI5 B6ITK3
B6ITL1 B6ITN1 B6ITR6 B6ITS0 B6ITS4 B6ITW5 B6IU54 B6IU92 B6IU97 B6IU98 B6IUC1
B6IUD9 B6IUP4 B6IUZ1 B6IV59 B6IV67 B6IVA4 B6IVA5 B6IVB4 B6IVC0 B6IVD0 B6IVF3
B6IVG1 B6IVG3 B6IVK5 B6IVL6 B6IVM2 B6IVT5 B6IVU4 B6IVU7 B6IVY6 B6IW12 B6IW31
B6IWC6 B6IWI0 B6IWI6 B6IWI7 B6IWP6 B6IWR7 B6IWS8 B6IWZ8 B6IX34 B6IX35 B6IXD4
B6IXI3 B6IXL9 B6IYC9 B6IYD3 B6IYP5 B6IYQ0 B6IYU2 B6IYU3 B6IYU4 B6IYU5 B6IYY4
B6IYZ9 B6IZ01 B6IZ09 B6IZ19 B6IZ24 B6IZ25 B6IZ59 B6IZ61 B6IZ62 B6IZ80 B6IZ81
B6IZ82 B6IZ97 B6IZA7 B6IZA9 B6IZD5 B6IZD8 B6IZE2 B6IZE4 B6IZF7 B6IZF9 B6IZG0
B6IZG2 B6IZH7 B6IZH9 B6IZI4 B6IZJ0 B6IZJ1 B6IZL4 B6IZL5 B6IZM3 B6IZM7 B6IZN1
B6IZN3 B6IZN7 B6IZS2 B6IZT4 B6IZT9 B6IZW5 B6IZY6 B6J016 B6J017 B6J031 B6J074
B6J083 B6J089 B6J0C7 B6J0G0 B6J0H0 B6J0J1 B6J0J6 B6J0K5 B6J0K9 B6J0P2 B6J0U7
B6J0V9 B6J0W0 B6J0W1 B6J119 B6J120 B6J130 B6J134 B6J185 B6J1A4 B6J1A6 B6J1B2
B6J1C6 B6J1D4 B6J1D9 B6J1E0 B6J1E8 B6J1I8 B6J1K1 B6J1L1 B6J1M3 B6J1N3 B6J1N4
B6J1N5 B6J1P6 B6J1Q3 B6J1Q4 B6J1S2 B6J1S7 B6J1S8 B6J1S9 B6J1W2 B6J1X5 B6J1X8
B6J1Y7 B6J1Z2 B6J202 B6J213 B6J216 B6J229 B6J237 B6J238 B6J239 B6J240 B6J241
B6J244 B6J245 B6J247 B6J248 B6J251 B6J252 B6J253 B6J255 B6J256 B6J257 B6J258
B6J259 B6J260 B6J261 B6J263 B6J266 B6J267 B6J270 B6J272 B6J273 B6J274 B6J275
B6J287 B6J289 B6J2A7 B6J2B1 B6J2B9 B6J2F3 B6J2H6 B6J2H7 B6J2H8 B6J2I0 B6J2J3
B6J2J7 B6J2L3 B6J2N7 B6J2Q1 B6J2Q2 B6J2Q5 B6J2R7 B6J2R9 B6J2S1 B6J2U5 B6J2V0
B6J2V1 B6J2X4 B6J307 B6J329 B6J330 B6J386 B6J390 B6J395 B6J3A6 B6J3B1 B6J3C2
B6J3D4 B6J3G6 B6J3H2 B6J3I0 B6J3I1 B6J3J2 B6J3K5 B6J3N7 B6J3P6 B6J3Q9 B6J3R0
B6J3R8 B6J3T6 B6J3T7 B6J3V5 B6J3V7 B6J3V8 B6J3W2 B6J3W7 B6J3X6 B6J433 B6J489
B6J495 B6J4A3 B6J4A4 B6J4B0 B6J4C4 B6J4E1 B6J4G5 B6J4H5 B6J4H6 B6J4I8 B6J4J7
B6J4J9 B6J4L4 B6J4Q5 B6J4R0 B6J4R1 B6J4T4 B6J4T6 B6J4T7 B6J4T8 B6J4U6 B6J4V8
B6J4W5 B6J4W8 B6J4X6 B6J4Z9 B6J503 B6J504 B6J505 B6J507 B6J535 B6J596 B6J5A0
B6J5B8 B6J5B9 B6J5C0 B6J5C1 B6J5C8 B6J5C9 B6J5D2 B6J5D4 B6J5D5 B6J5D6 B6J5D7
B6J5D8 B6J5D9 B6J5E0 B6J5E2 B6J5E3 B6J5E4 B6J5E7 B6J5E8 B6J5F0 B6J5F1 B6J5F4
B6J5F5 B6J5F6 B6J5F7 B6J5F8 B6J5G8 B6J5I5 B6J5K1 B6J5K2 B6J5K6 B6J5L7 B6J5L9

B6J5R6 B6J5S3 B6J5S6 B6J5S7 B6J5T7 B6J5U6 B6J5V1 B6J5W3 B6J5W6 B6J5Y2 B6J625
B6J630 B6J641 B6J646 B6J655 B6J667 B6J676 B6J6A9 B6J6B1 B6J6B2 B6J6H0 B6J6H1
B6J6H8 B6J6K0 B6J6K1 B6J6M2 B6J6M4 B6J6M5 B6J6N8 B6J6S3 B6J6S6 B6J6S7 B6J6T7
B6J6U2 B6J6W3 B6J725 B6J734 B6J738 B6J775 B6J7A4 B6J7C0 B6J7F6 B6J7H5 B6J7I2
B6J7K3 B6J7K7 B6J7L9 B6J7Q0 B6J7Q3 B6J7Q5 B6J7Q9 B6J7R3 B6J7S2 B6J7S3 B6J7U6
B6J7U7 B6J7V3 B6J7V9 B6J7W1 B6J7X5 B6J7X7 B6J7X8 B6J7Y0 B6J7Z1 B6J7Z2 B6J7Z4
B6J7Z9 B6J802 B6J836 B6J853 B6J855 B6J863 B6J870 B6J878 B6J887 B6J892 B6J893
B6J8A0 B6J8E7 B6J8F9 B6J8G8 B6J8H9 B6J8I9 B6J8J0 B6J8J1 B6J8K2 B6J8M4 B6J8M6
B6J8N7 B6J8Q7 B6J8Q8 B6J8Q9 B6J8R8 B6J8S3 B6J8S5 B6J8U0 B6J8U4 B6J8V2 B6J8W3
B6J8W4 B6J8W5 B6J928 B6J929 B6J938 B6J943 B6J9B6 B6J9D8 B6J9D9 B6J9H8 B6J9H9
B6J9J5 B6J9J9 B6J9K2 B6J9K3 B6J9Z3 B6JA06 B6JA22 B6JA27 B6JAE9 B6JAI4 B6JAJ1
B6JAK6 B6JAL0 B6JAL5 B6JAL6 B6JAP3 B6JAQ4 B6JAQ7 B6JAQ9 B6JAR2 B6JAT0 B6JAT9
B6JAU5 B6JAU6 B6JB38 B6JB45 B6JB67 B6JB74 B6JB83 B6JB87 B6JBS5 B6JBZ6 B6JC05
B6JC20 B6JCC3 B6JCF0 B6JCG4 B6JCG5 B6JCG7 B6JCH9 B6JCI0 B6JCI1 B6JCI3 B6JCI7
B6JCI8 B6JCI9 B6JCK5 B6JCK6 B6JCK7 B6JCN2 B6JCN5 B6JCN9 B6JCP1 B6JCP2 B6JCP3
B6JCQ3 B6JCQ7 B6JCR8 B6JCR9 B6JCS1 B6JCS7 B6JCT6 B6JCV5 B6JCW3 B6JD11 B6JD16
B6JD19 B6JD20 B6JD21 B6JD23 B6JD24 B6JD70 B6JDD7 B6JDF7 B6JDU6 B6JDY1 B6JDZ0
B6JE08 B6JE29 B6JE33 B6JEC6 B6JER5 B6JER6 B6JER7 B6JER8 B6JES1 B6JES8 B6JES9
B6JET0 B6JET1 B6JET2 B6JET3 B6JET4 B6JET6 B6JET7 B6JET9 B6JEU0 B6JEU2 B6JEU3
B6JEV8 B6JEX2 B6JEX6 B6JEX7 B6JEX8 B6JEX9 B6JEY0 B6JEY1 B6JEY2 B6JEY4 B6JEY9
B6JF54 B6JF93 B6JFC2 B6JFC5 B6JFI3 B6JFJ0 B6JFP0 B6JFP2 B6JFP3 B6JFV4 B6JFW0
B6JFW2 B6JFW6 B6JFZ2 B6JFZ6 B6JG28 B6JG29 B6JGA4 B6JGA6 B6JGF2 B6JGF6 B6JGG5
B6JGH2 B6JGH9 B6JGI0 B6JGI3 B6JGJ7 B6JGK9 B6JGL8 B6JGM4 B6JGN7 B6JGN8 B6JGR8
B6JGT5 B6JGT6 B6JGU6 B6JGU7 B6JGU8 B6JGW4 B6JGW6 B6JGY0 B6JH58 B6JH66 B6JH69
B6JH72 B6JH82 B6JH85 B6JHG1 B6JHG4 B6JHH8 B6JHM4 B6JHM9 B6JHT8 B6JI86 B6JIA0
B6JIL2 B6JIM2 B6JIP3 B6JIP4 B6JIV1 B6JIW1 B6JIZ6 B6JIZ7 B6JJ00 B6JJ31 B6JJ36
B6JJ39 B6JJ40 B6JJ68 B6JJF0 B6JJN6 B6JJN9 B6JJP1 B6JJP3 B6JJP4 B6JJP6 B6JJP7
B6JJP8 B6JJQ1 B6JJQ2 B6JJT0 B6JKD1 B6JKD3 B6JKE4 B6JKE5 B6JKE8 B6JKF2 B6JKG8
B6JKG9 B6JKH0 B6JKI4 B6JKI8 B6JKJ8 B6JKL0 B6JKL2 B6JKM3 B6JKM5 B6JKM6 B6JKN2
B6JKN5 B6JKP5 B6JKQ6 B6JKQ8 B6JKR0 B6JKT9 B6JKW1 B6JKW4 B6JKW5 B6JKX0 B6JKX5
B6JKX6 B6JKX7 B6JKX8 B6JKY3 B6JKY5 B6JL03 B6JL09 B6JL62 B6JL75 B6JL92 B6JL96
B6JL97 B6JL98 B6JLD2 B6JLD4 B6JLE4 B6JLE8 B6JLF2 B6JLF4 B6JLG3 B6JLI7 B6JLJ9
B6JLK1 B6JLL1 B6JLL3 B6JLM6 B6JLN1 B6JLN6 B6JLQ0 B6JLQ1 B6JLQ2 B6JLU2 B6JLW7
B6JLX1 B6JM01 B6JM02 B6JM09 B6JM24 B6JM26 B6JM29 B6JM32 B6JM52 B6JM54 B6JM61
B6JM63 B6JM65 B6JM81 B6JM82 B6JM89 B6JM90 B6JMA1 B6JMA7 B6JME6 B6JME7 B6JME9
B6JMF1 B6JMG2 B6JMG9 B6JMI0 B6JMJ2 B6JMM9 B6JMN5 B6JMP1 B6JMP3 B6JMP4 B6JMQ6
B6JMR9 B6JMS3 B6JMT1 B6JMT3 B6JMU2 B6JMY1 B6JMY7 B6JMZ1 B6JN00 B6JN06 B6JN17
B6JN18 B6JN30 B6JN34 B6JN35 B6JN36 B6JN38 B6JN39 B6JN40 B6JN41 B6JN44 B6JN53
B6JN68 B6JN78 B6JN80 B6JN84 B6JN86 B6JN89 B6JN96 B6JN97 B6JNB7 B6JNC0 B6JND2
B6JND4 B6JND5 B6JND6 B6JNE1 B6JNE2 B6JNE3 B6JNE4 B6JNE5 B6JNE6 B6JNE7 B6JNE8
B6JNE9 B6JNF0 B6JNF1 B6JNF2 B6JNF3 B6JNF4 B6JNF5 B6JNF6 B6JNF8 B6JNG1 B6JNH4
B6JNH5 B6JNI4 B6JNP9 B6JNQ5 B6JNQ7 B6JNR5 B6JNS0 B6JNS3 B6JNU0 B6JNU2 B6JNU5
B6JNX2 B6JNX3 B6JNX8 B6JNY0 B6JNZ3 B6JNZ4 B6JNZ5 B6JP11 B6JP23 B6JP37 B6JP47
B6JP48 B6JP67 B6JP76 B6JP91 B6JP92 B6JP93 B6JPA0 B6JPA1 B6JPA4 B6JPA5 B6JPA7

B6JPA8 B6JPC9 B6JPD0 B6JPH0 B6JPH1 B6JPH2 B6JPH3 B6JPH5 B6JPH6 B6JPH8 B6JPH9
B6JPI0 B6JPI5 B6JPI6 B6JPI8 B6JPL0 B6JPL1 B6JPM2 B6JPM4 B6JPM5 B6JPQ1 B6JPQ5
B6JPQ9 B6JPR7 B6JPS3 B6JPT1 B6JPT2 B6JPU4 B6JPU7 B6JUP5 B6JUV1 B6JVD0 B6JVP0
B6JWC1 B6JWF7 B6JX19 B6JXU1 B6JY16 B6JYB6 B6JYD2 B6JYI7 B6JZ70 B6K0X1 B6K180
B6K1G6 B6K1S2 B6K1V7 B6K1Y8 B6K286 B6K2L2 B6K2N0 B6K2P1 B6K4J2 B6K4Q2 B6K4Q8
B6K7R8 B6K8A0 B6KG46 B6LI37 B6LS00 B6MUN4 B6NK32 B6NXD5 B6PLN3 B6Q1K5 B6Q1N3
B6Q1T9 B6Q2W9 B6Q348 B6Q3B6 B6Q3C6 B6Q3G2 B6Q4Z5 B6Q5I3 B6Q5P5 B6Q702 B6Q8T5
B6QAR7 B6QAW7 B6QB86 B6QC06 B6QC56 B6QCA7 B6QD96 B6QEP5 B6QFH5 B6QFZ6 B6QG01
B6QGB9 B6QGS2 B6QGW6 B6QHL1 B6QHL4 B6QL24 B6QNA1 B6QP49 B6QQ13 B6QQH9 B6QRG1
B6QS64 B6QSZ4 B6QT52 B6QVD3 B6QWH9 B6RGY0 B6SCF3 B6SCF5 B6SCF6 B6SFA4 B6SHG7
B6SU46 B6SXF8 B6SXH2 B6SZW0 B6TB21 B6TCE8 B6TGN4 B6TN12 B6TNK6 B6TNW8 B6TPF2
B6TRH4 B6TTV8 B6TTW1 B6TZD1 B6U151 B6UFC7 B6UI56 B6UL32 B6V2N4 B6V2P4 B6VJS4
B6VP39 B6VQA1 B6VRE6 B6VRE8 B6YPZ6 B6YPZ8 B6YQ00 B6YQ01 B6YQ04 B6YQ08 B6YQ10
B6YQ14 B6YQ20 B6YQ21 B6YQ29 B6YQ30 B6YQ31 B6YQ32 B6YQ38 B6YQ41 B6YQ44 B6YQ59
B6YQ61 B6YQ62 B6YQ63 B6YQ67 B6YQ69 B6YQ71 B6YQ72 B6YQ74 B6YQ75 B6YQ78 B6YQ79
B6YQ80 B6YQ81 B6YQ82 B6YQ84 B6YQ85 B6YQ88 B6YQ89 B6YQ90 B6YQ92 B6YQ94 B6YQ95
B6YQA2 B6YQA4 B6YQA7 B6YQB6 B6YQB7 B6YQB8 B6YQB9 B6YQC2 B6YQC3 B6YQC6 B6YQC7
B6YQD4 B6YQE4 B6YQE9 B6YQF1 B6YQF6 B6YQI6 B6YQJ0 B6YQL2 B6YQL8 B6YQQ8 B6YQS6
B6YQS7 B6YQS8 B6YQT0 B6YQT1 B6YQT8 B6YQU1 B6YQU3 B6YQV1 B6YQY8 B6YQZ8 B6YR04
B6YR15 B6YR24 B6YR34 B6YR38 B6YR44 B6YR45 B6YR52 B6YR69 B6YRA4 B6YRA6 B6YRA7
B6YRB0 B6YRC6 B6YRD1 B6YRF7 B6YRH7 B6YRH8 B6YRI6 B6YRK7 B6YRL1 B6YRL2 B6YRL9
B6YRM5 B6YRN2 B6YRN4 B6YRP0 B6YRQ9 B6YRR7 B6YRR8 B6YRR9 B6YRS4 B6YRU5 B6YRU6
B6YRV5 B6YRV8 B6YS07 B6YS08 B6YS10 B6YS11 B6YS15 B6YS18 B6YS23 B6YS28 B6YS33
B6YS36 B6YS40 B6YS43 B6YS49 B6YSE7 B6YSH9 B6YSI2 B6YSJ5 B6YSL0 B6YSL3 B6YSL4
B6YSL5 B6YSL6 B6YSL7 B6YSL9 B6YSM0 B6YSM1 B6YSM2 B6YSM9 B6YSN0 B6YSN1 B6YSN3
B6YSN8 B6YSP0 B6YSP2 B6YSP4 B6YSQ1 B6YSQ2 B6YSQ4 B6YSQ7 B6YSQ8 B6YSR1 B6YST0
B6YST3 B6YST9 B6YSU2 B6YSU5 B6YSU8 B6YSW2 B6YSW7 B6YSX3 B6YSX7 B6YSX9 B6YSY5
B6YSY6 B6YSY7 B6YSZ4 B6YT00 B6YT11 B6YT12 B6YT14 B6YT17 B6YT21 B6YT34 B6YT35
B6YT36 B6YT45 B6YT50 B6YT59 B6YTB2 B6YTE3 B6YTE4 B6YTE8 B6YTF0 B6YTF2 B6YTG0
B6YTH4 B6YTH7 B6YTJ2 B6YTJ3 B6YTJ4 B6YTJ5 B6YTK7 B6YTM4 B6YTM6 B6YTN5 B6YTP6
B6YTR4 B6YTZ4 B6YTZ7 B6YTZ8 B6YU51 B6YU52 B6YU58 B6YU65 B6YUD7 B6YUD9 B6YUE9
B6YUF8 B6YUH1 B6YUJ5 B6YUL1 B6YUL3 B6YUN0 B6YUN2 B6YUQ3 B6YUQ8 B6YUR8 B6YUU0
B6YUU2 B6YUU5 B6YUU6 B6YUU9 B6YUV3 B6YUW9 B6YV04 B6YV39 B6YV61 B6YV97 B6YV98
B6YVB1 B6YVC3 B6YVF3 B6YVG1 B6YVG2 B6YVG5 B6YVN3 B6YVQ4 B6YVQ7 B6YVQ8 B6YVS9
B6YVT5 B6YVU5 B6YVU7 B6YVY6 B6YVZ1 B6YW35 B6YW36 B6YW69 B6YW70 B6YW91 B6YWA4
B6YWA8 B6YWA9 B6YWB8 B6YWC1 B6YWD6 B6YWD9 B6YWF5 B6YWG5 B6YWH0 B6YWH3 B6YWH7
B6YWH9 B6YWK6 B6YWL5 B6YWN8 B6YWS3 B6YWS6 B6YWT7 B6YWV3 B6YWW5 B6YWX4 B6YWX7
B6YWY1 B6YWY2 B6YWY3 B6YWY4 B6YX10 B6YX39 B6YX51 B6YXA5 B6YXA9 B6YXC8 B6YXE5
B6YXF3 B6YXH6 B6YXH7 B6YXI6 B6YXJ1 B6YXJ9 B6YXM1 B6YXM3 B6YXP0 B6YXP9 B6YXQ1
B6YXR2 B6YXT4 B6YXU4 B6YXV3 B6YXW4 B6YXW5 B6YXX2 B6YY21 B6ZK72 B6ZLK2 B7E321
B7E7M8 B7EA73 B7EIH2 B7EJ91 B7EZJ7 B7F7B9 B7F845 B7F8P5 B7F924 B7F958 B7F9I5
B7FAL5 B7FAS0 B7FAS6 B7FF06 B7FF07 B7FF08 B7FF09 B7FF12 B7FKW7 B7FM45 B7FNA9
B7FNU7 B7FRE8 B7FRU7 B7FTW3 B7FUM7 B7FX91 B7FXE8 B7G267 B7G3Y9 B7G5J1 B7G620
B7G6D3 B7G7Y7 B7GEY2 B7GF12 B7GF88 B7GFA1 B7GFA5 B7GFB6 B7GFF3 B7GFF9 B7GFG2

B7GFH0 B7GFH9 B7GFK8 B7GFL8 B7GFL9 B7GFM4 B7GFM6 B7GFN1 B7GFP2 B7GFP5 B7GFQ0
B7GFQ4 B7GFQ6 B7GFR0 B7GFR1 B7GFR5 B7GFR6 B7GFT5 B7GFT7 B7GFT8 B7GFU2 B7GFV1
B7GFV3 B7GFW3 B7GG21 B7GG69 B7GG70 B7GG75 B7GG79 B7GG89 B7GGC4 B7GGC6 B7GGC7
B7GGC8 B7GGD5 B7GGD6 B7GGD9 B7GGE3 B7GGE9 B7GGI1 B7GGJ0 B7GGJ1 B7GGJ8 B7GGK2
B7GGM5 B7GGN9 B7GGQ4 B7GGR2 B7GGR4 B7GGR5 B7GGS0 B7GGT1 B7GGT8 B7GGU7 B7GGU9
B7GGV1 B7GGV3 B7GGV9 B7GGW0 B7GGW8 B7GH12 B7GH18 B7GH19 B7GH21 B7GH22 B7GH24
B7GH25 B7GH28 B7GH35 B7GH39 B7GH71 B7GH78 B7GH81 B7GH89 B7GHD3 B7GHD6 B7GHE3
B7GHE5 B7GHF9 B7GHJ4 B7GHJ8 B7GHM4 B7GHM5 B7GHN1 B7GHP8 B7GHQ9 B7GHS8 B7GHT2
B7GHW7 B7GHW9 B7GI39 B7GI77 B7GI87 B7GIA1 B7GIA2 B7GIA3 B7GIB5 B7GIC0 B7GIC1
B7GIH1 B7GIK0 B7GIR0 B7GIR2 B7GIR9 B7GIU0 B7GIU6 B7GIU7 B7GIY9 B7GJ32 B7GJ46
B7GJ55 B7GJ57 B7GJ63 B7GJ64 B7GJ65 B7GJ68 B7GJ69 B7GJ71 B7GJ72 B7GJ73 B7GJ74
B7GJ76 B7GJ77 B7GJ78 B7GJ79 B7GJ80 B7GJ81 B7GJ84 B7GJ86 B7GJ91 B7GJ92 B7GJ93
B7GJ95 B7GJ99 B7GJH1 B7GJM6 B7GJN4 B7GJR4 B7GJU0 B7GJU8 B7GK04 B7GK53 B7GK78
B7GKC6 B7GKC7 B7GKC8 B7GKD0 B7GKG6 B7GKI1 B7GKQ7 B7GKQ8 B7GKT2 B7GKT8 B7GKU4
B7GKZ6 B7GKZ7 B7GL26 B7GL27 B7GL34 B7GL36 B7GL38 B7GL41 B7GL45 B7GL49 B7GLA4
B7GLD4 B7GLD5 B7GLD6 B7GLH3 B7GLK2 B7GLK6 B7GLS9 B7GLX3 B7GLZ2 B7GM02 B7GM51
B7GM52 B7GMG3 B7GMG4 B7GMH4 B7GMH5 B7GMJ3 B7GMK8 B7GML0 B7GMM1 B7GMU4 B7GMV9
B7GMW3 B7GMW8 B7GMX1 B7GMZ8 B7GN04 B7GN19 B7GN27 B7GN83 B7GN96 B7GNA1 B7GNA2
B7GNA3 B7GNB3 B7GNB5 B7GNB6 B7GNC1 B7GNC2 B7GNC3 B7GNC4 B7GNC5 B7GNC6 B7GNC7
B7GNC8 B7GNC9 B7GND0 B7GND1 B7GND3 B7GND4 B7GND5 B7GND6 B7GND7 B7GND8 B7GND9
B7GNE0 B7GNE1 B7GNE7 B7GNE8 B7GNF9 B7GNG8 B7GNG9 B7GNH2 B7GNH3 B7GNJ5 B7GNJ6
B7GNK0 B7GNK1 B7GNK2 B7GNK3 B7GNM0 B7GNM6 B7GNM7 B7GNR5 B7GNR6 B7GP38 B7GP48
B7GP73 B7GP74 B7GP85 B7GPC4 B7GPE3 B7GPE4 B7GPF7 B7GPR8 B7GPS9 B7GPT0 B7GPV4
B7GPV9 B7GPW0 B7GPW1 B7GPW2 B7GPZ1 B7GQ02 B7GQ05 B7GQ20 B7GQ24 B7GQ48 B7GQ51
B7GQ70 B7GQ76 B7GQ79 B7GQ87 B7GQA1 B7GQC3 B7GQC4 B7GQE1 B7GQG0 B7GQG1 B7GQI9
B7GQR8 B7GQR9 B7GQS1 B7GQS5 B7GQS6 B7GQU5 B7GQU6 B7GQU7 B7GQV7 B7GR13 B7GR15
B7GR16 B7GR17 B7GR18 B7GR20 B7GRM4 B7GRM5 B7GRN1 B7GRU8 B7GRV4 B7GRV8 B7GRY8
B7GS74 B7GS82 B7GS87 B7GS97 B7GSA0 B7GSB1 B7GSB6 B7GSF9 B7GSG2 B7GSP5 B7GSU4
B7GSV7 B7GSZ2 B7GT00 B7GT04 B7GT18 B7GT29 B7GT47 B7GT67 B7GT76 B7GT80 B7GTK5
B7GTL2 B7GTL3 B7GTM7 B7GTN2 B7GTN9 B7GTP0 B7GTP1 B7GTP2 B7GTP4 B7GTP9 B7GTS5
B7GTU6 B7GTU7 B7GTW0 B7GTZ5 B7GU04 B7GU46 B7GU48 B7GU49 B7GU64 B7GU81 B7GU90
B7GUB5 B7GUD5 B7GUE4 B7GUF2 B7GUF6 B7GUG7 B7GUH2 B7GUI8 B7GUN9 B7GUP3 B7GUP8
B7GUP9 B7GUQ0 B7GUQ1 B7GUR7 B7GUU5 B7GUU8 B7GUW2 B7GUW3 B7GUW9 B7GUX5 B7GUX7
B7GUY5 B7GUZ7 B7GUZ8 B7GV08 B7GV10 B7GV30 B7GV54 B7GV69 B7GV74 B7GV75 B7GV90
B7GVJ5 B7GVK1 B7GVN1 B7GVP5 B7GVQ9 B7GVR1 B7GVR8 B7GVS0 B7GVS1 B7GVV0 B7GVV1
B7GVW2 B7GVX2 B7GVX4 B7GW01 B7GW02 B7GW03 B7GW04 B7GW05 B7GW06 B7GW08 B7GW09
B7GW10 B7GW11 B7GW12 B7GW13 B7GW14 B7GW15 B7GW16 B7GW17 B7GW18 B7GW19 B7GW20
B7GW21 B7GW23 B7GW24 B7GW25 B7GW26 B7GW27 B7GW28 B7GW49 B7GW53 B7GW55 B7GW58
B7GW85 B7GWA1 B7GWH6 B7GWI2 B7GWK1 B7GWK9 B7GWL4 B7GWL7 B7GWN1 B7GWQ5 B7GWT8
B7GWV6 B7GWX1 B7GWX6 B7GX14 B7GX48 B7GX68 B7GXA3 B7GXC5 B7GXH8 B7GXH9 B7GXI2
B7GXK7 B7GXS7 B7GXS8 B7GXT4 B7GXU0 B7GXU2 B7GXU4 B7GXU5 B7GXW0 B7GXW8 B7GXX8
B7GY04 B7GY18 B7GY31 B7GY35 B7GY36 B7GY44 B7GY62 B7GY71 B7GY87 B7GY98 B7GYB5
B7GYB6 B7GYD0 B7GYE0 B7GYG4 B7GYG5 B7GYG6 B7GYK7 B7GYK9 B7GYL4 B7GYL6 B7GYM8
B7GYM9 B7GYN0 B7GYP8 B7GYQ4 B7GYQ7 B7GYQ9 B7GYR0 B7GYR1 B7GYS1 B7GYT0 B7GYV1

B7GYV2 B7GYZ5 B7GZ21 B7GZ27 B7GZ34 B7GZ38 B7GZ71 B7GZ77 B7GZ86 B7GZA8 B7GZI3
B7GZJ8 B7GZK6 B7GZN3 B7GZQ0 B7GZQ9 B7GZR6 B7GZV3 B7GZV4 B7GZZ4 B7GZZ9 B7H000
B7H005 B7H009 B7H010 B7H011 B7H018 B7H047 B7H051 B7H063 B7H065 B7H068 B7H069
B7H072 B7H073 B7H081 B7H092 B7H094 B7H096 B7H098 B7H0A1 B7H0A6 B7H0A7 B7H0I7
B7H0I9 B7H0J8 B7H0J9 B7H0K0 B7H0K8 B7H0L9 B7H0P4 B7H0P7 B7H0Q4 B7H0S9 B7H0T6
B7H0T7 B7H0U1 B7H0U3 B7H0W3 B7H0X1 B7H0Z1 B7H0Z2 B7H114 B7H115 B7H117 B7H141
B7H160 B7H1C7 B7H1F3 B7H1F6 B7H1G4 B7H1G7 B7H1J8 B7H1J9 B7H1K0 B7H1K1 B7H1K2
B7H1K5 B7H1L9 B7H1M6 B7H1N2 B7H1P9 B7H1Q6 B7H1U1 B7H1U3 B7H1U5 B7H1V4 B7H207
B7H211 B7H225 B7H2A9 B7H2E9 B7H2H4 B7H2J6 B7H2Y0 B7H2Y1 B7H2Y2 B7H2Y9 B7H316
B7H317 B7H343 B7H3A6 B7H3B2 B7H3B8 B7H3H0 B7H3H3 B7H3H4 B7H3I4 B7H3K6 B7H3L0
B7H3L4 B7H3N1 B7H3N8 B7H3Q0 B7H3Q6 B7H3S1 B7H3S4 B7H3U3 B7H3W7 B7H3X2 B7H4L8
B7H4L9 B7H4P7 B7H4Q3 B7H4Q6 B7H4Q7 B7H4Q8 B7H4T3 B7H4T5 B7H4T6 B7H4T8 B7H4U0
B7H4U8 B7H4V1 B7H4W1 B7H4W2 B7H4W3 B7H546 B7H560 B7H597 B7H5Z8 B7H636 B7H655
B7H673 B7H674 B7H6D1 B7H6K7 B7H6K9 B7H6L8 B7H6L9 B7H6M0 B7H6M1 B7H6M2 B7H6M4
B7H6M5 B7H6M7 B7H6N1 B7H6N5 B7H6P8 B7H6Q4 B7H6Q5 B7H6Q7 B7H6Q9 B7H6R5 B7H6S5
B7H6T6 B7H6U7 B7H6U8 B7H6V0 B7H6W3 B7H6W7 B7H6W8 B7H6Y4 B7H6Y5 B7H703 B7H711
B7H746 B7H758 B7H782 B7H798 B7H7A0 B7H7A5 B7H7A6 B7H7F6 B7H7H4 B7H7H5 B7H7K5
B7H7M2 B7H7P4 B7H841 B7H8F2 B7H8H3 B7H8H6 B7H8P4 B7H8Z2 B7H924 B7H927 B7H928
B7H929 B7H945 B7H948 B7H949 B7H965 B7H979 B7H9C3 B7H9C7 B7H9C8 B7H9Q9 B7H9R2
B7H9S1 B7H9S6 B7H9V0 B7H9V9 B7HA46 B7HAY0 B7HAY1 B7HAY4 B7HAY5 B7HAY7 B7HAZ0
B7HAZ1 B7HB02 B7HB04 B7HB12 B7HB35 B7HB48 B7HB50 B7HB51 B7HB52 B7HB53 B7HB68
B7HB70 B7HB75 B7HB98 B7HB99 B7HBA0 B7HBC4 B7HBC7 B7HBD4 B7HBG1 B7HBI0 B7HBI1
B7HBU7 B7HC95 B7HCC8 B7HCC9 B7HCD0 B7HCD1 B7HCI2 B7HCI3 B7HCQ3 B7HCQ9 B7HCR2
B7HCS0 B7HCS2 B7HCS7 B7HCT4 B7HCT8 B7HCT9 B7HCU0 B7HCU1 B7HCU2 B7HCU7 B7HCU8
B7HCV9 B7HCW1 B7HCZ6 B7HD14 B7HD62 B7HDF2 B7HDL9 B7HDP3 B7HDP4 B7HDP7 B7HDQ4
B7HDQ5 B7HDT0 B7HDT1 B7HDT4 B7HDT6 B7HDU0 B7HDU2 B7HDU7 B7HDU9 B7HDV0 B7HDV1
B7HDV2 B7HDV3 B7HDV5 B7HDV9 B7HDW0 B7HDW3 B7HDW4 B7HDW5 B7HDW7 B7HDX2 B7HDY1
B7HDY7 B7HDY9 B7HE08 B7HE15 B7HE21 B7HE22 B7HE37 B7HE40 B7HE42 B7HE51 B7HE52
B7HE54 B7HE55 B7HE61 B7HE73 B7HE75 B7HE77 B7HE82 B7HE86 B7HE87 B7HE93 B7HE96
B7HE98 B7HEA1 B7HEA4 B7HEA5 B7HEB3 B7HEB5 B7HEC5 B7HED4 B7HED5 B7HEF4 B7HEF6
B7HEG2 B7HEG4 B7HEK8 B7HEM8 B7HER4 B7HEV3 B7HEV9 B7HF33 B7HF56 B7HF67 B7HF77
B7HF86 B7HF87 B7HF94 B7HFA6 B7HFB3 B7HFC6 B7HFL2 B7HFL3 B7HFM8 B7HFM9 B7HFN6
B7HFN7 B7HFP5 B7HFR2 B7HFX0 B7HFZ0 B7HG04 B7HG19 B7HG46 B7HG50 B7HG60 B7HG67
B7HGB9 B7HGC9 B7HGD2 B7HGD4 B7HGE1 B7HGF9 B7HGP8 B7HGP9 B7HGR9 B7HGT7 B7HGV8
B7HGW3 B7HGY8 B7HGZ8 B7HGZ9 B7HH00 B7HH01 B7HH02 B7HH19 B7HH58 B7HH59 B7HH60
B7HH80 B7HH81 B7HH96 B7HH99 B7HHC6 B7HHE8 B7HHF7 B7HHF8 B7HHF9 B7HHG0 B7HHG2
B7HHG3 B7HHG4 B7HHG5 B7HHG6 B7HHG7 B7HHH6 B7HHH7 B7HHL7 B7HHQ0 B7HHQ2 B7HHQ7
B7HHR5 B7HHR7 B7HHS1 B7HHT6 B7HHT7 B7HHU0 B7HHU3 B7HHU4 B7HHU5 B7HHV4 B7HHW2
B7HHW3 B7HHX2 B7HID8 B7HIG1 B7HIH4 B7HIH7 B7HII3 B7HII4 B7HII5 B7HIJ2 B7HIJ3
B7HIJ8 B7HIK1 B7HIK9 B7HIL3 B7HIL6 B7HIP0 B7HIQ2 B7HIR1 B7HIT0 B7HJ05 B7HJ06
B7HJ16 B7HJ19 B7HJ25 B7HJ28 B7HJ35 B7HJ36 B7HJ37 B7HJ38 B7HJ40 B7HJ43 B7HJ44
B7HJ45 B7HJ46 B7HJ47 B7HJ48 B7HJ49 B7HJ50 B7HJ51 B7HJ52 B7HJ53 B7HJ55 B7HJ56
B7HJ57 B7HJ58 B7HJ59 B7HJ60 B7HJ61 B7HJ62 B7HJ64 B7HJ66 B7HJ67 B7HJ69 B7HJ72
B7HJ73 B7HJ74 B7HJ93 B7HJC1 B7HJE9 B7HJF3 B7HJI5 B7HJJ0 B7HJJ1 B7HJK3 B7HK47

B7HK48 B7HK63 B7HK66 B7HK95 B7HKB7 B7HKC6 B7HKC7 B7HKC8 B7HKC9 B7HKD1 B7HKD2
B7HKD3 B7HKD4 B7HKD5 B7HKD6 B7HKE5 B7HKE6 B7HKI9 B7HKJ0 B7HKJ1 B7HKJ2 B7HKR8
B7HKR9 B7HKU6 B7HKX2 B7HL06 B7HL09 B7HL14 B7HL21 B7HL23 B7HL27 B7HL30 B7HL46
B7HL47 B7HL50 B7HL53 B7HL54 B7HL55 B7HL64 B7HL73 B7HL74 B7HL82 B7HLA2 B7HLA3
B7HLA6 B7HLB5 B7HLE0 B7HLE1 B7HLE4 B7HLE6 B7HLF0 B7HLF2 B7HLF4 B7HLF7 B7HLF9
B7HLG0 B7HLG1 B7HLG2 B7HLG3 B7HLG5 B7HLG9 B7HLH0 B7HLH3 B7HLH4 B7HLH5 B7HLH7
B7HLI2 B7HLJ1 B7HLJ7 B7HLJ9 B7HLK3 B7HLK5 B7HLL7 B7HLL8 B7HLL9 B7HLM0 B7HLM2
B7HLM3 B7HLM5 B7HLM9 B7HM19 B7HM33 B7HM39 B7HM40 B7HM42 B7HM44 B7HMA5 B7HMA9
B7HMD4 B7HME5 B7HME6 B7HME8 B7HMG6 B7HMI7 B7HMM7 B7HMN9 B7HMP7 B7HMQ9 B7HMU7
B7HMV1 B7HMV2 B7HMX7 B7HN10 B7HN14 B7HN15 B7HN16 B7HN32 B7HN35 B7HN36 B7HN53
B7HN67 B7HNB7 B7HND7 B7HNI1 B7HNI5 B7HNM8 B7HNM9 B7HNN1 B7HNN4 B7HNP5 B7HNP7
B7HNP9 B7HNQ7 B7HNS7 B7HNT8 B7HNU0 B7HNU2 B7HNU3 B7HNU4 B7HNU5 B7HNW0 B7HNW2
B7HNW6 B7HNY9 B7HNZ0 B7HNZ1 B7HP29 B7HPD4 B7HPH8 B7HPI4 B7HPI8 B7HPJ5 B7HPJ7
B7HPK2 B7HPK8 B7HPL1 B7HPL2 B7HPL3 B7HPL4 B7HPL5 B7HPM0 B7HPM1 B7HPN1 B7HPN3
B7HPR7 B7HPS0 B7HPS6 B7HPS7 B7HPS8 B7HPT5 B7HPT6 B7HPU1 B7HPU4 B7HPV2 B7HPV6
B7HPV9 B7HPX7 B7HPX8 B7HPY8 B7HQ80 B7HQ94 B7HQD2 B7HQD7 B7HQD8 B7HQE4 B7HQE5
B7HQG0 B7HQG5 B7HQG7 B7HQH6 B7HQH7 B7HQH9 B7HQI0 B7HQI6 B7HQJ8 B7HQK0 B7HQK2
B7HQK7 B7HQL1 B7HQL2 B7HQM1 B7HQM4 B7HQM6 B7HQM9 B7HQN2 B7HQN3 B7HQP4 B7HQR4
B7HQS1 B7HQS4 B7HQT1 B7HQT2 B7HQT3 B7HQT4 B7HQT6 B7HQT9 B7HQU0 B7HQU1 B7HQU2
B7HQU3 B7HQU4 B7HQU5 B7HQU6 B7HQU7 B7HQU8 B7HQU9 B7HQV1 B7HQV2 B7HQV3 B7HQV4
B7HQV5 B7HQV6 B7HQV7 B7HQV8 B7HQW0 B7HQW2 B7HQW3 B7HQW5 B7HQW8 B7HQW9 B7HQX0
B7HQX2 B7HQX7 B7HQX8 B7HQZ0 B7HR31 B7HRD6 B7HRG3 B7HRI4 B7HRJ3 B7HRJ7 B7HRK2
B7HRK4 B7HRL2 B7HRL3 B7HRL9 B7HRN2 B7HRN9 B7HRQ4 B7HRS6 B7HRS7 B7HRZ5 B7HS01
B7HS04 B7HS05 B7HS29 B7HS31 B7HS32 B7HS34 B7HS36 B7HS46 B7HSI4 B7HSJ2 B7HSM3
B7HSM8 B7HSN5 B7HSR1 B7HSS7 B7HSV1 B7HSV6 B7HSV7 B7HSW5 B7HSW8 B7HSX9 B7HSY0
B7HSY1 B7HT69 B7HT70 B7HTE7 B7HTW3 B7HTX9 B7HTY6 B7HTZ9 B7HU51 B7HU54 B7HU62
B7HU66 B7HU74 B7HUD5 B7HUR5 B7HUS8 B7HUS9 B7HUW3 B7HUY1 B7HVH9 B7HVJ1 B7HVK6
B7HVS1 B7HVU3 B7HVU6 B7HVW2 B7HW45 B7HW54 B7HW55 B7HW81 B7HW83 B7HW89 B7HWF4
B7HWU2 B7HWY7 B7HWY9 B7HX62 B7HXH0 B7HXM4 B7HXM9 B7HY75 B7HY76 B7HY90 B7HY91
B7HY98 B7HY99 B7HYE4 B7HYE9 B7HYF9 B7HYG6 B7HYQ6 B7HYS8 B7HYU3 B7HYY0 B7HZ82
B7HZF6 B7HZF9 B7HZG1 B7HZG7 B7HZG8 B7HZH3 B7HZH4 B7HZH8 B7HZJ8 B7HZR4 B7HZR5
B7HZT6 B7HZV3 B7I056 B7I057 B7I063 B7I0B1 B7I0B6 B7I0E0 B7I0E9 B7I0F0 B7I0F1
B7I0F2 B7I0F3 B7I0H2 B7I0K9 B7I0L0 B7I0L1 B7I1E1 B7I1Q9 B7I1U9 B7I245 B7I251
B7I252 B7I268 B7I271 B7I289 B7I2B0 B7I2B2 B7I2C2 B7I2C3 B7I2K6 B7I2K7 B7I2M9
B7I2R7 B7I302 B7I309 B7I325 B7I331 B7I337 B7I338 B7I356 B7I357 B7I358 B7I359
B7I360 B7I3J1 B7I3K0 B7I3L8 B7I3Q5 B7I3R5 B7I3R7 B7I3R9 B7I3S0 B7I3U0 B7I3U1
B7I415 B7I420 B7I441 B7I451 B7I459 B7I499 B7I4A2 B7I4A8 B7I4B6 B7I4D5 B7I4D7
B7I4E1 B7I4E2 B7I4E8 B7I4G5 B7I4I4 B7I4I7 B7I4K2 B7I4K8 B7I4N4 B7I4T0 B7I4T7
B7I4U0 B7I4X1 B7I4X4 B7I4X6 B7I4X8 B7I4Y4 B7I4Z0 B7I501 B7I508 B7I526 B7I530
B7I534 B7I556 B7I557 B7I585 B7I595 B7I5A8 B7I5A9 B7I5C6 B7I5D6 B7I5E4 B7I5F5
B7I5G3 B7I5G4 B7I5G7 B7I5G8 B7I5H1 B7I5H3 B7I5I5 B7I5I9 B7I5N9 B7I5P9 B7I5Q0
B7I5Q3 B7I5V5 B7I5W1 B7I5W6 B7I5Y5 B7I5Z2 B7I610 B7I618 B7I619 B7I624 B7I636
B7I675 B7I682 B7I683 B7I684 B7I688 B7I693 B7I694 B7I698 B7I6C5 B7I6D7 B7I6M5
B7I6M6 B7I6M7 B7I6M8 B7I6T2 B7I6V5 B7I6V8 B7I6V9 B7I752 B7I774 B7I784 B7I790

B7I7A4 B7I7B4 B7I7B5 B7I7B6 B7I7N2 B7I7P3 B7I7P6 B7I7Q2 B7I7R9 B7I7S0 B7I7S1
B7I7S8 B7I7T5 B7I7T7 B7I7U2 B7I7U4 B7I842 B7I854 B7I894 B7I895 B7I896 B7I8C4
B7I8K2 B7I8M5 B7I8P7 B7I8Q3 B7I8T3 B7I8T4 B7I8T6 B7I8T8 B7I8U4 B7I8V1 B7I8W9
B7I920 B7I933 B7I936 B7I973 B7I976 B7I981 B7I989 B7I9B0 B7I9D2 B7I9D5 B7I9D7
B7I9E1 B7I9G6 B7I9K0 B7I9N8 B7I9U0 B7I9U9 B7I9V2 B7I9V4 B7I9W0 B7I9Y9 B7IA13
B7IA14 B7IA15 B7IA16 B7IA17 B7IA18 B7IA20 B7IA21 B7IA22 B7IA23 B7IA24 B7IA25
B7IA26 B7IA27 B7IA28 B7IA29 B7IA30 B7IA31 B7IA32 B7IA33 B7IA35 B7IA36 B7IA37
B7IA38 B7IA39 B7IA40 B7IA66 B7IA68 B7IA77 B7IA88 B7IA89 B7IAF3 B7IAG6 B7IAM8
B7IAP6 B7IAS9 B7IAT0 B7IAT2 B7IAU0 B7IAU2 B7IAV6 B7IAX1 B7IAZ8 B7IBA1 B7IBB3
B7IBC1 B7IBC2 B7IBC3 B7IBD1 B7IBD7 B7IBH5 B7IBH7 B7IBH8 B7IBK5 B7IBK6 B7IBN2
B7IBN8 B7IBR9 B7IBU3 B7IBV9 B7IBY8 B7IBY9 B7IC15 B7IC17 B7ICC6 B7ICC7 B7ICE3
B7ICE4 B7ICE9 B7ICG5 B7ICK4 B7ICM5 B7ICN0 B7ICN4 B7ICN6 B7ICR2 B7ICR3 B7ICR4
B7ICR5 B7ICT4 B7ICU2 B7ICX1 B7ICX7 B7ID00 B7ID01 B7ID25 B7ID34 B7ID58 B7ID76
B7ID82 B7ID85 B7ID86 B7ID87 B7IDB4 B7IDE2 B7IDG7 B7IDH3 B7IDI0 B7IDI7 B7IDK8
B7IDL6 B7IDQ7 B7IDR9 B7IDS3 B7IDS7 B7IDT4 B7IDT7 B7IDU1 B7IDU4 B7IDU7 B7IDX6
B7IDY7 B7IDZ1 B7IE21 B7IE23 B7IE26 B7IE29 B7IE30 B7IE32 B7IE34 B7IE38 B7IE73
B7IE81 B7IEE2 B7IEL8 B7IEM2 B7IEM8 B7IEQ3 B7IEQ5 B7IEQ6 B7IEQ8 B7IEU8 B7IEV2
B7IEV3 B7IEY2 B7IEY7 B7IEZ2 B7IEZ6 B7IF03 B7IF13 B7IF14 B7IF15 B7IF24 B7IF30
B7IF42 B7IF44 B7IF45 B7IF61 B7IF65 B7IFA6 B7IFA7 B7IFB4 B7IFB5 B7IFB6 B7IFB7
B7IFC4 B7IFD0 B7IFE0 B7IFG6 B7IFG8 B7IFM5 B7IFM6 B7IFN3 B7IFP5 B7IFP7 B7IFR8
B7IFS5 B7IFU1 B7IFU5 B7IFU6 B7IFU7 B7IFV2 B7IFV5 B7IFV7 B7IFV8 B7IFW4 B7IFW5
B7IFY1 B7IG15 B7IG18 B7IG20 B7IG29 B7IG50 B7IG99 B7IGA0 B7IGA5 B7IGB4 B7IGD0
B7IGK8 B7IGM6 B7IGP0 B7IGQ3 B7IGT1 B7IH57 B7IH59 B7IH68 B7IH86 B7IH88 B7IHA2
B7IHA8 B7IHE6 B7IHG0 B7IHH3 B7IHI2 B7IHK5 B7IHQ4 B7IHR3 B7IHS2 B7IHT2 B7IHT3
B7IHT4 B7IHU1 B7IHU2 B7IHU3 B7IHU4 B7IHU5 B7IHU6 B7IHU7 B7IHU9 B7IHV0 B7IHV1
B7IHV2 B7IHV3 B7IHV4 B7IHV5 B7IHV6 B7IHV7 B7IHV8 B7IHV9 B7IHW0 B7IHW1 B7IHW2
B7IHW3 B7IHW4 B7IHW5 B7IHW7 B7IHX0 B7IHX1 B7IHX2 B7IHX4 B7IHX7 B7II82 B7IIB8
B7IIM0 B7IIR4 B7IIR8 B7IIS0 B7IIS9 B7IIT0 B7IIT2 B7IIT3 B7IIT9 B7IIV2 B7IIV4
B7IIV6 B7IIW1 B7IIW5 B7IIW6 B7IIX2 B7IIX5 B7IIX7 B7IIY0 B7IIY3 B7IIY4 B7IIZ1
B7IIZ3 B7IJ18 B7IJ32 B7IJP9 B7IJU1 B7IJV1 B7IJW0 B7IJW2 B7IJX1 B7IJX2 B7IJX8
B7IJZ0 B7IJZ8 B7IK11 B7IK28 B7IK29 B7IK47 B7IK73 B7IK95 B7IKB1 B7IKC1 B7IKE2
B7IKP3 B7IKS0 B7IKV4 B7IKW0 B7IKW6 B7IKZ0 B7IL05 B7IL24 B7IL28 B7IL29 B7IL83
B7IL84 B7ILA4 B7ILC1 B7ILE3 B7ILE8 B7ILG9 B7ILI6 B7ILK1 B7ILK2 B7ILR6 B7IM37
B7IM63 B7IM73 B7IM74 B7IM75 B7IM76 B7IM77 B7IM94 B7IMD3 B7IMD4 B7IMD5 B7IMH6
B7IMR7 B7IMU0 B7IMV5 B7IMV6 B7IMZ7 B7IN18 B7IN19 B7IN35 B7IN38 B7IN64 B7IN87
B7IN95 B7IN97 B7IN98 B7INA0 B7INA1 B7INA2 B7INA3 B7INA4 B7INA5 B7INB4 B7INB5
B7INE4 B7INR1 B7INU9 B7IP12 B7IP22 B7IP23 B7IP41 B7IP75 B7IP77 B7IP82 B7IP89
B7IP91 B7IP95 B7IPB0 B7IPB1 B7IPB4 B7IPB7 B7IPB8 B7IPB9 B7IPC9 B7IPD7 B7IPD8
B7IPE6 B7IPP9 B7IPR7 B7IPR9 B7IPS5 B7IPS7 B7IPY9 B7IQN3 B7IQW8 B7IQW9 B7IQY3
B7IQY4 B7IQZ1 B7IQZ2 B7IR00 B7IR88 B7IRB0 B7IRQ5 B7IRR4 B7IRR8 B7IRS8 B7IRT5
B7IS03 B7IS13 B7IS16 B7IS18 B7IS20 B7IS23 B7IS29 B7IS30 B7IS31 B7IS38 B7IS39
B7IS44 B7IS56 B7IS63 B7IS80 B7ISF6 B7ISJ0 B7ISJ1 B7ISJ2 B7ISJ3 B7ISQ8 B7ISQ9
B7IST1 B7IST2 B7IST7 B7IST8 B7ISU0 B7ISU3 B7ISV1 B7ISV5 B7ISV8 B7ISX6 B7ISX7
B7ISY7 B7ISZ0 B7ISZ6 B7ISZ9 B7IT06 B7IT07 B7IT08 B7IT09 B7IT11 B7IT14 B7IT15

B7IT16 B7IT17 B7IT18 B7IT19 B7IT20 B7IT21 B7IT22 B7IT24 B7IT26 B7IT27 B7IT28
B7IT29 B7IT30 B7IT31 B7IT32 B7IT33 B7IT35 B7IT37 B7IT38 B7IT40 B7IT43 B7IT44
B7IT45 B7IT47 B7IT52 B7IT53 B7IT85 B7ITB1 B7ITF9 B7ITG3 B7ITJ6 B7ITM0 B7ITM1
B7ITM4 B7ITN1 B7ITN2 B7ITV9 B7IU36 B7IUA4 B7IUG5 B7IUG6 B7IUG9 B7IUH1 B7IUH5
B7IUH7 B7IUI3 B7IUI5 B7IUI6 B7IUI7 B7IUI8 B7IUI9 B7IUJ1 B7IUJ5 B7IUJ6 B7IUJ9
B7IUK0 B7IUK1 B7IUK3 B7IUK8 B7IUL7 B7IUM3 B7IUM5 B7IUM9 B7IUN1 B7IUP2 B7IUP3
B7IUP4 B7IUP5 B7IUP6 B7IUP8 B7IUP9 B7IUQ1 B7IUQ5 B7IUQ9 B7IUS2 B7IUS6 B7IUS9
B7IUT0 B7IUT1 B7IUV0 B7IUV2 B7IUV3 B7IUV5 B7IUV7 B7IUW5 B7IUW8 B7IUX8 B7IUX9
B7IUY0 B7IV32 B7IVD7 B7IVF3 B7IVF4 B7IVF6 B7IVF8 B7IVG6 B7IVH6 B7IVI6 B7IVJ7
B7IVJ8 B7IVK0 B7IVL3 B7IVL7 B7IVL8 B7IVP2 B7IVS8 B7IVV9 B7IW08 B7IW10 B7IW18
B7IW22 B7IW48 B7IWE5 B7IWE8 B7IWE9 B7IWF0 B7IWG6 B7IWG9 B7IWH0 B7IWI6 B7IWK1
B7IWM5 B7IWM6 B7IWM8 B7IWN1 B7IWN2 B7IWP4 B7IWQ5 B7IWR8 B7IWS7 B7IX20 B7IX37
B7IXF5 B7IXG8 B7IXH0 B7IXH1 B7IXH2 B7IXH3 B7IXI8 B7IXJ0 B7IXJ5 B7IXL2 B7IXL3
B7IXL4 B7IXM2 B7IXT0 B7IY02 B7IYD0 B7IYD6 B7IYD9 B7IYE7 B7IYE9 B7IYF4 B7IYG2
B7IYG5 B7IYG6 B7IYG7 B7IYG8 B7IYG9 B7IYH4 B7IYH5 B7IYI5 B7IYI7 B7IYM7 B7IYN4
B7IYP0 B7IYP1 B7IZN7 B7J0E6 B7J0E9 B7J0F0 B7J0G5 B7J0H6 B7J0I6 B7J0K1 B7J0L4
B7J0M3 B7J0M4 B7J0M6 B7J0M7 B7J0M8 B7J0N1 B7J0N2 B7J0N3 B7J0N4 B7J0P3 B7J0P5
B7J0P8 B7J0P9 B7J0Q0 B7J0Q1 B7J0Q4 B7J0R6 B7J0S6 B7J0S9 B7J0T5 B7J0T6 B7J0V1
B7J0V4 B7J0W2 B7J0W4 B7J0W5 B7J0W8 B7J0Y4 B7J0Y9 B7J0Z1 B7J0Z2 B7J0Z6 B7J100
B7J121 B7J132 B7J145 B7J146 B7J148 B7J154 B7J155 B7J156 B7J161 B7J164 B7J168
B7J183 B7J192 B7J1B0 B7J1B1 B7J1B2 B7J1B6 B7J1B7 B7J1C2 B7J1C3 B7J1C9 B7J1D2
B7J1E4 B7J1F4 B7J1F7 B7J1G1 B7J1G9 B7J1H5 B7J1H9 B7J1I4 B7J1M4 B7J1N4 B7J1P0
B7J1R3 B7J1R4 B7J1R6 B7J1S5 B7J1T7 B7J1T9 B7J1U2 B7J1U3 B7J1U8 B7J1V8 B7J1V9
B7J1W0 B7J1W1 B7J1W2 B7J1W3 B7J1W4 B7J1W5 B7J1W7 B7J1X2 B7J1Z1 B7J203 B7J206
B7J207 B7J213 B7J223 B7J228 B7J241 B7J242 B7J243 B7J244 B7J245 B7J246 B7J247
B7J248 B7J249 B7J250 B7J251 B7J252 B7J253 B7J254 B7J255 B7J256 B7J257 B7J258
B7J259 B7J262 B7J264 B7J265 B7J266 B7J268 B7J278 B7J282 B7J283 B7J2B3 B7J2E8
B7J2F0 B7J2F3 B7J2G1 B7J2G3 B7J2G8 B7J2H5 B7J2I0 B7J2K0 B7J2K5 B7J2L2 B7J2L3
B7J2N7 B7J2Q0 B7J3B2 B7J3C1 B7J3D6 B7J3F7 B7J3G0 B7J3G2 B7J3M4 B7J3Q3 B7J3R1
B7J3R3 B7J3R6 B7J3S7 B7J3V1 B7J3W0 B7J3W1 B7J3Y7 B7J403 B7J415 B7J427 B7J428
B7J429 B7J432 B7J437 B7J439 B7J440 B7J444 B7J455 B7J456 B7J457 B7J458 B7J460
B7J462 B7J463 B7J464 B7J466 B7J467 B7J468 B7J469 B7J470 B7J471 B7J472 B7J474
B7J475 B7J476 B7J477 B7J478 B7J480 B7J481 B7J482 B7J483 B7J484 B7J485 B7J486
B7J4A0 B7J4A1 B7J4A3 B7J4A5 B7J4A8 B7J4C0 B7J4C9 B7J4D2 B7J4D5 B7J4D7 B7J4D8
B7J4E2 B7J4G8 B7J4J4 B7J4K3 B7J4M2 B7J4R9 B7J4S8 B7J4S9 B7J4T0 B7J4X0 B7J4Y5
B7J4Y9 B7J502 B7J551 B7J561 B7J562 B7J583 B7J5B2 B7J5E0 B7J5H6 B7J5I9 B7J5P7
B7J5S3 B7J5S6 B7J5U0 B7J5V5 B7J5X3 B7J5X6 B7J629 B7J643 B7J648 B7J653 B7J6A7
B7J6C1 B7J6C4 B7J6R2 B7J6R6 B7J6V1 B7J781 B7J791 B7J7A3 B7J7A9 B7J7B0 B7J7C4
B7J7F3 B7J7N7 B7J7P8 B7J7S6 B7J7S7 B7J7S8 B7J7T0 B7J7U7 B7J7X4 B7J7X8 B7J7X9
B7J7Y0 B7J7Y4 B7J7Y5 B7J8G6 B7J8G8 B7J8H7 B7J8I7 B7J911 B7J913 B7J917 B7J921
B7J945 B7J948 B7J950 B7J9K3 B7J9K4 B7J9P2 B7J9P7 B7J9S1 B7J9S3 B7J9T2 B7J9Z4
B7JA12 B7JA13 B7JA16 B7JA18 B7JA19 B7JA24 B7JA81 B7JA99 B7JAG8 B7JAH6 B7JAI2
B7JAI4 B7JAM6 B7JAR2 B7JB30 B7JB95 B7JBC0 B7JBC1 B7JBC8 B7JBD6 B7JBE5 B7JBG5
B7JBH8 B7JBI1 B7JBI2 B7JBM7 B7JBM8 B7JBS5 B7JBX9 B7JC05 B7JC17 B7JC18 B7JC25

B7JC26 B7JC30 B7JC34 B7JC67 B7JCS0 B7JCU0 B7JCV9 B7JCX5 B7JD15 B7JDA2 B7JDC3
B7JDD6 B7JDF1 B7JDF6 B7JDH9 B7JDJ2 B7JDJ3 B7JDM3 B7JDM6 B7JDU3 B7JDU4 B7JDW4
B7JDY2 B7JE03 B7JEE3 B7JEJ3 B7JEP0 B7JEP5 B7JER8 B7JES7 B7JES8 B7JES9 B7JET1
B7JEU9 B7JF00 B7JF01 B7JF02 B7JF51 B7JFF5 B7JFG5 B7JFG6 B7JFI0 B7JFI2 B7JFI8
B7JFJ0 B7JFQ8 B7JFQ9 B7JFS4 B7JFS7 B7JFV5 B7JFX8 B7JFY5 B7JFY7 B7JFY8 B7JFY9
B7JFZ0 B7JFZ2 B7JFZ3 B7JFZ4 B7JFZ5 B7JFZ6 B7JFZ7 B7JG06 B7JG75 B7JGF0 B7JGP0
B7JGP1 B7JGQ4 B7JGU6 B7JGY2 B7JGY4 B7JGY9 B7JGZ6 B7JGZ8 B7JH02 B7JH05 B7JH17
B7JH18 B7JH21 B7JH71 B7JHF2 B7JHF3 B7JHG1 B7JHK6 B7JHL3 B7JHM3 B7JHN0 B7JHQ6
B7JHQ7 B7JHQ8 B7JHR7 B7JHS5 B7JHS6 B7JHT4 B7JI37 B7JI38 B7JI46 B7JI78 B7JI79
B7JI80 B7JI81 B7JIE0 B7JIE1 B7JIJ6 B7JIJ9 B7JIK1 B7JIK8 B7JIK9 B7JIL4 B7JIL5
B7JIX2 B7JJ20 B7JJ46 B7JJ47 B7JJ50 B7JJ59 B7JJ84 B7JJ85 B7JJ88 B7JJ95 B7JJ97
B7JJA2 B7JJA4 B7JJA5 B7JJA6 B7JJA7 B7JJA8 B7JJB0 B7JJB6 B7JJB7 B7JJC0 B7JJC6
B7JJC7 B7JJC8 B7JJD5 B7JJD6 B7JJE4 B7JJG8 B7JJI0 B7JJJ3 B7JJK4 B7JJS6 B7JJS7
B7JJT1 B7JJT6 B7JJU5 B7JJV1 B7JJV3 B7JJV7 B7JJV9 B7JJW9 B7JJX0 B7JJX1 B7JJX2
B7JJX3 B7JJX5 B7JJX6 B7JJX8 B7JJY2 B7JJY6 B7JK00 B7JK06 B7JK07 B7JK09 B7JK11
B7JK13 B7JK17 B7JK27 B7JK37 B7JK47 B7JK51 B7JK55 B7JK73 B7JK74 B7JK84 B7JK87
B7JK94 B7JK97 B7JKA5 B7JKA6 B7JKA7 B7JKA8 B7JKB4 B7JKB5 B7JKB6 B7JKB7 B7JKB8
B7JKB9 B7JKC0 B7JKC1 B7JKC2 B7JKC3 B7JKC4 B7JKC6 B7JKC7 B7JKC8 B7JKC9 B7JKD0
B7JKD1 B7JKD2 B7JKD3 B7JKD5 B7JKD7 B7JKD8 B7JKE1 B7JKE4 B7JKE5 B7JKE6 B7JKE8
B7JKF3 B7JKF4 B7JKG2 B7JKI8 B7JKM7 B7JKN1 B7JKT4 B7JKT5 B7JKT7 B7JKV0 B7JKV4
B7JKV5 B7JKY0 B7JL13 B7JL16 B7JL17 B7JL18 B7JL33 B7JL36 B7JL37 B7JL64 B7JLF2
B7JLG8 B7JLS6 B7JLU2 B7JLW6 B7JLW7 B7JLW9 B7JLX2 B7JLX3 B7JLY5 B7JLZ5 B7JM15
B7JM28 B7JM30 B7JM31 B7JM32 B7JM33 B7JM48 B7JM50 B7JM56 B7JM59 B7JM60 B7JM61
B7JM82 B7JM84 B7JM85 B7JM87 B7JM89 B7JM96 B7JM99 B7JMB1 B7JMB2 B7JMB3 B7JML4
B7JMS8 B7JMU9 B7JMV0 B7JMV1 B7JMV9 B7JN03 B7JN09 B7JN13 B7JN21 B7JN23 B7JN28
B7JN34 B7JN37 B7JN38 B7JN39 B7JN40 B7JN41 B7JN59 B7JN71 B7JN72 B7JNA2 B7JNE9
B7JNF2 B7JNG2 B7JNG6 B7JNT4 B7JNV3 B7JNV4 B7JNW4 B7JNW6 B7JP64 B7JP70 B7JP75
B7JPK2 B7JPM3 B7JPM4 B7JPX2 B7JPX3 B7JPY8 B7JPZ2 B7JPZ4 B7JQ03 B7JQ04 B7JQ14
B7JQ26 B7JQ28 B7JQ30 B7JQ35 B7JQ48 B7JQ54 B7JQ57 B7JQ59 B7JQ62 B7JQ65 B7JQ66
B7JQ74 B7JQF1 B7JQU1 B7JR24 B7JR48 B7JR57 B7JR67 B7JR69 B7JR80 B7JR81 B7JR87
B7JRB1 B7JRU3 B7JRW5 B7JRX2 B7JRY7 B7JS05 B7JS06 B7JS23 B7JS31 B7JS63 B7JS68
B7JS74 B7JSA1 B7JSD8 B7JSE3 B7JSQ5 B7JSV5 B7JSX2 B7JSY7 B7JT07 B7JT28 B7JT32
B7JT33 B7JTN6 B7JUE3 B7JUF5 B7JUH3 B7JUH4 B7JUI4 B7JUJ9 B7JUK2 B7JUP5 B7JUP7
B7JUP8 B7JUU6 B7JUZ9 B7JV16 B7JV38 B7JV66 B7JVC2 B7JVE4 B7JVE5 B7JVF9 B7JVG1
B7JVH4 B7JVH9 B7JVJ6 B7JVL5 B7JVP5 B7JVR6 B7JVR7 B7JVS6 B7JVV5 B7JVZ0 B7JVZ9
B7JW32 B7JW41 B7JW59 B7JW74 B7JW76 B7JWJ7 B7JWN0 B7JWQ8 B7JWS2 B7JWT5 B7JWT6
B7JWT7 B7JWT8 B7JWU1 B7JWV1 B7JWW6 B7JX16 B7JX26 B7JX32 B7JX56 B7JX64 B7JXB4
B7JXM2 B7JXN4 B7JXP0 B7JXS8 B7JY11 B7JY16 B7JY20 B7JY30 B7JY33 B7JY63 B7JY68
B7JYA9 B7JYF1 B7JYG9 B7JYJ9 B7JYK3 B7JYL4 B7JYM6 B7JYM8 B7JYN0 B7JYP4 B7JYV9
B7JYY6 B7JYZ1 B7JZ48 B7JZ83 B7JZA3 B7JZA4 B7JZB1 B7JZB2 B7JZG7 B7JZR4 B7JZS9
B7JZT6 B7JZW9 B7K034 B7K077 B7K079 B7K087 B7K0A2 B7K0A6 B7K0B0 B7K0H0 B7K0I0
B7K0I2 B7K0L9 B7K0S6 B7K0T5 B7K127 B7K165 B7K189 B7K1A8 B7K1E7 B7K1F0 B7K1G5
B7K1H5 B7K1K0 B7K1N1 B7K1R5 B7K1S0 B7K1T1 B7K1U3 B7K1Z3 B7K205 B7K213 B7K219
B7K221 B7K222 B7K223 B7K225 B7K226 B7K227 B7K231 B7K232 B7K233 B7K234 B7K235

B7K236 B7K238 B7K240 B7K241 B7K293 B7K2B2 B7K2B6 B7K2C0 B7K2E7 B7K2I1 B7K2J4
B7K2N3 B7K2S1 B7K318 B7K329 B7K330 B7K331 B7K332 B7K333 B7K334 B7K335 B7K339
B7K379 B7K399 B7K3A9 B7K3F6 B7K3J2 B7K3N8 B7K3R6 B7K400 B7K422 B7K468 B7K4A4
B7K4D0 B7K4F1 B7K4N4 B7K4P4 B7K4S8 B7K4S9 B7K4V4 B7K4V5 B7K4V8 B7K508 B7K5B1
B7K5E6 B7K5G2 B7K5G6 B7K5H5 B7K5I1 B7K5N3 B7K5R5 B7K5U5 B7K5U7 B7K5W1 B7K612
B7K639 B7K660 B7K6D3 B7K6S5 B7K6S6 B7K6Z1 B7K735 B7K739 B7K762 B7K765 B7K7E4
B7K7J4 B7K7P3 B7K7R4 B7K7T8 B7K7Y7 B7K825 B7K833 B7K834 B7K836 B7K837 B7K8F6
B7K8I7 B7K8M6 B7K8N1 B7K8N3 B7K8N5 B7K8P1 B7K8P2 B7K8R0 B7K8T7 B7K8U0 B7K8U6
B7K8X6 B7K906 B7K907 B7K911 B7K912 B7K961 B7K970 B7K991 B7K993 B7K9F0 B7K9I2
B7K9K1 B7K9M6 B7K9Q5 B7KA18 B7KA22 B7KAA8 B7KAD1 B7KAF7 B7KAT5 B7KAZ5 B7KB74
B7KB81 B7KB89 B7KBA7 B7KBE3 B7KBE4 B7KBH7 B7KBS1 B7KBV6 B7KBV9 B7KC04 B7KC08
B7KC96 B7KCB0 B7KCB8 B7KCC4 B7KCF3 B7KCF5 B7KCH0 B7KCI7 B7KCT7 B7KCX9 B7KCY0
B7KD38 B7KD42 B7KD52 B7KD70 B7KDA3 B7KDB6 B7KDB8 B7KDF3 B7KDJ9 B7KDM5 B7KDN8
B7KDQ8 B7KEB4 B7KEG3 B7KEI6 B7KEI7 B7KEJ8 B7KEM9 B7KEN6 B7KEP3 B7KEV7 B7KEW8
B7KF08 B7KF17 B7KF19 B7KF23 B7KF44 B7KF62 B7KFA6 B7KFE2 B7KFE3 B7KFE6 B7KFE8
B7KFF5 B7KFG7 B7KFJ9 B7KFK5 B7KFP0 B7KFR8 B7KFT1 B7KFV3 B7KFZ3 B7KG02 B7KG67
B7KG76 B7KGB7 B7KGB9 B7KGG1 B7KGG9 B7KGN0 B7KGN3 B7KGS0 B7KGT7 B7KGU8 B7KGX7
B7KH06 B7KH28 B7KH96 B7KHC2 B7KHD0 B7KHD2 B7KHE3 B7KHE4 B7KHE5 B7KHE9 B7KHF1
B7KHS7 B7KHU5 B7KHY7 B7KHY8 B7KHZ0 B7KHZ1 B7KHZ3 B7KHZ4 B7KHZ5 B7KHZ7 B7KHZ9
B7KI00 B7KI01 B7KI02 B7KI03 B7KI04 B7KI08 B7KI09 B7KI10 B7KI12 B7KI14 B7KI15
B7KI16 B7KI17 B7KI42 B7KI47 B7KIC1 B7KIC5 B7KID4 B7KIQ3 B7KIQ6 B7KIR2 B7KIR5
B7KIR6 B7KIR7 B7KIT8 B7KIU0 B7KIU2 B7KIU5 B7KJ74 B7KJ88 B7KJG8 B7KJH2 B7KJJ4
B7KJK8 B7KJM5 B7KJM8 B7KJQ2 B7KJX8 B7KJY0 B7KK78 B7KK85 B7KKJ9 B7KKQ8 B7KKT8
B7KKW4 B7KKX4 B7KKZ9 B7KL10 B7KL53 B7KL61 B7KL75 B7KLB5 B7KLD0 B7KLE7 B7KLG3
B7KLH9 B7KLL5 B7KN26 B7KN27 B7KN33 B7KN34 B7KN43 B7KN44 B7KN45 B7KN46 B7KN55
B7KN56 B7KN75 B7KN92 B7KNJ8 B7KNP6 B7KNQ3 B7KNS9 B7KNT0 B7KNT1 B7KNU7 B7KNU8
B7KNX9 B7KPG0 B7KPI6 B7KPK1 B7KPM4 B7KPM8 B7KPN8 B7KPU5 B7KPU6 B7KPV4 B7KPY4
B7KPY5 B7KPZ2 B7KQ28 B7KQF6 B7KQG2 B7KQJ7 B7KQK1 B7KQM2 B7KQP2 B7KR63 B7KRB3
B7KRC9 B7KRF3 B7KRK4 B7KRU8 B7KS47 B7KS48 B7KSC2 B7KSC5 B7KSC6 B7KSH0 B7KSH3
B7KSH4 B7KSJ4 B7KSK7 B7KSZ4 B7KSZ5 B7KT12 B7KT22 B7KT24 B7KT38 B7KT53 B7KTR5
B7KTS1 B7KU77 B7KU78 B7KUF7 B7KUK7 B7KUL9 B7KUS7 B7KUT4 B7KUU9 B7KUV9 B7KV55
B7KV98 B7KVA1 B7KVA2 B7KVA7 B7KVC8 B7KVD0 B7KVD3 B7KVE0 B7KVM8 B7KVN1 B7KVV7
B7KVW2 B7KVW3 B7KVW7 B7KVW8 B7KVX2 B7KW75 B7KW92 B7KW98 B7KWA9 B7KWB0 B7KWG7
B7KWJ1 B7KWN1 B7KWT3 B7KWT4 B7KWT5 B7KWT6 B7KWT7 B7KWY7 B7KWY8 B7KX11 B7KX68
B7KXC6 B7KXC8 B7KXI3 B7KY35 B7KYA2 B7KYC5 B7KYE7 B7KYG6 B7KYJ1 B7KYK4 B7KYZ3
B7KZ04 B7KZ47 B7KZ48 B7KZ56 B7KZ57 B7KZ66 B7KZD5 B7KZG0 B7KZG1 B7KZH3 B7KZL6
B7KZL8 B7KZZ9 B7L060 B7L065 B7L092 B7L097 B7L0A5 B7L0K9 B7L0L2 B7L0Q6 B7L0Q7
B7L0Q8 B7L0R0 B7L0R1 B7L0R2 B7L0R3 B7L0R4 B7L0R5 B7L0R6 B7L0R8 B7L0S0 B7L0S4
B7L0S5 B7L0S7 B7L0S8 B7L0S9 B7L0T1 B7L0T2 B7L0T3 B7L0T4 B7L0T5 B7L0T9 B7L0X9
B7L0Y0 B7L0Y4 B7L1H4 B7L246 B7L253 B7L255 B7L270 B7L273 B7L289 B7L2C1 B7L2C9
B7L2I7 B7L2K9 B7L2N9 B7L2P1 B7L2P2 B7L3Z1 B7L3Z2 B7L3Z4 B7L3Z5 B7L425 B7L427
B7L439 B7L441 B7L442 B7L492 B7L493 B7L4B3 B7L4B7 B7L4C3 B7L4C6 B7L4C8 B7L4D1
B7L4D9 B7L4E5 B7L4E8 B7L4F1 B7L4F2 B7L4F3 B7L4F8 B7L4G0 B7L4G1 B7L4G2 B7L4G4
B7L4G5 B7L4H2 B7L4H3 B7L4H4 B7L4H5 B7L4I0 B7L4I3 B7L4I4 B7L4J2 B7L4J3 B7L4J4

B7L4J5 B7L4J7 B7L4J8 B7L4J9 B7L4K0 B7L4K2 B7L4K3 B7L4K4 B7L4K5 B7L4K6 B7L4K8
B7L4K9 B7L4L0 B7L4L1 B7L4L6 B7L4L7 B7L4L8 B7L4L9 B7L4M1 B7L4M4 B7L4N1 B7L4P2
B7L4R5 B7L4S8 B7L4T1 B7L4T9 B7L4U0 B7L4U4 B7L4U8 B7L4U9 B7L4V4 B7L4W0 B7L4W1
B7L4W3 B7L4Z6 B7L503 B7L504 B7L505 B7L506 B7L507 B7L508 B7L534 B7L542 B7L543
B7L546 B7L548 B7L5B2 B7L5G2 B7L5H5 B7L5J0 B7L5J1 B7L5J3 B7L5J5 B7L5L1 B7L5N1
B7L5N2 B7L5P4 B7L5P5 B7L5S1 B7L5T4 B7L5X8 B7L603 B7L638 B7L639 B7L647 B7L649
B7L650 B7L654 B7L656 B7L657 B7L660 B7L673 B7L675 B7L681 B7L683 B7L6F9 B7L6H6
B7L6H9 B7L6I5 B7L6I7 B7L6I9 B7L6J0 B7L6J2 B7L6K7 B7L6L4 B7L6L6 B7L6L8 B7L6L9
B7L6M0 B7L6M1 B7L6M2 B7L6Q3 B7L6Q9 B7L6T9 B7L6U1 B7L6X1 B7L6X3 B7L6Y0 B7L6Z1
B7L708 B7L723 B7L735 B7L742 B7L745 B7L747 B7L757 B7L758 B7L759 B7L760 B7L761
B7L763 B7L766 B7L771 B7L773 B7L789 B7L794 B7L796 B7L797 B7L799 B7L7A0 B7L7A1
B7L7D0 B7L7D2 B7L7D8 B7L7F3 B7L7F6 B7L7H9 B7L7L9 B7L7M1 B7L7N3 B7L7N7 B7L7R3
B7L7R4 B7L7R6 B7L7R9 B7L7S3 B7L7S4 B7L7S7 B7L7S9 B7L7U5 B7L830 B7L831 B7L836
B7L842 B7L844 B7L846 B7L847 B7L851 B7L852 B7L853 B7L889 B7L890 B7L891 B7L894
B7L895 B7L896 B7L8A6 B7L8B3 B7L8B5 B7L8B6 B7L8C0 B7L8G3 B7L8J6 B7L8J7 B7L8K0
B7L8S5 B7L8T2 B7L8U9 B7L8W4 B7L8W6 B7L8Y4 B7L8Z1 B7L916 B7L959 B7L963 B7L968
B7L969 B7L996 B7L9A5 B7L9B9 B7L9D6 B7L9E7 B7L9F1 B7L9F3 B7L9F7 B7L9F8 B7L9G0
B7L9G1 B7L9H2 B7L9H4 B7L9H5 B7L9I0 B7L9I6 B7L9J7 B7L9K6 B7L9K8 B7L9L4 B7L9L7
B7L9P6 B7L9P8 B7L9Q1 B7L9Q2 B7L9T9 B7L9W7 B7L9X5 B7L9Y6 B7LA09 B7LA13 B7LA21
B7LA23 B7LA26 B7LA28 B7LA29 B7LA33 B7LA39 B7LA54 B7LA56 B7LA57 B7LA60 B7LA64
B7LA65 B7LA67 B7LA72 B7LA74 B7LA77 B7LA78 B7LA79 B7LA80 B7LA84 B7LA85 B7LA88
B7LA89 B7LA91 B7LA92 B7LAB9 B7LAC0 B7LAD4 B7LAF1 B7LAF6 B7LAF8 B7LAH4 B7LAI2
B7LAI5 B7LAJ5 B7LAK9 B7LAL0 B7LAL1 B7LAN4 B7LAP9 B7LAR4 B7LAR6 B7LAS7 B7LAS9
B7LAT0 B7LAT4 B7LAV1 B7LAV3 B7LAV8 B7LAW3 B7LAX0 B7LAY7 B7LB75 B7LBE5 B7LBE8
B7LBH2 B7LBH4 B7LBI3 B7LBI5 B7LBR9 B7LBS7 B7LBZ6 B7LC01 B7LC02 B7LC03 B7LC16
B7LC22 B7LC24 B7LC30 B7LC31 B7LC36 B7LC37 B7LC45 B7LC52 B7LC57 B7LC58 B7LC61
B7LC63 B7LC65 B7LC87 B7LC90 B7LC95 B7LCB8 B7LCE0 B7LCF6 B7LCG3 B7LCH1 B7LCH2
B7LCI0 B7LCI4 B7LCK7 B7LCL4 B7LCL6 B7LCN6 B7LCN8 B7LCN9 B7LCP7 B7LCP9 B7LCQ1
B7LCQ4 B7LCQ5 B7LCQ6 B7LCQ7 B7LCQ8 B7LCR2 B7LCR3 B7LCR4 B7LCS2 B7LCT1 B7LCU1
B7LCV3 B7LCV5 B7LCW4 B7LCX0 B7LD45 B7LD52 B7LD59 B7LD66 B7LD74 B7LD77 B7LD85
B7LD99 B7LDA0 B7LDA2 B7LDA4 B7LDA7 B7LDA9 B7LDB0 B7LDB5 B7LDB8 B7LDB9 B7LDC0
B7LDC2 B7LDC3 B7LDD1 B7LDD3 B7LDD4 B7LDE3 B7LDG0 B7LDG8 B7LDH2 B7LDH3 B7LDJ5
B7LDJ6 B7LDJ7 B7LDJ8 B7LDK2 B7LDK3 B7LDK6 B7LDK8 B7LDM7 B7LDX2 B7LDX6 B7LDY6
B7LDZ6 B7LE15 B7LE21 B7LE23 B7LE31 B7LE35 B7LE56 B7LE58 B7LE63 B7LE66 B7LE91
B7LEA4 B7LEA7 B7LEB0 B7LEB1 B7LEC2 B7LEE7 B7LEG1 B7LEG2 B7LEG3 B7LEG4 B7LEG5
B7LEG8 B7LEG9 B7LEH0 B7LEI1 B7LEI2 B7LEK0 B7LEK9 B7LEL6 B7LEM0 B7LEM7 B7LEM8
B7LEM9 B7LEN0 B7LEN2 B7LEN9 B7LEP0 B7LEP1 B7LEX4 B7LEX6 B7LEY2 B7LEY3 B7LEY4
B7LF04 B7LF07 B7LF08 B7LF20 B7LF59 B7LF76 B7LF83 B7LF85 B7LF89 B7LF90 B7LF91
B7LFA7 B7LFA8 B7LFA9 B7LFB3 B7LFB7 B7LFB8 B7LFB9 B7LFC0 B7LFC1 B7LFC2 B7LFE4
B7LFG6 B7LFG8 B7LFH2 B7LFH3 B7LFI3 B7LFJ6 B7LFK4 B7LFK7 B7LFL1 B7LFL2 B7LFL6
B7LFM5 B7LFU5 B7LFV1 B7LFV2 B7LFW1 B7LFX0 B7LFX2 B7LFY6 B7LFY7 B7LFY8 B7LFZ3
B7LFZ6 B7LG01 B7LG22 B7LG31 B7LG40 B7LG41 B7LG53 B7LGI1 B7LGI7 B7LGJ3 B7LGJ5
B7LGJ6 B7LGK8 B7LGL6 B7LGL8 B7LGM1 B7LGM2 B7LGM5 B7LGM7 B7LGN0 B7LGN1 B7LGN3
B7LGP5 B7LGQ2 B7LGR5 B7LGT0 B7LGT2 B7LGT4 B7LGT7 B7LGU9 B7LGW4 B7LGW9 B7LGX1

B7LGX2 B7LGY3 B7LGY7 B7LGZ2 B7LGZ9 B7LH01 B7LH04 B7LH08 B7LH20 B7LH44 B7LH62
B7LH67 B7LH72 B7LH86 B7LH87 B7LH88 B7LH93 B7LH99 B7LHA4 B7LHB5 B7LHB8 B7LHC0
B7LHF3 B7LHH6 B7LHJ3 B7LHN2 B7LHN3 B7LHN5 B7LHN6 B7LHN9 B7LHP2 B7LHP5 B7LHP6
B7LHP8 B7LHP9 B7LHQ9 B7LHT0 B7LHT2 B7LHT3 B7LHT7 B7LHT8 B7LHT9 B7LHU4 B7LHW6
B7LHW8 B7LHX9 B7LHY3 B7LHY6 B7LHY8 B7LHZ0 B7LHZ6 B7LHZ8 B7LHZ9 B7LI02 B7LI03
B7LI04 B7LI05 B7LJH5 B7LJI4 B7LJI7 B7LJQ3 B7LJR2 B7LJS7 B7LJS9 B7LJT0 B7LJT2
B7LJV2 B7LJV5 B7LJX9 B7LJY1 B7LJY3 B7LJY7 B7LJY8 B7LJZ2 B7LK02 B7LK04 B7LK09
B7LK20 B7LK29 B7LK34 B7LK42 B7LK45 B7LK46 B7LK49 B7LK50 B7LK85 B7LK89 B7LK90
B7LK93 B7LKA1 B7LKA4 B7LKA9 B7LKB1 B7LKB2 B7LKB3 B7LKB6 B7LKC0 B7LKC1 B7LKC3
B7LKC4 B7LKC7 B7LKD2 B7LKD4 B7LKE3 B7LKE4 B7LKE6 B7LKG3 B7LKG5 B7LKH0 B7LKJ3
B7LKJ9 B7LKK7 B7LKK8 B7LKL5 B7LKQ5 B7LKQ8 B7LKT3 B7LKT5 B7LKU3 B7LKU5 B7LKV4
B7LKZ8 B7LL66 B7LL73 B7LL88 B7LLD9 B7LLE1 B7LLF0 B7LLF2 B7LLH1 B7LLH4 B7LLH6
B7LLI1 B7LLI3 B7LLN2 B7LLP8 B7LLP9 B7LLQ1 B7LLS4 B7LLS5 B7LLS9 B7LLT9 B7LLU5
B7LLV2 B7LLV3 B7LLV4 B7LLV9 B7LLW0 B7LLX4 B7LLX8 B7LLX9 B7LLY0 B7LLY2 B7LLY3
B7LLY4 B7LLY5 B7LLZ8 B7LM66 B7LM67 B7LM69 B7LM80 B7LM95 B7LMB2 B7LMB6 B7LMD3
B7LMD5 B7LME1 B7LME3 B7LMG1 B7LMG4 B7LMG5 B7LMG7 B7LMH1 B7LMH2 B7LMH4 B7LMI1
B7LMI2 B7LMJ3 B7LMJ4 B7LMJ7 B7LMJ8 B7LMP0 B7LMQ3 B7LMR7 B7LMR8 B7LMS3 B7LMT2
B7LMT6 B7LMT7 B7LMU4 B7LMV1 B7LMV6 B7LMX3 B7LMZ9 B7LN16 B7LN23 B7LN30 B7LN38
B7LN49 B7LN53 B7LN61 B7LN70 B7LN72 B7LN74 B7LN79 B7LN84 B7LN86 B7LNF7 B7LNF8
B7LNG0 B7LNG1 B7LNI2 B7LNI5 B7LNK6 B7LNS1 B7LNS2 B7LNS3 B7LNS4 B7LNS6 B7LNT5
B7LNT7 B7LNU8 B7LNV2 B7LNW6 B7LNW8 B7LNX3 B7LNX6 B7LP01 B7LP19 B7LP27 B7LP46
B7LPA4 B7LPB2 B7LPB4 B7LPB7 B7LPB8 B7LPB9 B7LPC3 B7LPC8 B7LPC9 B7LPD5 B7LPE7
B7LPF5 B7LPH0 B7LPH3 B7LPH4 B7LPH8 B7LPI1 B7LPI3 B7LPI4 B7LPL0 B7LPL4 B7LPN4
B7LPN6 B7LPQ4 B7LPQ9 B7LPR1 B7LPR5 B7LPR6 B7LPS0 B7LPT2 B7LQ18 B7LQ20 B7LQ21
B7LQ44 B7LQ45 B7LQ46 B7LQ47 B7LQ48 B7LQ50 B7LQ52 B7LQ59 B7LQ71 B7LQ73 B7LQ74
B7LQ75 B7LQ77 B7LQ82 B7LQ85 B7LQ96 B7LQ97 B7LQA9 B7LQC3 B7LQC9 B7LQD4 B7LQD8
B7LQD9 B7LQE3 B7LQF8 B7LQG3 B7LQN0 B7LQN2 B7LQN3 B7LQQ4 B7LQX1 B7LR07 B7LR08
B7LR33 B7LR34 B7LR36 B7LR37 B7LR39 B7LR40 B7LR43 B7LR46 B7LR50 B7LR52 B7LR53
B7LR56 B7LR72 B7LR95 B7LRD0 B7LRD2 B7LRJ0 B7LRJ1 B7LRJ3 B7LRJ8 B7LRJ9 B7LRL0
B7LRL2 B7LRN4 B7LRN6 B7LRQ1 B7LRQ3 B7LRQ4 B7LRQ5 B7LRR1 B7LRR3 B7LRR4 B7LRR7
B7LRR8 B7LRS0 B7LRS1 B7LRS2 B7LRS3 B7LRS4 B7LRS5 B7LRS6 B7LRS7 B7LRS8 B7LRS9
B7LRT0 B7LRT1 B7LRT2 B7LRT3 B7LRT5 B7LRU9 B7LRZ1 B7LRZ3 B7LRZ7 B7LS01 B7LS19
B7LS20 B7LS30 B7LS40 B7LS41 B7LS46 B7LS47 B7LS48 B7LS49 B7LS50 B7LS51 B7LS54
B7LS61 B7LS75 B7LS90 B7LS91 B7LSA2 B7LSA4 B7LSB3 B7LSB5 B7LSB7 B7LSC1 B7LSC2
B7LSC7 B7LSE0 B7LSE1 B7LSE2 B7LSF2 B7LSH9 B7LSI0 B7LSI2 B7LSI6 B7LSJ5 B7LSK2
B7LSK4 B7LSK7 B7LSN5 B7LSP9 B7LST2 B7LSV6 B7LSX8 B7LSZ0 B7LT11 B7LT32 B7LT40
B7LT49 B7LT70 B7LT74 B7LT77 B7LT80 B7LT90 B7LTA1 B7LTC6 B7LTH3 B7LTH4 B7LTH9
B7LTI6 B7LTJ8 B7LTL6 B7LTN9 B7LTQ3 B7LTW5 B7LTX5 B7LTX6 B7LTX8 B7LTY8 B7LTZ5
B7LU01 B7LU25 B7LU45 B7LU47 B7LU51 B7LU53 B7LU75 B7LU76 B7LU79 B7LU82 B7LUF0
B7LUF2 B7LUF5 B7LUF6 B7LUJ5 B7LUJ6 B7LUK4 B7LUK5 B7LUK6 B7LUK7 B7LUK8 B7LUL2
B7LUL5 B7LUL6 B7LUL7 B7LUL8 B7LUL9 B7LUM3 B7LUM6 B7LUM8 B7LUN3 B7LUN5 B7LUN8
B7LUN9 B7LUR5 B7LUR8 B7LUS2 B7LUS3 B7LUS5 B7LUS6 B7LUS9 B7LUV4 B7LUV6 B7LUW4
B7LUW5 B7LUW6 B7LUW7 B7LUY5 B7LUY6 B7LUY9 B7LUZ7 B7LUZ9 B7LV13 B7LV14 B7LV29
B7LV30 B7LV66 B7LV75 B7LVA3 B7LVB3 B7LVB9 B7LVC3 B7LVC6 B7LVC9 B7LVD8 B7LVD9

B7LVE0 B7LVE1 B7LVE3 B7LVF3 B7LVF7 B7LVG2 B7LVH6 B7LVH8 B7LVJ5 B7LVJ6 B7LVJ7
B7LVJ8 B7LVJ9 B7LVK1 B7LVK2 B7LVK4 B7LVL1 B7LVN2 B7LVN5 B7LVN6 B7LVP5 B7LVP7
B7LVQ1 B7LVR1 B7LVS5 B7LVT0 B7LVT3 B7LVT4 B7LVU3 B7LVU4 B7LVU5 B7LVW4 B7LVW5
B7LVW9 B7LVY1 B7LVY2 B7LVY5 B7LVZ1 B7LW06 B7LW23 B7LW30 B7LW31 B7LW33 B7LW38
B7LW40 B7LW41 B7LW53 B7LW62 B7LW69 B7LW79 B7LW84 B7LW87 B7LW89 B7LW94 B7LW98
B7LWA5 B7LWA6 B7LWA8 B7LWA9 B7LWB1 B7LWB5 B7LWB7 B7LWC0 B7LWC1 B7LWC4 B7LWD9
B7LWE0 B7LWE2 B7LWF7 B7LWG6 B7LWG7 B7LWG9 B7LWH0 B7LWH8 B7LWJ1 B7LWJ2 B7LWK3
B7LWK4 B7LWK5 B7LWK8 B7LWK9 B7LWL0 B7LWM0 B7LWM1 B7LWM6 B7LWM8 B7LWM9 B7LWN0
B7LWN2 B7LWN3 B7LWN9 B7LWP0 B7LWP4 B7LWQ6 B7LWR1 B7LWR4 B7LX23 B7LX32 B7LX41
B7LX42 B7LX53 B7LX85 B7LX87 B7LX89 B7LX92 B7LXA3 B7LXB9 B7LXC3 B7LXC5 B7LXC6
B7LXE0 B7LXF5 B7LXF6 B7LXF7 B7LXF8 B7LXF9 B7LXG2 B7LXG3 B7LXG4 B7LXH5 B7LXH6
B7LXJ6 B7LXL0 B7LXL4 B7LXL8 B7LXM0 B7LXM2 B7LXN0 B7LXP0 B7LXS5 B7LXT2 B7LXT6
B7LXU3 B7LXU4 B7LXU5 B7LXU6 B7LXU8 B7LXV6 B7LXV9 B7LXY9 B7LY03 B7LY16 B7LY17
B7LY36 B7LY40 B7LY47 B7LY83 B7LY86 B7LY87 B7LY99 B7LYD8 B7LYF3 B7LYF5 B7LYG2
B7LYG4 B7LYG7 B7LYG8 B7LYG9 B7LYH3 B7LYL5 B7LYT1 B7LYT3 B7LYU4 B7LYW6 B7LYX2
B7LYX4 B7LYX8 B7LYX9 B7LYY3 B7LYZ2 B7LZ33 B7LZ73 B7LZC1 B7LZC3 B7LZJ1 B7LZJ8
B7LZK1 B7LZK6 B7LZL4 B7LZL5 B7LZL8 B7LZM2 B7LZZ6 B7M017 B7M032 B7M038 B7M045
B7M056 B7M057 B7M062 B7M063 B7M072 B7M073 B7M075 B7M076 B7M078 B7M079 B7M082
B7M085 B7M089 B7M091 B7M092 B7M095 B7M0A0 B7M0A3 B7M0B1 B7M0B9 B7M0C2 B7M0C5
B7M0C6 B7M0C7 B7M0D2 B7M0D4 B7M0D5 B7M0D6 B7M0D8 B7M0D9 B7M0E6 B7M0E7 B7M0E8
B7M0E9 B7M0F4 B7M0F7 B7M0F8 B7M0G6 B7M0H9 B7M0J4 B7M0J5 B7M0J7 B7M0J9 B7M0L5
B7M0N4 B7M0N5 B7M0P8 B7M0P9 B7M0S3 B7M0T5 B7M0T7 B7M0T8 B7M0U2 B7M0U3 B7M0U8
B7M0U9 B7M0X1 B7M0X3 B7M0Y7 B7M0Z0 B7M0Z2 B7M0Z3 B7M0Z4 B7M100 B7M102 B7M103
B7M106 B7M107 B7M116 B7M117 B7M119 B7M124 B7M125 B7M134 B7M143 B7M145 B7M160
B7M161 B7M162 B7M172 B7M174 B7M175 B7M187 B7M195 B7M197 B7M1A0 B7M1A1 B7M1A4
B7M1A5 B7M1A6 B7M1A9 B7M1B0 B7M1B1 B7M1C1 B7M1C3 B7M1C4 B7M1C5 B7M1C7 B7M1E5
B7M1F2 B7M1F4 B7M1F6 B7M1F7 B7M1F8 B7M1F9 B7M1G0 B7M1J2 B7M1J8 B7M1L8 B7M1L9
B7M1M0 B7M1M1 B7M1M2 B7M1M3 B7M1M4 B7M1M5 B7M1M6 B7M1M7 B7M1M8 B7M1M9 B7M1N0
B7M1N1 B7M1N3 B7M1N4 B7M1N5 B7M1P1 B7M1P2 B7M1P3 B7M1P4 B7M1P5 B7M1P6 B7M1P9
B7M1Q6 B7M1S1 B7M1U1 B7M1U2 B7M1V3 B7M1V7 B7M1W6 B7M1W8 B7M1X0 B7M1X4 B7M1Y6
B7M1Z3 B7M1Z9 B7M208 B7M211 B7M213 B7M252 B7M268 B7M269 B7M271 B7M272 B7M285
B7M287 B7M2A7 B7M2B1 B7M2F1 B7M2F2 B7M2F4 B7M2F7 B7M2G1 B7M2G2 B7M2G5 B7M2G7
B7M2H9 B7M2I0 B7M2I5 B7M2J2 B7M2J3 B7M2J4 B7M2N3 B7M2P4 B7M2Q7 B7M2U3 B7M2V5
B7M2V6 B7M2V7 B7M2Y9 B7M2Z5 B7M2Z6 B7M2Z7 B7M2Z8 B7M2Z9 B7M300 B7M332 B7M333
B7M340 B7M356 B7M363 B7M377 B7M392 B7M394 B7M3A5 B7M3B3 B7M3C7 B7M3D0 B7M3I1
B7M3I2 B7M3I8 B7M3J9 B7M3L6 B7M3M7 B7M3N0 B7M3N2 B7M3P4 B7M3P5 B7M3Q2 B7M3Q4
B7M3Q5 B7M3Q9 B7M3R1 B7M3R2 B7M3R5 B7M3S9 B7M3T1 B7M3T7 B7M3T9 B7M3W1 B7M3W3
B7M3W4 B7M3W6 B7M3W7 B7M3W8 B7M3Z8 B7M400 B7M403 B7M404 B7M449 B7M476 B7M477
B7M485 B7M496 B7M4A3 B7M4A5 B7M4B8 B7M4B9 B7M4C0 B7M4C1 B7M4C3 B7M4C5 B7M4C6
B7M4C7 B7M4D2 B7M4D4 B7M4F2 B7M4H5 B7M4H6 B7M4I1 B7M4I9 B7M4J1 B7M4K7 B7M4K9
B7M4L5 B7M4M9 B7M4N2 B7M4R1 B7M4U5 B7M4U6 B7M4U9 B7M4V9 B7M4Y8 B7M4Z9 B7M507
B7M510 B7M521 B7M535 B7M536 B7M537 B7M553 B7M555 B7M556 B7M559 B7M560 B7M596
B7M597 B7M5A0 B7M5A1 B7M5A2 B7M5A4 B7M5B2 B7M5B9 B7M5C2 B7M5C6 B7M5C7 B7M5F5
B7M5F7 B7M5F8 B7M5G3 B7M5G6 B7M5G9 B7M5H8 B7M5I6 B7M5I8 B7M5J6 B7M5J8 B7M5M7

B7M5M8 B7M5P1 B7M5Q1 B7M5R7 B7M5T3 B7M5U4 B7M5U6 B7M5U7 B7M5V1 B7M5X0 B7M5X3
B7M603 B7M605 B7M611 B7M613 B7M634 B7M638 B7M650 B7M664 B7M686 B7M693 B7M697
B7M6B3 B7M6B8 B7M6C0 B7M6D0 B7M6J9 B7M6K1 B7M6L0 B7M6L2 B7M6N5 B7M6P3 B7M6Q8
B7M6S1 B7M6S8 B7M6T6 B7M6T7 B7M6U5 B7M6V3 B7M6V4 B7M6V5 B7M6V6 B7M6V7 B7M6W7
B7M6X0 B7M6X7 B7M6X9 B7M6Y0 B7M6Y2 B7M6Y3 B7M6Y7 B7M6Z3 B7M710 B7M711 B7M713
B7M714 B7M718 B7M721 B7M723 B7M726 B7M730 B7M731 B7M732 B7M733 B7M734 B7M738
B7M747 B7M748 B7M751 B7M753 B7M755 B7M779 B7M782 B7M787 B7M7B0 B7M7C7 B7M7D4
B7M7E1 B7M7E9 B7M7H2 B7M7H9 B7M7I1 B7M7K0 B7M7K2 B7M7K3 B7M7L1 B7M7L3 B7M7L5
B7M7L8 B7M7L9 B7M7M1 B7M7M2 B7M7M6 B7M7M9 B7M7N0 B7M7N1 B7M7N3 B7M7N4 B7M7P2
B7M7P4 B7M7Q3 B7M7Q4 B7M7Q5 B7M7Q6 B7M7Q7 B7M7R5 B7M7R7 B7M7S1 B7M7S7 B7M7T4
B7M7V2 B7M801 B7M809 B7M813 B7M821 B7M835 B7M836 B7M839 B7M841 B7M846 B7M851
B7M853 B7M860 B7M864 B7M885 B7M887 B7M892 B7M895 B7M8A7 B7M8H8 B7M8I0 B7M8I9
B7M8J4 B7M8J5 B7M8P8 B7M8Q3 B7M8Q4 B7M8R0 B7M8S0 B7M8S4 B7M8T0 B7M8T1 B7M8T2
B7M8T7 B7M8T8 B7M8V1 B7M8V2 B7M8V6 B7M8Y1 B7M8Y2 B7M8Y3 B7M8Y8 B7M8Z2 B7M8Z3
B7M8Z4 B7M8Z5 B7M8Z6 B7M8Z7 B7M908 B7M930 B7M934 B7M937 B7M942 B7M975 B7M976
B7M977 B7M978 B7M983 B7M984 B7M987 B7M989 B7M9A2 B7M9B7 B7M9D0 B7M9D3 B7M9D5
B7M9D6 B7M9E8 B7M9F9 B7M9G0 B7M9G2 B7M9G3 B7M9G4 B7M9G5 B7M9H2 B7M9I1 B7M9J4
B7M9K4 B7M9K5 B7M9L3 B7M9L9 B7M9R6 B7M9R9 B7M9W3 B7M9X6 B7M9Z1 B7M9Z2 B7M9Z4
B7M9Z6 B7MA12 B7MA31 B7MA32 B7MAC3 B7MAC5 B7MAC8 B7MAD3 B7MAD6 B7MAE3 B7MAE6
B7MAE9 B7MAF0 B7MAF2 B7MAF8 B7MAG0 B7MAG1 B7MAG2 B7MAG4 B7MAG5 B7MAH4 B7MAH5
B7MAH6 B7MAI2 B7MAI5 B7MAI6 B7MAJ6 B7MAJ7 B7MAJ9 B7MAK4 B7MAK5 B7MAL4 B7MAM3
B7MAM4 B7MAM6 B7MAQ2 B7MAQ3 B7MAR4 B7MAR7 B7MAS3 B7MAS5 B7MAS6 B7MAS7 B7MAS9
B7MAU4 B7MAV1 B7MAV3 B7MAV5 B7MAV6 B7MAV7 B7MAV8 B7MAV9 B7MAY9 B7MB00 B7MB01
B7MB04 B7MB08 B7MB25 B7MB40 B7MB51 B7MB55 B7MB62 B7MB69 B7MB70 B7MB75 B7MB76
B7MB85 B7MB86 B7MB88 B7MB89 B7MB91 B7MB92 B7MB95 B7MBA2 B7MBA3 B7MBA4 B7MBB4
B7MBB6 B7MBB7 B7MBC9 B7MBD7 B7MBD9 B7MBE2 B7MBE3 B7MBE6 B7MBE7 B7MBE8 B7MBF0
B7MBF1 B7MBF3 B7MBG4 B7MBH1 B7MBH6 B7MBI5 B7MBI8 B7MBJ0 B7MBJ2 B7MBL7 B7MBL9
B7MBN8 B7MBP2 B7MBR9 B7MBS0 B7MBS2 B7MBS5 B7MBS9 B7MBT0 B7MBT3 B7MBT5 B7MBU8
B7MBV2 B7MBV4 B7MBV5 B7MBV8 B7MBX4 B7MBY4 B7MBY5 B7MBY7 B7MBY8 B7MBZ1 B7MBZ2
B7MBZ7 B7MBZ8 B7MC29 B7MC31 B7MC45 B7MC74 B7MC89 B7MC90 B7MC92 B7MC93 B7MCA6
B7MCD0 B7MCD1 B7MCD2 B7MCF4 B7MCH0 B7MCH6 B7MCJ3 B7MCK8 B7MCL0 B7MCM0 B7MCM7
B7MCQ0 B7MCQ2 B7MCQ3 B7MCQ4 B7MCR0 B7MCR2 B7MCR3 B7MCR6 B7MCR7 B7MCR9 B7MCS0
B7MCS1 B7MCS2 B7MCS3 B7MCS4 B7MCS5 B7MCS6 B7MCS7 B7MCS8 B7MCS9 B7MCT0 B7MCT1
B7MCT2 B7MCT4 B7MCT5 B7MCT6 B7MCV5 B7MCV6 B7MCV7 B7MCV8 B7MCV9 B7MCW0 B7MCW3
B7MCX0 B7MCY5 B7MD00 B7MD01 B7MD04 B7MD41 B7MD46 B7MD47 B7MD50 B7MD52 B7MD63
B7MD64 B7MD71 B7MD73 B7MD74 B7MD78 B7MD80 B7MD81 B7MD84 B7MD95 B7MD97 B7MDA3
B7MDA5 B7MDB7 B7MDH3 B7MDH5 B7MDH8 B7MDH9 B7MDM6 B7MDM9 B7MDN6 B7MDN8 B7MDP0
B7MDP4 B7MDP5 B7MDQ1 B7MDR4 B7MDR5 B7MDR6 B7MDW0 B7MDX1 B7MDZ1 B7MDZ3 B7MDZ4
B7MDZ6 B7MDZ7 B7MDZ8 B7ME26 B7ME28 B7ME34 B7ME49 B7ME52 B7ME78 B7MEF0 B7MEF1
B7MEF9 B7MEG9 B7MEJ2 B7MEM1 B7MER4 B7MER5 B7MES1 B7MEZ2 B7MF26 B7MF27 B7MF30
B7MF55 B7MF66 B7MF74 B7MF76 B7MF86 B7MFA0 B7MFA1 B7MFA2 B7MFC3 B7MFE1 B7MFF5
B7MFG0 B7MFH3 B7MFI0 B7MFI2 B7MFJ5 B7MFJ6 B7MFJ7 B7MFJ8 B7MFJ9 B7MFK1 B7MFK2
B7MFK3 B7MFK8 B7MFL0 B7MFN5 B7MFQ1 B7MFQ3 B7MFQ4 B7MFQ9 B7MFR2 B7MFR5 B7MFR7
B7MFS5 B7MFS7 B7MFT4 B7MFU7 B7MFX2 B7MFX3 B7MFZ8 B7MG11 B7MG14 B7MG18 B7MG29

B7MG31 B7MG32 B7MG36 B7MG55 B7MG58 B7MG82 B7MG84 B7MG93 B7MG95 B7MGA7 B7MGA8
B7MGB3 B7MGC1 B7MGC3 B7MGC4 B7MGC5 B7MGC8 B7MGC9 B7MGG0 B7MGG1 B7MGG4 B7MGG5
B7MGG6 B7MGG8 B7MGH6 B7MGI3 B7MGI6 B7MGI9 B7MGJ1 B7MGK7 B7MGL2 B7MGL4 B7MGM4
B7MGN3 B7MGN4 B7MGN7 B7MGN9 B7MGP1 B7MGR2 B7MGR5 B7MGS0 B7MGU2 B7MH26 B7MH34
B7MH48 B7MH66 B7MH68 B7MH73 B7MH75 B7MH94 B7MHC0 B7MHD2 B7MHE5 B7MHF6 B7MHG3
B7MHH1 B7MHH8 B7MHI7 B7MHJ7 B7MHK5 B7MHL6 B7MHL7 B7MHL9 B7MHM1 B7MHM6 B7MHN1
B7MHN3 B7MHP0 B7MHP4 B7MHR5 B7MHS2 B7MHT0 B7MHT7 B7MHU1 B7MHW4 B7MHW9 B7MHX7
B7MHX9 B7MHY0 B7MHY8 B7MHZ0 B7MHZ6 B7MHZ9 B7MI00 B7MI02 B7MI03 B7MI07 B7MI18
B7MI25 B7MI29 B7MI33 B7MI34 B7MI35 B7MI36 B7MI37 B7MI48 B7MI51 B7MI58 B7MI60
B7MI61 B7MI63 B7MI64 B7MI68 B7MI92 B7MI93 B7MI95 B7MI96 B7MIA0 B7MIA5 B7MIA7
B7MIA9 B7MIB2 B7MIB7 B7MIC0 B7MIE4 B7MIE5 B7MIE6 B7MIE9 B7MIF4 B7MIF5 B7MIF6
B7MIF7 B7MIF8 B7MIF9 B7MIH4 B7MIJ6 B7MIK0 B7MIK3 B7MIK8 B7MIL6 B7MIL7 B7MIL8
B7MIM0 B7MIM1 B7MIN5 B7MIQ0 B7MIQ2 B7MIR0 B7MIR4 B7MIR5 B7MIU4 B7MIU5 B7MIU6
B7MIU7 B7MIV1 B7MIV2 B7MIV5 B7MIV7 B7MIW5 B7MIW9 B7MIX0 B7MIX1 B7MIX2 B7MIX3
B7MIX6 B7MIY0 B7MIY1 B7MIY2 B7MIY3 B7MIY4 B7MIZ2 B7MIZ5 B7MIZ8 B7MJ04 B7MJ15
B7MJ29 B7MJ76 B7MJ85 B7MJ93 B7MJ94 B7MJ95 B7MJA6 B7MJB5 B7MJS0 B7MK69 B7MK71
B7MK73 B7MK76 B7MK86 B7MKA6 B7MKB0 B7MKB2 B7MKB3 B7MKE7 B7MKG2 B7MKG5 B7MKG7
B7MKG8 B7MKI0 B7MKK5 B7MKL7 B7MKL8 B7MKL9 B7MKM0 B7MKM1 B7MKM4 B7MKM5 B7MKM6
B7MKM9 B7MKN0 B7MKU2 B7MKU7 B7MKU8 B7MKV2 B7MKW2 B7MKW8 B7MKX4 B7MKX5 B7MKX6
B7MKY1 B7MKY5 B7ML00 B7ML76 B7ML77 B7MLA1 B7MLB5 B7MLB9 B7MLC3 B7MLC5 B7MLC7
B7MLH3 B7MLH6 B7MLH7 B7MLI3 B7MLJ6 B7MLJ7 B7MLK1 B7MLK2 B7MLK3 B7MLK5 B7MLK6
B7MLK7 B7MLK8 B7MLM2 B7MLN1 B7MLP2 B7MLQ2 B7MLQ3 B7MLQ9 B7MLR5 B7MLV3 B7MLV7
B7MLW3 B7MM21 B7MM49 B7MM60 B7MM76 B7MM78 B7MM85 B7MM87 B7MM89 B7MM90 B7MM91
B7MM95 B7MM99 B7MMA1 B7MMA9 B7MMC5 B7MMD2 B7MMD6 B7MMD7 B7MME1 B7MMH5 B7MMH6
B7MMH7 B7MMJ4 B7MML2 B7MML7 B7MMM2 B7MMS8 B7MMW1 B7MMY3 B7MMY5 B7MN09 B7MNB4
B7MNC1 B7MNI8 B7MNI9 B7MNJ0 B7MNJ1 B7MNJ3 B7MNK2 B7MNK3 B7MNK4 B7MNL4 B7MNM2
B7MNM8 B7MNN1 B7MNN4 B7MNN5 B7MNN7 B7MNP2 B7MNP4 B7MNP5 B7MNP6 B7MNP8 B7MNP9
B7MNQ8 B7MNQ9 B7MNR0 B7MNR1 B7MNR6 B7MNR9 B7MNS0 B7MNS9 B7MNT0 B7MNT2 B7MNU0
B7MNU1 B7MNV0 B7MNV9 B7MNW1 B7MNY1 B7MNY2 B7MNY3 B7MNZ3 B7MNZ5 B7MNZ6 B7MP08
B7MP16 B7MP18 B7MP21 B7MP22 B7MP25 B7MP27 B7MP29 B7MP30 B7MP32 B7MP43 B7MP50
B7MP55 B7MP64 B7MP71 B7MPB4 B7MPB5 B7MPB6 B7MPB7 B7MPB8 B7MPB9 B7MPE3 B7MPE8
B7MPE9 B7MPF2 B7MPF4 B7MPG6 B7MPG7 B7MPH5 B7MPH7 B7MPI3 B7MPI5 B7MPK9 B7MPL0
B7MPN4 B7MPN9 B7MPP1 B7MPQ2 B7MPT0 B7MPT1 B7MPT2 B7MQ18 B7MQ21 B7MQ23 B7MQ59
B7MQ75 B7MQ76 B7MQ78 B7MQ79 B7MQC8 B7MQD0 B7MQD1 B7MQD5 B7MQD7 B7MQD8 B7MQE1
B7MQF2 B7MQF4 B7MQG0 B7MQG2 B7MQI4 B7MQI6 B7MQI7 B7MQI9 B7MQJ0 B7MQJ1 B7MQK2
B7MQL5 B7MQL7 B7MQM2 B7MQM8 B7MQM9 B7MQN2 B7MQN5 B7MQN7 B7MQQ8 B7MQR1 B7MQR6
B7MQT8 B7MQV5 B7MQW3 B7MQX0 B7MQX7 B7MQY5 B7MQY8 B7MQZ1 B7MQZ3 B7MQZ4 B7MR01
B7MR18 B7MR21 B7MR29 B7MR48 B7MR51 B7MR55 B7MR65 B7MR70 B7MR71 B7MR83 B7MR93
B7MR95 B7MRA9 B7MRB2 B7MRB3 B7MRC0 B7MRC1 B7MRD0 B7MRD7 B7MRE6 B7MRF7 B7MRI9
B7MRJ2 B7MRL6 B7MRQ1 B7MRQ2 B7MRQ5 B7MRR5 B7MRR7 B7MRR8 B7MRS3 B7MRS9 B7MRT1
B7MRT8 B7MRU8 B7MRV6 B7MS22 B7MS23 B7MS25 B7MS27 B7MS35 B7MS40 B7MS42 B7MS49
B7MS53 B7MS67 B7MS69 B7MS74 B7MS77 B7MSF8 B7MSG3 B7MSG8 B7MSJ0 B7MSJ5 B7MSJ6
B7MSL0 B7MSL1 B7MSS8 B7MST0 B7MST2 B7MSU8 B7MSV9 B7MSW9 B7MSX2 B7MSX6 B7MSX9
B7MSY3 B7MSY4 B7MSY5 B7MSZ3 B7MT10 B7MT75 B7MT77 B7MT80 B7MTA1 B7MTB6 B7MTC0

B7MTC3 B7MTC8 B7MTC9 B7MTD2 B7MTE3 B7MTF1 B7MTF2 B7MTF3 B7MTF4 B7MTF5 B7MTG5
B7MTI7 B7MTJ0 B7MTJ3 B7MTJ8 B7MTL9 B7MTM8 B7MTN6 B7MTN7 B7MTN8 B7MTP9 B7MTQ8
B7MTV0 B7MTV2 B7MTV4 B7MTV7 B7MTW7 B7MTY8 B7MTZ2 B7MTZ4 B7MTZ5 B7MU19 B7MU35
B7MU99 B7MUA0 B7MUB9 B7MUC3 B7MUI7 B7MUN0 B7MUR3 B7MV59 B7MV86 B7MVA0 B7MVB5
B7MVB6 B7MVB8 B7MVC0 B7MVD4 B7MVF5 B7MVG8 B7MVG9 B7MVI2 B7MVI5 B7MVJ1 B7MVJ3
B7MVJ4 B7MVJ5 B7MVJ7 B7MVL2 B7MVL9 B7MVM1 B7MVM3 B7MVM4 B7MVM5 B7MVM6 B7MVM7
B7MVP7 B7MVQ9 B7MVR5 B7MVU0 B7MVU2 B7MVW1 B7MVW5 B7MVZ1 B7MVZ4 B7MVZ7 B7MW64
B7MW65 B7MW68 B7MW70 B7MW83 B7MWA0 B7MWA6 B7MWC6 B7MWE0 B7MWE2 B7MWF3 B7MWG3
B7MWP8 B7MWQ4 B7MWT8 B7MWU0 B7MWU2 B7MWU3 B7MWX9 B7MX24 B7MX26 B7MX36 B7MXE4
B7MXE7 B7MXF5 B7MXG6 B7MXH3 B7MXI6 B7MXK1 B7MXK2 B7MXK3 B7MXP1 B7MXR3 B7MXS6
B7MXS8 B7MXT2 B7MXU6 B7MXU7 B7MXV1 B7MXX0 B7MXZ5 B7MXZ7 B7MY06 B7MY08 B7MY19
B7MY25 B7MY33 B7MY46 B7MY64 B7MY71 B7MY79 B7MY86 B7MYA0 B7MYA1 B7MYA6 B7MYB5
B7MYB7 B7MYC3 B7MYC5 B7MYC7 B7MYD5 B7MYD7 B7MYE6 B7MYE9 B7MYF0 B7MYF2 B7MYF7
B7MYG1 B7MYG2 B7MYG3 B7MYG4 B7MYI0 B7MYJ3 B7MYJ8 B7MYJ9 B7MYK8 B7MYL1 B7MYL2
B7MYL3 B7MYN6 B7MYP6 B7MYP8 B7MYQ2 B7MYQ3 B7MYQ5 B7MYQ6 B7MYQ8 B7MYR0 B7MYR1
B7MYV7 B7MYW1 B7MYX8 B7MYY4 B7MYZ3 B7MYZ5 B7MZ27 B7MZ42 B7MZ44 B7MZ45 B7MZ46
B7MZ47 B7MZ51 B7MZ52 B7MZ53 B7MZ55 B7MZ56 B7MZ57 B7MZ67 B7MZ76 B7MZ90 B7MZ94
B7MZ98 B7MZA0 B7MZC6 B7MZD0 B7MZE3 B7MZI9 B7MZJ8 B7MZK6 B7MZL0 B7MZM0 B7MZN5
B7MZP3 B7MZP7 B7MZP8 B7MZQ2 B7MZR1 B7N052 B7N068 B7N095 B7N0B1 B7N0H8 B7N0J6
B7N0J8 B7N0K1 B7N0K6 B7N0L5 B7N0L6 B7N0L8 B7N0M1 B7N0M2 B7N0Q3 B7N0Q5 B7N0S2
B7N0S5 B7N0S6 B7N0S9 B7N0T3 B7N0T4 B7N0T9 B7N0U0 B7N0U2 B7N0U4 B7N0U9 B7N0V0
B7N0V1 B7N0V2 B7N0V3 B7N0V5 B7N0V6 B7N0V9 B7N0W0 B7N0W5 B7N0W8 B7N0X1 B7N0X6
B7N0X9 B7N0Z1 B7N0Z9 B7N102 B7N109 B7N133 B7N136 B7N143 B7N145 B7N147 B7N157
B7N171 B7N172 B7N176 B7N179 B7N181 B7N182 B7N185 B7N189 B7N190 B7N192 B7N193
B7N194 B7N196 B7N197 B7N198 B7N199 B7N1A0 B7N1A1 B7N1A3 B7N1A5 B7N1C2 B7N1C3
B7N1C5 B7N1C6 B7N1D6 B7N1F1 B7N1G6 B7N1J6 B7N1J7 B7N1L1 B7N1L9 B7N1M2 B7N1M3
B7N1Q8 B7N1R9 B7N1S0 B7N1S3 B7N1T5 B7N1U1 B7N1U3 B7N1V9 B7N1X4 B7N1Z1 B7N1Z6
B7N203 B7N204 B7N211 B7N212 B7N223 B7N237 B7N242 B7N247 B7N255 B7N260 B7N266
B7N269 B7N272 B7N276 B7N277 B7N279 B7N282 B7N287 B7N295 B7N297 B7N2A1 B7N2D2
B7N2E1 B7N2E7 B7N2E8 B7N2E9 B7N2F3 B7N2G7 B7N2H9 B7N2I0 B7N2I4 B7N2I5 B7N2I7
B7N2J2 B7N2J8 B7N2J9 B7N2K0 B7N2L5 B7N2M8 B7N2P3 B7N2P6 B7N2P7 B7N2Q7 B7N2R7
B7N2R9 B7N2S2 B7N2S7 B7N2U2 B7N2U6 B7N2V7 B7N2V8 B7N2W3 B7N2W4 B7N2X1 B7N2X2
B7N304 B7N362 B7N363 B7N381 B7N389 B7N391 B7N398 B7N3A4 B7N3B8 B7N3C1 B7N3C6
B7N3C9 B7N3F3 B7N3F4 B7N3F5 B7N3F9 B7N3G3 B7N3G4 B7N3G5 B7N3G6 B7N3G7 B7N3G8
B7N3I1 B7N3N7 B7N3W1 B7N3X6 B7N3X8 B7N3Y0 B7N3Y3 B7N3Z3 B7N415 B7N419 B7N421
B7N422 B7N444 B7N459 B7N473 B7N474 B7N493 B7N497 B7N4A3 B7N4F3 B7N4K1 B7N4P2
B7N4U4 B7N4U6 B7N515 B7N516 B7N528 B7N529 B7N541 B7N544 B7N550 B7N552 B7N553
B7N554 B7N556 B7N569 B7N576 B7N578 B7N580 B7N581 B7N582 B7N583 B7N584 B7N5B4
B7N5C2 B7N5C5 B7N5D5 B7N5E9 B7N5F0 B7N5F1 B7N5H2 B7N5J4 B7N5L0 B7N5L4 B7N5L6
B7N5M7 B7N5M9 B7N5N0 B7N5N4 B7N5Q3 B7N5Q6 B7N5T0 B7N5T2 B7N5U1 B7N5U3 B7N5W6
B7N5X4 B7N5Y9 B7N602 B7N609 B7N617 B7N618 B7N626 B7N630 B7N653 B7N658 B7N678
B7N680 B7N681 B7N691 B7N693 B7N699 B7N6A2 B7N6A3 B7N6A5 B7N6A6 B7N6B0 B7N6B3
B7N6B4 B7N6B5 B7N6B7 B7N6B8 B7N6C6 B7N6C8 B7N6C9 B7N6D8 B7N6F3 B7N6F5 B7N6G4
B7N6H0 B7N6H1 B7N6J2 B7N6J3 B7N6J4 B7N6J5 B7N6J9 B7N6K0 B7N6K3 B7N6K5 B7N6P3

B7N6R0 B7N6S3 B7N6S6 B7N6S8 B7N6S9 B7N6U1 B7N6W4 B7N6X3 B7N6X4 B7N6X5 B7N6X6
B7N6X7 B7N6Y0 B7N6Y1 B7N6Y2 B7N6Z6 B7N6Z7 B7N716 B7N728 B7N732 B7N736 B7N738
B7N740 B7N759 B7N762 B7N763 B7N775 B7N7B8 B7N7D3 B7N7D5 B7N7E2 B7N7E4 B7N7E6
B7N7E7 B7N7E8 B7N7F2 B7N7F6 B7N7G1 B7N7H3 B7N7I9 B7N7J5 B7N7J7 B7N7K1 B7N7K2
B7N7K6 B7N7L8 B7N7M8 B7N7N1 B7N7N9 B7N7P7 B7N7Q0 B7N7Q3 B7N7Q4 B7N7Q5 B7N7R0
B7N7R2 B7N7R3 B7N7R4 B7N7R6 B7N7R7 B7N7S4 B7N7S5 B7N7S6 B7N7S7 B7N7T3 B7N7T6
B7N7T7 B7N7U6 B7N7U7 B7N7U9 B7N7V4 B7N7V5 B7N7W4 B7N7X3 B7N7X5 B7N7Y9 B7N7Z0
B7N7Z1 B7N801 B7N803 B7N804 B7N815 B7N823 B7N825 B7N828 B7N829 B7N832 B7N833
B7N834 B7N836 B7N837 B7N839 B7N850 B7N857 B7N862 B7N871 B7N874 B7N876 B7N8A8
B7N8H0 B7N8H1 B7N8H3 B7N8H4 B7N8J2 B7N8J3 B7N8L3 B7N8L4 B7N8L5 B7N8P7 B7N8Q1
B7N8Q4 B7N8Q5 B7N8Q6 B7N8Q7 B7N8Q8 B7N8Q9 B7N8T4 B7N8T9 B7N8U0 B7N8U3 B7N8U5
B7N8V7 B7N8V8 B7N8W5 B7N8W7 B7N8W8 B7N8X3 B7N8X5 B7N8X6 B7N8X9 B7N8Z0 B7N8Z2
B7N8Z8 B7N900 B7N922 B7N924 B7N925 B7N927 B7N928 B7N929 B7N958 B7N960 B7N966
B7N980 B7N983 B7N9I5 B7N9K4 B7N9L9 B7N9M0 B7N9M3 B7N9N3 B7N9N6 B7N9N7 B7N9P2
B7N9P5 B7N9P8 B7N9Q7 B7N9R4 B7N9R6 B7N9S4 B7N9S6 B7N9V2 B7N9V3 B7N9W6 B7N9Z2
B7N9Z7 B7N9Z9 B7NA10 B7NA74 B7NA75 B7NA78 B7NA81 B7NA83 B7NAA4 B7NAA7 B7NAB2
B7NAI1 B7NAJ8 B7NAK5 B7NAL7 B7NAM4 B7NAN2 B7NAN6 B7NAP4 B7NAQ6 B7NAQ7 B7NAQ9
B7NAR1 B7NAR6 B7NAT1 B7NAT4 B7NAT7 B7NAU2 B7NAW5 B7NAX6 B7NAY4 B7NAY5 B7NAY6
B7NAZ7 B7NB63 B7NB76 B7NB91 B7NB92 B7NB94 B7NB96 B7NBB1 B7NBC9 B7NBF7 B7NBF9
B7NBH9 B7NBI3 B7NBL1 B7NBL2 B7NBL4 B7NBL7 B7NBM1 B7NBM2 B7NBM5 B7NBM7 B7NBP2
B7NBQ8 B7NBR4 B7NBU5 B7NBU7 B7NBV8 B7NBW5 B7NC25 B7NC59 B7NC61 B7NC64 B7NC65
B7NCA1 B7NCC5 B7NCC6 B7NCD4 B7NCE4 B7NCH2 B7NCI3 B7ND35 B7ND37 B7ND40 B7ND45
B7ND53 B7ND54 B7ND57 B7ND61 B7ND80 B7ND98 B7NDB2 B7NDB8 B7NDC5 B7NDD2 B7NDD3
B7NDD8 B7NDD9 B7NDF0 B7NDF1 B7NDF3 B7NDF4 B7NDF6 B7NDF7 B7NDG0 B7NDG3 B7NDG7
B7NDG9 B7NDH0 B7NDH3 B7NDI9 B7NDK1 B7NDK4 B7NDK8 B7NDK9 B7NDL4 B7NDL5 B7NDN7
B7NDN9 B7NDQ3 B7NDQ6 B7NDQ8 B7NDQ9 B7NDR0 B7NDR6 B7NDR8 B7NDR9 B7NDS2 B7NDS3
B7NDS5 B7NDS6 B7NDS7 B7NDS8 B7NDS9 B7NDT0 B7NDT1 B7NDT2 B7NDT3 B7NDT4 B7NDT5
B7NDT6 B7NDT7 B7NDT8 B7NDU0 B7NDU1 B7NDU2 B7NDU8 B7NDU9 B7NDV0 B7NDV1 B7NDV2
B7NDV3 B7NDV6 B7NDW3 B7NDX8 B7NDZ3 B7NDZ4 B7NE05 B7NE08 B7NE15 B7NE17 B7NE19
B7NE23 B7NE24 B7NE29 B7NE39 B7NE40 B7NE41 B7NE79 B7NEA8 B7NEC8 B7NEG2 B7NEL0
B7NEL1 B7NEL7 B7NEM8 B7NEP6 B7NEQ1 B7NER6 B7NES3 B7NES6 B7NET8 B7NET9 B7NEU0
B7NEU1 B7NEU2 B7NEU4 B7NEU5 B7NEU6 B7NEV1 B7NEV3 B7NEX9 B7NF03 B7NF04 B7NF09
B7NF17 B7NF19 B7NF20 B7NF21 B7NF24 B7NF25 B7NF28 B7NF56 B7NF57 B7NF60 B7NF61
B7NF62 B7NF64 B7NF72 B7NF78 B7NF81 B7NF84 B7NF85 B7NFA7 B7NFA9 B7NFB1 B7NFB3
B7NFD2 B7NFD6 B7NFE8 B7NFG1 B7NFI6 B7NFJ2 B7NFJ6 B7NFJ9 B7NFK0 B7NFK1 B7NFK2
B7NFK3 B7NFL3 B7NFL6 B7NFM3 B7NFM5 B7NFM8 B7NFN0 B7NFN1 B7NFN5 B7NFQ6 B7NFQ7
B7NFR0 B7NFR2 B7NFR5 B7NFR9 B7NFS3 B7NFS4 B7NFS5 B7NFS6 B7NFS7 B7NFT1 B7NFT5
B7NFT6 B7NFT7 B7NFT8 B7NFT9 B7NFU7 B7NFU9 B7NFV2 B7NFV7 B7NFW8 B7NFY3 B7NG75
B7NG80 B7NG81 B7NG85 B7NG94 B7NG98 B7NGA4 B7NGA5 B7NGA6 B7NGB1 B7NGB2 B7NGC5
B7NGC6 B7NGD0 B7NGD1 B7NGD2 B7NGD4 B7NGD5 B7NGD6 B7NGD7 B7NGE7 B7NGF6 B7NGG7
B7NGH7 B7NGH8 B7NGI6 B7NGJ2 B7NGT8 B7NGV6 B7NGX8 B7NGY3 B7NGY8 B7NGZ6 B7NH31
B7NH38 B7NH42 B7NH49 B7NH50 B7NH51 B7NH52 B7NH54 B7NH68 B7NH70 B7NH81 B7NHA4
B7NHA6 B7NHA9 B7NHB7 B7NHC8 B7NHD0 B7NHD3 B7NHD4 B7NHD5 B7NHD9 B7NHE1 B7NHE2
B7NHE3 B7NHE5 B7NHE6 B7NHF5 B7NHF6 B7NHF7 B7NHF8 B7NHG5 B7NHG8 B7NHG9 B7NHH9

B7NHI0 B7NHI2 B7NHI7 B7NHI8 B7NHJ7 B7NHK6 B7NHP6 B7NHU3 B7NHW2 B7NHW4 B7NHW5
B7NHW6 B7NHX0 B7NHX4 B7NHX5 B7NHY6 B7NI01 B7NI05 B7NI07 B7NI11 B7NI12 B7NI16
B7NI31 B7NI65 B7NI79 B7NI80 B7NI81 B7NI91 B7NI93 B7NI94 B7NIA9 B7NIB7 B7NIB9
B7NIC2 B7NIC3 B7NIC6 B7NIC7 B7NIC8 B7NID0 B7NID1 B7NID4 B7NIE5 B7NIF4 B7NIF5
B7NIF6 B7NIF8 B7NIF9 B7NIG1 B7NIJ1 B7NIJ4 B7NJ48 B7NJ50 B7NJ56 B7NJ58 B7NJ71
B7NJ74 B7NJ75 B7NJ77 B7NJ81 B7NJ82 B7NJ84 B7NJ96 B7NJ97 B7NJA9 B7NJB1 B7NJB4
B7NJB5 B7NJC0 B7NJF1 B7NJG3 B7NJG6 B7NJG8 B7NJH1 B7NJI4 B7NJQ5 B7NJR2 B7NJR5
B7NJR9 B7NJS7 B7NJS8 B7NJT2 B7NJT8 B7NJV5 B7NJX7 B7NJZ1 B7NK03 B7NK04 B7NK05
B7NK06 B7NK07 B7NK08 B7NK15 B7NK49 B7NK50 B7NK70 B7NK71 B7NK82 B7NK83 B7NK85
B7NK86 B7NKG0 B7NKH1 B7NKH2 B7NKI2 B7NKJ1 B7NKK3 B7NKL0 B7NKL7 B7NKL8 B7NKM3
B7NKM4 B7NKN3 B7NKN4 B7NKN6 B7NKN7 B7NKN9 B7NKP0 B7NKP3 B7NKP6 B7NKQ0 B7NKQ2
B7NKQ3 B7NKQ6 B7NKS2 B7NKT3 B7NKT4 B7NKT6 B7NKT7 B7NKU2 B7NKU3 B7NKU9 B7NKV0
B7NKV9 B7NKW4 B7NKW6 B7NKW9 B7NKX8 B7NKY7 B7NL20 B7NL23 B7NL67 B7NL72 B7NL75
B7NL80 B7NLA0 B7NLB4 B7NLB5 B7NLB6 B7NLB7 B7NLB8 B7NLB9 B7NLC1 B7NLC5 B7NLC6
B7NLC7 B7NLF0 B7NLF4 B7NLI5 B7NLI7 B7NLK0 B7NLK3 B7NLK6 B7NLK7 B7NLK8 B7NLL4
B7NLL6 B7NLL7 B7NLM0 B7NLM1 B7NLM3 B7NLM4 B7NLM5 B7NLM6 B7NLM7 B7NLM8 B7NLM9
B7NLN0 B7NLN1 B7NLN2 B7NLN3 B7NLN4 B7NLN5 B7NLN6 B7NLN8 B7NLN9 B7NLP0 B7NLP5
B7NLP6 B7NLP7 B7NLP8 B7NLP9 B7NLQ0 B7NLQ3 B7NLU0 B7NLX3 B7NLX4 B7NLX7 B7NLY7
B7NLY9 B7NLZ0 B7NLZ5 B7NLZ8 B7NM01 B7NM03 B7NM10 B7NM12 B7NM15 B7NM18 B7NM39
B7NM43 B7NM50 B7NM52 B7NM57 B7NM62 B7NM64 B7NM67 B7NM68 B7NM82 B7NM92 B7NM96
B7NMA8 B7NMB8 B7NMC2 B7NMD7 B7NMF2 B7NMF3 B7NMG4 B7NMG6 B7NMH5 B7NMH7 B7NMH9
B7NMI3 B7NMI4 B7NMI9 B7NMJ4 B7NMJ5 B7NMJ6 B7NMM9 B7NMN1 B7NMQ9 B7NMR0 B7NMS9
B7NMW2 B7NMX3 B7NMX6 B7NMY6 B7NN00 B7NN01 B7NN02 B7NN21 B7NN47 B7NN63 B7NN72
B7NN84 B7NN95 B7NNB9 B7NNF3 B7NNH7 B7NNH9 B7NNI9 B7NNK5 B7NNK6 B7NNK9 B7NNL1
B7NNL3 B7NNN5 B7NNN9 B7NNP4 B7NNR8 B7NNU1 B7NNU3 B7NNU4 B7NNU8 B7NNX0 B7NNX3
B7NNZ9 B7NP01 B7NP10 B7NP12 B7NP55 B7NP56 B7NP65 B7NP75 B7NP97 B7NPA3 B7NPB8
B7NPC5 B7NPC7 B7NPE0 B7NPE1 B7NPE2 B7NPE7 B7NPF5 B7NPG3 B7NPH0 B7NPL8 B7NPM9
B7NPN7 B7NPN8 B7NPQ8 B7NPS4 B7NPU7 B7NPV5 B7NPW3 B7NPW4 B7NPX2 B7NPX6 B7NPZ8
B7NPZ9 B7NQ01 B7NQ02 B7NQ04 B7NQ11 B7NQ13 B7NQC0 B7NQG5 B7NQG6 B7NQG9 B7NQH1
B7NQK9 B7NQL4 B7NQL6 B7NQN5 B7NQN7 B7NQN8 B7NQV3 B7NQV5 B7NQW0 B7NQW3 B7NQY6
B7NQY7 B7NQZ2 B7NR02 B7NR04 B7NR05 B7NR06 B7NR09 B7NR10 B7NR42 B7NR43 B7NR46
B7NR47 B7NR48 B7NR51 B7NR61 B7NR83 B7NRA8 B7NRB5 B7NRC7 B7NRC9 B7NRE7 B7NRG1
B7NRG4 B7NRG5 B7NRG7 B7NRG8 B7NRH2 B7NRH5 B7NRH6 B7NRH7 B7NRH9 B7NRI0 B7NRI8
B7NRJ0 B7NRJ1 B7NRK2 B7NRL8 B7NRM0 B7NRM8 B7NRN5 B7NRN6 B7NRR1 B7NRR2 B7NRR3
B7NRR4 B7NRR5 B7NRS3 B7NRS7 B7NRS8 B7NRS9 B7NRT0 B7NRT1 B7NRT9 B7NRU2 B7NRV3
B7NRZ0 B7NS03 B7NS11 B7NS27 B7NS42 B7NS44 B7NS47 B7NS48 B7NS52 B7NS55 B7NS57
B7NS58 B7NS84 B7NS88 B7NSA5 B7NSA6 B7NSA7 B7NSA8 B7NSB2 B7NSB3 B7NSB6 B7NSB8
B7NSE0 B7NSF9 B7NSH3 B7NSH6 B7NSH8 B7NSH9 B7NSJ1 B7NSW2 B7NSW4 B7NSZ2 B7NSZ8
B7NT29 B7NT30 B7NT31 B7NT32 B7NT33 B7NT35 B7NT37 B7NT44 B7NT58 B7NT60 B7NT61
B7NT62 B7NT64 B7NT79 B7NT87 B7NT88 B7NT89 B7NT90 B7NT91 B7NT94 B7NT95 B7NT96
B7NT99 B7NTA0 B7NTD1 B7NTD3 B7NTD8 B7NTE0 B7NTG2 B7NTG3 B7NTG7 B7NTJ6 B7NTK1
B7NTK2 B7NTK6 B7NTL6 B7NTM0 B7NTM7 B7NTM8 B7NTN3 B7NTN4 B7NTP7 B7NTP8 B7NTQ2
B7NTQ3 B7NTQ4 B7NTQ6 B7NTQ7 B7NTQ8 B7NTS5 B7NTS8 B7NTU3 B7NTU4 B7NTV6 B7NTV7
B7NTX5 B7NTZ0 B7NTZ2 B7NTZ4 B7NTZ5 B7NU11 B7NU26 B7NU32 B7NU33 B7NU35 B7NU38

B7NU40 B7NU43 B7NU44 B7NU69 B7NU73 B7NU74 B7NU76 B7NU79 B7NU81 B7NU88 B7NU91
B7NUA1 B7NUA2 B7NUA3 B7NUA6 B7NUA7 B7NUB4 B7NUB8 B7NUC3 B7NUD2 B7NUE9 B7NUG0
B7NUG1 B7NUG6 B7NUH4 B7NUM9 B7NUX0 B7NUX4 B7NUX6 B7NUX7 B7NUY4 B7NV06 B7NV19
B7NV33 B7NV51 B7NV70 B7NV92 B7NV97 B7NVB3 B7NVD2 B7NVD9 B7NVK2 B7NVL7 B7NVN0
B7NVN1 B7NVQ1 B7NVQ5 B7NVR1 B7NVU2 B7NVU6 B7NVV0 B7NVV2 B7NVV4 B7NVX2 B7NVX5
B7NVX7 B7NVY9 B7NW14 B7NW30 B7NW32 B7NW39 B7NW61 B7NW62 B7NW63 B7NW64 B7NW66
B7NW74 B7NW76 B7NZ88 B7NZC7 B7P877 B7PBI5 B7PDC5 B7PP17 B7PRF6 B7PS00 B7PY76
B7PZ18 B7Q1Q9 B7Q290 B7Q2M2 B7Q5K1 B7QDG3 B7QKS1 B7RR92 B7S4N1 B7SBD2 B7SNI3
B7SP66 B7SSM2 B7SSM6 B7STY1 B7STY2 B7SXM5 B7TWP7 B7U179 B7U9B5 B7UEU8 B7UFB2
B7UFB3 B7UFC1 B7UFD1 B7UFF9 B7UFG9 B7UFH5 B7UFH8 B7UFI8 B7UFK2 B7UFK3 B7UFK4
B7UFM4 B7UFP4 B7UFQ9 B7UFR3 B7UFS4 B7UFS6 B7UFS7 B7UFT1 B7UFU9 B7UFV2 B7UFX6
B7UFX8 B7UFY7 B7UFY9 B7UG84 B7UG97 B7UGB2 B7UGB9 B7UGC8 B7UGD5 B7UGM1 B7UGM6
B7UGN4 B7UGN6 B7UGN7 B7UGU5 B7UGU7 B7UGV7 B7UGW0 B7UGW1 B7UGW3 B7UGW4 B7UGW9
B7UGX2 B7UGX3 B7UGX4 B7UGX6 B7UGX7 B7UGZ1 B7UH07 B7UH15 B7UH18 B7UH19 B7UH55
B7UH56 B7UH57 B7UH58 B7UH62 B7UH63 B7UH66 B7UH68 B7UH78 B7UH94 B7UHB1 B7UHB4
B7UHB6 B7UHB7 B7UHC3 B7UHE9 B7UHG2 B7UHG3 B7UHG4 B7UHG5 B7UHG6 B7UHG9 B7UHH0
B7UHH1 B7UHH4 B7UHH5 B7UHJ6 B7UHL0 B7UHL4 B7UHL8 B7UHM0 B7UHM2 B7UHP4 B7UHP7
B7UHP9 B7UHR2 B7UHS3 B7UHT8 B7UHU0 B7UHU7 B7UHU9 B7UHV1 B7UHV2 B7UHV3 B7UHV7
B7UHW1 B7UHW2 B7UHW9 B7UHY3 B7UHY9 B7UHZ1 B7UHZ4 B7UHZ5 B7UHZ6 B7UI00 B7UI09
B7UI46 B7UI48 B7UI51 B7UI60 B7UI68 B7UI71 B7UI74 B7UI75 B7UI76 B7UI81 B7UI83
B7UI84 B7UI85 B7UI87 B7UI88 B7UI97 B7UI98 B7UI99 B7UIA5 B7UIA8 B7UIA9 B7UIC1
B7UIC2 B7UIC4 B7UID1 B7UID2 B7UIE1 B7UIF0 B7UIF1 B7UIF3 B7UIG7 B7UIG8 B7UIG9
B7UIH9 B7UII1 B7UII2 B7UIJ0 B7UIK1 B7UIK3 B7UIK6 B7UIK7 B7UIL0 B7UIL1 B7UIL2
B7UIL4 B7UIL5 B7UIL7 B7UIU7 B7UIV5 B7UIV7 B7UIW0 B7UIW4 B7UIX2 B7UIX3 B7UIY0
B7UIZ9 B7UJ14 B7UJ26 B7UJ30 B7UJ37 B7UJ44 B7UJ45 B7UJ49 B7UJ50 B7UJ59 B7UJ60
B7UJ62 B7UJ63 B7UJ65 B7UJ66 B7UJ69 B7UJ72 B7UJ76 B7UJ78 B7UJ79 B7UJ84 B7UJ91
B7UJ96 B7UJA5 B7UJA8 B7UJB0 B7UJB5 B7UJC7 B7UJC8 B7UJD0 B7UJD1 B7UJE2 B7UJE5
B7UJG4 B7UJG5 B7UJG6 B7UJI9 B7UJK6 B7UJL1 B7UJL2 B7UJL5 B7UJL7 B7UJM8 B7UJM9
B7UJN6 B7UJN8 B7UJN9 B7UJP3 B7UJP5 B7UJP6 B7UJP9 B7UJQ9 B7UJR1 B7UJR7 B7UJR9
B7UJS9 B7UJU5 B7UJV5 B7UJV6 B7UJV8 B7UJV9 B7UJW2 B7UJW3 B7UJW8 B7UJW9 B7UJZ0
B7UJZ2 B7UK05 B7UK08 B7UK10 B7UK11 B7UK12 B7UK18 B7UK20 B7UK21 B7UK25 B7UK26
B7UK28 B7UK29 B7UK30 B7UK32 B7UK33 B7UK34 B7UK35 B7UK36 B7UK37 B7UK38 B7UK39
B7UK40 B7UK41 B7UK43 B7UK44 B7UK45 B7UK50 B7UK51 B7UK52 B7UK53 B7UK54 B7UK55
B7UK58 B7UK65 B7UK79 B7UK94 B7UKA5 B7UKA7 B7UKB5 B7UKB7 B7UKB9 B7UKC3 B7UKC4
B7UKC9 B7UKD4 B7UKE9 B7UKF1 B7UKF2 B7UKF4 B7UKF5 B7UKF6 B7UKI1 B7UKI3 B7UKI9
B7UKK3 B7UKK6 B7UKM3 B7UKQ5 B7UKQ6 B7UKQ9 B7UKR9 B7UKS0 B7UKS1 B7UKS6 B7UKT2
B7UKT4 B7UKU2 B7UKV0 B7UKW1 B7UKY6 B7UKY7 B7UKY8 B7UL22 B7UL33 B7UL45 B7UL72
B7ULB9 B7ULC0 B7ULC6 B7ULD6 B7ULF0 B7ULF5 B7ULG5 B7ULH2 B7ULH4 B7ULI9 B7ULJ0
B7ULJ1 B7ULJ2 B7ULJ3 B7ULJ5 B7ULM3 B7ULM8 B7ULN0 B7ULP0 B7ULX2 B7ULX3 B7ULX6
B7ULX8 B7ULY0 B7UM00 B7UM03 B7UM08 B7UM46 B7UM47 B7UM49 B7UM50 B7UM69 B7UM71
B7UM74 B7UMF3 B7UMF4 B7UMG7 B7UMG9 B7UMH0 B7UMH1 B7UMH3 B7UMH5 B7UMK5 B7UMK6
B7UMK9 B7UML0 B7UML1 B7UML3 B7UMM1 B7UMM8 B7UMN1 B7UMN4 B7UMN5 B7UMU4 B7UMV1
B7UMV8 B7UMW5 B7UMX3 B7UMX6 B7UMY4 B7UMZ3 B7UMZ4 B7UMZ6 B7UMZ8 B7UN03 B7UN08
B7UN10 B7UN17 B7UN21 B7UN35 B7UN37 B7UN42 B7UN45 B7UN46 B7UNC2 B7UNC6 B7UNC8

B7UNG3 B7UNH5 B7UNI7 B7UNI8 B7UNK4 B7UNL0 B7UNL4 B7UNM2 B7UNM3 B7UNM4 B7UNM5
B7UNM6 B7UNN6 B7UNP6 B7UNP8 B7UNP9 B7UNQ1 B7UNQ2 B7UNQ6 B7UNQ8 B7UNT2 B7UNT3
B7UNT6 B7UNU0 B7UNU5 B7UNV0 B7UNY1 B7UNY3 B7UNY7 B7UNZ0 B7UNZ1 B7UNZ2 B7UNZ3
B7UNZ4 B7UNZ5 B7UP19 B7UP20 B7UP48 B7UP69 B7UP73 B7UP76 B7UP81 B7UPA5 B7UPB4
B7UPC2 B7UPC3 B7UPC4 B7UPD8 B7UPD9 B7UPE0 B7UPE1 B7UPE2 B7UPE5 B7UPE9 B7UPF0
B7UPF1 B7UPF2 B7UPF3 B7UPG1 B7UPG4 B7UPG7 B7UPH5 B7UPI6 B7UPK0 B7UPM7 B7UPW2
B7UPW3 B7UPW7 B7UPX7 B7UPY1 B7UPY3 B7UQ15 B7UQ36 B7UQ55 B7UQ59 B7UQ61 B7UQ64
B7UQ73 B7UQ92 B7UQ96 B7UQ98 B7UQ99 B7UQC5 B7UQD8 B7UQH9 B7UQI0 B7UQI1 B7UQI6
B7UQI7 B7UQK0 B7UQK1 B7UQK5 B7UQK6 B7UQK9 B7UQL0 B7UQL1 B7UQL2 B7UQM4 B7UQN3
B7UQP4 B7UQQ4 B7UQQ5 B7UQR7 B7UQW0 B7UQW3 B7UQX7 B7UQZ1 B7UR02 B7UR09 B7UR10
B7UR11 B7UR12 B7UR14 B7UR23 B7UR24 B7UR66 B7UR67 B7UR85 B7UR89 B7UR93 B7UR96
B7URE9 B7URJ0 B7URK1 B7URL9 B7URP6 B7URP8 B7URV0 B7URW2 B7URX8 B7URX9 B7URY1
B7URZ9 B7US18 B7US19 B7US31 B7US32 B7US42 B7US45 B7US95 B7US97 B7US99 B7USA0
B7USA1 B7USA2 B7USB4 B7USC1 B7USC3 B7USC5 B7USC6 B7USC7 B7USC8 B7USC9 B7USF8
B7USG4 B7USI6 B7USI8 B7USK8 B7USL2 B7USN7 B7USN8 B7USP0 B7USP3 B7USP7 B7USP8
B7USQ1 B7USQ3 B7USR7 B7UST2 B7UST8 B7USV1 B7USW6 B7USW8 B7USY4 B7UT20 B7UT45
B7UT56 B7UT58 B7UT61 B7UT62 B7UTA3 B7UTE8 B7UTE9 B7UUT0 B7UUU2 B7UUV7 B7UUY3
B7UUZ9 B7UV15 B7UV23 B7UV24 B7UV36 B7UV38 B7UV40 B7UV48 B7UV78 B7UV83 B7UVD3
B7UVE8 B7UVF9 B7UVG0 B7UVH9 B7UVI8 B7UVI9 B7UVJ3 B7UVK3 B7UVK4 B7UVR5 B7UVS7
B7UVT6 B7UVU3 B7UVU4 B7UVV3 B7UVV5 B7UW10 B7UW11 B7UW21 B7UW25 B7UW33 B7UW34
B7UWH1 B7UWH3 B7UWH7 B7UWI0 B7UWI1 B7UWI4 B7UWI8 B7UWI9 B7UWJ2 B7UWJ8 B7UWK0
B7UWK9 B7UWN0 B7UWP0 B7UWY4 B7UX01 B7UX27 B7UX51 B7UX89 B7UXE2 B7UXF2 B7UXJ1
B7UXJ9 B7UXM6 B7UXN9 B7UXT0 B7UXV6 B7UXW5 B7UXW6 B7UXW7 B7UXW8 B7UXW9 B7UXX2
B7UXX6 B7UXY9 B7UY29 B7UY33 B7UY35 B7UY63 B7UY78 B7UYA0 B7UYK0 B7UYK6 B7UYK7
B7UYK8 B7UYN4 B7UYP4 B7UYS5 B7UYW8 B7UYW9 B7UYX0 B7UYX2 B7UYZ2 B7UZ17 B7UZG3
B7UZG4 B7UZH3 B7UZI9 B7UZJ8 B7UZJ9 B7UZK2 B7UZK3 B7UZL0 B7UZL1 B7UZM0 B7UZU5
B7UZY2 B7UZY8 B7UZY9 B7V004 B7V012 B7V022 B7V023 B7V024 B7V055 B7V073 B7V0A0
B7V0A3 B7V0A6 B7V0A8 B7V0A9 B7V0B0 B7V0B1 B7V0C8 B7V0H2 B7V0J2 B7V0K1 B7V0K8
B7V0L1 B7V0L2 B7V0L5 B7V0L7 B7V0L8 B7V0N6 B7V0N8 B7V0P3 B7V0P4 B7V0Q3 B7V0Q9
B7V0S0 B7V0Z5 B7V109 B7V148 B7V159 B7V167 B7V188 B7V190 B7V1A6 B7V1D2 B7V1D3
B7V1D6 B7V1E2 B7V1E3 B7V1E4 B7V1F2 B7V1F3 B7V1F5 B7V1F6 B7V1F8 B7V1G0 B7V1G1
B7V1G4 B7V1H1 B7V1H2 B7V1H3 B7V1H4 B7V1H8 B7V1I0 B7V1I5 B7V1L9 B7V1R2 B7V1R6
B7V1S8 B7V1T0 B7V1T5 B7V1V4 B7V1V5 B7V1V6 B7V1Y3 B7V1Z5 B7V1Z7 B7V1Z8 B7V201
B7V202 B7V207 B7V208 B7V209 B7V265 B7V2C0 B7V2C1 B7V2K4 B7V2P3 B7V2P9 B7V2Q5
B7V2R6 B7V2S0 B7V2S6 B7V2T3 B7V2X1 B7V2Y0 B7V309 B7V310 B7V312 B7V341 B7V362
B7V388 B7V3B1 B7V3B6 B7V3C7 B7V3D1 B7V3D2 B7V3E2 B7V3E4 B7V3E6 B7V3E7 B7V3F6
B7V3F9 B7V3G0 B7V3H2 B7V3I5 B7V3J1 B7V3J3 B7V3K3 B7V3L1 B7V3M1 B7V3N3 B7V3N4
B7V3N7 B7V3P2 B7V3T9 B7V3U0 B7V3U4 B7V3V0 B7V3W7 B7V3X6 B7V3Y9 B7V3Z0 B7V3Z1
B7V3Z2 B7V419 B7V467 B7V485 B7V486 B7V489 B7V4D3 B7V4D9 B7V4G5 B7V4G6 B7V4H0
B7V4H3 B7V4H5 B7V4H6 B7V4H7 B7V4X6 B7V594 B7V5B5 B7V5D7 B7V5F0 B7V5F3 B7V5G9
B7V5H1 B7V5I9 B7V5K7 B7V5L0 B7V5L2 B7V5L4 B7V5M2 B7V5M5 B7V5M8 B7V5R3 B7V5R4
B7V5R5 B7V608 B7V609 B7V612 B7V620 B7V623 B7V628 B7V633 B7V634 B7V635 B7V636
B7V637 B7V639 B7V640 B7V643 B7V644 B7V645 B7V646 B7V647 B7V648 B7V649 B7V651
B7V652 B7V653 B7V654 B7V655 B7V656 B7V657 B7V658 B7V659 B7V660 B7V661 B7V662

B7V663 B7V665 B7V666 B7V667 B7V669 B7V6L2 B7V6T4 B7V6W0 B7V729 B7V732 B7V753
B7V776 B7V7A1 B7V7A2 B7V7A6 B7V7F5 B7V7F7 B7V7F8 B7V7G0 B7V7G3 B7V7J4 B7V7Q1
B7V7Q5 B7V7R1 B7V7R4 B7V7R6 B7V7T3 B7V7U6 B7V7V1 B7V7X2 B7V7X4 B7V7X6 B7V800
B7V8A3 B7V8A6 B7V8A7 B7V8A9 B7V8B5 B7V8B6 B7V8C1 B7V8C2 B7V8C3 B7V8C4 B7V8C8
B7V8D1 B7V8D2 B7V9C1 B7V9C9 B7V9D4 B7V9D9 B7V9E0 B7V9F5 B7V9F6 B7V9H3 B7V9J1
B7V9J5 B7V9J7 B7V9J9 B7V9K9 B7V9S6 B7V9Z4 B7VA51 B7VA96 B7VAB6 B7VAB7 B7VAB8
B7VAB9 B7VAL6 B7VAL8 B7VAP9 B7VAU6 B7VB28 B7VB54 B7VB62 B7VB64 B7VB75 B7VB77
B7VB83 B7VB84 B7VB85 B7VBA7 B7VBB0 B7VBC3 B7VBJ0 B7VBK5 B7VBP9 B7VBQ0 B7VBQ7
B7VGI0 B7VGI4 B7VGI6 B7VGK6 B7VGK7 B7VGL0 B7VGL9 B7VGS0 B7VGT7 B7VGU7 B7VGU8
B7VGZ1 B7VGZ7 B7VH15 B7VH16 B7VH19 B7VH29 B7VH32 B7VH34 B7VH36 B7VH49 B7VHB0
B7VHD5 B7VHE8 B7VHF2 B7VHH1 B7VHI0 B7VHJ7 B7VHJ8 B7VHK0 B7VHK2 B7VHK3 B7VHK4
B7VHK6 B7VHL5 B7VHM6 B7VHQ4 B7VHQ7 B7VHR3 B7VHR8 B7VHY7 B7VHZ1 B7VHZ9 B7VI00
B7VI01 B7VI24 B7VI35 B7VI46 B7VI54 B7VI55 B7VI65 B7VI66 B7VI67 B7VI70 B7VI73
B7VI85 B7VI86 B7VIC2 B7VIC9 B7VID0 B7VID3 B7VID4 B7VID5 B7VIE1 B7VIE2 B7VIE9
B7VIF2 B7VIH1 B7VIH2 B7VIH6 B7VII4 B7VII5 B7VII6 B7VIK3 B7VIL8 B7VIN4 B7VIP1
B7VIP5 B7VIQ1 B7VIQ4 B7VIR7 B7VIS0 B7VIS1 B7VIS8 B7VIT0 B7VIT2 B7VIT4 B7VIV4
B7VIX1 B7VIY3 B7VIY4 B7VIZ5 B7VJ01 B7VJ04 B7VJ13 B7VJ21 B7VJ36 B7VJ50 B7VJ56
B7VJ58 B7VJ74 B7VJ78 B7VJ84 B7VJ87 B7VJ90 B7VJ92 B7VJ96 B7VJ99 B7VJA1 B7VJA4
B7VJA9 B7VJB0 B7VJF8 B7VJG2 B7VJH3 B7VJH4 B7VJH5 B7VJH6 B7VJH7 B7VJH9 B7VJI1
B7VJI4 B7VJJ4 B7VJJ5 B7VJM5 B7VJN4 B7VJP8 B7VJR1 B7VJR7 B7VJR8 B7VJS5 B7VJS6
B7VJS9 B7VJT0 B7VJT3 B7VJT4 B7VJT5 B7VJT8 B7VJT9 B7VJU2 B7VJU8 B7VJW6 B7VJW7
B7VJW9 B7VJX0 B7VJZ6 B7VJZ8 B7VK03 B7VK04 B7VK24 B7VK26 B7VK27 B7VK28 B7VK29
B7VK42 B7VK59 B7VK62 B7VK66 B7VK69 B7VK72 B7VK77 B7VK79 B7VK90 B7VK95 B7VK98
B7VKB3 B7VKD0 B7VKD2 B7VKE6 B7VKE7 B7VKF3 B7VKF7 B7VKG1 B7VKG8 B7VKH0 B7VKH2
B7VKH3 B7VKP6 B7VKQ2 B7VKR3 B7VKR6 B7VKU1 B7VKV7 B7VKW1 B7VKW7 B7VKY2 B7VKY3
B7VL54 B7VL65 B7VL66 B7VL71 B7VL81 B7VL84 B7VL95 B7VL97 B7VL98 B7VLA3 B7VLC8
B7VLD1 B7VLD2 B7VLD3 B7VLD4 B7VLD6 B7VLD8 B7VLE0 B7VLE1 B7VLE2 B7VLE3 B7VLE5
B7VLE6 B7VLE8 B7VLF0 B7VLF1 B7VLF3 B7VLF4 B7VLF5 B7VLF6 B7VLF7 B7VLG1 B7VLG2
B7VLG9 B7VLH8 B7VLI4 B7VLJ6 B7VLL4 B7VLL5 B7VLM0 B7VLM2 B7VLM6 B7VLM7 B7VLN0
B7VLR2 B7VLU4 B7VLU6 B7VLW8 B7VLX0 B7VLX9 B7VM08 B7VM32 B7VM36 B7VM38 B7VM42
B7VM52 B7VM54 B7VM57 B7VM66 B7VM70 B7VM73 B7VM88 B7VM91 B7VMA4 B7VMD2 B7VMD4
B7VMD7 B7VMD9 B7VME9 B7VMF0 B7VMG4 B7VMG5 B7VMH0 B7VMH7 B7VMH8 B7VMI0 B7VMI1
B7VMI3 B7VMI4 B7VMK8 B7VMT0 B7VMU2 B7VMV3 B7VMV7 B7VMW2 B7VMW3 B7VMW8 B7VMX2
B7VMZ0 B7VN06 B7VN07 B7VN19 B7VN20 B7VN21 B7VNJ2 B7VNJ8 B7VNK3 B7VNL8 B7VNT9
B7VNU4 B7VPB2 B7VPD7 B7VPF1 B7VPF3 B7VPF6 B7VPK0 B7VPU4 B7VPY4 B7VQ27 B7VQ28
B7VQ39 B7VQ44 B7VQ85 B7VQM9 B7VQP0 B7VQP3 B7VQW0 B7VQW7 B7VQW9 B7VR44 B7VR52
B7VRZ7 B7VS36 B7VS66 B7VS90 B7VSB8 B7VSC8 B7VSJ2 B7VSK2 B7VSX1 B7VSY8 B7VSZ6
B7VT10 B7VTF0 B7VTF8 B7VTI0 B7VU31 B7WN72 B7X2B6 B7XDF1 B7XHC1 B7XHF3 B7XHM5
B7XHP6 B7XHQ4 B7XHS8 B7XHU4 B7XI98 B7XIA1 B7XIB8 B7XIV9 B7XIW9 B7XJI2 B7XJR9
B7XK16 B7XK66 B7XMD2 B7Z073 B7Z0I7 B7Z368 B7Z6K7 B7ZAP0 B7ZBB8 B7ZD04 B7ZFT1
B7ZMP1 B7ZNG4 B7ZQA9 B7ZRI2 B7ZRJ4 B7ZRT8 B7ZS37 B7ZS42 B7ZSG3 B7ZTW1 B7ZUF3
B7ZUL2 B7ZW38 B8A031 B8A4F4 B8A556 B8A5G9 B8A5I7 B8A8D2 B8A9B2 B8AA76 B8ADZ3
B8AE37 B8AE67 B8AEC1 B8AEH3 B8AI54 B8AIM8 B8AJL9 B8AK78 B8AL33 B8AM21 B8AMA8
B8AMB8 B8AMS4 B8ANW0 B8ANX9 B8APB5 B8APQ0 B8AR30 B8AR41 B8ARI7 B8ARW2 B8ASB2

B8ATT7 B8AU84 B8AVX5 B8AW67 B8AWM1 B8AX23 B8AX51 B8AX53 B8AXB6 B8AXU2 B8AY58
B8AZ14 B8AZ99 B8AZX3 B8B016 B8B0E2 B8B2G4 B8B406 B8B4H5 B8B5N8 B8B5U8 B8B624
B8B644 B8B7S5 B8B7X6 B8B8C5 B8B8I3 B8B9K6 B8BAI9 B8BBT7 B8BBZ7 B8BCZ8 B8BDK0
B8BDQ4 B8BDW1 B8BF91 B8BH95 B8BIU8 B8BJ39 B8BJV8 B8BK80 B8BKI7 B8BKI8 B8BLA3
B8BLT3 B8BM09 B8BM17 B8BMI1 B8BQY5 B8BVB6 B8BVM7 B8C4D7 B8C6S5 B8C6V3 B8CAZ5
B8CGT2 B8CGX4 B8CGY2 B8CGY4 B8CGY5 B8CH19 B8CH32 B8CH33 B8CH52 B8CH56 B8CH57
B8CH70 B8CH71 B8CH73 B8CH76 B8CH77 B8CH81 B8CHA2 B8CHB1 B8CHB5 B8CHB9 B8CHC9
B8CHJ7 B8CHM9 B8CHS0 B8CHY1 B8CHY3 B8CHZ1 B8CI06 B8CI18 B8CI19 B8CI29 B8CI99
B8CIB4 B8CID2 B8CID3 B8CIG1 B8CIL6 B8CIN2 B8CIP3 B8CIP5 B8CIQ1 B8CIQ2 B8CIQ3
B8CIQ4 B8CIQ9 B8CIT6 B8CIX3 B8CIX4 B8CIX6 B8CJ37 B8CJF0 B8CJF1 B8CJF9 B8CJG4
B8CJH9 B8CJM7 B8CJM8 B8CJN2 B8CJN3 B8CJP2 B8CJP8 B8CJP9 B8CJQ2 B8CJQ5 B8CJQ6
B8CJQ8 B8CJV9 B8CJW7 B8CJY1 B8CK16 B8CK17 B8CKA9 B8CKF0 B8CKF3 B8CKF4 B8CKF7
B8CKG5 B8CKG8 B8CKG9 B8CKH1 B8CKH3 B8CKH4 B8CKH5 B8CKH6 B8CKH8 B8CKI4 B8CKI5
B8CKL7 B8CKR1 B8CKR2 B8CKR5 B8CKS4 B8CL73 B8CLA1 B8CLC4 B8CLC5 B8CLG1 B8CLH4
B8CLI1 B8CLI5 B8CLJ4 B8CLM6 B8CM22 B8CM26 B8CM27 B8CM29 B8CM30 B8CM31 B8CM39
B8CM41 B8CM42 B8CM45 B8CM47 B8CM48 B8CM62 B8CME8 B8CMF1 B8CMG9 B8CMH2 B8CMH3
B8CMP0 B8CMS5 B8CMV8 B8CMW5 B8CMW9 B8CMX4 B8CMY5 B8CMY7 B8CN00 B8CN03 B8CN09
B8CN80 B8CN81 B8CN91 B8CNB7 B8CNC1 B8CNC2 B8CNC3 B8CNC4 B8CNC5 B8CNC6 B8CNC7
B8CNC8 B8CND3 B8CND4 B8CND5 B8CND6 B8CND7 B8CND8 B8CNE0 B8CNE1 B8CNE2 B8CNE3
B8CNE4 B8CNE5 B8CNE6 B8CNE7 B8CNE8 B8CNE9 B8CNF1 B8CNF2 B8CNF4 B8CNF5 B8CNF6
B8CNF7 B8CNF9 B8CNH3 B8CNH7 B8CNI0 B8CNI1 B8CNI9 B8CNK0 B8CNL2 B8CNP2 B8CNP8
B8CNQ6 B8CNR8 B8CNS0 B8CNS1 B8CNW1 B8CNX4 B8CNX5 B8CNX7 B8CNX9 B8CNY0 B8CNY1
B8CP82 B8CPA0 B8CPB0 B8CPE2 B8CPE7 B8CPG9 B8CPH7 B8CPI6 B8CPW5 B8CQ07 B8CQ71
B8CQ79 B8CQ82 B8CQ84 B8CQ85 B8CQG1 B8CQH4 B8CQI1 B8CQI2 B8CQI3 B8CQJ5 B8CQK7
B8CQL0 B8CQP2 B8CQQ1 B8CQR0 B8CQT6 B8CQV2 B8CQV3 B8CQV5 B8CQY2 B8CR00 B8CR48
B8CR53 B8CR54 B8CR57 B8CR59 B8CR70 B8CR73 B8CR83 B8CR95 B8CRF6 B8CRF7 B8CRF9
B8CRG0 B8CRT5 B8CS19 B8CS27 B8CS28 B8CS65 B8CS82 B8CS84 B8CSC1 B8CSC5 B8CSC8
B8CSD3 B8CSD4 B8CSF8 B8CSG9 B8CSH5 B8CSH6 B8CSH7 B8CSN6 B8CSV0 B8CSX5 B8CSX6
B8CSY3 B8CSY4 B8CSY5 B8CSY7 B8CSY8 B8CT15 B8CTC1 B8CTC6 B8CTC7 B8CTL3 B8CTW4
B8CTX5 B8CU18 B8CU29 B8CU40 B8CU41 B8CU43 B8CU98 B8CUC9 B8CUG6 B8CUS1 B8CUY8
B8CV32 B8CV40 B8CVC1 B8CVD1 B8CVD4 B8CVE5 B8CVJ1 B8CVJ3 B8CVL3 B8CVV5 B8CVV6
B8CVY1 B8CW05 B8CW13 B8CW18 B8CW20 B8CW21 B8CW53 B8CW54 B8CW56 B8CW61 B8CW65
B8CW69 B8CW72 B8CW73 B8CW77 B8CW78 B8CW79 B8CW89 B8CW97 B8CWA9 B8CWH0 B8CWH7
B8CWI5 B8CWI7 B8CWI8 B8CWK6 B8CWL4 B8CWM4 B8CWQ0 B8CWR7 B8CWS6 B8CWS7 B8CWU5
B8CWV6 B8CWV8 B8CWW0 B8CWY9 B8CX03 B8CX17 B8CX20 B8CX21 B8CX22 B8CX23 B8CX89
B8CX90 B8CX96 B8CX97 B8CX98 B8CXC3 B8CXE5 B8CXE6 B8CXE9 B8CXF1 B8CXG0 B8CXG1
B8CXG3 B8CXG4 B8CXG6 B8CXH4 B8CXH5 B8CXL0 B8CXL1 B8CXL9 B8CXR8 B8CXZ0 B8CXZ1
B8CY23 B8CY70 B8CY71 B8CY73 B8CY74 B8CY75 B8CYF2 B8CYG0 B8CYG6 B8CYG8 B8CYJ3
B8CYT9 B8CZ07 B8CZ30 B8CZ57 B8CZ59 B8CZN7 B8CZR7 B8CZT1 B8CZX0 B8CZZ9 B8D015
B8D067 B8D075 B8D093 B8D0A0 B8D0B2 B8D0B3 B8D0B4 B8D0B5 B8D0B9 B8D0C1 B8D0C4
B8D0C5 B8D0C7 B8D0C8 B8D0C9 B8D0D0 B8D0D1 B8D0D4 B8D0D5 B8D0D6 B8D0D7 B8D0D8
B8D0D9 B8D0E2 B8D0I2 B8D0L5 B8D0S4 B8D0S5 B8D0T3 B8D0T4 B8D0T6 B8D0T8 B8D0U9
B8D0W1 B8D0W3 B8D0Y6 B8D0Z3 B8D0Z4 B8D0Z5 B8D110 B8D111 B8D115 B8D116 B8D122
B8D124 B8D125 B8D151 B8D185 B8D1A3 B8D1C5 B8D1D5 B8D1D6 B8D1D7 B8D1G6 B8D1G8

B8D1H3 B8D1J4 B8D1K5 B8D1Z8 B8D274 B8D298 B8D2B0 B8D2C6 B8D2E3 B8D2E5 B8D2G0
B8D2H8 B8D2H9 B8D2I3 B8D2I6 B8D2J5 B8D355 B8D3V1 B8D3X7 B8D4I6 B8D4P0 B8D4W5
B8D4W8 B8D4Z8 B8D509 B8D537 B8D5I6 B8D5M3 B8D5N1 B8D5P9 B8D5Q6 B8D5Q7 B8D5T1
B8D5U3 B8D5U4 B8D5U8 B8D5U9 B8D5V1 B8D5V4 B8D5V5 B8D5V8 B8D5W2 B8D5W9 B8D610
B8D673 B8D680 B8D683 B8D687 B8D688 B8D691 B8D6B2 B8D6B7 B8D6C6 B8D6D0 B8D6D8
B8D6E8 B8D6F2 B8D6F5 B8D6G4 B8D6H2 B8D6L8 B8D6S0 B8D6T1 B8D6T2 B8D6T5 B8D6T6
B8D6T9 B8D6U7 B8D6V0 B8D6V1 B8D6V2 B8D6V3 B8D6V4 B8D6V9 B8D6W0 B8D6W1 B8D6W2
B8D6W5 B8D6W9 B8D6X0 B8D6X1 B8D6X3 B8D6X9 B8D6Z3 B8D6Z4 B8D6Z9 B8D701 B8D702
B8D703 B8D704 B8D705 B8D707 B8D708 B8D710 B8D715 B8D726 B8D730 B8D731 B8D732
B8D733 B8D736 B8D737 B8D738 B8D740 B8D741 B8D742 B8D743 B8D747 B8D751 B8D752
B8D754 B8D756 B8D757 B8D758 B8D775 B8D778 B8D787 B8D789 B8D797 B8D7A0 B8D7A1
B8D7A3 B8D7A8 B8D7B2 B8D7B3 B8D7B4 B8D7B5 B8D7B8 B8D7C1 B8D7C6 B8D7D2 B8D7D3
B8D7D5 B8D7D6 B8D7E1 B8D7E4 B8D7E7 B8D7F0 B8D7F1 B8D7F5 B8D7G3 B8D7G6 B8D7G7
B8D7H4 B8D7H6 B8D7I5 B8D7I7 B8D7J7 B8D7J8 B8D7K1 B8D7K3 B8D7K4 B8D7K5 B8D7K8
B8D7L7 B8D7L9 B8D7M3 B8D7N7 B8D7P8 B8D7Q2 B8D7Q4 B8D7Q5 B8D7Q6 B8D7Q7 B8D7R0
B8D7R1 B8D7R3 B8D7R4 B8D7R7 B8D7R9 B8D7S1 B8D7S2 B8D7S3 B8D7S4 B8D7S5 B8D7S6
B8D7S9 B8D7T2 B8D7T4 B8D7T5 B8D7T9 B8D7U0 B8D7U1 B8D7U4 B8D7U5 B8D7U6 B8D7U9
B8D7V3 B8D7V4 B8D7V6 B8D7V7 B8D7V9 B8D7W2 B8D7X0 B8D7X6 B8D7X7 B8D7Y0 B8D7Y8
B8D7Z2 B8D7Z3 B8D803 B8D805 B8D811 B8D813 B8D815 B8D816 B8D818 B8D820 B8D821
B8D822 B8D823 B8D825 B8D826 B8D827 B8D828 B8D830 B8D831 B8D833 B8D835 B8D837
B8D838 B8D839 B8D840 B8D841 B8D842 B8D843 B8D844 B8D845 B8D846 B8D847 B8D848
B8D849 B8D850 B8D851 B8D852 B8D853 B8D854 B8D855 B8D856 B8D857 B8D865 B8D866
B8D867 B8D868 B8D872 B8D875 B8D878 B8D879 B8D880 B8D881 B8D883 B8D884 B8D885
B8D887 B8D890 B8D891 B8D892 B8D894 B8D897 B8D899 B8D8A2 B8D8A3 B8D8A6 B8D8A8
B8D8A9 B8D8B0 B8D8B4 B8D8B5 B8D8B6 B8D8B8 B8D8B9 B8D8C1 B8D8C3 B8D8C4 B8D8C6
B8D8C7 B8D8C8 B8D8D0 B8D8D3 B8D8D4 B8D8D5 B8D8D7 B8D8E0 B8D8E2 B8D8E5 B8D8E6
B8D8E9 B8D8F1 B8D8F2 B8D8F3 B8D8F7 B8D8F8 B8D8F9 B8D8G1 B8D8G2 B8D8G4 B8D8G6
B8D8H7 B8D8H8 B8D8I1 B8D8I2 B8D8I5 B8D8J3 B8D8J6 B8D8J7 B8D8J8 B8D8J9 B8D8K0
B8D8K5 B8D8K6 B8D8K7 B8D8K8 B8D8L1 B8D8L5 B8D8L6 B8D8L7 B8D8L9 B8D8M5 B8D8N9
B8D8P0 B8D8P5 B8D8P7 B8D8P8 B8D8P9 B8D8Q0 B8D8Q1 B8D8Q3 B8D8Q4 B8D8Q6 B8D8R1
B8D8S2 B8D8S6 B8D8S7 B8D8S8 B8D8S9 B8D8T2 B8D8T3 B8D8T4 B8D8T6 B8D8T7 B8D8T8
B8D8T9 B8D8U3 B8D8U7 B8D8U8 B8D8V0 B8D8V2 B8D8V3 B8D8V4 B8D8X1 B8D8X4 B8D8Y4
B8D8Y9 B8D8Z2 B8D8Z5 B8D8Z6 B8D8Z8 B8D903 B8D907 B8D908 B8D909 B8D910 B8D913
B8D916 B8D922 B8D928 B8D929 B8D931 B8D932 B8D937 B8D940 B8D943 B8D946 B8D947
B8D951 B8D959 B8D962 B8D963 B8D970 B8D972 B8D981 B8D982 B8D983 B8D985 B8D995
B8D996 B8D999 B8D9A1 B8D9A2 B8D9A3 B8D9A6 B8D9B5 B8D9B7 B8D9C1 B8D9D5 B8D9E6
B8D9F0 B8D9F2 B8D9F3 B8D9F4 B8D9F5 B8D9F8 B8D9F9 B8D9G1 B8D9G2 B8D9G5 B8D9G7
B8D9G9 B8D9H0 B8D9H1 B8D9H2 B8D9H3 B8D9H4 B8D9H7 B8D9I0 B8D9I2 B8D9I3 B8D9I7
B8D9I8 B8D9I9 B8D9J2 B8D9J3 B8D9J4 B8D9J7 B8D9K1 B8D9K2 B8D9K4 B8D9K5 B8D9K7
B8D9L0 B8D9L8 B8D9M4 B8D9M5 B8D9M8 B8D9N6 B8D9P0 B8D9P1 B8D9Q1 B8D9Q3 B8D9Q9
B8D9R1 B8D9R3 B8D9R4 B8D9R6 B8D9R8 B8D9R9 B8D9S0 B8D9S1 B8D9S3 B8D9S4 B8D9S5
B8D9S6 B8D9S8 B8D9S9 B8D9T1 B8D9T3 B8D9T5 B8D9T6 B8D9T7 B8D9T8 B8D9T9 B8D9U0
B8D9U1 B8D9U2 B8D9U3 B8D9U4 B8D9U5 B8D9U6 B8D9U7 B8D9U8 B8D9U9 B8D9V0 B8D9V1
B8D9V2 B8D9V3 B8D9V4 B8D9V5 B8D9W3 B8D9W4 B8D9W5 B8D9W6 B8D9X0 B8D9X3 B8D9X6

B8D9X7 B8D9Y4 B8DA56 B8DA57 B8DA59 B8DA60 B8DA61 B8DA62 B8DA63 B8DA64 B8DA86
B8DA91 B8DAL0 B8DAL3 B8DAM0 B8DAM2 B8DAM3 B8DAQ5 B8DAQ9 B8DAR1 B8DAT8 B8DAT9
B8DAU8 B8DAY4 B8DAY5 B8DAY6 B8DAY7 B8DB07 B8DB08 B8DB09 B8DB10 B8DB12 B8DB13
B8DB15 B8DB16 B8DB17 B8DB18 B8DB19 B8DB20 B8DB21 B8DB22 B8DB23 B8DB24 B8DB26
B8DB27 B8DB29 B8DB31 B8DB32 B8DB33 B8DB34 B8DB35 B8DB42 B8DB43 B8DB44 B8DBE8
B8DBG1 B8DBG4 B8DBG6 B8DBH0 B8DBH1 B8DBM8 B8DBN0 B8DBN7 B8DBN9 B8DBP0 B8DBQ3
B8DBQ6 B8DBR2 B8DBR3 B8DBT6 B8DBU1 B8DBU2 B8DBU4 B8DBU5 B8DBU8 B8DBV5 B8DBV7
B8DBX2 B8DBX5 B8DBX6 B8DBY7 B8DBY9 B8DBZ3 B8DBZ5 B8DBZ7 B8DBZ8 B8DBZ9 B8DC01
B8DC03 B8DC05 B8DC19 B8DC20 B8DC21 B8DC24 B8DC25 B8DC26 B8DC39 B8DC56 B8DCC1
B8DCE2 B8DCE7 B8DCF0 B8DCF7 B8DCG2 B8DCI1 B8DCI6 B8DCJ3 B8DCJ5 B8DCT6 B8DCT7
B8DCT8 B8DCV5 B8DCW1 B8DCW2 B8DCW3 B8DD12 B8DD13 B8DD20 B8DD73 B8DD98 B8DDA8
B8DDD3 B8DDG0 B8DDL2 B8DDL4 B8DDL5 B8DDM5 B8DDM6 B8DDM8 B8DDP2 B8DDP3 B8DDP5
B8DDP6 B8DDR2 B8DDR4 B8DDR5 B8DDR7 B8DDR8 B8DDR9 B8DDS0 B8DDS6 B8DDS9 B8DDT6
B8DDU8 B8DDV6 B8DDW0 B8DDW1 B8DDW6 B8DDX3 B8DDX7 B8DDY5 B8DDY7 B8DDY8 B8DDZ0
B8DDZ2 B8DDZ8 B8DE07 B8DE09 B8DE13 B8DE15 B8DE17 B8DE23 B8DE31 B8DE36 B8DE37
B8DE38 B8DE39 B8DE40 B8DE42 B8DE46 B8DE51 B8DE52 B8DE53 B8DE62 B8DE74 B8DE89
B8DEA7 B8DEC3 B8DEC4 B8DEE7 B8DEG2 B8DEH8 B8DEK7 B8DEL2 B8DET0 B8DET2 B8DEV0
B8DEY8 B8DF04 B8DF05 B8DF06 B8DF07 B8DF15 B8DF19 B8DF43 B8DF44 B8DF87 B8DF98
B8DFA5 B8DFG5 B8DFG7 B8DFG8 B8DFG9 B8DFL4 B8DFM9 B8DFN5 B8DFN7 B8DFQ7 B8DFR1
B8DFR3 B8DFR4 B8DFS3 B8DFS4 B8DFS9 B8DFT0 B8DFU3 B8DFU4 B8DFV8 B8DFW1 B8DFW6
B8DFW8 B8DFW9 B8DFX3 B8DFX8 B8DFX9 B8DFY0 B8DFZ6 B8DFZ7 B8DG00 B8DG02 B8DG06
B8DG08 B8DG10 B8DG13 B8DG24 B8DG34 B8DG51 B8DG52 B8DG53 B8DG54 B8DG60 B8DG65
B8DG73 B8DGJ3 B8DGK3 B8DGK4 B8DGL2 B8DGL7 B8DGN4 B8DGN5 B8DGN7 B8DGQ7 B8DH02
B8DH16 B8DH17 B8DH27 B8DH28 B8DH58 B8DH59 B8DH75 B8DH77 B8DH80 B8DH87 B8DH88
B8DHB1 B8DHB2 B8DHB3 B8DHB4 B8DHB5 B8DHC7 B8DHD4 B8DHD7 B8DHE4 B8DHE5 B8DHE9
B8DHF0 B8DHF2 B8DHF4 B8DHH0 B8DHH7 B8DHI6 B8DHI9 B8DHJ0 B8DHJ1 B8DHJ2 B8DHJ5
B8DHJ9 B8DHK3 B8DHK6 B8DHK8 B8DHL0 B8DHL1 B8DHL5 B8DHL6 B8DHL7 B8DHL8 B8DHM4
B8DHM6 B8DHM9 B8DHN2 B8DHN7 B8DHN8 B8DHP2 B8DHP3 B8DHZ4 B8DHZ5 B8DHZ8 B8DI00
B8DI11 B8DIJ5 B8DIL5 B8DIU7 B8DJ34 B8DJ66 B8DJB8 B8DJC3 B8DJC4 B8DJF0 B8DJF1
B8DJF2 B8DJF7 B8DJG3 B8DJG5 B8DJI3 B8DJJ6 B8DJK7 B8DJL8 B8DJP7 B8DJQ9 B8DJR8
B8DJT5 B8DJT6 B8DJT9 B8DJU4 B8DKC9 B8DKD1 B8DKI5 B8DKL0 B8DKL3 B8DKL6 B8DKL7
B8DKM6 B8DKP2 B8DKS6 B8DKU0 B8DKV6 B8DKV9 B8DKW0 B8DKW2 B8DL62 B8DL94 B8DLK7
B8DLK8 B8DLL6 B8DLL9 B8DLM3 B8DLM4 B8DLM5 B8DLM7 B8DLN4 B8DLR0 B8DM45 B8DM91
B8DM92 B8DMA9 B8DMB1 B8DMG5 B8DMK7 B8DMM1 B8DMM2 B8DMV2 B8DN02 B8DN07 B8DN08
B8DN09 B8DN15 B8DN19 B8DN39 B8DN41 B8DN64 B8DN65 B8DN76 B8DN92 B8DN93 B8DN94
B8DN95 B8DN96 B8DN97 B8DN98 B8DN99 B8DNA0 B8DNA1 B8DNA2 B8DNA3 B8DNA4 B8DNA5
B8DNA6 B8DNA7 B8DNA8 B8DNA9 B8DNB0 B8DNB1 B8DNB2 B8DNB3 B8DNB4 B8DNB6 B8DNB9
B8DNF4 B8DNJ4 B8DNJ5 B8DNJ8 B8DNK1 B8DNK2 B8DNK5 B8DNK6 B8DNK7 B8DNK8 B8DNK9
B8DNL0 B8DNL1 B8DNL3 B8DNL4 B8DNM5 B8DNQ9 B8DNS7 B8DNV5 B8DNW9 B8DNX4 B8DP00
B8DP14 B8DP34 B8DP81 B8DP86 B8DP87 B8DP96 B8DP97 B8DPB3 B8DPD6 B8DPF1 B8DPG0
B8DPG1 B8DPG2 B8DPG3 B8DPH7 B8DPL7 B8DPL8 B8DPM3 B8DPM8 B8DPM9 B8DPN0 B8DPN1
B8DPN2 B8DPP9 B8DQ18 B8DQ42 B8DQP8 B8DQQ9 B8DQU5 B8DQW7 B8DQW8 B8DQX6 B8DQX7
B8DQY7 B8DR41 B8DRD6 B8DRD8 B8DRE0 B8DRI3 B8DRI7 B8DRL1 B8DRL2 B8DRL3 B8DRL4
B8DRN9 B8DRP0 B8DRP1 B8DRQ6 B8DRR1 B8DRR9 B8DRT6 B8DRU2 B8DRV3 B8DSA1 B8DSA5

B8DSA6 B8DSC3 B8DSC4 B8DSI2 B8DSI4 B8DSK3 B8DSQ0 B8DSQ1 B8DSS1 B8DSU7 B8DSV1
B8DSV5 B8DSW1 B8DSW4 B8DSX3 B8DSX7 B8DSY8 B8DT03 B8DT48 B8DT61 B8DT62 B8DT91
B8DTD1 B8DTI5 B8DTL4 B8DTM3 B8DTN1 B8DTP8 B8DTT6 B8DTV0 B8DTV4 B8DTV6 B8DTV7
B8DTW0 B8DTW1 B8DTW3 B8DTW7 B8DTW8 B8DTX1 B8DTX4 B8DTX8 B8DTZ2 B8DU04 B8DU32
B8DU51 B8DU54 B8DU56 B8DU64 B8DUA3 B8DUA4 B8DUA6 B8DUB2 B8DUC2 B8DUC3 B8DUE5
B8DUE8 B8DUE9 B8DUF0 B8DUF4 B8DUG2 B8DUH3 B8DUH4 B8DUJ6 B8DUK6 B8DUL7 B8DUM8
B8DUN4 B8DUQ4 B8DUR2 B8DUR4 B8DUT6 B8DUV2 B8DUY2 B8DV04 B8DV05 B8DV06 B8DV60
B8DVC2 B8DVE0 B8DVI9 B8DVK3 B8DVN0 B8DVN1 B8DVN7 B8DVP8 B8DVQ3 B8DVQ6 B8DVR7
B8DVS0 B8DVS6 B8DVT9 B8DVU0 B8DVU1 B8DVU2 B8DVU7 B8DVV7 B8DVV8 B8DVW3 B8DVY4
B8DVY5 B8DVZ4 B8DW05 B8DW06 B8DW12 B8DW14 B8DW15 B8DW16 B8DW17 B8DW18 B8DW19
B8DW20 B8DW22 B8DW23 B8DW24 B8DW26 B8DW27 B8DW28 B8DW29 B8DW30 B8DW37 B8DW39
B8DW40 B8DW43 B8DW52 B8DWC0 B8DWH9 B8DWI2 B8DWI3 B8DWJ5 B8DWK0 B8DWP6 B8DWR0
B8DWS0 B8DYH6 B8DYK8 B8DYM0 B8DYS1 B8DYS2 B8DZH0 B8DZH3 B8DZK4 B8DZP4 B8DZP8
B8DZS1 B8DZW2 B8DZX0 B8DZX8 B8DZY2 B8DZZ9 B8E001 B8E003 B8E004 B8E008 B8E009
B8E029 B8E032 B8E088 B8E089 B8E0B1 B8E0B2 B8E0B3 B8E0B4 B8E0D2 B8E0D5 B8E0D9
B8E0E2 B8E0E8 B8E0E9 B8E0F1 B8E0F9 B8E0G7 B8E0J0 B8E0J2 B8E0K0 B8E0K1 B8E0K2
B8E0K3 B8E0M0 B8E0N8 B8E0N9 B8E0R9 B8E0T1 B8E0T7 B8E0U2 B8E0X3 B8E0X6 B8E0X7
B8E0X9 B8E0Y1 B8E0Z3 B8E0Z4 B8E109 B8E120 B8E121 B8E181 B8E183 B8E191 B8E1A7
B8E1A9 B8E1B4 B8E1C0 B8E1C1 B8E1C2 B8E1C8 B8E1C9 B8E1D6 B8E1D7 B8E1D9 B8E1E3
B8E1E4 B8E1E5 B8E1E6 B8E1E7 B8E1E8 B8E1E9 B8E1F6 B8E1F7 B8E1F8 B8E1F9 B8E1G1
B8E1G5 B8E1G6 B8E1Q9 B8E200 B8E222 B8E227 B8E240 B8E246 B8E247 B8E257 B8E259
B8E280 B8E281 B8E291 B8E2A8 B8E2C2 B8E2C5 B8E2C7 B8E2C8 B8E2D1 B8E2D3 B8E2D8
B8E2E7 B8E2F5 B8E2G0 B8E2G3 B8E2G4 B8E2G7 B8E2H4 B8E2K1 B8E2N5 B8E2P3 B8E2P4
B8E2P6 B8E2R2 B8E2S1 B8E2S4 B8E2S5 B8E2S7 B8E2T7 B8E2U4 B8E2W8 B8E2W9 B8E2Y1
B8E2Y2 B8E2Y4 B8E2Z9 B8E324 B8E335 B8E358 B8E391 B8E395 B8E3F6 B8E3J4 B8E3K4
B8E3K7 B8E3K9 B8E3L5 B8E3L7 B8E3M2 B8E3P4 B8E3P6 B8E3Q5 B8E3Q6 B8E3R0 B8E3R4
B8E3S3 B8E3S8 B8E3W0 B8E3W7 B8E3X7 B8E465 B8E478 B8E4B8 B8E4C0 B8E4F6 B8E4I8
B8E4J0 B8E4J2 B8E4J3 B8E4J5 B8E4J6 B8E4J7 B8E4K4 B8E4K6 B8E4K7 B8E4K9 B8E4L1
B8E4L2 B8E4M8 B8E4P5 B8E4S1 B8E4S2 B8E4S9 B8E4U3 B8E4W4 B8E4W5 B8E4W6 B8E4X2
B8E4X4 B8E4X7 B8E520 B8E525 B8E544 B8E569 B8E599 B8E5A2 B8E5E8 B8E5E9 B8E5F0
B8E5F1 B8E5F2 B8E5Q1 B8E5Q5 B8E5T2 B8E5U4 B8E5U5 B8E5U7 B8E642 B8E665 B8E666
B8E670 B8E680 B8E682 B8E690 B8E699 B8E6B6 B8E6E3 B8E6E4 B8E6G9 B8E6H2 B8E6K8
B8E6L3 B8E6M4 B8E6N0 B8E6N2 B8E6N4 B8E6N5 B8E6N7 B8E6N9 B8E6P4 B8E6P5 B8E6P6
B8E6P7 B8E6Q7 B8E6W1 B8E6W2 B8E6W6 B8E6W7 B8E700 B8E702 B8E703 B8E718 B8E724
B8E732 B8E735 B8E744 B8E749 B8E759 B8E770 B8E786 B8E794 B8E7C8 B8E7F1 B8E7F3
B8E7F5 B8E7F7 B8E7F9 B8E7G0 B8E7G2 B8E7M2 B8E7N1 B8E7P4 B8E7Q2 B8E7R0 B8E7R3
B8E7R5 B8E7R6 B8E7T7 B8E801 B8E815 B8E816 B8E818 B8E835 B8E879 B8E8A9 B8E8D2
B8E8D7 B8E8I2 B8E8S1 B8E8S8 B8E8T0 B8E8T4 B8E8T6 B8E8T7 B8E8U0 B8E8U2 B8E8U6
B8E8Y9 B8E916 B8E918 B8E920 B8E927 B8E945 B8E950 B8E954 B8E958 B8E960 B8E961
B8E968 B8E975 B8E999 B8E9A0 B8E9A5 B8E9B6 B8E9D2 B8E9D5 B8E9D6 B8E9E1 B8E9F2
B8E9H2 B8E9K8 B8E9M3 B8E9M7 B8E9M8 B8E9N6 B8E9N7 B8E9N8 B8E9N9 B8E9P2 B8E9P5
B8E9P8 B8E9S4 B8E9S7 B8E9S8 B8E9T1 B8E9T2 B8E9T4 B8EA23 B8EA59 B8EA68 B8EA69
B8EA71 B8EA73 B8EA74 B8EA75 B8EA88 B8EA90 B8EA95 B8EA96 B8EA98 B8EAH9 B8EAI0
B8EAI2 B8EAI5 B8EAJ2 B8EAL3 B8EAL4 B8EAS7 B8EAT2 B8EAT7 B8EAU2 B8EAU7 B8EAU9

B8EAW7 B8EAX1 B8EAZ5 B8EB35 B8EB36 B8EB37 B8EB45 B8EB46 B8EB47 B8EB55 B8EB56
B8EB82 B8EBC2 B8EBD9 B8EBE0 B8EBF1 B8EBF9 B8EBG0 B8EBG2 B8EBG3 B8EBG6 B8EBH9
B8EBI1 B8EBI2 B8EBI3 B8EBI4 B8EBI6 B8EBI7 B8EBI8 B8EBI9 B8EBJ0 B8EBJ1 B8EBJ3
B8EBJ4 B8EBJ5 B8EBJ6 B8EBJ7 B8EBJ8 B8EBJ9 B8EBK0 B8EBK1 B8EBK2 B8EBK4 B8EBK5
B8EBK6 B8EBK9 B8EBL0 B8EBL1 B8EBL2 B8EBL3 B8EBL4 B8EBL5 B8EBL6 B8EBM0 B8EBM4
B8EBM5 B8EBM9 B8EBN0 B8EBN7 B8EBP0 B8EBP1 B8EBP2 B8EBP3 B8EBQ2 B8EBR5 B8EBR9
B8EBS0 B8EBS7 B8EBT5 B8EBT9 B8EBU3 B8EBU8 B8EBV5 B8EBV6 B8EC04 B8EC05 B8EC06
B8EC83 B8ECA3 B8ECA4 B8ECC0 B8ECC6 B8ECE1 B8ECJ0 B8ECN4 B8ECV6 B8EDD0 B8EDJ6
B8EDJ7 B8EDQ5 B8EDQ9 B8EDR6 B8EDS6 B8EDT1 B8EDT7 B8EDW0 B8EDW1 B8EDW7 B8EDW8
B8EDX3 B8EDX9 B8EE43 B8EE51 B8EE60 B8EE63 B8EEA5 B8EEA7 B8EEB4 B8EEB7 B8EEJ1
B8EEK5 B8EEN5 B8EES6 B8EES7 B8EES8 B8EES9 B8EET7 B8EEX7 B8EF24 B8EF35 B8EF44
B8EF62 B8EFC7 B8EFD0 B8EFD3 B8EFD7 B8EFD8 B8EI29 B8EI32 B8EI82 B8EIA0 B8EIA6
B8EIA7 B8EIB0 B8EIB7 B8EIL1 B8EIL3 B8EIP9 B8EIQ3 B8EIR0 B8EIS2 B8EIS6 B8EIT3
B8EIT4 B8EIT5 B8EIT6 B8EIU8 B8EIV1 B8EJ84 B8EJI4 B8EJI5 B8EJI6 B8EJL7 B8EJS1
B8EJS2 B8EJS8 B8EJT0 B8EJU1 B8EJY5 B8EJZ0 B8EJZ2 B8EK13 B8EK21 B8EK29 B8EK30
B8EK40 B8EK41 B8EK46 B8EKA0 B8EKA1 B8EKB9 B8EKE1 B8EKI5 B8EKP4 B8EKR5 B8EL92
B8ELB8 B8ELE2 B8ELE4 B8ELE6 B8ELE7 B8ELE8 B8ELE9 B8ELF2 B8ELF3 B8ELF4 B8ELF6
B8ELF7 B8ELF8 B8ELF9 B8ELG0 B8ELG2 B8ELG3 B8ELG4 B8ELG5 B8ELG6 B8ELG7 B8ELG8
B8ELG9 B8ELI3 B8ELJ0 B8ELJ6 B8ELJ7 B8ELV1 B8ELZ9 B8EM00 B8EM14 B8EM38 B8EM72
B8EM84 B8EM86 B8EM89 B8EM94 B8EMF2 B8EMJ5 B8EML2 B8EMN5 B8EMP4 B8EMP5 B8EMR7
B8EMR8 B8EMR9 B8EMS0 B8EMX4 B8EMY2 B8EMZ0 B8EMZ5 B8EN54 B8EN55 B8EN59 B8EN73
B8EN93 B8EN95 B8EN97 B8ENA6 B8ENB9 B8ENG6 B8ENG8 B8ENI5 B8ENI9 B8ENL8 B8ENN1
B8EP09 B8EP10 B8EP16 B8EP17 B8EP34 B8EP84 B8EPC0 B8EPC1 B8EPC2 B8EPC4 B8EPC5
B8EPE8 B8EPF2 B8EPG3 B8EPH9 B8EPI0 B8EPJ0 B8EPL9 B8EPM3 B8EPU6 B8EPU9 B8EPV0
B8EPV1 B8EPX5 B8EPX7 B8EQ21 B8EQB8 B8EQB9 B8EQC8 B8EQG6 B8EQH0 B8EQH4 B8EQH6
B8EQH7 B8EQM2 B8EQN3 B8EQP4 B8ER03 B8ER19 B8ER20 B8ER26 B8ER30 B8ER45 B8ERB9
B8ERC6 B8ERJ6 B8ERL9 B8ERN7 B8ERW8 B8ES21 B8ES25 B8ES44 B8ESC6 B8ESC7 B8ESH8
B8ESH9 B8ESI1 B8ESL2 B8ESQ6 B8ESQ9 B8ESV1 B8ESV2 B8ET39 B8ET77 B8ETE4 B8ETG9
B8ETL3 B8ETL4 B8ETM5 B8ETM9 B8ETN6 B8F314 B8F356 B8F382 B8F389 B8F392 B8F3A4
B8F3A7 B8F3A8 B8F3B7 B8F3B8 B8F3C7 B8F3C9 B8F3D3 B8F3D4 B8F3E3 B8F3E4 B8F3F6
B8F3F7 B8F3G3 B8F3G5 B8F3G6 B8F3G9 B8F3J3 B8F3K1 B8F3P6 B8F3Q0 B8F3R6 B8F3R7
B8F3R8 B8F3S8 B8F3T2 B8F3V1 B8F3W0 B8F3W5 B8F3W7 B8F3W8 B8F3Y5 B8F407 B8F412
B8F415 B8F420 B8F426 B8F454 B8F457 B8F472 B8F475 B8F480 B8F492 B8F4B1 B8F4B2
B8F4C0 B8F4C5 B8F4D0 B8F4P0 B8F4P3 B8F4P6 B8F4Q2 B8F4Q5 B8F4Q7 B8F4R5 B8F4R6
B8F4R9 B8F4T5 B8F4T7 B8F4U0 B8F4U8 B8F4V0 B8F4V1 B8F4V4 B8F4V8 B8F4W1 B8F4X2
B8F4X6 B8F4X7 B8F4Y1 B8F503 B8F506 B8F526 B8F531 B8F542 B8F557 B8F571 B8F586
B8F5E2 B8F5E6 B8F5F8 B8F5G6 B8F5G8 B8F5H9 B8F5J4 B8F5K3 B8F5K4 B8F5L9 B8F5M9
B8F5N0 B8F5N2 B8F5P3 B8F5Q2 B8F5Q3 B8F5Q7 B8F5Q8 B8F5Q9 B8F5R1 B8F5R2 B8F5T4
B8F5T5 B8F5T7 B8F5U5 B8F5X3 B8F5Z2 B8F5Z8 B8F605 B8F632 B8F634 B8F647 B8F662
B8F663 B8F667 B8F668 B8F671 B8F672 B8F678 B8F679 B8F689 B8F690 B8F6A3 B8F6C3
B8F6C4 B8F6C8 B8F6C9 B8F6D3 B8F6D8 B8F6E6 B8F6E9 B8F6G2 B8F6H5 B8F6H9 B8F6J5
B8F6J8 B8F6K3 B8F6K8 B8F6L5 B8F6M3 B8F6N0 B8F6N1 B8F6N2 B8F6N9 B8F6P5 B8F6P7
B8F6P8 B8F6Q2 B8F6Q3 B8F6Q5 B8F6Q6 B8F6Q7 B8F6Q9 B8F6R0 B8F6S2 B8F6T6 B8F6T7
B8F6T9 B8F6W0 B8F6X2 B8F6Y2 B8F6Y7 B8F704 B8F706 B8F711 B8F713 B8F726 B8F727

B8F738 B8F741 B8F744 B8F754 B8F755 B8F756 B8F758 B8F759 B8F760 B8F762 B8F763

B8F764 B8F777 B8F789 B8F790 B8F7C2 B8F7C5 B8F7C7 B8F7D1 B8F7D2 B8F7D3 B8F7G0

B8F7G7 B8F7H5 B8F7K7 B8F7P4 B8F7R8 B8F7S3 B8F7S6 B8F7T1 B8F7U6 B8F7W4 B8F7W5

B8F7W7 B8F7X3 B8F7X4 B8F7X5 B8F7Y7 B8F7Z1 B8F7Z2 B8F7Z3 B8F7Z4 B8F815 B8F822

B8F824 B8F853 B8F854 B8F858 B8F859 B8F864 B8F865 B8F870 B8F871 B8F874 B8F875

B8F877 B8F878 B8F882 B8F8E6 B8F8G9 B8F8H3 B8F8H4 B8F8H7 B8F8L6 B8F8N4 B8F8Q1

B8F8R7 B8F8W4 B8F9C9 B8F9F0 B8F9F2 B8F9J5 B8F9K1 B8F9W2 B8FA37 B8FA39 B8FA41

B8FA61 B8FA62 B8FA63 B8FAC4 B8FB10 B8FB52 B8FB71 B8FB83 B8FBA1 B8FBE8 B8FBE9

B8FBH4 B8FBL5 B8FBR7 B8FBS0 B8FBS6 B8FBT7 B8FC91 B8FCK9 B8FCL0 B8FCL2 B8FCY3

B8FCY5 B8FCY7 B8FCY9 B8FCZ0 B8FDS1 B8FEA2 B8FEA3 B8FEI1 B8FER0 B8FER1 B8FER2

B8FER6 B8FER7 B8FER9 B8FES0 B8FES2 B8FES3 B8FES5 B8FES7 B8FES8 B8FES9 B8FET0

B8FET1 B8FET2 B8FET3 B8FET4 B8FET5 B8FET6 B8FET8 B8FET9 B8FEU1 B8FEU2 B8FEU3

B8FEU4 B8FEU5 B8FEV1 B8FFL4 B8FFL8 B8FFQ6 B8FFR9 B8FFU1 B8FFU2 B8FFU3 B8FG16

B8FGA1 B8FGA3 B8FGE3 B8FGE6 B8FGS3 B8FGS4 B8FGS8 B8FGS9 B8FGU7 B8FH08 B8FH11

B8FH26 B8FH30 B8FH31 B8FH38 B8FHH0 B8FHH7 B8FHQ1 B8FHQ6 B8FHR2 B8FHU4 B8FI50

B8FI54 B8FI57 B8FI62 B8FIN2 B8FJ17 B8FJ67 B8FJ72 B8FJ73 B8FJ77 B8FJ78 B8FJ89

B8FJL5 B8FJP4 B8FJV3 B8FJW2 B8FJY6 B8FK13 B8FK51 B8FKD5 B8FKD8 B8FKD9 B8FKE1

B8FKS6 B8FKS7 B8FKV1 B8FL12 B8FL13 B8FL28 B8FL92 B8FLB8 B8FLC3 B8FLE5 B8FLE7

B8FLE8 B8FLF0 B8FLF4 B8FLI1 B8FLT9 B8FLU0 B8FLV5 B8FLW6 B8FM51 B8FM66 B8FM67

B8FM68 B8FM70 B8FM86 B8FM87 B8FMB1 B8FMM4 B8FMP4 B8FMU1 B8FMU9 B8FMV0 B8FMV2

B8FMW6 B8FN26 B8FN99 B8FNA6 B8FNF7 B8FNN4 B8FNN5 B8FNP1 B8FNP3 B8FNP6 B8FNT2

B8FNT6 B8FNT7 B8FP01 B8FP03 B8FP05 B8FP16 B8FP18 B8FP20 B8FP22 B8FP23 B8FP24

B8FP44 B8FP45 B8FPL1 B8FPN5 B8FPU4 B8FQ41 B8FQ45 B8FQ61 B8FQ76 B8FQ77 B8FQ92

B8FQB9 B8FQC1 B8FQR7 B8FQT8 B8FQU0 B8FQV2 B8FQV3 B8FQV5 B8FQV7 B8FQV9 B8FQZ3

B8FQZ6 B8FR00 B8FR05 B8FR10 B8FR28 B8FR31 B8FRA6 B8FRG2 B8FRG4 B8FRG5 B8FRG8

B8FRG9 B8FRH5 B8FRJ6 B8FRK8 B8FRL1 B8FRL6 B8FRL9 B8FRM1 B8FRT2 B8FS67 B8FS78

B8FS81 B8FS83 B8FSB9 B8FSC1 B8FSC3 B8FSD4 B8FSV8 B8FT13 B8FT31 B8FT33 B8FT35

B8FT58 B8FT64 B8FT65 B8FTK5 B8FTP6 B8FTR7 B8FTV6 B8FTV7 B8FTY8 B8FU83 B8FU92

B8FU95 B8FU98 B8FUB6 B8FUD2 B8FUD3 B8FUD4 B8FUI8 B8FUJ0 B8FUJ1 B8FUJ2 B8FUK8

B8FUK9 B8FUL3 B8FUM7 B8FUN2 B8FUN3 B8FUN4 B8FUN6 B8FUQ0 B8FUR3 B8FUR7 B8FUR9

B8FUS1 B8FUS3 B8FUT0 B8FVF5 B8FVF9 B8FVH0 B8FVH4 B8FVH5 B8FVI0 B8FW04 B8FW05

B8FW28 B8FWE3 B8FWE4 B8FWP2 B8FWT4 B8FWU2 B8FWU9 B8FWX2 B8FWX3 B8FXB6 B8FXD7

B8FXD8 B8FXS7 B8FXV6 B8FXW4 B8FXW8 B8FXX3 B8FY16 B8FY38 B8FY51 B8FYS8 B8FYW5

B8FZ68 B8FZ69 B8FZ76 B8FZ78 B8FZ81 B8FZ89 B8FZA5 B8FZD1 B8FZE0 B8FZE1 B8FZE5

B8FZK1 B8FZK2 B8FZM2 B8FZR3 B8FZR4 B8G089 B8G0H2 B8G0I5 B8G0I9 B8G0J4 B8G0K1

B8G0L2 B8G0L7 B8G0L8 B8G0N6 B8G0R5 B8G120 B8G177 B8G186 B8G187 B8G1D7 B8G1G1

B8G1G3 B8G1G5 B8G1I4 B8G1J0 B8G1S3 B8G1T9 B8G1U2 B8G1V0 B8G1V1 B8G1V3 B8G1V8

B8G1W1 B8G1W2 B8G1W3 B8G1W4 B8G1W5 B8G1W7 B8G1W8 B8G1W9 B8G1X0 B8G1X1 B8G1X2

B8G1X3 B8G1X4 B8G1X5 B8G1X6 B8G1X8 B8G1X9 B8G1Y0 B8G1Y1 B8G1Y2 B8G1Y3 B8G1Y4

B8G1Y5 B8G1Y7 B8G1Z0 B8G1Z1 B8G1Z2 B8G1Z3 B8G1Z4 B8G1Z5 B8G242 B8G289 B8G2K7

B8G2L6 B8G2M2 B8G2M3 B8G2P9 B8G2Q5 B8G2R0 B8G2S3 B8G2S7 B8G2V0 B8G2Z2 B8G358

B8G3A9 B8G3E6 B8G3F7 B8G3J6 B8G3M6 B8G3V5 B8G3X3 B8G3Z0 B8G3Z1 B8G418 B8G451

B8G458 B8G461 B8G4D0 B8G4H3 B8G4I7 B8G4I8 B8G4J9 B8G4R4 B8G4T8 B8G4U8 B8G4U9

B8G4V8 B8G4W8 B8G503 B8G513 B8G568 B8G5C5 B8G5F1 B8G5G4 B8G5H0 B8G5R5 B8G5S6

B8G5X1 B8G5X3 B8G5Y3 B8G5Y4 B8G607 B8G635 B8G643 B8G644 B8G663 B8G664 B8G6B1
B8G6F7 B8G6K6 B8G6M3 B8G6P2 B8G6P5 B8G6P6 B8G6P7 B8G6P9 B8G6Q1 B8G6Q2 B8G6Q6
B8G6Q7 B8G6Q8 B8G6Q9 B8G6R0 B8G6R1 B8G6R3 B8G6R4 B8G6R5 B8G6R7 B8G6R8 B8G6R9
B8G6S0 B8G6S1 B8G6S2 B8G6S5 B8G6S6 B8G6S9 B8G6T0 B8G6W9 B8G6X1 B8G6X2 B8G6Y1
B8G6Y3 B8G711 B8G736 B8G762 B8G782 B8G783 B8G7I4 B8G7I8 B8G7K1 B8G7M3 B8G7M4
B8G7S5 B8G7X1 B8G7X5 B8G7Y8 B8G806 B8G817 B8G821 B8G822 B8G845 B8G857 B8G8E0
B8G8K0 B8G8N0 B8G8R1 B8G8S1 B8G8T0 B8G8T1 B8G933 B8G943 B8G945 B8G974 B8G975
B8G988 B8G989 B8G990 B8G991 B8G9R5 B8G9R8 B8G9S2 B8G9S3 B8G9X1 B8G9X9 B8G9Z3
B8GA36 B8GA45 B8GA54 B8GAA4 B8GAC1 B8GAE1 B8GAE2 B8GAM6 B8GAQ5 B8GAQ9 B8GAY2
B8GAY7 B8GB21 B8GB37 B8GBB1 B8GBF3 B8GBF7 B8GBG7 B8GBK7 B8GBN6 B8GBV4 B8GBV5
B8GC03 B8GC39 B8GC51 B8GC73 B8GC95 B8GCC9 B8GCH4 B8GCI2 B8GCM2 B8GCQ8 B8GCS2
B8GCV1 B8GD12 B8GDA2 B8GDE2 B8GDE3 B8GDF2 B8GDH2 B8GDM7 B8GDM8 B8GDQ7 B8GDW0
B8GDZ3 B8GDZ6 B8GE19 B8GEA3 B8GEC0 B8GEC2 B8GEG1 B8GEG4 B8GEG7 B8GEI1 B8GET9
B8GEU2 B8GEU4 B8GEU5 B8GEU6 B8GEV3 B8GEX3 B8GEX4 B8GEZ3 B8GEZ4 B8GEZ7 B8GF03
B8GF04 B8GF12 B8GF42 B8GF55 B8GF70 B8GF73 B8GFB9 B8GFC3 B8GFC4 B8GFF8 B8GFL4
B8GFL8 B8GFN1 B8GFN4 B8GFN7 B8GFN8 B8GFS9 B8GFU5 B8GFW4 B8GG34 B8GG35 B8GG66
B8GG79 B8GG80 B8GGB8 B8GGN4 B8GGP6 B8GGQ0 B8GGQ3 B8GGS3 B8GGX9 B8GH05 B8GH15
B8GH19 B8GH25 B8GHE0 B8GHM1 B8GHN1 B8GHN2 B8GI76 B8GI94 B8GIA0 B8GIB0 B8GIB2
B8GIB3 B8GIC5 B8GIF8 B8GIQ3 B8GIQ4 B8GIR5 B8GIX8 B8GIY2 B8GJ30 B8GJF9 B8GJJ2
B8GJK8 B8GJL3 B8GJL7 B8GJL9 B8GJN8 B8GJQ7 B8GJQ9 B8GJR4 B8GJR6 B8GJW5 B8GJY2
B8GJY3 B8GKC0 B8GKC9 B8GKD3 B8GKD6 B8GKE2 B8GKE8 B8GKF2 B8GKF9 B8GKG7 B8GKG8
B8GKL6 B8GKX5 B8GKY4 B8GL18 B8GL19 B8GL29 B8GL94 B8GL95 B8GL96 B8GLA1 B8GLA7
B8GLD4 B8GLD8 B8GLE3 B8GLE4 B8GM40 B8GM72 B8GMA8 B8GMB3 B8GMG5 B8GMI2 B8GMK2
B8GMM0 B8GMM1 B8GMV6 B8GMX6 B8GMX9 B8GMY8 B8GN06 B8GN14 B8GN16 B8GN38 B8GN57
B8GN62 B8GN64 B8GN76 B8GN79 B8GNA4 B8GNA7 B8GNA8 B8GNB7 B8GNC6 B8GNC7 B8GND2
B8GND3 B8GNE2 B8GNF8 B8GNH0 B8GNH1 B8GNI1 B8GNS2 B8GNS3 B8GNS5 B8GNS7 B8GNT5
B8GNT6 B8GNT7 B8GNT9 B8GNX0 B8GNX1 B8GNX2 B8GNX3 B8GNX4 B8GNX7 B8GNX9 B8GNY2
B8GP00 B8GP02 B8GP03 B8GP05 B8GP06 B8GP32 B8GP39 B8GP44 B8GP63 B8GP96 B8GPA2
B8GPB2 B8GPB3 B8GPB9 B8GPC3 B8GPL4 B8GPM6 B8GPP4 B8GPR9 B8GPS1 B8GPS8 B8GPT9
B8GPU0 B8GPU3 B8GPV2 B8GPV3 B8GQ24 B8GQ25 B8GQ27 B8GQ39 B8GQ45 B8GQ48 B8GQ51
B8GQ53 B8GQ66 B8GQ73 B8GQ78 B8GQ80 B8GQ83 B8GQ85 B8GQ87 B8GQ90 B8GQ91 B8GQA2
B8GQC3 B8GQD6 B8GQD7 B8GQE5 B8GQN1 B8GQP8 B8GQQ1 B8GQQ2 B8GQQ3 B8GQQ4 B8GQQ5
B8GQR1 B8GQR4 B8GQT0 B8GQV3 B8GQV7 B8GR29 B8GR39 B8GR95 B8GRA4 B8GRC9 B8GRD3
B8GRD4 B8GRG3 B8GRI1 B8GRI2 B8GRI3 B8GRI4 B8GRI6 B8GRJ2 B8GRR4 B8GRS4 B8GRT8
B8GSD2 B8GSS3 B8GSU8 B8GSV2 B8GSX9 B8GT00 B8GT02 B8GT89 B8GTA0 B8GTA1 B8GTD2
B8GTE3 B8GTF1 B8GTF9 B8GTG2 B8GTH4 B8GTH6 B8GTL0 B8GTL3 B8GTN1 B8GTN4 B8GTN8
B8GTQ4 B8GTQ5 B8GTX6 B8GTY2 B8GU08 B8GU11 B8GU25 B8GU29 B8GU31 B8GU32 B8GU33
B8GU79 B8GU82 B8GUB6 B8GUD3 B8GUF9 B8GUJ4 B8GUJ5 B8GUJ7 B8GUK0 B8GUN6 B8GUQ6
B8GUR2 B8GUR8 B8GUV7 B8GV07 B8GV13 B8GV32 B8GV33 B8GV34 B8GV35 B8GV36 B8GV37
B8GV39 B8GV40 B8GV42 B8GV43 B8GV44 B8GV46 B8GV47 B8GV48 B8GV49 B8GV50 B8GV51
B8GV53 B8GV54 B8GV55 B8GV56 B8GV57 B8GV58 B8GV59 B8GV62 B8GV63 B8GV65 B8GV66
B8GV67 B8GV68 B8GV69 B8GVD0 B8GVD8 B8GVE1 B8GVE2 B8GVE7 B8GVF3 B8GVI7 B8GVL8
B8GVN4 B8GVN5 B8GVN6 B8GVS9 B8GVT2 B8GVW2 B8GVX4 B8GVY4 B8GVY5 B8GVZ0 B8GW06
B8GW12 B8GW15 B8GW16 B8GW20 B8GW27 B8GW30 B8GW31 B8GW32 B8GW33 B8GW34 B8GW40

B8GW41 B8GW43 B8GW64 B8GW65 B8GW82 B8GW85 B8GW92 B8GWL9 B8GWM3 B8GWN0 B8GWN2
B8GWN5 B8GWP8 B8GWP9 B8GWR7 B8GWS0 B8GWS2 B8GWS3 B8GWS6 B8GWT1 B8GWW0 B8GWW4
B8GWW5 B8GWW6 B8GWX0 B8GWX1 B8GWZ0 B8GWZ1 B8GWZ3 B8GX01 B8GX04 B8GX09 B8GX11
B8GX12 B8GX14 B8GX15 B8GX16 B8GX17 B8GX51 B8GX52 B8GX53 B8GX62 B8GX84 B8GXA8
B8GXC4 B8GXC6 B8GXF0 B8GXG1 B8GXH3 B8GXJ5 B8GXJ8 B8GXK7 B8GXL9 B8GXM4 B8GXN3
B8GXP2 B8GXP3 B8GXP4 B8GXP6 B8GXP9 B8GXQ0 B8GXU1 B8GXW2 B8GY23 B8GY65 B8GY66
B8GY92 B8GY96 B8GYB5 B8GYD9 B8GYE0 B8GYE3 B8GYE7 B8GYE8 B8GYF0 B8GYF1 B8GYF5
B8GYI6 B8GYI7 B8GYL6 B8GYM1 B8GYN6 B8GYP3 B8GYS2 B8GYT0 B8GYW7 B8GYX1 B8GYZ1
B8GZ28 B8GZ33 B8GZ34 B8GZ98 B8GZA6 B8GZB1 B8GZC1 B8GZD8 B8GZJ5 B8GZJ6 B8GZJ7
B8GZL9 B8GZM8 B8GZS0 B8GZS3 B8GZS7 B8GZU8 B8GZU9 B8GZV1 B8GZW1 B8GZW2 B8GZW7
B8GZX1 B8GZZ0 B8H053 B8H072 B8H085 B8H086 B8H096 B8H0A2 B8H0A3 B8H0P3 B8H0P4
B8H0R5 B8H0X4 B8H0Z4 B8H163 B8H164 B8H172 B8H173 B8H1A0 B8H1E7 B8H1E9 B8H1F0
B8H1H5 B8H1Q1 B8H1Q2 B8H1Q3 B8H1Q4 B8H1V7 B8H1Z0 B8H209 B8H292 B8H2F3 B8H2R9
B8H2S1 B8H2S2 B8H307 B8H308 B8H309 B8H312 B8H331 B8H332 B8H363 B8H367 B8H3C3
B8H3C9 B8H3F9 B8H3I7 B8H3S9 B8H3W7 B8H3X3 B8H3Z4 B8H414 B8H415 B8H416 B8H427
B8H428 B8H429 B8H454 B8H455 B8H462 B8H482 B8H4A1 B8H4A3 B8H4D3 B8H4D5 B8H4D6
B8H4D7 B8H4D8 B8H4E0 B8H4E1 B8H4E2 B8H4E3 B8H4E4 B8H4E5 B8H4E6 B8H4E8 B8H4E9
B8H4F0 B8H4F1 B8H4F2 B8H4F3 B8H4F5 B8H4F6 B8H4F7 B8H4F8 B8H4G0 B8H4J9 B8H4M5
B8H4M6 B8H4R9 B8H4S1 B8H4V4 B8H4V5 B8H4W0 B8H4W6 B8H4W7 B8H510 B8H516 B8H533
B8H535 B8H546 B8H554 B8H555 B8H5B6 B8H5C6 B8H5H3 B8H5P6 B8H5S2 B8H5U4 B8H5V4
B8H5V5 B8H5Y7 B8H607 B8H618 B8H621 B8H623 B8H627 B8H640 B8H661 B8H6A5 B8H6A7
B8H6A8 B8H6D3 B8H6E1 B8H6Q1 B8H6T9 B8H6U0 B8H6U1 B8H6U2 B8H717 B8H729 B8H775
B8H776 B8H7D1 B8H7K6 B8H7L9 B8H7M0 B8H7M2 B8H7M5 B8H7N3 B8H7S3 B8H7T9 B8H806
B8H838 B8H839 B8H841 B8H8I4 B8H8K8 B8H8L3 B8H8L6 B8H8L7 B8H8L8 B8H8Q9 B8H8R4
B8H8R9 B8H8S5 B8H8T1 B8H8U5 B8H8U9 B8H8V2 B8H8V3 B8H8V5 B8H8V7 B8H8Y9 B8H8Z3
B8H9C3 B8H9C7 B8H9D6 B8H9D7 B8H9D8 B8H9D9 B8H9E0 B8H9E8 B8H9H0 B8H9P2 B8H9Q1
B8H9W6 B8HA05 B8HA06 B8HA07 B8HA08 B8HA20 B8HA21 B8HA33 B8HA36 B8HA51 B8HA70
B8HA88 B8HA93 B8HA94 B8HA97 B8HAB3 B8HAB6 B8HAC4 B8HAC8 B8HAC9 B8HAS8 B8HAT7
B8HAX3 B8HAY3 B8HB03 B8HB09 B8HBH4 B8HBH8 B8HBR2 B8HBS2 B8HBV4 B8HCE1 B8HCE4
B8HCF8 B8HCH4 B8HCH9 B8HCI0 B8HCK2 B8HCL2 B8HCQ9 B8HCR6 B8HCR7 B8HCX3 B8HCX4
B8HCX5 B8HCX6 B8HCX7 B8HCY6 B8HCY9 B8HCZ2 B8HCZ3 B8HCZ4 B8HCZ5 B8HCZ6 B8HCZ9
B8HD00 B8HD01 B8HD02 B8HD03 B8HD04 B8HD06 B8HD07 B8HD11 B8HD12 B8HD13 B8HD14
B8HD15 B8HD21 B8HD24 B8HD25 B8HD40 B8HD44 B8HDC2 B8HDD2 B8HDH4 B8HEI6 B8HF09
B8HF37 B8HFE2 B8HFJ3 B8HFK0 B8HFK2 B8HFN5 B8HFN6 B8HFN8 B8HFR2 B8HFS1 B8HG44
B8HG52 B8HG54 B8HG55 B8HG66 B8HG67 B8HG89 B8HG92 B8HGA0 B8HGA8 B8HGC3 B8HGC5
B8HGC9 B8HGW1 B8HGW2 B8HGX6 B8HGY5 B8HGZ8 B8HH36 B8HH63 B8HIQ9 B8HJY4 B8HK54
B8HK56 B8HK77 B8HKC7 B8HKL1 B8HL79 B8HL86 B8HLD2 B8HLN1 B8HLW6 B8HM05 B8HM61
B8HMF0 B8HMJ9 B8HMK7 B8HMM0 B8HMN5 B8HMQ4 B8HMQ5 B8HMQ7 B8HMQ8 B8HMR1 B8HMR2
B8HMR4 B8HMR6 B8HMR7 B8HMR8 B8HMR9 B8HMS3 B8HMS5 B8HMS6 B8HMS7 B8HMS9 B8HMT1
B8HMX3 B8HMX6 B8HN58 B8HN65 B8HNE6 B8HNE7 B8HNG6 B8HNI6 B8HNJ0 B8HNJ7 B8HNK4
B8HNL6 B8HNM1 B8HNM9 B8HNW1 B8HP19 B8HP79 B8HP81 B8HPC6 B8HPG4 B8HPH2 B8HPI0
B8HPI6 B8HPI7 B8HPP0 B8HPS4 B8HPS9 B8HPV7 B8HPZ1 B8HQ19 B8HQ34 B8HQ99 B8HQA4
B8HQA9 B8HQE1 B8HQI6 B8HQP9 B8HQS5 B8HQZ8 B8HR03 B8HR50 B8HR51 B8HR52 B8HR57
B8HR59 B8HR63 B8HRF8 B8HRK7 B8HRK9 B8HRP1 B8HRT5 B8HS22 B8HS30 B8HS80 B8HSA9

B8HSI6 B8HSJ3 B8HSK3 B8HSK8 B8HSQ9 B8HST7 B8HSU2 B8HSX6 B8HT34 B8HTH5 B8HTI9
B8HTK3 B8HTT7 B8HTV6 B8HTV9 B8HTW2 B8HU20 B8HU22 B8HU41 B8HU54 B8HU56 B8HU67
B8HU74 B8HUA6 B8HUA9 B8HUL5 B8HUM7 B8HUQ3 B8HUR1 B8HUR2 B8HUV7 B8HUW5 B8HUY7
B8HUY8 B8HV17 B8HV20 B8HV47 B8HVB6 B8HVF7 B8HVI5 B8HVK1 B8HVK5 B8HVL2 B8HVL3
B8HVL5 B8HVN1 B8HVP2 B8HVR1 B8HVR7 B8HVR8 B8HVR9 B8HVS0 B8HVV6 B8HW13 B8HW27
B8HW50 B8HW51 B8HW52 B8HW91 B8HW95 B8HWD3 B8HWE3 B8HWL3 B8HWL8 B8HWM0 B8HWP9
B8HWS1 B8HWW4 B8HWY0 B8HWY2 B8HWY6 B8HX01 B8HX02 B8HX39 B8HXF0 B8HXK3 B8HXK4
B8HXN3 B8HXQ5 B8HXQ8 B8HXS0 B8HXS4 B8HXS5 B8HXT9 B8HXV0 B8HXV1 B8HY03 B8HY50
B8HY57 B8HY86 B8HY89 B8HY92 B8HYG3 B8HYJ8 B8HYK1 B8HYK3 B8HYL7 B8I005 B8I0I6
B8I0Q1 B8I0Q8 B8I0R1 B8I0R5 B8I0T8 B8I0U9 B8I0V0 B8I164 B8I168 B8I169 B8I176
B8I178 B8I179 B8I184 B8I1A0 B8I1A2 B8I1A3 B8I1A8 B8I1G1 B8I1M9 B8I1P0 B8I1T7
B8I1T8 B8I1Z8 B8I255 B8I259 B8I262 B8I298 B8I2A5 B8I2B5 B8I2N8 B8I2Q2 B8I2Q5
B8I2R5 B8I2R6 B8I2T0 B8I2T4 B8I2V4 B8I2Y2 B8I2Y3 B8I2Z2 B8I2Z3 B8I2Z6 B8I2Z7
B8I2Z9 B8I303 B8I304 B8I305 B8I307 B8I359 B8I363 B8I364 B8I385 B8I388 B8I3A2
B8I3A3 B8I3A9 B8I3B0 B8I3C7 B8I3F1 B8I3F5 B8I3F7 B8I3I1 B8I3I4 B8I3J4 B8I3J5
B8I3K7 B8I3R2 B8I3R5 B8I3S9 B8I3X2 B8I406 B8I442 B8I445 B8I447 B8I490 B8I492
B8I4B5 B8I4C1 B8I4D2 B8I4D3 B8I4G1 B8I4H0 B8I4L5 B8I4P0 B8I4S7 B8I4S8 B8I4X0
B8I4X3 B8I550 B8I551 B8I562 B8I567 B8I590 B8I592 B8I598 B8I5A9 B8I5B5 B8I5L3
B8I5M2 B8I5M9 B8I5N5 B8I5N6 B8I5N7 B8I5N8 B8I5R4 B8I5U8 B8I5U9 B8I5V1 B8I5V2
B8I5V5 B8I5V6 B8I5V9 B8I5W0 B8I601 B8I602 B8I6C2 B8I6D2 B8I6D9 B8I6E1 B8I6E3
B8I6E7 B8I6G5 B8I6G6 B8I6I0 B8I6M0 B8I6Q1 B8I6R1 B8I6T0 B8I754 B8I761 B8I769
B8I770 B8I784 B8I799 B8I7A3 B8I7D0 B8I7T2 B8I7T4 B8I7T5 B8I7T6 B8I7T9 B8I7X8
B8I7X9 B8I7Y0 B8I7Y1 B8I7Y2 B8I7Y3 B8I7Y6 B8I7Y7 B8I7Y8 B8I7Y9 B8I7Z1 B8I7Z2
B8I7Z3 B8I7Z4 B8I7Z5 B8I7Z7 B8I7Z8 B8I800 B8I804 B8I805 B8I807 B8I808 B8I809
B8I814 B8I815 B8I816 B8I894 B8I8F2 B8I8F4 B8I8F5 B8I8F6 B8I8G4 B8I8N7 B8I8V6
B8I944 B8I9H6 B8I9N8 B8I9T2 B8I9V8 B8IB09 B8IB34 B8IBI6 B8IBN8 B8IBP2 B8IBS4
B8IBT1 B8IBW1 B8IBW2 B8ICA2 B8ICE5 B8ICY0 B8ICZ4 B8ID24 B8IDD6 B8IDE7 B8IEB0
B8IEB4 B8IEC7 B8IEC9 B8IED1 B8IED2 B8IEL6 B8IEL7 B8IEL9 B8IEM1 B8IEM4 B8IER1
B8IEX1 B8IEX6 B8IF20 B8IFN4 B8IFN5 B8IFQ0 B8IFQ2 B8IFQ3 B8IFQ5 B8IFR7 B8IFW8
B8IGT3 B8IGW4 B8IGW5 B8IGX2 B8IGX3 B8IH88 B8IH95 B8IHL2 B8IHL3 B8IHL5 B8IHS8
B8IHY5 B8IHZ7 B8II06 B8II08 B8II09 B8II14 B8IIZ9 B8IJ00 B8IJ69 B8IJ81 B8IJ86
B8IJ89 B8IJB4 B8IJD1 B8IJD7 B8IJW7 B8IK26 B8IKN3 B8IL50 B8IL52 B8IL55 B8IL75
B8ILE2 B8ILM3 B8ILP3 B8ILR2 B8IM27 B8IM39 B8IMM7 B8IMT2 B8IMW6 B8IMX2 B8IN19
B8IN20 B8IN22 B8IN25 B8IN26 B8IN27 B8IN36 B8IN63 B8IN72 B8INF1 B8INF6 B8INK3
B8IP05 B8IP16 B8IP29 B8IP71 B8IP76 B8IP77 B8IP78 B8IP92 B8IP99 B8IPA1 B8IPG1
B8IPH2 B8IPH3 B8IPH4 B8IPI3 B8IPM0 B8IPM5 B8IPU4 B8IPV3 B8IPW1 B8IQG9 B8IQU9
B8IQW2 B8IQY5 B8IQY6 B8IRU5 B8IRX8 B8IRY2 B8IS49 B8IS51 B8IS61 B8IS72 B8IS73
B8IS75 B8IS76 B8IS78 B8IS79 B8IS80 B8IS81 B8IS82 B8IS84 B8IS85 B8IS86 B8IS87
B8IS88 B8IS89 B8IS90 B8IS92 B8IS94 B8IS96 B8ISA0 B8ISA1 B8ISA2 B8ISA3 B8ISA4
B8IST7 B8ISU9 B8ISV4 B8ISY7 B8IT13 B8IT21 B8IT23 B8IT32 B8IT33 B8IT34 B8IT35
B8IT36 B8ITG7 B8ITT7 B8ITV7 B8ITW0 B8ITX3 B8IU03 B8IU32 B8IU36 B8IU50 B8IUB0
B8IUB4 B8IUE9 B8IUF0 B8IUQ6 B8IUT2 B8IUW4 B8IYD5 B8IYG6 B8IYH2 B8IYH3 B8IYH4
B8IYH5 B8IYH6 B8IYH7 B8IYH8 B8IYH9 B8IYI2 B8IYI3 B8IYK3 B8IYK4 B8IYK5 B8IYK6
B8IYK7 B8IYK9 B8IYL0 B8IYL3 B8IYL4 B8IYL5 B8IYL6 B8IYL7 B8IYL8 B8IYM1 B8IYM2

B8IYS4 B8IYS8 B8IYW9 B8IZ52 B8IZ74 B8IZ95 B8IZ97 B8IZD1 B8IZD4 B8IZD9 B8IZF2
B8IZF5 B8IZG2 B8IZH6 B8IZJ2 B8IZM4 B8IZP2 B8IZP3 B8IZQ7 B8IZR8 B8IZS3 B8IZS6
B8IZT3 B8IZT9 B8IZU2 B8IZX7 B8IZX8 B8J008 B8J017 B8J025 B8J030 B8J031 B8J080
B8J0A8 B8J0B1 B8J0C2 B8J0C6 B8J0H2 B8J0K9 B8J0N0 B8J0P9 B8J0Q0 B8J0R0 B8J0R1
B8J0S2 B8J0S4 B8J0S7 B8J0S9 B8J0U1 B8J122 B8J123 B8J160 B8J162 B8J183 B8J189
B8J193 B8J194 B8J198 B8J1A0 B8J1A4 B8J1A5 B8J1A6 B8J1A7 B8J1A8 B8J1H5 B8J1K3
B8J1N0 B8J1Q4 B8J1Q5 B8J1S9 B8J1U2 B8J1X3 B8J1X9 B8J1Y0 B8J1Y4 B8J1Y7 B8J215
B8J216 B8J249 B8J283 B8J2A8 B8J2D5 B8J2E2 B8J2G0 B8J2G2 B8J2K3 B8J2Q8 B8J2T3
B8J2U6 B8J2X5 B8J2X6 B8J364 B8J369 B8J374 B8J3A3 B8J3A4 B8J3A8 B8J3A9 B8J3B0
B8J3C2 B8J3F6 B8J3F7 B8J3G1 B8J3G9 B8J3M5 B8J3M6 B8J3R3 B8J3V0 B8J3V1 B8J402
B8J405 B8J412 B8J457 B8J466 B8J468 B8J495 B8J4A8 B8J4C1 B8J4D3 B8J4E0 B8J4E1
B8J4E2 B8J4E4 B8J4F3 B8J4F9 B8J4G0 B8J4I9 B8J4L3 B8J4L4 B8J4L5 B8J4N1 B8J4Q1
B8J4Q3 B8J4Q5 B8J4R4 B8J4R6 B8J4S2 B8J4S7 B8J538 B8J561 B8J5G7 B8J5H5 B8J5I5
B8J637 B8J638 B8J6C6 B8J6S7 B8J6T5 B8J6V6 B8J6X9 B8J6Y3 B8J714 B8J731 B8J7C8
B8J7G1 B8J7G5 B8J7G7 B8J7H0 B8J7H2 B8J7P4 B8J7V5 B8J7X9 B8J7Y5 B8J801 B8J802
B8J804 B8J805 B8J807 B8J808 B8J817 B8J823 B8J825 B8J826 B8J829 B8J830 B8J831
B8J832 B8J836 B8J856 B8J857 B8J860 B8J861 B8J862 B8J863 B8J864 B8J865 B8J866
B8J867 B8J869 B8J870 B8J871 B8J872 B8J873 B8J874 B8J875 B8J876 B8J878 B8J879
B8J881 B8J883 B8J884 B8J886 B8J887 B8J888 B8J890 B8J893 B8J894 B8J8E0 B8J8E9
B8J8F0 B8J8F1 B8J8M9 B8J8R4 B8J8S6 B8J8T3 B8J8V3 B8J8W7 B8J8W8 B8J8W9 B8J8Z7
B8J916 B8J950 B8J9D7 B8J9E3 B8J9F5 B8J9L8 B8J9M0 B8J9N6 B8J9P3 B8J9V0 B8J9V9
B8J9Z2 B8JA66 B8JAC3 B8JAD8 B8JAE0 B8JAE8 B8JAE9 B8JAF5 B8JAJ1 B8JAL2 B8JAL5
B8JAM9 B8JAN4 B8JAN5 B8JAN8 B8JAR2 B8JAT1 B8JAT9 B8JAU0 B8JAU2 B8JB61 B8JB70
B8JB71 B8JB72 B8JB73 B8JB74 B8JB77 B8JBD5 B8JBE2 B8JBF4 B8JBF6 B8JBF9 B8JBM1
B8JBN6 B8JBP2 B8JBP3 B8JBP4 B8JBQ0 B8JBQ1 B8JBQ2 B8JBS8 B8JBU6 B8JBW0 B8JC48
B8JC53 B8JCT2 B8JCU6 B8JD56 B8JDD8 B8JDD9 B8JDE1 B8JDG6 B8JDH1 B8JDH9 B8JDI0
B8JDJ0 B8JDJ3 B8JDK2 B8JDK8 B8JDL0 B8JDL2 B8JDL4 B8JDQ9 B8JDR1 B8JDV1 B8JDY4
B8JDY7 B8JDY8 B8JDZ9 B8JEH9 B8JEI1 B8JEQ8 B8JES1 B8JES4 B8JEV0 B8JEW3 B8JEW4
B8JEW8 B8JEW9 B8JF03 B8JF04 B8JF05 B8JFS8 B8JFV3 B8JFW2 B8JFX1 B8JFY1 B8JFY3
B8JFY6 B8JFY8 B8JFZ0 B8JFZ1 B8JFZ2 B8JGW9 B8JH48 B8JH60 B8JH75 B8JH92 B8JHC9
B8JKD7 B8JKF4 B8JLQ0 B8JM82 B8JMH0 B8LBM8 B8LCD8 B8LIX8 B8LK84 B8LPG7 B8LT44
B8LTU5 B8LUH2 B8LVD6 B8LXM1 B8M072 B8M0Q1 B8M0U4 B8M0Z4 B8M2W9 B8M3A8 B8M3T8
B8M501 B8M6L1 B8M7D2 B8M7Q5 B8M990 B8M9J5 B8M9J9 B8M9K0 B8M9K3 B8M9K4 B8M9K5
B8M9K6 B8M9K8 B8M9L2 B8M9V7 B8M9W2 B8MBY6 B8MDC1 B8MDD1 B8MDN4 B8MDP8 B8MED8
B8MJJ5 B8MJJ8 B8MKR3 B8MLF7 B8MLU5 B8MNF2 B8MQZ4 B8MR69 B8MSM6 B8MWR8 B8MYL0
B8MYS5 B8MYS6 B8MZA3 B8MZI5 B8N0E6 B8N0E7 B8N0E8 B8N0F0 B8N0F2 B8N122 B8N195
B8N219 B8N3N5 B8N3R8 B8N4F5 B8N4G0 B8N4Q9 B8N513 B8N6M6 B8N8Q9 B8N8R0 B8N8R1
B8N8R2 B8N9H4 B8N9M2 B8N9M5 B8N9X2 B8NA06 B8NBX4 B8NC10 B8NDP1 B8NDQ2 B8NEI6
B8NEJ3 B8NG55 B8NG97 B8NGC8 B8NGT5 B8NHD6 B8NHD7 B8NHE0 B8NHE1 B8NHE2 B8NHE3
B8NHE4 B8NHF2 B8NHY4 B8NI18 B8NI19 B8NI20 B8NI22 B8NI23 B8NI24 B8NI26 B8NI28
B8NIM4 B8NJG3 B8NJG5 B8NJG9 B8NJH1 B8NJH3 B8NKS1 B8NL00 B8NLL0 B8NLM9 B8NLN6
B8NLU5 B8NM69 B8NM70 B8NM72 B8NM76 B8NM78 B8NM79 B8NQ08 B8NR69 B8NR70 B8NR71
B8NSD4 B8NSW2 B8NT06 B8NTI4 B8NTZ8 B8NTZ9 B8NU00 B8NUL8 B8NV37 B8NWW2 B8NWW5
B8NWW6 B8NXI4 B8NY85 B8NY88 B8NYX0 B8P4B0 B8PAR0 B8PBQ6 B8PBR2 B8PD44 B8PD53

B8PE34 B8PH83 B8PHE1 B8PIG4 B8PQ84 B8PZP3 B8RCD3 B8X8Z0 B8XA40 B8XA41 B8XIA5
B8Y466 B8Y6I0 B8Y7Y5 B8YB53 B8YB54 B8YB55 B8YB56 B8YB57 B8YB58 B8YB59 B8YB60
B8YB61 B8YB62 B8YB63 B8YB64 B8YB65 B8YEK4 B8YIE8 B8ZJH9 B8ZJI3 B8ZJI9 B8ZJJ5
B8ZJM7 B8ZJN0 B8ZJN3 B8ZJQ0 B8ZJQ9 B8ZJR8 B8ZJR9 B8ZJT5 B8ZJT9 B8ZJU1 B8ZJU2
B8ZJU9 B8ZJV6 B8ZJX3 B8ZJY0 B8ZK04 B8ZK08 B8ZK13 B8ZK23 B8ZKC9 B8ZKD0 B8ZKD4
B8ZKE4 B8ZKE6 B8ZKF6 B8ZKF7 B8ZKF8 B8ZKF9 B8ZKG0 B8ZKG1 B8ZKG2 B8ZKG3 B8ZKG4
B8ZKG5 B8ZKG6 B8ZKG7 B8ZKG8 B8ZKG9 B8ZKK3 B8ZKK4 B8ZKK9 B8ZKL1 B8ZKM0 B8ZKM1
B8ZKM3 B8ZKM4 B8ZKM5 B8ZKM6 B8ZKN7 B8ZKP4 B8ZKP5 B8ZKP6 B8ZKP8 B8ZKP9 B8ZKQ1
B8ZKQ3 B8ZKQ4 B8ZKQ6 B8ZKQ8 B8ZKS3 B8ZKS5 B8ZKT3 B8ZKT8 B8ZKT9 B8ZKU0 B8ZKX2
B8ZKX3 B8ZKX4 B8ZKX7 B8ZKZ5 B8ZL02 B8ZL13 B8ZL14 B8ZL19 B8ZL20 B8ZL21 B8ZL26
B8ZL30 B8ZL51 B8ZL58 B8ZL82 B8ZL83 B8ZL84 B8ZL95 B8ZLB8 B8ZLC2 B8ZLD4 B8ZLD9
B8ZLE1 B8ZLE9 B8ZLF5 B8ZLG6 B8ZLG8 B8ZLH1 B8ZLH4 B8ZLH8 B8ZLJ7 B8ZLJ9 B8ZLK0
B8ZLK1 B8ZLK2 B8ZLK3 B8ZLK5 B8ZLL0 B8ZLL9 B8ZLM2 B8ZLS6 B8ZLS8 B8ZLT0 B8ZLT1
B8ZLT8 B8ZLT9 B8ZLU4 B8ZLX1 B8ZLX5 B8ZLY7 B8ZLY8 B8ZLY9 B8ZM12 B8ZM16 B8ZM28
B8ZM32 B8ZM41 B8ZM52 B8ZM55 B8ZM59 B8ZM66 B8ZM67 B8ZM88 B8ZM89 B8ZM93 B8ZM94
B8ZMC0 B8ZMG1 B8ZMH4 B8ZMH8 B8ZMJ8 B8ZMK5 B8ZML8 B8ZML9 B8ZMP5 B8ZMQ1 B8ZMQ3
B8ZMQ6 B8ZMR0 B8ZMR7 B8ZMW2 B8ZMZ6 B8ZMZ9 B8ZN14 B8ZN21 B8ZN39 B8ZN40 B8ZN58
B8ZN59 B8ZN60 B8ZN61 B8ZN62 B8ZN87 B8ZNB3 B8ZNB7 B8ZNC8 B8ZND1 B8ZND2 B8ZNH8
B8ZNH9 B8ZNI0 B8ZNI1 B8ZNI5 B8ZNK7 B8ZNK8 B8ZNK9 B8ZNL0 B8ZNM3 B8ZNN8 B8ZNP0
B8ZNP1 B8ZNP4 B8ZNS6 B8ZNT8 B8ZNU0 B8ZNU7 B8ZNU8 B8ZNW6 B8ZNW7 B8ZNX4 B8ZNX6
B8ZNY9 B8ZNZ8 B8ZP15 B8ZP26 B8ZP34 B8ZP35 B8ZP37 B8ZP39 B8ZP43 B8ZP45 B8ZP47
B8ZP56 B8ZP57 B8ZP58 B8ZP64 B8ZPA3 B8ZPA4 B8ZPB8 B8ZPD4 B8ZPG1 B8ZPG2 B8ZPG7
B8ZPH1 B8ZPH5 B8ZPK0 B8ZPK2 B8ZPL8 B8ZPL9 B8ZPM2 B8ZPN7 B8ZPP6 B8ZPQ2 B8ZPS8
B8ZPT5 B8ZPT7 B8ZPU4 B8ZPU7 B8ZPU9 B8ZPV1 B8ZPV9 B8ZPW3 B8ZPW5 B8ZPX5 B8ZPY7
B8ZPY8 B8ZPZ6 B8ZQ14 B8ZQ22 B8ZQ24 B8ZQ31 B8ZQ32 B8ZQ37 B8ZQ39 B8ZQ48 B8ZQ59
B8ZQ64 B8ZQ65 B8ZQ66 B8ZQ75 B8ZQ76 B8ZQ77 B8ZQ84 B8ZQ95 B8ZQA5 B8ZQA6 B8ZQB8
B8ZQC9 B8ZQE9 B8ZQJ5 B8ZQJ6 B8ZQJ9 B8ZQK0 B8ZQK6 B8ZQK8 B8ZQM2 B8ZQP1 B8ZQP2
B8ZQP8 B8ZQQ1 B8ZQQ6 B8ZQT6 B8ZQU0 B8ZQU1 B8ZQU7 B8ZQW4 B8ZQW9 B8ZQX2 B8ZQY9
B8ZQZ1 B8ZR24 B8ZR28 B8ZR30 B8ZR31 B8ZR57 B8ZR59 B8ZR84 B8ZR85 B8ZR86 B8ZR90
B8ZRB0 B8ZRB1 B8ZRB4 B8ZRB5 B8ZRB9 B8ZRC1 B8ZRD9 B8ZRE4 B8ZRF0 B8ZRF3 B8ZRF4
B8ZRF5 B8ZRG8 B8ZRH2 B8ZRH6 B8ZRI2 B8ZRJ6 B8ZRJ7 B8ZRK2 B8ZRK7 B8ZRK9 B8ZRN0
B8ZRN2 B8ZRN3 B8ZRN4 B8ZRN5 B8ZRN7 B8ZRP1 B8ZRP5 B8ZRQ3 B8ZRQ6 B8ZRR3 B8ZRR4
B8ZRT2 B8ZRT3 B8ZRT4 B8ZRU5 B8ZRU7 B8ZRV0 B8ZRV3 B8ZRW3 B8ZRW5 B8ZRW7 B8ZRW9
B8ZRZ2 B8ZS00 B8ZS02 B8ZS07 B8ZS08 B8ZS11 B8ZS15 B8ZS22 B8ZS25 B8ZS27 B8ZS29
B8ZS49 B8ZS79 B8ZS96 B8ZSA0 B8ZSA1 B8ZSA2 B8ZSA4 B8ZSA5 B8ZSA6 B8ZSA7 B8ZSA8
B8ZSB2 B8ZSB3 B8ZSB4 B8ZSB5 B8ZSB6 B8ZSB7 B8ZSB8 B8ZSC0 B8ZSC1 B8ZSC2 B8ZSC3
B8ZSC4 B8ZSC5 B8ZSC6 B8ZSC7 B8ZSD0 B8ZSD5 B8ZSD6 B8ZSE2 B8ZSF6 B8ZSG6 B8ZSH2
B8ZSH3 B8ZSH5 B8ZSH7 B8ZSH8 B8ZSH9 B8ZSI0 B8ZSN3 B8ZSN5 B8ZSP0 B8ZSU0 B8ZSV5
B8ZSW0 B8ZSY2 B8ZT20 B8ZT53 B8ZT67 B8ZT86 B8ZT88 B8ZT90 B8ZT93 B8ZTB7 B8ZTC3
B8ZTF7 B8ZTI4 B8ZTI5 B8ZTJ5 B8ZTL4 B8ZTL5 B8ZTN3 B8ZTN6 B8ZTN7 B8ZTP0 B8ZTP1
B8ZTR5 B8ZTT3 B8ZTU2 B8ZTU5 B8ZTZ5 B8ZU00 B8ZU05 B8ZU08 B8ZU42 B8ZU50 B8ZU51
B8ZU59 B8ZU60 B8ZU62 B8ZU74 B8ZU80 B8ZU90 B8ZU92 B8ZU95 B8ZUA7 B8ZUA8 B8ZUA9
B8ZUB0 B8ZUB2 B8ZUB9 B8ZUC0 B8ZUC1 B8ZUD1 B8ZUE2 B8ZUG3 B8ZUG9 B8ZUH9 B8ZUI1
B8ZUI2 B8ZUJ3 B8ZUJ7 B8ZUK0 B8ZUK3 B8ZUK6 B8ZUK7 B8ZUL1 B8ZUL5 B8ZUM0 B8ZUM2

B8ZUM6 B8ZUN3 B8ZUN5 B8ZUN6 B8ZUN8 B8ZUP2 B8ZUQ2 B8ZUR8 B8ZUS6 B8ZUT0 B8ZUT3
B8ZUV0 B8ZUV3 B8ZUV4 B8ZUV9 B8ZUW8 B8ZUX8 B8ZV93 B8ZV94 B8ZYW0 B8ZYW1 B9A014
B9A1H3 B9DFA8 B9DFG3 B9DFZ0 B9DG24 B9DGG8 B9DGT7 B9DGU7 B9DHA6 B9DHD3 B9DHQ0
B9DI93 B9DI94 B9DI97 B9DIB8 B9DIC0 B9DID6 B9DIF1 B9DIJ2 B9DIJ5 B9DIP1 B9DIP2
B9DIP5 B9DIR3 B9DIS2 B9DIT7 B9DIU0 B9DIU3 B9DIU4 B9DIW9 B9DIX2 B9DIX8 B9DJ55
B9DJG9 B9DJH3 B9DJJ6 B9DJK4 B9DJK9 B9DJL1 B9DJL4 B9DJN0 B9DJN5 B9DJR3 B9DK09
B9DK21 B9DK28 B9DK35 B9DK67 B9DK71 B9DKF4 B9DKF5 B9DKF6 B9DKG1 B9DKJ7 B9DKM0
B9DKM7 B9DKN5 B9DKQ7 B9DKR5 B9DKS1 B9DKT0 B9DKT3 B9DKV0 B9DKV5 B9DKV6 B9DKV7
B9DKV8 B9DKW3 B9DKX1 B9DKX2 B9DKX7 B9DKX8 B9DL85 B9DLC1 B9DLD0 B9DLD5 B9DLD8
B9DLD9 B9DLF6 B9DLF7 B9DLK2 B9DLL5 B9DLM4 B9DLM7 B9DLR4 B9DLS6 B9DLW9 B9DLX8
B9DLX9 B9DLY0 B9DLY1 B9DM01 B9DM16 B9DM17 B9DM18 B9DM19 B9DM20 B9DM27 B9DM29
B9DM30 B9DM31 B9DM32 B9DM33 B9DM34 B9DM35 B9DM36 B9DM37 B9DM38 B9DM39 B9DM40
B9DM42 B9DM46 B9DM50 B9DM52 B9DM53 B9DM54 B9DM96 B9DM98 B9DMC3 B9DMC7 B9DMD1
B9DMD3 B9DME0 B9DMF2 B9DMF3 B9DMH1 B9DMI1 B9DMI9 B9DMJ2 B9DMJ5 B9DMJ6 B9DMJ8
B9DMK9 B9DML1 B9DMM2 B9DMM3 B9DMR8 B9DMT3 B9DMT7 B9DMT8 B9DMT9 B9DMU3 B9DMV6
B9DMW2 B9DMW9 B9DMX3 B9DMY5 B9DMZ6 B9DN00 B9DN09 B9DN19 B9DN20 B9DN29 B9DN51
B9DN57 B9DN65 B9DN75 B9DN77 B9DN91 B9DN93 B9DN94 B9DNA2 B9DNA8 B9DNB9 B9DNC0
B9DNC1 B9DNC4 B9DNC5 B9DNC6 B9DNC9 B9DND0 B9DND4 B9DND6 B9DND8 B9DNE1 B9DNE3
B9DNE4 B9DNE5 B9DNE7 B9DNE8 B9DNF0 B9DNF8 B9DNF9 B9DNG1 B9DNG3 B9DNG7 B9DNH1
B9DNH2 B9DNH6 B9DNI9 B9DNJ2 B9DNJ3 B9DNJ4 B9DNJ8 B9DNJ9 B9DNK0 B9DNK1 B9DNK2
B9DNK5 B9DNL1 B9DNL2 B9DNL6 B9DNM1 B9DNM2 B9DNM8 B9DNN6 B9DNN7 B9DNP9 B9DNQ0
B9DNQ4 B9DNS1 B9DNT1 B9DNT5 B9DNU7 B9DNV0 B9DNV1 B9DNV2 B9DNV3 B9DNV5 B9DNV9
B9DNW3 B9DNX1 B9DNX5 B9DNY9 B9DP23 B9DP24 B9DP25 B9DP27 B9DP50 B9DP52 B9DP67
B9DP91 B9DPA1 B9DPB9 B9DPC0 B9DPC5 B9DPD6 B9DPD9 B9DPE0 B9DPF1 B9DPF5 B9DPG0
B9DPG3 B9DPG4 B9DPG5 B9DPG6 B9DPG7 B9DPG8 B9DPH1 B9DPH3 B9DPI8 B9DPJ2 B9DPJ9
B9DPK1 B9DPK4 B9DPK8 B9DPL6 B9DPM5 B9DPM7 B9DPM9 B9DPN2 B9DPN4 B9DPP7 B9DPQ3
B9DPQ7 B9DPQ8 B9DPQ9 B9DPR3 B9DPT1 B9DPT7 B9DPU2 B9DPU4 B9DPU6 B9DPU8 B9DPV4
B9DPV5 B9DPW5 B9DPX1 B9DPX4 B9DPY3 B9DQ02 B9DQ10 B9DQ35 B9DQ38 B9DQ40 B9DQ42
B9DQ45 B9DQ86 B9DQ89 B9DQA0 B9DRE8 B9DRE9 B9DRF1 B9DRF3 B9DRF4 B9DRF7 B9DRH1
B9DRJ1 B9DRJ3 B9DRL9 B9DRN4 B9DRN6 B9DRP5 B9DRQ0 B9DRQ9 B9DRS2 B9DRS6 B9DRS8
B9DRU2 B9DRU9 B9DRV0 B9DRV3 B9DRV7 B9DRW3 B9DRW5 B9DRW6 B9DRX8 B9DRX9 B9DRZ0
B9DRZ6 B9DS19 B9DS21 B9DS39 B9DS40 B9DS45 B9DS48 B9DS53 B9DS62 B9DS69 B9DS79
B9DS83 B9DS93 B9DS97 B9DSA4 B9DSB9 B9DSE2 B9DSE8 B9DSH1 B9DSH7 B9DSI5 B9DSK8
B9DSL8 B9DSL9 B9DSM6 B9DSN2 B9DSN7 B9DSP2 B9DSR1 B9DSR4 B9DSR6 B9DSU1 B9DSU9
B9DSV0 B9DSV1 B9DSV3 B9DSV4 B9DSV5 B9DSV6 B9DSV7 B9DSV8 B9DSV9 B9DSW0 B9DSW1
B9DSW2 B9DSW3 B9DSW4 B9DSW5 B9DSW8 B9DSW9 B9DSX3 B9DSX4 B9DSX5 B9DSX7 B9DSY9
B9DSZ3 B9DSZ5 B9DSZ6 B9DT06 B9DT23 B9DT31 B9DTA9 B9DTB9 B9DTD1 B9DTD6 B9DTE2
B9DTE3 B9DTF3 B9DTJ0 B9DTJ5 B9DTJ7 B9DTK1 B9DTL0 B9DTN0 B9DTN4 B9DTP3 B9DTQ0
B9DTQ3 B9DTR1 B9DTR9 B9DTS3 B9DTS4 B9DTT7 B9DTU7 B9DTU8 B9DTV0 B9DTV7 B9DTW4
B9DTZ7 B9DU13 B9DU35 B9DU39 B9DU40 B9DU41 B9DU42 B9DU46 B9DU47 B9DU51 B9DU54
B9DU56 B9DU60 B9DU62 B9DU73 B9DU74 B9DU77 B9DU78 B9DU93 B9DU98 B9DUB2 B9DUD2
B9DUE6 B9DUH9 B9DUK0 B9DUK3 B9DUK4 B9DUK9 B9DUP2 B9DUP5 B9DUP8 B9DUQ2 B9DUR6
B9DUS0 B9DUS1 B9DUS6 B9DUS7 B9DUT8 B9DUT9 B9DUU0 B9DUV1 B9DUV7 B9DUW7 B9DUX3
B9DUX7 B9DUY3 B9DUY5 B9DUY9 B9DUZ0 B9DV19 B9DV29 B9DV30 B9DV45 B9DV47 B9DV61

B9DV62 B9DV63 B9DV66 B9DV78 B9DV79 B9DV80 B9DV90 B9DV92 B9DV93 B9DVB6 B9DVB7
B9DVC1 B9DVC2 B9DVD8 B9DVF2 B9DVF3 B9DVF5 B9DVG0 B9DVG5 B9DVG6 B9DVG7 B9DVG9
B9DVH6 B9DVI3 B9DVJ4 B9DVJ6 B9DVK2 B9DVK4 B9DVK7 B9DVL0 B9DVM1 B9DVN4 B9DVQ2
B9DVQ7 B9DVR0 B9DVS0 B9DVS2 B9DVS3 B9DVS4 B9DVT4 B9DVU8 B9DVU9 B9DVV7 B9DVV8
B9DVV9 B9DVY6 B9DVY7 B9DVZ9 B9DW16 B9DW28 B9DW29 B9DW41 B9DW42 B9DW54 B9DW61
B9DW62 B9DW63 B9DW65 B9DW66 B9DW68 B9DW70 B9DW73 B9DW76 B9DW80 B9DW81 B9DW82
B9DWC7 B9DWD0 B9DWD3 B9DWE6 B9DWF3 B9DWG4 B9DWL2 B9DWL7 B9DWL8 B9DWM9 B9DWN6
B9DWP3 B9DWV1 B9DWV2 B9DX75 B9DX85 B9DX96 B9DXE4 B9DXI3 B9DXI6 B9DXQ4 B9DXQ8
B9DXR0 B9DXR9 B9DXS6 B9DXS7 B9DXT9 B9DY32 B9DY39 B9DY44 B9DY52 B9DY69 B9DY72
B9DY74 B9DY75 B9DY81 B9DY87 B9DY88 B9DY98 B9DYA0 B9DYA1 B9DYA5 B9DYA6 B9DYA8
B9DYA9 B9DYB0 B9DYB1 B9DYB2 B9DYB3 B9DYB4 B9DYB5 B9DYB6 B9DYB7 B9DYB8 B9DYC1
B9DYC2 B9DYC3 B9DYC4 B9DYC7 B9DYC8 B9DYD0 B9DYD3 B9DYD4 B9DYD5 B9DYD7 B9DYD8
B9DYE3 B9DYE4 B9DYJ3 B9DYU5 B9DYW1 B9DYW7 B9DYY4 B9DYY5 B9DYY7 B9DZE4 B9DZF9
B9DZG0 B9DZK4 B9DZL6 B9DZL9 B9DZM2 B9DZP5 B9DZQ6 B9DZT0 B9DZV9 B9DZX5 B9E003
B9E004 B9E009 B9E018 B9E020 B9E021 B9E022 B9E033 B9E041 B9E043 B9E047 B9E052
B9E056 B9E076 B9E092 B9E0B1 B9E0B2 B9E0C6 B9E0C7 B9E0C8 B9E0D7 B9E0V5 B9E0W7
B9E100 B9E101 B9E102 B9E104 B9E106 B9E107 B9E115 B9E148 B9E149 B9E150 B9E151
B9E152 B9E156 B9E166 B9E168 B9E169 B9E171 B9E197 B9E198 B9E199 B9E1B6 B9E1C8
B9E1D6 B9E1D8 B9E1E8 B9E1F2 B9E1F8 B9E1G1 B9E1G8 B9E1H0 B9E1H4 B9E1H7 B9E1I7
B9E1I8 B9E1I9 B9E1J1 B9E1J4 B9E1J8 B9E1K3 B9E1K7 B9E1K8 B9E1Q9 B9E1W5 B9E1X0
B9E1X2 B9E1Y4 B9E1Z0 B9E1Z2 B9E1Z3 B9E234 B9E299 B9E2A2 B9E2H2 B9E2H5 B9E332
B9E333 B9E371 B9E377 B9E3M4 B9E3V3 B9E3V5 B9E4K6 B9E4K8 B9E4L0 B9E4Q5 B9E4U5
B9E4V7 B9E4X8 B9E560 B9E564 B9E5P2 B9E5P3 B9E5P8 B9E5Q6 B9E5Q7 B9E5Q8 B9E5R9
B9E5S1 B9E5U7 B9E5V8 B9E5V9 B9E5W0 B9E684 B9E685 B9E686 B9E695 B9E6A8 B9E6B4
B9E6E0 B9E6K7 B9E6K9 B9E6L5 B9E6L7 B9E6N3 B9E6P2 B9E6Q4 B9E6Q6 B9E6Q9 B9E6R3
B9E6R6 B9E6R9 B9E6S1 B9E6U1 B9E6U8 B9E6V6 B9E6V7 B9E6W1 B9E6W9 B9E6X0 B9E6X1
B9E6Y4 B9E6Y8 B9E6Y9 B9E706 B9E707 B9E709 B9E711 B9E716 B9E717 B9E718 B9E719
B9E724 B9E740 B9E743 B9E745 B9E746 B9E748 B9E764 B9E766 B9E782 B9E783 B9E784
B9E7A4 B9E7B2 B9E7B7 B9E7C5 B9E7C9 B9E7E0 B9E7E5 B9E7F6 B9E7G7 B9E7H4 B9E7J1
B9E7J8 B9E7M5 B9E7M7 B9E7Y3 B9E825 B9E826 B9E827 B9E830 B9E899 B9E8A0 B9E8C3
B9E8D0 B9E8D6 B9E8F4 B9E8F5 B9E8G1 B9E8G2 B9E8G3 B9E8H2 B9E8I3 B9E8I5 B9E8M1
B9E8N2 B9E8N6 B9E8Q0 B9E8Q1 B9E8Q2 B9E8Q3 B9E8Q5 B9E8Q7 B9E8Q8 B9E8Q9 B9E8T2
B9E8U9 B9E8V3 B9E8V7 B9E8V8 B9E8X2 B9E8X3 B9E8Y0 B9E8Y3 B9E8Z2 B9E8Z3 B9E8Z6
B9E8Z7 B9E903 B9E909 B9E913 B9E922 B9E9A2 B9E9B4 B9E9E3 B9E9F9 B9E9G8 B9E9G9
B9E9H0 B9E9J0 B9E9J1 B9E9J2 B9E9J3 B9E9J4 B9E9J5 B9E9J6 B9E9J8 B9E9K0 B9E9K1
B9E9K2 B9E9K3 B9E9K4 B9E9K5 B9E9K7 B9E9K9 B9E9L0 B9E9L4 B9E9L5 B9E9L6 B9E9L7
B9E9L8 B9E9M3 B9E9M4 B9E9W9 B9EA22 B9EAC1 B9EAC6 B9EAC8 B9EAD2 B9EAD7 B9EAF7
B9EAG2 B9EAG4 B9EAG5 B9EAG8 B9EAH5 B9EAJ9 B9EAM9 B9EAN2 B9EAQ8 B9EAS8 B9EAY2
B9EAY5 B9EAY8 B9EAZ0 B9EAZ2 B9EB04 B9EB18 B9EB42 B9EB46 B9EB47 B9EB51 B9EB57
B9EB60 B9EB64 B9EB66 B9EB67 B9EB69 B9EB94 B9EBB8 B9EBB9 B9EBC6 B9EBC7 B9EBD3
B9EBD5 B9EBD6 B9EBD7 B9EBD9 B9EBE2 B9EBE4 B9EBE6 B9EBE9 B9EBF4 B9EBF6 B9EBF7
B9EBG0 B9EBH1 B9EBI4 B9EBI5 B9EBI8 B9EBJ1 B9EBU2 B9EBZ6 B9EC41 B9EJ57 B9EJA2
B9EJR8 B9EJX3 B9EM04 B9EMY6 B9ENE7 B9ENM6 B9EQ30 B9EUM5 B9EXM2 B9EY52 B9EYZ1
B9F166 B9F1L8 B9F2L1 B9F2U5 B9F2Y7 B9F3B6 B9F4Q9 B9F7C8 B9FAF3 B9FDB8 B9FFA3

B9FI63 B9FJG3 B9FKM7 B9FL70 B9FS33 B9FS74 B9FSH5 B9FTR1 B9FUF9 B9FXV5 B9G2X9
B9G3M6 B9G434 B9G5N1 B9G8P1 B9GAG9 B9GBM3 B9GCH9 B9GE13 B9GHA6 B9GLX8 B9H5L9
B9HCR2 B9HQZ6 B9HRL7 B9HSR7 B9I2J6 B9I666 B9IJ21 B9INH0 B9IR64 B9IRA0 B9IRB4
B9IRF0 B9IRL0 B9IRU7 B9IRU8 B9IRW2 B9IRW3 B9IRX1 B9IRX2 B9IRY0 B9IS16 B9IS21
B9IS32 B9IS83 B9IS84 B9IS85 B9IS86 B9ISD5 B9ISF7 B9ISQ6 B9IT27 B9IT30 B9IT32
B9IT39 B9IT40 B9IT45 B9IT46 B9IT52 B9IT66 B9ITE8 B9ITE9 B9ITF9 B9ITG8 B9ITN8
B9ITX7 B9ITZ8 B9IU03 B9IU27 B9IU35 B9IU36 B9IU37 B9IU38 B9IU58 B9IUC0 B9IUC1
B9IUC7 B9IUG8 B9IUG9 B9IUH0 B9IUH1 B9IUM6 B9IUM7 B9IUP3 B9IUP4 B9IUP5 B9IUR5
B9IUR6 B9IUT0 B9IUT3 B9IUW2 B9IUY4 B9IUZ0 B9IUZ2 B9IUZ3 B9IUZ4 B9IUZ5 B9IUZ7
B9IUZ8 B9IUZ9 B9IV00 B9IV01 B9IV02 B9IV11 B9IV12 B9IV41 B9IV58 B9IV59 B9IV71
B9IV96 B9IV99 B9IVA1 B9IVA5 B9IVA8 B9IVB0 B9IVB5 B9IVB6 B9IVB7 B9IVB8 B9IVB9
B9IVC1 B9IVC5 B9IVC6 B9IVC9 B9IVD0 B9IVD1 B9IVD3 B9IVD8 B9IVE7 B9IVF3 B9IVF5
B9IVI6 B9IVL9 B9IVM2 B9IVM6 B9IVN3 B9IVN5 B9IVN9 B9IVP2 B9IVQ5 B9IVQ6 B9IVQ9
B9IVR2 B9IVR3 B9IVT0 B9IVT1 B9IVT8 B9IVU6 B9IVU8 B9IVW0 B9IVW1 B9IVW2 B9IVW3
B9IVW5 B9IVW6 B9IVW8 B9IVX2 B9IVX6 B9IVY9 B9IVZ5 B9IVZ6 B9IVZ9 B9IW02 B9IW07
B9IW18 B9IW28 B9IW40 B9IW43 B9IW56 B9IW60 B9IW61 B9IWL4 B9IWP6 B9IWP9 B9IWQ0
B9IWQ1 B9IWR7 B9IWS0 B9IWS1 B9IWT7 B9IWV1 B9IWX4 B9IWX5 B9IWX7 B9IWY0 B9IX36
B9IX72 B9IX93 B9IXA5 B9IXC5 B9IXC7 B9IXC9 B9IXD7 B9IXF8 B9IXG8 B9IXH2 B9IXH3
B9IXH4 B9IXH5 B9IXJ1 B9IXJ3 B9IXJ8 B9IXL7 B9IXL8 B9IXL9 B9IXM6 B9IXW1 B9IXY1
B9IY25 B9IY29 B9IY45 B9IY53 B9IY56 B9IY70 B9IY76 B9IY79 B9IY80 B9IY81 B9IY83
B9IY88 B9IY89 B9IY99 B9IYA1 B9IYE4 B9IYE5 B9IYF1 B9IYG1 B9IYG8 B9IYH1 B9IYH8
B9IYH9 B9IYI0 B9IYI8 B9IYI9 B9IYP4 B9IYX5 B9IYY0 B9IYY2 B9IYZ1 B9IYZ2 B9IYZ4
B9IZ04 B9IZ16 B9IZ18 B9IZ20 B9IZ25 B9IZ29 B9IZ30 B9IZ36 B9IZ39 B9IZ41 B9IZ44
B9IZ47 B9IZ48 B9IZ73 B9IZB3 B9IZC4 B9IZC8 B9IZD4 B9IZF0 B9IZG5 B9IZH1 B9IZH4
B9IZI2 B9IZI3 B9IZI4 B9IZI6 B9IZI9 B9IZJ0 B9IZJ1 B9IZJ2 B9IZJ3 B9IZJ4 B9IZJ5
B9IZJ6 B9IZJ7 B9IZJ8 B9IZJ9 B9IZK1 B9IZK3 B9IZK4 B9IZK5 B9IZK6 B9IZK8 B9IZL0
B9IZL2 B9IZL3 B9IZL5 B9IZL8 B9IZL9 B9IZM0 B9IZM2 B9IZM7 B9IZM8 B9IZT9 B9IZV2
B9IZZ7 B9J024 B9J044 B9J054 B9J063 B9J065 B9J073 B9J074 B9J092 B9J099 B9J0B4
B9J0E1 B9J0F1 B9J0P1 B9J0X9 B9J102 B9J151 B9J159 B9J189 B9J194 B9J1A0 B9J1C6
B9J1E1 B9J1G5 B9J1G9 B9J1H1 B9J1H2 B9J1K2 B9J1K4 B9J1K5 B9J1K7 B9J1K9 B9J1L6
B9J1L9 B9J1M9 B9J1N0 B9J1N1 B9J265 B9J266 B9J2F2 B9J2G5 B9J2G7 B9J2G8 B9J2L4
B9J2L5 B9J2P7 B9J2U7 B9J2V0 B9J2V8 B9J353 B9J3C5 B9J3H2 B9J3J0 B9J3N2 B9J3R0
B9J3S1 B9J484 B9J498 B9J499 B9J4E0 B9J4L7 B9J4M7 B9J4Q7 B9J4Q9 B9J4R5 B9J4V7
B9J523 B9J560 B9J587 B9J590 B9J5G3 B9J5N4 B9J6R3 B9J6R4 B9J6R6 B9J6R7 B9J6S5
B9J6V7 B9J6V8 B9J6V9 B9J738 B9J753 B9J795 B9J7B2 B9J7E0 B9J7E8 B9J7G1 B9J7G2
B9J7G4 B9J7L1 B9J7M8 B9J7P6 B9J7S8 B9J7T1 B9J7T9 B9J7U0 B9J7W1 B9J7X6 B9J885
B9J8C6 B9J8D0 B9J8D1 B9J8D8 B9J8E2 B9J8G8 B9J8H5 B9J8H7 B9J8H9 B9J8L8 B9J8M0
B9J8M3 B9J8M5 B9J8M6 B9J8M7 B9J8Q9 B9J8R3 B9J8R4 B9J8S0 B9J950 B9J966 B9J9F7
B9J9H0 B9J9I5 B9J9J0 B9J9J2 B9J9M4 B9J9P2 B9J9U0 B9J9Y1 B9J9Y2 B9JA11 B9JAA3
B9JAA4 B9JAF5 B9JAL7 B9JAL9 B9JAN3 B9JAQ2 B9JAQ3 B9JAR2 B9JAX6 B9JB75 B9JB78
B9JB95 B9JB96 B9JBA2 B9JBA3 B9JBE6 B9JBH4 B9JBS8 B9JBT6 B9JBW0 B9JC66 B9JC68
B9JC73 B9JC76 B9JCA5 B9JCB6 B9JCB8 B9JCB9 B9JCD8 B9JCF4 B9JCF5 B9JCG8 B9JCI9
B9JCJ2 B9JCL3 B9JCL9 B9JCM0 B9JCN5 B9JCQ5 B9JCT4 B9JCV0 B9JCV9 B9JCW7 B9JCX4
B9JCX5 B9JCY3 B9JCZ0 B9JD13 B9JD18 B9JDB9 B9JDC0 B9JDI1 B9JDI5 B9JDL2 B9JDM0

B9JDM7 B9JDM9 B9JDN2 B9JDP1 B9JDR7 B9JDR8 B9JDR9 B9JDS0 B9JDS1 B9JDS4 B9JDS5
B9JDS6 B9JDS9 B9JDT0 B9JDT1 B9JDT2 B9JDT4 B9JDT5 B9JDT6 B9JDT7 B9JDT8 B9JDT9
B9JDU0 B9JDU2 B9JDU3 B9JDU4 B9JDU9 B9JDV0 B9JDV1 B9JDV2 B9JDV3 B9JDW2 B9JDZ7
B9JE02 B9JEB9 B9JEG2 B9JEG3 B9JEH1 B9JEH2 B9JEI6 B9JEJ1 B9JEP8 B9JEQ7 B9JEQ8
B9JER1 B9JER2 B9JER3 B9JES1 B9JET5 B9JEU9 B9JEV4 B9JEW2 B9JEX0 B9JEX1 B9JEX3
B9JEZ6 B9JF01 B9JF08 B9JFA5 B9JFA8 B9JFD7 B9JFD8 B9JFG4 B9JFG9 B9JFK6 B9JFK7
B9JFP7 B9JFQ3 B9JFR4 B9JFR5 B9JFS7 B9JFX9 B9JFZ4 B9JG15 B9JG24 B9JG42 B9JG43
B9JG44 B9JG53 B9JG59 B9JG63 B9JG64 B9JG66 B9JG70 B9JG71 B9JG75 B9JGD3 B9JGH5
B9JGK2 B9JGS9 B9JGT0 B9JGT7 B9JGT9 B9JGW1 B9JGW2 B9JH03 B9JH26 B9JH27 B9JH32
B9JH39 B9JH49 B9JH59 B9JH60 B9JHB0 B9JHU6 B9JHZ9 B9JI94 B9JIT0 B9JJ30 B9JJB7
B9JJD5 B9JJD7 B9JJJ9 B9JJK5 B9JK65 B9JK73 B9JK75 B9JK80 B9JL06 B9JLF7 B9JLH9
B9JLT5 B9JLT6 B9JLT7 B9JLT8 B9JLT9 B9JML6 B9JML7 B9JMX8 B9JMY1 B9JN19 B9JN20
B9JNG1 B9JPK6 B9JQV3 B9JQV9 B9JQW1 B9JQW4 B9JQX0 B9JQX1 B9JQX3 B9JR40 B9JR44
B9JR47 B9JR74 B9JR75 B9JR81 B9JR84 B9JR86 B9JRA5 B9JRA7 B9JRD7 B9JRI5 B9JRI6
B9JRR8 B9JRX1 B9JRY5 B9JRY6 B9JS00 B9JS08 B9JS22 B9JS29 B9JS40 B9JS46 B9JSD2
B9JSD6 B9JSL2 B9JSL3 B9JSM6 B9JSN3 B9JSR6 B9JST0 B9JT13 B9JT39 B9JT47 B9JT86
B9JTJ2 B9JTL3 B9JTP4 B9JTR0 B9JTS8 B9JTS9 B9JU16 B9JU97 B9JUA0 B9JUB9 B9JUC4
B9JUC8 B9JUD6 B9JUE9 B9JUF7 B9JUH4 B9JUH7 B9JUH8 B9JUH9 B9JUI3 B9JUI4 B9JUM9
B9JUN3 B9JUN7 B9JUT9 B9JUU1 B9JUU2 B9JUV4 B9JUW1 B9JUY3 B9JV07 B9JV13 B9JV26
B9JV43 B9JV46 B9JV51 B9JV52 B9JV61 B9JV68 B9JV74 B9JV75 B9JV80 B9JV88 B9JVB9
B9JVC4 B9JVC5 B9JVD6 B9JVH6 B9JVI4 B9JVI8 B9JVJ0 B9JVJ1 B9JVJ7 B9JVK2 B9JVM4
B9JVM5 B9JVM6 B9JVM7 B9JVM8 B9JVN3 B9JVN4 B9JVN6 B9JVN7 B9JVN8 B9JVN9 B9JVP0
B9JVP1 B9JVP4 B9JVP5 B9JVP6 B9JVP7 B9JVP8 B9JVP9 B9JVQ1 B9JVQ2 B9JVQ3 B9JVQ4
B9JVQ5 B9JVQ6 B9JVQ8 B9JVQ9 B9JVR0 B9JVR1 B9JVR2 B9JVV3 B9JVW2 B9JW09 B9JW57
B9JW84 B9JW91 B9JWF1 B9JWF4 B9JWI2 B9JWI3 B9JWJ9 B9JWL2 B9JWU3 B9JWW7 B9JWX0
B9JWZ2 B9JX30 B9JX32 B9JX56 B9JX63 B9JX64 B9JX68 B9JXD7 B9JXD8 B9JXE6 B9JXE7
B9JXM6 B9JXN7 B9JXP7 B9JXQ5 B9JXS2 B9JXS3 B9JXT0 B9JXT9 B9JXV5 B9JXV6 B9JXV7
B9JXW4 B9JXW5 B9JXW8 B9JXX0 B9JXX1 B9JXX4 B9JXY6 B9JY41 B9JY52 B9JY53 B9JY62
B9JYI6 B9JYJ6 B9JYK3 B9JYK6 B9JYK7 B9JYK9 B9JYL0 B9JYM8 B9JYM9 B9JYN0 B9JYP7
B9JYP8 B9JYP9 B9JYQ2 B9JYQ3 B9JYQ5 B9JYQ6 B9JYR4 B9JYY6 B9JYZ7 B9JZ39 B9JZ61
B9JZ87 B9JZ89 B9JZ91 B9JZA3 B9JZC0 B9JZF4 B9JZF5 B9JZF6 B9JZG3 B9JZG5 B9JZH2
B9JZI2 B9JZJ4 B9JZK9 B9JZL3 B9JZL8 B9JZL9 B9JZM8 B9JZN5 B9JZQ5 B9JZR4 B9K0G2
B9K1J4 B9K1N4 B9K1W0 B9K1Y7 B9K1Y8 B9K201 B9K206 B9K253 B9K256 B9K286 B9K288
B9K2E5 B9K2I7 B9K2Z2 B9K4A6 B9K4G4 B9K4P2 B9K6M6 B9K6M8 B9K6N9 B9K6P5 B9K6P8
B9K6R1 B9K6R2 B9K6R4 B9K6R7 B9K6R9 B9K6S9 B9K6U6 B9K6Y6 B9K6Z3 B9K6Z6 B9K712
B9K734 B9K736 B9K748 B9K754 B9K767 B9K773 B9K795 B9K7B8 B9K7G3 B9K7G8 B9K7G9
B9K7H2 B9K7H3 B9K7I7 B9K7J6 B9K7K0 B9K7M5 B9K7M6 B9K7N7 B9K7P3 B9K7S4 B9K7S5
B9K7U6 B9K7W0 B9K7W1 B9K7W8 B9K7X0 B9K7Y7 B9K7Y8 B9K7Z1 B9K7Z6 B9K7Z8 B9K800
B9K819 B9K824 B9K844 B9K846 B9K852 B9K857 B9K861 B9K865 B9K866 B9K867 B9K877
B9K882 B9K883 B9K884 B9K885 B9K887 B9K888 B9K889 B9K890 B9K892 B9K893 B9K894
B9K895 B9K896 B9K897 B9K899 B9K8A0 B9K8A1 B9K8A2 B9K8A3 B9K8A4 B9K8A5 B9K8A7
B9K8B1 B9K8B2 B9K8B3 B9K8B5 B9K8C0 B9K8C2 B9K8D0 B9K8D3 B9K8E0 B9K8E2 B9K8E3
B9K8E5 B9K8F0 B9K8N7 B9K8P2 B9K8P4 B9K8P7 B9K8Q0 B9K8S3 B9K8S7 B9K8T5 B9K8U1
B9K8V0 B9K8V5 B9K8X3 B9K8Y9 B9K937 B9K942 B9K9C8 B9K9F6 B9K9K8 B9K9N3 B9K9R3

B9K9R7 B9K9R9 B9K9S0 B9K9S2 B9KA10 B9KA57 B9KA80 B9KA88 B9KA96 B9KAA5 B9KAB7
B9KAB8 B9KAB9 B9KAC4 B9KAE8 B9KAI3 B9KAN6 B9KAN7 B9KAP3 B9KAP7 B9KAQ6 B9KAQ7
B9KAQ8 B9KAS1 B9KAS5 B9KAT4 B9KAU2 B9KAY3 B9KAZ0 B9KAZ2 B9KB39 B9KB80 B9KB91
B9KB95 B9KB98 B9KBC2 B9KBC5 B9KBC9 B9KBD1 B9KBD7 B9KBI1 B9KBJ4 B9KBJ6 B9KBJ7
B9KBJ8 B9KBJ9 B9KBK3 B9KBQ2 B9KBR9 B9KBS2 B9KBT4 B9KBV1 B9KBV5 B9KBV6 B9KBW2
B9KBW7 B9KC39 B9KCG6 B9KCH0 B9KCH1 B9KCH8 B9KCI0 B9KCI3 B9KCI6 B9KCI8 B9KCJ4
B9KCK3 B9KCM6 B9KCN0 B9KCN6 B9KCQ6 B9KCR2 B9KCR7 B9KCR8 B9KCS8 B9KCT6 B9KCU7
B9KCW4 B9KCX0 B9KCX7 B9KD06 B9KD13 B9KD35 B9KD43 B9KD44 B9KD58 B9KD64 B9KD70
B9KD74 B9KD77 B9KD79 B9KDE1 B9KDF4 B9KDG5 B9KDH6 B9KDL1 B9KDL3 B9KDM7 B9KDM8
B9KDN6 B9KDP1 B9KDP2 B9KDP9 B9KDQ8 B9KDT6 B9KDU0 B9KDU7 B9KDV3 B9KDV7 B9KDZ6
B9KDZ7 B9KDZ8 B9KE09 B9KE24 B9KE25 B9KE50 B9KE56 B9KE62 B9KE78 B9KE83 B9KE87
B9KE89 B9KE98 B9KE99 B9KEA4 B9KEB1 B9KEB2 B9KEB7 B9KEB8 B9KED6 B9KED9 B9KEE1
B9KEE2 B9KEE3 B9KEE4 B9KEE5 B9KEE6 B9KEE7 B9KEE8 B9KEF0 B9KEF2 B9KEF3 B9KEF4
B9KEF5 B9KEF6 B9KEH2 B9KEH3 B9KEH4 B9KEH5 B9KEH7 B9KEH8 B9KEI4 B9KEJ4 B9KEK7
B9KEM3 B9KEN1 B9KEN8 B9KER1 B9KER2 B9KER3 B9KER4 B9KEU1 B9KEU8 B9KEV0 B9KEV2
B9KEV8 B9KEW8 B9KEX3 B9KEY7 B9KF91 B9KFA8 B9KFD9 B9KFF9 B9KFG3 B9KFG4 B9KFG5
B9KFG6 B9KFG7 B9KFG8 B9KFG9 B9KFH0 B9KFH1 B9KFJ1 B9KFJ8 B9KFK6 B9KFL4 B9KFM1
B9KFM9 B9KFN2 B9KFN5 B9KFN9 B9KFP0 B9KFP1 B9KFP6 B9KFQ2 B9KFR5 B9KFS8 B9KFT3
B9KFW0 B9KFX0 B9KFX4 B9KFX7 B9KFX8 B9KFY4 B9KFZ2 B9KFZ4 B9KG03 B9KG17 B9KG20
B9KG40 B9KG80 B9KG87 B9KG88 B9KG97 B9KG98 B9KG99 B9KGA9 B9KGB1 B9KGB2 B9KGL7
B9KGM2 B9KGT5 B9KGY6 B9KH06 B9KH18 B9KH59 B9KH69 B9KH73 B9KH76 B9KH92 B9KH95
B9KHB8 B9KHC6 B9KHC7 B9KHC9 B9KHF1 B9KHH1 B9KHH5 B9KHH6 B9KHI5 B9KHJ1 B9KHK6
B9KHK9 B9KHM7 B9KHN0 B9KHN2 B9KHN3 B9KHN8 B9KHP7 B9KHP8 B9KHQ5 B9KHQ6 B9KHS0
B9KHS3 B9KHV1 B9KHV2 B9KHV3 B9KHV8 B9KHW2 B9KHW8 B9KHZ4 B9KHZ5 B9KHZ8 B9KHZ9
B9KI16 B9KI57 B9KI71 B9KI83 B9KI89 B9KIA2 B9KIB9 B9KIE2 B9KIE3 B9KIG4 B9KIH0
B9KIJ4 B9KIJ7 B9KIK0 B9KIN8 B9KIP4 B9KIQ1 B9KIR2 B9KIR5 B9KIT5 B9KIU4 B9KIV1
B9KIV2 B9KIW9 B9KIX1 B9KIX9 B9KIY0 B9KJ19 B9KJ30 B9KJ41 B9KJ46 B9KJ49 B9KJ52
B9KJ53 B9KJ54 B9KJ56 B9KJ58 B9KJ60 B9KJ64 B9KJ65 B9KJ66 B9KJ67 B9KJ69 B9KJ71
B9KJ94 B9KJ95 B9KJ98 B9KJG2 B9KJP0 B9KJP1 B9KJP2 B9KJP3 B9KJP6 B9KJU5 B9KJU8
B9KJU9 B9KK15 B9KK24 B9KK25 B9KK27 B9KK42 B9KK43 B9KK46 B9KK50 B9KK51 B9KKA1
B9KKA7 B9KKB0 B9KKC7 B9KKD5 B9KKH3 B9KKI5 B9KKM5 B9KKM8 B9KKQ0 B9KKS1 B9KKV1
B9KKV4 B9KKV6 B9KKV7 B9KKW2 B9KKX4 B9KKZ0 B9KL02 B9KL48 B9KL51 B9KL69 B9KL75
B9KL79 B9KL80 B9KL82 B9KL83 B9KL87 B9KL88 B9KL90 B9KL92 B9KL93 B9KL94 B9KL95
B9KL96 B9KL97 B9KL98 B9KLA0 B9KLA1 B9KLA2 B9KLA3 B9KLA5 B9KLA7 B9KLA8 B9KLA9
B9KLB0 B9KLB3 B9KLB4 B9KLB5 B9KLB6 B9KLD3 B9KLE0 B9KLE1 B9KLE8 B9KLF5 B9KLF9
B9KLJ5 B9KLK2 B9KLK4 B9KLM3 B9KLQ5 B9KLR1 B9KLX3 B9KM05 B9KM14 B9KM22 B9KM35
B9KM37 B9KM57 B9KM96 B9KMA3 B9KMC3 B9KME7 B9KMG1 B9KMG3 B9KMG7 B9KMH8 B9KMJ2
B9KMK2 B9KMK4 B9KMK8 B9KML7 B9KMM4 B9KMR6 B9KMT6 B9KMT9 B9KMU9 B9KMV0 B9KMV1
B9KMW0 B9KN19 B9KN59 B9KN62 B9KNB3 B9KNB4 B9KNB6 B9KND1 B9KNE5 B9KNE8 B9KNI7
B9KNI8 B9KNJ5 B9KNK0 B9KNK2 B9KNS6 B9KNU3 B9KNX5 B9KNX6 B9KNX7 B9KNZ1 B9KNZ4
B9KP00 B9KP36 B9KP40 B9KP41 B9KP47 B9KP62 B9KPC7 B9KPC8 B9KPC9 B9KPD2 B9KPE1
B9KPF5 B9KPF7 B9KPF8 B9KPH3 B9KPH4 B9KPJ0 B9KPJ9 B9KPN3 B9KPP3 B9KPP7 B9KPP8
B9KPT5 B9KPU4 B9KPV7 B9KPW0 B9KQ02 B9KQ06 B9KQ40 B9KQ55 B9KQ95 B9KQC8 B9KQJ8
B9KQJ9 B9KQK1 B9KQL7 B9KQP1 B9KQP8 B9KQQ7 B9KQT7 B9KQU0 B9KQX9 B9KQZ1 B9KQZ2

B9KR64 B9KRA6 B9KRC9 B9KRE8 B9KRG9 B9KRH0 B9KRK2 B9KRK6 B9KRK8 B9KRL8 B9KRP4
B9KRR7 B9KRT2 B9KRT8 B9KRU0 B9KRU3 B9KRV8 B9KRX7 B9KSA1 B9KSA7 B9KSB0 B9KSB7
B9KSE2 B9KSE9 B9KSF0 B9KSH8 B9KSH9 B9KSJ9 B9KSS4 B9KST5 B9KST8 B9KSW5 B9KSW6
B9KSX6 B9KSY0 B9KSY5 B9KT07 B9KT27 B9KT32 B9KT42 B9KT43 B9KTA6 B9KTC8 B9KTD9
B9KTM3 B9KU51 B9KU92 B9KU94 B9KUI2 B9KUU6 B9KW01 B9KW04 B9KW06 B9KW45 B9KXB8
B9KXC8 B9KXJ0 B9KXJ4 B9KXJ6 B9KXL1 B9KXL4 B9KXM2 B9KXM5 B9KXM8 B9KXN0 B9KXN9
B9KXX3 B9KY70 B9KY71 B9KYD3 B9KYK0 B9KYQ3 B9KYR9 B9KYS0 B9KYS1 B9KYT2 B9KYT9
B9KYU7 B9KYV8 B9KYW4 B9KYX3 B9KYX4 B9KYX5 B9KZ04 B9KZ09 B9KZ23 B9KZ37 B9KZ38
B9KZ42 B9KZ43 B9KZ44 B9KZ59 B9KZ66 B9KZ87 B9KZB4 B9KZB5 B9KZD4 B9KZK4 B9KZP8
B9KZR0 B9KZS7 B9KZV9 B9KZW1 B9KZW2 B9KZW3 B9KZW4 B9KZW9 B9KZX2 B9KZX3 B9KZX6
B9KZX7 B9KZY0 B9KZY1 B9KZY2 B9KZY3 B9KZY4 B9KZY7 B9KZZ1 B9KZZ2 B9L003 B9L006
B9L044 B9L0C1 B9L0E3 B9L0E5 B9L0I4 B9L0J2 B9L0K8 B9L0L0 B9L0Q1 B9L0Q2 B9L0R1
B9L0T5 B9L0U6 B9L0W3 B9L0W9 B9L101 B9L111 B9L131 B9L146 B9L159 B9L1A9 B9L1E4
B9L1H7 B9L1J9 B9L1L6 B9L1M7 B9L1X1 B9L1Y8 B9L1Z5 B9L273 B9L274 B9L287 B9L2A7
B9L2B5 B9L2J0 B9L2J1 B9L2M3 B9L2M4 B9L4T8 B9L505 B9L5I6 B9L5K2 B9L5M5 B9L5S9
B9L5T1 B9L5U7 B9L5U8 B9L5V9 B9L5W5 B9L5Y7 B9L5Y8 B9L5Z7 B9L5Z8 B9L5Z9 B9L603
B9L605 B9L606 B9L607 B9L611 B9L615 B9L638 B9L698 B9L6B0 B9L6B5 B9L6E1 B9L6E6
B9L6I1 B9L6L8 B9L6L9 B9L6M0 B9L6M3 B9L6M6 B9L6M7 B9L6M8 B9L6M9 B9L6N0 B9L6N1
B9L6N4 B9L6S4 B9L6T1 B9L6T6 B9L6T8 B9L6T9 B9L6U0 B9L6X1 B9L6Y0 B9L6Y8 B9L718
B9L721 B9L729 B9L734 B9L735 B9L788 B9L789 B9L7B0 B9L7B6 B9L7E1 B9L7G2 B9L7G9
B9L7I8 B9L7J2 B9L7J3 B9L7J4 B9L7J5 B9L7J8 B9L7J9 B9L7K0 B9L7K6 B9L7Q2 B9L7R7
B9L7R8 B9L7R9 B9L7U2 B9L811 B9L822 B9L823 B9L851 B9L884 B9L885 B9L889 B9L8B2
B9L8B8 B9L8C3 B9L8E1 B9L8F4 B9L8G5 B9L8H3 B9L8K3 B9L8M1 B9L8Q2 B9L8R4 B9L8R5
B9L8T4 B9L8V5 B9L8Z0 B9L8Z4 B9L8Z9 B9L952 B9L985 B9L986 B9L987 B9L9A2 B9L9C1
B9L9C2 B9L9D4 B9L9G3 B9L9H5 B9L9H9 B9L9I3 B9L9I6 B9L9K0 B9L9K5 B9L9P0 B9L9T8
B9L9Y8 B9LA07 B9LA27 B9LA36 B9LA47 B9LA82 B9LA83 B9LA90 B9LA97 B9LAD4 B9LAE4
B9LAF4 B9LAF7 B9LAL2 B9LAL6 B9LAL7 B9LAM9 B9LAP2 B9LAT1 B9LB04 B9LB15 B9LB70
B9LB77 B9LB79 B9LBA2 B9LBJ2 B9LBJ3 B9LBJ9 B9LBK4 B9LBR4 B9LBS4 B9LBT1 B9LBT6
B9LBX1 B9LC05 B9LC30 B9LC55 B9LC61 B9LCC8 B9LCG3 B9LCM0 B9LCM1 B9LCN0 B9LCT8
B9LCT9 B9LCU6 B9LCU8 B9LD23 B9LD34 B9LD40 B9LD60 B9LD66 B9LD91 B9LDR9 B9LDT2
B9LDT6 B9LDY3 B9LE47 B9LE93 B9LE96 B9LED5 B9LEM0 B9LEM3 B9LEN6 B9LEQ9 B9LER8
B9LER9 B9LEV9 B9LF86 B9LFE4 B9LFG0 B9LFG6 B9LFJ4 B9LFL8 B9LFN2 B9LFP4 B9LFQ7
B9LFU4 B9LFU5 B9LGI0 B9LGM7 B9LGN1 B9LGY2 B9LGZ1 B9LH02 B9LH03 B9LH09 B9LH40
B9LH68 B9LHE7 B9LHI6 B9LHK4 B9LHR5 B9LI29 B9LI30 B9LI31 B9LI32 B9LI74 B9LI75
B9LI79 B9LI82 B9LIA5 B9LIB5 B9LIH3 B9LIH9 B9LII0 B9LIK3 B9LIK4 B9LIV1 B9LIV8
B9LIX9 B9LJ02 B9LJ32 B9LJ33 B9LJ65 B9LJ75 B9LJ76 B9LJ78 B9LJC7 B9LJC8 B9LJD1
B9LJD2 B9LJD5 B9LJD6 B9LJD7 B9LJD8 B9LJD9 B9LJE0 B9LJE2 B9LJE3 B9LJE4 B9LJE5
B9LJE6 B9LJE7 B9LJE8 B9LJE9 B9LJF0 B9LJF1 B9LJF5 B9LJF6 B9LJF8 B9LJG0 B9LJG1
B9LJG2 B9LJG5 B9LJI4 B9LK59 B9LK98 B9LKB2 B9LKB9 B9LKC1 B9LKC5 B9LKI9 B9LKK0
B9LKK2 B9LKK8 B9LKM2 B9LKM4 B9LKN1 B9LKQ0 B9LKR5 B9LKR9 B9LKS0 B9LKW0 B9LL90
B9LL91 B9LLE9 B9LLP6 B9LLY0 B9LLZ7 B9LM50 B9LM66 B9LM70 B9LM83 B9LMG2 B9LMH0
B9LMH2 B9LML7 B9LML9 B9LMM4 B9LMP6 B9LMQ1 B9LMX0 B9LMX4 B9LMX5 B9LND9 B9LNE4
B9LNJ8 B9LNK1 B9LNM9 B9LP76 B9LP84 B9LP86 B9LPA9 B9LPE3 B9LPF8 B9LPH6 B9LPT9
B9LPV1 B9LPV4 B9LPW4 B9LPW5 B9LPY2 B9LPY3 B9LPY5 B9LQ28 B9LQ33 B9LQ94 B9LQ96

B9LQ97 B9LQC4 B9LQE0 B9LQI7 B9LQJ1 B9LQK2 B9LQL7 B9LQR1 B9LQT0 B9LQZ7 B9LR02
B9LR22 B9LR51 B9LR58 B9LR64 B9LRB1 B9LRD9 B9LRF1 B9LRF2 B9LRR0 B9LRS2 B9LRS6
B9LS04 B9LS21 B9LS33 B9LS95 B9LSC8 B9LSD1 B9LSG7 B9LSH0 B9LSI7 B9LSK0 B9LSL4
B9LSM1 B9LSM2 B9LSM6 B9LSN4 B9LSQ5 B9LSQ8 B9LSR2 B9LSR4 B9LSR5 B9LSR7 B9LSS3
B9LSS4 B9LSS6 B9LSV1 B9LSX2 B9LSY2 B9LSZ3 B9LT08 B9LTA6 B9LTD6 B9LTF0 B9LTN3
B9LTP1 B9LTS6 B9LTT6 B9LTT8 B9LTX1 B9LTZ9 B9LU09 B9LU14 B9LU22 B9LU30 B9LU47
B9LU81 B9LUC6 B9LUC7 B9LUD2 B9LUG4 B9LUM0 B9LUM1 B9LUM9 B9LUS0 B9LUU4 B9LUU5
B9LUU8 B9LUU9 B9LUV0 B9LUV3 B9LUZ0 B9LUZ3 B9LUZ9 B9LV13 B9LV23 B9LV56 B9LW38
B9LWA7 B9LZ13 B9LZ17 B9LZ18 B9LZ19 B9LZ20 B9LZ35 B9LZ36 B9LZ47 B9LZ49 B9LZ53
B9LZ61 B9LZA9 B9LZC4 B9LZC5 B9LZC6 B9LZC7 B9LZG2 B9LZG4 B9LZI6 B9LZK1 B9LZK2
B9LZL6 B9LZU0 B9LZU2 B9LZV4 B9LZW2 B9LZW4 B9M0D5 B9M0D6 B9M0D7 B9M0D9 B9M0K9
B9M0L4 B9M0L7 B9M0L9 B9M0M0 B9M0M1 B9M0M2 B9M0M7 B9M0W5 B9M0X3 B9M0X4 B9M0Y0
B9M0Y1 B9M0Y2 B9M163 B9M164 B9M170 B9M173 B9M174 B9M197 B9M1C9 B9M1D2 B9M1E2
B9M1E6 B9M1E7 B9M1F7 B9M1G0 B9M1G1 B9M1G4 B9M1G5 B9M1G7 B9M1P4 B9M1R1 B9M1R2
B9M1X7 B9M1Y2 B9M1Y7 B9M211 B9M2A2 B9M2A3 B9M2C3 B9M2D5 B9M2J2 B9M2P7 B9M2R9
B9M2S2 B9M2S8 B9M2Y9 B9M305 B9M326 B9M357 B9M361 B9M363 B9M371 B9M374 B9M377
B9M380 B9M381 B9M384 B9M3D4 B9M3E3 B9M3I1 B9M3I5 B9M3M9 B9M3N0 B9M3Q9 B9M3W1
B9M3W2 B9M3W3 B9M3X6 B9M416 B9M417 B9M422 B9M4A8 B9M4F4 B9M4M5 B9M4M9 B9M4N1
B9M4Q6 B9M4S3 B9M4S4 B9M4U3 B9M4V3 B9M4Z1 B9M4Z4 B9M520 B9M521 B9M522 B9M543
B9M591 B9M593 B9M594 B9M596 B9M597 B9M598 B9M5A2 B9M5B1 B9M5B4 B9M5C0 B9M5C3
B9M5C5 B9M5C6 B9M5K2 B9M5K3 B9M5M0 B9M5M2 B9M5N6 B9M5P5 B9M5R7 B9M5S1 B9M5T8
B9M5U2 B9M5U4 B9M5U5 B9M5U8 B9M5U9 B9M5V1 B9M5V2 B9M5V4 B9M644 B9M699 B9M6F1
B9M6F3 B9M6F5 B9M6F7 B9M6F9 B9M6G0 B9M6G2 B9M6G3 B9M6G4 B9M6G5 B9M6G6 B9M6G7
B9M6G8 B9M6G9 B9M6H0 B9M6H1 B9M6H2 B9M6H3 B9M6H4 B9M6H6 B9M6U2 B9M6U4 B9M6U5
B9M6U6 B9M6U9 B9M6V0 B9M6V1 B9M6V2 B9M6V3 B9M6V4 B9M6V5 B9M6V6 B9M6W2 B9M6W3
B9M6W5 B9M769 B9M7D5 B9M7Q2 B9M7S1 B9M7S3 B9M7T4 B9M7V0 B9M861 B9M876 B9M8U3
B9M8U6 B9M8U7 B9M902 B9M905 B9M910 B9M914 B9M9F3 B9M9F6 B9M9G4 B9M9Q5 B9M9R6
B9M9W3 B9M9W4 B9M9X5 B9M9X6 B9MA02 B9MA32 B9MA35 B9MA54 B9MAC8 B9MAC9 B9MAD0
B9MAN2 B9MAU5 B9MAV8 B9MB32 B9MB68 B9MB69 B9MB70 B9MB72 B9MB73 B9MB74 B9MB76
B9MB77 B9MB78 B9MB80 B9MB81 B9MB82 B9MB93 B9MBD2 B9MBJ1 B9MBJ2 B9MBJ3 B9MBJ9
B9MBK7 B9MBL6 B9MBL7 B9MBN9 B9MBP0 B9MBR0 B9MBS3 B9MBS5 B9MBT7 B9MBU7 B9MBU8
B9MBU9 B9MBV1 B9MBV2 B9MBV3 B9MBV4 B9MBV5 B9MBV6 B9MBV8 B9MBV9 B9MBW0 B9MBW1
B9MBW2 B9MBW3 B9MC79 B9MC94 B9MC99 B9MCH0 B9MCH3 B9MCH5 B9MCI6 B9MCK1 B9MCK8
B9MCM3 B9MCM4 B9MCN0 B9MCT1 B9MCW1 B9MD32 B9MD45 B9MD52 B9MD53 B9MDC2 B9MDC5
B9MDC6 B9MDD1 B9MDD6 B9MDE4 B9MDJ2 B9MDJ3 B9MDJ5 B9MDJ6 B9MDJ7 B9MDJ8 B9MDK8
B9MDL1 B9MDL6 B9MDL9 B9MDP4 B9MDR1 B9MDS3 B9MDU0 B9MDV0 B9MDV1 B9MDV4 B9MDV5
B9MDV7 B9MDW3 B9MDW4 B9MDZ5 B9MDZ6 B9MDZ7 B9MDZ8 B9MDZ9 B9ME00 B9ME05 B9ME28
B9ME46 B9ME47 B9ME49 B9ME52 B9ME55 B9ME57 B9ME75 B9ME96 B9ME99 B9MEB7 B9MEB8
B9MEC5 B9MEG7 B9MEH2 B9MEI9 B9MEJ9 B9MEK5 B9MEK9 B9MEL0 B9MEQ7 B9MER6 B9MEU8
B9MEX0 B9MEY1 B9MEY7 B9MEY9 B9MEZ2 B9MF03 B9MF57 B9MF60 B9MF68 B9MFG4 B9MFH8
B9MFP8 B9MFQ0 B9MFQ3 B9MFR0 B9MFR1 B9MFS0 B9MFS1 B9MFS2 B9MFS3 B9MFT0 B9MFW6
B9MFX2 B9MFX7 B9MFY0 B9MFY1 B9MFY6 B9MFY7 B9MG13 B9MG14 B9MG15 B9MGH2 B9MGI1
B9MGI8 B9MGL7 B9MGM0 B9MGN1 B9MGN9 B9MGP2 B9MGQ0 B9MH46 B9MH47 B9MH48 B9MH49
B9MH50 B9MH51 B9MH63 B9MH67 B9MHC7 B9MHJ3 B9MHW0 B9MHX8 B9MHY0 B9MHY1 B9MHY2

B9MI07 B9MI46 B9MI80 B9MI82 B9MI87 B9MI90 B9MIA2 B9MIF6 B9MIH9 B9MIR7 B9MIT4
B9MIU6 B9MIW4 B9MJ02 B9MJ16 B9MJ26 B9MJ44 B9MJA6 B9MJB2 B9MJH4 B9MJK9 B9MJL1
B9MJP2 B9MJR4 B9MJR6 B9MJR9 B9MJS9 B9MJT3 B9MJT7 B9MJT8 B9MJU0 B9MJV0 B9MJW2
B9MJX0 B9MJX1 B9MJX4 B9MJX8 B9MJY9 B9MJZ0 B9MJZ1 B9MK54 B9MK78 B9MK80 B9MKA5
B9MKA6 B9MKC1 B9MKC6 B9MKD0 B9MKD4 B9MKD6 B9MKF2 B9MKF3 B9MKF4 B9MKF5 B9MKF6
B9MKF7 B9MKG1 B9MKG3 B9MKG4 B9MKG6 B9MKG7 B9MKG8 B9MKH0 B9MKH1 B9MKH2 B9MKH3
B9MKH4 B9MKH5 B9MKH7 B9MKH8 B9MKH9 B9MKI0 B9MKI1 B9MKI2 B9MKP8 B9MKQ0 B9MKQ1
B9MKR8 B9MKW9 B9MKY8 B9MKZ9 B9ML29 B9ML31 B9ML57 B9ML76 B9ML77 B9ML78 B9MLC6
B9MLF6 B9MLF7 B9MLK0 B9MLV3 B9MLV4 B9MLW8 B9MLX4 B9MLY0 B9MLY9 B9MLZ0 B9MM31
B9MM32 B9MM51 B9MM59 B9MMB0 B9MMM6 B9MMQ3 B9MMU5 B9MN03 B9MN04 B9MN06 B9MN44
B9MN52 B9MNC2 B9MND6 B9MND8 B9MNP4 B9MNR6 B9MNV1 B9MP50 B9MP52 B9MPB0 B9MPC0
B9MPC6 B9MPD7 B9MPF7 B9MPG8 B9MPK0 B9MPL7 B9MPM8 B9MPS5 B9MQ10 B9MQ16 B9MQ23
B9MQ24 B9MQ31 B9MQ33 B9MQ83 B9MQ85 B9MQ92 B9MQ93 B9MQ99 B9MQF1 B9MQF8 B9MQG5
B9MQG9 B9MQH0 B9MQH1 B9MQH7 B9MQJ8 B9MQK4 B9MQM8 B9MQN7 B9MQS8 B9MQW7 B9MQW9
B9MQX0 B9MQX2 B9MQX5 B9MQY2 B9MR36 B9MR45 B9MR49 B9MR53 B9MR54 B9MR57 B9MR65
B9MR68 B9MR74 B9MR75 B9MR76 B9MRB2 B9MRB3 B9MRB7 B9MRB8 B9MRD1 B9MRD4 B9MRE0
B9MRE7 B9MRJ1 B9MRJ7 B9MRK1 B9MRK2 B9MRK7 B9MRK8 B9MRM3 B9MRM7 B9MRP3 B9MRP4
B9MRP5 B9MRP6 B9MRQ5 B9MRR4 B9MRS0 B9MRS5 B9MRS9 B9MRU6 B9MRV5 B9MRW0 B9MRX1
B9MRX9 B9MRY0 B9MRZ1 B9MRZ5 B9MS19 B9MS21 B9MS22 B9MS23 B9MS26 B9MS28 B9MS46
B9MS55 B9MS76 B9MS82 B9MS89 B9MS91 B9N1F9 B9N2B0 B9NAN0 B9PVB9 B9Q0C2 B9R4E3
B9RAJ0 B9RBT0 B9RHP7 B9RHQ5 B9RHX7 B9RI00 B9RPM3 B9RR88 B9RRX2 B9RU15 B9RW49
B9RX57 B9RXW0 B9RXW4 B9S1N2 B9S9Z3 B9SBU9 B9SBV6 B9SCB6 B9SEZ6 B9SL58 B9SQI7
B9SQR9 B9SVG9 B9SZB6 B9T3R0 B9T536 B9T825 B9TAD0 B9U6P5 B9UYK5 B9UYK6 B9V973
B9V984 B9VJ80 B9VRJ2 B9VVJ6 B9VXI8 B9VXK4 B9VXM8 B9VXQ2 B9W4V8 B9W6S2 B9W790
B9W7H9 B9W7S3 B9W7X4 B9W892 B9W8A8 B9W8P6 B9W8S4 B9W8Y2 B9W900 B9W923 B9W9A9
B9W9T4 B9WAG5 B9WAM8 B9WAV2 B9WC12 B9WD30 B9WD86 B9WDB4 B9WEC5 B9WED1 B9WEV3
B9WFG1 B9WFS9 B9WFW5 B9WG98 B9WGW9 B9WHJ2 B9WHK1 B9WJ16 B9WJ47 B9WJ66 B9WJA2
B9WK98 B9WL71 B9WLA7 B9WLF5 B9WLQ5 B9WM88 B9WMA4 B9WN49 B9WYE6 B9WZ64 B9WZX0
B9WZX2 B9WZX3 B9WZX5 B9WZX7 B9XEI9 B9XEK4 B9XXL6 B9XZG7 C0BK17 C0CMQ7 C0CMQ8
C0H3P9 C0H3Q2 C0H3Q3 C0H3Q6 C0H3Q8 C0H3Q9 C0H3R0 C0H3R1 C0H3R3 C0H3R4 C0H3R5
C0H3R9 C0H3S3 C0H3S6 C0H3S8 C0H3S9 C0H3T1 C0H3T2 C0H3T5 C0H3T6 C0H3T9 C0H3U7
C0H3U9 C0H3V3 C0H3V5 C0H3V8 C0H3V9 C0H3W1 C0H3W2 C0H3W7 C0H3X2 C0H3X4 C0H3X8
C0H3X9 C0H3Y1 C0H3Y4 C0H3Y5 C0H3Y7 C0H3Y9 C0H3Z0 C0H3Z1 C0H3Z2 C0H3Z3 C0H3Z4
C0H3Z5 C0H3Z6 C0H3Z7 C0H3Z8 C0H3Z9 C0H400 C0H402 C0H403 C0H404 C0H405 C0H406
C0H408 C0H410 C0H411 C0H412 C0H413 C0H418 C0H419 C0H421 C0H424 C0H425 C0H429
C0H432 C0H434 C0H437 C0H440 C0H443 C0H444 C0H445 C0H446 C0H447 C0H450 C0H452
C0H454 C0H455 C0H458 C0H461 C0H462 C0H4K2 C0H4X5 C0H537 C0H559 C0H582 C0H677
C0H859 C0H8I2 C0H906 C0HAC0 C0HAV3 C0HBG1 C0HBT3 C0HFE5 C0HIA3 C0HJA4 C0HJA5
C0HJA6 C0HJA7 C0HJA9 C0HJB3 C0HJE0 C0HJE1 C0HJE2 C0HJE3 C0HJE4 C0HJE8 C0HJF2
C0HJF7 C0HJF9 C0HJG0 C0HJG2 C0HJG5 C0HJG7 C0HJG8 C0HJH8 C0HJL9 C0HJM6 C0HJQ3
C0HJQ9 C0HJR0 C0HJS0 C0HJS1 C0HJS2 C0HJS3 C0HJS5 C0HJS6 C0HJS7 C0HJS8 C0HJT6
C0HJT7 C0HJT8 C0HJU5 C0HJV0 C0HJV1 C0HJV2 C0HJW7 C0HJX1 C0HJX6 C0HJY1 C0HJY2
C0HJZ2 C0HJZ3 C0HJZ4 C0HJZ5 C0HK14 C0HK20 C0HK21 C0HK22 C0HK23 C0HK24 C0HK25
C0HK28 C0HK29 C0HK30 C0HK31 C0HK32 C0HK33 C0HK34 C0HK35 C0HK36 C0HK37 C0HK38

C0HK39 C0HK40 C0HK41 C0HK48 C0HK70 C0HK81 C0HK94 C0HK95 C0HK96 C0HKB4 C0HKB7
C0HKC0 C0HKC6 C0HKD6 C0HKD8 C0HKD9 C0HKE1 C0HKE2 C0HKE3 C0HKE4 C0HKE5 C0HKE6
C0HKE7 C0HKE8 C0HKE9 C0HKF5 C0HKF9 C0HKG2 C0HKG3 C0HKH0 C0HKH1 C0HKH2 C0HKH3
C0HKH4 C0HKH5 C0HKH6 C0HKH7 C0HKH8 C0HKI2 C0HKI3 C0HKI4 C0HKI5 C0HKI6 C0HKI7
C0HKI8 C0HKI9 C0HKJ0 C0HKJ1 C0HKJ2 C0HKJ3 C0HKJ4 C0HKJ5 C0HKJ6 C0HKJ7 C0HKJ8
C0HKJ9 C0HKK0 C0HKK1 C0HKK2 C0HKK3 C0HKK4 C0HKK5 C0HKQ3 C0HKQ4 C0HKQ9 C0HL14
C0HL15 C0HL16 C0HL20 C0HL21 C0HL22 C0HL23 C0HL24 C0HL25 C0HL26 C0HL27 C0HL28
C0HL29 C0HL30 C0HL31 C0HL32 C0HL33 C0HL34 C0HL35 C0HL36 C0HL37 C0HL66 C0HL67
C0HL89 C0HL90 C0HL94 C0HL99 C0HLK7 C0HLL6 C0HLL7 C0HLL8 C0HLM4 C0HLM5 C0HLM7
C0HLM8 C0HLM9 C0HLN3 C0HLN7 C0HLN8 C0HLN9 C0HLP0 C0HLP1 C0HLP2 C0HLP3 C0HLP4
C0HLP5 C0HLP6 C0HLP7 C0HLP8 C0HLQ0 C0HLQ1 C0HLQ2 C0HLQ3 C0HLQ6 C0HLR0 C0HLR1
C0HLR7 C0HLR9 C0HLS1 C0HLS6 C0HLS7 C0HLS8 C0HLS9 C0HLT0 C0HLT1 C0HLT2 C0HLT3
C0HLT4 C0HLT5 C0HLT6 C0HLU2 C0HLV0 C0HLV1 C0HLV3 C0HLV4 C0HLV5 C0HLV8 C0HLZ4
C0HLZ6 C0HM01 C0HM02 C0HM10 C0HM19 C0HM20 C0HM39 C0HM40 C0HM41 C0HM48 C0HM49
C0HM50 C0HM52 C0IN03 C0JRZ9 C0KTJ6 C0KWV3 C0KWV4 C0KWV5 C0KWV6 C0KWV7 C0LA90
C0LTL9 C0LTM1 C0LU16 C0LZJ1 C0M660 C0M691 C0M695 C0M6B4 C0M6C1 C0M6C7 C0M6F3
C0M6F4 C0M6G6 C0M6G7 C0M6I5 C0M6J7 C0M6L2 C0M6L7 C0M6X1 C0M6X2 C0M6X3 C0M6X5
C0M6X6 C0M712 C0M748 C0M749 C0M751 C0M756 C0M758 C0M760 C0M780 C0M782 C0M791
C0M792 C0M795 C0M7A8 C0M7B1 C0M7C0 C0M7C5 C0M7C6 C0M7C8 C0M7E1 C0M7E5 C0M7E7
C0M7E8 C0M7E9 C0M7F4 C0M7G8 C0M7H8 C0M7I0 C0M7I2 C0M7J9 C0M7L5 C0M7M2 C0M7M5
C0M7M6 C0M7M8 C0M7M9 C0M7N7 C0M7Q4 C0M7Q5 C0M7Q6 C0M7R0 C0M7R2 C0M7R3 C0M7R4
C0M7R5 C0M7S3 C0M7S4 C0M7T7 C0M7T9 C0M7V0 C0M7V3 C0M7X0 C0M7X5 C0M7X6 C0M7Z4
C0M809 C0M815 C0M821 C0M831 C0M852 C0M858 C0M890 C0M899 C0M8A9 C0M8D5 C0M8F7
C0M8H1 C0M8H8 C0M8K2 C0M8K6 C0M8K7 C0M8K8 C0M8L1 C0M8L4 C0M8M3 C0M8M4 C0M8N3
C0M8P2 C0M8P7 C0M8P8 C0M8Q6 C0M8Q7 C0M8R7 C0M8R8 C0M8S3 C0M8S4 C0M8S8 C0M8S9
C0M8T0 C0M8T1 C0M8T8 C0M8T9 C0M8U0 C0M8U4 C0M8V6 C0M8X3 C0M8Y3 C0M903 C0M913
C0M918 C0M919 C0M934 C0M935 C0M937 C0M940 C0M953 C0M970 C0M990 C0M9A7 C0M9B0
C0M9C4 C0M9D9 C0M9E1 C0M9E6 C0M9F6 C0M9G4 C0M9H1 C0M9H2 C0M9H3 C0M9J2 C0M9J5
C0M9J7 C0M9J8 C0M9M0 C0M9M7 C0M9N3 C0M9R7 C0M9U4 C0M9Z1 C0M9Z4 C0MA08 C0MA32
C0MA42 C0MA50 C0MA57 C0MA63 C0MA71 C0MA73 C0MAA1 C0MAE4 C0MAF0 C0MAG2 C0MAG4
C0MAG5 C0MAG7 C0MAH5 C0MAI3 C0MAM1 C0MAM2 C0MAM5 C0MAM9 C0MAN2 C0MAQ5 C0MAR3
C0MAR4 C0MAR6 C0MAR7 C0MAS2 C0MAS4 C0MAS5 C0MAS8 C0MAT4 C0MAV6 C0MB16 C0MB21
C0MB23 C0MB46 C0MB49 C0MB53 C0MB61 C0MB66 C0MB79 C0MB81 C0MBC1 C0MBC3 C0MBD6
C0MBD7 C0MBG1 C0MBH4 C0MBI1 C0MBI2 C0MBK0 C0MBK2 C0MBK4 C0MBK6 C0MBQ7 C0MBS1
C0MBS9 C0MBT1 C0MBT9 C0MBU0 C0MBW5 C0MBW6 C0MBY9 C0MBZ1 C0MBZ4 C0MC10 C0MC15
C0MC19 C0MC29 C0MC62 C0MC67 C0MC74 C0MC84 C0MC89 C0MCA1 C0MCA9 C0MCB0 C0MCB1
C0MCB2 C0MCB3 C0MCB4 C0MCB5 C0MCB6 C0MCB7 C0MCB8 C0MCB9 C0MCC0 C0MCC1 C0MCC2
C0MCC3 C0MCC4 C0MCC9 C0MCD1 C0MCD2 C0MCH3 C0MCK4 C0MCL4 C0MCM1 C0MCM8 C0MCN7
C0MCR4 C0MCU1 C0MCV8 C0MCW0 C0MCW5 C0MCW6 C0MCZ5 C0MCZ8 C0MD33 C0MD36 C0MD49
C0MD67 C0MD70 C0MD73 C0MD77 C0MD79 C0MDA9 C0MDB4 C0MDC6 C0MDJ6 C0MDJ9 C0MDK3
C0MDQ2 C0MDQ3 C0MDQ4 C0MDS5 C0MDT1 C0MDT8 C0MDV8 C0MDW7 C0MDX5 C0MDX6 C0MDY4
C0MDY5 C0MDZ0 C0ME07 C0ME20 C0ME21 C0ME23 C0ME24 C0ME26 C0ME28 C0ME29 C0ME30
C0ME32 C0ME44 C0ME45 C0ME50 C0ME57 C0ME74 C0ME76 C0ME81 C0ME83 C0ME86 C0ME88
C0ME92 C0ME96 C0ME97 C0ME98 C0MEA0 C0MEB7 C0MED0 C0MED6 C0MEE0 C0MEE3 C0MEE4

C0MEF7 C0MEG5 C0MEG6 C0MEH1 C0MEH6 C0MEI2 C0MEI3 C0MEL3 C0MEL9 C0MEM1 C0MEM8
C0MEP5 C0MEQ5 C0MER2 C0MES2 C0MES3 C0MET9 C0MEV8 C0MEW5 C0MEW6 C0MEX3 C0MEX9
C0MEY7 C0MEY9 C0MF02 C0MF03 C0MF07 C0MF11 C0MF12 C0MF22 C0MF25 C0MF26 C0MF27
C0MF36 C0MF49 C0MF82 C0MF91 C0MFA1 C0MFA6 C0MFA8 C0MFB1 C0MFB5 C0MFC7 C0MFD9
C0MFE9 C0MFF6 C0MFF9 C0MFG8 C0MFN0 C0MFQ1 C0MFQ4 C0MFQ7 C0MFQ8 C0MFQ9 C0MFR4
C0MFU5 C0MFY0 C0MFZ1 C0MG35 C0MG37 C0MG47 C0MG52 C0MG55 C0MG79 C0MGA5 C0MGB3
C0MGB4 C0MGB6 C0MGB7 C0MGC2 C0MGC4 C0MGC5 C0MGC7 C0MGC8 C0MGD3 C0MGD4 C0MGD6
C0MGE9 C0MGF5 C0MGG5 C0MGG9 C0MGH0 C0MGH1 C0MGI9 C0MGJ9 C0MGK8 C0MGL2 C0MGL3
C0MGL4 C0MGL8 C0MGL9 C0MGM0 C0MGM6 C0MGP5 C0MGP8 C0MGQ2 C0MGR5 C0MGS7 C0MGT4
C0MGT5 C0MGT7 C0MGT8 C0MGU1 C0MGU8 C0MGV5 C0MGV8 C0MGX1 C0MGX4 C0MGY6 C0MGY7
C0MH11 C0MH21 C0MH28 C0MH30 C0MH45 C0MH46 C0MH77 C0MH79 C0MHA6 C0MHB1 C0MHB4
C0MHD5 C0MHD7 C0MHG3 C0MHG4 C0MHH2 C0NBH5 C0NCM1 C0NDZ7 C0NE02 C0NE84 C0NF18
C0NFV9 C0NIF0 C0NIQ4 C0NIT8 C0NKW7 C0NL18 C0NL63 C0NN25 C0NN85 C0NNH7 C0NQL8
C0NQU6 C0NRC6 C0NRX4 C0NSF3 C0NTM8 C0NUP2 C0NV09 C0NWE3 C0NX86 C0NXP6 C0NYZ7
C0NZU2 C0NZV4 C0P0F6 C0P108 C0P9J6 C0PF72 C0PN26 C0PPM5 C0PPR1 C0PT35 C0PTH8
C0PVE1 C0PVG5 C0PVI8 C0PVJ3 C0PVN0 C0PVY6 C0PVZ1 C0PVZ2 C0PW16 C0PW17 C0PW19
C0PW29 C0PW62 C0PW71 C0PW78 C0PW90 C0PW97 C0PWA5 C0PWA7 C0PWD6 C0PWD7 C0PWE7
C0PWK6 C0PWL7 C0PWL9 C0PWM1 C0PWM2 C0PWN3 C0PWV2 C0PWW0 C0PWW2 C0PWX3 C0PWX5
C0PWX6 C0PWX8 C0PWX9 C0PWY2 C0PWY3 C0PWY7 C0PWZ0 C0PWZ2 C0PX17 C0PXA3 C0PXA5
C0PXA6 C0PXA7 C0PXB0 C0PXB1 C0PXC5 C0PXD6 C0PXF6 C0PXF9 C0PXG5 C0PXG7 C0PXG9
C0PXI9 C0PXJ2 C0PXJ3 C0PXK7 C0PXN2 C0PXN9 C0PXQ7 C0PXR3 C0PXS2 C0PXT0 C0PXU3
C0PXU4 C0PXU6 C0PXU8 C0PXV4 C0PXV9 C0PXW1 C0PXX2 C0PY11 C0PY12 C0PY21 C0PY23
C0PY26 C0PY27 C0PY28 C0PY33 C0PY37 C0PY38 C0PY46 C0PY54 C0PY69 C0PY73 C0PY74
C0PY89 C0PYB4 C0PYB6 C0PYJ5 C0PYK6 C0PYK7 C0PYK9 C0PYL0 C0PYL1 C0PYL4 C0PYM7
C0PYM8 C0PYN1 C0PYN4 C0PYP2 C0PYP4 C0PYQ6 C0PYQ8 C0PYR0 C0PYS1 C0PYS5 C0PYV8
C0PYW8 C0PYX3 C0PYX7 C0PYY1 C0PYY2 C0PYY5 C0PYZ1 C0PZ11 C0PZ22 C0PZ24 C0PZ33
C0PZ34 C0PZ38 C0PZ50 C0PZ51 C0PZ53 C0PZ54 C0PZ56 C0PZ59 C0PZ62 C0PZ83 C0PZ88
C0PZB4 C0PZD3 C0PZJ5 C0PZJ7 C0PZK2 C0PZL8 C0PZN4 C0PZN5 C0PZN9 C0PZP0 C0PZQ4
C0PZQ5 C0PZT1 C0PZV0 C0PZV1 C0PZV7 C0PZW0 C0PZW4 C0PZW5 C0PZW7 C0PZW8 C0PZX8
C0PZZ3 C0PZZ5 C0Q034 C0Q055 C0Q059 C0Q060 C0Q062 C0Q073 C0Q077 C0Q079 C0Q093
C0Q0A4 C0Q0A5 C0Q0A7 C0Q0A8 C0Q0A9 C0Q0B0 C0Q0B1 C0Q0B2 C0Q0B5 C0Q0B6 C0Q0B7
C0Q0C2 C0Q0C3 C0Q0C4 C0Q0C6 C0Q0C7 C0Q0D0 C0Q0D9 C0Q0F3 C0Q0G1 C0Q0H5 C0Q0I2
C0Q0I5 C0Q0I7 C0Q0J3 C0Q0K9 C0Q0L0 C0Q0L1 C0Q0R3 C0Q0R4 C0Q0R6 C0Q0S9 C0Q0T7
C0Q0U7 C0Q0V4 C0Q0Y1 C0Q0Y8 C0Q148 C0Q162 C0Q1B6 C0Q1B7 C0Q1C2 C0Q1C9 C0Q1E4
C0Q1G0 C0Q1G1 C0Q1J7 C0Q1J8 C0Q1K1 C0Q1K3 C0Q1L6 C0Q1L8 C0Q1P4 C0Q1R0 C0Q1U3
C0Q1V0 C0Q1V2 C0Q1W7 C0Q1W8 C0Q1W9 C0Q1X0 C0Q1X1 C0Q1X2 C0Q1X3 C0Q1X4 C0Q1X9
C0Q1Y1 C0Q241 C0Q263 C0Q287 C0Q2A0 C0Q2A7 C0Q2E3 C0Q2E4 C0Q2E8 C0Q2F1 C0Q2F4
C0Q2F5 C0Q2G8 C0Q2K7 C0Q2L0 C0Q2L4 C0Q2P1 C0Q2P4 C0Q2P5 C0Q2P8 C0Q2Q8 C0Q2R3
C0Q2R4 C0Q2R5 C0Q2R6 C0Q2R7 C0Q2S2 C0Q2S6 C0Q2S7 C0Q2S8 C0Q2S9 C0Q2T0 C0Q2T8
C0Q2U0 C0Q2U8 C0Q2V3 C0Q2V8 C0Q2V9 C0Q2Y5 C0Q2Y9 C0Q309 C0Q311 C0Q319 C0Q321
C0Q322 C0Q324 C0Q330 C0Q357 C0Q378 C0Q3B3 C0Q3B5 C0Q3E1 C0Q3F4 C0Q3G7 C0Q3G9
C0Q3J4 C0Q3K0 C0Q3K3 C0Q3L1 C0Q3L2 C0Q3L3 C0Q3L4 C0Q3N3 C0Q3N4 C0Q3Q9 C0Q3R1
C0Q3V6 C0Q414 C0Q436 C0Q438 C0Q439 C0Q441 C0Q442 C0Q446 C0Q452 C0Q463 C0Q472
C0Q475 C0Q476 C0Q478 C0Q481 C0Q486 C0Q489 C0Q494 C0Q4C6 C0Q4D8 C0Q4E1 C0Q4E3

C0Q4E5 C0Q4F3 C0Q4F4 C0Q4I2 C0Q4I9 C0Q4J2 C0Q4K5 C0Q4L1 C0Q4L2 C0Q4L3 C0Q4L5
C0Q4N4 C0Q4Z6 C0Q502 C0Q5E5 C0Q5E7 C0Q5F4 C0Q5H1 C0Q5H7 C0Q5H8 C0Q5I7 C0Q5J8
C0Q5K1 C0Q5M6 C0Q5M7 C0Q5P2 C0Q5P3 C0Q5P4 C0Q5R5 C0Q5S0 C0Q5S6 C0Q5T3 C0Q5T4
C0Q5T6 C0Q5T8 C0Q5V2 C0Q632 C0Q635 C0Q640 C0Q642 C0Q644 C0Q647 C0Q657 C0Q659
C0Q6A2 C0Q6A6 C0Q6B6 C0Q6C7 C0Q6C8 C0Q6C9 C0Q6D4 C0Q6E8 C0Q6F4 C0Q6F5 C0Q6F8
C0Q6F9 C0Q6G0 C0Q6G1 C0Q6G8 C0Q6H8 C0Q6I8 C0Q6J5 C0Q6K6 C0Q6L7 C0Q6M7 C0Q6N0
C0Q6N2 C0Q6T7 C0Q6T8 C0Q6U2 C0Q6X4 C0Q6X7 C0Q6X8 C0Q6X9 C0Q708 C0Q764 C0Q772
C0Q783 C0Q785 C0Q792 C0Q7B7 C0Q7B8 C0Q7C5 C0Q7D6 C0Q7I4 C0Q7K3 C0Q7L6 C0Q7M5
C0Q7M8 C0Q7R0 C0Q7R1 C0Q7R4 C0Q7S8 C0Q7S9 C0Q7U2 C0Q7U3 C0Q7U7 C0Q7U9 C0Q7V0
C0Q7V5 C0Q7V6 C0Q7X2 C0Q7X4 C0Q7Y0 C0Q7Z3 C0Q7Z4 C0Q807 C0Q809 C0Q810 C0Q812
C0Q840 C0Q845 C0Q871 C0Q886 C0Q893 C0Q894 C0Q8A0 C0Q8C5 C0Q8C6 C0Q8C9 C0Q8D4
C0Q8D6 C0Q8F3 C0Q8F4 C0Q8F5 C0Q8N4 C0Q8N5 C0Q8P4 C0Q8X7 C0Q9D3 C0Q9S7 C0Q9U6
C0Q9U7 C0Q9U8 C0Q9V0 C0Q9V1 C0Q9V4 C0Q9V5 C0Q9V7 C0Q9V8 C0Q9W1 C0Q9W2 C0Q9W3
C0Q9W4 C0Q9W5 C0Q9W6 C0Q9W7 C0Q9W8 C0Q9W9 C0Q9X1 C0Q9X2 C0Q9X3 C0Q9X6 C0Q9X7
C0Q9X9 C0Q9Y0 C0Q9Y2 C0Q9Y3 C0Q9Y7 C0Q9Z7 C0QA05 C0QA34 C0QAG9 C0QAP1 C0QAP3
C0QAT7 C0QAX9 C0QAY0 C0QAZ0 C0QB12 C0QB14 C0QB17 C0QB20 C0QB49 C0QBA3 C0QBU5
C0QBV1 C0QBV4 C0QBX7 C0QC86 C0QCR9 C0QD10 C0QDM7 C0QDY4 C0QET6 C0QFH9 C0QFI0
C0QFI1 C0QFJ2 C0QFJ4 C0QFK8 C0QFM5 C0QFS1 C0QFZ0 C0QGP2 C0QGP3 C0QGP6 C0QGS9
C0QGT0 C0QGV9 C0QH22 C0QHJ8 C0QHL6 C0QHL7 C0QHM0 C0QHM2 C0QHM3 C0QHM5 C0QHM6
C0QHN3 C0QHQ9 C0QHT9 C0QI20 C0QI21 C0QI24 C0QI26 C0QI33 C0QI55 C0QI73 C0QI75
C0QI90 C0QIA5 C0QIZ3 C0QIZ4 C0QJ00 C0QJ36 C0QJ54 C0QJ55 C0QJ67 C0QJ86 C0QJ89
C0QJ90 C0QJB0 C0QJG7 C0QJG8 C0QKG6 C0QKK8 C0QKM9 C0QKP4 C0QKP5 C0QKQ4 C0QKT7
C0QKT9 C0QKU4 C0QKW6 C0QKW7 C0QKX1 C0QKX3 C0QKX6 C0QKY2 C0QKZ5 C0QKZ6 C0QKZ8
C0QL01 C0QLA5 C0QLA8 C0QLD3 C0QLE7 C0QLE8 C0QLE9 C0QLF1 C0QLM0 C0QLV7 C0QM06
C0QM36 C0QMA1 C0QMG1 C0QMX0 C0QP68 C0QP76 C0QPF0 C0QPF6 C0QPI1 C0QPL9 C0QPQ6
C0QPU5 C0QPV2 C0QPW0 C0QPY2 C0QQ02 C0QQ35 C0QQ61 C0QQ98 C0QQA0 C0QQC2 C0QQC5
C0QQE4 C0QQH3 C0QQJ9 C0QQL2 C0QQL3 C0QQL5 C0QQL8 C0QQL9 C0QQM0 C0QQM2 C0QQM3
C0QQM6 C0QQM7 C0QQM9 C0QQN1 C0QQN3 C0QQN6 C0QQN7 C0QQN8 C0QQP0 C0QQP1 C0QQP7
C0QQQ0 C0QQQ2 C0QQW3 C0QQW6 C0QQX0 C0QQX4 C0QR28 C0QR62 C0QR68 C0QR76 C0QR78
C0QR82 C0QR95 C0QRE6 C0QRE8 C0QRF9 C0QRH2 C0QRH5 C0QRI1 C0QRI3 C0QRI5 C0QRI8
C0QRP9 C0QRQ2 C0QRW3 C0QRW6 C0QRY8 C0QS00 C0QS40 C0QS44 C0QS47 C0QS72 C0QSF6
C0QSH0 C0QSI8 C0QSL7 C0QSL9 C0QSS1 C0QST3 C0QSU0 C0QSY6 C0QT00 C0QT02 C0QT34
C0QT35 C0QT41 C0QT44 C0QT45 C0QT55 C0QT56 C0QT57 C0QT62 C0QT74 C0QTA7 C0QTB8
C0QTC4 C0QTD8 C0QTE2 C0QTG9 C0QTI3 C0QTI8 C0QTL3 C0QTL6 C0QTL9 C0QTM3 C0QTM4
C0QTN9 C0QTP7 C0QTV2 C0QTY9 C0QTZ1 C0QU23 C0QU24 C0QU54 C0QU74 C0QU76 C0QUB5
C0QUB7 C0QUC0 C0QUE5 C0QUE7 C0QUF7 C0QUG8 C0QUH9 C0QUI2 C0QUJ3 C0QUK5 C0QUK8
C0QUM2 C0QUM8 C0QV47 C0QVC0 C0QVC8 C0QVE5 C0QVG4 C0QVH2 C0QVI8 C0QVL4 C0QVL9
C0QVM0 C0QVU4 C0QVX1 C0QVZ2 C0QVZ4 C0QVZ5 C0QVZ6 C0QVZ7 C0QVZ8 C0QW00 C0QW01
C0QW06 C0QW08 C0QW11 C0QW13 C0QW91 C0QW93 C0QWA1 C0QWA9 C0QWB0 C0QWF7 C0QWJ2
C0QWM9 C0QWN8 C0QWR3 C0QWR6 C0QWW9 C0QWX2 C0QWX5 C0QWY0 C0QX38 C0QX49 C0QX60
C0QX75 C0QXF2 C0QXH9 C0QXK7 C0QXM0 C0QXU6 C0QY51 C0QY98 C0QYF1 C0QYH6 C0QYU7
C0QYU9 C0QYX7 C0QZ35 C0QZ40 C0QZ84 C0QZD0 C0QZD1 C0QZK3 C0QZQ2 C0QZT8 C0QZV6
C0QZV8 C0QZX1 C0R000 C0R046 C0R074 C0R078 C0R087 C0R090 C0R0B8 C0R0E0 C0R0J6
C0R0P3 C0R0Q0 C0R0R3 C0R0X6 C0R0Y0 C0R0Y4 C0R112 C0R175 C0R176 C0R1H5 C0R1J9

C0R1L5 C0R1L6 C0R1Q2 C0R1U9 C0R1X1 C0R1Z0 C0R1Z9 C0R214 C0R250 C0R292 C0R2B0
C0R2C9 C0R2E2 C0R2K4 C0R2K8 C0R2L2 C0R2L4 C0R2L5 C0R2M2 C0R2M3 C0R2N7 C0R2N9
C0R2Q3 C0R2Q4 C0R2Q8 C0R2R5 C0R2S2 C0R2S8 C0R2U2 C0R2U3 C0R2U5 C0R2U8 C0R2V3
C0R2V4 C0R2W3 C0R2X2 C0R2Y5 C0R2Y7 C0R2Y9 C0R2Z1 C0R2Z3 C0R2Z5 C0R2Z7 C0R2Z8
C0R2Z9 C0R300 C0R301 C0R302 C0R304 C0R305 C0R306 C0R308 C0R310 C0R392 C0R399
C0R3A3 C0R3A4 C0R3B8 C0R3K7 C0R3M3 C0R3M5 C0R3N5 C0R3P3 C0R3S5 C0R3S6 C0R3S7
C0R3T4 C0R3T5 C0R3U0 C0R3W7 C0R403 C0R405 C0R414 C0R430 C0R442 C0R443 C0R444
C0R449 C0R453 C0R462 C0R4C7 C0R4E1 C0R4F9 C0R4G6 C0R4G7 C0R4I6 C0R4J5 C0R4K8
C0R4L0 C0R4L3 C0R4M1 C0R4N2 C0R4N4 C0R4Q8 C0R4S1 C0R4S5 C0R4T5 C0R4W4 C0R4X2
C0R4X3 C0R4X5 C0R4X8 C0R4Y0 C0R4Z2 C0R4Z3 C0R512 C0R515 C0R541 C0R542 C0R543
C0R562 C0R573 C0R576 C0R579 C0R594 C0R595 C0R598 C0R5A2 C0R5B1 C0R5B6 C0R5B9
C0R5C2 C0R5C3 C0R5C9 C0R5D1 C0R5E5 C0R5F0 C0R5F9 C0R5G4 C0R5H4 C0R5H8 C0R5I4
C0R5J8 C0R5K8 C0R5L5 C0R5L9 C0R5M1 C0R5M6 C0R5N3 C0R5N5 C0R5P2 C0RE20 C0RE25
C0RE30 C0RE38 C0RE50 C0RE56 C0RE59 C0RE68 C0RE69 C0RE78 C0REB5 C0REC2 C0REE0
C0REH1 C0REH2 C0REI6 C0REM2 C0RES0 C0REU2 C0REU7 C0REV5 C0REW5 C0REW6 C0REW7
C0REX2 C0REX8 C0REX9 C0RF11 C0RF29 C0RF32 C0RF56 C0RF58 C0RF62 C0RF67 C0RF74
C0RF85 C0RF89 C0RF92 C0RF93 C0RF97 C0RF98 C0RFB5 C0RFB8 C0RFD1 C0RFD2 C0RFF1
C0RFF2 C0RFF8 C0RFF9 C0RFG8 C0RFH1 C0RFH2 C0RFH7 C0RFI1 C0RFL5 C0RFM2 C0RFM4
C0RFQ6 C0RFR9 C0RFS0 C0RFU8 C0RFV3 C0RFV5 C0RFV7 C0RFW0 C0RFW7 C0RFW8 C0RFW9
C0RFX2 C0RFX3 C0RFX4 C0RFX5 C0RFY3 C0RFZ2 C0RFZ4 C0RFZ5 C0RG03 C0RG04 C0RG06
C0RG11 C0RG13 C0RG38 C0RG44 C0RG48 C0RG50 C0RG51 C0RG65 C0RG67 C0RG90 C0RG91
C0RG96 C0RG97 C0RGD6 C0RGD9 C0RGE2 C0RGF9 C0RGH3 C0RGK0 C0RGK7 C0RGM6 C0RGM7
C0RGN1 C0RGP5 C0RGR3 C0RGX4 C0RGX7 C0RH03 C0RH21 C0RH45 C0RH46 C0RH89 C0RH97
C0RHB0 C0RHC3 C0RHD5 C0RHE3 C0RHF9 C0RHG1 C0RHG2 C0RHI3 C0RHI7 C0RHL3 C0RHQ0
C0RHQ3 C0RHQ4 C0RHY9 C0RHZ3 C0RHZ5 C0RI03 C0RI19 C0RI23 C0RI35 C0RI52 C0RI89
C0RI93 C0RIA2 C0RIA3 C0RIA7 C0RIB4 C0RIC3 C0RIC5 C0RIC6 C0RIF7 C0RIG4 C0RIG5
C0RIH1 C0RIH6 C0RIK0 C0RIL0 C0RIL8 C0RIL9 C0RIN3 C0RIN8 C0RIP1 C0RIT3 C0RIT4
C0RIW8 C0RJ23 C0RJ44 C0RJ47 C0RJ48 C0RJ65 C0RJ68 C0RJ73 C0RJ80 C0RJ81 C0RJ83
C0RJ95 C0RJA5 C0RJB0 C0RJB1 C0RJB2 C0RJC7 C0RJC9 C0RJD0 C0RJD8 C0RJF0 C0RJG2
C0RJG9 C0RJH5 C0RJH6 C0RJH7 C0RJH8 C0RJI0 C0RJI2 C0RJI3 C0RJI4 C0RJI5 C0RJI6
C0RJI7 C0RJI9 C0RJJ0 C0RJJ1 C0RJJ2 C0RJJ3 C0RJJ4 C0RJJ5 C0RJJ7 C0RJJ8 C0RJJ9
C0RJK0 C0RJK1 C0RJK2 C0RJK4 C0RJK5 C0RJK6 C0RJK9 C0RJL1 C0RJL2 C0RJL3 C0RJL4
C0RJL5 C0RJP4 C0RJP6 C0RJR0 C0RJS3 C0RK28 C0RK37 C0RK96 C0RKC7 C0RKC8 C0RKC9
C0RKD5 C0RKD6 C0RKF6 C0RKP1 C0RKP8 C0RKS9 C0RL20 C0RL24 C0RL26 C0RLB0 C0RLB5
C0RLB6 C0RLB8 C0RLC0 C0RLD0 C0RLJ5 C0RLN1 C0RLR7 C0RLT3 C0RLV7 C0RLW2 C0RM19
C0RM30 C0RM38 C0RM54 C0RM68 C0RM77 C0RM93 C0RMB1 C0RMG1 C0RMG2 C0RMG5 C0RMG8
C0RMH0 C0RMH2 C0RMH3 C0RMI7 C0RMJ1 C0RMK3 C0RMK4 C0RMK5 C0RMK6 C0RMR6 C0RWW9
C0RYV6 C0S0E2 C0S1C8 C0S3U2 C0S3Z3 C0S433 C0S4C4 C0S4M2 C0S4Z7 C0S8C7 C0S902
C0SC25 C0SCV1 C0SCV8 C0SDG9 C0SDW6 C0SG20 C0SGK2 C0SH35 C0SHQ0 C0SIM8 C0SIQ6
C0SP86 C0SP91 C0SP98 C0SPA0 C0SPA5 C0SPA8 C0SPB0 C0SPB1 C0SPB5 C0SPB6 C0SPB8
C0SPB9 C0SPC0 C0SPC1 C0SPC3 C0SQ89 C0STD9 C0SUW7 C0SV51 C0SVG5 C0SVM5 C0SVS4
C0SVV6 C0SVZ5 C0SVZ6 C0VHC9 C0XIJ3 C0Z2L5 C0Z361 C0Z4D6 C0Z4E4 C0Z4E5 C0Z4Q5
C0Z4V3 C0Z5A8 C0Z671 C0Z6L6 C0Z6N8 C0Z6P0 C0Z6P4 C0Z6P5 C0Z6Q2 C0Z6S0 C0Z6T0
C0Z6T1 C0Z6W4 C0Z787 C0Z7P4 C0Z7W7 C0Z7Z0 C0Z7Z5 C0Z818 C0Z828 C0Z832 C0Z838

C0Z887 C0Z8I6 C0Z8L5 C0Z8Y8 C0Z9B3 C0Z9B5 C0Z9B6 C0Z9C7 C0Z9C8 C0Z9F1 C0Z9I7
C0Z9W8 C0Z9X2 C0ZA28 C0ZA40 C0ZA42 C0ZA45 C0ZA47 C0ZA63 C0ZA70 C0ZA72 C0ZAF8
C0ZAG1 C0ZAG3 C0ZAG9 C0ZAH1 C0ZAH4 C0ZAJ4 C0ZAL3 C0ZAL7 C0ZAN2 C0ZAN3 C0ZAN4
C0ZAN8 C0ZAP9 C0ZAQ1 C0ZAS1 C0ZAS6 C0ZB07 C0ZB13 C0ZB21 C0ZB46 C0ZB48 C0ZB49
C0ZB50 C0ZB54 C0ZB72 C0ZB81 C0ZB86 C0ZB87 C0ZBP8 C0ZBR4 C0ZBS0 C0ZBW6 C0ZBW7
C0ZBW9 C0ZBY1 C0ZC05 C0ZC06 C0ZC10 C0ZC13 C0ZC50 C0ZC76 C0ZC77 C0ZCB6 C0ZCC7
C0ZCD6 C0ZCD8 C0ZCE2 C0ZCE4 C0ZCE7 C0ZCE9 C0ZCG5 C0ZCH6 C0ZCJ6 C0ZCK0 C0ZCX0
C0ZCX3 C0ZD92 C0ZDI3 C0ZDU5 C0ZDU6 C0ZDV1 C0ZDZ3 C0ZES4 C0ZEV2 C0ZEY6 C0ZEZ4
C0ZF00 C0ZF14 C0ZF43 C0ZF45 C0ZF49 C0ZF55 C0ZF57 C0ZF58 C0ZF60 C0ZF63 C0ZF65
C0ZF66 C0ZFA5 C0ZFA7 C0ZFA9 C0ZFB3 C0ZFN0 C0ZFN1 C0ZFN5 C0ZFQ2 C0ZFY2 C0ZG01
C0ZG03 C0ZG41 C0ZG45 C0ZG86 C0ZG90 C0ZGB3 C0ZGB4 C0ZGC8 C0ZGF5 C0ZGH7 C0ZGH9
C0ZGJ5 C0ZGL2 C0ZGV3 C0ZH32 C0ZH40 C0ZH53 C0ZH54 C0ZHA2 C0ZHB6 C0ZHC9 C0ZHD2
C0ZHD7 C0ZHH0 C0ZHK9 C0ZHP0 C0ZHT2 C0ZHY5 C0ZI91 C0ZIC7 C0ZIE6 C0ZIF7 C0ZIF8
C0ZIG5 C0ZIG6 C0ZIG8 C0ZIH0 C0ZIH3 C0ZIH4 C0ZIH5 C0ZIH6 C0ZIH9 C0ZII0 C0ZII1
C0ZII2 C0ZII3 C0ZII4 C0ZII5 C0ZII6 C0ZII7 C0ZII8 C0ZII9 C0ZIJ0 C0ZIJ1 C0ZIJ2
C0ZIJ3 C0ZIJ4 C0ZIJ5 C0ZIJ6 C0ZIJ8 C0ZIJ9 C0ZIK1 C0ZIK4 C0ZIK5 C0ZIK6 C0ZIK7
C0ZIK8 C0ZIL2 C0ZIL3 C0ZIL4 C0ZIM8 C0ZIX8 C0ZIY1 C0ZJ12 C0ZJ16 C0ZJF8 C0ZK51
C0ZKC3 C0ZKC5 C0ZKC8 C0ZKI1 C0ZKW2 C0ZL09 C0ZL15 C0ZL41 C0ZL73 C0ZLE1 C0ZLE4
C0ZM44 C0ZN46 C0ZNE8 C0ZNS6 C0ZP17 C0ZP44 C0ZPL4 C0ZPL5 C0ZPR7 C0ZPR8 C0ZPX0
C0ZPX1 C0ZRJ1 C0ZRP8 C0ZSV1 C0ZSW0 C0ZSY9 C0ZT83 C0ZT86 C0ZTG3 C0ZU49 C0ZU50
C0ZUD4 C0ZUQ6 C0ZUT0 C0ZUV4 C0ZUX9 C0ZV17 C0ZV24 C0ZV30 C0ZV31 C0ZV33 C0ZV42
C0ZV45 C0ZVK4 C0ZVK5 C0ZVL8 C0ZVP7 C0ZVP8 C0ZVQ6 C0ZVQ7 C0ZVT4 C0ZVT5 C0ZVT6
C0ZVT7 C0ZW24 C0ZW25 C0ZW27 C0ZW28 C0ZW29 C0ZW30 C0ZW31 C0ZW32 C0ZW33 C0ZW34
C0ZW35 C0ZW38 C0ZW40 C0ZW41 C0ZW43 C0ZW44 C0ZW45 C0ZW47 C0ZW50 C0ZW51 C0ZW52
C0ZW53 C0ZW55 C0ZW77 C0ZW78 C0ZW79 C0ZW96 C0ZWI9 C0ZWV2 C0ZX05 C0ZXD0 C0ZXG7
C0ZXH4 C0ZXK1 C0ZXK3 C0ZXK6 C0ZXK7 C0ZXL3 C0ZXM2 C0ZXM4 C0ZXN2 C0ZXN6 C0ZXN9
C0ZXQ0 C0ZXQ5 C0ZXR7 C0ZXR9 C0ZXT0 C0ZXT2 C0ZY15 C0ZY16 C0ZY18 C0ZY23 C0ZY77
C0ZY88 C0ZYA2 C0ZYA5 C0ZYA7 C0ZYB6 C0ZYB7 C0ZYM6 C0ZYN8 C0ZYQ1 C0ZYQ9 C0ZYU5
C0ZYV9 C0ZYX0 C0ZZ26 C0ZZ45 C0ZZ46 C0ZZ47 C0ZZ48 C0ZZ58 C0ZZA3 C0ZZA9 C0ZZB5
C0ZZB6 C0ZZC0 C0ZZC1 C0ZZC3 C0ZZD2 C0ZZD4 C0ZZD9 C0ZZE5 C0ZZE8 C0ZZE9 C0ZZF3
C0ZZU4 C0ZZU6 C0ZZU7 C0ZZV2 C0ZZV7 C0ZZV8 C0ZZW0 C0ZZW9 C0ZZZ0 C1A039 C1A051
C1A066 C1A067 C1A0L2 C1A0L6 C1A0L8 C1A0M3 C1A0S6 C1A0W9 C1A0X4 C1A0Y3 C1A0Y4
C1A138 C1A163 C1A1D6 C1A1F6 C1A1G4 C1A1H0 C1A1J0 C1A1L0 C1A1L5 C1A1L6 C1A1L7
C1A1M0 C1A1N6 C1A1N9 C1A1V2 C1A1X1 C1A1Z3 C1A1Z7 C1A2D8 C1A2N3 C1A2R3 C1A2V7
C1A2X7 C1A2Y4 C1A348 C1A349 C1A383 C1A399 C1A3C4 C1A3F2 C1A3S5 C1A3W7 C1A3Y1
C1A3Z4 C1A3Z5 C1A482 C1A492 C1A493 C1A494 C1A4A9 C1A4B1 C1A4E4 C1A4H0 C1A4H9
C1A4I5 C1A4I9 C1A4J3 C1A4J4 C1A4L9 C1A4M2 C1A5E9 C1A5H8 C1A601 C1A611 C1A615
C1A617 C1A6H2 C1A6J9 C1A6N3 C1A6N4 C1A6P1 C1A6P2 C1A6Q2 C1A6Q3 C1A6Q8 C1A6Q9
C1A6R1 C1A6R2 C1A6R3 C1A6R5 C1A6S0 C1A6S6 C1A7E5 C1A7F4 C1A7K7 C1A7N5 C1A7P9
C1A873 C1A8A2 C1A8F6 C1A8J7 C1A8K1 C1A8L2 C1A8L7 C1A8L8 C1A8M5 C1A8N5 C1A8N9
C1A8S7 C1A8T0 C1A8T3 C1A8T7 C1A8U3 C1A8W3 C1A8X3 C1A8X8 C1A8Z5 C1A930 C1A949
C1A9H6 C1A9T6 C1AA44 C1AA54 C1AAD9 C1AAX0 C1AB49 C1ABE1 C1ABL6 C1ACL9 C1AE25
C1AED7 C1AEQ6 C1AEQ8 C1AER2 C1AES0 C1AES4 C1AES9 C1AF01 C1AF02 C1AF05 C1AF08
C1AF23 C1AF48 C1AF49 C1AF61 C1AF62 C1AF63 C1AF74 C1AF75 C1AF77 C1AF78 C1AF85

C1AFD7 C1AFE1 C1AFF6 C1AFH2 C1AFH7 C1AFI5 C1AFI6 C1AFJ5 C1AFL5 C1AFL6 C1AFM7
C1AFN6 C1AFP7 C1AFP9 C1AFV2 C1AFV3 C1AFV6 C1AFW6 C1AFY4 C1AFY6 C1AFZ5 C1AFZ7
C1AG04 C1AG09 C1AG13 C1AG17 C1AG19 C1AG21 C1AG22 C1AG24 C1AG40 C1AG80 C1AG91
C1AG98 C1AG99 C1AGA2 C1AGA3 C1AGB0 C1AGC5 C1AGC7 C1AGC8 C1AGD0 C1AGE1 C1AGH0
C1AGI1 C1AGL9 C1AGW8 C1AH01 C1AH22 C1AH35 C1AH36 C1AH56 C1AHF8 C1AHH2 C1AHL0
C1AHN1 C1AHN2 C1AHQ0 C1AHQ1 C1AHQ2 C1AHR4 C1AHR6 C1AHR7 C1AHR8 C1AHR9 C1AHZ2
C1AHZ3 C1AI39 C1AI40 C1AI41 C1AI45 C1AI57 C1AI58 C1AI59 C1AI67 C1AIH0 C1AIM6
C1AIU0 C1AIV9 C1AIZ8 C1AJ00 C1AJ01 C1AJ35 C1AJ38 C1AJ64 C1AJ76 C1AJ79 C1AJD9
C1AJF3 C1AJG2 C1AJN3 C1AJN6 C1AJT1 C1AJX2 C1AJZ7 C1AJZ9 C1AK26 C1AK27 C1AK32
C1AK93 C1AK99 C1AKA5 C1AKA9 C1AKF6 C1AKG0 C1AKG4 C1AKG7 C1AKI9 C1AKJ0 C1AKK4
C1AKN3 C1AKN5 C1AKN8 C1AKP6 C1AKV1 C1AKX3 C1AKX4 C1AKY4 C1AL01 C1AL03 C1AL15
C1AL16 C1AL17 C1AL18 C1AL34 C1AL35 C1AL36 C1AL37 C1AL38 C1AL39 C1AL40 C1AL41
C1AL42 C1AL43 C1AL48 C1AL49 C1AL50 C1AL51 C1AL52 C1AL53 C1AL54 C1AL55 C1AL56
C1AL57 C1AL67 C1ALB6 C1ALD5 C1ALD8 C1ALF4 C1ALT2 C1ALT7 C1ALU3 C1ALW8 C1ALZ2
C1AM01 C1AM02 C1AM05 C1AM72 C1AM86 C1AM93 C1AMA1 C1AMA2 C1AMF4 C1AMG5 C1AMK7
C1AMN4 C1AMT6 C1AMU2 C1AMU3 C1AMX6 C1AMY3 C1AMY5 C1AN24 C1AN25 C1AN30 C1AN35
C1AN37 C1AN51 C1AN60 C1AN65 C1AN66 C1AN68 C1AN82 C1AN93 C1AND1 C1ANK7 C1ANL6
C1ANM2 C1ANM3 C1ANM5 C1ANM7 C1ANM8 C1ANN3 C1ANN5 C1ANR5 C1ANR6 C1ANS2 C1ANS5
C1ANS7 C1ANT0 C1ANT1 C1ANT2 C1ANW0 C1ANX1 C1ANY5 C1ANY6 C1AP18 C1APA3 C1APA9
C1APB4 C1APC5 C1APC6 C1APC7 C1APC8 C1APC9 C1APH7 C1API1 C1API8 C1AQ16 C1AQ25
C1AQ26 C1AQ31 C1AQ37 C1AQ38 C1AQ46 C1AQ68 C1AQ71 C1AQ81 C1AQ82 C1AQA7 C1AQC2
C1AQC8 C1AQD2 C1AQD3 C1AQS5 C1AQT7 C1AQV5 C1AQX2 C1AQX7 C1AQZ3 C1AQZ5 C1ARF7
C1ASP4 C1ASP6 C1ASP7 C1ASQ2 C1ASQ7 C1ASQ8 C1ASR0 C1ASS8 C1ASX3 C1ASX9 C1ASY3
C1ASZ5 C1ASZ6 C1ASZ8 C1AT07 C1AT08 C1AT55 C1AT85 C1ATA2 C1ATC6 C1ATN6 C1ATY6
C1ATY7 C1ATY9 C1ATZ4 C1ATZ5 C1AU49 C1AU54 C1AU63 C1AU64 C1AU95 C1AUB5 C1AUB9
C1AUI5 C1AUK2 C1AUK7 C1AUL1 C1AUP4 C1AV76 C1AVG2 C1AVG3 C1AVJ6 C1AVJ9 C1AVK1
C1AVK2 C1AVK4 C1AVQ3 C1AVQ6 C1AVW7 C1AW13 C1AW17 C1AW66 C1AWB6 C1AWU5 C1AWW0
C1AWW6 C1AX07 C1AX87 C1AXW8 C1AXY5 C1AXZ0 C1AXZ1 C1AXZ2 C1AXZ4 C1AY15 C1AY16
C1AYB3 C1AYB4 C1AYH1 C1AYI4 C1AYS3 C1AYS4 C1AYS5 C1AYS6 C1AYV8 C1AYV9 C1AYX3
C1AYX5 C1AYX6 C1AYY2 C1AYZ0 C1AZ30 C1AZ44 C1AZ61 C1AZ64 C1AZH2 C1AZL1 C1B011
C1B012 C1B014 C1B015 C1B016 C1B017 C1B018 C1B019 C1B020 C1B021 C1B022 C1B025
C1B026 C1B027 C1B029 C1B030 C1B031 C1B035 C1B038 C1B039 C1B040 C1B042 C1B056
C1B057 C1B058 C1B075 C1B0D9 C1B155 C1B1A6 C1B1F1 C1B1L3 C1B1N5 C1B1N8 C1B1P5
C1B1S2 C1B1S4 C1B1S8 C1B2L3 C1B2M1 C1B2M3 C1B2N5 C1B2N8 C1B2Q0 C1B2Q5 C1B2R7
C1B2R9 C1B2S4 C1B2S6 C1B2T3 C1B2U2 C1B2U3 C1B2U5 C1B2V0 C1B2V4 C1B2W9 C1B2Z4
C1B310 C1B313 C1B315 C1B323 C1B324 C1B381 C1B393 C1B3A6 C1B4A1 C1B4B5 C1B4C5
C1B4C8 C1B4C9 C1B4D0 C1B4D1 C1B4D6 C1B4E1 C1B4G8 C1B4H5 C1B4H9 C1B4I1 C1B4I4
C1B4I5 C1B4I7 C1B4J5 C1B4J7 C1B4K2 C1B4L2 C1B4L3 C1B4L5 C1B4L7 C1B5J0 C1B716
C1B719 C1B762 C1B7L0 C1B7L1 C1B7P4 C1B7S5 C1B7S6 C1B7S7 C1B7T0 C1B8H9 C1B8I0
C1B914 C1B997 C1B9N1 C1BA63 C1BA76 C1BA77 C1BAC2 C1BAC3 C1BAC4 C1BAJ3 C1BAJ4
C1BBD1 C1BE28 C1BE29 C1BH56 C1BJ98 C1BJB1 C1BJQ3 C1BKD1 C1BM18 C1BW56 C1BY38
C1BY64 C1BYA3 C1BZU2 C1C3R6 C1C488 C1C4M3 C1C4M8 C1C5C3 C1C5D6 C1C5D8 C1C5E1
C1C5E4 C1C5E8 C1C5G8 C1C5H0 C1C5H1 C1C5H2 C1C5H3 C1C5H4 C1C5H5 C1C5I0 C1C5L8
C1C5M3 C1C5N5 C1C5N6 C1C5N7 C1C5S3 C1C5S4 C1C5S8 C1C5S9 C1C5V0 C1C5V9 C1C5Y6

C1C5Z8 C1C5Z9 C1C625 C1C631 C1C636 C1C648 C1C651 C1C666 C1C673 C1C691 C1C692
C1C6A5 C1C6B0 C1C6B6 C1C6B9 C1C6C0 C1C6G3 C1C6G4 C1C6H0 C1C6H2 C1C6H3 C1C6H7
C1C6J3 C1C6L4 C1C6M6 C1C6M8 C1C6P0 C1C6Q3 C1C6R3 C1C6R4 C1C6R6 C1C6R8 C1C6S2
C1C6S4 C1C6S6 C1C6T6 C1C6T7 C1C6T8 C1C6U4 C1C6V3 C1C6Y4 C1C6Y5 C1C6Z0 C1C6Z5
C1C6Z9 C1C727 C1C785 C1C787 C1C794 C1C797 C1C799 C1C7A1 C1C7B1 C1C7B5 C1C7B7
C1C7C8 C1C7D0 C1C7G0 C1C7G1 C1C7G2 C1C7H5 C1C7H6 C1C7H7 C1C7I6 C1C7J4 C1C7L7
C1C7M6 C1C7M7 C1C7P9 C1C7Q3 C1C7Q5 C1C7Q6 C1C7R5 C1C7S2 C1C7S8 C1C7V4 C1C7V5
C1C7W1 C1C7W3 C1C7X2 C1C7X3 C1C7X5 C1C7X6 C1C7X7 C1C7X8 C1C7Z0 C1C803 C1C812
C1C813 C1C814 C1C818 C1C842 C1C851 C1C862 C1C863 C1C869 C1C870 C1C871 C1C881
C1C8A8 C1C8B2 C1C8C3 C1C8C8 C1C8D0 C1C8D9 C1C8E8 C1C8F1 C1C8F7 C1C8F9 C1C8G0
C1C8G7 C1C8G8 C1C8H3 C1C8I7 C1C8J2 C1C8L6 C1C8M1 C1C8N4 C1C8P5 C1C8P8 C1C8Q3
C1C8R1 C1C8S3 C1C8T6 C1C8U6 C1C8U9 C1C8X2 C1C8X4 C1C8X8 C1C922 C1C965 C1C966
C1C967 C1C968 C1C969 C1C982 C1C986 C1C992 C1C9B2 C1C9E1 C1C9E2 C1C9E7 C1C9H2
C1C9H3 C1C9H6 C1C9H7 C1C9J1 C1C9K8 C1C9K9 C1C9U8 C1C9V1 C1C9V4 C1C9Y5 C1C9Z1
C1CA05 C1CA06 C1CA14 C1CA16 C1CA29 C1CA38 C1CA87 C1CA89 C1CAA6 C1CAC7 C1CAE7
C1CAF1 C1CAF6 C1CAI5 C1CAI9 C1CAJ9 C1CAK0 C1CAK1 C1CAL1 C1CAL2 C1CAL3 C1CAL4
C1CAL5 C1CAL6 C1CAL7 C1CAL8 C1CAL9 C1CAM0 C1CAM1 C1CAM2 C1CAM3 C1CAM4 C1CAM6
C1CAM7 C1CAM8 C1CAN0 C1CAN1 C1CAN3 C1CAN5 C1CAN6 C1CAN7 C1CAN9 C1CAP1 C1CAQ5
C1CAQ7 C1CAS3 C1CAS4 C1CAS7 C1CAU1 C1CAV1 C1CAV6 C1CAV7 C1CAX1 C1CAX9 C1CAZ3
C1CAZ4 C1CB02 C1CB20 C1CB31 C1CB34 C1CB39 C1CB44 C1CB45 C1CB46 C1CB65 C1CB69
C1CBB2 C1CBC0 C1CBF0 C1CBF1 C1CBF2 C1CBG2 C1CBG4 C1CBI1 C1CBJ1 C1CBJ2 C1CBK4
C1CBL5 C1CBM0 C1CBM4 C1CBM7 C1CBQ4 C1CBR1 C1CBS9 C1CBU0 C1CBU5 C1CBV0 C1CBX8
C1CBX9 C1CBY3 C1CBZ3 C1CBZ4 C1CBZ5 C1CC05 C1CC06 C1CC07 C1CC08 C1CC09 C1CC10
C1CC11 C1CC12 C1CC13 C1CC14 C1CC15 C1CC16 C1CC17 C1CC18 C1CC20 C1CC21 C1CC22
C1CC24 C1CC25 C1CC27 C1CC29 C1CC30 C1CC31 C1CC33 C1CC35 C1CC47 C1CC50 C1CC55
C1CC60 C1CC61 C1CC62 C1CC81 C1CC86 C1CCA8 C1CCB4 C1CCD5 C1CCD6 C1CCD7 C1CCG5
C1CCG7 C1CCH0 C1CCH3 C1CCH7 C1CCJ6 C1CCJ8 C1CCJ9 C1CCK0 C1CCK1 C1CCK2 C1CCK4
C1CCK9 C1CCN9 C1CCP3 C1CCQ6 C1CCQ7 C1CCQ8 C1CCT3 C1CCT4 C1CCT8 C1CCT9 C1CCV6
C1CCW4 C1CCZ1 C1CD02 C1CD03 C1CD30 C1CD36 C1CD41 C1CD51 C1CD54 C1CD69 C1CD76
C1CD96 C1CD97 C1CDB0 C1CDC3 C1CDC6 C1CDC7 C1CDH6 C1CDH7 C1CDI3 C1CDI5 C1CDI6
C1CDJ0 C1CDN0 C1CDP2 C1CDP4 C1CDQ5 C1CDR9 C1CDS8 C1CDS9 C1CDT1 C1CDT3 C1CDT9
C1CDU1 C1CDU3 C1CDV4 C1CDV5 C1CDV6 C1CDW2 C1CDX1 C1CE02 C1CE03 C1CE08 C1CE12
C1CE16 C1CE75 C1CE77 C1CE84 C1CE90 C1CE92 C1CE94 C1CEA4 C1CEA8 C1CEB0 C1CEC3
C1CEC5 C1CED4 C1CEE7 C1CEF2 C1CEF3 C1CEF4 C1CEG9 C1CEH0 C1CEH1 C1CEH8 C1CEI6
C1CEJ5 C1CEK4 C1CEK5 C1CEM5 C1CEM9 C1CEN1 C1CEN2 C1CEP0 C1CEP8 C1CEQ4 C1CEU3
C1CEU4 C1CEU9 C1CEV1 C1CEW0 C1CEW1 C1CEW3 C1CEW4 C1CEW5 C1CEW6 C1CEX8 C1CEZ2
C1CF01 C1CF02 C1CF03 C1CF07 C1CF29 C1CF38 C1CF49 C1CF50 C1CF56 C1CF57 C1CF58
C1CF71 C1CFA2 C1CFA5 C1CFB5 C1CFC0 C1CFC2 C1CFD1 C1CFE0 C1CFE3 C1CFE9 C1CFF1
C1CFF2 C1CFF9 C1CFG6 C1CFH9 C1CFI5 C1CFJ7 C1CFK0 C1CFL4 C1CFM7 C1CFN0 C1CFN5
C1CFP2 C1CFP3 C1CFR7 C1CFT0 C1CFT4 C1CFV7 C1CFV9 C1CFW4 C1CG04 C1CG41 C1CG42
C1CG43 C1CG44 C1CG45 C1CG72 C1CG99 C1CGA0 C1CGA5 C1CGD2 C1CGD3 C1CGD7 C1CGD8
C1CGK9 C1CGM4 C1CGM5 C1CGP2 C1CGP3 C1CGP4 C1CGQ1 C1CGQ3 C1CGR5 C1CGS4 C1CGX2
C1CGX4 C1CGY8 C1CH06 C1CH08 C1CH24 C1CH25 C1CH28 C1CH44 C1CH52 C1CH59 C1CH67
C1CH81 C1CH85 C1CH91 C1CHE6 C1CHE7 C1CHF7 C1CHI6 C1CHI8 C1CHK3 C1CHL3 C1CHL4

C1CHM6 C1CHN8 C1CHV6 C1CHV9 C1CHW2 C1CHZ1 C1CHZ8 C1CI29 C1CI34 C1CI40 C1CI68
C1CI69 C1CI73 C1CI83 C1CI84 C1CI85 C1CI96 C1CI97 C1CI98 C1CI99 C1CIA0 C1CIA1
C1CIA2 C1CIA3 C1CIA4 C1CIA5 C1CIA6 C1CIA7 C1CIA8 C1CIA9 C1CIB1 C1CIB2 C1CIB3
C1CIB5 C1CIB6 C1CIB8 C1CIC0 C1CIC1 C1CIC2 C1CIC4 C1CIC6 C1CID7 C1CIE0 C1CIE5
C1CIF1 C1CIF2 C1CIF3 C1CIH2 C1CIH6 C1CIJ8 C1CIK6 C1CIN1 C1CIN2 C1CIN3 C1CIQ2
C1CIQ5 C1CIQ8 C1CIR2 C1CIT2 C1CIT4 C1CIT7 C1CIT8 C1CIT9 C1CIU0 C1CIU2 C1CIU7
C1CIY6 C1CIZ1 C1CJ04 C1CJ05 C1CJ06 C1CJ34 C1CJ35 C1CJ39 C1CJ40 C1CJ89 C1CJA5
C1CJA6 C1CJD0 C1CJD5 C1CJD7 C1CJE0 C1CJF2 C1CJF5 C1CJG8 C1CJH5 C1CJJ5 C1CJJ6
C1CJL0 C1CJL5 C1CJM1 C1CJM4 C1CJM5 C1CJR7 C1CJR8 C1CJS5 C1CJS7 C1CJS8 C1CJT2
C1CJW9 C1CJY2 C1CJY4 C1CK08 C1CK17 C1CK18 C1CK20 C1CK22 C1CK27 C1CK29 C1CK31
C1CK40 C1CK41 C1CK42 C1CK48 C1CK57 C1CK88 C1CK93 C1CK98 C1CKA2 C1CKF5 C1CKF7
C1CKG4 C1CKG7 C1CKG8 C1CKH0 C1CKI0 C1CKI5 C1CKI7 C1CKQ7 C1CKQ9 C1CKR8 C1CKT7
C1CKT8 C1CKT9 C1CKV2 C1CKV3 C1CKV4 C1CKW1 C1CKW9 C1CKX8 C1CKY7 C1CKY8 C1CL05
C1CL09 C1CL11 C1CL12 C1CL20 C1CL28 C1CL34 C1CL62 C1CL63 C1CL69 C1CL71 C1CL80
C1CL81 C1CL83 C1CL84 C1CL85 C1CL86 C1CL98 C1CLB2 C1CLC1 C1CLC2 C1CLC3 C1CLC7
C1CLE8 C1CLF5 C1CLG6 C1CLG7 C1CLH3 C1CLH4 C1CLH5 C1CLI6 C1CLL5 C1CLL9 C1CLN3
C1CLN8 C1CLP0 C1CLP7 C1CLQ6 C1CLQ9 C1CLR5 C1CLR7 C1CLR8 C1CLS5 C1CLS6 C1CLT1
C1CLU3 C1CLU8 C1CLW8 C1CLX1 C1CLY4 C1CLZ5 C1CLZ8 C1CM03 C1CM10 C1CM11 C1CM34
C1CM45 C1CM49 C1CM71 C1CM73 C1CM77 C1CMA0 C1CMD2 C1CMD3 C1CMD4 C1CMD5 C1CMD6
C1CMF7 C1CMG3 C1CMI5 C1CMI6 C1CMJ1 C1CML5 C1CML6 C1CML7 C1CML8 C1CMM9 C1CMP5
C1CMP6 C1CMQ6 C1CMQ7 C1CMQ8 C1CMR5 C1CMR7 C1CMT3 C1CMU0 C1CMY8 C1CMZ0 C1CN05
C1CN23 C1CN25 C1CN42 C1CN43 C1CN46 C1CN60 C1CN66 C1CN70 C1CN76 C1CNA0 C1CNA5
C1CNB3 C1CNC2 C1CND0 C1CNE9 C1CNF8 C1CNG1 C1CNH8 C1CNI7 C1CNI8 C1CNK0 C1CNL1
C1CNN8 C1CNP4 C1CNS2 C1CNS6 C1CNS9 C1CNV7 C1CNW3 C1CP03 C1CP08 C1CP14 C1CP43
C1CP44 C1CP48 C1CP74 C1CP75 C1CP76 C1CP87 C1CP88 C1CP89 C1CP90 C1CP91 C1CP92
C1CP93 C1CP94 C1CP95 C1CP96 C1CP97 C1CP98 C1CP99 C1CPA0 C1CPA2 C1CPA3 C1CPA4
C1CPA6 C1CPA7 C1CPA9 C1CPB1 C1CPB3 C1CPB5 C1CPB7 C1CPD0 C1CPD4 C1CPD8 C1CPE3
C1CPE4 C1CPE5 C1CPG4 C1CPG8 C1CPL0 C1CPL6 C1CPN7 C1CPN9 C1CPR1 C1CPR4 C1CPR7
C1CPS1 C1CPU1 C1CPU3 C1CPU6 C1CPU7 C1CPU8 C1CPU9 C1CPV0 C1CPV5 C1CQ00 C1CQ04
C1CQ16 C1CQ17 C1CQ18 C1CQ43 C1CQ44 C1CQ48 C1CQ49 C1CQ76 C1CQA2 C1CQB5 C1CQB6
C1CQE1 C1CQE7 C1CQE9 C1CQF2 C1CQG4 C1CQG7 C1CQI2 C1CQI9 C1CQK8 C1CQK9 C1CQM2
C1CQN4 C1CQP3 C1CQP4 C1CQP5 C1CQQ0 C1CQQ1 C1CQR2 C1CQS0 C1CQT7 C1CQU2 C1CQU3
C1CQU4 C1CQV3 C1CQW7 C1CQX8 C1CQX9 C1CQY0 C1CQY1 C1CQY3 C1CQY4 C1CQZ3 C1CQZ4
C1CR00 C1CR01 C1CR07 C1CR13 C1CR21 C1CR28 C1CR29 C1CR31 C1CR35 C1CR55 C1CR56
C1CR65 C1CR74 C1CR83 C1CR90 C1CR91 C1CR92 C1CRA4 C1CRA5 C1CRA6 C1CRC4 C1CRD1
C1CRD6 C1CRE0 C1CRE4 C1CRJ1 C1CRJ3 C1CRK0 C1CRK3 C1CRK5 C1CRK7 C1CRL7 C1CRM1
C1CRM3 C1CRN4 C1CRN6 C1CRP5 C1CRS4 C1CRT3 C1CRT9 C1CRU0 C1CRU1 C1CRV1 C1CRV3
C1CRV5 C1CRV7 C1CRV9 C1CRW1 C1CRW2 C1CRX2 C1CRY6 C1CRZ7 C1CRZ9 C1CS14 C1CS48
C1CS52 C1CS53 C1CS55 C1CS61 C1CS62 C1CSA6 C1CSA7 C1CSB0 C1CSD7 C1CSE1 C1CSF2
C1CSF7 C1CSF9 C1CSG8 C1CSH7 C1CSI0 C1CSI6 C1CSI8 C1CSJ6 C1CSJ8 C1CSK2 C1CSL6
C1CSM2 C1CSP2 C1CSP6 C1CSR0 C1CSS1 C1CSS4 C1CSS9 C1CST6 C1CST7 C1CSU8 C1CSV9
C1CSX0 C1CSX4 C1CSZ4 C1CSZ6 C1CT01 C1CT24 C1CT50 C1CT51 C1CT52 C1CT53 C1CT54
C1CT78 C1CTA5 C1CTA6 C1CTB1 C1CTD1 C1CTD2 C1CTD6 C1CTD7 C1CTF2 C1CTH6 C1CTH7
C1CTL2 C1CTL3 C1CTL4 C1CTM3 C1CTM5 C1CTN9 C1CTP8 C1CTU8 C1CTV0 C1CTW4 C1CTY2

C1CTY4 C1CTZ9 C1CU00 C1CU03 C1CU16 C1CU26 C1CU32 C1CU40 C1CU44 C1CU66 C1CU67
C1CU77 C1CU95 C1CUA3 C1CUA5 C1CUC1 C1CUD1 C1CUD2 C1CUE4 C1CUF5 C1CUI9 C1CUQ9
C1CUR2 C1CUT8 C1CUU1 C1CV18 C1CV26 C1CV29 C1CV34 C1CV35 C1CV60 C1CVC5 C1CVC7
C1CVD5 C1CVE0 C1CVF4 C1CVJ5 C1CVN8 C1CVN9 C1CVP1 C1CVP5 C1CVW7 C1CVW9 C1CVX7
C1CW06 C1CW51 C1CWA3 C1CWB4 C1CWB5 C1CWI8 C1CWJ1 C1CWJ7 C1CWM4 C1CWT7 C1CWW2
C1CWY8 C1CX26 C1CX71 C1CX80 C1CX98 C1CXB5 C1CXD3 C1CXD5 C1CXD6 C1CXD7 C1CXD8
C1CXE5 C1CXE7 C1CXE8 C1CXE9 C1CXF0 C1CXF1 C1CXF2 C1CXF3 C1CXF5 C1CXF6 C1CXF8
C1CXG0 C1CXG1 C1CXG2 C1CXG3 C1CXG4 C1CXG5 C1CXG7 C1CXH0 C1CXH1 C1CXJ1 C1CXJ9
C1CXK0 C1CXK2 C1CXN1 C1CXR4 C1CXR6 C1CXS0 C1CXS7 C1CXT2 C1CXX6 C1CXY3 C1CY24
C1CY25 C1CY73 C1CYB8 C1CYD7 C1CYD9 C1CYL1 C1CYP4 C1CYT5 C1CYT8 C1CYV3 C1CYX1
C1CZ29 C1CZ40 C1CZ84 C1CZ90 C1CZB4 C1CZB5 C1CZH9 C1CZJ9 C1CZK0 C1CZK3 C1CZP1
C1CZV9 C1CZX7 C1CZY8 C1CZZ4 C1D070 C1D087 C1D094 C1D0A7 C1D0A9 C1D0D5 C1D0F6
C1D0F7 C1D0J7 C1D0M2 C1D0N3 C1D0Q7 C1D0Q8 C1D0S3 C1D0S4 C1D0S5 C1D0S6 C1D0S9
C1D0V5 C1D0V9 C1D0W2 C1D0Z0 C1D0Z6 C1D0Z8 C1D141 C1D165 C1D166 C1D195 C1D1C3
C1D1F5 C1D1G6 C1D1L2 C1D1L3 C1D1L7 C1D1L8 C1D1Q9 C1D1R8 C1D1S1 C1D1U3 C1D1Y6
C1D3U5 C1D4B9 C1D4C1 C1D4D0 C1D4E5 C1D4H9 C1D4I5 C1D4J2 C1D4P1 C1D4P9 C1D4R5
C1D4R8 C1D4Z4 C1D4Z8 C1D504 C1D505 C1D538 C1D539 C1D543 C1D546 C1D549 C1D558
C1D559 C1D560 C1D5B2 C1D5E0 C1D5H2 C1D5H3 C1D5I3 C1D5L4 C1D5L5 C1D5M5 C1D5M8
C1D5N0 C1D5Q6 C1D5R7 C1D5S2 C1D5S5 C1D5T3 C1D5T4 C1D5W4 C1D5X9 C1D5Z6 C1D5Z8
C1D602 C1D603 C1D665 C1D672 C1D679 C1D687 C1D6C4 C1D6C9 C1D6E4 C1D6F1 C1D6F5
C1D6H7 C1D6I1 C1D6I2 C1D6I8 C1D6J0 C1D6J9 C1D6L0 C1D6U0 C1D6U5 C1D6V9 C1D6X4
C1D6Y0 C1D702 C1D7E9 C1D7F0 C1D7F5 C1D7F6 C1D7H3 C1D7H9 C1D7I2 C1D7J7 C1D7L6
C1D7L7 C1D7M1 C1D7P3 C1D7Y5 C1D7Z6 C1D890 C1D8A6 C1D8N3 C1D8Q0 C1D8R7 C1D8V1
C1D8V4 C1D8W8 C1D8W9 C1D8X2 C1D8X4 C1D8X7 C1D973 C1D9A7 C1D9B6 C1D9C2 C1D9F0
C1D9F3 C1D9F5 C1D9Q4 C1D9S8 C1D9T5 C1D9T9 C1D9W9 C1DA02 C1DA26 C1DA28 C1DA41
C1DAD5 C1DAG2 C1DAG4 C1DAI5 C1DAJ6 C1DAK1 C1DAK5 C1DAQ6 C1DAQ7 C1DAQ8 C1DAQ9
C1DAR2 C1DAR3 C1DAR4 C1DAR6 C1DAR7 C1DAR8 C1DAS0 C1DAS1 C1DAS2 C1DAS3 C1DAS4
C1DAS5 C1DAS7 C1DAS8 C1DAT1 C1DAT2 C1DAT3 C1DAT4 C1DAT5 C1DAT6 C1DAT9 C1DAU0
C1DAU1 C1DAU2 C1DAU3 C1DAU4 C1DAV4 C1DAW8 C1DAZ0 C1DB12 C1DB31 C1DB32 C1DB52
C1DB55 C1DB66 C1DB68 C1DB91 C1DBA1 C1DBA8 C1DBA9 C1DBB4 C1DBE8 C1DBF4 C1DBF8
C1DBG0 C1DBG1 C1DBS9 C1DBU8 C1DBV7 C1DBV9 C1DBX3 C1DBX7 C1DBY7 C1DC35 C1DC39
C1DC72 C1DC78 C1DC84 C1DCC0 C1DCC1 C1DCE3 C1DCF0 C1DCK5 C1DCM3 C1DCM4 C1DCN2
C1DCR1 C1DCR2 C1DCR4 C1DCT7 C1DCV3 C1DCV6 C1DCV9 C1DCW2 C1DCW5 C1DCX2 C1DD01
C1DD12 C1DD37 C1DD40 C1DD41 C1DD43 C1DD44 C1DD48 C1DD53 C1DD55 C1DD59 C1DD62
C1DD64 C1DD68 C1DD72 C1DD87 C1DD88 C1DDA2 C1DDA3 C1DDA4 C1DDA6 C1DDA7 C1DDB1
C1DDC6 C1DDV7 C1DDY6 C1DDY7 C1DDZ2 C1DE08 C1DE40 C1DE52 C1DE56 C1DE57 C1DE61
C1DE64 C1DE65 C1DE68 C1DE76 C1DE77 C1DE99 C1DEA4 C1DEA5 C1DEA6 C1DED8 C1DEF4
C1DEF6 C1DEH4 C1DEK7 C1DEK8 C1DEM7 C1DEN4 C1DEQ3 C1DEU0 C1DEU1 C1DEV0 C1DEV6
C1DEV7 C1DEW5 C1DEW6 C1DEW7 C1DEW8 C1DF10 C1DF11 C1DF42 C1DF44 C1DF45 C1DF46
C1DFA8 C1DFH7 C1DFI9 C1DFJ9 C1DFK0 C1DFK5 C1DFK6 C1DFK8 C1DFK9 C1DFL1 C1DFL3
C1DFL6 C1DFM1 C1DFM2 C1DFM4 C1DFN0 C1DFU2 C1DFU4 C1DFV1 C1DFV8 C1DFX8 C1DGB0
C1DGT9 C1DGW5 C1DH23 C1DH55 C1DH66 C1DH67 C1DHB9 C1DHC2 C1DHE5 C1DHE6 C1DHE7
C1DHE8 C1DHG2 C1DHH0 C1DHH2 C1DHP3 C1DHQ1 C1DHQ5 C1DHR5 C1DHR7 C1DHS0 C1DHS2
C1DHS5 C1DHS6 C1DHT6 C1DHU3 C1DHV0 C1DHW2 C1DHX9 C1DHY0 C1DHY4 C1DHY7 C1DHY8

C1DI45 C1DI48 C1DI51 C1DI53 C1DI55 C1DI57 C1DI61 C1DI67 C1DI68 C1DI75 C1DI77
C1DI94 C1DIB2 C1DIC2 C1DIK7 C1DIX0 C1DIX1 C1DIX4 C1DIX6 C1DIY3 C1DIY4 C1DIY8
C1DJ20 C1DJ48 C1DJ52 C1DJ56 C1DJ58 C1DJC2 C1DJC9 C1DJD2 C1DJD3 C1DJD9 C1DJE8
C1DJG1 C1DJH1 C1DJI2 C1DJJ0 C1DJM5 C1DJZ2 C1DJZ7 C1DK04 C1DK16 C1DK22 C1DKD3
C1DKE3 C1DKE9 C1DKJ6 C1DKK1 C1DKK2 C1DKK3 C1DKK4 C1DKK5 C1DKK7 C1DKK8 C1DKL2
C1DKL4 C1DKL5 C1DKL6 C1DKL7 C1DKM0 C1DKM1 C1DKM2 C1DKM4 C1DKM5 C1DKM9 C1DKN0
C1DKN1 C1DKN2 C1DKN4 C1DKN5 C1DKN6 C1DKN7 C1DKN9 C1DKY7 C1DKY8 C1DKZ4 C1DKZ8
C1DLA8 C1DLH3 C1DLH8 C1DLI3 C1DLJ1 C1DLJ6 C1DLP4 C1DLQ0 C1DLQ1 C1DLQ2 C1DLQ7
C1DLQ8 C1DLR2 C1DLR3 C1DLR5 C1DLS8 C1DLT0 C1DLW0 C1DM03 C1DM14 C1DM53 C1DM65
C1DM66 C1DMQ0 C1DMQ3 C1DMQ9 C1DMR0 C1DMR1 C1DMT1 C1DMT5 C1DMV9 C1DMW4 C1DMY5
C1DMY6 C1DMZ7 C1DN27 C1DN48 C1DN55 C1DN56 C1DNF9 C1DNG0 C1DNG2 C1DNG3 C1DNK5
C1DNK6 C1DNK9 C1DNN5 C1DNP5 C1DNP6 C1DNP9 C1DPC0 C1DPC3 C1DPE2 C1DPF6 C1DPG2
C1DPH5 C1DPH8 C1DPM9 C1DPN1 C1DPN9 C1DPP2 C1DPQ6 C1DPY6 C1DQ00 C1DQ36 C1DQ37
C1DQ38 C1DQ47 C1DQ50 C1DQ65 C1DQ66 C1DQ71 C1DQ77 C1DQ78 C1DQ79 C1DQ86 C1DQ90
C1DQ91 C1DQA0 C1DQB5 C1DQC1 C1DQD4 C1DQH9 C1DQL3 C1DQM6 C1DQM8 C1DQR0 C1DQS2
C1DQS7 C1DQS9 C1DQT0 C1DQV1 C1DQV2 C1DR10 C1DR20 C1DR31 C1DRE9 C1DRF8 C1DRG0
C1DRG2 C1DRG4 C1DRI2 C1DRI8 C1DRI9 C1DRJ0 C1DRK5 C1DRL5 C1DRM1 C1DRN1 C1DRQ3
C1DRQ7 C1DRQ9 C1DRR3 C1DRR5 C1DRW0 C1DSQ4 C1DSR1 C1DSR8 C1DSR9 C1DSS0 C1DSS8
C1DST2 C1DSU3 C1DSU5 C1DSU6 C1DSU8 C1DSX1 C1DSX5 C1DSZ9 C1DTV7 C1DTW6 C1DUY4
C1DW63 C1DW75 C1DXS9 C1E3X9 C1E9T6 C1E9Y5 C1EKC8 C1EKE4 C1EKG1 C1EKT5 C1EL61
C1EL70 C1EL88 C1ELB0 C1ELB5 C1ELD8 C1ELE7 C1ELE8 C1ELE9 C1ELF0 C1ELF1 C1ELG5
C1ELP8 C1EM10 C1EM11 C1EM12 C1EM34 C1EM49 C1EM52 C1EM80 C1EMA2 C1EMA9 C1EMB1
C1EMB2 C1EMB3 C1EME8 C1EMQ2 C1EMQ4 C1EMQ5 C1EMQ6 C1EMQ7 C1EMQ8 C1EMQ9 C1EMR8
C1EMR9 C1EMV9 C1EMZ4 C1EMZ6 C1EN01 C1EN08 C1EN10 C1EN14 C1EN17 C1EN29 C1EN30
C1EN33 C1EN36 C1EN37 C1EN38 C1EN48 C1EN49 C1EN57 C1EN91 C1EN92 C1EN93 C1EN94
C1ENE6 C1ENE7 C1ENI6 C1ENJ4 C1ENJ5 C1ENK3 C1ENW4 C1ENZ2 C1ENZ3 C1EP03 C1EP04
C1EP29 C1EP30 C1EP33 C1EP35 C1EP39 C1EP43 C1EP45 C1EP48 C1EP50 C1EP51 C1EP52
C1EP53 C1EP54 C1EP56 C1EP60 C1EP61 C1EP64 C1EP65 C1EP66 C1EP68 C1EP73 C1EP82
C1EP88 C1EP90 C1EP94 C1EP96 C1EPF1 C1EPH2 C1EPL6 C1EPN0 C1EPP7 C1EPP8 C1EPP9
C1EPQ0 C1EPQ2 C1EPQ3 C1EPQ5 C1EPQ9 C1EPR3 C1EPS6 C1EPT2 C1EPT3 C1EPT5 C1EPT7
C1EPT9 C1EPU3 C1EPV3 C1EPW3 C1EPX4 C1EPX5 C1EPX7 C1EPZ0 C1EPZ4 C1EPZ5 C1EQ20
C1EQ58 C1EQ68 C1EQ75 C1EQF9 C1EQH0 C1EQQ8 C1EQR1 C1EQR2 C1EQR3 C1EQS8 C1EQT1
C1EQT2 C1EQU8 C1EQW1 C1EQY5 C1EQY6 C1EQZ2 C1ER02 C1ER04 C1ER06 C1ER14 C1ER62
C1ER65 C1ER67 C1ER74 C1ER75 C1ER80 C1ER81 C1ER94 C1ERC2 C1ERH2 C1ERH6 C1ERL1
C1ERL6 C1ERL7 C1ERN7 C1ERP8 C1ERQ2 C1ERQ3 C1ERQ4 C1ERQ5 C1ERR9 C1ERS1 C1ERS6
C1ERU8 C1ERU9 C1ERV0 C1ERV8 C1ES01 C1ES07 C1ES08 C1ES11 C1ES17 C1ES18 C1ES19
C1ES26 C1ES27 C1ES32 C1ES35 C1ES77 C1ESI2 C1ESJ0 C1ESJ2 C1ESJ7 C1ESK3 C1ESK6
C1ESK7 C1ESK8 C1ESL0 C1ESL5 C1ESL6 C1ESM6 C1ESM8 C1ESR9 C1ESS0 C1ESS5 C1EST1
C1EST2 C1ESU7 C1ESV0 C1ESW1 C1ESW2 C1ESW4 C1ESW5 C1ESX0 C1ESX4 C1ESX8 C1ESZ6
C1ESZ7 C1ET10 C1ET16 C1ET19 C1ET26 C1ET27 C1ET28 C1ET29 C1ET31 C1ET34 C1ET35
C1ET36 C1ET37 C1ET38 C1ET39 C1ET40 C1ET41 C1ET42 C1ET43 C1ET44 C1ET46 C1ET47
C1ET48 C1ET49 C1ET50 C1ET51 C1ET52 C1ET53 C1ET55 C1ET57 C1ET58 C1ET60 C1ET63
C1ET64 C1ET65 C1ET67 C1ET72 C1ET73 C1ETF1 C1ETG6 C1ETM6 C1ETN7 C1ETN9 C1ETP1
C1ETP6 C1ETQ0 C1ETQ1 C1ETQ7 C1ETR0 C1ETR2 C1ETR5 C1ETR8 C1ETR9 C1ETS9 C1ETT1

C1ETV9 C1ETY1 C1ETZ0 C1ETZ4 C1EU00 C1EU02 C1EU11 C1EUA1 C1EUA7 C1EUB0 C1EUB1
C1EUB4 C1EUL0 C1EUQ5 C1EUR7 C1EUR8 C1EUS4 C1EUT6 C1EUU3 C1EUV8 C1EUX9 C1EUZ5
C1EV03 C1EV43 C1EV60 C1EV62 C1EV63 C1EV65 C1EV67 C1EV74 C1EV77 C1EV87 C1EV88
C1EV89 C1EVE5 C1EVE6 C1EVJ1 C1EVJ3 C1EVJ4 C1EVW0 C1EVW6 C1EVZ1 C1EW09 C1EW31
C1EW36 C1EW38 C1EWA3 C1EWD5 C1EWF1 C1EWF4 C1EWG2 C1EWG6 C1EWH5 C1EWI4 C1EWU0
C1EX54 C1EX85 C1EX98 C1EX99 C1EXK2 C1EXV7 C1EY56 C1EY77 C1EYD5 C1EYJ3 C1EYT0
C1EYX7 C1EYZ6 C1EZ15 C1EZ79 C1EZ92 C1EZA3 C1EZD4 C1EZD6 C1EZE2 C1EZE4 C1EZI8
C1EZL5 C1EZN7 C1EZP2 C1EZZ5 C1F026 C1F030 C1F0A7 C1F0N8 C1F0N9 C1F0Q7 C1F0Q8
C1F0R6 C1F0S5 C1F0Z2 C1F111 C1F121 C1F127 C1F130 C1F1B8 C1F1C7 C1F1E7 C1F1J0
C1F1L8 C1F1M9 C1F1S2 C1F1S6 C1F227 C1F2D2 C1F2F2 C1F2H6 C1F2J1 C1F354 C1F355
C1F3A9 C1F3B3 C1F3B5 C1F3C4 C1F3M3 C1F3P3 C1F3Q7 C1F3S1 C1F3T2 C1F3T7 C1F3Y0
C1F3Y1 C1F3Y2 C1F3Z9 C1F401 C1F407 C1F408 C1F425 C1F455 C1F460 C1F466 C1F467
C1F4B7 C1F4C3 C1F4E8 C1F4F1 C1F4F3 C1F4G5 C1F4K1 C1F4W9 C1F510 C1F513 C1F540
C1F541 C1F542 C1F548 C1F557 C1F613 C1F615 C1F617 C1F618 C1F623 C1F626 C1F627
C1F628 C1F630 C1F634 C1F635 C1F636 C1F638 C1F639 C1F641 C1F643 C1F645 C1F646
C1F647 C1F652 C1F658 C1F683 C1F695 C1F697 C1F6A5 C1F6H5 C1F6W4 C1F6Z5 C1F6Z9
C1F700 C1F735 C1F763 C1F792 C1F7F1 C1F7K8 C1F7R8 C1F7S7 C1F844 C1F857 C1F876
C1F8I0 C1F8M0 C1F8M1 C1F8M5 C1F8R9 C1F8U3 C1F8Z2 C1F902 C1F903 C1F904 C1F905
C1F911 C1F924 C1F933 C1F934 C1F935 C1F9K2 C1F9K5 C1FA53 C1FA54 C1FA63 C1FAD8
C1FAE8 C1FAF1 C1FI13 C1FJ20 C1FJY8 C1FK28 C1FKE5 C1FKE6 C1FKE9 C1FKF9 C1FKG0
C1FKG1 C1FKG2 C1FKG4 C1FKL4 C1FKM3 C1FKM7 C1FKM8 C1FKN0 C1FKN5 C1FKP7 C1FKS1
C1FKZ0 C1FKZ3 C1FL18 C1FL32 C1FLA5 C1FLA6 C1FLA7 C1FLB0 C1FLB3 C1FLB5 C1FLB6
C1FLD8 C1FLD9 C1FLE0 C1FLJ7 C1FLM3 C1FLS8 C1FLV2 C1FLV5 C1FLV6 C1FLW3 C1FLX0
C1FLY1 C1FM10 C1FM32 C1FM55 C1FM59 C1FME6 C1FMF8 C1FMJ6 C1FMJ8 C1FMK5 C1FMP0
C1FMR5 C1FMS1 C1FMS2 C1FMS4 C1FMS5 C1FMS6 C1FMT0 C1FMT2 C1FMT3 C1FMT5 C1FMT6
C1FMT7 C1FMT8 C1FMT9 C1FMU0 C1FMU1 C1FMU2 C1FMU3 C1FMU4 C1FMU5 C1FMU7 C1FMU8
C1FMU9 C1FMV0 C1FMV1 C1FMV2 C1FMV4 C1FMV5 C1FMV6 C1FMV9 C1FMW0 C1FMW2 C1FMW3
C1FMX3 C1FMX6 C1FMZ1 C1FN26 C1FN27 C1FN33 C1FN35 C1FN36 C1FN38 C1FN40 C1FN41
C1FN42 C1FN43 C1FN44 C1FNA5 C1FNC3 C1FNC6 C1FNC9 C1FND0 C1FNE6 C1FNF2 C1FNF4
C1FNF9 C1FNG8 C1FNH5 C1FNL3 C1FNL8 C1FNT6 C1FNT7 C1FNT8 C1FNT9 C1FNU7 C1FNY3
C1FNZ8 C1FP10 C1FP14 C1FP16 C1FP29 C1FP35 C1FP36 C1FPA5 C1FPA6 C1FPB1 C1FPB2
C1FPB3 C1FPC4 C1FPC5 C1FPC6 C1FPF1 C1FPG7 C1FPH3 C1FPH6 C1FPI7 C1FPJ5 C1FPJ6
C1FPJ9 C1FPN9 C1FQ26 C1FQ40 C1FQ79 C1FQ93 C1FQ94 C1FQ97 C1FQA3 C1FQK1 C1FQQ5
C1FQT8 C1FQU7 C1FQW3 C1FQW9 C1FQX0 C1FR28 C1FR34 C1FRL0 C1FS53 C1FS58 C1FS59
C1FS63 C1FS67 C1FS96 C1FS97 C1FSA8 C1FSB0 C1FSC1 C1FSC2 C1FSC4 C1FSH8 C1FSJ4
C1FSK2 C1FSK4 C1FSK5 C1FSK6 C1FSL0 C1FSL4 C1FSL5 C1FSL8 C1FSN0 C1FSR0 C1FSR3
C1FSR8 C1FSS3 C1FST3 C1FST5 C1FSU2 C1FSV2 C1FSW2 C1FSX6 C1FTB7 C1FTB8 C1FTB9
C1FTC4 C1FTE9 C1FTF1 C1FTL9 C1FTW3 C1FTW4 C1FU67 C1FU68 C1FUD5 C1FUD6 C1FUE9
C1FUW6 C1FUX4 C1FUY9 C1FV12 C1FV67 C1FV76 C1FV78 C1FV80 C1FVB2 C1FVB9 C1FVD5
C1FVJ1 C1FVP6 C1FVS9 C1FVT4 C1FVT8 C1FVU0 C1FVU8 C1FVV5 C1FVW3 C1FVW6 C1FVW8
C1FVX5 C1FVY2 C1FVY3 C1FW05 C1FW06 C1FW07 C1FW08 C1FXW2 C1FYF6 C1FYJ9 C1FYU3
C1FZ35 C1G198 C1G1P4 C1G2D1 C1G2L9 C1G3Q0 C1G788 C1G966 C1G9Q3 C1GAZ3 C1GB49
C1GD57 C1GEY4 C1GEZ1 C1GFM3 C1GG27 C1GGT8 C1GHS9 C1GIF2 C1GJU5 C1GLM4 C1GLQ3
C1GP53 C1GPD0 C1GR85 C1GTX5 C1GUB6 C1GUH5 C1GXI3 C1GYL0 C1H0K4 C1H3X3 C1H594

C1H765 C1H7M6 C1H861 C1H8L1 C1H8W8 C1H978 C1H986 C1H9Q9 C1HAB2 C1HAE1 C1HDW2
C1I201 C1I202 C1IWT1 C1JCT1 C1JCT2 C1K5M3 C1KEU0 C1KKR1 C1KV60 C1KV62 C1KV77
C1KV84 C1KV85 C1KV87 C1KV95 C1KV96 C1KV98 C1KVA3 C1KVA5 C1KVA6 C1KVA7 C1KVB2
C1KVB6 C1KVB8 C1KVB9 C1KVC0 C1KVC1 C1KVC2 C1KVC7 C1KVD5 C1KVE1 C1KVE3 C1KVE4
C1KVE8 C1KVE9 C1KVF0 C1KVF9 C1KVG6 C1KVG9 C1KVH1 C1KVH7 C1KVH8 C1KVH9 C1KVI0
C1KVI5 C1KVI7 C1KVI9 C1KVJ2 C1KVJ6 C1KVK0 C1KVK3 C1KVK4 C1KVK5 C1KVK6 C1KVK9
C1KVL8 C1KVM4 C1KVN6 C1KVN8 C1KVN9 C1KVP3 C1KVP5 C1KVQ2 C1KVQ5 C1KVS4 C1KVS5
C1KVS6 C1KVS7 C1KVS8 C1KVV3 C1KVV4 C1KVV7 C1KVV9 C1KVW3 C1KVY0 C1KVY4 C1KVY6
C1KVZ2 C1KW06 C1KW53 C1KW54 C1KW55 C1KW57 C1KW64 C1KW66 C1KW68 C1KW69 C1KW71
C1KW79 C1KW83 C1KW86 C1KW91 C1KW92 C1KW96 C1KWA4 C1KWB5 C1KWC2 C1KWC5 C1KWD1
C1KWD2 C1KWD3 C1KWD4 C1KWD6 C1KWD7 C1KWD9 C1KWF5 C1KWF6 C1KWF8 C1KWF9 C1KWH4
C1KWH6 C1KWH7 C1KWI4 C1KWI8 C1KWI9 C1KWJ1 C1KWK0 C1KWK1 C1KWK2 C1KWK5 C1KWK6
C1KWK7 C1KWM1 C1KWM3 C1KWM5 C1KWM7 C1KWM8 C1KWM9 C1KWN1 C1KWN3 C1KWN7 C1KWN9
C1KWQ1 C1KWQ2 C1KWQ5 C1KWR9 C1KWS1 C1KWS7 C1KWT0 C1KWT1 C1KWT3 C1KWT4 C1KWT9
C1KWW2 C1KWW3 C1KWW9 C1KWY3 C1KWZ4 C1KWZ5 C1KX02 C1KX05 C1KX07 C1KX21 C1KX22
C1KX25 C1KX28 C1KX43 C1KX44 C1KX53 C1KX54 C1KX66 C1KXE5 C1KXG0 C1KXG7 C1KXH5
C1KXH8 C1KXY3 C1KY36 C1KY63 C1KY87 C1KY97 C1KYC1 C1KYC3 C1KYC4 C1KYE1 C1KYE6
C1KYF4 C1KYF5 C1KYG4 C1KYG9 C1KYH2 C1KYI0 C1KYI1 C1KYI2 C1KYI3 C1KYJ3 C1KYN8
C1KYN9 C1KYP1 C1KYP8 C1KYQ0 C1KYV5 C1KYV6 C1KYW0 C1KYW2 C1KYW5 C1KYX7 C1KZ11
C1KZ12 C1KZ13 C1KZ18 C1KZ32 C1KZ34 C1KZ99 C1KZA0 C1KZA1 C1KZF4 C1KZF5 C1KZF6
C1KZF7 C1KZF9 C1KZG1 C1KZG2 C1KZG4 C1KZG5 C1KZG6 C1KZG7 C1KZG8 C1KZG9 C1KZH0
C1KZH1 C1KZH2 C1KZH3 C1KZH5 C1KZH6 C1KZH7 C1KZH8 C1KZH9 C1KZI0 C1KZI1 C1KZJ2
C1KZK6 C1KZK7 C1KZK8 C1KZK9 C1KZP7 C1KZQ7 C1KZQ8 C1KZZ3 C1KZZ5 C1L005 C1L051
C1L056 C1L082 C1L0C8 C1L0D5 C1L0D6 C1L0I0 C1L0I1 C1L0I2 C1L0I8 C1L0J0 C1L0J1
C1L0J2 C1L0J3 C1L0J4 C1L0J5 C1L0J7 C1L0J8 C1L199 C1L1F6 C1L1G4 C1L1G9 C1L1J5
C1L1K0 C1L1M8 C1L1P5 C1L1P6 C1L1Q9 C1L1T2 C1L1T3 C1L1U9 C1L1W7 C1L1W8 C1L1X2
C1L1X9 C1L1Y2 C1L1Y7 C1L207 C1L282 C1L299 C1L2B3 C1L2C6 C1L2D3 C1L2D7 C1L2D9
C1L2E2 C1L2E3 C1L2G7 C1L2G8 C1L2H5 C1L2H6 C1L2I1 C1L2I5 C1L2I6 C1L2I7 C1L2I8
C1L2K1 C1L2L0 C1L2M0 C1L2M3 C1L2M5 C1L2M7 C1L2N1 C1L2N3 C1L2N6 C1L2N7 C1L2Q4
C1L2Q5 C1L2Q6 C1L2R1 C1L2R5 C1L2R6 C1L2R7 C1L2R8 C1L2S1 C1L2S3 C1L2S6 C1L2T9
C1L2U0 C1L2V3 C1L2V4 C1L2V9 C1L2W0 C1L2X6 C1L2Z1 C1L2Z7 C1L2Z8 C1L302 C1MI97
C1MIE8 C1MR42 C1N138 C1N1Y2 C1P607 C1P608 C1P617 C2K1F0 C3JXF9 C3JXP7 C3JXX6
C3JXY0 C3JXY4 C3JXY6 C3JXY7 C3JY28 C3JY43 C3JY45 C3JY53 C3JY54 C3JY59 C3JY60
C3JY63 C3JY68 C3JYD3 C3JYI3 C3JYJ9 C3JYK0 C3JYK1 C3JYR1 C3JYR8 C3JYS7 C3JYS8
C3JYS9 C3JYT1 C3JYT7 C3JYU1 C3JZN1 C3JZN3 C3JZN4 C3JZN5 C3JZN6 C3JZN7 C3JZP4
C3JZR3 C3JZS5 C3K059 C3K071 C3K084 C3K0A7 C3K0A8 C3K0D3 C3K0D5 C3K0D7 C3K0G3
C3K0M9 C3K0N7 C3K0S7 C3K0T6 C3K0U6 C3K0W5 C3K0Y8 C3K0Z0 C3K102 C3K116 C3K148
C3K1A0 C3K1G4 C3K1G8 C3K1G9 C3K1H0 C3K1H1 C3K1H6 C3K1K0 C3K1K5 C3K1K7 C3K1L3
C3K1L4 C3K1L7 C3K1L8 C3K1M1 C3K1M2 C3K1M5 C3K1N5 C3K1P8 C3K1X7 C3K225 C3K238
C3K249 C3K255 C3K256 C3K258 C3K259 C3K261 C3K264 C3K267 C3K274 C3K275 C3K276
C3K282 C3K2F6 C3K2K2 C3K2K3 C3K2K8 C3K2L2 C3K2L8 C3K2L9 C3K2M0 C3K2M6 C3K2M9
C3K2P6 C3K2Q9 C3K2R1 C3K2R5 C3K2R8 C3K2V0 C3K2V2 C3K2V3 C3K2V4 C3K2V5 C3K2V7
C3K2V8 C3K2V9 C3K2W0 C3K2W1 C3K2W2 C3K2W4 C3K2W5 C3K2W6 C3K2W7 C3K2W8 C3K2W9
C3K2X1 C3K2X2 C3K2X3 C3K2X4 C3K2X5 C3K2X6 C3K2X7 C3K2X8 C3K2X9 C3K2Y0 C3K2Y1

C3K2Y2 C3K2Y3 C3K2Y4 C3K2Y5 C3K2Y6 C3K2Y7 C3K2Z1 C3K2Z6 C3K2Z7 C3K307 C3K308
C3K328 C3K329 C3K330 C3K332 C3K334 C3K338 C3K347 C3K348 C3K350 C3K3D1 C3K3D2
C3K3D3 C3K3F1 C3K3F5 C3K3F7 C3K3G6 C3K3I7 C3K3J9 C3K3K0 C3K3K2 C3K3L0 C3K3L6
C3K3L7 C3K3N4 C3K3N6 C3K3P0 C3K3Q8 C3K3R3 C3K3R7 C3K3V3 C3K407 C3K424 C3K426
C3K433 C3K438 C3K452 C3K469 C3K471 C3K473 C3K475 C3K476 C3K479 C3K482 C3K4G7
C3K4H5 C3K4I0 C3K4J4 C3K4J5 C3K4L3 C3K4L4 C3K4L5 C3K4Q4 C3K4W1 C3K568 C3K5A2
C3K5A5 C3K5A9 C3K5E4 C3K5E6 C3K5F2 C3K5F5 C3K5G4 C3K5G5 C3K5G7 C3K5H2 C3K5M3
C3K5U8 C3K5Z6 C3K5Z8 C3K601 C3K630 C3K6E0 C3K6E1 C3K6E2 C3K6G1 C3K6G5 C3K6H4
C3K6H8 C3K6J1 C3K6J5 C3K6J9 C3K6K2 C3K6L0 C3K6N0 C3K6N7 C3K6R3 C3K6T0 C3K6U5
C3K6U9 C3K6V2 C3K6V3 C3K6W5 C3K6Y3 C3K6Y4 C3K6Y5 C3K6Y7 C3K6Z0 C3K6Z3 C3K6Z4
C3K703 C3K704 C3K705 C3K706 C3K710 C3K732 C3K735 C3K758 C3K7C1 C3K7D0 C3K7D6
C3K7J3 C3K7L4 C3K7N8 C3K7T2 C3K803 C3K814 C3K839 C3K860 C3K863 C3K8E3 C3K8U0
C3K8U4 C3K8V3 C3K8W2 C3K8X4 C3K8X5 C3K902 C3K9E6 C3K9I8 C3K9Q0 C3K9S0 C3K9T0
C3KAH1 C3KAH3 C3KAH5 C3KAI3 C3KAJ8 C3KBB4 C3KBH3 C3KBM0 C3KBV5 C3KBY0 C3KBY5
C3KBZ0 C3KBZ2 C3KBZ7 C3KCR9 C3KCS0 C3KCS2 C3KCT1 C3KCT2 C3KDB8 C3KDC2 C3KDC5
C3KDC6 C3KDG0 C3KDG1 C3KDN7 C3KDP1 C3KDS6 C3KDU4 C3KDV4 C3KDW1 C3KDW2 C3KDW3
C3KDW7 C3KDW8 C3KDX1 C3KDX4 C3KDX7 C3KE01 C3KE40 C3KE41 C3KE47 C3KE61 C3KE67
C3KE68 C3KE70 C3KE92 C3KEA5 C3KEC6 C3KED7 C3KHG1 C3KJE6 C3KJF2 C3KSZ0 C3KT80
C3KTA3 C3KTB6 C3KTB7 C3KTC0 C3KTD0 C3KTD1 C3KTD2 C3KTD3 C3KTD5 C3KTI4 C3KTJ3
C3KTJ7 C3KTJ8 C3KTK0 C3KTK5 C3KTL7 C3KTV2 C3KTV5 C3KTY4 C3KU76 C3KU78 C3KU81
C3KU84 C3KU86 C3KU87 C3KU96 C3KU97 C3KU98 C3KUA1 C3KUC5 C3KUC8 C3KUC9 C3KUL4
C3KUP0 C3KUS0 C3KUS7 C3KUT7 C3KUV8 C3KUX9 C3KV06 C3KV10 C3KV11 C3KV13 C3KV20
C3KV90 C3KVA2 C3KVF0 C3KVJ0 C3KVL5 C3KVM1 C3KVM2 C3KVM4 C3KVM5 C3KVM6 C3KVN0
C3KVN2 C3KVN3 C3KVN5 C3KVN6 C3KVN7 C3KVN8 C3KVN9 C3KVP0 C3KVP1 C3KVP2 C3KVP3
C3KVP4 C3KVP5 C3KVP7 C3KVP8 C3KVP9 C3KVQ0 C3KVQ1 C3KVQ2 C3KVQ4 C3KVQ5 C3KVQ6
C3KVQ9 C3KVR0 C3KVR2 C3KVR3 C3KVS3 C3KVS6 C3KVT7 C3KVU0 C3KVU3 C3KVU4 C3KVW3
C3KVW4 C3KVW7 C3KVW9 C3KVX0 C3KVX2 C3KVX4 C3KVX5 C3KVX6 C3KVX7 C3KW44 C3KW96
C3KWA2 C3KWA4 C3KWA9 C3KWB8 C3KWC5 C3KWG3 C3KWG8 C3KWH4 C3KWH8 C3KWI0 C3KWJ3
C3KWJ9 C3KWK0 C3KX32 C3KX33 C3KX34 C3KX35 C3KX86 C3KXB9 C3KXC0 C3KXC5 C3KXC6
C3KXC7 C3KXD8 C3KXD9 C3KXE0 C3KXI0 C3KXJ8 C3KXP4 C3KXQ7 C3KXR0 C3KXS1 C3KXS9
C3KXT0 C3KXT3 C3KXW6 C3KXY4 C3KYF7 C3KYH1 C3KYH2 C3KYH5 C3KYI1 C3KYN9 C3KYP8
C3KYR4 C3KYY3 C3KZ28 C3KZ35 C3KZ52 C3KZ53 C3KZB6 C3KZE8 C3L033 C3L034 C3L035
C3L047 C3L049 C3L061 C3L062 C3L065 C3L0B0 C3L0B5 C3L0B6 C3L0C0 C3L0C4 C3L0C9
C3L0D2 C3L0D3 C3L0D4 C3L0D8 C3L0E2 C3L0E3 C3L0E6 C3L0F7 C3L0J1 C3L0J4 C3L0J9
C3L0K4 C3L0L4 C3L0L6 C3L0M1 C3L0Q0 C3L0R6 C3L117 C3L127 C3L140 C3L147 C3L148
C3L149 C3L179 C3L181 C3L195 C3L1M5 C3L1M6 C3L1U2 C3L1U3 C3L206 C3L207 C3L220
C3L254 C3L261 C3L2K4 C3L2L2 C3L2N7 C3L2U0 C3L2U9 C3L2V1 C3L2V3 C3L2Y4 C3L2Z3
C3L308 C3L357 C3L3A3 C3L3F6 C3L3G1 C3L3G5 C3L3G7 C3L3H5 C3L3I3 C3L3J1 C3L3J4
C3L3J6 C3L3K3 C3L3L0 C3L3L1 C3L3W1 C3L496 C3L503 C3L506 C3L507 C3L508 C3L529
C3L531 C3L532 C3L534 C3L536 C3L543 C3L546 C3L558 C3L559 C3L560 C3L5E9 C3L5F3
C3L5J1 C3L5M2 C3L5Q1 C3L5Q6 C3L5R2 C3L5R5 C3L5R6 C3L5R7 C3L5R9 C3L5S4 C3L5S5
C3L5T6 C3L5T8 C3L5Y0 C3L5Y1 C3L5Y5 C3L5Y6 C3L5Z2 C3L5Z3 C3L608 C3L611 C3L613
C3L626 C3L627 C3L658 C3L6A4 C3L6E8 C3L6F0 C3L6F2 C3L6F3 C3L6K8 C3L6L9 C3L6N3
C3L6P6 C3L6U1 C3L6U6 C3L6U7 C3L6U9 C3L6V0 C3L6V9 C3L6X4 C3L6X6 C3L6Y1 C3L6Y5

C3L6Y6 C3L6Z2 C3L6Z5 C3L6Z7 C3L700 C3L704 C3L705 C3L713 C3L726 C3L730 C3L735
C3L737 C3L738 C3L740 C3L741 C3L742 C3L743 C3L744 C3L755 C3L757 C3L761 C3L763
C3L769 C3L778 C3L783 C3L785 C3L786 C3L787 C3L790 C3L791 C3L795 C3L797 C3L798
C3L799 C3L7A0 C3L7A1 C3L7A3 C3L7A8 C3L7B0 C3L7B4 C3L7B6 C3L7B9 C3L7C0 C3L7E6
C3L7G5 C3L7T5 C3L7X5 C3L7Z8 C3L808 C3L822 C3L823 C3L855 C3L886 C3L887 C3L8M1
C3L8N0 C3L8N1 C3L8N9 C3L8P8 C3L8P9 C3L8Q0 C3L8Q3 C3L8Q6 C3L8Q7 C3L8R9 C3L8S2
C3L8S6 C3L8S8 C3L8T2 C3L8T8 C3L8U0 C3L8V1 C3L8V2 C3L8V8 C3L8X1 C3L8X8 C3L8Z4
C3L913 C3L981 C3L982 C3L983 C3L984 C3L9B6 C3L9C4 C3L9C5 C3L9F4 C3L9F9 C3L9G1
C3L9J6 C3L9N5 C3L9N6 C3L9P5 C3L9P6 C3L9P7 C3L9P8 C3L9P9 C3L9Q0 C3L9Q2 C3L9Q3
C3L9Q4 C3L9Q5 C3L9Q7 C3L9R4 C3L9T6 C3L9V4 C3L9W4 C3L9Z3 C3LA04 C3LA30 C3LA56
C3LAB5 C3LAH8 C3LAK7 C3LAL0 C3LAM8 C3LAQ2 C3LAQ3 C3LAQ4 C3LAU3 C3LAV7 C3LAV8
C3LAV9 C3LAW0 C3LAW1 C3LAX2 C3LAZ5 C3LAZ9 C3LB87 C3LBF0 C3LBP1 C3LBR5 C3LBS0
C3LBU2 C3LBW1 C3LBX0 C3LC26 C3LC57 C3LC72 C3LC73 C3LCA6 C3LCC7 C3LCL5 C3LCW8
C3LCY4 C3LCZ8 C3LD19 C3LD96 C3LDE9 C3LDI3 C3LDK1 C3LDU8 C3LDV2 C3LE50 C3LE56
C3LE93 C3LEC2 C3LEC4 C3LED0 C3LED2 C3LEJ7 C3LEQ2 C3LEW9 C3LEY8 C3LF06 C3LFA6
C3LFI9 C3LFJ0 C3LFK4 C3LFK5 C3LFL2 C3LFM1 C3LFR2 C3LFR7 C3LFS7 C3LFT4 C3LG25
C3LG97 C3LGL2 C3LGR6 C3LGS6 C3LGS9 C3LGT1 C3LGT8 C3LGT9 C3LGU4 C3LGU5 C3LH21
C3LH91 C3LHA0 C3LHA4 C3LHB3 C3LHB6 C3LHM9 C3LHY0 C3LHY1 C3LHY4 C3LHZ0 C3LHZ4
C3LI05 C3LI15 C3LI26 C3LI27 C3LI29 C3LI42 C3LI46 C3LI47 C3LI71 C3LIA3 C3LIA6
C3LIA7 C3LIA8 C3LIC2 C3LIC5 C3LIE5 C3LIG8 C3LIR9 C3LIS2 C3LIS3 C3LIU0 C3LIV3
C3LIX7 C3LIX8 C3LIY7 C3LIY8 C3LIY9 C3LIZ6 C3LIZ7 C3LJ02 C3LJ05 C3LJ13 C3LJ17
C3LJ21 C3LJ39 C3LJ40 C3LJ53 C3LJ59 C3LJ62 C3LJ69 C3LJ70 C3LJ71 C3LJ72 C3LJ74
C3LJ77 C3LJ78 C3LJ79 C3LJ80 C3LJ81 C3LJ82 C3LJ83 C3LJ84 C3LJ85 C3LJ86 C3LJ87
C3LJ89 C3LJ90 C3LJ91 C3LJ92 C3LJ93 C3LJ94 C3LJ95 C3LJ96 C3LJ98 C3LJA0 C3LJA1
C3LJA3 C3LJA6 C3LJH8 C3LJP5 C3LJQ5 C3LJQ7 C3LJQ9 C3LJR8 C3LJT8 C3LJU9 C3LJV1
C3LJV3 C3LJV4 C3LJV5 C3LJV6 C3LJW9 C3LJX0 C3LJX2 C3LJX7 C3LJX8 C3LJZ1 C3LK85
C3LKJ6 C3LKM5 C3LKM7 C3LKN2 C3LKQ2 C3LKQ3 C3LKQ4 C3LKR2 C3LKV7 C3LKW3 C3LKW7
C3LKX5 C3LL81 C3LLI5 C3LLJ0 C3LLK9 C3LLL0 C3LLL2 C3LLL5 C3LLL6 C3LLP9 C3LLQ0
C3LLQ1 C3LLQ4 C3LLQ5 C3LLR0 C3LLR5 C3LLR8 C3LLS2 C3LLS4 C3LLS7 C3LLT8 C3LLV3
C3LLY1 C3LM61 C3LM83 C3LMC8 C3LMI0 C3LMI5 C3LMJ0 C3LMN9 C3LMQ4 C3LMU1 C3LMW6
C3LN37 C3LN39 C3LN43 C3LN44 C3LN49 C3LN54 C3LN57 C3LN58 C3LNA0 C3LNA1 C3LND6
C3LNE8 C3LNE9 C3LNF0 C3LNF1 C3LNF5 C3LNI0 C3LNJ6 C3LNL5 C3LNL8 C3LNM5 C3LNM6
C3LNM7 C3LNR2 C3LNS3 C3LNS6 C3LNU9 C3LNW7 C3LNX6 C3LNZ0 C3LNZ1 C3LP35 C3LP59
C3LP65 C3LP68 C3LP79 C3LP80 C3LP87 C3LP93 C3LP97 C3LPA1 C3LPB8 C3LPC2 C3LPC8
C3LPC9 C3LPF7 C3LPG2 C3LPI4 C3LPI5 C3LPI7 C3LPI9 C3LPM9 C3LPN0 C3LPN8 C3LPP0
C3LPP2 C3LPP6 C3LPQ3 C3LPQ7 C3LPR5 C3LPS5 C3LPU2 C3LPV3 C3LPV6 C3LPW5 C3LPX0
C3LPZ2 C3LPZ5 C3LPZ8 C3LQ01 C3LQ18 C3LQ26 C3LQ29 C3LQ31 C3LQ32 C3LQ39 C3LQ47
C3LQ48 C3LQ53 C3LQ59 C3LQC0 C3LQC2 C3LQC7 C3LQD1 C3LQD2 C3LQD3 C3LQD5 C3LQF1
C3LQH9 C3LQI0 C3LQI1 C3LQI3 C3LQI4 C3LQK1 C3LQN2 C3LQP6 C3LQP9 C3LQQ3 C3LQR0
C3LQR1 C3LQT6 C3LQV4 C3LQX1 C3LQX2 C3LQY0 C3LQY9 C3LQZ1 C3LQZ5 C3LR02 C3LR04
C3LR15 C3LR19 C3LR22 C3LR32 C3LR34 C3LR35 C3LR42 C3LR47 C3LR49 C3LR52 C3LR55
C3LR56 C3LR57 C3LR58 C3LR59 C3LR63 C3LR68 C3LR77 C3LR78 C3LR79 C3LR84 C3LR91
C3LR92 C3LR98 C3LRA0 C3LRA1 C3LRA6 C3LRB7 C3LRB8 C3LRF4 C3LRG6 C3LRG7 C3LRH2
C3LRH3 C3LRJ5 C3LRK7 C3LRL2 C3LRM1 C3LRM2 C3LRN2 C3LRN3 C3LRN4 C3LRN5 C3LRN9

C3LRP0 C3LRP1 C3LRP2 C3LRP3 C3LRP4 C3LRP6 C3LRP7 C3LRP8 C3LRP9 C3LRQ0 C3LRQ1
C3LRQ2 C3LRQ3 C3LRQ4 C3LRQ5 C3LRQ6 C3LRQ7 C3LRQ8 C3LRQ9 C3LRR4 C3LRS3 C3LRS7
C3LRV2 C3LRV3 C3LRV4 C3LRV6 C3LRW4 C3LRW8 C3LRX0 C3LRX5 C3LRY3 C3LRY8 C3LRY9
C3LRZ9 C3LS10 C3LS11 C3LS19 C3LS20 C3LS21 C3LS22 C3LS25 C3LS34 C3LS38 C3LS51
C3LS52 C3LS53 C3LS54 C3LS60 C3LS61 C3LS70 C3LS81 C3LS82 C3LS89 C3LSA3 C3LSA4
C3LSA8 C3LSB1 C3LSB5 C3LSD6 C3LSE7 C3LSG5 C3LSG8 C3LSH8 C3LSJ9 C3LSK0 C3LSL0
C3LSN1 C3LSN2 C3LSP1 C3LSP4 C3LSP6 C3LSP8 C3LSQ0 C3LSQ1 C3LSQ2 C3LSR4 C3LSR6
C3LSS3 C3LST0 C3LST4 C3LST7 C3LSU7 C3LSV6 C3LSX1 C3LSY3 C3LSZ2 C3LSZ4 C3LT01
C3LT04 C3LT05 C3LT09 C3LT10 C3LT13 C3LT16 C3LT19 C3LT21 C3LT39 C3LT50 C3LT53
C3LT54 C3LT98 C3LTA3 C3LTA4 C3LTA5 C3LTA6 C3LTC2 C3LTC5 C3LTC8 C3LTD1 C3LTD9
C3LTE1 C3LTJ1 C3LTJ3 C3LTJ9 C3LTK8 C3LTL9 C3LTN3 C3LTN4 C3LTP5 C3LTP7 C3LTS1
C3LTS3 C3LTV0 C3LTV3 C3LTV5 C3LU07 C3LU31 C3LU38 C3LUB5 C3LUC2 C3LUI5 C3LUK2
C3LUL9 C3LUM7 C3LUU7 C3LUU8 C3LUV8 C3LUW0 C3LUW1 C3LVI5 C3LVI8 C3LVL9 C3LVM1
C3LVN8 C3LVX6 C3LW05 C3LW07 C3LW10 C3LW40 C3LW69 C3LW87 C3LW91 C3LWD9 C3LWF0
C3LWF8 C3LWJ3 C3LWL5 C3LWP9 C3LWT7 C3LWV8 C3LWX2 C3LWZ1 C3LX01 C3M8G8 C3M8L8
C3M8R6 C3M8S4 C3M8U4 C3M8W9 C3M8Y0 C3M930 C3M9A8 C3M9B0 C3M9B1 C3M9C2 C3M9C8
C3M9D7 C3M9F7 C3M9G8 C3M9I2 C3M9I9 C3M9J7 C3M9M8 C3M9S7 C3M9U0 C3M9V0 C3M9W6
C3M9X0 C3M9X5 C3M9X6 C3M9X9 C3M9Y0 C3M9Y4 C3M9Z5 C3M9Z9 C3MA29 C3MA32 C3MA33
C3MA45 C3MA88 C3MA97 C3MAA2 C3MAA3 C3MAA5 C3MAA7 C3MAB3 C3MAF7 C3MAG2 C3MAG3
C3MAI8 C3MAJ1 C3MAL6 C3MAR5 C3MAS0 C3MAW0 C3MAW7 C3MAW8 C3MAX1 C3MAX2 C3MAX5
C3MAX6 C3MAX7 C3MAX9 C3MAY1 C3MAY2 C3MAY3 C3MAY4 C3MAY6 C3MAY7 C3MAY8 C3MAY9
C3MAZ0 C3MAZ1 C3MAZ2 C3MAZ4 C3MAZ5 C3MAZ6 C3MAZ7 C3MB01 C3MB02 C3MB03 C3MB04
C3MB23 C3MB44 C3MB45 C3MB49 C3MB60 C3MB65 C3MB70 C3MB71 C3MB73 C3MB76 C3MB98
C3MB99 C3MBA0 C3MBB9 C3MBC0 C3MBC1 C3MBC2 C3MBC5 C3MBC6 C3MBC8 C3MBD2 C3MBH0
C3MBH2 C3MBH4 C3MBH5 C3MBM0 C3MBQ2 C3MBQ3 C3MBQ5 C3MBU2 C3MBU3 C3MBV9 C3MBW4
C3MBY3 C3MBY4 C3MBY7 C3MBY8 C3MC05 C3MC58 C3MC63 C3MC67 C3MC70 C3MC71 C3MC80
C3MCF1 C3MCF2 C3MCF3 C3MCH2 C3MCN8 C3MCP8 C3MCP9 C3MCQ0 C3MCQ9 C3MCS5 C3MCV0
C3MCY6 C3MCY7 C3MCY9 C3MCZ7 C3MD28 C3MD65 C3MD72 C3MD80 C3MD81 C3MD82 C3MD87
C3MD93 C3MDE7 C3MDF9 C3MDH9 C3MDR9 C3ME70 C3MEE8 C3MEJ1 C3MEJ2 C3MEL0 C3MEL6
C3MEL7 C3MEL9 C3MEM6 C3MEM7 C3MEM8 C3MEN7 C3MF10 C3MF25 C3MF26 C3MF35 C3MF41
C3MF47 C3MF51 C3MF80 C3MF81 C3MFB9 C3MFH1 C3MFM4 C3MFQ1 C3MFQ8 C3MFW1 C3MFX0
C3MFX7 C3MFZ6 C3MG06 C3MG23 C3MG45 C3MG57 C3MG60 C3MG71 C3MGH8 C3MGI0 C3MGI5
C3MGQ4 C3MGT3 C3MGT4 C3MGW0 C3MGX4 C3MGX6 C3MH72 C3MHC0 C3MHD5 C3MHI2 C3MHI8
C3MHK2 C3MHK7 C3MHQ9 C3MHU1 C3MHW1 C3MI00 C3MI14 C3MIE1 C3MIE4 C3MIE5 C3MIH6
C3MIH7 C3MIH9 C3MIJ2 C3MIK0 C3MIL5 C3MIM2 C3MIR3 C3MIS7 C3MJ04 C3MJ10 C3MJ14
C3MJ21 C3MJ22 C3MJ24 C3MJ28 C3MJ30 C3MJ34 C3MJ42 C3MJ44 C3MJ49 C3MJ86 C3MJ88
C3MJA1 C3MJA6 C3MJB1 C3MJB8 C3MJC4 C3MJC8 C3MJE5 C3MJI5 C3MJI6 C3MJN1 C3MJN7
C3MJP0 C3MJP1 C3MJP5 C3MJP7 C3MJQ1 C3MJQ3 C3MJQ6 C3MJW3 C3MJX9 C3MK52 C3MK60
C3MK63 C3MK79 C3MKC0 C3MKC1 C3MKC9 C3MKM4 C3MKR6 C3MKT5 C3MKT6 C3MMC3 C3MMK8
C3MN93 C3MP45 C3MP49 C3MPC7 C3MPG2 C3MPG3 C3MPG7 C3MPH2 C3MPH3 C3MPH4 C3MPH6
C3MPM7 C3MPM9 C3MPN0 C3MPN5 C3MPN7 C3MPN8 C3MPQ0 C3MPQ5 C3MPR5 C3MPR6 C3MPR8
C3MPS1 C3MPS2 C3MPT5 C3MPT6 C3MPU9 C3MPW2 C3MPZ8 C3MPZ9 C3MQ13 C3MQ16 C3MQ25
C3MQ26 C3MQ30 C3MQ36 C3MQ39 C3MQ42 C3MQ43 C3MQ47 C3MQ51 C3MQ53 C3MQ59 C3MQ60
C3MQ62 C3MQ71 C3MQ74 C3MQ76 C3MQ78 C3MQ80 C3MQ83 C3MQ85 C3MQ86 C3MQ89 C3MQ90

C3MQC7 C3MQE3 C3MQF3 C3MQG7 C3MQG8 C3MQH9 C3MQI3 C3MQI6 C3MQJ5 C3MQK4 C3MQK7
C3MQK8 C3MQL0 C3MQM6 C3MQN0 C3MQN2 C3MQN6 C3MQN7 C3MQV5 C3MQX1 C3MQX7 C3MQY1
C3MQY2 C3MQY4 C3MQZ8 C3MQZ9 C3MR04 C3MR05 C3MR07 C3MR14 C3MR15 C3MR21 C3MR62
C3MR64 C3MR75 C3MR76 C3MR77 C3MR78 C3MR79 C3MR80 C3MR83 C3MR85 C3MR87 C3MR91
C3MRB5 C3MRB7 C3MRB8 C3MRC8 C3MRD1 C3MRD7 C3MRE4 C3MRE5 C3MRF1 C3MRF5 C3MRH3
C3MRH5 C3MRI1 C3MRJ6 C3MRJ8 C3MRJ9 C3MRK1 C3MRK4 C3MRL0 C3MRR3 C3MRT0 C3MS47
C3MT45 C3MTF2 C3MTL3 C3MTN3 C3MTN8 C3MTP3 C3MTQ0 C3MTQ6 C3MTR0 C3MTS7 C3MTW8
C3MTW9 C3MU21 C3MU29 C3MU32 C3MU48 C3MU90 C3MU91 C3MU99 C3MUB4 C3MUC6 C3MUC7
C3MUU8 C3MUU9 C3MUV8 C3MUW1 C3MUZ1 C3MV89 C3MVB7 C3MVB8 C3MVD2 C3MVD5 C3MVE4
C3MVE5 C3MVE8 C3MVF4 C3MVF7 C3MVG0 C3MVG1 C3MVG5 C3MVG9 C3MVH1 C3MVH6 C3MVH8
C3MVH9 C3MVI1 C3MVJ0 C3MVJ3 C3MVJ5 C3MVJ7 C3MVJ9 C3MVK2 C3MVX7 C3MVX8 C3MVY1
C3MVY2 C3MW06 C3MW21 C3MW31 C3MW45 C3MW46 C3MW57 C3MW61 C3MW64 C3MW73 C3MW82
C3MW85 C3MW86 C3MW88 C3MW97 C3MWA4 C3MWN0 C3MWN2 C3MWN6 C3MWN7 C3MWU3 C3MWV9
C3MWW5 C3MWW9 C3MWX0 C3MWX2 C3MWY6 C3MWY7 C3MWZ2 C3MWZ3 C3MWZ5 C3MX02 C3MX03
C3MX60 C3MXC5 C3MXF1 C3MXF3 C3MXG4 C3MXG5 C3MXG6 C3MXG7 C3MXG8 C3MXG9 C3MXH2
C3MXH4 C3MXH6 C3MXI0 C3MXK4 C3MXK6 C3MXK7 C3MXL7 C3MXM0 C3MXY8 C3MY33 C3MY40
C3MY41 C3MY47 C3MY51 C3MY71 C3MY77 C3MY92 C3MY94 C3MY95 C3MY97 C3MYA0 C3MYA6
C3MYC2 C3MYC8 C3MYD2 C3MYD9 C3MYE0 C3MYE2 C3MYE7 C3MYF0 C3MYF1 C3MYF5 C3MYG0
C3MYG1 C3MYG2 C3MYG4 C3MYM1 C3MYM3 C3MYM4 C3MYM9 C3MYN1 C3MYN2 C3MYP4 C3MYP9
C3MYQ9 C3MYR0 C3MYR2 C3MYR5 C3MYR6 C3MYS2 C3MYS6 C3MYT4 C3MYT6 C3MYU1 C3MYX8
C3MYY0 C3MYY9 C3MYZ5 C3MYZ8 C3MYZ9 C3MZ03 C3MZ05 C3MZ09 C3MZ11 C3MZ14 C3MZ46
C3MZ47 C3MZ52 C3MZ53 C3MZ55 C3MZ62 C3MZ63 C3MZ69 C3MZ97 C3MZ99 C3MZB0 C3MZB1
C3MZB2 C3MZB3 C3MZB4 C3MZB5 C3MZB8 C3MZC0 C3MZC2 C3MZC6 C3MZF0 C3MZF2 C3MZF3
C3MZG3 C3MZG6 C3MZH2 C3MZH9 C3MZI0 C3MZI6 C3MZJ0 C3MZK0 C3MZK6 C3MZM1 C3MZM3
C3MZM4 C3MZM6 C3MZM9 C3MZN5 C3MZQ1 C3MZQ7 C3MZR1 C3MZR8 C3MZR9 C3MZS1 C3MZS5
C3MZS7 C3MZT1 C3MZT9 C3MZU1 C3MZU9 C3MZW9 C3MZX4 C3MZX9 C3MZY6 C3MZY8 C3N026
C3N028 C3N038 C3N044 C3N047 C3N048 C3N052 C3N054 C3N058 C3N060 C3N063 C3N0B6
C3N0C0 C3N0D7 C3N0H8 C3N0H9 C3N0N3 C3N0P1 C3N0Q3 C3N0R9 C3N120 C3N136 C3N178
C3N179 C3N187 C3N1A2 C3N1B4 C3N1B5 C3N1G8 C3N1X1 C3N2L8 C3N2M3 C3N323 C3N3Q3
C3N4K5 C3N509 C3N544 C3N545 C3N549 C3N554 C3N555 C3N556 C3N558 C3N5A3 C3N5A5
C3N5A6 C3N5B1 C3N5B3 C3N5B4 C3N5C6 C3N5D1 C3N5E1 C3N5E2 C3N5E4 C3N5E7 C3N5E8
C3N5G1 C3N5G2 C3N5H2 C3N5H5 C3N5I8 C3N5L6 C3N5L7 C3N5N1 C3N5N4 C3N5P3 C3N5P4
C3N5P7 C3N5Q3 C3N5Q6 C3N5Q9 C3N5R0 C3N5R4 C3N5R8 C3N5S0 C3N5S5 C3N5S7 C3N5S8
C3N5T0 C3N5T9 C3N5U2 C3N5U4 C3N5U6 C3N5U8 C3N5V1 C3N5V3 C3N5V4 C3N5V7 C3N5V8
C3N649 C3N664 C3N674 C3N688 C3N689 C3N6A0 C3N6A4 C3N6A7 C3N6B5 C3N6C4 C3N6C7
C3N6C8 C3N6D0 C3N6E6 C3N6F0 C3N6F2 C3N6F6 C3N6F7 C3N6L0 C3N6M6 C3N6N2 C3N6N6
C3N6N7 C3N6N9 C3N6Z8 C3N740 C3N746 C3N749 C3N750 C3N752 C3N766 C3N767 C3N772
C3N773 C3N775 C3N782 C3N783 C3N789 C3N792 C3N7B7 C3N7B9 C3N7D0 C3N7D1 C3N7D2
C3N7D3 C3N7D4 C3N7D5 C3N7D9 C3N7E1 C3N7E3 C3N7E8 C3N7H2 C3N7H4 C3N7H5 C3N7I5
C3N7I8 C3N7J4 C3N7K1 C3N7K2 C3N7K8 C3N7L2 C3N7M2 C3N7M8 C3N7P3 C3N7P5 C3N7P6
C3N7P8 C3N7Q1 C3N7Q8 C3N7V1 C3N7V6 C3N7W1 C3N7W8 C3N7X4 C3N7X8 C3N824 C3N830
C3N834 C3N841 C3N842 C3N844 C3N849 C3N851 C3N855 C3N864 C3N866 C3N871 C3N8A9
C3N8B1 C3N8C3 C3N8G6 C3N8G7 C3N8M1 C3N8M9 C3N8N2 C3N8Q2 C3N8Q8 C3N8R1 C3N8R2
C3N8R6 C3N8R8 C3N8S2 C3N8S4 C3N8S7 C3N8Y9 C3N905 C3N926 C3N968 C3N969 C3N977

C3N992 C3N9B3 C3N9B4 C3N9R7 C3NAL6 C3NAM2 C3NBY5 C3NCS3 C3NDK6 C3NDP2 C3NDP3
C3NDP7 C3NDQ2 C3NDQ3 C3NDQ4 C3NDQ6 C3NDV7 C3NDV9 C3NDW0 C3NDW5 C3NDW7 C3NDW8
C3NDY0 C3NDY5 C3NDZ5 C3NDZ6 C3NDZ8 C3NE01 C3NE02 C3NE25 C3NE26 C3NE39 C3NE52
C3NE80 C3NE81 C3NE95 C3NE98 C3NEA7 C3NEA8 C3NEB3 C3NEB9 C3NEC2 C3NEC5 C3NEC6
C3NED0 C3NED4 C3NED6 C3NEE3 C3NEE4 C3NEE6 C3NEF5 C3NEF8 C3NEG0 C3NEG2 C3NEG4
C3NEG7 C3NEG9 C3NEH0 C3NEH3 C3NEH4 C3NEK4 C3NEM0 C3NEN0 C3NEP5 C3NEP6 C3NEQ7
C3NER1 C3NER4 C3NES3 C3NET2 C3NET5 C3NET6 C3NET8 C3NEV4 C3NEV8 C3NEW0 C3NEW4
C3NEW5 C3NF41 C3NF46 C3NF48 C3NF56 C3NF60 C3NF62 C3NF67 C3NF69 C3NF70 C3NF77
C3NF81 C3NF87 C3NFE3 C3NFR5 C3NFS2 C3NFS5 C3NFS7 C3NFS8 C3NFT0 C3NFU5 C3NFV1
C3NFW2 C3NFW6 C3NFX2 C3NFX3 C3NFY0 C3NFY8 C3NFZ1 C3NG01 C3NG02 C3NG04 C3NG28
C3NG91 C3NG93 C3NG95 C3NG99 C3NGA0 C3NGA1 C3NGA2 C3NGA3 C3NGA4 C3NGB5 C3NGB7
C3NGE6 C3NGF2 C3NGF3 C3NGG0 C3NGG2 C3NGG8 C3NGG9 C3NGI3 C3NGI5 C3NGI6 C3NGJ0
C3NGJ6 C3NGL3 C3NGS9 C3NGT0 C3NGT4 C3NGT6 C3NGU0 C3NGV6 C3NGV8 C3NGV9 C3NGW2
C3NGX1 C3NGY0 C3NGY3 C3NGY7 C3NGZ8 C3NGZ9 C3NH14 C3NH23 C3NH39 C3NH77 C3NH78
C3NH82 C3NH83 C3NH85 C3NH88 C3NH89 C3NH90 C3NH92 C3NH94 C3NH97 C3NHA6 C3NHA8
C3NHA9 C3NHB6 C3NHB8 C3NHC2 C3NHC6 C3NHC7 C3NHD0 C3NHD3 C3NHD9 C3NHE1 C3NHE2
C3NHF1 C3NHF4 C3NHG8 C3NHG9 C3NHL1 C3NHM4 C3NHN7 C3NHN8 C3NHQ1 C3NHQ2 C3NHQ5
C3NHQ7 C3NHQ8 C3NHR8 C3NHS3 C3NHT5 C3NHT6 C3NHT8 C3NHU3 C3NHU4 C3NHU6 C3NHZ5
C3NHZ7 C3NHZ8 C3NHZ9 C3NI04 C3NI08 C3NI09 C3NJ57 C3NJ62 C3NJ67 C3NJ74 C3NJ80
C3NJ84 C3NJA0 C3NJQ5 C3NJQ6 C3NL04 C3NLR8 C3NM92 C3NMA0 C3NMA3 C3NMB9 C3NMG4
C3NMI0 C3NMP1 C3NMP4 C3NMP6 C3NMQ0 C3NMQ2 C3NMQ6 C3NMQ7 C3NMR0 C3NMR6 C3NMS5
C3NMS7 C3NMT7 C3NMT8 C3NMU5 C3NMW1 C3NMY1 C3NMY2 C3NQJ0 C3NZC1 C3NZD1 C3NZR4
C3NZS9 C3NZU4 C3P002 C3P025 C3P072 C3P0A6 C3P0C0 C3P0C2 C3P0C8 C3P0D0 C3P0E9
C3P0L4 C3P0R7 C3P0S0 C3P101 C3P178 C3P1G4 C3P1G5 C3P1J8 C3P1N3 C3P1N9 C3P1X2
C3P210 C3P292 C3P293 C3P2A0 C3P2A9 C3P2E4 C3P2E9 C3P2F8 C3P2G5 C3P2P9 C3P2S0
C3P2T3 C3P2U9 C3P341 C3P3E0 C3P3E3 C3P3E5 C3P3F2 C3P3F3 C3P3F8 C3P3F9 C3P3K3
C3P3L1 C3P3M8 C3P3Q0 C3P3Q5 C3P3S8 C3P3T7 C3P3T8 C3P3T9 C3P3U0 C3P3U1 C3P3V9
C3P449 C3P450 C3P458 C3P487 C3P4C3 C3P4C4 C3P4C5 C3P4E3 C3P4F6 C3P4F9 C3P4I7
C3P4M0 C3P4M1 C3P4M2 C3P4M3 C3P4Y3 C3P4Y4 C3P4Y9 C3P4Z6 C3P4Z8 C3P4Z9 C3P500
C3P501 C3P503 C3P504 C3P505 C3P506 C3P507 C3P516 C3P517 C3P552 C3P584 C3P586
C3P591 C3P598 C3P5A0 C3P5A4 C3P5A7 C3P5E3 C3P5H4 C3P5H5 C3P5I7 C3P5L0 C3P5L3
C3P5L5 C3P5L9 C3P5M1 C3P5M6 C3P5M8 C3P5M9 C3P5N0 C3P5N1 C3P5N2 C3P5N4 C3P5N7
C3P5N8 C3P5P1 C3P5P2 C3P5P3 C3P5P5 C3P5Q0 C3P5Q2 C3P5Q3 C3P5Q6 C3P5Q9 C3P5R0
C3P5R1 C3P5S0 C3P5S8 C3P5S9 C3P5T8 C3P629 C3P635 C3P637 C3P641 C3P643 C3P652
C3P653 C3P654 C3P655 C3P656 C3P658 C3P659 C3P661 C3P665 C3P669 C3P682 C3P688
C3P689 C3P691 C3P693 C3P6D6 C3P6N8 C3P6S2 C3P6T3 C3P6T7 C3P6U6 C3P6V6 C3P6W7
C3P6W8 C3P6X0 C3P6Y3 C3P6Y7 C3P6Y8 C3P710 C3P742 C3P745 C3P746 C3P747 C3P761
C3P764 C3P768 C3P778 C3P786 C3P7F5 C3P7H9 C3P7J3 C3P7L0 C3P7M3 C3P7P7 C3P7P8
C3P7Q4 C3P7R4 C3P7R6 C3P7R8 C3P7S4 C3P7U3 C3P7V4 C3P7V6 C3P7V8 C3P7V9 C3P7W0
C3P7W1 C3P7X5 C3P7X7 C3P7Z9 C3P803 C3P858 C3P8B4 C3P8D3 C3P8D4 C3P8D5 C3P8E3
C3P8I5 C3P8J1 C3P8J4 C3P8K2 C3P8K4 C3P8K9 C3P8L5 C3P8L8 C3P8L9 C3P8M0 C3P8M2
C3P8M7 C3P8M8 C3P8N9 C3P8P5 C3P8P8 C3P8Q3 C3P8Q4 C3P8Z9 C3P914 C3P964 C3P965
C3P968 C3P969 C3P975 C3P976 C3P990 C3P993 C3P995 C3P9A4 C3P9A5 C3P9A7 C3P9A8
C3P9B5 C3P9C8 C3P9D0 C3P9D5 C3P9D9 C3P9E0 C3P9E6 C3P9E9 C3P9F1 C3P9F4 C3P9F7

C3P9F8 C3P9G2 C3P9G9 C3P9H0 C3P9H5 C3P9H8 C3P9I6 C3P9J0 C3P9J4 C3P9L2 C3P9L3
C3P9M6 C3P9N2 C3P9N5 C3P9P2 C3P9P3 C3P9P4 C3P9P5 C3P9P7 C3P9Q0 C3P9Q1 C3P9Q2
C3P9Q3 C3P9Q4 C3P9Q5 C3P9Q6 C3P9Q7 C3P9Q8 C3P9Q9 C3P9R0 C3P9R2 C3P9R3 C3P9R4
C3P9R5 C3P9R6 C3P9R7 C3P9R8 C3P9R9 C3P9S1 C3P9S3 C3P9S4 C3P9S6 C3P9S9 C3P9T0
C3P9T1 C3P9T3 C3PA65 C3PA74 C3PAA9 C3PAD1 C3PAE0 C3PAE4 C3PAE9 C3PAF1 C3PAG2
C3PAG3 C3PAG9 C3PAI1 C3PAI8 C3PAK3 C3PAK4 C3PAL5 C3PAU2 C3PAU8 C3PAV0 C3PAV1
C3PAW8 C3PAZ0 C3PAZ3 C3PB71 C3PB77 C3PB96 C3PB97 C3PBB3 C3PBC1 C3PBF0 C3PBG1
C3PBI3 C3PBK0 C3PBL1 C3PBM7 C3PBM9 C3PBN0 C3PBN2 C3PBN4 C3PBP1 C3PBP4 C3PBQ3
C3PBQ4 C3PBQ5 C3PBW0 C3PBW1 C3PC50 C3PCC0 C3PCC4 C3PCC6 C3PCK3 C3PCL8 C3PCM9
C3PCX7 C3PCY0 C3PCY9 C3PCZ3 C3PD52 C3PDF7 C3PDH0 C3PDH1 C3PDK2 C3PDM2 C3PDZ9
C3PE72 C3PE74 C3PEA3 C3PEE7 C3PES4 C3PEY5 C3PF03 C3PF14 C3PF15 C3PF16 C3PF17
C3PFC1 C3PFD4 C3PFJ1 C3PFN8 C3PFP1 C3PFQ4 C3PFR8 C3PFT7 C3PFU5 C3PFU9 C3PFW7
C3PFX1 C3PFX5 C3PFZ5 C3PFZ9 C3PG15 C3PG24 C3PG27 C3PG33 C3PG35 C3PG36 C3PG40
C3PG89 C3PG92 C3PG93 C3PGA0 C3PGA3 C3PGI6 C3PGJ1 C3PGJ7 C3PGJ8 C3PGM1 C3PGM2
C3PGM7 C3PGN1 C3PGN5 C3PGQ6 C3PGQ7 C3PGR0 C3PGS1 C3PGV9 C3PGW4 C3PGX6 C3PGY9
C3PH04 C3PH05 C3PH06 C3PH09 C3PH10 C3PH13 C3PH18 C3PH19 C3PH22 C3PH25 C3PH30
C3PH48 C3PH50 C3PH66 C3PH72 C3PH76 C3PH91 C3PHA6 C3PHF5 C3PHG4 C3PHK3 C3PHK4
C3PHK5 C3PHK6 C3PHU5 C3PHU9 C3PHY3 C3PI04 C3PI05 C3PI20 C3PI25 C3PI36 C3PIP1
C3PIQ3 C3PIU4 C3PIW7 C3PJE8 C3PJJ2 C3PJK0 C3PJM6 C3PJW6 C3PK12 C3PKC8 C3PKD0
C3PKD8 C3PKD9 C3PKG1 C3PKG2 C3PKI4 C3PKL1 C3PKL7 C3PKL8 C3PKM7 C3PKM8 C3PKN2
C3PKN9 C3PKP0 C3PKP1 C3PKP2 C3PKQ1 C3PKQ2 C3PKQ3 C3PKQ4 C3PKQ5 C3PKQ6 C3PKQ7
C3PKQ8 C3PKQ9 C3PKR0 C3PKR1 C3PKU4 C3PKY4 C3PKY7 C3PL13 C3PL14 C3PL17 C3PL18
C3PL19 C3PL24 C3PL26 C3PL27 C3PL28 C3PL31 C3PL35 C3PL36 C3PL37 C3PL39 C3PL50
C3PL61 C3PLA3 C3PLD0 C3PLF4 C3PLF5 C3PLG0 C3PLG3 C3PLG9 C3PLH1 C3PLJ2 C3PLJ4
C3PLL5 C3PLL8 C3PLL9 C3PLM8 C3PLM9 C3PLP0 C3PLQ4 C3PLR3 C3PLU6 C3PLU8 C3PLX4
C3PLX7 C3PLY4 C3PLY5 C3PLZ3 C3PM02 C3PM12 C3PM18 C3PM20 C3PM21 C3PM22 C3PM25
C3PM28 C3PM29 C3PM56 C3PM61 C3PM69 C3PM73 C3PM74 C3PM75 C3PM93 C3PM94 C3PM95
C3PMA0 C3PMA3 C3PMA4 C3PMB0 C3PMB1 C3PMB3 C3PMC1 C3PMC2 C3PMC3 C3PMD1 C3PMD7
C3PMD9 C3PME7 C3PME8 C3PME9 C3PMF3 C3PMF5 C3PMG3 C3PMG8 C3PMG9 C3PMH0 C3PMH3
C3PMH4 C3PMH5 C3PMH6 C3PMH7 C3PMI0 C3PMI3 C3PMI6 C3PMI9 C3PMJ0 C3PMJ1 C3PMJ3
C3PMK2 C3PMK3 C3PMM6 C3PMM7 C3PMP2 C3PMP3 C3PMP4 C3PMP5 C3PMQ0 C3PMQ3 C3PMR4
C3PMR6 C3PMS9 C3PMT0 C3PMT3 C3PMT5 C3PMU4 C3PMU6 C3PMX2 C3PN02 C3PN11 C3PN13
C3PN29 C3PN30 C3PN35 C3PN45 C3PN57 C3PN64 C3PN66 C3PN86 C3PN92 C3PNA0 C3PNA1
C3PNA4 C3PNA6 C3PNB9 C3PNE1 C3PNE3 C3PNE7 C3PNF5 C3PNF6 C3PNF9 C3PNG6 C3PNG9
C3PNH0 C3PNI9 C3PNK2 C3PNK3 C3PNK4 C3PNN0 C3PNN8 C3PNP6 C3PNP9 C3PNQ8 C3PNS9
C3PNU6 C3PNU7 C3PNW8 C3PNX3 C3PNX7 C3PNZ9 C3PP00 C3PP03 C3PP13 C3PP14 C3PP41
C3PP46 C3PP47 C3PP50 C3PP51 C3PP52 C3PP58 C3PP60 C3PP72 C3PP75 C3PP76 C3PP81
C3PP82 C3PP83 C3PP84 C3PP85 C3PP86 C3PP88 C3PP89 C3PP90 C3PP91 C3PP92 C3PP93
C3PP95 C3PP96 C3PP97 C3PP98 C3PP99 C3PPA0 C3PPA1 C3PPA2 C3PPA3 C3PPA4 C3PPA5
C3PPA6 C3PPA7 C3PPA8 C3PPA9 C3PPC3 C3PPC7 C3PPD1 C3RSF5 C3SBS8 C3SBU4 C3SBU5
C3SBW0 C3UVB0 C3VEQ0 C3VEQ3 C3VEQ4 C3W7B0 C3W947 C3XR70 C3XRY1 C3XVM1 C3XVT5
C3Y3G4 C3YDS7 C3YEM5 C3YFB4 C3YN79 C3YZ51 C3Z1P5 C3Z724 C3ZAH2 C3ZBT0 C3ZDX5
C4B644 C4IZI7 C4JDR3 C4JE77 C4JF09 C4JFE4 C4JHR8 C4JJE8 C4JKS2 C4JMJ4 C4JMY4
C4JPW9 C4JSX6 C4JTA0 C4JTH6 C4JUN6 C4JWQ7 C4JXN2 C4JY72 C4JYB4 C4JYG6 C4JYP8

C4JZS6 C4K0B5 C4K0C5 C4K0C9 C4K0F2 C4K0F3 C4K0F4 C4K0H2 C4K0J2 C4K0J3 C4K0J6
C4K0K1 C4K0L7 C4K0N9 C4K0P8 C4K0Q1 C4K0Q4 C4K0Q5 C4K0R2 C4K0R3 C4K0R4 C4K0R5
C4K0R8 C4K0T1 C4K0T2 C4K0V3 C4K0V5 C4K0V7 C4K0V8 C4K0X3 C4K0X5 C4K0Y4 C4K0Y8
C4K0Z4 C4K0Z6 C4K0Z7 C4K110 C4K111 C4K124 C4K125 C4K126 C4K157 C4K159 C4K160
C4K161 C4K162 C4K175 C4K176 C4K183 C4K184 C4K188 C4K189 C4K199 C4K1A0 C4K1A1
C4K1B0 C4K1B7 C4K1B9 C4K1C8 C4K1C9 C4K1D2 C4K1D4 C4K1D6 C4K1E1 C4K1F5 C4K1G6
C4K1G7 C4K1H9 C4K1I0 C4K1I1 C4K1I8 C4K1J3 C4K1J4 C4K1K8 C4K1K9 C4K1L0 C4K1L1
C4K1L3 C4K1L4 C4K1L9 C4K1M0 C4K1N6 C4K1N9 C4K1P1 C4K1P2 C4K1P3 C4K1P6 C4K1P7
C4K1P8 C4K1Q3 C4K1R1 C4K1R3 C4K1R7 C4K1R9 C4K1W0 C4K1W2 C4K1X3 C4K1X4 C4K1X9
C4K1Z7 C4K208 C4K233 C4K236 C4K256 C4K258 C4K265 C4K280 C4K281 C4K287 C4K288
C4K297 C4K2A1 C4K2A3 C4K2C2 C4K2D4 C4K2D5 C4K2D7 C4K2D8 C4K2E0 C4K2E2 C4K2E4
C4K2F3 C4K2F4 C4K2F5 C4K2F6 C4K2F7 C4K2F8 C4K2G0 C4K2G1 C4K2G2 C4K2G3 C4K2G4
C4K2G5 C4K2G7 C4K2G8 C4K2G9 C4K2H0 C4K2H1 C4K2H2 C4K2H3 C4K2H4 C4K2H5 C4K2H6
C4K2H7 C4K2H8 C4K2H9 C4K2I0 C4K2I2 C4K2J5 C4K2K1 C4K2K3 C4K2K4 C4K2K8 C4K2L1
C4K2L5 C4K2L7 C4K2P9 C4K2Q1 C4K2Q9 C4K2S0 C4K2S8 C4K2T0 C4K2T9 C4K2U0 C4K2U2
C4K2U3 C4K2V3 C4K2V4 C4K2W2 C4K2W9 C4K2X8 C4K2Y1 C4K2Y3 C4K2Y5 C4K2Y8 C4K2Z1
C4K332 C4K338 C4K346 C4K359 C4K389 C4K392 C4K3B1 C4K3C2 C4K3E6 C4K3E7 C4K3E8
C4K3F0 C4K3F2 C4K3H0 C4K3H3 C4K3I3 C4K3I5 C4K3I6 C4K3I8 C4K3J5 C4K3J7 C4K3K4
C4K3L4 C4K3Q5 C4K3R2 C4K3R9 C4K3S0 C4K3T8 C4K3U1 C4K3V5 C4K3W7 C4K3W9 C4K3X0
C4K3X2 C4K3X3 C4K3X8 C4K3Z3 C4K3Z5 C4K405 C4K407 C4K415 C4K432 C4K434 C4K435
C4K450 C4K459 C4K460 C4K473 C4K477 C4K487 C4K491 C4K4A2 C4K4B6 C4K4C4 C4K4D1
C4K4D4 C4K4E2 C4K4E3 C4K4F1 C4K4F2 C4K4F3 C4K4F4 C4K4F5 C4K4F8 C4K4F9 C4K4G0
C4K4G1 C4K4G2 C4K4I4 C4K4I8 C4K4J2 C4K4J9 C4K4K0 C4K4K4 C4K4K8 C4K4K9 C4K4M5
C4K4M9 C4K4P2 C4K4P3 C4K4P6 C4K4R2 C4K4U5 C4K4U6 C4K4U7 C4K578 C4K5L7 C4K5S0
C4K5S1 C4K5T0 C4K5T1 C4K5T8 C4K5U1 C4K5W2 C4K5W3 C4K5W6 C4K5W8 C4K5W9 C4K6C1
C4K6F4 C4K6G4 C4K6G8 C4K6M5 C4K6M7 C4K6N8 C4K6P7 C4K6P9 C4K6Q1 C4K6Q9 C4K6R0
C4K6Y0 C4K6Y1 C4K6Y5 C4K6Z7 C4K708 C4K734 C4K735 C4K744 C4K745 C4K759 C4K764
C4K776 C4K777 C4K778 C4K779 C4K781 C4K789 C4K792 C4K794 C4K795 C4K796 C4K797
C4K799 C4K7A0 C4K7A2 C4K7A3 C4K7A4 C4K7A6 C4K7A7 C4K7A8 C4K7A9 C4K7B0 C4K7B1
C4K7B2 C4K7B3 C4K7B4 C4K7B5 C4K7B7 C4K7B8 C4K7D7 C4K7E1 C4K7E2 C4K7E4 C4K7E6
C4K7G0 C4K7G1 C4K7G3 C4K7H1 C4K7H4 C4K7I4 C4K7I5 C4K7I6 C4K7I7 C4K7J9 C4K7K5
C4K7K8 C4K7K9 C4K7L1 C4K7P4 C4K7P7 C4K7P8 C4K7R7 C4K7V2 C4K7W2 C4K7W8 C4K7X8
C4K7Y8 C4K7Z0 C4K7Z9 C4K832 C4K838 C4K841 C4K842 C4K846 C4K8F9 C4K8I1 C4K8I5
C4K8I6 C4K8I7 C4K8J6 C4K8L9 C4K8M3 C4K8M4 C4K8M8 C4K8N5 C4K8P2 C4K8Q4 C4K8X0
C4K8Y6 C4K8Z1 C4K8Z4 C4K900 C4K901 C4K911 C4KAR8 C4KBU2 C4KEG7 C4KEH1 C4KEU3
C4KF50 C4KG47 C4KGL2 C4KGP7 C4KGP8 C4KGQ2 C4KGQ7 C4KGQ8 C4KGQ9 C4KGR1 C4KGW9
C4KGX1 C4KGX2 C4KGX7 C4KGX9 C4KGY0 C4KGZ2 C4KGZ7 C4KH07 C4KH08 C4KH10 C4KH13
C4KH14 C4KH27 C4KH28 C4KH38 C4KH41 C4KH54 C4KH92 C4KH93 C4KHA7 C4KHB0 C4KHB9
C4KHC0 C4KHC6 C4KHD2 C4KHD5 C4KHD8 C4KHD9 C4KHE3 C4KHE7 C4KHE9 C4KHF6 C4KHF7
C4KHF9 C4KHG8 C4KHH1 C4KHH3 C4KHH5 C4KHH7 C4KHI0 C4KHI2 C4KHI3 C4KHI6 C4KHI7
C4KHL2 C4KHM7 C4KHN7 C4KHQ2 C4KHQ3 C4KHR4 C4KHR8 C4KHS1 C4KHT0 C4KHT9 C4KHU2
C4KHU3 C4KHU5 C4KHW1 C4KHW5 C4KHW7 C4KHX1 C4KHX2 C4KI81 C4KI98 C4KIA4 C4KIA7
C4KIA8 C4KIB0 C4KIC4 C4KIC5 C4KID0 C4KID1 C4KID3 C4KIE0 C4KIE1 C4KIE7 C4KIH5
C4KIH7 C4KII8 C4KII9 C4KIJ0 C4KIJ1 C4KIJ2 C4KIJ3 C4KIJ7 C4KIJ9 C4KIK1 C4KIK6

C4KIN0 C4KIN2 C4KIN3 C4KIP3 C4KIP6 C4KIQ2 C4KIQ9 C4KIR0 C4KIR6 C4KIS0 C4KIT0
C4KIT6 C4KIV1 C4KIV3 C4KIV4 C4KIV6 C4KIV9 C4KIW6 C4KIY2 C4KIY8 C4KIZ2 C4KIZ9
C4KJ00 C4KJ02 C4KJ07 C4KJ09 C4KJ13 C4KJ22 C4KJ24 C4KJ29 C4KJ66 C4KJ68 C4KJ77
C4KJ83 C4KJ86 C4KJ87 C4KJ91 C4KJ93 C4KJ97 C4KJ99 C4KJA2 C4KJC1 C4KJE6 C4KJF1
C4KJF8 C4KJG4 C4KJG8 C4KJI5 C4KJM8 C4KJM9 C4KJT0 C4KJU6 C4KK76 C4KK84 C4KK87
C4KKA3 C4KKE5 C4KKE6 C4KKJ8 C4KKT0 C4KKU5 C4KKV8 C4KKV9 C4KYT2 C4KYV2 C4KYV5
C4KYW2 C4KZ07 C4KZ14 C4KZ18 C4KZ51 C4KZ52 C4KZ55 C4KZ66 C4KZD1 C4KZE2 C4KZK4
C4KZL2 C4KZL3 C4KZL4 C4KZL8 C4KZM0 C4KZM1 C4KZM2 C4KZM5 C4KZM7 C4KZM8 C4KZN0
C4KZN2 C4KZN4 C4KZN5 C4KZN6 C4KZN7 C4KZN8 C4KZN9 C4KZP0 C4KZP1 C4KZP2 C4KZP3
C4KZP5 C4KZP6 C4KZP7 C4KZP9 C4KZQ0 C4KZQ2 C4KZQ4 C4KZQ9 C4KZR7 C4KZS5 C4KZT8
C4KZU9 C4KZV2 C4KZW9 C4KZX2 C4KZX3 C4KZX8 C4KZZ0 C4KZZ3 C4KZZ4 C4KZZ5 C4L001
C4L008 C4L012 C4L014 C4L022 C4L056 C4L088 C4L0B0 C4L0I9 C4L0S0 C4L0S1 C4L0S2
C4L0S9 C4L0T7 C4L0V1 C4L0V4 C4L0W8 C4L0Z5 C4L0Z7 C4L173 C4L176 C4L191 C4L192
C4L193 C4L1B9 C4L1C0 C4L1C9 C4L1E1 C4L1I8 C4L1L1 C4L1L2 C4L274 C4L296 C4L298
C4L299 C4L2A1 C4L2C1 C4L2C5 C4L2D2 C4L2D3 C4L2D4 C4L2E7 C4L2P3 C4L2S9 C4L2V3
C4L2Y4 C4L367 C4L379 C4L3C2 C4L3C9 C4L3F0 C4L3F5 C4L3F6 C4L3F7 C4L3G0 C4L3G2
C4L3K5 C4L3L1 C4L3U2 C4L3W2 C4L3Y8 C4L3Z5 C4L405 C4L408 C4L415 C4L423 C4L424
C4L425 C4L427 C4L432 C4L434 C4L450 C4L456 C4L461 C4L464 C4L466 C4L470 C4L493
C4L4A0 C4L4I7 C4L4J0 C4L4J1 C4L4J3 C4L4J5 C4L4J6 C4L4J9 C4L4K4 C4L4N1 C4L4N3
C4L4P1 C4L4Q4 C4L522 C4L523 C4L531 C4L533 C4L534 C4L535 C4L552 C4L559 C4L574
C4L5A2 C4L5G5 C4L5H7 C4L5I7 C4L5J5 C4L5K9 C4L5P2 C4L5P3 C4L5T1 C4L5T3 C4L5T5
C4L5T9 C4L5V1 C4L5W3 C4L5X2 C4L5Y4 C4L601 C4L602 C4L603 C4L604 C4L610 C4L614
C4L618 C4L650 C4L651 C4L656 C4L660 C4L690 C4L691 C4L6J5 C4L6J6 C4L6M0 C4L6N1
C4L6N3 C4L6N4 C4L6Q0 C4L6Q1 C4L6Q2 C4L755 C4L762 C4L763 C4L772 C4L7I7 C4L7J5
C4L7J8 C4L7K2 C4L7L3 C4L7L4 C4L7L6 C4L7Q5 C4L7R5 C4L7S3 C4L7S5 C4L7S8 C4L7S9
C4L7T0 C4L7T1 C4L7T2 C4L7T3 C4L7T4 C4L7T7 C4L7T8 C4L7T9 C4L7U0 C4L7U1 C4L7U2
C4L7U4 C4L7U5 C4L7U6 C4L7U7 C4L7U9 C4L7V2 C4L7V3 C4L7V4 C4L7V5 C4L7V6 C4L7W3
C4L7X3 C4L7Y3 C4L7Y4 C4L7Y7 C4L7Z1 C4L810 C4L812 C4L813 C4L814 C4L816 C4L818
C4L823 C4L858 C4L861 C4L863 C4L864 C4L868 C4L881 C4L887 C4L889 C4L8B5 C4L8C4
C4L8C6 C4L8D2 C4L8D4 C4L8E9 C4L8F7 C4L8H9 C4L8J2 C4L8J8 C4L8N9 C4L8P1 C4L8P4
C4L8U2 C4L8U4 C4L8U5 C4L8U7 C4L8V5 C4L8W1 C4L8X0 C4L8X1 C4L8X2 C4L8X3 C4L8X4
C4L8X6 C4L8Y1 C4L8Y4 C4L8Y5 C4L8Y7 C4L8Z0 C4L8Z2 C4L929 C4L930 C4L931 C4L932
C4L937 C4L944 C4L952 C4L968 C4L9A7 C4L9A9 C4L9J3 C4L9J8 C4L9L4 C4L9L6 C4L9M2
C4L9M3 C4L9M5 C4L9M9 C4L9N0 C4L9N1 C4L9W9 C4L9X0 C4L9Y1 C4L9Y9 C4L9Z0 C4L9Z3
C4LA04 C4LA08 C4LA10 C4LA13 C4LA16 C4LA17 C4LA26 C4LA34 C4LA41 C4LA51 C4LA78
C4LAA1 C4LAB3 C4LAB6 C4LAB7 C4LAB8 C4LAC1 C4LAC2 C4LAE0 C4LAE1 C4LAE5 C4LAE6
C4LAE7 C4LAF1 C4LAF3 C4LAG3 C4LAG5 C4LAG9 C4LAK4 C4LAP0 C4LAS5 C4LAV4 C4LAV5
C4LAV7 C4LAV8 C4LAY4 C4LAY8 C4LAY9 C4LB07 C4LB08 C4LB24 C4LB49 C4LB55 C4LB59
C4LB60 C4LBG4 C4LBH9 C4LBI8 C4LBK4 C4LBL2 C4LBL5 C4LBN0 C4LBN1 C4LBN2 C4LBN3
C4LBN4 C4LBN8 C4LBP7 C4LBQ0 C4LBQ6 C4LBQ9 C4LBR2 C4LBR5 C4LBS7 C4LBS8 C4LBU4
C4LBU5 C4LBU6 C4LBV1 C4LBV2 C4LBV3 C4LBV4 C4LBV5 C4LBV6 C4LBX2 C4LBX3 C4LBX6
C4LBX7 C4LC00 C4LC10 C4LC34 C4LC38 C4LC41 C4LC44 C4LC58 C4LC60 C4LC62 C4LC65
C4LC66 C4LC88 C4LC89 C4LC91 C4LC98 C4LCA2 C4LCA3 C4LCB6 C4LCC5 C4LCH1 C4LCK6
C4LCK7 C4LCM1 C4LCM6 C4LCN2 C4LCN4 C4LCN9 C4LCP0 C4LCT4 C4LCT8 C4LCV3 C4LCV5

C4LCY4 C4LCZ9 C4LD10 C4LD15 C4LD19 C4LD21 C4LD24 C4LD25 C4LD26 C4LD27 C4LD49
C4LD51 C4LD58 C4LDB2 C4LDB4 C4LDC0 C4LDD0 C4LDD2 C4LDE6 C4LDG0 C4LDK0 C4LDM6
C4LDS1 C4LDU8 C4LDV0 C4LDX1 C4LDX4 C4LE53 C4LE64 C4LEB8 C4LEG3 C4LEL9 C4LEQ5
C4LEV7 C4LEW9 C4LEX2 C4LEX3 C4LEX5 C4LEY6 C4LEY8 C4LEZ7 C4LF00 C4LF05 C4LF07
C4LF13 C4LF16 C4LF59 C4LF62 C4LF63 C4LF64 C4LF65 C4LFC1 C4LFE6 C4LFG7 C4LFG9
C4LFH0 C4LFH1 C4LFH3 C4LFH4 C4LFH8 C4LFI3 C4LFI4 C4LFK3 C4LFU8 C4LG01 C4LG04
C4LG69 C4LG87 C4LG97 C4LGF9 C4LGG1 C4LGJ8 C4LGQ1 C4LGQ3 C4LGT5 C4LGU9 C4LGV8
C4LH17 C4LH71 C4LHE0 C4LHF0 C4LHF2 C4LHF3 C4LHF4 C4LHF5 C4LHI6 C4LHT4 C4LHU9
C4LI04 C4LI41 C4LI42 C4LIA9 C4LID0 C4LID1 C4LID5 C4LIE1 C4LIF4 C4LIG1 C4LIK9
C4LIL2 C4LIL5 C4LIL9 C4LIR6 C4LIS3 C4LIT2 C4LIU1 C4LIU5 C4LIV4 C4LIW2 C4LIW5
C4LIX2 C4LIX8 C4LIY0 C4LJ12 C4LJ26 C4LJ33 C4LJ74 C4LJ75 C4LJ81 C4LJ85 C4LJA6
C4LJA8 C4LJA9 C4LJB9 C4LJC4 C4LJF0 C4LJH3 C4LJH6 C4LJI1 C4LJJ2 C4LJJ9 C4LJL1
C4LJM5 C4LJM9 C4LJR0 C4LJT1 C4LJU0 C4LJU5 C4LJU6 C4LJV6 C4LJV9 C4LJY5 C4LJY7
C4LJZ9 C4LK22 C4LK80 C4LKC1 C4LKV3 C4LKZ0 C4LKZ2 C4LKZ3 C4LKZ4 C4LL05 C4LL07
C4LL08 C4LL09 C4LL10 C4LL11 C4LL12 C4LL24 C4LL25 C4LL26 C4LL43 C4LL44 C4LL45
C4LL46 C4LL47 C4LL48 C4LL49 C4LL50 C4LL51 C4LL52 C4LL63 C4LL65 C4LL66 C4LL70
C4LL71 C4LL72 C4LL80 C4LL88 C4LL89 C4LL93 C4LLD4 C4LLD6 C4LSS0 C4LWQ8 C4LY96
C4LYI2 C4LZC2 C4LZV8 C4M1E4 C4M244 C4M2B8 C4M572 C4M6G8 C4M6T2 C4M7H7 C4MBZ5
C4NF90 C4NF91 C4NYP8 C4NYZ3 C4P6S0 C4PWA1 C4PYP8 C4PZQ3 C4Q0P6 C4QCD2 C4QGM3
C4QV17 C4QV62 C4QVB0 C4QVR9 C4QVX6 C4QXA5 C4QXH2 C4QXX8 C4QY17 C4QYF2 C4QYK0
C4QYL7 C4QYS6 C4QZ20 C4R0B5 C4R1K8 C4R1T9 C4R2K0 C4R2P3 C4R331 C4R3K5 C4R3R8
C4R492 C4R4R8 C4R591 C4R5N0 C4R5P7 C4R6H3 C4R6W8 C4R7D9 C4R7H4 C4R7Q5 C4R853
C4R8C9 C4R8E8 C4R8P2 C4R8S9 C4R970 C4RJF7 C4RJF8 C4SGN8 C4TP09 C4V6H6 C4V6K5
C4V6W1 C4V7I7 C4V7V1 C4V819 C4V847 C4V8K5 C4V8L1 C4V8Q1 C4V8R9 C4V922 C4V924
C4V943 C4V9G9 C4V9J0 C4V9L5 C4VA52 C4VAS3 C4VB43 C4VBI7 C4VBI8 C4WU37 C4WUT0
C4WXC1 C4XGC2 C4XGI2 C4XGI3 C4XGN5 C4XGN6 C4XGN7 C4XGU0 C4XGV9 C4XH03 C4XH04
C4XHA0 C4XHQ9 C4XHR1 C4XHR2 C4XHY9 C4XI54 C4XIH0 C4XIJ2 C4XIJ3 C4XIJ9 C4XIK0
C4XIK1 C4XIN9 C4XIP0 C4XIP2 C4XIP3 C4XIQ1 C4XIQ3 C4XIQ5 C4XIQ6 C4XIR3 C4XIR5
C4XIS0 C4XIS1 C4XJ76 C4XJA5 C4XJB6 C4XJB9 C4XJU6 C4XJX5 C4XK68 C4XK86 C4XK87
C4XKL7 C4XKM1 C4XL11 C4XL13 C4XL14 C4XL87 C4XLK0 C4XLK1 C4XLK2 C4XLK3 C4XLK4
C4XLR8 C4XLU8 C4XLW8 C4XLW9 C4XLX2 C4XLX3 C4XLX4 C4XLX5 C4XLX6 C4XLX7 C4XLX8
C4XLX9 C4XLY0 C4XLY1 C4XLY3 C4XLY4 C4XLY5 C4XLY6 C4XLY8 C4XLZ1 C4XLZ2 C4XLZ5
C4XMB3 C4XMR7 C4XMX9 C4XMY1 C4XMY2 C4XMZ1 C4XN25 C4XN27 C4XNJ4 C4XNQ1 C4XNQ2
C4XNQ9 C4XNW0 C4XNW1 C4XNX1 C4XNX3 C4XNY1 C4XPA8 C4XPD6 C4XPE9 C4XPF2 C4XPJ3
C4XPZ6 C4XPZ7 C4XPZ9 C4XQ09 C4XQ59 C4XQ61 C4XQ63 C4XQT5 C4XR49 C4XRE2 C4XRE5
C4XRH8 C4XRH9 C4XRK2 C4XS28 C4XS35 C4XS37 C4XS40 C4XS52 C4XS53 C4XS54 C4XS55
C4XS73 C4XSD3 C4XSE1 C4XSF4 C4XSI9 C4XSN4 C4XSN6 C4XSN8 C4XSQ4 C4XST3 C4XSW4
C4XSW5 C4XSX5 C4XSX9 C4XT34 C4XT36 C4XT58 C4XT92 C4XTC8 C4XTD0 C4XTP0 C4XTP4
C4XU39 C4XU70 C4XUD2 C4XWY8 C4XXB1 C4XXU2 C4XYD5 C4XYS2 C4XZU6 C4Y146 C4Y1F8
C4Y1G7 C4Y1P0 C4Y2C6 C4Y2I7 C4Y2J3 C4Y2K4 C4Y2M6 C4Y3M4 C4Y3N8 C4Y3W1 C4Y459
C4Y4A0 C4Y4V1 C4Y584 C4Y5P7 C4Y6U0 C4Y732 C4Y7Q2 C4Y8K5 C4Y9D5 C4Y9X0 C4YA37
C4YBE4 C4YBG0 C4YBJ8 C4YC37 C4YCB9 C4YCH0 C4YCU0 C4YD35 C4YD57 C4YDV1 C4YE90
C4YEN2 C4YF50 C4YFC7 C4YFD4 C4YFX2 C4YH38 C4YH95 C4YHS3 C4YIM0 C4YIU5 C4YJ78
C4YJE1 C4YJP2 C4YJQ8 C4YL88 C4YLC3 C4YLG6 C4YLK8 C4YLS2 C4YMA4 C4YMC1 C4YMW2

C4YN69 C4YNP4 C4YNR7 C4YNS4 C4YP90 C4YPI7 C4YPQ0 C4YPS3 C4YQD9 C4YR54 C4YRG6
C4YRQ5 C4YRY0 C4YSA9 C4YSG0 C4YSJ2 C4YSU5 C4YTS0 C4Z030 C4Z044 C4Z055 C4Z060
C4Z073 C4Z075 C4Z088 C4Z090 C4Z0A5 C4Z0B1 C4Z0C3 C4Z0C8 C4Z0D7 C4Z0E0 C4Z0F4
C4Z0F5 C4Z0G2 C4Z0G8 C4Z0G9 C4Z0J8 C4Z0N9 C4Z0P7 C4Z0Q0 C4Z0Q6 C4Z0T6 C4Z0V2
C4Z0V6 C4Z111 C4Z114 C4Z135 C4Z138 C4Z157 C4Z158 C4Z176 C4Z1C2 C4Z1D7 C4Z1I4
C4Z1J3 C4Z1J4 C4Z1K1 C4Z1Q7 C4Z1S4 C4Z1S7 C4Z1T3 C4Z1T5 C4Z1T6 C4Z1V6 C4Z1Z8
C4Z205 C4Z281 C4Z282 C4Z2R3 C4Z2R7 C4Z2R9 C4Z2S9 C4Z2T0 C4Z2T1 C4Z2T2 C4Z2T4
C4Z2T5 C4Z2T7 C4Z2T8 C4Z2T9 C4Z2U0 C4Z2U1 C4Z2U3 C4Z2U6 C4Z2V1 C4Z2V4 C4Z2V6
C4Z2V8 C4Z2W3 C4Z309 C4Z312 C4Z3J4 C4Z3J5 C4Z3J8 C4Z3N3 C4Z3N5 C4Z3P0 C4Z3R4
C4Z3X1 C4Z417 C4Z429 C4Z478 C4Z4C1 C4Z4C2 C4Z4L5 C4Z4L6 C4Z4L8 C4Z4L9 C4Z4Q1
C4Z4T8 C4Z4U3 C4Z4W1 C4Z4Y1 C4Z520 C4Z523 C4Z530 C4Z534 C4Z552 C4Z562 C4Z563
C4Z566 C4Z578 C4Z5E3 C4Z5K1 C4Z5N3 C4Z5N7 C4Z5N8 C4Z5P5 C4Z5P8 C4Z899 C4Z8N0
C4Z8V7 C4Z8W1 C4Z8W9 C4Z8Z2 C4Z900 C4Z911 C4Z916 C4Z917 C4Z929 C4Z940 C4Z987
C4Z9C9 C4Z9E6 C4Z9L7 C4Z9Q0 C4Z9Q2 C4Z9R8 C4Z9S1 C4Z9V4 C4Z9X7 C4Z9Y1 C4Z9Z6
C4Z9Z8 C4ZA36 C4ZA52 C4ZA59 C4ZA66 C4ZA70 C4ZA73 C4ZAS2 C4ZAV7 C4ZAW6 C4ZAX1
C4ZAY4 C4ZAY9 C4ZAZ2 C4ZB25 C4ZB57 C4ZB90 C4ZB99 C4ZBD3 C4ZBD4 C4ZBD5 C4ZBD8
C4ZBF3 C4ZBF8 C4ZBG1 C4ZBG2 C4ZBG7 C4ZBG8 C4ZBH1 C4ZBJ4 C4ZBL1 C4ZBS1 C4ZBS3
C4ZBS4 C4ZBS5 C4ZBS6 C4ZBS7 C4ZBS8 C4ZBS9 C4ZBT0 C4ZBT1 C4ZBT4 C4ZBT6 C4ZBT7
C4ZBT9 C4ZBU3 C4ZBU5 C4ZBV1 C4ZBV4 C4ZBV8 C4ZBW8 C4ZC24 C4ZCQ4 C4ZCY4 C4ZCZ0
C4ZCZ4 C4ZD15 C4ZD30 C4ZD46 C4ZD63 C4ZDB0 C4ZDL0 C4ZDP0 C4ZDU4 C4ZEG7 C4ZEG8
C4ZEK2 C4ZEV1 C4ZEV8 C4ZEV9 C4ZF43 C4ZF71 C4ZF78 C4ZFN0 C4ZFN9 C4ZG24 C4ZG66
C4ZGB4 C4ZGF3 C4ZGF4 C4ZGF5 C4ZGY6 C4ZH69 C4ZH75 C4ZHB9 C4ZHC0 C4ZHH4 C4ZHN1
C4ZI07 C4ZI12 C4ZI69 C4ZI84 C4ZI85 C4ZJM0 C4ZJU1 C4ZPS8 C4ZPT0 C4ZPT3 C4ZPU1
C4ZPU9 C4ZPV2 C4ZPV5 C4ZPV6 C4ZPV7 C4ZPW1 C4ZPW3 C4ZPW4 C4ZPW5 C4ZPW7 C4ZPW8
C4ZPX5 C4ZPX6 C4ZPX7 C4ZPX8 C4ZPY3 C4ZPY5 C4ZPY6 C4ZPZ4 C4ZPZ5 C4ZPZ7 C4ZQ03
C4ZQ04 C4ZQ11 C4ZQ19 C4ZQ33 C4ZQ34 C4ZQ36 C4ZQ38 C4ZQ43 C4ZQ48 C4ZQ50 C4ZQ57
C4ZQ61 C4ZQ82 C4ZQ84 C4ZQ89 C4ZQ92 C4ZQ93 C4ZQC6 C4ZQC7 C4ZQC8 C4ZQD2 C4ZQD5
C4ZQD6 C4ZQD7 C4ZQD8 C4ZQD9 C4ZQE0 C4ZQE4 C4ZQE5 C4ZQE7 C4ZQF0 C4ZQF4 C4ZQF5
C4ZQF8 C4ZQG0 C4ZQJ1 C4ZQJ7 C4ZQL1 C4ZQM6 C4ZQM8 C4ZQN7 C4ZQP5 C4ZQQ4 C4ZQR6
C4ZQV5 C4ZQV9 C4ZQW6 C4ZQW9 C4ZQX3 C4ZQY1 C4ZQY2 C4ZQY5 C4ZQY9 C4ZR08 C4ZR24
C4ZR38 C4ZR44 C4ZR47 C4ZR48 C4ZR49 C4ZR54 C4ZR55 C4ZR68 C4ZR69 C4ZR73 C4ZR74
C4ZR75 C4ZR77 C4ZR78 C4ZR79 C4ZR80 C4ZR85 C4ZR95 C4ZRA8 C4ZRB7 C4ZRB8 C4ZRD1
C4ZRH1 C4ZRI6 C4ZRJ5 C4ZRJ6 C4ZRJ8 C4ZRL2 C4ZRL3 C4ZRL4 C4ZRM4 C4ZRM6 C4ZRM7
C4ZRN7 C4ZRN9 C4ZRP7 C4ZRP9 C4ZRQ2 C4ZRQ3 C4ZRQ6 C4ZRQ8 C4ZRQ9 C4ZRR1 C4ZRR2
C4ZRR4 C4ZRS5 C4ZRT2 C4ZRT7 C4ZRU6 C4ZRU9 C4ZRV1 C4ZRV8 C4ZRX5 C4ZRZ5 C4ZRZ9
C4ZS02 C4ZS07 C4ZS29 C4ZS38 C4ZS46 C4ZS47 C4ZS48 C4ZS59 C4ZS67 C4ZS77 C4ZS88
C4ZS91 C4ZS93 C4ZSA8 C4ZSB0 C4ZSB3 C4ZSB4 C4ZSF3 C4ZSH9 C4ZSI0 C4ZSI8 C4ZSJ8
C4ZSM3 C4ZSN9 C4ZSP0 C4ZSP5 C4ZSP6 C4ZSQ5 C4ZSQ6 C4ZSQ8 C4ZSQ9 C4ZSR1 C4ZSR2
C4ZSR5 C4ZSR8 C4ZSS4 C4ZSS5 C4ZSS8 C4ZSU4 C4ZSW0 C4ZSW1 C4ZSW3 C4ZSW4 C4ZSW8
C4ZSW9 C4ZSX4 C4ZSX5 C4ZSZ5 C4ZSZ7 C4ZT10 C4ZT14 C4ZT19 C4ZT30 C4ZT45 C4ZT56
C4ZT63 C4ZT65 C4ZT66 C4ZT68 C4ZT75 C4ZT76 C4ZT77 C4ZT96 C4ZT97 C4ZT99 C4ZTA0
C4ZTB5 C4ZTE1 C4ZTE6 C4ZTE7 C4ZTF0 C4ZTF1 C4ZTG2 C4ZTG3 C4ZTH0 C4ZTH2 C4ZTH3
C4ZTH7 C4ZTH9 C4ZTI0 C4ZTI3 C4ZTJ3 C4ZTJ5 C4ZTK1 C4ZTK3 C4ZTM0 C4ZTM2 C4ZTN0
C4ZTP5 C4ZTP9 C4ZTQ1 C4ZTQ2 C4ZTS5 C4ZTU0 C4ZTV3 C4ZTX2 C4ZTX6 C4ZTY0 C4ZTY2

C4ZU03 C4ZU09 C4ZU12 C4ZU30 C4ZU31 C4ZU32 C4ZU53 C4ZU73 C4ZU87 C4ZU89 C4ZU91
C4ZU93 C4ZUA4 C4ZUA6 C4ZUA7 C4ZUB1 C4ZUD6 C4ZUD9 C4ZUE1 C4ZUE2 C4ZUE3 C4ZUE9
C4ZUF1 C4ZUF2 C4ZUF4 C4ZUF6 C4ZUF7 C4ZUF9 C4ZUG0 C4ZUG1 C4ZUG3 C4ZUG4 C4ZUG5
C4ZUG6 C4ZUG7 C4ZUG8 C4ZUG9 C4ZUH0 C4ZUH1 C4ZUH2 C4ZUH4 C4ZUH5 C4ZUH6 C4ZUJ5
C4ZUJ6 C4ZUJ7 C4ZUJ8 C4ZUJ9 C4ZUK0 C4ZUK3 C4ZUL0 C4ZUM4 C4ZUP4 C4ZUQ6 C4ZUQ8
C4ZUR5 C4ZUR7 C4ZUS3 C4ZUS5 C4ZUS6 C4ZUS8 C4ZUS9 C4ZUT0 C4ZUV3 C4ZUV5 C4ZUW1
C4ZUX4 C4ZUX7 C4ZV25 C4ZV92 C4ZVI3 C4ZVI4 C4ZVI7 C4ZVL1 C4ZVL3 C4ZVM2 C4ZVM4
C4ZVQ3 C4ZVR1 C4ZVS5 C4ZVU9 C4ZVV8 C4ZVV9 C4ZVW3 C4ZVX0 C4ZVX1 C4ZVX6 C4ZVX9
C4ZVY0 C4ZVY1 C4ZW19 C4ZW30 C4ZW44 C4ZW66 C4ZWA2 C4ZWA3 C4ZWA6 C4ZWB5 C4ZWB7
C4ZWB8 C4ZWC3 C4ZWC9 C4ZWD8 C4ZWE6 C4ZWF4 C4ZWF6 C4ZWI8 C4ZWI9 C4ZWK2 C4ZWN2
C4ZWP8 C4ZWT6 C4ZWT8 C4ZX09 C4ZX13 C4ZX46 C4ZX51 C4ZX71 C4ZX73 C4ZX74 C4ZX82
C4ZX84 C4ZX86 C4ZX89 C4ZX90 C4ZX92 C4ZX93 C4ZX98 C4ZXA1 C4ZXA2 C4ZXA3 C4ZXA5
C4ZXA6 C4ZXB4 C4ZXB6 C4ZXB7 C4ZXC6 C4ZXE9 C4ZXF0 C4ZXF6 C4ZXG6 C4ZXI2 C4ZXJ1
C4ZXK1 C4ZXK8 C4ZXL1 C4ZXM6 C4ZXM7 C4ZXM8 C4ZXM9 C4ZXN0 C4ZXN2 C4ZXN3 C4ZXN4
C4ZXN9 C4ZXP1 C4ZXQ7 C4ZXS1 C4ZXS6 C4ZXS8 C4ZXT8 C4ZXV1 C4ZXV3 C4ZXV5 C4ZXX6
C4ZXX9 C4ZXY4 C4ZY24 C4ZY31 C4ZY38 C4ZY45 C4ZY53 C4ZY73 C4ZY85 C4ZYA0 C4ZYA1
C4ZYA3 C4ZYA5 C4ZYC2 C4ZYE1 C4ZYE2 C4ZYF4 C4ZYF5 C4ZYG5 C4ZYG8 C4ZYH4 C4ZYH6
C4ZYH8 C4ZYH9 C4ZYI0 C4ZYI1 C4ZYJ0 C4ZYJ8 C4ZYK2 C4ZYK3 C4ZYM4 C4ZYM5 C4ZYM6
C4ZYM7 C4ZYN1 C4ZYN2 C4ZYN5 C4ZYN7 C4ZYR1 C4ZYS6 C4ZYT8 C4ZYU1 C4ZYU3 C4ZYU4
C4ZYV6 C4ZYW3 C4ZYW4 C4ZYW9 C4ZYX7 C4ZYX9 C4ZYY0 C4ZYY1 C4ZYY4 C4ZYY6 C4ZYY9
C4ZZ18 C4ZZ19 C4ZZ22 C4ZZ23 C4ZZ24 C4ZZ26 C4ZZ35 C4ZZ41 C4ZZ44 C4ZZ47 C4ZZ48
C4ZZ69 C4ZZ71 C4ZZ73 C4ZZ74 C4ZZ87 C4ZZ89 C4ZZ96 C4ZZ98 C4ZZA0 C4ZZA1 C4ZZA2
C4ZZA3 C4ZZA4 C4ZZD4 C4ZZE0 C4ZZG7 C4ZZG9 C4ZZI8 C4ZZJ2 C4ZZK6 C4ZZN7 C4ZZP7
C4ZZP8 C4ZZP9 C4ZZQ0 C4ZZQ1 C4ZZQ4 C4ZZQ5 C4ZZQ6 C4ZZR6 C4ZZR7 C4ZZT3 C4ZZU9
C4ZZV3 C4ZZV7 C4ZZV9 C4ZZW1 C4ZZX9 C4ZZY2 C4ZZY4 C5A009 C5A021 C5A033 C5A034
C5A055 C5A059 C5A063 C5A069 C5A070 C5A071 C5A072 C5A073 C5A074 C5A083 C5A093
C5A095 C5A096 C5A098 C5A099 C5A0A3 C5A0A5 C5A0C2 C5A0C3 C5A0C6 C5A0E6 C5A0G1
C5A0G3 C5A0H0 C5A0H2 C5A0H5 C5A0H6 C5A0H7 C5A0I1 C5A0I5 C5A0I6 C5A0J7 C5A0K6
C5A0L1 C5A0L3 C5A0L7 C5A0L8 C5A0M2 C5A0N1 C5A0Q8 C5A0Q9 C5A0R1 C5A0R4 C5A0R6
C5A0R9 C5A0S3 C5A0S4 C5A0S5 C5A0S6 C5A0S7 C5A0T1 C5A0T5 C5A0T6 C5A0T7 C5A0T8
C5A0T9 C5A0U7 C5A0V0 C5A0V3 C5A0V8 C5A0W2 C5A132 C5A1C9 C5A1D4 C5A1D5 C5A1D9
C5A1E9 C5A1F5 C5A1G1 C5A1H0 C5A1H1 C5A1H2 C5A1K2 C5A1K9 C5A1L0 C5A1L9 C5A1M3
C5A1N3 C5A1N9 C5A1P1 C5A1P4 C5A1S7 C5A1V0 C5A1V7 C5A1V9 C5A1W0 C5A1Y3 C5A203
C5A206 C5A207 C5A210 C5A216 C5A226 C5A241 C5A244 C5A250 C5A251 C5A255 C5A257
C5A259 C5A261 C5A266 C5A268 C5A269 C5A270 C5A278 C5A279 C5A280 C5A282 C5A283
C5A284 C5A285 C5A286 C5A293 C5A2A3 C5A2A8 C5A2B8 C5A2B9 C5A2C2 C5A2D5 C5A2D9
C5A2F3 C5A2F7 C5A2H1 C5A2H2 C5A2H8 C5A2X8 C5A355 C5A366 C5A3C2 C5A3C3 C5A3C9
C5A3G1 C5A3G3 C5A3H2 C5A3H8 C5A3K4 C5A3L5 C5A3N8 C5A3P0 C5A3Q1 C5A3R9 C5A3V9
C5A401 C5A403 C5A422 C5A426 C5A428 C5A430 C5A442 C5A443 C5A474 C5A475 C5A4B7
C5A4D6 C5A4G1 C5A4G9 C5A4H0 C5A4H7 C5A4I6 C5A4R5 C5A4R8 C5A4S1 C5A4U0 C5A4U6
C5A4Z4 C5A4Z9 C5A500 C5A545 C5A552 C5A558 C5A593 C5A594 C5A597 C5A5B0 C5A5D0
C5A5E2 C5A5J3 C5A5J8 C5A5L0 C5A5L9 C5A5M5 C5A5N6 C5A5P4 C5A5P5 C5A5R0 C5A5R8
C5A5S2 C5A5T8 C5A5V9 C5A5Y9 C5A5Z0 C5A628 C5A636 C5A639 C5A649 C5A697 C5A6A9
C5A6F6 C5A6G2 C5A6I4 C5A6L2 C5A6M8 C5A6N2 C5A6N7 C5A6P6 C5A6P8 C5A6R5 C5A6U0

C5A6X9 C5A6Y5 C5A6Z1 C5A739 C5A782 C5A784 C5A786 C5A797 C5A7A2 C5A7A4 C5A7C4
C5A7E9 C5A7F1 C5A7F5 C5A7H1 C5A7H2 C5A7H3 C5A7H4 C5A7J0 C5A7J1 C5A7L1 C5A7L6
C5APZ1 C5ATZ5 C5AZ74 C5B0U5 C5B0U6 C5B0U7 C5B0U8 C5B0U9 C5B135 C5B137 C5B138
C5B2R8 C5B6Z1 C5B714 C5B725 C5B747 C5B748 C5B754 C5B766 C5B780 C5B7A2 C5B7A4
C5B7H1 C5B7H2 C5B7H9 C5B7I2 C5B7K5 C5B7K7 C5B7L0 C5B7L7 C5B7L8 C5B7M4 C5B7M7
C5B7N2 C5B7N9 C5B7P0 C5B7R1 C5B7R2 C5B7R4 C5B7S2 C5B7T0 C5B7T3 C5B7W2 C5B806
C5B813 C5B814 C5B820 C5B825 C5B832 C5B835 C5B836 C5B846 C5B848 C5B849 C5B850
C5B852 C5B8B9 C5B8G5 C5B8J8 C5B8N0 C5B8V3 C5B8V6 C5B8X3 C5B8X6 C5B8Y2 C5B954
C5B965 C5B970 C5B995 C5B996 C5B9A2 C5B9B3 C5B9C9 C5B9D0 C5B9D1 C5B9D6 C5B9D7
C5B9D8 C5B9D9 C5B9E7 C5B9E8 C5B9G7 C5B9H0 C5B9I5 C5B9K4 C5B9K5 C5B9K6 C5B9S7
C5B9S9 C5B9T0 C5B9T2 C5B9V4 C5B9V6 C5B9W5 C5B9W9 C5BA64 C5BA73 C5BAA9 C5BAC7
C5BAC9 C5BAD8 C5BAF2 C5BAI6 C5BAK4 C5BAK8 C5BAL3 C5BAP4 C5BAP7 C5BAP9 C5BAR6
C5BAS5 C5BAT0 C5BAT1 C5BAT2 C5BAT6 C5BAU7 C5BAV1 C5BAV4 C5BAW4 C5BB03 C5BB22
C5BB78 C5BB79 C5BB81 C5BB97 C5BB99 C5BBA5 C5BBA8 C5BBD2 C5BBD4 C5BBH3 C5BBR3
C5BBR6 C5BBR9 C5BBS1 C5BBY4 C5BC01 C5BC06 C5BC26 C5BC34 C5BC48 C5BC57 C5BC58
C5BC59 C5BC62 C5BC63 C5BC71 C5BC84 C5BCA4 C5BCB6 C5BCC5 C5BCE8 C5BCF0 C5BCF2
C5BCF6 C5BCF7 C5BCH3 C5BCH5 C5BCH6 C5BCH9 C5BCI1 C5BCI2 C5BCJ0 C5BCJ5 C5BCJ6
C5BCK3 C5BCK8 C5BCL9 C5BCM3 C5BCM4 C5BCN2 C5BCP0 C5BCR7 C5BCS0 C5BCZ7 C5BCZ9
C5BD00 C5BD16 C5BD17 C5BD23 C5BD33 C5BD56 C5BD72 C5BD73 C5BDB6 C5BDB7 C5BDC2
C5BDE0 C5BDG5 C5BDG8 C5BDG9 C5BDK4 C5BDK5 C5BDL9 C5BDM9 C5BDP0 C5BDU0 C5BE32
C5BE34 C5BE36 C5BE45 C5BE90 C5BE98 C5BEA1 C5BEA9 C5BED3 C5BEJ2 C5BEL3 C5BEM7
C5BEP2 C5BER5 C5BER7 C5BES7 C5BET0 C5BET4 C5BET5 C5BEU1 C5BEU2 C5BEU3 C5BEU5
C5BEU6 C5BEV2 C5BEV5 C5BEW3 C5BEW8 C5BEX0 C5BF07 C5BF12 C5BF17 C5BF18 C5BF26
C5BF28 C5BF29 C5BF32 C5BF52 C5BF62 C5BF63 C5BF77 C5BF78 C5BF79 C5BF83 C5BF97
C5BF98 C5BFA2 C5BFB0 C5BFB3 C5BFB7 C5BFB8 C5BFC0 C5BFC1 C5BFL0 C5BFL5 C5BFM6
C5BFM7 C5BFN3 C5BFR3 C5BFT8 C5BG10 C5BG15 C5BG16 C5BG23 C5BG25 C5BG54 C5BG56
C5BGA4 C5BGA6 C5BGB7 C5BGC7 C5BGD2 C5BGD7 C5BGD9 C5BGF9 C5BGG0 C5BGG2 C5BGG7
C5BGH6 C5BGI4 C5BGI8 C5BGI9 C5BGJ0 C5BGJ4 C5BGK6 C5BGK7 C5BGK9 C5BGL0 C5BGL1
C5BGL3 C5BGL4 C5BGL5 C5BGL7 C5BGL8 C5BGL9 C5BGM2 C5BGM3 C5BGM4 C5BGM6 C5BGM8
C5BGM9 C5BGN0 C5BGP8 C5BGQ4 C5BGR8 C5BGS4 C5BGT3 C5BGT5 C5BGV9 C5BGW0 C5BGW5
C5BH21 C5BH54 C5BH76 C5BH86 C5BH95 C5BHA4 C5BHA5 C5BHB2 C5BHB3 C5BHB7 C5BHB9
C5BHC1 C5BHC5 C5BHC7 C5BHE0 C5BHE1 C5BHE2 C5BHE3 C5BHE7 C5BHE8 C5BHG0 C5BHG1
C5BHG5 C5BHH2 C5BHH8 C5BHI4 C5BHJ2 C5BHJ4 C5BHJ5 C5BHN8 C5BHP0 C5BHP3 C5BHQ4
C5BHQ5 C5BHX0 C5BI15 C5BI30 C5BIF4 C5BII1 C5BIJ7 C5BIJ8 C5BIK4 C5BIS1 C5BJ23
C5BJ25 C5BJC0 C5BJT4 C5BJZ8 C5BKL4 C5BKL9 C5BKM1 C5BKP0 C5BKQ0 C5BL09 C5BL65
C5BL66 C5BL71 C5BL73 C5BL83 C5BLF6 C5BLF9 C5BLG1 C5BLG3 C5BLG5 C5BLH1 C5BLJ8
C5BLL7 C5BLM4 C5BLP4 C5BLR8 C5BLU3 C5BLV6 C5BLW3 C5BM93 C5BM95 C5BM96 C5BMA0
C5BMA3 C5BMB1 C5BMC8 C5BMF2 C5BMF4 C5BMF5 C5BMG6 C5BMI0 C5BMM7 C5BMN7 C5BMQ2
C5BMR1 C5BMR5 C5BMZ8 C5BN45 C5BN46 C5BN48 C5BN51 C5BN52 C5BN73 C5BN92 C5BNE0
C5BNG6 C5BNH5 C5BNK6 C5BNN0 C5BNN2 C5BNR1 C5BNT8 C5BNW9 C5BNY8 C5BP01 C5BP08
C5BP09 C5BP33 C5BP36 C5BP42 C5BP43 C5BP49 C5BP61 C5BP64 C5BPA3 C5BPC0 C5BPC3
C5BPN9 C5BPP4 C5BPP8 C5BPQ2 C5BPQ3 C5BPQ5 C5BPQ9 C5BPR2 C5BPU7 C5BPV5 C5BPV6
C5BPV8 C5BPV9 C5BPW1 C5BQ00 C5BQ30 C5BQ31 C5BQ32 C5BQ33 C5BQ34 C5BQ37 C5BQ38
C5BQ39 C5BQ41 C5BQ42 C5BQ43 C5BQ44 C5BQ61 C5BQ62 C5BQ63 C5BQ64 C5BQ65 C5BQ66

C5BQ68 C5BQ69 C5BQ70 C5BQ71 C5BQ72 C5BQ73 C5BQ75 C5BQ76 C5BQ77 C5BQ79 C5BQ80
C5BQ83 C5BQ84 C5BQ85 C5BQ87 C5BQ92 C5BQB2 C5BQB3 C5BQB5 C5BQB7 C5BQD1 C5BQE9
C5BQF6 C5BQF7 C5BQF9 C5BQH7 C5BQN7 C5BQP1 C5BQT4 C5BQT5 C5BQT6 C5BQX2 C5BQZ6
C5BR43 C5BR49 C5BR54 C5BR59 C5BR73 C5BR76 C5BRB6 C5BRG2 C5BRH0 C5BRH1 C5BRH7
C5BRJ3 C5BRJ7 C5BRK5 C5BRK9 C5BRL0 C5BRL2 C5BRM0 C5BRP4 C5BRQ7 C5BS65 C5BS66
C5BS80 C5BS84 C5BS85 C5BS89 C5BS91 C5BSE0 C5BSJ0 C5BSJ5 C5BSJ7 C5BSK5 C5BSQ1
C5BSQ5 C5BSQ6 C5BSW3 C5BSW4 C5BSX6 C5BSY8 C5BSY9 C5BSZ6 C5BSZ7 C5BT03 C5BTB0
C5BTI9 C5BTJ0 C5BTK7 C5BTP7 C5BTQ4 C5BTX7 C5BTY4 C5BU55 C5BU64 C5BU70 C5BUK2
C5BUK5 C5BUL6 C5BUN7 C5BUN8 C5BUN9 C5BUP0 C5BUP6 C5BUY7 C5BV35 C5BV37 C5BV40
C5BV49 C5BV64 C5BV85 C5BV99 C5BVA2 C5BVA6 C5BVB1 C5BW20 C5BW32 C5BW33 C5BW67
C5BW68 C5BW79 C5BWI5 C5BWL5 C5BWQ6 C5BWR2 C5BWR3 C5BWS2 C5BWS3 C5BWS6 C5BWT4
C5BWT8 C5BWU0 C5BWU1 C5BWW3 C5BWW7 C5BWX1 C5BXA3 C5BXF8 C5BXG1 C5BY82 C5BY94
C5BYB6 C5BYK3 C5BYN4 C5BYT3 C5BZB0 C5BZT2 C5BZX2 C5BZY7 C5C020 C5C046 C5C0G0
C5C0G1 C5C0G2 C5C0G3 C5C0G4 C5C0H3 C5C0H4 C5C0H6 C5C0H8 C5C0I0 C5C0I1 C5C0I4
C5C0I5 C5C0I6 C5C0I7 C5C0I8 C5C0J2 C5C0J3 C5C0J4 C5C0J5 C5C0J6 C5C0K2 C5C0K5
C5C0K8 C5C0K9 C5C0N2 C5C0T0 C5C0V6 C5C1B3 C5C1C4 C5C1S7 C5C1V1 C5C226 C5C233
C5C246 C5C2D6 C5C2F6 C5C2I2 C5C2I9 C5C333 C5C3D1 C5C3F6 C5C3G5 C5C3P2 C5C3W5
C5C3W6 C5C430 C5C483 C5C491 C5C494 C5C4E1 C5C4P4 C5C4W6 C5C4Y6 C5C5B0 C5C5I0
C5C5Q3 C5C5Q7 C5C5S1 C5C5S5 C5C5T3 C5C659 C5C661 C5C680 C5C681 C5C683 C5C687
C5C692 C5C697 C5C6A2 C5C6N4 C5C6N8 C5C6N9 C5C6P4 C5C6R2 C5C6R3 C5C713 C5C7U8
C5C7U9 C5C7V1 C5C7X2 C5C7X3 C5C7X4 C5C7X6 C5C7X8 C5C7X9 C5C878 C5C8D9 C5C8H9
C5C8I6 C5C8U1 C5C8U2 C5C8U3 C5C8U4 C5C8U6 C5C8X5 C5C916 C5C958 C5C9D7 C5C9E5
C5C9I3 C5C9Q4 C5C9Q5 C5C9Q7 C5C9S6 C5C9T1 C5C9T2 C5C9T9 C5C9U0 C5CA18 C5CA24
C5CA38 C5CA39 C5CA60 C5CA81 C5CAA3 C5CAC9 C5CAE0 C5CAE3 C5CAG7 C5CAH0 C5CAH1
C5CAI7 C5CAM5 C5CAN3 C5CAP8 C5CAP9 C5CAX2 C5CAZ3 C5CAZ4 C5CB51 C5CB88 C5CB93
C5CBB6 C5CBI2 C5CBJ8 C5CBJ9 C5CBK0 C5CBK1 C5CBK2 C5CBK6 C5CBP2 C5CBP5 C5CBS3
C5CBT9 C5CBU4 C5CBV0 C5CBV8 C5CBZ4 C5CC02 C5CC09 C5CC33 C5CC37 C5CC38 C5CC39
C5CC42 C5CC44 C5CC45 C5CC48 C5CC49 C5CC50 C5CC51 C5CC52 C5CC53 C5CC55 C5CC56
C5CC57 C5CC58 C5CC59 C5CC60 C5CC63 C5CC66 C5CC67 C5CC68 C5CC69 C5CC70 C5CC72
C5CC73 C5CC74 C5CC75 C5CC85 C5CC88 C5CCC6 C5CCD6 C5CCF1 C5CCF5 C5CCG1 C5CCG8
C5CCH1 C5CCH2 C5CCH4 C5CCI2 C5CCI4 C5CCM4 C5CCN0 C5CD32 C5CD39 C5CD61 C5CD66
C5CD67 C5CD93 C5CDH9 C5CDI0 C5CDI2 C5CDZ1 C5CDZ4 C5CE09 C5CE10 C5CE41 C5CEA8
C5CEB3 C5CEB5 C5CEB8 C5CEC0 C5CEL1 C5CEP5 C5CEQ3 C5CEQ4 C5CER8 C5CEY4 C5CEY9
C5CF28 C5CF30 C5CF34 C5CF44 C5CF51 C5CFE0 C5CFP6 C5CFP9 C5CFQ6 C5CFQ7 C5CFR2
C5CFR8 C5CFR9 C5CFS3 C5CFU0 C5CFW1 C5CFW2 C5CG17 C5CG19 C5CG73 C5CGC3 C5CGD9
C5CGE0 C5CGE1 C5CGE2 C5CGE3 C5CGE8 C5CGF0 C5CGG7 C5CGG9 C5CGH2 C5CGH3 C5CGH5
C5CGH6 C5CGH7 C5CGH8 C5CGI1 C5CGI3 C5CGI4 C5CGI6 C5CGI7 C5CGI8 C5CGI9 C5CGJ0
C5CGM3 C5CGP2 C5CGQ3 C5CGQ4 C5CGQ5 C5CGQ6 C5CGQ7 C5CGQ8 C5CGR0 C5CGR1 C5CGR2
C5CGR3 C5CGR4 C5CGR5 C5CGR6 C5CGR8 C5CGR9 C5CGS3 C5CGS4 C5CGS8 C5CGT1 C5CGT4
C5CGU7 C5CGV0 C5CGW6 C5CH91 C5CHA1 C5CHA8 C5CHB8 C5CHD1 C5CHD5 C5CHN3 C5CHP2
C5CHW7 C5CHW9 C5CHX7 C5CHX9 C5CI63 C5CI64 C5CI78 C5CI88 C5CI89 C5CI92 C5CIH1
C5CIJ2 C5CIS7 C5CIU4 C5CIU9 C5CIV1 C5CJ15 C5CJ68 C5CJ75 C5CJ85 C5CJB5 C5CJH0
C5CJT3 C5CJT4 C5CJT5 C5CKE4 C5CKE9 C5CKF0 C5CKF1 C5CKF2 C5CKF4 C5CKN7 C5CKN8
C5CKS0 C5CKS1 C5CKU4 C5CKU7 C5CKV9 C5CKW1 C5CL68 C5CL71 C5CL84 C5CL95 C5CLI6

C5CLL8 C5CLM9 C5CLQ3 C5CLQ6 C5CLT2 C5CLT4 C5CLU5 C5CLW1 C5CLW3 C5CLW4 C5CM03
C5CM11 C5CM14 C5CM20 C5CM23 C5CM24 C5CM29 C5CM74 C5CM90 C5CMK6 C5CN79 C5CN80
C5CN81 C5CN82 C5CN84 C5CND4 C5CNE3 C5CNE5 C5CNE6 C5CNF5 C5CNG5 C5CNK9 C5CNM3
C5CNS8 C5CNS9 C5CNW9 C5CNX5 C5CP12 C5CP20 C5CP21 C5CP44 C5CP46 C5CP48 C5CP49
C5CP50 C5CP52 C5CP53 C5CP56 C5CP58 C5CP59 C5CP60 C5CP83 C5CPF1 C5CPN2 C5CPP8
C5CPP9 C5CPR2 C5CPT7 C5CPX0 C5CPY0 C5CPY1 C5CPZ2 C5CPZ9 C5CQ74 C5CQ75 C5CQ76
C5CQ77 C5CQ78 C5CQ79 C5CQ81 C5CQ83 C5CQ85 C5CQ86 C5CQ88 C5CQ89 C5CQ90 C5CQB2
C5CQC9 C5CQE1 C5CQF1 C5CQG3 C5CQI8 C5CQI9 C5CQJ5 C5CQK0 C5CQK2 C5CRA3 C5CRK1
C5CRM0 C5CRQ4 C5CRT0 C5CRW8 C5CS25 C5CS95 C5CSC4 C5CSE0 C5CSF5 C5CSF7 C5CSG3
C5CSH0 C5CSH4 C5CSH5 C5CSI5 C5CSI6 C5CSK7 C5CSM6 C5CSN3 C5CSQ9 C5CSV3 C5CT24
C5CT25 C5CT30 C5CT60 C5CTG9 C5CTL8 C5CU07 C5CU12 C5CU29 C5CUG9 C5CUK1 C5CUL9
C5CUN1 C5CUN7 C5CUP0 C5CUP2 C5CUP3 C5CUU2 C5CUU4 C5CUU5 C5CUU6 C5CUU9 C5CUX0
C5CV58 C5CVT2 C5CW10 C5CW30 C5CWN5 C5CWP0 C5CWV0 C5CWV9 C5CWX8 C5CWZ4 C5CX88
C5CXC5 C5CXD8 C5CXG2 C5CXH0 C5CXH1 C5CXH6 C5CXH7 C5CXR6 C5CXW3 C5CXW9 C5CXX0
C5CXX2 C5CXX4 C5CXX5 C5CXZ7 C5CXZ9 C5CY00 C5CY05 C5CY39 C5CY41 C5CYN9 C5CYP2
C5CYP5 C5CYY0 C5CYY5 C5CYY6 C5CYY8 C5CYZ1 C5CYZ4 C5CYZ6 C5CYZ7 C5CZC3 C5D2F9
C5D2I1 C5D2I2 C5D2K0 C5D2K6 C5D2N0 C5D2V1 C5D2V2 C5D2V7 C5D324 C5D327 C5D330
C5D337 C5D338 C5D340 C5D346 C5D347 C5D352 C5D355 C5D363 C5D367 C5D370 C5D373
C5D374 C5D391 C5D392 C5D3B1 C5D3B2 C5D3B3 C5D3B5 C5D3B8 C5D3D2 C5D3D3 C5D3D6
C5D3D7 C5D3E0 C5D3E1 C5D3E5 C5D3E7 C5D3E8 C5D3F4 C5D3G3 C5D3G6 C5D3L0 C5D3M9
C5D3N0 C5D3N8 C5D3P3 C5D3P6 C5D3Q4 C5D3Q5 C5D3Q6 C5D3R1 C5D3R2 C5D3R3 C5D3R4
C5D3R5 C5D3R6 C5D3R7 C5D3R8 C5D3R9 C5D3S0 C5D3S1 C5D3S2 C5D3S3 C5D3S4 C5D3S5
C5D3S6 C5D3S7 C5D3S8 C5D3S9 C5D3T0 C5D3T1 C5D3T3 C5D3T4 C5D3T5 C5D3T6 C5D3T8
C5D3U1 C5D3U2 C5D3U3 C5D3U5 C5D3U9 C5D3V0 C5D3V1 C5D403 C5D427 C5D428 C5D465
C5D467 C5D486 C5D492 C5D4A1 C5D4A2 C5D4D8 C5D4E6 C5D4E7 C5D4F3 C5D4F4 C5D4H9
C5D4I0 C5D4I2 C5D4J5 C5D4K0 C5D4K1 C5D4K5 C5D4K6 C5D4Q3 C5D4R0 C5D4R3 C5D4S1
C5D4S3 C5D4S8 C5D4T4 C5D4T6 C5D4U0 C5D4U1 C5D4U3 C5D4U8 C5D4V0 C5D4V7 C5D4W0
C5D4W2 C5D4X9 C5D4Y0 C5D4Y5 C5D4Z0 C5D4Z1 C5D4Z2 C5D509 C5D510 C5D512 C5D514
C5D571 C5D585 C5D586 C5D587 C5D5A6 C5D5A7 C5D5A9 C5D5E6 C5D5E8 C5D5E9 C5D5F1
C5D5F6 C5D5G7 C5D5G9 C5D5H7 C5D5H8 C5D5K7 C5D5L1 C5D5L4 C5D5L5 C5D5L7 C5D5L8
C5D5M0 C5D5M1 C5D5M7 C5D5M9 C5D5Q8 C5D5R2 C5D5S0 C5D5S3 C5D5T7 C5D5V2 C5D627
C5D630 C5D636 C5D637 C5D639 C5D643 C5D646 C5D654 C5D663 C5D672 C5D678 C5D679
C5D683 C5D693 C5D699 C5D6B2 C5D6B6 C5D6C2 C5D6D7 C5D6E9 C5D6I2 C5D6J1 C5D6K5
C5D6L2 C5D6L6 C5D6M5 C5D6M6 C5D6R2 C5D6U5 C5D720 C5D729 C5D742 C5D754 C5D764
C5D766 C5D781 C5D7A2 C5D7A6 C5D7C7 C5D7H5 C5D7J9 C5D7L1 C5D7L7 C5D7M4 C5D7M9
C5D7N1 C5D7N8 C5D7P0 C5D7P1 C5D7P3 C5D7P4 C5D7Q0 C5D7T4 C5D7T5 C5D7U5 C5D7U6
C5D7U9 C5D7V1 C5D7Y2 C5D802 C5D826 C5D827 C5D828 C5D830 C5D846 C5D8K0 C5D8K3
C5D8K7 C5D8K9 C5D8L4 C5D8L5 C5D8M2 C5D8N8 C5D8P2 C5D8P5 C5D8P7 C5D8P8 C5D8Q1
C5D8Q2 C5D8Q3 C5D8R0 C5D8R2 C5D8R6 C5D8S1 C5D8S5 C5D8T6 C5D8U1 C5D8U4 C5D8U6
C5D8U8 C5D8U9 C5D8V2 C5D8V8 C5D8V9 C5D8W0 C5D9B5 C5D9B6 C5D9B8 C5D9C1 C5D9C3
C5D9C5 C5D9C9 C5D9D1 C5D9D4 C5D9D5 C5D9G1 C5D9H5 C5D9H6 C5D9J0 C5D9K6 C5D9M4
C5D9N4 C5D9N8 C5D9P0 C5D9P4 C5D9R0 C5D9S1 C5D9S7 C5D9T3 C5D9W1 C5D9W9 C5D9X1
C5D9X4 C5D9X6 C5D9Y4 C5D9Y5 C5D9Z1 C5D9Z2 C5D9Z3 C5DA06 C5DA16 C5DA19 C5DA20
C5DAC8 C5DAE0 C5DAG6 C5DAN8 C5DAP4 C5DAQ1 C5DAR2 C5DAS5 C5DBR4 C5DBS6 C5DBV2

C5DCA2 C5DCN3 C5DCN9 C5DD02 C5DD31 C5DD42 C5DDJ3 C5DDM1 C5DDQ4 C5DDS4 C5DDT8
C5DE09 C5DE35 C5DE55 C5DEW2 C5DF10 C5DF48 C5DF79 C5DFG5 C5DFL1 C5DFM0 C5DFS5
C5DGD8 C5DGE2 C5DGG4 C5DGI8 C5DGQ7 C5DGT3 C5DGU9 C5DGZ5 C5DHJ1 C5DHJ9 C5DHL1
C5DHM2 C5DHP1 C5DHP6 C5DHU5 C5DIF1 C5DII3 C5DIR2 C5DJ44 C5DJG1 C5DJH5 C5DJH6
C5DJK6 C5DJL0 C5DJL7 C5DJV1 C5DKA0 C5DKJ7 C5DKM2 C5DLA7 C5DLH0 C5DLJ0 C5DMC1
C5DMF2 C5DMI3 C5DMP2 C5DMU9 C5DMW3 C5DN02 C5DN25 C5DP04 C5DP87 C5DPB7 C5DPS9
C5DQD7 C5DQW8 C5DRG2 C5DRI2 C5DRT0 C5DS11 C5DS56 C5DS65 C5DSC5 C5DSC9 C5DSD6
C5DSN0 C5DSV0 C5DT56 C5DT65 C5DTH5 C5DTI7 C5DTN0 C5DTR1 C5DU00 C5DU12 C5DUB3
C5DUI6 C5DUX2 C5DV23 C5DVG6 C5DWI2 C5DWK1 C5DX17 C5DX31 C5DXI0 C5DXN2 C5DXS6
C5DXZ1 C5DY07 C5DY19 C5DY20 C5DY36 C5DY61 C5DY71 C5DYE7 C5DYJ0 C5DYK5 C5DYR7
C5DYV8 C5DZ92 C5DZA9 C5DZE1 C5DZI5 C5DZT2 C5DZX2 C5E000 C5E042 C5E0B9 C5E0I9
C5E0N8 C5E1C0 C5E1C7 C5E1D0 C5E1J9 C5E268 C5E2K7 C5E2Q7 C5E2S6 C5E2U8 C5E3I1
C5E3P3 C5E3X9 C5E3Y4 C5E490 C5E4G1 C5E4I6 C5E4K0 C5E4V9 C5FC83 C5FD74 C5FEJ1
C5FEJ4 C5FF22 C5FF46 C5FFE1 C5FFQ7 C5FHR9 C5FHS6 C5FHU9 C5FIC5 C5FL48 C5FM57
C5FM58 C5FM60 C5FM61 C5FMD4 C5FNG9 C5FP02 C5FP68 C5FQ73 C5FQT8 C5FRS7 C5FS59
C5FSF3 C5FSR3 C5FSW4 C5FTM9 C5FTN1 C5FTN2 C5FTN3 C5FTT1 C5FUV0 C5FWH1 C5FY68
C5FZJ2 C5FZT5 C5G0N1 C5G6H8 C5G7E1 C5G874 C5G8P1 C5G8R4 C5G8V2 C5GA45 C5GAH0
C5GBK9 C5GCN0 C5GE44 C5GEF3 C5GF27 C5GFG2 C5GGA4 C5GGE5 C5GKT2 C5GLJ6 C5GLN0
C5GPA7 C5GPL2 C5GS07 C5GVJ9 C5GW39 C5GXZ4 C5GXZ9 C5GYL7 C5H881 C5H882 C5H883
C5H885 C5H8J1 C5I9X1 C5IY43 C5IY45 C5IY48 C5J049 C5J3R8 C5J697 C5J6H5 C5J7W8
C5JD40 C5JD82 C5JGS0 C5JIC2 C5JIS5 C5JJ91 C5JJW0 C5JK19 C5JKE6 C5JKJ6 C5JKQ2
C5JMX0 C5JQ04 C5JR12 C5JS82 C5JT39 C5JUE6 C5JVC1 C5JVG7 C5JW60 C5JYW1 C5JYZ5
C5JZ38 C5K0R2 C5K105 C5K110 C5K3T8 C5M1X2 C5M276 C5M2B8 C5M2G0 C5M2G8 C5M2I3
C5M2X8 C5M3P0 C5M3V9 C5M444 C5M5D9 C5M5Z6 C5M614 C5M685 C5M6H7 C5M799 C5M8M4
C5M9K8 C5M9N0 C5MA45 C5MAP5 C5MBA7 C5MBD4 C5MBF2 C5MCK5 C5MCY0 C5ME52 C5MED4
C5MFB4 C5MFG4 C5MG55 C5MH59 C5MHJ2 C5MHS7 C5MIB1 C5MJE8 C5MRT8 C5NU54 C5P263
C5P3I5 C5P441 C5P5Z4 C5P664 C5P7J2 C5P7K2 C5P7X2 C5P8M7 C5P8Q4 C5P9D1 C5PA09
C5PA86 C5PAD0 C5PAH2 C5PBM5 C5PBQ8 C5PDS9 C5PFU3 C5PFX0 C5PHH6 C5PHM7 C5PIC1
C5PIN8 C5VXY0 C5VZF1 C5VZR6 C5W331 C5W452 C5W7D8 C5W7S9 C5W865 C5W8B2 C5WBK5
C5WIJ4 C5WNV2 C5WPC2 C5WR30 C5WU23 C5WVW2 C5WWY0 C5WYI7 C5WZH0 C5X1F5 C5X2M4
C5XKZ1 C5XNN6 C5XX79 C5XYW4 C5XZK6 C5Y210 C5Y3V8 C5Y8Z8 C5Y9R0 C5YHH7 C5YHI2
C5YUK3 C5Z7K4 C5ZMD2 C5ZZZ3 C5ZZZ5 C6A011 C6A032 C6A048 C6A0F5 C6A0I0 C6A0I8
C6A0N1 C6A0R1 C6A0S1 C6A0S4 C6A0T3 C6A0Y7 C6A0Z6 C6A116 C6A117 C6A124 C6A159
C6A160 C6A161 C6A163 C6A165 C6A166 C6A168 C6A175 C6A177 C6A179 C6A184 C6A186
C6A187 C6A188 C6A1B1 C6A1B2 C6A1B6 C6A1F5 C6A1F7 C6A1F8 C6A1G2 C6A1I4 C6A1J2
C6A1J4 C6A1K8 C6A1K9 C6A1L0 C6A1L5 C6A1R1 C6A1R5 C6A1T2 C6A1U6 C6A1U9 C6A1V3
C6A1Y5 C6A1Z6 C6A222 C6A229 C6A236 C6A237 C6A238 C6A243 C6A264 C6A267 C6A270
C6A2A0 C6A2D7 C6A2E8 C6A2F6 C6A2L3 C6A2L5 C6A2P3 C6A2P7 C6A2Q1 C6A2R5 C6A2V9
C6A2Y3 C6A312 C6A317 C6A366 C6A382 C6A390 C6A3E1 C6A3F2 C6A3G4 C6A3M0 C6A3Q6
C6A3R1 C6A3U0 C6A3U9 C6A3X6 C6A429 C6A439 C6A452 C6A459 C6A460 C6A465 C6A4A3
C6A4A4 C6A4F1 C6A4G9 C6A4M0 C6A4N4 C6A4R7 C6A4T2 C6A4W3 C6A5A3 C6A5B5 C6A5C2
C6A5J5 C6A5J6 C6A7B2 C6A7B6 C6A8W4 C6A9K7 C6AEK6 C6AHX0 C6AHX4 C6AJH2 C6AKG2
C6AN26 C6ANQ9 C6AQR4 C6AQU6 C6AR33 C6AR38 C6AR50 C6ARU7 C6BAU4 C6BEJ1 C6BQP8
C6BRG8 C6BRG9 C6BRH1 C6BRT2 C6BRV2 C6BS14 C6BS23 C6BS43 C6BSC1 C6BSC2 C6BSD8

C6BSE1 C6BSE3 C6BSE9 C6BSF2 C6BSG3 C6BSH0 C6BSH4 C6BSL0 C6BSL6 C6BSW0 C6BSW3
C6BSW6 C6BSW8 C6BTR8 C6BTS9 C6BTT9 C6BTU7 C6BTU8 C6BTW3 C6BTW9 C6BU27 C6BU53
C6BU59 C6BU87 C6BUE5 C6BUE7 C6BUG4 C6BUG6 C6BUI7 C6BUK3 C6BUM7 C6BUS3 C6BUX8
C6BV55 C6BV58 C6BV66 C6BV68 C6BV69 C6BV77 C6BV89 C6BV99 C6BVA6 C6BVA9 C6BVB0
C6BVE8 C6BVJ0 C6BVJ9 C6BVN3 C6BVP4 C6BVQ1 C6BVR1 C6BVV4 C6BW12 C6BW21 C6BW35
C6BW42 C6BW69 C6BWN7 C6BX34 C6BXD5 C6BXE6 C6BXG0 C6BXG1 C6BXI4 C6BXJ6 C6BY79
C6BYB7 C6BYC7 C6BYD8 C6BYE0 C6BYG5 C6BYH4 C6BYH5 C6BYK5 C6BYK9 C6BYL8 C6BYN5
C6BYN6 C6BYQ4 C6BYQ6 C6BYX0 C6BYX7 C6BYY0 C6BZ93 C6BZ95 C6BZ96 C6BZQ4 C6BZY6
C6BZZ8 C6C000 C6C030 C6C060 C6C065 C6C067 C6C068 C6C0A2 C6C0A9 C6C0B2 C6C0C8
C6C0D6 C6C0E9 C6C0F0 C6C0F7 C6C0G0 C6C0I6 C6C0J4 C6C0L8 C6C0N2 C6C0P0 C6C0P1
C6C0T2 C6C0U6 C6C0U7 C6C0V0 C6C0W4 C6C125 C6C171 C6C175 C6C176 C6C177 C6C178
C6C179 C6C181 C6C182 C6C184 C6C185 C6C186 C6C187 C6C188 C6C189 C6C190 C6C191
C6C192 C6C193 C6C194 C6C195 C6C197 C6C198 C6C1A0 C6C1A2 C6C1A3 C6C1A4 C6C1A7
C6C1A8 C6C1A9 C6C1B0 C6C1B1 C6C1B2 C6C1D1 C6C1D3 C6C1D5 C6C1F4 C6C1J9 C6C1K0
C6C1K6 C6C1M7 C6C1T0 C6C1T1 C6C1T6 C6C1U5 C6C1U6 C6C1X6 C6C2Z3 C6C3U9 C6C4V4
C6C5A0 C6C608 C6C9K0 C6CAS5 C6CAW9 C6CB42 C6CBG9 C6CC07 C6CEU3 C6CFY1 C6CG41
C6CJW6 C6CL12 C6CLS0 C6CNL2 C6CVY9 C6CWS7 C6CXN6 C6D563 C6D8W9 C6D8X0 C6D8X1
C6D8Y4 C6D8Y7 C6D8Z5 C6D8Z6 C6D970 C6D972 C6D978 C6D9B8 C6D9C1 C6D9J8 C6D9M4
C6D9M5 C6D9P9 C6D9Q2 C6D9S0 C6DA17 C6DA49 C6DA52 C6DA54 C6DA76 C6DA85 C6DA87
C6DAC3 C6DAE5 C6DAE6 C6DAE9 C6DAG1 C6DAH1 C6DAH2 C6DAH4 C6DAH9 C6DAI1 C6DAI3
C6DAI4 C6DAI6 C6DAI7 C6DAJ7 C6DAK3 C6DAK6 C6DAM1 C6DAX0 C6DAX2 C6DAZ5 C6DB04
C6DB05 C6DB25 C6DB26 C6DB31 C6DB33 C6DB34 C6DB37 C6DB39 C6DB40 C6DB43 C6DB54
C6DB56 C6DB62 C6DB75 C6DB82 C6DB84 C6DB85 C6DB87 C6DBE0 C6DBG3 C6DBG5 C6DBH0
C6DBH3 C6DBH4 C6DBH8 C6DBI4 C6DBI7 C6DBI8 C6DBI9 C6DBJ1 C6DBJ2 C6DBN5 C6DBP2
C6DBP5 C6DBR0 C6DBR1 C6DBS2 C6DBV6 C6DBV7 C6DBV8 C6DBW3 C6DBW8 C6DBX6 C6DBY4
C6DBY6 C6DC00 C6DC08 C6DC10 C6DC11 C6DC27 C6DC32 C6DC42 C6DC47 C6DC48 C6DC49
C6DC58 C6DC59 C6DC61 C6DC65 C6DC67 C6DCC6 C6DCC7 C6DCC8 C6DCD6 C6DCF2 C6DCF6
C6DCH7 C6DCI4 C6DCI6 C6DCI7 C6DCP8 C6DCP9 C6DCQ0 C6DCQ1 C6DCQ5 C6DCQ8 C6DCR1
C6DCX4 C6DCX5 C6DCX9 C6DCY7 C6DCZ0 C6DCZ1 C6DCZ3 C6DCZ4 C6DCZ7 C6DD82 C6DDF6
C6DDF9 C6DDG0 C6DDG1 C6DDG2 C6DDH0 C6DDH2 C6DDH3 C6DDJ6 C6DDJ9 C6DDL3 C6DDL7
C6DDM6 C6DDN6 C6DDR1 C6DDS6 C6DDS7 C6DDU0 C6DE12 C6DE13 C6DE14 C6DE15 C6DE26
C6DE33 C6DE79 C6DED2 C6DEE2 C6DEF0 C6DEI9 C6DEJ0 C6DEJ2 C6DEK5 C6DEL1 C6DEM6
C6DEP1 C6DEP6 C6DEQ3 C6DEQ9 C6DET1 C6DET2 C6DEU1 C6DEU2 C6DEV1 C6DEV2 C6DEV8
C6DEW0 C6DEW1 C6DEY0 C6DEY6 C6DEY7 C6DEY8 C6DF00 C6DF01 C6DF04 C6DF06 C6DF09
C6DF10 C6DF16 C6DF18 C6DF26 C6DF27 C6DF34 C6DF35 C6DF39 C6DF43 C6DF46 C6DF54
C6DF64 C6DF65 C6DF66 C6DF68 C6DF71 C6DF72 C6DF75 C6DF77 C6DF85 C6DF89 C6DF90
C6DF99 C6DFA3 C6DFB1 C6DFB8 C6DFD0 C6DFD6 C6DFD9 C6DFE1 C6DFE2 C6DFE7 C6DFE9
C6DFF0 C6DFF1 C6DFF2 C6DFH7 C6DFH8 C6DFI3 C6DFI4 C6DFI8 C6DFI9 C6DFJ2 C6DFJ7
C6DFJ8 C6DFJ9 C6DFN0 C6DFN3 C6DFP1 C6DFP4 C6DFQ3 C6DFQ6 C6DFR0 C6DFR3 C6DFR5
C6DFR6 C6DFR7 C6DFS2 C6DFS4 C6DFS5 C6DFS6 C6DFT2 C6DFU6 C6DFW1 C6DFW7 C6DFY6
C6DFY7 C6DFY8 C6DFY9 C6DFZ0 C6DFZ2 C6DFZ6 C6DG29 C6DG36 C6DG38 C6DG39 C6DG41
C6DG55 C6DG58 C6DG59 C6DG60 C6DG62 C6DG63 C6DG64 C6DG65 C6DG66 C6DG67 C6DG68
C6DG69 C6DG70 C6DG71 C6DG73 C6DG74 C6DG75 C6DG80 C6DG81 C6DG82 C6DG83 C6DG84
C6DG85 C6DG94 C6DGA2 C6DGD4 C6DGE7 C6DGE8 C6DGH6 C6DGH7 C6DGH9 C6DGK1 C6DGL0

C6DGL1 C6DGR7 C6DGU9 C6DGV2 C6DGV8 C6DGW9 C6DGY2 C6DGZ4 C6DGZ5 C6DGZ7 C6DH19
C6DH20 C6DH28 C6DH49 C6DH68 C6DH73 C6DH76 C6DH77 C6DH78 C6DHB7 C6DHB9 C6DHC2
C6DHF5 C6DHF6 C6DHG3 C6DHH1 C6DHK2 C6DHM3 C6DHM4 C6DHM6 C6DHM7 C6DHN7 C6DHQ1
C6DHQ3 C6DHQ6 C6DHR1 C6DHR2 C6DHR3 C6DHR4 C6DHR5 C6DHS2 C6DHS7 C6DHS8 C6DHS9
C6DHT0 C6DHT5 C6DHY1 C6DHY2 C6DHY4 C6DHY7 C6DI64 C6DI71 C6DI77 C6DI79 C6DI80
C6DI81 C6DI82 C6DI84 C6DI86 C6DIA3 C6DIA7 C6DIA9 C6DIB6 C6DIB8 C6DIB9 C6DIC0
C6DIC1 C6DIC3 C6DIC4 C6DIC6 C6DIC7 C6DIG9 C6DIH5 C6DIH9 C6DII4 C6DII7 C6DII8
C6DIJ6 C6DIJ9 C6DIK5 C6DIM5 C6DIN0 C6DIP6 C6DIQ1 C6DIQ2 C6DIR5 C6DIR6 C6DIS1
C6DJ78 C6DJB2 C6DJC3 C6DJC5 C6DJD2 C6DJE0 C6DJE1 C6DJF3 C6DJF8 C6DJF9 C6DJG0
C6DJG3 C6DJG4 C6DJL2 C6DJL3 C6DJM6 C6DJN2 C6DJR0 C6DJR1 C6DJR2 C6DJR3 C6DJR4
C6DJR5 C6DJU6 C6DK05 C6DK13 C6DK25 C6DK28 C6DK30 C6DK36 C6DK50 C6DK97 C6DKA0
C6DKC6 C6DKC7 C6DKC8 C6DKE7 C6DKE8 C6DKG5 C6DKG8 C6DKG9 C6DKH0 C6DKH1 C6DKH6
C6DKH7 C6DKI3 C6DKI6 C6DKK3 C6DKK4 C6DKK6 C6DKK7 C6DKL5 C6DKL9 C6DKM0 C6DKM5
C6DKM6 C6DKP3 C6DKP5 C6DKQ4 C6DKR7 C6DKR8 C6DKS8 C6DKT7 C6DKU3 C6DKU9 C6DKX0
C6DKX3 C6DKX8 C6DKY4 C6DKY5 C6DKY6 C6DPT0 C6DPU6 C6DPV1 C6DPV2 C6DPW7 C6DQZ2
C6DT68 C6DUY7 C6DV63 C6DVC4 C6DW66 C6DY35 C6DY36 C6DY42 C6DY43 C6DYB7 C6DYE1
C6DYE3 C6DYJ1 C6DYJ2 C6DYJ3 C6DYJ5 C6DYM2 C6DYM5 C6DYR8 C6DYS3 C6DYS4 C6DYX1
C6DYX9 C6DYY7 C6DYY8 C6DZ51 C6DZ52 C6DZ53 C6DZ58 C6DZH8 C6DZJ7 C6DZJ8 C6DZK4
C6DZK7 C6DZK9 C6DZQ7 C6DZZ3 C6E082 C6E087 C6E091 C6E0B4 C6E0C4 C6E0H6 C6E0S2
C6E0V3 C6E0X3 C6E0Y2 C6E0Y6 C6E100 C6E105 C6E1L8 C6E1P3 C6E1S8 C6E1S9 C6E1T0
C6E1T1 C6E228 C6E229 C6E2B1 C6E2B2 C6E2B3 C6E2G6 C6E2G9 C6E2H4 C6E2L2 C6E2P3
C6E2P5 C6E2P6 C6E2P9 C6E2Q0 C6E2Q3 C6E2S9 C6E2T0 C6E2T1 C6E2U6 C6E2U7 C6E310
C6E348 C6E375 C6E3M1 C6E3M5 C6E3M6 C6E3N5 C6E3N6 C6E3V4 C6E3W7 C6E487 C6E496
C6E497 C6E4K4 C6E4K5 C6E4L2 C6E4M9 C6E4N1 C6E4N3 C6E4N4 C6E4N6 C6E4N8 C6E4N9
C6E4P1 C6E4P3 C6E4P4 C6E4P5 C6E4P6 C6E4P7 C6E4P8 C6E4P9 C6E4Q0 C6E4Q1 C6E4Q2
C6E4Q3 C6E4Q4 C6E4Q5 C6E4Q6 C6E4Q7 C6E4Q8 C6E4R1 C6E4R2 C6E4R3 C6E4R4 C6E4R5
C6E4R6 C6E4R8 C6E4S6 C6E4S8 C6E4S9 C6E4U6 C6E4W6 C6E500 C6E501 C6E503 C6E504
C6E506 C6E512 C6E513 C6E515 C6E518 C6E521 C6E543 C6E557 C6E559 C6E582 C6E585
C6E5H7 C6E5I4 C6E5I6 C6E5I8 C6E5J1 C6E5P5 C6E5P6 C6E5S6 C6E5Z3 C6E5Z5 C6E5Z6
C6E639 C6E643 C6E644 C6E657 C6E679 C6E689 C6E697 C6E6C1 C6E6C5 C6E6G4 C6E6I5
C6E6M9 C6E6N2 C6E6X2 C6E6X4 C6E6X6 C6E6Y1 C6E6Y6 C6E6Y9 C6E6Z1 C6E6Z2 C6E6Z3
C6E6Z4 C6E751 C6E770 C6E791 C6E7C1 C6E7E6 C6E7E7 C6E7F2 C6E7F4 C6E7F5 C6E7F7
C6E7G0 C6E7G6 C6E7L6 C6E7L8 C6E7L9 C6E7M0 C6E7P0 C6E7Q4 C6E7Q5 C6E7Q7 C6E7R9
C6E7U2 C6E7Z3 C6E805 C6E808 C6E870 C6E8C5 C6E8U1 C6E8U2 C6E8Y9 C6E916 C6E9G2
C6E9Q7 C6E9Q8 C6E9Q9 C6EHJ5 C6EHJ6 C6EHJ7 C6EHJ8 C6EHK0 C6EN00 C6FG12 C6FX52
C6GPU1 C6GW75 C6H178 C6H208 C6H2R1 C6H7R7 C6H861 C6HBB1 C6HBW5 C6HCG7 C6HGN3
C6HHV6 C6HI66 C6HLY2 C6HNY5 C6HQ17 C6HQ96 C6HQJ2 C6HRN2 C6HSY3 C6HSY8 C6HTE8
C6JS30 C6JWH0 C6KEM4 C6KI62 C6KRL6 C6KRN1 C6KSQ6 C6KSS5 C6KT50 C6KT68 C6KTD2
C6KYS2 C6L7U1 C6L7V8 C6L7V9 C6L862 C6LR75 C6LTG5 C6S3N3 C6SPT0 C6SXE1 C6T0L2
C6TAY1 C6TBN2 C6TEX6 C6TKZ5 C6UBI3 C6UCN7 C6UCT5 C6UFB8 C6UFC0 C6UFC1 C6UFC2
C6UFC3 C6UG39 C6UJK3 C6UM45 C6UMS6 C6UP45 C6UPM9 C6UPN1 C6UPN3 C6UPN4 C6UQ16
C6UQY1 C6USC2 C6V598 C6VS84 C6W9U8 C6WFL3 C6WIA0 C6WIC8 C6WID8 C6WID9 C6WIE4
C6WKX3 C6WPK3 C6WPR7 C6WX20 C6X2D2 C6X9I5 C6Y2G0 C6Y4A2 C6Y4A5 C6Y4B0 C6Y4B1
C6Y4B2 C6Y4B4 C6Y4B8 C6Y4B9 C6Y4C0 C6Y4C3 C6Y4C4 C6Y4C5 C6Y4D0 C6Y4D2 C6Y4D3

C6Y4D4 C6YCI2 C6ZJZ3 C7A278 C7A2A0 C7A639 C7AE94 C7AJA4 C7ASI9 C7ASJ5 C7AU21
C7BGY4 C7BKL0 C7BSD3 C7BZ94 C7BZQ3 C7C435 C7C436 C7C8V4 C7CFQ2 C7CM32 C7CM33
C7CM34 C7CM35 C7CM36 C7E3T4 C7E5V7 C7E5V8 C7E5V9 C7E5W0 C7EMF5 C7FF04 C7FF05
C7G046 C7G304 C7GIN5 C7GIQ7 C7GIZ9 C7GJ78 C7GJB0 C7GJG1 C7GJG9 C7GJX9 C7GJZ2
C7GK54 C7GKE1 C7GKF3 C7GKL8 C7GKY0 C7GL41 C7GL43 C7GL60 C7GL62 C7GLC0 C7GLG0
C7GLH8 C7GLQ0 C7GMC5 C7GMH9 C7GMI9 C7GML8 C7GMR4 C7GNV5 C7GP13 C7GP20 C7GP35
C7GPA7 C7GPC1 C7GPD7 C7GPE4 C7GPS8 C7GPZ8 C7GQ59 C7GQE4 C7GQI8 C7GQR4 C7GQV5
C7GQY3 C7GR11 C7GRF7 C7GRM0 C7GRM1 C7GRQ3 C7GRR3 C7GS66 C7GS77 C7GSF3 C7GSL4
C7GSL9 C7GTF9 C7GTL7 C7GTR2 C7GUH8 C7GUN6 C7GV13 C7GV42 C7GVJ8 C7GVL2 C7GVL7
C7GVQ9 C7GVS1 C7GVT8 C7GVY4 C7GVZ9 C7GWA2 C7GWA5 C7GWC1 C7GWG7 C7GWI3 C7GWJ7
C7GX58 C7GX69 C7GX75 C7GXS8 C7GXT0 C7GXX9 C7GYD5 C7GYE7 C7IVR4 C7IW64 C7J0A2
C7J2Z1 C7J8E5 C7JGP5 C7LJY3 C7LNQ3 C7LYP4 C7LYP7 C7LYP8 C7LYP9 C7M077 C7M0E4
C7MCY7 C7MCZ0 C7MCZ4 C7MGB3 C7MRC4 C7MSY6 C7MWV3 C7MWV6 C7MWV7 C7MWW2 C7MWX8
C7MY07 C7N8L9 C7NB67 C7NBY4 C7NCT6 C7NHQ9 C7NK43 C7NN78 C7NNA6 C7NQ99 C7NQP8
C7NST3 C7NTR1 C7NVQ4 C7NYH7 C7NZH8 C7P0W1 C7P2N1 C7P363 C7P4K0 C7P544 C7P592
C7P607 C7P6N4 C7P7L7 C7P7V6 C7P8V7 C7PC56 C7PDD8 C7PLM6 C7PLV2 C7PVU9 C7PVV2
C7PVV3 C7PVV4 C7PW26 C7Q0R3 C7Q4X5 C7Q4Y6 C7Q942 C7Q9B2 C7QJ42 C7QJN7 C7QKE2
C7QKE8 C7QKH8 C7R101 C7R1N6 C7R3Z4 C7R400 C7R403 C7R404 C7R405 C7RB40 C7RGH4
C7S340 C7SDN9 C7SG33 C7TEQ4 C7TMK0 C7U330 C7U331 C7UDU4 C7YHA8 C7YJ03 C7YK87
C7YKN8 C7YLQ3 C7YM38 C7YPY3 C7YQ16 C7YRJ1 C7YRR5 C7YRS7 C7YRT4 C7YS77 C7YS90
C7YSE7 C7YTD6 C7YTE3 C7YTF5 C7YTL6 C7YUB3 C7YVN8 C7YZ74 C7Z0Q8 C7Z125 C7Z147
C7Z193 C7Z508 C7Z527 C7Z622 C7Z638 C7Z6H2 C7Z7V7 C7Z837 C7Z8P6 C7Z9X4 C7Z9Z4
C7Z9Z7 C7ZBE4 C7ZBE5 C7ZC16 C7ZKL5 C7ZKL6 C7ZPG2 C8BKB7 C8BKC7 C8BKD0 C8NA81
C8TNB7 C8TNB9 C8TNC0 C8TNC1 C8TNC2 C8U5G9 C8U5H1 C8U5H2 C8U5H3 C8U5H4 C8UMM3
C8UMM5 C8UMM6 C8UMM7 C8UMM8 C8V1D1 C8V212 C8V213 C8V330 C8V3N0 C8V3W5 C8V3W6
C8V3Y7 C8V4D4 C8V4J5 C8V4J7 C8V7P4 C8V8H4 C8V8H5 C8V9B7 C8V9R7 C8V9Y5 C8VBH3
C8VBH4 C8VC05 C8VCM2 C8VDI1 C8VDQ3 C8VDQ4 C8VE82 C8VEJ5 C8VG90 C8VI80 C8VJ35
C8VJQ0 C8VJQ1 C8VJQ3 C8VJQ9 C8VJR6 C8VJR7 C8VP36 C8VP37 C8VP82 C8VPE5 C8VPS9
C8VPT2 C8VQ61 C8VQ62 C8VQ63 C8VQ71 C8VQ72 C8VQ92 C8VQ93 C8VSE8 C8VSZ2 C8VTR6
C8WAF3 C8WJZ5 C8WR67 C8WV58 C8X8S1 C8XA09 C8XA55 C8XAP5 C8XAQ3 C8XAQ4 C8XAR0
C8XBB4 C8XJN6 C8XPS0 C8YR32 C8Z3R4 C8Z3V4 C8Z430 C8Z4Y4 C8Z4Z5 C8Z543 C8Z5L6
C8Z5R9 C8Z779 C8Z792 C8Z7C7 C8Z7W7 C8Z7X8 C8Z8G8 C8Z944 C8Z9U3 C8Z9V6 C8Z9Z8
C8Z9Z9 C8ZA35 C8ZAF8 C8ZAJ0 C8ZAQ2 C8ZAR0 C8ZAU6 C8ZB86 C8ZBB1 C8ZBK1 C8ZBM7
C8ZBW6 C8ZC62 C8ZC77 C8ZC82 C8ZCB1 C8ZCJ2 C8ZCN3 C8ZCU0 C8ZD61 C8ZDA6 C8ZDL9
C8ZDR4 C8ZE17 C8ZE43 C8ZEF8 C8ZEK4 C8ZER4 C8ZER6 C8ZET4 C8ZET7 C8ZET8 C8ZEW0
C8ZF59 C8ZFE9 C8ZFN1 C8ZFY3 C8ZFZ7 C8ZG13 C8ZG19 C8ZG43 C8ZGE4 C8ZGV5 C8ZGX0
C8ZH19 C8ZH37 C8ZH56 C8ZHD6 C8ZHH9 C8ZHZ1 C8ZI12 C8ZI21 C8ZI80 C8ZIC3 C8ZIM0
C8ZIQ5 C8ZJB0 C8ZJD8 C8ZJE0 C8ZZN2 C9A1P5 C9CN91 C9DG08 C9DG80 C9DGK1 C9DQ21
C9DQ22 C9EIP1 C9J1S8 C9J202 C9J302 C9J3I9 C9J3V5 C9J6K1 C9J7I0 C9JBD0 C9JC47
C9JCN9 C9JE40 C9JFL3 C9JJ37 C9JLR9 C9JN71 C9JR72 C9JSJ3 C9JTQ0 C9K1X5 C9K1X6
C9K1X7 C9K203 C9K2Q2 C9K2Q3 C9NUM5 C9QZ64 C9QZ65 C9QZ66 C9QZ67 C9QZ69 C9RE37
C9REF5 C9REN7 C9REY1 C9RF94 C9RGE3 C9RH78 C9RLJ9 C9RPK4 C9RR52 C9RZ55 C9S0R2
C9S7Z6 C9S8A7 C9S8J9 C9S923 C9SB02 C9SB49 C9SDH0 C9SDK8 C9SE96 C9SEV5 C9SFZ6
C9SR45 C9STX5 C9SU87 C9SUS0 C9SV08 C9WPN6 C9X0M5 C9X1G5 C9X1G6 C9X1G7 C9XHG0

C9XIS5 C9XIS6 C9XIS7 C9XTA5 C9XUB3 C9XUE4 C9XVF1 C9XXI1 C9Y0Q0 C9Y0S2 C9Y0S4
C9Y0S5 C9Y0S6 C9Y0S7 C9Y0V8 C9Y5D5 C9YHW1 C9YHW2 C9YHW3 C9Z2Z3 C9Z319 C9Z3U4
C9Z4D0 C9Z4D1 C9Z4D4 C9ZBY0 C9ZFL1 C9ZH13 C9ZKW4 C9ZN16 C9ZUU9 D0A0G6 D0A6J7
D0A8W2 D0C6T7 D0C9N6 D0C9N8 D0CY60 D0E0C6 D0E0C7 D0E8I4 D0E8I5 D0EL35 D0EYG3
D0EZM8 D0ISG9 D0JFM7 D0JIM5 D0JZK4 D0K799 D0KC77 D0KC90 D0KG93 D0KN27 D0KQM8
D0KRX1 D0KTH0 D0KZ73 D0L1G3 D0L1T6 D0L3V5 D0L476 D0L658 D0LD40 D0LDR0 D0LDS6
D0LDT1 D0LDT2 D0LDT3 D0LHE3 D0LHE5 D0LI56 D0LI57 D0LI58 D0LID5 D0LID6 D0LV02
D0LWX4 D0MCQ4 D0MQ50 D0MY11 D0MY34 D0MYB4 D0N1U4 D0N381 D0N4E2 D0N761 D0NHA2
D0NK82 D0NLC2 D0NP70 D0NVH9 D0PX88 D0QMC3 D0QX26 D0TZF0 D0UFC7 D0UFC8 D0UZK2
D0V3Y4 D0VMR5 D0VMR6 D0VMR7 D0VMR8 D0VWM8 D0VWQ1 D0VWT0 D0VWY5 D0VX08 D0VX09
D0VX24 D0VYS2 D0WGK0 D0X4R4 D0Z5N4 D0Z8E6 D0Z9R0 D0ZE10 D0ZEW7 D0ZFT0 D0ZLR2
D0ZLR3 D0ZLR9 D0ZV07 D0ZVL4 D0ZW85 D1A2Q5 D1A2S5 D1A2S9 D1A2T0 D1A2T1 D1A3V2
D1A4F4 D1A4Q3 D1A7C3 D1A8M1 D1AB53 D1AFM9 D1AWS1 D1AZF3 D1BAA5 D1BD84 D1BE23
D1BHT7 D1BHT8 D1BHT9 D1BHU2 D1BHU9 D1BJD0 D1BMT0 D1BS17 D1BS23 D1BS26 D1BS27
D1BS28 D1BV84 D1BZ82 D1C883 D1CDA4 D1D8L6 D1FP53 D1FP57 D1FVF0 D1GUM7 D1GY43
D1I234 D1KF50 D1MF76 D1YUK8 D1YVK0 D1YZF2 D1Z042 D1Z199 D1Z1G0 D1Z4I6 D1Z670
D1Z6B1 D1ZA70 D1ZBF6 D1ZEB4 D1ZEM6 D1ZEN1 D1ZG64 D1ZHN6 D1ZHR9 D1ZJC1 D1ZK87
D1ZKF3 D1ZMW1 D1ZPB8 D1ZQ10 D1ZQL9 D1ZR99 D1ZRF1 D1ZRH0 D1ZRZ5 D1ZSG5 D1ZT73
D2A996 D2AC38 D2AC40 D2AC41 D2AC42 D2AC43 D2ATJ4 D2ATU8 D2ATV0 D2ATV1 D2ATX1
D2AU10 D2AV87 D2AVM1 D2B0T1 D2B129 D2B747 D2B7W7 D2B9F4 D2C357 D2CJS7 D2CSU4
D2CSU5 D2CVN6 D2CVN7 D2CVN8 D2D3B2 D2D3B6 D2DGW3 D2DRC4 D2DRC5 D2E9W9 D2E9X0
D2E9X1 D2EAC2 D2GVP7 D2GXM8 D2GXS7 D2GXY4 D2GZW6 D2H0H6 D2H0Y8 D2H163 D2H3M0
D2H526 D2H5P6 D2H8V8 D2HD32 D2HEW7 D2HNW6 D2HNY3 D2HQI1 D2HS90 D2HSB3 D2HWM5
D2HXI8 D2I1E3 D2I4M3 D2K759 D2K760 D2K805 D2KKB4 D2KQB0 D2N3G7 D2NGI4 D2NGI6
D2NGI7 D2NGI8 D2NGI9 D2NRH2 D2NS11 D2NT85 D2NT89 D2NT93 D2PDG6 D2PHL8 D2PK63
D2PN56 D2PPM7 D2PPM8 D2PPN2 D2PPT3 D2PT98 D2Q187 D2Q1C4 D2Q3K3 D2Q4G5 D2Q4H4
D2Q4H5 D2Q9C6 D2Q9D0 D2QN44 D2QPE6 D2QYP6 D2RES5 D2RFQ5 D2RGT4 D2RJU7 D2RQX3
D2RT08 D2RTP8 D2RUI0 D2RVT8 D2RW30 D2RWE9 D2RXF6 D2RXU5 D2S1F7 D2S4K7 D2S4P5
D2S6C9 D2S6D9 D2S6E2 D2S6E3 D2S6E4 D2S9Y0 D2S9Y8 D2SEH8 D2SMN4 D2T333 D2TDA7
D2TDP0 D2TMN6 D2TN58 D2TP58 D2TPR5 D2TPS5 D2TPS8 D2TS86 D2TT52 D2TUZ4 D2UYP1
D2V434 D2V9W6 D2VAA9 D2VN51 D2W6T1 D2WKD9 D2WL32 D2XV59 D2Y3X2 D2YW48 D2YZP9
D2Z024 D2Z025 D2Z026 D2Z027 D2Z028 D2Z030 D3AXF4 D3B3B7 D3BAV8 D3BH90 D3BMU4
D3BN56 D3BT31 D3BUN1 D3DFG8 D3DFP8 D3DFS4 D3DJ41 D3DJ42 D3DK66 D3DKC4 D3DTV9
D3DYU7 D3DZ18 D3E0A3 D3E0R7 D3E288 D3E2P3 D3E3B3 D3E3E4 D3E3M2 D3E3N9 D3E4V4
D3E4V5 D3EBC1 D3ENT3 D3ERF5 D3F5L6 D3FH24 D3FI09 D3FTV4 D3G0V0 D3G9N3 D3GKW6
D3GME4 D3H120 D3H122 D3H123 D3H124 D3H125 D3H5H5 D3HCJ2 D3HKY4 D3IUT5 D3J0Z1
D3JV03 D3JV05 D3K0N9 D3K5I7 D3K5L7 D3KCC4 D3KFX4 D3KVM5 D3KVM6 D3KYU2 D3KYU3
D3NXD9 D3P9D1 D3PW16 D3Q051 D3Q4A8 D3Q556 D3Q557 D3Q568 D3Q598 D3QFZ3 D3QLG0
D3QPK0 D3QPK2 D3QPK3 D3QPK4 D3QPK5 D3R4I7 D3RKJ0 D3RKL0 D3RKL1 D3RKL2 D3RKL3
D3RKL5 D3RNJ4 D3RNP0 D3RPB9 D3RPC0 D3RQU0 D3RW30 D3RX66 D3RZA9 D3RZQ1 D3S0H7
D3S3T0 D3S4A5 D3S534 D3S7P2 D3S8C7 D3S8M7 D3SN37 D3SQW0 D3SRA6 D3SSQ6 D3SY73
D3SYU6 D3SYZ8 D3T426 D3THI6 D3TLL6 D3TQJ5 D3U1D9 D3U715 D3U716 D3U717 D3UAG0
D3UAG1 D3UAG2 D3UAG3 D3UAG4 D3UAG5 D3UAG6 D3UAG7 D3UEC9 D3UEQ8 D3UET8 D3UEY7
D3UW26 D3UWU5 D3UX49 D3UZL8 D3V184 D3V9Q4 D3V9Q5 D3VBF7 D3VC10 D3VIS7 D3WAC0

D3WAC1 D3WAC2 D3WAC3 D3WAC4 D3WAC5 D3WAC7 D3WAC8 D3WAC9 D3WAD0 D3WAD2 D3WAD3
D3WZ85 D3WZ86 D3XDD3 D3YN49 D3YPD5 D3YT16 D3YU32 D3YU81 D3YUB6 D3YUG0 D3YV10
D3YV92 D3YWE6 D3YWJ0 D3YWP0 D3YY23 D3YY91 D3YYM0 D3YZI9 D3YZP9 D3Z070 D3Z0R2
D3Z1D3 D3Z2X2 D3Z3C6 D3Z3K2 D3Z423 D3Z4I3 D3Z4R1 D3Z4S3 D3Z5S8 D3Z6H8 D3Z6S9
D3Z750 D3Z7P3 D3Z8E6 D3Z8K2 D3Z8N2 D3Z8N4 D3Z8X7 D3Z8Y2 D3Z987 D3ZA12 D3ZA50
D3ZAP3 D3ZAQ5 D3ZBE5 D3ZBN0 D3ZBP4 D3ZCL3 D3ZCM9 D3ZDK2 D3ZDX9 D3ZEF4 D3ZEG1
D3ZF18 D3ZF42 D3ZG52 D3ZG83 D3ZGQ5 D3ZGX1 D3ZHP7 D3ZKB9 D3ZKD3 D3ZKV9 D3ZLB7
D3ZLZ7 D3ZMY7 D3ZND0 D3ZQD3 D3ZQF4 D3ZQG6 D3ZQL7 D3ZQX4 D3ZSC8 D3ZSK5 D3ZT37
D3ZTL1 D3ZTQ1 D3ZU57 D3ZUC4 D3ZUC6 D3ZUU2 D3ZV31 D3ZVF4 D3ZVK1 D3ZVM4 D3ZVP7
D3ZVR1 D3ZVV1 D3ZW55 D3ZW91 D3ZWK4 D3ZX08 D3ZXK7 D3ZXW3 D3ZY68 D3ZYB7 D3ZYW7
D3ZZC3 D3ZZW6 D4A0X3 D4A218 D4A2B7 D4A2K4 D4A3K5 D4A615 D4A666 D4A770 D4A7C0
D4A7U2 D4A7V9 D4A8G9 D4A8X0 D4A9E1 D4AAA5 D4AAT7 D4AAZ6 D4AB98 D4ABB4 D4ABH7
D4ABM4 D4ABP9 D4ACE5 D4ACP5 D4AE41 D4AE48 D4AEC2 D4AEP7 D4AIP9 D4AK44 D4AK46
D4AK47 D4AK48 D4AKN8 D4AL61 D4AM37 D4AMF9 D4AN26 D4AN96 D4AQY2 D4ARJ9 D4ARL8
D4AU26 D4AU28 D4AU29 D4AU30 D4AU31 D4AU56 D4AU57 D4AUV1 D4AVK1 D4AWG9 D4AWH0
D4AWH2 D4AWH3 D4AZ50 D4B0B2 D4B1Q8 D4B2G3 D4B3J8 D4CZZ2 D4CZZ3 D4CZZ5 D4CZZ6
D4D2Y6 D4D445 D4D447 D4D449 D4D4Q0 D4D6B8 D4D891 D4D8P3 D4D9R6 D4DB32 D4DDJ7
D4DE65 D4DG66 D4DHC4 D4FWG0 D4FZ53 D4G3R3 D4G8K6 D4GEU4 D4GEU5 D4GEU6 D4GEU7
D4GEU9 D4GGR2 D4GGV4 D4GIR2 D4GJ14 D4GKM1 D4GL26 D4GNJ8 D4GP28 D4GP31 D4GP33
D4GP40 D4GP41 D4GP42 D4GPG1 D4GPU7 D4GPU8 D4GQ08 D4GQ67 D4GQN4 D4GQN5 D4GQN6
D4GQN7 D4GQN8 D4GQN9 D4GQP0 D4GQP1 D4GR05 D4GS06 D4GS48 D4GSE6 D4GSF3 D4GSH6
D4GSS5 D4GST8 D4GSW7 D4GT97 D4GTA5 D4GTC1 D4GTL2 D4GTL3 D4GTL8 D4GTN6 D4GTS4
D4GU26 D4GU60 D4GU62 D4GU63 D4GU66 D4GU69 D4GU70 D4GU71 D4GU72 D4GU73 D4GU81
D4GU92 D4GUJ7 D4GUK0 D4GUK1 D4GV73 D4GVD5 D4GVD7 D4GVJ2 D4GVJ3 D4GVR1 D4GVW4
D4GVW6 D4GW05 D4GW48 D4GW50 D4GW73 D4GWB3 D4GWH7 D4GWJ7 D4GWY0 D4GWY2 D4GXC5
D4GY94 D4GY98 D4GYE0 D4GYE6 D4GYE7 D4GYE8 D4GYG5 D4GYH1 D4GYH5 D4GYI5 D4GYM0
D4GYV5 D4GYY1 D4GYZ1 D4GYZ4 D4GZ01 D4GZ42 D4GZ88 D4GZA2 D4GZB4 D4GZN2 D4GZW2
D4H071 D4H539 D4H6M0 D4H9Y2 D4HVG6 D4I2N1 D4ICL5 D4MUV9 D4N3P2 D4N3P3 D4N500
D4N501 D4N502 D4NUX0 D4NXE7 D4P095 D4P3R7 D4Q9Z4 D4Q9Z5 D4QF25 D4QFE7 D4YYG1
D4Z260 D4Z694 D4Z909 D4ZX34 D4ZX35 D5AKU6 D5AKX8 D5AKX9 D5AKY0 D5ANI3 D5ANI6
D5ANS3 D5AP15 D5AP78 D5AP79 D5AP83 D5AQI9 D5AR83 D5ARY6 D5ARY7 D5ARY8 D5ATX8
D5AUD5 D5AV13 D5AV14 D5AV20 D5AV94 D5BDP9 D5C1K8 D5CE05 D5CE32 D5CE33 D5CE34
D5CE35 D5CE37 D5CHP5 D5CIL6 D5CZG7 D5CZG9 D5CZH0 D5CZH1 D5CZH2 D5DIV9 D5DJL5
D5DKA5 D5DNA8 D5DVF4 D5E0A1 D5E3H2 D5E7G9 D5E7U7 D5E9R7 D5EAS6 D5EBS7 D5ECW5
D5EID4 D5FKJ3 D5GA77 D5GAC6 D5GBI7 D5GCI8 D5GDH4 D5GE59 D5GFR0 D5GHP2 D5GIH1
D5GN29 D5H0J3 D5HB86 D5IGG0 D5IGG4 D5IGG6 D5JGG0 D5JWB3 D5KXD2 D5KXJ0 D5L1S4
D5LHJ0 D5LXJ0 D5MCN2 D5MTF8 D5MTT1 D5SGZ8 D5SHR0 D5SJ87 D5SL78 D5SQS4 D5SRZ4
D5SXM0 D5T3F2 D5TAM8 D5TM67 D5TZL3 D5U396 D5UFA8 D5UJ42 D5UJR1 D5UPJ5 D5USX8
D5UU90 D5UVX0 D5UXA4 D5UYN8 D5V0N9 D5V2I8 D5V9A9 D5VAD8 D5VGV1 D5VGV2 D5VGV3
D5VGV4 D5VQA3 D5VQW1 D5VRB9 D5VRM1 D5VSX2 D5VT38 D5VUB0 D5W6H7 D5WQ56 D5WVB5
D5ZT17 D6C652 D6CFW3 D6GRK6 D6JLP0 D6R237 D6R3B6 D6RBM5 D6REC4 D6RGH6 D6RGX4
D6RIA3 D6RRG1 D6RRG7 D6XZ22 D6Y1N8 D6Y4C5 D6Y4U7 D6Y7M2 D6Z995 D6ZA79 D6ZE32
D6ZEJ5 D6ZFG9 D7A0Y0 D7A0Y1 D7A0Y2 D7A0Y3 D7A3L6 D7A989 D7AW65 D7AWZ7 D7B2S5
D7B412 D7B6L8 D7BAR0 D7BMJ2 D7BPX0 D7BQJ3 D7BSS4 D7C367 D7C376 D7CWZ0 D7DQH5

D7DQT8 D7DRZ8 D7DTT8 D7DV18 D7E8Z9 D7E9E0 D7E9E7 D7GCU5 D7GG24 D7KQR8 D7KWY6
D7PHY7 D7PHY9 D7PHZ0 D7PHZ1 D7PHZ2 D7PHZ3 D7PHZ4 D7PHZ5 D7PHZ7 D7PHZ8 D7PI00
D7PI10 D7PI11 D7PI12 D7PI14 D7PI15 D7PI16 D7PI17 D7PI18 D7PI19 D7REX8 D7REY3
D7REY4 D7REY5 D7SMN6 D7STK2 D7T737 D7TCD0 D7U6G6 D7UER1 D7UNT2 D7UQ41 D7UQ42
D7UQ43 D7UQ44 D7UQM5 D7URM0 D7URV0 D7UTD1 D7Y2H2 D7Y2H3 D7Y2H4 D7Y2H5 D8ADB5
D8I5S8 D8IJV0 D8IMC7 D8INT8 D8J0Q1 D8J4S4 D8J663 D8JA74 D8JBB7 D8JBB8 D8K235
D8K3Q1 D8KFT1 D8KIT5 D8KN09 D8MHV4 D8MKW4 D8MPW4 D8MUB2 D8PJU0 D8PR70 D8Q7V2
D8Q8R5 D8RLD3 D8RNZ9 D8S255 D8S5L4 D8UYN9 D8V070 D8V071 D8V073 D8V074 D8V076
D8V077 D8V078 D8VNC6 D9ICP2 D9IFD5 D9IL23 D9IVE5 D9J034 D9J0Z7 D9J0Z8 D9J100
D9J101 D9N129 D9PU00 D9PUH5 D9PUX5 D9PVI7 D9PVP5 D9PWM7 D9PX81 D9PXZ6 D9PYH9
D9PYS9 D9PZU4 D9Q2C4 D9Q7U8 D9R201 D9R4W7 D9RZ53 D9SEU0 D9SJ16 D9SM34 D9TQ02
D9TT09 D9TT10 D9UBG0 D9UBI3 D9UNB2 D9XD61 D9XDR8 D9XF45 E0CPH4 E0CSI1 E0CTF3
E0CTY1 E0CYV9 E0CZ16 E0D7H5 E0IWI3 E0J1Q3 E0J500 E0J5J4 E0J6W8 E0RQS7 E0RTD9
E0S6S0 E0SAK7 E0SAL0 E0SBQ8 E0SDG7 E0SGL7 E0SGL8 E0SIS1 E0SIS4 E0SLH6 E0SMM6
E0SNQ8 E0SPG5 E0SQX5 E0SRA9 E0T4V0 E0T5H9 E0TIQ1 E0TTS5 E0TU95 E0TVS5 E0TVT6
E0TXE1 E0TY11 E0TY12 E0TZF9 E0U070 E0U0H8 E0U0L7 E0U174 E0U4V7 E0UR96 E0V9D8
E0VI98 E0VLV0 E0VSF4 E0VSP9 E0VVF6 E0W481 E0WN94 E0X9N4 E0Y3X0 E0Y3X1 E0ZS45
E0ZS46 E0ZS47 E0ZS48 E1AB55 E1ACP6 E1ACQ0 E1ACQ4 E1ACQ7 E1ACQ8 E1ACQ9 E1ACR1
E1ACR2 E1ACR3 E1B7L7 E1B7Q7 E1B7X3 E1B8W3 E1B932 E1B9D8 E1BB03 E1BB50 E1BC15
E1BCH6 E1BD59 E1BEA8 E1BGH8 E1BGQ2 E1BH29 E1BI64 E1BJD1 E1BJL9 E1BJS7 E1BKF9
E1BKH1 E1BKK0 E1BLP6 E1BLZ4 E1BM70 E1BMN8 E1BMP7 E1BNQ4 E1BNS0 E1BNS6 E1BP36
E1BP74 E1BP92 E1BPH3 E1BPN0 E1BRE2 E1BSI0 E1BTG2 E1BUG7 E1BVR9 E1BXS0 E1BYJ2
E1BZ85 E1C1U1 E1C231 E1C2U2 E1C2V1 E1C2Z0 E1C3P4 E1C3S7 E1C6F0 E1C6Q1 E1C760
E1C762 E1C7T6 E1CHH8 E1F8H4 E1F8Q4 E1FU46 E1JH25 E1PL40 E1QBX4 E1QCT1 E1QU22
E1QUT8 E1QWZ1 E1R5X1 E1RHM5 E1SDF1 E1SHE4 E1SKR8 E1SM51 E1SSZ4 E1ST41 E1SVR5
E1U332 E1UV19 E1V4F9 E1V4J5 E1V4Y0 E1V7V7 E1V7V8 E1V7V9 E1V7W0 E1V7W1 E1V8I0
E1V8I1 E1V8J1 E1V931 E1VA04 E1VB09 E1VBK4 E1VBT6 E1W218 E1W6L8 E1W818 E1W823
E1W874 E1W9M1 E1WA38 E1WAB2 E1WAB4 E1WAB5 E1WAC0 E1WAC4 E1WAC5 E1WAQ4 E1WGF0
E1WGG8 E1WGG9 E1WGN1 E1WH94 E1WKT5 E1WNR6 E1WS50 E1WTS2 E1WTS4 E1X022 E1X791
E1XT70 E1XTG1 E1XTG6 E1XTG9 E1XTJ1 E1XTK3 E1XTK6 E2AB17 E2AXC7 E2D0Y9 E2E2N3
E2E2N4 E2E2N5 E2E2N7 E2E2N8 E2E2P0 E2E2P1 E2E2P2 E2FZM5 E2GIN1 E2IHE0 E2JA28
E2JA29 E2JA31 E2JA32 E2JFG1 E2JFG2 E2JKY7 E2NPN0 E2QSX5 E2QTD3 E2QUD0 E2QUI9
E2QV03 E2QXH7 E2R1I5 E2R222 E2R9X2 E2RBS6 E2RDV1 E2RDZ6 E2RGI3 E2RK33 E2RKN7
E2RTZ4 E2RU81 E2RYF8 E3E2E2 E3FKM2 E3G128 E3G1M3 E3G327 E3G381 E3G5B9 E3GCF1
E3GW42 E3GWN9 E3GWT5 E3GZ90 E3H7X6 E3H9T2 E3H9W1 E3HCT1 E3HDJ8 E3HQ85 E3HVU4
E3I7Z1 E3J0E1 E3JD18 E3JV98 E3K7C1 E3K7C3 E3KAB5 E3KGP2 E3KIY6 E3KWE1 E3KY53
E3L3Q8 E3L408 E3L9T1 E3LAN7 E3LAQ9 E3LB80 E3M3V1 E3MIE2 E3PRJ4 E3PRJ5 E3PRJ8
E3PRJ9 E3PRK0 E3PRK1 E3PTZ4 E3PV41 E3PVH7 E3PY95 E3PY96 E3PY97 E3PY98 E3PY99
E3Q897 E3QBC5 E3QBI3 E3QCU0 E3QDN5 E3QDQ4 E3QLU5 E3QRY8 E3QW41 E3QYP0 E3QZF5
E3RCY7 E3RD74 E3REX1 E3REX5 E3RNJ5 E3RP32 E3RPB1 E3RV84 E3RYW5 E3S405 E3S6N7
E3S7K9 E3SEW0 E3SXU4 E3SXU5 E3VWI2 E3VWI3 E3VWI4 E3VWI6 E3VWI7 E3VWI8 E3VWJ0
E3VWJ9 E3VWK0 E3VWK2 E3VWK3 E3VWK4 E3VXF2 E3W202 E3W203 E3W204 E3W205 E3W206
E3W207 E3W208 E3WDQ9 E3WPP8 E3WSB5 E3X5D6 E3XA68 E3XDZ8 E3XFR6 E4MYY0 E4N7E5
E4NQE5 E4PJF6 E4PMA5 E4QHI6 E4QJB8 E4QP00 E4QWH2 E4QWH3 E4RUV9 E4T0I2 E4TI44

E4UP58 E4UPA0 E4UQC3 E4USI8 E4V1Q7 E4V2N2 E4V2N3 E4V2N5 E4V2N6 E4V6L3 E4W8Y2
E4ZBM0 E4ZHV7 E4ZUJ1 E4ZYY3 E5A2Z3 E5A7D7 E5A7D9 E5A7E1 E5A7E2 E5A7E3 E5ABQ8
E5AD52 E5AE40 E5AE42 E5ASS2 E5AV36 E5AW43 E5AW45 E5DG73 E5EYS5 E5F146 E5FYH0
E5FYH1 E5KIB9 E5KIC0 E5L8F7 E5QZZ6 E5R278 E5R332 E5R3Z8 E5R4J3 E5R5D9 E5RG02
E5RJ46 E5RJM6 E5Y377 E5Y378 E5Y379 E5Y944 E5Y945 E5Y946 E6ELF6 E6ENP7 E6ENP9
E6ESW1 E6LHV2 E6LHV3 E6LHV4 E6LHV5 E6LHV6 E6LHV7 E6MUA0 E6MVV9 E6MVW0 E6MWF8
E6MXY4 E6MY99 E6N013 E6RHP9 E6SFG5 E6SFR5 E6SRF9 E6SRG2 E6TMM3 E6UB41 E6UBR9
E6UIS7 E6WEY6 E6WHK1 E6X4P3 E6Y2X0 E6YBW4 E6Z0R3 E6Z0R4 E6ZGB4 E6ZHJ8 E7BQV0
E7C196 E7CQW6 E7CY69 E7CY70 E7ETH6 E7EW31 E7EXT2 E7EZF3 E7F187 E7F1C4 E7F1H9
E7F3I6 E7F555 E7F5E1 E7F6F9 E7F7X0 E7F888 E7F9T0 E7FA21 E7FAM5 E7FAP1 E7FBU4
E7FCN8 E7FCP8 E7FDB3 E7FDW2 E7FDW8 E7FDX5 E7FFX1 E7FGY2 E7FH61 E7FHC4 E7FHF1
E7FHF8 E7FHN9 E7FHP1 E7FHU4 E7FHW8 E7FHX4 E7FHX6 E7FI44 E7FI45 E7KAR8 E7KB73
E7KBS9 E7KCH3 E7KDI6 E7KDM2 E7KDX7 E7KEM9 E7KEX7 E7KFK5 E7KFU6 E7KIF0 E7KIY3
E7KLI6 E7KM17 E7KMP1 E7KNF7 E7KPJ0 E7KPK0 E7KPZ3 E7KR23 E7KS00 E7KUA5 E7KV71
E7KVI3 E7LRT8 E7LSL5 E7LTG2 E7LUC9 E7LVH4 E7LVV3 E7LWL0 E7LWV5 E7LXI9 E7LXU7
E7LY32 E7M099 E7M1C7 E7NFN4 E7NG53 E7NGT6 E7NHM9 E7NIP0 E7NIP9 E7NK05 E7NKW8
E7NL30 E7NND9 E7NNK3 E7Q257 E7Q2L9 E7Q2X8 E7Q3U6 E7Q4T7 E7Q5V8 E7Q653 E7Q714
E7Q7C0 E7QA20 E7QAA9 E7QBZ1 E7QCS8 E7QDA1 E7QDX3 E7QFR8 E7QG80 E7QH16 E7QH95
E7QIA1 E7QIC5 E7QKN3 E7QLB7 E7QLJ6 E7RAM2 E7S4M0 E8LIC0 E8ME30 E8MF10 E8MF11
E8MF12 E8MF13 E8MGH8 E8MJ15 E8N877 E8NCH3 E8PLM2 E8R1E4 E8RMD3 E8T3K9 E8VHU4
E8W6C7 E8XYF5 E9AD19 E9AE57 E9AGH7 E9AIE8 E9AXF0 E9CTR7 E9CWP5 E9CY14 E9D0S5
E9DDK8 E9DV56 E9E2J4 E9E766 E9E9B2 E9EHG0 E9EHG1 E9EHG2 E9EK74 E9EMI7 E9EPV1
E9ES90 E9ES99 E9ET39 E9ET40 E9ET41 E9EUE6 E9EUE8 E9F5E7 E9F5E9 E9F5F0 E9F646
E9F647 E9F8L8 E9F8L9 E9F8M1 E9F8M2 E9F8M3 E9F8M4 E9F8R9 E9F9J8 E9FCP4 E9FCP6
E9FCP7 E9GD30 E9HCD7 E9JFX9 E9KMQ2 E9KMQ4 E9L7A5 E9P860 E9P886 E9P8D2 E9P974
E9PB15 E9PGG2 E9PI22 E9PIF3 E9PJ23 E9PJI5 E9PKD4 E9PQR5 E9PRG8 E9PT37 E9PT87
E9PU28 E9PV87 E9PVA8 E9PVB3 E9PVX6 E9PW05 E9PW74 E9PYH6 E9PYL2 E9PYQ0 E9PYY5
E9PZM4 E9PZZ1 E9Q0C6 E9Q173 E9Q1P8 E9Q1U1 E9Q286 E9Q2Z1 E9Q309 E9Q349 E9Q3T6
E9Q4F2 E9Q4F7 E9Q5F9 E9Q5G7 E9Q5M6 E9Q5Z5 E9Q6B2 E9Q6D6 E9Q6E9 E9Q6J5 E9Q6W4
E9Q6Z5 E9Q735 E9Q743 E9Q784 E9Q793 E9Q7F2 E9Q7R9 E9Q8D0 E9Q8G5 E9Q8I9 E9Q8T2
E9Q8T7 E9Q912 E9Q956 E9Q9D5 E9Q9M8 E9QA62 E9QAG8 E9QAU8 E9QI36 E9QJ05 E9QJ30
E9QMW4 E9QUT3 E9QYP0 E9RAH5 E9RD40 E9RFS9 E9RFT0 E9RFT1 E9RFT2 F0E1K6 F0JAI6
F0KFI5 F0KFI7 F0KKL2 F0LIT8 F0NBH8 F0NDL2 F0NID3 F0NJT3 F0Q2T3 F0QUB3 F0QYK7
F0SLK8 F0T7W2 F0ZDD0 F0ZHE2 F0ZK52 F0ZL92 F0ZLU4 F0ZPL2 F1A1D4 F1A1D6 F1A3V1
F1B281 F1C7I4 F1CKI6 F1CKI8 F1CKI9 F1CKJ1 F1CYZ5 F1DBA9 F1DBB2 F1DBB3 F1DBB4
F1LKH5 F1LKH6 F1LKH7 F1LKH8 F1LKH9 F1LMN3 F1LMZ8 F1LNJ2 F1LP64 F1LTE0 F1LU71
F1LV46 F1LWT0 F1LZ52 F1LZF0 F1M5F3 F1M5M3 F1M5N7 F1M8W4 F1MBP6 F1MDL2 F1MF21
F1MGG3 F1MH07 F1MHT9 F1MIW6 F1MJR8 F1MKX4 F1MLB4 F1MMV1 F1MNN4 F1MQW7 F1MRW8
F1MX48 F1N2N9 F1N7G5 F1N9S8 F1NB38 F1ND48 F1NEA7 F1NLH9 F1NW29 F1NZI4 F1PLN3
F1PT61 F1PZQ5 F1Q4R9 F1Q514 F1Q575 F1Q7Z7 F1Q8J0 F1Q8K0 F1Q8W0 F1QAJ4 F1QB54
F1QBY1 F1QCN0 F1QCV2 F1QCY8 F1QDF8 F1QDI9 F1QEB7 F1QFR9 F1QGC8 F1QGH9 F1QGW6
F1QH17 F1QHZ6 F1QJF4 F1QJX5 F1QLG5 F1QLY4 F1QMB9 F1QMV3 F1QMY1 F1QN54 F1QNW4
F1QNX7 F1QPR4 F1QQA8 F1QR98 F1QRC1 F1QRX7 F1QVR8 F1QWK4 F1QWM2 F1QXM5 F1QY75
F1QYJ6 F1QZ15 F1QZ88 F1R237 F1R2J1 F1R2X6 F1R345 F1R3W0 F1R4C4 F1R6N4 F1R7R1

F1R8Z9 F1R983 F1RBN2 F1RCE7 F1RCR6 F1RCY6 F1RET2 F1RKB1 F1RKQ4 F1RQM2 F1RRT2
F1SC07 F1SLM8 F1SRI0 F1SS88 F1SSF9 F1SWA0 F1SZ41 F1SZ42 F2EP55 F2G1U7 F2GA29
F2JTX5 F2JVT6 F2JWN1 F2JXJ2 F2L1X6 F2MMN7 F2MMN9 F2MMP0 F2MMP6 F2MTI0 F2MTI3
F2NV13 F2PWS5 F2PWS6 F2PWS8 F2PWS9 F2R776 F2RB77 F2RB78 F2RB79 F2RB80 F2RB81
F2RB83 F2RGQ4 F2S6Z9 F2S700 F2S702 F2S703 F2T0M0 F2T0M1 F2T0M3 F2T0M4 F2VYU4
F2XF92 F2XF93 F2XF94 F2XF95 F2XF96 F2XF97 F2XF98 F2XF99 F2XFA0 F2XFA1 F2XFA2
F2XFA4 F2XFA5 F2XFA6 F2XFA8 F2XFB2 F2XG53 F2Z266 F2Z398 F2Z3F1 F2Z3M2 F2Z3T4
F2Z461 F2Z5Z6 F2Z610 F2Z678 F2Z6D2 F2Z6F6 F3Y5P4 F4B8X5 F4B991 F4B9I5 F4BEN2
F4BT36 F4CQ77 F4CQ78 F4CUJ4 F4DAE8 F4FTW6 F4FYY6 F4G4E2 F4G5A4 F4HK38 F4HNU6
F4HNU8 F4HPA7 F4HPP7 F4HPY8 F4HQA1 F4HQE2 F4HQX1 F4HR03 F4HR11 F4HRC1 F4HRI2
F4HRJ4 F4HRV8 F4HSD4 F4HSD5 F4HSX9 F4HT49 F4HTL8 F4HTQ1 F4HTT6 F4HU58 F4HV65
F4HVA6 F4HVZ5 F4HW51 F4HWF9 F4HWL4 F4HWY6 F4HXP9 F4HXU3 F4HY56 F4HY61 F4HYD7
F4HYF3 F4HYJ7 F4HYV5 F4HZF0 F4HZG9 F4I096 F4I0K9 F4I0N3 F4I116 F4I169 F4I171
F4I1A6 F4I1N8 F4I1S7 F4I1T7 F4I1T9 F4I241 F4I274 F4I2H2 F4I2H7 F4I2J8 F4I2X9
F4I366 F4I3B3 F4I3Z5 F4I460 F4I4E1 F4I4P3 F4I4W2 F4I507 F4I532 F4I5N6 F4I5Q6
F4I6E6 F4I6M1 F4I6M4 F4I6V0 F4I7C7 F4I7I0 F4I7L1 F4I837 F4I878 F4I893 F4I8B9
F4I8F9 F4I8R6 F4I8U2 F4I8U3 F4I8W1 F4I907 F4I933 F4I9J7 F4I9Q5 F4I9R6 F4I9T0
F4IA73 F4IAE9 F4IAG2 F4IAR2 F4IAX0 F4IAX1 F4IBB2 F4IBE4 F4IBQ9 F4ICA0 F4ICD9
F4ICK8 F4ID16 F4IDI6 F4IDQ5 F4IDW9 F4IDY7 F4IE21 F4IE65 F4IE66 F4IEY4 F4IFC5
F4IFD0 F4IFF3 F4IG60 F4IG73 F4IGA5 F4IGL2 F4IGU3 F4IGZ2 F4IH25 F4IHL3 F4IIK6
F4IIS5 F4IIZ9 F4IJ79 F4IJK6 F4IJV4 F4IK01 F4IK92 F4IKM1 F4IL05 F4IL57 F4ILR7
F4IM84 F4IMK2 F4IMK4 F4IMQ0 F4IN32 F4IN78 F4INA9 F4INN2 F4INP9 F4IP06 F4IPE3
F4IPI4 F4IPK2 F4IPM3 F4IPY2 F4IQJ2 F4IQL7 F4IQQ9 F4IRQ5 F4IRR2 F4IRU3 F4IRW0
F4IS91 F4ISQ7 F4ISV6 F4ISY0 F4IUG9 F4IUX6 F4IUY8 F4IV16 F4IV45 F4IV66 F4IVR7
F4IVU1 F4IW10 F4IWB3 F4IXE4 F4IXE7 F4IXJ7 F4IY03 F4IY62 F4IYF8 F4IZM8 F4J077
F4J078 F4J0B7 F4J117 F4J1G1 F4J1H7 F4J1U4 F4J264 F4J284 F4J2J6 F4J2K4 F4J2M6
F4J2U9 F4J2Z7 F4J300 F4J394 F4J3D9 F4J3G5 F4J3P7 F4J3S1 F4J4A0 F4J4A1 F4J4P8
F4J4Y5 F4J5S1 F4J5U9 F4J738 F4J7T2 F4J7T3 F4J8G7 F4J8L3 F4J8R4 F4J9M5 F4J9Q6
F4JAA5 F4JBP3 F4JC97 F4JCB2 F4JCC1 F4JCN9 F4JCQ3 F4JCU0 F4JD14 F4JDI6 F4JEP5
F4JET1 F4JEX5 F4JF21 F4JFJ3 F4JFN3 F4JG10 F4JGB7 F4JGP4 F4JGZ1 F4JH53 F4JHA2
F4JHI7 F4JHT3 F4JI44 F4JIF5 F4JIJ7 F4JIU4 F4JJ23 F4JJE5 F4JJP1 F4JJR8 F4JJT9
F4JKB6 F4JKI3 F4JL28 F4JLC1 F4JLK2 F4JLM5 F4JLP5 F4JLS1 F4JM19 F4JMH9 F4JMJ3
F4JN26 F4JNY0 F4JP46 F4JP48 F4JPR7 F4JQ51 F4JQ95 F4JR57 F4JRA6 F4JRB0 F4JRF4
F4JRF5 F4JRJ6 F4JRP0 F4JRP8 F4JRR5 F4JSG3 F4JSU0 F4JSV3 F4JTE7 F4JTF6 F4JTI1
F4JTJ9 F4JTP5 F4JU69 F4JUI3 F4JUI9 F4JUL9 F4JUQ2 F4JUY5 F4JV59 F4JV80 F4JVH1
F4JVI3 F4JVJ1 F4JVN6 F4JW68 F4JWP9 F4JWR0 F4JWT7 F4JX00 F4JXF1 F4JXF9 F4JY12
F4JY24 F4JY37 F4JY84 F4JYE9 F4JYG0 F4JYJ1 F4JZ68 F4JZC2 F4JZJ2 F4K0A6 F4K0E8
F4K0J3 F4K128 F4K1B1 F4K1G2 F4K1J4 F4K1R2 F4K1Z0 F4K265 F4K295 F4K2A1 F4K2M8
F4K2R6 F4K3X8 F4K410 F4K460 F4K487 F4K4C5 F4K4E3 F4K4J0 F4K4L7 F4K4M0 F4K5J1
F4K5K6 F4K5R6 F4K5T4 F4K5X6 F4K657 F4K687 F4K6X0 F4K7D5 F4K7F6 F4K933 F4KAB8
F4KAS1 F4KB17 F4KBF5 F4KBW6 F4KBY0 F4KCE9 F4KD38 F4KDN0 F4KE63 F4KEA4 F4KEC6
F4KED2 F4KEM0 F4KEV7 F4KEY9 F4KF27 F4KFB3 F4KFC7 F4KFD8 F4KFG5 F4KFM8 F4KFS5
F4KFT6 F4KFT7 F4KFV7 F4KG50 F4KGE8 F4KGQ0 F4KGU4 F4KGY6 F4KH86 F4KH89 F4KHG6
F4KHG8 F4KI56 F4KIH4 F4KIN4 F4KIX1 F4KU78 F4LHM0 F4NUJ6 F4NYQ2 F4NZ38 F4P804

F4P9L8 F4Q7D4 F4QV02 F4ZCI3 F5A6E9 F5A894 F5BFC8 F5GYI3 F5H4B4 F5H8R6 F5H8S6
F5H991 F5H9F9 F5H9N4 F5HA06 F5HA10 F5HA93 F5HAM4 F5HAS7 F5HAU9 F5HAY6 F5HAZ3
F5HB71 F5HBC6 F5HBR4 F5HBX1 F5HC16 F5HC97 F5HCU8 F5HD03 F5HDC7 F5HDM3 F5HDQ6
F5HE12 F5HEN7 F5HEN8 F5HF23 F5HF44 F5HF49 F5HFA5 F5HFJ8 F5HG51 F5HG98 F5HGC2
F5HGG3 F5HGI9 F5HGJ4 F5HGT1 F5HH39 F5HHT4 F5HI68 F5HI85 F5HIG1 F5HIN9 F5HN73
F5HN75 F5HSE3 F5L9G2 F5RH07 F5WVJ2 F5Z7V5 F5ZEI8 F5ZTT9 F6BB32 F6BCS4 F6C6C1
F6CY50 F6D512 F6H2F8 F6H7K5 F6H9A9 F6HDM2 F6HDT7 F6HH45 F6HQ26 F6HS55 F6I457
F6IBC7 F6M8H4 F6M8H5 F6M8H6 F6M8H7 F6M8H8 F6M8I0 F6NSX9 F6P3G4 F6P6X0 F6PHZ6
F6QEU4 F6QXW0 F6R2G2 F6RCC2 F6RIX4 F6RJC2 F6RQL9 F6RRD7 F6S675 F6S9E6 F6SDF8
F6SNN2 F6SZT2 F6TFD9 F6TQD1 F6U5F9 F6UA42 F6UH96 F6ULY3 F6UMY3 F6V515 F6VSS6
F6W2R2 F6W3G8 F6WEQ6 F6WXT2 F6X2V8 F6XBL2 F6XLV1 F6XZJ7 F6Y089 F6Y9J3 F6YCR7
F6ZT52 F7A355 F7A3P2 F7AEX0 F7AQ22 F7ARS3 F7ASZ0 F7BHS0 F7BJB9 F7BLM1 F7CYY5
F7D4X9 F7DF15 F7DKV7 F7DP49 F7E540 F7E727 F7E7M3 F7EA39 F7EL49 F7EZ75 F7GSQ4
F7H9X2 F7J0M4 F7J0M6 F7J0N2 F7J5X9 F7V5D5 F7V998 F7V999 F7W4M2 F7XKY8 F7XQS2
F7YJG6 F7YQ08 F7ZDR3 F7ZKE9 F7ZLS5 F8AJL1 F8C4X8 F8DQP1 F8DT24 F8DT92 F8DT95
F8DVG5 F8DVL8 F8G0M4 F8GV06 F8GVD3 F8HWV7 F8I1D0 F8JJ25 F8JJ27 F8JX40 F8KEJ1
F8P1W3 F8P1W6 F8P2C8 F8P9P5 F8QQ74 F8SJR0 F8SJT5 F8UL80 F8UL81 F8VF07 F8VI19
F8VPN2 F8VPY8 F8VPZ5 F8VPZ9 F8VTS6 F8W1W9 F8WCM5 F8WFD2 F8WKW0 F8WKW1 F8WKW8
F8WLS6 F8WQ30 F8WQN2 F8WRK9 F9UL76 F9ULM5 F9UMS6 F9UN92 F9UP09 F9UPU7 F9US27
F9USS8 F9USS9 F9UST1 F9UST4 F9UT67 F9UT68 F9UU40 F9VEN7 F9VMT6 F9VN79 F9VND5
F9VPA4 F9VPG1 F9W301 F9WWF1 F9WZD2 F9X3D9 F9X9V1 F9X9V2 F9XLC1 F9XMW3 F9XMW5
F9XMW6 F9XMX6 F9YQP0 G0E1Z6 G0FS62 G0FS68 G0FUS0 G0GBS4 G0HQ31 G0HQ35 G0HQZ5
G0HQZ6 G0HV10 G0J2F8 G0J5N4 G0KYA7 G0KYB0 G0KYB1 G0KYB3 G0LD36 G0LES5 G0LET7
G0LET8 G0LET9 G0LEU0 G0LEU4 G0LFB0 G0LH85 G0LHZ8 G0R6S7 G0R6S8 G0R6S9 G0R6T2
G0R6T4 G0REX6 G0REX7 G0REX8 G0RH19 G0RNA2 G0RNN3 G0RS98 G0RYE0 G0S0Y3 G0S1B3
G0S215 G0S2G1 G0S2X1 G0S3J5 G0S3T2 G0S4X6 G0S5Q0 G0S616 G0S8E7 G0S8F1 G0S8G9
G0S902 G0S920 G0SAK8 G0SB20 G0SC29 G0SC91 G0SCK6 G0SEH7 G0SEQ5 G0SEV9 G0SF32
G0SF70 G0SFB5 G0SFM2 G0SGK0 G0SGU4 G0SHF3 G0WXL9 G0Y7D1 G0Y7D2 G0Y7D3 G0YF19
G1CWH0 G1CWH4 G1DGI7 G1ESZ9 G1JUH1 G1K3N4 G1NJB6 G1SRW8 G1UB44 G1UB67 G1UII1
G1X5Y2 G1XU02 G1XU04 G2HE57 G2HK15 G2IJ05 G2IN04 G2IN77 G2IQQ5 G2IQQ8 G2IQS7
G2IQS8 G2ITT5 G2JHL6 G2JZ14 G2JZ74 G2K042 G2K044 G2K045 G2K046 G2K047 G2K048
G2K265 G2K449 G2K4Q7 G2P5T1 G2Q9A5 G2Q9A7 G2SYN4 G2TRJ5 G2TRJ6 G2TRJ7 G2TRJ8
G2TRK1 G2TRK2 G2TRK3 G2TRK6 G2TRK8 G2TRL1 G2TRL2 G2TRL3 G2TRL4 G2TRL5 G2TRL6
G2TRL7 G2TRL8 G2TRM0 G2TRM2 G2TRM4 G2TRM5 G2TRM6 G2TRM7 G2TRM9 G2TRN0 G2TRN1
G2TRN4 G2TRN5 G2TRN6 G2TRN7 G2TRN9 G2TRP2 G2TRP3 G2TRP6 G2TRQ0 G2TRQ2 G2TRQ5
G2TRQ6 G2TRQ8 G2TRQ9 G2TRR1 G2TRR3 G2TRR5 G2TRR6 G2TRR7 G2TRR8 G2TRR9 G2TRS0
G2TRS1 G2TRS6 G2TRS9 G2TRT1 G2TRT2 G2TRT4 G2TRT5 G2TRU1 G2TRU3 G2TRU4 G2WJ80
G2WR64 G2X740 G2XQT1 G2XR75 G3CCC0 G3CCC1 G3E4M4 G3E4M6 G3ECR1 G3ECR2 G3ECR3
G3ECR4 G3F5K2 G3FFN4 G3FFN6 G3FFN8 G3FIN8 G3FIN9 G3FLZ7 G3FLZ8 G3GBU6 G3GBU7
G3GTP0 G3ISL7 G3JUI7 G3JUI8 G3KIM3 G3KIM4 G3KIM5 G3KIM6 G3KIM7 G3KIM8 G3KLH2
G3KLH3 G3KLH4 G3KLH5 G3KLH6 G3MWR8 G3MY25 G3MZQ6 G3N131 G3N1S4 G3S077 G3UW36
G3UW99 G3UWD5 G3UZ78 G3V211 G3V6S8 G3V6U9 G3V893 G3V8T1 G3V9M2 G3X982 G3X987
G3X9C2 G3X9G7 G3XCR8 G3XCU2 G3XCV0 G3XCV2 G3XCX3 G3XCY4 G3XCZ8 G3XD76 G3XD97
G3XGC1 G3XKQ9 G3XLL5 G3XMB7 G3XMB8 G3XMB9 G3XMC0 G3XMC4 G3XMC5 G3XMC6 G3XMD1

G3XNF3 G3XNF4 G3XRG4 G3XSI3 G3XSI5 G3XSI6 G3XUF0 G3Y415 G3Y419 G3Y422 G3Y423
G3Y424 G3YG17 G4FEF4 G4LTX4 G4MQX3 G4MUB2 G4MWZ7 G4MXJ9 G4MZI3 G4MZJ4 G4N134
G4N137 G4N1P8 G4N286 G4N290 G4N292 G4N296 G4N2A1 G4N2B2 G4N3L5 G4N708 G4N7S6
G4N7X0 G4NEB8 G4NEE4 G4NFB7 G4NID8 G4NYJ5 G4RES5 G4RJY5 G4RJY6 G4RJY8 G4RJY9
G4RJZ0 G4RJZ1 G4RJZ2 G4RJZ3 G4RJZ4 G4RJZ5 G4RK16 G4RT11 G4STG9 G4T4R7 G4V4T4
G4V4T5 G4V4T6 G4V4T7 G4VGH9 G4VGI0 G4VJD6 G4WJD3 G5CEW6 G5CV45 G5CV46 G5CV52
G5CV54 G5DDC2 G5DGA8 G5E869 G5E870 G5E872 G5E8A8 G5E8B9 G5E8F4 G5E8P0 G5E8P1
G5E9R7 G5EAZ2 G5EB19 G5EB20 G5EB37 G5EB87 G5EBD7 G5EBL3 G5EBM1 G5EBU4 G5EBV0
G5EBV6 G5EBX3 G5EBY0 G5EBZ4 G5EC36 G5EC37 G5EC44 G5EC77 G5EC89 G5ECG0 G5ECH5
G5ECH7 G5ECJ5 G5ECK3 G5ECN5 G5ECQ3 G5ECQ8 G5ECR9 G5ECT8 G5ECU7 G5ECW5 G5ED14
G5ED29 G5ED41 G5ED47 G5ED66 G5ED84 G5ED89 G5EDE9 G5EDF0 G5EDG2 G5EDJ0 G5EDM7
G5EDN3 G5EDN6 G5EDQ2 G5EDS1 G5EDT1 G5EDU6 G5EDW1 G5EDW7 G5EDZ2 G5EE18 G5EE86
G5EEA1 G5EED4 G5EEG1 G5EEG7 G5EEH5 G5EEJ1 G5EEL0 G5EEM0 G5EEM1 G5EEM5 G5EEN4
G5EER4 G5EES6 G5EEU2 G5EEV2 G5EEW6 G5EF51 G5EF68 G5EF76 G5EF84 G5EFA2 G5EFF1
G5EFF4 G5EFF7 G5EFJ4 G5EFL0 G5EFL9 G5EFQ5 G5EFS4 G5EFT4 G5EFV3 G5EFV5 G5EFV8
G5EFW7 G5EFY4 G5EFY5 G5EFZ1 G5EG14 G5EG17 G5EG38 G5EG44 G5EG51 G5EG86 G5EGA3
G5EGA6 G5EGC9 G5EGE9 G5EGF8 G5EGN8 G5EGQ2 G5EGQ5 G5EGS7 G5EGU9 G7CFI3 G7H7V7
G7ID19 G7J032 G7J1L1 G7JFU5 G7JMM0 G7JT50 G7L166 G7NY55 G7TAD9 G7WMS7 G7X8S6
G7XQ31 G7XTA8 G8B839 G8EWC8 G8FRC5 G8FRC6 G8GJ94 G8GJ95 G8GJ96 G8GTN7 G8GV69
G8H5M7 G8H5M8 G8H5M9 G8H5N0 G8H5N1 G8H5N2 G8H5N3 G8JVR4 G8Q5R9 G8QM63 G8T6H8
G8T8D0 G8XHD5 G8XHD7 G8XHD8 G8XR08 G9AIU0 G9EZR6 G9F1Y9 G9HXM2 G9M952 G9M973
G9M9M0 G9M9M1 G9M9M2 G9M9M3 G9M9M4 G9M9M5 G9MAN7 G9N4A1 G9N4A2 G9N4A3 G9N4A5
G9N4A6 G9N4A7 G9N4A9 G9N4B0 G9N4B1 G9N4B2 G9N4E3 G9NNY7 G9P9X3 H0SH23 H0UI37
H0UZ81 H0WFA5 H0Y7S4 H0YKK7 H0YM25 H1AAP2 H1UBK1 H1VN83 H1VQB1 H1ZV37 H1ZV38
H1ZZA4 H1ZZB0 H2AM11 H2AM13 H2AM14 H2DH19 H2DH23 H2DH24 H2E5X2 H2E7Q6 H2E7Q7
H2E7Q8 H2ELN1 H2FH31 H2I233 H2IFE7 H2IFX0 H2K885 H2K887 H2K888 H2K893 H2KY84
H2KY86 H2KY91 H2KYH3 H2KYH4 H2KYP0 H2KYQ5 H2KYW5 H2KZ49 H2KZK4 H2KZM9 H2KZZ6
H2L008 H2L056 H2L0H5 H2L0N3 H2L2B8 H2LBU8 H2LP95 H2MTR9 H2VFI5 H2VFR7 H3BKT1
H3BMG3 H3BNL1 H3BNL8 H3BPM6 H3BQW9 H3BRN8 H3BU77 H3BUK9 H3JQW0 H3JUC3 H3K2Y6
H3ZPL1 H3ZPU1 H5SQ95 H6AGY0 H6D7E6 H6LBB0 H6LBB1 H6LBS1 H6LGM6 H6LGM7 H6LGM8
H6QW39 H6VGI2 H6VGI3 H6VLG5 H6WZF2 H7BZ55 H7C794 H7C8I3 H8EVV4 H8F0D6 H8F0D7
H8F0N6 H8F1Z2 H8F3Q9 H8F3R6 H8L374 H8L901 H8L902 H8WR05 H8ZKS8 H8ZKW0 H8ZM70
H8ZM71 H8ZM73 H8ZPX2 H8ZW44 H9BFQ0 H9BFQ1 H9BFQ2 H9BFW7 H9C4A4 H9G301 H9IUR0
H9IWW7 H9JDT2 H9JU86 H9L427 H9L478 H9L9E5 H9N289 H9N290 H9N291 H9T8G7 H9T8I0
H9TB18 H9TB19 I0B6F4 I0DF35 I0HUK7 I0IUP3 I0IUP4 I0J062 I0J0E7 I0J1J1 I1C129
I1JLC8 I1K0K6 I1KEV6 I1L3T1 I1LCI8 I1LHU6 I1LPE9 I1M2G5 I1M2U5 I1MGE5 I1N2Z5
I1R9A8 I1R9A9 I1R9B0 I1R9B1 I1R9B2 I1R9B4 I1R9B5 I1R9B6 I1RAW4 I1RC73 I1RCN9
I1RF49 I1RF54 I1RF57 I1RFC6 I1RIV2 I1RL11 I1RL12 I1RL14 I1RL18 I1RMA2 I1RN13
I1RN14 I1RP53 I1RV17 I1RV18 I1RV20 I1RV21 I1RV22 I1RV23 I1RVD6 I1RVD8 I1RVD9
I1RW37 I1S0A8 I1S0E2 I1S0J7 I1S104 I1S160 I1S165 I1S166 I1S2J3 I1S2J4 I1S2J6
I1S2J8 I1S3K6 I1S3K7 I1S3K8 I1S3L0 I1S489 I1S490 I1S491 I1S5P3 I1SAJ7 I1SKR9
I1SSI5 I1WEI8 I1WFB9 I1WGA2 I1WGR6 I1WW99 I1YAP6 I1ZHA5 I2BA89 I2DDG2 I2HB52
I2N045 I3BN39 I3J7Q8 I3L0S3 I3L1E1 I3LHS8 I3LK75 I3LUP1 I3PB37 I3PFJ3 I3PFJ9
I3PLQ6 I3PLQ7 I3PLR2 I3PLR3 I3PLR4 I3R176 I3R584 I3R634 I3R636 I3R637 I3R7F1

I3R7F2 I3R7G3 I3R9K1 I3R9K3 I3R9Z2 I3R9Z3 I3R9Z4 I3RB26 I3V6A7 I3VE74 I3VE77
I3VRU1 I3X057 I3XHK1 I3ZNU9 I3ZRF1 I3ZRF2 I4AY86 I4DST8 I4DST9 I4EPA3 I6LDA6
I6LJ77 I6NXV7 I6PL68 I6QPS5 I6QSN0 I6R4V5 I6RAQ6 I6RE61 I6TCK3 I6UF18 I6UH61
I6WHP7 I6WJ72 I6WU39 I6WXS6 I6X235 I6X4W0 I6X7F9 I6XD65 I6XFZ8 I6Y0R5 I6Y276
I6Y4U9 I6Y8F7 I6Y8I5 I6Y9F7 I6Y9Q3 I6YBX3 I6YD99 I6YDK7 I6YEE1 I6YFP0 I6YG32
I6YGS0 I7A3T6 I7FA35 I7FJX8 I7GPX8 I7H0H9 I7H0I4 I7H727 I7H893 I7HHN4 I7HJS4
I7HUG0 I7K316 J3BZS6 J3F2B0 J3KSC0 J3QM76 J3QMY9 J3QPZ5 J3TRC5 J3TRD1 J4KLK4
J4KLU1 J4KMC1 J4UHQ6 J4UHQ8 J4VV61 J4VZG1 J5J930 J5JCC9 J5JPY5 J5JV76 J5K1E2
J7FCF0 J7FIX8 J7FK08 J7FQE8 J7GQ11 J7H5M4 J7H8Q3 J7HCX7 J7LH11 J7LJN5 J7LMP2
J7LP58 J7LQ09 J7M3T1 J7M799 J7MDG6 J7RC67 J7RUA5 J7SGD5 J7SHB8 J7SZ64 J7T0S1
J7T3V5 J7TF92 J8D1N4 J8GG30 J8H9C1 J8HQ06 J9MJK9 J9N5G7 J9PBR3 J9QS23 J9QS25
J9R1J8 J9R388 J9R393 J9R5V4 J9RLZ7 J9RYI6 J9VHE9 J9VHZ9 J9VI03 J9VI89 J9VL95
J9VND4 J9VP29 J9VQ03 J9VQB3 J9VQZ0 J9VR81 J9VS56 J9VT33 J9VT48 J9VT60 J9VVN9
J9VW97 J9VWH9 J9VX38 J9VXB3 J9VYP5 J9VZ70 J9W034 J9W0G9 J9ZCN8 K0BE10 K0DZ91
K0DZA0 K0E2F3 K0E2F6 K0E2G4 K0E3U5 K0E3V3 K0E4D7 K0E4E5 K0E678 K0E689 K0I210
K0I7Q2 K0I977 K0I986 K0ICR0 K0II72 K0J4Q8 K0K750 K0KPV8 K0KSN3 K0P2S0 K2PFJ6
K2QVI8 K2QXC4 K2QXP6 K2RGJ8 K2RLM6 K2RU68 K3UFY2 K3UIH8 K3VA96 K3VAW3 K3VAW8
K3VDP2 K3VDP7 K3VH30 K3VJJ1 K3VZ28 K4B6C9 K4BND8 K4BNG7 K4BW79 K4BWV2 K4C9E2
K4CA50 K4CJJ1 K4CWS6 K4D3U9 K4D3V7 K4D422 K4D9Y4 K4DEK0 K4DEY3 K4DF01 K4FF95
K4GGT4 K4GHR9 K4GHS2 K4GIP0 K4GKX2 K4GMD6 K4IAL8 K4ICC9 K4L7X3 K4L9M2 K4LMW2
K4LVZ1 K4N0V2 K4PW38 K4REZ6 K4RFM2 K5B7F3 K5BDL0 K5BJH8 K6UCV4 K7EIQ3 K7IM66
K7LFJ0 K7LSB9 K7MTW9 K7N5M5 K7N5M6 K7N6K2 K7NBW3 K7NCP5 K7NCV2 K7NCX6 K7QHS5
K7QKH1 K7QRJ5 K7QVW7 K7TLS0 K7TQE3 K7UPS5 K7UTH7 K7VCB9 K7W9N9 K7WCC7 K7WDC8
K7WDL7 K7WIZ6 K7WQ45 K8ERR8 K8ESC5 K9FWK0 K9GDC6 K9GKQ6 K9JHZ4 K9N4V0 K9N643
K9NBS6 K9NVA6 K9UV87 K9Y645 K9Y6N7 K9Y6Y9 K9Y7V9 L0BUM3 L0BWK3 L0E155 L0E168
L0E172 L0E1U3 L0E2P7 L0E2Q1 L0E2R0 L0E2T7 L0E2U2 L0E2U6 L0E2Y5 L0E2Z1 L0E301
L0E307 L0E4G3 L0E4H0 L0HAM7 L0HB77 L0PQJ9 L0R819 L0R8F8 L0T5V6 L0TAD5 L0TCG5
L0TDE1 L0TGF0 L7HT17 L7HUY5 L7HXG6 L7HZV6 L7I1W3 L7I518 L7IAU0 L7IBL2 L7IGD3
L7IK19 L7IKV1 L7IM20 L7N1X6 L7N5A6 L7N653 L7N689 L7N6B4 L7NCQ3 L7R9Z0 L7UZ85
L7WC64 L7WMF4 L7WMF9 L7WR46 L7WRQ4 L7WRR9 L7WRX9 L7WRZ7 L7WU80 L7X8G0 L7X8J4
L7XAV2 L8AXV5 L8B068 L8EBJ9 L8EUQ6 L8EYU3 L8FL04 L8FL29 L8FLZ2 L8FM21 L8FPI5
M0GNC6 M0HG00 M0QWB7 M0R6D8 M0R7T6 M0R7T9 M0R8U1 M0RD54 M0Y4P1 M1A3D5 M1BJF6
M1EAP5 M1ETA5 M1ETK3 M1GRN3 M1H607 M1J8U6 M1JJT8 M1RIR6 M1T9X3 M1UXG8 M1V4Y8
M1V9Q0 M1VDX3 M1VV65 M1VV66 M1VVW6 M1VWN2 M1VWN3 M1VWN4 M1W079 M1W0X8 M1W0X9
M1W0Y0 M1W268 M1W270 M1W271 M1W424 M1W426 M1W428 M1W5Z4 M1W5Z7 M1W600 M1W848
M1W850 M1W851 M1W853 M1W859 M1WA41 M1WA44 M1WCE9 M1WCF2 M1WCF5 M1WCF7 M1WCQ3
M1WEN5 M1WG89 M1WG92 M1WG94 M1WG96 M1YW29 M2PP75 M2SNN6 M2TF54 M2U578 M2V0Z8
M2V8C1 M2VWL5 M2WIS5 M2WJ74 M2WJF1 M2WKH6 M2XHU6 M2XHZ5 M2XJ44 M2XJV1 M2Y0N8
M2Y1A3 M2Y2F4 M2YIY2 M2YJ20 M2YJ38 M2YJK1 M2YJQ2 M2YKT1 M2YKT6 M2YMT1 M2YMU7
M2ZIX2 M2ZIX7 M3A326 M3ANL0 M3APK0 M3APK9 M3WHG5 M3XQV7 M4HXU6 M4HXW5 M4HY05
M4HY08 M4HYC6 M4HYC8 M4HYP3 M4I1C6 M4IQQ5 M4IQQ7 M4IQR7 M4IQS1 M4IRL4 M4IRL6
M4IRL9 M4IS88 M4IS90 M4IS92 M4ISH0 M4ISH1 M4ISH2 M4MR97 M4YFG7 M4YMP0 M5AW86
M5AWY0 M5EES5 M7U9B9 M9MRD5 M9MRI4 M9NE38 M9NEY8 M9SGV8 M9XB82 N0DKX5 N1P3Y5
N1PJ97 N4V6D4 N4WEA4 N4WHA7 N4WHE3 N4WQZ8 N4WR35 N4WW42 N4WYI1 N4XMB0 O00057

O00058 O00059 O00060 O00061 O00084 O00086 O00087 O00088 O00091 O00097 O00098
O00103 O00110 O00142 O00221 O00233 O00244 O00321 O00330 O00338 O00358 O00370
O00399 O00411 O00422 O00423 O00442 O00470 O00479 O00534 O00570 O00625 O00635
O00716 O00743 O00764 O00782 O00822 O00835 O00841 O00845 O00852 O00869 O00871
O00874 O00897 O00937 O00939 O00940 O01159 O01258 O01305 O01323 O01326 O01352
O01359 O01360 O01367 O01377 O01395 O01409 O01422 O01427 O01477 O01479 O01501
O01503 O01510 O01513 O01519 O01541 O01583 O01592 O01615 O01639 O01658 O01671
O01673 O01683 O01725 O01727 O01738 O01757 O01763 O01767 O01789 O01812 O01814
O01822 O01824 O01829 O01833 O01835 O01836 O01901 O01914 O01929 O01930 O01931
O01939 O01965 O01974 O01975 O01991 O01999 O02002 O02008 O02019 O02033 O02039
O02042 O02058 O02101 O02108 O02115 O02151 O02173 O02195 O02197 O02217 O02219
O02235 O02274 O02305 O02316 O02323 O02326 O02327 O02328 O02334 O02365 O02367
O02373 O02376 O02389 O02414 O02467 O02482 O02485 O02491 O02494 O02604 O02607
O02608 O02609 O02611 O02621 O02623 O02640 O02654 O02655 O02658 O02732 O02739
O02751 O02755 O02756 O02771 O02772 O02785 O02786 O02818 O02849 O03685 O03979
O03983 O03992 O04009 O04015 O04016 O04017 O04019 O04027 O04035 O04058 O04059
O04064 O04066 O04067 O04073 O04090 O04104 O04111 O04116 O04121 O04130 O04132
O04134 O04135 O04136 O04147 O04160 O04166 O04173 O04192 O04196 O04197 O04202
O04203 O04204 O04211 O04220 O04226 O04244 O04251 O04259 O04266 O04267 O04274
O04287 O04298 O04309 O04310 O04311 O04312 O04313 O04314 O04316 O04318 O04319
O04336 O04341 O04350 O04351 O04375 O04376 O04379 O04385 O04386 O04395 O04397
O04407 O04408 O04425 O04435 O04437 O04450 O04479 O04487 O04488 O04491 O04492
O04504 O04515 O04539 O04582 O04591 O04603 O04609 O04615 O04630 O04647 O04656
O04658 O04663 O04683 O04705 O04706 O04707 O04716 O04725 O04773 O04790 O04847
O04854 O04856 O04857 O04858 O04860 O04862 O04866 O04867 O04885 O04895 O04899
O04904 O04905 O04916 O04932 O04933 O04937 O04939 O04951 O04955 O04973 O04974
O04985 O04986 O04996 O04997 O05023 O05029 O05031 O05052 O05069 O05070 O05071
O05072 O05073 O05086 O05100 O05101 O05103 O05104 O05113 O05118 O05132 O05137
O05139 O05146 O05151 O05154 O05161 O05197 O05204 O05207 O05208 O05209 O05217
O05218 O05220 O05223 O05231 O05234 O05235 O05236 O05239 O05240 O05242 O05243
O05246 O05251 O05261 O05262 O05263 O05264 O05265 O05267 O05268 O05269 O05272
O05278 O05306 O05321 O05322 O05337 O05338 O05340 O05385 O05386 O05389 O05392
O05393 O05394 O05396 O05398 O05404 O05405 O05408 O05409 O05411 O05412 O05413
O05420 O05494 O05496 O05501 O05502 O05508 O05509 O05510 O05511 O05514 O05515
O05516 O05517 O05518 O05519 O05520 O05521 O05564 O05581 O05616 O05617 O05619
O05625 O05633 O05634 O05635 O05636 O05637 O05638 O05639 O05640 O05641 O05642
O05643 O05650 O05651 O05668 O05678 O05689 O05691 O05700 O05701 O05706 O05714
O05724 O05728 O05730 O05755 O05756 O05886 O05927 O05940 O05944 O05945 O05946
O05947 O05949 O05952 O05953 O05954 O05955 O05956 O05957 O05958 O05959 O05960
O05961 O05964 O05965 O05966 O05968 O05969 O05970 O05972 O05973 O05974 O05978
O05979 O05981 O05982 O06000 O06006 O06008 O06010 O06011 O06012 O06014 O06046
O06047 O06049 O06050 O06051 O06059 O06060 O06061 O06069 O06084 O06091 O06114
O06115 O06179 O06242 O06334 O06335 O06401 O06430 O06431 O06456 O06457 O06458
O06465 O06472 O06474 O06476 O06478 O06480 O06483 O06484 O06485 O06486 O06490

O06491 O06492 O06553 O06581 O06636 O06637 O06638 O06641 O06644 O06652 O06663
O06672 O06679 O06695 O06703 O06708 O06714 O06715 O06716 O06717 O06718 O06719
O06720 O06721 O06722 O06723 O06724 O06727 O06734 O06775 O06779 O06793 O06825
O06834 O06835 O06837 O06851 O06865 O06866 O06898 O06899 O06901 O06914 O06917
O06918 O06922 O06924 O06925 O06926 O06927 O06928 O06940 O06941 O06942 O06961
O06966 O06968 O06972 O06973 O06974 O06975 O06976 O06977 O06978 O06983 O06986
O06987 O06988 O06993 O06994 O06995 O06997 O06998 O07001 O07004 O07005 O07006
O07007 O07008 O07012 O07014 O07015 O07020 O07021 O07025 O07026 O07045 O07046
O07051 O07079 O07081 O07103 O07104 O07108 O07115 O07116 O07118 O07127 O07131
O07145 O07151 O07165 O07226 O07227 O07299 O07308 O07309 O07315 O07319 O07329
O07333 O07345 O07348 O07355 O07356 O07360 O07394 O07402 O07434 O07436 O07451
O07458 O07465 O07509 O07513 O07517 O07521 O07522 O07527 O07529 O07530 O07539
O07541 O07542 O07546 O07547 O07548 O07551 O07552 O07558 O07561 O07564 O07565
O07566 O07567 O07571 O07572 O07573 O07575 O07578 O07579 O07586 O07587 O07593
O07595 O07597 O07598 O07603 O07604 O07606 O07607 O07608 O07609 O07614 O07615
O07617 O07624 O07627 O07631 O07633 O07635 O07636 O07637 O07638 O07641 O07642
O07643 O07664 O07665 O07669 O07674 O07678 O07680 O07683 O07684 O07685 O07708
O07717 O07718 O07737 O07779 O07813 O07816 O07817 O07823 O07824 O07825 O07828
O07835 O07836 O07854 O07855 O07874 O07883 O07884 O07896 O07898 O07906 O07917
O07918 O07919 O07920 O07931 O07937 O07938 O07949 O08248 O08249 O08250 O08307
O08308 O08309 O08314 O08315 O08317 O08318 O08319 O08320 O08321 O08322 O08323
O08325 O08326 O08327 O08328 O08333 O08335 O08336 O08340 O08349 O08353 O08356
O08357 O08358 O08359 O08361 O08370 O08371 O08374 O08384 O08385 O08393 O08394
O08397 O08399 O08400 O08404 O08405 O08424 O08429 O08444 O08445 O08450 O08451
O08459 O08460 O08461 O08467 O08469 O08501 O08550 O08557 O08582 O08583 O08618
O08632 O08633 O08638 O08640 O08651 O08653 O08656 O08658 O08664 O08672 O08674
O08686 O08719 O08730 O08739 O08740 O08750 O08755 O08759 O08760 O08764 O08781
O08782 O08784 O08789 O08791 O08792 O08796 O08800 O08811 O08815 O08848 O08856
O08875 O08876 O08884 O08900 O08918 O08919 O08934 O08949 O08950 O08961 O08997
O09000 O09015 O09017 O09018 O09029 O09035 O09053 O09061 O09102 O09105 O09106
O09110 O09112 O09113 O09116 O09130 O09139 O09171 O09172 O09173 O09178 O09181
O09232 O09345 O09410 O09452 O09460 O09486 O09705 O09781 O09802 O10269 O10270
O10273 O10274 O10276 O10277 O10278 O10279 O10280 O10281 O10293 O10294 O10295
O10296 O10297 O10298 O10299 O10300 O10301 O10302 O10303 O10304 O10307 O10308
O10309 O10310 O10311 O10312 O10313 O10314 O10315 O10316 O10317 O10318 O10319
O10324 O10325 O10326 O10327 O10328 O10329 O10330 O10331 O10332 O10333 O10334
O10335 O10338 O10339 O10340 O10341 O10343 O10344 O10345 O10346 O10347 O10348
O10350 O10351 O10353 O10354 O10355 O10356 O10359 O10361 O10362 O10365 O10368
O10369 O10370 O10373 O10374 O10375 O10378 O10438 O10439 O10442 O10443 O11451
O11452 O11453 O11840 O12705 O12792 O12933 O12934 O12944 O12984 O12985 O12987
O12988 O12991 O12995 O13012 O13019 O13022 O13023 O13024 O13034 O13046 O13067
O13071 O13074 O13077 O13078 O13089 O13124 O13125 O13126 O13152 O13161 O13163
O13164 O13224 O13257 O13268 O13269 O13270 O13276 O13277 O13278 O13282 O13283
O13284 O13285 O13286 O13287 O13289 O13290 O13294 O13297 O13298 O13302 O13309

O13326 O13329 O13335 O13339 O13351 O13354 O13360 O13366 O13370 O13396 O13401
O13402 O13410 O13412 O13415 O13416 O13418 O13419 O13425 O13426 O13427 O13432
O13434 O13437 O13440 O13453 O13465 O13471 O13472 O13473 O13474 O13476 O13489
O13493 O13504 O13505 O13507 O13508 O13513 O13517 O13521 O13531 O13532 O13534
O13536 O13539 O13541 O13542 O13546 O13548 O13549 O13553 O13555 O13556 O13557
O13558 O13559 O13564 O13565 O13569 O13571 O13573 O13574 O13575 O13576 O13577
O13578 O13582 O13585 O13586 O13587 O13588 O13595 O13596 O13598 O13600 O13601
O13602 O13610 O13612 O13615 O13620 O13622 O13629 O13639 O13641 O13642 O13647
O13648 O13650 O13651 O13652 O13658 O13665 O13670 O13674 O13682 O13684 O13685
O13688 O13691 O13693 O13696 O13699 O13701 O13702 O13704 O13706 O13710 O13713
O13715 O13719 O13722 O13724 O13725 O13727 O13730 O13731 O13732 O13733 O13734
O13735 O13738 O13739 O13741 O13743 O13744 O13745 O13748 O13749 O13750 O13754
O13755 O13756 O13757 O13758 O13759 O13762 O13765 O13767 O13768 O13769 O13772
O13774 O13775 O13777 O13781 O13783 O13790 O13791 O13792 O13794 O13795 O13796
O13798 O13799 O13800 O13801 O13803 O13807 O13808 O13810 O13815 O13816 O13823
O13824 O13826 O13827 O13828 O13829 O13830 O13831 O13833 O13834 O13835 O13836
O13837 O13839 O13842 O13843 O13845 O13848 O13852 O13856 O13859 O13862 O13863
O13865 O13867 O13868 O13870 O13871 O13873 O13874 O13877 O13878 O13881 O13882
O13886 O13889 O13890 O13892 O13894 O13895 O13896 O13897 O13900 O13902 O13903
O13905 O13908 O13910 O13911 O13913 O13914 O13915 O13916 O13917 O13919 O13921
O13923 O13926 O13927 O13929 O13935 O13936 O13940 O13943 O13947 O13951 O13952
O13953 O13956 O13957 O13958 O13959 O13964 O13966 O13969 O13970 O13971 O13972
O13973 O13977 O13982 O13983 O13984 O13985 O13986 O13988 O13991 O13993 O13996
O13997 O13998 O14000 O14003 O14006 O14007 O14009 O14010 O14011 O14012 O14013
O14015 O14016 O14018 O14021 O14023 O14026 O14027 O14028 O14030 O14032 O14033
O14034 O14036 O14037 O14039 O14040 O14041 O14043 O14044 O14045 O14047 O14048
O14049 O14053 O14055 O14056 O14057 O14059 O14061 O14064 O14074 O14075 O14076
O14077 O14078 O14079 O14080 O14081 O14082 O14086 O14087 O14088 O14092 O14094
O14096 O14097 O14098 O14099 O14102 O14104 O14105 O14106 O14108 O14109 O14110
O14112 O14113 O14114 O14118 O14122 O14125 O14126 O14129 O14130 O14131 O14133
O14134 O14135 O14137 O14138 O14139 O14140 O14141 O14143 O14144 O14145 O14147
O14148 O14155 O14157 O14158 O14160 O14161 O14164 O14165 O14170 O14172 O14174
O14176 O14178 O14179 O14180 O14181 O14182 O14183 O14185 O14187 O14189 O14192
O14196 O14198 O14204 O14206 O14207 O14208 O14209 O14210 O14211 O14214 O14215
O14216 O14218 O14219 O14220 O14227 O14229 O14232 O14233 O14235 O14236 O14240
O14241 O14242 O14246 O14248 O14250 O14253 O14254 O14256 O14258 O14259 O14262
O14263 O14268 O14269 O14270 O14274 O14276 O14278 O14279 O14280 O14283 O14284
O14285 O14289 O14291 O14293 O14295 O14297 O14301 O14302 O14303 O14304 O14305
O14307 O14308 O14310 O14311 O14325 O14327 O14330 O14332 O14334 O14335 O14336
O14337 O14338 O14339 O14341 O14342 O14343 O14344 O14348 O14349 O14350 O14351
O14352 O14353 O14354 O14356 O14358 O14359 O14360 O14361 O14362 O14363 O14368
O14369 O14370 O14388 O14399 O14400 O14408 O14417 O14423 O14427 O14429 O14433
O14434 O14443 O14448 O14450 O14459 O14460 O14463 O14467 O14468 O14503 O14519
O14529 O14531 O14556 O14595 O14598 O14599 O14602 O14603 O14604 O14607 O14618

O14627 O14628 O14633 O14645 O14709 O14715 O14717 O14727 O14730 O14753 O14770
O14771 O14772 O14782 O14813 O14815 O14818 O14829 O14830 O14841 O14862 O14867
O14893 O14901 O14926 O14933 O14948 O14950 O14978 O15015 O15016 O15033 O15037
O15040 O15049 O15050 O15054 O15060 O15091 O15116 O15119 O15143 O15160 O15172
O15182 O15194 O15195 O15209 O15212 O15213 O15225 O15231 O15305 O15315 O15347
O15353 O15370 O15391 O15405 O15409 O15428 O15442 O15457 O15479 O15480 O15481
O15488 O15499 O15511 O15514 O15522 O15527 O15535 O15544 O15574 O15587 O15594
O15595 O15602 O15604 O15612 O15613 O15631 O15692 O15697 O15746 O15757 O15770
O15816 O15818 O15819 O15820 O15883 O15885 O15891 O15905 O15909 O15910 O15925
O15926 O15945 O15989 O15990 O15991 O15992 O15995 O15998 O16011 O16023 O16027
O16053 O16091 O16115 O16116 O16127 O16129 O16140 O16207 O16216 O16228 O16259
O16277 O16294 O16299 O16354 O16359 O16360 O16361 O16368 O16391 O16425 O16519
O16520 O16662 O16686 O16808 O16810 O16811 O16812 O16844 O16852 O16867 O16962
O16963 O16966 O16991 O16999 O17025 O17071 O17072 O17082 O17087 O17212 O17213
O17214 O17286 O17307 O17310 O17319 O17320 O17433 O17438 O17446 O17449 O17453
O17468 O17486 O17488 O17502 O17503 O17514 O17536 O17573 O17580 O17581 O17582
O17586 O17606 O17608 O17611 O17617 O17622 O17624 O17657 O17670 O17679 O17683
O17706 O17731 O17732 O17736 O17748 O17811 O17862 O17894 O17898 O17917 O17919
O17927 O17928 O17930 O17931 O17933 O17934 O17966 O18048 O18054 O18086 O18087
O18141 O18191 O18195 O18214 O18216 O18229 O18326 O18373 O18381 O18391 O18416
O18473 O18475 O18477 O18480 O18499 O18500 O18531 O18559 O18598 O18601 O18602
O18635 O18638 O18640 O18719 O18728 O18734 O18740 O18745 O18746 O18747 O18748
O18751 O18768 O18776 O18842 O18870 O18879 O18883 O18921 O18934 O19048 O19049
O19052 O19069 O19074 O19085 O19097 O19104 O19110 O19114 O19199 O19816 O19872
O19883 O19884 O19885 O19886 O19887 O19888 O19889 O19890 O19892 O19894 O19897
O19898 O19900 O19904 O19906 O19907 O19908 O19912 O19914 O19915 O19917 O19920
O19929 O19986 O19989 O19990 O19992 O19995 O19998 O20029 O20032 O20103 O20106
O20118 O20120 O20130 O20133 O20142 O20143 O20159 O20162 O20163 O20173 O20208
O20210 O20213 O20215 O20219 O20221 O20230 O20231 O20234 O20235 O20236 O20241
O20252 O20254 O20255 O20264 O20265 O20273 O20278 O20280 O20291 O20304 O20346
O20399 O21032 O21033 O21034 O21035 O21036 O21037 O21038 O21043 O21044 O21045
O21050 O21234 O21235 O21240 O21245 O21246 O21247 O21251 O21252 O21255 O21256
O21260 O21263 O21264 O21272 O21275 O21281 O21869 O21870 O21871 O21872 O21873
O21874 O21876 O21877 O21878 O21879 O21880 O21882 O21883 O21884 O21886 O21894
O21902 O21945 O21946 O21947 O21948 O21949 O21950 O21951 O21952 O21953 O21954
O21955 O21956 O21957 O21958 O21959 O21960 O21970 O22000 O22001 O22002 O22005
O22010 O22018 O22020 O22022 O22024 O22025 O22029 O22034 O22043 O22045 O22046
O22047 O22056 O22060 O22077 O22130 O22131 O22132 O22137 O22150 O22152 O22173
O22176 O22182 O22183 O22184 O22190 O22193 O22197 O22199 O22212 O22216 O22224
O22229 O22232 O22241 O22243 O22254 O22259 O22264 O22268 O22283 O22286 O22287
O22288 O22290 O22300 O22307 O22308 O22309 O22315 O22319 O22328 O22338 O22340
O22347 O22348 O22349 O22373 O22380 O22386 O22424 O22426 O22431 O22433 O22436
O22437 O22446 O22456 O22466 O22467 O22468 O22469 O22493 O22494 O22504 O22506
O22518 O22527 O22553 O22558 O22567 O22572 O22573 O22582 O22584 O22585 O22586

O22604 O22607 O22611 O22616 O22621 O22637 O22643 O22644 O22645 O22647 O22651
O22652 O22655 O22668 O22683 O22693 O22698 O22703 O22714 O22727 O22765 O22768
O22779 O22781 O22785 O22793 O22795 O22798 O22799 O22800 O22820 O22822 O22823
O22827 O22829 O22850 O22860 O22864 O22870 O22873 O22886 O22890 O22891 O22892
O22893 O22897 O22899 O22900 O22901 O22907 O22914 O22922 O22928 O22929 O22941
O22944 O22951 O22967 O22969 O23024 O23034 O23049 O23076 O23090 O23095 O23115
O23116 O23141 O23174 O23184 O23193 O23205 O23208 O23210 O23212 O23225 O23239
O23247 O23249 O23252 O23257 O23264 O23266 O23273 O23277 O23278 O23290 O23304
O23305 O23310 O23317 O23320 O23324 O23337 O23341 O23344 O23346 O23360 O23379
O23382 O23393 O23401 O23402 O23404 O23406 O23417 O23419 O23420 O23421 O23487
O23491 O23515 O23550 O23553 O23564 O23593 O23610 O23614 O23617 O23620 O23624
O23627 O23628 O23629 O23647 O23651 O23653 O23659 O23660 O23661 O23674 O23676
O23708 O23710 O23712 O23714 O23715 O23717 O23729 O23730 O23731 O23732 O23733
O23735 O23736 O23755 O23760 O23787 O23813 O23826 O23865 O23875 O23882 O23883
O23884 O23887 O23888 O23894 O23917 O23918 O23920 O23924 O23936 O23939 O23945
O23969 O23970 O23976 O23979 O24030 O24035 O24045 O24047 O24058 O24059 O24076
O24111 O24133 O24144 O24145 O24146 O24149 O24150 O24151 O24160 O24163 O24164
O24169 O24170 O24171 O24172 O24210 O24246 O24248 O24282 O24301 O24304 O24305
O24308 O24310 O24313 O24338 O24339 O24340 O24361 O24362 O24364 O24369 O24385
O24386 O24387 O24388 O24396 O24407 O24408 O24409 O24410 O24413 O24415 O24457
O24463 O24464 O24467 O24473 O24474 O24475 O24495 O24496 O24521 O24528 O24529
O24534 O24540 O24541 O24542 O24543 O24562 O24573 O24575 O24578 O24591 O24600
O24606 O24616 O24621 O24629 O24633 O24634 O24645 O24648 O24650 O24661 O24676
O24677 O24678 O24679 O24689 O24690 O24691 O24692 O24693 O24694 O24695 O24696
O24697 O24698 O24699 O24700 O24701 O24702 O24703 O24704 O24705 O24706 O24707
O24708 O24709 O24710 O24711 O24712 O24714 O24721 O24730 O24731 O24732 O24733
O24734 O24735 O24738 O24743 O24744 O24748 O24750 O24752 O24753 O24755 O24766
O24767 O24785 O24786 O24787 O24821 O24822 O24840 O24853 O24854 O24855 O24872
O24876 O24915 O24924 O24926 O24933 O24970 O24973 O24980 O24999 O25002 O25008
O25011 O25029 O25039 O25042 O25045 O25046 O25067 O25074 O25080 O25087 O25089
O25093 O25094 O25096 O25097 O25099 O25101 O25103 O25114 O25116 O25121 O25130
O25136 O25138 O25149 O25150 O25151 O25156 O25157 O25165 O25173 O25225 O25236
O25237 O25242 O25249 O25253 O25254 O25290 O25294 O25296 O25320 O25325 O25336
O25338 O25340 O25342 O25343 O25356 O25360 O25372 O25376 O25386 O25411 O25424
O25427 O25428 O25434 O25436 O25441 O25443 O25484 O25499 O25502 O25503 O25505
O25506 O25511 O25514 O25515 O25516 O25525 O25533 O25534 O25536 O25544 O25549
O25560 O25577 O25581 O25584 O25587 O25595 O25598 O25599 O25603 O25608 O25610
O25613 O25620 O25626 O25627 O25654 O25657 O25664 O25671 O25680 O25681 O25683
O25686 O25687 O25688 O25690 O25693 O25698 O25699 O25702 O25703 O25711 O25716
O25730 O25731 O25748 O25758 O25766 O25767 O25769 O25773 O25775 O25776 O25781
O25801 O25806 O25812 O25817 O25826 O25827 O25830 O25835 O25836 O25838 O25841
O25842 O25849 O25867 O25868 O25869 O25883 O25890 O25893 O25895 O25896 O25909
O25910 O25916 O25918 O25926 O25929 O25936 O25943 O25944 O25956 O25959 O25961
O25963 O25966 O25970 O25971 O25972 O25982 O25983 O25984 O25985 O25991 O26004

O26009 O26010 O26011 O26016 O26027 O26043 O26051 O26060 O26061 O26066 O26087
O26102 O26103 O26106 O26107 O26110 O26111 O26112 O26113 O26114 O26115 O26116
O26117 O26118 O26119 O26120 O26121 O26122 O26123 O26124 O26125 O26126 O26127
O26128 O26129 O26130 O26131 O26132 O26133 O26135 O26137 O26138 O26139 O26140
O26141 O26142 O26143 O26144 O26145 O26146 O26147 O26148 O26150 O26151 O26153
O26154 O26156 O26157 O26158 O26217 O26223 O26230 O26232 O26237 O26244 O26246
O26249 O26250 O26253 O26255 O26258 O26262 O26268 O26270 O26271 O26272 O26273
O26278 O26279 O26284 O26285 O26290 O26292 O26293 O26309 O26310 O26311 O26314
O26320 O26325 O26326 O26327 O26328 O26329 O26330 O26333 O26334 O26337 O26338
O26340 O26342 O26343 O26344 O26345 O26346 O26347 O26350 O26351 O26352 O26355
O26356 O26357 O26358 O26359 O26360 O26361 O26363 O26366 O26367 O26368 O26369
O26373 O26503 O26518 O26519 O26533 O26534 O26541 O26542 O26543 O26607 O26631
O26635 O26638 O26640 O26641 O26644 O26653 O26654 O26662 O26663 O26664 O26666
O26674 O26679 O26680 O26683 O26684 O26685 O26686 O26687 O26703 O26708 O26710
O26712 O26715 O26719 O26734 O26739 O26742 O26744 O26745 O26747 O26758 O26760
O26762 O26773 O26774 O26776 O26777 O26778 O26779 O26780 O26781 O26782 O26783
O26784 O26786 O26799 O26800 O26801 O26802 O26803 O26805 O26806 O26807 O26819
O26820 O26823 O26824 O26833 O26834 O26837 O26839 O26843 O26851 O26859 O26860
O26861 O26864 O26865 O26867 O26868 O26869 O26876 O26877 O26880 O26883 O26884
O26885 O26890 O26891 O26892 O26894 O26896 O26898 O26899 O26900 O26901 O26902
O26903 O26909 O26911 O26914 O26915 O26917 O26918 O26928 O26931 O26932 O26934
O26938 O26944 O26945 O26949 O26950 O26951 O26952 O26953 O26955 O26956 O26957
O26958 O26960 O26962 O26964 O26965 O26971 O26972 O26976 O26977 O26978 O26979
O26981 O27001 O27002 O27026 O27030 O27048 O27051 O27054 O27068 O27075 O27077
O27079 O27081 O27083 O27085 O27088 O27089 O27090 O27092 O27093 O27096 O27097
O27098 O27099 O27102 O27112 O27113 O27114 O27115 O27120 O27121 O27122 O27123
O27124 O27125 O27126 O27127 O27129 O27130 O27131 O27132 O27133 O27139 O27140
O27143 O27152 O27155 O27156 O27158 O27179 O27180 O27181 O27182 O27185 O27186
O27188 O27190 O27191 O27194 O27197 O27198 O27199 O27200 O27209 O27211 O27214
O27216 O27218 O27221 O27222 O27224 O27232 O27233 O27234 O27235 O27236 O27246
O27250 O27252 O27255 O27258 O27262 O27263 O27264 O27266 O27269 O27270 O27272
O27273 O27274 O27275 O27276 O27280 O27281 O27283 O27295 O27296 O27303 O27304
O27322 O27324 O27336 O27341 O27342 O27344 O27346 O27350 O27351 O27352 O27355
O27362 O27363 O27364 O27365 O27367 O27369 O27371 O27372 O27375 O27378 O27381
O27383 O27389 O27390 O27392 O27397 O27398 O27404 O27427 O27428 O27429 O27430
O27433 O27434 O27436 O27438 O27439 O27440 O27441 O27442 O27443 O27447 O27448
O27449 O27454 O27456 O27458 O27463 O27464 O27472 O27474 O27476 O27477 O27481
O27484 O27490 O27491 O27492 O27494 O27495 O27496 O27498 O27502 O27504 O27509
O27518 O27520 O27525 O27527 O27540 O27543 O27544 O27545 O27549 O27550 O27552
O27554 O27565 O27566 O27568 O27572 O27573 O27579 O27580 O27585 O27586 O27587
O27590 O27593 O27600 O27602 O27605 O27606 O27607 O27612 O27617 O27624 O27627
O27628 O27630 O27633 O27634 O27635 O27636 O27637 O27642 O27646 O27647 O27648
O27649 O27650 O27652 O27653 O27655 O27656 O27657 O27661 O27664 O27667 O27668
O27670 O27673 O27679 O27692 O27693 O27694 O27695 O27696 O27697 O27698 O27699

O27705 O27710 O27715 O27716 O27717 O27718 O27719 O27720 O27724 O27728 O27731
O27732 O27734 O27743 O27744 O27745 O27747 O27748 O27771 O27772 O27784 O27793
O27795 O27797 O27801 O27802 O27818 O27825 O27830 O27844 O27848 O27855 O27859
O27860 O27874 O27875 O27880 O27881 O27888 O27890 O27893 O27900 O27902 O27906
O27907 O27911 O27918 O27919 O27924 O27936 O27937 O27939 O27941 O27943 O27944
O27945 O27947 O27950 O27952 O27953 O27954 O27955 O27956 O27957 O27958 O27962
O27964 O27965 O27967 O27969 O27970 O27974 O27977 O27983 O27984 O27986 O27991
O27992 O27997 O27999 O28000 O28001 O28002 O28007 O28014 O28017 O28019 O28020
O28021 O28022 O28025 O28028 O28029 O28032 O28033 O28034 O28041 O28043 O28045
O28046 O28049 O28050 O28051 O28053 O28054 O28058 O28059 O28062 O28071 O28073
O28076 O28077 O28079 O28082 O28084 O28087 O28088 O28090 O28095 O28096 O28097
O28098 O28102 O28105 O28108 O28110 O28112 O28115 O28117 O28121 O28125 O28126
O28128 O28130 O28132 O28133 O28135 O28137 O28141 O28142 O28145 O28152 O28157
O28159 O28164 O28167 O28173 O28174 O28183 O28184 O28185 O28186 O28187 O28190
O28192 O28197 O28204 O28205 O28206 O28207 O28208 O28210 O28211 O28212 O28213
O28214 O28215 O28216 O28218 O28219 O28220 O28228 O28229 O28237 O28242 O28244
O28249 O28255 O28257 O28259 O28272 O28273 O28275 O28277 O28278 O28279 O28282
O28283 O28292 O28293 O28294 O28301 O28303 O28304 O28305 O28310 O28314 O28316
O28317 O28323 O28324 O28325 O28329 O28335 O28337 O28338 O28339 O28340 O28341
O28343 O28344 O28354 O28355 O28356 O28357 O28358 O28359 O28360 O28361 O28362
O28363 O28364 O28365 O28366 O28367 O28368 O28369 O28370 O28371 O28372 O28373
O28374 O28375 O28376 O28379 O28380 O28383 O28385 O28386 O28387 O28389 O28390
O28392 O28394 O28399 O28401 O28403 O28406 O28409 O28414 O28415 O28417 O28419
O28420 O28422 O28429 O28430 O28431 O28434 O28438 O28439 O28440 O28466 O28467
O28469 O28471 O28475 O28478 O28481 O28484 O28486 O28487 O28488 O28490 O28491
O28496 O28497 O28499 O28512 O28513 O28516 O28520 O28523 O28524 O28527 O28533
O28535 O28538 O28541 O28542 O28543 O28544 O28548 O28550 O28552 O28555 O28556
O28557 O28559 O28560 O28561 O28575 O28578 O28579 O28580 O28584 O28590 O28592
O28593 O28596 O28597 O28606 O28608 O28611 O28630 O28631 O28635 O28637 O28642
O28644 O28646 O28663 O28664 O28668 O28669 O28670 O28671 O28672 O28673 O28682
O28685 O28691 O28696 O28700 O28706 O28715 O28717 O28719 O28724 O28725 O28732
O28733 O28743 O28751 O28753 O28756 O28758 O28765 O28766 O28768 O28770 O28771
O28775 O28777 O28779 O28780 O28781 O28782 O28783 O28784 O28786 O28787 O28791
O28792 O28794 O28807 O28816 O28819 O28821 O28825 O28826 O28832 O28836 O28837
O28839 O28842 O28847 O28848 O28853 O28854 O28856 O28864 O28865 O28869 O28876
O28877 O28881 O28882 O28884 O28885 O28889 O28904 O28908 O28921 O28927 O28928
O28930 O28931 O28935 O28936 O28938 O28943 O28949 O28951 O28952 O28955 O28964
O28965 O28966 O28970 O28972 O28974 O28980 O28987 O28988 O28990 O28994 O28995
O28996 O28997 O28999 O29003 O29008 O29011 O29013 O29015 O29021 O29026 O29027
O29028 O29032 O29033 O29040 O29041 O29048 O29049 O29050 O29052 O29054 O29055
O29056 O29057 O29067 O29069 O29072 O29088 O29107 O29108 O29113 O29114 O29115
O29118 O29119 O29132 O29134 O29135 O29136 O29137 O29138 O29139 O29145 O29150
O29151 O29152 O29153 O29156 O29157 O29159 O29161 O29163 O29164 O29165 O29170
O29171 O29173 O29175 O29177 O29178 O29181 O29182 O29184 O29185 O29187 O29189

O29190 O29191 O29195 O29196 O29203 O29206 O29207 O29208 O29209 O29215 O29224
O29230 O29231 O29238 O29248 O29254 O29259 O29262 O29265 O29269 O29276 O29277
O29284 O29305 O29306 O29309 O29310 O29313 O29315 O29319 O29320 O29321 O29322
O29323 O29324 O29325 O29329 O29333 O29342 O29345 O29346 O29347 O29348 O29351
O29352 O29353 O29357 O29361 O29362 O29364 O29365 O29366 O29368 O29370 O29378
O29379 O29380 O29382 O29384 O29386 O29387 O29388 O29392 O29395 O29399 O29401
O29406 O29407 O29414 O29415 O29417 O29421 O29425 O29428 O29429 O29439 O29440
O29443 O29444 O29445 O29446 O29455 O29457 O29458 O29460 O29461 O29469 O29481
O29482 O29484 O29485 O29486 O29490 O29491 O29492 O29493 O29494 O29501 O29502
O29507 O29510 O29511 O29513 O29515 O29516 O29518 O29520 O29521 O29524 O29525
O29526 O29534 O29535 O29537 O29540 O29544 O29548 O29556 O29557 O29560 O29563
O29564 O29568 O29571 O29581 O29587 O29590 O29591 O29592 O29602 O29605 O29608
O29610 O29612 O29613 O29622 O29624 O29626 O29627 O29630 O29632 O29634 O29636
O29642 O29643 O29644 O29646 O29650 O29651 O29652 O29655 O29658 O29663 O29664
O29665 O29666 O29667 O29669 O29679 O29681 O29689 O29691 O29695 O29697 O29699
O29701 O29703 O29712 O29720 O29722 O29723 O29724 O29729 O29730 O29734 O29737
O29739 O29741 O29742 O29746 O29753 O29755 O29756 O29757 O29758 O29759 O29760
O29761 O29766 O29767 O29774 O29779 O29781 O29782 O29789 O29796 O29798 O29805
O29807 O29808 O29809 O29816 O29823 O29827 O29828 O29830 O29832 O29833 O29836
O29839 O29841 O29842 O29845 O29849 O29861 O29862 O29863 O29866 O29868 O29870
O29871 O29874 O29876 O29877 O29891 O29903 O29904 O29910 O29911 O29912 O29913
O29917 O29926 O29929 O29931 O29934 O29938 O29944 O29945 O29947 O29949 O29953
O29959 O29963 O29969 O29970 O29971 O29975 O29979 O29982 O29984 O29986 O29987
O29989 O29990 O29995 O29996 O29998 O29999 O30001 O30006 O30009 O30010 O30011
O30013 O30019 O30020 O30024 O30027 O30030 O30032 O30033 O30045 O30050 O30052
O30055 O30056 O30058 O30075 O30082 O30086 O30087 O30101 O30107 O30108 O30111
O30118 O30120 O30123 O30124 O30126 O30127 O30128 O30129 O30130 O30136 O30140
O30152 O30153 O30156 O30163 O30166 O30167 O30169 O30172 O30175 O30176 O30177
O30178 O30179 O30180 O30181 O30182 O30184 O30186 O30189 O30199 O30205 O30207
O30213 O30236 O30237 O30238 O30240 O30245 O30250 O30256 O30257 O30259 O30268
O30272 O30273 O30274 O30279 O30280 O30291 O30297 O30298 O30300 O30305 O30308
O30314 O30321 O30325 O30330 O30332 O30334 O30338 O30360 O30372 O30394 O30397
O30408 O30409 O30415 O30418 O30444 O30461 O30472 O30477 O30478 O30498 O30508
O30509 O30520 O30548 O30549 O30557 O30560 O30561 O30563 O30564 O30569 O30577
O30601 O30602 O30612 O30633 O30640 O30641 O30642 O30709 O30711 O30723 O30724
O30725 O30745 O30746 O30762 O30807 O30808 O30819 O30823 O30824 O30825 O30826
O30827 O30836 O30847 O30851 O30853 O30862 O30867 O30868 O30869 O30875 O30884
O30911 O30913 O30917 O30919 O30920 O30931 O30970 O30971 O30973 O30974 O30999
O31017 O31018 O31019 O31020 O31038 O31042 O31043 O31046 O31047 O31060 O31066
O31067 O31073 O31074 O31087 O31139 O31147 O31151 O31156 O31158 O31159 O31160
O31161 O31162 O31163 O31164 O31165 O31166 O31168 O31178 O31183 O31186 O31194
O31196 O31197 O31200 O31201 O31206 O31207 O31211 O31212 O31213 O31219 O31227
O31266 O31269 O31270 O31287 O31288 O31292 O31293 O31294 O31297 O31298 O31300
O31301 O31332 O31352 O31381 O31385 O31404 O31408 O31410 O31414 O31417 O31418

O31423 O31429 O31430 O31432 O31435 O31437 O31440 O31443 O31449 O31451 O31452
O31454 O31456 O31457 O31458 O31459 O31461 O31463 O31465 O31466 O31472 O31473
O31474 O31475 O31480 O31482 O31484 O31489 O31490 O31494 O31496 O31497 O31498
O31500 O31503 O31504 O31505 O31509 O31510 O31511 O31512 O31513 O31518 O31521
O31523 O31524 O31525 O31528 O31529 O31530 O31534 O31535 O31537 O31538 O31541
O31542 O31544 O31545 O31546 O31547 O31550 O31552 O31556 O31557 O31558 O31560
O31562 O31564 O31570 O31571 O31572 O31573 O31574 O31575 O31576 O31578 O31581
O31584 O31585 O31586 O31587 O31588 O31589 O31590 O31592 O31594 O31596 O31597
O31601 O31604 O31605 O31606 O31607 O31609 O31610 O31611 O31612 O31613 O31614
O31616 O31617 O31618 O31619 O31620 O31622 O31625 O31626 O31628 O31629 O31630
O31631 O31632 O31633 O31637 O31639 O31640 O31641 O31642 O31644 O31646 O31647
O31648 O31649 O31650 O31653 O31654 O31656 O31659 O31662 O31663 O31664 O31665
O31666 O31667 O31668 O31669 O31672 O31673 O31675 O31676 O31677 O31678 O31679
O31680 O31682 O31683 O31684 O31689 O31690 O31691 O31697 O31698 O31699 O31700
O31701 O31702 O31703 O31704 O31705 O31706 O31713 O31714 O31716 O31717 O31718
O31719 O31720 O31724 O31725 O31726 O31727 O31728 O31738 O31739 O31740 O31741
O31742 O31743 O31744 O31745 O31751 O31753 O31755 O31760 O31761 O31764 O31766
O31767 O31768 O31769 O31775 O31776 O31777 O31778 O31779 O31782 O31784 O31786
O31787 O31788 O31790 O31794 O31795 O31796 O31798 O31799 O31800 O31801 O31802
O31806 O31807 O31812 O31817 O31818 O31819 O31824 O31825 O31826 O31827 O31829
O31831 O31834 O31837 O31838 O31839 O31841 O31842 O31844 O31846 O31847 O31848
O31850 O31853 O31854 O31856 O31857 O31858 O31863 O31864 O31865 O31866 O31868
O31869 O31870 O31875 O31877 O31878 O31879 O31880 O31881 O31882 O31883 O31885
O31886 O31887 O31889 O31890 O31891 O31892 O31893 O31894 O31896 O31897 O31898
O31899 O31900 O31902 O31903 O31904 O31905 O31906 O31907 O31908 O31909 O31910
O31911 O31912 O31913 O31914 O31915 O31916 O31917 O31918 O31920 O31921 O31922
O31924 O31927 O31928 O31931 O31932 O31935 O31936 O31938 O31939 O31940 O31943
O31944 O31945 O31947 O31948 O31949 O31950 O31951 O31952 O31953 O31954 O31955
O31956 O31957 O31958 O31959 O31960 O31961 O31962 O31963 O31964 O31965 O31966
O31967 O31968 O31969 O31970 O31971 O31972 O31977 O31978 O31979 O31980 O31981
O31983 O31986 O31990 O31991 O31993 O31995 O31996 O31997 O31999 O32000 O32002
O32003 O32004 O32006 O32007 O32012 O32013 O32014 O32020 O32026 O32027 O32028
O32029 O32030 O32033 O32034 O32035 O32036 O32037 O32038 O32039 O32040 O32042
O32044 O32049 O32053 O32054 O32055 O32057 O32059 O32062 O32065 O32066 O32067
O32068 O32072 O32073 O32075 O32078 O32079 O32085 O32089 O32090 O32091 O32092
O32093 O32094 O32096 O32097 O32098 O32099 O32102 O32103 O32106 O32110 O32113
O32114 O32116 O32117 O32118 O32119 O32125 O32126 O32127 O32129 O32132 O32135
O32136 O32137 O32138 O32141 O32142 O32143 O32144 O32145 O32146 O32147 O32148
O32149 O32151 O32152 O32153 O32157 O32158 O32159 O32160 O32161 O32162 O32163
O32164 O32165 O32166 O32170 O32171 O32172 O32173 O32174 O32175 O32179 O32180
O32181 O32183 O32184 O32185 O32186 O32187 O32192 O32207 O32210 O32213 O32214
O32215 O32221 O32222 O32223 O32224 O32228 O32229 O32230 O32231 O32232 O32234
O32235 O32236 O32237 O32238 O32245 O32248 O32253 O32254 O32255 O32259 O32263
O32264 O32265 O32266 O32267 O32268 O32272 O32276 O32280 O32283 O32285 O32286

O32291 O32293 O32294 O32296 O32302 O32307 O32321 O32323 O32350 O32377 O32378
O32414 O32434 O32437 O32445 O32449 O32450 O32462 O32463 O32464 O32465 O32481
O32482 O32483 O32504 O32507 O32508 O32518 O32583 O32605 O32606 O32613 O32614
O32620 O32712 O32719 O32720 O32721 O32723 O32724 O32726 O32727 O32740 O32741
O32755 O32756 O32757 O32765 O32770 O32771 O32797 O32799 O32801 O32806 O32808
O32810 O32823 O32846 O32847 O32848 O32849 O32853 O32854 O32855 O32857 O32858
O32869 O32874 O32879 O32881 O32885 O32886 O32912 O32913 O32915 O32920 O32953
O32954 O32955 O32956 O32960 O32961 O32962 O32965 O32967 O32975 O32978 O32982
O32983 O32984 O32985 O32986 O32987 O32988 O32989 O32990 O32993 O32994 O32995
O32996 O32997 O32998 O32999 O33000 O33001 O33002 O33007 O33011 O33012 O33014
O33015 O33016 O33017 O33020 O33022 O33024 O33038 O33039 O33040 O33045 O33046
O33060 O33062 O33064 O33065 O33095 O33096 O33099 O33102 O33104 O33105 O33106
O33107 O33112 O33113 O33114 O33116 O33117 O33120 O33123 O33124 O33127 O33128
O33132 O33149 O33159 O33160 O33174 O33277 O33289 O33341 O33347 O33348 O33361
O33367 O33380 O33395 O33422 O33431 O33453 O33465 O33469 O33472 O33475 O33477
O33478 O33481 O33499 O33500 O33505 O33506 O33508 O33509 O33510 O33512 O33513
O33519 O33520 O33522 O33523 O33525 O33528 O33529 O33535 O33543 O33544 O33545
O33564 O33565 O33567 O33575 O33579 O33580 O33581 O33582 O33594 O33600 O33603
O33605 O33611 O33618 O33655 O33658 O33659 O33662 O33665 O33682 O33684 O33685
O33686 O33698 O33701 O33730 O33732 O33752 O33756 O33768 O33770 O33771 O33772
O33773 O33774 O33775 O33776 O33777 O33778 O33780 O33783 O33799 O33804 O33812
O33813 O33815 O33817 O33818 O33819 O33820 O33822 O33830 O33833 O33836 O33844
O33845 O33873 O33914 O33915 O33924 O33925 O33926 O33939 O33945 O33946 O33947
O33948 O33949 O33950 O33951 O33963 O33964 O33998 O34002 O34011 O34022 O34023
O34031 O34055 O34090 O34098 O34101 O34104 O34106 O34125 O34126 O34133 O34136
O34138 O34153 O34154 O34162 O34167 O34171 O34191 O34194 O34198 O34201 O34202
O34204 O34209 O34239 O34241 O34242 O34268 O34272 O34273 O34274 O34275 O34292
O34293 O34294 O34295 O34296 O34299 O34303 O34305 O34306 O34308 O34309 O34314
O34317 O34318 O34320 O34321 O34324 O34326 O34328 O34331 O34334 O34336 O34339
O34342 O34345 O34346 O34347 O34350 O34351 O34352 O34354 O34355 O34357 O34359
O34363 O34364 O34365 O34366 O34370 O34374 O34376 O34377 O34380 O34381 O34384
O34389 O34394 O34398 O34399 O34400 O34401 O34402 O34403 O34405 O34407 O34409
O34411 O34412 O34413 O34414 O34419 O34421 O34423 O34425 O34426 O34429 O34433
O34438 O34441 O34443 O34445 O34446 O34448 O34449 O34450 O34452 O34453 O34454
O34457 O34458 O34459 O34464 O34466 O34468 O34469 O34471 O34475 O34476 O34478
O34479 O34481 O34484 O34489 O34493 O34495 O34496 O34497 O34498 O34499 O34503
O34504 O34508 O34512 O34513 O34514 O34516 O34520 O34522 O34526 O34527 O34528
O34529 O34533 O34534 O34536 O34537 O34539 O34542 O34543 O34549 O34553 O34555
O34557 O34558 O34559 O34564 O34565 O34569 O34570 O34574 O34577 O34580 O34581
O34584 O34585 O34589 O34591 O34592 O34593 O34595 O34596 O34598 O34599 O34600
O34604 O34605 O34607 O34611 O34612 O34614 O34617 O34619 O34621 O34623 O34626
O34627 O34632 O34633 O34634 O34635 O34637 O34639 O34640 O34642 O34643 O34645
O34647 O34651 O34656 O34658 O34660 O34661 O34662 O34664 O34672 O34673 O34676
O34678 O34680 O34681 O34682 O34683 O34685 O34687 O34689 O34692 O34696 O34699

O34700 O34701 O34702 O34703 O34704 O34707 O34709 O34712 O34714 O34715 O34720
O34722 O34727 O34730 O34736 O34737 O34743 O34744 O34748 O34750 O34751 O34759
O34760 O34762 O34763 O34764 O34766 O34767 O34768 O34769 O34772 O34775 O34776
O34778 O34780 O34782 O34783 O34785 O34787 O34788 O34789 O34791 O34795 O34796
O34797 O34800 O34803 O34806 O34808 O34809 O34812 O34813 O34815 O34817 O34820
O34823 O34824 O34825 O34827 O34828 O34829 O34831 O34833 O34834 O34838 O34842
O34843 O34844 O34845 O34846 O34850 O34851 O34855 O34858 O34859 O34860 O34861
O34863 O34869 O34872 O34874 O34877 O34879 O34880 O34882 O34885 O34886 O34887
O34891 O34892 O34893 O34895 O34896 O34899 O34901 O34903 O34904 O34906 O34909
O34910 O34912 O34913 O34915 O34916 O34919 O34920 O34921 O34923 O34924 O34925
O34926 O34930 O34932 O34934 O34937 O34939 O34940 O34942 O34943 O34944 O34947
O34949 O34950 O34951 O34954 O34955 O34958 O34959 O34962 O34967 O34969 O34970
O34973 O34974 O34976 O34977 O34980 O34981 O34983 O34984 O34988 O34990 O34993
O34994 O34996 O35000 O35001 O35003 O35006 O35008 O35009 O35010 O35011 O35013
O35014 O35016 O35017 O35019 O35020 O35022 O35023 O35025 O35030 O35031 O35033
O35034 O35038 O35040 O35041 O35042 O35045 O35046 O35047 O35071 O35077 O35098
O35115 O35125 O35127 O35130 O35134 O35137 O35156 O35185 O35207 O35215 O35226
O35242 O35260 O35261 O35286 O35295 O35317 O35326 O35331 O35347 O35368 O35381
O35385 O35392 O35403 O35437 O35450 O35473 O35486 O35490 O35491 O35492 O35493
O35508 O35521 O35522 O35523 O35524 O35552 O35589 O35593 O35615 O35621 O35627
O35648 O35652 O35654 O35655 O35660 O35691 O35698 O35709 O35711 O35719 O35723
O35737 O35738 O35740 O35750 O35762 O35767 O35779 O35780 O35787 O35817 O35819
O35820 O35831 O35847 O35864 O35870 O35876 O35878 O35885 O35900 O35920 O35943
O35945 O35952 O35955 O35969 O35980 O35984 O35986 O36014 O36015 O36016 O36017
O36018 O36021 O36023 O36024 O36025 O36027 O36029 O36030 O36031 O36032 O36033
O36041 O36051 O36052 O36054 O36057 O36307 O36358 O36360 O36361 O36363 O36365
O36366 O36368 O36369 O36370 O36371 O36373 O36374 O36375 O36376 O36378 O36379
O36380 O36381 O36382 O36383 O36384 O36386 O36389 O36390 O36391 O36392 O36393
O36394 O36395 O36397 O36398 O36399 O36402 O36404 O36406 O36407 O36409 O36411
O36412 O36413 O36415 O36418 O36419 O36421 O36433 O36453 O36635 O36639 O36966
O37061 O37319 O39163 O39164 O39432 O39433 O39436 O39521 O39619 O39821 O39822
O39829 O39831 O39832 O39988 O40626 O40627 O40958 O40977 O40979 O40980 O40981
O40982 O40983 O40987 O41033 O41056 O41107 O41156 O41178 O41252 O41253 O41345
O41353 O41518 O41519 O42037 O42038 O42047 O42050 O42053 O42062 O42091 O42097
O42099 O42105 O42106 O42115 O42130 O42131 O42132 O42148 O42152 O42173 O42226
O42227 O42230 O42242 O42252 O42259 O42260 O42265 O42290 O42295 O42342 O42365
O42366 O42367 O42368 O42370 O42387 O42395 O42406 O42409 O42410 O42425 O42450
O42467 O42469 O42470 O42494 O42502 O42503 O42504 O42506 O42565 O42569 O42575
O42584 O42586 O42587 O42601 O42606 O42611 O42615 O42617 O42621 O42622 O42631
O42634 O42636 O42637 O42643 O42644 O42645 O42647 O42648 O42649 O42650 O42652
O42653 O42654 O42658 O42659 O42661 O42662 O42666 O42667 O42668 O42700 O42709
O42713 O42717 O42724 O42764 O42767 O42768 O42771 O42777 O42783 O42786 O42794
O42804 O42820 O42821 O42831 O42832 O42837 O42839 O42842 O42845 O42846 O42847
O42849 O42851 O42853 O42854 O42855 O42858 O42859 O42860 O42861 O42868 O42869

O42870 O42871 O42872 O42873 O42874 O42875 O42877 O42878 O42882 O42883 O42888
O42889 O42890 O42893 O42894 O42895 O42896 O42897 O42898 O42903 O42904 O42905
O42906 O42908 O42909 O42910 O42911 O42912 O42913 O42915 O42917 O42919 O42921
O42923 O42924 O42925 O42927 O42928 O42929 O42931 O42934 O42936 O42938 O42939
O42941 O42943 O42945 O42947 O42948 O42951 O42953 O42954 O42955 O42957 O42958
O42964 O42971 O42973 O42975 O42977 O42978 O42979 O42980 O42993 O42995 O42996
O42998 O43003 O43004 O43006 O43007 O43008 O43010 O43011 O43012 O43013 O43014
O43017 O43018 O43020 O43024 O43026 O43030 O43031 O43034 O43036 O43037 O43040
O43041 O43042 O43044 O43045 O43048 O43051 O43056 O43058 O43060 O43063 O43065
O43066 O43067 O43068 O43070 O43071 O43075 O43076 O43078 O43082 O43083 O43084
O43086 O43088 O43089 O43090 O43093 O43099 O43100 O43101 O43102 O43103 O43105
O43112 O43114 O43120 O43123 O43125 O43133 O43137 O43147 O43151 O43167 O43172
O43175 O43186 O43189 O43196 O43247 O43248 O43251 O43252 O43261 O43296 O43298
O43301 O43304 O43309 O43313 O43316 O43325 O43345 O43361 O43365 O43396 O43414
O43423 O43435 O43542 O43592 O43593 O43638 O43670 O43680 O43704 O43708 O43709
O43711 O43716 O43763 O43766 O43790 O43791 O43809 O43812 O43824 O43837 O43913
O43918 O43938 O43940 O43976 O43980 O43999 O44006 O44010 O44012 O44081 O44104
O44105 O44107 O44119 O44125 O44156 O44199 O44219 O44220 O44221 O44388 O44400
O44406 O44410 O44424 O44437 O44440 O44441 O44445 O44451 O44498 O44510 O44535
O44548 O44552 O44568 O44572 O44628 O44740 O44743 O44749 O44750 O44757 O44770
O44960 O45100 O45168 O45211 O45228 O45241 O45244 O45291 O45380 O45406 O45420
O45433 O45436 O45449 O45460 O45487 O45489 O45521 O45526 O45551 O45583 O45595
O45679 O45687 O45717 O45766 O45782 O45784 O45830 O45907 O45935 O45952 O46036
O46040 O46043 O46072 O46080 O46089 O46098 O46102 O46106 O46119 O46157 O46160
O46232 O46234 O46236 O46238 O46240 O46242 O46244 O46248 O46250 O46252 O46254
O46256 O46258 O46260 O46262 O46310 O46313 O46339 O46341 O46374 O46404 O46411
O46414 O46415 O46419 O46503 O46504 O46560 O46567 O46640 O46649 O46650 O46660
O46894 O46895 O46896 O46897 O46898 O46899 O46900 O46901 O46902 O46903 O46904
O46905 O46906 O46907 O46908 O46909 O46910 O46912 O46913 O46914 O46915 O46917
O46993 O46996 O47026 O47027 O47029 O47032 O47041 O47125 O47128 O47129 O47135
O47141 O47143 O47150 O47157 O47164 O47169 O47170 O48393 O48394 O48395 O48396
O48397 O48398 O48399 O48400 O48401 O48402 O48403 O48404 O48405 O48406 O48407
O48408 O48409 O48411 O48412 O48413 O48414 O48424 O48425 O48427 O48428 O48429
O48446 O48448 O48449 O48459 O48499 O48500 O48513 O48520 O48522 O48534 O48550
O48551 O48556 O48557 O48558 O48559 O48582 O48593 O48604 O48609 O48650 O48652
O48653 O48658 O48659 O48660 O48665 O48668 O48676 O48686 O48700 O48706 O48707
O48713 O48715 O48721 O48722 O48724 O48737 O48767 O48771 O48772 O48781 O48790
O48791 O48802 O48822 O48844 O48847 O48881 O48882 O48890 O48897 O48901 O48902
O48905 O48923 O48927 O48935 O48951 O48964 O48965 O49044 O49046 O49065 O49066
O49068 O49071 O49073 O49074 O49118 O49133 O49139 O49160 O49169 O49196 O49204
O49213 O49224 O49227 O49250 O49255 O49279 O49285 O49286 O49289 O49290 O49312
O49315 O49316 O49326 O49332 O49343 O49399 O49404 O49413 O49420 O49421 O49429
O49436 O49459 O49471 O49482 O49484 O49488 O49498 O49500 O49506 O49508 O49543
O49550 O49558 O49561 O49562 O49565 O49595 O49596 O49597 O49606 O49608 O49618

O49619 O49624 O49627 O49647 O49651 O49662 O49663 O49687 O49697 O49698 O49707
O49710 O49711 O49782 O49816 O49817 O49818 O49835 O49836 O49845 O49849 O49850
O49852 O49853 O49856 O49884 O49885 O49886 O49894 O49901 O49923 O49934 O49937
O49939 O49954 O49972 O50001 O50003 O50044 O50046 O50055 O50064 O50078 O50083
O50087 O50089 O50093 O50097 O50098 O50105 O50109 O50110 O50116 O50123 O50129
O50146 O50147 O50161 O50163 O50165 O50171 O50172 O50173 O50174 O50175 O50177
O50185 O50188 O50202 O50214 O50224 O50230 O50231 O50235 O50236 O50237 O50258
O50273 O50274 O50286 O50293 O50302 O50304 O50305 O50306 O50310 O50311 O50312
O50313 O50314 O50316 O50323 O50340 O50406 O50456 O50461 O50462 O50467 O50468
O50487 O50488 O50519 O50533 O50562 O50563 O50564 O50565 O50580 O50581 O50584
O50595 O50606 O50607 O50627 O50628 O50630 O50631 O50632 O50633 O50634 O50635
O50655 O50656 O50657 O50821 O50843 O50965 O50977 O50979 O50999 O51035 O51038
O51040 O51044 O51046 O51053 O51056 O51057 O51060 O51064 O51072 O51075 O51077
O51081 O51082 O51087 O51088 O51089 O51091 O51092 O51093 O51101 O51111 O51112
O51114 O51116 O51122 O51124 O51125 O51126 O51127 O51128 O51132 O51134 O51139
O51140 O51141 O51142 O51146 O51147 O51148 O51149 O51154 O51155 O51157 O51160
O51173 O51182 O51191 O51201 O51202 O51204 O51206 O51207 O51208 O51214 O51215
O51216 O51218 O51219 O51226 O51229 O51232 O51238 O51240 O51244 O51246 O51247
O51257 O51263 O51264 O51267 O51271 O51286 O51291 O51293 O51309 O51313 O51314
O51315 O51316 O51317 O51318 O51323 O51325 O51333 O51338 O51339 O51343 O51344
O51345 O51347 O51348 O51349 O51351 O51352 O51353 O51354 O51355 O51357 O51360
O51363 O51376 O51383 O51395 O51396 O51401 O51402 O51413 O51418 O51419 O51423
O51424 O51428 O51438 O51439 O51440 O51441 O51442 O51443 O51444 O51445 O51446
O51447 O51448 O51449 O51450 O51452 O51453 O51454 O51455 O51456 O51461 O51468
O51477 O51497 O51498 O51502 O51522 O51526 O51528 O51532 O51535 O51536 O51540
O51544 O51545 O51547 O51551 O51555 O51556 O51557 O51558 O51560 O51567 O51568
O51571 O51572 O51575 O51578 O51581 O51589 O51601 O51602 O51603 O51625 O51627
O51628 O51632 O51634 O51638 O51639 O51640 O51641 O51642 O51645 O51646 O51648
O51653 O51662 O51669 O51672 O51680 O51696 O51698 O51710 O51715 O51718 O51719
O51721 O51722 O51723 O51726 O51727 O51728 O51737 O51739 O51740 O51741 O51742
O51743 O51744 O51745 O51749 O51752 O51757 O51759 O51761 O51767 O51768 O51769
O51773 O51776 O51777 O51781 O51782 O51804 O51824 O51830 O51831 O51832 O51834
O51859 O51879 O51881 O51882 O51883 O51884 O51885 O51886 O51887 O51888 O51889
O51890 O51891 O51896 O51897 O51900 O51917 O51926 O51927 O51930 O51931 O51933
O51934 O51940 O51941 O51968 O51974 O52004 O52018 O52025 O52026 O52028 O52029
O52030 O52049 O52063 O52064 O52065 O52070 O52071 O52163 O52164 O52177 O52178
O52199 O52200 O52222 O52225 O52236 O52249 O52250 O52251 O52271 O52278 O52331
O52332 O52333 O52334 O52335 O52336 O52337 O52338 O52339 O52340 O52341 O52342
O52343 O52344 O52345 O52346 O52347 O52348 O52349 O52350 O52352 O52353 O52354
O52377 O52378 O52379 O52380 O52381 O52382 O52383 O52384 O52393 O52394 O52398
O52399 O52400 O52402 O52485 O52512 O52513 O52520 O52532 O52533 O52536 O52537
O52539 O52540 O52547 O52552 O52582 O52587 O52593 O52594 O52597 O52629 O52631
O52632 O52633 O52646 O52655 O52659 O52691 O52692 O52702 O52703 O52704 O52705
O52706 O52714 O52715 O52720 O52748 O52749 O52750 O52751 O52759 O52760 O52761

O52765 O52776 O52787 O52791 O52792 O52793 O52815 O52828 O52831 O52836 O52847
O52914 O52951 O52958 O52960 O52980 O53076 O53077 O53078 O53079 O53080 O53083
O53088 O53089 O53090 O53113 O53125 O53165 O53181 O53193 O53332 O53333 O53334
O53335 O53353 O53379 O53405 O53451 O53464 O53465 O53467 O53468 O53500 O53509
O53518 O53553 O53604 O53605 O53637 O53672 O53778 O53779 O53795 O53803 O53811
O53838 O54003 O54037 O54053 O54055 O54067 O54068 O54075 O54099 O54134 O54143
O54154 O54196 O54210 O54233 O54235 O54242 O54248 O54249 O54259 O54263 O54266
O54281 O54288 O54294 O54296 O54297 O54310 O54311 O54312 O54315 O54326 O54364
O54367 O54376 O54408 O54415 O54418 O54419 O54420 O54422 O54423 O54424 O54425
O54426 O54435 O54451 O54452 O54453 O54457 O54458 O54459 O54479 O54496 O54537
O54552 O54571 O54608 O54705 O54743 O54751 O54772 O54788 O54792 O54818 O54820
O54826 O54839 O54864 O54879 O54887 O54888 O54889 O54891 O54910 O54917 O54931
O54935 O54937 O54941 O54946 O54963 O54975 O54983 O54988 O55000 O55028 O55055
O55060 O55062 O55063 O55076 O55087 O55091 O55092 O55096 O55112 O55128 O55144
O55156 O55176 O55201 O55208 O55230 O55234 O55236 O55239 O55265 O55519 O55527
O55528 O55611 O55709 O55710 O55711 O55712 O55713 O55716 O55717 O55718 O55719
O55720 O55721 O55722 O55723 O55724 O55725 O55730 O55731 O55735 O55736 O55737
O55738 O55739 O55741 O55742 O55748 O55749 O55750 O55751 O55752 O55753 O55754
O55755 O55756 O55757 O55759 O55761 O55763 O55764 O55765 O55766 O55767 O55768
O55769 O55771 O55772 O55777 O55778 O55779 O56042 O56043 O56044 O56045 O56125
O56655 O56831 O56834 O56862 O57030 O57043 O57044 O57167 O57173 O57174 O57175
O57177 O57184 O57187 O57191 O57193 O57198 O57203 O57204 O57207 O57209 O57210
O57214 O57215 O57216 O57220 O57224 O57227 O57230 O57233 O57243 O57246 O57248
O57249 O57250 O57252 O57257 O57259 O57263 O57285 O57286 O57311 O57337 O57342
O57374 O57380 O57383 O57391 O57401 O57406 O57414 O57415 O57437 O57476 O57479
O57526 O57539 O57561 O57568 O57589 O57592 O57594 O57595 O57598 O57601 O57606
O57607 O57611 O57656 O57672 O57682 O57683 O57685 O57693 O57705 O57706 O57708
O57709 O57711 O57712 O57716 O57734 O57738 O57740 O57742 O57746 O57762 O57764
O57765 O57767 O57770 O57774 O57779 O57782 O57792 O57802 O57803 O57805 O57809
O57811 O57818 O57823 O57830 O57836 O57839 O57840 O57848 O57852 O57853 O57854
O57857 O57859 O57860 O57861 O57863 O57864 O57865 O57868 O57878 O57880 O57882
O57899 O57921 O57932 O57934 O57935 O57936 O57939 O57940 O57947 O57948 O57950
O57963 O57971 O57978 O57979 O57980 O57981 O57983 O58001 O58010 O58012 O58019
O58022 O58028 O58033 O58035 O58038 O58045 O58050 O58052 O58054 O58056 O58058
O58061 O58088 O58091 O58096 O58097 O58107 O58185 O58186 O58187 O58211 O58212
O58214 O58226 O58231 O58236 O58247 O58263 O58264 O58268 O58275 O58276 O58281
O58283 O58284 O58285 O58286 O58307 O58308 O58312 O58316 O58320 O58321 O58328
O58349 O58353 O58356 O58358 O58361 O58362 O58367 O58370 O58371 O58377 O58389
O58391 O58393 O58394 O58398 O58403 O58410 O58411 O58412 O58413 O58414 O58415
O58417 O58429 O58430 O58433 O58440 O58441 O58451 O58452 O58456 O58457 O58462
O58466 O58478 O58489 O58495 O58496 O58497 O58498 O58522 O58523 O58525 O58530
O58533 O58549 O58558 O58577 O58582 O58584 O58586 O58613 O58655 O58665 O58667
O58669 O58677 O58679 O58686 O58687 O58689 O58690 O58698 O58701 O58706 O58708
O58714 O58720 O58721 O58730 O58734 O58736 O58737 O58738 O58739 O58741 O58742

O58762 O58764 O58776 O58778 O58782 O58783 O58787 O58792 O58796 O58801 O58810
O58814 O58816 O58822 O58830 O58832 O58839 O58843 O58844 O58855 O58864 O58877
O58878 O58883 O58885 O58888 O58893 O58931 O58938 O58941 O58954 O58960 O58965
O58966 O58973 O58974 O58984 O58988 O58994 O58998 O59008 O59023 O59025 O59028
O59041 O59049 O59052 O59054 O59060 O59071 O59072 O59073 O59079 O59080 O59082
O59128 O59132 O59133 O59146 O59147 O59148 O59150 O59151 O59152 O59153 O59172
O59182 O59184 O59185 O59193 O59196 O59198 O59205 O59207 O59209 O59219 O59220
O59221 O59223 O59224 O59229 O59230 O59235 O59240 O59245 O59248 O59251 O59256
O59258 O59264 O59265 O59267 O59272 O59280 O59284 O59288 O59292 O59295 O59297
O59298 O59299 O59300 O59302 O59303 O59306 O59309 O59314 O59319 O59330 O59346
O59347 O59351 O59355 O59357 O59361 O59362 O59366 O59390 O59391 O59393 O59394
O59397 O59398 O59401 O59402 O59410 O59412 O59413 O59418 O59420 O59421 O59422
O59423 O59424 O59425 O59426 O59427 O59429 O59430 O59431 O59432 O59433 O59435
O59437 O59438 O59439 O59440 O59441 O59443 O59447 O59456 O59469 O59472 O59475
O59476 O59484 O59487 O59488 O59490 O59493 O59494 O59497 O59499 O59517 O59521
O59533 O59534 O59536 O59543 O59545 O59570 O59572 O59579 O59580 O59584 O59586
O59602 O59603 O59604 O59610 O59618 O59619 O59621 O59626 O59627 O59632 O59646
O59647 O59648 O59650 O59651 O59667 O59668 O59669 O59670 O59671 O59672 O59675
O59676 O59677 O59679 O59681 O59683 O59701 O59702 O59704 O59706 O59709 O59711
O59713 O59716 O59719 O59721 O59727 O59730 O59732 O59733 O59734 O59736 O59737
O59741 O59742 O59743 O59746 O59748 O59751 O59753 O59755 O59757 O59760 O59761
O59762 O59770 O59771 O59772 O59773 O59778 O59779 O59780 O59783 O59784 O59790
O59791 O59792 O59793 O59797 O59799 O59800 O59801 O59804 O59805 O59808 O59810
O59811 O59812 O59816 O59819 O59821 O59825 O59827 O59828 O59829 O59830 O59832
O59834 O59835 O59836 O59837 O59838 O59841 O59845 O59851 O59856 O59858 O59870
O59905 O59906 O59921 O59922 O59923 O59924 O59930 O59936 O59945 O59947 O59948
O59949 O59950 O59954 O59958 O60008 O60014 O60015 O60016 O60017 O60027 O60028
O60035 O60037 O60039 O60041 O60045 O60053 O60054 O60055 O60056 O60057 O60058
O60059 O60060 O60062 O60063 O60064 O60066 O60068 O60070 O60071 O60076 O60077
O60078 O60080 O60081 O60083 O60085 O60086 O60090 O60091 O60092 O60093 O60094
O60096 O60097 O60098 O60099 O60101 O60102 O60104 O60105 O60106 O60108 O60114
O60116 O60117 O60118 O60122 O60124 O60126 O60127 O60129 O60130 O60136 O60137
O60138 O60140 O60141 O60142 O60145 O60147 O60152 O60153 O60155 O60157 O60161
O60163 O60164 O60166 O60169 O60171 O60173 O60174 O60176 O60177 O60178 O60179
O60180 O60181 O60182 O60184 O60185 O60186 O60188 O60198 O60224 O60225 O60232
O60234 O60268 O60282 O60284 O60290 O60294 O60299 O60303 O60304 O60308 O60315
O60336 O60361 O60384 O60393 O60422 O60447 O60522 O60547 O60548 O60573 O60663
O60732 O60756 O60765 O60806 O60809 O60810 O60811 O60812 O60813 O60814 O60829
O60902 O60921 O60925 O60942 O60993 O60999 O61016 O61065 O61068 O61069 O61078
O61122 O61219 O61235 O61267 O61270 O61305 O61308 O61309 O61345 O61363 O61366
O61367 O61374 O61379 O61462 O61463 O61470 O61471 O61510 O61570 O61573 O61577
O61585 O61598 O61608 O61660 O61707 O61734 O61735 O61742 O61764 O61789 O61790
O61820 O61902 O61924 O61931 O61955 O62106 O62193 O62203 O62220 O62251 O62252
O62255 O62275 O62301 O62327 O62389 O62431 O62461 O62471 O62479 O62518 O62537

O62538 O62541 O62554 O62581 O62583 O62584 O62585 O62591 O62619 O62640 O62692
O62693 O62694 O62695 O62696 O62738 O62739 O62768 O62836 O62940 O62943 O62951
O62952 O62953 O62954 O62961 O62964 O62970 O62984 O62991 O62992 O63057 O63070
O63071 O63072 O63094 O63264 O63852 O63930 O64036 O64039 O64076 O64094 O64131
O64145 O64146 O64173 O64174 O64175 O64197 O64198 O64199 O64200 O64201 O64202
O64206 O64207 O64208 O64209 O64210 O64211 O64212 O64214 O64215 O64216 O64218
O64219 O64220 O64221 O64222 O64223 O64224 O64225 O64226 O64228 O64229 O64230
O64231 O64233 O64235 O64236 O64237 O64238 O64239 O64240 O64241 O64242 O64243
O64245 O64246 O64247 O64248 O64249 O64250 O64252 O64253 O64254 O64255 O64256
O64257 O64258 O64260 O64262 O64263 O64266 O64267 O64268 O64269 O64270 O64299
O64300 O64301 O64306 O64307 O64308 O64312 O64313 O64314 O64319 O64321 O64322
O64324 O64325 O64326 O64328 O64329 O64330 O64331 O64332 O64333 O64334 O64380
O64394 O64399 O64404 O64405 O64407 O64416 O64422 O64432 O64459 O64464 O64470
O64489 O64511 O64518 O64519 O64530 O64564 O64571 O64583 O64584 O64592 O64597
O64598 O64600 O64624 O64628 O64629 O64640 O64641 O64642 O64644 O64645 O64647
O64650 O64687 O64688 O64692 O64700 O64702 O64703 O64705 O64722 O64732 O64733
O64740 O64747 O64764 O64765 O64773 O64787 O64788 O64789 O64800 O64801 O64814
O64815 O64816 O64817 O64818 O64827 O64836 O64843 O64851 O64862 O64868 O64876
O64885 O64888 O64896 O64903 O64937 O64943 O64953 O64961 O64963 O64969 O64981
O64982 O64988 O65001 O65012 O65015 O65020 O65026 O65034 O65036 O65037 O65041
O65049 O65050 O65055 O65059 O65071 O65082 O65084 O65090 O65098 O65100 O65154
O65155 O65162 O65174 O65175 O65187 O65194 O65198 O65199 O65200 O65220 O65251
O65258 O65268 O65269 O65271 O65278 O65312 O65314 O65315 O65316 O65323 O65333
O65349 O65352 O65354 O65361 O65373 O65378 O65381 O65396 O65398 O65416 O65418
O65420 O65421 O65426 O65434 O65435 O65451 O65454 O65499 O65508 O65517 O65555
O65567 O65571 O65573 O65581 O65583 O65590 O65607 O65621 O65628 O65649 O65663
O65674 O65679 O65683 O65685 O65693 O65704 O65729 O65731 O65735 O65743 O65751
O65768 O65780 O65781 O65796 O65806 O65809 O65810 O65812 O65818 O65819 O65821
O65857 O65862 O65872 O65874 O65896 O65902 O65922 O65979 O65992 O66026 O66036
O66041 O66042 O66075 O66093 O66094 O66095 O66096 O66097 O66098 O66106 O66107
O66108 O66113 O66114 O66116 O66120 O66126 O66127 O66128 O66129 O66131 O66132
O66143 O66145 O66148 O66164 O66186 O66187 O66188 O66190 O66192 O66194 O66196
O66198 O66200 O66202 O66204 O66206 O66210 O66212 O66214 O66218 O66220 O66222
O66271 O66377 O66381 O66392 O66398 O66399 O66400 O66401 O66402 O66403 O66404
O66405 O66406 O66407 O66408 O66409 O66410 O66411 O66412 O66413 O66415 O66416
O66417 O66418 O66419 O66421 O66422 O66423 O66425 O66426 O66427 O66428 O66429
O66430 O66431 O66432 O66433 O66434 O66435 O66436 O66437 O66438 O66439 O66440
O66442 O66452 O66455 O66457 O66461 O66462 O66469 O66470 O66473 O66474 O66475
O66476 O66479 O66482 O66483 O66484 O66485 O66486 O66487 O66488 O66489 O66492
O66493 O66503 O66506 O66511 O66512 O66513 O66519 O66520 O66522 O66523 O66524
O66529 O66530 O66534 O66536 O66537 O66540 O66550 O66552 O66553 O66554 O66555
O66557 O66558 O66563 O66565 O66567 O66570 O66574 O66575 O66576 O66580 O66585
O66586 O66587 O66592 O66593 O66594 O66601 O66603 O66605 O66607 O66608 O66610
O66611 O66612 O66613 O66614 O66615 O66616 O66617 O66619 O66621 O66625 O66626

O66628 O66629 O66630 O66631 O66638 O66640 O66647 O66648 O66651 O66652 O66662
O66665 O66667 O66677 O66678 O66679 O66680 O66682 O66686 O66689 O66690 O66692
O66695 O66699 O66709 O66720 O66722 O66726 O66727 O66728 O66732 O66736 O66741
O66742 O66745 O66746 O66747 O66756 O66758 O66761 O66765 O66766 O66772 O66773
O66776 O66780 O66783 O66790 O66805 O66806 O66810 O66811 O66820 O66821 O66822
O66824 O66827 O66829 O66832 O66833 O66834 O66839 O66847 O66848 O66849 O66851
O66853 O66856 O66861 O66868 O66875 O66878 O66880 O66883 O66884 O66893 O66899
O66906 O66911 O66913 O66921 O66922 O66923 O66928 O66929 O66930 O66932 O66935
O66936 O66937 O66943 O66944 O66946 O66949 O66952 O66953 O66954 O66956 O66957
O66958 O66961 O66962 O66963 O66967 O66973 O66974 O66975 O66976 O66982 O66986
O66989 O66990 O66998 O67001 O67003 O67004 O67009 O67010 O67011 O67012 O67016
O67019 O67024 O67028 O67030 O67031 O67032 O67034 O67035 O67036 O67037 O67039
O67043 O67045 O67048 O67049 O67050 O67054 O67055 O67059 O67060 O67061 O67064
O67065 O67068 O67069 O67071 O67073 O67074 O67076 O67078 O67080 O67081 O67082
O67083 O67084 O67085 O67086 O67087 O67089 O67091 O67098 O67099 O67100 O67104
O67108 O67109 O67112 O67115 O67118 O67124 O67125 O67130 O67133 O67134 O67137
O67138 O67139 O67140 O67150 O67151 O67156 O67161 O67162 O67163 O67166 O67167
O67169 O67174 O67176 O67178 O67187 O67189 O67190 O67192 O67193 O67200 O67201
O67204 O67209 O67213 O67214 O67216 O67221 O67222 O67223 O67226 O67227 O67232
O67233 O67237 O67239 O67240 O67242 O67243 O67244 O67246 O67247 O67251 O67257
O67258 O67259 O67262 O67264 O67271 O67274 O67275 O67284 O67286 O67287 O67289
O67291 O67293 O67297 O67298 O67313 O67314 O67315 O67321 O67322 O67323 O67324
O67327 O67328 O67329 O67331 O67332 O67343 O67344 O67353 O67355 O67356 O67357
O67358 O67359 O67362 O67363 O67367 O67368 O67369 O67373 O67376 O67377 O67378
O67381 O67383 O67398 O67399 O67409 O67411 O67413 O67415 O67417 O67419 O67422
O67431 O67438 O67441 O67444 O67453 O67455 O67461 O67463 O67465 O67466 O67469
O67470 O67475 O67480 O67485 O67492 O67494 O67496 O67499 O67500 O67501 O67502
O67506 O67507 O67514 O67517 O67518 O67520 O67522 O67523 O67528 O67531 O67533
O67539 O67541 O67542 O67543 O67546 O67547 O67549 O67550 O67551 O67553 O67554
O67556 O67560 O67561 O67562 O67563 O67564 O67565 O67566 O67568 O67569 O67570
O67573 O67575 O67577 O67581 O67583 O67587 O67588 O67589 O67591 O67604 O67605
O67606 O67607 O67613 O67614 O67616 O67619 O67620 O67623 O67625 O67631 O67632
O67638 O67642 O67643 O67644 O67646 O67650 O67652 O67653 O67655 O67657 O67660
O67664 O67679 O67680 O67683 O67685 O67687 O67690 O67691 O67693 O67695 O67699
O67702 O67703 O67708 O67709 O67711 O67716 O67720 O67721 O67722 O67723 O67724
O67725 O67728 O67730 O67733 O67736 O67739 O67740 O67742 O67744 O67749 O67751
O67753 O67756 O67757 O67758 O67759 O67760 O67762 O67763 O67766 O67767 O67768
O67772 O67775 O67779 O67780 O67781 O67783 O67784 O67785 O67786 O67787 O67788
O67792 O67802 O67808 O67809 O67810 O67818 O67820 O67821 O67825 O67826 O67830
O67834 O67837 O67844 O67848 O67849 O67852 O67853 O67857 O67858 O67861 O67862
O67866 O67868 O67869 O67870 O67876 O67879 O67881 O67883 O67886 O67887 O67890
O67891 O67892 O67893 O67898 O67899 O67900 O67904 O67905 O67907 O67911 O67914
O67915 O67918 O67919 O67925 O67926 O67928 O67929 O67942 O67943 O67944 O67987
O67988 O68006 O68007 O68008 O68014 O68025 O68031 O68032 O68033 O68040 O68045

O68068 O68098 O68100 O68108 O68109 O68126 O68127 O68128 O68129 O68141 O68142
O68146 O68185 O68186 O68191 O68195 O68196 O68215 O68248 O68249 O68250 O68260
O68273 O68280 O68281 O68282 O68283 O68308 O68309 O68323 O68324 O68426 O68427
O68428 O68429 O68446 O68451 O68452 O68503 O68546 O68556 O68557 O68563 O68575
O68579 O68581 O68583 O68584 O68602 O68608 O68638 O68639 O68640 O68685 O68687
O68693 O68770 O68799 O68814 O68816 O68822 O68823 O68830 O68838 O68844 O68845
O68873 O68883 O68887 O68905 O68906 O68926 O68934 O68935 O68936 O68940 O68943
O68946 O68949 O68951 O68952 O68953 O68954 O68955 O68956 O68965 O68983 O68984
O68986 O68988 O68992 O69014 O69015 O69043 O69049 O69054 O69055 O69056 O69060
O69066 O69067 O69077 O69078 O69108 O69110 O69138 O69140 O69155 O69169 O69170
O69172 O69177 O69190 O69192 O69198 O69199 O69203 O69219 O69220 O69221 O69227
O69232 O69241 O69242 O69264 O69266 O69267 O69268 O69269 O69272 O69289 O69290
O69294 O69297 O69298 O69300 O69301 O69302 O69303 O69315 O69349 O69406 O69407
O69450 O69451 O69460 O69462 O69466 O69468 O69469 O69470 O69481 O69492 O69497
O69519 O69520 O69524 O69528 O69529 O69531 O69537 O69538 O69540 O69548 O69556
O69557 O69560 O69561 O69581 O69595 O69600 O69668 O69671 O69689 O69711 O69717
O69720 O69721 O69730 O69731 O69754 O69761 O69762 O69763 O69768 O69772 O69777
O69782 O69783 O69786 O69787 O69809 O69830 O69833 O69851 O69852 O69853 O69861
O69873 O69879 O69890 O69913 O69963 O69967 O69969 O69980 O69981 O69989 O70075
O70089 O70132 O70137 O70146 O70150 O70157 O70191 O70194 O70196 O70199 O70209
O70218 O70220 O70230 O70249 O70250 O70251 O70263 O70273 O70277 O70279 O70299
O70303 O70306 O70333 O70338 O70374 O70400 O70422 O70433 O70445 O70453 O70467
O70468 O70475 O70477 O70481 O70494 O70523 O70546 O70552 O70576 O70583 O70584
O70591 O70608 O70695 O70736 O70738 O70739 O70790 O70791 O71024 O71025 O71028
O71070 O71121 O71150 O71152 O71153 O71154 O71188 O71189 O71190 O71192 O71193
O71195 O71304 O72120 O72121 O72142 O72158 O72539 O72540 O72620 O72736 O72756
O72904 O72907 O72908 O72910 O73571 O73579 O73590 O73592 O73615 O73622 O73672
O73673 O73688 O73689 O73691 O73692 O73693 O73694 O73715 O73718 O73723 O73737
O73742 O73770 O73777 O73787 O73813 O73817 O73823 O73881 O73916 O73917 O73919
O73930 O73932 O73942 O73943 O73944 O73946 O73947 O73948 O73954 O73955 O73977
O73978 O73984 O73986 O74000 O74001 O74006 O74016 O74017 O74018 O74019 O74020
O74021 O74023 O74024 O74025 O74026 O74031 O74035 O74036 O74038 O74044 O74045
O74067 O74081 O74083 O74092 O74094 O74095 O74098 O74099 O74101 O74104 O74105
O74106 O74108 O74109 O74110 O74111 O74113 O74173 O74187 O74191 O74192 O74196
O74197 O74201 O74205 O74215 O74225 O74232 O74233 O74234 O74237 O74252 O74258
O74267 O74268 O74270 O74286 O74298 O74302 O74308 O74309 O74310 O74311 O74313
O74314 O74315 O74316 O74317 O74319 O74322 O74324 O74325 O74326 O74331 O74337
O74340 O74341 O74342 O74343 O74345 O74348 O74349 O74350 O74351 O74354 O74356
O74358 O74359 O74361 O74362 O74363 O74364 O74365 O74366 O74370 O74372 O74374
O74378 O74379 O74381 O74382 O74385 O74386 O74389 O74393 O74394 O74396 O74397
O74398 O74399 O74400 O74402 O74405 O74407 O74409 O74412 O74413 O74414 O74415
O74416 O74417 O74418 O74419 O74424 O74427 O74428 O74429 O74434 O74435 O74438
O74440 O74443 O74445 O74447 O74448 O74449 O74450 O74451 O74452 O74453 O74454
O74455 O74457 O74458 O74460 O74461 O74462 O74463 O74464 O74465 O74467 O74468

O74469 O74470 O74473 O74477 O74478 O74482 O74483 O74484 O74489 O74492 O74496
O74498 O74499 O74501 O74502 O74503 O74504 O74506 O74508 O74511 O74515 O74516
O74517 O74518 O74519 O74521 O74522 O74523 O74524 O74529 O74531 O74535 O74540
O74541 O74542 O74545 O74547 O74548 O74549 O74550 O74553 O74555 O74560 O74561
O74563 O74624 O74627 O74628 O74630 O74633 O74634 O74635 O74638 O74648 O74685
O74689 O74690 O74712 O74718 O74727 O74731 O74733 O74734 O74735 O74736 O74737
O74738 O74739 O74740 O74741 O74743 O74746 O74747 O74748 O74751 O74755 O74759
O74760 O74761 O74762 O74763 O74764 O74765 O74766 O74770 O74771 O74772 O74773
O74774 O74777 O74778 O74780 O74781 O74782 O74783 O74789 O74790 O74791 O74792
O74793 O74796 O74797 O74798 O74799 O74801 O74803 O74804 O74805 O74806 O74807
O74808 O74809 O74814 O74821 O74823 O74825 O74828 O74830 O74831 O74832 O74833
O74834 O74835 O74840 O74841 O74842 O74844 O74845 O74848 O74853 O74855 O74856
O74857 O74858 O74859 O74860 O74861 O74862 O74863 O74865 O74867 O74871 O74874
O74875 O74876 O74879 O74880 O74881 O74883 O74884 O74885 O74886 O74889 O74890
O74894 O74896 O74897 O74898 O74905 O74906 O74907 O74909 O74910 O74913 O74914
O74915 O74917 O74918 O74919 O74927 O74928 O74932 O74933 O74939 O74940 O74942
O74944 O74945 O74946 O74948 O74951 O74952 O74953 O74955 O74956 O74957 O74958
O74959 O74962 O74965 O74966 O74968 O74970 O74972 O74973 O74974 O74978 O74981
O74982 O74983 O74984 O74985 O74986 O74989 O74991 O74994 O74995 O74999 O75001
O75002 O75003 O75004 O75005 O75007 O75011 O75031 O75071 O75112 O75123 O75127
O75132 O75155 O75157 O75177 O75191 O75223 O75290 O75293 O75319 O75333 O75344
O75346 O75347 O75360 O75363 O75366 O75373 O75382 O75390 O75409 O75414 O75417
O75426 O75437 O75444 O75459 O75461 O75478 O75506 O75526 O75529 O75541 O75564
O75570 O75601 O75602 O75603 O75626 O75648 O75663 O75677 O75678 O75679 O75690
O75764 O75771 O75792 O75794 O75808 O75818 O75820 O75828 O75864 O75865 O75874
O75879 O75891 O75897 O75920 O75934 O75936 O75943 O75952 O75953 O75956 O76009
O76011 O76013 O76014 O76015 O76041 O76042 O76054 O76071 O76075 O76087 O76156
O76187 O76202 O76206 O76240 O76242 O76243 O76332 O76365 O76371 O76373 O76387
O76454 O76460 O76463 O76464 O76483 O76484 O76485 O76511 O76512 O76577 O76612
O76616 O76732 O76734 O76743 O76756 O76757 O76762 O76808 O76828 O76906 O76908
O76922 O76924 O76932 O76971 O77003 O77027 O77033 O77051 O77206 O77210 O77215
O77245 O77263 O77264 O77302 O77303 O77323 O77334 O77392 O77404 O77448 O77457
O77458 O77459 O77460 O77462 O77473 O77480 O77503 O77506 O77512 O77560 O77591
O77617 O77618 O77622 O77655 O77666 O77682 O77689 O77727 O77728 O77784 O77788
O77797 O78248 O78252 O78253 O78254 O78256 O78258 O78259 O78260 O78261 O78262
O78310 O78328 O78405 O78409 O78410 O78411 O78412 O78413 O78414 O78419 O78420
O78421 O78422 O78423 O78425 O78429 O78430 O78431 O78432 O78434 O78435 O78436
O78438 O78439 O78440 O78442 O78445 O78447 O78449 O78450 O78451 O78452 O78453
O78458 O78459 O78461 O78472 O78473 O78474 O78481 O78482 O78483 O78484 O78485
O78486 O78487 O78488 O78489 O78495 O78496 O78497 O78501 O78506 O78510 O78517
O78518 O78678 O79977 O80164 O80286 O80302 O80333 O80345 O80358 O80360 O80361
O80362 O80363 O80377 O80400 O80402 O80405 O80416 O80429 O80433 O80438 O80439
O80446 O80449 O80467 O80474 O80475 O80482 O80483 O80488 O80494 O80502 O80504
O80506 O80507 O80513 O80518 O80543 O80560 O80562 O80567 O80568 O80573 O80574

O80575 O80582 O80585 O80588 O80596 O80603 O80608 O80626 O80642 O80644 O80646
O80647 O80653 O80674 O80678 O80699 O80713 O80714 O80719 O80735 O80736 O80737
O80738 O80741 O80742 O80758 O80762 O80765 O80770 O80777 O80786 O80792 O80805
O80807 O80816 O80831 O80840 O80856 O80871 O80875 O80877 O80883 O80925 O80928
O80929 O80930 O80931 O80933 O80934 O80942 O80944 O80948 O80950 O80958 O80970
O80980 O80988 O80990 O80998 O81004 O81010 O81014 O81025 O81028 O81037 O81039
O81055 O81057 O81058 O81086 O81097 O81098 O81103 O81127 O81146 O81147 O81148
O81149 O81153 O81154 O81155 O81160 O81185 O81187 O81191 O81192 O81193 O81220
O81221 O81235 O81242 O81263 O81275 O81305 O81313 O81316 O81322 O81323 O81332
O81355 O81361 O81363 O81372 O81395 O81414 O81432 O81433 O81481 O81482 O81488
O81498 O81635 O81643 O81644 O81645 O81646 O81701 O81716 O81767 O81769 O81778
O81782 O81787 O81788 O81791 O81796 O81815 O81822 O81825 O81826 O81829 O81831
O81836 O81851 O81852 O81864 O81865 O81884 O81900 O81902 O81908 O81913 O81918
O81920 O81959 O81971 O81974 O81980 O81982 O81983 O82011 O82012 O82013 O82018
O82030 O82035 O82039 O82040 O82043 O82054 O82056 O82059 O82088 O82089 O82117
O82134 O82143 O82144 O82147 O82155 O82162 O82169 O82170 O82175 O82178 O82179
O82188 O82189 O82190 O82191 O82198 O82200 O82205 O82210 O82221 O82225 O82233
O82234 O82239 O82243 O82248 O82253 O82254 O82255 O82256 O82264 O82265 O82277
O82281 O82286 O82290 O82307 O82314 O82330 O82333 O82343 O82345 O82357 O82368
O82370 O82373 O82374 O82375 O82376 O82378 O82379 O82380 O82381 O82382 O82383
O82385 O82392 O82394 O82413 O82415 O82422 O82451 O82462 O82475 O82484 O82486
O82488 O82491 O82497 O82499 O82514 O82515 O82521 O82528 O82530 O82531 O82560
O82569 O82572 O82574 O82595 O82597 O82616 O82617 O82622 O82626 O82627 O82631
O82637 O82645 O82653 O82662 O82677 O82712 O82713 O82733 O82734 O82743 O82768
O82782 O82793 O82794 O82796 O82797 O82802 O82845 O82850 O82851 O82852 O82853
O82872 O82881 O82896 O82898 O82899 O82900 O82902 O82922 O83004 O83023 O83026
O83030 O83041 O83046 O83047 O83048 O83049 O83050 O83051 O83052 O83053 O83054
O83056 O83057 O83058 O83059 O83062 O83063 O83065 O83066 O83075 O83080 O83081
O83082 O83084 O83085 O83090 O83092 O83097 O83099 O83100 O83101 O83102 O83103
O83105 O83107 O83108 O83110 O83112 O83122 O83128 O83129 O83132 O83133 O83135
O83146 O83148 O83149 O83153 O83154 O83155 O83158 O83159 O83160 O83164 O83165
O83166 O83168 O83171 O83173 O83195 O83196 O83199 O83202 O83205 O83206 O83207
O83208 O83209 O83210 O83214 O83217 O83218 O83219 O83220 O83221 O83222 O83223
O83224 O83225 O83226 O83227 O83228 O83229 O83230 O83231 O83232 O83233 O83234
O83235 O83236 O83237 O83239 O83240 O83241 O83242 O83243 O83244 O83245 O83246
O83253 O83258 O83260 O83262 O83264 O83265 O83266 O83267 O83268 O83269 O83270
O83271 O83272 O83273 O83274 O83278 O83281 O83288 O83290 O83293 O83295 O83296
O83302 O83304 O83305 O83307 O83308 O83309 O83316 O83319 O83325 O83326 O83327
O83328 O83333 O83336 O83338 O83340 O83348 O83349 O83351 O83352 O83357 O83361
O83362 O83369 O83371 O83372 O83373 O83374 O83378 O83385 O83386 O83387 O83388
O83389 O83390 O83397 O83398 O83399 O83401 O83409 O83413 O83416 O83417 O83419
O83421 O83424 O83430 O83433 O83434 O83435 O83437 O83440 O83446 O83447 O83451
O83452 O83455 O83456 O83460 O83462 O83464 O83465 O83466 O83467 O83477 O83479
O83480 O83485 O83487 O83488 O83489 O83490 O83491 O83495 O83500 O83505 O83506

O83508 O83515 O83517 O83519 O83520 O83521 O83525 O83527 O83530 O83536 O83537
O83545 O83546 O83548 O83549 O83550 O83553 O83554 O83558 O83561 O83563 O83567
O83570 O83571 O83578 O83583 O83585 O83588 O83592 O83593 O83595 O83596 O83600
O83604 O83606 O83607 O83608 O83610 O83612 O83613 O83614 O83615 O83616 O83618
O83623 O83625 O83626 O83627 O83628 O83633 O83635 O83637 O83640 O83641 O83642
O83644 O83647 O83649 O83650 O83651 O83653 O83656 O83662 O83663 O83664 O83667
O83668 O83671 O83672 O83675 O83676 O83678 O83679 O83680 O83682 O83683 O83684
O83686 O83687 O83690 O83697 O83698 O83699 O83701 O83711 O83714 O83715 O83716
O83720 O83723 O83724 O83725 O83727 O83728 O83729 O83730 O83735 O83737 O83738
O83739 O83740 O83743 O83747 O83748 O83751 O83754 O83756 O83759 O83760 O83763
O83771 O83772 O83773 O83776 O83777 O83780 O83785 O83787 O83788 O83789 O83790
O83792 O83793 O83796 O83797 O83802 O83803 O83806 O83808 O83809 O83814 O83816
O83817 O83818 O83819 O83820 O83821 O83822 O83823 O83829 O83833 O83838 O83839
O83841 O83842 O83843 O83845 O83855 O83856 O83857 O83859 O83860 O83861 O83863
O83866 O83870 O83873 O83874 O83875 O83876 O83877 O83878 O83879 O83880 O83883
O83886 O83889 O83890 O83892 O83893 O83895 O83897 O83899 O83900 O83902 O83903
O83910 O83911 O83912 O83916 O83917 O83921 O83922 O83925 O83933 O83936 O83938
O83939 O83940 O83942 O83945 O83949 O83950 O83952 O83956 O83965 O83966 O83969
O83971 O83972 O83974 O83975 O83977 O83979 O83980 O83982 O83983 O83984 O83985
O83986 O83987 O83988 O83990 O83992 O83993 O83998 O84001 O84003 O84005 O84006
O84007 O84010 O84011 O84022 O84025 O84026 O84027 O84029 O84030 O84031 O84032
O84033 O84034 O84035 O84038 O84042 O84044 O84045 O84048 O84049 O84060 O84065
O84066 O84074 O84077 O84078 O84079 O84081 O84087 O84088 O84089 O84096 O84097
O84098 O84100 O84101 O84110 O84115 O84123 O84124 O84127 O84128 O84130 O84148
O84152 O84171 O84172 O84173 O84188 O84189 O84191 O84192 O84193 O84196 O84200
O84206 O84211 O84212 O84213 O84215 O84217 O84220 O84222 O84243 O84250 O84271
O84274 O84275 O84284 O84288 O84289 O84290 O84294 O84296 O84299 O84300 O84301
O84304 O84311 O84315 O84316 O84318 O84319 O84320 O84321 O84322 O84331 O84332
O84333 O84335 O84337 O84339 O84345 O84348 O84350 O84351 O84353 O84357 O84358
O84366 O84367 O84369 O84371 O84372 O84373 O84374 O84375 O84378 O84381 O84390
O84391 O84393 O84395 O84402 O84404 O84410 O84411 O84423 O84425 O84429 O84435
O84436 O84437 O84439 O84441 O84444 O84451 O84452 O84456 O84458 O84460 O84461
O84463 O84465 O84468 O84469 O84476 O84481 O84492 O84499 O84508 O84509 O84510
O84512 O84527 O84528 O84530 O84531 O84532 O84533 O84535 O84540 O84543 O84544
O84546 O84547 O84549 O84561 O84562 O84579 O84583 O84585 O84586 O84587 O84588
O84589 O84590 O84611 O84613 O84619 O84620 O84621 O84622 O84630 O84631 O84632
O84637 O84638 O84639 O84640 O84641 O84645 O84666 O84669 O84670 O84672 O84673
O84678 O84682 O84684 O84685 O84686 O84687 O84693 O84694 O84697 O84702 O84704
O84709 O84711 O84712 O84713 O84727 O84734 O84735 O84736 O84737 O84741 O84752
O84754 O84757 O84763 O84767 O84771 O84777 O84783 O84786 O84787 O84789 O84790
O84792 O84796 O84797 O84799 O84804 O84805 O84806 O84807 O84808 O84809 O84810
O84816 O84822 O84823 O84828 O84832 O84834 O84835 O84836 O84839 O84840 O84842
O84843 O84847 O84849 O84859 O84863 O84867 O84872 O84874 O84883 O84898 O84899
O84902 O84903 O84904 O84905 O84913 O84917 O84990 O84992 O85012 O85014 O85040

O85042 O85043 O85044 O85057 O85060 O85062 O85063 O85064 O85065 O85066 O85070
O85071 O85072 O85073 O85078 O85171 O85172 O85175 O85176 O85192 O85207 O85213
O85222 O85235 O85253 O85264 O85282 O85290 O85293 O85294 O85295 O85298 O85304
O85306 O85342 O85347 O85387 O85388 O85501 O85502 O85673 O85674 O85675 O85691
O85692 O85709 O85713 O85718 O85728 O85746 O85754 O85762 O85764 O85786 O85805
O85806 O85807 O85824 O85977 O85978 O85987 O86013 O86028 O86029 O86033 O86034
O86041 O86042 O86043 O86046 O86047 O86053 O86062 O86082 O86083 O86090 O86094
O86103 O86108 O86109 O86110 O86130 O86131 O86132 O86134 O86165 O86169 O86170
O86185 O86186 O86222 O86223 O86227 O86228 O86235 O86236 O86237 O86239 O86242
O86243 O86262 O86271 O86281 O86309 O86312 O86318 O86332 O86376 O86382 O86385
O86403 O86404 O86408 O86411 O86412 O86414 O86415 O86418 O86436 O86447 O86448
O86449 O86459 O86464 O86490 O86491 O86504 O86505 O86507 O86508 O86509 O86511
O86528 O86534 O86535 O86560 O86564 O86565 O86566 O86567 O86583 O86655 O86682
O86683 O86722 O86737 O86754 O86770 O86773 O86775 O86776 O86779 O86781 O86786
O86788 O86793 O86812 O86820 O86835 O86836 O86840 O86841 O86848 O86868 O86937
O86938 O86949 O86951 O86953 O86954 O86956 O86957 O86959 O86960 O87006 O87007
O87008 O87009 O87014 O87016 O87082 O87085 O87170 O87172 O87197 O87198 O87278
O87319 O87320 O87322 O87324 O87326 O87330 O87331 O87384 O87385 O87386 O87387
O87388 O87389 O87390 O87391 O87392 O87393 O87398 O87399 O87400 O87401 O87402
O87404 O87405 O87407 O87408 O87444 O87545 O87546 O87558 O87574 O87590 O87605
O87612 O87627 O87628 O87667 O87681 O87682 O87688 O87689 O87690 O87691 O87692
O87693 O87694 O87696 O87697 O87698 O87703 O87719 O87726 O87733 O87734 O87761
O87765 O87775 O87776 O87777 O87778 O87790 O87791 O87792 O87796 O87816 O87821
O87833 O87838 O87840 O87844 O87864 O87871 O87872 O87873 O87874 O87875 O87876
O87877 O87883 O87885 O87886 O87887 O87890 O87896 O87897 O87905 O87906 O87908
O87924 O87941 O87942 O87943 O87944 O87949 O87970 O87971 O88005 O88011 O88015
O88018 O88054 O88055 O88058 O88059 O88060 O88070 O88101 O88181 O88282 O88286
O88345 O88346 O88350 O88368 O88379 O88387 O88398 O88425 O88426 O88428 O88436
O88445 O88470 O88480 O88483 O88484 O88491 O88508 O88509 O88513 O88522 O88532
O88533 O88544 O88545 O88553 O88559 O88573 O88574 O88599 O88609 O88644 O88656
O88677 O88685 O88708 O88712 O88761 O88794 O88796 O88806 O88807 O88816 O88818
O88828 O88834 O88842 O88844 O88846 O88850 O88851 O88852 O88873 O88892 O88898
O88902 O88939 O88958 O88978 O88989 O88995 O89000 O89033 O89038 O89042 O89043
O89044 O89049 O89086 O89090 O89091 O89094 O89113 O89114 O89339 O89344 O90306
O91251 O91253 O91533 O91729 O91940 O92404 O92490 O92503 O92604 O92605 O93036
O93058 O93117 O93120 O93214 O93229 O93235 O93256 O93257 O93277 O93307 O93308
O93310 O93327 O93343 O93348 O93349 O93351 O93353 O93366 O93367 O93375 O93377
O93388 O93401 O93409 O93450 O93477 O93479 O93499 O93500 O93507 O93511 O93528
O93529 O93530 O93532 O93537 O93538 O93539 O93540 O93541 O93542 O93543 O93544
O93545 O93546 O93567 O93571 O93590 O93591 O93593 O93609 O93610 O93623 O93624
O93627 O93631 O93632 O93634 O93636 O93637 O93639 O93640 O93654 O93655 O93656
O93657 O93658 O93659 O93661 O93662 O93715 O93716 O93718 O93719 O93723 O93724
O93728 O93729 O93730 O93732 O93733 O93744 O93745 O93746 O93748 O93774 O93777
O93779 O93794 O93796 O93798 O93803 O93806 O93807 O93829 O93830 O93845 O93849

O93863 O93864 O93866 O93868 O93869 O93870 O93918 O93919 O93922 O93931 O93934
O93937 O93958 O93967 O94008 O94017 O94038 O94039 O94048 O94049 O94065 O94069
O94083 O94086 O94095 O94111 O94114 O94122 O94123 O94126 O94127 O94128 O94130
O94131 O94143 O94150 O94153 O94160 O94166 O94178 O94197 O94200 O94201 O94205
O94206 O94213 O94225 O94229 O94230 O94235 O94236 O94241 O94242 O94243 O94244
O94245 O94246 O94247 O94248 O94249 O94250 O94251 O94252 O94255 O94256 O94257
O94258 O94259 O94260 O94261 O94263 O94264 O94265 O94266 O94267 O94268 O94271
O94274 O94275 O94276 O94277 O94278 O94279 O94282 O94284 O94285 O94286 O94288
O94289 O94290 O94292 O94293 O94294 O94295 O94301 O94303 O94306 O94307 O94308
O94311 O94312 O94313 O94314 O94315 O94316 O94320 O94322 O94326 O94328 O94329
O94330 O94331 O94332 O94334 O94337 O94340 O94345 O94346 O94347 O94350 O94354
O94355 O94358 O94359 O94362 O94364 O94365 O94368 O94369 O94375 O94376 O94378
O94379 O94383 O94384 O94386 O94387 O94388 O94389 O94392 O94393 O94394 O94395
O94396 O94397 O94398 O94399 O94400 O94403 O94404 O94405 O94406 O94408 O94410
O94411 O94413 O94415 O94416 O94417 O94418 O94420 O94421 O94423 O94424 O94425
O94426 O94427 O94428 O94429 O94431 O94432 O94435 O94436 O94437 O94438 O94442
O94443 O94444 O94445 O94446 O94449 O94450 O94451 O94452 O94454 O94455 O94459
O94460 O94461 O94465 O94467 O94468 O94470 O94474 O94475 O94476 O94477 O94480
O94481 O94483 O94484 O94488 O94489 O94492 O94493 O94494 O94495 O94497 O94498
O94500 O94501 O94503 O94504 O94505 O94506 O94508 O94509 O94510 O94513 O94514
O94515 O94517 O94518 O94519 O94521 O94524 O94525 O94527 O94529 O94530 O94533
O94534 O94535 O94538 O94542 O94544 O94547 O94548 O94552 O94553 O94556 O94560
O94561 O94562 O94564 O94566 O94567 O94570 O94574 O94577 O94578 O94579 O94580
O94582 O94583 O94585 O94586 O94588 O94589 O94600 O94601 O94602 O94603 O94606
O94609 O94612 O94613 O94614 O94615 O94616 O94617 O94619 O94620 O94621 O94623
O94625 O94626 O94627 O94629 O94630 O94631 O94632 O94634 O94636 O94637 O94640
O94641 O94642 O94643 O94644 O94645 O94646 O94650 O94653 O94659 O94660 O94662
O94664 O94666 O94667 O94668 O94670 O94671 O94675 O94676 O94679 O94681 O94682
O94683 O94684 O94685 O94686 O94687 O94688 O94690 O94691 O94692 O94693 O94696
O94697 O94698 O94700 O94703 O94704 O94706 O94707 O94708 O94709 O94710 O94711
O94712 O94713 O94714 O94716 O94717 O94718 O94720 O94721 O94723 O94724 O94725
O94728 O94729 O94731 O94733 O94735 O94739 O94740 O94742 O94745 O94746 O94751
O94752 O94762 O94805 O94818 O94819 O94822 O94829 O94854 O94889 O94892 O94903
O94925 O94929 O94941 O94952 O95045 O95050 O95059 O95072 O95073 O95076 O95096
O95147 O95153 O95154 O95170 O95177 O95198 O95199 O95231 O95238 O95243 O95257
O95260 O95336 O95340 O95343 O95347 O95363 O95376 O95379 O95396 O95402 O95411
O95416 O95424 O95447 O95455 O95456 O95503 O95521 O95522 O95551 O95561 O95567
O95571 O95600 O95620 O95626 O95677 O95678 O95707 O95718 O95744 O95751 O95757
O95758 O95777 O95780 O95789 O95801 O95825 O95834 O95848 O95861 O95872 O95873
O95900 O95922 O95926 O95931 O95932 O95948 O95985 O95988 O96004 O96033 O96064
O96081 O96102 O96185 O96237 O96347 O96432 O96433 O96436 O96447 O96507 O96539
O96553 O96566 O96567 O96569 O96570 O96571 O96621 O96623 O96624 O96625 O96626
O96642 O96647 O96650 O96651 O96668 O96693 O96719 O96720 O96722 O96764 O96771
O96772 O96774 O96785 O96788 O96805 O96808 O96821 O96827 O96860 O96863 O96876

O96907 O96923 O96935 O96952 O97018 O97039 O97067 O97069 O97101 O97143 O97162
O97172 O97177 O97178 O97179 O97192 O97209 O97235 O97237 O97240 O97341 O97388
O97420 O97438 O97439 O97440 O97463 O97472 O97484 O97529 O97552 O97578 O97580
O97581 O97593 O97598 O97628 O97670 O97716 O97756 O97764 O97790 O97959 O97965
O97972 O97980 O98360 O98370 O98449 O98450 O98451 O98452 O98453 O98454 O98455
O98456 O98458 O98459 O98460 O98461 O98462 O98634 O98635 O98636 O98637 O98639
O98664 O98668 O98671 O98681 O98840 O98883 O98890 O98945 O98946 O98947 O98948
O98997 O99000 O99001 O99010 P00044 P00045 P00080 P00081 P00084 P00085 P00086
P00087 P00088 P00089 P00090 P00093 P00097 P00098 P00100 P00102 P00103 P00104
P00105 P00121 P00122 P00123 P00124 P00133 P00134 P00135 P00136 P00137 P00138
P00139 P00140 P00141 P00142 P00143 P00145 P00146 P00147 P00151 P00152 P00153
P00183 P00193 P00194 P00195 P00196 P00197 P00198 P00200 P00201 P00202 P00203
P00204 P00205 P00206 P00207 P00208 P00209 P00210 P00211 P00212 P00214 P00215
P00216 P00217 P00218 P00219 P00220 P00221 P00222 P00223 P00224 P00225 P00226
P00227 P00228 P00229 P00230 P00231 P00232 P00233 P00234 P00235 P00236 P00237
P00238 P00239 P00240 P00241 P00242 P00243 P00244 P00245 P00246 P00247 P00248
P00249 P00250 P00251 P00252 P00253 P00254 P00255 P00259 P00261 P00262 P00263
P00264 P00265 P00266 P00267 P00268 P00269 P00270 P00271 P00272 P00273 P00275
P00276 P00279 P00281 P00283 P00284 P00285 P00286 P00302 P00303 P00304 P00321
P00322 P00323 P00324 P00327 P00328 P00331 P00332 P00333 P00334 P00335 P00339
P00340 P00341 P00342 P00343 P00344 P00345 P00346 P00349 P00350 P00357 P00361
P00362 P00365 P00367 P00368 P00369 P00370 P00374 P00376 P00377 P00378 P00380
P00381 P00382 P00383 P00384 P00386 P00392 P00431 P00434 P00436 P00437 P00438
P00439 P00440 P00445 P00448 P00449 P00452 P00456 P00457 P00458 P00460 P00461
P00462 P00463 P00464 P00465 P00466 P00467 P00468 P00469 P00471 P00472 P00473
P00474 P00476 P00484 P00485 P00486 P00487 P00489 P00490 P00491 P00492 P00493
P00494 P00497 P00498 P00499 P00500 P00503 P00504 P00508 P00509 P00511 P00512
P00513 P00518 P00531 P00532 P00537 P00538 P00547 P00548 P00551 P00552 P00553
P00555 P00557 P00559 P00561 P00562 P00563 P00564 P00565 P00566 P00568 P00569
P00570 P00571 P00572 P00573 P00579 P00581 P00582 P00585 P00586 P00632 P00638
P00640 P00641 P00642 P00643 P00650 P00653 P00654 P00722 P00723 P00727 P00728
P00752 P00771 P00793 P00801 P00812 P00814 P00815 P00816 P00817 P00818 P00819
P00820 P00821 P00860 P00861 P00862 P00863 P00864 P00865 P00868 P00869 P00870
P00871 P00872 P00873 P00874 P00875 P00876 P00877 P00878 P00879 P00880 P00881
P00883 P00885 P00887 P00888 P00889 P00890 P00893 P00894 P00895 P00896 P00897
P00898 P00899 P00900 P00901 P00902 P00903 P00904 P00905 P00906 P00908 P00909
P00910 P00911 P00912 P00914 P00915 P00916 P00917 P00926 P00927 P00929 P00930
P00931 P00934 P00935 P00936 P00939 P00940 P00941 P00943 P00944 P00946 P00948
P00949 P00952 P00953 P00954 P00958 P00960 P00961 P00962 P00963 P00964 P00965
P00967 P00968 P00969 P00970 P00971 P00993 P01013 P01039 P01041 P01051 P01052
P01053 P01054 P01056 P01057 P01058 P01059 P01061 P01062 P01063 P01064 P01065
P01066 P01067 P01069 P01071 P01073 P01075 P01093 P01094 P01097 P01098 P01102
P01103 P01104 P01105 P01109 P01110 P01153 P01154 P01155 P01158 P01159 P01249
P01250 P01251 P01252 P01255 P01358 P01363 P01547 P01548 P01618 P01626 P01628

P01629 P01630 P01631 P01632 P01636 P01643 P01644 P01645 P01646 P01647 P01648
P01649 P01650 P01651 P01652 P01653 P01654 P01655 P01656 P01657 P01659 P01660
P01662 P01663 P01664 P01665 P01666 P01667 P01668 P01669 P01670 P01671 P01672
P01673 P01674 P01675 P01676 P01677 P01678 P01679 P01681 P01682 P01683 P01684
P01685 P01686 P01687 P01688 P01689 P01690 P01692 P01693 P01694 P01695 P01696
P01697 P01698 P01725 P01826 P01827 P01828 P01835 P01838 P01839 P01840 P01841
P01847 P01870 P01874 P01875 P01878 P01879 P01883 P01923 P01924 P01926 P01928
P01929 P01930 P01932 P01933 P01934 P01935 P01936 P01937 P01938 P01939 P01940
P01941 P01942 P01943 P01944 P01945 P01946 P01947 P01948 P01949 P01950 P01951
P01953 P01954 P01955 P01956 P01957 P01958 P01959 P01960 P01961 P01962 P01963
P01964 P01965 P01966 P01967 P01968 P01969 P01971 P01972 P01973 P01975 P01976
P01977 P01978 P01979 P01980 P01981 P01982 P01983 P01984 P01985 P01986 P01987
P01988 P01989 P01990 P01991 P01992 P01993 P01994 P01995 P01996 P01997 P01998
P01999 P02000 P02001 P02002 P02003 P02004 P02005 P02006 P02007 P02008 P02009
P02010 P02011 P02012 P02013 P02014 P02015 P02016 P02017 P02018 P02019 P02020
P02021 P02022 P02024 P02025 P02026 P02028 P02029 P02030 P02031 P02032 P02033
P02035 P02036 P02038 P02039 P02040 P02042 P02044 P02046 P02047 P02048 P02049
P02050 P02051 P02052 P02053 P02054 P02055 P02057 P02058 P02059 P02060 P02061
P02062 P02064 P02065 P02066 P02067 P02070 P02072 P02073 P02074 P02075 P02076
P02077 P02078 P02080 P02081 P02082 P02083 P02084 P02085 P02086 P02087 P02088
P02089 P02090 P02091 P02092 P02093 P02094 P02095 P02097 P02099 P02100 P02101
P02102 P02103 P02105 P02106 P02107 P02108 P02109 P02110 P02111 P02112 P02113
P02114 P02115 P02116 P02117 P02118 P02120 P02121 P02122 P02123 P02124 P02125
P02126 P02127 P02128 P02129 P02130 P02131 P02132 P02133 P02134 P02135 P02136
P02137 P02138 P02139 P02140 P02141 P02142 P02143 P02144 P02145 P02147 P02148
P02150 P02151 P02152 P02153 P02154 P02155 P02156 P02157 P02159 P02160 P02161
P02163 P02164 P02165 P02166 P02167 P02168 P02169 P02170 P02171 P02173 P02174
P02177 P02178 P02179 P02180 P02181 P02182 P02183 P02184 P02185 P02186 P02187
P02189 P02190 P02191 P02192 P02193 P02194 P02195 P02196 P02197 P02199 P02200
P02201 P02202 P02203 P02204 P02205 P02206 P02207 P02208 P02209 P02210 P02211
P02212 P02213 P02214 P02215 P02216 P02217 P02218 P02227 P02228 P02231 P02232
P02233 P02234 P02235 P02236 P02237 P02238 P02239 P02240 P02244 P02245 P02246
P02247 P02251 P02252 P02253 P02254 P02255 P02256 P02257 P02258 P02259 P02260
P02262 P02263 P02268 P02269 P02270 P02272 P02273 P02274 P02275 P02276 P02277
P02280 P02281 P02283 P02284 P02285 P02286 P02287 P02288 P02289 P02290 P02291
P02292 P02293 P02294 P02299 P02301 P02302 P02309 P02310 P02313 P02314 P02316
P02317 P02318 P02319 P02321 P02322 P02324 P02325 P02326 P02328 P02330 P02331
P02332 P02333 P02334 P02338 P02343 P02344 P02345 P02348 P02353 P02355 P02357
P02358 P02360 P02362 P02368 P02376 P02377 P02382 P02385 P02391 P02393 P02394
P02395 P02396 P02397 P02398 P02399 P02402 P02412 P02413 P02415 P02417 P02431
P02434 P02438 P02439 P02440 P02441 P02442 P02443 P02444 P02445 P02446 P02447
P02448 P02449 P02450 P02451 P02470 P02472 P02474 P02475 P02476 P02477 P02478
P02479 P02480 P02482 P02483 P02484 P02485 P02486 P02487 P02488 P02492 P02493
P02494 P02497 P02498 P02499 P02500 P02501 P02502 P02503 P02504 P02505 P02506

P02507 P02508 P02509 P02512 P02513 P02515 P02516 P02517 P02518 P02519 P02520
P02521 P02522 P02523 P02524 P02525 P02526 P02527 P02528 P02529 P02530 P02531
P02532 P02533 P02534 P02537 P02539 P02547 P02548 P02550 P02552 P02553 P02554
P02556 P02557 P02561 P02562 P02563 P02564 P02565 P02566 P02567 P02572 P02574
P02576 P02578 P02580 P02581 P02582 P02583 P02584 P02585 P02586 P02587 P02588
P02589 P02591 P02592 P02595 P02597 P02598 P02600 P02602 P02604 P02605 P02606
P02607 P02608 P02609 P02610 P02611 P02612 P02613 P02614 P02615 P02616 P02617
P02618 P02619 P02620 P02621 P02622 P02623 P02624 P02625 P02626 P02627 P02628
P02629 P02630 P02631 P02632 P02635 P02636 P02637 P02638 P02639 P02640 P02641
P02642 P02643 P02645 P02646 P02729 P02733 P02744 P02785 P02791 P02793 P02795
P02797 P02800 P02801 P02805 P02806 P02807 P02826 P02827 P02831 P02832 P02833
P02835 P02836 P02848 P02861 P02862 P02864 P02865 P02868 P02869 P02871 P02874
P02875 P02878 P02881 P02882 P02886 P02887 P02888 P02889 P02890 P02891 P02892
P02899 P02900 P02903 P02904 P02944 P02958 P02959 P02960 P02961 P02962 P02963
P02964 P02966 P02967 P02979 P02984 P02985 P02986 P02988 P02989 P02991 P02992
P02993 P03000 P03001 P03006 P03007 P03008 P03009 P03010 P03012 P03013 P03014
P03015 P03018 P03022 P03023 P03024 P03029 P03030 P03031 P03036 P03037 P03038
P03039 P03040 P03041 P03042 P03043 P03044 P03045 P03046 P03047 P03048 P03049
P03050 P03051 P03052 P03054 P03055 P03056 P03061 P03062 P03063 P03064 P03065
P03066 P03067 P03080 P03081 P03082 P03083 P03099 P03102 P03103 P03104 P03118
P03119 P03120 P03121 P03122 P03123 P03124 P03125 P03126 P03127 P03128 P03133
P03134 P03156 P03157 P03158 P03159 P03160 P03161 P03162 P03163 P03164 P03170
P03177 P03181 P03184 P03187 P03189 P03190 P03191 P03193 P03195 P03197 P03198
P03199 P03203 P03204 P03207 P03209 P03210 P03211 P03213 P03214 P03217 P03219
P03220 P03222 P03226 P03227 P03232 P03233 P03235 P03237 P03241 P03243 P03244
P03245 P03252 P03253 P03261 P03262 P03264 P03265 P03267 P03271 P03272 P03273
P03276 P03277 P03278 P03279 P03280 P03281 P03282 P03284 P03285 P03286 P03287
P03288 P03289 P03290 P03291 P03292 P03293 P03294 P03296 P03298 P03323 P03324
P03326 P03327 P03358 P03371 P03373 P03400 P03408 P03412 P03413 P03414 P03418
P03421 P03422 P03424 P03513 P03514 P03515 P03516 P03520 P03521 P03523 P03525
P03526 P03527 P03529 P03536 P03537 P03538 P03539 P03541 P03545 P03546 P03547
P03548 P03549 P03550 P03551 P03552 P03553 P03554 P03555 P03556 P03557 P03558
P03559 P03563 P03567 P03568 P03571 P03573 P03574 P03575 P03576 P03578 P03579
P03580 P03581 P03585 P03586 P03587 P03590 P03591 P03593 P03594 P03595 P03596
P03597 P03598 P03601 P03602 P03603 P03604 P03605 P03606 P03608 P03611 P03614
P03615 P03616 P03627 P03628 P03629 P03630 P03631 P03632 P03634 P03636 P03638
P03640 P03641 P03642 P03643 P03644 P03645 P03647 P03648 P03649 P03650 P03652
P03654 P03660 P03680 P03682 P03683 P03685 P03686 P03687 P03688 P03689 P03692
P03693 P03694 P03695 P03696 P03697 P03698 P03699 P03700 P03702 P03703 P03704
P03707 P03708 P03709 P03710 P03711 P03712 P03713 P03714 P03715 P03716 P03718
P03719 P03720 P03723 P03728 P03729 P03730 P03731 P03732 P03733 P03734 P03735
P03737 P03738 P03739 P03740 P03742 P03743 P03744 P03746 P03747 P03748 P03749
P03750 P03751 P03752 P03753 P03754 P03755 P03756 P03758 P03761 P03763 P03764
P03765 P03766 P03767 P03768 P03769 P03770 P03771 P03772 P03773 P03775 P03776

P03778 P03779 P03780 P03781 P03782 P03783 P03784 P03785 P03786 P03787 P03789
P03790 P03792 P03793 P03794 P03795 P03796 P03797 P03798 P03799 P03800 P03801
P03813 P03817 P03828 P03829 P03831 P03832 P03833 P03835 P03839 P03845 P03846
P03847 P03848 P03849 P03850 P03851 P03852 P03853 P03854 P03856 P03857 P03858
P03859 P03860 P03861 P03862 P03864 P03865 P03866 P03869 P03870 P03871 P03872
P03882 P03883 P03884 P03885 P03934 P03935 P03936 P03938 P03939 P03941 P03942
P03947 P03962 P03963 P03964 P03965 P03966 P03975 P03976 P03977 P03984 P04009
P04012 P04015 P04016 P04017 P04019 P04026 P04028 P04030 P04034 P04036 P04043
P04045 P04046 P04047 P04051 P04052 P04076 P04077 P04078 P04079 P04080 P04082
P04101 P04105 P04106 P04107 P04109 P04110 P04111 P04112 P04113 P04120 P04122
P04130 P04132 P04133 P04137 P04139 P04146 P04149 P04161 P04162 P04164 P04168
P04169 P04170 P04173 P04174 P04176 P04181 P04182 P04184 P04193 P04197 P04198
P04215 P04219 P04237 P04238 P04239 P04240 P04241 P04242 P04243 P04244 P04245
P04246 P04247 P04248 P04249 P04250 P04251 P04252 P04255 P04256 P04257 P04259
P04260 P04261 P04262 P04263 P04265 P04266 P04282 P04283 P04291 P04292 P04293
P04294 P04296 P04298 P04299 P04300 P04301 P04302 P04303 P04304 P04305 P04308
P04309 P04310 P04311 P04312 P04318 P04323 P04332 P04335 P04337 P04338 P04339
P04342 P04344 P04345 P04346 P04352 P04353 P04355 P04357 P04358 P04363 P04368
P04375 P04376 P04382 P04383 P04385 P04386 P04387 P04390 P04391 P04392 P04393
P04395 P04397 P04399 P04407 P04408 P04414 P04415 P04418 P04419 P04424 P04425
P04442 P04443 P04444 P04445 P04446 P04447 P04448 P04450 P04452 P04453 P04454
P04455 P04457 P04458 P04459 P04460 P04461 P04462 P04464 P04466 P04476 P04481
P04482 P04483 P04486 P04487 P04489 P04490 P04491 P04493 P04495 P04496 P04497
P04498 P04514 P04515 P04516 P04518 P04519 P04520 P04524 P04525 P04526 P04527
P04529 P04530 P04531 P04532 P04533 P04534 P04535 P04536 P04537 P04538 P04543
P04545 P04546 P04547 P04550 P04553 P04554 P04568 P04569 P04570 P04571 P04572
P04573 P04576 P04625 P04630 P04631 P04642 P04646 P04665 P04666 P04668 P04669
P04673 P04674 P04675 P04676 P04680 P04681 P04682 P04685 P04686 P04688 P04689
P04690 P04693 P04694 P04707 P04711 P04712 P04713 P04714 P04715 P04716 P04717
P04718 P04731 P04732 P04733 P04734 P04735 P04736 P04743 P04744 P04748 P04749
P04751 P04752 P04753 P04766 P04770 P04771 P04772 P04786 P04787 P04793 P04794
P04795 P04796 P04801 P04803 P04805 P04819 P04828 P04829 P04831 P04832 P04833
P04834 P04835 P04851 P04857 P04858 P04859 P04860 P04861 P04862 P04865 P04866
P04870 P04873 P04874 P04877 P04878 P04879 P04880 P04881 P04885 P04886 P04889
P04890 P04891 P04892 P04893 P04903 P04907 P04909 P04910 P04911 P04912 P04913
P04914 P04915 P04931 P04935 P04940 P04941 P04943 P04944 P04945 P04946 P04947
P04948 P04952 P04962 P04964 P04968 P04969 P04970 P04982 P04990 P04991 P04992
P04993 P04994 P04995 P05020 P05031 P05033 P05034 P05035 P05041 P05042 P05043
P05048 P05050 P05051 P05052 P05057 P05058 P05062 P05063 P05066 P05068 P05070
P05072 P05074 P05075 P05076 P05077 P05078 P05079 P05080 P05081 P05083 P05084
P05085 P05096 P05100 P05101 P05102 P05103 P05104 P05116 P05123 P05133 P05138
P05147 P05148 P05162 P05166 P05189 P05194 P05195 P05201 P05204 P05205 P05219
P05220 P05301 P05302 P05303 P05304 P05308 P05309 P05314 P05324 P05325 P05326
P05327 P05328 P05332 P05336 P05338 P05339 P05340 P05341 P05342 P05343 P05344

P05345 P05346 P05348 P05349 P05361 P05373 P05378 P05379 P05382 P05384 P05385
P05390 P05391 P05392 P05393 P05394 P05395 P05396 P05397 P05398 P05400 P05401
P05404 P05406 P05409 P05410 P05411 P05415 P05416 P05419 P05423 P05428 P05433
P05434 P05443 P05444 P05445 P05446 P05447 P05449 P05450 P05453 P05459 P05460
P05461 P05462 P05463 P05466 P05467 P05468 P05469 P05470 P05471 P05472 P05473
P05474 P05475 P05476 P05477 P05478 P05481 P05488 P05511 P05514 P05516 P05519
P05523 P05526 P05527 P05529 P05530 P05531 P05547 P05548 P05549 P05552 P05621
P05624 P05634 P05637 P05638 P05644 P05645 P05646 P05647 P05649 P05650 P05651
P05652 P05653 P05654 P05657 P05659 P05661 P05662 P05663 P05664 P05665 P05666
P05667 P05668 P05669 P05670 P05671 P05672 P05673 P05675 P05676 P05678 P05679
P05680 P05682 P05683 P05684 P05685 P05693 P05698 P05699 P05707 P05709 P05716
P05719 P05720 P05721 P05722 P05723 P05724 P05725 P05726 P05727 P05728 P05729
P05734 P05752 P05762 P05766 P05768 P05781 P05782 P05785 P05786 P05789 P05791
P05793 P05794 P05804 P05807 P05812 P05813 P05815 P05823 P05824 P05827 P05830
P05831 P05833 P05835 P05836 P05839 P05840 P05841 P05842 P05844 P05845 P05846
P05847 P05853 P05918 P05930 P05932 P05933 P05934 P05935 P05936 P05938 P05939
P05940 P05941 P05944 P05945 P05946 P05957 P05958 P05963 P05969 P05971 P05976
P05977 P05987 P05989 P05990 P05993 P05998 P06012 P06018 P06019 P06020 P06022
P06023 P06025 P06100 P06101 P06102 P06103 P06104 P06107 P06115 P06116 P06117
P06118 P06119 P06120 P06121 P06122 P06123 P06130 P06131 P06132 P06135 P06144
P06145 P06146 P06147 P06148 P06150 P06151 P06153 P06154 P06155 P06159 P06161
P06162 P06163 P06164 P06165 P06168 P06169 P06171 P06173 P06174 P06187 P06188
P06189 P06190 P06191 P06193 P06194 P06195 P06198 P06203 P06204 P06208 P06212
P06214 P06219 P06220 P06221 P06222 P06225 P06226 P06227 P06228 P06229 P06232
P06233 P06234 P06235 P06236 P06237 P06242 P06267 P06269 P06270 P06271 P06272
P06273 P06274 P06275 P06292 P06294 P06295 P06302 P06337 P06347 P06348 P06349
P06352 P06353 P06354 P06355 P06356 P06357 P06358 P06359 P06360 P06362 P06363
P06364 P06368 P06370 P06371 P06372 P06373 P06374 P06375 P06377 P06378 P06379
P06381 P06382 P06383 P06384 P06385 P06386 P06387 P06388 P06389 P06390 P06391
P06392 P06393 P06398 P06416 P06417 P06422 P06423 P06424 P06425 P06426 P06427
P06428 P06431 P06434 P06435 P06439 P06440 P06444 P06447 P06455 P06456 P06458
P06459 P06460 P06461 P06462 P06463 P06467 P06472 P06478 P06479 P06485 P06486
P06489 P06490 P06491 P06492 P06495 P06496 P06500 P06501 P06502 P06503 P06504
P06505 P06506 P06507 P06508 P06509 P06510 P06513 P06514 P06515 P06516 P06524
P06529 P06530 P06531 P06532 P06533 P06538 P06549 P06550 P06552 P06553 P06555
P06556 P06557 P06558 P06559 P06560 P06561 P06562 P06565 P06566 P06569 P06571
P06572 P06573 P06574 P06575 P06581 P06582 P06586 P06587 P06588 P06589 P06590
P06592 P06593 P06594 P06595 P06601 P06602 P06603 P06604 P06605 P06612 P06614
P06615 P06617 P06618 P06619 P06622 P06626 P06627 P06629 P06630 P06632 P06633
P06634 P06635 P06636 P06637 P06638 P06639 P06640 P06642 P06643 P06646 P06650
P06652 P06661 P06662 P06663 P06665 P06666 P06667 P06668 P06672 P06689 P06691
P06693 P06694 P06695 P06700 P06701 P06704 P06706 P06707 P06708 P06709 P06710
P06714 P06720 P06721 P06722 P06723 P06724 P06726 P06732 P06735 P06736 P06737
P06738 P06742 P06746 P06753 P06754 P06757 P06766 P06768 P06769 P06770 P06773

P06774 P06777 P06778 P06779 P06785 P06786 P06791 P06792 P06794 P06798 P06801
P06805 P06829 P06831 P06839 P06841 P06843 P06844 P06845 P06846 P06847 P06854
P06855 P06856 P06864 P06890 P06892 P06893 P06894 P06895 P06897 P06898 P06899
P06900 P06902 P06903 P06904 P06916 P06917 P06919 P06921 P06922 P06923 P06924
P06925 P06926 P06929 P06930 P06931 P06936 P06937 P06938 P06940 P06941 P06945
P06947 P06948 P06950 P06951 P06952 P06953 P06954 P06955 P06956 P06959 P06960
P06961 P06965 P06966 P06968 P06983 P06986 P06987 P06988 P06989 P06990 P06991
P06992 P06993 P06999 P07004 P07010 P07012 P07013 P07023 P07026 P07031 P07032
P07033 P07034 P07035 P07036 P07040 P07041 P07046 P07047 P07048 P07049 P07051
P07059 P07061 P07062 P07065 P07066 P07067 P07068 P07069 P07070 P07071 P07072
P07074 P07075 P07076 P07077 P07078 P07079 P07080 P07081 P07082 P07083 P07086
P07087 P07089 P07090 P07094 P07097 P07104 P07105 P07107 P07112 P07113 P07114
P07115 P07116 P07118 P07125 P07126 P07128 P07129 P07131 P07132 P07133 P07134
P07135 P07139 P07145 P07159 P07160 P07161 P07162 P07164 P07165 P07166 P07169
P07170 P07172 P07175 P07178 P07179 P07180 P07196 P07201 P07216 P07218 P07229
P07234 P07235 P07236 P07243 P07244 P07245 P07246 P07247 P07248 P07249 P07258
P07260 P07261 P07262 P07263 P07264 P07265 P07266 P07269 P07272 P07273 P07274
P07276 P07283 P07284 P07285 P07287 P07290 P07291 P07294 P07295 P07296 P07300
P07304 P07305 P07310 P07311 P07313 P07315 P07316 P07317 P07318 P07320 P07322
P07324 P07331 P07341 P07342 P07343 P07344 P07345 P07346 P07347 P07362 P07372
P07375 P07377 P07378 P07382 P07383 P07385 P07386 P07387 P07388 P07392 P07393
P07394 P07396 P07398 P07400 P07402 P07403 P07404 P07405 P07406 P07407 P07408
P07409 P07410 P07411 P07412 P07413 P07414 P07415 P07417 P07419 P07420 P07421
P07425 P07428 P07429 P07430 P07431 P07432 P07433 P07436 P07438 P07440 P07441
P07442 P07443 P07444 P07446 P07447 P07448 P07450 P07451 P07452 P07461 P07462
P07463 P07464 P07472 P07481 P07484 P07485 P07486 P07487 P07488 P07505 P07508
P07509 P07511 P07512 P07521 P07524 P07530 P07531 P07532 P07533 P07535 P07537
P07538 P07541 P07543 P07545 P07547 P07548 P07583 P07591 P07598 P07600 P07601
P07604 P07605 P07606 P07610 P07614 P07617 P07621 P07623 P07626 P07627 P07633
P07636 P07637 P07639 P07640 P07641 P07642 P07648 P07649 P07659 P07665 P07668
P07669 P07670 P07673 P07674 P07679 P07684 P07685 P07689 P07691 P07693 P07694
P07695 P07696 P07699 P07702 P07703 P07715 P07716 P07717 P07718 P07719 P07733
P07735 P07736 P07738 P07739 P07740 P07741 P07742 P07744 P07746 P07748 P07752
P07754 P07760 P07762 P07764 P07765 P07769 P07770 P07771 P07772 P07773 P07774
P07776 P07777 P07778 P07779 P07780 P07781 P07782 P07783 P07784 P07785 P07786
P07787 P07789 P07790 P07791 P07793 P07794 P07795 P07796 P07797 P07798 P07799
P07803 P07804 P07805 P07806 P07807 P07810 P07812 P07813 P07815 P07816 P07817
P07818 P07819 P07820 P07828 P07833 P07836 P07837 P07838 P07839 P07840 P07841
P07842 P07843 P07844 P07850 P07859 P07860 P07862 P07874 P07878 P07879 P07884
P07885 P07887 P07889 P07902 P07909 P07913 P07917 P07918 P07920 P07922 P07924
P07928 P07929 P07931 P07932 P07933 P07934 P07935 P07938 P07940 P07945 P07951
P07952 P07953 P07954 P07955 P07956 P07957 P07958 P07959 P07960 P07961 P07962
P07963 P07964 P07978 P07989 P07990 P07991 P07993 P07999 P08002 P08009 P08015
P08016 P08023 P08028 P08030 P08036 P08042 P08043 P08044 P08045 P08048 P08051

P08052 P08053 P08054 P08055 P08057 P08058 P08061 P08063 P08066 P08070 P08073
P08075 P08076 P08077 P08078 P08080 P08086 P08087 P08088 P08094 P08096 P08097
P08098 P08105 P08115 P08116 P08127 P08130 P08131 P08135 P08140 P08142 P08145
P08146 P08147 P08151 P08154 P08155 P08157 P08158 P08159 P08160 P08161 P08164
P08175 P08178 P08179 P08192 P08196 P08200 P08201 P08202 P08203 P08204 P08209
P08211 P08223 P08224 P08225 P08229 P08230 P08231 P08232 P08241 P08242 P08243
P08244 P08245 P08256 P08257 P08258 P08259 P08260 P08261 P08262 P08266 P08267
P08280 P08281 P08282 P08283 P08284 P08285 P08286 P08287 P08288 P08293 P08298
P08302 P08304 P08308 P08309 P08310 P08312 P08315 P08320 P08322 P08332 P08333
P08335 P08337 P08341 P08343 P08344 P08345 P08346 P08347 P08348 P08349 P08352
P08357 P08358 P08364 P08365 P08373 P08385 P08389 P08390 P08392 P08393 P08394
P08397 P08405 P08417 P08418 P08421 P08422 P08423 P08425 P08429 P08431 P08432
P08433 P08436 P08437 P08439 P08440 P08442 P08451 P08455 P08457 P08461 P08463
P08465 P08474 P08475 P08477 P08490 P08495 P08497 P08499 P08504 P08507 P08509
P08515 P08518 P08520 P08521 P08527 P08528 P08529 P08530 P08535 P08536 P08537
P08540 P08543 P08546 P08548 P08551 P08552 P08553 P08557 P08558 P08562 P08566
P08579 P08580 P08590 P08593 P08619 P08624 P08625 P08626 P08632 P08633 P08634
P08638 P08653 P08654 P08655 P08657 P08660 P08662 P08664 P08685 P08688 P08692
P08694 P08695 P08696 P08698 P08703 P08705 P08706 P08707 P08710 P08711 P08717
P08719 P08734 P08735 P08736 P08737 P08738 P08747 P08760 P08761 P08762 P08763
P08764 P08765 P08767 P08770 P08771 P08773 P08775 P08776 P08777 P08778 P08779
P08791 P08792 P08793 P08794 P08799 P08802 P08811 P08812 P08813 P08814 P08819
P08820 P08821 P08823 P08824 P08841 P08844 P08845 P08848 P08849 P08850 P08851
P08852 P08853 P08854 P08855 P08862 P08868 P08870 P08874 P08881 P08889 P08891
P08892 P08893 P08894 P08895 P08898 P08902 P08903 P08906 P08914 P08915 P08916
P08917 P08924 P08925 P08926 P08927 P08929 P08930 P08931 P08955 P08956 P08957
P08964 P08968 P08970 P08973 P08974 P08975 P08976 P08977 P08978 P08979 P08985
P08988 P08991 P08992 P08993 P08994 P08997 P09005 P09007 P09013 P09014 P09015
P09016 P09017 P09019 P09020 P09021 P09022 P09023 P09024 P09025 P09026 P09027
P09029 P09030 P09032 P09033 P09035 P09041 P09043 P09044 P09046 P09052 P09053
P09057 P09060 P09061 P09062 P09063 P09064 P09067 P09070 P09071 P09074 P09075
P09076 P09077 P09078 P09079 P09080 P09081 P09082 P09083 P09084 P09085 P09087
P09088 P09089 P09090 P09092 P09094 P09095 P09099 P09105 P09106 P09111 P09114
P09117 P09120 P09122 P09123 P09124 P09125 P09126 P09134 P09135 P09136 P09137
P09140 P09141 P09142 P09143 P09145 P09147 P09148 P09151 P09153 P09154 P09155
P09158 P09162 P09163 P09165 P09176 P09178 P09184 P09187 P09188 P09189 P09195
P09201 P09202 P09203 P09204 P09205 P09206 P09210 P09213 P09223 P09224 P09226
P09227 P09233 P09243 P09244 P09245 P09246 P09248 P09249 P09250 P09251 P09252
P09253 P09254 P09255 P09265 P09268 P09269 P09270 P09275 P09276 P09277 P09279
P09281 P09282 P09284 P09285 P09286 P09287 P09288 P09289 P09291 P09292 P09293
P09295 P09299 P09300 P09301 P09302 P09303 P09304 P09305 P09306 P09307 P09309
P09311 P09314 P09315 P09316 P09317 P09322 P09330 P09339 P09342 P09347 P09352
P09357 P09358 P09364 P09366 P09368 P09369 P09370 P09372 P09377 P09378 P09379
P09380 P09381 P09383 P09387 P09389 P09395 P09397 P09398 P09399 P09400 P09402

P09403 P09404 P09406 P09407 P09414 P09415 P09417 P09420 P09421 P09422 P09423
P09424 P09425 P09426 P09427 P09430 P09436 P09437 P09440 P09441 P09442 P09443
P09444 P09445 P09450 P09451 P09459 P09463 P09485 P09491 P09495 P09498 P09499
P09502 P09503 P09504 P09505 P09508 P09509 P09510 P09511 P09512 P09514 P09515
P09517 P09518 P09520 P09521 P09522 P09523 P09524 P09528 P09532 P09540 P09541
P09542 P09547 P09552 P09553 P09554 P09555 P09556 P09557 P09559 P09561 P09562
P09565 P09569 P09575 P09577 P09578 P09580 P09588 P09589 P09590 P09591 P09594
P09595 P09596 P09597 P09602 P09604 P09608 P09609 P09614 P09623 P09624 P09629
P09630 P09631 P09632 P09633 P09634 P09635 P09636 P09637 P09638 P09642 P09643
P09644 P09645 P09647 P09652 P09658 P09662 P09672 P09673 P09674 P09675 P09676
P09677 P09678 P09694 P09695 P09696 P09697 P09698 P09699 P09700 P09701 P09702
P09706 P09707 P09708 P09710 P09711 P09714 P09715 P09722 P09733 P09734 P09735
P09737 P09738 P09739 P09741 P09745 P09746 P09747 P09748 P09749 P09750 P09751
P09753 P09754 P09757 P09763 P09764 P09768 P09772 P09773 P09774 P09775 P09788
P09792 P09795 P09796 P09797 P09798 P09804 P09806 P09810 P09812 P09816 P09818
P09819 P09820 P09821 P09822 P09823 P09824 P09825 P09826 P09827 P09831 P09832
P09839 P09840 P09842 P09844 P09845 P09846 P09847 P09853 P09854 P09855 P09856
P09857 P09860 P09863 P09864 P09865 P09867 P09873 P09877 P09881 P09882 P09883
P09885 P09886 P09887 P09890 P09891 P09897 P09898 P09899 P09901 P09904 P09905
P09906 P09907 P09908 P09909 P09911 P09913 P09914 P09915 P09921 P09924 P09928
P09932 P09938 P09940 P09941 P09942 P09943 P09947 P09948 P09950 P09952 P09953
P09956 P09959 P09961 P09964 P09965 P09966 P09967 P09968 P09972 P09975 P09976
P09979 P09980 P09984 P09985 P09987 P09988 P09992 P09996 P0A004 P0A006 P0A009
P0A011 P0A012 P0A013 P0A014 P0A016 P0A017 P0A022 P0A023 P0A024 P0A025 P0A027
P0A028 P0A032 P0A033 P0A034 P0A036 P0A037 P0A038 P0A039 P0A040 P0A041 P0A043
P0A044 P0A045 P0A047 P0A048 P0A049 P0A050 P0A051 P0A052 P0A053 P0A054 P0A055
P0A061 P0A062 P0A078 P0A079 P0A080 P0A081 P0A082 P0A083 P0A084 P0A085 P0A086
P0A087 P0A088 P0A089 P0A091 P0A092 P0A094 P0A095 P0A097 P0A099 P0A0A1 P0A0A2
P0A0A3 P0A0A4 P0A0A5 P0A0A6 P0A0A8 P0A0A9 P0A0B0 P0A0B1 P0A0B2 P0A0B3 P0A0B5
P0A0B6 P0A0B7 P0A0B8 P0A0B9 P0A0C0 P0A0C1 P0A0C2 P0A0C3 P0A0C4 P0A0C5 P0A0C6
P0A0C7 P0A0C8 P0A0C9 P0A0D0 P0A0D2 P0A0E4 P0A0E5 P0A0E9 P0A0F0 P0A0F1 P0A0F3
P0A0F4 P0A0F5 P0A0F7 P0A0F8 P0A0F9 P0A0G1 P0A0G2 P0A0G3 P0A0G4 P0A0G5 P0A0G6
P0A0G7 P0A0G9 P0A0H0 P0A0H1 P0A0H3 P0A0H4 P0A0H5 P0A0H6 P0A0I8 P0A0I9 P0A0J0
P0A0J1 P0A0J2 P0A0J3 P0A0K0 P0A0K1 P0A0K2 P0A0K3 P0A0K4 P0A0K5 P0A0K6 P0A0K7
P0A0K8 P0A0K9 P0A0M5 P0A0M6 P0A0M7 P0A0M8 P0A0M9 P0A0N3 P0A0N4 P0A0N5 P0A0N6
P0A0N7 P0A0N9 P0A0P1 P0A0P2 P0A0P3 P0A0P4 P0A0P5 P0A0P6 P0A0P7 P0A0P8 P0A0P9
P0A0Q0 P0A0Q1 P0A0Q2 P0A0Q3 P0A0Q4 P0A0Q7 P0A0Q8 P0A0Q9 P0A0R0 P0A0R1 P0A0R2
P0A0R3 P0A0R4 P0A0R5 P0A0R6 P0A0R7 P0A0S7 P0A0S8 P0A0S9 P0A0T0 P0A0T1 P0A0T2
P0A0T3 P0A0T4 P0A0T5 P0A0T7 P0A0T8 P0A0T9 P0A0U0 P0A0U1 P0A0U2 P0A0U3 P0A0U4
P0A0U5 P0A0U6 P0A0U7 P0A0V4 P0A0V5 P0A0V6 P0A0V7 P0A0W0 P0A0W1 P0A0W2 P0A0W3
P0A0W4 P0A0W5 P0A0W6 P0A0W7 P0A0W8 P0A0W9 P0A0X0 P0A0X1 P0A0X2 P0A0X3 P0A0X4
P0A0X5 P0A0X6 P0A0X7 P0A0X8 P0A0X9 P0A0Y0 P0A0Y1 P0A0Y2 P0A0Y7 P0A0Y8 P0A0Y9
P0A0Z0 P0A0Z1 P0A0Z2 P0A0Z7 P0A0Z8 P0A0Z9 P0A100 P0A101 P0A102 P0A105 P0A106
P0A107 P0A108 P0A109 P0A110 P0A111 P0A112 P0A113 P0A114 P0A115 P0A116 P0A117

P0A118 P0A119 P0A120 P0A121 P0A122 P0A123 P0A124 P0A125 P0A126 P0A127 P0A128
P0A129 P0A132 P0A133 P0A134 P0A135 P0A136 P0A137 P0A142 P0A143 P0A144 P0A147
P0A148 P0A149 P0A150 P0A151 P0A152 P0A155 P0A156 P0A157 P0A158 P0A159 P0A160
P0A161 P0A162 P0A163 P0A164 P0A165 P0A166 P0A167 P0A168 P0A169 P0A170 P0A175
P0A176 P0A178 P0A183 P0A184 P0A185 P0A186 P0A194 P0A195 P0A196 P0A1A2 P0A1A3
P0A1A4 P0A1A5 P0A1A6 P0A1A7 P0A1A8 P0A1A9 P0A1B0 P0A1B1 P0A1B2 P0A1B3 P0A1B4
P0A1B5 P0A1B6 P0A1C8 P0A1C9 P0A1D0 P0A1D1 P0A1D2 P0A1D3 P0A1D4 P0A1D5 P0A1D6
P0A1D7 P0A1D8 P0A1D9 P0A1E0 P0A1E1 P0A1E2 P0A1E3 P0A1E4 P0A1F0 P0A1F1 P0A1F2
P0A1F3 P0A1F6 P0A1F7 P0A1F8 P0A1F9 P0A1G0 P0A1G1 P0A1G2 P0A1G3 P0A1G4 P0A1G5
P0A1G6 P0A1G7 P0A1G8 P0A1G9 P0A1H0 P0A1H3 P0A1H4 P0A1H5 P0A1H6 P0A1I7 P0A1I8
P0A1I9 P0A1J0 P0A1J1 P0A1J2 P0A1J3 P0A1J4 P0A1J7 P0A1J8 P0A1K7 P0A1K8 P0A1N0
P0A1N1 P0A1N6 P0A1N7 P0A1P0 P0A1P1 P0A1P7 P0A1P8 P0A1P9 P0A1Q1 P0A1Q2 P0A1Q3
P0A1Q4 P0A1Q5 P0A1Q6 P0A1Q7 P0A1Q8 P0A1Q9 P0A1R0 P0A1R1 P0A1R2 P0A1R3 P0A1R4
P0A1R5 P0A1R6 P0A1R7 P0A1R8 P0A1R9 P0A1S0 P0A1S1 P0A1S2 P0A1S3 P0A1S4 P0A1S5
P0A1S6 P0A1S7 P0A1S9 P0A1T2 P0A1T3 P0A1T4 P0A1T5 P0A1T8 P0A1T9 P0A1U4 P0A1U5
P0A1U6 P0A1U7 P0A1V0 P0A1V1 P0A1V2 P0A1V3 P0A1V4 P0A1V5 P0A1V6 P0A1V7 P0A1W0
P0A1W1 P0A1X6 P0A1X7 P0A1Y0 P0A1Y1 P0A1Y2 P0A1Y3 P0A1Z0 P0A1Z1 P0A1Z4 P0A1Z5
P0A1Z6 P0A1Z7 P0A206 P0A207 P0A208 P0A209 P0A210 P0A211 P0A212 P0A213 P0A214
P0A215 P0A229 P0A230 P0A239 P0A240 P0A241 P0A242 P0A245 P0A246 P0A248 P0A251
P0A252 P0A253 P0A254 P0A255 P0A256 P0A265 P0A266 P0A267 P0A268 P0A271 P0A272
P0A281 P0A282 P0A285 P0A286 P0A287 P0A288 P0A289 P0A290 P0A292 P0A293 P0A294
P0A295 P0A296 P0A297 P0A298 P0A299 P0A2A0 P0A2A1 P0A2A2 P0A2A3 P0A2A4 P0A2A5
P0A2A6 P0A2A7 P0A2A8 P0A2A9 P0A2B0 P0A2B1 P0A2B2 P0A2B3 P0A2B4 P0A2B5 P0A2B6
P0A2B7 P0A2B8 P0A2B9 P0A2C0 P0A2C1 P0A2C2 P0A2C3 P0A2C4 P0A2D1 P0A2D2 P0A2D7
P0A2D8 P0A2E1 P0A2E2 P0A2E3 P0A2E4 P0A2E6 P0A2E7 P0A2E8 P0A2E9 P0A2F2 P0A2F3
P0A2F4 P0A2F5 P0A2F6 P0A2F7 P0A2G1 P0A2G2 P0A2G7 P0A2G8 P0A2G9 P0A2H0 P0A2H1
P0A2H2 P0A2I1 P0A2I2 P0A2I3 P0A2I4 P0A2I5 P0A2I6 P0A2K1 P0A2K2 P0A2K3 P0A2K4
P0A2K5 P0A2K6 P0A2K9 P0A2L0 P0A2L1 P0A2L2 P0A2L4 P0A2L7 P0A2L8 P0A2L9 P0A2M0
P0A2M1 P0A2M2 P0A2M3 P0A2M4 P0A2M5 P0A2M6 P0A2M7 P0A2M8 P0A2N2 P0A2N3 P0A2N8
P0A2N9 P0A2P0 P0A2P1 P0A2P2 P0A2P3 P0A2P4 P0A2P5 P0A2P6 P0A2P7 P0A2P8 P0A2P9
P0A2Q0 P0A2Q1 P0A2Q2 P0A2Q3 P0A2Q4 P0A2Q5 P0A2Q6 P0A2Q7 P0A2Q8 P0A2Q9 P0A2R0
P0A2R1 P0A2R2 P0A2R3 P0A2R4 P0A2R5 P0A2R6 P0A2R7 P0A2S0 P0A2S3 P0A2S4 P0A2S5
P0A2S6 P0A2S9 P0A2T0 P0A2T2 P0A2T3 P0A2T4 P0A2T5 P0A2T6 P0A2T7 P0A2T8 P0A2T9
P0A2U0 P0A2U1 P0A2U2 P0A2U3 P0A2U5 P0A2W8 P0A2W9 P0A2X0 P0A2X1 P0A2X2 P0A2X3
P0A2X4 P0A2X5 P0A2X6 P0A2X7 P0A2X8 P0A2X9 P0A2Y0 P0A2Y1 P0A2Y2 P0A2Y3 P0A2Y4
P0A2Y5 P0A2Y6 P0A2Y7 P0A314 P0A315 P0A323 P0A324 P0A325 P0A326 P0A328 P0A329
P0A330 P0A335 P0A336 P0A337 P0A338 P0A339 P0A340 P0A341 P0A342 P0A343 P0A345
P0A346 P0A347 P0A348 P0A349 P0A351 P0A352 P0A353 P0A354 P0A355 P0A356 P0A357
P0A358 P0A361 P0A362 P0A363 P0A364 P0A366 P0A367 P0A368 P0A369 P0A370 P0A371
P0A372 P0A373 P0A374 P0A375 P0A376 P0A377 P0A378 P0A379 P0A380 P0A381 P0A390
P0A391 P0A392 P0A393 P0A394 P0A395 P0A396 P0A397 P0A398 P0A399 P0A3A2 P0A3A3
P0A3A4 P0A3A6 P0A3A7 P0A3A8 P0A3A9 P0A3B0 P0A3B1 P0A3B2 P0A3B3 P0A3B4 P0A3B5
P0A3B6 P0A3B7 P0A3B8 P0A3C7 P0A3C8 P0A3C9 P0A3D0 P0A3D1 P0A3D2 P0A3D3 P0A3D4
P0A3D5 P0A3D6 P0A3D7 P0A3D8 P0A3D9 P0A3E0 P0A3E2 P0A3E3 P0A3E4 P0A3E5 P0A3E6

P0A3E7 P0A3E9 P0A3F0 P0A3F1 P0A3F2 P0A3F3 P0A3F8 P0A3F9 P0A3G2 P0A3G3 P0A3G4
P0A3G7 P0A3G8 P0A3H0 P0A3H1 P0A3H2 P0A3H3 P0A3H4 P0A3H5 P0A3H6 P0A3H7 P0A3H8
P0A3I1 P0A3I2 P0A3I3 P0A3I4 P0A3I5 P0A3I6 P0A3I7 P0A3I8 P0A3I9 P0A3J0 P0A3J1
P0A3J2 P0A3J3 P0A3J4 P0A3J7 P0A3J8 P0A3J9 P0A3K0 P0A3K1 P0A3K2 P0A3K3 P0A3K4
P0A3K5 P0A3K6 P0A3K7 P0A3K8 P0A3K9 P0A3L0 P0A3L1 P0A3L2 P0A3L3 P0A3L4 P0A3L5
P0A3L6 P0A3L7 P0A3L8 P0A3L9 P0A3M0 P0A3M7 P0A3M8 P0A3M9 P0A3N0 P0A3N1 P0A3N2
P0A3N3 P0A3Q0 P0A3Q1 P0A3Q2 P0A3Q3 P0A3Q5 P0A3Q6 P0A3R1 P0A3R2 P0A3R3 P0A3R4
P0A3S0 P0A3S1 P0A3S2 P0A3T0 P0A3T1 P0A3T4 P0A3T5 P0A3T6 P0A3T7 P0A3T8 P0A3T9
P0A3U0 P0A3U1 P0A3U2 P0A3U3 P0A3U6 P0A3U7 P0A3V2 P0A3V3 P0A3V4 P0A3V5 P0A3X0
P0A3X1 P0A3X2 P0A3X3 P0A3X4 P0A3X5 P0A3X6 P0A3Y3 P0A3Y4 P0A3Y5 P0A3Y6 P0A3Y9
P0A3Z0 P0A3Z1 P0A3Z2 P0A3Z3 P0A3Z4 P0A3Z9 P0A400 P0A420 P0A421 P0A422 P0A440
P0A441 P0A442 P0A443 P0A448 P0A449 P0A450 P0A451 P0A452 P0A454 P0A455 P0A456
P0A457 P0A458 P0A459 P0A460 P0A461 P0A462 P0A463 P0A464 P0A465 P0A466 P0A467
P0A468 P0A469 P0A470 P0A471 P0A472 P0A473 P0A474 P0A475 P0A476 P0A477 P0A478
P0A479 P0A480 P0A481 P0A482 P0A483 P0A484 P0A485 P0A486 P0A487 P0A488 P0A489
P0A490 P0A491 P0A492 P0A493 P0A494 P0A495 P0A496 P0A497 P0A498 P0A499 P0A4A0
P0A4A2 P0A4A3 P0A4A4 P0A4A5 P0A4A6 P0A4A7 P0A4A8 P0A4A9 P0A4B0 P0A4B1 P0A4B2
P0A4B3 P0A4B4 P0A4B5 P0A4B6 P0A4B7 P0A4B8 P0A4C0 P0A4C1 P0A4C2 P0A4C3 P0A4C4
P0A4C5 P0A4C6 P0A4C7 P0A4C8 P0A4C9 P0A4D0 P0A4D1 P0A4D2 P0A4D3 P0A4D4 P0A4D5
P0A4D6 P0A4D7 P0A4D9 P0A4E0 P0A4E1 P0A4E2 P0A4E3 P0A4E4 P0A4E5 P0A4E6 P0A4E7
P0A4E8 P0A4E9 P0A4F0 P0A4F1 P0A4F2 P0A4F3 P0A4F4 P0A4F5 P0A4F6 P0A4F7 P0A4I1
P0A4I9 P0A4J0 P0A4J1 P0A4J2 P0A4J3 P0A4J4 P0A4J5 P0A4J6 P0A4J7 P0A4J8 P0A4J9
P0A4K0 P0A4K1 P0A4K2 P0A4L1 P0A4L2 P0A4L3 P0A4L4 P0A4L9 P0A4M0 P0A4M1 P0A4M2
P0A4M6 P0A4P1 P0A4P2 P0A4P3 P0A4P4 P0A4P5 P0A4P6 P0A4P7 P0A4P8 P0A4Q4 P0A4Q5
P0A4Q8 P0A4Q9 P0A4R2 P0A4R3 P0A4R4 P0A4R5 P0A4R6 P0A4R7 P0A4R8 P0A4R9 P0A4S0
P0A4S1 P0A4S2 P0A4S3 P0A4S4 P0A4S5 P0A4S6 P0A4S7 P0A4S8 P0A4S9 P0A4T0 P0A4T1
P0A4T2 P0A4T3 P0A4T4 P0A4T6 P0A4T7 P0A4T8 P0A4T9 P0A4U0 P0A4U1 P0A4U2 P0A4U3
P0A4U4 P0A4U5 P0A4U6 P0A4U7 P0A4U8 P0A4U9 P0A4V0 P0A4V1 P0A4X1 P0A4X3 P0A4X5
P0A4X7 P0A4Y5 P0A4Y7 P0A4Y9 P0A4Z1 P0A4Z3 P0A4Z5 P0A4Z7 P0A4Z9 P0A507 P0A509
P0A511 P0A515 P0A519 P0A523 P0A525 P0A527 P0A529 P0A531 P0A533 P0A535 P0A539
P0A541 P0A543 P0A545 P0A547 P0A549 P0A551 P0A553 P0A555 P0A557 P0A559 P0A573
P0A577 P0A579 P0A581 P0A583 P0A585 P0A587 P0A589 P0A591 P0A593 P0A594 P0A596
P0A598 P0A5A0 P0A5A2 P0A5A4 P0A5A6 P0A5A8 P0A5B0 P0A5B2 P0A5B4 P0A5B6 P0A5C0
P0A5C4 P0A5C8 P0A5D0 P0A5D2 P0A5D8 P0A5E0 P0A5E6 P0A5F0 P0A5F2 P0A5F6 P0A5F8
P0A5G0 P0A5G2 P0A5G4 P0A5G6 P0A5G8 P0A5H0 P0A5H4 P0A5H6 P0A5H7 P0A5I1 P0A5I3
P0A5I5 P0A5J3 P0A5J5 P0A5J7 P0A5J9 P0A5K5 P0A5K7 P0A5L1 P0A5L3 P0A5L5 P0A5L7
P0A5L9 P0A5M3 P0A5M5 P0A5M7 P0A5M9 P0A5N1 P0A5N3 P0A5N5 P0A5N7 P0A5N9 P0A5R1
P0A5R3 P0A5R5 P0A5R7 P0A5S1 P0A5S3 P0A5S7 P0A5S9 P0A5T1 P0A5T3 P0A5T5 P0A5T7
P0A5T9 P0A5U1 P0A5U3 P0A5U5 P0A5U7 P0A5U9 P0A5V1 P0A5V3 P0A5V5 P0A5V7 P0A5V9
P0A5W1 P0A5W3 P0A5W5 P0A5W7 P0A5W9 P0A5X1 P0A5X3 P0A5X5 P0A5X7 P0A5X9 P0A603
P0A605 P0A611 P0A613 P0A617 P0A621 P0A623 P0A633 P0A635 P0A637 P0A639 P0A641
P0A645 P0A647 P0A649 P0A653 P0A655 P0A657 P0A659 P0A661 P0A663 P0A665 P0A667
P0A673 P0A675 P0A677 P0A679 P0A681 P0A687 P0A688 P0A689 P0A691 P0A693 P0A697
P0A698 P0A699 P0A6A4 P0A6A5 P0A6A6 P0A6A7 P0A6B4 P0A6B5 P0A6B6 P0A6B7 P0A6B8

P0A6B9 P0A6C0 P0A6C1 P0A6C2 P0A6C5 P0A6C6 P0A6C7 P0A6C8 P0A6C9 P0A6D0 P0A6D1
P0A6D2 P0A6D3 P0A6D4 P0A6D5 P0A6D6 P0A6D7 P0A6D8 P0A6D9 P0A6E0 P0A6E1 P0A6E2
P0A6E3 P0A6E4 P0A6E5 P0A6E9 P0A6F0 P0A6F1 P0A6F2 P0A6F3 P0A6F4 P0A6F6 P0A6F7
P0A6F8 P0A6G0 P0A6G1 P0A6G2 P0A6G3 P0A6G4 P0A6G5 P0A6G6 P0A6G8 P0A6G9 P0A6H0
P0A6H2 P0A6H3 P0A6H4 P0A6H6 P0A6H7 P0A6I0 P0A6I1 P0A6I2 P0A6I3 P0A6I4 P0A6I5
P0A6I6 P0A6I7 P0A6I8 P0A6I9 P0A6J0 P0A6J1 P0A6J2 P0A6J8 P0A6J9 P0A6K0 P0A6K1
P0A6K2 P0A6K3 P0A6K4 P0A6K5 P0A6K6 P0A6K7 P0A6K8 P0A6K9 P0A6L1 P0A6L2 P0A6L3
P0A6L4 P0A6L5 P0A6L6 P0A6L7 P0A6L8 P0A6L9 P0A6M0 P0A6M1 P0A6M4 P0A6M5 P0A6M6
P0A6M7 P0A6M8 P0A6M9 P0A6N0 P0A6N4 P0A6N5 P0A6N6 P0A6N7 P0A6N8 P0A6N9 P0A6P0
P0A6P2 P0A6P3 P0A6P4 P0A6P5 P0A6P6 P0A6P7 P0A6P8 P0A6R3 P0A6R4 P0A6R5 P0A6R6
P0A6R7 P0A6R8 P0A6R9 P0A6T1 P0A6T2 P0A6T3 P0A6T4 P0A6T5 P0A6T6 P0A6T7 P0A6T8
P0A6T9 P0A6U0 P0A6U1 P0A6U2 P0A6U3 P0A6U4 P0A6U5 P0A6U6 P0A6U7 P0A6U8 P0A6U9
P0A6V0 P0A6V1 P0A6V2 P0A6V3 P0A6V4 P0A6V5 P0A6V6 P0A6V7 P0A6V8 P0A6V9 P0A6W0
P0A6W1 P0A6W2 P0A6W5 P0A6W6 P0A6W7 P0A6W8 P0A6W9 P0A6X0 P0A6X1 P0A6X2 P0A6X4
P0A6X5 P0A6X6 P0A6X7 P0A6X8 P0A6X9 P0A6Y0 P0A6Y1 P0A6Y2 P0A6Y3 P0A6Y4 P0A6Y5
P0A6Y6 P0A6Y7 P0A6Y9 P0A6Z0 P0A6Z1 P0A6Z2 P0A6Z4 P0A6Z5 P0A6Z6 P0A6Z7 P0A6Z8
P0A6Z9 P0A701 P0A702 P0A703 P0A704 P0A706 P0A708 P0A709 P0A717 P0A718 P0A719
P0A720 P0A721 P0A729 P0A730 P0A731 P0A732 P0A733 P0A735 P0A736 P0A737 P0A738
P0A739 P0A740 P0A741 P0A744 P0A745 P0A746 P0A747 P0A748 P0A749 P0A750 P0A751
P0A752 P0A753 P0A759 P0A760 P0A761 P0A762 P0A763 P0A764 P0A765 P0A772 P0A773
P0A774 P0A775 P0A776 P0A777 P0A778 P0A779 P0A780 P0A781 P0A782 P0A783 P0A784
P0A785 P0A787 P0A788 P0A789 P0A790 P0A791 P0A792 P0A793 P0A794 P0A795 P0A796
P0A797 P0A798 P0A799 P0A7A0 P0A7A1 P0A7A2 P0A7A3 P0A7A4 P0A7A5 P0A7A6 P0A7B0
P0A7B2 P0A7B3 P0A7B4 P0A7B6 P0A7B7 P0A7B9 P0A7C0 P0A7C1 P0A7C6 P0A7C7 P0A7D1
P0A7D2 P0A7D3 P0A7D5 P0A7D6 P0A7D8 P0A7D9 P0A7E0 P0A7E3 P0A7E4 P0A7E5 P0A7E6
P0A7E7 P0A7E8 P0A7F0 P0A7F1 P0A7F2 P0A7F3 P0A7F4 P0A7F5 P0A7F6 P0A7F7 P0A7F8
P0A7F9 P0A7G0 P0A7G1 P0A7G2 P0A7G3 P0A7G4 P0A7G5 P0A7G7 P0A7G8 P0A7G9 P0A7H0
P0A7H1 P0A7H2 P0A7H3 P0A7H4 P0A7H5 P0A7H6 P0A7H7 P0A7H8 P0A7H9 P0A7I0 P0A7I1
P0A7I2 P0A7I3 P0A7I4 P0A7I5 P0A7I6 P0A7I7 P0A7I8 P0A7I9 P0A7J1 P0A7J2 P0A7J3
P0A7J4 P0A7J5 P0A7J6 P0A7J7 P0A7J8 P0A7J9 P0A7K0 P0A7K1 P0A7K2 P0A7K3 P0A7K4
P0A7K5 P0A7K6 P0A7K7 P0A7K8 P0A7K9 P0A7L0 P0A7L1 P0A7L2 P0A7L3 P0A7L4 P0A7L5
P0A7L6 P0A7L7 P0A7L8 P0A7L9 P0A7M0 P0A7M1 P0A7M2 P0A7M3 P0A7M4 P0A7M5 P0A7M7
P0A7M8 P0A7M9 P0A7N0 P0A7N1 P0A7N2 P0A7N3 P0A7N4 P0A7N5 P0A7N6 P0A7N7 P0A7N8
P0A7N9 P0A7P0 P0A7P1 P0A7P2 P0A7P3 P0A7P4 P0A7P5 P0A7P6 P0A7P7 P0A7P8 P0A7P9
P0A7Q0 P0A7Q1 P0A7Q2 P0A7Q3 P0A7Q4 P0A7Q5 P0A7Q6 P0A7Q7 P0A7Q8 P0A7Q9 P0A7R0
P0A7R1 P0A7R2 P0A7R3 P0A7R4 P0A7R5 P0A7R6 P0A7R7 P0A7R8 P0A7R9 P0A7S0 P0A7S1
P0A7S2 P0A7S3 P0A7S4 P0A7S5 P0A7S6 P0A7S7 P0A7S8 P0A7S9 P0A7T0 P0A7T1 P0A7T2
P0A7T3 P0A7T4 P0A7T5 P0A7T6 P0A7T7 P0A7T8 P0A7T9 P0A7U0 P0A7U1 P0A7U2 P0A7U3
P0A7U4 P0A7U5 P0A7U6 P0A7U7 P0A7U8 P0A7U9 P0A7V0 P0A7V1 P0A7V2 P0A7V3 P0A7V4
P0A7V5 P0A7V6 P0A7V7 P0A7V8 P0A7V9 P0A7W0 P0A7W1 P0A7W2 P0A7W3 P0A7W4 P0A7W5
P0A7W6 P0A7W7 P0A7W8 P0A7W9 P0A7X0 P0A7X1 P0A7X2 P0A7X3 P0A7X4 P0A7X5 P0A7X6
P0A7X7 P0A7X8 P0A7X9 P0A7Y0 P0A7Y1 P0A7Y2 P0A7Y3 P0A7Y4 P0A7Y5 P0A7Y6 P0A7Y7
P0A7Y8 P0A7Y9 P0A7Z0 P0A7Z1 P0A7Z2 P0A7Z3 P0A7Z5 P0A7Z6 P0A7Z7 P0A7Z8 P0A7Z9
P0A800 P0A801 P0A802 P0A803 P0A804 P0A805 P0A806 P0A807 P0A808 P0A809 P0A810

P0A811 P0A812 P0A813 P0A814 P0A815 P0A816 P0A817 P0A818 P0A819 P0A820 P0A821
P0A822 P0A823 P0A824 P0A826 P0A827 P0A828 P0A829 P0A832 P0A833 P0A834 P0A835
P0A836 P0A837 P0A838 P0A839 P0A840 P0A841 P0A842 P0A848 P0A849 P0A851 P0A852
P0A854 P0A859 P0A860 P0A861 P0A863 P0A864 P0A865 P0A866 P0A867 P0A868 P0A869
P0A871 P0A872 P0A873 P0A874 P0A875 P0A876 P0A877 P0A878 P0A879 P0A880 P0A881
P0A882 P0A883 P0A884 P0A885 P0A886 P0A888 P0A889 P0A890 P0A891 P0A892 P0A893
P0A894 P0A895 P0A896 P0A897 P0A898 P0A899 P0A8A0 P0A8A1 P0A8A2 P0A8A3 P0A8A4
P0A8A5 P0A8A6 P0A8A7 P0A8A8 P0A8A9 P0A8B0 P0A8B1 P0A8B2 P0A8B3 P0A8B4 P0A8B5
P0A8B6 P0A8B7 P0A8B8 P0A8B9 P0A8C0 P0A8C1 P0A8C2 P0A8C3 P0A8C4 P0A8C5 P0A8C6
P0A8C7 P0A8D0 P0A8D1 P0A8D2 P0A8D3 P0A8D4 P0A8D5 P0A8D6 P0A8D7 P0A8D8 P0A8E1
P0A8E2 P0A8E3 P0A8E4 P0A8E5 P0A8E6 P0A8E7 P0A8E8 P0A8E9 P0A8F1 P0A8F2 P0A8F3
P0A8F4 P0A8F5 P0A8F6 P0A8F7 P0A8F8 P0A8F9 P0A8G0 P0A8G1 P0A8G2 P0A8G3 P0A8G4
P0A8G5 P0A8G7 P0A8G8 P0A8G9 P0A8H0 P0A8H1 P0A8H2 P0A8H6 P0A8H7 P0A8H8 P0A8H9
P0A8I0 P0A8I1 P0A8I2 P0A8I3 P0A8I4 P0A8I5 P0A8I6 P0A8I7 P0A8I8 P0A8I9 P0A8J0
P0A8J1 P0A8J2 P0A8J3 P0A8J4 P0A8J5 P0A8J6 P0A8J7 P0A8J8 P0A8J9 P0A8K0 P0A8K5
P0A8K6 P0A8K7 P0A8L1 P0A8L2 P0A8L3 P0A8L4 P0A8L5 P0A8L6 P0A8L7 P0A8L8 P0A8L9
P0A8M0 P0A8M1 P0A8M2 P0A8M3 P0A8M4 P0A8M5 P0A8M6 P0A8M7 P0A8M8 P0A8M9 P0A8N1
P0A8N2 P0A8N4 P0A8N6 P0A8N7 P0A8N8 P0A8N9 P0A8P0 P0A8P1 P0A8P2 P0A8P3 P0A8P4
P0A8P5 P0A8P6 P0A8P7 P0A8P8 P0A8P9 P0A8Q6 P0A8Q7 P0A8Q8 P0A8Q9 P0A8R0 P0A8R1
P0A8R2 P0A8R3 P0A8R4 P0A8R5 P0A8R6 P0A8R9 P0A8S0 P0A8S1 P0A8S2 P0A8S3 P0A8S4
P0A8S9 P0A8T0 P0A8T1 P0A8T2 P0A8T3 P0A8T4 P0A8T5 P0A8T6 P0A8T8 P0A8T9 P0A8U2
P0A8U3 P0A8U4 P0A8U5 P0A8U6 P0A8U7 P0A8U8 P0A8U9 P0A8V0 P0A8V1 P0A8V3 P0A8V4
P0A8V5 P0A8W0 P0A8W1 P0A8W2 P0A8W3 P0A8W4 P0A8W5 P0A8W6 P0A8W7 P0A8W8 P0A8W9
P0A8X0 P0A8X1 P0A8Y1 P0A8Y2 P0A8Y3 P0A8Y4 P0A8Y5 P0A8Y6 P0A8Y7 P0A8Y8 P0A8Y9
P0A8Z0 P0A8Z1 P0A8Z2 P0A8Z7 P0A8Z8 P0A8Z9 P0A900 P0A944 P0A945 P0A946 P0A947
P0A948 P0A949 P0A950 P0A951 P0A952 P0A956 P0A957 P0A958 P0A959 P0A960 P0A961
P0A968 P0A969 P0A970 P0A971 P0A972 P0A973 P0A974 P0A975 P0A976 P0A977 P0A978
P0A979 P0A980 P0A981 P0A982 P0A983 P0A984 P0A985 P0A986 P0A987 P0A988 P0A989
P0A990 P0A992 P0A993 P0A994 P0A995 P0A998 P0A999 P0A9A0 P0A9A1 P0A9A2 P0A9A3
P0A9A4 P0A9A5 P0A9B3 P0A9B4 P0A9B5 P0A9B6 P0A9B7 P0A9B8 P0A9B9 P0A9C3 P0A9C4
P0A9C6 P0A9C7 P0A9C8 P0A9C9 P0A9D0 P0A9D1 P0A9D2 P0A9D3 P0A9D4 P0A9D5 P0A9D6
P0A9D7 P0A9D8 P0A9D9 P0A9E0 P0A9E1 P0A9E2 P0A9E3 P0A9E4 P0A9E5 P0A9E6 P0A9E7
P0A9E8 P0A9E9 P0A9F0 P0A9F1 P0A9F2 P0A9F3 P0A9F4 P0A9F5 P0A9F6 P0A9F7 P0A9F8
P0A9F9 P0A9G0 P0A9G1 P0A9G2 P0A9G3 P0A9G4 P0A9G5 P0A9G6 P0A9G7 P0A9G8 P0A9G9
P0A9H0 P0A9H1 P0A9H2 P0A9H3 P0A9H4 P0A9H5 P0A9H6 P0A9I1 P0A9I2 P0A9I3 P0A9I4
P0A9I8 P0A9I9 P0A9J1 P0A9J2 P0A9J3 P0A9J4 P0A9J5 P0A9J6 P0A9J7 P0A9J8 P0A9J9
P0A9K0 P0A9K1 P0A9K2 P0A9K3 P0A9K4 P0A9K5 P0A9K6 P0A9L3 P0A9L4 P0A9L5 P0A9L6
P0A9L7 P0A9L8 P0A9L9 P0A9M0 P0A9M1 P0A9M2 P0A9M3 P0A9M4 P0A9M8 P0A9M9 P0A9N0
P0A9N1 P0A9N2 P0A9N3 P0A9N4 P0A9N5 P0A9N6 P0A9N7 P0A9N8 P0A9N9 P0A9P1 P0A9P2
P0A9P3 P0A9P5 P0A9P6 P0A9P7 P0A9P8 P0A9P9 P0A9Q0 P0A9Q8 P0A9R0 P0A9R1 P0A9R2
P0A9R3 P0A9R4 P0A9R5 P0A9R6 P0A9S1 P0A9S2 P0A9S3 P0A9S4 P0A9S5 P0A9S6 P0A9S8
P0A9S9 P0A9T0 P0A9T1 P0A9T2 P0A9T3 P0A9T4 P0A9T5 P0A9T6 P0A9T7 P0A9U3 P0A9U4
P0A9U5 P0A9U6 P0A9U7 P0A9U8 P0A9U9 P0A9V0 P0A9V5 P0A9V6 P0A9V7 P0A9V8 P0A9V9
P0A9W0 P0A9W1 P0A9W2 P0A9W3 P0A9W4 P0A9W5 P0A9W6 P0A9W7 P0A9W8 P0A9W9 P0A9X0

P0A9X9 P0A9Y0 P0A9Y1 P0A9Y2 P0A9Y3 P0A9Y4 P0A9Y5 P0A9Y7 P0A9Y8 P0A9Y9 P0A9Z0
P0A9Z2 P0A9Z3 P0A9Z4 P0A9Z5 P0A9Z6 P0AA01 P0AA02 P0AA03 P0AA10 P0AA11 P0AA12
P0AA13 P0AA14 P0AA15 P0AA25 P0AA26 P0AA27 P0AA28 P0AA29 P0AA30 P0AA31 P0AA32
P0AA33 P0AA34 P0AA35 P0AA36 P0AA37 P0AA38 P0AA39 P0AA40 P0AA41 P0AA42 P0AA43
P0AA44 P0AA45 P0AA46 P0AA90 P0AA97 P0AA98 P0AAB3 P0AAB8 P0AAB9 P0AAC0 P0AAC1
P0AAC2 P0AAC3 P0AAC8 P0AAC9 P0AAD0 P0AAD1 P0AAK1 P0AAK2 P0AAK3 P0AAK4 P0AAK5
P0AAK6 P0AAL1 P0AAL2 P0AAL6 P0AAL7 P0AAL8 P0AAL9 P0AAM0 P0AAM3 P0AAM4 P0AAM5
P0AAM6 P0AAM7 P0AAM8 P0AAM9 P0AAN0 P0AAN1 P0AAN2 P0AAN3 P0AAN4 P0AAN5 P0AAN6
P0AAN7 P0AAN8 P0AAN9 P0AAP0 P0AAP3 P0AAP4 P0AAP5 P0AAP6 P0AAQ3 P0AAQ4 P0AAQ5
P0AAQ6 P0AAQ7 P0AAQ8 P0AAQ9 P0AAR0 P0AAR1 P0AAR2 P0AAR4 P0AAS5 P0AAS6 P0AAS7
P0AAS8 P0AAS9 P0AAT0 P0AAT1 P0AAT2 P0AAT3 P0AAT6 P0AAT7 P0AAT8 P0AAT9 P0AAU0
P0AAU1 P0AAU7 P0AAU8 P0AAU9 P0AAV0 P0AAV1 P0AAV2 P0AAV3 P0AAV4 P0AAV5 P0AAV8
P0AAV9 P0AAW0 P0AAW1 P0AAW2 P0AAW3 P0AAW4 P0AAY1 P0AAY2 P0AAY3 P0AAY6 P0AAY7
P0AAY8 P0AAY9 P0AAZ5 P0AAZ6 P0AAZ7 P0AAZ8 P0AAZ9 P0AB00 P0AB03 P0AB04 P0AB05
P0AB14 P0AB15 P0AB16 P0AB17 P0AB18 P0AB19 P0AB20 P0AB21 P0AB22 P0AB23 P0AB28
P0AB29 P0AB30 P0AB33 P0AB34 P0AB43 P0AB44 P0AB45 P0AB49 P0AB50 P0AB51 P0AB52
P0AB53 P0AB54 P0AB55 P0AB56 P0AB57 P0AB61 P0AB62 P0AB63 P0AB64 P0AB65 P0AB66
P0AB71 P0AB72 P0AB73 P0AB74 P0AB75 P0AB76 P0AB77 P0AB78 P0AB79 P0AB80 P0AB81
P0AB82 P0AB83 P0AB84 P0AB87 P0AB88 P0AB89 P0AB90 P0AB91 P0AB92 P0AB96 P0AB97
P0ABD4 P0ABE3 P0ABE4 P0ABE9 P0ABF0 P0ABF2 P0ABF3 P0ABF4 P0ABF5 P0ABF6 P0ABF7
P0ABH7 P0ABH8 P0ABH9 P0ABI0 P0ABI1 P0ABK6 P0ABP9 P0ABQ0 P0ABQ1 P0ABQ2 P0ABQ3
P0ABQ4 P0ABQ5 P0ABQ6 P0ABQ7 P0ABQ8 P0ABR1 P0ABR2 P0ABR3 P0ABR4 P0ABR5 P0ABR6
P0ABR7 P0ABR8 P0ABR9 P0ABS0 P0ABS1 P0ABS2 P0ABS3 P0ABS4 P0ABS5 P0ABS6 P0ABS7
P0ABS8 P0ABS9 P0ABT0 P0ABT1 P0ABT3 P0ABT4 P0ABT5 P0ABT6 P0ABT7 P0ABU0 P0ABU1
P0ABU2 P0ABU3 P0ABU4 P0ABU5 P0ABU6 P0ABW0 P0ABW1 P0ABW2 P0ABW3 P0ABW4 P0ABW9
P0ABX0 P0ABX1 P0ABX2 P0ABX3 P0ABX4 P0ABX6 P0ABX7 P0ABY4 P0ABY5 P0ABY6 P0ABY7
P0ABY8 P0ABY9 P0ABZ0 P0AC00 P0AC01 P0AC11 P0AC12 P0AC13 P0AC14 P0AC15 P0AC16
P0AC17 P0AC18 P0AC19 P0AC20 P0AC21 P0AC22 P0AC28 P0AC29 P0AC33 P0AC34 P0AC35
P0AC36 P0AC37 P0AC39 P0AC40 P0AC51 P0AC52 P0AC53 P0AC54 P0AC59 P0AC60 P0AC61
P0AC62 P0AC63 P0AC64 P0AC65 P0AC66 P0AC67 P0AC68 P0AC69 P0AC70 P0AC71 P0AC72
P0AC73 P0AC74 P0AC81 P0AC82 P0AC83 P0AC84 P0AC85 P0AC86 P0AC87 P0AC88 P0AC89
P0AC90 P0AC91 P0AC92 P0AC93 P0ACA1 P0ACA2 P0ACA3 P0ACA4 P0ACA5 P0ACA6 P0ACA7
P0ACA8 P0ACA9 P0ACB0 P0ACB1 P0ACB2 P0ACB3 P0ACC1 P0ACC2 P0ACC3 P0ACC4 P0ACC5
P0ACC6 P0ACC9 P0ACD0 P0ACD1 P0ACD2 P0ACD3 P0ACD4 P0ACD5 P0ACD6 P0ACD7 P0ACE3
P0ACE4 P0ACE5 P0ACE6 P0ACE8 P0ACE9 P0ACF1 P0ACF2 P0ACF3 P0ACF4 P0ACF5 P0ACF6
P0ACF7 P0ACF9 P0ACG0 P0ACG1 P0ACG2 P0ACG3 P0ACG8 P0ACG9 P0ACH0 P0ACH1 P0ACH2
P0ACH3 P0ACH4 P0ACH5 P0ACH6 P0ACH7 P0ACH8 P0ACH9 P0ACI0 P0ACI1 P0ACI2 P0ACI3
P0ACI4 P0ACI5 P0ACI6 P0ACI7 P0ACI8 P0ACI9 P0ACJ0 P0ACJ1 P0ACJ2 P0ACJ3 P0ACJ4
P0ACJ5 P0ACJ6 P0ACJ7 P0ACJ8 P0ACJ9 P0ACK0 P0ACK1 P0ACK2 P0ACK3 P0ACK4 P0ACK5
P0ACK6 P0ACK7 P0ACK8 P0ACK9 P0ACL0 P0ACL1 P0ACL2 P0ACL3 P0ACL4 P0ACL5 P0ACL6
P0ACL7 P0ACL8 P0ACL9 P0ACM0 P0ACM1 P0ACM2 P0ACM3 P0ACM4 P0ACM5 P0ACM6 P0ACM7
P0ACM8 P0ACM9 P0ACN0 P0ACN1 P0ACN2 P0ACN3 P0ACN4 P0ACN5 P0ACN6 P0ACN7 P0ACN8
P0ACN9 P0ACP0 P0ACP1 P0ACP2 P0ACP3 P0ACP4 P0ACP5 P0ACP6 P0ACP7 P0ACP8 P0ACP9
P0ACQ0 P0ACQ1 P0ACQ2 P0ACQ3 P0ACQ4 P0ACQ5 P0ACQ6 P0ACQ7 P0ACQ8 P0ACQ9 P0ACR0

P0ACR1 P0ACR2 P0ACR3 P0ACR4 P0ACR5 P0ACR6 P0ACR7 P0ACR8 P0ACR9 P0ACS0 P0ACS1
P0ACS3 P0ACS4 P0ACS5 P0ACS6 P0ACS7 P0ACS8 P0ACS9 P0ACT0 P0ACT1 P0ACT2 P0ACT3
P0ACT4 P0ACT5 P0ACT6 P0ACT7 P0ACT8 P0ACT9 P0ACU0 P0ACU1 P0ACU2 P0ACU3 P0ACU4
P0ACU8 P0ACU9 P0ACV6 P0ACV7 P0ACV8 P0ACV9 P0ACW0 P0ACW1 P0ACW6 P0ACW7 P0ACW8
P0ACW9 P0ACX3 P0ACX4 P0ACX5 P0ACX6 P0ACX7 P0ACX8 P0ACX9 P0ACY0 P0ACY1 P0ACY2
P0ACY3 P0ACY4 P0ACY5 P0ACY9 P0ACZ0 P0ACZ1 P0ACZ2 P0ACZ3 P0AD05 P0AD06 P0AD07
P0AD08 P0AD09 P0AD10 P0AD11 P0AD21 P0AD22 P0AD23 P0AD24 P0AD25 P0AD26 P0AD33
P0AD34 P0AD35 P0AD36 P0AD37 P0AD38 P0AD39 P0AD40 P0AD41 P0AD49 P0AD50 P0AD51
P0AD52 P0AD53 P0AD54 P0AD55 P0AD56 P0AD57 P0AD58 P0AD62 P0AD72 P0AD73 P0AD74
P0AD75 P0AD76 P0AD77 P0AD78 P0AD79 P0AD80 P0AD81 P0AD82 P0AD83 P0AD84 P0AD85
P0AD86 P0AD87 P0AD88 P0AD89 P0AD90 P0AD91 P0AD92 P0AD93 P0AD94 P0AD95 P0ADC8
P0ADC9 P0ADD0 P0ADD1 P0ADD7 P0ADD8 P0ADD9 P0ADE0 P0ADE1 P0ADE2 P0ADE3 P0ADE6
P0ADE7 P0ADE8 P0ADE9 P0ADF0 P0ADF1 P0ADF2 P0ADF3 P0ADF4 P0ADF5 P0ADF6 P0ADF7
P0ADF8 P0ADF9 P0ADG0 P0ADG1 P0ADG2 P0ADG3 P0ADG5 P0ADG6 P0ADG8 P0ADG9 P0ADH1
P0ADH2 P0ADH4 P0ADH5 P0ADH6 P0ADH7 P0ADH8 P0ADH9 P0ADI0 P0ADI1 P0ADI2 P0ADI3
P0ADI5 P0ADI6 P0ADI7 P0ADI8 P0ADJ3 P0ADJ4 P0ADK0 P0ADK1 P0ADK2 P0ADK3 P0ADK6
P0ADK7 P0ADK8 P0ADK9 P0ADL0 P0ADL3 P0ADL4 P0ADL5 P0ADM8 P0ADM9 P0ADN1 P0ADN2
P0ADN3 P0ADN4 P0ADN5 P0ADP0 P0ADP1 P0ADP8 P0ADP9 P0ADQ0 P0ADQ1 P0ADQ5 P0ADQ6
P0ADR6 P0ADR7 P0ADR8 P0ADR9 P0ADS0 P0ADS1 P0ADS7 P0ADS8 P0ADS9 P0ADT0 P0ADT1
P0ADT3 P0ADT4 P0ADT5 P0ADT6 P0ADT7 P0ADU2 P0ADU3 P0ADU4 P0ADU7 P0ADU8 P0ADU9
P0ADV0 P0ADV5 P0ADV6 P0ADW6 P0ADW7 P0ADW8 P0ADW9 P0ADX0 P0ADX1 P0ADX2 P0ADX3
P0ADX4 P0ADX5 P0ADX6 P0ADX9 P0ADY0 P0ADY3 P0ADY4 P0ADY5 P0ADY6 P0ADY7 P0ADY8
P0ADY9 P0ADZ1 P0ADZ2 P0ADZ3 P0ADZ4 P0ADZ5 P0ADZ6 P0AE01 P0AE02 P0AE03 P0AE04
P0AE05 P0AE09 P0AE10 P0AE11 P0AE12 P0AE13 P0AE18 P0AE19 P0AE20 P0AE21 P0AE28
P0AE29 P0AE37 P0AE38 P0AE39 P0AE48 P0AE49 P0AE50 P0AE51 P0AE52 P0AE53 P0AE54
P0AE55 P0AE56 P0AE57 P0AE58 P0AE59 P0AE60 P0AE61 P0AE62 P0AE63 P0AE64 P0AE66
P0AE71 P0AE72 P0AE73 P0AE76 P0AE77 P0AE79 P0AE80 P0AE81 P0AEA8 P0AEA9 P0AEB7
P0AEB8 P0AEB9 P0AED1 P0AED2 P0AED3 P0AED4 P0AED7 P0AED8 P0AED9 P0AEE0 P0AEE8
P0AEE9 P0AEF0 P0AEF1 P0AEF2 P0AEF3 P0AEG8 P0AEG9 P0AEH0 P0AEH3 P0AEH4 P0AEI1
P0AEI2 P0AEI3 P0AEI4 P0AEI5 P0AEI6 P0AEI7 P0AEI8 P0AEI9 P0AEJ2 P0AEJ3 P0AEJ6
P0AEJ7 P0AEJ8 P0AEJ9 P0AEK0 P0AEK1 P0AEL3 P0AEL4 P0AEL5 P0AEM0 P0AEM1 P0AEM2
P0AEM3 P0AEM4 P0AEM5 P0AEM6 P0AEM7 P0AEM8 P0AEN1 P0AEN2 P0AEN3 P0AEN8 P0AEN9
P0AEP0 P0AEP3 P0AEP4 P0AEP5 P0AEP7 P0AEP8 P0AEQ1 P0AEQ2 P0AES0 P0AES1 P0AES2
P0AES3 P0AES5 P0AES6 P0AES7 P0AES8 P0AEV1 P0AEV4 P0AEV5 P0AEV6 P0AEV7 P0AEV8
P0AEV9 P0AEW0 P0AEW4 P0AEW5 P0AEW6 P0AEW7 P0AEW8 P0AEW9 P0AEX0 P0AEX1 P0AEX2
P0AEX5 P0AEX6 P0AEY3 P0AEY4 P0AEY5 P0AEY6 P0AEY7 P0AEZ1 P0AEZ2 P0AEZ9 P0AF00
P0AF03 P0AF04 P0AF05 P0AF08 P0AF09 P0AF10 P0AF11 P0AF12 P0AF13 P0AF14 P0AF15
P0AF18 P0AF19 P0AF20 P0AF21 P0AF22 P0AF23 P0AF24 P0AF25 P0AF36 P0AF37 P0AF38
P0AF39 P0AF48 P0AF49 P0AF50 P0AF51 P0AF59 P0AF60 P0AF62 P0AF63 P0AF64 P0AF65
P0AF66 P0AF67 P0AF68 P0AF69 P0AF76 P0AF77 P0AF90 P0AF91 P0AF92 P0AF94 P0AF95
P0AF96 P0AF97 P0AFB8 P0AFB9 P0AFC0 P0AFC1 P0AFC2 P0AFF6 P0AFF7 P0AFF8 P0AFF9
P0AFG0 P0AFG1 P0AFG2 P0AFG3 P0AFG4 P0AFG5 P0AFG6 P0AFG7 P0AFG9 P0AFH0 P0AFH1
P0AFI0 P0AFI1 P0AFI7 P0AFI8 P0AFJ1 P0AFJ2 P0AFJ3 P0AFJ4 P0AFJ5 P0AFJ6 P0AFK0
P0AFK1 P0AFP2 P0AFP3 P0AFP4 P0AFP5 P0AFP6 P0AFP7 P0AFP8 P0AFQ5 P0AFQ6 P0AFQ7

P0AFQ8 P0AFQ9 P0AFR4 P0AFR5 P0AFR6 P0AFS3 P0AFS4 P0AFT8 P0AFT9 P0AFU6 P0AFU7
P0AFU8 P0AFU9 P0AFW0 P0AFW1 P0AFW2 P0AFW3 P0AFW4 P0AFW5 P0AFW6 P0AFW7 P0AFW8
P0AFW9 P0AFX1 P0AFX2 P0AFX3 P0AFX4 P0AFX5 P0AFX6 P0AFY8 P0AFY9 P0AFZ0 P0AFZ1
P0AFZ2 P0AFZ4 P0AFZ6 P0AG03 P0AG04 P0AG05 P0AG06 P0AG07 P0AG08 P0AG09 P0AG10
P0AG16 P0AG17 P0AG18 P0AG19 P0AG21 P0AG22 P0AG23 P0AG25 P0AG26 P0AG27 P0AG28
P0AG29 P0AG31 P0AG32 P0AG33 P0AG40 P0AG41 P0AG42 P0AG43 P0AG44 P0AG45 P0AG46
P0AG47 P0AG48 P0AG49 P0AG50 P0AG51 P0AG52 P0AG53 P0AG54 P0AG55 P0AG56 P0AG57
P0AG58 P0AG59 P0AG60 P0AG61 P0AG62 P0AG63 P0AG64 P0AG65 P0AG66 P0AG68 P0AG69
P0AG70 P0AG71 P0AG72 P0AG73 P0AG74 P0AG75 P0AG76 P0AG77 P0AG84 P0AG85 P0AG87
P0AG88 P0AG89 P0AGB0 P0AGB1 P0AGB2 P0AGB4 P0AGB5 P0AGD4 P0AGD5 P0AGD6 P0AGE0
P0AGE1 P0AGE2 P0AGE3 P0AGE6 P0AGE7 P0AGE8 P0AGE9 P0AGF0 P0AGF1 P0AGF2 P0AGF3
P0AGF6 P0AGF7 P0AGF8 P0AGF9 P0AGG0 P0AGG1 P0AGG4 P0AGG5 P0AGG6 P0AGG7 P0AGG8
P0AGG9 P0AGH0 P0AGJ2 P0AGJ3 P0AGJ4 P0AGJ5 P0AGJ6 P0AGJ7 P0AGJ8 P0AGK0 P0AGK4
P0AGK5 P0AGK6 P0AGK7 P0AGK8 P0AGK9 P0AGL0 P0AGL1 P0AGL2 P0AGL3 P0AGL4 P0AGL5
P0AGL6 P0AGL7 P0AGL8 P0AGL9 P0AGM5 P0AGM6 P0C002 P0C003 P0C004 P0C012 P0C016
P0C017 P0C018 P0C019 P0C020 P0C021 P0C025 P0C026 P0C030 P0C031 P0C032 P0C036
P0C037 P0C038 P0C039 P0C040 P0C042 P0C043 P0C044 P0C045 P0C054 P0C055 P0C056
P0C057 P0C058 P0C059 P0C060 P0C061 P0C062 P0C063 P0C064 P0C065 P0C068 P0C073
P0C074 P0C076 P0C077 P0C078 P0C079 P0C080 P0C093 P0C094 P0C095 P0C096 P0C097
P0C098 P0C099 P0C0A0 P0C0A4 P0C0A5 P0C0A6 P0C0A7 P0C0A8 P0C0B0 P0C0B1 P0C0B2
P0C0B3 P0C0B4 P0C0B5 P0C0B6 P0C0B7 P0C0B8 P0C0C1 P0C0C2 P0C0C3 P0C0C4 P0C0C5
P0C0C6 P0C0C9 P0C0D0 P0C0D1 P0C0D3 P0C0D4 P0C0D5 P0C0D6 P0C0D7 P0C0D8 P0C0D9
P0C0E0 P0C0E1 P0C0E6 P0C0F0 P0C0F1 P0C0F2 P0C0F3 P0C0F4 P0C0F5 P0C0F8 P0C0F9
P0C0G0 P0C0G1 P0C0G2 P0C0G3 P0C0G4 P0C0G5 P0C0G6 P0C0G7 P0C0H2 P0C0H3 P0C0H4
P0C0H5 P0C0H6 P0C0H7 P0C0I8 P0C0I9 P0C0J2 P0C0J3 P0C0J4 P0C0J5 P0C0J6 P0C0J9
P0C0K0 P0C0K3 P0C0K4 P0C0K9 P0C0L0 P0C0L1 P0C0L2 P0C0L3 P0C0L9 P0C0M0 P0C0M1
P0C0M2 P0C0M3 P0C0M4 P0C0M5 P0C0M6 P0C0M7 P0C0M8 P0C0M9 P0C0N0 P0C0N1 P0C0N2
P0C0N3 P0C0N6 P0C0N7 P0C0N8 P0C0N9 P0C0P0 P0C0P1 P0C0P3 P0C0P4 P0C0Q6 P0C0Q7
P0C0Q8 P0C0Q9 P0C0R0 P0C0R2 P0C0R8 P0C0R9 P0C0S0 P0C0S4 P0C0S5 P0C0S6 P0C0S7
P0C0S9 P0C0T2 P0C0T3 P0C0T7 P0C0U4 P0C0U5 P0C0U6 P0C0U7 P0C0U8 P0C0U9 P0C0V2
P0C0V3 P0C0V4 P0C0V5 P0C0V6 P0C0V7 P0C0W2 P0C0W3 P0C0W4 P0C0W5 P0C0W6 P0C0W7
P0C0X1 P0C0X2 P0C0X3 P0C0X6 P0C0X7 P0C0X8 P0C0Y4 P0C0Y5 P0C0Y6 P0C0Z2 P0C0Z3
P0C0Z4 P0C0Z5 P0C0Z6 P0C0Z7 P0C0Z8 P0C100 P0C101 P0C103 P0C106 P0C107 P0C108
P0C113 P0C115 P0C116 P0C118 P0C119 P0C121 P0C122 P0C123 P0C124 P0C125 P0C126
P0C127 P0C128 P0C129 P0C130 P0C131 P0C132 P0C133 P0C134 P0C136 P0C138 P0C140
P0C141 P0C144 P0C145 P0C146 P0C151 P0C153 P0C154 P0C169 P0C170 P0C181 P0C189
P0C193 P0C196 P0C1A0 P0C1A1 P0C1B0 P0C1B2 P0C1B9 P0C1C4 P0C1C6 P0C1D3 P0C1D5
P0C1D8 P0C1D9 P0C1E0 P0C1E1 P0C1E2 P0C1E3 P0C1E4 P0C1E5 P0C1E6 P0C1E7 P0C1E8
P0C1E9 P0C1F9 P0C1G0 P0C1G2 P0C1G3 P0C1G4 P0C1G5 P0C1H3 P0C1H4 P0C1H5 P0C1H6
P0C1H7 P0C1H8 P0C1I1 P0C1I2 P0C1I3 P0C1I4 P0C1I5 P0C1I6 P0C1I7 P0C1I8 P0C1I9
P0C1J0 P0C1J1 P0C1J2 P0C1J3 P0C1J6 P0C1J7 P0C1J8 P0C1L0 P0C1P6 P0C1P8 P0C1P9
P0C1Q1 P0C1R5 P0C1R6 P0C1R9 P0C1S0 P0C1S1 P0C1S2 P0C1S3 P0C1S4 P0C1S7 P0C1T1
P0C1T2 P0C1T3 P0C1T4 P0C1U3 P0C1V0 P0C1V2 P0C1V3 P0C1V5 P0C1V6 P0C1Y0 P0C1Z1
P0C1Z2 P0C203 P0C218 P0C221 P0C224 P0C227 P0C230 P0C233 P0C237 P0C238 P0C239

P0C240 P0C242 P0C243 P0C263 P0C264 P0C265 P0C266 P0C267 P0C268 P0C269 P0C270
P0C271 P0C272 P0C275 P0C277 P0C281 P0C282 P0C290 P0C291 P0C2B4 P0C2B9 P0C2C0
P0C2C1 P0C2C2 P0C2C4 P0C2D0 P0C2D9 P0C2E1 P0C2E3 P0C2E4 P0C2E5 P0C2E6 P0C2E7
P0C2E8 P0C2F0 P0C2F1 P0C2F5 P0C2F6 P0C2F7 P0C2F8 P0C2F9 P0C2G0 P0C2G1 P0C2G2
P0C2G3 P0C2G4 P0C2G5 P0C2G6 P0C2G9 P0C2I4 P0C2I8 P0C2J2 P0C2J4 P0C2J6 P0C2J8
P0C2L9 P0C2M7 P0C2N0 P0C2N2 P0C2N3 P0C2N5 P0C2N6 P0C2N7 P0C2N8 P0C2P0 P0C2P1
P0C2P2 P0C2P3 P0C2P7 P0C2P8 P0C2P9 P0C2Q0 P0C2R1 P0C2R2 P0C2R3 P0C2R4 P0C2R7
P0C2R8 P0C2R9 P0C2S2 P0C2S3 P0C2T0 P0C2T3 P0C2T4 P0C2T6 P0C2T7 P0C2T9 P0C2U0
P0C2U1 P0C2U4 P0C2U5 P0C2V4 P0C2V5 P0C2V6 P0C2V7 P0C2V9 P0C2W6 P0C2W7 P0C2X1
P0C2X2 P0C2X3 P0C2X4 P0C2X5 P0C2X6 P0C2X8 P0C2X9 P0C2Y0 P0C2Y1 P0C304 P0C305
P0C306 P0C307 P0C308 P0C309 P0C310 P0C311 P0C312 P0C313 P0C314 P0C378 P0C379
P0C380 P0C381 P0C382 P0C383 P0C397 P0C398 P0C438 P0C439 P0C440 P0C441 P0C442
P0C443 P0C444 P0C445 P0C446 P0C447 P0C448 P0C449 P0C450 P0C451 P0C452 P0C453
P0C454 P0C455 P0C456 P0C457 P0C458 P0C459 P0C460 P0C461 P0C462 P0C463 P0C464
P0C465 P0C466 P0C467 P0C468 P0C469 P0C470 P0C471 P0C472 P0C473 P0C474 P0C475
P0C476 P0C477 P0C478 P0C479 P0C480 P0C481 P0C482 P0C483 P0C484 P0C485 P0C486
P0C487 P0C488 P0C489 P0C490 P0C491 P0C492 P0C493 P0C494 P0C495 P0C496 P0C497
P0C498 P0C499 P0C500 P0C501 P0C502 P0C503 P0C504 P0C505 P0C506 P0C507 P0C508
P0C509 P0C510 P0C511 P0C512 P0C525 P0C530 P0C534 P0C539 P0C540 P0C542 P0C543
P0C544 P0C545 P0C557 P0C558 P0C559 P0C561 P0C562 P0C563 P0C564 P0C565 P0C567
P0C568 P0C570 P0C580 P0C581 P0C585 P0C586 P0C587 P0C591 P0C592 P0C593 P0C594
P0C595 P0C596 P0C597 P0C598 P0C599 P0C5A0 P0C5A1 P0C5A2 P0C5A3 P0C5A4 P0C5A5
P0C5A7 P0C5B0 P0C5B1 P0C5B2 P0C5B3 P0C5B5 P0C5B7 P0C5C6 P0C5D3 P0C5D4 P0C5D7
P0C5D9 P0C5E1 P0C5E6 P0C5E9 P0C5H0 P0C5H5 P0C5H7 P0C5H8 P0C5I2 P0C5J1 P0C5K0
P0C5K1 P0C5K6 P0C5K7 P0C5K9 P0C5L0 P0C5L1 P0C5L3 P0C5L4 P0C5L5 P0C5L6 P0C5L7
P0C5L9 P0C5M0 P0C5M1 P0C5M4 P0C5M5 P0C5M8 P0C5M9 P0C5N1 P0C5N2 P0C5N3 P0C5N4
P0C5N5 P0C5N6 P0C5N7 P0C5P0 P0C5P1 P0C5P2 P0C5P4 P0C5P6 P0C5P7 P0C5P8 P0C5P9
P0C5Q0 P0C5Q1 P0C5Q2 P0C5Q5 P0C5Q6 P0C5Q7 P0C5Q8 P0C5Q9 P0C5R0 P0C5R2 P0C5R3
P0C5R4 P0C5R5 P0C5R6 P0C5S0 P0C5S1 P0C5S2 P0C5W1 P0C5W3 P0C5W5 P0C5Y0 P0C5Y1
P0C5Y2 P0C5Y3 P0C5Y4 P0C5Y5 P0C5Y6 P0C5Y7 P0C5Y9 P0C5Z0 P0C5Z1 P0C5Z2 P0C5Z3
P0C5Z4 P0C5Z5 P0C5Z6 P0C5Z7 P0C5Z9 P0C600 P0C601 P0C602 P0C606 P0C618 P0C619
P0C620 P0C621 P0C622 P0C624 P0C630 P0C631 P0C632 P0C634 P0C652 P0C654 P0C656
P0C657 P0C659 P0C660 P0C661 P0C664 P0C671 P0C674 P0C675 P0C676 P0C679 P0C688
P0C690 P0C691 P0C6A0 P0C6A2 P0C6C0 P0C6C1 P0C6C2 P0C6C8 P0C6D0 P0C6D1 P0C6D2
P0C6D3 P0C6D4 P0C6D6 P0C6D7 P0C6E1 P0C6E7 P0C6F1 P0C6F9 P0C6G0 P0C6G1 P0C6G2
P0C6G4 P0C6G5 P0C6G8 P0C6H0 P0C6H1 P0C6H9 P0C6I0 P0C6I7 P0C6K5 P0C6K6 P0C6K8
P0C6L1 P0C6L2 P0C6P0 P0C6P3 P0C6P6 P0C6P7 P0C6P8 P0C6P9 P0C6Q0 P0C6Q1 P0C6Q2
P0C6Q3 P0C6Q4 P0C6Q8 P0C6R2 P0C6R4 P0C6T1 P0C6Y7 P0C6Z0 P0C6Z2 P0C6Z8 P0C6Z9
P0C702 P0C703 P0C704 P0C705 P0C706 P0C710 P0C711 P0C712 P0C713 P0C717 P0C722
P0C723 P0C724 P0C725 P0C726 P0C727 P0C728 P0C731 P0C732 P0C733 P0C734 P0C740
P0C744 P0C749 P0C750 P0C752 P0C753 P0C754 P0C755 P0C756 P0C757 P0C758 P0C759
P0C760 P0C766 P0C768 P0C769 P0C774 P0C775 P0C777 P0C779 P0C780 P0C784 P0C785
P0C790 P0C7A0 P0C7A1 P0C7A2 P0C7B4 P0C7B5 P0C7B6 P0C7G9 P0C7H8 P0C7H9 P0C7I0
P0C7I3 P0C7I4 P0C7J1 P0C7J2 P0C7J3 P0C7K7 P0C7K8 P0C7K9 P0C7L0 P0C7L2 P0C7L3

P0C7L4 P0C7L5 P0C7L6 P0C7L7 P0C7L8 P0C7M6 P0C7N0 P0C7N2 P0C7N7 P0C7N9 P0C7P1
P0C7P2 P0C7P3 P0C7P7 P0C7Q1 P0C7Q7 P0C7Q8 P0C7Q9 P0C7R0 P0C7R1 P0C7R2 P0C7R3
P0C7R4 P0C7R5 P0C7T0 P0C7T1 P0C7T4 P0C7T6 P0C7T7 P0C7U6 P0C7V0 P0C7V1 P0C7V2
P0C7V4 P0C7V6 P0C7V8 P0C7V9 P0C7W0 P0C7W4 P0C7W6 P0C7W8 P0C7W9 P0C7X0 P0C7X2
P0C7X4 P0C7X8 P0C7X9 P0C7Y0 P0C7Y1 P0C7Y2 P0C7Y3 P0C7Y4 P0C7Y5 P0C7Y6 P0C7Y7
P0C7Y8 P0C7Y9 P0C7Z0 P0C7Z1 P0C7Z2 P0C7Z3 P0C7Z4 P0C7Z5 P0C7Z6 P0C7Z7 P0C7Z8
P0C7Z9 P0C800 P0C801 P0C802 P0C803 P0C804 P0C805 P0C806 P0C807 P0C808 P0C809
P0C810 P0C811 P0C812 P0C813 P0C814 P0C815 P0C816 P0C817 P0C818 P0C819 P0C820
P0C821 P0C822 P0C823 P0C824 P0C825 P0C826 P0C827 P0C835 P0C836 P0C841 P0C843
P0C853 P0C856 P0C860 P0C861 P0C864 P0C866 P0C870 P0C872 P0C875 P0C879 P0C880
P0C881 P0C883 P0C884 P0C885 P0C890 P0C894 P0C895 P0C896 P0C898 P0C8A0 P0C8B3
P0C8B4 P0C8B5 P0C8F2 P0C8F3 P0C8F4 P0C8F5 P0C8F6 P0C8F7 P0C8F8 P0C8F9 P0C8G0
P0C8G1 P0C8G2 P0C8G3 P0C8G4 P0C8G5 P0C8G8 P0C8G9 P0C8H0 P0C8H1 P0C8H2 P0C8H3
P0C8H4 P0C8H5 P0C8H6 P0C8H7 P0C8H8 P0C8H9 P0C8I0 P0C8I1 P0C8I2 P0C8I3 P0C8I4
P0C8I5 P0C8I6 P0C8J6 P0C8J7 P0C8J9 P0C8K1 P0C8K2 P0C8K3 P0C8K4 P0C8K5 P0C8K6
P0C8K8 P0C8L4 P0C8L5 P0C8L6 P0C8L7 P0C8L8 P0C8M4 P0C8M5 P0C8M8 P0C8M9 P0C8N9
P0C8P1 P0C8P2 P0C8P3 P0C8P4 P0C8P5 P0C8Q0 P0C8Q1 P0C8Q3 P0C8Q4 P0C8Q6 P0C8Q7
P0C8Q8 P0C8R0 P0C8R3 P0C8R4 P0C8R5 P0C8S1 P0C8S3 P0C8S4 P0C8Y1 P0C8Y3 P0C8Y9
P0C8Z0 P0C8Z1 P0C8Z4 P0C8Z5 P0C8Z6 P0C913 P0C914 P0C915 P0C916 P0C917 P0C919
P0C920 P0C921 P0C922 P0C934 P0C935 P0C936 P0C937 P0C938 P0C939 P0C945 P0C946
P0C949 P0C952 P0C953 P0C961 P0C962 P0C967 P0C968 P0C969 P0C970 P0C971 P0C972
P0C973 P0C974 P0C979 P0C980 P0C981 P0C982 P0C983 P0C984 P0C985 P0C986 P0C987
P0C988 P0C989 P0C990 P0C991 P0C992 P0C993 P0C994 P0C995 P0C996 P0C997 P0C998
P0C9A5 P0C9A6 P0C9A7 P0C9A8 P0C9A9 P0C9B0 P0C9B1 P0C9B2 P0C9B3 P0C9B4 P0C9B5
P0C9B6 P0C9B7 P0C9B8 P0C9B9 P0C9C0 P0C9C1 P0C9C2 P0C9C3 P0C9C4 P0C9C5 P0C9C6
P0C9C7 P0C9C8 P0C9C9 P0C9D0 P0C9D1 P0C9D2 P0C9D3 P0C9D4 P0C9D5 P0C9D6 P0C9D7
P0C9D8 P0C9D9 P0C9E3 P0C9E4 P0C9E5 P0C9E6 P0C9E7 P0C9E8 P0C9E9 P0C9F0 P0C9F1
P0C9F2 P0C9F3 P0C9F4 P0C9F5 P0C9F6 P0C9F7 P0C9F8 P0C9F9 P0C9G0 P0C9K8 P0C9K9
P0C9L0 P0C9L1 P0C9L3 P0C9L6 P0C9L8 P0C9L9 P0C9M0 P0C9M1 P0C9M2 P0C9M3 P0C9M4
P0C9M5 P0C9M6 P0C9M7 P0C9M8 P0C9M9 P0C9N0 P0C9N1 P0C9N2 P0C9N3 P0C9N4 P0C9N5
P0C9N6 P0C9N7 P0C9N8 P0C9N9 P0C9P0 P0C9P1 P0C9P2 P0C9P6 P0C9P7 P0C9P8 P0C9P9
P0C9Q0 P0C9Q1 P0C9Q2 P0C9Q3 P0C9Q4 P0C9Q5 P0C9Q6 P0C9Q7 P0C9Q8 P0C9Q9 P0C9R0
P0C9R1 P0C9R2 P0C9R3 P0C9R4 P0C9R5 P0C9R7 P0C9R8 P0C9R9 P0C9S0 P0C9S1 P0C9S2
P0C9S3 P0C9S4 P0C9S5 P0C9S6 P0C9S7 P0C9S8 P0C9S9 P0C9T0 P0C9T1 P0C9T2 P0C9T3
P0C9T4 P0C9T5 P0C9T6 P0C9T7 P0C9T8 P0C9T9 P0C9U0 P0C9U1 P0C9U2 P0C9U3 P0C9U4
P0C9U5 P0C9U6 P0C9U7 P0C9U8 P0C9U9 P0C9V0 P0C9V1 P0C9V2 P0C9V3 P0C9V4 P0C9V5
P0C9X1 P0C9X2 P0C9X3 P0C9X4 P0C9X5 P0C9X6 P0C9X7 P0C9X8 P0C9X9 P0C9Y0 P0C9Z1
P0C9Z2 P0C9Z3 P0C9Z4 P0CA09 P0CA10 P0CA11 P0CA12 P0CA16 P0CA17 P0CA18 P0CA22
P0CA23 P0CA24 P0CA25 P0CA30 P0CA31 P0CA32 P0CA33 P0CA34 P0CA35 P0CA36 P0CA37
P0CA38 P0CA39 P0CA40 P0CA41 P0CA42 P0CA43 P0CA44 P0CA45 P0CA46 P0CA47 P0CA48
P0CA49 P0CA50 P0CA51 P0CA73 P0CA74 P0CA75 P0CA76 P0CA77 P0CA78 P0CA79 P0CA80
P0CA81 P0CA82 P0CA83 P0CA84 P0CA85 P0CA86 P0CA87 P0CA88 P0CA89 P0CA90 P0CA91
P0CA92 P0CA93 P0CA97 P0CA98 P0CA99 P0CAA0 P0CAA4 P0CAA5 P0CAA6 P0CAA7 P0CAA8
P0CAA9 P0CAB3 P0CAB4 P0CAB5 P0CAB6 P0CAC1 P0CAC2 P0CAC3 P0CAC4 P0CAC5 P0CAC6

P0CAC7 P0CAC8 P0CAC9 P0CAD0 P0CAD1 P0CAD2 P0CAD3 P0CAD4 P0CAD5 P0CAD6 P0CAD7
P0CAD8 P0CAD9 P0CAE0 P0CAE1 P0CAE2 P0CAE7 P0CAE8 P0CAE9 P0CAF0 P0CAF1 P0CAF2
P0CAF3 P0CAF4 P0CAF5 P0CAF6 P0CAF7 P0CAF8 P0CAF9 P0CAG0 P0CAG1 P0CAG2 P0CAG3
P0CAG4 P0CAG5 P0CAG6 P0CAG7 P0CAG8 P0CAG9 P0CAH0 P0CAH1 P0CAH2 P0CAH3 P0CAH4
P0CAH5 P0CAH6 P0CAI0 P0CAI1 P0CAI2 P0CAI3 P0CAI4 P0CAI5 P0CAI6 P0CAI7 P0CAI8
P0CAI9 P0CAJ0 P0CAJ5 P0CAJ6 P0CAJ7 P0CAJ8 P0CAJ9 P0CAK0 P0CAK1 P0CAK2 P0CAK3
P0CAK4 P0CAK5 P0CAK6 P0CAK7 P0CAK8 P0CAK9 P0CAL1 P0CAL2 P0CAL3 P0CAM4 P0CAM5
P0CAM6 P0CAM7 P0CAM8 P0CAM9 P0CAN0 P0CAN1 P0CAN2 P0CAN3 P0CAN4 P0CAN5 P0CAN8
P0CAN9 P0CAP3 P0CAP4 P0CAP5 P0CAP8 P0CAT3 P0CAT7 P0CAT8 P0CAT9 P0CAU0 P0CAU1
P0CAU2 P0CAU3 P0CAU4 P0CAU5 P0CAV0 P0CAV1 P0CAV2 P0CAV3 P0CAV4 P0CAV5 P0CAV6
P0CAV7 P0CAV8 P0CAW0 P0CAW1 P0CAW2 P0CAW3 P0CAW4 P0CAW5 P0CAW6 P0CAW9 P0CAX0
P0CAX1 P0CAX2 P0CAX3 P0CAX8 P0CAZ0 P0CAZ1 P0CB00 P0CB01 P0CB02 P0CB03 P0CB04
P0CB16 P0CB19 P0CB21 P0CB23 P0CB24 P0CB25 P0CB26 P0CB27 P0CB33 P0CB34 P0CB35
P0CB36 P0CB37 P0CB38 P0CB41 P0CB42 P0CB47 P0CB48 P0CB49 P0CB50 P0CB52 P0CB53
P0CB57 P0CB58 P0CB61 P0CB65 P0CB66 P0CB75 P0CB76 P0CB77 P0CB78 P0CC03 P0CC06
P0CC07 P0CC09 P0CD65 P0CD66 P0CD71 P0CD72 P0CD73 P0CD77 P0CD78 P0CD80 P0CD81
P0CD82 P0CD83 P0CD84 P0CD85 P0CD86 P0CD87 P0CD88 P0CD96 P0CE01 P0CE02 P0CE03
P0CE04 P0CE05 P0CE06 P0CE07 P0CE08 P0CE09 P0CE10 P0CE13 P0CE15 P0CE16 P0CE17
P0CE18 P0CE19 P0CE20 P0CE21 P0CE22 P0CE23 P0CE41 P0CE49 P0CE50 P0CE51 P0CE52
P0CE53 P0CE54 P0CE55 P0CE56 P0CE57 P0CE58 P0CE59 P0CE60 P0CE61 P0CE62 P0CE63
P0CE64 P0CE65 P0CE67 P0CE71 P0CE72 P0CE86 P0CE87 P0CE95 P0CE96 P0CE97 P0CE98
P0CE99 P0CF00 P0CF06 P0CF07 P0CF08 P0CF09 P0CF10 P0CF11 P0CF12 P0CF13 P0CF18
P0CF21 P0CF22 P0CF23 P0CF24 P0CF25 P0CF26 P0CF27 P0CF28 P0CF29 P0CF30 P0CF31
P0CF35 P0CF36 P0CF40 P0CF41 P0CF42 P0CF43 P0CF44 P0CF45 P0CF46 P0CF47 P0CF48
P0CF49 P0CF50 P0CF52 P0CF53 P0CF54 P0CF55 P0CF56 P0CF57 P0CF58 P0CF59 P0CF60
P0CF61 P0CF62 P0CF63 P0CF64 P0CF65 P0CF66 P0CF67 P0CF68 P0CF69 P0CF70 P0CF71
P0CF72 P0CF73 P0CF75 P0CF79 P0CF80 P0CF81 P0CF82 P0CF83 P0CF84 P0CF85 P0CF86
P0CF87 P0CF88 P0CF89 P0CF90 P0CF91 P0CF92 P0CF93 P0CF95 P0CF96 P0CF97 P0CG00
P0CG10 P0CG11 P0CG13 P0CG14 P0CG15 P0CG18 P0CG19 P0CG20 P0CG21 P0CG23 P0CG24
P0CG25 P0CG28 P0CG29 P0CG30 P0CG31 P0CG32 P0CG34 P0CG35 P0CG38 P0CG39 P0CG40
P0CG42 P0CG43 P0CG70 P0CG71 P0CG72 P0CG73 P0CG74 P0CG75 P0CG76 P0CG82 P0CG83
P0CG84 P0CG85 P0CG86 P0CG87 P0CG89 P0CG94 P0CG96 P0CG97 P0CG98 P0CG99 P0CH00
P0CH04 P0CH05 P0CH06 P0CH07 P0CH10 P0CH11 P0CH25 P0CH26 P0CH27 P0CH30 P0CH31
P0CH32 P0CH33 P0CH34 P0CH35 P0CH36 P0CH37 P0CH62 P0CH63 P0CH64 P0CH65 P0CH66
P0CH67 P0CH68 P0CH92 P0CH93 P0CH94 P0CH95 P0CH96 P0CH97 P0CH98 P0CH99 P0CI00
P0CI01 P0CI02 P0CI25 P0CI26 P0CI29 P0CI31 P0CI32 P0CI34 P0CI35 P0CI61 P0CI62
P0CI63 P0CI64 P0CI65 P0CI66 P0CI67 P0CI68 P0CI69 P0CI72 P0CI73 P0CI74 P0CI75
P0CI76 P0CI77 P0CI78 P0CI79 P0CI80 P0CJ42 P0CJ43 P0CJ44 P0CJ47 P0CJ63 P0CJ64
P0CJ65 P0CJ66 P0CJ67 P0CJ78 P0CJ79 P0CJ80 P0CJ83 P0CJ84 P0CJ85 P0CJ86 P0CJ87
P0CJ88 P0CJ89 P0CJ90 P0CK18 P0CK19 P0CK24 P0CK31 P0CK32 P0CK35 P0CK36 P0CK39
P0CK40 P0CK53 P0CK54 P0CK55 P0CK56 P0CK57 P0CK58 P0CK59 P0CK63 P0CK64 P0CK65
P0CK66 P0CK67 P0CK68 P0CK69 P0CK70 P0CK71 P0CK72 P0CK73 P0CK74 P0CK75 P0CK76
P0CK77 P0CK78 P0CK79 P0CK80 P0CK81 P0CK82 P0CK83 P0CK84 P0CK85 P0CK86 P0CK87
P0CK88 P0CK89 P0CK90 P0CK91 P0CK92 P0CK98 P0CL01 P0CL02 P0CL03 P0CL05 P0CL06

P0CL07 P0CL08 P0CL09 P0CL10 P0CL11 P0CL12 P0CL16 P0CL17 P0CL18 P0CL19 P0CL20
P0CL21 P0CL22 P0CL23 P0CL24 P0CL25 P0CL26 P0CL44 P0CL45 P0CL50 P0CL51 P0CL53
P0CL54 P0CL56 P0CL57 P0CL58 P0CL60 P0CL61 P0CL62 P0CL64 P0CL69 P0CL72 P0CL73
P0CL74 P0CL75 P0CL76 P0CL77 P0CL78 P0CL79 P0CL80 P0CL81 P0CL82 P0CL83 P0CL84
P0CL85 P0CL86 P0CL87 P0CL88 P0CL89 P0CL90 P0CL91 P0CL96 P0CL97 P0CL98 P0CL99
P0CM00 P0CM01 P0CM02 P0CM03 P0CM04 P0CM05 P0CM06 P0CM07 P0CM08 P0CM09 P0CM12
P0CM13 P0CM14 P0CM15 P0CM18 P0CM19 P0CM20 P0CM21 P0CM22 P0CM23 P0CM26 P0CM27
P0CM44 P0CM45 P0CM46 P0CM47 P0CM49 P0CM52 P0CM53 P0CM54 P0CM55 P0CM58 P0CM59
P0CM62 P0CM63 P0CM64 P0CM65 P0CM66 P0CM67 P0CM70 P0CM71 P0CM74 P0CM75 P0CM76
P0CM77 P0CM78 P0CM79 P0CM86 P0CM87 P0CM88 P0CM89 P0CM90 P0CM91 P0CM94 P0CM95
P0CM96 P0CM97 P0CM98 P0CM99 P0CN00 P0CN01 P0CN02 P0CN03 P0CN04 P0CN05 P0CN06
P0CN07 P0CN08 P0CN09 P0CN10 P0CN11 P0CN12 P0CN13 P0CN14 P0CN15 P0CN18 P0CN19
P0CN20 P0CN21 P0CN22 P0CN23 P0CN24 P0CN25 P0CN26 P0CN27 P0CN28 P0CN29 P0CN30
P0CN31 P0CN32 P0CN33 P0CN34 P0CN35 P0CN38 P0CN39 P0CN40 P0CN41 P0CN42 P0CN43
P0CN44 P0CN45 P0CN46 P0CN47 P0CN48 P0CN49 P0CN50 P0CN51 P0CN52 P0CN53 P0CN54
P0CN55 P0CN56 P0CN57 P0CN58 P0CN59 P0CN60 P0CN61 P0CN64 P0CN65 P0CN68 P0CN69
P0CN74 P0CN75 P0CN76 P0CN77 P0CN78 P0CN79 P0CN80 P0CN81 P0CN82 P0CN83 P0CN84
P0CN85 P0CN86 P0CN87 P0CN88 P0CN89 P0CN94 P0CN95 P0CN98 P0CN99 P0CO02 P0CO03
P0CO04 P0CO05 P0CO06 P0CO07 P0CO08 P0CO09 P0CO10 P0CO11 P0CO12 P0CO13 P0CO14
P0CO15 P0CO16 P0CO17 P0CO18 P0CO19 P0CO20 P0CO21 P0CO22 P0CO23 P0CO24 P0CO25
P0CO26 P0CO27 P0CO28 P0CO29 P0CO30 P0CO31 P0CO34 P0CO35 P0CO36 P0CO37 P0CO38
P0CO39 P0CO40 P0CO41 P0CO42 P0CO43 P0CO44 P0CO45 P0CO46 P0CO47 P0CO50 P0CO51
P0CO52 P0CO53 P0CO56 P0CO57 P0CO58 P0CO59 P0CO62 P0CO63 P0CO64 P0CO65 P0CO72
P0CO73 P0CO74 P0CO75 P0CO76 P0CO77 P0CO84 P0CO85 P0CO86 P0CO87 P0CO88 P0CO89
P0CO90 P0CO91 P0CO92 P0CO93 P0CO94 P0CO95 P0CO96 P0CO97 P0CO98 P0CO99 P0CP02
P0CP03 P0CP04 P0CP05 P0CP10 P0CP11 P0CP12 P0CP13 P0CP16 P0CP17 P0CP18 P0CP19
P0CP20 P0CP21 P0CP22 P0CP23 P0CP24 P0CP25 P0CP26 P0CP27 P0CP34 P0CP35 P0CP38
P0CP39 P0CP40 P0CP41 P0CP42 P0CP43 P0CP44 P0CP45 P0CP46 P0CP47 P0CP50 P0CP51
P0CP54 P0CP55 P0CP56 P0CP57 P0CP58 P0CP59 P0CP60 P0CP61 P0CP67 P0CP72 P0CP73
P0CP80 P0CP81 P0CP83 P0CP84 P0CP85 P0CP86 P0CP87 P0CP88 P0CP89 P0CP90 P0CP91
P0CP92 P0CP93 P0CP94 P0CP95 P0CP98 P0CP99 P0CQ00 P0CQ01 P0CQ02 P0CQ03 P0CQ06
P0CQ07 P0CQ08 P0CQ09 P0CQ14 P0CQ15 P0CQ22 P0CQ23 P0CQ24 P0CQ25 P0CQ28 P0CQ29
P0CQ30 P0CQ31 P0CQ32 P0CQ33 P0CQ34 P0CQ35 P0CQ36 P0CQ37 P0CQ41 P0CQ44 P0CQ45
P0CQ48 P0CQ49 P0CQ50 P0CQ51 P0CQ52 P0CQ53 P0CQ54 P0CQ55 P0CQ56 P0CQ57 P0CQ58
P0CQ59 P0CQ62 P0CQ63 P0CQ64 P0CQ65 P0CQ66 P0CQ67 P0CQ68 P0CQ69 P0CQ70 P0CQ71
P0CQ72 P0CQ73 P0CQ74 P0CQ75 P0CQ76 P0CQ77 P0CQ78 P0CQ79 P0CQ80 P0CQ81 P0CQ82
P0CQ83 P0CQ88 P0CQ89 P0CQ90 P0CQ91 P0CQ92 P0CQ93 P0CQ94 P0CQ95 P0CQ96 P0CQ97
P0CQ98 P0CQ99 P0CR00 P0CR01 P0CR02 P0CR03 P0CR04 P0CR05 P0CR06 P0CR07 P0CR08
P0CR09 P0CR10 P0CR11 P0CR12 P0CR13 P0CR14 P0CR15 P0CR16 P0CR17 P0CR19 P0CR20
P0CR21 P0CR22 P0CR23 P0CR24 P0CR25 P0CR26 P0CR27 P0CR28 P0CR29 P0CR32 P0CR33
P0CR42 P0CR43 P0CR48 P0CR49 P0CR50 P0CR51 P0CR52 P0CR53 P0CR56 P0CR57 P0CR66
P0CR67 P0CR68 P0CR69 P0CR70 P0CR71 P0CR72 P0CR73 P0CR74 P0CR75 P0CR76 P0CR77
P0CR80 P0CR81 P0CR82 P0CR83 P0CR84 P0CR85 P0CR86 P0CR87 P0CR92 P0CR93 P0CS06
P0CS07 P0CS08 P0CS09 P0CS10 P0CS11 P0CS12 P0CS13 P0CS14 P0CS15 P0CS16 P0CS17

P0CS20 P0CS21 P0CS22 P0CS23 P0CS30 P0CS31 P0CS32 P0CS33 P0CS34 P0CS35 P0CS36
P0CS37 P0CS38 P0CS39 P0CS40 P0CS41 P0CS42 P0CS43 P0CS46 P0CS47 P0CS48 P0CS49
P0CS52 P0CS53 P0CS54 P0CS55 P0CS56 P0CS57 P0CS60 P0CS61 P0CS62 P0CS63 P0CS64
P0CS65 P0CS70 P0CS71 P0CS72 P0CS73 P0CS74 P0CS75 P0CS76 P0CS77 P0CS78 P0CS79
P0CS80 P0CS81 P0CS82 P0CS84 P0CS85 P0CS88 P0CS92 P0CS94 P0CS95 P0CS96 P0CS97
P0CS98 P0CS99 P0CT00 P0CT01 P0CT02 P0CT03 P0CT05 P0CT06 P0CT07 P0CT08 P0CT11
P0CT12 P0CT13 P0CT14 P0CT15 P0CT19 P0CT20 P0CT21 P0CT22 P0CT23 P0CT25 P0CT26
P0CT28 P0CT29 P0CT31 P0CT33 P0CT34 P0CT35 P0CT36 P0CT37 P0CT38 P0CT39 P0CT40
P0CT41 P0CT42 P0CT43 P0CT45 P0CT46 P0CT50 P0CT51 P0CT53 P0CT54 P0CT55 P0CT62
P0CT63 P0CT81 P0CT82 P0CT86 P0CT93 P0CU00 P0CU01 P0CU08 P0CU09 P0CU17 P0CU18
P0CU23 P0CU24 P0CU25 P0CU27 P0CU28 P0CU29 P0CU32 P0CU34 P0CU35 P0CU37 P0CU43
P0CU44 P0CU45 P0CU46 P0CU47 P0CU48 P0CU49 P0CU50 P0CU51 P0CU52 P0CU56 P0CU57
P0CU58 P0CU59 P0CU60 P0CU61 P0CU62 P0CU63 P0CU64 P0CU65 P0CU67 P0CU68 P0CU69
P0CU71 P0CU72 P0CU73 P0CU74 P0CU75 P0CU76 P0CU78 P0CU81 P0CU82 P0CU84 P0CU85
P0CV86 P0CV93 P0CV94 P0CV95 P0CV96 P0CV97 P0CV98 P0CV99 P0CW00 P0CW01 P0CW04
P0CW05 P0CW06 P0CW07 P0CW08 P0CW09 P0CW10 P0CW11 P0CW12 P0CW13 P0CW14 P0CW15
P0CW19 P0CW20 P0CW21 P0CW23 P0CW24 P0CW27 P0CW29 P0CW31 P0CW32 P0CW33 P0CW37
P0CW38 P0CW42 P0CW43 P0CW44 P0CW45 P0CW46 P0CW47 P0CW48 P0CW49 P0CW50 P0CW51
P0CW52 P0CW53 P0CW54 P0CW55 P0CW56 P0CW57 P0CW58 P0CW59 P0CW60 P0CW61 P0CW62
P0CW63 P0CW64 P0CW65 P0CW66 P0CW67 P0CW68 P0CW69 P0CW74 P0CW75 P0CW76 P0CW84
P0CW85 P0CW88 P0CW89 P0CW90 P0CW91 P0CW92 P0CW93 P0CW94 P0CW99 P0CX00 P0CX01
P0CX02 P0CX03 P0CX04 P0CX05 P0CX06 P0CX07 P0CX08 P0CX09 P0CX10 P0CX11 P0CX14
P0CX15 P0CX16 P0CX17 P0CX20 P0CX21 P0CX22 P0CX57 P0CX58 P0CX59 P0CX60 P0CX61
P0CX62 P0CX65 P0CX66 P0CX67 P0CX68 P0CX69 P0CX70 P0CX71 P0CX72 P0CX73 P0CX74
P0CX75 P0CX76 P0CX80 P0CX81 P0CX88 P0CX89 P0CX90 P0CX91 P0CY02 P0CY03 P0CY04
P0CY05 P0CY07 P0CY08 P0CY09 P0CY10 P0CY11 P0CY12 P0CY13 P0CY14 P0CY16 P0CY17
P0CY18 P0CY19 P0CY20 P0CY21 P0CY23 P0CY25 P0CY34 P0CY35 P0CY36 P0CY37 P0CY38
P0CY39 P0CY40 P0CY53 P0CY54 P0CY55 P0CY57 P0CY58 P0CY89 P0CY91 P0CY95 P0CY96
P0CY97 P0CY98 P0CY99 P0CZ02 P0CZ03 P0CZ12 P0CZ15 P0CZ16 P0CZ25 P0CZ46 P0CZ47
P0CZ52 P0CZ53 P0CZ54 P0CZ55 P0CZ56 P0CZ57 P0CZ58 P0CZ59 P0CZ60 P0CZ61 P0CZ62
P0CZ63 P0CZ64 P0CZ65 P0CZ66 P0CZ67 P0CZ68 P0CZ69 P0CZ70 P0CZ71 P0CZ72 P0CZ73
P0CZ74 P0CZ75 P0CZ76 P0CZ77 P0CZ78 P0CZ79 P0CZ80 P0CZ81 P0CZ82 P0CZ83 P0CZ84
P0CZ85 P0CZ86 P0CZ87 P0CZ88 P0CZ89 P0DA12 P0DA13 P0DA14 P0DA15 P0DA16 P0DA17
P0DA18 P0DA19 P0DA20 P0DA21 P0DA22 P0DA23 P0DA24 P0DA25 P0DA26 P0DA27 P0DA28
P0DA29 P0DA30 P0DA31 P0DA32 P0DA33 P0DA34 P0DA35 P0DA36 P0DA37 P0DA38 P0DA39
P0DA40 P0DA41 P0DA42 P0DA43 P0DA44 P0DA45 P0DA46 P0DA47 P0DA48 P0DA49 P0DA50
P0DA51 P0DA60 P0DA61 P0DA62 P0DA63 P0DA64 P0DA65 P0DA68 P0DA69 P0DA70 P0DA71
P0DA72 P0DA73 P0DA74 P0DA75 P0DA76 P0DA77 P0DA78 P0DA79 P0DA80 P0DA81 P0DA82
P0DA83 P0DA84 P0DA85 P0DA86 P0DA87 P0DA88 P0DA89 P0DA90 P0DA91 P0DA92 P0DA93
P0DB04 P0DB05 P0DB06 P0DB07 P0DB08 P0DB09 P0DB10 P0DB11 P0DB12 P0DB13 P0DB14
P0DB15 P0DB18 P0DB19 P0DB20 P0DB21 P0DB24 P0DB25 P0DB26 P0DB27 P0DB28 P0DB29
P0DB30 P0DB31 P0DB32 P0DB33 P0DB34 P0DB35 P0DB36 P0DB37 P0DB38 P0DB39 P0DB40
P0DB41 P0DB44 P0DB45 P0DB46 P0DB47 P0DB48 P0DB49 P0DB52 P0DB53 P0DB54 P0DB55
P0DB58 P0DB59 P0DB64 P0DB65 P0DB66 P0DB67 P0DB68 P0DB69 P0DB70 P0DB71 P0DB72

P0DB73 P0DB74 P0DB75 P0DB76 P0DB77 P0DB78 P0DB79 P0DB80 P0DB81 P0DB82 P0DB83
P0DB84 P0DB85 P0DB86 P0DB87 P0DB88 P0DB89 P0DB90 P0DB91 P0DB92 P0DB93 P0DB94
P0DB95 P0DB96 P0DB97 P0DB98 P0DB99 P0DC00 P0DC01 P0DC02 P0DC03 P0DC04 P0DC05
P0DC06 P0DC07 P0DC08 P0DC09 P0DC10 P0DC11 P0DC12 P0DC13 P0DC14 P0DC15 P0DC16
P0DC17 P0DC18 P0DC19 P0DC28 P0DC29 P0DC30 P0DC31 P0DC32 P0DC33 P0DC34 P0DC35
P0DC36 P0DC37 P0DC38 P0DC39 P0DC40 P0DC41 P0DC42 P0DC43 P0DC44 P0DC45 P0DC46
P0DC47 P0DC48 P0DC49 P0DC50 P0DC51 P0DC52 P0DC53 P0DC54 P0DC55 P0DC58 P0DC59
P0DC60 P0DC61 P0DC62 P0DC63 P0DC64 P0DC65 P0DC66 P0DC67 P0DC68 P0DC69 P0DC70
P0DC71 P0DC72 P0DC73 P0DC74 P0DC75 P0DC76 P0DC77 P0DC78 P0DC79 P0DC80 P0DC81
P0DC82 P0DC83 P0DC84 P0DC85 P0DC90 P0DC91 P0DC92 P0DC93 P0DC94 P0DC95 P0DC96
P0DC97 P0DC98 P0DC99 P0DD00 P0DD01 P0DD02 P0DD03 P0DD04 P0DD05 P0DD06 P0DD07
P0DD12 P0DD13 P0DD14 P0DD15 P0DD16 P0DD17 P0DD18 P0DD19 P0DD20 P0DD21 P0DD22
P0DD23 P0DD24 P0DD25 P0DD26 P0DD27 P0DD28 P0DD29 P0DD36 P0DD37 P0DD40 P0DD41
P0DD46 P0DD47 P0DD48 P0DD49 P0DD50 P0DD51 P0DD52 P0DD53 P0DD54 P0DD55 P0DD56
P0DD57 P0DD58 P0DD59 P0DD60 P0DD61 P0DD62 P0DD63 P0DD64 P0DD65 P0DD66 P0DD67
P0DD68 P0DD69 P0DD70 P0DD71 P0DD72 P0DD73 P0DD74 P0DD75 P0DD76 P0DD77 P0DD78
P0DD79 P0DD80 P0DD81 P0DD82 P0DD83 P0DD84 P0DD85 P0DD86 P0DD87 P0DD88 P0DD89
P0DD90 P0DD91 P0DD92 P0DD93 P0DD94 P0DD95 P0DD96 P0DD97 P0DD98 P0DD99 P0DE00
P0DE01 P0DE02 P0DE03 P0DE04 P0DE05 P0DE06 P0DE07 P0DE08 P0DE09 P0DE10 P0DE11
P0DE12 P0DE13 P0DE14 P0DE15 P0DE16 P0DE17 P0DE18 P0DE19 P0DE20 P0DE21 P0DE22
P0DE23 P0DE24 P0DE25 P0DE26 P0DE27 P0DE28 P0DE29 P0DE30 P0DE31 P0DE32 P0DE33
P0DE34 P0DE35 P0DE36 P0DE37 P0DE38 P0DE39 P0DE40 P0DE41 P0DE42 P0DE43 P0DE44
P0DE45 P0DE46 P0DE47 P0DE48 P0DE49 P0DE50 P0DE51 P0DE52 P0DE53 P0DE54 P0DE55
P0DE56 P0DE57 P0DE58 P0DE59 P0DE60 P0DE61 P0DE62 P0DE63 P0DE64 P0DE65 P0DE66
P0DE67 P0DE68 P0DE69 P0DE70 P0DE71 P0DE72 P0DE73 P0DE74 P0DE75 P0DE76 P0DE77
P0DE78 P0DE79 P0DE80 P0DE81 P0DE82 P0DE83 P0DE84 P0DE85 P0DE86 P0DE87 P0DE88
P0DE89 P0DE90 P0DE91 P0DE92 P0DE93 P0DE94 P0DE95 P0DE96 P0DE97 P0DE98 P0DE99
P0DF00 P0DF01 P0DF02 P0DF03 P0DF04 P0DF05 P0DF06 P0DF07 P0DF08 P0DF09 P0DF10
P0DF11 P0DF12 P0DF13 P0DF14 P0DF15 P0DF16 P0DF17 P0DF18 P0DF19 P0DF22 P0DF23
P0DF24 P0DF25 P0DF26 P0DF27 P0DF28 P0DF29 P0DF30 P0DF31 P0DF32 P0DF33 P0DF34
P0DF35 P0DF36 P0DF37 P0DF38 P0DF39 P0DF40 P0DF41 P0DF42 P0DF43 P0DF44 P0DF45
P0DF46 P0DF47 P0DF48 P0DF49 P0DF50 P0DF51 P0DF52 P0DF53 P0DF54 P0DF55 P0DF56
P0DF57 P0DF62 P0DF63 P0DF64 P0DF65 P0DF68 P0DF69 P0DF70 P0DF71 P0DF74 P0DF75
P0DF76 P0DF77 P0DF80 P0DF81 P0DF82 P0DF83 P0DF84 P0DF85 P0DF86 P0DF87 P0DF88
P0DF89 P0DF90 P0DF91 P0DF92 P0DF93 P0DF94 P0DF95 P0DF98 P0DF99 P0DG00 P0DG01
P0DG02 P0DG03 P0DG04 P0DG05 P0DG10 P0DG11 P0DG12 P0DG13 P0DG14 P0DG15 P0DG16
P0DG17 P0DG18 P0DG19 P0DG20 P0DG21 P0DG22 P0DG23 P0DG24 P0DG25 P0DG26 P0DG27
P0DG28 P0DG29 P0DG30 P0DG31 P0DG32 P0DG33 P0DG34 P0DG35 P0DG36 P0DG37 P0DG38
P0DG39 P0DG40 P0DG41 P0DG42 P0DG43 P0DG44 P0DG45 P0DG46 P0DG47 P0DG48 P0DG49
P0DG50 P0DG51 P0DG52 P0DG53 P0DG54 P0DG55 P0DG56 P0DG57 P0DG58 P0DG59 P0DG60
P0DG61 P0DG62 P0DG63 P0DG64 P0DG65 P0DG66 P0DG67 P0DG68 P0DG69 P0DG70 P0DG71
P0DG72 P0DG73 P0DG74 P0DG75 P0DG76 P0DG77 P0DG78 P0DG79 P0DG82 P0DG83 P0DG84
P0DG85 P0DG90 P0DG91 P0DG92 P0DG93 P0DG94 P0DG95 P0DG96 P0DG97 P0DG98 P0DG99
P0DH00 P0DH01 P0DH02 P0DH03 P0DH06 P0DH07 P0DH08 P0DH09 P0DH10 P0DH11 P0DH12

P0DH13 P0DH14 P0DH15 P0DH16 P0DH17 P0DH18 P0DH19 P0DH20 P0DH21 P0DH22 P0DH23
P0DH24 P0DH25 P0DH26 P0DH27 P0DH28 P0DH29 P0DH32 P0DH33 P0DH34 P0DH35 P0DH36
P0DH37 P0DH38 P0DH39 P0DH40 P0DH41 P0DH42 P0DH43 P0DH44 P0DH45 P0DH46 P0DH47
P0DH48 P0DH49 P0DH50 P0DH51 P0DH52 P0DH53 P0DH54 P0DH55 P0DH56 P0DH57 P0DH59
P0DH60 P0DH62 P0DH70 P0DH71 P0DH72 P0DH73 P0DH74 P0DH75 P0DH76 P0DH77 P0DH78
P0DH85 P0DH88 P0DH89 P0DH90 P0DH96 P0DH97 P0DH98 P0DI00 P0DI01 P0DI02 P0DI03
P0DI04 P0DI05 P0DI06 P0DI07 P0DI08 P0DI09 P0DI12 P0DI13 P0DI14 P0DI16 P0DI17
P0DI18 P0DI19 P0DI76 P0DI77 P0DI83 P0DJ09 P0DJ10 P0DJ13 P0DJ14 P0DJ15 P0DJ16
P0DJ17 P0DJ18 P0DJ19 P0DJ20 P0DJ21 P0DJ22 P0DJ23 P0DJ24 P0DJ25 P0DJ26 P0DJ27
P0DJ28 P0DJ51 P0DJ52 P0DJ53 P0DJ54 P0DJ55 P0DJ56 P0DJ57 P0DJ58 P0DJ59 P0DJ60
P0DJ61 P0DJA2 P0DJA5 P0DJA6 P0DJA7 P0DJA8 P0DJB1 P0DJD0 P0DJD1 P0DJD3 P0DJD4
P0DJF2 P0DJF8 P0DJF9 P0DJG4 P0DJG9 P0DJH0 P0DJH1 P0DJH6 P0DJH7 P0DJH9 P0DJJ0
P0DJL7 P0DJL8 P0DJL9 P0DJM1 P0DJM2 P0DJM3 P0DJM4 P0DJN9 P0DJO7 P0DJO8 P0DJO9
P0DJP0 P0DJP1 P0DJQ3 P0DJQ5 P0DJQ6 P0DJQ7 P0DJR0 P0DJR3 P0DJR4 P0DJR5 P0DJR6
P0DJR7 P0DJR8 P0DJR9 P0DJS0 P0DJS1 P0DJS2 P0DJS3 P0DJS4 P0DJS5 P0DJS6 P0DJS7
P0DJS8 P0DJS9 P0DJT0 P0DJT1 P0DJT2 P0DJT3 P0DJT4 P0DJT5 P0DJT6 P0DJT7 P0DJT8
P0DJT9 P0DJU0 P0DJU1 P0DJU2 P0DJU3 P0DJU4 P0DJU5 P0DJU6 P0DJU7 P0DJU8 P0DJU9
P0DJV0 P0DJV1 P0DJV2 P0DJV3 P0DJV4 P0DJV5 P0DJV6 P0DJV7 P0DJV8 P0DJV9 P0DJW0
P0DJW1 P0DJW2 P0DJW3 P0DJW4 P0DJW5 P0DJW6 P0DJW7 P0DJW8 P0DJW9 P0DJX0 P0DJX1
P0DJX2 P0DJX3 P0DJX4 P0DJX5 P0DJX6 P0DJX7 P0DJX8 P0DJY1 P0DJY2 P0DJY3 P0DJY4
P0DJY5 P0DJY6 P0DJY9 P0DJZ0 P0DJZ1 P0DJZ2 P0DJZ7 P0DKA0 P0DKA1 P0DKA2 P0DKA3
P0DKA4 P0DKA5 P0DKA6 P0DKA7 P0DKA8 P0DKA9 P0DKB0 P0DKB2 P0DKB4 P0DKC3 P0DKC4
P0DKC8 P0DKC9 P0DKD0 P0DKD1 P0DKD2 P0DKD3 P0DKD4 P0DKD5 P0DKD6 P0DKD7 P0DKD8
P0DKD9 P0DKE0 P0DKE1 P0DKE2 P0DKE3 P0DKE4 P0DKE5 P0DKE6 P0DKE7 P0DKE8 P0DKE9
P0DKF0 P0DKF1 P0DKF2 P0DKF3 P0DKF4 P0DKF5 P0DKF6 P0DKF7 P0DKF8 P0DKF9 P0DKG0
P0DKG1 P0DKG3 P0DKG4 P0DKG5 P0DKG6 P0DKG7 P0DKG8 P0DKH2 P0DKH3 P0DKH4 P0DKH7
P0DKH8 P0DKI3 P0DKI4 P0DKI5 P0DKI8 P0DKI9 P0DKJ0 P0DKJ1 P0DKJ2 P0DKJ7 P0DKJ8
P0DKJ9 P0DKK0 P0DKK1 P0DKK2 P0DKK3 P0DKK4 P0DKK7 P0DKK8 P0DKK9 P0DKL0 P0DKL1
P0DKL4 P0DKL6 P0DKL9 P0DKR1 P0DKR2 P0DKR7 P0DKR8 P0DKS2 P0DKS4 P0DKS5 P0DKS6
P0DKS7 P0DKS8 P0DKS9 P0DKT0 P0DKX0 P0DKX8 P0DKY4 P0DKY7 P0DKY8 P0DL09 P0DL13
P0DL28 P0DM32 P0DM33 P0DM35 P0DM41 P0DM42 P0DM58 P0DM63 P0DM65 P0DM81 P0DM85
P0DM86 P0DMB1 P0DMB2 P0DMC9 P0DMD0 P0DMG1 P0DMG2 P0DMG3 P0DMG4 P0DMI1 P0DMK2
P0DMK3 P0DMK4 P0DMK5 P0DMK8 P0DML0 P0DML1 P0DMM2 P0DMM3 P0DMM4 P0DMM5 P0DMN7
P0DMP2 P0DMP3 P0DMP4 P0DMP5 P0DMQ7 P0DMQ9 P0DMR1 P0DMR3 P0DMR5 P0DMR6 P0DMS0
P0DMU3 P0DMU7 P0DMU8 P0DMU9 P0DMV0 P0DMV1 P0DMV2 P0DMV5 P0DMV7 P0DN21 P0DN22
P0DN23 P0DN30 P0DN31 P0DN33 P0DN69 P0DN70 P0DN71 P0DN72 P0DN73 P0DN74 P0DN75
P0DN85 P0DN88 P0DN95 P0DO01 P0DO02 P0DO03 P0DO07 P0DO08 P0DO09 P0DO10 P0DO11
P0DO12 P0DO23 P0DO25 P0DO26 P0DO30 P0DO31 P0DO33 P0DO46 P0DO47 P0DO48 P0DO49
P0DO55 P0DO92 P0DO93 P0DO96 P0DO97 P0DO98 P0DOA5 P0DOB3 P0DOB5 P0DOB6 P0DOB7
P0DOB8 P0DOB9 P0DOC0 P0DOC6 P0DOC7 P0DOC8 P0DOD0 P0DOD1 P0DOD2 P0DOD3 P0DOD4
P0DOD5 P0DOD6 P0DOD7 P0DOD8 P0DOD9 P0DOE3 P0DOF3 P0DOG3 P0DOG4 P0DOG5 P0DOG6
P0DOG7 P0DOI8 P0DOI9 P0DOJ0 P0DOK3 P0DOK4 P0DOK5 P0DOK9 P0DOL0 P0DOL1 P0DOL2
P0DOM1 P0DOM2 P0DOM5 P0DOM6 P0DOM7 P0DOM8 P0DON3 P0DON4 P0DON5 P0DON6 P0DON9
P0DOO0 P0DOO1 P0DOO2 P0DOO3 P0DOO4 P0DOO5 P0DOO6 P0DOO7 P0DOO8 P0DOO9 P0DOP0

P0DOP3 P0DOP4 P0DOP5 P0DOP6 P0DOP7 P0DOP8 P0DOQ1 P0DOQ2 P0DOQ3 P0DOQ4 P0DOQ5
P0DOQ6 P0DOQ9 P0DOR0 P0DOR1 P0DOR2 P0DOR7 P0DOR8 P0DOR9 P0DOS0 P0DOS1 P0DOS2
P0DOS3 P0DOS4 P0DOS5 P0DOS6 P0DOS9 P0DOT0 P0DOT1 P0DOT2 P0DOT3 P0DOT4 P0DOT5
P0DOT6 P0DOT9 P0DOU0 P0DOU3 P0DOU4 P0DOU5 P0DOU6 P0DOU7 P0DOU8 P0DOV5 P0DOV6
P0DOV7 P0DOV8 P0DOV9 P0DOW0 P0DOW1 P0DOW4 P0DOW7 P0DOW8 P0DOW9 P0DOY1 P0DOY4
P0DOY6 P0DOY7 P0DOY8 P0DOY9 P0DP15 P0DP21 P0DP22 P0DP44 P0DP45 P0DP63 P0DP64
P0DP65 P0DP66 P0DP67 P0DP68 P0DP71 P0DP75 P0DP77 P0DP78 P0DP79 P0DP80 P0DP81
P0DP82 P0DP89 P0DP90 P0DP92 P0DPA1 P0DPA3 P0DPA5 P0DPA6 P0DPA8 P0DPA9 P0DPB0
P0DPB5 P0DPB6 P0DPB7 P0DPB8 P0DPC3 P0DPC5 P0DPC6 P0DPC7 P0DPC8 P0DPC9 P0DPD4
P0DPD5 P0DPD7 P0DPE0 P0DPE1 P0DPE4 P0DPE9 P0DPF0 P0DPF2 P0DPF3 P0DPF5 P0DPF6
P0DPH9 P0DPI2 P0DPI3 P0DPJ9 P0DPK2 P0DPK5 P0DPK6 P0DPM5 P0DPM6 P0DPM7 P0DPM8
P0DPM9 P0DPN2 P0DPN5 P0DPN6 P0DPN7 P0DPN8 P0DPN9 P0DPO0 P0DPO2 P0DPO3 P0DPO6
P0DPO7 P0DPO9 P0DPP1 P0DPP2 P0DPP3 P0DPP4 P0DPP5 P0DPP6 P0DPP8 P0DPP9 P0DPQ3
P0DPQ6 P0DPQ7 P0DPQ8 P0DPR8 P0DPS1 P0DPS8 P0DPS9 P0DQD8 P0DQP1 P0DQW0 P0DSE3
P0DSE4 P0DSE5 P0DSE6 P0DSE7 P0DSE8 P0DSE9 P0DSF0 P0DSF1 P0DSF2 P0DSF3 P0DSF4
P0DSF5 P0DSF6 P0DSF7 P0DSF8 P0DSF9 P0DSG0 P0DSG1 P0DSG2 P0DSG3 P0DSG4 P0DSG5
P0DSG6 P0DSG7 P0DSG8 P0DSG9 P0DSH0 P0DSH1 P0DSH2 P0DSH3 P0DSH4 P0DSH5 P0DSH6
P0DSH7 P0DSH8 P0DSH9 P0DSI0 P0DSM6 P0DSM7 P0DSM8 P0DSO1 P0DSO2 P0DSO3 P0DSP7
P0DSP8 P0DSP9 P0DSQ0 P0DSQ1 P0DSQ2 P0DSQ5 P0DSQ6 P0DSQ7 P0DSQ8 P0DSQ9 P0DSR0
P0DSR1 P0DSR2 P0DSR3 P0DSR4 P0DSR5 P0DSR6 P0DSR9 P0DSS0 P0DSS3 P0DSS4 P0DSS5
P0DSS6 P0DSS7 P0DSS8 P0DSS9 P0DST0 P0DST3 P0DST4 P0DST7 P0DST8 P0DST9 P0DSU0
P0DSU1 P0DSU2 P0DSU3 P0DSU4 P0DSU5 P0DSU6 P0DSU7 P0DSU8 P0DSU9 P0DSV0 P0DSV1
P0DSV2 P0DSV3 P0DSV4 P0DSV5 P0DSV6 P0DSV9 P0DSW0 P0DSW3 P0DSW4 P0DSW5 P0DSW6
P0DSW9 P0DSX0 P0DSX1 P0DSX2 P0DSX3 P0DSX4 P0DSX5 P0DSX6 P0DSX9 P0DSY0 P0DSZ1
P0DSZ2 P0DSZ5 P0DSZ6 P0DSZ7 P0DSZ8 P0DSZ9 P0DTA0 P0DTA1 P0DTA2 P0DTA3 P0DTA7
P0DTA8 P0DTA9 P0DTB0 P0DTB1 P0DTB2 P0DTD9 P0DTF1 P0DTF2 P0DTF3 P0DTF4 P0DTF5
P0DTF6 P0DTF7 P0DTF8 P0DTG0 P0DTG1 P0DTH5 P0DTH7 P0DTH8 P0DTI1 P0DTI3 P0DTI9
P0DTK3 P0DTK4 P0DTK5 P0DTK6 P0DTK7 P0DTK8 P0DTK9 P0DTL0 P0DTL1 P0DTL2 P0DTL6
P0DTL8 P0DTP6 P0DTP7 P0DTP8 P0DTP9 P0DTQ0 P0DTQ1 P0DTQ2 P0DTQ4 P0DTT0 P0DTV6
P0DTV7 P0DTV8 P0DTV9 P0DTW0 P0DTW1 P0DTX0 P0DTX1 P0DUB5 P0DUD1 P0DUD2 P0DUD3
P0DUD4 P0DUD5 P0DUD6 P0DUD9 P0DUE0 P0DUE2 P0DUE3 P0DUE4 P0DUE5 P0DUE9 P0DUF0
P0DUF1 P0DUF2 P0DUF3 P0DUF4 P0DUF5 P0DUF6 P0DUF7 P0DUF8 P0DUF9 P0DUG0 P0DUG1
P0DUH6 P0DUH7 P0DUK0 P0DUK1 P0DUK2 P0DUL1 P0DUL2 P0DUL5 P0DUL6 P0DUL7 P0DUL8
P0DUL9 P0DUM0 P0DUM1 P0DUM7 P0DUM8 P0DUM9 P0DUQ0 P0DUQ1 P0DUQ2 P0DUQ3 P0DUR5
P0DUR7 P0DUR8 P0DUR9 P0DUU5 P0DUV3 P0DUV4 P0DUV5 P0DUV6 P0DUV7 P0DUV8 P0DUV9
P0DUW0 P0DUW1 P0DUW2 P0DUW3 P0DUW4 P0DUX0 P0DUX1 P0DV14 P0DV19 P0DV20 P0DV21
P0DV22 P0DV23 P0DV29 P0DV33 P0DV34 P0DV35 P0DV41 P0DV54 P0DV55 P0DV56 P0DV57
P0DV58 P0DV59 P0DV60 P0DV65 P0DV66 P0DV73 P0DV74 P0DV75 P0DV76 P0DV77 P0DV79
P0DV86 P0DV87 P0DV88 P0DV89 P0DV90 P0DV91 P0DV92 P0DV93 P0DV94 P0DV98 P0DV99
P0DW04 P0DW05 P0DW06 P0DW07 P0DW08 P0DW11 P0DW12 P0DW13 P0DW14 P0DW16 P0DW27
P0DW28 P0DW32 P0DW33 P0DW34 P0DW35 P0DW36 P0DW37 P0DW38 P0DW39 P0DW40 P0DW41
P0DW42 P0DW43 P0DW44 P0DW46 P0DW47 P0DW48 P0DW49 P0DW52 P0DW53 P0DW79 P0DW80
P0DW81 P0DW82 P0DW85 P0DW87 P0DW89 P0DW91 P0DW92 P0DX00 P0DX01 P0DX02 P10018
P10020 P10021 P10022 P10023 P10024 P10025 P10027 P10030 P10031 P10032 P10035

P10036 P10037 P10042 P10043 P10044 P10045 P10050 P10051 P10052 P10053 P10054
P10055 P10057 P10058 P10059 P10060 P10061 P10062 P10065 P10066 P10067 P10068
P10069 P10072 P10073 P10074 P10075 P10076 P10077 P10078 P10080 P10083 P10085
P10088 P10091 P10094 P10095 P10097 P10104 P10105 P10112 P10118 P10119 P10123
P10125 P10127 P10128 P10129 P10130 P10131 P10132 P10133 P10134 P10135 P10136
P10137 P10138 P10139 P10140 P10141 P10142 P10143 P10151 P10156 P10166 P10169
P10172 P10173 P10178 P10179 P10180 P10181 P10182 P10183 P10186 P10187 P10188
P10189 P10190 P10191 P10192 P10193 P10200 P10201 P10202 P10205 P10208 P10209
P10212 P10216 P10217 P10221 P10225 P10234 P10236 P10239 P10242 P10243 P10245
P10246 P10248 P10249 P10251 P10255 P10265 P10266 P10270 P10277 P10283 P10284
P10290 P10299 P10303 P10304 P10306 P10308 P10309 P10310 P10311 P10312 P10326
P10331 P10336 P10337 P10340 P10341 P10343 P10350 P10351 P10356 P10357 P10359
P10363 P10365 P10367 P10368 P10369 P10370 P10371 P10372 P10381 P10382 P10383
P10390 P10392 P10394 P10395 P10396 P10399 P10400 P10401 P10403 P10405 P10420
P10421 P10426 P10427 P10428 P10429 P10430 P10431 P10432 P10433 P10434 P10435
P10436 P10437 P10442 P10449 P10462 P10463 P10468 P10470 P10472 P10473 P10482
P10484 P10485 P10486 P10487 P10488 P10489 P10490 P10497 P10498 P10505 P10511
P10512 P10514 P10518 P10525 P10528 P10533 P10537 P10538 P10539 P10540 P10545
P10546 P10550 P10551 P10559 P10560 P10563 P10565 P10567 P10570 P10571 P10572
P10575 P10576 P10577 P10579 P10580 P10581 P10582 P10583 P10584 P10585 P10587
P10593 P10618 P10621 P10623 P10624 P10626 P10627 P10628 P10629 P10630 P10647
P10650 P10651 P10652 P10653 P10654 P10655 P10656 P10658 P10659 P10691 P10711
P10713 P10723 P10724 P10725 P10726 P10727 P10728 P10733 P10734 P10735 P10738
P10739 P10746 P10748 P10751 P10755 P10759 P10760 P10768 P10770 P10771 P10772
P10773 P10775 P10777 P10778 P10780 P10781 P10782 P10783 P10784 P10785 P10786
P10788 P10789 P10790 P10791 P10792 P10796 P10797 P10798 P10801 P10802 P10806
P10807 P10816 P10819 P10821 P10828 P10831 P10832 P10835 P10836 P10841 P10847
P10848 P10849 P10850 P10853 P10854 P10856 P10857 P10860 P10862 P10868 P10869
P10871 P10872 P10873 P10874 P10875 P10876 P10878 P10880 P10881 P10883 P10885
P10887 P10892 P10893 P10902 P10914 P10916 P10922 P10928 P10929 P10930 P10932
P10937 P10938 P10941 P10943 P10944 P10952 P10961 P10963 P10964 P10967 P10969
P10971 P10978 P10979 P10981 P10982 P10984 P10987 P10988 P10990 P10992 P10993
P10994 P10995 P10996 P11003 P11014 P11018 P11020 P11025 P11035 P11037 P11038
P11039 P11040 P11043 P11044 P11045 P11053 P11054 P11055 P11065 P11066 P11067
P11068 P11071 P11079 P11080 P11081 P11084 P11086 P11088 P11094 P11102 P11107
P11108 P11109 P11110 P11111 P11112 P11115 P11118 P11120 P11121 P11122 P11123
P11124 P11125 P11126 P11131 P11138 P11141 P11143 P11144 P11145 P11154 P11156
P11157 P11158 P11175 P11178 P11179 P11180 P11181 P11182 P11183 P11186 P11189
P11190 P11191 P11192 P11202 P11203 P11204 P11205 P11207 P11208 P11210 P11211
P11212 P11213 P11217 P11231 P11237 P11245 P11246 P11248 P11250 P11251 P11252
P11256 P11257 P11260 P11272 P11289 P11292 P11299 P11300 P11301 P11302 P11304
P11314 P11315 P11319 P11320 P11321 P11322 P11324 P11325 P11326 P11329 P11330
P11331 P11333 P11334 P11335 P11336 P11337 P11338 P11340 P11341 P11342 P11343
P11347 P11348 P11353 P11368 P11369 P11370 P11375 P11376 P11386 P11405 P11406

P11408 P11409 P11410 P11411 P11412 P11414 P11415 P11419 P11421 P11422 P11424
P11425 P11426 P11428 P11435 P11436 P11437 P11444 P11446 P11447 P11451 P11452
P11455 P11458 P11459 P11468 P11469 P11470 P11475 P11480 P11481 P11482 P11500
P11503 P11504 P11512 P11513 P11514 P11517 P11519 P11521 P11522 P11523 P11524
P11534 P11536 P11538 P11539 P11540 P11544 P11546 P11553 P11559 P11561 P11563
P11564 P11565 P11566 P11567 P11568 P11569 P11570 P11573 P11596 P11600 P11605
P11612 P11621 P11624 P11626 P11632 P11633 P11635 P11637 P11638 P11639 P11640
P11641 P11642 P11650 P11651 P11652 P11653 P11660 P11663 P11664 P11671 P11673
P11675 P11689 P11690 P11703 P11704 P11705 P11708 P11709 P11720 P11724 P11726
P11727 P11728 P11731 P11732 P11733 P11734 P11735 P11736 P11737 P11738 P11739
P11741 P11742 P11745 P11746 P11747 P11748 P11749 P11750 P11751 P11752 P11753
P11754 P11755 P11756 P11757 P11758 P11759 P11765 P11778 P11794 P11795 P11796
P11804 P11805 P11806 P11807 P11817 P11818 P11819 P11820 P11821 P11822 P11823
P11824 P11825 P11826 P11832 P11833 P11837 P11842 P11843 P11844 P11857 P11860
P11861 P11863 P11864 P11865 P11866 P11868 P11870 P11871 P11872 P11873 P11875
P11880 P11886 P11887 P11888 P11890 P11891 P11892 P11893 P11894 P11896 P11897
P11899 P11901 P11904 P11905 P11906 P11907 P11908 P11917 P11918 P11922 P11926
P11927 P11929 P11931 P11932 P11934 P11935 P11938 P11954 P11956 P11957 P11959
P11960 P11961 P11964 P11966 P11974 P11975 P11977 P11978 P11979 P11980 P11982
P11983 P11988 P11989 P11998 P12000 P12002 P12004 P12008 P12010 P12011 P12013
P12015 P12026 P12031 P12033 P12035 P12036 P12037 P12039 P12040 P12041 P12042
P12043 P12044 P12045 P12046 P12047 P12048 P12049 P12052 P12053 P12054 P12055
P12057 P12059 P12063 P12066 P12070 P12073 P12076 P12078 P12081 P12114 P12115
P12116 P12134 P12136 P12139 P12142 P12149 P12151 P12153 P12173 P12174 P12176
P12195 P12196 P12198 P12208 P12210 P12227 P12230 P12232 P12233 P12243 P12248
P12249 P12250 P12254 P12256 P12258 P12261 P12262 P12269 P12271 P12274 P12275
P12281 P12282 P12288 P12289 P12290 P12291 P12294 P12295 P12297 P12298 P12299
P12300 P12304 P12305 P12306 P12307 P12308 P12309 P12310 P12311 P12320 P12321
P12322 P12323 P12325 P12326 P12335 P12339 P12340 P12341 P12343 P12347 P12355
P12356 P12364 P12366 P12378 P12380 P12381 P12383 P12385 P12393 P12395 P12400
P12411 P12413 P12414 P12415 P12418 P12419 P12421 P12423 P12424 P12426 P12427
P12431 P12432 P12433 P12434 P12435 P12438 P12455 P12456 P12457 P12458 P12459
P12460 P12461 P12462 P12464 P12465 P12466 P12468 P12472 P12475 P12500 P12501
P12521 P12523 P12524 P12525 P12529 P12531 P12533 P12535 P12536 P12537 P12538
P12539 P12540 P12543 P12551 P12552 P12576 P12577 P12579 P12580 P12609 P12611
P12613 P12614 P12620 P12622 P12623 P12628 P12629 P12639 P12646 P12653 P12659
P12660 P12675 P12677 P12678 P12679 P12680 P12689 P12692 P12693 P12694 P12695
P12712 P12715 P12716 P12717 P12723 P12726 P12731 P12734 P12741 P12742 P12743
P12744 P12745 P12746 P12747 P12753 P12754 P12758 P12759 P12762 P12768 P12769
P12770 P12779 P12780 P12781 P12782 P12783 P12792 P12793 P12794 P12795 P12806
P12807 P12808 P12809 P12810 P12811 P12812 P12817 P12818 P12819 P12826 P12828
P12829 P12832 P12833 P12834 P12835 P12839 P12844 P12845 P12847 P12848 P12851
P12854 P12858 P12859 P12860 P12863 P12864 P12867 P12873 P12874 P12875 P12877
P12878 P12879 P12880 P12881 P12882 P12883 P12884 P12885 P12886 P12887 P12888

P12894 P12898 P12899 P12900 P12902 P12917 P12918 P12923 P12927 P12929 P12932
P12933 P12940 P12944 P12945 P12948 P12949 P12951 P12952 P12954 P12955 P12957
P12959 P12962 P12963 P12964 P12965 P12971 P12972 P12973 P12974 P12977 P12979
P12980 P12994 P12995 P12996 P12997 P12998 P12999 P13000 P13001 P13004 P13005
P13007 P13009 P13015 P13018 P13019 P13022 P13024 P13026 P13027 P13029 P13035
P13039 P13041 P13043 P13044 P13045 P13054 P13057 P13058 P13059 P13060 P13062
P13064 P13065 P13066 P13069 P13070 P13071 P13075 P13077 P13078 P13079 P13081
P13082 P13086 P13088 P13089 P13091 P13092 P13093 P13094 P13095 P13096 P13097
P13098 P13099 P13100 P13104 P13105 P13106 P13110 P13111 P13121 P13122 P13123
P13125 P13154 P13157 P13158 P13159 P13161 P13179 P13188 P13191 P13192 P13193
P13203 P13212 P13215 P13218 P13222 P13226 P13227 P13228 P13230 P13239 P13242
P13243 P13246 P13249 P13250 P13251 P13254 P13255 P13256 P13257 P13258 P13262
P13263 P13264 P13266 P13267 P13268 P13269 P13270 P13273 P13274 P13275 P13278
P13279 P13280 P13281 P13282 P13283 P13286 P13288 P13295 P13298 P13299 P13300
P13302 P13303 P13305 P13306 P13307 P13308 P13309 P13310 P13311 P13312 P13313
P13314 P13315 P13316 P13317 P13318 P13319 P13320 P13321 P13322 P13323 P13324
P13325 P13326 P13327 P13328 P13329 P13330 P13331 P13335 P13336 P13337 P13338
P13339 P13340 P13341 P13342 P13349 P13350 P13351 P13360 P13361 P13363 P13364
P13365 P13367 P13375 P13376 P13378 P13382 P13390 P13392 P13393 P13394 P13397
P13398 P13411 P13412 P13413 P13416 P13417 P13418 P13419 P13420 P13431 P13433
P13434 P13439 P13440 P13441 P13442 P13443 P13444 P13445 P13446 P13448 P13449
P13453 P13456 P13458 P13459 P13460 P13461 P13462 P13463 P13468 P13469 P13474
P13476 P13477 P13478 P13479 P13483 P13484 P13486 P13491 P13496 P13499 P13503
P13505 P13513 P13520 P13522 P13523 P13524 P13525 P13526 P13528 P13533 P13535
P13537 P13538 P13539 P13540 P13541 P13542 P13543 P13544 P13545 P13549 P13550
P13551 P13552 P13553 P13556 P13557 P13558 P13561 P13564 P13566 P13567 P13570
P13571 P13575 P13576 P13577 P13601 P13602 P13603 P13604 P13606 P13615 P13622
P13623 P13629 P13630 P13634 P13640 P13649 P13651 P13653 P13657 P13658 P13659
P13660 P13665 P13669 P13675 P13676 P13681 P13682 P13694 P13697 P13699 P13702
P13703 P13705 P13706 P13707 P13708 P13712 P13714 P13715 P13716 P13722 P13733
P13736 P13737 P13740 P13741 P13742 P13743 P13769 P13770 P13771 P13772 P13774
P13775 P13776 P13777 P13778 P13779 P13780 P13781 P13782 P13783 P13784 P13785
P13786 P13787 P13788 P13789 P13792 P13803 P13804 P13805 P13809 P13813 P13822
P13824 P13832 P13833 P13834 P13836 P13840 P13846 P13848 P13849 P13853 P13857
P13859 P13863 P13867 P13870 P13877 P13884 P13885 P13894 P13895 P13903 P13904
P13906 P13911 P13912 P13913 P13914 P13920 P13921 P13922 P13925 P13927 P13934
P13939 P13940 P13946 P13949 P13951 P13952 P13955 P13956 P13957 P13959 P13960
P13961 P13962 P13963 P13964 P13965 P13966 P13967 P13968 P13971 P13972 P13973
P13975 P13976 P13978 P13982 P13985 P13988 P13989 P13991 P13995 P13997 P13998
P13999 P14001 P14002 P14004 P14010 P14011 P14012 P14016 P14017 P14019 P14021
P14022 P14023 P14024 P14025 P14026 P14027 P14029 P14030 P14031 P14032 P14033
P14034 P14035 P14036 P14037 P14038 P14039 P14040 P14041 P14042 P14044 P14057
P14062 P14064 P14065 P14070 P14071 P14072 P14073 P14081 P14083 P14088 P14107
P14108 P14109 P14110 P14111 P14112 P14113 P14116 P14117 P14119 P14125 P14128

P14129 P14134 P14140 P14141 P14145 P14149 P14150 P14152 P14155 P14156 P14157
P14158 P14159 P14161 P14164 P14165 P14166 P14167 P14168 P14169 P14173 P14179
P14188 P14193 P14194 P14195 P14196 P14197 P14199 P14201 P14202 P14203 P14205
P14208 P14218 P14219 P14228 P14229 P14230 P14232 P14233 P14235 P14239 P14240
P14241 P14242 P14243 P14244 P14245 P14247 P14248 P14249 P14251 P14252 P14253
P14254 P14259 P14260 P14261 P14265 P14266 P14267 P14268 P14269 P14284 P14288
P14289 P14291 P14294 P14295 P14298 P14299 P14300 P14301 P14303 P14307 P14309
P14310 P14311 P14312 P14313 P14316 P14318 P14321 P14322 P14325 P14326 P14327
P14328 P14330 P14332 P14333 P14345 P14346 P14347 P14348 P14352 P14355 P14356
P14357 P14358 P14360 P14361 P14362 P14363 P14364 P14365 P14366 P14368 P14369
P14370 P14371 P14372 P14379 P14380 P14381 P14385 P14386 P14387 P14388 P14389
P14390 P14391 P14392 P14393 P14394 P14395 P14396 P14397 P14398 P14399 P14402
P14405 P14407 P14408 P14412 P14425 P14485 P14486 P14487 P14490 P14491 P14492
P14493 P14498 P14499 P14500 P14501 P14503 P14504 P14505 P14506 P14508 P14509
P14510 P14511 P14520 P14521 P14522 P14523 P14524 P14525 P14526 P14527 P14528
P14533 P14540 P14547 P14549 P14552 P14561 P14563 P14564 P14565 P14567 P14568
P14577 P14582 P14583 P14586 P14587 P14588 P14589 P14590 P14591 P14592 P14594
P14597 P14607 P14610 P14611 P14620 P14621 P14633 P14634 P14636 P14637 P14638
P14640 P14641 P14642 P14643 P14647 P14649 P14651 P14652 P14653 P14654 P14655
P14656 P14657 P14670 P14673 P14674 P14675 P14678 P14685 P14686 P14696 P14697
P14705 P14706 P14707 P14708 P14710 P14711 P14714 P14717 P14718 P14720 P14721
P14723 P14724 P14725 P14726 P14734 P14736 P14737 P14739 P14741 P14742 P14743
P14746 P14747 P14748 P14751 P14752 P14755 P14757 P14758 P14759 P14760 P14761
P14762 P14766 P14776 P14785 P14795 P14799 P14802 P14805 P14807 P14808 P14809
P14810 P14814 P14815 P14816 P14817 P14820 P14821 P14822 P14826 P14827 P14828
P14829 P14830 P14831 P14832 P14836 P14837 P14838 P14839 P14840 P14843 P14845
P14846 P14848 P14849 P14850 P14858 P14864 P14865 P14870 P14871 P14874 P14875
P14881 P14883 P14885 P14886 P14887 P14888 P14889 P14890 P14898 P14899 P14900
P14902 P14903 P14905 P14908 P14909 P14912 P14913 P14922 P14928 P14930 P14932
P14934 P14935 P14936 P14937 P14938 P14939 P14940 P14941 P14942 P14945 P14949
P14951 P14952 P14953 P14957 P14958 P14959 P14960 P14961 P14962 P14963 P14964
P14965 P14967 P14969 P14972 P14975 P14977 P14980 P14982 P14990 P14991 P14994
P14995 P15000 P15001 P15005 P15006 P15008 P15009 P15010 P15019 P15020 P15024
P15025 P15026 P15032 P15033 P15034 P15038 P15042 P15043 P15047 P15052 P15057
P15058 P15059 P15062 P15063 P15066 P15067 P15075 P15076 P15081 P15082 P15093
P15094 P15095 P15096 P15099 P15100 P15102 P15109 P15110 P15111 P15112 P15113
P15114 P15115 P15124 P15132 P15135 P15137 P15138 P15139 P15142 P15143 P15158
P15159 P15160 P15161 P15162 P15163 P15164 P15165 P15166 P15170 P15171 P15172
P15173 P15178 P15179 P15180 P15181 P15183 P15184 P15187 P15188 P15189 P15190
P15197 P15201 P15204 P15206 P15214 P15234 P15235 P15236 P15237 P15241 P15244
P15245 P15249 P15250 P15251 P15254 P15255 P15256 P15259 P15262 P15266 P15268
P15274 P15275 P15276 P15280 P15281 P15288 P15305 P15308 P15314 P15315 P15316
P15317 P15326 P15327 P15329 P15332 P15333 P15334 P15335 P15339 P15340 P15341
P15342 P15343 P15345 P15348 P15349 P15353 P15355 P15356 P15360 P15361 P15370

P15373 P15375 P15394 P15395 P15397 P15398 P15401 P15402 P15403 P15406 P15412
P15419 P15424 P15426 P15434 P15435 P15436 P15443 P15444 P15446 P15447 P15448
P15449 P15454 P15465 P15466 P15469 P15474 P15480 P15482 P15489 P15494 P15505
P15512 P15518 P15523 P15524 P15525 P15526 P15527 P15528 P15534 P15556 P15557
P15558 P15562 P15563 P15564 P15567 P15587 P15593 P15594 P15596 P15597 P15598
P15599 P15602 P15603 P15604 P15605 P15606 P15607 P15608 P15609 P15610 P15611
P15612 P15613 P15614 P15615 P15616 P15617 P15618 P15619 P15620 P15621 P15622
P15623 P15624 P15625 P15628 P15629 P15630 P15631 P15632 P15633 P15634 P15635
P15639 P15640 P15646 P15648 P15649 P15653 P15687 P15699 P15700 P15705 P15714
P15717 P15719 P15723 P15730 P15732 P15733 P15734 P15735 P15736 P15738 P15744
P15755 P15757 P15758 P15759 P15760 P15761 P15762 P15764 P15765 P15766 P15767
P15768 P15769 P15770 P15771 P15772 P15786 P15787 P15788 P15789 P15790 P15792
P15796 P15799 P15801 P15804 P15807 P15808 P15820 P15821 P15824 P15825 P15826
P15837 P15840 P15844 P15846 P15850 P15854 P15856 P15857 P15858 P15859 P15860
P15861 P15862 P15864 P15865 P15866 P15867 P15868 P15869 P15870 P15871 P15872
P15873 P15874 P15875 P15881 P15885 P15886 P15888 P15889 P15890 P15894 P15895
P15904 P15905 P15908 P15912 P15913 P15916 P15925 P15931 P15934 P15935 P15937
P15938 P15950 P15960 P15962 P15964 P15965 P15966 P15972 P15973 P15974 P15975
P15976 P15977 P15984 P15986 P15987 P15990 P15991 P15992 P16006 P16007 P16010
P16011 P16015 P16016 P16018 P16019 P16020 P16023 P16024 P16025 P16031 P16032
P16034 P16037 P16040 P16048 P16053 P16058 P16062 P16063 P16064 P16072 P16073
P16075 P16076 P16078 P16079 P16080 P16083 P16084 P16088 P16089 P16091 P16093
P16095 P16096 P16097 P16098 P16099 P16100 P16101 P16103 P16107 P16108 P16113
P16114 P16115 P16117 P16118 P16120 P16121 P16122 P16123 P16124 P16126 P16128
P16129 P16130 P16131 P16132 P16133 P16134 P16135 P16136 P16137 P16138 P16142
P16143 P16146 P16147 P16148 P16149 P16165 P16166 P16167 P16171 P16172 P16181
P16184 P16222 P16238 P16241 P16243 P16245 P16246 P16247 P16249 P16250 P16251
P16252 P16253 P16263 P16266 P16267 P16268 P16269 P16272 P16273 P16274 P16275
P16276 P16280 P16281 P16285 P16289 P16290 P16304 P16306 P16309 P16320 P16323
P16329 P16331 P16332 P16340 P16343 P16346 P16347 P16348 P16349 P16350 P16351
P16352 P16355 P16356 P16370 P16372 P16373 P16374 P16375 P16379 P16380 P16384
P16385 P16387 P16394 P16400 P16405 P16407 P16409 P16411 P16412 P16415 P16417
P16418 P16419 P16420 P16421 P16423 P16424 P16425 P16426 P16431 P16433 P16437
P16440 P16443 P16447 P16448 P16451 P16453 P16456 P16458 P16461 P16463 P16464
P16467 P16468 P16470 P16472 P16480 P16485 P16486 P16489 P16490 P16491 P16494
P16495 P16515 P16516 P16518 P16521 P16522 P16523 P16524 P16525 P16528 P16533
P16538 P16539 P16540 P16541 P16542 P16543 P16544 P16550 P16551 P16553 P16556
P16557 P16558 P16559 P16560 P16577 P16578 P16594 P16595 P16596 P16597 P16600
P16605 P16606 P16607 P16608 P16609 P16616 P16618 P16625 P16626 P16631 P16633
P16634 P16635 P16639 P16640 P16641 P16642 P16643 P16649 P16650 P16653 P16654
P16656 P16659 P16666 P16667 P16668 P16678 P16679 P16681 P16684 P16685 P16686
P16687 P16688 P16689 P16690 P16691 P16692 P16703 P16706 P16709 P16712 P16713
P16714 P16715 P16716 P16722 P16723 P16725 P16726 P16728 P16729 P16730 P16731
P16732 P16734 P16735 P16736 P16737 P16738 P16746 P16747 P16748 P16749 P16752

P16755 P16759 P16760 P16761 P16762 P16763 P16764 P16765 P16766 P16767 P16769
P16770 P16771 P16773 P16775 P16776 P16777 P16779 P16781 P16782 P16783 P16784
P16786 P16787 P16788 P16789 P16790 P16792 P16793 P16796 P16797 P16798 P16799
P16801 P16802 P16803 P16804 P16805 P16806 P16807 P16810 P16811 P16812 P16813
P16814 P16816 P16817 P16818 P16819 P16820 P16821 P16822 P16824 P16825 P16826
P16827 P16828 P16829 P16830 P16835 P16836 P16838 P16839 P16840 P16842 P16847
P16855 P16856 P16857 P16865 P16866 P16867 P16868 P16878 P16883 P16884 P16886
P16887 P16888 P16889 P16890 P16891 P16895 P16898 P16900 P16901 P16902 P16904
P16905 P16909 P16911 P16913 P16914 P16916 P16917 P16918 P16919 P16923 P16924
P16925 P16928 P16929 P16931 P16932 P16937 P16938 P16939 P16940 P16942 P16943
P16944 P16953 P16954 P16961 P16965 P16966 P16971 P16972 P16992 P16993 P17007
P17009 P17010 P17012 P17014 P17017 P17020 P17021 P17022 P17023 P17024 P17025
P17026 P17027 P17030 P17031 P17032 P17035 P17036 P17038 P17039 P17040 P17041
P17044 P17051 P17052 P17054 P17055 P17056 P17058 P17059 P17060 P17063 P17067
P17070 P17072 P17086 P17087 P17088 P17089 P17090 P17091 P17092 P17093 P17094
P17095 P17100 P17103 P17104 P17105 P17106 P17109 P17115 P17117 P17119 P17120
P17122 P17123 P17126 P17128 P17130 P17133 P17136 P17138 P17141 P17142 P17143
P17144 P17145 P17147 P17148 P17149 P17158 P17160 P17161 P17162 P17163 P17166
P17167 P17169 P17170 P17171 P17172 P17173 P17174 P17192 P17193 P17196 P17197
P17198 P17199 P17202 P17204 P17208 P17209 P17210 P17211 P17212 P17214 P17217
P17218 P17220 P17221 P17222 P17223 P17224 P17225 P17239 P17240 P17241 P17245
P17248 P17257 P17262 P17264 P17265 P17268 P17271 P17276 P17277 P17278 P17279
P17289 P17290 P17291 P17292 P17293 P17294 P17295 P17296 P17297 P17303 P17304
P17305 P17306 P17307 P17308 P17309 P17310 P17311 P17312 P17313 P17319 P17329
P17330 P17331 P17337 P17338 P17339 P17346 P17353 P17355 P17356 P17358 P17359
P17360 P17361 P17362 P17365 P17366 P17367 P17368 P17369 P17370 P17371 P17372
P17373 P17374 P17377 P17381 P17383 P17384 P17385 P17386 P17388 P17393 P17394
P17395 P17396 P17402 P17410 P17411 P17418 P17419 P17420 P17421 P17422 P17423
P17424 P17429 P17430 P17433 P17434 P17435 P17436 P17440 P17441 P17445 P17446
P17449 P17456 P17457 P17459 P17460 P17461 P17462 P17467 P17468 P17469 P17470
P17473 P17474 P17478 P17479 P17481 P17482 P17483 P17486 P17487 P17488 P17492
P17493 P17499 P17500 P17502 P17505 P17506 P17507 P17508 P17509 P17511 P17512
P17518 P17521 P17522 P17525 P17526 P17529 P17530 P17536 P17537 P17545 P17546
P17548 P17550 P17552 P17556 P17557 P17560 P17562 P17565 P17569 P17570 P17571
P17574 P17575 P17576 P17578 P17584 P17585 P17586 P17587 P17588 P17589 P17591
P17597 P17603 P17606 P17611 P17613 P17615 P17616 P17617 P17618 P17620 P17622
P17624 P17625 P17628 P17629 P17631 P17639 P17641 P17642 P17648 P17649 P17652
P17653 P17657 P17663 P17667 P17671 P17672 P17673 P17676 P17678 P17679 P17688
P17689 P17695 P17699 P17703 P17707 P17708 P17714 P17718 P17719 P17720 P17721
P17724 P17725 P17726 P17729 P17730 P17731 P17734 P17735 P17736 P17738 P17743
P17744 P17746 P17750 P17751 P17752 P17761 P17768 P17770 P17771 P17772 P17776
P17779 P17780 P17782 P17783 P17784 P17786 P17788 P17789 P17802 P17808 P17814
P17817 P17818 P17819 P17820 P17828 P17829 P17830 P17831 P17832 P17833 P17834
P17839 P17842 P17843 P17845 P17846 P17847 P17853 P17854 P17856 P17857 P17863

P17867 P17868 P17869 P17871 P17874 P17877 P17878 P17882 P17885 P17887 P17888
P17889 P17890 P17893 P17894 P17903 P17904 P17906 P17907 P17908 P17909 P17910
P17915 P17916 P17917 P17919 P17920 P17921 P17922 P17923 P17924 P17926 P17928
P17930 P17932 P17933 P17937 P17938 P17941 P17947 P17950 P17952 P17957 P17958
P17959 P17960 P17961 P17962 P17965 P17966 P17969 P17976 P17978 P17983 P17984
P17985 P17991 P17992 P17994 P17998 P17999 P18000 P18002 P18005 P18007 P18016
P18017 P18018 P18019 P18020 P18021 P18022 P18023 P18025 P18026 P18027 P18028
P18029 P18030 P18032 P18036 P18037 P18046 P18047 P18048 P18051 P18053 P18055
P18056 P18057 P18058 P18060 P18061 P18062 P18063 P18076 P18079 P18086 P18087
P18088 P18100 P18105 P18111 P18115 P18117 P18119 P18120 P18128 P18133 P18134
P18138 P18139 P18140 P18147 P18148 P18150 P18154 P18155 P18157 P18158 P18162
P18174 P18175 P18178 P18179 P18182 P18183 P18184 P18185 P18186 P18188 P18204
P18207 P18208 P18240 P18241 P18243 P18244 P18248 P18249 P18250 P18253 P18255
P18256 P18258 P18261 P18264 P18267 P18268 P18270 P18271 P18272 P18279 P18281
P18283 P18284 P18285 P18286 P18288 P18294 P18296 P18297 P18298 P18299 P18300
P18304 P18305 P18310 P18311 P18314 P18315 P18316 P18317 P18318 P18319 P18320
P18321 P18322 P18324 P18325 P18326 P18327 P18333 P18335 P18338 P18339 P18346
P18349 P18351 P18352 P18353 P18355 P18358 P18377 P18378 P18379 P18380 P18383
P18385 P18386 P18387 P18388 P18393 P18394 P18407 P18408 P18412 P18416 P18420
P18421 P18425 P18426 P18432 P18435 P18436 P18437 P18444 P18451 P18452 P18455
P18457 P18459 P18474 P18478 P18480 P18483 P18485 P18486 P18487 P18488 P18491
P18492 P18494 P18499 P18500 P18501 P18514 P18515 P18516 P18517 P18520 P18544
P18547 P18548 P18549 P18550 P18555 P18561 P18562 P18566 P18568 P18571 P18576
P18579 P18591 P18592 P18593 P18600 P18601 P18602 P18603 P18605 P18606 P18608
P18612 P18616 P18617 P18622 P18623 P18625 P18629 P18638 P18642 P18644 P18645
P18655 P18661 P18663 P18664 P18665 P18666 P18667 P18668 P18670 P18671 P18672
P18673 P18674 P18675 P18676 P18677 P18679 P18681 P18682 P18683 P18685 P18686
P18689 P18690 P18691 P18699 P18700 P18707 P18712 P18713 P18714 P18715 P18716
P18717 P18718 P18719 P18720 P18721 P18722 P18723 P18724 P18725 P18726 P18727
P18728 P18729 P18730 P18731 P18732 P18733 P18734 P18735 P18736 P18737 P18738
P18739 P18740 P18741 P18742 P18743 P18744 P18745 P18746 P18747 P18748 P18749
P18750 P18751 P18752 P18753 P18755 P18756 P18758 P18769 P18771 P18778 P18779
P18781 P18782 P18785 P18786 P18787 P18789 P18797 P18807 P18808 P18809 P18811
P18816 P18818 P18819 P18820 P18821 P18831 P18834 P18843 P18847 P18853 P18854
P18855 P18863 P18864 P18865 P18866 P18867 P18868 P18869 P18896 P18898 P18899
P18904 P18905 P18906 P18911 P18912 P18921 P18922 P18925 P18949 P18953 P18955
P18957 P18960 P18961 P18969 P18970 P18971 P18972 P18973 P18974 P18975 P18976
P18977 P18978 P18979 P18980 P18981 P18982 P18983 P18984 P18985 P18986 P18987
P18988 P18989 P18990 P18991 P18992 P18993 P18994 P18995 P18996 P19002 P19012
P19013 P19014 P19015 P19016 P19018 P19026 P19028 P19029 P19031 P19039 P19055
P19059 P19060 P19061 P19062 P19064 P19066 P19067 P19068 P19071 P19076 P19077
P19078 P19079 P19080 P19088 P19089 P19102 P19107 P19109 P19113 P19115 P19117
P19118 P19119 P19123 P19126 P19127 P19128 P19130 P19132 P19133 P19135 P19140
P19141 P19142 P19143 P19148 P19149 P19151 P19160 P19161 P19162 P19163 P19164

P19165 P19166 P19167 P19168 P19169 P19170 P19175 P19176 P19177 P19178 P19179
P19183 P19187 P19188 P19189 P19191 P19192 P19193 P19194 P19195 P19198 P19200
P19201 P19205 P19206 P19210 P19211 P19212 P19213 P19216 P19219 P19220 P19229
P19230 P19231 P19237 P19239 P19241 P19242 P19243 P19246 P19248 P19251 P19252
P19257 P19262 P19263 P19264 P19265 P19267 P19268 P19270 P19271 P19272 P19273
P19274 P19275 P19276 P19277 P19278 P19279 P19280 P19281 P19282 P19283 P19284
P19285 P19286 P19287 P19288 P19289 P19290 P19291 P19292 P19293 P19294 P19295
P19296 P19297 P19298 P19299 P19300 P19301 P19302 P19303 P19304 P19305 P19306
P19307 P19308 P19309 P19310 P19311 P19312 P19314 P19315 P19335 P19348 P19351
P19352 P19356 P19358 P19359 P19360 P19363 P19364 P19368 P19371 P19374 P19375
P19376 P19379 P19382 P19385 P19386 P19387 P19388 P19389 P19392 P19414 P19415
P19416 P19417 P19418 P19421 P19422 P19429 P19432 P19433 P19434 P19435 P19436
P19441 P19446 P19452 P19454 P19457 P19458 P19459 P19460 P19461 P19463 P19464
P19465 P19466 P19471 P19475 P19480 P19486 P19496 P19497 P19498 P19499 P19500
P19509 P19510 P19514 P19520 P19522 P19527 P19529 P19537 P19538 P19541 P19543
P19563 P19572 P19574 P19582 P19583 P19587 P19592 P19593 P19595 P19601 P19617
P19623 P19624 P19625 P19626 P19629 P19631 P19632 P19636 P19639 P19645 P19646
P19650 P19653 P19654 P19655 P19662 P19663 P19664 P19665 P19666 P19668 P19669
P19670 P19672 P19679 P19680 P19682 P19683 P19684 P19685 P19688 P19689 P19691
P19692 P19693 P19704 P19705 P19709 P19710 P19715 P19717 P19721 P19722 P19723
P19724 P19726 P19727 P19728 P19729 P19730 P19731 P19732 P19733 P19734 P19735
P19736 P19737 P19738 P19740 P19745 P19747 P19748 P19749 P19753 P19755 P19757
P19759 P19760 P19763 P19765 P19766 P19767 P19768 P19769 P19770 P19771 P19772
P19775 P19778 P19779 P19780 P19782 P19786 P19789 P19791 P19800 P19808 P19810
P19812 P19815 P19818 P19821 P19822 P19824 P19831 P19832 P19834 P19839 P19840
P19848 P19852 P19853 P19858 P19860 P19862 P19864 P19865 P19866 P19867 P19868
P19870 P19873 P19881 P19885 P19886 P19887 P19888 P19891 P19893 P19894 P19895
P19896 P19897 P19898 P19899 P19900 P19904 P19907 P19908 P19910 P19911 P19912
P19913 P19914 P19915 P19916 P19917 P19918 P19919 P19920 P19921 P19922 P19923
P19924 P19930 P19931 P19932 P19936 P19937 P19938 P19940 P19943 P19946 P19947
P19948 P19950 P19951 P19952 P19953 P19954 P19955 P19962 P19966 P19971 P19975
P19976 P19977 P19978 P19979 P19980 P19981 P19982 P19983 P19984 P19985 P19986
P19991 P19993 P19994 P19997 P20000 P20004 P20009 P20011 P20012 P20015 P20016
P20018 P20019 P20024 P20025 P20028 P20030 P20031 P20032 P20035 P20044 P20045
P20046 P20047 P20049 P20050 P20051 P20053 P20054 P20065 P20074 P20077 P20080
P20081 P20083 P20085 P20086 P20087 P20088 P20089 P20090 P20095 P20098 P20102
P20104 P20105 P20110 P20111 P20115 P20119 P20123 P20124 P20125 P20126 P20127
P20129 P20130 P20131 P20134 P20135 P20137 P20139 P20140 P20141 P20144 P20154
P20158 P20159 P20161 P20167 P20170 P20173 P20174 P20177 P20178 P20179 P20180
P20182 P20183 P20184 P20185 P20186 P20187 P20188 P20189 P20190 P20191 P20193
P20194 P20195 P20196 P20197 P20198 P20199 P20200 P20201 P20202 P20203 P20204
P20205 P20206 P20207 P20208 P20209 P20210 P20211 P20212 P20213 P20214 P20215
P20216 P20217 P20218 P20219 P20220 P20221 P20222 P20224 P20226 P20227 P20228
P20230 P20232 P20234 P20235 P20238 P20243 P20244 P20245 P20246 P20247 P20263

P20266 P20267 P20268 P20269 P20270 P20271 P20275 P20276 P20277 P20278 P20279
P20280 P20281 P20282 P20284 P20287 P20296 P20299 P20300 P20301 P20302 P20304
P20307 P20308 P20311 P20312 P20313 P20314 P20315 P20316 P20317 P20318 P20319
P20320 P20321 P20322 P20323 P20324 P20325 P20326 P20327 P20328 P20329 P20330
P20342 P20343 P20345 P20348 P20349 P20351 P20356 P20359 P20360 P20364 P20365
P20368 P20369 P20370 P20371 P20372 P20373 P20376 P20378 P20384 P20385 P20389
P20397 P20399 P20400 P20402 P20403 P20405 P20406 P20409 P20412 P20413 P20418
P20421 P20425 P20427 P20429 P20430 P20432 P20433 P20434 P20435 P20436 P20437
P20438 P20439 P20441 P20442 P20445 P20447 P20448 P20455 P20457 P20458 P20459
P20461 P20463 P20464 P20467 P20470 P20472 P20473 P20477 P20478 P20479 P20480
P20482 P20483 P20486 P20492 P20493 P20495 P20498 P20499 P20502 P20503 P20504
P20505 P20506 P20509 P20510 P20511 P20512 P20513 P20514 P20515 P20516 P20517
P20518 P20519 P20520 P20523 P20524 P20525 P20526 P20527 P20530 P20531 P20534
P20536 P20537 P20538 P20542 P20543 P20546 P20547 P20548 P20549 P20550 P20551
P20552 P20553 P20554 P20555 P20556 P20557 P20558 P20559 P20560 P20561 P20562
P20563 P20564 P20565 P20566 P20567 P20568 P20569 P20570 P20571 P20574 P20575
P20577 P20578 P20579 P20583 P20586 P20587 P20588 P20589 P20590 P20593 P20596
P20605 P20608 P20615 P20617 P20618 P20619 P20620 P20621 P20622 P20623 P20624
P20627 P20628 P20629 P20633 P20634 P20635 P20636 P20637 P20639 P20640 P20641
P20642 P20643 P20654 P20655 P20658 P20659 P20664 P20665 P20666 P20667 P20668
P20669 P20689 P20690 P20691 P20692 P20703 P20704 P20706 P20707 P20708 P20709
P20710 P20711 P20712 P20714 P20717 P20719 P20721 P20728 P20731 P20732 P20733
P20734 P20743 P20744 P20745 P20746 P20748 P20751 P20758 P20763 P20768 P20771
P20772 P20780 P20784 P20801 P20802 P20805 P20811 P20819 P20821 P20822 P20824
P20825 P20828 P20829 P20830 P20831 P20833 P20835 P20836 P20837 P20839 P20841
P20842 P20847 P20854 P20855 P20856 P20857 P20860 P20884 P20894 P20901 P20902
P20903 P20904 P20906 P20907 P20911 P20912 P20913 P20914 P20930 P20941 P20942
P20945 P20951 P20952 P20955 P20958 P20960 P20964 P20965 P20967 P20969 P20970
P20971 P20972 P20973 P20979 P20980 P20981 P20982 P20985 P20986 P20989 P20992
P20994 P20995 P20997 P20998 P20999 P21000 P21001 P21007 P21008 P21009 P21010
P21011 P21012 P21013 P21016 P21017 P21019 P21020 P21021 P21022 P21026 P21027
P21028 P21031 P21033 P21034 P21036 P21037 P21038 P21039 P21040 P21042 P21043
P21044 P21045 P21046 P21047 P21052 P21054 P21060 P21066 P21067 P21068 P21069
P21073 P21074 P21075 P21076 P21078 P21079 P21080 P21081 P21082 P21084 P21085
P21087 P21089 P21095 P21096 P21098 P21099 P21100 P21101 P21102 P21103 P21104
P21105 P21106 P21107 P21108 P21110 P21111 P21112 P21113 P21114 P21117 P21119
P21120 P21121 P21122 P21123 P21132 P21133 P21134 P21139 P21148 P21149 P21152
P21153 P21154 P21156 P21157 P21159 P21161 P21162 P21164 P21165 P21169 P21170
P21172 P21173 P21174 P21179 P21182 P21186 P21187 P21189 P21190 P21192 P21193
P21197 P21198 P21199 P21200 P21201 P21203 P21204 P21207 P21212 P21213 P21216
P21219 P21220 P21222 P21225 P21227 P21228 P21232 P21233 P21234 P21236 P21238
P21239 P21240 P21241 P21244 P21248 P21249 P21251 P21254 P21256 P21258 P21260
P21263 P21264 P21265 P21266 P21269 P21270 P21277 P21286 P21287 P21288 P21290
P21293 P21294 P21295 P21296 P21297 P21298 P21299 P21302 P21304 P21305 P21308

P21310 P21311 P21312 P21314 P21315 P21316 P21318 P21320 P21321 P21322 P21323
P21324 P21325 P21328 P21329 P21330 P21331 P21332 P21335 P21336 P21337 P21340
P21341 P21342 P21343 P21344 P21348 P21357 P21358 P21360 P21361 P21362 P21363
P21368 P21369 P21370 P21373 P21374 P21375 P21376 P21379 P21380 P21394 P21399
P21402 P21403 P21406 P21417 P21418 P21421 P21422 P21423 P21424 P21434 P21437
P21438 P21442 P21455 P21461 P21464 P21465 P21466 P21467 P21468 P21469 P21470
P21471 P21472 P21473 P21475 P21476 P21477 P21478 P21479 P21481 P21499 P21505
P21506 P21507 P21508 P21509 P21510 P21511 P21512 P21514 P21517 P21518 P21519
P21522 P21523 P21524 P21525 P21528 P21532 P21538 P21541 P21542 P21547 P21553
P21557 P21558 P21561 P21568 P21569 P21572 P21574 P21577 P21578 P21593 P21594
P21595 P21596 P21597 P21598 P21599 P21601 P21603 P21604 P21605 P21606 P21607
P21609 P21610 P21613 P21615 P21622 P21623 P21631 P21632 P21633 P21635 P21636
P21637 P21638 P21639 P21640 P21643 P21650 P21651 P21656 P21657 P21659 P21660
P21666 P21667 P21668 P21670 P21672 P21673 P21678 P21679 P21680 P21681 P21682
P21683 P21684 P21686 P21688 P21689 P21690 P21691 P21693 P21695 P21696 P21698
P21699 P21700 P21701 P21703 P21705 P21711 P21725 P21726 P21732 P21733 P21735
P21738 P21739 P21740 P21752 P21753 P21762 P21763 P21766 P21767 P21768 P21769
P21770 P21772 P21777 P21778 P21787 P21788 P21794 P21795 P21797 P21798 P21799
P21800 P21816 P21820 P21829 P21833 P21834 P21843 P21846 P21852 P21861 P21866
P21867 P21868 P21869 P21871 P21872 P21873 P21874 P21878 P21879 P21880 P21881
P21882 P21883 P21885 P21886 P21887 P21888 P21889 P21893 P21894 P21895 P21896
P21897 P21898 P21899 P21900 P21901 P21907 P21908 P21909 P21920 P21921 P21922
P21923 P21924 P21925 P21937 P21938 P21939 P21940 P21943 P21947 P21951 P21952
P21954 P21955 P21966 P21967 P21968 P21969 P21970 P21971 P21972 P21973 P21974
P21975 P21977 P21983 P21984 P21985 P21986 P21987 P21988 P21989 P21990 P21991
P21992 P21996 P21999 P22008 P22011 P22012 P22013 P22014 P22022 P22025 P22026
P22031 P22033 P22034 P22035 P22038 P22039 P22040 P22041 P22042 P22043 P22044
P22047 P22048 P22049 P22050 P22051 P22058 P22061 P22065 P22066 P22069 P22073
P22075 P22076 P22080 P22081 P22082 P22095 P22096 P22097 P22098 P22099 P22100
P22101 P22102 P22103 P22106 P22108 P22114 P22115 P22116 P22117 P22118 P22119
P22120 P22121 P22130 P22131 P22132 P22133 P22134 P22136 P22138 P22139 P22143
P22144 P22149 P22151 P22155 P22156 P22157 P22158 P22159 P22162 P22163 P22166
P22168 P22169 P22172 P22173 P22174 P22175 P22176 P22177 P22178 P22186 P22188
P22190 P22191 P22192 P22193 P22194 P22197 P22198 P22200 P22209 P22216 P22220
P22221 P22225 P22227 P22229 P22230 P22231 P22232 P22233 P22238 P22239 P22240
P22241 P22244 P22245 P22246 P22248 P22249 P22250 P22253 P22259 P22262 P22271
P22275 P22276 P22281 P22291 P22293 P22294 P22296 P22299 P22302 P22305 P22308
P22317 P22319 P22320 P22321 P22322 P22326 P22333 P22335 P22336 P22339 P22341
P22344 P22346 P22347 P22348 P22349 P22350 P22351 P22358 P22359 P22360 P22367
P22368 P22369 P22370 P22371 P22372 P22373 P22374 P22375 P22376 P22377 P22385
P22392 P22399 P22400 P22401 P22402 P22403 P22404 P22405 P22406 P22416 P22418
P22420 P22421 P22422 P22424 P22431 P22433 P22434 P22438 P22439 P22441 P22442
P22446 P22447 P22448 P22450 P22452 P22464 P22468 P22469 P22470 P22472 P22473
P22474 P22486 P22488 P22492 P22494 P22495 P22496 P22498 P22499 P22501 P22502

P22505 P22508 P22509 P22510 P22515 P22516 P22517 P22524 P22527 P22537 P22544
P22547 P22548 P22555 P22556 P22558 P22562 P22564 P22566 P22567 P22572 P22573
P22574 P22577 P22578 P22579 P22580 P22581 P22582 P22583 P22584 P22585 P22587
P22588 P22589 P22590 P22591 P22592 P22601 P22602 P22605 P22611 P22613 P22615
P22616 P22623 P22624 P22625 P22634 P22635 P22636 P22639 P22643 P22644 P22645
P22647 P22649 P22654 P22657 P22658 P22659 P22660 P22661 P22665 P22672 P22674
P22675 P22676 P22677 P22678 P22679 P22696 P22697 P22698 P22701 P22702 P22703
P22704 P22705 P22706 P22707 P22711 P22713 P22714 P22715 P22737 P22738 P22740
P22741 P22742 P22743 P22744 P22745 P22746 P22753 P22754 P22755 P22759 P22761
P22764 P22765 P22766 P22768 P22769 P22772 P22773 P22780 P22781 P22782 P22787
P22793 P22795 P22797 P22798 P22799 P22800 P22805 P22806 P22807 P22808 P22809
P22810 P22811 P22813 P22814 P22816 P22818 P22822 P22823 P22824 P22835 P22836
P22837 P22838 P22839 P22840 P22841 P22842 P22843 P22844 P22845 P22846 P22849
P22850 P22851 P22852 P22853 P22857 P22859 P22860 P22862 P22866 P22867 P22868
P22871 P22872 P22873 P22874 P22875 P22876 P22877 P22878 P22879 P22880 P22884
P22886 P22896 P22898 P22899 P22900 P22901 P22902 P22903 P22904 P22906 P22907
P22914 P22915 P22916 P22917 P22919 P22920 P22921 P22924 P22925 P22926 P22927
P22928 P22931 P22936 P22939 P22944 P22945 P22946 P22948 P22949 P22950 P22955
P22958 P22959 P22960 P22962 P22970 P22971 P22972 P22973 P22974 P22975 P22976
P22978 P22979 P22980 P22983 P22984 P22988 P22989 P22991 P22993 P22994 P22995
P22996 P23001 P23002 P23003 P23005 P23007 P23009 P23010 P23011 P23012 P23013
P23014 P23015 P23016 P23017 P23018 P23019 P23020 P23021 P23023 P23029 P23032
P23033 P23034 P23038 P23046 P23047 P23048 P23050 P23051 P23053 P23055 P23056
P23057 P23059 P23060 P23065 P23070 P23071 P23072 P23076 P23078 P23081 P23084
P23085 P23086 P23091 P23092 P23098 P23099 P23100 P23101 P23102 P23103 P23105
P23106 P23107 P23109 P23112 P23116 P23119 P23120 P23121 P23122 P23123 P23124
P23125 P23128 P23129 P23130 P23133 P23138 P23139 P23140 P23145 P23146 P23147
P23149 P23153 P23154 P23155 P23156 P23157 P23158 P23159 P23160 P23161 P23162
P23163 P23164 P23165 P23166 P23167 P23168 P23170 P23171 P23172 P23177 P23179
P23180 P23181 P23182 P23183 P23184 P23185 P23186 P23189 P23190 P23191 P23192
P23198 P23201 P23202 P23205 P23208 P23209 P23210 P23211 P23212 P23216 P23225
P23226 P23234 P23236 P23237 P23238 P23241 P23244 P23247 P23249 P23251 P23252
P23254 P23255 P23256 P23257 P23261 P23262 P23263 P23277 P23278 P23279 P23281
P23283 P23286 P23287 P23293 P23295 P23296 P23300 P23301 P23302 P23303 P23305
P23307 P23308 P23309 P23310 P23311 P23312 P23315 P23320 P23324 P23325 P23326
P23330 P23331 P23333 P23334 P23335 P23337 P23339 P23341 P23342 P23343 P23344
P23345 P23346 P23349 P23350 P23351 P23353 P23354 P23357 P23361 P23364 P23367
P23370 P23371 P23374 P23375 P23376 P23379 P23386 P23390 P23391 P23393 P23394
P23395 P23397 P23398 P23400 P23401 P23402 P23403 P23404 P23405 P23406 P23407
P23408 P23409 P23410 P23412 P23417 P23418 P23419 P23428 P23429 P23430 P23434
P23437 P23441 P23444 P23445 P23446 P23449 P23451 P23454 P23455 P23456 P23457
P23459 P23460 P23461 P23463 P23474 P23476 P23477 P23478 P23479 P23481 P23487
P23489 P23490 P23492 P23493 P23494 P23495 P23496 P23509 P23511 P23512 P23522
P23524 P23526 P23527 P23535 P23538 P23541 P23542 P23543 P23544 P23548 P23552

P23553 P23554 P23555 P23556 P23557 P23558 P23565 P23566 P23569 P23570 P23571
P23572 P23575 P23579 P23580 P23581 P23583 P23587 P23589 P23591 P23594 P23595
P23599 P23600 P23601 P23602 P23607 P23608 P23611 P23612 P23617 P23618 P23619
P23620 P23623 P23626 P23627 P23628 P23629 P23630 P23632 P23635 P23636 P23637
P23647 P23651 P23652 P23656 P23657 P23660 P23666 P23668 P23669 P23670 P23672
P23673 P23679 P23681 P23682 P23683 P23686 P23687 P23688 P23689 P23690 P23691
P23692 P23693 P23696 P23706 P23708 P23712 P23715 P23721 P23722 P23724 P23725
P23729 P23730 P23731 P23733 P23734 P23736 P23737 P23738 P23740 P23741 P23742
P23744 P23748 P23750 P23751 P23753 P23754 P23755 P23756 P23757 P23758 P23759
P23760 P23762 P23767 P23768 P23770 P23773 P23774 P23777 P23778 P23779 P23788
P23789 P23790 P23792 P23793 P23796 P23799 P23801 P23803 P23808 P23812 P23813
P23822 P23823 P23824 P23826 P23828 P23829 P23832 P23835 P23839 P23840 P23841
P23845 P23848 P23862 P23869 P23871 P23872 P23873 P23874 P23880 P23881 P23882
P23883 P23893 P23899 P23901 P23906 P23908 P23909 P23914 P23915 P23916 P23917
P23918 P23919 P23920 P23921 P23922 P23923 P23926 P23939 P23940 P23941 P23949
P23955 P23966 P23970 P23971 P23973 P23974 P23981 P23983 P23984 P23985 P23986
P23987 P23989 P23991 P23992 P23995 P23996 P23999 P24002 P24005 P24007 P24015
P24018 P24022 P24024 P24030 P24032 P24033 P24034 P24035 P24042 P24044 P24056
P24057 P24058 P24060 P24061 P24064 P24066 P24068 P24069 P24070 P24074 P24075
P24076 P24078 P24079 P24080 P24081 P24087 P24094 P24096 P24098 P24099 P24111
P24112 P24113 P24114 P24116 P24117 P24118 P24124 P24125 P24130 P24131 P24134
P24146 P24147 P24150 P24154 P24157 P24163 P24164 P24166 P24168 P24169 P24174
P24175 P24178 P24186 P24188 P24192 P24193 P24197 P24200 P24202 P24203 P24211
P24215 P24218 P24220 P24221 P24222 P24226 P24230 P24238 P24240 P24242 P24252
P24260 P24261 P24263 P24264 P24265 P24267 P24269 P24273 P24274 P24276 P24277
P24278 P24279 P24281 P24283 P24286 P24290 P24291 P24292 P24295 P24297 P24298
P24299 P24300 P24301 P24304 P24309 P24312 P24313 P24314 P24315 P24316 P24319
P24320 P24321 P24322 P24323 P24324 P24325 P24329 P24339 P24340 P24341 P24342
P24343 P24344 P24349 P24350 P24352 P24353 P24354 P24355 P24356 P24357 P24359
P24360 P24362 P24371 P24372 P24375 P24378 P24381 P24382 P24384 P24395 P24397
P24399 P24401 P24402 P24403 P24404 P24417 P24418 P24421 P24423 P24424 P24425
P24427 P24428 P24429 P24431 P24432 P24434 P24435 P24436 P24437 P24438 P24439
P24440 P24441 P24442 P24443 P24444 P24445 P24446 P24447 P24466 P24467 P24468
P24472 P24473 P24475 P24477 P24479 P24480 P24481 P24482 P24486 P24488 P24493
P24494 P24495 P24496 P24497 P24500 P24501 P24502 P24508 P24509 P24517 P24519
P24525 P24528 P24531 P24532 P24536 P24537 P24540 P24542 P24543 P24546 P24547
P24554 P24555 P24558 P24562 P24566 P24567 P24569 P24575 P24576 P24577 P24580
P24581 P24582 P24589 P24590 P24598 P24599 P24600 P24601 P24602 P24607 P24610
P24613 P24617 P24618 P24619 P24620 P24621 P24622 P24623 P24624 P24629 P24630
P24631 P24632 P24633 P24634 P24635 P24637 P24641 P24642 P24646 P24647 P24648
P24649 P24650 P24651 P24652 P24654 P24655 P24656 P24657 P24658 P24659 P24660
P24661 P24664 P24669 P24670 P24671 P24672 P24673 P24674 P24675 P24676 P24677
P24678 P24679 P24680 P24681 P24682 P24683 P24685 P24694 P24696 P24698 P24699
P24700 P24701 P24703 P24704 P24705 P24706 P24707 P24708 P24709 P24710 P24711

P24712 P24713 P24714 P24715 P24716 P24718 P24719 P24720 P24722 P24730 P24731
P24732 P24733 P24734 P24743 P24744 P24745 P24748 P24750 P24751 P24753 P24757
P24760 P24762 P24768 P24769 P24772 P24773 P24782 P24783 P24784 P24785 P24788
P24789 P24790 P24793 P24795 P24796 P24804 P24805 P24808 P24809 P24810 P24812
P24819 P24820 P24824 P24825 P24826 P24827 P24828 P24830 P24831 P24832 P24833
P24834 P24835 P24836 P24838 P24842 P24844 P24845 P24846 P24847 P24850 P24852
P24861 P24862 P24863 P24864 P24865 P24868 P24869 P24870 P24871 P24898 P24899
P24900 P24902 P24907 P24909 P24910 P24911 P24912 P24913 P24914 P24915 P24920
P24922 P24923 P24924 P24925 P24927 P24929 P24931 P24932 P24933 P24936 P24938
P24939 P24940 P25008 P25009 P25010 P25011 P25012 P25013 P25015 P25016 P25017
P25018 P25019 P25022 P25026 P25027 P25030 P25032 P25034 P25038 P25039 P25040
P25042 P25044 P25046 P25052 P25053 P25055 P25066 P25070 P25071 P25077 P25078
P25079 P25080 P25081 P25082 P25084 P25096 P25097 P25113 P25114 P25124 P25125
P25126 P25127 P25128 P25132 P25133 P25134 P25135 P25136 P25137 P25139 P25141
P25143 P25144 P25145 P25148 P25150 P25151 P25154 P25156 P25158 P25161 P25165
P25166 P25167 P25170 P25171 P25172 P25177 P25183 P25186 P25201 P25202 P25203
P25204 P25206 P25207 P25208 P25209 P25210 P25211 P25214 P25215 P25217 P25219
P25220 P25229 P25233 P25237 P25238 P25239 P25240 P25241 P25242 P25243 P25244
P25245 P25246 P25247 P25251 P25257 P25258 P25259 P25260 P25261 P25262 P25263
P25264 P25265 P25266 P25267 P25269 P25270 P25277 P25278 P25279 P25280 P25281
P25282 P25283 P25287 P25293 P25294 P25298 P25301 P25302 P25306 P25309 P25313
P25314 P25315 P25316 P25317 P25319 P25320 P25323 P25324 P25325 P25327 P25328
P25332 P25335 P25336 P25339 P25345 P25347 P25350 P25352 P25355 P25357 P25359
P25361 P25362 P25364 P25366 P25367 P25368 P25369 P25370 P25372 P25373 P25374
P25375 P25377 P25379 P25382 P25383 P25387 P25393 P25405 P25406 P25407 P25408
P25409 P25413 P25414 P25415 P25420 P25426 P25436 P25437 P25438 P25439 P25453
P25454 P25456 P25457 P25458 P25459 P25460 P25461 P25462 P25469 P25470 P25475
P25476 P25477 P25478 P25479 P25480 P25482 P25483 P25484 P25486 P25499 P25502
P25503 P25516 P25524 P25531 P25536 P25539 P25544 P25545 P25550 P25552 P25553
P25554 P25555 P25558 P25559 P25561 P25562 P25563 P25566 P25567 P25569 P25571
P25572 P25575 P25576 P25579 P25582 P25583 P25586 P25588 P25593 P25600 P25601
P25602 P25603 P25605 P25608 P25611 P25612 P25614 P25615 P25616 P25617 P25620
P25629 P25630 P25631 P25632 P25635 P25637 P25638 P25640 P25644 P25646 P25649
P25650 P25651 P25652 P25654 P25655 P25657 P25659 P25665 P25666 P25685 P25690
P25691 P25692 P25698 P25699 P25700 P25704 P25719 P25720 P25721 P25724 P25728
P25734 P25735 P25738 P25745 P25748 P25749 P25750 P25756 P25762 P25767 P25768
P25769 P25770 P25771 P25772 P25791 P25795 P25800 P25801 P25806 P25808 P25810
P25811 P25812 P25813 P25814 P25816 P25817 P25820 P25821 P25822 P25823 P25824
P25825 P25826 P25827 P25828 P25829 P25830 P25831 P25832 P25833 P25834 P25835
P25836 P25837 P25838 P25839 P25840 P25843 P25846 P25847 P25851 P25853 P25855
P25860 P25861 P25862 P25863 P25864 P25865 P25866 P25867 P25869 P25872 P25873
P25875 P25879 P25888 P25889 P25890 P25891 P25892 P25906 P25907 P25908 P25915
P25919 P25920 P25921 P25923 P25924 P25932 P25933 P25934 P25938 P25939 P25947
P25948 P25949 P25950 P25951 P25952 P25969 P25970 P25971 P25972 P25973 P25976

P25977 P25979 P25980 P25981 P25983 P25984 P25985 P25986 P25987 P25988 P25991
P25993 P25994 P25995 P25996 P25997 P25998 P25999 P26000 P26001 P26002 P26003
P26004 P26005 P26014 P26017 P26018 P26026 P26027 P26029 P26030 P26033 P26035
P26036 P26037 P26144 P26145 P26146 P26154 P26156 P26157 P26158 P26159 P26160
P26161 P26162 P26163 P26164 P26165 P26166 P26167 P26168 P26169 P26172 P26173
P26174 P26175 P26177 P26179 P26180 P26181 P26182 P26183 P26184 P26186 P26187
P26188 P26189 P26190 P26191 P26192 P26194 P26195 P26197 P26198 P26199 P26200
P26206 P26208 P26209 P26210 P26211 P26212 P26223 P26226 P26227 P26228 P26236
P26238 P26239 P26242 P26247 P26248 P26249 P26250 P26251 P26252 P26254 P26256
P26257 P26258 P26259 P26263 P26267 P26268 P26269 P26270 P26275 P26277 P26281
P26283 P26284 P26285 P26297 P26298 P26300 P26301 P26302 P26306 P26307 P26311
P26315 P26317 P26318 P26323 P26325 P26337 P26340 P26341 P26343 P26344 P26346
P26348 P26350 P26351 P26352 P26353 P26354 P26355 P26356 P26357 P26359 P26369
P26371 P26372 P26375 P26388 P26409 P26410 P26411 P26412 P26413 P26416 P26420
P26421 P26422 P26423 P26424 P26425 P26444 P26448 P26449 P26462 P26464 P26474
P26475 P26477 P26480 P26481 P26482 P26483 P26484 P26485 P26486 P26488 P26490
P26491 P26492 P26494 P26496 P26497 P26498 P26499 P26500 P26503 P26505 P26506
P26507 P26508 P26512 P26516 P26517 P26518 P26519 P26520 P26521 P26535 P26536
P26537 P26541 P26545 P26546 P26547 P26548 P26549 P26550 P26551 P26552 P26553
P26554 P26555 P26556 P26563 P26564 P26565 P26566 P26567 P26568 P26569 P26571
P26573 P26574 P26575 P26576 P26577 P26578 P26585 P26586 P26589 P26590 P26593
P26596 P26600 P26608 P26609 P26612 P26613 P26616 P26620 P26624 P26625 P26630
P26632 P26633 P26634 P26635 P26637 P26639 P26640 P26642 P26643 P26646 P26649
P26658 P26659 P26667 P26669 P26673 P26676 P26683 P26685 P26686 P26688 P26690
P26692 P26693 P26698 P26699 P26700 P26701 P26711 P26712 P26713 P26719 P26720
P26721 P26722 P26723 P26724 P26745 P26746 P26747 P26748 P26749 P26750 P26751
P26752 P26754 P26755 P26764 P26767 P26768 P26772 P26777 P26778 P26788 P26791
P26793 P26795 P26796 P26797 P26799 P26800 P26801 P26802 P26811 P26812 P26813
P26815 P26816 P26820 P26821 P26822 P26823 P26825 P26826 P26830 P26832 P26833
P26834 P26836 P26837 P26838 P26839 P26840 P26841 P26859 P26860 P26861 P26862
P26863 P26864 P26865 P26866 P26867 P26868 P26869 P26870 P26871 P26872 P26873
P26874 P26875 P26879 P26880 P26881 P26899 P26901 P26902 P26908 P26909 P26910
P26911 P26915 P26916 P26919 P26920 P26921 P26922 P26923 P26924 P26925 P26929
P26930 P26931 P26935 P26938 P26939 P26940 P26941 P26942 P26943 P26944 P26945
P26946 P26947 P26950 P26958 P26959 P26960 P26961 P26962 P26963 P26964 P26969
P26970 P26971 P26977 P26978 P26983 P26984 P26985 P26987 P26988 P26993 P26995
P26996 P26997 P26998 P27000 P27001 P27002 P27003 P27004 P27009 P27010 P27011
P27012 P27013 P27014 P27015 P27016 P27017 P27018 P27027 P27030 P27031 P27047
P27048 P27055 P27059 P27063 P27064 P27065 P27066 P27067 P27068 P27069 P27070
P27071 P27072 P27073 P27074 P27075 P27076 P27077 P27078 P27079 P27082 P27084
P27086 P27087 P27095 P27096 P27097 P27098 P27099 P27100 P27101 P27102 P27107
P27111 P27116 P27117 P27118 P27119 P27120 P27121 P27124 P27130 P27131 P27132
P27134 P27136 P27137 P27138 P27141 P27142 P27143 P27144 P27145 P27150 P27151
P27152 P27153 P27154 P27155 P27156 P27157 P27158 P27159 P27161 P27163 P27164

P27165 P27166 P27172 P27176 P27186 P27187 P27188 P27189 P27190 P27192 P27193
P27194 P27199 P27203 P27204 P27205 P27207 P27208 P27209 P27210 P27211 P27213
P27214 P27217 P27222 P27223 P27224 P27225 P27226 P27227 P27228 P27229 P27232
P27233 P27238 P27245 P27248 P27253 P27257 P27258 P27259 P27260 P27264 P27265
P27266 P27269 P27272 P27276 P27279 P27280 P27281 P27294 P27296 P27297 P27302
P27305 P27306 P27309 P27310 P27311 P27314 P27316 P27317 P27318 P27319 P27320
P27322 P27325 P27326 P27327 P27328 P27329 P27330 P27334 P27335 P27336 P27342
P27344 P27345 P27346 P27347 P27356 P27362 P27363 P27368 P27369 P27371 P27375
P27394 P27396 P27397 P27398 P27403 P27404 P27405 P27406 P27407 P27408 P27409
P27410 P27411 P27412 P27413 P27417 P27418 P27419 P27421 P27422 P27431 P27438
P27439 P27440 P27442 P27443 P27454 P27456 P27459 P27462 P27464 P27472 P27473
P27476 P27482 P27484 P27486 P27488 P27499 P27500 P27501 P27503 P27504 P27505
P27506 P27507 P27508 P27509 P27510 P27515 P27526 P27527 P27532 P27536 P27538
P27541 P27543 P27546 P27550 P27552 P27553 P27554 P27555 P27557 P27560 P27568
P27569 P27570 P27574 P27577 P27578 P27579 P27581 P27585 P27586 P27588 P27592
P27598 P27602 P27603 P27604 P27605 P27608 P27609 P27610 P27616 P27622 P27623
P27624 P27625 P27626 P27627 P27629 P27630 P27632 P27633 P27634 P27637 P27639
P27640 P27642 P27643 P27649 P27651 P27652 P27653 P27660 P27663 P27672 P27673
P27679 P27683 P27684 P27686 P27687 P27692 P27696 P27699 P27705 P27709 P27710
P27711 P27714 P27718 P27722 P27723 P27725 P27726 P27727 P27730 P27735 P27736
P27737 P27738 P27739 P27744 P27745 P27746 P27747 P27749 P27750 P27751 P27753
P27754 P27756 P27763 P27764 P27765 P27766 P27768 P27770 P27771 P27774 P27775
P27777 P27782 P27783 P27785 P27787 P27788 P27789 P27790 P27792 P27793 P27795
P27800 P27802 P27803 P27804 P27805 P27806 P27807 P27818 P27819 P27838 P27840
P27848 P27859 P27862 P27864 P27865 P27866 P27868 P27871 P27872 P27873 P27874
P27875 P27876 P27879 P27880 P27882 P27887 P27888 P27891 P27892 P27893 P27894
P27895 P27897 P27898 P27901 P27903 P27905 P27921 P27923 P27926 P27927 P27928
P27942 P27943 P27946 P27947 P27948 P27949 P27950 P27962 P27964 P27966 P27976
P27984 P27988 P27989 P27990 P27991 P27992 P27993 P27994 P27995 P27996 P27997
P27998 P27999 P28000 P28002 P28003 P28004 P28005 P28006 P28007 P28009 P28010
P28011 P28012 P28013 P28014 P28015 P28016 P28019 P28020 P28021 P28022 P28024
P28025 P28032 P28034 P28037 P28038 P28040 P28042 P28043 P28044 P28045 P28046
P28054 P28055 P28061 P28062 P28063 P28064 P28065 P28069 P28072 P28073 P28074
P28075 P28076 P28077 P28079 P28080 P28147 P28148 P28149 P28150 P28151 P28152
P28153 P28154 P28155 P28156 P28157 P28166 P28167 P28173 P28174 P28181 P28183
P28191 P28193 P28194 P28196 P28197 P28199 P28200 P28201 P28202 P28203 P28204
P28205 P28206 P28207 P28208 P28209 P28210 P28211 P28212 P28213 P28214 P28215
P28216 P28217 P28218 P28219 P28227 P28237 P28241 P28243 P28247 P28248 P28256
P28258 P28259 P28260 P28261 P28262 P28263 P28266 P28267 P28268 P28269 P28270
P28274 P28275 P28277 P28279 P28280 P28281 P28284 P28287 P28295 P28298 P28299
P28304 P28305 P28318 P28322 P28329 P28332 P28337 P28338 P28342 P28343 P28348
P28349 P28350 P28354 P28355 P28356 P28357 P28358 P28359 P28361 P28362 P28363
P28364 P28365 P28367 P28368 P28369 P28371 P28372 P28373 P28376 P28377 P28378
P28379 P28380 P28381 P28382 P28383 P28384 P28385 P28386 P28387 P28388 P28389

P28390 P28391 P28392 P28393 P28394 P28395 P28396 P28397 P28398 P28399 P28400
P28401 P28402 P28403 P28404 P28405 P28406 P28407 P28408 P28409 P28410 P28411
P28412 P28413 P28414 P28415 P28416 P28417 P28418 P28419 P28420 P28421 P28422
P28423 P28424 P28425 P28426 P28427 P28428 P28429 P28430 P28431 P28432 P28433
P28434 P28435 P28436 P28437 P28438 P28439 P28440 P28441 P28442 P28443 P28444
P28445 P28446 P28447 P28448 P28449 P28450 P28451 P28452 P28453 P28454 P28455
P28456 P28457 P28458 P28459 P28460 P28461 P28462 P28463 P28464 P28467 P28468
P28469 P28475 P28478 P28480 P28483 P28484 P28485 P28486 P28487 P28488 P28492
P28515 P28516 P28519 P28520 P28521 P28522 P28523 P28524 P28528 P28547 P28551
P28562 P28567 P28574 P28575 P28577 P28578 P28579 P28580 P28586 P28587 P28588
P28593 P28594 P28595 P28598 P28599 P28600 P28601 P28602 P28603 P28604 P28605
P28606 P28609 P28610 P28613 P28615 P28616 P28617 P28618 P28619 P28624 P28629
P28630 P28631 P28632 P28634 P28638 P28639 P28640 P28641 P28644 P28651 P28659
P28661 P28668 P28669 P28670 P28671 P28697 P28698 P28701 P28702 P28704 P28705
P28706 P28707 P28709 P28710 P28711 P28718 P28719 P28720 P28723 P28724 P28725
P28727 P28734 P28735 P28736 P28738 P28739 P28742 P28743 P28748 P28750 P28752
P28753 P28755 P28757 P28758 P28759 P28760 P28761 P28762 P28763 P28764 P28765
P28766 P28767 P28768 P28769 P28776 P28777 P28779 P28780 P28781 P28786 P28787
P28789 P28796 P28803 P28804 P28805 P28807 P28808 P28809 P28811 P28812 P28813
P28814 P28815 P28816 P28819 P28820 P28821 P28822 P28823 P28830 P28831 P28832
P28833 P28834 P28836 P28839 P28844 P28846 P28848 P28851 P28852 P28853 P28854
P28855 P28857 P28858 P28859 P28861 P28866 P28870 P28875 P28887 P28888 P28889
P28890 P28892 P28893 P28894 P28896 P28897 P28899 P28901 P28903 P28904 P28905
P28911 P28912 P28916 P28917 P28918 P28919 P28920 P28921 P28925 P28926 P28927
P28928 P28932 P28934 P28935 P28938 P28939 P28942 P28943 P28944 P28945 P28946
P28947 P28949 P28950 P28953 P28956 P28957 P28962 P28964 P28965 P28970 P28972
P28974 P28975 P28976 P28978 P28979 P28983 P28985 P28988 P28989 P28990 P28996
P28997 P28998 P29010 P29012 P29013 P29035 P29036 P29037 P29038 P29044 P29045
P29046 P29047 P29048 P29049 P29051 P29052 P29054 P29055 P29056 P29065 P29069
P29078 P29079 P29082 P29083 P29084 P29087 P29088 P29090 P29091 P29093 P29094
P29097 P29098 P29102 P29107 P29115 P29116 P29123 P29124 P29128 P29129 P29133
P29134 P29137 P29139 P29140 P29144 P29151 P29153 P29156 P29157 P29158 P29159
P29160 P29161 P29164 P29166 P29170 P29174 P29176 P29177 P29184 P29185 P29188
P29190 P29192 P29193 P29194 P29195 P29196 P29197 P29208 P29214 P29215 P29217
P29219 P29220 P29221 P29222 P29223 P29224 P29225 P29226 P29227 P29231 P29237
P29238 P29239 P29246 P29247 P29248 P29249 P29251 P29252 P29253 P29257 P29259
P29260 P29261 P29262 P29263 P29264 P29265 P29266 P29269 P29271 P29272 P29277
P29278 P29280 P29281 P29283 P29284 P29286 P29287 P29289 P29290 P29291 P29298
P29299 P29300 P29303 P29304 P29305 P29306 P29307 P29308 P29309 P29315 P29324
P29331 P29333 P29338 P29340 P29341 P29342 P29343 P29344 P29345 P29346 P29347
P29356 P29363 P29364 P29365 P29367 P29368 P29372 P29378 P29379 P29381 P29382
P29383 P29384 P29385 P29386 P29389 P29390 P29391 P29393 P29394 P29395 P29396
P29397 P29398 P29405 P29406 P29407 P29408 P29409 P29410 P29411 P29412 P29422
P29427 P29428 P29429 P29433 P29434 P29435 P29436 P29437 P29438 P29439 P29440

P29441 P29442 P29443 P29444 P29445 P29446 P29447 P29448 P29449 P29450 P29454
P29458 P29461 P29467 P29468 P29469 P29471 P29472 P29480 P29489 P29490 P29495
P29496 P29497 P29499 P29500 P29501 P29502 P29504 P29506 P29507 P29509 P29514
P29516 P29520 P29521 P29522 P29535 P29537 P29538 P29539 P29540 P29541 P29542
P29543 P29544 P29545 P29546 P29548 P29549 P29550 P29551 P29552 P29555 P29556
P29557 P29562 P29564 P29565 P29566 P29567 P29568 P29569 P29570 P29571 P29572
P29573 P29574 P29575 P29576 P29577 P29578 P29579 P29580 P29581 P29582 P29583
P29584 P29585 P29586 P29587 P29588 P29589 P29595 P29596 P29602 P29603 P29604
P29613 P29616 P29620 P29623 P29624 P29625 P29626 P29627 P29628 P29629 P29673
P29674 P29675 P29680 P29683 P29684 P29686 P29691 P29693 P29694 P29696 P29702
P29703 P29711 P29714 P29715 P29718 P29726 P29727 P29729 P29730 P29734 P29735
P29736 P29737 P29738 P29741 P29745 P29746 P29747 P29748 P29749 P29758 P29763
P29764 P29766 P29770 P29771 P29772 P29773 P29774 P29775 P29776 P29778 P29779
P29780 P29781 P29782 P29783 P29784 P29785 P29792 P29802 P29804 P29806 P29807
P29808 P29809 P29810 P29811 P29814 P29815 P29818 P29819 P29820 P29830 P29836
P29840 P29847 P29848 P29849 P29852 P29882 P29883 P29884 P29886 P29887 P29888
P29890 P29891 P29892 P29893 P29901 P29902 P29903 P29906 P29908 P29909 P29910
P29927 P29928 P29929 P29930 P29931 P29932 P29933 P29934 P29935 P29937 P29943
P29944 P29946 P29950 P29951 P29952 P29953 P29954 P29962 P29964 P29976 P29978
P29979 P29980 P29985 P30003 P30007 P30008 P30012 P30014 P30015 P30020 P30024
P30028 P30038 P30046 P30051 P30052 P30053 P30055 P30056 P30057 P30058 P30059
P30060 P30061 P30062 P30063 P30065 P30066 P30067 P30068 P30069 P30070 P30071
P30072 P30073 P30074 P30075 P30076 P30077 P30078 P30079 P30080 P30081 P30085
P30102 P30103 P30104 P30106 P30107 P30108 P30109 P30110 P30111 P30112 P30113
P30114 P30116 P30117 P30118 P30119 P30124 P30125 P30126 P30128 P30131 P30136
P30137 P30138 P30139 P30140 P30147 P30151 P30155 P30156 P30157 P30161 P30162
P30163 P30164 P30165 P30167 P30168 P30169 P30170 P30171 P30172 P30173 P30174
P30175 P30176 P30177 P30178 P30180 P30181 P30182 P30183 P30184 P30185 P30186
P30187 P30188 P30189 P30190 P30192 P30196 P30197 P30202 P30211 P30212 P30218
P30219 P30220 P30221 P30222 P30226 P30227 P30228 P30231 P30232 P30235 P30241
P30242 P30243 P30244 P30251 P30252 P30253 P30254 P30255 P30256 P30257 P30258
P30259 P30261 P30265 P30266 P30267 P30268 P30271 P30274 P30278 P30283 P30284
P30286 P30287 P30290 P30297 P30300 P30303 P30304 P30308 P30309 P30310 P30311
P30313 P30314 P30315 P30316 P30317 P30318 P30319 P30320 P30321 P30322 P30325
P30326 P30328 P30331 P30332 P30333 P30334 P30338 P30339 P30340 P30341 P30342
P30346 P30347 P30350 P30352 P30359 P30360 P30362 P30363 P30364 P30366 P30373
P30389 P30390 P30397 P30401 P30402 P30416 P30417 P30418 P30428 P30435 P30436
P30520 P30521 P30522 P30523 P30524 P30525 P30526 P30527 P30528 P30529 P30534
P30537 P30538 P30539 P30541 P30562 P30563 P30564 P30565 P30566 P30568 P30569
P30570 P30571 P30610 P30611 P30613 P30614 P30615 P30616 P30620 P30621 P30627
P30629 P30630 P30631 P30634 P30635 P30636 P30637 P30640 P30641 P30642 P30643
P30644 P30646 P30647 P30648 P30651 P30653 P30655 P30660 P30661 P30664 P30665
P30666 P30668 P30672 P30674 P30693 P30694 P30707 P30711 P30713 P30717 P30719
P30720 P30721 P30722 P30724 P30725 P30726 P30727 P30732 P30733 P30734 P30735

P30736 P30738 P30739 P30741 P30744 P30745 P30748 P30749 P30753 P30755 P30756
P30757 P30759 P30760 P30761 P30762 P30763 P30764 P30765 P30766 P30767 P30768
P30769 P30770 P30771 P30773 P30774 P30775 P30776 P30778 P30779 P30780 P30781
P30786 P30787 P30788 P30791 P30792 P30794 P30795 P30797 P30798 P30799 P30806
P30812 P30813 P30817 P30819 P30820 P30821 P30827 P30828 P30829 P30830 P30840
P30841 P30842 P30850 P30851 P30852 P30864 P30866 P30871 P30877 P30883 P30890
P30892 P30893 P30900 P30901 P30903 P30918 P30924 P30925 P30927 P30928 P30929
P30949 P30950 P30956 P30958 P30967 P30979 P31002 P31003 P31004 P31005 P31006
P31008 P31011 P31012 P31013 P31014 P31015 P31017 P31018 P31019 P31020 P31023
P31026 P31027 P31028 P31029 P31031 P31032 P31033 P31034 P31035 P31036 P31037
P31046 P31047 P31050 P31051 P31052 P31061 P31062 P31068 P31072 P31073 P31074
P31076 P31078 P31082 P31089 P31101 P31102 P31103 P31104 P31108 P31111 P31112
P31113 P31114 P31115 P31116 P31117 P31118 P31120 P31131 P31142 P31153 P31155
P31156 P31158 P31160 P31161 P31162 P31163 P31164 P31165 P31166 P31169 P31170
P31171 P31177 P31179 P31180 P31181 P31182 P31183 P31184 P31186 P31188 P31189
P31190 P31191 P31192 P31193 P31194 P31195 P31196 P31197 P31198 P31199 P31200
P31201 P31202 P31203 P31204 P31205 P31209 P31212 P31225 P31232 P31233 P31237
P31238 P31239 P31244 P31246 P31247 P31249 P31251 P31252 P31254 P31255 P31256
P31257 P31258 P31259 P31260 P31261 P31262 P31263 P31264 P31267 P31269 P31270
P31271 P31272 P31273 P31274 P31275 P31276 P31277 P31278 P31279 P31280 P31281
P31282 P31293 P31294 P31295 P31296 P31300 P31301 P31307 P31308 P31310 P31311
P31313 P31314 P31315 P31316 P31318 P31319 P31320 P31325 P31326 P31330 P31331
P31332 P31333 P31335 P31337 P31352 P31353 P31354 P31357 P31360 P31363 P31364
P31365 P31367 P31368 P31369 P31370 P31373 P31376 P31380 P31383 P31384 P31385
P31386 P31393 P31396 P31425 P31426 P31427 P31433 P31434 P31441 P31443 P31447
P31449 P31450 P31453 P31455 P31459 P31460 P31463 P31467 P31469 P31470 P31471
P31473 P31475 P31485 P31490 P31491 P31494 P31495 P31496 P31503 P31504 P31505
P31506 P31507 P31508 P31509 P31516 P31521 P31525 P31526 P31528 P31531 P31533
P31534 P31535 P31536 P31537 P31538 P31540 P31541 P31542 P31543 P31551 P31552
P31558 P31561 P31568 P31569 P31570 P31572 P31574 P31592 P31593 P31605 P31610
P31611 P31612 P31617 P31624 P31630 P31638 P31653 P31654 P31655 P31656 P31657
P31658 P31660 P31663 P31665 P31667 P31668 P31670 P31671 P31672 P31673 P31674
P31677 P31678 P31681 P31682 P31683 P31684 P31685 P31686 P31687 P31688 P31718
P31730 P31732 P31752 P31753 P31754 P31756 P31758 P31759 P31764 P31765 P31766
P31767 P31768 P31769 P31770 P31771 P31773 P31774 P31777 P31778 P31781 P31782
P31784 P31787 P31790 P31792 P31795 P31798 P31799 P31806 P31808 P31811 P31812
P31813 P31814 P31815 P31816 P31817 P31822 P31823 P31824 P31825 P31832 P31834
P31843 P31849 P31850 P31851 P31852 P31858 P31860 P31861 P31863 P31864 P31865
P31866 P31867 P31870 P31874 P31878 P31879 P31880 P31881 P31882 P31888 P31893
P31895 P31900 P31901 P31902 P31903 P31904 P31905 P31906 P31907 P31915 P31916
P31918 P31919 P31920 P31921 P31922 P31923 P31925 P31926 P31927 P31928 P31937
P31941 P31950 P31954 P31957 P31958 P31959 P31960 P31961 P31965 P31967 P31968
P31970 P31972 P31974 P31991 P31992 P31998 P32000 P32001 P32002 P32003 P32005
P32006 P32008 P32009 P32012 P32021 P32024 P32027 P32028 P32029 P32030 P32031

P32033 P32034 P32035 P32039 P32043 P32044 P32046 P32047 P32048 P32053 P32055
P32056 P32057 P32060 P32067 P32068 P32069 P32070 P32072 P32073 P32081 P32083
P32084 P32085 P32086 P32087 P32088 P32090 P32091 P32092 P32094 P32096 P32097
P32098 P32101 P32102 P32103 P32104 P32109 P32110 P32111 P32112 P32114 P32115
P32119 P32125 P32131 P32134 P32138 P32139 P32140 P32141 P32143 P32144 P32151
P32152 P32153 P32156 P32157 P32162 P32169 P32170 P32171 P32173 P32177 P32178
P32179 P32180 P32183 P32185 P32186 P32190 P32192 P32197 P32206 P32208 P32209
P32216 P32217 P32218 P32219 P32220 P32221 P32222 P32223 P32224 P32226 P32227
P32228 P32230 P32231 P32233 P32234 P32235 P32242 P32243 P32255 P32256 P32257
P32260 P32263 P32264 P32265 P32267 P32268 P32270 P32271 P32273 P32274 P32275
P32276 P32277 P32278 P32279 P32280 P32281 P32282 P32283 P32284 P32285 P32286
P32287 P32289 P32290 P32292 P32293 P32294 P32295 P32296 P32315 P32316 P32318
P32320 P32321 P32324 P32325 P32326 P32327 P32330 P32338 P32345 P32347 P32348
P32349 P32350 P32351 P32356 P32362 P32364 P32373 P32375 P32381 P32382 P32385
P32389 P32390 P32393 P32395 P32396 P32398 P32400 P32410 P32411 P32419 P32425
P32426 P32427 P32428 P32429 P32430 P32431 P32432 P32437 P32438 P32440 P32441
P32442 P32445 P32447 P32448 P32449 P32450 P32451 P32452 P32453 P32454 P32460
P32461 P32469 P32471 P32473 P32475 P32477 P32479 P32480 P32481 P32483 P32488
P32489 P32492 P32493 P32495 P32496 P32497 P32501 P32502 P32503 P32504 P32508
P32509 P32511 P32512 P32516 P32517 P32518 P32523 P32524 P32526 P32527 P32528
P32529 P32530 P32531 P32532 P32533 P32534 P32535 P32538 P32539 P32542 P32552
P32553 P32555 P32558 P32561 P32565 P32566 P32569 P32570 P32574 P32575 P32576
P32579 P32580 P32582 P32583 P32586 P32588 P32589 P32590 P32591 P32593 P32597
P32598 P32604 P32605 P32606 P32607 P32608 P32613 P32617 P32622 P32626 P32628
P32630 P32634 P32635 P32636 P32637 P32638 P32639 P32641 P32642 P32644 P32645
P32651 P32652 P32656 P32657 P32662 P32664 P32669 P32674 P32675 P32677 P32680
P32681 P32684 P32685 P32693 P32694 P32695 P32700 P32704 P32718 P32719 P32725
P32726 P32727 P32728 P32729 P32730 P32731 P32732 P32733 P32734 P32738 P32740
P32742 P32743 P32744 P32761 P32764 P32769 P32770 P32771 P32772 P32773 P32774
P32775 P32776 P32780 P32782 P32783 P32785 P32786 P32787 P32788 P32793 P32794
P32806 P32807 P32809 P32810 P32811 P32812 P32813 P32814 P32815 P32816 P32817
P32818 P32828 P32829 P32831 P32832 P32833 P32835 P32836 P32838 P32841 P32848
P32849 P32859 P32860 P32862 P32868 P32872 P32875 P32877 P32888 P32893 P32895
P32896 P32899 P32910 P32911 P32914 P32920 P32921 P32922 P32924 P32925 P32928
P32931 P32932 P32933 P32934 P32935 P32941 P32943 P32944 P32953 P32960 P32961
P32963 P32964 P32966 P32968 P32975 P32976 P32985 P32986 P32987 P32989 P32990
P32999 P33006 P33007 P33008 P33009 P33010 P33012 P33014 P33018 P33020 P33022
P33025 P33029 P33030 P33035 P33036 P33037 P33038 P33044 P33045 P33052 P33054
P33062 P33071 P33073 P33074 P33075 P33076 P33077 P33078 P33079 P33080 P33081
P33082 P33083 P33084 P33086 P33095 P33097 P33098 P33099 P33100 P33101 P33102
P33103 P33104 P33105 P33106 P33107 P33109 P33110 P33111 P33115 P33117 P33118
P33119 P33122 P33127 P33144 P33149 P33156 P33158 P33159 P33160 P33161 P33162
P33163 P33164 P33165 P33166 P33167 P33168 P33169 P33170 P33171 P33174 P33177
P33178 P33179 P33180 P33181 P33182 P33187 P33188 P33189 P33190 P33191 P33192

P33194 P33195 P33197 P33198 P33200 P33201 P33203 P33204 P33205 P33206 P33211
P33214 P33216 P33217 P33221 P33222 P33224 P33227 P33228 P33229 P33230 P33234
P33238 P33245 P33246 P33248 P33249 P33271 P33275 P33279 P33280 P33283 P33284
P33285 P33286 P33287 P33288 P33293 P33294 P33297 P33298 P33299 P33301 P33304
P33306 P33309 P33312 P33313 P33315 P33317 P33318 P33319 P33320 P33321 P33322
P33323 P33324 P33327 P33330 P33331 P33334 P33339 P33345 P33346 P33347 P33348
P33349 P33352 P33353 P33355 P33358 P33368 P33371 P33380 P33381 P33401 P33403
P33404 P33405 P33411 P33416 P33417 P33418 P33419 P33420 P33422 P33423 P33424
P33427 P33428 P33429 P33437 P33441 P33444 P33447 P33451 P33452 P33453 P33454
P33458 P33459 P33462 P33471 P33472 P33473 P33474 P33477 P33479 P33483 P33485
P33486 P33494 P33499 P33514 P33520 P33521 P33529 P33530 P33536 P33537 P33538
P33539 P33540 P33541 P33542 P33544 P33552 P33561 P33562 P33563 P33564 P33565
P33567 P33569 P33570 P33571 P33585 P33586 P33596 P33609 P33610 P33611 P33623
P33624 P33625 P33626 P33627 P33628 P33629 P33630 P33631 P33632 P33634 P33637
P33640 P33642 P33643 P33644 P33647 P33651 P33652 P33653 P33654 P33655 P33656
P33657 P33658 P33660 P33661 P33663 P33664 P33667 P33670 P33672 P33674 P33676
P33677 P33678 P33683 P33696 P33697 P33700 P33701 P33728 P33732 P33734 P33744
P33746 P33748 P33749 P33751 P33752 P33753 P33757 P33761 P33762 P33763 P33768
P33769 P33770 P33771 P33776 P33777 P33778 P33779 P33785 P33787 P33789 P33791
P33794 P33796 P33800 P33801 P33802 P33812 P33813 P33825 P33842 P33859 P33861
P33862 P33863 P33864 P33868 P33876 P33880 P33881 P33884 P33885 P33887 P33888
P33889 P33892 P33893 P33895 P33904 P33905 P33906 P33907 P33909 P33910 P33912
P33919 P33920 P33953 P33954 P33959 P33960 P33963 P33967 P33968 P33973 P33974
P33975 P33976 P33977 P33983 P33984 P33986 P33987 P33988 P33989 P33997 P34002
P34003 P34012 P34018 P34028 P34029 P34030 P34031 P34032 P34035 P34037 P34038
P34039 P34040 P34041 P34047 P34048 P34050 P34051 P34052 P34053 P34055 P34060
P34061 P34064 P34065 P34066 P34067 P34072 P34081 P34085 P34087 P34089 P34091
P34093 P34097 P34101 P34102 P34105 P34106 P34107 P34108 P34111 P34114 P34117
P34118 P34119 P34120 P34122 P34123 P34124 P34127 P34138 P34153 P34154 P34155
P34156 P34159 P34161 P34167 P34169 P34172 P34173 P34175 P34176 P34177 P34178
P34183 P34184 P34185 P34203 P34204 P34205 P34206 P34214 P34215 P34217 P34218
P34219 P34220 P34221 P34228 P34232 P34233 P34234 P34238 P34241 P34243 P34245
P34246 P34247 P34249 P34252 P34253 P34254 P34256 P34257 P34258 P34259 P34263
P34264 P34266 P34268 P34274 P34275 P34277 P34278 P34279 P34280 P34281 P34282
P34284 P34286 P34287 P34291 P34292 P34293 P34294 P34300 P34301 P34303 P34304
P34305 P34306 P34307 P34308 P34310 P34318 P34320 P34321 P34324 P34325 P34326
P34327 P34328 P34333 P34335 P34337 P34338 P34339 P34340 P34341 P34342 P34344
P34345 P34346 P34347 P34348 P34350 P34357 P34360 P34361 P34362 P34363 P34365
P34366 P34368 P34369 P34371 P34375 P34376 P34378 P34384 P34385 P34388 P34390
P34391 P34392 P34394 P34395 P34396 P34397 P34398 P34399 P34401 P34402 P34404
P34406 P34409 P34412 P34415 P34416 P34418 P34419 P34420 P34423 P34424 P34425
P34427 P34428 P34429 P34430 P34431 P34433 P34436 P34437 P34438 P34440 P34441
P34442 P34447 P34450 P34451 P34452 P34456 P34457 P34458 P34459 P34460 P34466
P34467 P34468 P34469 P34470 P34471 P34473 P34475 P34476 P34477 P34478 P34480

P34481 P34482 P34483 P34484 P34487 P34491 P34492 P34493 P34494 P34495 P34496
P34497 P34498 P34507 P34509 P34510 P34511 P34514 P34515 P34517 P34518 P34520
P34522 P34523 P34524 P34526 P34527 P34529 P34531 P34534 P34536 P34537 P34538
P34540 P34541 P34544 P34545 P34549 P34550 P34553 P34555 P34556 P34558 P34560
P34562 P34563 P34564 P34565 P34566 P34567 P34568 P34569 P34575 P34580 P34582
P34583 P34584 P34587 P34590 P34592 P34593 P34594 P34596 P34600 P34601 P34603
P34604 P34606 P34607 P34611 P34613 P34614 P34615 P34618 P34619 P34621 P34622
P34625 P34626 P34627 P34629 P34630 P34636 P34638 P34640 P34645 P34646 P34647
P34648 P34649 P34650 P34651 P34653 P34654 P34656 P34657 P34658 P34659 P34661
P34663 P34665 P34667 P34668 P34670 P34671 P34673 P34674 P34677 P34680 P34681
P34683 P34685 P34686 P34687 P34689 P34690 P34691 P34694 P34695 P34696 P34699
P34700 P34701 P34705 P34706 P34707 P34708 P34715 P34716 P34717 P34718 P34719
P34720 P34721 P34723 P34725 P34733 P34734 P34735 P34736 P34737 P34738 P34739
P34742 P34743 P34748 P34751 P34761 P34762 P34763 P34764 P34765 P34766 P34767
P34768 P34769 P34770 P34771 P34772 P34774 P34775 P34776 P34777 P34778 P34779
P34780 P34781 P34782 P34783 P34785 P34786 P34787 P34789 P34793 P34794 P34795
P34796 P34797 P34800 P34801 P34802 P34803 P34804 P34806 P34807 P34808 P34809
P34811 P34813 P34816 P34817 P34818 P34823 P34824 P34825 P34826 P34827 P34828
P34829 P34830 P34831 P34832 P34877 P34878 P34879 P34880 P34881 P34882 P34883
P34887 P34890 P34893 P34894 P34895 P34896 P34898 P34899 P34905 P34906 P34915
P34916 P34917 P34918 P34919 P34920 P34921 P34922 P34923 P34924 P34926 P34933
P34936 P34937 P34938 P34939 P34940 P34945 P34946 P34948 P34949 P34952 P34961
P34967 P35006 P35007 P35014 P35017 P35018 P35019 P35020 P35021 P35023 P35024
P35025 P35026 P35027 P35028 P35029 P35055 P35057 P35059 P35060 P35061 P35062
P35064 P35065 P35066 P35067 P35068 P35069 P35072 P35074 P35076 P35079 P35080
P35081 P35082 P35083 P35084 P35085 P35086 P35087 P35100 P35112 P35128 P35129
P35130 P35131 P35132 P35133 P35134 P35135 P35136 P35137 P35138 P35139 P35140
P35141 P35142 P35143 P35144 P35145 P35146 P35147 P35148 P35151 P35153 P35154
P35155 P35156 P35159 P35163 P35168 P35170 P35172 P35173 P35174 P35175 P35177
P35178 P35184 P35187 P35189 P35190 P35191 P35192 P35194 P35195 P35201 P35202
P35203 P35207 P35208 P35209 P35217 P35218 P35219 P35244 P35249 P35250 P35258
P35259 P35262 P35263 P35266 P35267 P35270 P35297 P35298 P35299 P35300 P35301
P35302 P35303 P35304 P35306 P35307 P35308 P35309 P35310 P35311 P35312 P35313
P35314 P35319 P35320 P35327 P35328 P35340 P35341 P35385 P35394 P35415 P35416
P35417 P35421 P35422 P35423 P35424 P35425 P35428 P35429 P35430 P35432 P35433
P35450 P35452 P35453 P35458 P35467 P35469 P35470 P35471 P35473 P35474 P35478
P35479 P35480 P35484 P35485 P35486 P35487 P35488 P35489 P35491 P35492 P35493
P35494 P35497 P35510 P35511 P35512 P35513 P35514 P35516 P35517 P35540 P35545
P35547 P35549 P35550 P35551 P35552 P35553 P35554 P35557 P35567 P35571 P35573
P35574 P35584 P35608 P35614 P35615 P35616 P35617 P35622 P35623 P35627 P35630
P35631 P35632 P35635 P35638 P35639 P35640 P35642 P35643 P35644 P35649 P35660
P35661 P35662 P35663 P35664 P35667 P35668 P35669 P35673 P35675 P35677 P35678
P35681 P35682 P35683 P35684 P35685 P35686 P35687 P35691 P35692 P35693 P35700
P35704 P35707 P35710 P35711 P35712 P35713 P35718 P35719 P35722 P35725 P35727

P35728 P35732 P35733 P35734 P35736 P35744 P35745 P35748 P35749 P35756 P35758
P35767 P35768 P35769 P35770 P35773 P35774 P35788 P35789 P35791 P35798 P35805
P35806 P35810 P35812 P35816 P35817 P35820 P35838 P35840 P35850 P35851 P35852
P35853 P35854 P35856 P35857 P35861 P35862 P35863 P35864 P35866 P35868 P35870
P35871 P35872 P35873 P35879 P35880 P35881 P35883 P35884 P35885 P35886 P35889
P35890 P35892 P35900 P35901 P35902 P35904 P35907 P35912 P35919 P35920 P35921
P35923 P35924 P35925 P35926 P35927 P35928 P35929 P35931 P35932 P35933 P35934
P35935 P35939 P35940 P35941 P35942 P35943 P35944 P35945 P35946 P35948 P35958
P35970 P35972 P35974 P35975 P35977 P35978 P35982 P35983 P35984 P35985 P35986
P35987 P35993 P35995 P36001 P36008 P36009 P36010 P36011 P36013 P36022 P36023
P36024 P36025 P36030 P36031 P36038 P36040 P36041 P36042 P36047 P36048 P36049
P36050 P36052 P36053 P36054 P36056 P36058 P36059 P36061 P36065 P36067 P36069
P36070 P36071 P36072 P36073 P36074 P36075 P36076 P36077 P36079 P36080 P36082
P36083 P36084 P36085 P36086 P36087 P36088 P36089 P36093 P36097 P36099 P36100
P36102 P36103 P36104 P36106 P36109 P36111 P36113 P36114 P36118 P36119 P36120
P36121 P36123 P36124 P36127 P36128 P36129 P36131 P36132 P36133 P36134 P36136
P36138 P36140 P36141 P36143 P36144 P36145 P36146 P36150 P36151 P36152 P36153
P36154 P36155 P36156 P36157 P36158 P36159 P36160 P36162 P36167 P36168 P36174
P36176 P36177 P36179 P36186 P36187 P36188 P36197 P36200 P36201 P36203 P36205
P36206 P36207 P36208 P36211 P36214 P36215 P36216 P36220 P36221 P36223 P36224
P36225 P36226 P36228 P36229 P36230 P36231 P36232 P36234 P36236 P36237 P36238
P36239 P36243 P36244 P36245 P36247 P36248 P36249 P36250 P36251 P36252 P36254
P36255 P36256 P36257 P36258 P36260 P36261 P36262 P36263 P36264 P36265 P36266
P36272 P36273 P36274 P36275 P36276 P36279 P36281 P36282 P36283 P36284 P36285
P36286 P36287 P36288 P36289 P36292 P36293 P36294 P36296 P36297 P36303 P36304
P36309 P36311 P36314 P36315 P36322 P36324 P36326 P36333 P36341 P36345 P36351
P36352 P36353 P36354 P36358 P36384 P36385 P36387 P36388 P36389 P36390 P36391
P36392 P36393 P36394 P36395 P36396 P36398 P36399 P36413 P36414 P36415 P36416
P36417 P36418 P36419 P36420 P36422 P36424 P36426 P36427 P36428 P36429 P36430
P36431 P36432 P36433 P36434 P36435 P36436 P36437 P36439 P36440 P36441 P36443
P36444 P36447 P36451 P36454 P36456 P36459 P36461 P36464 P36465 P36466 P36467
P36470 P36472 P36473 P36476 P36477 P36478 P36479 P36480 P36481 P36482 P36483
P36484 P36485 P36486 P36487 P36488 P36489 P36490 P36493 P36496 P36505 P36508
P36515 P36522 P36530 P36547 P36553 P36556 P36562 P36566 P36568 P36569 P36570
P36571 P36572 P36577 P36580 P36589 P36590 P36592 P36594 P36595 P36597 P36598
P36599 P36600 P36601 P36602 P36603 P36605 P36607 P36609 P36611 P36612 P36613
P36614 P36617 P36618 P36620 P36621 P36622 P36623 P36624 P36625 P36626 P36627
P36629 P36630 P36631 P36632 P36644 P36645 P36659 P36661 P36662 P36673 P36674
P36675 P36677 P36680 P36683 P36684 P36686 P36687 P36688 P36692 P36693 P36695
P36697 P36698 P36700 P36701 P36702 P36704 P36706 P36708 P36709 P36710 P36711
P36712 P36713 P36714 P36715 P36716 P36717 P36731 P36732 P36733 P36734 P36735
P36736 P36737 P36738 P36740 P36741 P36742 P36743 P36767 P36771 P36772 P36773
P36774 P36778 P36779 P36780 P36781 P36782 P36783 P36784 P36785 P36786 P36787
P36788 P36789 P36790 P36791 P36792 P36793 P36794 P36795 P36796 P36797 P36798

P36799 P36800 P36801 P36802 P36803 P36804 P36805 P36806 P36807 P36808 P36809
P36810 P36811 P36812 P36813 P36814 P36815 P36838 P36839 P36841 P36842 P36843
P36844 P36845 P36847 P36848 P36849 P36850 P36851 P36852 P36853 P36854 P36855
P36856 P36858 P36859 P36866 P36867 P36868 P36869 P36870 P36872 P36875 P36876
P36882 P36883 P36889 P36891 P36892 P36906 P36916 P36922 P36923 P36928 P36929
P36930 P36932 P36933 P36934 P36935 P36936 P36944 P36945 P36946 P36951 P36954
P36958 P36959 P36960 P36961 P36962 P36972 P36973 P36979 P36981 P36982 P36995
P36999 P37000 P37003 P37005 P37006 P37007 P37008 P37012 P37014 P37024 P37025
P37029 P37030 P37031 P37032 P37033 P37046 P37047 P37051 P37052 P37057 P37061
P37062 P37063 P37075 P37077 P37078 P37079 P37084 P37092 P37094 P37095 P37097
P37098 P37099 P37100 P37101 P37102 P37111 P37112 P37113 P37119 P37120 P37121
P37122 P37123 P37124 P37125 P37127 P37128 P37133 P37135 P37137 P37138 P37142
P37143 P37144 P37145 P37146 P37155 P37164 P37165 P37167 P37168 P37171 P37174
P37182 P37197 P37200 P37202 P37209 P37210 P37213 P37215 P37216 P37218 P37219
P37220 P37221 P37222 P37223 P37224 P37225 P37226 P37227 P37228 P37229 P37233
P37234 P37245 P37247 P37248 P37249 P37251 P37252 P37253 P37254 P37262 P37263
P37269 P37270 P37272 P37273 P37276 P37282 P37283 P37292 P37293 P37294 P37304
P37306 P37309 P37317 P37323 P37326 P37330 P37332 P37333 P37334 P37335 P37337
P37342 P37348 P37349 P37351 P37352 P37353 P37355 P37356 P37357 P37358 P37363
P37365 P37366 P37369 P37370 P37371 P37372 P37374 P37375 P37376 P37380 P37382
P37383 P37384 P37391 P37392 P37393 P37394 P37395 P37397 P37398 P37402 P37408
P37411 P37412 P37413 P37414 P37415 P37416 P37417 P37418 P37419 P37424 P37425
P37426 P37427 P37430 P37431 P37437 P37440 P37444 P37449 P37450 P37454 P37455
P37459 P37462 P37463 P37464 P37465 P37467 P37468 P37469 P37470 P37471 P37472
P37473 P37474 P37477 P37479 P37480 P37481 P37485 P37486 P37487 P37490 P37495
P37496 P37497 P37499 P37500 P37501 P37502 P37503 P37504 P37506 P37507 P37509
P37510 P37512 P37515 P37516 P37517 P37518 P37521 P37522 P37523 P37524 P37525
P37526 P37527 P37528 P37529 P37530 P37531 P37532 P37533 P37534 P37535 P37536
P37537 P37539 P37540 P37541 P37542 P37543 P37544 P37545 P37547 P37549 P37550
P37551 P37552 P37553 P37554 P37556 P37557 P37560 P37561 P37562 P37563 P37564
P37565 P37567 P37568 P37569 P37570 P37571 P37572 P37574 P37578 P37584 P37586
P37587 P37588 P37595 P37596 P37609 P37610 P37613 P37615 P37623 P37627 P37629
P37631 P37634 P37635 P37639 P37640 P37641 P37646 P37647 P37657 P37664 P37671
P37672 P37673 P37677 P37678 P37679 P37680 P37682 P37685 P37686 P37689 P37693
P37694 P37695 P37705 P37706 P37707 P37708 P37709 P37712 P37713 P37714 P37715
P37719 P37722 P37724 P37725 P37728 P37733 P37743 P37752 P37754 P37756 P37757
P37762 P37765 P37766 P37767 P37769 P37771 P37775 P37787 P37788 P37789 P37790
P37793 P37794 P37800 P37801 P37803 P37804 P37805 P37806 P37807 P37818 P37820
P37821 P37822 P37823 P37824 P37825 P37829 P37830 P37832 P37833 P37836 P37838
P37839 P37842 P37843 P37844 P37845 P37846 P37847 P37848 P37849 P37850 P37851
P37852 P37853 P37854 P37855 P37856 P37857 P37858 P37859 P37860 P37861 P37862
P37863 P37864 P37865 P37866 P37867 P37870 P37871 P37872 P37874 P37875 P37876
P37877 P37878 P37880 P37882 P37883 P37893 P37895 P37898 P37899 P37900 P37903
P37906 P37928 P37929 P37934 P37935 P37936 P37937 P37938 P37939 P37940 P37941

P37942 P37944 P37945 P37946 P37954 P37955 P37956 P37958 P37959 P37960 P37963
P37964 P37967 P37968 P37969 P37970 P37971 P37977 P37980 P37981 P37982 P37983
P37984 P37985 P37986 P37987 P37988 P37991 P37992 P37993 P37995 P38001 P38002
P38003 P38005 P38007 P38014 P38015 P38016 P38017 P38018 P38019 P38020 P38021
P38023 P38024 P38025 P38032 P38033 P38036 P38037 P38038 P38039 P38042 P38048
P38049 P38051 P38056 P38062 P38063 P38066 P38067 P38070 P38073 P38074 P38075
P38076 P38081 P38082 P38086 P38088 P38089 P38092 P38095 P38096 P38098 P38099
P38100 P38103 P38104 P38105 P38110 P38111 P38112 P38113 P38115 P38117 P38118
P38121 P38122 P38123 P38126 P38128 P38129 P38132 P38140 P38141 P38144 P38145
P38148 P38149 P38151 P38160 P38161 P38162 P38165 P38170 P38171 P38174 P38177
P38178 P38180 P38183 P38184 P38188 P38191 P38193 P38194 P38195 P38197 P38198
P38200 P38201 P38202 P38203 P38204 P38205 P38207 P38208 P38209 P38210 P38213
P38214 P38218 P38219 P38222 P38223 P38230 P38231 P38232 P38235 P38236 P38238
P38239 P38241 P38243 P38246 P38249 P38251 P38254 P38255 P38256 P38257 P38260
P38262 P38265 P38274 P38275 P38277 P38278 P38281 P38282 P38283 P38284 P38285
P38289 P38290 P38291 P38293 P38294 P38296 P38299 P38302 P38303 P38304 P38305
P38306 P38309 P38313 P38314 P38315 P38319 P38321 P38322 P38324 P38326 P38328
P38331 P38333 P38334 P38336 P38337 P38338 P38340 P38344 P38345 P38346 P38347
P38348 P38349 P38350 P38351 P38352 P38354 P38364 P38365 P38366 P38372 P38373
P38393 P38394 P38413 P38420 P38421 P38423 P38424 P38425 P38426 P38427 P38429
P38430 P38431 P38434 P38437 P38439 P38442 P38443 P38448 P38450 P38451 P38452
P38455 P38456 P38457 P38458 P38463 P38464 P38465 P38466 P38467 P38468 P38469
P38470 P38471 P38472 P38473 P38474 P38475 P38476 P38478 P38479 P38481 P38483
P38485 P38487 P38488 P38491 P38492 P38493 P38494 P38496 P38497 P38500 P38502
P38505 P38506 P38508 P38509 P38510 P38511 P38512 P38513 P38514 P38515 P38516
P38517 P38518 P38519 P38520 P38521 P38524 P38525 P38526 P38527 P38529 P38530
P38531 P38532 P38533 P38534 P38541 P38542 P38543 P38546 P38547 P38548 P38550
P38552 P38554 P38557 P38559 P38560 P38561 P38562 P38563 P38564 P38569 P38576
P38581 P38583 P38584 P38585 P38587 P38588 P38589 P38590 P38609 P38611 P38613
P38618 P38619 P38620 P38621 P38622 P38625 P38627 P38628 P38632 P38635 P38636
P38637 P38639 P38640 P38641 P38642 P38643 P38644 P38645 P38652 P38653 P38654
P38656 P38662 P38664 P38665 P38667 P38668 P38669 P38671 P38672 P38673 P38674
P38675 P38677 P38678 P38681 P38684 P38689 P38690 P38692 P38694 P38697 P38698
P38699 P38705 P38707 P38708 P38709 P38712 P38714 P38715 P38716 P38719 P38720
P38722 P38730 P38732 P38737 P38738 P38741 P38744 P38746 P38747 P38749 P38751
P38758 P38763 P38764 P38765 P38766 P38768 P38769 P38771 P38772 P38773 P38774
P38777 P38779 P38780 P38781 P38782 P38786 P38787 P38788 P38789 P38790 P38791
P38792 P38793 P38794 P38795 P38796 P38797 P38798 P38801 P38803 P38804 P38805
P38806 P38808 P38809 P38811 P38814 P38816 P38818 P38820 P38821 P38823 P38824
P38826 P38827 P38828 P38829 P38830 P38831 P38833 P38835 P38839 P38840 P38850
P38852 P38854 P38858 P38859 P38863 P38864 P38867 P38874 P38876 P38877 P38882
P38883 P38884 P38885 P38886 P38887 P38889 P38890 P38891 P38893 P38898 P38899
P38900 P38902 P38903 P38907 P38910 P38912 P38913 P38915 P38918 P38920 P38922
P38927 P38930 P38937 P38940 P38941 P38942 P38944 P38945 P38946 P38947 P38948

P38949 P38950 P38953 P38956 P38957 P38960 P38961 P38972 P38974 P38975 P38980
P38981 P38984 P38986 P38987 P38989 P38991 P38996 P38997 P38999 P39014 P39015
P39016 P39017 P39022 P39040 P39043 P39044 P39047 P39050 P39051 P39057 P39058
P39065 P39066 P39067 P39070 P39071 P39072 P39075 P39076 P39077 P39078 P39079
P39081 P39095 P39096 P39097 P39110 P39112 P39113 P39115 P39118 P39119 P39120
P39121 P39122 P39123 P39124 P39125 P39126 P39127 P39130 P39131 P39132 P39133
P39134 P39135 P39138 P39140 P39142 P39143 P39144 P39145 P39146 P39147 P39148
P39149 P39151 P39152 P39153 P39155 P39156 P39157 P39158 P39160 P39161 P39163
P39173 P39177 P39197 P39198 P39199 P39200 P39204 P39206 P39207 P39208 P39211
P39212 P39213 P39220 P39221 P39222 P39223 P39224 P39225 P39226 P39228 P39229
P39231 P39232 P39236 P39237 P39238 P39239 P39240 P39241 P39242 P39243 P39244
P39245 P39247 P39248 P39249 P39250 P39251 P39252 P39253 P39254 P39255 P39256
P39257 P39258 P39259 P39260 P39261 P39262 P39267 P39274 P39279 P39280 P39283
P39286 P39288 P39293 P39295 P39298 P39300 P39304 P39305 P39306 P39309 P39315
P39317 P39332 P39333 P39334 P39336 P39337 P39342 P39343 P39346 P39347 P39351
P39353 P39354 P39356 P39358 P39359 P39360 P39361 P39362 P39364 P39366 P39368
P39369 P39374 P39375 P39376 P39377 P39380 P39383 P39384 P39389 P39390 P39391
P39393 P39394 P39399 P39400 P39404 P39406 P39408 P39409 P39410 P39411 P39413
P39415 P39416 P39418 P39419 P39420 P39421 P39423 P39424 P39425 P39426 P39427
P39430 P39431 P39432 P39433 P39439 P39440 P39450 P39451 P39452 P39458 P39460
P39461 P39462 P39463 P39466 P39467 P39468 P39469 P39470 P39471 P39472 P39473
P39474 P39475 P39476 P39477 P39478 P39482 P39483 P39484 P39485 P39487 P39488
P39489 P39490 P39491 P39492 P39493 P39494 P39495 P39496 P39497 P39498 P39499
P39500 P39501 P39502 P39505 P39506 P39507 P39508 P39510 P39512 P39514 P39520
P39521 P39522 P39523 P39526 P39529 P39533 P39534 P39550 P39564 P39566 P39567
P39572 P39573 P39574 P39575 P39576 P39581 P39583 P39585 P39586 P39587 P39588
P39592 P39593 P39594 P39598 P39601 P39605 P39606 P39609 P39610 P39614 P39615
P39616 P39617 P39620 P39621 P39623 P39624 P39625 P39626 P39629 P39630 P39631
P39633 P39635 P39638 P39639 P39640 P39641 P39643 P39644 P39645 P39646 P39647
P39648 P39650 P39651 P39657 P39661 P39662 P39666 P39667 P39669 P39671 P39673
P39674 P39676 P39678 P39679 P39682 P39683 P39692 P39693 P39696 P39697 P39698
P39701 P39703 P39706 P39708 P39710 P39711 P39714 P39715 P39720 P39721 P39724
P39725 P39726 P39728 P39729 P39730 P39731 P39732 P39735 P39736 P39737 P39739
P39740 P39744 P39746 P39749 P39750 P39752 P39753 P39756 P39757 P39758 P39759
P39761 P39762 P39768 P39769 P39770 P39771 P39772 P39773 P39776 P39778 P39780
P39781 P39782 P39783 P39784 P39785 P39786 P39787 P39788 P39789 P39791 P39792
P39795 P39796 P39797 P39801 P39805 P39806 P39807 P39808 P39809 P39812 P39813
P39814 P39815 P39820 P39821 P39825 P39829 P39831 P39839 P39840 P39841 P39842
P39845 P39846 P39847 P39849 P39852 P39856 P39857 P39858 P39859 P39860 P39861
P39862 P39863 P39864 P39865 P39866 P39867 P39868 P39869 P39870 P39871 P39875
P39881 P39882 P39883 P39884 P39885 P39887 P39888 P39889 P39890 P39896 P39897
P39902 P39907 P39908 P39909 P39911 P39912 P39914 P39915 P39916 P39918 P39919
P39921 P39922 P39927 P39933 P39935 P39936 P39937 P39943 P39946 P39947 P39949
P39950 P39954 P39955 P39959 P39961 P39964 P39965 P39966 P39969 P39970 P39971

P39972 P39973 P39974 P39975 P39976 P39977 P39978 P39979 P39983 P39984 P39985
P39987 P39988 P39990 P39992 P39995 P39998 P39999 P40000 P40001 P40002 P40003
P40005 P40007 P40010 P40011 P40013 P40014 P40016 P40019 P40020 P40021 P40022
P40023 P40024 P40025 P40026 P40028 P40029 P40032 P40034 P40036 P40037 P40038
P40040 P40041 P40042 P40045 P40047 P40048 P40050 P40052 P40053 P40054 P40055
P40057 P40058 P40059 P40060 P40062 P40063 P40065 P40068 P40070 P40072 P40076
P40079 P40080 P40081 P40082 P40083 P40084 P40089 P40090 P40091 P40093 P40095
P40096 P40099 P40101 P40102 P40103 P40104 P40105 P40108 P40110 P40111 P40112
P40114 P40115 P40117 P40118 P40119 P40127 P40128 P40129 P40130 P40134 P40149
P40151 P40154 P40156 P40157 P40160 P40161 P40162 P40164 P40165 P40166 P40167
P40168 P40170 P40171 P40173 P40174 P40175 P40176 P40177 P40178 P40181 P40182
P40183 P40185 P40186 P40187 P40188 P40191 P40192 P40193 P40194 P40196 P40201
P40204 P40205 P40206 P40208 P40210 P40211 P40214 P40217 P40218 P40227 P40228
P40229 P40231 P40261 P40262 P40263 P40264 P40265 P40266 P40267 P40268 P40269
P40270 P40273 P40274 P40275 P40276 P40277 P40278 P40279 P40280 P40281 P40282
P40283 P40284 P40285 P40286 P40287 P40288 P40291 P40294 P40296 P40301 P40304
P40306 P40307 P40320 P40323 P40325 P40326 P40328 P40329 P40333 P40334 P40339
P40340 P40342 P40347 P40348 P40352 P40354 P40355 P40359 P40360 P40361 P40362
P40363 P40366 P40369 P40370 P40371 P40373 P40374 P40375 P40376 P40377 P40379
P40380 P40381 P40382 P40383 P40384 P40385 P40386 P40387 P40389 P40390 P40391
P40396 P40397 P40398 P40399 P40402 P40405 P40408 P40412 P40413 P40414 P40418
P40421 P40422 P40426 P40427 P40428 P40431 P40432 P40433 P40434 P40446 P40447
P40448 P40449 P40450 P40454 P40455 P40456 P40459 P40460 P40461 P40462 P40463
P40465 P40466 P40467 P40468 P40469 P40470 P40473 P40479 P40480 P40481 P40483
P40486 P40487 P40488 P40489 P40490 P40491 P40492 P40493 P40494 P40495 P40498
P40503 P40505 P40506 P40507 P40510 P40511 P40512 P40513 P40516 P40518 P40521
P40522 P40523 P40529 P40530 P40535 P40536 P40537 P40539 P40541 P40542 P40543
P40545 P40546 P40547 P40551 P40553 P40554 P40555 P40558 P40560 P40561 P40562
P40563 P40565 P40567 P40568 P40571 P40572 P40573 P40574 P40575 P40577 P40579
P40580 P40584 P40586 P40588 P40590 P40591 P40592 P40593 P40594 P40595 P40596
P40597 P40598 P40599 P40607 P40609 P40610 P40611 P40612 P40613 P40615 P40618
P40619 P40620 P40621 P40622 P40623 P40625 P40626 P40628 P40629 P40630 P40631
P40632 P40633 P40634 P40635 P40636 P40637 P40638 P40639 P40640 P40641 P40642
P40643 P40644 P40645 P40646 P40647 P40649 P40650 P40651 P40652 P40653 P40654
P40655 P40656 P40657 P40665 P40666 P40667 P40668 P40669 P40670 P40671 P40676
P40677 P40683 P40685 P40687 P40688 P40690 P40691 P40693 P40699 P40713 P40714
P40720 P40731 P40732 P40733 P40737 P40738 P40740 P40744 P40745 P40746 P40747
P40757 P40761 P40762 P40764 P40765 P40778 P40780 P40781 P40783 P40784 P40788
P40796 P40798 P40801 P40802 P40804 P40805 P40806 P40807 P40808 P40810 P40811
P40813 P40814 P40815 P40816 P40817 P40819 P40820 P40821 P40824 P40825 P40828
P40829 P40830 P40831 P40832 P40833 P40834 P40847 P40850 P40851 P40852 P40853
P40854 P40856 P40859 P40861 P40865 P40870 P40871 P40872 P40873 P40874 P40875
P40880 P40882 P40883 P40884 P40889 P40891 P40892 P40893 P40895 P40896 P40898
P40899 P40903 P40904 P40905 P40906 P40909 P40910 P40911 P40914 P40917 P40918

P40920 P40921 P40923 P40924 P40927 P40935 P40936 P40937 P40938 P40942 P40947
P40950 P40952 P40956 P40957 P40958 P40960 P40962 P40963 P40965 P40968 P40969
P40971 P40974 P40976 P40978 P40981 P40982 P40983 P40984 P40987 P40991 P40992
P40993 P40997 P40998 P40999 P41000 P41001 P41003 P41004 P41005 P41007 P41008
P41016 P41018 P41019 P41020 P41021 P41022 P41023 P41024 P41030 P41035 P41039
P41040 P41041 P41044 P41045 P41046 P41048 P41049 P41054 P41055 P41059 P41060
P41061 P41062 P41063 P41064 P41065 P41067 P41068 P41070 P41072 P41078 P41079
P41080 P41081 P41082 P41084 P41087 P41089 P41090 P41091 P41095 P41096 P41098
P41099 P41101 P41102 P41103 P41107 P41110 P41111 P41112 P41113 P41114 P41115
P41116 P41117 P41118 P41119 P41120 P41121 P41122 P41127 P41128 P41129 P41133
P41138 P41139 P41141 P41142 P41150 P41152 P41153 P41154 P41156 P41157 P41158
P41161 P41163 P41164 P41165 P41174 P41175 P41176 P41177 P41178 P41179 P41184
P41185 P41186 P41187 P41188 P41189 P41190 P41191 P41192 P41193 P41197 P41198
P41199 P41200 P41201 P41202 P41203 P41204 P41205 P41206 P41207 P41209 P41210
P41213 P41224 P41226 P41228 P41230 P41249 P41255 P41256 P41257 P41258 P41259
P41260 P41261 P41262 P41267 P41268 P41269 P41270 P41320 P41326 P41327 P41328
P41329 P41330 P41331 P41332 P41338 P41339 P41340 P41341 P41347 P41348 P41349
P41351 P41352 P41353 P41354 P41357 P41359 P41368 P41369 P41370 P41371 P41372
P41374 P41375 P41376 P41378 P41379 P41380 P41381 P41382 P41383 P41384 P41385
P41386 P41387 P41388 P41389 P41390 P41392 P41393 P41394 P41395 P41396 P41397
P41398 P41399 P41400 P41401 P41402 P41403 P41404 P41407 P41409 P41411 P41412
P41415 P41417 P41418 P41420 P41421 P41422 P41423 P41424 P41425 P41426 P41427
P41429 P41430 P41431 P41432 P41433 P41434 P41435 P41436 P41437 P41446 P41447
P41448 P41449 P41450 P41452 P41454 P41455 P41456 P41457 P41458 P41459 P41460
P41461 P41462 P41463 P41464 P41465 P41466 P41467 P41469 P41470 P41471 P41472
P41473 P41474 P41476 P41477 P41478 P41479 P41480 P41481 P41482 P41495 P41500
P41501 P41504 P41505 P41506 P41508 P41511 P41512 P41513 P41515 P41516 P41545
P41548 P41549 P41550 P41551 P41552 P41553 P41554 P41556 P41557 P41558 P41559
P41560 P41561 P41562 P41563 P41564 P41565 P41566 P41568 P41569 P41570 P41571
P41572 P41573 P41574 P41575 P41576 P41577 P41578 P41579 P41580 P41581 P41582
P41583 P41605 P41606 P41607 P41609 P41610 P41612 P41621 P41629 P41630 P41631
P41632 P41633 P41634 P41635 P41636 P41637 P41638 P41641 P41645 P41646 P41648
P41651 P41652 P41653 P41654 P41655 P41657 P41658 P41659 P41660 P41661 P41662
P41664 P41665 P41666 P41667 P41668 P41669 P41670 P41671 P41673 P41674 P41676
P41677 P41678 P41679 P41680 P41681 P41686 P41691 P41695 P41696 P41699 P41700
P41703 P41704 P41706 P41707 P41708 P41709 P41710 P41711 P41712 P41714 P41716
P41719 P41720 P41724 P41726 P41727 P41728 P41729 P41730 P41737 P41741 P41742
P41745 P41747 P41751 P41752 P41753 P41754 P41755 P41756 P41757 P41758 P41759
P41760 P41762 P41764 P41765 P41766 P41767 P41768 P41769 P41772 P41777 P41778
P41779 P41784 P41785 P41790 P41791 P41792 P41793 P41795 P41798 P41799 P41812
P41813 P41814 P41816 P41817 P41819 P41824 P41825 P41826 P41827 P41828 P41829
P41831 P41833 P41835 P41836 P41838 P41842 P41843 P41844 P41845 P41846 P41847
P41848 P41853 P41877 P41878 P41880 P41882 P41883 P41884 P41886 P41888 P41889
P41890 P41891 P41892 P41894 P41895 P41896 P41900 P41902 P41905 P41906 P41907

P41908 P41910 P41911 P41913 P41914 P41915 P41917 P41918 P41919 P41921 P41923
P41926 P41927 P41928 P41929 P41931 P41935 P41936 P41937 P41939 P41940 P41949
P41952 P41953 P41954 P41959 P41960 P41961 P41962 P41971 P41972 P41973 P41976
P41977 P41978 P41979 P41980 P41981 P41982 P41988 P41991 P41992 P41993 P41995
P41998 P41999 P42000 P42001 P42016 P42017 P42019 P42021 P42022 P42023 P42030
P42033 P42034 P42035 P42036 P42037 P42039 P42040 P42041 P42044 P42045 P42048
P42058 P42059 P42060 P42066 P42068 P42073 P42074 P42075 P42076 P42077 P42078
P42079 P42080 P42083 P42084 P42085 P42088 P42089 P42090 P42091 P42092 P42093
P42095 P42096 P42097 P42100 P42101 P42103 P42105 P42106 P42113 P42115 P42119
P42120 P42121 P42122 P42128 P42129 P42131 P42132 P42136 P42137 P42138 P42139
P42142 P42143 P42145 P42147 P42148 P42152 P42170 P42172 P42173 P42179 P42180
P42183 P42193 P42195 P42198 P42206 P42207 P42209 P42212 P42217 P42218 P42219
P42220 P42222 P42223 P42233 P42234 P42235 P42236 P42238 P42239 P42240 P42242
P42244 P42246 P42253 P42265 P42267 P42268 P42269 P42270 P42271 P42275 P42281
P42282 P42283 P42284 P42285 P42286 P42294 P42295 P42298 P42299 P42300 P42301
P42302 P42305 P42310 P42313 P42315 P42316 P42317 P42318 P42319 P42320 P42321
P42322 P42327 P42328 P42329 P42332 P42340 P42341 P42342 P42343 P42344 P42349
P42350 P42352 P42353 P42355 P42357 P42358 P42359 P42365 P42366 P42369 P42370
P42371 P42373 P42374 P42375 P42376 P42377 P42378 P42379 P42380 P42381 P42382
P42383 P42386 P42387 P42388 P42389 P42390 P42391 P42392 P42393 P42395 P42398
P42403 P42404 P42405 P42407 P42408 P42409 P42410 P42412 P42413 P42414 P42415
P42416 P42418 P42419 P42420 P42425 P42427 P42428 P42430 P42433 P42434 P42435
P42436 P42437 P42438 P42439 P42441 P42442 P42443 P42444 P42445 P42446 P42447
P42449 P42450 P42451 P42452 P42453 P42454 P42455 P42456 P42457 P42458 P42460
P42463 P42471 P42472 P42473 P42474 P42475 P42476 P42477 P42478 P42479 P42480
P42481 P42482 P42484 P42485 P42486 P42487 P42488 P42489 P42490 P42491 P42492
P42493 P42494 P42495 P42496 P42497 P42498 P42499 P42500 P42502 P42503 P42504
P42505 P42506 P42507 P42508 P42509 P42510 P42511 P42515 P42516 P42518 P42521
P42524 P42527 P42529 P42530 P42534 P42535 P42537 P42538 P42539 P42540 P42541
P42542 P42543 P42544 P42545 P42546 P42547 P42556 P42570 P42571 P42577 P42580
P42581 P42582 P42583 P42584 P42585 P42586 P42587 P42588 P42589 P42596 P42597
P42599 P42604 P42620 P42624 P42625 P42626 P42630 P42636 P42637 P42638 P42639
P42648 P42649 P42650 P42653 P42654 P42661 P42663 P42669 P42671 P42672 P42673
P42675 P42678 P42691 P42698 P42700 P42709 P42710 P42711 P42712 P42713 P42719
P42720 P42721 P42722 P42723 P42724 P42725 P42727 P42728 P42729 P42730 P42732
P42733 P42735 P42738 P42739 P42740 P42743 P42744 P42745 P42746 P42748 P42749
P42750 P42751 P42752 P42754 P42755 P42756 P42757 P42760 P42764 P42769 P42770
P42772 P42773 P42774 P42775 P42776 P42777 P42778 P42786 P42791 P42793 P42797
P42798 P42799 P42805 P42806 P42807 P42808 P42809 P42810 P42811 P42816 P42818
P42821 P42822 P42823 P42826 P42827 P42828 P42832 P42841 P42842 P42843 P42845
P42846 P42849 P42850 P42851 P42855 P42856 P42860 P42861 P42862 P42863 P42865
P42868 P42869 P42870 P42871 P42873 P42874 P42875 P42876 P42877 P42878 P42879
P42880 P42882 P42883 P42884 P42885 P42886 P42887 P42888 P42894 P42895 P42896
P42897 P42898 P42906 P42912 P42919 P42920 P42921 P42923 P42924 P42926 P42927

P42928 P42929 P42931 P42932 P42933 P42935 P42936 P42937 P42938 P42940 P42941
P42942 P42943 P42944 P42945 P42948 P42950 P42955 P42957 P42958 P42961 P42962
P42963 P42966 P42968 P42972 P42973 P42976 P42977 P42979 P42980 P42981 P42982
P42984 P42985 P42992 P43011 P43012 P43013 P43014 P43015 P43019 P43038 P43039
P43041 P43043 P43044 P43048 P43049 P43050 P43051 P43052 P43053 P43055 P43056
P43060 P43061 P43062 P43064 P43065 P43066 P43067 P43072 P43073 P43074 P43078
P43084 P43085 P43086 P43087 P43099 P43100 P43101 P43102 P43112 P43122 P43123
P43124 P43126 P43127 P43128 P43129 P43130 P43132 P43135 P43136 P43139 P43152
P43160 P43161 P43165 P43166 P43167 P43168 P43169 P43176 P43177 P43178 P43179
P43180 P43183 P43184 P43185 P43186 P43187 P43188 P43209 P43210 P43211 P43222
P43223 P43224 P43225 P43226 P43227 P43228 P43229 P43230 P43239 P43241 P43242
P43247 P43248 P43267 P43273 P43274 P43275 P43276 P43277 P43278 P43279 P43280
P43281 P43282 P43283 P43284 P43290 P43299 P43300 P43301 P43305 P43309 P43310
P43311 P43312 P43313 P43314 P43316 P43320 P43321 P43329 P43332 P43333 P43334
P43335 P43336 P43337 P43338 P43340 P43343 P43344 P43345 P43346 P43347 P43348
P43349 P43351 P43352 P43357 P43358 P43359 P43360 P43361 P43362 P43363 P43364
P43365 P43366 P43386 P43387 P43388 P43389 P43390 P43392 P43393 P43394 P43396
P43398 P43399 P43400 P43401 P43402 P43408 P43409 P43410 P43411 P43412 P43413
P43414 P43415 P43417 P43418 P43419 P43420 P43421 P43422 P43423 P43424 P43429
P43433 P43437 P43448 P43449 P43450 P43456 P43458 P43459 P43460 P43461 P43462
P43463 P43464 P43465 P43467 P43468 P43469 P43471 P43472 P43473 P43479 P43484
P43485 P43486 P43491 P43492 P43493 P43494 P43495 P43496 P43503 P43504 P43506
P43518 P43520 P43521 P43522 P43523 P43525 P43533 P43534 P43535 P43538 P43539
P43540 P43541 P43544 P43545 P43546 P43547 P43550 P43554 P43567 P43568 P43569
P43572 P43574 P43576 P43577 P43579 P43580 P43582 P43583 P43586 P43587 P43589
P43596 P43597 P43598 P43599 P43603 P43605 P43609 P43610 P43612 P43615 P43616
P43618 P43619 P43622 P43623 P43625 P43634 P43638 P43639 P43640 P43641 P43642
P43643 P43645 P43646 P43651 P43659 P43677 P43678 P43680 P43684 P43687 P43688
P43689 P43690 P43691 P43692 P43693 P43694 P43695 P43696 P43697 P43698 P43699
P43700 P43701 P43703 P43704 P43705 P43706 P43707 P43708 P43722 P43723 P43724
P43725 P43726 P43727 P43730 P43731 P43732 P43733 P43734 P43735 P43736 P43737
P43738 P43739 P43740 P43741 P43743 P43744 P43745 P43746 P43747 P43748 P43749
P43750 P43751 P43752 P43753 P43754 P43755 P43757 P43758 P43759 P43760 P43761
P43762 P43763 P43764 P43766 P43767 P43770 P43771 P43772 P43773 P43774 P43775
P43776 P43777 P43779 P43780 P43783 P43784 P43785 P43786 P43788 P43790 P43791
P43792 P43793 P43795 P43796 P43797 P43802 P43807 P43809 P43811 P43812 P43813
P43814 P43815 P43816 P43817 P43818 P43819 P43820 P43821 P43822 P43823 P43824
P43825 P43826 P43827 P43828 P43829 P43830 P43831 P43832 P43833 P43834 P43835
P43836 P43837 P43841 P43844 P43845 P43846 P43847 P43848 P43849 P43850 P43851
P43852 P43853 P43854 P43855 P43856 P43857 P43858 P43860 P43861 P43862 P43863
P43864 P43865 P43866 P43867 P43868 P43869 P43870 P43871 P43875 P43876 P43877
P43878 P43879 P43880 P43881 P43882 P43885 P43886 P43890 P43891 P43892 P43893
P43894 P43895 P43896 P43898 P43899 P43900 P43901 P43902 P43903 P43904 P43905
P43906 P43907 P43908 P43909 P43910 P43911 P43912 P43913 P43914 P43915 P43916

P43917 P43918 P43919 P43920 P43923 P43924 P43925 P43926 P43927 P43928 P43929
P43930 P43933 P43934 P43935 P43936 P43941 P43942 P43944 P43948 P43954 P43955
P43964 P43965 P43966 P43967 P43968 P43971 P43974 P43976 P43977 P43980 P43981
P43985 P43987 P43990 P43992 P43994 P43995 P43996 P43998 P44000 P44001 P44002
P44003 P44004 P44011 P44013 P44014 P44015 P44017 P44021 P44025 P44027 P44029
P44030 P44031 P44033 P44034 P44036 P44037 P44039 P44040 P44041 P44044 P44045
P44047 P44048 P44050 P44052 P44053 P44055 P44059 P44061 P44062 P44064 P44065
P44066 P44068 P44072 P44073 P44074 P44075 P44076 P44077 P44078 P44083 P44085
P44086 P44088 P44093 P44094 P44096 P44098 P44099 P44101 P44104 P44105 P44107
P44111 P44116 P44117 P44118 P44119 P44120 P44121 P44123 P44124 P44126 P44134
P44138 P44140 P44143 P44144 P44148 P44150 P44151 P44152 P44153 P44154 P44156
P44158 P44159 P44160 P44161 P44162 P44164 P44166 P44167 P44168 P44169 P44171
P44172 P44173 P44174 P44175 P44176 P44177 P44178 P44179 P44180 P44181 P44183
P44184 P44187 P44189 P44190 P44191 P44192 P44193 P44194 P44195 P44197 P44198
P44199 P44204 P44208 P44209 P44210 P44211 P44212 P44213 P44214 P44215 P44216
P44218 P44221 P44223 P44224 P44225 P44226 P44227 P44228 P44230 P44231 P44232
P44233 P44234 P44235 P44238 P44239 P44240 P44241 P44242 P44243 P44244 P44246
P44247 P44248 P44251 P44254 P44255 P44256 P44258 P44259 P44260 P44262 P44264
P44267 P44268 P44271 P44280 P44281 P44283 P44286 P44294 P44295 P44297 P44298
P44299 P44303 P44304 P44305 P44306 P44307 P44309 P44310 P44311 P44312 P44313
P44314 P44315 P44316 P44317 P44318 P44319 P44320 P44321 P44322 P44323 P44324
P44325 P44326 P44327 P44328 P44329 P44330 P44331 P44334 P44335 P44336 P44337
P44338 P44339 P44340 P44341 P44342 P44343 P44344 P44345 P44346 P44347 P44348
P44349 P44350 P44351 P44353 P44354 P44355 P44356 P44357 P44358 P44359 P44360
P44361 P44362 P44363 P44364 P44365 P44366 P44367 P44368 P44369 P44371 P44372
P44373 P44374 P44375 P44376 P44377 P44379 P44380 P44381 P44382 P44383 P44386
P44387 P44388 P44389 P44390 P44391 P44392 P44393 P44394 P44395 P44396 P44397
P44398 P44399 P44400 P44401 P44402 P44403 P44404 P44405 P44406 P44409 P44410
P44411 P44413 P44414 P44416 P44417 P44418 P44420 P44421 P44422 P44423 P44425
P44426 P44427 P44428 P44429 P44430 P44431 P44433 P44434 P44435 P44436 P44437
P44438 P44439 P44440 P44441 P44442 P44444 P44445 P44447 P44449 P44452 P44453
P44455 P44458 P44459 P44460 P44461 P44462 P44463 P44464 P44465 P44470 P44471
P44479 P44480 P44481 P44482 P44487 P44488 P44489 P44492 P44494 P44495 P44496
P44497 P44499 P44500 P44501 P44502 P44503 P44504 P44505 P44506 P44507 P44509
P44514 P44515 P44516 P44521 P44524 P44527 P44529 P44533 P44534 P44536 P44537
P44538 P44539 P44540 P44541 P44551 P44552 P44554 P44556 P44557 P44558 P44561
P44562 P44564 P44565 P44568 P44571 P44580 P44583 P44584 P44586 P44588 P44589
P44593 P44594 P44595 P44598 P44605 P44606 P44609 P44611 P44612 P44613 P44617
P44618 P44619 P44622 P44624 P44627 P44628 P44630 P44631 P44632 P44633 P44634
P44635 P44638 P44639 P44641 P44642 P44643 P44644 P44645 P44647 P44648 P44649
P44650 P44656 P44658 P44659 P44665 P44667 P44668 P44669 P44670 P44672 P44675
P44676 P44679 P44680 P44681 P44682 P44683 P44684 P44685 P44686 P44687 P44688
P44689 P44690 P44694 P44697 P44700 P44701 P44702 P44703 P44708 P44709 P44710
P44711 P44712 P44718 P44719 P44722 P44723 P44725 P44726 P44728 P44732 P44733

P44734 P44738 P44740 P44743 P44744 P44746 P44748 P44749 P44751 P44754 P44755
P44756 P44757 P44758 P44759 P44760 P44761 P44762 P44763 P44765 P44766 P44769
P44770 P44771 P44773 P44774 P44777 P44778 P44779 P44780 P44781 P44782 P44784
P44786 P44787 P44788 P44793 P44794 P44797 P44801 P44803 P44804 P44805 P44807
P44808 P44811 P44812 P44813 P44814 P44815 P44817 P44818 P44819 P44821 P44822
P44827 P44830 P44831 P44834 P44837 P44838 P44839 P44841 P44842 P44844 P44851
P44853 P44854 P44859 P44861 P44862 P44864 P44865 P44866 P44868 P44869 P44876
P44877 P44878 P44879 P44880 P44881 P44882 P44886 P44887 P44889 P44895 P44896
P44897 P44899 P44900 P44901 P44902 P44904 P44905 P44906 P44907 P44909 P44910
P44915 P44916 P44918 P44920 P44921 P44922 P44923 P44931 P44932 P44940 P44941
P44945 P44946 P44948 P44951 P44952 P44953 P44954 P44955 P44956 P44957 P44959
P44960 P44961 P44965 P44966 P44967 P44968 P44976 P44978 P44979 P44987 P44988
P44989 P44990 P44991 P44995 P44996 P44997 P44999 P45000 P45008 P45018 P45025
P45026 P45040 P45043 P45049 P45055 P45056 P45057 P45060 P45061 P45063 P45066
P45071 P45076 P45077 P45078 P45080 P45083 P45085 P45097 P45099 P45100 P45101
P45102 P45103 P45104 P45105 P45106 P45107 P45110 P45112 P45113 P45116 P45118
P45119 P45120 P45121 P45124 P45125 P45127 P45128 P45131 P45134 P45135 P45138
P45141 P45142 P45148 P45149 P45150 P45153 P45154 P45157 P45158 P45162 P45164
P45167 P45172 P45173 P45175 P45176 P45177 P45178 P45179 P45180 P45181 P45182
P45183 P45184 P45185 P45186 P45187 P45188 P45189 P45194 P45197 P45198 P45199
P45200 P45201 P45202 P45204 P45205 P45206 P45207 P45208 P45209 P45210 P45211
P45213 P45215 P45241 P45242 P45244 P45245 P45248 P45249 P45250 P45252 P45253
P45255 P45256 P45257 P45260 P45261 P45262 P45265 P45266 P45267 P45269 P45271
P45273 P45277 P45281 P45283 P45284 P45291 P45292 P45293 P45294 P45297 P45298
P45300 P45302 P45303 P45305 P45308 P45309 P45310 P45311 P45312 P45313 P45314
P45324 P45331 P45334 P45337 P45339 P45340 P45344 P45347 P45349 P45350 P45353
P45358 P45359 P45360 P45362 P45363 P45366 P45367 P45368 P45369 P45370 P45371
P45372 P45373 P45374 P45375 P45377 P45378 P45379 P45381 P45382 P45383 P45388
P45410 P45413 P45414 P45415 P45416 P45418 P45421 P45423 P45424 P45425 P45429
P45430 P45431 P45438 P45439 P45440 P45441 P45443 P45444 P45446 P45447 P45450
P45451 P45454 P45455 P45461 P45462 P45463 P45465 P45469 P45470 P45472 P45475
P45480 P45486 P45487 P45488 P45489 P45491 P45493 P45494 P45496 P45497 P45505
P45511 P45512 P45513 P45514 P45515 P45516 P45517 P45521 P45527 P45530 P45531
P45532 P45541 P45543 P45544 P45545 P45548 P45549 P45550 P45551 P45552 P45554
P45555 P45558 P45563 P45568 P45573 P45574 P45575 P45579 P45581 P45583 P45584
P45585 P45586 P45587 P45588 P45600 P45601 P45602 P45603 P45605 P45606 P45607
P45612 P45614 P45616 P45619 P45620 P45621 P45622 P45623 P45624 P45625 P45626
P45627 P45631 P45632 P45633 P45647 P45648 P45651 P45671 P45672 P45674 P45679
P45680 P45681 P45683 P45684 P45686 P45687 P45688 P45689 P45690 P45693 P45694
P45702 P45703 P45707 P45718 P45719 P45720 P45721 P45722 P45724 P45725 P45726
P45727 P45728 P45729 P45730 P45731 P45732 P45733 P45734 P45735 P45736 P45737
P45738 P45739 P45740 P45742 P45743 P45744 P45745 P45748 P45749 P45750 P45753
P45771 P45785 P45786 P45787 P45788 P45792 P45795 P45799 P45805 P45807 P45809
P45811 P45812 P45813 P45815 P45818 P45821 P45831 P45832 P45833 P45834 P45835

P45836 P45837 P45840 P45841 P45842 P45855 P45857 P45858 P45859 P45863 P45864
P45868 P45870 P45871 P45872 P45873 P45875 P45876 P45882 P45883 P45885 P45886
P45887 P45888 P45889 P45890 P45891 P45898 P45899 P45902 P45903 P45904 P45905
P45906 P45907 P45908 P45909 P45910 P45911 P45912 P45913 P45914 P45915 P45916
P45917 P45918 P45920 P45921 P45922 P45923 P45924 P45925 P45926 P45927 P45929
P45930 P45931 P45932 P45933 P45934 P45935 P45936 P45937 P45938 P45939 P45940
P45941 P45943 P45944 P45945 P45947 P45948 P45949 P45950 P45951 P45956 P45959
P45960 P45961 P45962 P45964 P45965 P45966 P45967 P45968 P45969 P45970 P45972
P45975 P45976 P45978 P45981 P45982 P45986 P46003 P46012 P46015 P46016 P46017
P46033 P46034 P46035 P46036 P46037 P46038 P46039 P46040 P46041 P46042 P46043
P46044 P46045 P46046 P46047 P46048 P46049 P46050 P46051 P46052 P46053 P46054
P46056 P46057 P46058 P46064 P46067 P46068 P46070 P46072 P46079 P46080 P46082
P46088 P46099 P46103 P46107 P46117 P46118 P46126 P46142 P46144 P46148 P46151
P46154 P46155 P46172 P46173 P46174 P46175 P46176 P46177 P46178 P46179 P46180
P46181 P46182 P46183 P46184 P46185 P46190 P46191 P46195 P46198 P46199 P46200
P46204 P46205 P46207 P46208 P46209 P46211 P46212 P46213 P46214 P46215 P46216
P46217 P46218 P46219 P46220 P46221 P46224 P46225 P46226 P46227 P46228 P46230
P46232 P46234 P46235 P46240 P46241 P46243 P46244 P46245 P46246 P46248 P46251
P46252 P46254 P46255 P46256 P46257 P46258 P46259 P46263 P46264 P46265 P46266
P46267 P46272 P46275 P46276 P46277 P46278 P46279 P46280 P46281 P46283 P46284
P46285 P46286 P46289 P46290 P46291 P46292 P46293 P46295 P46296 P46297 P46298
P46299 P46300 P46301 P46302 P46303 P46307 P46308 P46309 P46314 P46315 P46320
P46321 P46323 P46325 P46326 P46327 P46328 P46329 P46330 P46331 P46334 P46336
P46337 P46343 P46347 P46350 P46351 P46352 P46353 P46354 P46355 P46356 P46357
P46358 P46361 P46362 P46363 P46364 P46365 P46366 P46367 P46368 P46369 P46370
P46371 P46372 P46373 P46374 P46376 P46377 P46378 P46380 P46381 P46382 P46385
P46386 P46387 P46388 P46389 P46390 P46391 P46393 P46394 P46395 P46396 P46397
P46398 P46399 P46409 P46413 P46415 P46416 P46417 P46418 P46419 P46420 P46423
P46426 P46427 P46428 P46429 P46430 P46431 P46432 P46433 P46434 P46436 P46437
P46439 P46440 P46441 P46446 P46449 P46451 P46453 P46456 P46457 P46465 P46466
P46468 P46470 P46471 P46472 P46479 P46484 P46485 P46486 P46487 P46488 P46489
P46493 P46494 P46495 P46496 P46500 P46502 P46505 P46507 P46509 P46510 P46511
P46512 P46513 P46514 P46515 P46516 P46517 P46518 P46519 P46520 P46521 P46522
P46523 P46525 P46526 P46532 P46534 P46535 P46536 P46537 P46538 P46539 P46541
P46542 P46543 P46544 P46545 P46547 P46548 P46550 P46551 P46553 P46554 P46562
P46563 P46575 P46577 P46581 P46582 P46583 P46585 P46586 P46588 P46589 P46595
P46600 P46601 P46602 P46603 P46604 P46605 P46606 P46608 P46609 P46610 P46618
P46635 P46637 P46639 P46640 P46643 P46646 P46655 P46660 P46664 P46665 P46666
P46667 P46668 P46669 P46670 P46672 P46674 P46675 P46676 P46677 P46678 P46679
P46680 P46681 P46683 P46684 P46692 P46696 P46697 P46700 P46701 P46702 P46703
P46705 P46707 P46708 P46710 P46711 P46712 P46713 P46714 P46715 P46716 P46718
P46723 P46724 P46725 P46726 P46728 P46729 P46730 P46731 P46732 P46736 P46737
P46740 P46741 P46742 P46743 P46744 P46745 P46746 P46747 P46748 P46749 P46750
P46751 P46752 P46753 P46754 P46755 P46756 P46757 P46758 P46759 P46760 P46761

P46762 P46763 P46764 P46765 P46766 P46767 P46768 P46770 P46771 P46772 P46773
P46775 P46780 P46786 P46787 P46788 P46789 P46790 P46792 P46794 P46795 P46796
P46797 P46799 P46801 P46805 P46806 P46807 P46808 P46809 P46810 P46811 P46813
P46814 P46815 P46817 P46818 P46819 P46820 P46828 P46831 P46833 P46834 P46835
P46836 P46837 P46843 P46844 P46846 P46849 P46850 P46852 P46853 P46854 P46855
P46856 P46857 P46859 P46861 P46862 P46864 P46865 P46867 P46869 P46870 P46871
P46872 P46873 P46874 P46875 P46879 P46881 P46887 P46888 P46891 P46893 P46894
P46897 P46898 P46899 P46903 P46905 P46906 P46908 P46910 P46914 P46915 P46916
P46917 P46924 P46926 P46936 P46942 P46945 P46946 P46948 P46949 P46952 P46954
P46955 P46957 P46958 P46959 P46962 P46963 P46969 P46973 P46974 P46979 P46980
P46983 P46984 P46987 P46988 P46994 P46995 P46997 P47004 P47005 P47006 P47007
P47009 P47011 P47012 P47015 P47016 P47017 P47018 P47019 P47021 P47023 P47027
P47035 P47036 P47037 P47038 P47039 P47042 P47043 P47044 P47047 P47049 P47050
P47051 P47053 P47058 P47059 P47063 P47066 P47068 P47072 P47073 P47074 P47076
P47077 P47078 P47079 P47083 P47084 P47085 P47086 P47087 P47089 P47092 P47093
P47095 P47096 P47099 P47101 P47103 P47105 P47107 P47108 P47110 P47112 P47115
P47118 P47119 P47120 P47122 P47123 P47125 P47126 P47128 P47129 P47130 P47132
P47134 P47136 P47137 P47138 P47143 P47146 P47149 P47151 P47152 P47156 P47158
P47162 P47163 P47164 P47165 P47166 P47167 P47168 P47169 P47171 P47172 P47173
P47174 P47175 P47176 P47177 P47181 P47182 P47183 P47184 P47188 P47199 P47200
P47201 P47202 P47206 P47207 P47208 P47209 P47227 P47228 P47229 P47230 P47231
P47232 P47233 P47235 P47236 P47237 P47238 P47239 P47240 P47242 P47243 P47246
P47247 P47248 P47249 P47250 P47251 P47252 P47253 P47254 P47255 P47256 P47257
P47258 P47259 P47264 P47265 P47266 P47267 P47268 P47269 P47270 P47271 P47272
P47273 P47275 P47276 P47277 P47278 P47280 P47281 P47282 P47283 P47284 P47292
P47293 P47295 P47296 P47297 P47298 P47299 P47300 P47302 P47303 P47304 P47305
P47306 P47311 P47312 P47316 P47319 P47320 P47327 P47328 P47329 P47330 P47331
P47333 P47334 P47335 P47336 P47337 P47338 P47339 P47340 P47342 P47343 P47345
P47346 P47347 P47348 P47349 P47350 P47352 P47353 P47354 P47356 P47357 P47358
P47359 P47361 P47362 P47363 P47364 P47368 P47370 P47371 P47372 P47373 P47374
P47378 P47380 P47382 P47385 P47386 P47387 P47388 P47389 P47391 P47394 P47396
P47397 P47398 P47399 P47400 P47401 P47402 P47403 P47404 P47405 P47406 P47407
P47408 P47409 P47410 P47411 P47412 P47413 P47414 P47415 P47418 P47419 P47420
P47421 P47422 P47423 P47424 P47428 P47429 P47433 P47436 P47437 P47438 P47439
P47440 P47441 P47442 P47443 P47444 P47445 P47447 P47448 P47449 P47450 P47451
P47453 P47455 P47456 P47457 P47458 P47459 P47461 P47463 P47464 P47465 P47469
P47470 P47471 P47472 P47473 P47474 P47476 P47477 P47478 P47479 P47480 P47481
P47482 P47486 P47487 P47488 P47490 P47491 P47492 P47493 P47494 P47495 P47496
P47499 P47500 P47505 P47506 P47507 P47508 P47510 P47512 P47513 P47514 P47515
P47516 P47518 P47520 P47524 P47525 P47526 P47527 P47530 P47534 P47537 P47538
P47540 P47541 P47542 P47543 P47547 P47553 P47566 P47567 P47571 P47572 P47574
P47575 P47576 P47577 P47579 P47581 P47582 P47583 P47584 P47585 P47587 P47588
P47589 P47591 P47592 P47593 P47595 P47596 P47597 P47599 P47603 P47604 P47605
P47606 P47607 P47609 P47610 P47612 P47613 P47614 P47615 P47616 P47617 P47618

P47619 P47620 P47621 P47622 P47623 P47624 P47628 P47631 P47632 P47633 P47634
P47636 P47648 P47649 P47656 P47657 P47658 P47660 P47661 P47663 P47664 P47665
P47667 P47669 P47670 P47672 P47673 P47682 P47683 P47684 P47686 P47687 P47691
P47692 P47693 P47696 P47698 P47699 P47700 P47701 P47703 P47704 P47705 P47706
P47707 P47714 P47715 P47716 P47717 P47718 P47719 P47720 P47721 P47722 P47723
P47724 P47728 P47729 P47732 P47734 P47737 P47738 P47762 P47763 P47765 P47767
P47768 P47771 P47772 P47773 P47788 P47792 P47794 P47808 P47813 P47814 P47815
P47816 P47822 P47823 P47825 P47826 P47827 P47828 P47829 P47830 P47831 P47832
P47834 P47837 P47838 P47839 P47840 P47846 P47848 P47849 P47850 P47855 P47856
P47859 P47897 P47902 P47903 P47904 P47905 P47906 P47907 P47908 P47909 P47910
P47916 P47917 P47919 P47920 P47921 P47922 P47923 P47924 P47928 P47938 P47943
P47944 P47945 P47947 P47948 P47949 P47950 P47952 P47956 P47957 P47958 P47959
P47961 P47967 P47976 P47977 P47979 P47980 P47988 P47990 P47995 P47996 P47999
P48000 P48001 P48002 P48003 P48004 P48006 P48007 P48008 P48009 P48011 P48012
P48013 P48015 P48024 P48026 P48028 P48031 P48045 P48046 P48053 P48054 P48063
P48064 P48069 P48070 P48072 P48073 P48074 P48075 P48077 P48099 P48100 P48101
P48110 P48118 P48119 P48120 P48124 P48125 P48126 P48127 P48128 P48129 P48130
P48131 P48132 P48133 P48134 P48135 P48136 P48137 P48138 P48139 P48140 P48151
P48153 P48155 P48161 P48162 P48163 P48190 P48200 P48204 P48205 P48206 P48207
P48208 P48209 P48210 P48212 P48213 P48214 P48215 P48216 P48219 P48220 P48223
P48224 P48226 P48228 P48229 P48232 P48234 P48235 P48237 P48238 P48239 P48240
P48246 P48247 P48254 P48255 P48260 P48261 P48262 P48267 P48270 P48271 P48273
P48274 P48275 P48276 P48277 P48279 P48282 P48286 P48288 P48289 P48291 P48292
P48293 P48294 P48295 P48296 P48308 P48311 P48312 P48319 P48322 P48323 P48324
P48326 P48327 P48328 P48329 P48330 P48332 P48334 P48335 P48336 P48337 P48338
P48339 P48340 P48341 P48342 P48343 P48344 P48345 P48350 P48351 P48352 P48354
P48358 P48361 P48362 P48363 P48364 P48365 P48367 P48368 P48369 P48370 P48371
P48372 P48378 P48379 P48380 P48381 P48383 P48384 P48385 P48386 P48387 P48388
P48389 P48390 P48391 P48392 P48393 P48394 P48395 P48396 P48397 P48398 P48399
P48400 P48401 P48402 P48403 P48404 P48406 P48407 P48408 P48409 P48416 P48418
P48419 P48420 P48424 P48425 P48427 P48428 P48429 P48430 P48433 P48435 P48437
P48438 P48443 P48445 P48446 P48447 P48451 P48456 P48458 P48459 P48460 P48461
P48462 P48463 P48464 P48465 P48466 P48467 P48479 P48480 P48481 P48482 P48483
P48484 P48485 P48486 P48487 P48488 P48489 P48490 P48492 P48493 P48494 P48495
P48496 P48497 P48498 P48499 P48500 P48501 P48507 P48508 P48510 P48511 P48512
P48513 P48514 P48515 P48516 P48523 P48525 P48526 P48527 P48528 P48529 P48534
P48535 P48536 P48539 P48556 P48557 P48568 P48570 P48571 P48572 P48573 P48574
P48575 P48576 P48577 P48578 P48579 P48580 P48581 P48584 P48585 P48586 P48587
P48590 P48591 P48592 P48596 P48597 P48598 P48599 P48600 P48601 P48603 P48605
P48606 P48609 P48611 P48633 P48635 P48636 P48637 P48638 P48639 P48640 P48641
P48642 P48646 P48647 P48648 P48649 P48668 P48671 P48672 P48673 P48674 P48677
P48682 P48683 P48684 P48685 P48686 P48687 P48688 P48689 P48690 P48691 P48692
P48693 P48694 P48695 P48696 P48697 P48698 P48699 P48700 P48701 P48702 P48703
P48704 P48705 P48706 P48707 P48708 P48709 P48710 P48711 P48712 P48713 P48714

P48715 P48716 P48717 P48718 P48719 P48720 P48722 P48724 P48726 P48728 P48735
P48742 P48743 P48752 P48753 P48769 P48774 P48775 P48776 P48778 P48779 P48780
P48781 P48782 P48783 P48784 P48785 P48786 P48787 P48788 P48790 P48791 P48793
P48795 P48797 P48809 P48810 P48812 P48814 P48815 P48817 P48822 P48826 P48828
P48840 P48843 P48844 P48847 P48848 P48849 P48850 P48851 P48852 P48853 P48854
P48855 P48856 P48857 P48858 P48859 P48860 P48861 P48862 P48864 P48865 P48883
P48936 P48938 P48939 P48940 P48941 P48943 P48944 P48945 P48946 P48948 P48949
P48950 P48951 P48952 P48953 P48955 P48956 P48957 P48958 P48959 P48963 P48964
P48965 P48966 P48967 P48968 P48971 P48972 P48976 P48977 P48983 P48984 P48985
P48987 P48988 P48989 P48991 P48992 P48993 P48996 P48997 P48998 P49004 P49016
P49026 P49030 P49031 P49032 P49033 P49034 P49035 P49036 P49037 P49039 P49040
P49050 P49055 P49057 P49058 P49067 P49068 P49077 P49078 P49079 P49080 P49085
P49088 P49089 P49090 P49091 P49092 P49093 P49094 P49095 P49102 P49105 P49114
P49115 P49117 P49119 P49120 P49128 P49140 P49148 P49149 P49152 P49153 P49155
P49156 P49162 P49163 P49164 P49165 P49168 P49169 P49170 P49172 P49193 P49198
P49199 P49200 P49202 P49205 P49206 P49208 P49209 P49210 P49211 P49212 P49213
P49214 P49215 P49216 P49221 P49225 P49226 P49228 P49230 P49231 P49232 P49233
P49234 P49235 P49239 P49240 P49248 P49249 P49251 P49261 P49264 P49292 P49293
P49295 P49298 P49299 P49304 P49305 P49306 P49307 P49309 P49310 P49311 P49313
P49314 P49315 P49316 P49317 P49318 P49320 P49322 P49323 P49324 P49330 P49332
P49335 P49336 P49342 P49348 P49357 P49358 P49359 P49360 P49361 P49362 P49363
P49364 P49365 P49367 P49372 P49373 P49378 P49379 P49381 P49383 P49384 P49385
P49386 P49387 P49388 P49389 P49391 P49393 P49394 P49395 P49396 P49397 P49398
P49399 P49401 P49402 P49403 P49404 P49408 P49409 P49412 P49413 P49417 P49420
P49421 P49422 P49423 P49425 P49428 P49431 P49432 P49433 P49434 P49435 P49440
P49448 P49449 P49451 P49455 P49459 P49462 P49463 P49464 P49465 P49466 P49467
P49468 P49469 P49490 P49491 P49492 P49493 P49494 P49495 P49496 P49497 P49498
P49499 P49500 P49501 P49502 P49503 P49504 P49505 P49506 P49507 P49518 P49519
P49520 P49521 P49522 P49530 P49532 P49533 P49534 P49535 P49536 P49537 P49540
P49541 P49544 P49545 P49546 P49547 P49548 P49549 P49550 P49551 P49552 P49553
P49554 P49555 P49556 P49557 P49558 P49559 P49560 P49561 P49562 P49563 P49564
P49565 P49566 P49567 P49568 P49569 P49570 P49572 P49574 P49576 P49577 P49592
P49594 P49595 P49596 P49601 P49604 P49607 P49608 P49609 P49611 P49612 P49613
P49617 P49622 P49623 P49624 P49627 P49628 P49629 P49632 P49633 P49636 P49637
P49639 P49640 P49644 P49645 P49665 P49667 P49668 P49669 P49671 P49672 P49673
P49677 P49678 P49679 P49680 P49690 P49692 P49693 P49694 P49695 P49696 P49700
P49704 P49707 P49709 P49714 P49716 P49717 P49718 P49720 P49722 P49723 P49724
P49725 P49726 P49730 P49731 P49735 P49739 P49740 P49741 P49742 P49743 P49749
P49760 P49762 P49770 P49774 P49775 P49776 P49777 P49778 P49811 P49812 P49814
P49818 P49819 P49823 P49824 P49826 P49827 P49828 P49829 P49830 P49831 P49832
P49833 P49834 P49835 P49836 P49837 P49838 P49839 P49843 P49844 P49846 P49847
P49849 P49850 P49851 P49852 P49854 P49855 P49859 P49860 P49861 P49865 P49867
P49868 P49869 P49870 P49871 P49872 P49880 P49881 P49882 P49883 P49889 P49890
P49900 P49902 P49905 P49906 P49910 P49911 P49914 P49917 P49925 P49943 P49944

P49945 P49946 P49947 P49948 P49949 P49953 P49954 P49955 P49956 P49957 P49958
P49960 P49973 P49974 P49975 P49979 P49981 P49982 P49983 P49985 P49986 P49987
P49988 P49990 P49991 P49996 P49997 P50016 P50018 P50019 P50020 P50021 P50022
P50023 P50025 P50026 P50027 P50028 P50037 P50038 P50044 P50045 P50046 P50047
P50049 P50053 P50056 P50059 P50061 P50062 P50063 P50064 P50065 P50066 P50067
P50068 P50069 P50070 P50071 P50074 P50075 P50084 P50086 P50089 P50090 P50094
P50096 P50097 P50100 P50104 P50105 P50106 P50107 P50109 P50111 P50113 P50120
P50121 P50124 P50125 P50134 P50135 P50136 P50138 P50139 P50141 P50143 P50157
P50158 P50159 P50160 P50161 P50162 P50163 P50164 P50165 P50166 P50167 P50173
P50174 P50176 P50177 P50178 P50179 P50180 P50181 P50182 P50183 P50184 P50185
P50186 P50187 P50188 P50189 P50190 P50191 P50192 P50193 P50194 P50195 P50196
P50199 P50201 P50202 P50203 P50204 P50205 P50206 P50207 P50208 P50209 P50210
P50211 P50212 P50213 P50214 P50215 P50216 P50217 P50218 P50219 P50220 P50221
P50237 P50238 P50239 P50240 P50241 P50242 P50243 P50244 P50245 P50246 P50247
P50249 P50250 P50251 P50252 P50254 P50255 P50256 P50257 P50258 P50259 P50260
P50261 P50262 P50263 P50264 P50265 P50266 P50275 P50277 P50278 P50286 P50288
P50292 P50293 P50294 P50295 P50297 P50298 P50299 P50300 P50301 P50302 P50303
P50304 P50305 P50306 P50307 P50308 P50310 P50311 P50312 P50313 P50314 P50315
P50317 P50319 P50320 P50321 P50322 P50323 P50324 P50326 P50327 P50328 P50329
P50330 P50331 P50337 P50338 P50344 P50345 P50346 P50347 P50348 P50349 P50352
P50353 P50354 P50355 P50358 P50359 P50360 P50361 P50362 P50371 P50372 P50373
P50374 P50375 P50376 P50377 P50378 P50379 P50380 P50381 P50382 P50383 P50384
P50386 P50387 P50388 P50389 P50410 P50430 P50431 P50432 P50433 P50434 P50435
P50436 P50437 P50439 P50444 P50445 P50446 P50452 P50455 P50456 P50457 P50465
P50471 P50472 P50474 P50475 P50476 P50480 P50481 P50483 P50484 P50485 P50486
P50503 P50504 P50506 P50507 P50508 P50509 P50510 P50511 P50512 P50513 P50514
P50521 P50524 P50525 P50529 P50531 P50532 P50533 P50534 P50536 P50539 P50540
P50541 P50543 P50546 P50547 P50549 P50553 P50554 P50557 P50558 P50559 P50561
P50562 P50563 P50564 P50565 P50566 P50567 P50569 P50574 P50575 P50576 P50577
P50581 P50583 P50584 P50587 P50589 P50590 P50594 P50597 P50602 P50603 P50610
P50611 P50614 P50615 P50618 P50619 P50620 P50621 P50623 P50624 P50625 P50626
P50628 P50639 P50641 P50642 P50643 P50646 P50647 P50648 P50649 P50650 P50651
P50652 P50719 P50727 P50728 P50729 P50734 P50735 P50736 P50740 P50746 P50747
P50751 P50752 P50753 P50754 P50755 P50756 P50766 P50767 P50768 P50769 P50770
P50771 P50772 P50773 P50774 P50775 P50776 P50777 P50786 P50787 P50788 P50789
P50790 P50791 P50792 P50793 P50802 P50803 P50804 P50805 P50806 P50807 P50809
P50810 P50812 P50813 P50814 P50815 P50816 P50817 P50818 P50819 P50820 P50821
P50822 P50823 P50824 P50825 P50826 P50830 P50831 P50832 P50833 P50834 P50835
P50838 P50840 P50842 P50843 P50844 P50845 P50846 P50848 P50849 P50853 P50854
P50855 P50856 P50857 P50858 P50861 P50862 P50863 P50866 P50867 P50868 P50869
P50870 P50871 P50872 P50874 P50875 P50879 P50883 P50884 P50885 P50886 P50888
P50889 P50891 P50892 P50893 P50894 P50896 P50898 P50901 P50902 P50906 P50908
P50909 P50910 P50912 P50913 P50917 P50918 P50919 P50921 P50922 P50923 P50924
P50926 P50932 P50933 P50934 P50935 P50936 P50937 P50938 P50947 P50970 P50971

P50972 P50977 P50978 P50979 P50986 P50987 P50988 P50991 P50998 P50999 P51007
P51008 P51009 P51010 P51011 P51012 P51013 P51014 P51015 P51016 P51017 P51018
P51019 P51020 P51023 P51024 P51025 P51026 P51031 P51032 P51033 P51034 P51035
P51036 P51037 P51038 P51039 P51040 P51041 P51042 P51043 P51044 P51045 P51053
P51056 P51058 P51059 P51060 P51061 P51062 P51063 P51064 P51066 P51067 P51069
P51070 P51071 P51072 P51073 P51074 P51075 P51076 P51077 P51078 P51079 P51080
P51081 P51082 P51083 P51084 P51085 P51086 P51087 P51088 P51089 P51090 P51091
P51092 P51093 P51094 P51101 P51103 P51104 P51105 P51106 P51107 P51108 P51109
P51110 P51115 P51117 P51118 P51119 P51120 P51121 P51122 P51126 P51129 P51137
P51138 P51139 P51145 P51163 P51181 P51182 P51185 P51187 P51188 P51189 P51190
P51191 P51192 P51201 P51202 P51203 P51204 P51205 P51207 P51208 P51209 P51210
P51211 P51213 P51214 P51216 P51217 P51218 P51219 P51223 P51224 P51225 P51226
P51227 P51228 P51229 P51230 P51231 P51233 P51234 P51235 P51237 P51238 P51240
P51241 P51248 P51249 P51250 P51251 P51252 P51253 P51254 P51255 P51256 P51257
P51266 P51267 P51268 P51269 P51270 P51271 P51272 P51273 P51277 P51278 P51281
P51282 P51286 P51287 P51288 P51289 P51290 P51291 P51292 P51293 P51294 P51295
P51296 P51298 P51299 P51300 P51301 P51302 P51303 P51304 P51305 P51306 P51307
P51308 P51309 P51310 P51311 P51312 P51313 P51314 P51319 P51320 P51329 P51331
P51332 P51333 P51334 P51335 P51336 P51337 P51338 P51339 P51342 P51344 P51345
P51346 P51348 P51349 P51350 P51351 P51352 P51354 P51355 P51359 P51361 P51362
P51366 P51372 P51373 P51375 P51376 P51379 P51380 P51382 P51383 P51384 P51386
P51394 P51400 P51406 P51407 P51408 P51409 P51411 P51413 P51414 P51415 P51416
P51417 P51418 P51419 P51420 P51421 P51422 P51423 P51425 P51426 P51427 P51428
P51429 P51431 P51434 P51438 P51440 P51441 P51442 P51443 P51452 P51464 P51465
P51469 P51501 P51502 P51503 P51504 P51505 P51506 P51507 P51508 P51509 P51514
P51521 P51522 P51530 P51534 P51535 P51537 P51540 P51541 P51544 P51545 P51546
P51549 P51550 P51551 P51552 P51553 P51554 P51555 P51558 P51560 P51561 P51562
P51565 P51566 P51567 P51568 P51583 P51585 P51591 P51592 P51594 P51595 P51596
P51597 P51598 P51599 P51600 P51601 P51602 P51603 P51604 P51605 P51606 P51607
P51615 P51616 P51618 P51634 P51649 P51650 P51652 P51666 P51667 P51668 P51687
P51691 P51695 P51696 P51702 P51703 P51704 P51705 P51706 P51707 P51711 P51713
P51714 P51715 P51716 P51717 P51718 P51719 P51720 P51721 P51722 P51723 P51724
P51725 P51726 P51730 P51731 P51732 P51733 P51734 P51735 P51736 P51737 P51738
P51739 P51754 P51755 P51767 P51768 P51772 P51774 P51775 P51776 P51777 P51778
P51781 P51783 P51785 P51786 P51808 P51814 P51815 P51820 P51826 P51827 P51837
P51838 P51844 P51845 P51846 P51850 P51851 P51852 P51853 P51854 P51855 P51856
P51861 P51873 P51874 P51879 P51889 P51892 P51893 P51894 P51896 P51897 P51900
P51902 P51903 P51908 P51911 P51913 P51945 P51947 P51948 P51949 P51950 P51951
P51953 P51954 P51956 P51958 P51959 P51960 P51961 P51962 P51963 P51967 P51968
P51971 P51974 P51978 P51979 P51981 P51984 P51985 P51986 P51987 P51988 P51989
P51990 P51992 P51994 P51997 P51999 P52000 P52001 P52006 P52007 P52009 P52012
P52017 P52021 P52022 P52023 P52024 P52025 P52026 P52027 P52028 P52029 P52030
P52031 P52034 P52035 P52037 P52039 P52040 P52043 P52044 P52045 P52048 P52052
P52053 P52055 P52056 P52057 P52060 P52061 P52062 P52063 P52064 P52069 P52070

P52086 P52087 P52089 P52093 P52095 P52096 P52097 P52102 P52111 P52116 P52119
P52123 P52124 P52125 P52126 P52127 P52128 P52129 P52130 P52131 P52132 P52133
P52139 P52140 P52141 P52144 P52147 P52148 P52150 P52151 P52152 P52153 P52154
P52155 P52156 P52157 P52158 P52160 P52161 P52162 P52164 P52167 P52168 P52172
P52179 P52184 P52197 P52200 P52201 P52207 P52208 P52209 P52210 P52213 P52215
P52216 P52217 P52227 P52228 P52230 P52231 P52232 P52233 P52270 P52271 P52273
P52274 P52275 P52276 P52277 P52278 P52279 P52283 P52284 P52285 P52288 P52297
P52299 P52304 P52308 P52309 P52310 P52311 P52313 P52316 P52317 P52318 P52322
P52323 P52324 P52325 P52326 P52327 P52328 P52329 P52331 P52334 P52337 P52338
P52339 P52341 P52342 P52343 P52344 P52345 P52346 P52347 P52348 P52349 P52354
P52355 P52356 P52357 P52359 P52360 P52363 P52364 P52365 P52367 P52374 P52375
P52376 P52377 P52378 P52379 P52384 P52385 P52386 P52387 P52388 P52389 P52390
P52391 P52392 P52393 P52415 P52416 P52417 P52418 P52419 P52420 P52421 P52422
P52423 P52424 P52425 P52426 P52427 P52428 P52432 P52434 P52435 P52437 P52438
P52439 P52440 P52441 P52442 P52446 P52447 P52448 P52450 P52451 P52452 P52453
P52454 P52455 P52456 P52457 P52458 P52462 P52463 P52464 P52467 P52468 P52469
P52470 P52471 P52472 P52475 P52478 P52480 P52482 P52483 P52484 P52486 P52487
P52489 P52491 P52492 P52493 P52494 P52495 P52496 P52499 P52502 P52506 P52507
P52516 P52517 P52518 P52519 P52520 P52521 P52522 P52523 P52524 P52525 P52527
P52528 P52529 P52530 P52531 P52532 P52533 P52534 P52535 P52536 P52537 P52538
P52539 P52540 P52541 P52544 P52545 P52548 P52550 P52551 P52552 P52553 P52554
P52557 P52558 P52559 P52560 P52561 P52562 P52563 P52570 P52574 P52575 P52576
P52578 P52579 P52580 P52581 P52595 P52596 P52598 P52601 P52602 P52603 P52604
P52605 P52606 P52608 P52609 P52611 P52614 P52617 P52618 P52619 P52620 P52621
P52622 P52623 P52624 P52627 P52629 P52640 P52641 P52643 P52644 P52647 P52651
P52652 P52653 P52654 P52655 P52656 P52657 P52658 P52659 P52660 P52661 P52662
P52665 P52666 P52667 P52669 P52670 P52671 P52672 P52673 P52674 P52675 P52676
P52677 P52678 P52679 P52680 P52681 P52683 P52684 P52685 P52686 P52689 P52690
P52691 P52692 P52693 P52694 P52695 P52696 P52697 P52698 P52704 P52705 P52709
P52713 P52720 P52721 P52722 P52723 P52724 P52725 P52726 P52727 P52728 P52729
P52730 P52733 P52736 P52737 P52738 P52740 P52741 P52742 P52743 P52744 P52746
P52747 P52761 P52762 P52763 P52767 P52771 P52772 P52776 P52777 P52778 P52779
P52780 P52782 P52783 P52788 P52790 P52807 P52809 P52810 P52811 P52812 P52813
P52816 P52817 P52818 P52820 P52827 P52831 P52832 P52833 P52835 P52836 P52837
P52838 P52839 P52845 P52847 P52851 P52852 P52854 P52855 P52856 P52857 P52858
P52860 P52861 P52863 P52864 P52865 P52866 P52868 P52874 P52877 P52878 P52888
P52889 P52890 P52892 P52893 P52894 P52898 P52899 P52900 P52901 P52902 P52903
P52904 P52906 P52907 P52908 P52909 P52910 P52913 P52914 P52920 P52922 P52944
P52949 P52950 P52951 P52952 P52954 P52955 P52957 P52958 P52959 P52960 P52964
P52965 P52967 P52968 P52969 P52971 P52972 P52973 P52974 P52978 P52979 P52980
P52981 P52982 P52983 P52984 P52985 P52986 P52987 P52989 P52990 P52991 P52992
P52993 P52994 P52995 P52996 P52997 P52998 P52999 P53000 P53001 P53003 P53009
P53010 P53013 P53014 P53015 P53016 P53017 P53018 P53019 P53020 P53021 P53022
P53023 P53027 P53028 P53031 P53032 P53033 P53034 P53035 P53036 P53038 P53040

P53043 P53050 P53051 P53054 P53056 P53060 P53061 P53063 P53064 P53065 P53066
P53068 P53070 P53071 P53072 P53076 P53077 P53078 P53080 P53081 P53082 P53084
P53085 P53086 P53088 P53090 P53091 P53092 P53094 P53095 P53096 P53097 P53098
P53100 P53101 P53102 P53103 P53105 P53110 P53111 P53115 P53119 P53123 P53124
P53125 P53126 P53127 P53128 P53131 P53132 P53133 P53135 P53136 P53137 P53138
P53143 P53144 P53145 P53147 P53150 P53156 P53158 P53162 P53165 P53166 P53167
P53168 P53169 P53170 P53172 P53175 P53177 P53183 P53184 P53185 P53186 P53187
P53188 P53196 P53200 P53201 P53202 P53204 P53205 P53207 P53210 P53211 P53212
P53213 P53215 P53216 P53218 P53222 P53225 P53227 P53228 P53231 P53232 P53233
P53234 P53235 P53236 P53238 P53240 P53243 P53246 P53250 P53251 P53254 P53255
P53256 P53258 P53261 P53263 P53265 P53267 P53270 P53274 P53275 P53276 P53277
P53278 P53280 P53284 P53286 P53287 P53289 P53291 P53294 P53295 P53297 P53298
P53300 P53302 P53303 P53304 P53307 P53312 P53313 P53314 P53315 P53316 P53317
P53321 P53323 P53327 P53329 P53330 P53331 P53332 P53333 P53335 P53336 P53340
P53348 P53352 P53358 P53360 P53361 P53362 P53363 P53364 P53370 P53371 P53372
P53373 P53374 P53375 P53376 P53378 P53381 P53382 P53383 P53385 P53395 P53397
P53398 P53399 P53400 P53401 P53405 P53406 P53407 P53408 P53409 P53410 P53411
P53412 P53413 P53414 P53415 P53416 P53419 P53421 P53423 P53424 P53429 P53430
P53433 P53434 P53435 P53437 P53440 P53441 P53442 P53444 P53445 P53446 P53447
P53448 P53449 P53451 P53455 P53456 P53457 P53458 P53459 P53460 P53461 P53463
P53464 P53465 P53466 P53467 P53468 P53469 P53470 P53471 P53472 P53473 P53474
P53475 P53476 P53479 P53480 P53482 P53483 P53486 P53487 P53488 P53489 P53490
P53491 P53494 P53496 P53497 P53498 P53499 P53500 P53502 P53503 P53504 P53506
P53515 P53523 P53524 P53527 P53528 P53529 P53531 P53532 P53533 P53534 P53535
P53536 P53537 P53538 P53540 P53544 P53545 P53546 P53547 P53549 P53551 P53552
P53553 P53554 P53555 P53556 P53557 P53558 P53559 P53560 P53568 P53569 P53570
P53571 P53572 P53573 P53574 P53575 P53576 P53577 P53578 P53579 P53580 P53581
P53582 P53583 P53587 P53588 P53589 P53590 P53591 P53592 P53593 P53594 P53595
P53596 P53597 P53598 P53603 P53607 P53608 P53609 P53610 P53612 P53617 P53623
P53627 P53628 P53630 P53632 P53638 P53639 P53641 P53642 P53643 P53644 P53645
P53646 P53648 P53650 P53651 P53652 P53653 P53655 P53656 P53658 P53661 P53662
P53672 P53673 P53674 P53681 P53685 P53686 P53687 P53688 P53689 P53693 P53695
P53696 P53699 P53704 P53705 P53709 P53716 P53718 P53719 P53720 P53725 P53727
P53728 P53729 P53731 P53734 P53737 P53738 P53741 P53742 P53743 P53749 P53750
P53754 P53755 P53757 P53758 P53759 P53764 P53765 P53766 P53768 P53769 P53770
P53771 P53772 P53773 P53774 P53775 P53776 P53780 P53781 P53783 P53784 P53787
P53795 P53797 P53803 P53805 P53806 P53819 P53820 P53821 P53823 P53824 P53829
P53830 P53831 P53833 P53834 P53837 P53839 P53840 P53841 P53846 P53849 P53850
P53851 P53852 P53853 P53854 P53858 P53859 P53861 P53863 P53864 P53866 P53869
P53870 P53871 P53873 P53876 P53877 P53883 P53886 P53887 P53888 P53889 P53890
P53891 P53892 P53893 P53897 P53899 P53900 P53902 P53904 P53905 P53907 P53909
P53910 P53911 P53912 P53913 P53914 P53915 P53919 P53922 P53923 P53924 P53927
P53929 P53930 P53934 P53935 P53936 P53940 P53941 P53942 P53944 P53946 P53948
P53949 P53952 P53957 P53958 P53960 P53962 P53963 P53967 P53968 P53970 P53971

P53972 P53973 P53974 P53977 P53978 P53980 P53981 P53982 P53984 P53995 P53996
P53997 P53998 P54000 P54004 P54005 P54006 P54007 P54008 P54009 P54010 P54011
P54012 P54013 P54014 P54015 P54016 P54017 P54018 P54019 P54020 P54021 P54022
P54023 P54024 P54025 P54026 P54027 P54028 P54029 P54030 P54031 P54032 P54033
P54034 P54035 P54036 P54037 P54038 P54039 P54040 P54041 P54042 P54043 P54044
P54045 P54046 P54047 P54048 P54049 P54050 P54051 P54052 P54053 P54054 P54055
P54056 P54057 P54058 P54059 P54060 P54061 P54062 P54063 P54064 P54065 P54066
P54067 P54073 P54075 P54076 P54077 P54081 P54084 P54087 P54088 P54089 P54090
P54092 P54093 P54094 P54095 P54096 P54098 P54109 P54110 P54113 P54114 P54115
P54117 P54118 P54119 P54120 P54121 P54122 P54123 P54124 P54125 P54137 P54138
P54139 P54149 P54150 P54151 P54152 P54153 P54154 P54155 P54157 P54160 P54161
P54162 P54164 P54165 P54167 P54168 P54169 P54171 P54174 P54177 P54182 P54184
P54186 P54187 P54188 P54199 P54202 P54203 P54205 P54206 P54207 P54208 P54213
P54214 P54216 P54217 P54220 P54221 P54222 P54224 P54225 P54226 P54231 P54233
P54234 P54235 P54236 P54237 P54238 P54239 P54240 P54241 P54242 P54243 P54260
P54261 P54262 P54263 P54264 P54269 P54270 P54271 P54272 P54273 P54275 P54277
P54280 P54293 P54294 P54296 P54298 P54303 P54304 P54306 P54307 P54308 P54309
P54322 P54324 P54325 P54326 P54328 P54329 P54330 P54331 P54332 P54333 P54334
P54335 P54336 P54337 P54338 P54339 P54340 P54341 P54342 P54343 P54344 P54345
P54346 P54347 P54348 P54353 P54354 P54357 P54359 P54361 P54363 P54364 P54365
P54366 P54372 P54374 P54375 P54376 P54377 P54378 P54379 P54380 P54381 P54382
P54383 P54384 P54385 P54386 P54387 P54388 P54389 P54390 P54391 P54394 P54395
P54398 P54401 P54402 P54403 P54404 P54405 P54407 P54408 P54409 P54410 P54411
P54412 P54413 P54414 P54415 P54416 P54418 P54419 P54420 P54426 P54429 P54430
P54431 P54432 P54433 P54435 P54436 P54441 P54443 P54446 P54448 P54452 P54453
P54454 P54455 P54456 P54457 P54458 P54459 P54460 P54461 P54462 P54464 P54468
P54470 P54471 P54472 P54473 P54474 P54475 P54476 P54477 P54479 P54480 P54482
P54489 P54490 P54491 P54492 P54494 P54495 P54497 P54499 P54501 P54503 P54504
P54508 P54509 P54511 P54512 P54517 P54518 P54519 P54520 P54521 P54522 P54523
P54524 P54525 P54528 P54530 P54531 P54532 P54533 P54540 P54541 P54542 P54543
P54545 P54547 P54548 P54549 P54550 P54551 P54552 P54553 P54555 P54556 P54557
P54560 P54561 P54562 P54563 P54564 P54565 P54566 P54568 P54569 P54570 P54572
P54573 P54574 P54575 P54579 P54584 P54586 P54588 P54590 P54591 P54592 P54597
P54601 P54604 P54605 P54606 P54607 P54608 P54610 P54613 P54621 P54622 P54635
P54636 P54638 P54649 P54653 P54655 P54656 P54658 P54659 P54664 P54665 P54666
P54667 P54669 P54670 P54671 P54680 P54685 P54686 P54688 P54689 P54691 P54692
P54693 P54694 P54696 P54697 P54698 P54705 P54712 P54714 P54716 P54717 P54721
P54722 P54724 P54728 P54731 P54733 P54734 P54735 P54737 P54739 P54740 P54741
P54742 P54746 P54752 P54765 P54766 P54767 P54768 P54769 P54770 P54771 P54772
P54775 P54776 P54778 P54779 P54784 P54785 P54786 P54787 P54788 P54789 P54791
P54794 P54795 P54796 P54797 P54798 P54799 P54800 P54805 P54806 P54807 P54808
P54809 P54810 P54814 P54816 P54819 P54821 P54822 P54824 P54835 P54836 P54838
P54840 P54841 P54842 P54843 P54844 P54845 P54846 P54847 P54858 P54866 P54875
P54877 P54879 P54882 P54885 P54887 P54888 P54889 P54890 P54891 P54892 P54893

P54894 P54895 P54896 P54897 P54898 P54899 P54901 P54902 P54903 P54904 P54905
P54906 P54907 P54918 P54919 P54922 P54923 P54933 P54934 P54935 P54938 P54943
P54944 P54947 P54948 P54949 P54950 P54951 P54955 P54956 P54959 P54964 P54966
P54967 P54972 P54973 P54974 P54975 P54976 P54977 P54978 P54979 P54980 P54981
P54983 P54984 P54985 P54987 P54988 P54989 P54990 P54991 P54992 P54993 P54994
P54995 P54996 P54997 P54998 P54999 P55004 P55005 P55022 P55023 P55035 P55037
P55038 P55044 P55045 P55049 P55070 P55071 P55080 P55094 P55120 P55121 P55124
P55132 P55133 P55134 P55135 P55136 P55137 P55138 P55141 P55142 P55143 P55149
P55150 P55164 P55165 P55166 P55168 P55169 P55170 P55171 P55172 P55173 P55174
P55175 P55176 P55177 P55178 P55179 P55180 P55181 P55185 P55186 P55192 P55193
P55195 P55216 P55217 P55218 P55219 P55222 P55224 P55228 P55229 P55230 P55231
P55232 P55233 P55234 P55235 P55236 P55238 P55239 P55240 P55241 P55242 P55243
P55248 P55250 P55251 P55255 P55267 P55271 P55272 P55273 P55275 P55299 P55300
P55301 P55303 P55304 P55305 P55306 P55308 P55310 P55311 P55316 P55318 P55323
P55324 P55326 P55327 P55339 P55341 P55342 P55343 P55347 P55349 P55350 P55351
P55353 P55354 P55355 P55356 P55357 P55358 P55359 P55360 P55361 P55362 P55363
P55364 P55365 P55366 P55370 P55371 P55373 P55378 P55379 P55380 P55381 P55382
P55383 P55385 P55387 P55388 P55389 P55390 P55391 P55392 P55393 P55394 P55395
P55398 P55399 P55400 P55407 P55408 P55409 P55411 P55412 P55413 P55415 P55418
P55419 P55420 P55422 P55423 P55424 P55425 P55426 P55429 P55434 P55435 P55437
P55438 P55440 P55444 P55445 P55446 P55451 P55455 P55457 P55458 P55459 P55461
P55462 P55463 P55464 P55465 P55466 P55467 P55468 P55470 P55472 P55473 P55474
P55477 P55479 P55481 P55482 P55485 P55486 P55487 P55489 P55494 P55495 P55496
P55500 P55501 P55502 P55503 P55504 P55507 P55508 P55509 P55510 P55511 P55512
P55514 P55515 P55517 P55521 P55524 P55525 P55526 P55528 P55529 P55530 P55531
P55532 P55533 P55534 P55536 P55537 P55538 P55539 P55540 P55541 P55542 P55543
P55544 P55547 P55548 P55549 P55550 P55551 P55552 P55555 P55558 P55559 P55560
P55563 P55564 P55565 P55566 P55567 P55573 P55575 P55576 P55577 P55579 P55582
P55584 P55586 P55587 P55588 P55589 P55590 P55597 P55598 P55599 P55605 P55606
P55607 P55608 P55609 P55611 P55612 P55613 P55614 P55615 P55616 P55617 P55618
P55619 P55620 P55623 P55624 P55625 P55626 P55627 P55628 P55629 P55630 P55631
P55632 P55634 P55635 P55636 P55637 P55638 P55639 P55640 P55641 P55643 P55645
P55646 P55648 P55650 P55651 P55653 P55654 P55655 P55656 P55658 P55663 P55664
P55665 P55666 P55667 P55668 P55670 P55673 P55674 P55675 P55676 P55677 P55678
P55680 P55681 P55683 P55684 P55685 P55686 P55687 P55688 P55689 P55690 P55693
P55694 P55695 P55696 P55697 P55700 P55706 P55707 P55708 P55709 P55710 P55713
P55715 P55716 P55717 P55718 P55719 P55729 P55730 P55731 P55733 P55736 P55742
P55743 P55744 P55747 P55748 P55750 P55757 P55758 P55759 P55767 P55768 P55769
P55770 P55771 P55784 P55785 P55790 P55792 P55793 P55800 P55804 P55805 P55810
P55811 P55812 P55813 P55814 P55818 P55819 P55822 P55823 P55825 P55826 P55827
P55829 P55832 P55833 P55834 P55835 P55836 P55837 P55838 P55839 P55840 P55842
P55843 P55844 P55852 P55857 P55858 P55859 P55861 P55862 P55863 P55864 P55866
P55871 P55872 P55873 P55874 P55875 P55876 P55877 P55878 P55879 P55880 P55881
P55882 P55883 P55886 P55888 P55895 P55897 P55900 P55907 P55909 P55911 P55912

P55913 P55914 P55915 P55923 P55924 P55925 P55932 P55933 P55938 P55940 P55941
P55942 P55943 P55945 P55946 P55947 P55948 P55949 P55950 P55951 P55952 P55953
P55959 P55960 P55961 P55964 P55965 P55968 P55970 P55971 P55972 P55973 P55975
P55976 P55977 P55979 P55982 P55983 P55984 P55985 P55986 P55990 P55991 P55992
P55993 P55994 P55995 P55996 P55997 P55998 P55999 P56000 P56001 P56002 P56003
P56004 P56006 P56007 P56008 P56009 P56010 P56011 P56013 P56018 P56020 P56021
P56022 P56023 P56026 P56027 P56028 P56030 P56031 P56032 P56033 P56034 P56035
P56036 P56038 P56039 P56040 P56041 P56042 P56043 P56044 P56045 P56046 P56047
P56049 P56050 P56052 P56054 P56058 P56061 P56062 P56063 P56064 P56065 P56066
P56067 P56068 P56069 P56070 P56071 P56072 P56073 P56074 P56075 P56076 P56077
P56078 P56080 P56081 P56088 P56089 P56090 P56091 P56094 P56099 P56102 P56103
P56104 P56105 P56106 P56107 P56108 P56109 P56110 P56111 P56114 P56115 P56116
P56118 P56119 P56120 P56121 P56122 P56123 P56124 P56125 P56126 P56127 P56128
P56129 P56130 P56131 P56132 P56133 P56137 P56138 P56139 P56140 P56141 P56142
P56143 P56144 P56145 P56146 P56148 P56149 P56153 P56154 P56155 P56156 P56157
P56158 P56160 P56162 P56163 P56167 P56168 P56170 P56172 P56175 P56176 P56178
P56179 P56183 P56184 P56185 P56186 P56187 P56188 P56189 P56191 P56194 P56195
P56196 P56197 P56200 P56204 P56206 P56208 P56209 P56210 P56212 P56213 P56214
P56216 P56218 P56220 P56221 P56222 P56223 P56224 P56225 P56250 P56251 P56252
P56255 P56256 P56257 P56259 P56260 P56262 P56263 P56265 P56268 P56272 P56275
P56276 P56277 P56278 P56281 P56285 P56286 P56287 P56288 P56289 P56290 P56291
P56292 P56298 P56299 P56300 P56302 P56303 P56304 P56317 P56329 P56330 P56331
P56333 P56335 P56336 P56337 P56344 P56345 P56346 P56347 P56350 P56351 P56352
P56353 P56354 P56355 P56356 P56357 P56358 P56359 P56360 P56361 P56362 P56363
P56364 P56365 P56366 P56367 P56368 P56372 P56374 P56375 P56376 P56380 P56387
P56389 P56390 P56394 P56396 P56397 P56398 P56407 P56408 P56414 P56415 P56417
P56418 P56419 P56420 P56421 P56422 P56428 P56429 P56431 P56452 P56453 P56454
P56455 P56456 P56457 P56458 P56459 P56460 P56461 P56462 P56463 P56465 P56466
P56467 P56468 P56471 P56472 P56474 P56503 P56506 P56507 P56511 P56512 P56517
P56518 P56519 P56521 P56523 P56530 P56531 P56532 P56533 P56535 P56536 P56538
P56540 P56544 P56545 P56546 P56547 P56553 P56555 P56561 P56562 P56565 P56567
P56568 P56570 P56571 P56572 P56573 P56574 P56575 P56576 P56582 P56584 P56585
P56586 P56596 P56597 P56598 P56599 P56600 P56607 P56613 P56614 P56616 P56623
P56624 P56625 P56634 P56635 P56647 P56648 P56649 P56653 P56659 P56660 P56661
P56662 P56663 P56664 P56665 P56666 P56667 P56668 P56669 P56670 P56671 P56672
P56679 P56681 P56683 P56689 P56690 P56691 P56692 P56701 P56702 P56707 P56709
P56717 P56721 P56723 P56724 P56725 P56727 P56728 P56737 P56739 P56740 P56741
P56744 P56762 P56763 P56764 P56772 P56784 P56786 P56791 P56792 P56793 P56794
P56795 P56796 P56797 P56798 P56799 P56801 P56802 P56804 P56805 P56806 P56807
P56808 P56809 P56810 P56811 P56813 P56815 P56816 P56820 P56821 P56822 P56823
P56824 P56825 P56826 P56835 P56838 P56839 P56840 P56841 P56842 P56844 P56845
P56848 P56849 P56850 P56858 P56859 P56860 P56861 P56862 P56863 P56864 P56866
P56868 P56869 P56870 P56871 P56872 P56873 P56875 P56876 P56878 P56879 P56881
P56883 P56884 P56885 P56886 P56887 P56888 P56889 P56890 P56891 P56892 P56893

P56894 P56898 P56899 P56900 P56901 P56903 P56904 P56905 P56906 P56916 P56925
P56926 P56927 P56928 P56930 P56931 P56932 P56933 P56934 P56935 P56936 P56944
P56946 P56947 P56948 P56953 P56954 P56955 P56956 P56957 P56958 P56960 P56961
P56965 P56968 P56969 P56970 P56977 P56978 P56979 P56980 P56981 P56985 P56986
P56987 P56989 P56990 P56991 P56992 P56993 P56994 P56995 P56996 P56997 P56998
P57000 P57001 P57002 P57006 P57007 P57008 P57009 P57014 P57015 P57017 P57018
P57020 P57028 P57029 P57035 P57036 P57040 P57053 P57055 P57064 P57065 P57071
P57072 P57074 P57076 P57077 P57079 P57082 P57084 P57085 P57086 P57089 P57090
P57098 P57099 P57100 P57101 P57102 P57108 P57109 P57111 P57112 P57113 P57114
P57115 P57116 P57117 P57126 P57127 P57128 P57129 P57130 P57132 P57133 P57134
P57135 P57136 P57138 P57140 P57142 P57143 P57144 P57145 P57146 P57147 P57148
P57149 P57150 P57151 P57153 P57154 P57155 P57156 P57157 P57158 P57159 P57160
P57161 P57162 P57163 P57164 P57165 P57166 P57167 P57168 P57169 P57171 P57172
P57173 P57174 P57177 P57180 P57181 P57187 P57188 P57190 P57191 P57192 P57194
P57195 P57196 P57197 P57198 P57199 P57200 P57201 P57202 P57203 P57204 P57205
P57206 P57207 P57208 P57209 P57210 P57211 P57212 P57219 P57220 P57221 P57222
P57224 P57225 P57226 P57227 P57228 P57229 P57230 P57231 P57232 P57233 P57235
P57236 P57237 P57238 P57241 P57242 P57243 P57244 P57245 P57246 P57247 P57249
P57250 P57251 P57265 P57266 P57267 P57268 P57269 P57270 P57271 P57272 P57273
P57274 P57275 P57276 P57277 P57278 P57279 P57280 P57281 P57282 P57284 P57285
P57286 P57287 P57288 P57289 P57290 P57291 P57292 P57293 P57294 P57295 P57298
P57299 P57300 P57301 P57302 P57303 P57304 P57305 P57306 P57307 P57310 P57313
P57315 P57316 P57319 P57320 P57321 P57323 P57324 P57325 P57326 P57327 P57328
P57329 P57332 P57333 P57336 P57337 P57338 P57340 P57341 P57342 P57343 P57346
P57349 P57351 P57354 P57356 P57357 P57358 P57359 P57360 P57362 P57365 P57366
P57367 P57369 P57371 P57372 P57373 P57374 P57375 P57376 P57377 P57378 P57379
P57380 P57381 P57384 P57385 P57386 P57387 P57388 P57389 P57390 P57391 P57393
P57394 P57395 P57396 P57397 P57398 P57399 P57400 P57401 P57404 P57405 P57407
P57409 P57410 P57412 P57413 P57416 P57417 P57419 P57420 P57421 P57422 P57423
P57424 P57427 P57430 P57431 P57434 P57435 P57436 P57438 P57439 P57441 P57442
P57444 P57445 P57446 P57447 P57448 P57449 P57450 P57451 P57452 P57453 P57454
P57455 P57456 P57457 P57458 P57459 P57461 P57463 P57464 P57465 P57466 P57467
P57468 P57469 P57470 P57471 P57472 P57474 P57475 P57476 P57477 P57478 P57479
P57480 P57481 P57483 P57484 P57485 P57486 P57487 P57488 P57489 P57490 P57491
P57493 P57494 P57495 P57497 P57498 P57499 P57500 P57501 P57502 P57503 P57504
P57506 P57507 P57508 P57509 P57510 P57511 P57512 P57513 P57515 P57516 P57517
P57518 P57519 P57520 P57521 P57522 P57523 P57525 P57526 P57528 P57529 P57530
P57532 P57533 P57534 P57535 P57536 P57537 P57539 P57545 P57546 P57547 P57548
P57549 P57553 P57554 P57555 P57556 P57557 P57558 P57559 P57561 P57562 P57563
P57564 P57565 P57566 P57567 P57568 P57569 P57570 P57572 P57573 P57574 P57575
P57576 P57577 P57578 P57579 P57580 P57581 P57582 P57583 P57584 P57585 P57586
P57587 P57588 P57589 P57590 P57591 P57592 P57593 P57594 P57595 P57596 P57597
P57598 P57599 P57600 P57602 P57603 P57604 P57605 P57606 P57607 P57608 P57609
P57610 P57611 P57612 P57613 P57614 P57615 P57616 P57617 P57618 P57619 P57620

P57621 P57622 P57623 P57625 P57626 P57627 P57628 P57629 P57632 P57633 P57634
P57636 P57637 P57639 P57640 P57641 P57642 P57643 P57644 P57645 P57646 P57649
P57650 P57651 P57652 P57653 P57654 P57655 P57656 P57657 P57658 P57659 P57660
P57661 P57662 P57665 P57666 P57667 P57668 P57670 P57671 P57673 P57674 P57676
P57677 P57690 P57691 P57692 P57693 P57694 P57696 P57697 P57702 P57703 P57704
P57705 P57706 P57707 P57708 P57710 P57711 P57712 P57713 P57714 P57715 P57717
P57718 P57719 P57720 P57721 P57722 P57723 P57741 P57743 P57749 P57750 P57753
P57755 P57761 P57762 P57763 P57765 P57766 P57767 P57775 P57776 P57777 P57778
P57779 P57781 P57782 P57783 P57784 P57790 P57793 P57794 P57795 P57797 P57798
P57799 P57801 P57802 P57803 P57805 P57807 P57808 P57809 P57810 P57811 P57812
P57813 P57815 P57818 P57819 P57820 P57821 P57822 P57823 P57824 P57825 P57826
P57827 P57828 P57829 P57830 P57831 P57832 P57833 P57836 P57837 P57838 P57840
P57841 P57842 P57843 P57844 P57845 P57847 P57848 P57849 P57850 P57851 P57852
P57854 P57855 P57856 P57857 P57859 P57860 P57861 P57862 P57863 P57865 P57866
P57867 P57868 P57869 P57870 P57871 P57872 P57873 P57874 P57875 P57876 P57877
P57878 P57879 P57880 P57881 P57882 P57884 P57885 P57886 P57887 P57888 P57889
P57890 P57891 P57892 P57893 P57894 P57895 P57896 P57897 P57898 P57899 P57900
P57902 P57903 P57904 P57905 P57906 P57907 P57908 P57909 P57910 P57911 P57912
P57913 P57914 P57915 P57916 P57918 P57919 P57920 P57921 P57922 P57923 P57924
P57925 P57926 P57927 P57928 P57930 P57931 P57932 P57933 P57934 P57935 P57936
P57937 P57938 P57939 P57940 P57941 P57942 P57943 P57944 P57945 P57946 P57947
P57948 P57949 P57951 P57952 P57953 P57954 P57955 P57956 P57957 P57958 P57960
P57961 P57962 P57963 P57964 P57965 P57966 P57967 P57968 P57969 P57971 P57972
P57973 P57974 P57977 P57978 P57979 P57980 P57981 P57982 P57983 P57984 P57985
P57986 P57987 P57988 P57989 P57990 P57991 P57992 P57993 P57994 P57995 P57997
P57998 P58001 P58002 P58003 P58005 P58006 P58009 P58011 P58013 P58014 P58015
P58016 P58017 P58018 P58023 P58024 P58025 P58026 P58032 P58033 P58036 P58037
P58040 P58052 P58054 P58055 P58056 P58057 P58058 P58060 P58063 P58065 P58066
P58067 P58072 P58074 P58075 P58077 P58078 P58079 P58080 P58082 P58083 P58084
P58085 P58086 P58088 P58091 P58092 P58093 P58094 P58095 P58097 P58100 P58101
P58102 P58103 P58105 P58106 P58109 P58110 P58111 P58112 P58113 P58114 P58115
P58116 P58117 P58121 P58122 P58123 P58125 P58127 P58129 P58131 P58132 P58133
P58134 P58135 P58136 P58138 P58139 P58140 P58145 P58146 P58147 P58148 P58149
P58150 P58151 P58152 P58153 P58156 P58157 P58159 P58160 P58161 P58167 P58168
P58171 P58174 P58176 P58177 P58178 P58179 P58183 P58189 P58190 P58192 P58193
P58201 P58202 P58207 P58208 P58209 P58210 P58211 P58213 P58217 P58218 P58221
P58222 P58223 P58226 P58227 P58228 P58230 P58231 P58232 P58233 P58235 P58236
P58237 P58238 P58243 P58245 P58249 P58250 P58251 P58254 P58256 P58258 P58259
P58262 P58263 P58264 P58267 P58268 P58269 P58270 P58271 P58272 P58275 P58276
P58277 P58278 P58280 P58283 P58284 P58285 P58286 P58288 P58289 P58290 P58291
P58292 P58293 P58299 P58301 P58302 P58304 P58306 P58313 P58314 P58315 P58316
P58317 P58318 P58319 P58322 P58326 P58329 P58331 P58332 P58333 P58334 P58336
P58338 P58339 P58346 P58347 P58348 P58349 P58350 P58364 P58372 P58374 P58375
P58376 P58378 P58379 P58382 P58384 P58388 P58389 P58393 P58394 P58395 P58407

P58408 P58409 P58413 P58415 P58416 P58422 P58423 P58424 P58429 P58430 P58431
P58432 P58433 P58434 P58435 P58436 P58437 P58438 P58439 P58441 P58442 P58445
P58446 P58447 P58448 P58449 P58451 P58452 P58453 P58456 P58458 P58462 P58463
P58465 P58466 P58467 P58468 P58469 P58470 P58471 P58472 P58473 P58474 P58475
P58477 P58478 P58479 P58480 P58481 P58482 P58484 P58485 P58486 P58487 P58493
P58494 P58495 P58496 P58497 P58500 P58501 P58503 P58505 P58506 P58507 P58508
P58509 P58510 P58512 P58513 P58515 P58516 P58517 P58518 P58520 P58521 P58522
P58523 P58524 P58525 P58526 P58527 P58528 P58532 P58533 P58534 P58535 P58536
P58538 P58539 P58540 P58541 P58542 P58543 P58545 P58546 P58552 P58554 P58555
P58557 P58559 P58561 P58562 P58563 P58567 P58571 P58572 P58574 P58578 P58579
P58580 P58581 P58582 P58584 P58585 P58586 P58587 P58588 P58589 P58590 P58591
P58592 P58600 P58602 P58614 P58615 P58616 P58617 P58618 P58619 P58620 P58621
P58626 P58627 P58628 P58629 P58630 P58631 P58632 P58633 P58634 P58635 P58636
P58637 P58638 P58639 P58640 P58641 P58642 P58643 P58644 P58652 P58660 P58661
P58665 P58666 P58667 P58668 P58669 P58670 P58671 P58672 P58673 P58674 P58675
P58676 P58677 P58678 P58679 P58680 P58685 P58687 P58688 P58692 P58693 P58694
P58695 P58696 P58697 P58698 P58699 P58700 P58703 P58704 P58708 P58709 P58711
P58713 P58714 P58715 P58716 P58717 P58718 P58719 P58723 P58724 P58726 P58728
P58729 P58730 P58731 P58733 P58736 P58737 P58739 P58744 P58745 P58746 P58747
P58748 P58750 P58758 P58759 P58760 P58761 P58768 P58772 P58773 P58774 P58775
P58776 P58777 P58788 P58789 P58790 P58791 P58792 P58793 P58794 P58795 P58796
P58797 P58798 P58799 P58800 P58812 P58813 P58814 P58815 P58816 P58817 P58818
P58824 P58827 P58828 P58829 P58830 P58831 P58832 P58833 P58834 P58835 P58836
P58837 P58838 P58839 P58847 P58850 P58851 P58852 P58853 P58854 P58855 P58856
P58858 P58859 P58860 P58861 P58862 P58863 P58865 P58866 P58867 P58868 P58869
P58876 P58877 P58878 P58879 P58880 P58881 P58882 P58883 P58884 P58885 P58886
P58887 P58888 P58889 P58890 P58891 P58892 P58893 P58894 P58895 P58896 P58897
P58898 P58899 P58900 P58901 P58902 P58903 P58904 P58907 P58909 P58929 P58939
P58940 P58941 P58942 P58943 P58944 P58945 P58946 P58947 P58948 P58949 P58963
P58964 P58965 P58966 P58967 P58968 P58969 P58970 P58971 P58972 P58973 P58974
P58977 P58978 P58979 P58980 P58981 P58982 P58983 P58984 P58987 P58988 P58991
P58992 P58993 P58994 P58995 P58998 P59001 P59002 P59003 P59004 P59005 P59008
P59009 P59010 P59011 P59013 P59014 P59019 P59020 P59021 P59022 P59023 P59027
P59028 P59029 P59030 P59031 P59032 P59033 P59036 P59037 P59038 P59039 P59040
P59045 P59048 P59049 P59050 P59051 P59052 P59054 P59055 P59056 P59057 P59058
P59059 P59060 P59061 P59063 P59064 P59065 P59075 P59076 P59077 P59078 P59079
P59081 P59082 P59083 P59085 P59086 P59090 P59091 P59098 P59099 P59100 P59101
P59102 P59103 P59106 P59107 P59109 P59117 P59118 P59119 P59121 P59122 P59123
P59124 P59125 P59126 P59127 P59128 P59129 P59130 P59131 P59132 P59133 P59134
P59135 P59136 P59137 P59138 P59139 P59140 P59141 P59142 P59143 P59144 P59145
P59146 P59147 P59148 P59149 P59150 P59151 P59152 P59153 P59154 P59155 P59156
P59157 P59159 P59160 P59161 P59162 P59163 P59164 P59165 P59166 P59167 P59168
P59173 P59174 P59175 P59176 P59177 P59178 P59179 P59180 P59181 P59182 P59183
P59184 P59186 P59187 P59188 P59189 P59191 P59192 P59193 P59196 P59197 P59199

P59200 P59203 P59204 P59207 P59209 P59210 P59211 P59212 P59217 P59223 P59224
P59225 P59230 P59231 P59232 P59233 P59234 P59236 P59237 P59238 P59244 P59245
P59271 P59272 P59273 P59274 P59278 P59280 P59282 P59283 P59284 P59285 P59286
P59287 P59288 P59289 P59291 P59292 P59293 P59294 P59295 P59296 P59297 P59298
P59299 P59300 P59301 P59302 P59303 P59304 P59305 P59306 P59307 P59308 P59309
P59310 P59311 P59312 P59313 P59314 P59315 P59316 P59317 P59318 P59319 P59320
P59321 P59322 P59323 P59324 P59326 P59327 P59328 P59329 P59330 P59331 P59332
P59336 P59337 P59348 P59349 P59350 P59351 P59352 P59353 P59361 P59362 P59369
P59370 P59371 P59372 P59373 P59374 P59375 P59377 P59378 P59379 P59380 P59381
P59385 P59386 P59388 P59389 P59390 P59391 P59392 P59393 P59394 P59395 P59396
P59397 P59398 P59399 P59400 P59401 P59403 P59404 P59405 P59406 P59407 P59408
P59409 P59410 P59411 P59412 P59413 P59414 P59415 P59416 P59417 P59418 P59419
P59420 P59421 P59422 P59423 P59425 P59426 P59427 P59428 P59429 P59430 P59431
P59432 P59433 P59434 P59435 P59437 P59439 P59441 P59442 P59443 P59444 P59445
P59446 P59447 P59448 P59450 P59451 P59452 P59453 P59454 P59455 P59456 P59457
P59458 P59459 P59460 P59461 P59462 P59463 P59464 P59465 P59466 P59467 P59468
P59469 P59470 P59472 P59473 P59474 P59476 P59477 P59478 P59480 P59482 P59483
P59484 P59485 P59487 P59488 P59489 P59490 P59491 P59492 P59493 P59495 P59496
P59497 P59498 P59499 P59500 P59501 P59502 P59503 P59504 P59505 P59506 P59507
P59508 P59512 P59513 P59514 P59515 P59516 P59517 P59518 P59519 P59520 P59521
P59522 P59524 P59525 P59526 P59527 P59553 P59554 P59555 P59556 P59557 P59558
P59560 P59561 P59562 P59563 P59564 P59565 P59567 P59568 P59569 P59573 P59574
P59575 P59576 P59577 P59578 P59579 P59580 P59581 P59582 P59583 P59584 P59586
P59587 P59591 P59597 P59599 P59600 P59601 P59602 P59603 P59604 P59605 P59606
P59607 P59608 P59609 P59610 P59611 P59612 P59613 P59614 P59615 P59616 P59617
P59618 P59619 P59620 P59621 P59623 P59624 P59625 P59626 P59627 P59628 P59629
P59630 P59631 P59642 P59643 P59650 P59651 P59655 P59656 P59657 P59658 P59659
P59660 P59661 P59663 P59664 P59667 P59669 P59671 P59672 P59673 P59674 P59675
P59677 P59678 P59679 P59680 P59681 P59682 P59683 P59686 P59687 P59688 P59689
P59690 P59691 P59692 P59696 P59697 P59698 P59700 P59701 P59707 P59708 P59714
P59716 P59717 P59718 P59719 P59720 P59721 P59722 P59723 P59724 P59725 P59726
P59727 P59728 P59730 P59731 P59732 P59733 P59734 P59736 P59739 P59740 P59742
P59743 P59744 P59745 P59746 P59747 P59748 P59749 P59752 P59753 P59754 P59755
P59756 P59758 P59759 P59774 P59775 P59776 P59778 P59779 P59781 P59782 P59785
P59787 P59790 P59793 P59794 P59795 P59797 P59799 P59800 P59801 P59805 P59806
P59812 P59815 P59816 P59817 P59818 P59819 P59820 P59821 P59825 P59829 P59830
P59831 P59834 P59835 P59838 P59840 P59842 P59843 P59846 P59860 P59871 P59872
P59873 P59874 P59876 P59877 P59878 P59879 P59880 P59881 P59882 P59883 P59884
P59885 P59892 P59893 P59895 P59910 P59911 P59912 P59916 P59918 P59919 P59920
P59921 P59923 P59927 P59928 P59930 P59931 P59932 P59933 P59935 P59937 P59942
P59945 P59946 P59948 P59951 P59952 P59953 P59954 P59955 P59956 P59957 P59958
P59959 P59964 P59965 P59966 P59967 P59968 P59969 P59971 P59973 P59974 P59975
P59978 P59979 P59980 P59982 P59986 P59989 P59990 P59991 P59992 P59993 P59997
P59998 P59999 P60001 P60002 P60003 P60004 P60005 P60006 P60007 P60008 P60009

P60010 P60011 P60012 P60013 P60014 P60024 P60027 P60038 P60039 P60040 P60046
P60047 P60048 P60049 P60050 P60053 P60054 P60055 P60056 P60070 P60071 P60072
P60073 P60074 P60075 P60077 P60078 P60079 P60080 P60082 P60083 P60086 P60087
P60089 P60090 P60091 P60092 P60093 P60094 P60095 P60096 P60097 P60098 P60099
P60100 P60101 P60102 P60103 P60104 P60105 P60106 P60107 P60109 P60110 P60111
P60119 P60121 P60166 P60167 P60168 P60169 P60175 P60180 P60184 P60185 P60186
P60187 P60188 P60189 P60190 P60193 P60194 P60195 P60196 P60197 P60199 P60200
P60204 P60205 P60206 P60216 P60217 P60218 P60219 P60220 P60221 P60223 P60224
P60225 P60226 P60227 P60230 P60231 P60232 P60238 P60239 P60240 P60241 P60246
P60279 P60280 P60281 P60282 P60283 P60284 P60286 P60287 P60288 P60289 P60290
P60291 P60292 P60293 P60294 P60295 P60296 P60297 P60299 P60312 P60313 P60314
P60315 P60316 P60317 P60318 P60319 P60320 P60321 P60322 P60323 P60324 P60325
P60326 P60327 P60328 P60329 P60330 P60331 P60332 P60333 P60335 P60336 P60338
P60339 P60340 P60341 P60342 P60343 P60344 P60345 P60346 P60347 P60348 P60349
P60350 P60351 P60352 P60353 P60354 P60355 P60356 P60357 P60358 P60360 P60362
P60364 P60365 P60366 P60367 P60368 P60369 P60370 P60371 P60372 P60373 P60374
P60375 P60376 P60377 P60378 P60380 P60381 P60382 P60383 P60384 P60385 P60386
P60387 P60388 P60389 P60390 P60391 P60393 P60394 P60395 P60396 P60397 P60398
P60399 P60400 P60401 P60402 P60403 P60404 P60405 P60406 P60407 P60408 P60409
P60410 P60411 P60412 P60413 P60414 P60415 P60416 P60417 P60418 P60419 P60420
P60422 P60423 P60424 P60425 P60426 P60427 P60428 P60429 P60430 P60431 P60433
P60434 P60435 P60436 P60437 P60438 P60439 P60440 P60441 P60443 P60444 P60445
P60446 P60447 P60448 P60450 P60452 P60453 P60454 P60455 P60456 P60457 P60458
P60470 P60471 P60476 P60477 P60480 P60481 P60482 P60485 P60486 P60488 P60489
P60490 P60491 P60492 P60493 P60494 P60495 P60499 P60500 P60501 P60502 P60504
P60515 P60516 P60523 P60524 P60525 P60526 P60527 P60529 P60530 P60531 P60532
P60533 P60534 P60535 P60536 P60537 P60538 P60539 P60540 P60541 P60542 P60543
P60544 P60545 P60546 P60547 P60548 P60549 P60550 P60551 P60552 P60553 P60554
P60555 P60556 P60557 P60558 P60559 P60560 P60561 P60562 P60564 P60565 P60566
P60567 P60575 P60576 P60577 P60579 P60580 P60581 P60582 P60583 P60584 P60585
P60586 P60587 P60593 P60594 P60595 P60596 P60597 P60598 P60599 P60600 P60601
P60617 P60619 P60622 P60624 P60625 P60626 P60628 P60629 P60630 P60631 P60651
P60652 P60653 P60654 P60655 P60656 P60657 P60658 P60659 P60661 P60662 P60663
P60664 P60665 P60666 P60667 P60668 P60700 P60702 P60704 P60705 P60714 P60715
P60716 P60717 P60718 P60719 P60720 P60721 P60722 P60723 P60724 P60725 P60726
P60727 P60728 P60730 P60731 P60732 P60733 P60734 P60736 P60738 P60739 P60740
P60741 P60742 P60743 P60744 P60745 P60746 P60757 P60758 P60760 P60776 P60780
P60781 P60795 P60797 P60799 P60800 P60801 P60802 P60803 P60804 P60805 P60806
P60807 P60809 P60816 P60817 P60818 P60819 P60820 P60821 P60822 P60824 P60825
P60826 P60828 P60829 P60831 P60832 P60833 P60834 P60835 P60836 P60837 P60838
P60840 P60841 P60843 P60846 P60847 P60848 P60849 P60850 P60851 P60854 P60858
P60859 P60860 P60861 P60862 P60863 P60864 P60865 P60875 P60876 P60884 P60885
P60886 P60887 P60888 P60889 P60890 P60891 P60892 P60896 P60897 P60898 P60899
P60900 P60901 P60906 P60907 P60908 P60909 P60911 P60912 P60913 P60914 P60915

P60916 P60917 P60918 P60919 P60920 P60921 P60922 P60923 P60954 P60983 P60984
P60995 P60996 P60997 P60998 P60999 P61000 P61001 P61002 P61003 P61004 P61005
P61024 P61025 P61029 P61030 P61031 P61054 P61055 P61056 P61057 P61058 P61060
P61061 P61062 P61063 P61064 P61065 P61066 P61067 P61068 P61069 P61070 P61071
P61074 P61075 P61080 P61081 P61082 P61083 P61084 P61090 P61091 P61093 P61108
P61111 P61112 P61113 P61114 P61115 P61116 P61118 P61120 P61122 P61123 P61124
P61127 P61128 P61129 P61133 P61136 P61151 P61154 P61155 P61157 P61159 P61161
P61167 P61173 P61174 P61175 P61176 P61177 P61178 P61179 P61180 P61182 P61183
P61184 P61185 P61186 P61187 P61188 P61189 P61190 P61191 P61192 P61193 P61194
P61196 P61197 P61198 P61199 P61200 P61202 P61203 P61215 P61216 P61217 P61219
P61220 P61234 P61235 P61236 P61237 P61238 P61239 P61240 P61241 P61242 P61243
P61245 P61246 P61253 P61258 P61259 P61272 P61282 P61286 P61288 P61290 P61291
P61292 P61293 P61295 P61296 P61297 P61300 P61301 P61302 P61303 P61304 P61305
P61306 P61307 P61308 P61309 P61310 P61311 P61323 P61324 P61325 P61326 P61330
P61331 P61332 P61333 P61334 P61335 P61336 P61337 P61338 P61339 P61340 P61341
P61342 P61343 P61344 P61345 P61346 P61347 P61348 P61349 P61350 P61360 P61361
P61362 P61363 P61367 P61368 P61369 P61370 P61371 P61372 P61373 P61374 P61375
P61376 P61379 P61396 P61397 P61400 P61401 P61404 P61405 P61406 P61407 P61410
P61411 P61412 P61413 P61414 P61422 P61423 P61424 P61425 P61426 P61427 P61428
P61430 P61431 P61432 P61433 P61434 P61435 P61436 P61437 P61438 P61439 P61440
P61441 P61442 P61443 P61444 P61445 P61446 P61447 P61448 P61449 P61456 P61457
P61458 P61459 P61461 P61462 P61463 P61476 P61477 P61478 P61479 P61480 P61486
P61487 P61488 P61489 P61490 P61491 P61492 P61493 P61494 P61495 P61496 P61497
P61498 P61499 P61500 P61501 P61502 P61503 P61511 P61512 P61515 P61517 P61518
P61519 P61520 P61521 P61522 P61523 P61524 P61525 P61526 P61527 P61528 P61529
P61530 P61531 P61532 P61533 P61534 P61535 P61536 P61537 P61538 P61539 P61540
P61548 P61580 P61581 P61582 P61583 P61590 P61591 P61592 P61593 P61594 P61595
P61597 P61600 P61603 P61615 P61616 P61617 P61618 P61623 P61624 P61650 P61651
P61652 P61658 P61659 P61660 P61661 P61662 P61663 P61664 P61665 P61666 P61667
P61668 P61669 P61670 P61671 P61672 P61673 P61675 P61676 P61677 P61678 P61679
P61680 P61681 P61682 P61683 P61684 P61685 P61686 P61687 P61688 P61689 P61690
P61691 P61692 P61693 P61694 P61695 P61696 P61697 P61698 P61699 P61700 P61701
P61702 P61703 P61704 P61705 P61706 P61707 P61708 P61709 P61710 P61711 P61712
P61713 P61714 P61715 P61717 P61718 P61719 P61720 P61721 P61723 P61724 P61725
P61726 P61727 P61728 P61729 P61730 P61731 P61732 P61733 P61734 P61735 P61736
P61760 P61772 P61773 P61774 P61775 P61778 P61779 P61780 P61781 P61782 P61783
P61786 P61787 P61788 P61798 P61799 P61800 P61801 P61802 P61816 P61817 P61818
P61819 P61820 P61825 P61827 P61830 P61831 P61832 P61833 P61834 P61835 P61836
P61841 P61842 P61845 P61846 P61847 P61848 P61851 P61852 P61853 P61854 P61857
P61858 P61859 P61860 P61861 P61865 P61867 P61868 P61873 P61874 P61875 P61876
P61877 P61878 P61879 P61880 P61881 P61882 P61883 P61884 P61885 P61886 P61890
P61891 P61892 P61894 P61895 P61896 P61897 P61906 P61907 P61908 P61909 P61910
P61911 P61912 P61913 P61919 P61920 P61921 P61922 P61925 P61926 P61929 P61930
P61931 P61932 P61933 P61934 P61935 P61936 P61937 P61938 P61940 P61941 P61942

P61945 P61946 P61947 P61948 P61949 P61950 P61951 P61960 P61961 P61962 P61963
P61964 P61965 P61968 P61969 P61973 P61974 P61975 P61976 P61977 P61979 P61980
P61984 P61985 P61986 P61987 P61988 P61989 P61990 P61991 P61992 P61993 P61994
P61995 P61996 P61997 P61998 P61999 P62000 P62001 P62002 P62003 P62004 P62005
P62006 P62007 P62008 P62009 P62010 P62011 P62012 P62013 P62014 P62015 P62025
P62026 P62028 P62029 P62030 P62031 P62032 P62033 P62034 P62035 P62036 P62037
P62038 P62039 P62040 P62041 P62042 P62043 P62044 P62045 P62046 P62047 P62048
P62049 P62050 P62051 P62052 P62053 P62054 P62055 P62056 P62057 P62058 P62059
P62060 P62061 P62062 P62063 P62064 P62065 P62066 P62067 P62085 P62086 P62087
P62088 P62089 P62117 P62118 P62119 P62120 P62121 P62122 P62123 P62124 P62125
P62126 P62127 P62128 P62129 P62131 P62132 P62134 P62135 P62139 P62142 P62143
P62145 P62146 P62147 P62148 P62153 P62154 P62162 P62163 P62164 P62165 P62169
P62170 P62171 P62175 P62176 P62177 P62178 P62179 P62180 P62181 P62182 P62183
P62184 P62185 P62186 P62187 P62188 P62189 P62196 P62198 P62199 P62200 P62201
P62202 P62203 P62207 P62208 P62209 P62210 P62211 P62212 P62213 P62214 P62215
P62216 P62217 P62218 P62219 P62220 P62221 P62222 P62223 P62224 P62225 P62226
P62227 P62228 P62229 P62230 P62231 P62232 P62233 P62234 P62235 P62236 P62237
P62238 P62239 P62240 P62247 P62251 P62252 P62253 P62254 P62255 P62256 P62257
P62262 P62265 P62279 P62283 P62284 P62285 P62286 P62287 P62288 P62289 P62290
P62291 P62292 P62293 P62294 P62295 P62296 P62297 P62299 P62300 P62302 P62303
P62304 P62305 P62306 P62307 P62308 P62309 P62310 P62311 P62312 P62313 P62314
P62315 P62317 P62318 P62320 P62321 P62322 P62323 P62324 P62325 P62326 P62327
P62329 P62334 P62335 P62338 P62339 P62340 P62346 P62347 P62348 P62349 P62350
P62351 P62352 P62353 P62354 P62355 P62356 P62357 P62358 P62359 P62360 P62363
P62364 P62365 P62366 P62367 P62368 P62369 P62370 P62371 P62372 P62373 P62374
P62378 P62379 P62380 P62381 P62382 P62386 P62387 P62391 P62392 P62393 P62399
P62400 P62401 P62402 P62404 P62405 P62406 P62409 P62410 P62411 P62412 P62413
P62414 P62415 P62416 P62417 P62418 P62419 P62420 P62421 P62422 P62423 P62426
P62427 P62428 P62429 P62430 P62431 P62432 P62433 P62434 P62435 P62436 P62437
P62438 P62440 P62441 P62442 P62443 P62444 P62445 P62446 P62448 P62449 P62450
P62451 P62452 P62453 P62454 P62455 P62456 P62457 P62458 P62459 P62460 P62468
P62469 P62470 P62471 P62472 P62473 P62474 P62475 P62476 P62477 P62478 P62479
P62485 P62486 P62487 P62488 P62489 P62495 P62496 P62497 P62498 P62499 P62500
P62501 P62505 P62506 P62507 P62508 P62509 P62510 P62511 P62515 P62516 P62521
P62522 P62523 P62524 P62525 P62526 P62527 P62528 P62529 P62534 P62535 P62536
P62538 P62539 P62550 P62552 P62553 P62554 P62555 P62556 P62557 P62558 P62559
P62572 P62573 P62574 P62577 P62578 P62579 P62580 P62584 P62585 P62588 P62589
P62590 P62591 P62592 P62597 P62599 P62600 P62601 P62602 P62603 P62604 P62611
P62612 P62613 P62615 P62616 P62617 P62618 P62619 P62620 P62621 P62622 P62623
P62624 P62625 P62627 P62628 P62633 P62634 P62642 P62649 P62650 P62652 P62653
P62654 P62655 P62656 P62657 P62658 P62659 P62660 P62661 P62662 P62663 P62664
P62665 P62666 P62667 P62668 P62669 P62672 P62673 P62674 P62675 P62676 P62677
P62678 P62679 P62680 P62681 P62682 P62696 P62697 P62698 P62699 P62700 P62704
P62705 P62707 P62708 P62709 P62710 P62711 P62712 P62713 P62726 P62727 P62728

P62729 P62731 P62732 P62733 P62734 P62735 P62739 P62740 P62741 P62742 P62752
P62757 P62765 P62766 P62767 P62768 P62769 P62770 P62771 P62774 P62776 P62777
P62778 P62779 P62780 P62781 P62782 P62783 P62784 P62785 P62786 P62787 P62788
P62789 P62790 P62791 P62792 P62793 P62794 P62795 P62807 P62808 P62809 P62819
P62840 P62842 P62848 P62860 P62862 P62863 P62864 P62865 P62866 P62867 P62868
P62869 P62870 P62875 P62876 P62883 P62884 P62885 P62886 P62887 P62904 P62905
P62915 P62916 P62920 P62921 P62923 P62924 P62925 P62932 P62946 P62947 P62948
P62949 P62958 P62959 P62962 P62963 P62964 P62972 P62975 P62976 P62978 P62980
P62981 P62985 P63002 P63003 P63006 P63007 P63008 P63013 P63014 P63020 P63021
P63022 P63023 P63028 P63029 P63054 P63055 P63074 P63083 P63084 P63105 P63106
P63107 P63108 P63109 P63110 P63111 P63112 P63113 P63114 P63119 P63120 P63121
P63122 P63123 P63124 P63125 P63127 P63129 P63131 P63132 P63133 P63135 P63136
P63139 P63140 P63150 P63151 P63156 P63157 P63162 P63163 P63164 P63169 P63177
P63178 P63179 P63180 P63187 P63188 P63189 P63191 P63193 P63194 P63195 P63196
P63198 P63199 P63200 P63201 P63202 P63203 P63205 P63206 P63207 P63237 P63238
P63248 P63249 P63254 P63255 P63256 P63262 P63263 P63264 P63265 P63266 P63267
P63268 P63269 P63270 P63271 P63272 P63274 P63275 P63280 P63281 P63282 P63283
P63285 P63286 P63287 P63301 P63302 P63303 P63312 P63313 P63314 P63315 P63317
P63335 P63336 P63337 P63339 P63346 P63358 P63381 P63382 P63389 P63390 P63396
P63402 P63406 P63410 P63411 P63412 P63413 P63414 P63415 P63417 P63418 P63419
P63420 P63421 P63422 P63423 P63424 P63426 P63428 P63430 P63432 P63433 P63434
P63435 P63436 P63453 P63462 P63463 P63475 P63476 P63477 P63478 P63479 P63481
P63482 P63483 P63485 P63486 P63487 P63488 P63491 P63493 P63495 P63497 P63499
P63501 P63503 P63505 P63507 P63508 P63510 P63511 P63512 P63513 P63515 P63517
P63518 P63525 P63526 P63532 P63534 P63536 P63537 P63539 P63541 P63542 P63543
P63544 P63545 P63546 P63548 P63549 P63550 P63552 P63553 P63555 P63556 P63557
P63558 P63559 P63560 P63561 P63563 P63564 P63565 P63566 P63567 P63569 P63570
P63572 P63573 P63574 P63575 P63576 P63578 P63579 P63581 P63582 P63583 P63584
P63586 P63587 P63588 P63589 P63590 P63592 P63593 P63594 P63595 P63596 P63597
P63599 P63600 P63601 P63602 P63603 P63604 P63605 P63606 P63607 P63608 P63609
P63610 P63612 P63613 P63614 P63615 P63616 P63617 P63618 P63621 P63622 P63623
P63624 P63626 P63630 P63631 P63632 P63633 P63634 P63636 P63637 P63639 P63640
P63641 P63643 P63644 P63645 P63646 P63647 P63649 P63651 P63698 P63699 P63700
P63703 P63704 P63705 P63706 P63708 P63712 P63716 P63718 P63720 P63722 P63724
P63729 P63730 P63731 P63732 P63733 P63734 P63735 P63737 P63738 P63739 P63741
P63742 P63743 P63746 P63747 P63748 P63750 P63766 P63767 P63768 P63769 P63770
P63772 P63773 P63774 P63776 P63778 P63784 P63785 P63786 P63787 P63788 P63789
P63791 P63792 P63793 P63795 P63796 P63804 P63805 P63806 P63807 P63808 P63809
P63811 P63812 P63813 P63814 P63815 P63816 P63817 P63818 P63819 P63820 P63821
P63823 P63824 P63825 P63827 P63828 P63829 P63830 P63832 P63834 P63836 P63840
P63842 P63843 P63845 P63846 P63847 P63849 P63850 P63851 P63870 P63872 P63874
P63875 P63876 P63877 P63878 P63889 P63890 P63891 P63893 P63894 P63895 P63896
P63898 P63901 P63902 P63903 P63904 P63905 P63906 P63907 P63908 P63909 P63910
P63913 P63914 P63915 P63916 P63917 P63918 P63919 P63920 P63921 P63922 P63923

P63924 P63925 P63926 P63927 P63930 P63932 P63933 P63934 P63936 P63938 P63939
P63940 P63943 P63944 P63946 P63947 P63948 P63949 P63951 P63952 P63953 P63954
P63955 P63956 P63963 P63964 P63965 P63967 P63968 P63969 P63970 P63972 P63974
P63976 P63978 P63979 P63980 P63981 P63983 P63984 P63986 P63988 P63989 P63990
P63991 P63992 P63996 P63997 P63998 P63999 P64001 P64004 P64005 P64006 P64007
P64008 P64009 P64013 P64022 P64023 P64024 P64025 P64026 P64028 P64029 P64030
P64031 P64032 P64033 P64035 P64037 P64038 P64039 P64040 P64041 P64042 P64043
P64044 P64045 P64046 P64047 P64048 P64049 P64050 P64051 P64052 P64053 P64054
P64055 P64056 P64058 P64059 P64061 P64062 P64063 P64064 P64065 P64066 P64067
P64068 P64069 P64070 P64072 P64073 P64095 P64096 P64098 P64119 P64120 P64121
P64123 P64124 P64126 P64127 P64128 P64129 P64130 P64131 P64132 P64133 P64135
P64136 P64137 P64138 P64140 P64141 P64142 P64144 P64145 P64146 P64147 P64148
P64149 P64151 P64152 P64153 P64154 P64155 P64157 P64159 P64172 P64179 P64180
P64181 P64183 P64185 P64193 P64194 P64195 P64196 P64197 P64198 P64200 P64201
P64202 P64203 P64204 P64206 P64208 P64209 P64210 P64211 P64212 P64213 P64215
P64216 P64217 P64219 P64221 P64222 P64223 P64224 P64226 P64227 P64229 P64231
P64232 P64233 P64234 P64236 P64237 P64238 P64239 P64240 P64242 P64243 P64244
P64246 P64248 P64250 P64251 P64252 P64253 P64254 P64262 P64264 P64267 P64268
P64269 P64271 P64272 P64273 P64274 P64275 P64276 P64277 P64278 P64280 P64281
P64282 P64283 P64284 P64285 P64287 P64289 P64290 P64291 P64294 P64295 P64296
P64297 P64298 P64299 P64300 P64302 P64304 P64305 P64306 P64308 P64309 P64310
P64311 P64312 P64315 P64318 P64321 P64323 P64324 P64327 P64329 P64330 P64331
P64332 P64333 P64335 P64337 P64340 P64342 P64343 P64344 P64345 P64346 P64347
P64348 P64349 P64350 P64351 P64352 P64355 P64356 P64359 P64360 P64361 P64362
P64363 P64364 P64365 P64366 P64367 P64369 P64370 P64371 P64372 P64373 P64374
P64375 P64377 P64378 P64379 P64380 P64381 P64382 P64383 P64386 P64387 P64388
P64389 P64390 P64391 P64392 P64393 P64394 P64395 P64396 P64397 P64399 P64400
P64401 P64402 P64403 P64404 P64405 P64406 P64407 P64408 P64409 P64410 P64412
P64413 P64414 P64415 P64417 P64419 P64420 P64421 P64422 P64443 P64444 P64448
P64449 P64450 P64455 P64456 P64457 P64458 P64462 P64463 P64464 P64465 P64466
P64467 P64468 P64469 P64470 P64474 P64475 P64476 P64477 P64478 P64479 P64480
P64483 P64484 P64488 P64489 P64490 P64491 P64492 P64493 P64494 P64495 P64496
P64497 P64498 P64503 P64504 P64505 P64509 P64510 P64511 P64517 P64518 P64519
P64520 P64521 P64522 P64523 P64524 P64525 P64526 P64527 P64529 P64530 P64531
P64532 P64533 P64540 P64541 P64554 P64555 P64556 P64557 P64558 P64560 P64561
P64562 P64563 P64568 P64569 P64570 P64571 P64572 P64573 P64574 P64575 P64576
P64577 P64578 P64579 P64580 P64588 P64589 P64594 P64595 P64599 P64600 P64601
P64610 P64611 P64612 P64613 P64616 P64617 P64618 P64624 P64625 P64626 P64631
P64632 P64633 P64636 P64637 P64638 P64639 P64649 P64650 P64657 P64658 P64659
P64660 P64663 P64664 P64665 P64666 P64668 P64670 P64672 P64674 P64676 P64678
P64692 P64694 P64702 P64704 P64706 P64708 P64710 P64714 P64720 P64726 P64728
P64734 P64740 P64742 P64746 P64748 P64750 P64754 P64762 P64764 P64766 P64768
P64770 P64774 P64778 P64780 P64786 P64790 P64796 P64798 P64800 P64804 P64806
P64810 P64814 P64820 P64822 P64826 P64828 P64834 P64836 P64844 P64846 P64852

P64854 P64860 P64862 P64864 P64868 P64872 P64874 P64878 P64880 P64882 P64886
P64888 P64898 P64900 P64902 P64908 P64910 P64912 P64914 P64916 P64918 P64920
P64922 P64924 P64926 P64930 P64936 P64937 P64938 P64942 P64944 P64956 P64960
P64964 P64966 P64968 P64974 P64976 P64982 P64984 P64986 P64988 P64990 P64992
P64994 P64996 P65002 P65004 P65006 P65010 P65014 P65018 P65028 P65032 P65034
P65036 P65040 P65042 P65044 P65046 P65050 P65052 P65054 P65056 P65058 P65060
P65064 P65066 P65068 P65070 P65074 P65076 P65078 P65080 P65086 P65088 P65090
P65092 P65094 P65096 P65098 P65099 P65100 P65101 P65102 P65103 P65105 P65106
P65107 P65108 P65109 P65110 P65111 P65112 P65113 P65114 P65115 P65116 P65117
P65118 P65120 P65121 P65122 P65123 P65125 P65126 P65127 P65128 P65129 P65130
P65132 P65133 P65136 P65137 P65138 P65139 P65141 P65142 P65143 P65144 P65145
P65146 P65148 P65150 P65151 P65152 P65153 P65155 P65156 P65157 P65158 P65159
P65160 P65162 P65164 P65166 P65168 P65169 P65171 P65173 P65176 P65179 P65180
P65181 P65182 P65184 P65185 P65186 P65191 P65192 P65193 P65194 P65199 P65200
P65201 P65202 P65203 P65204 P65205 P65217 P65218 P65219 P65220 P65221 P65223
P65225 P65226 P65227 P65228 P65229 P65230 P65233 P65234 P65236 P65238 P65239
P65240 P65241 P65242 P65243 P65245 P65246 P65247 P65248 P65249 P65250 P65251
P65252 P65255 P65257 P65258 P65259 P65261 P65275 P65276 P65278 P65279 P65280
P65281 P65282 P65284 P65285 P65287 P65328 P65337 P65341 P65342 P65343 P65344
P65345 P65347 P65349 P65350 P65351 P65353 P65354 P65355 P65356 P65358 P65359
P65360 P65361 P65362 P65364 P65365 P65366 P65367 P65368 P65369 P65382 P65383
P65385 P65386 P65387 P65388 P65389 P65393 P65394 P65395 P65396 P65397 P65398
P65399 P65400 P65401 P65403 P65404 P65406 P65407 P65409 P65413 P65414 P65415
P65416 P65417 P65418 P65420 P65421 P65423 P65424 P65426 P65427 P65428 P65430
P65431 P65433 P65434 P65435 P65437 P65438 P65440 P65442 P65443 P65444 P65445
P65447 P65448 P65449 P65450 P65451 P65452 P65453 P65454 P65455 P65456 P65458
P65459 P65461 P65462 P65464 P65465 P65466 P65467 P65468 P65469 P65470 P65471
P65473 P65474 P65476 P65478 P65479 P65486 P65488 P65489 P65490 P65491 P65493
P65494 P65495 P65498 P65500 P65501 P65502 P65503 P65504 P65505 P65506 P65507
P65508 P65509 P65512 P65513 P65514 P65515 P65516 P65517 P65518 P65519 P65520
P65521 P65522 P65523 P65529 P65531 P65532 P65533 P65534 P65535 P65536 P65537
P65538 P65539 P65546 P65547 P65549 P65551 P65552 P65553 P65554 P65555 P65556
P65557 P65558 P65559 P65560 P65577 P65579 P65580 P65581 P65582 P65583 P65585
P65590 P65591 P65592 P65595 P65596 P65598 P65599 P65600 P65601 P65603 P65604
P65605 P65606 P65607 P65608 P65610 P65611 P65612 P65614 P65615 P65616 P65617
P65619 P65620 P65634 P65635 P65639 P65640 P65641 P65642 P65643 P65644 P65646
P65649 P65654 P65655 P65657 P65658 P65661 P65662 P65663 P65664 P65665 P65666
P65669 P65671 P65672 P65673 P65676 P65677 P65678 P65679 P65680 P65681 P65683
P65685 P65687 P65689 P65691 P65692 P65693 P65694 P65695 P65696 P65697 P65698
P65699 P65701 P65702 P65703 P65704 P65705 P65708 P65709 P65711 P65717 P65722
P65723 P65725 P65744 P65745 P65747 P65748 P65749 P65750 P65751 P65752 P65754
P65755 P65756 P65757 P65759 P65761 P65763 P65769 P65770 P65771 P65772 P65773
P65774 P65775 P65776 P65778 P65779 P65780 P65781 P65783 P65787 P65789 P65790
P65791 P65792 P65793 P65794 P65795 P65796 P65798 P65799 P65800 P65802 P65803

P65804 P65805 P65806 P65807 P65808 P65809 P65827 P65828 P65830 P65831 P65832
P65833 P65834 P65835 P65836 P65837 P65840 P65841 P65843 P65844 P65845 P65847
P65848 P65849 P65850 P65851 P65852 P65853 P65854 P65856 P65857 P65858 P65859
P65861 P65862 P65863 P65864 P65866 P65867 P65868 P65869 P65870 P65871 P65872
P65878 P65879 P65881 P65882 P65883 P65884 P65885 P65886 P65887 P65888 P65889
P65890 P65891 P65892 P65894 P65895 P65897 P65899 P65900 P65903 P65904 P65905
P65907 P65910 P65911 P65914 P65915 P65916 P65917 P65918 P65920 P65921 P65922
P65923 P65924 P65925 P65927 P65928 P65930 P65931 P65932 P65933 P65934 P65935
P65937 P65938 P65940 P65942 P65943 P65945 P65946 P65947 P65948 P65949 P65950
P65952 P65954 P65955 P65956 P65957 P65958 P65959 P65960 P65961 P65962 P65963
P65965 P65966 P65968 P65969 P65970 P65971 P65973 P65974 P65975 P65976 P65977
P65978 P65979 P65980 P65981 P65982 P65984 P65985 P65986 P65987 P65989 P65991
P65992 P65993 P65994 P65996 P65998 P65999 P66000 P66001 P66002 P66005 P66007
P66008 P66009 P66010 P66011 P66012 P66013 P66015 P66017 P66018 P66019 P66020
P66021 P66023 P66024 P66025 P66027 P66029 P66030 P66031 P66032 P66033 P66035
P66036 P66037 P66038 P66039 P66040 P66041 P66042 P66043 P66045 P66046 P66047
P66048 P66049 P66050 P66051 P66052 P66053 P66054 P66055 P66057 P66058 P66059
P66061 P66062 P66063 P66066 P66067 P66068 P66070 P66071 P66072 P66073 P66074
P66075 P66077 P66078 P66079 P66081 P66082 P66084 P66086 P66087 P66088 P66091
P66092 P66093 P66094 P66095 P66096 P66097 P66098 P66099 P66100 P66101 P66102
P66103 P66104 P66106 P66107 P66109 P66110 P66111 P66112 P66113 P66114 P66116
P66118 P66119 P66120 P66122 P66123 P66124 P66125 P66126 P66128 P66129 P66130
P66131 P66132 P66134 P66135 P66136 P66137 P66139 P66140 P66141 P66142 P66143
P66144 P66145 P66149 P66150 P66151 P66152 P66153 P66154 P66155 P66156 P66157
P66159 P66160 P66161 P66162 P66163 P66164 P66165 P66166 P66167 P66168 P66169
P66170 P66171 P66172 P66174 P66175 P66176 P66178 P66179 P66180 P66182 P66183
P66184 P66185 P66186 P66188 P66189 P66190 P66191 P66192 P66193 P66194 P66195
P66197 P66198 P66199 P66200 P66201 P66202 P66204 P66205 P66206 P66207 P66208
P66209 P66211 P66213 P66214 P66215 P66216 P66217 P66218 P66219 P66220 P66221
P66222 P66225 P66226 P66227 P66228 P66229 P66230 P66232 P66233 P66234 P66235
P66237 P66238 P66239 P66240 P66241 P66242 P66243 P66244 P66245 P66246 P66247
P66248 P66249 P66250 P66251 P66252 P66253 P66254 P66255 P66256 P66257 P66258
P66260 P66261 P66262 P66263 P66264 P66265 P66266 P66267 P66268 P66269 P66270
P66272 P66273 P66274 P66275 P66276 P66277 P66278 P66279 P66280 P66282 P66283
P66284 P66285 P66286 P66287 P66288 P66289 P66290 P66291 P66292 P66293 P66294
P66295 P66296 P66297 P66298 P66299 P66300 P66301 P66302 P66303 P66305 P66306
P66307 P66308 P66309 P66310 P66312 P66313 P66314 P66316 P66317 P66319 P66320
P66321 P66322 P66324 P66325 P66326 P66327 P66328 P66329 P66330 P66331 P66332
P66333 P66335 P66336 P66337 P66338 P66339 P66340 P66341 P66343 P66344 P66345
P66346 P66347 P66348 P66349 P66350 P66351 P66352 P66353 P66354 P66355 P66356
P66358 P66359 P66360 P66362 P66363 P66364 P66365 P66366 P66367 P66368 P66369
P66370 P66371 P66372 P66373 P66374 P66375 P66376 P66378 P66379 P66380 P66381
P66382 P66383 P66384 P66385 P66386 P66387 P66389 P66390 P66391 P66392 P66393
P66394 P66396 P66397 P66398 P66399 P66400 P66401 P66402 P66406 P66407 P66408

P66409 P66410 P66411 P66412 P66413 P66414 P66415 P66416 P66417 P66418 P66419
P66420 P66421 P66423 P66424 P66426 P66427 P66428 P66430 P66431 P66432 P66433
P66434 P66436 P66437 P66438 P66439 P66441 P66442 P66443 P66444 P66445 P66446
P66448 P66449 P66450 P66451 P66452 P66453 P66454 P66455 P66456 P66457 P66458
P66459 P66460 P66461 P66462 P66466 P66467 P66469 P66470 P66471 P66472 P66473
P66474 P66475 P66477 P66478 P66479 P66480 P66481 P66482 P66483 P66484 P66485
P66486 P66487 P66488 P66489 P66490 P66491 P66492 P66493 P66495 P66496 P66498
P66499 P66500 P66501 P66502 P66503 P66504 P66506 P66507 P66508 P66509 P66511
P66512 P66513 P66514 P66515 P66516 P66517 P66518 P66520 P66522 P66523 P66524
P66525 P66526 P66528 P66529 P66530 P66531 P66532 P66533 P66534 P66535 P66536
P66538 P66539 P66541 P66542 P66543 P66545 P66546 P66547 P66548 P66549 P66550
P66551 P66552 P66554 P66555 P66557 P66558 P66559 P66560 P66561 P66562 P66564
P66565 P66566 P66567 P66569 P66570 P66571 P66572 P66573 P66575 P66576 P66577
P66578 P66580 P66581 P66582 P66583 P66585 P66586 P66587 P66592 P66593 P66594
P66595 P66596 P66597 P66598 P66599 P66601 P66602 P66603 P66604 P66605 P66606
P66607 P66608 P66609 P66610 P66611 P66612 P66613 P66614 P66615 P66617 P66618
P66619 P66620 P66621 P66622 P66623 P66624 P66626 P66627 P66628 P66629 P66631
P66632 P66633 P66634 P66636 P66637 P66638 P66640 P66641 P66642 P66643 P66644
P66645 P66647 P66648 P66650 P66651 P66652 P66654 P66655 P66657 P66661 P66663
P66664 P66665 P66667 P66669 P66670 P66672 P66673 P66674 P66675 P66676 P66677
P66678 P66679 P66680 P66681 P66682 P66683 P66684 P66685 P66686 P66687 P66688
P66689 P66691 P66692 P66693 P66694 P66695 P66696 P66697 P66699 P66700 P66702
P66703 P66704 P66705 P66707 P66708 P66709 P66710 P66711 P66712 P66713 P66714
P66716 P66717 P66718 P66719 P66722 P66723 P66724 P66725 P66727 P66728 P66729
P66730 P66731 P66732 P66733 P66735 P66736 P66737 P66738 P66739 P66740 P66741
P66742 P66743 P66745 P66746 P66747 P66748 P66749 P66750 P66751 P66754 P66755
P66756 P66757 P66759 P66761 P66762 P66763 P66764 P66765 P66766 P66768 P66770
P66774 P66775 P66776 P66778 P66780 P66782 P66784 P66793 P66794 P66795 P66802
P66803 P66804 P66805 P66806 P66808 P66810 P66814 P66815 P66816 P66817 P66818
P66819 P66820 P66821 P66822 P66823 P66828 P66829 P66830 P66832 P66833 P66834
P66835 P66836 P66837 P66839 P66840 P66841 P66846 P66847 P66848 P66849 P66850
P66851 P66852 P66854 P66855 P66856 P66857 P66858 P66859 P66860 P66861 P66862
P66863 P66864 P66865 P66866 P66867 P66868 P66869 P66870 P66871 P66872 P66874
P66876 P66878 P66879 P66880 P66881 P66882 P66886 P66898 P66899 P66900 P66901
P66903 P66904 P66906 P66907 P66908 P66910 P66912 P66914 P66915 P66917 P66918
P66920 P66921 P66922 P66927 P66928 P66929 P66931 P66932 P66933 P66934 P66935
P66936 P66938 P66941 P66942 P66943 P66947 P66953 P66954 P66955 P66956 P66957
P66958 P66959 P66961 P66962 P66963 P66969 P66970 P66971 P66972 P66973 P66974
P66975 P66977 P66978 P66979 P66981 P66982 P66983 P66985 P66986 P66987 P66988
P66989 P66990 P66991 P66993 P66994 P66995 P66996 P66997 P66998 P66999 P67000
P67002 P67003 P67004 P67005 P67006 P67007 P67008 P67009 P67010 P67012 P67013
P67014 P67016 P67018 P67021 P67022 P67023 P67024 P67025 P67026 P67027 P67028
P67029 P67030 P67031 P67033 P67034 P67035 P67036 P67037 P67038 P67039 P67040
P67042 P67043 P67045 P67046 P67047 P67048 P67049 P67050 P67051 P67055 P67056

P67061 P67063 P67064 P67065 P67068 P67070 P67072 P67073 P67074 P67075 P67076
P67077 P67078 P67079 P67080 P67081 P67082 P67084 P67086 P67087 P67088 P67089
P67090 P67091 P67092 P67093 P67094 P67095 P67096 P67097 P67100 P67101 P67102
P67103 P67105 P67107 P67108 P67110 P67111 P67112 P67113 P67120 P67122 P67124
P67126 P67132 P67133 P67135 P67136 P67137 P67138 P67139 P67140 P67142 P67152
P67160 P67172 P67173 P67174 P67175 P67176 P67178 P67179 P67180 P67181 P67183
P67184 P67185 P67186 P67187 P67188 P67190 P67191 P67192 P67193 P67194 P67195
P67196 P67197 P67198 P67199 P67200 P67201 P67203 P67212 P67213 P67215 P67216
P67217 P67218 P67219 P67220 P67221 P67223 P67227 P67228 P67229 P67231 P67233
P67234 P67235 P67237 P67239 P67241 P67243 P67246 P67247 P67257 P67258 P67259
P67260 P67261 P67262 P67263 P67264 P67265 P67266 P67267 P67268 P67269 P67270
P67271 P67272 P67273 P67274 P67276 P67286 P67287 P67299 P67310 P67311 P67312
P67314 P67315 P67316 P67317 P67318 P67319 P67320 P67322 P67323 P67324 P67325
P67326 P67327 P67328 P67329 P67330 P67331 P67332 P67333 P67334 P67335 P67336
P67337 P67338 P67339 P67340 P67341 P67342 P67343 P67344 P67349 P67350 P67351
P67352 P67353 P67354 P67356 P67357 P67358 P67359 P67360 P67361 P67362 P67363
P67365 P67366 P67367 P67381 P67382 P67383 P67395 P67397 P67398 P67399 P67400
P67402 P67403 P67405 P67406 P67407 P67408 P67409 P67410 P67412 P67413 P67414
P67415 P67418 P67419 P67421 P67423 P67424 P67427 P67428 P67429 P67431 P67433
P67435 P67437 P67441 P67443 P67448 P67449 P67450 P67451 P67452 P67454 P67455
P67457 P67458 P67459 P67460 P67461 P67462 P67463 P67464 P67465 P67466 P67467
P67469 P67472 P67474 P67476 P67477 P67478 P67479 P67480 P67481 P67482 P67484
P67485 P67486 P67488 P67489 P67490 P67491 P67492 P67493 P67495 P67496 P67497
P67499 P67500 P67501 P67502 P67503 P67504 P67506 P67507 P67508 P67511 P67512
P67514 P67515 P67516 P67517 P67518 P67519 P67520 P67521 P67522 P67523 P67524
P67525 P67526 P67527 P67528 P67529 P67530 P67531 P67532 P67534 P67535 P67536
P67537 P67538 P67539 P67540 P67542 P67543 P67545 P67546 P67549 P67550 P67551
P67552 P67562 P67563 P67564 P67565 P67566 P67567 P67568 P67570 P67571 P67573
P67574 P67575 P67577 P67578 P67580 P67581 P67583 P67584 P67586 P67587 P67588
P67589 P67591 P67592 P67594 P67595 P67596 P67598 P67600 P67603 P67604 P67605
P67606 P67608 P67609 P67612 P67613 P67614 P67615 P67616 P67617 P67618 P67622
P67623 P67624 P67625 P67626 P67627 P67629 P67630 P67631 P67632 P67633 P67634
P67637 P67640 P67641 P67642 P67648 P67649 P67650 P67651 P67652 P67655 P67657
P67660 P67661 P67662 P67663 P67664 P67666 P67668 P67670 P67672 P67673 P67674
P67675 P67676 P67677 P67678 P67679 P67680 P67681 P67683 P67684 P67685 P67688
P67689 P67690 P67691 P67692 P67693 P67694 P67695 P67696 P67698 P67699 P67700
P67701 P67702 P67703 P67705 P67706 P67707 P67708 P67709 P67710 P67711 P67714
P67715 P67716 P67718 P67719 P67720 P67721 P67724 P67726 P67727 P67728 P67732
P67734 P67735 P67736 P67738 P67740 P67742 P67743 P67746 P67748 P67756 P67758
P67759 P67760 P67761 P67762 P67763 P67764 P67766 P67767 P67769 P67771 P67773
P67804 P67805 P67815 P67816 P67817 P67818 P67819 P67820 P67821 P67822 P67823
P67824 P67825 P67828 P67829 P67830 P67831 P67832 P67833 P67834 P67835 P67836
P67837 P67838 P67839 P67840 P67841 P67842 P67843 P67844 P67845 P67846 P67847
P67848 P67849 P67850 P67851 P67852 P67853 P67854 P67855 P67856 P67869 P67872

P67873 P67884 P67897 P67898 P67899 P67900 P67901 P67902 P67903 P67904 P67905
P67906 P67916 P67917 P67925 P67937 P67963 P67982 P67983 P68011 P68012 P68013
P68014 P68015 P68016 P68017 P68018 P68019 P68020 P68021 P68022 P68023 P68024
P68025 P68026 P68027 P68028 P68029 P68030 P68031 P68033 P68034 P68035 P68036
P68037 P68038 P68041 P68044 P68045 P68046 P68047 P68048 P68049 P68050 P68051
P68052 P68053 P68054 P68055 P68056 P68057 P68058 P68059 P68060 P68061 P68062
P68063 P68064 P68065 P68067 P68068 P68069 P68070 P68071 P68077 P68078 P68079
P68080 P68081 P68082 P68083 P68084 P68085 P68086 P68087 P68101 P68133 P68134
P68135 P68136 P68137 P68138 P68139 P68140 P68158 P68163 P68164 P68165 P68166
P68168 P68169 P68170 P68171 P68172 P68173 P68176 P68177 P68189 P68190 P68191
P68192 P68193 P68194 P68197 P68200 P68202 P68203 P68205 P68206 P68207 P68208
P68210 P68211 P68222 P68223 P68224 P68225 P68226 P68227 P68228 P68229 P68230
P68231 P68232 P68234 P68235 P68236 P68237 P68239 P68240 P68246 P68247 P68251
P68252 P68253 P68256 P68257 P68258 P68261 P68262 P68263 P68264 P68271 P68272
P68276 P68277 P68278 P68279 P68280 P68281 P68282 P68283 P68284 P68285 P68286
P68287 P68288 P68289 P68301 P68302 P68303 P68304 P68305 P68306 P68311 P68312
P68313 P68314 P68315 P68316 P68317 P68318 P68319 P68321 P68322 P68335 P68336
P68337 P68339 P68340 P68341 P68342 P68345 P68349 P68351 P68352 P68353 P68354
P68355 P68356 P68357 P68358 P68359 P68360 P68361 P68365 P68367 P68395 P68396
P68397 P68398 P68399 P68405 P68406 P68427 P68428 P68429 P68430 P68432 P68433
P68438 P68439 P68442 P68443 P68448 P68449 P68450 P68451 P68454 P68455 P68456
P68457 P68464 P68465 P68466 P68467 P68472 P68473 P68474 P68475 P68476 P68477
P68478 P68479 P68480 P68481 P68482 P68483 P68484 P68485 P68486 P68487 P68488
P68489 P68490 P68491 P68492 P68493 P68494 P68495 P68497 P68498 P68499 P68501
P68502 P68503 P68504 P68505 P68506 P68507 P68508 P68522 P68523 P68524 P68525
P68535 P68536 P68544 P68545 P68546 P68547 P68550 P68552 P68555 P68556 P68559
P68560 P68561 P68562 P68563 P68564 P68565 P68566 P68567 P68568 P68573 P68574
P68575 P68576 P68581 P68582 P68583 P68584 P68585 P68586 P68587 P68590 P68591
P68595 P68596 P68601 P68602 P68603 P68608 P68609 P68610 P68611 P68612 P68613
P68614 P68620 P68621 P68624 P68625 P68626 P68627 P68628 P68629 P68630 P68631
P68634 P68635 P68644 P68645 P68646 P68647 P68648 P68649 P68650 P68651 P68652
P68653 P68654 P68655 P68656 P68657 P68658 P68659 P68660 P68661 P68662 P68663
P68664 P68665 P68666 P68669 P68671 P68673 P68675 P68679 P68680 P68681 P68682
P68683 P68684 P68685 P68686 P68687 P68688 P68689 P68690 P68691 P68692 P68693
P68694 P68695 P68696 P68697 P68698 P68707 P68708 P68709 P68710 P68713 P68714
P68715 P68716 P68717 P68718 P68719 P68720 P68728 P68729 P68730 P68731 P68733
P68737 P68738 P68739 P68740 P68741 P68742 P68743 P68746 P68747 P68748 P68749
P68750 P68751 P68752 P68753 P68754 P68763 P68764 P68765 P68766 P68767 P68768
P68770 P68771 P68773 P68775 P68777 P68778 P68780 P68781 P68785 P68786 P68787
P68788 P68790 P68791 P68792 P68793 P68794 P68797 P68807 P68809 P68810 P68811
P68812 P68814 P68815 P68816 P68818 P68819 P68820 P68821 P68822 P68823 P68824
P68825 P68826 P68828 P68829 P68830 P68831 P68834 P68835 P68837 P68839 P68840
P68842 P68843 P68845 P68846 P68847 P68848 P68849 P68850 P68852 P68861 P68862
P68863 P68864 P68865 P68867 P68868 P68869 P68870 P68871 P68872 P68873 P68876

P68878 P68886 P68890 P68892 P68893 P68895 P68896 P68897 P68898 P68899 P68900
P68901 P68902 P68903 P68905 P68906 P68907 P68908 P68910 P68912 P68916 P68918
P68919 P68922 P68923 P68924 P68925 P68926 P68927 P68928 P68929 P68930 P68931
P68932 P68933 P68934 P68935 P68936 P68937 P68938 P68941 P68942 P68944 P68945
P68946 P68947 P68948 P68949 P68958 P68959 P68960 P68961 P68962 P68963 P68966
P68967 P68968 P68969 P68970 P68971 P68972 P68973 P68974 P68975 P68977 P68982
P68983 P68984 P68985 P68986 P68993 P68994 P68995 P68996 P68997 P68998 P68999
P69000 P69001 P69002 P69003 P69004 P69005 P69006 P69007 P69008 P69009 P69010
P69011 P69012 P69013 P69014 P69015 P69037 P69038 P69039 P69040 P69041 P69042
P69050 P69053 P69060 P69061 P69064 P69065 P69066 P69069 P69070 P69071 P69072
P69073 P69074 P69075 P69076 P69077 P69078 P69079 P69080 P69081 P69082 P69083
P69084 P69085 P69086 P69090 P69091 P69097 P69098 P69099 P69100 P69103 P69104
P69110 P69111 P69112 P69113 P69114 P69115 P69116 P69117 P69118 P69119 P69120
P69121 P69122 P69123 P69124 P69125 P69126 P69127 P69139 P69140 P69141 P69142
P69149 P69150 P69151 P69152 P69153 P69154 P69163 P69164 P69170 P69171 P69172
P69173 P69174 P69175 P69176 P69177 P69180 P69181 P69182 P69183 P69184 P69185
P69186 P69189 P69190 P69191 P69196 P69197 P69198 P69199 P69200 P69201 P69203
P69204 P69205 P69206 P69207 P69209 P69222 P69223 P69224 P69225 P69226 P69227
P69231 P69242 P69243 P69244 P69245 P69246 P69248 P69249 P69250 P69251 P69280
P69281 P69282 P69283 P69284 P69285 P69286 P69287 P69288 P69309 P69310 P69313
P69315 P69317 P69322 P69325 P69326 P69329 P69330 P69331 P69341 P69342 P69346
P69347 P69348 P69349 P69365 P69366 P69376 P69377 P69406 P69413 P69418 P69439
P69442 P69466 P69467 P69470 P69471 P69472 P69473 P69474 P69475 P69484 P69485
P69486 P69487 P69489 P69503 P69504 P69505 P69506 P69507 P69508 P69509 P69510
P69514 P69515 P69516 P69517 P69545 P69546 P69547 P69548 P69549 P69567 P69568
P69569 P69570 P69571 P69572 P69573 P69574 P69575 P69576 P69577 P69578 P69579
P69580 P69581 P69582 P69583 P69584 P69585 P69586 P69587 P69588 P69589 P69590
P69591 P69592 P69593 P69594 P69595 P69608 P69614 P69615 P69620 P69621 P69622
P69623 P69624 P69625 P69626 P69627 P69628 P69629 P69630 P69631 P69632 P69633
P69634 P69635 P69636 P69637 P69638 P69639 P69640 P69641 P69642 P69643 P69644
P69645 P69646 P69647 P69648 P69649 P69650 P69651 P69652 P69653 P69654 P69655
P69656 P69659 P69660 P69661 P69662 P69663 P69664 P69665 P69666 P69667 P69668
P69679 P69683 P69684 P69687 P69688 P69689 P69701 P69702 P69715 P69716 P69717
P69737 P69738 P69772 P69773 P69774 P69782 P69815 P69834 P69848 P69850 P69851
P69852 P69859 P69882 P69883 P69884 P69886 P69887 P69889 P69890 P69891 P69892
P69893 P69895 P69898 P69899 P69900 P69901 P69902 P69903 P69906 P69907 P69909
P69911 P69912 P69913 P69914 P69915 P69916 P69917 P69918 P69920 P69921 P69922
P69923 P69924 P69925 P69932 P69933 P69934 P69935 P69936 P69946 P69948 P69952
P69953 P69954 P69957 P69958 P69959 P69960 P69961 P69962 P69963 P69964 P69967
P69968 P69970 P69971 P69974 P69975 P69976 P69977 P69978 P69979 P69988 P69989
P69990 P69991 P69992 P69993 P69994 P69995 P70005 P70010 P70011 P70030 P70032
P70033 P70039 P70040 P70047 P70052 P70054 P70060 P70061 P70062 P70063 P70064
P70066 P70076 P70080 P70082 P70094 P70112 P70120 P70121 P70122 P70178 P70195
P70202 P70210 P70213 P70217 P70255 P70257 P70265 P70266 P70270 P70284 P70288

P70298 P70303 P70318 P70321 P70323 P70324 P70325 P70331 P70333 P70338 P70353
P70354 P70368 P70372 P70378 P70388 P70390 P70396 P70397 P70399 P70402 P70404
P70429 P70434 P70436 P70447 P70459 P70475 P70501 P70512 P70531 P70537 P70541
P70563 P70583 P70595 P70645 P70660 P70661 P70662 P70671 P70675 P70696 P70697
P70698 P70700 P70708 P70712 P70715 P70718 P70727 P70728 P70730 P70736 P70742
P70752 P70753 P70754 P70755 P70782 P70785 P70787 P70788 P70789 P70790 P70792
P70794 P70795 P70796 P70814 P70826 P70827 P70828 P70830 P70838 P70842 P70843
P70844 P70865 P70870 P70877 P70878 P70881 P70882 P70889 P70910 P70911 P70917
P70918 P70920 P70936 P70937 P70938 P70947 P70949 P70955 P70958 P70962 P70964
P70965 P70973 P70974 P70976 P70981 P70985 P70986 P70993 P70994 P70996 P70998
P70999 P71000 P71002 P71003 P71004 P71015 P71016 P71017 P71020 P71021 P71025
P71031 P71032 P71035 P71036 P71037 P71039 P71043 P71046 P71047 P71049 P71053
P71054 P71055 P71057 P71058 P71059 P71063 P71065 P71066 P71073 P71080 P71084
P71086 P71088 P71089 P71103 P71104 P71119 P71121 P71128 P71145 P71146 P71147
P71148 P71153 P71154 P71156 P71165 P71169 P71198 P71237 P71239 P71242 P71243
P71296 P71297 P71298 P71301 P71311 P71317 P71318 P71319 P71331 P71333 P71335
P71339 P71340 P71344 P71346 P71348 P71349 P71350 P71351 P71353 P71357 P71358
P71359 P71362 P71363 P71365 P71366 P71373 P71374 P71376 P71378 P71379 P71381
P71384 P71385 P71387 P71388 P71389 P71390 P71394 P71398 P71404 P71405 P71422
P71426 P71447 P71478 P71479 P71481 P71496 P71498 P71499 P71500 P71512 P71513
P71517 P71592 P71615 P71622 P71635 P71650 P71668 P71716 P71820 P71839 P71889
P72020 P72021 P72065 P72077 P72131 P72138 P72139 P72154 P72156 P72157 P72158
P72170 P72171 P72173 P72174 P72180 P72183 P72188 P72190 P72191 P72192 P72208
P72220 P72227 P72230 P72231 P72232 P72233 P72235 P72238 P72241 P72255 P72263
P72270 P72273 P72294 P72299 P72300 P72314 P72315 P72324 P72329 P72332 P72338
P72339 P72349 P72357 P72360 P72366 P72394 P72396 P72403 P72449 P72469 P72478
P72483 P72484 P72495 P72524 P72533 P72539 P72540 P72541 P72542 P72546 P72578
P72579 P72580 P72581 P72583 P72584 P72587 P72588 P72622 P72623 P72641 P72642
P72644 P72648 P72652 P72657 P72663 P72666 P72667 P72680 P72687 P72689 P72699
P72702 P72703 P72711 P72720 P72721 P72723 P72725 P72731 P72736 P72743 P72745
P72749 P72753 P72757 P72758 P72759 P72760 P72761 P72772 P72776 P72777 P72785
P72794 P72795 P72796 P72797 P72800 P72811 P72817 P72818 P72821 P72828 P72830
P72835 P72848 P72849 P72851 P72854 P72856 P72861 P72866 P72871 P72872 P72873
P72874 P72913 P72925 P72931 P72933 P72934 P72940 P72943 P72946 P72955 P72965
P72970 P72979 P72980 P72983 P73002 P73003 P73014 P73042 P73047 P73053 P73055
P73056 P73057 P73058 P73061 P73063 P73067 P73068 P73071 P73079 P73090 P73092
P73097 P73098 P73121 P73123 P73125 P73127 P73133 P73136 P73138 P73141 P73161
P73162 P73163 P73177 P73191 P73201 P73211 P73212 P73245 P73246 P73248 P73257
P73263 P73269 P73270 P73274 P73289 P73290 P73292 P73293 P73294 P73295 P73296
P73297 P73298 P73299 P73300 P73301 P73302 P73303 P73304 P73305 P73306 P73307
P73308 P73309 P73310 P73311 P73312 P73313 P73314 P73315 P73316 P73317 P73318
P73319 P73320 P73321 P73326 P73327 P73334 P73335 P73336 P73340 P73341 P73345
P73346 P73349 P73367 P73374 P73376 P73389 P73390 P73392 P73406 P73407 P73408
P73410 P73411 P73412 P73418 P73421 P73423 P73426 P73436 P73440 P73441 P73443

P73455 P73460 P73467 P73469 P73471 P73473 P73475 P73481 P73492 P73493 P73505
P73511 P73515 P73526 P73527 P73530 P73532 P73534 P73538 P73554 P73558 P73560
P73571 P73572 P73577 P73580 P73585 P73599 P73602 P73617 P73618 P73623 P73625
P73632 P73634 P73636 P73638 P73639 P73646 P73648 P73651 P73655 P73658 P73660
P73662 P73664 P73667 P73668 P73672 P73690 P73709 P73715 P73720 P73728 P73735
P73737 P73747 P73754 P73755 P73761 P73769 P73789 P73790 P73796 P73807 P73810
P73820 P73821 P73825 P73826 P73832 P73833 P73839 P73841 P73844 P73846 P73849
P73851 P73859 P73860 P73881 P73882 P73883 P73889 P73890 P73891 P73903 P73910
P73911 P73922 P73925 P73935 P73942 P73955 P73960 P73971 P73997 P74002 P74003
P74007 P74008 P74034 P74035 P74038 P74040 P74041 P74045 P74060 P74061 P74070
P74071 P74081 P74089 P74095 P74101 P74104 P74106 P74117 P74120 P74122 P74125
P74130 P74132 P74133 P74134 P74142 P74143 P74148 P74163 P74177 P74178 P74192
P74193 P74207 P74208 P74215 P74217 P74218 P74220 P74221 P74226 P74227 P74228
P74229 P74230 P74232 P74240 P74241 P74246 P74257 P74260 P74261 P74263 P74266
P74267 P74268 P74269 P74287 P74290 P74292 P74296 P74297 P74299 P74301 P74304
P74306 P74309 P74312 P74313 P74323 P74324 P74327 P74328 P74330 P74331 P74334
P74342 P74346 P74352 P74355 P74360 P74361 P74368 P74371 P74373 P74374 P74383
P74384 P74386 P74391 P74395 P74397 P74410 P74415 P74416 P74421 P74422 P74423
P74430 P74432 P74435 P74438 P74442 P74456 P74457 P74458 P74459 P74466 P74467
P74476 P74494 P74497 P74503 P74507 P74514 P74516 P74518 P74519 P74521 P74523
P74528 P74529 P74533 P74557 P74561 P74562 P74565 P74572 P74576 P74578 P74582
P74587 P74591 P74592 P74593 P74594 P74596 P74606 P74607 P74618 P74641 P74644
P74645 P74646 P74650 P74662 P74667 P74676 P74689 P74690 P74696 P74706 P74709
P74724 P74733 P74737 P74739 P74741 P74745 P74750 P74755 P74759 P74762 P74764
P74766 P74770 P74786 P74794 P74811 P74836 P74838 P74839 P74840 P74855 P74858
P74861 P74879 P74881 P74883 P74892 P74893 P74897 P74909 P74910 P74918 P74921
P74925 P74926 P74933 P74935 P74936 P74941 P74945 P74946 P74956 P74963 P74965
P74984 P75010 P75012 P75022 P75030 P75033 P75035 P75036 P75037 P75038 P75042
P75043 P75044 P75045 P75046 P75049 P75050 P75051 P75052 P75053 P75055 P75064
P75065 P75066 P75067 P75068 P75069 P75070 P75072 P75073 P75074 P75075 P75076
P75077 P75078 P75079 P75080 P75081 P75082 P75084 P75085 P75086 P75087 P75088
P75089 P75090 P75091 P75092 P75093 P75096 P75097 P75098 P75099 P75100 P75103
P75104 P75105 P75106 P75107 P75110 P75111 P75113 P75114 P75115 P75116 P75117
P75118 P75119 P75122 P75123 P75124 P75128 P75129 P75131 P75132 P75133 P75135
P75136 P75138 P75139 P75140 P75141 P75142 P75143 P75144 P75148 P75159 P75161
P75162 P75163 P75164 P75165 P75169 P75170 P75171 P75172 P75173 P75175 P75176
P75177 P75178 P75179 P75180 P75181 P75182 P75183 P75187 P75188 P75194 P75196
P75197 P75199 P75200 P75202 P75203 P75205 P75206 P75209 P75211 P75214 P75215
P75216 P75217 P75218 P75219 P75220 P75221 P75222 P75223 P75224 P75225 P75226
P75227 P75228 P75229 P75230 P75231 P75233 P75234 P75235 P75236 P75237 P75238
P75239 P75240 P75241 P75242 P75243 P75244 P75245 P75246 P75247 P75248 P75249
P75250 P75252 P75253 P75254 P75256 P75257 P75258 P75259 P75260 P75261 P75264
P75266 P75267 P75268 P75269 P75270 P75271 P75272 P75273 P75274 P75275 P75276
P75277 P75278 P75279 P75281 P75282 P75283 P75285 P75286 P75287 P75288 P75289

P75290 P75293 P75294 P75297 P75298 P75299 P75300 P75301 P75303 P75304 P75305
P75306 P75308 P75309 P75310 P75311 P75313 P75315 P75317 P75319 P75320 P75321
P75325 P75326 P75333 P75335 P75338 P75340 P75341 P75344 P75345 P75347 P75348
P75349 P75350 P75351 P75353 P75354 P75359 P75360 P75361 P75364 P75365 P75368
P75370 P75372 P75374 P75375 P75380 P75381 P75382 P75386 P75388 P75389 P75394
P75395 P75396 P75398 P75399 P75400 P75401 P75402 P75403 P75404 P75406 P75407
P75410 P75411 P75413 P75414 P75415 P75416 P75417 P75419 P75420 P75423 P75424
P75425 P75426 P75427 P75429 P75430 P75431 P75432 P75433 P75434 P75435 P75436
P75437 P75438 P75442 P75444 P75446 P75447 P75448 P75451 P75452 P75453 P75454
P75455 P75456 P75457 P75458 P75460 P75461 P75465 P75466 P75467 P75469 P75470
P75473 P75475 P75476 P75477 P75478 P75481 P75482 P75483 P75485 P75486 P75487
P75488 P75490 P75491 P75492 P75494 P75495 P75497 P75499 P75500 P75502 P75504
P75508 P75509 P75510 P75511 P75512 P75517 P75518 P75519 P75521 P75522 P75523
P75524 P75525 P75526 P75527 P75529 P75530 P75531 P75532 P75533 P75534 P75536
P75537 P75539 P75540 P75541 P75542 P75543 P75544 P75545 P75546 P75548 P75549
P75550 P75558 P75560 P75561 P75562 P75563 P75564 P75565 P75566 P75567 P75568
P75570 P75571 P75572 P75575 P75576 P75577 P75578 P75579 P75580 P75581 P75584
P75587 P75589 P75590 P75591 P75592 P75593 P75594 P75598 P75599 P75600 P75602
P75604 P75605 P75612 P75617 P75620 P75672 P75675 P75676 P75677 P75678 P75679
P75680 P75682 P75684 P75688 P75691 P75692 P75697 P75704 P75711 P75713 P75717
P75718 P75719 P75726 P75741 P75745 P75746 P75764 P75767 P75779 P75782 P75791
P75792 P75793 P75794 P75809 P75811 P75817 P75821 P75823 P75824 P75825 P75828
P75829 P75836 P75838 P75843 P75849 P75862 P75863 P75864 P75867 P75869 P75874
P75876 P75893 P75894 P75895 P75897 P75898 P75914 P75915 P75931 P75936 P75937
P75938 P75942 P75946 P75948 P75949 P75959 P75960 P75966 P75967 P75969 P75970
P75972 P75973 P75975 P75976 P75977 P75978 P75980 P75981 P75987 P75988 P75989
P75990 P75994 P76008 P76010 P76012 P76014 P76015 P76016 P76023 P76034 P76035
P76038 P76041 P76043 P76044 P76046 P76049 P76052 P76053 P76055 P76056 P76057
P76062 P76064 P76065 P76066 P76068 P76069 P76071 P76072 P76073 P76077 P76078
P76079 P76080 P76081 P76083 P76084 P76085 P76086 P76096 P76097 P76100 P76102
P76104 P76106 P76112 P76113 P76114 P76117 P76118 P76119 P76121 P76122 P76123
P76127 P76134 P76135 P76138 P76141 P76145 P76148 P76149 P76150 P76154 P76155
P76156 P76157 P76159 P76160 P76161 P76162 P76163 P76164 P76165 P76168 P76187
P76192 P76194 P76196 P76201 P76204 P76205 P76210 P76213 P76214 P76215 P76216
P76217 P76222 P76231 P76234 P76235 P76241 P76243 P76245 P76250 P76251 P76254
P76257 P76268 P76270 P76273 P76275 P76280 P76290 P76291 P76296 P76308 P76316
P76318 P76319 P76322 P76323 P76329 P76346 P76349 P76359 P76362 P76364 P76369
P76393 P76394 P76395 P76396 P76402 P76403 P76406 P76418 P76419 P76421 P76422
P76423 P76440 P76469 P76481 P76483 P76484 P76485 P76486 P76491 P76505 P76508
P76509 P76510 P76512 P76513 P76514 P76515 P76516 P76524 P76535 P76536 P76539
P76540 P76541 P76542 P76543 P76544 P76545 P76546 P76547 P76548 P76550 P76551
P76553 P76555 P76556 P76558 P76561 P76562 P76569 P76573 P76575 P76584 P76585
P76586 P76594 P76611 P76612 P76613 P76616 P76632 P76633 P76641 P76692 P76938
P77044 P77072 P77087 P77129 P77136 P77147 P77148 P77150 P77154 P77161 P77163

P77169 P77170 P77171 P77174 P77181 P77182 P77183 P77187 P77206 P77212 P77213
P77215 P77216 P77218 P77231 P77234 P77237 P77243 P77245 P77247 P77252 P77256
P77258 P77260 P77277 P77280 P77295 P77296 P77297 P77300 P77301 P77309 P77316
P77319 P77324 P77326 P77329 P77334 P77337 P77357 P77359 P77360 P77364 P77366
P77376 P77378 P77379 P77381 P77390 P77393 P77396 P77397 P77402 P77414 P77425
P77427 P77432 P77433 P77434 P77444 P77445 P77453 P77454 P77455 P77460 P77467
P77475 P77488 P77489 P77493 P77495 P77499 P77503 P77515 P77522 P77526 P77527
P77528 P77536 P77539 P77541 P77544 P77546 P77551 P77552 P77555 P77559 P77561
P77564 P77567 P77569 P77570 P77580 P77581 P77585 P77596 P77598 P77601 P77607
P77608 P77609 P77615 P77624 P77625 P77626 P77627 P77634 P77649 P77650 P77656
P77657 P77658 P77667 P77668 P77671 P77674 P77688 P77689 P77692 P77695 P77698
P77699 P77700 P77704 P77712 P77713 P77714 P77718 P77721 P77730 P77731 P77732
P77735 P77736 P77739 P77743 P77744 P77746 P77748 P77756 P77759 P77766 P77768
P77775 P77779 P77781 P77788 P77790 P77791 P77806 P77808 P77809 P77810 P77814
P77819 P77828 P77829 P77832 P77834 P77835 P77836 P77837 P77839 P77844 P77845
P77866 P77872 P77873 P77874 P77875 P77877 P77879 P77880 P77882 P77883 P77884
P77885 P77886 P77887 P77888 P77889 P77891 P77892 P77913 P77914 P77915 P77917
P77918 P77919 P77923 P77925 P77929 P77930 P77933 P77935 P77938 P77941 P77947
P77949 P77951 P77952 P77953 P77958 P77960 P77962 P77965 P77966 P77967 P77968
P77969 P77971 P77973 P77978 P77980 P77983 P77984 P77985 P77986 P77989 P77992
P77993 P77994 P77998 P78003 P78004 P78005 P78006 P78008 P78009 P78010 P78011
P78012 P78013 P78014 P78015 P78016 P78017 P78018 P78019 P78020 P78021 P78022
P78023 P78024 P78025 P78026 P78027 P78028 P78029 P78030 P78032 P78033 P78034
P78035 P78055 P78061 P78218 P78271 P78285 P78330 P78358 P78367 P78385 P78386
P78396 P78411 P78412 P78413 P78414 P78415 P78417 P78424 P78426 P78540 P78543
P78545 P78567 P78568 P78569 P78571 P78574 P78578 P78586 P78587 P78590 P78594
P78599 P78600 P78605 P78612 P78621 P78688 P78697 P78698 P78704 P78706 P78711
P78712 P78716 P78717 P78724 P78736 P78748 P78750 P78753 P78773 P78774 P78780
P78790 P78794 P78795 P78804 P78811 P78812 P78814 P78825 P78827 P78833 P78847
P78859 P78860 P78870 P78871 P78875 P78890 P78917 P78920 P78921 P78926 P78929
P78937 P78954 P78955 P78956 P78957 P78958 P78962 P78963 P78965 P78972 P78974
P78978 P78983 P78985 P78987 P78992 P79005 P79007 P79008 P79009 P79011 P79015
P79016 P79023 P79024 P79051 P79060 P79063 P79064 P79065 P79075 P79081 P79083
P79089 P79122 P79145 P79149 P79183 P79226 P79269 P79280 P79288 P79290 P79293
P79303 P79324 P79334 P79364 P79376 P79377 P79378 P79379 P79380 P79384 P79387
P79398 P79431 P79457 P79522 P79621 P79677 P79678 P79679 P79680 P79686 P79703
P79724 P79732 P79733 P79741 P79742 P79745 P79746 P79764 P79769 P79772 P79774
P79775 P79777 P79778 P79779 P79782 P79797 P79888 P79896 P79912 P79913 P79920
P79926 P79937 P79942 P79944 P79955 P79958 P79987 P79990 P80001 P80002 P80008
P80011 P80017 P80018 P80019 P80026 P80028 P80037 P80039 P80040 P80041 P80042
P80043 P80044 P80045 P80046 P80048 P80050 P80052 P80053 P80064 P80072 P80073
P80074 P80077 P80078 P80079 P80080 P80081 P80089 P80092 P80094 P80099 P80145
P80147 P80148 P80153 P80158 P80159 P80160 P80161 P80164 P80166 P80168 P80173
P80175 P80176 P80187 P80193 P80194 P80196 P80197 P80205 P80206 P80210 P80211

P80212 P80213 P80216 P80220 P80222 P80230 P80231 P80232 P80233 P80237 P80238
P80239 P80240 P80241 P80242 P80243 P80246 P80247 P80248 P80249 P80251 P80252
P80254 P80255 P80256 P80258 P80270 P80271 P80272 P80288 P80289 P80290 P80291
P80292 P80293 P80294 P80297 P80298 P80304 P80305 P80306 P80312 P80313 P80314
P80315 P80316 P80317 P80318 P80319 P80320 P80321 P80322 P80338 P80339 P80340
P80341 P80343 P80346 P80347 P80348 P80349 P80350 P80354 P80355 P80357 P80358
P80359 P80360 P80361 P80363 P80364 P80367 P80371 P80372 P80373 P80374 P80375
P80376 P80377 P80378 P80379 P80380 P80381 P80382 P80389 P80394 P80402 P80403
P80404 P80405 P80408 P80409 P80410 P80411 P80412 P80413 P80414 P80416 P80424
P80427 P80428 P80435 P80436 P80437 P80438 P80448 P80458 P80459 P80461 P80462
P80464 P80465 P80466 P80467 P80469 P80472 P80473 P80474 P80483 P80484 P80485
P80486 P80487 P80488 P80490 P80491 P80492 P80501 P80502 P80503 P80505 P80506
P80507 P80509 P80510 P80512 P80514 P80515 P80516 P80517 P80518 P80519 P80521
P80522 P80523 P80524 P80525 P80526 P80527 P80528 P80529 P80530 P80532 P80533
P80536 P80537 P80538 P80539 P80540 P80541 P80542 P80543 P80545 P80546 P80550
P80551 P80552 P80554 P80562 P80563 P80564 P80565 P80567 P80568 P80572 P80573
P80574 P80575 P80576 P80577 P80578 P80579 P80580 P80581 P80582 P80584 P80585
P80593 P80594 P80602 P80605 P80606 P80608 P80609 P80610 P80611 P80612 P80613
P80614 P80615 P80616 P80617 P80618 P80619 P80620 P80621 P80622 P80623 P80624
P80625 P80626 P80627 P80628 P80629 P80630 P80631 P80632 P80633 P80634 P80635
P80636 P80637 P80638 P80639 P80640 P80641 P80642 P80644 P80645 P80647 P80649
P80657 P80659 P80668 P80673 P80674 P80675 P80676 P80679 P80680 P80681 P80682
P80683 P80684 P80685 P80692 P80698 P80699 P80700 P80701 P80702 P80704 P80705
P80706 P80707 P80709 P80717 P80718 P80721 P80722 P80725 P80726 P80727 P80728
P80732 P80733 P80734 P80735 P80736 P80737 P80738 P80739 P80740 P80741 P80750
P80857 P80858 P80859 P80860 P80861 P80862 P80863 P80864 P80865 P80866 P80867
P80868 P80869 P80870 P80872 P80873 P80874 P80875 P80876 P80878 P80879 P80880
P80885 P80886 P80887 P80889 P80890 P80891 P80892 P80893 P80894 P80895 P80900
P80901 P80902 P80903 P80904 P80905 P80906 P80907 P80908 P80909 P80910 P80911
P80912 P80924 P80925 P80933 P80934 P80935 P80936 P80938 P80939 P80940 P80941
P80944 P80945 P80946 P80951 P80959 P80968 P80969 P80994 P81006 P81007 P81008
P81009 P81010 P81011 P81012 P81021 P81023 P81024 P81041 P81042 P81043 P81044
P81045 P81046 P81056 P81063 P81064 P81065 P81066 P81067 P81068 P81069 P81072
P81073 P81074 P81078 P81079 P81080 P81081 P81082 P81083 P81084 P81085 P81086
P81087 P81095 P81096 P81100 P81101 P81102 P81104 P81106 P81107 P81108 P81109
P81110 P81123 P81124 P81135 P81136 P81138 P81140 P81149 P81151 P81152 P81153
P81154 P81161 P81163 P81164 P81171 P81178 P81180 P81181 P81183 P81192 P81193
P81195 P81196 P81197 P81198 P81199 P81200 P81201 P81202 P81205 P81207 P81221
P81225 P81227 P81228 P81229 P81235 P81246 P81248 P81266 P81269 P81270 P81271
P81281 P81284 P81285 P81287 P81288 P81289 P81290 P81291 P81292 P81293 P81296
P81302 P81303 P81307 P81308 P81312 P81315 P81319 P81321 P81325 P81326 P81330
P81332 P81333 P81334 P81335 P81336 P81338 P81339 P81340 P81341 P81342 P81344
P81345 P81346 P81347 P81348 P81349 P81350 P81351 P81352 P81353 P81354 P81355
P81356 P81358 P81359 P81360 P81361 P81362 P81364 P81365 P81366 P81371 P81372

P81373 P81390 P81391 P81392 P81393 P81394 P81395 P81396 P81406 P81407 P81409
P81412 P81413 P81414 P81415 P81419 P81420 P81421 P81422 P81424 P81430 P81431
P81432 P81433 P81434 P81435 P81440 P81445 P81456 P81457 P81460 P81461 P81472
P81473 P81477 P81483 P81484 P81493 P81517 P81527 P81528 P81530 P81531 P81532
P81533 P81535 P81536 P81539 P81540 P81541 P81542 P81543 P81551 P81552 P81554
P81559 P81572 P81580 P81581 P81582 P81583 P81584 P81585 P81586 P81587 P81588
P81589 P81590 P81591 P81593 P81600 P81601 P81621 P81622 P81624 P81625 P81628
P81629 P81637 P81638 P81639 P81643 P81650 P81660 P81664 P81666 P81667 P81669
P81670 P81671 P81672 P81673 P81674 P81675 P81695 P81701 P81705 P81706 P81708
P81709 P81710 P81712 P81713 P81728 P81729 P81747 P81754 P81758 P81760 P81762
P81764 P81766 P81778 P81784 P81786 P81795 P81799 P81800 P81801 P81802 P81826
P81831 P81833 P81834 P81859 P81860 P81870 P81874 P81875 P81876 P81886 P81893
P81898 P81901 P81902 P81903 P81904 P81905 P81906 P81907 P81925 P81926 P81929
P81930 P81942 P81943 P81947 P81949 P81954 P81955 P81956 P81958 P81959 P81987
P81988 P81989 P82004 P82005 P82006 P82010 P82024 P82025 P82111 P82112 P82113
P82116 P82118 P82119 P82120 P82121 P82122 P82125 P82129 P82130 P82137 P82139
P82147 P82150 P82151 P82159 P82160 P82162 P82163 P82165 P82166 P82167 P82168
P82169 P82170 P82171 P82174 P82175 P82177 P82180 P82185 P82186 P82190 P82191
P82192 P82193 P82194 P82195 P82197 P82202 P82204 P82205 P82206 P82207 P82208
P82210 P82212 P82217 P82218 P82219 P82221 P82222 P82230 P82231 P82244 P82245
P82248 P82249 P82257 P82258 P82259 P82264 P82266 P82277 P82278 P82291 P82315
P82316 P82322 P82323 P82324 P82325 P82326 P82327 P82328 P82329 P82330 P82331
P82332 P82333 P82335 P82336 P82337 P82338 P82340 P82341 P82342 P82343 P82345
P82371 P82381 P82385 P82403 P82411 P82412 P82413 P82452 P82453 P82454 P82456
P82457 P82458 P82469 P82473 P82474 P82530 P82531 P82533 P82536 P82537 P82595
P82596 P82598 P82602 P82604 P82605 P82606 P82607 P82608 P82610 P82611 P82612
P82648 P82655 P82657 P82671 P82678 P82679 P82680 P82681 P82682 P82694 P82695
P82696 P82697 P82708 P82736 P82796 P82799 P82802 P82804 P82808 P82809 P82810
P82816 P82853 P82856 P82858 P82865 P82866 P82886 P82887 P82888 P82890 P82897
P82898 P82899 P82902 P82903 P82936 P82937 P82938 P82939 P82944 P82946 P82947
P82954 P82955 P82956 P82957 P82961 P82964 P82968 P82973 P82974 P82976 P82979
P82980 P82989 P82990 P82992 P82993 P82996 P82997 P82998 P82999 P83000 P83001
P83003 P83004 P83038 P83049 P83054 P83061 P83062 P83063 P83064 P83065 P83067
P83071 P83072 P83073 P83075 P83076 P83077 P83090 P83092 P83098 P83102 P83104
P83113 P83114 P83115 P83116 P83117 P83123 P83124 P83126 P83130 P83131 P83132
P83133 P83134 P83135 P83139 P83140 P83141 P83142 P83143 P83145 P83146 P83147
P83148 P83155 P83156 P83157 P83160 P83161 P83170 P83171 P83179 P83181 P83183
P83184 P83186 P83187 P83194 P83199 P83200 P83201 P83211 P83212 P83213 P83214
P83215 P83216 P83217 P83221 P83233 P83241 P83245 P83246 P83252 P83264 P83265
P83266 P83267 P83268 P83270 P83271 P83272 P83273 P83282 P83283 P83284 P83289
P83298 P83299 P83309 P83310 P83311 P83315 P83323 P83324 P83325 P83328 P83329
P83331 P83340 P83341 P83342 P83344 P83352 P83353 P83354 P83355 P83356 P83357
P83358 P83359 P83360 P83361 P83363 P83364 P83365 P83366 P83367 P83368 P83369
P83372 P83373 P83375 P83378 P83399 P83401 P83402 P83410 P83422 P83423 P83424

P83426 P83438 P83439 P83440 P83442 P83447 P83448 P83449 P83450 P83453 P83454
P83465 P83466 P83468 P83474 P83477 P83478 P83479 P83481 P83488 P83491 P83497
P83501 P83511 P83516 P83518 P83520 P83522 P83523 P83524 P83525 P83526 P83527
P83529 P83530 P83531 P83532 P83533 P83534 P83535 P83536 P83537 P83538 P83539
P83540 P83549 P83550 P83564 P83565 P83571 P83572 P83573 P83574 P83575 P83576
P83581 P83582 P83583 P83584 P83585 P83596 P83597 P83598 P83599 P83600 P83601
P83602 P83611 P83612 P83613 P83614 P83623 P83624 P83625 P83630 P83631 P83643
P83645 P83647 P83657 P83662 P83664 P83665 P83666 P83672 P83678 P83682 P83684
P83688 P83690 P83691 P83692 P83695 P83700 P83707 P83708 P83709 P83710 P83711
P83712 P83713 P83715 P83721 P83722 P83723 P83724 P83725 P83726 P83727 P83728
P83729 P83730 P83733 P83736 P83737 P83752 P83753 P83757 P83758 P83761 P83763
P83764 P83765 P83766 P83767 P83770 P83772 P83773 P83775 P83776 P83777 P83778
P83780 P83783 P83788 P83790 P83811 P83813 P83827 P83829 P83830 P83834 P83835
P83836 P83837 P83838 P83839 P83840 P83842 P83843 P83844 P83847 P83851 P83855
P83856 P83857 P83863 P83864 P83865 P83870 P83871 P83889 P83890 P83891 P83898
P83899 P83907 P83908 P83914 P83918 P83937 P83938 P83944 P83946 P83949 P83950
P83955 P83956 P83960 P83961 P83966 P83967 P83968 P83969 P83986 P83987 P83988
P83989 P83990 P83991 P83992 P83993 P83994 P83995 P84040 P84041 P84042 P84043
P84044 P84045 P84046 P84047 P84048 P84049 P84050 P84051 P84052 P84053 P84054
P84055 P84056 P84057 P84059 P84061 P84064 P84066 P84071 P84072 P84073 P84090
P84101 P84102 P84104 P84107 P84117 P84138 P84139 P84140 P84142 P84148 P84149
P84152 P84155 P84158 P84169 P84172 P84177 P84178 P84179 P84182 P84183 P84184
P84185 P84187 P84188 P84193 P84199 P84200 P84203 P84204 P84205 P84206 P84208
P84209 P84216 P84217 P84235 P84236 P84237 P84238 P84239 P84244 P84245 P84246
P84247 P84248 P84250 P84251 P84252 P84253 P84254 P84282 P84283 P84291 P84308
P84311 P84312 P84313 P84314 P84315 P84316 P84317 P84318 P84319 P84320 P84321
P84322 P84323 P84324 P84325 P84326 P84327 P84328 P84330 P84331 P84335 P84339
P84342 P84390 P84394 P84395 P84398 P84401 P84402 P84403 P84404 P84405 P84445
P84446 P84447 P84450 P84456 P84457 P84458 P84459 P84460 P84463 P84468 P84469
P84476 P84477 P84478 P84517 P84520 P84522 P84525 P84526 P84528 P84529 P84530
P84531 P84532 P84533 P84534 P84537 P84538 P84539 P84540 P84541 P84542 P84543
P84544 P84545 P84547 P84548 P84549 P84550 P84551 P84553 P84554 P84555 P84556
P84557 P84558 P84560 P84561 P84562 P84564 P84565 P84575 P84576 P84577 P84579
P84580 P84581 P84583 P84584 P84585 P84587 P84588 P84589 P84604 P84609 P84610
P84611 P84612 P84613 P84624 P84625 P84627 P84633 P84635 P84636 P84637 P84638
P84639 P84640 P84641 P84642 P84643 P84644 P84645 P84647 P84648 P84649 P84650
P84652 P84653 P84675 P84699 P84702 P84708 P84712 P84714 P84717 P84719 P84721
P84722 P84723 P84724 P84725 P84726 P84729 P84730 P84731 P84733 P84734 P84748
P84749 P84754 P84755 P84760 P84761 P84762 P84763 P84764 P84765 P84766 P84767
P84768 P84769 P84775 P84778 P84779 P84780 P84781 P84782 P84783 P84784 P84785
P84786 P84790 P84791 P84792 P84793 P84794 P84795 P84796 P84797 P84798 P84799
P84811 P84812 P84813 P84814 P84819 P84820 P84847 P84848 P84849 P84850 P84852
P84853 P84854 P84856 P84863 P84865 P84866 P84867 P84868 P84869 P84870 P84871
P84872 P84873 P84874 P84875 P84881 P84882 P84884 P84890 P84910 P84911 P84920

P84961 P84963 P84964 P84965 P84966 P84967 P84968 P84969 P84970 P84971 P84972
P84973 P84974 P84975 P84976 P84977 P84978 P84979 P84980 P84982 P84983 P84984
P84985 P84986 P84994 P84995 P84997 P84998 P84999 P85002 P85003 P85004 P85026
P85028 P85035 P85036 P85051 P85052 P85063 P85064 P85076 P85077 P85078 P85080
P85081 P85082 P85083 P85084 P85085 P85086 P85087 P85089 P85094 P85096 P85100
P85101 P85102 P85111 P85112 P85119 P85121 P85126 P85127 P85128 P85129 P85130
P85131 P85132 P85133 P85134 P85135 P85156 P85157 P85172 P85175 P85177 P85178
P85179 P85180 P85181 P85182 P85183 P85184 P85185 P85186 P85187 P85188 P85191
P85192 P85193 P85198 P85199 P85200 P85203 P85217 P85225 P85227 P85228 P85229
P85230 P85231 P85232 P85233 P85234 P85238 P85239 P85240 P85241 P85242 P85244
P85245 P85246 P85248 P85249 P85250 P85251 P85252 P85260 P85278 P85280 P85281
P85282 P85283 P85284 P85285 P85286 P85287 P85288 P85289 P85290 P85291 P85292
P85299 P85303 P85304 P85305 P85310 P85311 P85312 P85313 P85328 P85329 P85330
P85343 P85363 P85364 P85365 P85366 P85367 P85368 P85369 P85370 P85371 P85372
P85373 P85374 P85375 P85376 P85377 P85378 P85379 P85380 P85381 P85382 P85383
P85384 P85385 P85386 P85387 P85388 P85389 P85390 P85391 P85392 P85393 P85394
P85395 P85396 P85397 P85398 P85399 P85400 P85401 P85402 P85403 P85404 P85405
P85406 P85407 P85408 P85409 P85415 P85416 P85417 P85418 P85419 P85421 P85422
P85423 P85425 P85426 P85427 P85440 P85441 P85442 P85444 P85498 P85500 P85501
P85502 P85503 P85508 P85509 P85510 P85512 P85516 P85517 P85524 P85525 P85526
P85801 P85814 P85825 P85829 P85834 P85835 P85836 P85837 P85838 P85839 P85863
P85866 P85888 P85896 P85898 P85899 P85900 P85901 P85902 P85903 P85904 P85905
P85906 P85907 P85908 P85910 P85911 P85912 P85913 P85914 P85915 P85916 P85917
P85918 P85919 P85920 P85921 P85922 P85923 P85926 P85927 P85929 P85930 P85931
P85932 P85934 P85935 P85936 P85937 P85938 P85939 P85940 P85941 P85942 P85943
P85944 P85945 P85946 P85947 P85948 P85949 P85950 P85951 P85952 P85953 P85954
P85955 P85956 P85957 P85958 P85959 P85960 P85961 P85962 P85963 P85964 P85965
P85971 P85973 P85977 P85978 P85984 P85987 P85988 P85989 P85990 P85991 P85993
P85997 P86005 P86029 P86030 P86041 P86049 P86052 P86054 P86055 P86056 P86057
P86058 P86059 P86060 P86061 P86062 P86063 P86064 P86065 P86066 P86067 P86068
P86069 P86070 P86071 P86072 P86073 P86074 P86076 P86077 P86079 P86080 P86081
P86082 P86083 P86084 P86086 P86087 P86088 P86089 P86090 P86101 P86102 P86172
P86173 P86174 P86175 P86178 P86179 P86181 P86184 P86195 P86196 P86197 P86199
P86200 P86204 P86206 P86208 P86209 P86211 P86214 P86216 P86219 P86221 P86222
P86225 P86226 P86231 P86234 P86235 P86237 P86238 P86240 P86241 P86245 P86246
P86247 P86249 P86251 P86252 P86254 P86265 P86277 P86287 P86288 P86292 P86293
P86317 P86318 P86319 P86320 P86321 P86322 P86323 P86324 P86330 P86331 P86345
P86346 P86347 P86349 P86352 P86353 P86355 P86363 P86364 P86365 P86366 P86367
P86384 P86388 P86390 P86391 P86409 P86413 P86415 P86416 P86431 P86432 P86434
P86448 P86449 P86451 P86452 P86478 P86479 P86480 P86481 P86496 P86498 P86499
P86518 P86519 P86521 P86549 P86600 P86601 P86624 P86626 P86699 P86700 P86701
P86703 P86704 P86708 P86712 P86722 P86723 P86739 P86740 P86741 P86742 P86743
P86744 P86745 P86746 P86747 P86748 P86749 P86750 P86751 P86752 P86753 P86754
P86755 P86756 P86757 P86758 P86759 P86760 P86761 P86762 P86763 P86764 P86765

P86766 P86767 P86768 P86769 P86770 P86771 P86772 P86773 P86774 P86775 P86776
P86777 P86778 P86779 P86781 P86782 P86784 P86786 P86787 P86788 P86794 P86807
P86808 P86830 P86831 P86832 P86833 P86834 P86837 P86839 P86843 P86844 P86845
P86846 P86847 P86848 P86849 P86850 P86851 P86852 P86873 P86874 P86876 P86879
P86880 P86881 P86882 P86883 P86884 P86885 P86887 P86888 P86890 P86893 P86895
P86896 P86897 P86898 P86909 P86910 P86912 P86916 P86919 P86922 P86924 P86925
P86926 P86927 P86930 P86931 P86932 P86933 P86934 P86937 P86938 P86939 P86971
P86978 P86979 P86980 P86981 P87000 P87012 P87017 P87025 P87027 P87039 P87047
P87048 P87051 P87052 P87053 P87054 P87057 P87060 P87064 P87066 P87072 P87078
P87090 P87102 P87112 P87114 P87115 P87118 P87120 P87121 P87123 P87125 P87126
P87127 P87132 P87133 P87137 P87138 P87142 P87143 P87144 P87151 P87152 P87153
P87154 P87156 P87159 P87163 P87164 P87167 P87168 P87169 P87171 P87172 P87173
P87174 P87175 P87176 P87177 P87183 P87185 P87186 P87187 P87191 P87197 P87206
P87208 P87214 P87215 P87216 P87218 P87219 P87227 P87228 P87229 P87231 P87232
P87233 P87235 P87237 P87238 P87240 P87241 P87242 P87243 P87244 P87250 P87256
P87257 P87261 P87263 P87268 P87269 P87271 P87282 P87283 P87286 P87288 P87292
P87294 P87295 P87297 P87299 P87303 P87304 P87305 P87306 P87307 P87308 P87309
P87310 P87311 P87312 P87313 P87314 P87315 P87318 P87324 P87347 P87362 P87393
P87394 P87497 P87503 P87505 P87552 P87553 P87563 P87568 P87577 P87604 P87607
P87617 P87630 P87724 P87744 P88812 P89028 P89029 P89036 P89037 P89079 P89083
P89105 P89114 P89116 P89201 P89202 P89216 P89263 P89429 P89430 P89431 P89432
P89437 P89439 P89440 P89441 P89442 P89444 P89446 P89447 P89448 P89451 P89452
P89453 P89455 P89456 P89458 P89461 P89469 P89471 P89474 P89476 P89478 P89479
P89495 P89522 P89657 P89658 P89659 P89676 P89677 P89678 P89681 P89682 P89684
P89886 P89888 P90211 P90212 P90245 P90335 P90463 P90493 P90513 P90516 P90518
P90519 P90520 P90521 P90543 P90548 P90587 P90597 P90648 P90666 P90682 P90689
P90697 P90702 P90703 P90707 P90727 P90740 P90761 P90783 P90795 P90815 P90829
P90846 P90866 P90897 P90900 P90916 P90917 P90925 P90947 P90953 P90971 P90978
P90986 P90990 P90993 P90994 P91001 P91127 P91134 P91136 P91167 P91252 P91253
P91254 P91277 P91278 P91285 P91309 P91318 P91341 P91343 P91349 P91351 P91353
P91375 P91399 P91408 P91416 P91420 P91424 P91427 P91430 P91443 P91455 P91477
P91478 P91499 P91529 P91599 P91607 P91613 P91622 P91623 P91641 P91664 P91686
P91697 P91698 P91705 P91711 P91716 P91753 P91754 P91759 P91791 P91798 P91800
P91809 P91829 P91849 P91851 P91853 P91856 P91859 P91868 P91869 P91870 P91875
P91882 P91883 P91890 P91895 P91902 P91910 P91917 P91918 P91919 P91931 P91938
P91958 P92119 P92120 P92176 P92179 P92182 P92186 P92188 P92189 P92203 P92204
P92209 P92220 P92225 P92306 P92389 P92392 P92397 P92401 P92406 P92407 P92418
P92429 P92439 P92445 P92463 P92510 P92512 P92513 P92515 P92516 P92517 P92518
P92519 P92520 P92521 P92522 P92523 P92524 P92525 P92526 P92529 P92530 P92531
P92532 P92534 P92535 P92537 P92538 P92539 P92540 P92542 P92543 P92544 P92548
P92551 P92552 P92554 P92555 P92557 P92560 P92561 P92562 P92563 P92566 P92567
P92812 P92892 P92918 P92948 P92953 P92965 P92966 P92969 P92973 P92974 P92979
P92980 P92981 P92983 P92985 P93006 P93008 P93017 P93028 P93031 P93032 P93033
P93045 P93047 P93052 P93082 P93083 P93087 P93099 P93101 P93105 P93107 P93111

P93119 P93134 P93143 P93162 P93171 P93176 P93203 P93227 P93231 P93243 P93253
P93254 P93255 P93256 P93257 P93258 P93259 P93262 P93263 P93275 P93276 P93277
P93281 P93282 P93287 P93290 P93291 P93292 P93293 P93294 P93295 P93297 P93299
P93300 P93301 P93302 P93304 P93305 P93308 P93309 P93310 P93311 P93315 P93317
P93318 P93324 P93330 P93333 P93338 P93340 P93348 P93351 P93354 P93371 P93372
P93373 P93374 P93375 P93376 P93377 P93394 P93395 P93407 P93411 P93422 P93431
P93438 P93447 P93526 P93528 P93530 P93531 P93541 P93554 P93568 P93584 P93585
P93586 P93587 P93594 P93629 P93665 P93673 P93692 P93693 P93711 P93732 P93736
P93738 P93740 P93748 P93751 P93755 P93757 P93762 P93768 P93771 P93789 P93804
P93805 P93820 P93821 P93824 P93830 P93831 P93832 P93836 P93841 P93847 P93848
P93849 P93869 P93880 P93890 P93906 P93913 P93936 P93956 P93960 P93962 P93964
P93968 P93974 P93988 P93993 P93998 P94026 P94029 P94044 P94063 P94092 P94102
P94126 P94132 P94135 P94136 P94147 P94160 P94164 P94165 P94166 P94169 P94188
P94190 P94200 P94210 P94211 P94212 P94252 P94266 P94267 P94268 P94269 P94270
P94271 P94272 P94273 P94281 P94282 P94283 P94284 P94298 P94300 P94316 P94317
P94319 P94323 P94325 P94326 P94327 P94328 P94335 P94338 P94340 P94352 P94358
P94359 P94368 P94370 P94374 P94377 P94378 P94382 P94387 P94388 P94389 P94390
P94393 P94396 P94398 P94400 P94403 P94404 P94405 P94407 P94409 P94413 P94415
P94417 P94423 P94424 P94425 P94426 P94427 P94428 P94433 P94437 P94443 P94447
P94453 P94459 P94461 P94462 P94463 P94464 P94465 P94474 P94475 P94478 P94479
P94480 P94481 P94482 P94483 P94484 P94487 P94490 P94494 P94495 P94496 P94498
P94500 P94501 P94502 P94504 P94512 P94514 P94521 P94523 P94524 P94525 P94526
P94531 P94533 P94536 P94537 P94538 P94541 P94544 P94545 P94550 P94551 P94552
P94556 P94558 P94559 P94560 P94562 P94563 P94565 P94568 P94569 P94573 P94578
P94583 P94586 P94588 P94591 P94592 P94593 P94598 P94601 P94604 P94605 P94612
P94613 P94614 P94616 P94631 P94632 P94661 P94666 P94667 P94668 P94669 P94678
P94688 P94689 P94692 P94693 P94694 P94695 P94702 P94777 P94797 P94798 P94799
P94804 P94819 P94820 P94844 P94846 P94868 P94869 P94870 P94871 P94872 P94876
P94890 P94891 P94892 P94899 P94907 P94915 P94919 P94929 P94939 P94948 P94951
P94954 P94965 P94966 P94967 P94968 P95004 P95006 P95007 P95038 P95225 P95231
P95257 P95262 P95276 P95296 P95297 P95329 P95331 P95333 P95334 P95337 P95340
P95354 P95361 P95364 P95368 P95370 P95389 P95399 P95400 P95405 P95412 P95413
P95414 P95415 P95442 P95453 P95454 P95458 P95459 P95463 P95468 P95469 P95474
P95477 P95479 P95480 P95481 P95482 P95483 P95506 P95508 P95509 P95516 P95519
P95524 P95525 P95526 P95539 P95544 P95547 P95575 P95576 P95578 P95596 P95598
P95607 P95608 P95609 P95629 P95631 P95645 P95646 P95647 P95648 P95649 P95650
P95651 P95653 P95669 P95676 P95677 P95678 P95689 P95690 P95691 P95694 P95713
P95721 P95724 P95727 P95734 P95735 P95744 P95746 P95747 P95748 P95756 P95757
P95758 P95765 P95778 P95780 P95781 P95800 P95801 P95803 P95829 P95830 P95836
P95837 P95846 P95847 P95855 P95862 P95867 P95871 P95872 P95879 P95895 P95896
P95907 P95914 P95928 P95931 P95937 P95949 P95957 P95958 P95960 P95961 P95962
P95963 P95967 P95968 P95970 P95979 P95980 P95982 P95986 P95987 P95988 P95989
P95990 P95991 P95992 P95993 P95994 P95997 P96010 P96022 P96026 P96036 P96037
P96038 P96039 P96040 P96041 P96045 P96050 P96051 P96053 P96060 P96061 P96072

P96074 P96076 P96077 P96078 P96079 P96081 P96082 P96084 P96086 P96095 P96105
P96108 P96110 P96111 P96112 P96113 P96115 P96117 P96121 P96122 P96123 P96128
P96129 P96130 P96131 P96132 P96133 P96134 P96136 P96137 P96142 P96155 P96158
P96166 P96172 P96174 P96175 P96177 P96178 P96185 P96188 P96189 P96190 P96192
P96193 P96194 P96195 P96196 P96197 P96198 P96199 P96274 P96284 P96314 P96315
P96318 P96329 P96343 P96349 P96379 P96426 P96440 P96457 P96486 P96488 P96489
P96495 P96502 P96551 P96556 P96574 P96575 P96576 P96578 P96579 P96582 P96583
P96590 P96591 P96595 P96596 P96597 P96598 P96602 P96605 P96608 P96609 P96611
P96612 P96613 P96621 P96622 P96623 P96624 P96625 P96628 P96629 P96630 P96631
P96632 P96633 P96636 P96637 P96638 P96642 P96643 P96651 P96652 P96653 P96654
P96655 P96658 P96659 P96660 P96662 P96663 P96666 P96669 P96670 P96671 P96673
P96676 P96677 P96679 P96681 P96682 P96683 P96684 P96688 P96690 P96692 P96693
P96695 P96698 P96699 P96700 P96701 P96703 P96705 P96707 P96708 P96711 P96714
P96717 P96719 P96720 P96721 P96723 P96724 P96725 P96726 P96728 P96738 P96741
P96744 P96749 P96750 P96763 P96769 P96771 P96789 P96794 P96797 P96800 P96801
P96803 P96847 P96886 P96955 P96963 P96972 P96974 P96975 P96977 P96981 P96985
P96993 P96994 P96995 P97014 P97028 P97030 P97032 P97043 P97051 P97052 P97084
P97086 P97089 P97115 P97117 P97195 P97227 P97247 P97273 P97302 P97304 P97310
P97311 P97328 P97329 P97334 P97346 P97355 P97357 P97358 P97360 P97364 P97366
P97367 P97368 P97371 P97372 P97384 P97386 P97427 P97434 P97436 P97443 P97457
P97458 P97459 P97473 P97480 P97481 P97489 P97493 P97496 P97499 P97500 P97503
P97516 P97532 P97536 P97587 P97608 P97609 P97615 P97616 P97760 P97762 P97764
P97765 P97767 P97770 P97779 P97801 P97807 P97817 P97830 P97832 P97861 P97863
P97868 P97875 P97876 P97888 P97891 P97930 P97931 P98080 P98081 P98168 P98169
P98170 P98179 P98182 P98185 P99012 P99026 P99031 P99032 P99085 P99123 P99172
P99503 P99506 P99507 P9WEN8 P9WEN9 P9WEP0 P9WEP2 P9WEP3 P9WEP6 P9WEP8 P9WEP9
P9WEQ0 P9WEQ2 P9WEQ5 P9WEQ8 P9WER0 P9WER1 P9WER3 P9WER6 P9WER8 P9WES1 P9WES2
P9WES4 P9WES5 P9WES6 P9WES8 P9WES9 P9WET0 P9WET1 P9WET2 P9WET3 P9WET4 P9WET5
P9WET6 P9WET7 P9WET8 P9WET9 P9WEU0 P9WEU5 P9WEU6 P9WEU7 P9WEU8 P9WEU9 P9WEV0
P9WEV1 P9WEV2 P9WEV6 P9WEV7 P9WEV8 P9WEW0 P9WEW1 P9WEW2 P9WEW8 P9WEW9 P9WEX0
P9WEX3 P9WEY4 P9WEY5 P9WEY6 P9WEY7 P9WEY9 P9WEZ0 P9WEZ2 P9WEZ3 P9WEZ4 P9WEZ5
P9WEZ7 P9WEZ8 P9WEZ9 P9WF00 P9WF01 P9WF02 P9WF08 P9WF09 P9WF10 P9WF11 P9WF12
P9WF13 P9WF14 P9WF16 P9WF17 P9WF18 P9WF19 P9WF20 P9WF21 P9WF22 P9WF23 P9WF24
P9WF25 P9WF26 P9WF27 P9WF28 P9WF29 P9WF30 P9WF31 P9WF32 P9WF33 P9WF34 P9WF36
P9WF37 P9WF38 P9WF40 P9WF42 P9WF43 P9WF44 P9WF46 P9WF47 P9WF48 P9WF49 P9WF50
P9WF51 P9WF52 P9WF54 P9WF55 P9WF56 P9WF57 P9WF58 P9WF62 P9WF64 P9WF65 P9WF66
P9WF67 P9WF68 P9WF70 P9WF71 P9WF72 P9WF73 P9WF74 P9WF75 P9WF76 P9WF77 P9WF78
P9WF79 P9WF80 P9WF81 P9WF84 P9WF85 P9WF86 P9WF88 P9WF89 P9WF90 P9WF91 P9WF92
P9WF94 P9WF95 P9WF96 P9WF97 P9WF98 P9WF99 P9WFA0 P9WFA2 P9WFA3 P9WFA4 P9WFA5
P9WFA6 P9WFA8 P9WFA9 P9WFB0 P9WFB1 P9WFB2 P9WFB3 P9WFB4 P9WFB5 P9WFB6 P9WFB7
P9WFB8 P9WFB9 P9WFC0 P9WFC1 P9WFC2 P9WFC3 P9WFC4 P9WFC6 P9WFC8 P9WFD0 P9WFD1
P9WFD2 P9WFD4 P9WFD6 P9WFD8 P9WFE0 P9WFE1 P9WFE2 P9WFE3 P9WFE4 P9WFE5 P9WFE6
P9WFE7 P9WFE8 P9WFE9 P9WFF0 P9WFF1 P9WFF2 P9WFG6 P9WFG8 P9WFH0 P9WFH1 P9WFH2
P9WFH3 P9WFH4 P9WFH6 P9WFH7 P9WFH8 P9WFI0 P9WFI1 P9WFI2 P9WFI4 P9WFI5 P9WFI6

P9WFI7 P9WFI8 P9WFJ0 P9WFJ1 P9WFJ2 P9WFJ8 P9WFK0 P9WFK2 P9WFK4 P9WFK8 P9WFL0
P9WFL1 P9WFL6 P9WFL7 P9WFM0 P9WFM1 P9WFM6 P9WFM7 P9WFM8 P9WFM9 P9WFN0 P9WFN4
P9WFN5 P9WFP4 P9WFP5 P9WFP6 P9WFP7 P9WFP8 P9WFP9 P9WFQ0 P9WFQ1 P9WFQ2 P9WFQ4
P9WFQ5 P9WFQ6 P9WFQ7 P9WFQ8 P9WFQ9 P9WFR0 P9WFR2 P9WFR8 P9WFR9 P9WFS0 P9WFS1
P9WFS6 P9WFS7 P9WFS8 P9WFT0 P9WFT2 P9WFT4 P9WFT6 P9WFT8 P9WFU0 P9WFU2 P9WFU4
P9WFU8 P9WFU9 P9WFV0 P9WFV2 P9WFV4 P9WFV6 P9WFV8 P9WFW0 P9WFW1 P9WFW2 P9WFW3
P9WFW4 P9WFW6 P9WFW8 P9WFW9 P9WFX0 P9WFX2 P9WFX4 P9WFX6 P9WFX8 P9WFX9 P9WFY0
P9WFY2 P9WFY3 P9WFY4 P9WFY5 P9WFY6 P9WFY7 P9WFY8 P9WFY9 P9WFZ0 P9WFZ4 P9WG24
P9WG32 P9WG33 P9WG34 P9WG35 P9WG36 P9WG38 P9WG39 P9WG40 P9WG41 P9WG42 P9WG44
P9WG46 P9WG48 P9WG52 P9WG53 P9WG54 P9WG56 P9WG57 P9WG58 P9WG66 P9WG68 P9WG70
P9WG71 P9WG72 P9WG73 P9WG74 P9WG75 P9WG76 P9WG77 P9WG78 P9WG80 P9WG82 P9WG83
P9WG94 P9WGA2 P9WGA3 P9WGA4 P9WGA5 P9WGA8 P9WGA9 P9WGB0 P9WGB1 P9WGB2 P9WGB4
P9WGB5 P9WGB6 P9WGB8 P9WGB9 P9WGC0 P9WGC2 P9WGC3 P9WGC4 P9WGC5 P9WGC6 P9WGC8
P9WGC9 P9WGD0 P9WGD1 P9WGD2 P9WGD3 P9WGD4 P9WGE0 P9WGE1 P9WGE2 P9WGE3 P9WGF2
P9WGF8 P9WGF9 P9WGG0 P9WGG1 P9WGG2 P9WGG3 P9WGG4 P9WGG6 P9WGG7 P9WGH0 P9WGH1
P9WGH2 P9WGH3 P9WGH8 P9WGH9 P9WGI0 P9WGI2 P9WGI4 P9WGI6 P9WGI8 P9WGJ0 P9WGJ2
P9WGJ4 P9WGJ6 P9WGJ7 P9WGK0 P9WGK1 P9WGK2 P9WGL9 P9WGN1 P9WGP6 P9WGP8 P9WGP9
P9WGQ0 P9WGQ2 P9WGQ3 P9WGQ4 P9WGQ6 P9WGQ8 P9WGR2 P9WGR3 P9WGR4 P9WGR6 P9WGR8
P9WGR9 P9WGS0 P9WGS2 P9WGS4 P9WGS5 P9WGS6 P9WGS7 P9WGS8 P9WGT4 P9WGV0 P9WGV2
P9WGV6 P9WGV7 P9WGV8 P9WGV9 P9WGW0 P9WGW1 P9WGW2 P9WGW4 P9WGW5 P9WGW6 P9WGW8
P9WGX0 P9WGX1 P9WGX2 P9WGX3 P9WGX6 P9WGX7 P9WGX8 P9WGX9 P9WGY0 P9WGY2 P9WGY3
P9WGY4 P9WGY5 P9WGY6 P9WGY8 P9WGZ0 P9WGZ2 P9WGZ3 P9WGZ4 P9WGZ5 P9WGZ6 P9WGZ8
P9WGZ9 P9WH00 P9WH01 P9WH02 P9WH03 P9WH04 P9WH05 P9WH06 P9WH07 P9WH08 P9WH09
P9WH10 P9WH11 P9WH12 P9WH14 P9WH16 P9WH18 P9WH19 P9WH20 P9WH21 P9WH24 P9WH26
P9WH27 P9WH28 P9WH30 P9WH32 P9WH34 P9WH36 P9WH38 P9WH40 P9WH41 P9WH42 P9WH44
P9WH46 P9WH48 P9WH49 P9WH50 P9WH52 P9WH54 P9WH56 P9WH57 P9WH58 P9WH59 P9WH60
P9WH62 P9WH63 P9WH64 P9WH66 P9WH70 P9WH71 P9WH72 P9WH73 P9WH74 P9WH76 P9WH78
P9WH80 P9WH82 P9WH84 P9WH86 P9WH88 P9WH89 P9WH90 P9WH91 P9WH92 P9WH93 P9WH94
P9WH95 P9WH96 P9WH97 P9WH98 P9WH99 P9WHA0 P9WHA1 P9WHA2 P9WHA3 P9WHA4 P9WHA6
P9WHA7 P9WHA8 P9WHA9 P9WHB0 P9WHB1 P9WHB2 P9WHB4 P9WHB5 P9WHB6 P9WHB7 P9WHB8
P9WHC0 P9WHC1 P9WHC2 P9WHC4 P9WHC6 P9WHC8 P9WHD0 P9WHD1 P9WHD2 P9WHD4 P9WHD6
P9WHD8 P9WHE0 P9WHE2 P9WHE4 P9WHE5 P9WHE6 P9WHE8 P9WHE9 P9WHF0 P9WHF1 P9WHF2
P9WHF4 P9WHF5 P9WHF6 P9WHF8 P9WHG0 P9WHG1 P9WHG2 P9WHG3 P9WHG4 P9WHG5 P9WHG6
P9WHG7 P9WHG8 P9WHH0 P9WHH1 P9WHH2 P9WHH3 P9WHH4 P9WHH6 P9WHH7 P9WHH8 P9WHI0
P9WHI2 P9WHI3 P9WHI4 P9WHI5 P9WHI6 P9WHI7 P9WHI8 P9WHJ0 P9WHJ2 P9WHJ3 P9WHJ4
P9WHJ5 P9WHJ6 P9WHJ7 P9WHJ8 P9WHJ9 P9WHK0 P9WHK1 P9WHK2 P9WHK3 P9WHK4 P9WHK6
P9WHK8 P9WHK9 P9WHL2 P9WHL3 P9WHL4 P9WHL6 P9WHL7 P9WHL8 P9WHM0 P9WHM1 P9WHM2
P9WHM3 P9WHM4 P9WHM5 P9WHM6 P9WHM8 P9WHM9 P9WHN0 P9WHN1 P9WHN2 P9WHN3 P9WHN4
P9WHN6 P9WHN8 P9WHN9 P9WHP0 P9WHP1 P9WHP2 P9WHP3 P9WHP6 P9WHP7 P9WHP8 P9WHP9
P9WHQ0 P9WHQ1 P9WHQ2 P9WHQ3 P9WHQ6 P9WHQ7 P9WHQ8 P9WHQ9 P9WHS0 P9WHS1 P9WHS8
P9WHS9 P9WHT0 P9WHT2 P9WHT4 P9WHT6 P9WHT7 P9WHT8 P9WHU0 P9WHU2 P9WHU3 P9WHU6
P9WHU8 P9WHU9 P9WHV0 P9WHV2 P9WHV3 P9WHV4 P9WHV5 P9WHV6 P9WHV7 P9WHV8 P9WHW0
P9WHW2 P9WHW6 P9WHW7 P9WHX2 P9WHX4 P9WHX8 P9WHX9 P9WHY0 P9WHY1 P9WHY2 P9WHY4
P9WHY5 P9WHY6 P9WHY7 P9WHY8 P9WHY9 P9WHZ2 P9WHZ3 P9WHZ4 P9WHZ5 P9WHZ6 P9WHZ7

P9WHZ8 P9WHZ9 P9WI02 P9WI03 P9WI04 P9WI05 P9WI06 P9WI07 P9WI08 P9WI09 P9WI10
P9WI11 P9WI13 P9WI14 P9WI15 P9WI16 P9WI17 P9WI18 P9WI19 P9WI24 P9WI25 P9WI32
P9WI33 P9WI36 P9WI37 P9WI44 P9WI45 P9WI50 P9WI51 P9WI54 P9WI56 P9WI84 P9WI88
P9WI90 P9WI91 P9WI92 P9WI93 P9WI98 P9WI99 P9WIA0 P9WIA2 P9WIC0 P9WIC2 P9WIC3
P9WIC4 P9WIC5 P9WIC6 P9WIC7 P9WIC8 P9WID0 P9WID2 P9WID3 P9WID4 P9WID6 P9WID8
P9WID9 P9WIE0 P9WIE2 P9WIE3 P9WIE4 P9WIE6 P9WIE7 P9WIE8 P9WIE9 P9WIF6 P9WIF7
P9WIF8 P9WIF9 P9WIG0 P9WIG1 P9WIG2 P9WIG3 P9WIG4 P9WIG5 P9WIH2 P9WIH4 P9WIH5
P9WIH6 P9WIH7 P9WIH8 P9WIH9 P9WII0 P9WII1 P9WII2 P9WII4 P9WII5 P9WII6 P9WII7
P9WII8 P9WIJ0 P9WIJ1 P9WIJ2 P9WIJ3 P9WIJ4 P9WIJ5 P9WIJ6 P9WIJ7 P9WIJ8 P9WIK0
P9WIK1 P9WIK2 P9WIK5 P9WIL0 P9WIL1 P9WIL2 P9WIL3 P9WIL4 P9WIL5 P9WIL6 P9WIL8
P9WIL9 P9WIM0 P9WIM2 P9WIM3 P9WIN0 P9WIN1 P9WIP4 P9WIP5 P9WIP6 P9WIP8 P9WIQ0
P9WIQ2 P9WIQ3 P9WIQ4 P9WIQ8 P9WIR4 P9WIR5 P9WIS0 P9WIS2 P9WIS3 P9WIS4 P9WIS6
P9WIS8 P9WIT0 P9WIT6 P9WIT8 P9WIT9 P9WIU0 P9WIU1 P9WIU2 P9WIU3 P9WIU6 P9WIU7
P9WIU8 P9WIV0 P9WIV1 P9WIV2 P9WIX4 P9WIX6 P9WIX7 P9WIX8 P9WIX9 P9WIY0 P9WIY1
P9WIY2 P9WIY3 P9WIY4 P9WIY5 P9WIZ0 P9WIZ1 P9WIZ2 P9WIZ3 P9WIZ6 P9WIZ8 P9WJ00
P9WJ01 P9WJ02 P9WJ03 P9WJ04 P9WJ05 P9WJ06 P9WJ07 P9WJ08 P9WJ09 P9WJ10 P9WJ11
P9WJ12 P9WJ13 P9WJ14 P9WJ15 P9WJ16 P9WJ17 P9WJ18 P9WJ19 P9WJ20 P9WJ21 P9WJ22
P9WJ23 P9WJ24 P9WJ25 P9WJ26 P9WJ27 P9WJ28 P9WJ29 P9WJ30 P9WJ31 P9WJ32 P9WJ34
P9WJ35 P9WJ36 P9WJ37 P9WJ38 P9WJ39 P9WJ40 P9WJ41 P9WJ42 P9WJ43 P9WJ44 P9WJ45
P9WJ46 P9WJ47 P9WJ48 P9WJ49 P9WJ50 P9WJ51 P9WJ52 P9WJ53 P9WJ54 P9WJ55 P9WJ56
P9WJ57 P9WJ58 P9WJ59 P9WJ60 P9WJ61 P9WJ68 P9WJ69 P9WJ72 P9WJ73 P9WJ74 P9WJ75
P9WJ78 P9WJ80 P9WJ81 P9WJ84 P9WJ85 P9WJ86 P9WJ87 P9WJ88 P9WJ90 P9WJ91 P9WJ94
P9WJ96 P9WJ97 P9WJA4 P9WJA5 P9WJA6 P9WJA7 P9WJA8 P9WJB6 P9WJB8 P9WJC0 P9WJC4
P9WJC6 P9WJC7 P9WJC8 P9WJC9 P9WJD0 P9WJF4 P9WJF6 P9WJF7 P9WJF8 P9WJF9 P9WJG0
P9WJG1 P9WJG2 P9WJG3 P9WJH6 P9WJH8 P9WJH9 P9WJI0 P9WJI4 P9WJI6 P9WJI8 P9WJI9
P9WJJ0 P9WJJ2 P9WJJ4 P9WJJ5 P9WJJ6 P9WJJ8 P9WJJ9 P9WJK0 P9WJK1 P9WJK4 P9WJK6
P9WJL0 P9WJL1 P9WJL2 P9WJL4 P9WJL5 P9WJL6 P9WJL7 P9WJL8 P9WJL9 P9WJM0 P9WJM1
P9WJM2 P9WJM3 P9WJM4 P9WJM5 P9WJM6 P9WJM7 P9WJM8 P9WJN0 P9WJN2 P9WJN6 P9WJN7
P9WJN8 P9WJN9 P9WJP0 P9WJP2 P9WJP3 P9WJP4 P9WJP6 P9WJP7 P9WJP8 P9WJQ4 P9WJQ5
P9WJQ6 P9WJQ8 P9WJR0 P9WJR1 P9WJR2 P9WJR3 P9WJR4 P9WJR5 P9WJR6 P9WJR7 P9WJR8
P9WJR9 P9WJS0 P9WJS1 P9WJS2 P9WJS4 P9WJS5 P9WJW0 P9WJW1 P9WJW2 P9WJW4 P9WJY8
P9WJZ0 P9WJZ2 P9WJZ3 P9WJZ4 P9WJZ5 P9WJZ6 P9WJZ8 P9WJZ9 P9WK00 P9WK01 P9WK02
P9WK03 P9WK04 P9WK05 P9WK06 P9WK08 P9WK09 P9WK10 P9WK12 P9WK14 P9WK15 P9WK16
P9WK18 P9WK19 P9WK20 P9WK22 P9WK23 P9WK24 P9WK25 P9WK26 P9WK27 P9WK28 P9WK34
P9WK35 P9WK82 P9WK86 P9WK87 P9WK90 P9WK91 P9WK94 P9WKD4 P9WKD5 P9WKD6 P9WKD7
P9WKD8 P9WKD9 P9WKE0 P9WKE1 P9WKE2 P9WKE4 P9WKE6 P9WKE7 P9WKE8 P9WKE9 P9WKF4
P9WKF8 P9WKF9 P9WKG0 P9WKG1 P9WKG2 P9WKG4 P9WKG5 P9WKG6 P9WKG7 P9WKG8 P9WKH2
P9WKH4 P9WKH6 P9WKH8 P9WKI0 P9WKI1 P9WKI2 P9WKI4 P9WKI6 P9WKI8 P9WKJ0 P9WKJ2
P9WKJ4 P9WKJ6 P9WKJ8 P9WKK0 P9WKK2 P9WKK4 P9WKK5 P9WKK6 P9WKK8 P9WKK9 P9WKL0
P9WKL4 P9WKL5 P9WKL8 P9WKM0 P9WKM2 P9WKM3 P9WKM4 P9WKM5 P9WKM6 P9WKM7 P9WKN0
P9WKN4 P9WKN5 P9WKN6 P9WKN8 P9WKN9 P9WKP0 P9WKP4 P9WKP5 P9WKP8 P9WKP9 P9WKQ2
P9WKQ3 P9WKQ6 P9WKQ7 P9WKR2 P9WKR3 P9WKS0 P9WKS2 P9WKS3 P9WKS4 P9WKS5 P9WKS8
P9WKT2 P9WKT4 P9WKT5 P9WKT6 P9WKT7 P9WKT8 P9WKT9 P9WKU4 P9WKU5 P9WKU6 P9WKU7
P9WKU8 P9WKU9 P9WKV0 P9WKV1 P9WKV2 P9WKV3 P9WKV4 P9WKW2 P9WKW3 P9WKW6 P9WKW7

P9WKX0 P9WKX1 P9WKX2 P9WKX4 P9WKX6 P9WKY0 P9WKY1 P9WKY2 P9WKY3 P9WKY8 P9WKY9
P9WKZ0 P9WKZ2 P9WKZ3 P9WKZ6 P9WKZ7 P9WL00 P9WL01 P9WL02 P9WL03 P9WL04 P9WL05
P9WL06 P9WL08 P9WL09 P9WL10 P9WL11 P9WL14 P9WL15 P9WL16 P9WL17 P9WL18 P9WL20
P9WL22 P9WL23 P9WL24 P9WL25 P9WL26 P9WL28 P9WL29 P9WL30 P9WL34 P9WL36 P9WL40
P9WL42 P9WL43 P9WL44 P9WL46 P9WL47 P9WL48 P9WL49 P9WL50 P9WL51 P9WL52 P9WL54
P9WL55 P9WL58 P9WL59 P9WL60 P9WL61 P9WL62 P9WL64 P9WL65 P9WL66 P9WL70 P9WL71
P9WL86 P9WL90 P9WL91 P9WL92 P9WL94 P9WL95 P9WL96 P9WL98 P9WL99 P9WLA2 P9WLA3
P9WLA4 P9WLA5 P9WLA6 P9WLA8 P9WLA9 P9WLB2 P9WLB3 P9WLB4 P9WLB5 P9WLB6 P9WLB7
P9WLB8 P9WLB9 P9WLC0 P9WLC1 P9WLC2 P9WLC3 P9WLC4 P9WLC5 P9WLD0 P9WLD2 P9WLD3
P9WLD4 P9WLD5 P9WLD6 P9WLE4 P9WLE5 P9WLE6 P9WLE7 P9WLE8 P9WLE9 P9WLF6 P9WLF7
P9WLF8 P9WLF9 P9WLG0 P9WLG1 P9WLG4 P9WLG8 P9WLG9 P9WLH2 P9WLH4 P9WLH5 P9WLH6
P9WLH7 P9WLH8 P9WLJ0 P9WLJ6 P9WLJ7 P9WLJ8 P9WLJ9 P9WLK0 P9WLK6 P9WLK7 P9WLK8
P9WLK9 P9WLL8 P9WLM0 P9WLM2 P9WLM3 P9WLM6 P9WLM7 P9WLM8 P9WLM9 P9WLN0 P9WLN4
P9WLN5 P9WLN6 P9WLN8 P9WLN9 P9WLP0 P9WLP2 P9WLP4 P9WLP6 P9WLP7 P9WLP8 P9WLP9
P9WLQ2 P9WLQ3 P9WLQ4 P9WLQ5 P9WLR0 P9WLR1 P9WLR2 P9WLR3 P9WLR4 P9WLS2 P9WLS6
P9WLS7 P9WLT0 P9WLT2 P9WLT3 P9WLT6 P9WLT7 P9WLT8 P9WLU0 P9WLU1 P9WLU2 P9WLU3
P9WLU4 P9WLU5 P9WLU6 P9WLU8 P9WLU9 P9WLV2 P9WLV3 P9WLV4 P9WLV6 P9WLV7 P9WLW2
P9WLW3 P9WLW4 P9WLW6 P9WLW7 P9WLW8 P9WLW9 P9WLX2 P9WLX3 P9WLX4 P9WLX6 P9WLY2
P9WLY3 P9WLY4 P9WLY5 P9WLZ0 P9WLZ2 P9WLZ4 P9WLZ5 P9WM04 P9WM05 P9WM08 P9WM09
P9WM10 P9WM12 P9WM16 P9WM17 P9WM24 P9WM25 P9WM26 P9WM27 P9WM32 P9WM33 P9WM36
P9WM37 P9WM40 P9WM48 P9WM50 P9WM51 P9WM52 P9WM53 P9WM54 P9WM55 P9WM56 P9WM57
P9WM58 P9WM72 P9WM78 P9WM79 P9WM80 P9WM81 P9WM82 P9WM83 P9WM84 P9WM85 P9WM90
P9WM92 P9WM93 P9WM94 P9WM95 P9WM96 P9WM97 P9WM98 P9WM99 P9WMA0 P9WMA1 P9WMA4
P9WMA5 P9WMA8 P9WMB0 P9WMB1 P9WMB2 P9WMB3 P9WMB6 P9WMB7 P9WMB8 P9WMB9 P9WMC0
P9WMC1 P9WMC2 P9WMC3 P9WMC4 P9WMC5 P9WMC6 P9WMC7 P9WMD0 P9WMD1 P9WMD2 P9WMD3
P9WMD4 P9WMD5 P9WMD6 P9WMD7 P9WMD8 P9WMD9 P9WME0 P9WME1 P9WME2 P9WME3 P9WME4
P9WME5 P9WME6 P9WME7 P9WME8 P9WME9 P9WMF0 P9WMF1 P9WMF2 P9WMF3 P9WMF4 P9WMF5
P9WMF6 P9WMF7 P9WMG0 P9WMG1 P9WMG2 P9WMG3 P9WMG4 P9WMG5 P9WMG6 P9WMG7 P9WMG8
P9WMG9 P9WMH0 P9WMH2 P9WMH4 P9WMH5 P9WMH7 P9WMH8 P9WMH9 P9WMI2 P9WMI3 P9WMI4
P9WMI5 P9WMI6 P9WMI8 P9WMI9 P9WMJ0 P9WMJ1 P9WMJ2 P9WMJ3 P9WMJ4 P9WMJ5 P9WMJ6
P9WMJ8 P9WMK2 P9WMK4 P9WMK5 P9WMK8 P9WML0 P9WML1 P9WML2 P9WML3 P9WML4 P9WML5
P9WML6 P9WML8 P9WML9 P9WMM0 P9WMM1 P9WMM2 P9WMM3 P9WMM4 P9WMM5 P9WMM6 P9WMM7
P9WMM8 P9WMM9 P9WMN0 P9WMN4 P9WMN8 P9WMP0 P9WMP1 P9WMP2 P9WMP4 P9WMP5 P9WMP6
P9WMP7 P9WMP8 P9WMP9 P9WMQ0 P9WMQ2 P9WMQ4 P9WMQ6 P9WMQ8 P9WMR0 P9WMR2 P9WMR3
P9WMR4 P9WMR6 P9WMR7 P9WMR8 P9WMS0 P9WMS2 P9WMS4 P9WMS5 P9WMS6 P9WMS7 P9WMT0
P9WMT2 P9WMT4 P9WMT5 P9WMT6 P9WMT7 P9WMT8 P9WMU0 P9WMU4 P9WMU5 P9WMV2 P9WMV3
P9WMV4 P9WMV6 P9WMV7 P9WMW0 P9WMW2 P9WMW3 P9WMW8 P9WMW9 P9WMX6 P9WMX7 P9WMX8
P9WMX9 P9WMY0 P9WMY1 P9WMY2 P9WMY6 P9WMY8 P9WMY9 P9WMZ0 P9WMZ1 P9WN08 P9WN10
P9WN12 P9WN14 P9WN16 P9WN17 P9WN18 P9WN24 P9WN25 P9WN28 P9WN30 P9WN32 P9WN34
P9WN36 P9WN38 P9WN40 P9WN41 P9WN42 P9WN43 P9WN44 P9WN46 P9WN48 P9WN50 P9WN51
P9WN52 P9WN54 P9WN55 P9WN56 P9WN57 P9WN58 P9WN59 P9WN60 P9WN62 P9WN63 P9WN64
P9WN65 P9WN66 P9WN68 P9WN70 P9WN72 P9WN78 P9WN80 P9WN82 P9WN84 P9WN86 P9WN87
P9WN92 P9WNB8 P9WNB9 P9WNC0 P9WNC1 P9WNC2 P9WNC4 P9WNC5 P9WNC6 P9WNC7 P9WNC8
P9WNC9 P9WND0 P9WND1 P9WND2 P9WND3 P9WND4 P9WND6 P9WNE0 P9WNE2 P9WNE4 P9WNE6

P9WNE7 P9WNF0 P9WNF1 P9WNF6 P9WNF7 P9WNG0 P9WNG1 P9WNG6 P9WNG8 P9WNH0 P9WNH1
P9WNH2 P9WNL2 P9WNM0 P9WNM2 P9WNM3 P9WNM4 P9WNM6 P9WNM8 P9WNN0 P9WNP4 P9WNR8
P9WNS0 P9WNS1 P9WNS2 P9WNS3 P9WNS4 P9WNS5 P9WNS6 P9WNS7 P9WNS8 P9WNS9 P9WNT0
P9WNT2 P9WNT4 P9WNT6 P9WNT8 P9WNT9 P9WNU0 P9WNU1 P9WNU2 P9WNU3 P9WNU4 P9WNU6
P9WNU8 P9WNU9 P9WNV0 P9WNV2 P9WNV4 P9WNV5 P9WNV6 P9WNV7 P9WNV8 P9WNW0 P9WNW2
P9WNW4 P9WNW5 P9WNW8 P9WNW9 P9WNX0 P9WNX1 P9WNX2 P9WNX4 P9WNX6 P9WNX8 P9WNY0
P9WNY2 P9WNY3 P9WNY4 P9WNY5 P9WNY6 P9WNY8 P9WNY9 P9WNZ0 P9WP00 P9WP01 P9WP02
P9WP03 P9WP16 P9WP17 P9WP18 P9WP20 P9WP22 P9WP24 P9WP26 P9WP27 P9WP30 P9WP32
P9WP33 P9WP48 P9WP50 P9WP52 P9WP53 P9WP54 P9WP55 P9WP58 P9WP59 P9WP72 P9WP73
P9WP74 P9WP75 P9WP76 P9WP78 P9WP80 P9WP81 P9WP82 P9WP83 P9WP84 P9WP85 P9WP86
P9WP87 P9WP88 P9WP89 P9WP94 P9WP95 P9WP96 P9WP98 P9WP99 P9WPA0 P9WPA2 P9WPA3
P9WPA4 P9WPA5 P9WPA6 P9WPA8 P9WPA9 P9WPB8 P9WPB9 P9WPC0 P9WPC1 P9WPC2 P9WPC4
P9WPC6 P9WPC7 P9WPC8 P9WPD0 P9WPD2 P9WPD3 P9WPD4 P9WPE0 P9WPE1 P9WPE2 P9WPE4
P9WPE8 P9WPF8 P9WPF9 P9WPH0 P9WPH2 P9WPH4 P9WPH5 P9WPI4 P9WPI5 P9WPI8 P9WPI9
P9WPJ0 P9WPJ1 P9WPJ2 P9WPJ3 P9WPJ4 P9WPJ5 P9WPJ6 P9WPJ8 P9WPJ9 P9WPK0 P9WPK1
P9WPK2 P9WPK4 P9WPK5 P9WPK8 P9WPK9 P9WPL0 P9WPL1 P9WPL2 P9WPL3 P9WPL8 P9WPL9
P9WPM0 P9WPM1 P9WPM2 P9WPM3 P9WPM4 P9WPM6 P9WPM7 P9WPM8 P9WPM9 P9WPN0 P9WPN1
P9WPN2 P9WPN3 P9WPN4 P9WPN5 P9WPN8 P9WPN9 P9WPP4 P9WPP6 P9WPP7 P9WPQ0 P9WPQ1
P9WPQ2 P9WPQ4 P9WPQ5 P9WPQ6 P9WPQ8 P9WPR0 P9WPR2 P9WPR3 P9WPR4 P9WPR5 P9WPW2
P9WPW3 P9WPW4 P9WPW5 P9WPW6 P9WPW8 P9WPX0 P9WPX2 P9WPX6 P9WPX7 P9WPX8 P9WPY0
P9WPY1 P9WPY2 P9WPY3 P9WPY4 P9WPY6 P9WPY7 P9WPY8 P9WPY9 P9WPZ0 P9WPZ1 P9WPZ2
P9WPZ3 P9WPZ4 P9WPZ5 P9WPZ6 P9WPZ7 P9WPZ8 P9WPZ9 P9WQ00 P9WQ02 P9WQ03 P9WQ04
P9WQ06 P9WQ08 P9WQ10 P9WQ12 P9WQ13 P9WQ14 P9WQ16 P9WQ18 P9WQ20 P9WQ21 P9WQ22
P9WQ23 P9WQ24 P9WQ25 P9WQ26 P9WQ38 P9WQ39 P9WQ40 P9WQ41 P9WQ66 P9WQ68 P9WQ70
P9WQ71 P9WQ72 P9WQ74 P9WQ75 P9WQ76 P9WQ77 P9WQ78 P9WQ80 P9WQ81 P9WQ82 P9WQ84
P9WQ85 P9WQ86 P9WQ88 P9WQ89 P9WQ90 P9WQ92 P9WQ93 P9WQ94 P9WQ96 P9WQ97 P9WQ98
P9WQ99 P9WQA0 P9WQA1 P9WQA2 P9WQA4 P9WQA6 P9WQA8 P9WQA9 P9WQB2 P9WQB4 P9WQB6
P9WQB8 P9WQB9 P9WQC0 P9WQC1 P9WQC2 P9WQC3 P9WQC4 P9WQC6 P9WQC8 P9WQC9 P9WQD0
P9WQF0 P9WQF1 P9WQF4 P9WQF5 P9WQF6 P9WQF8 P9WQF9 P9WQG0 P9WQG2 P9WQG4 P9WQG5
P9WQG6 P9WQG7 P9WQG8 P9WQH0 P9WQH1 P9WQH2 P9WQH3 P9WQH8 P9WQI0 P9WQI4 P9WQJ4
P9WQK0 P9WQK2 P9WQK6 P9WQL4 P9WQN0 P9WQN1 P9WQN4 P9WQP4 Q00002 Q00010 Q00011
Q00014 Q00016 Q00024 Q00028 Q00030 Q00033 Q00037 Q00040 Q00042 Q00043 Q00049
Q00050 Q00052 Q00053 Q00054 Q00055 Q00056 Q00063 Q00075 Q00080 Q00081 Q00092
Q00094 Q00095 Q00097 Q00098 Q00099 Q000A9 Q00100 Q00101 Q00103 Q00105 Q00106
Q00107 Q00108 Q00109 Q00110 Q00111 Q00112 Q00113 Q00114 Q00115 Q00116 Q00117
Q00118 Q00119 Q00120 Q00121 Q00122 Q00123 Q00124 Q00125 Q00126 Q00127 Q00128
Q00129 Q00130 Q00131 Q00132 Q00140 Q00142 Q00143 Q00144 Q00145 Q00146 Q00147
Q00148 Q00149 Q00150 Q00151 Q00152 Q00153 Q00154 Q00155 Q00156 Q00158 Q00159
Q00160 Q00161 Q00162 Q00163 Q00164 Q00165 Q00166 Q00167 Q00186 Q00187 Q00188
Q00189 Q00190 Q00191 Q00192 Q00203 Q00204 Q00213 Q00214 Q00215 Q00217 Q00234
Q00237 Q00239 Q00240 Q00241 Q00247 Q00251 Q00257 Q00258 Q00259 Q00260 Q00261
Q00264 Q00265 Q00266 Q00267 Q00268 Q00271 Q00272 Q00273 Q00277 Q00278 Q00285
Q00286 Q00288 Q002B5 Q00301 Q00302 Q00305 Q00312 Q00313 Q00314 Q00322 Q00332
Q00333 Q00334 Q00335 Q00338 Q00369 Q00379 Q00384 Q00388 Q00390 Q00391 Q00393

Q003J2 Q00400 Q00401 Q00402 Q00404 Q00405 Q00406 Q00407 Q00408 Q00409 Q00410
Q00412 Q00420 Q00422 Q00423 Q00436 Q00438 Q00441 Q00444 Q00445 Q00454 Q00455
Q00456 Q00457 Q00458 Q00459 Q00460 Q00461 Q00462 Q00464 Q00466 Q00467 Q00469
Q00472 Q00477 Q00479 Q00488 Q00492 Q00497 Q004B5 Q00520 Q00526 Q00531 Q00539
Q00547 Q00556 Q00566 Q00571 Q00572 Q00579 Q00584 Q00597 Q00616 Q00637 Q00640
Q00644 Q00648 Q00659 Q00664 Q00667 Q00669 Q00670 Q00671 Q00672 Q00674 Q00675
Q00678 Q00682 Q00683 Q00684 Q00686 Q00701 Q00703 Q00705 Q00706 Q00713 Q00714
Q00715 Q00717 Q00719 Q00721 Q00723 Q00727 Q00728 Q00730 Q00732 Q00733 Q00736
Q00737 Q00738 Q00739 Q00740 Q00742 Q00747 Q00753 Q00763 Q00764 Q00766 Q00767
Q00768 Q00769 Q00770 Q00771 Q00775 Q00777 Q00778 Q00791 Q00793 Q00794 Q007T1
Q00808 Q00813 Q00830 Q00834 Q00835 Q00840 Q00844 Q00846 Q00847 Q00852 Q00853
Q00856 Q00858 Q00859 Q00861 Q00862 Q00864 Q00865 Q00866 Q00868 Q00869 Q00872
Q00899 Q008S8 Q008X2 Q00909 Q00911 Q00916 Q00923 Q00924 Q00932 Q00937 Q00939
Q00942 Q00946 Q00957 Q00962 Q00965 Q00966 Q00967 Q00970 Q00972 Q00976 Q00985
Q00990 Q00997 Q00998 Q00999 Q00G37 Q00IB6 Q00LS8 Q00P32 Q00PI9 Q00PJ1 Q00PJ3
Q00PJ7 Q00PK1 Q00VB2 Q00VK4 Q00YU3 Q00ZY2 Q01001 Q01003 Q01004 Q01005 Q01006
Q01007 Q01008 Q01009 Q01010 Q01011 Q01012 Q01013 Q01014 Q01015 Q01019 Q01020
Q01021 Q01022 Q01023 Q01024 Q01026 Q01027 Q01028 Q01029 Q01030 Q01032 Q01033
Q01034 Q01036 Q01037 Q01040 Q01042 Q01043 Q01044 Q01046 Q01047 Q01050 Q01051
Q01052 Q01053 Q01055 Q01059 Q01060 Q01068 Q01069 Q01070 Q01071 Q01074 Q01075
Q01077 Q01080 Q01087 Q01089 Q01095 Q01096 Q01100 Q01101 Q01103 Q01108 Q01109
Q01116 Q01128 Q01137 Q01146 Q01159 Q01168 Q01170 Q01173 Q01179 Q01180 Q01181
Q01182 Q01195 Q01198 Q01202 Q01206 Q01207 Q01209 Q01210 Q01213 Q01214 Q01217
Q01219 Q01221 Q01222 Q01223 Q01225 Q01226 Q01229 Q01230 Q01234 Q01238 Q01239
Q01240 Q01241 Q01246 Q01247 Q01249 Q01250 Q01252 Q01253 Q01254 Q01256 Q01257
Q01259 Q01261 Q01262 Q01263 Q01264 Q01267 Q01268 Q01277 Q01284 Q01286 Q01287
Q01288 Q01289 Q01291 Q01292 Q01295 Q01297 Q01302 Q01306 Q01320 Q01322 Q01323
Q01324 Q01325 Q01326 Q01327 Q01329 Q01330 Q01331 Q01332 Q01333 Q01334 Q01335
Q01336 Q01346 Q01347 Q01348 Q01349 Q01350 Q01351 Q01352 Q01353 Q01357 Q01358
Q01360 Q01365 Q01367 Q01368 Q01369 Q01371 Q01372 Q01378 Q01386 Q01390 Q01397
Q01398 Q01399 Q013G9 Q01401 Q01402 Q01408 Q01410 Q01414 Q01415 Q01417 Q01434
Q01435 Q01436 Q01437 Q01438 Q01445 Q01448 Q01449 Q01454 Q01456 Q01457 Q01462
Q01463 Q01468 Q01470 Q01478 Q01511 Q01515 Q01516 Q01517 Q01520 Q01521 Q01529
Q01534 Q01538 Q01540 Q01544 Q01545 Q01546 Q01548 Q01549 Q01550 Q01551 Q01558
Q01560 Q01569 Q01572 Q01576 Q01577 Q01579 Q01582 Q01586 Q01587 Q01588 Q01592
Q01593 Q01597 Q015G2 Q01604 Q01607 Q01612 Q01620 Q01622 Q01624 Q01630 Q01637
Q01642 Q01643 Q01644 Q01645 Q01647 Q01649 Q01651 Q01655 Q01656 Q01657 Q01661
Q01662 Q01669 Q01670 Q01671 Q01684 Q01687 Q01688 Q01697 Q01698 Q016E7 Q01700
Q01702 Q01703 Q01704 Q01713 Q01714 Q01743 Q01750 Q01751 Q01752 Q01757 Q01761
Q01765 Q01766 Q01767 Q01769 Q01770 Q01777 Q01778 Q01779 Q01781 Q01782 Q01789
Q01790 Q01791 Q01792 Q01793 Q01795 Q01797 Q01798 Q01799 Q017T9 Q01800 Q01802
Q01808 Q01811 Q01820 Q01822 Q01832 Q01851 Q01861 Q01864 Q01870 Q01871 Q01872
Q01873 Q01874 Q01877 Q01879 Q01886 Q01892 Q01893 Q01899 Q01902 Q01904 Q01911
Q01912 Q01917 Q01923 Q01930 Q01936 Q01939 Q01941 Q01953 Q01966 Q01976 Q01980

Q01981 Q01982 Q01984 Q01987 Q01990 Q01992 Q01993 Q01994 Q01997 Q01998 Q01999
Q01CH5 Q01F03 Q01FJ6 Q01H90 Q01I07 Q01I23 Q01I35 Q01IJ3 Q01IK0 Q01IX6 Q01J96
Q01JG2 Q01K11 Q01K12 Q01K26 Q01NF9 Q01NI7 Q01NI9 Q01NJ5 Q01NV7 Q01NW2 Q01NW3
Q01NZ1 Q01NZ9 Q01P09 Q01PF1 Q01PF4 Q01PM8 Q01PT7 Q01PW6 Q01Q15 Q01Q19 Q01Q26
Q01Q40 Q01Q43 Q01Q46 Q01Q49 Q01Q91 Q01QC0 Q01QF9 Q01QM3 Q01QM5 Q01QQ9 Q01QR0
Q01QR3 Q01QR4 Q01QR5 Q01QR6 Q01QT2 Q01QW7 Q01QZ0 Q01RP1 Q01RQ2 Q01RT2 Q01RT6
Q01RU5 Q01RU9 Q01RW8 Q01RX8 Q01RY8 Q01S04 Q01S07 Q01SE4 Q01SP7 Q01SU0 Q01SW8
Q01SX1 Q01SX2 Q01SZ5 Q01T71 Q01T72 Q01TA4 Q01TU5 Q01TY0 Q01TY1 Q01U81 Q01UR1
Q01V13 Q01V55 Q01V56 Q01VA8 Q01VB0 Q01VB1 Q01VB2 Q01VJ5 Q01VN6 Q01VU1 Q01VX3
Q01W05 Q01W10 Q01W29 Q01W31 Q01W33 Q01W87 Q01W88 Q01W89 Q01W91 Q01W92 Q01W93
Q01W94 Q01W95 Q01W96 Q01W97 Q01W98 Q01W99 Q01WA0 Q01WA2 Q01WA3 Q01WA4 Q01WA6
Q01WA7 Q01WA8 Q01WA9 Q01WB0 Q01WB1 Q01WB6 Q01WB7 Q01WB8 Q01WB9 Q01WC0 Q01WC1
Q01WJ8 Q01WL4 Q01X27 Q01X76 Q01X96 Q01XA0 Q01XQ7 Q01Y56 Q01Y83 Q01YA2 Q01YD0
Q01YQ2 Q01Z80 Q01Z81 Q01ZM2 Q01ZN1 Q01ZT9 Q01ZU0 Q01ZU4 Q01ZU6 Q02000 Q02001
Q02002 Q02003 Q02007 Q02008 Q02009 Q02011 Q02025 Q02026 Q02027 Q02028 Q02029
Q02030 Q02031 Q02032 Q02033 Q02034 Q02035 Q02038 Q02045 Q02046 Q02047 Q02050
Q02051 Q02054 Q02055 Q02058 Q02059 Q02061 Q02062 Q02067 Q02068 Q02073 Q02078
Q02080 Q02084 Q02093 Q02099 Q02100 Q02102 Q02103 Q02106 Q02109 Q02119 Q02123
Q02129 Q02130 Q02131 Q02132 Q02133 Q02134 Q02135 Q02136 Q02137 Q02138 Q02139
Q02140 Q02141 Q02142 Q02143 Q02144 Q02145 Q02146 Q02147 Q02149 Q02150 Q02154
Q02155 Q02166 Q02169 Q02170 Q02171 Q02173 Q02174 Q02175 Q02177 Q02178 Q02184
Q02185 Q02186 Q02187 Q02188 Q02189 Q02190 Q02191 Q02196 Q02197 Q02198 Q021H8
Q021N5 Q021N6 Q021Y7 Q02202 Q02206 Q02208 Q02209 Q02210 Q02216 Q02225 Q02229
Q02230 Q02231 Q02232 Q02233 Q02234 Q02236 Q02237 Q02239 Q02240 Q02243 Q02245
Q02250 Q02252 Q02253 Q02254 Q02263 Q02264 Q02265 Q02266 Q02267 Q02268 Q02269
Q02270 Q02273 Q02274 Q02283 Q02285 Q02286 Q02287 Q02292 Q022D2 Q022E5 Q022F6
Q022G3 Q022G4 Q022G5 Q022S9 Q022T0 Q02308 Q02314 Q02322 Q02323 Q02328 Q02332
Q02333 Q02336 Q02346 Q02347 Q02348 Q02349 Q02354 Q02356 Q02358 Q02360 Q02361
Q02362 Q02382 Q02386 Q02394 Q02395 Q02398 Q023P9 Q023Q5 Q023V5 Q023V7 Q02400
Q02403 Q02404 Q02405 Q02406 Q02407 Q02408 Q02415 Q02418 Q02419 Q02425 Q02426
Q02427 Q02430 Q02431 Q02437 Q02438 Q02441 Q02452 Q02455 Q02457 Q02458 Q02479
Q02480 Q02484 Q02499 Q024L7 Q024M2 Q024T3 Q024T4 Q024V4 Q02513 Q02514 Q02515
Q02516 Q02518 Q02521 Q02522 Q02525 Q02526 Q02535 Q02539 Q02550 Q02554 Q02555
Q02556 Q02566 Q02570 Q02572 Q02574 Q02575 Q02576 Q02577 Q02578 Q02579 Q02582
Q02584 Q02586 Q02587 Q02588 Q02589 Q02590 Q02598 Q025A4 Q02603 Q02604 Q02605
Q02606 Q02607 Q02610 Q02614 Q02635 Q02636 Q02637 Q02638 Q02647 Q02648 Q02652
Q02659 Q02685 Q026Q1 Q026Q2 Q026Q4 Q026Q7 Q02710 Q02720 Q02721 Q02723 Q02724
Q02732 Q02733 Q02735 Q02736 Q02748 Q02749 Q02751 Q02754 Q02764 Q02770 Q02771
Q02773 Q02775 Q02777 Q02780 Q02781 Q02784 Q02787 Q02792 Q02793 Q02796 Q02799
Q027B2 Q027G1 Q027G2 Q027I0 Q027L3 Q027U4 Q02803 Q02814 Q02826 Q02831 Q02835
Q02842 Q02855 Q02856 Q02857 Q02861 Q02864 Q02866 Q02869 Q02872 Q02873 Q02875
Q02876 Q02879 Q02884 Q02885 Q02886 Q02890 Q02892 Q02895 Q02899 Q028J0 Q028J4
Q028M4 Q028S4 Q028Z8 Q02908 Q02909 Q02910 Q02912 Q02916 Q02918 Q02919 Q02926
Q02931 Q02939 Q02941 Q02943 Q02944 Q02953 Q02954 Q02957 Q02958 Q02961 Q02964

Q02965 Q02966 Q02967 Q02968 Q02970 Q02974 Q02979 Q02980 Q02981 Q02983 Q02984
Q02986 Q02987 Q02990 Q02991 Q02997 Q02998 Q02999 Q029I5 Q029J4 Q029K6 Q029R3
Q029R4 Q029T9 Q029U7 Q02A16 Q02A20 Q02A42 Q02A54 Q02A69 Q02AG6 Q02AG7 Q02AK4
Q02AK9 Q02AL1 Q02AL2 Q02AL3 Q02AL5 Q02AX2 Q02B43 Q02BG1 Q02BS4 Q02BT1 Q02BZ2
Q02BZ5 Q02C42 Q02C58 Q02CB5 Q02CM4 Q02CZ9 Q02D35 Q02D92 Q02DD2 Q02DD9 Q02DE1
Q02DE3 Q02DE4 Q02DG9 Q02DJ3 Q02DL4 Q02DL7 Q02DY8 Q02DY9 Q02DZ0 Q02E05 Q02E25
Q02E28 Q02E31 Q02E39 Q02E41 Q02E43 Q02E46 Q02E64 Q02E82 Q02E84 Q02EA0 Q02EA3
Q02EB7 Q02ED9 Q02EG0 Q02EG1 Q02EG7 Q02EH1 Q02EH2 Q02EL6 Q02EM2 Q02EM5 Q02EM6
Q02EN5 Q02EN8 Q02EP8 Q02EQ6 Q02ER6 Q02ER8 Q02ES4 Q02ET8 Q02EV0 Q02EV1 Q02EV4
Q02EW3 Q02EW4 Q02EW6 Q02EW8 Q02EX8 Q02EX9 Q02EY3 Q02EZ4 Q02EZ9 Q02F21 Q02F45
Q02F66 Q02F67 Q02F72 Q02F73 Q02F74 Q02F79 Q02F80 Q02F83 Q02F84 Q02F86 Q02F98
Q02FC5 Q02FC6 Q02FC7 Q02FE6 Q02FF2 Q02FF4 Q02FF5 Q02FG6 Q02FH0 Q02FL3 Q02FP9
Q02FQ4 Q02FQ6 Q02FR0 Q02FR1 Q02FR2 Q02FR3 Q02FS0 Q02FS3 Q02FS4 Q02FS6 Q02FS8
Q02FS9 Q02FT0 Q02FT1 Q02FT2 Q02FU0 Q02FU1 Q02FU2 Q02FU3 Q02FU9 Q02FV2 Q02FV3
Q02FX9 Q02G02 Q02G03 Q02G05 Q02G08 Q02G09 Q02G12 Q02G19 Q02G28 Q02G49 Q02G60
Q02G92 Q02GA9 Q02GB0 Q02GB2 Q02GB4 Q02GB7 Q02GC0 Q02GN4 Q02GQ9 Q02GS6 Q02GV7
Q02GV8 Q02GV9 Q02GX6 Q02GZ1 Q02GZ2 Q02GZ8 Q02H07 Q02H08 Q02H15 Q02H16 Q02H19
Q02H20 Q02H29 Q02H45 Q02H54 Q02H55 Q02H78 Q02HQ1 Q02HS2 Q02HS4 Q02HS5 Q02HS6
Q02HS7 Q02HZ4 Q02I17 Q02I31 Q02I59 Q02I61 Q02I63 Q02I65 Q02I76 Q02I83 Q02I93
Q02IA6 Q02IB8 Q02IC2 Q02IC5 Q02IC6 Q02IC7 Q02IC8 Q02IC9 Q02ID8 Q02IG9 Q02IH8
Q02IK5 Q02IL5 Q02IR7 Q02IV5 Q02IY2 Q02J09 Q02J28 Q02JB8 Q02JC0 Q02JD6 Q02JE0
Q02JF1 Q02JF2 Q02JM2 Q02JM4 Q02JZ7 Q02JZ8 Q02K12 Q02K16 Q02K17 Q02K26 Q02K46
Q02K47 Q02K58 Q02K73 Q02K83 Q02KC3 Q02KC8 Q02KG3 Q02KH1 Q02KH3 Q02KH4 Q02KH5
Q02KP7 Q02KR0 Q02KR1 Q02KR3 Q02KT5 Q02KT6 Q02KT7 Q02KU3 Q02KU5 Q02KV8 Q02KW6
Q02KY4 Q02KZ2 Q02L47 Q02L76 Q02LD2 Q02LD3 Q02LD4 Q02LD5 Q02LD6 Q02LF6 Q02LK0
Q02LM8 Q02LQ6 Q02LX5 Q02M20 Q02ML7 Q02ML8 Q02ML9 Q02MM0 Q02MM1 Q02MM2 Q02N15
Q02N70 Q02N77 Q02NA3 Q02NA4 Q02NA9 Q02NB0 Q02NB1 Q02NB3 Q02NB8 Q02NN5 Q02NN7
Q02NN8 Q02NN9 Q02NP1 Q02NV3 Q02NW3 Q02NZ0 Q02NZ5 Q02P14 Q02P38 Q02P70 Q02PC5
Q02PD3 Q02PE3 Q02PF5 Q02PG3 Q02PG5 Q02PG9 Q02PJ4 Q02PK7 Q02PQ1 Q02PS6 Q02PT3
Q02PT4 Q02PU7 Q02PW7 Q02PW9 Q02PX3 Q02PX5 Q02PX7 Q02PY1 Q02PZ9 Q02Q74 Q02Q76
Q02Q78 Q02Q95 Q02QB5 Q02QH1 Q02QJ1 Q02QK0 Q02QU2 Q02QU5 Q02QV2 Q02QW0 Q02QW9
Q02QX1 Q02QZ6 Q02QZ8 Q02QZ9 Q02R10 Q02R72 Q02R73 Q02R88 Q02R89 Q02R92 Q02R96
Q02R97 Q02R98 Q02R99 Q02RA5 Q02RA9 Q02RB4 Q02RC2 Q02RC5 Q02RC6 Q02RC7 Q02RC8
Q02RD0 Q02RF2 Q02RF8 Q02RH2 Q02RL5 Q02RL6 Q02RL7 Q02RL8 Q02RM4 Q02RS2 Q02RT0
Q02RV3 Q02RV6 Q02RV7 Q02RW0 Q02RW1 Q02RW4 Q02RW5 Q02RW8 Q02RX7 Q02RX8 Q02RY8
Q02RZ9 Q02S14 Q02S21 Q02S69 Q02S84 Q02SE4 Q02SE5 Q02SE8 Q02SF0 Q02SF5 Q02SG3
Q02SG4 Q02SG5 Q02SH1 Q02SH2 Q02SH3 Q02SH4 Q02SH7 Q02SJ2 Q02SK9 Q02SL1 Q02SL4
Q02SM4 Q02SV2 Q02T54 Q02T55 Q02T56 Q02T57 Q02T58 Q02T59 Q02T61 Q02T62 Q02T63
Q02T64 Q02T65 Q02T66 Q02T67 Q02T68 Q02T69 Q02T70 Q02T71 Q02T72 Q02T73 Q02T74
Q02T75 Q02T76 Q02T77 Q02T78 Q02T79 Q02T80 Q02T81 Q02T82 Q02T84 Q02T85 Q02T86
Q02T87 Q02T88 Q02T89 Q02T90 Q02T91 Q02T96 Q02TA1 Q02TA4 Q02TB2 Q02TB5 Q02TB6
Q02TH2 Q02TH3 Q02TH4 Q02TH6 Q02TH9 Q02TI3 Q02TI4 Q02TL3 Q02TL9 Q02TR2 Q02TR5
Q02TR6 Q02TT1 Q02TY0 Q02U07 Q02U08 Q02U09 Q02U10 Q02U23 Q02U31 Q02U32 Q02U51
Q02U58 Q02U62 Q02U73 Q02U79 Q02U86 Q02UC5 Q02UM5 Q02UM6 Q02UQ8 Q02UT0 Q02V26

Q02V45 Q02V57 Q02V59 Q02V63 Q02V72 Q02V73 Q02V78 Q02V80 Q02VB0 Q02VK8 Q02VK9
Q02VL7 Q02VM1 Q02VP7 Q02VQ6 Q02VR5 Q02VS3 Q02VS5 Q02VS6 Q02VS7 Q02VT9 Q02VU1
Q02VU7 Q02VW5 Q02VX0 Q02VX8 Q02VX9 Q02VZ1 Q02VZ7 Q02W18 Q02W24 Q02W25 Q02W26
Q02W27 Q02W28 Q02W29 Q02W30 Q02W31 Q02W32 Q02W33 Q02W34 Q02W35 Q02W36 Q02W37
Q02W39 Q02W40 Q02W41 Q02W42 Q02W43 Q02W44 Q02W46 Q02W47 Q02W48 Q02W49 Q02W50
Q02W51 Q02W52 Q02W76 Q02W78 Q02W92 Q02W94 Q02W95 Q02WA3 Q02WC4 Q02WC5 Q02WC9
Q02WD2 Q02WD3 Q02WH3 Q02WH8 Q02WI8 Q02WJ0 Q02WK6 Q02WK8 Q02WL3 Q02WM7 Q02WN1
Q02WN8 Q02WP2 Q02WP8 Q02WW4 Q02WY3 Q02WY9 Q02WZ5 Q02WZ9 Q02X03 Q02X09 Q02X10
Q02X21 Q02X30 Q02X44 Q02X53 Q02X59 Q02X60 Q02XB7 Q02XB8 Q02XC0 Q02XC2 Q02XD1
Q02XD2 Q02XD6 Q02XE9 Q02XF3 Q02XG6 Q02XH5 Q02XK6 Q02XL9 Q02XM1 Q02XM4 Q02XM8
Q02XT4 Q02XU7 Q02XV9 Q02XW0 Q02XW4 Q02XW6 Q02XX3 Q02XX9 Q02XY2 Q02Y08 Q02Y12
Q02Y14 Q02Y16 Q02Y20 Q02Y22 Q02Y43 Q02Y52 Q02Y55 Q02Y66 Q02Y75 Q02Y87 Q02YA8
Q02YB4 Q02YB7 Q02YB8 Q02YD2 Q02YD6 Q02YG5 Q02YJ2 Q02YJ4 Q02YJ6 Q02YK5 Q02YP2
Q02YQ3 Q02YQ7 Q02YQ8 Q02YQ9 Q02YR7 Q02YR8 Q02YS4 Q02YS6 Q02YT3 Q02YU5 Q02YW3
Q02YW4 Q02YW7 Q02YX0 Q02YX1 Q02YX4 Q02YX5 Q02YY1 Q02YY5 Q02YY8 Q02YZ3 Q02YZ4
Q02Z11 Q02Z13 Q02Z21 Q02Z22 Q02Z29 Q02Z38 Q02Z48 Q02Z51 Q02Z55 Q02Z86 Q02Z92
Q02Z93 Q02Z94 Q02ZA2 Q02ZA6 Q02ZA8 Q02ZC9 Q02ZD2 Q02ZM3 Q02ZM8 Q02ZN0 Q02ZN6
Q02ZN8 Q02ZQ7 Q02ZQ9 Q02ZR4 Q02ZS7 Q02ZS8 Q02ZT0 Q02ZT2 Q02ZX9 Q02ZY9 Q03000
Q03003 Q03004 Q03005 Q03006 Q03007 Q03009 Q03010 Q03012 Q03014 Q03016 Q03017
Q03018 Q03020 Q03021 Q03033 Q03034 Q03036 Q03039 Q03042 Q03046 Q03047 Q03049
Q03051 Q03052 Q03054 Q03055 Q03056 Q03057 Q03058 Q03059 Q03061 Q03062 Q03063
Q03065 Q03066 Q03067 Q03068 Q03071 Q03080 Q03081 Q03083 Q03088 Q03096 Q03099
Q030A1 Q030A3 Q030B4 Q030C8 Q030E7 Q030F9 Q030G2 Q030K1 Q030K2 Q030K6 Q030L6
Q030M1 Q030M2 Q030M6 Q030P1 Q030T6 Q030U0 Q030V9 Q030W3 Q030Z7 Q030Z9 Q03123
Q03124 Q03125 Q03131 Q03132 Q03133 Q03134 Q03144 Q03149 Q03154 Q03161 Q03162
Q03169 Q03172 Q03177 Q03182 Q03183 Q03184 Q03195 Q03196 Q031A0 Q031A3 Q031A8
Q031B7 Q031D5 Q031D7 Q031E4 Q031G7 Q031H9 Q031I3 Q031I5 Q031I6 Q031M8 Q031N1
Q031P2 Q031P9 Q031S0 Q031S3 Q031S8 Q031S9 Q031T7 Q031U1 Q031U8 Q031V6 Q031W1
Q031W5 Q031X0 Q031X8 Q031X9 Q031Y3 Q03205 Q03207 Q03208 Q03209 Q03210 Q03213
Q03214 Q03215 Q03216 Q03217 Q03219 Q03220 Q03221 Q03222 Q03223 Q03224 Q03231
Q03233 Q03237 Q03240 Q03241 Q03242 Q03243 Q03244 Q03245 Q03248 Q03249 Q03253
Q03254 Q03255 Q03256 Q03257 Q03262 Q03264 Q03266 Q03267 Q03269 Q03270 Q03271
Q03272 Q03273 Q03274 Q03275 Q03276 Q03277 Q03278 Q03279 Q03280 Q03282 Q03283
Q03284 Q03285 Q03286 Q03287 Q03290 Q03293 Q03294 Q03295 Q03296 Q03298 Q032D4
Q032G4 Q032G8 Q032G9 Q032K6 Q032K8 Q032L5 Q032L7 Q032L8 Q032Q6 Q032Q9 Q032R6
Q032T2 Q032T3 Q032T4 Q032T5 Q032U4 Q032W2 Q032W3 Q032W4 Q032W9 Q032X2 Q032X5
Q032X6 Q032Z1 Q03300 Q03301 Q03302 Q03304 Q03313 Q03314 Q03316 Q03319 Q03320
Q03321 Q03323 Q03326 Q03328 Q03330 Q03331 Q03332 Q03335 Q03337 Q03338 Q03339
Q03340 Q03341 Q03342 Q03346 Q03356 Q03357 Q03361 Q03362 Q03365 Q03369 Q03372
Q03373 Q03375 Q03377 Q03378 Q03381 Q03382 Q03384 Q03387 Q03388 Q03389 Q03392
Q03396 Q033A2 Q033A6 Q033B8 Q033B9 Q033E7 Q033G3 Q033H5 Q033I1 Q033I2 Q033I4
Q033K5 Q033L0 Q033L1 Q033M5 Q033M9 Q033U6 Q033U7 Q033W5 Q03406 Q03408 Q03413
Q03414 Q03415 Q03416 Q03417 Q03421 Q03425 Q03428 Q03432 Q03433 Q03435 Q03442
Q03443 Q03444 Q03447 Q03456 Q03459 Q03460 Q03466 Q03469 Q03470 Q03471 Q03474

Q03483 Q03484 Q03488 Q03489 Q03495 Q03496 Q034G9 Q034H8 Q034I2 Q034T0 Q034T9
Q034V1 Q034W5 Q034X0 Q034X1 Q034X6 Q034X7 Q034X8 Q034Y2 Q034Y3 Q034Y4 Q034Y5
Q034Y6 Q034Y7 Q034Y8 Q034Y9 Q034Z0 Q034Z1 Q034Z2 Q034Z3 Q034Z4 Q034Z5 Q034Z6
Q034Z7 Q034Z8 Q034Z9 Q03503 Q03505 Q03506 Q03508 Q03509 Q03511 Q03512 Q03513
Q03520 Q03522 Q03523 Q03525 Q03533 Q03544 Q03545 Q03546 Q03547 Q03548 Q03549
Q03550 Q03552 Q03553 Q03556 Q03557 Q03558 Q03559 Q03560 Q03562 Q03568 Q03569
Q03570 Q03571 Q03572 Q03573 Q03576 Q03577 Q03585 Q03586 Q03587 Q03588 Q03589
Q03590 Q03598 Q03599 Q035A0 Q035A1 Q035A5 Q035A6 Q035A7 Q035A8 Q035A9 Q035B0
Q035B5 Q035C3 Q035C4 Q035G1 Q035M3 Q035R5 Q035S0 Q035T6 Q035U1 Q035U8 Q035U9
Q035V5 Q035V6 Q035W7 Q035W9 Q035X2 Q035Y0 Q035Y8 Q035Y9 Q035Z2 Q035Z3 Q035Z4
Q03601 Q03604 Q03605 Q03606 Q03607 Q03609 Q03615 Q03629 Q03630 Q03631 Q03638
Q03652 Q03654 Q03660 Q03661 Q03662 Q03663 Q03664 Q03666 Q03667 Q03675 Q03683
Q03689 Q03690 Q03695 Q036B8 Q036H7 Q036P0 Q036S2 Q036S8 Q036T0 Q036X3 Q036Y4
Q036Y5 Q03703 Q03705 Q03708 Q03718 Q03722 Q03735 Q03740 Q03744 Q03745 Q03746
Q03748 Q03749 Q03750 Q03759 Q03761 Q03768 Q03771 Q03772 Q03774 Q03780 Q03782
Q03784 Q03786 Q03787 Q03792 Q03796 Q037J6 Q037K4 Q037L6 Q037M7 Q037P8 Q037S9
Q037T6 Q037U6 Q037U8 Q037U9 Q037V3 Q037W6 Q037X5 Q037Y3 Q037Y8 Q037Z3 Q037Z6
Q03828 Q03831 Q03834 Q03835 Q03856 Q03862 Q03864 Q03878 Q03879 Q03886 Q03888
Q03898 Q03899 Q038A1 Q038A2 Q038B5 Q038C3 Q038C7 Q038E3 Q038E8 Q038F8 Q038F9
Q038G0 Q038H0 Q038H3 Q038I0 Q038J7 Q038J9 Q038K0 Q038K1 Q038L1 Q038L2 Q038L3
Q038L4 Q038L5 Q038L9 Q038M0 Q038M1 Q038M5 Q038M6 Q038M8 Q038N1 Q038N3 Q038N5
Q038P1 Q038Q5 Q038Q9 Q038S0 Q038S1 Q038S5 Q038S6 Q038S9 Q038U2 Q038U3 Q038V3
Q038V6 Q038V7 Q038W1 Q038W5 Q038W6 Q038W7 Q038Y4 Q038Y9 Q038Z2 Q038Z3 Q03900
Q03902 Q03919 Q03920 Q03923 Q03924 Q03928 Q03935 Q03936 Q03937 Q03938 Q03940
Q03942 Q03954 Q03958 Q03964 Q03968 Q03973 Q03974 Q03975 Q03981 Q03983 Q03986
Q03989 Q039A9 Q039B0 Q039B2 Q039B3 Q039B4 Q039B5 Q039B6 Q039B7 Q039E0 Q039E1
Q039E2 Q039E6 Q039F0 Q039F7 Q039G0 Q039G4 Q039I0 Q039I6 Q039J0 Q039J3 Q039J7
Q039K2 Q039K4 Q039K9 Q039L3 Q039L4 Q039M3 Q039N7 Q039P0 Q039P7 Q039Q9 Q039R3
Q039R8 Q039S2 Q039U0 Q039U3 Q039Y8 Q03A25 Q03A26 Q03A29 Q03A55 Q03A64 Q03A83
Q03AB9 Q03AC0 Q03AC3 Q03AD2 Q03AF5 Q03AF8 Q03AI1 Q03AJ7 Q03AK5 Q03AK6 Q03AL0
Q03AL2 Q03AL4 Q03AL7 Q03AM8 Q03AQ1 Q03AR1 Q03AS0 Q03AS5 Q03AT8 Q03AU4 Q03AW0
Q03AZ2 Q03AZ5 Q03AZ8 Q03AZ9 Q03B00 Q03B02 Q03B17 Q03B18 Q03BB6 Q03BB8 Q03BD0
Q03C44 Q03CA5 Q03CB2 Q03CB3 Q03CD2 Q03CI6 Q03CJ9 Q03CK3 Q03CP2 Q03CR5 Q03CT6
Q03CU2 Q03CW1 Q03CW2 Q03CW3 Q03CY0 Q03CY1 Q03CY2 Q03CY3 Q03D45 Q03D47 Q03D52
Q03D55 Q03D56 Q03D57 Q03D59 Q03D60 Q03D61 Q03D70 Q03D71 Q03D72 Q03D86 Q03DF6
Q03DH7 Q03DK3 Q03DS0 Q03DS2 Q03DS8 Q03DT9 Q03DX9 Q03DY3 Q03DZ2 Q03DZ7 Q03DZ8
Q03E09 Q03E35 Q03E40 Q03E41 Q03E45 Q03E48 Q03E49 Q03E50 Q03E51 Q03E58 Q03E59
Q03E60 Q03E63 Q03E69 Q03E71 Q03E72 Q03E76 Q03E83 Q03EA5 Q03EA9 Q03EB0 Q03EB2
Q03EB3 Q03EB4 Q03EB5 Q03EB6 Q03EB7 Q03EB8 Q03EB9 Q03EC0 Q03EC1 Q03EC2 Q03EC3
Q03EC4 Q03EC6 Q03EC7 Q03EC8 Q03EC9 Q03ED0 Q03ED1 Q03ED2 Q03ED3 Q03ED4 Q03ED5
Q03ED8 Q03ED9 Q03EE0 Q03EE1 Q03EE2 Q03EE3 Q03EE8 Q03EE9 Q03EG0 Q03EG5 Q03EG6
Q03EI3 Q03EK4 Q03EK5 Q03EM9 Q03EN4 Q03EN8 Q03EQ4 Q03EQ7 Q03EQ8 Q03EQ9 Q03ER0
Q03ER1 Q03ER2 Q03ER8 Q03ER9 Q03ES0 Q03ES1 Q03ES2 Q03ES4 Q03ET4 Q03EU1 Q03EW2
Q03EW6 Q03EX6 Q03EX7 Q03EY1 Q03EY9 Q03EZ6 Q03F09 Q03F16 Q03F21 Q03F22 Q03F25

Q03F26 Q03F27 Q03F28 Q03F33 Q03F34 Q03F36 Q03F39 Q03F44 Q03F48 Q03F51 Q03F52
Q03F56 Q03F57 Q03F58 Q03F61 Q03F64 Q03F65 Q03F66 Q03F74 Q03F82 Q03F87 Q03F94
Q03F96 Q03FA3 Q03FJ5 Q03FK2 Q03FK3 Q03FM3 Q03FM6 Q03FN0 Q03FN1 Q03FN2 Q03FR6
Q03FR7 Q03FR8 Q03FR9 Q03FS1 Q03FS2 Q03FS3 Q03FS7 Q03FS8 Q03FS9 Q03FT3 Q03FT4
Q03FT5 Q03FT6 Q03FT8 Q03FU2 Q03FW2 Q03FW3 Q03FW4 Q03FW5 Q03FX0 Q03FX6 Q03FX9
Q03FY3 Q03FY6 Q03FZ3 Q03FZ4 Q03FZ5 Q03FZ6 Q03FZ9 Q03G01 Q03G75 Q03G83 Q03G94
Q03G99 Q03GA1 Q03GB0 Q03GB1 Q03GB7 Q03GC0 Q03GC2 Q03GC5 Q03GC9 Q03GD8 Q03GE2
Q03GG4 Q03GG8 Q03GI2 Q03GJ0 Q03GM4 Q03GM6 Q03GV0 Q03GV3 Q03GV8 Q03GW2 Q03GW6
Q03GW7 Q03GX1 Q03GX4 Q03GX6 Q03GY5 Q03H05 Q03H06 Q03H14 Q03H46 Q03H60 Q03H73
Q03H91 Q03HA9 Q03HB0 Q03HB2 Q03HB3 Q03HF4 Q03HF6 Q03HI6 Q03HJ9 Q03HL3 Q03HM9
Q03HN1 Q03HQ0 Q03HR3 Q03HS1 Q03HT0 Q03HT1 Q03HT2 Q03I50 Q03I52 Q03I57 Q03I60
Q03I69 Q03I76 Q03I88 Q03I89 Q03I91 Q03I94 Q03IA9 Q03IB0 Q03IB2 Q03IB4 Q03IC3
Q03IC4 Q03IC5 Q03IC6 Q03IE5 Q03IE9 Q03IF0 Q03IF1 Q03IF2 Q03IF4 Q03IF5 Q03IF6
Q03IF7 Q03IF8 Q03IF9 Q03IG0 Q03IG1 Q03IG2 Q03IG3 Q03IG4 Q03IG5 Q03IG6 Q03IG7
Q03IG8 Q03IG9 Q03IH0 Q03IH2 Q03IH3 Q03IH4 Q03IH5 Q03IH6 Q03IH7 Q03IH8 Q03IJ9
Q03IK0 Q03IK2 Q03IK3 Q03IL2 Q03IN0 Q03IN1 Q03IP3 Q03IP4 Q03IP7 Q03IP8 Q03IP9
Q03IQ0 Q03IQ3 Q03IQ5 Q03IR2 Q03IR9 Q03IS0 Q03IS1 Q03IS8 Q03IT3 Q03IT6 Q03IU1
Q03IU4 Q03IV3 Q03IV5 Q03IW4 Q03IW6 Q03IX8 Q03IZ7 Q03IZ8 Q03J02 Q03J03 Q03J09
Q03J17 Q03J44 Q03J54 Q03J75 Q03J80 Q03J81 Q03J82 Q03J93 Q03JA3 Q03JB0 Q03JB8
Q03JB9 Q03JC1 Q03JC2 Q03JC6 Q03JD8 Q03JF6 Q03JG3 Q03JH0 Q03JH6 Q03JI6 Q03JJ8
Q03JK4 Q03JL0 Q03JN0 Q03JN2 Q03JR1 Q03JS0 Q03JS1 Q03JS2 Q03JS8 Q03JW5 Q03K10
Q03K16 Q03K17 Q03K19 Q03K30 Q03K46 Q03K54 Q03K75 Q03K76 Q03K77 Q03K79 Q03K81
Q03K83 Q03K84 Q03K89 Q03K92 Q03K93 Q03K98 Q03K99 Q03KA9 Q03KB3 Q03KB4 Q03KB7
Q03KD6 Q03KD9 Q03KI4 Q03KI8 Q03KI9 Q03KJ0 Q03KJ1 Q03KJ8 Q03KJ9 Q03KK1 Q03KM6
Q03KP9 Q03KR8 Q03KS5 Q03KS6 Q03KT7 Q03KX0 Q03KX7 Q03L08 Q03L10 Q03L13 Q03L16
Q03L20 Q03L30 Q03L31 Q03L41 Q03L46 Q03L77 Q03L79 Q03L81 Q03L85 Q03L88 Q03L92
Q03L98 Q03LA9 Q03LD3 Q03LE7 Q03LE9 Q03LF7 Q03LG8 Q03LG9 Q03LH1 Q03LH3 Q03LH4
Q03LH5 Q03LH6 Q03LI8 Q03LJ1 Q03LJ5 Q03LJ7 Q03LK0 Q03LM9 Q03LN0 Q03LN3 Q03LP1
Q03LR5 Q03LS1 Q03LS2 Q03LT1 Q03LT2 Q03LT8 Q03LU0 Q03LU2 Q03LV6 Q03LV7 Q03LW8
Q03LW9 Q03LX0 Q03LY7 Q03LZ4 Q03LZ8 Q03M13 Q03M15 Q03M16 Q03M28 Q03M29 Q03M32
Q03M33 Q03M60 Q03M62 Q03M63 Q03M65 Q03M78 Q03M79 Q03M85 Q03M87 Q03M88 Q03M92
Q03MA3 Q03MB1 Q03MB4 Q03MB7 Q03MD9 Q03ME0 Q03ME1 Q03ME2 Q03ME3 Q03ME4 Q03ME5
Q03MF6 Q03MF7 Q03MG1 Q03MH4 Q03MH6 Q03MI1 Q03MI2 Q03MI3 Q03MI7 Q03MJ2 Q03MJ7
Q03MK3 Q03MK4 Q03MK9 Q03ML4 Q03ML5 Q03MM5 Q03MN5 Q03MP3 Q03MP6 Q03MP9 Q03MQ4
Q03MQ8 Q03MQ9 Q03MR4 Q03MR6 Q03MR7 Q03MR8 Q03MS2 Q03MU5 Q03MV9 Q03MW6 Q03MW7
Q03MX3 Q03MX5 Q03MX6 Q03MX8 Q03MY0 Q03MY4 Q03MY7 Q03MZ6 Q03MZ8 Q03N01 Q03N04
Q03N21 Q03N26 Q03N60 Q03N61 Q03N64 Q03N65 Q03NA6 Q03NA7 Q03NE0 Q03NE4 Q03NG2
Q03NP0 Q03NR9 Q03NS1 Q03NV0 Q03NY8 Q03NY9 Q03P20 Q03PA4 Q03PA8 Q03PB4 Q03PJ4
Q03PM5 Q03PP4 Q03PR5 Q03PU5 Q03PU9 Q03PV0 Q03PV2 Q03PV3 Q03PV4 Q03PV6 Q03PV7
Q03PV8 Q03PV9 Q03PW0 Q03PW1 Q03PW2 Q03PW3 Q03PW4 Q03PW5 Q03PW6 Q03PW7 Q03PW8
Q03PW9 Q03PX0 Q03PX1 Q03PX2 Q03PX3 Q03PX4 Q03PX5 Q03PX6 Q03PX9 Q03PY0 Q03PY1
Q03PY2 Q03PY3 Q03PY4 Q03PY8 Q03PY9 Q03PZ0 Q03PZ7 Q03PZ8 Q03Q10 Q03Q13 Q03Q16
Q03Q17 Q03Q37 Q03Q51 Q03Q52 Q03Q56 Q03Q74 Q03Q83 Q03QA0 Q03QA4 Q03QD4 Q03QD7
Q03QE7 Q03QF3 Q03QG9 Q03QH0 Q03QH9 Q03QI3 Q03QJ0 Q03QK0 Q03QL1 Q03QL3 Q03QM3

Q03QM5 Q03QN1 Q03QN2 Q03QN5 Q03QN6 Q03QN7 Q03QN8 Q03QP1 Q03QP2 Q03QP5 Q03QP8
Q03QR3 Q03QS1 Q03QS2 Q03QS3 Q03QS8 Q03QS9 Q03QT1 Q03QT5 Q03QT6 Q03QT7 Q03QU1
Q03QU2 Q03QV7 Q03QX7 Q03QY0 Q03QY1 Q03QZ9 Q03R06 Q03R08 Q03R12 Q03R28 Q03R29
Q03R33 Q03R34 Q03R35 Q03R36 Q03R37 Q03R43 Q03R44 Q03R45 Q03R46 Q03R49 Q03R52
Q03R67 Q03R97 Q03RH7 Q03RJ2 Q03RK0 Q03RK3 Q03RK5 Q03RK6 Q03RL4 Q03RL6 Q03RM5
Q03RM9 Q03RN9 Q03RP7 Q03RQ4 Q03RR0 Q03RR1 Q03RR3 Q03RS0 Q03RS3 Q03RT0 Q03RU4
Q03RU6 Q03RU7 Q03RU8 Q03S19 Q03S30 Q03S31 Q03S32 Q03S45 Q03S48 Q03S83 Q03S89
Q03S95 Q03S97 Q03SA1 Q03SC5 Q03SC6 Q03SC7 Q03SD0 Q03SD5 Q03SD6 Q03SE4 Q03SF0
Q03SF4 Q03SJ3 Q03SJ8 Q03SK6 Q03SK9 Q03SL6 Q03SL7 Q03SM3 Q03SM4 Q03SM6 Q03SM9
Q03SQ9 Q03SR0 Q03SR1 Q03SR5 Q03SS3 Q03SS5 Q03ST6 Q03ST7 Q03ST8 Q03ST9 Q03SU2
Q03SU6 Q03SU7 Q03SV2 Q03SX9 Q03SZ0 Q03SZ9 Q03T06 Q03T19 Q03T27 Q03T54 Q03T56
Q03TE7 Q03TS9 Q03TT0 Q03TX3 Q03U06 Q03U25 Q03U68 Q03UB1 Q03UC7 Q03UD6 Q03UD8
Q03UE1 Q03UE4 Q03UI8 Q03UI9 Q03UJ4 Q03UK8 Q03UL2 Q03UM2 Q03UM3 Q03UT3 Q03UT5
Q03UU4 Q03UV6 Q03UV9 Q03UX8 Q03V53 Q03V60 Q03V79 Q03VA1 Q03VB0 Q03VB1 Q03VC2
Q03VC3 Q03VF1 Q03VF5 Q03VG0 Q03VG4 Q03VG8 Q03VH8 Q03VI2 Q03VI4 Q03VI7 Q03VQ8
Q03VR7 Q03VS7 Q03VT9 Q03VU7 Q03VU8 Q03VV4 Q03VW0 Q03VW4 Q03VW5 Q03VX7 Q03VX9
Q03VY0 Q03VY2 Q03VY3 Q03W08 Q03W09 Q03W10 Q03W20 Q03W24 Q03W29 Q03W30 Q03W34
Q03W39 Q03W45 Q03W48 Q03W49 Q03W51 Q03W52 Q03W54 Q03W55 Q03W57 Q03W63 Q03W65
Q03W69 Q03W70 Q03W71 Q03W83 Q03W87 Q03W88 Q03WF7 Q03WG3 Q03WG8 Q03WG9 Q03WH0
Q03WH4 Q03WH5 Q03WI1 Q03WI2 Q03WL0 Q03WL7 Q03WM2 Q03WM8 Q03WN2 Q03WN4 Q03WU9
Q03WV9 Q03WW2 Q03WW4 Q03WW7 Q03WX5 Q03WX6 Q03WX7 Q03WX8 Q03WY0 Q03WY1 Q03WZ4
Q03WZ9 Q03X00 Q03X08 Q03X09 Q03X10 Q03XM8 Q03XW2 Q03XY9 Q03Y25 Q03Y28 Q03Y32
Q03Y35 Q03Y48 Q03Y50 Q03Y57 Q03Y61 Q03Y85 Q03Y89 Q03YA3 Q03YB1 Q03YC4 Q03YC9
Q03YD3 Q03YE0 Q03YE2 Q03YE3 Q03YE4 Q03YF0 Q03YF8 Q03YG0 Q03YH4 Q03YI2 Q03YJ2
Q03YJ6 Q03YJ8 Q03YK1 Q03YP0 Q03YP9 Q03YQ0 Q03YQ1 Q03YQ2 Q03YT6 Q03YU3 Q03YW0
Q03YX1 Q03YY4 Q03YZ6 Q03YZ8 Q03Z06 Q03Z41 Q03Z44 Q03Z52 Q03Z55 Q03Z94 Q03Z95
Q03ZA4 Q03ZE9 Q03ZF1 Q03ZG6 Q03ZG8 Q03ZH3 Q03ZH4 Q03ZH5 Q03ZH6 Q03ZH9 Q03ZI1
Q03ZL1 Q03ZL3 Q03ZL9 Q03ZM0 Q03ZM1 Q03ZM2 Q03ZM3 Q03ZM4 Q03ZM5 Q03ZM7 Q03ZM8
Q03ZM9 Q03ZN0 Q03ZN1 Q03ZN2 Q03ZN3 Q03ZN4 Q03ZN5 Q03ZN6 Q03ZN7 Q03ZN8 Q03ZN9
Q03ZP0 Q03ZP1 Q03ZP2 Q03ZP3 Q03ZP4 Q03ZP5 Q03ZP6 Q03ZQ2 Q03ZQ3 Q03ZQ4 Q03ZQ7
Q03ZQ9 Q03ZR0 Q03ZR1 Q04002 Q04003 Q04005 Q04019 Q04031 Q04047 Q04048 Q04049
Q04052 Q04062 Q04066 Q04067 Q04073 Q04081 Q04087 Q04088 Q04089 Q040B2 Q040E6
Q040F1 Q040F3 Q040F4 Q040J1 Q040L6 Q040U0 Q040Z7 Q04110 Q04116 Q04120 Q04149
Q04152 Q04158 Q04170 Q04173 Q04177 Q04178 Q04179 Q04183 Q04195 Q04199 Q041F5
Q041J1 Q041K2 Q041K5 Q041K6 Q041L3 Q041R2 Q041S2 Q041T4 Q041U1 Q041U3 Q041U6
Q041V2 Q041W0 Q041W5 Q041W8 Q041X4 Q041Y0 Q041Z0 Q041Z5 Q04201 Q04202 Q04208
Q04211 Q04212 Q04213 Q04215 Q04217 Q04221 Q04222 Q04223 Q04225 Q04226 Q04231
Q04235 Q04264 Q04276 Q04281 Q04285 Q04296 Q04299 Q042A7 Q042B4 Q042D2 Q042E2
Q042E5 Q042E7 Q042E8 Q042F2 Q042F3 Q042F7 Q042G6 Q042H3 Q042I3 Q042K0 Q042K5
Q042K8 Q042M7 Q042N0 Q042P3 Q042P4 Q042P8 Q042Q3 Q042R4 Q042S0 Q042S1 Q042S6
Q042T2 Q042T5 Q042T6 Q042T7 Q042T8 Q042X7 Q042Y1 Q042Y3 Q042Y5 Q04304 Q04305
Q04307 Q04314 Q04336 Q04341 Q04344 Q04351 Q04352 Q04353 Q04354 Q04368 Q04371
Q04373 Q04377 Q04383 Q04385 Q04399 Q043A0 Q043A6 Q043A9 Q043B0 Q043C5 Q043F6
Q043G5 Q043G6 Q043G9 Q043H4 Q043I0 Q043J4 Q043R3 Q043R6 Q043R9 Q043S2 Q043S5

Q043U2 Q043U6 Q043U7 Q043V2 Q043V6 Q043V9 Q043W1 Q043W2 Q043X4 Q043X5 Q043X8
Q04401 Q04406 Q04408 Q04409 Q04411 Q04412 Q04413 Q04416 Q04418 Q04429 Q04430
Q04432 Q04434 Q04436 Q04437 Q04438 Q04445 Q04446 Q04448 Q04449 Q04450 Q04451
Q04461 Q04462 Q04471 Q04474 Q04477 Q04489 Q04493 Q04499 Q044A1 Q044A4 Q044A8
Q044A9 Q044B0 Q044B1 Q044B5 Q044B6 Q044B7 Q044C1 Q044C3 Q044C7 Q044C8 Q044C9
Q044D0 Q044E2 Q044E4 Q044E5 Q044G3 Q044G6 Q044H0 Q044H2 Q044H8 Q044H9 Q044I0
Q04500 Q04502 Q04504 Q04505 Q04506 Q04511 Q04512 Q04513 Q04515 Q04518 Q04520
Q04522 Q04524 Q04526 Q04533 Q04535 Q04536 Q04537 Q04539 Q04540 Q04541 Q04542
Q04543 Q04545 Q04546 Q04550 Q04551 Q04554 Q04557 Q04558 Q04564 Q04570 Q04572
Q04574 Q04575 Q04580 Q04581 Q04583 Q04585 Q04590 Q04593 Q04595 Q045B8 Q045L5
Q045M0 Q045P1 Q045P3 Q045P4 Q045P5 Q045P6 Q045Q2 Q045Q3 Q045Q4 Q045Q5 Q045Q8
Q045Q9 Q045R4 Q045S7 Q045S8 Q045T6 Q045U2 Q045U7 Q045U8 Q045V5 Q045W0 Q045W4
Q045X8 Q045X9 Q045Z9 Q04600 Q04601 Q04603 Q04605 Q04607 Q04608 Q04610 Q04621
Q04624 Q04625 Q04628 Q04631 Q04633 Q04634 Q04636 Q04638 Q04644 Q04649 Q04650
Q04652 Q04655 Q04657 Q04659 Q04660 Q04661 Q04666 Q04667 Q04668 Q04673 Q04674
Q04688 Q04689 Q04693 Q04694 Q046A0 Q046A1 Q046A2 Q046A3 Q046A4 Q046A5 Q046A7
Q046A8 Q046A9 Q046B0 Q046B1 Q046B2 Q046B3 Q046B4 Q046B5 Q046B6 Q046B8 Q046B9
Q046C0 Q046C1 Q046C2 Q046C3 Q046C4 Q046C5 Q046C6 Q046C7 Q046C8 Q046C9 Q046D1
Q046D2 Q046D9 Q046E4 Q046E7 Q046F1 Q046F8 Q046I2 Q046I4 Q046V3 Q046Z0 Q04706
Q04708 Q04709 Q04710 Q04712 Q04713 Q04715 Q04716 Q04718 Q04720 Q04723 Q04728
Q04730 Q04731 Q04740 Q04741 Q04742 Q04743 Q04744 Q04747 Q04748 Q04750 Q04751
Q04752 Q04754 Q04755 Q04758 Q04763 Q04764 Q04765 Q04766 Q04768 Q04769 Q04770
Q04772 Q04773 Q04774 Q04775 Q04776 Q04777 Q04778 Q04779 Q04781 Q04786 Q04787
Q04788 Q04789 Q04792 Q04795 Q04796 Q04797 Q047E6 Q047E8 Q047F1 Q047F6 Q047F7
Q047F9 Q047G0 Q047I3 Q047M3 Q047M5 Q047M6 Q047M8 Q047N5 Q047Q7 Q047S1 Q047T2
Q04802 Q04805 Q04806 Q04809 Q04810 Q04811 Q04814 Q04820 Q04829 Q04830 Q04831
Q04832 Q04837 Q04841 Q04845 Q04848 Q04852 Q04855 Q04867 Q04868 Q04869 Q04870
Q04871 Q04890 Q04891 Q04892 Q04893 Q04894 Q04896 Q04897 Q04898 Q048F6 Q048G7
Q048S3 Q048S9 Q048T5 Q048T8 Q048T9 Q048U4 Q048U5 Q048W5 Q048W6 Q048W8 Q048X8
Q048Y2 Q048Y3 Q048Y4 Q048Y5 Q048Y6 Q048Y7 Q048Z3 Q048Z4 Q048Z5 Q048Z6 Q04902
Q04903 Q04907 Q04908 Q04913 Q04922 Q04923 Q04925 Q04930 Q04933 Q04937 Q04939
Q04942 Q04944 Q04945 Q04948 Q04949 Q04953 Q04957 Q04964 Q04965 Q04966 Q04971
Q04972 Q04973 Q04974 Q04975 Q04977 Q04978 Q04980 Q04983 Q04996 Q049B3 Q049B7
Q049D3 Q049E0 Q049E6 Q049E9 Q049F3 Q049F5 Q049G2 Q049G3 Q049G8 Q049I1 Q049I6
Q049K4 Q049L7 Q049L9 Q049M1 Q049M4 Q049M5 Q049N0 Q049N1 Q049N8 Q049P0 Q049Q0
Q049R6 Q049R7 Q049R8 Q049S3 Q049T2 Q049T6 Q049U2 Q049U3 Q049U4 Q049U5 Q049U9
Q049V1 Q049V5 Q049V6 Q049V7 Q049W4 Q049W5 Q049W6 Q049W7 Q049X1 Q049Z4 Q049Z7
Q049Z9 Q04A02 Q04A03 Q04A05 Q04A15 Q04A16 Q04A18 Q04A21 Q04A23 Q04A69 Q04AL4
Q04AM0 Q04AM5 Q04AM6 Q04AN7 Q04AN8 Q04AP3 Q04AV4 Q04AV5 Q04AV7 Q04AY6 Q04B07
Q04B09 Q04B11 Q04B12 Q04B17 Q04B18 Q04B29 Q04B34 Q04B36 Q04B37 Q04B40 Q04B41
Q04B51 Q04B52 Q04B58 Q04B65 Q04B68 Q04B73 Q04B77 Q04B78 Q04B89 Q04B92 Q04BB0
Q04BB3 Q04BD3 Q04BF6 Q04BG3 Q04BG7 Q04BH0 Q04BH1 Q04BH7 Q04BH8 Q04BI0 Q04BJ8
Q04BL3 Q04BM6 Q04BN0 Q04BQ8 Q04BV6 Q04BV7 Q04BV8 Q04BW0 Q04BY4 Q04BY8 Q04BY9
Q04BZ0 Q04BZ1 Q04BZ2 Q04BZ3 Q04BZ4 Q04BZ6 Q04BZ7 Q04BZ8 Q04BZ9 Q04C00 Q04C01

Q04C02 Q04C03 Q04C04 Q04C05 Q04C06 Q04C07 Q04C08 Q04C09 Q04C10 Q04C11 Q04C12
Q04C13 Q04C14 Q04C15 Q04C16 Q04C17 Q04C18 Q04C19 Q04C21 Q04C22 Q04C36 Q04C43
Q04C45 Q04C49 Q04C51 Q04C52 Q04C60 Q04CA4 Q04CA6 Q04CB0 Q04CC7 Q04CI2 Q04CJ7
Q04CM2 Q04CN9 Q04CQ7 Q04CW5 Q04CW7 Q04CX2 Q04CX5 Q04CX6 Q04CX7 Q04D20 Q04D35
Q04D45 Q04D49 Q04D98 Q04DA2 Q04DA3 Q04DD1 Q04DD2 Q04DK7 Q04DP1 Q04DQ8 Q04DR8
Q04DZ9 Q04E23 Q04E43 Q04E44 Q04E45 Q04E51 Q04E53 Q04E61 Q04E62 Q04E63 Q04E64
Q04E66 Q04E78 Q04E85 Q04E86 Q04ED4 Q04ED5 Q04ED6 Q04ED7 Q04EE0 Q04EE1 Q04EG2
Q04EG5 Q04EG6 Q04EH0 Q04EH1 Q04EH7 Q04EI0 Q04EI9 Q04EK0 Q04EL5 Q04EQ9 Q04ER0
Q04ER1 Q04ER2 Q04ES4 Q04ES5 Q04ES9 Q04ET4 Q04EU0 Q04EU6 Q04EU8 Q04EV5 Q04EX6
Q04EX9 Q04EY0 Q04EZ7 Q04EZ8 Q04EZ9 Q04F01 Q04F24 Q04F27 Q04F33 Q04F34 Q04F46
Q04F69 Q04F71 Q04F73 Q04F76 Q04F79 Q04F81 Q04F85 Q04F86 Q04F87 Q04F88 Q04FB0
Q04FB2 Q04FB3 Q04FB5 Q04FB7 Q04FC0 Q04FD8 Q04FE3 Q04FF4 Q04FF6 Q04FL2 Q04FL3
Q04FP5 Q04FP6 Q04FP8 Q04FQ4 Q04FR4 Q04FR7 Q04FS1 Q04FS2 Q04FS3 Q04FS5 Q04FS6
Q04FS7 Q04G05 Q04G13 Q04G28 Q04G36 Q04G42 Q04G43 Q04G44 Q04G51 Q04G52 Q04G54
Q04G59 Q04G60 Q04G61 Q04G62 Q04G63 Q04G64 Q04G65 Q04G67 Q04G68 Q04G69 Q04G70
Q04G71 Q04G72 Q04G73 Q04G74 Q04G75 Q04G76 Q04G77 Q04G78 Q04G79 Q04G80 Q04G81
Q04G82 Q04G83 Q04G84 Q04G85 Q04G86 Q04GA1 Q04GA2 Q04GA9 Q04GC3 Q04GD7 Q04GD9
Q04GL2 Q04GL9 Q04GM3 Q04GM9 Q04GN0 Q04GN4 Q04GN6 Q04GP2 Q04GX5 Q04GY7 Q04GY8
Q04H28 Q04H29 Q04H32 Q04H41 Q04HB6 Q04HB7 Q04HC1 Q04HC3 Q04HC7 Q04HG5 Q04HG9
Q04HN0 Q04HP9 Q04HQ3 Q04HQ5 Q04HQ7 Q04HR3 Q04HR6 Q04HS5 Q04HS9 Q04HU2 Q04HV1
Q04HW3 Q04HW4 Q04HX2 Q04HY8 Q04HZ0 Q04HZ7 Q04I02 Q04I16 Q04I25 Q04I36 Q04I37
Q04I44 Q04I50 Q04I59 Q04I74 Q04I77 Q04I78 Q04I94 Q04I96 Q04IA2 Q04IA3 Q04IB3
Q04IC8 Q04ID0 Q04IH6 Q04II4 Q04IJ8 Q04IK0 Q04IK6 Q04IK7 Q04IK8 Q04IM5 Q04IM6
Q04IN9 Q04IQ2 Q04IQ3 Q04IQ6 Q04IQ7 Q04IS7 Q04IT1 Q04IT2 Q04IT3 Q04IV9 Q04IY6
Q04IY7 Q04IY8 Q04IY9 Q04IZ0 Q04J12 Q04J33 Q04J38 Q04J40 Q04J42 Q04J60 Q04J64
Q04J74 Q04J97 Q04J98 Q04JA5 Q04JB0 Q04JB4 Q04JB8 Q04JC5 Q04JD7 Q04JE2 Q04JF3
Q04JF7 Q04JG7 Q04JH1 Q04JH2 Q04JH4 Q04JH9 Q04JI0 Q04JI2 Q04JI4 Q04JI5 Q04JI8
Q04JJ1 Q04JK0 Q04JK7 Q04JK8 Q04JL0 Q04JL4 Q04JL5 Q04JL8 Q04JM7 Q04JM8 Q04JM9
Q04JN5 Q04JN6 Q04JP7 Q04JQ8 Q04JT1 Q04JT4 Q04JT5 Q04JT6 Q04JT7 Q04JU6 Q04JV9
Q04JW5 Q04JX0 Q04JX1 Q04JX2 Q04JX3 Q04JX5 Q04JX6 Q04JY5 Q04JY7 Q04JZ3 Q04JZ4
Q04K26 Q04K32 Q04K39 Q04K45 Q04K46 Q04K48 Q04K52 Q04K71 Q04K72 Q04K81 Q04K90
Q04K99 Q04KA7 Q04KA9 Q04KB0 Q04KB1 Q04KC1 Q04KC2 Q04KC3 Q04KE2 Q04KF1 Q04KF4
Q04KF8 Q04KF9 Q04KG5 Q04KG7 Q04KG8 Q04KH2 Q04KI2 Q04KI4 Q04KI6 Q04KI9 Q04KJ6
Q04KJ8 Q04KK7 Q04KL8 Q04KR0 Q04KR4 Q04KR8 Q04KR9 Q04KS4 Q04KS5 Q04KV3 Q04KW1
Q04KW7 Q04KW8 Q04KW9 Q04KX8 Q04KY0 Q04KY1 Q04KY2 Q04KY8 Q04KZ0 Q04KZ2 Q04KZ3
Q04L01 Q04L24 Q04L26 Q04L36 Q04L55 Q04L69 Q04L73 Q04L74 Q04L76 Q04L81 Q04L82
Q04L84 Q04LC5 Q04LC6 Q04LD0 Q04LD5 Q04LE0 Q04LF3 Q04LF4 Q04LH2 Q04LH9 Q04LI2
Q04LJ2 Q04LJ3 Q04LK1 Q04LL3 Q04LL6 Q04LL8 Q04LM3 Q04LP6 Q04LP7 Q04LQ9 Q04LT4
Q04LT6 Q04LU1 Q04LV9 Q04LW0 Q04LW4 Q04LW5 Q04LY0 Q04LY1 Q04LY2 Q04LZ5 Q04LZ9
Q04M21 Q04M32 Q04M37 Q04M39 Q04M40 Q04M41 Q04M42 Q04M43 Q04M45 Q04M63 Q04M67
Q04M70 Q04M73 Q04M75 Q04M86 Q04M95 Q04M96 Q04M97 Q04MC1 Q04MC7 Q04MF5 Q04MF9
Q04MH6 Q04MH7 Q04MH8 Q04MH9 Q04MI4 Q04MI9 Q04MJ2 Q04ML0 Q04ML1 Q04ML2 Q04ML3
Q04ML4 Q04ML5 Q04ML7 Q04ML8 Q04ML9 Q04MM0 Q04MM1 Q04MM2 Q04MM3 Q04MM4 Q04MM5
Q04MM6 Q04MM7 Q04MM8 Q04MM9 Q04MN0 Q04MN1 Q04MN2 Q04MN3 Q04MN4 Q04MN5 Q04MN6

Q04MN7 Q04MP7 Q04MP8 Q04MP9 Q04MQ9 Q04MR3 Q04MR4 Q04MU2 Q04MU7 Q04MU9 Q04MV1
Q04MW6 Q04MW7 Q04MY5 Q04MZ3 Q04N19 Q04N22 Q04N25 Q04N32 Q04N48 Q04N53 Q04N59
Q04N63 Q04N79 Q04NB1 Q04NB4 Q04NC1 Q04NC2 Q04NC3 Q04NF4 Q04NG3 Q04NM2 Q04NM9
Q04NN6 Q04NU7 Q04NV4 Q04NX0 Q04NX9 Q04NY0 Q04P10 Q04P11 Q04P21 Q04P35 Q04P36
Q04P56 Q04P74 Q04P84 Q04P85 Q04P86 Q04PB3 Q04PD9 Q04PF0 Q04PH9 Q04PM6 Q04PM7
Q04PM8 Q04PM9 Q04PP1 Q04PP8 Q04PT6 Q04PT7 Q04PT8 Q04PT9 Q04PU0 Q04PU1 Q04PU2
Q04PU3 Q04PU4 Q04PU5 Q04PU6 Q04PU7 Q04PU8 Q04PV0 Q04PV1 Q04PV2 Q04PV3 Q04PV4
Q04PV5 Q04PV6 Q04PV7 Q04PV9 Q04PW0 Q04PW1 Q04PW2 Q04PW3 Q04PW4 Q04PW5 Q04PW6
Q04Q05 Q04Q48 Q04Q66 Q04Q78 Q04Q79 Q04Q87 Q04Q89 Q04Q90 Q04Q92 Q04Q93 Q04Q98
Q04QD0 Q04QG5 Q04QG6 Q04QG7 Q04QI5 Q04QI6 Q04QI7 Q04QI9 Q04QJ0 Q04QJ1 Q04QJ2
Q04QJ9 Q04QM3 Q04QQ5 Q04QS3 Q04QS4 Q04QS6 Q04QV4 Q04QV5 Q04QV8 Q04QW2 Q04QW8
Q04QY0 Q04R21 Q04R23 Q04R37 Q04R46 Q04R57 Q04R58 Q04R64 Q04R73 Q04RA5 Q04RD6
Q04RG9 Q04RH4 Q04RK6 Q04RK8 Q04RM0 Q04RM8 Q04RN4 Q04RN8 Q04RP4 Q04RQ6 Q04RS5
Q04RS7 Q04RS9 Q04RT2 Q04RT3 Q04RU2 Q04RU6 Q04RW4 Q04S01 Q04S02 Q04S66 Q04S67
Q04S98 Q04SA4 Q04SA5 Q04SA6 Q04SB0 Q04SB2 Q04SB3 Q04SB8 Q04SE4 Q04SK8 Q04SM4
Q04SV2 Q04SV8 Q04SW1 Q04SW3 Q04SW4 Q04SX0 Q04T16 Q04T19 Q04T45 Q04T48 Q04T56
Q04T59 Q04T63 Q04TA2 Q04TA4 Q04TB1 Q04TB2 Q04TD1 Q04TD5 Q04TD7 Q04TE9 Q04TF0
Q04TG1 Q04TG2 Q04TG5 Q04TH0 Q04TH1 Q04TH3 Q04TR3 Q04TR5 Q04TV4 Q04TV5 Q04TZ4
Q04U27 Q04U28 Q04U30 Q04U31 Q04U33 Q04U37 Q04U38 Q04U41 Q04U42 Q04U54 Q04U55
Q04U56 Q04U59 Q04U62 Q04U63 Q04U64 Q04U66 Q04U69 Q04U70 Q04U71 Q04U72 Q04U79
Q04U80 Q04UA3 Q04UA7 Q04UC6 Q04UG1 Q04UG6 Q04UI9 Q04UJ2 Q04UL2 Q04UL5 Q04UL7
Q04UM1 Q04UY8 Q04V34 Q04V47 Q04V49 Q04V79 Q04V89 Q04V91 Q04V97 Q04VC7 Q04VC8
Q04VC9 Q04VD0 Q04VF1 Q04VG7 Q04VH3 Q04VI6 Q04VK1 Q04VM3 Q04VP5 Q04VR0 Q04VV0
Q04W02 Q04W28 Q04W32 Q04W40 Q04W48 Q04W49 Q04W54 Q04W72 Q04W74 Q04W81 Q04W85
Q04W97 Q04WA6 Q04WC4 Q04WD4 Q04WE1 Q04WF5 Q04WF7 Q04WG0 Q04WJ1 Q04WJ4 Q04WK0
Q04WK1 Q04WK2 Q04WN5 Q04WP5 Q04WV5 Q04WW3 Q04WX0 Q04X34 Q04X52 Q04X60 Q04X69
Q04X87 Q04X97 Q04XA4 Q04XA6 Q04XB2 Q04XC3 Q04XC4 Q04XD2 Q04XE0 Q04XE4 Q04XH0
Q04XL9 Q04XR1 Q04XS7 Q04XV1 Q04XX5 Q04XY8 Q04Y01 Q04Y07 Q04Y23 Q04Y45 Q04Y46
Q04Y47 Q04Y48 Q04Y79 Q04Y85 Q04Y87 Q04Y97 Q04YC8 Q04YD0 Q04YE4 Q04YI9 Q04YV5
Q04YV8 Q04YW0 Q04YW2 Q04YY2 Q04YY5 Q04Z09 Q04Z14 Q04Z48 Q04Z61 Q04Z62 Q04Z65
Q04Z69 Q04Z75 Q04Z87 Q04ZD0 Q04ZD2 Q04ZE7 Q04ZF5 Q04ZG6 Q04ZG7 Q04ZG9 Q04ZH0
Q04ZH1 Q04ZI3 Q04ZI4 Q04ZI7 Q04ZI8 Q04ZJ3 Q04ZJ5 Q04ZJ6 Q04ZJ8 Q04ZJ9 Q04ZN1
Q04ZS0 Q04ZS1 Q04ZZ3 Q04ZZ4 Q05000 Q05006 Q05007 Q05015 Q05016 Q05021 Q05022
Q05024 Q05026 Q05027 Q05036 Q05043 Q05045 Q05046 Q05048 Q05053 Q05054 Q05055
Q05064 Q05065 Q05067 Q05068 Q05071 Q05072 Q05079 Q05092 Q05095 Q05099 Q050C4
Q050D0 Q050D1 Q050D2 Q050D6 Q050D8 Q050D9 Q050E4 Q050H2 Q050N5 Q050Q1 Q050X6
Q050Y2 Q050Y5 Q050Y7 Q050Y8 Q050Z4 Q05100 Q05101 Q05102 Q05103 Q05104 Q05105
Q05106 Q05107 Q05113 Q05114 Q05115 Q05118 Q05120 Q05121 Q05122 Q05123 Q05124
Q05125 Q05126 Q05127 Q05129 Q05136 Q05137 Q05138 Q05142 Q05145 Q05159 Q05174
Q05182 Q05183 Q05184 Q05185 Q05190 Q05191 Q05192 Q05196 Q051D8 Q051E1 Q051G7
Q051H0 Q051H8 Q051I1 Q051I5 Q051M4 Q051M6 Q051N3 Q051N4 Q051Q7 Q051S5 Q051S9
Q051T8 Q051T9 Q051U2 Q051U4 Q051U6 Q051W5 Q051X7 Q051Y3 Q051Y7 Q051Z3 Q05201
Q05212 Q05213 Q05214 Q05215 Q05218 Q05219 Q05220 Q05221 Q05222 Q05223 Q05224
Q05225 Q05226 Q05227 Q05228 Q05229 Q05230 Q05231 Q05232 Q05233 Q05234 Q05235

Q05236 Q05239 Q05240 Q05241 Q05242 Q05245 Q05246 Q05247 Q05248 Q05249 Q05250
Q05251 Q05252 Q05253 Q05254 Q05255 Q05256 Q05257 Q05258 Q05259 Q05261 Q05262
Q05263 Q05264 Q05265 Q05266 Q05267 Q05268 Q05269 Q05270 Q05271 Q05272 Q05273
Q05274 Q05275 Q05276 Q05277 Q05278 Q05279 Q05280 Q05281 Q05282 Q05283 Q05284
Q05286 Q05287 Q05288 Q05289 Q05290 Q05291 Q05292 Q05293 Q05294 Q05295 Q05297
Q05298 Q052A1 Q052B3 Q052B5 Q052F7 Q052G1 Q052G3 Q052H7 Q052H8 Q052J0 Q052J1
Q052J4 Q052J9 Q052K0 Q052K2 Q052U5 Q052U7 Q052X8 Q052X9 Q05300 Q05301 Q05302
Q05303 Q05304 Q05305 Q05309 Q05310 Q05311 Q05318 Q05323 Q05326 Q05327 Q05335
Q05338 Q05349 Q05353 Q05354 Q05355 Q05356 Q05357 Q05358 Q05361 Q05362 Q05363
Q05368 Q05370 Q05380 Q05381 Q053A4 Q053A8 Q053E2 Q053G0 Q053H5 Q053K4 Q053L4
Q053M2 Q053M5 Q053M6 Q053M7 Q053M9 Q053N2 Q053N3 Q053N4 Q053P2 Q053P3 Q053R1
Q053R5 Q053V3 Q053V5 Q053V6 Q053X4 Q05404 Q05423 Q05426 Q05427 Q05428 Q05437
Q05451 Q05459 Q05462 Q05466 Q05468 Q05470 Q05471 Q05473 Q05474 Q05481 Q05486
Q05488 Q05491 Q05492 Q05498 Q054A8 Q054D2 Q054D9 Q054E0 Q054E1 Q054E2 Q054E4
Q054E5 Q054E6 Q054G5 Q054G6 Q054G7 Q054K1 Q054N4 Q054N5 Q054P3 Q054P5 Q054P6
Q054P8 Q054P9 Q054Q4 Q054R6 Q054T5 Q054X8 Q05500 Q05501 Q05502 Q05506 Q05509
Q05510 Q05514 Q05515 Q05522 Q05526 Q05527 Q05528 Q05529 Q05530 Q05531 Q05534
Q05536 Q05541 Q05542 Q05543 Q05547 Q05549 Q05554 Q05557 Q05558 Q05559 Q05561
Q05562 Q05566 Q05569 Q05579 Q05580 Q05581 Q05582 Q05583 Q05584 Q05587 Q05590
Q05591 Q05592 Q05593 Q05597 Q05599 Q055B7 Q055B8 Q055B9 Q055C0 Q055C1 Q055C2
Q055C6 Q055C7 Q055D0 Q055D4 Q055D7 Q055D8 Q055D9 Q055E0 Q055E1 Q055E4 Q055E6
Q055M2 Q055M9 Q055P6 Q055P7 Q055P8 Q055P9 Q055U6 Q055X5 Q055Y6 Q05601 Q05603
Q05615 Q05620 Q05621 Q05623 Q05626 Q05627 Q05628 Q05631 Q05632 Q05634 Q05635
Q05636 Q05640 Q05646 Q05647 Q05650 Q05651 Q05670 Q05672 Q05681 Q056E8 Q056E9
Q056F0 Q056G0 Q056H8 Q056J8 Q056J9 Q056L2 Q056M2 Q056M3 Q056Q1 Q056Q2 Q056R5
Q056R9 Q056S6 Q056S8 Q056V0 Q056V2 Q056V5 Q056W3 Q056X2 Q056X3 Q056X4 Q056X6
Q056X7 Q056X8 Q056X9 Q056Y0 Q056Y6 Q056Z1 Q056Z3 Q056Z9 Q05709 Q05714 Q05728
Q05733 Q05736 Q05739 Q05741 Q05746 Q05749 Q05753 Q05755 Q05756 Q05758 Q05761
Q05762 Q05763 Q05766 Q05775 Q05778 Q05781 Q05783 Q05788 Q05791 Q05795 Q05796
Q05797 Q05798 Q05799 Q057A0 Q057A1 Q057A2 Q057A3 Q057A4 Q057A5 Q057A7 Q057A8
Q057A9 Q057B0 Q057B1 Q057B2 Q057B3 Q057B4 Q057B5 Q057B6 Q057B7 Q057B8 Q057C1
Q057C2 Q057C3 Q057C5 Q057C6 Q057C7 Q057C8 Q057C9 Q057D2 Q057D3 Q057D5 Q057D6
Q057D7 Q057D8 Q057D9 Q057E0 Q057F3 Q057G3 Q057G4 Q057G5 Q057G7 Q057H7 Q057H8
Q057I1 Q057I2 Q057I4 Q057I5 Q057I8 Q057J0 Q057J1 Q057J2 Q057J4 Q057J5 Q057K0
Q057K2 Q057K3 Q057L5 Q057M1 Q057M2 Q057M4 Q057N0 Q057N2 Q057N5 Q057N6 Q057N8
Q057N9 Q057P0 Q057P1 Q057P9 Q057Q1 Q057Q7 Q057Q9 Q057R1 Q057R4 Q057S1 Q057S8
Q057S9 Q057T0 Q057T1 Q057T6 Q057T9 Q057U7 Q057V0 Q057V1 Q057V4 Q057V8 Q057W1
Q057X7 Q057X8 Q057X9 Q057Y0 Q057Y8 Q057Z1 Q057Z2 Q05800 Q05801 Q05802 Q05803
Q05805 Q05815 Q05823 Q05831 Q05832 Q05840 Q05854 Q05856 Q05863 Q05865 Q05869
Q05870 Q05873 Q05874 Q05879 Q05880 Q05881 Q05885 Q05890 Q05892 Q05893 Q058A3
Q058A6 Q058B0 Q058B1 Q058B2 Q058B4 Q058B8 Q058B9 Q058C8 Q058C9 Q058D0 Q058D1
Q058D2 Q058D5 Q058D6 Q058E2 Q058E3 Q058E4 Q058E5 Q058E6 Q058E7 Q058F3 Q058F4
Q058F5 Q058F8 Q058F9 Q058G2 Q058N0 Q058P4 Q05900 Q05906 Q05907 Q05911 Q05913
Q05916 Q05917 Q05921 Q05924 Q05926 Q05931 Q05933 Q05935 Q05936 Q05937 Q05942

Q05943 Q05944 Q05945 Q05946 Q05947 Q05948 Q05949 Q05950 Q05954 Q05957 Q05959
Q05963 Q05964 Q05965 Q05966 Q05971 Q05972 Q05979 Q05980 Q05981 Q05983 Q05984
Q05985 Q05986 Q05987 Q05988 Q05989 Q05990 Q05991 Q05992 Q05993 Q05994 Q05995
Q059U7 Q05A36 Q05A62 Q05AH6 Q05AK5 Q05AK9 Q05AL1 Q05AL2 Q05AM5 Q05AQ8 Q05AT9
Q05AV0 Q05AV1 Q05AW7 Q05AW9 Q05AX3 Q05AY2 Q05B17 Q05B18 Q05B44 Q05B49 Q05B50
Q05B51 Q05B56 Q05B58 Q05B60 Q05B63 Q05B65 Q05B78 Q05B83 Q05B84 Q05B87 Q05B89
Q05B92 Q05BC3 Q05BQ5 Q05BU3 Q05BV3 Q05C16 Q05CL8 Q05D44 Q05D60 Q05DH4 Q05E21
Q05E29 Q05FH8 Q05FH9 Q05FI0 Q05FI1 Q05FI2 Q05FI3 Q05FI7 Q05FI8 Q05FJ0 Q05FJ1
Q05FJ3 Q05FJ6 Q05FJ9 Q05FK2 Q05FK3 Q05FK4 Q05FK5 Q05FN6 Q05FP5 Q05FQ0 Q05FR1
Q05FR8 Q05FS8 Q05FT9 Q05FX0 Q05FY8 Q05FY9 Q05HP6 Q05HS6 Q05HS7 Q05HZ7 Q05I08
Q05I49 Q05I79 Q05I90 Q06004 Q06005 Q06009 Q06012 Q06013 Q06014 Q06015 Q06016
Q06021 Q06022 Q06023 Q06024 Q06025 Q06030 Q06032 Q06036 Q06053 Q06060 Q06063
Q06069 Q06070 Q06071 Q06078 Q06085 Q06086 Q06087 Q06088 Q06091 Q06092 Q06095
Q06098 Q06099 Q06101 Q06102 Q06103 Q06106 Q06117 Q06118 Q06121 Q06125 Q06126
Q06128 Q06129 Q06132 Q06137 Q06139 Q06146 Q06149 Q06150 Q06151 Q06152 Q06156
Q06157 Q06160 Q06162 Q06163 Q06164 Q06168 Q06171 Q06172 Q06174 Q06176 Q06177
Q06178 Q06182 Q06183 Q06184 Q06188 Q06190 Q06191 Q06193 Q06196 Q06197 Q06198
Q06199 Q06201 Q06203 Q06204 Q06205 Q06206 Q06208 Q06211 Q06213 Q06214 Q06215
Q06216 Q06217 Q06218 Q06224 Q06234 Q06237 Q06238 Q06241 Q06244 Q06248 Q06250
Q06251 Q06253 Q06254 Q06255 Q06259 Q06260 Q06261 Q06262 Q06266 Q06267 Q06280
Q06281 Q06287 Q06309 Q06323 Q06331 Q06333 Q06337 Q06339 Q06340 Q06344 Q06348
Q06349 Q06355 Q06360 Q06364 Q06366 Q06372 Q06373 Q06374 Q06375 Q06378 Q06382
Q06383 Q06388 Q06390 Q06392 Q06398 Q06402 Q06406 Q06408 Q06411 Q06416 Q06419
Q06422 Q06423 Q06424 Q06425 Q06426 Q06427 Q06429 Q06433 Q06434 Q06436 Q06437
Q06440 Q06445 Q06447 Q06449 Q06453 Q06457 Q06458 Q06463 Q06464 Q06469 Q06479
Q06488 Q06489 Q06494 Q06504 Q06505 Q06506 Q06509 Q06511 Q06512 Q06523 Q06524
Q06525 Q06528 Q06539 Q06542 Q06543 Q06544 Q06546 Q06547 Q06549 Q06552 Q06554
Q06566 Q06568 Q06578 Q06579 Q06580 Q06581 Q06588 Q06592 Q06594 Q06597 Q06607
Q06608 Q06610 Q06615 Q06623 Q06624 Q06625 Q06629 Q06631 Q06632 Q06639 Q06640
Q06657 Q06659 Q06665 Q06666 Q06667 Q06672 Q06674 Q06675 Q06679 Q06680 Q06682
Q06683 Q06685 Q06687 Q06688 Q06691 Q06692 Q06695 Q06697 Q06698 Q06700 Q06706
Q06707 Q06709 Q06710 Q06712 Q06715 Q06716 Q06717 Q06718 Q06721 Q06725 Q06726
Q06727 Q06728 Q06730 Q06732 Q06734 Q06736 Q06741 Q06750 Q06752 Q06754 Q06755
Q06756 Q06774 Q06795 Q06796 Q06797 Q06798 Q06801 Q06803 Q06809 Q06811 Q06813
Q06817 Q06819 Q06820 Q06821 Q06822 Q06823 Q06827 Q06830 Q06834 Q06835 Q06841
Q06842 Q06847 Q06877 Q06879 Q06889 Q06891 Q06892 Q06903 Q06905 Q06909 Q06911
Q06923 Q06925 Q06929 Q06930 Q06931 Q06934 Q06938 Q06942 Q06943 Q06952 Q06965
Q06975 Q06996 Q06997 Q06998 Q06A28 Q06A37 Q06A98 Q06AA1 Q06AA3 Q06AA4 Q06AK7
Q06AN9 Q06AT9 Q06AU9 Q06AV1 Q06AV5 Q06BR1 Q06DK3 Q06DK7 Q06FK5 Q06FM0 Q06FM2
Q06FM3 Q06FM6 Q06FM7 Q06FM8 Q06FM9 Q06FN0 Q06FN1 Q06FN2 Q06FN4 Q06FN6 Q06FP3
Q06FP4 Q06FP5 Q06FP7 Q06FR2 Q06FV8 Q06FW2 Q06FW3 Q06FW9 Q06FX0 Q06FX1 Q06FX2
Q06FX9 Q06FY0 Q06GJ5 Q06GJ6 Q06GJ7 Q06GJ8 Q06GK0 Q06GK3 Q06GL2 Q06GL9 Q06GM0
Q06GM1 Q06GM2 Q06GM3 Q06GM4 Q06GM5 Q06GM6 Q06GM7 Q06GM8 Q06GN5 Q06GN6 Q06GN7
Q06GN8 Q06GN9 Q06GQ2 Q06GQ8 Q06GR2 Q06GR8 Q06GR9 Q06GS0 Q06GS1 Q06GS8 Q06GS9

Q06GT2 Q06GT3 Q06GT4 Q06GT5 Q06GT7 Q06GT9 Q06GU8 Q06GV6 Q06GV7 Q06GV8 Q06GV9
Q06GW0 Q06GW1 Q06GW2 Q06GW3 Q06GW4 Q06GW5 Q06GX2 Q06GX3 Q06GX4 Q06GX5 Q06GX6
Q06GY9 Q06GZ5 Q06GZ9 Q06H05 Q06H06 Q06H07 Q06H08 Q06H15 Q06H16 Q06J15 Q06J17
Q06J18 Q06J19 Q06J20 Q06J25 Q06J26 Q06J32 Q06J33 Q06J34 Q06J35 Q06J36 Q06J37
Q06J38 Q06J39 Q06J53 Q06J54 Q06J55 Q06J56 Q06J57 Q06J58 Q06J59 Q06J60 Q06J61
Q06J62 Q06J65 Q06R66 Q06R67 Q06R68 Q06R69 Q06R71 Q06R73 Q06R82 Q06R91 Q06R92
Q06R93 Q06R94 Q06R95 Q06R96 Q06R97 Q06R98 Q06R99 Q06RA0 Q06RA8 Q06RA9 Q06RB0
Q06RB1 Q06RC1 Q06RC7 Q06RD3 Q06RD9 Q06RE0 Q06RE1 Q06RE2 Q06RE9 Q06RF0 Q06S87
Q06SC7 Q06SD6 Q06SD8 Q06SE3 Q06SE5 Q06SE7 Q06SF2 Q06SF4 Q06SF5 Q06SF6 Q06SF7
Q06SF9 Q06SG0 Q06SG4 Q06SG5 Q06SG6 Q06SG9 Q06SH0 Q06SH3 Q06SH6 Q06SH7 Q06SH8
Q06SI3 Q06SI4 Q06SI6 Q06SJ3 Q06SJ4 Q06SJ5 Q06VC2 Q06VD3 Q06VD5 Q06VD6 Q06VD9
Q06VK9 Q06VN8 Q06VW1 Q06W39 Q06ZW3 Q07006 Q07010 Q07016 Q07017 Q07048 Q07049
Q07050 Q07051 Q07053 Q07064 Q07068 Q07083 Q07084 Q07088 Q07090 Q07097 Q07099
Q070J7 Q07100 Q07103 Q07120 Q07121 Q07123 Q07130 Q07133 Q07134 Q07135 Q07152
Q07158 Q07159 Q07160 Q07167 Q07177 Q07178 Q07179 Q07180 Q07182 Q07184 Q07187
Q07192 Q07197 Q071N0 Q07200 Q07201 Q07202 Q07204 Q07205 Q07209 Q07210 Q07211
Q07230 Q07231 Q07234 Q07251 Q07254 Q07262 Q07264 Q07271 Q07275 Q07281 Q07283
Q07286 Q07288 Q07291 Q07296 Q07297 Q07322 Q07340 Q07341 Q07342 Q07344 Q07346
Q07350 Q07351 Q07353 Q07362 Q07381 Q07384 Q07385 Q07395 Q07397 Q07398 Q07399
Q07400 Q07401 Q07402 Q07403 Q07410 Q07411 Q07412 Q07415 Q07422 Q07424 Q07427
Q07438 Q07442 Q07447 Q07448 Q07454 Q07457 Q07458 Q07463 Q07472 Q07474 Q07478
Q07500 Q07505 Q07506 Q07511 Q07512 Q07513 Q07517 Q07518 Q07521 Q07527 Q07535
Q07536 Q07538 Q07555 Q07559 Q07586 Q07587 Q07588 Q07589 Q07590 Q075B4 Q07601
Q07602 Q07605 Q07606 Q07607 Q07613 Q07617 Q07623 Q07626 Q07627 Q07630 Q07632
Q07635 Q07637 Q07648 Q07649 Q07655 Q07658 Q07661 Q07683 Q07684 Q07687 Q076A3
Q076A4 Q076A5 Q076A6 Q07703 Q07705 Q07729 Q07736 Q07738 Q07739 Q07740 Q07744
Q07746 Q07747 Q07748 Q07749 Q07750 Q07758 Q07759 Q07760 Q07761 Q07762 Q07765
Q07766 Q07785 Q07786 Q07796 Q07801 Q07802 Q07803 Q07805 Q07809 Q07818 Q07819
Q07821 Q07825 Q07834 Q07836 Q07838 Q07839 Q07840 Q07841 Q07843 Q07844 Q07845
Q07849 Q07850 Q07851 Q07852 Q07853 Q07854 Q07855 Q07856 Q07860 Q07861 Q07873
Q07874 Q07876 Q07886 Q07887 Q07888 Q07897 Q078T0 Q07903 Q07905 Q07906 Q07907
Q07908 Q07913 Q07915 Q07916 Q07921 Q07923 Q07927 Q07930 Q07933 Q07934 Q07935
Q07937 Q07938 Q07940 Q07942 Q07944 Q07945 Q07946 Q07947 Q07951 Q07953 Q07956
Q07957 Q07961 Q07963 Q07967 Q07970 Q07972 Q07975 Q07976 Q07978 Q07979 Q07980
Q07989 Q07993 Q07998 Q07DV1 Q07DV3 Q07DV7 Q07DW1 Q07DW4 Q07DX3 Q07DX4 Q07DX6
Q07DY0 Q07DY3 Q07DY4 Q07DY6 Q07DZ0 Q07DZ4 Q07DZ5 Q07DZ7 Q07E15 Q07E17 Q07E23
Q07E28 Q07E30 Q07E36 Q07E40 Q07E41 Q07E43 Q07E47 Q07FY3 Q07G10 Q07G17 Q07G42
Q07G43 Q07G87 Q07G98 Q07GU5 Q07GW8 Q07GY2 Q07H53 Q07H54 Q07H58 Q07H94 Q07H95
Q07H96 Q07H98 Q07HI5 Q07HS0 Q07HS5 Q07HT7 Q07HZ5 Q07I00 Q07I26 Q07I30 Q07I86
Q07I91 Q07IE7 Q07IG8 Q07IZ1 Q07J10 Q07J18 Q07J29 Q07J43 Q07J51 Q07J67 Q07JJ7
Q07K31 Q07K69 Q07K70 Q07K71 Q07K73 Q07K76 Q07K87 Q07KJ2 Q07KJ7 Q07KJ8 Q07KK5
Q07KK6 Q07KK7 Q07KK8 Q07KL3 Q07KL4 Q07KL5 Q07KL7 Q07KL8 Q07KL9 Q07KM1 Q07KM2
Q07KM3 Q07KM4 Q07KM5 Q07KM6 Q07KM7 Q07KM8 Q07KM9 Q07KN0 Q07KN1 Q07KN2 Q07KN3
Q07KN4 Q07KN5 Q07KN6 Q07KN7 Q07KP0 Q07KP1 Q07KP2 Q07KP3 Q07L14 Q07L15 Q07L16

Q07L24 Q07L39 Q07L51 Q07L69 Q07L79 Q07L84 Q07L90 Q07L94 Q07L96 Q07LA4 Q07LA7
Q07LD6 Q07LE1 Q07LE2 Q07LF4 Q07LG0 Q07LG4 Q07LH5 Q07LH9 Q07LX6 Q07LX8 Q07M57
Q07M58 Q07M65 Q07M67 Q07M70 Q07M79 Q07M86 Q07MA3 Q07MB1 Q07MG7 Q07MJ4 Q07ML3
Q07MN9 Q07MS0 Q07MT5 Q07MT8 Q07MT9 Q07MX7 Q07MY1 Q07MY5 Q07MZ2 Q07MZ5 Q07N39
Q07N42 Q07N44 Q07N47 Q07N61 Q07N65 Q07NE0 Q07NF1 Q07NF3 Q07NF5 Q07NF6 Q07NH8
Q07NI7 Q07NJ0 Q07NJ1 Q07NJ2 Q07NJ3 Q07NK2 Q07NN5 Q07NN6 Q07NN7 Q07P45 Q07PA9
Q07PD0 Q07PI3 Q07PI4 Q07PJ7 Q07PK4 Q07PP0 Q07PR9 Q07PS7 Q07PT0 Q07PT2 Q07PU1
Q07PW9 Q07PX1 Q07PZ6 Q07Q42 Q07Q53 Q07Q82 Q07QF5 Q07QK9 Q07QM9 Q07QP5 Q07QT9
Q07QZ3 Q07R54 Q07R72 Q07R87 Q07R90 Q07RK6 Q07RN1 Q07RP6 Q07RP7 Q07RR2 Q07RW5
Q07RX1 Q07RY2 Q07RY4 Q07RY5 Q07S20 Q07S91 Q07SD5 Q07SR2 Q07SR3 Q07SU0 Q07T66
Q07TA7 Q07TB7 Q07TF9 Q07TH7 Q07TH8 Q07TR7 Q07TW1 Q07TX0 Q07TX2 Q07TX3 Q07TX5
Q07TX9 Q07TY1 Q07TY4 Q07U69 Q07U70 Q07U75 Q07U76 Q07UC7 Q07UC8 Q07UF4 Q07UH0
Q07UH1 Q07UH2 Q07UH4 Q07UH9 Q07UI1 Q07UL7 Q07UN5 Q07UP0 Q07UP1 Q07UP2 Q07UP5
Q07UP9 Q07UQ3 Q07UQ5 Q07UQ8 Q07UQ9 Q07UR1 Q07UR7 Q07US4 Q07US6 Q07UT0 Q07UT2
Q07UT3 Q07UT6 Q07UU1 Q07UU3 Q07UU4 Q07UU6 Q07UU7 Q07UV4 Q07UX5 Q07UX6 Q07V02
Q07V06 Q07V09 Q07V29 Q07V68 Q07V70 Q07V74 Q07V75 Q07V78 Q07V79 Q07V81 Q07V82
Q07V91 Q07VA0 Q07VA9 Q07VB3 Q07VB6 Q07VC9 Q07VE4 Q07VK4 Q07VK8 Q07VM6 Q07VN5
Q07VN9 Q07VS2 Q07VS3 Q07VS4 Q07VS7 Q07VT3 Q07VT4 Q07VV2 Q07VW8 Q07VX0 Q07W23
Q07W34 Q07W36 Q07W45 Q07W50 Q07W60 Q07W63 Q07W69 Q07WC1 Q07WD2 Q07WF3 Q07WF8
Q07WF9 Q07WG1 Q07WG2 Q07WG3 Q07WG9 Q07WH1 Q07WH2 Q07WH4 Q07WH6 Q07WH7 Q07WI6
Q07WM2 Q07WM3 Q07WM5 Q07WM9 Q07WQ2 Q07WS1 Q07WS8 Q07WX6 Q07WX7 Q07X28 Q07X44
Q07XC8 Q07XD8 Q07XP8 Q07XR4 Q07XR8 Q07XR9 Q07XS1 Q07XS7 Q07XS9 Q07XT0 Q07XT3
Q07XW1 Q07XW5 Q07XW6 Q07XW7 Q07XY1 Q07XZ7 Q07XZ8 Q07Y01 Q07Y03 Q07Y38 Q07Y78
Q07Y86 Q07Y91 Q07Y94 Q07YA5 Q07YC7 Q07YC8 Q07YC9 Q07YH6 Q07YI6 Q07YI8 Q07YI9
Q07YJ0 Q07YJ6 Q07YJ7 Q07YJ8 Q07YJ9 Q07YK6 Q07YT2 Q07YU1 Q07YU5 Q07YU9 Q07YV2
Q07YV9 Q07YY0 Q07YY3 Q07YZ6 Q07Z05 Q07Z06 Q07Z07 Q07Z08 Q07Z24 Q07Z25 Q07Z29
Q07Z32 Q07Z51 Q07Z53 Q07Z62 Q07Z75 Q07ZA3 Q07ZD2 Q07ZD3 Q07ZD4 Q07ZD9 Q07ZE4
Q07ZE8 Q07ZF4 Q07ZH1 Q07ZL0 Q07ZM1 Q07ZM2 Q07ZQ7 Q07ZR3 Q07ZS6 Q07ZX5 Q07ZX6
Q07ZX7 Q07ZX8 Q07ZX9 Q08000 Q08014 Q08015 Q08016 Q08018 Q08021 Q08024 Q08032
Q08046 Q08047 Q08051 Q08052 Q08054 Q08066 Q08068 Q08069 Q08079 Q08080 Q08082
Q08091 Q08092 Q08093 Q08094 Q08096 Q08097 Q08099 Q080C3 Q080H8 Q080H9 Q080J1
Q080J4 Q080K9 Q080N2 Q080P6 Q080P9 Q080Q0 Q080Q2 Q080Q3 Q080R2 Q080R4 Q080R9
Q080T4 Q080X8 Q08104 Q08110 Q08111 Q08112 Q08113 Q08114 Q08115 Q08119 Q08121
Q08142 Q08143 Q08162 Q08168 Q08172 Q08182 Q08183 Q08184 Q08185 Q08186 Q08187
Q081C1 Q081F9 Q081G4 Q081G6 Q081G8 Q081H3 Q081H5 Q081H6 Q081H9 Q081I3 Q081J0
Q081K0 Q081L4 Q081M2 Q081N2 Q081N3 Q081N5 Q081N7 Q081N8 Q081N9 Q081Q0 Q081Q4
Q081Q7 Q081U0 Q081U4 Q081U5 Q081U7 Q081X4 Q08200 Q08204 Q08206 Q08208 Q08213
Q08216 Q08220 Q08222 Q08224 Q08225 Q08226 Q08230 Q08231 Q08235 Q08237 Q08241
Q08258 Q08259 Q08262 Q08263 Q08270 Q08274 Q08275 Q08276 Q08278 Q08282 Q08285
Q08287 Q08288 Q08290 Q08291 Q08293 Q08296 Q082E1 Q082E3 Q082E4 Q082E9 Q082N5
Q082Q5 Q082R1 Q082U2 Q082V3 Q08303 Q08304 Q08305 Q08306 Q08307 Q08308 Q08309
Q08310 Q08311 Q08312 Q08313 Q08314 Q08316 Q08317 Q08321 Q08324 Q08325 Q08327
Q08328 Q08329 Q08341 Q08346 Q08347 Q08350 Q08352 Q08356 Q08361 Q08364 Q08365
Q08368 Q08384 Q08385 Q08386 Q08387 Q08392 Q08393 Q083B6 Q083D7 Q083F0 Q083H5

Q083H7 Q083H8 Q083I0 Q083J5 Q083K0 Q083K1 Q083K5 Q083K7 Q083L3 Q083L6 Q083L8
Q083S7 Q083U9 Q08401 Q08402 Q08407 Q08411 Q08412 Q08415 Q08419 Q08421 Q08422
Q08425 Q08427 Q08432 Q08438 Q08444 Q08446 Q08449 Q08454 Q08455 Q08457 Q08465
Q08471 Q08473 Q08478 Q08479 Q08480 Q08486 Q08490 Q08491 Q08492 Q08496 Q084C2
Q084C4 Q084C6 Q084C7 Q084G9 Q084H1 Q084I8 Q084J4 Q084K7 Q084L0 Q084L1 Q084M1
Q084N6 Q084N8 Q084R8 Q084S1 Q084S3 Q084X6 Q084Y3 Q08503 Q08504 Q08506 Q08507
Q08508 Q08512 Q08513 Q08518 Q08519 Q08521 Q08522 Q08530 Q08531 Q08538 Q08539
Q08540 Q08541 Q08543 Q08544 Q08545 Q08553 Q08561 Q08562 Q08581 Q08582 Q08588
Q08590 Q08591 Q08593 Q08597 Q08598 Q085C8 Q085D6 Q085D9 Q085E0 Q085E1 Q085E2
Q085E5 Q085G4 Q085K0 Q085S1 Q085S4 Q085S6 Q085U2 Q085U5 Q085U6 Q085U9 Q085W8
Q085Z8 Q085Z9 Q08600 Q08601 Q08602 Q08603 Q08605 Q08620 Q08621 Q08623 Q08624
Q08628 Q08632 Q08634 Q08638 Q08639 Q08640 Q08641 Q08647 Q08649 Q08652 Q08653
Q08654 Q08655 Q08675 Q08681 Q08682 Q08683 Q08685 Q08686 Q08687 Q08689 Q08695
Q08696 Q08698 Q08699 Q086A0 Q086A1 Q086A4 Q086A5 Q086A7 Q086A8 Q086B1 Q086B4
Q086C3 Q086C4 Q086C8 Q086C9 Q086F7 Q086F8 Q086F9 Q086G0 Q086G5 Q086G7 Q086G9
Q086H0 Q086H1 Q086H2 Q086H4 Q086H6 Q086H7 Q086I0 Q086I8 Q086J2 Q086J3 Q086K1
Q086L3 Q086N8 Q086T6 Q086W4 Q086X9 Q08702 Q08704 Q08705 Q08713 Q08715 Q08716
Q08723 Q08726 Q08727 Q08728 Q08729 Q08742 Q08746 Q08747 Q08751 Q08753 Q08756
Q08759 Q08770 Q08773 Q08774 Q08783 Q08784 Q08787 Q08789 Q08792 Q087H5 Q087I4
Q087I5 Q087I7 Q087J1 Q087J4 Q087J6 Q087K2 Q087K5 Q087K7 Q087L3 Q087L4 Q087L5
Q087P4 Q087Q6 Q087Q7 Q087Q9 Q087R5 Q087U3 Q087W5 Q087X1 Q087X5 Q08810 Q08812
Q08815 Q08816 Q08818 Q08820 Q08821 Q08823 Q08831 Q08842 Q08854 Q08856 Q08862
Q08863 Q08864 Q08865 Q08872 Q08873 Q08874 Q08875 Q08876 Q08880 Q08882 Q08885
Q08887 Q08888 Q08889 Q08893 Q08897 Q088A4 Q088B4 Q088C3 Q088G0 Q088H1 Q088H8
Q088J2 Q088J8 Q088K0 Q088K8 Q088L3 Q088M9 Q088P1 Q088P6 Q088R4 Q088R5 Q088S6
Q088S7 Q088T7 Q088T8 Q088W8 Q088X0 Q088X9 Q08903 Q08904 Q08909 Q08911 Q08914
Q08916 Q08920 Q08923 Q08925 Q08932 Q08935 Q08937 Q08942 Q08949 Q08952 Q08953
Q08954 Q08956 Q08960 Q08961 Q08962 Q08963 Q08964 Q08965 Q08966 Q08968 Q08969
Q08971 Q08972 Q08975 Q08977 Q08979 Q08981 Q08984 Q08985 Q08990 Q08992 Q08993
Q08994 Q08995 Q08996 Q089C0 Q089F8 Q089K8 Q089L1 Q089L6 Q089M8 Q089M9 Q089N0
Q089N1 Q089N2 Q089N3 Q089N5 Q089N6 Q089N7 Q089N8 Q089N9 Q089P0 Q089P1 Q089P2
Q089P3 Q089P4 Q089P5 Q089P6 Q089P7 Q089P8 Q089P9 Q089Q0 Q089Q1 Q089Q2 Q089Q3
Q089Q4 Q089Q5 Q089Q6 Q089Q7 Q089Q8 Q089Q9 Q089R0 Q089R1 Q089R2 Q089R3 Q089R4
Q089R5 Q089R8 Q089R9 Q089S6 Q089S7 Q089T3 Q089V0 Q089V1 Q089W9 Q08A02 Q08A04
Q08A11 Q08A18 Q08A23 Q08A25 Q08A29 Q08A38 Q08A41 Q08A45 Q08A46 Q08A49 Q08A51
Q08A71 Q08AF3 Q08AG5 Q08AG7 Q08AI8 Q08AN1 Q08AV6 Q08AV8 Q08AW5 Q08AX9 Q08AY6
Q08AY9 Q08AZ1 Q08B09 Q08B12 Q08B20 Q08B36 Q08B53 Q08B72 Q08BA6 Q08BB1 Q08BB3
Q08BB5 Q08BC4 Q08BC6 Q08BD8 Q08BH5 Q08BH7 Q08BL7 Q08BM0 Q08BM8 Q08BN1 Q08BS4
Q08BU1 Q08BV2 Q08BW6 Q08BY0 Q08BY1 Q08BY5 Q08BZ4 Q08C33 Q08C53 Q08C57 Q08C69
Q08C72 Q08C76 Q08C81 Q08C84 Q08C92 Q08C99 Q08CA1 Q08CF3 Q08CG8 Q08CH3 Q08CH7
Q08CH8 Q08CK7 Q08CL3 Q08CL8 Q08CM4 Q08CN0 Q08CX1 Q08CX2 Q08CY1 Q08CY5 Q08CZ0
Q08CZ6 Q08D01 Q08D12 Q08D35 Q08D57 Q08D62 Q08D69 Q08D85 Q08D86 Q08D88 Q08D91
Q08DA0 Q08DA5 Q08DA7 Q08DA8 Q08DB0 Q08DF7 Q08DG4 Q08DG5 Q08DG7 Q08DG8 Q08DH3
Q08DH7 Q08DH8 Q08DI1 Q08DI8 Q08DJ0 Q08DJ7 Q08DJ8 Q08DK3 Q08DK9 Q08DL1 Q08DL5

Q08DM2 Q08DM7 Q08DM8 Q08DP0 Q08DQ2 Q08DR9 Q08DS0 Q08DU1 Q08DU9 Q08DV5 Q08DW2
Q08DX0 Q08DY0 Q08DY6 Q08DZ2 Q08DZ3 Q08DZ8 Q08E07 Q08E12 Q08E13 Q08E20 Q08E27
Q08E29 Q08E31 Q08E57 Q08E93 Q08ED0 Q08EJ0 Q08EN7 Q08ER8 Q08FE1 Q08FW1 Q08GK9
Q08GP4 Q08I43 Q08IB7 Q08IC0 Q08VU0 Q09003 Q09004 Q09005 Q09006 Q09007 Q09008
Q09009 Q09010 Q09011 Q09019 Q09020 Q09052 Q09053 Q09054 Q09063 Q09064 Q09065
Q09066 Q09067 Q09069 Q09094 Q09095 Q09097 Q09100 Q09102 Q09109 Q09110 Q09115
Q09119 Q09122 Q09123 Q09124 Q09125 Q09129 Q09130 Q09134 Q09135 Q09139 Q09142
Q09146 Q09148 Q09150 Q09158 Q09167 Q09171 Q09176 Q09177 Q09179 Q09181 Q09189
Q09190 Q09191 Q09196 Q09198 Q09214 Q09215 Q09217 Q09221 Q09227 Q09228 Q09230
Q09233 Q09234 Q09235 Q09237 Q09238 Q09239 Q09240 Q09242 Q09245 Q09246 Q09249
Q09250 Q09252 Q09253 Q09254 Q09255 Q09257 Q09259 Q09260 Q09261 Q09262 Q09264
Q09265 Q09266 Q09267 Q09268 Q09269 Q09270 Q09278 Q09279 Q09280 Q09281 Q09282
Q09283 Q09284 Q09287 Q09293 Q09294 Q09295 Q09296 Q09297 Q09298 Q09299 Q09301
Q09302 Q09305 Q09307 Q09309 Q09311 Q09312 Q09313 Q09314 Q09315 Q09316 Q09317
Q09318 Q09321 Q09329 Q09330 Q09336 Q09340 Q09345 Q09346 Q09348 Q09350 Q09354
Q09355 Q09356 Q09357 Q09358 Q09359 Q09360 Q09361 Q09362 Q09370 Q09373 Q09374
Q09375 Q09376 Q09377 Q09378 Q09380 Q09382 Q09383 Q09384 Q09385 Q09389 Q09390
Q09391 Q09392 Q09394 Q09399 Q09400 Q09407 Q09410 Q09411 Q09412 Q09413 Q09415
Q09420 Q09421 Q09424 Q09430 Q09433 Q09436 Q09437 Q09438 Q09439 Q09440 Q09441
Q09442 Q09443 Q09445 Q09446 Q09448 Q09449 Q09450 Q09451 Q09452 Q09454 Q09456
Q09458 Q09462 Q09464 Q09465 Q09474 Q09475 Q09477 Q09496 Q09501 Q09504 Q09505
Q09506 Q09507 Q09510 Q09511 Q09512 Q09522 Q09523 Q09524 Q09525 Q09526 Q09527
Q09528 Q09530 Q09535 Q09543 Q09549 Q09555 Q09556 Q09557 Q09560 Q09563 Q09564
Q09565 Q09568 Q09571 Q09575 Q09578 Q09580 Q09582 Q09583 Q09585 Q09587 Q09589
Q09591 Q09596 Q09599 Q09602 Q09604 Q09605 Q09607 Q09608 Q09609 Q09610 Q09618
Q09619 Q09622 Q09623 Q09629 Q09632 Q09637 Q09644 Q09646 Q09647 Q09651 Q09652
Q09655 Q09660 Q09663 Q09664 Q09665 Q09668 Q09669 Q09670 Q09671 Q09672 Q09673
Q09674 Q09675 Q09676 Q09677 Q09678 Q09682 Q09683 Q09685 Q09686 Q09687 Q09689
Q09692 Q09693 Q09694 Q09696 Q09698 Q09699 Q09700 Q09702 Q09704 Q09705 Q09708
Q09709 Q09713 Q09715 Q09716 Q09719 Q09720 Q09721 Q09722 Q09723 Q09724 Q09725
Q09727 Q09731 Q09732 Q09735 Q09737 Q09739 Q09740 Q09741 Q09742 Q09744 Q09745
Q09750 Q09751 Q09755 Q09756 Q09760 Q09761 Q09762 Q09763 Q09765 Q09768 Q09770
Q09771 Q09772 Q09773 Q09775 Q09776 Q09779 Q09784 Q09785 Q09786 Q09787 Q09789
Q09790 Q09791 Q09793 Q09794 Q09795 Q09797 Q09798 Q09799 Q09800 Q09801 Q09802
Q09804 Q09806 Q09807 Q09808 Q09810 Q09811 Q09813 Q09814 Q09815 Q09816 Q09817
Q09818 Q09819 Q09820 Q09824 Q09826 Q09828 Q09829 Q09830 Q09832 Q09833 Q09838
Q09839 Q09841 Q09842 Q09843 Q09844 Q09845 Q09847 Q09848 Q09855 Q09856 Q09858
Q09859 Q09860 Q09861 Q09862 Q09863 Q09865 Q09867 Q09868 Q09869 Q09870 Q09871
Q09872 Q09873 Q09878 Q09882 Q09884 Q09885 Q09886 Q09888 Q09890 Q09893 Q09894
Q09897 Q09899 Q09900 Q09902 Q09903 Q09908 Q09909 Q09911 Q09912 Q09913 Q09915
Q09918 Q09921 Q09922 Q09923 Q09924 Q09926 Q09927 Q09928 Q09929 Q09933 Q09936
Q09947 Q09949 Q09950 Q09956 Q09958 Q09961 Q09990 Q09994 Q09996 Q09EF7 Q09FC8
Q09FP6 Q09FP7 Q09FP8 Q09FP9 Q09FQ1 Q09FQ3 Q09FR2 Q09FS0 Q09FS1 Q09FS2 Q09FS3
Q09FS4 Q09FS5 Q09FS6 Q09FS7 Q09FS8 Q09FS9 Q09FT6 Q09FT7 Q09FT8 Q09FT9 Q09FU0

| | | | | | | | | | | |
|---|---|---|---|---|---|---|---|---|---|---|
| Q09FV9 | Q09FW3 | Q09FW9 | Q09FX0 | Q09FX1 | Q09FX2 | Q09FY0 | Q09FY3 | Q09FY4 | Q09FY5 | Q09FY9 |
| Q09FZ8 | Q09G05 | Q09G06 | Q09G07 | Q09G08 | Q09G09 | Q09G10 | Q09G11 | Q09G12 | Q09G13 | Q09G14 |
| Q09G21 | Q09G22 | Q09G23 | Q09G24 | Q09G25 | Q09G44 | Q09G48 | Q09G54 | Q09G55 | Q09G56 | Q09G57 |
| Q09G65 | Q09HN1 | Q09HN2 | Q09JK2 | Q09JT4 | Q09JW2 | Q09JZ4 | Q09KQ6 | Q09LL3 | Q09LX1 | Q09LY5 |
| Q09LZ8 | Q09M02 | Q09M05 | Q09MB1 | Q09MB2 | Q09MB3 | Q09MB4 | Q09MB5 | Q09MB7 | Q09MB9 | Q09MC8 |
| Q09MD9 | Q09ME0 | Q09ME1 | Q09ME2 | Q09ME3 | Q09ME4 | Q09ME5 | Q09MF2 | Q09MF3 | Q09MF5 | Q09MF6 |
| Q09MF7 | Q09MH0 | Q09MI0 | Q09MI6 | Q09MI7 | Q09MI8 | Q09MI9 | Q09MJ6 | Q09MJ7 | Q09MP3 | Q09MP4 |
| Q09MP5 | Q09WE7 | Q09WV5 | Q09WV6 | Q09WV7 | Q09WV9 | Q09WW1 | Q09WX0 | Q09WX7 | Q09WX8 | Q09WX9 |
| Q09WY0 | Q09WY1 | Q09WY2 | Q09WY3 | Q09WY4 | Q09WY5 | Q09WZ2 | Q09WZ3 | Q09WZ4 | Q09WZ5 | Q09WZ6 |
| Q09X09 | Q09X15 | Q09X19 | Q09X25 | Q09X26 | Q09X27 | Q09X28 | Q09X35 | Q09X36 | Q09XV5 | Q09YD1 |
| Q09YG9 | Q09YH1 | Q09YH6 | Q09YI0 | Q09YI1 | Q09YI3 | Q09YJ3 | Q09YJ5 | Q09YJ9 | Q09YK3 | Q09YK4 |
| Q09YK6 | Q09YL0 | Q09YL5 | Q09YL6 | Q09YM8 | Q09YN0 | Q09YN8 | Q0A4L3 | Q0A4L6 | Q0A4L7 | Q0A4L8 |
| Q0A4P0 | Q0A4Q5 | Q0A4R1 | Q0A4S2 | Q0A4T7 | Q0A4T8 | Q0A4X0 | Q0A4X2 | Q0A4Y2 | Q0A4Z6 | Q0A550 |
| Q0A551 | Q0A574 | Q0A577 | Q0A582 | Q0A583 | Q0A592 | Q0A598 | Q0A5A5 | Q0A5A6 | Q0A5B1 | Q0A5B4 |
| Q0A5B6 | Q0A5B9 | Q0A5C0 | Q0A5C9 | Q0A5D0 | Q0A5D2 | Q0A5D3 | Q0A5D4 | Q0A5H0 | Q0A5H6 | Q0A5J1 |
| Q0A5K0 | Q0A5M1 | Q0A5M7 | Q0A5M8 | Q0A5Q3 | Q0A5Q4 | Q0A5S1 | Q0A5T4 | Q0A5T5 | Q0A5T6 | Q0A5V1 |
| Q0A5V3 | Q0A5V4 | Q0A5V8 | Q0A5W1 | Q0A5W2 | Q0A653 | Q0A6A6 | Q0A6A7 | Q0A6A8 | Q0A6C9 | Q0A6D3 |
| Q0A6E7 | Q0A6F1 | Q0A6F3 | Q0A6F4 | Q0A6F5 | Q0A6G2 | Q0A6H8 | Q0A6H9 | Q0A6I9 | Q0A6J0 | Q0A6J3 |
| Q0A6J4 | Q0A6J7 | Q0A6K3 | Q0A6K4 | Q0A6K5 | Q0A6T1 | Q0A6U0 | Q0A706 | Q0A726 | Q0A738 | Q0A749 |
| Q0A751 | Q0A753 | Q0A757 | Q0A765 | Q0A769 | Q0A772 | Q0A773 | Q0A793 | Q0A795 | Q0A797 | Q0A798 |
| Q0A7A0 | Q0A7A1 | Q0A7D4 | Q0A7E2 | Q0A7E4 | Q0A7E5 | Q0A7F4 | Q0A7H5 | Q0A7H6 | Q0A7I0 | Q0A7I1 |
| Q0A7I2 | Q0A7I3 | Q0A7J3 | Q0A7K1 | Q0A7K2 | Q0A7K7 | Q0A7L4 | Q0A7L5 | Q0A7N2 | Q0A7N3 | Q0A7X7 |
| Q0A7X8 | Q0A7Z2 | Q0A812 | Q0A845 | Q0A879 | Q0A8A4 | Q0A8G6 | Q0A8H9 | Q0A8I0 | Q0A8I6 | Q0A8I9 |
| Q0A8J8 | Q0A8K3 | Q0A8K5 | Q0A8K7 | Q0A8K9 | Q0A8N7 | Q0A8P0 | Q0A8P2 | Q0A8P7 | Q0A8R1 | Q0A8R3 |
| Q0A8S2 | Q0A8V6 | Q0A8V7 | Q0A8X6 | Q0A8Y4 | Q0A8Y9 | Q0A8Z1 | Q0A911 | Q0A915 | Q0A923 | Q0A936 |
| Q0A939 | Q0A961 | Q0A977 | Q0A982 | Q0A986 | Q0A989 | Q0A990 | Q0A9A1 | Q0A9A9 | Q0A9B0 | Q0A9B2 |
| Q0A9C4 | Q0A9D7 | Q0A9E8 | Q0A9G3 | Q0A9G6 | Q0A9T9 | Q0A9W8 | Q0AA25 | Q0AA26 | Q0AA27 | Q0AA37 |
| Q0AA39 | Q0AA41 | Q0AA48 | Q0AA51 | Q0AA54 | Q0AA70 | Q0AA73 | Q0AA96 | Q0AA98 | Q0AAD1 | Q0AAD4 |
| Q0AAD6 | Q0AAD7 | Q0AAD8 | Q0AAG6 | Q0AAH7 | Q0AAL5 | Q0AAL7 | Q0AAL8 | Q0AAM1 | Q0AAU9 | Q0AAV2 |
| Q0AAV5 | Q0AAW0 | Q0AAX8 | Q0AB25 | Q0AB32 | Q0AB36 | Q0AB37 | Q0AB42 | Q0AB47 | Q0AB49 | Q0AB51 |
| Q0AB55 | Q0AB56 | Q0AB60 | Q0AB61 | Q0AB67 | Q0AB68 | Q0AB69 | Q0AB75 | Q0AB89 | Q0AB94 | Q0ABB1 |
| Q0ABB7 | Q0ABE6 | Q0ABE9 | Q0ABF0 | Q0ABF1 | Q0ABF2 | Q0ABF3 | Q0ABF4 | Q0ABF6 | Q0ABF7 | Q0ABF8 |
| Q0ABF9 | Q0ABG0 | Q0ABG1 | Q0ABG2 | Q0ABG3 | Q0ABG4 | Q0ABG5 | Q0ABG6 | Q0ABG7 | Q0ABG8 | Q0ABG9 |
| Q0ABH0 | Q0ABH1 | Q0ABH2 | Q0ABH3 | Q0ABH4 | Q0ABH5 | Q0ABH6 | Q0ABH7 | Q0ABH8 | Q0ABH9 | Q0ABI0 |
| Q0ABI1 | Q0ABI2 | Q0ABI3 | Q0ABI4 | Q0ABI5 | Q0ABI6 | Q0ABJ2 | Q0ABJ8 | Q0ABJ9 | Q0ABK6 | Q0ABM5 |
| Q0ABM7 | Q0ABN0 | Q0ABN3 | Q0ABN7 | Q0ABN9 | Q0ABQ3 | Q0ABQ4 | Q0ABQ8 | Q0ABQ9 | Q0ABR1 | Q0ABT6 |
| Q0ABU4 | Q0ABZ7 | Q0ABZ8 | Q0AC00 | Q0AC03 | Q0AC04 | Q0AC59 | Q0AC95 | Q0AC96 | Q0AC97 | Q0AC99 |
| Q0ACA0 | Q0ACA1 | Q0ACA3 | Q0ACB4 | Q0ACB5 | Q0ACB6 | Q0ACJ4 | Q0ACJ6 | Q0ACK7 | Q0ACK8 | Q0ACL3 |
| Q0ACL4 | Q0ACL9 | Q0ACP4 | Q0ACP5 | Q0ACP8 | Q0ACQ4 | Q0ACQ8 | Q0ACQ9 | Q0ACS7 | Q0AD35 | Q0AD36 |
| Q0AD37 | Q0AD60 | Q0AD61 | Q0AD66 | Q0AD92 | Q0ADD1 | Q0ADD6 | Q0ADD7 | Q0ADI4 | Q0ADI6 | Q0ADI8 |
| Q0ADI9 | Q0ADM4 | Q0ADM9 | Q0ADN4 | Q0ADN6 | Q0ADN7 | Q0ADP0 | Q0ADP1 | Q0ADP2 | Q0ADS2 | Q0ADS3 |
| Q0ADS5 | Q0ADS7 | Q0ADT8 | Q0ADU3 | Q0ADW4 | Q0ADX6 | Q0ADY0 | Q0AE32 | Q0AE34 | Q0AE38 | Q0AE39 |
| Q0AE42 | Q0AE43 | Q0AE46 | Q0AE53 | Q0AE54 | Q0AE55 | Q0AE58 | Q0AE59 | Q0AE64 | Q0AE72 | Q0AE73 |
| Q0AE87 | Q0AEA6 | Q0AEC9 | Q0AEE3 | Q0AEE4 | Q0AEG2 | Q0AEH4 | Q0AEH5 | Q0AEH6 | Q0AEH7 | Q0AEI0 |

Q0AEI5 Q0AEI7 Q0AEP5 Q0AEP6 Q0AEP7 Q0AEP8 Q0AEP9 Q0AEQ0 Q0AER1 Q0AES7 Q0AES9
Q0AET9 Q0AEU0 Q0AEU1 Q0AEU2 Q0AEU4 Q0AEU9 Q0AEV2 Q0AEV7 Q0AEV8 Q0AEY2 Q0AF46
Q0AF49 Q0AF50 Q0AF51 Q0AF53 Q0AF54 Q0AF59 Q0AF61 Q0AF70 Q0AF72 Q0AF74 Q0AFF1
Q0AFG8 Q0AFH8 Q0AFI1 Q0AFI5 Q0AFI6 Q0AFJ1 Q0AFJ3 Q0AFJ4 Q0AFK6 Q0AFM7 Q0AFN2
Q0AFN4 Q0AFP3 Q0AFP4 Q0AFQ9 Q0AFS9 Q0AFT2 Q0AFT6 Q0AFT7 Q0AFU6 Q0AFV0 Q0AFV1
Q0AFV7 Q0AFW3 Q0AFX7 Q0AFY5 Q0AFY6 Q0AG55 Q0AG56 Q0AG57 Q0AG72 Q0AG95 Q0AGI0
Q0AGI1 Q0AGJ0 Q0AGL1 Q0AGM0 Q0AGN1 Q0AGN2 Q0AGN5 Q0AGQ8 Q0AGS2 Q0AGU3 Q0AGV1
Q0AGV5 Q0AGX5 Q0AGX6 Q0AGX7 Q0AGY1 Q0AGY5 Q0AGY6 Q0AGY8 Q0AGZ0 Q0AGZ8 Q0AH80
Q0AH90 Q0AHA4 Q0AHC1 Q0AHC3 Q0AHC6 Q0AHF5 Q0AHF6 Q0AHG1 Q0AHG4 Q0AHG5 Q0AHG6
Q0AHG7 Q0AHH6 Q0AHI5 Q0AHK2 Q0AHQ0 Q0AHS7 Q0AHU1 Q0AHU2 Q0AHW1 Q0AHX2 Q0AHX4
Q0AHX8 Q0AHY0 Q0AHY1 Q0AHY2 Q0AHY7 Q0AHZ3 Q0AHZ4 Q0AHZ8 Q0AHZ9 Q0AI00 Q0AI04
Q0AI05 Q0AI06 Q0AI13 Q0AI14 Q0AI27 Q0AI30 Q0AI31 Q0AI71 Q0AIA2 Q0AIB2 Q0AID7
Q0AIE4 Q0AIE6 Q0AIF1 Q0AIF2 Q0AIF5 Q0AIG3 Q0AIG7 Q0AIH0 Q0AIH1 Q0AIH2 Q0AIH3
Q0AIH4 Q0AIH5 Q0AIH7 Q0AIH8 Q0AIH9 Q0AII2 Q0AII3 Q0AII4 Q0AII5 Q0AII6 Q0AII7
Q0AII8 Q0AII9 Q0AIJ0 Q0AIJ1 Q0AIJ2 Q0AIJ3 Q0AIJ4 Q0AIJ5 Q0AIJ6 Q0AIJ7 Q0AIJ8
Q0AIJ9 Q0AIK0 Q0AIL5 Q0AIM3 Q0AIP0 Q0AIP2 Q0AIP6 Q0AIP9 Q0AIS2 Q0AIU3 Q0AIU5
Q0AIU6 Q0AIU9 Q0AIV0 Q0AIV2 Q0AIY0 Q0AIY1 Q0AIY2 Q0AIZ1 Q0AIZ2 Q0AIZ4 Q0AIZ5
Q0AIZ8 Q0AJ02 Q0AJ07 Q0AJ12 Q0AJ13 Q0AJ18 Q0AJ41 Q0AJ67 Q0AJ68 Q0AJ79 Q0AJ90
Q0AJ91 Q0AJ94 Q0AJA3 Q0AJA4 Q0AJA5 Q0AJA6 Q0AJC4 Q0AJC5 Q0AJD2 Q0AJD3 Q0AJD9
Q0AJE2 Q0AJE3 Q0AJE4 Q0AJF3 Q0AJF4 Q0AJF8 Q0AJH8 Q0AJH9 Q0AJI1 Q0AJI2 Q0AJI3
Q0AJI9 Q0AJJ4 Q0AJP9 Q0AK27 Q0AK35 Q0AK44 Q0AK62 Q0AK63 Q0AK64 Q0AK66 Q0AK69
Q0AK72 Q0AK73 Q0AK77 Q0AK79 Q0AK91 Q0AK98 Q0AKA6 Q0AKB1 Q0AKB3 Q0AKC9 Q0AKD3
Q0AKD6 Q0AKD7 Q0AKD8 Q0AKD9 Q0AKE8 Q0AKE9 Q0AKF0 Q0AKF5 Q0AKH3 Q0AKJ6 Q0AKL3
Q0AKL5 Q0AKL6 Q0AKM4 Q0AKT3 Q0AKU3 Q0AKW7 Q0AKX2 Q0AKX6 Q0AKX7 Q0AL71 Q0AL88
Q0ALG1 Q0ALG8 Q0ALH6 Q0ALJ5 Q0ALN0 Q0ALP0 Q0ALQ9 Q0ALR7 Q0ALV0 Q0ALW1 Q0ALW2
Q0ALW8 Q0ALX7 Q0ALX8 Q0ALX9 Q0AM02 Q0AM03 Q0AM08 Q0AM20 Q0AM22 Q0AM30 Q0AMB1
Q0AME1 Q0AMI4 Q0AMI8 Q0AMJ2 Q0AMK3 Q0AML4 Q0AMQ6 Q0AMR3 Q0AMS2 Q0AMU7 Q0AMV9
Q0AMW8 Q0AMW9 Q0AMX1 Q0AMX9 Q0AN24 Q0AND9 Q0ANE1 Q0ANE2 Q0ANF3 Q0ANH8 Q0ANL8
Q0ANN1 Q0ANN9 Q0ANP0 Q0ANP1 Q0ANP2 Q0ANP3 Q0ANP4 Q0ANP5 Q0ANP6 Q0ANP7 Q0ANP9
Q0ANQ0 Q0ANQ1 Q0ANQ2 Q0ANQ3 Q0ANQ4 Q0ANQ5 Q0ANQ6 Q0ANQ7 Q0ANQ8 Q0ANQ9 Q0ANR0
Q0ANR1 Q0ANR2 Q0ANR3 Q0ANR4 Q0ANR5 Q0ANR6 Q0ANR7 Q0ANR8 Q0ANR9 Q0ANS1 Q0ANS2
Q0ANS3 Q0ANS4 Q0ANS5 Q0ANV0 Q0ANX4 Q0ANY2 Q0ANY8 Q0ANY9 Q0AP18 Q0AP20 Q0AP80
Q0AP83 Q0AP84 Q0AP91 Q0AP92 Q0AP94 Q0APB3 Q0APB9 Q0APC1 Q0APD6 Q0APF8 Q0APF9
Q0APG4 Q0APH3 Q0API1 Q0APM5 Q0APM9 Q0APP7 Q0APQ6 Q0APQ9 Q0APT3 Q0APT4 Q0APT7
Q0APU5 Q0APU9 Q0APW2 Q0APW3 Q0APW4 Q0APW5 Q0APX0 Q0APX5 Q0APZ9 Q0AQ06 Q0AQ07
Q0AQ08 Q0AQ60 Q0AQ67 Q0AQ76 Q0AQ78 Q0AQ81 Q0AQ87 Q0AQB0 Q0AQC2 Q0AQC3 Q0AQC4
Q0AQD2 Q0AQD3 Q0AQD4 Q0AQE4 Q0AQF4 Q0AQH3 Q0AQJ7 Q0AQL3 Q0AQL4 Q0AQL6 Q0AQL8
Q0AQP4 Q0AQT4 Q0AQU7 Q0AQW2 Q0AQX8 Q0AQY6 Q0AR08 Q0AR22 Q0ARD7 Q0ARD9 Q0ARE5
Q0ARG6 Q0ARM2 Q0ARN5 Q0ARN7 Q0ARN8 Q0ARQ1 Q0ARS7 Q0ARV1 Q0ARZ8 Q0AS39 Q0AS40
Q0AS86 Q0ASF3 Q0ASI7 Q0ASJ0 Q0ASJ5 Q0ASJ7 Q0ASK2 Q0ASK7 Q0ASK8 Q0AT09 Q0ATA0
Q0ATD8 Q0ATG2 Q0ATG3 Q0ATJ1 Q0ATJ4 Q0ATJ7 Q0ATN3 Q0ATP0 Q0ATQ2 Q0ATR3 Q0ATU1
Q0ATU2 Q0ATU5 Q0ATU6 Q0ATU7 Q0ATX6 Q0ATY3 Q0AU48 Q0AU94 Q0AU97 Q0AUA5 Q0AUB6
Q0AUB8 Q0AUC3 Q0AUE7 Q0AUF5 Q0AUF7 Q0AUG3 Q0AUG4 Q0AUG7 Q0AUG8 Q0AUH0 Q0AUH2
Q0AUH3 Q0AUH5 Q0AUH6 Q0AUH7 Q0AUH8 Q0AUI0 Q0AUI1 Q0AUI2 Q0AUI3 Q0AUI4 Q0AUI5

Q0AUI6 Q0AUI7 Q0AUI9 Q0AUJ0 Q0AUJ1 Q0AUJ2 Q0AUJ4 Q0AUJ5 Q0AUJ6 Q0AUJ7 Q0AUJ8
Q0AUJ9 Q0AUK1 Q0AUK4 Q0AUK5 Q0AUK6 Q0AUK7 Q0AUK8 Q0AUK9 Q0AUL0 Q0AUL4 Q0AUL5
Q0AUL6 Q0AUM4 Q0AV19 Q0AV47 Q0AV62 Q0AV72 Q0AVE3 Q0AVH7 Q0AVH8 Q0AVH9 Q0AVU0
Q0AVU6 Q0AVU7 Q0AVV0 Q0AVV1 Q0AW22 Q0AW29 Q0AW31 Q0AW37 Q0AW39 Q0AW40 Q0AW41
Q0AWE9 Q0AWF0 Q0AWF3 Q0AWF4 Q0AWG0 Q0AWG2 Q0AWG3 Q0AWH0 Q0AWJ0 Q0AWJ2 Q0AWJ4
Q0AWJ6 Q0AWK4 Q0AWL5 Q0AWM1 Q0AWM5 Q0AWM8 Q0AWN7 Q0AWP3 Q0AWT5 Q0AWT6 Q0AWT8
Q0AWU6 Q0AWV4 Q0AWV6 Q0AWV7 Q0AWV9 Q0AWX1 Q0AWX4 Q0AWX5 Q0AX10 Q0AX12 Q0AX14
Q0AX15 Q0AX16 Q0AX18 Q0AX38 Q0AX99 Q0AXC8 Q0AXG1 Q0AXG6 Q0AXG7 Q0AXH2 Q0AXI3
Q0AXI8 Q0AXK7 Q0AXK8 Q0AXK9 Q0AXL1 Q0AXL3 Q0AXL4 Q0AXL8 Q0AXM5 Q0AXN1 Q0AXT9
Q0AXU4 Q0AXU8 Q0AY00 Q0AY02 Q0AY03 Q0AY05 Q0AY75 Q0AYA9 Q0AYB0 Q0AYB2 Q0AYB5
Q0AYB7 Q0AYC7 Q0AYD0 Q0AYD5 Q0AYE0 Q0AYF8 Q0AYH7 Q0AYI2 Q0AYI3 Q0AYI5 Q0AYI7
Q0AYI8 Q0AYJ1 Q0AYJ4 Q0AYK1 Q0AYK2 Q0AYK3 Q0AYK4 Q0AYK9 Q0AYP1 Q0AYP2 Q0AYP4
Q0AYQ7 Q0AYR3 Q0AYR6 Q0AYS1 Q0AYS2 Q0AYS6 Q0AYV2 Q0AYV4 Q0AYW0 Q0AYW3 Q0AYW5
Q0AYW7 Q0AYX5 Q0AZ31 Q0AZ36 Q0AZ43 Q0AZ53 Q0AZ72 Q0AZ84 Q0AZ86 Q0AZ87 Q0AZD9
Q0AZE0 Q0AZE2 Q0AZF3 Q0AZF4 Q0AZF5 Q0AZF7 Q0AZH4 Q0AZJ3 Q0AZJ6 Q0AZP9 Q0AZQ0
Q0AZQ1 Q0AZQ5 Q0AZR6 Q0AZZ2 Q0AZZ3 Q0AZZ5 Q0B002 Q0B004 Q0B006 Q0B016 Q0B018
Q0B075 Q0B092 Q0B094 Q0B0A0 Q0B0F6 Q0B0F7 Q0B0N3 Q0B0N4 Q0B0N9 Q0B0P4 Q0B0P5
Q0B0P6 Q0B0S6 Q0B0S7 Q0B0T2 Q0B0W3 Q0B0Y5 Q0B0Z2 Q0B1N7 Q0B1U7 Q0B2K2 Q0B2Y2
Q0B3Y4 Q0B4C4 Q0B4U4 Q0B4V5 Q0B4Z6 Q0B569 Q0B5P1 Q0B5P6 Q0B5Q1 Q0B5Y5 Q0B602
Q0B628 Q0B6M7 Q0B711 Q0B712 Q0B713 Q0B744 Q0B746 Q0B7X8 Q0B880 Q0B8K0 Q0B8S0
Q0B950 Q0B953 Q0B9R9 Q0B9S2 Q0BA68 Q0BA70 Q0BAC2 Q0BAC6 Q0BAC8 Q0BAF9 Q0BAL6
Q0BAL7 Q0BAL8 Q0BAM0 Q0BAN2 Q0BAP9 Q0BAQ4 Q0BAT6 Q0BAU3 Q0BAV3 Q0BAV4 Q0BAV5
Q0BAW3 Q0BAX4 Q0BAX8 Q0BAX9 Q0BAY6 Q0BAY8 Q0BB33 Q0BB41 Q0BB90 Q0BBD4 Q0BBD5
Q0BBD6 Q0BBE2 Q0BBE9 Q0BBG3 Q0BBH9 Q0BBK5 Q0BBK8 Q0BBL1 Q0BBQ0 Q0BBQ3 Q0BBQ4
Q0BBQ6 Q0BBR2 Q0BBR3 Q0BBS3 Q0BBX9 Q0BBY4 Q0BBZ1 Q0BC07 Q0BC19 Q0BC21 Q0BC36
Q0BC71 Q0BC82 Q0BCE0 Q0BCE2 Q0BCF1 Q0BCG4 Q0BCG5 Q0BCG7 Q0BCH1 Q0BCH3 Q0BCK0
Q0BCK5 Q0BCK8 Q0BCL4 Q0BCL7 Q0BCL8 Q0BCL9 Q0BCQ2 Q0BCS3 Q0BCS7 Q0BCS8 Q0BCT3
Q0BCT7 Q0BCU9 Q0BD34 Q0BD35 Q0BD43 Q0BD59 Q0BD64 Q0BD80 Q0BD83 Q0BD85 Q0BD86
Q0BD88 Q0BDB6 Q0BDB9 Q0BDC5 Q0BDC6 Q0BDF6 Q0BDK6 Q0BDL0 Q0BDM3 Q0BDP0 Q0BDQ8
Q0BDR2 Q0BDS1 Q0BDU3 Q0BDV4 Q0BDX9 Q0BE03 Q0BE05 Q0BE10 Q0BE15 Q0BE16 Q0BE17
Q0BE18 Q0BE21 Q0BE29 Q0BE31 Q0BE35 Q0BE45 Q0BED6 Q0BED7 Q0BEF5 Q0BEF6 Q0BEF7
Q0BEH0 Q0BEH1 Q0BEN7 Q0BEQ6 Q0BEQ8 Q0BEQ9 Q0BEU6 Q0BEV3 Q0BEV4 Q0BEV7 Q0BEW2
Q0BEW4 Q0BEW5 Q0BEW8 Q0BEX1 Q0BEX2 Q0BEX7 Q0BEZ5 Q0BFF5 Q0BFQ6 Q0BFQ8 Q0BFQ9
Q0BFR5 Q0BFY2 Q0BFY3 Q0BFY5 Q0BG03 Q0BG06 Q0BG24 Q0BG57 Q0BG79 Q0BGI1 Q0BGI9
Q0BGJ2 Q0BGQ9 Q0BGU9 Q0BGV0 Q0BGZ5 Q0BGZ7 Q0BH19 Q0BH30 Q0BH38 Q0BH60 Q0BH66
Q0BH67 Q0BH68 Q0BH69 Q0BH74 Q0BH85 Q0BH90 Q0BH92 Q0BH93 Q0BH96 Q0BHA0 Q0BHD7
Q0BHF4 Q0BHF7 Q0BHG9 Q0BHH1 Q0BHH5 Q0BHJ2 Q0BHJ7 Q0BHJ8 Q0BHL9 Q0BHN2 Q0BHN3
Q0BHN4 Q0BHN5 Q0BHN6 Q0BHN7 Q0BHN8 Q0BHS4 Q0BHS6 Q0BHS7 Q0BHT1 Q0BHW3 Q0BHY0
Q0BI00 Q0BI05 Q0BI09 Q0BI15 Q0BI17 Q0BI18 Q0BI24 Q0BI25 Q0BI26 Q0BI48 Q0BI51
Q0BI73 Q0BI80 Q0BI81 Q0BI82 Q0BI83 Q0BI86 Q0BI88 Q0BI89 Q0BI90 Q0BI91 Q0BIB2
Q0BIB5 Q0BIB8 Q0BIF9 Q0BIH4 Q0BIH5 Q0BIH7 Q0BIH8 Q0BIH9 Q0BII0 Q0BII7 Q0BIJ9
Q0BIK0 Q0BIK3 Q0BIK9 Q0BIL0 Q0BIM7 Q0BIM8 Q0BIP9 Q0BIQ0 Q0BIW2 Q0BIW3 Q0BIW4
Q0BIW5 Q0BIW8 Q0BIX1 Q0BJ01 Q0BJ08 Q0BJ15 Q0BJ19 Q0BJ20 Q0BJ21 Q0BJ22 Q0BJ23
Q0BJ24 Q0BJ25 Q0BJ27 Q0BJ28 Q0BJ29 Q0BJ30 Q0BJ31 Q0BJ32 Q0BJ33 Q0BJ34 Q0BJ35

Q0BJ36 Q0BJ37 Q0BJ38 Q0BJ39 Q0BJ40 Q0BJ41 Q0BJ42 Q0BJ43 Q0BJ44 Q0BJ45 Q0BJ46
Q0BJ47 Q0BJ48 Q0BJ50 Q0BJ51 Q0BJ53 Q0BJ54 Q0BJ55 Q0BJ57 Q0BJ58 Q0BJB0 Q0BJI1
Q0BJM7 Q0BJM8 Q0BJP4 Q0BJQ3 Q0BJQ6 Q0BJW1 Q0BJW6 Q0BJY9 Q0BK01 Q0BK19 Q0BK24
Q0BK40 Q0BK41 Q0BK67 Q0BK68 Q0BK70 Q0BK71 Q0BK72 Q0BK98 Q0BK99 Q0BKA0 Q0BKB8
Q0BKC0 Q0BKC1 Q0BKC2 Q0BKC3 Q0BKC5 Q0BKC6 Q0BKC8 Q0BKD0 Q0BKD1 Q0BKD2 Q0BKD4
Q0BKE0 Q0BKE7 Q0BKE8 Q0BKF0 Q0BKF2 Q0BKF6 Q0BKF7 Q0BKI3 Q0BKJ6 Q0BKK1 Q0BKK2
Q0BKK9 Q0BKM8 Q0BKP4 Q0BKP7 Q0BKP9 Q0BKS4 Q0BKT2 Q0BKT3 Q0BKT7 Q0BKU0 Q0BKU1
Q0BKU2 Q0BKU5 Q0BKU7 Q0BKV5 Q0BKX7 Q0BKX8 Q0BKZ2 Q0BKZ4 Q0BL02 Q0BL03 Q0BL07
Q0BL35 Q0BL36 Q0BL41 Q0BL43 Q0BL44 Q0BL47 Q0BL54 Q0BL76 Q0BL82 Q0BL88 Q0BL95
Q0BLA3 Q0BLA4 Q0BLC3 Q0BLC5 Q0BLC7 Q0BLD2 Q0BLD5 Q0BLF0 Q0BLG2 Q0BLI0 Q0BLI1
Q0BLI7 Q0BLI9 Q0BLJ8 Q0BLK4 Q0BLK5 Q0BLK8 Q0BLL3 Q0BLL9 Q0BLP2 Q0BLP6 Q0BLS0
Q0BLS9 Q0BLU4 Q0BLU9 Q0BLV0 Q0BLV1 Q0BLW1 Q0BLW7 Q0BLX1 Q0BLY6 Q0BLY9 Q0BLZ2
Q0BM23 Q0BM40 Q0BM46 Q0BM50 Q0BM56 Q0BM63 Q0BM65 Q0BM66 Q0BM68 Q0BM81 Q0BM86
Q0BM87 Q0BM92 Q0BM93 Q0BM96 Q0BM98 Q0BMA6 Q0BMB7 Q0BMC7 Q0BMD2 Q0BME1 Q0BMF4
Q0BMF5 Q0BMG1 Q0BMI2 Q0BMI3 Q0BMJ8 Q0BMK6 Q0BMM0 Q0BMM2 Q0BMN1 Q0BMQ3 Q0BMQ5
Q0BMQ6 Q0BMQ7 Q0BMQ8 Q0BMS2 Q0BMT1 Q0BMU5 Q0BMU7 Q0BMV6 Q0BN06 Q0BN07 Q0BN25
Q0BN28 Q0BN37 Q0BN38 Q0BN41 Q0BN64 Q0BN65 Q0BN67 Q0BN71 Q0BN73 Q0BN93 Q0BNA8
Q0BNB5 Q0BNB8 Q0BND2 Q0BNE2 Q0BNF1 Q0BNG4 Q0BNH8 Q0BNJ1 Q0BNK1 Q0BNM7 Q0BNM8
Q0BNN3 Q0BNQ2 Q0BNQ3 Q0BNQ4 Q0BNQ5 Q0BNQ6 Q0BNQ8 Q0BNQ9 Q0BNR0 Q0BNR1 Q0BNR2
Q0BNR3 Q0BNR4 Q0BNR5 Q0BNR6 Q0BNR7 Q0BNR8 Q0BNR9 Q0BNS0 Q0BNS1 Q0BNS2 Q0BNS3
Q0BNS4 Q0BNS5 Q0BNS6 Q0BNS7 Q0BNS8 Q0BNS9 Q0BNT0 Q0BNT1 Q0BNT3 Q0BNT6 Q0BNT7
Q0BNT8 Q0BNT9 Q0BNU3 Q0BNX6 Q0BNX7 Q0BNY4 Q0BP04 Q0BP15 Q0BP45 Q0BP46 Q0BP50
Q0BP64 Q0BP66 Q0BP72 Q0BP79 Q0BPA2 Q0BPA5 Q0BPA8 Q0BPA9 Q0BPB6 Q0BPC9 Q0BPD5
Q0BPE2 Q0BPE6 Q0BPF5 Q0BPF6 Q0BPF9 Q0BPG2 Q0BPG3 Q0BPK2 Q0BPK3 Q0BPL3 Q0BPL4
Q0BPL8 Q0BPN7 Q0BPP0 Q0BPT7 Q0BPT8 Q0BPT9 Q0BPW9 Q0BPX1 Q0BPX3 Q0BPX5 Q0BPX7
Q0BPY8 Q0BPY9 Q0BPZ0 Q0BPZ9 Q0BQ24 Q0BQ38 Q0BQ41 Q0BQ42 Q0BQ43 Q0BQ87 Q0BQA2
Q0BQF8 Q0BQG5 Q0BQH9 Q0BQI0 Q0BQI4 Q0BQI7 Q0BQJ0 Q0BQJ9 Q0BQR6 Q0BQS6 Q0BQS8
Q0BQX7 Q0BQX8 Q0BQZ6 Q0BR06 Q0BR48 Q0BRE3 Q0BRE9 Q0BRF0 Q0BRF1 Q0BRH0 Q0BRH2
Q0BRH4 Q0BRH5 Q0BRI4 Q0BRP0 Q0BRW8 Q0BS00 Q0BS01 Q0BS04 Q0BS22 Q0BS26 Q0BS88
Q0BSB1 Q0BSD5 Q0BSF0 Q0BSG5 Q0BSJ5 Q0BSJ8 Q0BSL7 Q0BSM0 Q0BSM5 Q0BSM6 Q0BSQ2
Q0BSQ3 Q0BSS2 Q0BST5 Q0BSV8 Q0BSW6 Q0BT08 Q0BT18 Q0BT31 Q0BT42 Q0BT53 Q0BT56
Q0BT57 Q0BT69 Q0BTA0 Q0BTB0 Q0BTB7 Q0BTB8 Q0BTC4 Q0BTC5 Q0BTD5 Q0BTE2 Q0BTE6
Q0BTE7 Q0BTF1 Q0BTF8 Q0BTG0 Q0BTH9 Q0BTI5 Q0BTK1 Q0BTK2 Q0BTL9 Q0BTM0 Q0BTM1
Q0BTT2 Q0BTV7 Q0BTX6 Q0BTX8 Q0BTY4 Q0BTZ0 Q0BTZ5 Q0BU03 Q0BU46 Q0BU54 Q0BU70
Q0BU77 Q0BU81 Q0BUB3 Q0BUB7 Q0BUB8 Q0BUC5 Q0BUF0 Q0BUJ8 Q0BUK0 Q0BUL9 Q0BUM5
Q0BUM6 Q0BUM7 Q0BUM8 Q0BUN1 Q0BUN2 Q0BUN3 Q0BUN4 Q0BUN6 Q0BUN7 Q0BUN9 Q0BUP0
Q0BUP1 Q0BUP2 Q0BUP3 Q0BUP4 Q0BUP5 Q0BUP6 Q0BUP7 Q0BUP8 Q0BUP9 Q0BUQ0 Q0BUQ1
Q0BUQ2 Q0BUQ3 Q0BUQ4 Q0BUQ5 Q0BUQ6 Q0BUQ8 Q0BUR0 Q0BUR1 Q0BUS6 Q0BUU1 Q0BUV2
Q0BUV5 Q0BUV6 Q0BUX3 Q0BUX9 Q0BV08 Q0BV17 Q0BV26 Q0BV27 Q0BV28 Q0BV35 Q0BV38
Q0BV59 Q0BVB8 Q0BVE2 Q0BVF4 Q0BVG3 Q0BVG4 Q0BVH7 Q0BVJ1 Q0BVJ4 Q0BVJ5 Q0BVK2
Q0BVM8 Q0BVN0 Q0BVP4 Q0BVU4 Q0BVW2 Q0BVW4 Q0BVY6 Q0BVY8 Q0BW40 Q0BW57 Q0BW82
Q0BW92 Q0BW93 Q0BW94 Q0BW99 Q0BWA0 Q0BWA1 Q0BWA2 Q0BWA4 Q0BWA5 Q0BWA8 Q0BWA9
Q0BWB0 Q0BWB5 Q0BWC7 Q0BWH4 Q0BWI1 Q0BWJ4 Q0BWK6 Q0BWM8 Q0BWM9 Q0BWN3 Q0BWP1
Q0BWR4 Q0BWU4 Q0BWX0 Q0BWY9 Q0BWZ9 Q0BX02 Q0BX04 Q0BX22 Q0BX37 Q0BX67 Q0BX85

Q0BX92 Q0BXA0 Q0BXB5 Q0BXB7 Q0BXC2 Q0BXD3 Q0BXF6 Q0BXH7 Q0BXJ3 Q0BXT2 Q0BXT3
Q0BXT4 Q0BXU4 Q0BY00 Q0BY73 Q0BYA0 Q0BYA1 Q0BYA5 Q0BYA6 Q0BYA7 Q0BYA8 Q0BYA9
Q0BYB0 Q0BYB1 Q0BYB2 Q0BYB3 Q0BYB4 Q0BYB5 Q0BYB6 Q0BYB7 Q0BYB8 Q0BYC0 Q0BYC1
Q0BYC2 Q0BYC3 Q0BYC4 Q0BYC5 Q0BYC7 Q0BYC8 Q0BYC9 Q0BYD0 Q0BYD1 Q0BYD2 Q0BYD3
Q0BYD5 Q0BYD6 Q0BYD7 Q0BYD8 Q0BYD9 Q0BYG4 Q0BYG5 Q0BYJ5 Q0BYJ6 Q0BYK7 Q0BYK9
Q0BYM3 Q0BYP2 Q0BYP8 Q0BYQ0 Q0BYZ9 Q0BZ24 Q0BZ39 Q0BZ41 Q0BZ42 Q0BZB0 Q0BZK0
Q0BZK2 Q0BZK3 Q0BZT3 Q0BZU6 Q0BZU8 Q0BZU9 Q0BZV5 Q0BZW6 Q0BZX9 Q0BZZ1 Q0BZZ7
Q0C000 Q0C077 Q0C083 Q0C084 Q0C085 Q0C093 Q0C0C9 Q0C0G0 Q0C0H6 Q0C0I1 Q0C0I5
Q0C0I6 Q0C0J1 Q0C0L1 Q0C0L2 Q0C0N0 Q0C0N8 Q0C0R2 Q0C0T0 Q0C0T1 Q0C0T6 Q0C0U1
Q0C0W2 Q0C0W3 Q0C0Y0 Q0C137 Q0C154 Q0C161 Q0C187 Q0C190 Q0C195 Q0C197 Q0C1A1
Q0C1A2 Q0C1A6 Q0C1B8 Q0C1B9 Q0C1C0 Q0C1C1 Q0C1C5 Q0C1C9 Q0C1F4 Q0C1F7 Q0C1P4
Q0C1R0 Q0C2B5 Q0C2D0 Q0C2E7 Q0C2H1 Q0C2H2 Q0C2H9 Q0C2I2 Q0C2K0 Q0C347 Q0C348
Q0C3A9 Q0C3E2 Q0C3K3 Q0C3L9 Q0C3M1 Q0C3S3 Q0C3T7 Q0C3U2 Q0C3U4 Q0C3Z2 Q0C3Z3
Q0C406 Q0C427 Q0C429 Q0C438 Q0C441 Q0C444 Q0C451 Q0C454 Q0C4C9 Q0C4D8 Q0C4H1
Q0C4H3 Q0C4I8 Q0C4J3 Q0C4M2 Q0C4P9 Q0C4T2 Q0C4U8 Q0C4V1 Q0C504 Q0C511 Q0C516
Q0C530 Q0C532 Q0C533 Q0C535 Q0C545 Q0C548 Q0C551 Q0C566 Q0C570 Q0C577 Q0C592
Q0C593 Q0C5A1 Q0C5F3 Q0C5F9 Q0C5G6 Q0C5R4 Q0C5U1 Q0C5U8 Q0C5V2 Q0C5W2 Q0C5W4
Q0C5X3 Q0C5Z4 Q0C5Z5 Q0C5Z8 Q0C600 Q0C604 Q0C606 Q0C627 Q0C636 Q0C639 Q0C643
Q0C661 Q0C668 Q0C671 Q0C678 Q0C6A3 Q0C6A6 Q0C6A7 Q0C6A8 Q0C7C4 Q0C7E6 Q0C7M4
Q0C7Y1 Q0C807 Q0C838 Q0C898 Q0C899 Q0C8A0 Q0C8A2 Q0C8A3 Q0C8A4 Q0C8A5 Q0C8L3
Q0C8L6 Q0C8L8 Q0C8M0 Q0C8M1 Q0C8M2 Q0C8M3 Q0C9E6 Q0C9E8 Q0C9L6 Q0C9L7 Q0C9M9
Q0C9R2 Q0C9R3 Q0CA00 Q0CA25 Q0CA78 Q0CAS8 Q0CBD1 Q0CBD2 Q0CBE1 Q0CBN5 Q0CBS0
Q0CBX9 Q0CC19 Q0CCM5 Q0CCM6 Q0CCP0 Q0CCS0 Q0CCX4 Q0CCX5 Q0CCX8 Q0CCY0 Q0CCY1
Q0CCY2 Q0CCY3 Q0CCY4 Q0CCY5 Q0CD12 Q0CDB3 Q0CDB9 Q0CDP3 Q0CE14 Q0CE30 Q0CE91
Q0CEI2 Q0CEI3 Q0CEZ9 Q0CF43 Q0CF67 Q0CF68 Q0CF71 Q0CF73 Q0CF75 Q0CFD4 Q0CFY3
Q0CFZ0 Q0CG33 Q0CG34 Q0CG59 Q0CGJ9 Q0CH08 Q0CH39 Q0CH52 Q0CH56 Q0CH70 Q0CH76
Q0CHR0 Q0CHR1 Q0CI35 Q0CIC7 Q0CIL2 Q0CIL3 Q0CIQ3 Q0CJ13 Q0CJ54 Q0CJ56 Q0CJ57
Q0CJ59 Q0CJC3 Q0CJC5 Q0CJC6 Q0CJC7 Q0CJC9 Q0CJD1 Q0CJU7 Q0CK31 Q0CKB1 Q0CKB5
Q0CKU1 Q0CL13 Q0CL23 Q0CL68 Q0CL79 Q0CL94 Q0CLE0 Q0CLE8 Q0CLJ4 Q0CLJ6 Q0CLK1
Q0CLP3 Q0CLP9 Q0CLW8 Q0CLX0 Q0CM45 Q0CMB0 Q0CMM5 Q0CMM8 Q0CMY6 Q0CN46 Q0CNR3
Q0CNV1 Q0CNX1 Q0CNX5 Q0CP15 Q0CPA8 Q0CPB0 Q0CPF9 Q0CPP9 Q0CPV5 Q0CQ54 Q0CQF3
Q0CQF6 Q0CQH1 Q0CQK1 Q0CQK7 Q0CQL9 Q0CR17 Q0CR95 Q0CRF8 Q0CRP9 Q0CRQ4 Q0CRQ5
Q0CRW6 Q0CRX1 Q0CRX8 Q0CS61 Q0CS62 Q0CS64 Q0CS65 Q0CS66 Q0CS88 Q0CS89 Q0CS90
Q0CS91 Q0CS93 Q0CS94 Q0CS95 Q0CS96 Q0CS97 Q0CS99 Q0CSA0 Q0CSA2 Q0CSA3 Q0CSC0
Q0CSL5 Q0CSP9 Q0CSY3 Q0CT19 Q0CT27 Q0CT71 Q0CT94 Q0CTF3 Q0CTN3 Q0CTS1 Q0CTT3
Q0CU19 Q0CU52 Q0CU66 Q0CU99 Q0CUL0 Q0CUP6 Q0CV29 Q0CV52 Q0CVD6 Q0CVK8 Q0CVT0
Q0CW42 Q0CW83 Q0CWD0 Q0CWI2 Q0CWQ2 Q0CX45 Q0CX54 Q0CXD0 Q0CXH9 Q0CXN3 Q0CXS6
Q0CXW9 Q0CXX1 Q0CY06 Q0CY32 Q0CY48 Q0CYP4 Q0CYR5 Q0CZG5 Q0CZG8 Q0CZH1 Q0CZH2
Q0CZL3 Q0CZM6 Q0CZN5 Q0CZR3 Q0CZS8 Q0CZX6 Q0D034 Q0D0A1 Q0D0E8 Q0D0F3 Q0D0X6
Q0D131 Q0D154 Q0D155 Q0D158 Q0D159 Q0D1J4 Q0D1K3 Q0D1P3 Q0D1P4 Q0D1U6 Q0D205
Q0D252 Q0D2D2 Q0D2J5 Q0D2K2 Q0D2K3 Q0D2L3 Q0D3D2 Q0D3F2 Q0D3I7 Q0D3J9 Q0D3S3
Q0D4T4 Q0D4Z6 Q0D5B9 Q0D5G4 Q0D5W6 Q0D622 Q0D629 Q0D673 Q0D6F4 Q0D6M1 Q0D6V7
Q0D744 Q0D7I3 Q0D840 Q0D8D4 Q0D8N0 Q0D9C7 Q0D9R7 Q0DA50 Q0DAE4 Q0DB53 Q0DBD3
Q0DBS1 Q0DBU3 Q0DBU5 Q0DBW8 Q0DC89 Q0DDE3 Q0DDF6 Q0DDI1 Q0DEB8 Q0DEC8 Q0DEV5

Q0DFG8 Q0DFT7 Q0DG35 Q0DGG8 Q0DGS1 Q0DH40 Q0DHC4 Q0DHF6 Q0DI48 Q0DIQ5 Q0DIV0
Q0DJ94 Q0DJA3 Q0DJC7 Q0DJH7 Q0DJR9 Q0DJS1 Q0DJV6 Q0DK35 Q0DKA5 Q0DKE8 Q0DKF0
Q0DKF1 Q0DKM4 Q0DKN3 Q0DKP3 Q0DKP4 Q0DKY4 Q0DLA3 Q0DLB9 Q0DLD3 Q0DM51 Q0DNU1
Q0DQA9 Q0DR28 Q0DRV6 Q0DRX6 Q0DSH9 Q0DST9 Q0DSV9 Q0DTM7 Q0DUI1 Q0DUR2 Q0DV28
Q0DV32 Q0DV66 Q0DVU4 Q0DWF2 Q0DWH1 Q0DWH7 Q0DWQ7 Q0DWT8 Q0DX67 Q0DXB1 Q0DXS3
Q0DY72 Q0DY81 Q0DYB1 Q0DYP5 Q0DZE0 Q0DZP5 Q0DZT4 Q0E079 Q0E088 Q0E0A6 Q0E0Q3
Q0E0W7 Q0E0Y3 Q0E1D7 Q0E2L3 Q0E2P1 Q0E2Q3 Q0E2S4 Q0E2Y1 Q0E2Z7 Q0E342 Q0E3C3
Q0E3C8 Q0E3I9 Q0E3L5 Q0E3X4 Q0E474 Q0E4A8 Q0E517 Q0E559 Q0E592 Q0E7C4 Q0E7J7
Q0E7J8 Q0E908 Q0E940 Q0E959 Q0E9G3 Q0EAB8 Q0EDR2 Q0EF98 Q0FPQ4 Q0G819 Q0G825
Q0G838 Q0G9F5 Q0G9F6 Q0G9F7 Q0G9F8 Q0G9G2 Q0G9H1 Q0G9H8 Q0G9H9 Q0G9I0 Q0G9I1
Q0G9I2 Q0G9I3 Q0G9I4 Q0G9I5 Q0G9I6 Q0G9I7 Q0G9J4 Q0G9J5 Q0G9J6 Q0G9J7 Q0G9J8
Q0G9L7 Q0G9M1 Q0G9M7 Q0G9M8 Q0G9M9 Q0G9N0 Q0G9N7 Q0G9P9 Q0G9Q0 Q0G9Q1 Q0G9Q3
Q0G9Q5 Q0G9R4 Q0G9S1 Q0G9S2 Q0G9S3 Q0G9S4 Q0G9S5 Q0G9S6 Q0G9S7 Q0G9S8 Q0G9S9
Q0G9T0 Q0G9T7 Q0G9T8 Q0G9T9 Q0G9U0 Q0G9U1 Q0G9V4 Q0G9W0 Q0G9W4 Q0G9X0 Q0G9X1
Q0G9X2 Q0G9X3 Q0G9Y0 Q0G9Y1 Q0GA40 Q0GBX5 Q0GBX9 Q0GBY0 Q0GBY4 Q0GFF6 Q0GGX2
Q0GNE0 Q0GNE1 Q0GNN1 Q0GNQ5 Q0GP30 Q0GRC4 Q0GYU4 Q0GYU5 Q0GZS3 Q0H6L0 Q0H6L1
Q0H8C1 Q0H8C2 Q0H8C5 Q0H8D7 Q0H8X7 Q0H8X9 Q0H8Z0 Q0H8Z1 Q0H8Z2 Q0H8Z3 Q0HA25
Q0HD61 Q0HD62 Q0HD65 Q0HD68 Q0HD69 Q0HD92 Q0HD98 Q0HDA3 Q0HDB6 Q0HDC3 Q0HDC7
Q0HDH6 Q0HDK0 Q0HDN7 Q0HDU2 Q0HDU5 Q0HDU6 Q0HDU8 Q0HDU9 Q0HDV6 Q0HDV7 Q0HDW8
Q0HDW9 Q0HE08 Q0HE13 Q0HE39 Q0HE40 Q0HE49 Q0HE51 Q0HE53 Q0HE54 Q0HE56 Q0HE57
Q0HE58 Q0HE65 Q0HE67 Q0HE68 Q0HE70 Q0HE74 Q0HE75 Q0HE84 Q0HEA1 Q0HED0 Q0HED1
Q0HEF5 Q0HEF8 Q0HEI2 Q0HEJ5 Q0HEK0 Q0HEK4 Q0HEQ3 Q0HEQ4 Q0HEW1 Q0HEW2 Q0HEX0
Q0HEX1 Q0HEX2 Q0HEZ4 Q0HF12 Q0HF30 Q0HF31 Q0HF39 Q0HF41 Q0HF42 Q0HF52 Q0HF55
Q0HFA9 Q0HFC5 Q0HFC7 Q0HFC8 Q0HFE0 Q0HFL0 Q0HFL6 Q0HFL7 Q0HFM4 Q0HFM7 Q0HFR0
Q0HFS6 Q0HFX8 Q0HFX9 Q0HFY3 Q0HFY6 Q0HFY9 Q0HG41 Q0HG42 Q0HG49 Q0HG53 Q0HG57
Q0HG64 Q0HG72 Q0HG80 Q0HG81 Q0HG85 Q0HG98 Q0HGA8 Q0HGA9 Q0HGB0 Q0HGB1 Q0HGB4
Q0HGF3 Q0HGH0 Q0HGI8 Q0HGJ1 Q0HGL3 Q0HGL4 Q0HGL5 Q0HGM0 Q0HGM5 Q0HGN0 Q0HGN5
Q0HGT5 Q0HGV2 Q0HGV5 Q0HGV6 Q0HGV7 Q0HGV8 Q0HGW1 Q0HGW9 Q0HGX7 Q0HGZ0 Q0HGZ8
Q0HH98 Q0HH99 Q0HHA0 Q0HHA1 Q0HHA2 Q0HHF8 Q0HHH6 Q0HHI5 Q0HHJ4 Q0HHJ6 Q0HHL6
Q0HHL7 Q0HHM1 Q0HHN4 Q0HHN7 Q0HHP4 Q0HHQ6 Q0HHZ1 Q0HHZ4 Q0HHZ5 Q0HHZ7 Q0HI27
Q0HI69 Q0HI71 Q0HI72 Q0HI86 Q0HIB5 Q0HIB6 Q0HII5 Q0HIJ0 Q0HIK2 Q0HIK6 Q0HIL5
Q0HIL9 Q0HIM8 Q0HIR5 Q0HIS7 Q0HIT6 Q0HIT7 Q0HIT9 Q0HIW8 Q0HIX0 Q0HIX1 Q0HIX3
Q0HIX5 Q0HIZ1 Q0HIZ2 Q0HIZ3 Q0HIZ5 Q0HIZ7 Q0HIZ8 Q0HJ08 Q0HJ36 Q0HJ59 Q0HJ83
Q0HJ98 Q0HJ99 Q0HJD3 Q0HJE6 Q0HJE8 Q0HJF0 Q0HJF1 Q0HJF4 Q0HJG6 Q0HJG7 Q0HJM6
Q0HJN0 Q0HJN3 Q0HJN4 Q0HJN6 Q0HJP0 Q0HJP1 Q0HJP2 Q0HJP4 Q0HJQ4 Q0HJR6 Q0HJT7
Q0HJU9 Q0HJX9 Q0HJY0 Q0HJY2 Q0HJY5 Q0HJZ1 Q0HJZ9 Q0HK31 Q0HK48 Q0HK79 Q0HK81
Q0HKB3 Q0HKB6 Q0HKB8 Q0HKC3 Q0HKC5 Q0HKF7 Q0HKF8 Q0HKH4 Q0HKJ4 Q0HKK5 Q0HKU9
Q0HKV2 Q0HKV4 Q0HKV5 Q0HKV8 Q0HKV9 Q0HKW2 Q0HKY8 Q0HKZ7 Q0HL04 Q0HL06 Q0HL08
Q0HL58 Q0HL60 Q0HL62 Q0HL63 Q0HL66 Q0HL67 Q0HL69 Q0HL73 Q0HL77 Q0HL87 Q0HL88
Q0HL92 Q0HL93 Q0HLD7 Q0HLE7 Q0HLE8 Q0HLE9 Q0HLF0 Q0HLF4 Q0HLF6 Q0HLF8 Q0HLF9
Q0HLG0 Q0HLG1 Q0HLG3 Q0HLG5 Q0HLG6 Q0HLG9 Q0HLI0 Q0HLI3 Q0HLI5 Q0HLJ0 Q0HLJ3
Q0HLJ5 Q0HLK1 Q0HLK2 Q0HLK3 Q0HLL3 Q0HLL4 Q0HLM3 Q0HLM9 Q0HLN0 Q0HLP0 Q0HLQ7
Q0HLT2 Q0HLT3 Q0HLT4 Q0HLU0 Q0HLU1 Q0HLU2 Q0HLX7 Q0HLZ5 Q0HM10 Q0HM20 Q0HM44
Q0HM62 Q0HM63 Q0HM65 Q0HM78 Q0HMA6 Q0HMB1 Q0HMB2 Q0HMD9 Q0HMF0 Q0HMF2 Q0HMG7

Q0HMI1 Q0HMI8 Q0HMN9 Q0HMP0 Q0HMP1 Q0HMP6 Q0HMQ6 Q0HMR2 Q0HMS6 Q0HMU1 Q0HMX6
Q0HN48 Q0HN71 Q0HN72 Q0HN76 Q0HN86 Q0HN88 Q0HN93 Q0HN95 Q0HN97 Q0HN99 Q0HNC3
Q0HNG6 Q0HNG8 Q0HNL3 Q0HNM4 Q0HNR1 Q0HNR2 Q0HNR3 Q0HNR4 Q0HNR5 Q0HNR6 Q0HNR8
Q0HNR9 Q0HNS0 Q0HNS1 Q0HNS2 Q0HNS3 Q0HNS4 Q0HNS5 Q0HNS6 Q0HNS7 Q0HNS8 Q0HNS9
Q0HNT0 Q0HNT1 Q0HNT2 Q0HNT3 Q0HNT4 Q0HNT5 Q0HNT6 Q0HNT7 Q0HNT8 Q0HNT9 Q0HNU0
Q0HNU1 Q0HNU2 Q0HNU3 Q0HNU4 Q0HNU5 Q0HNU6 Q0HNU7 Q0HNU8 Q0HNV1 Q0HNV3 Q0HNV5
Q0HNV7 Q0HNV8 Q0HNW1 Q0HNW2 Q0HNY3 Q0HNY4 Q0HP01 Q0HP08 Q0HP10 Q0HP31 Q0HP37
Q0HP39 Q0HP46 Q0HP89 Q0HP91 Q0HP98 Q0HPA0 Q0HPA1 Q0HPA3 Q0HPA7 Q0HPB6 Q0HPC2
Q0HPC6 Q0HPC7 Q0HPD2 Q0HPD4 Q0HPE3 Q0HPE4 Q0HPE7 Q0HPF0 Q0HPF1 Q0HPH4 Q0HPI0
Q0HPI5 Q0HPJ5 Q0HPK4 Q0HPK8 Q0HPQ7 Q0HPU6 Q0HPU8 Q0HPY8 Q0HQ66 Q0HQ78 Q0HQC0
Q0HQC2 Q0HQG3 Q0HQI7 Q0HQI9 Q0HQJ2 Q0HQJ4 Q0HQJ9 Q0HQK1 Q0HQL1 Q0HQL5 Q0HQL6
Q0HQN5 Q0HQV5 Q0HQZ0 Q0HR05 Q0HR19 Q0HR25 Q0HR35 Q0HR40 Q0HR41 Q0HR42 Q0HRA0
Q0HRA7 Q0HRC1 Q0HRD6 Q0HRD8 Q0HRE9 Q0HRH5 Q0HRH6 Q0HRI1 Q0HRK8 Q0HRM1 Q0HRM3
Q0HRM4 Q0HRP2 Q0HRR5 Q0HRS5 Q0HRU1 Q0HRV9 Q0HRZ6 Q0HRZ7 Q0HRZ8 Q0HS04 Q0HS05
Q0HS06 Q0HS20 Q0HS71 Q0HS72 Q0HS76 Q0HS79 Q0HS82 Q0HSD4 Q0HSD5 Q0HSE2 Q0HSE6
Q0HSF0 Q0HSF7 Q0HSG5 Q0HSH3 Q0HSH4 Q0HSH8 Q0HSJ1 Q0HSK1 Q0HSK2 Q0HSK3 Q0HSK4
Q0HSK7 Q0HSP5 Q0HSR7 Q0HST5 Q0HST8 Q0HSW4 Q0HSW5 Q0HSW6 Q0HSX1 Q0HSX8 Q0HSY6
Q0HSZ1 Q0HT42 Q0HT59 Q0HT62 Q0HT63 Q0HT64 Q0HT65 Q0HT68 Q0HT76 Q0HT84 Q0HT97
Q0HTA5 Q0HTK4 Q0HTK5 Q0HTK6 Q0HTK7 Q0HTK8 Q0HTR2 Q0HTT0 Q0HTT9 Q0HTU8 Q0HTV0
Q0HTW8 Q0HTW9 Q0HTX3 Q0HTY6 Q0HTY9 Q0HTZ6 Q0HU08 Q0HU91 Q0HU94 Q0HU95 Q0HU97
Q0HUC4 Q0HUG5 Q0HUG7 Q0HUG8 Q0HUI2 Q0HUL2 Q0HUL3 Q0HUN4 Q0HUN5 Q0HUP0 Q0HUP8
Q0HUQ0 Q0HUQ2 Q0HUS4 Q0HUU6 Q0HUX4 Q0HUY4 Q0HUY5 Q0HUY7 Q0HUY9 Q0HUZ0 Q0HUZ1
Q0HV07 Q0HV09 Q0HV11 Q0HV12 Q0HV14 Q0HV45 Q0HV47 Q0HV48 Q0HV57 Q0HV71 Q0HVA7
Q0HVB6 Q0HVC0 Q0HVC9 Q0HVD3 Q0HVE5 Q0HVF0 Q0HVM5 Q0HVN6 Q0HVN8 Q0HVP0 Q0HVP1
Q0HVP4 Q0HVQ7 Q0HVQ8 Q0HVX2 Q0HVX6 Q0HVX9 Q0HVY0 Q0HVY2 Q0HVY6 Q0HVY7 Q0HVY8
Q0HVZ0 Q0HW00 Q0HW12 Q0HW33 Q0HW44 Q0HW75 Q0HW76 Q0HW78 Q0HW81 Q0HW92 Q0HWA0
Q0HWD2 Q0HWE9 Q0HWI1 Q0HWI3 Q0HWL5 Q0HWL8 Q0HWM0 Q0HWM5 Q0HWQ9 Q0HWR0 Q0HWS6
Q0HWU6 Q0HWV8 Q0HX47 Q0HX50 Q0HX52 Q0HX53 Q0HX56 Q0HX57 Q0HX60 Q0HX86 Q0HX95
Q0HXA2 Q0HXA4 Q0HXA6 Q0HXF6 Q0HXF8 Q0HXG0 Q0HXG1 Q0HXG4 Q0HXG5 Q0HXG7 Q0HXH1
Q0HXI0 Q0HXJ0 Q0HXJ1 Q0HXJ5 Q0HXJ6 Q0HXP1 Q0HXQ1 Q0HXQ2 Q0HXQ3 Q0HXQ4 Q0HXQ8
Q0HXR0 Q0HXR2 Q0HXR3 Q0HXR4 Q0HXR5 Q0HXR7 Q0HXR9 Q0HXS0 Q0HXS3 Q0HXT4 Q0HXT7
Q0HXT9 Q0HXU4 Q0HXU7 Q0HXU9 Q0HXV5 Q0HXV6 Q0HXV7 Q0HXZ4 Q0HXZ5 Q0HY04 Q0HY10
Q0HY11 Q0HY21 Q0HY38 Q0HY68 Q0HYA3 Q0HYA6 Q0HYB3 Q0HYB4 Q0HYC0 Q0HYJ5 Q0HYK8
Q0HYK9 Q0HYL1 Q0HYM7 Q0HYV7 Q0HYW0 Q0HYW3 Q0HYW4 Q0HYW6 Q0HYX4 Q0HYX5 Q0HYZ3
Q0HZ05 Q0HZ28 Q0HZ29 Q0HZ30 Q0HZ38 Q0HZ39 Q0HZ97 Q0HZ98 Q0HZE8 Q0HZF3 Q0HZF8
Q0HZH0 Q0HZJ3 Q0HZJ6 Q0HZL8 Q0HZL9 Q0HZP7 Q0HZR5 Q0HZS4 Q0HZS5 Q0HZS9 Q0HZT1
Q0HZT2 Q0HZT4 Q0HZU1 Q0HZU2 Q0HZU3 Q0HZU5 Q0HZU6 Q0HZU8 Q0HZV0 Q0HZX1 Q0HZX2
Q0HZZ6 Q0I001 Q0I040 Q0I041 Q0I052 Q0I053 Q0I060 Q0I061 Q0I063 Q0I064 Q0I067
Q0I079 Q0I080 Q0I081 Q0I082 Q0I083 Q0I084 Q0I086 Q0I087 Q0I088 Q0I089 Q0I090
Q0I091 Q0I092 Q0I093 Q0I094 Q0I095 Q0I096 Q0I097 Q0I098 Q0I099 Q0I0A0 Q0I0A1
Q0I0A2 Q0I0A3 Q0I0A4 Q0I0A5 Q0I0A6 Q0I0A7 Q0I0A8 Q0I0A9 Q0I0B0 Q0I0B1 Q0I0B2
Q0I0B3 Q0I0B4 Q0I0B5 Q0I0B6 Q0I0B9 Q0I0C1 Q0I0C3 Q0I0C5 Q0I0C6 Q0I0C9 Q0I0D0
Q0I0F1 Q0I0F2 Q0I0H0 Q0I0H7 Q0I0H9 Q0I0J9 Q0I0K5 Q0I0K7 Q0I0L4 Q0I0Q4 Q0I0Q6
Q0I0R4 Q0I0R6 Q0I0R7 Q0I0R9 Q0I0S3 Q0I0T2 Q0I0T6 Q0I0U0 Q0I0U1 Q0I0U6 Q0I0U8

Q0I0U9 Q0I0V0 Q0I0V1 Q0I0V2 Q0I0V8 Q0I0W6 Q0I0X6 Q0I0X7 Q0I0X8 Q0I0X9 Q0I0Y1
Q0I0Y5 Q0I0Y7 Q0I0Y8 Q0I0Y9 Q0I0Z2 Q0I101 Q0I127 Q0I129 Q0I131 Q0I132 Q0I136
Q0I137 Q0I138 Q0I139 Q0I140 Q0I141 Q0I143 Q0I144 Q0I145 Q0I146 Q0I147 Q0I148
Q0I149 Q0I150 Q0I151 Q0I152 Q0I154 Q0I155 Q0I156 Q0I157 Q0I158 Q0I159 Q0I160
Q0I161 Q0I162 Q0I163 Q0I164 Q0I178 Q0I181 Q0I182 Q0I195 Q0I1B3 Q0I1B4 Q0I1B6
Q0I1B7 Q0I1B8 Q0I1D1 Q0I1D2 Q0I1D8 Q0I1E1 Q0I1E2 Q0I1E6 Q0I1F1 Q0I1F8 Q0I1G1
Q0I1H1 Q0I1H7 Q0I1H8 Q0I1I3 Q0I1I7 Q0I1I8 Q0I1J7 Q0I1J8 Q0I1J9 Q0I1K0 Q0I1K5
Q0I1K8 Q0I1L2 Q0I1M1 Q0I1M3 Q0I1M4 Q0I1P5 Q0I1P7 Q0I1P8 Q0I1Q0 Q0I1Q1 Q0I1R2
Q0I1R4 Q0I1R5 Q0I1S1 Q0I1S4 Q0I1U0 Q0I1U8 Q0I1U9 Q0I1V8 Q0I1X0 Q0I1X3 Q0I1X5
Q0I1Y1 Q0I1Y6 Q0I1Y8 Q0I200 Q0I203 Q0I236 Q0I243 Q0I244 Q0I248 Q0I251 Q0I256
Q0I260 Q0I267 Q0I270 Q0I271 Q0I272 Q0I273 Q0I275 Q0I279 Q0I282 Q0I283 Q0I284
Q0I285 Q0I297 Q0I2A3 Q0I2D2 Q0I2D7 Q0I2E5 Q0I2E8 Q0I2E9 Q0I2G1 Q0I2G3 Q0I2G4
Q0I2G6 Q0I2G9 Q0I2H6 Q0I2K0 Q0I2M9 Q0I2N2 Q0I2N3 Q0I2N7 Q0I2P6 Q0I2Q5 Q0I2R1
Q0I2S8 Q0I2T0 Q0I2T8 Q0I2T9 Q0I2U1 Q0I2U5 Q0I2U9 Q0I2W9 Q0I2X4 Q0I2X6 Q0I2X8
Q0I2Y4 Q0I2Y5 Q0I2Y8 Q0I2Y9 Q0I316 Q0I320 Q0I322 Q0I355 Q0I368 Q0I370 Q0I371
Q0I378 Q0I379 Q0I390 Q0I3A1 Q0I3A8 Q0I3B0 Q0I3B5 Q0I3C7 Q0I3D1 Q0I3D2 Q0I3F6
Q0I3G1 Q0I3G4 Q0I3H7 Q0I3H9 Q0I3I0 Q0I3J0 Q0I3J1 Q0I3J3 Q0I3K1 Q0I3K2 Q0I3K3
Q0I3K4 Q0I3K7 Q0I3L5 Q0I3M2 Q0I3N1 Q0I3N5 Q0I3N6 Q0I3N9 Q0I3P2 Q0I3P3 Q0I3P4
Q0I3P5 Q0I3P7 Q0I3Q6 Q0I3Q7 Q0I3R1 Q0I3R3 Q0I3R5 Q0I3R7 Q0I3S3 Q0I3S4 Q0I3S5
Q0I3T0 Q0I3T3 Q0I3U1 Q0I3U5 Q0I3U8 Q0I3V1 Q0I3V2 Q0I3W7 Q0I3W8 Q0I3X9 Q0I3Y4
Q0I3Z1 Q0I414 Q0I426 Q0I432 Q0I441 Q0I442 Q0I447 Q0I456 Q0I457 Q0I458 Q0I462
Q0I463 Q0I471 Q0I476 Q0I490 Q0I491 Q0I495 Q0I4A3 Q0I4A5 Q0I4B4 Q0I4B6 Q0I4B9
Q0I4C6 Q0I4D8 Q0I4D9 Q0I4E5 Q0I4E6 Q0I4E7 Q0I4E8 Q0I4F0 Q0I4F2 Q0I4M1 Q0I4M6
Q0I4N3 Q0I4P2 Q0I4Q5 Q0I4Q7 Q0I4R7 Q0I4R8 Q0I4S4 Q0I4S5 Q0I4S8 Q0I4T0 Q0I4T7
Q0I4U1 Q0I4U4 Q0I4V7 Q0I4X5 Q0I4X6 Q0I4Y5 Q0I4Z6 Q0I508 Q0I511 Q0I514 Q0I516
Q0I525 Q0I534 Q0I535 Q0I536 Q0I537 Q0I544 Q0I554 Q0I555 Q0I557 Q0I559 Q0I560
Q0I563 Q0I569 Q0I572 Q0I586 Q0I591 Q0I593 Q0I5A5 Q0I5B0 Q0I5B7 Q0I5B8 Q0I5C3
Q0I5C4 Q0I5C5 Q0I5D1 Q0I5D2 Q0I5E0 Q0I5E1 Q0I5E2 Q0I5G1 Q0I5H4 Q0I5H7 Q0I5H8
Q0I5H9 Q0I5I6 Q0I5J0 Q0I5K4 Q0I5L8 Q0I5L9 Q0I5M2 Q0I5M5 Q0I5V1 Q0I5V7 Q0I5W2
Q0I5W4 Q0I5W5 Q0I5X6 Q0I5Y6 Q0I600 Q0I607 Q0I609 Q0I634 Q0I656 Q0I657 Q0I658
Q0I659 Q0I660 Q0I662 Q0I671 Q0I677 Q0I682 Q0I685 Q0I6C8 Q0I6D8 Q0I6E0 Q0I6E5
Q0I6G6 Q0I6J3 Q0I6K7 Q0I6K8 Q0I6K9 Q0I6L0 Q0I6M5 Q0I6N5 Q0I6R7 Q0I6S3 Q0I6T4
Q0I6T7 Q0I6V1 Q0I6Z1 Q0I6Z6 Q0I721 Q0I735 Q0I748 Q0I762 Q0I7A5 Q0I7A8 Q0I7I2
Q0I7K2 Q0I7K5 Q0I7L2 Q0I7L4 Q0I7L7 Q0I7L8 Q0I7L9 Q0I7M1 Q0I7M9 Q0I7U2 Q0I7U3
Q0I7V6 Q0I7X5 Q0I7Z0 Q0I7Z1 Q0I800 Q0I801 Q0I803 Q0I808 Q0I816 Q0I824 Q0I858
Q0I865 Q0I868 Q0I880 Q0I885 Q0I8C2 Q0I8C9 Q0I8D1 Q0I8D2 Q0I8G1 Q0I8H5 Q0I8I3
Q0I8J0 Q0I8J6 Q0I8J8 Q0I8L4 Q0I8L5 Q0I8L9 Q0I8P4 Q0I8P5 Q0I8P6 Q0I8Q2 Q0I8S8
Q0I8T6 Q0I8Y1 Q0I8Y2 Q0I8Z0 Q0I8Z9 Q0I902 Q0I9E5 Q0I9I2 Q0I9I3 Q0I9J4 Q0I9N0
Q0I9N6 Q0I9N9 Q0I9P0 Q0I9R7 Q0I9S3 Q0I9S4 Q0I9X0 Q0I9X3 Q0I9Y0 Q0I9Y8 Q0I9Z1
Q0IAA6 Q0IAB7 Q0IAE0 Q0IAF3 Q0IAF4 Q0IAH7 Q0IAH8 Q0IAJ5 Q0IAL0 Q0IAL3 Q0IAN8
Q0IAN9 Q0IAP2 Q0IAT7 Q0IAV3 Q0IB11 Q0IB24 Q0IBC2 Q0IBC8 Q0IBG0 Q0IBG9 Q0IBI3
Q0IBJ8 Q0IBN9 Q0IBR4 Q0IBR6 Q0IC13 Q0IC17 Q0IC27 Q0IC33 Q0IC41 Q0IC49 Q0IC57
Q0IC70 Q0IC71 Q0IC75 Q0IC80 Q0ICA6 Q0ICH2 Q0ICH5 Q0ICI5 Q0ICK0 Q0ICK8 Q0ICL5
Q0ICQ3 Q0ICR4 Q0ICR7 Q0ICV5 Q0ICW2 Q0ICW4 Q0ID00 Q0ID05 Q0ID06 Q0ID07 Q0ID08

Q0ID09 Q0ID10 Q0ID11 Q0ID12 Q0ID13 Q0ID14 Q0ID15 Q0ID16 Q0ID17 Q0ID18 Q0ID19
Q0ID20 Q0ID22 Q0ID24 Q0ID25 Q0ID26 Q0ID27 Q0ID28 Q0ID29 Q0ID31 Q0ID32 Q0ID33
Q0ID34 Q0ID56 Q0ID57 Q0ID58 Q0ID59 Q0ID60 Q0ID65 Q0ID68 Q0ID82 Q0ID96 Q0ID97
Q0IDA2 Q0IDA5 Q0IDB0 Q0IDB3 Q0IDC6 Q0IDD5 Q0IDD6 Q0IDD8 Q0IDE3 Q0IDE5 Q0IDE8
Q0IDF3 Q0IDG4 Q0IDH6 Q0IDK4 Q0IDP0 Q0IDV6 Q0IDW1 Q0IDW2 Q0IDW9 Q0IDX7 Q0IDZ4
Q0IDZ5 Q0IE03 Q0IE05 Q0IE10 Q0IE16 Q0IE18 Q0IE20 Q0IE25 Q0IE26 Q0IE37 Q0IE51
Q0IE55 Q0IE56 Q0IE57 Q0IE58 Q0IE70 Q0IE95 Q0IEG8 Q0IEH7 Q0IEK6 Q0IEN3 Q0IES7
Q0IES8 Q0IEW8 Q0IEY3 Q0IF90 Q0IFL2 Q0IFL5 Q0IG18 Q0IG34 Q0IGK1 Q0IGM7 Q0IH29
Q0IH40 Q0IH86 Q0IH88 Q0IH98 Q0IHA2 Q0IHB0 Q0IHB3 Q0IHB8 Q0IHF8 Q0IHH1 Q0IHH9
Q0IHI4 Q0IHI6 Q0IHI7 Q0IHJ3 Q0IHN5 Q0IHP3 Q0IHR1 Q0IHT1 Q0IHT3 Q0IHU5 Q0IHU9
Q0IHV2 Q0IHV9 Q0IHW3 Q0IHW6 Q0IHW8 Q0IHY4 Q0II04 Q0II25 Q0II26 Q0II29 Q0II48
Q0II50 Q0II59 Q0II68 Q0II70 Q0II83 Q0II87 Q0II90 Q0II91 Q0IIA6 Q0IIB1 Q0IIB6
Q0IIC2 Q0IID2 Q0IID7 Q0IIE6 Q0IIF0 Q0IIF6 Q0IIG6 Q0IIH4 Q0III0 Q0III6 Q0IIJ0
Q0IIJ2 Q0IIK5 Q0IIL7 Q0IIM1 Q0IIM3 Q0IIM8 Q0IIN9 Q0IIR9 Q0IIX4 Q0IIX9 Q0IIZ5
Q0IJ29 Q0IJ31 Q0ILZ4 Q0IMG5 Q0IMG9 Q0IMS9 Q0IMV4 Q0INC5 Q0ING3 Q0INT0 Q0INW1
Q0INY7 Q0INZ4 Q0IP69 Q0IQF7 Q0IQJ7 Q0IQN5 Q0IQN6 Q0IRB0 Q0ITS8 Q0ITU1 Q0ITW7
Q0IV63 Q0IWL9 Q0IWM5 Q0IXP9 Q0IZQ2 Q0IZS0 Q0J006 Q0J035 Q0J059 Q0J090 Q0J0H4
Q0J0S6 Q0J1C1 Q0J233 Q0J235 Q0J2C6 Q0J2L7 Q0J2R1 Q0J3H7 Q0J3L4 Q0J3Y6 Q0J3Y7
Q0J466 Q0J4D4 Q0J5F8 Q0J6N4 Q0J6P7 Q0J6T3 Q0J709 Q0J716 Q0J7J7 Q0J7N5 Q0J7P4
Q0J7U6 Q0J7V9 Q0J7Y8 Q0J8A4 Q0J8G4 Q0J8G8 Q0J8M2 Q0J8Q3 Q0J951 Q0J952 Q0J998
Q0J9J6 Q0J9V3 Q0J9V5 Q0J9W0 Q0JA75 Q0JA81 Q0JA82 Q0JA83 Q0JAA0 Q0JAD9 Q0JB88
Q0JB89 Q0JB92 Q0JBF0 Q0JBP5 Q0JBY9 Q0JBZ8 Q0JC10 Q0JC27 Q0JC44 Q0JCC3 Q0JCT1
Q0JCZ4 Q0JD07 Q0JDM0 Q0JDM4 Q0JE32 Q0JEB7 Q0JEF5 Q0JF02 Q0JF03 Q0JF48 Q0JF58
Q0JFN7 Q0JG89 Q0JG98 Q0JG99 Q0JGK4 Q0JGS5 Q0JGZ6 Q0JH50 Q0JHF1 Q0JHF8 Q0JHU7
Q0JHZ2 Q0JIC2 Q0JIF2 Q0JIL1 Q0JIZ1 Q0JJ01 Q0JJ05 Q0JJD1 Q0JJS8 Q0JKD0 Q0JKG7
Q0JKI9 Q0JKK6 Q0JKX1 Q0JKZ0 Q0JL44 Q0JL73 Q0JL75 Q0JLE4 Q0JLP9 Q0JM17 Q0JM76
Q0JMB2 Q0JMD4 Q0JMH0 Q0JMV4 Q0JMW0 Q0JMY1 Q0JMY8 Q0JNK6 Q0JNL3 Q0JNL7 Q0JNS6
Q0JP11 Q0JP62 Q0JP99 Q0JPA4 Q0JPT4 Q0JQ97 Q0JQF7 Q0JXE7 Q0JY21 Q0JYN7 Q0JZT9
Q0JZW8 Q0K0D3 Q0K0L8 Q0K0V6 Q0K0X8 Q0K181 Q0K1E0 Q0K1H0 Q0K292 Q0K2D1 Q0K2R3
Q0K391 Q0K3N1 Q0K3N2 Q0K3R9 Q0K3S5 Q0K3S6 Q0K4F5 Q0K4F6 Q0K4I4 Q0K4N8 Q0K4W7
Q0K572 Q0K5B8 Q0K5L6 Q0K5L7 Q0K5N1 Q0K5N7 Q0K5P5 Q0K5T4 Q0K5Z9 Q0K602 Q0K603
Q0K604 Q0K605 Q0K606 Q0K607 Q0K609 Q0K610 Q0K613 Q0K619 Q0K620 Q0K621 Q0K622
Q0K623 Q0K624 Q0K625 Q0K626 Q0K627 Q0K628 Q0K629 Q0K630 Q0K631 Q0K632 Q0K633
Q0K634 Q0K635 Q0K636 Q0K637 Q0K638 Q0K640 Q0K641 Q0K642 Q0K643 Q0K644 Q0K645
Q0K646 Q0K651 Q0K661 Q0K669 Q0K685 Q0K686 Q0K689 Q0K692 Q0K693 Q0K694 Q0K695
Q0K6G3 Q0K6H7 Q0K6I0 Q0K6I1 Q0K6L6 Q0K6M5 Q0K6M6 Q0K6N8 Q0K6P4 Q0K6P5 Q0K6P6
Q0K6P7 Q0K6P9 Q0K6Q0 Q0K6V7 Q0K6X3 Q0K6Y1 Q0K6Y5 Q0K6Y9 Q0K705 Q0K710 Q0K720
Q0K733 Q0K734 Q0K757 Q0K758 Q0K762 Q0K769 Q0K776 Q0K784 Q0K786 Q0K793 Q0K7A0
Q0K7B1 Q0K7B2 Q0K7B3 Q0K7D1 Q0K7F5 Q0K7H7 Q0K7I6 Q0K7K0 Q0K7L4 Q0K7M0 Q0K7N0
Q0K7N2 Q0K7S5 Q0K7T8 Q0K7T9 Q0K7V9 Q0K7W0 Q0K7W8 Q0K7X7 Q0K7X9 Q0K7Z0 Q0K808
Q0K823 Q0K844 Q0K845 Q0K846 Q0K848 Q0K860 Q0K861 Q0K862 Q0K889 Q0K8D7 Q0K8F5
Q0K8H3 Q0K8J6 Q0K8J9 Q0K8K6 Q0K8L7 Q0K8N3 Q0K8N6 Q0K8N9 Q0K8W6 Q0K8Y4 Q0K8Y7
Q0K8Z9 Q0K926 Q0K929 Q0K950 Q0K951 Q0K956 Q0K958 Q0K959 Q0K963 Q0K967 Q0K971
Q0K979 Q0K980 Q0K981 Q0K9B7 Q0K9B9 Q0K9C0 Q0K9E5 Q0K9E7 Q0K9E8 Q0K9I0 Q0K9J4

Q0K9U5 Q0K9V9 Q0KA02 Q0KA05 Q0KA11 Q0KA16 Q0KA17 Q0KA18 Q0KA19 Q0KA22 Q0KA30
Q0KA32 Q0KA36 Q0KA43 Q0KA59 Q0KAH8 Q0KAQ9 Q0KB67 Q0KBB2 Q0KBC0 Q0KBC7 Q0KBC8
Q0KBC9 Q0KBD1 Q0KBD2 Q0KBK3 Q0KBK4 Q0KBK5 Q0KBK6 Q0KBM9 Q0KBP1 Q0KBP2 Q0KBT2
Q0KBT3 Q0KBW5 Q0KBY9 Q0KBZ0 Q0KBZ1 Q0KBZ3 Q0KC14 Q0KCA9 Q0KCC0 Q0KCD1 Q0KCE5
Q0KCG3 Q0KCG7 Q0KCJ6 Q0KCP3 Q0KCP4 Q0KCP6 Q0KCP7 Q0KCT2 Q0KCT4 Q0KCT5 Q0KCT8
Q0KCU2 Q0KCW5 Q0KCY0 Q0KD22 Q0KD27 Q0KD59 Q0KD64 Q0KD76 Q0KD78 Q0KD79 Q0KD80
Q0KD81 Q0KDH0 Q0KDH7 Q0KDH9 Q0KDK9 Q0KDL6 Q0KDN9 Q0KDR7 Q0KDR8 Q0KE18 Q0KE20
Q0KE53 Q0KE68 Q0KE75 Q0KE78 Q0KE93 Q0KE95 Q0KEA7 Q0KEC0 Q0KEC1 Q0KEC3 Q0KEC4
Q0KED0 Q0KED6 Q0KED8 Q0KED9 Q0KEE2 Q0KEG8 Q0KEH6 Q0KEJ5 Q0KEM6 Q0KEN8 Q0KEP0
Q0KEP4 Q0KEP7 Q0KEP9 Q0KEQ0 Q0KEQ3 Q0KET5 Q0KET8 Q0KEU1 Q0KF09 Q0KF16 Q0KF25
Q0KF31 Q0KF34 Q0KF39 Q0KF42 Q0KF45 Q0KF58 Q0KF61 Q0KF69 Q0KF70 Q0KF86 Q0KF88
Q0KFB0 Q0KFE0 Q0KFE3 Q0KFE6 Q0KFE7 Q0KFF7 Q0KFF8 Q0KFG1 Q0KFG6 Q0KFI3 Q0KFI5
Q0KFM5 Q0KFR8 Q0KI28 Q0KIA2 Q0KIC3 Q0KIY0 Q0KIY1 Q0KIY2 Q0KIY3 Q0KIY5 Q0KIY6
Q0KIY7 Q0KIY9 Q0KJL4 Q0KK56 Q0MTC0 Q0MUU2 Q0MVN8 Q0MW30 Q0MXD0 Q0MXD1 Q0P140
Q0P3J3 Q0P3J7 Q0P3K4 Q0P3K9 Q0P3L0 Q0P3L1 Q0P3L2 Q0P3L3 Q0P3L4 Q0P3L5 Q0P3L6
Q0P3L7 Q0P3L8 Q0P3L9 Q0P3M0 Q0P3M3 Q0P3M4 Q0P3M5 Q0P3M6 Q0P3M7 Q0P3M8 Q0P3M9
Q0P3N0 Q0P3N2 Q0P3N5 Q0P3P4 Q0P3P5 Q0P3P7 Q0P3R5 Q0P3U3 Q0P3X7 Q0P3X8 Q0P3Z2
Q0P3Z8 Q0P424 Q0P427 Q0P448 Q0P451 Q0P457 Q0P464 Q0P466 Q0P485 Q0P499 Q0P4A6
Q0P4B9 Q0P4C4 Q0P4C5 Q0P4D1 Q0P4D6 Q0P4G8 Q0P4H0 Q0P4H6 Q0P4I1 Q0P4K0 Q0P4K8
Q0P4M4 Q0P4Q0 Q0P4R5 Q0P4R6 Q0P4S5 Q0P4U8 Q0P4V4 Q0P4W6 Q0P4W9 Q0P4X6 Q0P4Y1
Q0P565 Q0P571 Q0P574 Q0P593 Q0P5A3 Q0P5A4 Q0P5A5 Q0P5B2 Q0P5B3 Q0P5B4 Q0P5B9
Q0P5C2 Q0P5C3 Q0P5D1 Q0P5D2 Q0P5D8 Q0P5E2 Q0P5E7 Q0P5F2 Q0P5F9 Q0P5G1 Q0P5G4
Q0P5H2 Q0P5H7 Q0P5I0 Q0P5I5 Q0P5I6 Q0P5J4 Q0P5J6 Q0P5J7 Q0P5J9 Q0P5K1 Q0P5K3
Q0P5L0 Q0P5L8 Q0P5M0 Q0P5N2 Q0P5N6 Q0P5V2 Q0P5X1 Q0P5X5 Q0P641 Q0P678 Q0P6D6
Q0P7R8 Q0P7S6 Q0P7T4 Q0P7T9 Q0P7U6 Q0P7V6 Q0P823 Q0P832 Q0P891 Q0P897 Q0P899
Q0P8G1 Q0P8H3 Q0P8H7 Q0P8H8 Q0P8I7 Q0P8I9 Q0P8J1 Q0P8J6 Q0P8J7 Q0P8J8 Q0P8J9
Q0P8R5 Q0P8S7 Q0P8T0 Q0P8T1 Q0P8U0 Q0P8U2 Q0P8U3 Q0P8U4 Q0P8U5 Q0P8U6 Q0P8W3
Q0P8W4 Q0P903 Q0P9D3 Q0P9F1 Q0P9H9 Q0P9K4 Q0P9Q9 Q0P9S6 Q0P9V4 Q0PA13 Q0PA27
Q0PA35 Q0PA50 Q0PAS0 Q0PAS7 Q0PAT1 Q0PB48 Q0PB55 Q0PBB6 Q0PBW9 Q0PBX7 Q0PC20
Q0PC41 Q0PC43 Q0PCF4 Q0PCS3 Q0PDK6 Q0PDK7 Q0PF16 Q0PGJ6 Q0PGW2 Q0PHS9 Q0PHV7
Q0PHX9 Q0PHY0 Q0PI70 Q0PI71 Q0PKV7 Q0PNE2 Q0PUX0 Q0PUX7 Q0PXV9 Q0PXX8 Q0PXY6
Q0PZF7 Q0PZH5 Q0Q027 Q0Q043 Q0Q0E4 Q0Q4F0 Q0QEL7 Q0QFQ1 Q0QFQ2 Q0QFQ3 Q0QHL3
Q0QLE1 Q0QLE2 Q0QLE4 Q0QLE6 Q0QLE9 Q0QLF1 Q0QLF2 Q0QLF3 Q0QLF4 Q0QLF5 Q0QLF7
Q0QWS4 Q0QZR6 Q0R4F1 Q0RAP0 Q0RAR6 Q0RAR7 Q0RAW2 Q0RB64 Q0RB90 Q0RB93 Q0RB94
Q0RBA3 Q0RBB8 Q0RBC4 Q0RBL9 Q0RBM4 Q0RBN8 Q0RBQ7 Q0RBS5 Q0RC35 Q0RCD6 Q0RCJ1
Q0RCJ7 Q0RCR9 Q0RD46 Q0RD93 Q0RD99 Q0RDC6 Q0RDH6 Q0RDH8 Q0RDI8 Q0RDK7 Q0RDK8
Q0RDM7 Q0RDN6 Q0RDP2 Q0RDP3 Q0RDP4 Q0RDP5 Q0RDP7 Q0RDP9 Q0RDQ4 Q0RDQ5 Q0RDQ6
Q0RDQ8 Q0RDR2 Q0RDS4 Q0RDS6 Q0RDT0 Q0RDT1 Q0RDV0 Q0RDW5 Q0RDX2 Q0RDX5 Q0RE66
Q0RE94 Q0REL2 Q0RES2 Q0RF57 Q0RF65 Q0RF70 Q0RF77 Q0RF85 Q0RF89 Q0RF99 Q0RFA0
Q0RFA8 Q0RFB2 Q0RFD5 Q0RFE8 Q0RFF6 Q0RFI3 Q0RFX1 Q0RFX5 Q0RFX9 Q0RH01 Q0RH03
Q0RH06 Q0RH18 Q0RH38 Q0RH75 Q0RH96 Q0RHA0 Q0RI14 Q0RJ31 Q0RJ34 Q0RJF1 Q0RK33
Q0RKL9 Q0RKY6 Q0RLT2 Q0RLT6 Q0RLT7 Q0RLU1 Q0RLV2 Q0RM11 Q0RM12 Q0RNE2 Q0RNN4
Q0RNN9 Q0RNP9 Q0RNQ0 Q0RNU1 Q0RNU6 Q0RNU8 Q0RNU9 Q0RNV0 Q0RP74 Q0RP78 Q0RPE7
Q0RPF6 Q0RPF7 Q0RPG8 Q0RPH1 Q0RPH4 Q0RPW3 Q0RQ25 Q0RQL6 Q0RQR8 Q0RQR9 Q0RQS0

Q0RQS5 Q0RQV5 Q0RQV6 Q0RR00 Q0RRF4 Q0RRG0 Q0RRL9 Q0RRM3 Q0RRN7 Q0RRP2 Q0RRP3
Q0RRP4 Q0RRP5 Q0RRP6 Q0RRP7 Q0RRP8 Q0RRQ3 Q0RRQ4 Q0RRQ6 Q0RRQ7 Q0RRQ8 Q0RRQ9
Q0RRR0 Q0RRR1 Q0RRR3 Q0RRR4 Q0RRR5 Q0RRR6 Q0RRR7 Q0RRR8 Q0RRR9 Q0RRS1 Q0RRS2
Q0RRS3 Q0RRS4 Q0RRS5 Q0RRS6 Q0RRS9 Q0RRT0 Q0RRT2 Q0RRT3 Q0RRT5 Q0RRT6 Q0RRU1
Q0RRU6 Q0RRX8 Q0RS03 Q0RS17 Q0RS35 Q0RS49 Q0RS51 Q0RS64 Q0RT19 Q0RTD6 Q0RUF3
Q0RUP6 Q0RUR2 Q0RV72 Q0RV73 Q0RVH6 Q0RVH7 Q0RVJ9 Q0RWM7 Q0RWN1 Q0RXC3 Q0RXC4
Q0RZ64 Q0S005 Q0S006 Q0S032 Q0S0J3 Q0S0J4 Q0S0J6 Q0S0J8 Q0S0J9 Q0S0K2 Q0S0L4
Q0S0L6 Q0S0L7 Q0S0M4 Q0S0M6 Q0S0M7 Q0S0N1 Q0S0N2 Q0S0N6 Q0S0N7 Q0S0P3 Q0S1B6
Q0S1C1 Q0S1C6 Q0S1C7 Q0S1C8 Q0S1C9 Q0S1D2 Q0S1E2 Q0S1F6 Q0S1H1 Q0S1M4 Q0S1N7
Q0S1N9 Q0S1P3 Q0S1P9 Q0S208 Q0S209 Q0S212 Q0S217 Q0S219 Q0S222 Q0S235 Q0S251
Q0S258 Q0S275 Q0S277 Q0S282 Q0S283 Q0S284 Q0S285 Q0S2B2 Q0S2B9 Q0S2C1 Q0S2C5
Q0S2C6 Q0S2C8 Q0S2D6 Q0S2E0 Q0S2E4 Q0S2F6 Q0S2F9 Q0S2G8 Q0S2H1 Q0S2H3 Q0S2I1
Q0S2I8 Q0S2J2 Q0S2J4 Q0S2M1 Q0S2M9 Q0S2N2 Q0S2Q1 Q0S2Q6 Q0S2X1 Q0S319 Q0S365
Q0S375 Q0S392 Q0S3B9 Q0S3C0 Q0S3C4 Q0S3D6 Q0S3D9 Q0S3E6 Q0S3E7 Q0S3E8 Q0S3E9
Q0S3F0 Q0S3F1 Q0S3F2 Q0S3F3 Q0S3F7 Q0S3F8 Q0S3F9 Q0S3G0 Q0S3G1 Q0S3G2 Q0S3G3
Q0S3G5 Q0S3G6 Q0S3G7 Q0S3G8 Q0S3G9 Q0S3H0 Q0S3H1 Q0S3H2 Q0S3H3 Q0S3H4 Q0S3H6
Q0S3H7 Q0S3Y4 Q0S424 Q0S467 Q0S489 Q0S4A2 Q0S4F8 Q0S4F9 Q0S4R2 Q0S4R3 Q0S4S3
Q0S4S5 Q0S4S7 Q0S4S8 Q0S4S9 Q0S4T6 Q0S4V3 Q0S4V4 Q0S4Y6 Q0S4Y7 Q0S4Z0 Q0S4Z1
Q0S4Z2 Q0S533 Q0S539 Q0S555 Q0S5R4 Q0S5Y0 Q0S679 Q0S6R1 Q0S6R3 Q0S6X7 Q0S740
Q0S749 Q0S766 Q0S774 Q0S784 Q0S7Q9 Q0S7R0 Q0S7V2 Q0S815 Q0S816 Q0S868 Q0S888
Q0S889 Q0S890 Q0S8D3 Q0S8D4 Q0S8D5 Q0S8E7 Q0S8V5 Q0S8Y4 Q0S8Y5 Q0S962 Q0S9E8
Q0S9W9 Q0S9X0 Q0SAG3 Q0SAG4 Q0SAG7 Q0SAG8 Q0SAG9 Q0SAH7 Q0SAK3 Q0SAL1 Q0SAY4
Q0SB08 Q0SB51 Q0SB53 Q0SB54 Q0SBH9 Q0SED2 Q0SEI1 Q0SEI4 Q0SEQ7 Q0SET3 Q0SF06
Q0SF09 Q0SF26 Q0SF42 Q0SF82 Q0SF90 Q0SF91 Q0SF96 Q0SF97 Q0SF99 Q0SFB3 Q0SFB4
Q0SFF1 Q0SFF2 Q0SFF3 Q0SFF4 Q0SFL5 Q0SFL6 Q0SFQ8 Q0SFS2 Q0SG75 Q0SGH6 Q0SGJ6
Q0SGJ7 Q0SGN1 Q0SGN7 Q0SGN8 Q0SGQ9 Q0SGR9 Q0SGS7 Q0SGU7 Q0SGY9 Q0SGZ0 Q0SGZ1
Q0SGZ3 Q0SH07 Q0SH47 Q0SH49 Q0SH50 Q0SH51 Q0SH52 Q0SH55 Q0SH62 Q0SH75 Q0SH77
Q0SHB5 Q0SHC1 Q0SHD7 Q0SHJ0 Q0SHK6 Q0SHL0 Q0SHN1 Q0SHR2 Q0SHR3 Q0SHR9 Q0SHS2
Q0SHS7 Q0SHV6 Q0SHW6 Q0SHX9 Q0SHY0 Q0SHY5 Q0SHY7 Q0SHY8 Q0SHZ5 Q0SI46 Q0SI47
Q0SI54 Q0SI56 Q0SI58 Q0SI59 Q0SI65 Q0SI70 Q0SI74 Q0SI81 Q0SIC6 Q0SIE6 Q0SIE8
Q0SIE9 Q0SIF4 Q0SIF9 Q0SIG0 Q0SIG2 Q0SJ46 Q0SJ74 Q0SJD2 Q0SJD4 Q0SJD5 Q0SJK2
Q0SJK6 Q0SJK7 Q0SJK8 Q0SJK9 Q0SK44 Q0SL54 Q0SLI9 Q0SM07 Q0SM08 Q0SM15 Q0SM18
Q0SM30 Q0SM32 Q0SM34 Q0SM46 Q0SM47 Q0SM48 Q0SM49 Q0SM50 Q0SM52 Q0SM54 Q0SM64
Q0SM65 Q0SM66 Q0SM67 Q0SM71 Q0SM73 Q0SM74 Q0SM76 Q0SM77 Q0SM86 Q0SMA2 Q0SMB6
Q0SMC6 Q0SMD6 Q0SMF0 Q0SMF2 Q0SMF3 Q0SMF5 Q0SMF7 Q0SMF9 Q0SMG2 Q0SMG9 Q0SMH1
Q0SMJ5 Q0SMJ6 Q0SMK4 Q0SMK9 Q0SMP1 Q0SMP6 Q0SMP7 Q0SMQ0 Q0SMQ5 Q0SMQ8 Q0SMR4
Q0SMR8 Q0SMS1 Q0SMS3 Q0SMT3 Q0SMU9 Q0SMV7 Q0SMX0 Q0SMY2 Q0SMY9 Q0SMZ0 Q0SMZ4
Q0SMZ9 Q0SN04 Q0SN05 Q0SN06 Q0SN07 Q0SN08 Q0SN10 Q0SN11 Q0SN12 Q0SN13 Q0SN14
Q0SN15 Q0SN16 Q0SN17 Q0SN18 Q0SN19 Q0SN20 Q0SN21 Q0SN22 Q0SN23 Q0SN24 Q0SN25
Q0SN26 Q0SN27 Q0SN28 Q0SN29 Q0SN30 Q0SN31 Q0SN44 Q0SN50 Q0SN63 Q0SN68 Q0SN69
Q0SN72 Q0SN86 Q0SNA5 Q0SNB1 Q0SNB4 Q0SNB5 Q0SNB6 Q0SNB7 Q0SNB8 Q0SNB9 Q0SNC0
Q0SNC1 Q0SND7 Q0SNE1 Q0SNE2 Q0SNE3 Q0SNE8 Q0SNF0 Q0SNG2 Q0SNH0 Q0SNH1 Q0SNH3
Q0SNH4 Q0SNJ8 Q0SNK4 Q0SNL5 Q0SNQ5 Q0SNR0 Q0SNR4 Q0SNS0 Q0SNS8 Q0SNT2 Q0SNT6
Q0SNU2 Q0SNU8 Q0SNV1 Q0SNW2 Q0SNW3 Q0SNW8 Q0SNX3 Q0SNX4 Q0SNY0 Q0SNY1 Q0SNY5

Q0SNY6 Q0SNY7 Q0SNZ5 Q0SP13 Q0SP28 Q0SP32 Q0SP35 Q0SP41 Q0SP42 Q0SP43 Q0SP49
Q0SP51 Q0SP52 Q0SP57 Q0SP63 Q0SP65 Q0SP77 Q0SPA1 Q0SPA5 Q0SPA9 Q0SPB0 Q0SPB2
Q0SPB8 Q0SPD1 Q0SPD8 Q0SPD9 Q0SPE1 Q0SPE2 Q0SPE3 Q0SPE9 Q0SPF2 Q0SPP8 Q0SPP9
Q0SPQ3 Q0SPQ4 Q0SPQ5 Q0SPR6 Q0SPR8 Q0SPS2 Q0SPV0 Q0SPW1 Q0SPW9 Q0SPX8 Q0SPY0
Q0SPY4 Q0SPY8 Q0SPZ8 Q0SQ01 Q0SQ34 Q0SQ45 Q0SQ50 Q0SQ51 Q0SQ58 Q0SQ60 Q0SQ66
Q0SQ82 Q0SQ83 Q0SQ85 Q0SQ86 Q0SQ91 Q0SQA7 Q0SQB4 Q0SQB9 Q0SQC0 Q0SQC1 Q0SQC8
Q0SQC9 Q0SQD2 Q0SQD3 Q0SQD5 Q0SQD6 Q0SQD7 Q0SQD9 Q0SQE0 Q0SQE1 Q0SQE3 Q0SQE4
Q0SQE5 Q0SQE6 Q0SQE7 Q0SQE8 Q0SQE9 Q0SQF0 Q0SQF1 Q0SQF2 Q0SQF3 Q0SQF4 Q0SQF5
Q0SQF6 Q0SQF7 Q0SQF8 Q0SQF9 Q0SQG0 Q0SQG1 Q0SQG2 Q0SQG3 Q0SQG5 Q0SQG8 Q0SQG9
Q0SQH0 Q0SQH1 Q0SQH2 Q0SQH3 Q0SQH4 Q0SQH8 Q0SQH9 Q0SQI8 Q0SQK0 Q0SQL7 Q0SQM4
Q0SQN0 Q0SQP4 Q0SQQ6 Q0SQS9 Q0SQT0 Q0SQT6 Q0SQU1 Q0SQV5 Q0SQX1 Q0SQX5 Q0SQX8
Q0SQY1 Q0SQY5 Q0SR05 Q0SR12 Q0SR29 Q0SR36 Q0SR37 Q0SR42 Q0SR44 Q0SR51 Q0SR53
Q0SR54 Q0SR56 Q0SR63 Q0SR96 Q0SRC4 Q0SRC7 Q0SRD4 Q0SRD5 Q0SRE1 Q0SRE2 Q0SRE3
Q0SRE9 Q0SRF3 Q0SRF8 Q0SRG2 Q0SRH1 Q0SRK2 Q0SRK7 Q0SRK8 Q0SRM5 Q0SRN2 Q0SRN3
Q0SRN4 Q0SRN5 Q0SRP2 Q0SRP4 Q0SRP7 Q0SRP8 Q0SRQ2 Q0SRQ8 Q0SRS1 Q0SRS4 Q0SRS5
Q0SRS9 Q0SRT6 Q0SRT7 Q0SRU1 Q0SRU6 Q0SRV9 Q0SRX1 Q0SRX5 Q0SRY9 Q0SS00 Q0SS01
Q0SS02 Q0SS03 Q0SS05 Q0SS07 Q0SS18 Q0SS19 Q0SS30 Q0SS32 Q0SS39 Q0SS42 Q0SS43
Q0SS44 Q0SS45 Q0SS52 Q0SS62 Q0SS66 Q0SS72 Q0SS73 Q0SS74 Q0SS78 Q0SS81 Q0SS84
Q0SS87 Q0SS92 Q0SS95 Q0SS99 Q0SSA8 Q0SSB0 Q0SSB1 Q0SSB4 Q0SSB5 Q0SSB9 Q0SSC0
Q0SSC1 Q0SSC2 Q0SSC3 Q0SSC6 Q0SSD0 Q0SSD4 Q0SSD5 Q0SSD7 Q0SSD9 Q0SSE0 Q0SSE5
Q0SSE7 Q0SSI9 Q0SSK1 Q0SSL4 Q0SSM0 Q0SSN2 Q0SSQ2 Q0ST14 Q0ST16 Q0ST19 Q0ST25
Q0ST34 Q0ST36 Q0ST47 Q0ST52 Q0ST53 Q0ST54 Q0ST57 Q0ST77 Q0ST91 Q0ST92 Q0STA0
Q0STA1 Q0STA4 Q0STC3 Q0STD5 Q0STD6 Q0STD7 Q0STE4 Q0STE8 Q0STJ5 Q0STL9 Q0STN4
Q0STN6 Q0STN9 Q0STR1 Q0STR2 Q0STR3 Q0STR4 Q0STS2 Q0STS8 Q0STS9 Q0STT3 Q0STT4
Q0STW6 Q0STY9 Q0SU08 Q0SU11 Q0SU12 Q0SU18 Q0SUG8 Q0SUN0 Q0SUU2 Q0SUV5 Q0SV29
Q0SV31 Q0SV34 Q0SV35 Q0SV41 Q0SV48 Q0SV52 Q0SV55 Q0SV78 Q0SVA4 Q0SVB1 Q0SVG3
Q0SVI9 Q0SVK7 Q0SVK9 Q0SVU9 Q0SVV8 Q0SVZ5 Q0SW00 Q0SW34 Q0SW36 Q0SW37 Q0SW38
Q0SW58 Q0SW76 Q0SWB8 Q0SWI8 Q0SWI9 Q0SWK5 Q0SWK6 Q0SWM0 Q0SWN5 Q0SWR1 Q0SWS5
Q0SWS6 Q0SWS7 Q0SWT9 Q0SWV5 Q0SWW1 Q0SWW4 Q0SWW7 Q0SWW9 Q0SWX6 Q0SWY1 Q0SWZ1
Q0SWZ8 Q0SX03 Q0SX05 Q0SX08 Q0SX17 Q0SX19 Q0SX27 Q0SX28 Q0SX29 Q0SX30 Q0SX36
Q0SX40 Q0SX77 Q0SX83 Q0SX84 Q0SX85 Q0SX87 Q0SX88 Q0SX89 Q0SX94 Q0SXA4 Q0SXA5
Q0SXB0 Q0SXB1 Q0SXB2 Q0SXC1 Q0SXD0 Q0SXD6 Q0SXD7 Q0SXE2 Q0SXF6 Q0SXG5 Q0SXH4
Q0SXH6 Q0SXI5 Q0SXI6 Q0SXJ4 Q0SXK0 Q0SXM7 Q0SXN1 Q0SXP3 Q0SXQ6 Q0SXY7 Q0SXZ2
Q0SXZ4 Q0SY02 Q0SY03 Q0SY04 Q0SY05 Q0SY06 Q0SY07 Q0SY12 Q0SY13 Q0SY14 Q0SY15
Q0SY16 Q0SY17 Q0SY20 Q0SY24 Q0SY26 Q0SY34 Q0SY35 Q0SY37 Q0SY38 Q0SY48 Q0SY57
Q0SY58 Q0SY60 Q0SY61 Q0SY78 Q0SY79 Q0SYB5 Q0SYD5 Q0SYE5 Q0SYE8 Q0SYG2 Q0SYG3
Q0SYG4 Q0SYG5 Q0SYG7 Q0SYG8 Q0SYG9 Q0SYH0 Q0SYH4 Q0SYH7 Q0SYL1 Q0SYM9 Q0SYN0
Q0SYN3 Q0SYP0 Q0SYP3 Q0SYP6 Q0SYP7 Q0SYT0 Q0SYT1 Q0SYT2 Q0SYT5 Q0SYT6 Q0SYV0
Q0SYV7 Q0SYW3 Q0SYW9 Q0SYX0 Q0SYZ5 Q0SZ00 Q0SZ02 Q0SZ21 Q0SZ25 Q0SZ31 Q0SZ37
Q0SZ73 Q0SZ79 Q0SZ81 Q0SZ85 Q0SZ91 Q0SZ93 Q0SZ94 Q0SZ95 Q0SZ96 Q0SZA9 Q0SZB4
Q0SZG8 Q0SZJ2 Q0SZK3 Q0SZN2 Q0SZN3 Q0SZN4 Q0SZN5 Q0SZP1 Q0SZP7 Q0SZQ1 Q0SZQ8
Q0SZR5 Q0SZR7 Q0SZS9 Q0SZU8 Q0SZW2 Q0SZW9 Q0SZX2 Q0SZX3 Q0SZX4 Q0SZX5 Q0SZX6
Q0SZX7 Q0SZX8 Q0SZY2 Q0SZY3 Q0SZY4 Q0SZY6 Q0SZY7 Q0SZY8 Q0SZY9 Q0SZZ0 Q0SZZ1
Q0SZZ2 Q0SZZ3 Q0SZZ4 Q0SZZ5 Q0SZZ6 Q0SZZ7 Q0SZZ8 Q0SZZ9 Q0T000 Q0T001 Q0T003

Q0T004 Q0T005 Q0T006 Q0T007 Q0T008 Q0T014 Q0T015 Q0T016 Q0T018 Q0T019 Q0T020
Q0T029 Q0T031 Q0T051 Q0T052 Q0T058 Q0T065 Q0T067 Q0T068 Q0T079 Q0T095 Q0T098
Q0T099 Q0T0A1 Q0T0A5 Q0T0A8 Q0T0B0 Q0T0B1 Q0T0B3 Q0T0B4 Q0T0B5 Q0T0B6 Q0T0B7
Q0T0C6 Q0T0C7 Q0T0D2 Q0T0D3 Q0T0F8 Q0T0H6 Q0T0I4 Q0T0J2 Q0T0J4 Q0T0J8 Q0T0J9
Q0T0K7 Q0T0L1 Q0T0L7 Q0T0M1 Q0T0S5 Q0T0S8 Q0T0U3 Q0T0U8 Q0T0V0 Q0T0V2 Q0T0V3
Q0T0V7 Q0T0X3 Q0T0X4 Q0T0Y4 Q0T0Y5 Q0T0Z3 Q0T0Z4 Q0T0Z5 Q0T0Z9 Q0T106 Q0T107
Q0T121 Q0T133 Q0T134 Q0T137 Q0T154 Q0T156 Q0T157 Q0T161 Q0T174 Q0T175 Q0T176
Q0T181 Q0T182 Q0T186 Q0T1A7 Q0T1B9 Q0T1C2 Q0T1C3 Q0T1C4 Q0T1C5 Q0T1D5 Q0T1D6
Q0T1G0 Q0T1H4 Q0T1H5 Q0T1H6 Q0T1H7 Q0T1H8 Q0T1I1 Q0T1I2 Q0T1I3 Q0T1I6 Q0T1I7
Q0T1I8 Q0T1P8 Q0T1Q1 Q0T1R0 Q0T1S6 Q0T1S7 Q0T1T1 Q0T1T8 Q0T1U0 Q0T1W9 Q0T1X7
Q0T1X8 Q0T1X9 Q0T1Y0 Q0T1Y1 Q0T1Y7 Q0T1Y8 Q0T1Y9 Q0T1Z2 Q0T1Z3 Q0T1Z6 Q0T201
Q0T202 Q0T204 Q0T208 Q0T210 Q0T212 Q0T221 Q0T222 Q0T224 Q0T239 Q0T241 Q0T243
Q0T246 Q0T250 Q0T273 Q0T275 Q0T280 Q0T281 Q0T298 Q0T2A3 Q0T2B1 Q0T2C3 Q0T2D1
Q0T2F4 Q0T2F6 Q0T2G1 Q0T2G5 Q0T2G7 Q0T2J3 Q0T2J6 Q0T2L9 Q0T2M0 Q0T2N2 Q0T2P9
Q0T2T4 Q0T2T5 Q0T2V0 Q0T2W2 Q0T2W5 Q0T2X0 Q0T2X1 Q0T2Y0 Q0T324 Q0T334 Q0T335
Q0T342 Q0T343 Q0T366 Q0T3A2 Q0T3A3 Q0T3A6 Q0T3A8 Q0T3C4 Q0T3D7 Q0T3G3 Q0T3H9
Q0T3J2 Q0T3K8 Q0T3Q1 Q0T3Q3 Q0T3Q7 Q0T3Q8 Q0T3U0 Q0T3U3 Q0T3U5 Q0T3U6 Q0T3V7
Q0T3Z1 Q0T407 Q0T431 Q0T457 Q0T466 Q0T478 Q0T479 Q0T494 Q0T4B4 Q0T4C2 Q0T4D5
Q0T4D9 Q0T4E0 Q0T4F2 Q0T4F4 Q0T4G6 Q0T4H1 Q0T4K8 Q0T4L2 Q0T4L3 Q0T4L4 Q0T4R2
Q0T4S2 Q0T4S4 Q0T4S6 Q0T4S9 Q0T4U0 Q0T4U2 Q0T4U8 Q0T4V0 Q0T4V2 Q0T4V3 Q0T4V4
Q0T4X1 Q0T4Y5 Q0T504 Q0T506 Q0T524 Q0T529 Q0T5B1 Q0T5B2 Q0T5B6 Q0T5C0 Q0T5D5
Q0T5D6 Q0T5G4 Q0T5I4 Q0T5I5 Q0T5I7 Q0T5J1 Q0T5L2 Q0T5L9 Q0T5M1 Q0T5M9 Q0T5N1
Q0T5N8 Q0T5N9 Q0T5R1 Q0T5R7 Q0T5S8 Q0T5T0 Q0T5T8 Q0T5W5 Q0T5X0 Q0T5X2 Q0T5X6
Q0T5Z6 Q0T606 Q0T622 Q0T623 Q0T624 Q0T625 Q0T626 Q0T627 Q0T630 Q0T634 Q0T635
Q0T667 Q0T668 Q0T677 Q0T679 Q0T686 Q0T6A4 Q0T6B1 Q0T6B8 Q0T6C0 Q0T6C8 Q0T6F4
Q0T6F7 Q0T6G3 Q0T6H7 Q0T6I1 Q0T6I4 Q0T6I5 Q0T6J2 Q0T6N8 Q0T6P0 Q0T6P1 Q0T6P6
Q0T6P9 Q0T6Q2 Q0T6R1 Q0T6R8 Q0T6S0 Q0T6S6 Q0T6S8 Q0T6V3 Q0T6V4 Q0T6W6 Q0T6Y0
Q0T6Y5 Q0T6Y7 Q0T6Z9 Q0T702 Q0T703 Q0T746 Q0T773 Q0T776 Q0T788 Q0T790 Q0T7A9
Q0T7B0 Q0T7B1 Q0T7B2 Q0T7B3 Q0T7B5 Q0T7D7 Q0T7D9 Q0T7E5 Q0T7E6 Q0T7E7 Q0T7G3
Q0T7G6 Q0T7G7 Q0T7H3 Q0T7H6 Q0T7I3 Q0T7I4 Q0T7J7 Q0T7K0 Q0T7K8 Q0T7M8 Q0T7M9
Q0T7N0 Q0T7R0 Q0T7R1 Q0T7R5 Q0T7R6 Q0T802 Q0T805 Q0T815 Q0T820 Q0T826 Q0T837
Q0T838 Q0T839 Q0T840 Q0T842 Q0T843 Q0T844 Q0T846 Q0T847 Q0T850 Q0T852 Q0T860
Q0T867 Q0T868 Q0T870 Q0T879 Q0T880 Q0T894 Q0T896 Q0T897 Q0T8A6 Q0T8B5 Q0T8B6
Q0T8C4 Q0T8C6 Q0T8C7 Q0T8C8 Q0T8D4 Q0T8D5 Q0T8D8 Q0T8E3 Q0T8E4 Q0T8E5 Q0T8E6
Q0T8F3 Q0T8F5 Q0T8F6 Q0T8F7 Q0T8F9 Q0T8G4 Q0T8G5 Q0T8H0 Q0T8H5 Q0T8H6 Q0T8I2
Q0T8I5 Q0T8K2 Q0T8K3 Q0T8K8 Q0T8L9 Q0T8M9 Q0T8P1 Q0T8P2 Q0T8P4 Q0T8R6 Q0T8R8
Q0T8S7 Q0T8S9 Q0T8T0 Q0T8T2 Q0T8U3 Q0T8V0 Q0T8W8 Q0T8X6 Q0T8Y2 Q0T8Y5 Q0T953
Q0T964 Q0T9D1 Q0T9E0 Q0T9E3 Q0T9E4 Q0T9E7 Q0T9F5 Q0T9F7 Q0T9G6 Q0T9H5 Q0T9J0
Q0T9J1 Q0T9J2 Q0T9J5 Q0T9J6 Q0T9J7 Q0T9K1 Q0T9K2 Q0T9L5 Q0T9L6 Q0T9M1 Q0T9M2
Q0T9M3 Q0T9N4 Q0T9P4 Q0T9P8 Q0T9P9 Q0T9Q4 Q0TA23 Q0TA36 Q0TA47 Q0TA53 Q0TA56
Q0TA59 Q0TA67 Q0TA68 Q0TA69 Q0TA70 Q0TA71 Q0TA72 Q0TA75 Q0TA77 Q0TA78 Q0TA79
Q0TA80 Q0TA81 Q0TA82 Q0TA85 Q0TA86 Q0TA91 Q0TA96 Q0TAA0 Q0TAA1 Q0TAA3 Q0TAA4
Q0TAB7 Q0TAC6 Q0TAC8 Q0TAD2 Q0TAD3 Q0TAD5 Q0TAD6 Q0TAD8 Q0TAE5 Q0TAE8 Q0TAF7
Q0TAF8 Q0TAF9 Q0TAG0 Q0TAG1 Q0TAG2 Q0TAG5 Q0TAG8 Q0TAG9 Q0TAH5 Q0TAJ5 Q0TAJ6

Q0TAK9 Q0TAL4 Q0TAM1 Q0TAP6 Q0TAR4 Q0TAR6 Q0TAR9 Q0TAU3 Q0TAU6 Q0TAU9 Q0TAV4
Q0TAW1 Q0TAW3 Q0TAW4 Q0TAW5 Q0TAW8 Q0TAW9 Q0TB00 Q0TB01 Q0TB07 Q0TB15 Q0TB17
Q0TB20 Q0TB37 Q0TB41 Q0TB55 Q0TBG0 Q0TBG1 Q0TBG2 Q0TBG6 Q0TBG7 Q0TBG8 Q0TBG9
Q0TBH1 Q0TBH2 Q0TBH3 Q0TBH4 Q0TBH5 Q0TBI8 Q0TBJ0 Q0TBJ4 Q0TBJ7 Q0TBK7 Q0TBL4
Q0TBM6 Q0TBN7 Q0TBP3 Q0TBP4 Q0TBT4 Q0TBW6 Q0TBX7 Q0TBY8 Q0TC27 Q0TC28 Q0TC29
Q0TC30 Q0TC43 Q0TC48 Q0TC49 Q0TC53 Q0TC55 Q0TC57 Q0TC64 Q0TC78 Q0TC92 Q0TCA6
Q0TCB4 Q0TCB5 Q0TCB6 Q0TCB7 Q0TCB8 Q0TCB9 Q0TCC0 Q0TCE0 Q0TCE1 Q0TCE2 Q0TCE3
Q0TCE4 Q0TCE5 Q0TCE6 Q0TCE7 Q0TCE8 Q0TCE9 Q0TCF0 Q0TCF1 Q0TCF2 Q0TCF3 Q0TCF4
Q0TCF5 Q0TCF6 Q0TCF7 Q0TCF8 Q0TCF9 Q0TCG0 Q0TCG2 Q0TCG3 Q0TCG4 Q0TCG5 Q0TCG6
Q0TCG7 Q0TCH3 Q0TCH4 Q0TCH5 Q0TCH7 Q0TCH9 Q0TCI0 Q0TCJ5 Q0TCJ7 Q0TCL8 Q0TCM8
Q0TCN0 Q0TCN5 Q0TCN6 Q0TCP0 Q0TCP1 Q0TCP3 Q0TCP4 Q0TCQ4 Q0TCS1 Q0TCS4 Q0TCS5
Q0TCS7 Q0TCT1 Q0TCT4 Q0TCT8 Q0TCT9 Q0TCU1 Q0TCU2 Q0TCU3 Q0TCU4 Q0TCU5 Q0TCV4
Q0TCV5 Q0TCW0 Q0TCW1 Q0TCW8 Q0TCX5 Q0TCY1 Q0TCZ4 Q0TD14 Q0TD22 Q0TD34 Q0TD38
Q0TD41 Q0TD42 Q0TD44 Q0TD45 Q0TD46 Q0TD50 Q0TD55 Q0TD58 Q0TD60 Q0TD68 Q0TDI6
Q0TDN9 Q0TDP1 Q0TDP8 Q0TDQ2 Q0TDQ3 Q0TDR0 Q0TDR4 Q0TDS9 Q0TDT0 Q0TDT7 Q0TDT9
Q0TDU8 Q0TDU9 Q0TDV1 Q0TDV3 Q0TDW3 Q0TDX8 Q0TDY9 Q0TE01 Q0TE02 Q0TE05 Q0TE53
Q0TE55 Q0TE57 Q0TE61 Q0TE64 Q0TE74 Q0TE79 Q0TEA2 Q0TEA3 Q0TEA4 Q0TEA6 Q0TEA7
Q0TEA8 Q0TEB1 Q0TEB2 Q0TEB3 Q0TEB4 Q0TEC9 Q0TEG9 Q0TEH0 Q0TEI1 Q0TEI2 Q0TEI3
Q0TEI4 Q0TEI7 Q0TEK1 Q0TEL8 Q0TEM0 Q0TEM2 Q0TEM5 Q0TEM6 Q0TEN0 Q0TEN1 Q0TEN2
Q0TEN3 Q0TEQ8 Q0TEQ9 Q0TER3 Q0TES1 Q0TES3 Q0TET1 Q0TET8 Q0TEV3 Q0TEV4 Q0TEV5
Q0TEV7 Q0TEV8 Q0TEV9 Q0TEW2 Q0TEW7 Q0TEW8 Q0TEX0 Q0TEX1 Q0TEX4 Q0TEX9 Q0TEY1
Q0TEY9 Q0TEZ0 Q0TEZ2 Q0TF00 Q0TF05 Q0TF11 Q0TF29 Q0TF33 Q0TF35 Q0TF40 Q0TF48
Q0TF55 Q0TF64 Q0TF72 Q0TF87 Q0TF95 Q0TFB5 Q0TFB7 Q0TFC2 Q0TFC6 Q0TFC8 Q0TFF2
Q0TFF5 Q0TFH3 Q0TFH7 Q0TFH8 Q0TFI0 Q0TFJ1 Q0TFJ5 Q0TFJ7 Q0TFL0 Q0TFN1 Q0TFN3
Q0TFQ2 Q0TFQ3 Q0TFQ4 Q0TFR8 Q0TFS8 Q0TFT1 Q0TFT6 Q0TFT7 Q0TFU8 Q0TFX8 Q0TFY7
Q0TFY8 Q0TFZ6 Q0TFZ7 Q0TG25 Q0TG62 Q0TG63 Q0TG66 Q0TG68 Q0TG79 Q0TGD0 Q0TGE1
Q0TGH3 Q0TGJ4 Q0TGM8 Q0TGP0 Q0TGP2 Q0TGQ7 Q0TGU0 Q0TGU2 Q0TGV6 Q0TGV9 Q0TGW1
Q0TGW2 Q0TGW6 Q0TGW8 Q0TGW9 Q0TGX1 Q0TGX2 Q0TGY3 Q0TGZ7 Q0TH01 Q0TH20 Q0TH22
Q0TH44 Q0TH50 Q0TH64 Q0TH82 Q0TH83 Q0TH84 Q0TH85 Q0TH86 Q0TH88 Q0TH90 Q0TH97
Q0TH99 Q0THB0 Q0THB2 Q0THB3 Q0THB4 Q0THB6 Q0THC2 Q0THC5 Q0THD6 Q0THD7 Q0THE9
Q0THF0 Q0THG9 Q0THH6 Q0THI8 Q0THJ0 Q0THJ1 Q0THK2 Q0THK5 Q0THL8 Q0THM2 Q0THR7
Q0THR9 Q0THU0 Q0THU1 Q0THX6 Q0TI03 Q0TI19 Q0TI22 Q0TI78 Q0TI80 Q0TI84 Q0TI88
Q0TIA9 Q0TIB0 Q0TIC1 Q0TIC3 Q0TID8 Q0TIF9 Q0TIG0 Q0TIG2 Q0TIG6 Q0TII6 Q0TIJ4
Q0TIJ6 Q0TIK0 Q0TIK2 Q0TIK5 Q0TIK7 Q0TIL5 Q0TIR3 Q0TIT9 Q0TIU0 Q0TIU6 Q0TIV5
Q0TIW6 Q0TIW8 Q0TIX6 Q0TIY5 Q0TIY7 Q0TJ07 Q0TJ12 Q0TJ15 Q0TJ18 Q0TJ40 Q0TJ55
Q0TJ56 Q0TJ57 Q0TJ58 Q0TJ59 Q0TJ60 Q0TJ63 Q0TJ66 Q0TJ67 Q0TJ70 Q0TJ93 Q0TJ94
Q0TJA2 Q0TJA5 Q0TJB3 Q0TJB9 Q0TJC3 Q0TJC9 Q0TJD1 Q0TJD6 Q0TJE1 Q0TJE3 Q0TJE5
Q0TJE6 Q0TJF7 Q0TJG5 Q0TJG6 Q0TJG8 Q0TJH6 Q0TJI3 Q0TJI9 Q0TJJ6 Q0TJL3 Q0TJN6
Q0TJP1 Q0TJP4 Q0TJR5 Q0TJR9 Q0TJS2 Q0TJS3 Q0TJT3 Q0TJU4 Q0TJU6 Q0TJV1 Q0TJX8
Q0TJX9 Q0TK03 Q0TK13 Q0TK19 Q0TK21 Q0TK29 Q0TK31 Q0TK34 Q0TK37 Q0TK42 Q0TK43
Q0TK45 Q0TK55 Q0TK58 Q0TK59 Q0TK91 Q0TKB2 Q0TKB5 Q0TKD0 Q0TKD5 Q0TKD6 Q0TKD7
Q0TKD8 Q0TKG5 Q0TKG6 Q0TKG7 Q0TKG8 Q0TKG9 Q0TKH0 Q0TKH2 Q0TKJ5 Q0TKJ7 Q0TKK3
Q0TKK4 Q0TKK5 Q0TKL6 Q0TKL9 Q0TKM1 Q0TKM5 Q0TKM6 Q0TKM8 Q0TKN5 Q0TKN6 Q0TKP7
Q0TKP9 Q0TKQ3 Q0TKQ4 Q0TKQ9 Q0TKS6 Q0TKS7 Q0TKS8 Q0TKT1 Q0TKV9 Q0TKW0 Q0TKW1

Q0TKY8 Q0TKZ0 Q0TL73 Q0TL74 Q0TL76 Q0TL77 Q0TL85 Q0TL93 Q0TLC3 Q0TLC5 Q0TLD7
Q0TLE3 Q0TLF0 Q0TLG1 Q0TLG2 Q0TLG3 Q0TLG4 Q0TLG6 Q0TLG7 Q0TLG8 Q0TLH1 Q0TLH2
Q0TLH5 Q0TLH7 Q0TLI5 Q0TLJ8 Q0TLJ9 Q0TLK2 Q0TLL3 Q0TLL4 Q0TLL5 Q0TLN6 Q0TLN8
Q0TLN9 Q0TLP8 Q0TLQ1 Q0TLQ7 Q0TLQ8 Q0TLR5 Q0TLR7 Q0TLR8 Q0TLR9 Q0TLS7 Q0TLS8
Q0TLT1 Q0TLT6 Q0TLT7 Q0TLT8 Q0TLU7 Q0TLU8 Q0TLV0 Q0TLV1 Q0TLV2 Q0TLV3 Q0TLV4
Q0TLW0 Q0TLW1 Q0TLW2 Q0TLW5 Q0TLW7 Q0TLX5 Q0TLY2 Q0TLY6 Q0TLY8 Q0TLY9 Q0TLZ0
Q0TLZ4 Q0TLZ5 Q0TLZ7 Q0TM08 Q0TM10 Q0TM14 Q0TM49 Q0TM63 Q0TM73 Q0TM75 Q0TM79
Q0TM83 Q0TM95 Q0TMA0 Q0TMD6 Q0TME7 Q0TMF3 Q0TMG0 Q0TMG2 Q0TMG7 Q0TMI0 Q0TMI3
Q0TMI4 Q0TMI6 Q0TMI7 Q0TMJ3 Q0TML0 Q0TML8 Q0TMM2 Q0TMM3 Q0TMM4 Q0TMN0 Q0TMN1
Q0TMN4 Q0TMN5 Q0TMN7 Q0TMN8 Q0TMN9 Q0TMP1 Q0TMP2 Q0TMP3 Q0TMP5 Q0TMP6 Q0TMP7
Q0TMP8 Q0TMP9 Q0TMQ0 Q0TMQ1 Q0TMQ2 Q0TMQ3 Q0TMQ4 Q0TMQ5 Q0TMQ6 Q0TMQ7 Q0TMQ8
Q0TMQ9 Q0TMR0 Q0TMR1 Q0TMR2 Q0TMR3 Q0TMR4 Q0TMR5 Q0TMR7 Q0TMS0 Q0TMS1 Q0TMS2
Q0TMS3 Q0TMS4 Q0TMS5 Q0TMS6 Q0TMT1 Q0TMT2 Q0TMU2 Q0TMX1 Q0TMX8 Q0TMY4 Q0TN14
Q0TN26 Q0TN27 Q0TN51 Q0TN52 Q0TN59 Q0TN64 Q0TN80 Q0TNA0 Q0TNA4 Q0TNA7 Q0TNB0
Q0TNB4 Q0TND4 Q0TNE1 Q0TNG0 Q0TNG7 Q0TNG8 Q0TNH3 Q0TNH5 Q0TNI2 Q0TNI4 Q0TNI5
Q0TNI7 Q0TNJ4 Q0TNM7 Q0TNQ8 Q0TNR1 Q0TNR8 Q0TNR9 Q0TNS5 Q0TNS6 Q0TNS7 Q0TNT3
Q0TNT7 Q0TNU2 Q0TNU6 Q0TNV1 Q0TNY3 Q0TNY8 Q0TNY9 Q0TP06 Q0TP12 Q0TP13 Q0TP14
Q0TP15 Q0TP22 Q0TP24 Q0TP27 Q0TP28 Q0TP32 Q0TP38 Q0TP51 Q0TP54 Q0TP55 Q0TP59
Q0TP67 Q0TP68 Q0TP72 Q0TP77 Q0TP84 Q0TP92 Q0TPA4 Q0TPA8 Q0TPC2 Q0TPD3 Q0TPD4
Q0TPD5 Q0TPD6 Q0TPD8 Q0TPE0 Q0TPF1 Q0TPF2 Q0TPG4 Q0TPG6 Q0TPH2 Q0TPH4 Q0TPH5
Q0TPH6 Q0TPH7 Q0TPI4 Q0TPJ5 Q0TPJ9 Q0TPK5 Q0TPK6 Q0TPK7 Q0TPL4 Q0TPL7 Q0TPM0
Q0TPM5 Q0TPM8 Q0TPN2 Q0TPP1 Q0TPP3 Q0TPP4 Q0TPP5 Q0TPP7 Q0TPP8 Q0TPQ2 Q0TPQ3
Q0TPQ4 Q0TPQ5 Q0TPQ6 Q0TPQ9 Q0TPR3 Q0TPR7 Q0TPR8 Q0TPS0 Q0TPS2 Q0TPS3 Q0TPS8
Q0TPT0 Q0TPX4 Q0TPY5 Q0TPZ6 Q0TQ02 Q0TQ16 Q0TQ38 Q0TQ59 Q0TQ60 Q0TQA6 Q0TQG2
Q0TQG4 Q0TQG7 Q0TQH4 Q0TQI3 Q0TQI5 Q0TQJ7 Q0TQK1 Q0TQK2 Q0TQK3 Q0TQK6 Q0TQN4
Q0TQU4 Q0TQU5 Q0TQV3 Q0TQV4 Q0TQV7 Q0TQY7 Q0TQY8 Q0TQY9 Q0TQZ6 Q0TQZ9 Q0TR21
Q0TR61 Q0TR88 Q0TRA8 Q0TRB0 Q0TRB3 Q0TRD3 Q0TRD4 Q0TRD5 Q0TRD6 Q0TRE4 Q0TRE5
Q0TRF3 Q0TRF4 Q0TRJ4 Q0TRK5 Q0TRM0 Q0TRM9 Q0TRN3 Q0TRN4 Q0TRN9 Q0TSP9 Q0TSR5
Q0TSS9 Q0TSY6 Q0TSZ6 Q0TT93 Q0TT95 Q0TT99 Q0TTA0 Q0TTA4 Q0TTA5 Q0TTA6 Q0TTB0
Q0TTB2 Q0TTB4 Q0TTB7 Q0TTD0 Q0TTF4 Q0TTG1 Q0TTH1 Q0TTL3 Q0TTN7 Q0TTQ7 Q0TTQ9
Q0TU06 Q0TU16 Q0TU51 Q0TU55 Q0TU57 Q0TU86 Q0TU87 Q0TU88 Q0TU89 Q0TUB1 Q0TUD0
Q0TUH0 Q0TUK6 Q0TUP8 Q0TUP9 Q0TUR4 Q0TUR5 Q0TUR6 Q0TUT1 Q0TUU6 Q0TUX8 Q0TUZ0
Q0TUZ2 Q0TUZ4 Q0TV20 Q0TV21 Q0TV32 Q0TV42 Q0TV46 Q0TV47 Q0TV53 Q0TV61 Q0TV64
Q0TVK3 Q0TWJ7 Q0TXG3 Q0TXH0 Q0TXZ2 Q0TY47 Q0TYW1 Q0TZA1 Q0TZT0 Q0U086 Q0U0H7
Q0U0X1 Q0U154 Q0U194 Q0U1A0 Q0U1A1 Q0U1B1 Q0U1G2 Q0U2G5 Q0U324 Q0U397 Q0U3A4
Q0U3Y6 Q0U499 Q0U5F2 Q0U5H7 Q0U6E7 Q0U6E8 Q0U6G5 Q0U6J9 Q0U6Q5 Q0U6W8 Q0U6X2
Q0U6Z7 Q0U796 Q0U7C8 Q0U7S9 Q0U7W4 Q0U8L4 Q0U8U9 Q0U8V9 Q0U913 Q0U957 Q0U9D6
Q0U9X3 Q0UAL6 Q0UAM9 Q0UAT0 Q0UBQ5 Q0UBT2 Q0UCI9 Q0UDH1 Q0UE25 Q0UEB6 Q0UEM3
Q0UFP1 Q0UFP3 Q0UFY4 Q0UG00 Q0UG82 Q0UG91 Q0UHC0 Q0UHC4 Q0UHL8 Q0UHM7 Q0UI00
Q0UI01 Q0UI02 Q0UI04 Q0UI05 Q0UI06 Q0UI56 Q0UI93 Q0UIN2 Q0UIU6 Q0UJ25 Q0UJC3
Q0UJF3 Q0UJJ7 Q0UK12 Q0UK48 Q0UK50 Q0UK52 Q0UK53 Q0UK54 Q0UL22 Q0UM32 Q0UM75
Q0UMB6 Q0UMB9 Q0UME8 Q0UMR2 Q0UN56 Q0UN57 Q0UN58 Q0UNC6 Q0UP45 Q0UPL5 Q0UPV4
Q0UQJ8 Q0UQJ9 Q0UR48 Q0UR95 Q0USF2 Q0USG2 Q0USL4 Q0USP2 Q0USZ4 Q0UTA2 Q0UTE1
Q0UTI9 Q0UTL2 Q0UTQ5 Q0UTQ6 Q0UU86 Q0UUE2 Q0UUN6 Q0UVC3 Q0UVD1 Q0UVG7 Q0UVK8

| | | | | | | | | | | |
|---|---|---|---|---|---|---|---|---|---|---|
| Q0UWA6 | Q0UWC8 | Q0UWF1 | Q0UWF6 | Q0UWI9 | Q0UWT7 | Q0UXJ7 | Q0UXL8 | Q0UXP3 | Q0UY45 | Q0UY62 |
| Q0UY70 | Q0UYV9 | Q0UZ59 | Q0UZH1 | Q0UZK0 | Q0UZR3 | Q0UZV8 | Q0V090 | Q0V0I4 | Q0V147 | Q0V1B1 |
| Q0V1H5 | Q0V1Z7 | Q0V254 | Q0V2R4 | Q0V314 | Q0V320 | Q0V340 | Q0V389 | Q0V3D5 | Q0V3E0 | Q0V3H0 |
| Q0V4C4 | Q0V577 | Q0V5P6 | Q0V5W8 | Q0V640 | Q0V6H6 | Q0V6P9 | Q0V6Q1 | Q0V6Q2 | Q0V6Q3 | Q0V6Q4 |
| Q0V6Q6 | Q0V6Q7 | Q0V6U4 | Q0V6W0 | Q0V713 | Q0V786 | Q0V7M7 | Q0V7M8 | Q0V7P8 | Q0V7S0 | Q0V7S6 |
| Q0V7U7 | Q0V7W6 | Q0V7X4 | Q0V842 | Q0V865 | Q0V8E4 | Q0V8G2 | Q0V8G8 | Q0V8H6 | Q0V8M0 | Q0V8R7 |
| Q0V967 | Q0V984 | Q0V989 | Q0V9A3 | Q0V9A9 | Q0V9C9 | Q0V9D9 | Q0V9F1 | Q0V9F7 | Q0V9J5 | Q0V9K1 |
| Q0V9K5 | Q0V9L1 | Q0V9L3 | Q0V9N0 | Q0V9N5 | Q0V9P1 | Q0V9Q6 | Q0V9R0 | Q0V9R3 | Q0V9R4 | Q0V9S3 |
| Q0V9S9 | Q0V9T6 | Q0V9T8 | Q0V9U1 | Q0V9U8 | Q0V9V7 | Q0V9W6 | Q0V9X5 | Q0V9X7 | Q0V9Y8 | Q0VA03 |
| Q0VA04 | Q0VA16 | Q0VA20 | Q0VA40 | Q0VA52 | Q0VAA2 | Q0VAK6 | Q0VAV2 | Q0VAW7 | Q0VB26 | Q0VBB0 |
| Q0VBD2 | Q0VBK2 | Q0VBL1 | Q0VBV7 | Q0VBW2 | Q0VBY0 | Q0VBY1 | Q0VBY3 | Q0VBY7 | Q0VBY8 | Q0VBZ5 |
| Q0VBZ8 | Q0VC06 | Q0VC09 | Q0VC13 | Q0VC19 | Q0VC20 | Q0VC24 | Q0VC30 | Q0VC53 | Q0VC65 | Q0VC66 |
| Q0VC68 | Q0VC71 | Q0VC73 | Q0VC74 | Q0VC92 | Q0VCA1 | Q0VCA5 | Q0VCA8 | Q0VCB0 | Q0VCB2 | Q0VCC0 |
| Q0VCC5 | Q0VCD4 | Q0VCE2 | Q0VCE9 | Q0VCF3 | Q0VCF9 | Q0VCG0 | Q0VCG3 | Q0VCH3 | Q0VCI6 | Q0VCJ2 |
| Q0VCJ6 | Q0VCL3 | Q0VCL6 | Q0VCL9 | Q0VCM4 | Q0VCP1 | Q0VCQ1 | Q0VCQ4 | Q0VCQ7 | Q0VCR0 | Q0VCR1 |
| Q0VCR4 | Q0VCR7 | Q0VCR8 | Q0VCS4 | Q0VCS9 | Q0VCU0 | Q0VCU1 | Q0VCU8 | Q0VCW1 | Q0VCW3 | Q0VCW6 |
| Q0VCW8 | Q0VCX5 | Q0VCY7 | Q0VD00 | Q0VD01 | Q0VD18 | Q0VD22 | Q0VD23 | Q0VD24 | Q0VD26 | Q0VD27 |
| Q0VD31 | Q0VD34 | Q0VD35 | Q0VDD5 | Q0VDD8 | Q0VDG4 | Q0VDN7 | Q0VDQ9 | Q0VDT2 | Q0VEE6 | Q0VET5 |
| Q0VF22 | Q0VF49 | Q0VFA3 | Q0VFD6 | Q0VFG3 | Q0VFH3 | Q0VFH6 | Q0VFI5 | Q0VFL7 | Q0VFM6 | Q0VFN1 |
| Q0VFN8 | Q0VFP3 | Q0VFP5 | Q0VFP6 | Q0VFR9 | Q0VFT9 | Q0VFX4 | Q0VFX8 | Q0VFX9 | Q0VFZ6 | Q0VG06 |
| Q0VG62 | Q0VG73 | Q0VG85 | Q0VG99 | Q0VGB7 | Q0VGE8 | Q0VGK2 | Q0VGK3 | Q0VGY8 | Q0VH32 | Q0VH33 |
| Q0VHD6 | Q0VIA1 | Q0VKU5 | Q0VKU6 | Q0VKU8 | Q0VKW3 | Q0VKW4 | Q0VKX8 | Q0VKX9 | Q0VKY0 | Q0VKY1 |
| Q0VKY2 | Q0VKY4 | Q0VKY5 | Q0VKZ2 | Q0VKZ4 | Q0VL48 | Q0VL74 | Q0VL83 | Q0VL87 | Q0VL94 | Q0VLA6 |
| Q0VLA7 | Q0VLA8 | Q0VLB7 | Q0VLC7 | Q0VLQ4 | Q0VLR3 | Q0VLS6 | Q0VLY5 | Q0VM06 | Q0VM16 | Q0VM18 |
| Q0VM25 | Q0VM28 | Q0VM60 | Q0VM64 | Q0VM69 | Q0VM71 | Q0VM72 | Q0VM78 | Q0VM94 | Q0VMA1 | Q0VMA5 |
| Q0VMA6 | Q0VMA8 | Q0VMB0 | Q0VMC3 | Q0VMD0 | Q0VMD1 | Q0VMD3 | Q0VMD9 | Q0VME0 | Q0VME6 | Q0VME7 |
| Q0VMF3 | Q0VMF9 | Q0VMG1 | Q0VMG2 | Q0VMG4 | Q0VMH4 | Q0VMH5 | Q0VMI0 | Q0VMI4 | Q0VMK6 | Q0VML5 |
| Q0VML6 | Q0VMP7 | Q0VMU0 | Q0VMU1 | Q0VMU8 | Q0VMV2 | Q0VMX5 | Q0VMX9 | Q0VMY5 | Q0VN20 | Q0VN36 |
| Q0VN37 | Q0VN46 | Q0VN49 | Q0VN53 | Q0VN64 | Q0VN66 | Q0VND6 | Q0VND7 | Q0VNE0 | Q0VNE1 | Q0VNE4 |
| Q0VNE8 | Q0VNF0 | Q0VNG0 | Q0VNG1 | Q0VNG2 | Q0VNG4 | Q0VNJ1 | Q0VNJ7 | Q0VNK2 | Q0VNK4 | Q0VNK6 |
| Q0VNM6 | Q0VNM7 | Q0VNN3 | Q0VNQ9 | Q0VNR6 | Q0VNT0 | Q0VP19 | Q0VP22 | Q0VP26 | Q0VP51 | Q0VP64 |
| Q0VP73 | Q0VP77 | Q0VP88 | Q0VPA8 | Q0VPD2 | Q0VPE1 | Q0VPF7 | Q0VPG1 | Q0VPH7 | Q0VPI0 | Q0VPI1 |
| Q0VPI7 | Q0VPI8 | Q0VPJ0 | Q0VPK2 | Q0VPK3 | Q0VPK4 | Q0VPK5 | Q0VPK6 | Q0VPM2 | Q0VPW2 | Q0VPZ5 |
| Q0VQ05 | Q0VQ06 | Q0VQ12 | Q0VQ13 | Q0VQ14 | Q0VQ23 | Q0VQ25 | Q0VQ34 | Q0VQ52 | Q0VQ55 | Q0VQ76 |
| Q0VQ89 | Q0VQ90 | Q0VQ91 | Q0VQ92 | Q0VQC3 | Q0VQD1 | Q0VQD3 | Q0VQD8 | Q0VQE3 | Q0VQF4 | Q0VQF5 |
| Q0VQF6 | Q0VQF7 | Q0VQG3 | Q0VQG9 | Q0VQI2 | Q0VQM6 | Q0VQN4 | Q0VQN6 | Q0VQP1 | Q0VQR8 | Q0VQU5 |
| Q0VQU6 | Q0VQV5 | Q0VQX6 | Q0VQX7 | Q0VR01 | Q0VR14 | Q0VR70 | Q0VR89 | Q0VRB0 | Q0VRD0 | Q0VRE9 |
| Q0VRF0 | Q0VRF1 | Q0VRF2 | Q0VRH1 | Q0VRH4 | Q0VRI8 | Q0VRJ7 | Q0VRJ9 | Q0VRK0 | Q0VRK1 | Q0VRL1 |
| Q0VRM0 | Q0VRM2 | Q0VRM5 | Q0VRN6 | Q0VRR6 | Q0VRS0 | Q0VRV7 | Q0VRW6 | Q0VRW7 | Q0VS00 | Q0VS01 |
| Q0VS04 | Q0VS07 | Q0VS10 | Q0VS11 | Q0VS15 | Q0VS24 | Q0VS39 | Q0VS40 | Q0VS51 | Q0VS67 | Q0VS71 |
| Q0VS72 | Q0VS73 | Q0VS77 | Q0VS78 | Q0VS81 | Q0VS83 | Q0VS84 | Q0VSA0 | Q0VSA5 | Q0VSA8 | Q0VSB5 |
| Q0VSD8 | Q0VSE1 | Q0VSE4 | Q0VSE5 | Q0VSE6 | Q0VSE7 | Q0VSE8 | Q0VSH7 | Q0VSH8 | Q0VSH9 | Q0VSI0 |
| Q0VSI1 | Q0VSI2 | Q0VSI4 | Q0VSI5 | Q0VSI6 | Q0VSI7 | Q0VSI8 | Q0VSI9 | Q0VSJ0 | Q0VSJ1 | Q0VSJ2 |
| Q0VSJ3 | Q0VSJ4 | Q0VSJ5 | Q0VSJ6 | Q0VSJ7 | Q0VSJ8 | Q0VSJ9 | Q0VSK0 | Q0VSK1 | Q0VSK2 | Q0VSK3 |

Q0VSK4 Q0VSL7 Q0VSL8 Q0VSL9 Q0VSM0 Q0VSM1 Q0VSM2 Q0VSM3 Q0VSM4 Q0VSM5 Q0VSM6
Q0VSN1 Q0VSN8 Q0VSP5 Q0VSP6 Q0VSQ3 Q0VSR3 Q0VSR4 Q0VSR5 Q0VSR6 Q0VSR7 Q0VSR8
Q0VSS0 Q0VSS1 Q0VSS3 Q0VSS5 Q0VSS6 Q0VSS9 Q0VST5 Q0VST6 Q0VST7 Q0VSU3 Q0VSU5
Q0VSU8 Q0VSV6 Q0VSX4 Q0VSY7 Q0VT30 Q0VT56 Q0VT57 Q0VT58 Q0VT60 Q0VT62 Q0VT66
Q0VT94 Q0VT95 Q0VTA8 Q0VTA9 Q0VTB0 Q0VTD6 Q0VTD7 Q0VTD8 Q0VTE1 Q0VTE2 Q0VTG6
Q0VTI2 Q0VTI3 Q0VTM5 Q0VTN3 Q0VZ68 Q0VZ69 Q0VZ70 Q0W037 Q0W049 Q0W050 Q0W081
Q0W085 Q0W0C9 Q0W0H2 Q0W0J2 Q0W0J3 Q0W0L6 Q0W0P5 Q0W0T4 Q0W0X1 Q0W0X4 Q0W0X9
Q0W112 Q0W131 Q0W150 Q0W1B0 Q0W1C7 Q0W1F9 Q0W1L5 Q0W1L8 Q0W1L9 Q0W1M3 Q0W1N4
Q0W1V4 Q0W1V5 Q0W1V6 Q0W1W1 Q0W1W4 Q0W1W7 Q0W1W8 Q0W1W9 Q0W1X2 Q0W1X3 Q0W1X6
Q0W1X7 Q0W1X8 Q0W1X9 Q0W1Y2 Q0W1Y3 Q0W1Y4 Q0W1Y5 Q0W1Y6 Q0W1Y9 Q0W252 Q0W253
Q0W257 Q0W278 Q0W2C6 Q0W2D6 Q0W2E9 Q0W2G0 Q0W2J3 Q0W2J4 Q0W2K2 Q0W2L3 Q0W2P3
Q0W2P8 Q0W2Q3 Q0W2W0 Q0W2X4 Q0W2Y2 Q0W2Y3 Q0W2Y5 Q0W2Y7 Q0W2Z5 Q0W358 Q0W389
Q0W3B6 Q0W3K6 Q0W3L5 Q0W3P4 Q0W3U6 Q0W3X3 Q0W3X9 Q0W443 Q0W468 Q0W486 Q0W496
Q0W498 Q0W4B8 Q0W4I8 Q0W4J1 Q0W4J4 Q0W4L8 Q0W4N6 Q0W4U0 Q0W4U3 Q0W4W4 Q0W4Z8
Q0W5A6 Q0W5A8 Q0W5C7 Q0W5C8 Q0W5G9 Q0W5H1 Q0W5H2 Q0W5H4 Q0W5I2 Q0W5I3 Q0W5Q9
Q0W5R8 Q0W5T2 Q0W5T3 Q0W5T5 Q0W5U6 Q0W628 Q0W644 Q0W674 Q0W6F5 Q0W6F7 Q0W6G4
Q0W6H4 Q0W6I0 Q0W6K3 Q0W6L1 Q0W6L7 Q0W6M2 Q0W6M8 Q0W6P9 Q0W6Q2 Q0W6Q4 Q0W6Q8
Q0W6Q9 Q0W6S4 Q0W6V1 Q0W6V4 Q0W700 Q0W720 Q0W787 Q0W793 Q0W7A4 Q0W7E0 Q0W7K1
Q0W7V3 Q0W830 Q0W834 Q0W874 Q0W896 Q0W8A0 Q0W8A8 Q0W8B4 Q0W8C7 Q0W8D0 Q0W8E0
Q0W8E1 Q0W8G2 Q0W8G5 Q0W8L2 Q0W8M0 Q0W8N8 Q0W8P0 Q0W8P1 Q0W8R3 Q0W8R9 Q0W8S5
Q0W8U7 Q0W8U9 Q0W8W9 Q0W8X1 Q0W8X2 Q0W8Z4 Q0W932 Q0W933 Q0W938 Q0W941 Q0WAP4
Q0WB88 Q0WBC9 Q0WBD7 Q0WBF9 Q0WBQ9 Q0WD07 Q0WD32 Q0WDE1 Q0WDR2 Q0WEF0 Q0WGE8
Q0WHW5 Q0WJ78 Q0WJP7 Q0WJT1 Q0WJY2 Q0WKV3 Q0WKV8 Q0WKY2 Q0WKZ3 Q0WL52 Q0WL69
Q0WL81 Q0WLC6 Q0WLR1 Q0WM29 Q0WMN5 Q0WMR0 Q0WMV8 Q0WMY5 Q0WMZ6 Q0WN01 Q0WN60
Q0WN69 Q0WNB1 Q0WNN7 Q0WNR2 Q0WNR6 Q0WNX9 Q0WNZ5 Q0WP85 Q0WPF2 Q0WPH8 Q0WPJ7
Q0WPK4 Q0WPN0 Q0WPT7 Q0WPW4 Q0WPX7 Q0WPY0 Q0WPZ6 Q0WPZ7 Q0WQ91 Q0WQE7 Q0WQF7
Q0WQJ7 Q0WQM8 Q0WQN9 Q0WQQ1 Q0WQW5 Q0WQX9 Q0WR05 Q0WR66 Q0WRC9 Q0WRJ7 Q0WRU9
Q0WSK8 Q0WSN2 Q0WSY2 Q0WSZ8 Q0WT19 Q0WT24 Q0WTB4 Q0WTB8 Q0WTP1 Q0WTY4 Q0WUF6
Q0WUI9 Q0WUQ1 Q0WV25 Q0WV90 Q0WVD6 Q0WVE8 Q0WVF5 Q0WVF8 Q0WVK7 Q0WVL7 Q0WVU0
Q0WVU3 Q0WVV0 Q0WVW7 Q0WVX5 Q0WW21 Q0WW40 Q0WW55 Q0WW84 Q0WWT7 Q0WX00 Q0X0E2
Q0Z8B5 Q0ZA50 Q0ZB73 Q0ZB76 Q0ZB77 Q0ZB81 Q0ZCJ7 Q0ZFW8 Q0ZIH5 Q0ZIH7 Q0ZIV5
Q0ZIV6 Q0ZIV7 Q0ZIV9 Q0ZIW1 Q0ZIX0 Q0ZIX7 Q0ZIX8 Q0ZIY0 Q0ZIY1 Q0ZIY2 Q0ZIY3
Q0ZIY4 Q0ZIY5 Q0ZIY6 Q0ZIY7 Q0ZIZ4 Q0ZIZ5 Q0ZIZ7 Q0ZIZ8 Q0ZIZ9 Q0ZJ18 Q0ZJ22
Q0ZJ28 Q0ZJ29 Q0ZJ30 Q0ZJ31 Q0ZJ38 Q0ZJ39 Q0ZME2 Q0ZME6 Q0ZNK1 Q0ZPV7 Q0ZQ46
Q0ZQX0 Q0ZS27 Q0ZS46 Q10002 Q10003 Q10004 Q10006 Q10007 Q10008 Q10009 Q10012
Q10013 Q10020 Q10021 Q10026 Q10027 Q10030 Q10033 Q10034 Q10038 Q10040 Q10041
Q10043 Q10051 Q10055 Q10059 Q10060 Q10062 Q10064 Q10066 Q10067 Q10068 Q10069
Q10075 Q10076 Q10077 Q10078 Q10080 Q10081 Q10082 Q10083 Q10086 Q10089 Q10092
Q10093 Q10094 Q10095 Q10096 Q10098 Q10102 Q10103 Q10104 Q10105 Q10106 Q10107
Q10108 Q10110 Q10111 Q10112 Q10113 Q10115 Q10116 Q10121 Q10124 Q10126 Q10127
Q10129 Q10131 Q10134 Q10135 Q10136 Q10138 Q10139 Q10140 Q10143 Q10144 Q10145
Q10146 Q10147 Q10148 Q10154 Q10155 Q10156 Q10159 Q10160 Q10162 Q10163 Q10166
Q10167 Q10169 Q10170 Q10173 Q10174 Q10175 Q10178 Q10179 Q10180 Q10181 Q10183
Q10184 Q10186 Q10187 Q10189 Q10190 Q10191 Q10193 Q10195 Q10197 Q10198 Q10200

Q10202 Q10203 Q10204 Q10205 Q10206 Q10207 Q10208 Q10210 Q10212 Q10213 Q10214
Q10215 Q10216 Q10218 Q10220 Q10221 Q10222 Q10224 Q10225 Q10233 Q10234 Q10235
Q10236 Q10237 Q10238 Q10239 Q10241 Q10242 Q10247 Q10251 Q10253 Q10257 Q10258
Q10261 Q10262 Q10264 Q10267 Q10270 Q10271 Q10272 Q10273 Q10274 Q10277 Q10280
Q10290 Q10292 Q10295 Q10296 Q10298 Q10299 Q10300 Q10301 Q10302 Q10303 Q10304
Q10306 Q10307 Q10308 Q10310 Q10311 Q10312 Q10314 Q10315 Q10316 Q10317 Q10318
Q10319 Q10321 Q10325 Q10326 Q10327 Q10328 Q10329 Q10330 Q10332 Q10334 Q10335
Q10336 Q10337 Q10338 Q10339 Q10340 Q10342 Q10344 Q10347 Q10348 Q10349 Q10350
Q10352 Q10353 Q10355 Q10356 Q10360 Q10362 Q10368 Q10369 Q10370 Q10373 Q10407
Q10409 Q10414 Q10417 Q10420 Q10422 Q10423 Q10424 Q10425 Q10426 Q10427 Q10428
Q10429 Q10430 Q10432 Q10434 Q10435 Q10437 Q10438 Q10440 Q10441 Q10443 Q10445
Q10446 Q10447 Q10448 Q10451 Q10453 Q10454 Q10457 Q10474 Q10475 Q10477 Q10483
Q10486 Q10489 Q10490 Q10492 Q10494 Q10497 Q10498 Q10571 Q10574 Q10578 Q10579
Q10580 Q10581 Q10582 Q10653 Q10654 Q10655 Q10657 Q10660 Q10661 Q10663 Q10665
Q10667 Q10668 Q10669 Q10690 Q10712 Q10722 Q10725 Q10730 Q10732 Q10733 Q10736
Q10744 Q10752 Q10757 Q107X0 Q10891 Q108P1 Q108T9 Q108U1 Q108U5 Q108U9 Q10902
Q10905 Q10906 Q10912 Q10918 Q10919 Q10920 Q10921 Q10924 Q10926 Q10927 Q10928
Q10932 Q10933 Q10938 Q10942 Q10946 Q10948 Q10950 Q10958 Q10985 Q10986 Q10988
Q10989 Q10990 Q10992 Q10993 Q10994 Q10997 Q109B0 Q109G2 Q109R6 Q10A30 Q10AZ4
Q10AZ7 Q10B19 Q10B63 Q10B67 Q10B98 Q10BT5 Q10BX9 Q10CE7 Q10CE8 Q10CI8 Q10CQ1
Q10CQ8 Q10CT5 Q10D00 Q10D34 Q10D80 Q10DK7 Q10DU0 Q10DV5 Q10DV7 Q10E64 Q10EC6
Q10ED2 Q10EL1 Q10F03 Q10F25 Q10F39 Q10FF9 Q10FT0 Q10FZ7 Q10G39 Q10G56 Q10G81
Q10GB1 Q10GM3 Q10GM4 Q10GP0 Q10GX0 Q10HL3 Q10I26 Q10J20 Q10K62 Q10K98 Q10KF0
Q10KF5 Q10KL8 Q10L32 Q10L71 Q10LF7 Q10LG8 Q10LH0 Q10LI1 Q10LI6 Q10LJ0 Q10LR9
Q10LX4 Q10LZ1 Q10LZ9 Q10M00 Q10M12 Q10M18 Q10M29 Q10M41 Q10M74 Q10MB4 Q10MC0
Q10MH8 Q10MI4 Q10MI9 Q10MJ1 Q10MN5 Q10MN6 Q10MN8 Q10MQ2 Q10MR0 Q10MW3 Q10MX3
Q10N03 Q10N21 Q10N44 Q10N79 Q10NB9 Q10NJ6 Q10NQ3 Q10NQ9 Q10NY2 Q10P60 Q10P67
Q10P87 Q10PC5 Q10PQ9 Q10PS6 Q10PV9 Q10PZ4 Q10PZ6 Q10PZ9 Q10Q08 Q10Q26 Q10Q62
Q10Q63 Q10Q80 Q10QA2 Q10QF2 Q10QH1 Q10QL5 Q10QP3 Q10QR9 Q10QS7 Q10QS9 Q10R17
Q10R18 Q10R47 Q10RE2 Q10RE5 Q10RI7 Q10RZ1 Q10S32 Q10S44 Q10S55 Q10S58 Q10S65
Q10S72 Q10S83 Q10S93 Q10SM2 Q10SU5 Q10SX6 Q10UW6 Q10UX2 Q10UY0 Q10UY5 Q10V92
Q10V94 Q10VG1 Q10VG8 Q10VJ7 Q10VQ4 Q10VR1 Q10VR3 Q10VS0 Q10VU2 Q10VU5 Q10VV0
Q10VX5 Q10VZ5 Q10VZ7 Q10W70 Q10W80 Q10W95 Q10WA5 Q10WA6 Q10WA8 Q10WG3 Q10WH6
Q10WL4 Q10WP7 Q10WQ4 Q10WQ5 Q10WR6 Q10X06 Q10X25 Q10X41 Q10X46 Q10X61 Q10X90
Q10XA1 Q10XA5 Q10XB0 Q10XD8 Q10XE8 Q10XH7 Q10XJ9 Q10XL4 Q10XM0 Q10XM3 Q10XN7
Q10XP0 Q10XS2 Q10XT3 Q10XV0 Q10XW9 Q10XY9 Q10Y17 Q10Y19 Q10Y25 Q10Y33 Q10Y41
Q10Y48 Q10Y49 Q10Y54 Q10Y85 Q10Y87 Q10Y89 Q10Y90 Q10YG4 Q10YI1 Q10YI3 Q10YQ3
Q10YR2 Q10YV3 Q10YY1 Q10YY8 Q10Z39 Q10Z46 Q10Z79 Q10ZC3 Q10ZD0 Q10ZE2 Q10ZG6
Q10ZG7 Q10ZK3 Q10ZL5 Q10ZM7 Q10ZY2 Q11004 Q11010 Q11072 Q11074 Q11075 Q11076
Q11078 Q11081 Q11083 Q11084 Q11086 Q11089 Q11090 Q11091 Q11093 Q11094 Q11096
Q11097 Q110A7 Q110A8 Q110A9 Q110B0 Q110B1 Q110B2 Q110B3 Q110B4 Q110B5 Q110B6
Q110B7 Q110B8 Q110B9 Q110C0 Q110C1 Q110C2 Q110C4 Q110C6 Q110C7 Q110C8 Q110C9
Q110D0 Q110D1 Q110D2 Q110D4 Q110D6 Q110D7 Q110H1 Q110H2 Q110H3 Q110I1 Q110K2
Q110L1 Q110M9 Q110N1 Q110N5 Q110Q9 Q110R5 Q110R6 Q110U9 Q110V5 Q110Z7 Q110Z9

Q11100 Q11102 Q11103 Q11105 Q11107 Q11108 Q11109 Q11110 Q11111 Q11113 Q11115
Q11116 Q11118 Q11123 Q11125 Q11132 Q11136 Q11176 Q11178 Q11181 Q11183 Q11185
Q11187 Q11188 Q11189 Q11191 Q11192 Q11193 Q11194 Q111B4 Q111C2 Q111E0 Q111G3
Q111G4 Q111G7 Q111G9 Q111H1 Q111K4 Q111T6 Q111U2 Q111Z9 Q11212 Q112B6 Q112K0
Q112L2 Q112N2 Q112N5 Q112P2 Q112R6 Q112S1 Q112U2 Q112X4 Q113E3 Q113E7 Q113G4
Q113H3 Q113I7 Q113K0 Q113K5 Q113L8 Q113P2 Q113S1 Q113W0 Q114D6 Q114D7 Q114E1
Q114I1 Q114P5 Q114R3 Q114R5 Q114T6 Q114U6 Q114V5 Q115B3 Q115G0 Q115H2 Q115I0
Q115I9 Q115K2 Q115L1 Q115L3 Q115Q9 Q115S4 Q115Y7 Q115Z7 Q116D3 Q116D6 Q116J2
Q116J4 Q116L2 Q116P1 Q116Q2 Q116Q3 Q116Q9 Q116S1 Q116T4 Q116W4 Q117D0 Q117D2
Q117D6 Q117G4 Q117I4 Q117I5 Q117P0 Q117P8 Q117R7 Q117V9 Q117Z3 Q117Z4 Q117Z6
Q117Z7 Q117Z8 Q117Z9 Q118A6 Q118B0 Q118B3 Q118B4 Q118B7 Q118E6 Q118F4 Q118F5
Q118J2 Q118M1 Q118P4 Q118P6 Q118P8 Q118Y6 Q118Z1 Q118Z2 Q118Z3 Q118Z4 Q118Z5
Q119B9 Q119C3 Q119D0 Q119G2 Q119H9 Q119K2 Q119L1 Q119L7 Q119M3 Q119M4 Q119M5
Q119N9 Q119S1 Q119S3 Q119S6 Q119S7 Q119S8 Q119S9 Q11A09 Q11A13 Q11A17 Q11A19
Q11A96 Q11AD1 Q11AE3 Q11AV6 Q11AV8 Q11AV9 Q11AW7 Q11B05 Q11B27 Q11B39 Q11B41
Q11B42 Q11B48 Q11B64 Q11B66 Q11BC3 Q11BC4 Q11BC6 Q11BC7 Q11BC8 Q11BD1 Q11BD3
Q11BD7 Q11BD9 Q11BE3 Q11BH8 Q11C35 Q11C47 Q11C48 Q11C97 Q11CB2 Q11CB4 Q11CB5
Q11CD0 Q11CK2 Q11CK5 Q11CK6 Q11CK7 Q11CK8 Q11CL1 Q11CL2 Q11CL4 Q11CM1 Q11CM4
Q11CM5 Q11CM6 Q11CM8 Q11CN2 Q11CN3 Q11CN4 Q11CP3 Q11CP4 Q11CP6 Q11CP7 Q11CP8
Q11CQ1 Q11CQ6 Q11CQ9 Q11CR0 Q11CR1 Q11CR7 Q11CS8 Q11CT4 Q11CV8 Q11CV9 Q11CW3
Q11CY7 Q11D07 Q11D10 Q11D15 Q11D83 Q11D95 Q11DE1 Q11DE4 Q11DF0 Q11DG1 Q11DG3
Q11DG9 Q11DH4 Q11DH5 Q11DH7 Q11DI7 Q11DJ4 Q11DQ8 Q11DR9 Q11DS0 Q11DS3 Q11DS9
Q11DW5 Q11E01 Q11E04 Q11E17 Q11E18 Q11E20 Q11ED3 Q11EE0 Q11EH0 Q11EH9 Q11EJ9
Q11ES4 Q11EU7 Q11EW1 Q11EW2 Q11EW3 Q11EW4 Q11EW5 Q11EW7 Q11EW8 Q11EX1 Q11F81
Q11F82 Q11FK9 Q11G43 Q11G66 Q11G79 Q11GB4 Q11GC0 Q11GC5 Q11GC6 Q11GG5 Q11GR7
Q11GS0 Q11GS3 Q11GS6 Q11GS7 Q11GS8 Q11GT5 Q11GT9 Q11GV3 Q11GV7 Q11GW6 Q11GY5
Q11H14 Q11H15 Q11H16 Q11HA6 Q11HA9 Q11HB0 Q11HB1 Q11HB2 Q11HB3 Q11HB4 Q11HB8
Q11HC8 Q11HC9 Q11HD9 Q11HE1 Q11HG8 Q11HG9 Q11HH5 Q11HI0 Q11HI8 Q11HJ2 Q11HK9
Q11HL0 Q11HL3 Q11HL6 Q11HP7 Q11HP8 Q11HP9 Q11HQ1 Q11HQ2 Q11HQ3 Q11HQ4 Q11HQ5
Q11HQ6 Q11HQ8 Q11HQ9 Q11HR0 Q11HR1 Q11HR2 Q11HR3 Q11HR4 Q11HR5 Q11HR6 Q11HR7
Q11HR8 Q11HR9 Q11HS0 Q11HS1 Q11HS3 Q11HS4 Q11HS5 Q11HS6 Q11HS7 Q11HT5 Q11HU1
Q11HU2 Q11HU4 Q11HU6 Q11HV6 Q11HV9 Q11HW7 Q11HY2 Q11HZ9 Q11I44 Q11I48 Q11I83
Q11I86 Q11IA4 Q11IC3 Q11IE7 Q11IH4 Q11IH7 Q11IJ5 Q11IJ6 Q11IJ7 Q11IJ8 Q11IL4
Q11IL6 Q11IL7 Q11IL8 Q11IM0 Q11IM9 Q11IN8 Q11IU8 Q11IV0 Q11IV2 Q11IV8 Q11IV9
Q11IW3 Q11IW6 Q11J59 Q11J60 Q11J65 Q11J68 Q11J82 Q11J90 Q11J94 Q11JA2 Q11JA7
Q11JE6 Q11JI3 Q11JI8 Q11JM1 Q11JM4 Q11JM5 Q11JR3 Q11JS6 Q11JT3 Q11JT6 Q11JT7
Q11K42 Q11KC2 Q11KC5 Q11KC6 Q11KC7 Q11KE0 Q11KE5 Q11KF1 Q11KG2 Q11KG9 Q11KH2
Q11KI3 Q11KJ5 Q11KJ6 Q11KJ9 Q11KK1 Q11KK2 Q11KK8 Q11KK9 Q11KP9 Q11L51 Q11L77
Q11L79 Q11L80 Q11LA1 Q11LA2 Q11LC7 Q11LC8 Q11LD0 Q11LD2 Q11LD4 Q11LE1 Q11LE3
Q11LE6 Q11LE7 Q11LG4 Q11LK0 Q11LP6 Q11LR5 Q11LX8 Q11LY1 Q11LY3 Q11LY4 Q11LY8
Q11M18 Q11MA0 Q11NE6 Q11NE7 Q11NN5 Q11NN7 Q11NN8 Q11NR0 Q11NR3 Q11NS4 Q11NV4
Q11NY7 Q11NZ7 Q11P01 Q11PB4 Q11PC8 Q11PC9 Q11PK5 Q11PK7 Q11PK9 Q11PL9 Q11PM4
Q11PM7 Q11PN0 Q11PN9 Q11PP7 Q11PQ4 Q11PV4 Q11PY2 Q11PZ7 Q11Q02 Q11Q26 Q11Q45
Q11Q55 Q11Q84 Q11Q93 Q11Q97 Q11Q98 Q11QA1 Q11QA2 Q11QA3 Q11QA4 Q11QA5 Q11QA6

| | | | | | | | | | | |
|---|---|---|---|---|---|---|---|---|---|---|
| Q11QA8 | Q11QA9 | Q11QB0 | Q11QB1 | Q11QB2 | Q11QB3 | Q11QB5 | Q11QB6 | Q11QB7 | Q11QB8 | Q11QB9 |
| Q11QC0 | Q11QC1 | Q11QC2 | Q11QC3 | Q11QC5 | Q11QC6 | Q11QC7 | Q11QC9 | Q11QD0 | Q11QD1 | Q11QD4 |
| Q11QD5 | Q11QD6 | Q11QD7 | Q11QD8 | Q11QD9 | Q11QF0 | Q11QH3 | Q11QI8 | Q11QK1 | Q11QL8 | Q11QM8 |
| Q11QN5 | Q11QN6 | Q11QN7 | Q11QQ3 | Q11QT1 | Q11QT3 | Q11QU6 | Q11R22 | Q11R87 | Q11R91 | Q11RA1 |
| Q11RC7 | Q11RE0 | Q11RE2 | Q11RG5 | Q11RG6 | Q11RG9 | Q11RH3 | Q11RH6 | Q11RI7 | Q11RJ8 | Q11RK8 |
| Q11RM1 | Q11RM3 | Q11RN9 | Q11RP3 | Q11RP5 | Q11RT6 | Q11RT7 | Q11RW8 | Q11RW9 | Q11S24 | Q11S31 |
| Q11S34 | Q11S94 | Q11SS5 | Q11SV1 | Q11T17 | Q11T54 | Q11T61 | Q11TG8 | Q11TJ3 | Q11TJ6 | Q11TP2 |
| Q11TP3 | Q11TS0 | Q11TT4 | Q11TT6 | Q11TY6 | Q11TZ6 | Q11U13 | Q11U19 | Q11U26 | Q11U60 | Q11U61 |
| Q11U72 | Q11U74 | Q11U88 | Q11UA6 | Q11UD1 | Q11UE6 | Q11UK0 | Q11UK1 | Q11UK3 | Q11UL8 | Q11UM1 |
| Q11UN5 | Q11UP4 | Q11UQ9 | Q11UV2 | Q11UV9 | Q11V49 | Q11VI1 | Q11VI2 | Q11VI6 | Q11VK5 | Q11VK7 |
| Q11VM1 | Q11VM2 | Q11VM5 | Q11VN0 | Q11VN1 | Q11VN3 | Q11VT3 | Q11VU7 | Q11VZ3 | Q11W94 | Q11WG7 |
| Q11WK1 | Q11WK3 | Q11WK5 | Q11WK6 | Q11X05 | Q11X86 | Q11X87 | Q11X93 | Q11XA9 | Q11XC6 | Q11XD0 |
| Q11XD1 | Q11XI0 | Q11XI1 | Q11XK1 | Q11XK2 | Q11XT5 | Q11XW1 | Q11XW2 | Q11XX3 | Q11XX4 | Q11Y53 |
| Q11Y57 | Q11Y71 | Q11Y93 | Q11YI1 | Q11YP9 | Q11YS6 | Q11YV3 | Q11YV4 | Q11YV5 | Q11YV6 | Q11YW2 |
| Q11YX6 | Q11YY5 | Q11YY6 | Q11YY8 | Q11Z03 | Q11Z42 | Q11Z44 | Q11Z53 | Q11Z58 | Q11Z64 | Q12000 |
| Q12004 | Q12008 | Q12009 | Q12012 | Q12019 | Q12020 | Q12021 | Q12024 | Q12026 | Q12027 | Q12030 |
| Q12031 | Q12032 | Q12034 | Q12035 | Q12040 | Q12041 | Q12044 | Q12045 | Q12046 | Q12047 | Q12049 |
| Q12050 | Q12052 | Q12053 | Q12056 | Q12058 | Q12059 | Q12060 | Q12062 | Q12069 | Q12072 | Q12074 |
| Q12080 | Q12082 | Q12083 | Q12084 | Q12085 | Q12086 | Q12090 | Q12092 | Q12093 | Q12094 | Q12098 |
| Q12099 | Q120N9 | Q120P0 | Q12102 | Q12104 | Q12107 | Q12108 | Q12109 | Q12110 | Q12115 | Q12120 |
| Q12122 | Q12123 | Q12126 | Q12129 | Q12130 | Q12132 | Q12134 | Q12136 | Q12138 | Q12139 | Q12140 |
| Q12143 | Q12145 | Q12146 | Q12148 | Q12149 | Q12153 | Q12156 | Q12157 | Q12158 | Q12159 | Q12161 |
| Q12163 | Q12166 | Q12168 | Q12172 | Q12173 | Q12175 | Q12176 | Q12177 | Q12178 | Q12179 | Q12180 |
| Q12181 | Q12182 | Q12184 | Q12186 | Q12188 | Q12192 | Q12194 | Q12195 | Q12196 | Q12198 | Q121K8 |
| Q121L2 | Q121Q5 | Q12206 | Q12208 | Q12211 | Q12214 | Q12216 | Q12217 | Q12219 | Q12220 | Q12222 |
| Q12223 | Q12226 | Q12238 | Q12242 | Q12243 | Q12244 | Q12245 | Q12247 | Q12248 | Q12250 | Q12257 |
| Q12259 | Q12260 | Q12261 | Q12262 | Q12264 | Q12265 | Q12266 | Q12267 | Q12272 | Q12276 | Q12277 |
| Q12281 | Q12286 | Q12287 | Q12288 | Q12291 | Q12293 | Q12297 | Q12298 | Q122G2 | Q122H7 | Q122N0 |
| Q122P8 | Q122Q4 | Q122S5 | Q122U9 | Q12306 | Q12307 | Q12308 | Q12309 | Q12311 | Q12314 | Q12317 |
| Q12318 | Q12320 | Q12322 | Q12326 | Q12327 | Q12329 | Q12330 | Q12334 | Q12338 | Q12339 | Q12340 |
| Q12341 | Q12342 | Q12343 | Q12345 | Q12347 | Q12348 | Q12351 | Q12358 | Q12362 | Q12363 | Q12366 |
| Q12367 | Q12368 | Q12373 | Q12377 | Q12378 | Q12379 | Q12381 | Q12383 | Q12386 | Q12387 | Q12389 |
| Q12391 | Q12392 | Q12395 | Q12397 | Q12398 | Q123B6 | Q123D5 | Q123F3 | Q123F4 | Q123F6 | Q123F9 |
| Q123G0 | Q123G1 | Q123G2 | Q123G3 | Q123G4 | Q123S8 | Q123W3 | Q123X2 | Q123Y6 | Q12400 | Q12406 |
| Q12415 | Q12416 | Q12417 | Q12420 | Q12421 | Q12425 | Q12426 | Q12427 | Q12429 | Q12432 | Q12433 |
| Q12439 | Q12440 | Q12441 | Q12444 | Q12446 | Q12447 | Q12448 | Q12449 | Q12453 | Q12454 | Q12455 |
| Q12457 | Q12458 | Q12460 | Q12461 | Q12463 | Q12464 | Q12468 | Q12470 | Q12471 | Q12476 | Q12477 |
| Q12480 | Q12481 | Q12484 | Q12485 | Q12486 | Q12488 | Q12493 | Q12494 | Q12495 | Q12496 | Q12497 |
| Q12499 | Q124E7 | Q124G2 | Q124G8 | Q124N6 | Q124N7 | Q124P0 | Q124Q1 | Q124Q3 | Q124Q5 | Q124Q6 |
| Q124R0 | Q124V5 | Q12500 | Q12504 | Q12506 | Q12507 | Q12508 | Q12509 | Q12510 | Q12513 | Q12514 |
| Q12515 | Q12522 | Q12523 | Q12524 | Q12525 | Q12529 | Q12531 | Q12532 | Q12545 | Q12548 | Q12553 |
| Q12555 | Q12556 | Q12559 | Q12560 | Q12572 | Q12574 | Q12578 | Q12594 | Q12595 | Q12599 | Q125B6 |
| Q125Q0 | Q125U1 | Q125U2 | Q12600 | Q12604 | Q12608 | Q12609 | Q12611 | Q12613 | Q12627 | Q12629 |
| Q12630 | Q12634 | Q12640 | Q12641 | Q12645 | Q12650 | Q12657 | Q12658 | Q12659 | Q12663 | Q12669 |

Q12670 Q12676 Q12680 Q12698 Q126C9 Q126D2 Q126F5 Q126I4 Q126I6 Q126K5 Q126M0
Q126M1 Q126M3 Q126M8 Q126M9 Q126N9 Q126P7 Q126P9 Q126R4 Q126R8 Q126S5 Q12702
Q12704 Q12706 Q12709 Q12723 Q12724 Q12726 Q12731 Q12734 Q12740 Q12744 Q12748
Q12751 Q12754 Q12766 Q12788 Q12798 Q127P5 Q127U4 Q127W3 Q127W7 Q127W8 Q12800
Q12870 Q12882 Q12894 Q12899 Q128B2 Q128B6 Q128E7 Q128K1 Q128K2 Q128K3 Q128P8
Q128Q4 Q128Q7 Q128R7 Q128S1 Q128T0 Q128T5 Q128U1 Q128U3 Q128U4 Q12901 Q12946
Q12947 Q12950 Q12952 Q12988 Q129C6 Q129C8 Q129F0 Q129G8 Q129H3 Q129J0 Q129J1
Q129K2 Q129K3 Q129L2 Q129M0 Q129M4 Q129M7 Q129V0 Q129W6 Q12A06 Q12A25 Q12A31
Q12A32 Q12A33 Q12A34 Q12A45 Q12A51 Q12A85 Q12AB4 Q12AB5 Q12AC2 Q12AC3 Q12AC9
Q12AD4 Q12AE4 Q12AE5 Q12AK6 Q12AT7 Q12AU6 Q12AU7 Q12AU9 Q12AW5 Q12B25 Q12B26
Q12B36 Q12B43 Q12B44 Q12B47 Q12B50 Q12B88 Q12B97 Q12BQ7 Q12BR0 Q12BR1 Q12BR3
Q12BV1 Q12BV7 Q12BW0 Q12BY1 Q12BY3 Q12BZ9 Q12C18 Q12C38 Q12C39 Q12C42 Q12C45
Q12CG7 Q12CI6 Q12CL3 Q12CM2 Q12CN0 Q12CQ9 Q12CR1 Q12CT8 Q12CW3 Q12CW4 Q12CW5
Q12CW6 Q12D45 Q12D54 Q12D55 Q12D58 Q12D67 Q12D73 Q12D74 Q12D75 Q12DD9 Q12DF1
Q12DM9 Q12DT7 Q12DU1 Q12DU2 Q12DU3 Q12DU4 Q12DU5 Q12DU6 Q12DV7 Q12DV8 Q12DY9
Q12DZ0 Q12DZ2 Q12DZ5 Q12DZ8 Q12E00 Q12E01 Q12E50 Q12E97 Q12EE6 Q12EE8 Q12EE9
Q12EF7 Q12EH0 Q12EI9 Q12EK7 Q12EL3 Q12EL4 Q12EL7 Q12EM3 Q12EM4 Q12EQ4 Q12EY5
Q12EY8 Q12F12 Q12F35 Q12F38 Q12F39 Q12F40 Q12F41 Q12F47 Q12F85 Q12F86 Q12F97
Q12F98 Q12F99 Q12FA0 Q12FA3 Q12FC5 Q12FC6 Q12FC7 Q12FC8 Q12FD0 Q12FD4 Q12FF7
Q12FG5 Q12FG9 Q12FH6 Q12FH7 Q12FI4 Q12FK2 Q12FQ6 Q12FU8 Q12FW8 Q12G76 Q12G77
Q12G78 Q12G79 Q12G80 Q12G81 Q12G82 Q12G84 Q12G85 Q12G86 Q12G87 Q12G88 Q12G89
Q12G90 Q12G91 Q12G92 Q12G93 Q12GA2 Q12GB2 Q12GI7 Q12GR3 Q12GT8 Q12GW2 Q12GW3
Q12GW4 Q12GW5 Q12GW6 Q12GW7 Q12GW8 Q12GW9 Q12GX0 Q12GX1 Q12GX2 Q12GX4 Q12GX5
Q12GX6 Q12H06 Q12H34 Q12HF1 Q12HF2 Q12HI8 Q12HK7 Q12HM5 Q12HM6 Q12HM9 Q12HP0
Q12HP1 Q12HV5 Q12HV7 Q12I24 Q12I30 Q12I41 Q12IC3 Q12IH3 Q12IK4 Q12IL7 Q12IL8
Q12IN9 Q12IP4 Q12IQ6 Q12IT8 Q12IT9 Q12IU3 Q12IU6 Q12IW9 Q12J15 Q12J88 Q12J93
Q12J94 Q12J95 Q12JA9 Q12JC5 Q12JC6 Q12JD1 Q12JE5 Q12JH2 Q12JH7 Q12JN5 Q12JW2
Q12JX1 Q12JZ0 Q12K11 Q12K21 Q12K22 Q12K23 Q12K37 Q12K58 Q12K59 Q12K60 Q12KB6
Q12KB7 Q12KC4 Q12KC8 Q12KD2 Q12KD5 Q12KE6 Q12KG5 Q12KG8 Q12KI1 Q12KJ3 Q12KJ4
Q12KJ5 Q12KJ6 Q12KL2 Q12KL3 Q12KL8 Q12KM1 Q12KP8 Q12KQ0 Q12KQ6 Q12KR9 Q12KS3
Q12KS4 Q12KV7 Q12L03 Q12L09 Q12L24 Q12L25 Q12L26 Q12L31 Q12L35 Q12L39 Q12L53
Q12L63 Q12L82 Q12L98 Q12L99 Q12LA0 Q12LA1 Q12LA2 Q12LC6 Q12LE2 Q12LE4 Q12LJ8
Q12LK6 Q12LL4 Q12LL6 Q12LP2 Q12LR0 Q12LR1 Q12LR7 Q12LU7 Q12LU9 Q12LW4 Q12LW6
Q12LW9 Q12M48 Q12M63 Q12MB3 Q12ME4 Q12MF0 Q12MF1 Q12MI8 Q12MJ0 Q12MJ8 Q12MK0
Q12MM2 Q12MM4 Q12MM8 Q12MN3 Q12MQ1 Q12MQ7 Q12MT5 Q12MU6 Q12N03 Q12N04 Q12N06
Q12N08 Q12N09 Q12N10 Q12N51 Q12N55 Q12N57 Q12N59 Q12N64 Q12N78 Q12NA7 Q12NB8
Q12NC2 Q12ND8 Q12NE0 Q12NE1 Q12NE5 Q12NF7 Q12NI4 Q12NI7 Q12NL3 Q12NN4 Q12NR0
Q12NR2 Q12NR3 Q12NR5 Q12NS4 Q12NS9 Q12NT0 Q12NT8 Q12NU0 Q12NX3 Q12NY4 Q12NY5
Q12NY6 Q12NY7 Q12NZ0 Q12P04 Q12P28 Q12P38 Q12P55 Q12P58 Q12P60 Q12P69 Q12P79
Q12P80 Q12P83 Q12P94 Q12PA3 Q12PA8 Q12PB0 Q12PB2 Q12PB3 Q12PD9 Q12PE0 Q12PE3
Q12PR7 Q12PS0 Q12PS2 Q12PT0 Q12PT3 Q12PT4 Q12PT7 Q12PV1 Q12PX6 Q12PX7 Q12PY2
Q12PY3 Q12PY8 Q12PY9 Q12PZ1 Q12PZ2 Q12PZ5 Q12PZ8 Q12Q05 Q12Q07 Q12Q08 Q12Q42
Q12Q43 Q12Q47 Q12Q48 Q12Q96 Q12QA3 Q12QA4 Q12QA6 Q12QD2 Q12QG0 Q12QG1 Q12QG2
Q12QG7 Q12QG9 Q12QH8 Q12QH9 Q12QI0 Q12QI1 Q12QI3 Q12QI5 Q12QI6 Q12QI9 Q12QJ4

Q12QK6 Q12QM6 Q12QM9 Q12QQ5 Q12QR8 Q12QR9 Q12QS1 Q12QW7 Q12QX0 Q12QX1 Q12QY2
Q12R03 Q12R04 Q12R11 Q12R19 Q12R22 Q12R24 Q12R39 Q12R42 Q12R66 Q12R91 Q12RA0
Q12RC7 Q12RU3 Q12RU8 Q12RV3 Q12RW2 Q12RW6 Q12RW7 Q12RW9 Q12RX5 Q12RX7 Q12RX8
Q12RY2 Q12RZ8 Q12S16 Q12S21 Q12S23 Q12S35 Q12S38 Q12S40 Q12S61 Q12S62 Q12S90
Q12SC5 Q12SD4 Q12SD5 Q12SE4 Q12SE7 Q12SF3 Q12SF5 Q12SF7 Q12SF8 Q12SG0 Q12SG2
Q12SJ3 Q12SK8 Q12SK9 Q12SL8 Q12SL9 Q12SM6 Q12SM7 Q12SM9 Q12SN0 Q12SN4 Q12SP7
Q12SP8 Q12SQ2 Q12SR9 Q12SS1 Q12ST3 Q12ST4 Q12ST5 Q12ST6 Q12ST7 Q12ST8 Q12SU0
Q12SU1 Q12SU2 Q12SU3 Q12SU4 Q12SU5 Q12SU6 Q12SU7 Q12SU8 Q12SU9 Q12SV0 Q12SV1
Q12SV2 Q12SV3 Q12SV4 Q12SV5 Q12SV6 Q12SV7 Q12SV8 Q12SV9 Q12SW0 Q12SW1 Q12SW2
Q12SW3 Q12SW4 Q12SW5 Q12SW6 Q12SW7 Q12SW8 Q12SW9 Q12SX0 Q12SX3 Q12SX8 Q12T11
Q12T12 Q12T28 Q12T32 Q12T34 Q12T48 Q12T52 Q12T95 Q12T99 Q12TA0 Q12TA2 Q12TA6
Q12TB3 Q12TB6 Q12TC0 Q12TC1 Q12TC6 Q12TC8 Q12TF2 Q12TF9 Q12TG0 Q12TH8 Q12TI1
Q12TI2 Q12TI4 Q12TI6 Q12TL1 Q12TL2 Q12TM7 Q12TN0 Q12TN5 Q12TN7 Q12TP4 Q12TR2
Q12TR7 Q12TT2 Q12TT3 Q12TT4 Q12TU6 Q12TV0 Q12TW7 Q12TX1 Q12U08 Q12U18 Q12U19
Q12U23 Q12U25 Q12U30 Q12U31 Q12U80 Q12U93 Q12UA1 Q12UB6 Q12UF1 Q12UF8 Q12UF9
Q12UG6 Q12UG7 Q12UJ5 Q12UJ7 Q12UJ8 Q12UJ9 Q12UK2 Q12UK3 Q12UL5 Q12UL8 Q12UP8
Q12UP9 Q12UQ0 Q12UQ4 Q12UQ9 Q12UR3 Q12UT0 Q12UT1 Q12UU7 Q12UV0 Q12UV8 Q12UW5
Q12UX3 Q12UY1 Q12UY6 Q12V26 Q12V29 Q12V31 Q12V34 Q12V48 Q12V67 Q12V70 Q12V97
Q12V98 Q12VA2 Q12VA6 Q12VB0 Q12VC9 Q12VD1 Q12VF1 Q12VF4 Q12VF9 Q12VH0 Q12VH1
Q12VH2 Q12VH3 Q12VJ6 Q12VJ7 Q12VQ2 Q12VV9 Q12VZ8 Q12W02 Q12W06 Q12W15 Q12W26
Q12W27 Q12W31 Q12W43 Q12W53 Q12W64 Q12W69 Q12W71 Q12WA2 Q12WB4 Q12WC5 Q12WC6
Q12WC7 Q12WC8 Q12WD4 Q12WD5 Q12WD9 Q12WE6 Q12WF3 Q12WF5 Q12WF6 Q12WH5 Q12WH6
Q12WH8 Q12WI3 Q12WM5 Q12WM7 Q12WM8 Q12WN0 Q12WQ0 Q12WQ7 Q12WR3 Q12WS1 Q12WS3
Q12WT0 Q12WT3 Q12WT4 Q12WZ6 Q12X08 Q12X17 Q12X18 Q12X37 Q12X51 Q12X56 Q12X58
Q12X65 Q12X66 Q12X75 Q12X87 Q12XB4 Q12XB7 Q12XG0 Q12XG1 Q12XH1 Q12XH2 Q12XH4
Q12XH7 Q12XK2 Q12XK3 Q12XL6 Q12XL7 Q12XM4 Q12XP4 Q12XS2 Q12XS4 Q12XU7 Q12XX4
Q12XX5 Q12XX8 Q12Y03 Q12Y24 Q12Y46 Q12Y52 Q12YN5 Q12YN9 Q12YP2 Q12YR1 Q12YR6
Q12YU9 Q12YV7 Q12YV9 Q12Z05 Q12Z15 Q12Z63 Q12Z64 Q12Z92 Q12Z93 Q12Z94 Q12Z95
Q12ZB1 Q12ZB3 Q12ZB4 Q12ZB8 Q12ZC2 Q12ZC4 Q12ZC6 Q12ZD1 Q12ZG1 Q12ZG2 Q12ZG3
Q12ZI9 Q12ZJ1 Q12ZJ3 Q12ZJ5 Q12ZJ7 Q12ZL9 Q12ZM1 Q12ZN9 Q12ZP0 Q12ZP6 Q12ZP8
Q12ZQ0 Q12ZR4 Q12ZR5 Q12ZR6 Q12ZS5 Q12ZS7 Q12ZT0 Q12ZT1 Q12ZT2 Q12ZT5 Q12ZT6
Q12ZT7 Q12ZT8 Q12ZU0 Q12ZU1 Q12ZU2 Q12ZU3 Q12ZU5 Q12ZU6 Q12ZU7 Q12ZU8 Q12ZU9
Q12ZV0 Q12ZV1 Q12ZV2 Q12ZX1 Q13049 Q13064 Q13066 Q13069 Q13070 Q13099 Q130C0
Q130C9 Q130G7 Q130G8 Q130J1 Q130J4 Q130J6 Q130N3 Q130R0 Q130U9 Q130V1 Q130V3
Q130Z3 Q13106 Q13117 Q13118 Q13123 Q13126 Q13156 Q13166 Q131A2 Q131B9 Q131E0
Q131I8 Q131J1 Q131J2 Q131L9 Q131M0 Q131Q8 Q131U9 Q131V4 Q131X4 Q13202 Q13203
Q13207 Q13232 Q13278 Q13285 Q132J9 Q132N2 Q132N3 Q132N5 Q132P2 Q13347 Q13351
Q13360 Q13363 Q13394 Q13395 Q13398 Q133H2 Q133K4 Q133L1 Q133L2 Q133L3 Q133L6
Q133L9 Q133M2 Q133N2 Q133R5 Q133T3 Q133T5 Q133W3 Q133X2 Q133X3 Q133Y4 Q133Y8
Q13401 Q13434 Q13461 Q13472 Q13487 Q134B1 Q134B7 Q134D1 Q134L7 Q134L8 Q134R0
Q134R3 Q134R4 Q134R6 Q134R7 Q134R8 Q134R9 Q134S4 Q134S5 Q134S6 Q134S7 Q134S8
Q134S9 Q134T0 Q134T1 Q134T2 Q134T3 Q134T5 Q134T7 Q134T8 Q134T9 Q134U0 Q134U1
Q134U2 Q134U3 Q134U4 Q134U6 Q134U7 Q134V1 Q134V2 Q134V3 Q134V4 Q13503 Q13516
Q13536 Q135C1 Q135C2 Q135D0 Q135F0 Q135H6 Q135I5 Q135J0 Q135J5 Q135J7 Q135J9

Q135K5 Q135K8 Q135M5 Q135M7 Q135M8 Q135N9 Q135P2 Q135P8 Q135Q2 Q135R4 Q135R5
Q135W6 Q135W7 Q135W8 Q135Z6 Q13617 Q13630 Q13643 Q13670 Q13686 Q136A2 Q136A4
Q136A5 Q136A6 Q136A7 Q136B0 Q136B9 Q136D2 Q136D3 Q136D7 Q136E4 Q136G5 Q136H4
Q136H7 Q136H9 Q136I8 Q136L0 Q136Q2 Q136Q3 Q136S3 Q136T4 Q136T8 Q136V2 Q136V6
Q136W2 Q136W4 Q136W8 Q136X6 Q136X7 Q13772 Q137A8 Q137B2 Q137B6 Q137C3 Q137C6
Q137C7 Q137K5 Q137T4 Q137U7 Q13825 Q13867 Q13888 Q13889 Q138J5 Q138K0 Q138M7
Q138N7 Q138Z2 Q138Z3 Q13901 Q13939 Q13952 Q139A2 Q139A7 Q139D9 Q139E9 Q139F2
Q139H2 Q139H4 Q139H7 Q139V1 Q139X8 Q13A07 Q13A88 Q13AA1 Q13AB9 Q13AK7 Q13AN7
Q13AQ5 Q13AQ6 Q13B10 Q13B59 Q13B80 Q13B97 Q13BA4 Q13BB8 Q13BC6 Q13BD7 Q13BI7
Q13BJ4 Q13BM0 Q13BT8 Q13BU3 Q13BX8 Q13C64 Q13C78 Q13CA1 Q13CD3 Q13CJ6 Q13CL5
Q13CU0 Q13D88 Q13D92 Q13D98 Q13DA7 Q13DL2 Q13DL3 Q13DL7 Q13DM7 Q13DN0 Q13DN3
Q13DN7 Q13DR5 Q13DR6 Q13DT7 Q13DU2 Q13DU3 Q13DU5 Q13DU7 Q13DV3 Q13DY1 Q13DZ7
Q13E15 Q13E20 Q13E21 Q13E22 Q13E25 Q13E26 Q13E29 Q13E33 Q13E36 Q13E39 Q13E40
Q13E42 Q13E51 Q13E56 Q13E58 Q13E60 Q13E64 Q13E66 Q13E74 Q13E86 Q13EA6 Q13EA7
Q13ED0 Q13EE0 Q13EG6 Q13EK7 Q13EK9 Q13EL3 Q13EL5 Q13EL8 Q13EL9 Q13EM1 Q13EM2
Q13EM6 Q13EN7 Q13EN8 Q13EP1 Q13EQ1 Q13EQ2 Q13EQ3 Q13EQ5 Q13EQ8 Q13ER0 Q13ET3
Q13ET4 Q13ET5 Q13EZ9 Q13F02 Q13F06 Q13F13 Q13F14 Q13F18 Q13F19 Q13F20 Q13F22
Q13F24 Q13F27 Q13F29 Q13F98 Q13FB9 Q13IM9 Q13IY4 Q13J37 Q13JL5 Q13L77 Q13LW5
Q13LW8 Q13M01 Q13ME5 Q13MI0 Q13MI3 Q13MT2 Q13N12 Q13N17 Q13NE4 Q13NG5 Q13NG6
Q13NG7 Q13NQ1 Q13P97 Q13PC1 Q13PQ0 Q13Q00 Q13QH4 Q13QH6 Q13QH7 Q13QH9 Q13QI0
Q13QU8 Q13QV1 Q13R51 Q13RB8 Q13RW7 Q13RW9 Q13RX0 Q13RX1 Q13RY4 Q13S07 Q13S27
Q13S42 Q13SG2 Q13SH1 Q13SH2 Q13SH7 Q13SL2 Q13SL4 Q13SM2 Q13SP0 Q13SP1 Q13SQ6
Q13SR6 Q13SR7 Q13SW3 Q13SY1 Q13T36 Q13T51 Q13T64 Q13TF5 Q13TF8 Q13TF9 Q13TG0
Q13TG1 Q13TG2 Q13TG3 Q13TG5 Q13TG6 Q13TG7 Q13TG9 Q13TH0 Q13TH1 Q13TH2 Q13TH3
Q13TH4 Q13TH5 Q13TH6 Q13TH7 Q13TH8 Q13TH9 Q13TI0 Q13TI1 Q13TI2 Q13TI3 Q13TI4
Q13TI5 Q13TI6 Q13TI7 Q13TI8 Q13TI9 Q13TJ1 Q13TJ2 Q13TJ3 Q13TJ4 Q13TJ5 Q13TJ6
Q13TJ7 Q13TK1 Q13TK9 Q13TQ2 Q13TQ6 Q13TQ9 Q13TR0 Q13TR2 Q13TU2 Q13TU3 Q13TW4
Q13TW5 Q13TY3 Q13TY4 Q13TZ0 Q13TZ3 Q13TZ4 Q13U06 Q13U13 Q13U14 Q13U15 Q13U16
Q13U19 Q13U36 Q13U82 Q13U85 Q13UB2 Q13UB3 Q13UB4 Q13UB5 Q13UB7 Q13UC0 Q13UC1
Q13UC2 Q13UC4 Q13UD3 Q13UD5 Q13UE3 Q13UH9 Q13UI4 Q13UP0 Q13UP6 Q13UQ9 Q13UR0
Q13UR1 Q13UR4 Q13UR8 Q13US0 Q13US7 Q13UT1 Q13UU8 Q13UV5 Q13UV8 Q13UW2 Q13UX1
Q13UX2 Q13UX3 Q13V14 Q13V35 Q13V36 Q13V40 Q13V47 Q13V49 Q13V50 Q13V64 Q13V90
Q13VB4 Q13VC2 Q13VC3 Q13VC5 Q13VD0 Q13VE3 Q13VE8 Q13VE9 Q13VF0 Q13VF1 Q13VF6
Q13VF8 Q13VG0 Q13VL1 Q13VL3 Q13VN0 Q13VN6 Q13VN8 Q13VU0 Q13VU2 Q13VU3 Q13W47
Q13W50 Q13W97 Q13WE6 Q13WF7 Q13WF9 Q13WG0 Q13WM0 Q13WN4 Q13WU6 Q13WX0 Q13WZ0
Q13WZ5 Q13X05 Q13X25 Q13X26 Q13X29 Q13X32 Q13X33 Q13X39 Q13X40 Q13X51 Q13X60
Q13X76 Q13X79 Q13XA2 Q13XA4 Q13XB0 Q13XB6 Q13XB7 Q13XB8 Q13XB9 Q13XC2 Q13XD0
Q13XD2 Q13XD6 Q13XD8 Q13XE2 Q13Y97 Q13YI7 Q13YI8 Q13YL3 Q13YU3 Q13YY3 Q13YZ6
Q13Z07 Q13Z14 Q13Z15 Q13Z16 Q13Z30 Q13Z31 Q13Z56 Q13Z67 Q13Z72 Q13Z78 Q13Z79
Q13Z86 Q13ZG9 Q13ZH0 Q13ZH2 Q13ZJ7 Q13ZL9 Q13ZM1 Q13ZM2 Q13ZQ5 Q13ZW7 Q14061
Q14093 Q140U5 Q140U6 Q140U8 Q14117 Q14129 Q14139 Q14141 Q14159 Q14192 Q14194
Q141C9 Q141D1 Q141D2 Q141D3 Q141E6 Q141E8 Q141F5 Q14201 Q14206 Q14209 Q14240
Q14296 Q142D9 Q142H7 Q142H8 Q142I6 Q142L5 Q142L6 Q142L8 Q142M1 Q142M3 Q142P9
Q142Q7 Q142S4 Q142T1 Q142T5 Q14324 Q14353 Q14376 Q14390 Q14397 Q143B6 Q143B9

Q143C7 Q143C9 Q143D9 Q143F8 Q143I2 Q143I7 Q143J2 Q143J3 Q143P7 Q143P8 Q143Y9
Q14469 Q144D8 Q144E2 Q144E3 Q144E5 Q144E6 Q144E7 Q144P9 Q144Q1 Q144Q2 Q144Q6
Q14525 Q14532 Q14533 Q14541 Q14549 Q14558 Q14565 Q14568 Q14576 Q14582 Q14584
Q14585 Q14586 Q14587 Q14588 Q14590 Q14592 Q14593 Q145A5 Q145C5 Q145C6 Q145E8
Q145F0 Q145F1 Q145F3 Q145F5 Q145F6 Q145F7 Q145H9 Q145K0 Q145L1 Q145N3 Q145P0
Q145P6 Q145V4 Q145W4 Q145W5 Q145W8 Q145X1 Q145X3 Q145X4 Q145X7 Q14602 Q14657
Q14681 Q14691 Q14695 Q146B7 Q146F2 Q146G6 Q146I4 Q146J2 Q146K3 Q146K5 Q146L7
Q146Q2 Q146W0 Q146W2 Q146W9 Q146X0 Q146X4 Q146X7 Q146Y4 Q146Y5 Q146Y6 Q146Z6
Q14749 Q147A4 Q147F0 Q147G5 Q147U1 Q14863 Q14865 Q14894 Q14896 Q148B6 Q148C4
Q148C9 Q148D5 Q148E0 Q148F0 Q148F6 Q148G4 Q148G5 Q148G7 Q148H4 Q148H5 Q148H6
Q148H7 Q148H8 Q148I0 Q148I1 Q148I5 Q148J6 Q148K0 Q148L6 Q148L7 Q148M7 Q148M8
Q148N0 Q148W8 Q14919 Q14929 Q14990 Q14997 Q149C2 Q149F1 Q149F3 Q149G0 Q149M9
Q149S1 Q149T7 Q14AI0 Q14AI6 Q14AT2 Q14AW5 Q14AX6 Q14B48 Q14B70 Q14BI7 Q14BJ1
Q14BK3 Q14BP6 Q14BV6 Q14C37 Q14CH1 Q14CH7 Q14CN4 Q14CX7 Q14CZ7 Q14DL0 Q14DL3
Q14DQ1 Q14EA9 Q14F94 Q14F95 Q14F96 Q14F97 Q14F99 Q14FA1 Q14FB1 Q14FB6 Q14FB8
Q14FB9 Q14FC0 Q14FC1 Q14FC2 Q14FC3 Q14FC4 Q14FC5 Q14FD2 Q14FD4 Q14FD5 Q14FD6
Q14FE9 Q14FF5 Q14FF9 Q14FG5 Q14FG6 Q14FG7 Q14FG8 Q14FH5 Q14FN2 Q14FN3 Q14FN7
Q14FQ1 Q14FQ2 Q14FR3 Q14FU4 Q14FU5 Q14FU7 Q14FV3 Q14FW0 Q14FW2 Q14FX6 Q14G14
Q14G22 Q14G25 Q14G26 Q14G45 Q14G49 Q14G66 Q14G67 Q14GA7 Q14GB0 Q14GB1 Q14GD0
Q14GD9 Q14GG5 Q14GH2 Q14GJ0 Q14GJ6 Q14GK8 Q14GK9 Q14GL0 Q14GL1 Q14GL3 Q14GM5
Q14GM9 Q14GR3 Q14GR4 Q14GS7 Q14GS9 Q14GT5 Q14GU4 Q14GV0 Q14GV5 Q14GW0 Q14GW8
Q14GW9 Q14GX0 Q14GZ5 Q14H05 Q14H07 Q14H21 Q14H27 Q14H30 Q14H45 Q14H46 Q14H66
Q14H72 Q14H73 Q14H85 Q14H93 Q14HA0 Q14HB9 Q14HD0 Q14HF0 Q14HF1 Q14HF2 Q14HF3
Q14HF5 Q14HG2 Q14HG9 Q14HH3 Q14HH8 Q14HI8 Q14HJ0 Q14HJ1 Q14HJ6 Q14HK5 Q14HL3
Q14HL6 Q14HM0 Q14HM1 Q14HN8 Q14HQ0 Q14HR4 Q14HR7 Q14HS1 Q14HS3 Q14HS5 Q14HT2
Q14HT5 Q14HU9 Q14HV5 Q14HW8 Q14HY4 Q14I06 Q14I17 Q14I18 Q14I20 Q14I27 Q14I28
Q14I60 Q14I63 Q14I64 Q14I72 Q14I80 Q14IB6 Q14IC3 Q14IC5 Q14IC8 Q14IC9 Q14ID0
Q14ID3 Q14IE1 Q14IE8 Q14IF3 Q14IG1 Q14IH0 Q14IH1 Q14IH3 Q14II4 Q14II7 Q14IJ3
Q14IJ4 Q14IJ9 Q14IK0 Q14IK5 Q14IK7 Q14IL6 Q14IN1 Q14IN9 Q14IP6 Q14IT0 Q14IU7
Q14IU8 Q14IV5 Q14IY5 Q14IY7 Q14IZ0 Q14J05 Q14J34 Q14J36 Q14J40 Q14J42 Q14J59
Q14J73 Q14J74 Q14J81 Q14J90 Q14J94 Q14J95 Q14J96 Q14J97 Q14J98 Q14J99 Q14JA1
Q14JA2 Q14JA3 Q14JA4 Q14JA5 Q14JA6 Q14JA7 Q14JA8 Q14JA9 Q14JB0 Q14JB1 Q14JB2
Q14JB3 Q14JB4 Q14JB5 Q14JB6 Q14JB7 Q14JB8 Q14JB9 Q14JC0 Q14JC1 Q14JC2 Q14JC3
Q14JC4 Q14JC6 Q14JC9 Q14JD0 Q14JD1 Q14JD2 Q14JD8 Q14JH1 Q14JI3 Q14JK2 Q14JK5
Q14JL2 Q14JL3 Q14JN1 Q14JP9 Q14JR1 Q14JR2 Q14JR9 Q14JS6 Q14JS7 Q14JS8 Q14JS9
Q14JT0 Q14JT2 Q14JT4 Q14JT5 Q14JT6 Q14JT7 Q14JT8 Q14JT9 Q14JU2 Q14JU8 Q14JV7
Q14JV8 Q14JW3 Q14JW7 Q14JZ0 Q14JZ1 Q14JZ2 Q14K18 Q14K19 Q14K20 Q14K22 Q14K23
Q14K48 Q14K49 Q14KZ8 Q14L37 Q14LB9 Q14NV1 Q14NW7 Q14PJ7 Q14PJ9 Q14RS1 Q14RS2
Q14T92 Q14V87 Q15008 Q15024 Q15031 Q15034 Q15038 Q15048 Q15072 Q15118 Q15131
Q15257 Q15269 Q15274 Q152V3 Q15323 Q15369 Q15370 Q15386 Q15393 Q153Z0 Q15415
Q15427 Q15428 Q15475 Q15486 Q15528 Q15543 Q15560 Q15561 Q15569 Q15572 Q15573
Q15583 Q15631 Q15633 Q15695 Q15697 Q15699 Q156A1 Q15772 Q15777 Q15813 Q15819
Q15834 Q15843 Q15847 Q15906 Q15915 Q15916 Q15928 Q15935 Q15937 Q15940 Q15973
Q15BH7 Q15FB2 Q15FB3 Q15FB7 Q15GI3 Q15GI4 Q15JF5 Q15JF8 Q15JG1 Q15JG4 Q15JG5

Q15JG7 Q15MS4 Q15MS5 Q15MS8 Q15MS9 Q15MT3 Q15MT4 Q15MV7 Q15MV9 Q15MY4 Q15MY9
Q15N06 Q15N09 Q15N18 Q15N19 Q15N47 Q15N48 Q15N51 Q15N71 Q15NG5 Q15NG9 Q15NK1
Q15NK9 Q15NL2 Q15NL3 Q15NL4 Q15NN9 Q15NP1 Q15NP2 Q15NQ1 Q15NR0 Q15NR1 Q15NR8
Q15NS9 Q15NT8 Q15NV0 Q15NV5 Q15NV6 Q15P27 Q15P28 Q15P29 Q15P30 Q15P31 Q15P42
Q15P51 Q15PD3 Q15PD4 Q15PE1 Q15PE2 Q15PE8 Q15PE9 Q15PF0 Q15PG0 Q15PH6 Q15PI3
Q15PI4 Q15PJ1 Q15PJ2 Q15PM0 Q15PR3 Q15PR4 Q15PR6 Q15PS2 Q15PS5 Q15PU4 Q15PU5
Q15PX4 Q15Q03 Q15Q08 Q15Q09 Q15Q18 Q15Q19 Q15Q28 Q15Q55 Q15QB2 Q15QE8 Q15QF3
Q15QG8 Q15QK1 Q15QK3 Q15QK4 Q15QQ0 Q15QR2 Q15QR4 Q15QR7 Q15QR9 Q15QS3 Q15R01
Q15R04 Q15R07 Q15R35 Q15R43 Q15R45 Q15R46 Q15R47 Q15R52 Q15R53 Q15R57 Q15R60
Q15R61 Q15R63 Q15R69 Q15RG1 Q15RG5 Q15RG6 Q15RL9 Q15RN2 Q15RN4 Q15RN6 Q15RP8
Q15RR1 Q15RS7 Q15RT2 Q15RT5 Q15RT6 Q15RT7 Q15RU3 Q15RU5 Q15RU7 Q15RU8 Q15RZ5
Q15RZ7 Q15S25 Q15S48 Q15SN0 Q15SQ2 Q15SQ3 Q15SQ6 Q15SQ8 Q15SQ9 Q15SR5 Q15SS0
Q15SS1 Q15SU5 Q15SU8 Q15SX5 Q15SX6 Q15SX7 Q15SX9 Q15SY3 Q15SZ4 Q15SZ6 Q15T04
Q15T07 Q15T09 Q15T16 Q15T23 Q15T93 Q15T97 Q15T99 Q15TA0 Q15TA1 Q15TA7 Q15TB5
Q15TJ8 Q15TQ4 Q15TR2 Q15TR4 Q15TS0 Q15TS1 Q15TS5 Q15TS6 Q15TS8 Q15TT6 Q15TV2
Q15TZ2 Q15TZ4 Q15U36 Q15U56 Q15U72 Q15UB5 Q15UB6 Q15UC8 Q15UD2 Q15UD3 Q15UD4
Q15UF1 Q15UL9 Q15UM1 Q15UN9 Q15UT4 Q15UW6 Q15UY5 Q15UZ1 Q15V24 Q15V26 Q15V28
Q15V69 Q15V70 Q15V71 Q15V72 Q15V74 Q15VB5 Q15VB8 Q15VD0 Q15VD4 Q15VG2 Q15VI3
Q15VI6 Q15VI7 Q15VI8 Q15VI9 Q15VJ3 Q15VJ6 Q15VK0 Q15VK3 Q15VK5 Q15VL2 Q15VL3
Q15VL9 Q15VM5 Q15W53 Q15W54 Q15W93 Q15W94 Q15W97 Q15W98 Q15WA2 Q15WB3 Q15WD6
Q15WE9 Q15WG0 Q15WG1 Q15WG2 Q15WG3 Q15WH0 Q15WH1 Q15WI4 Q15WI5 Q15WQ1 Q15X11
Q15X21 Q15X31 Q15X40 Q15X43 Q15X47 Q15X48 Q15X49 Q15X50 Q15X51 Q15X52 Q15X54
Q15X55 Q15X56 Q15X57 Q15X58 Q15X59 Q15X60 Q15X62 Q15X63 Q15X77 Q15X80 Q15X83
Q15X84 Q15X88 Q15XD5 Q15XV2 Q15Y43 Q15Y44 Q15Y60 Q15YA7 Q15YA8 Q15YA9 Q15YB0
Q15YB1 Q15YB2 Q15YB3 Q15YB4 Q15YB5 Q15YC5 Q15YD8 Q15YF4 Q15YF9 Q15YG1 Q15YG5
Q15YH0 Q15YI8 Q15YL3 Q15YN0 Q15YN1 Q15YN2 Q15YN3 Q15YN4 Q15YN5 Q15YN6 Q15YN7
Q15YN8 Q15YN9 Q15YP0 Q15YP4 Q15YP8 Q15YR1 Q15YU1 Q15Z03 Q15Z09 Q15Z10 Q15Z11
Q15Z15 Q15Z55 Q15ZB1 Q15ZB4 Q15ZC9 Q15ZE1 Q15ZF7 Q15ZH0 Q15ZP6 Q15ZR4 Q15ZS0
Q15ZS7 Q15ZS8 Q15ZT1 Q15ZU0 Q15ZU3 Q15ZU4 Q15ZV3 Q15ZV5 Q15ZV7 Q15ZV8 Q15ZW0
Q15ZX4 Q15ZX7 Q15ZX9 Q15ZY3 Q15ZY9 Q15ZZ0 Q15ZZ5 Q16048 Q16082 Q160A3 Q160C0
Q160C1 Q160C2 Q160T4 Q160X3 Q160X4 Q160X5 Q160X6 Q160X7 Q160X8 Q160Y1 Q160Y2
Q160Y3 Q160Y4 Q16143 Q161G5 Q161G7 Q161H0 Q161H1 Q161H2 Q161H6 Q161H9 Q161I4
Q161I5 Q161I8 Q161S5 Q161S8 Q161T1 Q161T2 Q161T3 Q16254 Q162B6 Q162C5 Q162E2
Q162E3 Q162K5 Q162Q0 Q162Q1 Q162Q2 Q162Q6 Q162S3 Q162U5 Q16342 Q16352 Q16384
Q16385 Q163C5 Q163D2 Q163E9 Q163G0 Q163G3 Q163I0 Q163I6 Q163J1 Q163J3 Q163J4
Q163L1 Q163S6 Q163U2 Q163U3 Q163V1 Q163V5 Q163V6 Q163V9 Q163W3 Q163W4 Q163X3
Q163Y0 Q163Z1 Q16473 Q164A4 Q164B9 Q164C1 Q164D1 Q164D9 Q164E0 Q164E9 Q164F8
Q164G1 Q164G2 Q164H3 Q164J6 Q164K6 Q164M8 Q164N1 Q164N5 Q164N6 Q164N7 Q164R7
Q164S6 Q164T8 Q16517 Q16531 Q16534 Q16559 Q16560 Q16587 Q16589 Q16595 Q165D5
Q165E2 Q165E3 Q165E4 Q165F9 Q165G0 Q165G5 Q165J6 Q165K9 Q165M8 Q165N4 Q165P5
Q165P6 Q165Q6 Q165S4 Q165T7 Q165U5 Q165V7 Q165Y2 Q165Y3 Q165Y6 Q165Y7 Q165Y8
Q165Z3 Q165Z4 Q16600 Q16613 Q16621 Q16626 Q16633 Q16650 Q16654 Q16667 Q16670
Q16676 Q16695 Q166A7 Q166D0 Q166D3 Q166D6 Q166E0 Q166E3 Q166F5 Q166F6 Q166F7
Q166H0 Q166J5 Q166L9 Q16719 Q16762 Q16763 Q16773 Q16774 Q16775 Q16778 Q16798

Q167J3 Q167J5 Q167K4 Q167Q3 Q167R8 Q167V0 Q167Y9 Q16816 Q16828 Q16829 Q16831
Q16854 Q16877 Q16878 Q168A2 Q168A3 Q168B1 Q168C1 Q168E6 Q168G2 Q168H2 Q168I8
Q168I9 Q168J2 Q168J5 Q168K5 Q168N1 Q168N3 Q168P4 Q168R5 Q168S1 Q168S7 Q168U9
Q168V8 Q16959 Q16960 Q169A0 Q169C7 Q169E2 Q169F8 Q169H6 Q169N3 Q169N4 Q169P2
Q169Q9 Q169T2 Q169U9 Q169X5 Q16A32 Q16A96 Q16A99 Q16AA3 Q16AA4 Q16AB7 Q16AB8
Q16AB9 Q16AC0 Q16AC4 Q16AC6 Q16AC7 Q16AC8 Q16AC9 Q16AD0 Q16AD1 Q16AD2 Q16AD3
Q16AD4 Q16AD5 Q16AD6 Q16AE4 Q16AE5 Q16AE7 Q16AE8 Q16AF2 Q16AF3 Q16AF4 Q16AF5
Q16AH2 Q16AH6 Q16AI3 Q16AI6 Q16AJ0 Q16AJ3 Q16AJ6 Q16AK0 Q16AL2 Q16AN8 Q16AP0
Q16AP1 Q16AP3 Q16AP4 Q16AR3 Q16AR8 Q16AV0 Q16AV3 Q16AX0 Q16AX5 Q16AZ3 Q16B02
Q16B16 Q16B38 Q16BB4 Q16BB7 Q16BD5 Q16BG6 Q16BG8 Q16BG9 Q16BK4 Q16BL5 Q16BM5
Q16BN1 Q16BP2 Q16C40 Q16C60 Q16CE0 Q16CE5 Q16CE6 Q16CE8 Q16CF2 Q16CF6 Q16CH2
Q16CH7 Q16CI8 Q16CK7 Q16CN2 Q16CP0 Q16CW3 Q16CY0 Q16CY2 Q16CY7 Q16CZ0 Q16CZ1
Q16CZ5 Q16CZ6 Q16CZ7 Q16D02 Q16D05 Q16D10 Q16D32 Q16D35 Q16D38 Q16D44 Q16D45
Q16D64 Q16D84 Q16D98 Q16DB9 Q16DI8 Q16DK6 Q16DK7 Q16DL0 Q16DL1 Q16DM2 Q16DM5
Q16DM9 Q16DR2 Q16DR9 Q16DS0 Q16DS2 Q16DV7 Q16DW5 Q16DW6 Q16E06 Q16E16 Q16EE5
Q16FL6 Q16G71 Q16GH0 Q16H50 Q16HH9 Q16HW7 Q16I17 Q16IB4 Q16IR1 Q16IW3 Q16JA2
Q16JL4 Q16JS8 Q16K15 Q16K67 Q16KI5 Q16KN5 Q16LW2 Q16MK2 Q16ML2 Q16MW6 Q16MY0
Q16NS8 Q16NX0 Q16NZ8 Q16P20 Q16P87 Q16P90 Q16Q94 Q16QI1 Q16RF5 Q16RI1 Q16RL8
Q16RR7 Q16RX1 Q16RY9 Q16S14 Q16S21 Q16T79 Q16TA2 Q16TJ7 Q16U25 Q16U49 Q16UF8
Q16UG2 Q16UN6 Q16VC0 Q16VD3 Q16VW9 Q16W06 Q16W22 Q16W24 Q16WA6 Q16X15 Q16X92
Q16XV7 Q16Y28 Q16Y34 Q16YA8 Q16YB3 Q16ZR8 Q17015 Q17017 Q17045 Q170C2 Q17103
Q17107 Q17133 Q17134 Q17152 Q17172 Q171H3 Q171Y8 Q17237 Q17241 Q17285 Q17295
Q17299 Q172G3 Q172Y1 Q17308 Q17334 Q17335 Q17336 Q17339 Q17345 Q17348 Q17370
Q17381 Q17390 Q17391 Q17392 Q173M7 Q17413 Q17423 Q17424 Q17426 Q17427 Q17430
Q17435 Q17436 Q17439 Q17440 Q174D3 Q174R2 Q17533 Q17569 Q17574 Q175F8 Q17603
Q17604 Q17606 Q17607 Q17632 Q17635 Q17684 Q17693 Q17695 Q176V0 Q17745 Q17761
Q17768 Q17771 Q17784 Q177A7 Q177G4 Q177H0 Q177J8 Q17819 Q17820 Q17827 Q17828
Q17848 Q17856 Q17868 Q17886 Q17895 Q178A5 Q178E6 Q178L7 Q178M1 Q17902 Q17905
Q17949 Q179T2 Q17A58 Q17AF4 Q17AI4 Q17AN2 Q17BA4 Q17BB0 Q17BU3 Q17CA7 Q17CH9
Q17CJ5 Q17CQ8 Q17CS8 Q17D30 Q17DI7 Q17DK2 Q17DT0 Q17EN4 Q17FF6 Q17FP1 Q17FR9
Q17G65 Q17G75 Q17GM7 Q17GR9 Q17GW6 Q17GZ9 Q17H91 Q17I16 Q17IE8 Q17IN5 Q17J49
Q17JB7 Q17JT4 Q17L24 Q17LR9 Q17LZ2 Q17M80 Q17MI0 Q17MZ8 Q17N69 Q17N71 Q17NH9
Q17NP0 Q17P71 Q17P98 Q17PI0 Q17PP1 Q17PR1 Q17Q06 Q17Q32 Q17Q39 Q17Q89 Q17Q91
Q17Q97 Q17QA0 Q17QD3 Q17QE2 Q17QF0 Q17QF2 Q17QF6 Q17QH7 Q17QH8 Q17QH9 Q17QI1
Q17QI2 Q17QJ0 Q17QJ3 Q17QJ7 Q17QK1 Q17QK8 Q17QL1 Q17QM8 Q17QN0 Q17QN3 Q17QP0
Q17QQ2 Q17QQ4 Q17QQ9 Q17QR4 Q17QR5 Q17QR7 Q17QS0 Q17QT4 Q17QU3 Q17QU5 Q17QU6
Q17QV9 Q17QW4 Q17QX9 Q17QZ4 Q17QZ7 Q17R00 Q17R01 Q17R09 Q17R10 Q17R31 Q17R98
Q17RA5 Q17RB0 Q17RB8 Q17RC7 Q17RG1 Q17RH7 Q17RN3 Q17RP2 Q17RS7 Q17V54 Q17V60
Q17V73 Q17V74 Q17V90 Q17V95 Q17V96 Q17VA0 Q17VC6 Q17VC7 Q17VE9 Q17VF8 Q17VH2
Q17VI1 Q17VM4 Q17VM8 Q17VM9 Q17VN2 Q17VN3 Q17VN5 Q17VN6 Q17VN7 Q17VN8 Q17VN9
Q17VP5 Q17VS1 Q17VS2 Q17VS4 Q17VS7 Q17VS9 Q17VT0 Q17VT1 Q17VT4 Q17VT5 Q17VT7
Q17VU7 Q17VU8 Q17VU9 Q17VY3 Q17VY4 Q17VZ5 Q17VZ7 Q17W06 Q17W08 Q17W13 Q17W17
Q17W45 Q17W61 Q17W62 Q17W63 Q17W85 Q17W89 Q17W92 Q17WB0 Q17WB6 Q17WB9 Q17WC1
Q17WC3 Q17WC4 Q17WE3 Q17WH3 Q17WH6 Q17WI2 Q17WJ3 Q17WJ4 Q17WK1 Q17WL7 Q17WL9

Q17WM2 Q17WN8 Q17WP3 Q17WP4 Q17WP7 Q17WP8 Q17WQ8 Q17WQ9 Q17WR0 Q17WR1 Q17WS1
Q17WS3 Q17WU1 Q17WU7 Q17WW9 Q17WX2 Q17WX3 Q17WX5 Q17WX7 Q17WY3 Q17WZ1 Q17WZ6
Q17X03 Q17X26 Q17X32 Q17X37 Q17X40 Q17X54 Q17X65 Q17X66 Q17X68 Q17XC0 Q17XC9
Q17XD2 Q17XD3 Q17XD4 Q17XF1 Q17XG0 Q17XG1 Q17XG2 Q17XG8 Q17XH0 Q17XJ4 Q17XJ5
Q17XJ6 Q17XJ7 Q17XK9 Q17XL1 Q17XN2 Q17XN9 Q17XQ4 Q17XR6 Q17XU2 Q17XU9 Q17XV1
Q17XY7 Q17XZ1 Q17Y01 Q17Y32 Q17Y33 Q17Y58 Q17Y87 Q17Y93 Q17Y99 Q17YA0 Q17YA2
Q17YB5 Q17YC7 Q17YF5 Q17YG0 Q17YH9 Q17YI4 Q17YI6 Q17YK6 Q17YL7 Q17YM3 Q17YM7
Q17YP0 Q17YP1 Q17YQ4 Q17YQ6 Q17YQ7 Q17YR0 Q17YS0 Q17YS1 Q17YT0 Q17YT5 Q17YU3
Q17YU7 Q17YV1 Q17YZ0 Q17YZ7 Q17YZ8 Q17Z01 Q17Z05 Q17Z13 Q17Z16 Q17Z21 Q17Z25
Q17Z26 Q17Z29 Q17Z31 Q17Z34 Q17Z48 Q17Z49 Q17Z99 Q17ZA0 Q17ZA3 Q17ZA7 Q17ZB3
Q17ZB4 Q17ZB5 Q17ZB6 Q17ZB7 Q17ZB8 Q17ZC2 Q17ZC3 Q17ZC4 Q17ZC5 Q17ZC6 Q17ZC7
Q17ZC8 Q17ZC9 Q17ZD1 Q17ZD2 Q17ZD3 Q17ZD4 Q17ZD5 Q17ZD6 Q17ZD7 Q17ZD9 Q17ZE2
Q17ZF3 Q17ZF6 Q17ZF9 Q17ZG3 Q17ZK3 Q17ZL5 Q17ZL6 Q17ZN0 Q17ZN1 Q17ZN4 Q17ZN6
Q17ZP2 Q17ZQ6 Q17ZW5 Q17ZX2 Q17ZX7 Q17ZY4 Q18006 Q18008 Q18026 Q18034 Q18040
Q18053 Q18054 Q18086 Q180C9 Q180E4 Q180E5 Q180E8 Q180E9 Q180F3 Q180P4 Q180P5
Q180P8 Q180P9 Q180Q8 Q180R9 Q180S8 Q180T4 Q180T6 Q180X8 Q180Y2 Q180Y7 Q18161
Q18164 Q18169 Q18171 Q18192 Q18195 Q181A2 Q181B5 Q181B7 Q181C1 Q181F4 Q181F7
Q181G3 Q181G8 Q181Q1 Q181Q7 Q181R0 Q181R4 Q181R5 Q181S7 Q181S8 Q181T2 Q181T7
Q181T8 Q181Y7 Q181Z6 Q18212 Q18225 Q18232 Q18235 Q18237 Q18240 Q18248 Q18250
Q18262 Q18268 Q18273 Q18277 Q18299 Q182B8 Q182B9 Q182C0 Q182C6 Q182D5 Q182E7
Q182E8 Q182F0 Q182F1 Q182G1 Q182G9 Q182I1 Q182I5 Q182K8 Q182L1 Q182M7 Q182Q0
Q182S0 Q182S2 Q182S6 Q182S8 Q182T5 Q182V6 Q182W3 Q182X3 Q182Y7 Q182Z2 Q182Z8
Q18317 Q18336 Q183G0 Q183H5 Q183H9 Q183I0 Q183N8 Q183N9 Q183P3 Q183T0 Q18409
Q18412 Q18446 Q18452 Q18461 Q18490 Q18493 Q184M8 Q184N3 Q184N9 Q184Z6 Q18511
Q18533 Q18579 Q18590 Q185B2 Q185C5 Q185K3 Q185K4 Q185P4 Q185R8 Q185S5 Q185S7
Q185S8 Q185S9 Q18612 Q18616 Q18674 Q18677 Q18678 Q18680 Q18691 Q18694 Q186A6
Q186F4 Q186F8 Q186L5 Q186N3 Q186R0 Q186R1 Q186R7 Q186Z6 Q18726 Q18785 Q18786
Q18787 Q187C4 Q187C6 Q187D2 Q187E3 Q187M0 Q187M6 Q187N0 Q187N2 Q187N7 Q187Q1
Q187T2 Q187T5 Q187T6 Q187U6 Q18801 Q18885 Q188A2 Q188C8 Q188R7 Q18905 Q18919
Q18938 Q18953 Q18955 Q18964 Q18967 Q18990 Q189A4 Q189A6 Q189B8 Q189N0 Q189N4
Q189Q3 Q189Q7 Q189R2 Q189R8 Q189S4 Q189T1 Q189U8 Q189Y9 Q18A75 Q18A79 Q18A85
Q18A91 Q18A92 Q18AJ1 Q18AJ2 Q18AL0 Q18AN9 Q18AP0 Q18AR0 Q18AS6 Q18B06 Q18B07
Q18B11 Q18B13 Q18B16 Q18B22 Q18B24 Q18B27 Q18B36 Q18B37 Q18B38 Q18B55 Q18B61
Q18B66 Q18B68 Q18B70 Q18B86 Q18BA9 Q18BB6 Q18BB8 Q18BC1 Q18BC2 Q18BC9 Q18BD4
Q18BE2 Q18BE6 Q18BE7 Q18BF0 Q18BF1 Q18BG3 Q18BH3 Q18BH4 Q18BH6 Q18BH7 Q18BI0
Q18BI4 Q18BJ2 Q18BJ4 Q18BQ3 Q18BS8 Q18BW7 Q18C13 Q18C22 Q18C32 Q18C33 Q18C47
Q18C67 Q18C69 Q18C71 Q18C74 Q18C83 Q18C86 Q18CA2 Q18CA3 Q18CD1 Q18CD2 Q18CD3
Q18CD5 Q18CD6 Q18CD7 Q18CE3 Q18CE4 Q18CE5 Q18CE7 Q18CE8 Q18CE9 Q18CF1 Q18CF2
Q18CF3 Q18CF4 Q18CF5 Q18CF6 Q18CF8 Q18CG0 Q18CG1 Q18CG2 Q18CG3 Q18CG4 Q18CG5
Q18CG6 Q18CG7 Q18CG8 Q18CG9 Q18CH0 Q18CH1 Q18CH3 Q18CH4 Q18CH5 Q18CH7 Q18CH8
Q18CH9 Q18CI0 Q18CI1 Q18CI2 Q18CI5 Q18CI6 Q18CI7 Q18CI8 Q18CJ1 Q18CJ5 Q18CL0
Q18CL7 Q18CP0 Q18CP3 Q18CP4 Q18CP5 Q18CR5 Q18CS5 Q18CS6 Q18CS8 Q18CT5 Q18CT6
Q18CT8 Q18CW0 Q18CX3 Q18CX4 Q18D35 Q18DH0 Q18DI6 Q18DK8 Q18DL1 Q18DL2 Q18DP5
Q18DQ2 Q18DQ4 Q18DR1 Q18DR5 Q18DY6 Q18DY7 Q18E28 Q18E32 Q18E36 Q18E49 Q18E52

Q18E60 Q18E61 Q18E67 Q18E71 Q18E75 Q18E76 Q18EA0 Q18EB6 Q18EB7 Q18EB8 Q18EB9
Q18EC0 Q18ER3 Q18ES5 Q18EU1 Q18EU6 Q18EU7 Q18EU8 Q18EV0 Q18EV6 Q18EV7 Q18EV8
Q18EX1 Q18EX9 Q18EY0 Q18EY5 Q18EY6 Q18EZ9 Q18F03 Q18F09 Q18F10 Q18F17 Q18F19
Q18F28 Q18F41 Q18F42 Q18F47 Q18F55 Q18FC0 Q18FC1 Q18FC4 Q18FD1 Q18FF3 Q18FG3
Q18FI8 Q18FJ9 Q18FK7 Q18FK8 Q18FL9 Q18FM4 Q18FQ4 Q18FT0 Q18FU6 Q18FU7 Q18FY6
Q18FZ8 Q18G49 Q18G53 Q18G54 Q18G55 Q18G58 Q18G60 Q18G63 Q18G74 Q18G90 Q18G95
Q18G97 Q18GA1 Q18GA5 Q18GA8 Q18GA9 Q18GB1 Q18GB5 Q18GE9 Q18GF0 Q18GF1 Q18GF2
Q18GF3 Q18GF4 Q18GF5 Q18GF8 Q18GF9 Q18GG0 Q18GG2 Q18GG3 Q18GG4 Q18GG5 Q18GG8
Q18GG9 Q18GH0 Q18GH1 Q18GK0 Q18GK1 Q18GK9 Q18GL3 Q18GM4 Q18GQ9 Q18GR9 Q18GT4
Q18GW3 Q18GX1 Q18GX3 Q18GX5 Q18GX8 Q18GX9 Q18GY3 Q18GZ3 Q18GZ4 Q18H49 Q18H54
Q18H64 Q18H65 Q18H83 Q18HD7 Q18HD8 Q18HK0 Q18HQ3 Q18HQ4 Q18HQ7 Q18HT7 Q18I31
Q18IJ5 Q18J03 Q18J05 Q18J06 Q18J11 Q18J39 Q18J40 Q18J81 Q18J88 Q18JB5 Q18JE8
Q18JF4 Q18JG3 Q18JI2 Q18JI3 Q18JI5 Q18JK6 Q18JL4 Q18JS3 Q18JV4 Q18K02 Q18K08
Q18K60 Q18K99 Q18KA6 Q18KC5 Q18KD2 Q18KE9 Q18KI0 Q18KI1 Q18KI2 Q18KI6 Q18KJ5
Q18KL6 Q18KL7 Q18KM8 Q18KN3 Q18KN8 Q18KN9 Q18KP1 Q18KP5 Q18KP9 Q18KQ9 Q18KS0
Q18KS1 Q18LC9 Q18LD0 Q18LD1 Q18LD2 Q18LD3 Q18LD4 Q18LD5 Q18LD6 Q18LD7 Q18LD8
Q18LD9 Q18LE0 Q18LE1 Q18LE2 Q18LE3 Q18LE4 Q18LE6 Q18LE7 Q18LE8 Q18LF0 Q18LF2
Q18LF3 Q18LF4 Q18LF5 Q18LF7 Q18LF8 Q18LF9 Q18T09 Q18TV3 Q19000 Q19005 Q19013
Q19020 Q19052 Q19157 Q19191 Q19196 Q19203 Q19213 Q19264 Q19272 Q19286 Q19291
Q19294 Q19297 Q19311 Q19325 Q19326 Q19329 Q19336 Q19338 Q19341 Q19345 Q19366
Q19375 Q19404 Q19418 Q19503 Q19537 Q19541 Q19546 Q19555 Q19564 Q195N6 Q19614
Q19642 Q19683 Q19691 Q196T5 Q196T7 Q196T9 Q196U0 Q196U2 Q196U3 Q196U4 Q196U5
Q196U7 Q196U9 Q196V0 Q196V1 Q196V2 Q196V4 Q196V5 Q196V6 Q196V7 Q196V8 Q196V9
Q196W0 Q196W1 Q196W2 Q196W3 Q196W6 Q196W7 Q196W8 Q196W9 Q196X0 Q196X1 Q196X2
Q196X3 Q196X4 Q196X7 Q196X8 Q196X9 Q196Y0 Q196Y1 Q196Y2 Q196Y3 Q196Y4 Q196Y5
Q196Y6 Q196Y8 Q196Y9 Q196Z0 Q196Z1 Q196Z2 Q196Z3 Q196Z5 Q196Z6 Q196Z7 Q196Z8
Q196Z9 Q19706 Q19713 Q19720 Q19722 Q19743 Q19749 Q19753 Q19775 Q19782 Q19787
Q197A0 Q197A1 Q197A2 Q197A3 Q197A4 Q197A5 Q197A6 Q197A7 Q197A8 Q197B0 Q197B1
Q197B2 Q197B4 Q197B5 Q197B6 Q197B8 Q197B9 Q197C0 Q197C1 Q197C2 Q197C4 Q197C5
Q197C6 Q197C8 Q197C9 Q197D0 Q197D1 Q197D2 Q197D3 Q197D4 Q197D7 Q197D9 Q197E0
Q197E1 Q197E2 Q197E3 Q197E4 Q197E5 Q197E6 Q197E8 Q197E9 Q197F0 Q197F1 Q197F2
Q197F3 Q197F7 Q197F8 Q197F9 Q197W8 Q19802 Q19825 Q19826 Q19829 Q19832 Q19849
Q19857 Q19863 Q19872 Q19895 Q19905 Q19906 Q19910 Q19952 Q19980 Q19A30 Q19A40
Q19A41 Q19A54 Q19AV6 Q19KS6 Q19LA2 Q19NJ4 Q19PY3 Q19Q52 Q19QT9 Q19QU3 Q19UN5
Q19V52 Q19V53 Q19V64 Q19V71 Q19V72 Q19V74 Q19V75 Q19V76 Q19V89 Q19V92 Q19V98
Q19V99 Q19VA0 Q19VA1 Q19VA6 Q19VA8 Q19VA9 Q19VB0 Q19VB1 Q19VB2 Q19VB3 Q19VB4
Q19VB5 Q19VB6 Q19VC3 Q1A1A1 Q1A1A2 Q1A1A3 Q1A1A4 Q1A1A5 Q1A1A6 Q1A5X6 Q1A705
Q1A7B1 Q1A7B2 Q1A7B3 Q1ACD5 Q1ACD6 Q1ACD7 Q1ACD8 Q1ACE0 Q1ACE1 Q1ACF1 Q1ACF2
Q1ACF5 Q1ACF6 Q1ACF7 Q1ACF8 Q1ACF9 Q1ACG0 Q1ACG1 Q1ACG2 Q1ACG3 Q1ACG4 Q1ACG5
Q1ACG6 Q1ACG7 Q1ACG9 Q1ACH6 Q1ACH8 Q1ACH9 Q1ACI0 Q1ACI3 Q1ACK0 Q1ACK8 Q1ACK9
Q1ACL0 Q1ACL1 Q1ACL2 Q1ACL5 Q1ACM7 Q1ACN3 Q1ACN4 Q1ACN5 Q1ACN6 Q1ACP2 Q1ACP3
Q1AGS3 Q1AGS4 Q1AGS5 Q1AGS6 Q1AGS7 Q1AGS8 Q1AGS9 Q1AHB2 Q1AR58 Q1AR63 Q1AR71
Q1AR89 Q1AR90 Q1AR91 Q1ARE4 Q1ARU5 Q1AS21 Q1AS30 Q1AS32 Q1AS35 Q1AS42 Q1AS60
Q1AS71 Q1ASA8 Q1ASC4 Q1ASN4 Q1AST1 Q1AT03 Q1AT13 Q1AT14 Q1AT39 Q1AT41 Q1AT82

Q1ATE1 Q1ATH8 Q1ATM0 Q1ATN8 Q1ATQ4 Q1ATQ8 Q1AU02 Q1AU08 Q1AU09 Q1AU10 Q1AU11
Q1AU14 Q1AU15 Q1AU18 Q1AU19 Q1AU20 Q1AU21 Q1AU22 Q1AU23 Q1AU24 Q1AU25 Q1AU26
Q1AU28 Q1AU29 Q1AU30 Q1AU31 Q1AU32 Q1AU33 Q1AU34 Q1AU35 Q1AU36 Q1AU37 Q1AU38
Q1AU39 Q1AU40 Q1AU41 Q1AU42 Q1AU43 Q1AU44 Q1AU45 Q1AU46 Q1AU47 Q1AU48 Q1AU52
Q1AU53 Q1AU54 Q1AU55 Q1AU56 Q1AU57 Q1AU63 Q1AU64 Q1AU65 Q1AU89 Q1AU93 Q1AUE0
Q1AUF5 Q1AUH1 Q1AUH2 Q1AUH7 Q1AUH8 Q1AUK2 Q1AUK3 Q1AUK6 Q1AUT6 Q1AV95 Q1AVC5
Q1AVF5 Q1AVG0 Q1AVG3 Q1AVG5 Q1AVL3 Q1AVP8 Q1AVQ0 Q1AVS8 Q1AVS9 Q1AVT4 Q1AVT5
Q1AVT6 Q1AVU1 Q1AVU2 Q1AVU3 Q1AVU4 Q1AVU9 Q1AVV7 Q1AVW2 Q1AVW6 Q1AVX1 Q1AVX4
Q1AVY0 Q1AVY5 Q1AVY6 Q1AVY7 Q1AVY9 Q1AVZ3 Q1AVZ4 Q1AVZ9 Q1AW05 Q1AW08 Q1AW09
Q1AW18 Q1AW34 Q1AW38 Q1AW43 Q1AW48 Q1AW49 Q1AW53 Q1AW55 Q1AW57 Q1AW65 Q1AW66
Q1AW67 Q1AW68 Q1AW73 Q1AW75 Q1AW77 Q1AW89 Q1AW92 Q1AW97 Q1AWA3 Q1AWA5 Q1AWA6
Q1AWB1 Q1AWB4 Q1AWB5 Q1AWC4 Q1AWD5 Q1AWE0 Q1AWE1 Q1AWE2 Q1AWE6 Q1AWE7 Q1AWF5
Q1AWG1 Q1AWG2 Q1AWG4 Q1AWH4 Q1AWI8 Q1AWK3 Q1AWY5 Q1AX07 Q1AX09 Q1AX11 Q1AX12
Q1AX25 Q1AX26 Q1AX36 Q1AXB7 Q1AXE5 Q1AXF2 Q1AXG1 Q1AXL4 Q1AXL8 Q1AXL9 Q1AXQ5
Q1AXR0 Q1AXR1 Q1AXT0 Q1AXT1 Q1AXT3 Q1AXU6 Q1AXW6 Q1AXX5 Q1AXX6 Q1AXZ4 Q1AY33
Q1AY78 Q1AYB7 Q1AYD7 Q1AYE5 Q1AYH1 Q1AYH3 Q1AYP4 Q1AYR4 Q1AYW0 Q1AZ25 Q1AZ75
Q1AZ86 Q1AZC3 Q1AZC4 Q1AZV2 Q1B017 Q1B0P6 Q1B0P7 Q1B0S1 Q1B0S2 Q1B0S6 Q1B0U5
Q1B0W6 Q1B0W7 Q1B1D0 Q1B1L0 Q1B1N8 Q1B1S4 Q1B1U6 Q1B1V1 Q1B1Z8 Q1B276 Q1B277
Q1B2L8 Q1B2L9 Q1B2M0 Q1B2P3 Q1B2P4 Q1B2Q0 Q1B2X0 Q1B2X1 Q1B321 Q1B370 Q1B397
Q1B3A6 Q1B3F3 Q1B3G6 Q1B3H2 Q1B3Q8 Q1B3V2 Q1B3W0 Q1B3X7 Q1B416 Q1B425 Q1B487
Q1B4E6 Q1B4H2 Q1B4H4 Q1B4N5 Q1B4P5 Q1B4S9 Q1B4T4 Q1B4V6 Q1B510 Q1B531 Q1B535
Q1B541 Q1B542 Q1B568 Q1B571 Q1B585 Q1B5E2 Q1B5E3 Q1B5T7 Q1B5Z8 Q1B601 Q1B625
Q1B627 Q1B628 Q1B629 Q1B639 Q1B647 Q1B667 Q1B671 Q1B6B7 Q1B6C2 Q1B6C7 Q1B6D4
Q1B6P5 Q1B6R1 Q1B6R2 Q1B6R7 Q1B6T7 Q1B6W2 Q1B6W3 Q1B6X2 Q1B6X7 Q1B774 Q1B781
Q1B782 Q1B795 Q1B797 Q1B798 Q1B7F0 Q1B7F1 Q1B7F4 Q1B7F9 Q1B7G5 Q1B7G6 Q1B7G8
Q1B7H0 Q1B7H1 Q1B7H6 Q1B7P7 Q1B7P8 Q1B7Q7 Q1B7R1 Q1B7R2 Q1B7S1 Q1B7S2 Q1B7T3
Q1B832 Q1B860 Q1B870 Q1B871 Q1B872 Q1B874 Q1B8H0 Q1B8J4 Q1B959 Q1B968 Q1B9C0
Q1B9C1 Q1B9C8 Q1B9D0 Q1B9D1 Q1B9D5 Q1B9E3 Q1B9F3 Q1B9F6 Q1B9F9 Q1B9G3 Q1B9G7
Q1B9G8 Q1B9H4 Q1B9H9 Q1B9I0 Q1B9P7 Q1B9Q9 Q1B9R0 Q1B9R1 Q1B9S5 Q1B9S6 Q1B9T4
Q1B9W8 Q1BA06 Q1BA14 Q1BA19 Q1BA28 Q1BA29 Q1BA61 Q1BA64 Q1BA69 Q1BA76 Q1BA77
Q1BA81 Q1BA93 Q1BA94 Q1BA97 Q1BAA0 Q1BAA7 Q1BAC5 Q1BAG9 Q1BAH1 Q1BAH2 Q1BAI1
Q1BAI2 Q1BAI5 Q1BAJ0 Q1BAK3 Q1BAK5 Q1BAK8 Q1BAM3 Q1BAM5 Q1BAN7 Q1BAP2 Q1BAP6
Q1BAP9 Q1BAQ3 Q1BAQ4 Q1BAQ6 Q1BAR4 Q1BAR7 Q1BAT1 Q1BAU4 Q1BAU9 Q1BB09 Q1BB26
Q1BBA1 Q1BBA3 Q1BBA4 Q1BBL0 Q1BBL5 Q1BCB0 Q1BCD6 Q1BCE8 Q1BCI0 Q1BCN9 Q1BCR5
Q1BCR8 Q1BCW7 Q1BCX0 Q1BCY7 Q1BCZ0 Q1BD07 Q1BD08 Q1BD09 Q1BD10 Q1BD11 Q1BD12
Q1BD13 Q1BD73 Q1BD75 Q1BD76 Q1BD77 Q1BD87 Q1BD90 Q1BD91 Q1BD92 Q1BD93 Q1BD94
Q1BD95 Q1BD99 Q1BDA0 Q1BDA1 Q1BDA2 Q1BDA3 Q1BDA4 Q1BDA5 Q1BDA6 Q1BDA7 Q1BDA8
Q1BDD3 Q1BDD4 Q1BDD5 Q1BDD6 Q1BDE9 Q1BDF0 Q1BDF3 Q1BDJ5 Q1BDJ6 Q1BDK2 Q1BDY7
Q1BE01 Q1BE13 Q1BE73 Q1BE87 Q1BEA6 Q1BEB0 Q1BEI3 Q1BEJ0 Q1BEJ4 Q1BEL9 Q1BEM5
Q1BEP8 Q1BEV1 Q1BEY3 Q1BEZ2 Q1BF27 Q1BF44 Q1BFJ5 Q1BFN7 Q1BFP0 Q1BFX3 Q1BFX4
Q1BG58 Q1BG61 Q1BG90 Q1BGA7 Q1BGJ0 Q1BGJ2 Q1BGU5 Q1BGW5 Q1BGW9 Q1BGX0 Q1BGX3
Q1BGX6 Q1BGX7 Q1BGX9 Q1BGY4 Q1BGY7 Q1BGY8 Q1BGZ5 Q1BH33 Q1BH34 Q1BH36 Q1BH55
Q1BH70 Q1BH71 Q1BH79 Q1BH84 Q1BH85 Q1BH86 Q1BHA4 Q1BHA5 Q1BHF2 Q1BHG2 Q1BHG6
Q1BHG8 Q1BHH6 Q1BHH9 Q1BHI0 Q1BHI1 Q1BHI2 Q1BHI4 Q1BHI7 Q1BHJ3 Q1BHJ5 Q1BHL9

Q1BHP1 Q1BHQ1 Q1BHS2 Q1BHT1 Q1BHT5 Q1BHV3 Q1BHX0 Q1BHY1 Q1BHY5 Q1BIC1 Q1BIU1
Q1BJJ1 Q1BJL9 Q1BJT2 Q1BK24 Q1BK31 Q1BK36 Q1BK87 Q1BKI8 Q1BL32 Q1BL48 Q1BLH4
Q1BLX6 Q1BLY5 Q1BLY7 Q1BLY8 Q1BLY9 Q1BLZ1 Q1BM38 Q1BM55 Q1BM62 Q1BM66 Q1BNL1
Q1BNU0 Q1BP24 Q1BPK4 Q1BQE0 Q1BQE1 Q1BQE2 Q1BQP0 Q1BQX0 Q1BQX8 Q1BQZ3 Q1BR24
Q1BR97 Q1BR98 Q1BRE8 Q1BRE9 Q1BRL1 Q1BRP1 Q1BRT3 Q1BRT6 Q1BRT7 Q1BRT8 Q1BRT9
Q1BRU0 Q1BRU1 Q1BRU3 Q1BRU4 Q1BRU5 Q1BRU7 Q1BRU8 Q1BRU9 Q1BRV0 Q1BRV1 Q1BRV2
Q1BRV3 Q1BRV4 Q1BRV5 Q1BRV6 Q1BRV7 Q1BRV8 Q1BRV9 Q1BRW0 Q1BRW1 Q1BRW2 Q1BRW3
Q1BRW4 Q1BRW5 Q1BRW6 Q1BRW7 Q1BRW9 Q1BRX0 Q1BRX1 Q1BRX2 Q1BRX3 Q1BRX4 Q1BRX5
Q1BRX9 Q1BRY6 Q1BRZ3 Q1BS24 Q1BS27 Q1BS30 Q1BS31 Q1BS32 Q1BS33 Q1BSB1 Q1BSB3
Q1BSD2 Q1BSD3 Q1BSF4 Q1BSF9 Q1BSJ4 Q1BSK1 Q1BSL1 Q1BSL2 Q1BSL3 Q1BSM1 Q1BSM4
Q1BSM8 Q1BSM9 Q1BSN9 Q1BST9 Q1BSY9 Q1BT34 Q1BT35 Q1BT36 Q1BT42 Q1BT49 Q1BT63
Q1BT81 Q1BTB4 Q1BTG1 Q1BTG4 Q1BTG5 Q1BTG7 Q1BTH0 Q1BTH3 Q1BTH4 Q1BTI4 Q1BTJ3
Q1BTM9 Q1BTN4 Q1BTP1 Q1BTQ8 Q1BTS1 Q1BTS3 Q1BTT7 Q1BTW6 Q1BTX7 Q1BU34 Q1BU36
Q1BU45 Q1BU58 Q1BU59 Q1BU60 Q1BU63 Q1BU66 Q1BU68 Q1BU86 Q1BU94 Q1BU98 Q1BUA1
Q1BUA7 Q1BUB0 Q1BUB1 Q1BUB2 Q1BUE7 Q1BUH0 Q1BUH1 Q1BUH6 Q1BUI0 Q1BUI4 Q1BUR7
Q1BUR8 Q1BUS8 Q1BUU4 Q1BUU8 Q1BUW3 Q1BUW4 Q1BUW9 Q1BUX2 Q1BUZ9 Q1BV03 Q1BV08
Q1BV09 Q1BV39 Q1BV59 Q1BV70 Q1BVA7 Q1BVW7 Q1BWA2 Q1BWB8 Q1BWM0 Q1BWM2 Q1BWM3
Q1BWM9 Q1BWS6 Q1BWS7 Q1BWS9 Q1BWV0 Q1BWV1 Q1BWW8 Q1BX00 Q1BX19 Q1BXC7 Q1BXD0
Q1BXK3 Q1BXN9 Q1BXP0 Q1BXP1 Q1BXQ7 Q1BXT2 Q1BXT4 Q1BXT9 Q1BXV6 Q1BXV8 Q1BXW6
Q1BXX4 Q1BXZ3 Q1BXZ5 Q1BY01 Q1BY02 Q1BY03 Q1BY04 Q1BY09 Q1BY20 Q1BY25 Q1BY27
Q1BY28 Q1BY31 Q1BY35 Q1BY59 Q1BY72 Q1BY73 Q1BY75 Q1BY87 Q1BY89 Q1BYB1 Q1BYB6
Q1BYB7 Q1BYF4 Q1BYG9 Q1BYH0 Q1BYH1 Q1BYH2 Q1BYH3 Q1BYH4 Q1BYH5 Q1BYK8 Q1BYL0
Q1BYL1 Q1BYL5 Q1BYQ6 Q1BYR9 Q1BYW1 Q1BYW4 Q1BYX0 Q1BYX2 Q1BYX3 Q1BYX5 Q1BYX9
Q1BYY0 Q1BYY1 Q1BYZ9 Q1BZ02 Q1BZ27 Q1BZ33 Q1BZ34 Q1BZ35 Q1BZ36 Q1BZ40 Q1BZ42
Q1BZ43 Q1BZ44 Q1BZ45 Q1BZ67 Q1BZ70 Q1BZ73 Q1BZC1 Q1BZD6 Q1BZD7 Q1BZD9 Q1BZE0
Q1BZE1 Q1BZE2 Q1BZE9 Q1BZG1 Q1BZG2 Q1BZG5 Q1BZH1 Q1BZH2 Q1BZN5 Q1BZN8 Q1BZY8
Q1C086 Q1C087 Q1C0B3 Q1C0B6 Q1C0B7 Q1C0B8 Q1C0D3 Q1C0F6 Q1C0H5 Q1C0H6 Q1C0H7
Q1C0I8 Q1C0J0 Q1C0J3 Q1C0J5 Q1C0J8 Q1C0J9 Q1C0K7 Q1C0K9 Q1C0L5 Q1C0L6 Q1C0N0
Q1C0T7 Q1C0V2 Q1C0V7 Q1C0V8 Q1C0V9 Q1C0W1 Q1C0W2 Q1C0X5 Q1C0X9 Q1C0Y0 Q1C0Y3
Q1C0Y9 Q1C0Z2 Q1C0Z8 Q1C0Z9 Q1C100 Q1C105 Q1C106 Q1C133 Q1C134 Q1C142 Q1C153
Q1C156 Q1C165 Q1C166 Q1C1A9 Q1C1D9 Q1C1E0 Q1C1E1 Q1C1E2 Q1C1F7 Q1C1F8 Q1C1G9
Q1C1H0 Q1C1H6 Q1C1J3 Q1C1J5 Q1C1J6 Q1C1K6 Q1C1M9 Q1C1N7 Q1C1P4 Q1C1S2 Q1C1T4
Q1C1T7 Q1C1T8 Q1C1T9 Q1C1U0 Q1C1U1 Q1C1U2 Q1C1U5 Q1C1U8 Q1C1V0 Q1C1V1 Q1C1V3
Q1C1V9 Q1C1Y2 Q1C1Y8 Q1C1Z3 Q1C1Z4 Q1C1Z6 Q1C205 Q1C206 Q1C212 Q1C215 Q1C216
Q1C231 Q1C237 Q1C238 Q1C247 Q1C257 Q1C258 Q1C259 Q1C262 Q1C263 Q1C264 Q1C267
Q1C268 Q1C269 Q1C270 Q1C271 Q1C276 Q1C278 Q1C282 Q1C285 Q1C297 Q1C2A4 Q1C2B1
Q1C2C5 Q1C2D3 Q1C2L3 Q1C2L4 Q1C2L8 Q1C2M2 Q1C2M9 Q1C2N0 Q1C2P2 Q1C2P3 Q1C2P9
Q1C2R7 Q1C2T2 Q1C2T5 Q1C2T6 Q1C2T7 Q1C2T8 Q1C2T9 Q1C2U0 Q1C2U1 Q1C2U6 Q1C2U7
Q1C2U8 Q1C2U9 Q1C2V0 Q1C2V1 Q1C2V2 Q1C2V3 Q1C2V4 Q1C2V5 Q1C2V6 Q1C2V7 Q1C2V8
Q1C2V9 Q1C2W0 Q1C2W1 Q1C2W2 Q1C2W3 Q1C2W4 Q1C2W5 Q1C2W6 Q1C2W8 Q1C2W9 Q1C2X0
Q1C2X1 Q1C2X2 Q1C2X7 Q1C2X8 Q1C2X9 Q1C2Y1 Q1C2Y3 Q1C2Y4 Q1C300 Q1C310 Q1C365
Q1C366 Q1C370 Q1C374 Q1C377 Q1C380 Q1C3H9 Q1C3I0 Q1C3I5 Q1C3J5 Q1C3J8 Q1C3J9
Q1C3K9 Q1C3L0 Q1C3L8 Q1C3N3 Q1C3P1 Q1C3R8 Q1C3S7 Q1C3S8 Q1C3T0 Q1C3U4 Q1C3U6
Q1C3U7 Q1C3V6 Q1C3V7 Q1C3V8 Q1C3W4 Q1C3X1 Q1C3X3 Q1C3X7 Q1C3X8 Q1C3Y5 Q1C3Z0

Q1C3Z1 Q1C408 Q1C409 Q1C410 Q1C411 Q1C420 Q1C421 Q1C422 Q1C471 Q1C474 Q1C475
Q1C476 Q1C477 Q1C483 Q1C4C6 Q1C4D9 Q1C4E4 Q1C4E5 Q1C4E6 Q1C4E7 Q1C4F3 Q1C4F4
Q1C4F5 Q1C4H4 Q1C4I2 Q1C4I4 Q1C4I5 Q1C4I9 Q1C4J3 Q1C4J6 Q1C4K7 Q1C4K8 Q1C4K9
Q1C4L5 Q1C4L7 Q1C4N0 Q1C4N1 Q1C4P3 Q1C4P5 Q1C4P6 Q1C4P8 Q1C4P9 Q1C4Q0 Q1C4U1
Q1C4U4 Q1C507 Q1C508 Q1C509 Q1C516 Q1C518 Q1C521 Q1C529 Q1C531 Q1C537 Q1C540
Q1C553 Q1C555 Q1C564 Q1C569 Q1C570 Q1C571 Q1C575 Q1C596 Q1C5A9 Q1C5B1 Q1C5B2
Q1C5B3 Q1C5B5 Q1C5E3 Q1C5E7 Q1C5G0 Q1C5G8 Q1C5G9 Q1C5H2 Q1C5H3 Q1C5H7 Q1C5I4
Q1C5I5 Q1C5I8 Q1C5I9 Q1C5J2 Q1C5J4 Q1C5J6 Q1C5P2 Q1C5P3 Q1C5P5 Q1C5Q7 Q1C5R3
Q1C5T6 Q1C5T7 Q1C5T8 Q1C5U6 Q1C5U9 Q1C5X9 Q1C5Y4 Q1C5Z1 Q1C664 Q1C669 Q1C676
Q1C6A1 Q1C6A3 Q1C6A6 Q1C6D0 Q1C6D7 Q1C6D8 Q1C6E0 Q1C6E5 Q1C6F4 Q1C6F5 Q1C6L1
Q1C6L6 Q1C6M4 Q1C6M6 Q1C6N6 Q1C6Q0 Q1C6R0 Q1C6R3 Q1C6S0 Q1C6T8 Q1C6U2 Q1C6Y9
Q1C713 Q1C728 Q1C730 Q1C731 Q1C733 Q1C735 Q1C748 Q1C752 Q1C761 Q1C762 Q1C774
Q1C786 Q1C788 Q1C790 Q1C791 Q1C792 Q1C7C2 Q1C7D4 Q1C7J2 Q1C7J3 Q1C7L1 Q1C7L9
Q1C7P0 Q1C7P2 Q1C7P3 Q1C7S6 Q1C7U0 Q1C7U5 Q1C7V0 Q1C7Z0 Q1C7Z3 Q1C7Z4 Q1C7Z6
Q1C814 Q1C817 Q1C818 Q1C820 Q1C823 Q1C824 Q1C825 Q1C827 Q1C836 Q1C854 Q1C855
Q1C857 Q1C861 Q1C8A1 Q1C8A7 Q1C8A8 Q1C8A9 Q1C8B0 Q1C8B1 Q1C8L1 Q1C8L9 Q1C8V5
Q1C8V8 Q1C8V9 Q1C8W1 Q1C8Z9 Q1C930 Q1C931 Q1C932 Q1C938 Q1C942 Q1C946 Q1C947
Q1C949 Q1C960 Q1C964 Q1C982 Q1C9A5 Q1C9A6 Q1C9C3 Q1C9C5 Q1C9C6 Q1C9H5 Q1C9P9
Q1C9Q9 Q1C9R1 Q1C9R5 Q1C9R6 Q1C9T5 Q1C9T6 Q1C9T8 Q1C9U7 Q1C9U8 Q1CA17 Q1CA18
Q1CA21 Q1CA26 Q1CA41 Q1CA47 Q1CA55 Q1CA61 Q1CA70 Q1CA72 Q1CA73 Q1CA74 Q1CA85
Q1CA93 Q1CA94 Q1CA97 Q1CAA5 Q1CAC8 Q1CAE0 Q1CAE9 Q1CAG1 Q1CAG9 Q1CAH3 Q1CAH5
Q1CAJ5 Q1CAJ7 Q1CAK9 Q1CAL3 Q1CAM0 Q1CAN0 Q1CAN1 Q1CAN2 Q1CAN4 Q1CAN6 Q1CAN8
Q1CAN9 Q1CAP0 Q1CAP6 Q1CAP8 Q1CAQ2 Q1CAR0 Q1CAR9 Q1CAS2 Q1CAS3 Q1CB23 Q1CB34
Q1CB42 Q1CB43 Q1CB44 Q1CB48 Q1CB53 Q1CB61 Q1CB62 Q1CB69 Q1CB75 Q1CB76 Q1CB83
Q1CB91 Q1CB93 Q1CBE1 Q1CBE3 Q1CBE7 Q1CBE8 Q1CBF0 Q1CBG0 Q1CBG7 Q1CBM6 Q1CBM8
Q1CBN1 Q1CBQ4 Q1CBS1 Q1CBS9 Q1CBT5 Q1CBU2 Q1CBU5 Q1CBV4 Q1CBV5 Q1CBV8 Q1CBW4
Q1CBW5 Q1CBW6 Q1CBW7 Q1CBX2 Q1CBX8 Q1CBY3 Q1CBY6 Q1CBY7 Q1CBZ1 Q1CBZ2 Q1CBZ4
Q1CC00 Q1CC03 Q1CC05 Q1CC07 Q1CC08 Q1CC09 Q1CC10 Q1CC27 Q1CC34 Q1CC38 Q1CC43
Q1CC48 Q1CC49 Q1CC50 Q1CC63 Q1CCG6 Q1CCG7 Q1CCJ3 Q1CCJ6 Q1CCJ7 Q1CCJ8 Q1CCK7
Q1CCL5 Q1CCL9 Q1CCM6 Q1CCM7 Q1CCN9 Q1CCP0 Q1CCP6 Q1CCR5 Q1CCT0 Q1CCT3 Q1CCT4
Q1CCT5 Q1CCT6 Q1CCT7 Q1CCT8 Q1CCT9 Q1CCU3 Q1CCU4 Q1CCU5 Q1CCU6 Q1CCU7 Q1CCU8
Q1CCU9 Q1CCV0 Q1CCV1 Q1CCV2 Q1CCV3 Q1CCV4 Q1CCV5 Q1CCV6 Q1CCV7 Q1CCV8 Q1CCV9
Q1CCW0 Q1CCW1 Q1CCW2 Q1CCW3 Q1CCW5 Q1CCW6 Q1CCW7 Q1CCW8 Q1CCW9 Q1CCX4 Q1CCX5
Q1CCX6 Q1CCX8 Q1CCY0 Q1CCY1 Q1CCZ2 Q1CCZ3 Q1CCZ4 Q1CCZ7 Q1CCZ8 Q1CCZ9 Q1CD02
Q1CD03 Q1CD04 Q1CD05 Q1CD06 Q1CD11 Q1CD13 Q1CD17 Q1CD20 Q1CD33 Q1CD40 Q1CD47
Q1CD53 Q1CD55 Q1CD56 Q1CD60 Q1CD62 Q1CD74 Q1CD75 Q1CD85 Q1CD86 Q1CD89 Q1CD96
Q1CDA1 Q1CDA2 Q1CDB8 Q1CDD8 Q1CDI1 Q1CDL3 Q1CDL4 Q1CDL5 Q1CDL6 Q1CDQ8 Q1CDT5
Q1CDU2 Q1CDV0 Q1CDX2 Q1CDY2 Q1CDY3 Q1CDY5 Q1CE02 Q1CE08 Q1CE09 Q1CE20 Q1CE21
Q1CE37 Q1CE94 Q1CEA9 Q1CEB1 Q1CEB2 Q1CEB3 Q1CEB5 Q1CEB6 Q1CEC9 Q1CED3 Q1CED4
Q1CED7 Q1CEE4 Q1CEE7 Q1CEF3 Q1CEF4 Q1CEF5 Q1CEG0 Q1CEG1 Q1CEH0 Q1CEH1 Q1CEH2
Q1CEH3 Q1CEH8 Q1CEI4 Q1CEI9 Q1CEJ2 Q1CEJ3 Q1CEJ7 Q1CEJ8 Q1CEK0 Q1CEK6 Q1CEK9
Q1CEL1 Q1CEL3 Q1CEL4 Q1CEL5 Q1CEL6 Q1CEV2 Q1CEV4 Q1CEV6 Q1CEW2 Q1CEW9 Q1CEX0
Q1CEX6 Q1CEY2 Q1CEY3 Q1CEY8 Q1CEZ3 Q1CEZ7 Q1CEZ8 Q1CEZ9 Q1CF05 Q1CF16 Q1CFB1
Q1CFB2 Q1CFB5 Q1CFC4 Q1CFD2 Q1CFD3 Q1CFD5 Q1CFE1 Q1CFE3 Q1CFE4 Q1CFE6 Q1CFE7

Q1CFE9 Q1CFF0 Q1CFF1 Q1CFG1 Q1CFG8 Q1CFH2 Q1CFI4 Q1CFI6 Q1CFK6 Q1CFK8 Q1CFL2
Q1CFM0 Q1CFN2 Q1CFN6 Q1CFP0 Q1CFQ1 Q1CFQ3 Q1CFQ4 Q1CFQ8 Q1CFR2 Q1CFR7 Q1CFR8
Q1CFR9 Q1CFZ1 Q1CG37 Q1CG38 Q1CG40 Q1CG57 Q1CG58 Q1CG79 Q1CG96 Q1CGA8 Q1CGD1
Q1CGD9 Q1CGE2 Q1CGE3 Q1CGF1 Q1CGG5 Q1CGG6 Q1CGG8 Q1CGH4 Q1CGI0 Q1CGI7 Q1CGJ3
Q1CGK9 Q1CGL4 Q1CGL7 Q1CGL8 Q1CGL9 Q1CGT4 Q1CGT5 Q1CGU4 Q1CGU6 Q1CGU7 Q1CGW5
Q1CGW6 Q1CGX0 Q1CGX2 Q1CGY2 Q1CGZ2 Q1CH12 Q1CH51 Q1CH53 Q1CH54 Q1CH57 Q1CHC4
Q1CHD6 Q1CHE4 Q1CHK6 Q1CHL1 Q1CHL8 Q1CHP3 Q1CHP5 Q1CHP8 Q1CHS2 Q1CHS9 Q1CHT0
Q1CHT2 Q1CHT7 Q1CHU6 Q1CHU7 Q1CI03 Q1CI06 Q1CI09 Q1CI15 Q1CI17 Q1CI27 Q1CI38
Q1CI48 Q1CI51 Q1CI58 Q1CI76 Q1CI78 Q1CIC5 Q1CIE6 Q1CIG3 Q1CIG5 Q1CIG6 Q1CIG8
Q1CIH0 Q1CII3 Q1CII7 Q1CIJ6 Q1CIJ7 Q1CIL0 Q1CIM0 Q1CIM2 Q1CIM4 Q1CIM5 Q1CIM6
Q1CIQ7 Q1CIR9 Q1CIX8 Q1CIX9 Q1CIZ7 Q1CJ05 Q1CJ26 Q1CJ28 Q1CJ29 Q1CJ62 Q1CJ76
Q1CJ81 Q1CJ86 Q1CJE1 Q1CJE4 Q1CJE5 Q1CJE7 Q1CJG5 Q1CJG7 Q1CJG8 Q1CJH0 Q1CJH3
Q1CJH4 Q1CJH5 Q1CJH7 Q1CJI6 Q1CJK1 Q1CJK2 Q1CJK4 Q1CJK8 Q1CJP8 Q1CJU5 Q1CJV2
Q1CJV7 Q1CJY6 Q1CJY8 Q1CJZ6 Q1CJZ7 Q1CJZ8 Q1CK00 Q1CK18 Q1CK24 Q1CK37 Q1CK39
Q1CK40 Q1CK82 Q1CK84 Q1CK87 Q1CK90 Q1CK91 Q1CK94 Q1CK95 Q1CKA2 Q1CKA6 Q1CKA7
Q1CKB0 Q1CKB1 Q1CKB8 Q1CKD2 Q1CKD6 Q1CKE2 Q1CKE4 Q1CKF3 Q1CKF7 Q1CKF8 Q1CKF9
Q1CKG2 Q1CKI2 Q1CKJ5 Q1CKJ7 Q1CKJ8 Q1CKJ9 Q1CKK1 Q1CKN4 Q1CKN7 Q1CKP3 Q1CKP5
Q1CKQ4 Q1CKQ7 Q1CKQ9 Q1CKR4 Q1CKR5 Q1CKR6 Q1CKX8 Q1CKY1 Q1CL26 Q1CL27 Q1CL28
Q1CL30 Q1CL31 Q1CL33 Q1CL42 Q1CL43 Q1CL56 Q1CL58 Q1CL64 Q1CL65 Q1CL66 Q1CL83
Q1CL87 Q1CL91 Q1CL92 Q1CL94 Q1CLA2 Q1CLB8 Q1CLB9 Q1CLC0 Q1CLC6 Q1CLC7 Q1CLC8
Q1CLC9 Q1CLD4 Q1CLE7 Q1CLJ2 Q1CLJ3 Q1CLJ4 Q1CLJ5 Q1CLK8 Q1CLK9 Q1CLL0 Q1CLL1
Q1CLQ9 Q1CLR2 Q1CLR3 Q1CLR4 Q1CLR5 Q1CLR6 Q1CLS1 Q1CLS2 Q1CLS7 Q1CLS8 Q1CLT3
Q1CLU0 Q1CLU1 Q1CLU5 Q1CLU7 Q1CLV4 Q1CLW0 Q1CLW1 Q1CLW2 Q1CLX1 Q1CLX2 Q1CLX4
Q1CLY8 Q1CLZ0 Q1CLZ1 Q1CM01 Q1CM06 Q1CM28 Q1CM36 Q1CM51 Q1CMC8 Q1CMD1 Q1CMD4
Q1CMD8 Q1CME2 Q1CME3 Q1CMJ1 Q1CMJ9 Q1CMK0 Q1CMM5 Q1CMM6 Q1CMM9 Q1CMN5 Q1CMN6
Q1CMP5 Q1CMP7 Q1CMP8 Q1CMQ0 Q1CMQ6 Q1CMT2 Q1CMT3 Q1CMT4 Q1CMU5 Q1CMU8 Q1CMV1
Q1CMV3 Q1CMV6 Q1CMV7 Q1CMW4 Q1CMW6 Q1CMX2 Q1CMX3 Q1CMY7 Q1CMY8 Q1CMZ7 Q1CN00
Q1CN11 Q1CN19 Q1CN20 Q1CN42 Q1CN50 Q1CN51 Q1CN57 Q1CN60 Q1CN66 Q1CN68 Q1CN69
Q1CN71 Q1CN74 Q1CN77 Q1CN78 Q1CN79 Q1CN80 Q1CN81 Q1CN82 Q1CN86 Q1CN90 Q1CN98
Q1CNB4 Q1CNC1 Q1CNH8 Q1CNI0 Q1CNI3 Q1CNK3 Q1CNM0 Q1CNM8 Q1CNN4 Q1CNP1 Q1CNP4
Q1CNQ3 Q1CNQ4 Q1CNQ7 Q1CNS4 Q1CNT1 Q1CNT5 Q1CNU0 Q1CNU2 Q1CNU3 Q1CNU4 Q1CR24
Q1CR25 Q1CR39 Q1CR49 Q1CR51 Q1CR52 Q1CR59 Q1CR72 Q1CR79 Q1CRB6 Q1CRB7 Q1CRC2
Q1CRC4 Q1CRD7 Q1CRD8 Q1CRD9 Q1CRG8 Q1CRH7 Q1CRI1 Q1CRI2 Q1CRI5 Q1CRI6 Q1CRJ9
Q1CRK2 Q1CRK7 Q1CRL4 Q1CRL6 Q1CRM5 Q1CRN2 Q1CRQ2 Q1CRR1 Q1CRS0 Q1CRS1 Q1CRS3
Q1CRT7 Q1CRU0 Q1CRU2 Q1CRU3 Q1CRU4 Q1CRU5 Q1CRU6 Q1CRU7 Q1CRU8 Q1CRU9 Q1CRV0
Q1CRV1 Q1CRV2 Q1CRV3 Q1CRV4 Q1CRV5 Q1CRV6 Q1CRV7 Q1CRV8 Q1CRV9 Q1CRW2 Q1CRW3
Q1CRW4 Q1CRW5 Q1CRW6 Q1CRW7 Q1CRW8 Q1CRX0 Q1CRX3 Q1CS04 Q1CS05 Q1CS13 Q1CS16
Q1CS18 Q1CS22 Q1CS23 Q1CS25 Q1CS35 Q1CS51 Q1CS61 Q1CS64 Q1CS65 Q1CS66 Q1CS67
Q1CS68 Q1CS69 Q1CS70 Q1CS71 Q1CS74 Q1CS87 Q1CS88 Q1CSA0 Q1CSA6 Q1CSB2 Q1CSB5
Q1CSB7 Q1CSB9 Q1CSC6 Q1CSF2 Q1CSG4 Q1CSH3 Q1CSI9 Q1CSK1 Q1CSL3 Q1CSL4 Q1CSL6
Q1CSM2 Q1CSN0 Q1CSP9 Q1CSQ0 Q1CSQ2 Q1CSR1 Q1CSR7 Q1CSS2 Q1CST1 Q1CSU4 Q1CSU6
Q1CSU8 Q1CSU9 Q1CSY9 Q1CSZ2 Q1CSZ5 Q1CT06 Q1CT07 Q1CT10 Q1CT14 Q1CT15 Q1CT17
Q1CT25 Q1CT27 Q1CT35 Q1CT37 Q1CT39 Q1CT46 Q1CT48 Q1CT68 Q1CT72 Q1CT75 Q1CT76
Q1CT77 Q1CT93 Q1CT94 Q1CTA1 Q1CTA2 Q1CTC3 Q1CTD3 Q1CTD7 Q1CTG3 Q1CTK6 Q1CTK7

Q1CTK9 Q1CTL2 Q1CTM2 Q1CTM7 Q1CTN3 Q1CTP7 Q1CTP9 Q1CTQ6 Q1CTQ7 Q1CTQ9 Q1CTS0
Q1CTU5 Q1CTU9 Q1CTV4 Q1CTV6 Q1CTW0 Q1CTW4 Q1CTX6 Q1CTX8 Q1CU15 Q1CU16 Q1CU17
Q1CU21 Q1CU35 Q1CU38 Q1CU52 Q1CU72 Q1CU78 Q1CU96 Q1CU98 Q1CUA3 Q1CUA4 Q1CUA5
Q1CUA6 Q1CUB6 Q1CUB7 Q1CUC0 Q1CUC1 Q1CUC6 Q1CUE2 Q1CUF6 Q1CUG1 Q1CUG7 Q1CUG9
Q1CUH1 Q1CUH2 Q1CUI1 Q1CUI3 Q1CUJ7 Q1CUK0 Q1CUK6 Q1CUK7 Q1CUK9 Q1CUM0 Q1CUM2
Q1CUN4 Q1CUP4 Q1CUP8 Q1CUR3 Q1CUR4 Q1CUR5 Q1CUT8 Q1CUU1 Q1CUU2 Q1CUU4 Q1CUW1
Q1CUW3 Q1CUW4 Q1CUW7 Q1CUX5 Q1CUX6 Q1CUY2 Q1CUY8 Q1CUZ6 Q1CV00 Q1CV04 Q1CV30
Q1CV31 Q1CV33 Q1CV45 Q1CV46 Q1CV68 Q1CV70 Q1CV71 Q1CV77 Q1CV78 Q1CV79 Q1CV81
Q1CV82 Q1CV84 Q1CV86 Q1CV87 Q1CVC2 Q1CVC3 Q1CVE4 Q1CVE5 Q1CVE9 Q1CVF3 Q1CVF4
Q1CVF8 Q1CVG4 Q1CVH6 Q1CVH7 Q1CVR6 Q1CW44 Q1CW45 Q1CW46 Q1CW50 Q1CW51 Q1CWP8
Q1CX51 Q1CXC6 Q1CXF1 Q1CXI5 Q1CXV1 Q1CY37 Q1CY73 Q1CY77 Q1CY83 Q1CY84 Q1CYD8
Q1CYE1 Q1CYI2 Q1CZ49 Q1CZH4 Q1CZH6 Q1CZI7 Q1CZL3 Q1CZQ8 Q1CZV9 Q1CZW4 Q1D063
Q1D084 Q1D096 Q1D0P6 Q1D0P7 Q1D0S0 Q1D0S2 Q1D0S8 Q1D0T1 Q1D0T2 Q1D104 Q1D1F0
Q1D1H1 Q1D1H9 Q1D1I0 Q1D1I1 Q1D1I2 Q1D1W9 Q1D230 Q1D264 Q1D282 Q1D289 Q1D290
Q1D292 Q1D295 Q1D2J3 Q1D2J4 Q1D2J5 Q1D2J8 Q1D2L9 Q1D2Q7 Q1D2Q9 Q1D2S1 Q1D2S2
Q1D2S6 Q1D338 Q1D345 Q1D346 Q1D349 Q1D350 Q1D369 Q1D375 Q1D3A6 Q1D3G4 Q1D3Y5
Q1D3Y8 Q1D402 Q1D498 Q1D4A5 Q1D4L3 Q1D4M2 Q1D4M4 Q1D4N0 Q1D4R3 Q1D4U7 Q1D513
Q1D568 Q1D5J6 Q1D5V7 Q1D5X5 Q1D5X9 Q1D607 Q1D651 Q1D654 Q1D6D8 Q1D6E0 Q1D6E1
Q1D6E2 Q1D6E3 Q1D6H3 Q1D6H4 Q1D6H9 Q1D6I1 Q1D6I2 Q1D6I3 Q1D6I5 Q1D6I6 Q1D6I7
Q1D6I9 Q1D6K1 Q1D6L9 Q1D6N0 Q1D6P1 Q1D6R1 Q1D6W9 Q1D6Y8 Q1D706 Q1D739 Q1D747
Q1D748 Q1D749 Q1D750 Q1D751 Q1D752 Q1D753 Q1D756 Q1D757 Q1D758 Q1D759 Q1D760
Q1D761 Q1D762 Q1D764 Q1D765 Q1D766 Q1D767 Q1D768 Q1D769 Q1D770 Q1D771 Q1D772
Q1D773 Q1D774 Q1D775 Q1D776 Q1D777 Q1D778 Q1D779 Q1D7A6 Q1D7R9 Q1D7T4 Q1D7U2
Q1D7U3 Q1D7U4 Q1D7U5 Q1D7U6 Q1D7U7 Q1D7V0 Q1D7V1 Q1D7X3 Q1D7Y2 Q1D7Z0 Q1D801
Q1D802 Q1D804 Q1D841 Q1D898 Q1D8I9 Q1D8K2 Q1D8L9 Q1D8V5 Q1D8Y1 Q1D8Z8 Q1D8Z9
Q1D912 Q1D921 Q1D936 Q1D980 Q1D9P3 Q1D9P5 Q1D9V4 Q1D9W6 Q1D9W9 Q1D9X8 Q1D9Z7
Q1DA04 Q1DA76 Q1DAM1 Q1DAM2 Q1DAM4 Q1DAM6 Q1DAM9 Q1DAN7 Q1DAN8 Q1DAN9 Q1DAP2
Q1DAS2 Q1DAT0 Q1DAT1 Q1DAT4 Q1DAT6 Q1DAV2 Q1DAZ4 Q1DAZ8 Q1DAZ9 Q1DB04 Q1DBJ3
Q1DC47 Q1DC68 Q1DC88 Q1DC90 Q1DC95 Q1DC96 Q1DC97 Q1DCA2 Q1DCA3 Q1DCA4 Q1DCC8
Q1DCG7 Q1DCH2 Q1DCK9 Q1DCQ5 Q1DCS0 Q1DCS1 Q1DCU1 Q1DCV8 Q1DCV9 Q1DCX5 Q1DCX8
Q1DD74 Q1DDB7 Q1DDE5 Q1DDF6 Q1DDU5 Q1DDU6 Q1DDZ9 Q1DEJ2 Q1DEM0 Q1DES4 Q1DFA7
Q1DFP6 Q1DFW5 Q1DG16 Q1DGL5 Q1DGL6 Q1DHB2 Q1DHB6 Q1DHE1 Q1DHH9 Q1DI07 Q1DI97
Q1DJF0 Q1DJF7 Q1DJH5 Q1DJR2 Q1DK31 Q1DK38 Q1DKD8 Q1DKE7 Q1DKI1 Q1DKL9 Q1DLD2
Q1DLG1 Q1DLJ4 Q1DLM0 Q1DLP2 Q1DM35 Q1DME8 Q1DMX8 Q1DNB0 Q1DNB9 Q1DNE1 Q1DNW5
Q1DNY1 Q1DNY5 Q1DP56 Q1DP69 Q1DP77 Q1DPD9 Q1DPU4 Q1DQ20 Q1DQ73 Q1DQB2 Q1DQE9
Q1DQS5 Q1DQS9 Q1DQZ5 Q1DR06 Q1DR42 Q1DR50 Q1DR81 Q1DRB6 Q1DRC2 Q1DRC9 Q1DRN0
Q1DS65 Q1DSY6 Q1DTB3 Q1DTF9 Q1DTI6 Q1DTL7 Q1DTR6 Q1DU03 Q1DU75 Q1DUF5 Q1DUF9
Q1DVD1 Q1DW80 Q1DWF4 Q1DXC2 Q1DXH0 Q1DXS8 Q1DXU0 Q1DY43 Q1DY99 Q1DYR6 Q1DYR7
Q1DZ34 Q1DZA6 Q1DZB0 Q1DZK8 Q1DZL3 Q1E024 Q1E036 Q1E065 Q1E076 Q1E0A3 Q1E0G9
Q1E0Q7 Q1E0W9 Q1E0Z3 Q1E101 Q1E146 Q1E170 Q1E180 Q1E1H4 Q1E1N5 Q1E1Q5 Q1E1R2
Q1E1R7 Q1E225 Q1E2B2 Q1E2F4 Q1E2I3 Q1E2S2 Q1E306 Q1E3N5 Q1E3S4 Q1E3Z7 Q1E403
Q1E406 Q1E493 Q1E4I8 Q1E4L8 Q1E4Z3 Q1E554 Q1E556 Q1E5B0 Q1E5I0 Q1E5N0 Q1E5N1
Q1E5R1 Q1E5T3 Q1E6D3 Q1E6K9 Q1E6M1 Q1E6Q0 Q1E736 Q1E7S5 Q1E7Y4 Q1E803 Q1E873
Q1E891 Q1E9A1 Q1E9B7 Q1E9C0 Q1E9L2 Q1E9T9 Q1EA11 Q1EA54 Q1EAF9 Q1EAK2 Q1EAU8

Q1EAW8 Q1EB31 Q1EB38 Q1EBC4 Q1EBD3 Q1EBK0 Q1EBV4 Q1EBV6 Q1ECJ7 Q1ECS0 Q1ECV4
Q1ECV8 Q1ECW2 Q1ECW6 Q1ECY2 Q1ECZ2 Q1ECZ4 Q1ECZ9 Q1ED21 Q1EG27 Q1EG72 Q1EG91
Q1EHT7 Q1EHW4 Q1EIR0 Q1EMV2 Q1ENB6 Q1EPR4 Q1EPR5 Q1EQW7 Q1ERH9 Q1ERI0 Q1ERI1
Q1ERI2 Q1G391 Q1G3B8 Q1G3I2 Q1G3I7 Q1G3M2 Q1G3M3 Q1G3N6 Q1G3Q4 Q1G3U5 Q1G3U6
Q1G3U8 Q1G3Y4 Q1G3Z6 Q1G7G9 Q1G7Q5 Q1G7S0 Q1G7S5 Q1G7W2 Q1G7Z1 Q1G7Z2 Q1G7Z4
Q1G7Z5 Q1G817 Q1G825 Q1G864 Q1G866 Q1G867 Q1G889 Q1G8A5 Q1G8B5 Q1G8L9 Q1G8N2
Q1G8P4 Q1G8Y5 Q1G8Z0 Q1G8Z5 Q1G8Z8 Q1G8Z9 Q1G904 Q1G905 Q1G920 Q1G921 Q1G923
Q1G933 Q1G936 Q1G937 Q1G938 Q1G939 Q1G940 Q1G941 Q1G947 Q1G948 Q1G949 Q1G950
Q1G967 Q1G971 Q1G986 Q1G991 Q1G997 Q1G9A0 Q1G9A3 Q1G9A5 Q1G9B3 Q1G9B9 Q1G9C7
Q1G9D2 Q1G9D6 Q1G9E8 Q1G9F7 Q1G9F8 Q1G9G0 Q1G9G2 Q1G9G5 Q1G9H1 Q1G9H2 Q1G9H8
Q1G9H9 Q1G9J5 Q1G9L2 Q1G9L3 Q1G9L4 Q1G9L7 Q1G9M6 Q1G9N0 Q1G9N6 Q1G9N7 Q1G9N8
Q1G9N9 Q1G9P3 Q1G9P5 Q1G9P9 Q1G9Q0 Q1G9Q1 Q1G9R2 Q1G9R3 Q1G9R7 Q1G9U3 Q1G9U6
Q1G9U8 Q1G9V1 Q1G9V2 Q1G9V4 Q1G9W1 Q1G9W2 Q1G9W4 Q1G9W7 Q1G9Z4 Q1GA88 Q1GA94
Q1GA98 Q1GA99 Q1GAA9 Q1GAB0 Q1GAB5 Q1GAG8 Q1GAH0 Q1GAH2 Q1GAK5 Q1GAM0 Q1GAM2
Q1GAM3 Q1GAM4 Q1GAN0 Q1GAN1 Q1GAP2 Q1GAP7 Q1GAP9 Q1GAQ0 Q1GAQ3 Q1GAQ4 Q1GAR4
Q1GAR5 Q1GAS1 Q1GAS8 Q1GAT1 Q1GAT6 Q1GAU0 Q1GAU1 Q1GAV4 Q1GAX2 Q1GAX5 Q1GAY0
Q1GAZ3 Q1GB12 Q1GB18 Q1GB22 Q1GB25 Q1GB26 Q1GB31 Q1GB32 Q1GB34 Q1GB51 Q1GB65
Q1GB78 Q1GB82 Q1GBA8 Q1GBD4 Q1GBF3 Q1GBF4 Q1GBF5 Q1GBF7 Q1GBI6 Q1GBJ1 Q1GBJ2
Q1GBJ3 Q1GBJ4 Q1GBJ5 Q1GBJ6 Q1GBJ7 Q1GBJ9 Q1GBK0 Q1GBK1 Q1GBK2 Q1GBK3 Q1GBK4
Q1GBK5 Q1GBK6 Q1GBK7 Q1GBK8 Q1GBK9 Q1GBL0 Q1GBL1 Q1GBL2 Q1GBL3 Q1GBL4 Q1GBL5
Q1GBL6 Q1GBL7 Q1GBL8 Q1GBL9 Q1GBM0 Q1GBM1 Q1GBM2 Q1GBM4 Q1GBM5 Q1GBN7 Q1GBP4
Q1GBP6 Q1GBQ0 Q1GBQ2 Q1GBQ3 Q1GBR1 Q1GBU7 Q1GBU9 Q1GBV3 Q1GBV4 Q1GBW9 Q1GC33
Q1GC35 Q1GC40 Q1GC43 Q1GC53 Q1GC55 Q1GC56 Q1GC70 Q1GC80 Q1GC88 Q1GCG2 Q1GCG4
Q1GCH2 Q1GCH5 Q1GCH8 Q1GCK1 Q1GCK9 Q1GCL2 Q1GCL8 Q1GCL9 Q1GCM0 Q1GCM3 Q1GCM4
Q1GCM7 Q1GCN3 Q1GCN5 Q1GCQ5 Q1GCR5 Q1GCT5 Q1GCU3 Q1GCX7 Q1GD76 Q1GD84 Q1GDB7
Q1GDB8 Q1GDC3 Q1GDC5 Q1GDC7 Q1GDF7 Q1GDG1 Q1GDG5 Q1GDG8 Q1GDH8 Q1GDK0 Q1GDK6
Q1GDL4 Q1GDL5 Q1GDM0 Q1GDM1 Q1GDM3 Q1GDM5 Q1GDM6 Q1GDM9 Q1GDP8 Q1GDQ2 Q1GDR0
Q1GDR6 Q1GDU2 Q1GDU6 Q1GDV0 Q1GE00 Q1GE05 Q1GE13 Q1GE26 Q1GE28 Q1GE59 Q1GE74
Q1GE79 Q1GE81 Q1GEA9 Q1GEC6 Q1GEF8 Q1GEI7 Q1GEP8 Q1GES7 Q1GET3 Q1GEY0 Q1GEY7
Q1GEZ0 Q1GEZ2 Q1GEZ3 Q1GEZ5 Q1GF01 Q1GF19 Q1GF20 Q1GF45 Q1GF47 Q1GF61 Q1GF65
Q1GF74 Q1GF88 Q1GF96 Q1GFB3 Q1GFB4 Q1GFB5 Q1GFB8 Q1GFE0 Q1GFI1 Q1GFL4 Q1GFP1
Q1GFP2 Q1GFQ4 Q1GFQ6 Q1GFQ7 Q1GFV4 Q1GFW2 Q1GGA4 Q1GGF6 Q1GGF7 Q1GGG4 Q1GGK7
Q1GGM0 Q1GGQ1 Q1GGR8 Q1GGR9 Q1GGS0 Q1GGS3 Q1GGT2 Q1GGT5 Q1GGT9 Q1GGU2 Q1GGU4
Q1GGU8 Q1GGV4 Q1GGV8 Q1GGW2 Q1GGW9 Q1GGX7 Q1GH67 Q1GH90 Q1GHA1 Q1GHC2 Q1GHC3
Q1GHC8 Q1GHC9 Q1GHD0 Q1GHD2 Q1GHD3 Q1GHF1 Q1GHI0 Q1GHI5 Q1GHJ3 Q1GHJ4 Q1GHK0
Q1GHL0 Q1GHL1 Q1GHL5 Q1GHM4 Q1GHP0 Q1GHP9 Q1GHT0 Q1GHT8 Q1GHT9 Q1GHU0 Q1GHV0
Q1GHY9 Q1GHZ2 Q1GI16 Q1GI26 Q1GI38 Q1GI39 Q1GI42 Q1GI57 Q1GI59 Q1GI60 Q1GI67
Q1GI70 Q1GI80 Q1GI81 Q1GIA1 Q1GIB5 Q1GIC1 Q1GIL3 Q1GIL9 Q1GIM1 Q1GIT2 Q1GIU7
Q1GIU8 Q1GIV0 Q1GIV4 Q1GIW2 Q1GIW7 Q1GIY4 Q1GIZ8 Q1GJ36 Q1GJ37 Q1GJ55 Q1GJB3
Q1GJD0 Q1GJI4 Q1GJL4 Q1GJP5 Q1GJP6 Q1GJP7 Q1GJP8 Q1GJQ1 Q1GJQ2 Q1GJQ5 Q1GJU6
Q1GJX0 Q1GJX4 Q1GJX8 Q1GJY2 Q1GK03 Q1GK04 Q1GK05 Q1GK06 Q1GK09 Q1GK11 Q1GK12
Q1GK13 Q1GK14 Q1GK15 Q1GK16 Q1GK17 Q1GK18 Q1GK19 Q1GK20 Q1GK21 Q1GK27 Q1GK28
Q1GK29 Q1GK30 Q1GK31 Q1GK35 Q1GK36 Q1GK37 Q1GK38 Q1GK42 Q1GK43 Q1GK44 Q1GK48
Q1GK49 Q1GK50 Q1GK51 Q1GK53 Q1GK54 Q1GK67 Q1GK69 Q1GK77 Q1GK79 Q1GK81 Q1GK84

Q1GK88 Q1GK90 Q1GK93 Q1GK98 Q1GKB9 Q1GKF8 Q1GKJ2 Q1GKJ6 Q1GKJ7 Q1GKJ9 Q1GKK2
Q1GKK5 Q1GKM7 Q1GKN8 Q1GKQ4 Q1GKS3 Q1GKS4 Q1GKT2 Q1GL02 Q1GLD5 Q1GLE0 Q1GLE1
Q1GLQ5 Q1GLQ6 Q1GM98 Q1GMD1 Q1GMM6 Q1GN68 Q1GN74 Q1GN81 Q1GN82 Q1GNB4 Q1GNC0
Q1GNC2 Q1GNC3 Q1GNC9 Q1GND0 Q1GNH1 Q1GNH3 Q1GNM2 Q1GNP7 Q1GNR4 Q1GNR9 Q1GNT5
Q1GNW5 Q1GNW6 Q1GNX0 Q1GNX2 Q1GNX7 Q1GNX8 Q1GP30 Q1GP41 Q1GP42 Q1GP62 Q1GP63
Q1GP64 Q1GP74 Q1GP75 Q1GP76 Q1GP77 Q1GP88 Q1GP94 Q1GP95 Q1GP96 Q1GP97 Q1GP98
Q1GP99 Q1GPA0 Q1GPA1 Q1GPA2 Q1GPA3 Q1GPA4 Q1GPA5 Q1GPA6 Q1GPA7 Q1GPA8 Q1GPA9
Q1GPB0 Q1GPB1 Q1GPB2 Q1GPB3 Q1GPB4 Q1GPB5 Q1GPB6 Q1GPB7 Q1GPB8 Q1GPC0 Q1GPG1
Q1GPG3 Q1GPG7 Q1GPH4 Q1GPH5 Q1GPI7 Q1GPJ1 Q1GPJ7 Q1GPJ8 Q1GPK3 Q1GPS4 Q1GQ92
Q1GQI3 Q1GQI7 Q1GQJ3 Q1GQK9 Q1GQY3 Q1GQZ4 Q1GR07 Q1GR20 Q1GR22 Q1GR26 Q1GR54
Q1GR67 Q1GR82 Q1GR84 Q1GRD4 Q1GRE9 Q1GRG3 Q1GRH1 Q1GRH2 Q1GRI1 Q1GRI4 Q1GRI6
Q1GRJ2 Q1GRJ6 Q1GRK2 Q1GRQ0 Q1GRQ1 Q1GRQ2 Q1GRQ3 Q1GRR1 Q1GRR9 Q1GRU4 Q1GRU5
Q1GRV0 Q1GRV8 Q1GRV9 Q1GRW0 Q1GRX1 Q1GRX7 Q1GRY0 Q1GRY1 Q1GRY2 Q1GRZ2 Q1GS53
Q1GSA0 Q1GSA2 Q1GSC8 Q1GSC9 Q1GSD9 Q1GSF4 Q1GSF8 Q1GSF9 Q1GSG8 Q1GSH0 Q1GSH1
Q1GSN3 Q1GSP1 Q1GSP2 Q1GSQ7 Q1GSU5 Q1GSV9 Q1GSW9 Q1GT16 Q1GT23 Q1GT24 Q1GT31
Q1GT35 Q1GT37 Q1GT40 Q1GT67 Q1GT68 Q1GT81 Q1GT88 Q1GT91 Q1GTA2 Q1GTB8 Q1GTF3
Q1GTF8 Q1GTG6 Q1GTH6 Q1GTJ0 Q1GTJ4 Q1GTL4 Q1GTM7 Q1GTN0 Q1GTS7 Q1GTT5 Q1GTU1
Q1GTU3 Q1GTU9 Q1GTW7 Q1GTW9 Q1GTY6 Q1GTZ7 Q1GU73 Q1GU89 Q1GUB6 Q1GUG9 Q1GUJ1
Q1GUJ4 Q1GUM2 Q1GUM9 Q1GUN0 Q1GUW1 Q1GUW3 Q1GUW4 Q1GUW8 Q1GUX4 Q1GV11 Q1GV13
Q1GV16 Q1GV29 Q1GV37 Q1GV40 Q1GV79 Q1GV83 Q1GV98 Q1GVA9 Q1GVE2 Q1GVE7 Q1GVI8
Q1GVI9 Q1GVJ2 Q1GVL0 Q1GVL7 Q1GVM0 Q1GVM5 Q1GVN4 Q1GVN5 Q1GVN6 Q1GVN7 Q1GVQ2
Q1GVQ8 Q1GVQ9 Q1GVU1 Q1GVV2 Q1GVV6 Q1GVZ9 Q1GWA7 Q1GWB1 Q1GWB2 Q1GWB3 Q1GWB4
Q1GWB6 Q1GWI7 Q1GWJ5 Q1GWJ6 Q1GWJ9 Q1GWK0 Q1GWK1 Q1GWP7 Q1GWT3 Q1GWT5 Q1GX07
Q1GX86 Q1GXA1 Q1GXA4 Q1GXA9 Q1GXB2 Q1GXB4 Q1GXB8 Q1GXB9 Q1GXC7 Q1GXD2 Q1GXD3
Q1GXD4 Q1GXD5 Q1GXD6 Q1GXD8 Q1GXE6 Q1GXG2 Q1GXH3 Q1GXK1 Q1GXL1 Q1GXL3 Q1GXL7
Q1GXL9 Q1GXM0 Q1GXV8 Q1GXW0 Q1GXW1 Q1GXY7 Q1GXY8 Q1GYC1 Q1GYC2 Q1GYD2 Q1GYD3
Q1GYD4 Q1GYE8 Q1GYE9 Q1GYF5 Q1GYH7 Q1GYN8 Q1GYN9 Q1GYR5 Q1GYR6 Q1GYR7 Q1GYS6
Q1GYT5 Q1GYT6 Q1GYT7 Q1GYT8 Q1GYT9 Q1GYU4 Q1GYU5 Q1GYV1 Q1GYV3 Q1GYX9 Q1GYY7
Q1GYZ0 Q1GYZ2 Q1GYZ3 Q1GYZ9 Q1GZ02 Q1GZ04 Q1GZ11 Q1GZ15 Q1GZ16 Q1GZ23 Q1GZ28
Q1GZ51 Q1GZ52 Q1GZ53 Q1GZ54 Q1GZ59 Q1GZ66 Q1GZ82 Q1GZ85 Q1GZA1 Q1GZA6 Q1GZA7
Q1GZB2 Q1GZB5 Q1GZB7 Q1GZB8 Q1GZD3 Q1GZD6 Q1GZD7 Q1GZE7 Q1GZE9 Q1GZF5 Q1GZF7
Q1GZG8 Q1GZH0 Q1GZH1 Q1GZI2 Q1GZJ0 Q1GZK6 Q1GZR7 Q1GZR9 Q1GZS0 Q1GZS1 Q1GZS3
Q1GZY9 Q1H003 Q1H004 Q1H009 Q1H031 Q1H051 Q1H054 Q1H069 Q1H075 Q1H095 Q1H096
Q1H0B6 Q1H0D2 Q1H0E9 Q1H0F0 Q1H0F1 Q1H0F2 Q1H0F3 Q1H0F4 Q1H0F5 Q1H0G4 Q1H0I2
Q1H0J2 Q1H0L0 Q1H0L1 Q1H0L2 Q1H0L3 Q1H0L4 Q1H0L5 Q1H0L9 Q1H0M1 Q1H0N6 Q1H0Q5
Q1H0R4 Q1H0U5 Q1H0U6 Q1H0U9 Q1H0V0 Q1H0X3 Q1H0X9 Q1H0Y0 Q1H0Y5 Q1H0Y6 Q1H0Z5
Q1H139 Q1H140 Q1H141 Q1H142 Q1H145 Q1H153 Q1H156 Q1H160 Q1H169 Q1H188 Q1H190
Q1H1F9 Q1H1G0 Q1H1G1 Q1H1G6 Q1H1G9 Q1H1H9 Q1H1I4 Q1H1K1 Q1H1K2 Q1H1N8 Q1H1P0
Q1H1P1 Q1H1P3 Q1H1P4 Q1H1T1 Q1H1W4 Q1H274 Q1H280 Q1H290 Q1H2A2 Q1H2B1 Q1H2E4
Q1H2F4 Q1H2I8 Q1H2K1 Q1H2K2 Q1H2P1 Q1H361 Q1H365 Q1H372 Q1H378 Q1H384 Q1H387
Q1H396 Q1H397 Q1H3A0 Q1H3B7 Q1H3B8 Q1H3B9 Q1H3D2 Q1H3I3 Q1H3J0 Q1H3J2 Q1H3J3
Q1H3L5 Q1H3L6 Q1H3L8 Q1H3M2 Q1H3M3 Q1H3S0 Q1H3S1 Q1H3S3 Q1H3S5 Q1H3T1 Q1H3T4
Q1H3U7 Q1H3U9 Q1H3X8 Q1H403 Q1H417 Q1H424 Q1H425 Q1H450 Q1H497 Q1H4F1 Q1H4F2
Q1H4G7 Q1H4G9 Q1H4H0 Q1H4H6 Q1H4K1 Q1H4K3 Q1H4K4 Q1H4K5 Q1H4L1 Q1H4L2 Q1H4L3

Q1H4L4 Q1H4L5 Q1H4L6 Q1H4L8 Q1H4L9 Q1H4M0 Q1H4M1 Q1H4M2 Q1H4M3 Q1H4M4 Q1H4M5
Q1H4M6 Q1H4M7 Q1H4M8 Q1H4M9 Q1H4N0 Q1H4N1 Q1H4N2 Q1H4N3 Q1H4N4 Q1H4N5 Q1H4N6
Q1H4N7 Q1H4N8 Q1H4N9 Q1H4P0 Q1H4P1 Q1H4P2 Q1H4P3 Q1H4P4 Q1H4P5 Q1H4P6 Q1H4P7
Q1H4P8 Q1H4Q1 Q1H4R0 Q1H4R1 Q1H4R2 Q1H4R3 Q1H4R5 Q1H4R9 Q1H4S0 Q1H4T4 Q1H4T5
Q1H4T8 Q1H4U1 Q1H4U4 Q1H4W1 Q1H4W7 Q1H4X2 Q1H4X6 Q1H4Y0 Q1H501 Q1H521 Q1H537
Q1H595 Q1H5E9 Q1H9T6 Q1HCL7 Q1HDZ5 Q1HE00 Q1HE01 Q1HG60 Q1HGK7 Q1HPK6 Q1HPQ0
Q1HPS0 Q1HPS4 Q1HPU0 Q1HPU2 Q1HPW4 Q1HQ10 Q1HQK0 Q1HQN4 Q1HQY6 Q1HR24 Q1HR36
Q1HR47 Q1HR62 Q1HR65 Q1HR79 Q1HRR3 Q1HRT4 Q1HRV4 Q1HVB5 Q1HVB8 Q1HVC1 Q1HVC3
Q1HVD1 Q1HVD3 Q1HVE7 Q1HVF2 Q1HVF7 Q1HVF8 Q1HVG0 Q1HVG4 Q1HVH2 Q1HVH8 Q1HVI8
Q1I0U1 Q1I0U2 Q1I0U3 Q1I0U4 Q1I0U5 Q1I0U6 Q1I0U7 Q1I0U8 Q1I0U9 Q1I0V0 Q1I0V1
Q1I2G4 Q1I2G6 Q1I2G9 Q1I2H1 Q1I2H2 Q1I2H5 Q1I2H6 Q1I2H7 Q1I2K4 Q1I2L2 Q1I2L8
Q1I2R4 Q1I2S9 Q1I2T8 Q1I2T9 Q1I2U1 Q1I2U3 Q1I2U5 Q1I2V6 Q1I2V7 Q1I2W1 Q1I301
Q1I303 Q1I306 Q1I314 Q1I316 Q1I328 Q1I350 Q1I3B8 Q1I3C2 Q1I3G0 Q1I3G4 Q1I3G7
Q1I3G8 Q1I3L0 Q1I3N6 Q1I3N7 Q1I3Q0 Q1I3Q8 Q1I3R1 Q1I3R4 Q1I3S6 Q1I3S7 Q1I3T0
Q1I3T9 Q1I3U0 Q1I3U2 Q1I3U3 Q1I3U5 Q1I3W5 Q1I3X5 Q1I3X6 Q1I3Y0 Q1I411 Q1I419
Q1I439 Q1I440 Q1I447 Q1I448 Q1I449 Q1I453 Q1I454 Q1I461 Q1I462 Q1I464 Q1I469
Q1I490 Q1I491 Q1I4B1 Q1I4C1 Q1I4C5 Q1I4F0 Q1I4F1 Q1I4F3 Q1I4F9 Q1I4G0 Q1I4G1
Q1I4G7 Q1I4H2 Q1I4H5 Q1I4H6 Q1I4J8 Q1I4K4 Q1I4M5 Q1I4P0 Q1I4P1 Q1I4P2 Q1I4P8
Q1I4Q1 Q1I4R5 Q1I4S2 Q1I4S5 Q1I4S8 Q1I4U6 Q1I4X3 Q1I500 Q1I571 Q1I574 Q1I585
Q1I596 Q1I597 Q1I5A5 Q1I5A6 Q1I5A9 Q1I5B0 Q1I5B9 Q1I5C0 Q1I5D3 Q1I5E1 Q1I5E2
Q1I5F3 Q1I5F5 Q1I5F9 Q1I5K2 Q1I5K4 Q1I5K9 Q1I5M7 Q1I5M8 Q1I5P3 Q1I5T6 Q1I5V8
Q1I5W0 Q1I5W1 Q1I5Y3 Q1I5Y8 Q1I5Y9 Q1I5Z0 Q1I5Z1 Q1I614 Q1I624 Q1I626 Q1I627
Q1I628 Q1I629 Q1I640 Q1I644 Q1I648 Q1I652 Q1I653 Q1I654 Q1I655 Q1I659 Q1I6B7
Q1I6B8 Q1I6D1 Q1I6D5 Q1I6D7 Q1I6E0 Q1I6E5 Q1I6E7 Q1I6E8 Q1I6E9 Q1I6F8 Q1I6G9
Q1I6H8 Q1I6J5 Q1I6K3 Q1I6K6 Q1I6M7 Q1I6Y2 Q1I6Z5 Q1I6Z6 Q1I6Z8 Q1I749 Q1I7B1
Q1I7B2 Q1I7B3 Q1I7F0 Q1I7F9 Q1I7G7 Q1I7L3 Q1I7P9 Q1I7T2 Q1I7T4 Q1I7T9 Q1I7U0
Q1I833 Q1I898 Q1I8C2 Q1I8G2 Q1I8G3 Q1I926 Q1I984 Q1I9D2 Q1I9K4 Q1IA50 Q1IA73
Q1IAK7 Q1IAK8 Q1IAL0 Q1IAY6 Q1IB54 Q1IBC9 Q1IBD0 Q1IBD5 Q1IBD6 Q1IBD7 Q1IBE4
Q1IBF8 Q1IBN6 Q1IBN7 Q1IBN9 Q1IBP0 Q1IBP1 Q1IBP2 Q1IBP3 Q1IBP9 Q1IC09 Q1IC11
Q1IC12 Q1IC13 Q1IC14 Q1IC15 Q1IC21 Q1ICA8 Q1ICA9 Q1ICB3 Q1ICG2 Q1ICH9 Q1ICI0
Q1ICI1 Q1ICK0 Q1ICL2 Q1ICM4 Q1ICM7 Q1ICQ6 Q1ICS3 Q1ICT0 Q1ICV9 Q1ICW0 Q1ICW8
Q1ICY2 Q1ICZ0 Q1ID00 Q1ID16 Q1ID19 Q1ID35 Q1ID65 Q1ID82 Q1ID97 Q1ID99 Q1IDA2
Q1IDA4 Q1IDA6 Q1IDB3 Q1IDB9 Q1IDC3 Q1IDE2 Q1IDE4 Q1IDE6 Q1IDH6 Q1IDJ7 Q1IDJ9
Q1IDL2 Q1IDL9 Q1IDN2 Q1IDN4 Q1IDS5 Q1IDT9 Q1IDU5 Q1IDV1 Q1IDV2 Q1IDW1 Q1IDY5
Q1IDZ3 Q1IE03 Q1IE05 Q1IE06 Q1IE16 Q1IE31 Q1IE97 Q1IE99 Q1IEA0 Q1IEB5 Q1IEC3
Q1IED2 Q1IED3 Q1IED4 Q1IEF7 Q1IEH2 Q1IEH7 Q1IEI0 Q1IEI1 Q1IEI4 Q1IEI5 Q1IEI8
Q1IEI9 Q1IEJ2 Q1IEK2 Q1IEL0 Q1IEU2 Q1IEV9 Q1IEW1 Q1IEW8 Q1IEX1 Q1IEX2 Q1IEY5
Q1IEY7 Q1IEY8 Q1IF12 Q1IF13 Q1IF14 Q1IF15 Q1IF39 Q1IF40 Q1IF41 Q1IF42 Q1IF43
Q1IF45 Q1IF47 Q1IF48 Q1IF51 Q1IF57 Q1IF58 Q1IF59 Q1IF60 Q1IF64 Q1IF66 Q1IFE4
Q1IFI3 Q1IFK9 Q1IFL1 Q1IFL5 Q1IFL9 Q1IFM0 Q1IFM4 Q1IFP3 Q1IFU0 Q1IFU1 Q1IFU2
Q1IFU3 Q1IFU4 Q1IFU5 Q1IFU7 Q1IFU8 Q1IFU9 Q1IFV0 Q1IFV1 Q1IFV2 Q1IFV3 Q1IFV4
Q1IFV5 Q1IFV6 Q1IFV7 Q1IFV8 Q1IFV9 Q1IFW0 Q1IFW1 Q1IFW2 Q1IFW3 Q1IFW4 Q1IFW5
Q1IFW6 Q1IFW7 Q1IFW8 Q1IFX0 Q1IFX1 Q1IFX2 Q1IFX3 Q1IFX4 Q1IFX5 Q1IFX6 Q1IFX7
Q1IFY7 Q1IFY8 Q1IFZ9 Q1IG00 Q1IG21 Q1IG22 Q1IG23 Q1IG25 Q1IG27 Q1IG31 Q1IG32

Q1IG45 Q1IG46 Q1IG47 Q1IG48 Q1IG59 Q1IG62 Q1IG69 Q1IG70 Q1IG73 Q1IGA3 Q1IGA7
Q1IGA8 Q1IGB6 Q1IGB8 Q1IGC6 Q1IGD2 Q1IGD3 Q1IGF0 Q1IGF3 Q1IGF7 Q1IGK1 Q1IGK2
Q1IGK6 Q1IGL1 Q1IGL6 Q1IGY1 Q1IH20 Q1IH21 Q1IH28 Q1IH35 Q1IH41 Q1IH46 Q1IHA5
Q1IHA8 Q1IHC2 Q1IHD8 Q1IHG6 Q1IHG7 Q1IHH0 Q1IHH1 Q1IHH2 Q1IHH3 Q1IHH4 Q1IHH5
Q1IHJ4 Q1IHK6 Q1IHK7 Q1IHM1 Q1IHQ3 Q1IHT1 Q1IHU1 Q1IHV2 Q1IHV6 Q1IHW2 Q1IHW3
Q1IHW4 Q1IHW5 Q1IHZ8 Q1II13 Q1II25 Q1II28 Q1II29 Q1II76 Q1IIJ5 Q1IIL2 Q1IIS2
Q1IIT1 Q1IIT3 Q1IIT5 Q1IIU5 Q1IIY8 Q1IIZ0 Q1IIZ2 Q1IIZ3 Q1IJ06 Q1IJ49 Q1IJA9
Q1IJF8 Q1IJG5 Q1IJG7 Q1IJI6 Q1IJR5 Q1IJS5 Q1IJS7 Q1IJT2 Q1IJZ0 Q1IK18 Q1IK22
Q1IKB2 Q1IKB4 Q1IKB7 Q1IKC9 Q1IKF4 Q1IKG0 Q1IKG1 Q1IKG8 Q1IKH1 Q1IKI1 Q1IKI5
Q1IKJ1 Q1IKJ2 Q1IKQ3 Q1IKR2 Q1IL12 Q1ILA0 Q1ILA1 Q1ILG3 Q1ILN0 Q1ILY2 Q1ILZ0
Q1ILZ3 Q1IMB2 Q1IMI3 Q1IMI4 Q1IMM1 Q1IMM3 Q1IMM4 Q1IMM5 Q1IMM7 Q1IN52 Q1INB1
Q1INC3 Q1INE2 Q1INI5 Q1INI6 Q1INK7 Q1INP0 Q1INQ6 Q1INS6 Q1INT8 Q1INT9 Q1INU0
Q1IP68 Q1IPC9 Q1IPF0 Q1IPG4 Q1IPK2 Q1IPL8 Q1IPQ5 Q1IPQ9 Q1IPS4 Q1IPU1 Q1IPU7
Q1IPZ8 Q1IQ21 Q1IQ53 Q1IQ54 Q1IQ55 Q1IQ75 Q1IQ76 Q1IQG9 Q1IQH5 Q1IQH8 Q1IRA2
Q1IRD1 Q1IRD7 Q1IRD8 Q1IRS5 Q1IRT8 Q1IRZ2 Q1IS76 Q1IS80 Q1IS84 Q1IS92 Q1IS93
Q1IS94 Q1IS96 Q1IS97 Q1IS98 Q1IS99 Q1ISA3 Q1ISA4 Q1ISA5 Q1ISA6 Q1ISA7 Q1ISA8
Q1ISB1 Q1ISB2 Q1ISB3 Q1ISB5 Q1ISB6 Q1ISB7 Q1ISB8 Q1ISB9 Q1ISC0 Q1ISC1 Q1ISC2
Q1ISC3 Q1ISC5 Q1ISC6 Q1ISC7 Q1ISD3 Q1ISE0 Q1ISE2 Q1ISF4 Q1ISG1 Q1ISH2 Q1ISI9
Q1ISJ1 Q1ISQ0 Q1ISS4 Q1IST4 Q1ISV1 Q1ISX7 Q1IT15 Q1ITN7 Q1ITT0 Q1ITT1 Q1ITV2
Q1ITW5 Q1IU80 Q1IU82 Q1IU84 Q1IU89 Q1IUE4 Q1IUF1 Q1IUW9 Q1IV16 Q1IV19 Q1IV36
Q1IV55 Q1IV81 Q1IVA8 Q1IVB1 Q1IVD0 Q1IVF1 Q1IVF9 Q1IVL5 Q1IVP4 Q1IVQ3 Q1IVR7
Q1IVS1 Q1IVS5 Q1IVS6 Q1IVS7 Q1IVV5 Q1IVV7 Q1IW06 Q1IW58 Q1IW59 Q1IW70 Q1IW72
Q1IW73 Q1IW74 Q1IW81 Q1IW89 Q1IW96 Q1IWA0 Q1IWC9 Q1IWD2 Q1IWD8 Q1IWD9 Q1IWF2
Q1IWF4 Q1IWG2 Q1IWG9 Q1IWI7 Q1IWK0 Q1IWL4 Q1IWM0 Q1IWM3 Q1IWM5 Q1IWN0 Q1IWN1
Q1IWN4 Q1IWQ7 Q1IWV2 Q1IWZ8 Q1IX15 Q1IX16 Q1IX18 Q1IX33 Q1IX39 Q1IX43 Q1IX45
Q1IX66 Q1IX67 Q1IX68 Q1IX70 Q1IX71 Q1IX72 Q1IX73 Q1IX74 Q1IX75 Q1IX76 Q1IX77
Q1IX78 Q1IX79 Q1IX80 Q1IX81 Q1IX82 Q1IX83 Q1IX84 Q1IX85 Q1IX86 Q1IX87 Q1IX88
Q1IX89 Q1IX90 Q1IX91 Q1IX93 Q1IX94 Q1IX95 Q1IX96 Q1IX97 Q1IX98 Q1IX99 Q1IXA0
Q1IXD5 Q1IXJ0 Q1IXJ7 Q1IXK1 Q1IXK5 Q1IXK7 Q1IXK9 Q1IXM3 Q1IXM5 Q1IXQ0 Q1IXS5
Q1IXS9 Q1IXU8 Q1IXV7 Q1IXV8 Q1IXW9 Q1IXY5 Q1IY21 Q1IY22 Q1IY50 Q1IY56 Q1IY72
Q1IY76 Q1IY80 Q1IY82 Q1IYA3 Q1IYA4 Q1IYA6 Q1IYF7 Q1IYG7 Q1IYH9 Q1IYI1 Q1IYK3
Q1IYK9 Q1IYM0 Q1IYR1 Q1IYS0 Q1IYT4 Q1IYW2 Q1IYW5 Q1IYX1 Q1IYZ8 Q1IZ02 Q1IZ05
Q1IZ41 Q1IZ94 Q1IZA1 Q1IZA2 Q1IZA3 Q1IZB2 Q1IZB4 Q1IZB5 Q1IZC9 Q1IZE7 Q1IZH8
Q1IZI6 Q1IZI7 Q1IZI8 Q1IZI9 Q1IZK2 Q1IZL1 Q1IZM1 Q1IZN3 Q1IZP0 Q1IZP1 Q1IZP2
Q1IZP8 Q1IZQ3 Q1IZQ5 Q1IZU9 Q1IZV0 Q1J021 Q1J022 Q1J043 Q1J055 Q1J063 Q1J090
Q1J0A0 Q1J0B6 Q1J0B7 Q1J0C1 Q1J0C2 Q1J0F6 Q1J0F9 Q1J0I3 Q1J0I4 Q1J0I6 Q1J0L2
Q1J0N7 Q1J0P0 Q1J0P1 Q1J0P2 Q1J0P3 Q1J0P6 Q1J0P7 Q1J0P9 Q1J0R0 Q1J0S3 Q1J0T2
Q1J0T4 Q1J0V5 Q1J0V8 Q1J0V9 Q1J0W2 Q1J0W5 Q1J0Z7 Q1J115 Q1J118 Q1J119 Q1J123
Q1J129 Q1J141 Q1J147 Q1J148 Q1J154 Q1J162 Q1J163 Q1J188 Q1J199 Q1J1B1 Q1J1C0
Q1J1C7 Q1J1D0 Q1J1F4 Q1J1F6 Q1J1J3 Q1J1K4 Q1J1M8 Q1J1N6 Q1J1P4 Q1J1P5 Q1J1S9
Q1J1V8 Q1J1W0 Q1J1X6 Q1J200 Q1J201 Q1J215 Q1J217 Q1J218 Q1J237 Q1J245 Q1J288
Q1J2A2 Q1J2B4 Q1J2B6 Q1J2C9 Q1J2D0 Q1J2E1 Q1J2G7 Q1J391 Q1J3J2 Q1J3J3 Q1J434
Q1J443 Q1J455 Q1J457 Q1J459 Q1J462 Q1J478 Q1J479 Q1J486 Q1J489 Q1J492 Q1J494
Q1J495 Q1J497 Q1J498 Q1J499 Q1J4A6 Q1J4B3 Q1J4B4 Q1J4B6 Q1J4C0 Q1J4C4 Q1J4D1

Q1J4E0 Q1J4K0 Q1J4K7 Q1J4L9 Q1J4M0 Q1J4M3 Q1J4M6 Q1J4N2 Q1J4P6 Q1J4Q3 Q1J4Q7
Q1J4W4 Q1J4W7 Q1J4W9 Q1J4X1 Q1J4X4 Q1J4X9 Q1J4Y7 Q1J4Y8 Q1J4Z3 Q1J4Z8 Q1J501
Q1J509 Q1J513 Q1J524 Q1J530 Q1J532 Q1J545 Q1J564 Q1J569 Q1J570 Q1J576 Q1J578
Q1J594 Q1J5A9 Q1J5B0 Q1J5B4 Q1J5B5 Q1J5E0 Q1J5E1 Q1J5E2 Q1J5E4 Q1J5E9 Q1J5F5
Q1J5F6 Q1J5F8 Q1J5H3 Q1J5H4 Q1J5H5 Q1J5I6 Q1J5I7 Q1J5I9 Q1J5K6 Q1J5K7 Q1J5M4
Q1J5Q4 Q1J5Q6 Q1J5R0 Q1J5R2 Q1J5R7 Q1J5S5 Q1J5T2 Q1J5U5 Q1J5U7 Q1J5W1 Q1J5W6
Q1J5W7 Q1J5X1 Q1J5X5 Q1J5Y6 Q1J5Z4 Q1J626 Q1J633 Q1J634 Q1J652 Q1J665 Q1J667
Q1J694 Q1J6C1 Q1J6D8 Q1J6D9 Q1J6F7 Q1J6G5 Q1J6H1 Q1J6H8 Q1J6I6 Q1J6J1 Q1J6K1
Q1J6L1 Q1J6L7 Q1J6M1 Q1J6M6 Q1J6M7 Q1J6N4 Q1J6P0 Q1J6P2 Q1J6Q7 Q1J6U1 Q1J6U4
Q1J6W8 Q1J702 Q1J705 Q1J708 Q1J713 Q1J724 Q1J728 Q1J729 Q1J733 Q1J736 Q1J737
Q1J740 Q1J741 Q1J744 Q1J760 Q1J776 Q1J777 Q1J778 Q1J786 Q1J794 Q1J796 Q1J7A2
Q1J7A5 Q1J7B1 Q1J7B2 Q1J7B3 Q1J7B6 Q1J7B7 Q1J7B8 Q1J7B9 Q1J7C3 Q1J7E2 Q1J7E4
Q1J7E5 Q1J7G9 Q1J7J0 Q1J7J5 Q1J7K2 Q1J7K4 Q1J7K5 Q1J7N3 Q1J7N4 Q1J7Q8 Q1J7V3
Q1J7X1 Q1J7Y1 Q1J7Y3 Q1J7Y6 Q1J824 Q1J829 Q1J832 Q1J833 Q1J834 Q1J835 Q1J861
Q1J881 Q1J886 Q1J887 Q1J888 Q1J890 Q1J8A8 Q1J8A9 Q1J8B0 Q1J8B4 Q1J8C6 Q1J8C9
Q1J8D2 Q1J8D8 Q1J8E7 Q1J8E9 Q1J8I4 Q1J8I5 Q1J8I6 Q1J8J4 Q1J8K5 Q1J8K6 Q1J8K9
Q1J8L3 Q1J8N2 Q1J8P1 Q1J8Q8 Q1J8S0 Q1J8U9 Q1J8W0 Q1J8X0 Q1J8X1 Q1J8X3 Q1J8Y6
Q1J8Y7 Q1J8Y8 Q1J8Y9 Q1J8Z0 Q1J8Z1 Q1J8Z2 Q1J8Z4 Q1J8Z5 Q1J8Z6 Q1J8Z7 Q1J8Z8
Q1J8Z9 Q1J900 Q1J901 Q1J902 Q1J904 Q1J905 Q1J906 Q1J907 Q1J908 Q1J909 Q1J910
Q1J911 Q1J912 Q1J917 Q1J924 Q1J934 Q1J936 Q1J942 Q1J956 Q1J960 Q1J963 Q1J973
Q1J988 Q1J990 Q1J992 Q1J995 Q1J9B1 Q1J9B2 Q1J9B9 Q1J9C1 Q1J9C2 Q1J9C4 Q1J9C6
Q1J9C7 Q1J9C9 Q1J9D0 Q1J9D8 Q1J9E5 Q1J9E6 Q1J9E8 Q1J9F2 Q1J9F6 Q1J9G4 Q1J9H3
Q1J9P3 Q1J9R3 Q1J9R4 Q1J9R7 Q1J9S0 Q1J9S6 Q1J9U0 Q1J9U7 Q1J9U8 Q1J9V2 Q1J9W8
Q1J9X6 Q1J9X9 Q1J9Y4 Q1J9Y9 Q1J9Z2 Q1J9Z4 Q1J9Z9 Q1JA04 Q1JA07 Q1JA16 Q1JA20
Q1JA29 Q1JA31 Q1JA37 Q1JA39 Q1JA50 Q1JA74 Q1JA75 Q1JA81 Q1JA83 Q1JA99 Q1JAB6
Q1JAB7 Q1JAC1 Q1JAC2 Q1JAE8 Q1JAF0 Q1JAF2 Q1JAF7 Q1JAG2 Q1JAG3 Q1JAG5 Q1JAI0
Q1JAI1 Q1JAI2 Q1JAJ4 Q1JAJ5 Q1JAJ7 Q1JAL4 Q1JAL5 Q1JAN3 Q1JAR2 Q1JAR4 Q1JAR8
Q1JAS0 Q1JAS5 Q1JAT4 Q1JAU1 Q1JAV4 Q1JAV6 Q1JAX0 Q1JAX6 Q1JAX7 Q1JAY1 Q1JAY8
Q1JB00 Q1JB08 Q1JB36 Q1JB43 Q1JB44 Q1JBB7 Q1JBD1 Q1JBF7 Q1JBI5 Q1JBK1 Q1JBK2
Q1JBM0 Q1JBM8 Q1JBN4 Q1JBP0 Q1JBP9 Q1JBQ9 Q1JBR5 Q1JBR9 Q1JBS4 Q1JBS5 Q1JBT2
Q1JBT9 Q1JBU1 Q1JBV7 Q1JBZ4 Q1JC19 Q1JC56 Q1JC59 Q1JC62 Q1JC67 Q1JC76 Q1JC80
Q1JC81 Q1JC85 Q1JC88 Q1JC89 Q1JC92 Q1JC93 Q1JC96 Q1JCB2 Q1JCC9 Q1JCE0 Q1JCE8
Q1JCF0 Q1JCF7 Q1JCG0 Q1JCG5 Q1JCG6 Q1JCG7 Q1JCH0 Q1JCH1 Q1JCH2 Q1JCH3 Q1JCH7
Q1JCJ5 Q1JCJ8 Q1JCJ9 Q1JCM1 Q1JCP4 Q1JCP9 Q1JCQ4 Q1JCQ6 Q1JCQ7 Q1JCT5 Q1JCT6
Q1JCZ9 Q1JD10 Q1JD20 Q1JD22 Q1JD25 Q1JD48 Q1JD55 Q1JD56 Q1JD58 Q1JD59 Q1JD60
Q1JD61 Q1JD89 Q1JDA2 Q1JDA9 Q1JDD0 Q1JDD1 Q1JDD2 Q1JDD6 Q1JDE8 Q1JDF1 Q1JDF4
Q1JDG1 Q1JDG9 Q1JDH1 Q1JDK7 Q1JDK8 Q1JDL6 Q1JDM7 Q1JDM8 Q1JDN4 Q1JDQ8 Q1JDS0
Q1JDV2 Q1JDZ8 Q1JE09 Q1JE19 Q1JE20 Q1JE22 Q1JE32 Q1JE33 Q1JE34 Q1JE35 Q1JE36
Q1JE37 Q1JE38 Q1JE40 Q1JE41 Q1JE42 Q1JE43 Q1JE44 Q1JE45 Q1JE46 Q1JE47 Q1JE48
Q1JE49 Q1JE50 Q1JE51 Q1JE52 Q1JE53 Q1JE54 Q1JE55 Q1JE56 Q1JE59 Q1JE60 Q1JE69
Q1JE78 Q1JE80 Q1JE86 Q1JEA1 Q1JEA5 Q1JEB8 Q1JED4 Q1JED6 Q1JED8 Q1JEE1 Q1JEG0
Q1JEG1 Q1JEG8 Q1JEH0 Q1JEH4 Q1JEH6 Q1JEH7 Q1JEH9 Q1JEI0 Q1JEI1 Q1JEI8 Q1JEJ6
Q1JEJ7 Q1JEK0 Q1JEK4 Q1JEK8 Q1JEL5 Q1JEM3 Q1JET3 Q1JEU0 Q1JEV3 Q1JEV4 Q1JEV7
Q1JEW0 Q1JEW6 Q1JEY0 Q1JEY9 Q1JEZ3 Q1JF09 Q1JF12 Q1JF14 Q1JF16 Q1JF19 Q1JF32

Q1JF33 Q1JF38 Q1JF47 Q1JF50 Q1JF59 Q1JF63 Q1JF72 Q1JF74 Q1JF80 Q1JF82 Q1JF94
Q1JFB9 Q1JFC0 Q1JFC6 Q1JFC8 Q1JFE4 Q1JFF9 Q1JFG0 Q1JFG4 Q1JFG5 Q1JFJ0 Q1JFJ1
Q1JFJ2 Q1JFJ5 Q1JFK0 Q1JFK5 Q1JFK8 Q1JFM0 Q1JFM3 Q1JFM4 Q1JFM5 Q1JFN7 Q1JFN8
Q1JFP0 Q1JFQ7 Q1JFQ8 Q1JFS6 Q1JFY9 Q1JFZ1 Q1JFZ5 Q1JFZ7 Q1JG02 Q1JG10 Q1JG16
Q1JG29 Q1JG31 Q1JG44 Q1JG49 Q1JG50 Q1JG54 Q1JG57 Q1JG69 Q1JG77 Q1JGA1 Q1JGC1
Q1JGC2 Q1JGD5 Q1JGE8 Q1JGF0 Q1JGH6 Q1JGK3 Q1JGL9 Q1JGM0 Q1JGN8 Q1JGP6 Q1JGQ2
Q1JGR0 Q1JGR7 Q1JGS2 Q1JGT2 Q1JGU2 Q1JGU8 Q1JGV2 Q1JGV7 Q1JGV8 Q1JGW5 Q1JGX1
Q1JGX3 Q1JGY8 Q1JH22 Q1JH25 Q1JH49 Q1JH84 Q1JH87 Q1JH90 Q1JH95 Q1JHA5 Q1JHA9
Q1JHB0 Q1JHB7 Q1JHB8 Q1JHC1 Q1JHC2 Q1JHC5 Q1JHE0 Q1JHF5 Q1JHF6 Q1JHF7 Q1JHG4
Q1JHH2 Q1JHI2 Q1JHI4 Q1JHJ0 Q1JHJ1 Q1JHJ2 Q1JHJ5 Q1JHJ6 Q1JHJ7 Q1JHJ8 Q1JHK2
Q1JHL9 Q1JHM1 Q1JHM2 Q1JHP3 Q1JHR3 Q1JHR8 Q1JHS3 Q1JHS5 Q1JHS6 Q1JHV5 Q1JHV6
Q1JI19 Q1JI30 Q1JI40 Q1JI42 Q1JI45 Q1JI69 Q1JI74 Q1JI75 Q1JI77 Q1JI78 Q1JI79
Q1JI80 Q1JIB1 Q1JIC7 Q1JID3 Q1JID4 Q1JID6 Q1JID8 Q1JIF4 Q1JIF5 Q1JIF6 Q1JIG0
Q1JIH1 Q1JIH4 Q1JIH7 Q1JII3 Q1JIJ2 Q1JIJ4 Q1JIM6 Q1JIM7 Q1JIM8 Q1JIN6 Q1JIP7
Q1JIP8 Q1JIQ1 Q1JIQ6 Q1JIS1 Q1JIT0 Q1JIV9 Q1JIX0 Q1JJ01 Q1JJ12 Q1JJ22 Q1JJ23
Q1JJ25 Q1JJ34 Q1JJ35 Q1JJ36 Q1JJ37 Q1JJ38 Q1JJ39 Q1JJ40 Q1JJ43 Q1JJ44 Q1JJ45
Q1JJ46 Q1JJ47 Q1JJ48 Q1JJ49 Q1JJ50 Q1JJ51 Q1JJ52 Q1JJ53 Q1JJ54 Q1JJ55 Q1JJ56
Q1JJ57 Q1JJ58 Q1JJ59 Q1JJ60 Q1JJ61 Q1JJ62 Q1JJ63 Q1JJ71 Q1JJ80 Q1JJ82 Q1JJ89
Q1JJA3 Q1JJA7 Q1JJB0 Q1JJC0 Q1JJD5 Q1JJD7 Q1JJD9 Q1JJE2 Q1JJF9 Q1JJG0 Q1JJG7
Q1JJH0 Q1JJH1 Q1JJH3 Q1JJH5 Q1JJH6 Q1JJH8 Q1JJH9 Q1JJI0 Q1JJI7 Q1JJJ4 Q1JJJ5
Q1JJJ7 Q1JJK1 Q1JJK6 Q1JJL5 Q1JJM4 Q1JJU1 Q1JJU8 Q1JJW2 Q1JJW3 Q1JJW6 Q1JJW9
Q1JJX5 Q1JJY9 Q1JJZ6 Q1JJZ7 Q1JK01 Q1JK16 Q1JK21 Q1JK23 Q1JK27 Q1JK32 Q1JK37
Q1JK40 Q1JK41 Q1JK46 Q1JK51 Q1JK54 Q1JK63 Q1JK67 Q1JK76 Q1JK78 Q1JK85 Q1JK87
Q1JKA0 Q1JKC7 Q1JKC8 Q1JKD4 Q1JKD6 Q1JKF1 Q1JKG6 Q1JKG7 Q1JKH1 Q1JKH2 Q1JKJ9
Q1JKK0 Q1JKK1 Q1JKK3 Q1JKK8 Q1JKL3 Q1JKL4 Q1JKL7 Q1JKN2 Q1JKN3 Q1JKN4 Q1JKP6
Q1JKP7 Q1JKP9 Q1JKR7 Q1JKR8 Q1JKT5 Q1JKW3 Q1JKW5 Q1JKW9 Q1JKX1 Q1JKX6 Q1JKY4
Q1JKZ0 Q1JL04 Q1JL06 Q1JL20 Q1JL26 Q1JL27 Q1JL31 Q1JL34 Q1JL45 Q1JL53 Q1JL80
Q1JL87 Q1JL88 Q1JLA1 Q1JLB5 Q1JLB7 Q1JLD9 Q1JLG7 Q1JLI3 Q1JLI4 Q1JLK2 Q1JLL1
Q1JLL7 Q1JLM2 Q1JLN1 Q1JLP1 Q1JLP8 Q1JLQ2 Q1JLQ7 Q1JLQ8 Q1JLR5 Q1JLS1 Q1JLS3
Q1JLT8 Q1JLX3 Q1JLX6 Q1JM03 Q1JM38 Q1JM42 Q1JM45 Q1JM51 Q1JM60 Q1JM64 Q1JM65
Q1JM69 Q1JM72 Q1JM73 Q1JM76 Q1JM77 Q1JM80 Q1JM96 Q1JMB2 Q1JMB3 Q1JMB4 Q1JMC0
Q1JMC8 Q1JMD0 Q1JMD6 Q1JMD8 Q1JME3 Q1JME4 Q1JME5 Q1JME8 Q1JME9 Q1JMF0 Q1JMF1
Q1JMF5 Q1JMH3 Q1JMH5 Q1JMH6 Q1JMJ7 Q1JMM0 Q1JMM5 Q1JMN0 Q1JMN2 Q1JMN3 Q1JMR2
Q1JMR3 Q1JMX4 Q1JMY5 Q1JMZ5 Q1JMZ7 Q1JN00 Q1JN23 Q1JN28 Q1JN29 Q1JN31 Q1JN32
Q1JN33 Q1JN34 Q1JN60 Q1JN74 Q1JN82 Q1JN83 Q1JN85 Q1JN87 Q1JNA3 Q1JNA4 Q1JNA5
Q1JNA9 Q1JNC1 Q1JNC4 Q1JNC7 Q1JND4 Q1JNE3 Q1JNE5 Q1JNH7 Q1JNH8 Q1JNH9 Q1JNI8
Q1JNJ8 Q1JNJ9 Q1JNK3 Q1JNK7 Q1JNM4 Q1JNN4 Q1JNR2 Q1JNS3 Q1JNV5 Q1JNW6 Q1JNX6
Q1JNX8 Q1JNY0 Q1JNZ0 Q1JNZ1 Q1JNZ2 Q1JNZ3 Q1JNZ4 Q1JNZ5 Q1JNZ6 Q1JNZ8 Q1JNZ9
Q1JP00 Q1JP01 Q1JP02 Q1JP03 Q1JP04 Q1JP05 Q1JP06 Q1JP07 Q1JP08 Q1JP09 Q1JP10
Q1JP11 Q1JP12 Q1JP13 Q1JP14 Q1JP15 Q1JP16 Q1JP17 Q1JP18 Q1JP25 Q1JP34 Q1JP36
Q1JP42 Q1JP56 Q1JP60 Q1JP61 Q1JP73 Q1JP79 Q1JPA6 Q1JPD3 Q1JPD6 Q1JPG1 Q1JPH6
Q1JPJ0 Q1JPJ2 Q1JPJ8 Q1JPJ9 Q1JPL0 Q1JPN0 Q1JPP8 Q1JPS1 Q1JPT7 Q1JPX3 Q1JPX4
Q1JPY4 Q1JPY8 Q1JPZ7 Q1JQ73 Q1JQ92 Q1JQ94 Q1JQA1 Q1JQA8 Q1JQB5 Q1JQD2 Q1JQD4
Q1JQD6 Q1JQD7 Q1JQD9 Q1JQE0 Q1JQE2 Q1JQN1 Q1JU68 Q1JU72 Q1JUP4 Q1JUP5 Q1JUQ0

Q1JUQ1 Q1JZW3 Q1K4R6 Q1K615 Q1K6I4 Q1K6U0 Q1K7T5 Q1K8E7 Q1K8F6 Q1K8F7 Q1K8G0
Q1K9C2 Q1K9C4 Q1KKR6 Q1KKR7 Q1KKR8 Q1KKS7 Q1KKS8 Q1KKS9 Q1KKT0 Q1KKT1 Q1KKT2
Q1KKU6 Q1KKU7 Q1KKV1 Q1KKV2 Q1KKV3 Q1KKV4 Q1KKW8 Q1KKX0 Q1KKX1 Q1KKX5 Q1KKX7
Q1KKX9 Q1KKY0 Q1KKY1 Q1KKY2 Q1KKY3 Q1KKZ1 Q1KKZ2 Q1KKZ3 Q1KKZ4 Q1KKZ5 Q1KKZ6
Q1KL10 Q1KL11 Q1KL12 Q1KL13 Q1KL14 Q1KL16 Q1KL17 Q1KL22 Q1KLK0 Q1KLK1 Q1KLZ1
Q1KLZ2 Q1KMR4 Q1KS79 Q1KSC7 Q1KVR0 Q1KVR1 Q1KVR2 Q1KVR3 Q1KVR6 Q1KVR9 Q1KVS0
Q1KVS9 Q1KVT1 Q1KVT3 Q1KVT5 Q1KVT6 Q1KVT7 Q1KVT8 Q1KVT9 Q1KVU3 Q1KVU9 Q1KVV0
Q1KVV1 Q1KVV7 Q1KVV8 Q1KVV9 Q1KVW2 Q1KVW4 Q1KVX0 Q1KVX4 Q1KVX5 Q1KVX6 Q1KVX7
Q1KVX8 Q1KXP4 Q1KXP5 Q1KXP6 Q1KXP8 Q1KXP9 Q1KXQ1 Q1KXR0 Q1KXR8 Q1KXR9 Q1KXS0
Q1KXS1 Q1KXS2 Q1KXS3 Q1KXS4 Q1KXS5 Q1KXS6 Q1KXS7 Q1KXT4 Q1KXT6 Q1KXT7 Q1KXT8
Q1KXV7 Q1KXW1 Q1KXW9 Q1KXX0 Q1KXX1 Q1KXX2 Q1KXX7 Q1KXX8 Q1KXY0 Q1KYT0 Q1KZ55
Q1KZ57 Q1KZ58 Q1KZ59 Q1L5Z9 Q1L673 Q1L6A3 Q1L721 Q1L899 Q1L8G6 Q1L8G7 Q1L8H0
Q1L8I0 Q1L8J7 Q1L8L9 Q1L8T5 Q1L8W0 Q1L8X2 Q1L8Y6 Q1L981 Q1L987 Q1L994 Q1L9C7
Q1LAN7 Q1LB18 Q1LC96 Q1LCQ8 Q1LCQ9 Q1LCS4 Q1LCU1 Q1LFU5 Q1LFW3 Q1LH07 Q1LH12
Q1LH94 Q1LHJ8 Q1LHJ9 Q1LHL3 Q1LHL9 Q1LHM2 Q1LHM4 Q1LHS4 Q1LI13 Q1LI16 Q1LI17
Q1LI18 Q1LI19 Q1LI20 Q1LI21 Q1LI27 Q1LI28 Q1LI29 Q1LI31 Q1LI37 Q1LI38 Q1LI39
Q1LI40 Q1LI41 Q1LI42 Q1LI43 Q1LI44 Q1LI45 Q1LI46 Q1LI47 Q1LI48 Q1LI49 Q1LI50
Q1LI51 Q1LI52 Q1LI53 Q1LI54 Q1LI55 Q1LI56 Q1LI58 Q1LI59 Q1LI60 Q1LI61 Q1LI62
Q1LI63 Q1LI64 Q1LI69 Q1LI79 Q1LI88 Q1LIA4 Q1LIA5 Q1LIA8 Q1LIB1 Q1LIB2 Q1LIB3
Q1LIB4 Q1LIF3 Q1LIF4 Q1LIH0 Q1LIH4 Q1LIL8 Q1LIM4 Q1LIM7 Q1LIM8 Q1LIP2 Q1LIP8
Q1LIP9 Q1LIQ0 Q1LIQ1 Q1LIQ4 Q1LIS6 Q1LIT8 Q1LIU6 Q1LIV4 Q1LJ29 Q1LJ30 Q1LJ33
Q1LJ45 Q1LJ59 Q1LJ60 Q1LJ80 Q1LJ82 Q1LJ86 Q1LJ92 Q1LJ94 Q1LJ99 Q1LJA1 Q1LJA2
Q1LJA4 Q1LJB0 Q1LJB8 Q1LJD2 Q1LJD3 Q1LJF1 Q1LJJ6 Q1LJM9 Q1LJN4 Q1LJP8 Q1LJQ2
Q1LJR0 Q1LJR1 Q1LJR3 Q1LJU9 Q1LJV8 Q1LJV9 Q1LJX2 Q1LJY1 Q1LK00 Q1LK07 Q1LK22
Q1LK34 Q1LK35 Q1LK37 Q1LK40 Q1LK68 Q1LK78 Q1LK81 Q1LKE3 Q1LKG0 Q1LKH7 Q1LKI4
Q1LKJ4 Q1LKJ7 Q1LKK7 Q1LKL5 Q1LKL7 Q1LKN1 Q1LKN3 Q1LKN4 Q1LKN7 Q1LKN9 Q1LL27
Q1LL89 Q1LL91 Q1LLA8 Q1LLB1 Q1LLC3 Q1LLF7 Q1LLG0 Q1LLG9 Q1LLH0 Q1LLI5 Q1LLI7
Q1LLI8 Q1LLJ5 Q1LLJ6 Q1LLK0 Q1LLK1 Q1LLK8 Q1LLM6 Q1LLR4 Q1LLR6 Q1LLR7 Q1LLR8
Q1LLW8 Q1LLX0 Q1LLX1 Q1LLZ4 Q1LM60 Q1LM61 Q1LM62 Q1LM63 Q1LM68 Q1LM83 Q1LMB9
Q1LMC7 Q1LMQ4 Q1LN82 Q1LN92 Q1LND0 Q1LND7 Q1LNE1 Q1LNE4 Q1LNF2 Q1LNF5 Q1LNF6
Q1LNF7 Q1LNF8 Q1LNG0 Q1LNG5 Q1LNH0 Q1LNH3 Q1LNJ3 Q1LNM2 Q1LNN2 Q1LNS3 Q1LNS4
Q1LP24 Q1LP26 Q1LP48 Q1LP76 Q1LP77 Q1LP78 Q1LPA2 Q1LPE1 Q1LPG5 Q1LPH7 Q1LPI8
Q1LPK6 Q1LPL1 Q1LPN6 Q1LPN9 Q1LPR2 Q1LPS6 Q1LPS7 Q1LPT0 Q1LPT2 Q1LPT3 Q1LPW7
Q1LPW9 Q1LPX2 Q1LPX7 Q1LQ00 Q1LQ15 Q1LQ33 Q1LQ34 Q1LQ38 Q1LQ52 Q1LQ59 Q1LQ72
Q1LQ98 Q1LQA5 Q1LQB0 Q1LQB1 Q1LQD7 Q1LQD9 Q1LQE0 Q1LQE1 Q1LQG2 Q1LQG7 Q1LQG9
Q1LQJ5 Q1LQM3 Q1LQS4 Q1LQS5 Q1LR06 Q1LR08 Q1LR36 Q1LR54 Q1LR74 Q1LR87 Q1LR89
Q1LR92 Q1LRA1 Q1LRB3 Q1LRB4 Q1LRB6 Q1LRB7 Q1LRC2 Q1LRC5 Q1LRC6 Q1LRC9 Q1LRD0
Q1LRD4 Q1LRG2 Q1LRG9 Q1LRI7 Q1LRM9 Q1LRP1 Q1LRP3 Q1LRP7 Q1LRQ0 Q1LRQ2 Q1LRQ3
Q1LRQ5 Q1LRT6 Q1LRV9 Q1LRY0 Q1LS06 Q1LS11 Q1LS20 Q1LS25 Q1LS28 Q1LS34 Q1LS37
Q1LS40 Q1LS47 Q1LS50 Q1LS59 Q1LS73 Q1LS75 Q1LS79 Q1LS86 Q1LS94 Q1LSB6 Q1LSC6
Q1LSC9 Q1LSE0 Q1LSE1 Q1LSE4 Q1LSF5 Q1LSF7 Q1LSI9 Q1LSJ1 Q1LSJ4 Q1LSJ6 Q1LSJ7
Q1LSJ8 Q1LSJ9 Q1LSK1 Q1LSK2 Q1LSK5 Q1LSK8 Q1LSK9 Q1LSL0 Q1LSL1 Q1LSL2 Q1LSL6
Q1LSL7 Q1LSM0 Q1LSM1 Q1LSM6 Q1LSM8 Q1LSN4 Q1LSP0 Q1LSP1 Q1LSP4 Q1LSP5 Q1LSP7
Q1LSP8 Q1LSQ0 Q1LSQ2 Q1LSQ3 Q1LSQ5 Q1LSQ9 Q1LSR2 Q1LSR4 Q1LSR5 Q1LSR7 Q1LSS2

Q1LSS3 Q1LSS7 Q1LSS9 Q1LST1 Q1LST3 Q1LSU0 Q1LSU2 Q1LSV3 Q1LSV4 Q1LSV5 Q1LSV6
Q1LSV7 Q1LSV9 Q1LSW0 Q1LSW4 Q1LSW6 Q1LSX6 Q1LSX7 Q1LSX8 Q1LSX9 Q1LSY0 Q1LSY1
Q1LSY4 Q1LSY5 Q1LSY6 Q1LSY7 Q1LSZ3 Q1LSZ6 Q1LSZ8 Q1LT01 Q1LT02 Q1LT04 Q1LT06
Q1LT07 Q1LT18 Q1LT19 Q1LT27 Q1LT29 Q1LT35 Q1LT42 Q1LT51 Q1LT55 Q1LT56 Q1LT57
Q1LT59 Q1LT63 Q1LT66 Q1LT68 Q1LT69 Q1LT71 Q1LT72 Q1LT75 Q1LT76 Q1LT77 Q1LT82
Q1LT88 Q1LTA7 Q1LTA9 Q1LTB2 Q1LTB3 Q1LTB4 Q1LTB5 Q1LTB6 Q1LTB7 Q1LTB9 Q1LTC0
Q1LTC1 Q1LTC2 Q1LTC4 Q1LTC5 Q1LTC6 Q1LTC7 Q1LTC8 Q1LTC9 Q1LTD0 Q1LTD1 Q1LTD2
Q1LTD3 Q1LTD4 Q1LTD5 Q1LTD6 Q1LTD7 Q1LTD8 Q1LTD9 Q1LTE4 Q1LTE8 Q1LTE9 Q1LTF0
Q1LTF1 Q1LTF2 Q1LTF3 Q1LTG5 Q1LTG7 Q1LTG8 Q1LTH1 Q1LTH2 Q1LTH3 Q1LTH5 Q1LTH8
Q1LTI8 Q1LTI9 Q1LTJ8 Q1LTJ9 Q1LTK0 Q1LTK2 Q1LTK5 Q1LTK6 Q1LTK8 Q1LTL1 Q1LTL2
Q1LTL3 Q1LTL5 Q1LTL8 Q1LTM3 Q1LTM4 Q1LTM7 Q1LTM9 Q1LTN0 Q1LTN1 Q1LTN5 Q1LTN6
Q1LTN7 Q1LTP1 Q1LTP2 Q1LTP3 Q1LTP4 Q1LTP5 Q1LTP8 Q1LTQ0 Q1LTQ3 Q1LTQ4 Q1LTQ5
Q1LTQ6 Q1LTQ9 Q1LTR0 Q1LTR1 Q1LTR2 Q1LTS1 Q1LTS4 Q1LTS5 Q1LTS6 Q1LTT2 Q1LTT4
Q1LTT5 Q1LTT6 Q1LTT8 Q1LTU1 Q1LTU2 Q1LTU4 Q1LTU6 Q1LTU7 Q1LTV8 Q1LTW1 Q1LTW2
Q1LTX0 Q1LTX1 Q1LTX2 Q1LTX5 Q1LTX6 Q1LTX7 Q1LTY0 Q1LTY2 Q1LTY3 Q1LTZ4 Q1LU01
Q1LU04 Q1LU10 Q1LU15 Q1LU20 Q1LU21 Q1LU22 Q1LU25 Q1LU41 Q1LU45 Q1LU51 Q1LU53
Q1LU54 Q1LU57 Q1LU61 Q1LU62 Q1LU68 Q1LU70 Q1LU72 Q1LU73 Q1LU74 Q1LU77 Q1LU78
Q1LU81 Q1LU84 Q1LU93 Q1LUA8 Q1LUC1 Q1LUC3 Q1LUE5 Q1LUI2 Q1LUS5 Q1LUT1 Q1LV15
Q1LV17 Q1LV19 Q1LV22 Q1LVE8 Q1LVF3 Q1LVK9 Q1LVN8 Q1LVP6 Q1LVQ7 Q1LWB0 Q1LWG3
Q1LWH4 Q1LWJ6 Q1LWL6 Q1LWL8 Q1LWM5 Q1LWV4 Q1LWX3 Q1LWZ0 Q1LX49 Q1LXC9 Q1LXE6
Q1LXI5 Q1LXK4 Q1LXK5 Q1LXS2 Q1LXZ7 Q1LXZ9 Q1LY10 Q1LY51 Q1LY77 Q1LY84 Q1LYE3
Q1LYG0 Q1LYL8 Q1LYM6 Q1LZ08 Q1LZ50 Q1LZ53 Q1LZ74 Q1LZ78 Q1LZ79 Q1LZ80 Q1LZ83
Q1LZ87 Q1LZ94 Q1LZ96 Q1LZA1 Q1LZA3 Q1LZB4 Q1LZB6 Q1LZC0 Q1LZC9 Q1LZD3 Q1LZD9
Q1LZE0 Q1LZE1 Q1LZE2 Q1LZE8 Q1LZF1 Q1LZF2 Q1LZF3 Q1LZG6 Q1LZH0 Q1LZH5 Q1M0P5
Q1M3A7 Q1M3H2 Q1M4A0 Q1M4A1 Q1M4T3 Q1M5J2 Q1M5U3 Q1M5X7 Q1M667 Q1M7F3 Q1M7F4
Q1M813 Q1M901 Q1M964 Q1MA15 Q1MA18 Q1MA19 Q1MA20 Q1MA25 Q1MA35 Q1MA50 Q1MA52
Q1MA67 Q1MA73 Q1MA74 Q1MA75 Q1MA76 Q1MA80 Q1MA81 Q1MAC5 Q1MAD1 Q1MAE9 Q1MAF5
Q1MAI6 Q1MAJ9 Q1MAK2 Q1MAK3 Q1MAK4 Q1MAK5 Q1MAM8 Q1MAN2 Q1MAN9 Q1MAP9 Q1MAQ8
Q1MAW1 Q1MB26 Q1MB46 Q1MB56 Q1MB73 Q1MBA9 Q1MBB2 Q1MBD4 Q1MBE0 Q1MBK9 Q1MBL0
Q1MBL8 Q1MBM8 Q1MBR4 Q1MBS7 Q1MBS8 Q1MBS9 Q1MBW8 Q1MBX8 Q1MC25 Q1MC31 Q1MC52
Q1MC53 Q1MC56 Q1MC59 Q1MC77 Q1MC84 Q1MCF8 Q1MCQ8 Q1MCS2 Q1MCV4 Q1MCV5 Q1MCV6
Q1MCV9 Q1MCW2 Q1MCW3 Q1MCW4 Q1MDD6 Q1MDE8 Q1MDG4 Q1MDH2 Q1MDH4 Q1MDH6 Q1MDJ5
Q1MDK6 Q1MDL5 Q1MDR7 Q1MDT5 Q1MDX5 Q1ME24 Q1ME25 Q1ME31 Q1ME34 Q1ME35 Q1ME47
Q1ME53 Q1ME82 Q1ME89 Q1MED4 Q1MEZ8 Q1MFE8 Q1MFI6 Q1MFK4 Q1MFQ6 Q1MFT6 Q1MFT8
Q1MFZ6 Q1MFZ8 Q1MFZ9 Q1MG11 Q1MG12 Q1MG13 Q1MG23 Q1MG27 Q1MG28 Q1MG48 Q1MG62
Q1MG80 Q1MGA3 Q1MGA7 Q1MGC4 Q1MGE1 Q1MGE2 Q1MGI5 Q1MGM3 Q1MGM6 Q1MGZ1 Q1MH21
Q1MH26 Q1MH51 Q1MH52 Q1MH53 Q1MH54 Q1MHI1 Q1MHI4 Q1MHI9 Q1MHJ2 Q1MHJ5 Q1MHJ6
Q1MHJ7 Q1MHJ8 Q1MHK3 Q1MHK9 Q1MHL2 Q1MHM1 Q1MHM2 Q1MHN1 Q1MHV8 Q1MHW6 Q1MI52
Q1MIB2 Q1MIB6 Q1MIB7 Q1MIB8 Q1MIB9 Q1MIC2 Q1MIC3 Q1MIC4 Q1MIC5 Q1MIC6 Q1MIC7
Q1MIC8 Q1MIC9 Q1MID0 Q1MID1 Q1MID2 Q1MID3 Q1MID4 Q1MID5 Q1MID7 Q1MID8 Q1MID9
Q1MIE0 Q1MIE1 Q1MIE2 Q1MIE3 Q1MIE4 Q1MIE5 Q1MIE6 Q1MIE8 Q1MIE9 Q1MIF0 Q1MIF1
Q1MIF3 Q1MIF4 Q1MIH6 Q1MII4 Q1MIJ6 Q1MIM6 Q1MIM7 Q1MIP2 Q1MIP3 Q1MIP7 Q1MIS5
Q1MIT2 Q1MIU5 Q1MIV1 Q1MIW0 Q1MIW9 Q1MIX8 Q1MIY5 Q1MIZ4 Q1MJ01 Q1MJ02 Q1MJ10
Q1MJ11 Q1MJ13 Q1MJ22 Q1MJ50 Q1MJ54 Q1MJ60 Q1MJ62 Q1MJF2 Q1MJG9 Q1MJK8 Q1MJR4

Q1MJU3 Q1MJU4 Q1MJV3 Q1MK43 Q1MKH3 Q1MKH6 Q1MKH7 Q1MKH8 Q1MKJ4 Q1MKK1 Q1MKN4
Q1MKQ6 Q1MKS2 Q1MKS7 Q1MKU7 Q1MKX4 Q1MKX9 Q1MKY0 Q1ML15 Q1MLB6 Q1MLC7 Q1MLG6
Q1MLL4 Q1MLP1 Q1MLP2 Q1MLP3 Q1MLP5 Q1MLP8 Q1MLV9 Q1MM06 Q1MM09 Q1MM16 Q1MM57
Q1MM59 Q1MM66 Q1MM74 Q1MM75 Q1MM94 Q1MMA5 Q1MMB6 Q1MMB8 Q1MMC3 Q1MMC9 Q1MMD0
Q1MMD1 Q1MMD6 Q1MMD7 Q1MMD9 Q1MME0 Q1MMG6 Q1MMH6 Q1MMI1 Q1MMJ7 Q1MMN0 Q1MMP3
Q1MMP4 Q1MMP5 Q1MMV6 Q1MMV7 Q1MMV8 Q1MMY1 Q1MMY4 Q1MN11 Q1MN12 Q1MN15 Q1MN32
Q1MN36 Q1MN39 Q1MN40 Q1MN41 Q1MN43 Q1MN44 Q1MN56 Q1MNB0 Q1MNB3 Q1MNB5 Q1MNB6
Q1MNC1 Q1MNC6 Q1MND5 Q1MND6 Q1MND7 Q1MNF1 Q1MNF5 Q1MNF6 Q1MNF9 Q1MNG0 Q1MNG2
Q1MP36 Q1MP37 Q1MP42 Q1MP91 Q1MPA2 Q1MPA7 Q1MPB8 Q1MPC1 Q1MPC4 Q1MPC6 Q1MPD2
Q1MPD3 Q1MPF1 Q1MPG1 Q1MPH2 Q1MPH5 Q1MPK8 Q1MPL2 Q1MPL4 Q1MPL6 Q1MPL7 Q1MPN3
Q1MPN4 Q1MPP0 Q1MPP1 Q1MPP2 Q1MPP3 Q1MPP4 Q1MPP7 Q1MPP8 Q1MPP9 Q1MPQ0 Q1MPQ1
Q1MPQ2 Q1MPQ3 Q1MPQ4 Q1MPQ5 Q1MPQ6 Q1MPQ7 Q1MPQ8 Q1MPQ9 Q1MPR0 Q1MPR1 Q1MPR2
Q1MPR3 Q1MPR4 Q1MPS7 Q1MPS9 Q1MPT0 Q1MPT1 Q1MPT8 Q1MPU0 Q1MPU1 Q1MPU2 Q1MPU3
Q1MPU5 Q1MPW1 Q1MPW3 Q1MPW7 Q1MPW8 Q1MPW9 Q1MPX3 Q1MPX7 Q1MPY2 Q1MPY8 Q1MPZ0
Q1MPZ3 Q1MPZ8 Q1MPZ9 Q1MQ25 Q1MQ35 Q1MQ47 Q1MQ52 Q1MQ55 Q1MQ73 Q1MQ78 Q1MQ79
Q1MQ80 Q1MQ89 Q1MQB1 Q1MQB3 Q1MQD5 Q1MQD6 Q1MQD8 Q1MQE1 Q1MQG9 Q1MQH1 Q1MQH2
Q1MQJ1 Q1MQJ3 Q1MQJ5 Q1MQL6 Q1MQL7 Q1MQP8 Q1MQP9 Q1MQQ0 Q1MQQ1 Q1MQQ4 Q1MQQ6
Q1MQR5 Q1MQV0 Q1MQW3 Q1MQW4 Q1MQY8 Q1MQZ0 Q1MR01 Q1MR19 Q1MR44 Q1MR60 Q1MR63
Q1MR71 Q1MR76 Q1MR87 Q1MR90 Q1MR97 Q1MRA1 Q1MRB3 Q1MRD0 Q1MRD4 Q1MRD7 Q1MRD8
Q1MRE2 Q1MRE3 Q1MRE5 Q1MRF1 Q1MRF2 Q1MRG7 Q1MRG8 Q1MRI6 Q1MRI7 Q1MRJ6 Q1MRJ8
Q1MRK3 Q1MRK9 Q1MRL0 Q1MRL9 Q1MRM0 Q1MRM1 Q1MRM4 Q1MRN8 Q1MRN9 Q1MRP0 Q1MRQ0
Q1MRQ1 Q1MRQ2 Q1MRS4 Q1MRS7 Q1MRS8 Q1MRT4 Q1MRU1 Q1MRU5 Q1MRU7 Q1MRU8 Q1MRU9
Q1MRV5 Q1MRV6 Q1MRV8 Q1MRW7 Q1MRW8 Q1MRX8 Q1MS11 Q1MS15 Q1MS16 Q1MS17 Q1MS21
Q1MS56 Q1MS79 Q1MSA1 Q1MSA2 Q1MSD5 Q1MSE5 Q1MSG7 Q1MSG8 Q1MTD3 Q1MTI4 Q1MTM9
Q1MTN3 Q1MTN7 Q1MTN8 Q1MTN9 Q1MTP1 Q1MTQ0 Q1MTQ1 Q1MTQ7 Q1MTQ9 Q1MTR1 Q1MTR2
Q1MTR3 Q1MTR4 Q1MTS0 Q1MW86 Q1NAJ2 Q1NEI6 Q1NZ26 Q1PCB0 Q1PCB1 Q1PD52 Q1PD53
Q1PD54 Q1PD56 Q1PDC4 Q1PDC6 Q1PDC9 Q1PDD0 Q1PDI2 Q1PDL6 Q1PDP9 Q1PDT6 Q1PDV2
Q1PE04 Q1PE09 Q1PE10 Q1PE27 Q1PE49 Q1PE51 Q1PE57 Q1PE68 Q1PED2 Q1PED9 Q1PEF6
Q1PEL7 Q1PEN2 Q1PEN8 Q1PEP5 Q1PER6 Q1PES7 Q1PEU9 Q1PEW8 Q1PEY8 Q1PEZ8 Q1PF16
Q1PF17 Q1PF20 Q1PFA4 Q1PFC2 Q1PFC5 Q1PFD1 Q1PFG1 Q1PFH7 Q1PFI4 Q1PFK0 Q1PFN9
Q1PFQ9 Q1PFR7 Q1PFW3 Q1PRL4 Q1PS67 Q1PSI9 Q1PSW8 Q1Q7V4 Q1Q7V5 Q1Q7W4 Q1Q7X0
Q1Q803 Q1Q809 Q1Q833 Q1Q835 Q1Q845 Q1Q846 Q1Q883 Q1Q8A4 Q1Q8A5 Q1Q8A6 Q1Q8A7
Q1Q8B0 Q1Q8B2 Q1Q8B6 Q1Q8C5 Q1Q8C6 Q1Q8C9 Q1Q8F6 Q1Q8I2 Q1Q8I3 Q1Q8I4 Q1Q8I5
Q1Q8I7 Q1Q8J1 Q1Q8J2 Q1Q8J3 Q1Q8K9 Q1Q8L9 Q1Q8M8 Q1Q8N2 Q1Q8N9 Q1Q8P0 Q1Q8P1
Q1Q8P2 Q1Q8P5 Q1Q8P6 Q1Q8P7 Q1Q8P8 Q1Q8P9 Q1Q8Q0 Q1Q8Q8 Q1Q8Q9 Q1Q8S2 Q1Q8S6
Q1Q8V7 Q1Q8Y4 Q1Q929 Q1Q932 Q1Q945 Q1Q955 Q1Q962 Q1Q963 Q1Q972 Q1Q973 Q1Q974
Q1Q978 Q1Q986 Q1Q987 Q1Q994 Q1Q999 Q1Q9C6 Q1Q9E2 Q1Q9E3 Q1Q9E4 Q1Q9K1 Q1Q9K8
Q1Q9L2 Q1Q9M0 Q1Q9M6 Q1Q9X1 Q1Q9X2 Q1Q9X9 Q1Q9Y8 Q1QA11 Q1QA12 Q1QA13 Q1QA16
Q1QA23 Q1QA43 Q1QA44 Q1QA45 Q1QA46 Q1QA56 Q1QA59 Q1QA75 Q1QA78 Q1QA86 Q1QA88
Q1QA91 Q1QAD0 Q1QAD8 Q1QAF2 Q1QAF3 Q1QAF5 Q1QAF7 Q1QAG2 Q1QAL9 Q1QAN0 Q1QAP5
Q1QAQ4 Q1QAR3 Q1QAR8 Q1QAW3 Q1QAW5 Q1QAX7 Q1QAY1 Q1QB42 Q1QBB6 Q1QBB9 Q1QBF3
Q1QBI9 Q1QBM0 Q1QBM3 Q1QBM4 Q1QBT8 Q1QBW5 Q1QBY7 Q1QBZ0 Q1QBZ1 Q1QBZ2 Q1QC15
Q1QC16 Q1QC19 Q1QC33 Q1QC34 Q1QC36 Q1QC40 Q1QCD8 Q1QCE0 Q1QCE1 Q1QCE2 Q1QCE3
Q1QCH2 Q1QCI5 Q1QCJ2 Q1QCL3 Q1QCL5 Q1QCL7 Q1QCM0 Q1QCM1 Q1QCP2 Q1QCP3 Q1QCP8

Q1QCS6 Q1QCU5 Q1QCY5 Q1QCY6 Q1QCY9 Q1QCZ9 Q1QD10 Q1QD14 Q1QD17 Q1QD18 Q1QD22
Q1QD23 Q1QD25 Q1QD26 Q1QD30 Q1QD80 Q1QDA0 Q1QDA2 Q1QDA4 Q1QDC1 Q1QDC8 Q1QDC9
Q1QDD0 Q1QDD9 Q1QDE0 Q1QDE6 Q1QDF6 Q1QDG0 Q1QDG1 Q1QDG2 Q1QDG3 Q1QDG4 Q1QDG5
Q1QDG7 Q1QDG8 Q1QDG9 Q1QDH0 Q1QDH1 Q1QDH2 Q1QDH3 Q1QDH4 Q1QDH5 Q1QDH6 Q1QDH7
Q1QDH8 Q1QDH9 Q1QDI0 Q1QDI1 Q1QDI2 Q1QDI3 Q1QDI4 Q1QDI5 Q1QDI6 Q1QDI7 Q1QDI9
Q1QDJ4 Q1QDJ5 Q1QDK2 Q1QDK7 Q1QDK8 Q1QDL4 Q1QDM9 Q1QDN7 Q1QDP2 Q1QDP8 Q1QDQ5
Q1QDQ9 Q1QDT0 Q1QDT1 Q1QDU2 Q1QDV0 Q1QDW1 Q1QDW8 Q1QDX4 Q1QDZ7 Q1QE01 Q1QE02
Q1QE18 Q1QE21 Q1QE34 Q1QE46 Q1QE71 Q1QE74 Q1QE85 Q1QE88 Q1QE99 Q1QEB6 Q1QEC2
Q1QED0 Q1QED3 Q1QED5 Q1QED6 Q1QED7 Q1QEE8 Q1QEF7 Q1QEF9 Q1QEG9 Q1QEI9 Q1QEJ2
Q1QEJ3 Q1QEJ4 Q1QEK1 Q1QEK5 Q1QEM4 Q1QEM5 Q1QEM6 Q1QEN9 Q1QEP0 Q1QEP3 Q1QEP4
Q1QEP5 Q1QEP7 Q1QEP9 Q1QEQ3 Q1QEQ5 Q1QEQ7 Q1QEQ8 Q1QES5 Q1QET1 Q1QET5 Q1QEW1
Q1QEW7 Q1QFN5 Q1QFS0 Q1QGD4 Q1QGD7 Q1QGE2 Q1QH19 Q1QH22 Q1QH29 Q1QH30 Q1QH66
Q1QH69 Q1QH71 Q1QH73 Q1QH74 Q1QH76 Q1QH77 Q1QH78 Q1QHD2 Q1QHD7 Q1QHE0 Q1QHE3
Q1QHE6 Q1QHH8 Q1QHI4 Q1QHI6 Q1QHI7 Q1QHI8 Q1QHI9 Q1QHJ3 Q1QHK3 Q1QHL8 Q1QHM3
Q1QHP3 Q1QHP5 Q1QHP7 Q1QHU1 Q1QI02 Q1QI14 Q1QIH5 Q1QIJ1 Q1QIJ6 Q1QIL6 Q1QIM4
Q1QIP3 Q1QIR5 Q1QIS8 Q1QIV7 Q1QIV8 Q1QJ74 Q1QJ95 Q1QJC0 Q1QJD3 Q1QJD6 Q1QJF3
Q1QJL1 Q1QJR5 Q1QJT8 Q1QJU5 Q1QJU8 Q1QJV3 Q1QK71 Q1QKB6 Q1QKC3 Q1QKK3 Q1QKL6
Q1QKN4 Q1QKP2 Q1QKU9 Q1QKV0 Q1QKV7 Q1QKW0 Q1QKW6 Q1QL03 Q1QL23 Q1QL26 Q1QL28
Q1QL30 Q1QL34 Q1QL43 Q1QL45 Q1QL49 Q1QL57 Q1QL63 Q1QL75 Q1QL76 Q1QL77 Q1QLA8
Q1QLB2 Q1QLB8 Q1QLD8 Q1QLD9 Q1QLH8 Q1QLI2 Q1QLI8 Q1QLI9 Q1QLQ7 Q1QLT8 Q1QM39
Q1QM87 Q1QM92 Q1QM99 Q1QMA2 Q1QMA3 Q1QMB4 Q1QMB5 Q1QMB8 Q1QMB9 Q1QMC9 Q1QME1
Q1QME2 Q1QMG7 Q1QMH5 Q1QMH6 Q1QMJ0 Q1QMJ4 Q1QMJ8 Q1QMJ9 Q1QMM1 Q1QMM2 Q1QMN1
Q1QMN5 Q1QMN6 Q1QMN7 Q1QMP2 Q1QMP4 Q1QMP7 Q1QMS2 Q1QMT5 Q1QMU2 Q1QMU3 Q1QMV7
Q1QMW0 Q1QMW2 Q1QMY9 Q1QN05 Q1QN06 Q1QN07 Q1QN08 Q1QN09 Q1QN11 Q1QN12 Q1QN13
Q1QN14 Q1QN15 Q1QN16 Q1QN17 Q1QN18 Q1QN19 Q1QN20 Q1QN21 Q1QN22 Q1QN23 Q1QN24
Q1QN25 Q1QN26 Q1QN27 Q1QN28 Q1QN29 Q1QN30 Q1QN31 Q1QN32 Q1QN33 Q1QN34 Q1QN35
Q1QN43 Q1QN44 Q1QN45 Q1QN46 Q1QN47 Q1QN48 Q1QN61 Q1QN87 Q1QN95 Q1QN98 Q1QNB1
Q1QNH1 Q1QNQ1 Q1QNS3 Q1QNT2 Q1QNU0 Q1QNU2 Q1QNU5 Q1QNV1 Q1QNY6 Q1QNZ8 Q1QP05
Q1QP32 Q1QP77 Q1QP86 Q1QP91 Q1QPC1 Q1QPC2 Q1QPM0 Q1QPR8 Q1QPW6 Q1QQ39 Q1QQ40
Q1QQ46 Q1QQ51 Q1QQB5 Q1QQC7 Q1QQD3 Q1QQD5 Q1QQG6 Q1QQI4 Q1QQI9 Q1QQJ4 Q1QQJ9
Q1QQP5 Q1QQP6 Q1QQR2 Q1QQR3 Q1QQT4 Q1QQT7 Q1QQT9 Q1QQU0 Q1QQU2 Q1QQU3 Q1QQV5
Q1QQW6 Q1QQY7 Q1QQZ1 Q1QQZ3 Q1QQZ4 Q1QR12 Q1QR57 Q1QR91 Q1QRF8 Q1QRF9 Q1QRG2
Q1QRG4 Q1QRH1 Q1QRM1 Q1QRS7 Q1QRT1 Q1QRT2 Q1QRT7 Q1QRU1 Q1QRU4 Q1QRU5 Q1QRV0
Q1QRW8 Q1QRW9 Q1QRX0 Q1QRX1 Q1QRX4 Q1QRX5 Q1QRX6 Q1QRY0 Q1QRY3 Q1QRY4 Q1QRZ0
Q1QRZ1 Q1QRZ2 Q1QRZ8 Q1QS13 Q1QS20 Q1QS21 Q1QS22 Q1QS30 Q1QS32 Q1QS34 Q1QS44
Q1QS47 Q1QS58 Q1QS60 Q1QS61 Q1QS63 Q1QS64 Q1QS67 Q1QS68 Q1QS72 Q1QS74 Q1QS94
Q1QS96 Q1QS97 Q1QS99 Q1QSB0 Q1QSB5 Q1QSB9 Q1QSC0 Q1QSG3 Q1QSI0 Q1QSI1 Q1QSL0
Q1QSR0 Q1QSR9 Q1QSS9 Q1QSU7 Q1QSU9 Q1QSV1 Q1QSV4 Q1QSW3 Q1QSW9 Q1QSX0 Q1QSX1
Q1QSX7 Q1QSY2 Q1QSY5 Q1QSY6 Q1QSY8 Q1QSZ0 Q1QSZ1 Q1QSZ2 Q1QSZ3 Q1QSZ4 Q1QSZ7
Q1QSZ8 Q1QSZ9 Q1QT05 Q1QT13 Q1QT18 Q1QT39 Q1QT48 Q1QT49 Q1QT50 Q1QT51 Q1QT66
Q1QT72 Q1QT79 Q1QT83 Q1QT89 Q1QT91 Q1QT92 Q1QT93 Q1QTC8 Q1QTE5 Q1QTJ4 Q1QTJ5
Q1QTJ8 Q1QTJ9 Q1QTK0 Q1QTK4 Q1QTK7 Q1QTK8 Q1QTL1 Q1QTL2 Q1QTM9 Q1QTN0 Q1QTP6
Q1QTP9 Q1QTQ7 Q1QTQ8 Q1QU06 Q1QU16 Q1QU27 Q1QU47 Q1QU53 Q1QU73 Q1QU74 Q1QU75
Q1QU76 Q1QU77 Q1QU82 Q1QU94 Q1QUC5 Q1QUC9 Q1QUD2 Q1QUD9 Q1QUF2 Q1QUF8 Q1QUF9

Q1QUG3 Q1QUG4 Q1QUG5 Q1QUI0 Q1QUI7 Q1QUL4 Q1QUL6 Q1QUP5 Q1QUQ9 Q1QUR0 Q1QUR7
Q1QUS2 Q1QUS3 Q1QUS4 Q1QUU9 Q1QV15 Q1QV19 Q1QV22 Q1QV24 Q1QV38 Q1QV43 Q1QV50
Q1QV51 Q1QV52 Q1QV55 Q1QV64 Q1QV72 Q1QVB5 Q1QVB6 Q1QVC0 Q1QVC8 Q1QVE4 Q1QVF0
Q1QVF5 Q1QVF6 Q1QVF9 Q1QVG5 Q1QVG8 Q1QVG9 Q1QVJ7 Q1QVJ8 Q1QVK2 Q1QVK5 Q1QVL8
Q1QVM4 Q1QVM7 Q1QVM9 Q1QVN0 Q1QVN7 Q1QVP3 Q1QVP5 Q1QVQ3 Q1QVQ9 Q1QVR2 Q1QVU4
Q1QVV6 Q1QVV7 Q1QVV9 Q1QVW0 Q1QVW1 Q1QVW2 Q1QVW9 Q1QW62 Q1QW64 Q1QWA6 Q1QWC3
Q1QWF8 Q1QWG0 Q1QWG1 Q1QWG2 Q1QWG7 Q1QWJ4 Q1QWK1 Q1QWK3 Q1QWK4 Q1QWK5 Q1QWK7
Q1QWN5 Q1QWN6 Q1QWP0 Q1QWP1 Q1QWX0 Q1QX15 Q1QX18 Q1QX49 Q1QX57 Q1QX59 Q1QX67
Q1QX78 Q1QX86 Q1QXA4 Q1QXB1 Q1QXB3 Q1QXB4 Q1QXC6 Q1QXC7 Q1QXD0 Q1QXD2 Q1QXD3
Q1QXD7 Q1QXD9 Q1QXE1 Q1QXJ3 Q1QXJ5 Q1QXJ6 Q1QXV0 Q1QXV3 Q1QXV7 Q1QXV9 Q1QXW0
Q1QXW4 Q1QXY2 Q1QXY5 Q1QXZ6 Q1QY21 Q1QY22 Q1QY26 Q1QY27 Q1QY28 Q1QY34 Q1QY41
Q1QY43 Q1QY63 Q1QY92 Q1QYD5 Q1QYD6 Q1QYD9 Q1QYF4 Q1QYS2 Q1QYX7 Q1QYX8 Q1QZ10
Q1QZ16 Q1QZ19 Q1QZ29 Q1QZ30 Q1QZ31 Q1QZ38 Q1QZ60 Q1QZ62 Q1QZ63 Q1QZ69 Q1QZ76
Q1QZ91 Q1QZ94 Q1QZC2 Q1QZE1 Q1QZP3 Q1QZP4 Q1QZS5 Q1QZV1 Q1QZW9 Q1QZX1 Q1QZX2
Q1QZX3 Q1QZX5 Q1QZX9 Q1QZY3 Q1QZZ2 Q1QZZ4 Q1QZZ7 Q1R002 Q1R019 Q1R030 Q1R031
Q1R032 Q1R033 Q1R039 Q1R073 Q1R077 Q1R079 Q1R080 Q1R084 Q1R089 Q1R092 Q1R0B2
Q1R0B5 Q1R0B8 Q1R0B9 Q1R0C0 Q1R0C1 Q1R0C2 Q1R0E9 Q1R0F0 Q1R0F1 Q1R0F2 Q1R0F3
Q1R0F4 Q1R0F6 Q1R0F7 Q1R0F8 Q1R0F9 Q1R0G0 Q1R0G1 Q1R0G2 Q1R0G3 Q1R0G4 Q1R0G5
Q1R0G6 Q1R0G7 Q1R0G8 Q1R0G9 Q1R0H0 Q1R0H1 Q1R0H2 Q1R0H3 Q1R0H4 Q1R0H5 Q1R0H6
Q1R0H7 Q1R0H8 Q1R0H9 Q1R0I0 Q1R0I1 Q1R0I2 Q1R0I3 Q1R0I4 Q1R0I5 Q1R0I6 Q1R0I9
Q1R0L1 Q1R0L8 Q1R0Z3 Q1R153 Q1R183 Q1R185 Q1R1C8 Q1R1E5 Q1R1I5 Q1R1I6 Q1R1J5
Q1R1J7 Q1R1M6 Q1R1N5 Q1R1N7 Q1R1P0 Q1R1P2 Q1R1U5 Q1R1W4 Q1R1Y6 Q1R246 Q1R248
Q1R257 Q1R260 Q1R270 Q1R2D5 Q1R2E3 Q1R2G2 Q1R2J0 Q1R2J3 Q1R2M2 Q1R2Q6 Q1R2Z4
Q1R309 Q1R310 Q1R314 Q1R322 Q1R324 Q1R342 Q1R357 Q1R358 Q1R359 Q1R360 Q1R363
Q1R369 Q1R382 Q1R383 Q1R388 Q1R389 Q1R3A2 Q1R3B2 Q1R3B6 Q1R3B7 Q1R3C3 Q1R3P7
Q1R3R3 Q1R3S5 Q1R3T1 Q1R3T4 Q1R3U0 Q1R3U2 Q1R3U7 Q1R3V1 Q1R3V2 Q1R3V5 Q1R3X0
Q1R3Y5 Q1R3Y6 Q1R3Y9 Q1R3Z0 Q1R3Z9 Q1R413 Q1R414 Q1R415 Q1R418 Q1R420 Q1R424
Q1R425 Q1R431 Q1R447 Q1R448 Q1R465 Q1R477 Q1R4A4 Q1R4C3 Q1R4D1 Q1R4F9 Q1R4G0
Q1R4G3 Q1R4G6 Q1R4H3 Q1R4I4 Q1R4I6 Q1R4I7 Q1R4I8 Q1R4J1 Q1R4J2 Q1R4M8 Q1R4N2
Q1R4N3 Q1R4N5 Q1R4N7 Q1R4P4 Q1R4P6 Q1R4R6 Q1R4S0 Q1R4U2 Q1R4U3 Q1R4U5 Q1R4V2
Q1R4V4 Q1R4V5 Q1R4V6 Q1R4V8 Q1R4V9 Q1R4X2 Q1R4X4 Q1R4Y5 Q1R4Z6 Q1R503 Q1R519
Q1R536 Q1R537 Q1R584 Q1R5B9 Q1R5D7 Q1R5E9 Q1R5J4 Q1R5J5 Q1R5J6 Q1R5J7 Q1R5L5
Q1R5L6 Q1R5M0 Q1R5M2 Q1R5M4 Q1R5P8 Q1R5R3 Q1R5S8 Q1R5T5 Q1R5T8 Q1R5T9 Q1R5U0
Q1R5U1 Q1R5U2 Q1R5U3 Q1R5U4 Q1R5U7 Q1R5U9 Q1R5V0 Q1R5V1 Q1R5V3 Q1R5V4 Q1R5V9
Q1R5W1 Q1R5W2 Q1R5W3 Q1R5W4 Q1R5X1 Q1R5X9 Q1R5Y2 Q1R601 Q1R602 Q1R604 Q1R606
Q1R607 Q1R609 Q1R610 Q1R612 Q1R613 Q1R615 Q1R616 Q1R617 Q1R619 Q1R620 Q1R621
Q1R622 Q1R624 Q1R626 Q1R627 Q1R629 Q1R630 Q1R632 Q1R633 Q1R635 Q1R636 Q1R637
Q1R638 Q1R644 Q1R645 Q1R646 Q1R648 Q1R650 Q1R651 Q1R667 Q1R669 Q1R692 Q1R6A2
Q1R6A3 Q1R6A9 Q1R6B0 Q1R6B4 Q1R6B5 Q1R6C9 Q1R6E7 Q1R6F0 Q1R6F2 Q1R6F4 Q1R6F9
Q1R6G2 Q1R6G5 Q1R6G7 Q1R6H0 Q1R6H1 Q1R6H2 Q1R6H3 Q1R6H4 Q1R6I6 Q1R6I7 Q1R6J2
Q1R6J4 Q1R6K0 Q1R6K7 Q1R6L1 Q1R6M3 Q1R6P6 Q1R6Q7 Q1R6R1 Q1R6R5 Q1R6R7 Q1R6R9
Q1R6S0 Q1R6S1 Q1R6S5 Q1R6S9 Q1R6T2 Q1R6T4 Q1R6U6 Q1R763 Q1R766 Q1R778 Q1R779
Q1R783 Q1R788 Q1R791 Q1R7A6 Q1R7B4 Q1R7C6 Q1R7C8 Q1R7D0 Q1R7D2 Q1R7E2 Q1R7F8
Q1R7G9 Q1R7I1 Q1R7I3 Q1R7I6 Q1R7N4 Q1R7N6 Q1R7N8 Q1R7P3 Q1R7P8 Q1R7Q7 Q1R7R3

Q1R7T4 Q1R7T5 Q1R7T6 Q1R7T9 Q1R7U0 Q1R7U1 Q1R7U4 Q1R7U5 Q1R7U6 Q1R7U7 Q1R7U8
Q1R7W4 Q1R7Y9 Q1R7Z0 Q1R801 Q1R802 Q1R804 Q1R805 Q1R808 Q1R824 Q1R844 Q1R8A4
Q1R8A9 Q1R8B1 Q1R8B7 Q1R8B8 Q1R8B9 Q1R8F2 Q1R8F7 Q1R8G6 Q1R8G8 Q1R8H7 Q1R8I4
Q1R8K0 Q1R8K1 Q1R8K5 Q1R8K9 Q1R8L4 Q1R8L6 Q1R8L8 Q1R8L9 Q1R8M7 Q1R8M9 Q1R8N9
Q1R8P2 Q1R8Q6 Q1R8R1 Q1R8T1 Q1R8T7 Q1R8U4 Q1R8V2 Q1R8W1 Q1R8W9 Q1R8X8 Q1R8Z2
Q1R901 Q1R904 Q1R981 Q1R983 Q1R988 Q1R992 Q1R994 Q1R9C4 Q1R9E5 Q1R9E9 Q1R9F0
Q1R9F3 Q1R9G4 Q1R9G8 Q1R9H0 Q1R9I4 Q1R9K7 Q1R9L0 Q1R9N1 Q1R9N2 Q1R9P8 Q1R9R0
Q1R9R2 Q1R9R8 Q1R9R9 Q1R9T0 Q1R9V8 Q1R9W7 Q1R9W8 Q1R9X7 Q1R9X8 Q1RA05 Q1RA48
Q1RA49 Q1RA52 Q1RA54 Q1RA65 Q1RAB7 Q1RAE7 Q1RAG7 Q1RAH5 Q1RAI5 Q1RAI9 Q1RAM1
Q1RAM7 Q1RAP1 Q1RAP3 Q1RAQ8 Q1RAR0 Q1RAR3 Q1RAR4 Q1RAR8 Q1RAS0 Q1RAS1 Q1RAS3
Q1RAS4 Q1RAT6 Q1RAV2 Q1RAV6 Q1RAX9 Q1RAY1 Q1RB08 Q1RB45 Q1RB46 Q1RB47 Q1RB48
Q1RB49 Q1RB52 Q1RB54 Q1RB62 Q1RB64 Q1RB76 Q1RB78 Q1RB79 Q1RB83 Q1RB89 Q1RB92
Q1RBA3 Q1RBA4 Q1RBB6 Q1RBB7 Q1RBD7 Q1RBE3 Q1RBF5 Q1RBF7 Q1RBF8 Q1RBF9 Q1RBH1
Q1RBH5 Q1RBJ2 Q1RBP6 Q1RBP8 Q1RBT8 Q1RBT9 Q1RBX3 Q1RC02 Q1RC21 Q1RC77 Q1RC79
Q1RC84 Q1RCA6 Q1RCA7 Q1RCI6 Q1RCM7 Q1RCM8 Q1RCN0 Q1RCN4 Q1RCR7 Q1RCS3 Q1RCS7
Q1RCY4 Q1RD19 Q1RD20 Q1RD26 Q1RD35 Q1RD47 Q1RD48 Q1RD57 Q1RD68 Q1RD70 Q1RD90
Q1RD95 Q1RD98 Q1RDC7 Q1RDK5 Q1RDK6 Q1RDK7 Q1RDK8 Q1RDK9 Q1RDL0 Q1RDL3 Q1RDL6
Q1RDL7 Q1RDP5 Q1RDP6 Q1RDQ5 Q1RDQ8 Q1RDR6 Q1RDS2 Q1RDS7 Q1RDT3 Q1RDT5 Q1RDU1
Q1RDU6 Q1RDU8 Q1RDV0 Q1RDV1 Q1RE21 Q1RE29 Q1RE30 Q1RE32 Q1RE41 Q1RE52 Q1RE59
Q1RE66 Q1RE73 Q1RE90 Q1REB2 Q1REB8 Q1REC1 Q1REE5 Q1REF1 Q1REF4 Q1REF5 Q1REG3
Q1REH4 Q1REI1 Q1REK9 Q1REL0 Q1REN7 Q1REP9 Q1REQ5 Q1REQ7 Q1RER9 Q1RES6 Q1RET1
Q1RET2 Q1RET3 Q1REU3 Q1REU8 Q1REU9 Q1REY3 Q1RF04 Q1RF08 Q1RF23 Q1RF28 Q1RF29
Q1RF30 Q1RF31 Q1RF59 Q1RF60 Q1RF62 Q1RF63 Q1RF67 Q1RF89 Q1RF91 Q1RF97 Q1RF98
Q1RFA0 Q1RFB4 Q1RFB7 Q1RFB8 Q1RFC0 Q1RFC4 Q1RFC8 Q1RFD5 Q1RFD6 Q1RFF2 Q1RFF6
Q1RFG2 Q1RFI6 Q1RFI7 Q1RFI8 Q1RFJ2 Q1RFM1 Q1RFM3 Q1RFP9 Q1RFQ0 Q1RFQ3 Q1RFS7
Q1RFS8 Q1RFT0 Q1RFT1 Q1RFU0 Q1RFX9 Q1RFZ4 Q1RFZ9 Q1RG06 Q1RG17 Q1RG18 Q1RG19
Q1RG21 Q1RG23 Q1RG24 Q1RG25 Q1RG28 Q1RG29 Q1RG32 Q1RG34 Q1RG44 Q1RG56 Q1RG57
Q1RG60 Q1RG70 Q1RG71 Q1RG92 Q1RG94 Q1RG95 Q1RGA4 Q1RGA7 Q1RGB3 Q1RGC3 Q1RGC4
Q1RGC5 Q1RGC6 Q1RGC7 Q1RGD6 Q1RGD7 Q1RGE0 Q1RGE6 Q1RGE7 Q1RGE8 Q1RGF6 Q1RGF9
Q1RGG0 Q1RGG1 Q1RGG3 Q1RGH0 Q1RGH2 Q1RGH3 Q1RGH6 Q1RGH9 Q1RGI7 Q1RGI8 Q1RGJ7
Q1RGK1 Q1RGK3 Q1RGK4 Q1RGK5 Q1RGK7 Q1RGL3 Q1RGM2 Q1RGM6 Q1RGM7 Q1RGM8 Q1RGM9
Q1RGN9 Q1RGP3 Q1RGP6 Q1RGQ3 Q1RGQ4 Q1RGQ5 Q1RGR1 Q1RGS0 Q1RGS3 Q1RGS4 Q1RGT0
Q1RGT1 Q1RGT2 Q1RGT4 Q1RGU4 Q1RGV0 Q1RGV8 Q1RGV9 Q1RGX2 Q1RGX3 Q1RGX4 Q1RGX5
Q1RGX6 Q1RGY1 Q1RGY2 Q1RGY6 Q1RGY7 Q1RGY9 Q1RGZ7 Q1RH00 Q1RH01 Q1RH02 Q1RH04
Q1RH05 Q1RH06 Q1RH11 Q1RH23 Q1RH25 Q1RH40 Q1RH45 Q1RH48 Q1RH55 Q1RH56 Q1RH63
Q1RH65 Q1RH66 Q1RH68 Q1RH76 Q1RH78 Q1RH84 Q1RH88 Q1RH98 Q1RHA2 Q1RHA5 Q1RHB0
Q1RHB4 Q1RHC1 Q1RHC2 Q1RHC3 Q1RHC5 Q1RHC6 Q1RHC7 Q1RHC8 Q1RHC9 Q1RHD0 Q1RHD1
Q1RHD4 Q1RHE0 Q1RHE5 Q1RHE8 Q1RHE9 Q1RHF0 Q1RHF1 Q1RHF2 Q1RHG2 Q1RHG3 Q1RHG4
Q1RHG9 Q1RHH0 Q1RHH3 Q1RHI2 Q1RHI4 Q1RHI5 Q1RHI6 Q1RHI9 Q1RHJ0 Q1RHK6 Q1RHL6
Q1RHL9 Q1RHM0 Q1RHM1 Q1RHM2 Q1RHM3 Q1RHM4 Q1RHM5 Q1RHM6 Q1RHM7 Q1RHM8 Q1RHM9
Q1RHN0 Q1RHN1 Q1RHN2 Q1RHN3 Q1RHN4 Q1RHN5 Q1RHN6 Q1RHN7 Q1RHN8 Q1RHN9 Q1RHP0
Q1RHP3 Q1RHP4 Q1RHP5 Q1RHP6 Q1RHP7 Q1RHQ8 Q1RHR2 Q1RHR3 Q1RHT1 Q1RHT6 Q1RHT8
Q1RHT9 Q1RHU0 Q1RHU4 Q1RHU9 Q1RHV0 Q1RHV1 Q1RHV9 Q1RHW1 Q1RHX0 Q1RHX1 Q1RHX2
Q1RHY6 Q1RHZ6 Q1RHZ9 Q1RI04 Q1RI08 Q1RI12 Q1RI22 Q1RI24 Q1RI29 Q1RI31 Q1RI52

Q1RI57 Q1RI62 Q1RI66 Q1RI67 Q1RI68 Q1RI69 Q1RI71 Q1RI72 Q1RI78 Q1RI81 Q1RI82
Q1RI85 Q1RI92 Q1RI96 Q1RIB3 Q1RIB6 Q1RIB7 Q1RIC4 Q1RIC6 Q1RIC9 Q1RID0 Q1RID2
Q1RID6 Q1RID9 Q1RIE3 Q1RIF4 Q1RIG0 Q1RIG1 Q1RIG2 Q1RIG5 Q1RIG7 Q1RIH4 Q1RII8
Q1RIJ2 Q1RIJ3 Q1RIJ4 Q1RIJ6 Q1RIK2 Q1RIM1 Q1RIM2 Q1RIM4 Q1RIM7 Q1RIN4 Q1RIN6
Q1RIP8 Q1RIS7 Q1RIS8 Q1RIS9 Q1RIT1 Q1RIT9 Q1RIV0 Q1RIV1 Q1RIV2 Q1RIW0 Q1RIW9
Q1RIX0 Q1RIX2 Q1RIX6 Q1RIX7 Q1RIY1 Q1RIY2 Q1RIY5 Q1RIZ2 Q1RIZ3 Q1RIZ4 Q1RIZ8
Q1RIZ9 Q1RJ14 Q1RJ15 Q1RJ22 Q1RJ28 Q1RJ43 Q1RJ44 Q1RJ48 Q1RJ49 Q1RJ50 Q1RJ52
Q1RJ60 Q1RJ61 Q1RJ69 Q1RJ70 Q1RJ71 Q1RJ72 Q1RJ74 Q1RJ81 Q1RJ84 Q1RJ87 Q1RJ94
Q1RJC6 Q1RJC7 Q1RJC9 Q1RJD8 Q1RJH0 Q1RJH1 Q1RJH2 Q1RJJ3 Q1RJJ5 Q1RJL7 Q1RJM0
Q1RJM1 Q1RJM6 Q1RJN2 Q1RJN4 Q1RJN6 Q1RJN7 Q1RJQ1 Q1RJQ4 Q1RJQ6 Q1RJR2 Q1RJR4
Q1RJR6 Q1RJR7 Q1RJS2 Q1RJT5 Q1RJU1 Q1RJU4 Q1RJV5 Q1RJV8 Q1RJX2 Q1RJX3 Q1RJX7
Q1RJY5 Q1RJY6 Q1RJZ5 Q1RJZ6 Q1RK03 Q1RK09 Q1RK10 Q1RK13 Q1RK21 Q1RK25 Q1RK29
Q1RK40 Q1RK52 Q1RK56 Q1RK71 Q1RK72 Q1RK82 Q1RK83 Q1RK89 Q1RK90 Q1RK99 Q1RKA6
Q1RKB1 Q1RKB2 Q1RKB3 Q1RKB6 Q1RKC2 Q1RKD5 Q1RKF4 Q1RKF8 Q1RKF9 Q1RKG2 Q1RKH3
Q1RKH7 Q1RKH8 Q1RKI0 Q1RKI1 Q1RKI5 Q1RKJ6 Q1RKJ9 Q1RKK4 Q1RKK6 Q1RKM4 Q1RKN1
Q1RKN2 Q1RKN3 Q1RLC9 Q1RLU1 Q1RLU9 Q1RM03 Q1RM35 Q1RM55 Q1RMA6 Q1RMH9 Q1RMI3
Q1RMI8 Q1RMJ7 Q1RMK1 Q1RML1 Q1RML4 Q1RML7 Q1RMM0 Q1RMM1 Q1RMM5 Q1RMP0 Q1RMP7
Q1RMQ5 Q1RMR0 Q1RMR2 Q1RMR5 Q1RMS0 Q1RMS2 Q1RMS6 Q1RMS8 Q1RMT7 Q1RMU5 Q1RMW1
Q1RMX6 Q1RMX9 Q1RN00 Q1RNF8 Q1RP77 Q1RQI9 Q1RQJ0 Q1RQJ1 Q1RS89 Q1S053 Q1S9I9
Q1SF86 Q1SGF1 Q1SU99 Q1SWQ1 Q1T763 Q1T764 Q1T769 Q1T7B7 Q1T7B8 Q1T7C1 Q1T7F1
Q1W1G1 Q1W209 Q1W2L8 Q1W374 Q1W375 Q1W376 Q1W377 Q1W3E0 Q1W6C3 Q1WBT4 Q1WBV0
Q1WCC0 Q1WG82 Q1WIQ6 Q1WM15 Q1WM16 Q1WNZ7 Q1WQV0 Q1WQZ0 Q1WRE3 Q1WRE4 Q1WRF8
Q1WRF9 Q1WRG9 Q1WRH2 Q1WRH7 Q1WRQ9 Q1WRR3 Q1WRR5 Q1WRR6 Q1WRR9 Q1WRS0 Q1WRS8
Q1WRY7 Q1WRY8 Q1WRZ1 Q1WS60 Q1WS89 Q1WS90 Q1WS91 Q1WS92 Q1WS93 Q1WS94 Q1WS95
Q1WS96 Q1WS97 Q1WS98 Q1WS99 Q1WSA0 Q1WSA1 Q1WSA2 Q1WSA3 Q1WSA4 Q1WSA5 Q1WSA6
Q1WSA7 Q1WSA8 Q1WSA9 Q1WSB2 Q1WSB3 Q1WSB4 Q1WSB5 Q1WSB6 Q1WSB7 Q1WSC1 Q1WSC2
Q1WSC3 Q1WSC7 Q1WSF8 Q1WSG2 Q1WSG3 Q1WSH6 Q1WSH8 Q1WSH9 Q1WSI0 Q1WSL3 Q1WSL8
Q1WSM8 Q1WSM9 Q1WSN1 Q1WSR2 Q1WSS4 Q1WSS5 Q1WSS9 Q1WST2 Q1WST3 Q1WST4 Q1WST5
Q1WSU4 Q1WSU5 Q1WSU7 Q1WSU9 Q1WSV5 Q1WSV7 Q1WSV9 Q1WSW0 Q1WSX9 Q1WSY8 Q1WSZ5
Q1WSZ9 Q1WT04 Q1WT15 Q1WT16 Q1WT17 Q1WT18 Q1WT32 Q1WT33 Q1WT34 Q1WT35 Q1WT38
Q1WT45 Q1WT46 Q1WT50 Q1WT53 Q1WT60 Q1WT66 Q1WT71 Q1WT77 Q1WT80 Q1WT94 Q1WT95
Q1WT99 Q1WTA4 Q1WTA8 Q1WTA9 Q1WTB5 Q1WTD6 Q1WTI0 Q1WTI9 Q1WTJ0 Q1WTK0 Q1WTK3
Q1WTM0 Q1WTM2 Q1WTM3 Q1WTP0 Q1WTP1 Q1WTP7 Q1WTQ4 Q1WTR3 Q1WTS4 Q1WTT3 Q1WTT7
Q1WTU0 Q1WTU5 Q1WTU9 Q1WTV0 Q1WTV4 Q1WTW0 Q1WTW4 Q1WTX2 Q1WTX5 Q1WTX9 Q1WTY6
Q1WTY9 Q1WU04 Q1WU07 Q1WU09 Q1WU17 Q1WU19 Q1WU20 Q1WU25 Q1WU26 Q1WU33 Q1WU36
Q1WU48 Q1WU55 Q1WU57 Q1WU59 Q1WU60 Q1WU76 Q1WU81 Q1WU82 Q1WU83 Q1WU86 Q1WU87
Q1WU88 Q1WU89 Q1WU92 Q1WU93 Q1WU95 Q1WUA0 Q1WUA6 Q1WUB1 Q1WUB4 Q1WUB5 Q1WUD3
Q1WUD6 Q1WUE8 Q1WUF0 Q1WUF2 Q1WUF3 Q1WUF4 Q1WUF8 Q1WUF9 Q1WUG0 Q1WUG4 Q1WUG5
Q1WUH1 Q1WUH8 Q1WUI3 Q1WUJ2 Q1WUJ3 Q1WUJ4 Q1WUK1 Q1WUL4 Q1WUL5 Q1WUL9 Q1WUM1
Q1WUM9 Q1WUN0 Q1WUN2 Q1WUN5 Q1WUN7 Q1WUP1 Q1WUP5 Q1WUQ3 Q1WUQ7 Q1WUQ8 Q1WUR2
Q1WUT3 Q1WUT4 Q1WUT6 Q1WUV3 Q1WUZ2 Q1WUZ4 Q1WUZ8 Q1WV02 Q1WV14 Q1WV25 Q1WV30
Q1WV31 Q1WV48 Q1WV65 Q1WV66 Q1WV68 Q1WV73 Q1WV74 Q1WVA0 Q1WVA1 Q1WVA2 Q1WVA4
Q1WVA5 Q1WVA9 Q1WVB1 Q1WVB2 Q1WVB3 Q1WVE0 Q1WVH6 Q1WVH7 Q1WVJ0 Q1WVJ7 Q1WVN7
Q1WVP2 Q1WVQ8 Q1WVS1 Q1WVT3 Q1X8D7 Q1XBU5 Q1XBW5 Q1XD93 Q1XDA1 Q1XDA3 Q1XDA4

Q1XDA5 Q1XDA7 Q1XDA8 Q1XDB1 Q1XDB2 Q1XDB4 Q1XDB5 Q1XDC1 Q1XDC5 Q1XDC6 Q1XDC8
Q1XDD2 Q1XDD3 Q1XDD5 Q1XDD6 Q1XDD7 Q1XDD8 Q1XDD9 Q1XDE1 Q1XDE2 Q1XDE3 Q1XDE5
Q1XDE8 Q1XDE9 Q1XDF0 Q1XDF1 Q1XDF2 Q1XDF3 Q1XDF4 Q1XDF5 Q1XDF6 Q1XDF7 Q1XDF8
Q1XDG7 Q1XDG8 Q1XDH2 Q1XDH4 Q1XDH5 Q1XDH6 Q1XDH7 Q1XDH8 Q1XDH9 Q1XDI0 Q1XDI1
Q1XDI2 Q1XDI3 Q1XDI4 Q1XDI5 Q1XDI6 Q1XDI7 Q1XDI8 Q1XDI9 Q1XDJ0 Q1XDJ2 Q1XDJ3
Q1XDJ4 Q1XDJ5 Q1XDJ6 Q1XDJ7 Q1XDJ8 Q1XDJ9 Q1XDK0 Q1XDK1 Q1XDK2 Q1XDK8 Q1XDK9
Q1XDL3 Q1XDL4 Q1XDL5 Q1XDL6 Q1XDL7 Q1XDL8 Q1XDM0 Q1XDM1 Q1XDN0 Q1XDN1 Q1XDN2
Q1XDN3 Q1XDN4 Q1XDN5 Q1XDN6 Q1XDN7 Q1XDN8 Q1XDN9 Q1XDP6 Q1XDP7 Q1XDP9 Q1XDQ0
Q1XDQ2 Q1XDQ3 Q1XDQ4 Q1XDQ6 Q1XDQ7 Q1XDQ8 Q1XDQ9 Q1XDR3 Q1XDR4 Q1XDR8 Q1XDR9
Q1XDS0 Q1XDS2 Q1XDS3 Q1XDS6 Q1XDS7 Q1XDS8 Q1XDS9 Q1XDT0 Q1XDT2 Q1XDT3 Q1XDT4
Q1XDT5 Q1XDT6 Q1XDU5 Q1XDU6 Q1XDU7 Q1XDU8 Q1XDU9 Q1XDV0 Q1XFL1 Q1XGE2 Q1XH04
Q1XH05 Q1XH10 Q1XHT8 Q1XHU0 Q1XHV8 Q1XID0 Q1XIR9 Q1ZXB2 Q1ZXB9 Q1ZXC2 Q1ZXC6
Q1ZXC8 Q1ZXC9 Q1ZXD3 Q1ZXD5 Q1ZXE0 Q1ZXE6 Q1ZXF0 Q1ZXF7 Q1ZXG4 Q1ZXG8 Q1ZXH0
Q1ZXH2 Q1ZXI0 Q1ZXI2 Q1ZXI6 Q1ZXK1 Q1ZXK2 Q1ZXK3 Q1ZXL0 Q1ZXL6 Q1ZXQ1 Q1ZXQ8
Q1ZXU0 Q1ZZS1 Q1ZZU3 Q20010 Q20025 Q20052 Q20054 Q20057 Q20059 Q20060 Q20082
Q20117 Q20140 Q20201 Q20230 Q20255 Q20256 Q20257 Q20296 Q202I6 Q20318 Q20334
Q20347 Q20351 Q20363 Q20374 Q20443 Q20448 Q20480 Q20481 Q20497 Q20502 Q20561
Q20585 Q20588 Q20619 Q20624 Q20680 Q20681 Q20683 Q20689 Q20709 Q20716 Q20728
Q20733 Q20751 Q20765 Q20772 Q20819 Q20845 Q20848 Q20849 Q20870 Q20875 Q20893
Q20898 Q208S2 Q208S3 Q20932 Q20937 Q20938 Q20939 Q20970 Q20EU5 Q20EU9 Q20EV0
Q20EV1 Q20EV2 Q20EV4 Q20EV5 Q20EW0 Q20EW1 Q20EX0 Q20EX1 Q20EX2 Q20EX3 Q20EX7
Q20EX8 Q20EX9 Q20EY2 Q20F00 Q20F01 Q20F02 Q20F03 Q20F04 Q20F06 Q20F07 Q20F08
Q20F09 Q20F10 Q20F11 Q20F12 Q20F14 Q20F17 Q20HU6 Q20HU7 Q20WM1 Q20WN5 Q20WT9
Q20WV6 Q20WW7 Q20WW8 Q20X07 Q20X08 Q20X09 Q20X11 Q20X88 Q20X96 Q20XN0 Q20Y14
Q20Y19 Q20Y45 Q20Y47 Q20Y49 Q20Y76 Q20YA2 Q20YA7 Q20YB4 Q20YH9 Q20YI1 Q20YN2
Q20YV7 Q20YZ3 Q20YZ7 Q20Z21 Q20Z24 Q20Z33 Q20Z78 Q20ZC1 Q20ZD1 Q20ZE8 Q20ZR6
Q20ZU5 Q21006 Q21012 Q21018 Q21037 Q21045 Q21065 Q21067 Q21086 Q210E7 Q210F4
Q210F5 Q210F6 Q210F9 Q210G0 Q210G1 Q210H1 Q21102 Q21106 Q21122 Q21126 Q21127
Q21148 Q21187 Q21192 Q21193 Q211C4 Q211C5 Q211D7 Q211D8 Q211D9 Q211E3 Q211E4
Q211E5 Q211E6 Q211E7 Q211E8 Q211E9 Q211F0 Q211F1 Q211F2 Q211F3 Q211F4 Q211F5
Q211F6 Q211F7 Q211F8 Q211F9 Q211G0 Q211G1 Q211G2 Q211G3 Q211G4 Q211G5 Q211G6
Q211G7 Q211H0 Q211H1 Q211H2 Q211H3 Q211P3 Q211P4 Q211P6 Q211R2 Q211T9 Q211U0
Q211U3 Q211V0 Q211X8 Q211Y8 Q211Z6 Q21209 Q21215 Q21217 Q21227 Q21230 Q21244
Q21253 Q21263 Q21272 Q21276 Q21278 Q21279 Q21289 Q212A8 Q212A9 Q212F4 Q212H2
Q212N0 Q212N1 Q212Q9 Q212R3 Q212S9 Q21305 Q21307 Q21338 Q21346 Q21355 Q21361
Q21388 Q213I9 Q213J3 Q213K6 Q213M1 Q213P4 Q213W6 Q213Y7 Q21407 Q21412 Q21441
Q21446 Q21452 Q21454 Q21489 Q214D4 Q214E1 Q214G0 Q214H2 Q214H6 Q214H7 Q214I7
Q214J3 Q214J9 Q214K2 Q214K5 Q214L4 Q214L6 Q214P8 Q214Q4 Q214R1 Q214R4 Q214R5
Q214W1 Q214W4 Q214W6 Q214W8 Q214X7 Q214X8 Q21502 Q21554 Q21557 Q21568 Q21578
Q215A3 Q215B2 Q215B4 Q215B8 Q215B9 Q215D4 Q215E3 Q215E6 Q215E7 Q215E8 Q215E9
Q215F8 Q215J1 Q215J2 Q215J3 Q215P7 Q215P8 Q215Q6 Q215R4 Q215R8 Q215S4 Q215S7
Q215T6 Q215T7 Q215T9 Q215V6 Q215W3 Q215W8 Q215X2 Q215X7 Q215Y5 Q215Z0 Q215Z7
Q21625 Q21633 Q21657 Q21691 Q21693 Q216E0 Q216E6 Q216G8 Q216H0 Q216J6 Q216N4
Q216P6 Q216Q6 Q21701 Q21735 Q21738 Q21743 Q21755 Q21770 Q21773 Q21774 Q21775

Q21776 Q217M3 Q217T7 Q21806 Q21824 Q21827 Q21832 Q21836 Q21878 Q21890 Q218E3
Q218F9 Q218M1 Q218N3 Q218N7 Q218Q3 Q218T3 Q21902 Q21920 Q21924 Q21926 Q21939
Q21944 Q21950 Q21952 Q21986 Q21988 Q21993 Q219B6 Q219E2 Q219F8 Q219F9 Q219I5
Q219P7 Q219Q8 Q219R0 Q219R1 Q21A46 Q21A54 Q21A74 Q21A81 Q21A86 Q21AM9 Q21AW5
Q21AX3 Q21AX5 Q21B46 Q21B49 Q21B51 Q21B54 Q21B56 Q21B58 Q21B59 Q21B62 Q21B69
Q21B77 Q21B81 Q21BQ1 Q21BQ2 Q21BR2 Q21BW8 Q21BX8 Q21BZ6 Q21BZ9 Q21C03 Q21C12
Q21C20 Q21C27 Q21C28 Q21C30 Q21C31 Q21C34 Q21C37 Q21C41 Q21C43 Q21C62 Q21C63
Q21C64 Q21CB3 Q21CB6 Q21CB8 Q21CB9 Q21CD0 Q21CD4 Q21CE8 Q21CF0 Q21CH5 Q21CH6
Q21CH8 Q21CI1 Q21CI2 Q21CI5 Q21CI6 Q21CJ1 Q21CJ9 Q21CK1 Q21CK4 Q21CK7 Q21CK9
Q21CL3 Q21CL5 Q21CL6 Q21CL7 Q21CM0 Q21CM1 Q21CM2 Q21CM7 Q21CP5 Q21CS1 Q21CT1
Q21CT4 Q21CT6 Q21CT7 Q21CT8 Q21CT9 Q21CU6 Q21CW7 Q21CW8 Q21CZ3 Q21CZ7 Q21D00
Q21D03 Q21D04 Q21D07 Q21D08 Q21D57 Q21D94 Q21D98 Q21D99 Q21DB1 Q21DC2 Q21DC6
Q21DF6 Q21DF7 Q21DF8 Q21DG1 Q21DG2 Q21DG3 Q21E00 Q21E80 Q21EC8 Q21EC9 Q21EE1
Q21EE3 Q21EE4 Q21EE5 Q21EE6 Q21EE9 Q21EF2 Q21EF4 Q21EI0 Q21EI6 Q21EI7 Q21EN2
Q21EP1 Q21F15 Q21F19 Q21F20 Q21F24 Q21F87 Q21F93 Q21F95 Q21FA3 Q21FC9 Q21FD0
Q21FD2 Q21FD9 Q21FE3 Q21FG6 Q21FH2 Q21FH4 Q21FJ6 Q21FL6 Q21FL7 Q21FL9 Q21FM2
Q21FM3 Q21FP9 Q21FQ0 Q21FS7 Q21FS8 Q21FT3 Q21FU1 Q21FV2 Q21FV3 Q21FV8 Q21FV9
Q21FX6 Q21FY3 Q21FY4 Q21FY5 Q21FZ9 Q21G19 Q21G89 Q21GQ5 Q21GV1 Q21GW0 Q21H28
Q21H35 Q21H36 Q21H37 Q21H39 Q21H48 Q21H51 Q21H59 Q21H61 Q21H62 Q21H63 Q21H64
Q21H65 Q21H69 Q21H71 Q21H72 Q21H76 Q21H84 Q21H88 Q21H92 Q21HA1 Q21HA2 Q21HA7
Q21HA8 Q21HE7 Q21HF7 Q21HG1 Q21HG2 Q21HH2 Q21HH3 Q21HH4 Q21HH5 Q21HH8 Q21HI6
Q21HK3 Q21HK6 Q21HN0 Q21HN4 Q21HN5 Q21HN6 Q21HP4 Q21HQ0 Q21HQ5 Q21HU3 Q21HZ3
Q21HZ6 Q21I21 Q21I23 Q21I34 Q21I40 Q21I62 Q21IG1 Q21II4 Q21IJ4 Q21IN0 Q21IR9
Q21IS0 Q21IS4 Q21IS6 Q21IS8 Q21IS9 Q21IT4 Q21IW6 Q21IX0 Q21IX8 Q21IY3 Q21IY4
Q21IY5 Q21J16 Q21J37 Q21J46 Q21JE9 Q21JG6 Q21JI8 Q21JK7 Q21JL6 Q21JL7 Q21JS2
Q21JW7 Q21JW8 Q21JY6 Q21K21 Q21K22 Q21K28 Q21K29 Q21K31 Q21K42 Q21K60 Q21K91
Q21K93 Q21KA1 Q21KA8 Q21KA9 Q21KB0 Q21KC3 Q21KD4 Q21KD6 Q21KD7 Q21KD8 Q21KE0
Q21KG6 Q21KM1 Q21KP6 Q21KS9 Q21KT3 Q21KT6 Q21KT7 Q21KW1 Q21KW2 Q21KZ5 Q21L28
Q21L30 Q21L55 Q21L65 Q21L66 Q21L68 Q21L70 Q21L79 Q21L89 Q21L94 Q21LB1 Q21LB7
Q21LB9 Q21LC4 Q21LG1 Q21LG2 Q21LG3 Q21LG4 Q21LH6 Q21LJ5 Q21LK2 Q21LK8 Q21LL4
Q21LN7 Q21LT1 Q21LT5 Q21LV0 Q21LV5 Q21LV6 Q21LV8 Q21LV9 Q21LW0 Q21LY9 Q21LZ9
Q21M04 Q21M06 Q21M07 Q21M08 Q21M09 Q21M11 Q21M27 Q21M30 Q21M32 Q21M33 Q21M34
Q21M35 Q21M36 Q21M37 Q21M39 Q21M40 Q21M42 Q21M43 Q21M44 Q21M45 Q21M46 Q21M47
Q21M48 Q21M49 Q21M50 Q21M51 Q21M52 Q21M53 Q21M54 Q21M55 Q21M56 Q21M57 Q21M58
Q21M86 Q21M87 Q21M89 Q21M90 Q21M91 Q21M92 Q21M93 Q21M94 Q21M95 Q21M96 Q21M97
Q21MA1 Q21MA4 Q21MA8 Q21MC3 Q21MC6 Q21MD3 Q21MF7 Q21MG7 Q21MG8 Q21MH1 Q21MH7
Q21MH8 Q21MI1 Q21MI4 Q21MI9 Q21MJ0 Q21MK4 Q21MK7 Q21MP3 Q21MR4 Q21MS9 Q21MT0
Q21MT2 Q21MT5 Q21MU7 Q21MU8 Q21MZ8 Q21N29 Q21NB9 Q21ND2 Q21NG2 Q21NG8 Q21NH1
Q21NH5 Q21NH6 Q21NH8 Q21NJ1 Q21NJ5 Q21NP8 Q21NU6 Q21NV7 Q21NW5 Q21NW8 Q21NX6
Q21NX7 Q21NX9 Q21P11 Q21P75 Q21P78 Q21P92 Q21P93 Q21P94 Q21P95 Q21P96 Q21P97
Q21PP0 Q21PT9 Q21PU0 Q21PU1 Q21PU3 Q21PV3 Q21PV5 Q21PW4 Q21Q74 Q21QL4 Q21QL5
Q21QM1 Q21QM5 Q21QM7 Q21QN3 Q21QN4 Q21QN5 Q21QN6 Q21QN7 Q21QN8 Q21QN9 Q21QP0
Q21QP1 Q21QP2 Q21QP4 Q21QP5 Q21QP6 Q21QP7 Q21QP8 Q21QP9 Q21QQ0 Q21R10 Q21R65
Q21RF9 Q21RG9 Q21RH8 Q21RH9 Q21RI0 Q21RM4 Q21RN1 Q21RP5 Q21RR4 Q21RV3 Q21RV4

Q21RV5 Q21RV6 Q21RV7 Q21RV8 Q21RV9 Q21RW0 Q21RW1 Q21RW2 Q21RW3 Q21RW4 Q21RW5
Q21RW6 Q21RW7 Q21RX7 Q21RZ7 Q21S04 Q21S14 Q21S63 Q21S75 Q21SD6 Q21SE7 Q21SE8
Q21SE9 Q21SF0 Q21SF3 Q21SF4 Q21SF5 Q21SF6 Q21SF7 Q21SF8 Q21SJ0 Q21SJ8 Q21SM4
Q21SV0 Q21SV8 Q21SW1 Q21SW7 Q21SX0 Q21SX1 Q21SX7 Q21SY6 Q21SY7 Q21SY8 Q21SZ2
Q21SZ4 Q21SZ5 Q21T18 Q21T29 Q21T70 Q21T97 Q21TA1 Q21TB9 Q21TE2 Q21TF1 Q21TK4
Q21TL3 Q21TL4 Q21TN3 Q21TN6 Q21TN8 Q21TP6 Q21TP7 Q21U05 Q21U08 Q21U09 Q21U10
Q21U27 Q21U33 Q21U56 Q21U89 Q21U90 Q21U91 Q21U92 Q21U94 Q21U97 Q21U98 Q21UC5
Q21UD7 Q21UF7 Q21UG7 Q21UL3 Q21UQ9 Q21UR3 Q21UT1 Q21UY4 Q21UZ5 Q21UZ9 Q21V06
Q21V17 Q21V18 Q21V28 Q21V29 Q21V39 Q21V41 Q21V75 Q21VE0 Q21VH4 Q21VL4 Q21VL5
Q21VM4 Q21VS7 Q21VT1 Q21VT3 Q21VU1 Q21VV5 Q21VW4 Q21VW6 Q21VY7 Q21VY8 Q21VZ1
Q21W24 Q21W25 Q21W32 Q21W33 Q21W38 Q21W39 Q21W43 Q21W48 Q21W76 Q21W87 Q21W89
Q21W90 Q21WC0 Q21WC3 Q21WC8 Q21WD3 Q21WD4 Q21WD6 Q21WD9 Q21WG7 Q21WJ4 Q21WJ5
Q21WJ7 Q21WK7 Q21WL3 Q21WM7 Q21WM8 Q21WN2 Q21WP8 Q21WQ0 Q21WQ3 Q21WR0 Q21WR9
Q21WT2 Q21WT4 Q21WW5 Q21WW8 Q21WX6 Q21WY6 Q21WY7 Q21WY8 Q21WY9 Q21X03 Q21X08
Q21X09 Q21X17 Q21X23 Q21X56 Q21X72 Q21X81 Q21XE3 Q21XH1 Q21XI0 Q21XI1 Q21XI4
Q21XI6 Q21XI7 Q21XM2 Q21XM3 Q21XN1 Q21XP6 Q21XP8 Q21XQ9 Q21XR4 Q21XR6 Q21XW1
Q21XW2 Q21XW4 Q21XW7 Q21XX0 Q21XX2 Q21XX3 Q21XX5 Q21XZ3 Q21Y51 Q21Y60 Q21Y66
Q21Y67 Q21Y68 Q21Y86 Q21YC9 Q21YD1 Q21YD2 Q21YF3 Q21YF6 Q21YF8 Q21YG1 Q21YG7
Q21YL5 Q21YL6 Q21YL7 Q21YL8 Q21YM7 Q21YM9 Q21YN9 Q21YP1 Q21YP2 Q21YQ0 Q21YS0
Q21YS7 Q21YS9 Q21YT0 Q21YT1 Q21YT7 Q21YU0 Q21YV7 Q21YW0 Q21YX2 Q21YY8 Q21YY9
Q21YZ5 Q21YZ6 Q21YZ8 Q21YZ9 Q21Z00 Q21Z01 Q21ZD0 Q21ZD1 Q21ZD5 Q21ZD6 Q22002
Q22004 Q22006 Q22017 Q22018 Q22019 Q22024 Q22031 Q22037 Q22053 Q22058 Q22067
Q22071 Q22099 Q220A0 Q220D5 Q220I2 Q220I3 Q220I8 Q220J4 Q220N0 Q220P2 Q220P5
Q220R4 Q220R6 Q220R7 Q220S3 Q220S7 Q220S8 Q220T1 Q220T2 Q22100 Q22111 Q22122
Q22127 Q22156 Q221G2 Q221I2 Q221N8 Q221N9 Q221R3 Q221U6 Q221U7 Q22230 Q22236
Q22258 Q22289 Q22292 Q222U2 Q22306 Q223E6 Q223G5 Q223N7 Q22431 Q22438 Q22453
Q22460 Q22510 Q22544 Q22555 Q22557 Q22616 Q22618 Q22639 Q22640 Q22670 Q22678
Q22695 Q22707 Q22708 Q22717 Q22756 Q22799 Q22811 Q22812 Q22813 Q22830 Q22835
Q22836 Q22866 Q22885 Q22888 Q22908 Q22909 Q22910 Q22915 Q22918 Q22992 Q229Z6
Q22A30 Q22AX5 Q22DH9 Q22RG7 Q22SV3 Q22X38 Q22XZ3 Q23045 Q23058 Q23088 Q230X8
Q23116 Q23120 Q23121 Q23122 Q23126 Q23175 Q23192 Q23202 Q23223 Q23232 Q23236
Q23237 Q23243 Q23246 Q23247 Q23255 Q23256 Q23261 Q23270 Q23280 Q23294 Q23295
Q23314 Q23357 Q23381 Q23400 Q23402 Q23405 Q23449 Q23450 Q23462 Q23489 Q23495
Q23500 Q23519 Q23523 Q23525 Q23541 Q23551 Q23579 Q23588 Q23590 Q235M8 Q23622
Q23623 Q23629 Q23639 Q23646 Q23647 Q23651 Q23652 Q23670 Q23679 Q23688 Q23695
Q23696 Q23697 Q236T0 Q23711 Q23716 Q23758 Q23794 Q23884 Q23885 Q23886 Q23887
Q23888 Q23889 Q23919 Q23921 Q23925 Q23935 Q23939 Q23946 Q23949 Q23972 Q23973
Q23976 Q23989 Q23AS2 Q23BV5 Q23DE3 Q23F13 Q23F79 Q23FE2 Q23G98 Q23H79 Q23K29
Q23MT7 Q23SI8 Q23TC2 Q23TC8 Q24009 Q24020 Q24039 Q24040 Q24087 Q24117 Q24118
Q24133 Q24134 Q24141 Q24142 Q24143 Q24152 Q24167 Q24168 Q24169 Q24174 Q24178
Q24186 Q24206 Q24207 Q24208 Q24214 Q24239 Q24248 Q24255 Q24256 Q24266 Q24306
Q24307 Q24317 Q24318 Q24320 Q24325 Q24337 Q24338 Q24368 Q24371 Q24399 Q24423
Q24432 Q24434 Q24437 Q24459 Q24472 Q24478 Q24491 Q24492 Q24524 Q24533 Q24535
Q24536 Q24537 Q24558 Q24560 Q24562 Q24572 Q24573 Q245F2 Q24617 Q24618 Q24621

Q24625 Q24629 Q24630 Q24640 Q24641 Q24645 Q24648 Q24654 Q24656 Q24669 Q24695
Q24703 Q24704 Q24705 Q24708 Q24731 Q24732 Q24733 Q24740 Q24741 Q24742 Q24747
Q24753 Q24756 Q24761 Q24762 Q24766 Q24789 Q24799 Q24800 Q24803 Q24829 Q24849
Q24857 Q24902 Q24956 Q24957 Q24C27 Q24DE2 Q24F59 Q24FH8 Q24JJ9 Q24JK4 Q24JL3
Q24JP3 Q24JP4 Q24JV4 Q24JX8 Q24JX9 Q24JY1 Q24JY3 Q24JY8 Q24K03 Q24K06 Q24K12
Q24K21 Q24LI1 Q24LI4 Q24LJ4 Q24M93 Q24M98 Q24M99 Q24MA0 Q24MA9 Q24MB7 Q24MB9
Q24MC2 Q24MC6 Q24MD8 Q24MI3 Q24MK1 Q24MK7 Q24MK9 Q24ML5 Q24ML8 Q24ML9 Q24MM6
Q24MM7 Q24MV8 Q24MW0 Q24MW8 Q24NA9 Q24NB7 Q24NW7 Q24PE3 Q24PG6 Q24PK7 Q24PK8
Q24PT9 Q24PX9 Q24Q31 Q24Q41 Q24Q90 Q24QC9 Q24QE2 Q24QE3 Q24QH2 Q24QH4 Q24QH6
Q24QI7 Q24QI9 Q24QJ1 Q24QJ2 Q24QJ3 Q24QJ4 Q24QJ5 Q24QL5 Q24QL6 Q24R48 Q24R88
Q24RN9 Q24RW2 Q24S01 Q24S27 Q24S49 Q24SG7 Q24SG9 Q24SJ8 Q24SJ9 Q24SK4 Q24SK5
Q24SK9 Q24SM2 Q24SM3 Q24SN0 Q24SN7 Q24SN8 Q24SP0 Q24SP2 Q24SP5 Q24SP6 Q24SP7
Q24SP8 Q24SQ8 Q24SS4 Q24SS5 Q24ST0 Q24SU4 Q24SU9 Q24SV0 Q24SW7 Q24SW9 Q24SX0
Q24SX2 Q24T28 Q24T29 Q24T63 Q24T84 Q24TC7 Q24TD7 Q24TD8 Q24TE4 Q24TG6 Q24TH1
Q24TH3 Q24TH5 Q24TJ1 Q24TX3 Q24TX4 Q24TX5 Q24U06 Q24U09 Q24U12 Q24U24 Q24U35
Q24U37 Q24U48 Q24UA0 Q24UA2 Q24UA5 Q24UA8 Q24UA9 Q24UB1 Q24UB4 Q24UC6 Q24UF0
Q24UF3 Q24UF7 Q24UF8 Q24UF9 Q24UG0 Q24UG4 Q24UG7 Q24UI2 Q24UI6 Q24UI7 Q24UJ1
Q24UJ4 Q24UJ8 Q24UK0 Q24UN3 Q24UN5 Q24UN6 Q24UN7 Q24UN9 Q24UQ1 Q24UQ3 Q24US1
Q24US9 Q24UT2 Q24UT3 Q24UT5 Q24UV9 Q24UW1 Q24UX5 Q24UX6 Q24UY9 Q24UZ7 Q24V00
Q24V01 Q24V05 Q24V09 Q24V41 Q24V65 Q24V91 Q24V96 Q24VA2 Q24VA6 Q24VC7 Q24VC8
Q24VD4 Q24VE4 Q24VT7 Q24VU3 Q24VU8 Q24VW3 Q24VW5 Q24VW9 Q24W00 Q24W34 Q24W37
Q24W96 Q24WC0 Q24WI0 Q24WR0 Q24WW7 Q24WW8 Q24WY8 Q24X58 Q24X59 Q24X61 Q24X66
Q24X67 Q24XQ0 Q24XQ1 Q24XR6 Q24XR7 Q24XT2 Q24XT4 Q24XT5 Q24XU1 Q24XW2 Q24XZ0
Q24XZ4 Q24YA3 Q24YJ4 Q24YU1 Q24ZG1 Q24ZG7 Q24ZG8 Q24ZJ1 Q24ZL1 Q24ZM5 Q24ZM6
Q24ZT5 Q24ZZ6 Q24ZZ8 Q25008 Q25009 Q25010 Q25019 Q25020 Q250C6 Q250I5 Q250K3
Q250K4 Q250K5 Q250K6 Q250K7 Q250K8 Q250K9 Q250L1 Q250L3 Q250L4 Q250L5 Q250L6
Q250L7 Q250L8 Q250L9 Q250M0 Q250M1 Q250M2 Q250M3 Q250M4 Q250M5 Q250M6 Q250M7
Q250M8 Q250M9 Q250N0 Q250N1 Q250N2 Q250N3 Q250N4 Q250N5 Q250N6 Q250N7 Q250N9
Q250P0 Q250P5 Q250P7 Q250P8 Q250Q1 Q250Q5 Q250Q6 Q250Q8 Q250Q9 Q250S5 Q25109
Q25113 Q25132 Q25145 Q25189 Q251D2 Q251F6 Q251G4 Q251I2 Q251I4 Q251I5 Q251N7
Q251P1 Q251P2 Q251P8 Q251Q1 Q251S4 Q251V5 Q251W8 Q251Z0 Q25206 Q25214 Q25221
Q25225 Q25264 Q25278 Q252J5 Q252K0 Q252M0 Q252M1 Q252N9 Q252Q4 Q252Q5 Q252Q6
Q252Q7 Q252S3 Q252T8 Q252V3 Q252V4 Q252V5 Q252V6 Q252V7 Q252V8 Q252V9 Q252W0
Q252W1 Q252W2 Q252W3 Q252W4 Q252W5 Q252W6 Q252W7 Q252W8 Q252W9 Q252X0 Q252X2
Q252X3 Q252X4 Q252X5 Q252X9 Q252Y0 Q252Y1 Q252Y3 Q252Y9 Q252Z2 Q252Z7 Q25306
Q25330 Q25337 Q25379 Q25380 Q25381 Q25386 Q25390 Q253C0 Q253C1 Q253C8 Q253D1
Q253D3 Q253E2 Q253E9 Q253F0 Q253F1 Q253F2 Q253F4 Q253F6 Q253F7 Q253F8 Q253G3
Q253H1 Q253I4 Q253I5 Q253I8 Q253J5 Q253J9 Q253K1 Q253K2 Q253K3 Q253K4 Q253K9
Q253M6 Q253Q1 Q253Q2 Q253Q7 Q253R6 Q253R7 Q253R9 Q253S0 Q253S1 Q253S4 Q253S7
Q253S9 Q253T0 Q253T5 Q253T6 Q253U4 Q253V2 Q253V7 Q253W3 Q253Y0 Q253Y4 Q25417
Q25435 Q25451 Q25456 Q25457 Q25469 Q25472 Q25478 Q254B3 Q254B4 Q254B9 Q254D2
Q254F0 Q254F3 Q254F5 Q254F7 Q254G4 Q254H2 Q254H3 Q254H4 Q254J2 Q254L6 Q254M8
Q254N4 Q254N9 Q254P0 Q254P2 Q254Q0 Q254Q1 Q254S6 Q254S7 Q254S9 Q254T1 Q254T8
Q254U7 Q254V9 Q254W0 Q254X5 Q254X9 Q254Z2 Q254Z3 Q254Z5 Q25514 Q25515 Q25520

Q25537 Q25538 Q25544 Q25555 Q25563 Q25566 Q255A0 Q255A1 Q255A6 Q255A8 Q255A9
Q255B0 Q255B1 Q255B2 Q255B5 Q255B6 Q255D2 Q255D3 Q255E4 Q255E5 Q255E6 Q255E8
Q255E9 Q255F0 Q255F3 Q255F4 Q255G0 Q255G3 Q255G8 Q255H0 Q255H3 Q255H5 Q255H6
Q255I0 Q255I4 Q255I6 Q255J4 Q255L3 Q255L4 Q255L6 Q255M0 Q255M1 Q255M2 Q255M5
Q255M7 Q255N7 Q255P2 Q255Q9 Q255R7 Q255R8 Q255R9 Q255S0 Q255S1 Q255S2 Q255S3
Q255T1 Q255T2 Q255T5 Q255T6 Q255T7 Q255T8 Q255U1 Q255U2 Q255V8 Q255W7 Q255X6
Q255Z6 Q255Z9 Q25604 Q25626 Q25632 Q256A6 Q256B8 Q256C2 Q256C3 Q256C5 Q256D0
Q256D6 Q256F3 Q256F9 Q256I5 Q256I7 Q256J8 Q256K7 Q256S2 Q256S4 Q256S6 Q256S7
Q25802 Q25820 Q25861 Q258Y5 Q25998 Q259G4 Q25A68 Q25AA3 Q25BG0 Q25BG2 Q25BG4
Q25BG5 Q25BG6 Q25BH3 Q25BH6 Q25BH7 Q25BH9 Q25BI0 Q25BI1 Q25BI3 Q25BI6 Q25BI7
Q25BI8 Q25BI9 Q25BJ0 Q25BJ1 Q25BJ2 Q25BJ3 Q25BJ4 Q25BN1 Q25BW4 Q25BW5 Q25C93
Q25QU7 Q25QX6 Q26065 Q26228 Q26231 Q26240 Q26255 Q26287 Q26288 Q26289 Q26327
Q26366 Q26417 Q26430 Q26454 Q26457 Q26458 Q26486 Q26487 Q26496 Q26497 Q26503
Q26516 Q26519 Q26537 Q26540 Q26544 Q26547 Q26548 Q26565 Q26571 Q26600 Q26601
Q26602 Q26603 Q26604 Q26605 Q26609 Q26612 Q26613 Q26619 Q26622 Q26626 Q26630
Q26635 Q26648 Q26656 Q26657 Q26671 Q26677 Q26678 Q26680 Q26695 Q26696 Q26734
Q26768 Q26789 Q26891 Q26997 Q26998 Q27001 Q27021 Q27031 Q27032 Q27052 Q27128
Q27135 Q27139 Q27140 Q27152 Q27171 Q27177 Q27178 Q27179 Q27200 Q27203 Q27245
Q27249 Q27268 Q27272 Q27274 Q27287 Q27294 Q27297 Q27302 Q27319 Q27350 Q27352
Q27355 Q27365 Q27368 Q27371 Q27380 Q27403 Q27404 Q27415 Q27421 Q27442 Q27443
Q27450 Q27464 Q27474 Q27483 Q27484 Q27488 Q27489 Q27490 Q27497 Q27511 Q27513
Q27514 Q27515 Q27516 Q27517 Q27518 Q27519 Q27520 Q27521 Q27522 Q27527 Q27532
Q27535 Q27536 Q27539 Q27541 Q27543 Q27546 Q27549 Q27556 Q27563 Q27564 Q27567
Q27575 Q27580 Q27601 Q27607 Q27638 Q27640 Q27650 Q27651 Q27652 Q27655 Q27665
Q27666 Q27683 Q27684 Q27685 Q27686 Q27705 Q27707 Q27709 Q27710 Q27713 Q27740
Q27743 Q27746 Q27749 Q27750 Q27765 Q27772 Q27775 Q27778 Q27783 Q27788 Q27793
Q27797 Q27828 Q27850 Q27869 Q27873 Q27876 Q27877 Q27883 Q27884 Q27888 Q27889
Q27894 Q27896 Q27934 Q27940 Q27949 Q27957 Q27958 Q27976 Q27991 Q27995 Q27GK7
Q27GV2 Q27I66 Q27IK6 Q27IK7 Q27RY7 Q27RZ0 Q27S09 Q27S24 Q27S25 Q27S52 Q27U48
Q27W01 Q27W02 Q27YE1 Q27YE3 Q27YE5 Q27YG7 Q27YG8 Q27YH0 Q27YU7 Q27Z07 Q28007
Q28009 Q28037 Q28055 Q28068 Q28088 Q28092 Q28115 Q28151 Q28161 Q28165 Q28220
Q28221 Q28222 Q28283 Q28314 Q28333 Q28337 Q28338 Q28346 Q28350 Q28356 Q28372
Q28399 Q283L0 Q283L3 Q283N4 Q283Q6 Q28447 Q28452 Q28466 Q28479 Q28480 Q28488
Q28496 Q28503 Q28507 Q28521 Q28555 Q28566 Q28570 Q28571 Q28580 Q28598 Q28599
Q285L8 Q28600 Q28601 Q28641 Q28658 Q28670 Q28690 Q28706 Q28717 Q28772 Q28775
Q28778 Q28779 Q28783 Q28798 Q28810 Q28811 Q288N6 Q288N7 Q28901 Q28931 Q28932
Q28943 Q28955 Q28970 Q28986 Q28987 Q28988 Q28999 Q28BK4 Q28BL7 Q28BR5 Q28BS0
Q28BS5 Q28BT7 Q28BT8 Q28BU7 Q28BZ1 Q28BZ2 Q28BZ7 Q28C22 Q28C26 Q28C33 Q28C44
Q28C59 Q28C61 Q28C65 Q28C69 Q28C89 Q28C91 Q28C98 Q28CA1 Q28CB1 Q28CH2 Q28CH3
Q28CI6 Q28CJ6 Q28CK1 Q28CM3 Q28CQ4 Q28CQ7 Q28CR0 Q28CU0 Q28CW2 Q28CX0 Q28CY2
Q28CZ7 Q28CZ9 Q28D40 Q28D67 Q28D79 Q28D84 Q28D99 Q28DB1 Q28DB3 Q28DB5 Q28DE0
Q28DE6 Q28DE7 Q28DF4 Q28DG7 Q28DH0 Q28DH2 Q28DK1 Q28DP6 Q28DR7 Q28DS0 Q28DT7
Q28DT9 Q28DV7 Q28DW0 Q28DZ0 Q28DZ1 Q28E28 Q28E41 Q28E45 Q28E54 Q28E95 Q28EE8
Q28EG9 Q28EM1 Q28EM3 Q28EM7 Q28EN2 Q28ER3 Q28ES8 Q28ET4 Q28ET8 Q28EW4 Q28EX9

Q28F19 Q28F43 Q28F51 Q28F55 Q28F67 Q28F72 Q28F89 Q28FA8 Q28FB9 Q28FC1 Q28FE0
Q28FE2 Q28FE4 Q28FE5 Q28FE8 Q28FH2 Q28FI7 Q28FJ0 Q28FK7 Q28FN6 Q28FQ5 Q28FR6
Q28FT4 Q28FX0 Q28FY0 Q28FY7 Q28G02 Q28G05 Q28G12 Q28G67 Q28G71 Q28G87 Q28G88
Q28G94 Q28GB0 Q28GD1 Q28GD4 Q28GD5 Q28GD9 Q28GE1 Q28GG4 Q28GG8 Q28GH3 Q28GJ2
Q28GK6 Q28GL6 Q28GP0 Q28GQ3 Q28GR1 Q28GU6 Q28GV1 Q28H12 Q28H21 Q28H30 Q28H51
Q28H63 Q28H65 Q28H71 Q28H76 Q28H87 Q28HA8 Q28HC7 Q28HC9 Q28HD5 Q28HE9 Q28HL3
Q28HM1 Q28HN4 Q28HT3 Q28HX0 Q28HX4 Q28HY7 Q28I03 Q28I19 Q28I29 Q28I42 Q28I85
Q28IB1 Q28IB3 Q28IC1 Q28ID3 Q28IE5 Q28IG6 Q28IG8 Q28IH8 Q28IN4 Q28IN9 Q28IQ9
Q28IU6 Q28IV3 Q28IV6 Q28IV8 Q28IW8 Q28J15 Q28J59 Q28J76 Q28J83 Q28J92 Q28JB2
Q28JI3 Q28JI5 Q28JW5 Q28K03 Q28K09 Q28K11 Q28K15 Q28K17 Q28K48 Q28K56 Q28KC6
Q28KE3 Q28KE9 Q28KF4 Q28KG7 Q28KJ9 Q28KN6 Q28KP2 Q28L59 Q28L73 Q28L99 Q28LA3
Q28LA6 Q28LA8 Q28LB0 Q28LB1 Q28LN3 Q28LS1 Q28LS3 Q28LT2 Q28LW5 Q28LY6 Q28LY7
Q28M21 Q28M40 Q28M91 Q28MB2 Q28MD3 Q28MG4 Q28MG6 Q28MH0 Q28MH9 Q28MK0 Q28MN0
Q28MN1 Q28MN2 Q28N04 Q28N58 Q28NA9 Q28NB3 Q28NC3 Q28NC5 Q28NF0 Q28NI7 Q28NI8
Q28NJ9 Q28NK0 Q28NK3 Q28NK7 Q28NM6 Q28NN2 Q28NN6 Q28NP1 Q28NP2 Q28NS0 Q28NS8
Q28NX0 Q28P47 Q28P75 Q28PA0 Q28PA2 Q28PC5 Q28PD8 Q28PE1 Q28PE4 Q28PE7 Q28PF8
Q28PH0 Q28PI4 Q28PI5 Q28PI7 Q28PJ0 Q28PK5 Q28PU4 Q28PV7 Q28PV9 Q28PY7 Q28PY9
Q28PZ0 Q28PZ5 Q28Q10 Q28Q11 Q28Q12 Q28Q14 Q28Q15 Q28Q51 Q28Q60 Q28QB4 Q28QC2
Q28QC6 Q28QC7 Q28QS5 Q28QV8 Q28QW4 Q28QX4 Q28QY7 Q28R10 Q28R34 Q28R42 Q28R66
Q28R69 Q28R79 Q28R88 Q28RA3 Q28RA7 Q28RA9 Q28RC3 Q28RD6 Q28RD9 Q28RE3 Q28RF2
Q28RF9 Q28RG2 Q28RI9 Q28RJ0 Q28RJ3 Q28RJ8 Q28RK0 Q28RK1 Q28RK3 Q28RK4 Q28RK6
Q28RL2 Q28RM8 Q28RQ1 Q28RX3 Q28RX4 Q28RX5 Q28RY2 Q28S09 Q28S88 Q28S90 Q28SE8
Q28SW8 Q28SZ3 Q28T07 Q28T20 Q28T23 Q28T24 Q28T47 Q28TH5 Q28TH7 Q28TH8 Q28TI5
Q28TK3 Q28TL1 Q28TL3 Q28TN0 Q28TR2 Q28TR4 Q28TS9 Q28TT0 Q28TU1 Q28TV0 Q28U74
Q28U77 Q28U96 Q28UD5 Q28UE6 Q28UE7 Q28UF1 Q28UF3 Q28UF4 Q28UI7 Q28UI8 Q28UI9
Q28UJ2 Q28UJ8 Q28UR1 Q28UR2 Q28US7 Q28US8 Q28US9 Q28UT0 Q28UT2 Q28UT4 Q28UT6
Q28UT7 Q28UT8 Q28UT9 Q28UU0 Q28UU1 Q28UU2 Q28UU3 Q28UU4 Q28UU5 Q28UU6 Q28UV3
Q28UV4 Q28UV5 Q28UV6 Q28UV7 Q28UW0 Q28UW1 Q28UW2 Q28UW3 Q28UW7 Q28UW8 Q28UW9
Q28UX0 Q28UX6 Q28UX7 Q28UX8 Q28UX9 Q28UY1 Q28UY2 Q28V06 Q28V18 Q28V23 Q28V29
Q28V32 Q28V43 Q28V47 Q28V48 Q28V59 Q28V76 Q28V93 Q28V94 Q28V95 Q28VA1 Q28VA7
Q28VC1 Q28VD4 Q28VF4 Q28VG7 Q28VJ1 Q28VJ6 Q28VK7 Q28VL4 Q28VP7 Q28VR1 Q28VS6
Q28VY3 Q28VY4 Q28VY6 Q28VZ4 Q28VZ5 Q28VZ6 Q28VZ8 Q28VZ9 Q28W00 Q28W02 Q28W08
Q28W10 Q28W25 Q28W27 Q28W34 Q28W35 Q28W37 Q28W60 Q28W67 Q28W72 Q28WA7 Q28WC8
Q28WF4 Q28WF7 Q28WG2 Q28WG4 Q28WI0 Q28WI9 Q28WL0 Q28WQ1 Q28WQ8 Q28X18 Q28X44
Q28X71 Q28XA5 Q28XE2 Q28XF0 Q28XK6 Q28XT3 Q28XY0 Q28Y69 Q28YD9 Q28YQ7 Q28YX9
Q28YY2 Q28Z18 Q28Z41 Q28ZA9 Q28ZC1 Q28ZG0 Q28ZT4 Q28ZX3 Q29013 Q29024 Q29029
Q29031 Q29058 Q29068 Q29076 Q29077 Q29087 Q29090 Q29099 Q290A8 Q290F0 Q290L5
Q290L7 Q290P4 Q290S5 Q290X3 Q290Z2 Q29131 Q29187 Q29197 Q291A0 Q291E4 Q291H4
Q291J5 Q291K8 Q291L4 Q291L9 Q29218 Q29221 Q29228 Q29236 Q29261 Q29290 Q29293
Q292B6 Q292D0 Q292E8 Q292F0 Q292F9 Q292G3 Q292R5 Q292U5 Q29303 Q29308 Q29315
Q29329 Q29333 Q29350 Q29371 Q29373 Q29375 Q29384 Q29387 Q293Q2 Q293V2 Q293V6
Q293Y0 Q29407 Q29415 Q29419 Q29425 Q29426 Q29438 Q29490 Q29491 Q29495 Q294B9
Q294E0 Q294G7 Q294Y7 Q29503 Q29504 Q29513 Q29536 Q29555 Q29558 Q29561 Q29563
Q29576 Q295E6 Q295G5 Q295I4 Q295P6 Q295S1 Q295U7 Q295Y7 Q296B0 Q296H8 Q296J6

Q296Q5 Q296V2 Q297A7 Q297F5 Q297G3 Q297L2 Q297N8 Q297U0 Q297U9 Q297V5 Q298C0
Q298E0 Q299D3 Q299F9 Q299L3 Q299P8 Q29A33 Q29AI1 Q29AK9 Q29AP0 Q29AP1 Q29B63
Q29B72 Q29BB3 Q29BD5 Q29BL3 Q29BR3 Q29BT3 Q29CT2 Q29CU2 Q29CV2 Q29CW0 Q29CW2
Q29DB7 Q29DI0 Q29DV9 Q29DY1 Q29E01 Q29EE9 Q29EQ3 Q29EZ6 Q29FC1 Q29FE1 Q29FF7
Q29FS3 Q29FV5 Q29G73 Q29G82 Q29GM0 Q29GU0 Q29H55 Q29H56 Q29HB2 Q29HG9 Q29HT0
Q29HV4 Q29HZ1 Q29I03 Q29I16 Q29I93 Q29IG6 Q29IK4 Q29IK8 Q29IM3 Q29IM7 Q29IN4
Q29J14 Q29JI9 Q29JQ0 Q29JT7 Q29K06 Q29K59 Q29KF5 Q29KH2 Q29KL8 Q29KQ0 Q29L19
Q29L39 Q29L43 Q29L50 Q29L58 Q29L80 Q29LT4 Q29LW1 Q29MA5 Q29MB2 Q29MT7 Q29MZ0
Q29NB4 Q29NF3 Q29NG1 Q29NT9 Q29NV5 Q29NZ8 Q29P70 Q29P71 Q29PG4 Q29PG5 Q29PZ1
Q29PZ2 Q29Q81 Q29R99 Q29RA3 Q29RH2 Q29RH4 Q29RI9 Q29RJ0 Q29RJ1 Q29RJ9 Q29RK0
Q29RK1 Q29RK4 Q29RL0 Q29RL1 Q29RL9 Q29RM2 Q29RM4 Q29RM5 Q29RP8 Q29RP9 Q29RR1
Q29RS0 Q29RS4 Q29RS8 Q29RT6 Q29RU3 Q29RU8 Q29RV0 Q29RW1 Q29RY4 Q29RY9 Q29RZ2
Q29RZ4 Q29RZ5 Q29RZ9 Q29S05 Q29S07 Q29S11 Q29S16 Q29S17 Q29S19 Q29S20 Q29S21
Q29S22 Q29U70 Q29VN1 Q29VN2 Q29VN3 Q29W20 Q29W37 Q29W39 Q29ZJ1 Q2A068 Q2A167
Q2A182 Q2A1A1 Q2A1B4 Q2A1D8 Q2A1D9 Q2A1G5 Q2A1G6 Q2A1G8 Q2A1G9 Q2A1H0 Q2A1J5
Q2A1J6 Q2A1K3 Q2A1M0 Q2A1M3 Q2A1M4 Q2A1M5 Q2A1M6 Q2A1M7 Q2A1M8 Q2A1N0 Q2A1N2
Q2A1N3 Q2A1N4 Q2A1N5 Q2A1N6 Q2A1P2 Q2A1P9 Q2A1Q0 Q2A1Q2 Q2A1Q4 Q2A1T7 Q2A1V2
Q2A1V7 Q2A1V8 Q2A1W9 Q2A1Z9 Q2A205 Q2A215 Q2A218 Q2A220 Q2A253 Q2A260 Q2A261
Q2A265 Q2A268 Q2A269 Q2A270 Q2A273 Q2A275 Q2A282 Q2A285 Q2A2B0 Q2A2B1 Q2A2C5
Q2A2C7 Q2A2E4 Q2A2E5 Q2A2E9 Q2A2I5 Q2A2I6 Q2A2J0 Q2A2J2 Q2A2J3 Q2A2J7 Q2A2K6
Q2A2N1 Q2A2N7 Q2A2P9 Q2A2Q6 Q2A2S0 Q2A2S1 Q2A2S8 Q2A2U4 Q2A2U6 Q2A2U8 Q2A2V6
Q2A2V9 Q2A2X3 Q2A2Y6 Q2A303 Q2A304 Q2A307 Q2A311 Q2A322 Q2A327 Q2A328 Q2A329
Q2A332 Q2A337 Q2A342 Q2A369 Q2A373 Q2A397 Q2A3A9 Q2A3C8 Q2A3D3 Q2A3D4 Q2A3D7
Q2A3E7 Q2A3F5 Q2A3F9 Q2A3H3 Q2A3H5 Q2A3H6 Q2A3H7 Q2A3H9 Q2A3K7 Q2A3L2 Q2A3M7
Q2A3N5 Q2A3P1 Q2A3P9 Q2A3Q7 Q2A3Q9 Q2A3R0 Q2A3R2 Q2A3S7 Q2A3T5 Q2A3T6 Q2A3U1
Q2A3U2 Q2A3U3 Q2A3U6 Q2A3U8 Q2A3X6 Q2A3Y7 Q2A3Z7 Q2A406 Q2A418 Q2A419 Q2A425
Q2A446 Q2A447 Q2A464 Q2A467 Q2A472 Q2A486 Q2A488 Q2A498 Q2A4C4 Q2A4C6 Q2A4C7
Q2A4C8 Q2A4C9 Q2A4E2 Q2A4E9 Q2A4H0 Q2A4H7 Q2A4N1 Q2A4N2 Q2A4P9 Q2A4Q4 Q2A4R1
Q2A4R2 Q2A4R5 Q2A4U1 Q2A4U4 Q2A4U5 Q2A4U7 Q2A4V1 Q2A4V3 Q2A4X4 Q2A4Y6 Q2A4Z4
Q2A4Z7 Q2A515 Q2A524 Q2A535 Q2A549 Q2A562 Q2A587 Q2A5B0 Q2A5B5 Q2A5B6 Q2A5B9
Q2A5D5 Q2A5E0 Q2A5E4 Q2A5E5 Q2A5E6 Q2A5E7 Q2A5E8 Q2A5E9 Q2A5F1 Q2A5F2 Q2A5F3
Q2A5F4 Q2A5F5 Q2A5F6 Q2A5F7 Q2A5F8 Q2A5F9 Q2A5G0 Q2A5G1 Q2A5G2 Q2A5G3 Q2A5G4
Q2A5G5 Q2A5G6 Q2A5G7 Q2A5G8 Q2A5G9 Q2A5H0 Q2A5H1 Q2A5H2 Q2A5H3 Q2A5H4 Q2A5H6
Q2A5H9 Q2A5I0 Q2A5I1 Q2A5I2 Q2A5I8 Q2A5M3 Q2A5M4 Q2A5N1 Q2A5N4 Q2A5Q2 Q2A5R3
Q2A5V3 Q2A5V4 Q2A5V8 Q2A5X5 Q2A5X7 Q2A5Y4 Q2A5Z4 Q2A622 Q2A629 Q2A630 Q2A640
Q2ABE5 Q2ABP6 Q2BN77 Q2CEE2 Q2CEE3 Q2CIN0 Q2EEP9 Q2EEQ3 Q2EEQ8 Q2EER5 Q2EES0
Q2EES3 Q2EES9 Q2EET0 Q2EEU2 Q2EEV4 Q2EEV7 Q2EEV8 Q2EEV9 Q2EEW0 Q2EEW1 Q2EEW2
Q2EEW3 Q2EEW4 Q2EEW5 Q2EEW6 Q2EEW7 Q2EEW8 Q2EEW9 Q2EEX0 Q2EEX1 Q2EEX2 Q2EEX3
Q2EEX8 Q2EEX9 Q2EF73 Q2EGB9 Q2EGP9 Q2EI20 Q2EI21 Q2EN76 Q2F5J3 Q2F5L3 Q2F5R8
Q2F637 Q2F7Z4 Q2F8F7 Q2F969 Q2FAY8 Q2FDE6 Q2FDE7 Q2FDE8 Q2FDE9 Q2FDF0 Q2FDH1
Q2FDH2 Q2FDI1 Q2FDI2 Q2FDI5 Q2FDI7 Q2FDI8 Q2FDM7 Q2FDM8 Q2FDN2 Q2FDP9 Q2FDQ4
Q2FDQ7 Q2FDR0 Q2FDR1 Q2FDR2 Q2FDS6 Q2FDS9 Q2FDU3 Q2FDU5 Q2FDU6 Q2FDU9 Q2FDV3
Q2FDV8 Q2FDY2 Q2FDY9 Q2FE21 Q2FE72 Q2FE74 Q2FE81 Q2FEA5 Q2FEB2 Q2FEC4 Q2FED3
Q2FEF1 Q2FEF3 Q2FEG2 Q2FEG3 Q2FEG6 Q2FEI5 Q2FEI9 Q2FEJ8 Q2FEJ9 Q2FEK0 Q2FEK1

Q2FEK2 Q2FEK3 Q2FEK4 Q2FEK5 Q2FEL2 Q2FEL8 Q2FEM3 Q2FEM4 Q2FEN5 Q2FEN8 Q2FEN9
Q2FEP0 Q2FEP1 Q2FEP2 Q2FEP3 Q2FEP4 Q2FEP5 Q2FEP6 Q2FEP8 Q2FEP9 Q2FEQ0 Q2FEQ1
Q2FEQ2 Q2FEQ3 Q2FEQ4 Q2FEQ5 Q2FEQ6 Q2FEQ7 Q2FEQ8 Q2FER1 Q2FER2 Q2FER3 Q2FER4
Q2FER5 Q2FER6 Q2FES0 Q2FES1 Q2FES2 Q2FES9 Q2FET7 Q2FET8 Q2FET9 Q2FEU0 Q2FEU3
Q2FEV0 Q2FEV9 Q2FEW1 Q2FEX1 Q2FEX4 Q2FEZ3 Q2FEZ8 Q2FF00 Q2FF01 Q2FF06 Q2FF08
Q2FF09 Q2FF10 Q2FF13 Q2FF14 Q2FF15 Q2FF16 Q2FF32 Q2FF34 Q2FF35 Q2FF43 Q2FF45
Q2FF55 Q2FF56 Q2FF59 Q2FF62 Q2FF63 Q2FF64 Q2FF65 Q2FF66 Q2FF67 Q2FF68 Q2FF71
Q2FF75 Q2FF78 Q2FF94 Q2FF95 Q2FFH4 Q2FFH6 Q2FFI3 Q2FFI7 Q2FFJ1 Q2FFJ4 Q2FFJ5
Q2FFJ6 Q2FFK2 Q2FFL5 Q2FFM2 Q2FFM8 Q2FFN1 Q2FFN4 Q2FFQ0 Q2FFR2 Q2FFU9 Q2FFV5
Q2FFV6 Q2FFW3 Q2FFW6 Q2FFX3 Q2FFY0 Q2FFY7 Q2FFZ0 Q2FFZ5 Q2FFZ8 Q2FG06 Q2FG09
Q2FG18 Q2FG23 Q2FG27 Q2FG28 Q2FG29 Q2FG30 Q2FG31 Q2FG40 Q2FG49 Q2FG51 Q2FG54
Q2FG56 Q2FG57 Q2FG58 Q2FG61 Q2FG62 Q2FG63 Q2FG64 Q2FG66 Q2FG69 Q2FG72 Q2FG75
Q2FG80 Q2FG82 Q2FG83 Q2FG84 Q2FG85 Q2FG86 Q2FG87 Q2FG88 Q2FG92 Q2FG94 Q2FG96
Q2FG97 Q2FGA1 Q2FGA5 Q2FGA8 Q2FGA9 Q2FGB0 Q2FGB1 Q2FGB5 Q2FGB6 Q2FGC1 Q2FGC5
Q2FGC8 Q2FGD0 Q2FGD8 Q2FGE1 Q2FGE2 Q2FGE3 Q2FGE4 Q2FGE5 Q2FGE8 Q2FGF3 Q2FGF7
Q2FGF8 Q2FGG0 Q2FGG6 Q2FGH0 Q2FGH2 Q2FGI5 Q2FGI6 Q2FGI7 Q2FGJ3 Q2FGK9 Q2FGL0
Q2FGL1 Q2FGL3 Q2FGL9 Q2FGM9 Q2FGN7 Q2FGN8 Q2FGW7 Q2FGX3 Q2FGX4 Q2FGX5 Q2FGX6
Q2FGX8 Q2FGY5 Q2FGY6 Q2FGZ1 Q2FGZ3 Q2FGZ4 Q2FH00 Q2FH01 Q2FH09 Q2FH10 Q2FH11
Q2FH15 Q2FH17 Q2FH25 Q2FH26 Q2FH29 Q2FH32 Q2FH34 Q2FH36 Q2FH40 Q2FH41 Q2FH42
Q2FH43 Q2FH47 Q2FH63 Q2FH64 Q2FH65 Q2FH66 Q2FH67 Q2FH71 Q2FH78 Q2FH83 Q2FH88
Q2FH89 Q2FH96 Q2FH97 Q2FH98 Q2FH99 Q2FHA3 Q2FHD6 Q2FHD8 Q2FHD9 Q2FHE2 Q2FHE6
Q2FHG3 Q2FHG4 Q2FHG5 Q2FHG7 Q2FHG8 Q2FHG9 Q2FHH3 Q2FHH4 Q2FHH5 Q2FHH9 Q2FHI0
Q2FHI1 Q2FHI2 Q2FHI3 Q2FHI4 Q2FHI5 Q2FHI6 Q2FHI7 Q2FHI8 Q2FHJ3 Q2FHJ4 Q2FHJ7
Q2FHJ8 Q2FHJ9 Q2FHK0 Q2FHK5 Q2FHL2 Q2FHL4 Q2FHM0 Q2FHM2 Q2FHM8 Q2FHM9 Q2FHN4
Q2FHN5 Q2FHN8 Q2FHP0 Q2FHP4 Q2FHP8 Q2FHQ4 Q2FHQ8 Q2FHQ9 Q2FHR0 Q2FHR7 Q2FHT1
Q2FHT5 Q2FHT6 Q2FHT7 Q2FHU1 Q2FHU2 Q2FHU3 Q2FHU5 Q2FHV3 Q2FHV5 Q2FHV6 Q2FHV9
Q2FHW8 Q2FHX4 Q2FHY3 Q2FHZ0 Q2FHZ1 Q2FI05 Q2FI07 Q2FI09 Q2FI10 Q2FI12 Q2FI15
Q2FI23 Q2FI34 Q2FI57 Q2FI59 Q2FI62 Q2FI70 Q2FI72 Q2FI75 Q2FI79 Q2FIA5 Q2FIA7
Q2FIA8 Q2FIA9 Q2FIB3 Q2FIB4 Q2FIB5 Q2FIC1 Q2FID6 Q2FIE3 Q2FIE5 Q2FIE9 Q2FIF3
Q2FIF6 Q2FIG2 Q2FII7 Q2FIJ1 Q2FIJ2 Q2FIL2 Q2FIL9 Q2FIM0 Q2FIM4 Q2FIM5 Q2FIM6
Q2FIM8 Q2FIN3 Q2FIN9 Q2FIP8 Q2FIQ3 Q2FIR0 Q2FIR2 Q2FIR7 Q2FIS8 Q2FIV8 Q2FIX0
Q2FIX1 Q2FJ20 Q2FJ29 Q2FJ31 Q2FJ50 Q2FJ56 Q2FJ62 Q2FJ70 Q2FJ71 Q2FJ74 Q2FJ80
Q2FJ87 Q2FJ88 Q2FJ89 Q2FJ92 Q2FJ93 Q2FJ94 Q2FJ95 Q2FJ96 Q2FJ97 Q2FJ98 Q2FJA0
Q2FJA1 Q2FJA2 Q2FJA3 Q2FJB0 Q2FJB2 Q2FJB5 Q2FJB6 Q2FJB8 Q2FJC0 Q2FJC1 Q2FJC3
Q2FJC9 Q2FJD8 Q2FJD9 Q2FJE0 Q2FJE4 Q2FJE7 Q2FJE9 Q2FJF4 Q2FJG0 Q2FJG2 Q2FJG3
Q2FJM5 Q2FJM6 Q2FJM8 Q2FJN3 Q2FJN4 Q2FJP6 Q2FJP8 Q2FJQ7 Q2FJQ9 Q2FJU6 Q2FJU9
Q2FK01 Q2FK11 Q2FK15 Q2FK29 Q2FK43 Q2FK44 Q2FK58 Q2FK71 Q2FK78 Q2FK80 Q2FK94
Q2FK96 Q2FKC0 Q2FKC6 Q2FKE3 Q2FKF1 Q2FKJ8 Q2FKN2 Q2FKP1 Q2FKP7 Q2FKP8 Q2FKQ0
Q2FKQ1 Q2FKQ2 Q2FKQ5 Q2FKX5 Q2FL30 Q2FL78 Q2FL79 Q2FLB2 Q2FLD3 Q2FLD8 Q2FLD9
Q2FLE8 Q2FLH1 Q2FLH5 Q2FLL0 Q2FLL1 Q2FLN5 Q2FLY5 Q2FM20 Q2FM37 Q2FM63 Q2FMT0
Q2FMZ1 Q2FMZ3 Q2FN10 Q2FN12 Q2FN13 Q2FN14 Q2FN18 Q2FN43 Q2FN79 Q2FNA1 Q2FNA2
Q2FNB7 Q2FNC9 Q2FNE1 Q2FNJ0 Q2FNQ9 Q2FNR0 Q2FNR1 Q2FNT7 Q2FNX1 Q2FNX6 Q2FP80
Q2FPA4 Q2FPC6 Q2FPC8 Q2FPD1 Q2FPD4 Q2FPD8 Q2FPJ4 Q2FPJ8 Q2FPK3 Q2FPM5 Q2FPN5
Q2FPP7 Q2FPQ2 Q2FPV1 Q2FPW6 Q2FPX3 Q2FPZ8 Q2FQ31 Q2FQ32 Q2FQ35 Q2FQ45 Q2FQ51

Q2FQ54 Q2FQ56 Q2FQ74 Q2FQ93 Q2FQ95 Q2FQD6 Q2FQH4 Q2FQJ1 Q2FQL8 Q2FQM1 Q2FQM7
Q2FQM9 Q2FQN9 Q2FQQ2 Q2FQQ4 Q2FQR4 Q2FQT9 Q2FQV3 Q2FQY7 Q2FR07 Q2FR16 Q2FR18
Q2FR24 Q2FR36 Q2FR55 Q2FRE6 Q2FRI1 Q2FRI3 Q2FRP5 Q2FRR2 Q2FRU5 Q2FRU6 Q2FRU9
Q2FRW3 Q2FRY8 Q2FS27 Q2FS30 Q2FS35 Q2FS38 Q2FS43 Q2FS46 Q2FS65 Q2FS66 Q2FS67
Q2FS69 Q2FS70 Q2FS90 Q2FS92 Q2FS96 Q2FS97 Q2FS98 Q2FS99 Q2FSA9 Q2FSC3 Q2FSC7
Q2FSD2 Q2FSD3 Q2FSE6 Q2FSF4 Q2FSF9 Q2FSG1 Q2FSG2 Q2FSG3 Q2FSG4 Q2FSG5 Q2FSH0
Q2FSK9 Q2FSN8 Q2FSR2 Q2FSV9 Q2FT21 Q2FT22 Q2FT32 Q2FT33 Q2FT34 Q2FT35 Q2FT36
Q2FT37 Q2FT38 Q2FT39 Q2FT43 Q2FT51 Q2FT55 Q2FT56 Q2FT57 Q2FT79 Q2FT96 Q2FT97
Q2FT98 Q2FTA2 Q2FTC2 Q2FTC5 Q2FTC7 Q2FTC8 Q2FTF1 Q2FTH0 Q2FTH1 Q2FTH4 Q2FTH6
Q2FTJ7 Q2FTK3 Q2FTK5 Q2FTL0 Q2FTM0 Q2FTM6 Q2FTN4 Q2FTN5 Q2FTN7 Q2FTS5 Q2FTU1
Q2FTV8 Q2FTY4 Q2FTZ8 Q2FU12 Q2FU45 Q2FU48 Q2FU50 Q2FU56 Q2FU77 Q2FU89 Q2FU91
Q2FU92 Q2FU93 Q2FU94 Q2FUA5 Q2FUA9 Q2FUB2 Q2FUB7 Q2FUQ0 Q2FUQ1 Q2FUQ2 Q2FUQ3
Q2FUQ4 Q2FUS6 Q2FUS7 Q2FUT6 Q2FUT7 Q2FUU0 Q2FUU2 Q2FUW3 Q2FUW4 Q2FUW5 Q2FUX7
Q2FUY1 Q2FV11 Q2FV17 Q2FV21 Q2FV22 Q2FV23 Q2FV31 Q2FV39 Q2FV42 Q2FV57 Q2FV59
Q2FV60 Q2FV63 Q2FV74 Q2FVA4 Q2FVB2 Q2FVD5 Q2FVJ6 Q2FVJ7 Q2FVJ9 Q2FVK8 Q2FVM6
Q2FVN3 Q2FVP8 Q2FVQ7 Q2FVQ9 Q2FVR9 Q2FVS1 Q2FVT2 Q2FVT3 Q2FVT6 Q2FVV9 Q2FVW4
Q2FVW5 Q2FVX3 Q2FVX9 Q2FVY4 Q2FVY9 Q2FW03 Q2FW06 Q2FW07 Q2FW08 Q2FW10 Q2FW11
Q2FW12 Q2FW13 Q2FW14 Q2FW15 Q2FW16 Q2FW17 Q2FW18 Q2FW19 Q2FW20 Q2FW21 Q2FW22
Q2FW23 Q2FW28 Q2FW29 Q2FW30 Q2FW31 Q2FW32 Q2FW33 Q2FW37 Q2FW38 Q2FW39 Q2FW52
Q2FW66 Q2FW70 Q2FW72 Q2FW79 Q2FW81 Q2FW96 Q2FWC1 Q2FWC7 Q2FWD0 Q2FWD1 Q2FWD6
Q2FWD8 Q2FWE0 Q2FWE1 Q2FWE3 Q2FWE4 Q2FWE5 Q2FWE6 Q2FWG0 Q2FWG2 Q2FWG3 Q2FWH3
Q2FWH5 Q2FWH6 Q2FWI7 Q2FWI8 Q2FWJ3 Q2FWJ6 Q2FWJ9 Q2FWK0 Q2FWK1 Q2FWK2 Q2FWK3
Q2FWK4 Q2FWK7 Q2FWL2 Q2FWL6 Q2FWN3 Q2FWN4 Q2FWX8 Q2FWX9 Q2FWY1 Q2FWY8 Q2FWY9
Q2FWZ0 Q2FWZ8 Q2FX13 Q2FX77 Q2FX88 Q2FXA0 Q2FXA3 Q2FXA4 Q2FXA5 Q2FXF2 Q2FXF5
Q2FXG2 Q2FXH2 Q2FXH9 Q2FXI2 Q2FXI7 Q2FXJ0 Q2FXJ5 Q2FXK6 Q2FXL1 Q2FXL5 Q2FXL6
Q2FXL7 Q2FXL9 Q2FXM0 Q2FXM8 Q2FXM9 Q2FXP3 Q2FXP7 Q2FXP9 Q2FXQ0 Q2FXQ1 Q2FXQ6
Q2FXQ7 Q2FXQ8 Q2FXQ9 Q2FXR3 Q2FXR4 Q2FXR8 Q2FXS8 Q2FXT0 Q2FXT1 Q2FXT2 Q2FXT3
Q2FXT4 Q2FXT5 Q2FXT6 Q2FXU0 Q2FXU2 Q2FXU4 Q2FXU5 Q2FXV1 Q2FXV6 Q2FXV9 Q2FXW0
Q2FXW1 Q2FXW2 Q2FXW6 Q2FXW7 Q2FXX2 Q2FXX8 Q2FXY1 Q2FXY3 Q2FXY6 Q2FXZ0 Q2FXZ1
Q2FXZ2 Q2FXZ3 Q2FXZ4 Q2FXZ7 Q2FY02 Q2FY08 Q2FY10 Q2FY15 Q2FY16 Q2FY19 Q2FY22
Q2FY32 Q2FY33 Q2FY34 Q2FY35 Q2FY37 Q2FY41 Q2FY45 Q2FY46 Q2FY47 Q2FY49 Q2FY55
Q2FY67 Q2FY76 Q2FY77 Q2FY79 Q2FYG0 Q2FYG7 Q2FYG9 Q2FYH0 Q2FYH3 Q2FYH5 Q2FYH6
Q2FYI4 Q2FYI5 Q2FYJ2 Q2FYJ3 Q2FYK3 Q2FYK4 Q2FYK5 Q2FYL1 Q2FYM1 Q2FYM2 Q2FYM7
Q2FYM9 Q2FYN2 Q2FYN6 Q2FYN7 Q2FYP3 Q2FYR3 Q2FYR4 Q2FYR5 Q2FYR6 Q2FYR7 Q2FYS1
Q2FYS5 Q2FYS7 Q2FYT3 Q2FYT4 Q2FYU4 Q2FYU5 Q2FYU6 Q2FYU7 Q2FYV2 Q2FYZ2 Q2FYZ4
Q2FYZ5 Q2FYZ9 Q2FZ02 Q2FZ19 Q2FZ20 Q2FZ21 Q2FZ22 Q2FZ23 Q2FZ25 Q2FZ28 Q2FZ29
Q2FZ30 Q2FZ31 Q2FZ32 Q2FZ37 Q2FZ38 Q2FZ42 Q2FZ43 Q2FZ44 Q2FZ45 Q2FZ50 Q2FZ58
Q2FZ60 Q2FZ66 Q2FZ68 Q2FZ71 Q2FZ72 Q2FZ75 Q2FZ77 Q2FZ82 Q2FZ86 Q2FZ92 Q2FZ97
Q2FZ98 Q2FZA9 Q2FZC6 Q2FZD0 Q2FZD2 Q2FZD3 Q2FZD7 Q2FZD9 Q2FZE2 Q2FZF1 Q2FZF4
Q2FZF5 Q2FZF8 Q2FZG8 Q2FZG9 Q2FZI6 Q2FZI8 Q2FZJ0 Q2FZJ1 Q2FZJ3 Q2FZJ6 Q2FZL7
Q2FZP4 Q2FZP6 Q2FZP9 Q2FZT4 Q2FZT5 Q2FZT6 Q2FZU0 Q2FZU1 Q2FZU2 Q2FZU9 Q2FZV9
Q2FZW6 Q2FZX0 Q2FZX4 Q2FZX9 Q2FZY3 Q2FZY9 Q2FZZ8 Q2G002 Q2G023 Q2G030 Q2G031
Q2G035 Q2G036 Q2G037 Q2G039 Q2G045 Q2G055 Q2G064 Q2G069 Q2G079 Q2G081 Q2G087
Q2G0B1 Q2G0B6 Q2G0D0 Q2G0D1 Q2G0F8 Q2G0G1 Q2G0I5 Q2G0I9 Q2G0J1 Q2G0J7 Q2G0K7

Q2G0K8 Q2G0L1 Q2G0L7 Q2G0M5 Q2G0M6 Q2G0M7 Q2G0N0 Q2G0N1 Q2G0N5 Q2G0N9 Q2G0P0
Q2G0P2 Q2G0P5 Q2G0P6 Q2G0P7 Q2G0P8 Q2G0Q0 Q2G0Q1 Q2G0Q3 Q2G0Q9 Q2G0R8 Q2G0R9
Q2G0S5 Q2G0S8 Q2G0T0 Q2G0T1 Q2G0T3 Q2G0T4 Q2G0Y6 Q2G0Y7 Q2G0Y9 Q2G0Z5 Q2G111
Q2G113 Q2G122 Q2G124 Q2G157 Q2G160 Q2G178 Q2G1A5 Q2G1B9 Q2G1C0 Q2G1C4 Q2G1D7
Q2G1D8 Q2G1F2 Q2G1G6 Q2G1H4 Q2G1H6 Q2G1J0 Q2G1J2 Q2G1M5 Q2G1M6 Q2G1M8 Q2G1M9
Q2G1N1 Q2G1N2 Q2G1N3 Q2G1N7 Q2G1P1 Q2G1T3 Q2G1T7 Q2G1T9 Q2G1U0 Q2G1U5 Q2G1U6
Q2G1W2 Q2G1W4 Q2G1X6 Q2G1Y0 Q2G1Y5 Q2G1Z4 Q2G1Z8 Q2G1Z9 Q2G200 Q2G203 Q2G218
Q2G227 Q2G230 Q2G236 Q2G241 Q2G252 Q2G261 Q2G272 Q2G273 Q2G275 Q2G282 Q2G283
Q2G296 Q2G297 Q2G2A2 Q2G2B1 Q2G2C3 Q2G2C4 Q2G2D0 Q2G2D3 Q2G2D5 Q2G2G2 Q2G2G7
Q2G2G9 Q2G2J8 Q2G2K5 Q2G2K6 Q2G2K7 Q2G2K8 Q2G2M4 Q2G2M6 Q2G2M8 Q2G2N4 Q2G2P7
Q2G2P8 Q2G2Q0 Q2G2Q1 Q2G2Q3 Q2G2Q4 Q2G2R0 Q2G2R4 Q2G2S0 Q2G2T3 Q2G2U9 Q2G2W7
Q2G2X2 Q2G2Z1 Q2G2Z4 Q2G300 Q2G319 Q2G338 Q2G341 Q2G348 Q2G365 Q2G373 Q2G3C3
Q2G3F0 Q2G3G8 Q2G3J8 Q2G3K1 Q2G3S7 Q2G3S9 Q2G3T3 Q2G3T4 Q2G467 Q2G469 Q2G480
Q2G484 Q2G491 Q2G492 Q2G493 Q2G4F2 Q2G4F4 Q2G4F5 Q2G4F6 Q2G4K2 Q2G4V4 Q2G4W0
Q2G4X5 Q2G512 Q2G552 Q2G586 Q2G5A8 Q2G5A9 Q2G5B0 Q2G5B1 Q2G5E4 Q2G5E7 Q2G5E9
Q2G5F0 Q2G5F3 Q2G5F4 Q2G5F5 Q2G5P7 Q2G5Q6 Q2G5R9 Q2G601 Q2G623 Q2G626 Q2G630
Q2G632 Q2G636 Q2G639 Q2G641 Q2G644 Q2G646 Q2G656 Q2G669 Q2G6G8 Q2G6G9 Q2G6I4
Q2G6M4 Q2G6M5 Q2G6N0 Q2G6P1 Q2G6Q3 Q2G6Q7 Q2G6R0 Q2G6R1 Q2G6R7 Q2G6T1 Q2G6V6
Q2G6W8 Q2G6X5 Q2G6Y3 Q2G708 Q2G712 Q2G718 Q2G733 Q2G781 Q2G794 Q2G798 Q2G7A9
Q2G7C4 Q2G7E9 Q2G7F1 Q2G7H8 Q2G7J5 Q2G7S5 Q2G7S6 Q2G7S7 Q2G8D5 Q2G8D6 Q2G8E3
Q2G8G3 Q2G8G4 Q2G8G5 Q2G8H1 Q2G8H2 Q2G8H3 Q2G8H4 Q2G8J9 Q2G8K7 Q2G8K8 Q2G8K9
Q2G8L0 Q2G8P1 Q2G8Q3 Q2G8S2 Q2G8S7 Q2G8S8 Q2G8V0 Q2G8W1 Q2G8W2 Q2G8W3 Q2G8W4
Q2G8W5 Q2G8W6 Q2G8W7 Q2G8W8 Q2G8W9 Q2G8X0 Q2G8X1 Q2G8X3 Q2G8X4 Q2G8X5 Q2G8X6
Q2G8X7 Q2G8X8 Q2G8X9 Q2G8Y0 Q2G8Y1 Q2G8Y2 Q2G8Y3 Q2G8Y4 Q2G8Y5 Q2G933 Q2G946
Q2G955 Q2G958 Q2G982 Q2G983 Q2G992 Q2G993 Q2G994 Q2G997 Q2G9A3 Q2G9B0 Q2G9C7
Q2G9D9 Q2G9E3 Q2G9G8 Q2G9H0 Q2G9H1 Q2G9I7 Q2G9J6 Q2G9L7 Q2G9L9 Q2G9M2 Q2G9M5
Q2G9P6 Q2G9P8 Q2G9R1 Q2G9T9 Q2G9U3 Q2G9U8 Q2G9U9 Q2G9W6 Q2G9X2 Q2G9X9 Q2G9Y4
Q2G9Z2 Q2G9Z6 Q2GA15 Q2GA28 Q2GA32 Q2GA44 Q2GA45 Q2GA52 Q2GAB9 Q2GAF7 Q2GAF9
Q2GAI1 Q2GAI7 Q2GAI8 Q2GAJ0 Q2GAJ1 Q2GAJ2 Q2GAJ3 Q2GAJ4 Q2GAK1 Q2GAT0 Q2GAU0
Q2GAU5 Q2GAU7 Q2GAV1 Q2GB44 Q2GB53 Q2GB76 Q2GBA2 Q2GBA3 Q2GBA4 Q2GBA6 Q2GBJ7
Q2GBK2 Q2GBM0 Q2GBN6 Q2GBN7 Q2GBP1 Q2GBV3 Q2GBX1 Q2GC13 Q2GC28 Q2GC31 Q2GC33
Q2GC37 Q2GC38 Q2GC39 Q2GC56 Q2GC57 Q2GC58 Q2GC60 Q2GC80 Q2GC97 Q2GCB4 Q2GCB7
Q2GCC6 Q2GCC7 Q2GCD1 Q2GCD2 Q2GCD7 Q2GCD8 Q2GCF2 Q2GCG4 Q2GCG8 Q2GCH5 Q2GCH6
Q2GCH7 Q2GCI4 Q2GCI5 Q2GCI6 Q2GCI7 Q2GCL0 Q2GCL1 Q2GCN1 Q2GCN2 Q2GCN3 Q2GCN4
Q2GCP9 Q2GCQ2 Q2GCS3 Q2GCS7 Q2GCT8 Q2GCU4 Q2GCU9 Q2GCV3 Q2GCW3 Q2GCX9 Q2GCZ0
Q2GCZ2 Q2GD18 Q2GD19 Q2GD33 Q2GD44 Q2GD50 Q2GD53 Q2GD63 Q2GD69 Q2GD80 Q2GD81
Q2GD82 Q2GD83 Q2GD87 Q2GD90 Q2GD91 Q2GD99 Q2GDA1 Q2GDC5 Q2GDC6 Q2GDC7 Q2GDD5
Q2GDE9 Q2GDF4 Q2GDF7 Q2GDG2 Q2GDI3 Q2GDJ0 Q2GDJ1 Q2GDJ5 Q2GDK4 Q2GDL6 Q2GDL7
Q2GDP4 Q2GDQ6 Q2GDR2 Q2GDR7 Q2GDS9 Q2GDU3 Q2GDV1 Q2GDW7 Q2GDX3 Q2GE03 Q2GE16
Q2GE40 Q2GE45 Q2GE48 Q2GE55 Q2GE60 Q2GE74 Q2GE82 Q2GE84 Q2GE85 Q2GE99 Q2GEA0
Q2GEB1 Q2GEB5 Q2GEB6 Q2GEB7 Q2GEB8 Q2GEB9 Q2GEC2 Q2GEC5 Q2GEC6 Q2GEC7 Q2GEC8
Q2GEC9 Q2GED2 Q2GED3 Q2GED4 Q2GED5 Q2GED6 Q2GED8 Q2GEE0 Q2GEE6 Q2GEE7 Q2GEE8
Q2GEF1 Q2GEF3 Q2GEG6 Q2GEH7 Q2GEI3 Q2GEI6 Q2GEI7 Q2GEM3 Q2GEM5 Q2GEP2 Q2GER2
Q2GES2 Q2GES3 Q2GEU6 Q2GEW1 Q2GEY4 Q2GEY5 Q2GEY7 Q2GEY8 Q2GEZ6 Q2GF15 Q2GF33

Q2GF34 Q2GF44 Q2GF82 Q2GF90 Q2GFE4 Q2GFI0 Q2GFK0 Q2GFL5 Q2GFL7 Q2GFN3 Q2GFN4
Q2GFN5 Q2GFN6 Q2GFP0 Q2GFP2 Q2GFP4 Q2GFP8 Q2GFQ3 Q2GFQ7 Q2GFR2 Q2GFT0 Q2GFT7
Q2GFT8 Q2GFT9 Q2GFU1 Q2GFU2 Q2GFU3 Q2GFV4 Q2GFW6 Q2GFY5 Q2GFZ4 Q2GG00 Q2GG09
Q2GG13 Q2GG24 Q2GG27 Q2GG30 Q2GG34 Q2GG37 Q2GG39 Q2GG41 Q2GG43 Q2GG44 Q2GG58
Q2GG67 Q2GG78 Q2GG92 Q2GGA2 Q2GGA3 Q2GGA4 Q2GGC4 Q2GGC5 Q2GGC7 Q2GGE6 Q2GGG8
Q2GGH5 Q2GGH6 Q2GGH7 Q2GGH9 Q2GGI2 Q2GGJ4 Q2GGK1 Q2GGK2 Q2GGL8 Q2GGM5 Q2GGM7
Q2GGM8 Q2GGN4 Q2GGQ8 Q2GGR4 Q2GGS4 Q2GGS5 Q2GGS7 Q2GGS8 Q2GGT3 Q2GGT4 Q2GGT6
Q2GGV3 Q2GGV4 Q2GGV6 Q2GGW3 Q2GGX7 Q2GGY3 Q2GH05 Q2GH06 Q2GH19 Q2GH22 Q2GH25
Q2GH32 Q2GH33 Q2GH34 Q2GH35 Q2GH38 Q2GH39 Q2GH40 Q2GH42 Q2GH43 Q2GH44 Q2GH45
Q2GH46 Q2GH47 Q2GH48 Q2GH49 Q2GH50 Q2GH51 Q2GH52 Q2GH53 Q2GH55 Q2GH57 Q2GH90
Q2GH92 Q2GH98 Q2GH99 Q2GHA4 Q2GHB1 Q2GHC9 Q2GHD8 Q2GHE0 Q2GHE2 Q2GHE3 Q2GHF1
Q2GHF2 Q2GHF3 Q2GHF4 Q2GHG1 Q2GHH1 Q2GHH2 Q2GHI5 Q2GHJ5 Q2GHJ6 Q2GHJ9 Q2GHM9
Q2GHN3 Q2GHQ1 Q2GHR1 Q2GHR2 Q2GHR3 Q2GHR5 Q2GHR7 Q2GHS3 Q2GHT7 Q2GHU0 Q2GHV0
Q2GHW1 Q2GHW3 Q2GHW4 Q2GHX0 Q2GHY2 Q2GI11 Q2GI13 Q2GI32 Q2GI42 Q2GI72 Q2GI75
Q2GI91 Q2GI92 Q2GI94 Q2GI95 Q2GI96 Q2GIC8 Q2GID5 Q2GIG5 Q2GIH0 Q2GII5 Q2GII6
Q2GIJ7 Q2GIJ8 Q2GIK8 Q2GIL4 Q2GIL5 Q2GIN1 Q2GIQ3 Q2GIW0 Q2GIZ5 Q2GJ25 Q2GJ36
Q2GJ42 Q2GJ58 Q2GJ59 Q2GJ60 Q2GJ61 Q2GJ65 Q2GJ67 Q2GJ68 Q2GJ69 Q2GJ70 Q2GJ76
Q2GJ82 Q2GJ99 Q2GJA0 Q2GJB3 Q2GJB4 Q2GJB5 Q2GJB8 Q2GJC1 Q2GJC2 Q2GJE2 Q2GJF3
Q2GJG8 Q2GJI3 Q2GJJ1 Q2GJJ5 Q2GJL7 Q2GJT4 Q2GJV3 Q2GJV4 Q2GJV5 Q2GJX4 Q2GJY5
Q2GJY7 Q2GK20 Q2GK21 Q2GK22 Q2GK25 Q2GK29 Q2GK70 Q2GK85 Q2GK88 Q2GK91 Q2GKB1
Q2GKD6 Q2GKF4 Q2GKG2 Q2GKJ2 Q2GKJ8 Q2GKK2 Q2GKK3 Q2GKK4 Q2GKM2 Q2GKN0 Q2GKQ2
Q2GKQ8 Q2GKR7 Q2GKS1 Q2GKS7 Q2GKS8 Q2GKT8 Q2GKU8 Q2GKU9 Q2GKV0 Q2GKV7 Q2GKX7
Q2GKY5 Q2GL11 Q2GL24 Q2GL30 Q2GL35 Q2GL36 Q2GL37 Q2GL38 Q2GL42 Q2GL43 Q2GL45
Q2GL46 Q2GL47 Q2GL48 Q2GL49 Q2GL50 Q2GL52 Q2GL53 Q2GL54 Q2GL55 Q2GL56 Q2GL58
Q2GL60 Q2GL89 Q2GL91 Q2GL93 Q2GL94 Q2GLA8 Q2GLB4 Q2GLD8 Q2GLF2 Q2GLF7 Q2GLG0
Q2GLG1 Q2GLH3 Q2GLH4 Q2GLH5 Q2GLH6 Q2GLI8 Q2GLJ4 Q2GLK4 Q2GLL2 Q2GLM6 Q2GLN0
Q2GLP1 Q2GLP6 Q2GLQ7 Q2GLQ8 Q2GLR4 Q2GLR6 Q2GLS0 Q2GLS9 Q2GLU6 Q2GLU9 Q2GM53
Q2GM68 Q2GMF6 Q2GMX1 Q2GN26 Q2GNN3 Q2GNS7 Q2GNY7 Q2GNZ6 Q2GP45 Q2GPK5 Q2GQ93
Q2GQS2 Q2GRJ3 Q2GS86 Q2GSA4 Q2GSJ7 Q2GSJ9 Q2GSV2 Q2GSX8 Q2GT28 Q2GTG9 Q2GTM8
Q2GUI4 Q2GUW6 Q2GV19 Q2GV49 Q2GV75 Q2GVC2 Q2GVK1 Q2GVK6 Q2GVR6 Q2GW22 Q2GW51
Q2GW61 Q2GWF3 Q2GWJ5 Q2GWJ7 Q2GWZ4 Q2GXM5 Q2GXN6 Q2GXT0 Q2GXY3 Q2GXZ7 Q2GY21
Q2GYA8 Q2GYB6 Q2GYV9 Q2GZK0 Q2GZN9 Q2GZQ4 Q2GZU7 Q2H080 Q2H085 Q2H0C0 Q2H0G2
Q2H0I3 Q2H0R2 Q2H0S9 Q2H0U6 Q2H132 Q2H137 Q2H1L0 Q2H1Q1 Q2H1Q8 Q2H2D7 Q2H2I4
Q2H2J1 Q2H317 Q2H334 Q2H3D0 Q2H3R6 Q2H401 Q2H454 Q2H4D0 Q2H5T8 Q2H5Z7 Q2H679
Q2H683 Q2H6A9 Q2H6C5 Q2H6G4 Q2H6N9 Q2H6R9 Q2H720 Q2H731 Q2H7G2 Q2H7U1 Q2H8D5
Q2H8S0 Q2H8T2 Q2H988 Q2H991 Q2H9E8 Q2H9L1 Q2H9N4 Q2H9P7 Q2HA12 Q2HA26 Q2HAD8
Q2HAJ4 Q2HAN0 Q2HAX7 Q2HBE7 Q2HBI0 Q2HBN0 Q2HBN1 Q2HBN2 Q2HBN3 Q2HBN4 Q2HBN5
Q2HBN6 Q2HBQ2 Q2HBX6 Q2HCM6 Q2HCV1 Q2HCV7 Q2HD63 Q2HDD6 Q2HDI2 Q2HDP2 Q2HDQ7
Q2HDR6 Q2HDU8 Q2HDW0 Q2HE65 Q2HE69 Q2HE71 Q2HE94 Q2HEB0 Q2HEF1 Q2HEK0 Q2HEW4
Q2HEW5 Q2HEW6 Q2HEW7 Q2HFA6 Q2HFD4 Q2HFP1 Q2HFZ9 Q2HG76 Q2HGJ1 Q2HGJ2 Q2HGY8
Q2HH37 Q2HH38 Q2HH48 Q2HHB4 Q2HHH2 Q2HHS4 Q2HIJ8 Q2HIV9 Q2HJ27 Q2HJ38 Q2HJ41
Q2HJ46 Q2HJ47 Q2HJ48 Q2HJ51 Q2HJ52 Q2HJ56 Q2HJ58 Q2HJ67 Q2HJ73 Q2HJ78 Q2HJ84
Q2HJ86 Q2HJ87 Q2HJ88 Q2HJ90 Q2HJ91 Q2HJ92 Q2HJ93 Q2HJ98 Q2HJA2 Q2HJA9 Q2HJB0
Q2HJC9 Q2HJD5 Q2HJD7 Q2HJF2 Q2HJF3 Q2HJF5 Q2HJG2 Q2HJG4 Q2HJG7 Q2HJG8 Q2HJH0

Q2HJH1 Q2HJH3 Q2HJH6 Q2HJH8 Q2HJI3 Q2HJJ0 Q2HJN4 Q2HJN6 Q2HJN8 Q2HJN9 Q2HNT1
Q2HNT2 Q2HR95 Q2HU65 Q2HU68 Q2HVD6 Q2HW56 Q2HXI5 Q2HXI6 Q2HXI7 Q2HXL9 Q2HY40
Q2HZ26 Q2HZ33 Q2I0M6 Q2I327 Q2I689 Q2I8V6 Q2IA00 Q2IBA2 Q2IBA3 Q2IBA7 Q2IBB1
Q2IBB2 Q2IBB4 Q2IBC3 Q2IBD4 Q2IBE3 Q2IBE6 Q2IBE7 Q2IBF5 Q2IBF7 Q2IBF8 Q2IBG0
Q2IBH0 Q2IE39 Q2IEC3 Q2IEW0 Q2IEW1 Q2IEX1 Q2IEX2 Q2IEX8 Q2IF61 Q2IF62 Q2IFJ5
Q2IFK0 Q2IFM3 Q2IFP3 Q2IFP4 Q2IFQ4 Q2IFT1 Q2IFT5 Q2IFT9 Q2IFU2 Q2IFU3 Q2IFU7
Q2IG17 Q2IG19 Q2IG29 Q2IG30 Q2IG33 Q2IG34 Q2IG40 Q2IG43 Q2IG55 Q2IG97 Q2IGB1
Q2IGK0 Q2IGK6 Q2IGM2 Q2IGM4 Q2IGT4 Q2IGV6 Q2IGV9 Q2IGW6 Q2IGX0 Q2IGX5 Q2IGX8
Q2IH01 Q2IH17 Q2IH69 Q2IH81 Q2IH83 Q2IH84 Q2IH91 Q2IH93 Q2IH94 Q2IH95 Q2IH96
Q2IHC1 Q2IHD1 Q2IHD7 Q2IHM1 Q2IHN8 Q2IHP8 Q2IHR2 Q2IHR4 Q2IHV2 Q2IHX6 Q2IHZ5
Q2IHZ6 Q2II09 Q2II14 Q2II25 Q2II77 Q2II78 Q2II80 Q2II81 Q2II82 Q2II85 Q2II86
Q2II87 Q2IIA8 Q2IIB0 Q2IIB1 Q2IIB9 Q2IIC1 Q2IIC2 Q2IIC5 Q2IID8 Q2IIJ3 Q2IIK1
Q2IIL0 Q2IIL9 Q2IIM3 Q2IIV4 Q2IIW6 Q2IJ48 Q2IJ53 Q2IJ54 Q2IJ55 Q2IJ57 Q2IJ58
Q2IJ60 Q2IJ62 Q2IJ63 Q2IJ64 Q2IJ65 Q2IJ66 Q2IJ67 Q2IJ68 Q2IJ70 Q2IJ71 Q2IJ72
Q2IJ73 Q2IJ74 Q2IJ76 Q2IJ77 Q2IJ78 Q2IJ79 Q2IJ80 Q2IJ81 Q2IJ82 Q2IJ83 Q2IJ85
Q2IJ86 Q2IJ87 Q2IJ88 Q2IJ91 Q2IJ92 Q2IJ93 Q2IJA7 Q2IJB1 Q2IJB3 Q2IJB7 Q2IJB8
Q2IJB9 Q2IJC0 Q2IJC2 Q2IJC3 Q2IJC6 Q2IJD0 Q2IJD4 Q2IJI0 Q2IJJ5 Q2IJK7 Q2IJL8
Q2IJN4 Q2IJX6 Q2IK16 Q2IK81 Q2IKE9 Q2IKH9 Q2IKI2 Q2IKJ8 Q2IKN1 Q2IKQ7 Q2IKT3
Q2IKU4 Q2IKW1 Q2IKX7 Q2IKY3 Q2IL38 Q2IL45 Q2ILB9 Q2ILC5 Q2ILD4 Q2ILE0 Q2ILG1
Q2ILG4 Q2ILH7 Q2ILH8 Q2ILI1 Q2ILI2 Q2ILJ1 Q2ILU8 Q2ILX2 Q2IM14 Q2IM25 Q2IM40
Q2IM41 Q2IM42 Q2IM61 Q2IM62 Q2IM64 Q2IM65 Q2IM67 Q2IM68 Q2IM81 Q2IM96 Q2IM98
Q2IMB4 Q2IMC5 Q2IMC7 Q2IMC8 Q2IMD4 Q2IMF3 Q2IMG0 Q2IMI1 Q2IML9 Q2IMM0 Q2IMM1
Q2IMM2 Q2IMM4 Q2IMQ8 Q2IMV5 Q2IMV6 Q2IMW2 Q2IN94 Q2INH7 Q2INI7 Q2INJ4 Q2INT4
Q2INT8 Q2INT9 Q2INV5 Q2INW0 Q2INW2 Q2INZ4 Q2INZ6 Q2IP38 Q2IP67 Q2IPE3 Q2IPF0
Q2IPJ5 Q2IPJ6 Q2IPJ7 Q2IPT3 Q2IPT7 Q2IPW2 Q2IPX5 Q2IPY2 Q2IPY3 Q2IPZ0 Q2IPZ2
Q2IPZ5 Q2IPZ7 Q2IQ00 Q2IQ01 Q2IQ02 Q2IQC1 Q2IQC5 Q2IQD6 Q2IQE1 Q2IQE7 Q2IQF7
Q2IQF8 Q2IQG7 Q2IQG8 Q2IQH0 Q2IQJ5 Q2IQJ6 Q2IQJ8 Q2IQR1 Q2IQW5 Q2IQY5 Q2IRD5
Q2IRH1 Q2IRH5 Q2IRI6 Q2IRL7 Q2IRL8 Q2IRX1 Q2IRX4 Q2IRX7 Q2IRY0 Q2IS06 Q2IS49
Q2IS53 Q2IS68 Q2IS70 Q2IS91 Q2ISE2 Q2ISF7 Q2ISF8 Q2ISK2 Q2ISQ1 Q2ISV3 Q2ISY6
Q2ISY7 Q2ISY9 Q2ISZ6 Q2IT03 Q2ITA9 Q2ITC6 Q2ITM9 Q2ITP9 Q2ITQ2 Q2ITR6 Q2ITZ4
Q2IU23 Q2IU52 Q2IUB0 Q2IUE0 Q2IUI7 Q2IUJ4 Q2IUM8 Q2IUP0 Q2IUR8 Q2IUS3 Q2IUT3
Q2IUT4 Q2IV18 Q2IV30 Q2IV63 Q2IV68 Q2IV75 Q2IVL9 Q2IVN3 Q2IVP0 Q2IVR1 Q2IVV2
Q2IW19 Q2IW20 Q2IW23 Q2IW30 Q2IW34 Q2IW68 Q2IW74 Q2IW80 Q2IW81 Q2IW82 Q2IW83
Q2IW86 Q2IW96 Q2IWB0 Q2IWB1 Q2IWB8 Q2IWC9 Q2IWF7 Q2IWF8 Q2IWG6 Q2IWG9 Q2IWH1
Q2IWI0 Q2IWJ9 Q2IWN6 Q2IWN7 Q2IWQ7 Q2IWR9 Q2IWS3 Q2IWS4 Q2IWT7 Q2IWU2 Q2IWU8
Q2IWV0 Q2IWV3 Q2IWW0 Q2IWZ3 Q2IWZ4 Q2IWZ5 Q2IX59 Q2IX60 Q2IX63 Q2IX70 Q2IX74
Q2IX80 Q2IX83 Q2IX91 Q2IX92 Q2IX94 Q2IXA7 Q2IXB0 Q2IXB5 Q2IXB7 Q2IXC1 Q2IXC6
Q2IXD1 Q2IXE1 Q2IXG7 Q2IXI4 Q2IXI7 Q2IXN5 Q2IXN6 Q2IXN7 Q2IXN8 Q2IXP1 Q2IXP2
Q2IXP3 Q2IXP4 Q2IXP5 Q2IXP6 Q2IXP7 Q2IXP8 Q2IXP9 Q2IXQ0 Q2IXQ1 Q2IXQ2 Q2IXQ3
Q2IXQ4 Q2IXQ5 Q2IXQ6 Q2IXQ7 Q2IXQ8 Q2IXQ9 Q2IXR0 Q2IXR1 Q2IXR2 Q2IXR3 Q2IXR4
Q2IXR5 Q2IXR9 Q2IXS0 Q2IXS1 Q2IXS2 Q2IXS7 Q2IXS8 Q2IY19 Q2IY20 Q2IYB5 Q2IYG1
Q2IYG7 Q2IYJ4 Q2IYK4 Q2IYK6 Q2IYL0 Q2IYL6 Q2IYP6 Q2IYP8 Q2IYT0 Q2IYX0 Q2IYY0
Q2IYY4 Q2IYY5 Q2IZ07 Q2IZ16 Q2IZ49 Q2IZ51 Q2IZ52 Q2IZ53 Q2IZ60 Q2IZ89 Q2IZP7
Q2IZR1 Q2IZR2 Q2IZZ7 Q2J055 Q2J074 Q2J082 Q2J097 Q2J0A5 Q2J0B6 Q2J0F8 Q2J0F9

Q2J0H0 Q2J0J3 Q2J0J8 Q2J0T8 Q2J0U5 Q2J0X3 Q2J164 Q2J169 Q2J1G7 Q2J1I1 Q2J1J9
Q2J1K3 Q2J1P2 Q2J1V2 Q2J1X2 Q2J299 Q2J2B4 Q2J2B7 Q2J2B9 Q2J2C3 Q2J2C4 Q2J2C5
Q2J2C6 Q2J2D2 Q2J2D3 Q2J2F6 Q2J2F8 Q2J2G1 Q2J2G3 Q2J2G4 Q2J2G5 Q2J2H6 Q2J2H9
Q2J2I0 Q2J2I9 Q2J2J4 Q2J2J5 Q2J2J8 Q2J2J9 Q2J2K2 Q2J2K3 Q2J2K7 Q2J2K9 Q2J2Q3
Q2J2S1 Q2J2S7 Q2J2T5 Q2J2X6 Q2J2Y8 Q2J302 Q2J307 Q2J308 Q2J314 Q2J316 Q2J319
Q2J320 Q2J322 Q2J323 Q2J324 Q2J329 Q2J338 Q2J340 Q2J341 Q2J344 Q2J346 Q2J350
Q2J352 Q2J353 Q2J354 Q2J357 Q2J358 Q2J359 Q2J364 Q2J390 Q2J397 Q2J398 Q2J399
Q2J3A0 Q2J3A1 Q2J3A7 Q2J3B4 Q2J3D4 Q2J3F5 Q2J3G8 Q2J3G9 Q2J3I9 Q2J3J3 Q2J3J6
Q2J3J7 Q2J3J8 Q2J3K2 Q2J3K3 Q2J3Y9 Q2J3Z0 Q2J3Z9 Q2J405 Q2J419 Q2J423 Q2J429
Q2J454 Q2J455 Q2J497 Q2J498 Q2J4A4 Q2J4C6 Q2J4C7 Q2J4D1 Q2J4F8 Q2J4G6 Q2J4M9
Q2J4N0 Q2J4P8 Q2J4R7 Q2J4S0 Q2J4S8 Q2J528 Q2J529 Q2J542 Q2J543 Q2J545 Q2J5A7
Q2J5Z0 Q2J5Z1 Q2J641 Q2J645 Q2J678 Q2J6D7 Q2J6H8 Q2J6H9 Q2J6L9 Q2J6M0 Q2J6P5
Q2J6Q9 Q2J6U1 Q2J6U3 Q2J6U6 Q2J6U8 Q2J6V2 Q2J6V8 Q2J6W9 Q2J6X0 Q2J6X4 Q2J6Y1
Q2J6Y2 Q2J6Y5 Q2J6Y7 Q2J6Z2 Q2J6Z6 Q2J6Z8 Q2J6Z9 Q2J700 Q2J702 Q2J704 Q2J710
Q2J711 Q2J713 Q2J717 Q2J725 Q2J728 Q2J730 Q2J734 Q2J735 Q2J750 Q2J752 Q2J764
Q2J771 Q2J773 Q2J781 Q2J7C9 Q2J7E7 Q2J7H0 Q2J7K4 Q2J7K8 Q2J7L5 Q2J816 Q2J821
Q2J822 Q2J826 Q2J827 Q2J829 Q2J830 Q2J833 Q2J838 Q2J840 Q2J841 Q2J843 Q2J845
Q2J850 Q2J854 Q2J855 Q2J860 Q2J862 Q2J866 Q2J867 Q2J868 Q2J878 Q2J886 Q2J897
Q2J8F9 Q2J8K9 Q2J8L0 Q2J8L2 Q2J8L3 Q2J8P2 Q2J8X3 Q2J9A8 Q2J9A9 Q2J9H6 Q2J9L3
Q2J9L7 Q2J9L8 Q2J9N8 Q2J9Q0 Q2J9Q3 Q2J9Q4 Q2J9R8 Q2J9S7 Q2JA12 Q2JAZ7 Q2JB35
Q2JBP8 Q2JCE9 Q2JCG8 Q2JCH8 Q2JCI0 Q2JCI2 Q2JCI3 Q2JCJ1 Q2JCX4 Q2JD44 Q2JD48
Q2JD52 Q2JD58 Q2JD59 Q2JD90 Q2JD94 Q2JD96 Q2JD97 Q2JD98 Q2JDD1 Q2JDD9 Q2JDJ2
Q2JDJ6 Q2JDK0 Q2JDK3 Q2JDN2 Q2JDN7 Q2JDP1 Q2JDP2 Q2JDP3 Q2JDP4 Q2JDQ4 Q2JDQ7
Q2JDQ8 Q2JDQ9 Q2JDR0 Q2JE00 Q2JEN8 Q2JES4 Q2JES5 Q2JES6 Q2JES8 Q2JEV7 Q2JEW7
Q2JF65 Q2JFA7 Q2JFC5 Q2JFC9 Q2JFD6 Q2JFE2 Q2JFE4 Q2JFE6 Q2JFE7 Q2JFE8 Q2JFE9
Q2JFF0 Q2JFF1 Q2JFF2 Q2JFF3 Q2JFF7 Q2JFF8 Q2JFF9 Q2JFG0 Q2JFG1 Q2JFG2 Q2JFG3
Q2JFG4 Q2JFG5 Q2JFG6 Q2JFG7 Q2JFG9 Q2JFH0 Q2JFH1 Q2JFH2 Q2JFH3 Q2JFH4 Q2JFH5
Q2JFH6 Q2JFH7 Q2JFH8 Q2JFH9 Q2JFI0 Q2JFI1 Q2JFI4 Q2JFI5 Q2JFI6 Q2JFI7 Q2JFI8
Q2JFI9 Q2JFJ5 Q2JFK0 Q2JFM8 Q2JFQ1 Q2JFS1 Q2JFV0 Q2JFV4 Q2JFW4 Q2JG22 Q2JGD6
Q2JGG1 Q2JGW9 Q2JH03 Q2JH49 Q2JH51 Q2JH53 Q2JH54 Q2JHA7 Q2JHC7 Q2JHF7 Q2JHM1
Q2JHN6 Q2JHP5 Q2JHQ5 Q2JHS1 Q2JHS2 Q2JHS3 Q2JHU0 Q2JHU3 Q2JHW2 Q2JHW3 Q2JHX0
Q2JHZ4 Q2JI02 Q2JI03 Q2JI25 Q2JI26 Q2JI31 Q2JI33 Q2JI36 Q2JI39 Q2JI46 Q2JI48
Q2JI70 Q2JIA6 Q2JIB8 Q2JIE6 Q2JIE9 Q2JIJ2 Q2JIK6 Q2JIK8 Q2JIL0 Q2JIL2 Q2JIL3
Q2JIL4 Q2JIL5 Q2JIL6 Q2JIL7 Q2JIL8 Q2JIL9 Q2JIM0 Q2JIM1 Q2JIM2 Q2JIM3 Q2JIM4
Q2JIM5 Q2JIM6 Q2JIM7 Q2JIN3 Q2JIP1 Q2JIP3 Q2JIQ7 Q2JIR9 Q2JIV0 Q2JJ17 Q2JJ18
Q2JJ19 Q2JJ22 Q2JJ30 Q2JJ72 Q2JJ73 Q2JJ74 Q2JJ90 Q2JJC6 Q2JJE2 Q2JJG8 Q2JJG9
Q2JJI0 Q2JJK5 Q2JJM6 Q2JJM9 Q2JJN4 Q2JJP4 Q2JJQ0 Q2JJQ1 Q2JJQ9 Q2JJU2 Q2JJV2
Q2JJY4 Q2JK09 Q2JK10 Q2JK17 Q2JK54 Q2JK57 Q2JK60 Q2JK61 Q2JK64 Q2JK68 Q2JK82
Q2JK90 Q2JK96 Q2JKB7 Q2JKC1 Q2JKD4 Q2JKE8 Q2JKH3 Q2JKK9 Q2JKL2 Q2JKL5 Q2JKN4
Q2JKN5 Q2JKP0 Q2JKQ8 Q2JKS2 Q2JKS4 Q2JKS7 Q2JKT3 Q2JKT7 Q2JKU0 Q2JKV7 Q2JKV8
Q2JKX6 Q2JKY0 Q2JKY3 Q2JKZ9 Q2JL07 Q2JL27 Q2JL28 Q2JL30 Q2JL42 Q2JL43 Q2JL51
Q2JL68 Q2JL69 Q2JL70 Q2JL71 Q2JL73 Q2JL74 Q2JL75 Q2JL76 Q2JL77 Q2JL88 Q2JL97
Q2JLB2 Q2JLB3 Q2JLC4 Q2JLD4 Q2JLD5 Q2JLD7 Q2JLG7 Q2JLJ5 Q2JLK3 Q2JLL9 Q2JLM3
Q2JLP6 Q2JLQ2 Q2JLR0 Q2JLR9 Q2JLT0 Q2JLT1 Q2JLU2 Q2JLU3 Q2JLV1 Q2JLV2 Q2JLX1

Q2JM08 Q2JM09 Q2JM21 Q2JM39 Q2JM43 Q2JM48 Q2JM49 Q2JM50 Q2JM51 Q2JM55 Q2JM78
Q2JMA3 Q2JMA8 Q2JMD7 Q2JMD8 Q2JMG5 Q2JMH0 Q2JMI4 Q2JMK8 Q2JMN2 Q2JMP1 Q2JMP3
Q2JMP5 Q2JMP7 Q2JMQ3 Q2JMR2 Q2JMR8 Q2JMX6 Q2JMX7 Q2JMX8 Q2JMX9 Q2JMY0 Q2JMZ3
Q2JN07 Q2JN09 Q2JN19 Q2JN62 Q2JN71 Q2JN87 Q2JNA9 Q2JNE1 Q2JNE2 Q2JNF7 Q2JNH2
Q2JNH3 Q2JNH4 Q2JNI1 Q2JNK3 Q2JNM6 Q2JNW7 Q2JNW8 Q2JNZ0 Q2JP15 Q2JP46 Q2JP53
Q2JP65 Q2JP68 Q2JP78 Q2JPE6 Q2JPF1 Q2JPG2 Q2JPG6 Q2JPH1 Q2JPJ7 Q2JPK0 Q2JPK6
Q2JPL4 Q2JPM4 Q2JPQ5 Q2JPS6 Q2JPT2 Q2JPT8 Q2JPU2 Q2JPU4 Q2JPX4 Q2JPZ9 Q2JQ25
Q2JQ26 Q2JQ33 Q2JQ36 Q2JQ57 Q2JQ59 Q2JQ60 Q2JQ64 Q2JQ65 Q2JQ67 Q2JQ88 Q2JQB4
Q2JQB9 Q2JQC0 Q2JQD7 Q2JQF4 Q2JQF8 Q2JQG8 Q2JQK3 Q2JQL3 Q2JQL5 Q2JQL7 Q2JQN1
Q2JQN2 Q2JQN3 Q2JQN4 Q2JQQ0 Q2JQS1 Q2JQT4 Q2JQT5 Q2JQT6 Q2JQU3 Q2JQU4 Q2JQW9
Q2JQX0 Q2JQX5 Q2JQX6 Q2JQY9 Q2JQZ7 Q2JR37 Q2JR49 Q2JR50 Q2JR54 Q2JR69 Q2JR73
Q2JRC3 Q2JRC6 Q2JRD6 Q2JRF6 Q2JRF7 Q2JRG2 Q2JRG5 Q2JRH1 Q2JRH2 Q2JRH9 Q2JRI4
Q2JRJ4 Q2JRJ6 Q2JRJ9 Q2JRL0 Q2JRM5 Q2JRN8 Q2JRN9 Q2JRQ8 Q2JRQ9 Q2JRR3 Q2JRU4
Q2JRV8 Q2JRX6 Q2JRY1 Q2JS04 Q2JS42 Q2JS69 Q2JS70 Q2JS72 Q2JS74 Q2JS78 Q2JSB6
Q2JSB7 Q2JSC8 Q2JSC9 Q2JSD0 Q2JSF0 Q2JSF1 Q2JSG0 Q2JSG7 Q2JSH9 Q2JSI0 Q2JSQ2
Q2JSU8 Q2JSV1 Q2JSX8 Q2JSZ9 Q2JT08 Q2JT18 Q2JT19 Q2JT26 Q2JT35 Q2JT47 Q2JT50
Q2JT83 Q2JT84 Q2JTF4 Q2JTG5 Q2JTG8 Q2JTH3 Q2JTH4 Q2JTJ1 Q2JTK2 Q2JTL7 Q2JTM7
Q2JTN8 Q2JTP2 Q2JTQ2 Q2JTQ3 Q2JTQ5 Q2JTQ6 Q2JTR0 Q2JTU8 Q2JTV2 Q2JTW6 Q2JTX2
Q2JTX5 Q2JTX6 Q2JTY2 Q2JTZ2 Q2JTZ8 Q2JU21 Q2JU22 Q2JU28 Q2JU40 Q2JU45 Q2JU97
Q2JUA5 Q2JUC0 Q2JUD0 Q2JUD2 Q2JUD6 Q2JUE5 Q2JUH1 Q2JUJ3 Q2JUJ4 Q2JUJ5 Q2JUJ6
Q2JUJ8 Q2JUJ9 Q2JUK0 Q2JUK1 Q2JUM6 Q2JUN7 Q2JUN8 Q2JUS2 Q2JUS3 Q2JUT8 Q2JUV6
Q2JUX3 Q2JUX4 Q2JUX5 Q2JUX6 Q2JUX7 Q2JUZ8 Q2JV04 Q2JV13 Q2JV16 Q2JV31 Q2JV45
Q2JV46 Q2JV67 Q2JV68 Q2JV74 Q2JV81 Q2JV82 Q2JV83 Q2JV84 Q2JV85 Q2JV86 Q2JV87
Q2JV88 Q2JV89 Q2JV90 Q2JV91 Q2JV92 Q2JV94 Q2JV96 Q2JV98 Q2JVC0 Q2JVC3 Q2JVE0
Q2JVG1 Q2JVI1 Q2JVK5 Q2JVL6 Q2JVR0 Q2JVR1 Q2JVR3 Q2JVR9 Q2JVS9 Q2JVU4 Q2JVV4
Q2JVW2 Q2JVX9 Q2JW12 Q2JW45 Q2JW63 Q2JW64 Q2JW78 Q2JW85 Q2JW90 Q2JWA7 Q2JWE1
Q2JWE3 Q2JWE4 Q2JWH1 Q2JWI9 Q2JWJ1 Q2JWJ9 Q2JWK2 Q2JWL1 Q2JWL5 Q2JWL7 Q2JWN6
Q2JWN7 Q2JWN9 Q2JWP0 Q2JWS5 Q2JWU6 Q2JWU7 Q2JWZ1 Q2JWZ2 Q2JX00 Q2JX13 Q2JX22
Q2JX64 Q2JX67 Q2JX75 Q2JX86 Q2JX97 Q2JXB3 Q2JXB4 Q2JXC9 Q2JXD0 Q2JXD4 Q2JXD5
Q2JXF0 Q2JXF5 Q2JXF6 Q2JXG4 Q2JXG8 Q2JXG9 Q2JXJ4 Q2JXL1 Q2JXL2 Q2JXL8 Q2JXP8
Q2JXQ9 Q2JXR2 Q2JXT8 Q2JXY2 Q2JXY9 Q2JY05 Q2JY06 Q2JY34 Q2JY68 Q2JYW6 Q2JZ21
Q2JZH2 Q2JZQ0 Q2JZQ2 Q2JZT8 Q2JZU0 Q2JZU1 Q2JZU4 Q2JZV0 Q2K0S6 Q2K0X8 Q2K196
Q2K206 Q2K2S0 Q2K2S1 Q2K2S2 Q2K2S7 Q2K2T1 Q2K2T6 Q2K2V0 Q2K2V2 Q2K2W7 Q2K2X3
Q2K2X4 Q2K2X5 Q2K2X6 Q2K2Y0 Q2K2Y1 Q2K302 Q2K327 Q2K333 Q2K364 Q2K376 Q2K379
Q2K380 Q2K381 Q2K382 Q2K386 Q2K3A7 Q2K3B1 Q2K3B7 Q2K3C8 Q2K3D7 Q2K3E9 Q2K3F0
Q2K3K0 Q2K3L3 Q2K3M2 Q2K3M6 Q2K3P2 Q2K3S8 Q2K3T1 Q2K3W4 Q2K3X0 Q2K414 Q2K424
Q2K425 Q2K433 Q2K443 Q2K474 Q2K485 Q2K486 Q2K487 Q2K4C4 Q2K4D4 Q2K4E7 Q2K4H4
Q2K4J8 Q2K4J9 Q2K4K2 Q2K4K5 Q2K4M3 Q2K4M8 Q2K4Q9 Q2K4W8 Q2K4X9 Q2K511 Q2K512
Q2K514 Q2K517 Q2K521 Q2K522 Q2K5I1 Q2K5L2 Q2K5M6 Q2K5N6 Q2K5P2 Q2K5P8 Q2K5Q0
Q2K5S3 Q2K5T4 Q2K5U1 Q2K5U4 Q2K600 Q2K618 Q2K619 Q2K665 Q2K6B3 Q2K6B9 Q2K6C2
Q2K6C3 Q2K6D3 Q2K6E0 Q2K6G8 Q2K6H4 Q2K6M2 Q2K7B5 Q2K7G6 Q2K7L4 Q2K7M6 Q2K7M8
Q2K7T3 Q2K7T4 Q2K7U4 Q2K7U6 Q2K7X0 Q2K7X5 Q2K7Z9 Q2K813 Q2K814 Q2K827 Q2K833
Q2K853 Q2K859 Q2K869 Q2K871 Q2K879 Q2K880 Q2K8D6 Q2K8D9 Q2K8U6 Q2K8V5 Q2K8V9
Q2K8Y4 Q2K8Y5 Q2K8Y6 Q2K8Y7 Q2K927 Q2K933 Q2K936 Q2K939 Q2K940 Q2K941 Q2K942

Q2K946 Q2K952 Q2K957 Q2K966 Q2K967 Q2K975 Q2K976 Q2K9C7 Q2K9E5 Q2K9I7 Q2K9J1
Q2K9J2 Q2K9J3 Q2K9J4 Q2K9J5 Q2K9J7 Q2K9J8 Q2K9J9 Q2K9K0 Q2K9K1 Q2K9K2 Q2K9K3
Q2K9K4 Q2K9K5 Q2K9K6 Q2K9K7 Q2K9K8 Q2K9K9 Q2K9L0 Q2K9L1 Q2K9L2 Q2K9L3 Q2K9L4
Q2K9L5 Q2K9L6 Q2K9L7 Q2K9L8 Q2K9L9 Q2K9M0 Q2K9M1 Q2K9M3 Q2K9M4 Q2K9M5 Q2K9M6
Q2K9M7 Q2K9M8 Q2K9N2 Q2K9P5 Q2K9Q3 Q2K9R4 Q2K9U6 Q2K9U7 Q2K9W3 Q2K9W4 Q2K9W7
Q2KA00 Q2KA07 Q2KA24 Q2KA25 Q2KA31 Q2KA45 Q2KA54 Q2KA61 Q2KA77 Q2KA84 Q2KA85
Q2KA93 Q2KA94 Q2KA96 Q2KAA8 Q2KAB9 Q2KAD9 Q2KAE3 Q2KAE9 Q2KAF1 Q2KAR0 Q2KAR9
Q2KAU2 Q2KAV8 Q2KAZ5 Q2KB10 Q2KB42 Q2KB43 Q2KB48 Q2KBK9 Q2KBL2 Q2KBL3 Q2KBN4
Q2KBP0 Q2KBQ9 Q2KBR2 Q2KBT5 Q2KBU1 Q2KBV2 Q2KBV7 Q2KBX7 Q2KBZ7 Q2KC02 Q2KC03
Q2KC36 Q2KCC6 Q2KCD7 Q2KCG0 Q2KCH6 Q2KCN3 Q2KCN4 Q2KCN5 Q2KCN7 Q2KCP0 Q2KCU6
Q2KCY8 Q2KCZ1 Q2KCZ6 Q2KD13 Q2KD25 Q2KD28 Q2KD36 Q2KD46 Q2KD47 Q2KD50 Q2KD65
Q2KD76 Q2KD88 Q2KD90 Q2KD95 Q2KD99 Q2KDA1 Q2KDA7 Q2KDA9 Q2KDB0 Q2KDD4 Q2KDE4
Q2KDF0 Q2KDG7 Q2KDI7 Q2KDK0 Q2KDK1 Q2KDK2 Q2KDR4 Q2KDR5 Q2KDT0 Q2KDT8 Q2KDT9
Q2KDW6 Q2KDW7 Q2KDX0 Q2KDY8 Q2KDZ2 Q2KDZ5 Q2KDZ6 Q2KDZ7 Q2KDZ9 Q2KE00 Q2KE51
Q2KE54 Q2KE55 Q2KE56 Q2KE60 Q2KE62 Q2KE63 Q2KE66 Q2KE72 Q2KE81 Q2KE82 Q2KE83
Q2KE96 Q2KE98 Q2KE99 Q2KEA0 Q2KEA1 Q2KF83 Q2KFH6 Q2KHI9 Q2KHM9 Q2KHN1 Q2KHP8
Q2KHT8 Q2KHU0 Q2KHU3 Q2KHU8 Q2KHU9 Q2KHV5 Q2KHW8 Q2KHX6 Q2KHX8 Q2KHX9 Q2KHY5
Q2KHY7 Q2KHZ2 Q2KHZ4 Q2KHZ9 Q2KI00 Q2KI04 Q2KI06 Q2KI09 Q2KI14 Q2KI15 Q2KI19
Q2KI24 Q2KI39 Q2KI42 Q2KI45 Q2KI54 Q2KI56 Q2KI58 Q2KI75 Q2KI83 Q2KI84 Q2KI85
Q2KI89 Q2KI95 Q2KIA0 Q2KIA1 Q2KIA2 Q2KIA3 Q2KIA6 Q2KIA9 Q2KIB0 Q2KIB9 Q2KIC0
Q2KIC2 Q2KIC5 Q2KID2 Q2KID4 Q2KID8 Q2KIE4 Q2KIF1 Q2KIF8 Q2KIG1 Q2KIG2 Q2KIG4
Q2KIH1 Q2KIH7 Q2KIH8 Q2KII1 Q2KII4 Q2KII6 Q2KIJ1 Q2KIJ9 Q2KIK0 Q2KIL1 Q2KIL7
Q2KIL8 Q2KIM1 Q2KIN0 Q2KIN3 Q2KIN4 Q2KIN5 Q2KIN8 Q2KIP3 Q2KIQ2 Q2KIQ5 Q2KIR0
Q2KIR1 Q2KIR4 Q2KIR7 Q2KIR8 Q2KIS1 Q2KIS3 Q2KIS6 Q2KIT1 Q2KIT6 Q2KIU7 Q2KIV0
Q2KIW0 Q2KIW3 Q2KIW9 Q2KIX1 Q2KIY0 Q2KIY3 Q2KIY7 Q2KIZ3 Q2KIZ9 Q2KJ16 Q2KJ18
Q2KJ21 Q2KJ24 Q2KJ25 Q2KJ29 Q2KJ34 Q2KJ36 Q2KJ38 Q2KJ44 Q2KJ46 Q2KJ58 Q2KJ61
Q2KJ64 Q2KJ69 Q2KJA4 Q2KJA6 Q2KJB1 Q2KJB9 Q2KJC6 Q2KJD3 Q2KJD7 Q2KJD8 Q2KJE0
Q2KJE1 Q2KJE4 Q2KJE5 Q2KJF7 Q2KJF9 Q2KJG2 Q2KJG8 Q2KJH5 Q2KJH9 Q2KJI1 Q2KJJ0
Q2KJJ5 Q2KJY2 Q2KMM2 Q2KN93 Q2KN94 Q2KN95 Q2KN96 Q2KN97 Q2KN98 Q2KN99 Q2KNA0
Q2KNA1 Q2KNB9 Q2KNL5 Q2KNL6 Q2KPA3 Q2KPA4 Q2KS03 Q2KS10 Q2KTC2 Q2KTI2 Q2KTI4
Q2KTI7 Q2KTI8 Q2KTI9 Q2KTJ9 Q2KTM3 Q2KTN9 Q2KTQ5 Q2KTS9 Q2KTT0 Q2KTT1 Q2KTT2
Q2KTT4 Q2KTT7 Q2KTT8 Q2KU14 Q2KU24 Q2KU63 Q2KU70 Q2KU71 Q2KU72 Q2KU73 Q2KU83
Q2KU87 Q2KU88 Q2KU92 Q2KU93 Q2KU98 Q2KU99 Q2KUD7 Q2KUE4 Q2KUG1 Q2KUI1 Q2KUI9
Q2KUK2 Q2KUK7 Q2KUK9 Q2KUM5 Q2KUS4 Q2KUS6 Q2KUT1 Q2KUT2 Q2KUU3 Q2KUU5 Q2KUV4
Q2KUX6 Q2KUY9 Q2KUZ3 Q2KV01 Q2KV03 Q2KV05 Q2KV08 Q2KV10 Q2KV13 Q2KV15 Q2KV16
Q2KV56 Q2KV81 Q2KV83 Q2KVA0 Q2KVB0 Q2KVE4 Q2KVF5 Q2KVG1 Q2KVG4 Q2KVJ9 Q2KVQ6
Q2KVR0 Q2KVR5 Q2KVT3 Q2KVU4 Q2KVU6 Q2KVV8 Q2KVW5 Q2KVW8 Q2KVX0 Q2KVY2 Q2KVY6
Q2KVY8 Q2KW23 Q2KW46 Q2KW47 Q2KW92 Q2KW99 Q2KWA2 Q2KWA4 Q2KWB0 Q2KWB3 Q2KWD6
Q2KWE0 Q2KWF1 Q2KWF5 Q2KWG1 Q2KWH2 Q2KWH7 Q2KWI4 Q2KWI6 Q2KWN1 Q2KWR7 Q2KWX0
Q2KWY0 Q2KWZ4 Q2KX12 Q2KX20 Q2KX23 Q2KX26 Q2KX38 Q2KX74 Q2KXA0 Q2KXA2 Q2KXB6
Q2KXE0 Q2KXE5 Q2KXH7 Q2KXH8 Q2KXM4 Q2KXM7 Q2KXM9 Q2KXN2 Q2KXN6 Q2KXU5 Q2KXU7
Q2KXU8 Q2KXW3 Q2KXW4 Q2KXY5 Q2KXY7 Q2KXY8 Q2KXY9 Q2KXZ0 Q2KXZ3 Q2KY06 Q2KY19
Q2KY40 Q2KY48 Q2KY72 Q2KY79 Q2KY81 Q2KY83 Q2KY86 Q2KY87 Q2KY92 Q2KY99 Q2KYA0
Q2KYA2 Q2KYA6 Q2KYA7 Q2KYB0 Q2KYB7 Q2KYC8 Q2KYE8 Q2KYE9 Q2KYF3 Q2KYF7 Q2KYJ8

Q2KYK7 Q2KYL4 Q2KYL5 Q2KYL7 Q2KYL9 Q2KYM0 Q2KYM1 Q2KYM2 Q2KYQ1 Q2KYQ6 Q2KYR0
Q2KYS4 Q2KYS7 Q2KYV3 Q2KYV4 Q2KYV6 Q2KYW0 Q2KYW5 Q2KYY2 Q2KZ15 Q2KZ16 Q2KZ18
Q2KZ21 Q2KZ23 Q2KZ46 Q2KZ71 Q2KZD0 Q2KZD7 Q2KZE4 Q2KZE7 Q2KZF3 Q2KZF8 Q2KZI1
Q2KZL7 Q2KZL8 Q2KZM3 Q2KZT1 Q2KZT7 Q2KZZ3 Q2KZZ5 Q2L002 Q2L006 Q2L008 Q2L016
Q2L048 Q2L058 Q2L060 Q2L062 Q2L066 Q2L067 Q2L068 Q2L083 Q2L090 Q2L0A6 Q2L0A9
Q2L0C4 Q2L0K7 Q2L0Y2 Q2L136 Q2L142 Q2L146 Q2L155 Q2L159 Q2L160 Q2L161 Q2L162
Q2L165 Q2L182 Q2L1C6 Q2L1D0 Q2L1D8 Q2L1E2 Q2L1E5 Q2L1G0 Q2L1G9 Q2L1H6 Q2L1J6
Q2L1K6 Q2L1L0 Q2L1P5 Q2L1U0 Q2L1W2 Q2L1Z2 Q2L1Z5 Q2L201 Q2L224 Q2L235 Q2L236
Q2L238 Q2L242 Q2L244 Q2L245 Q2L248 Q2L250 Q2L254 Q2L255 Q2L256 Q2L258 Q2L260
Q2L263 Q2L266 Q2L268 Q2L270 Q2L273 Q2L279 Q2L282 Q2L284 Q2L2B2 Q2L2B5 Q2L2B8
Q2L2C0 Q2L2D7 Q2L2E3 Q2L2E7 Q2L2F1 Q2L2F4 Q2L2F7 Q2L2G0 Q2L2G3 Q2L2G6 Q2L2H1
Q2L2H3 Q2L2H8 Q2L2J5 Q2L2J6 Q2L2K5 Q2L2L3 Q2L2L5 Q2L2M3 Q2L2M5 Q2L2M6 Q2L2N3
Q2L2N9 Q2L2P5 Q2L2Q1 Q2L2R8 Q2L2S3 Q2L2S7 Q2L2T1 Q2L2T3 Q2L2T5 Q2L4H3 Q2L4T2
Q2L4X1 Q2L6E3 Q2L875 Q2L8Y5 Q2L8Y7 Q2L8Z4 Q2L8Z5 Q2L8Z6 Q2L8Z9 Q2L904 Q2L908
Q2L926 Q2L927 Q2L928 Q2L929 Q2L936 Q2L937 Q2L938 Q2L940 Q2L941 Q2L942 Q2L943
Q2L944 Q2L945 Q2L946 Q2L948 Q2L949 Q2L954 Q2L961 Q2L964 Q2LAE1 Q2LAJ3 Q2LAP6
Q2LCP0 Q2LCP1 Q2LCP2 Q2LCP4 Q2LCQ0 Q2LCQ5 Q2LCR7 Q2LCR8 Q2LCR9 Q2LD53 Q2LE08
Q2LE78 Q2LEC2 Q2LKV5 Q2LKX9 Q2LPJ8 Q2LPL3 Q2LPL5 Q2LPL7 Q2LPM2 Q2LPM7 Q2LPW5
Q2LPY6 Q2LQ62 Q2LQ68 Q2LQ82 Q2LQ83 Q2LQ85 Q2LQ86 Q2LQ87 Q2LQ88 Q2LQ89 Q2LQ90
Q2LQ91 Q2LQ94 Q2LQ95 Q2LQ96 Q2LQ97 Q2LQ98 Q2LQ99 Q2LQA0 Q2LQA1 Q2LQA2 Q2LQA3
Q2LQA4 Q2LQA6 Q2LQA7 Q2LQA8 Q2LQA9 Q2LQB0 Q2LQB1 Q2LQB2 Q2LQB3 Q2LQB5 Q2LQB8
Q2LQB9 Q2LQC0 Q2LQC1 Q2LQC2 Q2LQC8 Q2LQC9 Q2LQD0 Q2LQD1 Q2LQD2 Q2LQK4 Q2LQK5
Q2LQK7 Q2LQK8 Q2LQL0 Q2LQM2 Q2LQM6 Q2LQM8 Q2LQN1 Q2LQN4 Q2LQN9 Q2LQP0 Q2LQU2
Q2LQU3 Q2LR03 Q2LR20 Q2LR24 Q2LR26 Q2LR28 Q2LR39 Q2LR40 Q2LR42 Q2LR47 Q2LR50
Q2LR56 Q2LR71 Q2LR75 Q2LR76 Q2LR77 Q2LR78 Q2LR98 Q2LRA8 Q2LRA9 Q2LRB1 Q2LRC1
Q2LRD0 Q2LRK2 Q2LRN9 Q2LRX3 Q2LRX4 Q2LS66 Q2LSD6 Q2LSF6 Q2LSG2 Q2LSI8 Q2LSK9
Q2LSL4 Q2LSN7 Q2LSP8 Q2LSQ4 Q2LSQ5 Q2LSQ6 Q2LST8 Q2LSU0 Q2LT32 Q2LT33 Q2LT41
Q2LT92 Q2LT94 Q2LT95 Q2LT96 Q2LT97 Q2LT98 Q2LTA0 Q2LTA1 Q2LTB7 Q2LTB9 Q2LTE0
Q2LTE1 Q2LTG7 Q2LTI8 Q2LTJ0 Q2LTJ5 Q2LTK5 Q2LTK6 Q2LTL4 Q2LTM9 Q2LTN8 Q2LTQ4
Q2LTQ5 Q2LTQ6 Q2LTQ7 Q2LTR7 Q2LTS1 Q2LTS3 Q2LTS7 Q2LTT5 Q2LTU0 Q2LU42 Q2LU46
Q2LU82 Q2LU85 Q2LU86 Q2LUA0 Q2LUA7 Q2LUB1 Q2LUC0 Q2LUC8 Q2LUD6 Q2LUD8 Q2LUE0
Q2LUH4 Q2LUH6 Q2LUH7 Q2LUJ5 Q2LUJ7 Q2LUJ9 Q2LUK5 Q2LUK6 Q2LUL3 Q2LUL6 Q2LUM5
Q2LUP0 Q2LUP6 Q2LUP9 Q2LUQ6 Q2LUR0 Q2LUR5 Q2LUS9 Q2LUT1 Q2LUT2 Q2LUT4 Q2LUY2
Q2LVE9 Q2LVF5 Q2LVG1 Q2LVG6 Q2LVI0 Q2LVI1 Q2LVI3 Q2LVI4 Q2LVI9 Q2LVL3 Q2LVR6
Q2LVS9 Q2LVT7 Q2LVU2 Q2LVU4 Q2LVU5 Q2LVU6 Q2LVU9 Q2LVV5 Q2LVW6 Q2LVW9 Q2LVX4
Q2LW77 Q2LWA6 Q2LWC5 Q2LWJ1 Q2LWJ2 Q2LWJ3 Q2LWS7 Q2LWT4 Q2LWT5 Q2LWT9 Q2LWU1
Q2LWU2 Q2LWU4 Q2LWU6 Q2LWU8 Q2LWU9 Q2LWV1 Q2LWX9 Q2LWY9 Q2LX49 Q2LXE3 Q2LXM9
Q2LXN0 Q2LXN1 Q2LXN4 Q2LXP6 Q2LXR2 Q2LXR3 Q2LXU2 Q2LXU8 Q2LXZ9 Q2LY09 Q2LY85
Q2LYF5 Q2LYK1 Q2LYK3 Q2LYX9 Q2LYY4 Q2LZ59 Q2LZ79 Q2LZI9 Q2M021 Q2M0G4 Q2M0J9
Q2M0S3 Q2M197 Q2M1D1 Q2M1K5 Q2M1V0 Q2M218 Q2M238 Q2M243 Q2M296 Q2M2D7 Q2M2I5
Q2M2N2 Q2M2S5 Q2M2S7 Q2M2T1 Q2M2T2 Q2M2U3 Q2M2U4 Q2M2U5 Q2M2Z5 Q2M329 Q2M3A8
Q2M3C7 Q2M3D2 Q2M3W8 Q2M3X9 Q2M5Q2 Q2M7R5 Q2M7X4 Q2MEY3 Q2MF12 Q2MF16 Q2MF17
Q2MF22 Q2MF66 Q2MF70 Q2MF71 Q2MF72 Q2MFP1 Q2MFP2 Q2MFP3 Q2MFU6 Q2MGI2 Q2MGL3
Q2MH31 Q2MHH1 Q2MHH2 Q2MHH3 Q2MHH4 Q2MHN1 Q2MHN2 Q2MHN3 Q2MHQ9 Q2MHT0 Q2MI39

Q2MI43 Q2MI54 Q2MI58 Q2MI59 Q2MI60 Q2MI61 Q2MI62 Q2MI63 Q2MI64 Q2MI65 Q2MI67
Q2MI68 Q2MI69 Q2MI76 Q2MI77 Q2MI78 Q2MI79 Q2MI98 Q2MIA2 Q2MIA8 Q2MIA9 Q2MIB0
Q2MIB1 Q2MIB8 Q2MIB9 Q2MIC2 Q2MIC5 Q2MID0 Q2MID9 Q2MIE1 Q2MIE3 Q2MIE5 Q2MIE6
Q2MIE7 Q2MIE8 Q2MIE9 Q2MIF0 Q2MIF1 Q2MIF2 Q2MIF4 Q2MIF5 Q2MIF6 Q2MIG3 Q2MIG4
Q2MIG5 Q2MIG6 Q2MIH9 Q2MII5 Q2MII9 Q2MIJ5 Q2MIJ6 Q2MIJ7 Q2MIJ8 Q2MIK5 Q2MIK6
Q2MIK8 Q2MJB4 Q2MJN4 Q2MJS2 Q2MJT0 Q2MJU7 Q2N2K6 Q2N5P5 Q2N5Q3 Q2N5Q5 Q2N5Q7
Q2N5R4 Q2N5R5 Q2N5R9 Q2N5U2 Q2N5U6 Q2N5U7 Q2N5V5 Q2N5V9 Q2N5X0 Q2N5X1 Q2N612
Q2N631 Q2N635 Q2N659 Q2N669 Q2N682 Q2N695 Q2N6A6 Q2N6B2 Q2N6F2 Q2N6F4 Q2N6I7
Q2N6I8 Q2N6I9 Q2N6J8 Q2N6J9 Q2N6Q7 Q2N6R1 Q2N6U5 Q2N6X5 Q2N758 Q2N784 Q2N7E1
Q2N7E3 Q2N7F4 Q2N7G6 Q2N7Q3 Q2N7Q6 Q2N7R4 Q2N7X0 Q2N7X8 Q2N7Y7 Q2N7Y8 Q2N7Z9
Q2N806 Q2N810 Q2N811 Q2N814 Q2N843 Q2N850 Q2N886 Q2N8D9 Q2N8I5 Q2N8I7 Q2N8I9
Q2N8K3 Q2N8R7 Q2N8W9 Q2N8X0 Q2N8X7 Q2N8X9 Q2N908 Q2N923 Q2N927 Q2N948 Q2N950
Q2N970 Q2N9A5 Q2N9A6 Q2N9A7 Q2N9A8 Q2N9B1 Q2N9B2 Q2N9B4 Q2N9B5 Q2N9B6 Q2N9B7
Q2N9B8 Q2N9B9 Q2N9C0 Q2N9C1 Q2N9C2 Q2N9C4 Q2N9C5 Q2N9C6 Q2N9C7 Q2N9C8 Q2N9C9
Q2N9D1 Q2N9D3 Q2N9D5 Q2N9D6 Q2N9D7 Q2N9D8 Q2N9H9 Q2N9J2 Q2N9J7 Q2N9K0 Q2N9K5
Q2N9N0 Q2N9N2 Q2N9N4 Q2N9P7 Q2N9Q5 Q2N9R0 Q2N9S1 Q2N9S6 Q2N9V1 Q2N9V6 Q2N9Y3
Q2NA15 Q2NA81 Q2NA91 Q2NA95 Q2NA97 Q2NAA7 Q2NAC8 Q2NAD7 Q2NAE1 Q2NAH8 Q2NAJ5
Q2NAJ6 Q2NAJ7 Q2NAK2 Q2NAK3 Q2NAM2 Q2NAN1 Q2NAN2 Q2NAP7 Q2NAR9 Q2NAS1 Q2NAS6
Q2NAT1 Q2NAT8 Q2NAU5 Q2NAU8 Q2NAY0 Q2NAZ6 Q2NB00 Q2NB52 Q2NB65 Q2NB77 Q2NB79
Q2NB98 Q2NBL8 Q2NBQ6 Q2NBR6 Q2NBU1 Q2NBU2 Q2NBU5 Q2NBU6 Q2NBZ2 Q2NBZ3 Q2NC10
Q2NC14 Q2NC39 Q2NC49 Q2NCA3 Q2NCC9 Q2NCD6 Q2NCE3 Q2NCF5 Q2NCK2 Q2NCK7 Q2NCP5
Q2NCS5 Q2NCT3 Q2NCT4 Q2NCT7 Q2NCV0 Q2NCX8 Q2NCY7 Q2NCY8 Q2NCY9 Q2NCZ8 Q2ND39
Q2ND71 Q2ND76 Q2ND78 Q2NDC1 Q2NDC7 Q2NDE3 Q2NDM7 Q2NDM8 Q2NDN5 Q2NDR7 Q2NDY5
Q2NDY6 Q2NE10 Q2NE12 Q2NE13 Q2NE42 Q2NE44 Q2NE45 Q2NE55 Q2NE59 Q2NE61 Q2NE66
Q2NE86 Q2NE95 Q2NEA8 Q2NEB3 Q2NEB6 Q2NEC4 Q2NEH1 Q2NEH4 Q2NEH8 Q2NEI3 Q2NEI6
Q2NEK1 Q2NEK6 Q2NEK8 Q2NEK9 Q2NEL0 Q2NEL1 Q2NEL2 Q2NEL3 Q2NEL4 Q2NEL6 Q2NEM0
Q2NEM4 Q2NEN0 Q2NEP1 Q2NEP2 Q2NEP6 Q2NEP8 Q2NEQ0 Q2NEQ4 Q2NER5 Q2NER8 Q2NES6
Q2NES7 Q2NET3 Q2NET4 Q2NEU6 Q2NEU9 Q2NEV4 Q2NEW0 Q2NEW1 Q2NEW2 Q2NEW4 Q2NEX3
Q2NEX6 Q2NEY3 Q2NEZ5 Q2NEZ9 Q2NF29 Q2NF42 Q2NF45 Q2NF48 Q2NF51 Q2NF52 Q2NF55
Q2NF63 Q2NF79 Q2NFC3 Q2NFD4 Q2NFD9 Q2NFE4 Q2NFG2 Q2NFH1 Q2NFI3 Q2NFJ6 Q2NFJ8
Q2NFK9 Q2NFL1 Q2NFL9 Q2NFM3 Q2NFM9 Q2NFU1 Q2NFU2 Q2NFU9 Q2NFV6 Q2NFV7 Q2NFV8
Q2NFV9 Q2NFW0 Q2NFW1 Q2NFW2 Q2NFW4 Q2NFW5 Q2NFW6 Q2NFW7 Q2NFW8 Q2NFW9 Q2NFX0
Q2NFX1 Q2NFX2 Q2NFX3 Q2NFX6 Q2NFX7 Q2NFX8 Q2NFX9 Q2NFY3 Q2NFY4 Q2NFZ3 Q2NFZ4
Q2NFZ5 Q2NFZ6 Q2NFZ8 Q2NFZ9 Q2NG04 Q2NG79 Q2NGG1 Q2NGH3 Q2NGH9 Q2NGI3 Q2NGI4
Q2NGK6 Q2NGK8 Q2NGM2 Q2NGM5 Q2NGM6 Q2NGM7 Q2NGN3 Q2NGN4 Q2NGN5 Q2NGN7 Q2NGQ5
Q2NGQ7 Q2NGR0 Q2NGR3 Q2NGS6 Q2NGS8 Q2NGT1 Q2NGT2 Q2NGV2 Q2NGV4 Q2NGX9 Q2NGY3
Q2NGY4 Q2NGY8 Q2NH05 Q2NH50 Q2NH88 Q2NH89 Q2NHA5 Q2NHB2 Q2NHB3 Q2NHB4 Q2NHD1
Q2NHD6 Q2NHD9 Q2NHE1 Q2NHE3 Q2NHE8 Q2NHF0 Q2NHF9 Q2NHG2 Q2NHG6 Q2NHG8 Q2NHH7
Q2NHI8 Q2NHL6 Q2NHL8 Q2NHN5 Q2NHN9 Q2NHQ1 Q2NHQ2 Q2NHQ3 Q2NHT0 Q2NHT6 Q2NHU7
Q2NHW0 Q2NHW2 Q2NHW6 Q2NHX5 Q2NHY7 Q2NHZ0 Q2NHZ6 Q2NI00 Q2NI23 Q2NI31 Q2NI42
Q2NI56 Q2NI68 Q2NI73 Q2NI83 Q2NI84 Q2NI86 Q2NI95 Q2NI97 Q2NIA4 Q2NIA9 Q2NIC0
Q2NIC5 Q2NIF5 Q2NIG1 Q2NIG3 Q2NIG5 Q2NIG6 Q2NIH1 Q2NIH4 Q2NIH7 Q2NIH8 Q2NIJ3
Q2NIJ4 Q2NIJ6 Q2NIJ7 Q2NIJ9 Q2NIK0 Q2NIM1 Q2NIM2 Q2NIM3 Q2NIM6 Q2NIM9 Q2NIN2
Q2NIN3 Q2NIN4 Q2NIP1 Q2NIP5 Q2NIP6 Q2NIP7 Q2NIQ6 Q2NIQ9 Q2NIR6 Q2NIS2 Q2NIS3

Q2NIS4 Q2NIS5 Q2NIS7 Q2NIT8 Q2NIT9 Q2NIV1 Q2NIV2 Q2NIV4 Q2NIV5 Q2NIV6 Q2NIV7
Q2NIV8 Q2NIV9 Q2NIW0 Q2NIW1 Q2NIW3 Q2NIW4 Q2NIW5 Q2NIW6 Q2NIW7 Q2NIW8 Q2NIW9
Q2NIX0 Q2NIX1 Q2NIX2 Q2NIX3 Q2NIX5 Q2NIX7 Q2NIX8 Q2NIX9 Q2NIY0 Q2NIY1 Q2NIY2
Q2NIY3 Q2NIY4 Q2NIZ2 Q2NIZ4 Q2NJ01 Q2NJ02 Q2NJ06 Q2NJ11 Q2NJ12 Q2NJ15 Q2NJ16
Q2NJ17 Q2NJ18 Q2NJ19 Q2NJ20 Q2NJ22 Q2NJ23 Q2NJ30 Q2NJ31 Q2NJ38 Q2NJ40 Q2NJ46
Q2NJ58 Q2NJ60 Q2NJ77 Q2NJ78 Q2NJE2 Q2NJE3 Q2NJF5 Q2NJF9 Q2NJG7 Q2NJG8 Q2NJG9
Q2NJH0 Q2NJM5 Q2NJN5 Q2NJY3 Q2NJZ2 Q2NJZ3 Q2NJZ8 Q2NK04 Q2NK14 Q2NK15 Q2NK66
Q2NK67 Q2NKC0 Q2NKC1 Q2NKC3 Q2NKC5 Q2NKI2 Q2NKR3 Q2NKS3 Q2NKS9 Q2NKT2 Q2NKU0
Q2NKU6 Q2NKV2 Q2NKX9 Q2NKY8 Q2NKZ1 Q2NKZ4 Q2NL00 Q2NL05 Q2NL08 Q2NL11 Q2NL13
Q2NL16 Q2NL18 Q2NL22 Q2NL24 Q2NL26 Q2NL30 Q2NL31 Q2NL37 Q2NL67 Q2NL68 Q2NL82
Q2NL98 Q2NLA8 Q2NLB0 Q2NNB6 Q2NNU1 Q2NQ68 Q2NQ69 Q2NQ72 Q2NQ94 Q2NQ95 Q2NQA6
Q2NQA9 Q2NQB1 Q2NQB2 Q2NQF4 Q2NQH2 Q2NQH5 Q2NQH6 Q2NQI3 Q2NQI4 Q2NQJ1 Q2NQJ2
Q2NQK2 Q2NQK8 Q2NQL1 Q2NQL2 Q2NQL3 Q2NQL4 Q2NQL5 Q2NQL6 Q2NQL7 Q2NQM1 Q2NQM2
Q2NQM3 Q2NQM4 Q2NQM5 Q2NQM6 Q2NQM7 Q2NQM8 Q2NQM9 Q2NQN0 Q2NQN1 Q2NQN2 Q2NQN3
Q2NQN4 Q2NQN5 Q2NQN6 Q2NQN7 Q2NQN8 Q2NQN9 Q2NQP0 Q2NQP1 Q2NQP3 Q2NQP4 Q2NQP5
Q2NQP6 Q2NQP7 Q2NQP8 Q2NQQ2 Q2NQQ3 Q2NQQ4 Q2NQQ5 Q2NQQ7 Q2NQQ8 Q2NQR6 Q2NQS5
Q2NQS8 Q2NQS9 Q2NQT7 Q2NQT9 Q2NQU0 Q2NQU2 Q2NQU3 Q2NQU4 Q2NQU5 Q2NQV7 Q2NQW5
Q2NQX2 Q2NQX4 Q2NQX9 Q2NQY0 Q2NQY1 Q2NQY2 Q2NQY5 Q2NQY6 Q2NQZ0 Q2NQZ1 Q2NQZ3
Q2NQZ6 Q2NQZ7 Q2NQZ8 Q2NR04 Q2NR06 Q2NR28 Q2NR34 Q2NR35 Q2NR38 Q2NR41 Q2NRB4
Q2NRB6 Q2NRC0 Q2NRC6 Q2NRC7 Q2NRD1 Q2NRD3 Q2NRD5 Q2NRD9 Q2NRE0 Q2NRE8 Q2NRE9
Q2NRF0 Q2NRF3 Q2NRG1 Q2NRH0 Q2NRH3 Q2NRJ6 Q2NRJ8 Q2NRK0 Q2NRK3 Q2NRK4 Q2NRK5
Q2NRK7 Q2NRK8 Q2NRK9 Q2NRL0 Q2NRL1 Q2NRM1 Q2NRN0 Q2NRR9 Q2NRS1 Q2NRS3 Q2NRS4
Q2NRT4 Q2NRZ6 Q2NRZ8 Q2NS01 Q2NS02 Q2NS03 Q2NS13 Q2NS15 Q2NS22 Q2NS25 Q2NS29
Q2NS30 Q2NS36 Q2NS37 Q2NS40 Q2NS41 Q2NS44 Q2NS49 Q2NS50 Q2NS52 Q2NS68 Q2NS69
Q2NS74 Q2NS76 Q2NS82 Q2NS86 Q2NS87 Q2NS88 Q2NS89 Q2NSC2 Q2NSC4 Q2NSH3 Q2NSH9
Q2NSJ4 Q2NSL7 Q2NSM6 Q2NSM8 Q2NST6 Q2NSV5 Q2NSV8 Q2NT02 Q2NT03 Q2NT05 Q2NT07
Q2NT14 Q2NT22 Q2NT26 Q2NT27 Q2NT28 Q2NT29 Q2NT30 Q2NT31 Q2NT36 Q2NT50 Q2NT52
Q2NT53 Q2NT85 Q2NTA6 Q2NTA8 Q2NTB3 Q2NTB4 Q2NTB6 Q2NTC0 Q2NTD5 Q2NTH8 Q2NTI8
Q2NTI9 Q2NTJ0 Q2NTJ1 Q2NTJ3 Q2NTJ7 Q2NTJ8 Q2NTJ9 Q2NTK2 Q2NTK9 Q2NTL2 Q2NTS9
Q2NTW6 Q2NTX0 Q2NTX2 Q2NTX3 Q2NTX5 Q2NTX6 Q2NTY9 Q2NTZ7 Q2NTZ9 Q2NU15 Q2NU35
Q2NU44 Q2NU46 Q2NU52 Q2NU62 Q2NU63 Q2NU66 Q2NU68 Q2NU80 Q2NU84 Q2NU85 Q2NU97
Q2NUA3 Q2NUA6 Q2NUA8 Q2NUA9 Q2NUB0 Q2NUC6 Q2NUC7 Q2NUD0 Q2NUD2 Q2NUD3 Q2NUE3
Q2NUH6 Q2NUI3 Q2NUJ1 Q2NUJ6 Q2NUJ7 Q2NUK0 Q2NUK1 Q2NUK6 Q2NUL1 Q2NUM2 Q2NUP0
Q2NUT9 Q2NUU1 Q2NUU7 Q2NUV0 Q2NUV2 Q2NUV6 Q2NUV7 Q2NUV8 Q2NV44 Q2NV45 Q2NV56
Q2NV57 Q2NV58 Q2NV59 Q2NV60 Q2NV62 Q2NV67 Q2NV74 Q2NV78 Q2NV79 Q2NV80 Q2NV91
Q2NV92 Q2NV94 Q2NV97 Q2NV98 Q2NVA0 Q2NVA4 Q2NVA5 Q2NVB3 Q2NVB6 Q2NVE9 Q2NVF0
Q2NVF5 Q2NVF9 Q2NVG1 Q2NVK1 Q2NVK2 Q2NVK3 Q2NVK4 Q2NVK9 Q2NVL1 Q2NVL2 Q2NVL3
Q2NVL9 Q2NVM1 Q2NVM2 Q2NVM3 Q2NVM4 Q2NVM8 Q2NVM9 Q2NVN0 Q2NVN2 Q2NVN3 Q2NVN4
Q2NVN8 Q2NVP7 Q2NVP9 Q2NVQ0 Q2NVQ7 Q2NVR2 Q2NVR3 Q2NVR4 Q2NVS1 Q2NVS2 Q2NVT9
Q2NVU1 Q2NVV0 Q2NVV3 Q2NVV6 Q2NVV9 Q2NVW0 Q2NVW3 Q2NVW4 Q2NVW5 Q2NVW6 Q2NVX0
Q2NVX5 Q2NVX6 Q2NVX7 Q2NVX8 Q2NVY2 Q2NVY3 Q2NVY6 Q2NVY8 Q2NVZ0 Q2NVZ1 Q2NVZ3
Q2NVZ5 Q2NVZ9 Q2NW04 Q2NW05 Q2NW08 Q2NW11 Q2NW13 Q2NW19 Q2NW20 Q2NW21 Q2NW22
Q2NW23 Q2NW25 Q2NW27 Q2NW30 Q2NW34 Q2NW36 Q2NW37 Q2NW41 Q2NW44 Q2NW52 Q2NW53
Q2NW55 Q2NW58 Q2NW63 Q2NW64 Q2NW86 Q2NW87 Q2NW90 Q2NW91 Q2NW94 Q2NW95 Q2NWC7

Q2NWC8 Q2NWD5 Q2NWD7 Q2NWD9 Q2NWE2 Q2NWE6 Q2NWE7 Q2NWI3 Q2NWI4 Q2NWI9 Q2NWK4
Q2NWN5 Q2NWP6 Q2NWP9 Q2NWQ0 Q2NWQ7 Q2NWR1 Q2NWR2 Q2NWR3 Q2NWR5 Q2NWR6 Q2NWR7
Q2NWR8 Q2NWR9 Q2NWS0 Q2NWS4 Q2NWT2 Q2NWV8 Q2NWX1 Q2NWZ6 Q2NX28 Q2NX35 Q2NX38
Q2NX39 Q2NX47 Q2NX49 Q2NX50 Q2NX51 Q2NX54 Q2NXC6 Q2NXC7 Q2NXD0 Q2NXE2 Q2NXQ2
Q2NXR2 Q2NXR3 Q2NXR4 Q2NXS9 Q2NY21 Q2NY28 Q2NY29 Q2NY49 Q2NY70 Q2NY76 Q2NYA6
Q2NYB8 Q2NYD6 Q2NYD7 Q2NYE0 Q2NYE3 Q2NYE4 Q2NYE5 Q2NYH0 Q2NYH1 Q2NYM2 Q2NYM7
Q2NYP3 Q2NYU4 Q2NYV5 Q2NYV8 Q2NYV9 Q2NYW4 Q2NYZ0 Q2NYZ1 Q2NYZ2 Q2NZ41 Q2NZ59
Q2NZ83 Q2NZ85 Q2NZ91 Q2NZ92 Q2NZA5 Q2NZB0 Q2NZB1 Q2NZB4 Q2NZB9 Q2NZE3 Q2NZE4
Q2NZF1 Q2NZH9 Q2NZI2 Q2NZI3 Q2NZI4 Q2NZK8 Q2NZL7 Q2NZP2 Q2NZR7 Q2NZS1 Q2NZV7
Q2NZW2 Q2NZW3 Q2NZW6 Q2NZW8 Q2NZW9 Q2NZX1 Q2NZX4 Q2NZX5 Q2NZX6 Q2NZX7 Q2NZX8
Q2NZX9 Q2NZY0 Q2NZY1 Q2NZY2 Q2NZY4 Q2NZY5 Q2NZY6 Q2NZY7 Q2NZY8 Q2NZY9 Q2NZZ0
Q2NZZ1 Q2NZZ2 Q2NZZ3 Q2NZZ4 Q2NZZ5 Q2NZZ6 Q2NZZ7 Q2NZZ8 Q2NZZ9 Q2P000 Q2P001
Q2P002 Q2P003 Q2P004 Q2P006 Q2P007 Q2P008 Q2P009 Q2P010 Q2P021 Q2P034 Q2P036
Q2P047 Q2P0F7 Q2P0H1 Q2P0H2 Q2P0H4 Q2P0J4 Q2P0J5 Q2P0N0 Q2P0N7 Q2P0P1 Q2P0P6
Q2P0Q1 Q2P0R9 Q2P0S4 Q2P0T3 Q2P0T9 Q2P0U0 Q2P0U2 Q2P0U3 Q2P0U5 Q2P0V3 Q2P0W9
Q2P0X1 Q2P0X2 Q2P0X3 Q2P0X4 Q2P0X5 Q2P0Z3 Q2P0Z5 Q2P0Z7 Q2P0Z8 Q2P0Z9 Q2P100
Q2P101 Q2P135 Q2P192 Q2P1A2 Q2P1A3 Q2P1K5 Q2P1L0 Q2P1L1 Q2P1L2 Q2P1L4 Q2P1L5
Q2P1M0 Q2P1M3 Q2P1M4 Q2P1M9 Q2P1N0 Q2P1N1 Q2P1N2 Q2P1Q3 Q2P1R7 Q2P1U6 Q2P1V9
Q2P241 Q2P263 Q2P266 Q2P290 Q2P295 Q2P2A5 Q2P2A9 Q2P2B0 Q2P2C1 Q2P2C4 Q2P2E8
Q2P2F6 Q2P2F7 Q2P2G0 Q2P2G1 Q2P2Q7 Q2P2Q9 Q2P2R1 Q2P2R2 Q2P2S9 Q2P2T0 Q2P2T5
Q2P2U8 Q2P2V0 Q2P2V7 Q2P2X0 Q2P309 Q2P310 Q2P316 Q2P317 Q2P320 Q2P326 Q2P328
Q2P329 Q2P330 Q2P334 Q2P337 Q2P345 Q2P354 Q2P356 Q2P374 Q2P377 Q2P378 Q2P379
Q2P380 Q2P3D0 Q2P3F2 Q2P3J8 Q2P3J9 Q2P3K0 Q2P3K1 Q2P3K2 Q2P3K3 Q2P3K4 Q2P3K8
Q2P3M1 Q2P3M7 Q2P3P2 Q2P3P5 Q2P3Q9 Q2P3R1 Q2P3S1 Q2P3S3 Q2P3S4 Q2P3W4 Q2P3W5
Q2P3W6 Q2P3X9 Q2P3Y4 Q2P3Z2 Q2P425 Q2P458 Q2P459 Q2P460 Q2P468 Q2P472 Q2P492
Q2P497 Q2P4A5 Q2P4A6 Q2P4A8 Q2P4A9 Q2P4B2 Q2P4B9 Q2P4E9 Q2P4L0 Q2P4L7 Q2P4L9
Q2P4Q8 Q2P4S4 Q2P4S5 Q2P4Y6 Q2P4Y7 Q2P4Y8 Q2P4Y9 Q2P553 Q2P575 Q2P577 Q2P578
Q2P579 Q2P582 Q2P5A8 Q2P5B0 Q2P5B3 Q2P5B4 Q2P5B5 Q2P5B6 Q2P5C8 Q2P5E6 Q2P5F7
Q2P5H9 Q2P5M8 Q2P5R0 Q2P5R1 Q2P5V2 Q2P5W1 Q2P5W2 Q2P5W3 Q2P631 Q2P644 Q2P650
Q2P651 Q2P652 Q2P653 Q2P674 Q2P682 Q2P692 Q2P693 Q2P6C7 Q2P6C8 Q2P6C9 Q2P6D1
Q2P6E4 Q2P6G8 Q2P6M2 Q2P6M7 Q2P6N0 Q2P6N2 Q2P6N8 Q2P6P4 Q2P6P9 Q2P6S9 Q2P6V6
Q2P6V7 Q2P6W1 Q2P6Y2 Q2P6Y4 Q2P6Y9 Q2P6Z0 Q2P6Z1 Q2P736 Q2P743 Q2P755 Q2P757
Q2P761 Q2P762 Q2P764 Q2P765 Q2P782 Q2P7C0 Q2P7C2 Q2P7G2 Q2P7G5 Q2P7I5 Q2P7M5
Q2P7M9 Q2P7S5 Q2P7U1 Q2P814 Q2P825 Q2P829 Q2P833 Q2P849 Q2P856 Q2P866 Q2P884
Q2P892 Q2P894 Q2P898 Q2P8A8 Q2P8B0 Q2P8B1 Q2P8E4 Q2P8F2 Q2P8F3 Q2P8H9 Q2P8I1
Q2P8K0 Q2P8V2 Q2P8W4 Q2P8Z6 Q2P8Z8 Q2P902 Q2P907 Q2P915 Q2P928 Q2P949 Q2P9F6
Q2P9F7 Q2P9K6 Q2P9L0 Q2P9L9 Q2P9M1 Q2PBY4 Q2PBY5 Q2PC20 Q2PC95 Q2PDF6 Q2PE14
Q2PF16 Q2PFM2 Q2PFS4 Q2PFU1 Q2PFX0 Q2PFX4 Q2PFX5 Q2PFZ3 Q2PG37 Q2PG46 Q2PG52
Q2PG53 Q2PG84 Q2PGG3 Q2PIU8 Q2PMM3 Q2PMM4 Q2PMM5 Q2PMN0 Q2PMN9 Q2PMP7 Q2PMP8
Q2PMP9 Q2PMQ0 Q2PMQ2 Q2PMQ3 Q2PMR0 Q2PMR2 Q2PMR3 Q2PMS5 Q2PMT2 Q2PMU1 Q2PMU5
Q2PQH8 Q2PQM1 Q2PQN9 Q2PS26 Q2PS27 Q2PT27 Q2PUJ9 Q2PW47 Q2PWU9 Q2PYN8 Q2Q493
Q2Q5Y8 Q2Q5Y9 Q2QAV0 Q2QCI8 Q2QD07 Q2QD12 Q2QD30 Q2QD42 Q2QD64 Q2QD68 Q2QDA6
Q2QDA7 Q2QGD7 Q2QKB3 Q2QKB4 Q2QKL5 Q2QL78 Q2QL82 Q2QL84 Q2QL92 Q2QLA1 Q2QLA2
Q2QLA4 Q2QLB2 Q2QLB3 Q2QLB5 Q2QLB9 Q2QLC3 Q2QLC6 Q2QLD1 Q2QLF4 Q2QLF8 Q2QLG0

Q2QLG4 Q2QLG9 Q2QLH1 Q2QLH9 Q2QLI6 Q2QLT8 Q2QLW3 Q2QLY4 Q2QLY5 Q2QM00 Q2QM17
Q2QM47 Q2QM55 Q2QM59 Q2QM62 Q2QM84 Q2QMB3 Q2QMG2 Q2QMH1 Q2QMT2 Q2QMT5 Q2QMT6
Q2QMT7 Q2QMW0 Q2QMW1 Q2QN26 Q2QN36 Q2QNC3 Q2QNE8 Q2QNF7 Q2QNM1 Q2QNM3 Q2QNU0
Q2QP54 Q2QP86 Q2QPW1 Q2QPW2 Q2QQ32 Q2QQ96 Q2QQ99 Q2QQA3 Q2QQJ5 Q2QQS5 Q2QQX6
Q2QRK9 Q2QRX6 Q2QS13 Q2QS14 Q2QS71 Q2QSB9 Q2QSL4 Q2QT65 Q2QTC2 Q2QTL0 Q2QTY2
Q2QVG9 Q2QVI1 Q2QW44 Q2QW53 Q2QW55 Q2QWE3 Q2QWE9 Q2QWK1 Q2QWK9 Q2QWT4 Q2QWU2
Q2QX01 Q2QXB2 Q2QXB3 Q2QXL0 Q2QXR8 Q2QXZ2 Q2QY10 Q2QYF3 Q2QYR5 Q2QYW1 Q2QYW5
Q2QZ37 Q2QZF1 Q2QZF2 Q2QZJ8 Q2QZL4 Q2R073 Q2R075 Q2R076 Q2R0J4 Q2R0K3 Q2R0Q1
Q2R0W8 Q2R0Z3 Q2R114 Q2R185 Q2R1J3 Q2R1M8 Q2R1U4 Q2R1U8 Q2R1V8 Q2R1Z5 Q2R2B4
Q2R2P7 Q2R2W1 Q2R2Z0 Q2R374 Q2R3A1 Q2R3B4 Q2R3F5 Q2R3K5 Q2R480 Q2R482 Q2R483
Q2R4J4 Q2R4Z4 Q2R4Z5 Q2R4Z7 Q2R4Z8 Q2R637 Q2R837 Q2R8L1 Q2R8P2 Q2R8Z5 Q2R9D2
Q2RAC5 Q2RAD9 Q2RAK2 Q2RAQ5 Q2RB28 Q2RB33 Q2RB59 Q2RBB1 Q2RBJ4 Q2RBJ6 Q2RBP2
Q2RFI8 Q2RFI9 Q2RFJ0 Q2RFJ5 Q2RFK3 Q2RFM0 Q2RFM1 Q2RFM3 Q2RFM4 Q2RFN2 Q2RFN5
Q2RFN6 Q2RFN7 Q2RFN8 Q2RFN9 Q2RFP0 Q2RFP2 Q2RFP3 Q2RFP4 Q2RFP5 Q2RFP6 Q2RFP7
Q2RFP8 Q2RFP9 Q2RFQ0 Q2RFQ1 Q2RFQ2 Q2RFQ3 Q2RFQ4 Q2RFQ5 Q2RFQ6 Q2RFQ7 Q2RFQ8
Q2RFQ9 Q2RFR0 Q2RFR1 Q2RFR2 Q2RFR3 Q2RFR4 Q2RFR5 Q2RFR6 Q2RFR8 Q2RFS1 Q2RFS2
Q2RFS3 Q2RFS4 Q2RFS5 Q2RFS6 Q2RFS7 Q2RFT1 Q2RFT2 Q2RFT3 Q2RFU1 Q2RFU7 Q2RFU8
Q2RFV4 Q2RFV8 Q2RFW0 Q2RFW1 Q2RFW7 Q2RG43 Q2RG62 Q2RG64 Q2RG67 Q2RG68 Q2RG86
Q2RG93 Q2RG98 Q2RG99 Q2RGA0 Q2RGA6 Q2RGG9 Q2RGI2 Q2RGI8 Q2RGJ3 Q2RGL2 Q2RGL8
Q2RGM7 Q2RGP2 Q2RGU4 Q2RGU5 Q2RGU7 Q2RGU9 Q2RGV6 Q2RGV7 Q2RGV8 Q2RGV9 Q2RGW0
Q2RGW1 Q2RGW2 Q2RGX6 Q2RGY0 Q2RGY3 Q2RGY4 Q2RGY5 Q2RH46 Q2RH48 Q2RH52 Q2RH55
Q2RH96 Q2RHB8 Q2RHC4 Q2RHE9 Q2RHF1 Q2RHG3 Q2RHH3 Q2RHL2 Q2RHL3 Q2RHM6 Q2RHN1
Q2RHN2 Q2RHN7 Q2RHN8 Q2RHR1 Q2RHR2 Q2RHR4 Q2RHR5 Q2RHS8 Q2RHT3 Q2RHT5 Q2RHT6
Q2RHT7 Q2RHU2 Q2RHU3 Q2RHV8 Q2RHW0 Q2RHW1 Q2RHX4 Q2RHY2 Q2RHY7 Q2RHZ3 Q2RHZ4
Q2RHZ7 Q2RI40 Q2RI41 Q2RI42 Q2RI72 Q2RI73 Q2RI74 Q2RI90 Q2RI92 Q2RIA9 Q2RIB4
Q2RIB5 Q2RIB6 Q2RIB9 Q2RIC1 Q2RIC2 Q2RID0 Q2RIL2 Q2RIM5 Q2RIR6 Q2RIS6 Q2RIT6
Q2RIT8 Q2RIU8 Q2RIV4 Q2RIW8 Q2RIX1 Q2RIZ3 Q2RJ00 Q2RJ25 Q2RJ26 Q2RJ31 Q2RJ46
Q2RJ59 Q2RJD1 Q2RJD5 Q2RJE3 Q2RJF1 Q2RJF9 Q2RJG0 Q2RJG2 Q2RJG3 Q2RJH7 Q2RJI7
Q2RJK1 Q2RJK7 Q2RJL2 Q2RJL9 Q2RJM0 Q2RJM2 Q2RJM4 Q2RJM5 Q2RJM9 Q2RJN1 Q2RJN4
Q2RJP0 Q2RJP1 Q2RJP2 Q2RJP3 Q2RJP6 Q2RJP8 Q2RJT6 Q2RJU8 Q2RJV3 Q2RJV4 Q2RJV5
Q2RJV7 Q2RJW1 Q2RJX2 Q2RJY1 Q2RJY3 Q2RK02 Q2RK04 Q2RK09 Q2RK16 Q2RK24 Q2RK25
Q2RK28 Q2RK30 Q2RK31 Q2RK33 Q2RK34 Q2RK39 Q2RK43 Q2RK59 Q2RK63 Q2RK77 Q2RK78
Q2RK81 Q2RK87 Q2RK88 Q2RKA1 Q2RKC1 Q2RKH2 Q2RKI6 Q2RKK1 Q2RKL6 Q2RKL8 Q2RKS7
Q2RKU0 Q2RKV3 Q2RKV5 Q2RKV6 Q2RKV8 Q2RKW4 Q2RKW9 Q2RKX4 Q2RKX6 Q2RKY2 Q2RKY6
Q2RKZ1 Q2RKZ5 Q2RKZ6 Q2RKZ7 Q2RKZ8 Q2RL02 Q2RL13 Q2RL16 Q2RL23 Q2RL24 Q2RL30
Q2RL31 Q2RL32 Q2RL40 Q2RL44 Q2RL66 Q2RL67 Q2RL77 Q2RL78 Q2RL95 Q2RLC7 Q2RLC8
Q2RLS9 Q2RLT9 Q2RLU0 Q2RLU1 Q2RLU4 Q2RLU6 Q2RLU8 Q2RM06 Q2RM40 Q2RM49 Q2RM55
Q2RM59 Q2RM60 Q2RM63 Q2RM68 Q2RM70 Q2RM72 Q2RM79 Q2RM91 Q2RM94 Q2RMC1 Q2RMC2
Q2RMC8 Q2RMD6 Q2RME8 Q2RMH0 Q2RMH1 Q2RMH8 Q2RMI9 Q2RMJ4 Q2RMP9 Q2RMQ2 Q2RMR6
Q2RMR7 Q2RMR8 Q2RMR9 Q2RMS0 Q2RMS3 Q2RMS4 Q2RMS5 Q2RMS9 Q2RMT1 Q2RMV0 Q2RMV4
Q2RMV5 Q2RMW0 Q2RMW1 Q2RN09 Q2RN46 Q2RN60 Q2RN61 Q2RN70 Q2RN75 Q2RN76 Q2RN77
Q2RN85 Q2RN86 Q2RN87 Q2RN88 Q2RN89 Q2RN91 Q2RNA1 Q2RNA4 Q2RNA5 Q2RNA6 Q2RNA7
Q2RNA8 Q2RNC0 Q2RND2 Q2RNE6 Q2RNG0 Q2RNG2 Q2RNG5 Q2RNH7 Q2RNH8 Q2RNH9 Q2RNI0
Q2RNI5 Q2RNI6 Q2RNL4 Q2RNM1 Q2RNM2 Q2RNM9 Q2RNN0 Q2RNQ3 Q2RNR2 Q2RNS6 Q2RNS7

Q2RNU4 Q2RNZ9 Q2RP10 Q2RP11 Q2RP15 Q2RP22 Q2RP30 Q2RP31 Q2RP80 Q2RP84 Q2RP86
Q2RPE1 Q2RPE5 Q2RPG0 Q2RPH2 Q2RPH3 Q2RPH4 Q2RPH5 Q2RPU1 Q2RPU6 Q2RPU7 Q2RPV0
Q2RPV1 Q2RPW9 Q2RQ43 Q2RQ44 Q2RQ51 Q2RQ52 Q2RQ67 Q2RQB1 Q2RQH9 Q2RQI0 Q2RQI5
Q2RQI7 Q2RQK4 Q2RQM6 Q2RQM7 Q2RQM8 Q2RQN0 Q2RQN9 Q2RQU6 Q2RQU9 Q2RQV0 Q2RQV1
Q2RQV2 Q2RQV3 Q2RQV4 Q2RQV5 Q2RQV6 Q2RQV7 Q2RQV8 Q2RQV9 Q2RQW0 Q2RQW1 Q2RQW2
Q2RQW3 Q2RQW4 Q2RQW5 Q2RQW6 Q2RQW7 Q2RQW8 Q2RQW9 Q2RQX0 Q2RQX1 Q2RQX2 Q2RQX3
Q2RQX4 Q2RQX5 Q2RQX6 Q2RQX7 Q2RQX8 Q2RQX9 Q2RQY2 Q2RQY3 Q2RQY4 Q2RQY5 Q2RQZ4
Q2RR21 Q2RR25 Q2RR29 Q2RR43 Q2RR72 Q2RRH8 Q2RRI3 Q2RRK9 Q2RRL3 Q2RRM4 Q2RRP2
Q2RRP5 Q2RRT3 Q2RRV2 Q2RRZ2 Q2RS06 Q2RS27 Q2RS49 Q2RS50 Q2RS75 Q2RS98 Q2RSC6
Q2RSC8 Q2RSE3 Q2RSE4 Q2RSF1 Q2RSM3 Q2RSR3 Q2RSR4 Q2RSS1 Q2RST6 Q2RST7 Q2RSU5
Q2RSU7 Q2RSU9 Q2RSY6 Q2RSY8 Q2RT17 Q2RT48 Q2RT49 Q2RT58 Q2RT68 Q2RT81 Q2RT88
Q2RT91 Q2RT96 Q2RTA0 Q2RTB8 Q2RTB9 Q2RTC4 Q2RTF9 Q2RTG6 Q2RTH5 Q2RTH6 Q2RTH7
Q2RTI8 Q2RTK2 Q2RTR1 Q2RTS1 Q2RTS7 Q2RTT0 Q2RTY3 Q2RTZ1 Q2RTZ4 Q2RU07 Q2RU08
Q2RU09 Q2RU19 Q2RU25 Q2RU44 Q2RU45 Q2RU46 Q2RUI7 Q2RUN7 Q2RUU0 Q2RUU1 Q2RUU3
Q2RV02 Q2RV03 Q2RV04 Q2RV05 Q2RV06 Q2RV09 Q2RV14 Q2RV33 Q2RV34 Q2RV50 Q2RV53
Q2RV54 Q2RV55 Q2RV56 Q2RV57 Q2RV58 Q2RV59 Q2RV90 Q2RV91 Q2RVA2 Q2RVD8 Q2RVD9
Q2RVE4 Q2RVF5 Q2RVF6 Q2RVF7 Q2RVF8 Q2RVG7 Q2RVH2 Q2RVK1 Q2RVS0 Q2RVS1 Q2RVT5
Q2RVT6 Q2RVU2 Q2RVU5 Q2RVU6 Q2RVU7 Q2RVV5 Q2RVX7 Q2RW60 Q2RW61 Q2RWC5 Q2RWE0
Q2RWE1 Q2RWE2 Q2RWE3 Q2RWE4 Q2RWE9 Q2RWF6 Q2RWI2 Q2RWI3 Q2RWJ0 Q2RWK2 Q2RWK3
Q2RWM9 Q2RWR8 Q2RWR9 Q2RWS1 Q2RWT4 Q2RWV4 Q2RWY4 Q2RX71 Q2RX74 Q2RX88 Q2RXA9
Q2RXB1 Q2RXC6 Q2RXC7 Q2RXC9 Q2RXD1 Q2RXE3 Q2RXG7 Q2RXH9 Q2RXI0 Q2RXI1 Q2RXI5
Q2RXP5 Q2RXP6 Q2RXP8 Q2RXQ9 Q2RXS7 Q2RXU5 Q2RXY0 Q2RY11 Q2RY12 Q2RY21 Q2RY28
Q2RY67 Q2RY87 Q2RYA4 Q2RYB8 Q2RYD6 Q2RYT3 Q2RYT4 Q2RYT5 Q2RYT6 Q2RYU8 Q2RYY4
Q2RZ59 Q2RZ60 Q2RZ65 Q2RZ66 Q2RZ77 Q2RZF8 Q2RZH4 Q2RZH8 Q2RZI1 Q2RZI8 Q2RZJ2
Q2RZN9 Q2RZV7 Q2RZX7 Q2S006 Q2S008 Q2S009 Q2S026 Q2S030 Q2S042 Q2S0A0 Q2S0I2
Q2S0L2 Q2S0M6 Q2S0M8 Q2S0M9 Q2S0P5 Q2S0P9 Q2S0Q3 Q2S0T3 Q2S0U3 Q2S0U5 Q2S0U9
Q2S0V0 Q2S0V3 Q2S0V7 Q2S0V8 Q2S0W2 Q2S158 Q2S176 Q2S184 Q2S1B9 Q2S1G3 Q2S1G4
Q2S1H6 Q2S1H7 Q2S1I0 Q2S1I3 Q2S1J1 Q2S1J6 Q2S1L8 Q2S1M6 Q2S1N5 Q2S1N7 Q2S1N8
Q2S1P0 Q2S1P1 Q2S1P8 Q2S1Q1 Q2S1Q2 Q2S1Q3 Q2S1Q4 Q2S1Q5 Q2S1Q6 Q2S1S3 Q2S1V4
Q2S1W0 Q2S1Z2 Q2S1Z4 Q2S1Z5 Q2S1Z6 Q2S204 Q2S210 Q2S211 Q2S232 Q2S233 Q2S234
Q2S235 Q2S237 Q2S238 Q2S254 Q2S280 Q2S289 Q2S295 Q2S296 Q2S299 Q2S2A1 Q2S2A3
Q2S2A5 Q2S2A8 Q2S2B5 Q2S2B6 Q2S2D3 Q2S2F5 Q2S2F9 Q2S2G5 Q2S2I3 Q2S2L8 Q2S2N7
Q2S2N8 Q2S2P5 Q2S2P6 Q2S2U0 Q2S2U1 Q2S2U3 Q2S2V6 Q2S2X3 Q2S304 Q2S307 Q2S316
Q2S320 Q2S325 Q2S338 Q2S354 Q2S355 Q2S371 Q2S3A7 Q2S3C0 Q2S3C9 Q2S3D4 Q2S3D8
Q2S3E3 Q2S3F9 Q2S3H9 Q2S3M1 Q2S3M6 Q2S3N1 Q2S3N5 Q2S3N8 Q2S3N9 Q2S3P0 Q2S3P1
Q2S3P2 Q2S3P3 Q2S3P6 Q2S3P7 Q2S3P8 Q2S3P9 Q2S3Q0 Q2S3Q1 Q2S3Q2 Q2S3Q3 Q2S3Q4
Q2S3Q5 Q2S3Q7 Q2S3Q8 Q2S3Q9 Q2S3R0 Q2S3R1 Q2S3R3 Q2S3R4 Q2S3R5 Q2S3R7 Q2S3R8
Q2S3R9 Q2S3T1 Q2S3T5 Q2S3W3 Q2S413 Q2S415 Q2S440 Q2S459 Q2S498 Q2S4A0 Q2S4B9
Q2S4C3 Q2S4D1 Q2S4D2 Q2S4G9 Q2S4I1 Q2S4I8 Q2S4K5 Q2S4L5 Q2S4L6 Q2S4Q4 Q2S4R2
Q2S4R9 Q2S4S2 Q2S4S9 Q2S4T0 Q2S527 Q2S530 Q2S535 Q2S538 Q2S544 Q2S560 Q2S567
Q2S580 Q2S591 Q2S5A6 Q2S5C8 Q2S5G4 Q2S5Z3 Q2S5Z8 Q2S6G1 Q2S6G3 Q2S6G7 Q2S6G8
Q2S6H2 Q2S6I6 Q2S6I8 Q2S6J0 Q2S6J1 Q2S6J2 Q2S6J3 Q2S6J5 Q2S6K1 Q2S6K2 Q2S6K8
Q2S6M6 Q2S6M9 Q2S6N0 Q2S6U2 Q2S6X1 Q2S6X2 Q2S6X5 Q2S6Z8 Q2S7U9 Q2S8B2 Q2S8D3
Q2S8F7 Q2S8L4 Q2S8M0 Q2S8N8 Q2S8P0 Q2S8R2 Q2S8R3 Q2S8V6 Q2S8V7 Q2S8W1 Q2S8W2

Q2S8W3 Q2S8X2 Q2S8X9 Q2S8Y5 Q2S8Y7 Q2S8Y9 Q2S8Z5 Q2S8Z8 Q2S901 Q2S902 Q2S903
Q2S904 Q2S905 Q2S906 Q2S907 Q2S908 Q2S909 Q2S911 Q2S912 Q2S913 Q2S914 Q2S915
Q2S916 Q2S917 Q2S918 Q2S919 Q2S920 Q2S921 Q2S922 Q2S923 Q2S924 Q2S925 Q2S926
Q2S927 Q2S928 Q2S929 Q2S930 Q2S931 Q2S933 Q2S934 Q2S935 Q2S936 Q2S937 Q2S938
Q2S990 Q2S999 Q2S9C3 Q2S9D5 Q2S9L7 Q2S9M0 Q2S9N9 Q2S9P3 Q2S9P4 Q2S9P7 Q2S9Q5
Q2S9R0 Q2S9R4 Q2S9R5 Q2S9R9 Q2S9S0 Q2S9T1 Q2S9T2 Q2S9T3 Q2S9T4 Q2S9T5 Q2S9T9
Q2S9U2 Q2S9V9 Q2S9X0 Q2S9X2 Q2S9X3 Q2S9Y0 Q2S9Y4 Q2S9Z0 Q2S9Z3 Q2SA07 Q2SA08
Q2SA17 Q2SA20 Q2SA27 Q2SA28 Q2SA30 Q2SA36 Q2SA37 Q2SA38 Q2SAE6 Q2SB21 Q2SBA6
Q2SBB3 Q2SBB4 Q2SBC1 Q2SBC2 Q2SBC7 Q2SBC8 Q2SBD4 Q2SBD5 Q2SBD7 Q2SBE7 Q2SBF2
Q2SBF4 Q2SBG0 Q2SBG3 Q2SBG4 Q2SBG5 Q2SBH1 Q2SBH8 Q2SBJ4 Q2SBJ5 Q2SBJ7 Q2SBL9
Q2SBP0 Q2SBP7 Q2SBP8 Q2SBP9 Q2SBQ0 Q2SBQ3 Q2SBR2 Q2SBR5 Q2SBR7 Q2SBT9 Q2SBU2
Q2SC73 Q2SCE2 Q2SCE3 Q2SCE5 Q2SCE8 Q2SCF2 Q2SCF7 Q2SCG0 Q2SCH0 Q2SCH1 Q2SCJ1
Q2SCJ3 Q2SCK8 Q2SCK9 Q2SCL1 Q2SCL2 Q2SCL3 Q2SCM1 Q2SCM9 Q2SCX8 Q2SD07 Q2SD35
Q2SD47 Q2SD50 Q2SDF4 Q2SDF9 Q2SDG0 Q2SDI1 Q2SDJ2 Q2SDJ4 Q2SDJ5 Q2SDJ6 Q2SDJ7
Q2SDJ8 Q2SDN9 Q2SDP4 Q2SDP8 Q2SDP9 Q2SDQ0 Q2SDQ1 Q2SDQ2 Q2SDQ3 Q2SDQ4 Q2SDT2
Q2SDV0 Q2SDW0 Q2SDW1 Q2SDW4 Q2SDW8 Q2SE61 Q2SE63 Q2SEP6 Q2SEP7 Q2SEP8 Q2SF17
Q2SF51 Q2SFI6 Q2SFI7 Q2SFU4 Q2SFX1 Q2SGY7 Q2SHA6 Q2SHM3 Q2SHM4 Q2SHN2 Q2SHN3
Q2SHX7 Q2SIN3 Q2SIP2 Q2SIQ1 Q2SIW7 Q2SIW9 Q2SIX0 Q2SJ45 Q2SJ47 Q2SJB7 Q2SJB8
Q2SJD2 Q2SJD6 Q2SJD7 Q2SJD8 Q2SJE4 Q2SJK8 Q2SJK9 Q2SJL1 Q2SJL8 Q2SJP9 Q2SJQ9
Q2SJS1 Q2SJV9 Q2SJW0 Q2SJW1 Q2SJX0 Q2SJX3 Q2SJZ6 Q2SJZ8 Q2SK05 Q2SK22 Q2SK35
Q2SK36 Q2SK37 Q2SK39 Q2SK40 Q2SK46 Q2SK53 Q2SK56 Q2SK70 Q2SK73 Q2SK75 Q2SK84
Q2SKD0 Q2SKK6 Q2SKP0 Q2SKP1 Q2SKR0 Q2SKT3 Q2SKU2 Q2SKW3 Q2SKW4 Q2SKW6 Q2SKW7
Q2SKX2 Q2SKZ0 Q2SKZ1 Q2SKZ2 Q2SL17 Q2SL21 Q2SL23 Q2SL27 Q2SL30 Q2SL32 Q2SL47
Q2SL55 Q2SL56 Q2SL57 Q2SL58 Q2SL86 Q2SL95 Q2SL96 Q2SLA0 Q2SLA2 Q2SLA3 Q2SLB1
Q2SLB3 Q2SLB4 Q2SLB5 Q2SLC0 Q2SLS9 Q2SLT3 Q2SLT4 Q2SLV0 Q2SLV8 Q2SLW3 Q2SM69
Q2SM94 Q2SMA3 Q2SMA7 Q2SMB2 Q2SMB3 Q2SMB5 Q2SMB8 Q2SMF5 Q2SMK9 Q2SML0 Q2SML1
Q2SML2 Q2SML3 Q2SML5 Q2SML7 Q2SML8 Q2SMM1 Q2SMM6 Q2SMM7 Q2SMM8 Q2SMN3 Q2SMN6
Q2SMS4 Q2SMY0 Q2SN06 Q2SN12 Q2SN16 Q2SN31 Q2SN34 Q2SN53 Q2SN65 Q2SN67 Q2SN69
Q2SN70 Q2SN71 Q2SN75 Q2SN79 Q2SNC4 Q2SNS0 Q2SNV0 Q2SPF3 Q2SPF8 Q2SPF9 Q2SPG7
Q2SPG9 Q2SPH0 Q2SPJ2 Q2SPJ7 Q2SPM9 Q2SPQ2 Q2SPT0 Q2SQ64 Q2SQ67 Q2SQ70 Q2SQB9
Q2SQE7 Q2SQL5 Q2SQL8 Q2SQM5 Q2SQW5 Q2SQW8 Q2SQX1 Q2SQX2 Q2SQY0 Q2SQZ4 Q2SQZ9
Q2SR15 Q2SR32 Q2SR35 Q2SR42 Q2SR45 Q2SR47 Q2SR48 Q2SR96 Q2SRA2 Q2SRA7 Q2SRB2
Q2SRE1 Q2SRE4 Q2SRE7 Q2SRH5 Q2SRH7 Q2SRH8 Q2SRH9 Q2SRI4 Q2SRI6 Q2SRI9 Q2SRJ4
Q2SRJ6 Q2SRK6 Q2SRN2 Q2SRP7 Q2SRR6 Q2SRR7 Q2SRU4 Q2SRU5 Q2SRU7 Q2SRU8 Q2SRV0
Q2SRV9 Q2SRW8 Q2SRX1 Q2SRX4 Q2SRY2 Q2SRY4 Q2SRY5 Q2SRZ3 Q2SRZ8 Q2SS02 Q2SS11
Q2SS13 Q2SS24 Q2SS50 Q2SS60 Q2SS69 Q2SS74 Q2SS76 Q2SS95 Q2SS96 Q2SS97 Q2SS99
Q2SSA0 Q2SSA3 Q2SSA4 Q2SSA5 Q2SSA7 Q2SSA8 Q2SSB0 Q2SSE2 Q2SSE6 Q2SSE8 Q2SSF2
Q2SSF4 Q2SSG0 Q2SSM4 Q2SSM5 Q2SSN1 Q2SSN9 Q2SSQ6 Q2SSQ8 Q2SSR1 Q2SSR8 Q2SSS1
Q2SSS2 Q2SSS3 Q2SSW4 Q2SSW8 Q2SSW9 Q2SSX0 Q2SSX1 Q2SSX9 Q2SSZ1 Q2ST11 Q2ST15
Q2ST16 Q2ST19 Q2ST23 Q2ST29 Q2ST42 Q2ST43 Q2ST47 Q2ST48 Q2ST50 Q2ST51 Q2ST52
Q2ST53 Q2ST57 Q2ST89 Q2ST91 Q2ST93 Q2STA4 Q2STD7 Q2STD8 Q2STF3 Q2STK2 Q2STL6
Q2STL7 Q2STM2 Q2STR0 Q2STS7 Q2STU0 Q2STV7 Q2SU15 Q2SU16 Q2SU17 Q2SU18 Q2SU19
Q2SU20 Q2SU22 Q2SU23 Q2SU24 Q2SU25 Q2SU26 Q2SU27 Q2SU28 Q2SU29 Q2SU30 Q2SU31
Q2SU32 Q2SU33 Q2SU34 Q2SU35 Q2SU36 Q2SU37 Q2SU38 Q2SU39 Q2SU40 Q2SU41 Q2SU42

Q2SU43 Q2SU44 Q2SU45 Q2SU46 Q2SU48 Q2SU49 Q2SU50 Q2SU51 Q2SU52 Q2SU53 Q2SU54
Q2SU58 Q2SU65 Q2SU71 Q2SU89 Q2SUA1 Q2SUA4 Q2SUA7 Q2SUA8 Q2SUB0 Q2SUG2 Q2SUG3
Q2SUH7 Q2SUI0 Q2SUI1 Q2SUP3 Q2SUW0 Q2SUW3 Q2SV73 Q2SV78 Q2SVC0 Q2SVD8 Q2SVE0
Q2SVE1 Q2SVE2 Q2SVE3 Q2SVG6 Q2SVG8 Q2SVG9 Q2SVM0 Q2SVM1 Q2SVM3 Q2SVM8 Q2SVY2
Q2SW40 Q2SWB9 Q2SWD3 Q2SWD4 Q2SWD9 Q2SWE3 Q2SWE6 Q2SWE7 Q2SWE9 Q2SWF4 Q2SWF5
Q2SWF7 Q2SWF8 Q2SWG5 Q2SWJ6 Q2SWJ7 Q2SWJ9 Q2SWM4 Q2SWN8 Q2SWN9 Q2SWP6 Q2SWQ5
Q2SWQ6 Q2SWQ8 Q2SWT5 Q2SWT6 Q2SWX6 Q2SWY2 Q2SWY4 Q2SWZ2 Q2SWZ5 Q2SWZ6 Q2SWZ7
Q2SWZ8 Q2SX00 Q2SX03 Q2SX11 Q2SX33 Q2SX36 Q2SX78 Q2SXA6 Q2SXC7 Q2SXD6 Q2SXE0
Q2SXE1 Q2SXF8 Q2SXI2 Q2SXJ3 Q2SXN7 Q2SXN8 Q2SXN9 Q2SXQ4 Q2SXS5 Q2SXS7 Q2SXT2
Q2SXU9 Q2SXV1 Q2SXZ8 Q2SXZ9 Q2SY00 Q2SY01 Q2SY06 Q2SY18 Q2SY23 Q2SY25 Q2SY32
Q2SY55 Q2SY57 Q2SY71 Q2SY72 Q2SY74 Q2SY86 Q2SY88 Q2SY90 Q2SYB2 Q2SYC5 Q2SYE0
Q2SYF4 Q2SYF5 Q2SYF6 Q2SYF7 Q2SYF8 Q2SYG0 Q2SYI5 Q2SYI8 Q2SYJ0 Q2SYJ1 Q2SYJ5
Q2SYQ6 Q2SYS3 Q2SYS4 Q2SYY3 Q2SYY5 Q2SYZ1 Q2SYZ3 Q2SYZ4 Q2SYZ6 Q2SYZ9 Q2SZ00
Q2SZ01 Q2SZ20 Q2SZ45 Q2SZ52 Q2SZ53 Q2SZ54 Q2SZ55 Q2SZ60 Q2SZ62 Q2SZ63 Q2SZ67
Q2SZ68 Q2SZ69 Q2SZ88 Q2SZ91 Q2SZ94 Q2SZA4 Q2SZE1 Q2SZF6 Q2SZF7 Q2SZF9 Q2SZG0
Q2SZG1 Q2SZG2 Q2SZG8 Q2SZG9 Q2SZI1 Q2SZI2 Q2SZI5 Q2SZJ1 Q2SZJ2 Q2SZK6 Q2SZN9
Q2SZP0 Q2SZP8 Q2SZS7 Q2SZS9 Q2SZT0 Q2SZT5 Q2SZT6 Q2SZW2 Q2SZW6 Q2SZY3 Q2SZY9
Q2SZZ0 Q2T071 Q2T079 Q2T083 Q2T087 Q2T095 Q2T096 Q2T0B6 Q2T0C0 Q2T0G2 Q2T0G3
Q2T0G4 Q2T0G7 Q2T0H3 Q2T0H6 Q2T0I1 Q2T0I7 Q2T0K2 Q2T0K4 Q2T0K8 Q2T0L2 Q2T0L3
Q2T0M6 Q2T0N2 Q2T0U7 Q2T0Z3 Q2T101 Q2T103 Q2T114 Q2T132 Q2T139 Q2T144 Q2T199
Q2T1A9 Q2T1B0 Q2T1B3 Q2T1B6 Q2T1B8 Q2T1B9 Q2T1C2 Q2T1H2 Q2T1H5 Q2T1K5 Q2T1K6
Q2T1M2 Q2T1N2 Q2T1N6 Q2T1P5 Q2T1Q2 Q2T1Q3 Q2T1Q4 Q2T1V1 Q2T1W2 Q2T213 Q2T267
Q2T269 Q2T277 Q2T281 Q2T284 Q2T292 Q2T293 Q2T294 Q2T2A4 Q2T2B1 Q2T2K1 Q2T2K3
Q2T321 Q2T3J4 Q2T3K6 Q2T3N9 Q2T3W7 Q2T437 Q2T4M4 Q2T4N0 Q2T4N1 Q2T4N2 Q2T4P1
Q2T4T8 Q2T559 Q2T5T2 Q2T602 Q2T6A1 Q2T6C8 Q2T6D0 Q2T6K1 Q2T6L2 Q2T6R2 Q2T6R3
Q2T7G9 Q2T7H3 Q2T7H6 Q2T7H7 Q2T7H8 Q2T7J2 Q2T7N1 Q2T7N3 Q2T7N4 Q2T7N5 Q2T7P9
Q2T7S9 Q2T7T0 Q2T8Q2 Q2T8Y7 Q2T9E1 Q2T9I5 Q2T9I9 Q2T9K2 Q2T9K6 Q2T9L4 Q2T9L8
Q2T9L9 Q2T9M0 Q2T9M4 Q2T9M9 Q2T9N0 Q2T9N1 Q2T9N3 Q2T9P0 Q2T9P9 Q2T9Q2 Q2T9Q3
Q2T9Q6 Q2T9Q7 Q2T9R6 Q2T9S3 Q2T9S7 Q2T9S8 Q2T9T3 Q2T9T5 Q2T9T9 Q2T9U2 Q2T9U5
Q2T9U9 Q2T9V2 Q2T9V5 Q2T9V7 Q2T9V8 Q2T9V9 Q2T9W2 Q2T9W3 Q2T9W4 Q2T9W6 Q2T9W8
Q2T9W9 Q2T9X2 Q2T9X3 Q2T9X5 Q2T9X7 Q2T9X8 Q2T9Y0 Q2T9Y1 Q2T9Y6 Q2T9Z0 Q2T9Z1
Q2T9Z2 Q2T9Z5 Q2T9Z6 Q2T9Z7 Q2TA00 Q2TA05 Q2TA10 Q2TA11 Q2TA16 Q2TA17 Q2TA20
Q2TA38 Q2TA39 Q2TA42 Q2TA43 Q2TA45 Q2TA51 Q2TAA8 Q2TAC2 Q2TAC6 Q2TAD9 Q2TAF3
Q2TAF4 Q2TAG6 Q2TAK8 Q2TAL5 Q2TAM9 Q2TAP8 Q2TAP9 Q2TAQ1 Q2TAQ8 Q2TAR3 Q2TAW0
Q2TAY7 Q2TB02 Q2TB18 Q2TBA0 Q2TBC4 Q2TBG7 Q2TBH0 Q2TBH6 Q2TBH8 Q2TBI1 Q2TBI2
Q2TBI7 Q2TBI8 Q2TBJ0 Q2TBK0 Q2TBK7 Q2TBL4 Q2TBL6 Q2TBL8 Q2TBN1 Q2TBN5 Q2TBN6
Q2TBN7 Q2TBN9 Q2TBP0 Q2TBP2 Q2TBP4 Q2TBP7 Q2TBP8 Q2TBQ0 Q2TBQ3 Q2TBQ6 Q2TBQ8
Q2TBQ9 Q2TBR0 Q2TBR5 Q2TBR6 Q2TBR9 Q2TBS3 Q2TBS4 Q2TBS9 Q2TBT5 Q2TBT8 Q2TBU5
Q2TBU8 Q2TBV1 Q2TBV3 Q2TBV5 Q2TBW0 Q2TBW5 Q2TBX0 Q2TBX2 Q2TBX6 Q2TBX7 Q2TBY0
Q2TDY4 Q2TE74 Q2TFN9 Q2TGZ2 Q2TL60 Q2TN88 Q2TPA8 Q2TSC7 Q2TUM3 Q2TVY9 Q2TVZ2
Q2TW34 Q2TWN3 Q2TX99 Q2TXF1 Q2TXF3 Q2TXF5 Q2TXF6 Q2TXG0 Q2TXG2 Q2TXG3 Q2TXG4
Q2TYB1 Q2TZ06 Q2TZ19 Q2TZ33 Q2TZB2 Q2TZG4 Q2TZH2 Q2TZH4 Q2TZM9 Q2TZN9 Q2U002
Q2U038 Q2U041 Q2U059 Q2U070 Q2U0C4 Q2U0E0 Q2U0J4 Q2U0J5 Q2U0J7 Q2U0J9 Q2U0M4
Q2U0N3 Q2U0Q9 Q2U100 Q2U256 Q2U2J1 Q2U2J6 Q2U2U3 Q2U2Z2 Q2U316 Q2U3T8 Q2U3V3

Q2U3V4 Q2U3W4 Q2U469 Q2U4D6 Q2U4H2 Q2U4W2 Q2U513 Q2U561 Q2U587 Q2U5A2 Q2U5A8
Q2U5A9 Q2U5H9 Q2U5I0 Q2U5K2 Q2U5W8 Q2U600 Q2U616 Q2U639 Q2U664 Q2U667 Q2U685
Q2U696 Q2U6A1 Q2U6C4 Q2U6D4 Q2U6E8 Q2U6H2 Q2U6P7 Q2U6Q1 Q2U6U0 Q2U7A3 Q2U7R8
Q2U7S5 Q2U852 Q2U913 Q2U988 Q2U9B0 Q2U9B5 Q2U9C3 Q2U9C6 Q2U9E2 Q2U9E7 Q2U9G3
Q2U9H6 Q2U9L6 Q2U9X7 Q2UA27 Q2UA42 Q2UA71 Q2UAK1 Q2UAX9 Q2UAZ9 Q2UB00 Q2UB17
Q2UB88 Q2UBF1 Q2UBH0 Q2UBH8 Q2UBM4 Q2UBU2 Q2UBW0 Q2UBZ5 Q2UBZ9 Q2UC04 Q2UC58
Q2UCB7 Q2UCC9 Q2UCL6 Q2UCP5 Q2UCP6 Q2UCQ0 Q2UD93 Q2UDB2 Q2UDE2 Q2UDW7 Q2UDZ9
Q2UE04 Q2UE66 Q2UEA0 Q2UEA6 Q2UEF7 Q2UEK2 Q2UEK4 Q2UEK6 Q2UEK8 Q2UEL0 Q2UEL1
Q2UEN6 Q2UES9 Q2UEX1 Q2UF33 Q2UF93 Q2UF95 Q2UFL0 Q2UFN0 Q2UFN8 Q2UG94 Q2UGF7
Q2UGG5 Q2UGK1 Q2UGK3 Q2UGN1 Q2UGP9 Q2UGQ2 Q2UGQ8 Q2UGT1 Q2UGU1 Q2UGU3 Q2UH00
Q2UH11 Q2UH56 Q2UHB7 Q2UHC1 Q2UHT9 Q2UHU3 Q2UHU9 Q2UIM3 Q2UJ16 Q2UJ66 Q2UJE8
Q2UJJ4 Q2UJU5 Q2UJY8 Q2UK72 Q2UKD0 Q2UKG6 Q2UKH7 Q2UKS0 Q2UKS9 Q2UKV0 Q2UKV4
Q2UKV7 Q2UKV8 Q2UKX3 Q2UKX8 Q2UL45 Q2UL89 Q2ULA2 Q2ULT1 Q2ULU6 Q2UM19 Q2UM42
Q2UM46 Q2UMB1 Q2UMB8 Q2UMC5 Q2UMG1 Q2UMH3 Q2UMH8 Q2UMJ1 Q2UMM2 Q2UMM4 Q2UMQ5
Q2UMQ8 Q2UMY7 Q2UN27 Q2UN37 Q2UN81 Q2UNA2 Q2UNB7 Q2UNX4 Q2UNX5 Q2UNX8 Q2UPA6
Q2UPA9 Q2UPB1 Q2UPB2 Q2UPB3 Q2UPB4 Q2UPB5 Q2UPB6 Q2UPB7 Q2UPB9 Q2UPC2 Q2UPC3
Q2UPC5 Q2UPG7 Q2UPH5 Q2UPI0 Q2UPK0 Q2UPL9 Q2UPM0 Q2UPS5 Q2UPU5 Q2UPY3 Q2UPZ7
Q2UQ34 Q2UQ97 Q2UQC6 Q2UQH9 Q2UQI6 Q2UQJ5 Q2UQQ2 Q2UQW3 Q2UQY2 Q2UQZ5 Q2UR29
Q2URB7 Q2URI6 Q2URJ0 Q2URJ3 Q2URJ5 Q2USD7 Q2USL9 Q2UST1 Q2UT70 Q2UTI8 Q2UTN6
Q2UTN7 Q2UTQ9 Q2UU13 Q2UU72 Q2UUH5 Q2UUI3 Q2UUM2 Q2UUT4 Q2V057 Q2V0C9 Q2V0P1
Q2V0V2 Q2V0W1 Q2V0Z5 Q2V2M9 Q2V2P0 Q2V2P1 Q2V2P7 Q2V2P8 Q2V2Q0 Q2V2Q3 Q2V2T9
Q2V2V3 Q2V3B2 Q2V3H0 Q2V3L6 Q2V3N5 Q2V414 Q2V6J9 Q2V6K0 Q2V6K1 Q2V8X7 Q2V9B0
Q2VA69 Q2VB19 Q2VEB7 Q2VEC4 Q2VED2 Q2VED6 Q2VED8 Q2VEE1 Q2VEE2 Q2VEE3 Q2VEE4
Q2VEE5 Q2VEE6 Q2VEE7 Q2VEE8 Q2VEF5 Q2VEF6 Q2VEF7 Q2VEH5 Q2VEI4 Q2VEI5 Q2VEI6
Q2VEI7 Q2VEQ7 Q2VF18 Q2VF19 Q2VIR3 Q2VIS4 Q2VL50 Q2VL51 Q2VL52 Q2VL53 Q2VL54
Q2VL56 Q2VL57 Q2VL58 Q2VL59 Q2VL60 Q2VL61 Q2VL62 Q2VL76 Q2VL77 Q2VL78 Q2VL79
Q2VL80 Q2VL82 Q2VL83 Q2VL85 Q2VL86 Q2VLB9 Q2VLC6 Q2VLK6 Q2VNJ2 Q2VPA6 Q2VPG3
Q2VPH1 Q2VPJ9 Q2VPL9 Q2VPM4 Q2VPM9 Q2VPQ9 Q2VUH7 Q2VWA4 Q2VWH6 Q2VWQ3 Q2VY69
Q2VYH1 Q2VYH2 Q2VYH3 Q2VYH4 Q2VYH5 Q2VYH7 Q2VYI5 Q2VYI7 Q2VYI9 Q2VYJ0 Q2VYJ1
Q2VYL8 Q2VYM5 Q2VYN2 Q2VYQ1 Q2VYQ2 Q2VYQ6 Q2VYQ8 Q2VYT0 Q2VYT1 Q2VYT6 Q2VYV2
Q2VYV5 Q2VYV7 Q2VYX5 Q2VYY7 Q2VYY8 Q2VYZ1 Q2VYZ7 Q2VZ00 Q2VZ01 Q2VZ17 Q2VZ18
Q2VZ23 Q2VZ24 Q2VZ28 Q2VZA8 Q2VZA9 Q2VZE7 Q2VZP8 Q2VZQ4 Q2VZQ5 Q2VZR6 Q2VZR7
Q2VZT7 Q2VZT9 Q2VZU1 Q2VZU2 Q2VZU3 Q2VZU4 Q2VZU7 Q2VZV0 Q2VZV1 Q2VZV2 Q2VZV3
Q2VZV4 Q2VZV5 Q2VZV6 Q2VZV7 Q2VZV8 Q2VZZ9 Q2W000 Q2W010 Q2W019 Q2W020 Q2W047
Q2W063 Q2W064 Q2W095 Q2W0C9 Q2W0E7 Q2W0E8 Q2W0G3 Q2W0H0 Q2W0H1 Q2W0H2 Q2W0H5
Q2W0I1 Q2W0M1 Q2W0P2 Q2W0V3 Q2W136 Q2W141 Q2W156 Q2W1A2 Q2W1B4 Q2W1D9 Q2W1G2
Q2W1G5 Q2W1X8 Q2W1Y0 Q2W1Y2 Q2W1Y3 Q2W2A2 Q2W2A3 Q2W2A5 Q2W2C2 Q2W2D0 Q2W2D1
Q2W2H3 Q2W2H8 Q2W2H9 Q2W2I0 Q2W2I1 Q2W2I2 Q2W2I6 Q2W2I7 Q2W2I8 Q2W2J0 Q2W2J1
Q2W2J2 Q2W2J3 Q2W2J4 Q2W2J5 Q2W2J6 Q2W2J7 Q2W2J8 Q2W2K0 Q2W2K1 Q2W2K2 Q2W2K3
Q2W2K4 Q2W2K5 Q2W2K6 Q2W2K7 Q2W2K8 Q2W2K9 Q2W2L1 Q2W2L2 Q2W2L3 Q2W2L4 Q2W2Q6
Q2W2S4 Q2W2T3 Q2W2T6 Q2W2T7 Q2W2V2 Q2W367 Q2W369 Q2W3A9 Q2W3C2 Q2W3D3 Q2W3D5
Q2W3F1 Q2W3F8 Q2W3G8 Q2W3H2 Q2W3H8 Q2W3H9 Q2W3I0 Q2W3J9 Q2W3K9 Q2W3L2 Q2W3L4
Q2W3N1 Q2W3R9 Q2W3S5 Q2W3W0 Q2W3X1 Q2W3X9 Q2W402 Q2W440 Q2W477 Q2W485 Q2W492
Q2W4A0 Q2W4A1 Q2W4A2 Q2W4C3 Q2W4C4 Q2W4C5 Q2W4C6 Q2W4D9 Q2W4P9 Q2W4Q8 Q2W4R6

Q2W4R9 Q2W4S5 Q2W4S6 Q2W4T1 Q2W4T2 Q2W4W4 Q2W4X7 Q2W519 Q2W5E3 Q2W5F9 Q2W5G0
Q2W5G1 Q2W5G8 Q2W5H0 Q2W5H3 Q2W5J7 Q2W5N3 Q2W5T9 Q2W5W8 Q2W6A0 Q2W6V3 Q2W6W0
Q2W6W7 Q2W6Y1 Q2W6Y3 Q2W6Y4 Q2W6Y6 Q2W6Y7 Q2W744 Q2W7M7 Q2W7P4 Q2W7T6 Q2W7Y5
Q2W7Y6 Q2W896 Q2W897 Q2W899 Q2W8V2 Q2W8Y5 Q2W992 Q2W993 Q2W9A3 Q2W9A4 Q2W9A8
Q2W9A9 Q2W9C3 Q2W9F0 Q2W9N6 Q2W9P9 Q2WAI1 Q2WAN4 Q2WAS9 Q2WAT0 Q2WAV0 Q2WAW8
Q2WB09 Q2WBD9 Q2WBE4 Q2WBG8 Q2WBG9 Q2WBH0 Q2WBN3 Q2WF59 Q2WG76 Q2WG77 Q2WG78
Q2WG79 Q2WG80 Q2WGC1 Q2WGF1 Q2WGI1 Q2WGJ6 Q2XPP4 Q2XQU3 Q2XQV4 Q2XQY7 Q2XSC4
Q2XSC5 Q2XSC6 Q2Y0E2 Q2Y0E3 Q2Y0E4 Q2Y0E5 Q2Y0E6 Q2Y0E7 Q2Y0E8 Q2Y0E9 Q2Y0F0
Q2Y2L8 Q2Y2M1 Q2Y2M2 Q2Y2M5 Q2Y2M6 Q2Y5A5 Q2Y5A9 Q2Y5B4 Q2Y5B5 Q2Y5F0 Q2Y5G2
Q2Y5G4 Q2Y5G5 Q2Y5H4 Q2Y5H5 Q2Y5H6 Q2Y5I7 Q2Y5J2 Q2Y5P2 Q2Y5P3 Q2Y5Q1 Q2Y5Q5
Q2Y5Q6 Q2Y5R7 Q2Y5S2 Q2Y5W9 Q2Y5X8 Q2Y5X9 Q2Y5Y2 Q2Y5Y6 Q2Y5Z0 Q2Y5Z1 Q2Y629
Q2Y630 Q2Y636 Q2Y639 Q2Y640 Q2Y641 Q2Y651 Q2Y653 Q2Y654 Q2Y676 Q2Y681 Q2Y6A2
Q2Y6A3 Q2Y6A9 Q2Y6B0 Q2Y6B3 Q2Y6B8 Q2Y6F2 Q2Y6F3 Q2Y6F9 Q2Y6G0 Q2Y6G2 Q2Y6I5
Q2Y6I6 Q2Y6I9 Q2Y6J0 Q2Y6J1 Q2Y6K2 Q2Y6L7 Q2Y6P4 Q2Y6R0 Q2Y6S0 Q2Y6T2 Q2Y6T6
Q2Y6T9 Q2Y6U0 Q2Y6W3 Q2Y6Y6 Q2Y6Z3 Q2Y720 Q2Y738 Q2Y739 Q2Y740 Q2Y742 Q2Y748
Q2Y751 Q2Y752 Q2Y763 Q2Y791 Q2Y796 Q2Y797 Q2Y7A9 Q2Y7B1 Q2Y7B6 Q2Y7B7 Q2Y7I8
Q2Y7I9 Q2Y7J7 Q2Y7L1 Q2Y7L8 Q2Y7M5 Q2Y7M7 Q2Y7M8 Q2Y7P0 Q2Y7P9 Q2Y7Q8 Q2Y7R3
Q2Y7R4 Q2Y7R5 Q2Y7U5 Q2Y7W0 Q2Y7W2 Q2Y7W3 Q2Y7W4 Q2Y7X0 Q2Y7X4 Q2Y7X5 Q2Y7Y3
Q2Y807 Q2Y808 Q2Y809 Q2Y863 Q2Y865 Q2Y867 Q2Y8F8 Q2Y8K1 Q2Y8L4 Q2Y8L6 Q2Y8M6
Q2Y8S0 Q2Y8S8 Q2Y8V8 Q2Y981 Q2Y9J7 Q2Y9K0 Q2Y9K4 Q2Y9M4 Q2Y9M5 Q2Y9M6 Q2Y9M7
Q2Y9M8 Q2Y9M9 Q2Y9N0 Q2Y9P1 Q2Y9R5 Q2Y9R9 Q2Y9Y5 Q2Y9Y6 Q2Y9Y7 Q2Y9Y8 Q2Y9Y9
Q2Y9Z4 Q2Y9Z7 Q2YA02 Q2YA09 Q2YA48 Q2YA50 Q2YA74 Q2YA75 Q2YA81 Q2YA91 Q2YAB2
Q2YAC1 Q2YAC5 Q2YAI3 Q2YAN2 Q2YAN9 Q2YAP0 Q2YAQ4 Q2YAU4 Q2YAU5 Q2YAU6 Q2YAU7
Q2YAU8 Q2YAU9 Q2YAV0 Q2YAV1 Q2YAV2 Q2YAX1 Q2YAX2 Q2YAX3 Q2YAX4 Q2YAX5 Q2YAX6
Q2YAX8 Q2YAX9 Q2YAY0 Q2YAY1 Q2YAY2 Q2YAY3 Q2YAY4 Q2YAY5 Q2YAY6 Q2YAY7 Q2YAY8
Q2YAZ0 Q2YAZ1 Q2YAZ2 Q2YAZ3 Q2YAZ4 Q2YAZ5 Q2YAZ6 Q2YAZ7 Q2YAZ8 Q2YAZ9 Q2YB00
Q2YB01 Q2YB02 Q2YB04 Q2YB05 Q2YB06 Q2YB07 Q2YB08 Q2YB09 Q2YB18 Q2YB22 Q2YB30
Q2YB46 Q2YB47 Q2YB48 Q2YB52 Q2YB53 Q2YB59 Q2YB61 Q2YB78 Q2YB94 Q2YB96 Q2YBA1
Q2YBA4 Q2YBA5 Q2YBA6 Q2YBA7 Q2YBB2 Q2YBB7 Q2YBC0 Q2YBG1 Q2YBH2 Q2YBH3 Q2YBH5
Q2YBI1 Q2YBJ1 Q2YBJ2 Q2YBJ3 Q2YBJ4 Q2YBJ8 Q2YBK2 Q2YBK3 Q2YBK4 Q2YBK7 Q2YBK8
Q2YBL2 Q2YBM5 Q2YBP5 Q2YBP9 Q2YBQ2 Q2YBQ4 Q2YBR1 Q2YBR7 Q2YBR8 Q2YBS0 Q2YBS1
Q2YBS2 Q2YBS3 Q2YBS4 Q2YBS6 Q2YBS8 Q2YBT1 Q2YBT6 Q2YBU1 Q2YBU4 Q2YBW0 Q2YBW3
Q2YBW4 Q2YBW6 Q2YBX1 Q2YBX2 Q2YBX7 Q2YBZ1 Q2YC20 Q2YC27 Q2YC32 Q2YC50 Q2YC51
Q2YC56 Q2YC67 Q2YC90 Q2YC91 Q2YC92 Q2YCH6 Q2YCH7 Q2YCI8 Q2YCJ2 Q2YCK0 Q2YCK1
Q2YCK3 Q2YCK7 Q2YCL8 Q2YCN3 Q2YCR1 Q2YCX0 Q2YCY1 Q2YCZ2 Q2YCZ3 Q2YCZ6 Q2YD10
Q2YD11 Q2YD40 Q2YD43 Q2YD50 Q2YD51 Q2YD57 Q2YD58 Q2YD61 Q2YDC9 Q2YDD1 Q2YDD2
Q2YDE4 Q2YDE5 Q2YDE7 Q2YDE8 Q2YDF2 Q2YDF9 Q2YDG2 Q2YDG3 Q2YDH1 Q2YDH8 Q2YDH9
Q2YDI2 Q2YDI3 Q2YDI7 Q2YDI9 Q2YDJ0 Q2YDJ4 Q2YDJ9 Q2YDK0 Q2YDK1 Q2YDK4 Q2YDL7
Q2YDM2 Q2YDM5 Q2YDM7 Q2YDN4 Q2YDP3 Q2YDP6 Q2YDP8 Q2YDQ5 Q2YDS1 Q2YDU6 Q2YEI7
Q2YEJ1 Q2YEM8 Q2YEM9 Q2YEN0 Q2YEN2 Q2YIH2 Q2YIH8 Q2YIH9 Q2YIJ2 Q2YIJ3 Q2YIJ9
Q2YIL0 Q2YIL5 Q2YIL6 Q2YIL8 Q2YIN9 Q2YIQ1 Q2YIQ2 Q2YIQ3 Q2YIQ4 Q2YIQ6 Q2YIQ9
Q2YIS9 Q2YIT5 Q2YIT8 Q2YIU9 Q2YIW0 Q2YJ11 Q2YJ35 Q2YJ50 Q2YJ71 Q2YJ82 Q2YJG3
Q2YJM1 Q2YJM4 Q2YJN6 Q2YJN7 Q2YJN9 Q2YJP0 Q2YJQ3 Q2YJQ4 Q2YJQ6 Q2YJQ7 Q2YJR0
Q2YJR1 Q2YJR4 Q2YJT9 Q2YJZ2 Q2YK00 Q2YK17 Q2YK38 Q2YK44 Q2YKB2 Q2YKB4 Q2YKB6

Q2YKB8 Q2YKI5 Q2YKK8 Q2YKL3 Q2YKL4 Q2YKL5 Q2YKL6 Q2YKL8 Q2YKM8 Q2YKR3 Q2YKU4
Q2YKV1 Q2YKW2 Q2YKX9 Q2YKY0 Q2YL15 Q2YL33 Q2YL51 Q2YL58 Q2YL59 Q2YLB4 Q2YLD3
Q2YLF3 Q2YLG4 Q2YLH0 Q2YLH5 Q2YLH9 Q2YLI1 Q2YLI7 Q2YLI8 Q2YLJ0 Q2YLJ4 Q2YLK6
Q2YLL7 Q2YLL8 Q2YLM2 Q2YLM3 Q2YLM4 Q2YLM5 Q2YLN5 Q2YLN8 Q2YLP6 Q2YLP7 Q2YLR4
Q2YLR9 Q2YLS2 Q2YLT0 Q2YLT1 Q2YLU8 Q2YLX2 Q2YLX5 Q2YLX6 Q2YLY6 Q2YLY7 Q2YLZ6
Q2YLZ8 Q2YLZ9 Q2YM00 Q2YM02 Q2YM03 Q2YM04 Q2YM05 Q2YM06 Q2YM07 Q2YM08 Q2YM11
Q2YM12 Q2YM13 Q2YM14 Q2YM15 Q2YM16 Q2YM22 Q2YM28 Q2YM57 Q2YM63 Q2YM69 Q2YM98
Q2YMB5 Q2YMC3 Q2YMD1 Q2YME4 Q2YMF0 Q2YMF8 Q2YMG9 Q2YMH0 Q2YMH2 Q2YMH5 Q2YMI3
Q2YMI4 Q2YMJ2 Q2YMK2 Q2YML2 Q2YMM3 Q2YMP8 Q2YMQ2 Q2YMS6 Q2YMY0 Q2YMZ9 Q2YN02
Q2YN10 Q2YN11 Q2YN12 Q2YN15 Q2YN22 Q2YN35 Q2YN40 Q2YN59 Q2YN92 Q2YN94 Q2YN95
Q2YNA4 Q2YNA7 Q2YNB8 Q2YNC5 Q2YNC6 Q2YND8 Q2YND9 Q2YNE1 Q2YNE3 Q2YNG1 Q2YNG3
Q2YNG6 Q2YNH1 Q2YNH2 Q2YNJ9 Q2YNK0 Q2YNK7 Q2YNL2 Q2YNL3 Q2YNL7 Q2YNM5 Q2YNM6
Q2YNP1 Q2YNP4 Q2YNQ9 Q2YNR0 Q2YNR6 Q2YNT7 Q2YNV0 Q2YNV3 Q2YNV6 Q2YNW9 Q2YNZ0
Q2YNZ1 Q2YP18 Q2YP29 Q2YP33 Q2YP46 Q2YP58 Q2YPB6 Q2YPB8 Q2YPB9 Q2YPC1 Q2YPC2
Q2YPC3 Q2YPD2 Q2YPD3 Q2YPD5 Q2YPD6 Q2YPD7 Q2YPD8 Q2YPF3 Q2YPH0 Q2YPH2 Q2YPI1
Q2YPI2 Q2YPK6 Q2YPM3 Q2YPM9 Q2YPN0 Q2YPN5 Q2YPN6 Q2YPQ8 Q2YPS7 Q2YPT3 Q2YPU8
Q2YPV8 Q2YPW1 Q2YPW9 Q2YPX1 Q2YPX2 Q2YPX3 Q2YPX5 Q2YPX7 Q2YPY4 Q2YPY7 Q2YQ02
Q2YQ03 Q2YQ07 Q2YQ23 Q2YQ56 Q2YQB2 Q2YQD5 Q2YQD6 Q2YQD7 Q2YQD8 Q2YQD9 Q2YQE0
Q2YQH6 Q2YQH8 Q2YQI5 Q2YQK2 Q2YQL5 Q2YQM7 Q2YQN2 Q2YQP6 Q2YQP8 Q2YQR3 Q2YQR5
Q2YQR6 Q2YQR7 Q2YQS0 Q2YQS1 Q2YQS6 Q2YQS8 Q2YQU9 Q2YQV1 Q2YQV4 Q2YQV5 Q2YQV7
Q2YQV8 Q2YQW4 Q2YQW5 Q2YQW7 Q2YQX8 Q2YQY6 Q2YQY7 Q2YQY8 Q2YQY9 Q2YQZ0 Q2YQZ2
Q2YQZ3 Q2YQZ4 Q2YR03 Q2YR04 Q2YR05 Q2YR06 Q2YR07 Q2YR11 Q2YR12 Q2YR13 Q2YR40
Q2YR42 Q2YR43 Q2YR56 Q2YR81 Q2YR87 Q2YR89 Q2YR96 Q2YRA0 Q2YRA1 Q2YRA3 Q2YRA4
Q2YRA5 Q2YRA6 Q2YRA7 Q2YRA8 Q2YRA9 Q2YRB0 Q2YRC4 Q2YRD0 Q2YRD1 Q2YRD2 Q2YRF2
Q2YRF3 Q2YRF9 Q2YRG1 Q2YRG4 Q2YRJ1 Q2YRK0 Q2YRK5 Q2YRN8 Q2YRP5 Q2YRP7 Q2YRQ9
Q2YRR3 Q2YRR4 Q2YRR5 Q2YRT2 Q2YRT3 Q2YRT4 Q2YRT5 Q2YRT7 Q2YRT8 Q2YRT9 Q2YRU0
Q2YRU1 Q2YRU7 Q2YRU8 Q2YRV4 Q2YRW5 Q2YRY7 Q2YS10 Q2YS22 Q2YS31 Q2YS71 Q2YS77
Q2YS83 Q2YS94 Q2YS95 Q2YS98 Q2YSA2 Q2YSA8 Q2YSA9 Q2YSB0 Q2YSB3 Q2YSB4 Q2YSB5
Q2YSB6 Q2YSB8 Q2YSB9 Q2YSC2 Q2YSC3 Q2YSC4 Q2YSC7 Q2YSD1 Q2YSD3 Q2YSD6 Q2YSD7
Q2YSD9 Q2YSE1 Q2YSF0 Q2YSF1 Q2YSF6 Q2YSF8 Q2YSG0 Q2YSG2 Q2YSG8 Q2YSH5 Q2YSH7
Q2YSI3 Q2YSI6 Q2YSJ1 Q2YSJ9 Q2YSK2 Q2YSL9 Q2YSQ0 Q2YSR2 Q2YSR3 Q2YSV9 Q2YSW8
Q2YSX0 Q2YSX9 Q2YSZ2 Q2YSZ4 Q2YT00 Q2YT04 Q2YT09 Q2YT13 Q2YT14 Q2YT16 Q2YT22
Q2YT26 Q2YT29 Q2YT32 Q2YT34 Q2YT41 Q2YT45 Q2YT46 Q2YT47 Q2YT48 Q2YT49 Q2YT57
Q2YT60 Q2YT61 Q2YT62 Q2YT63 Q2YT67 Q2YT68 Q2YT73 Q2YT75 Q2YT78 Q2YT83 Q2YT85
Q2YT86 Q2YT87 Q2YT88 Q2YT89 Q2YT90 Q2YT91 Q2YT95 Q2YT97 Q2YT99 Q2YTA0 Q2YTA2
Q2YTA4 Q2YTA7 Q2YTA9 Q2YTB0 Q2YTB1 Q2YTB4 Q2YTB5 Q2YTB6 Q2YTB7 Q2YTB9 Q2YTC2
Q2YTC9 Q2YTD0 Q2YTD1 Q2YTD2 Q2YTD3 Q2YTE2 Q2YTE3 Q2YTF0 Q2YTF6 Q2YTG1 Q2YTH0
Q2YTH9 Q2YTI5 Q2YTI8 Q2YTJ3 Q2YTK0 Q2YTK1 Q2YTL0 Q2YTL3 Q2YTM0 Q2YTP0 Q2YTZ2
Q2YU05 Q2YU14 Q2YU19 Q2YU22 Q2YU25 Q2YU27 Q2YU32 Q2YU34 Q2YU47 Q2YU51 Q2YU53
Q2YU60 Q2YU66 Q2YU70 Q2YU75 Q2YU76 Q2YUD7 Q2YUD8 Q2YUE5 Q2YUE8 Q2YUE9 Q2YUF0
Q2YUF1 Q2YUF2 Q2YUF3 Q2YUF6 Q2YUG0 Q2YUH1 Q2YUH3 Q2YUI4 Q2YUI5 Q2YUI8 Q2YUJ0
Q2YUJ1 Q2YUJ2 Q2YUK4 Q2YUL0 Q2YUL1 Q2YUL2 Q2YUL3 Q2YUL6 Q2YUL7 Q2YUM3 Q2YUM5
Q2YUM6 Q2YUM9 Q2YUN1 Q2YUN3 Q2YUN4 Q2YUN5 Q2YUN8 Q2YUP1 Q2YUP7 Q2YUQ6 Q2YUR2
Q2YUR3 Q2YUS3 Q2YUU0 Q2YUU3 Q2YUU4 Q2YUU9 Q2YUY9 Q2YUZ7 Q2YV11 Q2YV20 Q2YV38

Q2YV52 Q2YV53 Q2YV69 Q2YV73 Q2YV76 Q2YVA4 Q2YVA8 Q2YVB1 Q2YVF5 Q2YVH7 Q2YVJ0
Q2YVJ2 Q2YVK2 Q2YVL5 Q2YVL6 Q2YVL8 Q2YVM4 Q2YVU7 Q2YVV0 Q2YVV2 Q2YVV7 Q2YVV9
Q2YVW3 Q2YVW5 Q2YVW6 Q2YVW8 Q2YVX3 Q2YVY2 Q2YVY3 Q2YVY4 Q2YVY6 Q2YVY8 Q2YVZ6
Q2YW87 Q2YW94 Q2YWA6 Q2YWB1 Q2YWC8 Q2YWD0 Q2YWE5 Q2YWE7 Q2YWE8 Q2YWF3 Q2YWF6
Q2YWF9 Q2YWG0 Q2YWG1 Q2YWH4 Q2YWH5 Q2YWJ5 Q2YWJ9 Q2YWK0 Q2YWK9 Q2YWN2 Q2YWN8
Q2YWP4 Q2YWQ1 Q2YWQ8 Q2YWR1 Q2YWR5 Q2YWS3 Q2YWS4 Q2YWS5 Q2YWT2 Q2YWU2 Q2YWW1
Q2YWW3 Q2YWW4 Q2YWW8 Q2YWX6 Q2YWX8 Q2YWY1 Q2YWY4 Q2YWY9 Q2YX08 Q2YX11 Q2YX30
Q2YX35 Q2YX50 Q2YX52 Q2YX54 Q2YX55 Q2YX57 Q2YX59 Q2YX63 Q2YX68 Q2YX76 Q2YX86
Q2YX87 Q2YX89 Q2YX97 Q2YX99 Q2YXA0 Q2YXA3 Q2YXC5 Q2YXC9 Q2YXD0 Q2YXD1 Q2YXD9
Q2YXE3 Q2YXE4 Q2YXE5 Q2YXE6 Q2YXF2 Q2YXF5 Q2YXF8 Q2YXF9 Q2YXG4 Q2YXG5 Q2YXH0
Q2YXH4 Q2YXH8 Q2YXI3 Q2YXJ0 Q2YXJ2 Q2YXJ8 Q2YXK0 Q2YXK2 Q2YXK3 Q2YXK4 Q2YXK5
Q2YXK9 Q2YXL0 Q2YXL1 Q2YXL2 Q2YXL3 Q2YXL4 Q2YXL5 Q2YXL6 Q2YXL7 Q2YXL8 Q2YXM1
Q2YXM2 Q2YXM4 Q2YXM5 Q2YXN2 Q2YXP1 Q2YXP2 Q2YXP3 Q2YXP5 Q2YXP6 Q2YXP7 Q2YXR3
Q2YXR5 Q2YXR6 Q2YXR9 Q2YXS2 Q2YXS9 Q2YXT0 Q2YXT1 Q2YXT2 Q2YXT6 Q2YXU8 Q2YXU9
Q2YXV4 Q2YXW0 Q2YXW5 Q2YXX0 Q2YXX1 Q2YXX2 Q2YXX3 Q2YXX4 Q2YXX8 Q2YXZ4 Q2YXZ5
Q2YXZ6 Q2YXZ7 Q2YXZ8 Q2YY05 Q2YY06 Q2YY09 Q2YY12 Q2YY14 Q2YY16 Q2YY21 Q2YY38
Q2YY39 Q2YY40 Q2YY44 Q2YY49 Q2YY51 Q2YY52 Q2YY57 Q2YY59 Q2YY60 Q2YY66 Q2YY67
Q2YY79 Q2YY85 Q2YY87 Q2YY88 Q2YY89 Q2YY90 Q2YYA2 Q2YYB7 Q2YYB8 Q2YYB9 Q2YYC1
Q2YYC7 Q2YYE6 Q2YYE8 Q2YYG3 Q2YYH2 Q2YYH4 Q2YYJ3 Q2YYJ4 Q2YYJ5 Q2YYJ6 Q2YYJ9
Q2YYK6 Q2YYK7 Q2YYK8 Q2YYK9 Q2YYL0 Q2YYL1 Q2YYL2 Q2YYL3 Q2YYL4 Q2YYL5 Q2YYL6
Q2YYL9 Q2YYM0 Q2YYM1 Q2YYM2 Q2YYM3 Q2YYM7 Q2YYM8 Q2YYM9 Q2YYN0 Q2YYP5 Q2YYP6
Q2YYP7 Q2YYP8 Q2YYP9 Q2YYQ0 Q2YYQ1 Q2YYQ2 Q2YYQ3 Q2YYQ5 Q2YYQ6 Q2YYQ7 Q2YYQ8
Q2YYR6 Q2YYS2 Q2YYS7 Q2YYS8 Q2YYS9 Q2YYT1 Q2YYT9 Q2YYV0 Q2YYV1 Q2YYV2 Q2YYV4
Q2YYV8 Q2YYY9 Q2YYZ2 Q2YZ12 Q2YZ22 Q2YZ41 Q2YZ47 Q2YZ66 Q2YZ69 Q2YZ70 Q2YZ71
Q2YZ91 Q2YZA2 Q2YZA3 Q2YZB2 Q2YZB3 Q2YZB4 Q2YZB6 Q2YZB7 Q2YZB8 Q2YZB9 Q2YZC0
Q2Z1R2 Q2Z2E9 Q2Z2F4 Q2Z2F6 Q307Y9 Q308N0 Q309B1 Q30BK9 Q30DN6 Q30DW5 Q30DW8
Q30DW9 Q30NQ3 Q30NQ5 Q30NR4 Q30NR6 Q30NR7 Q30NS7 Q30NT6 Q30NT9 Q30NU0 Q30NW0
Q30NW2 Q30NW3 Q30NW8 Q30NX0 Q30NX1 Q30NY4 Q30NY7 Q30NY8 Q30NZ0 Q30P05 Q30P36
Q30P60 Q30P84 Q30P92 Q30P97 Q30PA6 Q30PA9 Q30PD7 Q30PF0 Q30PH2 Q30PJ5 Q30PK5
Q30PK6 Q30PL1 Q30PL3 Q30PM2 Q30PN0 Q30PN2 Q30PN4 Q30PQ2 Q30PQ4 Q30PQ5 Q30PR1
Q30PS0 Q30PV8 Q30PX5 Q30PX6 Q30PY0 Q30PY1 Q30PY7 Q30PZ2 Q30PZ4 Q30PZ7 Q30Q10
Q30Q11 Q30Q40 Q30Q52 Q30Q82 Q30Q92 Q30QB8 Q30QC1 Q30QG7 Q30QG8 Q30QK7 Q30QK9
Q30QN8 Q30QN9 Q30QP0 Q30QP2 Q30QR5 Q30QS8 Q30QS9 Q30QT1 Q30QV9 Q30QX0 Q30R29
Q30R39 Q30R53 Q30R77 Q30R91 Q30R94 Q30R95 Q30RD1 Q30RE1 Q30RE2 Q30RH3 Q30RH7
Q30RI5 Q30RJ0 Q30RJ4 Q30RK1 Q30RK2 Q30RL0 Q30RL7 Q30RL8 Q30RM3 Q30RN2 Q30RN3
Q30RP0 Q30RP4 Q30RQ3 Q30RQ5 Q30RS1 Q30RT3 Q30RT5 Q30RX6 Q30S35 Q30S45 Q30S46
Q30S57 Q30S58 Q30S64 Q30S65 Q30S81 Q30S82 Q30S83 Q30S84 Q30S87 Q30S90 Q30S93
Q30S94 Q30S99 Q30SA5 Q30SC1 Q30SF5 Q30SG0 Q30SG7 Q30SH2 Q30SH9 Q30SM1 Q30SM6
Q30SN2 Q30SN5 Q30SP2 Q30SR5 Q30SS1 Q30SS4 Q30SS6 Q30SS8 Q30SW8 Q30SX0 Q30SX1
Q30T22 Q30T29 Q30T46 Q30T53 Q30T55 Q30T61 Q30T74 Q30T75 Q30T80 Q30T90 Q30TA4
Q30TB0 Q30TB2 Q30TC5 Q30TC9 Q30TD4 Q30TD5 Q30TD9 Q30TE4 Q30TE5 Q30TE6 Q30TE7
Q30TE9 Q30TF1 Q30TF9 Q30TG0 Q30TG2 Q30TH2 Q30TH8 Q30TI2 Q30TI8 Q30TJ7 Q30TJ8
Q30TK8 Q30TL4 Q30TL7 Q30TM4 Q30TN9 Q30TP0 Q30TP3 Q30TP4 Q30TP5 Q30TP6 Q30TP7
Q30TP8 Q30TP9 Q30TQ0 Q30TQ1 Q30TQ4 Q30TQ5 Q30TR6 Q30TS2 Q30TS3 Q30TS4 Q30TS5

Q30TS6 Q30TS7 Q30TU3 Q30TU7 Q30TU8 Q30TV0 Q30TV1 Q30TV2 Q30TV3 Q30TV4 Q30TV5
Q30TV7 Q30TV8 Q30TV9 Q30TW0 Q30TW1 Q30TW3 Q30TW5 Q30U33 Q30U43 Q30U44 Q30U81
Q30U93 Q30UA5 Q30UB9 Q30UF7 Q30UF8 Q30UH5 Q30UH9 Q30UI3 Q30UI5 Q30UJ4 Q30UL1
Q30UL7 Q30UM8 Q30UP6 Q30UP9 Q30US7 Q30UT5 Q30UZ9 Q30V54 Q30V89 Q30VA9 Q30VD6
Q30VD9 Q30VM1 Q30W70 Q30W71 Q30WB0 Q30WD0 Q30WD2 Q30WD3 Q30WE0 Q30WF9 Q30WH2
Q30WH3 Q30WI5 Q30WI6 Q30WI7 Q30WJ0 Q30WJ2 Q30WK2 Q30WK3 Q30WL8 Q30WM4 Q30WZ8
Q30X04 Q30X05 Q30X07 Q30X08 Q30X09 Q30X10 Q30X13 Q30X18 Q30X20 Q30X35 Q30X38
Q30X45 Q30X46 Q30X61 Q30XF7 Q30XN3 Q30XS2 Q30XS4 Q30XT0 Q30XT1 Q30XT2 Q30XT6
Q30XT7 Q30XU3 Q30XV8 Q30XV9 Q30XW0 Q30XW1 Q30XZ5 Q30Y02 Q30Y11 Q30Y13 Q30Y14
Q30Y15 Q30Y16 Q30Y29 Q30Y32 Q30Y35 Q30Y42 Q30Y43 Q30Y72 Q30Y73 Q30YB7 Q30YB8
Q30YB9 Q30YH0 Q30YH5 Q30YH6 Q30YK6 Q30YL0 Q30YL1 Q30YL2 Q30YL7 Q30YM4 Q30YP7
Q30YQ3 Q30YQ7 Q30YS1 Q30YS5 Q30YS6 Q30YV9 Q30YW1 Q30YX3 Q30YX4 Q30YX5 Q30YX6
Q30YX7 Q30YX9 Q30Z04 Q30Z06 Q30Z08 Q30Z13 Q30Z14 Q30Z36 Q30Z37 Q30Z38 Q30Z41
Q30Z42 Q30Z43 Q30Z44 Q30Z45 Q30Z46 Q30Z47 Q30Z48 Q30Z49 Q30Z50 Q30Z51 Q30Z52
Q30Z53 Q30Z54 Q30Z55 Q30Z56 Q30Z57 Q30Z58 Q30Z59 Q30Z60 Q30Z61 Q30Z65 Q30Z66
Q30Z67 Q30Z68 Q30Z69 Q30Z77 Q30Z79 Q30Z80 Q30Z93 Q30Z97 Q30Z99 Q30ZD3 Q30ZF7
Q30ZH1 Q30ZH2 Q30ZH4 Q30ZH5 Q30ZH8 Q30ZJ9 Q30ZK1 Q30ZK3 Q30ZK6 Q30ZK7 Q30ZM2
Q30ZN0 Q30ZN1 Q30ZP0 Q30ZP3 Q30ZQ6 Q30ZR4 Q30ZR5 Q30ZR6 Q30ZR8 Q30ZS4 Q30ZT4
Q30ZT5 Q30ZW0 Q30ZX0 Q30ZX3 Q30ZX9 Q30ZZ6 Q30ZZ9 Q310P0 Q310Q2 Q310R5 Q310R6
Q310S5 Q310T4 Q310X3 Q310Z1 Q311A6 Q311A7 Q311C7 Q311E8 Q311G6 Q311P4 Q311T0
Q311U7 Q311U9 Q311V1 Q311X7 Q311Y0 Q311Y2 Q311Z4 Q311Z5 Q311Z9 Q312A8 Q312C5
Q312D5 Q312D6 Q312D7 Q312G9 Q312I9 Q312S2 Q312S3 Q313E8 Q313G2 Q313G3 Q313G6
Q313G7 Q313H2 Q313J6 Q313J8 Q313J9 Q313K0 Q313K1 Q313K2 Q313K7 Q313P5 Q313Q1
Q313Q2 Q313Q5 Q313R1 Q313R2 Q313S2 Q313S4 Q313S5 Q313S6 Q313T1 Q313T7 Q314N8
Q315F5 Q315M7 Q315P9 Q315T1 Q315T9 Q31655 Q31669 Q31672 Q31673 Q31674 Q31680
Q316D5 Q316F5 Q316H9 Q316L3 Q316L4 Q316N1 Q316P9 Q316Q1 Q316U6 Q316U7 Q316U8
Q316W7 Q316Z9 Q31708 Q31715 Q31730 Q31738 Q31750 Q31795 Q317B9 Q317E4 Q317F9
Q317H9 Q317J6 Q317L9 Q317N3 Q317R3 Q317R4 Q317T4 Q317T5 Q317T6 Q317V0 Q317V2
Q317V3 Q317V6 Q317W7 Q317W9 Q317X3 Q317X8 Q317Y0 Q317Y1 Q317Y5 Q317Y6 Q317Y7
Q317Y9 Q317Z7 Q317Z9 Q31809 Q31827 Q31859 Q31886 Q318A5 Q318A6 Q318B6 Q318C3
Q318C5 Q318C6 Q318D0 Q318D6 Q318E9 Q318F0 Q318F2 Q318F3 Q318G0 Q318G7 Q318I0
Q318I4 Q318I5 Q318I6 Q318I7 Q318I8 Q318I9 Q318J0 Q318J1 Q318J2 Q318J3 Q318J4
Q318J5 Q318J6 Q318J7 Q318J8 Q318J9 Q318K0 Q318K1 Q318K4 Q318K5 Q318K6 Q318K7
Q318K8 Q318K9 Q318L0 Q318L1 Q318L2 Q318L3 Q318N2 Q318N3 Q318N4 Q318N5 Q318N6
Q318P1 Q318P3 Q318P8 Q318Q1 Q318Q3 Q318Q4 Q318Q5 Q318Q7 Q318Q8 Q318Q9 Q318R1
Q318R7 Q318T1 Q318V5 Q318V6 Q318W1 Q31946 Q31948 Q31951 Q31983 Q31992 Q319C4
Q319D4 Q319D8 Q319E0 Q319E2 Q319E3 Q319E4 Q319F5 Q319F8 Q319F9 Q319H3 Q319H4
Q319H9 Q319I3 Q319I4 Q319I9 Q319J2 Q319J8 Q319K4 Q319K9 Q319L1 Q319M0 Q319M1
Q319M2 Q319M8 Q319U4 Q319U5 Q319V1 Q319V2 Q319X4 Q319X7 Q319X8 Q319Z8 Q319Z9
Q31AA3 Q31AA9 Q31AB8 Q31AC8 Q31AD1 Q31AD2 Q31AF6 Q31AF9 Q31AG5 Q31AH1 Q31AM1
Q31AP2 Q31AR4 Q31AS6 Q31AS9 Q31AT8 Q31AV3 Q31AV4 Q31AV5 Q31AW7 Q31AW9 Q31AX4
Q31AX5 Q31AX7 Q31AY7 Q31AZ2 Q31B18 Q31B19 Q31B21 Q31B28 Q31B29 Q31B32 Q31B38
Q31B49 Q31B50 Q31B51 Q31B52 Q31B53 Q31B54 Q31B55 Q31B81 Q31B86 Q31B89 Q31B91
Q31B97 Q31BA3 Q31BA9 Q31BB0 Q31BC3 Q31BC4 Q31BD0 Q31BD5 Q31BK2 Q31BL9 Q31BR1

Q31BS1 Q31BS5 Q31BS7 Q31BT7 Q31BT9 Q31BU4 Q31BV0 Q31BV1 Q31BW9 Q31BX4 Q31BX5
Q31BY4 Q31BY9 Q31BZ0 Q31BZ1 Q31C09 Q31C12 Q31C13 Q31C15 Q31C25 Q31C27 Q31C34
Q31C38 Q31C42 Q31C50 Q31C53 Q31C59 Q31C61 Q31C64 Q31C67 Q31C80 Q31C83 Q31C96
Q31CA5 Q31CB9 Q31CC9 Q31CD3 Q31CD7 Q31CE3 Q31CE7 Q31CK0 Q31CK8 Q31CL5 Q31CL7
Q31CQ1 Q31CR1 Q31CR6 Q31CR8 Q31CS0 Q31CS4 Q31CS5 Q31CS6 Q31CT2 Q31CU1 Q31CU2
Q31CU4 Q31CU8 Q31CV7 Q31CV9 Q31CW3 Q31CX7 Q31CX8 Q31CX9 Q31CY0 Q31CY6 Q31CZ2
Q31CZ4 Q31CZ9 Q31D08 Q31D10 Q31D17 Q31D25 Q31D51 Q31D56 Q31D68 Q31D78 Q31DB4
Q31DD4 Q31DD5 Q31DD7 Q31DD8 Q31DD9 Q31DE7 Q31DE8 Q31DF8 Q31DG4 Q31DG8 Q31DH2
Q31DI1 Q31DI6 Q31DJ0 Q31DJ1 Q31DJ4 Q31DK8 Q31DK9 Q31DN1 Q31DP2 Q31DP7 Q31DP8
Q31DP9 Q31DQ3 Q31DQ5 Q31DX2 Q31DX4 Q31DX5 Q31E14 Q31E38 Q31E41 Q31E43 Q31E54
Q31E55 Q31E63 Q31E64 Q31E65 Q31E66 Q31E67 Q31E77 Q31E80 Q31E84 Q31E86 Q31EA3
Q31EB2 Q31EB4 Q31EB5 Q31EB6 Q31EC7 Q31ED1 Q31ED4 Q31EF5 Q31EF8 Q31EG9 Q31EI1
Q31EJ2 Q31EK2 Q31EK3 Q31EK4 Q31EL0 Q31EM1 Q31EM2 Q31ES7 Q31ET3 Q31EW3 Q31EY6
Q31F14 Q31F16 Q31F17 Q31F18 Q31F19 Q31F20 Q31F24 Q31F31 Q31F42 Q31F43 Q31F52
Q31F53 Q31F54 Q31F66 Q31F68 Q31F72 Q31F87 Q31F89 Q31F91 Q31F93 Q31FB1 Q31FB3
Q31FD3 Q31FE3 Q31FE7 Q31FF5 Q31FF6 Q31FF9 Q31FG4 Q31FH4 Q31FI6 Q31FN8 Q31FR9
Q31FS6 Q31FS7 Q31FT1 Q31FT2 Q31FV4 Q31FW4 Q31FW5 Q31FW7 Q31FY5 Q31FZ4 Q31FZ5
Q31G14 Q31G32 Q31G37 Q31G41 Q31G43 Q31G44 Q31G45 Q31G46 Q31G56 Q31G58 Q31G65
Q31G66 Q31G71 Q31G89 Q31GB2 Q31GC7 Q31GD2 Q31GD4 Q31GD8 Q31GE9 Q31GF1 Q31GF2
Q31GG2 Q31GG8 Q31GI3 Q31GJ9 Q31GK0 Q31GK3 Q31GK4 Q31GK5 Q31GK7 Q31GL4 Q31GL5
Q31GL7 Q31GM4 Q31GM9 Q31GN4 Q31GN5 Q31GP2 Q31GP3 Q31GQ3 Q31GT2 Q31GU1 Q31GV5
Q31GW8 Q31GX8 Q31GY9 Q31GZ0 Q31GZ1 Q31H08 Q31H19 Q31H49 Q31H51 Q31H54 Q31H63
Q31H76 Q31H83 Q31H84 Q31H85 Q31H87 Q31H97 Q31HA6 Q31HA7 Q31HA8 Q31HB0 Q31HC6
Q31HD1 Q31HD2 Q31HD3 Q31HD4 Q31HD5 Q31HD6 Q31HD8 Q31HD9 Q31HG2 Q31HG3 Q31HG6
Q31HH3 Q31HH6 Q31HI0 Q31HI2 Q31HK9 Q31HP0 Q31HQ4 Q31HR9 Q31HS1 Q31HX7 Q31HX8
Q31HX9 Q31HY0 Q31HZ1 Q31HZ4 Q31HZ6 Q31I05 Q31I07 Q31I08 Q31I16 Q31I19 Q31I28
Q31I33 Q31I36 Q31I42 Q31I43 Q31I55 Q31I61 Q31I68 Q31I69 Q31I95 Q31I96 Q31IB5
Q31IB6 Q31IB9 Q31IC2 Q31IC8 Q31ID5 Q31IE1 Q31IE3 Q31IE5 Q31IE8 Q31IF2 Q31IF4
Q31IG5 Q31IG6 Q31IH6 Q31II2 Q31II5 Q31II8 Q31IJ9 Q31IK0 Q31IK3 Q31IL7 Q31IM5
Q31IN3 Q31IN4 Q31IN7 Q31IN8 Q31IR4 Q31IS6 Q31IS8 Q31IT1 Q31IT2 Q31IT3 Q31IT4
Q31IT5 Q31IT6 Q31IV7 Q31IV8 Q31IV9 Q31IW0 Q31IW1 Q31IW3 Q31IW4 Q31IW5 Q31IW6
Q31IW7 Q31IW8 Q31IW9 Q31IX0 Q31IX1 Q31IX2 Q31IX3 Q31IX5 Q31IX6 Q31IX7 Q31IX8
Q31IX9 Q31IY0 Q31IY1 Q31IY2 Q31IY3 Q31IY4 Q31IY5 Q31IY6 Q31IY7 Q31IY8 Q31IY9
Q31IZ0 Q31IZ1 Q31IZ2 Q31IZ3 Q31J09 Q31J37 Q31J39 Q31J60 Q31J61 Q31J79 Q31J84
Q31J85 Q31JA4 Q31JB0 Q31JB9 Q31JC4 Q31JC9 Q31JD2 Q31JE7 Q31JF2 Q31JF4 Q31JF6
Q31JF8 Q31JG5 Q31JG7 Q31JG9 Q31JJ7 Q31JJ8 Q31JM9 Q31JQ7 Q31JS3 Q31JS5 Q31JY3
Q31K19 Q31K20 Q31K30 Q31K36 Q31K48 Q31K58 Q31K59 Q31KC5 Q31KC6 Q31KC7 Q31KD2
Q31KE9 Q31KG6 Q31KH4 Q31KI0 Q31KJ1 Q31KK8 Q31KL4 Q31KL5 Q31KM4 Q31KN6 Q31KN8
Q31KP1 Q31KP9 Q31KT1 Q31KU0 Q31KU1 Q31KX4 Q31KX6 Q31KY0 Q31KY9 Q31KZ2 Q31KZ5
Q31L04 Q31L07 Q31L08 Q31L09 Q31L10 Q31L11 Q31L12 Q31L13 Q31L14 Q31L15 Q31L16
Q31L17 Q31L18 Q31L19 Q31L20 Q31L21 Q31L22 Q31L23 Q31L26 Q31L27 Q31L28 Q31L29
Q31L30 Q31L31 Q31L33 Q31L35 Q31L36 Q31L42 Q31LA3 Q31LA7 Q31LB6 Q31LB7 Q31LC2
Q31LC9 Q31LF9 Q31LH6 Q31LI0 Q31LJ3 Q31LJ9 Q31LL0 Q31LL6 Q31LL9 Q31LP5 Q31LP7
Q31LT0 Q31LU4 Q31LW2 Q31LW9 Q31LY2 Q31LZ1 Q31LZ7 Q31M42 Q31M81 Q31M94 Q31MB5

Q31MB7 Q31MD1 Q31ME3 Q31ME5 Q31MH4 Q31MH9 Q31MM2 Q31MM7 Q31MN3 Q31MQ1 Q31MS4
Q31MS5 Q31MY7 Q31N04 Q31N15 Q31N17 Q31N19 Q31N26 Q31N27 Q31N30 Q31N34 Q31N39
Q31N52 Q31N55 Q31N58 Q31N99 Q31NA4 Q31NA6 Q31NB2 Q31NB3 Q31ND6 Q31NH0 Q31NK6
Q31NM4 Q31NM6 Q31NR1 Q31NR2 Q31P19 Q31P38 Q31P66 Q31P67 Q31P73 Q31P90 Q31P93
Q31P96 Q31PB6 Q31PF9 Q31PG2 Q31PH2 Q31PJ2 Q31PK5 Q31PL4 Q31PM2 Q31PN0 Q31PU2
Q31PU5 Q31PU7 Q31PV3 Q31PV4 Q31PV6 Q31PW0 Q31PW3 Q31PX4 Q31PY6 Q31PY7 Q31Q03
Q31Q05 Q31Q18 Q31Q39 Q31Q54 Q31Q60 Q31QC4 Q31QE2 Q31QE7 Q31QF1 Q31QF6 Q31QG5
Q31QG7 Q31QG8 Q31QJ2 Q31QK3 Q31QK4 Q31QK5 Q31QK6 Q31QK9 Q31QL1 Q31QM1 Q31QM9
Q31QN4 Q31QP9 Q31QR1 Q31QR8 Q31QS4 Q31QS8 Q31QW7 Q31QY0 Q31QY1 Q31QY2 Q31QY3
Q31R15 Q31R18 Q31R68 Q31R76 Q31R79 Q31R85 Q31R91 Q31RB6 Q31RD2 Q31RD9 Q31RH5
Q31RH6 Q31RH7 Q31RJ2 Q31RK2 Q31RK3 Q31RK5 Q31RK7 Q31RL0 Q31RL4 Q31RM0 Q31RQ5
Q31RR0 Q31RS5 Q31RS6 Q31RS8 Q31RU7 Q31S00 Q31S12 Q31S19 Q31S64 Q31S95 Q31SA6
Q31SC2 Q31SC5 Q31SC8 Q31SU3 Q31SU5 Q31SV2 Q31SV5 Q31SV6 Q31SV7 Q31SV8 Q31SW4
Q31SW8 Q31SX4 Q31SY8 Q31T16 Q31T20 Q31T77 Q31T78 Q31T82 Q31TA0 Q31TA1 Q31TA2
Q31TA7 Q31TA8 Q31TC2 Q31TC3 Q31TC7 Q31TC8 Q31TD1 Q31TD2 Q31TD3 Q31TD4 Q31TE9
Q31TF9 Q31TG8 Q31TH0 Q31TI0 Q31TI1 Q31TJ4 Q31TK0 Q31TK2 Q31TU7 Q31TW1 Q31TX1
Q31TY2 Q31TY7 Q31TY9 Q31TZ1 Q31U00 Q31U01 Q31U02 Q31U03 Q31U04 Q31U06 Q31U09
Q31U10 Q31U11 Q31U12 Q31U13 Q31U14 Q31U19 Q31U21 Q31U23 Q31U28 Q31U29 Q31U31
Q31U32 Q31U45 Q31U49 Q31U54 Q31U58 Q31U59 Q31U61 Q31U62 Q31U64 Q31U71 Q31U74
Q31U83 Q31U84 Q31U85 Q31U86 Q31U87 Q31U88 Q31U98 Q31UA2 Q31UA6 Q31UC9 Q31UD0
Q31UE2 Q31UF3 Q31UF9 Q31UH7 Q31UI1 Q31UK5 Q31UK6 Q31UL0 Q31UL3 Q31UM0 Q31UM5
Q31UN9 Q31UP0 Q31UP3 Q31UP5 Q31UQ6 Q31UQ7 Q31US3 Q31US8 Q31UU6 Q31UV3 Q31UV5
Q31UV6 Q31UV7 Q31UW0 Q31UW5 Q31UY3 Q31UY4 Q31UY6 Q31UY8 Q31UY9 Q31UZ0 Q31UZ1
Q31UZ2 Q31V04 Q31V06 Q31V09 Q31V12 Q31V27 Q31V29 Q31V36 Q31V43 Q31V53 Q31V61
Q31V62 Q31VA6 Q31VC8 Q31VE5 Q31VF6 Q31VJ1 Q31VJ2 Q31VJ3 Q31VJ4 Q31VK8 Q31VL3
Q31VL4 Q31VL8 Q31VM0 Q31VM2 Q31VN1 Q31VN3 Q31VP4 Q31VP5 Q31VP9 Q31VS0 Q31VT6
Q31VU2 Q31VU5 Q31VU6 Q31VU7 Q31VU8 Q31VU9 Q31VV0 Q31VV5 Q31VV6 Q31VV7 Q31VV8
Q31VV9 Q31VW0 Q31VW1 Q31VW2 Q31VW3 Q31VW4 Q31VW5 Q31VW6 Q31VW7 Q31VW8 Q31VW9
Q31VX0 Q31VX1 Q31VX2 Q31VX3 Q31VX4 Q31VX5 Q31VX7 Q31VX8 Q31VX9 Q31VY0 Q31VY1
Q31VY2 Q31VY8 Q31VY9 Q31VZ0 Q31VZ2 Q31VZ4 Q31VZ5 Q31W07 Q31W09 Q31W27 Q31W28
Q31W33 Q31W34 Q31W43 Q31W44 Q31W45 Q31W46 Q31W47 Q31W49 Q31W50 Q31W53 Q31W56
Q31W60 Q31W62 Q31W63 Q31W66 Q31W81 Q31W92 Q31W94 Q31W95 Q31W98 Q31W99 Q31WA4
Q31WA5 Q31WB6 Q31WE3 Q31WF1 Q31WF2 Q31WG0 Q31WG2 Q31WG4 Q31WG5 Q31WG6 Q31WH4
Q31WH5 Q31WI5 Q31WI6 Q31WK0 Q31WK2 Q31WK4 Q31WK6 Q31WL0 Q31WL1 Q31WM2 Q31WM4
Q31WQ5 Q31WS0 Q31WU2 Q31WU9 Q31WW1 Q31WW5 Q31WW9 Q31WX0 Q31WX2 Q31WX3 Q31WX9
Q31WY3 Q31WZ1 Q31WZ5 Q31X44 Q31X58 Q31X61 Q31X62 Q31X63 Q31X64 Q31X74 Q31X75
Q31X95 Q31XA5 Q31XA6 Q31XA7 Q31XA8 Q31XA9 Q31XB2 Q31XB3 Q31XB4 Q31XC6 Q31XD1
Q31XD2 Q31XD6 Q31XD7 Q31XD8 Q31XD9 Q31XE5 Q31XF6 Q31XF7 Q31XG0 Q31XI5 Q31XI7
Q31XJ2 Q31XJ6 Q31XK5 Q31XL0 Q31XM4 Q31XM5 Q31XM6 Q31XQ4 Q31XQ5 Q31XR1 Q31XS0
Q31XS2 Q31XS3 Q31XT0 Q31XT6 Q31XU9 Q31XV0 Q31XV1 Q31XV2 Q31XV8 Q31XV9 Q31XW2
Q31XW3 Q31XW4 Q31XW7 Q31XX2 Q31XX3 Q31XX5 Q31XX6 Q31XY1 Q31XY3 Q31XZ3 Q31XZ6
Q31Y13 Q31Y15 Q31Y20 Q31Y24 Q31Y44 Q31Y46 Q31Y51 Q31Y52 Q31Y68 Q31Y73 Q31Y83
Q31Y97 Q31YA5 Q31YC7 Q31YC9 Q31YD4 Q31YD8 Q31YE0 Q31YJ3 Q31YJ4 Q31YJ6 Q31YL1
Q31YL9 Q31YM1 Q31YM9 Q31YN0 Q31YN1 Q31YN2 Q31YQ9 Q31YR0 Q31YR1 Q31YR6 Q31YR7

Q31YS6 Q31YS8 Q31YT3 Q31YT8 Q31YU0 Q31YU3 Q31YU4 Q31YW1 Q31YW2 Q31YW6 Q31YW9
Q31YX9 Q31YZ2 Q31YZ3 Q31YZ4 Q31Z33 Q31Z65 Q31Z78 Q31Z80 Q31Z90 Q31ZB5 Q31ZB7
Q31ZC2 Q31ZE2 Q31ZE4 Q31ZF3 Q31ZG3 Q31ZG4 Q31ZH6 Q31ZK3 Q31ZK9 Q31ZL0 Q31ZL8
Q31ZL9 Q31ZM1 Q31ZN0 Q31ZP7 Q31ZQ1 Q31ZQ3 Q31ZQ4 Q31ZS3 Q31ZS8 Q31ZV3 Q31ZV4
Q31ZX1 Q31ZX5 Q31ZY2 Q32001 Q32016 Q32026 Q32040 Q32041 Q32042 Q32063 Q32065
Q32067 Q320D9 Q320H8 Q320R8 Q320R9 Q320T0 Q320W2 Q320W6 Q320Y0 Q320Y2 Q320Z3
Q320Z4 Q320Z5 Q320Z7 Q321A7 Q321B7 Q321D8 Q321D9 Q321F0 Q321G0 Q321G3 Q321K4
Q321K5 Q321K6 Q321K7 Q321K8 Q321L0 Q321N2 Q321N4 Q321N6 Q321N7 Q321N8 Q321N9
Q321P0 Q321Q8 Q321R4 Q321S2 Q321V7 Q321V9 Q321Y0 Q321Y4 Q32214 Q32223 Q32255
Q32256 Q32271 Q32283 Q322B0 Q322D4 Q322D6 Q322D7 Q322E5 Q322E6 Q322F8 Q322G6
Q322I9 Q322J0 Q322J3 Q322J5 Q322K5 Q322P1 Q322Q2 Q322Q3 Q322V3 Q322W7 Q32303
Q32344 Q32345 Q32360 Q32397 Q323B4 Q323B8 Q323C6 Q323E8 Q323I7 Q323I8 Q323I9
Q323J1 Q323J2 Q323J3 Q323L0 Q323L8 Q323L9 Q323M1 Q323M9 Q323N6 Q323P2 Q323P9
Q323T3 Q323V7 Q323Y1 Q323Y7 Q323Z0 Q323Z6 Q32450 Q324A9 Q324B1 Q324B3 Q324B6
Q324B7 Q324C4 Q324G2 Q324G4 Q324G9 Q324I4 Q324J6 Q324J7 Q324M4 Q324M6 Q324N4
Q324N6 Q324P5 Q324Q2 Q324Q5 Q324Q7 Q324R2 Q324R3 Q324R5 Q324S4 Q324W2 Q324Y3
Q324Y6 Q324Z1 Q324Z4 Q324Z5 Q325B0 Q325C0 Q325C1 Q325C2 Q325C3 Q325C4 Q325C5
Q325C7 Q325F5 Q325F7 Q325G3 Q325G4 Q325G5 Q325H5 Q325H8 Q325H9 Q325I1 Q325I5
Q325I6 Q325I8 Q325J5 Q325J6 Q325K5 Q325K7 Q325L0 Q325L1 Q325L6 Q325P3 Q325P4
Q325P6 Q325P7 Q325R5 Q325R6 Q325S4 Q325T2 Q325T4 Q325T7 Q325U6 Q325V1 Q325V7
Q325W8 Q325W9 Q325X0 Q325X1 Q325X3 Q325X4 Q325X6 Q325X9 Q325Y0 Q325Y3 Q325Y5
Q325Z3 Q32616 Q32619 Q32620 Q32625 Q32663 Q32668 Q32686 Q32699 Q326A3 Q326A4
Q326A6 Q326B5 Q326B6 Q326D1 Q326D2 Q326D3 Q326D4 Q326E4 Q326E7 Q326F3 Q326F4
Q326G1 Q326G2 Q326G3 Q326G4 Q326G6 Q326H2 Q326H3 Q326H6 Q326I1 Q326I2 Q326I3
Q326I4 Q326J2 Q326J3 Q326J4 Q326J7 Q326K0 Q326K6 Q326K7 Q326L3 Q326L6 Q326L9
Q32701 Q32742 Q32759 Q32766 Q32770 Q327K0 Q327K2 Q327K9 Q327L2 Q327L3 Q327L4
Q327L5 Q327M2 Q327M6 Q327N4 Q327U6 Q327W0 Q32836 Q32890 Q328B8 Q328C4 Q328C5
Q328D1 Q328E3 Q328E9 Q328F0 Q328F1 Q328F6 Q328F7 Q328H8 Q328I9 Q328J6 Q328J7
Q328J8 Q328J9 Q328K0 Q328K1 Q328K5 Q328K6 Q328L9 Q328M0 Q328N3 Q328N5 Q328S1
Q328S3 Q328S8 Q328U0 Q328U1 Q328V2 Q328V4 Q328X7 Q328Y9 Q328Z9 Q32908 Q329B1
Q329B4 Q329B5 Q329B6 Q329B8 Q329C3 Q329C6 Q329F0 Q329H3 Q329K7 Q329K8 Q329L3
Q329L4 Q329L7 Q329L9 Q329M0 Q329M1 Q329M2 Q329M3 Q329N6 Q329N8 Q329P2 Q329P5
Q329S9 Q329T0 Q329T3 Q329T4 Q329T5 Q329U3 Q329V0 Q329V3 Q329V7 Q329V8 Q329Y2
Q329Y7 Q32A07 Q32A11 Q32A22 Q32A36 Q32A57 Q32A61 Q32A65 Q32A67 Q32A68 Q32A70
Q32A71 Q32A80 Q32A84 Q32A92 Q32A94 Q32AA2 Q32AA4 Q32AA5 Q32AA9 Q32AB5 Q32AB6
Q32AB8 Q32AB9 Q32AD6 Q32AE5 Q32AE8 Q32AF0 Q32AF5 Q32AF6 Q32AF7 Q32AF8 Q32AF9
Q32AG0 Q32AG3 Q32AG6 Q32AG7 Q32AG8 Q32AG9 Q32AH0 Q32AH1 Q32AH9 Q32AJ7 Q32AK1
Q32AK2 Q32AL3 Q32AL7 Q32AM4 Q32AM6 Q32AM7 Q32AN4 Q32AN7 Q32AQ0 Q32AR4 Q32AV3
Q32AV4 Q32AV5 Q32AV6 Q32AW6 Q32AZ8 Q32B11 Q32B18 Q32B21 Q32B22 Q32B23 Q32B24
Q32B25 Q32B26 Q32B27 Q32B30 Q32B31 Q32B32 Q32B33 Q32B34 Q32B35 Q32B36 Q32B37
Q32B38 Q32B39 Q32B40 Q32B41 Q32B42 Q32B43 Q32B44 Q32B45 Q32B46 Q32B47 Q32B48
Q32B49 Q32B50 Q32B52 Q32B53 Q32B54 Q32B55 Q32B56 Q32B57 Q32B61 Q32B62 Q32B63
Q32B65 Q32B67 Q32B68 Q32B77 Q32B79 Q32B92 Q32BA2 Q32BA3 Q32BA8 Q32BA9 Q32BB3
Q32BB4 Q32BB6 Q32BC8 Q32BE4 Q32BE7 Q32BE8 Q32BF0 Q32BF4 Q32BF7 Q32BG2 Q32BG3

Q32BG5 Q32BG6 Q32BG7 Q32BG8 Q32BG9 Q32BH8 Q32BH9 Q32BI3 Q32BI4 Q32BJ0 Q32BK8
Q32BN8 Q32BP9 Q32BQ2 Q32BQ3 Q32BQ8 Q32BR2 Q32BR9 Q32BS1 Q32BU3 Q32BV2 Q32BV3
Q32BW1 Q32BW4 Q32BW5 Q32BW6 Q32BW7 Q32BX5 Q32BX8 Q32BY6 Q32BY7 Q32C02 Q32C08
Q32C11 Q32C13 Q32C17 Q32C18 Q32C29 Q32C31 Q32C90 Q32C91 Q32C94 Q32CB2 Q32CB4
Q32CB6 Q32CC0 Q32CC4 Q32CD0 Q32CD5 Q32CG3 Q32CG4 Q32CG5 Q32CH8 Q32CH9 Q32CI0
Q32CI3 Q32CI4 Q32CI5 Q32CI6 Q32CI7 Q32CJ6 Q32CL7 Q32CL8 Q32CM9 Q32CN0 Q32CN1
Q32CN2 Q32CN5 Q32CQ1 Q32CT9 Q32CU0 Q32CU6 Q32CV4 Q32CV6 Q32CV7 Q32CW9 Q32CX4
Q32CX5 Q32CX9 Q32CY0 Q32CY1 Q32CY2 Q32D15 Q32D21 Q32D32 Q32D33 Q32D36 Q32D37
Q32D38 Q32D41 Q32D44 Q32D45 Q32D47 Q32D48 Q32D53 Q32D55 Q32D69 Q32D72 Q32D87
Q32D90 Q32D98 Q32DA4 Q32DB3 Q32DB7 Q32DB9 Q32DC6 Q32DC7 Q32DD5 Q32DE2 Q32DE6
Q32DF7 Q32DG9 Q32DH8 Q32DK6 Q32DK7 Q32DK8 Q32DL2 Q32DL6 Q32DL8 Q32DP8 Q32DS4
Q32DS5 Q32DS7 Q32DU2 Q32DV8 Q32DY4 Q32DZ1 Q32E01 Q32E03 Q32E04 Q32E12 Q32E24
Q32E25 Q32E30 Q32E32 Q32E37 Q32E42 Q32E44 Q32E51 Q32E52 Q32E73 Q32E86 Q32E99
Q32EA0 Q32EA9 Q32ED0 Q32EE8 Q32EE9 Q32EF0 Q32EF2 Q32EF3 Q32EF4 Q32EH3 Q32EI8
Q32EM3 Q32EN8 Q32EQ3 Q32EQ7 Q32ER0 Q32ER8 Q32ES3 Q32ES9 Q32EU9 Q32EV8 Q32EW6
Q32EW7 Q32EX9 Q32EY5 Q32EZ1 Q32EZ2 Q32EZ5 Q32EZ9 Q32F19 Q32F26 Q32F89 Q32FB3
Q32FB8 Q32FD3 Q32FD5 Q32FD6 Q32FD7 Q32FE8 Q32FF0 Q32FG8 Q32FI0 Q32FI2 Q32FI3
Q32FI5 Q32FI6 Q32FI7 Q32FJ7 Q32FL5 Q32FQ5 Q32FR6 Q32FR8 Q32G00 Q32G05 Q32G72
Q32G79 Q32G84 Q32G86 Q32G87 Q32G88 Q32G89 Q32GA8 Q32GC7 Q32GE1 Q32GE2 Q32GI6
Q32GP4 Q32GP7 Q32GQ2 Q32GQ6 Q32GS9 Q32GT0 Q32GW8 Q32GX3 Q32GZ5 Q32GZ6 Q32GZ7
Q32GZ9 Q32H03 Q32H27 Q32H41 Q32H45 Q32H47 Q32H58 Q32H60 Q32H69 Q32H71 Q32H78
Q32H79 Q32H95 Q32H97 Q32H98 Q32HA0 Q32HA1 Q32HB3 Q32HE0 Q32HG9 Q32HI7 Q32HI9
Q32HN6 Q32HQ0 Q32HQ1 Q32HQ2 Q32HQ3 Q32HQ4 Q32HQ6 Q32HR2 Q32HR4 Q32HT7 Q32HT9
Q32HU0 Q32HV1 Q32HV8 Q32HW0 Q32HX1 Q32HY5 Q32HY6 Q32HY7 Q32I11 Q32I44 Q32I45
Q32I48 Q32I50 Q32I52 Q32I80 Q32I85 Q32I91 Q32ID0 Q32ID7 Q32IG8 Q32IH0 Q32IH8
Q32IJ3 Q32IJ5 Q32IK3 Q32IL9 Q32IM0 Q32IN7 Q32IQ0 Q32IQ2 Q32IR1 Q32IR3 Q32IT9
Q32IU2 Q32IU4 Q32IU9 Q32IV0 Q32IV2 Q32IX2 Q32J06 Q32J17 Q32J22 Q32J23 Q32J26
Q32J27 Q32J49 Q32J51 Q32J52 Q32J53 Q32J54 Q32J55 Q32J95 Q32J96 Q32J98 Q32JB1
Q32JB6 Q32JD0 Q32JD6 Q32JD7 Q32JE0 Q32JE7 Q32JF9 Q32JG0 Q32JG7 Q32JG9 Q32JH0
Q32JH8 Q32JI0 Q32JI1 Q32JI4 Q32JJ2 Q32JJ3 Q32JJ4 Q32JJ5 Q32JK2 Q32JK7 Q32JL0
Q32JN6 Q32JP9 Q32JQ1 Q32JQ4 Q32JR3 Q32JR8 Q32JS6 Q32JT7 Q32JT8 Q32JT9 Q32JU0
Q32JU2 Q32JU3 Q32JU4 Q32JU9 Q32JV2 Q32JV4 Q32JW1 Q32JW8 Q32JW9 Q32JY8 Q32JY9
Q32JZ0 Q32JZ1 Q32K01 Q32K04 Q32K10 Q32K11 Q32K20 Q32K21 Q32K22 Q32K23 Q32K24
Q32K30 Q32K31 Q32K36 Q32K42 Q32K43 Q32K44 Q32K45 Q32K55 Q32K58 Q32K59 Q32K60
Q32K62 Q32K66 Q32K67 Q32K68 Q32K71 Q32K74 Q32K78 Q32K90 Q32K91 Q32KA4 Q32KA5
Q32KB0 Q32KB2 Q32KB5 Q32KB7 Q32KD2 Q32KG4 Q32KL2 Q32KL4 Q32KL8 Q32KM1 Q32KM2
Q32KM6 Q32KN0 Q32KN5 Q32KN7 Q32KN8 Q32KP0 Q32KP2 Q32KP7 Q32KQ1 Q32KQ2 Q32KQ7
Q32KR8 Q32KS0 Q32KS2 Q32KS7 Q32KT0 Q32KT5 Q32KT7 Q32KU3 Q32KU4 Q32KV0 Q32KV2
Q32KV4 Q32KV8 Q32KW0 Q32KW2 Q32KW7 Q32KX0 Q32KX1 Q32KX5 Q32KX9 Q32KY1 Q32KY3
Q32KY6 Q32KY7 Q32KY8 Q32KY9 Q32KZ1 Q32KZ2 Q32KZ5 Q32KZ9 Q32L00 Q32L03 Q32L04
Q32L08 Q32L17 Q32L18 Q32L19 Q32L22 Q32L26 Q32L27 Q32L31 Q32L34 Q32L37 Q32L40
Q32L48 Q32L49 Q32L52 Q32L62 Q32L67 Q32L72 Q32L75 Q32L78 Q32L89 Q32L91 Q32L92
Q32L93 Q32L96 Q32LA4 Q32LA7 Q32LB0 Q32LB1 Q32LB6 Q32LC1 Q32LC2 Q32LD1 Q32LD2
Q32LD4 Q32LE3 Q32LE4 Q32LE5 Q32LE6 Q32LF5 Q32LH1 Q32LH3 Q32LH4 Q32LH7 Q32LI1

Q32LI3 Q32LI5 Q32LJ3 Q32LJ7 Q32LK1 Q32LK9 Q32LL5 Q32LL6 Q32LL9 Q32LM4 Q32LM5
Q32LM7 Q32LN0 Q32LN1 Q32LN3 Q32LN4 Q32LN5 Q32LN7 Q32LP0 Q32LP9 Q32LQ1 Q32LQ3
Q32LQ4 Q32LR5 Q32LU7 Q32M02 Q32M07 Q32M08 Q32M78 Q32M84 Q32M88 Q32M92 Q32MG2
Q32MH5 Q32MK0 Q32MQ0 Q32MW3 Q32N19 Q32N22 Q32N55 Q32N66 Q32N87 Q32N90 Q32N92
Q32N93 Q32NA2 Q32NC0 Q32NG0 Q32NH9 Q32NI9 Q32NJ2 Q32NJ6 Q32NK7 Q32NM1 Q32NN2
Q32NP7 Q32NP8 Q32NQ7 Q32NQ8 Q32NR3 Q32NR4 Q32NS2 Q32NS4 Q32NT4 Q32NU8 Q32NV1
Q32NW2 Q32P12 Q32P41 Q32P51 Q32P59 Q32P66 Q32P67 Q32P68 Q32P71 Q32P73 Q32P76
Q32P78 Q32P79 Q32P85 Q32P96 Q32P97 Q32PA0 Q32PA2 Q32PA5 Q32PA8 Q32PB2 Q32PB4
Q32PB5 Q32PD0 Q32PD7 Q32PE0 Q32PE4 Q32PE7 Q32PE9 Q32PF6 Q32PF7 Q32PG3 Q32PG5
Q32PG9 Q32PH0 Q32PH7 Q32PI0 Q32PJ3 Q32PJ6 Q32PJ8 Q32PL1 Q32PL8 Q32PP3 Q32PP9
Q32PV0 Q32PV5 Q32PW3 Q32PX2 Q32PX7 Q32PY6 Q32PY9 Q32PZ3 Q32PZ9 Q32RF8 Q32RG0
Q32RG1 Q32RG2 Q32RH2 Q32RH3 Q32RJ9 Q32RK4 Q32RK6 Q32RK7 Q32RL3 Q32RL5 Q32RM2
Q32RM8 Q32RM9 Q32RN0 Q32RN1 Q32RN2 Q32RN3 Q32RN4 Q32RN5 Q32RN6 Q32RN7 Q32RN8
Q32RN9 Q32RP7 Q32RP8 Q32RP9 Q32RQ0 Q32RQ1 Q32RQ2 Q32RQ7 Q32RQ8 Q32RQ9 Q32RS0
Q32RS1 Q32RS2 Q32RS5 Q32RT7 Q32RT8 Q32RT9 Q32RU0 Q32RU7 Q32RU8 Q32RU9 Q32RV0
Q32RV1 Q32RV2 Q32RV3 Q32RV4 Q32RV5 Q32RV6 Q32RV7 Q32RV8 Q32RW0 Q32RW5 Q32RX0
Q32RY1 Q32RY2 Q32RY3 Q32RY4 Q32RY7 Q32RZ3 Q32RZ7 Q32RZ8 Q32S00 Q32S07 Q32S12
Q32SG3 Q32SG4 Q32SG5 Q32SG6 Q32TF8 Q32TG3 Q32ZM1 Q33015 Q33050 Q33062 Q33076
Q33078 Q33093 Q330M9 Q33101 Q33119 Q33134 Q33142 Q33162 Q331Q7 Q331R1 Q331T6
Q331T7 Q331T8 Q33245 Q33274 Q332R5 Q332R6 Q332R7 Q332R9 Q332S0 Q332S9 Q332T6
Q332T7 Q332T8 Q332T9 Q332U0 Q332U1 Q332U2 Q332U3 Q332U4 Q332U5 Q332V2 Q332V3
Q332V5 Q332V6 Q332V7 Q332X6 Q332Y0 Q332Z6 Q332Z7 Q33301 Q33323 Q33339 Q33369
Q33383 Q33406 Q33438 Q33443 Q33446 Q33451 Q33528 Q33569 Q33584 Q33600 Q33619
Q336M2 Q336M7 Q336N8 Q336R3 Q336R9 Q336V5 Q336V8 Q337A5 Q337Y2 Q338B9 Q338C0
Q338G6 Q338N2 Q338P6 Q338P8 Q338X7 Q338Z7 Q339D2 Q339W7 Q33BP8 Q33BV3 Q33BV4
Q33BW5 Q33BX4 Q33BY2 Q33BY9 Q33BZ0 Q33BZ1 Q33BZ2 Q33BZ3 Q33BZ5 Q33BZ6 Q33BZ7
Q33BZ8 Q33BZ9 Q33C00 Q33C07 Q33C08 Q33C09 Q33C10 Q33C11 Q33C26 Q33C33 Q33C38
Q33C46 Q33C47 Q33C48 Q33C49 Q33C57 Q33DK1 Q33DL3 Q33DV3 Q33DX5 Q33E94 Q34806
Q34807 Q34940 Q35811 Q35905 Q36570 Q36580 Q36610 Q36665 Q36800 Q36863 Q36907
Q36996 Q36997 Q37001 Q37066 Q37069 Q37082 Q37106 Q37192 Q37227 Q37282 Q37328
Q37683 Q37757 Q37761 Q37869 Q37870 Q37871 Q37873 Q37874 Q37882 Q37883 Q37884
Q37885 Q37886 Q37887 Q37888 Q37889 Q37891 Q37892 Q37893 Q37894 Q37897 Q37898
Q37907 Q37928 Q37989 Q37993 Q37BZ5 Q37D32 Q37G91 Q37M24 Q37M27 Q37M36 Q38036
Q38037 Q38038 Q38039 Q38040 Q38041 Q38042 Q38055 Q38067 Q38087 Q38100 Q38156
Q38167 Q38168 Q38169 Q38199 Q381A3 Q381F9 Q381M1 Q381V6 Q38200 Q38278 Q382V8
Q38300 Q38305 Q38361 Q38362 Q38394 Q383A1 Q383H9 Q383Q5 Q38419 Q38423 Q38424
Q38433 Q38439 Q38440 Q38441 Q38442 Q38477 Q38479 Q38480 Q38482 Q38488 Q38490
Q38492 Q38494 Q384Y0 Q384Y1 Q38503 Q38504 Q38563 Q38580 Q38581 Q38582 Q38584
Q38617 Q38621 Q38625 Q38661 Q38662 Q38663 Q38664 Q38665 Q38666 Q38667 Q38668
Q38669 Q38692 Q38693 Q38694 Q386E7 Q386N1 Q38707 Q38709 Q38710 Q38732 Q38771
Q38772 Q38773 Q38774 Q38775 Q38786 Q38793 Q38796 Q38799 Q387H6 Q38802 Q38803
Q38804 Q38805 Q38813 Q38819 Q38821 Q38822 Q38824 Q38825 Q38826 Q38827 Q38828
Q38829 Q38830 Q38831 Q38832 Q38836 Q38837 Q38838 Q38840 Q38841 Q38847 Q38850
Q38851 Q38854 Q38859 Q38876 Q38884 Q38895 Q38896 Q38897 Q38899 Q388A7 Q38905

Q38914 Q38920 Q38929 Q38931 Q38933 Q38942 Q38945 Q38950 Q38953 Q38960 Q38961
Q38A34 Q38AH0 Q38C92 Q38CE9 Q38E83 Q38F76 Q38FI2 Q38HM4 Q38HX2 Q38HX3 Q38HX4
Q38IY3 Q38J50 Q38JA7 Q38JH8 Q38JU2 Q38M35 Q38UE3 Q38UE4 Q38UE9 Q38UF0 Q38UH2
Q38UM4 Q38UP9 Q38UQ2 Q38UQ3 Q38UQ7 Q38UQ8 Q38UQ9 Q38UR1 Q38UR2 Q38UR3 Q38UR4
Q38UR5 Q38UR6 Q38UR7 Q38UR8 Q38UR9 Q38US0 Q38US1 Q38US2 Q38US3 Q38US4 Q38US5
Q38US6 Q38US7 Q38US8 Q38US9 Q38UT0 Q38UT2 Q38UT3 Q38UT4 Q38UT5 Q38UT6 Q38UT7
Q38UT8 Q38UV3 Q38UV4 Q38UV5 Q38UZ6 Q38V01 Q38V05 Q38V06 Q38V07 Q38V10 Q38V11
Q38V12 Q38V22 Q38V25 Q38V36 Q38V48 Q38V49 Q38V50 Q38V64 Q38V71 Q38V72 Q38V81
Q38V83 Q38VA7 Q38VB9 Q38VC5 Q38VC9 Q38VD0 Q38VK6 Q38VL2 Q38VN3 Q38VN8 Q38VP5
Q38VQ9 Q38VS0 Q38VS5 Q38VS6 Q38VS7 Q38VT0 Q38VT3 Q38VT4 Q38VU3 Q38VU9 Q38VV3
Q38VV4 Q38VV6 Q38VX2 Q38VX5 Q38VY5 Q38VZ2 Q38VZ6 Q38W03 Q38W22 Q38W49 Q38W53
Q38W55 Q38W63 Q38W64 Q38W65 Q38W66 Q38W67 Q38W72 Q38W73 Q38W81 Q38W82 Q38W87
Q38W91 Q38W92 Q38W93 Q38W94 Q38WA3 Q38WF1 Q38WH3 Q38WI8 Q38WJ3 Q38WJ4 Q38WJ7
Q38WJ8 Q38WK8 Q38WN0 Q38WN9 Q38WP3 Q38WQ2 Q38WQ5 Q38WQ7 Q38WR3 Q38WR4 Q38WR7
Q38WR8 Q38WS0 Q38WS5 Q38WT4 Q38WT6 Q38WU0 Q38WV4 Q38WW3 Q38WX5 Q38WX8 Q38WY3
Q38WY6 Q38WZ1 Q38X21 Q38X22 Q38X24 Q38X26 Q38X27 Q38X29 Q38X32 Q38X59 Q38X60
Q38X61 Q38X65 Q38X69 Q38X70 Q38X74 Q38X86 Q38X88 Q38X91 Q38X92 Q38X99 Q38XA0
Q38XA1 Q38XA4 Q38XA7 Q38XA8 Q38XB2 Q38XB4 Q38XB6 Q38XB7 Q38XC8 Q38XD1 Q38XD3
Q38XD6 Q38XD7 Q38XI0 Q38XI1 Q38XI2 Q38XI3 Q38XK7 Q38XM0 Q38XM4 Q38XM9 Q38XN3
Q38XN4 Q38XN8 Q38XP2 Q38XQ6 Q38XQ7 Q38XQ8 Q38XR0 Q38XR8 Q38XS9 Q38XT2 Q38XT6
Q38XT9 Q38XU0 Q38XU7 Q38XU8 Q38XU9 Q38XV0 Q38XV5 Q38XW3 Q38XW7 Q38XW8 Q38XX0
Q38XX6 Q38XY8 Q38XZ3 Q38Y03 Q38Y19 Q38Y20 Q38Y85 Q38Y96 Q38Y99 Q38YA1 Q38YA4
Q38YB1 Q38YD9 Q38YF0 Q38YF8 Q38YJ1 Q38YK9 Q38YM1 Q38YM3 Q38YN3 Q38YN6 Q38YN7
Q38YN8 Q38YN9 Q38YP9 Q38YQ6 Q38YQ8 Q38YQ9 Q38YR0 Q38YR2 Q38YR4 Q38YR7 Q38YR8
Q38YS5 Q38YT3 Q38YT6 Q38YT8 Q38YY5 Q38YY6 Q38YZ5 Q38Z09 Q38Z74 Q38Z79 Q38ZC4
Q38ZE1 Q38ZE2 Q38ZF1 Q38ZH8 Q38ZJ0 Q38ZJ4 Q38ZL1 Q38ZL5 Q38ZL9 Q38ZM1 Q38ZM8
Q38ZR1 Q38ZR4 Q38ZR6 Q38ZR8 Q38ZS1 Q38ZS4 Q38ZS8 Q38ZT5 Q38ZU0 Q39009 Q39013
Q39030 Q39048 Q39054 Q39055 Q39056 Q39061 Q39067 Q39068 Q39069 Q39070 Q39071
Q39072 Q39081 Q39083 Q39084 Q39088 Q39089 Q39090 Q39092 Q39097 Q390D0 Q390P5
Q390P6 Q390Q5 Q390T2 Q390V3 Q390Z5 Q39103 Q39110 Q39111 Q39112 Q39117 Q39129
Q39138 Q39140 Q39147 Q39160 Q39161 Q39162 Q39163 Q39189 Q39190 Q39199 Q391E0
Q391K1 Q391N3 Q391Y9 Q39211 Q39212 Q39218 Q39224 Q39230 Q39234 Q39236 Q39237
Q39239 Q39242 Q39244 Q39247 Q39250 Q39251 Q39255 Q39256 Q39261 Q39262 Q39263
Q39265 Q39266 Q39269 Q39290 Q39295 Q392C9 Q392V6 Q39315 Q39336 Q39362 Q39366
Q39371 Q39375 Q393M6 Q393P2 Q393P4 Q393P5 Q393P6 Q393P8 Q393V1 Q393X2 Q393X4
Q393X8 Q393Y2 Q39411 Q39419 Q39445 Q39458 Q39459 Q39469 Q39471 Q39472 Q39490
Q39491 Q39493 Q394J5 Q394V3 Q39511 Q39516 Q39522 Q39535 Q39548 Q39557 Q39565
Q39566 Q39575 Q39578 Q39579 Q39580 Q39582 Q39584 Q39586 Q39591 Q39592 Q39596
Q395F2 Q39604 Q39610 Q39613 Q39617 Q39618 Q39635 Q39644 Q39664 Q39676 Q39677
Q39685 Q39697 Q396C5 Q396N9 Q396P0 Q396P7 Q396Y3 Q396Y9 Q39705 Q39724 Q39727
Q39732 Q39734 Q39735 Q39743 Q39744 Q39745 Q39746 Q39747 Q39748 Q39749 Q39752
Q39757 Q39758 Q39760 Q39761 Q39769 Q39779 Q39785 Q397E7 Q397Q9 Q397R0 Q397X6
Q397Y5 Q39818 Q39827 Q39836 Q39839 Q39846 Q39891 Q398D7 Q398V5 Q39967 Q39978
Q39979 Q39980 Q39982 Q399J3 Q39A02 Q39A07 Q39A26 Q39A42 Q39A43 Q39A44 Q39AM3

Q39AN3 Q39AP8 Q39AS8 Q39B00 Q39BA7 Q39BD4 Q39BG3 Q39BP9 Q39BQ4 Q39BS5 Q39BU5
Q39BV2 Q39BW2 Q39BW3 Q39BW4 Q39BX2 Q39BX5 Q39BY0 Q39BY1 Q39BZ1 Q39C10 Q39C36
Q39C44 Q39C93 Q39CE4 Q39CE5 Q39CE6 Q39CF3 Q39CG0 Q39CH4 Q39CJ1 Q39CN6 Q39CN9
Q39CT5 Q39CT7 Q39CT8 Q39CT9 Q39CU1 Q39CU4 Q39CU7 Q39CU8 Q39CV8 Q39CW7 Q39D07
Q39D13 Q39D20 Q39D37 Q39D49 Q39D51 Q39D93 Q39D96 Q39DA3 Q39DG2 Q39DG4 Q39DH2
Q39DI5 Q39DI6 Q39DI7 Q39DI9 Q39DJ3 Q39DJ5 Q39DL2 Q39DM1 Q39DM5 Q39DM8 Q39DN4
Q39DN7 Q39DN8 Q39DN9 Q39DS5 Q39DS9 Q39DU8 Q39DU9 Q39DV4 Q39DV8 Q39DW2 Q39E36
Q39E37 Q39E45 Q39E61 Q39E66 Q39E95 Q39E96 Q39E98 Q39E99 Q39EA1 Q39EA2 Q39EA4
Q39ED2 Q39ED5 Q39EE0 Q39EE1 Q39EH1 Q39EK2 Q39EM7 Q39EP0 Q39EP4 Q39EQ6 Q39ES6
Q39EU4 Q39EV7 Q39EX8 Q39EY8 Q39F07 Q39F31 Q39F33 Q39F38 Q39F41 Q39F43 Q39F44
Q39F45 Q39F46 Q39F49 Q39F57 Q39F59 Q39F63 Q39F73 Q39FB8 Q39FB9 Q39FE7 Q39FE8
Q39FE9 Q39FF4 Q39FG2 Q39FG3 Q39FH9 Q39FJ8 Q39FK0 Q39FK1 Q39FN8 Q39FP5 Q39FP6
Q39FP8 Q39FP9 Q39FQ4 Q39FQ6 Q39FQ7 Q39FR0 Q39FR3 Q39FR4 Q39FR9 Q39FS0 Q39FU0
Q39GJ4 Q39GL2 Q39GX1 Q39GX3 Q39GX4 Q39GY0 Q39H28 Q39H29 Q39H30 Q39H32 Q39H50
Q39H51 Q39H52 Q39H53 Q39H55 Q39H65 Q39H98 Q39HB7 Q39HM0 Q39HM9 Q39HN2 Q39HY9
Q39I23 Q39I24 Q39I25 Q39I40 Q39I64 Q39I66 Q39I70 Q39I71 Q39I88 Q39I90 Q39I98
Q39IC5 Q39IC7 Q39ID3 Q39ID4 Q39ID5 Q39ID6 Q39IE1 Q39IF3 Q39IF8 Q39IG0 Q39IG1
Q39IG4 Q39IG8 Q39IJ4 Q39IL1 Q39IL2 Q39IL4 Q39IN0 Q39IN2 Q39IN7 Q39IQ4 Q39IR3
Q39IU7 Q39IW0 Q39IW1 Q39IW2 Q39IW3 Q39IW4 Q39IW5 Q39IW6 Q39J02 Q39J04 Q39J05
Q39J12 Q39J54 Q39J71 Q39J72 Q39JB5 Q39JB9 Q39JC5 Q39JC7 Q39JC8 Q39JD0 Q39JD2
Q39JD3 Q39JF1 Q39JF4 Q39JI1 Q39JI8 Q39JI9 Q39JJ0 Q39JJ1 Q39JJ5 Q39JJ6 Q39JJ7
Q39JJ8 Q39JJ9 Q39JK0 Q39JM2 Q39JM5 Q39JM8 Q39JS9 Q39JU3 Q39JU4 Q39JU6 Q39JU7
Q39JU8 Q39JU9 Q39JV5 Q39JV6 Q39JW0 Q39JW9 Q39JX2 Q39JX8 Q39JX9 Q39JZ6 Q39JZ7
Q39K00 Q39K17 Q39K18 Q39K83 Q39K84 Q39K85 Q39K86 Q39K87 Q39K89 Q39K90 Q39K91
Q39K92 Q39K93 Q39KA4 Q39KC4 Q39KC5 Q39KC9 Q39KD6 Q39KE0 Q39KE1 Q39KE2 Q39KE3
Q39KE4 Q39KE5 Q39KE6 Q39KE8 Q39KE9 Q39KF0 Q39KF1 Q39KF2 Q39KF3 Q39KF4 Q39KF5
Q39KF6 Q39KF7 Q39KF8 Q39KF9 Q39KG0 Q39KG1 Q39KG2 Q39KG3 Q39KG4 Q39KG5 Q39KG6
Q39KG7 Q39KG8 Q39KH0 Q39KH1 Q39KH2 Q39KH4 Q39KH5 Q39KH6 Q39KH7 Q39KH8 Q39KH9
Q39KI2 Q39KM3 Q39KQ2 Q39KT9 Q39KU1 Q39KY8 Q39KY9 Q39L06 Q39L14 Q39L16 Q39L82
Q39LF7 Q39LF8 Q39LH7 Q39LH8 Q39LI1 Q39LI8 Q39M02 Q39M27 Q39M29 Q39M92 Q39MM8
Q39N21 Q39N89 Q39NL8 Q39PA6 Q39PN9 Q39PQ5 Q39PQ9 Q39PR0 Q39PR1 Q39PS5 Q39PT6
Q39PT7 Q39PU1 Q39Q42 Q39Q58 Q39Q76 Q39Q78 Q39QA6 Q39QC2 Q39QC5 Q39QK1 Q39QK2
Q39QM7 Q39QM8 Q39QN3 Q39QP7 Q39QQ2 Q39QQ5 Q39QQ6 Q39QQ9 Q39QR1 Q39QR2 Q39QR3
Q39QR4 Q39QR5 Q39QR6 Q39QR8 Q39QT6 Q39QU2 Q39QV5 Q39QW1 Q39R35 Q39R36 Q39R75
Q39R76 Q39R83 Q39RA1 Q39RG3 Q39RG4 Q39RG5 Q39RH2 Q39RJ6 Q39RJ8 Q39RJ9 Q39RK1
Q39RN5 Q39RN7 Q39RN8 Q39RP0 Q39RP1 Q39RP2 Q39RP7 Q39RQ7 Q39RQ9 Q39RR0 Q39RR2
Q39RR3 Q39RR5 Q39RS2 Q39RT4 Q39RX3 Q39RY8 Q39S09 Q39S27 Q39S28 Q39S70 Q39S71
Q39SM1 Q39SM2 Q39SN2 Q39SQ3 Q39SQ6 Q39SQ9 Q39SS2 Q39SU1 Q39SY7 Q39T29 Q39T72
Q39T73 Q39T76 Q39T82 Q39T85 Q39T99 Q39TA2 Q39TA3 Q39TA7 Q39TV7 Q39TZ3 Q39U59
Q39U97 Q39U98 Q39UA9 Q39UB1 Q39UC4 Q39UD2 Q39UE8 Q39UF9 Q39UG0 Q39UG9 Q39UH1
Q39UH2 Q39UH3 Q39UJ7 Q39UK1 Q39UK4 Q39UK8 Q39UR5 Q39US3 Q39US4 Q39UT1 Q39UT4
Q39UV7 Q39UW8 Q39V17 Q39V40 Q39V47 Q39V50 Q39V51 Q39V65 Q39V66 Q39V70 Q39V87
Q39V95 Q39V99 Q39VA1 Q39VA2 Q39VA3 Q39VA5 Q39VA6 Q39VA9 Q39VB0 Q39VQ6 Q39VR6
Q39VR8 Q39VS0 Q39VS1 Q39VS4 Q39VS5 Q39VS6 Q39VS7 Q39VS8 Q39VU8 Q39VY0 Q39VY3

Q39VY5 Q39VY6 Q39W34 Q39W35 Q39W48 Q39W52 Q39W62 Q39W70 Q39W76 Q39W79 Q39W82
Q39W85 Q39W86 Q39W87 Q39W88 Q39WD1 Q39WF4 Q39WH4 Q39WK9 Q39WQ8 Q39WU7 Q39WV0
Q39X01 Q39X04 Q39X05 Q39X06 Q39X41 Q39X47 Q39X60 Q39X62 Q39X66 Q39X78 Q39X87
Q39X89 Q39XA9 Q39XB6 Q39XB8 Q39XD1 Q39XE0 Q39XL1 Q39XN6 Q39XN7 Q39XN8 Q39XN9
Q39XS5 Q39XX8 Q39XX9 Q39XY0 Q39XY1 Q39XY2 Q39XY3 Q39XY7 Q39XY8 Q39XY9 Q39XZ0
Q39XZ1 Q39XZ2 Q39XZ3 Q39XZ4 Q39XZ5 Q39XZ6 Q39XZ7 Q39XZ8 Q39XZ9 Q39Y00 Q39Y01
Q39Y02 Q39Y03 Q39Y04 Q39Y05 Q39Y06 Q39Y07 Q39Y08 Q39Y09 Q39Y10 Q39Y11 Q39Y12
Q39Y13 Q39Y14 Q39Y15 Q39Y16 Q39Y17 Q39Y23 Q39Y25 Q39Y29 Q39Y84 Q39YD4 Q39YD7
Q39YF3 Q39YG0 Q39YG4 Q39YG6 Q39YJ0 Q39YJ1 Q39YL7 Q39YL8 Q39YM1 Q39YM4 Q39YM7
Q39YM8 Q39YP0 Q39YP1 Q39YP2 Q39YP3 Q39YP4 Q39YP5 Q39YP6 Q39YP7 Q39YP8 Q39YP9
Q39YQ1 Q39YQ4 Q39Z11 Q39Z17 Q39Z32 Q39Z65 Q39Z66 Q39Z67 Q39Z69 Q39Z72 Q39Z73
Q39Z75 Q39Z77 Q39Z80 Q39Z82 Q39Z84 Q39Z86 Q39ZJ9 Q39ZK0 Q39ZK2 Q39ZK3 Q39ZL4
Q39ZL5 Q39ZL6 Q39ZL8 Q39ZN3 Q39ZP5 Q39ZP6 Q39ZQ5 Q39ZQ8 Q39ZS1 Q39ZS3 Q39ZS6
Q39ZT0 Q39ZT1 Q39ZT2 Q39ZX2 Q39ZZ2 Q3A002 Q3A008 Q3A009 Q3A090 Q3A091 Q3A0B8
Q3A0G9 Q3A0V1 Q3A0V2 Q3A0Y0 Q3A127 Q3A129 Q3A131 Q3A132 Q3A133 Q3A134 Q3A135
Q3A136 Q3A137 Q3A187 Q3A1D3 Q3A1D6 Q3A1D7 Q3A1D8 Q3A1D9 Q3A1E0 Q3A1E1 Q3A1E3
Q3A1E5 Q3A1Q7 Q3A1R3 Q3A1T0 Q3A1U5 Q3A1U6 Q3A1U7 Q3A1V0 Q3A1V1 Q3A1V3 Q3A1V5
Q3A1V6 Q3A1V7 Q3A1W0 Q3A1W1 Q3A1W5 Q3A1W7 Q3A201 Q3A209 Q3A220 Q3A221 Q3A230
Q3A231 Q3A232 Q3A233 Q3A241 Q3A246 Q3A247 Q3A248 Q3A253 Q3A258 Q3A282 Q3A286
Q3A287 Q3A2A9 Q3A2B3 Q3A2C7 Q3A2D2 Q3A2D6 Q3A2E4 Q3A2E6 Q3A2E7 Q3A2E9 Q3A2F0
Q3A2F1 Q3A2F4 Q3A2F5 Q3A2F7 Q3A2F8 Q3A2G4 Q3A2G7 Q3A2G8 Q3A2G9 Q3A2I0 Q3A2J0
Q3A2J4 Q3A2J8 Q3A2J9 Q3A2K0 Q3A2K1 Q3A2N5 Q3A2N6 Q3A2P8 Q3A2U5 Q3A2Y7 Q3A2Z2
Q3A2Z3 Q3A2Z4 Q3A311 Q3A313 Q3A314 Q3A317 Q3A318 Q3A320 Q3A334 Q3A346 Q3A371
Q3A381 Q3A388 Q3A392 Q3A395 Q3A396 Q3A397 Q3A398 Q3A3A5 Q3A3B2 Q3A3B5 Q3A3B6
Q3A3B7 Q3A3B8 Q3A3C1 Q3A3D1 Q3A3D2 Q3A3D3 Q3A3F2 Q3A3F8 Q3A3G6 Q3A3G8 Q3A3H7
Q3A3I6 Q3A3I7 Q3A3I8 Q3A3J2 Q3A3N8 Q3A3P5 Q3A3X3 Q3A3X4 Q3A3X5 Q3A3X6 Q3A3Z3
Q3A3Z6 Q3A3Z7 Q3A3Z8 Q3A413 Q3A423 Q3A448 Q3A449 Q3A450 Q3A452 Q3A499 Q3A4A1
Q3A4A2 Q3A4A3 Q3A4A5 Q3A4A7 Q3A4B0 Q3A4F1 Q3A4K3 Q3A4L2 Q3A4L3 Q3A4L7 Q3A4L9
Q3A4M7 Q3A4N0 Q3A4N4 Q3A4N5 Q3A4N8 Q3A4N9 Q3A4P0 Q3A4P1 Q3A4P3 Q3A4P5 Q3A4P8
Q3A4P9 Q3A4Q5 Q3A4Q8 Q3A4X0 Q3A4X5 Q3A503 Q3A504 Q3A508 Q3A515 Q3A519 Q3A522
Q3A523 Q3A526 Q3A538 Q3A575 Q3A587 Q3A590 Q3A594 Q3A5A7 Q3A5D3 Q3A5G2 Q3A5Q4
Q3A5Q6 Q3A5R6 Q3A5R7 Q3A5S0 Q3A5T0 Q3A5V4 Q3A5V5 Q3A668 Q3A6G9 Q3A6H9 Q3A6I7
Q3A6I8 Q3A6J1 Q3A6K6 Q3A6L6 Q3A6L9 Q3A6M0 Q3A6M1 Q3A6M2 Q3A6M3 Q3A6M4 Q3A6M6
Q3A6M8 Q3A6M9 Q3A6N0 Q3A6N1 Q3A6N2 Q3A6N3 Q3A6N4 Q3A6N5 Q3A6N6 Q3A6N7 Q3A6N8
Q3A6N9 Q3A6P0 Q3A6P1 Q3A6P2 Q3A6P3 Q3A6P4 Q3A6P5 Q3A6P6 Q3A6P7 Q3A6P8 Q3A6P9
Q3A6Q0 Q3A6Q1 Q3A6Q2 Q3A6Q3 Q3A6Q4 Q3A6Q5 Q3A6Q6 Q3A6Q7 Q3A6Q8 Q3A6R1 Q3A6R2
Q3A6R4 Q3A6R6 Q3A6S7 Q3A6Y1 Q3A6Y9 Q3A701 Q3A709 Q3A734 Q3A798 Q3A7A3 Q3A7A9
Q3A7E2 Q3A7G2 Q3A7H9 Q3A7I0 Q3A7K8 Q3A7N4 Q3A7P5 Q3A7P7 Q3A7Q8 Q3A7R3 Q3A7T7
Q3A7V6 Q3A7W5 Q3A7Y3 Q3A7Y4 Q3A827 Q3A828 Q3A829 Q3A831 Q3A834 Q3A8C2 Q3A8C3
Q3A8C6 Q3A8C7 Q3A8C9 Q3A8D0 Q3A8E8 Q3A8E9 Q3A8G4 Q3A8J5 Q3A8J6 Q3A8K5 Q3A8M6
Q3A8N5 Q3A8N6 Q3A8N7 Q3A8P5 Q3A8Q0 Q3A8Q5 Q3A8R0 Q3A8S2 Q3A8S5 Q3A8X5 Q3A926
Q3A928 Q3A934 Q3A949 Q3A9E0 Q3A9J4 Q3A9J5 Q3A9K2 Q3A9L0 Q3A9L1 Q3A9L2 Q3A9L7
Q3A9M2 Q3A9N7 Q3A9N8 Q3A9N9 Q3A9P1 Q3A9P8 Q3A9P9 Q3A9Q2 Q3A9Q3 Q3A9Q4 Q3A9Q5
Q3A9Q7 Q3A9Q8 Q3A9R0 Q3A9R1 Q3A9R2 Q3A9R3 Q3A9R5 Q3A9R6 Q3A9R7 Q3A9R8 Q3A9R9

Q3A9S0 Q3A9S1 Q3A9S2 Q3A9S3 Q3A9S4 Q3A9S5 Q3A9S6 Q3A9S7 Q3A9S8 Q3A9S9 Q3A9T0
Q3A9T1 Q3A9T2 Q3A9T3 Q3A9T4 Q3A9T5 Q3A9T9 Q3A9U0 Q3A9U1 Q3A9U2 Q3A9U3 Q3A9U4
Q3A9U5 Q3A9V1 Q3A9V2 Q3A9W1 Q3A9W2 Q3A9W4 Q3A9W5 Q3A9W6 Q3A9Z9 Q3AA16 Q3AA17
Q3AA24 Q3AA34 Q3AA35 Q3AAC8 Q3AAD6 Q3AAD7 Q3AAE4 Q3AAE7 Q3AAE8 Q3AAE9 Q3AAF7
Q3AAG3 Q3AAH3 Q3AAI8 Q3AAK3 Q3AAM1 Q3AAM5 Q3AAM6 Q3AAN0 Q3AAN2 Q3AAN4 Q3AAQ2
Q3AAS0 Q3AAT6 Q3AAT7 Q3AAT8 Q3AAU6 Q3AAV2 Q3AAV5 Q3AAW2 Q3AAW3 Q3AAY6 Q3AAZ0
Q3AAZ2 Q3AB25 Q3AB53 Q3AB61 Q3AB71 Q3AB73 Q3AB74 Q3AB77 Q3AB78 Q3AB79 Q3AB80
Q3AB86 Q3AB89 Q3AB94 Q3AB98 Q3AB99 Q3ABA1 Q3ABA4 Q3ABA6 Q3ABA7 Q3ABL2 Q3ABL4
Q3ABL6 Q3ABS0 Q3ABS1 Q3ABS4 Q3ABS6 Q3ABS8 Q3ABS9 Q3ABT4 Q3ABT5 Q3ABT8 Q3ABU1
Q3ABU4 Q3ABU6 Q3ABV0 Q3ABX6 Q3ABX7 Q3ABX8 Q3ABX9 Q3ABY0 Q3ABY6 Q3ABZ3 Q3AC00
Q3AC01 Q3AC05 Q3AC06 Q3AC07 Q3AC09 Q3AC10 Q3AC13 Q3AC14 Q3AC15 Q3AC18 Q3AC19
Q3AC22 Q3AC25 Q3AC30 Q3AC32 Q3AC33 Q3AC43 Q3AC47 Q3AC49 Q3AC58 Q3AC64 Q3AC69
Q3AC71 Q3AC72 Q3AC74 Q3AC76 Q3AC88 Q3ACA4 Q3ACA5 Q3ACA6 Q3ACA8 Q3ACA9 Q3ACF5
Q3ACG0 Q3ACM0 Q3ACM1 Q3ACS9 Q3ACT3 Q3ACT4 Q3ACU3 Q3ACX5 Q3ACY8 Q3AD35 Q3AD36
Q3AD37 Q3AD38 Q3AD48 Q3AD49 Q3AD50 Q3AD51 Q3AD52 Q3AD53 Q3AD54 Q3AD55 Q3AD62
Q3AD64 Q3AD65 Q3AD70 Q3AD96 Q3ADC2 Q3ADG3 Q3ADI0 Q3ADI2 Q3ADI5 Q3ADL7 Q3ADP5
Q3ADR7 Q3ADX3 Q3ADX7 Q3ADX8 Q3AE11 Q3AE32 Q3AE33 Q3AE55 Q3AE81 Q3AE90 Q3AEF4
Q3AEF5 Q3AEJ6 Q3AEN3 Q3AEN4 Q3AEN6 Q3AEN8 Q3AEP2 Q3AEQ2 Q3AEQ5 Q3AEQ7 Q3AER0
Q3AET1 Q3AET5 Q3AET6 Q3AET7 Q3AEU2 Q3AEU3 Q3AEV6 Q3AEW1 Q3AEY1 Q3AEY5 Q3AEZ6
Q3AF02 Q3AF06 Q3AF07 Q3AF08 Q3AF09 Q3AF11 Q3AF17 Q3AF30 Q3AF34 Q3AF39 Q3AF41
Q3AF51 Q3AF53 Q3AF54 Q3AF72 Q3AF80 Q3AF81 Q3AF82 Q3AF87 Q3AF96 Q3AF97 Q3AFA0
Q3AFA7 Q3AFA8 Q3AFC2 Q3AFD0 Q3AFD1 Q3AFD7 Q3AFE0 Q3AFE5 Q3AFF4 Q3AFH3 Q3AFJ7
Q3AFL7 Q3AFM4 Q3AFS4 Q3AFT3 Q3AFT7 Q3AFT8 Q3AFV0 Q3AFZ5 Q3AG04 Q3AG12 Q3AG23
Q3AG25 Q3AG27 Q3AG54 Q3AG55 Q3AG56 Q3AG60 Q3AG61 Q3AG69 Q3AG78 Q3AG80 Q3AGB7
Q3AGD0 Q3AGD1 Q3AGD2 Q3AGD3 Q3AGD6 Q3AGF3 Q3AGF7 Q3AGF9 Q3AGK2 Q3AGK9 Q3AGL2
Q3AGL5 Q3AGN4 Q3AGP0 Q3AGQ3 Q3AGR0 Q3AGS9 Q3AGT0 Q3AGT1 Q3AGT2 Q3AGU1 Q3AGU7
Q3AGU9 Q3AGV6 Q3AGW5 Q3AGW9 Q3AGY2 Q3AH20 Q3AH24 Q3AH50 Q3AH55 Q3AH63 Q3AH76
Q3AH85 Q3AHC4 Q3AHI8 Q3AHM3 Q3AHM4 Q3AHN5 Q3AHP0 Q3AHP8 Q3AHR1 Q3AHR5 Q3AHR6
Q3AHS3 Q3AHT4 Q3AHU0 Q3AHU2 Q3AHV1 Q3AHW1 Q3AHW4 Q3AHX1 Q3AHX3 Q3AHX5 Q3AHX6
Q3AHX7 Q3AHX9 Q3AHY7 Q3AI13 Q3AI17 Q3AI20 Q3AI26 Q3AI29 Q3AI33 Q3AI71 Q3AI74
Q3AI88 Q3AI95 Q3AIA2 Q3AIB3 Q3AIC0 Q3AIG3 Q3AIG9 Q3AIH2 Q3AIH4 Q3AII4 Q3AIK3
Q3AIK8 Q3AIZ7 Q3AJ26 Q3AJ44 Q3AJ48 Q3AJ51 Q3AJ96 Q3AJA6 Q3AJM8 Q3AJN8 Q3AJP8
Q3AJS3 Q3AJS6 Q3AJT5 Q3AJT9 Q3AJU7 Q3AJU8 Q3AJW3 Q3AJW5 Q3AJY9 Q3AJZ0 Q3AJZ1
Q3AK18 Q3AK21 Q3AK30 Q3AK46 Q3AK67 Q3AK73 Q3AK74 Q3AKA4 Q3AKA5 Q3AKB1 Q3AKD9
Q3AKE0 Q3AKE4 Q3AKG6 Q3AKG7 Q3AKH0 Q3AKJ1 Q3AKJ4 Q3AKJ6 Q3AKK1 Q3AKK3 Q3AKR1
Q3AKU0 Q3AKU3 Q3AKU8 Q3AKV6 Q3AKV7 Q3AKY8 Q3AL07 Q3AL08 Q3AL65 Q3AL68 Q3AL81
Q3AL87 Q3AL94 Q3AL98 Q3ALB4 Q3ALB5 Q3ALB9 Q3ALC4 Q3ALC5 Q3ALF2 Q3ALF6 Q3ALG7
Q3ALH7 Q3ALJ5 Q3ALJ7 Q3ALL1 Q3ALL6 Q3ALL7 Q3ALL9 Q3ALP2 Q3ALP7 Q3ALP8 Q3ALR5
Q3ALR7 Q3ALS4 Q3ALS9 Q3ALU9 Q3ALV6 Q3ALW5 Q3ALW7 Q3ALX3 Q3ALX6 Q3ALY8 Q3ALZ3
Q3AM28 Q3AM35 Q3AM37 Q3AM38 Q3AM91 Q3AM97 Q3AMA6 Q3AMB7 Q3AMB9 Q3AMC8 Q3AMD9
Q3AME2 Q3AMI3 Q3AMJ0 Q3AMJ2 Q3AMK3 Q3AML2 Q3AMM5 Q3AMN0 Q3AMN1 Q3AMN2 Q3AMN3
Q3AMN4 Q3AMN5 Q3AMN6 Q3AMN7 Q3AMN8 Q3AMN9 Q3AMP0 Q3AMP1 Q3AMP2 Q3AMP3 Q3AMP4
Q3AMP5 Q3AMP6 Q3AMP7 Q3AMQ0 Q3AMQ1 Q3AMQ2 Q3AMQ3 Q3AMQ4 Q3AMQ5 Q3AMQ6 Q3AMQ7
Q3AMQ8 Q3AMQ9 Q3AMT3 Q3AMT4 Q3AMT5 Q3AMT6 Q3AMT7 Q3AMU2 Q3AMU5 Q3AMV2 Q3AMW4

Q3AMW9 Q3AMX6 Q3AMX9 Q3AMY6 Q3AMZ1 Q3AN00 Q3AN01 Q3AN03 Q3AN08 Q3AN10 Q3AN13
Q3AN18 Q3AN25 Q3AN36 Q3AN75 Q3ANC6 Q3AND3 Q3AND8 Q3AND9 Q3ANE6 Q3ANF4 Q3ANG7
Q3ANG8 Q3ANH5 Q3ANH6 Q3ANI5 Q3ANI7 Q3ANI8 Q3ANI9 Q3ANJ5 Q3ANJ6 Q3ANJ9 Q3ANK7
Q3ANK8 Q3ANM0 Q3ANM4 Q3ANM5 Q3ANM6 Q3ANM7 Q3ANN0 Q3ANN7 Q3ANP3 Q3ANP7 Q3ANQ3
Q3ANQ6 Q3ANR2 Q3ANR3 Q3ANR5 Q3ANS5 Q3ANU2 Q3ANV2 Q3ANV5 Q3ANW1 Q3ANX4 Q3ANX8
Q3ANY2 Q3ANY3 Q3ANY4 Q3ANZ4 Q3AP21 Q3AP34 Q3AP35 Q3AP59 Q3APB9 Q3APC3 Q3APC4
Q3APD2 Q3APF2 Q3APF4 Q3APG8 Q3APG9 Q3APH0 Q3APH1 Q3APH2 Q3APH3 Q3APH4 Q3APH5
Q3APH6 Q3APH7 Q3APH8 Q3APH9 Q3API0 Q3API2 Q3API3 Q3API4 Q3API5 Q3API6 Q3API7
Q3API8 Q3API9 Q3APJ0 Q3APJ1 Q3APJ2 Q3APJ5 Q3APJ6 Q3APJ7 Q3APJ8 Q3APJ9 Q3APK0
Q3APK1 Q3APK2 Q3APK6 Q3APK7 Q3APK8 Q3APK9 Q3APL0 Q3APL1 Q3APL6 Q3APL9 Q3APM2
Q3APN5 Q3APP2 Q3APQ7 Q3APR0 Q3APR5 Q3APT0 Q3APT1 Q3APT3 Q3APU0 Q3APU1 Q3APU2
Q3APV9 Q3APW0 Q3APW1 Q3APW2 Q3APW3 Q3APW7 Q3APX4 Q3APY4 Q3APY5 Q3AQ07 Q3AQ14
Q3AQ15 Q3AQ17 Q3AQ22 Q3AQ23 Q3AQ24 Q3AQ25 Q3AQ26 Q3AQ27 Q3AQ40 Q3AQ41 Q3AQ69
Q3AQ75 Q3AQ83 Q3AQC1 Q3AQC2 Q3AQC3 Q3AQC6 Q3AQD7 Q3AQF1 Q3AQF4 Q3AQF5 Q3AQF7
Q3AQK2 Q3AQK5 Q3AQK7 Q3AQK8 Q3AQK9 Q3AQL1 Q3AQL3 Q3AQL9 Q3AQM4 Q3AQM9 Q3AQN3
Q3AQP3 Q3AQP5 Q3AQR4 Q3AQS3 Q3AQS4 Q3AQT7 Q3AQX0 Q3AQX4 Q3AQY1 Q3AQY2 Q3AQZ8
Q3AR00 Q3AR02 Q3AR09 Q3AR25 Q3AR26 Q3AR32 Q3AR47 Q3AR70 Q3AR93 Q3AR94 Q3AR95
Q3ARA7 Q3ARC0 Q3ARC3 Q3ARC4 Q3ARD7 Q3ARE0 Q3ARF1 Q3ARF2 Q3ARF7 Q3ARK3 Q3ARL1
Q3ARM4 Q3ARM7 Q3ARP2 Q3ARP5 Q3ARQ5 Q3ARU4 Q3ARV1 Q3ARZ7 Q3AS18 Q3AS25 Q3AS30
Q3AS33 Q3AS37 Q3AS53 Q3AS55 Q3AS74 Q3AS96 Q3AS97 Q3AS98 Q3ASD2 Q3ASD4 Q3ASF5
Q3ASF6 Q3ASF7 Q3ASF8 Q3ASH7 Q3ASI0 Q3ASI2 Q3ASI3 Q3ASI5 Q3ASI6 Q3ASM4 Q3ASP1
Q3ASP3 Q3ASQ9 Q3ASZ0 Q3AT00 Q3AT13 Q3AT23 Q3AT37 Q3AT43 Q3AT62 Q3AT63 Q3AT74
Q3ATA5 Q3ATA7 Q3ATB5 Q3ATB8 Q3ATD7 Q3ATE5 Q3ATE6 Q3ATG0 Q3ATI5 Q3ATL0 Q3ATL3
Q3ATL6 Q3ATM0 Q3ATN0 Q3ATN3 Q3ATP1 Q3ATP4 Q3ATP5 Q3ATP6 Q3ATP7 Q3ATP8 Q3ATP9
Q3ATR4 Q3ATS5 Q3ATS6 Q3ATT3 Q3ATT4 Q3ATT8 Q3ATV7 Q3ATV8 Q3ATW8 Q3ATX4 Q3ATZ1
Q3ATZ5 Q3AU08 Q3AU11 Q3AU36 Q3AU39 Q3AU43 Q3AU60 Q3AU73 Q3AU82 Q3AU83 Q3AUA2
Q3AUA4 Q3AUB5 Q3AUB6 Q3AUB7 Q3AUD8 Q3AUE2 Q3AUE4 Q3AUF2 Q3AUG1 Q3AUG4 Q3AUG8
Q3AUG9 Q3AUH7 Q3AUJ5 Q3AUJ6 Q3AUJ7 Q3AUJ8 Q3AUL9 Q3AUM0 Q3AUM3 Q3AUM7 Q3AUN6
Q3AUQ0 Q3AUR5 Q3AUR6 Q3AUU4 Q3AUU5 Q3AUU9 Q3AUV1 Q3AUV9 Q3AUW0 Q3AUW1 Q3AUW2
Q3AUW3 Q3AUW7 Q3AUW8 Q3AUW9 Q3AUX0 Q3AUX8 Q3AUZ9 Q3AV08 Q3AV13 Q3AV41 Q3AV42
Q3AV44 Q3AV45 Q3AV47 Q3AV73 Q3AV87 Q3AV89 Q3AV90 Q3AV97 Q3AV98 Q3AVB7 Q3AVC2
Q3AVC3 Q3AVE5 Q3AVF6 Q3AVF9 Q3AVH3 Q3AVJ1 Q3AVJ4 Q3AVJ5 Q3AVK0 Q3AVK1 Q3AVK4
Q3AVL9 Q3AVM1 Q3AVP2 Q3AVP9 Q3AVR1 Q3AVR2 Q3AVR3 Q3AVR4 Q3AVR7 Q3AVS9 Q3AVT0
Q3AVV3 Q3AVV5 Q3AVV7 Q3AVV9 Q3AVX6 Q3AVY3 Q3AVY5 Q3AVZ5 Q3AW00 Q3AW11 Q3AW12
Q3AW18 Q3AW20 Q3AW21 Q3AW32 Q3AW35 Q3AW42 Q3AW44 Q3AW47 Q3AW52 Q3AW53 Q3AW54
Q3AW55 Q3AW56 Q3AW64 Q3AW65 Q3AW66 Q3AW67 Q3AW68 Q3AW69 Q3AW70 Q3AW71 Q3AW72
Q3AW73 Q3AW74 Q3AW76 Q3AW77 Q3AW78 Q3AW79 Q3AW80 Q3AW81 Q3AW82 Q3AW83 Q3AW84
Q3AW85 Q3AWA8 Q3AWE6 Q3AWE9 Q3AWF3 Q3AWF5 Q3AWG6 Q3AWK9 Q3AWM1 Q3AWM4 Q3AWM6
Q3AWM9 Q3AWN7 Q3AWP4 Q3AWQ4 Q3AWR0 Q3AWR7 Q3AWR9 Q3AWT4 Q3AWT5 Q3AWT6 Q3AWU1
Q3AWW1 Q3AWW5 Q3AWX1 Q3AWY4 Q3AWZ4 Q3AX02 Q3AX63 Q3AX77 Q3AX81 Q3AX82 Q3AXB7
Q3AXB9 Q3AXD5 Q3AXE9 Q3AXF2 Q3AXF3 Q3AXF4 Q3AXI3 Q3AXI9 Q3AXJ0 Q3AXK3 Q3AXK4
Q3AXL0 Q3AXM7 Q3AXP3 Q3AXQ4 Q3AXR9 Q3AXU8 Q3AXV0 Q3AXV7 Q3AXV8 Q3AXW8 Q3AXX8
Q3AXY2 Q3AXZ4 Q3AY04 Q3AY98 Q3AYC6 Q3AYC9 Q3AYD4 Q3AYE2 Q3AYE3 Q3AYG5 Q3AYG9
Q3AYH5 Q3AYI1 Q3AYI2 Q3AYN1 Q3AYN5 Q3AYP0 Q3AYR1 Q3AYS1 Q3AYS3 Q3AYS6 Q3AYT2

Q3AYW4 Q3AYX2 Q3AYY2 Q3AYZ0 Q3AYZ7 Q3AZ17 Q3AZ20 Q3AZ46 Q3AZ50 Q3AZ58 Q3AZ61
Q3AZ65 Q3AZ92 Q3AZA0 Q3AZA2 Q3AZA3 Q3AZA4 Q3AZA6 Q3AZA8 Q3AZB4 Q3AZB7 Q3AZD9
Q3AZE2 Q3AZE8 Q3AZF6 Q3AZG3 Q3AZG4 Q3AZG7 Q3AZH7 Q3AZI5 Q3AZJ0 Q3AZK3 Q3AZK4
Q3AZN8 Q3AZU8 Q3AZX9 Q3AZZ0 Q3B002 Q3B009 Q3B014 Q3B039 Q3B043 Q3B078 Q3B080
Q3B083 Q3B088 Q3B095 Q3B0A6 Q3B0D3 Q3B0H7 Q3B0I6 Q3B0J1 Q3B0J2 Q3B0J9 Q3B0K7
Q3B0M1 Q3B0M2 Q3B0Q0 Q3B0S2 Q3B0T8 Q3B0U0 Q3B0U1 Q3B0U2 Q3B0U8 Q3B0U9 Q3B0V2
Q3B0W1 Q3B0W2 Q3B0X4 Q3B0X8 Q3B0Y0 Q3B0Y4 Q3B0Z0 Q3B100 Q3B107 Q3B116 Q3B120
Q3B121 Q3B127 Q3B130 Q3B146 Q3B147 Q3B151 Q3B169 Q3B172 Q3B189 Q3B192 Q3B1A1
Q3B1A2 Q3B1B4 Q3B1E7 Q3B1F0 Q3B1F7 Q3B1F8 Q3B1H3 Q3B1H4 Q3B1H5 Q3B1H6 Q3B1H7
Q3B1H8 Q3B1I7 Q3B1J3 Q3B1L6 Q3B1Z1 Q3B1Z6 Q3B1Z8 Q3B1Z9 Q3B200 Q3B202 Q3B204
Q3B208 Q3B213 Q3B218 Q3B219 Q3B228 Q3B241 Q3B250 Q3B256 Q3B278 Q3B281 Q3B286
Q3B287 Q3B298 Q3B2C7 Q3B2D4 Q3B2D5 Q3B2E1 Q3B2E2 Q3B2E3 Q3B2F0 Q3B2F5 Q3B2F8
Q3B2G2 Q3B2G3 Q3B2G4 Q3B2H0 Q3B2H1 Q3B2H3 Q3B2H9 Q3B2I6 Q3B2I7 Q3B2P6 Q3B2Q0
Q3B2Q6 Q3B2Q8 Q3B2R4 Q3B2R5 Q3B2R6 Q3B2S6 Q3B2T0 Q3B2T3 Q3B2T4 Q3B2T5 Q3B2U3
Q3B2U9 Q3B2V0 Q3B2V8 Q3B2V9 Q3B2W3 Q3B2Y6 Q3B2Z5 Q3B2Z6 Q3B304 Q3B314 Q3B315
Q3B316 Q3B317 Q3B327 Q3B365 Q3B368 Q3B373 Q3B380 Q3B391 Q3B394 Q3B397 Q3B3A1
Q3B3A7 Q3B3B1 Q3B3D4 Q3B3D6 Q3B3D7 Q3B3E6 Q3B3J7 Q3B3L3 Q3B3L6 Q3B3M5 Q3B3M9
Q3B3N3 Q3B3N9 Q3B3P0 Q3B3P4 Q3B3Q8 Q3B3Y0 Q3B3Y5 Q3B422 Q3B424 Q3B425 Q3B426
Q3B4A0 Q3B4A1 Q3B4A2 Q3B4A3 Q3B4B1 Q3B4B5 Q3B4B8 Q3B4C0 Q3B4D9 Q3B4E2 Q3B4Y6
Q3B4Z2 Q3B531 Q3B532 Q3B533 Q3B561 Q3B577 Q3B589 Q3B594 Q3B595 Q3B5A0 Q3B5A4
Q3B5A5 Q3B5C9 Q3B5D0 Q3B5D1 Q3B5D4 Q3B5D7 Q3B5E2 Q3B5E3 Q3B5F2 Q3B5F4 Q3B5F5
Q3B5F6 Q3B5I8 Q3B5J2 Q3B5J6 Q3B5K1 Q3B5K8 Q3B5N2 Q3B5N7 Q3B5P3 Q3B5P7 Q3B5R1
Q3B5R5 Q3B5T8 Q3B5T9 Q3B5U0 Q3B5U1 Q3B5U2 Q3B5U3 Q3B5U7 Q3B5V8 Q3B5Y2 Q3B5Y3
Q3B5Z4 Q3B5Z5 Q3B605 Q3B606 Q3B612 Q3B619 Q3B622 Q3B625 Q3B659 Q3B660 Q3B6A2
Q3B6A6 Q3B6B4 Q3B6C3 Q3B6C6 Q3B6C7 Q3B6C8 Q3B6D0 Q3B6D1 Q3B6D2 Q3B6D4 Q3B6D5
Q3B6D6 Q3B6D7 Q3B6D8 Q3B6D9 Q3B6E0 Q3B6E3 Q3B6E4 Q3B6E5 Q3B6E6 Q3B6E7 Q3B6E8
Q3B6E9 Q3B6F0 Q3B6F1 Q3B6F2 Q3B6F3 Q3B6F5 Q3B6F6 Q3B6F7 Q3B6F8 Q3B6F9 Q3B6G0
Q3B6G1 Q3B6G2 Q3B6G3 Q3B6G4 Q3B6G5 Q3B6G6 Q3B6H6 Q3B6H8 Q3B6I3 Q3B6K6 Q3B6L3
Q3B6L9 Q3B6M0 Q3B6M1 Q3B6M2 Q3B6M4 Q3B6M5 Q3B6M7 Q3B6N5 Q3B6P1 Q3B6R4 Q3B6R6
Q3B6X8 Q3B6Y3 Q3B6Y7 Q3B725 Q3B748 Q3B752 Q3B7D1 Q3B7D9 Q3B7G7 Q3B7I2 Q3B7I8
Q3B7K9 Q3B7L6 Q3B7L8 Q3B7L9 Q3B7M1 Q3B7M2 Q3B7M4 Q3B7M5 Q3B7M7 Q3B7M9 Q3B7N1
Q3B7Q7 Q3B7T1 Q3B7T8 Q3B7U2 Q3B7U4 Q3B807 Q3B820 Q3B8C3 Q3B8D4 Q3B8D5 Q3B8D7
Q3B8E3 Q3B8E5 Q3B8E8 Q3B8E9 Q3B8G0 Q3B8G8 Q3B8H4 Q3B8I4 Q3B8K3 Q3B8M3 Q3B8N0
Q3B8N2 Q3B8N5 Q3B8N7 Q3B8N8 Q3B8Q1 Q3B8Q2 Q3B8S0 Q3BAH0 Q3BAH2 Q3BAH3 Q3BAH4
Q3BAH6 Q3BAI0 Q3BAI2 Q3BAJ4 Q3BAJ7 Q3BAJ8 Q3BAJ9 Q3BAK0 Q3BAK1 Q3BAK2 Q3BAK3
Q3BAK4 Q3BAK5 Q3BAL3 Q3BAL6 Q3BAL8 Q3BAL9 Q3BAN4 Q3BAN8 Q3BAP2 Q3BAP8 Q3BAQ1
Q3BAQ2 Q3BAQ3 Q3BAR0 Q3BAR2 Q3BAR5 Q3BBV0 Q3BBV2 Q3BCQ8 Q3BCR0 Q3BCR1 Q3BCR2
Q3BCR3 Q3BCR4 Q3BCR5 Q3BCR6 Q3BCR8 Q3BCR9 Q3BJS0 Q3BJS1 Q3BJS3 Q3BJY7 Q3BLZ5
Q3BLZ6 Q3BLZ9 Q3BM80 Q3BM92 Q3BM98 Q3BMA2 Q3BMA4 Q3BMC5 Q3BMF2 Q3BMF4 Q3BMH1
Q3BML8 Q3BMM5 Q3BMM6 Q3BMT4 Q3BMW1 Q3BN06 Q3BN23 Q3BN25 Q3BN93 Q3BNA3 Q3BNA4
Q3BNA6 Q3BNB6 Q3BNC0 Q3BNC2 Q3BNE1 Q3BNE2 Q3BNG9 Q3BNH4 Q3BNH8 Q3BNI2 Q3BNJ9
Q3BNK3 Q3BNK7 Q3BNL9 Q3BNP7 Q3BNX3 Q3BNX4 Q3BNZ1 Q3BP42 Q3BP46 Q3BP90 Q3BP98
Q3BPA1 Q3BPD5 Q3BPG1 Q3BPJ6 Q3BPJ7 Q3BPJ8 Q3BPJ9 Q3BPK3 Q3BPK7 Q3BPM9 Q3BPP7
Q3BPQ4 Q3BPQ9 Q3BPR2 Q3BPR8 Q3BPS0 Q3BPS1 Q3BPS6 Q3BPW7 Q3BPY3 Q3BPY4 Q3BPY6

Q3BQ14 Q3BQ15 Q3BQ65 Q3BQ76 Q3BQ83 Q3BQB5 Q3BQB7 Q3BQE6 Q3BQF0 Q3BQF1 Q3BQF2
Q3BQF5 Q3BQG4 Q3BQI4 Q3BQI5 Q3BQI6 Q3BQI7 Q3BQN6 Q3BQN7 Q3BQQ3 Q3BQR9 Q3BQS1
Q3BQS2 Q3BQS3 Q3BQW0 Q3BQW1 Q3BQX2 Q3BQX8 Q3BQZ6 Q3BR19 Q3BR20 Q3BR21 Q3BR29
Q3BR52 Q3BR53 Q3BR75 Q3BRB4 Q3BRE4 Q3BRF1 Q3BRF4 Q3BRG1 Q3BRG7 Q3BRJ0 Q3BRJ5
Q3BRK5 Q3BRL0 Q3BRL1 Q3BRL3 Q3BRL7 Q3BRL9 Q3BRM7 Q3BRP3 Q3BRP5 Q3BRP6 Q3BRP7
Q3BRP8 Q3BRP9 Q3BRT7 Q3BRT9 Q3BRU0 Q3BRU1 Q3BRU2 Q3BRU3 Q3BRW8 Q3BRX9 Q3BS21
Q3BS23 Q3BS29 Q3BS36 Q3BS37 Q3BS67 Q3BS90 Q3BS94 Q3BSA6 Q3BSC5 Q3BSD0 Q3BSE3
Q3BSE4 Q3BSF5 Q3BSF8 Q3BSH9 Q3BSI7 Q3BSI8 Q3BSJ0 Q3BSJ1 Q3BSK1 Q3BSN4 Q3BSN5
Q3BSP5 Q3BSP7 Q3BSV3 Q3BSV4 Q3BSW1 Q3BTC3 Q3BTE6 Q3BTH9 Q3BTI2 Q3BTU8 Q3BTW0
Q3BTW5 Q3BTW6 Q3BTY0 Q3BTY1 Q3BTY6 Q3BUA1 Q3BUB6 Q3BUB7 Q3BUC6 Q3BUD7 Q3BUE6
Q3BUE7 Q3BUE8 Q3BUF2 Q3BUF3 Q3BUF4 Q3BUF5 Q3BUF6 Q3BUF7 Q3BUF8 Q3BUG0 Q3BUJ8
Q3BUK3 Q3BUL3 Q3BUL4 Q3BUL5 Q3BUL6 Q3BUL9 Q3BUN6 Q3BUQ6 Q3BUQ7 Q3BUQ8 Q3BUQ9
Q3BUR4 Q3BUR5 Q3BUR8 Q3BUS3 Q3BUS4 Q3BUS6 Q3BUS7 Q3BUS8 Q3BUT3 Q3BUY7 Q3BUY8
Q3BUZ1 Q3BUZ3 Q3BV03 Q3BV06 Q3BV11 Q3BV14 Q3BV19 Q3BV20 Q3BV21 Q3BV23 Q3BV37
Q3BV40 Q3BV41 Q3BV46 Q3BV47 Q3BV71 Q3BV87 Q3BVB7 Q3BVB8 Q3BVB9 Q3BVC0 Q3BVC6
Q3BVI6 Q3BVJ3 Q3BVJ6 Q3BVK3 Q3BVK4 Q3BVK6 Q3BVK7 Q3BVL0 Q3BVP1 Q3BVU6 Q3BVV1
Q3BVV3 Q3BVV6 Q3BVY0 Q3BVY2 Q3BVY7 Q3BVY8 Q3BVY9 Q3BVZ0 Q3BW40 Q3BW42 Q3BW45
Q3BW46 Q3BW47 Q3BW48 Q3BW59 Q3BW63 Q3BW74 Q3BW76 Q3BW89 Q3BWA6 Q3BWF4 Q3BWJ0
Q3BWK3 Q3BWK4 Q3BWM5 Q3BWP3 Q3BWP5 Q3BWQ0 Q3BWQ1 Q3BWQ2 Q3BWT0 Q3BWU8 Q3BWV8
Q3BWV9 Q3BWW0 Q3BWW1 Q3BWW2 Q3BWW4 Q3BWW5 Q3BWW6 Q3BWW7 Q3BWW8 Q3BWW9 Q3BWX0
Q3BWX1 Q3BWX2 Q3BWX3 Q3BWX4 Q3BWX5 Q3BWX6 Q3BWX7 Q3BWX8 Q3BWX9 Q3BWY0 Q3BWY1
Q3BWY2 Q3BWY3 Q3BWY4 Q3BWY6 Q3BWY7 Q3BWY8 Q3BWY9 Q3BWZ0 Q3BWZ1 Q3BWZ2 Q3BWZ3
Q3BWZ4 Q3BWZ5 Q3BX00 Q3BX01 Q3BX03 Q3BX06 Q3BX07 Q3BX12 Q3BX43 Q3BX45 Q3BX52
Q3BX58 Q3BX82 Q3BX83 Q3BX84 Q3BX87 Q3BXB7 Q3BXC4 Q3BXC6 Q3BXF1 Q3BXF9 Q3BXG0
Q3BXG6 Q3BXH9 Q3BXI0 Q3BXI6 Q3BXI8 Q3BXJ3 Q3BXK1 Q3BXK7 Q3BXK8 Q3BXK9 Q3BXQ3
Q3BXQ4 Q3BXT9 Q3BXU0 Q3BXU3 Q3BXV3 Q3BY34 Q3BY37 Q3BY45 Q3BY61 Q3BY62 Q3BY66
Q3BY72 Q3BY85 Q3BY98 Q3BYA7 Q3BYA8 Q3BYA9 Q3BYB2 Q3BYB5 Q3BYB6 Q3BYD2 Q3BYE8
Q3BYF4 Q3BYI0 Q3BYI1 Q3BYM9 Q3BYN0 Q3BYP1 Q3BYR0 Q3BYS5 Q3BZ76 Q3BZD0 Q3BZE1
Q3BZI4 Q3BZR9 Q3BZS9 Q3BZT1 Q3C167 Q3C1A6 Q3C1D2 Q3C1D4 Q3C1D5 Q3C1E0 Q3C1E1
Q3C1E2 Q3C1E3 Q3C1E9 Q3C1G1 Q3C1G7 Q3C1G8 Q3C1G9 Q3C1H0 Q3C1H5 Q3C1I0 Q3C1J4
Q3C1K4 Q3C1K5 Q3C1K6 Q3C1K7 Q3C1K8 Q3C1L5 Q3C1L6 Q3C1L8 Q3C1L9 Q3C1M0 Q3C1M1
Q3C1M2 Q3C1M3 Q3C1M8 Q3C1N3 Q3C1N4 Q3C1N5 Q3C1N6 Q3C1N7 Q3C1N8 Q3C1P7 Q3C1V1
Q3C1V8 Q3C1V9 Q3C251 Q3C2C4 Q3C2E5 Q3E6P4 Q3E6Q1 Q3E6S8 Q3E6S9 Q3E6X9 Q3E705
Q3E731 Q3E732 Q3E736 Q3E737 Q3E740 Q3E741 Q3E742 Q3E744 Q3E746 Q3E747 Q3E758
Q3E762 Q3E764 Q3E766 Q3E767 Q3E769 Q3E771 Q3E772 Q3E774 Q3E775 Q3E778 Q3E781
Q3E784 Q3E785 Q3E786 Q3E787 Q3E789 Q3E791 Q3E793 Q3E794 Q3E795 Q3E7A0 Q3E7A2
Q3E7A5 Q3E7A6 Q3E7A8 Q3E7A9 Q3E7B3 Q3E7B4 Q3E7B5 Q3E7B7 Q3E7C1 Q3E7D1 Q3E7I5
Q3E7J7 Q3E7K4 Q3E7K7 Q3E7K8 Q3E7L7 Q3E7N5 Q3E7S2 Q3E7T8 Q3E7X8 Q3E7Y4 Q3E7Y5
Q3E7Y6 Q3E7Y7 Q3E7Y8 Q3E7Y9 Q3E7Z0 Q3E7Z1 Q3E7Z2 Q3E7Z3 Q3E7Z5 Q3E7Z6 Q3E7Z7
Q3E7Z9 Q3E801 Q3E802 Q3E804 Q3E805 Q3E806 Q3E807 Q3E808 Q3E809 Q3E811 Q3E813
Q3E816 Q3E817 Q3E818 Q3E819 Q3E825 Q3E826 Q3E827 Q3E829 Q3E832 Q3E833 Q3E835
Q3E838 Q3E840 Q3E841 Q3E843 Q3E846 Q3E8A8 Q3E8H7 Q3E8J4 Q3E8K6 Q3E8L3 Q3E8L4
Q3E8L5 Q3E8Y5 Q3E8Y7 Q3E902 Q3E906 Q3E911 Q3E922 Q3E944 Q3E960 Q3E984 Q3E9B4
Q3E9C3 Q3E9D5 Q3E9F0 Q3E9F5 Q3E9F6 Q3E9F7 Q3E9H1 Q3E9N1 Q3E9N2 Q3E9T2 Q3E9Z8

Q3EA00 Q3EA33 Q3EA38 Q3EAE5 Q3EAE7 Q3EAF8 Q3EAF9 Q3EAI1 Q3EAL1 Q3EAR3 Q3EAS6
Q3EAZ3 Q3EB08 Q3EBC8 Q3EBD3 Q3EBI7 Q3EBJ8 Q3EBP0 Q3EBP1 Q3EBQ3 Q3EBY8 Q3EBZ2
Q3EC60 Q3EC97 Q3ECB8 Q3ECD0 Q3ECE2 Q3ECH0 Q3ECH5 Q3ECK2 Q3ECP0 Q3ECQ9 Q3ECR3
Q3ECS5 Q3ECU8 Q3ECW2 Q3ECY6 Q3ED38 Q3ED65 Q3ED68 Q3ED78 Q3ED92 Q3ED93 Q3EDA9
Q3EDF8 Q3EDH0 Q3EDJ0 Q3HRN7 Q3HRN8 Q3HRP1 Q3HRP2 Q3HT77 Q3HVN1 Q3I3X0 Q3I409
Q3I410 Q3I411 Q3I5G5 Q3I5I5 Q3IBU5 Q3IBW7 Q3IBX5 Q3IBX6 Q3IC07 Q3IC08 Q3IC33
Q3IC38 Q3IC91 Q3ICC5 Q3ICD4 Q3ICF0 Q3ICF4 Q3ICF9 Q3ICG0 Q3ICG2 Q3ICJ5 Q3ICL0
Q3ICQ7 Q3ICS0 Q3ICU8 Q3ICU9 Q3ICY1 Q3ICZ5 Q3ICZ6 Q3ICZ9 Q3ID04 Q3ID09 Q3ID15
Q3ID16 Q3ID31 Q3IDC8 Q3IDD2 Q3IDD3 Q3IDD5 Q3IDE6 Q3IDE8 Q3IDH7 Q3IDH9 Q3IDI2
Q3IDI3 Q3IDI6 Q3IDK9 Q3IDL0 Q3IDL2 Q3IDL9 Q3IDM3 Q3IDM6 Q3IDN1 Q3IDP4 Q3IDP9
Q3IDQ1 Q3IDQ4 Q3IDQ5 Q3IDQ6 Q3IDT2 Q3IDT7 Q3IDT9 Q3IDU0 Q3IE34 Q3IE36 Q3IE45
Q3IE60 Q3IE61 Q3IE64 Q3IE99 Q3IEA2 Q3IEA5 Q3IEB7 Q3IEC6 Q3IEC7 Q3IEC8 Q3IEC9
Q3IEF7 Q3IEN6 Q3IEQ9 Q3IES7 Q3IEW0 Q3IEW3 Q3IEY7 Q3IEY8 Q3IEZ3 Q3IF18 Q3IF19
Q3IF20 Q3IF21 Q3IF22 Q3IF23 Q3IF24 Q3IF25 Q3IF26 Q3IF34 Q3IF36 Q3IF40 Q3IF46
Q3IF51 Q3IF52 Q3IF53 Q3IF68 Q3IF86 Q3IF87 Q3IF89 Q3IFA0 Q3IFA1 Q3IFD1 Q3IFD2
Q3IFE5 Q3IFE7 Q3IFF4 Q3IFF5 Q3IFF6 Q3IFG5 Q3IFH4 Q3IFH5 Q3IFI3 Q3IFM1 Q3IFM3
Q3IFM4 Q3IFM6 Q3IFN6 Q3IFN9 Q3IFP4 Q3IFP5 Q3IFQ5 Q3IFT4 Q3IFT7 Q3IFV9 Q3IFW0
Q3IFW1 Q3IFY1 Q3IFY2 Q3IFZ5 Q3IFZ6 Q3IG02 Q3IG11 Q3IG20 Q3IG26 Q3IG30 Q3IG44
Q3IG47 Q3IG50 Q3IG57 Q3IG75 Q3IG80 Q3IG97 Q3IGA3 Q3IGA7 Q3IGS6 Q3IGS9 Q3IGT7
Q3IGU4 Q3IGV1 Q3IGV3 Q3IGW5 Q3IGW8 Q3IGX7 Q3IGZ2 Q3IGZ3 Q3IH16 Q3IH21 Q3IH34
Q3IH36 Q3IH61 Q3IH67 Q3IH68 Q3IH69 Q3IH70 Q3IH71 Q3IH90 Q3IHA0 Q3IHC1 Q3IHD0
Q3IHD5 Q3IHD7 Q3IHD9 Q3IHG8 Q3IHI4 Q3IHI5 Q3IHJ1 Q3IHJ4 Q3IHK6 Q3IHK8 Q3IHM8
Q3IHQ0 Q3IHQ6 Q3IHR0 Q3IHS1 Q3IHS5 Q3IHT8 Q3IHU4 Q3IHW0 Q3IHW1 Q3IHW4 Q3IHX1
Q3IHX7 Q3IHY7 Q3IHZ2 Q3II09 Q3II11 Q3II23 Q3II24 Q3II28 Q3II29 Q3II35 Q3II43
Q3II62 Q3II73 Q3II81 Q3II82 Q3IIA2 Q3IIA6 Q3IIB6 Q3IIC0 Q3IIC3 Q3IID1 Q3IID3
Q3IIE1 Q3IIE4 Q3III5 Q3IIJ0 Q3IIJ1 Q3IIJ2 Q3IIL2 Q3IIL3 Q3IIM0 Q3IIN7 Q3IIQ2
Q3IIQ3 Q3IIQ4 Q3IIQ6 Q3IIR9 Q3IIS1 Q3IIT0 Q3IIW9 Q3IIX4 Q3IIX6 Q3IIZ3 Q3IIZ6
Q3IIZ7 Q3IJ06 Q3IJ21 Q3IJ25 Q3IJ28 Q3IJ32 Q3IJ42 Q3IJ50 Q3IJ52 Q3IJ53 Q3IJ55
Q3IJ73 Q3IJ74 Q3IJ75 Q3IJ79 Q3IJ80 Q3IJ81 Q3IJ93 Q3IJ98 Q3IJA2 Q3IJB4 Q3IJC5
Q3IJC7 Q3IJC9 Q3IJD2 Q3IJD5 Q3IJG3 Q3IJH1 Q3IJH7 Q3IJH8 Q3IJI0 Q3IJI1 Q3IJI6
Q3IJI7 Q3IJJ5 Q3IJJ6 Q3IJJ7 Q3IJJ8 Q3IJJ9 Q3IJK0 Q3IJK1 Q3IJK2 Q3IJK3 Q3IJK4
Q3IJK6 Q3IJK7 Q3IJL9 Q3IJQ5 Q3IJS3 Q3IJS4 Q3IJS5 Q3IJT8 Q3IJU2 Q3IJU8 Q3IJV0
Q3IJV1 Q3IJV7 Q3IJW1 Q3IJW9 Q3IK32 Q3IK39 Q3IK40 Q3IK51 Q3IK52 Q3IK56 Q3IK75
Q3IK77 Q3IK85 Q3IK86 Q3IK95 Q3IK96 Q3IK98 Q3IKB2 Q3IKB8 Q3IKE8 Q3IKF4 Q3IKH4
Q3IKJ0 Q3IKN2 Q3IKN3 Q3IKN5 Q3IKQ1 Q3IKQ2 Q3IKQ7 Q3IKR2 Q3IKR4 Q3IKS6 Q3IKY4
Q3IKY9 Q3IL08 Q3IL10 Q3IL20 Q3IL56 Q3IL78 Q3IL79 Q3IL80 Q3IL99 Q3ILA2 Q3ILA3
Q3ILA5 Q3ILB8 Q3ILB9 Q3ILD8 Q3ILF1 Q3ILF3 Q3ILI9 Q3ILJ1 Q3ILJ2 Q3ILJ5 Q3ILJ7
Q3ILJ9 Q3ILK1 Q3ILK5 Q3ILL1 Q3ILL7 Q3ILL8 Q3ILM2 Q3ILP2 Q3ILP3 Q3ILP4 Q3ILP5
Q3ILP6 Q3ILP7 Q3ILP8 Q3ILP9 Q3ILQ0 Q3ILQ2 Q3ILQ3 Q3ILQ4 Q3ILQ9 Q3ILR0 Q3IM07
Q3IMD8 Q3IME1 Q3IME2 Q3IMF0 Q3IMF1 Q3IMF3 Q3IMI0 Q3IMI4 Q3IMJ0 Q3IMK8 Q3IMM5
Q3IMM9 Q3IMN0 Q3IMN2 Q3IMP5 Q3IMP6 Q3IMR2 Q3IMS0 Q3IMS5 Q3IMV2 Q3IMV8 Q3IMW0
Q3IMW6 Q3IMW7 Q3IMW8 Q3IMW9 Q3IMX0 Q3IMX2 Q3IMX3 Q3IMX4 Q3IMX5 Q3IMX6 Q3IMX7
Q3IMX8 Q3IMY1 Q3IMY2 Q3IMY3 Q3IMY4 Q3IMY5 Q3IMY6 Q3IMY7 Q3IMY8 Q3IN28 Q3IN29
Q3IN38 Q3INE3 Q3INE8 Q3INF0 Q3ING1 Q3INI5 Q3INI6 Q3INI7 Q3INK4 Q3INN3 Q3INN9

Q3INP4 Q3INQ4 Q3INQ6 Q3INU3 Q3INV7 Q3INY0 Q3INY2 Q3INZ0 Q3INZ6 Q3IP06 Q3IP08
Q3IP34 Q3IP35 Q3IP66 Q3IP72 Q3IP73 Q3IP82 Q3IPG8 Q3IPH3 Q3IPI5 Q3IPI7 Q3IPJ1
Q3IPJ2 Q3IPJ7 Q3IPL2 Q3IPL6 Q3IPL7 Q3IPM0 Q3IPM6 Q3IPM8 Q3IPM9 Q3IPU7 Q3IPU8
Q3IPW6 Q3IPW7 Q3IPX0 Q3IPX1 Q3IPX9 Q3IQ04 Q3IQ21 Q3IQ37 Q3IQ38 Q3IQ95 Q3IQA0
Q3IQA3 Q3IQA5 Q3IQC4 Q3IQF5 Q3IQG6 Q3IQL3 Q3IQR5 Q3IQS9 Q3IQT2 Q3IQT4 Q3IQT6
Q3IQT7 Q3IQT8 Q3IQT9 Q3IQU8 Q3IQZ0 Q3IQZ9 Q3IR53 Q3IR95 Q3IR96 Q3IR98 Q3IRB3
Q3IRE7 Q3IRF2 Q3IRF6 Q3IRJ0 Q3IRL6 Q3IRL7 Q3IRM1 Q3IRM3 Q3IRM5 Q3IRN3 Q3IRP7
Q3IRQ2 Q3IRQ4 Q3IRT0 Q3IRT1 Q3IRT2 Q3IRT5 Q3IRU8 Q3IRX3 Q3IRX5 Q3IRY6 Q3IRZ1
Q3IRZ2 Q3IRZ3 Q3IRZ4 Q3IRZ5 Q3IRZ6 Q3IS04 Q3IS14 Q3IS15 Q3IS17 Q3IS55 Q3IS99
Q3ISA5 Q3ISB2 Q3ISB3 Q3ISC7 Q3ISC9 Q3ISI7 Q3ISI8 Q3ISJ9 Q3ISN0 Q3ISP9 Q3ISQ0
Q3ISR6 Q3IST1 Q3ISU0 Q3ISW0 Q3ISY4 Q3IT10 Q3IT14 Q3IT15 Q3IT20 Q3IT24 Q3IT33
Q3IT46 Q3IT47 Q3IT59 Q3IT64 Q3ITA0 Q3ITB1 Q3ITB2 Q3ITB6 Q3ITD5 Q3ITD6 Q3ITF7
Q3ITJ2 Q3ITK1 Q3ITN8 Q3ITN9 Q3ITP9 Q3ITQ4 Q3ITQ5 Q3ITR7 Q3ITS0 Q3ITS6 Q3ITT7
Q3ITU5 Q3ITY2 Q3ITY5 Q3ITY6 Q3ITY7 Q3IU04 Q3IU11 Q3IU15 Q3IU43 Q3IU46 Q3IU54
Q3IU57 Q3IU60 Q3IU70 Q3IU82 Q3IU96 Q3IUB0 Q3IUB3 Q3IUB7 Q3IUD7 Q3IUD8 Q3IUE5
Q3IUE8 Q3IUG3 Q3IUG6 Q3IUH3 Q3IUI0 Q3IUI1 Q3IUJ2 Q3IUJ6 Q3IUJ7 Q3IUM4 Q3IUM7
Q3IUM8 Q3IUP2 Q3IUP8 Q3IUQ3 Q3IUQ6 Q3IUS0 Q3IUT3 Q3IVU8 Q3IW19 Q3IW60 Q3IW81
Q3IW83 Q3IW86 Q3IW88 Q3IW89 Q3IWB0 Q3IWD2 Q3IWG2 Q3IWV4 Q3IX44 Q3IX55 Q3IX60
Q3IX78 Q3IXC0 Q3IXK6 Q3IXP4 Q3IXX7 Q3IXX8 Q3IXX9 Q3IXY1 Q3IXY2 Q3IY00 Q3IY06
Q3IY15 Q3IY22 Q3IY46 Q3IY61 Q3IY62 Q3IY64 Q3IY65 Q3IYB9 Q3IYC2 Q3IYC6 Q3IYG6
Q3IYG8 Q3IYG9 Q3IYH0 Q3IYH3 Q3IYH4 Q3IYH5 Q3IYI4 Q3IYI5 Q3IYJ1 Q3IYM4 Q3IYM5
Q3IYM7 Q3IYM8 Q3IYN5 Q3IYN8 Q3IYR6 Q3IYR9 Q3IYS2 Q3IYT7 Q3IYT8 Q3IYU1 Q3IYU3
Q3IYU4 Q3IYU8 Q3IYV6 Q3IYX6 Q3IYY2 Q3IYY4 Q3IYY8 Q3IYZ1 Q3IZ02 Q3IZ03 Q3IZ05
Q3IZ06 Q3IZ07 Q3IZ49 Q3IZ52 Q3IZ66 Q3IZ78 Q3IZ83 Q3IZ84 Q3IZ90 Q3IZ91 Q3IZB6
Q3IZB8 Q3IZC1 Q3IZF4 Q3IZG8 Q3IZH6 Q3IZI5 Q3IZI8 Q3IZJ0 Q3IZJ3 Q3IZJ9 Q3IZL4
Q3IZM8 Q3IZN0 Q3IZN1 Q3IZN2 Q3IZP1 Q3IZP4 Q3IZQ8 Q3IZR0 Q3IZR5 Q3IZS9 Q3IZX0
Q3IZY2 Q3IZZ0 Q3IZZ6 Q3J047 Q3J049 Q3J055 Q3J061 Q3J077 Q3J079 Q3J080 Q3J085
Q3J086 Q3J0B2 Q3J0C4 Q3J0D9 Q3J0F0 Q3J0F5 Q3J0F6 Q3J0F8 Q3J0G1 Q3J0H1 Q3J0I4
Q3J0K8 Q3J0N0 Q3J0R1 Q3J0S2 Q3J0T3 Q3J0W6 Q3J0W9 Q3J0X4 Q3J0Z1 Q3J0Z3 Q3J101
Q3J149 Q3J151 Q3J154 Q3J156 Q3J157 Q3J158 Q3J1A0 Q3J1A1 Q3J1A8 Q3J1G6 Q3J1G7
Q3J1H1 Q3J1L8 Q3J1L9 Q3J1M0 Q3J1M1 Q3J1S6 Q3J1Z4 Q3J1Z5 Q3J205 Q3J210 Q3J252
Q3J263 Q3J273 Q3J276 Q3J277 Q3J281 Q3J289 Q3J290 Q3J2A5 Q3J2B6 Q3J2B7 Q3J2B9
Q3J2D0 Q3J2H4 Q3J2I9 Q3J2J0 Q3J2K8 Q3J2K9 Q3J2L7 Q3J2N4 Q3J2N5 Q3J2P4 Q3J2Q5
Q3J2U4 Q3J2V7 Q3J2X1 Q3J2X4 Q3J2X5 Q3J2X6 Q3J2Y1 Q3J2Z5 Q3J317 Q3J340 Q3J349
Q3J355 Q3J358 Q3J363 Q3J366 Q3J3D0 Q3J3D1 Q3J3E2 Q3J3I2 Q3J3I3 Q3J3I6 Q3J3L4
Q3J3M6 Q3J3M8 Q3J3N4 Q3J3P6 Q3J3U4 Q3J3V1 Q3J3Y3 Q3J3Y8 Q3J419 Q3J420 Q3J436
Q3J445 Q3J446 Q3J459 Q3J460 Q3J462 Q3J474 Q3J4D4 Q3J4D7 Q3J4E3 Q3J4E6 Q3J4E9
Q3J4H6 Q3J4K2 Q3J4L8 Q3J4L9 Q3J4M1 Q3J4M6 Q3J4N3 Q3J4N4 Q3J4W8 Q3J4X5 Q3J4Z5
Q3J4Z8 Q3J517 Q3J555 Q3J561 Q3J5A0 Q3J5A6 Q3J5C0 Q3J5C2 Q3J5C7 Q3J5D3 Q3J5E0
Q3J5E8 Q3J5E9 Q3J5H3 Q3J5H6 Q3J5K7 Q3J5L0 Q3J5L6 Q3J5M2 Q3J5M3 Q3J5M4 Q3J5M6
Q3J5N0 Q3J5N1 Q3J5P7 Q3J5P8 Q3J5P9 Q3J5Q0 Q3J5Q3 Q3J5Q4 Q3J5Q5 Q3J5Q6 Q3J5Q7
Q3J5Q8 Q3J5Q9 Q3J5R0 Q3J5R1 Q3J5R2 Q3J5R3 Q3J5R5 Q3J5R6 Q3J5R7 Q3J5R8 Q3J5R9
Q3J5S0 Q3J5S1 Q3J5S2 Q3J5S3 Q3J5S4 Q3J5S5 Q3J5S6 Q3J5S7 Q3J5S9 Q3J5T0 Q3J5T1
Q3J5T2 Q3J5T3 Q3J5T4 Q3J5V2 Q3J5V6 Q3J5V8 Q3J5W1 Q3J5W4 Q3J5W7 Q3J5X0 Q3J6B1

Q3J6B2 Q3J6D2 Q3J6F2 Q3J6H3 Q3J6K8 Q3J6K9 Q3J6L0 Q3J6L6 Q3J6L7 Q3J6L9 Q3J6M0
Q3J6M1 Q3J6P9 Q3J6Q0 Q3J6Q1 Q3J6Q2 Q3J6Q3 Q3J6Q4 Q3J6R8 Q3J6R9 Q3J6S0 Q3J6S1
Q3J6S2 Q3J6T9 Q3J6U0 Q3J6U3 Q3J6U6 Q3J6V4 Q3J6V6 Q3J6V9 Q3J6W1 Q3J713 Q3J716
Q3J728 Q3J729 Q3J767 Q3J768 Q3J769 Q3J770 Q3J771 Q3J772 Q3J773 Q3J780 Q3J781
Q3J787 Q3J790 Q3J791 Q3J792 Q3J7A0 Q3J7A3 Q3J7B4 Q3J7C1 Q3J7C4 Q3J7D1 Q3J7D4
Q3J7D7 Q3J7D8 Q3J7D9 Q3J7E0 Q3J7E3 Q3J7E7 Q3J7F2 Q3J7G9 Q3J7H0 Q3J7H1 Q3J7H2
Q3J7M5 Q3J7P0 Q3J7R5 Q3J7R6 Q3J7S6 Q3J7S8 Q3J7T1 Q3J7T4 Q3J7T5 Q3J7U5 Q3J7V2
Q3J7V3 Q3J7V7 Q3J7V8 Q3J7W3 Q3J7W4 Q3J7W7 Q3J7X3 Q3J7X9 Q3J7Y0 Q3J7Z5 Q3J7Z6
Q3J7Z7 Q3J7Z8 Q3J7Z9 Q3J802 Q3J803 Q3J807 Q3J808 Q3J818 Q3J822 Q3J823 Q3J826
Q3J827 Q3J846 Q3J869 Q3J871 Q3J872 Q3J875 Q3J877 Q3J897 Q3J8A0 Q3J8D5 Q3J8F0
Q3J8G0 Q3J8H5 Q3J8N5 Q3J8P8 Q3J8Q0 Q3J8Q3 Q3J8Q4 Q3J8Q5 Q3J8Q6 Q3J8Q7 Q3J8Q8
Q3J8Q9 Q3J8R0 Q3J8R1 Q3J8R3 Q3J8R4 Q3J8R5 Q3J8R6 Q3J8R7 Q3J8R8 Q3J8R9 Q3J8S0
Q3J8S1 Q3J8S2 Q3J8S3 Q3J8S4 Q3J8S5 Q3J8S6 Q3J8S7 Q3J8S8 Q3J8S9 Q3J8T0 Q3J8T1
Q3J8T2 Q3J8T4 Q3J8T5 Q3J8T6 Q3J8T7 Q3J8T8 Q3J8T9 Q3J8U2 Q3J8U5 Q3J8V9 Q3J8W1
Q3J8X0 Q3J8X8 Q3J8X9 Q3J8Y0 Q3J8Y1 Q3J8Y4 Q3J8Y8 Q3J8Y9 Q3J8Z0 Q3J910 Q3J953
Q3J9B0 Q3J9B1 Q3J9B4 Q3J9B6 Q3J9B7 Q3J9B8 Q3J9B9 Q3J9C0 Q3J9C8 Q3J9D5 Q3J9D6
Q3J9G2 Q3J9J6 Q3J9J7 Q3J9K8 Q3J9K9 Q3J9L3 Q3J9L6 Q3J9L7 Q3J9M1 Q3J9N3 Q3JAC4
Q3JAD1 Q3JAD5 Q3JAF3 Q3JAF4 Q3JAF6 Q3JAH8 Q3JAI1 Q3JAJ7 Q3JAJ8 Q3JAJ9 Q3JAK5
Q3JAM9 Q3JAN0 Q3JAP1 Q3JAS2 Q3JAW3 Q3JAW6 Q3JAW7 Q3JB93 Q3JBA8 Q3JBB6 Q3JBE4
Q3JBF6 Q3JBJ1 Q3JBT2 Q3JBT3 Q3JBU6 Q3JBV4 Q3JBW2 Q3JBW4 Q3JBZ6 Q3JBZ7 Q3JBZ8
Q3JBZ9 Q3JC00 Q3JC02 Q3JC03 Q3JC50 Q3JC80 Q3JC85 Q3JC88 Q3JC90 Q3JC91 Q3JC94
Q3JCA2 Q3JCB0 Q3JCB2 Q3JCB7 Q3JCB9 Q3JCC0 Q3JCC4 Q3JCC5 Q3JCG5 Q3JCI4 Q3JCK6
Q3JCK8 Q3JCL5 Q3JCM3 Q3JCM5 Q3JCM7 Q3JCM9 Q3JCN0 Q3JCN1 Q3JCN4 Q3JCN5 Q3JCN7
Q3JCP6 Q3JCP7 Q3JCP8 Q3JCP9 Q3JCS0 Q3JCS7 Q3JCS8 Q3JCS9 Q3JCT3 Q3JCW1 Q3JCW9
Q3JCX2 Q3JCX3 Q3JCX4 Q3JCX5 Q3JCX7 Q3JCY0 Q3JCY2 Q3JCY8 Q3JCY9 Q3JCZ3 Q3JCZ4
Q3JD12 Q3JD71 Q3JD89 Q3JDG3 Q3JDH3 Q3JDH7 Q3JDL8 Q3JDM3 Q3JDQ7 Q3JDQ8 Q3JDR0
Q3JDR3 Q3JDR4 Q3JDR7 Q3JDS2 Q3JDS3 Q3JE31 Q3JE52 Q3JE53 Q3JE54 Q3JE55 Q3JE65
Q3JE87 Q3JEA2 Q3JEA9 Q3JEB6 Q3JEC8 Q3JEF2 Q3JEG3 Q3JEG4 Q3JEH3 Q3JEH7 Q3JEH9
Q3JEI0 Q3JEI2 Q3JEJ2 Q3JEJ4 Q3JEJ5 Q3JEJ6 Q3JEM4 Q3JEN4 Q3JEN6 Q3JEN8 Q3JEP4
Q3JER4 Q3JES3 Q3JES4 Q3JES5 Q3JES6 Q3JEW3 Q3JEY0 Q3JF12 Q3JF13 Q3JF20 Q3JF23
Q3JF27 Q3JF33 Q3JF36 Q3JF39 Q3JFK2 Q3JGP5 Q3JGX2 Q3JH62 Q3JH97 Q3JHA9 Q3JIC1
Q3JID4 Q3JJ17 Q3JJ76 Q3JJ81 Q3JK54 Q3JK55 Q3JK86 Q3JKA3 Q3JKA4 Q3JKA5 Q3JKA9
Q3JKE9 Q3JKG6 Q3JKG8 Q3JKG9 Q3JKH3 Q3JL79 Q3JLJ5 Q3JLK5 Q3JLL7 Q3JLL8 Q3JLL9
Q3JLQ0 Q3JLY2 Q3JM44 Q3JME7 Q3JMP6 Q3JMP9 Q3JMQ0 Q3JMQ1 Q3JMQ2 Q3JMQ3 Q3JMQ4
Q3JMQ7 Q3JMQ8 Q3JMR0 Q3JMR2 Q3JMR3 Q3JMR4 Q3JMR5 Q3JMR6 Q3JMR7 Q3JMR8 Q3JMR9
Q3JMS0 Q3JMS1 Q3JMS2 Q3JMS3 Q3JMS4 Q3JMS5 Q3JMS6 Q3JMS7 Q3JMS8 Q3JMS9 Q3JMT0
Q3JMT1 Q3JMT2 Q3JMT4 Q3JMT5 Q3JMT6 Q3JMT7 Q3JMT8 Q3JMT9 Q3JMU0 Q3JMU4 Q3JMV4
Q3JMW0 Q3JMW1 Q3JMY2 Q3JMZ4 Q3JMZ5 Q3JMZ6 Q3JMZ7 Q3JMZ8 Q3JN00 Q3JN01 Q3JN02
Q3JN03 Q3JN04 Q3JN85 Q3JN88 Q3JNA7 Q3JNB0 Q3JNB1 Q3JND0 Q3JND6 Q3JND9 Q3JNE1
Q3JNE9 Q3JNF2 Q3JNF3 Q3JNF9 Q3JNG0 Q3JNG1 Q3JNG3 Q3JNG4 Q3JNI0 Q3JNM7 Q3JNN1
Q3JNN5 Q3JNR1 Q3JNR2 Q3JNR3 Q3JNR7 Q3JNS0 Q3JNS1 Q3JNS5 Q3JNS6 Q3JNS7 Q3JNS8
Q3JNT6 Q3JNW2 Q3JNW6 Q3JP04 Q3JP05 Q3JP06 Q3JP08 Q3JP10 Q3JP12 Q3JP15 Q3JP21
Q3JP23 Q3JP41 Q3JP81 Q3JP82 Q3JPA3 Q3JPF6 Q3JPG1 Q3JPG2 Q3JPG4 Q3JPG7 Q3JPJ3
Q3JPJ4 Q3JPJ5 Q3JPJ6 Q3JPJ7 Q3JPJ8 Q3JPJ9 Q3JPL2 Q3JPN1 Q3JPQ8 Q3JPR1 Q3JPR4

Q3JPS9 Q3JPT0 Q3JPU4 Q3JPU6 Q3JPX1 Q3JPY2 Q3JPY8 Q3JQ00 Q3JQ06 Q3JQ07 Q3JQ08
Q3JQ09 Q3JQ14 Q3JQ61 Q3JQ63 Q3JQ79 Q3JQ80 Q3JQ86 Q3JQ88 Q3JQA6 Q3JQC6 Q3JQC7
Q3JQC8 Q3JQH7 Q3JQJ0 Q3JQL8 Q3JQN8 Q3JQN9 Q3JQP3 Q3JQQ4 Q3JQS6 Q3JQT6 Q3JQY8
Q3JQZ3 Q3JR15 Q3JR19 Q3JR23 Q3JR26 Q3JR28 Q3JR29 Q3JR30 Q3JR31 Q3JR34 Q3JR43
Q3JR45 Q3JR53 Q3JR65 Q3JR99 Q3JRA0 Q3JRC7 Q3JRC9 Q3JRE2 Q3JRF0 Q3JRF1 Q3JRG5
Q3JRJ2 Q3JRJ4 Q3JRJ5 Q3JRM8 Q3JRN6 Q3JRN7 Q3JRP0 Q3JRP1 Q3JRP8 Q3JRQ0 Q3JRQ1
Q3JRQ3 Q3JRQ5 Q3JRR3 Q3JRR4 Q3JRT1 Q3JRV4 Q3JSR8 Q3JSS7 Q3JST3 Q3JSY7 Q3JSY8
Q3JSY9 Q3JSZ1 Q3JT07 Q3JT08 Q3JT10 Q3JT11 Q3JT13 Q3JT35 Q3JT80 Q3JT87 Q3JTG5
Q3JTT2 Q3JTT5 Q3JU20 Q3JU77 Q3JUB3 Q3JUB4 Q3JUB9 Q3JUC2 Q3JUF3 Q3JUF6 Q3JUF8
Q3JUG4 Q3JUG5 Q3JUI8 Q3JUJ4 Q3JUL7 Q3JUM4 Q3JUM5 Q3JUN4 Q3JUP8 Q3JUW3 Q3JUY1
Q3JUY8 Q3JUY9 Q3JV12 Q3JV16 Q3JV52 Q3JV54 Q3JV55 Q3JV72 Q3JV77 Q3JV86 Q3JVA9
Q3JVB1 Q3JVB6 Q3JVB9 Q3JVC0 Q3JVD0 Q3JVD6 Q3JVD7 Q3JVD8 Q3JVL5 Q3JVV1 Q3JVV3
Q3JVW6 Q3JVY8 Q3JVZ6 Q3JW01 Q3JW68 Q3JW78 Q3JW79 Q3JW82 Q3JW85 Q3JW87 Q3JW88
Q3JW89 Q3JW91 Q3JWH4 Q3JWH7 Q3JWL1 Q3JWL3 Q3JWP7 Q3JWQ8 Q3JWR6 Q3JWR7 Q3JWR8
Q3JWX0 Q3JWY8 Q3JX94 Q3JXA4 Q3JXA9 Q3JXB2 Q3JXC0 Q3JXC1 Q3JXC2 Q3JXD3 Q3JXE1
Q3JXI0 Q3JXI6 Q3JXS0 Q3JXS3 Q3JXT0 Q3JXU7 Q3JXU8 Q3JY08 Q3JY47 Q3JY54 Q3JY64
Q3JY79 Q3JY95 Q3JYD2 Q3JYE2 Q3JYE9 Q3JYG1 Q3JYG3 Q3JYG5 Q3JYG8 Q3JYL4 Q3JYL5
Q3JYL6 Q3JYL7 Q3JYM1 Q3JYM3 Q3JYM6 Q3JYM8 Q3JYN0 Q3JYN3 Q3JYN4 Q3JYN5 Q3JYP3
Q3JYQ0 Q3JYQ4 Q3JYQ5 Q3JYR6 Q3JYS3 Q3JYX9 Q3JZ09 Q3JZ18 Q3JZ19 Q3JZ20 Q3JZ37
Q3JZ45 Q3JZ54 Q3JZ55 Q3JZ62 Q3JZ68 Q3JZ91 Q3JZ94 Q3JZA5 Q3JZB3 Q3JZB4 Q3JZB5
Q3JZB8 Q3JZC4 Q3JZC7 Q3JZC9 Q3JZD6 Q3JZF9 Q3JZG9 Q3JZH3 Q3JZH6 Q3JZI0 Q3JZI2
Q3JZJ4 Q3JZJ6 Q3JZK5 Q3JZL4 Q3JZL8 Q3JZM0 Q3JZM1 Q3JZM2 Q3JZN3 Q3JZP3 Q3JZQ7
Q3JZR3 Q3JZR6 Q3JZS0 Q3JZS3 Q3JZT5 Q3JZT9 Q3JZU0 Q3JZU1 Q3JZU9 Q3JZV0 Q3JZV1
Q3JZV8 Q3JZW0 Q3K003 Q3K004 Q3K010 Q3K011 Q3K013 Q3K014 Q3K020 Q3K028 Q3K029
Q3K035 Q3K046 Q3K051 Q3K076 Q3K081 Q3K093 Q3K0B5 Q3K0C1 Q3K0C3 Q3K0C7 Q3K0C8
Q3K0C9 Q3K0D0 Q3K0D3 Q3K0D4 Q3K0D5 Q3K0D6 Q3K0E0 Q3K0E2 Q3K0E8 Q3K0F4 Q3K0F7
Q3K0F8 Q3K0F9 Q3K0J7 Q3K0K0 Q3K0K1 Q3K0K8 Q3K0K9 Q3K0N2 Q3K0P1 Q3K0P4 Q3K0P8
Q3K0S0 Q3K0S1 Q3K0S8 Q3K0X8 Q3K0Y1 Q3K0Y5 Q3K0Z6 Q3K0Z7 Q3K103 Q3K110 Q3K111
Q3K118 Q3K119 Q3K122 Q3K138 Q3K145 Q3K146 Q3K148 Q3K150 Q3K175 Q3K189 Q3K192
Q3K1A9 Q3K1B8 Q3K1C0 Q3K1C8 Q3K1D6 Q3K1D7 Q3K1E6 Q3K1F5 Q3K1G0 Q3K1H1 Q3K1I2
Q3K1I4 Q3K1I5 Q3K1I7 Q3K1I9 Q3K1J2 Q3K1K2 Q3K1K4 Q3K1K5 Q3K1K8 Q3K1L5 Q3K1L6
Q3K1L7 Q3K1M7 Q3K1N1 Q3K1P7 Q3K1Q7 Q3K1Q9 Q3K1S8 Q3K1T4 Q3K1T9 Q3K1U0 Q3K1U2
Q3K1U3 Q3K1U4 Q3K1U6 Q3K1V5 Q3K1V6 Q3K1W9 Q3K1X2 Q3K1Y2 Q3K1Z4 Q3K203 Q3K204
Q3K2B0 Q3K2D0 Q3K2D4 Q3K2D6 Q3K2D9 Q3K2E1 Q3K2E5 Q3K2F4 Q3K2F6 Q3K2G4 Q3K2M3
Q3K2M4 Q3K2M6 Q3K2N4 Q3K2N8 Q3K2P4 Q3K2Q5 Q3K2R2 Q3K2R6 Q3K2R9 Q3K2S4 Q3K2X2
Q3K2X3 Q3K2X6 Q3K2Y2 Q3K2Y7 Q3K301 Q3K302 Q3K306 Q3K307 Q3K326 Q3K365 Q3K367
Q3K368 Q3K378 Q3K379 Q3K380 Q3K383 Q3K394 Q3K395 Q3K396 Q3K3A6 Q3K3A8 Q3K3A9
Q3K3B0 Q3K3B1 Q3K3F8 Q3K3G5 Q3K3G9 Q3K3H0 Q3K3K1 Q3K3L1 Q3K3L2 Q3K3L4 Q3K3L9
Q3K3N3 Q3K3Q0 Q3K3Q3 Q3K3Q4 Q3K3R1 Q3K3R5 Q3K3S2 Q3K3S3 Q3K3S4 Q3K3S6 Q3K3S9
Q3K3T1 Q3K3T2 Q3K3T3 Q3K3T4 Q3K3U3 Q3K3U4 Q3K3U5 Q3K3U6 Q3K3U7 Q3K3U8 Q3K3V0
Q3K3V1 Q3K3V2 Q3K3V3 Q3K3V4 Q3K3V5 Q3K3V6 Q3K3V7 Q3K3V8 Q3K3V9 Q3K3W0 Q3K3W1
Q3K3W2 Q3K3W3 Q3K3W4 Q3K3W5 Q3K3W6 Q3K3W8 Q3K3W9 Q3K3X0 Q3K3X8 Q3K3Z4 Q3K3Z7
Q3K400 Q3K403 Q3K407 Q3K419 Q3K425 Q3K426 Q3K429 Q3K430 Q3K431 Q3K4B0 Q3K4B8
Q3K4G4 Q3K4L7 Q3K4L8 Q3K4M0 Q3K4M3 Q3K4M4 Q3K4M6 Q3K4M8 Q3K4N0 Q3K4P7 Q3K4R6

Q3K4R8 Q3K4S1 Q3K4S8 Q3K4T0 Q3K4U8 Q3K4Y2 Q3K4Z1 Q3K4Z8 Q3K535 Q3K539 Q3K559
Q3K566 Q3K569 Q3K587 Q3K588 Q3K594 Q3K595 Q3K5A3 Q3K5A5 Q3K5A6 Q3K5B6 Q3K5B7
Q3K5C0 Q3K5D7 Q3K5E7 Q3K5E8 Q3K5F2 Q3K5H3 Q3K5H4 Q3K5H5 Q3K5K9 Q3K5N4 Q3K5P1
Q3K5P2 Q3K5Q7 Q3K5Q9 Q3K5R0 Q3K5S0 Q3K5S1 Q3K5S5 Q3K5S7 Q3K5S9 Q3K5T0 Q3K5T1
Q3K5T2 Q3K5U8 Q3K5V3 Q3K5V4 Q3K5V8 Q3K5W5 Q3K5W6 Q3K5W7 Q3K5W9 Q3K5X2 Q3K5X4
Q3K5X7 Q3K5X8 Q3K5X9 Q3K5Y0 Q3K5Y1 Q3K5Y2 Q3K5Y3 Q3K5Y4 Q3K5Y5 Q3K5Y7 Q3K5Y8
Q3K5Y9 Q3K5Z0 Q3K5Z1 Q3K5Z2 Q3K5Z3 Q3K5Z4 Q3K5Z5 Q3K5Z6 Q3K5Z7 Q3K5Z8 Q3K5Z9
Q3K600 Q3K602 Q3K603 Q3K604 Q3K605 Q3K606 Q3K607 Q3K609 Q3K610 Q3K611 Q3K612
Q3K613 Q3K647 Q3K652 Q3K660 Q3K662 Q3K675 Q3K692 Q3K693 Q3K696 Q3K6A2 Q3K6A3
Q3K6A4 Q3K6B0 Q3K6B5 Q3K6B7 Q3K6C0 Q3K6C1 Q3K6G9 Q3K6H7 Q3K6J0 Q3K6K7 Q3K6K8
Q3K6K9 Q3K6L0 Q3K6L2 Q3K6L5 Q3K6L8 Q3K6P3 Q3K6P4 Q3K6Q4 Q3K6Q5 Q3K6Q6 Q3K6R5
Q3K6T1 Q3K6W2 Q3K6W3 Q3K6W5 Q3K6W8 Q3K6W9 Q3K6Y7 Q3K6Y9 Q3K705 Q3K723 Q3K724
Q3K731 Q3K732 Q3K735 Q3K736 Q3K742 Q3K745 Q3K746 Q3K795 Q3K7A5 Q3K7A8 Q3K7A9
Q3K7B2 Q3K7B3 Q3K7B6 Q3K7B7 Q3K7C0 Q3K7C3 Q3K7C5 Q3K7D0 Q3K7D4 Q3K7D5 Q3K7I5
Q3K7J7 Q3K7J8 Q3K7K0 Q3K7L0 Q3K7L5 Q3K7L6 Q3K7N4 Q3K7U0 Q3K7U6 Q3K7V1 Q3K7V6
Q3K7V7 Q3K7V8 Q3K7V9 Q3K7W0 Q3K7W9 Q3K7X0 Q3K7X5 Q3K885 Q3K886 Q3K888 Q3K892
Q3K8D3 Q3K8D5 Q3K8E9 Q3K8H1 Q3K8J3 Q3K8J6 Q3K8K5 Q3K8K7 Q3K8L5 Q3K8R4 Q3K8R9
Q3K8S7 Q3K8T6 Q3K8T8 Q3K8U2 Q3K8U4 Q3K8V7 Q3K918 Q3K932 Q3K935 Q3K955 Q3K965
Q3K999 Q3K9A2 Q3K9C8 Q3K9C9 Q3K9D1 Q3K9F5 Q3K9G4 Q3K9H0 Q3K9R2 Q3K9V5 Q3K9W7
Q3K9W8 Q3K9W9 Q3K9X0 Q3K9Y2 Q3K9Y3 Q3K9Y9 Q3KA28 Q3KA70 Q3KA77 Q3KA78 Q3KA79
Q3KA84 Q3KA85 Q3KAS8 Q3KAU3 Q3KBG7 Q3KBL2 Q3KBQ0 Q3KC83 Q3KC85 Q3KCB2 Q3KCJ2
Q3KD08 Q3KD65 Q3KD78 Q3KDV1 Q3KDW0 Q3KE62 Q3KE68 Q3KE77 Q3KE79 Q3KEL1 Q3KEX6
Q3KEX7 Q3KEX8 Q3KEX9 Q3KEY0 Q3KEY2 Q3KEY9 Q3KF16 Q3KF21 Q3KF35 Q3KF38 Q3KFA3
Q3KFA5 Q3KFA7 Q3KFB2 Q3KFC5 Q3KFE1 Q3KFE4 Q3KFG8 Q3KFI7 Q3KFI8 Q3KFI9 Q3KFJ1
Q3KFK9 Q3KFL0 Q3KFL8 Q3KFM0 Q3KFM3 Q3KFP5 Q3KFR5 Q3KFR7 Q3KFT0 Q3KFT8 Q3KFU6
Q3KG28 Q3KG65 Q3KGD1 Q3KGE4 Q3KGF8 Q3KGH1 Q3KGH4 Q3KGI9 Q3KGL4 Q3KGV8 Q3KGY4
Q3KGZ5 Q3KH11 Q3KH16 Q3KH18 Q3KH22 Q3KH38 Q3KH79 Q3KH83 Q3KH84 Q3KH85 Q3KH86
Q3KH90 Q3KH94 Q3KH95 Q3KH98 Q3KHA6 Q3KHA9 Q3KHB0 Q3KHB1 Q3KHB2 Q3KHB4 Q3KHC5
Q3KHD6 Q3KHE3 Q3KHE4 Q3KHF3 Q3KHJ1 Q3KHJ2 Q3KHJ3 Q3KHJ4 Q3KHJ9 Q3KHL4 Q3KHL6
Q3KHL7 Q3KHL9 Q3KHM4 Q3KHM8 Q3KHN0 Q3KHQ2 Q3KHV3 Q3KHY6 Q3KHZ1 Q3KHZ2 Q3KHZ3
Q3KHZ4 Q3KI09 Q3KI17 Q3KI22 Q3KI26 Q3KI27 Q3KI28 Q3KI36 Q3KI56 Q3KI64 Q3KI80
Q3KI81 Q3KI82 Q3KI83 Q3KI84 Q3KI86 Q3KI88 Q3KI89 Q3KI92 Q3KI98 Q3KIA0 Q3KIA1
Q3KIA5 Q3KIA8 Q3KIF2 Q3KIJ1 Q3KIP5 Q3KIR0 Q3KIS7 Q3KIS8 Q3KIT1 Q3KIT2 Q3KIU3
Q3KIU9 Q3KIV0 Q3KIV1 Q3KIW4 Q3KIW5 Q3KIX5 Q3KIX7 Q3KIX8 Q3KIY4 Q3KIY5 Q3KIY9
Q3KIZ0 Q3KIZ7 Q3KIZ8 Q3KJ20 Q3KJ28 Q3KJ68 Q3KJ97 Q3KJA1 Q3KJA2 Q3KJB1 Q3KJB3
Q3KJB6 Q3KJC5 Q3KJC8 Q3KJC9 Q3KJE0 Q3KJE5 Q3KJE6 Q3KJG0 Q3KJG6 Q3KJH6 Q3KJH9
Q3KJI5 Q3KJI6 Q3KJI8 Q3KJI9 Q3KJJ3 Q3KJK7 Q3KJK8 Q3KJP4 Q3KJP5 Q3KJQ0 Q3KJQ5
Q3KK58 Q3KK59 Q3KKA7 Q3KKE0 Q3KKE2 Q3KKE6 Q3KKF2 Q3KKF3 Q3KKF9 Q3KKG1 Q3KKH0
Q3KKH8 Q3KKJ6 Q3KKJ7 Q3KKJ9 Q3KKK3 Q3KKK4 Q3KKK5 Q3KKK7 Q3KKK8 Q3KKL0 Q3KKL8
Q3KKM5 Q3KKN1 Q3KKN7 Q3KKN8 Q3KKN9 Q3KKP0 Q3KKP1 Q3KKP2 Q3KKP3 Q3KKQ0 Q3KKQ1
Q3KKQ5 Q3KKQ6 Q3KKQ7 Q3KKQ8 Q3KKR0 Q3KKR1 Q3KKS0 Q3KKS6 Q3KKT4 Q3KKU4 Q3KKU6
Q3KKW2 Q3KKW5 Q3KKX2 Q3KKY7 Q3KKY8 Q3KKY9 Q3KKZ1 Q3KKZ6 Q3KL01 Q3KL14 Q3KL15
Q3KL16 Q3KL17 Q3KL32 Q3KL45 Q3KL61 Q3KL68 Q3KL69 Q3KL70 Q3KL78 Q3KL88 Q3KL90
Q3KLB1 Q3KLB6 Q3KLC2 Q3KLD9 Q3KLF4 Q3KLF5 Q3KLG7 Q3KLG9 Q3KLH0 Q3KLH1 Q3KLH2

Q3KLH3 Q3KLH4 Q3KLH5 Q3KLH6 Q3KLH8 Q3KLH9 Q3KLI0 Q3KLI1 Q3KLI2 Q3KLI3 Q3KLI4
Q3KLI5 Q3KLI6 Q3KLI8 Q3KLI9 Q3KLJ0 Q3KLJ1 Q3KLJ5 Q3KLJ6 Q3KLJ7 Q3KLJ9 Q3KLK5
Q3KLL2 Q3KLM3 Q3KLM8 Q3KLN5 Q3KLN6 Q3KLN9 Q3KLP1 Q3KLP3 Q3KLP4 Q3KLP6 Q3KLQ4
Q3KLR1 Q3KLR2 Q3KLR3 Q3KLR4 Q3KLR6 Q3KLR8 Q3KLR9 Q3KLS0 Q3KLS2 Q3KLS9 Q3KLT3
Q3KLT4 Q3KLT5 Q3KLU7 Q3KLV7 Q3KLV8 Q3KLV9 Q3KLW0 Q3KLW3 Q3KLW5 Q3KLX6 Q3KLX8
Q3KLY3 Q3KLY7 Q3KLY8 Q3KLZ0 Q3KLZ2 Q3KLZ5 Q3KM04 Q3KM05 Q3KM09 Q3KM11 Q3KM12
Q3KM16 Q3KM17 Q3KM24 Q3KM28 Q3KM30 Q3KM31 Q3KM32 Q3KM33 Q3KM39 Q3KM40 Q3KM43
Q3KM44 Q3KM45 Q3KM46 Q3KM47 Q3KM48 Q3KM49 Q3KM60 Q3KM61 Q3KM65 Q3KM70 Q3KM75
Q3KM80 Q3KM90 Q3KM93 Q3KMC2 Q3KME4 Q3KME9 Q3KMF1 Q3KMF2 Q3KMF3 Q3KMG5 Q3KMG9
Q3KMH4 Q3KMH8 Q3KMH9 Q3KMJ6 Q3KML9 Q3KMM3 Q3KMP1 Q3KMP3 Q3KMP4 Q3KMQ8 Q3KMQ9
Q3KMS4 Q3KMS5 Q3KMT5 Q3KMU3 Q3KMU5 Q3KMU7 Q3KMV0 Q3KMV9 Q3KMW4 Q3KMX6 Q3KMY1
Q3KMY3 Q3KMZ0 Q3KMZ2 Q3KMZ3 Q3KMZ4 Q3KMZ5 Q3KMZ6 Q3KMZ9 Q3KN00 Q3KN03 Q3KN14
Q3KN18 Q3KN19 Q3KN20 Q3KN66 Q3KNI6 Q3KNJ2 Q3KNL4 Q3KNS6 Q3KNT7 Q3KNV8 Q3KNW1
Q3KNY5 Q3KP31 Q3KP44 Q3KPK4 Q3KPL3 Q3KPN2 Q3KPR1 Q3KPR5 Q3KPT0 Q3KPT3 Q3KPT5
Q3KPU0 Q3KPU6 Q3KPU8 Q3KPW1 Q3KPW4 Q3KPZ2 Q3KQ10 Q3KQ35 Q3KQ45 Q3KQ62 Q3KQ77
Q3KQ80 Q3KQ85 Q3KQB6 Q3KQD1 Q3KQF0 Q3KQF4 Q3KQP7 Q3KQV3 Q3KQV9 Q3KQW6 Q3KQW7
Q3KR53 Q3KR54 Q3KR64 Q3KR66 Q3KR73 Q3KRA6 Q3KRC5 Q3KRD0 Q3KRD8 Q3KRE6 Q3KRF3
Q3KRG3 Q3KSM3 Q3KSN9 Q3KSP1 Q3KSP3 Q3KSP4 Q3KSP5 Q3KSQ2 Q3KSR3 Q3KSR5 Q3KSR8
Q3KSR9 Q3KSS1 Q3KSS4 Q3KSS5 Q3KSS7 Q3KST0 Q3KST1 Q3KST2 Q3KST3 Q3KST5 Q3KST7
Q3KSU1 Q3KSU3 Q3KSU5 Q3KSU7 Q3KSV2 Q3KSV5 Q3KVN1 Q3L181 Q3L1C9 Q3L6L4 Q3L7F5
Q3LFX6 Q3LH63 Q3LHL9 Q3LHN0 Q3LHN1 Q3LHN2 Q3LI54 Q3LI58 Q3LI59 Q3LI60 Q3LI61
Q3LI62 Q3LI63 Q3LI64 Q3LI66 Q3LI67 Q3LI68 Q3LI70 Q3LI72 Q3LI73 Q3LI76 Q3LI77
Q3LI81 Q3LI83 Q3LIE5 Q3LRV4 Q3LSM4 Q3LSN4 Q3LSN5 Q3LSS0 Q3LT96 Q3LTE0 Q3LU38
Q3LU39 Q3LU40 Q3LU41 Q3LUD3 Q3LXA3 Q3LXA7 Q3M1B1 Q3M1N6 Q3M244 Q3M356 Q3M3A3
Q3M3B9 Q3M3F2 Q3M3F3 Q3M3H8 Q3M3I0 Q3M3M2 Q3M3N7 Q3M3N9 Q3M3Q5 Q3M3R4 Q3M3Y3
Q3M438 Q3M446 Q3M450 Q3M454 Q3M455 Q3M473 Q3M482 Q3M4F6 Q3M4J7 Q3M4K7 Q3M4P0
Q3M4P1 Q3M4S2 Q3M4T0 Q3M4U9 Q3M4V2 Q3M4Y1 Q3M4Y3 Q3M4Z2 Q3M502 Q3M504 Q3M534
Q3M542 Q3M546 Q3M5C8 Q3M5C9 Q3M5D0 Q3M5D2 Q3M5D3 Q3M5D7 Q3M5M3 Q3M5Q6 Q3M5R7
Q3M5W4 Q3M5Z3 Q3M614 Q3M629 Q3M644 Q3M672 Q3M674 Q3M687 Q3M691 Q3M698 Q3M6A1
Q3M6B5 Q3M6C3 Q3M6G3 Q3M6H4 Q3M6H7 Q3M6L1 Q3M6L2 Q3M6L5 Q3M6M1 Q3M6N4 Q3M6R3
Q3M6S3 Q3M6T2 Q3M6T3 Q3M6Z4 Q3M704 Q3M708 Q3M709 Q3M710 Q3M712 Q3M717 Q3M725
Q3M726 Q3M727 Q3M745 Q3M769 Q3M782 Q3M785 Q3M790 Q3M7E8 Q3M7F0 Q3M7G3 Q3M7I1
Q3M7I2 Q3M7J6 Q3M7J7 Q3M7K5 Q3M7N0 Q3M7N8 Q3M7P3 Q3M7Q7 Q3M7Y0 Q3M7Y8 Q3M7Z9
Q3M859 Q3M882 Q3M890 Q3M892 Q3M8C8 Q3M8F0 Q3M8H2 Q3M8N0 Q3M8N2 Q3M8N9 Q3M8Q4
Q3M8R1 Q3M8T9 Q3M8X6 Q3M925 Q3M929 Q3M998 Q3M9A3 Q3M9A4 Q3M9B9 Q3M9C9 Q3M9E1
Q3M9G1 Q3M9G2 Q3M9J6 Q3M9K5 Q3M9L4 Q3M9M0 Q3M9Q2 Q3M9Q8 Q3M9U1 Q3MA16 Q3MA17
Q3MA18 Q3MA19 Q3MA22 Q3MA33 Q3MA53 Q3MA58 Q3MA60 Q3MA89 Q3MAA1 Q3MAB0 Q3MAF5
Q3MAI6 Q3MAK6 Q3MAL4 Q3MAN6 Q3MAN8 Q3MAP3 Q3MAP6 Q3MAP7 Q3MAP8 Q3MAP9 Q3MAQ3
Q3MAQ7 Q3MAV1 Q3MAV8 Q3MAV9 Q3MAW5 Q3MAX6 Q3MAZ7 Q3MB19 Q3MB82 Q3MB98 Q3MB99
Q3MBA3 Q3MBD2 Q3MBD6 Q3MBD8 Q3MBE2 Q3MBE6 Q3MBF3 Q3MBF4 Q3MBG0 Q3MBG1 Q3MBH6
Q3MBJ4 Q3MBJ9 Q3MBL3 Q3MBM3 Q3MBM5 Q3MBQ2 Q3MBU3 Q3MBV3 Q3MBZ3 Q3MBZ7 Q3MC34
Q3MC52 Q3MC53 Q3MC57 Q3MC63 Q3MC76 Q3MC79 Q3MCB5 Q3MCD0 Q3MCD2 Q3MCE0 Q3MCG0
Q3MCI4 Q3MCK2 Q3MCR5 Q3MCS7 Q3MCT9 Q3MCZ7 Q3MCZ8 Q3MD34 Q3MD38 Q3MD39 Q3MD46
Q3MD78 Q3MD91 Q3MDJ1 Q3MDJ2 Q3MDK7 Q3MDL2 Q3MDM2 Q3MDM3 Q3MDM4 Q3MDM5 Q3MDM6

Q3MDN0 Q3MDN5 Q3MDN9 Q3MDP1 Q3MDT4 Q3MDY8 Q3ME15 Q3ME71 Q3MEA0 Q3MEA8 Q3MEA9
Q3MED4 Q3MEE5 Q3MEF4 Q3MEF5 Q3MEH7 Q3MEJ6 Q3MEJ8 Q3MEM9 Q3MEN7 Q3MEN8 Q3MEV6
Q3MEV7 Q3MEX6 Q3MEZ3 Q3MF19 Q3MF32 Q3MF37 Q3MF40 Q3MF82 Q3MF83 Q3MF89 Q3MF91
Q3MF92 Q3MF93 Q3MF94 Q3MF96 Q3MF97 Q3MF98 Q3MF99 Q3MFA0 Q3MFA1 Q3MFA4 Q3MFA5
Q3MFA6 Q3MFA7 Q3MFA8 Q3MFA9 Q3MFB0 Q3MFB1 Q3MFB2 Q3MFB3 Q3MFB4 Q3MFB5 Q3MFB6
Q3MFB7 Q3MFB8 Q3MFB9 Q3MFC0 Q3MFC1 Q3MFH1 Q3MFI4 Q3MFM3 Q3MFQ5 Q3MFQ8 Q3MFQ9
Q3MFR5 Q3MFS2 Q3MFW5 Q3MFX2 Q3MFY8 Q3MG27 Q3MG30 Q3MG40 Q3MG53 Q3MG63 Q3MG79
Q3MG81 Q3MG83 Q3MGA0 Q3MGA3 Q3MGD1 Q3MGD2 Q3MGE7 Q3MGH8 Q3MGI4 Q3MGJ7 Q3MGL3
Q3MGL8 Q3MGN4 Q3MGR4 Q3MGT8 Q3MGW8 Q3MGX7 Q3MGZ5 Q3MH17 Q3MH53 Q3MHA9 Q3MHC2
Q3MHE2 Q3MHE4 Q3MHF7 Q3MHF8 Q3MHG6 Q3MHH0 Q3MHH1 Q3MHH2 Q3MHH3 Q3MHH4 Q3MHH5
Q3MHI7 Q3MHI8 Q3MHJ0 Q3MHL3 Q3MHL4 Q3MHL6 Q3MHL7 Q3MHL8 Q3MHL9 Q3MHN0 Q3MHN7
Q3MHN8 Q3MHP0 Q3MHP1 Q3MHP3 Q3MHP5 Q3MHQ0 Q3MHR2 Q3MHR5 Q3MHS2 Q3MHT3 Q3MHT4
Q3MHU3 Q3MHV6 Q3MHW0 Q3MHX1 Q3MHX3 Q3MHY6 Q3MHY8 Q3MHZ2 Q3MI00 Q3MI03 Q3MIF2
Q3MIF4 Q3MIS6 Q3MIV0 Q3MJ40 Q3MJ62 Q3MJK5 Q3MK94 Q3MKA1 Q3MKB3 Q3MKB9 Q3MKM6
Q3MKP9 Q3MNT0 Q3MSM3 Q3MSM4 Q3MSQ8 Q3MSU9 Q3S2T9 Q3S2U0 Q3S2U1 Q3S2U2 Q3S2U4
Q3S2U6 Q3S3T7 Q3S405 Q3S4B7 Q3S664 Q3S8P6 Q3S8P9 Q3S8Q4 Q3S8R0 Q3SA46 Q3SA47
Q3SA48 Q3SA49 Q3SB08 Q3SB09 Q3SB10 Q3SB11 Q3SB12 Q3SB13 Q3SB14 Q3SB15 Q3SEK0
Q3SEK2 Q3SER0 Q3SEU5 Q3SEU6 Q3SEU7 Q3SEU8 Q3SEU9 Q3SEX4 Q3SEY2 Q3SEZ6 Q3SF18
Q3SF19 Q3SF20 Q3SF21 Q3SF39 Q3SF55 Q3SF56 Q3SF80 Q3SFE8 Q3SFF1 Q3SFF3 Q3SFG3
Q3SFH1 Q3SFR3 Q3SFR4 Q3SFR5 Q3SFR7 Q3SFS1 Q3SFS3 Q3SFS4 Q3SFS6 Q3SFT5 Q3SFU6
Q3SFU7 Q3SFW2 Q3SFY2 Q3SFZ9 Q3SG10 Q3SG11 Q3SG32 Q3SG43 Q3SG58 Q3SG61 Q3SG64
Q3SG65 Q3SG76 Q3SG95 Q3SGA3 Q3SGB2 Q3SGB3 Q3SGC1 Q3SGC2 Q3SGC3 Q3SGD0 Q3SGD1
Q3SGD2 Q3SGD3 Q3SGE9 Q3SGF0 Q3SGF7 Q3SGJ3 Q3SGK8 Q3SGL1 Q3SGR3 Q3SGR5 Q3SGS1
Q3SGS2 Q3SGS9 Q3SGT0 Q3SGT1 Q3SGT3 Q3SGT8 Q3SGU8 Q3SGU9 Q3SGX4 Q3SGX5 Q3SGZ3
Q3SH10 Q3SH11 Q3SH24 Q3SH26 Q3SH29 Q3SH31 Q3SH33 Q3SH34 Q3SH36 Q3SH50 Q3SH52
Q3SH53 Q3SH61 Q3SH62 Q3SH73 Q3SH75 Q3SH78 Q3SH79 Q3SH81 Q3SH82 Q3SH83 Q3SH93
Q3SH94 Q3SH96 Q3SHA3 Q3SHA4 Q3SHA5 Q3SHE4 Q3SHE7 Q3SHI2 Q3SHJ2 Q3SHL1 Q3SHL2
Q3SHL3 Q3SHL7 Q3SHL8 Q3SHL9 Q3SHM0 Q3SHN1 Q3SHP2 Q3SHP3 Q3SHS3 Q3SHS6 Q3SHT1
Q3SHT2 Q3SHT4 Q3SHT9 Q3SHU1 Q3SI16 Q3SI24 Q3SI29 Q3SI46 Q3SI67 Q3SI68 Q3SI69
Q3SI97 Q3SI98 Q3SI99 Q3SIB0 Q3SIB2 Q3SID4 Q3SIF9 Q3SIH8 Q3SIK7 Q3SIL8 Q3SIM2
Q3SIM9 Q3SIN3 Q3SIN4 Q3SIN5 Q3SIN9 Q3SIQ5 Q3SIR0 Q3SIR1 Q3SIR6 Q3SIR8 Q3SIV9
Q3SJE0 Q3SJH5 Q3SJH9 Q3SJI1 Q3SJI4 Q3SJP0 Q3SJQ7 Q3SJR2 Q3SJR5 Q3SJS2 Q3SJS7
Q3SJT0 Q3SJU1 Q3SJU8 Q3SJW8 Q3SK28 Q3SK29 Q3SK30 Q3SK31 Q3SK32 Q3SK34 Q3SK37
Q3SK38 Q3SK65 Q3SK67 Q3SK69 Q3SK74 Q3SK77 Q3SK83 Q3SK85 Q3SK88 Q3SK91 Q3SKD9
Q3SKF1 Q3SKF2 Q3SKG2 Q3SKG3 Q3SKG4 Q3SKG8 Q3SKG9 Q3SKH1 Q3SKH7 Q3SKJ2 Q3SKJ3
Q3SKM1 Q3SKM7 Q3SKN6 Q3SKN7 Q3SKN8 Q3SKN9 Q3SKP1 Q3SKP9 Q3SKU2 Q3SKW8 Q3SKW9
Q3SKX1 Q3SKX2 Q3SKX3 Q3SKX4 Q3SKX5 Q3SKX9 Q3SKY5 Q3SL02 Q3SL03 Q3SL40 Q3SL45
Q3SL47 Q3SL57 Q3SL58 Q3SL66 Q3SL69 Q3SL73 Q3SL74 Q3SL86 Q3SL94 Q3SLA9 Q3SLD3
Q3SLE3 Q3SLF4 Q3SLI4 Q3SLI5 Q3SLJ3 Q3SLJ6 Q3SLJ8 Q3SLK0 Q3SLK1 Q3SLL2 Q3SLM3
Q3SLM4 Q3SLM5 Q3SLM6 Q3SLM7 Q3SLM8 Q3SLN0 Q3SLN1 Q3SLN2 Q3SLN3 Q3SLN4 Q3SLN5
Q3SLN6 Q3SLN7 Q3SLN8 Q3SLN9 Q3SLP0 Q3SLP1 Q3SLP2 Q3SLP3 Q3SLP4 Q3SLP5 Q3SLP6
Q3SLP7 Q3SLP8 Q3SLP9 Q3SLQ0 Q3SLQ1 Q3SLQ2 Q3SLQ3 Q3SLQ4 Q3SLQ5 Q3SLQ6 Q3SLQ7
Q3SLQ8 Q3SLQ9 Q3SLR3 Q3SLR4 Q3SLR6 Q3SLR9 Q3SLS2 Q3SLX9 Q3SLY0 Q3SLY3 Q3SLY4
Q3SM23 Q3SM28 Q3SM39 Q3SM40 Q3SM42 Q3SM51 Q3SM68 Q3SM81 Q3SM89 Q3SMA6 Q3SMB4

Q3SMB7 Q3SMB8 Q3SMC1 Q3SMC6 Q3SMD2 Q3SMH0 Q3SMH1 Q3SMH2 Q3SMH5 Q3SMI1 Q3SMI2
Q3SMI5 Q3SMK0 Q3SMK1 Q3SMP9 Q3SMQ9 Q3SMR4 Q3SMS3 Q3SMS6 Q3SMT7 Q3SMV0 Q3SMV1
Q3SN20 Q3SN22 Q3SN24 Q3SN25 Q3SN27 Q3SN28 Q3SN92 Q3SN93 Q3SNB4 Q3SNB8 Q3SNC6
Q3SNG2 Q3SNK1 Q3SNS6 Q3SNS9 Q3SNT2 Q3SNT5 Q3SNU3 Q3SNU7 Q3SNV3 Q3SNV5 Q3SNV6
Q3SNV7 Q3SNV8 Q3SNX3 Q3SNY6 Q3SNZ1 Q3SP09 Q3SP12 Q3SP13 Q3SP14 Q3SP44 Q3SP50
Q3SP51 Q3SPC5 Q3SPE0 Q3SPE5 Q3SPG4 Q3SPH2 Q3SPJ0 Q3SPJ6 Q3SPL2 Q3SPM6 Q3SPM7
Q3SPQ7 Q3SPS0 Q3SPS2 Q3SPT8 Q3SPW7 Q3SPZ6 Q3SQ36 Q3SQ43 Q3SQ46 Q3SQ49 Q3SQJ5
Q3SQN5 Q3SQN7 Q3SQP4 Q3SQS3 Q3SR08 Q3SR12 Q3SR25 Q3SR28 Q3SR29 Q3SR31 Q3SR35
Q3SR45 Q3SRA4 Q3SRA8 Q3SRB0 Q3SRB3 Q3SRB6 Q3SRC0 Q3SRD1 Q3SRD2 Q3SRD3 Q3SRG6
Q3SRG9 Q3SRH2 Q3SRH3 Q3SRH4 Q3SRH5 Q3SRH8 Q3SRI7 Q3SRI8 Q3SRJ3 Q3SRJ4 Q3SRJ8
Q3SRK3 Q3SRM3 Q3SRN1 Q3SRN4 Q3SRQ5 Q3SRQ7 Q3SRR6 Q3SRS3 Q3SRS9 Q3SRU5 Q3SRU9
Q3SRV3 Q3SRV4 Q3SRV7 Q3SRV8 Q3SRX8 Q3SRY6 Q3SRY8 Q3SRY9 Q3SRZ7 Q3SRZ8 Q3SS04
Q3SS12 Q3SS13 Q3SS37 Q3SS79 Q3SS83 Q3SS86 Q3SS92 Q3SSH7 Q3SSL6 Q3SSM7 Q3SSN1
Q3SSN8 Q3SSP1 Q3SSP2 Q3SSP4 Q3SSP7 Q3SSQ8 Q3SSS2 Q3SSU1 Q3SSU2 Q3SSU3 Q3SSU4
Q3SSU5 Q3SSU7 Q3SSU8 Q3SSU9 Q3SSV0 Q3SSV1 Q3SSV2 Q3SSV3 Q3SSV4 Q3SSV5 Q3SSV6
Q3SSV7 Q3SSV8 Q3SSV9 Q3SSW0 Q3SSW1 Q3SSW2 Q3SSW3 Q3SSW4 Q3SSW5 Q3SSW6 Q3SSW7
Q3SSW8 Q3SSW9 Q3SSX0 Q3SSX1 Q3SSX9 Q3SSY0 Q3SSY1 Q3SSY2 Q3SSY3 Q3SSY4 Q3ST00
Q3ST14 Q3ST22 Q3ST24 Q3ST43 Q3ST46 Q3ST50 Q3STA1 Q3STA8 Q3STB0 Q3STC2 Q3STP0
Q3STQ8 Q3STR7 Q3STS5 Q3STS7 Q3STT0 Q3STT6 Q3STY0 Q3STZ0 Q3STZ6 Q3SU01 Q3SU08
Q3SU34 Q3SUR0 Q3SUS4 Q3SUZ0 Q3SUZ1 Q3SUZ5 Q3SV02 Q3SV21 Q3SV33 Q3SV39 Q3SV41
Q3SVB0 Q3SVC7 Q3SVD0 Q3SVD4 Q3SVE4 Q3SVF3 Q3SVF4 Q3SVH3 Q3SVH7 Q3SVI0 Q3SVI1
Q3SVI2 Q3SVI4 Q3SVI5 Q3SVJ8 Q3SVK4 Q3SVK5 Q3SVL8 Q3SVM8 Q3SVM9 Q3SVN6 Q3SVP3
Q3SVP7 Q3SVQ1 Q3SVQ6 Q3SVQ7 Q3SVS4 Q3SVW2 Q3SVZ7 Q3SW25 Q3SW26 Q3SW27 Q3SW30
Q3SW60 Q3SW64 Q3SW65 Q3SW71 Q3SW72 Q3SW76 Q3SW78 Q3SW79 Q3SW80 Q3SWE0 Q3SWE4
Q3SWE6 Q3SWE7 Q3SWE9 Q3SWF0 Q3SWF1 Q3SWF2 Q3SWF3 Q3SWF7 Q3SWG7 Q3SWG9 Q3SWH0
Q3SWH1 Q3SWH5 Q3SWH6 Q3SWH7 Q3SWI2 Q3SWJ8 Q3SWK5 Q3SWK6 Q3SWK7 Q3SWK8 Q3SWL5
Q3SWL7 Q3SWL9 Q3SWM9 Q3SWN3 Q3SWP3 Q3SWP5 Q3SWP6 Q3SWP7 Q3SWP8 Q3SWP9 Q3SWQ2
Q3SWQ3 Q3SWQ7 Q3SWS8 Q3SWT1 Q3SWT4 Q3SWT6 Q3SWU0 Q3SWU3 Q3SWU4 Q3SWX2 Q3SWX7
Q3SWX8 Q3SWX9 Q3SWY1 Q3SWY3 Q3SWY6 Q3SWZ4 Q3SWZ6 Q3SWZ7 Q3SWZ8 Q3SX04 Q3SX07
Q3SX13 Q3SX14 Q3SX21 Q3SX22 Q3SX24 Q3SX30 Q3SX40 Q3SX41 Q3SX43 Q3SX44 Q3SXD3
Q3SXM0 Q3SXR2 Q3SXZ3 Q3SXZ7 Q3SY00 Q3SY05 Q3SY46 Q3SY52 Q3SY56 Q3SY69 Q3SY84
Q3SYA9 Q3SYB3 Q3SYF9 Q3SYK4 Q3SYR3 Q3SYS4 Q3SYS7 Q3SYS9 Q3SYT1 Q3SYT7 Q3SYU1
Q3SYU2 Q3SYU6 Q3SYV7 Q3SYW0 Q3SYW5 Q3SYW6 Q3SYW9 Q3SYX3 Q3SYX6 Q3SYX9 Q3SYY7
Q3SYZ6 Q3SYZ8 Q3SYZ9 Q3SZ07 Q3SZ12 Q3SZ18 Q3SZ19 Q3SZ21 Q3SZ25 Q3SZ31 Q3SZ40
Q3SZ43 Q3SZ44 Q3SZ52 Q3SZ54 Q3SZ55 Q3SZ60 Q3SZ62 Q3SZ63 Q3SZ65 Q3SZ68 Q3SZ70
Q3SZ71 Q3SZ76 Q3SZ84 Q3SZ85 Q3SZA0 Q3SZA5 Q3SZB8 Q3SZC0 Q3SZC2 Q3SZD4 Q3SZD5
Q3SZE2 Q3SZE5 Q3SZE9 Q3SZF0 Q3SZF3 Q3SZF8 Q3SZG8 Q3SZH6 Q3SZI0 Q3SZI8 Q3SZJ0
Q3SZJ5 Q3SZJ9 Q3SZK4 Q3SZL6 Q3SZL8 Q3SZM1 Q3SZM5 Q3SZM6 Q3SZM9 Q3SZN0 Q3SZN5
Q3SZP0 Q3SZP7 Q3SZP8 Q3SZR0 Q3SZR5 Q3SZR8 Q3SZT7 Q3SZU0 Q3SZU4 Q3SZV0 Q3SZV3
Q3SZW1 Q3SZW6 Q3SZX4 Q3SZX7 Q3SZX9 Q3SZY7 Q3SZY9 Q3SZZ0 Q3T014 Q3T016 Q3T020
Q3T022 Q3T024 Q3T025 Q3T026 Q3T028 Q3T035 Q3T042 Q3T044 Q3T056 Q3T058 Q3T062
Q3T063 Q3T064 Q3T067 Q3T071 Q3T077 Q3T078 Q3T084 Q3T086 Q3T090 Q3T0A6 Q3T0B2
Q3T0B9 Q3T0C7 Q3T0C8 Q3T0D0 Q3T0D3 Q3T0D6 Q3T0E0 Q3T0E8 Q3T0F2 Q3T0F7 Q3T0G1
Q3T0G5 Q3T0H0 Q3T0I4 Q3T0J1 Q3T0J9 Q3T0K2 Q3T0K9 Q3T0L7 Q3T0M3 Q3T0Q0 Q3T0Q3

Q3T0Q4 Q3T0Q6 Q3T0Q8 Q3T0S5 Q3T0T1 Q3T0T5 Q3T0U7 Q3T0V9 Q3T0W3 Q3T0W4 Q3T0X5
Q3T0X7 Q3T0X9 Q3T0Y3 Q3T0Y5 Q3T0Z3 Q3T0Z5 Q3T0Z7 Q3T0Z8 Q3T102 Q3T105 Q3T106
Q3T108 Q3T112 Q3T122 Q3T123 Q3T135 Q3T136 Q3T138 Q3T140 Q3T145 Q3T147 Q3T148
Q3T168 Q3T174 Q3T183 Q3T191 Q3T1H5 Q3T1H6 Q3T1I3 Q3T1I4 Q3T1I9 Q3T1J8 Q3T1L0
Q3T5N4 Q3T6E2 Q3T7C9 Q3T8Q6 Q3TC33 Q3TC72 Q3TC93 Q3TCH7 Q3TCJ1 Q3TCX3 Q3TDD9
Q3TDE8 Q3TDK6 Q3TDX8 Q3TEA8 Q3TEI4 Q3TFK5 Q3TFQ1 Q3TGF2 Q3THE2 Q3THG9 Q3THK7
Q3THS6 Q3THW5 Q3TIR1 Q3TIU4 Q3TIV5 Q3TIX9 Q3TJD7 Q3TJZ6 Q3TKT4 Q3TKY6 Q3TL26
Q3TL54 Q3TLD5 Q3TLH4 Q3TLS3 Q3TMH2 Q3TMV7 Q3TMW1 Q3TNA1 Q3TNU4 Q3TPE9 Q3TPJ7
Q3TQ03 Q3TQF0 Q3TQI7 Q3TQQ9 Q3TRJ4 Q3TRM8 Q3TRR0 Q3TSG4 Q3TT38 Q3TTC2 Q3TTI8
Q3TTJ4 Q3TTL0 Q3TTY5 Q3TUF7 Q3TUU5 Q3TV65 Q3TV70 Q3TVC7 Q3TVI4 Q3TW96 Q3TWF6
Q3TWW8 Q3TX08 Q3TX51 Q3TXS7 Q3TXT3 Q3TXU5 Q3TXZ1 Q3TY60 Q3TY65 Q3TY92 Q3TYG6
Q3TYL0 Q3TYR5 Q3TYX3 Q3TYX8 Q3TZ65 Q3TZA2 Q3TZI6 Q3TZW7 Q3TZX8 Q3U034 Q3U0J8
Q3U0K8 Q3U0L2 Q3U0M1 Q3U0P1 Q3U0V1 Q3U108 Q3U129 Q3U133 Q3U155 Q3U186 Q3U1C6
Q3U1D0 Q3U1D9 Q3U1G5 Q3U1J1 Q3U1J4 Q3U1T3 Q3U1U7 Q3U269 Q3U276 Q3U285 Q3U288
Q3U2A8 Q3U2E2 Q3U2I3 Q3U2J5 Q3U2K0 Q3U2K5 Q3U2U7 Q3U2V3 Q3U308 Q3U381 Q3U3C9
Q3U3E2 Q3U3I9 Q3U3N0 Q3U3Q1 Q3U3T8 Q3U3V8 Q3U3W5 Q3U410 Q3U487 Q3U4G0 Q3U4H6
Q3U595 Q3U5Q7 Q3U6N9 Q3U6U5 Q3U7U3 Q3U821 Q3U829 Q3U898 Q3U8K7 Q3U9F6 Q3UA06
Q3UAW9 Q3UB74 Q3UBZ5 Q3UC65 Q3UD01 Q3UD82 Q3UDE2 Q3UDK1 Q3UDP0 Q3UE17 Q3UE37
Q3UEB3 Q3UEG6 Q3UEN2 Q3UFB2 Q3UFM5 Q3UFS0 Q3UFS4 Q3UFT3 Q3UFY4 Q3UFY7 Q3UFY8
Q3UG20 Q3UG98 Q3UGC7 Q3UGM2 Q3UGP9 Q3UGR5 Q3UGS4 Q3UH06 Q3UH60 Q3UH68 Q3UHB1
Q3UHB8 Q3UHC0 Q3UHD3 Q3UHF3 Q3UHG7 Q3UHH1 Q3UHI0 Q3UHI4 Q3UHJ8 Q3UHR0 Q3UHV4
Q3UHX0 Q3UHX8 Q3UHX9 Q3UHZ5 Q3UI43 Q3UI66 Q3UIK4 Q3UIL6 Q3UIW8 Q3UIZ8 Q3UJB3
Q3UJB9 Q3UJC8 Q3UJK4 Q3UJV1 Q3UJZ3 Q3UK37 Q3UK78 Q3UKC1 Q3UKJ7 Q3UKZ7 Q3UL36
Q3UL97 Q3ULD5 Q3ULG3 Q3ULM6 Q3ULZ2 Q3UM18 Q3UM45 Q3UM89 Q3UMB5 Q3UMC0 Q3UMF0
Q3UMG5 Q3UMM4 Q3UMQ8 Q3UMU9 Q3UMY5 Q3UN58 Q3UN90 Q3UNA4 Q3UNU4 Q3UNZ8 Q3UP44
Q3UPC7 Q3UPE3 Q3UPH1 Q3UPL5 Q3UQ44 Q3UQ84 Q3UQA7 Q3UQS8 Q3UQU0 Q3UQV5 Q3UR50
Q3URF8 Q3URK1 Q3URQ0 Q3URQ7 Q3URR0 Q3URR7 Q3URS2 Q3URY0 Q3URY2 Q3URY6 Q3US16
Q3US17 Q3US41 Q3USH1 Q3USH5 Q3USL1 Q3USS3 Q3USZ2 Q3UTA9 Q3UTB7 Q3UTQ7 Q3UUE9
Q3UUF8 Q3UUJ4 Q3UUX7 Q3UV17 Q3UV70 Q3UVG3 Q3UVV9 Q3UW68 Q3UWA4 Q3UWM4 Q3UWQ3
Q3UWR2 Q3UWS6 Q3UWW6 Q3UWY1 Q3UWZ0 Q3UX61 Q3UX62 Q3UX83 Q3UXL4 Q3UXZ6 Q3UXZ9
Q3UY23 Q3UY96 Q3UYG1 Q3UYG8 Q3UYK3 Q3UYR4 Q3UYV8 Q3UZ01 Q3UZ18 Q3UZ45 Q3UZ57
Q3UZA1 Q3UZB0 Q3UZD5 Q3UZD7 Q3UZW7 Q3V036 Q3V037 Q3V038 Q3V079 Q3V080 Q3V089
Q3V096 Q3V0A6 Q3V0C1 Q3V0C3 Q3V0E1 Q3V0F0 Q3V0J4 Q3V0L5 Q3V0M2 Q3V0P1 Q3V0Q1
Q3V0Y1 Q3V124 Q3V125 Q3V129 Q3V1D3 Q3V1H1 Q3V1H9 Q3V1L4 Q3V1L6 Q3V1U8 Q3V1V3
Q3V2C1 Q3V2J0 Q3V2K1 Q3V2Q8 Q3V300 Q3V3A1 Q3V3E1 Q3V3Q4 Q3V3R1 Q3V4B5 Q3V4Q0
Q3V4Q1 Q3V4Q2 Q3V4Q3 Q3V4Q4 Q3V4Q6 Q3V4Q7 Q3V4Q8 Q3V4Q9 Q3V4R0 Q3V4R1 Q3V4R2
Q3V4R3 Q3V4R4 Q3V4R5 Q3V4R6 Q3V4R7 Q3V4R8 Q3V4R9 Q3V4S0 Q3V4S1 Q3V4S2 Q3V4S3
Q3V4S4 Q3V4S5 Q3V4S6 Q3V4S8 Q3V4S9 Q3V4T0 Q3V4T1 Q3V4T2 Q3V4T3 Q3V4T4 Q3V4T5
Q3V4T6 Q3V4T7 Q3V4T8 Q3V4T9 Q3V4U1 Q3V4U2 Q3V4U4 Q3V4U6 Q3V4U7 Q3V4U8 Q3V4U9
Q3V4V1 Q3V4V2 Q3V4V3 Q3V4V5 Q3V4V7 Q3V4V8 Q3V4V9 Q3V4W0 Q3V4W2 Q3V4W3 Q3V4W6
Q3V4W7 Q3V4W8 Q3V4W9 Q3V4X0 Q3V4X1 Q3V4X2 Q3V4X7 Q3V4Y6 Q3V4Z0 Q3V4Z3 Q3V4Z4
Q3V4Z5 Q3V4Z6 Q3V4Z7 Q3V4Z8 Q3V4Z9 Q3V500 Q3V501 Q3V502 Q3V509 Q3V510 Q3V512
Q3V513 Q3V514 Q3V526 Q3V532 Q3V536 Q3V542 Q3V543 Q3V544 Q3V545 Q3V5W6 Q3V5Z9
Q3V7F6 Q3V7G1 Q3V7G6 Q3V7I4 Q3V7I5 Q3V7I6 Q3V7J3 Q3V7K1 Q3V7K9 Q3V7M2 Q3V7N2

Q3V7N3 Q3V7P0 Q3V7Q4 Q3V7R4 Q3V7R9 Q3V7S0 Q3V7S1 Q3V7S9 Q3V7T4 Q3V7T5 Q3V7Z9
Q3V812 Q3V814 Q3V818 Q3V870 Q3V891 Q3V8A2 Q3V8A3 Q3V8B2 Q3V8B3 Q3V8C9 Q3XZZ9
Q3Y452 Q3Y4E1 Q3Y4F4 Q3Y598 Q3Y8L7 Q3Y8M6 Q3YAT3 Q3YAT5 Q3YFL6 Q3YLA6 Q3YN09
Q3YTZ9 Q3YU01 Q3YU08 Q3YU09 Q3YU10 Q3YU11 Q3YU12 Q3YU19 Q3YU23 Q3YU31 Q3YU43
Q3YU59 Q3YU89 Q3YU91 Q3YU98 Q3YUA6 Q3YUA7 Q3YUB7 Q3YUB9 Q3YUC8 Q3YUD8 Q3YUE4
Q3YUE5 Q3YUE6 Q3YUE7 Q3YUF0 Q3YUF1 Q3YUF2 Q3YUF5 Q3YUF6 Q3YUG5 Q3YUG6 Q3YUH1
Q3YUH2 Q3YUH3 Q3YUJ2 Q3YUJ7 Q3YUJ8 Q3YUK3 Q3YUU5 Q3YUW0 Q3YUW9 Q3YUX6 Q3YUX8
Q3YUY7 Q3YUY8 Q3YUY9 Q3YUZ0 Q3YUZ1 Q3YUZ4 Q3YUZ6 Q3YUZ7 Q3YUZ8 Q3YUZ9 Q3YV00
Q3YV01 Q3YV04 Q3YV06 Q3YV08 Q3YV11 Q3YV16 Q3YV17 Q3YV19 Q3YV20 Q3YV31 Q3YV35
Q3YV42 Q3YV46 Q3YV47 Q3YV49 Q3YV50 Q3YV59 Q3YV62 Q3YV71 Q3YV72 Q3YV73 Q3YV74
Q3YV75 Q3YV81 Q3YV85 Q3YV89 Q3YVA7 Q3YVA8 Q3YVC0 Q3YVD2 Q3YVD7 Q3YVF5 Q3YVG0
Q3YVI4 Q3YVI5 Q3YVJ0 Q3YVJ3 Q3YVK2 Q3YVK9 Q3YVP4 Q3YVP5 Q3YVP8 Q3YVP9 Q3YVQ0
Q3YVU0 Q3YVU1 Q3YVV0 Q3YVV3 Q3YVW2 Q3YVW4 Q3YVX4 Q3YVX7 Q3YVY1 Q3YVY3 Q3YVZ6
Q3YVZ7 Q3YVZ8 Q3YVZ9 Q3YW02 Q3YW04 Q3YW05 Q3YW09 Q3YW11 Q3YW12 Q3YW34 Q3YW47
Q3YW58 Q3YW93 Q3YW94 Q3YW95 Q3YW96 Q3YWA3 Q3YWA7 Q3YWB0 Q3YWB1 Q3YWB2 Q3YWB4
Q3YWC2 Q3YWC5 Q3YWE3 Q3YWE7 Q3YWJ3 Q3YWK2 Q3YWK6 Q3YWK7 Q3YWL1 Q3YWL3 Q3YWL5
Q3YWM2 Q3YWM3 Q3YWN3 Q3YWN4 Q3YWN8 Q3YWQ3 Q3YWR7 Q3YWS4 Q3YWS7 Q3YWS8 Q3YWS9
Q3YWT0 Q3YWT1 Q3YWT2 Q3YWT3 Q3YWT8 Q3YWT9 Q3YWU0 Q3YWU1 Q3YWU2 Q3YWU3 Q3YWU4
Q3YWU5 Q3YWU6 Q3YWU7 Q3YWU8 Q3YWU9 Q3YWV0 Q3YWV1 Q3YWV2 Q3YWV3 Q3YWV4 Q3YWV5
Q3YWV6 Q3YWV7 Q3YWV8 Q3YWW0 Q3YWW1 Q3YWW2 Q3YWW3 Q3YWW4 Q3YWW5 Q3YWX1 Q3YWX2
Q3YWX3 Q3YWX5 Q3YWX6 Q3YWX7 Q3YWY8 Q3YWZ0 Q3YX01 Q3YX10 Q3YX11 Q3YX17 Q3YX18
Q3YX22 Q3YX23 Q3YX25 Q3YX36 Q3YX52 Q3YX55 Q3YX56 Q3YX58 Q3YX62 Q3YX65 Q3YX68
Q3YX71 Q3YX73 Q3YX74 Q3YX75 Q3YX76 Q3YX77 Q3YX86 Q3YX87 Q3YX92 Q3YX93 Q3YXA5
Q3YXB9 Q3YXD9 Q3YXE7 Q3YXE9 Q3YXH0 Q3YXH4 Q3YXH8 Q3YXH9 Q3YXI1 Q3YXI2 Q3YXI6
Q3YXJ0 Q3YXJ3 Q3YXK0 Q3YXK4 Q3YXQ6 Q3YXS2 Q3YXS3 Q3YXS7 Q3YXS9 Q3YXT2 Q3YXT4
Q3YXU5 Q3YXU6 Q3YXV6 Q3YXV7 Q3YXW5 Q3YXW6 Q3YXW7 Q3YXX0 Q3YXX2 Q3YXY0 Q3YY15
Q3YY29 Q3YY30 Q3YY33 Q3YY51 Q3YY53 Q3YY58 Q3YY62 Q3YY71 Q3YY76 Q3YY94 Q3YY95
Q3YY96 Q3YYB0 Q3YYB1 Q3YYB2 Q3YYB5 Q3YYB6 Q3YYB7 Q3YYB8 Q3YYB9 Q3YYC9 Q3YYF3
Q3YYF4 Q3YYG4 Q3YYG5 Q3YYG6 Q3YYG7 Q3YYH0 Q3YYM4 Q3YYM5 Q3YYM9 Q3YYN0 Q3YYN1
Q3YYN2 Q3YYQ2 Q3YYT5 Q3YYT6 Q3YYU0 Q3YYU9 Q3YYV1 Q3YYV2 Q3YYZ8 Q3YZ04 Q3YZ11
Q3YZ12 Q3YZ13 Q3YZ14 Q3YZ15 Q3YZ20 Q3YZ21 Q3YZ24 Q3YZ25 Q3YZ26 Q3YZ29 Q3YZ34
Q3YZ35 Q3YZ37 Q3YZ38 Q3YZ43 Q3YZ45 Q3YZ53 Q3YZ56 Q3YZ74 Q3YZ76 Q3YZ81 Q3YZ86
Q3YZ89 Q3YZA3 Q3YZA7 Q3YZA9 Q3YZB4 Q3YZB5 Q3YZC3 Q3YZD1 Q3YZD6 Q3YZE4 Q3YZF6
Q3YZG4 Q3YZH8 Q3YZN0 Q3YZN2 Q3YZN7 Q3YZP1 Q3YZP3 Q3YZR9 Q3YZU2 Q3YZU3 Q3YZU5
Q3YZV5 Q3YZV9 Q3YZW1 Q3YZX6 Q3YZZ7 Q3Z020 Q3Z021 Q3Z022 Q3Z035 Q3Z044 Q3Z047
Q3Z052 Q3Z053 Q3Z062 Q3Z096 Q3Z0A5 Q3Z0A6 Q3Z0B4 Q3Z0D8 Q3Z0F9 Q3Z0G0 Q3Z0G1
Q3Z0G2 Q3Z0G4 Q3Z0G5 Q3Z0G6 Q3Z0K4 Q3Z0K8 Q3Z0L5 Q3Z0M5 Q3Z0N4 Q3Z0N6 Q3Z0Q1
Q3Z0S2 Q3Z0S6 Q3Z0S8 Q3Z0S9 Q3Z0U8 Q3Z0V3 Q3Z108 Q3Z109 Q3Z127 Q3Z131 Q3Z135
Q3Z146 Q3Z194 Q3Z1D9 Q3Z1H6 Q3Z1K4 Q3Z1M2 Q3Z1M3 Q3Z1N3 Q3Z1Q7 Q3Z1V9 Q3Z1W3
Q3Z1X8 Q3Z1Y0 Q3Z1Z2 Q3Z1Z3 Q3Z1Z4 Q3Z1Z6 Q3Z208 Q3Z213 Q3Z232 Q3Z233 Q3Z246
Q3Z247 Q3Z250 Q3Z253 Q3Z260 Q3Z261 Q3Z262 Q3Z264 Q3Z265 Q3Z267 Q3Z278 Q3Z280
Q3Z287 Q3Z289 Q3Z292 Q3Z293 Q3Z294 Q3Z295 Q3Z2B0 Q3Z2B6 Q3Z2F2 Q3Z2F4 Q3Z2H0
Q3Z2H4 Q3Z2K0 Q3Z2L8 Q3Z2L9 Q3Z2M1 Q3Z2M2 Q3Z2M4 Q3Z2P6 Q3Z2P7 Q3Z2Q2 Q3Z2Q4
Q3Z2S0 Q3Z2S4 Q3Z2T7 Q3Z2U3 Q3Z2V9 Q3Z2X0 Q3Z2X3 Q3Z2X5 Q3Z2X6 Q3Z2Y2 Q3Z2Y5

Q3Z2Y6 Q3Z2Z2 Q3Z2Z8 Q3Z310 Q3Z311 Q3Z319 Q3Z328 Q3Z330 Q3Z349 Q3Z354 Q3Z357
Q3Z361 Q3Z392 Q3Z3A3 Q3Z3A4 Q3Z3A5 Q3Z3A6 Q3Z3B7 Q3Z3C3 Q3Z3C4 Q3Z3F2 Q3Z3F4
Q3Z3G3 Q3Z3G5 Q3Z3H3 Q3Z3I9 Q3Z3J0 Q3Z3J7 Q3Z3J9 Q3Z3K4 Q3Z3K9 Q3Z3L1 Q3Z3L4
Q3Z3L5 Q3Z3M7 Q3Z3N5 Q3Z3N6 Q3Z3N8 Q3Z3R0 Q3Z3R7 Q3Z3S5 Q3Z3T2 Q3Z3U9 Q3Z3X3
Q3Z3X8 Q3Z3Y1 Q3Z3Y7 Q3Z401 Q3Z403 Q3Z405 Q3Z408 Q3Z409 Q3Z422 Q3Z453 Q3Z455
Q3Z476 Q3Z489 Q3Z490 Q3Z4C0 Q3Z4C2 Q3Z4D1 Q3Z4D3 Q3Z4E1 Q3Z4F0 Q3Z4F3 Q3Z4F6
Q3Z4G1 Q3Z4G2 Q3Z4G4 Q3Z4H6 Q3Z4H9 Q3Z4I0 Q3Z4J6 Q3Z4L2 Q3Z4P5 Q3Z4P8 Q3Z4Q0
Q3Z4S3 Q3Z4S4 Q3Z4S5 Q3Z4S6 Q3Z4S7 Q3Z4S8 Q3Z4T0 Q3Z4V2 Q3Z4V4 Q3Z4W5 Q3Z4W6
Q3Z4W7 Q3Z4Y3 Q3Z4Y6 Q3Z4Y7 Q3Z4Y9 Q3Z4Z3 Q3Z4Z4 Q3Z4Z6 Q3Z503 Q3Z504 Q3Z515
Q3Z519 Q3Z522 Q3Z523 Q3Z529 Q3Z549 Q3Z550 Q3Z551 Q3Z554 Q3Z555 Q3Z556 Q3Z583
Q3Z585 Q3Z586 Q3Z594 Q3Z595 Q3Z597 Q3Z598 Q3Z5A4 Q3Z5B4 Q3Z5E9 Q3Z5F1 Q3Z5F4
Q3Z5G3 Q3Z5G7 Q3Z5H5 Q3Z5I6 Q3Z5I7 Q3Z5I8 Q3Z5I9 Q3Z5J1 Q3Z5J2 Q3Z5J4 Q3Z5J8
Q3Z5K1 Q3Z5K3 Q3Z5L1 Q3Z5M6 Q3Z5M7 Q3Z5M9 Q3Z5N9 Q3Z5P0 Q3Z5P1 Q3Z5Q5 Q3Z5Q6
Q3Z5Q7 Q3Z5Q8 Q3Z5R8 Q3Z5S1 Q3Z5S7 Q3Z5S8 Q3Z5T6 Q3Z5T7 Q3Z5T8 Q3Z5T9 Q3Z5U2
Q3Z5U8 Q3Z5U9 Q3Z5V2 Q3Z5V7 Q3Z5V8 Q3Z5V9 Q3Z5W0 Q3Z5W6 Q3Z5W9 Q3Z5X2 Q3Z5X4
Q3Z5X9 Q3Z5Y0 Q3Z5Y1 Q3Z5Y4 Q3Z5Y7 Q3Z600 Q3Z601 Q3Z606 Q3Z608 Q3Z611 Q3Z613
Q3Z614 Q3Z617 Q3Z629 Q3Z637 Q3Z663 Q3Z6A1 Q3Z6E3 Q3Z6G1 Q3Z6G4 Q3Z6G5 Q3Z6G6
Q3Z6J2 Q3Z6L1 Q3Z6L3 Q3Z6L5 Q3Z6N1 Q3Z6N2 Q3Z6N9 Q3Z6P1 Q3Z6Q4 Q3Z6R8 Q3Z6S3
Q3Z6U4 Q3Z6V3 Q3Z6V4 Q3Z6V7 Q3Z6W1 Q3Z6W2 Q3Z6W3 Q3Z6W5 Q3Z6X0 Q3Z6Z5 Q3Z6Z8
Q3Z6Z9 Q3Z705 Q3Z709 Q3Z712 Q3Z717 Q3Z726 Q3Z727 Q3Z730 Q3Z731 Q3Z767 Q3Z770
Q3Z776 Q3Z782 Q3Z787 Q3Z788 Q3Z799 Q3Z7C7 Q3Z7F2 Q3Z7F5 Q3Z7F6 Q3Z7G4 Q3Z7N7
Q3Z7N9 Q3Z7P1 Q3Z7P5 Q3Z7Q6 Q3Z7Q8 Q3Z7S9 Q3Z7T0 Q3Z7T3 Q3Z7T4 Q3Z7T5 Q3Z7T6
Q3Z7U3 Q3Z7U4 Q3Z7U5 Q3Z7V0 Q3Z7V3 Q3Z7V5 Q3Z7V6 Q3Z7V7 Q3Z7Y8 Q3Z876 Q3Z878
Q3Z879 Q3Z880 Q3Z886 Q3Z888 Q3Z891 Q3Z896 Q3Z8B3 Q3Z8B4 Q3Z8B5 Q3Z8C0 Q3Z8C1
Q3Z8C6 Q3Z8D6 Q3Z8D7 Q3Z8D8 Q3Z8D9 Q3Z8E1 Q3Z8E2 Q3Z8E7 Q3Z8F6 Q3Z8G2 Q3Z8G4
Q3Z8G5 Q3Z8G9 Q3Z8H2 Q3Z8H3 Q3Z8H5 Q3Z8J8 Q3Z8J9 Q3Z8K0 Q3Z8K3 Q3Z8K6 Q3Z8V0
Q3Z8V1 Q3Z8V3 Q3Z8V4 Q3Z8V6 Q3Z8V9 Q3Z8W1 Q3Z8W2 Q3Z8X4 Q3Z8X7 Q3Z8X9 Q3Z8Y0
Q3Z8Y8 Q3Z8Z0 Q3Z914 Q3Z937 Q3Z943 Q3Z949 Q3Z950 Q3Z952 Q3Z953 Q3Z954 Q3Z955
Q3Z956 Q3Z957 Q3Z958 Q3Z960 Q3Z962 Q3Z963 Q3Z964 Q3Z965 Q3Z966 Q3Z967 Q3Z968
Q3Z969 Q3Z970 Q3Z971 Q3Z972 Q3Z973 Q3Z974 Q3Z975 Q3Z976 Q3Z977 Q3Z978 Q3Z979
Q3Z980 Q3Z981 Q3Z982 Q3Z983 Q3Z984 Q3Z985 Q3Z988 Q3Z989 Q3Z990 Q3Z991 Q3Z992
Q3Z993 Q3Z997 Q3Z9A4 Q3Z9B1 Q3Z9B2 Q3Z9B3 Q3Z9B4 Q3Z9B9 Q3Z9C5 Q3Z9C7 Q3Z9E9
Q3Z9F0 Q3Z9G0 Q3Z9G6 Q3Z9H3 Q3Z9H5 Q3Z9H6 Q3Z9H8 Q3Z9H9 Q3Z9I5 Q3Z9I7 Q3Z9I8
Q3Z9K2 Q3Z9K8 Q3Z9L1 Q3Z9L2 Q3Z9L3 Q3Z9L4 Q3Z9Y3 Q3ZA07 Q3ZA11 Q3ZA71 Q3ZAD5
Q3ZAD6 Q3ZAD7 Q3ZAE2 Q3ZAE3 Q3ZAE4 Q3ZAF8 Q3ZAF9 Q3ZAH5 Q3ZAI1 Q3ZAI3 Q3ZAI8
Q3ZAJ1 Q3ZAJ2 Q3ZAV0 Q3ZAV8 Q3ZB90 Q3ZB98 Q3ZBA7 Q3ZBC0 Q3ZBC2 Q3ZBD0 Q3ZBD2
Q3ZBD3 Q3ZBD4 Q3ZBE0 Q3ZBE7 Q3ZBF0 Q3ZBG0 Q3ZBG4 Q3ZBH0 Q3ZBI3 Q3ZBI6 Q3ZBJ0
Q3ZBJ1 Q3ZBK1 Q3ZBK5 Q3ZBK6 Q3ZBL0 Q3ZBL5 Q3ZBL9 Q3ZBM1 Q3ZBM6 Q3ZBM7 Q3ZBN0
Q3ZBP0 Q3ZBP3 Q3ZBP4 Q3ZBP8 Q3ZBQ0 Q3ZBQ1 Q3ZBQ4 Q3ZBR5 Q3ZBR6 Q3ZBR9 Q3ZBS0
Q3ZBS9 Q3ZBT0 Q3ZBT2 Q3ZBU3 Q3ZBV2 Q3ZBV3 Q3ZBV4 Q3ZBV8 Q3ZBW4 Q3ZBW7 Q3ZBX9
Q3ZBY2 Q3ZBY3 Q3ZC01 Q3ZC07 Q3ZC09 Q3ZC10 Q3ZC12 Q3ZC15 Q3ZC21 Q3ZC22 Q3ZC34
Q3ZC52 Q3ZC61 Q3ZC62 Q3ZC66 Q3ZC67 Q3ZC82 Q3ZC84 Q3ZC89 Q3ZCA1 Q3ZCA2 Q3ZCB2
Q3ZCB6 Q3ZCC4 Q3ZCC5 Q3ZCC8 Q3ZCE0 Q3ZCE8 Q3ZCF2 Q3ZCF5 Q3ZCH9 Q3ZCI3 Q3ZCI6
Q3ZCI9 Q3ZCJ7 Q3ZCK1 Q3ZCK3 Q3ZCK5 Q3ZCK8 Q3ZCK9 Q3ZCT1 Q3ZCT8 Q3ZCU0 Q3ZCV2

Q3ZCX4 Q3ZE13 Q3ZEE5 Q3ZFI7 Q3ZIZ0 Q3ZIZ5 Q3ZJ05 Q3ZJ12 Q3ZJ23 Q3ZJ24 Q3ZJ37
Q3ZJ38 Q3ZJ39 Q3ZJ40 Q3ZJ41 Q3ZJ47 Q3ZJ48 Q3ZJ74 Q3ZJ77 Q3ZJ78 Q3ZJ79 Q3ZJ80
Q3ZJ81 Q3ZJ82 Q3ZJ83 Q3ZJ84 Q3ZJ85 Q3ZJ86 Q3ZJ87 Q3ZJ88 Q3ZJ89 Q3ZJ90 Q3ZJ92
Q3ZJ93 Q3ZK55 Q3ZK56 Q3ZK57 Q3ZK61 Q3ZK62 Q3ZK63 Q3ZK65 Q3ZLR7 Q3ZM63 Q3ZN05
Q3ZPN4 Q3ZUZ8 Q3ZUZ9 Q3ZV00 Q3ZW41 Q3ZW59 Q3ZW73 Q3ZW88 Q3ZW89 Q3ZWA4 Q3ZWB0
Q3ZWB2 Q3ZWB6 Q3ZWC1 Q3ZWC2 Q3ZWC3 Q3ZWD7 Q3ZWD8 Q3ZWE1 Q3ZWG4 Q3ZWG5 Q3ZWJ9
Q3ZWQ1 Q3ZWQ5 Q3ZWU0 Q3ZWX3 Q3ZWX7 Q3ZWX9 Q3ZWY0 Q3ZWY7 Q3ZWY8 Q3ZWZ9 Q3ZX00
Q3ZX01 Q3ZX03 Q3ZX06 Q3ZX08 Q3ZX09 Q3ZX40 Q3ZX43 Q3ZX54 Q3ZX82 Q3ZX83 Q3ZX84
Q3ZXA9 Q3ZXB5 Q3ZXB7 Q3ZXB8 Q3ZXC2 Q3ZXC5 Q3ZXC6 Q3ZXC8 Q3ZXD5 Q3ZXD6 Q3ZXD7
Q3ZXD8 Q3ZXE0 Q3ZXE1 Q3ZXF9 Q3ZXG0 Q3ZXG1 Q3ZXG5 Q3ZXG6 Q3ZXH1 Q3ZXH7 Q3ZXH9
Q3ZXI2 Q3ZXI7 Q3ZXL5 Q3ZXL7 Q3ZXL8 Q3ZXL9 Q3ZXQ8 Q3ZXU3 Q3ZXU4 Q3ZXU5 Q3ZXV0
Q3ZXV3 Q3ZXV5 Q3ZXV6 Q3ZXV7 Q3ZXX3 Q3ZXX4 Q3ZXX7 Q3ZXX8 Q3ZXX9 Q3ZXY0 Q3ZXY7
Q3ZXY9 Q3ZY12 Q3ZY14 Q3ZY16 Q3ZY20 Q3ZY29 Q3ZY38 Q3ZY50 Q3ZY75 Q3ZY85 Q3ZY97
Q3ZYA0 Q3ZYA3 Q3ZYA9 Q3ZYB5 Q3ZYF0 Q3ZYF9 Q3ZYG0 Q3ZYG3 Q3ZYG4 Q3ZYG8 Q3ZYH5
Q3ZYH8 Q3ZYH9 Q3ZYI5 Q3ZYI9 Q3ZYJ9 Q3ZYL6 Q3ZYM5 Q3ZYM6 Q3ZYM9 Q3ZYN3 Q3ZYN4
Q3ZYN5 Q3ZYN7 Q3ZYP4 Q3ZYS0 Q3ZYT5 Q3ZYU1 Q3ZYU9 Q3ZYV1 Q3ZYW6 Q3ZYW8 Q3ZYX0
Q3ZYX5 Q3ZYZ2 Q3ZZ10 Q3ZZ13 Q3ZZ14 Q3ZZ15 Q3ZZ31 Q3ZZ95 Q3ZZA1 Q3ZZA4 Q3ZZA6
Q3ZZA7 Q3ZZB2 Q3ZZB8 Q3ZZC1 Q3ZZC3 Q3ZZC4 Q3ZZC9 Q3ZZD0 Q3ZZD2 Q3ZZD3 Q3ZZE5
Q3ZZE6 Q3ZZF6 Q3ZZG3 Q3ZZG9 Q3ZZH1 Q3ZZI5 Q3ZZI6 Q3ZZJ0 Q3ZZJ7 Q3ZZK4 Q3ZZK5
Q3ZZK6 Q3ZZK7 Q3ZZK9 Q3ZZL0 Q3ZZL1 Q3ZZL2 Q3ZZL3 Q3ZZL4 Q3ZZL5 Q3ZZL6 Q3ZZL7
Q3ZZL8 Q3ZZL9 Q3ZZM0 Q3ZZM1 Q3ZZM2 Q3ZZM3 Q3ZZM4 Q3ZZM5 Q3ZZM6 Q3ZZM7 Q3ZZM8
Q3ZZN1 Q3ZZN2 Q3ZZN3 Q3ZZN4 Q3ZZN7 Q3ZZP2 Q3ZZP3 Q3ZZP5 Q3ZZP6 Q3ZZP7 Q3ZZP8
Q3ZZP9 Q3ZZQ0 Q3ZZQ1 Q3ZZQ3 Q3ZZQ5 Q3ZZQ6 Q3ZZQ7 Q3ZZQ8 Q3ZZT1 Q3ZZT5 Q3ZZV8
Q3ZZW0 Q3ZZW3 Q3ZZX5 Q40004 Q40034 Q40061 Q40062 Q40066 Q40073 Q40082 Q400C7
Q400C8 Q400C9 Q400G9 Q400K2 Q400K3 Q400K4 Q40106 Q40134 Q40147 Q40152 Q40153
Q40157 Q40158 Q40168 Q40170 Q40250 Q40255 Q40256 Q40280 Q40281 Q40284 Q40285
Q40286 Q40287 Q40288 Q40289 Q402B2 Q402E1 Q402E2 Q40302 Q40313 Q40316 Q40345
Q40353 Q40359 Q40396 Q40410 Q40450 Q40460 Q40465 Q40466 Q40467 Q40468 Q40469
Q40470 Q40471 Q40504 Q40505 Q40545 Q40546 Q40554 Q40565 Q40577 Q40592 Q40606
Q40665 Q40677 Q40680 Q40682 Q40691 Q40702 Q40703 Q40704 Q40742 Q40762 Q40784
Q40831 Q40832 Q40854 Q40863 Q40864 Q40872 Q40884 Q40885 Q40910 Q40941 Q40976
Q40977 Q40983 Q41005 Q41011 Q41014 Q41046 Q41058 Q41059 Q41060 Q41086 Q41112
Q41140 Q41141 Q41219 Q41247 Q41249 Q41261 Q41274 Q41276 Q41328 Q41330 Q41344
Q41351 Q41365 Q41373 Q41382 Q41389 Q41391 Q41407 Q41420 Q41436 Q41452 Q41558
Q41560 Q41575 Q41578 Q41583 Q41593 Q41594 Q41595 Q41607 Q41608 Q41639 Q41651
Q41657 Q41669 Q41670 Q41709 Q41720 Q41738 Q41739 Q41741 Q41761 Q41764 Q41768
Q41769 Q41771 Q41782 Q41783 Q41784 Q41785 Q41803 Q41807 Q41808 Q41811 Q41819
Q41822 Q41852 Q41853 Q41864 Q41874 Q41969 Q42112 Q42134 Q42182 Q42202 Q42340
Q42344 Q42351 Q42362 Q42368 Q42376 Q42379 Q42384 Q42404 Q42410 Q42429 Q42430
Q42435 Q42440 Q42450 Q42453 Q42463 Q42464 Q42472 Q42480 Q42485 Q42498 Q42516
Q42521 Q42522 Q42523 Q42524 Q42527 Q42529 Q42538 Q42539 Q42551 Q42553 Q42565
Q42575 Q42581 Q42583 Q42586 Q42588 Q42592 Q42598 Q42601 Q42609 Q42611 Q42612
Q42613 Q42624 Q42649 Q42652 Q42653 Q42654 Q42662 Q42663 Q42665 Q42667 Q42669
Q42671 Q42675 Q42676 Q42677 Q42679 Q42681 Q42682 Q42684 Q42686 Q42688 Q42689

Q42690 Q42693 Q42694 Q42695 Q42698 Q42699 Q42707 Q42711 Q42726 Q42736 Q42777
Q42796 Q42800 Q42805 Q42808 Q42822 Q42823 Q42829 Q42831 Q42836 Q42840 Q42843
Q42850 Q42855 Q42857 Q42858 Q42876 Q42877 Q42881 Q42882 Q42884 Q42885 Q42891
Q42896 Q42899 Q42908 Q42910 Q42915 Q42919 Q42942 Q42945 Q42946 Q42948 Q42954
Q42961 Q42962 Q42963 Q42966 Q42967 Q42968 Q42971 Q42972 Q42982 Q42997 Q43008
Q43011 Q43036 Q43046 Q43047 Q43052 Q43056 Q43058 Q43064 Q43066 Q43070 Q43072
Q43075 Q43082 Q43083 Q43086 Q43087 Q43088 Q43090 Q43092 Q43093 Q43095 Q43117
Q43124 Q43130 Q43134 Q43137 Q43138 Q43139 Q43153 Q43154 Q43155 Q43161 Q43163
Q43175 Q43187 Q43188 Q43199 Q43207 Q43208 Q43209 Q43210 Q43213 Q43214 Q43215
Q43216 Q43217 Q43236 Q43237 Q43239 Q43247 Q43260 Q43261 Q43266 Q43272 Q43291
Q43292 Q43295 Q43296 Q43298 Q43299 Q43309 Q43312 Q43314 Q43317 Q43321 Q43349
Q43364 Q43383 Q43385 Q43415 Q43467 Q43470 Q43472 Q43473 Q43484 Q43492 Q43497
Q43503 Q43512 Q43513 Q43515 Q43517 Q43560 Q43578 Q43582 Q43585 Q43592 Q43594
Q43609 Q43621 Q43636 Q43641 Q43654 Q43694 Q43695 Q43697 Q43704 Q43714 Q43716
Q43725 Q43743 Q43744 Q43754 Q43768 Q43772 Q43776 Q43779 Q43784 Q43785 Q43792
Q43794 Q43814 Q43820 Q43831 Q43832 Q43845 Q43846 Q43847 Q43870 Q43876 Q43883
Q43884 Q43885 Q43895 Q43899 Q43907 Q43908 Q43909 Q43914 Q43922 Q43925 Q43926
Q43931 Q43932 Q43935 Q43947 Q43948 Q43949 Q43950 Q43951 Q43954 Q43955 Q43957
Q43958 Q43959 Q43963 Q43966 Q43971 Q43973 Q43974 Q43984 Q43989 Q43990 Q43992
Q44004 Q44015 Q44020 Q44022 Q44023 Q44037 Q44044 Q44045 Q44057 Q44065 Q44078
Q44080 Q44081 Q44088 Q44093 Q44104 Q44115 Q44116 Q44118 Q44141 Q44144 Q44145
Q44148 Q44149 Q44151 Q44160 Q44161 Q44176 Q44177 Q44178 Q44184 Q44185 Q44212
Q44217 Q44233 Q44238 Q44243 Q44244 Q44245 Q44247 Q44249 Q44256 Q44257 Q44258
Q44261 Q44262 Q44270 Q44271 Q44272 Q44273 Q44274 Q44275 Q44276 Q44277 Q44278
Q44290 Q44297 Q44311 Q44315 Q44316 Q44317 Q44333 Q44335 Q44336 Q44337 Q44338
Q44347 Q44348 Q44349 Q44361 Q44362 Q44363 Q44406 Q44407 Q44408 Q44427 Q44428
Q44433 Q44452 Q44467 Q44471 Q44472 Q44473 Q44482 Q44484 Q44491 Q44498 Q44501
Q44507 Q44510 Q44524 Q44532 Q44540 Q44554 Q44556 Q44557 Q44560 Q44571 Q44581
Q44582 Q44592 Q44593 Q44598 Q44602 Q44603 Q44604 Q44607 Q44625 Q44633 Q44634
Q44636 Q44643 Q44653 Q44654 Q44678 Q44679 Q44680 Q44681 Q44683 Q44685 Q44686
Q44687 Q44688 Q44689 Q44691 Q44695 Q44696 Q44697 Q44730 Q44737 Q44740 Q44754
Q44767 Q44772 Q44773 Q44777 Q44843 Q44904 Q44951 Q45032 Q45056 Q45057 Q45058
Q45060 Q45061 Q45062 Q45063 Q45065 Q45069 Q45072 Q45087 Q45096 Q45097 Q45099
Q45100 Q45101 Q45119 Q45120 Q45129 Q45131 Q45133 Q45218 Q45219 Q45220 Q45221
Q45222 Q45231 Q45251 Q45252 Q45256 Q45257 Q45263 Q45264 Q45269 Q45288 Q45291
Q45292 Q45296 Q45305 Q45338 Q45345 Q45347 Q45352 Q45388 Q45401 Q45410 Q45421
Q45423 Q45477 Q45478 Q45480 Q45486 Q45488 Q45489 Q45494 Q45495 Q45497 Q45498
Q45500 Q45515 Q45535 Q45536 Q45537 Q45538 Q45550 Q45551 Q45552 Q45557 Q45560
Q45581 Q45582 Q45583 Q45584 Q45591 Q45595 Q45596 Q45597 Q45598 Q45599 Q45600
Q45601 Q45603 Q45604 Q45618 Q45619 Q45620 Q45633 Q45692 Q45694 Q45695 Q45696
Q45697 Q45704 Q45705 Q45706 Q45707 Q45708 Q45709 Q45710 Q45712 Q45715 Q45716
Q45718 Q45722 Q45729 Q45730 Q45733 Q45738 Q45739 Q45743 Q45744 Q45746 Q45747
Q45748 Q45752 Q45753 Q45754 Q45755 Q45758 Q45760 Q45774 Q45791 Q45811 Q45812
Q45825 Q45826 Q45831 Q45881 Q45914 Q45918 Q45920 Q45952 Q45966 Q45968 Q45972

Q45978 Q45982 Q45983 Q45EK7 Q45FF9 Q45FY6 Q45GQ7 Q45GW3 Q45KJ4 Q45KJ6 Q45QI7
Q45U48 Q45UF0 Q45UF4 Q45UF6 Q45UF7 Q45UF9 Q45UG0 Q45VV3 Q45XH3 Q45XH5 Q45XH6
Q45XH7 Q45XH8 Q45XI4 Q45XI5 Q45XI6 Q45XI7 Q45XI8 Q45XI9 Q45XJ0 Q46051 Q46078
Q46082 Q46083 Q46084 Q46087 Q460N3 Q46104 Q46106 Q46108 Q46121 Q46124 Q46127
Q46130 Q46135 Q46139 Q46158 Q46159 Q46171 Q46172 Q46175 Q46185 Q46189 Q46204
Q46205 Q46206 Q46213 Q46228 Q46229 Q46244 Q46257 Q46258 Q46259 Q46260 Q46261
Q46262 Q46263 Q46264 Q46266 Q46267 Q46289 Q46293 Q462Q7 Q462R2 Q46306 Q46323
Q46336 Q46337 Q46338 Q46339 Q46363 Q46367 Q46370 Q46372 Q46373 Q46381 Q46383
Q46386 Q46389 Q46393 Q46397 Q46398 Q46403 Q46404 Q46435 Q46436 Q46437 Q46438
Q46439 Q46440 Q46441 Q46442 Q46451 Q46455 Q46463 Q46464 Q46466 Q46473 Q46480
Q46482 Q46490 Q46491 Q46495 Q46497 Q464W3 Q464Y2 Q464Y4 Q464Z1 Q464Z2 Q464Z3
Q464Z4 Q464Z5 Q46501 Q46505 Q46506 Q46507 Q46508 Q46509 Q46526 Q46550 Q46558
Q46560 Q46561 Q46565 Q46577 Q46582 Q465C4 Q465F3 Q465F4 Q465G4 Q465J4 Q465K9
Q465N2 Q465Q4 Q465Q9 Q465R0 Q465R3 Q465R4 Q465S1 Q465S2 Q465U3 Q465U8 Q465V4
Q465X9 Q465Y5 Q465Y6 Q46604 Q46630 Q46634 Q46635 Q46637 Q46638 Q46664 Q46666
Q466A1 Q466A2 Q466A3 Q466A4 Q466D1 Q466D4 Q466D5 Q466D6 Q466H2 Q466J4 Q466J5
Q466Q1 Q466Q9 Q466S6 Q466S9 Q466T6 Q466V2 Q466V4 Q466W7 Q466X0 Q466X1 Q466X7
Q466Y5 Q46731 Q46751 Q46786 Q46787 Q46789 Q46793 Q46795 Q46796 Q46799 Q467J7
Q467K0 Q467V2 Q467Z2 Q467Z4 Q46800 Q46801 Q46802 Q46803 Q46806 Q46807 Q46808
Q46809 Q46810 Q46811 Q46812 Q46814 Q46819 Q46820 Q46822 Q46829 Q46840 Q46843
Q46844 Q46845 Q46851 Q46855 Q46856 Q46857 Q46861 Q46864 Q46865 Q46866 Q46871
Q46877 Q46888 Q46889 Q46890 Q46891 Q46893 Q46896 Q46897 Q46898 Q46899 Q468I5
Q468I6 Q468N4 Q468N6 Q468N7 Q468S9 Q46901 Q46904 Q46905 Q46906 Q46907 Q46911
Q46915 Q46919 Q46920 Q46925 Q46938 Q46939 Q46941 Q46948 Q46953 Q46967 Q46968
Q46978 Q46995 Q46996 Q469A3 Q469F5 Q469I9 Q469J2 Q469M0 Q469M4 Q469M6 Q469M7
Q469M8 Q469W0 Q469Y2 Q469Z7 Q469Z8 Q46A30 Q46A31 Q46A50 Q46A52 Q46A53 Q46A65
Q46AE4 Q46AH3 Q46AH5 Q46AI3 Q46AJ8 Q46AN2 Q46AN4 Q46AN5 Q46AR1 Q46AR9 Q46AS7
Q46AS9 Q46AT6 Q46AU2 Q46AU8 Q46AW0 Q46AY7 Q46AZ4 Q46B07 Q46B32 Q46B88 Q46B95
Q46BA3 Q46BC9 Q46BD6 Q46BD7 Q46BE1 Q46BF1 Q46BI1 Q46BJ4 Q46BK8 Q46BL0 Q46BQ2
Q46BQ5 Q46BZ5 Q46BZ6 Q46C04 Q46C09 Q46C10 Q46C14 Q46C16 Q46C63 Q46C69 Q46C74
Q46C77 Q46C79 Q46CA1 Q46CA2 Q46CB8 Q46CC5 Q46CC9 Q46CD2 Q46CE2 Q46CE4 Q46CG5
Q46CG8 Q46CH0 Q46CH1 Q46CH3 Q46CH4 Q46CH5 Q46CH6 Q46CH7 Q46CJ6 Q46CK6 Q46CK8
Q46CL0 Q46CL1 Q46CL4 Q46CQ6 Q46CR0 Q46CS0 Q46CU6 Q46CU8 Q46CW6 Q46CW7 Q46CW8
Q46CW9 Q46D07 Q46D27 Q46D52 Q46D63 Q46D71 Q46D82 Q46D91 Q46D93 Q46D96 Q46DA4
Q46DA6 Q46DA7 Q46DC3 Q46DC4 Q46DC5 Q46DH3 Q46DH6 Q46DI6 Q46DR6 Q46DT9 Q46DU3
Q46DU5 Q46DV8 Q46DV9 Q46DW1 Q46DY5 Q46DY6 Q46DZ6 Q46DZ7 Q46DZ8 Q46E00 Q46E16
Q46E19 Q46E28 Q46E34 Q46E35 Q46E38 Q46E39 Q46E46 Q46E56 Q46E58 Q46E62 Q46E72
Q46E77 Q46E78 Q46E79 Q46EB1 Q46EC7 Q46EF4 Q46EG2 Q46EH1 Q46EH3 Q46EH4 Q46EH7
Q46EK8 Q46EL9 Q46EN9 Q46EU3 Q46EU7 Q46EU8 Q46EU9 Q46EW4 Q46EW5 Q46F10 Q46F14
Q46F18 Q46F20 Q46F25 Q46F27 Q46F30 Q46F76 Q46F78 Q46F79 Q46F99 Q46FA1 Q46FA3
Q46FA5 Q46FF6 Q46FK9 Q46FL0 Q46FL1 Q46FL2 Q46FL3 Q46FR6 Q46FS5 Q46FS6 Q46FS7
Q46FS9 Q46FV0 Q46FV2 Q46FV4 Q46FY8 Q46FZ5 Q46G00 Q46G04 Q46G05 Q46G12 Q46G14
Q46G28 Q46G29 Q46G70 Q46G95 Q46G96 Q46G97 Q46G98 Q46G99 Q46GA0 Q46GA1 Q46GA2
Q46GA3 Q46GA4 Q46GA5 Q46GA6 Q46GA7 Q46GA8 Q46GA9 Q46GB0 Q46GB1 Q46GB4 Q46GB5

Q46GB6 Q46GB7 Q46GB9 Q46GC2 Q46GC3 Q46GC7 Q46GC8 Q46GC9 Q46GE2 Q46GJ5 Q46GJ6
Q46GM0 Q46GM1 Q46GN1 Q46GN4 Q46GN5 Q46GN6 Q46GQ1 Q46GQ4 Q46GQ5 Q46GQ7 Q46GR7
Q46GR9 Q46GS3 Q46GS7 Q46GT1 Q46GT9 Q46GU2 Q46GU7 Q46GU9 Q46GV3 Q46GV6 Q46GW4
Q46GW7 Q46GY9 Q46GZ7 Q46H06 Q46H10 Q46H16 Q46H20 Q46H25 Q46H41 Q46H47 Q46H54
Q46H57 Q46H93 Q46H95 Q46HA8 Q46HB0 Q46HB2 Q46HB6 Q46HB7 Q46HB8 Q46HC4 Q46HD3
Q46HD5 Q46HD8 Q46HE0 Q46HE7 Q46HE9 Q46HF4 Q46HG9 Q46HH0 Q46HH1 Q46HH2 Q46HH3
Q46HH8 Q46HI4 Q46HI6 Q46HJ0 Q46HJ9 Q46HK1 Q46HK9 Q46HL7 Q46HP4 Q46HP8 Q46HR0
Q46HR4 Q46HU0 Q46HV9 Q46I00 Q46I07 Q46I08 Q46I09 Q46I11 Q46I13 Q46I26 Q46I27
Q46I36 Q46I40 Q46I41 Q46I42 Q46I43 Q46I46 Q46I50 Q46I55 Q46I59 Q46I72 Q46I73
Q46I76 Q46I95 Q46I97 Q46I98 Q46I99 Q46IA2 Q46IA7 Q46IA9 Q46IB4 Q46IB6 Q46IC0
Q46IC1 Q46IC6 Q46IC8 Q46IC9 Q46ID3 Q46ID4 Q46ID5 Q46ID7 Q46ID9 Q46IE1 Q46IE5
Q46IE7 Q46IH3 Q46IH4 Q46IJ5 Q46IJ6 Q46IK0 Q46IK7 Q46IM0 Q46IM1 Q46IM3 Q46IM4
Q46IN4 Q46IQ5 Q46IQ9 Q46IR0 Q46IR1 Q46IR2 Q46IR3 Q46IR4 Q46IR5 Q46IR6 Q46IR7
Q46IR8 Q46IR9 Q46IS0 Q46IS1 Q46IS2 Q46IS3 Q46IS4 Q46IS5 Q46IS6 Q46IS8 Q46IS9
Q46IT0 Q46IT1 Q46IT2 Q46IT3 Q46IT5 Q46IT7 Q46IT8 Q46IW1 Q46IW2 Q46IW3 Q46IW4
Q46IW5 Q46IX0 Q46IX2 Q46IX8 Q46IY3 Q46IY4 Q46IY5 Q46IY6 Q46IY9 Q46J13 Q46J16
Q46J18 Q46J19 Q46J20 Q46J22 Q46J23 Q46J24 Q46J26 Q46J33 Q46J47 Q46J69 Q46J70
Q46J72 Q46J74 Q46JA8 Q46JB7 Q46JB9 Q46JC1 Q46JC2 Q46JC3 Q46JC7 Q46JD7 Q46JD8
Q46JF6 Q46JF7 Q46JG1 Q46JG5 Q46JG6 Q46JH1 Q46JH4 Q46JH9 Q46JI5 Q46JK0 Q46JK1
Q46JK2 Q46JM7 Q46JM8 Q46JN0 Q46JN4 Q46JN5 Q46JQ9 Q46JR2 Q46JR6 Q46JR8 Q46JT6
Q46JT8 Q46JY8 Q46JZ5 Q46K01 Q46K05 Q46K32 Q46K33 Q46K45 Q46K52 Q46KI4 Q46KV4
Q46KV7 Q46KW1 Q46KX0 Q46KY2 Q46KZ6 Q46KZ7 Q46KZ8 Q46L08 Q46L10 Q46L14 Q46L15
Q46L16 Q46L18 Q46L21 Q46L29 Q46L31 Q46L36 Q46L44 Q46L57 Q46L58 Q46L60 Q46L68
Q46L69 Q46L72 Q46L78 Q46LD2 Q46LD7 Q46LE0 Q46LE2 Q46LE9 Q46LF6 Q46LG2 Q46LG3
Q46LH8 Q46LH9 Q46LM0 Q46LN9 Q46LS5 Q46LT2 Q46LT5 Q46LT7 Q46LU1 Q46LU3 Q46LV3
Q46LV5 Q46LW0 Q46LW6 Q46LW7 Q46LX8 Q46LZ6 Q46LZ9 Q46M54 Q46M57 Q46N53 Q46NS0
Q46NW7 Q46NW8 Q46P49 Q46P61 Q46PH8 Q46PI5 Q46PN1 Q46PQ4 Q46PS3 Q46PT7 Q46QA5
Q46QB2 Q46QJ0 Q46QQ0 Q46QQ8 Q46QU8 Q46QZ0 Q46RR4 Q46S32 Q46S75 Q46SS9 Q46TF2
Q46TH1 Q46TZ9 Q46UA4 Q46UT0 Q46UV8 Q46VL3 Q46VM0 Q46VT8 Q46VW9 Q46VX0 Q46VY4
Q46VZ0 Q46VZ5 Q46VZ7 Q46W52 Q46W60 Q46WC7 Q46WD0 Q46WD1 Q46WD2 Q46WD3 Q46WD4
Q46WD5 Q46WD8 Q46WD9 Q46WE0 Q46WE2 Q46WE4 Q46WE5 Q46WE6 Q46WE7 Q46WE8 Q46WE9
Q46WF0 Q46WF1 Q46WF2 Q46WF3 Q46WF4 Q46WF5 Q46WF6 Q46WF7 Q46WF8 Q46WF9 Q46WG0
Q46WG1 Q46WG2 Q46WG3 Q46WG5 Q46WG6 Q46WG7 Q46WG8 Q46WG9 Q46WH0 Q46WH5 Q46WI6
Q46WJ3 Q46WJ4 Q46WL0 Q46WL1 Q46WL2 Q46WL3 Q46WL4 Q46WL6 Q46WL7 Q46WL8 Q46WL9
Q46WM0 Q46WS9 Q46WT0 Q46WU4 Q46WU7 Q46WU8 Q46WY5 Q46WY6 Q46WZ2 Q46WZ5 Q46WZ6
Q46X04 Q46X09 Q46X10 Q46X15 Q46X16 Q46X17 Q46X18 Q46X20 Q46X21 Q46X58 Q46X89
Q46X95 Q46XA5 Q46XB3 Q46XB7 Q46XD7 Q46XD8 Q46XE1 Q46XF1 Q46XG4 Q46XG5 Q46XI7
Q46XI8 Q46XJ2 Q46XJ9 Q46XK2 Q46XK7 Q46XK9 Q46XL0 Q46XL1 Q46XL2 Q46XL9 Q46XM7
Q46XN8 Q46XN9 Q46XP0 Q46XQ9 Q46XT9 Q46XW5 Q46XW6 Q46XZ7 Q46Y42 Q46Y48 Q46Y50
Q46Y52 Q46Y54 Q46Y59 Q46Y82 Q46Y83 Q46Y84 Q46Y85 Q46Y87 Q46Y94 Q46Y95 Q46YA0
Q46YB8 Q46YE8 Q46YE9 Q46YH2 Q46YH7 Q46YI9 Q46YJ3 Q46YS9 Q46YU4 Q46YV7 Q46YV9
Q46YW0 Q46YW3 Q46YW7 Q46YX7 Q46YY3 Q46YZ3 Q46Z01 Q46Z03 Q46Z18 Q46Z20 Q46Z21
Q46Z24 Q46Z26 Q46Z37 Q46Z41 Q46Z81 Q46Z82 Q46Z98 Q46ZA1 Q46ZB3 Q46ZE6 Q46ZE9
Q46ZF8 Q46ZF9 Q46ZH1 Q46ZH2 Q46ZH7 Q46ZH9 Q46ZI0 Q46ZI4 Q46ZI7 Q46ZI8 Q46ZJ2

Q46ZJ3 Q46ZK0 Q46ZK1 Q46ZK2 Q46ZL2 Q46ZN9 Q46ZP1 Q46ZP2 Q46ZP3 Q46ZR3 Q46ZR5
Q46ZR6 Q46ZU3 Q46ZV6 Q47005 Q47013 Q47015 Q47036 Q47037 Q47077 Q47083 Q47098
Q470C3 Q470C6 Q470D2 Q470D5 Q470D7 Q470D8 Q470D9 Q470E0 Q470E3 Q470F1 Q470F3
Q470F7 Q470F9 Q470G4 Q470L8 Q470S7 Q47112 Q47129 Q47132 Q47133 Q47138 Q47140
Q47141 Q47147 Q47149 Q47150 Q47152 Q47154 Q47155 Q47156 Q47157 Q47158 Q47163
Q47189 Q471B8 Q471D2 Q471H6 Q471I0 Q471I1 Q471I2 Q471X2 Q471X9 Q47266 Q47268
Q47269 Q47270 Q47274 Q47281 Q47282 Q472B7 Q472C7 Q472D2 Q472D3 Q472D4 Q472D5
Q472F1 Q472F2 Q472G1 Q472J4 Q472J5 Q472L4 Q472N3 Q472N4 Q472N5 Q472N6 Q472N7
Q472Q5 Q47309 Q47316 Q47318 Q47319 Q47329 Q47334 Q473D3 Q473E4 Q473F6 Q473G7
Q473I5 Q473J2 Q473K3 Q473L4 Q473L7 Q473L9 Q473P0 Q473Q6 Q473Q7 Q473Q8 Q473Q9
Q473R0 Q473R2 Q473R3 Q473U5 Q473U7 Q473U8 Q473V1 Q473V5 Q473Z4 Q47400 Q47417
Q47418 Q47427 Q474C0 Q474C1 Q474C2 Q474D4 Q474E1 Q474F4 Q474I9 Q474J5 Q474K5
Q474L3 Q474L4 Q474N4 Q474N5 Q474P5 Q474U3 Q474U5 Q474U6 Q474V6 Q474W3 Q474X7
Q474Y3 Q474Y6 Q47505 Q47511 Q47526 Q47527 Q47536 Q47588 Q475B2 Q475F5 Q475F9
Q475G1 Q475J5 Q475L1 Q475M1 Q475M4 Q475N7 Q475N9 Q475Q1 Q475Q2 Q475R4 Q475R5
Q475R7 Q475R8 Q475S4 Q475S5 Q475S7 Q475T0 Q475T2 Q475T3 Q475T7 Q475W2 Q475X0
Q475Y4 Q475Y9 Q47608 Q47679 Q47684 Q47685 Q47688 Q47690 Q476D8 Q476E9 Q476F1
Q476F5 Q476F8 Q476G0 Q476G1 Q476G3 Q476J4 Q476J7 Q476K0 Q476M7 Q476N0 Q476N1
Q476S4 Q476S9 Q476T8 Q476U3 Q476U6 Q476V0 Q476V3 Q476W5 Q476X1 Q476Y3 Q476Y4
Q47710 Q47718 Q47719 Q47747 Q47748 Q47749 Q47758 Q47776 Q477A1 Q477A3 Q477A8
Q477B5 Q477C4 Q477E8 Q477F9 Q477G2 Q477G5 Q477G6 Q477H7 Q477I0 Q477I9 Q477Q1
Q477Q5 Q477V3 Q477W2 Q477X0 Q47810 Q47823 Q47824 Q47826 Q47827 Q47829 Q47836
Q47839 Q47840 Q47851 Q47860 Q47861 Q47862 Q47878 Q47881 Q47884 Q47887 Q478A1
Q478A2 Q478D8 Q478E2 Q478E3 Q478E5 Q478G3 Q478G7 Q478M5 Q478R6 Q478S1 Q478T2
Q478U6 Q478V1 Q478V2 Q478W0 Q478W1 Q478W3 Q478W6 Q478Y7 Q478Z2 Q478Z3 Q478Z5
Q47907 Q47914 Q47917 Q47921 Q47922 Q47944 Q47948 Q47950 Q47951 Q47953 Q47956
Q479A2 Q479A8 Q479A9 Q479B1 Q479B7 Q479E5 Q479E6 Q479G9 Q479H0 Q479H7 Q479I3
Q479J6 Q479L3 Q479M0 Q479M1 Q479M3 Q479M6 Q479P3 Q479P4 Q479P5 Q479R3 Q479R5
Q479S1 Q479S8 Q479T3 Q479U6 Q479W2 Q479W5 Q479X4 Q47A08 Q47A13 Q47A19 Q47A50
Q47A67 Q47A73 Q47A75 Q47A95 Q47A96 Q47AA2 Q47AA5 Q47AA6 Q47AB5 Q47AB8 Q47AC6
Q47AC7 Q47AC9 Q47AD0 Q47AD1 Q47AD2 Q47AF2 Q47AK9 Q47AL4 Q47AL5 Q47AL9 Q47AM0
Q47AM1 Q47AM2 Q47AM3 Q47AM4 Q47AM5 Q47AN8 Q47AR4 Q47AR7 Q47AU8 Q47AW0 Q47AX3
Q47AX4 Q47AZ9 Q47B12 Q47B13 Q47B44 Q47B46 Q47B64 Q47B65 Q47B66 Q47B76 Q47B77
Q47B80 Q47BA7 Q47BA8 Q47BA9 Q47BB1 Q47BD0 Q47BF5 Q47BF7 Q47BG6 Q47BG9 Q47BH8
Q47BI0 Q47BI5 Q47BI6 Q47BI7 Q47BI8 Q47BI9 Q47BJ0 Q47BJ2 Q47BK5 Q47BK8 Q47BL6
Q47BL9 Q47BM0 Q47BM2 Q47BM3 Q47BQ2 Q47BR2 Q47BR3 Q47BR6 Q47BR7 Q47BS0 Q47BS1
Q47BS6 Q47BS7 Q47C02 Q47C04 Q47C08 Q47C22 Q47C26 Q47C34 Q47C51 Q47C53 Q47CF0
Q47CF8 Q47CH1 Q47CM4 Q47CM6 Q47CM7 Q47CM8 Q47CM9 Q47CN0 Q47CW9 Q47D24 Q47D25
Q47D77 Q47D81 Q47D82 Q47D83 Q47D92 Q47D94 Q47D95 Q47D96 Q47D97 Q47D98 Q47DH9
Q47DJ8 Q47DJ9 Q47DK0 Q47DK3 Q47DK8 Q47DZ5 Q47E74 Q47EE8 Q47EF1 Q47EF4 Q47EF5
Q47EF7 Q47EH3 Q47EH5 Q47EI2 Q47EL1 Q47EL2 Q47EN5 Q47EP3 Q47ER3 Q47EU6 Q47F12
Q47F76 Q47F78 Q47F86 Q47F90 Q47F91 Q47F92 Q47F95 Q47FA0 Q47FA8 Q47FB0 Q47FB5
Q47FB6 Q47FB7 Q47FE4 Q47FN7 Q47FN9 Q47G40 Q47G52 Q47G53 Q47G54 Q47G55 Q47G56
Q47G57 Q47G58 Q47G67 Q47G87 Q47G91 Q47GC1 Q47GC6 Q47GC7 Q47GC8 Q47GC9 Q47GF4

Q47GG0 Q47GH9 Q47GJ3 Q47GJ8 Q47GP2 Q47GP8 Q47GQ6 Q47GQ9 Q47GS9 Q47GT0 Q47GU0
Q47GW8 Q47GX8 Q47GY0 Q47GZ8 Q47H37 Q47H71 Q47HH8 Q47HH9 Q47HI2 Q47HI9 Q47HJ4
Q47HJ5 Q47HJ6 Q47HK1 Q47HK2 Q47HK6 Q47HL4 Q47HL8 Q47HL9 Q47HM0 Q47HM4 Q47HQ4
Q47HQ5 Q47HQ6 Q47HQ7 Q47HR1 Q47HR2 Q47HR4 Q47HR7 Q47HS4 Q47HS9 Q47HW7 Q47HW8
Q47HZ8 Q47I44 Q47IC1 Q47IC5 Q47IC9 Q47ID0 Q47IE0 Q47IE5 Q47IE9 Q47IF6 Q47IG1
Q47IH0 Q47IH1 Q47II1 Q47II8 Q47II9 Q47IJ3 Q47IJ4 Q47IJ7 Q47IK0 Q47IN0 Q47IN4
Q47IP4 Q47IP5 Q47IR5 Q47IS1 Q47IS3 Q47IT5 Q47IX1 Q47IX6 Q47IZ2 Q47IZ8 Q47IZ9
Q47J76 Q47J77 Q47J78 Q47J79 Q47J80 Q47J81 Q47J82 Q47J84 Q47J85 Q47J86 Q47J87
Q47J88 Q47J89 Q47J90 Q47J91 Q47J92 Q47J93 Q47J94 Q47J95 Q47J96 Q47J97 Q47J98
Q47J99 Q47JA0 Q47JA1 Q47JA2 Q47JA3 Q47JA4 Q47JA5 Q47JA6 Q47JA7 Q47JA8 Q47JA9
Q47JB0 Q47JB1 Q47JB2 Q47JB3 Q47JB4 Q47JC0 Q47JD4 Q47JD5 Q47JE5 Q47JK6 Q47JK7
Q47JL4 Q47JL8 Q47JL9 Q47JN0 Q47JN4 Q47JQ0 Q47JQ1 Q47JQ3 Q47JS4 Q47JS8 Q47JU2
Q47JV2 Q47JV4 Q47JV5 Q47K02 Q47K28 Q47K40 Q47K50 Q47K67 Q47K71 Q47K72 Q47K73
Q47K77 Q47K94 Q47K96 Q47K97 Q47KH1 Q47KH3 Q47KH4 Q47KH7 Q47KQ7 Q47KR7 Q47KS6
Q47KS8 Q47KT3 Q47KU2 Q47KU5 Q47KV5 Q47KV7 Q47KW7 Q47L17 Q47LA0 Q47LA6 Q47LA7
Q47LB3 Q47LE2 Q47LG7 Q47LH2 Q47LH3 Q47LH5 Q47LI1 Q47LI2 Q47LI3 Q47LI4 Q47LI5
Q47LI6 Q47LI8 Q47LI9 Q47LJ0 Q47LJ1 Q47LJ2 Q47LJ3 Q47LJ4 Q47LJ6 Q47LJ7 Q47LJ8
Q47LJ9 Q47LK0 Q47LK1 Q47LK2 Q47LK3 Q47LK4 Q47LK5 Q47LK6 Q47LK7 Q47LK8 Q47LK9
Q47LL0 Q47LL1 Q47LL2 Q47LL4 Q47LL7 Q47LL8 Q47LL9 Q47LM0 Q47LM1 Q47LM2 Q47LM5
Q47LM6 Q47LM7 Q47LM9 Q47LN3 Q47LN7 Q47LP1 Q47LQ0 Q47LR8 Q47LS4 Q47LV0 Q47LX2
Q47M13 Q47M69 Q47M70 Q47MA0 Q47MD6 Q47MD7 Q47MS7 Q47MU1 Q47MU2 Q47MU3 Q47MU8
Q47MV4 Q47MV5 Q47MV6 Q47MV7 Q47MW1 Q47MY0 Q47MY6 Q47MZ1 Q47N28 Q47N39 Q47N40
Q47N41 Q47N42 Q47N43 Q47N45 Q47N50 Q47N66 Q47N67 Q47N71 Q47N72 Q47N75 Q47N76
Q47N79 Q47N80 Q47N81 Q47N84 Q47N85 Q47N93 Q47NA3 Q47NB5 Q47NB6 Q47NB8 Q47ND3
Q47NE4 Q47NK0 Q47NK1 Q47NL9 Q47NN7 Q47NS4 Q47NV4 Q47NX7 Q47NZ4 Q47NZ5 Q47NZ6
Q47PQ7 Q47PU3 Q47Q20 Q47Q21 Q47Q90 Q47QN2 Q47QN5 Q47QN9 Q47QR3 Q47QR5 Q47QS0
Q47QS3 Q47QS4 Q47QS5 Q47QS7 Q47QS8 Q47QS9 Q47QV9 Q47QW3 Q47QW8 Q47QX1 Q47QX7
Q47QX8 Q47QY5 Q47QY7 Q47QY8 Q47QY9 Q47QZ8 Q47R04 Q47R14 Q47R16 Q47R17 Q47R19
Q47R25 Q47R33 Q47R37 Q47R55 Q47R66 Q47R87 Q47R88 Q47R98 Q47RL8 Q47RM6 Q47RM7
Q47RM9 Q47RN4 Q47RP1 Q47RQ2 Q47RR3 Q47RR4 Q47RR8 Q47RS5 Q47RT4 Q47RU3 Q47RU5
Q47RU6 Q47RU8 Q47RU9 Q47RV1 Q47RV4 Q47RV7 Q47RX0 Q47RY0 Q47S46 Q47S50 Q47S51
Q47S52 Q47S53 Q47S60 Q47S63 Q47S64 Q47S65 Q47S67 Q47S76 Q47S77 Q47S78 Q47S79
Q47S81 Q47S84 Q47SA2 Q47SA3 Q47SA6 Q47SB4 Q47SB6 Q47SC0 Q47SC3 Q47SC4 Q47SC6
Q47SD2 Q47SE7 Q47SP3 Q47SR1 Q47SS3 Q47SW2 Q47SW3 Q47SX2 Q47SX4 Q47SZ0 Q47T68
Q47T77 Q47TB0 Q47TD1 Q47TE8 Q47TG4 Q47TI0 Q47TK1 Q47TL8 Q47TT4 Q47TY1 Q47TY2
Q47TZ5 Q47U20 Q47U23 Q47U25 Q47U32 Q47U36 Q47U38 Q47U39 Q47UB2 Q47UB3 Q47UN7
Q47UP4 Q47UP5 Q47UQ0 Q47UR1 Q47UT5 Q47UU9 Q47UV5 Q47UV6 Q47UV7 Q47UV8 Q47UV9
Q47UW0 Q47UW1 Q47UW2 Q47UW3 Q47UW8 Q47UY2 Q47UY4 Q47UY5 Q47UZ9 Q47V01 Q47V03
Q47V69 Q47V96 Q47VA3 Q47VB1 Q47VD0 Q47VD1 Q47VG7 Q47VH8 Q47VJ4 Q47VJ7 Q47VJ8
Q47VJ9 Q47VK9 Q47VL0 Q47VL2 Q47VL3 Q47VL4 Q47VM6 Q47VN0 Q47VQ0 Q47VQ1 Q47VQ7
Q47VR0 Q47VR1 Q47VS2 Q47VS4 Q47VT1 Q47VT2 Q47VU0 Q47VU1 Q47VY2 Q47VY5 Q47VZ4
Q47VZ9 Q47W03 Q47W08 Q47W35 Q47W36 Q47W53 Q47W59 Q47W60 Q47W65 Q47W67 Q47WB0
Q47WB6 Q47WB7 Q47WC0 Q47WC1 Q47WC5 Q47WD1 Q47WD4 Q47WG1 Q47WG2 Q47WG3 Q47WJ3
Q47WK1 Q47WL9 Q47WM2 Q47WN0 Q47WN3 Q47WN5 Q47WN6 Q47WP5 Q47WP8 Q47WQ1 Q47WQ9

Q47WR3 Q47WR5 Q47WT4 Q47WT5 Q47WU8 Q47WU9 Q47WV0 Q47WV1 Q47WV5 Q47WW8 Q47WY2
Q47XA7 Q47XA8 Q47XB3 Q47XB4 Q47XB5 Q47XB6 Q47XB7 Q47XB8 Q47XD4 Q47XD5 Q47XG2
Q47XG4 Q47XH8 Q47XI3 Q47XI4 Q47XI6 Q47XI7 Q47XJ9 Q47XK1 Q47XK4 Q47XL5 Q47XL6
Q47XL7 Q47XL8 Q47XL9 Q47XN8 Q47XQ8 Q47XR0 Q47XU3 Q47XW2 Q47XX0 Q47XX7 Q47Y27
Q47Y31 Q47Y68 Q47Y78 Q47Y80 Q47Y87 Q47Y88 Q47Y90 Q47Y92 Q47Y94 Q47YA5 Q47YA8
Q47YB3 Q47YC2 Q47YC9 Q47YI0 Q47YI6 Q47YJ3 Q47YJ7 Q47YJ8 Q47YL9 Q47Z09 Q47Z13
Q47Z18 Q47Z40 Q47Z50 Q47Z58 Q47Z78 Q47Z95 Q47ZC3 Q47ZE0 Q47ZQ7 Q47ZS4 Q47ZS5
Q47ZS6 Q47ZU5 Q47ZU7 Q47ZY2 Q48082 Q480A9 Q480B1 Q480B2 Q480B3 Q480B8 Q480C4
Q480C7 Q480C9 Q480D3 Q480H5 Q480H7 Q480P2 Q480P3 Q480P4 Q480Q2 Q48154 Q481E4
Q481G0 Q481G3 Q481H1 Q481H3 Q481S6 Q481U4 Q48214 Q48215 Q48230 Q48255 Q48260
Q48295 Q48296 Q482F9 Q482G2 Q482G4 Q482G5 Q482G8 Q482J5 Q482K3 Q482K5 Q482M9
Q482P4 Q482P5 Q482S1 Q482T5 Q482T6 Q482T7 Q482T8 Q482T9 Q482U1 Q482Z9 Q48303
Q48311 Q48325 Q48328 Q48335 Q48389 Q483A0 Q483B4 Q483C4 Q483C5 Q483C6 Q483C7
Q483C9 Q483D0 Q483D3 Q483H8 Q483Q8 Q483R0 Q483R5 Q483R6 Q483W3 Q48409 Q48413
Q48433 Q48436 Q48439 Q48440 Q48441 Q48447 Q48448 Q48449 Q48451 Q48453 Q48454
Q48455 Q48458 Q48459 Q48462 Q48463 Q48464 Q48468 Q48493 Q484F1 Q484J7 Q484Q5
Q484R1 Q484R7 Q484R8 Q484U8 Q48514 Q48543 Q48544 Q48558 Q48585 Q485E3 Q485F4
Q485G4 Q485G7 Q485G8 Q485G9 Q485H0 Q485H4 Q485H5 Q485H8 Q485J1 Q485J2 Q485Q8
Q485R8 Q485S0 Q48660 Q48661 Q48662 Q48665 Q48677 Q486F9 Q486G0 Q486J6 Q486J8
Q486L6 Q486M0 Q486M1 Q486R1 Q486R8 Q486U0 Q486U3 Q486Y6 Q486Y7 Q486Z0 Q48708
Q48709 Q48712 Q48722 Q48725 Q48726 Q48727 Q48733 Q48734 Q48735 Q48739 Q48745
Q48759 Q48761 Q48762 Q48791 Q48793 Q48796 Q487A3 Q487A4 Q487B0 Q487B1 Q487C3
Q487C7 Q487D1 Q487D3 Q487E5 Q487E6 Q487E7 Q487E8 Q487E9 Q487G8 Q487H4 Q487H5
Q487H6 Q487J6 Q487J9 Q487K8 Q487P4 Q487Q9 Q487R0 Q487R8 Q487V5 Q487Z1 Q487Z2
Q487Z3 Q487Z4 Q487Z5 Q487Z6 Q487Z7 Q487Z8 Q487Z9 Q48806 Q48833 Q48835 Q48838
Q48841 Q48846 Q48847 Q48864 Q48878 Q48879 Q48883 Q48884 Q488A0 Q488A1 Q488A2
Q488A4 Q488J6 Q488J7 Q488M4 Q488M6 Q488N5 Q488N6 Q488Q3 Q488Y0 Q488Y7 Q488Y8
Q488Y9 Q488Z0 Q488Z1 Q488Z2 Q488Z4 Q488Z5 Q488Z6 Q488Z7 Q488Z8 Q488Z9 Q48900
Q48907 Q48924 Q48925 Q48927 Q48935 Q48943 Q48949 Q48952 Q48960 Q48965 Q48972
Q48979 Q48980 Q489A0 Q489A1 Q489A2 Q489A3 Q489A4 Q489F4 Q489G6 Q489N3 Q489N4
Q489P3 Q489P5 Q489Q8 Q489T8 Q489T9 Q489U1 Q48A12 Q48A14 Q48A17 Q48A22 Q48A23
Q48A24 Q48A63 Q48A85 Q48A93 Q48A96 Q48AC6 Q48AD4 Q48AD6 Q48AD7 Q48AG4 Q48AK7
Q48AM0 Q48AM4 Q48AN2 Q48AN3 Q48AN9 Q48AR8 Q48AS2 Q48AS5 Q48AS7 Q48AS9 Q48AT5
Q48AT8 Q48AU1 Q48AU5 Q48AV5 Q48AW7 Q48AW8 Q48BE9 Q48BF0 Q48BF3 Q48BF4 Q48BF5
Q48BL6 Q48BQ6 Q48BY7 Q48BY8 Q48BZ2 Q48C02 Q48C04 Q48C72 Q48C77 Q48C78 Q48C80
Q48C81 Q48C87 Q48C90 Q48CC1 Q48CC6 Q48CD3 Q48CE0 Q48CH3 Q48CH7 Q48CJ2 Q48CK1
Q48CL9 Q48CM0 Q48CM5 Q48CM6 Q48CM7 Q48CP3 Q48CS1 Q48CS2 Q48CT3 Q48CT4 Q48CT5
Q48CT6 Q48CT7 Q48D01 Q48D21 Q48D25 Q48D26 Q48D27 Q48D28 Q48D29 Q48D30 Q48D31
Q48D32 Q48D33 Q48D34 Q48D35 Q48D36 Q48D37 Q48D38 Q48D39 Q48D40 Q48D41 Q48D42
Q48D43 Q48D44 Q48D45 Q48D46 Q48D47 Q48D48 Q48D49 Q48D50 Q48D51 Q48D52 Q48D53
Q48D54 Q48D55 Q48D57 Q48D58 Q48D59 Q48D60 Q48D61 Q48D62 Q48DB7 Q48DC2 Q48DC7
Q48DC8 Q48DC9 Q48DE6 Q48DE7 Q48DF0 Q48DF1 Q48DG3 Q48DI0 Q48DI3 Q48DL2 Q48DL4
Q48DL7 Q48DM5 Q48DM6 Q48DN1 Q48DN6 Q48DN8 Q48DP1 Q48DP2 Q48DQ3 Q48DT0 Q48DT7
Q48DU7 Q48DZ2 Q48E20 Q48E42 Q48E54 Q48E57 Q48E61 Q48E62 Q48E63 Q48E64 Q48E69

Q48E72 Q48E73 Q48E75 Q48E77 Q48E78 Q48E79 Q48E80 Q48E81 Q48E91 Q48E92 Q48E93
Q48E97 Q48EB2 Q48EC7 Q48EC8 Q48EC9 Q48ED0 Q48ED3 Q48EE0 Q48EE1 Q48EE6 Q48EE9
Q48EF0 Q48EF6 Q48EF9 Q48EG7 Q48EH2 Q48EI4 Q48EI5 Q48ET5 Q48EU3 Q48EV3 Q48EV5
Q48EV6 Q48EV7 Q48EV8 Q48F01 Q48F11 Q48F12 Q48F14 Q48F26 Q48F34 Q48F36 Q48F42
Q48F49 Q48F57 Q48F59 Q48F60 Q48F61 Q48F62 Q48F65 Q48F73 Q48F77 Q48F81 Q48F85
Q48F86 Q48F87 Q48F88 Q48F92 Q48F94 Q48F95 Q48FA7 Q48FB2 Q48FB4 Q48FB7 Q48FC1
Q48FC2 Q48FC3 Q48FC4 Q48FC5 Q48FD4 Q48FG7 Q48FG8 Q48FH7 Q48FI4 Q48FJ7 Q48FJ9
Q48FL6 Q48FM4 Q48FM6 Q48FN1 Q48FN3 Q48FP6 Q48FR2 Q48FT7 Q48G05 Q48G09 Q48G19
Q48G20 Q48G22 Q48G25 Q48GB5 Q48GE1 Q48GH3 Q48GK0 Q48GK6 Q48GS7 Q48GY3 Q48H03
Q48H61 Q48H65 Q48H67 Q48H68 Q48H69 Q48H74 Q48H75 Q48HJ1 Q48HN1 Q48HR7 Q48HV8
Q48IB1 Q48IE2 Q48IG5 Q48IH8 Q48IK9 Q48J73 Q48J96 Q48JG9 Q48JI0 Q48JK4 Q48JR7
Q48JR8 Q48JR9 Q48JS0 Q48JS1 Q48JS3 Q48JU8 Q48JX8 Q48JX9 Q48JZ3 Q48JZ6 Q48K64
Q48K68 Q48K97 Q48K98 Q48K99 Q48KA5 Q48KA6 Q48KH5 Q48KI2 Q48KI8 Q48KM6 Q48KM7
Q48KM8 Q48KN0 Q48KP5 Q48KQ0 Q48KQ3 Q48KQ4 Q48KQ5 Q48KR2 Q48KS4 Q48KS5 Q48KS9
Q48KX6 Q48KX8 Q48KY9 Q48KZ0 Q48KZ1 Q48KZ8 Q48L06 Q48L25 Q48L36 Q48L44 Q48L46
Q48L51 Q48L53 Q48L73 Q48L75 Q48L80 Q48LA4 Q48LC2 Q48LC5 Q48LC8 Q48LD2 Q48LD5
Q48LF6 Q48LG4 Q48LP5 Q48LQ7 Q48LR7 Q48LT4 Q48LT7 Q48LV7 Q48LV8 Q48LV9 Q48LW0
Q48LW5 Q48LX3 Q48LY0 Q48LY3 Q48LY5 Q48LY6 Q48LZ0 Q48LZ3 Q48LZ4 Q48LZ7 Q48LZ8
Q48M01 Q48M05 Q48M15 Q48ME3 Q48MF2 Q48MF3 Q48MF9 Q48MI2 Q48MI5 Q48MQ4 Q48MQ5
Q48MQ6 Q48MQ7 Q48MQ8 Q48MR4 Q48MT3 Q48MT5 Q48MT7 Q48MV4 Q48MV5 Q48MV8 Q48MW0
Q48MW1 Q48N21 Q48N66 Q48N77 Q48N86 Q48N87 Q48N88 Q48NC2 Q48ND8 Q48NK3 Q48NK6
Q48NK9 Q48NL1 Q48NL2 Q48NL3 Q48NL4 Q48NL9 Q48NN3 Q48NN5 Q48NN6 Q48NN7 Q48NP3
Q48NP7 Q48NP8 Q48NS2 Q48NT7 Q48NT9 Q48NU0 Q48NU2 Q48NU4 Q48NU8 Q48NU9 Q48NX0
Q48NY1 Q48NY2 Q48NZ0 Q48NZ2 Q48NZ3 Q48NZ7 Q48NZ8 Q48P02 Q48P03 Q48P10 Q48P11
Q48P32 Q48P39 Q48P81 Q48P90 Q48P91 Q48P92 Q48P93 Q48PA1 Q48PA6 Q48PB0 Q48PB5
Q48PD1 Q48PD2 Q48PG2 Q48PH0 Q48PH1 Q48PI0 Q48PI2 Q48PI4 Q48PI5 Q48PJ4 Q48PJ7
Q48PJ8 Q48PK8 Q48PL3 Q48PR0 Q48PV6 Q48PX1 Q48PY9 Q48PZ1 Q48Q02 Q48Q04 Q48Q06
Q48Q08 Q48Q10 Q48Q15 Q48Q16 Q48Q55 Q48QC3 Q48QD3 Q48QG6 Q48QG7 Q48QH6 Q48QH8
Q48QI2 Q48QJ1 Q48QJ8 Q48QK0 Q48QK3 Q48QL2 Q48QM8 Q48QN0 Q48QN2 Q48QN5 Q48QS5
Q48QS6 Q48QS7 Q48QS8 Q48QT3 Q48QT6 Q48QW4 Q48QW6 Q48QW7 Q48QW9 Q48QX0 Q48QX1
Q48QX8 Q48QY5 Q48QY6 Q48QY8 Q48QZ2 Q48QZ6 Q48R03 Q48R12 Q48R70 Q48R77 Q48R89
Q48R90 Q48R93 Q48R96 Q48RA2 Q48RA7 Q48RB6 Q48RC2 Q48RC3 Q48RC4 Q48RC8 Q48RE4
Q48RE9 Q48RF1 Q48RF4 Q48RF9 Q48RG4 Q48RG7 Q48RG8 Q48RH2 Q48RI2 Q48RI5 Q48RJ4
Q48RJ8 Q48RK8 Q48RL0 Q48RL6 Q48RL8 Q48RM8 Q48RQ4 Q48RQ5 Q48RR1 Q48RR3 Q48RS8
Q48RU3 Q48RU4 Q48RU8 Q48RU9 Q48RV9 Q48RW0 Q48RX2 Q48RX3 Q48RX4 Q48RX6 Q48RY1
Q48RY6 Q48RY7 Q48RZ0 Q48S05 Q48S06 Q48S07 Q48S18 Q48S19 Q48S21 Q48S38 Q48S39
Q48S56 Q48S61 Q48SA4 Q48SI7 Q48SJ3 Q48SJ5 Q48SK0 Q48SK8 Q48SL4 Q48SL8 Q48SM7
Q48SM9 Q48SP2 Q48SP7 Q48SP8 Q48SQ1 Q48SQ4 Q48SR6 Q48SS4 Q48SU7 Q48SU8 Q48SV4
Q48SX1 Q48SX8 Q48SZ1 Q48SZ3 Q48T86 Q48TA2 Q48TB3 Q48TC9 Q48TD0 Q48TE8 Q48TF6
Q48TG2 Q48TG9 Q48TH6 Q48TI1 Q48TJ0 Q48TK0 Q48TK6 Q48TL0 Q48TL1 Q48TL5 Q48TL6
Q48TM3 Q48TM9 Q48TN1 Q48TP5 Q48TS3 Q48TS4 Q48TS5 Q48TS8 Q48TV1 Q48TY5 Q48TY8
Q48TZ1 Q48TZ6 Q48U05 Q48U09 Q48U10 Q48U17 Q48U18 Q48U21 Q48U22 Q48U25 Q48U28
Q48U40 Q48U54 Q48U55 Q48U56 Q48U63 Q48U71 Q48U73 Q48U79 Q48U81 Q48U87 Q48U88
Q48U89 Q48U92 Q48U93 Q48U94 Q48U95 Q48U99 Q48UB6 Q48UB8 Q48UB9 Q48UC5 Q48UD0

Q48UE1 Q48UG3 Q48UG8 Q48UH3 Q48UH5 Q48UH6 Q48UK4 Q48UK5 Q48UR6 Q48US7 Q48UT7
Q48UT9 Q48UU2 Q48UU5 Q48UU6 Q48UW8 Q48UX3 Q48UX4 Q48UX6 Q48UX7 Q48UX8 Q48UX9
Q48V07 Q48V20 Q48V24 Q48V25 Q48V26 Q48V28 Q48V41 Q48V44 Q48V45 Q48V46 Q48V50
Q48V60 Q48V63 Q48V66 Q48V72 Q48V81 Q48V83 Q48VB6 Q48VB7 Q48VB8 Q48VC6 Q48VD6
Q48VD7 Q48VE1 Q48VE5 Q48VF9 Q48VG9 Q48VJ7 Q48VK8 Q48VN9 Q48VQ0 Q48VR0 Q48VR1
Q48VR3 Q48VS2 Q48VS3 Q48VS4 Q48VS5 Q48VS6 Q48VS7 Q48VS8 Q48VT0 Q48VT1 Q48VT2
Q48VT3 Q48VT4 Q48VT5 Q48VT6 Q48VT7 Q48VT8 Q48VT9 Q48VU0 Q48VU1 Q48VU2 Q48VU3
Q48VU4 Q48VU5 Q48VU6 Q48VU7 Q48VU8 Q48VU9 Q48VV0 Q48VW8 Q48VX2 Q48VX3 Q48VX9
Q48VY1 Q48VY9 Q48WA5 Q48XR0 Q49006 Q49025 Q49091 Q49092 Q49093 Q490D8 Q49110
Q49112 Q49113 Q49115 Q49117 Q49118 Q49127 Q49129 Q49131 Q49136 Q49148 Q49149
Q49150 Q49155 Q49156 Q49157 Q49158 Q49160 Q49161 Q49162 Q49163 Q49164 Q49166
Q49168 Q49171 Q49172 Q49173 Q49174 Q49178 Q49179 Q49180 Q49184 Q491U9 Q491V0
Q491V4 Q491V7 Q491W2 Q491W4 Q491W5 Q491W8 Q491X0 Q491X1 Q491X2 Q491X9 Q491Y3
Q491Y5 Q491Z0 Q491Z2 Q491Z6 Q491Z7 Q492A3 Q492A4 Q492A5 Q492A9 Q492B0 Q492B1
Q492B2 Q492B5 Q492B6 Q492B7 Q492B8 Q492B9 Q492C0 Q492C6 Q492C7 Q492D1 Q492D2
Q492D5 Q492D9 Q492E0 Q492E1 Q492E3 Q492E5 Q492E6 Q492E7 Q492E8 Q492E9 Q492F0
Q492F1 Q492F3 Q492F5 Q492F7 Q492G5 Q492G6 Q492G9 Q492H2 Q492H3 Q492J5 Q492J6
Q492J7 Q492J8 Q492K0 Q492K2 Q492K3 Q492K4 Q492K6 Q492L0 Q492L1 Q492M3 Q492N1
Q492N6 Q492N9 Q492P3 Q492P5 Q492Q2 Q492Q6 Q492R5 Q492S0 Q492S4 Q492S5 Q492S6
Q492T1 Q492T7 Q492T8 Q492T9 Q492U0 Q492U9 Q492V0 Q492V1 Q492V2 Q492V3 Q492V8
Q492W0 Q492W5 Q492W6 Q492Z0 Q492Z3 Q49329 Q49373 Q49374 Q49375 Q49396 Q49398
Q493A0 Q493A1 Q493A3 Q493A5 Q493B0 Q493B4 Q493B5 Q493B8 Q493C8 Q493C9 Q493D0
Q493D1 Q493D2 Q493D4 Q493D9 Q493E1 Q493E3 Q493E8 Q493F0 Q493F2 Q493F3 Q493F4
Q493F6 Q493F8 Q493G5 Q493G7 Q493G9 Q493H3 Q493H7 Q493H8 Q493H9 Q493I0 Q493I1
Q493I2 Q493I3 Q493I4 Q493I5 Q493I6 Q493I7 Q493I8 Q493J0 Q493J1 Q493J2 Q493J4
Q493J5 Q493J6 Q493J7 Q493J8 Q493J9 Q493K1 Q493K2 Q493K3 Q493K4 Q493K5 Q493K6
Q493K7 Q493K8 Q493K9 Q493L3 Q493M0 Q493M1 Q493M2 Q493M3 Q493M5 Q493M6 Q493M8
Q493N1 Q493N3 Q493N6 Q493N7 Q493P4 Q493Q0 Q493Q3 Q493Q9 Q493R0 Q493R1 Q493R2
Q493R3 Q493R7 Q493S0 Q493S1 Q493S3 Q493S5 Q493S6 Q493S7 Q493S9 Q493T3 Q493T4
Q493T5 Q493T6 Q493T7 Q493U0 Q493U3 Q493U4 Q493U5 Q493U6 Q493U7 Q493V3 Q493V4
Q493V5 Q493V9 Q493W0 Q493W6 Q493W7 Q493W8 Q493X3 Q493X4 Q493X8 Q493X9 Q493Y5
Q493Y6 Q493Y9 Q493Z9 Q49400 Q49404 Q49405 Q49406 Q49407 Q49408 Q49415 Q49416
Q49418 Q49419 Q49420 Q49421 Q49422 Q49423 Q49424 Q49425 Q49426 Q49428 Q49429
Q49434 Q49435 Q49436 Q49467 Q49492 Q494B7 Q494B8 Q494C0 Q494D6 Q494D7 Q494D8
Q494E6 Q494E8 Q494F3 Q494F8 Q494Q1 Q494R0 Q494R4 Q494V2 Q494X3 Q49539 Q49564
Q49573 Q49593 Q49594 Q495C1 Q495D7 Q495X7 Q495Y7 Q495Y8 Q495Z4 Q49605 Q49608
Q49610 Q49611 Q49616 Q49617 Q49622 Q49640 Q49645 Q49649 Q49670 Q49682 Q49689
Q49690 Q496A3 Q496M5 Q496Y0 Q496Z0 Q496Z9 Q49700 Q49721 Q49723 Q49729 Q49734
Q49736 Q49741 Q49742 Q49744 Q49751 Q49752 Q49755 Q49757 Q49762 Q49775 Q49776
Q49780 Q49784 Q49786 Q497B0 Q497B8 Q497C3 Q497H0 Q497I4 Q497K5 Q497M3 Q497N6
Q497N7 Q497P3 Q497Q6 Q497V5 Q497V6 Q49808 Q49823 Q49842 Q49844 Q49864 Q49869
Q49884 Q49885 Q49890 Q49896 Q49897 Q498C5 Q498D1 Q498D9 Q498E2 Q498E6 Q498E7
Q498G2 Q498H0 Q498J7 Q498J9 Q498K9 Q498L0 Q498L2 Q498L9 Q498M4 Q498M5 Q498P2
Q498R1 Q498R7 Q498S6 Q498T2 Q498T3 Q498T4 Q498U4 Q498Z6 Q49900 Q49901 Q49945

Q49946 Q49988 Q49992 Q499A3 Q499B3 Q499D0 Q499E5 Q499E6 Q499M4 Q499M5 Q499N3
Q499N6 Q499R0 Q499T7 Q499U2 Q499U4 Q499U6 Q499V3 Q499V6 Q499W2 Q499X9 Q499Y3
Q499Z3 Q499Z4 Q49A33 Q49A92 Q49AG3 Q49AM1 Q49AM3 Q49AR2 Q49AS3 Q49CA2 Q49CB8
Q49CC1 Q49I55 Q49I57 Q49KF8 Q49KF9 Q49KG0 Q49KL9 Q49KT4 Q49KT5 Q49KT6 Q49KT8
Q49KT9 Q49KU1 Q49KV0 Q49KV8 Q49KV9 Q49KW0 Q49KW1 Q49KW2 Q49KW3 Q49KW4 Q49KW5
Q49KW6 Q49KX3 Q49KX4 Q49KX6 Q49KX7 Q49KX8 Q49KZ0 Q49KZ6 Q49L00 Q49L06 Q49L07
Q49L08 Q49L09 Q49L16 Q49L17 Q49PZ0 Q49SP3 Q49SP4 Q49SP5 Q49SP6 Q49SP7 Q49U13
Q49UF2 Q49UH1 Q49UI2 Q49UI4 Q49UI5 Q49UI6 Q49UL7 Q49UL9 Q49UM4 Q49UM5 Q49UQ0
Q49UQ2 Q49UT8 Q49UU0 Q49UU5 Q49UU6 Q49UU8 Q49UU9 Q49UX2 Q49UY8 Q49UY9 Q49UZ1
Q49UZ5 Q49UZ7 Q49UZ9 Q49V02 Q49V04 Q49V07 Q49V09 Q49V10 Q49V11 Q49V12 Q49V21
Q49V26 Q49V28 Q49V29 Q49V31 Q49V33 Q49V34 Q49V37 Q49V39 Q49V41 Q49V44 Q49V47
Q49V48 Q49V49 Q49V50 Q49V52 Q49V53 Q49V55 Q49V56 Q49V57 Q49V58 Q49V87 Q49VA1
Q49VB1 Q49VB3 Q49VB6 Q49VB7 Q49VD1 Q49VE0 Q49VE6 Q49VF3 Q49VL1 Q49VM7 Q49VN8
Q49VQ7 Q49VS0 Q49VS6 Q49VS8 Q49VT7 Q49VU2 Q49VV1 Q49VV3 Q49VV6 Q49VV8 Q49VW3
Q49VW5 Q49VY9 Q49VZ2 Q49VZ4 Q49W00 Q49W01 Q49W02 Q49W09 Q49W17 Q49W40 Q49W41
Q49W45 Q49W57 Q49W60 Q49W64 Q49W68 Q49W78 Q49W93 Q49W96 Q49W99 Q49WA0 Q49WA1
Q49WA5 Q49WA6 Q49WA8 Q49WB0 Q49WC6 Q49WC7 Q49WD1 Q49WD4 Q49WD6 Q49WE4 Q49WE9
Q49WG0 Q49WG6 Q49WG8 Q49WI7 Q49WI9 Q49WJ0 Q49WJ2 Q49WJ3 Q49WJ5 Q49WJ7 Q49WL3
Q49WL4 Q49WM2 Q49WN1 Q49WN7 Q49WP9 Q49WQ0 Q49WQ2 Q49WQ5 Q49WQ6 Q49WQ7 Q49WR1
Q49WR2 Q49WR3 Q49WR7 Q49WS7 Q49WS9 Q49WT0 Q49WT1 Q49WT7 Q49WV8 Q49WV9 Q49WW0
Q49WW1 Q49WW5 Q49WW9 Q49WX1 Q49WX5 Q49WX9 Q49WY1 Q49WY3 Q49WY4 Q49WY5 Q49WY6
Q49WZ1 Q49WZ2 Q49WZ9 Q49X02 Q49X05 Q49X07 Q49X17 Q49X24 Q49X25 Q49X26 Q49X27
Q49X31 Q49X32 Q49X35 Q49X36 Q49X37 Q49X38 Q49X39 Q49X40 Q49X41 Q49X42 Q49X43
Q49X44 Q49X45 Q49X48 Q49X49 Q49X50 Q49X54 Q49X58 Q49X59 Q49X61 Q49X62 Q49X63
Q49X73 Q49X85 Q49X88 Q49X89 Q49X93 Q49X94 Q49X96 Q49XB7 Q49XC1 Q49XD0 Q49XD1
Q49XD2 Q49XE0 Q49XE1 Q49XE6 Q49XF1 Q49XF7 Q49XG3 Q49XG5 Q49XH5 Q49XH6 Q49XH8
Q49XH9 Q49XJ2 Q49XJ7 Q49XJ8 Q49XJ9 Q49XK0 Q49XK3 Q49XK7 Q49XL1 Q49XM4 Q49XM5
Q49XN2 Q49XN4 Q49XN8 Q49XN9 Q49XP0 Q49XQ3 Q49XQ5 Q49XR0 Q49XR1 Q49XR3 Q49XS0
Q49XS1 Q49XS2 Q49XS3 Q49XS5 Q49XS9 Q49XT0 Q49XT1 Q49XU2 Q49XU3 Q49XU5 Q49XV0
Q49XV1 Q49XV9 Q49XW5 Q49XW8 Q49XW9 Q49XX3 Q49XX9 Q49XY0 Q49XY1 Q49XY2 Q49XZ4
Q49XZ6 Q49Y00 Q49Y06 Q49Y08 Q49Y09 Q49Y13 Q49Y18 Q49Y20 Q49Y21 Q49Y22 Q49Y23
Q49Y24 Q49Y27 Q49Y35 Q49Y37 Q49Y40 Q49Y42 Q49Y48 Q49Y49 Q49Y53 Q49Y54 Q49Y55
Q49Y56 Q49Y59 Q49Y63 Q49Y68 Q49Y69 Q49Y71 Q49Y73 Q49Y78 Q49Y79 Q49Y80 Q49Y81
Q49Y82 Q49Y83 Q49Y85 Q49Y92 Q49Y96 Q49Y99 Q49YA2 Q49YA4 Q49YA5 Q49YA6 Q49YA7
Q49YB0 Q49YB1 Q49YB2 Q49YB4 Q49YB7 Q49YB9 Q49YC7 Q49YD6 Q49YD7 Q49YD9 Q49YE0
Q49YE2 Q49YE4 Q49YE8 Q49YF2 Q49YG2 Q49YH1 Q49YH4 Q49YH9 Q49YI2 Q49YI8 Q49YJ3
Q49YJ6 Q49YJ7 Q49YL6 Q49YL7 Q49YM6 Q49YM7 Q49YN8 Q49YP8 Q49YQ6 Q49YQ9 Q49YR2
Q49YR8 Q49YS0 Q49YS5 Q49YS6 Q49YT7 Q49YT9 Q49YU3 Q49YU4 Q49YU5 Q49YU8 Q49YV3
Q49YV6 Q49YW3 Q49YW6 Q49YY5 Q49YY6 Q49Z00 Q49Z04 Q49Z08 Q49Z11 Q49Z12 Q49Z13
Q49Z14 Q49Z15 Q49Z16 Q49Z17 Q49Z22 Q49Z23 Q49Z24 Q49Z29 Q49Z31 Q49Z39 Q49Z40
Q49Z42 Q49Z47 Q49Z59 Q49Z60 Q49Z62 Q49Z65 Q49Z66 Q49Z67 Q49Z69 Q49Z71 Q49Z73
Q49Z74 Q49Z76 Q49Z82 Q49Z84 Q49ZA6 Q49ZA7 Q49ZB5 Q49ZB7 Q49ZB8 Q49ZC7 Q49ZD5
Q49ZD6 Q49ZD7 Q49ZE1 Q49ZE2 Q49ZE3 Q49ZE4 Q49ZE5 Q49ZE6 Q49ZE9 Q49ZF0 Q49ZF1
Q49ZF2 Q49ZF3 Q49ZF4 Q49ZF5 Q49ZF6 Q49ZF7 Q49ZF8 Q49ZF9 Q49ZG0 Q49ZG1 Q49ZG2

Q49ZG3 Q49ZG4 Q49ZG5 Q49ZG6 Q49ZG7 Q49ZG8 Q49ZG9 Q49ZH2 Q49ZI6 Q49ZI7 Q49ZJ2
Q49ZK0 Q49ZK2 Q49ZM5 Q49ZN0 Q49ZN6 Q49ZQ6 Q49ZQ7 Q49ZR6 Q49ZR8 Q49ZS3 Q49ZU9
Q49ZX3 Q49ZZ2 Q4A002 Q4A040 Q4A041 Q4A043 Q4A044 Q4A046 Q4A047 Q4A048 Q4A049
Q4A054 Q4A071 Q4A087 Q4A094 Q4A0A8 Q4A0E7 Q4A0F0 Q4A0G2 Q4A0J3 Q4A0J4 Q4A0J5
Q4A0J6 Q4A0J7 Q4A0J8 Q4A0J9 Q4A0M9 Q4A0N0 Q4A0N1 Q4A0N2 Q4A0Q1 Q4A0R6 Q4A0S3
Q4A0T9 Q4A0U0 Q4A0W6 Q4A127 Q4A155 Q4A164 Q4A166 Q4A172 Q4A173 Q4A177 Q4A180
Q4A548 Q4A577 Q4A578 Q4A587 Q4A589 Q4A597 Q4A5A8 Q4A5B4 Q4A5C0 Q4A5C1 Q4A5C2
Q4A5C4 Q4A5C5 Q4A5C6 Q4A5C7 Q4A5C8 Q4A5C9 Q4A5D0 Q4A5D1 Q4A5D2 Q4A5D3 Q4A5D4
Q4A5D5 Q4A5D6 Q4A5D7 Q4A5D8 Q4A5D9 Q4A5E0 Q4A5E1 Q4A5E2 Q4A5E8 Q4A5F3 Q4A5F4
Q4A5F6 Q4A5F7 Q4A5G1 Q4A5G4 Q4A5I6 Q4A5I7 Q4A5I8 Q4A5I9 Q4A5J0 Q4A5J1 Q4A5J6
Q4A5K1 Q4A5K5 Q4A5L2 Q4A5L3 Q4A5M9 Q4A5N1 Q4A5P7 Q4A5P8 Q4A5Q7 Q4A5S0 Q4A5S6
Q4A5S7 Q4A5S8 Q4A5S9 Q4A5T2 Q4A5T4 Q4A5T5 Q4A5V9 Q4A5W6 Q4A5X2 Q4A5X8 Q4A5Y3
Q4A5Y8 Q4A5Y9 Q4A5Z5 Q4A5Z6 Q4A611 Q4A620 Q4A632 Q4A633 Q4A634 Q4A636 Q4A645
Q4A647 Q4A656 Q4A658 Q4A666 Q4A667 Q4A696 Q4A6A2 Q4A6A3 Q4A6B0 Q4A6B1 Q4A6L9
Q4A6N4 Q4A6N6 Q4A6N7 Q4A6N8 Q4A6N9 Q4A6Q0 Q4A6Q8 Q4A6S5 Q4A6S6 Q4A6S7 Q4A6T7
Q4A6V0 Q4A6V9 Q4A6W2 Q4A6X3 Q4A6X4 Q4A6X9 Q4A6Y4 Q4A6Z8 Q4A701 Q4A702 Q4A703
Q4A724 Q4A734 Q4A737 Q4A738 Q4A739 Q4A743 Q4A749 Q4A750 Q4A755 Q4A759 Q4A769
Q4A772 Q4A775 Q4A776 Q4A777 Q4A782 Q4A786 Q4A789 Q4A792 Q4A7A7 Q4A7A8 Q4A7A9
Q4A7B0 Q4A7B8 Q4A7D1 Q4A7E2 Q4A7F0 Q4A7I8 Q4A7J2 Q4A7J6 Q4A7K0 Q4A7P8 Q4A7Q7
Q4A7R3 Q4A7R7 Q4A7S3 Q4A7U1 Q4A7U4 Q4A7W4 Q4A7W5 Q4A7W9 Q4A7X8 Q4A7X9 Q4A7Y3
Q4A870 Q4A872 Q4A879 Q4A880 Q4A881 Q4A890 Q4A891 Q4A898 Q4A899 Q4A8A6 Q4A8C0
Q4A8C3 Q4A8D5 Q4A8D9 Q4A8E1 Q4A8F6 Q4A8F8 Q4A8G1 Q4A8G3 Q4A8G4 Q4A8H0 Q4A8H1
Q4A8H2 Q4A8H3 Q4A8H4 Q4A8H5 Q4A8H6 Q4A8H7 Q4A8H8 Q4A8I0 Q4A8I1 Q4A8I2 Q4A8I3
Q4A8I4 Q4A8I5 Q4A8I7 Q4A8I8 Q4A8I9 Q4A8J3 Q4A8J4 Q4A8J5 Q4A8J6 Q4A8J7 Q4A8J8
Q4A8K7 Q4A8L2 Q4A8L3 Q4A8L6 Q4A8M4 Q4A8M6 Q4A8M8 Q4A8N3 Q4A8N4 Q4A8P0 Q4A8P1
Q4A8P2 Q4A8Q0 Q4A8Q3 Q4A8Q9 Q4A8R0 Q4A8T5 Q4A8T6 Q4A8T7 Q4A8U3 Q4A8U5 Q4A8U9
Q4A8V0 Q4A8V6 Q4A8V7 Q4A8X1 Q4A8X3 Q4A8Y0 Q4A900 Q4A902 Q4A908 Q4A909 Q4A912
Q4A913 Q4A917 Q4A921 Q4A930 Q4A933 Q4A936 Q4A937 Q4A938 Q4A943 Q4A947 Q4A952
Q4A967 Q4A968 Q4A969 Q4A970 Q4A977 Q4A992 Q4A9A2 Q4A9B0 Q4A9E9 Q4A9F3 Q4A9F7
Q4A9G1 Q4A9L4 Q4A9M3 Q4A9M9 Q4A9N3 Q4A9N9 Q4A9Q7 Q4A9R0 Q4A9R6 Q4A9R7 Q4A9S1
Q4A9T0 Q4A9T1 Q4A9V2 Q4AA43 Q4AA45 Q4AA52 Q4AA53 Q4AA54 Q4AA63 Q4AA64 Q4AA71
Q4AA72 Q4AA79 Q4AA93 Q4AA96 Q4AAA6 Q4AAB0 Q4AAB2 Q4AAC5 Q4AAC7 Q4AAD0 Q4AAD2
Q4AAD3 Q4AAD9 Q4AAE0 Q4AAE1 Q4AAE2 Q4AAE3 Q4AAE4 Q4AAE5 Q4AAE6 Q4AAE7 Q4AAE9
Q4AAF0 Q4AAF1 Q4AAF2 Q4AAF3 Q4AAF4 Q4AAF6 Q4AAF7 Q4AAF8 Q4AAG2 Q4AAG3 Q4AAG4
Q4AAG5 Q4AAG6 Q4AAG7 Q4AAH6 Q4AAI1 Q4AAI2 Q4AAI5 Q4AAJ3 Q4AAJ5 Q4AAK2 Q4AAK9
Q4AAL0 Q4AAL1 Q4AAL9 Q4AAM2 Q4AAM8 Q4AAQ4 Q4AAQ5 Q4AAQ6 Q4AAR2 Q4AAR4 Q4AAS1
Q4AAT7 Q4AAT9 Q4AAU5 Q4AAU6 Q4AAV4 Q4AAV6 Q4AAW7 Q4AAW8 Q4AAX3 Q4AAX4 Q4AAX7
Q4AAY2 Q4AC99 Q4ADK4 Q4ADK7 Q4AE11 Q4AE12 Q4AE28 Q4AE62 Q4AEH7 Q4AEH8 Q4AEH9
Q4AEI0 Q4CNL4 Q4CQJ5 Q4CTY5 Q4CWX1 Q4CY36 Q4D248 Q4D2R9 Q4D3B7 Q4D3W2 Q4D4Y5
Q4D6B2 Q4D7L5 Q4DBW3 Q4DBX5 Q4DC43 Q4DCH3 Q4DER6 Q4DKQ5 Q4DLT6 Q4DLX9 Q4DPN8
Q4DRX4 Q4DTX9 Q4DW94 Q4DYL7 Q4DZ41 Q4DZP2 Q4E097 Q4E123 Q4E162 Q4F6N6 Q4F8H9
Q4F8I0 Q4F8Z9 Q4FCZ5 Q4FE45 Q4FE47 Q4FG66 Q4FG71 Q4FG72 Q4FG73 Q4FK66 Q4FL49
Q4FL89 Q4FLB9 Q4FLD7 Q4FLI7 Q4FLJ0 Q4FLJ4 Q4FLJ5 Q4FLJ7 Q4FLK0 Q4FLK5 Q4FLK8
Q4FLK9 Q4FLL0 Q4FLL1 Q4FLL2 Q4FLL3 Q4FLL4 Q4FLL5 Q4FLL6 Q4FLL7 Q4FLL8 Q4FLL9

Q4FLM0 Q4FLM1 Q4FLM2 Q4FLM3 Q4FLM4 Q4FLM5 Q4FLM6 Q4FLM7 Q4FLM8 Q4FLM9 Q4FLN0
Q4FLN1 Q4FLN2 Q4FLN4 Q4FLN5 Q4FLN6 Q4FLN8 Q4FLN9 Q4FLP0 Q4FLP1 Q4FLP2 Q4FLP4
Q4FLP6 Q4FLQ4 Q4FLQ5 Q4FLQ6 Q4FLQ7 Q4FLQ8 Q4FLR0 Q4FLS1 Q4FLS2 Q4FLS6 Q4FLS9
Q4FLT4 Q4FLT5 Q4FLU5 Q4FLZ4 Q4FM02 Q4FM10 Q4FM11 Q4FM19 Q4FM20 Q4FM31 Q4FM35
Q4FM39 Q4FM53 Q4FM56 Q4FM64 Q4FM68 Q4FM73 Q4FM76 Q4FM94 Q4FM95 Q4FME4 Q4FMH0
Q4FMI9 Q4FMN4 Q4FMN5 Q4FMP0 Q4FMP8 Q4FMR0 Q4FMR1 Q4FMR5 Q4FMR6 Q4FMR7 Q4FMS8
Q4FMU2 Q4FMV1 Q4FMV2 Q4FMV9 Q4FMW6 Q4FMW8 Q4FMX1 Q4FMX6 Q4FMZ1 Q4FN05 Q4FN06
Q4FN07 Q4FN13 Q4FN15 Q4FN26 Q4FN35 Q4FN36 Q4FN38 Q4FN40 Q4FN45 Q4FN99 Q4FNA0
Q4FNA2 Q4FNA8 Q4FND0 Q4FND9 Q4FNE1 Q4FNE2 Q4FNE4 Q4FNE5 Q4FNF2 Q4FNF4 Q4FNF5
Q4FNF9 Q4FNG0 Q4FNG1 Q4FNG5 Q4FNH4 Q4FNH5 Q4FNH6 Q4FNH7 Q4FNH9 Q4FNK3 Q4FNK9
Q4FNM6 Q4FNM7 Q4FNM8 Q4FNM9 Q4FNN1 Q4FNN3 Q4FNN5 Q4FNP9 Q4FNQ0 Q4FNQ1 Q4FNR5
Q4FNR6 Q4FNR7 Q4FNS1 Q4FNS3 Q4FNS4 Q4FNS5 Q4FNS8 Q4FNT4 Q4FNT5 Q4FNT6 Q4FNT7
Q4FNV6 Q4FNW1 Q4FNX4 Q4FP10 Q4FP12 Q4FP13 Q4FP14 Q4FP15 Q4FP16 Q4FP17 Q4FP21
Q4FP28 Q4FP29 Q4FP40 Q4FP43 Q4FP45 Q4FP46 Q4FP47 Q4FP48 Q4FP49 Q4FP51 Q4FP52
Q4FP74 Q4FP80 Q4FPA5 Q4FPA6 Q4FPA9 Q4FPF7 Q4FPG0 Q4FPG2 Q4FPG9 Q4FPH0 Q4FPI1
Q4FPK6 Q4FPK7 Q4FPK8 Q4FPL3 Q4FPN1 Q4FPN3 Q4FPN5 Q4FPQ6 Q4FPR4 Q4FPR8 Q4FPR9
Q4FPS7 Q4FPS9 Q4FPT7 Q4FPU1 Q4FPV8 Q4FPW3 Q4FPZ1 Q4FPZ3 Q4FQ04 Q4FQ05 Q4FQ21
Q4FQ23 Q4FQ25 Q4FQ42 Q4FQ43 Q4FQ44 Q4FQ45 Q4FQ48 Q4FQ54 Q4FQ63 Q4FQ64 Q4FQ65
Q4FQ66 Q4FQ67 Q4FQ90 Q4FQ91 Q4FQB0 Q4FQB1 Q4FQB2 Q4FQB4 Q4FQB8 Q4FQB9 Q4FQC0
Q4FQD6 Q4FQE4 Q4FQF2 Q4FQF6 Q4FQF7 Q4FQF8 Q4FQF9 Q4FQG3 Q4FQG4 Q4FQG5 Q4FQG6
Q4FQG9 Q4FQH0 Q4FQH1 Q4FQH2 Q4FQH3 Q4FQH4 Q4FQI2 Q4FQI3 Q4FQJ5 Q4FQJ9 Q4FQK7
Q4FQL0 Q4FQM6 Q4FQQ0 Q4FQU4 Q4FQU7 Q4FQW0 Q4FQX1 Q4FQX4 Q4FQX8 Q4FQX9 Q4FQY6
Q4FQY7 Q4FQY8 Q4FQZ1 Q4FQZ9 Q4FR00 Q4FR06 Q4FR09 Q4FR14 Q4FR30 Q4FR69 Q4FR76
Q4FR79 Q4FR86 Q4FR92 Q4FRD7 Q4FRD8 Q4FRE5 Q4FRF4 Q4FRH4 Q4FRH5 Q4FRH6 Q4FRH9
Q4FRI6 Q4FRK6 Q4FRK7 Q4FRK8 Q4FRK9 Q4FRL9 Q4FRM3 Q4FRN8 Q4FRP0 Q4FRP5 Q4FRP8
Q4FRQ0 Q4FRQ5 Q4FRS8 Q4FRT5 Q4FRU7 Q4FRU8 Q4FRV0 Q4FRV1 Q4FRV6 Q4FRX3 Q4FRZ9
Q4FS01 Q4FS02 Q4FS19 Q4FS20 Q4FS21 Q4FS24 Q4FS37 Q4FS54 Q4FS63 Q4FSB4 Q4FSB5
Q4FSB8 Q4FSF4 Q4FSF9 Q4FSH2 Q4FSQ4 Q4FST7 Q4FSU1 Q4FSV2 Q4FSV3 Q4FT44 Q4FT48
Q4FT57 Q4FT67 Q4FT85 Q4FT93 Q4FTH7 Q4FTI5 Q4FTJ2 Q4FTK5 Q4FTK7 Q4FTK9 Q4FTL1
Q4FTL2 Q4FTN3 Q4FTN4 Q4FTN9 Q4FTQ5 Q4FTR7 Q4FTT9 Q4FTU0 Q4FTU8 Q4FTV0 Q4FTW3
Q4FTW6 Q4FTW7 Q4FTX0 Q4FTX1 Q4FTX2 Q4FTX3 Q4FTX6 Q4FU23 Q4FU67 Q4FU68 Q4FU71
Q4FU86 Q4FU93 Q4FU94 Q4FU95 Q4FUA5 Q4FUA6 Q4FUB6 Q4FUC6 Q4FUD0 Q4FUD1 Q4FUD2
Q4FUD3 Q4FUD4 Q4FUD5 Q4FUD7 Q4FUD8 Q4FUD9 Q4FUE0 Q4FUE1 Q4FUE2 Q4FUE3 Q4FUE4
Q4FUE5 Q4FUE6 Q4FUE7 Q4FUE8 Q4FUE9 Q4FUF0 Q4FUF1 Q4FUF2 Q4FUF3 Q4FUF4 Q4FUF5
Q4FUF6 Q4FUF7 Q4FUF9 Q4FUG4 Q4FUG5 Q4FUL2 Q4FUL6 Q4FUL7 Q4FUP5 Q4FUQ0 Q4FUQ3
Q4FUQ9 Q4FUR3 Q4FUR4 Q4FUR9 Q4FUS5 Q4FUT8 Q4FUT9 Q4FUU5 Q4FUU7 Q4FUV3 Q4FUV6
Q4FUW4 Q4FUW9 Q4FUX4 Q4FUZ5 Q4FUZ8 Q4FUZ9 Q4FV14 Q4FV16 Q4FV29 Q4FV41 Q4FV61
Q4FV64 Q4FV71 Q4FV74 Q4FV85 Q4FVA1 Q4FVA9 Q4FVB2 Q4FVB4 Q4FVB5 Q4FVB6 Q4FVC4
Q4FVD4 Q4FVD6 Q4FVE5 Q4FVF3 Q4FVG3 Q4FVG6 Q4FVG7 Q4FVG8 Q4FVH5 Q4FVH9 Q4FVJ6
Q4FVJ7 Q4FVJ8 Q4FVL0 Q4FVL1 Q4FVL3 Q4FVL4 Q4FVL5 Q4FVL7 Q4FVL9 Q4FVM3 Q4FVM5
Q4FVM7 Q4FVM8 Q4FVP8 Q4FVQ4 Q4FVQ6 Q4FVQ7 Q4FVS5 Q4FVS6 Q4FX73 Q4FXY8 Q4FYU7
Q4FZB7 Q4FZD7 Q4FZF3 Q4FZG9 Q4FZP2 Q4FZQ6 Q4FZR5 Q4FZT2 Q4FZT6 Q4FZT8 Q4FZT9
Q4FZU2 Q4FZU3 Q4FZV3 Q4FZX0 Q4FZX5 Q4G008 Q4G009 Q4G033 Q4G055 Q4G056 Q4G059
Q4G061 Q4G069 Q4G073 Q4G074 Q4G078 Q4G0F8 Q4G0G2 Q4G0N4 Q4G0N7 Q4G0P3 Q4G0S7

Q4G0U5 Q4G0W2 Q4G0X4 Q4G0X9 Q4G0Z9 Q4G112 Q4G163 Q4G2I2 Q4G2Y1 Q4G338 Q4G341
Q4G342 Q4G343 Q4G344 Q4G345 Q4G346 Q4G347 Q4G348 Q4G349 Q4G350 Q4G352 Q4G353
Q4G354 Q4G355 Q4G356 Q4G357 Q4G358 Q4G359 Q4G360 Q4G361 Q4G362 Q4G363 Q4G364
Q4G366 Q4G384 Q4G386 Q4G387 Q4G391 Q4G392 Q4G393 Q4G3A3 Q4G3A4 Q4G3A5 Q4G3A6
Q4G3A7 Q4G3B1 Q4G3D5 Q4G3D9 Q4G3E0 Q4G3E8 Q4G3E9 Q4G3F4 Q4G3H4 Q4G4B1 Q4GX86
Q4GX87 Q4GXG7 Q4GXH5 Q4GXH6 Q4GXH7 Q4GXP2 Q4GXP3 Q4GXU6 Q4GZD1 Q4H0T5 Q4H174
Q4H178 Q4H186 Q4H192 Q4H195 Q4H1A1 Q4H1A5 Q4H1B9 Q4H1G3 Q4H1G4 Q4H2R2 Q4H3N8
Q4H3P5 Q4H424 Q4H4E3 Q4H4E4 Q4H4E5 Q4H4E7 Q4H4E9 Q4H4F0 Q4H4F3 Q4H4F5 Q4H4F7
Q4H4F8 Q4HTS9 Q4HTT1 Q4HTT2 Q4HVQ9 Q4HVW2 Q4HW67 Q4HW93 Q4HWB3 Q4HWE2 Q4HWQ2
Q4HXF0 Q4HXF6 Q4HY90 Q4HYB8 Q4HYC1 Q4HYI0 Q4HYQ4 Q4HYR2 Q4HZ34 Q4HZ35 Q4HZ42
Q4HZ47 Q4HZ68 Q4HZB8 Q4HZI0 Q4HZK7 Q4HZQ1 Q4HZS8 Q4I061 Q4I0C5 Q4I0J6 Q4I0M9
Q4I174 Q4I1M9 Q4I1N3 Q4I1Q6 Q4I1Y1 Q4I1Y5 Q4I2D7 Q4I2X5 Q4I327 Q4I5I4 Q4I5M4
Q4I5R3 Q4I5U9 Q4I5V2 Q4I5V3 Q4I5W5 Q4I5Z5 Q4I613 Q4I624 Q4I650 Q4I662 Q4I665
Q4I6A4 Q4I7F9 Q4I7G0 Q4I7K4 Q4I7L0 Q4I7N9 Q4I7S1 Q4I7X1 Q4I830 Q4I877 Q4I8C9
Q4I8S2 Q4I8X0 Q4I948 Q4I963 Q4I9B7 Q4I9U7 Q4IA56 Q4IAA0 Q4IAJ1 Q4IAK7 Q4IAS1
Q4IB50 Q4IB70 Q4IB89 Q4IB96 Q4IBK5 Q4IBL8 Q4IBN1 Q4IBR4 Q4IBS2 Q4IBS9 Q4IBU5
Q4IBX0 Q4IC11 Q4ICA8 Q4ICG8 Q4ICI6 Q4ICM0 Q4ID21 Q4IE79 Q4IEA5 Q4IER0 Q4IER8
Q4IEV4 Q4IEV5 Q4IF44 Q4IF76 Q4IFH8 Q4IFI0 Q4IJ11 Q4IJ56 Q4IJ84 Q4IJH1 Q4IJK2
Q4IJM4 Q4IJP1 Q4IJT5 Q4IJU0 Q4IL82 Q4ILI9 Q4IMJ1 Q4IMM0 Q4IMN9 Q4IN52 Q4IN63
Q4ING3 Q4INI2 Q4INS6 Q4INW6 Q4INZ9 Q4IP34 Q4IP51 Q4IPA4 Q4IPB3 Q4IPB8 Q4IPC8
Q4IPH4 Q4IPI1 Q4IPI4 Q4IPN9 Q4IQ72 Q4IQC1 Q4IQK7 Q4IQM6 Q4IQT8 Q4IQW0 Q4IQX3
Q4IR08 Q4IR09 Q4IR18 Q4IR87 Q4IRI9 Q4J6A5 Q4J6D8 Q4J6D9 Q4J6F2 Q4J6G3 Q4J6H1
Q4J6J6 Q4J6M3 Q4J6M5 Q4J6M6 Q4J6Z4 Q4J781 Q4J7A3 Q4J7W0 Q4J860 Q4J894 Q4J8E7
Q4J8E8 Q4J8E9 Q4J8F0 Q4J8F1 Q4J8F8 Q4J8G0 Q4J8G7 Q4J8I4 Q4J8I5 Q4J8I7 Q4J8I8
Q4J8I9 Q4J8J1 Q4J8J2 Q4J8J7 Q4J8J9 Q4J8K9 Q4J8L2 Q4J8L3 Q4J8L4 Q4J8M3 Q4J8P1
Q4J8P2 Q4J8P4 Q4J8P8 Q4J8P9 Q4J8Q0 Q4J8Q2 Q4J8S2 Q4J8T0 Q4J8U0 Q4J8X1 Q4J8X2
Q4J8X3 Q4J8X6 Q4J8X7 Q4J8X8 Q4J8X9 Q4J8Y4 Q4J8Z2 Q4J904 Q4J912 Q4J915 Q4J924
Q4J929 Q4J932 Q4J933 Q4J937 Q4J939 Q4J940 Q4J945 Q4J946 Q4J947 Q4J955 Q4J963
Q4J965 Q4J966 Q4J973 Q4J978 Q4J985 Q4J986 Q4J989 Q4J9A0 Q4J9A3 Q4J9A4 Q4J9A6
Q4J9A8 Q4J9B0 Q4J9B1 Q4J9C4 Q4J9C6 Q4J9D2 Q4J9D4 Q4J9H1 Q4J9L3 Q4J9R1 Q4J9S8
Q4JA46 Q4JA54 Q4JA57 Q4JA60 Q4JA70 Q4JA78 Q4JAA8 Q4JAB0 Q4JAB1 Q4JAB2 Q4JAB4
Q4JAC4 Q4JAC6 Q4JAE7 Q4JAE8 Q4JAF4 Q4JAF8 Q4JAF9 Q4JAG0 Q4JAG1 Q4JAH5 Q4JAH6
Q4JAI1 Q4JAI2 Q4JAI3 Q4JAI4 Q4JAI6 Q4JAI7 Q4JAJ2 Q4JAJ6 Q4JAJ9 Q4JAK2 Q4JAK8
Q4JAL1 Q4JAL7 Q4JAM0 Q4JAM1 Q4JAM7 Q4JAM9 Q4JAN1 Q4JAN4 Q4JAP5 Q4JAP7 Q4JAP8
Q4JAP9 Q4JAQ0 Q4JAQ2 Q4JAQ3 Q4JAS8 Q4JAT0 Q4JAT5 Q4JAU3 Q4JAU9 Q4JAV0 Q4JAV1
Q4JAV2 Q4JAV3 Q4JAX5 Q4JAY2 Q4JAY3 Q4JAY4 Q4JAY6 Q4JAZ3 Q4JAZ7 Q4JAZ8 Q4JAZ9
Q4JB11 Q4JB12 Q4JB16 Q4JB17 Q4JB20 Q4JB24 Q4JB26 Q4JB27 Q4JB32 Q4JB40 Q4JB41
Q4JB42 Q4JB43 Q4JB44 Q4JB45 Q4JB46 Q4JB47 Q4JB49 Q4JB50 Q4JB72 Q4JB73 Q4JB78
Q4JB80 Q4JB89 Q4JB92 Q4JBF2 Q4JBG0 Q4JBG6 Q4JBG7 Q4JBJ3 Q4JBL6 Q4JBL7 Q4JBN2
Q4JBN3 Q4JBP0 Q4JBR3 Q4JBR4 Q4JBU5 Q4JBV4 Q4JBW3 Q4JBY3 Q4JBY6 Q4JC00 Q4JC09
Q4JC10 Q4JC35 Q4JC46 Q4JC60 Q4JC68 Q4JC71 Q4JC74 Q4JC75 Q4JC76 Q4JC84 Q4JCA6
Q4JCA7 Q4JCB9 Q4JCD8 Q4JCG3 Q4JCH6 Q4JCH7 Q4JCI0 Q4JCI5 Q4JCJ1 Q4JCK7 Q4JCL6
Q4JCM0 Q4JCM9 Q4JCN8 Q4JCP0 Q4JCP8 Q4JF75 Q4JFH9 Q4JFW8 Q4JG17 Q4JHE0 Q4JHG3
Q4JIJ2 Q4JIJ3 Q4JJB1 Q4JK59 Q4JM28 Q4JM65 Q4JQJ5 Q4JQS7 Q4JQS9 Q4JQT7 Q4JQU0

Q4JQU3 Q4JQU6 Q4JQU7 Q4JQV6 Q4JQW2 Q4JQW5 Q4JQW7 Q4JQW9 Q4JQX1 Q4JQX2 Q4JZ13
Q4K0R3 Q4K0S9 Q4K2S1 Q4K2U1 Q4K394 Q4K396 Q4K399 Q4K3A0 Q4K3E4 Q4K3E8 Q4K3F6
Q4K3M0 Q4K3R3 Q4K3R4 Q4K3R6 Q4K3R9 Q4K3S0 Q4K3S2 Q4K3S4 Q4K3S6 Q4K3T9 Q4K3U3
Q4K3W0 Q4K3W5 Q4K3X1 Q4K3X3 Q4K3Y6 Q4K406 Q4K416 Q4K418 Q4K423 Q4K434 Q4K469
Q4K475 Q4K498 Q4K4A5 Q4K4A7 Q4K4C5 Q4K4C6 Q4K4C7 Q4K4D3 Q4K4D4 Q4K4E2 Q4K4E3
Q4K4E4 Q4K4E5 Q4K4F6 Q4K4F7 Q4K4G0 Q4K4H7 Q4K4I6 Q4K4I7 Q4K4J1 Q4K4K7 Q4K4K8
Q4K4K9 Q4K4P6 Q4K4S5 Q4K4T2 Q4K4T3 Q4K4U8 Q4K4U9 Q4K4W3 Q4K4W4 Q4K4W8 Q4K4X0
Q4K4X2 Q4K4X3 Q4K4X4 Q4K4X5 Q4K4Z4 Q4K502 Q4K503 Q4K514 Q4K515 Q4K517 Q4K519
Q4K522 Q4K523 Q4K524 Q4K525 Q4K526 Q4K527 Q4K528 Q4K529 Q4K530 Q4K532 Q4K533
Q4K534 Q4K535 Q4K536 Q4K537 Q4K538 Q4K539 Q4K540 Q4K541 Q4K542 Q4K543 Q4K544
Q4K545 Q4K546 Q4K547 Q4K548 Q4K549 Q4K550 Q4K551 Q4K552 Q4K554 Q4K555 Q4K556
Q4K557 Q4K558 Q4K592 Q4K597 Q4K5A5 Q4K5A7 Q4K5E1 Q4K5F9 Q4K5G0 Q4K5G2 Q4K5G9
Q4K5H6 Q4K5I1 Q4K5I3 Q4K5I6 Q4K5I7 Q4K5P2 Q4K5Q4 Q4K5R9 Q4K5T6 Q4K5T7 Q4K5T8
Q4K5T9 Q4K5U1 Q4K5U4 Q4K5U7 Q4K5X0 Q4K5X1 Q4K5Y1 Q4K5Y3 Q4K5Y4 Q4K5Z0 Q4K5Z3
Q4K608 Q4K629 Q4K688 Q4K689 Q4K691 Q4K694 Q4K695 Q4K6B3 Q4K6B5 Q4K6D2 Q4K6H2
Q4K6H3 Q4K6I1 Q4K6I4 Q4K6I5 Q4K6J1 Q4K6J4 Q4K6M3 Q4K6S8 Q4K6T8 Q4K6U2 Q4K6U5
Q4K6U6 Q4K6U9 Q4K6V0 Q4K6V3 Q4K6V7 Q4K6V9 Q4K6W4 Q4K6X6 Q4K6X7 Q4K706 Q4K734
Q4K746 Q4K747 Q4K749 Q4K758 Q4K763 Q4K764 Q4K779 Q4K7C2 Q4K7C7 Q4K7D1 Q4K7D5
Q4K7D6 Q4K7D7 Q4K7D8 Q4K7D9 Q4K7E8 Q4K7H5 Q4K7Q8 Q4K837 Q4K838 Q4K840 Q4K843
Q4K880 Q4K882 Q4K898 Q4K8A5 Q4K8B9 Q4K8C1 Q4K8D9 Q4K8G0 Q4K8H2 Q4K8H3 Q4K8J3
Q4K8J5 Q4K8K5 Q4K8K9 Q4K8L6 Q4K8M4 Q4K8M6 Q4K8N0 Q4K8N2 Q4K8S7 Q4K8U2 Q4K8U5
Q4K8Z0 Q4K9I2 Q4K9J4 Q4K9J5 Q4K9J6 Q4K9L7 Q4K9L8 Q4K9M3 Q4K9S1 Q4K9U2 Q4K9U7
Q4K9U9 Q4K9V0 Q4K9V1 Q4K9V7 Q4K9V8 Q4KA27 Q4KAD4 Q4KAH1 Q4KAR9 Q4KB60 Q4KBD3
Q4KBH9 Q4KBI1 Q4KBJ2 Q4KBL2 Q4KBW8 Q4KC17 Q4KCP0 Q4KCQ3 Q4KCY5 Q4KCY6 Q4KCZ0
Q4KD86 Q4KDI5 Q4KDX1 Q4KE22 Q4KEB8 Q4KEE7 Q4KEN7 Q4KEV8 Q4KEV9 Q4KEW0 Q4KEW1
Q4KEW2 Q4KEW3 Q4KEW9 Q4KEZ9 Q4KF05 Q4KF07 Q4KF08 Q4KF16 Q4KF19 Q4KF31 Q4KF90
Q4KF97 Q4KFA6 Q4KFD1 Q4KFD3 Q4KFD4 Q4KFF5 Q4KFF7 Q4KFF9 Q4KFH1 Q4KFI6 Q4KFJ3
Q4KFJ6 Q4KFR2 Q4KFS0 Q4KFS2 Q4KFT1 Q4KFT4 Q4KFV3 Q4KFX8 Q4KFY6 Q4KG70 Q4KG97
Q4KGG0 Q4KGH6 Q4KGJ3 Q4KGK7 Q4KGL0 Q4KGL6 Q4KGP5 Q4KGR0 Q4KGS0 Q4KGT8 Q4KGU2
Q4KH47 Q4KH71 Q4KH98 Q4KHA5 Q4KHC0 Q4KHC1 Q4KHE3 Q4KHE7 Q4KHE8 Q4KHE9 Q4KHF0
Q4KHF4 Q4KHF8 Q4KHG2 Q4KHH0 Q4KHH3 Q4KHH4 Q4KHH5 Q4KHH6 Q4KHH8 Q4KHI7 Q4KHJ7
Q4KHK4 Q4KHK5 Q4KHL4 Q4KHQ5 Q4KHQ6 Q4KHQ7 Q4KHQ8 Q4KHR3 Q4KHS6 Q4KHS8 Q4KHS9
Q4KHT1 Q4KHT9 Q4KHU0 Q4KHU2 Q4KHX7 Q4KI35 Q4KI67 Q4KI72 Q4KI73 Q4KI74 Q4KI75
Q4KI90 Q4KI98 Q4KIA3 Q4KIA7 Q4KIA8 Q4KIA9 Q4KIC9 Q4KID6 Q4KIF2 Q4KIF3 Q4KIF4
Q4KIF5 Q4KIF6 Q4KIF8 Q4KIF9 Q4KIG0 Q4KIG3 Q4KIG9 Q4KIH0 Q4KIH1 Q4KIH2 Q4KIH6
Q4KIH9 Q4KIP3 Q4KIP4 Q4KIS2 Q4KIX1 Q4KIX5 Q4KIY6 Q4KJ05 Q4KJ06 Q4KJ09 Q4KJ10
Q4KJ26 Q4KJ32 Q4KJ33 Q4KJ34 Q4KJ59 Q4KJ61 Q4KJ62 Q4KJ68 Q4KJ69 Q4KJ73 Q4KJ74
Q4KJ81 Q4KJ82 Q4KJA1 Q4KJA9 Q4KJF3 Q4KJI4 Q4KJI8 Q4KJI9 Q4KJJ8 Q4KJK0 Q4KJK2
Q4KJK3 Q4KJL5 Q4KJL6 Q4KJM7 Q4KJN0 Q4KJP6 Q4KJR5 Q4KJR8 Q4KJS4 Q4KJS5 Q4KJS7
Q4KJS8 Q4KJT2 Q4KJT9 Q4KJX9 Q4KK03 Q4KK04 Q4KK09 Q4KK14 Q4KKJ8 Q4KKP4 Q4KKP5
Q4KKQ4 Q4KKQ6 Q4KKR0 Q4KKR7 Q4KKR8 Q4KKS8 Q4KKT0 Q4KKX4 Q4KKZ1 Q4KL13 Q4KL35
Q4KL78 Q4KL92 Q4KL94 Q4KL97 Q4KLA0 Q4KLB0 Q4KLC4 Q4KLE6 Q4KLF3 Q4KLF5 Q4KLF8
Q4KLG2 Q4KLH4 Q4KLH5 Q4KLI1 Q4KLI2 Q4KLJ8 Q4KLK8 Q4KLK9 Q4KLL0 Q4KLL5 Q4KLM5
Q4KLN4 Q4KLN6 Q4KLN8 Q4KLP0 Q4KLP8 Q4KLQ5 Q4KLS8 Q4KLT3 Q4KLT6 Q4KLU2 Q4KLU7

Q4KLV2 Q4KLV6 Q4KLV7 Q4KLY2 Q4KLY4 Q4KLY6 Q4KLZ4 Q4KLZ6 Q4KM14 Q4KM32 Q4KM35
Q4KM45 Q4KM47 Q4KM51 Q4KM65 Q4KM73 Q4KM79 Q4KM86 Q4KM92 Q4KM95 Q4KMA0 Q4KMA2
Q4KMC4 Q4KMC8 Q4KMC9 Q4KMD3 Q4KMD7 Q4KMJ1 Q4KMK0 Q4KML2 Q4KML4 Q4KMM3 Q4KMX7
Q4KMZ1 Q4KSH9 Q4KTA7 Q4KTC0 Q4KTD9 Q4KTE2 Q4KTG9 Q4KTY1 Q4KYM5 Q4L1M7 Q4L2X9
Q4L2Y9 Q4L2Z0 Q4L2Z1 Q4L2Z2 Q4L2Z3 Q4L2Z4 Q4L320 Q4L321 Q4L330 Q4L332 Q4L340
Q4L342 Q4L376 Q4L378 Q4L382 Q4L383 Q4L385 Q4L386 Q4L3D5 Q4L3D6 Q4L3D9 Q4L3E5
Q4L3E7 Q4L3F0 Q4L3F2 Q4L3F5 Q4L3F7 Q4L3F8 Q4L3F9 Q4L3G0 Q4L3H0 Q4L3H5 Q4L3H8
Q4L3H9 Q4L3I1 Q4L3I3 Q4L3I4 Q4L3I7 Q4L3I9 Q4L3J1 Q4L3J6 Q4L3J7 Q4L3K0 Q4L3K1
Q4L3K3 Q4L3K4 Q4L3K6 Q4L3K7 Q4L3K8 Q4L3K9 Q4L3L8 Q4L3N6 Q4L3P5 Q4L3P8 Q4L3Q1
Q4L3Q2 Q4L3Q5 Q4L3Q7 Q4L3R3 Q4L3R9 Q4L425 Q4L491 Q4L4A4 Q4L4D0 Q4L4E7 Q4L4F1
Q4L4F3 Q4L4G3 Q4L4G8 Q4L4H7 Q4L4H9 Q4L4I5 Q4L4I7 Q4L4J2 Q4L4J4 Q4L4J5 Q4L4J7
Q4L4K4 Q4L4K5 Q4L4K6 Q4L4L2 Q4L4R0 Q4L4R1 Q4L4R5 Q4L4S3 Q4L4T1 Q4L4T4 Q4L4T8
Q4L4U2 Q4L4U5 Q4L4V4 Q4L4W9 Q4L4X6 Q4L4X7 Q4L4X8 Q4L4Y2 Q4L4Y3 Q4L4Y4 Q4L4Y7
Q4L505 Q4L506 Q4L510 Q4L513 Q4L515 Q4L522 Q4L525 Q4L527 Q4L547 Q4L549 Q4L572
Q4L574 Q4L575 Q4L577 Q4L578 Q4L580 Q4L582 Q4L5A3 Q4L5A4 Q4L5B1 Q4L5C0 Q4L5C6
Q4L5D8 Q4L5D9 Q4L5E1 Q4L5E3 Q4L5E4 Q4L5E5 Q4L5E9 Q4L5F0 Q4L5F1 Q4L5F5 Q4L5M7
Q4L5M8 Q4L5M9 Q4L5N0 Q4L5N4 Q4L5N8 Q4L5P0 Q4L5P4 Q4L5Q0 Q4L5Q2 Q4L5Q4 Q4L5Q5
Q4L5Q6 Q4L5Q7 Q4L5R1 Q4L5R2 Q4L5R9 Q4L5S2 Q4L5S5 Q4L5S7 Q4L5T5 Q4L5U0 Q4L5U1
Q4L5U2 Q4L5U3 Q4L5U7 Q4L5U8 Q4L5V2 Q4L5V3 Q4L5V4 Q4L5V5 Q4L5V6 Q4L5V7 Q4L5V8
Q4L5V9 Q4L5W0 Q4L5W1 Q4L5W2 Q4L5W5 Q4L5W6 Q4L5W7 Q4L5X1 Q4L5X2 Q4L5X4 Q4L5X6
Q4L5X7 Q4L5X8 Q4L5Y8 Q4L5Z6 Q4L5Z9 Q4L605 Q4L607 Q4L608 Q4L610 Q4L639 Q4L643
Q4L644 Q4L645 Q4L646 Q4L654 Q4L655 Q4L660 Q4L665 Q4L670 Q4L672 Q4L676 Q4L677
Q4L678 Q4L680 Q4L695 Q4L6A0 Q4L6A1 Q4L6A2 Q4L6A3 Q4L6A7 Q4L6A9 Q4L6B1 Q4L6B6
Q4L6C3 Q4L6C4 Q4L6D1 Q4L6D3 Q4L6D7 Q4L6D8 Q4L6D9 Q4L6F0 Q4L6F1 Q4L6F8 Q4L6F9
Q4L6G1 Q4L6G8 Q4L6G9 Q4L6H0 Q4L6H1 Q4L6H3 Q4L6I0 Q4L6I1 Q4L6I2 Q4L6J4 Q4L6J5
Q4L6J7 Q4L6K3 Q4L6K4 Q4L6L5 Q4L6M1 Q4L6M3 Q4L6M4 Q4L6M5 Q4L6M9 Q4L6N5 Q4L6N6
Q4L6N7 Q4L6N8 Q4L6Q1 Q4L6Q3 Q4L6Q7 Q4L6R3 Q4L6R5 Q4L6R6 Q4L6S0 Q4L6S5 Q4L6S8
Q4L6T0 Q4L6T1 Q4L6T2 Q4L6T5 Q4L6U5 Q4L6U7 Q4L6V0 Q4L6V2 Q4L6V7 Q4L6V8 Q4L6W2
Q4L6W3 Q4L6W4 Q4L6W5 Q4L6W8 Q4L6X5 Q4L6X6 Q4L6X8 Q4L6Y0 Q4L6Y4 Q4L6Y5 Q4L6Y6
Q4L6Y7 Q4L6Y8 Q4L6Y9 Q4L6Z1 Q4L6Z3 Q4L700 Q4L703 Q4L706 Q4L709 Q4L711 Q4L712
Q4L715 Q4L716 Q4L719 Q4L720 Q4L727 Q4L730 Q4L739 Q4L748 Q4L749 Q4L750 Q4L751
Q4L752 Q4L754 Q4L756 Q4L760 Q4L774 Q4L776 Q4L784 Q4L787 Q4L792 Q4L795 Q4L7A3
Q4L7B0 Q4L7B1 Q4L7C7 Q4L7C9 Q4L7D8 Q4L7E9 Q4L7F5 Q4L7G4 Q4L7G9 Q4L7H2 Q4L7H9
Q4L7I2 Q4L7J0 Q4L7J1 Q4L7K6 Q4L7K8 Q4L7L3 Q4L7L4 Q4L7L5 Q4L7L9 Q4L7M3 Q4L7M6
Q4L7N2 Q4L7N4 Q4L7R4 Q4L7R5 Q4L7T0 Q4L7T2 Q4L7T6 Q4L7T9 Q4L7U0 Q4L7U1 Q4L7U2
Q4L7U3 Q4L7U4 Q4L7V3 Q4L7V4 Q4L7W0 Q4L7W2 Q4L7X3 Q4L7X4 Q4L7X6 Q4L7Y1 Q4L7Z3
Q4L7Z4 Q4L7Z5 Q4L7Z6 Q4L7Z9 Q4L800 Q4L801 Q4L803 Q4L805 Q4L808 Q4L811 Q4L817
Q4L818 Q4L819 Q4L837 Q4L846 Q4L848 Q4L849 Q4L860 Q4L868 Q4L871 Q4L872 Q4L873
Q4L874 Q4L880 Q4L881 Q4L882 Q4L887 Q4L888 Q4L889 Q4L890 Q4L891 Q4L892 Q4L895
Q4L896 Q4L897 Q4L898 Q4L899 Q4L8A0 Q4L8A1 Q4L8A2 Q4L8A3 Q4L8A4 Q4L8A5 Q4L8A6
Q4L8A7 Q4L8A8 Q4L8A9 Q4L8B0 Q4L8B1 Q4L8B2 Q4L8B3 Q4L8B4 Q4L8B5 Q4L8B8 Q4L8D2
Q4L8D3 Q4L8D8 Q4L8E4 Q4L8G4 Q4L8G8 Q4L8H4 Q4L8J3 Q4L8J6 Q4L8K2 Q4L8K4 Q4L8N5
Q4L8Q3 Q4L8Q7 Q4L8T0 Q4L8T8 Q4L8X9 Q4L8Y1 Q4L904 Q4L907 Q4L919 Q4L920 Q4L941
Q4L966 Q4L967 Q4L971 Q4L975 Q4L976 Q4L978 Q4L9B6 Q4L9C4 Q4L9D7 Q4L9E6 Q4L9E7

Q4L9F1 Q4L9F3 Q4L9I0 Q4L9J7 Q4L9J9 Q4L9R7 Q4L9S3 Q4L9T4 Q4L9T7 Q4L9T8 Q4L9U2
Q4L9U7 Q4L9V7 Q4L9V8 Q4L9Y4 Q4LA33 Q4LA34 Q4LA35 Q4LA40 Q4LA96 Q4LAA8 Q4LAB0
Q4LAB5 Q4LAJ3 Q4LAK0 Q4LAK2 Q4LAK8 Q4LAL2 Q4LAL5 Q4LAL6 Q4LAM6 Q4LAY1 Q4LB21
Q4LB22 Q4LB23 Q4LB35 Q4LBC6 Q4LBY8 Q4LDG9 Q4LDQ3 Q4LEZ3 Q4MKK8 Q4MPR6 Q4MQ55
Q4MQ58 Q4MTG0 Q4MVY3 Q4MY95 Q4MYA4 Q4MYA8 Q4MYY1 Q4MYY2 Q4MYZ4 Q4MZD7 Q4N1J1
Q4N1R8 Q4N3F4 Q4N3P0 Q4N3S6 Q4N438 Q4N4B9 Q4N6K2 Q4N756 Q4N8L0 Q4N921 Q4N927
Q4N938 Q4P0G6 Q4P0H7 Q4P0K3 Q4P0N6 Q4P0P0 Q4P0P3 Q4P0Y5 Q4P101 Q4P112 Q4P144
Q4P149 Q4P184 Q4P1Q7 Q4P1U2 Q4P1V1 Q4P209 Q4P219 Q4P235 Q4P257 Q4P2E8 Q4P2E9
Q4P2Q5 Q4P2Q7 Q4P2U5 Q4P2W8 Q4P2Y1 Q4P308 Q4P328 Q4P331 Q4P358 Q4P384 Q4P3D8
Q4P3F1 Q4P3S3 Q4P3U9 Q4P3W3 Q4P3X7 Q4P400 Q4P426 Q4P447 Q4P456 Q4P460 Q4P4C1
Q4P4C2 Q4P4G2 Q4P4G8 Q4P4R3 Q4P4W3 Q4P4X6 Q4P4Y2 Q4P521 Q4P525 Q4P553 Q4P555
Q4P5F5 Q4P5F9 Q4P5P0 Q4P5R8 Q4P5U1 Q4P5U4 Q4P5V5 Q4P5V8 Q4P647 Q4P652 Q4P6D5
Q4P6E2 Q4P6E9 Q4P6G4 Q4P6G5 Q4P6I3 Q4P6I8 Q4P6N7 Q4P6R3 Q4P733 Q4P763 Q4P7A4
Q4P7C7 Q4P7F2 Q4P7G1 Q4P7H2 Q4P7J7 Q4P7L6 Q4P7M1 Q4P7R2 Q4P7S1 Q4P7X6 Q4P804
Q4P821 Q4P827 Q4P8F6 Q4P8G2 Q4P8R9 Q4P8S7 Q4P8T5 Q4P914 Q4P965 Q4P985 Q4P9E5
Q4P9P3 Q4P9P9 Q4P9T2 Q4P9T3 Q4P9X4 Q4P9Z3 Q4PA25 Q4PA27 Q4PA45 Q4PA50 Q4PA86
Q4PAB2 Q4PAB8 Q4PAR1 Q4PB04 Q4PB36 Q4PB37 Q4PB52 Q4PB56 Q4PB95 Q4PBE6 Q4PBF2
Q4PBL3 Q4PBP5 Q4PBP6 Q4PBU8 Q4PBY6 Q4PBZ9 Q4PC17 Q4PC84 Q4PC85 Q4PCH8 Q4PCI7
Q4PCR0 Q4PCS5 Q4PCV8 Q4PCY0 Q4PD06 Q4PD40 Q4PD66 Q4PD88 Q4PDA7 Q4PDF6 Q4PDT1
Q4PEB3 Q4PEF8 Q4PEF9 Q4PEJ3 Q4PEM9 Q4PEN0 Q4PEN1 Q4PEQ5 Q4PER6 Q4PEV8 Q4PEX7
Q4PEZ0 Q4PEZ2 Q4PF30 Q4PF67 Q4PF70 Q4PF85 Q4PFA5 Q4PFD9 Q4PFI5 Q4PFU7 Q4PFV5
Q4PG15 Q4PG71 Q4PGG5 Q4PGL2 Q4PGM6 Q4PGT8 Q4PGU6 Q4PH43 Q4PHA6 Q4PHA8 Q4PHN4
Q4PHU4 Q4PHU9 Q4PHV3 Q4PHW2 Q4PI64 Q4PI72 Q4PI84 Q4PI88 Q4PI89 Q4PIC4 Q4PII9
Q4PIL0 Q4PIM8 Q4PIN7 Q4PIR3 Q4PJT6 Q4PJW2 Q4PJX5 Q4PLJ0 Q4PLZ3 Q4PM04 Q4PM12
Q4PM15 Q4PM31 Q4PM47 Q4PM54 Q4PM64 Q4PMB3 Q4PMC9 Q4PMD1 Q4PNS1 Q4PNY0 Q4PS85
Q4PSA3 Q4PSD1 Q4PSE2 Q4PSE4 Q4PSE7 Q4PSE8 Q4PSI6 Q4PSK1 Q4PSK3 Q4PSL7 Q4PSN6
Q4PSN8 Q4PSR7 Q4PSU4 Q4PSY4 Q4PT00 Q4PT02 Q4Q089 Q4Q0M0 Q4Q0Q0 Q4Q0R9 Q4Q0V1
Q4Q1E7 Q4Q2Y6 Q4Q882 Q4Q9W0 Q4Q9X8 Q4QAU9 Q4QB33 Q4QCU9 Q4QDI6 Q4QDX3 Q4QEY9
Q4QFD3 Q4QFD4 Q4QFF0 Q4QFL5 Q4QFY1 Q4QG35 Q4QG98 Q4QH39 Q4QHM7 Q4QIS1 Q4QJK3
Q4QJK4 Q4QJK7 Q4QJL3 Q4QJL5 Q4QJL8 Q4QJM0 Q4QJM4 Q4QJM6 Q4QJM8 Q4QJM9 Q4QJN7
Q4QJR9 Q4QJS0 Q4QJT7 Q4QJU4 Q4QJU5 Q4QJU8 Q4QJV1 Q4QJV5 Q4QJV7 Q4QJW2 Q4QJW4
Q4QJW5 Q4QJW6 Q4QJX4 Q4QJY8 Q4QJZ3 Q4QJZ5 Q4QK35 Q4QK37 Q4QK42 Q4QK43 Q4QK45
Q4QK52 Q4QK58 Q4QK60 Q4QK61 Q4QK64 Q4QK65 Q4QK67 Q4QK69 Q4QK70 Q4QK76 Q4QK77
Q4QK81 Q4QK86 Q4QKA1 Q4QKA4 Q4QKA6 Q4QKF3 Q4QKF5 Q4QKF6 Q4QKG0 Q4QKG3 Q4QKG4
Q4QKG6 Q4QKG9 Q4QKH0 Q4QKH3 Q4QKH4 Q4QKH7 Q4QKI9 Q4QKJ2 Q4QKJ8 Q4QKK3 Q4QKK6
Q4QKK7 Q4QKM2 Q4QKM3 Q4QKM4 Q4QKM7 Q4QKN1 Q4QKN2 Q4QKN8 Q4QKP3 Q4QKP6 Q4QKR3
Q4QKS1 Q4QL07 Q4QL12 Q4QL19 Q4QL43 Q4QL45 Q4QL46 Q4QL66 Q4QL68 Q4QL69 Q4QL70
Q4QL71 Q4QL79 Q4QL80 Q4QL83 Q4QL88 Q4QL89 Q4QL90 Q4QL95 Q4QLA3 Q4QLA4 Q4QLA8
Q4QLB7 Q4QLC5 Q4QLC9 Q4QLD1 Q4QLD2 Q4QLD6 Q4QLE5 Q4QLF0 Q4QLF6 Q4QLF7 Q4QLG0
Q4QLG3 Q4QLG6 Q4QLG7 Q4QLG9 Q4QLH6 Q4QLH8 Q4QLI0 Q4QLI2 Q4QLJ0 Q4QLK7 Q4QLL2
Q4QLM2 Q4QLM7 Q4QLP7 Q4QLQ0 Q4QLQ5 Q4QLQ9 Q4QLR2 Q4QLR3 Q4QLR9 Q4QLS1 Q4QLS2
Q4QLS3 Q4QLS4 Q4QLS6 Q4QLS8 Q4QLS9 Q4QLT0 Q4QLT2 Q4QLT8 Q4QLU2 Q4QLU5 Q4QLU8
Q4QLU9 Q4QLV1 Q4QLV2 Q4QLV3 Q4QLV5 Q4QLV6 Q4QLV7 Q4QLV9 Q4QLW0 Q4QLW2 Q4QLW5
Q4QLY1 Q4QLY5 Q4QLY8 Q4QLZ5 Q4QLZ6 Q4QM04 Q4QM07 Q4QM08 Q4QM17 Q4QM18 Q4QM19

Q4QM28 Q4QM29 Q4QM31 Q4QM32 Q4QM33 Q4QM38 Q4QM44 Q4QM46 Q4QM52 Q4QM53 Q4QM57
Q4QM60 Q4QM70 Q4QM83 Q4QM86 Q4QM87 Q4QM89 Q4QM90 Q4QM91 Q4QM92 Q4QM93 Q4QM96
Q4QM97 Q4QM98 Q4QM99 Q4QMA0 Q4QMA2 Q4QMA3 Q4QMA4 Q4QMA5 Q4QMA6 Q4QMA7 Q4QMA8
Q4QMA9 Q4QMB0 Q4QMB1 Q4QMB3 Q4QMB4 Q4QMB5 Q4QMB6 Q4QMB7 Q4QMB8 Q4QMB9 Q4QMC0
Q4QMC1 Q4QMC2 Q4QMC3 Q4QMD5 Q4QMD9 Q4QME1 Q4QME2 Q4QME6 Q4QME8 Q4QMF4 Q4QMF8
Q4QMG5 Q4QMG7 Q4QMH2 Q4QMI0 Q4QMI4 Q4QMI7 Q4QMJ8 Q4QMK5 Q4QMK6 Q4QML1 Q4QML2
Q4QML4 Q4QML8 Q4QML9 Q4QMM7 Q4QMN4 Q4QMN7 Q4QMN8 Q4QMP0 Q4QMP4 Q4QMP5 Q4QMP6
Q4QMP8 Q4QMR0 Q4QMR1 Q4QMR6 Q4QMS6 Q4QMS7 Q4QMT5 Q4QMT6 Q4QMT7 Q4QMT8 Q4QMT9
Q4QMU5 Q4QMV1 Q4QMV2 Q4QMV6 Q4QMV7 Q4QMW1 Q4QMW6 Q4QMX2 Q4QMY0 Q4QMY5 Q4QMZ7
Q4QMZ9 Q4QN00 Q4QN01 Q4QN02 Q4QN03 Q4QN04 Q4QN05 Q4QN06 Q4QN07 Q4QN08 Q4QN09
Q4QN10 Q4QN11 Q4QN12 Q4QN13 Q4QN17 Q4QN19 Q4QN23 Q4QN30 Q4QN31 Q4QN32 Q4QN33
Q4QN34 Q4QN38 Q4QN45 Q4QN49 Q4QN50 Q4QN56 Q4QN69 Q4QN70 Q4QN71 Q4QN72 Q4QN73
Q4QN74 Q4QN75 Q4QN79 Q4QN84 Q4QNA2 Q4QNB2 Q4QNB3 Q4QNC1 Q4QNC6 Q4QNC8 Q4QND2
Q4QND8 Q4QND9 Q4QNE5 Q4QNE7 Q4QNE8 Q4QNG9 Q4QNH3 Q4QNH4 Q4QNH7 Q4QNJ1 Q4QNK0
Q4QNK7 Q4QNK8 Q4QNL2 Q4QNL5 Q4QNL7 Q4QNL8 Q4QNL9 Q4QNM1 Q4QNM3 Q4QNM4 Q4QNM5
Q4QNM6 Q4QNN8 Q4QNN9 Q4QNP9 Q4QNQ7 Q4QNR2 Q4QNR5 Q4QNR6 Q4QNR7 Q4QNS0 Q4QNT3
Q4QNU2 Q4QNU3 Q4QNV7 Q4QNW4 Q4QNX8 Q4QNY2 Q4QNY3 Q4QNY6 Q4QNY7 Q4QNY8 Q4QNY9
Q4QNZ0 Q4QNZ4 Q4QP08 Q4QP16 Q4QP24 Q4QP29 Q4QP43 Q4QP46 Q4QP49 Q4QP54 Q4QP55
Q4QP66 Q4QP70 Q4QP81 Q4QP83 Q4QP96 Q4QPG7 Q4QPH3 Q4QPH6 Q4QPH7 Q4QPI7 Q4QPI9
Q4QPJ6 Q4QPL1 Q4QPL6 Q4QPL7 Q4QPL8 Q4QPM0 Q4QPM5 Q4QPM6 Q4QPM7 Q4QPN0 Q4QPN2
Q4QPN5 Q4QPN9 Q4QPP0 Q4QQN5 Q4QQQ4 Q4QQQ7 Q4QQS2 Q4QQT0 Q4QQT2 Q4QQT3 Q4QQT5
Q4QQT6 Q4QQU1 Q4QQU7 Q4QQW4 Q4QQW5 Q4QQZ4 Q4QR00 Q4QR03 Q4QR06 Q4QR29 Q4QR58
Q4QR75 Q4QR76 Q4QR77 Q4QR80 Q4QR85 Q4QR87 Q4QR88 Q4QRB2 Q4QRC6 Q4QRD7 Q4QRJ7
Q4QRK0 Q4QSC8 Q4QSN4 Q4QXM3 Q4QXS7 Q4QZZ9 Q4R0H9 Q4R0I0 Q4R0J7 Q4R0W1 Q4R0W2
Q4R0W3 Q4R101 Q4R102 Q4R180 Q4R1E8 Q4R1I9 Q4R1J6 Q4R1R9 Q4R1S7 Q4R2Y2 Q4R2Z0
Q4R2Z6 Q4R2Z8 Q4R304 Q4R312 Q4R317 Q4R318 Q4R319 Q4R323 Q4R327 Q4R333 Q4R347
Q4R350 Q4R353 Q4R361 Q4R363 Q4R366 Q4R372 Q4R374 Q4R380 Q4R381 Q4R383 Q4R3A0
Q4R3A6 Q4R3B7 Q4R3D4 Q4R3D6 Q4R3E0 Q4R3F0 Q4R3G2 Q4R3G4 Q4R3H2 Q4R3I5 Q4R3I8
Q4R3J0 Q4R3J5 Q4R3J7 Q4R3K4 Q4R3K5 Q4R3M1 Q4R3P6 Q4R3Q1 Q4R3Q6 Q4R3Q7 Q4R3R9
Q4R3S3 Q4R3T1 Q4R3U8 Q4R3V2 Q4R3W5 Q4R3X1 Q4R3X5 Q4R3X7 Q4R3X9 Q4R3Z4 Q4R407
Q4R4C7 Q4R4E4 Q4R4F1 Q4R4F5 Q4R4F8 Q4R4H5 Q4R4H7 Q4R4I1 Q4R4I2 Q4R4J0 Q4R4J1
Q4R4K0 Q4R4K2 Q4R4K5 Q4R4K6 Q4R4M6 Q4R4N1 Q4R4P3 Q4R4P5 Q4R4P6 Q4R4P8 Q4R4Q6
Q4R4R0 Q4R4R7 Q4R4S8 Q4R4T9 Q4R4U1 Q4R4U3 Q4R4V3 Q4R4V8 Q4R4X5 Q4R4X8 Q4R4X9
Q4R4Y2 Q4R4Y9 Q4R4Z1 Q4R4Z6 Q4R502 Q4R506 Q4R517 Q4R526 Q4R531 Q4R535 Q4R537
Q4R538 Q4R539 Q4R543 Q4R550 Q4R555 Q4R568 Q4R572 Q4R578 Q4R579 Q4R584 Q4R588
Q4R594 Q4R596 Q4R599 Q4R5A4 Q4R5A5 Q4R5B3 Q4R5C6 Q4R5D0 Q4R5D4 Q4R5E4 Q4R5E6
Q4R5F0 Q4R5F2 Q4R5F5 Q4R5F6 Q4R5G1 Q4R5G2 Q4R5H0 Q4R5H5 Q4R5H6 Q4R5H7 Q4R5H8
Q4R5I3 Q4R5J0 Q4R5J4 Q4R5K7 Q4R5L1 Q4R5M3 Q4R5N1 Q4R5P3 Q4R5P6 Q4R5P9 Q4R5Q0
Q4R5Q4 Q4R5S7 Q4R5S9 Q4R5T4 Q4R5U5 Q4R5U8 Q4R5V1 Q4R5X9 Q4R5Y0 Q4R5Y2 Q4R5Y4
Q4R5Y8 Q4R5Z4 Q4R616 Q4R626 Q4R627 Q4R628 Q4R630 Q4R632 Q4R642 Q4R646 Q4R657
Q4R676 Q4R678 Q4R684 Q4R698 Q4R6C1 Q4R6C2 Q4R6C5 Q4R6C7 Q4R6D2 Q4R6F6 Q4R6F8
Q4R6G8 Q4R6I1 Q4R6I5 Q4R6J2 Q4R6J4 Q4R6J9 Q4R6L7 Q4R6M4 Q4R6M8 Q4R6N3 Q4R6N4
Q4R6P2 Q4R6P7 Q4R6Q3 Q4R6Q9 Q4R6R4 Q4R6T3 Q4R6T7 Q4R6U7 Q4R6V2 Q4R6V9 Q4R6W3
Q4R6W4 Q4R6W7 Q4R6W9 Q4R6X4 Q4R6X5 Q4R6X9 Q4R6Y6 Q4R6Y7 Q4R6Y8 Q4R712 Q4R713

Q4R723 Q4R729 Q4R731 Q4R736 Q4R739 Q4R741 Q4R744 Q4R747 Q4R755 Q4R760 Q4R767
Q4R779 Q4R786 Q4R793 Q4R796 Q4R7A4 Q4R7A8 Q4R7B1 Q4R7B9 Q4R7D0 Q4R7D3 Q4R7D5
Q4R7D9 Q4R7E8 Q4R7G2 Q4R7G7 Q4R7H0 Q4R7H3 Q4R7H5 Q4R7I0 Q4R7I4 Q4R7I8 Q4R7J8
Q4R7K7 Q4R7L3 Q4R7L5 Q4R7M0 Q4R7M4 Q4R7N3 Q4R7Q1 Q4R7Q7 Q4R7R3 Q4R7R9 Q4R7T5
Q4R7T7 Q4R7T8 Q4R7U4 Q4R7U8 Q4R7V1 Q4R7V3 Q4R7X8 Q4R7Y2 Q4R7Y5 Q4R7Y8 Q4R7Z4
Q4R7Z7 Q4R7Z9 Q4R802 Q4R803 Q4R804 Q4R809 Q4R812 Q4R813 Q4R816 Q4R821 Q4R826
Q4R828 Q4R836 Q4R837 Q4R842 Q4R846 Q4R848 Q4R856 Q4R871 Q4R881 Q4R882 Q4R888
Q4R898 Q4R899 Q4R8B6 Q4R8C3 Q4R8C7 Q4R8D2 Q4R8E0 Q4R8E6 Q4R8E7 Q4R8E8 Q4R8G4
Q4R8G6 Q4R8H1 Q4R8H9 Q4R8K5 Q4R8L2 Q4R8M9 Q4R8N2 Q4R8R3 Q4R8R4 Q4R8S8 Q4R8T1
Q4R8T4 Q4R8T9 Q4R8V8 Q4R8V9 Q4R8X4 Q4R8Y5 Q4R8Y8 Q4R8Z2 Q4R907 Q4R912 Q4R918
Q4R924 Q4R930 Q4R932 Q4R941 Q4R945 Q4R947 Q4R956 Q4R959 Q4R963 Q4R964 Q4R970
Q4R979 Q4R989 Q4R992 Q4R998 Q4R9A9 Q4R9C4 Q4R9D1 Q4R9E9 Q4R9F7 Q4R9N3 Q4RGM4
Q4RJN5 Q4RLT3 Q4RQD3 Q4RSW7 Q4RWX9 Q4S2L4 Q4S5X1 Q4S6U8 Q4SBY6 Q4SFA0 Q4SK88
Q4SPU8 Q4SQ06 Q4SRI6 Q4SS66 Q4SUE2 Q4T2X8 Q4T8S6 Q4TT88 Q4TU14 Q4TVV3 Q4TWH7
Q4U0F3 Q4U0G1 Q4U0S5 Q4U1U2 Q4U2V3 Q4U331 Q4U3E8 Q4U3F6 Q4U3F7 Q4U3L0 Q4U3P8
Q4U3U3 Q4U3U5 Q4U3U9 Q4U3V0 Q4U3Y2 Q4U4N4 Q4U4S6 Q4U5R3 Q4U5S1 Q4U8U5 Q4U923
Q4U925 Q4UAQ0 Q4UB74 Q4UB86 Q4UBI4 Q4UF75 Q4UFD5 Q4UFP0 Q4UFS9 Q4UGL5 Q4UGN5
Q4UH15 Q4UIF8 Q4UJ14 Q4UJ75 Q4UJB0 Q4UJB1 Q4UJC0 Q4UJC4 Q4UJJ2 Q4UJK6 Q4UJK7
Q4UJM2 Q4UJM3 Q4UJN4 Q4UJN5 Q4UJN8 Q4UJN9 Q4UJP2 Q4UJQ1 Q4UJQ2 Q4UJQ3 Q4UJQ5
Q4UJR2 Q4UJS3 Q4UJS8 Q4UJT0 Q4UJT1 Q4UJT2 Q4UJT5 Q4UJU1 Q4UJU2 Q4UJU4 Q4UJV1
Q4UJV2 Q4UJV4 Q4UJV5 Q4UJV6 Q4UJW3 Q4UJW4 Q4UJW9 Q4UJX3 Q4UJY2 Q4UJZ3 Q4UJZ5
Q4UJZ6 Q4UK06 Q4UK20 Q4UK35 Q4UK39 Q4UK40 Q4UK48 Q4UK55 Q4UK63 Q4UK70 Q4UK84
Q4UK85 Q4UK86 Q4UK93 Q4UK94 Q4UK95 Q4UK96 Q4UK97 Q4UK98 Q4UK99 Q4UKA4 Q4UKA8
Q4UKB1 Q4UKB5 Q4UKC5 Q4UKC6 Q4UKC9 Q4UKD0 Q4UKD1 Q4UKD2 Q4UKD3 Q4UKD4 Q4UKD5
Q4UKD7 Q4UKE2 Q4UKE7 Q4UKF0 Q4UKF1 Q4UKF2 Q4UKF3 Q4UKF4 Q4UKG1 Q4UKG2 Q4UKG8
Q4UKH2 Q4UKI1 Q4UKI6 Q4UKI7 Q4UKI8 Q4UKJ1 Q4UKJ3 Q4UKJ5 Q4UKK3 Q4UKL2 Q4UKL3
Q4UKL4 Q4UKL5 Q4UKL6 Q4UKL7 Q4UKM1 Q4UKM5 Q4UKM7 Q4UKN9 Q4UKP0 Q4UKP1 Q4UKQ3
Q4UKQ7 Q4UKR1 Q4UKR7 Q4UKR8 Q4UKS1 Q4UKS5 Q4UKS7 Q4UKS8 Q4UKT4 Q4UKT5 Q4UKU3
Q4UKW0 Q4UKW2 Q4UKX2 Q4UKX5 Q4UKX7 Q4UKX8 Q4UKY3 Q4UKZ9 Q4UL00 Q4UL10 Q4UL17
Q4UL18 Q4UL43 Q4UL45 Q4UL49 Q4UL51 Q4UL54 Q4UL56 Q4UL59 Q4UL62 Q4UL64 Q4UL77
Q4UL98 Q4ULA5 Q4ULC2 Q4ULC3 Q4ULD7 Q4ULD9 Q4ULE0 Q4ULF0 Q4ULF1 Q4ULF2 Q4ULF3
Q4ULF7 Q4ULG0 Q4ULG2 Q4ULI1 Q4ULI7 Q4ULJ9 Q4ULK0 Q4ULK1 Q4ULK8 Q4ULM2 Q4ULM3
Q4ULN0 Q4ULN5 Q4ULP1 Q4ULP7 Q4ULQ0 Q4ULQ3 Q4ULQ7 Q4ULQ8 Q4ULR1 Q4ULR2 Q4ULR3
Q4ULR6 Q4ULS1 Q4ULS3 Q4ULS4 Q4ULS5 Q4ULS7 Q4ULT8 Q4ULU8 Q4ULV2 Q4ULV5 Q4ULV6
Q4ULW6 Q4ULW8 Q4ULX2 Q4ULX3 Q4ULX9 Q4ULY2 Q4ULZ0 Q4ULZ2 Q4UM01 Q4UM12 Q4UM15
Q4UM17 Q4UM23 Q4UM29 Q4UM33 Q4UM37 Q4UM42 Q4UM43 Q4UM45 Q4UM50 Q4UM53 Q4UM54
Q4UM57 Q4UM71 Q4UM73 Q4UM78 Q4UM83 Q4UM86 Q4UM95 Q4UMA9 Q4UMB0 Q4UMD5 Q4UMD8
Q4UME7 Q4UME8 Q4UMF2 Q4UMF3 Q4UMH6 Q4UMI6 Q4UMI9 Q4UMJ0 Q4UMJ3 Q4UMK4 Q4UMK5
Q4UMK7 Q4UMK8 Q4UMK9 Q4UML0 Q4UML8 Q4UML9 Q4UMM0 Q4UMM7 Q4UMP3 Q4UMQ3 Q4UMQ4
Q4UMQ5 Q4UMQ6 Q4UMQ7 Q4UMQ8 Q4UMR0 Q4UMR1 Q4UMR2 Q4UMR3 Q4UMR4 Q4UMR5 Q4UMR6
Q4UMR7 Q4UMR8 Q4UMR9 Q4UMS0 Q4UMS1 Q4UMS2 Q4UMS3 Q4UMS4 Q4UMS5 Q4UMS6 Q4UMS7
Q4UMS8 Q4UMS9 Q4UMT0 Q4UMT4 Q4UMT9 Q4UMU1 Q4UMU5 Q4UMV1 Q4UMV4 Q4UMV9 Q4UMW4
Q4UMW5 Q4UMX2 Q4UMX3 Q4UMX8 Q4UMY2 Q4UMY8 Q4UMZ0 Q4UN06 Q4UN15 Q4UN16 Q4UN27
Q4UN30 Q4UN31 Q4UN57 Q4UN61 Q4UN64 Q4UN65 Q4UN66 Q4UN67 Q4UN71 Q4UN87 Q4UN88

Q4UN89 Q4UN91 Q4UN97 Q4UN98 Q4UNA7 Q4UNA8 Q4UNB0 Q4UNB1 Q4UNB2 Q4UNB3 Q4UNC2
Q4UNC3 Q4UNC4 Q4UNC8 Q4UND3 Q4UND7 Q4UND8 Q4UND9 Q4UNE4 Q4UNF0 Q4UNF2 Q4UNF3
Q4UNG8 Q4UNH2 Q4UNH4 Q4UNJ0 Q4UNJ4 Q4UNJ5 Q4UNK3 Q4UNK5 Q4UNK6 Q4UNK7 Q4UNL0
Q4UNT9 Q4UNU6 Q4UNV0 Q4UNV7 Q4UNV9 Q4UNX6 Q4UNZ4 Q4UNZ5 Q4UNZ6 Q4UP07 Q4UP08
Q4UP35 Q4UP54 Q4UP61 Q4UP62 Q4UP76 Q4UP98 Q4UPE3 Q4UPF9 Q4UPG4 Q4UPN0 Q4UPN9
Q4UPP0 Q4UPP2 Q4UPQ3 Q4UPQ7 Q4UPQ9 Q4UPU4 Q4UPU5 Q4UPX6 Q4UPY1 Q4UPY5 Q4UPY9
Q4UQ07 Q4UQ11 Q4UQ15 Q4UQ26 Q4UQB4 Q4UQB5 Q4UQD0 Q4UQI2 Q4UQI6 Q4UQN5 Q4UQP3
Q4UQP6 Q4UQS8 Q4UQT6 Q4UQT7 Q4UQU3 Q4UQU4 Q4UQV6 Q4UQW2 Q4UQW3 Q4UQW6 Q4UQX1
Q4UQZ8 Q4UR00 Q4UR08 Q4UR35 Q4UR38 Q4UR39 Q4UR40 Q4UR65 Q4UR70 Q4UR77 Q4UR79
Q4URB1 Q4URB6 Q4URB7 Q4URC0 Q4URC2 Q4URC3 Q4URC5 Q4URC8 Q4URC9 Q4URD0 Q4URD1
Q4URD2 Q4URD3 Q4URD4 Q4URD5 Q4URD6 Q4URD7 Q4URD8 Q4URD9 Q4URE0 Q4URE1 Q4URE2
Q4URE3 Q4URE4 Q4URE5 Q4URE6 Q4URE7 Q4URE8 Q4URE9 Q4URF0 Q4URF1 Q4URF2 Q4URF3
Q4URF4 Q4URF5 Q4URF6 Q4URF7 Q4URF8 Q4URG0 Q4URG1 Q4URG2 Q4URG3 Q4URG4 Q4URH2
Q4URI9 Q4URL3 Q4URL4 Q4URL5 Q4URM1 Q4URM3 Q4URN0 Q4URN2 Q4URN6 Q4URN7 Q4URP7
Q4URP9 Q4URT6 Q4URY1 Q4URZ4 Q4US05 Q4US07 Q4US21 Q4US26 Q4US34 Q4US35 Q4US36
Q4US37 Q4US40 Q4US42 Q4US82 Q4US83 Q4US84 Q4US85 Q4US88 Q4US90 Q4US93 Q4USF7
Q4USF9 Q4USG1 Q4USG7 Q4USM5 Q4USP6 Q4USQ4 Q4USQ7 Q4USQ8 Q4USR1 Q4USR2 Q4USS0
Q4USS4 Q4UST1 Q4UT03 Q4UT11 Q4UT12 Q4UT42 Q4UT60 Q4UT85 Q4UT86 Q4UT92 Q4UT93
Q4UT95 Q4UTA1 Q4UTA3 Q4UTA4 Q4UTA5 Q4UTB0 Q4UTB3 Q4UTB9 Q4UTC0 Q4UTC9 Q4UTD1
Q4UTD4 Q4UTN9 Q4UTP4 Q4UTP5 Q4UTP6 Q4UTP8 Q4UTP9 Q4UTQ4 Q4UTQ7 Q4UTQ8 Q4UTR4
Q4UTR5 Q4UTR6 Q4UTR7 Q4UTT5 Q4UTU6 Q4UTV2 Q4UTV5 Q4UTV6 Q4UTV7 Q4UTV8 Q4UTW8
Q4UTX4 Q4UU32 Q4UU37 Q4UU39 Q4UU40 Q4UU41 Q4UU42 Q4UU43 Q4UU44 Q4UU45 Q4UU49
Q4UU50 Q4UU51 Q4UU61 Q4UU71 Q4UU84 Q4UU98 Q4UUJ7 Q4UUJ9 Q4UUK1 Q4UUK2 Q4UUL4
Q4UUL5 Q4UUM0 Q4UUW4 Q4UUW5 Q4UUX2 Q4UV60 Q4UV74 Q4UV84 Q4UVA7 Q4UVB0 Q4UVC5
Q4UVC7 Q4UVD5 Q4UVD7 Q4UVD8 Q4UVH1 Q4UVI0 Q4UVI1 Q4UVI4 Q4UVI5 Q4UVJ3 Q4UVK6
Q4UVL3 Q4UVL4 Q4UVL7 Q4UVM8 Q4UVM9 Q4UVN3 Q4UVN8 Q4UVP3 Q4UVP8 Q4UVR0 Q4UVS2
Q4UVS5 Q4UVU9 Q4UVX2 Q4UVX3 Q4UVY0 Q4UVY5 Q4UVY7 Q4UW14 Q4UW29 Q4UW51 Q4UW52
Q4UW53 Q4UW54 Q4UW55 Q4UW57 Q4UW79 Q4UW98 Q4UW99 Q4UWA0 Q4UWA1 Q4UWA2 Q4UWA4
Q4UWC0 Q4UWC8 Q4UWD0 Q4UWD2 Q4UWD4 Q4UWD5 Q4UWE0 Q4UWF3 Q4UWF9 Q4UWI7 Q4UWI8
Q4UWJ4 Q4UWJ9 Q4UWK2 Q4UWK8 Q4UWM5 Q4UWS6 Q4UWV1 Q4UWV2 Q4UWX9 Q4UWY7 Q4UWY8
Q4UWY9 Q4UX14 Q4UX30 Q4UX36 Q4UX46 Q4UX47 Q4UX84 Q4UX85 Q4UX86 Q4UX89 Q4UXA5
Q4UXC1 Q4UXC2 Q4UXF8 Q4UXI1 Q4UXI2 Q4UXI3 Q4UXI4 Q4UXK8 Q4UXL7 Q4UXM0 Q4UXM1
Q4UXM2 Q4UXM5 Q4UXQ4 Q4UXR7 Q4UXU0 Q4UXU8 Q4UXV5 Q4UXY4 Q4UXY5 Q4UY06 Q4UY08
Q4UY09 Q4UY23 Q4UY33 Q4UY37 Q4UY39 Q4UY41 Q4UY42 Q4UY45 Q4UY54 Q4UY95 Q4UYA0
Q4UYA4 Q4UYA7 Q4UYC6 Q4UYF2 Q4UYF7 Q4UYG1 Q4UYG2 Q4UYG4 Q4UYG5 Q4UYJ6 Q4UYL9
Q4UYM8 Q4UYN4 Q4UYN5 Q4UYN8 Q4UYT2 Q4UYT3 Q4UYT4 Q4UYX6 Q4UYX7 Q4UYY0 Q4UYY2
Q4UYZ4 Q4UZ66 Q4UZ69 Q4UZ74 Q4UZA7 Q4UZA8 Q4UZB2 Q4UZB8 Q4UZD2 Q4UZE2 Q4UZF0
Q4UZF1 Q4UZF2 Q4UZF5 Q4UZF8 Q4UZF9 Q4UZH5 Q4UZI9 Q4UZJ5 Q4UZL7 Q4UZL8 Q4UZN8
Q4UZN9 Q4UZP1 Q4UZP9 Q4UZR8 Q4UZT2 Q4UZX0 Q4V073 Q4V0C6 Q4V0E2 Q4V0I4 Q4V0R7
Q4V0S6 Q4V0S8 Q4V182 Q4V1F4 Q4V1F5 Q4V1F6 Q4V1F7 Q4V2J8 Q4V321 Q4V326 Q4V339
Q4V348 Q4V399 Q4V3C1 Q4V3D3 Q4V3E2 Q4V516 Q4V5A3 Q4V5I9 Q4V6M7 Q4V7A0 Q4V7A4
Q4V7A6 Q4V7A9 Q4V7B1 Q4V7B2 Q4V7B4 Q4V7B5 Q4V7B8 Q4V7C4 Q4V7C6 Q4V7C8 Q4V7C9
Q4V7D2 Q4V7D6 Q4V7D7 Q4V7D8 Q4V7E0 Q4V7E1 Q4V7E5 Q4V7E8 Q4V7F0 Q4V7F3 Q4V7F5
Q4V7H1 Q4V7H8 Q4V7J0 Q4V7K4 Q4V7L5 Q4V7N3 Q4V7N4 Q4V7N8 Q4V7P7 Q4V7Q1 Q4V7Q6

Q4V7Q7 Q4V7R4 Q4V7R8 Q4V7T1 Q4V7T5 Q4V7T8 Q4V7V8 Q4V7W2 Q4V7W3 Q4V7W5 Q4V7W8
Q4V7X9 Q4V7Y4 Q4V7Y7 Q4V7Z0 Q4V7Z1 Q4V817 Q4V831 Q4V838 Q4V842 Q4V847 Q4V860
Q4V862 Q4V863 Q4V869 Q4V872 Q4V890 Q4V891 Q4V893 Q4V8A0 Q4V8A2 Q4V8A8 Q4V8B0
Q4V8B3 Q4V8B5 Q4V8B7 Q4V8B8 Q4V8C3 Q4V8C4 Q4V8C7 Q4V8C9 Q4V8D6 Q4V8D7 Q4V8D9
Q4V8E4 Q4V8E6 Q4V8E9 Q4V8F1 Q4V8F5 Q4V8F7 Q4V8F9 Q4V8G0 Q4V8G2 Q4V8G4 Q4V8G5
Q4V8I2 Q4V8I4 Q4V8I5 Q4V8J2 Q4V8J7 Q4V8K3 Q4V8P4 Q4V8R6 Q4V8R7 Q4V8T0 Q4V8V1
Q4V8V2 Q4V8W7 Q4V8X4 Q4V8Y7 Q4V908 Q4V909 Q4V9H5 Q4V9P6 Q4V9P9 Q4V9W2 Q4VA36
Q4VA44 Q4VA45 Q4VA53 Q4VA55 Q4VA72 Q4VAB4 Q4VAC9 Q4VBD9 Q4VBE8 Q4VBF2 Q4VBH4
Q4VBJ9 Q4VBS4 Q4VBT5 Q4VBV9 Q4VC12 Q4VC31 Q4VDN5 Q4VGL6 Q4VID0 Q4VK78 Q4VKB4
Q4VKU9 Q4VKV0 Q4VKV1 Q4VKV2 Q4VKV6 Q4VKV7 Q4VKV8 Q4VNK0 Q4VPH7 Q4VPH8 Q4VPP0
Q4VPP5 Q4VPP7 Q4VPP9 Q4VPQ3 Q4VPQ4 Q4VQY1 Q4VQY3 Q4VR96 Q4VR99 Q4VRV8 Q4VUF1
Q4VWF8 Q4VWW3 Q4VX62 Q4VXU2 Q4VYA0 Q4VYC8 Q4VYR7 Q4VYS1 Q4VZH3 Q4VZJ2 Q4VZJ4
Q4VZJ5 Q4VZK0 Q4VZK1 Q4VZK2 Q4VZK3 Q4VZK5 Q4VZK6 Q4VZK9 Q4VZL1 Q4VZM8 Q4VZM9
Q4VZN0 Q4VZN1 Q4VZN5 Q4VZP1 Q4VZP2 Q4VZP3 Q4VZP4 Q4W382 Q4W4Y0 Q4W4Z2 Q4W557
Q4W568 Q4W580 Q4W5G0 Q4W5Z4 Q4W7G7 Q4W7G8 Q4W7G9 Q4W7H0 Q4W7H1 Q4W7T3 Q4W8D0
Q4W944 Q4W945 Q4W946 Q4W9B7 Q4W9B8 Q4W9B9 Q4W9F7 Q4W9G3 Q4W9H1 Q4W9M3 Q4W9M7
Q4W9R0 Q4W9R2 Q4W9S8 Q4W9V0 Q4W9Z9 Q4WA21 Q4WA22 Q4WA58 Q4WA59 Q4WA61 Q4WA62
Q4WA70 Q4WAQ9 Q4WAR8 Q4WAS9 Q4WAU7 Q4WAW3 Q4WAW6 Q4WAW7 Q4WAW9 Q4WAX1 Q4WAY7
Q4WAY8 Q4WAZ0 Q4WAZ8 Q4WAZ9 Q4WB00 Q4WB01 Q4WB03 Q4WB07 Q4WBL6 Q4WBR0 Q4WCF1
Q4WCH5 Q4WCJ7 Q4WCL1 Q4WCP8 Q4WCR3 Q4WCV5 Q4WCW2 Q4WD42 Q4WD43 Q4WD45 Q4WD47
Q4WD48 Q4WDA4 Q4WDD0 Q4WDD7 Q4WDE1 Q4WDE9 Q4WDF1 Q4WDF7 Q4WDG1 Q4WDH3 Q4WDH9
Q4WDK5 Q4WE58 Q4WE62 Q4WEB4 Q4WED6 Q4WED7 Q4WED9 Q4WEH7 Q4WEI5 Q4WEL6 Q4WEN1
Q4WER3 Q4WEU2 Q4WEY4 Q4WF29 Q4WF30 Q4WF53 Q4WF56 Q4WF61 Q4WFF8 Q4WFT3 Q4WG40
Q4WG69 Q4WGB7 Q4WGK6 Q4WGS4 Q4WH83 Q4WH96 Q4WHA3 Q4WHB7 Q4WHD1 Q4WHD2 Q4WHF8
Q4WHG0 Q4WHG1 Q4WHG5 Q4WHK3 Q4WHP6 Q4WHT7 Q4WHT8 Q4WHT9 Q4WHU0 Q4WHU1 Q4WHW0
Q4WHW1 Q4WHY5 Q4WHZ9 Q4WI01 Q4WI37 Q4WI38 Q4WI71 Q4WI96 Q4WID9 Q4WII3 Q4WIK9
Q4WJ02 Q4WJ21 Q4WJ31 Q4WJ81 Q4WJ90 Q4WJE9 Q4WJH6 Q4WJI7 Q4WJJ0 Q4WJM6 Q4WJQ1
Q4WJR4 Q4WJS2 Q4WJT7 Q4WJT9 Q4WJX0 Q4WJX7 Q4WK03 Q4WK35 Q4WK56 Q4WKB2 Q4WKB4
Q4WKB5 Q4WKB9 Q4WKD9 Q4WKH9 Q4WKL7 Q4WKW9 Q4WKX0 Q4WKX1 Q4WLA7 Q4WLA8 Q4WLC9
Q4WLD4 Q4WLD5 Q4WLG1 Q4WLG9 Q4WLI5 Q4WLM7 Q4WLM9 Q4WLN1 Q4WLU1 Q4WLV6 Q4WLW5
Q4WLW7 Q4WLW8 Q4WM60 Q4WM95 Q4WMB6 Q4WMF7 Q4WMG1 Q4WMI1 Q4WMJ1 Q4WMJ5 Q4WMJ7
Q4WMJ8 Q4WMJ9 Q4WMK0 Q4WMK2 Q4WMS3 Q4WMU1 Q4WMU5 Q4WMV5 Q4WMX7 Q4WN25 Q4WN42
Q4WN99 Q4WNA1 Q4WNC1 Q4WNC6 Q4WND3 Q4WNG7 Q4WNH8 Q4WNI0 Q4WNI1 Q4WNQ6 Q4WNT9
Q4WNW3 Q4WNX1 Q4WNY4 Q4WP02 Q4WP03 Q4WP10 Q4WP13 Q4WP38 Q4WP54 Q4WP57 Q4WP65
Q4WP83 Q4WP96 Q4WPE6 Q4WPE9 Q4WPF5 Q4WPF7 Q4WPF8 Q4WPL7 Q4WPM6 Q4WPM8 Q4WPN0
Q4WPU8 Q4WPU9 Q4WPV8 Q4WPW2 Q4WPW8 Q4WPX2 Q4WQ54 Q4WQ87 Q4WQB9 Q4WQG5 Q4WQH4
Q4WQL4 Q4WQM6 Q4WQN1 Q4WQY4 Q4WQY5 Q4WQY7 Q4WQY9 Q4WQZ0 Q4WQZ1 Q4WQZ2 Q4WQZ3
Q4WQZ4 Q4WQZ5 Q4WQZ6 Q4WQZ7 Q4WR21 Q4WR23 Q4WR24 Q4WR82 Q4WRB8 Q4WRC2 Q4WRE2
Q4WRE4 Q4WRE8 Q4WRH5 Q4WRP2 Q4WRQ7 Q4WRU4 Q4WRV2 Q4WRV9 Q4WRX4 Q4WRY5 Q4WS57
Q4WS76 Q4WSA8 Q4WSI0 Q4WSM6 Q4WT34 Q4WT40 Q4WT66 Q4WT84 Q4WT99 Q4WTA6 Q4WTC4
Q4WTH0 Q4WTH1 Q4WTI3 Q4WTI4 Q4WTL0 Q4WTM9 Q4WTN5 Q4WTN9 Q4WTQ4 Q4WTT2 Q4WTV7
Q4WU00 Q4WU07 Q4WU09 Q4WU59 Q4WUA0 Q4WUD3 Q4WUE0 Q4WUG4 Q4WUJ6 Q4WUL6 Q4WUR1
Q4WUT7 Q4WUV8 Q4WUV9 Q4WV19 Q4WV66 Q4WV71 Q4WV91 Q4WVA0 Q4WVA5 Q4WVD1 Q4WVE3
Q4WVE5 Q4WVF4 Q4WVG8 Q4WVH3 Q4WVH4 Q4WVI6 Q4WVK2 Q4WVK5 Q4WVM1 Q4WVM7 Q4WVQ7

Q4WVS2 Q4WVU5 Q4WVZ0 Q4WW81 Q4WWC5 Q4WWC6 Q4WWD3 Q4WWE9 Q4WWH6 Q4WWJ1 Q4WWL0
Q4WWN8 Q4WWP1 Q4WWR2 Q4WWW5 Q4WWW9 Q4WX30 Q4WX43 Q4WX78 Q4WX89 Q4WX90 Q4WX99
Q4WXA1 Q4WXD3 Q4WXE9 Q4WXH8 Q4WXK4 Q4WXP0 Q4WXQ7 Q4WXV2 Q4WXV6 Q4WXW1 Q4WXX4
Q4WXX5 Q4WXX9 Q4WXY0 Q4WXY3 Q4WY08 Q4WY16 Q4WY33 Q4WY53 Q4WY67 Q4WYA1 Q4WYE5
Q4WYF1 Q4WYG2 Q4WYG3 Q4WYJ7 Q4WYK1 Q4WYK9 Q4WYN5 Q4WYN6 Q4WYP0 Q4WYR6 Q4WYS1
Q4WYS7 Q4WYU4 Q4WYX0 Q4WZ01 Q4WZ44 Q4WZ60 Q4WZ63 Q4WZ64 Q4WZ66 Q4WZ67 Q4WZ69
Q4WZ70 Q4WZB3 Q4WZC3 Q4WZJ0 Q4WZJ6 Q4WZL3 Q4WZP2 Q4WZS2 Q4WZT9 Q4WZV6 Q4X054
Q4X095 Q4X099 Q4X0A9 Q4X0C2 Q4X0I5 Q4X0I7 Q4X0N1 Q4X0S7 Q4X0T4 Q4X0W8 Q4X0X6
Q4X0Z3 Q4X0Z5 Q4X122 Q4X156 Q4X161 Q4X195 Q4X1A4 Q4X1D2 Q4X1D4 Q4X1D7 Q4X1F8
Q4X1I3 Q4X1Q4 Q4X1R1 Q4X1V0 Q4X1W0 Q4X1W8 Q4X1X0 Q4X1Y0 Q4X212 Q4X228 Q4X267
Q4X2S8 Q4XFJ8 Q4XRM7 Q4XS21 Q4XUG7 Q4XWV5 Q4XXP8 Q4XZN2 Q4XZZ5 Q4Y4B1 Q4YBG7
Q4YU52 Q4YU79 Q4YUG2 Q4YUZ9 Q4YZC6 Q4YZZ7 Q4Z015 Q4Z1L3 Q4Z694 Q4Z6R1 Q4Z971
Q4Z9F2 Q4Z9G3 Q4ZGE2 Q4ZHW1 Q4ZL08 Q4ZL12 Q4ZL14 Q4ZL75 Q4ZLB5 Q4ZLL3 Q4ZLP3
Q4ZLP8 Q4ZLP9 Q4ZLQ1 Q4ZLQ2 Q4ZLR0 Q4ZLR3 Q4ZLU8 Q4ZLV2 Q4ZLW0 Q4ZLW7 Q4ZLX9
Q4ZLZ9 Q4ZM01 Q4ZM05 Q4ZM24 Q4ZM34 Q4ZM37 Q4ZM55 Q4ZM56 Q4ZM61 Q4ZM62 Q4ZM63
Q4ZM83 Q4ZMA8 Q4ZMA9 Q4ZMC1 Q4ZMC2 Q4ZMC3 Q4ZMC4 Q4ZMC5 Q4ZMF3 Q4ZMF4 Q4ZMF8
Q4ZMG0 Q4ZMG2 Q4ZMG3 Q4ZMG5 Q4ZMI2 Q4ZML2 Q4ZML7 Q4ZMM3 Q4ZMM4 Q4ZMM5 Q4ZMM7
Q4ZMM9 Q4ZMN3 Q4ZMN4 Q4ZMN5 Q4ZMN6 Q4ZMN7 Q4ZMN8 Q4ZMN9 Q4ZMP0 Q4ZMP1 Q4ZMP2
Q4ZMP3 Q4ZMP4 Q4ZMP5 Q4ZMP6 Q4ZMP7 Q4ZMP8 Q4ZMP9 Q4ZMQ0 Q4ZMQ1 Q4ZMQ2 Q4ZMQ3
Q4ZMQ4 Q4ZMQ5 Q4ZMQ6 Q4ZMQ7 Q4ZMQ8 Q4ZMQ9 Q4ZMR0 Q4ZMR1 Q4ZMR2 Q4ZMR3 Q4ZMR5
Q4ZMR6 Q4ZMR7 Q4ZMR8 Q4ZMR9 Q4ZMV1 Q4ZMX9 Q4ZMY4 Q4ZMY9 Q4ZMZ1 Q4ZN06 Q4ZN07
Q4ZN10 Q4ZN11 Q4ZN23 Q4ZN40 Q4ZN43 Q4ZN72 Q4ZN73 Q4ZN74 Q4ZN76 Q4ZN82 Q4ZN83
Q4ZN84 Q4ZN90 Q4ZN95 Q4ZN97 Q4ZNA0 Q4ZNA1 Q4ZNF5 Q4ZNG3 Q4ZNH2 Q4ZNI9 Q4ZNL5
Q4ZNN4 Q4ZNN9 Q4ZNP2 Q4ZNP6 Q4ZNP7 Q4ZNP8 Q4ZNP9 Q4ZNQ4 Q4ZNR2 Q4ZNR3 Q4ZNR4
Q4ZNR5 Q4ZNR6 Q4ZNS6 Q4ZNS7 Q4ZNS8 Q4ZNT2 Q4ZNU2 Q4ZNV7 Q4ZNV8 Q4ZNV9 Q4ZNW0
Q4ZNW4 Q4ZNX2 Q4ZNX3 Q4ZNX7 Q4ZNX8 Q4ZNY1 Q4ZNY2 Q4ZNY8 Q4ZNZ1 Q4ZNZ9 Q4ZP05
Q4ZP19 Q4ZP20 Q4ZPC2 Q4ZPD1 Q4ZPE1 Q4ZPE3 Q4ZPE4 Q4ZPE5 Q4ZPE6 Q4ZPH2 Q4ZPI7
Q4ZPJ7 Q4ZPJ8 Q4ZPK0 Q4ZPK6 Q4ZPK9 Q4ZPM9 Q4ZPN9 Q4ZPQ0 Q4ZPZ9 Q4ZQ34 Q4ZQ35
Q4ZQ44 Q4ZQ51 Q4ZQ64 Q4ZQ66 Q4ZQ82 Q4ZQ90 Q4ZQ92 Q4ZQ94 Q4ZQ97 Q4ZQ99 Q4ZQB2
Q4ZQC3 Q4ZQC6 Q4ZQG4 Q4ZQG8 Q4ZQH8 Q4ZQH9 Q4ZQI1 Q4ZQI4 Q4ZQQ7 Q4ZQW4 Q4ZQY7
Q4ZQZ4 Q4ZR64 Q4ZRC2 Q4ZRE3 Q4ZRK0 Q4ZRK6 Q4ZRK8 Q4ZRK9 Q4ZRL0 Q4ZRL5 Q4ZRL6
Q4ZS10 Q4ZS35 Q4ZS46 Q4ZSC0 Q4ZSF5 Q4ZTG0 Q4ZTJ2 Q4ZTL7 Q4ZTN0 Q4ZTR4 Q4ZU05
Q4ZU37 Q4ZU60 Q4ZUA4 Q4ZUC9 Q4ZUD1 Q4ZUD2 Q4ZUD4 Q4ZUG1 Q4ZUG2 Q4ZUG3 Q4ZUG4
Q4ZUG5 Q4ZUG7 Q4ZUH3 Q4ZUJ4 Q4ZUM1 Q4ZUN7 Q4ZUP0 Q4ZUW2 Q4ZUW7 Q4ZUZ4 Q4ZUZ5
Q4ZUZ6 Q4ZV02 Q4ZV05 Q4ZV64 Q4ZV71 Q4ZV77 Q4ZVB8 Q4ZVB9 Q4ZVC0 Q4ZVC2 Q4ZVD9
Q4ZVE4 Q4ZVE7 Q4ZVE8 Q4ZVE9 Q4ZVF6 Q4ZVG8 Q4ZVG9 Q4ZVH4 Q4ZVL1 Q4ZVL3 Q4ZVM6
Q4ZVM7 Q4ZVM8 Q4ZVN1 Q4ZVN9 Q4ZVT9 Q4ZVW1 Q4ZVX2 Q4ZVY0 Q4ZVY2 Q4ZVY7 Q4ZW11
Q4ZW13 Q4ZW15 Q4ZW36 Q4ZW57 Q4ZW60 Q4ZW63 Q4ZW72 Q4ZW75 Q4ZW81 Q4ZWK2 Q4ZWL1
Q4ZWL2 Q4ZWL3 Q4ZWL4 Q4ZWL5 Q4ZWL9 Q4ZWM2 Q4ZWM4 Q4ZWM9 Q4ZWP2 Q4ZWP3 Q4ZWP5
Q4ZWP9 Q4ZWQ0 Q4ZWQ1 Q4ZWQ2 Q4ZWQ6 Q4ZWR0 Q4ZWR4 Q4ZWS2 Q4ZWS5 Q4ZWS6 Q4ZWS7
Q4ZWS8 Q4ZWT0 Q4ZWU0 Q4ZWU6 Q4ZWV2 Q4ZWV3 Q4ZWW0 Q4ZWY6 Q4ZWY7 Q4ZWY8 Q4ZWY9
Q4ZWZ4 Q4ZX02 Q4ZX09 Q4ZX12 Q4ZX14 Q4ZX15 Q4ZX19 Q4ZX22 Q4ZX23 Q4ZX26 Q4ZX27
Q4ZX30 Q4ZX34 Q4ZXA2 Q4ZXH2 Q4ZXH7 Q4ZXI0 Q4ZXI1 Q4ZXI3 Q4ZXJ9 Q4ZXK2 Q4ZXR7
Q4ZXR8 Q4ZXR9 Q4ZXS0 Q4ZXS1 Q4ZXT0 Q4ZXU7 Q4ZXU9 Q4ZXW7 Q4ZXW8 Q4ZXX1 Q4ZXX3

Q4ZXX4 Q4ZY26 Q4ZY66 Q4ZY77 Q4ZY80 Q4ZY86 Q4ZY87 Q4ZY88 Q4ZYB9 Q4ZYC0 Q4ZYD2
Q4ZYJ1 Q4ZYJ4 Q4ZYJ7 Q4ZYK0 Q4ZYK1 Q4ZYK2 Q4ZYK3 Q4ZYK5 Q4ZYP1 Q4ZYU6 Q4ZYU8
Q4ZYV7 Q4ZYV8 Q4ZYW8 Q4ZYX0 Q4ZYX1 Q4ZYX5 Q4ZYX6 Q4ZYY0 Q4ZYY1 Q4ZYY8 Q4ZYY9
Q4ZZ08 Q4ZZ15 Q4ZZ54 Q4ZZ58 Q4ZZ67 Q4ZZ68 Q4ZZ69 Q4ZZ70 Q4ZZ78 Q4ZZ83 Q4ZZ92
Q4ZZA8 Q4ZZA9 Q4ZZD8 Q4ZZE3 Q4ZZE4 Q4ZZF3 Q4ZZF5 Q4ZZF8 Q4ZZG3 Q4ZZG6 Q4ZZG7
Q4ZZH6 Q4ZZI0 Q4ZZT0 Q4ZZU4 Q4ZZU5 Q4ZZW2 Q4ZZW4 Q4ZZX5 Q4ZZX7 Q4ZZX9 Q4ZZY1
Q4ZZY3 Q4ZZY6 Q4ZZY8 Q4ZZY9 Q50000 Q50008 Q50022 Q50023 Q50025 Q50028 Q50036
Q50049 Q50068 Q500B4 Q500B6 Q500B9 Q500C6 Q500D0 Q500D1 Q500M2 Q500N3 Q500N6
Q500R4 Q500R5 Q500S4 Q500S6 Q500T0 Q500T7 Q500U1 Q500U5 Q500U7 Q500U8 Q500V5
Q500V9 Q500Y9 Q500Z7 Q50140 Q50144 Q50167 Q50178 Q50191 Q50192 Q501D1 Q501D5
Q501E9 Q501G5 Q501J6 Q501L1 Q501L3 Q501Q9 Q501R9 Q501S4 Q501V0 Q501X6 Q501Z5
Q50201 Q50203 Q50212 Q50220 Q50228 Q50229 Q50234 Q50235 Q50239 Q50255 Q50264
Q50265 Q50266 Q50267 Q50270 Q50271 Q50284 Q50286 Q50287 Q50290 Q50291 Q50295
Q50296 Q50297 Q50298 Q502A3 Q502B0 Q502C1 Q502D1 Q502G5 Q502H1 Q502I3 Q502I6
Q502J7 Q502K2 Q502K3 Q502M5 Q502M6 Q502M8 Q502P0 Q502P7 Q502W6 Q502W7 Q502X0
Q50300 Q50301 Q50302 Q50303 Q50304 Q50305 Q50306 Q50307 Q50308 Q50309 Q50310
Q50312 Q50313 Q50314 Q50315 Q50316 Q50319 Q50336 Q50338 Q50355 Q50362 Q50363
Q50373 Q50382 Q50383 Q50388 Q503C8 Q503D6 Q503E1 Q503F2 Q503H1 Q503I8 Q503I9
Q503J2 Q503L9 Q503N9 Q503P2 Q503R4 Q503S1 Q503T5 Q503U0 Q503U1 Q503V9 Q503X2
Q503Y4 Q503Y7 Q503Y8 Q50412 Q50422 Q50423 Q50428 Q50429 Q50436 Q50437 Q50438
Q50441 Q50485 Q50497 Q504A5 Q504E7 Q504H8 Q504L7 Q504L8 Q504N7 Q504Q3 Q504T8
Q504U0 Q504Y3 Q50500 Q50501 Q50504 Q50519 Q50521 Q50533 Q50538 Q50539 Q50543
Q50562 Q50566 Q50570 Q505B7 Q505B8 Q505F5 Q505G4 Q505G8 Q505H7 Q505I4 Q505I5
Q505J8 Q505K2 Q50744 Q50746 Q50754 Q50756 Q50757 Q50758 Q50759 Q50770 Q50777
Q50781 Q50782 Q50783 Q50784 Q50785 Q50786 Q50787 Q50788 Q507P6 Q507Q9 Q507R6
Q507T3 Q507Y2 Q50826 Q50827 Q50834 Q50837 Q50840 Q50841 Q50852 Q50864 Q50899
Q50900 Q50939 Q50945 Q50961 Q50962 Q50968 Q50973 Q50974 Q50975 Q50976 Q50983
Q50D79 Q50EJ9 Q50EK2 Q50EL0 Q50EW2 Q50EX7 Q50H32 Q50H63 Q50HP3 Q50HP4 Q50HR5
Q50HR6 Q50I20 Q50KA8 Q50L36 Q50L39 Q50L96 Q50LE4 Q50LE5 Q50LE6 Q51003 Q51008
Q51062 Q51080 Q51083 Q51151 Q51152 Q51161 Q51238 Q51296 Q51320 Q51334 Q51342
Q51344 Q51363 Q51375 Q51382 Q51383 Q51384 Q51385 Q51390 Q51391 Q51404 Q51422
Q51424 Q51425 Q51426 Q51440 Q51462 Q51470 Q51472 Q51473 Q51480 Q51481 Q51484
Q51493 Q51494 Q51495 Q51506 Q51507 Q51508 Q51517 Q51518 Q51519 Q51521 Q51551
Q51553 Q51561 Q51564 Q51567 Q51576 Q51577 Q51601 Q51602 Q51603 Q51645 Q51647
Q51649 Q51651 Q51660 Q51664 Q51669 Q51676 Q51687 Q51697 Q51701 Q51703 Q51725
Q51732 Q51739 Q51741 Q51742 Q51758 Q51761 Q51762 Q51772 Q51773 Q51782 Q51785
Q51786 Q51787 Q51788 Q51789 Q51790 Q51791 Q51792 Q51793 Q51795 Q51796 Q51798
Q51799 Q51800 Q51801 Q51802 Q51803 Q51804 Q51805 Q51813 Q51831 Q51832 Q51834
Q51841 Q51843 Q51853 Q51854 Q51856 Q51857 Q51858 Q51872 Q51879 Q51892 Q51896
Q51929 Q51945 Q51947 Q51948 Q51956 Q51973 Q51974 Q51975 Q51978 Q51983 Q51D88
Q51LA6 Q51LS1 Q51NJ3 Q51TF7 Q52008 Q52011 Q52028 Q52036 Q52039 Q52040 Q52042
Q52043 Q52060 Q52069 Q52070 Q52071 Q52078 Q52083 Q52086 Q52087 Q52093 Q52094
Q52109 Q52110 Q52118 Q52126 Q52149 Q52154 Q52185 Q52186 Q521C0 Q52219 Q52221
Q52243 Q52277 Q52278 Q52279 Q52281 Q52282 Q52287 Q52312 Q52313 Q52328 Q52329

Q52331 Q52346 Q52347 Q52353 Q52357 Q52358 Q52360 Q52366 Q52377 Q52400 Q52419
Q52424 Q52428 Q52438 Q52439 Q52440 Q52448 Q52449 Q52462 Q52472 Q524Z5 Q52500
Q52521 Q52522 Q52523 Q52524 Q52525 Q52528 Q52532 Q52556 Q52561 Q52562 Q52563
Q52620 Q52658 Q52671 Q52673 Q52698 Q52701 Q52702 Q52710 Q52720 Q52725 Q52728
Q52741 Q52742 Q52772 Q52779 Q52784 Q527H0 Q52811 Q52815 Q52828 Q52838 Q52839
Q52845 Q52846 Q52871 Q52873 Q52892 Q52905 Q52928 Q52944 Q52945 Q52946 Q52953
Q52955 Q52964 Q52967 Q52975 Q52984 Q52985 Q52986 Q52987 Q52990 Q52991 Q52992
Q52993 Q52994 Q52996 Q52997 Q52998 Q52B63 Q52B93 Q52BY4 Q52C47 Q52CS0 Q52DF3
Q52DM9 Q52E49 Q52E73 Q52F10 Q52FM1 Q52G60 Q52JK6 Q52K82 Q52K88 Q52KB5 Q52KD8
Q52KE7 Q52KG4 Q52KI8 Q52KJ8 Q52KK4 Q52KN3 Q52KN9 Q52KR3 Q52KS4 Q52KV5 Q52L34
Q52L45 Q52LG2 Q52LR7 Q52M58 Q52M75 Q52M93 Q52M98 Q52MB2 Q52NW7 Q52NY7 Q52PJ5
Q52PV9 Q52QH4 Q52QT8 Q52QW5 Q52S23 Q52TG1 Q52TG6 Q52TG9 Q52US9 Q52UT0 Q52UT1
Q52V16 Q52ZT5 Q52ZX4 Q53005 Q53009 Q53017 Q53034 Q53044 Q53046 Q53062 Q53079
Q53080 Q53081 Q53083 Q53084 Q53091 Q53107 Q53117 Q53118 Q53120 Q53121 Q53122
Q53123 Q53124 Q53138 Q53139 Q53158 Q53195 Q53196 Q53197 Q53198 Q53199 Q53200
Q53201 Q53202 Q53203 Q53204 Q53205 Q53207 Q53208 Q53209 Q53210 Q53211 Q53212
Q53213 Q53215 Q53216 Q53217 Q53218 Q53222 Q53226 Q53238 Q53245 Q53252 Q53308
Q53353 Q53389 Q53391 Q53450 Q53462 Q53464 Q53479 Q53514 Q53515 Q53526 Q53538
Q53539 Q53546 Q53547 Q53552 Q53554 Q53560 Q53562 Q53563 Q53586 Q53589 Q53595
Q53596 Q53597 Q535K8 Q53608 Q53609 Q53612 Q53634 Q53638 Q53685 Q53692 Q53700
Q53727 Q53750 Q53770 Q53778 Q537H7 Q53816 Q53844 Q53865 Q53867 Q53871 Q53872
Q53874 Q53875 Q53876 Q53877 Q53878 Q53882 Q538Z0 Q53915 Q53932 Q53979 Q53984
Q53B70 Q53B74 Q53B75 Q53DU6 Q53E76 Q53EK2 Q53EQ6 Q53EV4 Q53FA7 Q53FD0 Q53FE4
Q53G44 Q53GG5 Q53GI3 Q53H54 Q53H64 Q53H82 Q53H96 Q53HC5 Q53HV7 Q53HY2 Q53IP3
Q53IZ1 Q53JF7 Q53JI9 Q53K16 Q53K52 Q53KK6 Q53KS8 Q53KS9 Q53KW9 Q53LY0 Q53M52
Q53MB7 Q53MD1 Q53MD3 Q53N47 Q53N87 Q53N90 Q53NF7 Q53NI2 Q53P09 Q53P49 Q53PH2
Q53PP5 Q53Q11 Q53Q31 Q53Q32 Q53Q70 Q53QI0 Q53QK0 Q53QV2 Q53QW1 Q53R41 Q53RE8
Q53RK8 Q53RM0 Q53S08 Q53SZ7 Q53T94 Q53TQ3 Q53TZ2 Q53U08 Q53U10 Q53U11 Q53U14
Q53U15 Q53U18 Q53U19 Q53U20 Q53U21 Q53UH4 Q53UH5 Q53VB8 Q53VV5 Q53VV6 Q53VY0
Q53VY1 Q53VY2 Q53W05 Q53W17 Q53W19 Q53W21 Q53W34 Q53W62 Q53W63 Q53W83 Q53W92
Q53WD3 Q53WG6 Q53WG8 Q53WG9 Q53WH9 Q53WI0 Q53WJ1 Q53ZE5 Q54087 Q54088 Q54177
Q54215 Q54225 Q54235 Q54238 Q54240 Q54243 Q54258 Q54264 Q54271 Q54277 Q54304
Q54305 Q54316 Q54318 Q54340 Q54367 Q54387 Q54428 Q54430 Q54433 Q54441 Q54465
Q54506 Q54513 Q54518 Q54527 Q54528 Q54530 Q54699 Q54709 Q54728 Q54735 Q54751
Q54757 Q54760 Q54764 Q54765 Q54769 Q54775 Q54796 Q54800 Q54801 Q54807 Q54808
Q54809 Q54818 Q54823 Q54869 Q54900 Q54944 Q54970 Q54975 Q54986 Q54AK5 Q54AK6
Q54AU8 Q54AV4 Q54AX0 Q54AX5 Q54B10 Q54B20 Q54B23 Q54B33 Q54B37 Q54B46 Q54B48
Q54B59 Q54B60 Q54B64 Q54B65 Q54B68 Q54B75 Q54B82 Q54B97 Q54BA2 Q54BA6 Q54BB3
Q54BC2 Q54BC6 Q54BC8 Q54BC9 Q54BD1 Q54BD5 Q54BD6 Q54BD8 Q54BD9 Q54BE5 Q54BF3
Q54BF4 Q54BH8 Q54BI5 Q54BK0 Q54BK1 Q54BL2 Q54BL9 Q54BM1 Q54BM2 Q54BM5 Q54BM8
Q54BN3 Q54BT9 Q54BU6 Q54BU9 Q54BV6 Q54BW1 Q54BW5 Q54BW6 Q54BX3 Q54BX4 Q54BX5
Q54BX6 Q54BZ0 Q54BZ3 Q54BZ4 Q54BZ5 Q54BZ6 Q54BZ7 Q54BZ8 Q54C00 Q54C02 Q54C19
Q54C22 Q54C25 Q54C29 Q54C30 Q54C38 Q54C45 Q54C49 Q54C51 Q54C62 Q54C70 Q54C85
Q54C92 Q54CC5 Q54CC7 Q54CD6 Q54CD8 Q54CE0 Q54CE4 Q54CF6 Q54CG9 Q54CH6 Q54CI5

Q54CI8 Q54CJ3 Q54CJ8 Q54CK9 Q54CL0 Q54CL2 Q54CN7 Q54CN8 Q54CP1 Q54CP4 Q54CQ8
Q54CR9 Q54CS2 Q54CS7 Q54CS9 Q54CT1 Q54CT4 Q54CT8 Q54CU7 Q54CU8 Q54CU9 Q54CV5
Q54CW2 Q54CW7 Q54CX6 Q54CY2 Q54CY3 Q54CY5 Q54CY9 Q54CZ6 Q54D00 Q54D04 Q54D05
Q54D06 Q54D13 Q54D37 Q54D44 Q54D58 Q54D67 Q54D73 Q54D75 Q54D79 Q54D80 Q54D81
Q54D93 Q54D99 Q54DA1 Q54DA5 Q54DA8 Q54DB0 Q54DC8 Q54DC9 Q54DD0 Q54DD2 Q54DD3
Q54DD4 Q54DF9 Q54DG3 Q54DH7 Q54DH8 Q54DI8 Q54DJ0 Q54DJ6 Q54DK1 Q54DK4 Q54DL6
Q54DL7 Q54DM1 Q54DM7 Q54DN6 Q54DP0 Q54DP1 Q54DQ3 Q54DU5 Q54DU8 Q54DV0 Q54DV3
Q54DY0 Q54DY1 Q54DY3 Q54DZ6 Q54E17 Q54E21 Q54E29 Q54E33 Q54E34 Q54E43 Q54E49
Q54E53 Q54E63 Q54E71 Q54E83 Q54E86 Q54E95 Q54E98 Q54EC2 Q54ED2 Q54ED4 Q54ED5
Q54ED6 Q54ED7 Q54EF2 Q54EF7 Q54EH2 Q54EH6 Q54EJ5 Q54EJ7 Q54EK4 Q54EP0 Q54EP1
Q54EP3 Q54EQ1 Q54EQ9 Q54ER0 Q54ER4 Q54ER5 Q54ER6 Q54ER7 Q54ES6 Q54ES9 Q54ET3
Q54ET5 Q54ET6 Q54EW1 Q54EW8 Q54EW9 Q54EX7 Q54EY1 Q54EY2 Q54F05 Q54F07 Q54F11
Q54F23 Q54F30 Q54F33 Q54F38 Q54F40 Q54F46 Q54F60 Q54F84 Q54F87 Q54F89 Q54F93
Q54F99 Q54FA8 Q54FB7 Q54FD2 Q54FD6 Q54FD7 Q54FE8 Q54FE9 Q54FF0 Q54FF1 Q54FF2
Q54FG7 Q54FI0 Q54FI7 Q54FJ7 Q54FJ9 Q54FK1 Q54FK2 Q54FK4 Q54FL0 Q54FL3 Q54FM1
Q54FM3 Q54FM4 Q54FN2 Q54FN9 Q54FP4 Q54FP8 Q54FQ1 Q54FQ2 Q54FQ3 Q54FQ8 Q54FS1
Q54FS2 Q54FU2 Q54FU3 Q54FV1 Q54FW9 Q54FX2 Q54FX8 Q54G05 Q54G06 Q54G07 Q54G14
Q54G19 Q54G21 Q54G30 Q54G38 Q54G70 Q54G78 Q54G80 Q54G94 Q54G96 Q54GB2 Q54GC8
Q54GC9 Q54GE5 Q54GE6 Q54GF2 Q54GG8 Q54GG9 Q54GH0 Q54GH1 Q54GH2 Q54GH4 Q54GJ2
Q54GJ7 Q54GK6 Q54GL7 Q54GM5 Q54GN2 Q54GN5 Q54GP1 Q54GP3 Q54GP8 Q54GP9 Q54GQ8
Q54GQ9 Q54GR1 Q54GR2 Q54GR3 Q54GS8 Q54GT3 Q54GT6 Q54GU4 Q54GU5 Q54GV0 Q54GV2
Q54GV7 Q54GW3 Q54GZ0 Q54H10 Q54H11 Q54H36 Q54H44 Q54H55 Q54H69 Q54H71 Q54H97
Q54HA4 Q54HB2 Q54HB4 Q54HE7 Q54HE9 Q54HF0 Q54HF1 Q54HF6 Q54HG1 Q54HG4 Q54HG5
Q54HG7 Q54HH2 Q54HH8 Q54HI0 Q54HJ2 Q54HJ9 Q54HL6 Q54HL8 Q54HM1 Q54HN4 Q54HP1
Q54HP3 Q54HR6 Q54HS1 Q54HS3 Q54HT5 Q54HT7 Q54HV7 Q54HV9 Q54HW1 Q54HW2 Q54HW5
Q54HX0 Q54HX2 Q54HX6 Q54HY7 Q54I08 Q54I10 Q54I30 Q54I39 Q54I43 Q54I51 Q54I56
Q54I57 Q54I58 Q54I60 Q54I63 Q54I65 Q54I66 Q54I78 Q54I79 Q54I80 Q54I81 Q54I82
Q54I83 Q54I89 Q54I91 Q54IA0 Q54IA7 Q54IB0 Q54ID0 Q54ID4 Q54IE7 Q54IF0 Q54IF9
Q54IG2 Q54IG9 Q54II4 Q54II8 Q54IJ0 Q54IJ3 Q54IJ9 Q54IK1 Q54IK8 Q54IL2 Q54IL7
Q54IM4 Q54IN6 Q54IP0 Q54IR1 Q54IR4 Q54IS0 Q54IS6 Q54IT1 Q54IT3 Q54IU7 Q54IU8
Q54IU9 Q54IV3 Q54IV4 Q54IV8 Q54IW9 Q54IX3 Q54IY3 Q54IY5 Q54IZ9 Q54J05 Q54J07
Q54J11 Q54J14 Q54J26 Q54J34 Q54J44 Q54J47 Q54J48 Q54J55 Q54J58 Q54J66 Q54J68
Q54J73 Q54J75 Q54J83 Q54J88 Q54J90 Q54JB0 Q54JC0 Q54JC2 Q54JC8 Q54JD0 Q54JD2
Q54JD4 Q54JD9 Q54JE3 Q54JE4 Q54JH6 Q54JH8 Q54JI1 Q54JI2 Q54JI7 Q54JK1 Q54JK5
Q54JL0 Q54JL3 Q54JL4 Q54JM1 Q54JM2 Q54JM3 Q54JM5 Q54JM9 Q54JN0 Q54JP5 Q54JQ1
Q54JQ2 Q54JR4 Q54JR5 Q54JR9 Q54JS0 Q54JU0 Q54JU1 Q54JU3 Q54JU6 Q54JV5 Q54JV8
Q54JW0 Q54JW9 Q54JY2 Q54JZ1 Q54K00 Q54K19 Q54K21 Q54K27 Q54K33 Q54K37 Q54K39
Q54K47 Q54K57 Q54K66 Q54K67 Q54K76 Q54K78 Q54K91 Q54K93 Q54K95 Q54K96 Q54KA1
Q54KB7 Q54KB8 Q54KC7 Q54KD0 Q54KD2 Q54KD8 Q54KD9 Q54KE6 Q54KF1 Q54KF2 Q54KF3
Q54KG1 Q54KG6 Q54KH0 Q54KH3 Q54KI0 Q54KI3 Q54KJ1 Q54KJ8 Q54KK8 Q54KL0 Q54KL5
Q54KL8 Q54KM3 Q54KM6 Q54KM7 Q54KN2 Q54KN7 Q54KP9 Q54KQ1 Q54KQ5 Q54KR1 Q54KR4
Q54KR5 Q54KR9 Q54KS4 Q54KS5 Q54KS8 Q54KT2 Q54KT7 Q54KU3 Q54KU5 Q54KU7 Q54KU8
Q54KU9 Q54KV4 Q54KW6 Q54KW9 Q54KX0 Q54KX4 Q54KY8 Q54KZ7 Q54KZ8 Q54L22 Q54L25
Q54L28 Q54L35 Q54L40 Q54L43 Q54L48 Q54L50 Q54L54 Q54L63 Q54L68 Q54L71 Q54L72

Q54L80 Q54L82 Q54L83 Q54L92 Q54LA1 Q54LA3 Q54LA5 Q54LB3 Q54LD5 Q54LD8 Q54LF0
Q54LF2 Q54LH4 Q54LH5 Q54LJ8 Q54LK8 Q54LN9 Q54LP2 Q54LP8 Q54LQ6 Q54LR0 Q54LR3
Q54LS0 Q54LS2 Q54LS3 Q54LT4 Q54LT8 Q54LU5 Q54LU9 Q54LV0 Q54LV1 Q54LV3 Q54LV6
Q54LV7 Q54LV8 Q54LW0 Q54LW3 Q54LW6 Q54LY5 Q54LY8 Q54LZ2 Q54LZ8 Q54M18 Q54M20
Q54M22 Q54M23 Q54M29 Q54M64 Q54M70 Q54M71 Q54M74 Q54MA5 Q54MA9 Q54MB1 Q54MB4
Q54MB5 Q54MB8 Q54MC8 Q54MC9 Q54MD4 Q54MF2 Q54MG1 Q54MG7 Q54MH0 Q54MH3 Q54MH6
Q54MH9 Q54MI6 Q54MI7 Q54MJ7 Q54ML1 Q54ML2 Q54ML4 Q54ML6 Q54ML8 Q54ML9 Q54MM1
Q54MP2 Q54MP8 Q54MP9 Q54MQ7 Q54MQ9 Q54MR1 Q54MR3 Q54MR6 Q54MT0 Q54MT2 Q54MT6
Q54MZ3 Q54MZ8 Q54MZ9 Q54N00 Q54N03 Q54N08 Q54N12 Q54N19 Q54N41 Q54N44 Q54N45
Q54N47 Q54N48 Q54N54 Q54N57 Q54N65 Q54N68 Q54N70 Q54N72 Q54N83 Q54N87 Q54N95
Q54N97 Q54NA6 Q54NB6 Q54NC0 Q54NC5 Q54NC6 Q54NC8 Q54ND1 Q54ND3 Q54NE6 Q54NF3
Q54NG9 Q54NI1 Q54NI7 Q54NJ4 Q54NJ8 Q54NK1 Q54NK2 Q54NK3 Q54NK8 Q54NM5 Q54NM6
Q54NM8 Q54NP8 Q54NQ0 Q54NQ8 Q54NS1 Q54NS3 Q54NS8 Q54NU0 Q54NU5 Q54NU9 Q54NW0
Q54NW4 Q54NW6 Q54NX0 Q54NX2 Q54NX9 Q54NY7 Q54NZ1 Q54NZ5 Q54NZ6 Q54NZ7 Q54NZ8
Q54P04 Q54P14 Q54P25 Q54P26 Q54P40 Q54P49 Q54P56 Q54P62 Q54P65 Q54P70 Q54P71
Q54P72 Q54P77 Q54P78 Q54P79 Q54P86 Q54P92 Q54P93 Q54P94 Q54PA0 Q54PA9 Q54PB2
Q54PB4 Q54PB8 Q54PC2 Q54PD9 Q54PE0 Q54PE1 Q54PE6 Q54PF1 Q54PF3 Q54PG1 Q54PH5
Q54PI5 Q54PI9 Q54PJ1 Q54PJ7 Q54PK4 Q54PL2 Q54PM6 Q54PM7 Q54PN3 Q54PN5 Q54PN7
Q54PN9 Q54PP6 Q54PP7 Q54PT1 Q54PT6 Q54PU1 Q54PU2 Q54PU6 Q54PV7 Q54PV8 Q54PV9
Q54PW5 Q54PX0 Q54PX7 Q54PY1 Q54PY5 Q54PY9 Q54PZ2 Q54Q02 Q54Q03 Q54Q04 Q54Q05
Q54Q24 Q54Q37 Q54Q38 Q54Q39 Q54Q45 Q54Q48 Q54Q71 Q54Q72 Q54Q80 Q54Q89 Q54Q90
Q54Q94 Q54Q99 Q54QA0 Q54QA6 Q54QC0 Q54QC1 Q54QD1 Q54QD3 Q54QD5 Q54QE4 Q54QE6
Q54QG0 Q54QG4 Q54QG5 Q54QG9 Q54QI2 Q54QI4 Q54QJ8 Q54QJ9 Q54QK1 Q54QK4 Q54QK7
Q54QL5 Q54QM4 Q54QM6 Q54QN0 Q54QP0 Q54QP4 Q54QP5 Q54QP8 Q54QQ0 Q54QQ2 Q54QQ6
Q54QR1 Q54QR2 Q54QS3 Q54QT7 Q54QT8 Q54QT9 Q54QU0 Q54QU3 Q54QU5 Q54QU9 Q54QV7
Q54QW1 Q54QW5 Q54QW9 Q54QX0 Q54QX3 Q54QY7 Q54QZ7 Q54QZ8 Q54QZ9 Q54R00 Q54R01
Q54R03 Q54R14 Q54R16 Q54R21 Q54R42 Q54R47 Q54R59 Q54R60 Q54R62 Q54R65 Q54R66
Q54R69 Q54R73 Q54R75 Q54R98 Q54RA2 Q54RA4 Q54RB2 Q54RC3 Q54RC7 Q54RD4 Q54RF0
Q54RF3 Q54RF4 Q54RF5 Q54RG7 Q54RI9 Q54RK5 Q54RK6 Q54RM0 Q54RM2 Q54RM7 Q54RP0
Q54RP7 Q54RS2 Q54RT9 Q54RU3 Q54RV6 Q54RW9 Q54RX0 Q54RX1 Q54RX6 Q54RY6 Q54RY8
Q54RZ9 Q54S00 Q54S02 Q54S03 Q54S04 Q54S08 Q54S20 Q54S29 Q54S38 Q54S43 Q54S59
Q54S60 Q54S79 Q54S83 Q54S94 Q54S95 Q54S98 Q54SA5 Q54SA7 Q54SC4 Q54SD4 Q54SD7
Q54SE2 Q54SF4 Q54SF5 Q54SF7 Q54SF9 Q54SG7 Q54SH0 Q54SI1 Q54SJ5 Q54SK3 Q54SM3
Q54SN4 Q54SP1 Q54SP2 Q54SP5 Q54SP6 Q54SP7 Q54SP8 Q54SP9 Q54SR0 Q54SR6 Q54ST0
Q54ST2 Q54ST3 Q54ST4 Q54SU1 Q54SU3 Q54SV3 Q54SV8 Q54SY2 Q54SY5 Q54SZ6 Q54T05
Q54T16 Q54T17 Q54T27 Q54T36 Q54T40 Q54T43 Q54T44 Q54T46 Q54T58 Q54T76 Q54T81
Q54T85 Q54TA0 Q54TA5 Q54TA7 Q54TB4 Q54TB7 Q54TB9 Q54TC2 Q54TC5 Q54TC6 Q54TD3
Q54TD7 Q54TD8 Q54TD9 Q54TE3 Q54TE7 Q54TF6 Q54TF8 Q54TH8 Q54TI3 Q54TI6 Q54TJ4
Q54TK0 Q54TM6 Q54TM7 Q54TN2 Q54TN3 Q54TN4 Q54TP6 Q54TR4 Q54TR6 Q54TU3 Q54TU5
Q54TW4 Q54TW8 Q54TY5 Q54TY7 Q54U16 Q54U49 Q54U55 Q54U64 Q54U94 Q54UA9 Q54UB1
Q54UB5 Q54UC4 Q54UD0 Q54UE3 Q54UF3 Q54UF7 Q54UG2 Q54UG3 Q54UG4 Q54UG8 Q54UH8
Q54UI3 Q54UI4 Q54UJ0 Q54UJ5 Q54UK3 Q54UK6 Q54UK7 Q54UK8 Q54UK9 Q54UL0 Q54UL2
Q54UL3 Q54UL4 Q54UL5 Q54UL6 Q54UL8 Q54UM1 Q54UM6 Q54UM7 Q54UM8 Q54UN0 Q54UN1
Q54UN2 Q54UN3 Q54UN4 Q54UN8 Q54UP4 Q54UP6 Q54UQ2 Q54UQ4 Q54UR0 Q54UR4 Q54UR6

Q54UR9 Q54US7 Q54UT2 Q54UU2 Q54UU6 Q54UU8 Q54UV7 Q54UV8 Q54UV9 Q54UW0 Q54UW1
Q54UW4 Q54UW5 Q54V15 Q54V32 Q54V36 Q54V37 Q54V44 Q54V55 Q54V60 Q54V71 Q54V77
Q54V82 Q54V85 Q54VA2 Q54VA8 Q54VA9 Q54VB4 Q54VB7 Q54VD7 Q54VD8 Q54VE8 Q54VF1
Q54VF4 Q54VG0 Q54VH6 Q54VI0 Q54VI1 Q54VI3 Q54VI4 Q54VI5 Q54VJ3 Q54VJ9 Q54VK3
Q54VK6 Q54VL5 Q54VM2 Q54VM3 Q54VM4 Q54VN0 Q54VP3 Q54VP4 Q54VP7 Q54VQ0 Q54VQ5
Q54VQ6 Q54VQ7 Q54VR8 Q54VS1 Q54VT4 Q54VT8 Q54VU7 Q54VV5 Q54VV7 Q54VV8 Q54VY3
Q54VZ6 Q54VZ9 Q54W04 Q54W06 Q54W07 Q54W16 Q54W43 Q54W52 Q54W62 Q54W64 Q54W83
Q54W96 Q54WA4 Q54WA5 Q54WB6 Q54WB8 Q54WD6 Q54WD9 Q54WE5 Q54WF3 Q54WF4 Q54WG0
Q54WG6 Q54WH5 Q54WH6 Q54WH7 Q54WI4 Q54WI8 Q54WK1 Q54WM4 Q54WN5 Q54WN7 Q54WN8
Q54WP3 Q54WR1 Q54WR2 Q54WR8 Q54WR9 Q54WS6 Q54WS9 Q54WT4 Q54WT6 Q54WT9 Q54WU0
Q54WU3 Q54WU6 Q54WV0 Q54WV4 Q54WW4 Q54WX4 Q54WY0 Q54WY5 Q54WZ0 Q54WZ8 Q54WZ9
Q54X01 Q54X03 Q54X04 Q54X08 Q54X15 Q54X16 Q54X22 Q54X24 Q54X31 Q54X41 Q54X49
Q54X65 Q54X73 Q54X75 Q54X77 Q54X95 Q54X97 Q54XB0 Q54XC1 Q54XD8 Q54XE6 Q54XF2
Q54XF7 Q54XG0 Q54XH6 Q54XH9 Q54XI9 Q54XJ4 Q54XK0 Q54XK9 Q54XL1 Q54XL6 Q54XM7
Q54XM9 Q54XN7 Q54XP2 Q54XP3 Q54XP4 Q54XP6 Q54XQ8 Q54XR2 Q54XS0 Q54XS1 Q54XS2
Q54XS6 Q54XT8 Q54XV3 Q54XX5 Q54Y03 Q54Y06 Q54Y09 Q54Y18 Q54Y20 Q54Y25 Q54Y27
Q54Y51 Q54Y73 Q54Y76 Q54Y81 Q54Y85 Q54Y96 Q54Y98 Q54YA5 Q54YA7 Q54YB7 Q54YD4
Q54YD9 Q54YE1 Q54YE2 Q54YG5 Q54YG9 Q54YI1 Q54YJ6 Q54YJ7 Q54YL2 Q54YL3 Q54YN2
Q54YS0 Q54YS6 Q54YU6 Q54YW4 Q54YW8 Q54YX3 Q54YX7 Q54YY1 Q54YZ0 Q54YZ4 Q54YZ5
Q54Z01 Q54Z07 Q54Z08 Q54Z20 Q54Z23 Q54Z24 Q54Z26 Q54Z27 Q54Z40 Q54Z54 Q54Z60
Q54ZD0 Q54ZI9 Q54ZN3 Q54ZP5 Q54ZP8 Q54ZS8 Q54ZT0 Q54ZX8 Q54ZY3 Q55002 Q55004
Q55005 Q55012 Q55028 Q55029 Q55030 Q55031 Q55032 Q55038 Q55039 Q55041 Q55053
Q55054 Q55055 Q55056 Q55057 Q55058 Q55060 Q55074 Q55075 Q55078 Q55080 Q55081
Q55085 Q55087 Q55088 Q550D2 Q550D5 Q550E3 Q550E5 Q550E9 Q550I8 Q550K8 Q550L8
Q550R2 Q55116 Q55117 Q55119 Q55121 Q55128 Q55132 Q55135 Q55141 Q55142 Q55154
Q55156 Q55158 Q55163 Q55167 Q55170 Q55185 Q55187 Q551A3 Q551F8 Q551G1 Q551H0
Q551H4 Q551M2 Q551M3 Q551S9 Q55214 Q55215 Q55216 Q55217 Q55233 Q55235 Q55244
Q55267 Q55276 Q552D9 Q552J0 Q552K9 Q552L5 Q552P9 Q552Q3 Q552Q7 Q552R1 Q552S7
Q552U1 Q552X2 Q552Y8 Q552Z2 Q552Z6 Q55309 Q55317 Q55336 Q55338 Q55346 Q55348
Q55367 Q55368 Q55369 Q55370 Q55371 Q55373 Q55382 Q55383 Q55385 Q55389 Q55393
Q55397 Q553B1 Q553B6 Q553D3 Q553E0 Q553E8 Q553E9 Q553F3 Q553J1 Q553K1 Q553L4
Q553R3 Q553S1 Q553S5 Q553V1 Q553V2 Q553X7 Q55404 Q55418 Q55421 Q55422 Q55423
Q55424 Q55428 Q55433 Q55434 Q55435 Q55450 Q55451 Q55458 Q55459 Q55467 Q55468
Q55469 Q55470 Q55480 Q55482 Q55484 Q55486 Q55497 Q55498 Q55499 Q554C5 Q554C9
Q554D9 Q554G3 Q554J3 Q554L3 Q554M9 Q554P0 Q554S6 Q554U9 Q55503 Q55505 Q55506
Q55508 Q55510 Q55511 Q55512 Q55513 Q55515 Q55522 Q55535 Q55550 Q55563 Q55566
Q55569 Q55574 Q55575 Q55578 Q55585 Q55587 Q55593 Q55595 Q555C8 Q555D2 Q555H8
Q555L5 Q555L9 Q555N6 Q555Q9 Q555V7 Q55607 Q55613 Q55621 Q55622 Q55624 Q55625
Q55626 Q55637 Q55638 Q55643 Q55650 Q55653 Q55663 Q55664 Q55665 Q55670 Q55673
Q55674 Q55680 Q55681 Q55682 Q55687 Q55688 Q55690 Q55692 Q55694 Q556F2 Q556G3
Q556G4 Q556H2 Q556H5 Q556I2 Q556J0 Q556J6 Q556K3 Q556K4 Q556L8 Q556P3 Q556Q0
Q556Q3 Q556R9 Q556U5 Q556V9 Q556Y1 Q556Y2 Q556Y8 Q556Y9 Q556Z0 Q556Z3 Q55702
Q55710 Q55716 Q55724 Q55725 Q55729 Q55731 Q55738 Q55749 Q55750 Q55752 Q55756
Q55758 Q55759 Q55760 Q55761 Q55766 Q55777 Q55778 Q55781 Q55786 Q55787 Q55790

Q55791 Q55792 Q55793 Q55797 Q55798 Q557A2 Q557B6 Q557C9 Q557D1 Q557D2 Q557E4
Q557F6 Q557F7 Q557G1 Q557G3 Q557H1 Q557H3 Q557H7 Q557I1 Q557J3 Q557J5 Q557J8
Q557M8 Q55801 Q55803 Q55805 Q55806 Q55810 Q55819 Q55823 Q55828 Q55837 Q55842
Q55843 Q55848 Q55849 Q55850 Q55854 Q55855 Q55860 Q55863 Q55874 Q55875 Q55880
Q55884 Q55891 Q55894 Q55898 Q558U1 Q558W0 Q558W4 Q558X5 Q558Y7 Q558Z0 Q558Z3
Q55901 Q55905 Q55914 Q55921 Q55928 Q55940 Q55946 Q55963 Q55971 Q55978 Q55982
Q55983 Q55988 Q559A2 Q559A9 Q559G8 Q559M9 Q559R1 Q55A45 Q55AB6 Q55AF0 Q55AI5
Q55AL7 Q55AL8 Q55AQ2 Q55AR3 Q55AR8 Q55AS5 Q55AX2 Q55B51 Q55B72 Q55BA4 Q55BA5
Q55BD5 Q55BF2 Q55BF5 Q55BF8 Q55BG3 Q55BG7 Q55BI0 Q55BI2 Q55BI8 Q55BJ9 Q55BK0
Q55BK3 Q55BM0 Q55BM1 Q55BM4 Q55BM5 Q55BN0 Q55BN8 Q55BN9 Q55BP0 Q55BP5 Q55BQ3
Q55BR6 Q55BR9 Q55BT1 Q55BT2 Q55BU0 Q55BU1 Q55BU7 Q55BV5 Q55BV7 Q55BW2 Q55BX3
Q55BX7 Q55BY9 Q55C16 Q55C21 Q55C24 Q55C49 Q55C50 Q55C52 Q55C62 Q55C72 Q55C75
Q55C77 Q55C80 Q55C82 Q55C84 Q55C91 Q55C99 Q55CA6 Q55CB6 Q55CD9 Q55CE0 Q55CJ3
Q55CN0 Q55CP6 Q55CQ6 Q55CS7 Q55CS8 Q55CT8 Q55CU2 Q55CU7 Q55CV0 Q55CV9 Q55CW0
Q55CW3 Q55CX9 Q55CY3 Q55CZ6 Q55D12 Q55D13 Q55D15 Q55D16 Q55D17 Q55D51 Q55D61
Q55D66 Q55DA2 Q55DA3 Q55DA6 Q55DB0 Q55DB4 Q55DB6 Q55DC6 Q55DE2 Q55DE7 Q55DF5
Q55DH6 Q55DH8 Q55DJ3 Q55DJ5 Q55DJ8 Q55DJ9 Q55DK2 Q55DL0 Q55DL2 Q55DM7 Q55DP8
Q55DP9 Q55DQ4 Q55DQ5 Q55DS4 Q55DT8 Q55DU1 Q55DU3 Q55DU4 Q55DV9 Q55DY7 Q55DZ8
Q55E06 Q55E13 Q55E21 Q55E23 Q55E34 Q55E54 Q55E60 Q55E65 Q55E68 Q55E83 Q55E93
Q55E94 Q55EA2 Q55ED0 Q55ED1 Q55ED4 Q55EE0 Q55EI0 Q55EI6 Q55EJ3 Q55EK9 Q55EL2
Q55EL3 Q55EN5 Q55EP5 Q55EP9 Q55EQ0 Q55ER5 Q55ER8 Q55EU4 Q55EU6 Q55EX5 Q55EX6
Q55EX7 Q55EX9 Q55EY1 Q55EY8 Q55EZ4 Q55F19 Q55F27 Q55F34 Q55F37 Q55F45 Q55F67
Q55F71 Q55F75 Q55F78 Q55F83 Q55F95 Q55FA4 Q55FB9 Q55FC2 Q55FC3 Q55FC4 Q55FC5
Q55FC6 Q55FC7 Q55FD6 Q55FE0 Q55FE3 Q55FF3 Q55FI1 Q55FI3 Q55FI4 Q55FI7 Q55FJ0
Q55FJ1 Q55FJ3 Q55FJ5 Q55FL3 Q55FM1 Q55FM5 Q55FM6 Q55FN5 Q55FN7 Q55FP1 Q55FQ0
Q55FS0 Q55FS8 Q55FT1 Q55FT4 Q55FT8 Q55FT9 Q55FU5 Q55FV1 Q55FV2 Q55FV3 Q55FV5
Q55FV7 Q55FW7 Q55FW8 Q55G04 Q55G10 Q55G14 Q55G16 Q55G18 Q55G20 Q55G22 Q55G28
Q55G34 Q55G59 Q55G75 Q55G81 Q55G83 Q55G84 Q55G88 Q55G91 Q55G93 Q55G94 Q55G97
Q55G98 Q55GA4 Q55GB8 Q55GC2 Q55GD2 Q55GD5 Q55GD6 Q55GE7 Q55GF9 Q55GG6 Q55GH4
Q55GH8 Q55GI5 Q55GJ6 Q55GK0 Q55GK1 Q55GK3 Q55GM5 Q55GM9 Q55GN3 Q55GN8 Q55GQ5
Q55GQ6 Q55GR4 Q55GR5 Q55GS1 Q55GS2 Q55GS4 Q55GS6 Q55GU1 Q55GU2 Q55GU9 Q55GV9
Q55GW2 Q55GW8 Q55H65 Q56002 Q56016 Q56017 Q56022 Q56037 Q56056 Q56062 Q56063
Q56066 Q56067 Q56073 Q56083 Q56093 Q56106 Q56108 Q56109 Q56112 Q56114 Q56115
Q56121 Q56143 Q56148 Q56178 Q56184 Q56185 Q56194 Q56195 Q56198 Q561K4 Q561L1
Q561L5 Q561L6 Q561M0 Q561P3 Q561P5 Q561Q8 Q561R2 Q561R3 Q561R7 Q561R9 Q561X3
Q561Y0 Q56200 Q56201 Q56206 Q56207 Q56208 Q56209 Q56210 Q56211 Q56213 Q56214
Q56215 Q56216 Q56230 Q56231 Q56232 Q56235 Q56236 Q56237 Q56239 Q56242 Q56243
Q56251 Q56252 Q56253 Q56254 Q56259 Q56260 Q56268 Q56273 Q56275 Q562A2 Q562B4
Q562D5 Q562D6 Q562E2 Q56301 Q56303 Q56304 Q56307 Q56308 Q56313 Q56316 Q56317
Q56318 Q56319 Q56320 Q56326 Q56346 Q56353 Q56362 Q56366 Q563C3 Q56405 Q56415
Q56416 Q56419 Q56425 Q56435 Q56462 Q56464 Q564K3 Q564U4 Q564X7 Q56559 Q56560
Q56561 Q56562 Q56564 Q56573 Q56575 Q56581 Q56585 Q56586 Q56598 Q565D8 Q56632
Q56647 Q56648 Q56660 Q56691 Q56693 Q56694 Q566H4 Q566I1 Q566L4 Q566L7 Q566L8
Q566M1 Q566N9 Q566Q8 Q566R3 Q566T0 Q566U4 Q566V0 Q566V3 Q566X6 Q566X8 Q566Y1

Q566Y8 Q566Z6 Q56704 Q56715 Q56731 Q56732 Q56733 Q56734 Q56738 Q56741 Q56743
Q56764 Q56774 Q567C3 Q567C6 Q567G2 Q567I9 Q567J8 Q567V6 Q567W6 Q567Y6 Q567Z6
Q56815 Q56829 Q56837 Q56839 Q56840 Q56841 Q56847 Q56855 Q56893 Q56894 Q56895
Q56897 Q568A0 Q568B7 Q568D2 Q568D5 Q568E2 Q568F6 Q568G4 Q568K2 Q568K9 Q568M3
Q568P1 Q568P9 Q568Q0 Q568R1 Q568V3 Q568W0 Q568Y2 Q568Z1 Q568Z9 Q56922 Q56928
Q56978 Q56981 Q56990 Q56998 Q569B5 Q569B7 Q569C4 Q569E7 Q569G3 Q569H4 Q569K4
Q569K6 Q569M2 Q569M3 Q569R5 Q569Z1 Q56A04 Q56A08 Q56A10 Q56A18 Q56A24 Q56A35
Q56A40 Q56A73 Q56B59 Q56E62 Q56FC8 Q56FG8 Q56FG9 Q56FH2 Q56FL2 Q56JW4 Q56JX3
Q56JZ5 Q56JZ7 Q56JZ9 Q56K04 Q56K12 Q56K13 Q56P10 Q56P11 Q56P12 Q56P13 Q56P27
Q56R04 Q56R14 Q56RZ3 Q56RZ7 Q56S04 Q56SA8 Q56TU3 Q56TU4 Q56UQ5 Q56V19 Q56VL3
Q56W59 Q56W64 Q56WH1 Q56WH4 Q56WL5 Q56X05 Q56X76 Q56XE4 Q56XJ7 Q56XM9 Q56XR0
Q56XR6 Q56XS8 Q56XU4 Q56XV8 Q56XW8 Q56XX3 Q56XY2 Q56Y42 Q56Y52 Q56Y85 Q56YF8
Q56YH2 Q56YJ8 Q56YN3 Q56YT7 Q56YU0 Q56YW9 Q57004 Q57006 Q57014 Q57022 Q57039
Q57045 Q57051 Q57060 Q57065 Q57066 Q57070 Q57074 Q57083 Q57089 Q570C0 Q570N1
Q570P7 Q570R7 Q570U6 Q570Y6 Q57105 Q57109 Q57118 Q57120 Q57122 Q57125 Q57133
Q57140 Q57144 Q57146 Q57151 Q57152 Q57153 Q57154 Q57160 Q57163 Q57168 Q57179
Q57184 Q57190 Q57194 Q57195 Q57199 Q571F5 Q571F8 Q571G4 Q571H0 Q571I9 Q571J5
Q571K4 Q57209 Q57213 Q57220 Q57231 Q57233 Q57239 Q57240 Q57242 Q57252 Q57261
Q57267 Q57270 Q57273 Q57278 Q57287 Q57290 Q57300 Q57301 Q57302 Q57337 Q57346
Q57354 Q57374 Q57390 Q57394 Q57396 Q573B8 Q573B9 Q573C1 Q573C3 Q573C4 Q573C5
Q573C8 Q573C9 Q573D0 Q573D1 Q573D2 Q573D4 Q573D5 Q573D6 Q573D7 Q573D8 Q573D9
Q573E2 Q573E3 Q573E4 Q573E5 Q573E6 Q573E7 Q573E8 Q573E9 Q573F2 Q573F3 Q573F4
Q573F5 Q573F6 Q573F7 Q573F8 Q573F9 Q573G0 Q573G1 Q573G2 Q573G3 Q573G4 Q573G5
Q573G6 Q573G7 Q573G8 Q57409 Q57423 Q57431 Q57434 Q57458 Q57466 Q57471 Q57475
Q57476 Q57498 Q57501 Q57517 Q57525 Q57527 Q57532 Q57534 Q57544 Q57549 Q57551
Q57553 Q57555 Q57557 Q57558 Q57559 Q57560 Q57562 Q57563 Q57564 Q57566 Q57567
Q57568 Q57569 Q57572 Q57576 Q57577 Q57578 Q57579 Q57580 Q57581 Q57582 Q57583
Q57584 Q57585 Q57586 Q57587 Q57588 Q57589 Q57590 Q57591 Q57592 Q57594 Q57596
Q57597 Q57598 Q57599 Q575S8 Q575S9 Q575T0 Q57600 Q57602 Q57605 Q57606 Q57607
Q57608 Q57609 Q57610 Q57611 Q57612 Q57614 Q57615 Q57616 Q57617 Q57619 Q57620
Q57621 Q57622 Q57623 Q57624 Q57625 Q57626 Q57628 Q57630 Q57631 Q57632 Q57635
Q57636 Q57637 Q57638 Q57639 Q57640 Q57641 Q57642 Q57645 Q57646 Q57647 Q57648
Q57649 Q57650 Q57651 Q57652 Q57653 Q57655 Q57656 Q57657 Q57658 Q57659 Q57660
Q57661 Q57662 Q57663 Q57664 Q57665 Q57666 Q57667 Q57668 Q57676 Q57677 Q57678
Q57679 Q57680 Q57681 Q57682 Q57684 Q57686 Q57687 Q57688 Q57689 Q57690 Q57691
Q57692 Q57693 Q57694 Q57695 Q57696 Q57697 Q57698 Q57699 Q576I9 Q576P8 Q576P9
Q576Q0 Q576R3 Q576R4 Q576T0 Q576T1 Q576T3 Q576T5 Q576T8 Q576T9 Q576U2 Q576X0
Q57700 Q57701 Q57702 Q57703 Q57705 Q57706 Q57707 Q57709 Q57710 Q57711 Q57712
Q57713 Q57714 Q57715 Q57716 Q57717 Q57720 Q57721 Q57722 Q57724 Q57725 Q57726
Q57728 Q57729 Q57730 Q57731 Q57732 Q57733 Q57734 Q57735 Q57738 Q57741 Q57742
Q57743 Q57744 Q57745 Q57746 Q57747 Q57748 Q57749 Q57750 Q57751 Q57752 Q57754
Q57755 Q57756 Q57757 Q57758 Q57759 Q57760 Q57761 Q57762 Q57763 Q57764 Q57765
Q57766 Q57767 Q57768 Q57770 Q57771 Q57773 Q57774 Q57775 Q57776 Q57777 Q57778
Q57779 Q57780 Q57781 Q57782 Q57783 Q57784 Q57785 Q57786 Q57787 Q57788 Q57790

Q57791 Q57792 Q57793 Q57794 Q57795 Q57796 Q57797 Q57798 Q577C7 Q577D5 Q577F0
Q577H0 Q577P3 Q577P6 Q577P8 Q577W6 Q577Y1 Q577Y8 Q577Z3 Q577Z4 Q577Z5 Q577Z6
Q577Z8 Q57800 Q57801 Q57802 Q57803 Q57804 Q57805 Q57807 Q57808 Q57809 Q57810
Q57811 Q57812 Q57813 Q57814 Q57815 Q57818 Q57819 Q57820 Q57821 Q57822 Q57823
Q57824 Q57825 Q57826 Q57827 Q57828 Q57829 Q57830 Q57831 Q57832 Q57833 Q57834
Q57835 Q57836 Q57838 Q57839 Q57840 Q57841 Q57842 Q57843 Q57844 Q57845 Q57846
Q57847 Q57849 Q57850 Q57851 Q57854 Q57855 Q57857 Q57859 Q57860 Q57864 Q57865
Q57866 Q57867 Q57868 Q57869 Q57870 Q57871 Q57872 Q57874 Q57875 Q57876 Q57877
Q57878 Q57880 Q57881 Q57884 Q57885 Q57886 Q57887 Q57888 Q57889 Q57890 Q57892
Q57893 Q57894 Q57895 Q57896 Q57897 Q57899 Q578A8 Q578E4 Q578H4 Q578I2 Q578J2
Q578P4 Q578R0 Q578T1 Q578Z3 Q57900 Q57901 Q57902 Q57903 Q57905 Q57906 Q57907
Q57908 Q57909 Q57910 Q57911 Q57912 Q57913 Q57914 Q57916 Q57917 Q57918 Q57919
Q57920 Q57921 Q57923 Q57924 Q57925 Q57926 Q57927 Q57929 Q57930 Q57931 Q57932
Q57933 Q57934 Q57935 Q57936 Q57937 Q57949 Q57950 Q57951 Q57953 Q57954 Q57955
Q57956 Q57957 Q57958 Q57959 Q57960 Q57961 Q57962 Q57963 Q57964 Q57968 Q57969
Q57970 Q57971 Q57972 Q57973 Q57975 Q57976 Q57977 Q57978 Q57979 Q57980 Q57981
Q57982 Q57983 Q57984 Q57987 Q57988 Q57989 Q57990 Q57991 Q57992 Q57993 Q57995
Q57997 Q57998 Q57999 Q579A4 Q579B1 Q579B2 Q579C9 Q579E2 Q579E7 Q579E8 Q579F0
Q579G4 Q579I3 Q579M7 Q579Q6 Q579Q7 Q579Q8 Q579U0 Q579W9 Q579Z7 Q57A81 Q57A83
Q57A96 Q57A98 Q57A99 Q57AA0 Q57AA3 Q57AA4 Q57AA9 Q57AB1 Q57AD4 Q57AD6 Q57AD7
Q57AD9 Q57AE0 Q57AE2 Q57AE3 Q57AE9 Q57AF0 Q57AF2 Q57AG3 Q57AH1 Q57AH2 Q57AH3
Q57AH4 Q57AH5 Q57AH7 Q57AH8 Q57AH9 Q57AI8 Q57AI9 Q57AJ0 Q57AJ1 Q57AJ2 Q57AJ6
Q57AJ7 Q57AJ8 Q57AM8 Q57AM9 Q57AP2 Q57AR7 Q57AS3 Q57AS5 Q57AT2 Q57AV8 Q57AW2
Q57AW6 Q57AX1 Q57AX2 Q57AX5 Q57AY3 Q57AY4 Q57AY9 Q57AZ0 Q57B07 Q57B08 Q57B22
Q57B45 Q57B46 Q57B47 Q57B48 Q57B50 Q57B54 Q57B65 Q57B71 Q57B76 Q57B80 Q57B82
Q57B94 Q57BA5 Q57BA8 Q57BC9 Q57BD5 Q57BG3 Q57BG4 Q57BH6 Q57BH7 Q57BH8 Q57BI8
Q57BJ0 Q57BK5 Q57BM8 Q57BT1 Q57BW7 Q57BX7 Q57BY2 Q57C09 Q57C28 Q57C34 Q57C70
Q57C76 Q57C79 Q57C80 Q57C89 Q57C92 Q57C98 Q57CA9 Q57CB7 Q57CC2 Q57CC7 Q57CE4
Q57CE5 Q57CE6 Q57CE8 Q57CE9 Q57CF0 Q57CI7 Q57CK0 Q57CK2 Q57CL3 Q57CL5 Q57CP4
Q57CP5 Q57CP8 Q57CP9 Q57CQ3 Q57CQ4 Q57CQ5 Q57CQ7 Q57CQ8 Q57CQ9 Q57CR0 Q57CR1
Q57CR2 Q57CR3 Q57CR4 Q57CR6 Q57CR7 Q57CR8 Q57CR9 Q57CS0 Q57CS1 Q57CS2 Q57CS3
Q57CS4 Q57CS5 Q57CS6 Q57CS7 Q57CS9 Q57CT0 Q57CT1 Q57CT2 Q57CT3 Q57CU0 Q57CU6
Q57CV7 Q57CX7 Q57CX8 Q57CX9 Q57CY0 Q57CY1 Q57CZ2 Q57CZ7 Q57CZ8 Q57CZ9 Q57D01
Q57D05 Q57D15 Q57D18 Q57D38 Q57D42 Q57D45 Q57D60 Q57D61 Q57D65 Q57D82 Q57DB3
Q57DG3 Q57DG4 Q57DI4 Q57DI5 Q57DK4 Q57DK8 Q57DK9 Q57DL3 Q57DM1 Q57DM2 Q57DN0
Q57DN1 Q57DN9 Q57DR3 Q57DR5 Q57DR8 Q57DS4 Q57DS5 Q57DT1 Q57DW1 Q57DW2 Q57DW4
Q57DX3 Q57DY0 Q57DY1 Q57DY4 Q57DY5 Q57DZ4 Q57DZ8 Q57E30 Q57E41 Q57E46 Q57E58
Q57E65 Q57E78 Q57E81 Q57E89 Q57E90 Q57E91 Q57E95 Q57E96 Q57EH9 Q57EI0 Q57EI4
Q57EL6 Q57EP1 Q57EP4 Q57EP5 Q57EP6 Q57EQ3 Q57ER2 Q57ER3 Q57ER5 Q57ET1 Q57ET8
Q57EU9 Q57EW2 Q57EW9 Q57EX6 Q57EY6 Q57F22 Q57F29 Q57F30 Q57F54 Q57F56 Q57F73
Q57F88 Q57F97 Q57F99 Q57FA0 Q57FA3 Q57FG5 Q57FI5 Q57FJ9 Q57FK3 Q57FM3 Q57FM9
Q57FN0 Q57FQ9 Q57FS2 Q57FT6 Q57FT9 Q57FU2 Q57FU7 Q57FY1 Q57FY2 Q57FY7 Q57G08
Q57G10 Q57G26 Q57G27 Q57G28 Q57G36 Q57G38 Q57G39 Q57G40 Q57G48 Q57G51 Q57G60
Q57G77 Q57G99 Q57GA1 Q57GC3 Q57GC5 Q57GD5 Q57GE1 Q57GE2 Q57GG5 Q57GG7 Q57GH3

Q57GI1 Q57GI8 Q57GI9 Q57GJ0 Q57GJ1 Q57GJ4 Q57GJ5 Q57GJ6 Q57GK0 Q57GK1 Q57GL3
Q57GL4 Q57GL9 Q57GM0 Q57GM1 Q57GM7 Q57GN3 Q57GP7 Q57GQ4 Q57GZ4 Q57H06 Q57H37
Q57H42 Q57H45 Q57H57 Q57H58 Q57H59 Q57H60 Q57H61 Q57H62 Q57H64 Q57H68 Q57H69
Q57H70 Q57H71 Q57H72 Q57H73 Q57H76 Q57H79 Q57H81 Q57H86 Q57H88 Q57H91 Q57H93
Q57H94 Q57H96 Q57H97 Q57HA5 Q57HA8 Q57HB6 Q57HC1 Q57HC5 Q57HC6 Q57HC8 Q57HC9
Q57HD1 Q57HD6 Q57HD8 Q57HD9 Q57HE3 Q57HE4 Q57HF3 Q57HG7 Q57HG8 Q57HG9 Q57HH0
Q57HH1 Q57HH3 Q57HH9 Q57HI2 Q57HI8 Q57HL1 Q57HL3 Q57HL7 Q57HM5 Q57HN8 Q57HP2
Q57HR5 Q57HT6 Q57HT7 Q57HU2 Q57HU7 Q57HV1 Q57HW2 Q57HW4 Q57HW5 Q57HW6 Q57HW9
Q57HX0 Q57HZ6 Q57HZ9 Q57I00 Q57I02 Q57I11 Q57I17 Q57I24 Q57I25 Q57I32 Q57I62
Q57I91 Q57I92 Q57I94 Q57I99 Q57IA0 Q57IA1 Q57IA2 Q57IA4 Q57IA5 Q57IA6 Q57IA7
Q57IA8 Q57IC3 Q57IC5 Q57IC9 Q57ID2 Q57IE4 Q57IE9 Q57IG0 Q57IG5 Q57IG6 Q57IL9
Q57IN4 Q57IP3 Q57IP9 Q57IT8 Q57IT9 Q57IU0 Q57IU1 Q57IV0 Q57IV9 Q57IW3 Q57IW5
Q57IW7 Q57IX4 Q57IX6 Q57IY7 Q57IY8 Q57IZ2 Q57IZ5 Q57J09 Q57J18 Q57J21 Q57J22
Q57J23 Q57J24 Q57J25 Q57J26 Q57J31 Q57J32 Q57J33 Q57J34 Q57J35 Q57J36 Q57J38
Q57J39 Q57J40 Q57J41 Q57J42 Q57J43 Q57J44 Q57J45 Q57J46 Q57J47 Q57J48 Q57J49
Q57J50 Q57J51 Q57J53 Q57J54 Q57J55 Q57J56 Q57J57 Q57J62 Q57J63 Q57J64 Q57J66
Q57J68 Q57J69 Q57J83 Q57J85 Q57J97 Q57JA8 Q57JA9 Q57JC3 Q57JC4 Q57JC8 Q57JC9
Q57JE5 Q57JG1 Q57JG5 Q57JG7 Q57JH1 Q57JH3 Q57JH7 Q57JH9 Q57JI0 Q57JI1 Q57JI2
Q57JI3 Q57JJ4 Q57JJ5 Q57JJ9 Q57JK8 Q57JK9 Q57JL0 Q57JM2 Q57JN9 Q57JP1 Q57JP7
Q57JQ0 Q57JQ1 Q57JQ5 Q57JQ9 Q57JR3 Q57JR5 Q57JS5 Q57JX8 Q57JY0 Q57JY6 Q57K04
Q57K06 Q57K19 Q57K20 Q57K24 Q57K26 Q57K30 Q57K36 Q57K44 Q57K45 Q57K51 Q57K52
Q57K60 Q57K61 Q57K65 Q57K67 Q57K69 Q57K76 Q57K77 Q57K94 Q57K99 Q57KB2 Q57KB3
Q57KB6 Q57KD6 Q57KD8 Q57KE0 Q57KE5 Q57KE9 Q57KF9 Q57KG9 Q57KH7 Q57KH8 Q57KH9
Q57KI9 Q57KJ0 Q57KJ1 Q57KJ4 Q57KJ5 Q57KJ6 Q57KJ7 Q57KJ8 Q57KL5 Q57KT2 Q57KT3
Q57KU4 Q57KU5 Q57KU6 Q57KU7 Q57KV3 Q57KW6 Q57L18 Q57L20 Q57L24 Q57L29 Q57L30
Q57L31 Q57L32 Q57L54 Q57L55 Q57L59 Q57LD0 Q57LD2 Q57LD3 Q57LE0 Q57LE6 Q57LF5
Q57LF7 Q57LG7 Q57LG8 Q57LG9 Q57LH1 Q57LH2 Q57LH5 Q57LI3 Q57LI4 Q57LI6 Q57LI7
Q57LJ0 Q57LJ8 Q57LL0 Q57LL1 Q57LL3 Q57LM4 Q57LM8 Q57LN3 Q57LN7 Q57LQ6 Q57LQ8
Q57LS3 Q57LT1 Q57LU0 Q57LV0 Q57LX0 Q57LX5 Q57LY4 Q57LY6 Q57M20 Q57M23 Q57M43
Q57M47 Q57M48 Q57M50 Q57M59 Q57M62 Q57M64 Q57M77 Q57MB1 Q57MB2 Q57MB4 Q57MC8
Q57MD6 Q57ME6 Q57MF5 Q57MI5 Q57MJ3 Q57MN1 Q57MN2 Q57MR8 Q57MR9 Q57MS0 Q57MS2
Q57MS3 Q57MS4 Q57MT7 Q57MW5 Q57MX8 Q57N02 Q57N15 Q57N17 Q57N33 Q57N48 Q57N56
Q57N70 Q57N89 Q57N91 Q57N92 Q57N96 Q57N98 Q57N99 Q57NA2 Q57NA3 Q57NB7 Q57NF9
Q57NG3 Q57NI6 Q57NI8 Q57NJ6 Q57NJ8 Q57NJ9 Q57NK9 Q57NM8 Q57NN2 Q57NN4 Q57NN5
Q57NQ5 Q57NR0 Q57NR9 Q57NT2 Q57NT3 Q57NU9 Q57NV3 Q57NV7 Q57NV9 Q57P06 Q57P18
Q57P61 Q57P88 Q57P89 Q57PB3 Q57PG8 Q57PH6 Q57PI7 Q57PI8 Q57PI9 Q57PJ1 Q57PK2
Q57PK6 Q57PR1 Q57PR2 Q57PS7 Q57PS8 Q57PT8 Q57PU1 Q57PU2 Q57PU7 Q57PU8 Q57PU9
Q57PV0 Q57PV1 Q57PV3 Q57PW5 Q57PW7 Q57PX4 Q57PX7 Q57PX9 Q57Q09 Q57QC0 Q57QC7
Q57QD5 Q57QE5 Q57QE6 Q57QE7 Q57QF5 Q57QG6 Q57QG8 Q57QI9 Q57QJ4 Q57QK2 Q57QM1
Q57QP2 Q57QP3 Q57QP5 Q57QP9 Q57QR9 Q57QS3 Q57QS4 Q57QS7 Q57QT2 Q57QT4 Q57QU1
Q57QZ3 Q57QZ8 Q57R05 Q57R11 Q57R19 Q57R21 Q57R23 Q57R24 Q57R38 Q57R46 Q57R48
Q57R56 Q57R63 Q57R70 Q57R80 Q57R87 Q57RA8 Q57RC9 Q57RD7 Q57RF3 Q57RF9 Q57RG2
Q57RG3 Q57RG6 Q57RG7 Q57RG9 Q57RH0 Q57RH2 Q57RI3 Q57RI5 Q57RK0 Q57RL3 Q57RM1
Q57RM2 Q57RP8 Q57RQ0 Q57RQ7 Q57RQ9 Q57RR6 Q57RS7 Q57RT0 Q57RT4 Q57RU1 Q57RU3

| | | | | | | | | | | |
|---|---|---|---|---|---|---|---|---|---|---|
| Q57RV7 | Q57RZ1 | Q57S24 | Q57S29 | Q57S43 | Q57S48 | Q57S50 | Q57S51 | Q57S73 | Q57S74 | Q57S76 |
| Q57S77 | Q57S78 | Q57S79 | Q57S81 | Q57S93 | Q57S94 | Q57SA6 | Q57SA8 | Q57SB4 | Q57SB5 | Q57SB6 |
| Q57SC9 | Q57SD2 | Q57SD3 | Q57SD4 | Q57SD9 | Q57SE0 | Q57SE2 | Q57SE6 | Q57SE7 | Q57SE9 | Q57SF8 |
| Q57SF9 | Q57SH3 | Q57SH6 | Q57SH7 | Q57SI1 | Q57SN0 | Q57SN1 | Q57ST2 | Q57ST3 | Q57ST5 | Q57ST6 |
| Q57SU1 | Q57SU4 | Q57SZ6 | Q57SZ8 | Q57T01 | Q57T14 | Q57T17 | Q57T19 | Q57T25 | Q57T35 | Q57T36 |
| Q57T37 | Q57T38 | Q57T39 | Q57T41 | Q57T42 | Q57T44 | Q57T47 | Q57T48 | Q57T51 | Q57T53 | Q57T68 |
| Q57T70 | Q57T73 | Q57T74 | Q57T75 | Q57T89 | Q57T90 | Q57TB5 | Q57TB6 | Q57TB7 | Q57TB8 | Q57TC9 |
| Q57TD2 | Q57TD5 | Q57TD8 | Q57TD9 | Q57TE6 | Q57TE7 | Q57TE8 | Q57TE9 | Q57TF2 | Q57TF9 | Q57TG4 |
| Q57TH0 | Q57TH1 | Q57TH2 | Q57TI8 | Q57TI9 | Q57TJ0 | Q57TJ1 | Q57TJ2 | Q57TJ7 | Q57TJ8 | Q57TL0 |
| Q57TL2 | Q57TL5 | Q57TL8 | Q57TP2 | Q57TP3 | Q57TP8 | Q57TQ0 | Q57TQ2 | Q57TQ4 | Q57U83 | Q57UZ7 |
| Q57V41 | Q57VB1 | Q57VU6 | Q57W26 | Q57WH1 | Q57WW6 | Q57X81 | Q57XV5 | Q57XX3 | Q57Y88 | Q57YG0 |
| Q57YG2 | Q57YZ9 | Q57ZL2 | Q57ZP2 | Q57ZQ7 | Q58000 | Q58001 | Q58002 | Q58003 | Q58006 | Q58008 |
| Q58009 | Q58010 | Q58011 | Q58012 | Q58013 | Q58014 | Q58015 | Q58016 | Q58017 | Q58018 | Q58019 |
| Q58020 | Q58021 | Q58022 | Q58024 | Q58025 | Q58027 | Q58029 | Q58030 | Q58031 | Q58033 | Q58034 |
| Q58035 | Q58036 | Q58037 | Q58038 | Q58040 | Q58041 | Q58042 | Q58043 | Q58044 | Q58045 | Q58046 |
| Q58048 | Q58049 | Q58050 | Q58051 | Q58052 | Q58053 | Q58054 | Q58055 | Q58056 | Q58057 | Q58058 |
| Q58059 | Q58060 | Q58062 | Q58063 | Q58064 | Q58065 | Q58067 | Q58068 | Q58069 | Q58070 | Q58071 |
| Q58073 | Q58074 | Q58075 | Q58077 | Q58078 | Q58079 | Q58080 | Q58081 | Q58082 | Q58083 | Q58084 |
| Q58085 | Q58087 | Q58088 | Q58089 | Q58090 | Q58092 | Q58093 | Q58094 | Q58095 | Q58096 | Q58097 |
| Q58099 | Q580P9 | Q580W5 | Q580X6 | Q58100 | Q58101 | Q58102 | Q58103 | Q58104 | Q58105 | Q58108 |
| Q58109 | Q58110 | Q58112 | Q58113 | Q58114 | Q58116 | Q58118 | Q58120 | Q58123 | Q58124 | Q58125 |
| Q58126 | Q58127 | Q58129 | Q58130 | Q58131 | Q58132 | Q58133 | Q58134 | Q58135 | Q58136 | Q58137 |
| Q58138 | Q58139 | Q58140 | Q58141 | Q58142 | Q58143 | Q58144 | Q58145 | Q58146 | Q58147 | Q58148 |
| Q58149 | Q58150 | Q58151 | Q58152 | Q58153 | Q58154 | Q58155 | Q58156 | Q58157 | Q58158 | Q58161 |
| Q58162 | Q58164 | Q58167 | Q58168 | Q58170 | Q58171 | Q58173 | Q58174 | Q58175 | Q58176 | Q58177 |
| Q58178 | Q58179 | Q58180 | Q58181 | Q58182 | Q58184 | Q58185 | Q58187 | Q58188 | Q58192 | Q58193 |
| Q58194 | Q58195 | Q58196 | Q58197 | Q58198 | Q58199 | Q581T1 | Q58201 | Q58208 | Q58209 | Q58210 |
| Q58211 | Q58212 | Q58214 | Q58215 | Q58216 | Q58217 | Q58218 | Q58219 | Q58220 | Q58221 | Q58222 |
| Q58223 | Q58224 | Q58225 | Q58226 | Q58228 | Q58229 | Q58230 | Q58231 | Q58234 | Q58235 | Q58238 |
| Q58239 | Q58240 | Q58241 | Q58242 | Q58243 | Q58244 | Q58246 | Q58247 | Q58248 | Q58249 | Q58250 |
| Q58251 | Q58252 | Q58253 | Q58254 | Q58255 | Q58256 | Q58264 | Q58265 | Q58266 | Q58267 | Q58268 |
| Q58269 | Q58270 | Q58271 | Q58272 | Q58273 | Q58274 | Q58275 | Q58276 | Q58277 | Q58278 | Q58280 |
| Q58284 | Q58285 | Q58289 | Q58291 | Q58292 | Q58293 | Q58294 | Q58295 | Q58296 | Q58297 | Q58298 |
| Q58299 | Q582A7 | Q582G4 | Q58300 | Q58301 | Q58302 | Q58303 | Q58304 | Q58305 | Q58306 | Q58309 |
| Q58315 | Q58316 | Q58317 | Q58318 | Q58319 | Q58320 | Q58321 | Q58322 | Q58323 | Q58324 | Q58325 |
| Q58326 | Q58327 | Q58328 | Q58329 | Q58330 | Q58331 | Q58332 | Q58335 | Q58336 | Q58337 | Q58338 |
| Q58339 | Q58340 | Q58341 | Q58342 | Q58344 | Q58345 | Q58346 | Q58347 | Q58349 | Q58350 | Q58352 |
| Q58354 | Q58356 | Q58357 | Q58358 | Q58359 | Q58360 | Q58361 | Q58362 | Q58365 | Q58366 | Q58367 |
| Q58369 | Q58370 | Q58371 | Q58372 | Q58373 | Q58374 | Q58375 | Q58376 | Q58377 | Q58378 | Q58379 |
| Q58380 | Q58383 | Q58384 | Q58385 | Q58387 | Q58388 | Q58390 | Q58391 | Q58392 | Q58393 | Q58394 |
| Q58395 | Q58396 | Q58397 | Q58398 | Q58399 | Q58400 | Q58401 | Q58403 | Q58405 | Q58406 | Q58407 |
| Q58409 | Q58410 | Q58411 | Q58412 | Q58413 | Q58414 | Q58416 | Q58417 | Q58422 | Q58424 | Q58425 |
| Q58426 | Q58427 | Q58428 | Q58429 | Q58431 | Q58432 | Q58433 | Q58434 | Q58435 | Q58436 | Q58441 |
| Q58443 | Q58444 | Q58445 | Q58446 | Q58447 | Q58448 | Q58449 | Q58450 | Q58451 | Q58452 | Q58453 |

Q58454 Q58455 Q58456 Q58457 Q58458 Q58459 Q58460 Q58462 Q58463 Q58464 Q58465
Q58466 Q58469 Q58471 Q58473 Q58475 Q58476 Q58477 Q58481 Q58482 Q58483 Q58484
Q58485 Q58486 Q58493 Q58494 Q58495 Q58496 Q58497 Q58498 Q58499 Q584D5 Q584U2
Q58500 Q58501 Q58502 Q58503 Q58504 Q58506 Q58508 Q58509 Q58510 Q58511 Q58514
Q58515 Q58516 Q58517 Q58518 Q58519 Q58520 Q58521 Q58522 Q58523 Q58524 Q58525
Q58526 Q58527 Q58529 Q58530 Q58531 Q58532 Q58533 Q58534 Q58535 Q58536 Q58539
Q58540 Q58544 Q58546 Q58548 Q58549 Q58550 Q58551 Q58552 Q58553 Q58554 Q58555
Q58556 Q58557 Q58558 Q58559 Q58560 Q58562 Q58563 Q58564 Q58565 Q58566 Q58567
Q58568 Q58569 Q58570 Q58571 Q58572 Q58573 Q58574 Q58575 Q58576 Q58577 Q58579
Q58581 Q58582 Q58583 Q58584 Q58585 Q58587 Q58588 Q58591 Q58592 Q58593 Q58595
Q58597 Q58598 Q58599 Q585F3 Q585P3 Q58600 Q58601 Q58602 Q58603 Q58604 Q58605
Q58606 Q58608 Q58610 Q58611 Q58612 Q58613 Q58614 Q58615 Q58617 Q58619 Q58621
Q58622 Q58624 Q58625 Q58626 Q58627 Q58628 Q58629 Q58630 Q58631 Q58632 Q58633
Q58634 Q58635 Q58636 Q58637 Q58638 Q58639 Q58640 Q58641 Q58642 Q58643 Q58644
Q58645 Q58646 Q58648 Q58649 Q58652 Q58653 Q58654 Q58655 Q58656 Q58657 Q58658
Q58659 Q58660 Q58661 Q58664 Q58667 Q58668 Q58669 Q58670 Q58672 Q58673 Q58674
Q58675 Q58677 Q58678 Q58683 Q58684 Q58688 Q58690 Q58691 Q58692 Q58693 Q58694
Q58695 Q58696 Q58697 Q58698 Q58699 Q58700 Q58701 Q58707 Q58708 Q58709 Q58711
Q58712 Q58716 Q58717 Q58718 Q58719 Q58720 Q58721 Q58722 Q58723 Q58724 Q58725
Q58727 Q58728 Q58729 Q58730 Q58731 Q58732 Q58733 Q58734 Q58735 Q58736 Q58737
Q58738 Q58740 Q58741 Q58742 Q58743 Q58744 Q58745 Q58746 Q58747 Q58750 Q58751
Q58754 Q58755 Q58758 Q58759 Q58761 Q58764 Q58765 Q58766 Q58767 Q58769 Q58771
Q58772 Q58773 Q58774 Q58775 Q58776 Q58777 Q58779 Q58780 Q58782 Q58783 Q58784
Q58785 Q58786 Q58787 Q58793 Q58794 Q58796 Q58799 Q587E3 Q587J7 Q587J8 Q587K4
Q587N6 Q587N7 Q58800 Q58801 Q58802 Q58803 Q58804 Q58805 Q58806 Q58807 Q58808
Q58809 Q58810 Q58811 Q58813 Q58814 Q58815 Q58816 Q58817 Q58818 Q58819 Q58820
Q58821 Q58822 Q58823 Q58824 Q58825 Q58826 Q58827 Q58829 Q58830 Q58831 Q58832
Q58834 Q58835 Q58836 Q58839 Q58840 Q58841 Q58842 Q58843 Q58844 Q58845 Q58846
Q58847 Q58848 Q58849 Q58850 Q58851 Q58853 Q58854 Q58855 Q58856 Q58857 Q58858
Q58859 Q58860 Q58861 Q58862 Q58868 Q58869 Q58870 Q58871 Q58873 Q58874 Q58875
Q58876 Q58877 Q58879 Q58881 Q58882 Q58883 Q58884 Q58885 Q58888 Q58889 Q58891
Q58892 Q58893 Q58894 Q58895 Q58897 Q58898 Q58899 Q588U8 Q588V7 Q58900 Q58904
Q58905 Q58906 Q58907 Q58908 Q58909 Q58910 Q58912 Q58913 Q58914 Q58915 Q58917
Q58919 Q58921 Q58923 Q58924 Q58925 Q58926 Q58927 Q58928 Q58929 Q58931 Q58932
Q58933 Q58934 Q58935 Q58936 Q58937 Q58938 Q58939 Q58940 Q58941 Q58942 Q58943
Q58944 Q58945 Q58946 Q58947 Q58950 Q58952 Q58953 Q58954 Q58958 Q58959 Q58960
Q58962 Q58963 Q58968 Q58969 Q58970 Q58973 Q58974 Q58976 Q58977 Q58978 Q58979
Q58980 Q58981 Q58982 Q58987 Q58988 Q58989 Q58991 Q58992 Q58993 Q58994 Q58996
Q58997 Q58998 Q58999 Q589A4 Q589A6 Q589A7 Q589A9 Q589B0 Q589B8 Q589B9 Q589C0
Q589R5 Q589Y0 Q58A44 Q58A45 Q58CM8 Q58CN7 Q58CP0 Q58CQ2 Q58CQ5 Q58CR1 Q58CR3
Q58CU0 Q58CU3 Q58CU6 Q58CV6 Q58CW6 Q58CX2 Q58CX9 Q58CZ0 Q58CZ9 Q58D00 Q58D05
Q58D08 Q58D13 Q58D15 Q58D20 Q58D49 Q58D56 Q58D65 Q58DA0 Q58DB0 Q58DC0 Q58DC2
Q58DC5 Q58DC7 Q58DC8 Q58DD9 Q58DE2 Q58DF3 Q58DF7 Q58DG1 Q58DG5 Q58DG6 Q58DH1
Q58DH8 Q58DK4 Q58DK5 Q58DK8 Q58DL1 Q58DL7 Q58DM4 Q58DN4 Q58DP0 Q58DR0 Q58DR8

Q58DT0 Q58DT4 Q58DU0 Q58DU8 Q58DV7 Q58DW4 Q58DZ6 Q58E77 Q58E95 Q58EA2 Q58EB4
Q58ED9 Q58EE9 Q58EK5 Q58EL0 Q58EL7 Q58EM4 Q58FA4 Q58FG0 Q58FK9 Q58FL6 Q58FS2
Q58FS3 Q58FV0 Q58FV5 Q58FY4 Q58FZ0 Q58G53 Q58G87 Q58GE8 Q58H57 Q58HI1 Q58HK4
Q58L83 Q58MU6 Q58NQ4 Q58PK7 Q58VP7 Q58VT1 Q58Y43 Q58Y44 Q58Z64 Q59000 Q59001
Q59002 Q59003 Q59004 Q59005 Q59006 Q59007 Q59008 Q59009 Q59010 Q59011 Q59013
Q59014 Q59015 Q59017 Q59018 Q59019 Q59021 Q59022 Q59023 Q59024 Q59025 Q59026
Q59027 Q59028 Q59029 Q59030 Q59032 Q59033 Q59034 Q59035 Q59036 Q59037 Q59039
Q59040 Q59041 Q59042 Q59043 Q59044 Q59045 Q59046 Q59047 Q59048 Q59049 Q59050
Q59051 Q59052 Q59053 Q59054 Q59055 Q59056 Q59057 Q59058 Q59059 Q59060 Q59061
Q59062 Q59063 Q59064 Q59065 Q59066 Q59067 Q59068 Q59069 Q59070 Q59072 Q59073
Q59074 Q59075 Q59081 Q59083 Q59087 Q59088 Q59091 Q59092 Q59093 Q59097 Q59098
Q59100 Q59101 Q59102 Q59104 Q59106 Q59111 Q59112 Q59118 Q59119 Q59126 Q59136
Q59140 Q59149 Q59156 Q59159 Q59160 Q59168 Q59169 Q59175 Q59176 Q59177 Q59178
Q59179 Q59180 Q59181 Q59182 Q59185 Q59189 Q59190 Q59191 Q59192 Q59196 Q59198
Q59199 Q59200 Q59202 Q59203 Q59204 Q59205 Q59212 Q59214 Q59219 Q59224 Q59226
Q59228 Q59229 Q59240 Q59244 Q59258 Q59263 Q59266 Q59268 Q59270 Q59279 Q59280
Q59281 Q59282 Q59283 Q59284 Q59291 Q59292 Q59293 Q59294 Q59295 Q59296 Q59309
Q59321 Q59322 Q59324 Q59330 Q59331 Q59334 Q59335 Q59337 Q59342 Q59397 Q59408
Q59418 Q59435 Q59437 Q59447 Q59450 Q59458 Q59459 Q59460 Q59461 Q59462 Q59477
Q59482 Q59485 Q59486 Q59487 Q59490 Q59491 Q59495 Q59498 Q59499 Q59500 Q59509
Q59516 Q59519 Q59523 Q59526 Q59533 Q59536 Q59545 Q59547 Q59549 Q59556 Q59560
Q59561 Q59565 Q59579 Q59587 Q59591 Q59592 Q59594 Q59595 Q59596 Q59597 Q59599
Q595W7 Q59602 Q59603 Q59605 Q59606 Q59613 Q59614 Q59615 Q59619 Q59622 Q59627
Q59628 Q59629 Q59633 Q59637 Q59638 Q59642 Q59643 Q59645 Q59649 Q59650 Q59653
Q59654 Q59662 Q59666 Q59676 Q59677 Q59678 Q59679 Q59683 Q59684 Q59685 Q59686
Q59687 Q59695 Q59699 Q59701 Q59702 Q59711 Q59712 Q59714 Q59717 Q59723 Q59726
Q59727 Q59728 Q59730 Q59732 Q59734 Q59735 Q59738 Q59741 Q59742 Q59745 Q59747
Q59748 Q59750 Q59752 Q59753 Q59754 Q59757 Q59758 Q59759 Q59767 Q59768 Q59771
Q59776 Q59778 Q59782 Q59787 Q59800 Q59803 Q59818 Q59821 Q59827 Q59828 Q59829
Q59831 Q59832 Q59833 Q59838 Q59905 Q59906 Q59914 Q59918 Q59919 Q59920 Q59925
Q59928 Q59931 Q59934 Q59935 Q59938 Q59939 Q59940 Q59943 Q59948 Q59961 Q59966
Q59967 Q59968 Q59969 Q59971 Q59975 Q59977 Q59978 Q59984 Q59985 Q59987 Q59988
Q59989 Q59990 Q59992 Q59993 Q59994 Q59996 Q59A28 Q59A32 Q59DX8 Q59E36 Q59E55
Q59GN2 Q59H18 Q59HN8 Q59I44 Q59I47 Q59JU3 Q59K07 Q59K86 Q59KF3 Q59KI4 Q59KL6
Q59KN8 Q59L13 Q59L89 Q59LC9 Q59LF9 Q59LN9 Q59LQ5 Q59LR2 Q59LU0 Q59LV8 Q59LX5
Q59LY1 Q59M69 Q59MA3 Q59MA9 Q59MB6 Q59MD2 Q59MG1 Q59MP1 Q59MU1 Q59MV1 Q59MV9
Q59MW2 Q59N20 Q59N29 Q59N80 Q59NB8 Q59NG5 Q59NN8 Q59NQ9 Q59NW5 Q59NX5 Q59NX9
Q59P39 Q59P44 Q59P50 Q59P87 Q59P96 Q59PD3 Q59PD6 Q59PE7 Q59PG6 Q59PL9 Q59PP0
Q59PP6 Q59PR3 Q59PR9 Q59PT6 Q59PZ7 Q59Q38 Q59Q39 Q59Q46 Q59QC2 Q59QC4 Q59QD6
Q59QH0 Q59QH6 Q59QW5 Q59R32 Q59R35 Q59RA0 Q59RH5 Q59RN6 Q59RP7 Q59RR0 Q59RR3
Q59RW5 Q59S06 Q59S27 Q59S42 Q59S43 Q59S45 Q59S50 Q59S59 Q59S63 Q59S66 Q59SJ9
Q59SN0 Q59ST1 Q59ST6 Q59ST8 Q59SU5 Q59T36 Q59TB2 Q59TD3 Q59TN1 Q59U10 Q59U11
Q59U67 Q59U73 Q59U81 Q59UG4 Q59UH5 Q59UY7 Q59V88 Q59VC6 Q59VF4 Q59VF9 Q59VM4
Q59VM7 Q59VN2 Q59VP1 Q59VP2 Q59VP7 Q59VQ3 Q59VQ8 Q59VR1 Q59VR3 Q59VX8 Q59VX9

Q59W04 Q59W50 Q59W52 Q59WB9 Q59WD5 Q59WG0 Q59WH0 Q59WH7 Q59WI7 Q59WJ4 Q59WJ5
Q59WK2 Q59X26 Q59X38 Q59X40 Q59X67 Q59X94 Q59XQ1 Q59XV0 Q59XY0 Q59XY9 Q59Y20
Q59Y41 Q59Y46 Q59YD8 Q59YD9 Q59YH3 Q59YL9 Q59YU1 Q59Z14 Q59Z17 Q59Z29 Q59ZB1
Q59ZC8 Q59ZG8 Q59ZH9 Q59ZU1 Q59ZV4 Q59ZV5 Q59ZW4 Q59ZY9 Q59ZZ6 Q5A013 Q5A0A9
Q5A0E2 Q5A0E5 Q5A0J9 Q5A0W6 Q5A0W7 Q5A0W9 Q5A0Y2 Q5A0Y8 Q5A119 Q5A1A0 Q5A1D5
Q5A1E3 Q5A1M3 Q5A1M4 Q5A1N6 Q5A1V3 Q5A1W9 Q5A201 Q5A216 Q5A218 Q5A220 Q5A246
Q5A2A2 Q5A2B9 Q5A2K0 Q5A2T0 Q5A2W2 Q5A2Z1 Q5A301 Q5A309 Q5A310 Q5A387 Q5A3J1
Q5A3K2 Q5A3Q0 Q5A3V6 Q5A3Y5 Q5A3Z5 Q5A416 Q5A449 Q5A4E2 Q5A4F3 Q5A4G2 Q5A4G9
Q5A4H4 Q5A4H9 Q5A4K7 Q5A4N5 Q5A4P9 Q5A4Q1 Q5A4W8 Q5A4X0 Q5A519 Q5A5N5 Q5A5Q6
Q5A644 Q5A660 Q5A692 Q5A6A4 Q5A6B6 Q5A6K8 Q5A6Q4 Q5A6Q7 Q5A6T5 Q5A6T8 Q5A761
Q5A7P3 Q5A7Q2 Q5A7Q3 Q5A7Q6 Q5A7S5 Q5A847 Q5A860 Q5A884 Q5A893 Q5A8H7 Q5A8K2
Q5A8W9 Q5A8X7 Q5A8X9 Q5A917 Q5A922 Q5A931 Q5A940 Q5A960 Q5A961 Q5A985 Q5A9Z1
Q5A9Z6 Q5AA26 Q5AAF4 Q5AAG1 Q5AAI8 Q5AAJ7 Q5AAJ8 Q5AAR0 Q5AAT0 Q5AAV3 Q5AB15
Q5ABA2 Q5ABE2 Q5ABE5 Q5ABG1 Q5ABV4 Q5ABX0 Q5ABZ2 Q5AC37 Q5AC48 Q5ACI8 Q5ACK7
Q5ACL9 Q5ACM4 Q5ACM9 Q5ACU4 Q5ACU6 Q5ACW2 Q5ACW8 Q5ACX1 Q5AD13 Q5AD27 Q5AD49
Q5AD56 Q5AD59 Q5AD67 Q5ADL0 Q5ADL4 Q5ADL8 Q5ADL9 Q5ADP6 Q5ADP9 Q5ADQ0 Q5ADS3
Q5ADT1 Q5ADT9 Q5ADW3 Q5ADX2 Q5ADX5 Q5AED6 Q5AED9 Q5AEF0 Q5AEM5 Q5AEN1 Q5AEN6
Q5AF03 Q5AF56 Q5AF95 Q5AFG1 Q5AFK0 Q5AFK3 Q5AFP3 Q5AFT3 Q5AFV3 Q5AFX2 Q5AG31
Q5AG73 Q5AG86 Q5AG97 Q5AGC1 Q5AGC7 Q5AGE5 Q5AGM0 Q5AGV4 Q5AGZ9 Q5AH25 Q5AH38
Q5AH60 Q5AH87 Q5AHB8 Q5AHH4 Q5AHI7 Q5AHJ5 Q5AHK2 Q5AHZ7 Q5AI15 Q5AI37 Q5AI58
Q5AI80 Q5AI86 Q5AI90 Q5AI97 Q5AJ71 Q5AJ77 Q5AJ85 Q5AJC1 Q5AJF1 Q5AJS6 Q5AK10
Q5AK25 Q5AK42 Q5AK51 Q5AK64 Q5AK73 Q5AKA5 Q5AL10 Q5AL13 Q5AL29 Q5AL36 Q5AL45
Q5AL63 Q5ALL3 Q5ALL8 Q5ALP1 Q5ALR8 Q5ALU2 Q5ALW7 Q5ALX3 Q5ALX8 Q5ALY0 Q5AM44
Q5AM50 Q5AM80 Q5AM84 Q5AME2 Q5AMG5 Q5AMH6 Q5AML1 Q5AML2 Q5AML6 Q5AMM4 Q5AMQ6
Q5ANA8 Q5ANB1 Q5ANB2 Q5ANB7 Q5ANC8 Q5ANL6 Q5AP59 Q5AP80 Q5AP87 Q5AP89 Q5AP95
Q5AP97 Q5APB6 Q5APC8 Q5APF0 Q5APF2 Q5APG6 Q5APK0 Q5APK7 Q5APM7 Q5APS5 Q5APT8
Q5AQ12 Q5AQ33 Q5AQ47 Q5AQ57 Q5AQ62 Q5AQA3 Q5AQE4 Q5AQJ2 Q5AQL1 Q5AQN0 Q5AR06
Q5AR28 Q5AR31 Q5AR33 Q5AR34 Q5AR44 Q5AR45 Q5AR47 Q5AR48 Q5AR53 Q5AR54 Q5AR55
Q5AR56 Q5AR57 Q5ARI5 Q5ARK3 Q5ARL6 Q5AS60 Q5AS80 Q5ASA5 Q5ASA6 Q5ASE0 Q5ASE3
Q5ASK8 Q5ASK9 Q5ASN8 Q5ASQ0 Q5ASR2 Q5AT14 Q5AT95 Q5ATG5 Q5ATG6 Q5ATG8 Q5ATH0
Q5ATH1 Q5ATJ7 Q5ATV8 Q5AU18 Q5AU50 Q5AU62 Q5AUG9 Q5AUI1 Q5AUL4 Q5AUW6 Q5AUW7
Q5AUW9 Q5AUX1 Q5AUX7 Q5AUX9 Q5AUY1 Q5AUY5 Q5AUZ6 Q5AV02 Q5AV04 Q5AV05 Q5AV06
Q5AV07 Q5AV81 Q5AVC5 Q5AVC7 Q5AVF0 Q5AVF8 Q5AVK6 Q5AVS0 Q5AVZ0 Q5AW04 Q5AW32
Q5AW46 Q5AW61 Q5AW62 Q5AW69 Q5AW75 Q5AW89 Q5AW92 Q5AWA2 Q5AWA6 Q5AWM5 Q5AWS1
Q5AWS6 Q5AWZ6 Q5AX35 Q5AX75 Q5AXA9 Q5AXB0 Q5AXH1 Q5AXH3 Q5AXT5 Q5AXT6 Q5AXW5
Q5AY46 Q5AY89 Q5AYE3 Q5AYI6 Q5AYI7 Q5AYN6 Q5AYR1 Q5AYR3 Q5AYR9 Q5AYS4 Q5AYT1
Q5AYT7 Q5AYW6 Q5AYZ1 Q5AZ32 Q5AZA6 Q5AZJ0 Q5AZJ7 Q5AZQ6 Q5AZQ7 Q5AZR6 Q5AZS0
Q5AZX0 Q5AZY1 Q5AZY3 Q5B020 Q5B058 Q5B0C9 Q5B0D0 Q5B0D6 Q5B0F5 Q5B0H6 Q5B0H8
Q5B0I1 Q5B0J9 Q5B0M9 Q5B0N3 Q5B0R3 Q5B0Y2 Q5B0Y5 Q5B0Z5 Q5B122 Q5B128 Q5B135
Q5B136 Q5B1J4 Q5B1L4 Q5B1X8 Q5B1Z0 Q5B288 Q5B2E9 Q5B2Q1 Q5B2Q3 Q5B2V2 Q5B2V8
Q5B2X8 Q5B323 Q5B367 Q5B3B1 Q5B3C8 Q5B3G6 Q5B3H2 Q5B3H7 Q5B3I8 Q5B3I9 Q5B3J5
Q5B3U2 Q5B3U7 Q5B3Y9 Q5B423 Q5B4D1 Q5B4H8 Q5B4J6 Q5B4M1 Q5B4Q8 Q5B4R1 Q5B4R9
Q5B4S6 Q5B4T5 Q5B4U6 Q5B4W9 Q5B4X6 Q5B4Z3 Q5B503 Q5B590 Q5B5C1 Q5B5E7 Q5B5I6
Q5B5W1 Q5B601 Q5B630 Q5B6C5 Q5B6D6 Q5B6H7 Q5B6J2 Q5B6J8 Q5B6K3 Q5B6P3 Q5B6T1

Q5B6U3 Q5B714 Q5B748 Q5B778 Q5B787 Q5B7I4 Q5B7I5 Q5B7I6 Q5B7I7 Q5B7I8 Q5B7J0
Q5B7Q7 Q5B7T4 Q5B7U4 Q5B8A0 Q5B8A3 Q5B8A6 Q5B8A8 Q5B8F4 Q5B8F8 Q5B8G3 Q5B8K7
Q5B8L2 Q5B8S5 Q5B8U0 Q5B8X0 Q5B8X6 Q5B8Y3 Q5B8Y4 Q5B912 Q5B948 Q5B973 Q5B995
Q5B998 Q5B9G5 Q5B9L6 Q5B9N7 Q5BA81 Q5BA83 Q5BAB7 Q5BAD0 Q5BAD5 Q5BAE1 Q5BAH2
Q5BAH7 Q5BAJ7 Q5BAZ5 Q5BB35 Q5BB37 Q5BB41 Q5BB47 Q5BB99 Q5BBA8 Q5BBC5 Q5BBF1
Q5BBF4 Q5BBF7 Q5BBH7 Q5BBL5 Q5BBM1 Q5BBP5 Q5BBP8 Q5BBV4 Q5BBV9 Q5BBX9 Q5BC69
Q5BC71 Q5BC73 Q5BCC5 Q5BCG1 Q5BCI0 Q5BCL0 Q5BCN2 Q5BCU6 Q5BD77 Q5BD89 Q5BDC8
Q5BDG0 Q5BDG7 Q5BDI1 Q5BDI8 Q5BDJ5 Q5BDL3 Q5BDL9 Q5BDM6 Q5BDU4 Q5BDW0 Q5BDW4
Q5BDX1 Q5BDY8 Q5BE03 Q5BE04 Q5BE22 Q5BEG5 Q5BEJ4 Q5BEJ6 Q5BEJ8 Q5BEK1 Q5BEN5
Q5BEP0 Q5BER4 Q5BET6 Q5BEY7 Q5BF48 Q5BF62 Q5BFB1 Q5BFG0 Q5BFG8 Q5BFH3 Q5BFM4
Q5BFN9 Q5BFU1 Q5BFU7 Q5BG07 Q5BGA7 Q5BGA9 Q5BGC4 Q5BGE2 Q5BGG1 Q5BGH1 Q5BGK3
Q5BGN2 Q5BGN7 Q5BGP0 Q5BGP1 Q5BGW9 Q5BGX6 Q5BH14 Q5BH19 Q5BH30 Q5BH31 Q5BH32
Q5BH33 Q5BH34 Q5BH52 Q5BH63 Q5BH69 Q5BH83 Q5BH88 Q5BH89 Q5BHB8 Q5BHD6 Q5BHE2
Q5BI31 Q5BIM1 Q5BIM2 Q5BIM8 Q5BIN2 Q5BIN5 Q5BIP7 Q5BIP8 Q5BIR5 Q5BIR6 Q5BIV3
Q5BIX7 Q5BJ10 Q5BJ29 Q5BJ39 Q5BJ41 Q5BJ48 Q5BJ52 Q5BJ53 Q5BJ65 Q5BJ78 Q5BJ90
Q5BJ92 Q5BJA5 Q5BJB7 Q5BJC4 Q5BJE1 Q5BJI6 Q5BJJ5 Q5BJJ6 Q5BJJ7 Q5BJJ8 Q5BJL5
Q5BJN5 Q5BJN8 Q5BJP2 Q5BJP3 Q5BJP6 Q5BJP9 Q5BJQ6 Q5BJQ7 Q5BJR4 Q5BJR8 Q5BJS0
Q5BJS9 Q5BJT0 Q5BJT1 Q5BJT6 Q5BJU3 Q5BJV9 Q5BJX0 Q5BJX1 Q5BJX5 Q5BJY3 Q5BJY6
Q5BJZ6 Q5BK06 Q5BK07 Q5BK10 Q5BK18 Q5BK22 Q5BK24 Q5BK30 Q5BK43 Q5BK46 Q5BK60
Q5BK64 Q5BK67 Q5BK68 Q5BK82 Q5BKC6 Q5BKE0 Q5BKF8 Q5BKG8 Q5BKH4 Q5BKH5 Q5BKI6
Q5BKL1 Q5BKL8 Q5BKM6 Q5BKN5 Q5BKS4 Q5BKU9 Q5BKW9 Q5BKY1 Q5BKY6 Q5BL31 Q5BL35
Q5BL41 Q5BL56 Q5BL73 Q5BL87 Q5BLD8 Q5BLF0 Q5BLK4 Q5BLX8 Q5BMD4 Q5BN22 Q5BN23
Q5BN31 Q5BN45 Q5BN46 Q5BP74 Q5BPG5 Q5BPM6 Q5BPN1 Q5BPP4 Q5BPS0 Q5BPS3 Q5BPT7
Q5BQN5 Q5BQN8 Q5BU09 Q5BUB4 Q5C8T6 Q5C8T8 Q5C8U1 Q5C8U3 Q5C8U4 Q5C9I9 Q5C9L6
Q5C9L7 Q5CCK4 Q5CD17 Q5CD77 Q5CD84 Q5CD95 Q5CD96 Q5CD97 Q5CG95 Q5CKJ0 Q5CP76
Q5CPC9 Q5CPU3 Q5CT62 Q5CWQ7 Q5CX58 Q5CZ79 Q5CZA5 Q5CZC0 Q5CZI8 Q5CZL1 Q5D013
Q5D014 Q5D016 Q5D018 Q5D0E6 Q5D0W8 Q5D0X4 Q5D144 Q5D189 Q5D1B9 Q5D1C1 Q5D1Z3
Q5D525 Q5D6D3 Q5D6D7 Q5D704 Q5D7H7 Q5D7H8 Q5D7I0 Q5D7I1 Q5D7I2 Q5D7I3 Q5D7I5
Q5D7I6 Q5D7I9 Q5D7J0 Q5D7J1 Q5D7J2 Q5D862 Q5D869 Q5D873 Q5D892 Q5D8V5 Q5DA90
Q5DAA3 Q5DC69 Q5DD96 Q5DFU0 Q5DGJ1 Q5DHJ5 Q5DHT6 Q5DJT8 Q5DJU3 Q5DM57 Q5DNB1
Q5DP50 Q5DTH5 Q5DTI6 Q5DTM8 Q5DTN8 Q5DTT1 Q5DTT3 Q5DTT4 Q5DTT8 Q5DTV4 Q5DTW7
Q5DU09 Q5DU37 Q5DU99 Q5DUX5 Q5DUX7 Q5DUY3 Q5DVH6 Q5DW34 Q5DWI3 Q5DWN0 Q5DX34
Q5DYB5 Q5DYC3 Q5DYF2 Q5DYV8 Q5DZ63 Q5DZ79 Q5DZB8 Q5DZC0 Q5DZC1 Q5DZD2 Q5DZF2
Q5DZH9 Q5DZI1 Q5DZM3 Q5DZM4 Q5DZN1 Q5DZP0 Q5DZP1 Q5DZU8 Q5DZX0 Q5DZX2 Q5DZX4
Q5E006 Q5E022 Q5E084 Q5E0C3 Q5E0C4 Q5E0C6 Q5E0F0 Q5E0F9 Q5E0G4 Q5E0H4 Q5E0M1
Q5E0Q5 Q5E0T7 Q5E0X3 Q5E0Z0 Q5E176 Q5E1G0 Q5E1G4 Q5E1M6 Q5E1M7 Q5E1P2 Q5E1Q7
Q5E1Q8 Q5E1R1 Q5E1R3 Q5E1R4 Q5E1R5 Q5E1R6 Q5E1S3 Q5E1W5 Q5E1W7 Q5E1X3 Q5E1X4
Q5E1Z0 Q5E203 Q5E206 Q5E212 Q5E215 Q5E218 Q5E225 Q5E232 Q5E234 Q5E236 Q5E238
Q5E239 Q5E249 Q5E257 Q5E260 Q5E263 Q5E279 Q5E294 Q5E2A2 Q5E2B1 Q5E2B3 Q5E2C2
Q5E2C3 Q5E2C6 Q5E2C7 Q5E2C8 Q5E2D3 Q5E2D6 Q5E2D9 Q5E2E0 Q5E2E1 Q5E2E3 Q5E2E4
Q5E2E5 Q5E2E6 Q5E2E7 Q5E2E8 Q5E2F9 Q5E2G0 Q5E2G4 Q5E2G5 Q5E2G9 Q5E2H3 Q5E2H8
Q5E2K1 Q5E2K7 Q5E2L7 Q5E2M9 Q5E2N0 Q5E2N9 Q5E2P2 Q5E2P8 Q5E2Q1 Q5E2R0 Q5E2R1
Q5E2R2 Q5E2T1 Q5E2T2 Q5E2T7 Q5E2U7 Q5E2W4 Q5E2W6 Q5E2W7 Q5E2X3 Q5E2Y4 Q5E2Y6
Q5E2Y8 Q5E2Z2 Q5E2Z6 Q5E2Z9 Q5E304 Q5E305 Q5E314 Q5E316 Q5E317 Q5E320 Q5E325

Q5E328 Q5E329 Q5E330 Q5E331 Q5E332 Q5E345 Q5E399 Q5E3A4 Q5E3A5 Q5E3A7 Q5E3A8
Q5E3C5 Q5E3D1 Q5E3D2 Q5E3D6 Q5E3D9 Q5E3E0 Q5E3E1 Q5E3E2 Q5E3E5 Q5E3F3 Q5E3G3
Q5E3G5 Q5E3H3 Q5E3I3 Q5E3I7 Q5E3L1 Q5E3L4 Q5E3U4 Q5E3U5 Q5E3U6 Q5E3V7 Q5E3V8
Q5E3W7 Q5E3W9 Q5E3X0 Q5E3Y1 Q5E3Y2 Q5E3Y5 Q5E3Y7 Q5E3Y9 Q5E3Z0 Q5E3Z1 Q5E3Z3
Q5E3Z6 Q5E3Z7 Q5E3Z9 Q5E405 Q5E407 Q5E416 Q5E424 Q5E430 Q5E445 Q5E447 Q5E448
Q5E450 Q5E452 Q5E455 Q5E463 Q5E465 Q5E466 Q5E469 Q5E480 Q5E484 Q5E491 Q5E4A2
Q5E4C5 Q5E4D0 Q5E4E4 Q5E4F1 Q5E4F2 Q5E4H1 Q5E4H8 Q5E4H9 Q5E4I0 Q5E4J3 Q5E4N9
Q5E4P6 Q5E4P9 Q5E4Q3 Q5E4R4 Q5E4R6 Q5E4T4 Q5E4V7 Q5E4Z3 Q5E4Z6 Q5E524 Q5E531
Q5E590 Q5E5B4 Q5E5B5 Q5E5C2 Q5E5C7 Q5E5C9 Q5E5G1 Q5E5G4 Q5E5G5 Q5E5G6 Q5E5I3
Q5E5I4 Q5E5J8 Q5E5L2 Q5E5P2 Q5E5T7 Q5E5Y7 Q5E623 Q5E624 Q5E633 Q5E634 Q5E635
Q5E637 Q5E638 Q5E639 Q5E699 Q5E6A0 Q5E6A1 Q5E6A2 Q5E6A3 Q5E6A4 Q5E6B0 Q5E6B5
Q5E6E3 Q5E6F2 Q5E6F5 Q5E6F9 Q5E6I2 Q5E6I9 Q5E6L2 Q5E6L4 Q5E6M6 Q5E6N6 Q5E6P2
Q5E6Q4 Q5E6Q5 Q5E6Q6 Q5E6Q7 Q5E6Q8 Q5E6Q9 Q5E6T2 Q5E6T3 Q5E6T4 Q5E6T6 Q5E6U2
Q5E6U4 Q5E6U8 Q5E6V2 Q5E6V8 Q5E6V9 Q5E6W0 Q5E6W1 Q5E6X8 Q5E6Y5 Q5E6Y8 Q5E6Z0
Q5E6Z7 Q5E6Z8 Q5E702 Q5E706 Q5E719 Q5E725 Q5E726 Q5E727 Q5E728 Q5E729 Q5E730
Q5E731 Q5E735 Q5E739 Q5E743 Q5E749 Q5E760 Q5E763 Q5E765 Q5E768 Q5E771 Q5E772
Q5E775 Q5E776 Q5E780 Q5E781 Q5E7A0 Q5E7A3 Q5E7A7 Q5E7A8 Q5E7B6 Q5E7E1 Q5E7E2
Q5E7E9 Q5E7F0 Q5E7F1 Q5E7F2 Q5E7F8 Q5E7G3 Q5E7G5 Q5E7G6 Q5E7G7 Q5E7H1 Q5E7H2
Q5E7J4 Q5E7J5 Q5E7J6 Q5E7J7 Q5E7K1 Q5E7L1 Q5E7L2 Q5E7L3 Q5E7L4 Q5E7L5 Q5E7L7
Q5E7M0 Q5E7M3 Q5E7N0 Q5E7N1 Q5E7N4 Q5E7P1 Q5E7P5 Q5E7P8 Q5E7P9 Q5E7Q6 Q5E7Q9
Q5E7R0 Q5E7R1 Q5E7R2 Q5E7R7 Q5E7R8 Q5E7S8 Q5E7T0 Q5E7T8 Q5E7U0 Q5E7U3 Q5E7U4
Q5E7U5 Q5E7U6 Q5E7U8 Q5E7V0 Q5E7W7 Q5E7X1 Q5E7X6 Q5E814 Q5E830 Q5E831 Q5E839
Q5E840 Q5E841 Q5E847 Q5E856 Q5E857 Q5E858 Q5E859 Q5E865 Q5E866 Q5E871 Q5E872
Q5E873 Q5E875 Q5E876 Q5E881 Q5E887 Q5E888 Q5E889 Q5E890 Q5E891 Q5E892 Q5E893
Q5E895 Q5E896 Q5E897 Q5E898 Q5E899 Q5E8A0 Q5E8A1 Q5E8A2 Q5E8A3 Q5E8A5 Q5E8A6
Q5E8A7 Q5E8A8 Q5E8A9 Q5E8B0 Q5E8B1 Q5E8B2 Q5E8B3 Q5E8B4 Q5E8B7 Q5E8B9 Q5E8C0
Q5E8C1 Q5E8C2 Q5E8C3 Q5E8C7 Q5E8E1 Q5E8E5 Q5E8E6 Q5E8E7 Q5E8E9 Q5E8J9 Q5E8L9
Q5E8M1 Q5E8M3 Q5E8M4 Q5E8M5 Q5E8M6 Q5E8M7 Q5E8N3 Q5E8P0 Q5E8P5 Q5E8P6 Q5E8Q0
Q5E8Q8 Q5E8R9 Q5E8S3 Q5E8S6 Q5E8T5 Q5E8U5 Q5E8U7 Q5E8W6 Q5E8W9 Q5E8Y2 Q5E8Y5
Q5E8Y6 Q5E8Z0 Q5E8Z2 Q5E8Z6 Q5E8Z7 Q5E8Z9 Q5E913 Q5E915 Q5E924 Q5E938 Q5E947
Q5E948 Q5E950 Q5E951 Q5E953 Q5E956 Q5E959 Q5E963 Q5E964 Q5E966 Q5E977 Q5E981
Q5E982 Q5E983 Q5E984 Q5E987 Q5E992 Q5E996 Q5E997 Q5E999 Q5E9A0 Q5E9A3 Q5E9A4
Q5E9A5 Q5E9A9 Q5E9B3 Q5E9B4 Q5E9B5 Q5E9C4 Q5E9D0 Q5E9D6 Q5E9E1 Q5E9E2 Q5E9F5
Q5E9F6 Q5E9F8 Q5E9G0 Q5E9G1 Q5E9G3 Q5E9G4 Q5E9H2 Q5E9H9 Q5E9I1 Q5E9I7 Q5E9I8
Q5E9J1 Q5E9J3 Q5E9J4 Q5E9J7 Q5E9J9 Q5E9K0 Q5E9K1 Q5E9K2 Q5E9K3 Q5E9K7 Q5E9L3
Q5E9L4 Q5E9L5 Q5E9M5 Q5E9N2 Q5E9N3 Q5E9N4 Q5E9N9 Q5E9P1 Q5E9P9 Q5E9S2 Q5E9S3
Q5E9S4 Q5E9S8 Q5E9T1 Q5E9T4 Q5E9T7 Q5E9U0 Q5E9U4 Q5E9V3 Q5E9V5 Q5E9V6 Q5E9X1
Q5E9X6 Q5E9X8 Q5E9Y5 Q5E9Y6 Q5E9Y9 Q5E9Z1 Q5E9Z7 Q5E9Z8 Q5EA11 Q5EA15 Q5EA19
Q5EA24 Q5EA28 Q5EA33 Q5EA36 Q5EA37 Q5EA53 Q5EA76 Q5EA79 Q5EA80 Q5EA83 Q5EA88
Q5EA89 Q5EA95 Q5EA98 Q5EAB2 Q5EAC2 Q5EAC5 Q5EAC6 Q5EAC7 Q5EAC9 Q5EAD2 Q5EAD3
Q5EAD5 Q5EAD8 Q5EAE2 Q5EAE3 Q5EAE6 Q5EAF6 Q5EAK6 Q5EAP5 Q5EAQ1 Q5EAR5 Q5EAV6
Q5EAW4 Q5EAW9 Q5EAY2 Q5EB20 Q5EB25 Q5EB28 Q5EB30 Q5EB39 Q5EB59 Q5EB68 Q5EB92
Q5EB96 Q5EB97 Q5EBA1 Q5EBA3 Q5EBD9 Q5EBE1 Q5EBE8 Q5EBF1 Q5EBF2 Q5EBG2 Q5EBG8
Q5EBJ4 Q5EBL2 Q5EBM0 Q5EBM4 Q5EBN2 Q5EDG0 Q5EE01 Q5EE38 Q5EEL8 Q5EF31 Q5EF38

Q5EF39 Q5EF40 Q5EF74 Q5EFR0 Q5EFR1 Q5EG05 Q5EG55 Q5EIC4 Q5ENZ0 Q5EU40 Q5EU41
Q5EXK1 Q5EXX3 Q5EY87 Q5F1R7 Q5F201 Q5F204 Q5F293 Q5F2C3 Q5F2E7 Q5F310 Q5F359
Q5F361 Q5F368 Q5F371 Q5F3B1 Q5F3B2 Q5F3D1 Q5F3D7 Q5F3F2 Q5F3G6 Q5F3I0 Q5F3J5
Q5F3K0 Q5F3K4 Q5F3L4 Q5F3M0 Q5F3N1 Q5F3N9 Q5F3P8 Q5F3R2 Q5F3R7 Q5F3T7 Q5F3T9
Q5F3U0 Q5F3U9 Q5F3V0 Q5F3V3 Q5F3W5 Q5F3X4 Q5F3Z1 Q5F3Z3 Q5F3Z5 Q5F3Z7 Q5F3Z9
Q5F408 Q5F414 Q5F415 Q5F416 Q5F418 Q5F428 Q5F450 Q5F457 Q5F471 Q5F476 Q5F478
Q5F479 Q5F485 Q5F489 Q5F4A1 Q5F4A3 Q5F4A9 Q5F4B3 Q5F4K8 Q5F4M7 Q5F4W2 Q5F4W3
Q5F4W9 Q5F4X1 Q5F4X9 Q5F4Y7 Q5F4Y8 Q5F506 Q5F513 Q5F529 Q5F536 Q5F541 Q5F542
Q5F551 Q5F553 Q5F554 Q5F556 Q5F584 Q5F585 Q5F588 Q5F5A4 Q5F5A5 Q5F5A9 Q5F5B2
Q5F5B4 Q5F5C2 Q5F5C4 Q5F5C8 Q5F5D0 Q5F5D6 Q5F5E2 Q5F5E3 Q5F5E9 Q5F5F3 Q5F5F4
Q5F5F5 Q5F5G5 Q5F5H5 Q5F5H9 Q5F5I5 Q5F5J4 Q5F5J8 Q5F5K5 Q5F5K6 Q5F5M1 Q5F5P0
Q5F5P1 Q5F5P6 Q5F5P7 Q5F5Q3 Q5F5Q8 Q5F5R1 Q5F5R2 Q5F5R3 Q5F5R4 Q5F5R5 Q5F5R6
Q5F5S1 Q5F5S2 Q5F5S3 Q5F5S5 Q5F5S7 Q5F5S8 Q5F5S9 Q5F5T0 Q5F5T1 Q5F5T2 Q5F5T3
Q5F5T4 Q5F5T5 Q5F5T6 Q5F5T7 Q5F5T8 Q5F5T9 Q5F5U0 Q5F5U1 Q5F5U2 Q5F5U3 Q5F5U4
Q5F5U5 Q5F5U6 Q5F5U8 Q5F5U9 Q5F5V0 Q5F5V1 Q5F5V2 Q5F5V3 Q5F5V4 Q5F5V5 Q5F5V8
Q5F5W0 Q5F5W1 Q5F5X0 Q5F5X2 Q5F5X3 Q5F5X7 Q5F5X9 Q5F5Y0 Q5F5Y7 Q5F5Z0 Q5F632
Q5F633 Q5F636 Q5F641 Q5F645 Q5F678 Q5F682 Q5F683 Q5F685 Q5F686 Q5F690 Q5F691
Q5F694 Q5F695 Q5F6A6 Q5F6E8 Q5F6E9 Q5F6F7 Q5F6G1 Q5F6H5 Q5F6I7 Q5F6K6 Q5F6K7
Q5F6L6 Q5F6L9 Q5F6M0 Q5F6P1 Q5F6P9 Q5F6Q6 Q5F6R3 Q5F6R6 Q5F6T1 Q5F6U2 Q5F6W5
Q5F6W8 Q5F6X1 Q5F6Y0 Q5F6Y8 Q5F732 Q5F746 Q5F750 Q5F752 Q5F753 Q5F756 Q5F757
Q5F761 Q5F774 Q5F775 Q5F776 Q5F783 Q5F786 Q5F788 Q5F792 Q5F797 Q5F799 Q5F7A0
Q5F7B7 Q5F7C0 Q5F7C4 Q5F7D6 Q5F7D7 Q5F7D8 Q5F7E0 Q5F7E5 Q5F7E6 Q5F7G0 Q5F7G4
Q5F7G6 Q5F7H5 Q5F7I3 Q5F7I8 Q5F7J0 Q5F7J4 Q5F7K9 Q5F7N6 Q5F7V9 Q5F7W3 Q5F7W4
Q5F812 Q5F816 Q5F829 Q5F830 Q5F831 Q5F835 Q5F836 Q5F840 Q5F849 Q5F852 Q5F856
Q5F858 Q5F860 Q5F861 Q5F863 Q5F878 Q5F881 Q5F882 Q5F889 Q5F8B4 Q5F8C0 Q5F8C3
Q5F8D3 Q5F8D4 Q5F8D8 Q5F8E8 Q5F8F6 Q5F8G2 Q5F8G6 Q5F8G9 Q5F8H5 Q5F8I0 Q5F8I1
Q5F8I2 Q5F8I7 Q5F8K5 Q5F8K9 Q5F8L2 Q5F8M6 Q5F8M8 Q5F8M9 Q5F8N0 Q5F8N2 Q5F8P8
Q5F8Q0 Q5F8S9 Q5F8T1 Q5F8T3 Q5F8T6 Q5F8U7 Q5F8U8 Q5F8V0 Q5F8V2 Q5F8V5 Q5F8W5
Q5F8W6 Q5F8W8 Q5F8X4 Q5F8Y9 Q5F8Z9 Q5F904 Q5F911 Q5F913 Q5F914 Q5F915 Q5F922
Q5F923 Q5F924 Q5F925 Q5F928 Q5F929 Q5F931 Q5F935 Q5F936 Q5F938 Q5F956 Q5F973
Q5F9D8 Q5F9E0 Q5F9F4 Q5F9F6 Q5F9F8 Q5F9F9 Q5F9G8 Q5F9H3 Q5F9I5 Q5F9J0 Q5F9J2
Q5F9J3 Q5F9J5 Q5F9J6 Q5F9J9 Q5F9K3 Q5F9K6 Q5F9L0 Q5F9L3 Q5F9L4 Q5F9N1 Q5F9P0
Q5F9Q7 Q5F9R5 Q5F9R9 Q5F9S5 Q5F9S6 Q5F9T1 Q5F9T5 Q5F9T6 Q5F9U0 Q5F9U1 Q5F9U2
Q5F9U3 Q5F9U4 Q5F9U5 Q5F9U6 Q5F9W3 Q5F9W4 Q5F9X5 Q5F9X7 Q5F9X9 Q5F9Y0 Q5F9Y1
Q5F9Y6 Q5F9Z0 Q5F9Z5 Q5F9Z9 Q5FA03 Q5FA15 Q5FA21 Q5FA22 Q5FA23 Q5FA24 Q5FA25
Q5FA32 Q5FA41 Q5FA47 Q5FA52 Q5FA57 Q5FA58 Q5FA59 Q5FA60 Q5FA76 Q5FA84 Q5FA94
Q5FA99 Q5FAA9 Q5FAB1 Q5FAC0 Q5FAC2 Q5FAC6 Q5FAD3 Q5FAD4 Q5FAF2 Q5FAF5 Q5FAF7
Q5FAG1 Q5FAH3 Q5FAH7 Q5FAH8 Q5FAH9 Q5FAI1 Q5FAI2 Q5FAI3 Q5FAJ0 Q5FAJ2 Q5FAJ3
Q5FAK5 Q5FB23 Q5FB24 Q5FB30 Q5FB34 Q5FB93 Q5FBR4 Q5FC18 Q5FC62 Q5FCW3 Q5FEH6
Q5FF10 Q5FF19 Q5FF20 Q5FF38 Q5FF40 Q5FF55 Q5FF69 Q5FF72 Q5FF75 Q5FF92 Q5FFA0
Q5FFA2 Q5FFA3 Q5FFA6 Q5FFA8 Q5FFB2 Q5FFC3 Q5FFD0 Q5FFD4 Q5FFD8 Q5FFD9 Q5FFE2
Q5FFE6 Q5FFE7 Q5FFE8 Q5FFE9 Q5FFG2 Q5FFG3 Q5FFG4 Q5FFG6 Q5FFG7 Q5FFG8 Q5FFJ0
Q5FFJ4 Q5FFK4 Q5FFL5 Q5FFL8 Q5FFM4 Q5FFP1 Q5FFQ2 Q5FFQ3 Q5FFR4 Q5FFR5 Q5FFR6
Q5FFR9 Q5FFS0 Q5FFS1 Q5FFS2 Q5FFS3 Q5FFS7 Q5FFT0 Q5FFT3 Q5FFT9 Q5FFU1 Q5FFU3

Q5FFU4 Q5FFU5 Q5FFU6 Q5FFU7 Q5FFU8 Q5FFU9 Q5FFV0 Q5FFV1 Q5FFV2 Q5FFV3 Q5FFV4
Q5FFY2 Q5FFY7 Q5FFZ0 Q5FFZ1 Q5FFZ5 Q5FFZ7 Q5FFZ9 Q5FG00 Q5FG09 Q5FG20 Q5FG27
Q5FG28 Q5FG29 Q5FG30 Q5FG35 Q5FG39 Q5FG40 Q5FG57 Q5FG65 Q5FG66 Q5FG73 Q5FG88
Q5FG91 Q5FG94 Q5FGB6 Q5FGD0 Q5FGD5 Q5FGH0 Q5FGJ5 Q5FGN1 Q5FGP0 Q5FGP1 Q5FGP4
Q5FGP5 Q5FGP6 Q5FGP9 Q5FGQ8 Q5FGS1 Q5FGT9 Q5FGU3 Q5FGU6 Q5FGV7 Q5FGW4 Q5FGW9
Q5FGX3 Q5FGY8 Q5FGZ2 Q5FGZ3 Q5FGZ7 Q5FGZ8 Q5FGZ9 Q5FH15 Q5FH27 Q5FH30 Q5FH31
Q5FH38 Q5FH45 Q5FH54 Q5FH56 Q5FH68 Q5FH69 Q5FH70 Q5FH81 Q5FH83 Q5FH87 Q5FH91
Q5FH93 Q5FH94 Q5FH96 Q5FH97 Q5FHA4 Q5FHA6 Q5FHC4 Q5FHD1 Q5FHD2 Q5FHE8 Q5FHG1
Q5FHG5 Q5FHG7 Q5FHG9 Q5FHH0 Q5FHH4 Q5FHH8 Q5FHI9 Q5FHJ7 Q5FHK5 Q5FHK6 Q5FHK7
Q5FHL5 Q5FHM5 Q5FHN7 Q5FHN8 Q5FHQ2 Q5FHQ3 Q5FHQ5 Q5FHQ6 Q5FHT1 Q5FHY0 Q5FHY3
Q5FHY4 Q5FHZ6 Q5FI29 Q5FI43 Q5FI54 Q5FIF8 Q5FIJ4 Q5FIN5 Q5FIN8 Q5FIP3 Q5FIQ9
Q5FIR6 Q5FIS5 Q5FIS8 Q5FIU5 Q5FIV0 Q5FIV1 Q5FIV3 Q5FIV5 Q5FIV8 Q5FIV9 Q5FIW0
Q5FIW3 Q5FIW8 Q5FIW9 Q5FIX8 Q5FIY6 Q5FIY7 Q5FIZ7 Q5FJ02 Q5FJ20 Q5FJ41 Q5FJB3
Q5FJB5 Q5FJB7 Q5FJB8 Q5FJC0 Q5FJF9 Q5FJG0 Q5FJG2 Q5FJG5 Q5FJG6 Q5FJG7 Q5FJH2
Q5FJH3 Q5FJH5 Q5FJH9 Q5FJI3 Q5FJJ7 Q5FJK5 Q5FJK6 Q5FJK7 Q5FJK8 Q5FJL5 Q5FJM3
Q5FJM4 Q5FJM5 Q5FJM7 Q5FJN0 Q5FJN2 Q5FJN6 Q5FJN7 Q5FJN8 Q5FJP9 Q5FJS7 Q5FJT2
Q5FJT3 Q5FJT6 Q5FJT8 Q5FJU0 Q5FJW9 Q5FJX0 Q5FJX7 Q5FJX8 Q5FJY2 Q5FK05 Q5FK38
Q5FKD8 Q5FKE1 Q5FKE4 Q5FKE7 Q5FKF0 Q5FKF4 Q5FKG6 Q5FKH0 Q5FKH3 Q5FKH5 Q5FKH6
Q5FKI4 Q5FKI5 Q5FKI6 Q5FKI8 Q5FKK2 Q5FKK7 Q5FKL6 Q5FKQ8 Q5FKQ9 Q5FKR0 Q5FKR1
Q5FKR5 Q5FKR7 Q5FKR8 Q5FKS2 Q5FKS7 Q5FKT3 Q5FKU0 Q5FKU5 Q5FKU8 Q5FKV3 Q5FKV7
Q5FKW5 Q5FKW9 Q5FKX2 Q5FKZ0 Q5FKZ1 Q5FL04 Q5FL22 Q5FL35 Q5FL42 Q5FL46 Q5FL49
Q5FL50 Q5FL55 Q5FL56 Q5FL58 Q5FL61 Q5FL65 Q5FL67 Q5FL68 Q5FL72 Q5FL82 Q5FL98
Q5FLA9 Q5FLF6 Q5FLL0 Q5FLL1 Q5FLL2 Q5FLL4 Q5FLT4 Q5FLU4 Q5FLV7 Q5FLV8 Q5FLW2
Q5FLW3 Q5FLW4 Q5FLW5 Q5FLW6 Q5FLX2 Q5FLX3 Q5FLX4 Q5FLX5 Q5FLY5 Q5FLZ2 Q5FLZ3
Q5FM02 Q5FM04 Q5FM05 Q5FM12 Q5FM13 Q5FM22 Q5FM23 Q5FM34 Q5FM35 Q5FM59 Q5FM60
Q5FM64 Q5FM65 Q5FM66 Q5FM67 Q5FM68 Q5FM69 Q5FM70 Q5FM72 Q5FM73 Q5FM74 Q5FM75
Q5FM76 Q5FM77 Q5FM78 Q5FM79 Q5FM80 Q5FM81 Q5FM82 Q5FM83 Q5FM84 Q5FM85 Q5FM86
Q5FM87 Q5FM88 Q5FM89 Q5FM90 Q5FM91 Q5FM92 Q5FM93 Q5FM94 Q5FM96 Q5FM97 Q5FMA2
Q5FMA9 Q5FMB0 Q5FMB2 Q5FMB6 Q5FMC0 Q5FMD6 Q5FMD9 Q5FME5 Q5FME6 Q5FME7 Q5FMG3
Q5FMI6 Q5FMJ2 Q5FMJ3 Q5FMN6 Q5FMR0 Q5FMT1 Q5FMT8 Q5FMX8 Q5FN05 Q5FN07 Q5FN09
Q5FN12 Q5FN15 Q5FN16 Q5FN17 Q5FN20 Q5FN26 Q5FN28 Q5FNA1 Q5FNA2 Q5FNB8 Q5FND1
Q5FND2 Q5FND3 Q5FND4 Q5FNK4 Q5FNL0 Q5FNL1 Q5FNL4 Q5FNM1 Q5FNM7 Q5FNN2 Q5FNN7
Q5FNP1 Q5FNQ1 Q5FNS2 Q5FNS3 Q5FNS6 Q5FNS8 Q5FNT5 Q5FNT7 Q5FP70 Q5FP90 Q5FP94
Q5FP95 Q5FPC9 Q5FPD9 Q5FPE5 Q5FPF1 Q5FPF3 Q5FPG9 Q5FPH2 Q5FPK0 Q5FPK1 Q5FPK5
Q5FPL0 Q5FPN1 Q5FPQ6 Q5FPS3 Q5FPS6 Q5FPT6 Q5FPX2 Q5FPX3 Q5FPX5 Q5FPX6 Q5FPX8
Q5FPX9 Q5FPY1 Q5FPY2 Q5FPZ1 Q5FPZ4 Q5FPZ5 Q5FQ21 Q5FQ27 Q5FQ31 Q5FQ35 Q5FQ48
Q5FQ54 Q5FQ60 Q5FQ87 Q5FQ88 Q5FQ96 Q5FQ97 Q5FQ98 Q5FQA6 Q5FQB4 Q5FQB6 Q5FQC4
Q5FQC5 Q5FQC6 Q5FQC7 Q5FQD0 Q5FQD6 Q5FQE8 Q5FQK0 Q5FQL9 Q5FQM0 Q5FQM1 Q5FQM2
Q5FQM3 Q5FQM6 Q5FQP6 Q5FQR2 Q5FQT4 Q5FQY8 Q5FQZ4 Q5FQZ9 Q5FR00 Q5FR06 Q5FR18
Q5FR39 Q5FR73 Q5FR80 Q5FRF2 Q5FRF4 Q5FRG7 Q5FRG8 Q5FRI6 Q5FRJ1 Q5FRJ2 Q5FRJ3
Q5FRJ7 Q5FRN3 Q5FRR4 Q5FRR8 Q5FRS2 Q5FRS9 Q5FRT1 Q5FRT2 Q5FRT7 Q5FRT8 Q5FRY0
Q5FRY8 Q5FRZ1 Q5FS11 Q5FS12 Q5FS13 Q5FS37 Q5FS38 Q5FS51 Q5FS52 Q5FS71 Q5FS72
Q5FS83 Q5FS84 Q5FS85 Q5FSH9 Q5FSJ7 Q5FSJ9 Q5FSK6 Q5FSK8 Q5FSL0 Q5FSL4 Q5FSL5
Q5FSM1 Q5FSM2 Q5FSM8 Q5FSN7 Q5FSP9 Q5FSQ7 Q5FST0 Q5FSV8 Q5FSY7 Q5FT39 Q5FT82

Q5FT88 Q5FTB0 Q5FTB5 Q5FTB6 Q5FTC0 Q5FTF9 Q5FTL1 Q5FTN2 Q5FTN3 Q5FTN5 Q5FTN7
Q5FTU6 Q5FTV3 Q5FTW1 Q5FTX3 Q5FTX4 Q5FTX5 Q5FTX6 Q5FTX7 Q5FTX8 Q5FTX9 Q5FTY0
Q5FTY1 Q5FTY2 Q5FTY3 Q5FTY4 Q5FTY5 Q5FTY6 Q5FTY7 Q5FTY8 Q5FTY9 Q5FTZ0 Q5FTZ1
Q5FTZ2 Q5FTZ3 Q5FTZ4 Q5FTZ5 Q5FTZ6 Q5FTZ7 Q5FTZ8 Q5FTZ9 Q5FU00 Q5FU01 Q5FU02
Q5FU05 Q5FU06 Q5FU07 Q5FU08 Q5FU18 Q5FU29 Q5FU33 Q5FU57 Q5FU58 Q5FU59 Q5FU61
Q5FU70 Q5FU99 Q5FUA0 Q5FUA1 Q5FUA2 Q5FUA8 Q5FUA9 Q5FUB1 Q5FUB3 Q5FUC7 Q5FUE6
Q5FUF4 Q5FUG0 Q5FUG1 Q5FUG2 Q5FUG3 Q5FUG4 Q5FUG5 Q5FUH7 Q5FUH8 Q5FUH9 Q5FUI5
Q5FUJ2 Q5FUJ3 Q5FUJ4 Q5FUJ7 Q5FUK3 Q5FUK4 Q5FUK8 Q5FUL1 Q5FUL2 Q5FUL3 Q5FUL4
Q5FUM8 Q5FUN5 Q5FUP5 Q5FUP6 Q5FUR2 Q5FUR3 Q5FUR4 Q5FUR7 Q5FUR8 Q5FUR9 Q5FUU1
Q5FUU6 Q5FUU9 Q5FUV0 Q5FUV3 Q5FUV7 Q5FUV8 Q5FUW9 Q5FUX0 Q5FUX5 Q5FUY5 Q5FVA9
Q5FVB0 Q5FVB2 Q5FVB6 Q5FVD6 Q5FVF1 Q5FVG6 Q5FVG8 Q5FVH7 Q5FVI9 Q5FVJ5 Q5FVJ7
Q5FVJ8 Q5FVL4 Q5FVL7 Q5FVM4 Q5FVN6 Q5FVN8 Q5FVP2 Q5FVP5 Q5FVQ9 Q5FVR2 Q5FVR4
Q5FVR6 Q5FVR8 Q5FVY2 Q5FW05 Q5FW06 Q5FW12 Q5FW17 Q5FW25 Q5FW37 Q5FW38 Q5FW46
Q5FW48 Q5FW53 Q5FW57 Q5FWF6 Q5FWF7 Q5FWH2 Q5FWH6 Q5FWL0 Q5FWL3 Q5FWM2 Q5FWM3
Q5FWN8 Q5FWN9 Q5FWP2 Q5FWP4 Q5FWP8 Q5FWP9 Q5FWQ0 Q5FWQ6 Q5FWR0 Q5FWS6 Q5FWT1
Q5FWT5 Q5FWT7 Q5FWT8 Q5FWT9 Q5FWU5 Q5FWY3 Q5FWY4 Q5FX67 Q5G1S8 Q5G1T1 Q5G5C4
Q5G5T6 Q5GA22 Q5GA85 Q5GA88 Q5GA89 Q5GA90 Q5GA98 Q5GAA8 Q5GAA9 Q5GAB0 Q5GF59
Q5GF60 Q5GF62 Q5GF63 Q5GF67 Q5GF70 Q5GF76 Q5GFD8 Q5GFD9 Q5GGR6 Q5GGS4 Q5GHF7
Q5GIQ4 Q5GIQ5 Q5GIR6 Q5GIZ6 Q5GJ59 Q5GJ60 Q5GLH2 Q5GLZ8 Q5GM68 Q5GN51 Q5GRH8
Q5GRH9 Q5GRI0 Q5GRI1 Q5GRI5 Q5GRJ9 Q5GRK0 Q5GRK1 Q5GRK4 Q5GRK8 Q5GRL3 Q5GRM3
Q5GRP0 Q5GRP6 Q5GRQ2 Q5GRR4 Q5GRR7 Q5GRR9 Q5GRT2 Q5GRT5 Q5GRT9 Q5GRU0 Q5GRU5
Q5GRV1 Q5GRV2 Q5GRV6 Q5GRW6 Q5GRX3 Q5GRX8 Q5GRX9 Q5GRY3 Q5GRY6 Q5GRY9 Q5GRZ4
Q5GS00 Q5GS03 Q5GS04 Q5GS20 Q5GS25 Q5GS31 Q5GS35 Q5GS47 Q5GS56 Q5GS59 Q5GS62
Q5GS66 Q5GS68 Q5GS82 Q5GS83 Q5GS84 Q5GS87 Q5GS88 Q5GS95 Q5GS99 Q5GSA3 Q5GSA5
Q5GSB6 Q5GSB8 Q5GSC8 Q5GSD4 Q5GSD5 Q5GSD6 Q5GSD9 Q5GSE1 Q5GSG4 Q5GSH2 Q5GSH5
Q5GSI3 Q5GSI7 Q5GSJ0 Q5GSJ1 Q5GSJ3 Q5GSK9 Q5GSM4 Q5GSM7 Q5GSM8 Q5GSM9 Q5GSN3
Q5GSP0 Q5GSP9 Q5GSQ1 Q5GSR4 Q5GSS2 Q5GSS3 Q5GSS5 Q5GSS8 Q5GSS9 Q5GST1 Q5GST5
Q5GST6 Q5GST9 Q5GSU0 Q5GSU1 Q5GSU2 Q5GSU5 Q5GSU7 Q5GSU8 Q5GSU9 Q5GSV0 Q5GSV1
Q5GSV3 Q5GSV4 Q5GSV5 Q5GSV6 Q5GSV7 Q5GSV8 Q5GSW0 Q5GSW1 Q5GSW2 Q5GSW4 Q5GSW5
Q5GSW6 Q5GSW7 Q5GSW8 Q5GSY2 Q5GSY3 Q5GSZ3 Q5GT09 Q5GT12 Q5GT26 Q5GT31 Q5GT33
Q5GT34 Q5GT37 Q5GT39 Q5GT41 Q5GT50 Q5GT54 Q5GT55 Q5GT56 Q5GT66 Q5GT78 Q5GT91
Q5GT94 Q5GTA4 Q5GTA6 Q5GTB0 Q5GTB4 Q5GTC8 Q5GTD2 Q5GTD5 Q5GTE5 Q5GTF2 Q5GTF3
Q5GTG3 Q5GTG5 Q5GTG9 Q5GTH5 Q5GTI3 Q5GTI9 Q5GTJ0 Q5GTJ5 Q5GTK2 Q5GTK5 Q5GTK9
Q5GTN6 Q5GTP4 Q5GTQ0 Q5GTR2 Q5GTR3 Q5GTS0 Q5GTS7 Q5GTT1 Q5GTT5 Q5GU11 Q5GU13
Q5GUF2 Q5GUF3 Q5GUF4 Q5GUR8 Q5GUS2 Q5GUT1 Q5GUV2 Q5GUY0 Q5GUY8 Q5GV22 Q5GV34
Q5GV62 Q5GV66 Q5GV67 Q5GV68 Q5GV94 Q5GVE4 Q5GVE9 Q5GVG5 Q5GVL2 Q5GVM7 Q5GVR6
Q5GVR7 Q5GVY6 Q5GW07 Q5GW09 Q5GW16 Q5GW28 Q5GW33 Q5GW34 Q5GW37 Q5GW42 Q5GW71
Q5GW76 Q5GWB5 Q5GWB7 Q5GWB8 Q5GWB9 Q5GWE6 Q5GWF4 Q5GWI3 Q5GWL8 Q5GWQ4 Q5GWQ9
Q5GWR0 Q5GWR3 Q5GWR5 Q5GWR6 Q5GWR8 Q5GWS1 Q5GWS3 Q5GWS4 Q5GWS5 Q5GWS6 Q5GWS8
Q5GWS9 Q5GWT2 Q5GWT3 Q5GWT5 Q5GWT6 Q5GWT7 Q5GWT8 Q5GWT9 Q5GWU0 Q5GWU1 Q5GWU2
Q5GWU3 Q5GWU4 Q5GWU6 Q5GWU7 Q5GWU8 Q5GWU9 Q5GWV0 Q5GWV1 Q5GWV3 Q5GWV4 Q5GWV6
Q5GWV8 Q5GWV9 Q5GWW0 Q5GWY5 Q5GWZ8 Q5GX78 Q5GXA2 Q5GXE4 Q5GXE5 Q5GXI8 Q5GXJ5
Q5GXK1 Q5GXL3 Q5GXL9 Q5GXN9 Q5GXP5 Q5GXQ6 Q5GXR2 Q5GXR3 Q5GXR6 Q5GXR8 Q5GXU6
Q5GXU9 Q5GXV0 Q5GXV1 Q5GXV2 Q5GXV3 Q5GXX4 Q5GXY0 Q5GXY2 Q5GXY3 Q5GXY4 Q5GXY5

Q5GXY6 Q5GY22 Q5GYA0 Q5GYK0 Q5GYK6 Q5GYK7 Q5GYK8 Q5GYL1 Q5GYL2 Q5GYL7 Q5GYM0
Q5GYM1 Q5GYM8 Q5GYM9 Q5GYQ7 Q5GYR2 Q5GYV9 Q5GZ27 Q5GZ48 Q5GZ51 Q5GZ75 Q5GZ80
Q5GZ85 Q5GZ90 Q5GZ95 Q5GZ99 Q5GZA0 Q5GZB5 Q5GZD9 Q5GZF1 Q5GZF2 Q5GZR1 Q5GZR3
Q5GZR5 Q5GZR6 Q5GZT2 Q5GZT3 Q5GZV7 Q5GZV9 Q5GZY2 Q5GZY3 Q5H029 Q5H038 Q5H039
Q5H043 Q5H049 Q5H051 Q5H052 Q5H053 Q5H054 Q5H057 Q5H060 Q5H071 Q5H079 Q5H081
Q5H0A0 Q5H0A3 Q5H0A4 Q5H0A6 Q5H0A7 Q5H0F2 Q5H0G1 Q5H0H5 Q5H0K6 Q5H0K7 Q5H0K8
Q5H0K9 Q5H0L0 Q5H0L1 Q5H0L2 Q5H0L6 Q5H0N1 Q5H0N8 Q5H0Q5 Q5H0Q8 Q5H0S2 Q5H0S4
Q5H0T8 Q5H0X8 Q5H0X9 Q5H0Y0 Q5H0Z4 Q5H124 Q5H185 Q5H186 Q5H187 Q5H188 Q5H194
Q5H1A0 Q5H1C8 Q5H1D3 Q5H1E0 Q5H1E1 Q5H1E3 Q5H1E4 Q5H1E7 Q5H1F4 Q5H1J3 Q5H1Q0
Q5H1Q8 Q5H1R0 Q5H1R2 Q5H1V2 Q5H1X2 Q5H1X3 Q5H232 Q5H233 Q5H234 Q5H288 Q5H2A4
Q5H2A6 Q5H2A7 Q5H2A8 Q5H2B1 Q5H2C5 Q5H2D9 Q5H2E1 Q5H2E4 Q5H2E5 Q5H2E6 Q5H2E7
Q5H2F6 Q5H2H4 Q5H2I4 Q5H2R3 Q5H2V7 Q5H2Y1 Q5H2Z0 Q5H2Z1 Q5H2Z2 Q5H389 Q5H390
Q5H391 Q5H392 Q5H3C0 Q5H3C7 Q5H3D6 Q5H3D7 Q5H3H2 Q5H3H3 Q5H3H4 Q5H3H5 Q5H3I5
Q5H3K7 Q5H3R4 Q5H3R9 Q5H3S2 Q5H3S6 Q5H3T0 Q5H3U0 Q5H3X2 Q5H401 Q5H402 Q5H426
Q5H432 Q5H433 Q5H434 Q5H435 Q5H496 Q5H4A4 Q5H4C0 Q5H4C1 Q5H4C6 Q5H4C7 Q5H4D0
Q5H4D1 Q5H4F2 Q5H4J1 Q5H4N2 Q5H4N5 Q5H4Q7 Q5H505 Q5H522 Q5H5B9 Q5H5D0 Q5H5D4
Q5H5D8 Q5H5H7 Q5H5J1 Q5H5K1 Q5H5K4 Q5H5L9 Q5H5M1 Q5H5M2 Q5H5Q2 Q5H5R0 Q5H5R1
Q5H5T5 Q5H5T7 Q5H667 Q5H680 Q5H6B9 Q5H6C1 Q5H6C5 Q5H6D1 Q5H6D9 Q5H6E7 Q5H6H2
Q5H6Q3 Q5H6Q4 Q5H701 Q5H704 Q5H713 Q5H715 Q5H737 Q5H7N8 Q5H7P5 Q5H861 Q5H873
Q5H9B9 Q5H9I0 Q5H9K5 Q5H9L2 Q5H9L4 Q5H9M0 Q5H9N3 Q5H9T9 Q5H9U9 Q5H9Z3 Q5H9Z4
Q5H9Z6 Q5H9Z7 Q5H9Z8 Q5HA01 Q5HA25 Q5HA31 Q5HA74 Q5HA98 Q5HAB3 Q5HAC1 Q5HAD6
Q5HAD9 Q5HAF8 Q5HAF9 Q5HAG4 Q5HAI0 Q5HAI7 Q5HAJ4 Q5HAJ5 Q5HAJ6 Q5HAJ7 Q5HAK2
Q5HAK4 Q5HAK5 Q5HAK7 Q5HAK8 Q5HAL5 Q5HAN0 Q5HAN4 Q5HAP2 Q5HAP4 Q5HAS0 Q5HAS1
Q5HAS3 Q5HAS5 Q5HAS6 Q5HAS7 Q5HAS8 Q5HAS9 Q5HAT0 Q5HAT1 Q5HAT2 Q5HAT3 Q5HAT4
Q5HAT5 Q5HAT6 Q5HAT8 Q5HAT9 Q5HAU0 Q5HAU3 Q5HAU4 Q5HAU5 Q5HAU6 Q5HAU7 Q5HAV1
Q5HAV4 Q5HAV7 Q5HAW7 Q5HAW8 Q5HAY1 Q5HAZ6 Q5HB02 Q5HB13 Q5HB16 Q5HB21 Q5HB22
Q5HB23 Q5HB39 Q5HB43 Q5HB44 Q5HB45 Q5HB47 Q5HB48 Q5HB55 Q5HB57 Q5HB61 Q5HB76
Q5HB80 Q5HB81 Q5HB82 Q5HB92 Q5HB98 Q5HBA3 Q5HBA4 Q5HBC0 Q5HBC3 Q5HBC6 Q5HBC7
Q5HBD4 Q5HBF2 Q5HBG3 Q5HBG4 Q5HBH6 Q5HBH7 Q5HBH8 Q5HBI6 Q5HBJ6 Q5HBK5 Q5HBL2
Q5HBM5 Q5HBM8 Q5HBN0 Q5HBN2 Q5HBN3 Q5HBN7 Q5HBP0 Q5HBP2 Q5HBQ7 Q5HBR5 Q5HBT2
Q5HBV2 Q5HBV6 Q5HBX0 Q5HBX1 Q5HBX2 Q5HBX4 Q5HBX5 Q5HBX6 Q5HBY1 Q5HBZ2 Q5HBZ4
Q5HBZ7 Q5HC01 Q5HC04 Q5HC05 Q5HC08 Q5HC12 Q5HC13 Q5HC14 Q5HC25 Q5HC27 Q5HC38
Q5HC39 Q5HC40 Q5HC42 Q5HC44 Q5HC48 Q5HC60 Q5HC74 Q5HC82 Q5HC84 Q5HC85 Q5HC98
Q5HC99 Q5HCA2 Q5HCB3 Q5HCB5 Q5HCC9 Q5HCD7 Q5HCG1 Q5HCG4 Q5HCI1 Q5HCI2 Q5HCI3
Q5HCI4 Q5HCI5 Q5HCK7 Q5HCK8 Q5HCL7 Q5HCL8 Q5HCL9 Q5HCM1 Q5HCM2 Q5HCM3 Q5HCM4
Q5HCM5 Q5HCN2 Q5HCR2 Q5HCR3 Q5HCR5 Q5HCR6 Q5HCS0 Q5HCU1 Q5HCU5 Q5HCU6 Q5HCV0
Q5HCV3 Q5HCV4 Q5HCV5 Q5HCW9 Q5HCX3 Q5HCX7 Q5HCY6 Q5HCY8 Q5HCY9 Q5HCZ2 Q5HCZ5
Q5HCZ6 Q5HD02 Q5HD29 Q5HD30 Q5HD37 Q5HD55 Q5HD56 Q5HD73 Q5HDC9 Q5HDD1 Q5HDD9
Q5HDG4 Q5HDG9 Q5HDI3 Q5HDJ0 Q5HDJ2 Q5HDL0 Q5HDL2 Q5HDM1 Q5HDM3 Q5HDM4 Q5HDM6
Q5HDM7 Q5HDP9 Q5HDQ4 Q5HDR3 Q5HDR4 Q5HDR5 Q5HDR6 Q5HDR7 Q5HDR8 Q5HDR9 Q5HDS0
Q5HDS7 Q5HDT2 Q5HDT3 Q5HDT4 Q5HDT6 Q5HDT8 Q5HDT9 Q5HDU2 Q5HDU4 Q5HDV4 Q5HDV7
Q5HDV8 Q5HDV9 Q5HDW0 Q5HDW1 Q5HDW2 Q5HDW3 Q5HDW4 Q5HDW5 Q5HDW6 Q5HDW7 Q5HDW8
Q5HDW9 Q5HDX0 Q5HDX1 Q5HDX2 Q5HDX3 Q5HDX4 Q5HDX5 Q5HDX6 Q5HDX7 Q5HDX9 Q5HDY0
Q5HDY1 Q5HDY2 Q5HDY3 Q5HDY4 Q5HDY5 Q5HDY9 Q5HDZ0 Q5HDZ1 Q5HE00 Q5HE07 Q5HE08

Q5HE10 Q5HE11 Q5HE12 Q5HE13 Q5HE16 Q5HE19 Q5HE23 Q5HE32 Q5HE34 Q5HE43 Q5HE49
Q5HE63 Q5HE64 Q5HE70 Q5HE72 Q5HE73 Q5HE75 Q5HE76 Q5HE78 Q5HE80 Q5HE81 Q5HE82
Q5HE85 Q5HE86 Q5HE87 Q5HE88 Q5HEA0 Q5HEA5 Q5HEA7 Q5HEA8 Q5HEB7 Q5HEB9 Q5HED1
Q5HED2 Q5HED3 Q5HED6 Q5HED9 Q5HEE0 Q5HEE1 Q5HEE2 Q5HEE3 Q5HEE4 Q5HEE5 Q5HEE8
Q5HEF2 Q5HEF5 Q5HEH1 Q5HEH2 Q5HEJ8 Q5HEK1 Q5HEK7 Q5HEK9 Q5HEL4 Q5HEL7 Q5HEL8
Q5HEM1 Q5HEM2 Q5HEM3 Q5HEM5 Q5HEM7 Q5HEN0 Q5HEN6 Q5HEP4 Q5HEP9 Q5HEQ1 Q5HEQ7
Q5HER0 Q5HER3 Q5HES4 Q5HET0 Q5HEU2 Q5HEU3 Q5HEY2 Q5HEY8 Q5HEY9 Q5HEZ8 Q5HF01
Q5HF07 Q5HF08 Q5HF12 Q5HF16 Q5HF23 Q5HF24 Q5HF31 Q5HF34 Q5HF38 Q5HF39 Q5HF42
Q5HF45 Q5HF54 Q5HF59 Q5HF61 Q5HF63 Q5HF64 Q5HF65 Q5HF67 Q5HF71 Q5HF75 Q5HF76
Q5HF79 Q5HF85 Q5HF86 Q5HF87 Q5HF90 Q5HF92 Q5HF93 Q5HF94 Q5HF97 Q5HF98 Q5HF99
Q5HFA0 Q5HFA2 Q5HFA4 Q5HFA5 Q5HFA8 Q5HFB1 Q5HFB6 Q5HFB8 Q5HFB9 Q5HFC0 Q5HFC1
Q5HFC2 Q5HFC3 Q5HFC4 Q5HFC8 Q5HFD0 Q5HFD2 Q5HFD3 Q5HFD7 Q5HFE1 Q5HFE4 Q5HFE5
Q5HFE6 Q5HFE7 Q5HFF1 Q5HFF2 Q5HFF7 Q5HFG2 Q5HFG5 Q5HFG7 Q5HFH5 Q5HFH8 Q5HFH9
Q5HFI0 Q5HFI1 Q5HFI2 Q5HFI5 Q5HFJ0 Q5HFJ4 Q5HFJ5 Q5HFJ7 Q5HFJ8 Q5HFJ9 Q5HFK1
Q5HFK3 Q5HFK6 Q5HFK7 Q5HFK9 Q5HFM1 Q5HFM2 Q5HFM3 Q5HFM4 Q5HFN0 Q5HFP7 Q5HFP8
Q5HFP9 Q5HFQ1 Q5HFQ8 Q5HFR2 Q5HFR8 Q5HFR9 Q5HFS5 Q5HFS7 Q5HFS8 Q5HFU6 Q5HFU7
Q5HFU8 Q5HFV0 Q5HFV2 Q5HFV4 Q5HFV7 Q5HFV8 Q5HFV9 Q5HFW1 Q5HFW6 Q5HFW8 Q5HFW9
Q5HFX7 Q5HFX8 Q5HFY4 Q5HFY5 Q5HFZ4 Q5HFZ5 Q5HFZ6 Q5HFZ7 Q5HG00 Q5HG06 Q5HG07
Q5HG10 Q5HG14 Q5HG16 Q5HG18 Q5HG20 Q5HG21 Q5HG22 Q5HG23 Q5HG24 Q5HG25 Q5HG29
Q5HG44 Q5HG45 Q5HG46 Q5HG47 Q5HG48 Q5HG54 Q5HG56 Q5HG58 Q5HG62 Q5HG65 Q5HG67
Q5HG69 Q5HG72 Q5HG73 Q5HG77 Q5HG78 Q5HG83 Q5HG84 Q5HG85 Q5HG86 Q5HG90 Q5HGC3
Q5HGC7 Q5HGC9 Q5HGD1 Q5HGD2 Q5HGD5 Q5HGD6 Q5HGD9 Q5HGE6 Q5HGF6 Q5HGF7 Q5HGF8
Q5HGG0 Q5HGG1 Q5HGG2 Q5HGG6 Q5HGG7 Q5HGG8 Q5HGH1 Q5HGH2 Q5HGH3 Q5HGH4 Q5HGH6
Q5HGH7 Q5HGH8 Q5HGH9 Q5HGI0 Q5HGI1 Q5HGI2 Q5HGI6 Q5HGI7 Q5HGI8 Q5HGJ1 Q5HGJ2
Q5HGJ3 Q5HGJ4 Q5HGJ9 Q5HGK6 Q5HGK7 Q5HGK9 Q5HGL4 Q5HGL6 Q5HGL7 Q5HGM2 Q5HGM3
Q5HGM7 Q5HGM8 Q5HGM9 Q5HGN0 Q5HGN1 Q5HGN2 Q5HGN4 Q5HGN8 Q5HGP2 Q5HGP4 Q5HGP8
Q5HGQ2 Q5HGQ3 Q5HGQ4 Q5HGQ5 Q5HGR2 Q5HGR3 Q5HGT2 Q5HGT3 Q5HGT7 Q5HGT9 Q5HGU0
Q5HGU4 Q5HGU5 Q5HGU6 Q5HGU8 Q5HGV6 Q5HGV8 Q5HGV9 Q5HGW2 Q5HGX2 Q5HGX8 Q5HGY7
Q5HGY9 Q5HGZ0 Q5HGZ1 Q5HGZ3 Q5HGZ4 Q5HGZ5 Q5HH10 Q5HH11 Q5HH12 Q5HH13 Q5HH14
Q5HH15 Q5HH16 Q5HH18 Q5HH19 Q5HH21 Q5HH30 Q5HH37 Q5HH38 Q5HH40 Q5HH71 Q5HH78
Q5HH80 Q5HH83 Q5HH86 Q5HH87 Q5HH88 Q5HHB0 Q5HHB4 Q5HHB6 Q5HHB7 Q5HHB8 Q5HHC2
Q5HHC4 Q5HHC5 Q5HHC7 Q5HHC8 Q5HHD1 Q5HHE6 Q5HHF2 Q5HHF6 Q5HHG0 Q5HHG4 Q5HHG8
Q5HHH0 Q5HHH4 Q5HHK8 Q5HHL2 Q5HHL3 Q5HHN6 Q5HHP2 Q5HHP3 Q5HHP4 Q5HHP5 Q5HHP9
Q5HHQ0 Q5HHQ1 Q5HHQ3 Q5HHQ4 Q5HHQ6 Q5HHQ8 Q5HHQ9 Q5HHR0 Q5HHR5 Q5HHR8 Q5HHS7
Q5HHT2 Q5HHU1 Q5HHU4 Q5HHU8 Q5HHU9 Q5HHV8 Q5HHY2 Q5HHY7 Q5HHZ9 Q5HI00 Q5HI28
Q5HI51 Q5HI60 Q5HI63 Q5HI83 Q5HI88 Q5HI89 Q5HI95 Q5HI96 Q5HI97 Q5HI99 Q5HIA0
Q5HIA5 Q5HIA6 Q5HIA9 Q5HIB5 Q5HIC1 Q5HIC2 Q5HIC3 Q5HIC4 Q5HIC5 Q5HIC7 Q5HIC8
Q5HIC9 Q5HID0 Q5HID1 Q5HID2 Q5HID3 Q5HID5 Q5HID6 Q5HID7 Q5HID8 Q5HID9 Q5HIE3
Q5HIE5 Q5HIE6 Q5HIE7 Q5HIF0 Q5HIF2 Q5HIF4 Q5HIF5 Q5HIF7 Q5HIG0 Q5HIG1 Q5HIG2
Q5HIG3 Q5HIG5 Q5HIG6 Q5HIH2 Q5HIH3 Q5HIH4 Q5HIH5 Q5HIH8 Q5HII1 Q5HII3 Q5HII6
Q5HII9 Q5HIJ1 Q5HIJ5 Q5HIJ7 Q5HIJ8 Q5HIQ6 Q5HIQ7 Q5HIQ9 Q5HIR0 Q5HIR4 Q5HIR5
Q5HIR6 Q5HIS7 Q5HIS8 Q5HIS9 Q5HIT8 Q5HIU0 Q5HIW9 Q5HJ26 Q5HJ50 Q5HJ53 Q5HJA7
Q5HJB7 Q5HJC1 Q5HJC5 Q5HJD7 Q5HJF3 Q5HJF4 Q5HJG8 Q5HJH8 Q5HJI1 Q5HJI8 Q5HJI9
Q5HJJ0 Q5HJJ1 Q5HJK3 Q5HJK5 Q5HJM9 Q5HJN0 Q5HJN5 Q5HJP2 Q5HJQ3 Q5HJQ7 Q5HJR1

Q5HJR8 Q5HJT5 Q5HJX8 Q5HJY1 Q5HJY7 Q5HJY8 Q5HJZ0 Q5HJZ1 Q5HJZ2 Q5HJZ4 Q5HJZ5
Q5HJZ6 Q5HJZ9 Q5HK00 Q5HK02 Q5HK03 Q5HK04 Q5HK07 Q5HK14 Q5HK16 Q5HK23 Q5HK32
Q5HK76 Q5HKB7 Q5HKD5 Q5HKD6 Q5HKF0 Q5HKF3 Q5HKG6 Q5HKH9 Q5HKI0 Q5HKI1 Q5HKJ3
Q5HKJ6 Q5HKJ7 Q5HKM8 Q5HKN2 Q5HKN6 Q5HKN7 Q5HKN8 Q5HKN9 Q5HKP0 Q5HKP1 Q5HKP2
Q5HKP3 Q5HKQ1 Q5HKS7 Q5HKU2 Q5HKU6 Q5HKZ3 Q5HKZ5 Q5HKZ7 Q5HL01 Q5HL02 Q5HL11
Q5HL18 Q5HL19 Q5HL21 Q5HL31 Q5HL35 Q5HL36 Q5HL37 Q5HL55 Q5HL59 Q5HL86 Q5HL88
Q5HLA0 Q5HLA1 Q5HLA3 Q5HLB1 Q5HLB4 Q5HLD8 Q5HLE0 Q5HLI0 Q5HLK5 Q5HLK8 Q5HLN0
Q5HLP8 Q5HLQ3 Q5HLQ6 Q5HLQ9 Q5HLR4 Q5HLT3 Q5HLU4 Q5HLV7 Q5HLV8 Q5HLV9 Q5HLW0
Q5HLW1 Q5HLW2 Q5HLW3 Q5HLX4 Q5HLX5 Q5HLX6 Q5HLX8 Q5HLY0 Q5HLY1 Q5HLZ4 Q5HLZ8
Q5HLZ9 Q5HM00 Q5HM01 Q5HM02 Q5HM03 Q5HM04 Q5HM05 Q5HM06 Q5HM07 Q5HM08 Q5HM09
Q5HM10 Q5HM11 Q5HM12 Q5HM13 Q5HM14 Q5HM15 Q5HM16 Q5HM17 Q5HM18 Q5HM20 Q5HM21
Q5HM22 Q5HM23 Q5HM24 Q5HM25 Q5HM26 Q5HM30 Q5HM31 Q5HM32 Q5HM33 Q5HM34 Q5HM35
Q5HM36 Q5HM37 Q5HM38 Q5HM41 Q5HM44 Q5HM47 Q5HM56 Q5HM57 Q5HM59 Q5HM67 Q5HM69
Q5HM84 Q5HM85 Q5HM86 Q5HM92 Q5HM94 Q5HM95 Q5HM97 Q5HM98 Q5HMA0 Q5HMA2 Q5HMA3
Q5HMA4 Q5HMA8 Q5HMB0 Q5HMB1 Q5HMC2 Q5HMC7 Q5HMC9 Q5HMD0 Q5HMD8 Q5HME0 Q5HME5
Q5HME6 Q5HME7 Q5HME9 Q5HMF0 Q5HMF3 Q5HMF5 Q5HMF6 Q5HMF7 Q5HMF8 Q5HMF9 Q5HMG0
Q5HMG3 Q5HMG7 Q5HMH1 Q5HMP3 Q5HMP6 Q5HMV4 Q5HMZ0 Q5HMZ1 Q5HN15 Q5HN17 Q5HN23
Q5HN26 Q5HN29 Q5HN30 Q5HN33 Q5HN34 Q5HN35 Q5HN37 Q5HN39 Q5HN41 Q5HN46 Q5HN53
Q5HN54 Q5HN59 Q5HN61 Q5HN67 Q5HN71 Q5HN74 Q5HN85 Q5HN91 Q5HNA4 Q5HNA5 Q5HNB2
Q5HNB7 Q5HNB8 Q5HNE3 Q5HNE4 Q5HNF1 Q5HNF9 Q5HNG0 Q5HNG2 Q5HNG6 Q5HNG9 Q5HNH3
Q5HNH4 Q5HNH6 Q5HNI5 Q5HNJ0 Q5HNJ2 Q5HNJ4 Q5HNJ5 Q5HNJ6 Q5HNJ7 Q5HNJ8 Q5HNK2
Q5HNK6 Q5HNK7 Q5HNL1 Q5HNL6 Q5HNL7 Q5HNL8 Q5HNM1 Q5HNM3 Q5HNM4 Q5HNM5 Q5HNM8
Q5HNM9 Q5HNN0 Q5HNN1 Q5HNN3 Q5HNN5 Q5HNN6 Q5HNN9 Q5HNQ4 Q5HNQ6 Q5HNQ7 Q5HNQ8
Q5HNQ9 Q5HNR0 Q5HNR1 Q5HNR2 Q5HNR6 Q5HNR8 Q5HNR9 Q5HNS1 Q5HNS2 Q5HNS9 Q5HNT2
Q5HNT3 Q5HNT4 Q5HNT5 Q5HNT9 Q5HNU0 Q5HNU6 Q5HNU8 Q5HNV1 Q5HNV3 Q5HNW1 Q5HNW4
Q5HNW5 Q5HNW6 Q5HNW7 Q5HNW8 Q5HNX1 Q5HNX7 Q5HNY1 Q5HNY2 Q5HNY5 Q5HNY6 Q5HNY7
Q5HNY9 Q5HNZ1 Q5HNZ4 Q5HNZ5 Q5HNZ8 Q5HP11 Q5HP12 Q5HP13 Q5HP14 Q5HP16 Q5HP20
Q5HP21 Q5HP22 Q5HP23 Q5HP24 Q5HP28 Q5HP29 Q5HP30 Q5HP32 Q5HP38 Q5HP42 Q5HP47
Q5HP48 Q5HP53 Q5HP55 Q5HP56 Q5HP68 Q5HP69 Q5HP70 Q5HP74 Q5HP76 Q5HP77 Q5HP78
Q5HP79 Q5HP81 Q5HP86 Q5HP88 Q5HP89 Q5HP94 Q5HP96 Q5HP97 Q5HPA7 Q5HPB1 Q5HPB4
Q5HPB6 Q5HPC6 Q5HPC7 Q5HPD3 Q5HPD6 Q5HPE0 Q5HPE4 Q5HPE5 Q5HPE6 Q5HPE7 Q5HPF1
Q5HPG5 Q5HPG6 Q5HPG9 Q5HPH0 Q5HPH1 Q5HPH2 Q5HPH3 Q5HPH6 Q5HPH8 Q5HPI3 Q5HPI6
Q5HPI8 Q5HPJ0 Q5HPJ3 Q5HPJ4 Q5HPJ9 Q5HPK0 Q5HPK5 Q5HPK6 Q5HPK7 Q5HPK8 Q5HPL2
Q5HPN2 Q5HPN8 Q5HPP0 Q5HPP1 Q5HPP4 Q5HPP5 Q5HPP8 Q5HPQ6 Q5HPR6 Q5HPR7 Q5HPR8
Q5HPS0 Q5HPS1 Q5HPS2 Q5HPS6 Q5HPS7 Q5HPS8 Q5HPT1 Q5HPT2 Q5HPT3 Q5HPT4 Q5HPT5
Q5HPT7 Q5HPT8 Q5HPT9 Q5HPU0 Q5HPU1 Q5HPU2 Q5HPU4 Q5HPU5 Q5HPU6 Q5HPV0 Q5HPV1
Q5HPV2 Q5HPV3 Q5HPV8 Q5HPW4 Q5HPW5 Q5HPW7 Q5HPX0 Q5HPX3 Q5HPX5 Q5HPX6 Q5HPY0
Q5HPY1 Q5HPY6 Q5HPY7 Q5HPY8 Q5HPY9 Q5HPZ0 Q5HPZ1 Q5HPZ3 Q5HPZ9 Q5HQ03 Q5HQ05
Q5HQ09 Q5HQ13 Q5HQ14 Q5HQ15 Q5HQ16 Q5HQ23 Q5HQ24 Q5HQ28 Q5HQ29 Q5HQ30 Q5HQ34
Q5HQ35 Q5HQ36 Q5HQ39 Q5HQ41 Q5HQ42 Q5HQ45 Q5HQ56 Q5HQ63 Q5HQ72 Q5HQ74 Q5HQ75
Q5HQ76 Q5HQ78 Q5HQ79 Q5HQ80 Q5HQ96 Q5HQ97 Q5HQ98 Q5HQ99 Q5HQA0 Q5HQA1 Q5HQA2
Q5HQA4 Q5HQA5 Q5HQA7 Q5HQC3 Q5HQC5 Q5HQE4 Q5HQE6 Q5HQE9 Q5HQG2 Q5HQG4 Q5HQG8
Q5HQH1 Q5HQH3 Q5HQH4 Q5HQI5 Q5HQI9 Q5HQJ3 Q5HQJ4 Q5HQJ5 Q5HQJ9 Q5HQK0 Q5HQK1
Q5HQK4 Q5HQK8 Q5HQM3 Q5HQN0 Q5HQN3 Q5HQN7 Q5HQP8 Q5HQQ0 Q5HQQ4 Q5HQR3 Q5HQR7

Q5HQR8 Q5HQU5 Q5HQV1 Q5HQV2 Q5HQV3 Q5HQV4 Q5HQV8 Q5HQW0 Q5HQW1 Q5HQW3 Q5HQW4
Q5HQW8 Q5HQW9 Q5HQX0 Q5HQX5 Q5HQX7 Q5HQY6 Q5HQZ1 Q5HR00 Q5HR02 Q5HR07 Q5HR08
Q5HR20 Q5HR46 Q5HR57 Q5HR69 Q5HR71 Q5HRD1 Q5HRD6 Q5HRE0 Q5HRF2 Q5HRF6 Q5HRF7
Q5HRF8 Q5HRG5 Q5HRG7 Q5HRG9 Q5HRH2 Q5HRH3 Q5HRH7 Q5HRH8 Q5HRI1 Q5HRI4 Q5HRI7
Q5HRJ7 Q5HRJ8 Q5HRJ9 Q5HRK4 Q5HRK5 Q5HRK6 Q5HRK7 Q5HRK9 Q5HRL0 Q5HRL2 Q5HRL3
Q5HRL4 Q5HRL5 Q5HRL6 Q5HRL8 Q5HRM1 Q5HRM3 Q5HRM4 Q5HRM5 Q5HRM8 Q5HRM9 Q5HRN1
Q5HRN4 Q5HRN5 Q5HRN7 Q5HRN9 Q5HRP0 Q5HRP1 Q5HRP2 Q5HRP4 Q5HRP5 Q5HRQ2 Q5HRQ3
Q5HRQ4 Q5HRQ5 Q5HRQ7 Q5HRR0 Q5HRR2 Q5HRR5 Q5HRR7 Q5HRR9 Q5HRS3 Q5HRS7 Q5HRS8
Q5HRX1 Q5HRX2 Q5HRX4 Q5HRX5 Q5HRX9 Q5HRY1 Q5HRY2 Q5HRZ3 Q5HRZ5 Q5HS05 Q5HS07
Q5HS34 Q5HS35 Q5HS36 Q5HS37 Q5HS38 Q5HS68 Q5HS71 Q5HS73 Q5HS76 Q5HS77 Q5HS78
Q5HS79 Q5HS83 Q5HS85 Q5HS89 Q5HS91 Q5HS92 Q5HS93 Q5HS94 Q5HS95 Q5HS96 Q5HS97
Q5HS98 Q5HS99 Q5HSA0 Q5HSA1 Q5HSA2 Q5HSA3 Q5HSA4 Q5HSA5 Q5HSA6 Q5HSA7 Q5HSB2
Q5HSB7 Q5HSC0 Q5HSF7 Q5HSF8 Q5HSF9 Q5HSH9 Q5HSI0 Q5HSI4 Q5HSI8 Q5HSJ0 Q5HSJ1
Q5HSJ2 Q5HSJ3 Q5HSJ4 Q5HSJ5 Q5HSJ6 Q5HSJ7 Q5HSJ8 Q5HSJ9 Q5HSK0 Q5HSK1 Q5HSP5
Q5HSQ5 Q5HSQ6 Q5HSS7 Q5HST3 Q5HSV2 Q5HSX3 Q5HSX6 Q5HSX7 Q5HSX8 Q5HT13 Q5HT15
Q5HT16 Q5HT33 Q5HT36 Q5HT48 Q5HT53 Q5HT61 Q5HT65 Q5HT67 Q5HT90 Q5HTB5 Q5HTB9
Q5HTC1 Q5HTI5 Q5HTI9 Q5HTJ0 Q5HTJ2 Q5HTJ5 Q5HTK0 Q5HTK3 Q5HTK8 Q5HTL1 Q5HTL3
Q5HTL6 Q5HTL9 Q5HTM4 Q5HTM5 Q5HTP2 Q5HTP3 Q5HTR3 Q5HTR7 Q5HTS6 Q5HTT2 Q5HTT3
Q5HTT7 Q5HTT9 Q5HTU2 Q5HTW1 Q5HTY7 Q5HTZ5 Q5HTZ7 Q5HTZ9 Q5HU00 Q5HU02 Q5HU06
Q5HU13 Q5HU23 Q5HU29 Q5HU32 Q5HU34 Q5HU41 Q5HU43 Q5HU47 Q5HU51 Q5HU53 Q5HU56
Q5HU66 Q5HUF2 Q5HUF4 Q5HUG5 Q5HUG8 Q5HUG9 Q5HUI4 Q5HUJ8 Q5HUJ9 Q5HUK3 Q5HUK4
Q5HUK6 Q5HUM7 Q5HUM8 Q5HUN2 Q5HUP6 Q5HUQ3 Q5HUR1 Q5HUR2 Q5HUR3 Q5HUR4 Q5HUS3
Q5HUT8 Q5HUU1 Q5HUU3 Q5HUV1 Q5HUV6 Q5HUW5 Q5HUW9 Q5HUX2 Q5HUY2 Q5HUY6 Q5HUZ0
Q5HUZ3 Q5HUZ4 Q5HV17 Q5HV25 Q5HV26 Q5HV27 Q5HV33 Q5HV34 Q5HV67 Q5HV68 Q5HV69
Q5HV71 Q5HV77 Q5HV88 Q5HV92 Q5HV95 Q5HVA1 Q5HVA9 Q5HVB0 Q5HVB1 Q5HVB2 Q5HVC1
Q5HVD0 Q5HVD1 Q5HVD2 Q5HVD4 Q5HVD6 Q5HVD9 Q5HVI0 Q5HVI4 Q5HVI9 Q5HVJ3 Q5HVL5
Q5HVM1 Q5HVM3 Q5HVM4 Q5HVN3 Q5HVP5 Q5HVQ1 Q5HVQ6 Q5HVQ7 Q5HVR0 Q5HVR3 Q5HVX6
Q5HVX7 Q5HVX8 Q5HVY8 Q5HVY9 Q5HVZ0 Q5HVZ1 Q5HVZ2 Q5HVZ3 Q5HVZ6 Q5HVZ7 Q5HW09
Q5HW14 Q5HW17 Q5HW32 Q5HW34 Q5HW65 Q5HW66 Q5HW69 Q5HW73 Q5HW78 Q5HW79 Q5HW81
Q5HW84 Q5HW85 Q5HW86 Q5HW93 Q5HW97 Q5HWA7 Q5HWB8 Q5HWB9 Q5HWC0 Q5HWD9 Q5HWE1
Q5HWE5 Q5HWE6 Q5HWE7 Q5HWF0 Q5HWF4 Q5HWF9 Q5HWG0 Q5HWG3 Q5HWH3 Q5HWH4 Q5HWH5
Q5HWH7 Q5HWI0 Q5HWI6 Q5HWL3 Q5HWL9 Q5HWM0 Q5HWM8 Q5HWM9 Q5HWN6 Q5HWN8 Q5HWW2
Q5HWW3 Q5HWX1 Q5HWX4 Q5HWX5 Q5HWX6 Q5HX00 Q5HX04 Q5HX10 Q5HX17 Q5HX28 Q5HX29
Q5HX30 Q5HX32 Q5HX40 Q5HX45 Q5HX68 Q5HX70 Q5HX71 Q5HX72 Q5HX80 Q5HX97 Q5HXB2
Q5HXD1 Q5HXD2 Q5HXE2 Q5HXE4 Q5HXE7 Q5HXF5 Q5HXU1 Q5HY64 Q5HY92 Q5HY98 Q5HYC2
Q5HYK9 Q5HYM0 Q5HYN5 Q5HYR2 Q5HYW2 Q5HYW3 Q5HZ00 Q5HZ05 Q5HZ09 Q5HZ54 Q5HZ60
Q5HZ68 Q5HZ97 Q5HZA3 Q5HZA4 Q5HZB6 Q5HZE4 Q5HZF2 Q5HZH2 Q5HZJ0 Q5HZK1 Q5HZL1
Q5HZL4 Q5HZL9 Q5HZM1 Q5HZM3 Q5HZM6 Q5HZN1 Q5HZN8 Q5HZN9 Q5HZP1 Q5HZQ8 Q5HZT0
Q5HZV9 Q5HZZ6 Q5I019 Q5I033 Q5I050 Q5I082 Q5I085 Q5I0A0 Q5I0B4 Q5I0B5 Q5I0C3
Q5I0C5 Q5I0D1 Q5I0D5 Q5I0D7 Q5I0E2 Q5I0E3 Q5I0E6 Q5I0E8 Q5I0F1 Q5I0G3 Q5I0H5
Q5I0I5 Q5I0J8 Q5I0J9 Q5I0K3 Q5I0K5 Q5I0K8 Q5I0L3 Q5I0L4 Q5I0L6 Q5I0L7 Q5I0M0
Q5I0M2 Q5I0P2 Q5I0R4 Q5I0S8 Q5I0X4 Q5I0X7 Q5I123 Q5I124 Q5I125 Q5I126 Q5I127
Q5I128 Q5I129 Q5I131 Q5I132 Q5I133 Q5I134 Q5I135 Q5I136 Q5I137 Q5I138 Q5I139
Q5I140 Q5I141 Q5I142 Q5I143 Q5I144 Q5I145 Q5I147 Q5I148 Q5I149 Q5I150 Q5I151

Q5I152 Q5I153 Q5I154 Q5I155 Q5I156 Q5I157 Q5I158 Q5I159 Q5I160 Q5I162 Q5I163
Q5I1X5 Q5I2A0 Q5I2D0 Q5I2J3 Q5I2P5 Q5I4H3 Q5I597 Q5I598 Q5I5J1 Q5I6K1 Q5I6K6
Q5I7K3 Q5I7K9 Q5IBC5 Q5IBH6 Q5IBH7 Q5ICL9 Q5IFJ7 Q5IFK1 Q5IFN2 Q5IH13 Q5IH14
Q5IH81 Q5IR70 Q5IS44 Q5IS55 Q5IS58 Q5IS68 Q5IS79 Q5IS89 Q5ISC6 Q5ISW3 Q5ISW5
Q5IW36 Q5IW40 Q5J1R2 Q5J1R4 Q5J2D0 Q5J2U8 Q5J2V4 Q5J2V8 Q5J2W0 Q5J2W2 Q5J2W5
Q5J2X4 Q5J2X6 Q5J2Z9 Q5J300 Q5J306 Q5J308 Q5J309 Q5J3Q5 Q5J3W6 Q5J907 Q5JBK6
Q5JCX1 Q5JCX3 Q5JCY1 Q5JCY6 Q5JCY8 Q5JCZ1 Q5JD03 Q5JD05 Q5JD25 Q5JD26 Q5JD29
Q5JD31 Q5JD38 Q5JD44 Q5JD45 Q5JD67 Q5JDB0 Q5JDB6 Q5JDB7 Q5JDB8 Q5JDC5 Q5JDC7
Q5JDD5 Q5JDG3 Q5JDG5 Q5JDG6 Q5JDG7 Q5JDG9 Q5JDH1 Q5JDH2 Q5JDH3 Q5JDH4 Q5JDH5
Q5JDH6 Q5JDH7 Q5JDH8 Q5JDH9 Q5JDI0 Q5JDI6 Q5JDJ0 Q5JDJ1 Q5JDJ4 Q5JDJ7 Q5JDJ9
Q5JDK8 Q5JDL3 Q5JDM5 Q5JDM6 Q5JDM8 Q5JDN6 Q5JDS7 Q5JDS9 Q5JDU3 Q5JDU6 Q5JDW8
Q5JDW9 Q5JDZ7 Q5JE04 Q5JE15 Q5JE20 Q5JE31 Q5JE34 Q5JE35 Q5JE38 Q5JE39 Q5JE48
Q5JE49 Q5JE50 Q5JE63 Q5JE64 Q5JE70 Q5JE75 Q5JE78 Q5JE80 Q5JE88 Q5JEC9 Q5JED2
Q5JEF8 Q5JEG5 Q5JEK6 Q5JEK8 Q5JEL5 Q5JEP3 Q5JEQ6 Q5JER0 Q5JER1 Q5JER9 Q5JES6
Q5JET0 Q5JET2 Q5JET4 Q5JET9 Q5JEW1 Q5JEW3 Q5JEW5 Q5JEX8 Q5JEY0 Q5JEZ2 Q5JF06
Q5JF09 Q5JF17 Q5JF22 Q5JF28 Q5JF30 Q5JF32 Q5JF37 Q5JF39 Q5JF42 Q5JF45 Q5JF49
Q5JF63 Q5JF83 Q5JF85 Q5JF89 Q5JFD3 Q5JFE1 Q5JFE7 Q5JFG9 Q5JFH5 Q5JFI4 Q5JFJ9
Q5JFK6 Q5JFK7 Q5JFL6 Q5JFL7 Q5JFL8 Q5JFM1 Q5JFM4 Q5JFM9 Q5JFN1 Q5JFN5 Q5JFN7
Q5JFN8 Q5JFP1 Q5JFP4 Q5JFR1 Q5JFR2 Q5JFR5 Q5JFR6 Q5JFT1 Q5JFT2 Q5JFT3 Q5JFT4
Q5JFU6 Q5JFU8 Q5JFU9 Q5JFV0 Q5JFV1 Q5JFV6 Q5JFV7 Q5JFW1 Q5JFW3 Q5JFW4 Q5JFX1
Q5JFX3 Q5JFX9 Q5JFZ3 Q5JFZ4 Q5JFZ5 Q5JFZ7 Q5JG00 Q5JG26 Q5JG28 Q5JG47 Q5JGA9
Q5JGB4 Q5JGB7 Q5JGC2 Q5JGC8 Q5JGD1 Q5JGD6 Q5JGD9 Q5JGE1 Q5JGE6 Q5JGF0 Q5JGF1
Q5JGF3 Q5JGG6 Q5JGH7 Q5JGJ4 Q5JGJ7 Q5JGK6 Q5JGK8 Q5JGM4 Q5JGM7 Q5JGN0 Q5JGN1
Q5JGN3 Q5JGN7 Q5JGN9 Q5JGP4 Q5JGQ6 Q5JGR3 Q5JGR4 Q5JGR5 Q5JGR7 Q5JGR9 Q5JGT3
Q5JGT4 Q5JGT5 Q5JGT6 Q5JGT7 Q5JGV3 Q5JGV4 Q5JGV6 Q5JGX5 Q5JGX6 Q5JH10 Q5JH13
Q5JH14 Q5JH16 Q5JH21 Q5JH26 Q5JH27 Q5JH30 Q5JH34 Q5JH35 Q5JH36 Q5JH43 Q5JH47
Q5JH51 Q5JH53 Q5JH68 Q5JH71 Q5JH72 Q5JH75 Q5JH78 Q5JH81 Q5JH82 Q5JH90 Q5JHA3
Q5JHB2 Q5JHB5 Q5JHC0 Q5JHC1 Q5JHE8 Q5JHF1 Q5JHF4 Q5JHG1 Q5JHG3 Q5JHG5 Q5JHG7
Q5JHH2 Q5JHL2 Q5JHL8 Q5JHM9 Q5JHN0 Q5JHN1 Q5JHN3 Q5JHN5 Q5JHP1 Q5JHP2 Q5JHP5
Q5JHQ3 Q5JHR7 Q5JHS5 Q5JHT1 Q5JHT4 Q5JHU5 Q5JI02 Q5JI16 Q5JI17 Q5JI21 Q5JI38
Q5JI42 Q5JI44 Q5JI47 Q5JI51 Q5JI54 Q5JI69 Q5JI74 Q5JI77 Q5JI79 Q5JI92 Q5JI98
Q5JIB3 Q5JIB9 Q5JID8 Q5JIE0 Q5JIE1 Q5JIE3 Q5JIE6 Q5JIG7 Q5JIH3 Q5JIH9 Q5JIK2
Q5JIL7 Q5JIM2 Q5JIQ0 Q5JIR5 Q5JIR6 Q5JIR7 Q5JIT0 Q5JIU9 Q5JIX0 Q5JIY3 Q5JIY7
Q5JIY8 Q5JIY9 Q5JIZ3 Q5JIZ8 Q5JJ31 Q5JJ32 Q5JJ38 Q5JJ62 Q5JJ64 Q5JJ65 Q5JJ72
Q5JJ73 Q5JJ82 Q5JJ90 Q5JJ91 Q5JJB4 Q5JJB8 Q5JJC3 Q5JJC9 Q5JJD0 Q5JJD2 Q5JJD3
Q5JJE2 Q5JJE3 Q5JJE4 Q5JJE8 Q5JJF1 Q5JJF2 Q5JJF3 Q5JJF4 Q5JJF5 Q5JJF6 Q5JJF7
Q5JJF8 Q5JJF9 Q5JJG0 Q5JJG1 Q5JJG2 Q5JJG3 Q5JJG4 Q5JJG5 Q5JJG7 Q5JJG8 Q5JJG9
Q5JJH0 Q5JJH2 Q5JJY4 Q5JK20 Q5JK24 Q5JK52 Q5JK68 Q5JK84 Q5JK90 Q5JKB0 Q5JKF2
Q5JKN2 Q5JKW1 Q5JL96 Q5JLA7 Q5JLB5 Q5JLR7 Q5JM82 Q5JME8 Q5JMF2 Q5JMK6 Q5JMM1
Q5JMR9 Q5JN07 Q5JNA1 Q5JNS0 Q5JNT2 Q5JNT6 Q5JNZ3 Q5JNZ5 Q5JPB2 Q5JPF3 Q5JPH6
Q5JPI3 Q5JQC4 Q5JQF7 Q5JQF8 Q5JQS6 Q5JR12 Q5JR59 Q5JR98 Q5JRC9 Q5JRK9 Q5JRX3
Q5JS54 Q5JSQ8 Q5JSS6 Q5JST6 Q5JT82 Q5JTD7 Q5JTN6 Q5JTY5 Q5JTZ5 Q5JTZ9 Q5JU00
Q5JUK2 Q5JUK9 Q5JUQ0 Q5JUR7 Q5JUW0 Q5JUX0 Q5JVF3 Q5JVG2 Q5JVG8 Q5JVL4 Q5JVX7
Q5JWF8 Q5JXB2 Q5JY77 Q5JYT7 Q5K037 Q5K130 Q5K131 Q5K2C1 Q5K2C4 Q5K2K3 Q5K2K6

Q5K2K7 Q5K2N3 Q5K2N9 Q5K2P2 Q5K2P4 Q5K2P6 Q5K471 Q5K472 Q5K473 Q5K474 Q5K486
Q5K4R0 Q5K5B6 Q5K5C1 Q5K654 Q5K8L4 Q5KBP2 Q5KC16 Q5KD76 Q5KEG8 Q5KEQ1 Q5KJ55
Q5KMX3 Q5KMX8 Q5KP44 Q5KPF3 Q5KPJ5 Q5KPU2 Q5KQA1 Q5KQI4 Q5KQI6 Q5KQL9 Q5KR05
Q5KR47 Q5KR48 Q5KR49 Q5KSN4 Q5KSN5 Q5KSV0 Q5KTM9 Q5KTN0 Q5KTS7 Q5KU05 Q5KU53
Q5KU54 Q5KU57 Q5KU58 Q5KU59 Q5KU60 Q5KU69 Q5KU71 Q5KU74 Q5KU76 Q5KU84 Q5KUC1
Q5KUC3 Q5KUD0 Q5KUD5 Q5KUD7 Q5KUD9 Q5KUF0 Q5KUG0 Q5KUG2 Q5KUG6 Q5KUG7 Q5KUH0
Q5KUH1 Q5KUH3 Q5KUI1 Q5KUI2 Q5KUI3 Q5KUL0 Q5KUX2 Q5KV67 Q5KV68 Q5KV89 Q5KVB2
Q5KVB5 Q5KVB9 Q5KVC4 Q5KVC5 Q5KVC6 Q5KVC7 Q5KVC8 Q5KVC9 Q5KVD0 Q5KVD5 Q5KVD7
Q5KVD9 Q5KVE4 Q5KVE5 Q5KVE6 Q5KVF8 Q5KVI4 Q5KVJ7 Q5KVM7 Q5KVP3 Q5KVP4 Q5KVP7
Q5KVS7 Q5KVX6 Q5KVX9 Q5KW01 Q5KW02 Q5KW03 Q5KW09 Q5KW28 Q5KW33 Q5KW36 Q5KW48
Q5KW49 Q5KW56 Q5KW59 Q5KW66 Q5KW94 Q5KW95 Q5KWA1 Q5KWB1 Q5KWB7 Q5KWC3 Q5KWC4
Q5KWC6 Q5KWC8 Q5KWD2 Q5KWD4 Q5KWD5 Q5KWD9 Q5KWE4 Q5KWE5 Q5KWF4 Q5KWF8 Q5KWH6
Q5KWI3 Q5KWI5 Q5KWI6 Q5KWJ2 Q5KWJ3 Q5KWJ4 Q5KWJ5 Q5KWJ6 Q5KWJ8 Q5KWJ9 Q5KWK2
Q5KWK4 Q5KWK6 Q5KWK9 Q5KWL3 Q5KWN2 Q5KWN3 Q5KWN7 Q5KWP1 Q5KWP3 Q5KWP5 Q5KWP6
Q5KWQ2 Q5KWQ7 Q5KWQ9 Q5KWR0 Q5KWR3 Q5KWR4 Q5KWS2 Q5KWS4 Q5KWS8 Q5KWS9 Q5KWT8
Q5KWU5 Q5KWU6 Q5KWU7 Q5KWU8 Q5KWV3 Q5KWV4 Q5KWV8 Q5KWV9 Q5KWX7 Q5KWX9 Q5KWY9
Q5KWZ5 Q5KWZ6 Q5KWZ7 Q5KWZ8 Q5KWZ9 Q5KX02 Q5KX04 Q5KX10 Q5KX15 Q5KX17 Q5KX24
Q5KX27 Q5KX35 Q5KX51 Q5KX76 Q5KX77 Q5KX81 Q5KX85 Q5KX91 Q5KXA4 Q5KXB1 Q5KXD3
Q5KXG8 Q5KXG9 Q5KXI7 Q5KXJ2 Q5KXK7 Q5KXL1 Q5KXS4 Q5KXT0 Q5KXT7 Q5KXU0 Q5KXU2
Q5KXU4 Q5KXU5 Q5KXU8 Q5KXU9 Q5KXV2 Q5KXV3 Q5KXV5 Q5KXW6 Q5KXW7 Q5KXX0 Q5KXX2
Q5KXX3 Q5KXX4 Q5KXY4 Q5KY22 Q5KY23 Q5KY26 Q5KY48 Q5KY53 Q5KY94 Q5KY95 Q5KYA2
Q5KYA3 Q5KYA5 Q5KYI8 Q5KYK0 Q5KYK6 Q5KYL9 Q5KYM0 Q5KYM1 Q5KYP6 Q5KYP7 Q5KYR0
Q5KYR1 Q5KYR2 Q5KYR3 Q5KYR4 Q5KYS6 Q5KZ00 Q5KZ06 Q5KZ14 Q5KZ20 Q5KZ25 Q5KZ39
Q5KZ63 Q5KZ91 Q5KZE8 Q5KZM1 Q5KZN0 Q5KZN1 Q5KZP0 Q5KZP1 Q5KZR0 Q5KZR8 Q5KZT7
Q5KZU0 Q5KZU2 Q5KZU9 Q5L010 Q5L014 Q5L025 Q5L041 Q5L046 Q5L051 Q5L055 Q5L078
Q5L083 Q5L084 Q5L086 Q5L091 Q5L0A5 Q5L0A7 Q5L0C0 Q5L0C2 Q5L0D8 Q5L0D9 Q5L0E4
Q5L0E5 Q5L0E8 Q5L0F7 Q5L0I2 Q5L0I3 Q5L0I5 Q5L0I6 Q5L0I8 Q5L0J2 Q5L0J4 Q5L0J6
Q5L0J8 Q5L0J9 Q5L0K0 Q5L0K1 Q5L0K2 Q5L0K6 Q5L0N0 Q5L0N1 Q5L0N2 Q5L0N7 Q5L0P0
Q5L0P3 Q5L0P4 Q5L0P5 Q5L0P8 Q5L0Q3 Q5L0R4 Q5L0R8 Q5L0S1 Q5L0S3 Q5L0S7 Q5L0S8
Q5L0S9 Q5L0U0 Q5L0U1 Q5L0U2 Q5L0U5 Q5L0U6 Q5L0U8 Q5L0V9 Q5L0W3 Q5L0X7 Q5L0Y0
Q5L0Y4 Q5L0Y5 Q5L0Z0 Q5L0Z2 Q5L0Z6 Q5L0Z9 Q5L118 Q5L141 Q5L144 Q5L145 Q5L146
Q5L172 Q5L186 Q5L1B7 Q5L1C0 Q5L1D9 Q5L1E0 Q5L1E1 Q5L1E2 Q5L1E5 Q5L1E6 Q5L1G9
Q5L1H7 Q5L1N1 Q5L1R3 Q5L1R5 Q5L1R7 Q5L1S1 Q5L1T7 Q5L1V3 Q5L1V4 Q5L1V5 Q5L203
Q5L263 Q5L264 Q5L283 Q5L284 Q5L293 Q5L296 Q5L299 Q5L2A5 Q5L2C0 Q5L2C2 Q5L2C9
Q5L2G3 Q5L2S0 Q5L2S4 Q5L2S7 Q5L2T4 Q5L2U0 Q5L2U3 Q5L2V9 Q5L2X7 Q5L2X9 Q5L2Y0
Q5L2Y4 Q5L300 Q5L301 Q5L302 Q5L310 Q5L325 Q5L3B2 Q5L3B3 Q5L3B4 Q5L3B9 Q5L3D0
Q5L3D2 Q5L3D3 Q5L3E1 Q5L3E6 Q5L3E7 Q5L3F3 Q5L3F4 Q5L3F6 Q5L3G0 Q5L3G4 Q5L3G9
Q5L3H2 Q5L3K8 Q5L3P0 Q5L3P1 Q5L3Q5 Q5L3Q6 Q5L3Q7 Q5L3R1 Q5L3R2 Q5L3R3 Q5L3R4
Q5L3R5 Q5L3R6 Q5L3R8 Q5L3S0 Q5L3S1 Q5L3S2 Q5L3S3 Q5L3S9 Q5L3T0 Q5L3T3 Q5L3U7
Q5L3U8 Q5L3V1 Q5L3V4 Q5L3V8 Q5L3W9 Q5L3X4 Q5L3X5 Q5L3Y0 Q5L3Y1 Q5L3Y2 Q5L3Y9
Q5L3Z2 Q5L3Z3 Q5L3Z4 Q5L3Z5 Q5L3Z6 Q5L3Z7 Q5L3Z8 Q5L3Z9 Q5L400 Q5L401 Q5L402
Q5L403 Q5L404 Q5L405 Q5L407 Q5L408 Q5L409 Q5L410 Q5L411 Q5L412 Q5L413 Q5L414
Q5L415 Q5L416 Q5L417 Q5L418 Q5L419 Q5L420 Q5L421 Q5L424 Q5L429 Q5L431 Q5L432
Q5L433 Q5L437 Q5L4N6 Q5L4P3 Q5L4Q4 Q5L4Q6 Q5L4S6 Q5L4T1 Q5L4T7 Q5L4V4 Q5L4W0

Q5L4W4 Q5L4W6 Q5L4W7 Q5L4W8 Q5L4X1 Q5L4Y3 Q5L4Z0 Q5L4Z3 Q5L510 Q5L512 Q5L521
Q5L530 Q5L545 Q5L546 Q5L548 Q5L549 Q5L550 Q5L553 Q5L554 Q5L562 Q5L563 Q5L564
Q5L565 Q5L566 Q5L567 Q5L568 Q5L574 Q5L588 Q5L589 Q5L593 Q5L5A2 Q5L5A3 Q5L5A6
Q5L5A7 Q5L5A8 Q5L5B2 Q5L5B4 Q5L5B5 Q5L5D3 Q5L5E1 Q5L5E3 Q5L5E7 Q5L5F1 Q5L5F2
Q5L5F4 Q5L5F7 Q5L5F9 Q5L5G4 Q5L5G7 Q5L5H5 Q5L5H6 Q5L5H9 Q5L5I0 Q5L5I1 Q5L5I2
Q5L5I3 Q5L5I4 Q5L5I5 Q5L5J6 Q5L5J7 Q5L5L0 Q5L5L3 Q5L5L4 Q5L5L7 Q5L5L8 Q5L5L9
Q5L5M0 Q5L5M1 Q5L5M3 Q5L5N7 Q5L5N9 Q5L5P0 Q5L5Q2 Q5L5Q6 Q5L5S0 Q5L5S1 Q5L5T1
Q5L5V3 Q5L5W0 Q5L5W2 Q5L5W3 Q5L5W6 Q5L5W7 Q5L5X3 Q5L5Y4 Q5L609 Q5L627 Q5L628
Q5L629 Q5L637 Q5L644 Q5L646 Q5L648 Q5L651 Q5L661 Q5L669 Q5L681 Q5L686 Q5L687
Q5L6B2 Q5L6B5 Q5L6D1 Q5L6D8 Q5L6E3 Q5L6F1 Q5L6F7 Q5L6F8 Q5L6G2 Q5L6G3 Q5L6G5
Q5L6G8 Q5L6H0 Q5L6H1 Q5L6H2 Q5L6H4 Q5L6H5 Q5L6I4 Q5L6I8 Q5L6I9 Q5L6K4 Q5L6M0
Q5L6M2 Q5L6M5 Q5L6N0 Q5L6N4 Q5L6P3 Q5L6P4 Q5L6Q6 Q5L6R4 Q5L6R5 Q5L6R9 Q5L6S0
Q5L6S1 Q5L6S2 Q5L6S4 Q5L6S5 Q5L6S6 Q5L6S7 Q5L6T4 Q5L6U0 Q5L6U2 Q5L6U4 Q5L6U5
Q5L6U7 Q5L6U9 Q5L6V2 Q5L6V3 Q5L6W2 Q5L6W7 Q5L6X6 Q5L6Y4 Q5L6Z0 Q5L6Z2 Q5L6Z3
Q5L6Z4 Q5L6Z8 Q5L6Z9 Q5L700 Q5L701 Q5L703 Q5L704 Q5L705 Q5L706 Q5L707 Q5L708
Q5L709 Q5L710 Q5L711 Q5L712 Q5L713 Q5L714 Q5L715 Q5L716 Q5L717 Q5L718 Q5L719
Q5L720 Q5L722 Q5L734 Q5L735 Q5L748 Q5L763 Q5L764 Q5L765 Q5L766 Q5L782 Q5L798
Q5L799 Q5L7A0 Q5L7B4 Q5L7B7 Q5L7F1 Q5L7I8 Q5L7M0 Q5L7N5 Q5L7Q8 Q5L7W2 Q5L7W4
Q5L7Z7 Q5L883 Q5L887 Q5L890 Q5L893 Q5L894 Q5L896 Q5L897 Q5L898 Q5L8A5 Q5L8A6
Q5L8A7 Q5L8A8 Q5L8A9 Q5L8B0 Q5L8B2 Q5L8B3 Q5L8B4 Q5L8B5 Q5L8B6 Q5L8B7 Q5L8B8
Q5L8B9 Q5L8C0 Q5L8C1 Q5L8C2 Q5L8C3 Q5L8C4 Q5L8C5 Q5L8C6 Q5L8C7 Q5L8C8 Q5L8D1
Q5L8D2 Q5L8D3 Q5L8D4 Q5L8D5 Q5L8D6 Q5L8D7 Q5L8E4 Q5L8K8 Q5L8L6 Q5L8L7 Q5L8M5
Q5L8V7 Q5L8W2 Q5L8W6 Q5L8W7 Q5L8W8 Q5L8X0 Q5L8X2 Q5L8X6 Q5L8Y2 Q5L8Z8 Q5L911
Q5L912 Q5L917 Q5L923 Q5L925 Q5L975 Q5L9B5 Q5L9B6 Q5L9B7 Q5L9D0 Q5L9D9 Q5L9E0
Q5L9E3 Q5L9E5 Q5L9G0 Q5L9H9 Q5L9I8 Q5L9K3 Q5L9K4 Q5L9L6 Q5L9N8 Q5L9N9 Q5L9P3
Q5L9P4 Q5L9P6 Q5L9P9 Q5L9Q0 Q5L9Q6 Q5L9Q7 Q5L9Y1 Q5LA67 Q5LAB1 Q5LAB4 Q5LAD5
Q5LAE0 Q5LAF5 Q5LAF6 Q5LAN8 Q5LAQ8 Q5LAS4 Q5LAS7 Q5LAT7 Q5LAU6 Q5LAX0 Q5LAY7
Q5LAZ7 Q5LAZ8 Q5LAZ9 Q5LB00 Q5LBA7 Q5LBD5 Q5LBD6 Q5LBD7 Q5LBG5 Q5LBU1 Q5LBZ2
Q5LBZ5 Q5LBZ8 Q5LC03 Q5LC24 Q5LC42 Q5LC56 Q5LC76 Q5LC77 Q5LCA5 Q5LCA6 Q5LCA7
Q5LCE1 Q5LCE6 Q5LCF0 Q5LCF3 Q5LCF4 Q5LCF5 Q5LCF6 Q5LCF8 Q5LCH4 Q5LCI0 Q5LCI1
Q5LCI2 Q5LCK2 Q5LCS1 Q5LCT3 Q5LCU8 Q5LCV9 Q5LCW4 Q5LD54 Q5LD55 Q5LD58 Q5LD60
Q5LD90 Q5LD96 Q5LDB2 Q5LDK5 Q5LDK8 Q5LDL2 Q5LDM3 Q5LDN0 Q5LDN2 Q5LDN9 Q5LE70
Q5LED3 Q5LED4 Q5LEQ1 Q5LEQ3 Q5LEQ4 Q5LEQ5 Q5LEQ6 Q5LEQ9 Q5LER0 Q5LES3 Q5LES4
Q5LES5 Q5LEW2 Q5LEY1 Q5LFI4 Q5LFI5 Q5LFI6 Q5LFJ5 Q5LFN5 Q5LFS5 Q5LFT6 Q5LFT7
Q5LG28 Q5LG30 Q5LG50 Q5LG55 Q5LG56 Q5LG80 Q5LGG9 Q5LGH0 Q5LGK5 Q5LGL2 Q5LGN1
Q5LGN7 Q5LGQ7 Q5LGR5 Q5LGR6 Q5LGS0 Q5LGS1 Q5LGU0 Q5LGU6 Q5LGV6 Q5LGW6 Q5LGW9
Q5LGZ6 Q5LH03 Q5LH04 Q5LH11 Q5LH12 Q5LH18 Q5LH44 Q5LH66 Q5LH68 Q5LHC9 Q5LHD7
Q5LHE6 Q5LHG9 Q5LHJ8 Q5LHK5 Q5LHL2 Q5LHL3 Q5LHP5 Q5LHS7 Q5LHT1 Q5LHW2 Q5LHY8
Q5LHZ1 Q5LI10 Q5LI12 Q5LI24 Q5LI31 Q5LI34 Q5LI35 Q5LI41 Q5LI65 Q5LI89 Q5LIH9
Q5LII9 Q5LIJ0 Q5LIJ3 Q5LIJ5 Q5LIK3 Q5LIK4 Q5LIL0 Q5LIL2 Q5LIM4 Q5LIN1 Q5LIN3
Q5LIN8 Q5LIQ2 Q5LIQ3 Q5LIQ7 Q5LIR8 Q5LIS2 Q5LIS5 Q5LIT7 Q5LIV6 Q5LIX0 Q5LIY0
Q5LIY5 Q5LIY6 Q5LIY7 Q5LIY8 Q5LJ16 Q5LJ19 Q5LJ26 Q5LJ28 Q5LJ29 Q5LJ70 Q5LJP9
Q5LJZ2 Q5LKH2 Q5LKW3 Q5LL27 Q5LL69 Q5LL90 Q5LLE7 Q5LLE9 Q5LLF0 Q5LLF5 Q5LLG6
Q5LLK8 Q5LLL2 Q5LLL4 Q5LLM0 Q5LLM7 Q5LLN0 Q5LLN2 Q5LLN4 Q5LLP0 Q5LLP2 Q5LLR7

Q5LLT1 Q5LLT4 Q5LLT6 Q5LLT7 Q5LLU4 Q5LLU5 Q5LLU6 Q5LLU7 Q5LLU8 Q5LLW1 Q5LLW5
Q5LLW6 Q5LLW7 Q5LM03 Q5LM10 Q5LM13 Q5LM18 Q5LM31 Q5LMA4 Q5LMA7 Q5LMB5 Q5LMC7
Q5LMG4 Q5LMG5 Q5LMG9 Q5LMH8 Q5LMJ4 Q5LMK7 Q5LMP9 Q5LMQ0 Q5LMQ3 Q5LMQ4 Q5LMQ5
Q5LMQ6 Q5LMR2 Q5LMR3 Q5LMR4 Q5LMR5 Q5LMS3 Q5LMW0 Q5LMY0 Q5LMZ1 Q5LN23 Q5LN38
Q5LN53 Q5LN66 Q5LN74 Q5LN75 Q5LN80 Q5LN98 Q5LNC8 Q5LND3 Q5LNE6 Q5LNE8 Q5LNE9
Q5LNF0 Q5LNF2 Q5LNI8 Q5LNJ2 Q5LNJ7 Q5LNK0 Q5LNK5 Q5LNK7 Q5LNM2 Q5LNM6 Q5LNN5
Q5LNR0 Q5LNR7 Q5LNS5 Q5LNT6 Q5LNT7 Q5LNT8 Q5LNU7 Q5LNV0 Q5LNV2 Q5LNY7 Q5LP30
Q5LP58 Q5LP74 Q5LP79 Q5LP80 Q5LP83 Q5LPA8 Q5LPG8 Q5LPK1 Q5LPT8 Q5LPV9 Q5LQ08
Q5LQ09 Q5LQ42 Q5LQ43 Q5LQ58 Q5LQ75 Q5LQ80 Q5LQ99 Q5LQC3 Q5LQI6 Q5LQJ4 Q5LQJ7
Q5LQK5 Q5LQK6 Q5LQK9 Q5LQM7 Q5LQN0 Q5LQP4 Q5LQY2 Q5LR04 Q5LR07 Q5LR43 Q5LR48
Q5LR49 Q5LR50 Q5LR59 Q5LR94 Q5LRA3 Q5LRC0 Q5LRC9 Q5LRD8 Q5LRD9 Q5LRF9 Q5LRH2
Q5LRH8 Q5LRI7 Q5LRJ6 Q5LRL6 Q5LRN5 Q5LRP4 Q5LRS9 Q5LRU0 Q5LRU1 Q5LRV2 Q5LRY0
Q5LRY1 Q5LRY4 Q5LRY5 Q5LRY6 Q5LRZ4 Q5LRZ5 Q5LRZ7 Q5LS25 Q5LS56 Q5LS57 Q5LS78
Q5LS81 Q5LS92 Q5LS93 Q5LS98 Q5LSB0 Q5LSJ4 Q5LSL4 Q5LSN6 Q5LSN8 Q5LSP9 Q5LSQ0
Q5LSQ1 Q5LSQ2 Q5LSQ3 Q5LSQ4 Q5LSQ5 Q5LSS5 Q5LST0 Q5LST2 Q5LST6 Q5LST9 Q5LSU9
Q5LSV2 Q5LSV4 Q5LSV5 Q5LSX9 Q5LSY0 Q5LT18 Q5LT54 Q5LTB0 Q5LTB1 Q5LTB2 Q5LTB3
Q5LTB4 Q5LTH8 Q5LTJ1 Q5LTL1 Q5LTM4 Q5LTN6 Q5LTP7 Q5LTP9 Q5LTR1 Q5LTR9 Q5LTT5
Q5LTU4 Q5LTV0 Q5LTX8 Q5LTY7 Q5LU04 Q5LU15 Q5LU57 Q5LU58 Q5LU62 Q5LU71 Q5LU79
Q5LU80 Q5LU92 Q5LU93 Q5LU97 Q5LU99 Q5LUA0 Q5LUD3 Q5LUE3 Q5LUI2 Q5LUP4 Q5LUP9
Q5LUQ0 Q5LUS2 Q5LUS9 Q5LUT9 Q5LUV8 Q5LUX6 Q5LUY8 Q5LV15 Q5LV16 Q5LV46 Q5LV87
Q5LV90 Q5LV91 Q5LV94 Q5LV98 Q5LVI8 Q5LVJ2 Q5LVN0 Q5LVN3 Q5LVV1 Q5LW05 Q5LW06
Q5LW09 Q5LW13 Q5LW17 Q5LW31 Q5LW32 Q5LW33 Q5LW34 Q5LW38 Q5LW40 Q5LW41 Q5LW42
Q5LW43 Q5LW44 Q5LW45 Q5LW46 Q5LW47 Q5LW48 Q5LW49 Q5LW50 Q5LW55 Q5LW56 Q5LW57
Q5LW58 Q5LW59 Q5LW89 Q5LW93 Q5LWB2 Q5LWD3 Q5LWE2 Q5LWE5 Q5LWF2 Q5LWF3 Q5LWF6
Q5LWF7 Q5LWG3 Q5LWH0 Q5LWJ1 Q5LWJ5 Q5LWJ6 Q5LWK3 Q5LWL1 Q5LWL4 Q5LWL6 Q5LWM6
Q5LWN8 Q5LWP5 Q5LWQ9 Q5LWR1 Q5LWR2 Q5LWT6 Q5LWT9 Q5LWV4 Q5LWV5 Q5LWZ5 Q5LX06
Q5LX07 Q5LX16 Q5LX20 Q5LX29 Q5LX40 Q5LX42 Q5LX60 Q5LX75 Q5LX84 Q5LX98 Q5LXA4
Q5LXC0 Q5LXE1 Q5LXG9 Q5LXI0 Q5LXI7 Q5LXJ9 Q5LXK0 Q5LXK2 Q5LXK5 Q5LXM5 Q5LXM6
Q5LXM9 Q5LXN1 Q5LXN3 Q5LXN9 Q5LXP0 Q5LXP1 Q5LXP2 Q5LXQ5 Q5LXQ9 Q5LXR0 Q5LXR1
Q5LXR2 Q5LXR4 Q5LXR5 Q5LXR6 Q5LXR7 Q5LXR8 Q5LXR9 Q5LXS0 Q5LXS1 Q5LXS2 Q5LXS3
Q5LXS4 Q5LXS5 Q5LXS6 Q5LXS7 Q5LXS8 Q5LXS9 Q5LXT0 Q5LXT2 Q5LXT3 Q5LXT4 Q5LXT5
Q5LXT6 Q5LXT7 Q5LXT8 Q5LXU2 Q5LXV0 Q5LXV1 Q5LXV3 Q5LXV4 Q5LXY2 Q5LXY3 Q5LXZ5
Q5LXZ6 Q5LXZ7 Q5LXZ8 Q5LXZ9 Q5LY00 Q5LY03 Q5LY05 Q5LY12 Q5LY19 Q5LY20 Q5LY21
Q5LY23 Q5LY29 Q5LY32 Q5LY37 Q5LY40 Q5LY48 Q5LY50 Q5LY60 Q5LY62 Q5LY71 Q5LY79
Q5LY80 Q5LY83 Q5LY84 Q5LY91 Q5LY98 Q5LYA9 Q5LYC2 Q5LYE2 Q5LYE9 Q5LYF0 Q5LYG0
Q5LYG9 Q5LYH1 Q5LYH7 Q5LYI4 Q5LYI5 Q5LYI7 Q5LYI8 Q5LYJ2 Q5LYK1 Q5LYL3 Q5LYL9
Q5LYM4 Q5LYM8 Q5LYN3 Q5LYP0 Q5LYR4 Q5LYS1 Q5LYS8 Q5LYU6 Q5LYV4 Q5LYX1 Q5LYX2
Q5LYX4 Q5LYY1 Q5LYY2 Q5LYY3 Q5LYY5 Q5LYY7 Q5LYY8 Q5LZ08 Q5LZ62 Q5LZ68 Q5LZ70
Q5LZ72 Q5LZ83 Q5LZ98 Q5LZA7 Q5LZD1 Q5LZD4 Q5LZD5 Q5LZD9 Q5LZE0 Q5LZF1 Q5LZF3
Q5LZF4 Q5LZF5 Q5LZH5 Q5LZH6 Q5LZI0 Q5LZI4 Q5LZL2 Q5LZL6 Q5LZL7 Q5LZL8 Q5LZL9
Q5LZM1 Q5LZM5 Q5LZM6 Q5LZQ6 Q5LZR6 Q5LZT9 Q5LZU9 Q5LZV2 Q5LZW3 Q5LZW8 Q5LZW9
Q5M011 Q5M014 Q5M021 Q5M029 Q5M056 Q5M058 Q5M061 Q5M064 Q5M067 Q5M079 Q5M080
Q5M083 Q5M0B4 Q5M0B7 Q5M0B8 Q5M0C3 Q5M0C5 Q5M0C9 Q5M0D5 Q5M0E6 Q5M0J7 Q5M0J9
Q5M0L5 Q5M0L7 Q5M0L9 Q5M0M0 Q5M0M2 Q5M0N4 Q5M0N7 Q5M0N9 Q5M0P0 Q5M0P3 Q5M0P4

Q5M0P7 Q5M0S3 Q5M0S4 Q5M0S7 Q5M0T3 Q5M0V5 Q5M0W1 Q5M0W4 Q5M0W5 Q5M0W9 Q5M0X1
Q5M0X3 Q5M0Y4 Q5M0Y6 Q5M0Z9 Q5M100 Q5M101 Q5M111 Q5M117 Q5M122 Q5M123 Q5M126
Q5M138 Q5M140 Q5M141 Q5M153 Q5M154 Q5M158 Q5M159 Q5M189 Q5M191 Q5M192 Q5M194
Q5M1A8 Q5M1A9 Q5M1B4 Q5M1B8 Q5M1B9 Q5M1C3 Q5M1D9 Q5M1E2 Q5M1E6 Q5M1G2 Q5M1G3
Q5M1G4 Q5M1G5 Q5M1G6 Q5M1G7 Q5M1G8 Q5M1H9 Q5M1I0 Q5M1I1 Q5M1I4 Q5M1I5 Q5M1J6
Q5M1J9 Q5M1K5 Q5M1K6 Q5M1K7 Q5M1L1 Q5M1L6 Q5M1M3 Q5M1M9 Q5M1N3 Q5M1N6 Q5M1N8
Q5M1N9 Q5M1P9 Q5M1Q8 Q5M1R6 Q5M1R9 Q5M1S1 Q5M1S5 Q5M1S7 Q5M1S8 Q5M1S9 Q5M1T6
Q5M1T7 Q5M1T8 Q5M1U0 Q5M1V6 Q5M1W6 Q5M1W9 Q5M1X4 Q5M1X5 Q5M1Y0 Q5M1Y2 Q5M1Y3
Q5M1Y4 Q5M1Y6 Q5M1Z0 Q5M1Z3 Q5M203 Q5M206 Q5M209 Q5M216 Q5M217 Q5M222 Q5M226
Q5M229 Q5M237 Q5M249 Q5M250 Q5M252 Q5M255 Q5M277 Q5M278 Q5M281 Q5M283 Q5M285
Q5M291 Q5M292 Q5M293 Q5M294 Q5M2A7 Q5M2B1 Q5M2B2 Q5M2B3 Q5M2B4 Q5M2B6 Q5M2B7
Q5M2B8 Q5M2B9 Q5M2C0 Q5M2C1 Q5M2C2 Q5M2C3 Q5M2C4 Q5M2C5 Q5M2C6 Q5M2C7 Q5M2C8
Q5M2C9 Q5M2D0 Q5M2D1 Q5M2D2 Q5M2D4 Q5M2D5 Q5M2D6 Q5M2D7 Q5M2D8 Q5M2D9 Q5M2E0
Q5M2E4 Q5M2F2 Q5M2F3 Q5M2F5 Q5M2F6 Q5M2I4 Q5M2I5 Q5M2J9 Q5M2K0 Q5M2K1 Q5M2K2
Q5M2K3 Q5M2K4 Q5M2K7 Q5M2K9 Q5M2L6 Q5M2M4 Q5M2M5 Q5M2M6 Q5M2N4 Q5M2N7 Q5M2P2
Q5M2P5 Q5M2Q3 Q5M2Q5 Q5M2R5 Q5M2R7 Q5M2S7 Q5M2T6 Q5M2T7 Q5M2U0 Q5M2U1 Q5M2U6
Q5M2V3 Q5M2X3 Q5M2Y5 Q5M305 Q5M312 Q5M313 Q5M323 Q5M332 Q5M334 Q5M340 Q5M347
Q5M348 Q5M350 Q5M351 Q5M355 Q5M364 Q5M376 Q5M382 Q5M387 Q5M391 Q5M396 Q5M3A4
Q5M3B3 Q5M3C8 Q5M3D5 Q5M3E1 Q5M3F9 Q5M3G7 Q5M3I4 Q5M3I5 Q5M3I7 Q5M3J4 Q5M3J5
Q5M3J6 Q5M3J8 Q5M3K1 Q5M3M1 Q5M3R7 Q5M3S3 Q5M3S5 Q5M3S7 Q5M3T6 Q5M3V0 Q5M3V8
Q5M3Y3 Q5M3Y6 Q5M3Y7 Q5M3Z1 Q5M3Z2 Q5M405 Q5M406 Q5M407 Q5M427 Q5M430 Q5M434
Q5M439 Q5M465 Q5M469 Q5M470 Q5M471 Q5M472 Q5M474 Q5M481 Q5M482 Q5M4C1 Q5M4D1
Q5M4E9 Q5M4F9 Q5M4G2 Q5M4H3 Q5M4H9 Q5M4I0 Q5M4M4 Q5M4M7 Q5M4N6 Q5M4P4 Q5M4R7
Q5M4R9 Q5M4S2 Q5M4S5 Q5M4S8 Q5M4T8 Q5M4T9 Q5M4U2 Q5M4W1 Q5M4W4 Q5M4W5 Q5M4W6
Q5M4X0 Q5M4X2 Q5M4X6 Q5M4Y2 Q5M4Z3 Q5M533 Q5M535 Q5M551 Q5M553 Q5M555 Q5M556
Q5M558 Q5M570 Q5M573 Q5M575 Q5M576 Q5M579 Q5M580 Q5M583 Q5M5A9 Q5M5B0 Q5M5B3
Q5M5B9 Q5M5E1 Q5M5E8 Q5M5F1 Q5M5F2 Q5M5F6 Q5M5F8 Q5M5H1 Q5M5H3 Q5M5I6 Q5M5I8
Q5M5J8 Q5M5K5 Q5M5L0 Q5M5L1 Q5M5L5 Q5M5M7 Q5M5M9 Q5M5N0 Q5M5P1 Q5M5P2 Q5M5P6
Q5M5P7 Q5M5S6 Q5M5S8 Q5M5S9 Q5M5T1 Q5M5U4 Q5M5U5 Q5M5V0 Q5M5V4 Q5M5V5 Q5M5V9
Q5M5X5 Q5M5X8 Q5M5Y2 Q5M603 Q5M604 Q5M605 Q5M606 Q5M607 Q5M608 Q5M609 Q5M621
Q5M622 Q5M623 Q5M626 Q5M627 Q5M638 Q5M641 Q5M646 Q5M647 Q5M648 Q5M652 Q5M657
Q5M664 Q5M670 Q5M674 Q5M677 Q5M679 Q5M690 Q5M699 Q5M6A7 Q5M6B0 Q5M6B2 Q5M6B7
Q5M6C0 Q5M6C1 Q5M6C2 Q5M6C9 Q5M6D0 Q5M6D1 Q5M6D3 Q5M6E6 Q5M6F6 Q5M6F9 Q5M6G4
Q5M6G5 Q5M6H0 Q5M6H2 Q5M6H3 Q5M6H5 Q5M6H7 Q5M6I1 Q5M6I4 Q5M6J5 Q5M6J8 Q5M6K1
Q5M6K8 Q5M6K9 Q5M6L4 Q5M6L8 Q5M6W3 Q5M721 Q5M729 Q5M731 Q5M750 Q5M759 Q5M782
Q5M786 Q5M7C3 Q5M7C8 Q5M7D1 Q5M7D6 Q5M7D8 Q5M7E3 Q5M7F6 Q5M7F8 Q5M7G4 Q5M7K4
Q5M7L2 Q5M7L9 Q5M7N6 Q5M7N8 Q5M7P5 Q5M7P8 Q5M7T1 Q5M7T2 Q5M7T3 Q5M7T9 Q5M7U6
Q5M7V7 Q5M7W1 Q5M7W5 Q5M7W7 Q5M818 Q5M819 Q5M820 Q5M823 Q5M827 Q5M831 Q5M840
Q5M845 Q5M864 Q5M865 Q5M870 Q5M882 Q5M886 Q5M887 Q5M888 Q5M8E6 Q5M8G6 Q5M8I4
Q5M8J0 Q5M8K8 Q5M8M2 Q5M8N4 Q5M8Q2 Q5M8S7 Q5M8V0 Q5M8V2 Q5M8V6 Q5M8V8 Q5M8W9
Q5M8X5 Q5M8Y2 Q5M8Y7 Q5M8Z0 Q5M8Z2 Q5M8Z6 Q5M905 Q5M929 Q5M934 Q5M939 Q5M947
Q5M948 Q5M951 Q5M954 Q5M956 Q5M965 Q5M969 Q5M974 Q5M985 Q5M990 Q5M9B1 Q5M9F0
Q5M9F1 Q5M9F5 Q5M9F8 Q5M9G0 Q5M9G5 Q5M9G6 Q5M9G7 Q5M9G8 Q5M9G9 Q5M9H0 Q5M9H1
Q5M9I6 Q5M9N0 Q5M9Q1 Q5MAI5 Q5MAV4 Q5MAV5 Q5MBG2 Q5MBR3 Q5MBR5 Q5MBR6 Q5MCW4

Q5MFV5 Q5MFW3 Q5MGD1 Q5MGJ4 Q5MGL3 Q5MGM6 Q5MI53 Q5MIB5 Q5MIB6 Q5MIM5 Q5MIP6
Q5MIR6 Q5MJ07 Q5MJ08 Q5MJ09 Q5MJ10 Q5MJ12 Q5MJ68 Q5MJ70 Q5MJ86 Q5MK11 Q5MKM1
Q5MKM7 Q5MNH1 Q5MNH2 Q5MNH3 Q5MNH4 Q5MNH6 Q5MNH7 Q5MNH8 Q5MNH9 Q5MNI0 Q5MNI2
Q5MNI4 Q5MNI5 Q5MNI7 Q5MNV8 Q5MQ85 Q5MQC5 Q5MQC9 Q5MYA4 Q5MYW4 Q5MZ21 Q5MZ22
Q5MZ23 Q5MZ25 Q5MZ26 Q5MZ36 Q5MZ40 Q5MZ45 Q5MZ48 Q5MZ57 Q5MZ63 Q5MZ83 Q5MZ99
Q5MZA4 Q5MZA8 Q5MZE9 Q5MZF4 Q5MZH9 Q5MZI0 Q5MZI3 Q5MZJ6 Q5MZL0 Q5MZL2 Q5MZN2
Q5MZP5 Q5MZR5 Q5MZS4 Q5MZT0 Q5MZT3 Q5MZU1 Q5MZV1 Q5MZX7 Q5MZY1 Q5MZY3 Q5MZZ2
Q5N006 Q5N013 Q5N019 Q5N031 Q5N040 Q5N044 Q5N076 Q5N078 Q5N092 Q5N0A5 Q5N0A8
Q5N0B4 Q5N0C6 Q5N0D2 Q5N0D3 Q5N0E6 Q5N0F0 Q5N0G0 Q5N0G1 Q5N0G7 Q5N0H4 Q5N0J1
Q5N0J8 Q5N0K5 Q5N0K8 Q5N0L0 Q5N0L1 Q5N0M0 Q5N0M4 Q5N0T0 Q5N0T6 Q5N0T7 Q5N0T9
Q5N0U8 Q5N0W7 Q5N0X6 Q5N0X9 Q5N0Y2 Q5N0Z1 Q5N0Z5 Q5N0Z7 Q5N127 Q5N128 Q5N129
Q5N133 Q5N136 Q5N163 Q5N167 Q5N172 Q5N175 Q5N180 Q5N192 Q5N199 Q5N1A0 Q5N1A6
Q5N1B8 Q5N1C3 Q5N1D4 Q5N1D9 Q5N1E7 Q5N1G1 Q5N1G4 Q5N1I0 Q5N1I3 Q5N1I9 Q5N1J0
Q5N1J4 Q5N1J7 Q5N1K1 Q5N1L4 Q5N1M1 Q5N1N0 Q5N1N6 Q5N1N8 Q5N1P1 Q5N1P5 Q5N1P6
Q5N1P7 Q5N1Q0 Q5N1Q1 Q5N1Q7 Q5N1R1 Q5N1S9 Q5N1T5 Q5N1T9 Q5N1U0 Q5N1U7 Q5N1V0
Q5N1X6 Q5N1Y6 Q5N1Z2 Q5N1Z5 Q5N1Z8 Q5N223 Q5N249 Q5N294 Q5N2A1 Q5N2B2 Q5N2F8
Q5N2F9 Q5N2H7 Q5N2H9 Q5N2I0 Q5N2J4 Q5N2J9 Q5N2N4 Q5N2P0 Q5N2P4 Q5N2P7 Q5N2P9
Q5N2R3 Q5N2S7 Q5N2S8 Q5N2W5 Q5N2X2 Q5N2X5 Q5N2Y8 Q5N317 Q5N322 Q5N324 Q5N332
Q5N376 Q5N381 Q5N384 Q5N3B4 Q5N3B5 Q5N3B6 Q5N3B7 Q5N3D4 Q5N3G7 Q5N3H1 Q5N3I3
Q5N3J1 Q5N3J4 Q5N3J5 Q5N3K9 Q5N3L4 Q5N3M2 Q5N3N2 Q5N3N4 Q5N3N6 Q5N3N7 Q5N3N8
Q5N3N9 Q5N3S0 Q5N3S1 Q5N3S5 Q5N3T2 Q5N3T6 Q5N3T7 Q5N3U1 Q5N3W3 Q5N421 Q5N426
Q5N440 Q5N453 Q5N459 Q5N474 Q5N490 Q5N492 Q5N4A4 Q5N4B5 Q5N4B8 Q5N4C2 Q5N4D1
Q5N4D3 Q5N4D6 Q5N4F9 Q5N4J6 Q5N4L0 Q5N4L8 Q5N4L9 Q5N4N2 Q5N4Q0 Q5N4R0 Q5N4R3
Q5N4V3 Q5N4W5 Q5N4W6 Q5N4X3 Q5N4X6 Q5N4X9 Q5N4Z6 Q5N502 Q5N503 Q5N517 Q5N530
Q5N549 Q5N568 Q5N569 Q5N570 Q5N575 Q5N595 Q5N597 Q5N598 Q5N5A0 Q5N5A3 Q5N5F3
Q5N5F4 Q5N5F7 Q5N5G8 Q5N5I5 Q5N5I8 Q5N5J0 Q5N5K7 Q5N5P2 Q5N5S6 Q5N5U1 Q5N5U2
Q5N5V5 Q5N5V7 Q5N5W2 Q5N5W8 Q5N605 Q5N606 Q5N623 Q5N636 Q5N638 Q5N643 Q5N665
Q5N667 Q5N6V0 Q5N712 Q5N725 Q5N749 Q5N7C7 Q5N7W4 Q5N829 Q5N863 Q5N870 Q5N897
Q5N8G1 Q5N8J3 Q5N8Q4 Q5N8Z0 Q5N9Q7 Q5NAI7 Q5NAN5 Q5NAV3 Q5NAW2 Q5NAY4 Q5NAZ7
Q5NB25 Q5NB82 Q5NB83 Q5NB85 Q5NBI2 Q5NBJ3 Q5NBM0 Q5NBM8 Q5NBN9 Q5NBT9 Q5NBU5
Q5NBU8 Q5NBY9 Q5NC05 Q5NC57 Q5NC83 Q5NCC9 Q5NCF2 Q5NCI0 Q5NCQ5 Q5NCR9 Q5NCX5
Q5NCY0 Q5NCY3 Q5ND04 Q5ND52 Q5NDM2 Q5NDM8 Q5NDN1 Q5NE79 Q5NE80 Q5NE84 Q5NE98
Q5NE99 Q5NEB0 Q5NEE1 Q5NEE2 Q5NEE4 Q5NEF0 Q5NEF7 Q5NEF9 Q5NEH3 Q5NEL0 Q5NEL1
Q5NEL9 Q5NEM2 Q5NEM3 Q5NEP2 Q5NEP6 Q5NEQ9 Q5NER3 Q5NER4 Q5NEV4 Q5NEV7 Q5NEV8
Q5NEW3 Q5NEX0 Q5NEX7 Q5NEY6 Q5NF12 Q5NF19 Q5NF37 Q5NF43 Q5NF55 Q5NF56 Q5NF57
Q5NF58 Q5NF60 Q5NF72 Q5NF76 Q5NFB0 Q5NFB1 Q5NFC4 Q5NFC6 Q5NFD2 Q5NFD4 Q5NFE1
Q5NFE7 Q5NFF3 Q5NFF8 Q5NFG6 Q5NFG7 Q5NFG8 Q5NFJ3 Q5NFK3 Q5NFK5 Q5NFL9 Q5NFM5
Q5NFM8 Q5NFN4 Q5NFP3 Q5NFP4 Q5NFR4 Q5NFS0 Q5NFS1 Q5NFS4 Q5NFT3 Q5NFU1 Q5NFU8
Q5NFW7 Q5NFX8 Q5NFZ8 Q5NFZ9 Q5NG00 Q5NG01 Q5NG03 Q5NG10 Q5NG17 Q5NG21 Q5NG26
Q5NG36 Q5NG38 Q5NG39 Q5NG44 Q5NG53 Q5NG54 Q5NG61 Q5NG64 Q5NG68 Q5NG69 Q5NG86
Q5NG98 Q5NGB2 Q5NGB5 Q5NGB9 Q5NGC1 Q5NGC3 Q5NGD0 Q5NGD3 Q5NGE7 Q5NGF3 Q5NGG6
Q5NGI2 Q5NGK4 Q5NGL5 Q5NGL6 Q5NGL8 Q5NGM5 Q5NGM6 Q5NGQ8 Q5NGR1 Q5NGR2 Q5NGS0
Q5NGS8 Q5NGW1 Q5NGW4 Q5NGX1 Q5NGX3 Q5NGX6 Q5NGX7 Q5NGX8 Q5NGY1 Q5NGY3 Q5NGY5
Q5NGY9 Q5NGZ6 Q5NH01 Q5NH09 Q5NH17 Q5NH18 Q5NH19 Q5NH21 Q5NH32 Q5NH35 Q5NH41

Q5NH42 Q5NH46 Q5NH47 Q5NH48 Q5NH50 Q5NH53 Q5NH55 Q5NH64 Q5NH87 Q5NH94 Q5NHE5
Q5NHE6 Q5NHF3 Q5NHI3 Q5NHI5 Q5NHI8 Q5NHJ8 Q5NHK3 Q5NHL7 Q5NHN2 Q5NHN4 Q5NHN8
Q5NHP0 Q5NHQ7 Q5NHQ9 Q5NHS1 Q5NHS2 Q5NHS9 Q5NHT8 Q5NHU2 Q5NHU3 Q5NHU4 Q5NHU5
Q5NHU6 Q5NHU7 Q5NHU9 Q5NHV0 Q5NHV1 Q5NHV2 Q5NHV3 Q5NHV4 Q5NHV5 Q5NHV6 Q5NHV7
Q5NHV8 Q5NHV9 Q5NHW0 Q5NHW1 Q5NHW2 Q5NHW3 Q5NHW4 Q5NHW5 Q5NHW6 Q5NHW7 Q5NHW8
Q5NHW9 Q5NHX0 Q5NHX1 Q5NHX2 Q5NHX4 Q5NHX7 Q5NHX8 Q5NHX9 Q5NHY0 Q5NHY6 Q5NHZ4
Q5NI19 Q5NI31 Q5NI50 Q5NI53 Q5NI60 Q5NI61 Q5NI78 Q5NI96 Q5NIA7 Q5NIA8 Q5NIA9
Q5NIB6 Q5NIC3 Q5NIC4 Q5NIC5 Q5NIC6 Q5NIC7 Q5NIC9 Q5NID1 Q5NID2 Q5NID3 Q5NID4
Q5NID5 Q5NID6 Q5NID9 Q5NIE5 Q5NIF4 Q5NIF5 Q5NIG0 Q5NIG4 Q5NII7 Q5NII8 Q5NII9
Q5NIL5 Q5NIL6 Q5NIL7 Q5NIL9 Q5NIM0 Q5NIP5 Q5NIP6 Q5NIQ4 Q5NIQ8 Q5NKZ8 Q5NL05
Q5NL06 Q5NL15 Q5NL16 Q5NL22 Q5NL34 Q5NL75 Q5NL76 Q5NL79 Q5NL80 Q5NL82 Q5NLA8
Q5NLB9 Q5NLF0 Q5NLG3 Q5NLJ0 Q5NLJ4 Q5NLR1 Q5NLR2 Q5NLR6 Q5NLS3 Q5NLS8 Q5NLU3
Q5NLU9 Q5NLX2 Q5NLX3 Q5NLX5 Q5NLX8 Q5NLZ3 Q5NLZ4 Q5NM04 Q5NM38 Q5NMA4 Q5NMC0
Q5NMC1 Q5NMC2 Q5NMC3 Q5NMC7 Q5NMD3 Q5NMD4 Q5NMD5 Q5NMD6 Q5NMD7 Q5NME2 Q5NMF5
Q5NMJ1 Q5NMK1 Q5NMV7 Q5NMW1 Q5NMX2 Q5NMX3 Q5NMY4 Q5NN19 Q5NN40 Q5NN52 Q5NN59
Q5NN60 Q5NN61 Q5NN84 Q5NN85 Q5NNB4 Q5NNE1 Q5NNK7 Q5NNK8 Q5NNK9 Q5NNL0 Q5NNL4
Q5NNN4 Q5NNQ0 Q5NNQ4 Q5NNR3 Q5NNS2 Q5NNS3 Q5NNS5 Q5NNT2 Q5NNT4 Q5NNV0 Q5NNV1
Q5NNW5 Q5NNY7 Q5NNY8 Q5NNZ0 Q5NP09 Q5NP13 Q5NP22 Q5NP36 Q5NP61 Q5NP65 Q5NP67
Q5NP84 Q5NPA7 Q5NPC4 Q5NPC5 Q5NPC9 Q5NPE5 Q5NPF1 Q5NPF3 Q5NPF4 Q5NPG7 Q5NPH0
Q5NPH1 Q5NPJ8 Q5NPK4 Q5NPK5 Q5NPK7 Q5NPK9 Q5NPL0 Q5NPL1 Q5NPL5 Q5NPL6 Q5NPM1
Q5NPM2 Q5NPM9 Q5NPP5 Q5NPQ9 Q5NPR4 Q5NPS5 Q5NPS6 Q5NPY7 Q5NPY8 Q5NPZ5 Q5NPZ7
Q5NQ24 Q5NQ27 Q5NQ28 Q5NQ29 Q5NQ30 Q5NQ31 Q5NQ32 Q5NQ36 Q5NQ39 Q5NQ40 Q5NQ41
Q5NQ42 Q5NQ45 Q5NQ47 Q5NQ48 Q5NQ49 Q5NQ50 Q5NQ51 Q5NQ52 Q5NQ53 Q5NQ54 Q5NQ55
Q5NQ56 Q5NQ57 Q5NQ58 Q5NQ59 Q5NQ60 Q5NQ61 Q5NQ62 Q5NQ63 Q5NQ64 Q5NQ65 Q5NQ66
Q5NQ67 Q5NQA8 Q5NQB9 Q5NQC7 Q5NQE8 Q5NQF0 Q5NQH8 Q5NQN1 Q5NQQ2 Q5NQQ4 Q5NQQ7
Q5NQV7 Q5NQY1 Q5NQY4 Q5NR05 Q5NR19 Q5NR21 Q5NR23 Q5NR24 Q5NR44 Q5NR48 Q5NR50
Q5NR58 Q5NR72 Q5NR75 Q5NR76 Q5NR77 Q5NR81 Q5NRB3 Q5NRC4 Q5NRC5 Q5NRD6 Q5NRF0
Q5NRF3 Q5NRF6 Q5NRH4 Q5NRI7 Q5NRJ0 Q5NRL4 Q5NRL5 Q5NRL6 Q5NRL7 Q5NRM1 Q5NRM2
Q5NRM3 Q5NRM4 Q5NRM5 Q5NSW5 Q5NSW9 Q5NTH0 Q5NTH3 Q5NTH4 Q5NU13 Q5NU14 Q5NUA6
Q5NUF3 Q5NUF4 Q5NVA2 Q5NVA9 Q5NVB2 Q5NVC2 Q5NVC8 Q5NVD0 Q5NVD7 Q5NVE1 Q5NVE2
Q5NVE6 Q5NVF5 Q5NVF9 Q5NVH8 Q5NVI3 Q5NVJ8 Q5NVK0 Q5NVK5 Q5NVK7 Q5NVL7 Q5NVM0
Q5NVM3 Q5NVM5 Q5NVM8 Q5NVN0 Q5NVN7 Q5NVN8 Q5NVN9 Q5NVP2 Q5NVP8 Q5NVR0 Q5NVR2
Q5NVR5 Q5NVS0 Q5NVT1 Q5NVT2 Q5NWG5 Q5NXL5 Q5NXM1 Q5NXM4 Q5NXM5 Q5NXM6 Q5NXM7
Q5NXN2 Q5NXP4 Q5NXQ2 Q5NXQ3 Q5NXR0 Q5NXR4 Q5NXU8 Q5NXV7 Q5NXZ4 Q5NXZ7 Q5NY25
Q5NY30 Q5NY31 Q5NY33 Q5NY71 Q5NYA7 Q5NYA9 Q5NYD9 Q5NYE0 Q5NYE5 Q5NYJ9 Q5NYK0
Q5NYP6 Q5NYP9 Q5NYT0 Q5NYT1 Q5NYT7 Q5NYT9 Q5NYU3 Q5NYY1 Q5NZ71 Q5NZ92 Q5NZ93
Q5NZF3 Q5NZF5 Q5NZF6 Q5NZF8 Q5NZG1 Q5NZG9 Q5NZH2 Q5NZH3 Q5NZH4 Q5NZH5 Q5NZH8
Q5NZR7 Q5NZR8 Q5NZR9 Q5NZS0 Q5NZS1 Q5NZS3 Q5NZS9 Q5NZU2 Q5NZX8 Q5NZY0 Q5NZY1
Q5P011 Q5P080 Q5P081 Q5P083 Q5P084 Q5P086 Q5P0D8 Q5P0E0 Q5P0E7 Q5P0T2 Q5P0T4
Q5P0T5 Q5P0Z4 Q5P0Z5 Q5P0Z7 Q5P0Z8 Q5P102 Q5P106 Q5P119 Q5P160 Q5P161 Q5P162
Q5P1A7 Q5P1A8 Q5P1C5 Q5P1F7 Q5P1G0 Q5P1G6 Q5P1G9 Q5P1H1 Q5P1H4 Q5P1H5 Q5P1H7
Q5P1H8 Q5P1I9 Q5P1J2 Q5P1J6 Q5P1J7 Q5P1J8 Q5P1K1 Q5P1L3 Q5P1Y4 Q5P1Z7 Q5P1Z8
Q5P201 Q5P203 Q5P212 Q5P221 Q5P224 Q5P228 Q5P229 Q5P233 Q5P236 Q5P239 Q5P252
Q5P255 Q5P261 Q5P262 Q5P280 Q5P286 Q5P2G1 Q5P2G2 Q5P2I9 Q5P2J9 Q5P2L1 Q5P2L4

Q5P2L5 Q5P2M2 Q5P2M5 Q5P2P8 Q5P2R4 Q5P2S1 Q5P2U2 Q5P2U3 Q5P2U6 Q5P2U7 Q5P2V2
Q5P2V5 Q5P2Y9 Q5P306 Q5P307 Q5P308 Q5P309 Q5P310 Q5P311 Q5P313 Q5P314 Q5P315
Q5P316 Q5P317 Q5P318 Q5P319 Q5P320 Q5P321 Q5P322 Q5P323 Q5P324 Q5P325 Q5P326
Q5P327 Q5P328 Q5P329 Q5P330 Q5P331 Q5P332 Q5P333 Q5P334 Q5P335 Q5P336 Q5P337
Q5P338 Q5P339 Q5P340 Q5P341 Q5P342 Q5P343 Q5P3B0 Q5P3B7 Q5P3C5 Q5P3C8 Q5P3E7
Q5P3H3 Q5P3H4 Q5P3K0 Q5P3L9 Q5P3M0 Q5P3M1 Q5P3R2 Q5P3R8 Q5P3S4 Q5P3T0 Q5P3T7
Q5P409 Q5P4A4 Q5P4B8 Q5P4B9 Q5P4D0 Q5P4D1 Q5P4G4 Q5P4H2 Q5P4H6 Q5P4J6 Q5P4J7
Q5P4L8 Q5P4P0 Q5P4P1 Q5P4P2 Q5P4P5 Q5P4Q4 Q5P4R1 Q5P4Z2 Q5P502 Q5P503 Q5P5G2
Q5P5G3 Q5P5G4 Q5P5G5 Q5P5G6 Q5P5I4 Q5P5J1 Q5P5J5 Q5P5J6 Q5P5L5 Q5P5L9 Q5P5M2
Q5P5P2 Q5P5P3 Q5P6B7 Q5P6C0 Q5P6F0 Q5P6F1 Q5P6G9 Q5P6I2 Q5P6I4 Q5P6I6 Q5P6I7
Q5P6J5 Q5P6J6 Q5P6J9 Q5P6L3 Q5P6P9 Q5P6Y3 Q5P6Y5 Q5P6Y8 Q5P6Y9 Q5P6Z4 Q5P6Z7
Q5P6Z8 Q5P706 Q5P714 Q5P720 Q5P721 Q5P722 Q5P723 Q5P725 Q5P728 Q5P730 Q5P760
Q5P762 Q5P766 Q5P787 Q5P789 Q5P790 Q5P791 Q5P792 Q5P793 Q5P794 Q5P795 Q5P796
Q5P797 Q5P7A7 Q5P7A9 Q5P7B0 Q5P7B3 Q5P7B4 Q5P7B7 Q5P7B8 Q5P7C3 Q5P7C4 Q5P7F0
Q5P7F1 Q5P7F8 Q5P7G1 Q5P7G2 Q5P7G6 Q5P7G7 Q5P7H3 Q5P7H5 Q5P7I0 Q5P7J0 Q5P7J1
Q5P7J7 Q5P7K8 Q5P7L9 Q5P7M2 Q5P7M6 Q5P7N1 Q5P7N4 Q5P7P0 Q5P7P1 Q5P7P8 Q5P7Q3
Q5P7R7 Q5P7T1 Q5P7T2 Q5P7T7 Q5P7T8 Q5P7T9 Q5P7U0 Q5P7U2 Q5P7U3 Q5P7U7 Q5P7X5
Q5P7X6 Q5P7X7 Q5P7X8 Q5P7X9 Q5P7Y0 Q5P7Y1 Q5P7Z3 Q5P7Z4 Q5P7Z5 Q5P7Z6 Q5P7Z7
Q5P7Z9 Q5P801 Q5P811 Q5P832 Q5P833 Q5P838 Q5P841 Q5P8D0 Q5P8D4 Q5P8J4 Q5P8W0
Q5P905 Q5P922 Q5P951 Q5PBZ2 Q5PBZ5 Q5PC05 Q5PC07 Q5PC09 Q5PC10 Q5PC25 Q5PC26
Q5PC27 Q5PC28 Q5PC30 Q5PC31 Q5PC32 Q5PC39 Q5PC40 Q5PC41 Q5PC60 Q5PC77 Q5PC82
Q5PC86 Q5PC89 Q5PC93 Q5PC95 Q5PCB2 Q5PCB8 Q5PCE2 Q5PCE5 Q5PCF4 Q5PCF5 Q5PCG0
Q5PCH1 Q5PCH2 Q5PCH6 Q5PCH8 Q5PCK6 Q5PCL7 Q5PCM7 Q5PCN5 Q5PCQ1 Q5PCQ9 Q5PCR0
Q5PCR2 Q5PCR7 Q5PCT4 Q5PCT5 Q5PCT8 Q5PCU0 Q5PCU1 Q5PCU4 Q5PCV6 Q5PCV8 Q5PCW7
Q5PCX2 Q5PCX4 Q5PCY1 Q5PCY8 Q5PCZ6 Q5PCZ8 Q5PD06 Q5PD17 Q5PD31 Q5PD33 Q5PD43
Q5PD46 Q5PD49 Q5PD51 Q5PD54 Q5PD56 Q5PD57 Q5PD59 Q5PD60 Q5PD61 Q5PD62 Q5PD63
Q5PD71 Q5PD76 Q5PD80 Q5PD84 Q5PD96 Q5PDA0 Q5PDA1 Q5PDB8 Q5PDB9 Q5PDC2 Q5PDC6
Q5PDC7 Q5PDD7 Q5PDD8 Q5PDD9 Q5PDE0 Q5PDE1 Q5PDF0 Q5PDF1 Q5PDF2 Q5PDG1 Q5PDG2
Q5PDG3 Q5PDG4 Q5PDG5 Q5PDH1 Q5PDH4 Q5PDH5 Q5PDH8 Q5PDH9 Q5PDJ4 Q5PDJ5 Q5PDL9
Q5PDM2 Q5PDM3 Q5PDM5 Q5PDM6 Q5PDM7 Q5PDN6 Q5PDP2 Q5PDP3 Q5PDP4 Q5PDP5 Q5PDP6
Q5PDP7 Q5PDQ9 Q5PDT5 Q5PDU3 Q5PDU9 Q5PDV8 Q5PDX5 Q5PDX6 Q5PE20 Q5PE22 Q5PE23
Q5PE39 Q5PE50 Q5PE52 Q5PE68 Q5PEA8 Q5PEE6 Q5PEE9 Q5PEG1 Q5PEG2 Q5PEG3 Q5PEG4
Q5PEH1 Q5PEH2 Q5PEH3 Q5PEH7 Q5PEH8 Q5PEH9 Q5PEJ0 Q5PEJ4 Q5PEK0 Q5PEK5 Q5PEK7
Q5PEK9 Q5PEL3 Q5PEN2 Q5PEN3 Q5PEN6 Q5PEP3 Q5PEQ5 Q5PES4 Q5PEY3 Q5PF04 Q5PF10
Q5PF15 Q5PF16 Q5PF17 Q5PF18 Q5PF19 Q5PF36 Q5PF65 Q5PF66 Q5PF68 Q5PF69 Q5PF73
Q5PF76 Q5PFD6 Q5PFD8 Q5PFF6 Q5PFF7 Q5PFF8 Q5PFF9 Q5PFG2 Q5PFG7 Q5PFG9 Q5PFJ1
Q5PFJ2 Q5PFK3 Q5PFK5 Q5PFK6 Q5PFK7 Q5PFK8 Q5PFL9 Q5PFM0 Q5PFM4 Q5PFN3 Q5PFN4
Q5PFN5 Q5PFP1 Q5PFQ2 Q5PFQ4 Q5PFQ9 Q5PFR0 Q5PFR1 Q5PFR6 Q5PFR8 Q5PFS6 Q5PFS8
Q5PFS9 Q5PFT5 Q5PFT6 Q5PFU0 Q5PFV0 Q5PFV4 Q5PFV5 Q5PG12 Q5PG19 Q5PG24 Q5PG37
Q5PG39 Q5PG48 Q5PG49 Q5PG52 Q5PG56 Q5PG58 Q5PG59 Q5PG61 Q5PG75 Q5PG77 Q5PG85
Q5PG91 Q5PG96 Q5PGA0 Q5PGA1 Q5PGA2 Q5PGB1 Q5PGB5 Q5PGB6 Q5PGB8 Q5PGB9 Q5PGC6
Q5PGD4 Q5PGD6 Q5PGD8 Q5PGE3 Q5PGF0 Q5PGF4 Q5PGF9 Q5PGG1 Q5PGG5 Q5PGG7 Q5PGG9
Q5PGI6 Q5PGJ4 Q5PGJ6 Q5PGK0 Q5PGK4 Q5PGL4 Q5PGM5 Q5PGN2 Q5PGP7 Q5PGR8 Q5PGR9
Q5PGS9 Q5PGT0 Q5PGT4 Q5PGT7 Q5PGT9 Q5PGW5 Q5PGW9 Q5PGX0 Q5PGZ4 Q5PH01 Q5PH07

Q5PH15 Q5PH69 Q5PH77 Q5PH79 Q5PH83 Q5PH84 Q5PH85 Q5PH86 Q5PH88 Q5PH90 Q5PH91
Q5PH92 Q5PH96 Q5PHA6 Q5PHB3 Q5PHB6 Q5PHB9 Q5PHC0 Q5PHC1 Q5PHC2 Q5PHC8 Q5PHE4
Q5PHN5 Q5PHS1 Q5PHS8 Q5PHV8 Q5PHY6 Q5PHY7 Q5PHZ3 Q5PI07 Q5PI10 Q5PI26 Q5PI28
Q5PI32 Q5PI50 Q5PI52 Q5PI54 Q5PI83 Q5PI85 Q5PI88 Q5PIA0 Q5PIA7 Q5PIB8 Q5PIC3
Q5PIE1 Q5PIE3 Q5PIF2 Q5PIG3 Q5PIG5 Q5PIG8 Q5PIH6 Q5PII3 Q5PII8 Q5PIK4 Q5PIK6
Q5PIK7 Q5PIK8 Q5PIK9 Q5PIL0 Q5PIL1 Q5PIL2 Q5PIL9 Q5PIM9 Q5PIN3 Q5PIN4 Q5PIN6
Q5PIP0 Q5PIR6 Q5PIR9 Q5PIS0 Q5PIS1 Q5PIS3 Q5PIS8 Q5PIS9 Q5PIT6 Q5PIT7 Q5PIT8
Q5PIU0 Q5PIU1 Q5PIU2 Q5PIU3 Q5PIU4 Q5PIU5 Q5PIU6 Q5PIU7 Q5PIU8 Q5PIU9 Q5PIV0
Q5PIV1 Q5PIV2 Q5PIV3 Q5PIV4 Q5PIV5 Q5PIV6 Q5PIV7 Q5PIV8 Q5PIW0 Q5PIW1 Q5PIW2
Q5PIW3 Q5PIW4 Q5PJ43 Q5PJ49 Q5PJ50 Q5PJ55 Q5PJ56 Q5PJ57 Q5PJ58 Q5PJ61 Q5PJ62
Q5PJ63 Q5PJ71 Q5PJ74 Q5PJ81 Q5PJ90 Q5PJB2 Q5PJB3 Q5PJB8 Q5PJD4 Q5PJE0 Q5PJE2
Q5PJE3 Q5PJE8 Q5PJE9 Q5PJF2 Q5PJF4 Q5PJF5 Q5PJG1 Q5PJG4 Q5PJG5 Q5PJH0 Q5PJH1
Q5PJH7 Q5PJI5 Q5PJI6 Q5PJJ0 Q5PJJ2 Q5PJJ7 Q5PJJ8 Q5PJN1 Q5PJN7 Q5PJN8 Q5PJS5
Q5PJS6 Q5PJS9 Q5PJU3 Q5PJV0 Q5PJW3 Q5PJW5 Q5PJX1 Q5PJX2 Q5PJX6 Q5PJX8 Q5PJX9
Q5PJY0 Q5PJY9 Q5PJZ0 Q5PJZ5 Q5PK00 Q5PK01 Q5PK02 Q5PK03 Q5PK05 Q5PK06 Q5PK07
Q5PK08 Q5PK09 Q5PK10 Q5PK12 Q5PK16 Q5PK20 Q5PK21 Q5PK22 Q5PK29 Q5PK30 Q5PK31
Q5PK44 Q5PK45 Q5PK48 Q5PK51 Q5PK55 Q5PK58 Q5PK60 Q5PK68 Q5PK71 Q5PK73 Q5PK74
Q5PK75 Q5PK76 Q5PK78 Q5PK79 Q5PK80 Q5PK81 Q5PK87 Q5PK92 Q5PK93 Q5PK94 Q5PK95
Q5PK96 Q5PKA2 Q5PKA3 Q5PKA4 Q5PKA5 Q5PKA6 Q5PKA7 Q5PKA9 Q5PKB8 Q5PKC6 Q5PKE2
Q5PKF1 Q5PKG0 Q5PKG1 Q5PKG2 Q5PKG3 Q5PKG4 Q5PKG5 Q5PKG7 Q5PKI2 Q5PKI4 Q5PKJ2
Q5PKJ6 Q5PKK4 Q5PKL7 Q5PKM6 Q5PKM9 Q5PKP4 Q5PKP9 Q5PKQ1 Q5PKR3 Q5PKS5 Q5PKS6
Q5PKT4 Q5PKU1 Q5PKU4 Q5PKU5 Q5PKU6 Q5PKU8 Q5PKV5 Q5PKV9 Q5PKX5 Q5PKX8 Q5PKX9
Q5PKY6 Q5PKZ6 Q5PL07 Q5PL18 Q5PL24 Q5PL30 Q5PL31 Q5PL41 Q5PL47 Q5PL48 Q5PL49
Q5PL57 Q5PL58 Q5PL62 Q5PL63 Q5PL69 Q5PL77 Q5PL82 Q5PL86 Q5PL88 Q5PL92 Q5PL93
Q5PL97 Q5PLA5 Q5PLA8 Q5PLB0 Q5PLB1 Q5PLB2 Q5PLB3 Q5PLB4 Q5PLB5 Q5PLB7 Q5PLC1
Q5PLC4 Q5PLD1 Q5PLD3 Q5PLE9 Q5PLF1 Q5PLF2 Q5PLG7 Q5PLH2 Q5PLH7 Q5PLH8 Q5PLI3
Q5PLI6 Q5PLM6 Q5PLN1 Q5PLN2 Q5PLN6 Q5PLQ2 Q5PLR2 Q5PLR3 Q5PLR5 Q5PLR9 Q5PLS1
Q5PLS8 Q5PLT0 Q5PLV3 Q5PLV8 Q5PLX1 Q5PLX2 Q5PLX6 Q5PLX8 Q5PLY0 Q5PLY6 Q5PLY8
Q5PLZ0 Q5PLZ7 Q5PM00 Q5PM06 Q5PM07 Q5PM08 Q5PM09 Q5PM73 Q5PM78 Q5PM81 Q5PM85
Q5PM88 Q5PM97 Q5PM99 Q5PMA0 Q5PMA7 Q5PMB1 Q5PMF1 Q5PMJ4 Q5PMK2 Q5PMK3 Q5PMK9
Q5PML0 Q5PMM1 Q5PMN6 Q5PMP2 Q5PMP8 Q5PMT8 Q5PMU5 Q5PMV9 Q5PMW8 Q5PMY8 Q5PMY9
Q5PMZ0 Q5PMZ2 Q5PMZ4 Q5PMZ6 Q5PN00 Q5PN01 Q5PN20 Q5PN21 Q5PN45 Q5PN69 Q5PN73
Q5PN75 Q5PN76 Q5PN90 Q5PNB8 Q5PNC8 Q5PNE2 Q5PNE5 Q5PNF2 Q5PNG1 Q5PNG2 Q5PNG3
Q5PNG5 Q5PNH1 Q5PNH2 Q5PNH7 Q5PNI1 Q5PNI2 Q5PNI4 Q5PNI6 Q5PNJ7 Q5PNJ8 Q5PNL2
Q5PNL7 Q5PNL9 Q5PNM0 Q5PNN8 Q5PNP6 Q5PNS0 Q5PNU4 Q5PNY6 Q5PP38 Q5PP62 Q5PP65
Q5PP70 Q5PPG4 Q5PPG6 Q5PPG7 Q5PPH0 Q5PPH4 Q5PPI1 Q5PPI3 Q5PPI9 Q5PPJ2 Q5PPJ4
Q5PPL1 Q5PPL2 Q5PPL8 Q5PPM5 Q5PPN2 Q5PPN4 Q5PPN5 Q5PPP3 Q5PPP4 Q5PPR6 Q5PPS5
Q5PPU8 Q5PPV2 Q5PPV3 Q5PPV4 Q5PPV5 Q5PPX0 Q5PPY1 Q5PPY6 Q5PPZ5 Q5PQ04 Q5PQ23
Q5PQ25 Q5PQ30 Q5PQ44 Q5PQ59 Q5PQ71 Q5PQ76 Q5PQ89 Q5PQ90 Q5PQ92 Q5PQA5 Q5PQJ6
Q5PQJ7 Q5PQJ9 Q5PQK1 Q5PQK4 Q5PQK8 Q5PQL2 Q5PQL4 Q5PQL8 Q5PQM2 Q5PQM4 Q5PQM8
Q5PQN1 Q5PQN4 Q5PQN5 Q5PQN6 Q5PQN7 Q5PQN9 Q5PQP0 Q5PQP1 Q5PQP5 Q5PQQ1 Q5PQQ2
Q5PQQ6 Q5PQQ7 Q5PQR3 Q5PQR4 Q5PQR5 Q5PQR6 Q5PQR8 Q5PQS2 Q5PQS6 Q5PQS7 Q5PQT3
Q5PR01 Q5PR19 Q5PR55 Q5PR66 Q5PR67 Q5PR69 Q5PRA2 Q5PRB3 Q5PRC7 Q5PRE5 Q5PRF0
Q5PSV4 Q5PU89 Q5PWZ8 Q5PXE9 Q5PXQ6 Q5PXT2 Q5PYI0 Q5PZ43 Q5Q0B3 Q5Q0E6 Q5Q0U5

Q5QA71 Q5QA72 Q5QA74 Q5QA75 Q5QA76 Q5QA83 Q5QCP2 Q5QD03 Q5QE78 Q5QE79 Q5QE80
Q5QGS0 Q5QGZ8 Q5QIT3 Q5QJ38 Q5QJ74 Q5QJC3 Q5QJC4 Q5QJF4 Q5QJJ0 Q5QJV7 Q5QKR7
Q5QKR8 Q5QL03 Q5QLR2 Q5QLR5 Q5QLS7 Q5QM06 Q5QM27 Q5QMG3 Q5QMM3 Q5QMN3 Q5QMN4
Q5QMN5 Q5QMP3 Q5QN75 Q5QNA6 Q5QNI5 Q5QNV8 Q5QNW6 Q5QT59 Q5QTW7 Q5QTY6 Q5QTY8
Q5QTY9 Q5QTZ0 Q5QTZ1 Q5QTZ3 Q5QU03 Q5QU05 Q5QU19 Q5QU21 Q5QU38 Q5QUB2 Q5QUB6
Q5QUB8 Q5QUC3 Q5QUC4 Q5QUC5 Q5QUC6 Q5QUD0 Q5QUE0 Q5QUE2 Q5QUE5 Q5QUF4 Q5QUG6
Q5QUK1 Q5QUK3 Q5QUN4 Q5QUN6 Q5QUN8 Q5QUP2 Q5QUP9 Q5QUS3 Q5QUS6 Q5QUT9 Q5QUU1
Q5QUU8 Q5QUU9 Q5QUV0 Q5QUV7 Q5QUV8 Q5QUZ5 Q5QUZ6 Q5QUZ8 Q5QV03 Q5QV04 Q5QV06
Q5QV08 Q5QV29 Q5QV30 Q5QV32 Q5QV35 Q5QV36 Q5QV38 Q5QV40 Q5QV44 Q5QV46 Q5QV48
Q5QV55 Q5QV59 Q5QVA8 Q5QVB3 Q5QVD9 Q5QVE5 Q5QVE8 Q5QVF0 Q5QVF3 Q5QVL1 Q5QVL2
Q5QVL5 Q5QVL6 Q5QVM3 Q5QVM4 Q5QVM7 Q5QVN7 Q5QVP2 Q5QVP3 Q5QVP5 Q5QVP9 Q5QVQ4
Q5QVQ6 Q5QVQ8 Q5QVR4 Q5QVR5 Q5QVR9 Q5QVT1 Q5QVT3 Q5QVT4 Q5QVT8 Q5QVT9 Q5QVU2
Q5QVW2 Q5QVW3 Q5QVW5 Q5QVX3 Q5QVZ8 Q5QW04 Q5QW08 Q5QW15 Q5QW31 Q5QW40 Q5QW47
Q5QW88 Q5QW89 Q5QW95 Q5QW99 Q5QWA0 Q5QWA3 Q5QWA4 Q5QWA5 Q5QWA6 Q5QWA7 Q5QWA8
Q5QWA9 Q5QWB4 Q5QWB5 Q5QWB6 Q5QWC6 Q5QWC7 Q5QWE2 Q5QWH6 Q5QWI1 Q5QWI9 Q5QWP6
Q5QWP8 Q5QWQ5 Q5QWQ6 Q5QWQ7 Q5QWQ9 Q5QWR0 Q5QWR1 Q5QWR2 Q5QWR3 Q5QWS1 Q5QWW0
Q5QWW5 Q5QWY0 Q5QWY8 Q5QWZ3 Q5QWZ9 Q5QX02 Q5QX59 Q5QX63 Q5QX64 Q5QX79 Q5QX82
Q5QX98 Q5QXB8 Q5QXC3 Q5QXE0 Q5QXH5 Q5QXH9 Q5QXI3 Q5QXI5 Q5QXI6 Q5QXJ0 Q5QXJ3
Q5QXJ4 Q5QXJ5 Q5QXK8 Q5QXL1 Q5QXL2 Q5QXL5 Q5QXL7 Q5QXL8 Q5QXL9 Q5QXM6 Q5QXN7
Q5QXN8 Q5QXN9 Q5QXP4 Q5QXQ2 Q5QXS0 Q5QXS1 Q5QXS2 Q5QXS3 Q5QXS9 Q5QXT0 Q5QXT4
Q5QXT5 Q5QXT8 Q5QXT9 Q5QXU1 Q5QXU4 Q5QXV0 Q5QXV4 Q5QXV5 Q5QXV6 Q5QXV7 Q5QXV8
Q5QXV9 Q5QXW0 Q5QXW1 Q5QXW2 Q5QXW3 Q5QXW4 Q5QXW8 Q5QXW9 Q5QXX0 Q5QXX1 Q5QXX2
Q5QXX3 Q5QXX9 Q5QXY0 Q5QXY1 Q5QXY2 Q5QXY3 Q5QXY4 Q5QXY9 Q5QXZ0 Q5QXZ1 Q5QY01
Q5QY03 Q5QY04 Q5QY05 Q5QY14 Q5QY15 Q5QY23 Q5QY35 Q5QY37 Q5QY39 Q5QY46 Q5QY51
Q5QY57 Q5QY58 Q5QY61 Q5QY68 Q5QY69 Q5QY76 Q5QY78 Q5QYA9 Q5QYB0 Q5QYB3 Q5QYB9
Q5QYC0 Q5QYC3 Q5QYC5 Q5QYC8 Q5QYD1 Q5QYD9 Q5QYE1 Q5QYE5 Q5QYE6 Q5QYE7 Q5QYF8
Q5QYG2 Q5QYH9 Q5QYI0 Q5QYL5 Q5QYN4 Q5QYN5 Q5QYN6 Q5QYN7 Q5QYN9 Q5QYP3 Q5QYU4
Q5QYU5 Q5QYV0 Q5QYV2 Q5QYV3 Q5QYV9 Q5QYY1 Q5QYY9 Q5QYZ7 Q5QZ01 Q5QZ06 Q5QZ13
Q5QZ16 Q5QZ19 Q5QZ32 Q5QZ35 Q5QZ37 Q5QZ42 Q5QZ46 Q5QZ48 Q5QZ49 Q5QZ50 Q5QZ53
Q5QZA7 Q5QZA9 Q5QZB3 Q5QZB4 Q5QZB6 Q5QZB7 Q5QZC0 Q5QZC1 Q5QZC4 Q5QZC8 Q5QZH5
Q5QZI8 Q5QZI9 Q5QZJ5 Q5QZK0 Q5QZK1 Q5QZK7 Q5QZL0 Q5QZL3 Q5QZR7 Q5QZS3 Q5QZS6
Q5R027 Q5R030 Q5R033 Q5R039 Q5R041 Q5R049 Q5R053 Q5R059 Q5R064 Q5R080 Q5R0A6
Q5R0B0 Q5R0B5 Q5R0C3 Q5R0C5 Q5R0F4 Q5R0G6 Q5R0H4 Q5R0I2 Q5R0J5 Q5R0K5 Q5R0K6
Q5R0L0 Q5R0L7 Q5R0L8 Q5R0M0 Q5R0M3 Q5R0N1 Q5R0N3 Q5R0N4 Q5R0N5 Q5R0Q5 Q5R0Q6
Q5R0Q7 Q5R0R2 Q5R0R3 Q5R0V3 Q5R0Z7 Q5R106 Q5R108 Q5R109 Q5R111 Q5R142 Q5R146
Q5R191 Q5R192 Q5R1S9 Q5R1T0 Q5R1U3 Q5R1W1 Q5R1W4 Q5R1W5 Q5R1W9 Q5R1X0 Q5R1X4
Q5R2J2 Q5R3Z9 Q5R405 Q5R408 Q5R413 Q5R414 Q5R421 Q5R431 Q5R432 Q5R434 Q5R444
Q5R462 Q5R467 Q5R469 Q5R481 Q5R483 Q5R486 Q5R490 Q5R493 Q5R498 Q5R4A0 Q5R4A6
Q5R4B1 Q5R4B3 Q5R4B4 Q5R4B7 Q5R4C1 Q5R4C4 Q5R4C7 Q5R4D6 Q5R4E0 Q5R4E2 Q5R4E9
Q5R4F3 Q5R4F5 Q5R4F9 Q5R4H1 Q5R4H3 Q5R4H5 Q5R4I2 Q5R4I8 Q5R4J1 Q5R4J9 Q5R4K0
Q5R4K5 Q5R4K8 Q5R4L5 Q5R4L6 Q5R4M1 Q5R4M2 Q5R4N5 Q5R4N9 Q5R4P6 Q5R4P8 Q5R4Q5
Q5R4Q8 Q5R4R2 Q5R4R3 Q5R4R4 Q5R4S3 Q5R4S6 Q5R4S7 Q5R4T3 Q5R4T7 Q5R4T8 Q5R4T9
Q5R4U0 Q5R4U2 Q5R4U3 Q5R4V3 Q5R4V4 Q5R4V9 Q5R4W0 Q5R4W3 Q5R4W4 Q5R4W7 Q5R4W8
Q5R4X0 Q5R4X1 Q5R4X5 Q5R4Y2 Q5R4Z1 Q5R4Z6 Q5R503 Q5R507 Q5R514 Q5R516 Q5R522

Q5R526 Q5R538 Q5R539 Q5R558 Q5R559 Q5R561 Q5R562 Q5R565 Q5R573 Q5R578 Q5R580
Q5R581 Q5R585 Q5R587 Q5R590 Q5R592 Q5R596 Q5R5C2 Q5R5C8 Q5R5C9 Q5R5D0 Q5R5D8
Q5R5E5 Q5R5E7 Q5R5E9 Q5R5F0 Q5R5F1 Q5R5F5 Q5R5F8 Q5R5G8 Q5R5H1 Q5R5H8 Q5R5I4
Q5R5J3 Q5R5J5 Q5R5J6 Q5R5J7 Q5R5K6 Q5R5M1 Q5R5M3 Q5R5M6 Q5R5N4 Q5R5N8 Q5R5N9
Q5R5P0 Q5R5P3 Q5R5P4 Q5R5Q2 Q5R5Q6 Q5R5R2 Q5R5R3 Q5R5R4 Q5R5R5 Q5R5S1 Q5R5S4
Q5R5S6 Q5R5S9 Q5R5T0 Q5R5T1 Q5R5T5 Q5R5U3 Q5R5V0 Q5R5V3 Q5R5V9 Q5R5W0 Q5R5W2
Q5R5W9 Q5R5X0 Q5R5X3 Q5R5X9 Q5R5Y2 Q5R5Y7 Q5R5Z0 Q5R5Z5 Q5R600 Q5R606 Q5R607
Q5R610 Q5R614 Q5R623 Q5R628 Q5R629 Q5R631 Q5R633 Q5R636 Q5R637 Q5R638 Q5R648
Q5R649 Q5R650 Q5R652 Q5R654 Q5R655 Q5R663 Q5R667 Q5R669 Q5R670 Q5R673 Q5R675
Q5R678 Q5R679 Q5R680 Q5R691 Q5R694 Q5R698 Q5R6B4 Q5R6C7 Q5R6C9 Q5R6D0 Q5R6D6
Q5R6E0 Q5R6E1 Q5R6F3 Q5R6G1 Q5R6G3 Q5R6H6 Q5R6H7 Q5R6I0 Q5R6I2 Q5R6I3 Q5R6J0
Q5R6J8 Q5R6L1 Q5R6L3 Q5R6L5 Q5R6P2 Q5R6P5 Q5R6P6 Q5R6Q5 Q5R6R2 Q5R6R5 Q5R6R7
Q5R6R8 Q5R6S3 Q5R6U8 Q5R6W3 Q5R6W9 Q5R6X9 Q5R6Y0 Q5R6Y5 Q5R6Y9 Q5R6Z6 Q5R6Z7
Q5R6Z8 Q5R706 Q5R715 Q5R723 Q5R724 Q5R732 Q5R737 Q5R740 Q5R741 Q5R746 Q5R754
Q5R760 Q5R761 Q5R762 Q5R774 Q5R776 Q5R782 Q5R788 Q5R789 Q5R790 Q5R796 Q5R797
Q5R7A8 Q5R7B0 Q5R7B3 Q5R7B8 Q5R7C3 Q5R7C4 Q5R7D1 Q5R7D3 Q5R7D8 Q5R7F2 Q5R7F7
Q5R7F8 Q5R7F9 Q5R7G1 Q5R7G6 Q5R7G9 Q5R7H0 Q5R7H2 Q5R7H5 Q5R7H8 Q5R7I3 Q5R7I8
Q5R7J6 Q5R7J9 Q5R7K0 Q5R7K1 Q5R7K3 Q5R7K4 Q5R7L2 Q5R7L3 Q5R7L4 Q5R7L9 Q5R7M2
Q5R7N3 Q5R7N8 Q5R7P2 Q5R7P3 Q5R7P6 Q5R7P7 Q5R7Q4 Q5R7Q6 Q5R7R2 Q5R7R9 Q5R7S6
Q5R7S7 Q5R7T2 Q5R7T5 Q5R7T9 Q5R7U0 Q5R7U1 Q5R7U7 Q5R7U8 Q5R7V5 Q5R7W0 Q5R7W3
Q5R7X0 Q5R7X1 Q5R7X2 Q5R7X9 Q5R7Y1 Q5R7Y8 Q5R7Z5 Q5R7Z8 Q5R802 Q5R810 Q5R811
Q5R814 Q5R815 Q5R816 Q5R824 Q5R829 Q5R830 Q5R837 Q5R840 Q5R844 Q5R846 Q5R863
Q5R864 Q5R869 Q5R886 Q5R887 Q5R889 Q5R893 Q5R895 Q5R896 Q5R899 Q5R8A2 Q5R8A4
Q5R8B2 Q5R8C2 Q5R8C4 Q5R8C5 Q5R8D0 Q5R8D1 Q5R8D4 Q5R8D7 Q5R8D8 Q5R8E2 Q5R8E4
Q5R8F7 Q5R8F8 Q5R8G6 Q5R8G9 Q5R8H0 Q5R8H5 Q5R8I2 Q5R8J5 Q5R8J6 Q5R8J7 Q5R8K2
Q5R8K3 Q5R8K4 Q5R8K6 Q5R8K9 Q5R8L6 Q5R8M4 Q5R8M9 Q5R8N8 Q5R8P0 Q5R8P3 Q5R8Q7
Q5R8Q8 Q5R8R1 Q5R8R4 Q5R8S0 Q5R8S2 Q5R8S9 Q5R8T6 Q5R8T8 Q5R8U1 Q5R8U3 Q5R8V2
Q5R8W1 Q5R8W6 Q5R8W8 Q5R8X0 Q5R8X1 Q5R8X8 Q5R8X9 Q5R8Y3 Q5R8Y8 Q5R8Z3 Q5R8Z4
Q5R903 Q5R905 Q5R919 Q5R924 Q5R925 Q5R928 Q5R931 Q5R938 Q5R939 Q5R943 Q5R947
Q5R952 Q5R962 Q5R966 Q5R971 Q5R974 Q5R978 Q5R979 Q5R995 Q5R9B3 Q5R9B8 Q5R9C1
Q5R9C3 Q5R9C5 Q5R9C9 Q5R9D1 Q5R9D2 Q5R9E0 Q5R9E5 Q5R9F0 Q5R9G0 Q5R9G3 Q5R9H0
Q5R9H2 Q5R9H4 Q5R9I5 Q5R9I6 Q5R9J1 Q5R9J2 Q5R9J5 Q5R9K8 Q5R9L2 Q5R9L3 Q5R9L4
Q5R9L6 Q5R9L8 Q5R9M8 Q5R9N0 Q5R9N7 Q5R9P4 Q5R9P5 Q5R9P9 Q5R9Q5 Q5R9Q6 Q5R9Q7
Q5R9R1 Q5R9R4 Q5R9S0 Q5R9S5 Q5R9T6 Q5R9T9 Q5R9U6 Q5R9U8 Q5R9U9 Q5R9V1 Q5R9V6
Q5R9W0 Q5R9W6 Q5R9W8 Q5R9W9 Q5R9X6 Q5R9Y3 Q5R9Y9 Q5R9Z6 Q5R9Z7 Q5RA02 Q5RA07
Q5RA17 Q5RA20 Q5RA22 Q5RA23 Q5RA29 Q5RA30 Q5RA37 Q5RA38 Q5RA42 Q5RA50 Q5RA52
Q5RA61 Q5RA63 Q5RA67 Q5RA68 Q5RA72 Q5RA75 Q5RA78 Q5RA80 Q5RA87 Q5RA89 Q5RA91
Q5RA93 Q5RA95 Q5RA96 Q5RAA0 Q5RAA9 Q5RAB0 Q5RAD5 Q5RAE0 Q5RAE6 Q5RAF2 Q5RAG0
Q5RAG3 Q5RAG9 Q5RAH3 Q5RAH6 Q5RAH8 Q5RAH9 Q5RAI7 Q5RAI8 Q5RAJ5 Q5RAJ7 Q5RAK6
Q5RAK7 Q5RAM2 Q5RAM7 Q5RAM8 Q5RAM9 Q5RAN1 Q5RAN2 Q5RAP0 Q5RAP1 Q5RAQ2 Q5RAQ3
Q5RAQ5 Q5RAQ7 Q5RAQ8 Q5RAR0 Q5RAR5 Q5RAR8 Q5RAS1 Q5RAS2 Q5RAS7 Q5RAT4 Q5RAT5
Q5RAT8 Q5RAU7 Q5RAV3 Q5RAV7 Q5RAV9 Q5RAW7 Q5RAW8 Q5RAX0 Q5RAX6 Q5RAX7 Q5RAX9
Q5RAY0 Q5RAY5 Q5RAY7 Q5RAZ2 Q5RAZ4 Q5RAZ5 Q5RAZ9 Q5RB07 Q5RB16 Q5RB30 Q5RB32
Q5RB33 Q5RB36 Q5RB44 Q5RB52 Q5RB58 Q5RB59 Q5RB63 Q5RB68 Q5RB69 Q5RB71 Q5RB73

Q5RB75 Q5RB77 Q5RB79 Q5RB81 Q5RB83 Q5RB90 Q5RB99 Q5RBA1 Q5RBA3 Q5RBA4 Q5RBA5
Q5RBA8 Q5RBB3 Q5RBB5 Q5RBB8 Q5RBB9 Q5RBD3 Q5RBD4 Q5RBD5 Q5RBD6 Q5RBD9 Q5RBE4
Q5RBE5 Q5RBF3 Q5RBF9 Q5RBG2 Q5RBH2 Q5RBI3 Q5RBI5 Q5RBI7 Q5RBJ3 Q5RBJ6 Q5RBK0
Q5RBK1 Q5RBK9 Q5RBM6 Q5RBM7 Q5RBM8 Q5RBN6 Q5RBN8 Q5RBN9 Q5RBP7 Q5RBQ3 Q5RBQ7
Q5RBR0 Q5RBR4 Q5RBR6 Q5RBR8 Q5RBR9 Q5RBS5 Q5RBS7 Q5RBT0 Q5RBT2 Q5RBT4 Q5RBU1
Q5RBU2 Q5RBU3 Q5RBU4 Q5RBV4 Q5RBV9 Q5RBW0 Q5RBW7 Q5RBW9 Q5RBX0 Q5RBY6 Q5RBY8
Q5RBY9 Q5RBZ2 Q5RBZ3 Q5RBZ4 Q5RC01 Q5RC02 Q5RC03 Q5RC05 Q5RC14 Q5RC20 Q5RC21
Q5RC28 Q5RC31 Q5RC32 Q5RC34 Q5RC36 Q5RC37 Q5RC41 Q5RC42 Q5RC43 Q5RC51 Q5RC53
Q5RC59 Q5RC62 Q5RC63 Q5RC65 Q5RC67 Q5RC69 Q5RC79 Q5RC80 Q5RC87 Q5RC94 Q5RCA3
Q5RCB7 Q5RCC2 Q5RCC9 Q5RCD2 Q5RCD9 Q5RCE0 Q5RCE4 Q5RCE5 Q5RCE6 Q5RCF3 Q5RCG0
Q5RCG9 Q5RCH1 Q5RCH4 Q5RCI5 Q5RCJ0 Q5RCJ2 Q5RCJ3 Q5RCJ6 Q5RCM2 Q5RCP1 Q5RCP3
Q5RCP7 Q5RCP8 Q5RCQ0 Q5RCQ2 Q5RCQ9 Q5RCR3 Q5RCR9 Q5RCS6 Q5RCS9 Q5RCT1 Q5RCU0
Q5RCU2 Q5RCU4 Q5RCV7 Q5RCW7 Q5RCX2 Q5RCX3 Q5RCX4 Q5RCX5 Q5RCY1 Q5RCY2 Q5RCY5
Q5RCZ5 Q5RCZ7 Q5RD03 Q5RD06 Q5RD08 Q5RD13 Q5RD18 Q5RD21 Q5RD26 Q5RD40 Q5RD48
Q5RD50 Q5RD56 Q5RD58 Q5RD60 Q5RD68 Q5RD71 Q5RD73 Q5RD75 Q5RD78 Q5RD86 Q5RD88
Q5RD94 Q5RD97 Q5RDA1 Q5RDA3 Q5RDA9 Q5RDB9 Q5RDC3 Q5RDC7 Q5RDD2 Q5RDE1 Q5RDE3
Q5RDE7 Q5RDF9 Q5RDG2 Q5RDG3 Q5RDG7 Q5RDH2 Q5RDH8 Q5RDI0 Q5RDI2 Q5RDI7 Q5RDJ2
Q5RDK0 Q5RDK1 Q5RDK5 Q5RDL2 Q5RDL5 Q5RDL6 Q5RDL7 Q5RDN3 Q5RDN9 Q5RDP3 Q5RDP4
Q5RDQ0 Q5RDQ3 Q5RDQ4 Q5RDQ6 Q5RDR2 Q5RDS2 Q5RDT5 Q5RDU4 Q5RDU7 Q5RDU9 Q5RDV0
Q5RDV1 Q5RDV8 Q5RDW0 Q5RDW3 Q5RDW9 Q5RDX1 Q5RDX4 Q5RDY1 Q5RDY3 Q5RDZ0 Q5RDZ1
Q5RDZ2 Q5RE03 Q5RE06 Q5RE08 Q5RE09 Q5RE12 Q5RE14 Q5RE15 Q5RE16 Q5RE18 Q5RE21
Q5RE25 Q5RE30 Q5RE46 Q5RE47 Q5RE50 Q5RE52 Q5RE62 Q5RE63 Q5RE65 Q5RE70 Q5RE79
Q5RE82 Q5RE88 Q5RE95 Q5REA0 Q5REA8 Q5REB1 Q5REC0 Q5REC6 Q5RED5 Q5REE0 Q5REE1
Q5REE6 Q5REE8 Q5REF1 Q5REF4 Q5REF9 Q5REG1 Q5REG4 Q5REG6 Q5REG8 Q5REH3 Q5REH5
Q5REH8 Q5REI6 Q5REI7 Q5REJ0 Q5REJ1 Q5REJ2 Q5REJ9 Q5REK1 Q5REK3 Q5REK4 Q5REL3
Q5REL6 Q5REM2 Q5REM3 Q5REN1 Q5REN2 Q5REN4 Q5REP4 Q5REP9 Q5REQ8 Q5REQ9 Q5RER5
Q5RER6 Q5RER9 Q5RES2 Q5RES3 Q5RES4 Q5RES7 Q5RES8 Q5RET3 Q5RET9 Q5REU2 Q5REW0
Q5REW7 Q5REX3 Q5REX5 Q5REX9 Q5REY0 Q5REY2 Q5REY3 Q5REY4 Q5REY7 Q5REY8 Q5REY9
Q5REZ5 Q5RF00 Q5RF02 Q5RF04 Q5RF05 Q5RF14 Q5RF18 Q5RF24 Q5RF28 Q5RF31 Q5RF32
Q5RF36 Q5RF43 Q5RF51 Q5RF54 Q5RF63 Q5RF69 Q5RF70 Q5RF77 Q5RF83 Q5RF88 Q5RF92
Q5RF97 Q5RF99 Q5RFA0 Q5RFA2 Q5RFA3 Q5RFA6 Q5RFA9 Q5RFB0 Q5RFB3 Q5RFB8 Q5RFD3
Q5RFE6 Q5RFE9 Q5RFF4 Q5RFF7 Q5RFF8 Q5RFG2 Q5RFG5 Q5RFG8 Q5RFG9 Q5RFH1 Q5RFH3
Q5RFH5 Q5RFI3 Q5RFI6 Q5RFI8 Q5RFJ0 Q5RFJ1 Q5RFJ7 Q5RFJ8 Q5RFK0 Q5RFK5 Q5RFL8
Q5RFL9 Q5RFM3 Q5RFM7 Q5RFM9 Q5RFN0 Q5RFN2 Q5RFN4 Q5RFN6 Q5RFP0 Q5RFP4 Q5RFP5
Q5RFQ0 Q5RFQ1 Q5RFQ3 Q5RFR0 Q5RFR7 Q5RFS0 Q5RFS7 Q5RFT7 Q5RFT9 Q5RFV8 Q5RFZ7
Q5RG44 Q5RG82 Q5RGA4 Q5RGB8 Q5RGE4 Q5RGE5 Q5RGJ5 Q5RGJ6 Q5RGP9 Q5RGS7 Q5RGT6
Q5RGV1 Q5RGZ1 Q5RH01 Q5RHD1 Q5RHH4 Q5RHI5 Q5RHP9 Q5RHQ8 Q5RHU7 Q5RHX6 Q5RHY1
Q5RHZ6 Q5RI75 Q5RIA9 Q5RIC0 Q5RII9 Q5RIM0 Q5RIV4 Q5RIW8 Q5RJ20 Q5RJA1 Q5RJB0
Q5RJC5 Q5RJG1 Q5RJG7 Q5RJH6 Q5RJK8 Q5RJL0 Q5RJL2 Q5RJM0 Q5RJN7 Q5RJP9 Q5RJQ7
Q5RJR4 Q5RJS9 Q5RJT0 Q5RJT2 Q5RJU3 Q5RJV1 Q5RJW4 Q5RJW5 Q5RJX8 Q5RK00 Q5RK03
Q5RK09 Q5RK28 Q5RK30 Q5RKG1 Q5RKG3 Q5RKG6 Q5RKH0 Q5RKH1 Q5RKH3 Q5RKI1 Q5RKI3
Q5RKI9 Q5RKJ0 Q5RKN3 Q5RKN4 Q5RKQ0 Q5RKV6 Q5RKZ7 Q5RL51 Q5RL73 Q5RLY7 Q5RM09
Q5RT64 Q5S003 Q5S1S4 Q5S1U1 Q5S260 Q5S2C3 Q5S3I1 Q5S3I2 Q5S3I3 Q5S3I4 Q5S4V7
Q5S6T3 Q5SBK7 Q5SBL1 Q5SBL4 Q5SBL6 Q5SBP0 Q5SBP1 Q5SBP2 Q5SBP3 Q5SBP4 Q5SBP5

Q5SBP6 Q5SBP7 Q5SCB5 Q5SCV9 Q5SCW3 Q5SCW6 Q5SCW7 Q5SCX2 Q5SCX8 Q5SCX9 Q5SCY1
Q5SCY6 Q5SCY8 Q5SCY9 Q5SD00 Q5SD01 Q5SD08 Q5SD10 Q5SD12 Q5SD13 Q5SD14 Q5SD15
Q5SD16 Q5SD17 Q5SD18 Q5SD19 Q5SD20 Q5SD21 Q5SD25 Q5SD26 Q5SD27 Q5SD28 Q5SD32
Q5SD36 Q5SD37 Q5SF07 Q5SFA6 Q5SFD1 Q5SFM8 Q5SGK3 Q5SGV8 Q5SGX1 Q5SGX2 Q5SGY0
Q5SGY6 Q5SGZ3 Q5SH02 Q5SH03 Q5SH15 Q5SH22 Q5SH23 Q5SH28 Q5SH88 Q5SHA1 Q5SHB1
Q5SHB2 Q5SHD4 Q5SHD6 Q5SHE9 Q5SHF0 Q5SHF5 Q5SHH3 Q5SHH5 Q5SHI8 Q5SHL6 Q5SHN3
Q5SHN5 Q5SHN6 Q5SHN7 Q5SHN8 Q5SHN9 Q5SHP0 Q5SHP2 Q5SHP3 Q5SHP6 Q5SHP8 Q5SHP9
Q5SHQ0 Q5SHQ4 Q5SHQ5 Q5SHQ6 Q5SHQ7 Q5SHQ9 Q5SHR1 Q5SHR2 Q5SHR6 Q5SHS3 Q5SHS6
Q5SHT5 Q5SHT7 Q5SHT8 Q5SHU9 Q5SHW0 Q5SHW1 Q5SHX3 Q5SHY0 Q5SHY1 Q5SHZ1 Q5SHZ2
Q5SHZ3 Q5SHZ4 Q5SHZ8 Q5SI12 Q5SI14 Q5SI16 Q5SI18 Q5SI21 Q5SI26 Q5SI28 Q5SI32
Q5SI38 Q5SI49 Q5SI56 Q5SI57 Q5SI68 Q5SIA8 Q5SIC2 Q5SID2 Q5SID6 Q5SID7 Q5SIE3
Q5SIH0 Q5SIH3 Q5SIH5 Q5SIH7 Q5SII2 Q5SII4 Q5SII5 Q5SIJ1 Q5SIJ5 Q5SIL0 Q5SIL4
Q5SIL5 Q5SIM2 Q5SIP0 Q5SIP7 Q5SIQ7 Q5SIR5 Q5SIT1 Q5SIV8 Q5SIW2 Q5SIW6 Q5SIY0
Q5SIY4 Q5SJ03 Q5SJ18 Q5SJ26 Q5SJ28 Q5SJ35 Q5SJ39 Q5SJ45 Q5SJ47 Q5SJ49 Q5SJ58
Q5SJ65 Q5SJ75 Q5SJ76 Q5SJ85 Q5SJ93 Q5SJ94 Q5SJA1 Q5SJB8 Q5SJB9 Q5SJC0 Q5SJC4
Q5SJC8 Q5SJD2 Q5SJD8 Q5SJD9 Q5SJE1 Q5SJE2 Q5SJE8 Q5SJF8 Q5SJG3 Q5SJG6 Q5SJG7
Q5SJH3 Q5SJH5 Q5SJH6 Q5SJM4 Q5SJP7 Q5SJP8 Q5SJQ1 Q5SJQ9 Q5SJR1 Q5SJS4 Q5SJS7
Q5SJS9 Q5SJV1 Q5SJV8 Q5SJV9 Q5SJX7 Q5SJX8 Q5SJZ6 Q5SK01 Q5SK28 Q5SK31 Q5SK48
Q5SK49 Q5SK65 Q5SK66 Q5SK67 Q5SK88 Q5SK89 Q5SKC1 Q5SKC5 Q5SKD2 Q5SKD6 Q5SKD9
Q5SKG5 Q5SKG7 Q5SKG9 Q5SKL7 Q5SKM7 Q5SKM8 Q5SKM9 Q5SKN1 Q5SKN4 Q5SKN6 Q5SKN7
Q5SKP2 Q5SKR5 Q5SKS1 Q5SKS4 Q5SKS6 Q5SKT7 Q5SKU1 Q5SKU2 Q5SKV7 Q5SKW7 Q5SKW9
Q5SKX0 Q5SL28 Q5SL35 Q5SL36 Q5SL64 Q5SL86 Q5SL87 Q5SLC1 Q5SLC9 Q5SLE8 Q5SLF7
Q5SLH2 Q5SLH3 Q5SLH4 Q5SLI7 Q5SLI8 Q5SLL0 Q5SLL3 Q5SLL5 Q5SLL6 Q5SLM2 Q5SLM3
Q5SLN5 Q5SLP1 Q5SLP6 Q5SLP7 Q5SLP8 Q5SLP9 Q5SLQ0 Q5SLQ1 Q5SLQ3 Q5SLQ4 Q5SLR3
Q5SLR4 Q5SLR5 Q5SLS1 Q5SLS6 Q5SLU5 Q5SLV5 Q5SLW9 Q5SLX2 Q5SM01 Q5SM03 Q5SM09
Q5SM14 Q5SM15 Q5SM16 Q5SM25 Q5SM27 Q5SM28 Q5SM45 Q5SM60 Q5SMC1 Q5SMC5 Q5SMC7
Q5SMC8 Q5SMD7 Q5SME9 Q5SMF6 Q5SMH0 Q5SMH2 Q5SMH8 Q5SMI4 Q5SMK6 Q5SMM6 Q5SMM8
Q5SMQ0 Q5SMY5 Q5SN75 Q5SND2 Q5SNL7 Q5SNN4 Q5SNV9 Q5SP50 Q5SP85 Q5SP90 Q5SPB6
Q5SPD7 Q5SPH9 Q5SPJ8 Q5SPL2 Q5SPR8 Q5SPV6 Q5SPX1 Q5SQ80 Q5SQH8 Q5SQM0 Q5SQP1
Q5SQQ9 Q5SQS7 Q5SQS8 Q5SQX6 Q5SQY2 Q5SR53 Q5SRE7 Q5SRH9 Q5SRT8 Q5SRY7 Q5SS00
Q5SS90 Q5SSF7 Q5SSK3 Q5SSQ6 Q5SSW2 Q5ST30 Q5SU73 Q5SUD9 Q5SUE7 Q5SUE8 Q5SUF2
Q5SUR0 Q5SUS0 Q5SUV1 Q5SUV5 Q5SV06 Q5SV42 Q5SV77 Q5SVD0 Q5SVJ3 Q5SVQ0 Q5SVQ8
Q5SVT3 Q5SVZ6 Q5SW19 Q5SW24 Q5SW25 Q5SW75 Q5SWA1 Q5SWD9 Q5SWL7 Q5SWL8 Q5SWW4
Q5SWW7 Q5SX39 Q5SX40 Q5SXH7 Q5SXM1 Q5SXM8 Q5SY13 Q5SY68 Q5SY85 Q5SYL1 Q5SZ99
Q5SZB4 Q5SZD1 Q5SZJ8 Q5SZL2 Q5SZQ8 Q5SZT7 Q5T013 Q5T035 Q5T036 Q5T089 Q5T0J3
Q5T0J7 Q5T0U0 Q5T124 Q5T160 Q5T1B0 Q5T1J5 Q5T1J6 Q5T1N1 Q5T280 Q5T2L2 Q5T2N8
Q5T2S8 Q5T319 Q5T3Y7 Q5T440 Q5T447 Q5T481 Q5T4H9 Q5T4I8 Q5T4T6 Q5T5A8 Q5T5B0
Q5T5D7 Q5T5F5 Q5T5J6 Q5T5M9 Q5T5N4 Q5T5P2 Q5T5S1 Q5T619 Q5T655 Q5T681 Q5T686
Q5T6C5 Q5T6F0 Q5T6J7 Q5T6M2 Q5T6S3 Q5T6V5 Q5T750 Q5T751 Q5T752 Q5T753 Q5T754
Q5T764 Q5T7N3 Q5T7N8 Q5T7P2 Q5T7P3 Q5T7V8 Q5T7W0 Q5T7W7 Q5T870 Q5T871 Q5T890
Q5T8A7 Q5T8I3 Q5T8I9 Q5T8R8 Q5T953 Q5T9C2 Q5T9S5 Q5TA31 Q5TA45 Q5TA76 Q5TA77
Q5TA78 Q5TA79 Q5TA81 Q5TA82 Q5TA89 Q5TAA0 Q5TAB7 Q5TAG4 Q5TAP6 Q5TAX3 Q5TBB1
Q5TBC7 Q5TBE3 Q5TBK1 Q5TC04 Q5TC12 Q5TC63 Q5TC79 Q5TC82 Q5TCM9 Q5TD94 Q5TD97
Q5TEA3 Q5TEC3 Q5TEC6 Q5TEZ4 Q5TEZ5 Q5TF58 Q5TFE4 Q5TFG8 Q5TG53 Q5TGI4 Q5TGJ6

Q5TGL8 Q5TGP6 Q5TGS1 Q5THK1 Q5THR3 Q5TI25 Q5TIA1 Q5TID7 Q5TIS6 Q5TJ03 Q5TJ55
Q5TJ56 Q5TJ65 Q5TJE2 Q5TJE6 Q5TJE7 Q5TJF7 Q5TKD8 Q5TKP8 Q5TKR9 Q5TM17 Q5TM19
Q5TM26 Q5TM48 Q5TM49 Q5TM50 Q5TM51 Q5TM52 Q5TM55 Q5TM57 Q5TM59 Q5TM61 Q5TM64
Q5TM66 Q5TM74 Q5TM83 Q5TM84 Q5TNH5 Q5TP13 Q5TT27 Q5TTP0 Q5TU55 Q5TUE9 Q5TUF1
Q5TW90 Q5TXB2 Q5TYM7 Q5TYP4 Q5TYQ1 Q5TYQ3 Q5TYQ8 Q5TYS0 Q5TYS5 Q5TYV4 Q5TYW1
Q5TYW6 Q5TYX0 Q5TZ80 Q5TZE1 Q5TZF3 Q5TZK3 Q5U1W2 Q5U1W5 Q5U1X0 Q5U1X1 Q5U1Y4
Q5U1Y9 Q5U203 Q5U206 Q5U208 Q5U209 Q5U216 Q5U231 Q5U236 Q5U241 Q5U243 Q5U245
Q5U259 Q5U263 Q5U274 Q5U2M7 Q5U2N0 Q5U2N8 Q5U2P3 Q5U2Q3 Q5U2Q7 Q5U2R0 Q5U2R2
Q5U2R4 Q5U2R6 Q5U2S0 Q5U2T3 Q5U2T7 Q5U2U0 Q5U2W5 Q5U2X1 Q5U2X2 Q5U2Y8 Q5U2Y9
Q5U2Z0 Q5U309 Q5U311 Q5U374 Q5U378 Q5U379 Q5U385 Q5U3A8 Q5U3F2 Q5U3F8 Q5U3G4
Q5U3H2 Q5U3J8 Q5U3K5 Q5U3N5 Q5U3N7 Q5U3P0 Q5U3S1 Q5U3U0 Q5U3U4 Q5U3U6 Q5U3W0
Q5U3W3 Q5U3Y0 Q5U3Z0 Q5U3Z3 Q5U3Z6 Q5U4C1 Q5U4C3 Q5U4D9 Q5U4E8 Q5U4F3 Q5U4F6
Q5U4I3 Q5U4M5 Q5U4N7 Q5U4Q0 Q5U4Q9 Q5U4T7 Q5U4T8 Q5U4U4 Q5U4U6 Q5U4V2 Q5U4W1
Q5U4W6 Q5U4X3 Q5U4Z3 Q5U4Z8 Q5U504 Q5U508 Q5U526 Q5U528 Q5U538 Q5U550 Q5U560
Q5U575 Q5U580 Q5U581 Q5U593 Q5U597 Q5U5A6 Q5U5D0 Q5U5E5 Q5U5Q9 Q5U5R9 Q5U5V2
Q5U5X8 Q5U5Z8 Q5U623 Q5U649 Q5U752 Q5U8S9 Q5U8Z7 Q5U907 Q5U923 Q5U924 Q5U925
Q5UAK0 Q5UAL3 Q5UAM9 Q5UAP0 Q5UAP4 Q5UAP8 Q5UAS1 Q5UAS2 Q5UB07 Q5UBX3 Q5UBY2
Q5UE94 Q5UG13 Q5UL10 Q5ULZ4 Q5UNS7 Q5UNS8 Q5UNS9 Q5UNT0 Q5UNT1 Q5UNT3 Q5UNT4
Q5UNT5 Q5UNT6 Q5UNT7 Q5UNT8 Q5UNT9 Q5UNU0 Q5UNU1 Q5UNU2 Q5UNU3 Q5UNU4 Q5UNU5
Q5UNU9 Q5UNV0 Q5UNV1 Q5UNV2 Q5UNV3 Q5UNV4 Q5UNV6 Q5UNV7 Q5UNV8 Q5UNV9 Q5UNW2
Q5UNW3 Q5UNW4 Q5UNW5 Q5UNW6 Q5UNW7 Q5UNW8 Q5UNW9 Q5UNX1 Q5UNX2 Q5UNX3 Q5UNX4
Q5UNX5 Q5UNX6 Q5UNX7 Q5UNX8 Q5UNX9 Q5UNY1 Q5UNY2 Q5UNY3 Q5UNY4 Q5UNY5 Q5UNY6
Q5UNY8 Q5UNY9 Q5UNZ0 Q5UNZ2 Q5UNZ3 Q5UNZ4 Q5UNZ5 Q5UNZ6 Q5UNZ7 Q5UNZ8 Q5UNZ9
Q5UP00 Q5UP01 Q5UP02 Q5UP03 Q5UP04 Q5UP05 Q5UP06 Q5UP07 Q5UP08 Q5UP09 Q5UP10
Q5UP11 Q5UP12 Q5UP13 Q5UP14 Q5UP15 Q5UP16 Q5UP17 Q5UP18 Q5UP19 Q5UP20 Q5UP21
Q5UP22 Q5UP23 Q5UP24 Q5UP25 Q5UP26 Q5UP27 Q5UP29 Q5UP30 Q5UP31 Q5UP32 Q5UP34
Q5UP35 Q5UP36 Q5UP37 Q5UP38 Q5UP39 Q5UP40 Q5UP41 Q5UP42 Q5UP43 Q5UP44 Q5UP45
Q5UP46 Q5UP47 Q5UP48 Q5UP49 Q5UP50 Q5UP51 Q5UP52 Q5UP53 Q5UP54 Q5UP56 Q5UP57
Q5UP58 Q5UP59 Q5UP60 Q5UP61 Q5UP62 Q5UP63 Q5UP64 Q5UP65 Q5UP66 Q5UP67 Q5UP68
Q5UP69 Q5UP71 Q5UP72 Q5UP74 Q5UP75 Q5UP76 Q5UP77 Q5UP78 Q5UP79 Q5UP80 Q5UP81
Q5UP82 Q5UP83 Q5UP84 Q5UP85 Q5UP86 Q5UP87 Q5UP88 Q5UP89 Q5UP91 Q5UP92 Q5UP93
Q5UP94 Q5UP95 Q5UP96 Q5UP97 Q5UP98 Q5UP99 Q5UPA0 Q5UPA1 Q5UPA2 Q5UPA3 Q5UPA4
Q5UPA5 Q5UPA6 Q5UPA7 Q5UPA8 Q5UPA9 Q5UPB0 Q5UPB1 Q5UPB2 Q5UPB3 Q5UPB4 Q5UPB5
Q5UPB6 Q5UPB7 Q5UPB8 Q5UPB9 Q5UPC0 Q5UPC2 Q5UPC3 Q5UPC4 Q5UPC5 Q5UPC6 Q5UPC7
Q5UPC8 Q5UPC9 Q5UPD0 Q5UPD1 Q5UPD2 Q5UPD3 Q5UPD4 Q5UPD5 Q5UPD6 Q5UPD7 Q5UPD8
Q5UPD9 Q5UPE1 Q5UPE2 Q5UPE3 Q5UPE4 Q5UPE5 Q5UPE6 Q5UPE8 Q5UPE9 Q5UPF0 Q5UPF1
Q5UPF2 Q5UPF3 Q5UPF4 Q5UPF5 Q5UPF6 Q5UPF7 Q5UPF8 Q5UPF9 Q5UPG0 Q5UPG1 Q5UPG2
Q5UPG3 Q5UPG4 Q5UPG5 Q5UPG6 Q5UPG7 Q5UPG8 Q5UPG9 Q5UPH0 Q5UPH1 Q5UPH2 Q5UPH3
Q5UPH4 Q5UPH5 Q5UPH6 Q5UPH7 Q5UPH8 Q5UPH9 Q5UPI0 Q5UPI1 Q5UPI2 Q5UPI3 Q5UPI4
Q5UPI5 Q5UPI6 Q5UPI7 Q5UPI8 Q5UPI9 Q5UPJ0 Q5UPJ2 Q5UPJ3 Q5UPJ4 Q5UPJ6 Q5UPJ7
Q5UPJ8 Q5UPJ9 Q5UPK0 Q5UPK1 Q5UPK2 Q5UPK3 Q5UPK4 Q5UPK5 Q5UPK6 Q5UPK8 Q5UPK9
Q5UPL0 Q5UPL1 Q5UPL3 Q5UPL4 Q5UPL5 Q5UPL6 Q5UPL8 Q5UPL9 Q5UPM0 Q5UPM1 Q5UPM2
Q5UPM3 Q5UPM4 Q5UPM5 Q5UPM6 Q5UPM7 Q5UPM8 Q5UPM9 Q5UPN0 Q5UPN1 Q5UPN2 Q5UPN3
Q5UPN4 Q5UPN5 Q5UPN6 Q5UPN7 Q5UPN8 Q5UPN9 Q5UPP0 Q5UPP1 Q5UPP2 Q5UPP3 Q5UPP4

Q5UPP5 Q5UPP6 Q5UPP7 Q5UPP8 Q5UPP9 Q5UPQ0 Q5UPQ1 Q5UPQ2 Q5UPQ3 Q5UPQ4 Q5UPQ5
Q5UPQ6 Q5UPQ7 Q5UPQ8 Q5UPQ9 Q5UPR0 Q5UPR1 Q5UPR3 Q5UPR4 Q5UPR6 Q5UPR7 Q5UPR8
Q5UPR9 Q5UPS0 Q5UPS1 Q5UPS2 Q5UPS3 Q5UPS4 Q5UPS5 Q5UPS6 Q5UPS7 Q5UPS8 Q5UPS9
Q5UPT0 Q5UPT1 Q5UPT2 Q5UPT4 Q5UPT5 Q5UPT6 Q5UPT7 Q5UPT8 Q5UPT9 Q5UPU0 Q5UPU1
Q5UPU2 Q5UPU3 Q5UPU4 Q5UPU5 Q5UPU6 Q5UPU7 Q5UPU9 Q5UPV0 Q5UPV1 Q5UPV2 Q5UPV3
Q5UPV4 Q5UPV5 Q5UPV6 Q5UPV7 Q5UPV9 Q5UPW1 Q5UPW2 Q5UPW3 Q5UPW4 Q5UPW6 Q5UPW8
Q5UPW9 Q5UPX0 Q5UPX1 Q5UPX3 Q5UPX4 Q5UPX5 Q5UPX6 Q5UPX7 Q5UPX8 Q5UPX9 Q5UPY0
Q5UPY1 Q5UPY2 Q5UPY3 Q5UPY4 Q5UPY5 Q5UPY6 Q5UPY7 Q5UPY9 Q5UPZ0 Q5UPZ2 Q5UPZ3
Q5UPZ6 Q5UPZ7 Q5UPZ8 Q5UPZ9 Q5UQ00 Q5UQ01 Q5UQ02 Q5UQ03 Q5UQ04 Q5UQ05 Q5UQ06
Q5UQ07 Q5UQ08 Q5UQ09 Q5UQ10 Q5UQ11 Q5UQ12 Q5UQ13 Q5UQ14 Q5UQ15 Q5UQ16 Q5UQ17
Q5UQ18 Q5UQ19 Q5UQ20 Q5UQ21 Q5UQ22 Q5UQ24 Q5UQ25 Q5UQ26 Q5UQ29 Q5UQ31 Q5UQ32
Q5UQ34 Q5UQ35 Q5UQ36 Q5UQ38 Q5UQ39 Q5UQ40 Q5UQ41 Q5UQ42 Q5UQ43 Q5UQ45 Q5UQ46
Q5UQ47 Q5UQ48 Q5UQ49 Q5UQ50 Q5UQ51 Q5UQ52 Q5UQ53 Q5UQ54 Q5UQ55 Q5UQ57 Q5UQ58
Q5UQ59 Q5UQ60 Q5UQ61 Q5UQ62 Q5UQ64 Q5UQ65 Q5UQ66 Q5UQ67 Q5UQ68 Q5UQ69 Q5UQ70
Q5UQ71 Q5UQ72 Q5UQ73 Q5UQ74 Q5UQ75 Q5UQ76 Q5UQ78 Q5UQ79 Q5UQ80 Q5UQ81 Q5UQ83
Q5UQ84 Q5UQ85 Q5UQ87 Q5UQ89 Q5UQ91 Q5UQ92 Q5UQ93 Q5UQ94 Q5UQ95 Q5UQ96 Q5UQ97
Q5UQ98 Q5UQ99 Q5UQA1 Q5UQA2 Q5UQA3 Q5UQA4 Q5UQA5 Q5UQA6 Q5UQA7 Q5UQA8 Q5UQA9
Q5UQB0 Q5UQB1 Q5UQB2 Q5UQB3 Q5UQB4 Q5UQB5 Q5UQB6 Q5UQB8 Q5UQB9 Q5UQC0 Q5UQC1
Q5UQC2 Q5UQC3 Q5UQC4 Q5UQC5 Q5UQC6 Q5UQC7 Q5UQC8 Q5UQC9 Q5UQD0 Q5UQD1 Q5UQD2
Q5UQD4 Q5UQD5 Q5UQD6 Q5UQD7 Q5UQD8 Q5UQD9 Q5UQE0 Q5UQE1 Q5UQE5 Q5UQE6 Q5UQE7
Q5UQE8 Q5UQE9 Q5UQF0 Q5UQF1 Q5UQF2 Q5UQF3 Q5UQF4 Q5UQF5 Q5UQF6 Q5UQF8 Q5UQG1
Q5UQG2 Q5UQG3 Q5UQG4 Q5UQG6 Q5UQG9 Q5UQH0 Q5UQH1 Q5UQH2 Q5UQH3 Q5UQH5 Q5UQH6
Q5UQH7 Q5UQH8 Q5UQI0 Q5UQI1 Q5UQI6 Q5UQI7 Q5UQI8 Q5UQJ0 Q5UQJ1 Q5UQJ2 Q5UQJ3
Q5UQJ4 Q5UQJ6 Q5UQJ7 Q5UQJ9 Q5UQK0 Q5UQK1 Q5UQK2 Q5UQK3 Q5UQK5 Q5UQK6 Q5UQK7
Q5UQK8 Q5UQK9 Q5UQL0 Q5UQL1 Q5UQL2 Q5UQL3 Q5UQL4 Q5UQL6 Q5UQL7 Q5UQL8 Q5UQL9
Q5UQM0 Q5UQM1 Q5UQM2 Q5UQM3 Q5UQM4 Q5UQM5 Q5UQM6 Q5UQM7 Q5UQM8 Q5UQM9 Q5UQN0
Q5UQN1 Q5UQN3 Q5UQN4 Q5UQN6 Q5UQN7 Q5UQN9 Q5UQP1 Q5UQP3 Q5UQP4 Q5UQP5 Q5UQP6
Q5UQP8 Q5UQP9 Q5UQQ0 Q5UQQ1 Q5UQQ2 Q5UQQ3 Q5UQQ4 Q5UQQ5 Q5UQQ6 Q5UQQ7 Q5UQQ8
Q5UQQ9 Q5UQR0 Q5UQR1 Q5UQR2 Q5UQR4 Q5UQR5 Q5UQR7 Q5UQS0 Q5UQS1 Q5UQS2 Q5UQS4
Q5UQS5 Q5UQS6 Q5UQS7 Q5UQS8 Q5UQS9 Q5UQT0 Q5UQT1 Q5UQT3 Q5UQT4 Q5UQT5 Q5UQT6
Q5UQT7 Q5UQT9 Q5UQU0 Q5UQU1 Q5UQU2 Q5UQU3 Q5UQU4 Q5UQU5 Q5UQU6 Q5UQU7 Q5UQU8
Q5UQU9 Q5UQV0 Q5UQV1 Q5UQV3 Q5UQV4 Q5UQV5 Q5UQV7 Q5UQV8 Q5UQV9 Q5UQW0 Q5UQW1
Q5UQW2 Q5UQW5 Q5UQW7 Q5UQW8 Q5UQW9 Q5UQX0 Q5UQX1 Q5UQX2 Q5UQX3 Q5UQX4 Q5UQX5
Q5UQX6 Q5UQX8 Q5UQY0 Q5UQY1 Q5UQY2 Q5UQY4 Q5UQY5 Q5UQY6 Q5UQY7 Q5UQY8 Q5UQY9
Q5UQZ0 Q5UQZ2 Q5UQZ3 Q5UQZ4 Q5UQZ5 Q5UQZ6 Q5UQZ7 Q5UQZ9 Q5UR01 Q5UR03 Q5UR04
Q5UR05 Q5UR06 Q5UR07 Q5UR08 Q5UR09 Q5UR10 Q5UR12 Q5UR13 Q5UR14 Q5UR17 Q5UR18
Q5UR20 Q5UR21 Q5UR22 Q5UR23 Q5UR24 Q5UR25 Q5UR26 Q5UR27 Q5UR28 Q5UR29 Q5UR30
Q5UR32 Q5UR33 Q5UR35 Q5UR36 Q5UR37 Q5UR38 Q5UR39 Q5UR40 Q5UR41 Q5UR43 Q5UR44
Q5UR45 Q5UR47 Q5UR48 Q5UR49 Q5UR50 Q5UR51 Q5UR52 Q5UR53 Q5UR54 Q5UR55 Q5UR56
Q5UR57 Q5UR58 Q5UR59 Q5UR60 Q5UR62 Q5UR64 Q5UR66 Q5UR67 Q5UR68 Q5UR69 Q5UR70
Q5UR71 Q5UR72 Q5UR73 Q5UR74 Q5UR75 Q5UR76 Q5UR79 Q5UR82 Q5UR83 Q5UR84 Q5UR85
Q5UR86 Q5UR87 Q5UR89 Q5UR90 Q5UR91 Q5UR92 Q5UR93 Q5UR94 Q5UR95 Q5UR96 Q5UR97
Q5UR98 Q5URA0 Q5URA1 Q5URA2 Q5URA3 Q5URA4 Q5URA5 Q5URA6 Q5URA8 Q5URA9 Q5URB0
Q5URB1 Q5URB2 Q5URB3 Q5URB4 Q5URB5 Q5URB6 Q5URB7 Q5URB8 Q5URB9 Q5UT56 Q5UU75

Q5UWH2 Q5UWT2 Q5UWY4 Q5UX13 Q5UX21 Q5UX22 Q5UX23 Q5UX45 Q5UX56 Q5UX57 Q5UX65
Q5UX95 Q5UXB3 Q5UXB8 Q5UXB9 Q5UXD1 Q5UXG4 Q5UXG8 Q5UXH0 Q5UXH1 Q5UXH9 Q5UXK3
Q5UXM3 Q5UXM7 Q5UXN0 Q5UXN1 Q5UXP9 Q5UXQ3 Q5UXT4 Q5UXU0 Q5UXU6 Q5UXW0 Q5UXW4
Q5UXW7 Q5UXX3 Q5UXX4 Q5UXX8 Q5UXY3 Q5UY26 Q5UY41 Q5UY62 Q5UY71 Q5UY84 Q5UY89
Q5UY95 Q5UYC8 Q5UYF3 Q5UYF6 Q5UYG6 Q5UYI1 Q5UYM0 Q5UYM1 Q5UYM9 Q5UYP1 Q5UYP9
Q5UYQ8 Q5UYR3 Q5UYR5 Q5UYR8 Q5UYS2 Q5UYU5 Q5UYV3 Q5UYV8 Q5UZ24 Q5UZ40 Q5UZ46
Q5UZ47 Q5UZ50 Q5UZ63 Q5UZ65 Q5UZ84 Q5UZ90 Q5UZD3 Q5UZD7 Q5UZE5 Q5UZE6 Q5UZF0
Q5UZG9 Q5UZH1 Q5UZH6 Q5UZI1 Q5UZJ9 Q5UZM2 Q5UZP9 Q5UZQ7 Q5UZR5 Q5UZR8 Q5UZS7
Q5UZW0 Q5UZW6 Q5UZW7 Q5UZX1 Q5V030 Q5V059 Q5V061 Q5V0C3 Q5V0E6 Q5V0G6 Q5V0I9
Q5V0J7 Q5V0S5 Q5V0S8 Q5V0X0 Q5V0Z6 Q5V103 Q5V117 Q5V125 Q5V138 Q5V140 Q5V143
Q5V145 Q5V1B3 Q5V1B4 Q5V1C1 Q5V1D1 Q5V1D3 Q5V1D4 Q5V1D5 Q5V1E6 Q5V1F7 Q5V1G7
Q5V1H0 Q5V1H7 Q5V1I1 Q5V1I8 Q5V1I9 Q5V1K0 Q5V1L3 Q5V1M2 Q5V1N2 Q5V1P6 Q5V1R7
Q5V1W6 Q5V211 Q5V212 Q5V213 Q5V214 Q5V216 Q5V230 Q5V231 Q5V296 Q5V2B7 Q5V2C7
Q5V2C9 Q5V2D3 Q5V2G6 Q5V2M3 Q5V2N1 Q5V2P8 Q5V2S1 Q5V2S5 Q5V2T3 Q5V2T4 Q5V2X0
Q5V2X5 Q5V2X7 Q5V2X8 Q5V2Y4 Q5V2Y6 Q5V311 Q5V323 Q5V325 Q5V341 Q5V375 Q5V385
Q5V3D7 Q5V3D9 Q5V3F0 Q5V3G6 Q5V3K1 Q5V3L1 Q5V3P6 Q5V3Q5 Q5V3Q6 Q5V3R0 Q5V3R2
Q5V3R4 Q5V3T2 Q5V3Z1 Q5V448 Q5V450 Q5V452 Q5V463 Q5V464 Q5V465 Q5V466 Q5V467
Q5V468 Q5V470 Q5V474 Q5V4I4 Q5V4J1 Q5V4J7 Q5V4K3 Q5V4L7 Q5V4Q4 Q5V4Q5 Q5V4R0
Q5V4R4 Q5V4R5 Q5V4S6 Q5V4S7 Q5V518 Q5V520 Q5V545 Q5V547 Q5V554 Q5V574 Q5V584
Q5V588 Q5V5C9 Q5V5D4 Q5V5D5 Q5V5E2 Q5V5F7 Q5V5H0 Q5V5K5 Q5V5M1 Q5V5M5 Q5V5N9
Q5V5R5 Q5V5R9 Q5V5T7 Q5V5U5 Q5V5Y2 Q5V600 Q5V630 Q5V631 Q5V632 Q5V6G0 Q5V7B0
Q5V7E8 Q5V7I2 Q5V883 Q5V9E9 Q5V9F0 Q5VCS6 Q5VH41 Q5VH43 Q5VH50 Q5VIY5 Q5VKP2
Q5VKP6 Q5VM82 Q5VMI0 Q5VMJ3 Q5VMP0 Q5VMQ5 Q5VN06 Q5VNK1 Q5VNM3 Q5VNN5 Q5VPE3
Q5VPE5 Q5VQ09 Q5VQG4 Q5VQG8 Q5VQL1 Q5VR07 Q5VRD1 Q5VRG3 Q5VRH4 Q5VRH9 Q5VRL3
Q5VRM0 Q5VRN0 Q5VRR0 Q5VRW2 Q5VRY0 Q5VRY1 Q5VS55 Q5VS72 Q5VSA8 Q5VSB7 Q5VSD8
Q5VSR9 Q5VT03 Q5VT28 Q5VT33 Q5VT40 Q5VT79 Q5VT98 Q5VTA0 Q5VTB9 Q5VTE0 Q5VTE6
Q5VTH9 Q5VTM2 Q5VTT5 Q5VU43 Q5VU57 Q5VU69 Q5VU92 Q5VUE5 Q5VUJ5 Q5VUJ6 Q5VUJ9
Q5VUM1 Q5VUR7 Q5VV11 Q5VV16 Q5VVC0 Q5VVH5 Q5VVM6 Q5VVS0 Q5VVX9 Q5VVY1 Q5VW00
Q5VW22 Q5VWI1 Q5VWK0 Q5VWM3 Q5VWM4 Q5VWM6 Q5VWP2 Q5VX52 Q5VXD3 Q5VXH4 Q5VXH5
Q5VXU9 Q5VY09 Q5VYM1 Q5VYV0 Q5VYV7 Q5VYY1 Q5VZ03 Q5VZ18 Q5VZ52 Q5VZ66 Q5VZB9
Q5VZF2 Q5VZP5 Q5VZQ5 Q5VZR2 Q5VZV1 Q5W041 Q5W0A0 Q5W0Q7 Q5W0V3 Q5W111 Q5W150
Q5W1J5 Q5W1J6 Q5W251 Q5W252 Q5W259 Q5W262 Q5W263 Q5W265 Q5W266 Q5W267 Q5W268
Q5W269 Q5W271 Q5W283 Q5W416 Q5W5U4 Q5W5X9 Q5W670 Q5W6D6 Q5W6F9 Q5W6L9 Q5W726
Q5W770 Q5W794 Q5W7C3 Q5W7C6 Q5W915 Q5W9E3 Q5W9E7 Q5W9R9 Q5WA76 Q5WAC9 Q5WAD6
Q5WAD7 Q5WAE0 Q5WAF1 Q5WAF9 Q5WAG0 Q5WAG3 Q5WAG4 Q5WAG5 Q5WAG6 Q5WAH3 Q5WAH5
Q5WAH8 Q5WAI0 Q5WAI7 Q5WAR0 Q5WAV8 Q5WAX6 Q5WAZ6 Q5WAZ9 Q5WB17 Q5WB31 Q5WB34
Q5WB42 Q5WB43 Q5WB47 Q5WB48 Q5WB51 Q5WB52 Q5WB55 Q5WB66 Q5WB67 Q5WB82 Q5WB95
Q5WBF8 Q5WBJ6 Q5WC30 Q5WCF2 Q5WCI9 Q5WCM4 Q5WCP7 Q5WCP8 Q5WCR7 Q5WD10 Q5WD17
Q5WD36 Q5WDE9 Q5WDF0 Q5WDG7 Q5WDH1 Q5WDH6 Q5WDH7 Q5WDH8 Q5WDH9 Q5WDI0 Q5WDI1
Q5WDI2 Q5WDJ0 Q5WDJ2 Q5WDK0 Q5WDK6 Q5WDK7 Q5WDK8 Q5WDM4 Q5WDS1 Q5WDS6 Q5WDT6
Q5WDU0 Q5WDV0 Q5WDV1 Q5WDW4 Q5WDW9 Q5WDX0 Q5WDX3 Q5WDZ7 Q5WDZ8 Q5WE04 Q5WE05
Q5WE80 Q5WE98 Q5WEA5 Q5WEA8 Q5WEB8 Q5WEB9 Q5WEC6 Q5WED5 Q5WED8 Q5WED9 Q5WEE4
Q5WEF6 Q5WEG2 Q5WEG7 Q5WEG9 Q5WEH1 Q5WEH3 Q5WEI6 Q5WEI7 Q5WEI8 Q5WEJ3 Q5WEJ5
Q5WEJ6 Q5WEJ7 Q5WEK0 Q5WEK8 Q5WEL9 Q5WEM1 Q5WEM3 Q5WEM4 Q5WEM9 Q5WEN2 Q5WEN3

Q5WEN4 Q5WEN5 Q5WEN6 Q5WEN8 Q5WEN9 Q5WEP2 Q5WEP3 Q5WEP5 Q5WEP8 Q5WEQ4 Q5WEQ9
Q5WER0 Q5WER4 Q5WES3 Q5WES5 Q5WEW7 Q5WEW8 Q5WEW9 Q5WEY6 Q5WEY9 Q5WEZ7 Q5WF00
Q5WF04 Q5WF06 Q5WF30 Q5WF31 Q5WF32 Q5WF38 Q5WF43 Q5WF59 Q5WF60 Q5WF61 Q5WF63
Q5WF65 Q5WF77 Q5WF92 Q5WF94 Q5WF95 Q5WFA1 Q5WFA2 Q5WFC0 Q5WFC9 Q5WFE5 Q5WFE8
Q5WFF2 Q5WFF4 Q5WFF6 Q5WFF9 Q5WFG0 Q5WFG1 Q5WFG8 Q5WFH9 Q5WFI3 Q5WFI9 Q5WFJ0
Q5WFJ3 Q5WFJ4 Q5WFJ5 Q5WFK2 Q5WFK3 Q5WFK4 Q5WFK7 Q5WFL1 Q5WFL4 Q5WFM6 Q5WFM7
Q5WFN2 Q5WFN5 Q5WFN6 Q5WFN8 Q5WFP1 Q5WFP4 Q5WFP8 Q5WFQ0 Q5WFQ1 Q5WFQ2 Q5WFS5
Q5WFS8 Q5WFS9 Q5WFT0 Q5WFT1 Q5WFT2 Q5WFT4 Q5WFT6 Q5WFT8 Q5WFU2 Q5WFU4 Q5WFU5
Q5WFU7 Q5WFU8 Q5WFW9 Q5WFX1 Q5WFY0 Q5WFY3 Q5WFY4 Q5WFY6 Q5WFY7 Q5WG05 Q5WG41
Q5WG44 Q5WG56 Q5WGA1 Q5WGA3 Q5WGA4 Q5WGA5 Q5WGA8 Q5WGB1 Q5WGB6 Q5WGB8 Q5WGC3
Q5WGC5 Q5WGF4 Q5WGF5 Q5WGL4 Q5WGL6 Q5WGN4 Q5WGQ3 Q5WGQ4 Q5WGR5 Q5WGR7 Q5WGR9
Q5WGS0 Q5WGS1 Q5WGS2 Q5WGS3 Q5WGS4 Q5WGS7 Q5WGS8 Q5WGT0 Q5WGT4 Q5WGU8 Q5WH04
Q5WH07 Q5WH08 Q5WH11 Q5WH26 Q5WH37 Q5WH45 Q5WH53 Q5WH54 Q5WH73 Q5WH79 Q5WH85
Q5WH87 Q5WH99 Q5WHB2 Q5WHC2 Q5WHC6 Q5WHD2 Q5WHD3 Q5WHD5 Q5WHD6 Q5WHD7 Q5WHE2
Q5WHF1 Q5WHF4 Q5WHF9 Q5WHG0 Q5WHG1 Q5WHG2 Q5WHG3 Q5WHG9 Q5WHI0 Q5WHI2 Q5WHL7
Q5WHM1 Q5WHM5 Q5WHM6 Q5WHM7 Q5WHM8 Q5WHN4 Q5WHP3 Q5WHP4 Q5WHP9 Q5WHQ1 Q5WHR2
Q5WHR3 Q5WHR5 Q5WHR6 Q5WHS1 Q5WHS7 Q5WHS8 Q5WHT1 Q5WHT9 Q5WHU1 Q5WHU5 Q5WHV1
Q5WHX3 Q5WHY0 Q5WHY3 Q5WHZ3 Q5WI27 Q5WIE9 Q5WIF2 Q5WIG0 Q5WIU9 Q5WIY8 Q5WJ19
Q5WJ20 Q5WJ21 Q5WJ30 Q5WJ39 Q5WJ42 Q5WJ82 Q5WJ84 Q5WJ86 Q5WJ87 Q5WJ89 Q5WJC1
Q5WJC2 Q5WJD9 Q5WJE0 Q5WJE1 Q5WJE6 Q5WJI0 Q5WJL4 Q5WJM5 Q5WJM9 Q5WJN4 Q5WJN5
Q5WJP3 Q5WJP5 Q5WJS9 Q5WJU6 Q5WK52 Q5WK54 Q5WKB5 Q5WKB6 Q5WKC8 Q5WKD5 Q5WKE7
Q5WKF4 Q5WKF5 Q5WKJ3 Q5WKM9 Q5WKW1 Q5WKW2 Q5WKY5 Q5WKY6 Q5WKY7 Q5WKY8 Q5WKY9
Q5WKZ0 Q5WKZ1 Q5WL05 Q5WL06 Q5WL37 Q5WL39 Q5WL64 Q5WL77 Q5WL78 Q5WL79 Q5WL84
Q5WLA2 Q5WLE9 Q5WLG7 Q5WLG9 Q5WLM8 Q5WLM9 Q5WLN0 Q5WLN4 Q5WLN5 Q5WLN6 Q5WLN7
Q5WLN8 Q5WLN9 Q5WLP1 Q5WLP3 Q5WLP4 Q5WLP5 Q5WLP6 Q5WLP7 Q5WLP8 Q5WLP9 Q5WLQ0
Q5WLQ1 Q5WLQ2 Q5WLQ3 Q5WLQ4 Q5WLQ5 Q5WLQ6 Q5WLQ7 Q5WLQ8 Q5WLQ9 Q5WLR0 Q5WLR1
Q5WLR2 Q5WLR3 Q5WLR4 Q5WLR5 Q5WLR6 Q5WLR7 Q5WLR9 Q5WLS0 Q5WLS2 Q5WLS3 Q5WLS4
Q5WLS5 Q5WLS8 Q5WLT3 Q5WLT5 Q5WLT6 Q5WLT7 Q5WLU2 Q5WLU5 Q5WLV4 Q5WLV5 Q5WLV8
Q5WLW2 Q5WLX8 Q5WLY2 Q5WLZ3 Q5WLZ4 Q5WLZ7 Q5WM19 Q5WM28 Q5WM31 Q5WM45 Q5WN60
Q5WNE3 Q5WNI9 Q5WPZ8 Q5WPZ9 Q5WQ00 Q5WQ01 Q5WR10 Q5WRL1 Q5WRN3 Q5WSE6 Q5WSE7
Q5WSF0 Q5WSJ3 Q5WSM2 Q5WSM3 Q5WSM5 Q5WSQ8 Q5WSS3 Q5WSS4 Q5WSS7 Q5WST1 Q5WSU1
Q5WSU5 Q5WSX5 Q5WSX6 Q5WSX8 Q5WSX9 Q5WSY8 Q5WSZ9 Q5WT02 Q5WT13 Q5WT16 Q5WT18
Q5WT34 Q5WT36 Q5WT37 Q5WT38 Q5WT39 Q5WT40 Q5WT52 Q5WT53 Q5WT58 Q5WT59 Q5WT65
Q5WT78 Q5WT82 Q5WT83 Q5WT84 Q5WT85 Q5WT86 Q5WT87 Q5WT88 Q5WT90 Q5WTA1 Q5WTD6
Q5WTD7 Q5WTE7 Q5WTE8 Q5WTE9 Q5WTF0 Q5WTF1 Q5WTF5 Q5WTG3 Q5WTG8 Q5WTH5 Q5WTI5
Q5WTI7 Q5WTI8 Q5WTI9 Q5WTK7 Q5WTW4 Q5WTW6 Q5WTW9 Q5WTX7 Q5WU58 Q5WU88 Q5WU89
Q5WU94 Q5WUB1 Q5WUB2 Q5WUB9 Q5WUD4 Q5WUE1 Q5WUE6 Q5WUG0 Q5WUK0 Q5WUR6 Q5WUY8
Q5WV03 Q5WV09 Q5WV12 Q5WV14 Q5WV15 Q5WV16 Q5WV18 Q5WV28 Q5WV30 Q5WV31 Q5WV37
Q5WV42 Q5WV43 Q5WVC4 Q5WVD1 Q5WVE0 Q5WVG7 Q5WVI4 Q5WVI9 Q5WVJ0 Q5WVJ1 Q5WVK6
Q5WVL3 Q5WVP6 Q5WVQ0 Q5WVQ1 Q5WVQ2 Q5WVS0 Q5WVS7 Q5WVS9 Q5WVV9 Q5WVW0 Q5WVX1
Q5WVX4 Q5WVX6 Q5WVY7 Q5WVY8 Q5WVY9 Q5WVZ0 Q5WVZ3 Q5WVZ4 Q5WW20 Q5WW59 Q5WW81
Q5WW92 Q5WW97 Q5WW99 Q5WWA0 Q5WWA3 Q5WWD9 Q5WWF5 Q5WWF9 Q5WWG1 Q5WWG8 Q5WWH1
Q5WWH4 Q5WWH5 Q5WWJ5 Q5WWK4 Q5WWK6 Q5WWL7 Q5WWL9 Q5WWM0 Q5WWN7 Q5WWQ2 Q5WWS3
Q5WWS9 Q5WWT0 Q5WWT7 Q5WWU5 Q5WWU9 Q5WWV7 Q5WWV9 Q5WWW0 Q5WWW5 Q5WWW8 Q5WWX2

Q5WWX5 Q5WWX9 Q5WWY2 Q5WWZ8 Q5WX01 Q5WX12 Q5WX29 Q5WX31 Q5WX32 Q5WX33 Q5WX34
Q5WX38 Q5WX41 Q5WX47 Q5WX48 Q5WX49 Q5WX50 Q5WX53 Q5WX75 Q5WX86 Q5WX91 Q5WX92
Q5WX93 Q5WX94 Q5WX95 Q5WXA8 Q5WXC3 Q5WXV3 Q5WXW5 Q5WXX1 Q5WXX5 Q5WXX6 Q5WXZ0
Q5WXZ4 Q5WXZ5 Q5WXZ6 Q5WY16 Q5WY20 Q5WY21 Q5WY34 Q5WY41 Q5WY61 Q5WY74 Q5WY75
Q5WY82 Q5WY89 Q5WYB4 Q5WYB9 Q5WYE0 Q5WYE9 Q5WYF5 Q5WYG1 Q5WYH2 Q5WYH3 Q5WYH4
Q5WYI0 Q5WYI8 Q5WYJ0 Q5WYJ7 Q5WYK1 Q5WYK5 Q5WYL2 Q5WYL3 Q5WYL5 Q5WYQ0 Q5WYQ8
Q5WYQ9 Q5WYR0 Q5WYR3 Q5WYT2 Q5WYT4 Q5WYW3 Q5WYW4 Q5WYW5 Q5WYX3 Q5WYX4 Q5WYY6
Q5WYY8 Q5WYZ4 Q5WZ02 Q5WZ04 Q5WZ31 Q5WZ43 Q5WZ49 Q5WZ50 Q5WZ56 Q5WZ61 Q5WZ62
Q5WZ63 Q5WZ76 Q5WZ79 Q5WZB4 Q5WZC5 Q5WZE0 Q5WZE1 Q5WZE2 Q5WZE3 Q5WZH5 Q5WZI6
Q5WZI7 Q5WZI8 Q5WZI9 Q5WZJ0 Q5WZJ1 Q5WZJ3 Q5WZJ4 Q5WZJ5 Q5WZJ6 Q5WZJ7 Q5WZJ8
Q5WZJ9 Q5WZK0 Q5WZK1 Q5WZK2 Q5WZK3 Q5WZK4 Q5WZK5 Q5WZK6 Q5WZK7 Q5WZK8 Q5WZK9
Q5WZL0 Q5WZL1 Q5WZL2 Q5WZL3 Q5WZL4 Q5WZL5 Q5WZL6 Q5WZL7 Q5WZL8 Q5WZL9 Q5WZM0
Q5WZM1 Q5WZM2 Q5WZM3 Q5WZP1 Q5WZT6 Q5WZW1 Q5WZX8 Q5X084 Q5X090 Q5X097 Q5X0A4
Q5X0A5 Q5X0A6 Q5X0A8 Q5X0C7 Q5X0D2 Q5X0E6 Q5X0J6 Q5X0L0 Q5X0L4 Q5X0L8 Q5X0L9
Q5X0M0 Q5X0M3 Q5X0N1 Q5X0R9 Q5X0V0 Q5X0V1 Q5X0V3 Q5X0X6 Q5X0Z7 Q5X0Z8 Q5X101
Q5X105 Q5X115 Q5X119 Q5X148 Q5X149 Q5X153 Q5X154 Q5X168 Q5X170 Q5X186 Q5X189
Q5X1A0 Q5X1A3 Q5X1A5 Q5X1C1 Q5X1C3 Q5X1C4 Q5X1C5 Q5X1C6 Q5X1C7 Q5X1D9 Q5X1E0
Q5X1E5 Q5X1E6 Q5X1F2 Q5X1G7 Q5X1H3 Q5X1H4 Q5X1H5 Q5X1H6 Q5X1H7 Q5X1H8 Q5X1H9
Q5X1I1 Q5X1J2 Q5X1M7 Q5X1M8 Q5X1N7 Q5X1N8 Q5X1N9 Q5X1P0 Q5X1P1 Q5X1P5 Q5X1Q4
Q5X1Q9 Q5X1R6 Q5X1S4 Q5X1S6 Q5X1S7 Q5X1S8 Q5X1U6 Q5X240 Q5X242 Q5X245 Q5X252
Q5X2D9 Q5X2M8 Q5X2T0 Q5X2T1 Q5X2T6 Q5X2V3 Q5X2V4 Q5X2W0 Q5X2Y1 Q5X2Y6 Q5X300
Q5X340 Q5X3A8 Q5X3K0 Q5X3L5 Q5X3M1 Q5X3M4 Q5X3M6 Q5X3M7 Q5X3M8 Q5X3N0 Q5X3P0
Q5X3P2 Q5X3P3 Q5X3P9 Q5X3Q4 Q5X3Q5 Q5X3X9 Q5X3Y6 Q5X3Z5 Q5X427 Q5X446 Q5X450
Q5X451 Q5X452 Q5X472 Q5X479 Q5X4B2 Q5X4B5 Q5X4B6 Q5X4B7 Q5X4D5 Q5X4E2 Q5X4E4
Q5X4H4 Q5X4H5 Q5X4I6 Q5X4I9 Q5X4J1 Q5X4J7 Q5X4J8 Q5X4J9 Q5X4K0 Q5X4K3 Q5X4K4
Q5X4M9 Q5X4T2 Q5X4U9 Q5X4X0 Q5X4X1 Q5X4X3 Q5X4Y4 Q5X4Y6 Q5X4Z5 Q5X515 Q5X516
Q5X519 Q5X522 Q5X529 Q5X531 Q5X535 Q5X551 Q5X586 Q5X589 Q5X590 Q5X592 Q5X597
Q5X5A8 Q5X5C5 Q5X5E0 Q5X5E6 Q5X5E7 Q5X5F4 Q5X5G2 Q5X5G6 Q5X5H4 Q5X5H6 Q5X5H7
Q5X5I2 Q5X5I5 Q5X5I9 Q5X5J3 Q5X5J7 Q5X5K0 Q5X5L8 Q5X5M1 Q5X5N2 Q5X5P9 Q5X5Q1
Q5X5Q2 Q5X5Q3 Q5X5Q4 Q5X5Q9 Q5X5R2 Q5X5R9 Q5X5S0 Q5X5S1 Q5X5S2 Q5X5S5 Q5X5U7
Q5X5W4 Q5X5W9 Q5X5X0 Q5X5X1 Q5X5X2 Q5X5X3 Q5X5Y6 Q5X601 Q5X6E6 Q5X6G0 Q5X6G6
Q5X6H0 Q5X6H1 Q5X6I6 Q5X6J0 Q5X6J1 Q5X6J2 Q5X6L6 Q5X6L7 Q5X6N0 Q5X6N6 Q5X6Q6
Q5X6R9 Q5X6S0 Q5X6S7 Q5X6T6 Q5X6W4 Q5X6W9 Q5X6Y8 Q5X6Z7 Q5X703 Q5X709 Q5X720
Q5X721 Q5X722 Q5X728 Q5X739 Q5X741 Q5X748 Q5X752 Q5X756 Q5X762 Q5X763 Q5X765
Q5X7A1 Q5X7B0 Q5X7B1 Q5X7B2 Q5X7B5 Q5X7D5 Q5X7D7 Q5X7G7 Q5X7G8 Q5X7G9 Q5X7H7
Q5X7I9 Q5X7J1 Q5X7J6 Q5X7K4 Q5X7K6 Q5X7N0 Q5X7P2 Q5X7P8 Q5X7P9 Q5X7Q5 Q5X7R0
Q5X7R1 Q5X7R2 Q5X7S5 Q5X7S8 Q5X7W3 Q5X7X4 Q5X7Y9 Q5X7Z0 Q5X7Z1 Q5X7Z2 Q5X822
Q5X833 Q5X834 Q5X835 Q5X836 Q5X837 Q5X838 Q5X840 Q5X841 Q5X842 Q5X843 Q5X844
Q5X845 Q5X846 Q5X847 Q5X848 Q5X849 Q5X850 Q5X851 Q5X852 Q5X853 Q5X854 Q5X855
Q5X856 Q5X857 Q5X858 Q5X859 Q5X860 Q5X862 Q5X863 Q5X864 Q5X865 Q5X866 Q5X867
Q5X868 Q5X869 Q5X870 Q5X873 Q5X888 Q5X8E2 Q5X8H6 Q5X8J5 Q5X8U1 Q5X8U7 Q5X8V4
Q5X8W1 Q5X8W2 Q5X8W3 Q5X8W5 Q5X8Y4 Q5X8Y9 Q5X906 Q5X969 Q5X982 Q5X986 Q5X990
Q5X994 Q5X9A2 Q5X9A3 Q5X9A5 Q5X9A7 Q5X9A8 Q5X9C1 Q5X9C2 Q5X9C4 Q5X9C7 Q5X9E3
Q5X9E4 Q5X9E5 Q5X9E6 Q5X9F1 Q5X9F2 Q5X9F3 Q5X9H5 Q5X9H7 Q5X9H9 Q5X9I0 Q5X9I2

Q5X9I3 Q5X9I4 Q5X9J1 Q5X9J8 Q5X9J9 Q5X9K2 Q5X9K4 Q5X9K7 Q5X9L1 Q5X9L4 Q5X9L7
Q5X9L8 Q5X9M0 Q5X9M6 Q5X9R3 Q5X9S8 Q5X9T2 Q5X9T5 Q5X9U4 Q5X9U7 Q5X9U8 Q5X9V1
Q5X9V4 Q5X9W0 Q5X9W5 Q5X9X4 Q5X9Y1 Q5X9Y2 Q5X9Y3 Q5X9Y4 Q5X9Y7 Q5XA03 Q5XA06
Q5XA08 Q5XA09 Q5XA10 Q5XA12 Q5XA13 Q5XA14 Q5XA18 Q5XA23 Q5XA26 Q5XA27 Q5XA31
Q5XA33 Q5XA37 Q5XA40 Q5XA48 Q5XA52 Q5XA61 Q5XA62 Q5XA63 Q5XA86 Q5XA92 Q5XA93
Q5XA94 Q5XAA4 Q5XAC3 Q5XAC6 Q5XAC7 Q5XAC8 Q5XAD4 Q5XAD5 Q5XAD6 Q5XAD7 Q5XAF1
Q5XAG6 Q5XAG7 Q5XAH1 Q5XAH2 Q5XAI1 Q5XAI2 Q5XAI4 Q5XAJ5 Q5XAJ6 Q5XAJ7 Q5XAJ8
Q5XAJ9 Q5XAK0 Q5XAK4 Q5XAK5 Q5XAK9 Q5XAL0 Q5XAL3 Q5XAM5 Q5XAM8 Q5XAM9 Q5XAN0
Q5XAP1 Q5XAP2 Q5XAP4 Q5XAP6 Q5XAQ1 Q5XAQ3 Q5XAQ7 Q5XAR1 Q5XAU1 Q5XAV7 Q5XAW2
Q5XAW8 Q5XAX5 Q5XAY0 Q5XAY2 Q5XAY3 Q5XAY4 Q5XAY8 Q5XAZ0 Q5XAZ6 Q5XB04 Q5XB10
Q5XB14 Q5XB24 Q5XB25 Q5XB26 Q5XB35 Q5XB88 Q5XB92 Q5XB93 Q5XB97 Q5XBA0 Q5XBG3
Q5XBG4 Q5XBH1 Q5XBI8 Q5XBJ6 Q5XBJ7 Q5XBK5 Q5XBK8 Q5XBL1 Q5XBM1 Q5XBN4 Q5XBN5
Q5XBV1 Q5XBV2 Q5XBV6 Q5XBW6 Q5XBW7 Q5XBX7 Q5XBX9 Q5XBY0 Q5XBZ3 Q5XBZ4 Q5XC09
Q5XC12 Q5XC20 Q5XC26 Q5XC31 Q5XC34 Q5XC41 Q5XC46 Q5XC49 Q5XC58 Q5XC59 Q5XC60
Q5XC63 Q5XC65 Q5XC69 Q5XC70 Q5XC71 Q5XC74 Q5XC75 Q5XC82 Q5XC85 Q5XC88 Q5XC90
Q5XCA5 Q5XCA6 Q5XCA7 Q5XCA8 Q5XCA9 Q5XCB4 Q5XCB5 Q5XCB7 Q5XCB9 Q5XCC0 Q5XCD0
Q5XCE0 Q5XCG7 Q5XCH0 Q5XCH4 Q5XCH7 Q5XCI0 Q5XCI5 Q5XCK2 Q5XCK3 Q5XCK7 Q5XCK8
Q5XCL5 Q5XCL6 Q5XCL7 Q5XCL9 Q5XCM0 Q5XCM3 Q5XCM6 Q5XCN7 Q5XCQ0 Q5XCQ2 Q5XCQ3
Q5XCQ4 Q5XCR1 Q5XCR6 Q5XCR8 Q5XCR9 Q5XCS1 Q5XCS5 Q5XCS7 Q5XCT0 Q5XCT5 Q5XCT6
Q5XCT7 Q5XCU0 Q5XCU1 Q5XCU2 Q5XCU3 Q5XCU7 Q5XCW4 Q5XCW6 Q5XCW7 Q5XCX3 Q5XCX4
Q5XCX7 Q5XCY8 Q5XD05 Q5XD07 Q5XD12 Q5XD16 Q5XD17 Q5XD18 Q5XD19 Q5XD20 Q5XD24
Q5XD45 Q5XD48 Q5XD49 Q5XD52 Q5XD62 Q5XD71 Q5XD73 Q5XD99 Q5XDA6 Q5XDC4 Q5XDD4
Q5XDD6 Q5XDD9 Q5XDE3 Q5XDE4 Q5XDG5 Q5XDH0 Q5XDH1 Q5XDH3 Q5XDH4 Q5XDH6 Q5XDH7
Q5XDK5 Q5XDK8 Q5XDL5 Q5XDL7 Q5XDL8 Q5XDM2 Q5XDM3 Q5XDM4 Q5XDM5 Q5XDM7 Q5XDN3
Q5XDP3 Q5XDP4 Q5XDP5 Q5XDP8 Q5XDP9 Q5XDQ7 Q5XDR0 Q5XDR3 Q5XDR6 Q5XDS2 Q5XDT1
Q5XDT3 Q5XDT7 Q5XDV7 Q5XDW3 Q5XDW4 Q5XDW5 Q5XDW6 Q5XDX3 Q5XDX4 Q5XDY5 Q5XDY6
Q5XDZ0 Q5XDZ4 Q5XDZ5 Q5XDZ7 Q5XE04 Q5XE09 Q5XE14 Q5XE18 Q5XE35 Q5XE43 Q5XE45
Q5XE74 Q5XE77 Q5XE86 Q5XE96 Q5XE97 Q5XE99 Q5XEA3 Q5XEA8 Q5XEA9 Q5XEB0 Q5XEB1
Q5XEB2 Q5XEB3 Q5XEB4 Q5XEB6 Q5XEB7 Q5XEB8 Q5XEB9 Q5XEC0 Q5XEC1 Q5XEC2 Q5XEC3
Q5XEC4 Q5XEC5 Q5XEC6 Q5XEC7 Q5XEC8 Q5XEC9 Q5XED0 Q5XED1 Q5XED2 Q5XED3 Q5XED4
Q5XED5 Q5XED6 Q5XEE4 Q5XEF0 Q5XEF2 Q5XEF4 Q5XEF7 Q5XEK9 Q5XEL0 Q5XEL6 Q5XEL7
Q5XEM3 Q5XEM7 Q5XEM9 Q5XEN5 Q5XET4 Q5XET5 Q5XET6 Q5XEY7 Q5XEZ8 Q5XF07 Q5XF11
Q5XF20 Q5XF33 Q5XF57 Q5XF59 Q5XF75 Q5XF78 Q5XF82 Q5XFQ6 Q5XFR0 Q5XFW2 Q5XFW6
Q5XFX0 Q5XFY4 Q5XFZ0 Q5XG16 Q5XG21 Q5XG24 Q5XG48 Q5XG50 Q5XG58 Q5XG85 Q5XGA9
Q5XGB9 Q5XGC0 Q5XGC5 Q5XGC7 Q5XGC9 Q5XGD9 Q5XGE2 Q5XGE4 Q5XGG2 Q5XGG3 Q5XGI0
Q5XGI3 Q5XGI5 Q5XGL1 Q5XGL6 Q5XGL8 Q5XGM3 Q5XGM5 Q5XGP7 Q5XGR8 Q5XGS8 Q5XGT8
Q5XGU5 Q5XGW7 Q5XGY0 Q5XGY1 Q5XGY9 Q5XGZ8 Q5XH07 Q5XH09 Q5XH12 Q5XH16 Q5XH17
Q5XH28 Q5XH29 Q5XH30 Q5XH48 Q5XH61 Q5XH73 Q5XH91 Q5XHA1 Q5XHA8 Q5XHC1 Q5XHF8
Q5XHI2 Q5XHI4 Q5XHI9 Q5XHJ5 Q5XHK2 Q5XHX2 Q5XHX6 Q5XHX8 Q5XHX9 Q5XHY3 Q5XHY5
Q5XHY8 Q5XHZ2 Q5XHZ9 Q5XI01 Q5XI03 Q5XI06 Q5XI13 Q5XI14 Q5XI20 Q5XI29 Q5XI37
Q5XI46 Q5XI53 Q5XI54 Q5XI56 Q5XI57 Q5XI58 Q5XI63 Q5XI65 Q5XI67 Q5XI69 Q5XI70
Q5XI72 Q5XI79 Q5XI94 Q5XI95 Q5XI96 Q5XI97 Q5XIA2 Q5XIA3 Q5XIA4 Q5XIA9 Q5XIB1
Q5XIB2 Q5XIB3 Q5XIB4 Q5XIB5 Q5XIB9 Q5XIC1 Q5XIC3 Q5XIC4 Q5XIC7 Q5XIC8 Q5XID1
Q5XIE2 Q5XIE3 Q5XIE6 Q5XIE9 Q5XIG4 Q5XIG6 Q5XIG8 Q5XIH4 Q5XIH6 Q5XIH9 Q5XII5

Q5XII9 Q5XIJ1 Q5XIJ6 Q5XIJ8 Q5XIJ9 Q5XIK1 Q5XIK5 Q5XIK6 Q5XIK7 Q5XIK8 Q5XIL3
Q5XIL5 Q5XIL8 Q5XIM5 Q5XIM9 Q5XIN3 Q5XIN5 Q5XIN9 Q5XIP4 Q5XIP6 Q5XIQ5 Q5XIR6
Q5XIR8 Q5XIR9 Q5XIS1 Q5XIT0 Q5XIT1 Q5XIT9 Q5XIU1 Q5XIU2 Q5XIU5 Q5XIU7 Q5XIV0
Q5XIW8 Q5XIY4 Q5XIZ6 Q5XJ10 Q5XJ13 Q5XJ34 Q5XJ54 Q5XJ56 Q5XJ57 Q5XJ97 Q5XJA0
Q5XJA2 Q5XJA3 Q5XJB9 Q5XJD3 Q5XJE5 Q5XJK9 Q5XJN6 Q5XJP1 Q5XJQ5 Q5XJQ7 Q5XJS5
Q5XJV7 Q5XJX1 Q5XJY6 Q5XK67 Q5XK72 Q5XK83 Q5XK84 Q5XK85 Q5XKE0 Q5XKE5 Q5XKK7
Q5XKL5 Q5XKR4 Q5XKR9 Q5XL24 Q5XLA6 Q5XLD3 Q5XLE5 Q5XLR0 Q5XLR1 Q5XLY0 Q5XNP0
Q5XNS0 Q5XPJ6 Q5XPJ9 Q5XPK0 Q5XQN5 Q5XQP3 Q5XQP9 Q5XTN8 Q5XTY7 Q5XUN4 Q5XUX0
Q5XUX6 Q5XV31 Q5XVA8 Q5XVC4 Q5XVC7 Q5XVG3 Q5XVH0 Q5XVH5 Q5XVJ4 Q5XW77 Q5XWD9
Q5XWE1 Q5XX01 Q5XX03 Q5XX07 Q5XX08 Q5XX13 Q5XXA7 Q5XXB5 Q5XZD9 Q5Y223 Q5Y2C3
Q5Y9A9 Q5YB04 Q5YCU7 Q5YCU8 Q5YD48 Q5YDB5 Q5YJU0 Q5YJU1 Q5YJU6 Q5YJU7 Q5YJV6
Q5YJV9 Q5YJX3 Q5YJX4 Q5YJX5 Q5YJY5 Q5YK00 Q5YK01 Q5YK03 Q5YK05 Q5YK33 Q5YK47
Q5YK50 Q5YK53 Q5YLB4 Q5YLB5 Q5YLH5 Q5YMR6 Q5YMR7 Q5YMS1 Q5YMT5 Q5YN14 Q5YN15
Q5YN17 Q5YN65 Q5YN78 Q5YNA5 Q5YNB0 Q5YNG0 Q5YNI0 Q5YNI2 Q5YNL6 Q5YNN4 Q5YNP4
Q5YNP9 Q5YNQ2 Q5YNS3 Q5YNU2 Q5YNY7 Q5YP02 Q5YP36 Q5YP42 Q5YP47 Q5YP50 Q5YP69
Q5YP70 Q5YP79 Q5YPA6 Q5YPB0 Q5YPB9 Q5YPC0 Q5YPC4 Q5YPC5 Q5YPC7 Q5YPC8 Q5YPE0
Q5YPE1 Q5YPG1 Q5YPG2 Q5YPG3 Q5YPG4 Q5YPL7 Q5YPP1 Q5YPP8 Q5YPR1 Q5YPR2 Q5YPR3
Q5YPR4 Q5YPR6 Q5YPR7 Q5YPY6 Q5YPY8 Q5YPZ5 Q5YPZ6 Q5YQ00 Q5YQ21 Q5YQ72 Q5YQ76
Q5YQ91 Q5YQ92 Q5YQ94 Q5YQC8 Q5YQD2 Q5YQD7 Q5YQG3 Q5YQP4 Q5YQR9 Q5YQS0 Q5YQS7
Q5YQW3 Q5YQZ4 Q5YQZ5 Q5YQZ6 Q5YR52 Q5YR60 Q5YR63 Q5YRN0 Q5YRS5 Q5YRS7 Q5YRT3
Q5YRT4 Q5YRV4 Q5YRV8 Q5YRW2 Q5YRX2 Q5YRX6 Q5YRY0 Q5YRY1 Q5YRY5 Q5YRY7 Q5YRZ2
Q5YRZ6 Q5YS03 Q5YS07 Q5YS08 Q5YS09 Q5YS19 Q5YS35 Q5YS37 Q5YS38 Q5YS45 Q5YS47
Q5YS49 Q5YS61 Q5YS62 Q5YS63 Q5YS64 Q5YS67 Q5YS72 Q5YS74 Q5YS78 Q5YSA4 Q5YSB9
Q5YSC3 Q5YSC6 Q5YSC8 Q5YSE0 Q5YSV5 Q5YSV6 Q5YSW2 Q5YT00 Q5YT08 Q5YT12 Q5YT45
Q5YT53 Q5YT90 Q5YTA2 Q5YTA5 Q5YTD8 Q5YTE0 Q5YTE1 Q5YTE6 Q5YTE7 Q5YTE8 Q5YTF5
Q5YTH9 Q5YTI1 Q5YTJ4 Q5YTJ9 Q5YTK0 Q5YTK6 Q5YTK8 Q5YTL1 Q5YTL2 Q5YTM0 Q5YTM7
Q5YTM8 Q5YTN0 Q5YTN5 Q5YTP3 Q5YTP9 Q5YTQ0 Q5YTQ2 Q5YTQ5 Q5YTQ6 Q5YTV5 Q5YTY9
Q5YU18 Q5YU63 Q5YUA1 Q5YUH4 Q5YUH5 Q5YUS0 Q5YUV8 Q5YUV9 Q5YUW4 Q5YUX2 Q5YUX3
Q5YUX4 Q5YV93 Q5YV94 Q5YVA5 Q5YVJ4 Q5YVL0 Q5YVN7 Q5YW64 Q5YW75 Q5YW76 Q5YW77
Q5YW78 Q5YWQ4 Q5YWR8 Q5YWR9 Q5YWS0 Q5YWU1 Q5YWV1 Q5YWV4 Q5YWV9 Q5YX61 Q5YXN4
Q5YXQ1 Q5YY71 Q5YY83 Q5YY90 Q5YY98 Q5YYA4 Q5YYA5 Q5YYB6 Q5YYD2 Q5YYD3 Q5YYF4
Q5YYF7 Q5YYF8 Q5YYG7 Q5YYH6 Q5YYH7 Q5YYH9 Q5YYI0 Q5YYJ6 Q5YYN3 Q5YYP1 Q5YYP3
Q5YYP5 Q5YYP7 Q5YYP8 Q5YYP9 Q5YYQ0 Q5YYR3 Q5YYS5 Q5YYW9 Q5YYX3 Q5YYX8 Q5YYY1
Q5YYY7 Q5YYY8 Q5YZ21 Q5YZ49 Q5YZ53 Q5YZ58 Q5YZ59 Q5YZ95 Q5YZH4 Q5YZS7 Q5YZT7
Q5YZU4 Q5YZV0 Q5YZW3 Q5YZX0 Q5YZX1 Q5Z000 Q5Z025 Q5Z040 Q5Z041 Q5Z042 Q5Z043
Q5Z045 Q5Z048 Q5Z057 Q5Z061 Q5Z062 Q5Z063 Q5Z064 Q5Z066 Q5Z0A9 Q5Z0G1 Q5Z0G2
Q5Z0M2 Q5Z0M4 Q5Z0V0 Q5Z0V1 Q5Z0W0 Q5Z0W8 Q5Z0X7 Q5Z0Z2 Q5Z0Z3 Q5Z0Z5 Q5Z0Z9
Q5Z110 Q5Z154 Q5Z175 Q5Z187 Q5Z1C4 Q5Z1D1 Q5Z1E9 Q5Z1F9 Q5Z1G0 Q5Z1G2 Q5Z1H8
Q5Z1H9 Q5Z1I0 Q5Z1K7 Q5Z1K8 Q5Z1K9 Q5Z1L0 Q5Z1L1 Q5Z1L2 Q5Z1L3 Q5Z1L4 Q5Z1P0
Q5Z1Q1 Q5Z1Q3 Q5Z1Q4 Q5Z1Q5 Q5Z1Q6 Q5Z1Q7 Q5Z1Q8 Q5Z1R9 Q5Z1S0 Q5Z1S1 Q5Z1S2
Q5Z1U0 Q5Z1V4 Q5Z1V5 Q5Z1V6 Q5Z1V7 Q5Z1V8 Q5Z1V9 Q5Z1W0 Q5Z1W1 Q5Z1W2 Q5Z1W3
Q5Z1W4 Q5Z236 Q5Z297 Q5Z2A5 Q5Z2C3 Q5Z2C8 Q5Z2N2 Q5Z2N3 Q5Z2R1 Q5Z2R2 Q5Z2R3
Q5Z2U1 Q5Z2U2 Q5Z2U3 Q5Z2W1 Q5Z354 Q5Z355 Q5Z380 Q5Z3C0 Q5Z3K9 Q5Z3N4 Q5Z3P5
Q5Z3Z6 Q5Z3Z8 Q5Z4U5 Q5Z5B2 Q5Z5I4 Q5Z5Q3 Q5Z627 Q5Z676 Q5Z6B6 Q5Z6F6 Q5Z6G5

Q5Z6N9 Q5Z749 Q5Z754 Q5Z807 Q5Z856 Q5Z8K3 Q5Z8L1 Q5Z8P2 Q5Z8R1 Q5Z8S0 Q5Z8T3
Q5Z8V7 Q5Z8V9 Q5Z8Y4 Q5Z8Z7 Q5Z987 Q5Z9E5 Q5Z9Z3 Q5ZA07 Q5ZA21 Q5ZAQ2 Q5ZBH5
Q5ZC82 Q5ZC88 Q5ZCK5 Q5ZCL8 Q5ZD81 Q5ZDJ6 Q5ZEF1 Q5ZEP9 Q5ZHK9 Q5ZHN5 Q5ZHP3
Q5ZHP7 Q5ZHP8 Q5ZHQ2 Q5ZHS0 Q5ZHS1 Q5ZHS3 Q5ZHV2 Q5ZHV5 Q5ZHV7 Q5ZHV8 Q5ZHW7
Q5ZHX5 Q5ZHX9 Q5ZHY5 Q5ZHZ0 Q5ZHZ4 Q5ZI08 Q5ZI11 Q5ZI16 Q5ZI23 Q5ZI33 Q5ZI34
Q5ZI40 Q5ZI42 Q5ZI49 Q5ZI58 Q5ZI66 Q5ZI67 Q5ZI72 Q5ZI74 Q5ZI78 Q5ZI89 Q5ZIA0
Q5ZIA2 Q5ZIB9 Q5ZIC8 Q5ZID0 Q5ZID1 Q5ZID2 Q5ZID6 Q5ZIE6 Q5ZIE8 Q5ZIF3 Q5ZIF8
Q5ZIG2 Q5ZIH0 Q5ZIH9 Q5ZII9 Q5ZIJ0 Q5ZIJ2 Q5ZIJ8 Q5ZIJ9 Q5ZIK6 Q5ZIL4 Q5ZIL9
Q5ZIM5 Q5ZIM6 Q5ZIN2 Q5ZIP3 Q5ZIP4 Q5ZIQ3 Q5ZIR5 Q5ZIT8 Q5ZIU1 Q5ZIU8 Q5ZIW1
Q5ZIW2 Q5ZIX3 Q5ZIX8 Q5ZIY4 Q5ZIY5 Q5ZIZ2 Q5ZIZ4 Q5ZJ07 Q5ZJ20 Q5ZJ23 Q5ZJ24
Q5ZJ26 Q5ZJ27 Q5ZJ32 Q5ZJ37 Q5ZJ39 Q5ZJ40 Q5ZJ54 Q5ZJ60 Q5ZJ64 Q5ZJ65 Q5ZJ66
Q5ZJ69 Q5ZJ74 Q5ZJ85 Q5ZJ97 Q5ZJA2 Q5ZJA3 Q5ZJA4 Q5ZJB4 Q5ZJC8 Q5ZJD3 Q5ZJF3
Q5ZJF6 Q5ZJG3 Q5ZJH5 Q5ZJH6 Q5ZJH9 Q5ZJI6 Q5ZJI9 Q5ZJJ1 Q5ZJJ2 Q5ZJJ8 Q5ZJK5
Q5ZJK8 Q5ZJL4 Q5ZJL5 Q5ZJL7 Q5ZJL9 Q5ZJM3 Q5ZJM7 Q5ZJN0 Q5ZJN1 Q5ZJP9 Q5ZJQ2
Q5ZJQ3 Q5ZJQ7 Q5ZJR9 Q5ZJS6 Q5ZJT0 Q5ZJT1 Q5ZJU2 Q5ZJU3 Q5ZJU4 Q5ZJU5 Q5ZJU8
Q5ZJV4 Q5ZJV6 Q5ZJV7 Q5ZJV9 Q5ZJX4 Q5ZJY5 Q5ZJZ6 Q5ZK13 Q5ZK14 Q5ZK17 Q5ZK21
Q5ZK25 Q5ZK28 Q5ZK35 Q5ZK36 Q5ZK40 Q5ZK47 Q5ZK63 Q5ZK69 Q5ZK74 Q5ZK77 Q5ZK88
Q5ZK95 Q5ZKA2 Q5ZKA3 Q5ZKA4 Q5ZKA5 Q5ZKB6 Q5ZKB8 Q5ZKC1 Q5ZKD4 Q5ZKD7 Q5ZKD9
Q5ZKE7 Q5ZKF4 Q5ZKF6 Q5ZKG1 Q5ZKG3 Q5ZKH0 Q5ZKH3 Q5ZKH6 Q5ZKI0 Q5ZKI2 Q5ZKI7
Q5ZKJ4 Q5ZKK1 Q5ZKK2 Q5ZKK3 Q5ZKK5 Q5ZKL9 Q5ZKM1 Q5ZKN1 Q5ZKP2 Q5ZKP4 Q5ZKQ3
Q5ZKQ6 Q5ZKQ7 Q5ZKR5 Q5ZKT1 Q5ZKT6 Q5ZKT8 Q5ZKT9 Q5ZKU0 Q5ZKU1 Q5ZKV4 Q5ZKV8
Q5ZKW0 Q5ZKW2 Q5ZKW5 Q5ZKW8 Q5ZKW9 Q5ZKX6 Q5ZKY2 Q5ZKY4 Q5ZKY9 Q5ZL12 Q5ZL26
Q5ZL33 Q5ZL34 Q5ZL35 Q5ZL38 Q5ZL42 Q5ZL54 Q5ZL79 Q5ZL91 Q5ZL95 Q5ZL98 Q5ZLA5
Q5ZLC1 Q5ZLC6 Q5ZLC8 Q5ZLD3 Q5ZLE6 Q5ZLE9 Q5ZLF0 Q5ZLG3 Q5ZLG4 Q5ZLH0 Q5ZLH9
Q5ZLK1 Q5ZLK6 Q5ZLL9 Q5ZLM2 Q5ZLM8 Q5ZLN0 Q5ZLN1 Q5ZLN4 Q5ZLN5 Q5ZLN7 Q5ZLN8
Q5ZLP2 Q5ZLP8 Q5ZLQ4 Q5ZLR2 Q5ZLR4 Q5ZLS2 Q5ZLS3 Q5ZLS7 Q5ZLS8 Q5ZLT3 Q5ZLT5
Q5ZLT7 Q5ZLV2 Q5ZLX2 Q5ZLX5 Q5ZLY0 Q5ZLY2 Q5ZLY8 Q5ZM11 Q5ZM16 Q5ZM19 Q5ZM20
Q5ZM30 Q5ZM33 Q5ZM35 Q5ZM36 Q5ZM41 Q5ZM46 Q5ZM55 Q5ZM71 Q5ZM74 Q5ZM86 Q5ZM98
Q5ZMA3 Q5ZMA6 Q5ZMB2 Q5ZMB7 Q5ZMC0 Q5ZMC3 Q5ZMC6 Q5ZMD2 Q5ZMD6 Q5ZME2 Q5ZME8
Q5ZMF3 Q5ZMG0 Q5ZMG5 Q5ZMJ4 Q5ZMJ9 Q5ZMK7 Q5ZML0 Q5ZML3 Q5ZML4 Q5ZML5 Q5ZML9
Q5ZMM1 Q5ZMM5 Q5ZMN2 Q5ZMS3 Q5ZMS4 Q5ZMS6 Q5ZMU0 Q5ZMU6 Q5ZMV2 Q5ZMV7 Q5ZMW0
Q5ZMW6 Q5ZNC8 Q5ZND2 Q5ZPI9 Q5ZPV8 Q5ZQU2 Q5ZR76 Q5ZR78 Q5ZR82 Q5ZR89 Q5ZRC6
Q5ZRF3 Q5ZRF4 Q5ZRF6 Q5ZRH9 Q5ZRJ1 Q5ZRJ2 Q5ZRJ5 Q5ZRJ9 Q5ZRK9 Q5ZRL3 Q5ZRP1
Q5ZRP2 Q5ZRP4 Q5ZRP5 Q5ZRQ7 Q5ZRQ9 Q5ZRS2 Q5ZRT1 Q5ZRT4 Q5ZRT6 Q5ZRV2 Q5ZRV4
Q5ZRV5 Q5ZRV6 Q5ZRV7 Q5ZRV8 Q5ZRX0 Q5ZRX1 Q5ZRX9 Q5ZRY0 Q5ZRY6 Q5ZS01 Q5ZS05
Q5ZS07 Q5ZS08 Q5ZS09 Q5ZS10 Q5ZS12 Q5ZS23 Q5ZS57 Q5ZS58 Q5ZS66 Q5ZS67 Q5ZS68
Q5ZS69 Q5ZS70 Q5ZS74 Q5ZS83 Q5ZS95 Q5ZSA3 Q5ZSA5 Q5ZSA6 Q5ZSA7 Q5ZSC5 Q5ZSM8
Q5ZSN0 Q5ZSN3 Q5ZSP1 Q5ZSQ2 Q5ZSX7 Q5ZT07 Q5ZT08 Q5ZT13 Q5ZT29 Q5ZT30 Q5ZT34
Q5ZT36 Q5ZT51 Q5ZT57 Q5ZT62 Q5ZT75 Q5ZT84 Q5ZTB9 Q5ZTI6 Q5ZTK6 Q5ZTV7 Q5ZTX2
Q5ZTX7 Q5ZTY0 Q5ZTY2 Q5ZTY3 Q5ZTY4 Q5ZTY6 Q5ZTZ6 Q5ZTZ8 Q5ZU04 Q5ZU09 Q5ZU10
Q5ZU79 Q5ZU80 Q5ZU87 Q5ZU96 Q5ZUC1 Q5ZUD5 Q5ZUD8 Q5ZUD9 Q5ZUE0 Q5ZUF4 Q5ZUG1
Q5ZUJ3 Q5ZUJ7 Q5ZUJ9 Q5ZUL6 Q5ZUM3 Q5ZUM5 Q5ZUQ6 Q5ZUQ7 Q5ZUS0 Q5ZUS2 Q5ZUS8
Q5ZUS9 Q5ZUT0 Q5ZUT1 Q5ZUT4 Q5ZUT5 Q5ZV19 Q5ZV33 Q5ZV48 Q5ZV49 Q5ZV51 Q5ZV64
Q5ZV75 Q5ZV92 Q5ZV93 Q5ZV96 Q5ZV99 Q5ZVA6 Q5ZVA8 Q5ZVB2 Q5ZVC9 Q5ZVG2 Q5ZVG5

Q5ZVG6 Q5ZVG8 Q5ZVH3 Q5ZVI4 Q5ZVK0 Q5ZVL5 Q5ZVM1 Q5ZVM2 Q5ZVN7 Q5ZVP1 Q5ZVP9
Q5ZVQ1 Q5ZVQ2 Q5ZVQ7 Q5ZVR0 Q5ZVR4 Q5ZVR7 Q5ZVS1 Q5ZVS4 Q5ZVU2 Q5ZVU5 Q5ZVV6
Q5ZVY1 Q5ZVY3 Q5ZVY4 Q5ZVY5 Q5ZVY6 Q5ZVZ1 Q5ZVZ3 Q5ZW00 Q5ZW01 Q5ZW02 Q5ZW72
Q5ZW83 Q5ZW87 Q5ZW88 Q5ZW89 Q5ZW90 Q5ZW91 Q5ZWA4 Q5ZWB8 Q5ZWR3 Q5ZWX6 Q5ZWZ0
Q5ZWZ6 Q5ZX00 Q5ZX01 Q5ZX15 Q5ZX19 Q5ZX20 Q5ZX22 Q5ZX45 Q5ZX46 Q5ZX60 Q5ZX66
Q5ZX85 Q5ZX98 Q5ZX99 Q5ZXA6 Q5ZXB5 Q5ZXD1 Q5ZXE5 Q5ZXH2 Q5ZXH8 Q5ZXH9 Q5ZXI0
Q5ZXI1 Q5ZXI6 Q5ZXJ2 Q5ZXK5 Q5ZXK6 Q5ZXL2 Q5ZXM0 Q5ZXM2 Q5ZXM9 Q5ZXN3 Q5ZXN7
Q5ZXP4 Q5ZXP6 Q5ZXT1 Q5ZXU0 Q5ZXU1 Q5ZXU2 Q5ZXU5 Q5ZXZ2 Q5ZXZ3 Q5ZXZ4 Q5ZY07
Q5ZY18 Q5ZY20 Q5ZY26 Q5ZY37 Q5ZY59 Q5ZY71 Q5ZY77 Q5ZY78 Q5ZY86 Q5ZY91 Q5ZY93
Q5ZY94 Q5ZYA8 Q5ZYB1 Q5ZYE7 Q5ZYF8 Q5ZYH3 Q5ZYH5 Q5ZYH6 Q5ZYH7 Q5ZYK6 Q5ZYL7
Q5ZYL8 Q5ZYL9 Q5ZYM0 Q5ZYM1 Q5ZYM2 Q5ZYM4 Q5ZYM5 Q5ZYM6 Q5ZYM7 Q5ZYM8 Q5ZYM9
Q5ZYN0 Q5ZYN1 Q5ZYN2 Q5ZYN3 Q5ZYN4 Q5ZYN5 Q5ZYN6 Q5ZYN7 Q5ZYN8 Q5ZYP0 Q5ZYP1
Q5ZYP2 Q5ZYP3 Q5ZYP4 Q5ZYP5 Q5ZYP6 Q5ZYP7 Q5ZYP8 Q5ZYP9 Q5ZYQ0 Q5ZYQ1 Q5ZYQ2
Q5ZYQ3 Q5ZYQ4 Q5ZYS4 Q5ZYX0 Q5ZYX7 Q5ZYZ5 Q5ZZ14 Q5ZZ74 Q5ZZ80 Q5ZZ86 Q5ZZ93
Q5ZZ94 Q5ZZ95 Q5ZZ97 Q5ZZB7 Q5ZZC2 Q5ZZD9 Q5ZZI8 Q5ZZK1 Q5ZZK4 Q5ZZK8 Q5ZZK9
Q5ZZL4 Q5ZZL8 Q5ZZM8 Q5ZZN1 Q5ZZN4 Q5ZZN5 Q5ZZN6 Q5ZZP2 Q5ZZP6 Q5ZZQ1 Q5ZZR9
Q5ZZS0 Q5ZZS1 Q5ZZS2 Q5ZZT0 Q5ZZU2 Q5ZZV6 Q5ZZW4 Q60006 Q60009 Q60012 Q60013
Q60022 Q60023 Q60024 Q60028 Q60030 Q60034 Q60036 Q60040 Q60049 Q60050 Q60080
Q60099 Q600A4 Q600A8 Q600B2 Q600B5 Q600B6 Q600J2 Q600J7 Q600K1 Q600K7 Q600M2
Q600M5 Q600N3 Q600N9 Q600P9 Q600Q0 Q600Q7 Q600Z6 Q600Z8 Q60101 Q60103 Q60107
Q60112 Q60132 Q60151 Q60169 Q60172 Q60174 Q60176 Q60177 Q60178 Q60179 Q60180
Q60181 Q60182 Q601D3 Q601D6 Q601E3 Q601E4 Q601E5 Q601F2 Q601F3 Q601F9 Q601G1
Q601G9 Q601H2 Q601H3 Q601H8 Q601I8 Q601I9 Q601J0 Q601J1 Q601J2 Q601J6 Q601J7
Q601J8 Q601K0 Q601K1 Q601K2 Q601K3 Q601K4 Q601K5 Q601K7 Q601K8 Q601K9 Q601L1
Q601L2 Q601L3 Q601L4 Q601L5 Q601M2 Q601M3 Q601M5 Q601M6 Q601N0 Q601P7 Q601P9
Q601Q3 Q601R3 Q601R6 Q601T1 Q601T8 Q601T9 Q601U1 Q601U6 Q601U7 Q601V5 Q601V6
Q601V7 Q601W6 Q601W7 Q601W8 Q601X7 Q601X8 Q601Y5 Q601Y6 Q601Z2 Q601Z3 Q60216
Q60228 Q60252 Q60257 Q60258 Q60259 Q60260 Q60261 Q60262 Q60264 Q60265 Q60266
Q60267 Q60268 Q60269 Q60270 Q60271 Q60272 Q60273 Q60274 Q60275 Q60276 Q60277
Q60278 Q60279 Q60280 Q60281 Q60282 Q60283 Q60284 Q60285 Q60286 Q60287 Q60288
Q60289 Q60290 Q60291 Q60292 Q60295 Q60296 Q60297 Q60298 Q60299 Q602A7 Q602A9
Q602B7 Q602D8 Q602E0 Q602E6 Q602E7 Q602H4 Q602J1 Q602J5 Q602K3 Q602L8 Q602M5
Q602M8 Q602M9 Q602N0 Q602N2 Q602N9 Q602P0 Q602R3 Q602S0 Q602S1 Q602T2 Q602T6
Q60300 Q60301 Q60302 Q60303 Q60305 Q60306 Q60307 Q60308 Q60310 Q60311 Q60312
Q60313 Q60314 Q60315 Q60316 Q60317 Q60319 Q60320 Q60321 Q60322 Q60323 Q60325
Q60326 Q60327 Q60328 Q60329 Q60330 Q60331 Q60332 Q60334 Q60335 Q60337 Q60338
Q60339 Q60340 Q60341 Q60342 Q60343 Q60344 Q60345 Q60346 Q60347 Q60348 Q60349
Q60350 Q60351 Q60352 Q60353 Q60354 Q60355 Q60356 Q60357 Q60358 Q60359 Q60360
Q60361 Q60363 Q60364 Q60365 Q60367 Q60368 Q60369 Q60370 Q60372 Q60373 Q60374
Q60375 Q60376 Q60377 Q60378 Q60379 Q60380 Q60381 Q60382 Q60383 Q60384 Q60385
Q60386 Q60387 Q60388 Q60389 Q60390 Q60391 Q603B5 Q603B8 Q603C0 Q603C1 Q603C7
Q603G2 Q603G6 Q603G8 Q603I8 Q603K0 Q603K1 Q603K2 Q603K3 Q603K4 Q603K5 Q603L1
Q603L4 Q603L5 Q603M1 Q603M3 Q603Q7 Q603S0 Q603S2 Q603T9 Q60420 Q60435 Q60436
Q60443 Q60446 Q60462 Q60493 Q604B7 Q604C6 Q604D9 Q604E1 Q604F8 Q604K2 Q604M0

Q604M1 Q604M2 Q604M6 Q604P1 Q604P3 Q604P4 Q604Q5 Q604U3 Q604U5 Q604U6 Q604U8
Q604U9 Q604V0 Q604V5 Q604V8 Q604V9 Q604W0 Q604W8 Q60525 Q60539 Q60542 Q60554
Q60561 Q60564 Q60584 Q60585 Q60595 Q60596 Q605A7 Q605A8 Q605A9 Q605B0 Q605B1
Q605B2 Q605B3 Q605B4 Q605B5 Q605B6 Q605B7 Q605B8 Q605B9 Q605C0 Q605C1 Q605C2
Q605C3 Q605C4 Q605C5 Q605C6 Q605C7 Q605C8 Q605C9 Q605D0 Q605D1 Q605D3 Q605D4
Q605D5 Q605D6 Q605D7 Q605D8 Q605E4 Q605E5 Q605E8 Q605E9 Q605F1 Q605G6 Q605G8
Q605G9 Q605H0 Q605H3 Q605I0 Q605I7 Q605K7 Q605K9 Q605M0 Q605M7 Q605N0 Q605N1
Q605N2 Q605N3 Q605N4 Q605Q9 Q605S7 Q605W1 Q60605 Q60632 Q60636 Q60665 Q60668
Q60688 Q60692 Q606B9 Q606C1 Q606C5 Q606C7 Q606C8 Q606C9 Q606D2 Q606D6 Q606D8
Q606F1 Q606F2 Q606F3 Q606F4 Q606H1 Q606H8 Q606H9 Q606I1 Q606J4 Q606J5 Q606J9
Q606K1 Q606K2 Q606K3 Q606K7 Q606K9 Q606L1 Q606M0 Q606M6 Q606N2 Q606N3 Q606N7
Q606N8 Q606P2 Q606P5 Q606Q0 Q606Q1 Q606Q2 Q606W5 Q606Y1 Q606Y2 Q606Y5 Q60707
Q60722 Q60756 Q60764 Q60765 Q60772 Q60773 Q60775 Q60793 Q607A2 Q607A3 Q607A4
Q607A5 Q607A6 Q607A7 Q607B0 Q607B2 Q607B4 Q607C3 Q607C7 Q607D1 Q607E5 Q607E7
Q607F5 Q607G8 Q607H0 Q607H2 Q607H5 Q607I0 Q607I4 Q607L0 Q607L6 Q607M3 Q607M5
Q607P2 Q607Q3 Q607R4 Q607S4 Q607S6 Q607T4 Q607T5 Q607U4 Q607U5 Q607V0 Q607X6
Q60806 Q60809 Q60821 Q60843 Q60848 Q60854 Q60862 Q60866 Q60872 Q60899 Q608B0
Q608D9 Q608F1 Q608G0 Q608K0 Q608L4 Q608L5 Q608L6 Q608M7 Q608M9 Q608N1 Q608N3
Q608N7 Q608P0 Q608P1 Q608P3 Q608P4 Q608S3 Q608S5 Q608S6 Q608S8 Q60900 Q60925
Q60929 Q60945 Q60948 Q60949 Q60954 Q60969 Q60972 Q60973 Q60976 Q60980 Q60987
Q60988 Q60990 Q609A0 Q609A8 Q609A9 Q609B5 Q609B8 Q609C0 Q609C1 Q609C2 Q609C5
Q609C6 Q609F9 Q609G2 Q609I3 Q609I6 Q609I7 Q609J9 Q609K0 Q609L0 Q609L1 Q609L2
Q609L3 Q609M1 Q609M6 Q609U9 Q609V0 Q609V1 Q609V2 Q609W4 Q609X7 Q609Y2 Q609Z5
Q60A05 Q60A06 Q60A07 Q60A08 Q60A09 Q60A10 Q60A14 Q60A15 Q60A17 Q60A20 Q60A21
Q60A25 Q60A28 Q60AF7 Q60AF8 Q60AJ7 Q60AK3 Q60AM9 Q60AN1 Q60AN2 Q60AP7 Q60AP8
Q60AP9 Q60AQ2 Q60AU5 Q60AU6 Q60AV5 Q60AV6 Q60AW8 Q60AX2 Q60AX4 Q60AX5 Q60AY0
Q60AY5 Q60AY8 Q60AY9 Q60AZ0 Q60AZ1 Q60AZ2 Q60AZ3 Q60AZ8 Q60B10 Q60B13 Q60B48
Q60B71 Q60B77 Q60B95 Q60BA1 Q60BA4 Q60BA7 Q60BA8 Q60BA9 Q60BB0 Q60BB2 Q60BD3
Q60BD8 Q60BE7 Q60BE8 Q60BF6 Q60BF9 Q60BG7 Q60BG8 Q60BH8 Q60BJ2 Q60BQ2 Q60BS0
Q60BS1 Q60BS2 Q60BS3 Q60BS4 Q60BS6 Q60BS8 Q60BT7 Q60BT8 Q60BU2 Q60BV9 Q60BW3
Q60BW4 Q60BW5 Q60BX0 Q60BY1 Q60BY2 Q60BY3 Q60BZ1 Q60C21 Q60C56 Q60C59 Q60C60
Q60C62 Q60C64 Q60C67 Q60C68 Q60C69 Q60CC4 Q60CC5 Q60CE9 Q60CG7 Q60CG8 Q60CJ6
Q60CJ7 Q60CK7 Q60CK8 Q60CK9 Q60CL4 Q60CM4 Q60CN9 Q60CQ4 Q60CQ5 Q60CR0 Q60CS4
Q60CS5 Q60CT7 Q60CU8 Q60CV0 Q60CV1 Q60CV4 Q60DG4 Q60DW3 Q60DX1 Q60E60 Q60E61
Q60EJ6 Q60EQ4 Q60EX6 Q60EX7 Q60EY1 Q60F88 Q60FC2 Q60FC6 Q60FS1 Q60FX6 Q60FX9
Q60FY0 Q60GC1 Q60GU1 Q60HC2 Q60HC3 Q60HC7 Q60HC9 Q60HD0 Q60HD3 Q60HD6 Q60HD7
Q60HD8 Q60HD9 Q60HE5 Q60HE6 Q60HF9 Q60HG3 Q60HH2 Q60HH3 Q60HH9 Q60J38 Q60JJ0
Q60K16 Q60LV4 Q60LV7 Q60LW7 Q60M48 Q60P76 Q60P81 Q60PP8 Q60Q85 Q60Q90 Q60QD1
Q60QR6 Q60R52 Q60SG7 Q60SM4 Q60TI7 Q60UW4 Q60UZ5 Q60V43 Q60V67 Q60V73 Q60W34
Q60WM1 Q60YA8 Q60YF6 Q60YI3 Q60YJ7 Q60YN5 Q60YT5 Q60YU0 Q60ZM2 Q60ZS1 Q61024
Q61035 Q61039 Q61048 Q61060 Q61080 Q61116 Q61122 Q61127 Q61133 Q61136 Q61164
Q61171 Q61179 Q61188 Q611X0 Q61211 Q61216 Q61239 Q612G6 Q612Q9 Q612R3 Q612X6
Q61312 Q61313 Q61321 Q61333 Q61345 Q61390 Q613L4 Q61412 Q61414 Q61456 Q61457
Q61458 Q61467 Q61474 Q61493 Q61496 Q614K9 Q61539 Q61550 Q61574 Q61575 Q61585

Q61597 Q615B0 Q61624 Q61645 Q61655 Q61657 Q61658 Q61660 Q61663 Q61666 Q61670
Q61699 Q616T6 Q61749 Q61751 Q61753 Q61765 Q61781 Q61806 Q61820 Q61827 Q61845
Q61850 Q61858 Q61880 Q61881 Q61884 Q61897 Q618K0 Q61908 Q61909 Q61967 Q61976
Q61979 Q61990 Q61999 Q619T5 Q61A42 Q61A92 Q61B10 Q61BU1 Q61CW2 Q61CX7 Q61D43
Q61D44 Q61DA8 Q61DH4 Q61DI9 Q61DP2 Q61DP3 Q61E22 Q61E36 Q61E63 Q61FA3 Q61FQ3
Q61FW2 Q61GH1 Q61IR6 Q61IX2 Q61J08 Q61J97 Q61J98 Q61JF9 Q61JI9 Q61JK7 Q61JN8
Q61JR3 Q61JT8 Q61JZ2 Q61K76 Q61KX5 Q61L47 Q61LA1 Q61LN7 Q61MR2 Q61QN4 Q61QT3
Q61R02 Q61RF0 Q61S97 Q61SD1 Q61SE7 Q61SK8 Q61SU7 Q61SU8 Q61T02 Q61T16 Q61UC4
Q61WP7 Q61WR2 Q61WW9 Q61XD3 Q61XH2 Q61XX9 Q61Y48 Q61YQ1 Q61YZ4 Q61Z20 Q62018
Q62066 Q62070 Q62082 Q62084 Q62093 Q62095 Q62100 Q62136 Q62168 Q62188 Q62189
Q62193 Q621J7 Q621Q3 Q621Z7 Q62203 Q62220 Q62231 Q62232 Q62233 Q62234 Q62240
Q62241 Q62247 Q62249 Q62255 Q62276 Q62282 Q622B7 Q622K8 Q62311 Q62348 Q62376
Q62377 Q62383 Q62384 Q62392 Q62393 Q62394 Q62396 Q623D4 Q623U2 Q62407 Q62424
Q62426 Q62445 Q62447 Q62448 Q62481 Q62504 Q62511 Q62513 Q62559 Q62563 Q62565
Q62599 Q62630 Q62649 Q62655 Q62671 Q626C1 Q626F4 Q626I0 Q626S1 Q62722 Q62724
Q62725 Q62728 Q62751 Q62764 Q62782 Q62785 Q62792 Q62798 Q62806 Q62818 Q62826
Q62834 Q62857 Q62880 Q628A6 Q62901 Q62924 Q62947 Q62950 Q62951 Q62952 Q62981
Q629Z7 Q62A02 Q62AD1 Q62AG8 Q62AI6 Q62AI8 Q62AI9 Q62AJ2 Q62AJ6 Q62BG3 Q62BI3
Q62CD5 Q62CE3 Q62CH6 Q62CH7 Q62CH8 Q62CJ3 Q62CK6 Q62CM5 Q62CU8 Q62DC9 Q62DD0
Q62DD3 Q62DF2 Q62DF3 Q62DG4 Q62DI5 Q62DT5 Q62DT7 Q62DU0 Q62DU1 Q62DU3 Q62DV9
Q62EC5 Q62EK3 Q62EL5 Q62EM1 Q62EM2 Q62EM6 Q62EN9 Q62EQ4 Q62EU1 Q62EU9 Q62EX3
Q62EZ1 Q62EZ3 Q62EZ9 Q62F00 Q62F04 Q62F07 Q62F43 Q62F46 Q62F99 Q62FB8 Q62FC0
Q62FC1 Q62FC2 Q62FC4 Q62FC7 Q62FD0 Q62FD1 Q62FE9 Q62FI7 Q62FM9 Q62FN1 Q62FS7
Q62FS8 Q62FU4 Q62FV3 Q62FV6 Q62G11 Q62G22 Q62G58 Q62GA2 Q62GA9 Q62GB0 Q62GC6
Q62GD7 Q62GD8 Q62GD9 Q62GE0 Q62GE1 Q62GE3 Q62GE4 Q62GE5 Q62GE6 Q62GE7 Q62GJ3
Q62GJ4 Q62GJ5 Q62GJ6 Q62GJ7 Q62GK0 Q62GK1 Q62GK2 Q62GK3 Q62GK4 Q62GK5 Q62GK6
Q62GK7 Q62GK8 Q62GK9 Q62GL0 Q62GL1 Q62GL2 Q62GL3 Q62GL4 Q62GL5 Q62GL6 Q62GL7
Q62GL8 Q62GL9 Q62GM0 Q62GM1 Q62GM2 Q62GM3 Q62GM4 Q62GM6 Q62GM7 Q62GM8 Q62GM9
Q62GN0 Q62GN1 Q62GN2 Q62GN6 Q62GR8 Q62GR9 Q62GS5 Q62GS8 Q62GS9 Q62GT9 Q62GU2
Q62GU3 Q62GV2 Q62GV3 Q62GV4 Q62GV5 Q62GV7 Q62GV8 Q62GX2 Q62H08 Q62H14 Q62H17
Q62H20 Q62H23 Q62H74 Q62H77 Q62H98 Q62HA6 Q62HA7 Q62HA8 Q62HA9 Q62HB3 Q62HB4
Q62HB5 Q62HB8 Q62HB9 Q62HC0 Q62HC9 Q62HD0 Q62HD1 Q62HD3 Q62HD5 Q62HD6 Q62HD8
Q62HE4 Q62HE6 Q62HH7 Q62HI0 Q62HI3 Q62HK4 Q62HL1 Q62HL4 Q62HM0 Q62HM4 Q62HM5
Q62HM6 Q62HM8 Q62HQ3 Q62HR7 Q62HR8 Q62HR9 Q62HS0 Q62HS1 Q62HS2 Q62HS3 Q62HT7
Q62HV3 Q62HV4 Q62HV8 Q62HY1 Q62HY5 Q62HY7 Q62HZ7 Q62HZ8 Q62I00 Q62I01 Q62I16
Q62I17 Q62I34 Q62I82 Q62I86 Q62I87 Q62I89 Q62ID1 Q62ID2 Q62ID7 Q62IF4 Q62IG0
Q62IG1 Q62II4 Q62II5 Q62II8 Q62IJ0 Q62IJ1 Q62IJ3 Q62IM0 Q62IN0 Q62IN1 Q62IU9
Q62IX5 Q62IZ4 Q62IZ5 Q62IZ9 Q62J00 Q62J08 Q62J26 Q62J37 Q62J40 Q62J82 Q62J86
Q62JA4 Q62JB1 Q62JB3 Q62JB9 Q62JC2 Q62JC4 Q62JC5 Q62JC6 Q62JC7 Q62JD0 Q62JD8
Q62JE0 Q62JE7 Q62JF6 Q62JH0 Q62JH1 Q62JI5 Q62JI6 Q62JK6 Q62JK7 Q62JK8 Q62JL8
Q62JM6 Q62JM7 Q62JN9 Q62JQ9 Q62JR1 Q62JR2 Q62JU1 Q62JU9 Q62JV0 Q62JV2 Q62JV3
Q62JV9 Q62JW1 Q62JW2 Q62JW4 Q62JW5 Q62JW9 Q62JX4 Q62JX6 Q62JZ1 Q62K38 Q62K46
Q62K50 Q62KC5 Q62KD8 Q62KD9 Q62KI2 Q62KI4 Q62KI5 Q62KI6 Q62KI7 Q62KK7 Q62KK9
Q62KL0 Q62KL1 Q62KU7 Q62KW5 Q62KY7 Q62KZ3 Q62L77 Q62L80 Q62LC0 Q62LC1 Q62LD1

Q62LD5 Q62LE0 Q62LE7 Q62LE8 Q62LF1 Q62LG7 Q62LH2 Q62LJ2 Q62LJ6 Q62LN3 Q62LN4
Q62LN5 Q62LP6 Q62LS1 Q62LS3 Q62LS7 Q62LU4 Q62LU6 Q62LW1 Q62LZ0 Q62LZ2 Q62M18
Q62M26 Q62M28 Q62M34 Q62M46 Q62M50 Q62M51 Q62M52 Q62M53 Q62M59 Q62M61 Q62M74
Q62M76 Q62M77 Q62M78 Q62M91 Q62M97 Q62M98 Q62M99 Q62MG7 Q62MH0 Q62MK8 Q62ML1
Q62MM2 Q62MN2 Q62MP0 Q62MP4 Q62MP8 Q62MR3 Q62MR4 Q62MR5 Q62MS6 Q62MT4 Q62MW9
Q62MX0 Q62MX1 Q62MX7 Q62MY7 Q62MZ8 Q62N15 Q62N16 Q62Q36 Q62ZD8 Q63003 Q63042
Q63065 Q63073 Q63087 Q630B4 Q630B5 Q630B8 Q630B9 Q630C0 Q630C1 Q630C8 Q630D0
Q630D3 Q630D5 Q630E3 Q630E8 Q630J1 Q630J5 Q630K2 Q630L2 Q630L7 Q630N6 Q630Q9
Q630R0 Q630R4 Q630R7 Q630R8 Q630R9 Q630T2 Q630T3 Q630T4 Q630W1 Q63117 Q63150
Q63164 Q63170 Q63185 Q63186 Q63187 Q631C7 Q631E1 Q631H3 Q631I6 Q631J0 Q631J2
Q631J8 Q631K0 Q631K2 Q631L9 Q631M0 Q631M1 Q631M9 Q631P8 Q631P9 Q631X2 Q631X9
Q631Z7 Q63228 Q63244 Q63245 Q63246 Q63247 Q63248 Q63249 Q63250 Q63251 Q632C6
Q632C7 Q632D6 Q632E1 Q632E8 Q632G4 Q632G8 Q632H1 Q632H2 Q632H4 Q632I2 Q632I5
Q632S4 Q632S5 Q632S8 Q632V2 Q632W7 Q632Z1 Q632Z7 Q63302 Q63340 Q63342 Q63347
Q63379 Q63398 Q633A2 Q633D2 Q633E0 Q633E8 Q633F3 Q633F8 Q633G4 Q633G5 Q633I4
Q633J8 Q633K5 Q633L1 Q633L3 Q633L5 Q633L6 Q633M1 Q633M2 Q633M3 Q633N2 Q633N4
Q633N5 Q633N6 Q633N8 Q633P2 Q633Q1 Q633S1 Q633V2 Q633V4 Q633V5 Q633X1 Q633X2
Q633X5 Q633X8 Q633Y0 Q633Y3 Q633Y6 Q633Y9 Q633Z0 Q633Z4 Q633Z9 Q63410 Q63413
Q63468 Q63486 Q63487 Q634A1 Q634A3 Q634B5 Q634C3 Q634C4 Q634C6 Q634C7 Q634D6
Q634D8 Q634E2 Q634E3 Q634E9 Q634F6 Q634F7 Q634F8 Q634F9 Q634G4 Q634G5 Q634G9
Q634H0 Q634K8 Q634L0 Q634L9 Q634M0 Q634M5 Q634M6 Q634M7 Q634M8 Q634M9 Q634N2
Q634N8 Q634P2 Q634P4 Q634Q0 Q634Q3 Q634Q9 Q634V6 Q634V7 Q634V8 Q634Y0 Q634Y5
Q634Y7 Q63505 Q63518 Q63528 Q63530 Q63562 Q63569 Q63570 Q63572 Q63598 Q635A2
Q635A3 Q635A4 Q635A5 Q635A7 Q635A9 Q635C1 Q635D0 Q635E0 Q635E2 Q635F1 Q635F2
Q635F4 Q635G3 Q635G4 Q635G7 Q635G9 Q635H0 Q635J3 Q635K7 Q635M2 Q635M3 Q635M6
Q635P0 Q635P1 Q635P2 Q635P3 Q635P6 Q635P7 Q635S6 Q635U7 Q635U8 Q635V2 Q635W3
Q635W6 Q635X7 Q635Y7 Q635Z4 Q635Z7 Q63610 Q63618 Q63623 Q63624 Q63625 Q63627
Q63630 Q63638 Q63679 Q636A1 Q636A3 Q636A5 Q636A7 Q636A8 Q636B4 Q636B7 Q636C7
Q636D1 Q636D5 Q636D7 Q636D8 Q636D9 Q636E0 Q636E1 Q636E2 Q636E3 Q636F4 Q636F5
Q636F6 Q636G0 Q636G2 Q636G8 Q636H7 Q636I2 Q636I4 Q636I5 Q636I6 Q636I9 Q636J0
Q636J4 Q636J6 Q636J7 Q636J8 Q636J9 Q636K0 Q636K2 Q636K5 Q636K7 Q636K9 Q636L3
Q636L5 Q636L6 Q636L8 Q636L9 Q636P5 Q636Q4 Q636Q7 Q636Q8 Q636T5 Q636W2 Q636W3
Q63716 Q63737 Q63750 Q63754 Q63755 Q63797 Q63798 Q63799 Q637D9 Q637H7 Q637H8
Q637H9 Q637I0 Q637L7 Q637V9 Q63801 Q63829 Q63862 Q63871 Q638M6 Q638Q6 Q638R8
Q638W1 Q638W9 Q638X2 Q63945 Q63955 Q639D2 Q639M2 Q639M5 Q639M6 Q639P8 Q639W8
Q639W9 Q639Y8 Q63A04 Q63A05 Q63A07 Q63A51 Q63AC7 Q63AF7 Q63AG0 Q63AJ0 Q63AJ1
Q63AJ2 Q63AN7 Q63B62 Q63B72 Q63B73 Q63B74 Q63B75 Q63B92 Q63BB2 Q63BL0 Q63BP5
Q63BQ9 Q63BV2 Q63BV5 Q63BZ8 Q63C01 Q63C03 Q63C61 Q63CC5 Q63CC9 Q63CG2 Q63CI8
Q63CL4 Q63CN1 Q63CP4 Q63CR4 Q63CR9 Q63CS8 Q63CU1 Q63CV3 Q63CV4 Q63D03 Q63D23
Q63DG7 Q63DH5 Q63DH6 Q63DI3 Q63DJ1 Q63DJ2 Q63DJ3 Q63DJ6 Q63DJ9 Q63DK0 Q63DL2
Q63DL4 Q63DL5 Q63DL6 Q63DL7 Q63DL9 Q63DM1 Q63DM8 Q63DN3 Q63DN5 Q63DR9 Q63DV8
Q63DV9 Q63DW6 Q63DW7 Q63DW8 Q63DW9 Q63DX0 Q63DX1 Q63DX2 Q63DX3 Q63DX4 Q63DX5
Q63DX6 Q63DX7 Q63DX8 Q63DX9 Q63DY5 Q63E02 Q63E28 Q63E31 Q63E45 Q63E46 Q63E65
Q63E66 Q63E67 Q63EB1 Q63EC6 Q63EC7 Q63EC8 Q63EC9 Q63ED0 Q63ED9 Q63EG2 Q63EG5

Q63EG7 Q63EH0 Q63EI7 Q63EK2 Q63EK7 Q63EM2 Q63EM3 Q63ES4 Q63ES5 Q63EV6 Q63EX2
Q63EZ0 Q63F05 Q63F14 Q63F37 Q63FA5 Q63FD4 Q63FE7 Q63FJ0 Q63FR5 Q63FX6 Q63FY0
Q63G33 Q63G97 Q63GA5 Q63GB4 Q63GB8 Q63GC6 Q63GF1 Q63GH2 Q63GI2 Q63GK3 Q63GK4
Q63GK5 Q63GN3 Q63GQ9 Q63GR0 Q63GR1 Q63GS0 Q63GS3 Q63GS9 Q63GT1 Q63GT3 Q63GT4
Q63GT6 Q63GV4 Q63GV7 Q63GV8 Q63GW0 Q63GW2 Q63GW6 Q63GX5 Q63H01 Q63H45 Q63H57
Q63H58 Q63H59 Q63H63 Q63H64 Q63H65 Q63H66 Q63H67 Q63H69 Q63H71 Q63H72 Q63H73
Q63H74 Q63H75 Q63H76 Q63H77 Q63H78 Q63H79 Q63H80 Q63H81 Q63H83 Q63H84 Q63H85
Q63H86 Q63H87 Q63H88 Q63H89 Q63H90 Q63H91 Q63H92 Q63H93 Q63H94 Q63H95 Q63H97
Q63H98 Q63HA0 Q63HA1 Q63HA2 Q63HA3 Q63HB0 Q63HB2 Q63HB3 Q63HB4 Q63HB9 Q63HC2
Q63HD2 Q63HD3 Q63HD6 Q63HE8 Q63HE9 Q63HF6 Q63HF7 Q63HF8 Q63HG4 Q63HG7 Q63HH3
Q63HH5 Q63HH8 Q63HI2 Q63HI5 Q63HI8 Q63HJ1 Q63HK3 Q63HM1 Q63HN1 Q63IA7 Q63IF8
Q63IV6 Q63JB0 Q63JB1 Q63JD9 Q63JF2 Q63JF4 Q63JF5 Q63JF6 Q63JF8 Q63JK9 Q63JL1
Q63JL2 Q63JL6 Q63JM0 Q63K83 Q63KI8 Q63KJ7 Q63KK7 Q63KK8 Q63KK9 Q63KN7 Q63KP6
Q63KV4 Q63L05 Q63L73 Q63LV9 Q63MT0 Q63MV1 Q63N20 Q63NC4 Q63NE3 Q63NF6 Q63NG7
Q63NL2 Q63NV1 Q63PC7 Q63PD9 Q63PF1 Q63PG8 Q63PG9 Q63PI5 Q63PL2 Q63PL4 Q63PS1
Q63PT2 Q63PU8 Q63PZ6 Q63PZ9 Q63Q00 Q63Q01 Q63Q02 Q63Q03 Q63Q04 Q63Q06 Q63Q07
Q63Q08 Q63Q10 Q63Q11 Q63Q12 Q63Q13 Q63Q14 Q63Q15 Q63Q16 Q63Q17 Q63Q18 Q63Q19
Q63Q20 Q63Q21 Q63Q22 Q63Q23 Q63Q24 Q63Q25 Q63Q26 Q63Q27 Q63Q28 Q63Q29 Q63Q30
Q63Q32 Q63Q33 Q63Q34 Q63Q35 Q63Q36 Q63Q37 Q63Q38 Q63Q42 Q63Q50 Q63Q56 Q63Q57
Q63Q73 Q63Q84 Q63Q85 Q63Q86 Q63Q87 Q63Q88 Q63Q90 Q63Q91 Q63Q92 Q63Q93 Q63Q94
Q63QE9 Q63QF0 Q63QF1 Q63QG7 Q63QH0 Q63QH1 Q63QI8 Q63QI9 Q63QJ5 Q63QJ8 Q63QJ9
Q63QK7 Q63QL1 Q63QM0 Q63QM1 Q63QM4 Q63QM5 Q63QN9 Q63QS4 Q63QS9 Q63QT3 Q63QT6
Q63QT9 Q63QW3 Q63QW4 Q63QW5 Q63QW9 Q63QX0 Q63QX1 Q63QX5 Q63QX6 Q63QX7 Q63QX8
Q63QY5 Q63R06 Q63R14 Q63R41 Q63R42 Q63R43 Q63R45 Q63R47 Q63R49 Q63R55 Q63R58
Q63RB4 Q63RB5 Q63RD3 Q63RH9 Q63RI0 Q63RI2 Q63RL0 Q63RL1 Q63RL2 Q63RL3 Q63RL4
Q63RL5 Q63RL6 Q63RN0 Q63RP4 Q63RP5 Q63RP8 Q63RS1 Q63RS4 Q63RS6 Q63RV4 Q63RV5
Q63RV7 Q63RV8 Q63RX4 Q63RX6 Q63RZ8 Q63S05 Q63S07 Q63S25 Q63S30 Q63S31 Q63S32
Q63S33 Q63S39 Q63S41 Q63S63 Q63S79 Q63S81 Q63S98 Q63SA2 Q63SA4 Q63SC2 Q63SD7
Q63SD8 Q63SD9 Q63SH6 Q63SI0 Q63SJ4 Q63SL6 Q63SN4 Q63SN5 Q63SN9 Q63SP0 Q63SP8
Q63SR9 Q63SS8 Q63ST1 Q63SX4 Q63SX7 Q63SZ9 Q63T00 Q63T02 Q63T07 Q63T10 Q63T12
Q63T13 Q63T14 Q63T15 Q63T18 Q63T26 Q63T28 Q63T34 Q63T42 Q63T53 Q63T70 Q63T71
Q63T91 Q63T93 Q63TF9 Q63TG5 Q63TI8 Q63TK6 Q63TM2 Q63TM4 Q63TM5 Q63TM6 Q63TM7
Q63TM8 Q63TP6 Q63TP8 Q63TP9 Q63TQ0 Q63TU6 Q63TV2 Q63TV3 Q63TV5 Q63U59 Q63U72
Q63U73 Q63U95 Q63UQ8 Q63US2 Q63US3 Q63US8 Q63US9 Q63UT2 Q63UT3 Q63UT5 Q63UT6
Q63UT8 Q63UU3 Q63UU4 Q63UU6 Q63UU7 Q63UV5 Q63UY3 Q63UY4 Q63UY6 Q63V08 Q63V22
Q63V23 Q63V31 Q63V40 Q63V41 Q63V42 Q63V83 Q63V86 Q63VB7 Q63VF5 Q63VK4 Q63VN7
Q63VN8 Q63VP3 Q63VP6 Q63VS6 Q63VS8 Q63VS9 Q63VT1 Q63VT4 Q63VT5 Q63VX9 Q63VY0
Q63VY4 Q63W00 Q63W06 Q63W07 Q63W31 Q63W73 Q63W75 Q63W86 Q63W90 Q63W97 Q63W98
Q63WA1 Q63WB9 Q63WC3 Q63WH0 Q63WH3 Q63WH4 Q63WH7 Q63WI3 Q63WI6 Q63WJ1 Q63WJ7
Q63WL4 Q63WL6 Q63WM0 Q63WM2 Q63WM3 Q63WM4 Q63WN0 Q63WN7 Q63WP3 Q63WP4 Q63WP5
Q63WW2 Q63WX9 Q63X50 Q63X54 Q63X63 Q63X65 Q63X76 Q63X85 Q63X93 Q63XA0 Q63XA4
Q63XK0 Q63XL0 Q63XL1 Q63XL4 Q63XL7 Q63XL8 Q63XL9 Q63XM0 Q63XM1 Q63XM3 Q63XR8
Q63XU4 Q63XU7 Q63XX6 Q63XZ2 Q63Y06 Q63Y16 Q63Y23 Q63Y24 Q63Y25 Q63Y74 Q63YC3
Q63YD1 Q63YH5 Q63YH7 Q63YI3 Q63YI4 Q63YI7 Q63YJ0 Q63YJ8 Q63YJ9 Q63YL0 Q63YR6

Q63YV9 Q63YW3 Q63YW4 Q63YW5 Q63Z34 Q63Z36 Q63ZG9 Q63ZH2 Q63ZH8 Q63ZJ3 Q63ZK1
Q63ZL2 Q63ZM7 Q63ZM9 Q63ZP1 Q63ZP7 Q63ZS0 Q63ZT8 Q63ZV0 Q63ZV7 Q63ZY6 Q63ZY7
Q64028 Q64029 Q64060 Q64092 Q640A2 Q640B4 Q640C5 Q640E9 Q640F2 Q640G7 Q640I9
Q640J6 Q640K6 Q640L5 Q640M1 Q640P7 Q640Q5 Q640S2 Q640T1 Q640V2 Q640V3 Q640V9
Q640W2 Q640W6 Q640Z1 Q640Z9 Q64104 Q64105 Q64112 Q64119 Q64122 Q64124 Q641A1
Q641B2 Q641B9 Q641C9 Q641F0 Q641F1 Q641F2 Q641F8 Q641G3 Q641G4 Q641G7 Q641I1
Q641J8 Q641P0 Q641R9 Q641S2 Q641W2 Q641W4 Q641W7 Q641W9 Q641X1 Q641X2 Q641X6
Q641X7 Q641X9 Q641Y6 Q641Y7 Q641Y8 Q641Y9 Q641Z1 Q641Z4 Q641Z5 Q64201 Q64202
Q64204 Q64205 Q64210 Q64214 Q64249 Q64256 Q64261 Q64279 Q64282 Q64291 Q642A0
Q642A4 Q642A5 Q642B2 Q642B6 Q642B9 Q642H3 Q642J4 Q642J5 Q642M9 Q642Q1 Q642Q3
Q642T7 Q64305 Q64311 Q64318 Q64327 Q64336 Q64345 Q64346 Q64350 Q64361 Q64368
Q64380 Q64399 Q643C1 Q643C8 Q64413 Q64414 Q64415 Q64422 Q64427 Q64433 Q64467
Q64471 Q64475 Q64478 Q64507 Q64511 Q64514 Q64520 Q64522 Q64523 Q64525 Q64531
Q64536 Q64560 Q64565 Q64598 Q645M6 Q64602 Q64611 Q64620 Q64666 Q64674 Q64693
Q646L1 Q646M7 Q646P3 Q646R8 Q64707 Q64709 Q64731 Q64732 Q64733 Q64744 Q64745
Q64746 Q64748 Q64749 Q64750 Q64751 Q64753 Q64754 Q64755 Q64756 Q64759 Q64760
Q64761 Q64762 Q64763 Q64765 Q64766 Q64767 Q64769 Q64770 Q647I9 Q64846 Q64858
Q64865 Q64898 Q64903 Q64904 Q64905 Q64909 Q64912 Q64913 Q64914 Q64917 Q64919
Q64929 Q64962 Q64963 Q64966 Q64BR7 Q64ET8 Q64EV9 Q64EW0 Q64EW1 Q64EW2 Q64EW3
Q64EW6 Q64FJ6 Q64FN2 Q64FN9 Q64FP0 Q64G17 Q64GL0 Q64HY3 Q64HZ8 Q64HZ9 Q64I00
Q64I01 Q64L94 Q64LC9 Q64LD2 Q64LH0 Q64LH1 Q64LH2 Q64M78 Q64MA1 Q64MA3 Q64MG2
Q64MG4 Q64MG7 Q64MK4 Q64MP7 Q64MT2 Q64MT6 Q64MV4 Q64MX8 Q64N34 Q64N36 Q64N73
Q64NF0 Q64NF1 Q64NI3 Q64NI7 Q64NJ3 Q64NJ4 Q64NJ6 Q64NJ7 Q64NJ8 Q64NK4 Q64NK5
Q64NK6 Q64NK7 Q64NK8 Q64NK9 Q64NL1 Q64NL2 Q64NL3 Q64NL4 Q64NL5 Q64NL6 Q64NL7
Q64NL8 Q64NL9 Q64NM0 Q64NM1 Q64NM2 Q64NM3 Q64NM4 Q64NM5 Q64NM6 Q64NM7 Q64NN0
Q64NN1 Q64NN2 Q64NN3 Q64NN4 Q64NN5 Q64NN6 Q64NP4 Q64NV0 Q64NV3 Q64NW2 Q64NW3
Q64NX1 Q64P19 Q64P24 Q64P28 Q64P29 Q64P30 Q64P34 Q64P39 Q64P45 Q64P62 Q64P72
Q64P77 Q64P83 Q64P85 Q64PD6 Q64PH7 Q64PH8 Q64PH9 Q64PL4 Q64PM3 Q64PM4 Q64PM7
Q64PM9 Q64PP7 Q64PR7 Q64PS6 Q64PU1 Q64PU2 Q64PV5 Q64PY0 Q64PY1 Q64PY6 Q64PY7
Q64PY9 Q64PZ2 Q64PZ3 Q64PZ9 Q64Q87 Q64QI5 Q64QP1 Q64QR7 Q64QS2 Q64QU1 Q64QU2
Q64R36 Q64R56 Q64R71 Q64R85 Q64R94 Q64RB7 Q64RD6 Q64RE6 Q64RE7 Q64RE8 Q64RE9
Q64RQ5 Q64RT0 Q64RT1 Q64RT2 Q64RZ8 Q64SR3 Q64SX3 Q64SX6 Q64SX9 Q64SY6 Q64SY7
Q64T09 Q64T27 Q64T40 Q64T65 Q64T66 Q64T99 Q64TA0 Q64TA1 Q64TD9 Q64TE4 Q64TJ6
Q64TJ9 Q64TK0 Q64TK1 Q64TK2 Q64TK5 Q64TM7 Q64TM8 Q64TN0 Q64TQ1 Q64TX7 Q64TY9
Q64U07 Q64U20 Q64U25 Q64U74 Q64U75 Q64U78 Q64U80 Q64UB2 Q64UB9 Q64UD5 Q64UM6
Q64UM9 Q64UN4 Q64UP8 Q64UQ5 Q64UQ7 Q64UR4 Q64VB2 Q64VI6 Q64VI7 Q64VN7 Q64VN9
Q64VP0 Q64VP1 Q64VP2 Q64VP5 Q64VP6 Q64VQ9 Q64VR0 Q64VR1 Q64VV4 Q64VX4 Q64VX6
Q64WA8 Q64WA9 Q64WB0 Q64WC2 Q64WF9 Q64WG8 Q64WR0 Q64WS3 Q64WS4 Q64WY5 Q64X01
Q64X24 Q64X29 Q64X30 Q64X55 Q64X95 Q64XA5 Q64XA6 Q64XE8 Q64XF4 Q64XH6 Q64XI2
Q64XK5 Q64XL6 Q64XL7 Q64XM2 Q64XM3 Q64XP2 Q64XQ0 Q64XQ7 Q64XR8 Q64XS0 Q64XS1
Q64XV0 Q64XV7 Q64XV8 Q64XW5 Q64XW6 Q64XX2 Q64Y02 Q64Y97 Q64YA5 Q64YB4 Q64YD8
Q64YH3 Q64YI2 Q64YJ0 Q64YJ1 Q64YL9 Q64YS9 Q64YT3 Q64YW4 Q64YZ2 Q64YZ5 Q64Z15
Q64Z17 Q64Z29 Q64Z37 Q64Z40 Q64Z48 Q64Z72 Q64ZA1 Q64ZK3 Q64ZL3 Q64ZL4 Q64ZL7
Q64ZL9 Q64ZM7 Q64ZM9 Q64ZN5 Q64ZN7 Q64ZQ7 Q64ZR4 Q64ZR6 Q64ZS4 Q64ZT7 Q64ZT8

Q64ZU2 Q64ZV5 Q64ZV9 Q64ZW2 Q64ZY1 Q65030 Q650A0 Q650B4 Q650C1 Q650H5 Q650H6
Q650H7 Q650H8 Q650K4 Q650K7 Q650L4 Q650L7 Q650L8 Q650P7 Q650T9 Q65108 Q65109
Q65111 Q65114 Q65121 Q65122 Q65123 Q65124 Q65125 Q65126 Q65127 Q65128 Q65129
Q65132 Q65133 Q65137 Q65141 Q65143 Q65144 Q65146 Q65147 Q65148 Q65151 Q65152
Q65153 Q65154 Q65155 Q65156 Q65157 Q65159 Q65160 Q65161 Q65162 Q65163 Q65168
Q65169 Q65170 Q65171 Q65173 Q65174 Q65175 Q65177 Q65180 Q65181 Q65182 Q65183
Q65184 Q65185 Q65186 Q65187 Q65189 Q65190 Q65191 Q65192 Q65193 Q65195 Q65196
Q65199 Q651A1 Q651B6 Q651E8 Q651U1 Q651Z5 Q651Z7 Q65202 Q65203 Q65204 Q65205
Q65206 Q65207 Q65208 Q65209 Q65210 Q65211 Q65212 Q65213 Q65215 Q65216 Q65217
Q65218 Q65219 Q65221 Q65224 Q65225 Q65227 Q65228 Q65229 Q65230 Q65231 Q65232
Q65234 Q65235 Q65236 Q65237 Q65238 Q65239 Q65243 Q65246 Q65248 Q65249 Q65258
Q65259 Q65269 Q652A1 Q652A8 Q652K4 Q652K6 Q652L2 Q652L6 Q652Q8 Q652Z7 Q65329
Q65359 Q65361 Q65362 Q65365 Q65366 Q65370 Q65371 Q65372 Q65386 Q65388 Q65389
Q653H7 Q653N3 Q653S3 Q653T6 Q653U3 Q65474 Q65477 Q65478 Q65479 Q654B3 Q654M1
Q654V6 Q65526 Q655R6 Q655V4 Q655V5 Q655X0 Q65652 Q65657 Q65667 Q65670 Q65674
Q65677 Q65695 Q65696 Q656A5 Q65701 Q65707 Q65708 Q65709 Q65729 Q65748 Q65749
Q65750 Q657B3 Q657G7 Q658K8 Q658L1 Q658T7 Q65914 Q65942 Q65943 Q65945 Q65946
Q65949 Q65952 Q65955 Q65956 Q65959 Q65961 Q65962 Q65982 Q65983 Q659V3 Q65CC1
Q65CC7 Q65CJ7 Q65CL1 Q65CM7 Q65CN1 Q65CN2 Q65CN3 Q65CN4 Q65CP3 Q65CP5 Q65CQ3
Q65CQ5 Q65CR5 Q65CS8 Q65CX4 Q65CZ5 Q65D00 Q65D01 Q65D02 Q65D03 Q65D04 Q65D06
Q65D08 Q65D09 Q65D22 Q65D74 Q65DF9 Q65DH1 Q65DJ9 Q65DK0 Q65DN9 Q65DQ3 Q65DR6
Q65DS9 Q65DT6 Q65DT7 Q65DU2 Q65DU5 Q65DU6 Q65DU9 Q65DV1 Q65DW4 Q65DW5 Q65DW6
Q65DX8 Q65DY4 Q65DY5 Q65E10 Q65E18 Q65E52 Q65E56 Q65E69 Q65EB6 Q65EB7 Q65EC2
Q65ED8 Q65EF0 Q65EF7 Q65EF8 Q65EF9 Q65EG0 Q65EG1 Q65EG2 Q65EG3 Q65EH0 Q65EH2
Q65EI5 Q65EM0 Q65EM1 Q65EM9 Q65EN0 Q65EN1 Q65EQ3 Q65EW5 Q65EW6 Q65EX3 Q65F21
Q65F75 Q65FC4 Q65FD4 Q65FD6 Q65FE0 Q65FE6 Q65FL6 Q65FN2 Q65FR5 Q65FS4 Q65FS5
Q65FS7 Q65FT2 Q65FT5 Q65FU0 Q65FV7 Q65FV8 Q65FX8 Q65G12 Q65G19 Q65G22 Q65G33
Q65G35 Q65G42 Q65G52 Q65G58 Q65G65 Q65G67 Q65G68 Q65G70 Q65G82 Q65G89 Q65G93
Q65G95 Q65G98 Q65GA0 Q65GB4 Q65GB5 Q65GC0 Q65GC1 Q65GD0 Q65GD2 Q65GD3 Q65GD8
Q65GE2 Q65GF0 Q65GG2 Q65GG3 Q65GI7 Q65GI8 Q65GI9 Q65GJ0 Q65GJ1 Q65GJ3 Q65GJ4
Q65GJ7 Q65GJ9 Q65GK1 Q65GK4 Q65GK8 Q65GL2 Q65GL3 Q65GM3 Q65GM5 Q65GM7 Q65GM8
Q65GN4 Q65GN7 Q65GP2 Q65GP5 Q65GP6 Q65GP8 Q65GP9 Q65GQ8 Q65GR0 Q65GR3 Q65GR4
Q65GR9 Q65GS5 Q65GS6 Q65GS7 Q65GS8 Q65GT3 Q65GT4 Q65GT8 Q65GT9 Q65GY3 Q65H34
Q65H36 Q65H46 Q65H47 Q65H52 Q65H53 Q65H54 Q65H55 Q65H56 Q65H59 Q65H61 Q65H70
Q65H74 Q65H75 Q65H78 Q65H79 Q65H97 Q65HA0 Q65HA9 Q65HC5 Q65HF9 Q65HG0 Q65HG1
Q65HG7 Q65HH2 Q65HH4 Q65HI7 Q65HI8 Q65HI9 Q65HJ0 Q65HJ2 Q65HL2 Q65HN6 Q65HN7
Q65HN9 Q65HT9 Q65HU3 Q65HW7 Q65HX1 Q65HX2 Q65I08 Q65I10 Q65I15 Q65I24 Q65I26
Q65I28 Q65I29 Q65I32 Q65I33 Q65I34 Q65I35 Q65I36 Q65I37 Q65I39 Q65I41 Q65I50
Q65I51 Q65I54 Q65I56 Q65I57 Q65I58 Q65I63 Q65I68 Q65I79 Q65I80 Q65I86 Q65IA5
Q65IA6 Q65IB0 Q65IB1 Q65ID1 Q65ID2 Q65IE2 Q65IE9 Q65IG4 Q65IH4 Q65II8 Q65IJ4
Q65IS8 Q65IS9 Q65IV4 Q65IW0 Q65IX1 Q65J10 Q65J23 Q65J24 Q65J28 Q65J29 Q65J39
Q65J41 Q65J44 Q65JA6 Q65JA8 Q65JA9 Q65JC2 Q65JC5 Q65JE1 Q65JE2 Q65JE5 Q65JE7
Q65JF2 Q65JF3 Q65JH5 Q65JH6 Q65JH8 Q65JH9 Q65JI1 Q65JI5 Q65JJ1 Q65JJ3 Q65JJ6
Q65JJ7 Q65JJ8 Q65JJ9 Q65JK2 Q65JN2 Q65JN3 Q65JN4 Q65JN5 Q65JN6 Q65JP0 Q65JP3

Q65JP4 Q65JP5 Q65JP6 Q65JP7 Q65JQ0 Q65JQ5 Q65JR6 Q65JS3 Q65JS5 Q65JS9 Q65JT0
Q65JT1 Q65JT8 Q65JT9 Q65JU3 Q65JU4 Q65JU6 Q65JU9 Q65JV0 Q65JV7 Q65JW1 Q65JX9
Q65JY2 Q65JY4 Q65JY8 Q65JY9 Q65JZ0 Q65JZ2 Q65JZ4 Q65JZ6 Q65JZ9 Q65K01 Q65K17
Q65K35 Q65K49 Q65K51 Q65K78 Q65K83 Q65K84 Q65K85 Q65KI3 Q65KI6 Q65KJ5 Q65KJ6
Q65KJ7 Q65KJ8 Q65KK1 Q65KK2 Q65KT8 Q65KU7 Q65KU8 Q65KX8 Q65L04 Q65L57 Q65L74
Q65L94 Q65LA0 Q65LA2 Q65LA4 Q65LA8 Q65LC6 Q65LE8 Q65LE9 Q65LF0 Q65LF6 Q65LG2
Q65LJ9 Q65LK0 Q65LN0 Q65LS8 Q65LT6 Q65LU8 Q65M01 Q65M11 Q65M53 Q65M61 Q65M65
Q65M66 Q65M82 Q65M90 Q65MA1 Q65MA3 Q65MB1 Q65MG3 Q65MJ1 Q65MK9 Q65ML0 Q65ML1
Q65ML3 Q65ML7 Q65MM5 Q65MP7 Q65MP8 Q65MP9 Q65MR2 Q65MS6 Q65MS8 Q65MS9 Q65MU0
Q65MW0 Q65MZ8 Q65MZ9 Q65N04 Q65N05 Q65N07 Q65N43 Q65N49 Q65N62 Q65N65 Q65N90
Q65NA1 Q65NA4 Q65NA8 Q65NB0 Q65NN2 Q65NN5 Q65NN6 Q65NP0 Q65NX1 Q65P46 Q65P47
Q65P54 Q65P72 Q65P73 Q65P74 Q65P78 Q65P79 Q65P80 Q65P81 Q65P82 Q65P83 Q65P89
Q65P90 Q65P91 Q65P92 Q65P93 Q65P94 Q65P95 Q65P96 Q65P97 Q65P98 Q65P99 Q65PA0
Q65PA1 Q65PA2 Q65PA3 Q65PA4 Q65PA5 Q65PA6 Q65PA7 Q65PA8 Q65PA9 Q65PB0 Q65PB1
Q65PB2 Q65PB4 Q65PB5 Q65PB7 Q65PB8 Q65PB9 Q65PC0 Q65PC3 Q65PC8 Q65PC9 Q65PD0
Q65PD1 Q65PD2 Q65PD7 Q65PF0 Q65PF1 Q65PF4 Q65PG8 Q65PG9 Q65PH2 Q65PH5 Q65PH9
Q65PI6 Q65PI8 Q65PJ3 Q65PJ9 Q65PK0 Q65PL0 Q65PL1 Q65PL2 Q65PL9 Q65PM2 Q65PY0
Q65PY5 Q65PY6 Q65PZ8 Q65PZ9 Q65Q00 Q65Q15 Q65Q22 Q65Q24 Q65Q25 Q65Q26 Q65Q29
Q65Q30 Q65Q31 Q65Q32 Q65Q41 Q65Q60 Q65Q78 Q65Q79 Q65Q80 Q65Q81 Q65Q91 Q65QA0
Q65QA6 Q65QA9 Q65QB0 Q65QB1 Q65QB3 Q65QC1 Q65QC6 Q65QD4 Q65QD9 Q65QE8 Q65QF1
Q65QF2 Q65QG5 Q65QG6 Q65QH0 Q65QS1 Q65QT9 Q65QU0 Q65QU6 Q65QV4 Q65QV5 Q65QV6
Q65QV7 Q65QV8 Q65QV9 Q65QW0 Q65QW1 Q65QW2 Q65QW3 Q65QW4 Q65QW6 Q65QW7 Q65QW8
Q65QW9 Q65QX0 Q65QX1 Q65QX2 Q65QX3 Q65QX4 Q65QX6 Q65QX7 Q65QX8 Q65QX9 Q65QY0
Q65QY1 Q65R06 Q65R16 Q65R34 Q65R35 Q65R47 Q65R50 Q65R52 Q65R57 Q65R59 Q65R60
Q65R61 Q65R66 Q65R67 Q65R70 Q65R71 Q65R73 Q65R74 Q65R75 Q65R83 Q65R84 Q65R87
Q65R92 Q65R93 Q65RA4 Q65RB0 Q65RB1 Q65RB2 Q65RB3 Q65RB5 Q65RB8 Q65RC0 Q65RC1
Q65RC3 Q65RD4 Q65RE1 Q65RE8 Q65RF1 Q65RF4 Q65RF6 Q65RF7 Q65RG4 Q65RG7 Q65RG8
Q65RH0 Q65RH6 Q65RH7 Q65RJ7 Q65RK3 Q65RK9 Q65RL3 Q65RL9 Q65RN7 Q65RN9 Q65RP0
Q65RP1 Q65RP4 Q65RP7 Q65RQ2 Q65RQ4 Q65RQ6 Q65RQ9 Q65RR5 Q65RR6 Q65RS7 Q65RT1
Q65RT2 Q65RT5 Q65RU6 Q65RV1 Q65RV7 Q65RW3 Q65RX0 Q65RX3 Q65RX7 Q65RX8 Q65RY4
Q65RY7 Q65RY9 Q65S20 Q65S22 Q65S32 Q65S38 Q65S49 Q65S54 Q65S55 Q65S59 Q65S61
Q65S65 Q65S71 Q65S72 Q65S78 Q65S79 Q65S80 Q65S81 Q65S82 Q65S87 Q65S94 Q65S95
Q65S97 Q65SA3 Q65SB0 Q65SB6 Q65SB8 Q65SC0 Q65SC6 Q65SD0 Q65SD5 Q65SD6 Q65SD7
Q65SE1 Q65SE2 Q65SE6 Q65SE7 Q65SE8 Q65SF0 Q65SH2 Q65SH4 Q65SH6 Q65SH8 Q65SI1
Q65SI5 Q65SK3 Q65SK7 Q65SK9 Q65SL1 Q65SL2 Q65SS8 Q65ST2 Q65SV1 Q65SV7 Q65SX3
Q65SX9 Q65SZ6 Q65SZ8 Q65T08 Q65T14 Q65T20 Q65T21 Q65T36 Q65T37 Q65T41 Q65T49
Q65T53 Q65T54 Q65T71 Q65T73 Q65TB1 Q65TC1 Q65TC2 Q65TC6 Q65TC8 Q65TE9 Q65TF0
Q65TF2 Q65TG8 Q65TI0 Q65TI2 Q65TI3 Q65TI7 Q65TL1 Q65TL2 Q65TL3 Q65TL4 Q65TL6
Q65TL7 Q65TL8 Q65TL9 Q65TM9 Q65TN0 Q65TN4 Q65TP0 Q65TP1 Q65TP2 Q65TP4 Q65TP7
Q65TP8 Q65TQ0 Q65TQ6 Q65TR0 Q65TR1 Q65TT1 Q65TT2 Q65TT7 Q65TU7 Q65TV4 Q65TW5
Q65TX7 Q65TX8 Q65TY2 Q65TY6 Q65TZ0 Q65U07 Q65U16 Q65U19 Q65U29 Q65U43 Q65U54
Q65U55 Q65U59 Q65U60 Q65U83 Q65U87 Q65U90 Q65UA8 Q65UB4 Q65UE8 Q65UE9 Q65UF7
Q65UH7 Q65UI1 Q65UJ6 Q65UJ9 Q65UK6 Q65UL1 Q65UP0 Q65UP1 Q65UP2 Q65UP3 Q65UP5
Q65UP7 Q65UQ4 Q65US2 Q65US7 Q65US9 Q65UT4 Q65UT5 Q65UU9 Q65UV5 Q65UX2 Q65UX3

Q65UX5 Q65UX7 Q65UY1 Q65V05 Q65V07 Q65V08 Q65V34 Q65V45 Q65V47 Q65V59 Q65V66
Q65V70 Q65V72 Q65V80 Q65V95 Q65VB0 Q65VB3 Q65VB6 Q65VB8 Q65VB9 Q65VC0 Q65VC3
Q65VC6 Q65VD1 Q65VD2 Q65VD4 Q65VE4 Q65VF5 Q65VG0 Q65VG1 Q65VG2 Q65VG3 Q65VI0
Q65VI3 Q65VJ3 Q65VJ5 Q65VJ8 Q65VK7 Q65VK8 Q65VM6 Q65VM9 Q65VN0 Q65VN1 Q65VN5
Q65VP4 Q65VP5 Q65VQ5 Q65VQ6 Q65VR5 Q65VR9 Q65VS0 Q65VT1 Q65VT5 Q65VT7 Q65VU4
Q65VV2 Q65VW2 Q65VX7 Q65VY1 Q65VY2 Q65VZ2 Q65VZ3 Q65VZ8 Q65W01 Q65W02 Q65W08
Q65W13 Q65W14 Q65W16 Q65W17 Q65W23 Q65W40 Q65W41 Q65W43 Q65W44 Q65W45 Q65W46
Q65W50 Q65W70 Q65W71 Q65W80 Q65W81 Q65W89 Q65W90 Q65W91 Q65W97 Q65WA1 Q65WA7
Q65WC1 Q65WJ5 Q65WJ9 Q65WK8 Q65WL7 Q65WM2 Q65WM5 Q65WT0 Q65WX1 Q65X70 Q65X92
Q65XA0 Q65XF2 Q65XG6 Q65XK0 Q65XK1 Q65XK7 Q65XL5 Q65XV2 Q65XV7 Q65XV8 Q65XX2
Q65XX7 Q65YR7 Q65YR8 Q65YU6 Q65YU8 Q65YU9 Q65YV0 Q65YV1 Q65YW9 Q65YX0 Q65Z23
Q65Z54 Q65Z55 Q65Z56 Q65Z57 Q65Z58 Q65Z95 Q65ZA6 Q65ZG8 Q65ZT0 Q65ZT1 Q65ZT8
Q65ZU1 Q65ZV3 Q65ZV5 Q65ZV7 Q65ZW4 Q65ZW8 Q65ZW9 Q65ZX0 Q65ZX1 Q65ZX2 Q65ZX4
Q65ZX6 Q65ZY6 Q65ZY7 Q65ZY8 Q65ZY9 Q65ZZ2 Q65ZZ3 Q65ZZ4 Q65ZZ6 Q65ZZ7 Q66011
Q66012 Q66095 Q66096 Q66098 Q660A6 Q660C0 Q660D3 Q660D6 Q660E3 Q660F3 Q660G7
Q660G9 Q660H0 Q660H2 Q660H3 Q660H4 Q660H6 Q660H9 Q660I6 Q660I8 Q660L2 Q660M1
Q660M6 Q660Q0 Q660Q7 Q660R1 Q660R2 Q660R3 Q660R6 Q660S1 Q660S3 Q660S4 Q660S8
Q660T2 Q660T5 Q660U9 Q660W4 Q660X2 Q660X7 Q660Y4 Q660Z6 Q66104 Q66112 Q66114
Q66117 Q66119 Q66120 Q66134 Q66135 Q66136 Q66137 Q66138 Q66139 Q66140 Q66141
Q66142 Q66143 Q66145 Q66154 Q661A2 Q661A3 Q661A4 Q661A7 Q661B2 Q661B7 Q661B8
Q661B9 Q661C0 Q661C1 Q661C3 Q661C4 Q661C5 Q661C6 Q661C7 Q661C8 Q661C9 Q661D0
Q661D1 Q661D2 Q661D3 Q661D4 Q661D5 Q661D6 Q661D7 Q661D8 Q661D9 Q661E0 Q661E2
Q661E3 Q661E4 Q661E5 Q661E8 Q661G2 Q661H4 Q661I0 Q661I1 Q661I4 Q661J3 Q661J6
Q661L6 Q661M2 Q661M5 Q661M6 Q661M7 Q661M9 Q661N0 Q661N1 Q661N2 Q661P3 Q661P7
Q661P8 Q661P9 Q661Q3 Q661Q5 Q661R7 Q661S5 Q661S6 Q661S8 Q661S9 Q661V2 Q661V4
Q661V8 Q661W9 Q66220 Q66221 Q66222 Q66283 Q66284 Q662A9 Q662B4 Q662B8 Q662C3
Q662D1 Q662D5 Q662D8 Q662E4 Q662F0 Q662F3 Q662G3 Q662G4 Q662G9 Q662H4 Q662H5
Q662H6 Q662I0 Q662I1 Q662I5 Q662I6 Q662I7 Q662J5 Q662L3 Q662M6 Q662N0 Q662N3
Q662N9 Q662P0 Q662P1 Q662P7 Q662P9 Q662Q0 Q662Q5 Q662R1 Q662R3 Q662S5 Q662V0
Q662V4 Q662V8 Q662V9 Q662W1 Q662W7 Q662X9 Q662Y6 Q662Y7 Q662Y9 Q662Z0 Q662Z1
Q662Z7 Q663A0 Q663I2 Q663J7 Q663P9 Q663Q0 Q663R1 Q663S6 Q663S9 Q663T0 Q663T2
Q663T4 Q663U3 Q663U4 Q663U8 Q663V5 Q663V8 Q663V9 Q663W9 Q663X0 Q663Y3 Q66403
Q66453 Q664B7 Q664B8 Q664F4 Q664I2 Q664I3 Q664I4 Q664I5 Q664J2 Q664J3 Q664J6
Q664J8 Q664K0 Q664K7 Q664K8 Q664L9 Q664M0 Q664M4 Q664M6 Q664P4 Q664Q8 Q664R1
Q664R2 Q664R3 Q664R4 Q664R5 Q664R6 Q664R7 Q664S0 Q664S6 Q664S7 Q664S8 Q664S9
Q664T0 Q664T1 Q664T2 Q664T3 Q664T4 Q664T5 Q664T6 Q664T7 Q664T8 Q664T9 Q664U0
Q664U2 Q664U3 Q664U4 Q664U5 Q664U6 Q664U7 Q664V2 Q664V3 Q664V4 Q664V6 Q664V8
Q664V9 Q664W1 Q664W4 Q664W9 Q665B4 Q665E1 Q665E3 Q665E8 Q665F6 Q665H6 Q665I5
Q665I6 Q665I8 Q665K4 Q665K9 Q665L0 Q665M1 Q665M2 Q665N8 Q665P6 Q665U4 Q665U5
Q665U9 Q665V3 Q665V6 Q665V9 Q66610 Q66611 Q66612 Q66616 Q66617 Q66619 Q66621
Q66622 Q66623 Q66624 Q66626 Q66627 Q66628 Q66629 Q66632 Q66633 Q66634 Q66635
Q66636 Q66638 Q66639 Q66640 Q66641 Q66643 Q66645 Q66646 Q66647 Q66648 Q66650
Q66651 Q66652 Q66654 Q66656 Q66658 Q66659 Q66661 Q66663 Q66664 Q66665 Q66667
Q66668 Q66670 Q66671 Q66674 Q666M3 Q666M5 Q666N1 Q666N2 Q666N7 Q666N9 Q666P0

Q666P4 Q666P5 Q666Q0 Q666Q1 Q666Q6 Q666Q7 Q666R1 Q666R5 Q666R6 Q666R7 Q666S2
Q666S3 Q666T2 Q666T3 Q667F5 Q667F6 Q667F9 Q667G9 Q667H7 Q667H8 Q667I0 Q667I5
Q667I6 Q667I7 Q667I9 Q667J0 Q667J1 Q667J2 Q667J3 Q667K3 Q667K7 Q667L0 Q667L4
Q667M5 Q667M7 Q667N6 Q667R4 Q667T3 Q667T6 Q667T7 Q667T8 Q667U2 Q667V1 Q667V3
Q667V7 Q667W1 Q667X1 Q667X9 Q667Y0 Q667Y3 Q667Y4 Q667Y5 Q667Y8 Q667Z5 Q667Z6
Q667Z9 Q66802 Q668A0 Q668A3 Q668A6 Q668A8 Q668E5 Q668E6 Q668E8 Q668F7 Q668F9
Q668G5 Q668I3 Q668I5 Q668I6 Q668I7 Q668J7 Q668J9 Q668M6 Q668N0 Q668N7 Q668V5
Q668W0 Q668W7 Q668W9 Q668Z0 Q668Z3 Q668Z7 Q66914 Q66915 Q66916 Q66929 Q669C1
Q669C7 Q669C8 Q669D0 Q669D4 Q669E1 Q669E2 Q669J6 Q669J9 Q669K2 Q669K7 Q669K9
Q669L8 Q669N5 Q669N6 Q669P5 Q669P6 Q669P7 Q669Q4 Q669R9 Q669S1 Q669S5 Q669V5
Q669X6 Q669Z0 Q669Z1 Q669Z2 Q669Z3 Q669Z5 Q669Z6 Q66A10 Q66A11 Q66A13 Q66A22
Q66A23 Q66A32 Q66A36 Q66A46 Q66A48 Q66A49 Q66A50 Q66A78 Q66A90 Q66AE3 Q66AF0
Q66AF7 Q66AF8 Q66AH3 Q66AI1 Q66AI6 Q66AK2 Q66AK4 Q66AK5 Q66AM0 Q66AM8 Q66AN2
Q66AP0 Q66AP6 Q66AQ1 Q66AQ6 Q66AR5 Q66AR8 Q66AR9 Q66AS1 Q66AT9 Q66AU0 Q66AU1
Q66AU2 Q66AU4 Q66AU7 Q66AU8 Q66AU9 Q66AV1 Q66AV6 Q66AW1 Q66AX5 Q66AX6 Q66AX8
Q66AY1 Q66B14 Q66B17 Q66B18 Q66B19 Q66B20 Q66B21 Q66B88 Q66BN5 Q66BP3 Q66BQ4
Q66BW2 Q66BW5 Q66BW6 Q66BW8 Q66C05 Q66C31 Q66C39 Q66C48 Q66C49 Q66C50 Q66C51
Q66C52 Q66C53 Q66C54 Q66C69 Q66C70 Q66C72 Q66C79 Q66C80 Q66CD8 Q66CD9 Q66CE0
Q66CE1 Q66CE4 Q66CE9 Q66CF1 Q66CG1 Q66CG6 Q66CG7 Q66CH3 Q66CH4 Q66CH5 Q66CH9
Q66CI5 Q66CI7 Q66CI8 Q66CI9 Q66CJ9 Q66CK7 Q66CK8 Q66CL0 Q66CL7 Q66CN7 Q66CP8
Q66CR5 Q66CS8 Q66CT6 Q66CT7 Q66CV3 Q66CV4 Q66CV5 Q66CZ4 Q66D52 Q66D53 Q66D54
Q66D58 Q66D62 Q66D66 Q66D67 Q66D69 Q66D80 Q66D83 Q66D87 Q66DA0 Q66DA8 Q66DA9
Q66DB4 Q66DC5 Q66DC7 Q66DD3 Q66DD5 Q66DE3 Q66DE6 Q66DE8 Q66DF3 Q66DF4 Q66DF5
Q66DK7 Q66DK9 Q66DP7 Q66DP8 Q66DP9 Q66DQ0 Q66DQ2 Q66DR1 Q66DR2 Q66DS5 Q66DS7
Q66DT3 Q66DT4 Q66DT5 Q66DU7 Q66DV0 Q66DV2 Q66DV4 Q66DV7 Q66DV8 Q66DW0 Q66DW5
Q66DW6 Q66DY1 Q66DY2 Q66DY7 Q66DY8 Q66DY9 Q66DZ0 Q66DZ1 Q66DZ7 Q66E06 Q66E15
Q66E16 Q66E17 Q66E18 Q66E19 Q66E56 Q66E57 Q66E58 Q66E59 Q66E64 Q66E67 Q66E68
Q66E69 Q66E70 Q66EB5 Q66EB9 Q66EC0 Q66EC1 Q66EC2 Q66EC3 Q66EC7 Q66EC8 Q66EC9
Q66ED2 Q66ED3 Q66ED4 Q66ED9 Q66EE5 Q66EE6 Q66EE9 Q66EF1 Q66EF8 Q66EG0 Q66EG3
Q66EG4 Q66EG5 Q66EH3 Q66EH4 Q66EH6 Q66EJ0 Q66EJ1 Q66EJ2 Q66EJ3 Q66EK3 Q66EK4
Q66EK7 Q66EL3 Q66EL4 Q66EM1 Q66EM2 Q66EM3 Q66EM4 Q66EM6 Q66EN4 Q66EQ8 Q66EQ9
Q66ER0 Q66ER8 Q66ER9 Q66ES1 Q66ES4 Q66ES6 Q66ES9 Q66ET0 Q66ET5 Q66ET6 Q66ET8
Q66EU3 Q66EU4 Q66EU5 Q66EV7 Q66EV8 Q66EV9 Q66EW0 Q66EW6 Q66EW9 Q66EY7 Q66EZ3
Q66EZ4 Q66F09 Q66F11 Q66F14 Q66F34 Q66F42 Q66F56 Q66F57 Q66F58 Q66F59 Q66F60
Q66F62 Q66F64 Q66F67 Q66F73 Q66F75 Q66F76 Q66F81 Q66F84 Q66F89 Q66F95 Q66F99
Q66FA0 Q66FA1 Q66FA2 Q66FA9 Q66FB0 Q66FB5 Q66FB6 Q66FB7 Q66FC2 Q66FC3 Q66FC6
Q66FD2 Q66FD5 Q66FD6 Q66FE0 Q66FF1 Q66FF2 Q66FF3 Q66FF4 Q66FF5 Q66FF7 Q66FH1
Q66FM8 Q66FN5 Q66FP0 Q66FP1 Q66FP3 Q66FP4 Q66FP5 Q66FP6 Q66FP9 Q66FQ1 Q66FQ2
Q66FQ3 Q66FQ4 Q66FQ5 Q66FQ6 Q66FQ9 Q66FR1 Q66FR3 Q66FR7 Q66FT0 Q66FT7 Q66FX1
Q66FZ5 Q66FZ7 Q66G00 Q66G19 Q66G20 Q66G37 Q66G45 Q66G51 Q66G56 Q66G58 Q66G61
Q66G70 Q66G71 Q66G72 Q66G73 Q66G74 Q66G78 Q66G80 Q66G84 Q66G85 Q66G87 Q66G93
Q66GA1 Q66GA8 Q66GB9 Q66GC2 Q66GC5 Q66GC7 Q66GD2 Q66GD3 Q66GD4 Q66GD5 Q66GD6
Q66GD8 Q66GD9 Q66GE0 Q66GE1 Q66GE4 Q66GE5 Q66GE6 Q66GF4 Q66GG2 Q66GG3 Q66GH4
Q66GH6 Q66GH7 Q66GH8 Q66GI4 Q66GN9 Q66GP4 Q66GQ2 Q66GQ6 Q66GR3 Q66GV6 Q66H03

Q66H04 Q66H07 Q66H10 Q66H16 Q66H17 Q66H19 Q66H20 Q66H24 Q66H30 Q66H33 Q66H34

Q66H40 Q66H45 Q66H47 Q66H53 Q66H54 Q66H56 Q66H58 Q66H59 Q66H60 Q66H61 Q66H63

Q66H65 Q66H68 Q66H73 Q66H85 Q66H99 Q66HA1 Q66HA4 Q66HA8 Q66HB3 Q66HB5 Q66HB6

Q66HC0 Q66HC1 Q66HC4 Q66HC7 Q66HC8 Q66HD2 Q66HD5 Q66HD6 Q66HD8 Q66HE1 Q66HE7

Q66HF9 Q66HG1 Q66HG4 Q66HG6 Q66HG7 Q66HG8 Q66HH3 Q66HP6 Q66HS7 Q66HX8 Q66HY7

Q66HZ2 Q66I06 Q66I14 Q66I21 Q66I22 Q66I24 Q66I59 Q66I68 Q66I71 Q66I73 Q66I74

Q66I75 Q66I84 Q66I85 Q66IB8 Q66IC8 Q66IC9 Q66IG8 Q66IH2 Q66IH9 Q66II5 Q66II8

Q66IK1 Q66IK8 Q66IM5 Q66IN2 Q66IQ6 Q66IT9 Q66IV5 Q66IW8 Q66IY8 Q66IZ5 Q66IZ7

Q66J36 Q66J51 Q66J63 Q66J69 Q66J74 Q66J77 Q66J85 Q66J90 Q66J96 Q66J97 Q66J99

Q66JA8 Q66JB7 Q66JC5 Q66JD1 Q66JD7 Q66JD9 Q66JE4 Q66JF1 Q66JF8 Q66JG1 Q66JG3

Q66JG5 Q66JG8 Q66JJ3 Q66JJ4 Q66JK1 Q66JK4 Q66JL1 Q66JS6 Q66JT5 Q66JV4 Q66JV7

Q66JY2 Q66JZ4 Q66K41 Q66K80 Q66K94 Q66K99 Q66KB0 Q66KB7 Q66KC4 Q66KC9 Q66KD5

Q66KD6 Q66KD9 Q66KE8 Q66KE9 Q66KF6 Q66KG5 Q66KH8 Q66KH9 Q66KI6 Q66KI9 Q66KJ5

Q66KJ9 Q66KK3 Q66KK8 Q66KL9 Q66KM5 Q66KT3 Q66KU4 Q66KX0 Q66KX4 Q66KY0 Q66KZ3

Q66L17 Q66L33 Q66L42 Q66LE6 Q66LM5 Q66LM6 Q66MI6 Q66PF3 Q66PF4 Q66PF5 Q66QC7

Q66T60 Q66T65 Q66T82 Q66UV4 Q66UW5 Q66V57 Q66V72 Q66V81 Q66WF5 Q66WM4 Q66WU1

Q66WV0 Q66X01 Q66X03 Q66X52 Q66X92 Q672F7 Q672H9 Q672I0 Q672I1 Q672I7 Q672V5

Q672V6 Q673H1 Q673V5 Q67472 Q67473 Q67475 Q67480 Q674M7 Q67591 Q675L0 Q675L4

Q675L5 Q67616 Q67618 Q67620 Q67652 Q67653 Q67669 Q67683 Q67685 Q67704 Q67705

Q67706 Q67707 Q67708 Q67724 Q67815 Q67863 Q67878 Q67897 Q67925 Q679P3 Q67BD8

Q67BS9 Q67ER4 Q67EU8 Q67EX6 Q67EX7 Q67FW5 Q67FY2 Q67FY3 Q67IA4 Q67IB0 Q67IB6

Q67IC5 Q67ID1 Q67IE0 Q67IE3 Q67IG9 Q67II8 Q67IL8 Q67IP2 Q67J15 Q67J17 Q67J28

Q67J29 Q67J33 Q67J34 Q67J35 Q67J48 Q67J50 Q67J52 Q67J55 Q67J65 Q67J66 Q67JB0

Q67JB9 Q67JC2 Q67JC7 Q67JD0 Q67JG9 Q67JH4 Q67JH5 Q67JH7 Q67JH9 Q67JI4 Q67JI5

Q67JK8 Q67JN4 Q67JP5 Q67JP6 Q67JP9 Q67JQ0 Q67JQ1 Q67JS5 Q67JS8 Q67JS9 Q67JT1

Q67JT2 Q67JT3 Q67JT4 Q67JT8 Q67JT9 Q67JU0 Q67JU1 Q67JU2 Q67JU3 Q67JU4 Q67JU5

Q67JU6 Q67JU7 Q67JU8 Q67JU9 Q67JV0 Q67JV1 Q67JV2 Q67JV3 Q67JV4 Q67JV5 Q67JV6

Q67JV7 Q67JV8 Q67JV9 Q67JW0 Q67JW1 Q67JW2 Q67JW4 Q67JW6 Q67JW7 Q67JW8 Q67JW9

Q67JX0 Q67JX1 Q67JX2 Q67JX6 Q67JY2 Q67JZ8 Q67K79 Q67K93 Q67K94 Q67K97 Q67KA0

Q67KB7 Q67KB8 Q67KC3 Q67KC5 Q67KD3 Q67KE1 Q67KE6 Q67KE7 Q67KF7 Q67KF8 Q67KG0

Q67KG3 Q67KH4 Q67KH5 Q67KH6 Q67KH7 Q67KH8 Q67KH9 Q67KI0 Q67KI2 Q67KI8 Q67KJ1

Q67KJ2 Q67KJ3 Q67KM3 Q67KQ0 Q67KQ1 Q67KX6 Q67KZ3 Q67L72 Q67L76 Q67LA3 Q67LA5

Q67LB8 Q67LC1 Q67LC2 Q67LD9 Q67LE6 Q67LM3 Q67LN1 Q67LN9 Q67LP0 Q67LP5 Q67LR6

Q67LS2 Q67M16 Q67M29 Q67MD4 Q67MD5 Q67MF5 Q67MJ2 Q67MJ3 Q67MP0 Q67MS9 Q67MT5

Q67MV8 Q67MW0 Q67MW4 Q67N06 Q67N08 Q67N09 Q67N10 Q67N11 Q67N36 Q67N37 Q67N40

Q67N41 Q67N73 Q67N85 Q67N86 Q67N94 Q67NA9 Q67NB1 Q67NB3 Q67NB6 Q67NC1 Q67NC2

Q67ND8 Q67NF6 Q67NG0 Q67NI4 Q67NJ9 Q67NK1 Q67NL0 Q67NL1 Q67NL2 Q67NP4 Q67NP7

Q67NP8 Q67NQ5 Q67NR4 Q67NS3 Q67NS9 Q67NT4 Q67NU0 Q67NU3 Q67NW2 Q67NW5 Q67NX5

Q67P26 Q67P59 Q67P64 Q67P69 Q67P77 Q67P80 Q67P83 Q67P85 Q67P86 Q67P90 Q67PA6

Q67PA7 Q67PA9 Q67PB4 Q67PB5 Q67PB6 Q67PB7 Q67PC6 Q67PD3 Q67PD4 Q67PD7 Q67PD8

Q67PD9 Q67PE2 Q67PF5 Q67PG5 Q67PG9 Q67PJ3 Q67PJ5 Q67PJ9 Q67PP9 Q67PQ0 Q67PQ7

Q67PR4 Q67PR5 Q67PR8 Q67PR9 Q67PS0 Q67PZ3 Q67Q01 Q67Q27 Q67Q30 Q67Q46 Q67Q47

Q67Q54 Q67Q55 Q67Q57 Q67Q58 Q67Q63 Q67Q92 Q67Q93 Q67Q97 Q67Q98 Q67Q99 Q67QB5

Q67QD6 Q67QE3 Q67QF2 Q67QF3 Q67QG5 Q67QG6 Q67QM6 Q67QM7 Q67QQ0 Q67QQ3 Q67QQ4

Q67QU3 Q67QV9 Q67QZ4 Q67QZ8 Q67R57 Q67R59 Q67RD6 Q67RL6 Q67RV4 Q67RX3 Q67RZ0
Q67RZ3 Q67S27 Q67S44 Q67S51 Q67S53 Q67S54 Q67S56 Q67S57 Q67S72 Q67S82 Q67S99
Q67SB4 Q67SC4 Q67SC6 Q67SC7 Q67SC9 Q67SD7 Q67SI1 Q67SI3 Q67SI7 Q67SI8 Q67SJ2
Q67SJ6 Q67SJ9 Q67SK0 Q67SK1 Q67SL4 Q67SL5 Q67SL6 Q67SL8 Q67SW0 Q67SW4 Q67SW6
Q67T12 Q67T14 Q67T19 Q67T22 Q67T37 Q67T48 Q67T51 Q67T57 Q67T86 Q67T90 Q67T95
Q67TC9 Q67TD0 Q67TD5 Q67TD7 Q67TE2 Q67TG8 Q67TI9 Q67TJ2 Q67TJ3 Q67TJ8 Q67TK4
Q67TK7 Q67U26 Q67U69 Q67UE2 Q67UK9 Q67UP9 Q67UR2 Q67UU0 Q67UX0 Q67UX6 Q67V61
Q67V78 Q67V81 Q67VC8 Q67VL7 Q67VR7 Q67VW6 Q67W29 Q67W65 Q67W82 Q67WN8 Q67WR2
Q67WR5 Q67X83 Q67X99 Q67XC9 Q67XD9 Q67XG0 Q67XJ2 Q67XL4 Q67XN8 Q67XT3 Q67XV2
Q67XX3 Q67Y55 Q67Y69 Q67Y93 Q67Y99 Q67YC0 Q67YE6 Q67YG7 Q67Z52 Q67Z93 Q67ZB3
Q67ZB6 Q67ZM4 Q680A6 Q680B9 Q680D9 Q680I0 Q680K8 Q680Q4 Q680U9 Q680Z7 Q68101
Q681N2 Q681X4 Q681Y3 Q681Z2 Q682D3 Q682T9 Q682V0 Q683D5 Q683I9 Q684M4 Q687E1
Q688C0 Q688C4 Q688D0 Q688Q9 Q688R3 Q688T8 Q688U3 Q688X9 Q688Y7 Q689D6 Q68A21
Q68A65 Q68AP4 Q68BJ6 Q68BJ7 Q68CJ6 Q68CL5 Q68CM3 Q68CQ4 Q68D06 Q68D20 Q68D51
Q68D86 Q68DE3 Q68DI1 Q68DL7 Q68DN1 Q68DQ2 Q68DY1 Q68DY9 Q68EA5 Q68EH7 Q68EH8
Q68EH9 Q68EI0 Q68EI3 Q68EJ0 Q68EK9 Q68EL2 Q68EL6 Q68EN5 Q68EP2 Q68EP9 Q68EQ1
Q68EQ8 Q68ER9 Q68EV5 Q68EV6 Q68EV9 Q68EW0 Q68EW7 Q68EY3 Q68EY7 Q68EY9 Q68EZ2
Q68EZ3 Q68F20 Q68F45 Q68F47 Q68F50 Q68F53 Q68F54 Q68F55 Q68F60 Q68F64 Q68F70
Q68F77 Q68FA2 Q68FA4 Q68FB1 Q68FE8 Q68FE9 Q68FF0 Q68FF7 Q68FG3 Q68FH4 Q68FI1
Q68FI3 Q68FI4 Q68FJ4 Q68FJ8 Q68FJ9 Q68FK4 Q68FL6 Q68FN9 Q68FP8 Q68FQ0 Q68FQ4
Q68FQ7 Q68FQ8 Q68FR3 Q68FR6 Q68FR7 Q68FR9 Q68FS1 Q68FS2 Q68FS3 Q68FS6 Q68FT5
Q68FU3 Q68FU4 Q68FU5 Q68FV5 Q68FV8 Q68FW2 Q68FW6 Q68FW7 Q68FX0 Q68FX1 Q68FX5
Q68G30 Q68G74 Q68G84 Q68HB4 Q68J44 Q68LP1 Q68RJ5 Q68RU2 Q68RU3 Q68RU4 Q68RU5
Q68RU7 Q68RU9 Q68RV8 Q68RW6 Q68RW7 Q68RW8 Q68RW9 Q68RX0 Q68RX1 Q68RX2 Q68RX3
Q68RX4 Q68RX5 Q68RY2 Q68RY3 Q68RY4 Q68RY5 Q68RZ8 Q68S04 Q68S08 Q68S14 Q68S15
Q68S16 Q68S17 Q68S24 Q68S28 Q68UI8 Q68US1 Q68US6 Q68UT5 Q68VM8 Q68VM9 Q68VN1
Q68VN4 Q68VN5 Q68VN6 Q68VN7 Q68VN9 Q68VP6 Q68VQ2 Q68VQ4 Q68VQ5 Q68VQ6 Q68VQ7
Q68VR3 Q68VR4 Q68VR6 Q68VS1 Q68VS2 Q68VS3 Q68VT2 Q68VT4 Q68VT5 Q68VU1 Q68VV0
Q68VW6 Q68VX0 Q68VX1 Q68VX6 Q68VY3 Q68VY8 Q68VZ0 Q68VZ7 Q68VZ8 Q68VZ9 Q68W04
Q68W05 Q68W06 Q68W07 Q68W08 Q68W09 Q68W10 Q68W16 Q68W17 Q68W20 Q68W26 Q68W27
Q68W43 Q68W45 Q68W48 Q68W50 Q68W53 Q68W54 Q68W57 Q68W59 Q68W63 Q68W66 Q68W69
Q68W74 Q68W77 Q68W78 Q68W79 Q68W80 Q68W81 Q68W82 Q68W83 Q68W84 Q68W85 Q68W86
Q68W87 Q68W88 Q68W89 Q68W90 Q68W91 Q68W92 Q68W93 Q68W94 Q68W95 Q68W96 Q68W97
Q68W99 Q68WA0 Q68WA1 Q68WA2 Q68WA3 Q68WA4 Q68WA8 Q68WB4 Q68WB5 Q68WC2 Q68WC4
Q68WC8 Q68WC9 Q68WD0 Q68WD1 Q68WD3 Q68WD4 Q68WD8 Q68WE2 Q68WE9 Q68WF1 Q68WF6
Q68WF7 Q68WF8 Q68WG4 Q68WG6 Q68WG7 Q68WH9 Q68WI2 Q68WI4 Q68WI7 Q68WI9 Q68WJ1
Q68WJ2 Q68WJ4 Q68WK0 Q68WK5 Q68WK7 Q68WL0 Q68WL3 Q68WL4 Q68WL5 Q68WL9 Q68WM0
Q68WM5 Q68WN1 Q68WN2 Q68WN3 Q68WP2 Q68WP6 Q68WR2 Q68WR3 Q68WS3 Q68WS4 Q68WS6
Q68WS8 Q68WT1 Q68WT3 Q68WT8 Q68WU0 Q68WU3 Q68WU5 Q68WV4 Q68WV8 Q68WW0 Q68WW1
Q68WW2 Q68WW4 Q68WW9 Q68WX4 Q68WX8 Q68WY1 Q68WY2 Q68WY7 Q68WZ0 Q68WZ1 Q68WZ3
Q68WZ4 Q68WZ6 Q68WZ8 Q68X04 Q68X22 Q68X23 Q68X25 Q68X27 Q68X33 Q68X41 Q68X45
Q68X46 Q68X49 Q68X52 Q68X53 Q68X56 Q68X64 Q68X66 Q68X70 Q68X72 Q68X73 Q68X96
Q68X97 Q68XA5 Q68XA8 Q68XB2 Q68XC0 Q68XC1 Q68XC2 Q68XD1 Q68XD3 Q68XD4 Q68XD5
Q68XE1 Q68XE8 Q68XE9 Q68XF0 Q68XF1 Q68XF6 Q68XG1 Q68XG2 Q68XG4 Q68XG5 Q68XG6

Q68XG7 Q68XG8 Q68XH5 Q68XI2 Q68XI3 Q68XI5 Q68XI7 Q68XI8 Q68XI9 Q68XJ6 Q68XK5
Q68XK6 Q68XL3 Q68XL4 Q68XL5 Q68XL6 Q68XL7 Q68XM0 Q68XM3 Q68XM6 Q68XM7 Q68XM9
Q68XN0 Q68XN1 Q68XN2 Q68XN3 Q68XN6 Q68XN7 Q68XP2 Q68XP6 Q68XQ6 Q68XQ8 Q68XR2
Q68XR3 Q68XR5 Q68XR6 Q68XR7 Q68XR8 Q68XS0 Q68XS9 Q68XT0 Q68XT1 Q68XT3 Q68XT8
Q68XT9 Q68XU3 Q68XU4 Q68XU6 Q68XV4 Q68XV5 Q68XV6 Q68XV9 Q68XW3 Q68XW7 Q68XW8
Q68XW9 Q68XX7 Q68XX8 Q68XX9 Q68XY0 Q68XY5 Q68XY7 Q68XZ1 Q68XZ3 Q68Y00 Q68Y01
Q68Y02 Q68Y04 Q68Y08 Q68Y09 Q68Y11 Q68Y12 Q68Y28 Q68Y29 Q68Y30 Q68Y31 Q68Y48
Q69014 Q69015 Q69016 Q69022 Q69025 Q69028 Q69066 Q69138 Q69140 Q69151 Q69153
Q69154 Q692V3 Q693B1 Q694A3 Q694B3 Q694B4 Q694B5 Q694B6 Q694B7 Q694B8 Q694B9
Q694C0 Q694C1 Q694C2 Q694C4 Q694C5 Q69535 Q69545 Q69546 Q69548 Q69549 Q69551
Q69554 Q69560 Q69561 Q69562 Q69567 Q695L6 Q69602 Q69605 Q696W0 Q698V5 Q69AA9
Q69AB1 Q69AB2 Q69BZ7 Q69CJ9 Q69DB9 Q69FB1 Q69GZ5 Q69HT9 Q69IL4 Q69J40 Q69JW2
Q69JZ7 Q69K07 Q69K55 Q69KJ0 Q69KL9 Q69KN0 Q69KP0 Q69LD8 Q69LE0 Q69LE7 Q69LX2
Q69NG5 Q69NK8 Q69NX5 Q69P88 Q69PS6 Q69Q02 Q69QA6 Q69QB5 Q69QB8 Q69QZ0 Q69R21
Q69RG7 Q69RJ0 Q69S43 Q69S81 Q69T21 Q69T31 Q69T99 Q69TB1 Q69TF4 Q69TG5 Q69TU6
Q69TW5 Q69TY4 Q69TY5 Q69U54 Q69UD7 Q69UE6 Q69UI1 Q69UI2 Q69UP6 Q69UP7 Q69V10
Q69V36 Q69VD5 Q69VD9 Q69VE0 Q69WS3 Q69XJ4 Q69XM7 Q69XQ3 Q69XQ6 Q69YN2 Q69YU3
Q69Z23 Q69Z36 Q69Z37 Q69Z61 Q69Z66 Q69Z69 Q69Z99 Q69ZB0 Q69ZB3 Q69ZB8 Q69ZC8
Q69ZF8 Q69ZI1 Q69ZK6 Q69ZK7 Q69ZL1 Q69ZQ2 Q69ZS7 Q69ZT1 Q69ZX6 Q69ZX8 Q6A000
Q6A009 Q6A025 Q6A037 Q6A039 Q6A065 Q6A070 Q6A085 Q6A098 Q6A0A2 Q6A162 Q6A163
Q6A1B7 Q6A1I3 Q6A2H0 Q6A2H1 Q6A2H2 Q6A331 Q6A332 Q6A333 Q6A3P9 Q6A555 Q6A5A1
Q6A5B2 Q6A5D0 Q6A5D2 Q6A5H2 Q6A5L3 Q6A5M9 Q6A5N1 Q6A5N2 Q6A5P0 Q6A5S3 Q6A5T5
Q6A5T8 Q6A5U5 Q6A5U6 Q6A5U7 Q6A5U8 Q6A5U9 Q6A5X5 Q6A5X8 Q6A5Z1 Q6A651 Q6A655
Q6A661 Q6A662 Q6A6A0 Q6A6A3 Q6A6B0 Q6A6C0 Q6A6J5 Q6A6J6 Q6A6J8 Q6A6K2 Q6A6K3
Q6A6K4 Q6A6K5 Q6A6K6 Q6A6K7 Q6A6L2 Q6A6L4 Q6A6L5 Q6A6L7 Q6A6M1 Q6A6M4 Q6A6M5
Q6A6M6 Q6A6M7 Q6A6M8 Q6A6M9 Q6A6N0 Q6A6N1 Q6A6N2 Q6A6N3 Q6A6N4 Q6A6N5 Q6A6N6
Q6A6N7 Q6A6N8 Q6A6N9 Q6A6P0 Q6A6P1 Q6A6P2 Q6A6P3 Q6A6P4 Q6A6P5 Q6A6Q4 Q6A6Q6
Q6A6Q7 Q6A6Q8 Q6A6Q9 Q6A6R0 Q6A6R1 Q6A6R2 Q6A6S8 Q6A6S9 Q6A6T3 Q6A6T4 Q6A6T5
Q6A6T7 Q6A6V2 Q6A6W1 Q6A6W2 Q6A6X1 Q6A6Y0 Q6A6Y6 Q6A6Y8 Q6A6Z2 Q6A6Z5 Q6A707
Q6A725 Q6A759 Q6A760 Q6A761 Q6A763 Q6A780 Q6A786 Q6A7A2 Q6A7E7 Q6A7E9 Q6A7F0
Q6A7F1 Q6A7F4 Q6A7J7 Q6A7J8 Q6A7J9 Q6A7K0 Q6A7K4 Q6A7K8 Q6A7L2 Q6A7L9 Q6A7M3
Q6A7M5 Q6A7M9 Q6A7P0 Q6A7P5 Q6A7P6 Q6A7P7 Q6A7Q4 Q6A7S4 Q6A7S6 Q6A7S7 Q6A7T1
Q6A7T3 Q6A7T5 Q6A7T9 Q6A7U7 Q6A7V2 Q6A7V3 Q6A7V5 Q6A7V6 Q6A7V7 Q6A7W0 Q6A7W9
Q6A7X4 Q6A7Y1 Q6A7Y8 Q6A7Z2 Q6A808 Q6A811 Q6A814 Q6A816 Q6A8B2 Q6A8B4 Q6A8F1
Q6A8F7 Q6A8H1 Q6A8H3 Q6A8H5 Q6A8H6 Q6A8H8 Q6A8I2 Q6A8I4 Q6A8I7 Q6A8I8 Q6A8J1
Q6A8J7 Q6A8K3 Q6A8K7 Q6A8K8 Q6A8K9 Q6A8L0 Q6A8L1 Q6A8L2 Q6A8L3 Q6A8L4 Q6A8L5
Q6A8M7 Q6A8P2 Q6A8P4 Q6A8P5 Q6A8Q1 Q6A8Q7 Q6A8U3 Q6A8V3 Q6A8W1 Q6A8W6 Q6A8Y9
Q6A8Z5 Q6A8Z7 Q6A908 Q6A910 Q6A911 Q6A912 Q6A914 Q6A917 Q6A947 Q6A948 Q6A964
Q6A969 Q6A976 Q6A997 Q6A998 Q6A9B0 Q6A9B3 Q6A9C3 Q6A9C5 Q6A9H6 Q6A9H7 Q6A9H8
Q6A9I2 Q6A9I3 Q6A9I7 Q6A9J2 Q6A9J3 Q6A9J5 Q6A9J7 Q6A9J8 Q6A9K7 Q6A9M5 Q6A9P7
Q6A9Q1 Q6A9Q4 Q6A9R0 Q6A9R1 Q6A9R6 Q6A9R8 Q6A9W5 Q6A9W7 Q6A9X0 Q6AA18 Q6AA20
Q6AA30 Q6AA77 Q6AA78 Q6AA89 Q6AA93 Q6AAC7 Q6AAC8 Q6AAD6 Q6AAD7 Q6AAE4 Q6AAI8
Q6AAP6 Q6AAR3 Q6AAT0 Q6AAU3 Q6AAU8 Q6AAV7 Q6AAV8 Q6AAW4 Q6AAW5 Q6AAZ9 Q6AB02
Q6AB05 Q6AB08 Q6AB12 Q6AB26 Q6AB49 Q6AB62 Q6AB88 Q6AB89 Q6AB99 Q6ABA0 Q6ABL2

Q6ABL5 Q6ABP6 Q6ABS5 Q6ABU3 Q6ABV3 Q6ABV7 Q6ABW7 Q6ABW9 Q6ABX0 Q6ABX1 Q6ABX6
Q6ABX8 Q6ABX9 Q6ABY4 Q6ABY5 Q6ABY6 Q6AC27 Q6AC71 Q6AC73 Q6AC76 Q6AC77 Q6ACB0
Q6ACC3 Q6ACD4 Q6ACE5 Q6ACE6 Q6ACL1 Q6ACP7 Q6ACP8 Q6ACQ1 Q6ACQ8 Q6ACX5 Q6ACX6
Q6ACX8 Q6ACY7 Q6ACY8 Q6ACY9 Q6ACZ0 Q6ACZ4 Q6ACZ5 Q6ACZ6 Q6ACZ7 Q6ACZ8 Q6ACZ9
Q6AD00 Q6AD01 Q6AD02 Q6AD04 Q6AD05 Q6AD06 Q6AD07 Q6AD08 Q6AD09 Q6AD10 Q6AD11
Q6AD12 Q6AD13 Q6AD15 Q6AD17 Q6AD18 Q6AD19 Q6AD20 Q6AD21 Q6AD22 Q6AD25 Q6AD27
Q6AD28 Q6AD31 Q6AD32 Q6AD38 Q6AD41 Q6AD51 Q6AD57 Q6AD62 Q6ADB9 Q6ADC8 Q6ADC9
Q6ADF0 Q6ADF1 Q6ADH0 Q6ADH3 Q6ADI0 Q6ADI1 Q6ADJ8 Q6ADK2 Q6ADP1 Q6ADP2 Q6ADQ7
Q6ADQ8 Q6ADR4 Q6ADU8 Q6ADU9 Q6ADV0 Q6AE00 Q6AE03 Q6AE04 Q6AE05 Q6AE13 Q6AE14
Q6AE15 Q6AE23 Q6AE55 Q6AE56 Q6AE59 Q6AE62 Q6AE65 Q6AE69 Q6AE76 Q6AE77 Q6AE97
Q6AE98 Q6AE99 Q6AEA0 Q6AEA3 Q6AEA5 Q6AEA8 Q6AEB3 Q6AEB9 Q6AEC0 Q6AEC4 Q6AED3
Q6AED5 Q6AEE4 Q6AEE5 Q6AEE8 Q6AEE9 Q6AEG5 Q6AEK8 Q6AEK9 Q6AEL0 Q6AEL7 Q6AEN9
Q6AEP2 Q6AEP6 Q6AEQ1 Q6AER9 Q6AEU4 Q6AEV4 Q6AEV5 Q6AEV6 Q6AEV7 Q6AEW2 Q6AEX8
Q6AEX9 Q6AEY1 Q6AF19 Q6AF21 Q6AF36 Q6AF43 Q6AF44 Q6AF47 Q6AF48 Q6AF49 Q6AF52
Q6AF53 Q6AF66 Q6AF67 Q6AF68 Q6AF72 Q6AF73 Q6AF75 Q6AF77 Q6AF79 Q6AF81 Q6AF82
Q6AF87 Q6AF88 Q6AF89 Q6AF91 Q6AF92 Q6AF95 Q6AF97 Q6AFA0 Q6AFA1 Q6AFA4 Q6AFB4
Q6AFB5 Q6AFB6 Q6AFB7 Q6AFB8 Q6AFB9 Q6AFC1 Q6AFC3 Q6AFD5 Q6AFE0 Q6AFG2 Q6AFG6
Q6AFG7 Q6AFH4 Q6AFI0 Q6AFI7 Q6AFJ3 Q6AFJ4 Q6AFJ5 Q6AFJ7 Q6AFK6 Q6AFK7 Q6AFQ2
Q6AFQ4 Q6AFR0 Q6AFX7 Q6AFX9 Q6AFY0 Q6AFY1 Q6AFY2 Q6AFY3 Q6AFZ0 Q6AFZ2 Q6AFZ6
Q6AFZ7 Q6AFZ8 Q6AG00 Q6AG41 Q6AG48 Q6AG49 Q6AG71 Q6AG73 Q6AG81 Q6AG94 Q6AG99
Q6AGC2 Q6AGC6 Q6AGC8 Q6AGC9 Q6AGD1 Q6AGD3 Q6AGD6 Q6AGD7 Q6AGF6 Q6AGF7 Q6AGG5
Q6AGG7 Q6AGN3 Q6AGQ7 Q6AGR0 Q6AGR7 Q6AGS4 Q6AGT5 Q6AGY2 Q6AGY5 Q6AGY7 Q6AH10
Q6AH11 Q6AH25 Q6AH26 Q6AH30 Q6AH32 Q6AH61 Q6AHB4 Q6AHC2 Q6AHE8 Q6AHF1 Q6AHF2
Q6AHF5 Q6AHF6 Q6AHF7 Q6AHH7 Q6AHN3 Q6AHN6 Q6AHZ1 Q6AI12 Q6AIL0 Q6AIL1 Q6AIM2
Q6AIP3 Q6AIT0 Q6AIT8 Q6AIZ3 Q6AIZ4 Q6AIZ6 Q6AJ19 Q6AJ23 Q6AJ42 Q6AJ43 Q6AJ44
Q6AJ46 Q6AJ47 Q6AJ52 Q6AJ54 Q6AJ55 Q6AJ82 Q6AJ90 Q6AJ91 Q6AJA8 Q6AJD2 Q6AJE2
Q6AJE4 Q6AJE5 Q6AJE6 Q6AJE8 Q6AJF1 Q6AJF4 Q6AJG5 Q6AJH7 Q6AJJ9 Q6AJL0 Q6AJL4
Q6AJL6 Q6AJL7 Q6AJL8 Q6AJL9 Q6AJM2 Q6AJM4 Q6AJM5 Q6AJM7 Q6AJN1 Q6AJQ1 Q6AJQ3
Q6AJQ6 Q6AJQ9 Q6AJR0 Q6AJV8 Q6AJW0 Q6AJY2 Q6AJY4 Q6AJY5 Q6AJY7 Q6AJY8 Q6AJY9
Q6AJZ0 Q6AJZ3 Q6AJZ7 Q6AJZ9 Q6AK01 Q6AK02 Q6AK05 Q6AK06 Q6AK07 Q6AK08 Q6AK09
Q6AK10 Q6AK45 Q6AK48 Q6AK58 Q6AK59 Q6AK60 Q6AK61 Q6AK63 Q6AK68 Q6AK95 Q6AK99
Q6AKB8 Q6AKK4 Q6AKP0 Q6AKP3 Q6AKP5 Q6AKW8 Q6AL12 Q6AL19 Q6AL28 Q6AL30 Q6AL31
Q6AL48 Q6AL57 Q6AL71 Q6AL73 Q6AL80 Q6AL81 Q6ALS1 Q6ALS3 Q6ALS7 Q6ALU4 Q6ALU8
Q6ALV3 Q6ALV4 Q6ALW1 Q6ALW3 Q6ALW6 Q6ALW9 Q6ALZ3 Q6ALZ5 Q6AM21 Q6AMA0 Q6AMD2
Q6AMF7 Q6AMG3 Q6AMI5 Q6AMK5 Q6AML3 Q6AMN0 Q6AMQ3 Q6AMQ5 Q6AMR4 Q6AMR6 Q6AMS4
Q6AMS5 Q6AMS8 Q6AMT5 Q6AN63 Q6AN65 Q6ANB6 Q6ANB9 Q6ANC0 Q6ANC1 Q6ANC2 Q6ANH1
Q6ANH2 Q6ANH6 Q6ANL3 Q6ANL4 Q6ANL7 Q6ANL8 Q6ANL9 Q6ANM0 Q6ANM1 Q6ANP1 Q6ANQ6
Q6ANR1 Q6ANR2 Q6ANR3 Q6ANR5 Q6ANS2 Q6ANV0 Q6ANV2 Q6ANW2 Q6AP31 Q6AP32 Q6AP35
Q6AP38 Q6AP39 Q6AP40 Q6AP41 Q6AP42 Q6AP43 Q6AP44 Q6AP45 Q6AP46 Q6AP47 Q6AP48
Q6AP51 Q6AP52 Q6AP53 Q6AP54 Q6AP58 Q6AP59 Q6AP60 Q6AP61 Q6AP62 Q6AP64 Q6AP65
Q6AP66 Q6AP67 Q6AP68 Q6AP69 Q6AP70 Q6AP71 Q6AP72 Q6AP73 Q6AP74 Q6AP75 Q6AP76
Q6AP77 Q6AP78 Q6AP79 Q6AP80 Q6AP81 Q6AP82 Q6AP86 Q6AP93 Q6AP94 Q6APB0 Q6API9
Q6APT2 Q6APU2 Q6APV7 Q6APY7 Q6APZ0 Q6APZ2 Q6APZ9 Q6AQ04 Q6AQ16 Q6AQ27 Q6AQ32
Q6AQ43 Q6AQ48 Q6AQ78 Q6AQ96 Q6AQ97 Q6AQ98 Q6AQA5 Q6AQD0 Q6AQD7 Q6AQE3 Q6AQF1

Q6AQH2 Q6AQH6 Q6AQI3 Q6AQK0 Q6AQK1 Q6AQK2 Q6AQQ3 Q6AQS3 Q6AQS4 Q6AQT4 Q6AQT5
Q6AQW7 Q6AQX3 Q6AQZ6 Q6AR56 Q6AR58 Q6AR59 Q6AR60 Q6AR63 Q6AR64 Q6AR67 Q6ARD7
Q6ARJ7 Q6ARJ9 Q6ARK0 Q6ARL2 Q6ARL8 Q6ARM0 Q6ARM1 Q6ARN9 Q6ARP3 Q6ARP5 Q6ARS6
Q6ARV5 Q6ARV6 Q6ARX8 Q6ARX9 Q6AS16 Q6AS26 Q6AS28 Q6AS31 Q6AS40 Q6AS41 Q6AS42
Q6AS45 Q6AS54 Q6ASC7 Q6ASD8 Q6ASE4 Q6ASN4 Q6ASQ0 Q6ASS2 Q6ASW7 Q6AT10 Q6AT12
Q6AT25 Q6AT33 Q6AT41 Q6AT90 Q6ATB2 Q6ATB4 Q6ATG6 Q6ATJ8 Q6ATR0 Q6ATS0 Q6AU07
Q6AU80 Q6AUF3 Q6AUG0 Q6AUK5 Q6AUK6 Q6AUQ4 Q6AUQ7 Q6AUR2 Q6AUV1 Q6AUW3 Q6AV21
Q6AV22 Q6AV34 Q6AVA8 Q6AVG6 Q6AVI0 Q6AVI1 Q6AVK1 Q6AVN2 Q6AVT2 Q6AVZ9 Q6AW06
Q6AW21 Q6AW23 Q6AW86 Q6AWC8 Q6AWG9 Q6AWU6 Q6AWV1 Q6AWW4 Q6AX33 Q6AX50 Q6AX58
Q6AX59 Q6AX81 Q6AXA6 Q6AXB1 Q6AXB2 Q6AXC6 Q6AXJ2 Q6AXJ3 Q6AXJ7 Q6AXJ9 Q6AXK4
Q6AXL5 Q6AXL8 Q6AXM7 Q6AXM9 Q6AXN0 Q6AXN9 Q6AXP1 Q6AXP3 Q6AXP4 Q6AXQ0 Q6AXQ2
Q6AXQ5 Q6AXQ7 Q6AXQ8 Q6AXQ9 Q6AXS8 Q6AXS9 Q6AXT1 Q6AXT4 Q6AXT7 Q6AXT8 Q6AXU1
Q6AXU3 Q6AXU4 Q6AXU6 Q6AXU8 Q6AXU9 Q6AXV2 Q6AXV6 Q6AXV9 Q6AXW1 Q6AXW4 Q6AXX0
Q6AXX1 Q6AXX2 Q6AXX3 Q6AXX4 Q6AXX9 Q6AXY0 Q6AXY2 Q6AXY4 Q6AXY7 Q6AXY9 Q6AXZ0
Q6AXZ2 Q6AXZ3 Q6AXZ4 Q6AY08 Q6AY09 Q6AY13 Q6AY14 Q6AY16 Q6AY22 Q6AY23 Q6AY26
Q6AY27 Q6AY31 Q6AY34 Q6AY37 Q6AY43 Q6AY45 Q6AY46 Q6AY52 Q6AY53 Q6AY55 Q6AY59
Q6AY63 Q6AY70 Q6AY72 Q6AY75 Q6AY79 Q6AY80 Q6AY87 Q6AY91 Q6AY96 Q6AYA1 Q6AYA8
Q6AYB3 Q6AYC8 Q6AYD1 Q6AYD2 Q6AYD6 Q6AYE3 Q6AYF5 Q6AYG3 Q6AYG7 Q6AYH0 Q6AYH2
Q6AYH4 Q6AYH9 Q6AYI2 Q6AYI4 Q6AYJ1 Q6AYJ2 Q6AYJ3 Q6AYJ4 Q6AYJ5 Q6AYJ8 Q6AYK1
Q6AYK4 Q6AYK5 Q6AYK6 Q6AYK8 Q6AYK9 Q6AYL4 Q6AYL5 Q6AYL8 Q6AYM0 Q6AYM1 Q6AYM2
Q6AYM4 Q6AYN3 Q6AYN4 Q6AYN9 Q6AYP0 Q6AYP3 Q6AYP4 Q6AYP6 Q6AYP7 Q6AYQ0 Q6AYQ3
Q6AYQ6 Q6AYQ8 Q6AYR6 Q6AYR8 Q6AYS7 Q6AYT0 Q6AYT4 Q6AYT5 Q6AYU0 Q6AYU1 Q6AYU3
Q6AZ54 Q6AZB3 Q6AZB8 Q6AZD3 Q6AZF8 Q6AZG6 Q6AZJ9 Q6AZL2 Q6AZN0 Q6AZN4 Q6AZR2
Q6AZS2 Q6AZT4 Q6AZT7 Q6AZV7 Q6AZZ1 Q6B0B8 Q6B0I6 Q6B0K8 Q6B0K9 Q6B0R7 Q6B0V7
Q6B0W2 Q6B0X1 Q6B0Y7 Q6B0Y8 Q6B108 Q6B113 Q6B2U8 Q6B345 Q6B3N0 Q6B458 Q6B4J5
Q6B4U7 Q6B4U9 Q6B4Z3 Q6B4Z5 Q6B6N4 Q6B856 Q6B857 Q6B858 Q6B8I0 Q6B8L1 Q6B8L2
Q6B8L5 Q6B8L8 Q6B8N4 Q6B8N5 Q6B8P0 Q6B8P2 Q6B8Q1 Q6B8R4 Q6B8R5 Q6B8R6 Q6B8R7
Q6B8R8 Q6B8R9 Q6B8S0 Q6B8S1 Q6B8S2 Q6B8T1 Q6B8T4 Q6B8T5 Q6B8U1 Q6B8U8 Q6B8V2
Q6B8V3 Q6B8V4 Q6B8V5 Q6B8V6 Q6B8V7 Q6B8V8 Q6B8V9 Q6B8W0 Q6B8W1 Q6B8W2 Q6B8W3
Q6B8W4 Q6B8W5 Q6B8W6 Q6B8W7 Q6B8W8 Q6B8W9 Q6B8X1 Q6B8X2 Q6B8X3 Q6B8X4 Q6B8X5
Q6B8X6 Q6B8X7 Q6B8X8 Q6B8X9 Q6B8Y0 Q6B8Y1 Q6B8Z0 Q6B8Z3 Q6B8Z8 Q6B905 Q6B906
Q6B907 Q6B908 Q6B909 Q6B910 Q6B911 Q6B913 Q6B914 Q6B922 Q6B924 Q6B931 Q6B932
Q6B934 Q6B936 Q6B940 Q6B949 Q6B950 Q6B951 Q6B952 Q6B966 Q6B9Y8 Q6BBI8 Q6BDG1
Q6BDG8 Q6BDH0 Q6BDH1 Q6BDH3 Q6BDH5 Q6BDI8 Q6BDJ4 Q6BDR8 Q6BDS1 Q6BDZ9 Q6BEB4
Q6BEH3 Q6BER5 Q6BER6 Q6BET5 Q6BEV5 Q6BF16 Q6BF17 Q6BFF6 Q6BGR7 Q6BGR9 Q6BGU0
Q6BGU8 Q6BGV5 Q6BGV9 Q6BGW1 Q6BGW3 Q6BGW8 Q6BGY8 Q6BH05 Q6BH13 Q6BH22 Q6BH34
Q6BH66 Q6BH91 Q6BHA5 Q6BHA8 Q6BHK3 Q6BHL8 Q6BHS5 Q6BHU0 Q6BHV8 Q6BHX4 Q6BHZ2
Q6BI19 Q6BI20 Q6BI21 Q6BI59 Q6BI60 Q6BI69 Q6BI82 Q6BI90 Q6BI95 Q6BIB1 Q6BIC4
Q6BIE4 Q6BIF0 Q6BIF5 Q6BIH2 Q6BIH5 Q6BIH7 Q6BII5 Q6BII9 Q6BIJ0 Q6BIJ5 Q6BIK6
Q6BIN3 Q6BIP2 Q6BIP4 Q6BIR7 Q6BIR9 Q6BIS1 Q6BIT7 Q6BIV1 Q6BIX5 Q6BIX6 Q6BIY6
Q6BJ25 Q6BJ48 Q6BJ51 Q6BJ73 Q6BJ75 Q6BJ82 Q6BJ87 Q6BJD4 Q6BJD8 Q6BJE2 Q6BJF4
Q6BJF7 Q6BJG4 Q6BJH5 Q6BJM5 Q6BJW8 Q6BJX6 Q6BJY4 Q6BJY5 Q6BK07 Q6BK34 Q6BK42
Q6BK66 Q6BKA3 Q6BKA6 Q6BKB1 Q6BKB5 Q6BKC2 Q6BKD4 Q6BKH3 Q6BKH9 Q6BKI1 Q6BKI2
Q6BKI8 Q6BKK7 Q6BKL0 Q6BKL7 Q6BKM0 Q6BKN1 Q6BKN2 Q6BKP5 Q6BKT4 Q6BKU3 Q6BKV2

Q6BKW1 Q6BKW7 Q6BKW8 Q6BKX6 Q6BKY9 Q6BKZ4 Q6BL01 Q6BL27 Q6BL34 Q6BL58 Q6BL68
Q6BL86 Q6BLA0 Q6BLA1 Q6BLC3 Q6BLC4 Q6BLF4 Q6BLG4 Q6BLH9 Q6BLJ9 Q6BLK0 Q6BLM3
Q6BLM5 Q6BLN9 Q6BLQ3 Q6BLS3 Q6BLS5 Q6BLT2 Q6BLT3 Q6BLT5 Q6BLU6 Q6BLU8 Q6BLU9
Q6BLY5 Q6BLZ2 Q6BLZ8 Q6BM04 Q6BM23 Q6BM25 Q6BM45 Q6BM53 Q6BM74 Q6BM98 Q6BMB3
Q6BMB8 Q6BMD1 Q6BME5 Q6BMF6 Q6BMH4 Q6BMI2 Q6BMK0 Q6BMK7 Q6BML1 Q6BML9 Q6BMM5
Q6BMN7 Q6BMP5 Q6BMQ3 Q6BMT2 Q6BMU4 Q6BMU6 Q6BMW3 Q6BMZ5 Q6BN05 Q6BN12 Q6BN19
Q6BN80 Q6BNC2 Q6BNC5 Q6BND2 Q6BNE1 Q6BNF3 Q6BNG2 Q6BNI6 Q6BNL6 Q6BNN6 Q6BNP4
Q6BNQ8 Q6BNS9 Q6BNU4 Q6BNU7 Q6BNW4 Q6BNX0 Q6BNX8 Q6BNY5 Q6BP03 Q6BP09 Q6BP10
Q6BP45 Q6BP48 Q6BP70 Q6BP94 Q6BP96 Q6BPA9 Q6BPC1 Q6BPD3 Q6BPF6 Q6BPG7 Q6BPI1
Q6BPI5 Q6BPI8 Q6BPK1 Q6BPL7 Q6BPN4 Q6BPT6 Q6BPT8 Q6BPU5 Q6BPW7 Q6BPZ9 Q6BQ02
Q6BQ18 Q6BQ20 Q6BQ23 Q6BQ34 Q6BQ49 Q6BQ61 Q6BQ63 Q6BQ83 Q6BQD2 Q6BQF2 Q6BQG5
Q6BQK3 Q6BQP3 Q6BQR8 Q6BQV7 Q6BQZ1 Q6BR77 Q6BR90 Q6BRB3 Q6BRB4 Q6BRC6 Q6BRF0
Q6BRG2 Q6BRG3 Q6BRG6 Q6BRH3 Q6BRL3 Q6BRN4 Q6BRN7 Q6BRP6 Q6BRU4 Q6BRV5 Q6BRY1
Q6BRY2 Q6BRY4 Q6BRZ5 Q6BS00 Q6BS60 Q6BS64 Q6BS75 Q6BSB8 Q6BSD9 Q6BSE2 Q6BSE4
Q6BSE7 Q6BSI7 Q6BSL1 Q6BSL7 Q6BSM3 Q6BSP4 Q6BSP7 Q6BSQ1 Q6BSS5 Q6BST1 Q6BSU5
Q6BSV3 Q6BSY1 Q6BSY5 Q6BSZ4 Q6BSZ8 Q6BT10 Q6BT11 Q6BT27 Q6BT30 Q6BT38 Q6BT42
Q6BT51 Q6BT58 Q6BT60 Q6BTA4 Q6BTB1 Q6BTC0 Q6BTC8 Q6BTL5 Q6BTS9 Q6BTT1 Q6BTW5
Q6BTZ6 Q6BU18 Q6BU36 Q6BU46 Q6BU51 Q6BU54 Q6BU68 Q6BU76 Q6BU84 Q6BU94 Q6BU95
Q6BU97 Q6BU98 Q6BUA6 Q6BUE3 Q6BUG5 Q6BUH8 Q6BUI3 Q6BUI4 Q6BUJ2 Q6BUM6 Q6BUN9
Q6BUP7 Q6BUP9 Q6BUQ1 Q6BUQ2 Q6BUR3 Q6BUT3 Q6BUV9 Q6BV06 Q6BV14 Q6BV58 Q6BV76
Q6BV91 Q6BV94 Q6BVA0 Q6BVB8 Q6BVE0 Q6BVE1 Q6BVF9 Q6BVH4 Q6BVJ8 Q6BVY4 Q6BVZ3
Q6BVZ7 Q6BW07 Q6BW21 Q6BW39 Q6BW41 Q6BW57 Q6BW83 Q6BWA5 Q6BWB8 Q6BWB9 Q6BWC0
Q6BWC2 Q6BWF6 Q6BWG3 Q6BWH6 Q6BWJ5 Q6BWK1 Q6BWK6 Q6BWL0 Q6BWQ9 Q6BWR0 Q6BWS4
Q6BWS8 Q6BWW4 Q6BWW6 Q6BWX4 Q6BWZ7 Q6BX14 Q6BX32 Q6BX45 Q6BX71 Q6BX78 Q6BXB6
Q6BXC3 Q6BXC7 Q6BXE5 Q6BXF6 Q6BXG0 Q6BXG1 Q6BXG5 Q6BXG6 Q6BXH8 Q6BXJ4 Q6BXL7
Q6BXL9 Q6BXM5 Q6BXN0 Q6BXN1 Q6BXN9 Q6BXQ8 Q6BXX1 Q6BXX5 Q6BXX6 Q6BXZ7 Q6BY20
Q6BY27 Q6BY37 Q6BY69 Q6BY86 Q6BYC4 Q6BYD9 Q6BYE5 Q6BYF0 Q6BYF8 Q6BYI4 Q6BYJ1
Q6BYK1 Q6BYP2 Q6BYP7 Q6BYP9 Q6BYU0 Q6BYU4 Q6BYZ4 Q6BZ28 Q6BZ43 Q6BZ44 Q6BZ77
Q6BZ78 Q6BZD9 Q6BZF6 Q6BZF8 Q6BZF9 Q6BZG0 Q6BZG3 Q6BZG7 Q6BZH5 Q6BZP5 Q6BZQ9
Q6BZR4 Q6BZT5 Q6BZW8 Q6BZX4 Q6C007 Q6C022 Q6C024 Q6C030 Q6C036 Q6C061 Q6C070
Q6C093 Q6C097 Q6C0B6 Q6C0C4 Q6C0E6 Q6C0G3 Q6C0I0 Q6C0K1 Q6C0K9 Q6C0M4 Q6C0M9
Q6C0P9 Q6C0Q2 Q6C0S8 Q6C0U2 Q6C0U8 Q6C0W7 Q6C0X2 Q6C0Z6 Q6C100 Q6C101 Q6C116
Q6C141 Q6C161 Q6C169 Q6C186 Q6C187 Q6C193 Q6C195 Q6C1B1 Q6C1B5 Q6C1E0 Q6C1F3
Q6C1H8 Q6C1I3 Q6C1L3 Q6C1L4 Q6C1N1 Q6C1T4 Q6C1V5 Q6C1V6 Q6C1W9 Q6C1X5 Q6C202
Q6C229 Q6C233 Q6C239 Q6C265 Q6C2A3 Q6C2E2 Q6C2F4 Q6C2H4 Q6C2Q7 Q6C2R8 Q6C2R9
Q6C2S3 Q6C2T8 Q6C2T9 Q6C2U0 Q6C2Y6 Q6C2Z7 Q6C347 Q6C3E0 Q6C3J2 Q6C3J3 Q6C3L4
Q6C3P1 Q6C3P4 Q6C3R3 Q6C3S1 Q6C3T0 Q6C3V4 Q6C3X6 Q6C3Z9 Q6C417 Q6C423 Q6C435
Q6C443 Q6C452 Q6C462 Q6C482 Q6C4A3 Q6C4C9 Q6C4D4 Q6C4F8 Q6C4G1 Q6C4H1 Q6C4I6
Q6C4I7 Q6C4I9 Q6C4J0 Q6C4K3 Q6C4R1 Q6C4R3 Q6C4U6 Q6C4U7 Q6C4X9 Q6C4Y5 Q6C519
Q6C531 Q6C553 Q6C5D0 Q6C5G3 Q6C5G5 Q6C5H4 Q6C5K2 Q6C5K4 Q6C5P0 Q6C5P9 Q6C5R2
Q6C5U8 Q6C5V7 Q6C603 Q6C626 Q6C627 Q6C656 Q6C662 Q6C668 Q6C673 Q6C6C6 Q6C6D6
Q6C6D9 Q6C6I6 Q6C6M5 Q6C6Q7 Q6C6T1 Q6C6V8 Q6C6X6 Q6C6Y4 Q6C703 Q6C705 Q6C709
Q6C710 Q6C712 Q6C715 Q6C725 Q6C740 Q6C747 Q6C764 Q6C783 Q6C784 Q6C791 Q6C793
Q6C799 Q6C7A6 Q6C7B9 Q6C7C3 Q6C7C9 Q6C7D0 Q6C7D7 Q6C7G0 Q6C7G2 Q6C7G8 Q6C7H6

Q6C7K5 Q6C7K8 Q6C7L4 Q6C7Q4 Q6C7S6 Q6C7U1 Q6C7U7 Q6C7V4 Q6C7X8 Q6C804 Q6C818
Q6C830 Q6C835 Q6C842 Q6C846 Q6C855 Q6C872 Q6C882 Q6C8A3 Q6C8C5 Q6C8D1 Q6C8F5
Q6C8H2 Q6C8L8 Q6C8R5 Q6C8T3 Q6C8V1 Q6C8Y7 Q6C922 Q6C931 Q6C953 Q6C961 Q6C982
Q6C997 Q6C9C4 Q6C9E7 Q6C9G0 Q6C9G2 Q6C9H2 Q6C9I6 Q6C9I7 Q6C9M6 Q6C9M9 Q6C9N0
Q6C9N2 Q6C9P3 Q6C9Q1 Q6C9Q9 Q6C9T0 Q6C9W0 Q6C9Y4 Q6C9Y6 Q6C9Z2 Q6CA08 Q6CA26
Q6CA53 Q6CA67 Q6CA79 Q6CA87 Q6CAB5 Q6CAC7 Q6CAC9 Q6CAD1 Q6CAD2 Q6CAE9 Q6CAH5
Q6CAR6 Q6CAU9 Q6CAX1 Q6CAX3 Q6CAX5 Q6CAY6 Q6CB23 Q6CB30 Q6CB51 Q6CB52 Q6CB69
Q6CBA2 Q6CBB0 Q6CBB5 Q6CBC3 Q6CBI0 Q6CBI8 Q6CBK1 Q6CBN5 Q6CBS7 Q6CBW0 Q6CBX2
Q6CBZ0 Q6CC11 Q6CC39 Q6CC70 Q6CC77 Q6CC79 Q6CC82 Q6CC84 Q6CC99 Q6CCB0 Q6CCH0
Q6CCK2 Q6CCT1 Q6CCU3 Q6CCU7 Q6CCV4 Q6CCW0 Q6CCY2 Q6CCZ1 Q6CCZ5 Q6CD24 Q6CDG5
Q6CDH3 Q6CDK0 Q6CDK1 Q6CDL0 Q6CDL9 Q6CDM0 Q6CDN5 Q6CDR5 Q6CDS6 Q6CDV4 Q6CDX0
Q6CE12 Q6CE38 Q6CE60 Q6CE69 Q6CE80 Q6CE96 Q6CEB8 Q6CEC9 Q6CEE1 Q6CEE9 Q6CEF3
Q6CEJ6 Q6CEK8 Q6CER3 Q6CER9 Q6CES3 Q6CEU8 Q6CEV5 Q6CEW7 Q6CEW9 Q6CEY7 Q6CF20
Q6CF24 Q6CF35 Q6CF40 Q6CF41 Q6CF49 Q6CF50 Q6CF69 Q6CF78 Q6CFD2 Q6CFE7 Q6CFG6
Q6CFJ2 Q6CFK6 Q6CFQ0 Q6CFR3 Q6CFS7 Q6CFT1 Q6CFU2 Q6CFV3 Q6CFW4 Q6CFX1 Q6CFX5
Q6CG02 Q6CG37 Q6CG46 Q6CG48 Q6CG56 Q6CG57 Q6CGB2 Q6CGD1 Q6CGD3 Q6CGE1 Q6CGE6
Q6CGE7 Q6CGJ5 Q6CGK4 Q6CGM0 Q6CGP9 Q6CGR3 Q6CGS4 Q6CGV4 Q6CH03 Q6CH58 Q6CH63
Q6CH90 Q6CHA2 Q6CHE4 Q6CHE5 Q6CHE9 Q6CHI1 Q6CHJ7 Q6CHJ8 Q6CHK0 Q6CHN5 Q6CHP9
Q6CHS6 Q6CI08 Q6CI62 Q6CIB4 Q6CIC9 Q6CID6 Q6CIF0 Q6CIG2 Q6CIH2 Q6CIH3 Q6CII8
Q6CII9 Q6CIJ2 Q6CIL8 Q6CIN0 Q6CIN8 Q6CIP4 Q6CIQ1 Q6CIR0 Q6CIS0 Q6CIT4 Q6CIT6
Q6CIT9 Q6CIU0 Q6CIV2 Q6CIV8 Q6CIZ1 Q6CJ06 Q6CJ08 Q6CJ30 Q6CJ38 Q6CJ45 Q6CJ48
Q6CJ49 Q6CJ57 Q6CJ60 Q6CJ61 Q6CJ62 Q6CJ65 Q6CJ86 Q6CJA3 Q6CJB5 Q6CJC4 Q6CJD7
Q6CJE6 Q6CJF4 Q6CJG5 Q6CJI0 Q6CJI3 Q6CJK2 Q6CJM4 Q6CJQ3 Q6CJR7 Q6CJS0 Q6CJS8
Q6CJT2 Q6CJU4 Q6CJV1 Q6CJY0 Q6CJZ2 Q6CJZ6 Q6CK06 Q6CK11 Q6CK29 Q6CK32 Q6CK33
Q6CK38 Q6CK52 Q6CK57 Q6CK59 Q6CK60 Q6CK62 Q6CK68 Q6CK85 Q6CKB6 Q6CKE8 Q6CKE9
Q6CKF0 Q6CKF3 Q6CKH1 Q6CKH5 Q6CKH7 Q6CKI0 Q6CKI1 Q6CKI5 Q6CKL8 Q6CKR8 Q6CKS3
Q6CKU1 Q6CKU6 Q6CKV6 Q6CKX0 Q6CKX3 Q6CKY3 Q6CKZ4 Q6CKZ5 Q6CL13 Q6CL21 Q6CL23
Q6CL78 Q6CL82 Q6CL84 Q6CL99 Q6CLA4 Q6CLC9 Q6CLD3 Q6CLF6 Q6CLH3 Q6CLH4 Q6CLJ6
Q6CLJ7 Q6CLM5 Q6CLP9 Q6CLQ5 Q6CLR2 Q6CLR3 Q6CLR6 Q6CLS0 Q6CLS2 Q6CLS8 Q6CLY8
Q6CLZ6 Q6CM00 Q6CM04 Q6CM12 Q6CM22 Q6CM42 Q6CM46 Q6CM69 Q6CM87 Q6CM95 Q6CM99
Q6CMA2 Q6CMA8 Q6CMB5 Q6CMB6 Q6CMB8 Q6CMC0 Q6CMC2 Q6CMD3 Q6CME1 Q6CMF5 Q6CMG4
Q6CMJ8 Q6CMK4 Q6CMK8 Q6CMK9 Q6CMM2 Q6CMN0 Q6CMN5 Q6CMT3 Q6CMV8 Q6CMZ9 Q6CN12
Q6CN28 Q6CN40 Q6CN45 Q6CN53 Q6CN57 Q6CN59 Q6CN69 Q6CN92 Q6CN95 Q6CN96 Q6CNA0
Q6CNC2 Q6CNC8 Q6CND0 Q6CND9 Q6CNF3 Q6CNF8 Q6CNG7 Q6CNG8 Q6CNI5 Q6CNI6 Q6CNJ1
Q6CNM6 Q6CNM7 Q6CNQ2 Q6CNQ3 Q6CNQ9 Q6CNR2 Q6CNS3 Q6CNT6 Q6CNU6 Q6CNW3 Q6CNY4
Q6CNY5 Q6CP05 Q6CP16 Q6CP18 Q6CP20 Q6CP23 Q6CP44 Q6CP50 Q6CP69 Q6CP71 Q6CP76
Q6CP80 Q6CP83 Q6CPA5 Q6CPA6 Q6CPB1 Q6CPC2 Q6CPC3 Q6CPD5 Q6CPE2 Q6CPF9 Q6CPG3
Q6CPH8 Q6CPI4 Q6CPK5 Q6CPK7 Q6CPN1 Q6CPQ8 Q6CPQ9 Q6CPR4 Q6CPS7 Q6CPU0 Q6CPV6
Q6CPW2 Q6CPW4 Q6CQ02 Q6CQ06 Q6CQ07 Q6CQ22 Q6CQ49 Q6CQ59 Q6CQ61 Q6CQ65 Q6CQ76
Q6CQ86 Q6CQ91 Q6CQ94 Q6CQA1 Q6CQA6 Q6CQA9 Q6CQB7 Q6CQC4 Q6CQC6 Q6CQE5 Q6CQH2
Q6CQI2 Q6CQI9 Q6CQJ9 Q6CQL7 Q6CQM2 Q6CQN1 Q6CQN6 Q6CQN7 Q6CQP3 Q6CQQ0 Q6CQQ1
Q6CQQ8 Q6CQR6 Q6CQU2 Q6CQU4 Q6CQV4 Q6CQV5 Q6CQW3 Q6CQX2 Q6CQX4 Q6CQX5 Q6CQY2
Q6CQZ5 Q6CR42 Q6CR51 Q6CR57 Q6CR69 Q6CR76 Q6CR96 Q6CR99 Q6CRA2 Q6CRA9 Q6CRE9
Q6CRF4 Q6CRH2 Q6CRI9 Q6CRJ7 Q6CRJ8 Q6CRK4 Q6CRM3 Q6CRN7 Q6CRQ9 Q6CRS3 Q6CRT5

Q6CRT7 Q6CRT8 Q6CRV9 Q6CRY0 Q6CRY2 Q6CRY5 Q6CRY6 Q6CS47 Q6CS52 Q6CS90 Q6CSB9
Q6CSD1 Q6CSF8 Q6CSG8 Q6CSH0 Q6CSI1 Q6CSN1 Q6CSN3 Q6CSP8 Q6CSP9 Q6CSR1 Q6CSR7
Q6CSR8 Q6CSS3 Q6CSS6 Q6CST6 Q6CSV3 Q6CSW1 Q6CSW8 Q6CSY9 Q6CSZ0 Q6CSZ7 Q6CT00
Q6CT09 Q6CT22 Q6CT29 Q6CT34 Q6CT43 Q6CT46 Q6CT49 Q6CT50 Q6CT57 Q6CT60 Q6CT61
Q6CT72 Q6CT73 Q6CT76 Q6CT85 Q6CT90 Q6CTA7 Q6CTB9 Q6CTC3 Q6CTD5 Q6CTE4 Q6CTF5
Q6CTH3 Q6CTH5 Q6CTH8 Q6CTI2 Q6CTL5 Q6CTM7 Q6CTN7 Q6CTQ9 Q6CTR7 Q6CTS4 Q6CTS8
Q6CTT2 Q6CTT6 Q6CTT7 Q6CTU5 Q6CTU8 Q6CTY9 Q6CTZ8 Q6CU12 Q6CU41 Q6CU48 Q6CU52
Q6CU55 Q6CU59 Q6CU61 Q6CU63 Q6CU65 Q6CU66 Q6CU71 Q6CU86 Q6CUA3 Q6CUA5 Q6CUB4
Q6CUC6 Q6CUD2 Q6CUD4 Q6CUD9 Q6CUE1 Q6CUE5 Q6CUE9 Q6CUI0 Q6CUI5 Q6CUI6 Q6CUM6
Q6CUQ5 Q6CUQ9 Q6CUR6 Q6CUS7 Q6CUS9 Q6CUU2 Q6CUV0 Q6CUV9 Q6CUY0 Q6CUZ3 Q6CV12
Q6CV81 Q6CV82 Q6CV83 Q6CV86 Q6CVA2 Q6CVE5 Q6CVE9 Q6CVF1 Q6CVF3 Q6CVG0 Q6CVG8
Q6CVH3 Q6CVK1 Q6CVK3 Q6CVL1 Q6CVN2 Q6CVN3 Q6CVQ9 Q6CVR3 Q6CVS2 Q6CVS3 Q6CVT0
Q6CVU5 Q6CVU7 Q6CVX9 Q6CVZ8 Q6CW21 Q6CW22 Q6CW24 Q6CW35 Q6CW43 Q6CW48 Q6CW60
Q6CW75 Q6CWB2 Q6CWC1 Q6CWC4 Q6CWD0 Q6CWD7 Q6CWE8 Q6CWH3 Q6CWH8 Q6CWI4 Q6CWJ7
Q6CWK2 Q6CWM0 Q6CWM4 Q6CWM9 Q6CWN7 Q6CWQ4 Q6CWQ5 Q6CWQ8 Q6CWR0 Q6CWU8 Q6CWV0
Q6CWV1 Q6CWW0 Q6CWW6 Q6CWW9 Q6CWX1 Q6CWX4 Q6CWX6 Q6CWX7 Q6CWY0 Q6CX23 Q6CX30
Q6CX31 Q6CX42 Q6CX53 Q6CX73 Q6CX91 Q6CX98 Q6CXA6 Q6CXB9 Q6CXH7 Q6CXH8 Q6CXI8
Q6CXI9 Q6CXL5 Q6CXN0 Q6CXN5 Q6CXP5 Q6CXR1 Q6CXR4 Q6CXS0 Q6CXT4 Q6CXT6 Q6CXV0
Q6CXW0 Q6CXW3 Q6CXW4 Q6CXX3 Q6CXY0 Q6CY29 Q6CY34 Q6CY47 Q6CY84 Q6CYA2 Q6CYE2
Q6CYH1 Q6CYI6 Q6CYI7 Q6CYJ9 Q6CYM1 Q6CYQ9 Q6CYR1 Q6CYR2 Q6CYR4 Q6CYR6 Q6CYT9
Q6CYU4 Q6CYV2 Q6CYV3 Q6CZ00 Q6CZ23 Q6CZ24 Q6CZ25 Q6CZ26 Q6CZ46 Q6CZ80 Q6CZ81
Q6CZ88 Q6CZ89 Q6CZ91 Q6CZ92 Q6CZ96 Q6CZA0 Q6CZB1 Q6CZB4 Q6CZB6 Q6CZC7 Q6CZD1
Q6CZD8 Q6CZD9 Q6CZG3 Q6CZG6 Q6CZG8 Q6CZK0 Q6CZK1 Q6CZK2 Q6CZK3 Q6CZL2 Q6CZL7
Q6CZL9 Q6CZP5 Q6CZP6 Q6CZQ8 Q6CZQ9 Q6CZR3 Q6CZS0 Q6CZU0 Q6CZV1 Q6CZV5 Q6CZW0
Q6CZW1 Q6CZW2 Q6CZW3 Q6CZW4 Q6CZW5 Q6CZW6 Q6CZW9 Q6CZX0 Q6CZX1 Q6CZX2 Q6CZX3
Q6CZX4 Q6CZX5 Q6CZX6 Q6CZX7 Q6CZX8 Q6CZX9 Q6CZY0 Q6CZY1 Q6CZY2 Q6CZY3 Q6CZY4
Q6CZY5 Q6CZY6 Q6CZY7 Q6CZY8 Q6CZY9 Q6CZZ1 Q6CZZ2 Q6CZZ3 Q6CZZ4 Q6CZZ5 Q6CZZ6
Q6D000 Q6D001 Q6D002 Q6D004 Q6D006 Q6D009 Q6D012 Q6D022 Q6D025 Q6D033 Q6D036
Q6D037 Q6D065 Q6D066 Q6D067 Q6D072 Q6D074 Q6D075 Q6D076 Q6D080 Q6D081 Q6D086
Q6D087 Q6D092 Q6D093 Q6D0A0 Q6D0A1 Q6D0A9 Q6D0B1 Q6D0B2 Q6D0B7 Q6D0B8 Q6D0C1
Q6D0C3 Q6D0C6 Q6D0C7 Q6D0C8 Q6D0D8 Q6D0D9 Q6D0E0 Q6D0E6 Q6D0F8 Q6D0G5 Q6D0G6
Q6D0G7 Q6D0G8 Q6D0H4 Q6D0H5 Q6D0H8 Q6D0I1 Q6D0I4 Q6D0I5 Q6D0J4 Q6D0J5 Q6D0J6
Q6D0M1 Q6D0N7 Q6D0N8 Q6D0W5 Q6D0W6 Q6D0W7 Q6D0W8 Q6D0W9 Q6D0X0 Q6D117 Q6D118
Q6D125 Q6D127 Q6D135 Q6D136 Q6D137 Q6D138 Q6D142 Q6D151 Q6D160 Q6D164 Q6D165
Q6D178 Q6D181 Q6D1A1 Q6D1A2 Q6D1A3 Q6D1A4 Q6D1A5 Q6D1B3 Q6D1B4 Q6D1B5 Q6D1B6
Q6D1B9 Q6D1G1 Q6D1G2 Q6D1G3 Q6D1H0 Q6D1H3 Q6D1H8 Q6D1I1 Q6D1I2 Q6D1I3 Q6D1I4
Q6D1S8 Q6D1S9 Q6D1T0 Q6D1T1 Q6D1T4 Q6D1T8 Q6D1T9 Q6D1U0 Q6D1U1 Q6D1V5 Q6D1V7
Q6D1W1 Q6D1W3 Q6D1W4 Q6D1X4 Q6D1X5 Q6D1X6 Q6D1X9 Q6D1Y1 Q6D1Z0 Q6D1Z1 Q6D1Z4
Q6D206 Q6D208 Q6D209 Q6D211 Q6D219 Q6D221 Q6D234 Q6D238 Q6D245 Q6D246 Q6D257
Q6D258 Q6D259 Q6D261 Q6D262 Q6D263 Q6D266 Q6D272 Q6D273 Q6D276 Q6D277 Q6D280
Q6D286 Q6D288 Q6D299 Q6D2E4 Q6D2E6 Q6D2L3 Q6D2M4 Q6D2M6 Q6D2N1 Q6D2N5 Q6D2N7
Q6D2Q6 Q6D2Q8 Q6D2R1 Q6D2U2 Q6D2V8 Q6D2Y8 Q6D363 Q6D3B3 Q6D3B4 Q6D3B9 Q6D3C0
Q6D3C1 Q6D3C4 Q6D3C7 Q6D3F5 Q6D3H7 Q6D3J6 Q6D3J7 Q6D3J9 Q6D3L2 Q6D3L8 Q6D3N3
Q6D3N6 Q6D3R0 Q6D3R5 Q6D3S2 Q6D3S8 Q6D3T2 Q6D3T7 Q6D3U1 Q6D3U2 Q6D3V0 Q6D400

Q6D401 Q6D402 Q6D404 Q6D406 Q6D407 Q6D408 Q6D409 Q6D410 Q6D411 Q6D412 Q6D421
Q6D425 Q6D434 Q6D439 Q6D445 Q6D446 Q6D447 Q6D453 Q6D454 Q6D459 Q6D465 Q6D471
Q6D474 Q6D476 Q6D482 Q6D483 Q6D497 Q6D499 Q6D4A0 Q6D4A1 Q6D4A2 Q6D4B6 Q6D4C8
Q6D4E3 Q6D4E9 Q6D4G8 Q6D4G9 Q6D4H0 Q6D4H1 Q6D4H2 Q6D4H3 Q6D4H4 Q6D4H8 Q6D4L1
Q6D4L3 Q6D4M0 Q6D4M2 Q6D4M3 Q6D4M5 Q6D4N9 Q6D4P1 Q6D4P7 Q6D4Q8 Q6D4R3 Q6D4S1
Q6D4T9 Q6D4U0 Q6D4U2 Q6D4W5 Q6D4W6 Q6D4X4 Q6D4Z5 Q6D551 Q6D552 Q6D554 Q6D557
Q6D5N6 Q6D5P6 Q6D5S7 Q6D5T3 Q6D5T7 Q6D5T8 Q6D5U5 Q6D5V1 Q6D5V2 Q6D5V3 Q6D5V5
Q6D5V7 Q6D5W0 Q6D5W3 Q6D5X9 Q6D624 Q6D625 Q6D636 Q6D646 Q6D662 Q6D673 Q6D674
Q6D683 Q6D692 Q6D694 Q6D698 Q6D6A4 Q6D6C3 Q6D6C4 Q6D6C5 Q6D6C8 Q6D6D3 Q6D6D9
Q6D6E0 Q6D6E1 Q6D6F8 Q6D6J5 Q6D6J8 Q6D6K0 Q6D6Y6 Q6D6Y7 Q6D736 Q6D796 Q6D797
Q6D7B3 Q6D7B4 Q6D7B6 Q6D7C2 Q6D7C4 Q6D7E3 Q6D7E7 Q6D7G2 Q6D7H0 Q6D7H1 Q6D7J7
Q6D7J9 Q6D7K6 Q6D7K8 Q6D7L6 Q6D7L9 Q6D7M1 Q6D7M6 Q6D7M7 Q6D7M8 Q6D7N4 Q6D7Q9
Q6D7R1 Q6D7R8 Q6D7S0 Q6D7S1 Q6D7U0 Q6D7W1 Q6D7W3 Q6D7W4 Q6D7X5 Q6D7Z4 Q6D7Z5
Q6D7Z6 Q6D7Z7 Q6D7Z8 Q6D800 Q6D807 Q6D808 Q6D820 Q6D826 Q6D827 Q6D828 Q6D838
Q6D841 Q6D842 Q6D844 Q6D847 Q6D848 Q6D850 Q6D855 Q6D856 Q6D871 Q6D872 Q6D882
Q6D8C1 Q6D8C5 Q6D8C9 Q6D8D9 Q6D8E0 Q6D8E1 Q6D8E2 Q6D8E3 Q6D8E5 Q6D8E6 Q6D8E7
Q6D8F4 Q6D8F6 Q6D8F7 Q6D8H5 Q6D8I6 Q6D8I9 Q6D8J0 Q6D8J7 Q6D8J9 Q6D8S0 Q6D8S1
Q6D8S3 Q6D8U7 Q6D8U8 Q6D8U9 Q6D8V3 Q6D8V4 Q6D8V5 Q6D8V6 Q6D8V9 Q6D8W0 Q6D8X4
Q6D8X9 Q6D8Y0 Q6D8Y5 Q6D945 Q6D946 Q6D958 Q6D960 Q6D961 Q6D974 Q6D975 Q6D976
Q6D984 Q6D989 Q6D990 Q6D991 Q6D994 Q6D9A1 Q6D9A2 Q6D9A3 Q6D9A4 Q6D9A5 Q6D9A7
Q6D9B6 Q6D9B9 Q6D9C3 Q6D9C5 Q6D9C6 Q6D9D1 Q6D9D2 Q6D9D3 Q6D9D4 Q6D9D7 Q6D9G1
Q6D9G9 Q6D9H0 Q6D9I4 Q6D9I9 Q6D9J0 Q6D9J1 Q6D9J5 Q6D9W6 Q6D9Z5 Q6DA21 Q6DA22
Q6DA23 Q6DA24 Q6DA25 Q6DA26 Q6DA52 Q6DA54 Q6DA62 Q6DA71 Q6DA73 Q6DA74 Q6DAC6
Q6DAD4 Q6DAE6 Q6DAE7 Q6DAF2 Q6DAG8 Q6DAH2 Q6DAI3 Q6DAJ1 Q6DAJ5 Q6DAJ8 Q6DAL2
Q6DAL7 Q6DAL8 Q6DAL9 Q6DAM0 Q6DAM1 Q6DAM4 Q6DAM9 Q6DAN0 Q6DAN1 Q6DAN2 Q6DAN3
Q6DAN4 Q6DAN8 Q6DAP0 Q6DAP4 Q6DAQ1 Q6DAQ7 Q6DAQ9 Q6DAR0 Q6DAR1 Q6DAR2 Q6DAR4
Q6DAR6 Q6DAS2 Q6DAT1 Q6DAT5 Q6DAT7 Q6DAV2 Q6DAV4 Q6DAV5 Q6DAV6 Q6DAV7 Q6DAV9
Q6DAW0 Q6DAW2 Q6DAZ8 Q6DB05 Q6DB09 Q6DB14 Q6DB17 Q6DB18 Q6DB47 Q6DB58 Q6DB60
Q6DB61 Q6DB68 Q6DB75 Q6DB76 Q6DB88 Q6DB93 Q6DB94 Q6DB95 Q6DBA5 Q6DBN1 Q6DBN5
Q6DBN6 Q6DBP8 Q6DBQ1 Q6DBR0 Q6DBR4 Q6DBT3 Q6DBU1 Q6DBW0 Q6DBX1 Q6DBY2 Q6DBY5
Q6DC02 Q6DC03 Q6DC17 Q6DC37 Q6DC53 Q6DC60 Q6DC61 Q6DC64 Q6DCB7 Q6DCC6 Q6DCD0
Q6DCD1 Q6DCD7 Q6DCF2 Q6DCP3 Q6DCP5 Q6DCQ0 Q6DCQ5 Q6DCS1 Q6DCU7 Q6DCV0 Q6DCW1
Q6DCX2 Q6DCX3 Q6DCY9 Q6DCZ9 Q6DD06 Q6DD09 Q6DD17 Q6DD19 Q6DD21 Q6DD24 Q6DD27
Q6DD30 Q6DD33 Q6DD39 Q6DD45 Q6DD51 Q6DDA3 Q6DDA4 Q6DDB9 Q6DDF0 Q6DDH0 Q6DDH2
Q6DDH6 Q6DDI6 Q6DDJ3 Q6DDJ5 Q6DDK1 Q6DDQ5 Q6DDR8 Q6DDT5 Q6DDT6 Q6DDU5 Q6DDU8
Q6DDU9 Q6DDV1 Q6DDV8 Q6DDW4 Q6DDW7 Q6DDX3 Q6DDX7 Q6DDX8 Q6DDY6 Q6DDY9 Q6DE00
Q6DE02 Q6DE05 Q6DE08 Q6DE14 Q6DE18 Q6DE73 Q6DE84 Q6DE87 Q6DE94 Q6DE96 Q6DE97
Q6DEB1 Q6DEB4 Q6DEE4 Q6DEE7 Q6DEF4 Q6DEF6 Q6DEH3 Q6DEI8 Q6DEJ5 Q6DEL7 Q6DEN0
Q6DER1 Q6DER6 Q6DES0 Q6DET9 Q6DEU9 Q6DEV3 Q6DEV7 Q6DEW4 Q6DEW6 Q6DEY7 Q6DEY8
Q6DEZ7 Q6DF11 Q6DF30 Q6DF57 Q6DF62 Q6DF67 Q6DF78 Q6DF94 Q6DFA8 Q6DFB7 Q6DFB8
Q6DFC2 Q6DFC6 Q6DFF4 Q6DFF6 Q6DFF9 Q6DFH5 Q6DFH9 Q6DFJ6 Q6DFJ9 Q6DFL0 Q6DFL2
Q6DFM1 Q6DFN5 Q6DFN7 Q6DFP9 Q6DFQ2 Q6DFS2 Q6DFT3 Q6DFU2 Q6DFV5 Q6DFV7 Q6DFW4
Q6DFZ6 Q6DG03 Q6DG43 Q6DG50 Q6DG52 Q6DG60 Q6DG88 Q6DG91 Q6DG99 Q6DGA6 Q6DGB6
Q6DGD3 Q6DGE9 Q6DGF6 Q6DGG1 Q6DGH9 Q6DGI3 Q6DGJ1 Q6DGK2 Q6DGK9 Q6DGL6 Q6DGL8

Q6DGM1 Q6DGM3 Q6DGM8 Q6DGN0 Q6DGN6 Q6DGP2 Q6DGP7 Q6DGQ0 Q6DGQ4 Q6DGR4 Q6DGS3
Q6DGT7 Q6DGV1 Q6DGX3 Q6DGY7 Q6DGZ0 Q6DGZ3 Q6DH02 Q6DH13 Q6DH26 Q6DH42 Q6DH44
Q6DH65 Q6DH74 Q6DH86 Q6DHB1 Q6DHF7 Q6DHF9 Q6DHG0 Q6DHH7 Q6DHI2 Q6DHI5 Q6DHJ1
Q6DHJ6 Q6DHL5 Q6DHL7 Q6DHP6 Q6DHP9 Q6DHU4 Q6DHU8 Q6DHV7 Q6DI35 Q6DI37 Q6DI40
Q6DI51 Q6DI86 Q6DI92 Q6DIA2 Q6DIA7 Q6DIA9 Q6DIB4 Q6DID3 Q6DID5 Q6DIE2 Q6DIF3
Q6DIH3 Q6DIH5 Q6DII2 Q6DIJ5 Q6DIL6 Q6DIM3 Q6DIN3 Q6DIN8 Q6DIP1 Q6DIP3 Q6DIP5
Q6DIQ3 Q6DIR5 Q6DIS2 Q6DIS8 Q6DIT8 Q6DIV2 Q6DIV4 Q6DIW6 Q6DIX6 Q6DIY3 Q6DIY4
Q6DIY9 Q6DIZ0 Q6DJ03 Q6DJ05 Q6DJ06 Q6DJ13 Q6DJ25 Q6DJ48 Q6DJ55 Q6DJ66 Q6DJ71
Q6DJ78 Q6DJ95 Q6DJA3 Q6DJB3 Q6DJB9 Q6DJC2 Q6DJD3 Q6DJE4 Q6DJE5 Q6DJF8 Q6DJG6
Q6DJH2 Q6DJH5 Q6DJI4 Q6DJI5 Q6DJI8 Q6DJI9 Q6DJK9 Q6DJL7 Q6DJM6 Q6DJN3 Q6DJQ6
Q6DJQ9 Q6DJR9 Q6DJS0 Q6DK85 Q6DK91 Q6DK93 Q6DK98 Q6DK99 Q6DKA1 Q6DKB0 Q6DKE0
Q6DKE6 Q6DKE9 Q6DKG0 Q6DKI1 Q6DKI2 Q6DKJ4 Q6DKN2 Q6DKP5 Q6DL03 Q6DLH9 Q6DLR9
Q6DLV9 Q6DMN8 Q6DN03 Q6DN56 Q6DN58 Q6DN64 Q6DN65 Q6DN66 Q6DN67 Q6DQL1 Q6DQW3
Q6DR13 Q6DR20 Q6DR24 Q6DRA6 Q6DRB1 Q6DRC3 Q6DRC4 Q6DRC5 Q6DRD3 Q6DRF9 Q6DRG1
Q6DRH5 Q6DRI1 Q6DRI4 Q6DRI7 Q6DRJ4 Q6DRL4 Q6DRL5 Q6DRL8 Q6DRP2 Q6DRP4 Q6DSS2
Q6DT37 Q6DT83 Q6DT84 Q6DTN4 Q6DTY7 Q6DUB7 Q6DUX3 Q6DUZ9 Q6DV01 Q6DV14 Q6DYE4
Q6E0U3 Q6E0W6 Q6E0W9 Q6E0X0 Q6E0X1 Q6E244 Q6E2N3 Q6E3C7 Q6E3C9 Q6E3D0 Q6E3D2
Q6E3D4 Q6E3D5 Q6E3F0 Q6E3F2 Q6E3F3 Q6E434 Q6E4Q0 Q6E4Q3 Q6E4Q8 Q6E593 Q6E6B4
Q6E6J3 Q6E6S7 Q6E7D7 Q6E7H0 Q6EAS5 Q6ED65 Q6EE22 Q6EE31 Q6EEF3 Q6EEV4 Q6EEV6
Q6EGX7 Q6EI05 Q6EI06 Q6EI12 Q6EIC1 Q6EIG9 Q6EIH3 Q6EIJ2 Q6EIJ3 Q6EIJ4 Q6EIJ6
Q6EIJ9 Q6EIK1 Q6EIK6 Q6EIZ1 Q6EJ97 Q6EJ98 Q6EJB6 Q6EJB7 Q6EKJ0 Q6EMI9 Q6EN80
Q6EN83 Q6EN90 Q6EN94 Q6ENA9 Q6ENB2 Q6ENB7 Q6ENC0 Q6ENC1 Q6ENC8 Q6ENC9 Q6END4
Q6END5 Q6END6 Q6END7 Q6END8 Q6END9 Q6ENE0 Q6ENE1 Q6ENE2 Q6ENE9 Q6ENF0 Q6ENF1
Q6ENF2 Q6ENG6 Q6ENH2 Q6ENH6 Q6ENI1 Q6ENI2 Q6ENI3 Q6ENI4 Q6ENI8 Q6ENJ5 Q6ENJ6
Q6ENP9 Q6ENQ1 Q6ENQ5 Q6ENQ6 Q6ENQ9 Q6ENR2 Q6ENR3 Q6ENR9 Q6ENS0 Q6ENS1 Q6ENS3
Q6ENS4 Q6ENS5 Q6ENS6 Q6ENS7 Q6ENS8 Q6ENS9 Q6ENT0 Q6ENT1 Q6ENT2 Q6ENT9 Q6ENU0
Q6ENU1 Q6ENU2 Q6ENU3 Q6ENV5 Q6ENW1 Q6ENW5 Q6ENX0 Q6ENX1 Q6ENX2 Q6ENX3 Q6ENX7
Q6ENX8 Q6ENY4 Q6EP49 Q6EP77 Q6EPG8 Q6EPN6 Q6EPW7 Q6EPZ0 Q6EQG2 Q6EQX3 Q6ER21
Q6ER22 Q6ER94 Q6ERW5 Q6ERW7 Q6ERW9 Q6ERX1 Q6ES10 Q6ES52 Q6ESI7 Q6ESZ9 Q6ET36
Q6ET49 Q6ETK3 Q6ETL2 Q6ETN3 Q6ETX0 Q6EU10 Q6EU14 Q6EU30 Q6EU39 Q6EUF1 Q6EUH7
Q6EUK7 Q6EUP4 Q6EV04 Q6EV23 Q6EV97 Q6EVK8 Q6EVY5 Q6EVY6 Q6EVY7 Q6EVY8 Q6EVZ0
Q6EVZ3 Q6EW02 Q6EW11 Q6EW12 Q6EW13 Q6EW14 Q6EW15 Q6EW16 Q6EW17 Q6EW18 Q6EW19
Q6EW20 Q6EW27 Q6EW28 Q6EW30 Q6EW31 Q6EW32 Q6EW46 Q6EW50 Q6EW56 Q6EW57 Q6EW58
Q6EW59 Q6EW66 Q6EW67 Q6EW71 Q6EWG8 Q6EWQ6 Q6EWX1 Q6EZQ4 Q6F047 Q6F050 Q6F054
Q6F059 Q6F0D4 Q6F0D5 Q6F0E1 Q6F0E4 Q6F0E6 Q6F0F5 Q6F0G6 Q6F0G9 Q6F0H8 Q6F0H9
Q6F0I2 Q6F0I3 Q6F0I4 Q6F0J2 Q6F0J3 Q6F0J4 Q6F0J5 Q6F0K3 Q6F0K7 Q6F0K8 Q6F0L3
Q6F0L4 Q6F0L7 Q6F0L8 Q6F0L9 Q6F0M8 Q6F0M9 Q6F0Q4 Q6F0Q5 Q6F0Q9 Q6F0R0 Q6F0R1
Q6F0R9 Q6F0S3 Q6F0S4 Q6F0S5 Q6F0S6 Q6F0T3 Q6F0T6 Q6F0U3 Q6F0U4 Q6F0W3 Q6F0X2
Q6F0X3 Q6F0X5 Q6F0Y1 Q6F0Y5 Q6F0Y7 Q6F0Y9 Q6F101 Q6F123 Q6F125 Q6F145 Q6F147
Q6F148 Q6F149 Q6F150 Q6F152 Q6F155 Q6F157 Q6F159 Q6F165 Q6F166 Q6F168 Q6F169
Q6F170 Q6F175 Q6F184 Q6F196 Q6F199 Q6F1A0 Q6F1A9 Q6F1B1 Q6F1C4 Q6F1D3 Q6F1D5
Q6F1H1 Q6F1H5 Q6F1I2 Q6F1I4 Q6F1J0 Q6F1J8 Q6F1J9 Q6F1K1 Q6F1L2 Q6F1M4 Q6F1M6
Q6F1N5 Q6F1N9 Q6F1P8 Q6F1Q6 Q6F1R0 Q6F1R7 Q6F1R8 Q6F1S1 Q6F1S6 Q6F1S7 Q6F1S8
Q6F1U6 Q6F1V0 Q6F1V4 Q6F1V7 Q6F1W2 Q6F1W6 Q6F1W7 Q6F1W8 Q6F1W9 Q6F1X0 Q6F1X1

Q6F1X6 Q6F1X7 Q6F1X8 Q6F1X9 Q6F1Y0 Q6F1Y1 Q6F1Y2 Q6F1Y3 Q6F1Y4 Q6F1Y5 Q6F1Y7
Q6F1Y9 Q6F1Z0 Q6F1Z1 Q6F1Z2 Q6F1Z3 Q6F1Z4 Q6F1Z5 Q6F1Z6 Q6F210 Q6F211 Q6F215
Q6F216 Q6F228 Q6F230 Q6F234 Q6F235 Q6F237 Q6F240 Q6F243 Q6F265 Q6F271 Q6F273
Q6F2A0 Q6F2A3 Q6F2A9 Q6F2B4 Q6F2B5 Q6F2E2 Q6F2E3 Q6F2E4 Q6F2F0 Q6F2N0 Q6F2U9
Q6F2Y7 Q6F332 Q6F334 Q6F353 Q6F362 Q6F368 Q6F3F0 Q6F3F1 Q6F3F3 Q6F442 Q6F444
Q6F450 Q6F473 Q6F477 Q6F482 Q6F4M7 Q6F4M8 Q6F4M9 Q6F4N1 Q6F598 Q6F5E1 Q6F5E2
Q6F5E6 Q6F5E7 Q6F6A2 Q6F6B3 Q6F6K7 Q6F6K8 Q6F6L1 Q6F6N3 Q6F6N4 Q6F6P9 Q6F6Q0
Q6F6Q6 Q6F6R1 Q6F6R2 Q6F6U0 Q6F6U8 Q6F6V2 Q6F6V3 Q6F6W0 Q6F6W1 Q6F6Y1 Q6F6Y2
Q6F6Z6 Q6F704 Q6F705 Q6F717 Q6F718 Q6F799 Q6F7A5 Q6F7A7 Q6F7A8 Q6F7C8 Q6F7D1
Q6F7E4 Q6F7E5 Q6F7G9 Q6F7H0 Q6F7H9 Q6F7I0 Q6F7I2 Q6F7I5 Q6F7L8 Q6F7N0 Q6F7N4
Q6F7N5 Q6F7Q4 Q6F7R1 Q6F7R2 Q6F7R3 Q6F7R4 Q6F7R5 Q6F7R6 Q6F7R7 Q6F7R8 Q6F7R9
Q6F7S0 Q6F7S1 Q6F7S2 Q6F7S3 Q6F7S4 Q6F7S5 Q6F7S6 Q6F7S7 Q6F7S8 Q6F7S9 Q6F7T0
Q6F7T1 Q6F7T3 Q6F7T4 Q6F7T5 Q6F7T6 Q6F7T7 Q6F7T8 Q6F7U1 Q6F7W2 Q6F7W6 Q6F7W8
Q6F7X1 Q6F7Y7 Q6F7Y9 Q6F7Z8 Q6F801 Q6F805 Q6F817 Q6F821 Q6F823 Q6F847 Q6F854
Q6F855 Q6F856 Q6F860 Q6F861 Q6F867 Q6F868 Q6F872 Q6F873 Q6F874 Q6F880 Q6F898
Q6F8A0 Q6F8F0 Q6F8F4 Q6F8G1 Q6F8G2 Q6F8G5 Q6F8G6 Q6F8H3 Q6F8H8 Q6F8I3 Q6F8I6
Q6F8I7 Q6F8I8 Q6F8J0 Q6F8J3 Q6F8J9 Q6F8K0 Q6F8L4 Q6F8M7 Q6F8N1 Q6F8N7 Q6F8P2
Q6F8P5 Q6F8P6 Q6F8Q3 Q6F943 Q6F950 Q6F961 Q6F963 Q6F973 Q6F982 Q6F991 Q6F992
Q6F9A3 Q6F9A5 Q6F9B5 Q6F9C2 Q6F9D2 Q6F9D8 Q6F9D9 Q6F9E6 Q6F9F4 Q6F9F5 Q6F9F6
Q6F9F7 Q6F9G0 Q6F9H7 Q6F9H8 Q6F9I9 Q6F9J0 Q6F9J1 Q6F9J2 Q6F9J3 Q6F9M1 Q6F9N7
Q6F9P9 Q6F9Q1 Q6F9Q8 Q6F9Q9 Q6F9R0 Q6F9R8 Q6F9S2 Q6F9U4 Q6F9U5 Q6F9W1 Q6F9W7
Q6F9Y2 Q6F9Y6 Q6F9Z3 Q6FA18 Q6FA20 Q6FA38 Q6FA43 Q6FA53 Q6FA54 Q6FA61 Q6FA66
Q6FA79 Q6FA87 Q6FA99 Q6FAA5 Q6FAA9 Q6FAK7 Q6FAL8 Q6FAM5 Q6FAM6 Q6FAP5 Q6FAP9
Q6FAQ2 Q6FAQ6 Q6FAR2 Q6FAS9 Q6FAT7 Q6FAU1 Q6FAU4 Q6FAX7 Q6FAZ1 Q6FB05 Q6FB08
Q6FB19 Q6FB57 Q6FBI2 Q6FBI3 Q6FBS0 Q6FBW1 Q6FC54 Q6FC71 Q6FCE5 Q6FCE6 Q6FCF4
Q6FCF6 Q6FCG1 Q6FCG9 Q6FCH2 Q6FCH3 Q6FCH4 Q6FCH8 Q6FCI2 Q6FCI4 Q6FCI6 Q6FCK9
Q6FCL0 Q6FCN1 Q6FCN6 Q6FCP8 Q6FCP9 Q6FCQ0 Q6FCR5 Q6FCR7 Q6FCR9 Q6FCS7 Q6FCS8
Q6FCT2 Q6FCT4 Q6FCT7 Q6FCU0 Q6FCV6 Q6FCW2 Q6FCX7 Q6FCY1 Q6FD29 Q6FD45 Q6FD71
Q6FD79 Q6FD81 Q6FD83 Q6FD84 Q6FD85 Q6FD86 Q6FDA0 Q6FDB8 Q6FDC1 Q6FDF8 Q6FDF9
Q6FDG0 Q6FDH0 Q6FDN0 Q6FDN3 Q6FDQ8 Q6FDR0 Q6FDR5 Q6FDR7 Q6FDS1 Q6FDS2 Q6FDS6
Q6FDS7 Q6FDS8 Q6FDU0 Q6FDV0 Q6FDW0 Q6FDW7 Q6FDX8 Q6FDY3 Q6FDY4 Q6FE14 Q6FE23
Q6FE29 Q6FE36 Q6FE58 Q6FE77 Q6FE90 Q6FE96 Q6FEB3 Q6FEC7 Q6FEC8 Q6FEC9 Q6FED0
Q6FED8 Q6FEE6 Q6FEF1 Q6FEH6 Q6FEJ3 Q6FEJ4 Q6FEM2 Q6FEM3 Q6FEM6 Q6FEM7 Q6FEN5
Q6FEP2 Q6FEP7 Q6FEP8 Q6FEP9 Q6FEQ1 Q6FEQ3 Q6FEQ6 Q6FER7 Q6FES9 Q6FET0 Q6FEV2
Q6FEV4 Q6FEV6 Q6FEV8 Q6FEW0 Q6FEW7 Q6FEZ1 Q6FEZ6 Q6FEZ7 Q6FF12 Q6FF13 Q6FF39
Q6FF40 Q6FF42 Q6FF44 Q6FF45 Q6FF48 Q6FF50 Q6FF51 Q6FF54 Q6FF55 Q6FF64 Q6FF82
Q6FF89 Q6FF90 Q6FF91 Q6FF92 Q6FF93 Q6FF94 Q6FF97 Q6FFA9 Q6FFB2 Q6FFB3 Q6FFB6
Q6FFC0 Q6FFC2 Q6FFD5 Q6FFD8 Q6FFE5 Q6FFG4 Q6FFH9 Q6FFL8 Q6FFM0 Q6FFN2 Q6FFQ0
Q6FFQ1 Q6FFQ2 Q6FFR5 Q6FFX5 Q6FFY1 Q6FFZ4 Q6FFZ5 Q6FFZ6 Q6FFZ7 Q6FFZ9 Q6FG02
Q6FG10 Q6FG11 Q6FG12 Q6FG19 Q6FG21 Q6FIF0 Q6FIJ6 Q6FIK5 Q6FIN2 Q6FIN4 Q6FIP2
Q6FIP7 Q6FIQ6 Q6FIS9 Q6FIT7 Q6FIT9 Q6FIU0 Q6FIU7 Q6FIV4 Q6FIX7 Q6FIZ7 Q6FJ00
Q6FJ04 Q6FJ14 Q6FJ19 Q6FJ22 Q6FJ28 Q6FJ36 Q6FJ46 Q6FJ48 Q6FJ54 Q6FJ55 Q6FJ73
Q6FJ85 Q6FJ97 Q6FJA3 Q6FJA5 Q6FJB0 Q6FJC4 Q6FJC7 Q6FJE8 Q6FJG1 Q6FJH3 Q6FJI3
Q6FJI9 Q6FJJ1 Q6FJJ8 Q6FJK6 Q6FJL9 Q6FJM0 Q6FJN0 Q6FJP1 Q6FJQ5 Q6FJQ6 Q6FJQ9

Q6FJR2 Q6FJS0 Q6FJV0 Q6FJV4 Q6FJV8 Q6FJW4 Q6FJX4 Q6FJX6 Q6FJY3 Q6FJZ9 Q6FK00
Q6FK07 Q6FK11 Q6FK21 Q6FK47 Q6FK48 Q6FK59 Q6FK61 Q6FK63 Q6FK71 Q6FK87 Q6FK91
Q6FKA3 Q6FKA5 Q6FKB1 Q6FKB6 Q6FKC2 Q6FKC6 Q6FKD6 Q6FKE0 Q6FKE7 Q6FKG4 Q6FKH1
Q6FKH3 Q6FKH7 Q6FKI2 Q6FKJ9 Q6FKK0 Q6FKK3 Q6FKM8 Q6FKN4 Q6FKN6 Q6FKP2 Q6FKQ4
Q6FKQ6 Q6FKS1 Q6FKS5 Q6FKS8 Q6FKT4 Q6FKY1 Q6FKY3 Q6FKZ2 Q6FL15 Q6FL17 Q6FL25
Q6FL35 Q6FL44 Q6FL63 Q6FL71 Q6FL77 Q6FL80 Q6FL81 Q6FL84 Q6FLA4 Q6FLA8 Q6FLB1
Q6FLB2 Q6FLB8 Q6FLC3 Q6FLD4 Q6FLD5 Q6FLE5 Q6FLF3 Q6FLF8 Q6FLG1 Q6FLI1 Q6FLI2
Q6FLK5 Q6FLL9 Q6FLP2 Q6FLQ1 Q6FLQ5 Q6FLQ6 Q6FLR5 Q6FLT7 Q6FLU4 Q6FLZ0 Q6FM01
Q6FM02 Q6FM05 Q6FM20 Q6FM28 Q6FM30 Q6FM31 Q6FM32 Q6FM43 Q6FM49 Q6FM51 Q6FM81
Q6FM84 Q6FM86 Q6FM92 Q6FM94 Q6FM98 Q6FMA3 Q6FMA5 Q6FMB1 Q6FMB3 Q6FMB5 Q6FMC0
Q6FMC8 Q6FMD8 Q6FME9 Q6FMF8 Q6FMG7 Q6FMH9 Q6FMI2 Q6FMI5 Q6FMJ3 Q6FML0 Q6FML4
Q6FML5 Q6FML7 Q6FML9 Q6FMN0 Q6FMP0 Q6FMP4 Q6FMR7 Q6FMT2 Q6FMT6 Q6FMU7 Q6FMU9
Q6FMV8 Q6FMW9 Q6FMX1 Q6FN12 Q6FN16 Q6FN18 Q6FN48 Q6FN65 Q6FN68 Q6FN88 Q6FN96
Q6FNA2 Q6FNC2 Q6FNE9 Q6FNF9 Q6FNH1 Q6FNH8 Q6FNI6 Q6FNJ1 Q6FNK3 Q6FNK8 Q6FNL3
Q6FNL6 Q6FNM5 Q6FNM7 Q6FNN7 Q6FNP9 Q6FNR0 Q6FNR1 Q6FNR4 Q6FNR8 Q6FNU6 Q6FNU7
Q6FNU9 Q6FNV1 Q6FNX0 Q6FNX7 Q6FNY7 Q6FNZ4 Q6FP02 Q6FP10 Q6FP17 Q6FP19 Q6FP38
Q6FP41 Q6FP43 Q6FP72 Q6FP74 Q6FP78 Q6FP96 Q6FPD9 Q6FPF3 Q6FPH9 Q6FPI4 Q6FPJ5
Q6FPK1 Q6FPL6 Q6FPN5 Q6FPN7 Q6FPP7 Q6FPQ3 Q6FPQ4 Q6FPT7 Q6FPU0 Q6FPV6 Q6FPW3
Q6FPX3 Q6FPX5 Q6FPX9 Q6FPY0 Q6FPZ6 Q6FQ21 Q6FQ26 Q6FQ30 Q6FQ31 Q6FQ33 Q6FQ36
Q6FQ44 Q6FQ55 Q6FQ75 Q6FQ83 Q6FQ87 Q6FQ93 Q6FQA0 Q6FQA6 Q6FQB2 Q6FQE3 Q6FQG2
Q6FQH3 Q6FQJ4 Q6FQJ5 Q6FQL7 Q6FQQ6 Q6FQT2 Q6FQU2 Q6FQV5 Q6FQX5 Q6FQY1 Q6FQY4
Q6FQZ0 Q6FR01 Q6FR30 Q6FR35 Q6FR37 Q6FR39 Q6FR44 Q6FR48 Q6FR54 Q6FR56 Q6FR65
Q6FR76 Q6FR78 Q6FRB7 Q6FRC6 Q6FRE7 Q6FRF5 Q6FRG0 Q6FRG6 Q6FRH1 Q6FRI3 Q6FRJ6
Q6FRL4 Q6FRM1 Q6FRM6 Q6FRN1 Q6FRN2 Q6FRP0 Q6FRQ9 Q6FRR0 Q6FRS1 Q6FRS9 Q6FRT2
Q6FRU8 Q6FRV0 Q6FRV2 Q6FRW1 Q6FRX6 Q6FRY2 Q6FRY5 Q6FRZ5 Q6FRZ6 Q6FRZ9 Q6FS03
Q6FS31 Q6FS36 Q6FS48 Q6FS54 Q6FS55 Q6FS63 Q6FS70 Q6FS75 Q6FS76 Q6FS81 Q6FSB0
Q6FSB1 Q6FSD2 Q6FSD4 Q6FSD7 Q6FSF1 Q6FSH0 Q6FSH5 Q6FSJ4 Q6FSK8 Q6FSM4 Q6FSM8
Q6FSN5 Q6FSN6 Q6FSP5 Q6FSQ0 Q6FSQ6 Q6FSS0 Q6FST5 Q6FSU0 Q6FSU1 Q6FSW2 Q6FSZ8
Q6FT36 Q6FT39 Q6FT66 Q6FT85 Q6FTE0 Q6FTG1 Q6FTG6 Q6FTI5 Q6FTJ2 Q6FTJ8 Q6FTK3
Q6FTL0 Q6FTQ8 Q6FTR8 Q6FTT0 Q6FTT8 Q6FTU2 Q6FTV1 Q6FTV3 Q6FTW0 Q6FTW6 Q6FTX0
Q6FTZ3 Q6FTZ5 Q6FU05 Q6FU23 Q6FU41 Q6FU44 Q6FU45 Q6FU57 Q6FU69 Q6FU75 Q6FU78
Q6FU81 Q6FU86 Q6FU96 Q6FUA5 Q6FUA6 Q6FUC3 Q6FUD0 Q6FUD8 Q6FUD9 Q6FUF3 Q6FUF9
Q6FUG1 Q6FUH3 Q6FUI5 Q6FUI7 Q6FUJ0 Q6FUJ7 Q6FUM3 Q6FUM5 Q6FUP6 Q6FUQ5 Q6FUQ6
Q6FUS6 Q6FUV0 Q6FUV2 Q6FUV3 Q6FUX0 Q6FUZ2 Q6FV04 Q6FV05 Q6FV07 Q6FV12 Q6FV34
Q6FV36 Q6FV37 Q6FV72 Q6FV93 Q6FV94 Q6FVC6 Q6FVC7 Q6FVD3 Q6FVD8 Q6FVG5 Q6FVI8
Q6FVJ1 Q6FVL2 Q6FVL4 Q6FVM9 Q6FVN6 Q6FVQ3 Q6FVQ8 Q6FVR6 Q6FVV1 Q6FVV7 Q6FVX3
Q6FVY2 Q6FVY4 Q6FVY5 Q6FVZ6 Q6FVZ7 Q6FVZ8 Q6FW12 Q6FW13 Q6FW42 Q6FW54 Q6FW56
Q6FW60 Q6FW61 Q6FW62 Q6FW66 Q6FW76 Q6FW77 Q6FW79 Q6FWA7 Q6FWB9 Q6FWE1 Q6FWE3
Q6FWF3 Q6FWF4 Q6FWI1 Q6FWI7 Q6FWJ7 Q6FWL5 Q6FWM1 Q6FWM4 Q6FWM7 Q6FWM8 Q6FWQ2
Q6FWQ3 Q6FWS1 Q6FWS2 Q6FWT4 Q6FWT9 Q6FWU4 Q6FWW0 Q6FWW3 Q6FWX8 Q6FWY5 Q6FX19
Q6FX31 Q6FX42 Q6FX50 Q6FX63 Q6FXA4 Q6FXA5 Q6FXD0 Q6FXE3 Q6FXI2 Q6FXI5 Q6FXI8
Q6FXJ8 Q6FXK9 Q6FXM4 Q6FXN3 Q6FXP4 Q6FXQ4 Q6FXQ8 Q6FXR9 Q6FXS6 Q6FXX1 Q6FXY9
Q6FXZ0 Q6FXZ4 Q6FY22 Q6FY24 Q6FY25 Q6FY30 Q6FY35 Q6FY61 Q6FY67 Q6FY76 Q6FY89
Q6FYA6 Q6FYA7 Q6FYB6 Q6FYB8 Q6FYB9 Q6FYC0 Q6FYC3 Q6FYC7 Q6FYD0 Q6FYD1 Q6FYD4

Q6FYF1 Q6FYF9 Q6FYG4 Q6FYG6 Q6FYH3 Q6FYH4 Q6FYH5 Q6FYL4 Q6FYL6 Q6FYL8 Q6FYM8
Q6FYN4 Q6FYN6 Q6FYN9 Q6FYP4 Q6FYP5 Q6FYP6 Q6FYQ2 Q6FYQ5 Q6FYQ7 Q6FYR2 Q6FYT1
Q6FYT6 Q6FYU2 Q6FYU5 Q6FYU9 Q6FYZ3 Q6FYZ6 Q6FYZ7 Q6FZ00 Q6FZ12 Q6FZ13 Q6FZ15
Q6FZ17 Q6FZ18 Q6FZ21 Q6FZ30 Q6FZ46 Q6FZ57 Q6FZ66 Q6FZ67 Q6FZ74 Q6FZ81 Q6FZ91
Q6FZ96 Q6FZ98 Q6FZB1 Q6FZB7 Q6FZB8 Q6FZB9 Q6FZC0 Q6FZC1 Q6FZC2 Q6FZC3 Q6FZC4
Q6FZC5 Q6FZC6 Q6FZC7 Q6FZC8 Q6FZC9 Q6FZD0 Q6FZD1 Q6FZD2 Q6FZD3 Q6FZD4 Q6FZD5
Q6FZD6 Q6FZD7 Q6FZD8 Q6FZD9 Q6FZE0 Q6FZE1 Q6FZE3 Q6FZE4 Q6FZE5 Q6FZE6 Q6FZE7
Q6FZF0 Q6FZF1 Q6FZG1 Q6FZH0 Q6FZH1 Q6FZH6 Q6FZI4 Q6FZI9 Q6FZJ0 Q6FZJ2 Q6FZK1
Q6FZK6 Q6FZK7 Q6FZL2 Q6FZL5 Q6FZL6 Q6FZL7 Q6FZL8 Q6FZM0 Q6FZM5 Q6FZN0 Q6FZN2
Q6FZN3 Q6FZN9 Q6FZP1 Q6FZP6 Q6FZP9 Q6FZQ5 Q6FZQ6 Q6FZQ9 Q6FZR9 Q6FZS5 Q6FZS7
Q6FZS8 Q6FZS9 Q6FZU1 Q6FZU3 Q6FZU4 Q6FZU8 Q6FZW4 Q6FZX5 Q6FZZ4 Q6FZZ7 Q6FZZ9
Q6G004 Q6G005 Q6G008 Q6G009 Q6G014 Q6G027 Q6G029 Q6G036 Q6G041 Q6G043 Q6G052
Q6G053 Q6G060 Q6G061 Q6G063 Q6G073 Q6G078 Q6G079 Q6G080 Q6G082 Q6G086 Q6G088
Q6G090 Q6G091 Q6G099 Q6G0A2 Q6G0C7 Q6G0C8 Q6G0C9 Q6G0D4 Q6G0D5 Q6G0D7 Q6G0E0
Q6G0E6 Q6G0F8 Q6G0F9 Q6G0G3 Q6G0H5 Q6G0I1 Q6G0I9 Q6G0J4 Q6G0K6 Q6G0L3 Q6G0M7
Q6G0M8 Q6G0N1 Q6G0N3 Q6G0N8 Q6G0N9 Q6G0P2 Q6G0P3 Q6G0P4 Q6G0P6 Q6G0P7 Q6G0P9
Q6G0Q0 Q6G0Q6 Q6G0R0 Q6G0R4 Q6G0S2 Q6G0S6 Q6G0S7 Q6G0S8 Q6G0S9 Q6G0T7 Q6G0T8
Q6G0T9 Q6G0U2 Q6G0V6 Q6G0V7 Q6G0W9 Q6G0X1 Q6G0X3 Q6G0X4 Q6G0X7 Q6G0Y3 Q6G0Y4
Q6G104 Q6G107 Q6G116 Q6G122 Q6G125 Q6G126 Q6G132 Q6G164 Q6G166 Q6G177 Q6G178
Q6G197 Q6G1A4 Q6G1B3 Q6G1B4 Q6G1B5 Q6G1B6 Q6G1C0 Q6G1D6 Q6G1E4 Q6G1E5 Q6G1E6
Q6G1F4 Q6G1F7 Q6G1F8 Q6G1F9 Q6G1G3 Q6G1G6 Q6G1G7 Q6G1G8 Q6G1G9 Q6G1H4 Q6G1H5
Q6G1H9 Q6G1I2 Q6G1I3 Q6G1K8 Q6G1K9 Q6G1L0 Q6G1L3 Q6G1L7 Q6G1M0 Q6G1M1 Q6G1N8
Q6G1N9 Q6G1Q5 Q6G1R3 Q6G1R8 Q6G1R9 Q6G1S0 Q6G1W0 Q6G1W2 Q6G1W4 Q6G1X4 Q6G1X6
Q6G1Y5 Q6G1Z4 Q6G1Z7 Q6G202 Q6G226 Q6G234 Q6G252 Q6G253 Q6G289 Q6G2E7 Q6G2F0
Q6G2F1 Q6G2G3 Q6G2G5 Q6G2G7 Q6G2G8 Q6G2H1 Q6G2I3 Q6G2J9 Q6G2L1 Q6G2L2 Q6G2M7
Q6G2P7 Q6G2Q3 Q6G2Q6 Q6G2Q7 Q6G2Q8 Q6G2R5 Q6G2S8 Q6G2T3 Q6G2T6 Q6G2V3 Q6G2W1
Q6G2W4 Q6G2W5 Q6G2W6 Q6G2W7 Q6G2W8 Q6G2W9 Q6G2X0 Q6G2X1 Q6G2X2 Q6G2X3 Q6G2X4
Q6G2X5 Q6G2X6 Q6G2X7 Q6G2X8 Q6G2X9 Q6G2Y0 Q6G2Y1 Q6G2Y2 Q6G2Y3 Q6G2Y4 Q6G2Y6
Q6G2Y7 Q6G2Y8 Q6G2Z0 Q6G2Z3 Q6G2Z4 Q6G304 Q6G316 Q6G317 Q6G322 Q6G327 Q6G329
Q6G330 Q6G335 Q6G336 Q6G338 Q6G3A4 Q6G3B6 Q6G3D5 Q6G3D7 Q6G3E9 Q6G3F0 Q6G3F1
Q6G3F3 Q6G3F9 Q6G3I3 Q6G3I6 Q6G3I7 Q6G3J8 Q6G3K1 Q6G3K3 Q6G3K8 Q6G3L2 Q6G3L3
Q6G3L8 Q6G3U8 Q6G3V1 Q6G3V6 Q6G3V8 Q6G3X4 Q6G3X6 Q6G3X7 Q6G3X8 Q6G3X9 Q6G3Y7
Q6G3Y8 Q6G3Y9 Q6G3Z2 Q6G3Z3 Q6G3Z8 Q6G401 Q6G410 Q6G419 Q6G424 Q6G426 Q6G438
Q6G439 Q6G446 Q6G447 Q6G449 Q6G456 Q6G457 Q6G459 Q6G463 Q6G465 Q6G467 Q6G468
Q6G477 Q6G480 Q6G4C1 Q6G4C3 Q6G4C4 Q6G4C5 Q6G4D1 Q6G4D2 Q6G4D4 Q6G4D7 Q6G4E3
Q6G4E4 Q6G4F1 Q6G4G5 Q6G4R4 Q6G4S5 Q6G4T2 Q6G4V1 Q6G4V2 Q6G4V6 Q6G4V8 Q6G4W3
Q6G4W4 Q6G4W7 Q6G4W8 Q6G4Y4 Q6G4Y7 Q6G4Z0 Q6G4Z1 Q6G4Z2 Q6G4Z3 Q6G507 Q6G508
Q6G512 Q6G523 Q6G526 Q6G527 Q6G540 Q6G542 Q6G543 Q6G545 Q6G546 Q6G547 Q6G549
Q6G552 Q6G553 Q6G554 Q6G563 Q6G564 Q6G565 Q6G567 Q6G570 Q6G577 Q6G584 Q6G592
Q6G5A8 Q6G5A9 Q6G5B0 Q6G5B1 Q6G5C6 Q6G5C7 Q6G5C8 Q6G5C9 Q6G5D4 Q6G5D6 Q6G5E0
Q6G5E1 Q6G5E2 Q6G5E3 Q6G5E4 Q6G5E5 Q6G5F0 Q6G5F1 Q6G5F5 Q6G5F7 Q6G5F8 Q6G5G0
Q6G5G1 Q6G5G9 Q6G5H3 Q6G5H6 Q6G5I2 Q6G5I3 Q6G5J3 Q6G5J7 Q6G5K3 Q6G5M0 Q6G5M1
Q6G5N5 Q6G5P2 Q6G5P5 Q6G5P7 Q6G5Q9 Q6G5R0 Q6G5R1 Q6G5R9 Q6G5S5 Q6G5S7 Q6G5T1
Q6G5T6 Q6G5T7 Q6G5W2 Q6G5W3 Q6G5W4 Q6G5W5 Q6G5W6 Q6G5Y4 Q6G5Y5 Q6G5Z5 Q6G5Z6

Q6G5Z7 Q6G5Z9 Q6G600 Q6G601 Q6G602 Q6G603 Q6G609 Q6G639 Q6G640 Q6G642 Q6G643
Q6G664 Q6G669 Q6G670 Q6G674 Q6G677 Q6G678 Q6G679 Q6G693 Q6G696 Q6G6A0 Q6G6B0
Q6G6B2 Q6G6B3 Q6G6B6 Q6G6B9 Q6G6C0 Q6G6C6 Q6G6F1 Q6G6F2 Q6G6F9 Q6G6H6 Q6G6H7
Q6G6J1 Q6G6P6 Q6G6P8 Q6G6Q5 Q6G6S9 Q6G6T5 Q6G6U7 Q6G6V5 Q6G6V7 Q6G6X4 Q6G6X6
Q6G6Y5 Q6G6Y7 Q6G6Y9 Q6G6Z0 Q6G711 Q6G716 Q6G727 Q6G728 Q6G729 Q6G730 Q6G731
Q6G732 Q6G733 Q6G734 Q6G742 Q6G747 Q6G748 Q6G749 Q6G751 Q6G753 Q6G754 Q6G755
Q6G757 Q6G767 Q6G770 Q6G771 Q6G772 Q6G773 Q6G774 Q6G775 Q6G776 Q6G777 Q6G778
Q6G779 Q6G780 Q6G781 Q6G782 Q6G783 Q6G784 Q6G785 Q6G786 Q6G787 Q6G788 Q6G789
Q6G790 Q6G792 Q6G793 Q6G794 Q6G795 Q6G796 Q6G797 Q6G798 Q6G7A2 Q6G7A3 Q6G7A4
Q6G7B7 Q6G7B8 Q6G7B9 Q6G7C0 Q6G7C1 Q6G7C2 Q6G7C5 Q6G7C8 Q6G7D2 Q6G7E1 Q6G7E3
Q6G7E9 Q6G7F2 Q6G7F3 Q6G7F8 Q6G7H3 Q6G7H4 Q6G7I0 Q6G7I2 Q6G7I3 Q6G7I5 Q6G7I6
Q6G7I8 Q6G7J0 Q6G7J1 Q6G7J2 Q6G7J5 Q6G7J6 Q6G7J7 Q6G7J8 Q6G7L0 Q6G7L6 Q6G7L7
Q6G7L8 Q6G7L9 Q6G7M7 Q6G7M9 Q6G7N9 Q6G7P0 Q6G7P1 Q6G7P3 Q6G7P4 Q6G7P7 Q6G7P8
Q6G7P9 Q6G7Q0 Q6G7Q1 Q6G7Q2 Q6G7Q4 Q6G7Q8 Q6G7R1 Q6G7S7 Q6G7S8 Q6G7T6 Q6G7V6
Q6G811 Q6G813 Q6G818 Q6G820 Q6G825 Q6G828 Q6G829 Q6G832 Q6G833 Q6G834 Q6G836
Q6G838 Q6G840 Q6G846 Q6G854 Q6G859 Q6G861 Q6G867 Q6G870 Q6G873 Q6G884 Q6G890
Q6G8A2 Q6G8A3 Q6G8D6 Q6G8E2 Q6G8E3 Q6G8F3 Q6G8G1 Q6G8G2 Q6G8G5 Q6G8G9 Q6G8H6
Q6G8H7 Q6G8H9 Q6G8I4 Q6G8I7 Q6G8J1 Q6G8J2 Q6G8J5 Q6G8J9 Q6G8K8 Q6G8L2 Q6G8L4
Q6G8L6 Q6G8L7 Q6G8L8 Q6G8L9 Q6G8M0 Q6G8M4 Q6G8M8 Q6G8M9 Q6G8N2 Q6G8N8 Q6G8N9
Q6G8P0 Q6G8P3 Q6G8P5 Q6G8P6 Q6G8P7 Q6G8Q0 Q6G8Q1 Q6G8Q2 Q6G8Q3 Q6G8Q5 Q6G8Q7
Q6G8Q8 Q6G8R2 Q6G8R5 Q6G8S2 Q6G8S4 Q6G8S5 Q6G8S6 Q6G8S7 Q6G8S8 Q6G8S9 Q6G8T0
Q6G8T4 Q6G8T5 Q6G8T6 Q6G8T8 Q6G8T9 Q6G8U4 Q6G8U8 Q6G8V1 Q6G8V2 Q6G8V3 Q6G8V4
Q6G8V8 Q6G8V9 Q6G8W4 Q6G8W9 Q6G8X2 Q6G8X4 Q6G8Y2 Q6G8Y5 Q6G8Y6 Q6G8Y7 Q6G8Y8
Q6G8Y9 Q6G8Z2 Q6G8Z7 Q6G901 Q6G902 Q6G903 Q6G904 Q6G905 Q6G907 Q6G909 Q6G912
Q6G913 Q6G915 Q6G928 Q6G929 Q6G930 Q6G931 Q6G937 Q6G941 Q6G942 Q6G943 Q6G945
Q6G951 Q6G954 Q6G960 Q6G961 Q6G967 Q6G969 Q6G970 Q6G986 Q6G987 Q6G988 Q6G990
Q6G992 Q6G994 Q6G996 Q6G997 Q6G998 Q6G9A0 Q6G9A5 Q6G9A7 Q6G9A8 Q6G9B3 Q6G9B6
Q6G9B7 Q6G9C3 Q6G9C4 Q6G9D2 Q6G9D3 Q6G9D4 Q6G9D5 Q6G9D7 Q6G9D8 Q6G9E0 Q6G9E8
Q6G9E9 Q6G9F2 Q6G9F5 Q6G9F7 Q6G9F9 Q6G9G3 Q6G9G4 Q6G9G5 Q6G9G6 Q6G9H0 Q6G9I4
Q6G9I5 Q6G9I6 Q6G9I7 Q6G9I8 Q6G9I9 Q6G9J0 Q6G9J4 Q6G9J6 Q6G9J8 Q6G9K2 Q6G9K5
Q6G9K7 Q6G9K9 Q6G9L2 Q6G9L3 Q6G9L6 Q6G9L7 Q6G9M1 Q6G9M2 Q6G9M3 Q6G9M4 Q6G9M8
Q6G9Q4 Q6G9Q8 Q6G9R0 Q6G9R2 Q6G9R3 Q6G9R7 Q6G9R8 Q6G9S1 Q6G9S8 Q6G9T8 Q6G9T9
Q6G9U0 Q6G9U2 Q6G9U3 Q6G9U4 Q6G9U8 Q6G9U9 Q6G9V0 Q6G9V3 Q6G9V4 Q6G9V5 Q6G9V6
Q6G9V7 Q6G9V8 Q6G9V9 Q6G9W0 Q6G9W1 Q6G9W2 Q6G9W3 Q6G9W7 Q6G9W8 Q6G9W9 Q6G9X2
Q6G9X3 Q6G9X4 Q6G9X5 Q6G9Y0 Q6G9Y6 Q6G9Y7 Q6G9Y9 Q6G9Z2 Q6G9Z5 Q6G9Z7 Q6G9Z8
Q6GA03 Q6GA04 Q6GA08 Q6GA09 Q6GA10 Q6GA11 Q6GA12 Q6GA13 Q6GA15 Q6GA19 Q6GA23
Q6GA25 Q6GA29 Q6GA33 Q6GA34 Q6GA35 Q6GA36 Q6GA44 Q6GA45 Q6GA63 Q6GA64 Q6GA68
Q6GA69 Q6GA70 Q6GA74 Q6GA75 Q6GA76 Q6GA79 Q6GA87 Q6GA89 Q6GA90 Q6GA93 Q6GAA2
Q6GAA8 Q6GAB7 Q6GAB9 Q6GAC0 Q6GAC1 Q6GAC3 Q6GAC4 Q6GAC5 Q6GAD9 Q6GAE0 Q6GAE1
Q6GAE2 Q6GAE3 Q6GAE4 Q6GAE5 Q6GAE7 Q6GAE8 Q6GAF0 Q6GAF9 Q6GAG6 Q6GAG7 Q6GAG9
Q6GAQ7 Q6GAQ9 Q6GAR2 Q6GAS0 Q6GAS2 Q6GAS5 Q6GAS8 Q6GAS9 Q6GAT0 Q6GAV1 Q6GAV6
Q6GAV8 Q6GAV9 Q6GAW0 Q6GAW4 Q6GAW5 Q6GAW6 Q6GAW8 Q6GAW9 Q6GAX2 Q6GAY7 Q6GAZ4
Q6GAZ7 Q6GB01 Q6GB05 Q6GB09 Q6GB11 Q6GB15 Q6GB23 Q6GB27 Q6GB28 Q6GB49 Q6GB55
Q6GB56 Q6GB57 Q6GB58 Q6GB61 Q6GB62 Q6GB63 Q6GB65 Q6GB66 Q6GB68 Q6GB70 Q6GB71

Q6GB72 Q6GB76 Q6GB78 Q6GB87 Q6GB92 Q6GBA0 Q6GBA2 Q6GBA5 Q6GBA6 Q6GBB8 Q6GBE4
Q6GBE9 Q6GBG1 Q6GBG2 Q6GBJ0 Q6GBL2 Q6GBM1 Q6GBM4 Q6GBP3 Q6GBP8 Q6GBP9 Q6GBQ6
Q6GBQ8 Q6GBR0 Q6GBR1 Q6GBR7 Q6GBR8 Q6GBS1 Q6GBS7 Q6GBT3 Q6GBT4 Q6GBT5 Q6GBT6
Q6GBT7 Q6GBT9 Q6GBU0 Q6GBU1 Q6GBU2 Q6GBU3 Q6GBU4 Q6GBU5 Q6GBU7 Q6GBU8 Q6GBU9
Q6GBV0 Q6GBV1 Q6GBV3 Q6GBV6 Q6GBV8 Q6GBV9 Q6GBW0 Q6GBW3 Q6GBW4 Q6GBW6 Q6GBW8
Q6GBW9 Q6GBX1 Q6GBX3 Q6GBX4 Q6GBX5 Q6GBX6 Q6GBX8 Q6GBX9 Q6GBY5 Q6GBY6 Q6GBY7
Q6GBY8 Q6GBZ0 Q6GBZ3 Q6GBZ5 Q6GBZ8 Q6GC00 Q6GC02 Q6GC06 Q6GC08 Q6GC09 Q6GC81
Q6GC82 Q6GC84 Q6GC85 Q6GC90 Q6GC91 Q6GC92 Q6GCA5 Q6GCA6 Q6GCA7 Q6GCB6 Q6GCB8
Q6GCE6 Q6GCF3 Q6GCF6 Q6GCK3 Q6GCL3 Q6GCL7 Q6GCM3 Q6GCN4 Q6GCP9 Q6GCQ0 Q6GCR4
Q6GCT1 Q6GCT5 Q6GCU1 Q6GCU2 Q6GCU3 Q6GCU4 Q6GCV9 Q6GCW1 Q6GCY5 Q6GCY6 Q6GCZ1
Q6GCZ8 Q6GD09 Q6GD13 Q6GD21 Q6GD38 Q6GD66 Q6GD73 Q6GD75 Q6GD81 Q6GD82 Q6GD84
Q6GD85 Q6GD86 Q6GD88 Q6GD89 Q6GD90 Q6GD91 Q6GD92 Q6GD93 Q6GD94 Q6GDB4 Q6GDB5
Q6GDC4 Q6GDC5 Q6GDC6 Q6GDC8 Q6GDC9 Q6GDD0 Q6GDD1 Q6GDD2 Q6GDD9 Q6GDG7 Q6GDG8
Q6GDH0 Q6GDH1 Q6GDJ1 Q6GDJ6 Q6GDJ7 Q6GDK1 Q6GDK4 Q6GDK5 Q6GDK6 Q6GDM0 Q6GDM3
Q6GDM7 Q6GDN5 Q6GDN7 Q6GDP0 Q6GDP3 Q6GDP4 Q6GDQ0 Q6GDS2 Q6GDS9 Q6GDV6 Q6GE08
Q6GE10 Q6GE17 Q6GE41 Q6GE46 Q6GE59 Q6GE66 Q6GE71 Q6GE88 Q6GE90 Q6GE99 Q6GEA1
Q6GEA2 Q6GEA4 Q6GEA5 Q6GEC4 Q6GEC9 Q6GED9 Q6GEE0 Q6GEE1 Q6GEE2 Q6GEE3 Q6GEE4
Q6GEE5 Q6GEE6 Q6GEF4 Q6GEF9 Q6GEG0 Q6GEG1 Q6GEG3 Q6GEG5 Q6GEG6 Q6GEG7 Q6GEG9
Q6GEH9 Q6GEI2 Q6GEI3 Q6GEI4 Q6GEI5 Q6GEI6 Q6GEI7 Q6GEI8 Q6GEI9 Q6GEJ0 Q6GEJ1
Q6GEJ2 Q6GEJ3 Q6GEJ4 Q6GEJ5 Q6GEJ6 Q6GEJ7 Q6GEJ8 Q6GEJ9 Q6GEK0 Q6GEK1 Q6GEK2
Q6GEK4 Q6GEK5 Q6GEK6 Q6GEK7 Q6GEK8 Q6GEK9 Q6GEL0 Q6GEL6 Q6GEL7 Q6GEL8 Q6GEM5
Q6GEN2 Q6GEN3 Q6GEN4 Q6GEN5 Q6GEN6 Q6GEN7 Q6GEP0 Q6GEP3 Q6GEP7 Q6GEQ6 Q6GEQ8
Q6GER3 Q6GER6 Q6GER8 Q6GES3 Q6GET8 Q6GET9 Q6GEU5 Q6GEU7 Q6GEU8 Q6GEV0 Q6GEV1
Q6GEV3 Q6GEV5 Q6GEV6 Q6GEV7 Q6GEW0 Q6GEW1 Q6GEW2 Q6GEW3 Q6GEX5 Q6GEY1 Q6GEY2
Q6GEY3 Q6GEY4 Q6GEZ1 Q6GEZ3 Q6GF03 Q6GF04 Q6GF05 Q6GF07 Q6GF08 Q6GF11 Q6GF13
Q6GF14 Q6GF15 Q6GF16 Q6GF17 Q6GF19 Q6GF23 Q6GF26 Q6GF42 Q6GF43 Q6GF48 Q6GF73
Q6GFD5 Q6GFD7 Q6GFE2 Q6GFE4 Q6GFE9 Q6GFF2 Q6GFF3 Q6GFF6 Q6GFF7 Q6GFF8 Q6GFG0
Q6GFG2 Q6GFG4 Q6GFG9 Q6GFH7 Q6GFI2 Q6GFI4 Q6GFJ0 Q6GFJ3 Q6GFJ6 Q6GFK5 Q6GFL1
Q6GFM3 Q6GFM4 Q6GFR0 Q6GFR5 Q6GFR6 Q6GFS7 Q6GFT5 Q6GFT6 Q6GFT9 Q6GFU3 Q6GFV0
Q6GFV1 Q6GFV3 Q6GFW5 Q6GFW8 Q6GFX2 Q6GFX3 Q6GFX6 Q6GFX9 Q6GFY8 Q6GFZ2 Q6GFZ4
Q6GFZ6 Q6GFZ7 Q6GFZ8 Q6GFZ9 Q6GG00 Q6GG04 Q6GG08 Q6GG09 Q6GG12 Q6GG18 Q6GG19
Q6GG20 Q6GG23 Q6GG25 Q6GG26 Q6GG27 Q6GG30 Q6GG31 Q6GG32 Q6GG33 Q6GG35 Q6GG37
Q6GG38 Q6GG42 Q6GG57 Q6GG59 Q6GG60 Q6GG61 Q6GG62 Q6GG63 Q6GG64 Q6GG65 Q6GG69
Q6GG70 Q6GG71 Q6GG72 Q6GG73 Q6GG78 Q6GG82 Q6GG85 Q6GG86 Q6GG87 Q6GG88 Q6GG92
Q6GG93 Q6GG98 Q6GGA2 Q6GGA5 Q6GGA7 Q6GGB5 Q6GGB8 Q6GGB9 Q6GGC0 Q6GGC1 Q6GGC2
Q6GGC5 Q6GGD0 Q6GGD4 Q6GGD5 Q6GGD6 Q6GGD7 Q6GGD8 Q6GGE0 Q6GGE2 Q6GGE5 Q6GGE6
Q6GGE8 Q6GGG1 Q6GGG2 Q6GGG3 Q6GGG4 Q6GGH0 Q6GGH4 Q6GGH5 Q6GGH6 Q6GGH8 Q6GGI4
Q6GGI7 Q6GGJ4 Q6GGJ5 Q6GGK1 Q6GGK3 Q6GGK4 Q6GGT4 Q6GGT5 Q6GGT6 Q6GGT8 Q6GGU0
Q6GGU2 Q6GGU3 Q6GGU4 Q6GGU5 Q6GGU7 Q6GGV2 Q6GGV4 Q6GGV5 Q6GGW0 Q6GGW3 Q6GGW4
Q6GGW9 Q6GGX0 Q6GGX8 Q6GGX9 Q6GGY0 Q6GGY1 Q6GGY3 Q6GGY4 Q6GGY6 Q6GGZ5 Q6GGZ6
Q6GGZ9 Q6GH02 Q6GH04 Q6GH06 Q6GH10 Q6GH11 Q6GH12 Q6GH13 Q6GH17 Q6GH30 Q6GH31
Q6GH32 Q6GH33 Q6GH34 Q6GH35 Q6GH39 Q6GH41 Q6GH44 Q6GH48 Q6GH51 Q6GH53 Q6GH55
Q6GH60 Q6GH61 Q6GH64 Q6GH65 Q6GH69 Q6GH70 Q6GH71 Q6GH72 Q6GH76 Q6GHC6 Q6GHD0
Q6GHD2 Q6GHD4 Q6GHD5 Q6GHD9 Q6GHE0 Q6GHE3 Q6GHF0 Q6GHG0 Q6GHG1 Q6GHG2 Q6GHG4

Q6GHG5 Q6GHG6 Q6GHH0 Q6GHH1 Q6GHH2 Q6GHH5 Q6GHH6 Q6GHH7 Q6GHH8 Q6GHH9 Q6GHI0
Q6GHI1 Q6GHI2 Q6GHI3 Q6GHI4 Q6GHI5 Q6GHI9 Q6GHJ0 Q6GHJ1 Q6GHJ4 Q6GHJ5 Q6GHJ6
Q6GHJ7 Q6GHK2 Q6GHK8 Q6GHK9 Q6GHL1 Q6GHL4 Q6GHL7 Q6GHL9 Q6GHM0 Q6GHM5 Q6GHM6
Q6GHN0 Q6GHN1 Q6GHN2 Q6GHN3 Q6GHN4 Q6GHN5 Q6GHN7 Q6GHP2 Q6GHP6 Q6GHP8 Q6GHQ2
Q6GHQ6 Q6GHQ7 Q6GHQ8 Q6GHQ9 Q6GHR6 Q6GHR7 Q6GHT4 Q6GHT5 Q6GHT9 Q6GHU0 Q6GHU1
Q6GHU5 Q6GHU6 Q6GHU7 Q6GHV0 Q6GHV8 Q6GHW0 Q6GHW1 Q6GHW4 Q6GHX3 Q6GHX9 Q6GHY8
Q6GHZ0 Q6GHZ1 Q6GHZ2 Q6GHZ4 Q6GHZ5 Q6GHZ6 Q6GI10 Q6GI11 Q6GI12 Q6GI13 Q6GI14
Q6GI15 Q6GI16 Q6GI18 Q6GI19 Q6GI21 Q6GI30 Q6GI36 Q6GI37 Q6GI39 Q6GI64 Q6GI66
Q6GI69 Q6GI79 Q6GI81 Q6GI84 Q6GI87 Q6GI88 Q6GI89 Q6GIB2 Q6GIB7 Q6GIC0 Q6GIC1
Q6GIC2 Q6GIC6 Q6GIC7 Q6GIC8 Q6GID0 Q6GID1 Q6GID4 Q6GIE9 Q6GIF6 Q6GIF9 Q6GIG3
Q6GIG7 Q6GIH0 Q6GIH2 Q6GIH6 Q6GII3 Q6GII7 Q6GII8 Q6GIK9 Q6GIL5 Q6GIL6 Q6GIL7
Q6GIL8 Q6GIM1 Q6GIM3 Q6GIM4 Q6GIM6 Q6GIM7 Q6GIM9 Q6GIN1 Q6GIN2 Q6GIN3 Q6GIN7
Q6GIN9 Q6GIP8 Q6GIQ3 Q6GIR1 Q6GIR3 Q6GIR7 Q6GIR8 Q6GIT0 Q6GIV6 Q6GIW1 Q6GIX3
Q6GIZ3 Q6GJ02 Q6GJ03 Q6GJ30 Q6GJ52 Q6GJ61 Q6GJ63 Q6GJ75 Q6GJ80 Q6GJ81 Q6GJ88
Q6GJ90 Q6GJ92 Q6GJ93 Q6GJ99 Q6GJA0 Q6GJA3 Q6GJA8 Q6GJB4 Q6GJB5 Q6GJB6 Q6GJB7
Q6GJB8 Q6GJC0 Q6GJC1 Q6GJC2 Q6GJC3 Q6GJC4 Q6GJC5 Q6GJC6 Q6GJC8 Q6GJC9 Q6GJD0
Q6GJD1 Q6GJD2 Q6GJD4 Q6GJD7 Q6GJD9 Q6GJE0 Q6GJE1 Q6GJE4 Q6GJE5 Q6GJE7 Q6GJE9
Q6GJF0 Q6GJF4 Q6GJF6 Q6GJF7 Q6GJF8 Q6GJF9 Q6GJG1 Q6GJG2 Q6GJG8 Q6GJG9 Q6GJH0
Q6GJH1 Q6GJH3 Q6GJH6 Q6GJH8 Q6GJI1 Q6GJI3 Q6GJI5 Q6GJI9 Q6GJJ3 Q6GJJ4 Q6GJQ6
Q6GJQ7 Q6GJQ9 Q6GJR0 Q6GJR6 Q6GJR7 Q6GJR8 Q6GJV1 Q6GJV3 Q6GJW2 Q6GJW4 Q6GJZ1
Q6GJZ8 Q6GK01 Q6GK26 Q6GK42 Q6GK53 Q6GK57 Q6GK63 Q6GK73 Q6GK89 Q6GK90 Q6GKA4
Q6GKB1 Q6GKB5 Q6GKC1 Q6GKC2 Q6GKC3 Q6GKC4 Q6GKD8 Q6GKE0 Q6GKG6 Q6GKG7 Q6GKH2
Q6GKH9 Q6GKI9 Q6GKJ3 Q6GKK1 Q6GKK9 Q6GKL1 Q6GKM5 Q6GKQ0 Q6GKR3 Q6GKS1 Q6GKS8
Q6GKT0 Q6GKT6 Q6GKT7 Q6GKT9 Q6GKU0 Q6GKU1 Q6GKU3 Q6GKU4 Q6GKX5 Q6GL23 Q6GL38
Q6GL39 Q6GL41 Q6GL52 Q6GL62 Q6GL65 Q6GL68 Q6GL69 Q6GL72 Q6GL74 Q6GL75 Q6GL77
Q6GL87 Q6GL89 Q6GL92 Q6GLA1 Q6GLB0 Q6GLB5 Q6GLB9 Q6GLD8 Q6GLD9 Q6GLE8 Q6GLH8
Q6GLI9 Q6GLJ6 Q6GLM9 Q6GLP4 Q6GLP7 Q6GLQ4 Q6GLT5 Q6GLT8 Q6GLV4 Q6GLW1 Q6GLX3
Q6GLX7 Q6GLY8 Q6GLZ0 Q6GLZ1 Q6GLZ5 Q6GLZ8 Q6GM07 Q6GM16 Q6GM53 Q6GM71 Q6GM74
Q6GM78 Q6GM82 Q6GMB0 Q6GMD2 Q6GME1 Q6GME2 Q6GMF2 Q6GMG5 Q6GMH0 Q6GMH3 Q6GMI7
Q6GMK7 Q6GMK8 Q6GML1 Q6GML7 Q6GMV2 Q6GMV3 Q6GMY9 Q6GN11 Q6GN15 Q6GN21 Q6GN86
Q6GN91 Q6GN98 Q6GNB7 Q6GNG8 Q6GNI0 Q6GNI4 Q6GNJ8 Q6GNL7 Q6GNM3 Q6GNM6 Q6GNP2
Q6GNS3 Q6GNT9 Q6GNU1 Q6GNV6 Q6GNW0 Q6GNX1 Q6GNY1 Q6GP04 Q6GP15 Q6GP17 Q6GP25
Q6GP48 Q6GP53 Q6GP60 Q6GP65 Q6GP80 Q6GP89 Q6GP95 Q6GPA7 Q6GPB5 Q6GPB6 Q6GPB8
Q6GPB9 Q6GPC6 Q6GPD9 Q6GPE5 Q6GPE9 Q6GPH4 Q6GPJ4 Q6GPJ5 Q6GPJ8 Q6GPK9 Q6GPL3
Q6GPL9 Q6GPM1 Q6GPM3 Q6GPM4 Q6GPN6 Q6GPP0 Q6GPQ4 Q6GPQ5 Q6GPQ6 Q6GPR5 Q6GPR9
Q6GPT7 Q6GPV5 Q6GPW6 Q6GPY6 Q6GPZ1 Q6GQ33 Q6GQ34 Q6GQ37 Q6GQ44 Q6GQ48 Q6GQ53
Q6GQ63 Q6GQ66 Q6GQ67 Q6GQ68 Q6GQ71 Q6GQ73 Q6GQ80 Q6GQ82 Q6GQ95 Q6GQA1 Q6GQA6
Q6GQD0 Q6GQD1 Q6GQD3 Q6GQD5 Q6GQD7 Q6GQD9 Q6GQG3 Q6GQI5 Q6GQJ2 Q6GQJ7 Q6GQJ8
Q6GQL1 Q6GQL2 Q6GQN4 Q6GQN5 Q6GQN7 Q6GQR8 Q6GQT0 Q6GQU0 Q6GQU2 Q6GQV0 Q6GQV1
Q6GQV7 Q6GQX2 Q6GQX6 Q6GR00 Q6GR08 Q6GR09 Q6GR16 Q6GR25 Q6GR30 Q6GR31 Q6GR34
Q6GR35 Q6GR37 Q6GR45 Q6GSS7 Q6GV10 Q6GV29 Q6GVH4 Q6GVH5 Q6GVM3 Q6GVM5 Q6GVM6
Q6GVM7 Q6GWS6 Q6GWX0 Q6GX84 Q6GX86 Q6GZ04 Q6GZM7 Q6GZM8 Q6GZM9 Q6GZN0 Q6GZN1
Q6GZN2 Q6GZN3 Q6GZN6 Q6GZN7 Q6GZN8 Q6GZN9 Q6GZP0 Q6GZP1 Q6GZP4 Q6GZP5 Q6GZP6
Q6GZP7 Q6GZP8 Q6GZP9 Q6GZQ1 Q6GZQ2 Q6GZQ3 Q6GZQ4 Q6GZQ5 Q6GZQ7 Q6GZQ8 Q6GZQ9

Q6GZR0 Q6GZR1 Q6GZR2 Q6GZR3 Q6GZR4 Q6GZR5 Q6GZR6 Q6GZR7 Q6GZR8 Q6GZR9 Q6GZS1
Q6GZS2 Q6GZS4 Q6GZS5 Q6GZS6 Q6GZS7 Q6GZS8 Q6GZS9 Q6GZT0 Q6GZT1 Q6GZT2 Q6GZT4
Q6GZT5 Q6GZT7 Q6GZT8 Q6GZT9 Q6GZU0 Q6GZU1 Q6GZU4 Q6GZU5 Q6GZU6 Q6GZU7 Q6GZU8
Q6GZU9 Q6GZV2 Q6GZV3 Q6GZV4 Q6GZV6 Q6GZV8 Q6GZW0 Q6GZW1 Q6GZW2 Q6GZW3 Q6GZW6
Q6GZW7 Q6GZW8 Q6GZW9 Q6GZX0 Q6GZX4 Q6H001 Q6H090 Q6H1U7 Q6H236 Q6H3Y7 Q6H434
Q6H442 Q6H4D6 Q6H500 Q6H543 Q6H595 Q6H5L4 Q6H601 Q6H612 Q6H628 Q6H638 Q6H647
Q6H668 Q6H6C3 Q6H6E6 Q6H6N5 Q6H6S3 Q6H6V4 Q6H711 Q6H754 Q6H795 Q6H7E4 Q6H7J3
Q6H7J4 Q6H7J5 Q6H7M1 Q6H7P8 Q6H7S2 Q6H7U2 Q6H849 Q6H874 Q6H8A9 Q6H8Q1 Q6H8R9
Q6H9K2 Q6H9K4 Q6H9K5 Q6H9K7 Q6H9K8 Q6H9L0 Q6H9L1 Q6H9L2 Q6H9L3 Q6H9L5 Q6H9L6
Q6HA23 Q6HA24 Q6HAE9 Q6HAF2 Q6HAF3 Q6HAF4 Q6HAF5 Q6HAG2 Q6HAG4 Q6HAG7 Q6HAG9
Q6HAH7 Q6HAI0 Q6HAM6 Q6HAN0 Q6HAN7 Q6HAP7 Q6HAQ2 Q6HAS1 Q6HAT6 Q6HAU4 Q6HAU5
Q6HAU9 Q6HAV2 Q6HAV3 Q6HAV4 Q6HAW8 Q6HAW9 Q6HAX0 Q6HAZ8 Q6HB59 Q6HBA8 Q6HBC2
Q6HBC6 Q6HBC8 Q6HBD4 Q6HBD6 Q6HBD8 Q6HBF0 Q6HBF1 Q6HBF2 Q6HBG1 Q6HBI9 Q6HBJ0
Q6HBM0 Q6HBQ8 Q6HBR6 Q6HBT4 Q6HBW7 Q6HBW8 Q6HBX6 Q6HBY2 Q6HBZ1 Q6HC08 Q6HC14
Q6HC15 Q6HC16 Q6HC18 Q6HC26 Q6HC29 Q6HCB3 Q6HCB4 Q6HCB8 Q6HCE2 Q6HCF7 Q6HCI1
Q6HCI7 Q6HCJ2 Q6HCM0 Q6HCM2 Q6HCN0 Q6HCN5 Q6HCP0 Q6HCP7 Q6HCP8 Q6HCR8 Q6HCT3
Q6HCU0 Q6HCU5 Q6HCU7 Q6HCV0 Q6HCV6 Q6HCV7 Q6HCW5 Q6HCW7 Q6HCW8 Q6HCW9 Q6HCX2
Q6HCX6 Q6HCY5 Q6HD05 Q6HD42 Q6HD43 Q6HD53 Q6HD54 Q6HD57 Q6HD60 Q6HD62 Q6HD65
Q6HD68 Q6HD71 Q6HD72 Q6HD76 Q6HD81 Q6HD83 Q6HD85 Q6HD97 Q6HDA5 Q6HDA6 Q6HDA8
Q6HDA9 Q6HDB8 Q6HDC0 Q6HDC2 Q6HDC3 Q6HDD0 Q6HDD7 Q6HDD8 Q6HDD9 Q6HDE0 Q6HDE5
Q6HDE6 Q6HDF0 Q6HDF1 Q6HDI8 Q6HDJ0 Q6HDJ9 Q6HDK0 Q6HDK5 Q6HDK6 Q6HDK7 Q6HDK8
Q6HDK9 Q6HDL2 Q6HDL8 Q6HDM2 Q6HDM4 Q6HDN0 Q6HDN3 Q6HDN9 Q6HDT6 Q6HDT7 Q6HDT8
Q6HDW1 Q6HDW6 Q6HDW8 Q6HDY3 Q6HDY4 Q6HDY5 Q6HDY6 Q6HDY8 Q6HDZ0 Q6HE00 Q6HE08
Q6HE20 Q6HE28 Q6HE29 Q6HE30 Q6HE32 Q6HE47 Q6HE48 Q6HE51 Q6HE53 Q6HE54 Q6HE79
Q6HE93 Q6HEA8 Q6HEA9 Q6HEB2 Q6HEC6 Q6HEC7 Q6HEC8 Q6HEC9 Q6HED2 Q6HED3 Q6HEG3
Q6HEI4 Q6HEI5 Q6HEI9 Q6HEJ9 Q6HEK2 Q6HEL5 Q6HEM5 Q6HEN2 Q6HEN5 Q6HEN9 Q6HEP1
Q6HEP3 Q6HEP5 Q6HEP6 Q6HEQ2 Q6HEQ5 Q6HER5 Q6HER9 Q6HES3 Q6HES5 Q6HES6 Q6HES7
Q6HES8 Q6HES9 Q6HET0 Q6HET1 Q6HET2 Q6HEU3 Q6HEU4 Q6HEU5 Q6HEU9 Q6HEV1 Q6HEV7
Q6HEW6 Q6HEX1 Q6HEX3 Q6HEX4 Q6HEX5 Q6HEX8 Q6HEX9 Q6HEY3 Q6HEY5 Q6HEY6 Q6HEY7
Q6HEY8 Q6HEY9 Q6HEZ1 Q6HEZ2 Q6HEZ4 Q6HEZ6 Q6HEZ8 Q6HF02 Q6HF04 Q6HF05 Q6HF07
Q6HF08 Q6HF34 Q6HF43 Q6HF46 Q6HF47 Q6HF68 Q6HF97 Q6HF98 Q6HFB2 Q6HFE8 Q6HFE9
Q6HFF0 Q6HFF1 Q6HFG9 Q6HFI9 Q6HFT0 Q6HFZ7 Q6HG28 Q6HG59 Q6HG86 Q6HGE0 Q6HGG0
Q6HGI0 Q6HGP0 Q6HH05 Q6HH08 Q6HH09 Q6HH29 Q6HH75 Q6HHC1 Q6HHC2 Q6HHE0 Q6HHF5
Q6HHF6 Q6HHF8 Q6HHK0 Q6HHR6 Q6HHU6 Q6HHU9 Q6HHX7 Q6HHX8 Q6HHX9 Q6HI24 Q6HIJ3
Q6HIK1 Q6HIK2 Q6HIK3 Q6HIK4 Q6HIL9 Q6HIP4 Q6HJ19 Q6HJ64 Q6HJ78 Q6HJC2 Q6HJC7
Q6HJF2 Q6HJF5 Q6HJF7 Q6HJK9 Q6HJT7 Q6HJU1 Q6HJW8 Q6HJZ3 Q6HK19 Q6HK31 Q6HK44
Q6HK57 Q6HK62 Q6HK75 Q6HK87 Q6HKA0 Q6HKA1 Q6HKD8 Q6HKG3 Q6HKZ0 Q6HKZ8 Q6HKZ9
Q6HL06 Q6HL14 Q6HL15 Q6HL16 Q6HL19 Q6HL22 Q6HL23 Q6HL35 Q6HL37 Q6HL38 Q6HL39
Q6HL40 Q6HL42 Q6HL44 Q6HL51 Q6HL56 Q6HL58 Q6HL92 Q6HLA6 Q6HLD3 Q6HLD4 Q6HLE1
Q6HLE2 Q6HLE3 Q6HLE4 Q6HLE5 Q6HLE6 Q6HLE7 Q6HLE8 Q6HLE9 Q6HLF0 Q6HLF1 Q6HLF2
Q6HLF3 Q6HLF4 Q6HLG0 Q6HLH7 Q6HLK3 Q6HLK6 Q6HLM0 Q6HLM1 Q6HLP0 Q6HLP1 Q6HLP2
Q6HLS8 Q6HLU3 Q6HLU4 Q6HLU5 Q6HLU6 Q6HLU7 Q6HLV6 Q6HLX9 Q6HLY2 Q6HLY4 Q6HLY7
Q6HM04 Q6HM20 Q6HM28 Q6HM43 Q6HM44 Q6HM97 Q6HM98 Q6HMB8 Q6HMD5 Q6HME9 Q6HMG1
Q6HMH0 Q6HMH4 Q6HMS8 Q6HMS9 Q6HMV4 Q6HMV9 Q6HMZ5 Q6HN08 Q6HN84 Q6HNE4 Q6HNE8

Q6HNK1 Q6HNR1 Q6HNR9 Q6HNS8 Q6HNT2 Q6HNU0 Q6HNW5 Q6HNY7 Q6HNZ7 Q6HP17 Q6HP18
Q6HP19 Q6HP28 Q6HP29 Q6HP30 Q6HP49 Q6HP54 Q6HP80 Q6HP81 Q6HP82 Q6HP91 Q6HP94
Q6HPA0 Q6HPA2 Q6HPA4 Q6HPA5 Q6HPA7 Q6HPC6 Q6HPC7 Q6HPC8 Q6HPD1 Q6HPD3 Q6HPD7
Q6HPE6 Q6HPH4 Q6HPL2 Q6HPL3 Q6HPM5 Q6HPM6 Q6HPM7 Q6HPN1 Q6HPN2 Q6HPN3 Q6HPN4
Q6HPN5 Q6HPN7 Q6HPN9 Q6HPP0 Q6HPP1 Q6HPP2 Q6HPP3 Q6HPP4 Q6HPP5 Q6HPP6 Q6HPP7
Q6HPP8 Q6HPP9 Q6HPQ1 Q6HPQ2 Q6HPQ3 Q6HPQ4 Q6HPQ5 Q6HPQ6 Q6HPQ7 Q6HPQ8 Q6HPQ9
Q6HPR0 Q6HPR1 Q6HPR2 Q6HPR3 Q6HPR5 Q6HPR6 Q6HPR8 Q6HPR9 Q6HPS0 Q6HPS1 Q6HPS8
Q6HPT0 Q6HPT1 Q6HPT2 Q6HPT7 Q6HPU0 Q6HPV0 Q6HPV1 Q6HPV4 Q6HPW6 Q6HPW9 Q6HPX2
Q6HPX5 Q6HPY1 Q6HPY3 Q6HPY6 Q6HPZ0 Q6HPZ1 Q6HPZ8 Q6HQ00 Q6HQ03 Q6HQ04 Q6HQ05
Q6HQY9 Q6HSF4 Q6HSG2 Q6HUG3 Q6HX62 Q6HYJ1 Q6I3T8 Q6I4E9 Q6I4Q2 Q6I511 Q6I535
Q6I581 Q6I588 Q6I5Y0 Q6I6D4 Q6I6F4 Q6I7B0 Q6I7B6 Q6I7C0 Q6I7C4 Q6I7R3 Q6I7R5
Q6I9Y2 Q6IA69 Q6IA86 Q6IB77 Q6IC83 Q6ICC9 Q6ICG8 Q6ICL3 Q6ICX4 Q6ICX6 Q6ID70
Q6ID77 Q6IDB0 Q6IDB6 Q6IDS6 Q6IE24 Q6IE70 Q6IE77 Q6IE82 Q6IEG0 Q6IEI0 Q6IFU7
Q6IFU8 Q6IFV1 Q6IFV3 Q6IFV4 Q6IFW2 Q6IFW3 Q6IFW5 Q6IFW8 Q6IFX0 Q6IFX1 Q6IFX2
Q6IFX3 Q6IFX4 Q6IFZ6 Q6IFZ9 Q6IG00 Q6IG01 Q6IG02 Q6IG03 Q6IG04 Q6IG05 Q6IG12
Q6IM78 Q6IMA7 Q6IME9 Q6IMF0 Q6IMF1 Q6IMG9 Q6IMH0 Q6IMI4 Q6IMK0 Q6IMK3 Q6IMM1
Q6IMN8 Q6IMX7 Q6IMY8 Q6IMZ0 Q6IN36 Q6IN84 Q6INB0 Q6IND4 Q6IND6 Q6IND7 Q6INF3
Q6ING2 Q6ING7 Q6ING9 Q6INH0 Q6INH1 Q6INH5 Q6INI5 Q6INI7 Q6INL2 Q6INM2 Q6INN7
Q6INN8 Q6INP8 Q6INR1 Q6INS1 Q6INS5 Q6INU2 Q6INU3 Q6INU8 Q6INV3 Q6INV8 Q6INX4
Q6IP06 Q6IP09 Q6IP18 Q6IP48 Q6IP50 Q6IP57 Q6IP65 Q6IP67 Q6IP73 Q6IP91 Q6IPR1
Q6IPR3 Q6IPT2 Q6IPW1 Q6IPX3 Q6IQ16 Q6IQ19 Q6IQ21 Q6IQ63 Q6IQ86 Q6IQ97 Q6IQE0
Q6IQE9 Q6IQS6 Q6IQS9 Q6IQT4 Q6IQU4 Q6IQU6 Q6IQU7 Q6IQX0 Q6IQX8 Q6IR42 Q6IR52
Q6IR61 Q6IR68 Q6IR70 Q6IR75 Q6IR85 Q6IRA8 Q6IRB1 Q6IRB2 Q6IRB8 Q6IRC5 Q6IRC7
Q6IRL9 Q6IRM9 Q6IRN0 Q6IRN2 Q6IRN6 Q6IRP0 Q6IRQ5 Q6IRQ9 Q6IRS2 Q6IRT3 Q6IRU2
Q6IT00 Q6ITZ3 Q6IUF8 Q6IUP1 Q6IUP3 Q6IUR4 Q6IV56 Q6IV68 Q6IV72 Q6IVC2 Q6IVC3
Q6IVW0 Q6IVY4 Q6IZ48 Q6J171 Q6J1H4 Q6J1I7 Q6J1T3 Q6J1Z5 Q6J1Z6 Q6J212 Q6J256
Q6J272 Q6J2E6 Q6J2K6 Q6J2U6 Q6J362 Q6J3N5 Q6J3Q7 Q6J5J3 Q6J6I6 Q6J8G2 Q6J9Z4
Q6JAH0 Q6JB10 Q6JB11 Q6JD69 Q6JD70 Q6JD71 Q6JD72 Q6JD73 Q6JE37 Q6JE38 Q6JEL2
Q6JEL3 Q6JIY4 Q6JIY5 Q6JJ61 Q6JL78 Q6JLC9 Q6JM09 Q6JPI3 Q6JQN1 Q6JTB2 Q6JV40
Q6JVN2 Q6JXI5 Q6JZK2 Q6K1L7 Q6K1S6 Q6K1U4 Q6K1Z1 Q6K1Z6 Q6K223 Q6K271 Q6K2E1
Q6K2E8 Q6K332 Q6K3B1 Q6K3B2 Q6K3R9 Q6K471 Q6K498 Q6K4N0 Q6K4V3 Q6K537 Q6K5F8
Q6K5G8 Q6K5I0 Q6K5K2 Q6K609 Q6K638 Q6K641 Q6K669 Q6K678 Q6K687 Q6K6K7 Q6K6N7
Q6K762 Q6K765 Q6K7E6 Q6K7E9 Q6K7H2 Q6K7R9 Q6K7V6 Q6K833 Q6K846 Q6K881 Q6K8J4
Q6K8S0 Q6K8S1 Q6K8S3 Q6K8S5 Q6K8Z4 Q6K908 Q6K953 Q6K972 Q6K977 Q6K990 Q6K9A2
Q6K9C1 Q6K9C3 Q6K9N6 Q6K9U2 Q6K9X3 Q6KAE5 Q6KAI0 Q6KAJ4 Q6KAJ8 Q6KAQ7 Q6KAS7
Q6KAU7 Q6KB66 Q6KBA8 Q6KC51 Q6KCA4 Q6KCJ3 Q6KCJ5 Q6KCJ6 Q6KCK0 Q6KCK3 Q6KGH8
Q6KGI8 Q6KGV8 Q6KGW1 Q6KGW4 Q6KGW7 Q6KGX3 Q6KGX6 Q6KGX9 Q6KGY2 Q6KH13 Q6KH14
Q6KH17 Q6KH18 Q6KH30 Q6KH54 Q6KH72 Q6KH77 Q6KH80 Q6KH82 Q6KH87 Q6KH90 Q6KH93
Q6KH95 Q6KH96 Q6KH97 Q6KHA3 Q6KHA6 Q6KHA7 Q6KHB0 Q6KHB1 Q6KHC4 Q6KHC8 Q6KHC9
Q6KHD7 Q6KHE2 Q6KHE8 Q6KHF9 Q6KHG7 Q6KHG8 Q6KHH3 Q6KHH5 Q6KHH8 Q6KHK4 Q6KHK5
Q6KHK6 Q6KHL4 Q6KHL5 Q6KHL6 Q6KHM2 Q6KHM5 Q6KHM9 Q6KHN3 Q6KHN7 Q6KHP2 Q6KHP5
Q6KHR2 Q6KHR3 Q6KHR6 Q6KHS0 Q6KHS3 Q6KHS4 Q6KHS5 Q6KHT6 Q6KHT7 Q6KHU1 Q6KHU4
Q6KHV1 Q6KHV9 Q6KHX0 Q6KHY5 Q6KHY6 Q6KI08 Q6KI09 Q6KI17 Q6KI22 Q6KI26 Q6KI28
Q6KI29 Q6KI30 Q6KI31 Q6KI32 Q6KI33 Q6KI35 Q6KI37 Q6KI38 Q6KI39 Q6KI41 Q6KI42

Q6KI43 Q6KI44 Q6KI45 Q6KI46 Q6KI47 Q6KI48 Q6KI49 Q6KI50 Q6KI51 Q6KI52 Q6KI53
Q6KI54 Q6KI55 Q6KI56 Q6KI60 Q6KI61 Q6KI65 Q6KI66 Q6KI92 Q6KI95 Q6KI99 Q6KIA0
Q6KIB3 Q6KIB4 Q6KIB8 Q6KID0 Q6KID2 Q6KID5 Q6KID6 Q6KID8 Q6KIE1 Q6KIE2 Q6KIE6
Q6KIF0 Q6KIF1 Q6KIG0 Q6KIG2 Q6KIG6 Q6KIH7 Q6KIH8 Q6KIH9 Q6KIJ8 Q6KIK5 Q6KIL4
Q6KIL6 Q6KIM1 Q6KIN5 Q6KIN6 Q6KIP6 Q6KIP9 Q6KIQ6 Q6KIT3 Q6KMS8 Q6KNA9 Q6KSL4
Q6KYU3 Q6KYV2 Q6KYV3 Q6KYZ5 Q6KZ11 Q6KZ23 Q6KZ24 Q6KZ25 Q6KZ30 Q6KZ38 Q6KZ41
Q6KZ42 Q6KZ54 Q6KZ89 Q6KZA3 Q6KZB1 Q6KZB3 Q6KZC5 Q6KZD1 Q6KZD3 Q6KZF1 Q6KZF9
Q6KZH9 Q6KZI1 Q6KZI4 Q6KZI5 Q6KZI6 Q6KZI7 Q6KZI8 Q6KZJ4 Q6KZJ6 Q6KZK4 Q6KZK7
Q6KZL0 Q6KZL5 Q6KZM3 Q6KZM5 Q6KZN5 Q6KZP5 Q6KZP6 Q6KZP7 Q6KZP8 Q6KZQ3 Q6KZQ5
Q6KZR2 Q6KZR7 Q6KZS5 Q6KZV3 Q6KZW2 Q6KZZ4 Q6KZZ9 Q6L016 Q6L032 Q6L046 Q6L047
Q6L080 Q6L0A7 Q6L0A8 Q6L0C3 Q6L0D1 Q6L0D3 Q6L0E6 Q6L0E8 Q6L0F1 Q6L0F2 Q6L0F3
Q6L0J9 Q6L0K5 Q6L0L2 Q6L0N9 Q6L0P2 Q6L0P4 Q6L0P7 Q6L0P8 Q6L0P9 Q6L0R3 Q6L0R4
Q6L0R6 Q6L0S6 Q6L0S7 Q6L0S8 Q6L0U7 Q6L0V5 Q6L0V9 Q6L0W3 Q6L0W5 Q6L0Y7 Q6L123
Q6L125 Q6L126 Q6L135 Q6L140 Q6L150 Q6L152 Q6L153 Q6L171 Q6L178 Q6L181 Q6L195
Q6L1A0 Q6L1A6 Q6L1A7 Q6L1A8 Q6L1B0 Q6L1B1 Q6L1B2 Q6L1B3 Q6L1B4 Q6L1B5 Q6L1B6
Q6L1B7 Q6L1B8 Q6L1B9 Q6L1C0 Q6L1C1 Q6L1C2 Q6L1C3 Q6L1C4 Q6L1C5 Q6L1C7 Q6L1C8
Q6L1E2 Q6L1E3 Q6L1F0 Q6L1F4 Q6L1F5 Q6L1F6 Q6L1G5 Q6L1G8 Q6L1H6 Q6L1H9 Q6L1I0
Q6L1J7 Q6L1K2 Q6L1K7 Q6L1K9 Q6L1L0 Q6L1L7 Q6L1M2 Q6L1M4 Q6L1M5 Q6L1N2 Q6L1N3
Q6L1N5 Q6L1N7 Q6L1P9 Q6L1Q0 Q6L1Q1 Q6L1Q5 Q6L1R1 Q6L1R2 Q6L1R3 Q6L1S1 Q6L1T2
Q6L1T6 Q6L1U1 Q6L1V1 Q6L1V2 Q6L1W3 Q6L1X8 Q6L1X9 Q6L1Y0 Q6L1Y6 Q6L1Z4 Q6L200
Q6L201 Q6L202 Q6L204 Q6L206 Q6L222 Q6L225 Q6L227 Q6L228 Q6L233 Q6L240 Q6L243
Q6L270 Q6L271 Q6L273 Q6L277 Q6L285 Q6L292 Q6L293 Q6L294 Q6L295 Q6L2D8 Q6L2G4
Q6L2G8 Q6L2G9 Q6L2H3 Q6L2I0 Q6L2I9 Q6L2J1 Q6L2J8 Q6L2J9 Q6L2K9 Q6L2L0 Q6L2L1
Q6L2L2 Q6L2L3 Q6L2M3 Q6L2M4 Q6L2M8 Q6L2S4 Q6L2T8 Q6L2V8 Q6L2W1 Q6L2Z8 Q6L2Z9
Q6L366 Q6L367 Q6L368 Q6L369 Q6L370 Q6L377 Q6L378 Q6L379 Q6L380 Q6L391 Q6L397
Q6L3A0 Q6L3A4 Q6L3A5 Q6L3A6 Q6L3A7 Q6L3B4 Q6L3B6 Q6L3B8 Q6L3B9 Q6L3C0 Q6L3C4
Q6L3C6 Q6L3C7 Q6L3C8 Q6L3C9 Q6L3D3 Q6L3E2 Q6L3F1 Q6L3F9 Q6L3G0 Q6L3G1 Q6L3G2
Q6L482 Q6L4D4 Q6L4H4 Q6L4L4 Q6L4N4 Q6L4R7 Q6L4S0 Q6L4V0 Q6L500 Q6L506 Q6L512
Q6L534 Q6L545 Q6L589 Q6L5C4 Q6L5E5 Q6L5F4 Q6L5G1 Q6L5H6 Q6L5Q5 Q6L5S6 Q6L612
Q6L732 Q6L733 Q6L738 Q6L739 Q6L741 Q6L743 Q6L8G4 Q6L8G5 Q6L8G8 Q6L8G9 Q6L8H1
Q6L8H2 Q6L8H4 Q6L8J9 Q6L8K0 Q6L8K1 Q6L8K7 Q6L8L1 Q6L8L2 Q6L8L3 Q6L8L5 Q6L8L6
Q6L8L7 Q6LA04 Q6LA06 Q6LA17 Q6LA20 Q6LA53 Q6LA56 Q6LAF1 Q6LAF3 Q6LAN3 Q6LAN4
Q6LAQ8 Q6LBA6 Q6LBE3 Q6LBF0 Q6LCE4 Q6LCK1 Q6LCW8 Q6LD29 Q6LDD9 Q6LDH0 Q6LDH1
Q6LED0 Q6LFD5 Q6LFH8 Q6LFN2 Q6LFR3 Q6LFS0 Q6LFW5 Q6LG09 Q6LG16 Q6LG35 Q6LG56
Q6LGH5 Q6LGK4 Q6LGY8 Q6LH12 Q6LH19 Q6LH20 Q6LH46 Q6LHF5 Q6LHK5 Q6LHN1 Q6LHN5
Q6LHN7 Q6LHY2 Q6LI45 Q6LIL7 Q6LIU7 Q6LIW0 Q6LIW6 Q6LJ30 Q6LJI1 Q6LJR6 Q6LK43
Q6LKA1 Q6LKA2 Q6LKE5 Q6LKM4 Q6LKQ1 Q6LL16 Q6LL68 Q6LLA7 Q6LLD4 Q6LLF0 Q6LLF7
Q6LLF8 Q6LLH3 Q6LLH7 Q6LLJ2 Q6LLJ6 Q6LLJ9 Q6LLK0 Q6LLK1 Q6LLL4 Q6LLL5 Q6LLM4
Q6LLN3 Q6LLN5 Q6LLQ5 Q6LLQ8 Q6LLR9 Q6LLS2 Q6LLS3 Q6LLT9 Q6LLU2 Q6LLV2 Q6LLV5
Q6LLV9 Q6LLW0 Q6LLW1 Q6LLW2 Q6LLW3 Q6LLW5 Q6LLW6 Q6LLX8 Q6LLY3 Q6LLY5 Q6LLZ0
Q6LLZ1 Q6LM05 Q6LM06 Q6LM07 Q6LM11 Q6LM16 Q6LM23 Q6LM29 Q6LM35 Q6LM37 Q6LM38
Q6LM40 Q6LM41 Q6LM42 Q6LM43 Q6LM44 Q6LM51 Q6LM58 Q6LM59 Q6LM60 Q6LM67 Q6LM68
Q6LM69 Q6LMA4 Q6LMB3 Q6LMC9 Q6LMD4 Q6LME3 Q6LMF4 Q6LMF7 Q6LMG5 Q6LMG6 Q6LMG7
Q6LMI4 Q6LMJ5 Q6LMJ6 Q6LMJ9 Q6LMK1 Q6LMK7 Q6LML0 Q6LML1 Q6LML4 Q6LMM2 Q6LMM3

Q6LMM8 Q6LMN0 Q6LMN6 Q6LMP0 Q6LMR1 Q6LMR2 Q6LMS2 Q6LMS4 Q6LMS7 Q6LMT0 Q6LMT3
Q6LMT4 Q6LMT5 Q6LMT6 Q6LMT7 Q6LMU0 Q6LMU2 Q6LMU3 Q6LMU5 Q6LMV1 Q6LMV7 Q6LMV8
Q6LMV9 Q6LMW0 Q6LMY0 Q6LMZ6 Q6LN05 Q6LN06 Q6LN10 Q6LN21 Q6LN22 Q6LN24 Q6LN25
Q6LN26 Q6LN27 Q6LN30 Q6LN38 Q6LN41 Q6LN45 Q6LN48 Q6LN63 Q6LN73 Q6LN74 Q6LN85
Q6LN87 Q6LN88 Q6LN94 Q6LN98 Q6LNA3 Q6LNA5 Q6LNA9 Q6LNB1 Q6LNB3 Q6LNB4 Q6LNF6
Q6LNI6 Q6LNJ0 Q6LNN4 Q6LNT9 Q6LNU2 Q6LNU5 Q6LNV7 Q6LNV9 Q6LNW0 Q6LNW1 Q6LNX0
Q6LNX3 Q6LP05 Q6LP06 Q6LP11 Q6LP12 Q6LP25 Q6LP50 Q6LP62 Q6LPA3 Q6LPA4 Q6LPA6
Q6LPD7 Q6LPD9 Q6LPE1 Q6LPE2 Q6LPE4 Q6LPE7 Q6LPI7 Q6LPK8 Q6LPL4 Q6LPQ4 Q6LPQ6
Q6LPR2 Q6LPR3 Q6LPR5 Q6LPV0 Q6LPV5 Q6LPX9 Q6LQ03 Q6LQ36 Q6LQ38 Q6LQ56 Q6LQ57
Q6LQ58 Q6LQ59 Q6LQ71 Q6LQ72 Q6LQ73 Q6LQB9 Q6LQG2 Q6LQJ9 Q6LQK3 Q6LQL4 Q6LQM6
Q6LQR1 Q6LQR4 Q6LQU5 Q6LQV0 Q6LQV7 Q6LQY0 Q6LR24 Q6LR31 Q6LR94 Q6LRA0 Q6LRA1
Q6LRA6 Q6LRB9 Q6LRC2 Q6LRD1 Q6LRD9 Q6LRE4 Q6LRI0 Q6LRI4 Q6LRI5 Q6LRY1 Q6LRY9
Q6LRZ3 Q6LS21 Q6LS31 Q6LSD6 Q6LSS4 Q6LST1 Q6LST5 Q6LSW5 Q6LSX4 Q6LSX6 Q6LSY3
Q6LSZ7 Q6LT00 Q6LT10 Q6LT12 Q6LT14 Q6LT18 Q6LT40 Q6LT47 Q6LT48 Q6LT49 Q6LT51
Q6LT52 Q6LT53 Q6LT55 Q6LT56 Q6LT67 Q6LT75 Q6LT81 Q6LT85 Q6LT86 Q6LT88 Q6LTA1
Q6LTB5 Q6LTC6 Q6LTD1 Q6LTE0 Q6LTE1 Q6LTE2 Q6LTE6 Q6LTE9 Q6LTJ7 Q6LTT8 Q6LTU5
Q6LTW8 Q6LTX2 Q6LTX3 Q6LTX4 Q6LTX5 Q6LTZ3 Q6LU02 Q6LU05 Q6LU07 Q6LU11 Q6LU12
Q6LU16 Q6LU17 Q6LU24 Q6LU31 Q6LU33 Q6LU45 Q6LU49 Q6LU52 Q6LU53 Q6LU58 Q6LU59
Q6LU62 Q6LU63 Q6LU69 Q6LU70 Q6LU78 Q6LUA6 Q6LUA7 Q6LUA8 Q6LUA9 Q6LUB4 Q6LUG1
Q6LUG2 Q6LUH1 Q6LUH2 Q6LUH3 Q6LUH4 Q6LUI8 Q6LUI9 Q6LUJ0 Q6LUJ2 Q6LUJ4 Q6LUJ6
Q6LUJ9 Q6LUK6 Q6LUK7 Q6LUK8 Q6LUL1 Q6LUL5 Q6LUM1 Q6LUM5 Q6LUM6 Q6LUN2 Q6LUN9
Q6LUP1 Q6LUP5 Q6LUP6 Q6LUP9 Q6LUQ1 Q6LUQ3 Q6LUR4 Q6LUR9 Q6LUS1 Q6LUS3 Q6LUS5
Q6LUW0 Q6LUW1 Q6LUW5 Q6LUW9 Q6LUX0 Q6LUX1 Q6LUX5 Q6LUY7 Q6LUZ5 Q6LV05 Q6LV10
Q6LV11 Q6LV24 Q6LV25 Q6LV26 Q6LV27 Q6LV34 Q6LV41 Q6LV42 Q6LV43 Q6LV46 Q6LV48
Q6LV49 Q6LV53 Q6LV59 Q6LV65 Q6LV74 Q6LV90 Q6LV91 Q6LV92 Q6LV93 Q6LV94 Q6LV95
Q6LV97 Q6LV98 Q6LV99 Q6LVA0 Q6LVA1 Q6LVA2 Q6LVA3 Q6LVA4 Q6LVA5 Q6LVA6 Q6LVA7
Q6LVA8 Q6LVA9 Q6LVB0 Q6LVB1 Q6LVB2 Q6LVB3 Q6LVB4 Q6LVB5 Q6LVB6 Q6LVB7 Q6LVC0
Q6LVC1 Q6LVC2 Q6LVC3 Q6LVC6 Q6LVD0 Q6LVE5 Q6LVF1 Q6LVF5 Q6LVF6 Q6LVG8 Q6LVG9
Q6LVH1 Q6LVH8 Q6LVI2 Q6LVI3 Q6LVI5 Q6LVI6 Q6LVK2 Q6LVL0 Q6LVL2 Q6LVM8 Q6LVN0
Q6LVN2 Q6LVN3 Q6LVN4 Q6LVN6 Q6LVN7 Q6LVN8 Q6LVP5 Q6LVP6 Q6LVQ7 Q6LVQ9 Q6LVS4
Q6LVT0 Q6LVX7 Q6LVY0 Q6LVZ5 Q6LW10 Q6LW15 Q6LW50 Q6LW54 Q6LW56 Q6LWJ8 Q6LWK3
Q6LWK4 Q6LWL2 Q6LWL3 Q6LWL8 Q6LWM0 Q6LWM3 Q6LWM9 Q6LWN0 Q6LWQ0 Q6LWR0 Q6LWU0
Q6LWW5 Q6LWX1 Q6LWX2 Q6LWX5 Q6LWX6 Q6LWX9 Q6LWY7 Q6LX07 Q6LX08 Q6LX09 Q6LX10
Q6LX26 Q6LX40 Q6LX41 Q6LX43 Q6LX56 Q6LX60 Q6LX61 Q6LX62 Q6LX63 Q6LX65 Q6LX78
Q6LX80 Q6LX81 Q6LX82 Q6LX84 Q6LXA7 Q6LXB3 Q6LXC0 Q6LXC1 Q6LXC5 Q6LXC6 Q6LXC7
Q6LXD1 Q6LXD2 Q6LXD3 Q6LXD4 Q6LXD6 Q6LXD7 Q6LXD8 Q6LXD9 Q6LXE0 Q6LXE2 Q6LXE3
Q6LXE4 Q6LXE5 Q6LXE6 Q6LXE7 Q6LXF0 Q6LXF3 Q6LXF7 Q6LXF9 Q6LXH7 Q6LXH8 Q6LXI0
Q6LXI1 Q6LXI2 Q6LXI3 Q6LXI4 Q6LXI6 Q6LXI7 Q6LXJ8 Q6LXL7 Q6LXM4 Q6LXM5 Q6LXM6
Q6LXM7 Q6LXM9 Q6LXN0 Q6LXN1 Q6LXN9 Q6LXP1 Q6LXQ4 Q6LXQ9 Q6LXR0 Q6LXR3 Q6LXR7
Q6LXT2 Q6LXU2 Q6LXU3 Q6LXV6 Q6LXV8 Q6LXW0 Q6LXX9 Q6LXY3 Q6LXY6 Q6LXY7 Q6LXY8
Q6LY02 Q6LY04 Q6LY06 Q6LY19 Q6LY38 Q6LY39 Q6LY46 Q6LY53 Q6LY59 Q6LY87 Q6LYB4
Q6LYB5 Q6LYC4 Q6LYD1 Q6LYD6 Q6LYD7 Q6LYD8 Q6LYE0 Q6LYF4 Q6LYF7 Q6LYG0 Q6LYG5
Q6LYI0 Q6LYI4 Q6LYI7 Q6LYI9 Q6LYK1 Q6LYK4 Q6LYM1 Q6LYN8 Q6LYP1 Q6LYP5 Q6LYQ1
Q6LYQ2 Q6LYQ3 Q6LYR0 Q6LYR4 Q6LYR5 Q6LYR8 Q6LYS1 Q6LYT2 Q6LYU3 Q6LYU4 Q6LYV9

Q6LYX3 Q6LYX4 Q6LYY1 Q6LZ92 Q6LZA1 Q6LZA2 Q6LZC1 Q6LZC3 Q6LZC4 Q6LZC5 Q6LZC7
Q6LZD2 Q6LZD3 Q6LZD4 Q6LZE1 Q6LZE3 Q6LZE9 Q6LZF1 Q6LZG0 Q6LZG2 Q6LZG8 Q6LZH1
Q6LZH4 Q6LZI3 Q6LZI6 Q6LZI8 Q6LZK0 Q6LZK1 Q6LZK2 Q6LZK8 Q6LZL0 Q6LZL5 Q6LZL7
Q6LZL8 Q6LZM1 Q6LZM2 Q6LZM7 Q6LZM9 Q6LZN6 Q6LZP0 Q6LZP7 Q6LZP8 Q6LZQ3 Q6LZS1
Q6M029 Q6M030 Q6M044 Q6M056 Q6M057 Q6M060 Q6M066 Q6M067 Q6M079 Q6M086 Q6M099
Q6M0B4 Q6M0B9 Q6M0E4 Q6M0E5 Q6M0E6 Q6M0E9 Q6M0F1 Q6M0F3 Q6M0G7 Q6M0H0 Q6M0H3
Q6M0H5 Q6M0I0 Q6M0I4 Q6M0I6 Q6M0I7 Q6M0K2 Q6M0K7 Q6M0K9 Q6M0L0 Q6M0L1 Q6M0L3
Q6M0L9 Q6M0M1 Q6M0M3 Q6M0M5 Q6M0P3 Q6M0P5 Q6M0Q7 Q6M0T4 Q6M0U0 Q6M0U8 Q6M0V9
Q6M0W1 Q6M0X3 Q6M0Y7 Q6M0Z1 Q6M0Z5 Q6M101 Q6M102 Q6M115 Q6M116 Q6M125 Q6M130
Q6M132 Q6M133 Q6M139 Q6M146 Q6M154 Q6M155 Q6M156 Q6M157 Q6M160 Q6M161 Q6M174
Q6M176 Q6M181 Q6M183 Q6M191 Q6M197 Q6M1A3 Q6M1B0 Q6M1B4 Q6M1C1 Q6M259 Q6M2E8
Q6M4B7 Q6M6E7 Q6M7C6 Q6M8X7 Q6M923 Q6M981 Q6M9L0 Q6M9M7 Q6M9Z8 Q6M9Z9 Q6MA36
Q6MA55 Q6MA75 Q6MA76 Q6MAA3 Q6MAB0 Q6MAB4 Q6MAB5 Q6MAB7 Q6MAC0 Q6MAC7 Q6MAF1
Q6MAG7 Q6MAG8 Q6MAG9 Q6MAH4 Q6MAI4 Q6MAM1 Q6MAN0 Q6MAQ1 Q6MAR9 Q6MAS0 Q6MAS1
Q6MAS9 Q6MAT1 Q6MAT3 Q6MAT4 Q6MAT5 Q6MAT6 Q6MAW8 Q6MAY1 Q6MAZ6 Q6MB06 Q6MB12
Q6MB26 Q6MB27 Q6MB28 Q6MB45 Q6MBE8 Q6MBE9 Q6MBF0 Q6MBF8 Q6MBG3 Q6MBJ7 Q6MBK1
Q6MBL0 Q6MBL8 Q6MBM7 Q6MBP0 Q6MBR6 Q6MBS4 Q6MBS8 Q6MBT2 Q6MBU0 Q6MBU9 Q6MBV4
Q6MBZ5 Q6MC00 Q6MC06 Q6MC67 Q6MC72 Q6MC93 Q6MC97 Q6MCA2 Q6MCA7 Q6MCB5 Q6MCC1
Q6MCC8 Q6MCL7 Q6MCT3 Q6MCT6 Q6MCU1 Q6MCW1 Q6MCW9 Q6MD01 Q6MD02 Q6MD07 Q6MD15
Q6MD24 Q6MD31 Q6MD44 Q6MD56 Q6MD64 Q6MD65 Q6MD66 Q6MD85 Q6MD97 Q6MDC0 Q6MDC1
Q6MDC2 Q6MDC8 Q6MDD0 Q6MDD9 Q6MDE0 Q6MDE3 Q6MDE6 Q6MDF5 Q6MDF6 Q6MDF9 Q6MDG0
Q6MDG3 Q6MDG7 Q6MDG8 Q6MDG9 Q6MDH0 Q6MDH3 Q6MDH4 Q6MDI1 Q6MDI2 Q6MDI6 Q6MDJ3
Q6MDK2 Q6MDK5 Q6MDK6 Q6MDK7 Q6MDL5 Q6MDM0 Q6MDM1 Q6MDM2 Q6MDM3 Q6MDM4 Q6MDM5
Q6MDN0 Q6MDN1 Q6MDY1 Q6MDZ8 Q6ME07 Q6ME08 Q6ME13 Q6ME16 Q6ME32 Q6ME41 Q6ME42
Q6ME43 Q6ME44 Q6ME46 Q6ME47 Q6ME48 Q6ME49 Q6ME50 Q6ME51 Q6ME52 Q6ME53 Q6ME54
Q6ME55 Q6ME56 Q6ME57 Q6ME58 Q6ME59 Q6ME60 Q6ME63 Q6ME71 Q6ME80 Q6ME90 Q6ME91
Q6ME97 Q6MEA6 Q6MEA7 Q6MEA9 Q6MEB9 Q6MEC8 Q6MEC9 Q6MED0 Q6MED6 Q6MEE7 Q6MEE8
Q6MEF6 Q6MEG4 Q6MEG8 Q6MEJ2 Q6MEK1 Q6MEK7 Q6MEL5 Q6MEM7 Q6MEN1 Q6MEN7 Q6MEN8
Q6MEP0 Q6MEP4 Q6MEP6 Q6MEQ3 Q6MEQ6 Q6MEQ7 Q6MEQ8 Q6MER6 Q6MER7 Q6MER8 Q6MER9
Q6MEV1 Q6MEW4 Q6MEX3 Q6MEY8 Q6MEY9 Q6MEZ3 Q6MF40 Q6MF52 Q6MF95 Q6MFA3 Q6MFA5
Q6MFA6 Q6MFC0 Q6MFP4 Q6MFS1 Q6MFY5 Q6MFY6 Q6MFY8 Q6MFZ5 Q6MG06 Q6MG07 Q6MG08
Q6MG12 Q6MG21 Q6MG48 Q6MG51 Q6MG60 Q6MG62 Q6MG81 Q6MGL5 Q6MGL6 Q6MGL7 Q6MGP8
Q6MGQ3 Q6MGR8 Q6MGS3 Q6MGS4 Q6MGS5 Q6MGS7 Q6MGT1 Q6MGU3 Q6MGU4 Q6MGW1 Q6MGX6
Q6MGY5 Q6MGY6 Q6MH01 Q6MH10 Q6MH11 Q6MH12 Q6MH30 Q6MHH9 Q6MHI0 Q6MHI2 Q6MHI3
Q6MHJ5 Q6MHP7 Q6MHQ3 Q6MHQ6 Q6MHT3 Q6MHT9 Q6MHW4 Q6MHW5 Q6MI44 Q6MI59 Q6MI74
Q6MID2 Q6MIF8 Q6MII5 Q6MIK8 Q6MIP0 Q6MIV0 Q6MIW2 Q6MJ00 Q6MJ04 Q6MJ05 Q6MJ06
Q6MJ09 Q6MJ10 Q6MJ11 Q6MJ12 Q6MJ13 Q6MJ14 Q6MJ15 Q6MJ16 Q6MJ17 Q6MJ18 Q6MJ19
Q6MJ20 Q6MJ21 Q6MJ22 Q6MJ23 Q6MJ24 Q6MJ25 Q6MJ26 Q6MJ27 Q6MJ28 Q6MJ29 Q6MJ30
Q6MJ31 Q6MJ33 Q6MJ34 Q6MJ35 Q6MJ36 Q6MJ37 Q6MJE4 Q6MJE9 Q6MJH6 Q6MJJ2 Q6MJL7
Q6MJM2 Q6MJP3 Q6MJP9 Q6MJQ2 Q6MJR3 Q6MJR4 Q6MJR8 Q6MJR9 Q6MJY0 Q6MK42 Q6MK43
Q6MK45 Q6MK56 Q6MKB4 Q6MKB6 Q6MKB7 Q6MKE8 Q6MKE9 Q6MKR6 Q6MKR7 Q6MKS2 Q6MKW4
Q6MKX0 Q6MKZ5 Q6ML19 Q6ML45 Q6ML73 Q6ML86 Q6ML96 Q6ML97 Q6ML99 Q6MLA0 Q6MLC6
Q6MLD2 Q6MLE7 Q6MLF5 Q6MLI2 Q6MLJ4 Q6MLK1 Q6MLR5 Q6MLR7 Q6MLS4 Q6MLS8 Q6MM37
Q6MMA3 Q6MMJ0 Q6MMJ1 Q6MMJ2 Q6MMJ3 Q6MMK2 Q6MMK3 Q6MMK4 Q6MMK8 Q6MML0 Q6MMQ7

Q6MMQ9 Q6MMS2 Q6MMS3 Q6MMS4 Q6MMS5 Q6MMS6 Q6MMS8 Q6MMY3 Q6MMZ9 Q6MN06 Q6MN40
Q6MNC0 Q6MNF1 Q6MNF8 Q6MNG0 Q6MNG7 Q6MNN2 Q6MNP6 Q6MNQ2 Q6MNT9 Q6MNX8 Q6MNX9
Q6MNY2 Q6MP09 Q6MP18 Q6MP28 Q6MP77 Q6MPB7 Q6MPB8 Q6MPK2 Q6MPK7 Q6MPP0 Q6MPP2
Q6MPS6 Q6MPV7 Q6MPZ6 Q6MQ03 Q6MQ46 Q6MQ47 Q6MQ48 Q6MQ60 Q6MQ94 Q6MQB7 Q6MQC7
Q6MQD1 Q6MQH5 Q6MQH6 Q6MQJ7 Q6MQK8 Q6MQS6 Q6MQU0 Q6MQU3 Q6MQW5 Q6MQY9 Q6MR60
Q6MR96 Q6MRI1 Q6MRJ2 Q6MRK1 Q6MRK7 Q6MRK9 Q6MRL0 Q6MRL7 Q6MRL9 Q6MRM6 Q6MRM8
Q6MRM9 Q6MRP1 Q6MRP3 Q6MRS1 Q6MRS2 Q6MRS3 Q6MRU5 Q6MRX5 Q6MRX6 Q6MRX7 Q6MRX8
Q6MRY7 Q6MS23 Q6MS27 Q6MS28 Q6MS38 Q6MS67 Q6MS85 Q6MS86 Q6MSC1 Q6MSC4 Q6MSC7
Q6MSE7 Q6MSF0 Q6MSF2 Q6MSF4 Q6MSK8 Q6MSL3 Q6MSM5 Q6MSM6 Q6MSM7 Q6MSM8 Q6MSM9
Q6MSN0 Q6MSN1 Q6MSN2 Q6MSN4 Q6MSN5 Q6MSN6 Q6MSN8 Q6MSN9 Q6MSP0 Q6MSP1 Q6MSP2
Q6MSP3 Q6MSP5 Q6MSP7 Q6MSP8 Q6MSP9 Q6MSQ0 Q6MSQ4 Q6MSQ6 Q6MSR0 Q6MSR6 Q6MSR8
Q6MST3 Q6MT00 Q6MT04 Q6MT05 Q6MT06 Q6MT07 Q6MT14 Q6MT17 Q6MT18 Q6MT20 Q6MT21
Q6MT28 Q6MT48 Q6MT50 Q6MT55 Q6MT68 Q6MTA3 Q6MTB4 Q6MTB6 Q6MTC5 Q6MTC9 Q6MTD4
Q6MTE4 Q6MTE5 Q6MTE7 Q6MTF4 Q6MTF5 Q6MTF7 Q6MTF8 Q6MTG1 Q6MTG9 Q6MTH8 Q6MTI0
Q6MTI1 Q6MTI4 Q6MTJ6 Q6MTJ7 Q6MTM6 Q6MTP6 Q6MTQ0 Q6MTQ2 Q6MTR0 Q6MTR3 Q6MTR9
Q6MTU2 Q6MTU8 Q6MTU9 Q6MTV5 Q6MTW7 Q6MTX3 Q6MTX5 Q6MTY7 Q6MTZ0 Q6MTZ7 Q6MU20
Q6MU21 Q6MU22 Q6MU25 Q6MU75 Q6MU81 Q6MU82 Q6MU88 Q6MU89 Q6MUA3 Q6MUA6 Q6MUC7
Q6MUD1 Q6MUE6 Q6MUE7 Q6MUF2 Q6MUG2 Q6MUG5 Q6MUI1 Q6MUI5 Q6MUI9 Q6MUK2 Q6MUK4
Q6MUK6 Q6MUM4 Q6MUM5 Q6MUM7 Q6MVH7 Q6MVL2 Q6MVL6 Q6MW04 Q6MW50 Q6MW51 Q6MWS8
Q6MWV1 Q6MWY2 Q6MWZ7 Q6MX01 Q6MX07 Q6MX40 Q6MX50 Q6MX60 Q6MY48 Q6MY57 Q6MY91
Q6MYX3 Q6MZN7 Q6MZZ7 Q6N063 Q6N069 Q6N0C7 Q6N0J0 Q6N0J2 Q6N0K6 Q6N0L9 Q6N0N8
Q6N0T9 Q6N0W5 Q6N0W9 Q6N103 Q6N140 Q6N143 Q6N176 Q6N178 Q6N181 Q6N1B4 Q6N1B7
Q6N1G8 Q6N1H2 Q6N1I5 Q6N1I7 Q6N1M8 Q6N1N4 Q6N1P8 Q6N1P9 Q6N1S9 Q6N1T6 Q6N1V9
Q6N1Y1 Q6N1Y2 Q6N1Y3 Q6N223 Q6N233 Q6N234 Q6N239 Q6N253 Q6N260 Q6N274 Q6N2A3
Q6N2H8 Q6N2N5 Q6N2R5 Q6N2U4 Q6N344 Q6N376 Q6N3F1 Q6N3F2 Q6N3F3 Q6N3G0 Q6N3I8
Q6N3M9 Q6N3N0 Q6N3N1 Q6N3N3 Q6N3N5 Q6N3P9 Q6N3T8 Q6N3W7 Q6N3Y0 Q6N409 Q6N423
Q6N447 Q6N453 Q6N461 Q6N4D3 Q6N4E7 Q6N4Q4 Q6N4R5 Q6N4R7 Q6N4R8 Q6N4R9 Q6N4S0
Q6N4T2 Q6N4T3 Q6N4T4 Q6N4T6 Q6N4T7 Q6N4T8 Q6N4T9 Q6N4U0 Q6N4U1 Q6N4U2 Q6N4U3
Q6N4U4 Q6N4U5 Q6N4U6 Q6N4U7 Q6N4U8 Q6N4U9 Q6N4V0 Q6N4V1 Q6N4V2 Q6N4V5 Q6N4V6
Q6N4V7 Q6N4V8 Q6N509 Q6N510 Q6N514 Q6N528 Q6N535 Q6N554 Q6N565 Q6N569 Q6N571
Q6N575 Q6N583 Q6N586 Q6N5A1 Q6N5A3 Q6N5A4 Q6N5B4 Q6N5B9 Q6N5C3 Q6N5C8 Q6N5D5
Q6N5D6 Q6N5K3 Q6N5L2 Q6N5L3 Q6N5L4 Q6N5P6 Q6N5Q2 Q6N5Q3 Q6N5Q6 Q6N5R5 Q6N5T0
Q6N5T4 Q6N5U5 Q6N5W6 Q6N5X5 Q6N5X7 Q6N5Z1 Q6N608 Q6N622 Q6N643 Q6N650 Q6N651
Q6N676 Q6N688 Q6N692 Q6N693 Q6N6A7 Q6N6B8 Q6N6C1 Q6N6C5 Q6N6C9 Q6N6D6 Q6N6E2
Q6N6K3 Q6N6L0 Q6N6L4 Q6N6L8 Q6N6M5 Q6N6M8 Q6N6N1 Q6N6U0 Q6N6V1 Q6N702 Q6N703
Q6N726 Q6N734 Q6N765 Q6N7A6 Q6N7Q9 Q6N7R5 Q6N7T7 Q6N7U8 Q6N858 Q6N859 Q6N869
Q6N874 Q6N8A7 Q6N8C3 Q6N8F3 Q6N8F4 Q6N8F5 Q6N8F6 Q6N8F8 Q6N8N2 Q6N8V0 Q6N976
Q6N9F2 Q6N9G0 Q6N9J0 Q6N9J7 Q6N9K0 Q6N9K2 Q6N9K3 Q6N9S5 Q6NA55 Q6NA79 Q6NAI2
Q6NAI4 Q6NAI9 Q6NAL3 Q6NAM1 Q6NAQ6 Q6NAT0 Q6NAT1 Q6NAZ1 Q6NAZ6 Q6NB57 Q6NB75
Q6NB76 Q6NB80 Q6NBB0 Q6NBB5 Q6NBB9 Q6NBF3 Q6NBY8 Q6NC39 Q6NC42 Q6NC44 Q6NC47
Q6NC49 Q6NC50 Q6NC51 Q6NC55 Q6NC56 Q6NC62 Q6NC66 Q6NC69 Q6NCE6 Q6NCF3 Q6NCG8
Q6NCG9 Q6NCH9 Q6NCM4 Q6NCM6 Q6NCM9 Q6NCN2 Q6NCN5 Q6NCN6 Q6NCN7 Q6NCN8 Q6NCP1
Q6NCQ6 Q6NCS3 Q6NCS7 Q6NCT7 Q6NCT8 Q6NCU3 Q6NCX0 Q6NCX7 Q6NCX8 Q6NCY0 Q6NCY3
Q6NCY4 Q6NCY6 Q6NCY7 Q6NCY8 Q6NCY9 Q6NCZ4 Q6NCZ5 Q6NCZ6 Q6ND01 Q6ND03 Q6ND07

Q6ND09 Q6ND10 Q6ND11 Q6ND14 Q6ND15 Q6ND16 Q6ND22 Q6ND40 Q6ND47 Q6ND54 Q6ND59
Q6ND66 Q6ND67 Q6ND68 Q6ND69 Q6ND74 Q6ND82 Q6NDB7 Q6NDE1 Q6NDE4 Q6NDE5 Q6NDE6
Q6NDE9 Q6NDF0 Q6NDF6 Q6NDI4 Q6NDM1 Q6NDM2 Q6NDN5 Q6NDN6 Q6NDN7 Q6NDN9 Q6NDP1
Q6NDP2 Q6NDP4 Q6NDR5 Q6NDR6 Q6NDR7 Q6NDR9 Q6NDV3 Q6NE95 Q6NE96 Q6NE98 Q6NEB5
Q6NEB7 Q6NEB8 Q6NEC1 Q6NEC9 Q6NEF5 Q6NEH3 Q6NEI3 Q6NEI5 Q6NEL2 Q6NEN2 Q6NET4
Q6NET5 Q6NEW7 Q6NEY9 Q6NEZ0 Q6NEZ1 Q6NF05 Q6NF38 Q6NF47 Q6NF59 Q6NF74 Q6NF90
Q6NFC1 Q6NFC2 Q6NFC3 Q6NFG4 Q6NFH6 Q6NFH9 Q6NFK1 Q6NFL5 Q6NFN0 Q6NFN4 Q6NFN6
Q6NFN7 Q6NFU3 Q6NFU4 Q6NFU5 Q6NFU7 Q6NFV0 Q6NFV3 Q6NFV5 Q6NFV6 Q6NFV7 Q6NFV9
Q6NFW0 Q6NFX0 Q6NG13 Q6NG14 Q6NG17 Q6NG19 Q6NG30 Q6NG33 Q6NG70 Q6NG87 Q6NG90
Q6NGC0 Q6NGC6 Q6NGD7 Q6NGD8 Q6NGH3 Q6NGH4 Q6NGH6 Q6NGI4 Q6NGI5 Q6NGI8 Q6NGJ8
Q6NGK0 Q6NGK5 Q6NGK6 Q6NGK9 Q6NGL1 Q6NGL3 Q6NGL6 Q6NGL9 Q6NGM7 Q6NGM9 Q6NGN2
Q6NGN3 Q6NGP0 Q6NGP1 Q6NGP2 Q6NGP3 Q6NGP4 Q6NGQ9 Q6NGR0 Q6NGR5 Q6NGS9 Q6NGT4
Q6NGV2 Q6NGV3 Q6NGW2 Q6NGW9 Q6NGX3 Q6NGX4 Q6NGY0 Q6NGZ4 Q6NGZ8 Q6NH02 Q6NH03
Q6NH04 Q6NH05 Q6NH07 Q6NH08 Q6NH12 Q6NH13 Q6NH15 Q6NH23 Q6NH31 Q6NH32 Q6NH34
Q6NH37 Q6NH60 Q6NH62 Q6NH63 Q6NH66 Q6NH81 Q6NH87 Q6NH92 Q6NH95 Q6NHD4 Q6NHE2
Q6NHE6 Q6NHF7 Q6NHG2 Q6NHG4 Q6NHG5 Q6NHG6 Q6NHG8 Q6NHH0 Q6NHH1 Q6NHH2 Q6NHH5
Q6NHH6 Q6NHI3 Q6NHI5 Q6NHJ8 Q6NHK6 Q6NHK9 Q6NHL0 Q6NHL3 Q6NHM1 Q6NHM7 Q6NHN2
Q6NHN6 Q6NHQ1 Q6NHQ2 Q6NHQ4 Q6NHQ7 Q6NHR6 Q6NHS6 Q6NHU0 Q6NHU3 Q6NHU6 Q6NHY8
Q6NI09 Q6NI36 Q6NI37 Q6NI38 Q6NI47 Q6NI48 Q6NI56 Q6NI77 Q6NI78 Q6NIA1 Q6NIA2
Q6NIA5 Q6NIC4 Q6NIC5 Q6NIC6 Q6NIC7 Q6NIC8 Q6NIC9 Q6NID8 Q6NIE5 Q6NIF2 Q6NIL8
Q6NIM2 Q6NIW2 Q6NIX4 Q6NIZ3 Q6NIZ6 Q6NJ04 Q6NJ37 Q6NJ38 Q6NJ39 Q6NJ50 Q6NJ52
Q6NJ62 Q6NJ63 Q6NJ64 Q6NJ65 Q6NJ66 Q6NJ71 Q6NJ83 Q6NJ84 Q6NJ85 Q6NJ86 Q6NJ87
Q6NJ88 Q6NJ91 Q6NJC1 Q6NJC2 Q6NJC6 Q6NJC7 Q6NJC8 Q6NJC9 Q6NJD0 Q6NJD1 Q6NJD2
Q6NJD3 Q6NJD5 Q6NJD6 Q6NJD7 Q6NJD8 Q6NJF6 Q6NJG5 Q6NJG6 Q6NJG9 Q6NJH8 Q6NJJ2
Q6NJJ4 Q6NJK0 Q6NJK1 Q6NJL2 Q6NJL3 Q6NJX0 Q6NJY0 Q6NJY1 Q6NJY9 Q6NK10 Q6NKC7
Q6NKI3 Q6NKI6 Q6NKL5 Q6NKL7 Q6NKN8 Q6NKN9 Q6NKS1 Q6NKW7 Q6NKX1 Q6NKX3 Q6NKX5
Q6NL02 Q6NLB0 Q6NLB1 Q6NLC8 Q6NLE0 Q6NLG3 Q6NLG7 Q6NLH0 Q6NLL1 Q6NLP2 Q6NLQ7
Q6NLQ8 Q6NLS8 Q6NLU2 Q6NLU6 Q6NLV0 Q6NLV4 Q6NLW5 Q6NLY8 Q6NM26 Q6NMB0 Q6NMC6
Q6NME7 Q6NMK6 Q6NML5 Q6NMR8 Q6NN02 Q6NN40 Q6NN85 Q6NNI8 Q6NNL9 Q6NNM0 Q6NPC9
Q6NPD7 Q6NPF4 Q6NPF9 Q6NPG7 Q6NPL9 Q6NPM8 Q6NPN3 Q6NPN5 Q6NPR6 Q6NPS8 Q6NPT8
Q6NQ54 Q6NQ64 Q6NQ72 Q6NQ79 Q6NQ81 Q6NQ83 Q6NQ88 Q6NQD9 Q6NQH9 Q6NQJ7 Q6NQJ8
Q6NQL4 Q6NQP5 Q6NR09 Q6NRB0 Q6NRB5 Q6NRC7 Q6NRC9 Q6NRD3 Q6NRD5 Q6NRE4 Q6NRE8
Q6NRF1 Q6NRF4 Q6NRF9 Q6NRG0 Q6NRG5 Q6NRG6 Q6NRH0 Q6NRH3 Q6NRH7 Q6NRI0 Q6NRI5
Q6NRI8 Q6NRJ5 Q6NRJ7 Q6NRK1 Q6NRK3 Q6NRK9 Q6NRL0 Q6NRL1 Q6NRM0 Q6NRM6 Q6NRM8
Q6NRP2 Q6NRP6 Q6NRQ7 Q6NRT0 Q6NRT3 Q6NRT5 Q6NRV0 Q6NRV8 Q6NRW0 Q6NRW2 Q6NRW3
Q6NRW5 Q6NRX3 Q6NRY2 Q6NRY9 Q6NRZ0 Q6NS15 Q6NS21 Q6NS23 Q6NS38 Q6NS45 Q6NS46
Q6NS57 Q6NS60 Q6NSI1 Q6NSI4 Q6NSI8 Q6NSJ2 Q6NSN1 Q6NSN5 Q6NSQ7 Q6NSR8 Q6NSV7
Q6NSW3 Q6NSW5 Q6NSZ9 Q6NT04 Q6NT46 Q6NT89 Q6NT99 Q6NTA4 Q6NTF7 Q6NTJ3 Q6NTM6
Q6NTP2 Q6NTR1 Q6NTS3 Q6NTS7 Q6NTV6 Q6NTV8 Q6NTV9 Q6NTW1 Q6NTW6 Q6NTY6 Q6NTY8
Q6NU04 Q6NU05 Q6NU07 Q6NU08 Q6NU12 Q6NU13 Q6NU14 Q6NU18 Q6NU19 Q6NU21 Q6NU27
Q6NU28 Q6NU29 Q6NU40 Q6NU47 Q6NU53 Q6NU56 Q6NU60 Q6NU94 Q6NU95 Q6NU98 Q6NUA0
Q6NUA1 Q6NUB2 Q6NUC1 Q6NUC2 Q6NUC6 Q6NUD0 Q6NUD4 Q6NUD7 Q6NUF4 Q6NUH0 Q6NUI1
Q6NUJ5 Q6NUM6 Q6NUN7 Q6NUN9 Q6NUP7 Q6NUR6 Q6NUW9 Q6NV18 Q6NV26 Q6NV31 Q6NV32
Q6NV66 Q6NV72 Q6NV74 Q6NV83 Q6NVB0 Q6NVE3 Q6NVE9 Q6NVF9 Q6NVH7 Q6NVI7 Q6NVJ5

Q6NVK9 Q6NVL5 Q6NVL6 Q6NVM0 Q6NVM2 Q6NVM6 Q6NVM8 Q6NVN0 Q6NVN3 Q6NVP7 Q6NVR1
Q6NVR5 Q6NVR8 Q6NVS5 Q6NVT7 Q6NVU6 Q6NVV0 Q6NVV1 Q6NVV7 Q6NVY1 Q6NW52 Q6NW59
Q6NW85 Q6NW95 Q6NW96 Q6NWA8 Q6NWC6 Q6NWE0 Q6NWE1 Q6NWF6 Q6NWG4 Q6NWH0 Q6NWH1
Q6NWI1 Q6NWI4 Q6NWJ0 Q6NWJ4 Q6NWL4 Q6NWX7 Q6NWZ9 Q6NX08 Q6NX30 Q6NX31 Q6NX45
Q6NX49 Q6NXA4 Q6NXA8 Q6NXA9 Q6NXE6 Q6NXG1 Q6NXH8 Q6NXH9 Q6NXI6 Q6NXI8 Q6NXK2
Q6NXK5 Q6NXL6 Q6NXM2 Q6NXN1 Q6NXP0 Q6NXP2 Q6NXP6 Q6NXR4 Q6NXT2 Q6NXW6 Q6NXY1
Q6NXY9 Q6NXZ1 Q6NY15 Q6NY19 Q6NY34 Q6NY64 Q6NY74 Q6NY77 Q6NY82 Q6NY89 Q6NY98
Q6NYA0 Q6NYB4 Q6NYD4 Q6NYD7 Q6NYF0 Q6NYG6 Q6NYG8 Q6NYH1 Q6NYL5 Q6NYM1 Q6NYP8
Q6NYR8 Q6NYT1 Q6NYT3 Q6NYU2 Q6NYU6 Q6NYU7 Q6NYV9 Q6NYX8 Q6NYY4 Q6NZ03 Q6NZ04
Q6NZ06 Q6NZ18 Q6NZ22 Q6NZ36 Q6NZA9 Q6NZE7 Q6NZF1 Q6NZG4 Q6NZH6 Q6NZJ6 Q6NZK5
Q6NZK8 Q6NZL6 Q6NZN0 Q6NZN1 Q6NZP1 Q6NZP2 Q6NZQ0 Q6NZQ6 Q6NZR2 Q6NZS4 Q6NZT2
Q6NZT6 Q6NZW0 Q6NZZ9 Q6P047 Q6P050 Q6P087 Q6P0B1 Q6P0D0 Q6P0D5 Q6P0D9 Q6P0E1
Q6P0E7 Q6P0F4 Q6P0F9 Q6P0G6 Q6P0G8 Q6P0H6 Q6P0I6 Q6P0I8 Q6P0U0 Q6P0U3 Q6P0X2
Q6P104 Q6P116 Q6P132 Q6P1B1 Q6P1C1 Q6P1D7 Q6P1E7 Q6P1E8 Q6P1F6 Q6P1G0 Q6P1G2
Q6P1G6 Q6P1K8 Q6P1L6 Q6P1M9 Q6P1N9 Q6P1R4 Q6P1S4 Q6P1S6 Q6P1U0 Q6P1U3 Q6P1U7
Q6P1V4 Q6P1V8 Q6P1W5 Q6P1X6 Q6P1Y1 Q6P1Z5 Q6P205 Q6P255 Q6P256 Q6P268 Q6P280
Q6P286 Q6P298 Q6P2A1 Q6P2C0 Q6P2C8 Q6P2D0 Q6P2D8 Q6P2I3 Q6P2K3 Q6P2K6 Q6P2L6
Q6P2M8 Q6P2P2 Q6P2S7 Q6P2T7 Q6P2U9 Q6P2W5 Q6P2Y1 Q6P2Y2 Q6P2Y3 Q6P2Z0 Q6P2Z3
Q6P2Z6 Q6P315 Q6P320 Q6P381 Q6P387 Q6P3B2 Q6P3B9 Q6P3D7 Q6P3E7 Q6P3G4 Q6P3H4
Q6P3H7 Q6P3K3 Q6P3N6 Q6P3P1 Q6P3P4 Q6P3R1 Q6P3R8 Q6P3V2 Q6P3V7 Q6P3W2 Q6P3W6
Q6P3X3 Q6P3X8 Q6P3Y5 Q6P3Z3 Q6P423 Q6P435 Q6P4D5 Q6P4F2 Q6P4I2 Q6P4J6 Q6P4J8
Q6P4K1 Q6P4K5 Q6P4K6 Q6P4K7 Q6P4K8 Q6P4L6 Q6P4L9 Q6P4T2 Q6P4W0 Q6P4W5 Q6P4W8
Q6P4Y1 Q6P4Z6 Q6P4Z9 Q6P502 Q6P539 Q6P542 Q6P560 Q6P566 Q6P575 Q6P582 Q6P587
Q6P589 Q6P5C5 Q6P5C7 Q6P5D3 Q6P5D8 Q6P5G3 Q6P5G6 Q6P5H2 Q6P5H6 Q6P5J6 Q6P5L7
Q6P5L8 Q6P5M2 Q6P5P3 Q6P5Q4 Q6P5U7 Q6P5U8 Q6P5X5 Q6P616 Q6P618 Q6P632 Q6P635
Q6P637 Q6P640 Q6P642 Q6P643 Q6P656 Q6P6B1 Q6P6E6 Q6P6G7 Q6P6J0 Q6P6J4 Q6P6M7
Q6P6P7 Q6P6Q6 Q6P6T4 Q6P6Z0 Q6P6Z4 Q6P6Z8 Q6P747 Q6P773 Q6P788 Q6P798 Q6P7A2
Q6P7B0 Q6P7D3 Q6P7E4 Q6P7G6 Q6P7H4 Q6P7I0 Q6P7I6 Q6P7K0 Q6P7L0 Q6P7L9 Q6P7N1
Q6P7N4 Q6P7P5 Q6P7Q0 Q6P7Q1 Q6P7W0 Q6P7W5 Q6P7Y3 Q6P809 Q6P819 Q6P823 Q6P824
Q6P833 Q6P839 Q6P848 Q6P859 Q6P861 Q6P864 Q6P870 Q6P875 Q6P878 Q6P881 Q6P882
Q6P886 Q6P8A1 Q6P8A3 Q6P8A7 Q6P8B3 Q6P8C4 Q6P8D9 Q6P8E9 Q6P8G0 Q6P8G1 Q6P8G3
Q6P8J2 Q6P8M1 Q6P8W9 Q6P8Y0 Q6P8Y1 Q6P8Y7 Q6P8Z1 Q6P902 Q6P926 Q6P949 Q6P950
Q6P963 Q6P964 Q6P9A1 Q6P9A3 Q6P9F5 Q6P9G0 Q6P9G9 Q6P9H5 Q6P9I7 Q6P9J5 Q6P9L6
Q6P9P6 Q6P9P8 Q6P9R1 Q6P9S1 Q6P9U1 Q6P9U3 Q6P9U4 Q6P9U8 Q6P9Y7 Q6P9Z4 Q6P9Z8
Q6PA14 Q6PA15 Q6PA41 Q6PA43 Q6PA48 Q6PA69 Q6PA72 Q6PA79 Q6PA97 Q6PAB3 Q6PAC3
Q6PAC9 Q6PAD2 Q6PAE6 Q6PAF4 Q6PAF6 Q6PAJ3 Q6PAL7 Q6PAM1 Q6PAN7 Q6PAQ4 Q6PAR0
Q6PAT0 Q6PAV2 Q6PAV5 Q6PAV8 Q6PAW0 Q6PAY6 Q6PAY8 Q6PB04 Q6PB06 Q6PB19 Q6PB51
Q6PB60 Q6PB97 Q6PBA2 Q6PBA8 Q6PBC1 Q6PBC3 Q6PBC9 Q6PBD4 Q6PBD6 Q6PBE2 Q6PBF6
Q6PBI2 Q6PBI4 Q6PBJ2 Q6PBK1 Q6PBM1 Q6PBM7 Q6PBM8 Q6PBP3 Q6PBR7 Q6PBT5 Q6PBT9
Q6PBU0 Q6PBU5 Q6PBV4 Q6PBV6 Q6PBY3 Q6PBY7 Q6PC19 Q6PC30 Q6PC40 Q6PC45 Q6PC55
Q6PC60 Q6PC62 Q6PC91 Q6PCD5 Q6PCF7 Q6PCG0 Q6PCG7 Q6PCI6 Q6PCI7 Q6PCJ1 Q6PCL0
Q6PCM1 Q6PCM2 Q6PCN7 Q6PCR6 Q6PCR7 Q6PCT3 Q6PCT5 Q6PCW6 Q6PD05 Q6PD10 Q6PD29
Q6PD83 Q6PDB4 Q6PDC0 Q6PDH4 Q6PDJ6 Q6PDK2 Q6PDM4 Q6PDQ2 Q6PDU1 Q6PDW6 Q6PDX6
Q6PDY0 Q6PDY2 Q6PDZ2 Q6PE01 Q6PE15 Q6PE25 Q6PE54 Q6PE65 Q6PE87 Q6PEB4 Q6PEB9

Q6PEC0 Q6PEC1 Q6PEE2 Q6PEE3 Q6PEH4 Q6PEH5 Q6PEH8 Q6PEI7 Q6PEM6 Q6PER3 Q6PEV3
Q6PEV8 Q6PEW1 Q6PEX3 Q6PF04 Q6PF05 Q6PF15 Q6PF18 Q6PF21 Q6PF24 Q6PF35 Q6PF39
Q6PF42 Q6PFE3 Q6PFG8 Q6PFH3 Q6PFJ1 Q6PFJ9 Q6PFK1 Q6PFL0 Q6PFL6 Q6PFM9 Q6PFP3
Q6PFP6 Q6PFQ2 Q6PFS2 Q6PFX2 Q6PFX7 Q6PFX8 Q6PFY8 Q6PG04 Q6PG16 Q6PG31 Q6PG37
Q6PG95 Q6PGA0 Q6PGB6 Q6PGC1 Q6PGE4 Q6PGF2 Q6PGF3 Q6PGG6 Q6PGH0 Q6PGH4 Q6PGP7
Q6PGQ1 Q6PGT0 Q6PGU2 Q6PGY5 Q6PGZ3 Q6PH11 Q6PH18 Q6PH37 Q6PH52 Q6PH85 Q6PH90
Q6PHE8 Q6PHG4 Q6PHG8 Q6PHK6 Q6PHN1 Q6PHR2 Q6PI20 Q6PI26 Q6PI47 Q6PI48 Q6PI53
Q6PI77 Q6PI79 Q6PI97 Q6PI98 Q6PID8 Q6PIF2 Q6PIF6 Q6PII3 Q6PII5 Q6PIJ4 Q6PIP5
Q6PIU9 Q6PIV2 Q6PIY5 Q6PJ21 Q6PJ69 Q6PJN8 Q6PJP8 Q6PJQ5 Q6PK57 Q6PK81 Q6PKU1
Q6PL11 Q6PL24 Q6PL31 Q6PLS1 Q6PMI5 Q6PNC0 Q6PP78 Q6PP79 Q6PQJ9 Q6PQQ4 Q6PR31
Q6PR32 Q6PR33 Q6PRD7 Q6PS57 Q6PSB9 Q6PSC4 Q6PSC6 Q6PSE2 Q6PTT0 Q6PUA2 Q6PUF8
Q6PUR6 Q6PUR7 Q6PUV5 Q6PV61 Q6PVZ1 Q6PW23 Q6PWT7 Q6PWU2 Q6PY58 Q6PZ02 Q6Q0M6
Q6Q0N1 Q6Q0N2 Q6Q0N3 Q6Q137 Q6Q151 Q6Q152 Q6Q271 Q6Q272 Q6Q2A8 Q6Q2B2 Q6Q305
Q6Q311 Q6Q3H0 Q6Q3H3 Q6Q4D0 Q6Q546 Q6Q547 Q6Q560 Q6Q571 Q6Q572 Q6Q5F3 Q6Q5H1
Q6Q5K6 Q6Q5P6 Q6Q6R3 Q6Q6R5 Q6Q6X0 Q6Q759 Q6Q783 Q6Q795 Q6Q7J5 Q6Q7X3 Q6Q7X9
Q6Q870 Q6Q873 Q6Q874 Q6Q875 Q6Q877 Q6Q878 Q6Q880 Q6Q882 Q6Q883 Q6Q886 Q6Q887
Q6Q888 Q6Q8A5 Q6Q9H6 Q6Q9I2 Q6QA27 Q6QB00 Q6QDC9 Q6QEF8 Q6QEJ7 Q6QGC0 Q6QGD2
Q6QGD3 Q6QGD4 Q6QGD5 Q6QGD6 Q6QGD7 Q6QGD8 Q6QGD9 Q6QGE0 Q6QGE1 Q6QGE2 Q6QGE3
Q6QGE6 Q6QGE8 Q6QGE9 Q6QGF0 Q6QGG9 Q6QGH9 Q6QGJ4 Q6QGJ5 Q6QGL2 Q6QGM3 Q6QGM6
Q6QGP4 Q6QGS2 Q6QGT3 Q6QGW5 Q6QHD9 Q6QHE0 Q6QHK0 Q6QHK4 Q6QI06 Q6QI44 Q6QI89
Q6QIA5 Q6QIB0 Q6QIB5 Q6QJ72 Q6QLN1 Q6QM28 Q6QME8 Q6QMY5 Q6QMZ4 Q6QMZ7 Q6QN04
Q6QN05 Q6QNF8 Q6QNL9 Q6QNM1 Q6QPJ6 Q6QPM2 Q6QUK9 Q6QUL7 Q6QUW1 Q6QUY7 Q6QUY8
Q6QUY9 Q6QUZ0 Q6QVU4 Q6QW47 Q6QWR1 Q6QZN6 Q6QZQ4 Q6R053 Q6R0C4 Q6R0E3 Q6R0G4
Q6R0H0 Q6R0H1 Q6R1L1 Q6R2V6 Q6R2W3 Q6R311 Q6R3M4 Q6R3Q6 Q6R4B4 Q6R4R6 Q6R4R7
Q6R4R8 Q6R4R9 Q6R4S3 Q6R4S6 Q6R518 Q6R645 Q6R648 Q6R649 Q6R650 Q6R795 Q6R796
Q6R797 Q6R798 Q6R7A0 Q6R7A1 Q6R7A2 Q6R7A3 Q6R7B1 Q6R7B2 Q6R7B3 Q6R7B4 Q6R7B5
Q6R7B6 Q6R7B7 Q6R7B8 Q6R7C0 Q6R7C1 Q6R7C2 Q6R7C3 Q6R7C4 Q6R7C5 Q6R7C7 Q6R7C8
Q6R7C9 Q6R7D0 Q6R7D1 Q6R7D2 Q6R7D3 Q6R7D4 Q6R7D5 Q6R7D6 Q6R7D7 Q6R7D8 Q6R7D9
Q6R7E0 Q6R7E2 Q6R7E3 Q6R7E4 Q6R7E7 Q6R7E8 Q6R7F0 Q6R7F1 Q6R7F3 Q6R7F4 Q6R7F7
Q6R7F8 Q6R7F9 Q6R7G1 Q6R7G3 Q6R7G4 Q6R7G5 Q6R7G7 Q6R7G9 Q6R7H0 Q6R7H2 Q6R7H3
Q6R7H4 Q6R7H5 Q6R7H6 Q6R7H7 Q6R7H8 Q6R7H9 Q6R7I0 Q6R7I1 Q6R7I2 Q6R7I4 Q6R7I6
Q6R7I7 Q6R7I9 Q6R7J0 Q6R7J1 Q6R7J2 Q6R7J3 Q6R7J4 Q6R7J5 Q6R7J6 Q6R7J7 Q6R7J9
Q6R7K0 Q6R7K2 Q6R7K3 Q6R7K4 Q6R7K5 Q6R7K6 Q6R7K7 Q6R7K9 Q6R7L1 Q6R7L3 Q6R7L4
Q6R7L5 Q6R7L6 Q6R7L7 Q6RCE1 Q6REU5 Q6REU8 Q6RF66 Q6RFH8 Q6RFL5 Q6RFY2 Q6RG04
Q6RG77 Q6RH25 Q6RH26 Q6RHW0 Q6RI63 Q6RI85 Q6RJS2 Q6RSH7 Q6RSN5 Q6RSS2 Q6RUG5
Q6RUI8 Q6RUR1 Q6RUT8 Q6RVD6 Q6RVD7 Q6RWG0 Q6RWK4 Q6RX33 Q6RXC1 Q6RYS3 Q6RYT0
Q6RZB9 Q6RZD4 Q6RZD7 Q6RZE1 Q6RZF4 Q6RZF8 Q6RZF9 Q6RZG4 Q6RZJ4 Q6RZM1 Q6RZN2
Q6RZN3 Q6RZP5 Q6RZR1 Q6RZR2 Q6RZS3 Q6RZZ9 Q6S000 Q6S001 Q6S002 Q6S003 Q6S004
Q6S343 Q6S3I3 Q6S4N8 Q6S4P4 Q6S545 Q6S591 Q6S592 Q6S5H5 Q6S5I3 Q6S6N9 Q6S6P0
Q6S6P1 Q6S6P9 Q6S6R7 Q6S6R8 Q6S6S2 Q6S6S5 Q6S6S9 Q6S6T1 Q6S6T4 Q6S6T5 Q6S6U0
Q6S6U7 Q6S6V0 Q6S6W5 Q6S7B0 Q6S8J3 Q6S8J7 Q6S9C5 Q6S9E0 Q6S9V5 Q6S9V6 Q6S9V7
Q6S9V8 Q6S9V9 Q6S9Z5 Q6SEH4 Q6SEH5 Q6SJ82 Q6SJ84 Q6SJ91 Q6SJ93 Q6SJ94 Q6SJ95
Q6SJ96 Q6SJQ8 Q6SJR1 Q6SKR2 Q6SKR5 Q6SP92 Q6SQH4 Q6SR68 Q6SR69 Q6SSJ3 Q6SSJ6
Q6STH5 Q6STM1 Q6SVX2 Q6SW03 Q6SW04 Q6SW29 Q6SW46 Q6SW48 Q6SW50 Q6SW55 Q6SW63

Q6SW65 Q6SW66 Q6SW70 Q6SW73 Q6SW77 Q6SW79 Q6SW82 Q6SW85 Q6SW87 Q6SW89 Q6SWA3
Q6SWA4 Q6SWC5 Q6SWD3 Q6SWD5 Q6SWN9 Q6SWP7 Q6SXP0 Q6SYB9 Q6SYV5 Q6SZ65 Q6T1F5
Q6T1F6 Q6T1W6 Q6T1W7 Q6T1W8 Q6T1X6 Q6T267 Q6T308 Q6T311 Q6T367 Q6T3B0 Q6T412
Q6T486 Q6T5B7 Q6T6K9 Q6T723 Q6T724 Q6T7E8 Q6T893 Q6TA25 Q6TAS3 Q6TBX7 Q6TC27
Q6TC28 Q6TC30 Q6TC31 Q6TC32 Q6TC33 Q6TC34 Q6TC36 Q6TC37 Q6TC38 Q6TC39 Q6TC40
Q6TC41 Q6TC43 Q6TC44 Q6TC45 Q6TC46 Q6TC50 Q6TCF2 Q6TDP3 Q6TDP4 Q6TDU1 Q6TDU7
Q6TDU8 Q6TEC1 Q6TEL1 Q6TEM9 Q6TEP1 Q6TFL3 Q6TFL4 Q6TGC6 Q6TGJ8 Q6TGV7 Q6TGZ4
Q6TH01 Q6TH15 Q6TK73 Q6TKS7 Q6TLF6 Q6TLH3 Q6TM71 Q6TM72 Q6TMA3 Q6TMG4 Q6TMK2
Q6TMK3 Q6TMK8 Q6TN15 Q6TNA5 Q6TNA6 Q6TND0 Q6TND4 Q6TNU3 Q6TNU4 Q6TQE1 Q6TRW4
Q6TS43 Q6TU48 Q6TUF2 Q6TUI4 Q6TUQ9 Q6TUY0 Q6TUZ4 Q6TV19 Q6TVP7 Q6TWC4 Q6TXD2
Q6TXG9 Q6TY21 Q6TYB5 Q6U2P6 Q6U5S4 Q6U6G5 Q6U7Q0 Q6U949 Q6UAQ8 Q6UB28 Q6UB98
Q6UBI3 Q6UBL8 Q6UBQ3 Q6UDF2 Q6UDF3 Q6UDG0 Q6UDG1 Q6UDG2 Q6UDG6 Q6UDG7 Q6UDG8
Q6UDG9 Q6UDH0 Q6UDH1 Q6UDH2 Q6UDH3 Q6UDH6 Q6UDH8 Q6UDH9 Q6UDI0 Q6UDI1 Q6UDI2
Q6UDI3 Q6UDI4 Q6UDI6 Q6UDI9 Q6UDJ0 Q6UDJ2 Q6UDJ3 Q6UDJ4 Q6UDJ6 Q6UDJ8 Q6UDJ9
Q6UDK1 Q6UDK2 Q6UDK3 Q6UDK6 Q6UDK7 Q6UDK8 Q6UDK9 Q6UDL1 Q6UDL2 Q6UDL3 Q6UDL4
Q6UDL5 Q6UDL8 Q6UDM0 Q6UDM1 Q6UDM2 Q6UDM4 Q6UEF0 Q6UEF1 Q6UEF2 Q6UEF3 Q6UEG2
Q6UEG5 Q6UEG8 Q6UEH5 Q6UFS5 Q6UFZ2 Q6UFZ3 Q6UFZ8 Q6UFZ9 Q6UGN0 Q6UIL3 Q6UIM1
Q6UIM2 Q6UK63 Q6UL01 Q6UP17 Q6UQ76 Q6UR06 Q6UR32 Q6UR35 Q6UR41 Q6UR44 Q6UR50
Q6UR53 Q6UR59 Q6UR62 Q6UR67 Q6UR70 Q6UR75 Q6UR78 Q6UR83 Q6UR86 Q6URB0 Q6URC2
Q6URW6 Q6USK1 Q6UY61 Q6UY63 Q6UY65 Q6UY68 Q6UY69 Q6UY70 Q6UYE1 Q6UZF7 Q6V1Q2
Q6V1Q9 Q6V1X1 Q6V289 Q6V291 Q6V3V8 Q6V3W0 Q6V3W6 Q6V3X0 Q6V4H0 Q6V595 Q6V5K9
Q6V702 Q6V9I2 Q6V9I6 Q6V9R5 Q6V9S9 Q6VAA3 Q6VAA4 Q6VAA5 Q6VAA6 Q6VAA7 Q6VAA8
Q6VAA9 Q6VAB0 Q6VAB1 Q6VAB2 Q6VAB3 Q6VAB4 Q6VAF4 Q6VAF5 Q6VAF6 Q6VAF7 Q6VAF8
Q6VAF9 Q6VAG0 Q6VAG1 Q6VAM4 Q6VAY5 Q6VB84 Q6VBQ6 Q6VBQ8 Q6VE94 Q6VEU1 Q6VEU3
Q6VFT5 Q6VFT6 Q6VFT7 Q6VH22 Q6VMN7 Q6VMV8 Q6VMV9 Q6VMW0 Q6VMW1 Q6VMW2 Q6VN46
Q6VNZ9 Q6VQA8 Q6VQA9 Q6VTH5 Q6VUC0 Q6VUC1 Q6VUP9 Q6VV72 Q6VVD7 Q6VWJ6 Q6VY05
Q6VZ17 Q6VZL8 Q6VZQ4 Q6W0C5 Q6W102 Q6W163 Q6W1D5 Q6W210 Q6W2G5 Q6W2J0 Q6W349
Q6W3C0 Q6W3Z9 Q6W4T2 Q6W4T3 Q6W8Q3 Q6W8W3 Q6W990 Q6WAT6 Q6WAU0 Q6WAU1 Q6WB93
Q6WB96 Q6WB97 Q6WBA1 Q6WBX7 Q6WBX8 Q6WFW1 Q6WG17 Q6WG18 Q6WHE1 Q6WHE6 Q6WHE8
Q6WHG9 Q6WHH0 Q6WHI1 Q6WI70 Q6WI75 Q6WI88 Q6WIT9 Q6WKW9 Q6WKZ8 Q6WLH3 Q6WLH4
Q6WN20 Q6WN21 Q6WN22 Q6WN25 Q6WN26 Q6WN27 Q6WN28 Q6WN29 Q6WNK7 Q6WNP1 Q6WNP2
Q6WNP3 Q6WNV6 Q6WNV7 Q6WNV8 Q6WNW3 Q6WNW5 Q6WQI6 Q6WRH6 Q6WRT6 Q6WRX3 Q6WRX6
Q6WRX7 Q6WRX8 Q6WRX9 Q6WS85 Q6WUC1 Q6WUC2 Q6WV66 Q6WV67 Q6WV69 Q6WV72 Q6WV73
Q6WV74 Q6WV88 Q6WV90 Q6WVH5 Q6WVP7 Q6WY08 Q6WZ17 Q6WZ18 Q6WZ19 Q6WZ20 Q6WZ83
Q6X1C0 Q6X1D4 Q6X1D8 Q6X1E1 Q6X2E1 Q6X2E2 Q6X2E3 Q6X4M3 Q6X4T0 Q6X7J3 Q6X7J5
Q6X7K0 Q6X7K1 Q6X7S9 Q6X9E4 Q6X9Z5 Q6XBH1 Q6XBJ3 Q6XCG6 Q6XD66 Q6XD76 Q6XDB5
Q6XDK7 Q6XDK8 Q6XDT4 Q6XDT6 Q6XE24 Q6XHI1 Q6XHX5 Q6XI08 Q6XI71 Q6XIM7 Q6XIM8
Q6XIN1 Q6XIP0 Q6XIR2 Q6XJ13 Q6XJU9 Q6XK03 Q6XKE6 Q6XL52 Q6XLA1 Q6XLJ0 Q6XMH6
Q6XMH7 Q6XMI3 Q6XNL5 Q6XNP7 Q6XPT5 Q6XQ67 Q6XQB2 Q6XQC4 Q6XQH2 Q6XQH7 Q6XQI4
Q6XQI9 Q6XQN1 Q6XSH4 Q6XUW5 Q6XUZ5 Q6XXL8 Q6XXM1 Q6XXM7 Q6XXX2 Q6XYB7 Q6XYU1
Q6XYX0 Q6XYX2 Q6XYX3 Q6XYX4 Q6XYX5 Q6XYX6 Q6XYY0 Q6XYY1 Q6XYY2 Q6XZ05 Q6XZ07
Q6XZ78 Q6XZ79 Q6XZR0 Q6Y0Z3 Q6Y1R6 Q6Y1Z4 Q6Y237 Q6Y239 Q6Y263 Q6Y7S3 Q6Y8Q8
Q6YBV1 Q6YCH1 Q6YCH2 Q6YDN7 Q6YFE4 Q6YFE5 Q6YFE6 Q6YHU8 Q6YI30 Q6YI57 Q6YI93
Q6YIA3 Q6YJI5 Q6YLT2 Q6YLW2 Q6YLX4 Q6YNC8 Q6YNI5 Q6YNQ3 Q6YNQ4 Q6YNQ5 Q6YNQ6

Q6YNQ7 Q6YNR6 Q6YP15 Q6YP18 Q6YP19 Q6YP21 Q6YPD0 Q6YPG5 Q6YPH3 Q6YPI0 Q6YPI4
Q6YPI5 Q6YPJ3 Q6YPJ4 Q6YPL9 Q6YPM0 Q6YPM1 Q6YPM2 Q6YPV1 Q6YPV2 Q6YPW4 Q6YPW7
Q6YPX7 Q6YPX9 Q6YPY6 Q6YPY9 Q6YPZ6 Q6YQA4 Q6YQA5 Q6YQA6 Q6YQC1 Q6YQC6 Q6YQD6
Q6YQE1 Q6YQR8 Q6YQR9 Q6YQT0 Q6YQT8 Q6YQU0 Q6YQU7 Q6YQU8 Q6YQV5 Q6YQV6 Q6YQV8
Q6YQV9 Q6YQW0 Q6YQW1 Q6YQW2 Q6YQW3 Q6YQW5 Q6YQW6 Q6YQX2 Q6YQX5 Q6YQX6 Q6YQX7
Q6YQY4 Q6YQY6 Q6YQZ4 Q6YQZ5 Q6YQZ6 Q6YQZ7 Q6YQZ8 Q6YR01 Q6YR04 Q6YR05 Q6YR06
Q6YR07 Q6YR08 Q6YR09 Q6YR10 Q6YR11 Q6YR12 Q6YR14 Q6YR15 Q6YR16 Q6YR17 Q6YR18
Q6YR19 Q6YR20 Q6YR34 Q6YR36 Q6YR37 Q6YR48 Q6YR50 Q6YR51 Q6YR57 Q6YR64 Q6YR66
Q6YR70 Q6YR75 Q6YR76 Q6YR77 Q6YR80 Q6YR87 Q6YR88 Q6YR90 Q6YR94 Q6YRA9 Q6YRB0
Q6YRB1 Q6YRC6 Q6YRC7 Q6YRC9 Q6YRD0 Q6YRD2 Q6YRD4 Q6YRJ6 Q6YRK0 Q6YRK2 Q6YRK3
Q6YRK5 Q6YRL3 Q6YRW7 Q6YS30 Q6YSF3 Q6YSJ5 Q6YSY5 Q6YT00 Q6YTI2 Q6YTI3 Q6YTQ4
Q6YTW6 Q6YTY3 Q6YU51 Q6YU88 Q6YUA8 Q6YUB8 Q6YV04 Q6YV23 Q6YVJ0 Q6YVX4 Q6YVX9
Q6YVY0 Q6YW46 Q6YW64 Q6YWP9 Q6YWR4 Q6YXH5 Q6YXH8 Q6YXJ7 Q6YXJ8 Q6YXJ9 Q6YXK5
Q6YXK6 Q6YXK7 Q6YXK8 Q6YXK9 Q6YXL0 Q6YXL1 Q6YXL2 Q6YXM3 Q6YXM4 Q6YXM5 Q6YXM6
Q6YXM7 Q6YXN5 Q6YXN6 Q6YXP3 Q6YXP4 Q6YXP6 Q6YXQ4 Q6YXQ5 Q6YXQ7 Q6YXS0 Q6YXW6
Q6YXY2 Q6YXZ4 Q6YXZ7 Q6YY75 Q6YYA3 Q6YYB0 Q6YYC0 Q6YYX9 Q6YYZ1 Q6YYZ2 Q6YZ49
Q6YZ52 Q6YZ54 Q6YZE2 Q6YZI0 Q6YZM6 Q6YZW0 Q6YZW2 Q6YZX6 Q6Z0D2 Q6Z0D9 Q6Z1C0
Q6Z1G7 Q6Z1Y6 Q6Z1Z3 Q6Z234 Q6Z244 Q6Z248 Q6Z256 Q6Z2G9 Q6Z2K1 Q6Z2L5 Q6Z2T6
Q6Z2T8 Q6Z2W3 Q6Z2X3 Q6Z2Z4 Q6Z312 Q6Z345 Q6Z348 Q6Z351 Q6Z358 Q6Z398 Q6Z3A3
Q6Z3L4 Q6Z3U3 Q6Z433 Q6Z4A7 Q6Z4F1 Q6Z4G3 Q6Z4H0 Q6Z4I3 Q6Z4K6 Q6Z4N3 Q6Z4N4
Q6Z4N6 Q6Z4T3 Q6Z4T5 Q6Z4U2 Q6Z517 Q6Z528 Q6Z541 Q6Z5I0 Q6Z5J6 Q6Z5M0 Q6Z5N4
Q6Z690 Q6Z6G5 Q6Z6G6 Q6Z6L5 Q6Z6S1 Q6Z6W2 Q6Z746 Q6Z784 Q6Z7F4 Q6Z7H3 Q6Z7L3
Q6Z7V2 Q6Z836 Q6Z869 Q6Z8B9 Q6Z9A3 Q6Z9C3 Q6Z9D2 Q6Z9U7 Q6ZA74 Q6ZAC1 Q6ZAG3
Q6ZAL2 Q6ZB90 Q6ZBH6 Q6ZBP3 Q6ZBS8 Q6ZC69 Q6ZCF0 Q6ZD63 Q6ZD89 Q6ZD95 Q6ZDJ7
Q6ZDQ1 Q6ZEA5 Q6ZEC6 Q6ZEI1 Q6ZEM6 Q6ZEZ5 Q6ZFT5 Q6ZFU6 Q6ZG77 Q6ZGP8 Q6ZH29
Q6ZH94 Q6ZHC8 Q6ZHE5 Q6ZHJ5 Q6ZHS4 Q6ZHZ1 Q6ZI17 Q6ZIK0 Q6ZIK7 Q6ZJ05 Q6ZJ08
Q6ZJ48 Q6ZJK7 Q6ZJM9 Q6ZJS7 Q6ZJW8 Q6ZJW9 Q6ZJX0 Q6ZK48 Q6ZK57 Q6ZKL8 Q6ZKM2
Q6ZKV8 Q6ZL42 Q6ZL43 Q6ZL57 Q6ZL94 Q6ZLA3 Q6ZLA7 Q6ZLB0 Q6ZLC4 Q6ZM19 Q6ZM63
Q6ZMN7 Q6ZMN8 Q6ZMR3 Q6ZMS4 Q6ZMS7 Q6ZMT4 Q6ZMU1 Q6ZMV5 Q6ZMV7 Q6ZMV8 Q6ZMV9
Q6ZMW2 Q6ZMW3 Q6ZMY3 Q6ZMY6 Q6ZMY9 Q6ZN01 Q6ZN03 Q6ZN06 Q6ZN08 Q6ZN11 Q6ZN19
Q6ZN28 Q6ZN32 Q6ZN54 Q6ZN57 Q6ZN66 Q6ZN79 Q6ZN84 Q6ZN92 Q6ZNA1 Q6ZNB5 Q6ZNC4
Q6ZNE9 Q6ZNG0 Q6ZNG2 Q6ZNG9 Q6ZNH5 Q6ZNK6 Q6ZNM6 Q6ZNW5 Q6ZNX1 Q6ZP01 Q6ZP65
Q6ZP68 Q6ZP82 Q6ZPA2 Q6ZPG2 Q6ZPK0 Q6ZPL9 Q6ZPR1 Q6ZPS2 Q6ZPT1 Q6ZPU9 Q6ZPV2
Q6ZPY2 Q6ZPY5 Q6ZPY7 Q6ZPZ3 Q6ZQ03 Q6ZQ08 Q6ZQ38 Q6ZQ58 Q6ZQ73 Q6ZQB6 Q6ZQH8
Q6ZQJ5 Q6ZQK0 Q6ZQL4 Q6ZQN5 Q6ZQQ6 Q6ZQR2 Q6ZQT0 Q6ZQT7 Q6ZQV5 Q6ZQW0 Q6ZQX7
Q6ZQY2 Q6ZQY3 Q6ZQY7 Q6ZR03 Q6ZR08 Q6ZR52 Q6ZR54 Q6ZR62 Q6ZR85 Q6ZRC1 Q6ZRF7
Q6ZRG5 Q6ZRI6 Q6ZRK6 Q6ZRM9 Q6ZRN7 Q6ZRP0 Q6ZRP5 Q6ZRQ5 Q6ZRR7 Q6ZRS2 Q6ZRS4
Q6ZRT6 Q6ZRV3 Q6ZRX8 Q6ZRY4 Q6ZS02 Q6ZS11 Q6ZS27 Q6ZS46 Q6ZS49 Q6ZS52 Q6ZS86
Q6ZS92 Q6ZSA8 Q6ZSB3 Q6ZSB9 Q6ZSC3 Q6ZSG1 Q6ZSG2 Q6ZSI9 Q6ZSJ8 Q6ZSK4 Q6ZSN1
Q6ZSR3 Q6ZSR6 Q6ZSR9 Q6ZSS3 Q6ZST2 Q6ZST4 Q6ZSV7 Q6ZT07 Q6ZT52 Q6ZT77 Q6ZT98
Q6ZTB9 Q6ZTC4 Q6ZTI0 Q6ZTI6 Q6ZTK2 Q6ZTR5 Q6ZTR6 Q6ZTU2 Q6ZTW0 Q6ZTY9 Q6ZTZ1
Q6ZU15 Q6ZU52 Q6ZU67 Q6ZU80 Q6ZUA9 Q6ZUF6 Q6ZUG5 Q6ZUI0 Q6ZUJ4 Q6ZUL3 Q6ZUS5
Q6ZUS6 Q6ZUT4 Q6ZUT6 Q6ZUU3 Q6ZUX3 Q6ZV50 Q6ZV60 Q6ZV65 Q6ZV77 Q6ZV80 Q6ZV89
Q6ZVC0 Q6ZVH6 Q6ZVH7 Q6ZVK8 Q6ZVL8 Q6ZVN7 Q6ZVQ6 Q6ZVS7 Q6ZVT0 Q6ZVT6 Q6ZVU0

Q6ZVW7 Q6ZVX7 Q6ZW13 Q6ZW61 Q6ZW76 Q6ZWC4 Q6ZWH5 Q6ZWI9 Q6ZWM4 Q6ZWS8 Q6ZWX6
Q6ZWY8 Q6ZWY9 Q6ZWZ2 Q6ZXC1 Q6ZXC2 Q6ZXX3 Q6ZY51 Q6ZYA7 Q6ZYL4 Q6ZZC6 Q6ZZF4
Q6ZZF5 Q700C7 Q700D0 Q700D9 Q700E3 Q700E4 Q700X4 Q701N2 Q701N4 Q701N9 Q702N8
Q703G9 Q703I1 Q703I2 Q703W7 Q704B2 Q704D0 Q704D1 Q704D2 Q704V3 Q705D2 Q705F3
Q705F6 Q706S1 Q707Y0 Q707Y3 Q707Y6 Q707Y7 Q707Y8 Q708A1 Q708A7 Q708T3 Q708W1
Q708W2 Q708Y0 Q709N3 Q70AC7 Q70C33 Q70CP7 Q70CZ6 Q70D04 Q70D24 Q70D30 Q70D37
Q70D38 Q70D40 Q70D41 Q70EK8 Q70FD1 Q70FG7 Q70G58 Q70GH6 Q70GK9 Q70I53 Q70IY1
Q70JN8 Q70JS2 Q70JY7 Q70JZ1 Q70KD0 Q70KD4 Q70KF0 Q70KH9 Q70KY4 Q70LB1 Q70LB4
Q70LB5 Q70LB6 Q70LB7 Q70LB9 Q70LC0 Q70LC1 Q70LC3 Q70LC4 Q70LC5 Q70LC8 Q70LD0
Q70LD1 Q70LD3 Q70LD4 Q70LD5 Q70LD6 Q70LD7 Q70LD8 Q70LD9 Q70LE0 Q70LE1 Q70LE2
Q70LE3 Q70LE4 Q70LE5 Q70LE6 Q70LE7 Q70LE8 Q70LE9 Q70LF0 Q70LM4 Q70LM5 Q70LM6
Q70LM7 Q70LM8 Q70PP2 Q70PR7 Q70Q35 Q70SJ2 Q70UN9 Q70UP5 Q70UP6 Q70US9 Q70UV1
Q70WY6 Q70WY9 Q70XU7 Q70XV3 Q70XV4 Q70XV7 Q70XW5 Q70XX0 Q70XX1 Q70XX2 Q70XX3
Q70XX4 Q70XX5 Q70XX6 Q70XY2 Q70XY3 Q70XY4 Q70XY5 Q70XZ5 Q70Y01 Q70Y05 Q70Y11
Q70Y15 Q70Y16 Q70YC4 Q70YC5 Q70YI5 Q710C4 Q710D7 Q710E8 Q711A6 Q711G1 Q711M0
Q711Q0 Q711T2 Q711T7 Q711T9 Q712G9 Q712K3 Q712U6 Q714U9 Q715L4 Q715S9 Q715T0
Q716F7 Q717B2 Q717B4 Q717R8 Q717X1 Q717X5 Q717Z2 Q719H9 Q719I0 Q71A37 Q71A38
Q71A39 Q71AW2 Q71EW5 Q71F54 Q71F78 Q71FD7 Q71G51 Q71H53 Q71H73 Q71HN5 Q71HP2
Q71L16 Q71L19 Q71L24 Q71L33 Q71L39 Q71LE2 Q71LX6 Q71ML1 Q71ML5 Q71N41 Q71PD7
Q71RI9 Q71SY5 Q71T09 Q71T45 Q71T50 Q71TD6 Q71TD7 Q71TF8 Q71TL0 Q71U11 Q71U53
Q71U98 Q71UE8 Q71UF8 Q71UF9 Q71UG0 Q71UG1 Q71UG2 Q71UG3 Q71UG4 Q71UG5 Q71UG6
Q71UG7 Q71UG8 Q71UG9 Q71UH5 Q71UI9 Q71V09 Q71V10 Q71V11 Q71V12 Q71V13 Q71V14
Q71V16 Q71V17 Q71V18 Q71V19 Q71V20 Q71V22 Q71V39 Q71V89 Q71VG4 Q71VG5 Q71VG7
Q71VG8 Q71VG9 Q71VH0 Q71VP4 Q71VP5 Q71VP6 Q71VP7 Q71VP8 Q71VP9 Q71VQ6 Q71VQ7
Q71VR3 Q71VR4 Q71VR5 Q71VR6 Q71VT6 Q71VV0 Q71VV1 Q71VW1 Q71VW2 Q71VW9 Q71VZ1
Q71W03 Q71W17 Q71W21 Q71W25 Q71W68 Q71W69 Q71W79 Q71WB6 Q71WB7 Q71WB8 Q71WB9
Q71WD4 Q71WE5 Q71WE6 Q71WE7 Q71WE8 Q71WE9 Q71WF0 Q71WF1 Q71WF2 Q71WF3 Q71WF4
Q71WF5 Q71WF6 Q71WF7 Q71WF8 Q71WF9 Q71WG0 Q71WG1 Q71WG2 Q71WG3 Q71WG4 Q71WG5
Q71WG7 Q71WG8 Q71WG9 Q71WH0 Q71WH1 Q71WH2 Q71WH3 Q71WI0 Q71WI1 Q71WI2 Q71WJ5
Q71WL5 Q71WL8 Q71WL9 Q71WM0 Q71WM7 Q71WN0 Q71WN3 Q71WN4 Q71WN5 Q71WN9 Q71WP0
Q71WQ2 Q71WR9 Q71WT9 Q71WU0 Q71WU4 Q71WU6 Q71WV3 Q71WV5 Q71WV6 Q71WV9 Q71WW8
Q71WW9 Q71WX0 Q71WX8 Q71X03 Q71X33 Q71X35 Q71X61 Q71X98 Q71X99 Q71XE0 Q71XE2
Q71XE5 Q71XF3 Q71XF4 Q71XF9 Q71XG0 Q71XG7 Q71XH4 Q71XH5 Q71XK8 Q71XP5 Q71XQ0
Q71XR2 Q71XR3 Q71XS3 Q71XS4 Q71XT8 Q71XU1 Q71XU5 Q71XU6 Q71XW0 Q71XW2 Q71XW5
Q71XW7 Q71XX1 Q71XX2 Q71XX5 Q71XX7 Q71XY3 Q71XZ5 Q71Y01 Q71Y02 Q71Y27 Q71Y32
Q71Y33 Q71Y34 Q71Y35 Q71Y36 Q71Y38 Q71Y44 Q71Y46 Q71Y53 Q71Y59 Q71Y60 Q71Y63
Q71Y64 Q71Y76 Q71Y78 Q71Y82 Q71Y84 Q71Y86 Q71Y87 Q71Y88 Q71Y90 Q71Y92 Q71Y94
Q71YA8 Q71YA9 Q71YB0 Q71YB3 Q71YB4 Q71YB5 Q71YB9 Q71YC4 Q71YC6 Q71YC7 Q71YD1
Q71YD4 Q71YD8 Q71YD9 Q71YE1 Q71YF0 Q71YF6 Q71YF7 Q71YF9 Q71YG0 Q71YH6 Q71YH8
Q71YH9 Q71YI0 Q71YI1 Q71YI2 Q71YI3 Q71YI4 Q71YI5 Q71YI9 Q71YJ0 Q71YJ3 Q71YJ7
Q71YK0 Q71YL2 Q71YL9 Q71YM0 Q71YM3 Q71YM4 Q71YM9 Q71YN3 Q71YN4 Q71YN5 Q71YN8
Q71YP6 Q71YP8 Q71YP9 Q71YQ1 Q71YQ3 Q71YQ4 Q71YR0 Q71YR2 Q71YR3 Q71YR4 Q71YR7
Q71YS6 Q71YW0 Q71YW4 Q71YX4 Q71YY0 Q71YY2 Q71YY6 Q71YZ2 Q71YZ5 Q71Z03 Q71Z07
Q71Z09 Q71Z11 Q71Z12 Q71Z37 Q71Z38 Q71Z39 Q71Z40 Q71Z41 Q71Z49 Q71Z52 Q71Z59

Q71Z62 Q71Z70 Q71Z71 Q71Z75 Q71Z76 Q71Z77 Q71Z78 Q71Z79 Q71Z80 Q71Z81 Q71Z84
Q71Z86 Q71Z90 Q71Z96 Q71ZA3 Q71ZA4 Q71ZA5 Q71ZA9 Q71ZB0 Q71ZB1 Q71ZB2 Q71ZB5
Q71ZB6 Q71ZC0 Q71ZC5 Q71ZC8 Q71ZD0 Q71ZD2 Q71ZD3 Q71ZD5 Q71ZD6 Q71ZD7 Q71ZD8
Q71ZD9 Q71ZE0 Q71ZE6 Q71ZE8 Q71ZF0 Q71ZF1 Q71ZF8 Q71ZG6 Q71ZG7 Q71ZG8 Q71ZG9
Q71ZH3 Q71ZH4 Q71ZH6 Q71ZI2 Q71ZJ0 Q71ZJ5 Q71ZJ6 Q71ZJ7 Q71ZJ8 Q71ZJ9 Q71ZK1
Q71ZK5 Q71ZL0 Q71ZL1 Q71ZL2 Q71ZL3 Q71ZL4 Q71ZL9 Q71ZM1 Q71ZM2 Q71ZM9 Q71ZN5
Q71ZQ0 Q71ZR3 Q71ZR6 Q71ZR7 Q71ZS2 Q71ZS3 Q71ZT6 Q71ZT7 Q71ZT9 Q71ZV2 Q71ZV5
Q71ZV7 Q71ZW0 Q71ZW1 Q71ZW2 Q71ZW3 Q71ZW7 Q71ZX2 Q71ZX3 Q71ZX4 Q71ZY7 Q71ZZ0
Q71ZZ1 Q71ZZ2 Q71ZZ4 Q71ZZ5 Q71ZZ7 Q720A1 Q720A2 Q720A3 Q720A5 Q720A7 Q720A8
Q720A9 Q720B7 Q720B8 Q720C8 Q720E1 Q720E2 Q720E3 Q720E4 Q720E5 Q720E8 Q720F3
Q720F5 Q720G2 Q720G3 Q720J0 Q720J1 Q720J2 Q720J5 Q720J7 Q720K1 Q720K2 Q720K7
Q720K8 Q720M1 Q720N5 Q720N8 Q720Q2 Q720R2 Q720X7 Q720Y0 Q720Y7 Q721A0 Q721A5
Q721A8 Q721B5 Q721B9 Q721C0 Q721D8 Q721F4 Q721F5 Q721H8 Q721J0 Q721J8 Q721K2
Q721K9 Q721N0 Q721N6 Q721P1 Q721R7 Q721S2 Q721S9 Q721W2 Q721Y7 Q722D2 Q722H1
Q722I5 Q722M0 Q722T9 Q722V6 Q722Y1 Q722Y2 Q722Y3 Q722Y4 Q722Y5 Q722Y6 Q722Y7
Q722Y8 Q722Y9 Q723B1 Q723B6 Q723B7 Q723E1 Q723F3 Q723F8 Q723F9 Q723G3 Q723G7
Q723G9 Q723S4 Q723S8 Q723S9 Q723T0 Q723T1 Q723U1 Q723V9 Q723W2 Q723Y9 Q723Z1
Q724B0 Q724E9 Q724F0 Q724G1 Q724G2 Q724G3 Q724G4 Q724G8 Q724H3 Q724H5 Q724H6
Q724H7 Q724I1 Q724I4 Q724J1 Q724J2 Q724J4 Q724K0 Q724K2 Q724K3 Q724K5 Q724L4
Q724L6 Q724L7 Q724M2 Q724M3 Q724M5 Q724N6 Q724P3 Q724P5 Q725A8 Q725B1 Q725B8
Q725C0 Q725C1 Q725C4 Q725C6 Q725C8 Q725G6 Q725H0 Q725I1 Q725I2 Q725K9 Q725L9
Q725M4 Q725M6 Q725M7 Q725Q1 Q725Q3 Q725Q7 Q725Q9 Q725S3 Q725W6 Q725W7 Q725Z4
Q726D3 Q726D5 Q726F4 Q726F7 Q726H4 Q726J1 Q726J4 Q726N6 Q726S6 Q726S7 Q726X1
Q726X3 Q726X4 Q726Y4 Q727C0 Q727C6 Q727C7 Q727C8 Q727C9 Q727D0 Q727D4 Q727D5
Q727D9 Q727E1 Q727E3 Q727F1 Q727F4 Q727G3 Q727G4 Q727N0 Q727N1 Q727P4 Q728E4
Q728E5 Q728G0 Q728P4 Q728P5 Q728Q1 Q728Q5 Q728R5 Q728R6 Q728R7 Q728R8 Q728R9
Q728S0 Q728T3 Q728T4 Q728T9 Q728U6 Q728V0 Q729A3 Q729A4 Q729A6 Q729I7 Q729K0
Q729L7 Q729M2 Q729Q7 Q729T9 Q729U1 Q729U3 Q729U4 Q729U6 Q729V4 Q729V5 Q729Z3
Q72A55 Q72A57 Q72AL1 Q72AL5 Q72AL6 Q72AN9 Q72AR0 Q72AR5 Q72AU1 Q72AU2 Q72AU3
Q72AU4 Q72AV0 Q72AV1 Q72AV2 Q72AV5 Q72AV7 Q72AW6 Q72AX2 Q72AZ1 Q72AZ9 Q72B00
Q72B11 Q72B14 Q72B41 Q72B46 Q72B63 Q72BE3 Q72BF9 Q72BH0 Q72BH2 Q72BI1 Q72BK4
Q72BL2 Q72BL6 Q72BM6 Q72BM7 Q72BN0 Q72BN2 Q72BN4 Q72BQ5 Q72BQ7 Q72BQ8 Q72BR0
Q72BR1 Q72BV1 Q72BV2 Q72C17 Q72C18 Q72C23 Q72C25 Q72C30 Q72C32 Q72C44 Q72C58
Q72C59 Q72CA6 Q72CB8 Q72CD3 Q72CD5 Q72CD8 Q72CD9 Q72CE6 Q72CE7 Q72CE8 Q72CE9
Q72CF1 Q72CF3 Q72CF4 Q72CF5 Q72CF6 Q72CF7 Q72CF8 Q72CG1 Q72CG2 Q72CG3 Q72CG4
Q72CG5 Q72CG6 Q72CG7 Q72CG8 Q72CG9 Q72CH0 Q72CH1 Q72CH2 Q72CH3 Q72CH4 Q72CH5
Q72CH6 Q72CH7 Q72CH8 Q72CH9 Q72CI0 Q72CI1 Q72CI3 Q72CI4 Q72CI5 Q72CK1 Q72CM9
Q72CN3 Q72CN5 Q72CQ9 Q72CS2 Q72CS4 Q72CT0 Q72CT6 Q72CT7 Q72CU1 Q72CW9 Q72D35
Q72D36 Q72D51 Q72D55 Q72D86 Q72DA0 Q72DA1 Q72DA4 Q72DE5 Q72DG6 Q72DH1 Q72DH2
Q72DH3 Q72DH4 Q72DH6 Q72DJ9 Q72DK0 Q72DK1 Q72DK2 Q72DK5 Q72DL8 Q72DM3 Q72DM4
Q72DM8 Q72DM9 Q72DN0 Q72DN5 Q72DN7 Q72DQ5 Q72DQ6 Q72DQ8 Q72DQ9 Q72DR3 Q72DS4
Q72DS5 Q72DU2 Q72DU3 Q72DU4 Q72DU5 Q72DU6 Q72DV1 Q72DW3 Q72DW6 Q72DW7 Q72DW8
Q72DX0 Q72DX1 Q72DX2 Q72DX8 Q72E47 Q72E53 Q72E56 Q72E89 Q72EH1 Q72EK2 Q72EP2
Q72EP7 Q72EP8 Q72EQ8 Q72ER1 Q72ER4 Q72ER5 Q72ER6 Q72ET7 Q72EU7 Q72EX7 Q72EX8

Q72F03 Q72F80 Q72F91 Q72F93 Q72FC5 Q72FH6 Q72FX9 Q72G08 Q72G53 Q72G65 Q72G68
Q72G72 Q72G84 Q72GC7 Q72GE2 Q72GF9 Q72GG0 Q72GI1 Q72GI8 Q72GJ4 Q72GN2 Q72GN3
Q72GN6 Q72GQ0 Q72GR0 Q72GS1 Q72GS2 Q72GS9 Q72GT4 Q72GU0 Q72GU1 Q72GU2 Q72GV1
Q72GV2 Q72GV5 Q72GV9 Q72GW3 Q72GW4 Q72GX1 Q72GY6 Q72GY7 Q72GY9 Q72GZ0 Q72GZ4
Q72GZ9 Q72H00 Q72H18 Q72H19 Q72H31 Q72H32 Q72H33 Q72H57 Q72H70 Q72H75 Q72H81
Q72H87 Q72H88 Q72H89 Q72H90 Q72H91 Q72H98 Q72HA0 Q72HA1 Q72HA9 Q72HB4 Q72HB8
Q72HC1 Q72HF4 Q72HH1 Q72HI4 Q72HJ6 Q72HK9 Q72HM5 Q72HM6 Q72HN2 Q72HP1 Q72HP3
Q72HP8 Q72HR2 Q72HR3 Q72HR4 Q72HR5 Q72HS0 Q72HY0 Q72I01 Q72I04 Q72I05 Q72I06
Q72I07 Q72I09 Q72I11 Q72I12 Q72I14 Q72I15 Q72I16 Q72I19 Q72I20 Q72I22 Q72I23
Q72I25 Q72I28 Q72I32 Q72I33 Q72I39 Q72I42 Q72I52 Q72I53 Q72I65 Q72I77 Q72I89
Q72I96 Q72I97 Q72IA7 Q72IA8 Q72IA9 Q72IB8 Q72ID1 Q72ID5 Q72ID6 Q72ID8 Q72IE1
Q72IE5 Q72IF0 Q72IF6 Q72IG5 Q72IH2 Q72IH5 Q72IH7 Q72IH9 Q72IJ0 Q72IJ9 Q72IK5
Q72IK6 Q72IK9 Q72IN0 Q72IN1 Q72IQ5 Q72IQ9 Q72IV3 Q72IV5 Q72IW2 Q72IW3 Q72IW9
Q72IX3 Q72IZ2 Q72IZ3 Q72IZ9 Q72J02 Q72J22 Q72J23 Q72J35 Q72J39 Q72J47 Q72J62
Q72J87 Q72J91 Q72J95 Q72JA8 Q72JB2 Q72JB3 Q72JC8 Q72JE7 Q72JE9 Q72JF9 Q72JG1
Q72JG7 Q72JJ7 Q72JJ8 Q72JK8 Q72JL2 Q72JL6 Q72JL7 Q72JM6 Q72JN8 Q72JN9 Q72JP3
Q72JP7 Q72JQ1 Q72JQ8 Q72JR0 Q72JR1 Q72JR6 Q72JR7 Q72JS5 Q72JS7 Q72JT0 Q72JT5
Q72JT8 Q72JT9 Q72JU7 Q72JU8 Q72JU9 Q72JV2 Q72JZ8 Q72K02 Q72K16 Q72K27 Q72K35
Q72K37 Q72K55 Q72K66 Q72K83 Q72K85 Q72K87 Q72K90 Q72K91 Q72KA1 Q72KA9 Q72KB0
Q72KD8 Q72KD9 Q72KE0 Q72KE2 Q72KE8 Q72KF7 Q72KF8 Q72KG1 Q72KG3 Q72KH6 Q72KH8
Q72KL7 Q72KL8 Q72KN9 Q72KP6 Q72KS4 Q72KT9 Q72KU0 Q72L05 Q72L14 Q72L15 Q72L16
Q72L21 Q72L22 Q72L31 Q72L32 Q72L49 Q72L53 Q72L58 Q72L59 Q72L62 Q72L64 Q72L76
Q72L77 Q72L85 Q72L95 Q72LB0 Q72LB1 Q72LF3 Q72LF4 Q72LG3 Q72LH0 Q72LH1 Q72LI2
Q72LI9 Q72LL6 Q72LM0 Q72LQ3 Q72LR0 Q72LR3 Q72LS1 Q72LS5 Q72LS6 Q72LY0 Q72LY8
Q72LY9 Q72LZ4 Q72M00 Q72M35 Q72M46 Q72M65 Q72M66 Q72M80 Q72M93 Q72MA6 Q72MG4
Q72MG8 Q72MJ5 Q72MJ6 Q72MK0 Q72ML5 Q72MN0 Q72MQ5 Q72MR4 Q72MT7 Q72MU5 Q72MV4
Q72MY7 Q72N01 Q72N73 Q72N87 Q72N95 Q72NF1 Q72NF9 Q72NG0 Q72NG1 Q72NG2 Q72NG3
Q72NG4 Q72NG5 Q72NG6 Q72NG7 Q72NG8 Q72NG9 Q72NH0 Q72NH1 Q72NH2 Q72NH3 Q72NH4
Q72NH5 Q72NH6 Q72NH7 Q72NH8 Q72NH9 Q72NI0 Q72NI2 Q72NI4 Q72NI5 Q72NI6 Q72NI7
Q72NI8 Q72NI9 Q72NJ3 Q72NJ5 Q72NL9 Q72NN2 Q72NP0 Q72NP4 Q72NP6 Q72NP7 Q72NQ4
Q72NQ6 Q72NQ7 Q72NQ8 Q72NQ9 Q72NR0 Q72NR4 Q72NU6 Q72NW0 Q72NX1 Q72NX3 Q72NX4
Q72NX5 Q72NX6 Q72NX7 Q72NZ6 Q72NZ8 Q72P39 Q72P59 Q72PA2 Q72PA4 Q72PA7 Q72PC8
Q72PD0 Q72PF1 Q72PF7 Q72PG3 Q72PJ7 Q72PK5 Q72PK6 Q72PK8 Q72PK9 Q72PL0 Q72PL7
Q72PM1 Q72PM5 Q72PQ0 Q72PQ1 Q72PR5 Q72PR7 Q72PW7 Q72PX0 Q72PY2 Q72Q12 Q72Q44
Q72Q66 Q72Q67 Q72QA1 Q72QK0 Q72QK2 Q72QK3 Q72QK5 Q72QK8 Q72QL1 Q72QL2 Q72QL9
Q72QN9 Q72QP1 Q72QP5 Q72QT1 Q72QT3 Q72QT4 Q72QU2 Q72R01 Q72R21 Q72R38 Q72R59
Q72R81 Q72R93 Q72R95 Q72RA8 Q72RB6 Q72RB7 Q72RC1 Q72RC4 Q72RC5 Q72RD1 Q72RD4
Q72RG6 Q72RG8 Q72RH2 Q72RH7 Q72RH8 Q72RI5 Q72RI9 Q72RJ9 Q72RL9 Q72RM8 Q72RN8
Q72RP1 Q72RP6 Q72RR0 Q72RS7 Q72RT5 Q72RT8 Q72RU2 Q72RY7 Q72RY8 Q72RY9 Q72RZ0
Q72RZ7 Q72RZ8 Q72S00 Q72S23 Q72S27 Q72S28 Q72S29 Q72S30 Q72S34 Q72S40 Q72S46
Q72S57 Q72S65 Q72S74 Q72SA5 Q72SC3 Q72SC4 Q72SD0 Q72SG5 Q72SG6 Q72SG7 Q72SJ7
Q72SM5 Q72T26 Q72T46 Q72T73 Q72T98 Q72TC0 Q72TI2 Q72TL3 Q72TP1 Q72TR2 Q72U01
Q72U04 Q72U05 Q72U06 Q72U08 Q72U10 Q72U11 Q72U12 Q72U13 Q72U14 Q72U21 Q72U22
Q72U64 Q72U97 Q72U99 Q72UA5 Q72UA6 Q72UA7 Q72UA8 Q72UA9 Q72UB0 Q72UB1 Q72UC6

Q72UH6 Q72UH8 Q72UK9 Q72UP7 Q72UP9 Q72UU3 Q72UV8 Q72UY6 Q72V30 Q72V31 Q72V33
Q72V44 Q72V55 Q72V57 Q72V72 Q72V73 Q72V75 Q72V97 Q72VB8 Q72VI2 Q72VI6 Q72VI7
Q72VI8 Q72VK0 Q72VK9 Q72VM5 Q72VP7 Q72VS5 Q72VT0 Q72VY3 Q72VY6 Q72VZ0 Q72W01
Q72W10 Q72W11 Q72W15 Q72W38 Q72W44 Q72W54 Q72W63 Q72W72 Q72WC1 Q72WD4 Q72WD6
Q72WF4 Q72WT9 Q72WU0 Q72WU3 Q72WU4 Q72WU5 Q72WU6 Q72WV3 Q72WV5 Q72WV8 Q72WW1
Q72WX0 Q72WY4 Q72X39 Q72X44 Q72X51 Q72X61 Q72X65 Q72X78 Q72X85 Q72XA0 Q72XA8
Q72XB0 Q72XB5 Q72XC0 Q72XC1 Q72XC2 Q72XD6 Q72XD7 Q72XD8 Q72XG6 Q72XT9 Q72XV1
Q72XV8 Q72XW4 Q72XW6 Q72XW9 Q72XY3 Q72XY4 Q72XZ2 Q72Y09 Q72Y10 Q72Y44 Q72Y91
Q72YB2 Q72YE6 Q72YE7 Q72YF6 Q72YG1 Q72YG8 Q72YI4 Q72YJ3 Q72YJ4 Q72YJ6 Q72YK6
Q72YK9 Q72YV4 Q72YV6 Q72YW2 Q72YY8 Q72Z05 Q72Z28 Q72Z34 Q72Z73 Q72Z76 Q72Z85
Q72Z91 Q72Z96 Q72ZA4 Q72ZC1 Q72ZD8 Q72ZE5 Q72ZF1 Q72ZF3 Q72ZF6 Q72ZF7 Q72ZG2
Q72ZG3 Q72ZG4 Q72ZH9 Q72ZI1 Q72ZI2 Q72ZI3 Q72ZI6 Q72ZJ0 Q72ZK2 Q72ZM1 Q72ZT1
Q72ZT2 Q72ZV3 Q72ZV4 Q72ZV7 Q72ZW0 Q72ZW2 Q72ZW5 Q72ZW8 Q72ZX1 Q72ZX2 Q72ZY1
Q72ZY3 Q72ZY5 Q730A4 Q730B1 Q730B3 Q730B4 Q730C4 Q730C6 Q730D1 Q730D7 Q730E6
Q730E7 Q730E8 Q730F4 Q730F5 Q730F9 Q730G0 Q730J2 Q730K1 Q730K3 Q730L3 Q730L4
Q730L9 Q730M0 Q730M1 Q730M2 Q730M3 Q730M6 Q730N3 Q730N8 Q730P0 Q730P8 Q730Q8
Q730S0 Q730W1 Q730W2 Q730Y9 Q730Z4 Q730Z6 Q731B1 Q731B2 Q731B3 Q731B4 Q731B7
Q731B9 Q731D0 Q731D9 Q731F6 Q731G3 Q731G5 Q731G7 Q731H9 Q731I2 Q731I5 Q731L0
Q731M3 Q731P0 Q731P1 Q731P4 Q731Q9 Q731R0 Q731R1 Q731R2 Q731R6 Q731R7 Q731W2
Q731Y5 Q731Y6 Q731Z0 Q732A6 Q732A7 Q732B8 Q732D5 Q732D8 Q732E3 Q732E5 Q732E7
Q732E9 Q732F6 Q732F9 Q732H0 Q732H4 Q732H8 Q732I0 Q732I1 Q732I2 Q732I3 Q732I4
Q732I5 Q732I6 Q732I7 Q732J9 Q732K0 Q732K6 Q732K9 Q732L2 Q732M1 Q732M8 Q732M9
Q732N0 Q732N3 Q732N4 Q732N8 Q732P2 Q732P3 Q732P5 Q732P6 Q732P8 Q732Q0 Q732Q5
Q732Q9 Q732R1 Q732R2 Q732R4 Q732R5 Q732U5 Q732V4 Q732V8 Q732Y5 Q732Z5 Q733B9
Q733C0 Q733H7 Q733H8 Q733H9 Q733I0 Q733J4 Q733J5 Q733J6 Q733J7 Q733J9 Q733K0
Q733K3 Q733L9 Q733M2 Q733M9 Q733N0 Q733Y5 Q734S7 Q734W9 Q735E5 Q735N3 Q735N6
Q735N7 Q735Q6 Q735X2 Q735X3 Q735Z0 Q736A4 Q736A5 Q736A7 Q736G1 Q736P6 Q736S8
Q736T1 Q736W3 Q736W4 Q736W5 Q737C1 Q737X5 Q737Z7 Q738L2 Q738Q8 Q738S2 Q738X4
Q738X7 Q739A1 Q739A9 Q739D2 Q739F4 Q739I4 Q739I5 Q739J0 Q739M9 Q739N4 Q739R5
Q739T5 Q739W8 Q739Z4 Q73A11 Q73A25 Q73A47 Q73AC5 Q73AH3 Q73AS5 Q73AT3 Q73AT4
Q73AU3 Q73AV2 Q73AV3 Q73AV4 Q73AV7 Q73AW0 Q73AW1 Q73AX5 Q73AX7 Q73AX8 Q73AX9
Q73AY0 Q73AY2 Q73AY4 Q73AZ1 Q73AZ6 Q73B32 Q73B49 Q73B75 Q73B76 Q73B97 Q73B98
Q73B99 Q73BA0 Q73BA1 Q73BA6 Q73BD2 Q73BF7 Q73BG0 Q73BH8 Q73BH9 Q73BK0 Q73BK1
Q73BK2 Q73BN8 Q73BQ6 Q73BQ7 Q73BQ8 Q73BQ9 Q73BR0 Q73BS0 Q73BU4 Q73BU7 Q73BU9
Q73BV2 Q73BX2 Q73BY1 Q73BZ5 Q73C08 Q73C23 Q73C24 Q73C98 Q73C99 Q73CB7 Q73CE0
Q73CF4 Q73CH0 Q73CI0 Q73CI4 Q73CJ5 Q73CR9 Q73CS0 Q73CV5 Q73CW1 Q73D01 Q73DB1
Q73DH4 Q73DH8 Q73DN1 Q73DV4 Q73DX4 Q73DX8 Q73DY6 Q73DY9 Q73E07 Q73E17 Q73E18
Q73E40 Q73E50 Q73E73 Q73E74 Q73E86 Q73E87 Q73E88 Q73EA6 Q73EJ5 Q73EK7 Q73EK8
Q73EK9 Q73EM3 Q73EN1 Q73EN3 Q73EN5 Q73EN6 Q73EN8 Q73ER7 Q73ER9 Q73ES0 Q73ES3
Q73ES6 Q73ET1 Q73EU1 Q73EY2 Q73F49 Q73F50 Q73F62 Q73F63 Q73F64 Q73F68 Q73F69
Q73F70 Q73F71 Q73F72 Q73F75 Q73F77 Q73F78 Q73F79 Q73F80 Q73F81 Q73F82 Q73F83
Q73F84 Q73F85 Q73F86 Q73F87 Q73F88 Q73F89 Q73F90 Q73F91 Q73F92 Q73F93 Q73F94
Q73F95 Q73F96 Q73F97 Q73F98 Q73F99 Q73FA0 Q73FA1 Q73FA3 Q73FA4 Q73FA6 Q73FA7
Q73FA8 Q73FB2 Q73FB7 Q73FB9 Q73FC0 Q73FC1 Q73FC6 Q73FD1 Q73FE1 Q73FE2 Q73FE5

Q73FF7 Q73FG0 Q73FG3 Q73FG7 Q73FH3 Q73FH5 Q73FH8 Q73FI4 Q73FI5 Q73FJ3 Q73FJ4
Q73FJ5 Q73FK2 Q73FK5 Q73FL0 Q73FM5 Q73FP4 Q73FP5 Q73FQ1 Q73FR5 Q73FR9 Q73FS2
Q73FT3 Q73FW3 Q73FX5 Q73FX6 Q73FY6 Q73FY9 Q73FZ1 Q73G09 Q73G10 Q73G18 Q73G24
Q73G26 Q73G32 Q73G44 Q73G50 Q73G51 Q73G53 Q73G60 Q73G71 Q73G84 Q73G93 Q73GC3
Q73GC9 Q73GD4 Q73GD5 Q73GD6 Q73GE2 Q73GE7 Q73GG3 Q73GG7 Q73GH2 Q73GH3 Q73GH6
Q73GL5 Q73GL7 Q73GL9 Q73GN4 Q73GN5 Q73GP1 Q73GR4 Q73GR6 Q73GR7 Q73GR8 Q73GT3
Q73GV7 Q73GW6 Q73GX9 Q73GZ0 Q73GZ5 Q73GZ6 Q73H00 Q73H01 Q73H02 Q73H09 Q73H48
Q73H59 Q73H71 Q73H85 Q73H86 Q73H89 Q73H90 Q73H91 Q73H92 Q73H93 Q73H94 Q73H95
Q73H96 Q73H97 Q73H98 Q73H99 Q73HA0 Q73HA2 Q73HA3 Q73HA4 Q73HA6 Q73HA7 Q73HA8
Q73HA9 Q73HB0 Q73HK1 Q73HK3 Q73HL4 Q73HM1 Q73HM2 Q73HM7 Q73HP2 Q73HQ3 Q73HS5
Q73HS9 Q73HV1 Q73HV5 Q73HW5 Q73HW7 Q73HW8 Q73HX6 Q73HY6 Q73HZ4 Q73HZ5 Q73HZ8
Q73I07 Q73I13 Q73I23 Q73I40 Q73I41 Q73I44 Q73I47 Q73I58 Q73I59 Q73I60 Q73I70
Q73I71 Q73I74 Q73IC2 Q73IC3 Q73IE4 Q73IE9 Q73IG5 Q73IH1 Q73IH9 Q73II8 Q73II9
Q73IJ6 Q73IK6 Q73IL1 Q73IL4 Q73IL5 Q73IM3 Q73IM5 Q73IP1 Q73IQ0 Q73IQ8 Q73IR2
Q73IR3 Q73IS9 Q73IT2 Q73IT7 Q73IU2 Q73IV4 Q73IW5 Q73IW9 Q73IX2 Q73IX6 Q73IX7
Q73IX8 Q73IX9 Q73IZ0 Q73J26 Q73J84 Q73JB2 Q73JF6 Q73JG6 Q73JJ0 Q73JJ4 Q73JJ5
Q73JJ6 Q73JJ7 Q73JJ8 Q73JL8 Q73JM9 Q73JN2 Q73JP8 Q73JQ7 Q73JR1 Q73JR4 Q73JT1
Q73JU6 Q73JU8 Q73JV7 Q73K17 Q73K18 Q73K21 Q73K67 Q73K68 Q73K74 Q73K75 Q73K76
Q73K78 Q73K81 Q73K92 Q73KB2 Q73KB6 Q73KC6 Q73KE1 Q73KE3 Q73KE7 Q73KH5 Q73KI2
Q73KJ8 Q73KL7 Q73KM3 Q73KN7 Q73KR2 Q73KR3 Q73KR6 Q73KS3 Q73KU4 Q73KZ3 Q73L97
Q73LB4 Q73LC2 Q73LE9 Q73LF2 Q73LF4 Q73LJ8 Q73LJ9 Q73LW1 Q73LW3 Q73LW4 Q73LZ4
Q73M02 Q73M14 Q73M27 Q73M32 Q73M34 Q73M35 Q73M37 Q73M38 Q73M39 Q73M53 Q73M65
Q73M68 Q73M71 Q73M76 Q73M83 Q73MB8 Q73MC2 Q73MC7 Q73MD3 Q73MF9 Q73MG1 Q73MH0
Q73MK4 Q73MP4 Q73MQ8 Q73MR1 Q73MV2 Q73MY1 Q73MY7 Q73MZ2 Q73N15 Q73N16 Q73N17
Q73N18 Q73N26 Q73N77 Q73N90 Q73NB3 Q73NB9 Q73NE3 Q73NE4 Q73NF6 Q73NH7 Q73NM0
Q73NM2 Q73NP0 Q73NP4 Q73NP6 Q73NP7 Q73NP8 Q73NQ6 Q73NQ7 Q73NR2 Q73NS2 Q73NT9
Q73NV3 Q73NV4 Q73NV5 Q73NW1 Q73NW2 Q73NX5 Q73NX7 Q73P09 Q73P19 Q73P28 Q73P31
Q73P36 Q73P65 Q73P66 Q73P70 Q73PB4 Q73PB5 Q73PB6 Q73PD8 Q73PE7 Q73PH1 Q73PH3
Q73PH4 Q73PK2 Q73PK7 Q73PK8 Q73PK9 Q73PL0 Q73PL1 Q73PL3 Q73PL4 Q73PL5 Q73PL6
Q73PL7 Q73PL8 Q73PL9 Q73PM0 Q73PM1 Q73PM2 Q73PM3 Q73PM4 Q73PM5 Q73PM6 Q73PM7
Q73PM8 Q73PM9 Q73PN0 Q73PN1 Q73PN2 Q73PN3 Q73PQ2 Q73PV6 Q73PV9 Q73Q01 Q73Q03
Q73Q15 Q73Q16 Q73Q17 Q73Q56 Q73Q68 Q73QI7 Q73QJ2 Q73QJ5 Q73QK5 Q73QV3 Q73QV6
Q73QW4 Q73QY2 Q73QZ3 Q73R08 Q73R77 Q73R87 Q73RB6 Q73RG3 Q73RJ5 Q73RJ6 Q73RN1
Q73RN8 Q73RR5 Q73RR6 Q73RS2 Q73RS4 Q73S13 Q73S23 Q73S29 Q73S30 Q73S31 Q73S41
Q73S42 Q73S43 Q73S44 Q73S45 Q73S46 Q73S47 Q73S77 Q73S79 Q73S85 Q73S86 Q73S87
Q73S90 Q73S91 Q73S92 Q73S93 Q73S94 Q73S95 Q73S96 Q73S97 Q73S98 Q73SA5 Q73SA6
Q73SA7 Q73SA8 Q73SA9 Q73SB0 Q73SB1 Q73SB2 Q73SB3 Q73SB4 Q73SC5 Q73SC8 Q73SD1
Q73SD2 Q73SD3 Q73SD4 Q73SE3 Q73SE4 Q73SE8 Q73SE9 Q73SG1 Q73SG5 Q73SG7 Q73SG8
Q73SJ4 Q73SJ5 Q73SK9 Q73SL0 Q73SL8 Q73SM1 Q73SQ3 Q73SS0 Q73SS1 Q73SS7 Q73SU2
Q73SU4 Q73SU8 Q73SV2 Q73T00 Q73T03 Q73T20 Q73T33 Q73T38 Q73T50 Q73T66 Q73T77
Q73T78 Q73T98 Q73TE1 Q73TE7 Q73TE9 Q73TF0 Q73TF1 Q73TP5 Q73TQ5 Q73TS2 Q73TS5
Q73TT7 Q73U04 Q73U05 Q73U14 Q73U39 Q73U79 Q73U90 Q73U92 Q73UA5 Q73UB0 Q73UD0
Q73UD7 Q73UG7 Q73UI3 Q73UJ3 Q73UJ4 Q73UK6 Q73UN2 Q73V46 Q73V62 Q73V97 Q73VB3
Q73VF7 Q73VG8 Q73VG9 Q73VH7 Q73VI1 Q73VI4 Q73VI7 Q73VI8 Q73VJ1 Q73VJ2 Q73VJ4

Q73VJ7 Q73VK6 Q73VL1 Q73VL8 Q73VL9 Q73VM2 Q73VM3 Q73VN8 Q73VN9 Q73VP1 Q73VP3
Q73VP7 Q73VQ7 Q73VR6 Q73VR8 Q73VS1 Q73VS3 Q73VS8 Q73VU0 Q73VU8 Q73VV1 Q73VV4
Q73VV5 Q73VW3 Q73VW9 Q73VX0 Q73VY3 Q73VZ2 Q73VZ6 Q73VZ7 Q73W15 Q73W20 Q73W21
Q73W26 Q73W27 Q73W57 Q73W61 Q73WB6 Q73WE4 Q73WF0 Q73WF2 Q73WG0 Q73WG1 Q73WH3
Q73WH4 Q73WH6 Q73WJ9 Q73WM7 Q73WP7 Q73WP9 Q73WU1 Q73WU6 Q73WV2 Q73WX4 Q73X40
Q73X44 Q73X47 Q73X48 Q73X63 Q73X75 Q73X80 Q73X85 Q73X87 Q73X89 Q73XM4 Q73XM7
Q73XM8 Q73XM9 Q73XN1 Q73XN4 Q73XN8 Q73XP1 Q73XP3 Q73XP4 Q73XP5 Q73XP6 Q73XR2
Q73XR5 Q73XT5 Q73XT9 Q73XV0 Q73XV3 Q73XZ5 Q73XZ6 Q73XZ9 Q73Y16 Q73Y21 Q73Y26
Q73Y81 Q73Y99 Q73YC1 Q73YI6 Q73YJ7 Q73YK2 Q73YK4 Q73YK7 Q73YM4 Q73YP9 Q73YQ3
Q73YQ6 Q73YR4 Q73YS9 Q73YW8 Q73YW9 Q73YX0 Q73ZD5 Q73ZH8 Q73ZI0 Q73ZI1 Q73ZL4
Q73ZM8 Q73ZP6 Q73ZP7 Q73ZP8 Q73ZP9 Q73ZQ2 Q73ZQ6 Q73ZR2 Q740D6 Q740D7 Q740E5
Q740E9 Q740F5 Q740F6 Q740I3 Q740I4 Q740I5 Q740I7 Q740I9 Q740J0 Q740J1 Q740J6
Q740J7 Q740M4 Q740P2 Q740P4 Q740Q3 Q740Q9 Q740R7 Q740T9 Q740U6 Q740Y1 Q74123
Q741B5 Q741C5 Q741E2 Q741E4 Q741E5 Q741F1 Q741F8 Q741G5 Q741G7 Q741H0 Q741H2
Q741H5 Q741H6 Q741J2 Q741J3 Q741J6 Q741J7 Q741J8 Q741J9 Q741L2 Q741M7 Q741P3
Q741Q1 Q741Q2 Q741V8 Q741V9 Q741W1 Q741W2 Q742C2 Q742E4 Q742F3 Q742K7 Q742L2
Q742Q2 Q742Z0 Q742Z1 Q743C2 Q743D5 Q743E2 Q743F0 Q743F9 Q743L8 Q743Q8 Q743Q9
Q743V5 Q743W2 Q743W3 Q743W4 Q743W5 Q743W9 Q743Y3 Q743Y4 Q743Y5 Q743Z2 Q744A3
Q744M4 Q744M5 Q744T3 Q744V3 Q744V6 Q744X5 Q744X6 Q745I2 Q745I8 Q745J2 Q745R0
Q745W0 Q746B8 Q746C2 Q746F4 Q746G0 Q746H1 Q746H2 Q746I8 Q746J8 Q746M5 Q746Q3
Q746Q4 Q746Q5 Q746R5 Q746Y5 Q746Y7 Q746Y8 Q746Z9 Q747A0 Q747A2 Q747B9 Q747C7
Q747C8 Q747F5 Q747F9 Q747I1 Q747I2 Q747I7 Q747K3 Q747K8 Q747M9 Q747N0 Q747Q2
Q747Q3 Q747Q4 Q747Q5 Q747Q7 Q747Q8 Q747R0 Q747R1 Q747R4 Q747V4 Q747W9 Q747X9
Q748A8 Q748A9 Q748B1 Q748B2 Q748B3 Q748B4 Q748C9 Q748D2 Q748D4 Q748D8 Q748G1
Q748G2 Q748I9 Q748J3 Q748M1 Q748M4 Q748X2 Q748X4 Q748X6 Q748X8 Q748Y3 Q748Y4
Q748Y5 Q748Y6 Q748Y7 Q748Y8 Q748Z0 Q748Z1 Q748Z2 Q748Z3 Q748Z4 Q748Z5 Q748Z6
Q748Z7 Q748Z8 Q748Z9 Q749A0 Q749A1 Q749A2 Q749A3 Q749A4 Q749A5 Q749A6 Q749A8
Q749B0 Q749B1 Q749B2 Q749B3 Q749B4 Q749F2 Q749I1 Q749N5 Q749N8 Q749W3 Q749W5
Q749X2 Q749Y6 Q749Y7 Q749Y8 Q74A22 Q74A44 Q74A46 Q74A61 Q74AC8 Q74AE2 Q74AH4
Q74AH7 Q74AI3 Q74AR8 Q74AW2 Q74AW5 Q74AX0 Q74AX1 Q74AX4 Q74AZ0 Q74AZ3 Q74B44
Q74B57 Q74BE1 Q74BF0 Q74BF4 Q74BF6 Q74BH0 Q74BH6 Q74BJ6 Q74BL4 Q74BL5 Q74BL6
Q74BL9 Q74BP0 Q74BP1 Q74BU2 Q74BW1 Q74BW2 Q74BW4 Q74BW7 Q74BW9 Q74BX5 Q74BY3
Q74BZ3 Q74C11 Q74C12 Q74C57 Q74C63 Q74C65 Q74C70 Q74C76 Q74C79 Q74C80 Q74C81
Q74C82 Q74C83 Q74CA7 Q74CA8 Q74CB0 Q74CB6 Q74CB9 Q74CC2 Q74CC6 Q74CF3 Q74CG4
Q74CG7 Q74CG8 Q74CI1 Q74CI6 Q74CM2 Q74CR0 Q74CR5 Q74CS3 Q74CS9 Q74CT0 Q74CT2
Q74CT3 Q74CT6 Q74CT7 Q74CT9 Q74CU0 Q74CZ2 Q74CZ3 Q74CZ5 Q74CZ6 Q74CZ9 Q74D00
Q74D01 Q74D02 Q74D03 Q74D04 Q74D29 Q74D53 Q74D56 Q74D58 Q74D59 Q74D60 Q74D67
Q74DC4 Q74DI7 Q74DI8 Q74DK5 Q74DP4 Q74DP5 Q74DP6 Q74DR9 Q74DS2 Q74DU6 Q74DU7
Q74E08 Q74E20 Q74E25 Q74E52 Q74E53 Q74E54 Q74E87 Q74E88 Q74EA5 Q74EB5 Q74ED9
Q74EG5 Q74EM9 Q74EQ6 Q74ER9 Q74ES0 Q74EU2 Q74EU7 Q74F05 Q74F83 Q74FC3 Q74FD5
Q74FE1 Q74FE3 Q74FE4 Q74FE6 Q74FE7 Q74FE9 Q74FF1 Q74FF9 Q74FG0 Q74FG1 Q74FG3
Q74FG4 Q74FJ9 Q74FK1 Q74FK2 Q74FL9 Q74FM7 Q74FM8 Q74FM9 Q74FS7 Q74FT1 Q74FT5
Q74FU2 Q74FW6 Q74FW8 Q74FW9 Q74FY7 Q74G02 Q74G05 Q74G62 Q74G73 Q74GA9 Q74GB4
Q74GH6 Q74GL3 Q74GN1 Q74GS4 Q74GT3 Q74GT5 Q74GT6 Q74GT9 Q74GU2 Q74GU4 Q74GV1

Q74GV3 Q74GV6 Q74GW4 Q74GW7 Q74GZ7 Q74H35 Q74H36 Q74H55 Q74H58 Q74H59 Q74H60
Q74H61 Q74H71 Q74H90 Q74H92 Q74H94 Q74H95 Q74HC2 Q74HC3 Q74HC4 Q74HC8 Q74HE9
Q74HK4 Q74HL7 Q74HL8 Q74HM3 Q74HT0 Q74HT9 Q74HZ5 Q74I36 Q74I49 Q74I51 Q74I52
Q74I68 Q74I89 Q74I98 Q74IA4 Q74IB5 Q74IB6 Q74IB7 Q74IC0 Q74IC4 Q74IC5 Q74IC6
Q74ID3 Q74ID8 Q74IE1 Q74IE2 Q74IE7 Q74IF3 Q74IG2 Q74IG3 Q74IG4 Q74IG8 Q74IL4
Q74IL5 Q74IL7 Q74IM0 Q74IM1 Q74IM2 Q74IM7 Q74IM9 Q74IN3 Q74IN6 Q74IP8 Q74IQ5
Q74IQ6 Q74IQ7 Q74IR7 Q74IR8 Q74IR9 Q74IS2 Q74IS4 Q74IS8 Q74IS9 Q74IT0 Q74IT4
Q74IT5 Q74IT6 Q74IT7 Q74IU0 Q74IU3 Q74IX0 Q74IX3 Q74IX4 Q74IY4 Q74IY5 Q74IY8
Q74IZ0 Q74IZ2 Q74J23 Q74J27 Q74J28 Q74J29 Q74J31 Q74J32 Q74J34 Q74J95 Q74J96
Q74JA5 Q74JA6 Q74JA9 Q74JB4 Q74JC1 Q74JD5 Q74JJ5 Q74JJ8 Q74JK1 Q74JK4 Q74JK7
Q74JL6 Q74JL8 Q74JM2 Q74JM3 Q74JM8 Q74JN2 Q74JN5 Q74JN7 Q74JN8 Q74JU4 Q74JU9
Q74JV0 Q74JV6 Q74JW2 Q74JW3 Q74JW9 Q74JX6 Q74JY0 Q74JY5 Q74JZ0 Q74JZ8 Q74K02
Q74K05 Q74K25 Q74K26 Q74K31 Q74K59 Q74K66 Q74K70 Q74K75 Q74K79 Q74K84 Q74K85
Q74K87 Q74K90 Q74KC7 Q74KF3 Q74KI9 Q74KR5 Q74KS1 Q74KS4 Q74KS6 Q74KU4 Q74KU5
Q74KU8 Q74KV0 Q74KV1 Q74KV2 Q74KV9 Q74KW0 Q74KX0 Q74KY3 Q74KY8 Q74KZ4 Q74KZ7
Q74KZ9 Q74L00 Q74L08 Q74L09 Q74L13 Q74L17 Q74L26 Q74L27 Q74L45 Q74L58 Q74L59
Q74L63 Q74L64 Q74L65 Q74L66 Q74L67 Q74L69 Q74L71 Q74L72 Q74L73 Q74L74 Q74L75
Q74L76 Q74L78 Q74L79 Q74L80 Q74L81 Q74L82 Q74L83 Q74L84 Q74L85 Q74L86 Q74L87
Q74L88 Q74L89 Q74L90 Q74L91 Q74L92 Q74L94 Q74L95 Q74L99 Q74LA3 Q74LA8 Q74LB6
Q74LC1 Q74LF7 Q74LF9 Q74LG1 Q74LG2 Q74LG3 Q74LI0 Q74LL0 Q74LL9 Q74LQ9 Q74LS5
Q74LU5 Q74LW2 Q74LY0 Q74LZ9 Q74M23 Q74M26 Q74M28 Q74M31 Q74M34 Q74M50 Q74M51
Q74M52 Q74M58 Q74M64 Q74M65 Q74M72 Q74M83 Q74M84 Q74M95 Q74M98 Q74MA1 Q74MA7
Q74MB1 Q74MB5 Q74MB7 Q74MB8 Q74MD3 Q74MD4 Q74MH2 Q74MH3 Q74MH8 Q74MI1 Q74MI5
Q74MI6 Q74MJ0 Q74MJ4 Q74MJ5 Q74MN8 Q74MN9 Q74MP3 Q74MP4 Q74MP5 Q74MQ6 Q74MR5
Q74MR6 Q74MS7 Q74MU5 Q74MV1 Q74MV8 Q74MW8 Q74MX2 Q74MX9 Q74MY6 Q74MZ1 Q74MZ3
Q74MZ4 Q74MZ5 Q74N11 Q74N14 Q74N21 Q74N24 Q74N29 Q74N45 Q74N46 Q74N55 Q74N58
Q74N73 Q74N77 Q74N79 Q74N81 Q74N89 Q74N90 Q74NA6 Q74NA9 Q74NC6 Q74ND5 Q74ND7
Q74NE4 Q74NE6 Q74NF3 Q74NF7 Q74NG1 Q74NH9 Q74NJ4 Q74NK1 Q74NK2 Q74NT6 Q74P24
Q74P25 Q74P74 Q74PY8 Q74R94 Q74RQ9 Q74S44 Q74SF4 Q74SR9 Q74TH2 Q74VT6 Q74X11
Q74XD3 Q74XX5 Q74Y23 Q74Y45 Q74Y93 Q74Z08 Q74Z24 Q74Z26 Q74Z27 Q74Z32 Q74Z37
Q74Z47 Q74Z51 Q74Z52 Q74Z75 Q74ZA0 Q74ZA4 Q74ZA9 Q74ZB0 Q74ZB5 Q74ZC0 Q74ZC1
Q74ZC9 Q74ZD8 Q74ZF0 Q74ZF1 Q74ZF6 Q74ZG4 Q74ZH6 Q74ZH9 Q74ZJ0 Q74ZJ6 Q74ZK3
Q74ZK4 Q74ZK6 Q74ZK8 Q74ZL1 Q74ZL4 Q74ZL5 Q74ZL7 Q74ZM2 Q74ZM9 Q74ZN0 Q74ZN9
Q74ZQ0 Q74ZQ8 Q74ZR2 Q74ZS6 Q74ZU6 Q74ZV8 Q74ZW4 Q74ZX6 Q74ZZ1 Q750A4 Q750A5
Q750B5 Q750C2 Q750D5 Q750E0 Q750E9 Q750F0 Q750F4 Q750F5 Q750F8 Q750G3 Q750J3
Q750K9 Q750N3 Q750P5 Q750P7 Q750Q4 Q750Q9 Q750R1 Q750S2 Q750S3 Q750S5 Q750T7
Q750U1 Q750U8 Q750V7 Q750W3 Q750X2 Q750X3 Q750Y3 Q750Y8 Q751A7 Q751B2 Q751D1
Q751D2 Q751E7 Q751F5 Q751F9 Q751G1 Q751G9 Q751H1 Q751I6 Q751I7 Q751J3 Q751K7
Q751K8 Q751L1 Q751L7 Q751L8 Q751N0 Q751P0 Q751P4 Q751Q9 Q751S5 Q751T1 Q751U1
Q751X1 Q751Y8 Q751Z6 Q752A3 Q752B6 Q752B8 Q752D7 Q752E4 Q752H4 Q752J5 Q752J8
Q752L7 Q752N3 Q752N9 Q752P7 Q752Q3 Q752S4 Q752S6 Q752T7 Q752U2 Q752U3 Q752U5
Q752U6 Q752Y2 Q752Z8 Q752Z9 Q753A0 Q753A3 Q753C4 Q753C6 Q753D3 Q753D4 Q753F1
Q753F4 Q753F5 Q753I0 Q753I4 Q753K4 Q753L9 Q753M0 Q753M3 Q753N1 Q753P9 Q753Q4
Q753S8 Q753U4 Q753V5 Q753W8 Q753W9 Q753Y0 Q753Y2 Q753Y3 Q753Y5 Q753Z8 Q754C8

Q754C9 Q754E7 Q754G5 Q754H6 Q754J2 Q754J3 Q754J8 Q754K8 Q754N8 Q754P2 Q754Q2
Q754R8 Q754T3 Q754T8 Q754U2 Q754U4 Q754U7 Q754V2 Q754W7 Q754X1 Q754X6 Q754Z1
Q755A0 Q755A2 Q755A3 Q755A5 Q755A7 Q755B3 Q755C4 Q755C5 Q755C8 Q755D0 Q755D2
Q755D9 Q755E7 Q755F7 Q755G5 Q755J5 Q755L7 Q755M2 Q755M3 Q755M5 Q755M8 Q755N4
Q755P0 Q755Q2 Q755R1 Q755R3 Q755R9 Q755T1 Q755T2 Q755T5 Q755T8 Q755V6 Q755V9
Q755W0 Q755Z1 Q755Z9 Q756A7 Q756B7 Q756C2 Q756C3 Q756C7 Q756D0 Q756E1 Q756E2
Q756F4 Q756F6 Q756G2 Q756G5 Q756H2 Q756J5 Q756J6 Q756K3 Q756K6 Q756K8 Q756K9
Q756L0 Q756L2 Q756L3 Q756P0 Q756Q4 Q756Q5 Q756R8 Q756V1 Q756X3 Q756X6 Q756Y3
Q756Y8 Q756Z2 Q757B6 Q757B7 Q757C7 Q757D4 Q757D7 Q757D9 Q757E3 Q757G1 Q757G5
Q757G7 Q757H5 Q757H6 Q757H7 Q757I0 Q757I2 Q757I6 Q757I9 Q757J8 Q757J9 Q757K0
Q757K3 Q757L1 Q757L4 Q757M0 Q757M3 Q757M7 Q757N1 Q757N6 Q757Q0 Q757Q6 Q757R4
Q757S1 Q757T0 Q757T2 Q757U8 Q757V8 Q757W2 Q757Y8 Q758A1 Q758A4 Q758D8 Q758E4
Q758F0 Q758H6 Q758K0 Q758L0 Q758L3 Q758M1 Q758M2 Q758N7 Q758R2 Q758R8 Q758R9
Q758S2 Q758S3 Q758S6 Q758T2 Q758T3 Q758T8 Q758V1 Q758V8 Q758W7 Q758X9 Q758Y4
Q758Z4 Q759A9 Q759B6 Q759B7 Q759D1 Q759F6 Q759G5 Q759G7 Q759I2 Q759K3 Q759L2
Q759L6 Q759L8 Q759M1 Q759Q5 Q759Q6 Q759R7 Q759S0 Q759T0 Q759T3 Q759T4 Q759T6
Q759T9 Q759U5 Q759U7 Q759V4 Q759W1 Q759Y1 Q759Y5 Q759Y6 Q759Y8 Q75A03 Q75A06
Q75A07 Q75A17 Q75A34 Q75A36 Q75A37 Q75A43 Q75A44 Q75A47 Q75A58 Q75A59 Q75A64
Q75A74 Q75A83 Q75AA5 Q75AA7 Q75AA8 Q75AB5 Q75AB6 Q75AC2 Q75AC3 Q75AD1 Q75AD3
Q75AE1 Q75AF2 Q75AF9 Q75AG5 Q75AH3 Q75AH9 Q75AI1 Q75AK8 Q75AL5 Q75AP1 Q75AP6
Q75AP7 Q75AQ9 Q75AR4 Q75AS0 Q75AS3 Q75AT3 Q75AV2 Q75AV4 Q75AV8 Q75AW1 Q75AW4
Q75AX1 Q75AX7 Q75AX9 Q75AY1 Q75AZ2 Q75AZ9 Q75B32 Q75B37 Q75B43 Q75B47 Q75B48
Q75B50 Q75B51 Q75B70 Q75B78 Q75B82 Q75B88 Q75B89 Q75B93 Q75B95 Q75BA8 Q75BB0
Q75BB3 Q75BD8 Q75BE7 Q75BF1 Q75BI3 Q75BI6 Q75BJ7 Q75BK0 Q75BK9 Q75BL6 Q75BL8
Q75BN0 Q75BP2 Q75BQ6 Q75BR7 Q75BS7 Q75BT5 Q75BT9 Q75BU2 Q75BU9 Q75BV8 Q75BW4
Q75BX3 Q75BY1 Q75BY2 Q75BY3 Q75C07 Q75C17 Q75C22 Q75C26 Q75C29 Q75C70 Q75C76
Q75C80 Q75C90 Q75C99 Q75CA4 Q75CC5 Q75CD6 Q75CE0 Q75CF3 Q75CK5 Q75CM4 Q75CM8
Q75CN1 Q75CP5 Q75CP6 Q75CQ9 Q75CZ2 Q75CZ5 Q75CZ6 Q75D00 Q75D20 Q75D21 Q75D29
Q75D46 Q75D53 Q75D58 Q75D66 Q75D77 Q75D81 Q75D84 Q75D88 Q75D94 Q75DA3 Q75DA4
Q75DB8 Q75DC2 Q75DC5 Q75DC6 Q75DD1 Q75DE4 Q75DE8 Q75DE9 Q75DG4 Q75DJ1 Q75DJ3
Q75DK9 Q75DP8 Q75DR2 Q75DR9 Q75DS1 Q75DS4 Q75DS8 Q75DT8 Q75DU5 Q75DU9 Q75DV4
Q75DV9 Q75DX7 Q75DX9 Q75DY0 Q75DY2 Q75DZ2 Q75E14 Q75E19 Q75E22 Q75E32 Q75E35
Q75E45 Q75E56 Q75E60 Q75E62 Q75E67 Q75E72 Q75E79 Q75E92 Q75E95 Q75EA5 Q75EB3
Q75EB8 Q75EC0 Q75ED7 Q75EE0 Q75EF0 Q75EG6 Q75EH2 Q75EI1 Q75EI4 Q75EI5 Q75EJ1
Q75EK2 Q75EM1 Q75EN0 Q75EP8 Q75EQ0 Q75ET5 Q75ET6 Q75EV6 Q75EV9 Q75EW4 Q75EW5
Q75EW9 Q75EX7 Q75EY2 Q75EY3 Q75EY8 Q75EZ2 Q75EZ5 Q75EZ7 Q75F02 Q75F10 Q75F23
Q75F26 Q75F44 Q75F65 Q75F95 Q75FC8 Q75FF2 Q75FF4 Q75FG2 Q75FG3 Q75FL1 Q75FP0
Q75FQ8 Q75FT0 Q75FU1 Q75FU8 Q75FW3 Q75FW5 Q75G04 Q75G41 Q75G68 Q75G87 Q75GB1
Q75GK0 Q75GK1 Q75GM1 Q75GR5 Q75GS4 Q75GT3 Q75H81 Q75HA6 Q75HC2 Q75HE5 Q75HE6
Q75HJ0 Q75HV0 Q75HV1 Q75HX5 Q75HY5 Q75HZ9 Q75I13 Q75I54 Q75IM9 Q75IP6 Q75IR6
Q75IS2 Q75IV7 Q75IZ7 Q75J90 Q75JB3 Q75JC1 Q75JC2 Q75JC6 Q75JC7 Q75JC8 Q75JD5
Q75JD8 Q75JF1 Q75JF2 Q75JF4 Q75JF9 Q75JG8 Q75JL2 Q75JM4 Q75JN1 Q75JN8 Q75JP5
Q75JP7 Q75JQ1 Q75JQ6 Q75JQ9 Q75JR2 Q75JR3 Q75JU2 Q75JW3 Q75JW9 Q75JX0 Q75JZ0
Q75JZ1 Q75K12 Q75K24 Q75K28 Q75K81 Q75KE5 Q75KP8 Q75KR1 Q75KV9 Q75L11 Q75L30

Q75L34 Q75LI2 Q75LJ3 Q75LL2 Q75LL6 Q75LR2 Q75LS8 Q75LU8 Q75LV5 Q75LX7 Q75LX9
Q75M35 Q75MW2 Q75N62 Q75NG9 Q75NR7 Q75NT5 Q75NZ0 Q75PQ8 Q75QI0 Q75QN2 Q75R64
Q75SP7 Q75SY5 Q75TB7 Q75TB9 Q75TC0 Q75TC1 Q75TC2 Q75TC3 Q75TC9 Q75TH5 Q75UN0
Q75UN1 Q75UQ2 Q75UV1 Q75VA5 Q75VA8 Q75VA9 Q75VB1 Q75VB2 Q75VB3 Q75VB4 Q75VB5
Q75VN3 Q75VW3 Q75VX8 Q75W16 Q75W17 Q75W54 Q75W95 Q75WB5 Q75WD2 Q75WE7 Q75WM6
Q75WS4 Q75WV3 Q75ZP8 Q75ZQ2 Q75ZQ3 Q75ZQ4 Q75ZQ5 Q75ZQ6 Q75ZX4 Q75ZZ6 Q760R4
Q760R5 Q760R7 Q760R8 Q760R9 Q760S4 Q760S6 Q760S7 Q760T4 Q760T5 Q761Z9 Q762H5
Q763T4 Q764B9 Q764D2 Q764M6 Q764S0 Q764S1 Q765S0 Q765S2 Q76637 Q766U5 Q766Z3
Q767C8 Q767F1 Q767K6 Q767K8 Q767K9 Q767L0 Q767L1 Q767M0 Q767M3 Q768S8 Q768T2
Q768T3 Q768T4 Q768T5 Q76B40 Q76BS5 Q76BS7 Q76C99 Q76CY5 Q76CY8 Q76CZ1 Q76DY9
Q76E23 Q76EJ0 Q76EV4 Q76EZ2 Q76F73 Q76FD7 Q76FD9 Q76FE5 Q76FE7 Q76FE9 Q76FF1
Q76FF3 Q76FF5 Q76FS2 Q76FS3 Q76FS5 Q76G09 Q76G15 Q76G20 Q76H85 Q76HI7 Q76HM5
Q76HM7 Q76HN4 Q76I90 Q76IC5 Q76IH4 Q76IH5 Q76IQ1 Q76IQ2 Q76IQ3 Q76IQ9 Q76K71
Q76KA2 Q76KA3 Q76KA7 Q76KA8 Q76KC2 Q76KD6 Q76KF9 Q76KJ5 Q76KK0 Q76KX0 Q76KX8
Q76L36 Q76L37 Q76L39 Q76L87 Q76LA0 Q76LC1 Q76LC6 Q76LL6 Q76LL9 Q76LM0 Q76LW3
Q76MH8 Q76MH9 Q76MR7 Q76MT0 Q76MT4 Q76MU7 Q76MV0 Q76MX2 Q76MY1 Q76N23 Q76N24
Q76N32 Q76N59 Q76NT0 Q76NT9 Q76NU3 Q76NU5 Q76NV1 Q76NV5 Q76NV7 Q76NW2 Q76NW7
Q76NZ7 Q76P00 Q76PC4 Q76PC7 Q76PD2 Q76PD3 Q76QK5 Q76QY7 Q76QZ8 Q76QZ9 Q76R05
Q76RA8 Q76RB3 Q76RD8 Q76RE7 Q76VG7 Q76VG8 Q76VG9 Q76VH0 Q76VH1 Q76VH2 Q76VV3
Q76ZJ3 Q76ZL3 Q76ZN5 Q76ZN8 Q76ZP7 Q76ZQ8 Q76ZR2 Q76ZS0 Q76ZS7 Q76ZT1 Q76ZV3
Q76ZW9 Q76ZX2 Q774I1 Q775C3 Q775D6 Q775N4 Q775N7 Q775Q3 Q775Q4 Q775T6 Q775U0
Q775W3 Q775X4 Q775Z7 Q776A6 Q779J8 Q77DJ5 Q77DS0 Q77E03 Q77FQ3 Q77FQ5 Q77HP2
Q77IS8 Q77J49 Q77K03 Q77KZ6 Q77M43 Q77MP8 Q77MP9 Q77MQ0 Q77MQ9 Q77MR1 Q77MR2
Q77MR3 Q77MR5 Q77MR9 Q77MS1 Q77MS2 Q77MS6 Q77MS7 Q77MT0 Q77MT3 Q77NC1 Q77NC3
Q77NN7 Q77NN9 Q77PU6 Q77SJ8 Q77TF6 Q77TG7 Q77TG9 Q77TH1 Q77TH6 Q77TH8 Q77TL8
Q77TL9 Q77WY2 Q77Z83 Q77ZF7 Q77ZF8 Q789F3 Q78E60 Q78E65 Q78J03 Q78JE5 Q78JW9
Q78T81 Q78WH7 Q78WZ7 Q78XR0 Q78Y63 Q78ZR5 Q791B0 Q791N7 Q792H5 Q792Q4 Q793I8
Q794E4 Q794H2 Q795J3 Q795M6 Q795R8 Q795U4 Q795Y4 Q796A8 Q796K9 Q796Q6 Q796S4
Q796V8 Q796Y5 Q796Y8 Q796Z1 Q797E6 Q79AF6 Q79C01 Q79CE8 Q79E92 Q79EC5 Q79EM7
Q79EM8 Q79F77 Q79FH3 Q79FK6 Q79FP3 Q79FU2 Q79FU3 Q79FV4 Q79G84 Q79GC6 Q79PF6
Q79V15 Q79V60 Q79V61 Q79V62 Q79VI4 Q79VI7 Q79VJ0 Q79VJ1 Q79VJ2 Q79VJ3 Q79VJ4
Q79ZV4 Q79ZX9 Q79ZY4 Q79ZZ6 Q7A004 Q7A005 Q7A018 Q7A028 Q7A057 Q7A064 Q7A065
Q7A071 Q7A075 Q7A078 Q7A079 Q7A080 Q7A081 Q7A083 Q7A084 Q7A085 Q7A087 Q7A089
Q7A098 Q7A0A0 Q7A0B9 Q7A0C8 Q7A0D2 Q7A0D6 Q7A0D7 Q7A0G9 Q7A0H1 Q7A0H5 Q7A0I3
Q7A0J0 Q7A0J4 Q7A0K2 Q7A0L8 Q7A0M7 Q7A0N0 Q7A0N4 Q7A0P4 Q7A0Q2 Q7A0Q3 Q7A0R0
Q7A0R2 Q7A0R5 Q7A0R9 Q7A0T9 Q7A0U1 Q7A0U9 Q7A0V5 Q7A0V6 Q7A0W1 Q7A0W2 Q7A0W3
Q7A0W8 Q7A0X1 Q7A0X3 Q7A0X6 Q7A0Y7 Q7A0Y9 Q7A105 Q7A110 Q7A115 Q7A116 Q7A120
Q7A133 Q7A135 Q7A138 Q7A154 Q7A161 Q7A171 Q7A189 Q7A198 Q7A1B6 Q7A1B9 Q7A1C0
Q7A1C9 Q7A1D4 Q7A1E0 Q7A1E3 Q7A1F7 Q7A1G0 Q7A1G5 Q7A1H9 Q7A1I5 Q7A1J9 Q7A1K6
Q7A1N5 Q7A1P1 Q7A1P6 Q7A1P8 Q7A1P9 Q7A1Q4 Q7A1Q7 Q7A1Q9 Q7A1R4 Q7A1R6 Q7A1S4
Q7A1T6 Q7A1V6 Q7A1W0 Q7A1W8 Q7A1W9 Q7A1Y2 Q7A1Y7 Q7A1Y8 Q7A1Z6 Q7A1Z8 Q7A224
Q7A225 Q7A2K5 Q7A2K9 Q7A2M3 Q7A2N3 Q7A2R1 Q7A2R8 Q7A2S0 Q7A2U9 Q7A2W5 Q7A2W7
Q7A2W8 Q7A2W9 Q7A2Y8 Q7A352 Q7A3C7 Q7A3E1 Q7A3K1 Q7A3V0 Q7A3W9 Q7A428 Q7A441
Q7A442 Q7A455 Q7A4D4 Q7A4E1 Q7A4E2 Q7A4G8 Q7A4H2 Q7A4Q5 Q7A4Q9 Q7A4T2 Q7A4T8

Q7A505 Q7A511 Q7A5K2 Q7A5L0 Q7A5M8 Q7A5N7 Q7A5P1 Q7A5P3 Q7A5P6 Q7A5P7 Q7A5R7
Q7A5S2 Q7A5S7 Q7A617 Q7A621 Q7A647 Q7A649 Q7A658 Q7A668 Q7A6A8 Q7A6B1 Q7A6E2
Q7A6H5 Q7A6J6 Q7A6K4 Q7A6Q6 Q7A6Q9 Q7A6T4 Q7A6U7 Q7A6Y6 Q7A754 Q7A767 Q7A768
Q7A777 Q7A7A6 Q7A7C3 Q7A7J0 Q7A7T6 Q7A805 Q7A875 Q7A882 Q7A978 Q7A9L5 Q7A9X4
Q7ABA6 Q7ABI1 Q7ABK6 Q7ADE6 Q7ADE7 Q7ADF8 Q7ADI4 Q7ADI5 Q7AFV7 Q7AH91 Q7AHT0
Q7AJA5 Q7AJA9 Q7AKE4 Q7AKF0 Q7AKG8 Q7AKG9 Q7AKI6 Q7AKI9 Q7AKN0 Q7AKQ5 Q7AKQ6
Q7AMH5 Q7ANK1 Q7ANU3 Q7ANU4 Q7ANU5 Q7ANU6 Q7ANU7 Q7ANU8 Q7ANW8 Q7ANY4 Q7ANZ1
Q7AP11 Q7AP29 Q7AP36 Q7ARC3 Q7ARI8 Q7ATH7 Q7ATH9 Q7B8C3 Q7B8C5 Q7B9Y6 Q7B9Y9
Q7BEF3 Q7BI34 Q7BQ71 Q7BS31 Q7BS32 Q7BSH1 Q7BSX2 Q7BUE1 Q7BVT7 Q7BZ80 Q7BZ90
Q7C093 Q7C0D0 Q7C0F9 Q7C1E6 Q7C254 Q7C397 Q7C3P5 Q7C3R3 Q7C3S2 Q7C637 Q7CCK6
Q7CEG6 Q7CFS8 Q7CFT2 Q7CFT4 Q7CFT7 Q7CFU4 Q7CFX5 Q7CG46 Q7CG47 Q7CG56 Q7CGH1
Q7CGS4 Q7CGT3 Q7CH07 Q7CH25 Q7CH40 Q7CH65 Q7CH66 Q7CH67 Q7CHK5 Q7CHK7 Q7CHL5
Q7CHM0 Q7CHM1 Q7CHU1 Q7CHW2 Q7CHW4 Q7CIA2 Q7CIB6 Q7CIB7 Q7CID0 Q7CIP8 Q7CIR8
Q7CIS2 Q7CIZ3 Q7CJ65 Q7CJ72 Q7CJ75 Q7CJ76 Q7CJ96 Q7CJF3 Q7CJI9 Q7CJM9 Q7CJN8
Q7CJY7 Q7CK25 Q7CK59 Q7CKD4 Q7CKG8 Q7CKJ6 Q7CKM4 Q7CKT7 Q7CLB2 Q7CLC9 Q7CLE5
Q7CLU4 Q7CLU5 Q7CLU6 Q7CLV7 Q7CMP0 Q7CMQ8 Q7CMR3 Q7CMS1 Q7CMS5 Q7CMT4 Q7CMY7
Q7CMZ3 Q7CMZ7 Q7CN13 Q7CN70 Q7CN84 Q7CNB0 Q7CNC3 Q7CNH7 Q7CNK9 Q7CNN9 Q7CNP1
Q7CNP2 Q7CNP3 Q7CNP4 Q7CNP5 Q7CNP7 Q7CNP8 Q7CNP9 Q7CNQ0 Q7CNQ1 Q7CNQ9 Q7CNT2
Q7CNT3 Q7CNT4 Q7CP66 Q7CP77 Q7CP78 Q7CP92 Q7CP93 Q7CPA2 Q7CPB2 Q7CPC4 Q7CPD4
Q7CPD8 Q7CPF1 Q7CPG1 Q7CPH8 Q7CPJ9 Q7CPK5 Q7CPL3 Q7CPL4 Q7CPL5 Q7CPL6 Q7CPL7
Q7CPP7 Q7CPP9 Q7CPQ7 Q7CPQ8 Q7CPR7 Q7CPS1 Q7CPU5 Q7CPV1 Q7CPV2 Q7CPV3 Q7CPW1
Q7CPZ4 Q7CQ01 Q7CQ05 Q7CQ08 Q7CQ10 Q7CQ12 Q7CQ21 Q7CQ25 Q7CQ56 Q7CQ70 Q7CQ71
Q7CQC4 Q7CQE0 Q7CQF3 Q7CQJ0 Q7CQK5 Q7CQK7 Q7CQM8 Q7CQN5 Q7CQR2 Q7CQR9 Q7CQS3
Q7CQS8 Q7CQS9 Q7CQT0 Q7CQT7 Q7CQV5 Q7CQV9 Q7CQY1 Q7CQZ6 Q7CR08 Q7CR17 Q7CR22
Q7CR30 Q7CR46 Q7CR52 Q7CR62 Q7CR64 Q7CR66 Q7CR91 Q7CRA3 Q7CRQ0 Q7CS13 Q7CTU0
Q7CU92 Q7CUX4 Q7CV90 Q7CVH1 Q7CVK1 Q7CW97 Q7CWE8 Q7CWI1 Q7CWK7 Q7CWR3 Q7CWX2
Q7CWX3 Q7CX57 Q7CXF0 Q7CXF9 Q7CXI0 Q7CXU0 Q7CYD2 Q7CYR2 Q7D0L8 Q7D0R6 Q7D1A4
Q7D1M2 Q7D255 Q7D3B2 Q7D3U0 Q7D513 Q7D515 Q7D5F9 Q7D5T7 Q7D7P7 Q7DAF6 Q7DBA7
Q7DC27 Q7DD49 Q7DDI1 Q7DDL2 Q7DDM0 Q7DDM4 Q7DDM9 Q7DDN1 Q7DDR4 Q7DDR9 Q7DDS9
Q7DDT0 Q7DDT3 Q7DDT4 Q7DF80 Q7DF83 Q7DFU6 Q7DFV4 Q7DJ53 Q7DKW4 Q7DLR9 Q7DLS1
Q7DMN9 Q7DMP0 Q7EAG4 Q7EYV7 Q7EZ44 Q7EZ57 Q7EZD5 Q7EZT1 Q7F0J0 Q7F0R1 Q7F1M0
Q7F1X5 Q7F2E4 Q7F2Z1 Q7F7I7 Q7F830 Q7F8R0 Q7F8S5 Q7F8T6 Q7F9I1 Q7F9U3 Q7F9W2
Q7FAD5 Q7FAH2 Q7FAT5 Q7FAY6 Q7FGZ2 Q7FNR4 Q7FNR5 Q7FNS1 Q7FNS5 Q7FNS6 Q7FNT2
Q7G065 Q7G188 Q7G191 Q7G192 Q7G193 Q7G1L2 Q7G4G7 Q7G737 Q7G764 Q7G765 Q7G7J6
Q7G8Y3 Q7G8Y5 Q7G959 Q7G9E7 Q7G9P4 Q7GBC8 Q7GBF7 Q7GBF8 Q7GBK0 Q7GD79 Q7GDL5
Q7GDW5 Q7GED0 Q7GED9 Q7GUC7 Q7GUC8 Q7GUD0 Q7GUD1 Q7H6F6 Q7H6G1 Q7H6I5 Q7H6J0
Q7H6J1 Q7H6J2 Q7HEK3 Q7HKI2 Q7HKI3 Q7HKW9 Q7HKX4 Q7HTB5 Q7HTB8 Q7HUK6 Q7I9M7
Q7ICQ5 Q7ICQ6 Q7IJL0 Q7IS11 Q7IZ70 Q7J1C8 Q7J5Z8 Q7J5Z9 Q7J602 Q7JD61 Q7JEV7
Q7JEV9 Q7JEW0 Q7JEW6 Q7JEW7 Q7JF34 Q7JF70 Q7JFN6 Q7JFR7 Q7JGF7 Q7JGF9 Q7JGP2
Q7JGR2 Q7JH04 Q7JH05 Q7JH06 Q7JH07 Q7JHM8 Q7JHM9 Q7JIX8 Q7JIX9 Q7JIY0 Q7JJU6
Q7JK39 Q7JKP6 Q7JM44 Q7JNS1 Q7JPQ6 Q7JQ07 Q7JQD3 Q7JQW6 Q7JRJ1 Q7JRP4 Q7JUR5
Q7JUR6 Q7JUX9 Q7JUY7 Q7JVA5 Q7JVI3 Q7JVL3 Q7JVP4 Q7JVR7 Q7JWG9 Q7JWR9 Q7JWW5
Q7JXF5 Q7JXG9 Q7JYV7 Q7JZB4 Q7JZM8 Q7K0F0 Q7K0S9 Q7K0Y1 Q7K1Y4 Q7K1Z5 Q7K284
Q7K2B0 Q7K2Y9 Q7K3B9 Q7K3D8 Q7K480 Q7K486 Q7K4B3 Q7K4B4 Q7K4H4 Q7K4M4 Q7K4N3

Q7K4Q5 Q7K4V4 Q7K550 Q7K5K3 Q7K734 Q7K738 Q7K7A9 Q7K8C0 Q7KBL8 Q7KF89 Q7KF90
Q7KHG2 Q7KKH3 Q7KLV9 Q7KM13 Q7KMJ6 Q7KN79 Q7KN90 Q7KNF2 Q7KNS3 Q7KPA5 Q7KQD1
Q7KQJ9 Q7KQL5 Q7KQL8 Q7KQM0 Q7KQM1 Q7KQM6 Q7KQZ4 Q7KR04 Q7KRI2 Q7KRR5 Q7KRW8
Q7KS38 Q7KSD8 Q7KTH8 Q7KTX8 Q7KU24 Q7KU86 Q7KV12 Q7KV13 Q7KV14 Q7KV15 Q7KV19
Q7KV22 Q7KV23 Q7KVH9 Q7KVQ0 Q7KVS9 Q7KW09 Q7KW14 Q7KW39 Q7KWK5 Q7KWL3 Q7KWM9
Q7KWN3 Q7KWN4 Q7KWN5 Q7KWN6 Q7KWN7 Q7KWN8 Q7KWP3 Q7KWP9 Q7KWQ2 Q7KWR7 Q7KWS1
Q7KWT8 Q7KWU0 Q7KWX9 Q7KWY4 Q7L0R7 Q7L0X2 Q7L0Y3 Q7L1V2 Q7L266 Q7L2K0 Q7L2R6
Q7L3S4 Q7L3T8 Q7L4P6 Q7L592 Q7L5A3 Q7L5Y1 Q7L5Y6 Q7L7V1 Q7L8L6 Q7L8W6 Q7L945
Q7LG56 Q7LIF1 Q7LIF2 Q7LJF8 Q7LJR6 Q7LKT3 Q7LKV1 Q7LKY2 Q7LKZ5 Q7LL04 Q7LL14
Q7LL15 Q7LL22 Q7LLZ8 Q7LML9 Q7LWY0 Q7LX22 Q7LXK4 Q7LXL5 Q7LXP6 Q7LXU0 Q7LYI9
Q7LYJ6 Q7LYW4 Q7LYW5 Q7LZB2 Q7LZB6 Q7LZB9 Q7LZC1 Q7LZC2 Q7LZC3 Q7LZM1 Q7LZM2
Q7LZM3 Q7LZM4 Q7LZM5 Q7LZT1 Q7M0E7 Q7M0R2 Q7M0Z3 Q7M181 Q7M1B9 Q7M1G9 Q7M1L6
Q7M1L7 Q7M1R5 Q7M1Z2 Q7M258 Q7M2N1 Q7M2P1 Q7M2V4 Q7M2Y4 Q7M2Y5 Q7M316 Q7M317
Q7M3B6 Q7M3B8 Q7M3C1 Q7M3C2 Q7M3K2 Q7M3M6 Q7M3M8 Q7M3S9 Q7M3Y0 Q7M3Y3 Q7M3Y8
Q7M3Z5 Q7M400 Q7M416 Q7M424 Q7M443 Q7M4E8 Q7M4E9 Q7M4F0 Q7M4F1 Q7M4F2 Q7M4F3
Q7M4F4 Q7M4G7 Q7M4L6 Q7M4S9 Q7M4T5 Q7M4T6 Q7M523 Q7M537 Q7M6N6 Q7M6R1 Q7M6U3
Q7M6Y5 Q7M6Y6 Q7M6Z4 Q7M6Z5 Q7M757 Q7M767 Q7M7F1 Q7M7H7 Q7M7H8 Q7M7I1 Q7M7I2
Q7M7I6 Q7M7I7 Q7M7I8 Q7M7J0 Q7M7J4 Q7M7K0 Q7M7K5 Q7M7L1 Q7M7L6 Q7M7L9 Q7M7M2
Q7M7M3 Q7M7M4 Q7M7P6 Q7M7P9 Q7M7Q5 Q7M7Q6 Q7M7Q9 Q7M7R0 Q7M7R1 Q7M7S2 Q7M7T6
Q7M7U1 Q7M7U5 Q7M7U7 Q7M7U8 Q7M7V2 Q7M7V3 Q7M7V7 Q7M7V8 Q7M7W8 Q7M7W9 Q7M7X5
Q7M7X6 Q7M7Y2 Q7M7Y6 Q7M7Z0 Q7M7Z2 Q7M7Z3 Q7M800 Q7M806 Q7M818 Q7M831 Q7M840
Q7M841 Q7M842 Q7M847 Q7M851 Q7M877 Q7M878 Q7M879 Q7M886 Q7M887 Q7M8A0 Q7M8A5
Q7M8A6 Q7M8B4 Q7M8B5 Q7M8C4 Q7M8D4 Q7M8D5 Q7M8D6 Q7M8D7 Q7M8D8 Q7M8D9 Q7M8E0
Q7M8E1 Q7M8E2 Q7M8E3 Q7M8E4 Q7M8E5 Q7M8E6 Q7M8E7 Q7M8E8 Q7M8E9 Q7M8F0 Q7M8F3
Q7M8F4 Q7M8F5 Q7M8F6 Q7M8F7 Q7M8F8 Q7M8F9 Q7M8G1 Q7M8H4 Q7M8H6 Q7M8H7 Q7M8I0
Q7M8I1 Q7M8I9 Q7M8J0 Q7M8K0 Q7M8K2 Q7M8K5 Q7M8K9 Q7M8L1 Q7M8L2 Q7M8N8 Q7M8N9
Q7M8P7 Q7M8Q1 Q7M8S6 Q7M8U5 Q7M8U8 Q7M8W6 Q7M8W7 Q7M8W8 Q7M8X1 Q7M8Y6 Q7M8Y7
Q7M8Y8 Q7M8Y9 Q7M8Z2 Q7M8Z4 Q7M8Z5 Q7M8Z6 Q7M901 Q7M904 Q7M926 Q7M933 Q7M936
Q7M938 Q7M9A0 Q7M9A6 Q7M9A7 Q7M9B0 Q7M9C1 Q7M9C2 Q7M9F6 Q7M9I4 Q7M9J0 Q7M9J1
Q7M9J5 Q7M9J8 Q7M9K2 Q7M9K3 Q7M9L1 Q7M9L5 Q7M9L7 Q7M9L8 Q7M9L9 Q7M9M2 Q7M9M3
Q7M9M4 Q7M9M5 Q7M9M7 Q7M9N0 Q7M9N1 Q7M9N2 Q7M9N9 Q7M9R5 Q7M9S1 Q7M9T3 Q7M9U0
Q7M9U1 Q7M9U3 Q7M9W4 Q7M9W9 Q7M9X0 Q7M9X1 Q7M9X2 Q7M9X4 Q7M9X5 Q7M9X7 Q7M9Y2
Q7M9Y9 Q7M9Z9 Q7MA00 Q7MA03 Q7MA04 Q7MA26 Q7MA27 Q7MA28 Q7MA29 Q7MA30 Q7MA34
Q7MA35 Q7MA53 Q7MA54 Q7MA55 Q7MA56 Q7MA57 Q7MA58 Q7MA59 Q7MA63 Q7MA68 Q7MA70
Q7MA71 Q7MA77 Q7MAA5 Q7MAC1 Q7MAC4 Q7MAC6 Q7MAC9 Q7MAD1 Q7MAD3 Q7MAD7 Q7MAD8
Q7MAE0 Q7MAE2 Q7MAE3 Q7MAE6 Q7MAE7 Q7MAE8 Q7MAE9 Q7MAF0 Q7MAG3 Q7MAH6 Q7MAH8
Q7MAI3 Q7MAI9 Q7MAJ5 Q7MAK0 Q7MAK1 Q7MAK5 Q7MAK6 Q7MAL0 Q7MAN4 Q7MAP3 Q7MAP4
Q7MAP8 Q7MAQ8 Q7MAR0 Q7MAR1 Q7MAS1 Q7MAS4 Q7MAV4 Q7MAW4 Q7MAW6 Q7MAW8 Q7MAW9
Q7MAX1 Q7MAX2 Q7MAX3 Q7MAX5 Q7MAX6 Q7MAX8 Q7MAX9 Q7MAY1 Q7MAZ1 Q7MAZ6 Q7MAZ7
Q7MAZ9 Q7MB01 Q7MB17 Q7MB22 Q7MB41 Q7MB61 Q7MB63 Q7MB64 Q7MB85 Q7MB99 Q7MBA6
Q7MBB2 Q7MBB4 Q7MBB8 Q7MBB9 Q7MBC1 Q7MBC7 Q7MBC8 Q7MBC9 Q7MBD4 Q7MBD6 Q7MBE1
Q7MBE2 Q7MBE8 Q7MBF4 Q7MBF7 Q7MBG5 Q7MBG6 Q7MBG9 Q7MBH2 Q7MBQ4 Q7MBS3 Q7MBT9
Q7MBZ3 Q7MBZ9 Q7MCF1 Q7MCN7 Q7MCQ2 Q7MCR1 Q7MCS5 Q7MD09 Q7MD11 Q7MD31 Q7MD32
Q7MD57 Q7MDD4 Q7MDD5 Q7MDE5 Q7MDF5 Q7MDL6 Q7MDN2 Q7ME60 Q7ME78 Q7ME85 Q7ME92

Q7MEE8 Q7MEE9 Q7MEH7 Q7MEH8 Q7MEH9 Q7MEL2 Q7MEL4 Q7MEP0 Q7MEV2 Q7MEW6 Q7MF05
Q7MF12 Q7MF13 Q7MF14 Q7MF44 Q7MF46 Q7MF74 Q7MFB0 Q7MFB3 Q7MFG6 Q7MFL5 Q7MFN1
Q7MFN7 Q7MFR0 Q7MFU0 Q7MFY7 Q7MFY8 Q7MG04 Q7MG84 Q7MG88 Q7MG90 Q7MG94 Q7MGA2
Q7MGA4 Q7MGC2 Q7MGE1 Q7MGF9 Q7MGG3 Q7MGG9 Q7MGH0 Q7MGI8 Q7MGK4 Q7MGK5 Q7MGK6
Q7MGK9 Q7MGL1 Q7MGL2 Q7MGL3 Q7MGL4 Q7MGL5 Q7MGM3 Q7MGM7 Q7MGN5 Q7MGP6 Q7MGP7
Q7MGQ8 Q7MGQ9 Q7MGR1 Q7MGR5 Q7MGR6 Q7MGR7 Q7MGR8 Q7MGS0 Q7MGS7 Q7MGT4 Q7MGT5
Q7MGT8 Q7MGU2 Q7MGU3 Q7MGW6 Q7MGX2 Q7MGX9 Q7MGY8 Q7MGZ2 Q7MGZ6 Q7MGZ7 Q7MH01
Q7MH02 Q7MH07 Q7MH09 Q7MH10 Q7MH13 Q7MH14 Q7MH15 Q7MH19 Q7MH21 Q7MH24 Q7MH33
Q7MH37 Q7MH38 Q7MH40 Q7MH41 Q7MH42 Q7MH43 Q7MH47 Q7MH53 Q7MH54 Q7MH56 Q7MH57
Q7MH61 Q7MH63 Q7MH68 Q7MH69 Q7MH70 Q7MH71 Q7MH72 Q7MH73 Q7MH83 Q7MH84 Q7MH88
Q7MH89 Q7MH90 Q7MH91 Q7MH94 Q7MH97 Q7MHA5 Q7MHA6 Q7MHA7 Q7MHB1 Q7MHB5 Q7MHE0
Q7MHE6 Q7MHF0 Q7MHF1 Q7MHF2 Q7MHF3 Q7MHG1 Q7MHG4 Q7MHH4 Q7MHI4 Q7MHI6 Q7MHI7
Q7MHJ0 Q7MHJ2 Q7MHJ8 Q7MHK0 Q7MHK1 Q7MHK4 Q7MHK6 Q7MHK7 Q7MHL0 Q7MHL1 Q7MHL5
Q7MHL9 Q7MHM2 Q7MHM7 Q7MHM8 Q7MHN8 Q7MHP0 Q7MHP1 Q7MHP7 Q7MHQ0 Q7MHQ4 Q7MHQ5
Q7MHQ6 Q7MHQ7 Q7MHQ8 Q7MHR2 Q7MHR4 Q7MHR5 Q7MHR6 Q7MHR8 Q7MHS5 Q7MHT1 Q7MHT2
Q7MHT3 Q7MHT4 Q7MHT5 Q7MHV3 Q7MHV4 Q7MHV7 Q7MHV9 Q7MHW8 Q7MHY6 Q7MHY9 Q7MHZ0
Q7MHZ4 Q7MI01 Q7MI04 Q7MI07 Q7MI09 Q7MI10 Q7MI11 Q7MI12 Q7MI20 Q7MI34 Q7MI38
Q7MI39 Q7MI40 Q7MI41 Q7MI49 Q7MI80 Q7MI93 Q7MIA6 Q7MIC5 Q7MIC7 Q7MID6 Q7MIE3
Q7MIE5 Q7MIE7 Q7MIE8 Q7MIF1 Q7MIF7 Q7MIF8 Q7MIG0 Q7MIG1 Q7MIG2 Q7MIG3 Q7MIG6
Q7MIH3 Q7MIH8 Q7MII1 Q7MII4 Q7MII6 Q7MIK1 Q7MIK2 Q7MIL2 Q7MIL6 Q7MIS9 Q7MIT1
Q7MIT5 Q7MIT6 Q7MIV1 Q7MIV3 Q7MIV4 Q7MIW1 Q7MIX2 Q7MIZ4 Q7MJ09 Q7MJ13 Q7MJ16
Q7MJ40 Q7MJ42 Q7MJ43 Q7MJ45 Q7MJ49 Q7MJ50 Q7MJ57 Q7MJ58 Q7MJ62 Q7MJ63 Q7MJ64
Q7MJ71 Q7MJ72 Q7MJ74 Q7MJ75 Q7MJ77 Q7MJ78 Q7MJ90 Q7MJC0 Q7MJI2 Q7MJL8 Q7MJM6
Q7MJM8 Q7MJS0 Q7MK12 Q7MK24 Q7MK39 Q7MK40 Q7MK41 Q7MK44 Q7MK46 Q7MK47 Q7MK48
Q7MK58 Q7MK59 Q7MK61 Q7MK65 Q7MK67 Q7MK68 Q7MKQ9 Q7MKR2 Q7MKT7 Q7MKU7 Q7MKW4
Q7MKX1 Q7MKY1 Q7MKY6 Q7MKZ1 Q7ML77 Q7ML80 Q7ML86 Q7MLC1 Q7MLD5 Q7MLD9 Q7MLE8
Q7MLF3 Q7MLF9 Q7MLG3 Q7MLH6 Q7MLK7 Q7MLM1 Q7MLR4 Q7MLR5 Q7MLR9 Q7MLS0 Q7MLS1
Q7MLS2 Q7MLS3 Q7MLS4 Q7MLS5 Q7MLS6 Q7MLS7 Q7MLT4 Q7MLU7 Q7MLU9 Q7MLV0 Q7MLV2
Q7MLX2 Q7MLX5 Q7MLX7 Q7MLY4 Q7MLZ6 Q7MM02 Q7MM04 Q7MM12 Q7MM14 Q7MM27 Q7MM39
Q7MM44 Q7MM45 Q7MM54 Q7MM56 Q7MM57 Q7MM73 Q7MM75 Q7MM80 Q7MM91 Q7MMA6 Q7MMC4
Q7MMF6 Q7MMF9 Q7MMG6 Q7MMG7 Q7MMG8 Q7MMI2 Q7MMK9 Q7MML0 Q7MML2 Q7MMM3 Q7MMN3
Q7MMQ4 Q7MMR4 Q7MMR5 Q7MMR7 Q7MMT5 Q7MMW8 Q7MMY6 Q7MMY7 Q7MMZ0 Q7MMZ1 Q7MMZ2
Q7MN00 Q7MN02 Q7MN06 Q7MN10 Q7MN15 Q7MN16 Q7MN17 Q7MN19 Q7MN30 Q7MN34 Q7MN36
Q7MN44 Q7MN47 Q7MN49 Q7MN52 Q7MN53 Q7MN57 Q7MN58 Q7MN59 Q7MN60 Q7MN61 Q7MN70
Q7MN84 Q7MN85 Q7MN92 Q7MN93 Q7MN97 Q7MN99 Q7MND7 Q7MNE1 Q7MNE3 Q7MNE7 Q7MNF0
Q7MNF1 Q7MNF3 Q7MNF4 Q7MNF5 Q7MNF8 Q7MNF9 Q7MNG2 Q7MNG3 Q7MNH1 Q7MNH2 Q7MNH9
Q7MNJ0 Q7MNJ5 Q7MNK1 Q7MNK7 Q7MNK8 Q7MNL7 Q7MNM5 Q7MNM8 Q7MNN1 Q7MNN6 Q7MNP0
Q7MNP1 Q7MNP6 Q7MNQ0 Q7MNQ4 Q7MNQ6 Q7MNR0 Q7MNR1 Q7MNR2 Q7MNR4 Q7MNR7 Q7MNT0
Q7MNU0 Q7MNU1 Q7MNU2 Q7MNV0 Q7MNV3 Q7MNV6 Q7MNV9 Q7MNW2 Q7MNX0 Q7MNX1 Q7MNZ9
Q7MP00 Q7MP37 Q7MP60 Q7MP62 Q7MP77 Q7MP78 Q7MP79 Q7MP80 Q7MP85 Q7MP86 Q7MP87
Q7MP88 Q7MP92 Q7MP93 Q7MP94 Q7MP97 Q7MP98 Q7MPA3 Q7MPB9 Q7MPC4 Q7MPD0 Q7MPD2
Q7MPF0 Q7MPF2 Q7MPF3 Q7MPG3 Q7MPG4 Q7MPG5 Q7MPG6 Q7MPG7 Q7MPG9 Q7MPH0 Q7MPH1
Q7MPH2 Q7MPH3 Q7MPH4 Q7MPH5 Q7MPH6 Q7MPH7 Q7MPH8 Q7MPH9 Q7MPI0 Q7MPI1 Q7MPI2
Q7MPI3 Q7MPI4 Q7MPI5 Q7MPI6 Q7MPI7 Q7MPI8 Q7MPI9 Q7MPK7 Q7MPK8 Q7MPP3 Q7MPP4

Q7MPS0 Q7MPS3 Q7MPS5 Q7MPS6 Q7MPS7 Q7MPS9 Q7MPT0 Q7MPT2 Q7MPT3 Q7MPW9 Q7MPX0
Q7MPX6 Q7MPY0 Q7MPY4 Q7MQ02 Q7MQ03 Q7MQ06 Q7MQ19 Q7MQ23 Q7MQ33 Q7MQ40 Q7MQ55
Q7MQ79 Q7MQ83 Q7MQ88 Q7MQ91 Q7MQB9 Q7MQC1 Q7MQC4 Q7MQC7 Q7MQD7 Q7MQH3 Q7MQI4
Q7MQI7 Q7MQI8 Q7MQJ5 Q7MQJ7 Q7MQK3 Q7MQK4 Q7MQK6 Q7MQK8 Q7MQS0 Q7MQU0 Q7MQU1
Q7MQV6 Q7MQW9 Q7MQZ4 Q7MQZ8 Q7MQZ9 Q7MR19 Q7MR25 Q7MR31 Q7MR38 Q7MR59 Q7MR67
Q7MRV3 Q7MRV4 Q7MS14 Q7MS24 Q7MS38 Q7MS42 Q7MS89 Q7MSA5 Q7MSC2 Q7MSC3 Q7MSD5
Q7MSD7 Q7MSD9 Q7MSG7 Q7MSH1 Q7MSI8 Q7MSL3 Q7MSQ0 Q7MSS6 Q7MSS9 Q7MST2 Q7MST9
Q7MSU1 Q7MSW1 Q7MSY1 Q7MSY2 Q7MSZ3 Q7MT06 Q7MT07 Q7MT32 Q7MT44 Q7MT48 Q7MT50
Q7MT66 Q7MT71 Q7MT73 Q7MT83 Q7MT92 Q7MT94 Q7MT97 Q7MTB0 Q7MTB3 Q7MTB5 Q7MTC4
Q7MTE3 Q7MTF5 Q7MTG7 Q7MTG8 Q7MTG9 Q7MTJ4 Q7MTJ5 Q7MTK9 Q7MTL0 Q7MTL1 Q7MTL2
Q7MTL3 Q7MTL4 Q7MTL6 Q7MTL7 Q7MTL8 Q7MTL9 Q7MTM0 Q7MTM1 Q7MTM2 Q7MTM3 Q7MTM4
Q7MTM5 Q7MTM6 Q7MTM7 Q7MTM8 Q7MTM9 Q7MTN0 Q7MTN1 Q7MTN2 Q7MTN5 Q7MTN6 Q7MTN7
Q7MTN8 Q7MTN9 Q7MTP0 Q7MTP1 Q7MTP5 Q7MTP6 Q7MTQ0 Q7MTR6 Q7MTS0 Q7MU00 Q7MU04
Q7MU16 Q7MU19 Q7MU42 Q7MU43 Q7MU50 Q7MU77 Q7MU78 Q7MUB2 Q7MUC6 Q7MUD3 Q7MUF7
Q7MUG4 Q7MUH3 Q7MUI9 Q7MUQ9 Q7MUR5 Q7MUV7 Q7MUV9 Q7MUW1 Q7MUW2 Q7MUW3 Q7MUX4
Q7MUX8 Q7MUY1 Q7MUY2 Q7MUZ6 Q7MUZ8 Q7MV04 Q7MV08 Q7MV10 Q7MV30 Q7MV32 Q7MV45
Q7MV47 Q7MV54 Q7MV70 Q7MVD7 Q7MVE5 Q7MVE7 Q7MVE9 Q7MVF1 Q7MVG0 Q7MVG5 Q7MVG9
Q7MVI0 Q7MVJ6 Q7MVQ5 Q7MVQ6 Q7MVQ7 Q7MVQ8 Q7MVR7 Q7MVR8 Q7MVS7 Q7MVS9 Q7MVT6
Q7MVT7 Q7MVT9 Q7MVU8 Q7MVU9 Q7MVV7 Q7MVW8 Q7MVX9 Q7MVY0 Q7MVZ2 Q7MW36 Q7MW43
Q7MW49 Q7MW52 Q7MW54 Q7MW55 Q7MW79 Q7MWA2 Q7MWB6 Q7MWC6 Q7MWG0 Q7MWG1 Q7MWI5
Q7MWI7 Q7MWK6 Q7MWK7 Q7MWK9 Q7MWL1 Q7MWL2 Q7MWL3 Q7MWL6 Q7MWL9 Q7MWM0 Q7MWM5
Q7MWM7 Q7MWN0 Q7MWN4 Q7MWR7 Q7MWS7 Q7MWS9 Q7MWT4 Q7MWT5 Q7MWU9 Q7MWV3 Q7MWV8
Q7MWW8 Q7MWY2 Q7MX09 Q7MX10 Q7MX15 Q7MX24 Q7MX26 Q7MX27 Q7MX28 Q7MX30 Q7MX31
Q7MX34 Q7MX40 Q7MX41 Q7MX42 Q7MX47 Q7MX57 Q7MX58 Q7MX68 Q7MX71 Q7MX82 Q7MX94
Q7MX95 Q7MXD5 Q7MXD8 Q7MXE4 Q7MXE6 Q7MXF6 Q7MXG0 Q7MXI4 Q7MXJ4 Q7MXK7 Q7MXK8
Q7MXK9 Q7MXM0 Q7MXM3 Q7MXN7 Q7MXP1 Q7MXP4 Q7MXR4 Q7MXR6 Q7MXR7 Q7MXT5 Q7MXT6
Q7MXU9 Q7MXW0 Q7MXW3 Q7MXX0 Q7MXX4 Q7MXZ1 Q7MXZ2 Q7MY02 Q7MY03 Q7MY11 Q7MY25
Q7MY26 Q7MY28 Q7MY29 Q7MY30 Q7MY36 Q7MY37 Q7MY48 Q7MY53 Q7MY63 Q7MYB3 Q7MYB6
Q7MYB8 Q7MYB9 Q7MYC1 Q7MYC2 Q7MYC6 Q7MYC7 Q7MYD5 Q7MYD6 Q7MYD7 Q7MYD8 Q7MYD9
Q7MYE4 Q7MYE5 Q7MYE7 Q7MYE8 Q7MYF1 Q7MYF2 Q7MYF3 Q7MYF4 Q7MYF5 Q7MYF6 Q7MYF7
Q7MYF8 Q7MYF9 Q7MYG0 Q7MYG1 Q7MYG2 Q7MYG3 Q7MYG4 Q7MYG5 Q7MYG6 Q7MYG7 Q7MYG8
Q7MYG9 Q7MYH0 Q7MYH2 Q7MYH3 Q7MYH4 Q7MYH5 Q7MYH6 Q7MYI0 Q7MYI1 Q7MYI2 Q7MYI5
Q7MYI6 Q7MYJ5 Q7MYK9 Q7MYL1 Q7MYL2 Q7MYN1 Q7MYN3 Q7MYN5 Q7MYS7 Q7MYS8 Q7MYT9
Q7MYU0 Q7MYU4 Q7MYU6 Q7MYU7 Q7MYU8 Q7MYU9 Q7MYW1 Q7MYW6 Q7MYW8 Q7MYW9 Q7MYX4
Q7MYX5 Q7MYX6 Q7MYY0 Q7MYY3 Q7MYY5 Q7MYY7 Q7MYY8 Q7MYY9 Q7MYZ0 Q7MZ08 Q7MZ10
Q7MZ14 Q7MZ16 Q7MZ18 Q7MZ25 Q7MZ27 Q7MZ74 Q7MZ76 Q7MZ81 Q7MZ93 Q7MZ97 Q7MZA0
Q7MZB4 Q7MZB5 Q7MZD8 Q7MZD9 Q7MZE1 Q7MZH1 Q7MZI3 Q7MZN0 Q7MZN3 Q7MZX9 Q7MZY6
Q7MZZ5 Q7N015 Q7N030 Q7N035 Q7N042 Q7N053 Q7N068 Q7N078 Q7N079 Q7N089 Q7N090
Q7N0A6 Q7N0B6 Q7N0C8 Q7N0D4 Q7N0G0 Q7N0I3 Q7N0I7 Q7N0I9 Q7N0J0 Q7N0J7 Q7N0J8
Q7N0J9 Q7N0K1 Q7N0L2 Q7N0L3 Q7N0L4 Q7N0M0 Q7N0N9 Q7N0P1 Q7N0P2 Q7N0P4 Q7N0P5
Q7N0P6 Q7N0S5 Q7N118 Q7N119 Q7N121 Q7N122 Q7N124 Q7N125 Q7N126 Q7N127 Q7N128
Q7N129 Q7N138 Q7N139 Q7N142 Q7N145 Q7N148 Q7N149 Q7N157 Q7N158 Q7N179 Q7N185
Q7N189 Q7N193 Q7N197 Q7N198 Q7N199 Q7N1C0 Q7N1C1 Q7N1C8 Q7N1I2 Q7N1I3 Q7N1U1
Q7N1U3 Q7N1U6 Q7N1U7 Q7N1U8 Q7N1V4 Q7N1W7 Q7N1X5 Q7N1X7 Q7N1X8 Q7N1Z4 Q7N215

Q7N216 Q7N222 Q7N223 Q7N224 Q7N226 Q7N227 Q7N228 Q7N231 Q7N278 Q7N290 Q7N291
Q7N295 Q7N296 Q7N299 Q7N2A3 Q7N2B2 Q7N2D5 Q7N2D6 Q7N2E0 Q7N2E1 Q7N2G7 Q7N2G8
Q7N2G9 Q7N2H0 Q7N2H1 Q7N2I1 Q7N2I4 Q7N2K3 Q7N2K4 Q7N2K6 Q7N2M5 Q7N2S8 Q7N2T7
Q7N2U4 Q7N2V3 Q7N351 Q7N352 Q7N360 Q7N364 Q7N376 Q7N380 Q7N382 Q7N391 Q7N397
Q7N3D8 Q7N3F7 Q7N3F8 Q7N3G5 Q7N3H3 Q7N3J4 Q7N3K2 Q7N3K4 Q7N3M0 Q7N3M2 Q7N3N2
Q7N3P6 Q7N3P8 Q7N3P9 Q7N3Q0 Q7N3Q1 Q7N3Q2 Q7N3T4 Q7N3U5 Q7N3U6 Q7N3V8 Q7N3W0
Q7N3W4 Q7N3W5 Q7N3W6 Q7N3W7 Q7N3Y7 Q7N3Z6 Q7N400 Q7N406 Q7N451 Q7N457 Q7N466
Q7N485 Q7N486 Q7N488 Q7N4C1 Q7N4F5 Q7N4H5 Q7N4H8 Q7N4H9 Q7N4I0 Q7N4P7 Q7N4V5
Q7N4V6 Q7N4V7 Q7N4V8 Q7N4V9 Q7N4W0 Q7N4X5 Q7N4Y3 Q7N4Y4 Q7N4Y5 Q7N4Y7 Q7N4Y8
Q7N4Y9 Q7N511 Q7N519 Q7N521 Q7N522 Q7N524 Q7N547 Q7N548 Q7N549 Q7N550 Q7N555
Q7N556 Q7N557 Q7N561 Q7N565 Q7N574 Q7N575 Q7N586 Q7N587 Q7N589 Q7N590 Q7N5A1
Q7N5C7 Q7N5H9 Q7N5I5 Q7N5J9 Q7N5W0 Q7N5W3 Q7N5Z1 Q7N5Z2 Q7N5Z3 Q7N5Z7 Q7N601
Q7N603 Q7N612 Q7N621 Q7N622 Q7N6B7 Q7N6B8 Q7N6B9 Q7N6C3 Q7N6D2 Q7N6D4 Q7N6D5
Q7N6D6 Q7N6E7 Q7N6F5 Q7N6F7 Q7N6G6 Q7N6I0 Q7N6I1 Q7N6I2 Q7N6I3 Q7N6I4 Q7N6I5
Q7N6I6 Q7N6J3 Q7N6J4 Q7N6J6 Q7N6K3 Q7N6K4 Q7N6K7 Q7N6P5 Q7N6P6 Q7N6Q1 Q7N6Q6
Q7N6Q7 Q7N6R1 Q7N6S0 Q7N6S3 Q7N6V5 Q7N6Y0 Q7N6Y2 Q7N6Z8 Q7N6Z9 Q7N702 Q7N705
Q7N706 Q7N709 Q7N710 Q7N743 Q7N748 Q7N755 Q7N760 Q7N765 Q7N766 Q7N767 Q7N772
Q7N788 Q7N798 Q7N799 Q7N7A0 Q7N7A4 Q7N7A5 Q7N7A6 Q7N7B0 Q7N7B1 Q7N7B2 Q7N7B3
Q7N7B6 Q7N7F4 Q7N7G6 Q7N7G7 Q7N7H2 Q7N7H7 Q7N7H9 Q7N7I1 Q7N7Z5 Q7N7Z6 Q7N807
Q7N809 Q7N836 Q7N839 Q7N841 Q7N843 Q7N845 Q7N864 Q7N866 Q7N869 Q7N870 Q7N871
Q7N892 Q7N893 Q7N8B1 Q7N8C9 Q7N8D1 Q7N8D4 Q7N8K0 Q7N8K3 Q7N8K4 Q7N8K5 Q7N8K6
Q7N8K7 Q7N8L0 Q7N8L1 Q7N8L2 Q7N8L5 Q7N8L6 Q7N8M0 Q7N8M7 Q7N8N0 Q7N8N3 Q7N8P3
Q7N8P4 Q7N8P5 Q7N8P6 Q7N8P7 Q7N8P9 Q7N8Q0 Q7N8Q1 Q7N8Q7 Q7N8Q9 Q7N8R4 Q7N8S4
Q7N8U4 Q7N8U7 Q7N8V1 Q7N8V7 Q7N8V8 Q7N8V9 Q7N8W2 Q7N8W3 Q7N8W9 Q7N8X2 Q7N8X4
Q7N8Y3 Q7N8Y4 Q7N8Z1 Q7N8Z3 Q7N8Z5 Q7N900 Q7N901 Q7N902 Q7N930 Q7N931 Q7N932
Q7N954 Q7N959 Q7N960 Q7N961 Q7N963 Q7N964 Q7N966 Q7N976 Q7N9A3 Q7N9A4 Q7N9A5
Q7N9A6 Q7N9A7 Q7N9B1 Q7N9B2 Q7N9B3 Q7N9B4 Q7N9B5 Q7N9B6 Q7N9B7 Q7N9C0 Q7N9D8
Q7N9E2 Q7N9E5 Q7N9E9 Q7N9M5 Q7N9M7 Q7N9N4 Q7N9P2 Q7N9P3 Q7N9S2 Q7N9T4 Q7N9V4
Q7N9V7 Q7N9W0 Q7N9W2 Q7N9W3 Q7N9X5 Q7N9Y1 Q7NA24 Q7NA33 Q7NA45 Q7NA46 Q7NA55
Q7NA56 Q7NA60 Q7NA61 Q7NA64 Q7NA80 Q7NA81 Q7NA82 Q7NA83 Q7NA86 Q7NA87 Q7NA97
Q7NA98 Q7NA99 Q7NAD1 Q7NAD3 Q7NAD6 Q7NAD7 Q7NAD9 Q7NAE2 Q7NAE4 Q7NAE5 Q7NAF8
Q7NAH7 Q7NAI9 Q7NAK6 Q7NAK8 Q7NAM0 Q7NAM1 Q7NAM3 Q7NAM4 Q7NAN0 Q7NAN2 Q7NAN6
Q7NAQ0 Q7NAQ4 Q7NAQ5 Q7NAQ9 Q7NAR6 Q7NAR7 Q7NAT8 Q7NAU0 Q7NAU6 Q7NAV0 Q7NAV1
Q7NAV2 Q7NAV3 Q7NAV5 Q7NAW1 Q7NAW2 Q7NAX4 Q7NAZ2 Q7NAZ3 Q7NAZ6 Q7NB15 Q7NB30
Q7NB34 Q7NB37 Q7NB47 Q7NB76 Q7NB77 Q7NB78 Q7NB79 Q7NB85 Q7NB88 Q7NBA6 Q7NBA9
Q7NBB0 Q7NBB2 Q7NBB7 Q7NBB8 Q7NBC0 Q7NBC1 Q7NBC4 Q7NBC5 Q7NBE4 Q7NBE9 Q7NBG0
Q7NBG2 Q7NBG4 Q7NBG7 Q7NBH2 Q7NBH3 Q7NBH4 Q7NBH8 Q7NBH9 Q7NBM9 Q7NBP5 Q7NBS1
Q7NBS4 Q7NBS9 Q7NBT5 Q7NBU2 Q7NBV1 Q7NBV2 Q7NBW0 Q7NBW2 Q7NBW3 Q7NBW4 Q7NBX3
Q7NBX7 Q7NBY3 Q7NBY7 Q7NBY8 Q7NBZ0 Q7NBZ3 Q7NBZ4 Q7NC17 Q7NC19 Q7NC20 Q7NC21
Q7NC27 Q7NC34 Q7NC56 Q7NC65 Q7NC67 Q7NC69 Q7NC71 Q7NCB3 Q7NCB9 Q7NCC0 Q7NCE2
Q7NCE3 Q7NCF5 Q7NCG8 Q7NCI5 Q7NCK4 Q7NCP5 Q7NCQ1 Q7NCQ2 Q7NCQ9 Q7NCU0 Q7NCU2
Q7NCV1 Q7NCY1 Q7ND16 Q7ND17 Q7ND21 Q7ND22 Q7ND61 Q7ND67 Q7NDA8 Q7NDB8 Q7NDC1
Q7NDF6 Q7NDF7 Q7NDF8 Q7NDG8 Q7NDH1 Q7NDJ2 Q7NDK7 Q7NDN9 Q7NDP0 Q7NDP1 Q7NDQ2
Q7NDR9 Q7NDU3 Q7NDW3 Q7NDX4 Q7NE01 Q7NE10 Q7NE46 Q7NE48 Q7NE54 Q7NE64 Q7NE65

Q7NE68 Q7NE70 Q7NE87 Q7NEE9 Q7NEF1 Q7NEF2 Q7NEF3 Q7NEF4 Q7NEF6 Q7NEF7 Q7NEF8
Q7NEF9 Q7NEG0 Q7NEG1 Q7NEG2 Q7NEG3 Q7NEG4 Q7NEG5 Q7NEG6 Q7NEG7 Q7NEG8 Q7NEG9
Q7NEH0 Q7NEK2 Q7NEK3 Q7NEP1 Q7NEP3 Q7NEW1 Q7NEW2 Q7NEX4 Q7NEY1 Q7NEZ5 Q7NF32
Q7NF44 Q7NF75 Q7NFA1 Q7NFA4 Q7NFC3 Q7NFD3 Q7NFD4 Q7NFD9 Q7NFE9 Q7NFF1 Q7NFF2
Q7NFF3 Q7NFF4 Q7NFF5 Q7NFF6 Q7NFH4 Q7NFH8 Q7NFI7 Q7NFJ4 Q7NFJ5 Q7NFJ9 Q7NFK0
Q7NFL5 Q7NFM5 Q7NFM6 Q7NFN2 Q7NFN6 Q7NFV7 Q7NFV8 Q7NFW7 Q7NFY1 Q7NFZ1 Q7NG18
Q7NG31 Q7NG34 Q7NG49 Q7NG51 Q7NG73 Q7NG74 Q7NG91 Q7NG93 Q7NGF7 Q7NGF9 Q7NGI6
Q7NGJ2 Q7NGK1 Q7NGL0 Q7NGM0 Q7NGN4 Q7NGN9 Q7NGP0 Q7NGP3 Q7NGP9 Q7NGR7 Q7NGU2
Q7NGU6 Q7NGV2 Q7NGV3 Q7NGW6 Q7NGX9 Q7NGZ0 Q7NH22 Q7NH27 Q7NH80 Q7NH85 Q7NH89
Q7NH99 Q7NHA1 Q7NHA5 Q7NHC2 Q7NHC5 Q7NHC6 Q7NHE8 Q7NHG0 Q7NHH9 Q7NHI0 Q7NHJ3
Q7NHK6 Q7NHL0 Q7NHM2 Q7NHM8 Q7NHP4 Q7NHT3 Q7NHU3 Q7NHU4 Q7NHW1 Q7NHX6 Q7NHY0
Q7NHZ4 Q7NI06 Q7NI14 Q7NI15 Q7NI36 Q7NI42 Q7NI44 Q7NI66 Q7NI67 Q7NI70 Q7NI89
Q7NI93 Q7NIA0 Q7NIA2 Q7NIB2 Q7NID1 Q7NIE6 Q7NIF5 Q7NIG8 Q7NIM7 Q7NIP7 Q7NIP9
Q7NIR9 Q7NIT2 Q7NIT8 Q7NIV3 Q7NJ40 Q7NJ41 Q7NJA4 Q7NJB6 Q7NJE8 Q7NJF7 Q7NJH7
Q7NJK1 Q7NJK2 Q7NJK3 Q7NJM5 Q7NJM6 Q7NJR0 Q7NJS6 Q7NJS7 Q7NJT1 Q7NJU8 Q7NJV3
Q7NJY2 Q7NJY9 Q7NK22 Q7NK24 Q7NK35 Q7NK51 Q7NK59 Q7NK60 Q7NK76 Q7NK77 Q7NK78
Q7NK81 Q7NK82 Q7NK98 Q7NKE0 Q7NKF0 Q7NKF4 Q7NKH3 Q7NKI2 Q7NKI3 Q7NKK3 Q7NKL2
Q7NKL3 Q7NKN1 Q7NKT5 Q7NKX2 Q7NKY9 Q7NKZ0 Q7NL01 Q7NL02 Q7NL03 Q7NL20 Q7NL41
Q7NL65 Q7NL66 Q7NL83 Q7NLA8 Q7NLB5 Q7NLC0 Q7NLD3 Q7NLE2 Q7NLJ1 Q7NLJ6 Q7NLK5
Q7NLN7 Q7NLS0 Q7NLS3 Q7NLS9 Q7NLT1 Q7NLT3 Q7NLT7 Q7NLT8 Q7NLU2 Q7NLW9 Q7NLX2
Q7NLZ1 Q7NM37 Q7NM43 Q7NM64 Q7NM65 Q7NM67 Q7NM86 Q7NMB5 Q7NMB9 Q7NMC7 Q7NMD1
Q7NMD8 Q7NME1 Q7NME2 Q7NMG4 Q7NMH4 Q7NMI9 Q7NMJ6 Q7NMK3 Q7NMK9 Q7NML1 Q7NML9
Q7NMN8 Q7NMP5 Q7NMP9 Q7NMQ5 Q7NMQ7 Q7NMQ9 Q7NMR2 Q7NMT7 Q7NN18 Q7NN66 Q7NN71
Q7NN75 Q7NN81 Q7NND8 Q7NNG7 Q7NNH3 Q7NNH7 Q7NNJ7 Q7NNJ8 Q7NNK4 Q7NNK8 Q7NNL0
Q7NNL2 Q7NNL4 Q7NNW1 Q7NNX0 Q7NP43 Q7NP62 Q7NP63 Q7NP97 Q7NPB2 Q7NPE7 Q7NPF2
Q7NPF3 Q7NPG8 Q7NPG9 Q7NPH0 Q7NPH1 Q7NPI4 Q7NPI6 Q7NPL5 Q7NPM5 Q7NPQ5 Q7NPQ6
Q7NPQ7 Q7NPS3 Q7NPT6 Q7NPT9 Q7NPV7 Q7NPW1 Q7NPW2 Q7NPW5 Q7NPY6 Q7NPY7 Q7NPY8
Q7NPZ0 Q7NPZ1 Q7NPZ4 Q7NPZ7 Q7NQ00 Q7NQ01 Q7NQ07 Q7NQ20 Q7NQ37 Q7NQ51 Q7NQ65
Q7NQ72 Q7NQ75 Q7NQ76 Q7NQ92 Q7NQA9 Q7NQB5 Q7NQB7 Q7NQC4 Q7NQC5 Q7NQC8 Q7NQE3
Q7NQE4 Q7NQE5 Q7NQE6 Q7NQE7 Q7NQE8 Q7NQE9 Q7NQF0 Q7NQF1 Q7NQF2 Q7NQF3 Q7NQF4
Q7NQF5 Q7NQF6 Q7NQF7 Q7NQF8 Q7NQF9 Q7NQG0 Q7NQG1 Q7NQG2 Q7NQG3 Q7NQG4 Q7NQG5
Q7NQG6 Q7NQG7 Q7NQG8 Q7NQG9 Q7NQH0 Q7NQH1 Q7NQH3 Q7NQH4 Q7NQH5 Q7NQH6 Q7NQH7
Q7NQH8 Q7NQH9 Q7NQI5 Q7NQI8 Q7NQK6 Q7NQK7 Q7NQS0 Q7NQS5 Q7NQS8 Q7NQS9 Q7NQT0
Q7NQT1 Q7NQX1 Q7NR63 Q7NR74 Q7NR75 Q7NR81 Q7NRF7 Q7NRF8 Q7NRI3 Q7NRI6 Q7NRJ5
Q7NRJ9 Q7NRK0 Q7NRK9 Q7NRL1 Q7NRL2 Q7NRL4 Q7NRL5 Q7NRN0 Q7NRN3 Q7NRP0 Q7NRQ7
Q7NRS0 Q7NRS9 Q7NRT0 Q7NRT1 Q7NRT2 Q7NRT3 Q7NRT4 Q7NRT5 Q7NRT6 Q7NRV0 Q7NRV1
Q7NRV5 Q7NRV6 Q7NRV7 Q7NRW1 Q7NRW7 Q7NRY7 Q7NRY9 Q7NRZ0 Q7NS06 Q7NS12 Q7NS23
Q7NS40 Q7NS57 Q7NS59 Q7NS67 Q7NS72 Q7NS84 Q7NS85 Q7NS88 Q7NS89 Q7NS92 Q7NS93
Q7NS97 Q7NS98 Q7NSA6 Q7NSB3 Q7NSB9 Q7NSD8 Q7NSG1 Q7NSG2 Q7NSG4 Q7NSG9 Q7NSH0
Q7NSH1 Q7NSI9 Q7NSJ1 Q7NSJ3 Q7NSJ4 Q7NSJ5 Q7NSK2 Q7NSK3 Q7NSK5 Q7NSP3 Q7NSP5
Q7NSR4 Q7NSR8 Q7NSS5 Q7NSS7 Q7NST4 Q7NST5 Q7NSY7 Q7NT31 Q7NT63 Q7NT72 Q7NT84
Q7NT87 Q7NTA7 Q7NTD5 Q7NTF9 Q7NTG0 Q7NTG1 Q7NTH5 Q7NTI2 Q7NTK7 Q7NTK9 Q7NTL2
Q7NTN5 Q7NTY1 Q7NTY6 Q7NTY9 Q7NTZ8 Q7NTZ9 Q7NU26 Q7NU27 Q7NU77 Q7NUB6 Q7NUB8
Q7NUC2 Q7NUD6 Q7NUD8 Q7NUD9 Q7NUE3 Q7NUI6 Q7NUK5 Q7NUK7 Q7NUM8 Q7NUP9 Q7NUY8

Q7NUY9 Q7NUZ0 Q7NV18 Q7NV65 Q7NV67 Q7NV72 Q7NVB6 Q7NVB8 Q7NVF2 Q7NVF3 Q7NVF7
Q7NVH1 Q7NVH4 Q7NVH5 Q7NVH7 Q7NVK0 Q7NVL7 Q7NVM0 Q7NVN5 Q7NVN6 Q7NVP1 Q7NVT9
Q7NVY8 Q7NVZ0 Q7NVZ1 Q7NVZ2 Q7NVZ3 Q7NVZ4 Q7NW07 Q7NW10 Q7NW17 Q7NW22 Q7NWB7
Q7NWC0 Q7NWC1 Q7NWC4 Q7NWE3 Q7NWH3 Q7NWI9 Q7NWJ5 Q7NWJ6 Q7NWM1 Q7NWN5 Q7NWN7
Q7NWN9 Q7NWP4 Q7NWP9 Q7NWQ6 Q7NWR6 Q7NWT1 Q7NWU9 Q7NWY0 Q7NX05 Q7NX30 Q7NX31
Q7NX34 Q7NX61 Q7NX81 Q7NX82 Q7NX86 Q7NX89 Q7NX94 Q7NXD4 Q7NXH5 Q7NXI1 Q7NXI3
Q7NXI4 Q7NXI6 Q7NXI9 Q7NXJ0 Q7NXJ1 Q7NXK2 Q7NXL4 Q7NXL5 Q7NXL8 Q7NXL9 Q7NXM0
Q7NXM2 Q7NXN4 Q7NXP1 Q7NXQ0 Q7NXR0 Q7NXU5 Q7NXX6 Q7NXX7 Q7NY10 Q7NY11 Q7NY12
Q7NY13 Q7NY15 Q7NY19 Q7NY61 Q7NY96 Q7NY99 Q7NYA1 Q7NYC0 Q7NYC1 Q7NYC2 Q7NYC3
Q7NYC4 Q7NYC5 Q7NYC6 Q7NYC7 Q7NYF6 Q7NYG9 Q7NYI7 Q7NYI8 Q7NYJ7 Q7NYL4 Q7NYL5
Q7NYL6 Q7NYL8 Q7NYM3 Q7NYM4 Q7NYR7 Q7NYT5 Q7NYV9 Q7NYX3 Q7NYZ2 Q7NYZ5 Q7NZ00
Q7NZ02 Q7NZ10 Q7NZ19 Q7NZ31 Q7NZ33 Q7NZ47 Q7NZ60 Q7NZ62 Q7NZ91 Q7NZ95 Q7NZB2
Q7NZD3 Q7NZD4 Q7NZD5 Q7NZF7 Q7NZF9 Q7NZI3 Q7NZJ6 Q7NZS1 Q7NZS2 Q7NZS3 Q7NZU3
Q7NZU4 Q7NZV7 Q7NZY3 Q7P012 Q7P071 Q7P0A5 Q7P0A6 Q7P0B3 Q7P0E6 Q7P0E7 Q7P0E9
Q7P0F0 Q7P0F1 Q7P0F3 Q7P0F4 Q7P0F6 Q7P0F7 Q7P0F8 Q7P0H2 Q7P0H9 Q7P0J3 Q7P0L7
Q7P0M1 Q7P0P7 Q7P0P8 Q7P0P9 Q7P0R1 Q7P0S3 Q7P0S6 Q7P0S9 Q7P0T8 Q7P0V2 Q7P0W0
Q7P0W5 Q7P0X5 Q7P112 Q7P113 Q7P144 Q7P145 Q7P188 Q7P190 Q7P192 Q7P1B9 Q7P1G2
Q7P1H8 Q7P1I1 Q7P1M4 Q7P1P8 Q7P1R3 Q7P1R4 Q7P1R6 Q7P1T8 Q7P1U1 Q7P1U7 Q7P1Y2
Q7P1Y3 Q7P1Z5 Q7P203 Q7P206 Q7P207 Q7P221 Q7P259 Q7P9E2 Q7PAL9 Q7PBU9 Q7PC63
Q7PCC4 Q7PCJ6 Q7PCJ8 Q7PCJ9 Q7PCK7 Q7PGE8 Q7PHR1 Q7PK92 Q7PKS9 Q7PLI2 Q7PLK0
Q7PLS8 Q7PMS9 Q7PMT1 Q7PMU8 Q7PN68 Q7PNC0 Q7PND3 Q7PNM6 Q7PP77 Q7PQ25 Q7PR38
Q7PR68 Q7PRA4 Q7PRB8 Q7PRG3 Q7PRH1 Q7PS09 Q7PS24 Q7PSY2 Q7PTC5 Q7PTL0 Q7PUD7
Q7PV10 Q7PVQ9 Q7PVR3 Q7PVX8 Q7PVZ2 Q7PWB1 Q7PWB8 Q7PWQ6 Q7PWT9 Q7PX35 Q7PXE2
Q7PXG4 Q7PXU6 Q7PXY4 Q7PY41 Q7PYH1 Q7PYI4 Q7PYQ5 Q7PYU6 Q7PZ36 Q7PZ66 Q7PZ81
Q7PZ96 Q7PZD5 Q7PZK5 Q7PZY1 Q7Q068 Q7Q0G5 Q7Q0Q1 Q7Q0U1 Q7Q107 Q7Q161 Q7Q1K8
Q7Q2B7 Q7Q2P7 Q7Q323 Q7Q3I6 Q7Q412 Q7Q4F5 Q7Q547 Q7Q566 Q7Q5R5 Q7Q5Y8 Q7Q5Z3
Q7Q6B5 Q7Q6D6 Q7Q6D9 Q7Q6I6 Q7Q6S8 Q7Q6Y4 Q7Q727 Q7Q968 Q7Q9C0 Q7Q9F6 Q7Q9I4
Q7Q9N9 Q7QAD7 Q7QB13 Q7QB45 Q7QBJ0 Q7QBW3 Q7QBX2 Q7QC84 Q7QCB6 Q7QCJ9 Q7QCK2
Q7QCW2 Q7QD36 Q7QEH1 Q7QEI1 Q7QFL7 Q7QG28 Q7QGK4 Q7QGL0 Q7QGL9 Q7QGS3 Q7QGX9
Q7QH62 Q7QH64 Q7QHX4 Q7QIL2 Q7QIL8 Q7QIX1 Q7QJ33 Q7QJM5 Q7QJV0 Q7QJW7 Q7QJX9
Q7R7N4 Q7RAV5 Q7RBT0 Q7REH6 Q7RI54 Q7RJF9 Q7RJY1 Q7RKK8 Q7RM49 Q7RM92 Q7RME1
Q7RME3 Q7RNM5 Q7RPB4 Q7RT90 Q7RTN6 Q7RTS3 Q7RTT3 Q7RTT4 Q7RTT5 Q7RTT6 Q7RTU0
Q7RTU1 Q7RTU3 Q7RTU4 Q7RTU5 Q7RTU7 Q7RTV0 Q7RTV2 Q7RTV3 Q7RTV5 Q7RTY3 Q7RUA4
Q7RUX3 Q7RV75 Q7RV85 Q7RV88 Q7RVA8 Q7RVF7 Q7RVI1 Q7RVL0 Q7RVN2 Q7RVR8 Q7RVS9
Q7RVT0 Q7RVY5 Q7RW48 Q7RWR8 Q7RWT1 Q7RWV9 Q7RWW0 Q7RX33 Q7RX39 Q7RX73 Q7RX84
Q7RXA6 Q7RXH4 Q7RXK7 Q7RXQ1 Q7RXQ8 Q7RXR1 Q7RXR3 Q7RXX8 Q7RXY1 Q7RXY2 Q7RY30
Q7RY59 Q7RY68 Q7RY78 Q7RY84 Q7RYE7 Q7RYH7 Q7RYI3 Q7RYL4 Q7RYL6 Q7RYN6 Q7RYR4
Q7RYU8 Q7RYV5 Q7RYW6 Q7RYY4 Q7RYZ1 Q7RYZ7 Q7RZ35 Q7RZA5 Q7RZC1 Q7RZH4 Q7RZI0
Q7RZK5 Q7RZK7 Q7RZM8 Q7RZP7 Q7RZQ3 Q7RZS0 Q7RZT9 Q7RZU4 Q7RZV2 Q7S045 Q7S0I1
Q7S0P8 Q7S0S2 Q7S133 Q7S154 Q7S1H9 Q7S1P9 Q7S1X9 Q7S1Y0 Q7S232 Q7S2A9 Q7S2D4
Q7S2I2 Q7S2L7 Q7S2R9 Q7S2V0 Q7S2W6 Q7S2X0 Q7S2Y8 Q7S2Z9 Q7S302 Q7S304 Q7S332
Q7S3C4 Q7S3C8 Q7S432 Q7S438 Q7S454 Q7S464 Q7S4F7 Q7S4G7 Q7S4H6 Q7S4I9 Q7S4N5
Q7S4P1 Q7S519 Q7S532 Q7S565 Q7S5C0 Q7S5D9 Q7S5H0 Q7S5N8 Q7S5R1 Q7S604 Q7S634
Q7S6F3 Q7S6M5 Q7S6N6 Q7S6P8 Q7S6U4 Q7S6W5 Q7S6X4 Q7S6X6 Q7S724 Q7S743 Q7S745

Q7S747 Q7S7C0 Q7S7F1 Q7S7G7 Q7S7L4 Q7S7L7 Q7S7N3 Q7S7R6 Q7S860 Q7S873 Q7S891
Q7S8C2 Q7S8C8 Q7S8J7 Q7S8M2 Q7S8R3 Q7S8R8 Q7S8V3 Q7S8W7 Q7S8Z0 Q7S906 Q7S910
Q7S931 Q7S949 Q7S986 Q7S9B6 Q7S9B7 Q7S9D2 Q7S9H0 Q7S9J4 Q7S9L2 Q7S9P4 Q7S9Y9
Q7SA04 Q7SA44 Q7SA95 Q7SAA9 Q7SAB7 Q7SAF6 Q7SAK5 Q7SAK9 Q7SAN9 Q7SAP7 Q7SAX8
Q7SB49 Q7SB53 Q7SB62 Q7SB74 Q7SBC6 Q7SBD4 Q7SBD5 Q7SBJ9 Q7SBR1 Q7SBS7 Q7SBU6
Q7SBU7 Q7SBX8 Q7SC06 Q7SC15 Q7SC91 Q7SC98 Q7SCC1 Q7SCN0 Q7SCP4 Q7SCW9 Q7SD11
Q7SD49 Q7SD63 Q7SDA6 Q7SDF3 Q7SDH9 Q7SDM8 Q7SDP4 Q7SDU2 Q7SDV9 Q7SDW6 Q7SDX3
Q7SDY3 Q7SDY6 Q7SDZ1 Q7SE17 Q7SE75 Q7SE80 Q7SEF9 Q7SEI7 Q7SEK1 Q7SEK8 Q7SEL0
Q7SEM3 Q7SEW2 Q7SEY2 Q7SEZ0 Q7SF46 Q7SF55 Q7SF72 Q7SF81 Q7SF84 Q7SFC2 Q7SFC8
Q7SFJ3 Q7SFJ9 Q7SFQ6 Q7SFX6 Q7SFX7 Q7SFY0 Q7SG01 Q7SG09 Q7SGD2 Q7SGG0 Q7SGG8
Q7SGL3 Q7SGQ0 Q7SGR2 Q7SGS1 Q7SGW5 Q7SGZ5 Q7SH14 Q7SH17 Q7SH33 Q7SHE8 Q7SHG9
Q7SHH9 Q7SHI0 Q7SHI1 Q7SHI3 Q7SHI4 Q7SHI5 Q7SHI6 Q7SHI7 Q7SHR0 Q7SHR8 Q7SHT5
Q7SHU9 Q7SHY8 Q7SHZ2 Q7SI01 Q7SI09 Q7SI58 Q7SI59 Q7SI68 Q7SI91 Q7SI94 Q7SI95
Q7SI97 Q7SIA8 Q7SIB1 Q7SIB7 Q7SIB9 Q7SIC0 Q7SIC2 Q7SIC3 Q7SIC7 Q7SID0 Q7SID2
Q7SID3 Q7SID7 Q7SIE0 Q7SIF4 Q7SIG4 Q7SIG5 Q7SIG7 Q7SQ98 Q7SX95 Q7SX99 Q7SXA5
Q7SXA6 Q7SXA9 Q7SXB5 Q7SXC6 Q7SXE8 Q7SXF2 Q7SXG4 Q7SXI6 Q7SXK5 Q7SXL3 Q7SXL7
Q7SXM0 Q7SXN4 Q7SXP2 Q7SXP4 Q7SXP8 Q7SXR1 Q7SXS7 Q7SXS8 Q7SXT7 Q7SXU0 Q7SXV1
Q7SXV2 Q7SXW2 Q7SXW6 Q7SXY4 Q7SY07 Q7SY21 Q7SY23 Q7SY48 Q7SY53 Q7SY54 Q7SY65
Q7SYB3 Q7SYB5 Q7SYD9 Q7SYF8 Q7SYJ9 Q7SYL3 Q7SYM0 Q7SYN4 Q7SYN5 Q7SYR1 Q7SYS2
Q7SYS9 Q7SYU0 Q7SYV0 Q7SYV1 Q7SZ07 Q7SZ28 Q7SZ73 Q7SZ78 Q7SZB4 Q7SZE8 Q7SZF1
Q7SZK7 Q7SZL5 Q7SZM9 Q7SZP2 Q7SZS1 Q7SZS2 Q7SZT7 Q7T010 Q7T012 Q7T037 Q7T044
Q7T076 Q7T0B0 Q7T0B1 Q7T0L7 Q7T0M3 Q7T0M6 Q7T0P4 Q7T0P6 Q7T0Q0 Q7T0Q1 Q7T0Q3
Q7T0Q5 Q7T0S2 Q7T0S7 Q7T0U5 Q7T0W1 Q7T0W5 Q7T0Y4 Q7T0Z0 Q7T105 Q7T1H5 Q7T1K4
Q7T1R4 Q7T287 Q7T291 Q7T293 Q7T2A3 Q7T2A5 Q7T2B3 Q7T2B9 Q7T2D0 Q7T2G1 Q7T2N4
Q7T2N5 Q7T2P0 Q7T2T1 Q7T2U9 Q7T2V3 Q7T308 Q7T320 Q7T322 Q7T330 Q7T338 Q7T346
Q7T347 Q7T364 Q7T370 Q7T3B0 Q7T3C6 Q7T3E4 Q7T3E5 Q7T3E8 Q7T3F7 Q7T3H2 Q7T3H7
Q7T3H9 Q7T3I7 Q7T3M9 Q7T3N1 Q7T3N2 Q7T3N3 Q7T3N4 Q7T3N6 Q7T3N7 Q7T3N8 Q7T3N9
Q7T3P0 Q7T3P1 Q7T3P8 Q7T3T8 Q7T3T9 Q7T3U0 Q7T3X9 Q7T3Y0 Q7T6T2 Q7T6X0 Q7T6X1
Q7T6X3 Q7T6X4 Q7T6X5 Q7T6X6 Q7T6X7 Q7T6X8 Q7T6X9 Q7T6Y0 Q7T6Y1 Q7T6Y3 Q7T6Y4
Q7T6Y5 Q7T6Y8 Q7T6Y9 Q7TBL3 Q7TD08 Q7TD10 Q7TD11 Q7TD12 Q7TD13 Q7TDB4 Q7TDB5
Q7TDB6 Q7TF74 Q7TF75 Q7TLC4 Q7TLC7 Q7TM95 Q7TM96 Q7TMA2 Q7TMC8 Q7TMD5 Q7TME2
Q7TMF2 Q7TMI3 Q7TMV3 Q7TMW6 Q7TMX5 Q7TMY4 Q7TN02 Q7TN08 Q7TN33 Q7TNB2 Q7TNB4
Q7TNB8 Q7TNC4 Q7TNC6 Q7TND5 Q7TNE1 Q7TNF8 Q7TNF9 Q7TNG5 Q7TNK1 Q7TNK3 Q7TNK6
Q7TNL3 Q7TNL4 Q7TNM2 Q7TNS8 Q7TNU6 Q7TNZ6 Q7TP52 Q7TP98 Q7TPB1 Q7TPD0 Q7TPD2
Q7TPD6 Q7TPE5 Q7TPM1 Q7TPM3 Q7TPM5 Q7TPM6 Q7TPN9 Q7TPQ3 Q7TPV2 Q7TPX8 Q7TPX9
Q7TPY9 Q7TQ07 Q7TQ20 Q7TQ40 Q7TQ82 Q7TQ84 Q7TQ94 Q7TQC7 Q7TQD2 Q7TQE7 Q7TQF2
Q7TQG0 Q7TQI7 Q7TQI8 Q7TQK1 Q7TQK4 Q7TQM5 Q7TS63 Q7TS68 Q7TS73 Q7TS74 Q7TS99
Q7TSA6 Q7TSC3 Q7TSD4 Q7TSF4 Q7TSG3 Q7TSH3 Q7TSH6 Q7TSH9 Q7TSI0 Q7TSI3 Q7TSP2
Q7TSQ8 Q7TSS2 Q7TST9 Q7TSU0 Q7TSV4 Q7TSV6 Q7TSV9 Q7TSX9 Q7TSY6 Q7TT00 Q7TT23
Q7TT28 Q7TT37 Q7TT52 Q7TT79 Q7TT91 Q7TTF9 Q7TTP0 Q7TTP2 Q7TTQ1 Q7TTR0 Q7TTR4
Q7TTR7 Q7TTS1 Q7TTS3 Q7TTS6 Q7TTS8 Q7TTT1 Q7TTT6 Q7TTT8 Q7TTU0 Q7TTU3 Q7TTU4
Q7TTU5 Q7TTU6 Q7TTU8 Q7TTU9 Q7TTV4 Q7TTW1 Q7TTW4 Q7TTW6 Q7TTW9 Q7TTX0 Q7TTX1
Q7TTX2 Q7TTX6 Q7TTX7 Q7TTX9 Q7TTY1 Q7TTY8 Q7TTY9 Q7TTZ0 Q7TTZ4 Q7TTZ5 Q7TU07
Q7TU09 Q7TU18 Q7TU19 Q7TU20 Q7TU24 Q7TU25 Q7TU27 Q7TU28 Q7TU29 Q7TU30 Q7TU31

Q7TU38 Q7TU43 Q7TU44 Q7TU56 Q7TU60 Q7TU64 Q7TU65 Q7TU67 Q7TU75 Q7TU81 Q7TU88
Q7TU90 Q7TU92 Q7TU96 Q7TU99 Q7TUA0 Q7TUA3 Q7TUA7 Q7TUA9 Q7TUC7 Q7TUC8 Q7TUD1
Q7TUG0 Q7TUG1 Q7TUG2 Q7TUG9 Q7TUH0 Q7TUI1 Q7TUI6 Q7TUI9 Q7TUJ1 Q7TUJ2 Q7TUJ9
Q7TUK1 Q7TUK9 Q7TUL2 Q7TUM6 Q7TUM7 Q7TUN0 Q7TUN3 Q7TUN4 Q7TUP1 Q7TUP2 Q7TUP3
Q7TUP4 Q7TUP5 Q7TUR3 Q7TUR7 Q7TUS3 Q7TUS4 Q7TUS7 Q7TUT0 Q7TUT1 Q7TUT2 Q7TUT7
Q7TUU2 Q7TUU4 Q7TUV5 Q7TUV8 Q7TUW5 Q7TUX0 Q7TUZ8 Q7TV01 Q7TV03 Q7TV05 Q7TV07
Q7TV13 Q7TV16 Q7TV18 Q7TV26 Q7TV28 Q7TV34 Q7TV42 Q7TV43 Q7TV44 Q7TV48 Q7TV75
Q7TV76 Q7TV80 Q7TV86 Q7TV89 Q7TV92 Q7TV93 Q7TVA1 Q7TVA3 Q7TVA4 Q7TVA6 Q7TVB0
Q7TVI1 Q7TVJ6 Q7TVN7 Q7TVQ0 Q7TVQ7 Q7TVV6 Q7TW42 Q7TW52 Q7TW54 Q7TW55 Q7TW97
Q7TWE1 Q7TWH9 Q7TWJ2 Q7TWJ6 Q7TWL7 Q7TWL9 Q7TWN0 Q7TWT6 Q7TWT7 Q7TWV3 Q7TWV4
Q7TWW7 Q7TWY4 Q7TX14 Q7TX80 Q7TX84 Q7TXA9 Q7TXG2 Q7TXH6 Q7TXH8 Q7TXH9 Q7TXI6
Q7TXJ4 Q7TXJ8 Q7TXK6 Q7TXM7 Q7TXN0 Q7TXN6 Q7TXQ5 Q7TXW0 Q7TXX0 Q7TY17 Q7TY47
Q7TY92 Q7TYA6 Q7TYB1 Q7TYC8 Q7TYI9 Q7TYJ1 Q7TYK5 Q7TYM1 Q7TYN1 Q7TYP6 Q7TYQ1
Q7TYZ5 Q7TZ13 Q7TZ14 Q7TZA8 Q7TZB9 Q7TZC0 Q7TZG8 Q7TZN1 Q7TZP3 Q7TZP5 Q7U040
Q7U051 Q7U054 Q7U055 Q7U057 Q7U076 Q7U078 Q7U0A6 Q7U0D0 Q7U0E1 Q7U0E7 Q7U0G9
Q7U0H2 Q7U0N5 Q7U0N6 Q7U0P8 Q7U0Y2 Q7U0Y5 Q7U0Z0 Q7U0Z1 Q7U140 Q7U163 Q7U173
Q7U178 Q7U184 Q7U1B9 Q7U1E6 Q7U1E7 Q7U1E9 Q7U1K2 Q7U1K4 Q7U1L7 Q7U1S4 Q7U206
Q7U272 Q7U297 Q7U2D4 Q7U2I0 Q7U2J0 Q7U2R2 Q7U2R3 Q7U2S7 Q7U2U1 Q7U2X3 Q7U313
Q7U314 Q7U317 Q7U318 Q7U320 Q7U321 Q7U323 Q7U326 Q7U328 Q7U330 Q7U331 Q7U335
Q7U338 Q7U339 Q7U342 Q7U344 Q7U345 Q7U347 Q7U348 Q7U349 Q7U350 Q7U353 Q7U356
Q7U358 Q7U359 Q7U361 Q7U363 Q7U365 Q7U366 Q7U367 Q7U370 Q7U372 Q7U374 Q7U375
Q7U376 Q7U381 Q7U383 Q7U384 Q7U386 Q7U387 Q7U391 Q7U3A4 Q7U3B3 Q7U3B9 Q7U3C1
Q7U3C4 Q7U3D7 Q7U3E2 Q7U3H2 Q7U3I3 Q7U3I4 Q7U3I5 Q7U3I6 Q7U3I7 Q7U3I9 Q7U3K1
Q7U3K4 Q7U3K6 Q7U3L0 Q7U3N4 Q7U3P1 Q7U3P8 Q7U3Q2 Q7U3Q5 Q7U3R9 Q7U3S0 Q7U3S1
Q7U3S3 Q7U3S5 Q7U3S6 Q7U3T3 Q7U3T6 Q7U3T7 Q7U3T8 Q7U3T9 Q7U3V0 Q7U3V6 Q7U3V8
Q7U3W4 Q7U3W6 Q7U3W8 Q7U3X6 Q7U3X9 Q7U3Z7 Q7U406 Q7U437 Q7U442 Q7U469 Q7U477
Q7U490 Q7U4A1 Q7U4C3 Q7U4C6 Q7U4D0 Q7U4D1 Q7U4D2 Q7U4D3 Q7U4D4 Q7U4D9 Q7U4H2
Q7U4H3 Q7U4H4 Q7U4H5 Q7U4H6 Q7U4H7 Q7U4H8 Q7U4H9 Q7U4I0 Q7U4I1 Q7U4I3 Q7U4I4
Q7U4I5 Q7U4I6 Q7U4I7 Q7U4I8 Q7U4I9 Q7U4J0 Q7U4J1 Q7U4J2 Q7U4J3 Q7U4J4 Q7U4J5
Q7U4J6 Q7U4J7 Q7U4J8 Q7U4J9 Q7U4K0 Q7U4K5 Q7U4L7 Q7U4L8 Q7U4M0 Q7U4M7 Q7U4N8
Q7U4P0 Q7U4P6 Q7U4P7 Q7U4S6 Q7U4S9 Q7U4U1 Q7U4U9 Q7U4V1 Q7U4V2 Q7U4X5 Q7U4Y5
Q7U4Z8 Q7U4Z9 Q7U515 Q7U516 Q7U518 Q7U524 Q7U525 Q7U559 Q7U570 Q7U573 Q7U581
Q7U583 Q7U591 Q7U598 Q7U5B3 Q7U5C2 Q7U5C7 Q7U5D4 Q7U5D6 Q7U5E8 Q7U5F4 Q7U5F5
Q7U5G0 Q7U5G1 Q7U5I2 Q7U5I3 Q7U5I4 Q7U5I8 Q7U5J9 Q7U5K1 Q7U5K9 Q7U5L3 Q7U5L7
Q7U5M8 Q7U5N2 Q7U5Q1 Q7U5Q2 Q7U5Q9 Q7U5R0 Q7U5R5 Q7U5R6 Q7U5U1 Q7U5U4 Q7U5U5
Q7U5V5 Q7U5Z8 Q7U5Z9 Q7U605 Q7U618 Q7U635 Q7U637 Q7U645 Q7U646 Q7U654 Q7U659
Q7U662 Q7U697 Q7U6F4 Q7U6M8 Q7U6N7 Q7U6P6 Q7U6Q8 Q7U6R1 Q7U6R7 Q7U6S0 Q7U6S4
Q7U6T0 Q7U6T1 Q7U6T8 Q7U6U0 Q7U6V8 Q7U6V9 Q7U6W0 Q7U6W1 Q7U6X6 Q7U6Y8 Q7U6Z1
Q7U712 Q7U733 Q7U763 Q7U769 Q7U770 Q7U794 Q7U795 Q7U7A1 Q7U7D2 Q7U7D3 Q7U7D4
Q7U7D7 Q7U7F0 Q7U7F1 Q7U7F4 Q7U7F9 Q7U7H6 Q7U7H8 Q7U7I2 Q7U7I5 Q7U7L5 Q7U7P7
Q7U7P8 Q7U7P9 Q7U7Q2 Q7U7Q3 Q7U7S1 Q7U7T2 Q7U7W2 Q7U804 Q7U807 Q7U831 Q7U840
Q7U842 Q7U845 Q7U851 Q7U892 Q7U899 Q7U8A6 Q7U8C0 Q7U8C3 Q7U8E8 Q7U8F2 Q7U8F8
Q7U8G2 Q7U8G6 Q7U8I1 Q7U8J1 Q7U8K0 Q7U8K2 Q7U8K3 Q7U8K4 Q7U8K6 Q7U8K7 Q7U8K8
Q7U8L5 Q7U8L9 Q7U8M8 Q7U8P3 Q7U8Q1 Q7U8Q9 Q7U8R3 Q7U8R4 Q7U8R5 Q7U8R6 Q7U8R7

Q7U8R9 Q7U8S4 Q7U8T1 Q7U8T6 Q7U8U2 Q7U8Z6 Q7U923 Q7U924 Q7U9D4 Q7U9F0 Q7U9G2
Q7U9G6 Q7U9G9 Q7U9H3 Q7U9H6 Q7U9I1 Q7U9I5 Q7U9J4 Q7U9J5 Q7U9J7 Q7U9K2 Q7U9K4
Q7U9M0 Q7U9M8 Q7U9Q6 Q7U9V4 Q7U9W2 Q7U9W7 Q7U9W8 Q7U9X5 Q7U9Y3 Q7UA08 Q7UA19
Q7UA23 Q7UA24 Q7UA26 Q7UA33 Q7UA35 Q7UA36 Q7UA37 Q7UA42 Q7UA43 Q7UA46 Q7UA53
Q7UA54 Q7UA67 Q7UA71 Q7UA72 Q7UA73 Q7UA74 Q7UA76 Q7UA77 Q7UA92 Q7UAE6 Q7UAE7
Q7UAH4 Q7UAV7 Q7UB34 Q7UB84 Q7UBC5 Q7UBD3 Q7UBJ0 Q7UBT0 Q7UBU3 Q7UBW5 Q7UC03
Q7UC91 Q7UCI6 Q7UCI7 Q7UCT6 Q7UCU1 Q7UCW4 Q7UD30 Q7UDD0 Q7UDF6 Q7UDL2 Q7UDQ6
Q7UDT4 Q7UDT5 Q7UDT8 Q7UDU1 Q7UEP2 Q7UEU9 Q7UEV3 Q7UEW6 Q7UEX1 Q7UEY4 Q7UEY5
Q7UF53 Q7UF84 Q7UF86 Q7UF90 Q7UFG2 Q7UFG7 Q7UFH9 Q7UFN3 Q7UFN6 Q7UFQ9 Q7UFR1
Q7UFR2 Q7UFS3 Q7UFS5 Q7UFU6 Q7UFW2 Q7UFW4 Q7UFW7 Q7UFX8 Q7UFY2 Q7UFY6 Q7UFZ7
Q7UG04 Q7UG12 Q7UG38 Q7UG70 Q7UG74 Q7UG92 Q7UGJ6 Q7UGJ7 Q7UGM3 Q7UGV0 Q7UGV1
Q7UGZ7 Q7UH14 Q7UH37 Q7UH38 Q7UHE1 Q7UHE2 Q7UHU7 Q7UHW3 Q7UHZ5 Q7UHZ6 Q7UI03
Q7UI12 Q7UI13 Q7UI46 Q7UI51 Q7UI59 Q7UIA4 Q7UIA6 Q7UIA7 Q7UIC4 Q7UIC5 Q7UIC6
Q7UIC7 Q7UID0 Q7UIE1 Q7UIM1 Q7UIR4 Q7UIY1 Q7UJ17 Q7UJ19 Q7UJ52 Q7UJ58 Q7UJ62
Q7UJ69 Q7UJ86 Q7UJA8 Q7UJB1 Q7UJC4 Q7UJD7 Q7UJI2 Q7UJI3 Q7UJJ0 Q7UJJ7 Q7UJK0
Q7UJK7 Q7UJL3 Q7UK08 Q7UK39 Q7UK56 Q7UK64 Q7UK66 Q7UK67 Q7UKD1 Q7UKG6 Q7UKG9
Q7UKH3 Q7UKH4 Q7UKI3 Q7UKJ7 Q7UKJ8 Q7UKJ9 Q7UKQ9 Q7UKT5 Q7UKT6 Q7UKU7 Q7UKU9
Q7UKV0 Q7UKV1 Q7UKV4 Q7UKY0 Q7UKZ2 Q7UKZ8 Q7ULJ4 Q7ULM9 Q7ULP3 Q7ULT2 Q7ULT3
Q7ULT4 Q7ULU1 Q7ULV7 Q7ULX4 Q7UM14 Q7UM15 Q7UM23 Q7UM31 Q7UM33 Q7UM39 Q7UM42
Q7UM95 Q7UM96 Q7UM97 Q7UM98 Q7UM99 Q7UMC0 Q7UMN0 Q7UMW0 Q7UMW2 Q7UMY7 Q7UMZ0
Q7UMZ2 Q7UN00 Q7UN02 Q7UN03 Q7UN04 Q7UN05 Q7UN06 Q7UN07 Q7UN08 Q7UN09 Q7UN10
Q7UN11 Q7UN12 Q7UN13 Q7UN14 Q7UN15 Q7UN16 Q7UN17 Q7UN18 Q7UN19 Q7UN20 Q7UN22
Q7UN31 Q7UN32 Q7UN36 Q7UNC2 Q7UNC3 Q7UNC6 Q7UNE1 Q7UNF9 Q7UNG9 Q7UNH1 Q7UNL6
Q7UNN9 Q7UNQ2 Q7UNR1 Q7UNR2 Q7UNR3 Q7UNR5 Q7UNV2 Q7UNZ2 Q7UP05 Q7UP26 Q7UP45
Q7UP72 Q7UP74 Q7UP75 Q7UP77 Q7UP89 Q7UP92 Q7UP93 Q7UPC9 Q7UPG1 Q7UPG4 Q7UPM4
Q7UPM9 Q7UPN0 Q7UPN5 Q7UPP0 Q7UPR5 Q7UPS1 Q7UPT7 Q7UPY2 Q7UPZ8 Q7UQ02 Q7UQ43
Q7UQ68 Q7UQA4 Q7UQB3 Q7UQH1 Q7UQL3 Q7UQN2 Q7UQU4 Q7UQW3 Q7UQZ6 Q7UQZ8 Q7UR74
Q7UR95 Q7UR97 Q7URC7 Q7URE2 Q7URF1 Q7URG0 Q7URG5 Q7URJ8 Q7URL0 Q7URL1 Q7URM5
Q7URN0 Q7URP3 Q7URR0 Q7URR1 Q7URR4 Q7URU7 Q7URV2 Q7URW4 Q7URW6 Q7US70 Q7US71
Q7US84 Q7USA1 Q7USA9 Q7USB0 Q7USC2 Q7USC9 Q7USI0 Q7USI7 Q7USY7 Q7UT33 Q7UT69
Q7UTH0 Q7UTH1 Q7UTP4 Q7UTQ8 Q7UTS2 Q7UTZ9 Q7UU60 Q7UU71 Q7UU80 Q7UU94 Q7UUJ7
Q7UUW5 Q7UUW9 Q7UUZ6 Q7UVA0 Q7UVA1 Q7UVF9 Q7UVG2 Q7UVG8 Q7UVG9 Q7UVH1 Q7UVH5
Q7UVH6 Q7UVL4 Q7UVM5 Q7UW59 Q7UWB5 Q7UWB7 Q7UWB8 Q7UWC8 Q7UWK9 Q7UWN8 Q7UWS5
Q7UWY8 Q7UX39 Q7UX42 Q7UXF5 Q7UXH9 Q7UXK9 Q7UXM6 Q7UXN7 Q7UXX6 Q7UXZ8 Q7UYE5
Q7UYK5 Q7UYS5 Q7UYX5 Q7UYZ6 Q7UZ14 Q7UZ20 Q7UZG0 Q7UZG1 Q7UZG2 Q7UZG3 Q7UZH5
Q7UZH7 Q7UZH8 Q7UZI0 Q7UZI3 Q7UZJ1 Q7UZJ9 Q7UZK1 Q7UZK2 Q7UZK6 Q7UZK7 Q7UZK8
Q7UZK9 Q7UZL1 Q7UZL3 Q7UZL6 Q7UZL7 Q7UZN3 Q7UZP0 Q7UZP2 Q7UZP3 Q7UZP7 Q7UZQ3
Q7UZR6 Q7UZR7 Q7UZS3 Q7UZT0 Q7UZU3 Q7UZU7 Q7UZU8 Q7UZU9 Q7UZV0 Q7UZV1 Q7UZV2
Q7UZV3 Q7UZV4 Q7UZV5 Q7UZV6 Q7UZV7 Q7UZV8 Q7UZV9 Q7UZW0 Q7UZW1 Q7UZW3 Q7UZW4
Q7UZW5 Q7UZW6 Q7UZW7 Q7UZW8 Q7UZW9 Q7UZX0 Q7UZX1 Q7UZY1 Q7UZY4 Q7UZY5 Q7UZY6
Q7UZY7 Q7UZY8 Q7UZZ2 Q7UZZ3 Q7UZZ9 Q7V002 Q7V003 Q7V004 Q7V006 Q7V007 Q7V008
Q7V010 Q7V016 Q7V024 Q7V027 Q7V048 Q7V051 Q7V054 Q7V0A5 Q7V0B9 Q7V0C1 Q7V0C2
Q7V0C3 Q7V0C4 Q7V0C7 Q7V0D4 Q7V0D7 Q7V0D8 Q7V0F0 Q7V0F1 Q7V0F6 Q7V0F7 Q7V0G0
Q7V0G1 Q7V0I3 Q7V0I6 Q7V0I7 Q7V0I8 Q7V0J3 Q7V0R5 Q7V0R7 Q7V0S1 Q7V0T8 Q7V0U2

Q7V0U3 Q7V0W0 Q7V0W1 Q7V0X5 Q7V0Y7 Q7V0Z7 Q7V0Z8 Q7V100 Q7V101 Q7V102 Q7V111
Q7V121 Q7V127 Q7V132 Q7V179 Q7V1B0 Q7V1B1 Q7V1B2 Q7V1B3 Q7V1B4 Q7V1B5 Q7V1B6
Q7V1D0 Q7V1D3 Q7V1D9 Q7V1E1 Q7V1E2 Q7V1G6 Q7V1H0 Q7V1H4 Q7V1H9 Q7V1I1 Q7V1I2
Q7V1I7 Q7V1I9 Q7V1J8 Q7V1L2 Q7V1M1 Q7V1N1 Q7V1N4 Q7V1R4 Q7V1R7 Q7V1R9 Q7V1S5
Q7V1T1 Q7V1T7 Q7V1T8 Q7V1V0 Q7V1W0 Q7V209 Q7V215 Q7V226 Q7V263 Q7V272 Q7V275
Q7V277 Q7V286 Q7V287 Q7V289 Q7V293 Q7V299 Q7V2A0 Q7V2A3 Q7V2C0 Q7V2C5 Q7V2C6
Q7V2C7 Q7V2C9 Q7V2D0 Q7V2D1 Q7V2D3 Q7V2D5 Q7V2D6 Q7V2D7 Q7V2F5 Q7V2F8 Q7V2F9
Q7V2G1 Q7V2H0 Q7V2H1 Q7V2I1 Q7V2I5 Q7V2J3 Q7V2J5 Q7V2K1 Q7V2K3 Q7V2K6 Q7V2K9
Q7V2M1 Q7V2M3 Q7V2P2 Q7V2Q2 Q7V2Q8 Q7V2R1 Q7V2R3 Q7V2R8 Q7V2S2 Q7V2S6 Q7V2S9
Q7V2U0 Q7V2X4 Q7V2X8 Q7V2Y0 Q7V2Y6 Q7V2Y8 Q7V2Z2 Q7V2Z3 Q7V314 Q7V316 Q7V329
Q7V331 Q7V335 Q7V336 Q7V342 Q7V349 Q7V350 Q7V352 Q7V354 Q7V355 Q7V364 Q7V368
Q7V381 Q7V382 Q7V383 Q7V384 Q7V390 Q7V395 Q7V396 Q7V3A1 Q7V3A9 Q7V3B1 Q7V3C2
Q7V3E6 Q7V3F1 Q7V3G3 Q7V3H3 Q7V3K7 Q7V3M5 Q7V3M6 Q7V3M7 Q7V3M8 Q7V3M9 Q7V3N0
Q7V3N7 Q7V3N8 Q7V3P5 Q7V3P8 Q7V3P9 Q7V3Q0 Q7V3Q3 Q7V3Q4 Q7V3R5 Q7V3S4 Q7V3S9
Q7V3T1 Q7V3T5 Q7V3U5 Q7V3V2 Q7V3V3 Q7V3V4 Q7V3V5 Q7V3V8 Q7V3W6 Q7V3W8 Q7V3X1
Q7V3X3 Q7V3Y6 Q7V3Y9 Q7V3Z9 Q7V406 Q7V411 Q7V419 Q7V425 Q7V435 Q7V436 Q7V461
Q7V476 Q7V477 Q7V478 Q7V479 Q7V491 Q7V493 Q7V498 Q7V4A7 Q7V4B2 Q7V4D3 Q7V4D8
Q7V4G2 Q7V4G6 Q7V4I8 Q7V4K0 Q7V4P3 Q7V4R6 Q7V4S3 Q7V4S7 Q7V4T7 Q7V4T9 Q7V4U3
Q7V4U5 Q7V4V4 Q7V4V8 Q7V4W4 Q7V4W7 Q7V4X7 Q7V4Y9 Q7V4Z3 Q7V4Z5 Q7V4Z9 Q7V500
Q7V501 Q7V502 Q7V503 Q7V507 Q7V518 Q7V519 Q7V520 Q7V521 Q7V522 Q7V523 Q7V524
Q7V525 Q7V526 Q7V528 Q7V529 Q7V530 Q7V531 Q7V532 Q7V533 Q7V534 Q7V535 Q7V536
Q7V537 Q7V538 Q7V539 Q7V540 Q7V541 Q7V542 Q7V547 Q7V558 Q7V559 Q7V561 Q7V568
Q7V581 Q7V583 Q7V585 Q7V586 Q7V5A2 Q7V5A5 Q7V5B5 Q7V5B7 Q7V5D4 Q7V5F9 Q7V5L6
Q7V5M4 Q7V5M7 Q7V5N8 Q7V5N9 Q7V5P1 Q7V5P2 Q7V5P3 Q7V5P5 Q7V5Q2 Q7V5R2 Q7V5R6
Q7V5U0 Q7V5U5 Q7V5U8 Q7V5V5 Q7V5W1 Q7V5W3 Q7V5W4 Q7V5W5 Q7V5W6 Q7V5W7 Q7V5X1
Q7V5X8 Q7V5Y2 Q7V606 Q7V608 Q7V612 Q7V613 Q7V643 Q7V647 Q7V660 Q7V668 Q7V677
Q7V689 Q7V697 Q7V6A8 Q7V6B8 Q7V6C1 Q7V6C2 Q7V6C6 Q7V6D4 Q7V6E7 Q7V6E8 Q7V6E9
Q7V6F7 Q7V6F8 Q7V6F9 Q7V6G1 Q7V6G4 Q7V6G8 Q7V6G9 Q7V6H4 Q7V6H6 Q7V6H8 Q7V6J8
Q7V6K0 Q7V6L9 Q7V6M5 Q7V6Q5 Q7V6R9 Q7V6S0 Q7V6S2 Q7V6T7 Q7V6T8 Q7V6T9 Q7V7D2
Q7V7D4 Q7V7D8 Q7V7G9 Q7V7H4 Q7V7I8 Q7V7K4 Q7V7K5 Q7V7K6 Q7V7L7 Q7V7L9 Q7V7M5
Q7V7M6 Q7V7N1 Q7V7N4 Q7V7N7 Q7V7P3 Q7V7Q3 Q7V7R2 Q7V7S1 Q7V7S6 Q7V7T6 Q7V7T9
Q7V7V1 Q7V7V8 Q7V7W0 Q7V7W1 Q7V7Y8 Q7V7Z4 Q7V808 Q7V809 Q7V834 Q7V842 Q7V844
Q7V847 Q7V8B1 Q7V8B4 Q7V8B5 Q7V8C3 Q7V8D0 Q7V8D2 Q7V8E7 Q7V8F0 Q7V8F4 Q7V8F6
Q7V8G6 Q7V8G9 Q7V8H9 Q7V8I0 Q7V8I3 Q7V8L0 Q7V8L1 Q7V8L8 Q7V8L9 Q7V8M4 Q7V8M5
Q7V8P9 Q7V8Q1 Q7V8Q8 Q7V8S6 Q7V8U6 Q7V8U9 Q7V8V2 Q7V8V9 Q7V8W2 Q7V8W6 Q7V8X0
Q7V8X5 Q7V8X7 Q7V8Y5 Q7V8Y7 Q7V8Z5 Q7V904 Q7V906 Q7V910 Q7V911 Q7V926 Q7V958
Q7V959 Q7V977 Q7V978 Q7V987 Q7V990 Q7V991 Q7V992 Q7V993 Q7V998 Q7V999 Q7V9A2
Q7V9A9 Q7V9B0 Q7V9B9 Q7V9C3 Q7V9C4 Q7V9C5 Q7V9C6 Q7V9C8 Q7V9C9 Q7V9E5 Q7V9F8
Q7V9F9 Q7V9G2 Q7V9H8 Q7V9H9 Q7V9I0 Q7V9I1 Q7V9I2 Q7V9I5 Q7V9I9 Q7V9J2 Q7V9J7
Q7V9J9 Q7V9K3 Q7V9K4 Q7V9K9 Q7V9L1 Q7V9L2 Q7V9L5 Q7V9L6 Q7V9L7 Q7V9L9 Q7V9M1
Q7V9M5 Q7V9M7 Q7V9N2 Q7V9P3 Q7V9Q0 Q7V9Q3 Q7V9Q4 Q7V9Q8 Q7V9R4 Q7V9S8 Q7V9S9
Q7V9T7 Q7V9U4 Q7V9U7 Q7V9V8 Q7V9W2 Q7V9W3 Q7V9W4 Q7V9W5 Q7V9W6 Q7V9W7 Q7V9W8
Q7V9W9 Q7V9X0 Q7V9X1 Q7V9X2 Q7V9X3 Q7V9X4 Q7V9X5 Q7V9X6 Q7V9X7 Q7V9X8 Q7V9X9
Q7V9Y1 Q7V9Y2 Q7V9Y3 Q7V9Y4 Q7V9Y5 Q7V9Y6 Q7V9Y7 Q7V9Y8 Q7V9Y9 Q7V9Z0 Q7V9Z2

Q7V9Z9 Q7VA02 Q7VA03 Q7VA04 Q7VA05 Q7VA06 Q7VA11 Q7VA14 Q7VA20 Q7VA23 Q7VA25
Q7VA26 Q7VA27 Q7VA29 Q7VA30 Q7VA32 Q7VA38 Q7VA47 Q7VA53 Q7VA74 Q7VA78 Q7VA81
Q7VAC5 Q7VAL5 Q7VAM1 Q7VAM5 Q7VAN0 Q7VAN2 Q7VAN3 Q7VAN4 Q7VAN5 Q7VAN8 Q7VAP4
Q7VAP7 Q7VAP8 Q7VAR8 Q7VAR9 Q7VAS4 Q7VAS5 Q7VAS9 Q7VAT0 Q7VAT3 Q7VAT9 Q7VAU5
Q7VAV5 Q7VAV8 Q7VAV9 Q7VAW0 Q7VAW6 Q7VAZ5 Q7VAZ6 Q7VAZ8 Q7VB02 Q7VB29 Q7VB32
Q7VB41 Q7VB43 Q7VB62 Q7VB63 Q7VBB3 Q7VBE8 Q7VBF5 Q7VBG1 Q7VBH4 Q7VBI9 Q7VBJ0
Q7VBJ1 Q7VBJ3 Q7VBK3 Q7VBL0 Q7VBL3 Q7VBL4 Q7VBM1 Q7VBM6 Q7VBM8 Q7VBM9 Q7VBN8
Q7VBP0 Q7VBP6 Q7VBR5 Q7VBS7 Q7VBT3 Q7VBU5 Q7VBV0 Q7VBW1 Q7VBX4 Q7VBX5 Q7VBX6
Q7VBX7 Q7VBY8 Q7VBZ0 Q7VBZ5 Q7VBZ6 Q7VBZ8 Q7VC01 Q7VC04 Q7VC06 Q7VC09 Q7VC14
Q7VC22 Q7VC38 Q7VC41 Q7VC70 Q7VC75 Q7VC78 Q7VC80 Q7VC88 Q7VC95 Q7VCA1 Q7VCB5
Q7VCB6 Q7VCC2 Q7VCG1 Q7VCH6 Q7VCH7 Q7VCH9 Q7VCI6 Q7VCI7 Q7VCJ0 Q7VCJ6 Q7VCZ9
Q7VD04 Q7VD07 Q7VD15 Q7VD22 Q7VD23 Q7VD33 Q7VD37 Q7VD38 Q7VD39 Q7VD58 Q7VD61
Q7VD62 Q7VD64 Q7VD75 Q7VD77 Q7VD84 Q7VD89 Q7VD93 Q7VDA1 Q7VDA4 Q7VDB0 Q7VDB2
Q7VDB5 Q7VDB6 Q7VDC7 Q7VDF1 Q7VDF8 Q7VDG4 Q7VDG7 Q7VDH4 Q7VDH8 Q7VDI4 Q7VDI8
Q7VDJ2 Q7VDK6 Q7VDL2 Q7VDL4 Q7VDL7 Q7VDM4 Q7VDM7 Q7VDN2 Q7VDN6 Q7VDQ5 Q7VDQ7
Q7VDR1 Q7VDR5 Q7VDS0 Q7VDS2 Q7VDS4 Q7VDS8 Q7VDS9 Q7VDT0 Q7VDT7 Q7VDU7 Q7VDU8
Q7VDV0 Q7VDV3 Q7VDV6 Q7VDW9 Q7VDY5 Q7VDY6 Q7VDY7 Q7VDY8 Q7VDY9 Q7VDZ4 Q7VE01
Q7VE03 Q7VE08 Q7VE16 Q7VE18 Q7VE24 Q7VE26 Q7VE34 Q7VE61 Q7VE69 Q7VE81 Q7VE85
Q7VE92 Q7VEB3 Q7VED2 Q7VEF9 Q7VEG0 Q7VEG1 Q7VEG3 Q7VEG4 Q7VEG5 Q7VEH5 Q7VEH6
Q7VEI7 Q7VEI9 Q7VEJ1 Q7VEJ2 Q7VEJ3 Q7VEJ4 Q7VEJ6 Q7VEJ7 Q7VEK0 Q7VEK9 Q7VEN0
Q7VEN4 Q7VEN5 Q7VEP7 Q7VET6 Q7VET8 Q7VEU0 Q7VEV3 Q7VEV4 Q7VEW8 Q7VEZ0 Q7VF06
Q7VF14 Q7VF27 Q7VF28 Q7VF36 Q7VF43 Q7VF52 Q7VF68 Q7VF72 Q7VF74 Q7VF76 Q7VF77
Q7VF98 Q7VF99 Q7VFA0 Q7VFA6 Q7VFC6 Q7VFD3 Q7VFD7 Q7VFE5 Q7VFE6 Q7VFE9 Q7VFF0
Q7VFF1 Q7VFF5 Q7VFF6 Q7VFF8 Q7VFG4 Q7VFH6 Q7VFL0 Q7VFL1 Q7VFP9 Q7VFQ4 Q7VFQ6
Q7VFS1 Q7VFU3 Q7VFU4 Q7VFV4 Q7VFW0 Q7VFX7 Q7VFY5 Q7VFY6 Q7VFY9 Q7VFZ4 Q7VG06
Q7VG29 Q7VG30 Q7VG34 Q7VG37 Q7VG38 Q7VG78 Q7VGA7 Q7VGA8 Q7VGB3 Q7VGB8 Q7VGB9
Q7VGC0 Q7VGC1 Q7VGC2 Q7VGC3 Q7VGC7 Q7VGC8 Q7VGC9 Q7VGD0 Q7VGD2 Q7VGD3 Q7VGD4
Q7VGD5 Q7VGD6 Q7VGD7 Q7VGD8 Q7VGD9 Q7VGE0 Q7VGE1 Q7VGE2 Q7VGE3 Q7VGE4 Q7VGE5
Q7VGF0 Q7VGF2 Q7VGF6 Q7VGF9 Q7VGG0 Q7VGH1 Q7VGH7 Q7VGI1 Q7VGJ2 Q7VGJ6 Q7VGK6
Q7VGL0 Q7VGL3 Q7VGM5 Q7VGN1 Q7VGP1 Q7VGP6 Q7VGR5 Q7VGS1 Q7VGS3 Q7VGT3 Q7VGT8
Q7VGU1 Q7VGU6 Q7VGU9 Q7VGW6 Q7VGX8 Q7VGY5 Q7VGY9 Q7VGZ1 Q7VGZ3 Q7VGZ8 Q7VH30
Q7VH31 Q7VH32 Q7VH33 Q7VH56 Q7VH57 Q7VH74 Q7VH95 Q7VH96 Q7VHA5 Q7VHF4 Q7VHF6
Q7VHF9 Q7VHH1 Q7VHJ0 Q7VHJ1 Q7VHK3 Q7VHL1 Q7VHL2 Q7VHM1 Q7VHM7 Q7VHN0 Q7VHN2
Q7VHN3 Q7VHN8 Q7VHS5 Q7VHT1 Q7VHV1 Q7VHV4 Q7VHW3 Q7VHX0 Q7VHX2 Q7VHX9 Q7VHY0
Q7VHY1 Q7VHY5 Q7VHY7 Q7VI02 Q7VI03 Q7VI04 Q7VI05 Q7VI13 Q7VI24 Q7VI31 Q7VI68
Q7VI69 Q7VI99 Q7VIA1 Q7VIA7 Q7VIA8 Q7VIA9 Q7VIC7 Q7VIC9 Q7VIE2 Q7VIE3 Q7VIE6
Q7VIH2 Q7VIH7 Q7VIJ1 Q7VIJ3 Q7VIJ7 Q7VIL0 Q7VIM2 Q7VIM8 Q7VIN2 Q7VIN5 Q7VIN7
Q7VIN8 Q7VIP6 Q7VIR3 Q7VIR6 Q7VIR8 Q7VIS3 Q7VIT0 Q7VIT3 Q7VIT7 Q7VIU0 Q7VIU7
Q7VIU8 Q7VIV5 Q7VIV6 Q7VIW7 Q7VIW9 Q7VIY7 Q7VJ02 Q7VJ06 Q7VJ07 Q7VJ08 Q7VJ09
Q7VJ10 Q7VJ12 Q7VJ31 Q7VJ35 Q7VJ36 Q7VJ74 Q7VJ75 Q7VJ78 Q7VJ79 Q7VJ80 Q7VJ81
Q7VJ82 Q7VJ83 Q7VJ84 Q7VJ85 Q7VJ91 Q7VJ92 Q7VJA8 Q7VJB1 Q7VJB2 Q7VJB6 Q7VJB8
Q7VJB9 Q7VJD2 Q7VJD3 Q7VJN5 Q7VJQ0 Q7VJR4 Q7VJT9 Q7VJU0 Q7VJU3 Q7VJU4 Q7VJV4
Q7VJV5 Q7VJW2 Q7VJW4 Q7VJW9 Q7VJX1 Q7VJX6 Q7VJX7 Q7VJY2 Q7VJY3 Q7VJY4 Q7VK40
Q7VK48 Q7VK54 Q7VK55 Q7VK57 Q7VK64 Q7VK65 Q7VK73 Q7VK79 Q7VK82 Q7VK83 Q7VK84

Q7VK98 Q7VKA1 Q7VKA3 Q7VKA8 Q7VKB0 Q7VKB1 Q7VKB3 Q7VKB4 Q7VKB6 Q7VKC2 Q7VKC4
Q7VKC6 Q7VKC9 Q7VKD1 Q7VKD2 Q7VKD3 Q7VKD4 Q7VKD5 Q7VKD6 Q7VKD7 Q7VKD8 Q7VKD9
Q7VKE0 Q7VKE3 Q7VKE4 Q7VKE5 Q7VKE6 Q7VKE7 Q7VKE8 Q7VKE9 Q7VKF0 Q7VKF1 Q7VKF2
Q7VKF4 Q7VKF5 Q7VKF6 Q7VKF7 Q7VKF8 Q7VKF9 Q7VKG0 Q7VKG1 Q7VKG2 Q7VKG5 Q7VKG6
Q7VKG8 Q7VKH4 Q7VKH9 Q7VKI0 Q7VKI2 Q7VKI5 Q7VKJ0 Q7VKJ3 Q7VKJ6 Q7VKJ9 Q7VKK0
Q7VKK2 Q7VKK4 Q7VKK9 Q7VKL4 Q7VKL5 Q7VKL7 Q7VKL8 Q7VKM9 Q7VKN0 Q7VKN1 Q7VKP3
Q7VKP5 Q7VKQ4 Q7VKR0 Q7VKR5 Q7VKR6 Q7VKR7 Q7VKR8 Q7VKR9 Q7VKS0 Q7VKS1 Q7VKS2
Q7VKS6 Q7VKS7 Q7VKU5 Q7VKU9 Q7VKV0 Q7VKV3 Q7VKV4 Q7VKV5 Q7VKW2 Q7VKX3 Q7VKY7
Q7VKZ5 Q7VKZ7 Q7VL00 Q7VL09 Q7VL11 Q7VL12 Q7VL13 Q7VL22 Q7VL26 Q7VL27 Q7VL28
Q7VL44 Q7VL50 Q7VL54 Q7VL55 Q7VL75 Q7VL77 Q7VL79 Q7VL80 Q7VL82 Q7VL83 Q7VL86
Q7VL87 Q7VL91 Q7VL94 Q7VL95 Q7VL96 Q7VLE3 Q7VLE9 Q7VLF2 Q7VLF6 Q7VLF7 Q7VLG2
Q7VLG3 Q7VLG4 Q7VLG6 Q7VLH2 Q7VLH3 Q7VLI0 Q7VLI2 Q7VLI3 Q7VLI5 Q7VLK0 Q7VLK1
Q7VLK4 Q7VLL3 Q7VLL7 Q7VLM1 Q7VLM3 Q7VLM6 Q7VLM7 Q7VLN1 Q7VLN2 Q7VLN4 Q7VLN5
Q7VLN6 Q7VLN8 Q7VLN9 Q7VLP0 Q7VLP5 Q7VLP8 Q7VLP9 Q7VLQ4 Q7VLR0 Q7VLR3 Q7VLR5
Q7VLR8 Q7VLS0 Q7VLS2 Q7VLS3 Q7VLT3 Q7VLT5 Q7VLU0 Q7VLU5 Q7VLW2 Q7VLW6 Q7VLX2
Q7VLX4 Q7VLX6 Q7VLX8 Q7VLX9 Q7VLY1 Q7VLY6 Q7VLZ2 Q7VLZ5 Q7VM15 Q7VM19 Q7VM27
Q7VM29 Q7VM31 Q7VM33 Q7VM34 Q7VM42 Q7VM44 Q7VM59 Q7VM62 Q7VM64 Q7VM65 Q7VM66
Q7VM69 Q7VM70 Q7VM84 Q7VM86 Q7VM87 Q7VM90 Q7VMA0 Q7VMA4 Q7VMA6 Q7VMA9 Q7VMB7
Q7VMC0 Q7VMC2 Q7VMC4 Q7VMC9 Q7VMD0 Q7VMD5 Q7VMD6 Q7VMD7 Q7VME1 Q7VME2 Q7VME5
Q7VMF1 Q7VMF2 Q7VMG8 Q7VMH0 Q7VMH2 Q7VMH6 Q7VMI1 Q7VMI2 Q7VMJ1 Q7VMJ6 Q7VMS5
Q7VMS6 Q7VMS8 Q7VMS9 Q7VMU2 Q7VMV0 Q7VMV9 Q7VMW1 Q7VMW3 Q7VMW7 Q7VMX4 Q7VMX7
Q7VMY0 Q7VMY1 Q7VMY2 Q7VMZ9 Q7VN15 Q7VN18 Q7VN20 Q7VN27 Q7VN28 Q7VN33 Q7VN34
Q7VN41 Q7VN43 Q7VN44 Q7VN48 Q7VN49 Q7VN51 Q7VN52 Q7VN53 Q7VN67 Q7VN74 Q7VN81
Q7VN90 Q7VN93 Q7VN98 Q7VN99 Q7VNA2 Q7VNB3 Q7VNB5 Q7VNC2 Q7VNC4 Q7VNC9 Q7VND8
Q7VNF0 Q7VNF3 Q7VNF5 Q7VNG0 Q7VNG1 Q7VNG7 Q7VNH1 Q7VNH6 Q7VNH9 Q7VNI4 Q7VNI7
Q7VNI8 Q7VNM5 Q7VNN7 Q7VNN8 Q7VNP1 Q7VNP2 Q7VNP7 Q7VNQ2 Q7VNR4 Q7VNR5 Q7VNR9
Q7VNS2 Q7VNS3 Q7VNS6 Q7VNS7 Q7VNU9 Q7VNV0 Q7VNV1 Q7VNV2 Q7VNV5 Q7VNW5 Q7VNW8
Q7VNX4 Q7VNY0 Q7VNY1 Q7VNY2 Q7VNZ0 Q7VNZ1 Q7VNZ3 Q7VNZ4 Q7VP02 Q7VP04 Q7VP10
Q7VP34 Q7VP36 Q7VP37 Q7VP38 Q7VP41 Q7VP42 Q7VP51 Q7VP56 Q7VP59 Q7VP62 Q7VP66
Q7VP67 Q7VP68 Q7VP74 Q7VP78 Q7VP79 Q7VP83 Q7VP85 Q7VP87 Q7VP91 Q7VP92 Q7VP94
Q7VP95 Q7VPB1 Q7VPB2 Q7VPB7 Q7VPC0 Q7VPC3 Q7VPJ2 Q7VPK3 Q7VPK4 Q7VPK9 Q7VPM9
Q7VPN0 Q7VPN5 Q7VPN7 Q7VPN8 Q7VPP8 Q7VPP9 Q7VQB1 Q7VQB2 Q7VQB6 Q7VQB7 Q7VQB8
Q7VQB9 Q7VQC0 Q7VQC1 Q7VQC2 Q7VQC3 Q7VQC4 Q7VQC5 Q7VQC6 Q7VQC7 Q7VQC9 Q7VQD0
Q7VQD1 Q7VQD2 Q7VQD4 Q7VQD5 Q7VQD6 Q7VQD7 Q7VQD8 Q7VQD9 Q7VQE1 Q7VQE2 Q7VQE3
Q7VQE4 Q7VQE5 Q7VQE6 Q7VQE7 Q7VQE8 Q7VQE9 Q7VQF2 Q7VQF3 Q7VQF8 Q7VQF9 Q7VQG0
Q7VQG2 Q7VQG3 Q7VQG9 Q7VQH1 Q7VQH2 Q7VQH4 Q7VQH5 Q7VQI7 Q7VQI9 Q7VQJ2 Q7VQJ5
Q7VQJ6 Q7VQJ7 Q7VQJ8 Q7VQJ9 Q7VQK3 Q7VQK8 Q7VQL0 Q7VQL2 Q7VQL3 Q7VQL4 Q7VQL6
Q7VQM0 Q7VQM1 Q7VQM2 Q7VQM3 Q7VQM5 Q7VQM8 Q7VQN1 Q7VQN2 Q7VQN7 Q7VQN8 Q7VQN9
Q7VQP0 Q7VQP1 Q7VQP7 Q7VQQ0 Q7VQQ4 Q7VQQ6 Q7VQQ9 Q7VQR0 Q7VQR6 Q7VQS0 Q7VQT0
Q7VQT2 Q7VQT3 Q7VQV0 Q7VQV1 Q7VQV3 Q7VQW3 Q7VQW4 Q7VQW5 Q7VQW6 Q7VQW7 Q7VQW8
Q7VQW9 Q7VQX0 Q7VQX1 Q7VQX3 Q7VQX7 Q7VQX8 Q7VQY9 Q7VQZ1 Q7VQZ6 Q7VQZ9 Q7VR00
Q7VR05 Q7VR09 Q7VR10 Q7VR16 Q7VR23 Q7VR35 Q7VR39 Q7VR40 Q7VR41 Q7VR42 Q7VR46
Q7VR50 Q7VR52 Q7VR53 Q7VR54 Q7VR55 Q7VR65 Q7VR66 Q7VR67 Q7VR68 Q7VR69 Q7VR70
Q7VR74 Q7VR76 Q7VR80 Q7VR81 Q7VR99 Q7VRA3 Q7VRA6 Q7VRB3 Q7VRB4 Q7VRB6 Q7VRB8

Q7VRC5 Q7VRC8 Q7VRC9 Q7VRD2 Q7VRE2 Q7VRE3 Q7VRE4 Q7VRE5 Q7VRE6 Q7VRE8 Q7VRF3
Q7VRF5 Q7VRF7 Q7VRG1 Q7VRG3 Q7VRG4 Q7VRG5 Q7VRG6 Q7VRG7 Q7VRG9 Q7VRH0 Q7VRH1
Q7VRH9 Q7VRI1 Q7VRI2 Q7VRI4 Q7VRI8 Q7VRJ1 Q7VRJ7 Q7VRK0 Q7VRK1 Q7VRK2 Q7VRK9
Q7VRL0 Q7VRL2 Q7VRL8 Q7VRM0 Q7VRM4 Q7VRM5 Q7VRN1 Q7VRN2 Q7VRN3 Q7VRN5 Q7VRN7
Q7VRN8 Q7VRN9 Q7VRP0 Q7VRP3 Q7VRP4 Q7VRP5 Q7VRP6 Q7VRP7 Q7VRP8 Q7VRQ5 Q7VRQ6
Q7VRQ7 Q7VRR0 Q7VRR1 Q7VRR4 Q7VRR8 Q7VRS0 Q7VRS2 Q7VRS3 Q7VRS4 Q7VRS5 Q7VRS6
Q7VRS7 Q7VRS8 Q7VRS9 Q7VRT1 Q7VRT2 Q7VRT4 Q7VRU2 Q7VRU3 Q7VRU6 Q7VRU9 Q7VRX1
Q7VRX2 Q7VRX3 Q7VRX4 Q7VRX7 Q7VRX8 Q7VRY3 Q7VRZ3 Q7VS43 Q7VS46 Q7VS51 Q7VS76
Q7VS88 Q7VS89 Q7VS95 Q7VSA0 Q7VSB4 Q7VSD8 Q7VSD9 Q7VSE0 Q7VSF4 Q7VSG7 Q7VSH3
Q7VSM2 Q7VSN1 Q7VSN2 Q7VSN3 Q7VSN4 Q7VSR5 Q7VST8 Q7VSU7 Q7VSV1 Q7VSV5 Q7VSW3
Q7VSY4 Q7VSY5 Q7VSY6 Q7VSY7 Q7VSY8 Q7VSY9 Q7VSZ0 Q7VSZ1 Q7VSZ2 Q7VSZ3 Q7VT61
Q7VT71 Q7VT92 Q7VT93 Q7VT94 Q7VT96 Q7VTA1 Q7VTA6 Q7VTA7 Q7VTA8 Q7VTA9 Q7VTB0
Q7VTB2 Q7VTB3 Q7VTB4 Q7VTB5 Q7VTB6 Q7VTB7 Q7VTB8 Q7VTB9 Q7VTC0 Q7VTC1 Q7VTC4
Q7VTC5 Q7VTC6 Q7VTC7 Q7VTC8 Q7VTC9 Q7VTD0 Q7VTD1 Q7VTD2 Q7VTD3 Q7VTD4 Q7VTD5
Q7VTD6 Q7VTD7 Q7VTE5 Q7VTF0 Q7VTF1 Q7VTF7 Q7VTJ7 Q7VTJ8 Q7VTJ9 Q7VTK4 Q7VTM9
Q7VTN3 Q7VTN4 Q7VTP4 Q7VTR8 Q7VTT6 Q7VTT9 Q7VTU0 Q7VTZ7 Q7VTZ8 Q7VU11 Q7VU17
Q7VU32 Q7VU41 Q7VU56 Q7VU61 Q7VU66 Q7VU67 Q7VU72 Q7VU91 Q7VU93 Q7VU94 Q7VUD0
Q7VUD2 Q7VUD3 Q7VUD4 Q7VUD5 Q7VUE3 Q7VUF4 Q7VUG2 Q7VUG7 Q7VUH0 Q7VUH1 Q7VUH2
Q7VUH3 Q7VUJ9 Q7VUK0 Q7VUL5 Q7VUL8 Q7VUM8 Q7VUN6 Q7VUP5 Q7VUP6 Q7VUQ1 Q7VUQ4
Q7VUQ5 Q7VUS1 Q7VUS7 Q7VUU5 Q7VUV5 Q7VUV8 Q7VUV9 Q7VUW0 Q7VUW2 Q7VUW3 Q7VUW5
Q7VUW7 Q7VUW8 Q7VUZ0 Q7VUZ9 Q7VV53 Q7VV85 Q7VV87 Q7VV88 Q7VV89 Q7VV91 Q7VV92
Q7VVA6 Q7VVB1 Q7VVB6 Q7VVC1 Q7VVF1 Q7VVF8 Q7VVG6 Q7VVH7 Q7VVH8 Q7VVJ8 Q7VVM4
Q7VVR1 Q7VVR2 Q7VVR3 Q7VVR4 Q7VVR5 Q7VVR6 Q7VVS6 Q7VVU3 Q7VVU5 Q7VVU9 Q7VVW3
Q7VVX6 Q7VVX7 Q7VVX8 Q7VVY0 Q7VVY2 Q7VVY3 Q7VW08 Q7VW21 Q7VW25 Q7VW27 Q7VW39
Q7VW48 Q7VW76 Q7VW97 Q7VWC1 Q7VWC4 Q7VWE3 Q7VWE4 Q7VWE6 Q7VWE7 Q7VWE9 Q7VWH8
Q7VWK6 Q7VWK7 Q7VWK8 Q7VWL0 Q7VWL1 Q7VWL4 Q7VWL5 Q7VWL6 Q7VWM0 Q7VWM1 Q7VWM3
Q7VWP1 Q7VWV5 Q7VWV7 Q7VWV8 Q7VWW0 Q7VWY1 Q7VWY2 Q7VWY7 Q7VWZ0 Q7VWZ5 Q7VX16
Q7VX46 Q7VX83 Q7VX92 Q7VX96 Q7VXB5 Q7VXD2 Q7VXD7 Q7VXD8 Q7VXE1 Q7VXF8 Q7VXG7
Q7VXG8 Q7VXH4 Q7VXI6 Q7VXI7 Q7VXI8 Q7VXJ5 Q7VXJ6 Q7VXK3 Q7VXL5 Q7VXM6 Q7VXN2
Q7VXN4 Q7VXX8 Q7VXY3 Q7VXZ3 Q7VXZ4 Q7VXZ8 Q7VY01 Q7VY12 Q7VY13 Q7VY42 Q7VY62
Q7VY63 Q7VY64 Q7VY68 Q7VY73 Q7VY74 Q7VY75 Q7VY88 Q7VY92 Q7VY99 Q7VYA8 Q7VYB4
Q7VYB6 Q7VYC4 Q7VYC6 Q7VYC7 Q7VYC8 Q7VYC9 Q7VYD0 Q7VYD2 Q7VYI1 Q7VYK0 Q7VYK4
Q7VYQ1 Q7VYR1 Q7VYR2 Q7VYR4 Q7VYS2 Q7VYS4 Q7VYS5 Q7VYS6 Q7VYS7 Q7VYV4 Q7VYV6
Q7VZ05 Q7VZ35 Q7VZ37 Q7VZ47 Q7VZ56 Q7VZ59 Q7VZ77 Q7VZ79 Q7VZ81 Q7VZ86 Q7VZ88
Q7VZA1 Q7VZA3 Q7VZB8 Q7VZC8 Q7VZD6 Q7VZE0 Q7VZG1 Q7VZG4 Q7VZG7 Q7VZM5 Q7VZM9
Q7VZN1 Q7VZN2 Q7VZT5 Q7VZU0 Q7VZU1 Q7VZU4 Q7VZX4 Q7VZX5 Q7VZX6 Q7VZX7 Q7W001
Q7W006 Q7W014 Q7W022 Q7W023 Q7W041 Q7W044 Q7W045 Q7W046 Q7W049 Q7W050 Q7W066
Q7W069 Q7W089 Q7W0A2 Q7W0A6 Q7W0A7 Q7W0D0 Q7W0D1 Q7W0D3 Q7W0D7 Q7W0E3 Q7W0E4
Q7W0E5 Q7W0F2 Q7W0F7 Q7W0F8 Q7W0G3 Q7W0H1 Q7W0K0 Q7W0K8 Q7W0K9 Q7W0L0 Q7W0M8
Q7W0P6 Q7W0Q0 Q7W0Q6 Q7W0R8 Q7W0R9 Q7W0S0 Q7W0S1 Q7W0S2 Q7W0S3 Q7W0S9 Q7W0T0
Q7W0T4 Q7W134 Q7W138 Q7W142 Q7W143 Q7W154 Q7W179 Q7W180 Q7W181 Q7W182 Q7W185
Q7W189 Q7W197 Q7W1A9 Q7W1C4 Q7W1C6 Q7W1D3 Q7W1D8 Q7W1D9 Q7W1E4 Q7W1I6 Q7W1P0
Q7W1P1 Q7W1P2 Q7W1P3 Q7W1Q0 Q7W1Q6 Q7W1Q9 Q7W1R5 Q7W1T9 Q7W1U6 Q7W1V2 Q7W1V3
Q7W1Y7 Q7W1Z7 Q7W200 Q7W215 Q7W216 Q7W222 Q7W223 Q7W224 Q7W275 Q7W294 Q7W2B5

Q7W2B6 Q7W2B7 Q7W2B9 Q7W2C4 Q7W2D0 Q7W2D1 Q7W2D2 Q7W2D3 Q7W2D4 Q7W2D6 Q7W2D7
Q7W2D8 Q7W2D9 Q7W2E0 Q7W2E1 Q7W2E2 Q7W2E3 Q7W2E4 Q7W2E5 Q7W2E7 Q7W2E8 Q7W2E9
Q7W2F0 Q7W2F1 Q7W2F2 Q7W2F3 Q7W2F4 Q7W2F5 Q7W2F6 Q7W2F7 Q7W2F8 Q7W2F9 Q7W2G0
Q7W2G8 Q7W2G9 Q7W2H0 Q7W2H1 Q7W2H2 Q7W2H3 Q7W2I0 Q7W2I1 Q7W2J0 Q7W2K3 Q7W2K4
Q7W2K5 Q7W2L1 Q7W2M1 Q7W2N4 Q7W2N5 Q7W2N9 Q7W2P4 Q7W2Q2 Q7W2S3 Q7W2T7 Q7W2X7
Q7W2X8 Q7W2X9 Q7W2Y0 Q7W2Y1 Q7W2Y2 Q7W2Y3 Q7W2Y4 Q7W2Y5 Q7W2Y6 Q7W334 Q7W387
Q7W388 Q7W393 Q7W398 Q7W3B3 Q7W3E7 Q7W3F2 Q7W3F8 Q7W3H5 Q7W3H6 Q7W3H7 Q7W3H8
Q7W3I6 Q7W3J1 Q7W3J5 Q7W3K5 Q7W3L2 Q7W3M5 Q7W3N6 Q7W3P5 Q7W3Q2 Q7W3R1 Q7W3R4
Q7W3R8 Q7W3S6 Q7W3U1 Q7W3U2 Q7W3U5 Q7W3U6 Q7W3U7 Q7W3V0 Q7W3V3 Q7W3V9 Q7W3W2
Q7W3X9 Q7W3Y8 Q7W3Z1 Q7W3Z2 Q7W3Z3 Q7W3Z5 Q7W3Z6 Q7W3Z8 Q7W400 Q7W401 Q7W417
Q7W418 Q7W419 Q7W455 Q7W481 Q7W482 Q7W488 Q7W497 Q7W4A6 Q7W4A7 Q7W4B2 Q7W4B5
Q7W4B6 Q7W4D9 Q7W4J0 Q7W4J6 Q7W4K0 Q7W4N4 Q7W4N5 Q7W4N9 Q7W4Q1 Q7W4S7 Q7W4S9
Q7W4T0 Q7W4T6 Q7W4T9 Q7W4U0 Q7W4W7 Q7W4W8 Q7W4Z7 Q7W509 Q7W510 Q7W513 Q7W514
Q7W515 Q7W517 Q7W519 Q7W520 Q7W545 Q7W561 Q7W566 Q7W569 Q7W570 Q7W575 Q7W5C9
Q7W5D0 Q7W5G3 Q7W5G8 Q7W5G9 Q7W5I0 Q7W5I2 Q7W5J6 Q7W5K6 Q7W5N6 Q7W5Q8 Q7W5T6
Q7W5V2 Q7W5Z6 Q7W5Z9 Q7W602 Q7W603 Q7W605 Q7W641 Q7W668 Q7W670 Q7W671 Q7W676
Q7W690 Q7W6B1 Q7W6B2 Q7W6C1 Q7W6C6 Q7W6K7 Q7W6L4 Q7W6L5 Q7W6N4 Q7W6N6 Q7W6N7
Q7W6P3 Q7W6P4 Q7W6P5 Q7W6P6 Q7W6P7 Q7W6Q0 Q7W6Q1 Q7W6Q2 Q7W6Q6 Q7W6Q7 Q7W6Q9
Q7W749 Q7W785 Q7W789 Q7W791 Q7W7A7 Q7W7C5 Q7W7C6 Q7W7C7 Q7W7C8 Q7W7C9 Q7W7E8
Q7W7F4 Q7W7F8 Q7W7G0 Q7W7G3 Q7W7H6 Q7W7H7 Q7W7H8 Q7W7I7 Q7W7N2 Q7W7P5 Q7W7P6
Q7W7P8 Q7W7P9 Q7W7Q0 Q7W7Q2 Q7W7R1 Q7W7T8 Q7W7U0 Q7W7U1 Q7W7U3 Q7W7U4 Q7W7W9
Q7W850 Q7W859 Q7W862 Q7W880 Q7W8H4 Q7W8R3 Q7W8R6 Q7W8T4 Q7W8T6 Q7W8U9 Q7W8V0
Q7W8V6 Q7W8X1 Q7W8X2 Q7W8X3 Q7W8Y2 Q7W8Y3 Q7W8Z2 Q7W8Z8 Q7W910 Q7W911 Q7W912
Q7W923 Q7W929 Q7W930 Q7W931 Q7W950 Q7W954 Q7W972 Q7W977 Q7W978 Q7W979 Q7W980
Q7W9A4 Q7W9A5 Q7W9A7 Q7W9B4 Q7W9B6 Q7W9B7 Q7W9B8 Q7W9E3 Q7W9E6 Q7W9I4 Q7W9J1
Q7W9J4 Q7W9J5 Q7W9J7 Q7W9L9 Q7W9M0 Q7W9M4 Q7W9M7 Q7W9N2 Q7W9Q2 Q7W9Q5 Q7W9T3
Q7W9T4 Q7W9T9 Q7W9W1 Q7WA31 Q7WA41 Q7WA44 Q7WA46 Q7WA54 Q7WA56 Q7WA57 Q7WA58
Q7WA59 Q7WA60 Q7WA62 Q7WAB3 Q7WAC0 Q7WAC9 Q7WAE4 Q7WAK8 Q7WAV6 Q7WAY2 Q7WAY7
Q7WAY8 Q7WB15 Q7WB25 Q7WB43 Q7WB57 Q7WB58 Q7WB60 Q7WB66 Q7WB68 Q7WB85 Q7WB92
Q7WBA1 Q7WBC1 Q7WBD0 Q7WBF2 Q7WBG5 Q7WC23 Q7WC32 Q7WC84 Q7WC96 Q7WC98 Q7WCA6
Q7WCA8 Q7WCA9 Q7WCE3 Q7WCE4 Q7WCJ7 Q7WCJ8 Q7WCM7 Q7WCP1 Q7WCP6 Q7WCP9 Q7WCQ0
Q7WCQ5 Q7WCW3 Q7WCW4 Q7WCZ8 Q7WD04 Q7WD05 Q7WD16 Q7WD18 Q7WD32 Q7WD42 Q7WD72
Q7WD94 Q7WDI4 Q7WDJ7 Q7WDJ8 Q7WDJ9 Q7WDK7 Q7WDL7 Q7WDN0 Q7WDN1 Q7WDN7 Q7WDP2
Q7WDQ0 Q7WDS1 Q7WDT5 Q7WDX7 Q7WDX8 Q7WDX9 Q7WDY0 Q7WDY1 Q7WDY2 Q7WDY3 Q7WDY4
Q7WDY5 Q7WDY6 Q7WE36 Q7WE57 Q7WE96 Q7WE97 Q7WEA9 Q7WEK6 Q7WEK7 Q7WEL1 Q7WEL6
Q7WEN2 Q7WER7 Q7WES2 Q7WES8 Q7WEU9 Q7WEV0 Q7WEV1 Q7WEV2 Q7WEW1 Q7WEW6 Q7WEX0
Q7WEY0 Q7WEY7 Q7WF01 Q7WF12 Q7WF21 Q7WF29 Q7WF38 Q7WF42 Q7WF46 Q7WF54 Q7WF71
Q7WF72 Q7WF75 Q7WF76 Q7WF77 Q7WF80 Q7WF83 Q7WF89 Q7WF92 Q7WFA9 Q7WFC0 Q7WFC3
Q7WFC4 Q7WFC5 Q7WFC7 Q7WFC8 Q7WFD0 Q7WFD2 Q7WFD3 Q7WFF2 Q7WFF3 Q7WFL2 Q7WFN9
Q7WFP0 Q7WFP6 Q7WFQ5 Q7WFR4 Q7WFR5 Q7WFS0 Q7WFS3 Q7WFS4 Q7WFU7 Q7WG14 Q7WG20
Q7WG25 Q7WG60 Q7WG61 Q7WG65 Q7WG77 Q7WGA3 Q7WGA5 Q7WGA6 Q7WGB3 Q7WGB6 Q7WGB7
Q7WGE5 Q7WGE6 Q7WGH4 Q7WGH5 Q7WGH8 Q7WGH9 Q7WGI0 Q7WGI2 Q7WGI4 Q7WGI5 Q7WGI7
Q7WGK3 Q7WGT9 Q7WGU2 Q7WGW8 Q7WGX3 Q7WGX4 Q7WH03 Q7WH06 Q7WH25 Q7WH30 Q7WH33
Q7WH38 Q7WH39 Q7WH63 Q7WH65 Q7WH66 Q7WH69 Q7WH76 Q7WHB6 Q7WHB9 Q7WHE7 Q7WHE8

Q7WHE9 Q7WHF0 Q7WHF2 Q7WHG0 Q7WHG2 Q7WHG3 Q7WHI7 Q7WHI8 Q7WHI9 Q7WHJ0 Q7WHJ6
Q7WHL6 Q7WHL9 Q7WHM0 Q7WHM6 Q7WHM7 Q7WHM8 Q7WHN0 Q7WHN1 Q7WHN4 Q7WHN5 Q7WHN6
Q7WHP0 Q7WHP1 Q7WHP3 Q7WHW5 Q7WHW8 Q7WI07 Q7WI34 Q7WI36 Q7WI37 Q7WI42 Q7WI59
Q7WI79 Q7WI80 Q7WI90 Q7WI95 Q7WII1 Q7WII7 Q7WIJ0 Q7WIN3 Q7WJ55 Q7WJ63 Q7WJ73
Q7WJ78 Q7WJ80 Q7WJ88 Q7WJ90 Q7WJ91 Q7WJ92 Q7WJ93 Q7WJ94 Q7WJ96 Q7WJG1 Q7WJH0
Q7WJI7 Q7WJP7 Q7WJQ9 Q7WK13 Q7WK44 Q7WK46 Q7WK59 Q7WK60 Q7WK66 Q7WK82 Q7WK83
Q7WK84 Q7WKC6 Q7WKD5 Q7WKE3 Q7WKF5 Q7WKF6 Q7WKF7 Q7WKG8 Q7WKH4 Q7WKH5 Q7WKH6
Q7WKJ5 Q7WKJ9 Q7WKM1 Q7WKM5 Q7WKM7 Q7WKP4 Q7WKR3 Q7WKR4 Q7WKR5 Q7WKR6 Q7WKR7
Q7WKT7 Q7WKU2 Q7WKU6 Q7WKU8 Q7WKV2 Q7WKW5 Q7WKW6 Q7WKW7 Q7WKX5 Q7WL20 Q7WL32
Q7WL33 Q7WL35 Q7WL36 Q7WL37 Q7WL39 Q7WL48 Q7WL78 Q7WL80 Q7WL81 Q7WL83 Q7WL84
Q7WLA9 Q7WLE5 Q7WLJ8 Q7WLK7 Q7WLL0 Q7WLT5 Q7WM36 Q7WMD0 Q7WMD3 Q7WME6 Q7WMF1
Q7WMF2 Q7WMI1 Q7WMJ1 Q7WMK9 Q7WMM4 Q7WMM5 Q7WMM7 Q7WMN3 Q7WMN5 Q7WMQ3 Q7WMR0
Q7WMS0 Q7WMU0 Q7WMU9 Q7WMX3 Q7WMY6 Q7WNK2 Q7WNM3 Q7WNS4 Q7WNS8 Q7WNT2 Q7WNT3
Q7WNU4 Q7WNY2 Q7WNY3 Q7WNY4 Q7WNY5 Q7WNZ3 Q7WP01 Q7WP13 Q7WP29 Q7WP31 Q7WP40
Q7WP45 Q7WP46 Q7WP51 Q7WPH6 Q7WQ22 Q7WQ31 Q7WQ88 Q7WQA0 Q7WQA2 Q7WQB0 Q7WQB2
Q7WQB3 Q7WQE9 Q7WQF0 Q7WQL6 Q7WQL7 Q7WQL8 Q7WQL9 Q7WQM6 Q7WQN2 Q7WQN5 Q7WQP1
Q7WQR6 Q7WQS2 Q7WQS8 Q7WQS9 Q7WQW5 Q7WQX5 Q7WQX8 Q7WQZ3 Q7WQZ4 Q7WR00 Q7WR01
Q7WR02 Q7WR42 Q7WR61 Q7WR83 Q7WR84 Q7WR85 Q7WR87 Q7WR92 Q7WR98 Q7WR99 Q7WRA0
Q7WRA1 Q7WRA2 Q7WRA4 Q7WRA5 Q7WRA6 Q7WRA7 Q7WRA8 Q7WRA9 Q7WRB0 Q7WRB1 Q7WRB2
Q7WRB3 Q7WRB6 Q7WRB7 Q7WRB8 Q7WRB9 Q7WRC0 Q7WRC1 Q7WRC2 Q7WRC3 Q7WRC4 Q7WRC5
Q7WRC6 Q7WRC7 Q7WRC8 Q7WRC9 Q7WRD8 Q7WRD9 Q7WRE0 Q7WRE1 Q7WRE2 Q7WRE3 Q7WRF0
Q7WRF1 Q7WS85 Q7WSH2 Q7WSJ4 Q7WSY8 Q7WT72 Q7WTB0 Q7WTB1 Q7WTB3 Q7WTB4 Q7WTJ2
Q7WTJ6 Q7WTT9 Q7WTU1 Q7WUL3 Q7WUL4 Q7WUM3 Q7WUM8 Q7WVY0 Q7WY56 Q7WY57 Q7WY58
Q7WY59 Q7WY61 Q7WY62 Q7WY63 Q7WY65 Q7WY66 Q7WY67 Q7WY68 Q7WY69 Q7WY72 Q7WY73
Q7WY75 Q7WY76 Q7WY77 Q7WYG5 Q7WYN1 Q7WZ28 Q7WZ32 Q7WZE4 Q7WZE5 Q7WZY5 Q7X143
Q7X146 Q7X149 Q7X222 Q7X279 Q7X281 Q7X284 Q7X2C7 Q7X2H8 Q7X2N3 Q7X2N6 Q7X2P1
Q7X2S2 Q7X2S8 Q7X3G5 Q7X3T2 Q7X3X5 Q7X4S1 Q7X5C9 Q7X5H2 Q7X5K6 Q7X5K8 Q7X5K9
Q7X5L0 Q7X5L1 Q7X5L2 Q7X5L3 Q7X5L4 Q7X5L5 Q7X5L6 Q7X5L7 Q7X6A5 Q7X6Y7 Q7X742
Q7X745 Q7X7A4 Q7X7A9 Q7X7C9 Q7X7E9 Q7X7H4 Q7X7H9 Q7X7L3 Q7X7N2 Q7X7N3 Q7X8D4
Q7X8R5 Q7X8Z8 Q7X923 Q7X993 Q7X999 Q7X9A0 Q7X9A8 Q7X9B9 Q7X9H9 Q7X9I0 Q7X9V3
Q7XA39 Q7XA40 Q7XA42 Q7XA48 Q7XA65 Q7XA66 Q7XA67 Q7XA77 Q7XAK4 Q7XAM0 Q7XAP4
Q7XAQ6 Q7XAS7 Q7XB08 Q7XB10 Q7XB11 Q7XBA5 Q7XBQ9 Q7XBY6 Q7XC35 Q7XC54 Q7XC57
Q7XD82 Q7XD96 Q7XDB8 Q7XDC8 Q7XDH8 Q7XDP2 Q7XDY9 Q7XE48 Q7XEJ5 Q7XEY9 Q7XFU9
Q7XGM6 Q7XH05 Q7XH85 Q7XHC4 Q7XHI5 Q7XHI7 Q7XHI9 Q7XHL3 Q7XHN8 Q7XHW4 Q7XI08
Q7XI14 Q7XI40 Q7XI41 Q7XI73 Q7XI75 Q7XI85 Q7XI96 Q7XIM7 Q7XIZ1 Q7XJ02 Q7XJ53
Q7XJ60 Q7XJ81 Q7XJ96 Q7XJE5 Q7XJE6 Q7XJK5 Q7XJK6 Q7XJK8 Q7XJM8 Q7XJN0 Q7XJS0
Q7XJU1 Q7XJU2 Q7XK25 Q7XKC0 Q7XKC4 Q7XKC5 Q7XKD0 Q7XKF3 Q7XKI7 Q7XKK6 Q7XKQ8
Q7XKR9 Q7XKU5 Q7XL03 Q7XL73 Q7XLP4 Q7XM13 Q7XM16 Q7XM21 Q7XM31 Q7XMI0 Q7XMJ2
Q7XMK0 Q7XMK1 Q7XMK8 Q7XN11 Q7XN12 Q7XNS7 Q7XP01 Q7XP65 Q7XPE8 Q7XPJ0 Q7XPK1
Q7XPK7 Q7XPL2 Q7XPT0 Q7XPW5 Q7XPW7 Q7XPW8 Q7XPY5 Q7XQ45 Q7XQ83 Q7XQ85 Q7XQ97
Q7XQN1 Q7XQQ2 Q7XQR9 Q7XQU7 Q7XQZ6 Q7XR02 Q7XR06 Q7XR47 Q7XR60 Q7XR61 Q7XR62
Q7XR80 Q7XR83 Q7XR84 Q7XRA1 Q7XRF1 Q7XRV0 Q7XRX1 Q7XS74 Q7XS75 Q7XSA2 Q7XSB2
Q7XSJ6 Q7XSN8 Q7XSN9 Q7XSQ7 Q7XSS9 Q7XSV4 Q7XT07 Q7XT34 Q7XT97 Q7XT99 Q7XTC7
Q7XTG7 Q7XTJ3 Q7XTK3 Q7XTS3 Q7XTS4 Q7XTT4 Q7XTV7 Q7XTY9 Q7XU27 Q7XU84 Q7XUC5

Q7XUF4 Q7XUJ5 Q7XUN2 Q7XUP6 Q7XUP7 Q7XUR2 Q7XUS2 Q7XUW3 Q7XVF9 Q7XVM8 Q7XW27
Q7XWU3 Q7XXD4 Q7XXI9 Q7XXL2 Q7XXN4 Q7XXN5 Q7XXP3 Q7XYA7 Q7XYB0 Q7XYP4 Q7XYS8
Q7XYS9 Q7XYT0 Q7XYY2 Q7XYZ0 Q7XZF7 Q7XZQ6 Q7XZQ7 Q7XZQ8 Q7XZR1 Q7XZT8 Q7XZZ1
Q7XZZ3 Q7Y001 Q7Y021 Q7Y040 Q7Y052 Q7Y0E8 Q7Y0F2 Q7Y0F6 Q7Y0S0 Q7Y0Y6 Q7Y0Y8
Q7Y138 Q7Y180 Q7Y183 Q7Y1B6 Q7Y1C4 Q7Y1C5 Q7Y1C8 Q7Y1H8 Q7Y1I7 Q7Y1L9 Q7Y1W5
Q7Y1W9 Q7Y214 Q7Y219 Q7Y232 Q7Y2C9 Q7Y2D3 Q7Y2D5 Q7Y2D9 Q7Y4Y9 Q7Y5B1 Q7Y5D9
Q7Y5N4 Q7Y5P0 Q7Y618 Q7Y667 Q7Y677 Q7Y6G9 Q7Y7N3 Q7YEV0 Q7YJG1 Q7YJG2 Q7YJH1
Q7YJS7 Q7YJT5 Q7YJT7 Q7YJT8 Q7YJT9 Q7YJU0 Q7YJU1 Q7YJU2 Q7YJU3 Q7YJU4 Q7YJU5
Q7YJU6 Q7YJV2 Q7YJV3 Q7YJV4 Q7YJV5 Q7YJW4 Q7YJX0 Q7YJX4 Q7YJX8 Q7YJX9 Q7YJY0
Q7YJY1 Q7YJY6 Q7YJY7 Q7YJZ1 Q7YKL5 Q7YKM5 Q7YKQ2 Q7YKQ3 Q7YKQ5 Q7YKQ9 Q7YKW4
Q7YKX3 Q7YKY4 Q7YKY5 Q7YM14 Q7YM28 Q7YM30 Q7YM31 Q7YMF8 Q7YMG2 Q7YMV1 Q7YMV6
Q7YMV7 Q7YMX0 Q7YMX3 Q7YN57 Q7YN68 Q7YN74 Q7YN79 Q7YN81 Q7YN83 Q7YQJ3 Q7YQJ4
Q7YQJ9 Q7YQK8 Q7YQL3 Q7YQM1 Q7YQM2 Q7YR23 Q7YR24 Q7YR25 Q7YR26 Q7YR30 Q7YR31
Q7YR32 Q7YR33 Q7YR34 Q7YR35 Q7YR36 Q7YR37 Q7YR38 Q7YR39 Q7YR46 Q7YR48 Q7YR49
Q7YR76 Q7YRA3 Q7YRC1 Q7YRC3 Q7YRC6 Q7YRD0 Q7YRG5 Q7YRL2 Q7YRU4 Q7YRX0 Q7YS61
Q7YS80 Q7YS81 Q7YS82 Q7YS91 Q7YSW8 Q7YTB0 Q7YTC2 Q7YU81 Q7YWN4 Q7YWU9 Q7YZA2
Q7YZH1 Q7YZN9 Q7YZT0 Q7YZT6 Q7Z0G7 Q7Z0T8 Q7Z142 Q7Z144 Q7Z2B8 Q7Z2F6 Q7Z2T5
Q7Z2V1 Q7Z2X7 Q7Z2Y8 Q7Z2Z2 Q7Z340 Q7Z353 Q7Z392 Q7Z398 Q7Z3E2 Q7Z3H0 Q7Z3H4
Q7Z3I7 Q7Z3J3 Q7Z3V4 Q7Z3V5 Q7Z3Y7 Q7Z3Y8 Q7Z3Y9 Q7Z3Z0 Q7Z3Z3 Q7Z3Z4 Q7Z465
Q7Z4G1 Q7Z4G4 Q7Z4H3 Q7Z4H9 Q7Z4K8 Q7Z4L9 Q7Z4M0 Q7Z4Q2 Q7Z4T9 Q7Z4V0 Q7Z4W3
Q7Z572 Q7Z5D8 Q7Z5Q1 Q7Z5Q5 Q7Z5U6 Q7Z5W3 Q7Z5Y7 Q7Z624 Q7Z6I5 Q7Z6I8 Q7Z6J8
Q7Z6K1 Q7Z6K3 Q7Z6K5 Q7Z6M2 Q7Z6M4 Q7Z6R9 Q7Z6V5 Q7Z6W7 Q7Z745 Q7Z794 Q7Z7E8
Q7Z7J5 Q7Z7K0 Q7Z7K2 Q7Z7L1 Q7Z7L7 Q7Z7L9 Q7Z7S3 Q7Z7W5 Q7Z7W6 Q7Z891 Q7Z892
Q7Z893 Q7Z894 Q7Z895 Q7Z8E7 Q7Z8L4 Q7Z8M6 Q7Z8P9 Q7Z991 Q7Z995 Q7Z996 Q7Z9I2
Q7Z9I3 Q7Z9I4 Q7Z9J4 Q7Z9Z2 Q7ZA43 Q7ZA45 Q7ZA46 Q7ZAG7 Q7ZAI9 Q7ZAJ0 Q7ZAJ2
Q7ZAJ3 Q7ZAJ4 Q7ZAJ5 Q7ZAJ8 Q7ZAK0 Q7ZAK1 Q7ZAK3 Q7ZAK7 Q7ZAK8 Q7ZAK9 Q7ZAL1
Q7ZAL2 Q7ZAL5 Q7ZAL6 Q7ZAL9 Q7ZAM0 Q7ZAM1 Q7ZAM3 Q7ZAM4 Q7ZAM5 Q7ZAM7 Q7ZAM8
Q7ZAN1 Q7ZAN4 Q7ZAN6 Q7ZAN7 Q7ZAP1 Q7ZAP3 Q7ZSX3 Q7ZSY6 Q7ZT01 Q7ZT11 Q7ZT42
Q7ZT46 Q7ZT47 Q7ZT48 Q7ZT82 Q7ZTI3 Q7ZTI5 Q7ZTK4 Q7ZTM9 Q7ZTN8 Q7ZTP2 Q7ZTQ5
Q7ZTS4 Q7ZTU9 Q7ZTY4 Q7ZTY6 Q7ZTZ2 Q7ZU45 Q7ZU90 Q7ZU92 Q7ZUB2 Q7ZUF2 Q7ZUG0
Q7ZUH1 Q7ZUH5 Q7ZUK0 Q7ZUK7 Q7ZUP0 Q7ZUP1 Q7ZUV2 Q7ZUV7 Q7ZUX1 Q7ZUX6 Q7ZUZ0
Q7ZV20 Q7ZV22 Q7ZV26 Q7ZV34 Q7ZV35 Q7ZV55 Q7ZV79 Q7ZV84 Q7ZV90 Q7ZVA0 Q7ZVA6
Q7ZVB2 Q7ZVC2 Q7ZVC9 Q7ZVE0 Q7ZVF0 Q7ZVG6 Q7ZVJ8 Q7ZVK3 Q7ZVM9 Q7ZVN7 Q7ZVP1
Q7ZVQ8 Q7ZVR1 Q7ZVR8 Q7ZVS8 Q7ZVS9 Q7ZVT3 Q7ZVT5 Q7ZVU1 Q7ZVW9 Q7ZVX0 Q7ZVX6
Q7ZVY5 Q7ZVZ6 Q7ZW16 Q7ZW33 Q7ZW41 Q7ZW57 Q7ZW78 Q7ZW86 Q7ZWC3 Q7ZWD4 Q7ZWE3
Q7ZWE7 Q7ZWE9 Q7ZWF0 Q7ZWI4 Q7ZWL6 Q7ZWM3 Q7ZWM4 Q7ZWM8 Q7ZWN4 Q7ZWN5 Q7ZWN6
Q7ZWR2 Q7ZWS1 Q7ZWS2 Q7ZWU1 Q7ZWU9 Q7ZWV3 Q7ZWV9 Q7ZWX9 Q7ZWY2 Q7ZWZ4 Q7ZWZ5
Q7ZX03 Q7ZX06 Q7ZX20 Q7ZX23 Q7ZX31 Q7ZX36 Q7ZX41 Q7ZX42 Q7ZX59 Q7ZX60 Q7ZX64
Q7ZX83 Q7ZXB1 Q7ZXB5 Q7ZXB8 Q7ZXC9 Q7ZXD5 Q7ZXE2 Q7ZXE9 Q7ZXF3 Q7ZXG4 Q7ZXG7
Q7ZXH2 Q7ZXH3 Q7ZXH9 Q7ZXK9 Q7ZXP0 Q7ZXP1 Q7ZXP9 Q7ZXR3 Q7ZXS3 Q7ZXT3 Q7ZXV3
Q7ZXV6 Q7ZXV8 Q7ZXW9 Q7ZXX2 Q7ZXX9 Q7ZXY1 Q7ZXY4 Q7ZXZ0 Q7ZXZ2 Q7ZY08 Q7ZY11
Q7ZY13 Q7ZY18 Q7ZY29 Q7ZY31 Q7ZY35 Q7ZY47 Q7ZY56 Q7ZY60 Q7ZY69 Q7ZY73 Q7ZY78
Q7ZY87 Q7ZYA5 Q7ZYA6 Q7ZYB4 Q7ZYF1 Q7ZYF2 Q7ZYG4 Q7ZYG5 Q7ZYH1 Q7ZYI9 Q7ZYJ0
Q7ZYJ9 Q7ZYL5 Q7ZYM8 Q7ZYP0 Q7ZYP6 Q7ZYQ0 Q7ZYQ6 Q7ZYR8 Q7ZYS1 Q7ZYS8 Q7ZYU8

Q7ZYV9 Q7ZYZ7 Q7ZZ00 Q7ZZ03 Q7ZZC8 Q7ZZH7 Q7ZZP9 Q800D3 Q800L3 Q800L6 Q800Q5
Q801E0 Q801E1 Q801E2 Q801F8 Q801N6 Q801P0 Q801P1 Q801P7 Q801Q7 Q801R4 Q801R7
Q801R8 Q801S3 Q801S4 Q801X6 Q802A6 Q802G6 Q802R8 Q802R9 Q802W2 Q802W4 Q802Y8
Q803A0 Q803A6 Q803A7 Q803C1 Q803E1 Q803H0 Q803I4 Q803J8 Q803K4 Q803M0 Q803M3
Q803M5 Q803N9 Q803P1 Q803S3 Q803U7 Q803V3 Q803V5 Q803W1 Q803X1 Q803X4 Q803Y4
Q804A5 Q804A9 Q804H7 Q804Q5 Q804R0 Q804S6 Q804T6 Q804W2 Q805A1 Q805B4 Q805F9
Q805H0 Q805H2 Q805K3 Q805K5 Q805M1 Q805P6 Q805P9 Q805Q9 Q806Y6 Q80872 Q80903
Q80907 Q80910 Q80916 Q80920 Q80923 Q80930 Q80937 Q80941 Q80944 Q80948 Q80951
Q80955 Q80958 Q80961 Q809J9 Q809K1 Q80A33 Q80B88 Q80DP8 Q80DS7 Q80DT0 Q80DT4
Q80DV1 Q80DV6 Q80DX6 Q80E01 Q80E12 Q80H93 Q80HT6 Q80HT9 Q80HU0 Q80HU3 Q80HU9
Q80HV0 Q80HV2 Q80HV3 Q80HV7 Q80HW9 Q80HX0 Q80HX3 Q80HX4 Q80HX7 Q80IQ5 Q80IU4
Q80IU7 Q80KJ6 Q80LT4 Q80QL5 Q80ST9 Q80SU3 Q80SU7 Q80SW5 Q80SX8 Q80SY5 Q80SY6
Q80T18 Q80T74 Q80T85 Q80TA9 Q80TB7 Q80TC5 Q80TC6 Q80TE0 Q80TF4 Q80TG1 Q80TI1
Q80TL0 Q80TM6 Q80TP3 Q80TR8 Q80TS5 Q80TT8 Q80TY4 Q80U12 Q80U19 Q80U22 Q80U35
Q80U38 Q80U49 Q80U59 Q80U70 Q80U78 Q80U95 Q80UF7 Q80UG5 Q80UG8 Q80UJ9 Q80UK7
Q80UN9 Q80US4 Q80US8 Q80UU1 Q80UU2 Q80UW3 Q80UW5 Q80UW8 Q80UY1 Q80UZ2 Q80V23
Q80V24 Q80V31 Q80V62 Q80V63 Q80V85 Q80V86 Q80V91 Q80VC6 Q80VC9 Q80VF6 Q80VG1
Q80VH0 Q80VI1 Q80VJ3 Q80VJ4 Q80VJ6 Q80VK6 Q80VL1 Q80VM3 Q80VM4 Q80VM7 Q80VM8
Q80VN0 Q80VP2 Q80VP5 Q80VR3 Q80VV3 Q80VX4 Q80VY2 Q80VY9 Q80W00 Q80W03 Q80W14
Q80W21 Q80W22 Q80W31 Q80W32 Q80W49 Q80W69 Q80W90 Q80W93 Q80W96 Q80WB0 Q80WB5
Q80WC1 Q80WC7 Q80WE4 Q80WG7 Q80WL2 Q80WP8 Q80WQ4 Q80WQ8 Q80WQ9 Q80WR5 Q80WS1
Q80WS3 Q80WT2 Q80WV7 Q80WY3 Q80X32 Q80X44 Q80X50 Q80X53 Q80X56 Q80X60 Q80X76
Q80X91 Q80X95 Q80XA6 Q80XC2 Q80XC6 Q80XI1 Q80XI3 Q80XI6 Q80XJ2 Q80XJ3 Q80XS6
Q80XU3 Q80Y14 Q80Y20 Q80Y39 Q80Y44 Q80Y55 Q80Y73 Q80Y75 Q80Y81 Q80Y84 Q80YA7
Q80YD1 Q80YD6 Q80YE2 Q80YG3 Q80YN3 Q80YP0 Q80YP6 Q80YQ2 Q80YQ8 Q80YR2 Q80YR3
Q80YR4 Q80YR7 Q80YR9 Q80YS5 Q80YS9 Q80YT7 Q80YV4 Q80YY7 Q80Z32 Q80Z36 Q80Z37
Q80ZI1 Q80ZJ6 Q80ZJ7 Q80ZJ8 Q80ZK9 Q80ZM3 Q80ZM7 Q80ZQ3 Q80ZQ9 Q80ZU5 Q80ZV0
Q80ZW0 Q80ZX8 Q81018 Q81021 Q81022 Q810A1 Q810A3 Q810A5 Q810A7 Q810B3 Q810D6
Q810F8 Q810H6 Q810I1 Q810L3 Q810M1 Q810M6 Q810N5 Q810N6 Q810N9 Q810P2 Q810P3
Q810Q5 Q810T5 Q810U0 Q810V0 Q810Y8 Q81165 Q811B5 Q811C2 Q811C4 Q811F1 Q811I0
Q811J3 Q811P6 Q811Q1 Q811R2 Q811S7 Q811S9 Q811T9 Q811V6 Q811W0 Q811W1 Q811X5
Q811X6 Q812A3 Q812A5 Q812B1 Q812D3 Q812E0 Q812E8 Q812H1 Q812I1 Q812I4 Q812J6
Q812J8 Q812N6 Q812P6 Q812Q3 Q812Q9 Q812R6 Q812R8 Q812R9 Q812S0 Q812S1 Q812T1
Q812T2 Q812U1 Q812U6 Q812V9 Q812W2 Q812W8 Q812X2 Q812X3 Q812X4 Q812X5 Q812X7
Q812X9 Q812Y0 Q812Y4 Q812Y7 Q813A2 Q813A5 Q813D0 Q813J0 Q813M3 Q813Q5 Q813S8
Q813T6 Q813V3 Q813W6 Q813Z5 Q813Z9 Q814A6 Q814B0 Q814B1 Q814C2 Q814C3 Q814C4
Q814C5 Q814C9 Q814F2 Q814F3 Q814F6 Q814F7 Q814F8 Q814F9 Q814G5 Q814G6 Q814G7
Q814H0 Q814H1 Q814I1 Q814I2 Q814I6 Q814J1 Q814L8 Q814M3 Q814M9 Q814N7 Q814N9
Q814P5 Q814Q1 Q814Q2 Q814Q8 Q814T1 Q814T2 Q814T6 Q814T9 Q814U0 Q814U1 Q814V1
Q814V2 Q814V3 Q814X9 Q815D5 Q815E7 Q815E8 Q815H3 Q815I3 Q815I8 Q815J4 Q815J6
Q815K7 Q815L5 Q815R9 Q815S0 Q815U6 Q815X8 Q815Y3 Q816A0 Q816D0 Q816D1 Q816D9
Q816E3 Q816G0 Q816G3 Q816G6 Q816G7 Q816G8 Q816H3 Q816H7 Q816I1 Q816Q7 Q816Q8
Q816R0 Q816R1 Q816T0 Q816U4 Q816X6 Q817A2 Q817A4 Q817B1 Q817B5 Q817C0 Q817C6
Q817C7 Q817E2 Q817F3 Q817F9 Q817G5 Q817G7 Q817G9 Q817H0 Q817H6 Q817H7 Q817I4

Q817I6 Q817I7 Q817I8 Q817J0 Q817L9 Q817P3 Q817P4 Q817Q2 Q817Q5 Q817Q8 Q817R0
Q817R3 Q817R6 Q817R9 Q817T7 Q817U2 Q817U4 Q817U6 Q817V8 Q817W3 Q817W4 Q817W5
Q817W6 Q817X3 Q817X5 Q817X7 Q817X8 Q817Z0 Q817Z5 Q817Z6 Q81862 Q818A1 Q818C5
Q818D0 Q818D2 Q818E1 Q818E2 Q818E7 Q818E8 Q818E9 Q818F0 Q818F1 Q818G3 Q818H2
Q818H3 Q818H8 Q818I1 Q818J0 Q818M3 Q818M4 Q818M5 Q818P0 Q818P5 Q818P8 Q818R4
Q818R5 Q818R6 Q818R7 Q818R9 Q818S1 Q818T1 Q818U0 Q818U9 Q818V3 Q818W0 Q818W1
Q818W2 Q818W3 Q818W9 Q818X0 Q818X2 Q818X3 Q818X5 Q818X6 Q818Z9 Q819B3 Q819C8
Q819D1 Q819E5 Q819E6 Q819E7 Q819E8 Q819F1 Q819F2 Q819H7 Q819I5 Q819J5 Q819J6
Q819K0 Q819K2 Q819L3 Q819L4 Q819L5 Q819L6 Q819M7 Q819P1 Q819P4 Q819P6 Q819P8
Q819Q0 Q819Q2 Q819Q4 Q819R4 Q819R8 Q819S0 Q819S1 Q819S2 Q819S3 Q819S4 Q819S5
Q819S6 Q819S7 Q819T5 Q819T6 Q819T7 Q819U0 Q819U1 Q819U3 Q819U6 Q819U9 Q819V8
Q819W3 Q819W5 Q819W6 Q819W8 Q819W9 Q819X0 Q819X4 Q819X6 Q819X7 Q819X9 Q819Y0
Q819Y1 Q819Y3 Q819Y4 Q819Y5 Q819Y8 Q819Z0 Q819Z1 Q81A15 Q81A16 Q81A25 Q81A26
Q81A48 Q81A74 Q81AC5 Q81AC6 Q81AC7 Q81AC8 Q81AD9 Q81AF2 Q81AF4 Q81AF5 Q81AG1
Q81AG2 Q81AU5 Q81AZ5 Q81B01 Q81B10 Q81B21 Q81B49 Q81B71 Q81BC0 Q81BF4 Q81BK3
Q81BN7 Q81BT0 Q81BT3 Q81BT4 Q81BV3 Q81BX6 Q81C10 Q81C25 Q81C42 Q81C43 Q81C45
Q81C81 Q81CB2 Q81CE5 Q81CG8 Q81CH1 Q81CK0 Q81CK1 Q81D33 Q81DD0 Q81DD2 Q81DF9
Q81DH0 Q81DK5 Q81DR5 Q81DW0 Q81E02 Q81E05 Q81E30 Q81E36 Q81E85 Q81E87 Q81EF2
Q81EF6 Q81EI2 Q81EK4 Q81EK9 Q81EN2 Q81EP4 Q81EX6 Q81EY8 Q81EZ2 Q81F03 Q81F14
Q81F26 Q81F27 Q81F55 Q81F73 Q81F85 Q81FB5 Q81FL2 Q81FL9 Q81FM6 Q81FN5 Q81FN6
Q81FN7 Q81FP0 Q81FP2 Q81FP3 Q81FP4 Q81FQ1 Q81FQ2 Q81FQ3 Q81FQ4 Q81FQ6 Q81FQ8
Q81FR5 Q81FS0 Q81FS2 Q81FS9 Q81FV5 Q81FW4 Q81FZ0 Q81FZ1 Q81G00 Q81G01 Q81G02
Q81G03 Q81G04 Q81G05 Q81G06 Q81G07 Q81G08 Q81G09 Q81G10 Q81G11 Q81G13 Q81G17
Q81G39 Q81G66 Q81G67 Q81G69 Q81G81 Q81G82 Q81GA3 Q81GA4 Q81GA5 Q81GF2 Q81GG4
Q81GG5 Q81GG6 Q81GG7 Q81GG8 Q81GH5 Q81GJ7 Q81GJ9 Q81GK1 Q81GK5 Q81GK7 Q81GL8
Q81GM5 Q81GN2 Q81GN7 Q81GP2 Q81GP9 Q81GQ0 Q81GQ6 Q81GW5 Q81GW6 Q81GZ2 Q81H06
Q81H14 Q81H23 Q81H26 Q81H98 Q81HB0 Q81HB1 Q81HD0 Q81HD5 Q81HH1 Q81HQ5 Q81HW6
Q81HW9 Q81I05 Q81I11 Q81I66 Q81I72 Q81I85 Q81I89 Q81I94 Q81I97 Q81IB3 Q81ID9
Q81IE9 Q81IG8 Q81IG9 Q81IH1 Q81II0 Q81II1 Q81II2 Q81IK7 Q81IL5 Q81IN2 Q81IN3
Q81IN4 Q81IP2 Q81IP9 Q81IQ1 Q81IQ3 Q81IQ4 Q81IQ6 Q81IS3 Q81IS6 Q81IS8 Q81IT2
Q81IT9 Q81IU1 Q81IX0 Q81J01 Q81J03 Q81J12 Q81J13 Q81J17 Q81J18 Q81J19 Q81J20
Q81J22 Q81J23 Q81J24 Q81J25 Q81J26 Q81J27 Q81J28 Q81J29 Q81J30 Q81J31 Q81J32
Q81J33 Q81J34 Q81J35 Q81J36 Q81J37 Q81J38 Q81J39 Q81J40 Q81J41 Q81J42 Q81J43
Q81J44 Q81J45 Q81J47 Q81J48 Q81J50 Q81J51 Q81J52 Q81J53 Q81J58 Q81J59 Q81J61
Q81J62 Q81J63 Q81J67 Q81J70 Q81J80 Q81J81 Q81J84 Q81J96 Q81J97 Q81J99 Q81JA2
Q81JA5 Q81JA6 Q81JA8 Q81JB2 Q81JB5 Q81JB9 Q81JC4 Q81JC5 Q81JC6 Q81JD2 Q81JD9
Q81JE5 Q81JG1 Q81JG7 Q81JG8 Q81JG9 Q81JH0 Q81JH3 Q81JH4 Q81JH5 Q81JI2 Q81JI3
Q81JI4 Q81JI7 Q81JI9 Q81JJ9 Q81JK8 Q81JP2 Q81JQ4 Q81JR3 Q81JR5 Q81JR8 Q81JS3
Q81JT0 Q81JT1 Q81JT7 Q81JV2 Q81JW0 Q81JW1 Q81JW5 Q81JW9 Q81JX0 Q81JX1 Q81JX2
Q81JY3 Q81JY4 Q81JY5 Q81K13 Q81K75 Q81K80 Q81K82 Q81K83 Q81K85 Q81K90 Q81K94
Q81K97 Q81KH8 Q81KI0 Q81KI3 Q81KI4 Q81KK6 Q81KM1 Q81KP9 Q81KQ2 Q81KS9 Q81KT2
Q81KU0 Q81KU5 Q81KV0 Q81KV7 Q81KV8 Q81KX6 Q81KZ0 Q81L04 Q81L09 Q81L10 Q81L15
Q81L16 Q81L17 Q81L28 Q81L30 Q81L31 Q81L32 Q81L36 Q81L40 Q81L50 Q81L74 Q81LA9
Q81LB0 Q81LB8 Q81LB9 Q81LC2 Q81LC5 Q81LC7 Q81LD0 Q81LD3 Q81LD6 Q81LD7 Q81LE1

Q81LE6 Q81LE8 Q81LF0 Q81LG1 Q81LG8 Q81LG9 Q81LH1 Q81LH2 Q81LI1 Q81LI3 Q81LI6
Q81LI7 Q81LK0 Q81LK1 Q81LK2 Q81LK3 Q81LK8 Q81LK9 Q81LL3 Q81LL4 Q81LP3 Q81LR4
Q81LR5 Q81LS0 Q81LS1 Q81LS2 Q81LS3 Q81LS4 Q81LS7 Q81LT4 Q81LT9 Q81LU1 Q81LU9
Q81LV1 Q81LV7 Q81LW0 Q81LW9 Q81M06 Q81M07 Q81M08 Q81M24 Q81M27 Q81M32 Q81M49
Q81M50 Q81M51 Q81M52 Q81M54 Q81M56 Q81M67 Q81M74 Q81M86 Q81M88 Q81M95 Q81M96
Q81M97 Q81M98 Q81M99 Q81MA9 Q81MB0 Q81MB2 Q81MB3 Q81MB5 Q81MB6 Q81ME0 Q81MF4
Q81MH0 Q81MH2 Q81MH5 Q81MI9 Q81MJ0 Q81MJ1 Q81MJ2 Q81MJ5 Q81MJ6 Q81MN9 Q81MQ2
Q81MQ3 Q81MQ7 Q81MQ9 Q81MS1 Q81MS2 Q81MS3 Q81MS4 Q81MT5 Q81MW4 Q81MX1 Q81MY9
Q81N10 Q81N41 Q81N49 Q81ND5 Q81NJ6 Q81NN0 Q81NT0 Q81NT5 Q81NT6 Q81NV9 Q81P26
Q81P27 Q81P44 Q81P61 Q81P62 Q81P64 Q81PB2 Q81PE9 Q81PH1 Q81PH9 Q81PL1 Q81PL4
Q81PP8 Q81PP9 Q81PQ0 Q81PU5 Q81Q29 Q81QA7 Q81QB5 Q81QB6 Q81QB7 Q81QD4 Q81QG0
Q81QK2 Q81QR5 Q81QW3 Q81QX8 Q81R20 Q81R23 Q81R75 Q81R81 Q81R83 Q81RE1 Q81RK2
Q81RK6 Q81RP3 Q81RQ4 Q81RQ5 Q81RQ7 Q81RQ8 Q81RS0 Q81RU6 Q81RW4 Q81RX7 Q81RY8
Q81RZ3 Q81S00 Q81S14 Q81S26 Q81S27 Q81S64 Q81S85 Q81SA0 Q81SD1 Q81SQ5 Q81SR3
Q81SR4 Q81SS2 Q81ST1 Q81ST2 Q81ST3 Q81ST6 Q81ST7 Q81ST8 Q81ST9 Q81SU0 Q81SV3
Q81SV5 Q81SV6 Q81SV7 Q81SV8 Q81SW0 Q81SW2 Q81SW4 Q81SW9 Q81SX4 Q81SX6 Q81SY4
Q81T09 Q81T23 Q81T48 Q81T49 Q81T58 Q81T59 Q81T60 Q81T61 Q81T62 Q81T63 Q81T64
Q81T65 Q81T66 Q81T67 Q81T68 Q81T69 Q81T74 Q81T97 Q81TC4 Q81TC7 Q81TE0 Q81TE1
Q81TG0 Q81TG1 Q81TG2 Q81TK1 Q81TL7 Q81TL8 Q81TL9 Q81TM0 Q81TM1 Q81TM6 Q81TN0
Q81TQ0 Q81TQ3 Q81TQ5 Q81TQ8 Q81TR3 Q81TR6 Q81TS6 Q81TS7 Q81TT4 Q81TU4 Q81TU9
Q81TV3 Q81TW1 Q81TW2 Q81TW8 Q81U22 Q81U23 Q81U58 Q81U72 Q81U83 Q81U95 Q81UB2
Q81UH1 Q81UJ9 Q81UK5 Q81UP1 Q81UX4 Q81V33 Q81V37 Q81V80 Q81V87 Q81VE0 Q81VE1
Q81VE2 Q81VE5 Q81VF1 Q81VG0 Q81VJ1 Q81VN5 Q81VN7 Q81VP8 Q81VP9 Q81VQ3 Q81VQ4
Q81VQ5 Q81VQ6 Q81VQ7 Q81VQ9 Q81VR1 Q81VR2 Q81VR3 Q81VR4 Q81VR5 Q81VR6 Q81VR7
Q81VR8 Q81VR9 Q81VS0 Q81VS1 Q81VS2 Q81VS3 Q81VS4 Q81VS5 Q81VS6 Q81VS7 Q81VS8
Q81VS9 Q81VT0 Q81VT1 Q81VT2 Q81VT3 Q81VT4 Q81VT5 Q81VT8 Q81VU0 Q81VU1 Q81VU2
Q81VU3 Q81VU6 Q81VV1 Q81VV3 Q81VV4 Q81VV5 Q81VW0 Q81VW3 Q81VX3 Q81VX4 Q81VX7
Q81VY9 Q81VZ0 Q81VZ2 Q81VZ6 Q81W00 Q81W01 Q81W03 Q81W06 Q81W11 Q81W16 Q81W17
Q81W25 Q81W26 Q81W27 Q81W35 Q81W41 Q81W43 Q81W47 Q81W73 Q81WC0 Q81WC2 Q81WC3
Q81WC7 Q81WC9 Q81WD1 Q81WE0 Q81WE4 Q81WE7 Q81WE9 Q81WF0 Q81WF1 Q81WF2 Q81WF3
Q81WF4 Q81WF5 Q81WF6 Q81WG6 Q81WG7 Q81WG8 Q81WH1 Q81WH2 Q81WH4 Q81WH7 Q81WI0
Q81WI8 Q81WJ3 Q81WJ5 Q81WJ6 Q81WJ9 Q81WK0 Q81WK3 Q81WK5 Q81WK6 Q81WK7 Q81WK8
Q81WK9 Q81WL0 Q81WL1 Q81WL2 Q81WL4 Q81WL6 Q81WL9 Q81WM3 Q81WM5 Q81WM7 Q81WM8
Q81WQ3 Q81WR0 Q81WR3 Q81WR4 Q81WT0 Q81WU4 Q81WX1 Q81WY7 Q81X04 Q81X05 Q81X47
Q81X51 Q81X59 Q81X61 Q81X63 Q81X75 Q81X76 Q81X77 Q81XA8 Q81XC5 Q81XC6 Q81XF9
Q81XJ7 Q81XK8 Q81XM8 Q81XR1 Q81XR2 Q81XS0 Q81XS5 Q81XT3 Q81Y08 Q81Y44 Q81Y45
Q81Y46 Q81Y47 Q81Y79 Q81Y87 Q81Y88 Q81YI0 Q81YT1 Q81YT7 Q81YU1 Q81YV0 Q81YV4
Q81YW1 Q81YW4 Q81YY0 Q81Z48 Q81Z66 Q81Z76 Q81Z95 Q81Z96 Q81Z97 Q81ZB7 Q81ZC2
Q81ZD6 Q81ZE7 Q81ZE8 Q81ZE9 Q81ZF8 Q81ZG1 Q81ZG8 Q81ZH0 Q81ZH2 Q81ZH3 Q81ZH5
Q81ZK5 Q81ZL2 Q81ZS3 Q81ZS7 Q81ZU6 Q81ZV1 Q81ZV5 Q81ZX2 Q81ZZ2 Q81ZZ4 Q81ZZ5
Q81ZZ8 Q81ZZ9 Q82037 Q82041 Q82044 Q82045 Q82049 Q82050 Q82051 Q82052 Q82053
Q82054 Q82080 Q820A2 Q820A9 Q820B5 Q820C5 Q820D2 Q820D4 Q820D5 Q820D6 Q820D7
Q820D8 Q820D9 Q820E1 Q820E4 Q820E9 Q820F2 Q820F3 Q820F4 Q820F5 Q820F8 Q820G1
Q820G3 Q820I0 Q820I7 Q820J1 Q820J5 Q820J6 Q820J7 Q820K1 Q820K3 Q820K4 Q820M0

Q820M4 Q820M5 Q820M6 Q820M7 Q820N1 Q820N3 Q820N6 Q820P2 Q820P8 Q820Q0 Q820Q1
Q820Q4 Q820Q7 Q820Q8 Q820Q9 Q820R0 Q820R1 Q820S0 Q820S1 Q820S3 Q820S5 Q820T0
Q820U0 Q820U1 Q820U3 Q820V9 Q820W2 Q820W3 Q820X1 Q820X8 Q820X9 Q820Y0 Q820Y1
Q820Y5 Q820Y7 Q820Y9 Q820Z0 Q820Z2 Q820Z5 Q820Z6 Q820Z9 Q82171 Q821A5 Q821A6
Q821B0 Q821B6 Q821C1 Q821C2 Q821D9 Q821E2 Q821E6 Q821F7 Q821F9 Q821H8 Q821H9
Q821I3 Q821I9 Q821K6 Q821L2 Q821L7 Q821L9 Q821M0 Q821M4 Q821N6 Q821P2 Q821P5
Q821Q5 Q821R0 Q821R2 Q821R5 Q821S1 Q821S4 Q821T4 Q821U6 Q821U7 Q821V0 Q821V1
Q821V2 Q821V5 Q821V6 Q821W4 Q821W5 Q821W6 Q821W7 Q821W8 Q821W9 Q821X0 Q821X7
Q821Z6 Q821Z7 Q822A1 Q822B0 Q822B2 Q822B3 Q822B4 Q822B5 Q822B9 Q822C1 Q822C2
Q822D5 Q822D9 Q822E7 Q822E9 Q822F3 Q822F7 Q822F8 Q822G0 Q822G7 Q822H0 Q822I3
Q822I4 Q822I7 Q822I8 Q822I9 Q822J0 Q822J1 Q822J2 Q822J3 Q822K4 Q822K5 Q822L8
Q822M1 Q822M2 Q822M5 Q822M6 Q822M7 Q822M8 Q822N0 Q822P7 Q822P9 Q822Q0 Q822Q4
Q822R1 Q822R2 Q822R7 Q822T1 Q822T2 Q822U8 Q822V6 Q822W2 Q822W3 Q822W4 Q822W6
Q822W7 Q822W9 Q822X6 Q822Y3 Q822Y4 Q822Z1 Q822Z3 Q822Z4 Q822Z9 Q823A0 Q823A6
Q823A7 Q823B7 Q823D7 Q823E6 Q823F2 Q823F3 Q823F7 Q823G2 Q823G4 Q823G6 Q823G9
Q823H9 Q823I7 Q823J9 Q823K0 Q823K4 Q823K6 Q823N2 Q823N6 Q823P0 Q823Q9 Q823R4
Q823S6 Q823T2 Q823T3 Q823T8 Q823T9 Q823U1 Q823U4 Q823U7 Q823U8 Q823U9 Q823V1
Q823V2 Q823W2 Q823W6 Q823Y5 Q82446 Q82461 Q82462 Q82484 Q824A4 Q824A6 Q824A9
Q824B0 Q824B1 Q824B2 Q824B4 Q824B8 Q824C7 Q824C8 Q824E0 Q824E8 Q824E9 Q824F3
Q824F4 Q824F5 Q824F7 Q824F9 Q824G0 Q824G1 Q824G2 Q824H0 Q824H2 Q824H4 Q824H7
Q824I4 Q824J8 Q824K8 Q824L3 Q824L6 Q824M2 Q824M4 Q824M5 Q824M6 Q824N0 Q824N1
Q824N2 Q824N4 Q824N5 Q824N6 Q824N7 Q824N8 Q824N9 Q824P0 Q824P1 Q824P2 Q824P3
Q824P4 Q824P5 Q824P6 Q824P7 Q824P8 Q824P9 Q824Q0 Q824Q1 Q824Q3 Q824R3 Q824S8
Q824U3 Q824U4 Q824U5 Q824U6 Q824W1 Q824W6 Q824X6 Q824X7 Q824X8 Q824Y6 Q825I8
Q825Q8 Q825U9 Q82630 Q82635 Q82676 Q82680 Q82681 Q82684 Q82685 Q826D9 Q826E8
Q826F2 Q826F5 Q826F6 Q826J2 Q826P6 Q826Q0 Q826R6 Q826R7 Q826R9 Q826S0 Q826S1
Q826S4 Q826T2 Q826T4 Q826W3 Q826Z7 Q82707 Q82708 Q827B3 Q827D7 Q827G1 Q827H4
Q827I4 Q827I5 Q827L7 Q827L8 Q827L9 Q827P7 Q827Q0 Q827Q2 Q827Q3 Q827Q7 Q827Q8
Q827R1 Q827R4 Q827R5 Q827R8 Q827R9 Q827S0 Q827S3 Q827S4 Q827S7 Q827T0 Q827T5
Q827T9 Q827U0 Q827U8 Q827Z0 Q827Z1 Q82852 Q828A2 Q828A3 Q828A4 Q828A5 Q828A6
Q828C6 Q828C7 Q828D0 Q828D1 Q828D2 Q828I4 Q828I7 Q828I8 Q828J2 Q828K5 Q828K8
Q828K9 Q828M2 Q828S5 Q828W8 Q828Y7 Q828Y9 Q82994 Q829A7 Q829B4 Q829C5 Q829F2
Q829J5 Q829L3 Q829L4 Q829L5 Q829L6 Q829P9 Q829U5 Q829V7 Q829W1 Q829W2 Q829W3
Q829W5 Q829W7 Q829W8 Q829X3 Q829X9 Q829Y6 Q82A24 Q82A81 Q82A82 Q82A84 Q82A89
Q82A99 Q82AA1 Q82AA2 Q82AA4 Q82AA5 Q82AA6 Q82AC9 Q82AD2 Q82AD8 Q82AE1 Q82AE4
Q82AF9 Q82AG5 Q82AG6 Q82AK1 Q82AM7 Q82AN2 Q82AP6 Q82AP7 Q82AW2 Q82BN0 Q82BR9
Q82BS5 Q82BS6 Q82BV3 Q82BX0 Q82BX4 Q82BX8 Q82BY4 Q82BY5 Q82BZ4 Q82C66 Q82C81
Q82C83 Q82C85 Q82C86 Q82C87 Q82CA0 Q82CA3 Q82CA5 Q82CA6 Q82CA7 Q82CC3 Q82CI4
Q82CM8 Q82CS8 Q82CX8 Q82D15 Q82D17 Q82D28 Q82D55 Q82D77 Q82D82 Q82DC9 Q82DE2
Q82DE3 Q82DI5 Q82DK2 Q82DL5 Q82DL7 Q82DL8 Q82DL9 Q82DM1 Q82DM2 Q82DM3 Q82DM5
Q82DM7 Q82DM8 Q82DM9 Q82DN0 Q82DN1 Q82DN2 Q82DN3 Q82DN4 Q82DN5 Q82DN6 Q82DN7
Q82DN8 Q82DN9 Q82DP0 Q82DP1 Q82DP2 Q82DP4 Q82DP5 Q82DQ0 Q82DQ1 Q82DQ2 Q82DQ4
Q82DQ5 Q82DQ6 Q82DQ7 Q82DQ8 Q82DQ9 Q82DR2 Q82DR4 Q82DR8 Q82DS2 Q82E21 Q82E68
Q82E76 Q82E77 Q82E78 Q82E91 Q82E99 Q82EA9 Q82EC5 Q82EC8 Q82EE8 Q82EF1 Q82EF7

Q82EG9 Q82EH1 Q82EH2 Q82EH4 Q82EL5 Q82EQ7 Q82EQ8 Q82ER6 Q82EU1 Q82EU9 Q82EX7
Q82EX8 Q82EX9 Q82EZ9 Q82F60 Q82F68 Q82F74 Q82FB5 Q82FD4 Q82FD5 Q82FD8 Q82FD9
Q82FE0 Q82FE3 Q82FG4 Q82FG5 Q82FG6 Q82FI7 Q82FJ1 Q82FK8 Q82FS4 Q82FV6 Q82FW3
Q82FW4 Q82G40 Q82G68 Q82G71 Q82G74 Q82G92 Q82GB8 Q82GC8 Q82GC9 Q82GD0 Q82GG6
Q82GH3 Q82GN9 Q82GP1 Q82GV2 Q82H75 Q82HC3 Q82HD9 Q82HE3 Q82HE5 Q82HE6 Q82HE7
Q82HL2 Q82HL5 Q82HM9 Q82HR6 Q82HS2 Q82HU1 Q82I23 Q82I33 Q82I51 Q82I57 Q82I71
Q82IC1 Q82IC5 Q82ID7 Q82IE6 Q82IE7 Q82IE8 Q82IJ5 Q82IU1 Q82IV0 Q82IW0 Q82IY4
Q82IY7 Q82IY8 Q82IY9 Q82IZ1 Q82IZ2 Q82J66 Q82J69 Q82J70 Q82J98 Q82JD1 Q82JD2
Q82JE3 Q82JF0 Q82JI0 Q82JI1 Q82JI2 Q82JK2 Q82JK7 Q82JK9 Q82JL0 Q82JL2 Q82JN6
Q82JN8 Q82JN9 Q82JP0 Q82JR3 Q82JR8 Q82JR9 Q82JS2 Q82JS5 Q82JT5 Q82JT8 Q82JT9
Q82JU0 Q82JU5 Q82JV8 Q82JW0 Q82JW2 Q82JW3 Q82JW4 Q82JX1 Q82JX7 Q82JX8 Q82JX9
Q82JY0 Q82JY3 Q82K09 Q82K41 Q82K43 Q82K46 Q82K50 Q82K53 Q82K55 Q82K56 Q82K79
Q82K80 Q82K82 Q82K86 Q82K95 Q82KB1 Q82KC4 Q82KC8 Q82KE1 Q82KE2 Q82KF0 Q82KI9
Q82KJ6 Q82KK4 Q82KP4 Q82KQ4 Q82KS8 Q82KW8 Q82KY4 Q82KZ4 Q82L07 Q82LC4 Q82LG4
Q82LL4 Q82LL5 Q82LU9 Q82M02 Q82M23 Q82M40 Q82M83 Q82M90 Q82M93 Q82MF1 Q82MH6
Q82MI6 Q82MJ7 Q82ML3 Q82ML4 Q82MN2 Q82MT8 Q82MX6 Q82N59 Q82ND8 Q82NI9 Q82NM9
Q82NQ8 Q82NZ3 Q82NZ4 Q82P45 Q82P95 Q82P98 Q82PC4 Q82PM3 Q82PP4 Q82PX1 Q82PY3
Q82PZ0 Q82QG3 Q82QK2 Q82QN1 Q82QR5 Q82RR7 Q82S07 Q82S08 Q82S12 Q82S73 Q82S78
Q82S79 Q82S90 Q82S93 Q82S94 Q82S98 Q82SA7 Q82SD3 Q82SD7 Q82SD8 Q82SG6 Q82SH1
Q82SI5 Q82SI6 Q82SI7 Q82SI9 Q82SJ7 Q82SJ8 Q82SL7 Q82SL8 Q82SM5 Q82SM6 Q82SP3
Q82SP4 Q82SP6 Q82SP7 Q82SQ2 Q82SQ3 Q82SQ5 Q82SQ9 Q82ST4 Q82ST5 Q82SU1 Q82SU4
Q82SV5 Q82SW5 Q82T00 Q82T07 Q82T49 Q82T56 Q82T57 Q82T68 Q82T69 Q82T70 Q82T71
Q82T72 Q82T73 Q82T74 Q82T75 Q82T76 Q82T88 Q82TB9 Q82TC0 Q82TC8 Q82TD1 Q82TD4
Q82TD5 Q82TD7 Q82TD8 Q82TF6 Q82TF8 Q82TG6 Q82TH3 Q82TH4 Q82TK9 Q82TL0 Q82TM5
Q82TM7 Q82TN0 Q82TN2 Q82TN5 Q82TP3 Q82TP9 Q82TQ1 Q82TQ3 Q82TQ4 Q82TQ5 Q82TQ6
Q82TU0 Q82TU1 Q82TV8 Q82TW4 Q82TW6 Q82TX6 Q82TY3 Q82TZ6 Q82TZ7 Q82TZ8 Q82TZ9
Q82U01 Q82U06 Q82U08 Q82U36 Q82U37 Q82U38 Q82U41 Q82U42 Q82U44 Q82U62 Q82U64
Q82U74 Q82U77 Q82U82 Q82U84 Q82UC1 Q82UI9 Q82UJ4 Q82UJ5 Q82UJ6 Q82UJ7 Q82UK0
Q82UK1 Q82UK2 Q82UK6 Q82UK7 Q82UK8 Q82UL3 Q82UL7 Q82UL8 Q82UL9 Q82UM1 Q82UP3
Q82UP4 Q82UP5 Q82UP6 Q82UP9 Q82UQ0 Q82UQ1 Q82UQ3 Q82UQ4 Q82UQ7 Q82UQ8 Q82UR4
Q82UR9 Q82US7 Q82UZ3 Q82UZ8 Q82UZ9 Q82V16 Q82V18 Q82V19 Q82V20 Q82V23 Q82V28
Q82V29 Q82VB0 Q82VC2 Q82VD1 Q82VD3 Q82VD5 Q82VF0 Q82VF1 Q82VN1 Q82VN4 Q82VP4
Q82VS0 Q82VS1 Q82VS2 Q82VS5 Q82VS8 Q82VT1 Q82VT2 Q82VV0 Q82VV1 Q82VV3 Q82VV4
Q82VV5 Q82VV6 Q82VV7 Q82VV9 Q82VW0 Q82VY8 Q82W02 Q82W15 Q82W26 Q82W27 Q82W32
Q82W45 Q82W51 Q82WA8 Q82WA9 Q82WB9 Q82WC9 Q82WD0 Q82WD1 Q82WE9 Q82WF3 Q82WG1
Q82WG7 Q82WI1 Q82WI2 Q82WI3 Q82WI6 Q82WI7 Q82WI9 Q82WL1 Q82WL2 Q82WL9 Q82WM1
Q82WM3 Q82WM4 Q82WM5 Q82WM6 Q82WM7 Q82WP0 Q82WP9 Q82WQ0 Q82WQ3 Q82WQ4 Q82WQ5
Q82WQ6 Q82WR3 Q82WR4 Q82WS2 Q82WS3 Q82WU3 Q82WX6 Q82WY3 Q82WZ5 Q82X50 Q82X51
Q82X54 Q82X61 Q82X65 Q82X68 Q82X69 Q82X70 Q82X71 Q82X73 Q82X74 Q82X75 Q82X77
Q82X78 Q82X79 Q82X80 Q82X81 Q82X82 Q82X83 Q82X84 Q82X85 Q82X86 Q82X87 Q82X88
Q82X89 Q82X93 Q82X98 Q82X99 Q82XA1 Q82XC1 Q82XC3 Q82XC7 Q82XD1 Q82XD4 Q82XE0
Q82XE8 Q82XF2 Q82XI7 Q82XJ3 Q82XJ4 Q82XN1 Q82XN2 Q82XN5 Q82XP4 Q82XP5 Q82XP6
Q82XQ6 Q82XQ7 Q82XQ8 Q82XQ9 Q82XS2 Q82XS4 Q82XS9 Q82XT0 Q82XU6 Q82XU9 Q82XV0
Q82XV4 Q82XV5 Q82XV8 Q82XW0 Q82XW4 Q82XY0 Q82XY2 Q82XY4 Q82XY7 Q82XZ6 Q82Y02

Q82Y12 Q82Y13 Q82Y15 Q82Y17 Q82Y18 Q82Y30 Q82Y45 Q82Y50 Q82Y56 Q82Y57 Q82Y58
Q82Y60 Q82Y61 Q82Y74 Q82Y84 Q82YU9 Q82YV0 Q82YW1 Q82YX0 Q82YY1 Q82YY9 Q82Z12
Q82Z40 Q82Z41 Q82Z47 Q82Z70 Q82Z98 Q82ZA2 Q82ZA4 Q82ZA5 Q82ZC9 Q82ZD5 Q82ZD6
Q82ZD8 Q82ZE1 Q82ZE4 Q82ZG1 Q82ZG8 Q82ZH9 Q82ZI6 Q82ZJ0 Q82ZJ1 Q82ZJ2 Q82ZQ1
Q82ZT7 Q82ZW3 Q82ZW4 Q82ZW6 Q82ZX1 Q82ZX4 Q82ZZ8 Q83044 Q83045 Q83047 Q83048
Q83049 Q83050 Q83051 Q83096 Q830B9 Q830C2 Q830C3 Q830C9 Q830D0 Q830K2 Q830K4
Q830K5 Q830L4 Q830L6 Q830M0 Q830P3 Q830Q5 Q830Q6 Q830Q7 Q830Q8 Q830R6 Q830R7
Q830S9 Q830U2 Q830U3 Q830U4 Q830V0 Q830V2 Q830Y0 Q830Y9 Q831A8 Q831B2 Q831F5
Q831F6 Q831F7 Q831F9 Q831G0 Q831K9 Q831N0 Q831N1 Q831N9 Q831P3 Q831P7 Q831P9
Q831Q7 Q831R4 Q831S1 Q831T0 Q831T1 Q831T3 Q831T7 Q831U0 Q831U2 Q831U3 Q831U6
Q831U9 Q831V0 Q831V1 Q831V2 Q831W7 Q831X4 Q831Y0 Q831Y7 Q83250 Q83251 Q83252
Q83253 Q83262 Q83265 Q83268 Q83269 Q83271 Q832G5 Q832Z5 Q833B6 Q833G5 Q833H0
Q833H5 Q833H9 Q833I1 Q833I2 Q833I5 Q833I9 Q833J0 Q833M7 Q833M8 Q833N6 Q833P4
Q833P5 Q833S5 Q833S6 Q833S7 Q833U3 Q833U4 Q833U9 Q833W8 Q833W9 Q833Y3 Q833Y6
Q833Y8 Q833Y9 Q833Z1 Q833Z3 Q83417 Q83424 Q83444 Q83453 Q83457 Q83482 Q83485
Q83486 Q834C7 Q834D0 Q834D6 Q834E0 Q834E2 Q834E3 Q834F9 Q834G2 Q834G6 Q834H2
Q834J7 Q834K1 Q834K3 Q834K4 Q834L8 Q834M4 Q834N3 Q834Q9 Q834R1 Q834R3 Q834S1
Q834S2 Q834S5 Q834S6 Q834S9 Q834T4 Q834T6 Q834U3 Q834U4 Q834V4 Q834W6 Q834X4
Q835G1 Q835H1 Q835H3 Q835J5 Q835J8 Q835K6 Q835R2 Q835R5 Q835R7 Q835R8 Q835R9
Q835U7 Q835U8 Q835V2 Q835V5 Q835Z6 Q836B1 Q836B2 Q836B3 Q836D1 Q836D5 Q836E3
Q836E5 Q836E9 Q836G3 Q836G5 Q836H7 Q836H8 Q836J0 Q836J5 Q836J6 Q836J8 Q836J9
Q836N8 Q836P0 Q836R0 Q836R3 Q836T1 Q836V1 Q836V4 Q836W5 Q836W6 Q836W7 Q836W8
Q836X4 Q836X6 Q836Z5 Q83730 Q837A4 Q837A7 Q837A8 Q837B3 Q837C7 Q837C8 Q837C9
Q837E2 Q837E7 Q837G1 Q837G2 Q837G3 Q837H2 Q837J5 Q837K0 Q837L6 Q837N5 Q837P9
Q837R0 Q837R3 Q837R5 Q837R9 Q837T2 Q837U5 Q837U7 Q837V0 Q837V2 Q837V3 Q837V4
Q837V6 Q837W3 Q837X4 Q837X8 Q837X9 Q837Z3 Q837Z7 Q83884 Q83885 Q838A4 Q838A9
Q838C9 Q838D6 Q838I3 Q838I4 Q838I8 Q838J7 Q838J9 Q838K1 Q838L0 Q838L1 Q838L2
Q838L3 Q838L9 Q838N8 Q838Z5 Q83906 Q83924 Q83927 Q83928 Q83929 Q83944 Q83948
Q83949 Q83951 Q83956 Q83957 Q839A8 Q839B0 Q839B2 Q839B3 Q839C0 Q839C1 Q839C6
Q839D8 Q839D9 Q839E0 Q839E1 Q839E2 Q839E3 Q839E5 Q839E6 Q839E7 Q839E8 Q839E9
Q839F0 Q839F1 Q839F2 Q839F3 Q839F4 Q839F5 Q839F6 Q839F7 Q839F8 Q839F9 Q839G0
Q839G1 Q839G2 Q839G3 Q839G4 Q839G5 Q839G8 Q839G9 Q839H0 Q839H1 Q839H2 Q839H4
Q839I2 Q839J1 Q839J4 Q839J7 Q839J8 Q839Q5 Q839Q8 Q839T5 Q839T6 Q839T8 Q839U9
Q839V4 Q839V5 Q839V7 Q839V8 Q839W2 Q839W3 Q839Y4 Q839Y6 Q839Y8 Q839Y9 Q839Z0
Q839Z1 Q839Z2 Q839Z5 Q839Z9 Q83A16 Q83A17 Q83A18 Q83A19 Q83A28 Q83A36 Q83A37
Q83A42 Q83A62 Q83A77 Q83A78 Q83A83 Q83A90 Q83A94 Q83A95 Q83A98 Q83AA2 Q83AA6
Q83AA8 Q83AB0 Q83AB7 Q83AC1 Q83AC2 Q83AD4 Q83AD8 Q83AH2 Q83AJ2 Q83AJ3 Q83AK3
Q83AK6 Q83AK7 Q83AK9 Q83AL0 Q83AP0 Q83AP1 Q83AQ1 Q83AR4 Q83AR8 Q83AU6 Q83AV6
Q83AX3 Q83AX8 Q83AX9 Q83AY0 Q83AY4 Q83AY5 Q83AZ0 Q83AZ6 Q83B06 Q83B07 Q83B08
Q83B09 Q83B24 Q83B31 Q83B36 Q83B60 Q83BB0 Q83BB6 Q83BB9 Q83BE0 Q83BE2 Q83BE3
Q83BE4 Q83BE5 Q83BF8 Q83BG2 Q83BH2 Q83BI9 Q83BJ9 Q83BL1 Q83BL6 Q83BL7 Q83BM5
Q83BM8 Q83BM9 Q83BQ3 Q83BR9 Q83BS0 Q83BS1 Q83BS2 Q83BT3 Q83BT4 Q83BT5 Q83BV3
Q83BV4 Q83BY4 Q83BY7 Q83BZ6 Q83BZ8 Q83BZ9 Q83C10 Q83C11 Q83C12 Q83C13 Q83C14
Q83C15 Q83C16 Q83C29 Q83C31 Q83C38 Q83C41 Q83C63 Q83C64 Q83C71 Q83C77 Q83C80

Q83C83 Q83C87 Q83C88 Q83C93 Q83CA6 Q83CA8 Q83CC6 Q83CD1 Q83CD5 Q83CD9 Q83CG2
Q83CJ1 Q83CJ2 Q83CL8 Q83CM5 Q83CM9 Q83CP2 Q83CP8 Q83CP9 Q83CQ2 Q83CQ4 Q83CQ6
Q83CQ8 Q83CU5 Q83CU8 Q83CV4 Q83CV9 Q83CW4 Q83CW6 Q83CY8 Q83D06 Q83D14 Q83D18
Q83D39 Q83D67 Q83D70 Q83D71 Q83D73 Q83D75 Q83D76 Q83D86 Q83D87 Q83D88 Q83D91
Q83DC7 Q83DD0 Q83DE3 Q83DF6 Q83DI0 Q83DJ1 Q83DJ2 Q83DJ3 Q83DN1 Q83DN2 Q83DN9
Q83DP0 Q83DP1 Q83DP8 Q83DV5 Q83DX3 Q83DX5 Q83DY1 Q83DY4 Q83DZ6 Q83E06 Q83E09
Q83E10 Q83E11 Q83E12 Q83E18 Q83E35 Q83E41 Q83E63 Q83E64 Q83E75 Q83E83 Q83E84
Q83E85 Q83E97 Q83EA2 Q83EA3 Q83EA4 Q83EC9 Q83ED6 Q83ED8 Q83ED9 Q83EE0 Q83EH9
Q83EI7 Q83EJ0 Q83EK3 Q83EK7 Q83EL0 Q83EL6 Q83EL7 Q83EL9 Q83EM4 Q83EM7 Q83EP0
Q83EP4 Q83EQ1 Q83EQ2 Q83EQ3 Q83EQ4 Q83EQ7 Q83EQ8 Q83EQ9 Q83ER0 Q83ER1 Q83ER2
Q83ER4 Q83ER5 Q83ER6 Q83ER7 Q83ER8 Q83ER9 Q83ES0 Q83ES1 Q83ES2 Q83ES3 Q83ES4
Q83ES5 Q83ES6 Q83ES7 Q83ES8 Q83ES9 Q83ET0 Q83ET2 Q83ET3 Q83ET4 Q83EV3 Q83EV9
Q83EX7 Q83EX9 Q83F01 Q83F02 Q83F03 Q83F16 Q83F17 Q83F20 Q83F28 Q83F35 Q83F36
Q83F37 Q83F39 Q83F55 Q83F67 Q83F94 Q83FA3 Q83FA4 Q83FA5 Q83FB4 Q83FB6 Q83FD0
Q83FD6 Q83FD9 Q83FE8 Q83FF0 Q83FF5 Q83FI1 Q83FI4 Q83FI9 Q83FJ0 Q83FJ2 Q83FJ5
Q83FJ7 Q83FK2 Q83FK5 Q83FK8 Q83FL0 Q83FL1 Q83FM3 Q83FM6 Q83FN1 Q83FN9 Q83FP0
Q83FP1 Q83FQ7 Q83FQ8 Q83FR1 Q83FR7 Q83FS2 Q83FT1 Q83FT2 Q83FT9 Q83FU2 Q83FU3
Q83FV5 Q83FX9 Q83FY1 Q83FY2 Q83FY5 Q83FY6 Q83FY7 Q83FY9 Q83FZ0 Q83FZ1 Q83FZ2
Q83FZ3 Q83FZ4 Q83FZ5 Q83FZ6 Q83FZ7 Q83FZ8 Q83FZ9 Q83G00 Q83G01 Q83G02 Q83G03
Q83G04 Q83G05 Q83G07 Q83G08 Q83G10 Q83G11 Q83G21 Q83G28 Q83G32 Q83G43 Q83G46
Q83G48 Q83G49 Q83G50 Q83G51 Q83G52 Q83G55 Q83G56 Q83G58 Q83G63 Q83G64 Q83G66
Q83G68 Q83G71 Q83G72 Q83G73 Q83G93 Q83G94 Q83G99 Q83GA1 Q83GA2 Q83GA7 Q83GA8
Q83GC6 Q83GC9 Q83GD1 Q83GD2 Q83GD5 Q83GD7 Q83GE3 Q83GE4 Q83GE9 Q83GF1 Q83GG2
Q83GH3 Q83GH5 Q83GH8 Q83GI1 Q83GI2 Q83GJ0 Q83GJ1 Q83GJ5 Q83GK0 Q83GK3 Q83GK5
Q83GK7 Q83GM3 Q83GN2 Q83GN3 Q83GN7 Q83GP4 Q83GP6 Q83GP9 Q83GR4 Q83GS7 Q83GS8
Q83GS9 Q83GT0 Q83GT1 Q83GT2 Q83GT8 Q83GU5 Q83GU6 Q83GV1 Q83GW1 Q83GW6 Q83GW7
Q83GW8 Q83GX1 Q83GX2 Q83GX3 Q83GX4 Q83GX6 Q83GX8 Q83GY8 Q83GZ6 Q83H08 Q83H66
Q83H67 Q83H71 Q83H72 Q83H88 Q83H91 Q83H92 Q83H94 Q83H98 Q83HA1 Q83HA5 Q83HA7
Q83HA9 Q83HB1 Q83HB2 Q83HB8 Q83HB9 Q83HC3 Q83HC7 Q83HC8 Q83HD5 Q83HD6 Q83HD8
Q83HE6 Q83HF0 Q83HF6 Q83HF7 Q83HF9 Q83HG3 Q83HG7 Q83HH0 Q83HH1 Q83HH2 Q83HH3
Q83HH4 Q83HH5 Q83HI1 Q83HI6 Q83HI9 Q83HJ1 Q83HJ5 Q83HJ6 Q83HJ9 Q83HK0 Q83HK9
Q83HM1 Q83HM3 Q83HM5 Q83HM6 Q83HN0 Q83HN3 Q83HN7 Q83HN8 Q83HP5 Q83HP8 Q83HP9
Q83HQ3 Q83HQ6 Q83HQ8 Q83HR7 Q83HS8 Q83HT0 Q83HT7 Q83HT8 Q83HU4 Q83HU6 Q83HU9
Q83HV0 Q83HV1 Q83HV5 Q83HW8 Q83HW9 Q83HX1 Q83HX2 Q83HX4 Q83HX5 Q83HX8 Q83HZ5
Q83HZ6 Q83HZ7 Q83I01 Q83I04 Q83I06 Q83I07 Q83I10 Q83I11 Q83I12 Q83I16 Q83I17
Q83I18 Q83I19 Q83I20 Q83I22 Q83I32 Q83I36 Q83I43 Q83I55 Q83I57 Q83I58 Q83I60
Q83I61 Q83I62 Q83I63 Q83I64 Q83I65 Q83I66 Q83I67 Q83I68 Q83I69 Q83I70 Q83I71
Q83I72 Q83I73 Q83I74 Q83I75 Q83I77 Q83I78 Q83I79 Q83I82 Q83I83 Q83I84 Q83I86
Q83I94 Q83I95 Q83IA0 Q83IA1 Q83IA3 Q83IA7 Q83IB4 Q83IB5 Q83IB6 Q83IB9 Q83IC0
Q83IC1 Q83IC2 Q83IC3 Q83IC4 Q83IC8 Q83ID3 Q83ID9 Q83II5 Q83II8 Q83IJ0 Q83IJ9
Q83IK0 Q83IL8 Q83IM2 Q83IM5 Q83IN9 Q83IP6 Q83IR6 Q83IR9 Q83IS3 Q83IS7 Q83IT3
Q83IU3 Q83IU4 Q83IU5 Q83IV7 Q83IW0 Q83IW8 Q83IX2 Q83IY2 Q83IY4 Q83IY9 Q83IZ5
Q83J15 Q83J28 Q83J38 Q83J60 Q83J89 Q83J92 Q83J94 Q83J96 Q83J98 Q83JA5 Q83JB1
Q83JB2 Q83JB3 Q83JC0 Q83JC1 Q83JC2 Q83JC3 Q83JC4 Q83JD1 Q83JD2 Q83JE1 Q83JF9

Q83JG0 Q83JG5 Q83JG9 Q83JH9 Q83JI4 Q83JJ8 Q83JL2 Q83JS0 Q83JS1 Q83JS4 Q83JS6
Q83JS7 Q83JU1 Q83JU2 Q83JU6 Q83JW6 Q83JW7 Q83JW9 Q83JX8 Q83JX9 Q83JY2 Q83JY3
Q83JZ0 Q83K04 Q83K06 Q83K21 Q83K38 Q83K39 Q83K42 Q83K43 Q83K44 Q83K58 Q83K75
Q83K78 Q83K81 Q83K84 Q83K86 Q83KB9 Q83KC1 Q83KD7 Q83KD8 Q83KE6 Q83KF1 Q83KH0
Q83KH2 Q83KH7 Q83KI3 Q83KJ5 Q83KJ6 Q83KJ7 Q83KJ8 Q83KK3 Q83KK8 Q83KN1 Q83KQ3
Q83KQ5 Q83KR4 Q83KR5 Q83KS0 Q83KS7 Q83KV7 Q83KW2 Q83KX8 Q83KY0 Q83KY1 Q83KZ1
Q83KZ2 Q83L15 Q83L16 Q83L29 Q83L33 Q83L36 Q83L37 Q83L38 Q83L49 Q83L52 Q83L66
Q83L75 Q83L76 Q83LB6 Q83LB9 Q83LD1 Q83LD8 Q83LE1 Q83LF2 Q83LF3 Q83LF6 Q83LF7
Q83LH9 Q83LI5 Q83LI7 Q83LJ0 Q83LK1 Q83LK6 Q83LK7 Q83LK8 Q83LK9 Q83LL9 Q83LM0
Q83LN3 Q83LP1 Q83LP3 Q83LP7 Q83LS2 Q83LT1 Q83LU4 Q83LU7 Q83LU8 Q83LX5 Q83LY1
Q83LY3 Q83LY4 Q83LZ7 Q83M01 Q83M07 Q83M14 Q83M27 Q83M36 Q83M39 Q83M40 Q83M42
Q83M49 Q83M51 Q83M63 Q83M66 Q83M67 Q83M81 Q83M86 Q83MA8 Q83MC1 Q83MC8 Q83MD3
Q83MD4 Q83MD6 Q83ME2 Q83ME5 Q83MF0 Q83MF4 Q83MF7 Q83MF8 Q83MF9 Q83MG0 Q83MG1
Q83MG5 Q83MG8 Q83MG9 Q83MH0 Q83MH2 Q83MH5 Q83MH9 Q83MI1 Q83MI6 Q83MJ0 Q83MJ8
Q83MJ9 Q83MK2 Q83MK8 Q83ML6 Q83ML8 Q83MN1 Q83MQ1 Q83MQ8 Q83MR9 Q83MS7 Q83MT2
Q83MU8 Q83MU9 Q83MV5 Q83MW6 Q83MX3 Q83MY5 Q83MY6 Q83MZ4 Q83MZ8 Q83MZ9 Q83N09
Q83N13 Q83N18 Q83N20 Q83N34 Q83N51 Q83N52 Q83N75 Q83N78 Q83N83 Q83NA0 Q83NB2
Q83ND3 Q83NE4 Q83NF6 Q83NI2 Q83NI7 Q83NI9 Q83NK3 Q83NK8 Q83NN0 Q83NN3 Q83NN8
Q83NP1 Q83NS5 Q83NT9 Q83NU2 Q83NZ4 Q83NZ5 Q83P00 Q83P01 Q83P02 Q83P06 Q83P19
Q83P26 Q83P27 Q83P30 Q83P33 Q83P35 Q83P43 Q83P53 Q83P61 Q83P64 Q83P77 Q83PB5
Q83PB7 Q83PB8 Q83PB9 Q83PC0 Q83PC2 Q83PC3 Q83PC5 Q83PC7 Q83PD4 Q83PD9 Q83PE0
Q83PE1 Q83PE5 Q83PE8 Q83PF0 Q83PF4 Q83PG0 Q83PH4 Q83PI3 Q83PI6 Q83PJ1 Q83PJ3
Q83PJ4 Q83PJ6 Q83PJ7 Q83PL3 Q83PM8 Q83PM9 Q83PN2 Q83PN3 Q83PQ0 Q83PQ6 Q83PR0
Q83PR1 Q83PS8 Q83PV3 Q83PV4 Q83PW0 Q83PX0 Q83PX1 Q83PX4 Q83PY1 Q83PY4 Q83PY5
Q83PY6 Q83PY7 Q83PY8 Q83PY9 Q83PZ0 Q83PZ1 Q83PZ3 Q83PZ5 Q83Q04 Q83Q07 Q83Q08
Q83Q14 Q83Q15 Q83Q42 Q83Q43 Q83Q91 Q83Q93 Q83Q96 Q83QA2 Q83QB2 Q83QC1 Q83QC7
Q83QD5 Q83QD9 Q83QE0 Q83QE3 Q83QE9 Q83QI2 Q83QI6 Q83QJ8 Q83QK3 Q83QK4 Q83QK5
Q83QL0 Q83QL1 Q83QL4 Q83QM8 Q83QN1 Q83QN4 Q83QQ5 Q83QQ8 Q83QR1 Q83QR3 Q83QS1
Q83QS4 Q83QT4 Q83QT6 Q83QW6 Q83QY5 Q83QZ5 Q83R03 Q83R04 Q83R05 Q83R06 Q83R14
Q83R35 Q83R57 Q83R81 Q83R90 Q83RA1 Q83RA2 Q83RC0 Q83RC9 Q83RE0 Q83RE1 Q83RE2
Q83RE8 Q83RF3 Q83RF4 Q83RG1 Q83RG5 Q83RI9 Q83RJ2 Q83RJ3 Q83RK8 Q83RM1 Q83RP1
Q83RR6 Q83RR8 Q83RR9 Q83RT8 Q83RT9 Q83RV3 Q83RV4 Q83RX1 Q83RX5 Q83RY8 Q83S05
Q83S14 Q83S18 Q83S40 Q83S42 Q83S45 Q83S46 Q83S80 Q83S86 Q83SC8 Q83SD5 Q83SE1
Q83SE5 Q83SE6 Q83SG1 Q83SG2 Q83SH9 Q83SI1 Q83SL4 Q83SL9 Q83SM0 Q83SM9 Q83SN7
Q83SP0 Q83SP1 Q83SP3 Q83SQ1 Q83SQ2 Q83SQ6 Q83SQ7 Q83SQ8 Q83SQ9 Q83SR1 Q83SW5
Q83TH2 Q83TH3 Q83TH4 Q83TK1 Q83U21 Q83U59 Q83V26 Q83VZ3 Q83VZ4 Q83WC3 Q83WC4
Q83WC8 Q83WF2 Q83WF4 Q83WF5 Q83WI8 Q83WI9 Q83WJ5 Q83WL3 Q83WL4 Q83WS0 Q83WU4
Q83XD3 Q83XM3 Q83YI7 Q83YR8 Q84122 Q84123 Q84132 Q84133 Q84135 Q84136 Q84169
Q84173 Q84176 Q84177 Q841V6 Q84240 Q84291 Q84294 Q84295 Q84296 Q84361 Q84362
Q84424 Q84513 Q845S7 Q845S8 Q845S9 Q845U5 Q845U6 Q845U7 Q845U8 Q845U9 Q845V1
Q845V2 Q845V3 Q845V4 Q845W3 Q845X9 Q84687 Q84688 Q846B4 Q846V6 Q84709 Q84711
Q847G1 Q847R1 Q847S1 Q84842 Q84843 Q848I4 Q848I5 Q848R7 Q848W2 Q84939 Q84940
Q849P2 Q84AP0 Q84AP1 Q84B70 Q84BK9 Q84BL4 Q84BP4 Q84BQ9 Q84BU3 Q84BU4 Q84BU5
Q84BU6 Q84BV2 Q84BZ0 Q84BZ1 Q84BZ2 Q84BZ3 Q84BZ4 Q84CG1 Q84CH1 Q84CP9 Q84CZ9

Q84D00 Q84DC3 Q84DC4 Q84F14 Q84F15 Q84F16 Q84F85 Q84F86 Q84F94 Q84FD3 Q84FH6
Q84FI0 Q84FI1 Q84FY8 Q84G06 Q84GJ3 Q84GJ9 Q84H41 Q84H44 Q84HB6 Q84HC5 Q84HC6
Q84HC8 Q84HF4 Q84HF5 Q84HF6 Q84HH6 Q84HH8 Q84I51 Q84I52 Q84I53 Q84I54 Q84I55
Q84I56 Q84IF4 Q84IF5 Q84IL8 Q84J39 Q84J46 Q84J50 Q84J62 Q84J70 Q84J71 Q84J78
Q84JC0 Q84JC2 Q84JE8 Q84JF0 Q84JF4 Q84JF7 Q84JG2 Q84JH5 Q84JH7 Q84JI6 Q84JK2
Q84JL2 Q84JL3 Q84JL5 Q84JM4 Q84JN6 Q84JP1 Q84JQ8 Q84JR3 Q84JR9 Q84JS6 Q84JT6
Q84JT7 Q84JZ6 Q84JZ8 Q84K34 Q84K35 Q84K52 Q84K79 Q84K90 Q84K96 Q84KG5 Q84KI6
Q84KJ3 Q84KJ5 Q84KK4 Q84KK5 Q84KK6 Q84KL3 Q84KL4 Q84KL5 Q84KL6 Q84KP0 Q84L14
Q84L30 Q84L31 Q84L32 Q84L33 Q84LB2 Q84LB6 Q84LF0 Q84LF1 Q84LF2 Q84LG0 Q84LG2
Q84LH3 Q84LK7 Q84LM4 Q84M85 Q84M91 Q84M92 Q84M94 Q84M96 Q84M98 Q84MA1 Q84MA2
Q84MB2 Q84MB3 Q84MB6 Q84MC1 Q84MD6 Q84MD8 Q84MH1 Q84MM2 Q84MM9 Q84MN0 Q84MP7
Q84MQ9 Q84N28 Q84N37 Q84N38 Q84N48 Q84N49 Q84NC2 Q84NC5 Q84NC8 Q84NC9 Q84ND0
Q84ND9 Q84NJ4 Q84NN4 Q84P52 Q84P53 Q84P54 Q84PB3 Q84PD8 Q84PP7 Q84PW3 Q84Q72
Q84Q77 Q84Q92 Q84QC1 Q84QM3 Q84R11 Q84R27 Q84R36 Q84RD0 Q84RD1 Q84RE1 Q84RF2
Q84RK2 Q84RK6 Q84RQ7 Q84RQ8 Q84RQ9 Q84RR0 Q84RR2 Q84SC3 Q84SL2 Q84SM8 Q84T03
Q84T08 Q84T65 Q84T68 Q84T92 Q84TA3 Q84TB3 Q84TB6 Q84TC2 Q84TD6 Q84TD8 Q84TE9
Q84TF0 Q84TF4 Q84TF5 Q84TG1 Q84TG3 Q84TG9 Q84TH4 Q84TI3 Q84TI6 Q84TK0 Q84TK1
Q84TQ7 Q84TV4 Q84TX2 Q84U21 Q84U86 Q84UB4 Q84UB5 Q84UI6 Q84UL5 Q84UQ1 Q84UQ3
Q84UR0 Q84UU4 Q84UU9 Q84UV0 Q84V03 Q84V15 Q84V25 Q84V36 Q84V37 Q84V83 Q84VA7
Q84VG0 Q84VG6 Q84VG7 Q84VT7 Q84VV6 Q84VW5 Q84VW9 Q84VX0 Q84VX3 Q84VX4 Q84VZ1
Q84W55 Q84W56 Q84W65 Q84W80 Q84W89 Q84W91 Q84W92 Q84WD3 Q84WI0 Q84WJ0 Q84WJ2
Q84WJ9 Q84WK0 Q84WK5 Q84WL6 Q84WM0 Q84WP3 Q84WP6 Q84WQ5 Q84WS8 Q84WU4 Q84WU6
Q84WU9 Q84WV0 Q84WV1 Q84WV8 Q84WW1 Q84WW6 Q84WZ8 Q84X02 Q84X07 Q84X17 Q84X40
Q84X43 Q84X53 Q84XA3 Q84XV2 Q84XW5 Q84Y18 Q84YL3 Q84Z02 Q84ZA1 Q84ZD2 Q84ZT0
Q84ZT9 Q84ZX8 Q85014 Q85036 Q85055 Q85056 Q851C7 Q851Q6 Q851R2 Q851R5 Q851S7
Q851S8 Q851V1 Q851V5 Q851W4 Q851W5 Q851Y7 Q85228 Q85232 Q85280 Q85297 Q852K3
Q852K5 Q852K6 Q852K8 Q852M0 Q852M1 Q852M2 Q852N2 Q852S5 Q852U6 Q85318 Q85365
Q853F7 Q853W0 Q85424 Q85428 Q85431 Q85433 Q85434 Q85435 Q85436 Q85437 Q85438
Q85440 Q85442 Q85443 Q85446 Q85447 Q85448 Q85450 Q85672 Q856K7 Q857H2 Q858H1
Q85967 Q859K5 Q859K6 Q859P9 Q859Q5 Q859W3 Q859W7 Q85A01 Q85A69 Q85A72 Q85A82
Q85AC2 Q85AI6 Q85AL7 Q85AM2 Q85AT7 Q85AW7 Q85AX4 Q85B41 Q85B60 Q85B65 Q85B74
Q85BJ7 Q85BK3 Q85BR7 Q85BU5 Q85BW1 Q85BW6 Q85BX6 Q85BZ1 Q85C28 Q85C49 Q85C71
Q85C77 Q85CF2 Q85CH6 Q85CL6 Q85CS9 Q85CT4 Q85CU9 Q85CV5 Q85FG3 Q85FG4 Q85FG5
Q85FG6 Q85FG8 Q85FH7 Q85FH8 Q85FI0 Q85FI1 Q85FI2 Q85FI3 Q85FI4 Q85FI5 Q85FI6
Q85FI7 Q85FI8 Q85FI9 Q85FJ0 Q85FJ1 Q85FJ8 Q85FJ9 Q85FK0 Q85FK1 Q85FL8 Q85FM1
Q85FM7 Q85FM8 Q85FM9 Q85FN0 Q85FN8 Q85FN9 Q85FP0 Q85FR4 Q85FR5 Q85FR6 Q85FR7
Q85FR8 Q85FR9 Q85FT7 Q85FT8 Q85FT9 Q85FU0 Q85FU3 Q85FU4 Q85FU5 Q85FU7 Q85FU8
Q85FV0 Q85FV1 Q85FV2 Q85FV3 Q85FV4 Q85FV6 Q85FV7 Q85FV8 Q85FV9 Q85FW0 Q85FW2
Q85FW3 Q85FW4 Q85FW5 Q85FX1 Q85FX4 Q85FY4 Q85FY5 Q85G10 Q85G11 Q85G12 Q85G22
Q85G24 Q85G29 Q85G31 Q85G45 Q85G60 Q85G62 Q85G72 Q85G77 Q85G81 Q85G82 Q85G85
Q85UK6 Q85UY0 Q85UZ5 Q85UZ8 Q85V01 Q85V90 Q85V93 Q85V94 Q85WB7 Q85WL0 Q85WL2
Q85WS5 Q85WS6 Q85WS8 Q85WT6 Q85WU5 Q85WU8 Q85WV0 Q85WV2 Q85WV5 Q85WV9 Q85WW7
Q85WX8 Q85WY6 Q85WY8 Q85WY9 Q85WZ0 Q85WZ6 Q85WZ7 Q85WZ9 Q85X01 Q85X35 Q85X37
Q85X43 Q85X49 Q85X50 Q85X54 Q85X62 Q85X64 Q85X76 Q85XY6 Q85XY8 Q85ZM2 Q85ZS6

Q85ZS9 Q85ZU4 Q85ZU5 Q85ZV6 Q86108 Q86112 Q86117 Q86119 Q86132 Q86134 Q86194
Q86195 Q86197 Q86199 Q861Q9 Q861R7 Q861U7 Q861U8 Q861U9 Q861V0 Q861W0 Q861Y6
Q86217 Q86218 Q86281 Q86282 Q86284 Q86285 Q86286 Q86287 Q862A7 Q862Z4 Q862Z5
Q86305 Q863A2 Q863A4 Q863A7 Q863B6 Q863B7 Q863C0 Q863C2 Q863D4 Q863D9 Q863Z0
Q863Z4 Q86484 Q864P3 Q864P4 Q864P5 Q864P6 Q864P7 Q864P8 Q864P9 Q864Q0 Q864Q1
Q864Q2 Q864Q3 Q864Q4 Q864Q5 Q864Q6 Q864Q7 Q864Q8 Q864Q9 Q864R0 Q864R1 Q864R2
Q86504 Q86505 Q86517 Q86523 Q86563 Q86564 Q865B6 Q865B7 Q865C5 Q865F8 Q865U8
Q865V0 Q865W2 Q86606 Q86609 Q86625 Q866P6 Q86780 Q86783 Q867T7 Q867X0 Q867Z4
Q868H8 Q86993 Q869E1 Q869K3 Q869L3 Q869L4 Q869N6 Q869N9 Q869P0 Q869Q0 Q869Q2
Q869Q3 Q869R8 Q869R9 Q869S7 Q869S8 Q869S9 Q869T1 Q869T2 Q869T7 Q869W0 Q869W6
Q869W9 Q869X2 Q869X7 Q869Y7 Q86A05 Q86A14 Q86A17 Q86A21 Q86A24 Q86A67 Q86A68
Q86A84 Q86A85 Q86A88 Q86A90 Q86A92 Q86A97 Q86AA1 Q86AD2 Q86AD5 Q86AD6 Q86AD7
Q86AD9 Q86AE7 Q86AF2 Q86AF3 Q86AG8 Q86AH1 Q86AH2 Q86AH4 Q86AQ5 Q86AS0 Q86AS5
Q86AS6 Q86AU1 Q86AV6 Q86AW9 Q86AX3 Q86B00 Q86B05 Q86B06 Q86B09 Q86B11 Q86B14
Q86B20 Q86B47 Q86B79 Q86B87 Q86BL4 Q86BP6 Q86BS6 Q86BY9 Q86CR9 Q86CT3 Q86D25
Q86DC6 Q86DS1 Q86E67 Q86FP7 Q86H28 Q86H36 Q86H39 Q86H43 Q86H45 Q86H49 Q86H59
Q86H60 Q86H62 Q86H78 Q86H86 Q86H87 Q86H91 Q86H93 Q86H94 Q86HA0 Q86HB7 Q86HD3
Q86HF4 Q86HG0 Q86HG3 Q86HG4 Q86HG9 Q86HL1 Q86HN0 Q86HN7 Q86HQ8 Q86HS5 Q86HS8
Q86HS9 Q86HT3 Q86HV3 Q86HW0 Q86HW6 Q86HX0 Q86HX1 Q86HX2 Q86HX9 Q86I06 Q86I14
Q86I22 Q86I25 Q86I32 Q86I40 Q86I44 Q86I75 Q86I79 Q86I85 Q86I89 Q86I91 Q86I92
Q86IA1 Q86IA8 Q86IB5 Q86IC8 Q86IC9 Q86IG2 Q86IG7 Q86IH1 Q86II4 Q86II5 Q86IJ1
Q86IJ5 Q86IJ6 Q86IK1 Q86IL5 Q86IM3 Q86IV7 Q86IV8 Q86IW1 Q86IW8 Q86IX3 Q86IY1
Q86IY6 Q86IZ3 Q86IZ9 Q86J11 Q86J12 Q86J13 Q86J23 Q86J25 Q86JA1 Q86JB0 Q86JD9
Q86JF8 Q86JG2 Q86JI5 Q86JJ0 Q86JJ5 Q86JL6 Q86JM3 Q86JM4 Q86K32 Q86K56 Q86K64
Q86K86 Q86K91 Q86KA2 Q86KD0 Q86KD1 Q86KD3 Q86KE7 Q86KF5 Q86KH4 Q86KK2 Q86KL1
Q86KL4 Q86KR9 Q86KS1 Q86KT9 Q86KU0 Q86KU2 Q86KU6 Q86KZ2 Q86KZ5 Q86L14 Q86LA2
Q86MA2 Q86MV5 Q86MW9 Q86NP2 Q86P48 Q86QH9 Q86QI0 Q86QS6 Q86QW7 Q86RN8 Q86S05
Q86SE5 Q86SE9 Q86SG2 Q86SG3 Q86SG4 Q86SG5 Q86SH2 Q86SX3 Q86SX6 Q86SY8 Q86SZ2
Q86T23 Q86T29 Q86T65 Q86T75 Q86TA4 Q86TI4 Q86TJ2 Q86TJ5 Q86TP1 Q86TU6 Q86TU7
Q86TZ1 Q86U28 Q86U37 Q86UA1 Q86UA6 Q86UB2 Q86UC2 Q86UD4 Q86UD7 Q86UE3 Q86UF4
Q86UN6 Q86UP3 Q86UP8 Q86UQ0 Q86UQ8 Q86UT8 Q86UU5 Q86UV6 Q86UV7 Q86UY8 Q86UZ6
Q86V20 Q86V42 Q86V59 Q86V71 Q86V86 Q86V87 Q86VD1 Q86VE3 Q86VG3 Q86VH2 Q86VI1
Q86VK4 Q86VQ0 Q86VQ3 Q86VQ6 Q86VS3 Q86VV4 Q86VV8 Q86VX2 Q86VY4 Q86VZ2 Q86W11
Q86W24 Q86W25 Q86W28 Q86W34 Q86W42 Q86W50 Q86W54 Q86W67 Q86WC6 Q86WQ0 Q86WR6
Q86WS4 Q86WT1 Q86WT6 Q86WV5 Q86WZ0 Q86WZ6 Q86X02 Q86X45 Q86X51 Q86X59 Q86X67
Q86X76 Q86X83 Q86XA0 Q86XA9 Q86XD8 Q86XE0 Q86XE5 Q86XF7 Q86XG9 Q86XH1 Q86XI8
Q86XN6 Q86XN7 Q86XN8 Q86XU0 Q86XW9 Q86Y13 Q86Y25 Q86Y26 Q86Y33 Q86Y37 Q86Y46
Q86Y56 Q86Y79 Q86YD7 Q86YE8 Q86YG4 Q86YH2 Q86YN6 Q86YV6 Q86Z42 Q86Z65 Q86Z98
Q86ZC1 Q86ZD9 Q86ZF1 Q86ZF9 Q86ZM3 Q86ZN7 Q86ZP5 Q86ZR7 Q86ZU6 Q86ZV0 Q87010
Q87011 Q87012 Q87014 Q87019 Q87022 Q87026 Q870A3 Q870B9 Q870G1 Q870I4 Q870Q1
Q870S2 Q870W1 Q870Y1 Q871A4 Q871G3 Q871M9 Q871N9 Q871U2 Q871W6 Q871Z4 Q872F4
Q872I5 Q872T7 Q872U8 Q873B7 Q873B8 Q873C7 Q873E4 Q873E8 Q873H9 Q873W8 Q873X0
Q873Y3 Q874A5 Q874B3 Q874B9 Q874J6 Q874L7 Q874M2 Q874N0 Q874N1 Q874P0 Q874R3
Q874T6 Q874T7 Q874W3 Q874Z5 Q875B6 Q875B7 Q875B8 Q875D4 Q875I3 Q875L2 Q875L3

Q875M9 Q875N2 Q875P5 Q875Q8 Q875S0 Q875S4 Q875V0 Q875X3 Q875X4 Q875Z2 Q875Z9
Q876B2 Q876B4 Q876G9 Q876L8 Q877B5 Q877B6 Q877G5 Q877H6 Q877I0 Q877L9 Q877P8
Q877T5 Q878E3 Q879S2 Q879S5 Q879Y6 Q879Z0 Q87A16 Q87A17 Q87A18 Q87A21 Q87A22
Q87A23 Q87A24 Q87A27 Q87A28 Q87A29 Q87A30 Q87A31 Q87A32 Q87A33 Q87A34 Q87A35
Q87A39 Q87A46 Q87A51 Q87A52 Q87A63 Q87A70 Q87A73 Q87A76 Q87A82 Q87A84 Q87A85
Q87A88 Q87A92 Q87A98 Q87AA7 Q87AB0 Q87AB2 Q87AB3 Q87AB4 Q87AB5 Q87AB6 Q87AB7
Q87AC0 Q87AC5 Q87AC9 Q87AD1 Q87AD5 Q87AD6 Q87AE8 Q87AE9 Q87AF1 Q87AF2 Q87AF5
Q87AG0 Q87AG1 Q87AH7 Q87AH8 Q87AI0 Q87AI5 Q87AJ1 Q87AJ9 Q87AK0 Q87AK1 Q87AK5
Q87AK6 Q87AP2 Q87AP4 Q87AQ5 Q87AQ6 Q87AQ7 Q87AR0 Q87AR1 Q87AR7 Q87AR8 Q87AS0
Q87AS2 Q87AS3 Q87AS7 Q87AS8 Q87AT3 Q87AT6 Q87AV1 Q87AV9 Q87AX5 Q87AX6 Q87AX8
Q87AY2 Q87AY6 Q87AY7 Q87AZ9 Q87B10 Q87B16 Q87B17 Q87B25 Q87B33 Q87B35 Q87B36
Q87B41 Q87B49 Q87B68 Q87B70 Q87B72 Q87B73 Q87B82 Q87B84 Q87B85 Q87B89 Q87B94
Q87BA5 Q87BC0 Q87BC5 Q87BE2 Q87BE3 Q87BG0 Q87BG1 Q87BG5 Q87BG6 Q87BG9 Q87BI2
Q87BI4 Q87BJ5 Q87BJ6 Q87BJ8 Q87BK4 Q87BK6 Q87BK9 Q87BL0 Q87BL1 Q87BL2 Q87BL5
Q87BP7 Q87BP9 Q87BQ0 Q87BQ1 Q87BR3 Q87BR5 Q87BR7 Q87BS0 Q87BS2 Q87BS5 Q87BS6
Q87BS7 Q87BS8 Q87BS9 Q87BT4 Q87BU0 Q87BU2 Q87BV3 Q87BW3 Q87BX2 Q87BX8 Q87BY1
Q87BY8 Q87BZ2 Q87C00 Q87C03 Q87C04 Q87C06 Q87C18 Q87C20 Q87C22 Q87C24 Q87C26
Q87C28 Q87C29 Q87C30 Q87C31 Q87C32 Q87C33 Q87C34 Q87C35 Q87C36 Q87C37 Q87C38
Q87C43 Q87C45 Q87C62 Q87C65 Q87C74 Q87C76 Q87C83 Q87C84 Q87C85 Q87C86 Q87CA2
Q87CB7 Q87CC9 Q87CI8 Q87CJ5 Q87CK2 Q87CK7 Q87CK8 Q87CL6 Q87CL8 Q87CM2 Q87CU3
Q87CW0 Q87CW1 Q87CY6 Q87CZ0 Q87CZ1 Q87D00 Q87D02 Q87D03 Q87D04 Q87D06 Q87D16
Q87D18 Q87D19 Q87D24 Q87D27 Q87D42 Q87D45 Q87D46 Q87D47 Q87D58 Q87D59 Q87D63
Q87D65 Q87D69 Q87D75 Q87D84 Q87D89 Q87DA0 Q87DA5 Q87DB3 Q87DC2 Q87DD1 Q87DD3
Q87DD4 Q87DD6 Q87DF7 Q87DF8 Q87DG2 Q87DG3 Q87DG5 Q87DG6 Q87DG7 Q87DG8 Q87DH0
Q87DH1 Q87DH6 Q87DI1 Q87DI2 Q87DI4 Q87DJ2 Q87DJ6 Q87DK2 Q87DL5 Q87DL6 Q87DL8
Q87DM0 Q87DM6 Q87DM8 Q87DN0 Q87DN2 Q87DN8 Q87DP7 Q87DP8 Q87DQ5 Q87DR1 Q87DR8
Q87DR9 Q87DS0 Q87DS1 Q87DS4 Q87DS5 Q87DS9 Q87DT2 Q87DT3 Q87DT7 Q87DU4 Q87DU6
Q87DU7 Q87DU8 Q87DU9 Q87DV0 Q87DW2 Q87DW7 Q87DY3 Q87DY4 Q87DY8 Q87DZ7 Q87DZ8
Q87DZ9 Q87E04 Q87E07 Q87E35 Q87E48 Q87E50 Q87E51 Q87E52 Q87E59 Q87E60 Q87E61
Q87E63 Q87E64 Q87E65 Q87E66 Q87E67 Q87E68 Q87E69 Q87E70 Q87E71 Q87E72 Q87E73
Q87E74 Q87E75 Q87E76 Q87E77 Q87E78 Q87E79 Q87E80 Q87E81 Q87E82 Q87E99 Q87EA3
Q87EA7 Q87EB3 Q87EB7 Q87EB8 Q87EB9 Q87EC0 Q87EC4 Q87EF7 Q87EG6 Q87EG7 Q87EH1
Q87EH6 Q87EH9 Q87EI6 Q87EI8 Q87EJ8 Q87EK9 Q87EL0 Q87EL1 Q87EL2 Q87EL4 Q87EL7
Q87EM8 Q87EN0 Q87EQ7 Q87ES6 Q87EV0 Q87EV1 Q87EV2 Q87EV3 Q87EV4 Q87EV6 Q87EV7
Q87EV8 Q87EV9 Q87EW0 Q87EW6 Q87EX2 Q87EX3 Q87EX9 Q87EY0 Q87EZ6 Q87EZ7 Q87F03
Q87F06 Q87F08 Q87F16 Q87F19 Q87F21 Q87F22 Q87F25 Q87F26 Q87F28 Q87F32 Q87F36
Q87F43 Q87F44 Q87F49 Q87F51 Q87F53 Q87F54 Q87F55 Q87F56 Q87F63 Q87F66 Q87F70
Q87F71 Q87F85 Q87F88 Q87F90 Q87F94 Q87FA3 Q87FB2 Q87FC4 Q87FC6 Q87FD6 Q87FE9
Q87FH3 Q87FH9 Q87FK3 Q87FK5 Q87FP6 Q87FP9 Q87FQ5 Q87FR0 Q87FT1 Q87FZ3 Q87G42
Q87GA3 Q87GM3 Q87GR5 Q87GT9 Q87GU2 Q87GU3 Q87GX2 Q87GY4 Q87GY5 Q87GZ6 Q87H06
Q87H51 Q87H52 Q87H78 Q87HN1 Q87HP7 Q87HS0 Q87HS9 Q87HX2 Q87HX3 Q87I03 Q87I04
Q87I05 Q87I22 Q87I38 Q87ID3 Q87IJ5 Q87IJ9 Q87IL9 Q87IM1 Q87IM9 Q87IV1 Q87J02
Q87J41 Q87J46 Q87J80 Q87J85 Q87JD1 Q87JE4 Q87JG6 Q87JG7 Q87JH5 Q87JL4 Q87JL6
Q87JP5 Q87JQ6 Q87JS5 Q87JS8 Q87JV8 Q87JW7 Q87JW8 Q87JZ5 Q87K03 Q87K41 Q87K60

Q87K87 Q87K98 Q87K99 Q87KB6 Q87KD3 Q87KD4 Q87KD5 Q87KD8 Q87KE0 Q87KE1 Q87KE2
Q87KE3 Q87KE4 Q87KF0 Q87KF4 Q87KG2 Q87KH4 Q87KH5 Q87KI9 Q87KJ1 Q87KJ4 Q87KJ6
Q87KM5 Q87KM8 Q87KN5 Q87KN8 Q87KP1 Q87KP5 Q87KP9 Q87KQ0 Q87KQ1 Q87KQ2 Q87KQ3
Q87KQ4 Q87KQ5 Q87KQ7 Q87KR0 Q87KS8 Q87KT0 Q87KU1 Q87KU2 Q87KU6 Q87KU7 Q87KX0
Q87KX3 Q87KX9 Q87KY6 Q87KZ3 Q87KZ5 Q87L00 Q87L01 Q87L05 Q87L06 Q87L07 Q87L11
Q87L12 Q87L13 Q87L20 Q87L22 Q87L25 Q87L36 Q87L40 Q87L41 Q87L43 Q87L44 Q87L45
Q87L49 Q87L54 Q87L55 Q87L56 Q87L67 Q87L68 Q87L72 Q87L73 Q87L75 Q87L78 Q87L81
Q87L85 Q87L90 Q87L91 Q87L92 Q87L95 Q87LA0 Q87LA3 Q87LC4 Q87LD0 Q87LD3 Q87LD5
Q87LD9 Q87LF4 Q87LF7 Q87LF8 Q87LF9 Q87LG0 Q87LG6 Q87LG8 Q87LH9 Q87LI5 Q87LI7
Q87LI9 Q87LJ1 Q87LJ3 Q87LJ9 Q87LK0 Q87LK1 Q87LK4 Q87LK6 Q87LK8 Q87LL0 Q87LL1
Q87LL6 Q87LL9 Q87LM3 Q87LM8 Q87LM9 Q87LN9 Q87LP1 Q87LP2 Q87LP5 Q87LP9 Q87LQ2
Q87LQ3 Q87LQ4 Q87LQ5 Q87LQ6 Q87LQ9 Q87LR1 Q87LR2 Q87LR3 Q87LR5 Q87LS2 Q87LS8
Q87LS9 Q87LT0 Q87LT1 Q87LT2 Q87LT3 Q87LT4 Q87LV1 Q87LV2 Q87LV6 Q87LV9 Q87LW4
Q87LY1 Q87LY3 Q87LY4 Q87LY8 Q87LZ2 Q87LZ4 Q87LZ7 Q87M00 Q87M02 Q87M03 Q87M04
Q87M05 Q87M06 Q87M18 Q87M22 Q87M23 Q87M24 Q87M25 Q87M30 Q87M60 Q87M72 Q87M87
Q87MA4 Q87MA6 Q87MB4 Q87MC1 Q87MC3 Q87MC5 Q87MC8 Q87MD4 Q87MD5 Q87MD6 Q87MD8
Q87MD9 Q87ME0 Q87ME3 Q87MF1 Q87MF4 Q87MF7 Q87MG0 Q87MG2 Q87MH0 Q87MH1 Q87MI6
Q87MM7 Q87MM9 Q87MN6 Q87MN8 Q87MP1 Q87MQ1 Q87MQ3 Q87MQ4 Q87MQ8 Q87MR2 Q87MS5
Q87MV2 Q87MV3 Q87MX6 Q87MY0 Q87MY2 Q87MZ4 Q87N01 Q87N07 Q87N09 Q87N15 Q87N16
Q87N18 Q87N26 Q87N39 Q87N44 Q87N46 Q87N49 Q87N95 Q87NA1 Q87NA8 Q87NB7 Q87NB9
Q87NC0 Q87NC4 Q87ND5 Q87NE7 Q87NH5 Q87NU5 Q87NW7 Q87NZ6 Q87NZ7 Q87NZ8 Q87P80
Q87P89 Q87P91 Q87PA5 Q87PB0 Q87PC0 Q87PC8 Q87PG9 Q87PH8 Q87PK5 Q87PK8 Q87PM1
Q87PP9 Q87Q20 Q87Q26 Q87Q30 Q87Q40 Q87Q46 Q87Q52 Q87Q53 Q87Q54 Q87Q56 Q87Q59
Q87Q60 Q87Q67 Q87Q69 Q87Q70 Q87Q74 Q87Q75 Q87Q76 Q87Q77 Q87Q87 Q87Q92 Q87QA2
Q87QA3 Q87QD4 Q87QD9 Q87QG0 Q87QJ8 Q87QJ9 Q87QK4 Q87QK5 Q87QK6 Q87QK7 Q87QK8
Q87QK9 Q87QL0 Q87QL1 Q87QL2 Q87QL9 Q87QN3 Q87QN5 Q87QN6 Q87QP1 Q87QT6 Q87QU7
Q87QU8 Q87QV1 Q87QV2 Q87QV4 Q87QV5 Q87QW2 Q87QW3 Q87QW4 Q87QW8 Q87QW9 Q87QX5
Q87QX6 Q87QX9 Q87QY1 Q87QY2 Q87QY5 Q87QY8 Q87R08 Q87R12 Q87R29 Q87R69 Q87R72
Q87R73 Q87R79 Q87R80 Q87R81 Q87R88 Q87RB8 Q87RC2 Q87RC3 Q87RC5 Q87RD1 Q87RD6
Q87RE8 Q87RG4 Q87RH3 Q87RH4 Q87RH5 Q87RJ4 Q87RL6 Q87RN4 Q87RN5 Q87RN7 Q87RN8
Q87RN9 Q87RP4 Q87RP6 Q87RQ0 Q87RQ4 Q87RQ9 Q87RR0 Q87RR1 Q87RR2 Q87RS6 Q87RT0
Q87RT2 Q87RT6 Q87RT8 Q87RU0 Q87RU3 Q87RU4 Q87RU8 Q87RU9 Q87RV0 Q87RV1 Q87RV2
Q87RW0 Q87RX2 Q87RX3 Q87RX5 Q87RX6 Q87RY0 Q87RY2 Q87S07 Q87S09 Q87S12 Q87S15
Q87S16 Q87S19 Q87S20 Q87S21 Q87S24 Q87S25 Q87S28 Q87S29 Q87S36 Q87S37 Q87S45
Q87S51 Q87S56 Q87S63 Q87S65 Q87S70 Q87S71 Q87S80 Q87S81 Q87S87 Q87S90 Q87S93
Q87SA0 Q87SA4 Q87SA6 Q87SB1 Q87SB8 Q87SC0 Q87SC4 Q87SC6 Q87SC7 Q87SC9 Q87SD2
Q87SE5 Q87SF3 Q87SF4 Q87SF5 Q87SG3 Q87SG6 Q87SG9 Q87SH5 Q87SI4 Q87SI5 Q87SK9
Q87SL5 Q87SQ3 Q87SR3 Q87SS7 Q87SS8 Q87SS9 Q87ST0 Q87ST5 Q87ST6 Q87ST7 Q87ST8
Q87SU2 Q87SU3 Q87SU4 Q87SU7 Q87SU8 Q87SW0 Q87SW1 Q87SW6 Q87SX6 Q87SX9 Q87SY0
Q87SY9 Q87SZ0 Q87SZ1 Q87SZ2 Q87SZ4 Q87SZ5 Q87SZ6 Q87SZ7 Q87SZ8 Q87SZ9 Q87T00
Q87T01 Q87T02 Q87T03 Q87T04 Q87T05 Q87T06 Q87T07 Q87T08 Q87T09 Q87T10 Q87T11
Q87T12 Q87T13 Q87T15 Q87T20 Q87T21 Q87T23 Q87T24 Q87T31 Q87T80 Q87T81 Q87T84
Q87T85 Q87T86 Q87T89 Q87T90 Q87T92 Q87T93 Q87TA9 Q87TB0 Q87TB8 Q87TC2 Q87TC4
Q87TE0 Q87TE1 Q87TE4 Q87TF3 Q87TF6 Q87TG0 Q87TH4 Q87TJ6 Q87TN4 Q87TP4 Q87TP7

Q87TP8 Q87TQ5 Q87TQ7 Q87TR3 Q87TR4 Q87TR6 Q87TR8 Q87TR9 Q87TS2 Q87TS3 Q87TS4
Q87TT8 Q87TZ6 Q87U40 Q87UB2 Q87UC3 Q87UC6 Q87UC7 Q87UF5 Q87UG0 Q87UG1 Q87UG3
Q87UG4 Q87UH3 Q87UK7 Q87UL1 Q87UL4 Q87UL9 Q87UM1 Q87UM6 Q87UP6 Q87US2 Q87UX4
Q87UX9 Q87UY8 Q87UY9 Q87UZ2 Q87V00 Q87V01 Q87V03 Q87V05 Q87V14 Q87V15 Q87V23
Q87V55 Q87V56 Q87V59 Q87V73 Q87V83 Q87V88 Q87V90 Q87V99 Q87VA0 Q87VA1 Q87VA2
Q87VA4 Q87VB1 Q87VF4 Q87VG1 Q87VI5 Q87VI6 Q87VJ2 Q87VJ3 Q87VJ4 Q87VJ8 Q87VJ9
Q87VK2 Q87VK3 Q87VK4 Q87VK6 Q87VK8 Q87VL4 Q87VL5 Q87VM6 Q87VM7 Q87VN8 Q87VP0
Q87VP1 Q87VP4 Q87VP5 Q87VQ6 Q87VR8 Q87VR9 Q87VS1 Q87VS3 Q87VS6 Q87VU3 Q87VV5
Q87VV6 Q87VV7 Q87VV9 Q87VW5 Q87VW6 Q87VW7 Q87VX3 Q87VX9 Q87VY1 Q87VY5 Q87VY6
Q87VZ5 Q87W79 Q87WA0 Q87WA9 Q87WC1 Q87WD6 Q87WH1 Q87WJ7 Q87WL6 Q87WN1 Q87WN4
Q87WN9 Q87WP0 Q87WP1 Q87WP2 Q87WP3 Q87WP4 Q87WP5 Q87WP7 Q87WQ0 Q87WQ1 Q87WQ3
Q87WQ5 Q87WQ6 Q87WQ7 Q87WQ8 Q87WR8 Q87WR9 Q87WS0 Q87WS4 Q87WS9 Q87WT7 Q87WV2
Q87WV4 Q87WV5 Q87WV6 Q87WW0 Q87WW4 Q87WW7 Q87WW8 Q87WX3 Q87WX6 Q87WX7 Q87WY0
Q87WY3 Q87WY6 Q87WY7 Q87WZ4 Q87X00 Q87X13 Q87X14 Q87XE2 Q87XF1 Q87XG1 Q87XG3
Q87XG4 Q87XG5 Q87XG6 Q87XJ5 Q87XJ6 Q87XJ9 Q87XL0 Q87XL4 Q87XM0 Q87XM1 Q87XM3
Q87XN0 Q87XN2 Q87XP9 Q87XR0 Q87XS2 Q87XW6 Q87XY7 Q87XY8 Q87Y01 Q87Y22 Q87Y24
Q87Y27 Q87Y31 Q87Y32 Q87Y33 Q87Y34 Q87Y35 Q87Y44 Q87Y50 Q87Y59 Q87Y79 Q87Y82
Q87Y85 Q87YA4 Q87YC5 Q87YC6 Q87YC8 Q87YF6 Q87YF8 Q87YG2 Q87YG4 Q87YH1 Q87YI1
Q87YK2 Q87YQ1 Q87YQ2 Q87YR0 Q87YR5 Q87YR6 Q87YR7 Q87YS8 Q87YT0 Q87YW7 Q87YX3
Q87YX4 Q87YY6 Q87YZ4 Q87Z00 Q87Z65 Q87Z72 Q87Z79 Q87ZC3 Q87ZD5 Q87ZG3 Q87ZG4
Q87ZR6 Q87ZS0 Q87ZS2 Q87ZS3 Q87ZS4 Q87ZS9 Q87ZT0 Q880J5 Q880L8 Q880M9 Q880X7
Q880Z4 Q881F7 Q881H6 Q881T7 Q881X0 Q881Z4 Q882A9 Q882E2 Q882G0 Q882K4 Q882K8
Q882P7 Q882V6 Q882Y4 Q88305 Q883B5 Q883F1 Q883F2 Q883F3 Q883F5 Q883H6 Q883H7
Q883H8 Q883H9 Q883K6 Q883N9 Q883Q6 Q883Q9 Q883R9 Q883T5 Q883Y9 Q883Z4 Q88419
Q88420 Q88423 Q88434 Q88435 Q88469 Q884C0 Q884C1 Q884C2 Q884C6 Q884C8 Q884C9
Q884G7 Q884H4 Q884I1 Q884I6 Q884N3 Q884P1 Q884P2 Q884P3 Q884P5 Q884Q3 Q884R0
Q884R6 Q884R9 Q884S0 Q884S1 Q884U6 Q884X9 Q885A6 Q885J0 Q885J3 Q885J4 Q885J6
Q885J7 Q885K0 Q885K7 Q885L1 Q885Q1 Q885Q4 Q885R6 Q885T0 Q885T2 Q885T5 Q885T7
Q885T9 Q885U7 Q885V0 Q885W6 Q885W8 Q885X7 Q885Y1 Q885Y8 Q885Z7 Q885Z8 Q886D9
Q886E0 Q886E9 Q886F0 Q886I1 Q886I3 Q886L4 Q886L5 Q886L6 Q886L7 Q886M1 Q886M5
Q886M6 Q886M9 Q886N7 Q886P0 Q886P1 Q886P2 Q886P3 Q886P5 Q886Q4 Q886R2 Q886R8
Q886R9 Q886S6 Q886U9 Q886V0 Q886V1 Q886V2 Q886V7 Q886W6 Q886X5 Q886X9 Q886Y1
Q886Y2 Q886Y6 Q886Y9 Q886Z0 Q886Z3 Q886Z4 Q886Z7 Q886Z8 Q887A1 Q887B0 Q887D9
Q887L4 Q887L5 Q887M0 Q887M3 Q887M4 Q887M6 Q887N8 Q887P4 Q887P7 Q887P8 Q887Q0
Q887Q9 Q887V1 Q887Y4 Q887Z4 Q88887 Q88888 Q88890 Q88894 Q88897 Q88898 Q88899
Q888A0 Q888A2 Q888A4 Q888B8 Q888C1 Q888C2 Q888C5 Q888C6 Q888C7 Q888C8 Q888H1
Q888I3 Q888N4 Q888P6 Q888P9 Q888Q5 Q888Q6 Q888Q7 Q888U3 Q888U4 Q888V7 Q88918
Q88920 Q88921 Q88922 Q88936 Q88939 Q889C2 Q889E1 Q889E4 Q889E8 Q889F0 Q889F2
Q889F3 Q889F6 Q889F7 Q889J2 Q889M3 Q889M7 Q889P9 Q889Q1 Q889Q3 Q889Q5 Q889Q6
Q889R0 Q889U1 Q889U5 Q889U6 Q889U7 Q889U8 Q889U9 Q889V2 Q889V3 Q889V4 Q889V5
Q889V6 Q889V7 Q889V8 Q889V9 Q889W0 Q889W1 Q889W2 Q889W3 Q889W4 Q889W5 Q889W6
Q889W7 Q889W8 Q889W9 Q889X0 Q889X1 Q889X2 Q889X3 Q889X4 Q889X5 Q889X6 Q889X7
Q889X8 Q889X9 Q889Y0 Q889Y1 Q889Y2 Q889Y6 Q889Y8 Q889Z1 Q889Z2 Q889Z3 Q889Z9
Q88A03 Q88A04 Q88A13 Q88A30 Q88A45 Q88A46 Q88A47 Q88A48 Q88A49 Q88A51 Q88A53

Q88A57 Q88A58 Q88A79 Q88A80 Q88A81 Q88A82 Q88A83 Q88A84 Q88A97 Q88A98 Q88AB2
Q88AD1 Q88AE7 Q88AE9 Q88AF0 Q88AF5 Q88AF6 Q88AH3 Q88AH6 Q88AI5 Q88AK3 Q88AK7
Q88AK8 Q88AM9 Q88AP4 Q88AR1 Q88AR2 Q88AR9 Q88AS0 Q88AZ4 Q88AZ5 Q88AZ9 Q88B06
Q88B09 Q88B11 Q88B35 Q88B36 Q88B41 Q88B42 Q88B43 Q88B46 Q88B49 Q88B60 Q88B61
Q88B91 Q88B94 Q88BA5 Q88BB6 Q88BC7 Q88BC8 Q88BD1 Q88BD3 Q88BD5 Q88BD7 Q88BE0
Q88BE2 Q88BE3 Q88BK1 Q88BK3 Q88BX8 Q88BY5 Q88C26 Q88C66 Q88C82 Q88C87 Q88C89
Q88C92 Q88C95 Q88C97 Q88C99 Q88CA0 Q88CB1 Q88CB2 Q88CB6 Q88CF1 Q88CF3 Q88CF6
Q88CG0 Q88CG4 Q88CI0 Q88CI7 Q88CI8 Q88CI9 Q88CN0 Q88CN4 Q88CN9 Q88CQ2 Q88CQ5
Q88CQ7 Q88CS6 Q88CS7 Q88CT0 Q88CT3 Q88CU1 Q88CU3 Q88CV1 Q88CV2 Q88CV6 Q88CW2
Q88CW6 Q88CW7 Q88CX4 Q88CX7 Q88CY4 Q88CY9 Q88CZ6 Q88CZ7 Q88CZ9 Q88D02 Q88D14
Q88D15 Q88D17 Q88D27 Q88D28 Q88D30 Q88D32 Q88D33 Q88D35 Q88D59 Q88D60 Q88D64
Q88D93 Q88DA5 Q88DC4 Q88DC5 Q88DD1 Q88DD2 Q88DD3 Q88DD7 Q88DD8 Q88DE7 Q88DE8
Q88DE9 Q88DF1 Q88DF3 Q88DF6 Q88DF7 Q88DH7 Q88DH8 Q88DJ4 Q88DK2 Q88DK3 Q88DK5
Q88DK7 Q88DL4 Q88DL5 Q88DL7 Q88DM3 Q88DM4 Q88DM5 Q88DN1 Q88DN6 Q88DP0 Q88DP1
Q88DT6 Q88DU1 Q88DU2 Q88DU3 Q88DU4 Q88DU5 Q88DU6 Q88DU7 Q88DV0 Q88DV3 Q88DV4
Q88DV5 Q88DV7 Q88DV8 Q88DV9 Q88DW0 Q88DW6 Q88DW7 Q88DW8 Q88DW9 Q88DX4 Q88DX7
Q88DZ0 Q88E14 Q88E20 Q88E47 Q88E69 Q88E76 Q88E79 Q88E89 Q88EA1 Q88EA5 Q88EA7
Q88EC8 Q88EH2 Q88EH6 Q88EI4 Q88EI5 Q88EI7 Q88EI8 Q88EI9 Q88EM0 Q88EQ6 Q88ET3
Q88EV6 Q88F25 Q88F31 Q88F32 Q88F33 Q88F51 Q88FB2 Q88FB9 Q88FF3 Q88FF5 Q88FF8
Q88FI4 Q88FJ7 Q88FN1 Q88FN9 Q88FQ4 Q88FQ8 Q88FR9 Q88FS4 Q88FS6 Q88FS7 Q88FT2
Q88FT3 Q88FX7 Q88FX9 Q88FY0 Q88FY1 Q88FY3 Q88FY4 Q88FY5 Q88G93 Q88GG4 Q88GQ0
Q88GW8 Q88H32 Q88HA3 Q88HH7 Q88HL5 Q88HM9 Q88HV0 Q88I82 Q88IS4 Q88IS9 Q88IU0
Q88IU4 Q88IU5 Q88IX6 Q88IY3 Q88J00 Q88J01 Q88J03 Q88J04 Q88J05 Q88J06 Q88J07
Q88J44 Q88J85 Q88JD1 Q88JG8 Q88JU3 Q88JU7 Q88JX5 Q88JX7 Q88JX8 Q88JX9 Q88JY0
Q88JY1 Q88K22 Q88K23 Q88K24 Q88K25 Q88K26 Q88K27 Q88K33 Q88K50 Q88K57 Q88K58
Q88K84 Q88K93 Q88KB2 Q88KH5 Q88KI9 Q88KJ0 Q88KJ1 Q88KM5 Q88KT0 Q88KT1 Q88KV1
Q88KX0 Q88KX1 Q88KX9 Q88KY8 Q88L20 Q88L21 Q88L23 Q88L39 Q88L51 Q88L54 Q88L76
Q88LC3 Q88LE0 Q88LE5 Q88LE7 Q88LE8 Q88LF2 Q88LF6 Q88LF9 Q88LI7 Q88LK1 Q88LL9
Q88LM1 Q88LM5 Q88LM8 Q88LQ6 Q88LQ9 Q88LS0 Q88LS1 Q88LU7 Q88LV4 Q88LW2 Q88LW9
Q88M04 Q88M07 Q88M09 Q88M10 Q88M11 Q88M17 Q88M20 Q88M24 Q88M28 Q88M41 Q88M43
Q88M44 Q88M72 Q88M95 Q88M97 Q88MA9 Q88MB9 Q88MD2 Q88MD4 Q88ME3 Q88ME4 Q88ME7
Q88MF0 Q88MF1 Q88MF2 Q88MF3 Q88MF7 Q88MG1 Q88MG3 Q88MG6 Q88MH4 Q88MH7 Q88MH8
Q88MH9 Q88MI0 Q88MI2 Q88MP1 Q88MP5 Q88MQ7 Q88MR4 Q88MS3 Q88MS8 Q88MV0 Q88MV3
Q88MV4 Q88MV5 Q88MV6 Q88MW1 Q88MX7 Q88MX8 Q88MY2 Q88MY3 Q88MY5 Q88MZ4 Q88N05
Q88N27 Q88N29 Q88N55 Q88N56 Q88N74 Q88N75 Q88N78 Q88N81 Q88N84 Q88N85 Q88N88
Q88N96 Q88N97 Q88NA1 Q88NA9 Q88NB7 Q88NC0 Q88NC3 Q88NC4 Q88NC6 Q88ND5 Q88NF3
Q88NF9 Q88NG9 Q88NI3 Q88NJ0 Q88NJ1 Q88NJ2 Q88NJ3 Q88NJ4 Q88NJ9 Q88NK2 Q88NK4
Q88NL0 Q88NN6 Q88NR0 Q88NS5 Q88NT5 Q88NV4 Q88NV7 Q88NW7 Q88NW8 Q88NX4 Q88NX8
Q88P16 Q88P21 Q88P26 Q88P28 Q88P32 Q88P52 Q88P53 Q88P61 Q88P64 Q88P65 Q88P73
Q88P76 Q88P77 Q88P80 Q88P81 Q88P86 Q88P87 Q88P88 Q88PA1 Q88PA9 Q88PB4 Q88PB8
Q88PB9 Q88PC0 Q88PD5 Q88PH8 Q88PI8 Q88PJ3 Q88PJ6 Q88PJ7 Q88PK0 Q88PK1 Q88PK4
Q88PK5 Q88PK8 Q88PL7 Q88PL8 Q88PM7 Q88PP0 Q88PS4 Q88PT7 Q88PU7 Q88PV2 Q88PV4
Q88PW2 Q88PW5 Q88PW6 Q88PX5 Q88PX6 Q88PX7 Q88PX8 Q88Q07 Q88Q08 Q88Q09 Q88Q10
Q88Q27 Q88Q44 Q88Q65 Q88Q66 Q88Q71 Q88Q83 Q88Q89 Q88Q92 Q88Q95 Q88QA3 Q88QC7

Q88QD4 Q88QF2 Q88QF6 Q88QG4 Q88QG5 Q88QG7 Q88QH1 Q88QH5 Q88QH6 Q88QI0 Q88QJ8
Q88QJ9 Q88QK5 Q88QK7 Q88QL0 Q88QL1 Q88QL2 Q88QL3 Q88QL6 Q88QL7 Q88QL8 Q88QL9
Q88QM0 Q88QM1 Q88QM2 Q88QM3 Q88QM4 Q88QM5 Q88QM6 Q88QM7 Q88QM8 Q88QM9 Q88QN0
Q88QN1 Q88QN2 Q88QN3 Q88QN4 Q88QN5 Q88QN6 Q88QN7 Q88QN8 Q88QN9 Q88QP0 Q88QP1
Q88QP2 Q88QP3 Q88QP4 Q88QP5 Q88QP8 Q88QQ0 Q88QQ2 Q88QQ4 Q88QQ5 Q88QQ6 Q88QR6
Q88QR7 Q88QS2 Q88QT2 Q88QT3 Q88QT5 Q88QT6 Q88QT7 Q88QT8 Q88QT9 Q88QU2 Q88QU6
Q88QV3 Q88QV4 Q88QV5 Q88QV6 Q88QV7 Q88QV8 Q88QX1 Q88QX2 Q88QX8 Q88QZ8 Q88R12
Q88R32 Q88R41 Q88R42 Q88R44 Q88R45 Q88R49 Q88R68 Q88R81 Q88R85 Q88R90 Q88R95
Q88R97 Q88RA3 Q88RB9 Q88RC0 Q88RE5 Q88RK5 Q88RP6 Q88RP7 Q88RQ6 Q88RQ9 Q88RR1
Q88RR2 Q88RR3 Q88RR8 Q88RR9 Q88RW7 Q88RW8 Q88RX0 Q88RX5 Q88RX6 Q88RZ2 Q88S33
Q88S37 Q88S38 Q88S49 Q88S50 Q88S51 Q88S52 Q88S84 Q88S87 Q88S98 Q88SB0 Q88SC3
Q88SD8 Q88SE8 Q88SF0 Q88SI6 Q88SK6 Q88SR1 Q88ST4 Q88SV5 Q88SV6 Q88SY9 Q88T09
Q88T35 Q88TV5 Q88TY1 Q88U22 Q88U24 Q88U25 Q88U29 Q88U30 Q88U40 Q88U41 Q88U67
Q88UC0 Q88UD8 Q88UD9 Q88UE0 Q88UE1 Q88UE2 Q88UE3 Q88UE4 Q88UE5 Q88UE6 Q88UH8
Q88UH9 Q88UI4 Q88UT1 Q88UT2 Q88UT5 Q88UU5 Q88UV8 Q88UW5 Q88UW9 Q88UX0 Q88UX4
Q88UX7 Q88UZ3 Q88UZ4 Q88UZ7 Q88UZ8 Q88V02 Q88V03 Q88V04 Q88V05 Q88V07 Q88V10
Q88V12 Q88V13 Q88V14 Q88V16 Q88V19 Q88V20 Q88V23 Q88V24 Q88V33 Q88V41 Q88V55
Q88V56 Q88V61 Q88V67 Q88V75 Q88V76 Q88V80 Q88V85 Q88V89 Q88V90 Q88V96 Q88VA5
Q88VB0 Q88VB2 Q88VB7 Q88VC6 Q88VC8 Q88VD4 Q88VD5 Q88VE0 Q88VE1 Q88VE2 Q88VE3
Q88VE9 Q88VG4 Q88VG6 Q88VH0 Q88VH3 Q88VI7 Q88VJ1 Q88VJ2 Q88VJ4 Q88VJ5 Q88VJ6
Q88VJ7 Q88VK1 Q88VK2 Q88VK3 Q88VK7 Q88VK8 Q88VL0 Q88VL2 Q88VL5 Q88VL8 Q88VL9
Q88VM0 Q88VM1 Q88VM6 Q88VP8 Q88VP9 Q88VQ3 Q88VQ7 Q88VQ8 Q88VR2 Q88VR4 Q88VR5
Q88VR8 Q88VS1 Q88VS2 Q88VS3 Q88VX7 Q88VY1 Q88VY7 Q88VY8 Q88VZ4 Q88VZ6 Q88W01
Q88W02 Q88W05 Q88W10 Q88W20 Q88W23 Q88W25 Q88W26 Q88W32 Q88W33 Q88W46 Q88W76
Q88W95 Q88W96 Q88W97 Q88WA8 Q88WB6 Q88WD0 Q88WD3 Q88WF3 Q88WH9 Q88WI0 Q88WI1
Q88WI2 Q88WI3 Q88WJ1 Q88WJ2 Q88WJ3 Q88WJ5 Q88WJ9 Q88WK0 Q88WK6 Q88WK9 Q88WL3
Q88WL6 Q88WL7 Q88WM6 Q88WM7 Q88WM8 Q88WM9 Q88WN1 Q88WN3 Q88WN5 Q88WP5 Q88WQ9
Q88WR0 Q88WR2 Q88WR3 Q88WR7 Q88WS9 Q88WT1 Q88WT5 Q88WU6 Q88WU7 Q88WU8 Q88WU9
Q88WV2 Q88WV3 Q88WV4 Q88WW9 Q88WY1 Q88WZ5 Q88WZ9 Q88X16 Q88X33 Q88X36 Q88X53
Q88X60 Q88X61 Q88X63 Q88XA9 Q88XB8 Q88XC3 Q88XF3 Q88XG4 Q88XG5 Q88XJ7 Q88XP4
Q88XP6 Q88XP7 Q88XP8 Q88XQ0 Q88XQ4 Q88XS8 Q88XT4 Q88XU0 Q88XU7 Q88XU8 Q88XU9
Q88XV9 Q88XW0 Q88XW1 Q88XW2 Q88XW3 Q88XW4 Q88XW5 Q88XW7 Q88XW8 Q88XW9 Q88XX0
Q88XX1 Q88XX2 Q88XX3 Q88XX4 Q88XX5 Q88XX6 Q88XX7 Q88XX8 Q88XX9 Q88XY0 Q88XY1
Q88XY2 Q88XY3 Q88XY4 Q88XY5 Q88XY6 Q88XY7 Q88XY8 Q88XY9 Q88XZ0 Q88XZ2 Q88XZ3
Q88XZ6 Q88XZ7 Q88Y23 Q88Y39 Q88Y40 Q88Y41 Q88Y74 Q88Y75 Q88YC0 Q88YD2 Q88YD9
Q88YE7 Q88YE8 Q88YF4 Q88YG1 Q88YG7 Q88YH4 Q88YH5 Q88YH9 Q88YI2 Q88YI4 Q88YI8
Q88YK3 Q88YL5 Q88YM5 Q88YM6 Q88YM8 Q88YN0 Q88YN8 Q88YN9 Q88YP3 Q88YP6 Q88YP8
Q88YP9 Q88YW7 Q88YW8 Q88YW9 Q88YX0 Q88YX4 Q88YX9 Q88YY0 Q88YY5 Q88YY8 Q88Z02
Q88Z08 Q88Z10 Q88Z14 Q88Z28 Q88Z30 Q88Z33 Q88Z39 Q88Z52 Q88Z54 Q88Z60 Q88Z64
Q88Z76 Q88Z77 Q88Z84 Q88Z91 Q88Z93 Q88Z97 Q88ZA0 Q88ZF1 Q88ZS1 Q88ZS6 Q88ZS9
Q88ZU5 Q890C0 Q890C2 Q890D7 Q890D8 Q890E7 Q890H4 Q890I8 Q890J0 Q890J1 Q890J4
Q890J5 Q890J8 Q890K0 Q890K1 Q890K2 Q890K5 Q890K8 Q890L4 Q890L6 Q890M1 Q890M2
Q890M4 Q890M5 Q890M6 Q890N1 Q890N2 Q890N3 Q890N4 Q890N5 Q890N6 Q890N7 Q890N8
Q890N9 Q890P0 Q890P1 Q890P2 Q890P3 Q890P5 Q890P6 Q890P7 Q890P8 Q890P9 Q890Q0

Q890Q1 Q890Q3 Q890Q5 Q890Q7 Q890Q8 Q890Q9 Q890R0 Q890R1 Q890R5 Q890R6 Q890R7
Q890S0 Q890U1 Q890U2 Q890X1 Q890X8 Q890X9 Q890Y5 Q890Y6 Q890Y7 Q890Y9 Q890Z2
Q89121 Q89181 Q89183 Q891A0 Q891B8 Q891C7 Q891D3 Q891E7 Q891F1 Q891G4 Q891G7
Q891H2 Q891H3 Q891I0 Q891I1 Q891I2 Q891I4 Q891I6 Q891I9 Q891J1 Q891J4 Q891J6
Q891J7 Q891J8 Q891L4 Q891M0 Q891N2 Q891P9 Q891Q1 Q891Q9 Q891R3 Q891R5 Q891S2
Q891S3 Q891S6 Q891S7 Q891T0 Q891T1 Q891T2 Q891T7 Q891T8 Q891U1 Q891W4 Q891Y7
Q891Z5 Q891Z7 Q891Z8 Q891Z9 Q89249 Q89250 Q89251 Q89268 Q89269 Q89270 Q89273
Q892A0 Q892A7 Q892A9 Q892B2 Q892B5 Q892D0 Q892J3 Q892J5 Q892M0 Q892M1 Q892M6
Q892N4 Q892N6 Q892N8 Q892N9 Q892Q8 Q892Q9 Q892R0 Q892R1 Q892R2 Q892R9 Q892S8
Q892U0 Q892U4 Q892X1 Q892X3 Q892X7 Q892Z2 Q893F8 Q893G4 Q893H3 Q893K6 Q893M2
Q893Q3 Q893Q9 Q893R0 Q893R2 Q893S5 Q893T7 Q89420 Q89425 Q89445 Q89462 Q89485
Q89496 Q894B3 Q894B4 Q894B7 Q894C6 Q894C9 Q894D2 Q894F6 Q894G5 Q894G6 Q894G7
Q894H0 Q894H2 Q894H3 Q894L2 Q894L3 Q894M8 Q894N6 Q894P6 Q894S7 Q894Z2 Q894Z3
Q89508 Q89525 Q89544 Q89554 Q89565 Q89576 Q89581 Q89584 Q89592 Q895A7 Q895G1
Q895G2 Q895G3 Q895H1 Q895H2 Q895H3 Q895H4 Q895I5 Q895I7 Q895J3 Q895J5 Q895J7
Q895J8 Q895K1 Q895K2 Q895K3 Q895K5 Q895K8 Q895K9 Q895L0 Q895L1 Q895L2 Q895L3
Q895L7 Q895L8 Q895M1 Q895M2 Q895M3 Q895M9 Q895N2 Q895N4 Q895N8 Q895P5 Q895P8
Q895Q1 Q895Q2 Q895Q5 Q895R1 Q895U7 Q895W4 Q895X8 Q895Y6 Q89613 Q89621 Q89635
Q89642 Q89662 Q89667 Q89675 Q89681 Q89682 Q89698 Q896E3 Q896E8 Q896F1 Q896F2
Q896F5 Q896G3 Q896G4 Q896H4 Q896J4 Q896M5 Q896N5 Q896T7 Q896W8 Q896X8 Q89702
Q89703 Q89707 Q89709 Q89717 Q89740 Q89743 Q89746 Q89768 Q89769 Q89777 Q89785
Q89786 Q897B2 Q897C2 Q897E6 Q897F7 Q897K1 Q897K3 Q897K4 Q897K9 Q897L4 Q897P5
Q897P8 Q897Q7 Q897W7 Q89808 Q89814 Q89822 Q89828 Q89869 Q89882 Q89893 Q898A3
Q898C7 Q898E6 Q898E8 Q898F0 Q898L3 Q898N5 Q898P3 Q898Q7 Q898R1 Q898R2 Q898R3
Q898T5 Q898V2 Q898W7 Q898X3 Q898X9 Q898Y5 Q898Y8 Q898Z5 Q89907 Q89924 Q89925
Q89926 Q89933 Q89940 Q899A2 Q899C3 Q899D9 Q899E9 Q899F3 Q899G3 Q899G5 Q899G6
Q899G7 Q899H1 Q899H5 Q899I4 Q899I8 Q899J1 Q899J5 Q899K4 Q899K8 Q899M1 Q899M3
Q899M5 Q899M9 Q899P8 Q899Q4 Q899Q8 Q899R2 Q899S0 Q899S1 Q899S2 Q899S7 Q899T6
Q899U2 Q899U3 Q899U6 Q89A14 Q89A15 Q89A16 Q89A17 Q89A18 Q89A19 Q89A20 Q89A21
Q89A22 Q89A25 Q89A26 Q89A27 Q89A28 Q89A29 Q89A30 Q89A31 Q89A32 Q89A34 Q89A35
Q89A37 Q89A38 Q89A41 Q89A42 Q89A43 Q89A44 Q89A45 Q89A46 Q89A47 Q89A48 Q89A49
Q89A50 Q89A51 Q89A52 Q89A53 Q89A54 Q89A55 Q89A56 Q89A57 Q89A58 Q89A59 Q89A61
Q89A62 Q89A63 Q89A64 Q89A65 Q89A66 Q89A67 Q89A68 Q89A69 Q89A70 Q89A71 Q89A72
Q89A73 Q89A74 Q89A75 Q89A76 Q89A77 Q89A78 Q89A79 Q89A80 Q89A81 Q89A82 Q89A83
Q89A84 Q89A86 Q89A87 Q89A88 Q89A89 Q89A90 Q89A91 Q89A92 Q89A93 Q89A94 Q89A95
Q89A99 Q89AA0 Q89AA1 Q89AA2 Q89AA8 Q89AB0 Q89AB1 Q89AB2 Q89AB3 Q89AB4 Q89AB6
Q89AB8 Q89AB9 Q89AC0 Q89AC1 Q89AC2 Q89AC3 Q89AC4 Q89AC5 Q89AC6 Q89AC8 Q89AC9
Q89AD0 Q89AD1 Q89AD3 Q89AD4 Q89AD5 Q89AD6 Q89AD7 Q89AD8 Q89AD9 Q89AE0 Q89AE1
Q89AE2 Q89AE3 Q89AE5 Q89AE6 Q89AE7 Q89AE8 Q89AE9 Q89AF0 Q89AF1 Q89AF3 Q89AF4
Q89AF5 Q89AF6 Q89AF7 Q89AF8 Q89AF9 Q89AG0 Q89AG1 Q89AG2 Q89AG3 Q89AG4 Q89AG5
Q89AG7 Q89AG8 Q89AH1 Q89AH2 Q89AH4 Q89AH7 Q89AH8 Q89AH9 Q89AI0 Q89AI2 Q89AI4
Q89AI5 Q89AI7 Q89AI8 Q89AJ2 Q89AJ3 Q89AJ4 Q89AJ6 Q89AJ7 Q89AJ8 Q89AJ9 Q89AK0
Q89AK1 Q89AK2 Q89AK3 Q89AK4 Q89AK5 Q89AK6 Q89AK7 Q89AK8 Q89AK9 Q89AL1 Q89AL4
Q89AL6 Q89AL7 Q89AL8 Q89AM0 Q89AM3 Q89AM8 Q89AM9 Q89AN0 Q89AN1 Q89AN3 Q89AN4

Q89AN7 Q89AN8 Q89AP1 Q89AP2 Q89AP3 Q89AP4 Q89AP7 Q89AP8 Q89AQ1 Q89AQ2 Q89AQ6
Q89AQ7 Q89AQ8 Q89AQ9 Q89AR0 Q89AR3 Q89AR4 Q89AR5 Q89AR6 Q89AR7 Q89AR8 Q89AR9
Q89AS0 Q89AS1 Q89AS2 Q89AS3 Q89AS4 Q89AS5 Q89AS6 Q89AS7 Q89AS8 Q89AS9 Q89AT0
Q89AT2 Q89AU7 Q89AU8 Q89AU9 Q89AV1 Q89AV2 Q89AV3 Q89AV5 Q89AV6 Q89AV8 Q89AV9
Q89AW0 Q89AW2 Q89AW3 Q89AW4 Q89AX1 Q89AX3 Q89AX4 Q89AX5 Q89AX6 Q89AX7 Q89AX8
Q89AX9 Q89AY0 Q89AY1 Q89AY2 Q89AY3 Q89AY5 Q89AY6 Q89AY8 Q89AY9 Q89AZ5 Q89AZ8
Q89B01 Q89B02 Q89B06 Q89B07 Q89B08 Q89B09 Q89B10 Q89B11 Q89B12 Q89B13 Q89B15
Q89B16 Q89B17 Q89B18 Q89B19 Q89B20 Q89B21 Q89B22 Q89B23 Q89B24 Q89B25 Q89B27
Q89B29 Q89B30 Q89B31 Q89B32 Q89B33 Q89B35 Q89B36 Q89B37 Q89B46 Q89B47 Q89BK7
Q89BL4 Q89BN3 Q89BN4 Q89BN9 Q89BP0 Q89BP1 Q89BP2 Q89BP4 Q89BP7 Q89BP9 Q89BQ2
Q89C25 Q89CF5 Q89CK8 Q89CN4 Q89D74 Q89D92 Q89DA6 Q89DB4 Q89DD2 Q89DE6 Q89DF8
Q89DG5 Q89DG9 Q89DI2 Q89DI3 Q89DI7 Q89DJ1 Q89DJ8 Q89DJ9 Q89DN1 Q89DR1 Q89DR8
Q89DX7 Q89E06 Q89E48 Q89EA6 Q89EF0 Q89EG8 Q89EM1 Q89EM9 Q89EP4 Q89ER2 Q89ES7
Q89F27 Q89F47 Q89FD3 Q89FG1 Q89FG2 Q89FG3 Q89FG4 Q89FG5 Q89FH0 Q89FP0 Q89FP2
Q89FT9 Q89FU2 Q89FU5 Q89FU8 Q89FU9 Q89FV8 Q89FW1 Q89FY1 Q89G35 Q89G43 Q89G50
Q89G86 Q89G87 Q89GA7 Q89GB0 Q89GE3 Q89GF9 Q89GR1 Q89GR3 Q89GV2 Q89GV3 Q89GV4
Q89GX0 Q89GX9 Q89H51 Q89H53 Q89HA2 Q89HP5 Q89HP6 Q89HR3 Q89HV6 Q89HW8 Q89HZ2
Q89I27 Q89I84 Q89I86 Q89I87 Q89I89 Q89IB0 Q89IB6 Q89IC0 Q89ID3 Q89IK8 Q89IW2
Q89IW3 Q89J69 Q89J70 Q89J72 Q89J73 Q89J74 Q89J75 Q89J79 Q89J80 Q89J81 Q89J82
Q89J83 Q89J84 Q89J85 Q89J86 Q89J87 Q89J88 Q89J89 Q89J90 Q89J91 Q89J92 Q89J93
Q89J94 Q89J95 Q89J96 Q89J97 Q89J98 Q89J99 Q89JA0 Q89JA1 Q89JA2 Q89JA3 Q89JA6
Q89JA7 Q89JA8 Q89JL7 Q89JL8 Q89JM6 Q89JV7 Q89JW5 Q89K02 Q89K13 Q89K19 Q89K24
Q89K28 Q89K29 Q89K31 Q89K72 Q89K77 Q89K80 Q89K81 Q89KE4 Q89KE5 Q89KG0 Q89KG1
Q89KG2 Q89KM8 Q89KP3 Q89KP4 Q89KP5 Q89KP6 Q89KP7 Q89KP9 Q89KR5 Q89KR6 Q89KU3
Q89KU5 Q89KV2 Q89KV4 Q89KY5 Q89KZ0 Q89L00 Q89L02 Q89L21 Q89L46 Q89L50 Q89L55
Q89L59 Q89LB1 Q89LG2 Q89LG9 Q89LH6 Q89LK8 Q89LM5 Q89LP8 Q89LQ2 Q89LQ8 Q89LR6
Q89LR7 Q89LR8 Q89LY4 Q89M07 Q89M62 Q89M63 Q89MA1 Q89MA7 Q89MB9 Q89MI2 Q89MN4
Q89MP9 Q89MQ0 Q89MS3 Q89MT4 Q89MT9 Q89MU0 Q89MV4 Q89MW3 Q89MW4 Q89MW6 Q89MZ0
Q89MZ4 Q89N37 Q89N53 Q89N72 Q89N76 Q89NA7 Q89ND7 Q89NF3 Q89NF8 Q89NP8 Q89NS0
Q89NS9 Q89NT2 Q89NW6 Q89P00 Q89P49 Q89PS5 Q89PT3 Q89PW2 Q89Q06 Q89Q54 Q89Q71
Q89QG3 Q89QH2 Q89QK7 Q89QM1 Q89QU8 Q89QW7 Q89QX6 Q89R17 Q89R51 Q89RB7 Q89RJ4
Q89RW0 Q89RW1 Q89RW4 Q89RX6 Q89RY1 Q89RY8 Q89S69 Q89S70 Q89S79 Q89S84 Q89S87
Q89SB5 Q89SC0 Q89SC2 Q89SJ4 Q89SJ7 Q89SJ8 Q89SK1 Q89SS3 Q89T99 Q89TD1 Q89TZ6
Q89U80 Q89U81 Q89U82 Q89U83 Q89U95 Q89UB0 Q89UC1 Q89UC5 Q89UE3 Q89UE4 Q89UF7
Q89UF8 Q89UG0 Q89UG2 Q89UG3 Q89UG4 Q89UH1 Q89UK6 Q89UL2 Q89UL7 Q89UL9 Q89UU3
Q89UU4 Q89UU5 Q89UZ9 Q89V00 Q89V02 Q89V30 Q89VC7 Q89VE6 Q89VE8 Q89VE9 Q89VS2
Q89VT5 Q89VT6 Q89VT8 Q89VU7 Q89VV9 Q89VX1 Q89VX9 Q89W41 Q89W61 Q89W71 Q89W81
Q89W91 Q89W97 Q89W99 Q89WA4 Q89WA6 Q89WA9 Q89WB0 Q89WB1 Q89WB2 Q89WB3 Q89WB4
Q89WC9 Q89WD0 Q89WE4 Q89WE5 Q89WE6 Q89WE8 Q89WF1 Q89WF2 Q89WH7 Q89WH9 Q89WI0
Q89WI1 Q89WI2 Q89WI8 Q89WK1 Q89WK2 Q89WK4 Q89WK5 Q89WK9 Q89WL0 Q89WM2 Q89WM3
Q89WM4 Q89WM5 Q89WM6 Q89WM8 Q89WM9 Q89WN2 Q89WN7 Q89WN9 Q89WP0 Q89WP1 Q89WP4
Q89WP5 Q89WP6 Q89WQ1 Q89WR7 Q89WS6 Q89WU7 Q89WV5 Q89WW1 Q89WX9 Q89WY0 Q89WZ6
Q89X03 Q89X19 Q89X27 Q89X34 Q89X35 Q89X36 Q89X39 Q89X40 Q89X45 Q89X59 Q89X60
Q89X68 Q89X78 Q89X82 Q89X84 Q89X85 Q89X86 Q89X89 Q89X93 Q89X94 Q89X98 Q89XA0

Q89XE9 Q89XH1 Q89XM2 Q89XM4 Q89XN9 Q89XP2 Q89XR6 Q89XT5 Q89XT8 Q89XT9 Q89XU2
Q89XW5 Q89XW7 Q89XW8 Q89XZ3 Q89YN0 Q89YP0 Q89YS5 Q89YT0 Q89YW6 Q89YY3 Q89YY6
Q89YZ1 Q89YZ2 Q89YZ6 Q89YZ7 Q89Z26 Q89Z59 Q89ZC3 Q89ZF2 Q89ZG1 Q89ZI8 Q89ZI9
Q89ZJ0 Q89ZK3 Q89ZK5 Q89ZL1 Q89ZM4 Q89ZN1 Q89ZQ3 Q89ZR2 Q89ZR3 Q89ZR7 Q89ZR8
Q89ZV6 Q89ZW6 Q89ZW8 Q89ZW9 Q89ZY4 Q8A004 Q8A005 Q8A011 Q8A012 Q8A018 Q8A029
Q8A044 Q8A094 Q8A0B5 Q8A0C2 Q8A0H8 Q8A0L8 Q8A0M6 Q8A0S6 Q8A0U0 Q8A0U2 Q8A0U8
Q8A0V3 Q8A0V4 Q8A0W0 Q8A0X7 Q8A0Y3 Q8A0Y7 Q8A0Y9 Q8A0Z3 Q8A0Z4 Q8A0Z5 Q8A0Z8
Q8A102 Q8A120 Q8A128 Q8A129 Q8A135 Q8A138 Q8A155 Q8A1A0 Q8A1A2 Q8A1A3 Q8A1A4
Q8A1A7 Q8A1D3 Q8A1D5 Q8A1D7 Q8A1E0 Q8A1E7 Q8A1E8 Q8A1E9 Q8A1F3 Q8A1F6 Q8A1F7
Q8A1G8 Q8A245 Q8A250 Q8A251 Q8A254 Q8A256 Q8A275 Q8A276 Q8A287 Q8A293 Q8A295
Q8A2A1 Q8A2A3 Q8A2B0 Q8A2B1 Q8A2B2 Q8A2E4 Q8A2E8 Q8A2E9 Q8A2F1 Q8A2F4 Q8A2F6
Q8A2H2 Q8A2M0 Q8A2M2 Q8A2N7 Q8A2N8 Q8A2N9 Q8A2P0 Q8A2S4 Q8A2S5 Q8A2T6 Q8A2U2
Q8A2W0 Q8A327 Q8A329 Q8A334 Q8A3C0 Q8A3H9 Q8A3M1 Q8A3X5 Q8A407 Q8A414 Q8A418
Q8A455 Q8A463 Q8A465 Q8A466 Q8A468 Q8A469 Q8A470 Q8A472 Q8A473 Q8A474 Q8A475
Q8A476 Q8A477 Q8A479 Q8A480 Q8A481 Q8A482 Q8A483 Q8A484 Q8A485 Q8A486 Q8A487
Q8A488 Q8A489 Q8A490 Q8A491 Q8A492 Q8A493 Q8A494 Q8A495 Q8A498 Q8A499 Q8A4A0
Q8A4A1 Q8A4A2 Q8A4A3 Q8A4B1 Q8A4M8 Q8A4N6 Q8A4P5 Q8A4S8 Q8A4T0 Q8A4V8 Q8A525
Q8A528 Q8A531 Q8A5G4 Q8A5I9 Q8A5J0 Q8A5J7 Q8A5K3 Q8A5S5 Q8A5S6 Q8A5S7 Q8A5V6
Q8A5V7 Q8A5W2 Q8A5W4 Q8A602 Q8A606 Q8A608 Q8A624 Q8A625 Q8A626 Q8A639 Q8A675
Q8A677 Q8A681 Q8A682 Q8A684 Q8A6A7 Q8A6L7 Q8A6M0 Q8A6N4 Q8A6N7 Q8A6P7 Q8A6P8
Q8A712 Q8A733 Q8A749 Q8A753 Q8A756 Q8A765 Q8A767 Q8A7B8 Q8A7C8 Q8A7D1 Q8A7E7
Q8A7F8 Q8A7G0 Q8A7G2 Q8A7M7 Q8A7S9 Q8A7T2 Q8A7U2 Q8A7Z4 Q8A7Z5 Q8A7Z6 Q8A7Z7
Q8A800 Q8A806 Q8A8C3 Q8A8C4 Q8A8D9 Q8A8H7 Q8A8L4 Q8A8M0 Q8A8M2 Q8A8R2 Q8A8R5
Q8A8S8 Q8A8T4 Q8A8T6 Q8A8T9 Q8A8Y4 Q8A947 Q8A988 Q8A997 Q8A999 Q8A9A0 Q8A9A2
Q8A9B8 Q8A9C1 Q8A9C2 Q8A9C3 Q8A9E3 Q8A9H7 Q8A9J0 Q8A9J2 Q8A9K9 Q8A9M2 Q8A9M7
Q8A9S3 Q8A9S4 Q8A9S7 Q8A9S8 Q8A9W1 Q8A9W7 Q8A9X8 Q8AA13 Q8AA14 Q8AA15 Q8AA22
Q8AA38 Q8AA39 Q8AA41 Q8AA75 Q8AA95 Q8AAB1 Q8AAB7 Q8AAB8 Q8AAD2 Q8AAD5 Q8AAD6
Q8AAD8 Q8AAI7 Q8AAN7 Q8AAN9 Q8AAP0 Q8AAP1 Q8AAP2 Q8AAP4 Q8AAP5 Q8AAP9 Q8AAQ0
Q8AAQ1 Q8AAW1 Q8AAZ8 Q8AB53 Q8AB57 Q8ABA7 Q8ABA8 Q8ABA9 Q8ABB0 Q8ABD1 Q8ABF5
Q8ABH7 Q8ABV5 Q8AIH9 Q8AV19 Q8AV66 Q8AVC1 Q8AVD3 Q8AVF5 Q8AVG9 Q8AVH1 Q8AVH4
Q8AVJ0 Q8AVJ1 Q8AVJ2 Q8AVK2 Q8AVL0 Q8AVM7 Q8AVN9 Q8AVP1 Q8AVQ3 Q8AVQ6 Q8AVR2
Q8AVS4 Q8AVS6 Q8AVS9 Q8AVT9 Q8AVU0 Q8AVU2 Q8AVU4 Q8AVV6 Q8AVY1 Q8AW52 Q8AW92
Q8AW93 Q8AWA7 Q8AWD2 Q8AWF2 Q8AWF4 Q8AWF5 Q8AWG1 Q8AWG2 Q8AWG6 Q8AWH1 Q8AWH2
Q8AWH3 Q8AWW7 Q8AWY2 Q8AWY9 Q8AWZ0 Q8AWZ2 Q8AX00 Q8AX01 Q8AX02 Q8AX98 Q8AXB4
Q8AXF4 Q8AXL1 Q8AXQ4 Q8AXV5 Q8AXV6 Q8AXW8 Q8AXX2 Q8AXX8 Q8AY65 Q8AY73 Q8AY75
Q8AYB8 Q8AYC1 Q8AYC2 Q8AYG3 Q8AYI2 Q8AZ66 Q8AZJ3 Q8AZJ5 Q8AZK7 Q8AZL8 Q8AZL9
Q8AZM0 Q8B0H0 Q8B0H3 Q8B0H4 Q8B0H5 Q8B0H8 Q8B0H9 Q8B0I0 Q8B0I3 Q8B0U2 Q8B6J5
Q8B6J9 Q8B9B2 Q8B9L3 Q8B9Q8 Q8BB04 Q8BB12 Q8BB22 Q8BB47 Q8BCV3 Q8BCV4 Q8BCV5
Q8BCV6 Q8BCV7 Q8BCV8 Q8BCV9 Q8BD28 Q8BD31 Q8BDD6 Q8BDD9 Q8BDE4 Q8BDE5 Q8BDE6
Q8BDG4 Q8BDG7 Q8BFQ4 Q8BFQ9 Q8BFR1 Q8BFR6 Q8BFS9 Q8BFT6 Q8BFU2 Q8BFU3 Q8BFV2
Q8BFW3 Q8BFW4 Q8BFY7 Q8BFZ4 Q8BG02 Q8BG15 Q8BG17 Q8BG18 Q8BG30 Q8BG32 Q8BG34
Q8BG36 Q8BG41 Q8BG60 Q8BG71 Q8BG79 Q8BG80 Q8BG81 Q8BG89 Q8BG93 Q8BG94 Q8BG99
Q8BGA5 Q8BGA7 Q8BGB7 Q8BGC0 Q8BGC1 Q8BGD0 Q8BGD7 Q8BGD9 Q8BGE5 Q8BGE7 Q8BGF3
Q8BGF6 Q8BGF7 Q8BGI4 Q8BGJ3 Q8BGK2 Q8BGL1 Q8BGN2 Q8BGQ2 Q8BGQ6 Q8BGQ7 Q8BGR6

Q8BGR8 Q8BGR9 Q8BGS2 Q8BGS3 Q8BGT5 Q8BGT8 Q8BGV0 Q8BGV4 Q8BGW0 Q8BGW3 Q8BGW4
Q8BGW5 Q8BGW8 Q8BGX1 Q8BGX4 Q8BGX7 Q8BGY4 Q8BGZ2 Q8BGZ3 Q8BGZ4 Q8BGZ7 Q8BH00
Q8BH05 Q8BH15 Q8BH50 Q8BH53 Q8BH55 Q8BH58 Q8BH70 Q8BH75 Q8BH83 Q8BH86 Q8BH93
Q8BHA0 Q8BHA3 Q8BHB7 Q8BHC4 Q8BHD1 Q8BHD7 Q8BHE0 Q8BHE1 Q8BHG2 Q8BHI4 Q8BHK0
Q8BHL8 Q8BHN5 Q8BHN7 Q8BHP2 Q8BHR8 Q8BHS3 Q8BHW6 Q8BHW9 Q8BHZ4 Q8BHZ5 Q8BI17
Q8BI22 Q8BI66 Q8BI67 Q8BI69 Q8BI71 Q8BI73 Q8BI79 Q8BI99 Q8BIA3 Q8BIA4 Q8BID6
Q8BID8 Q8BIE6 Q8BIF2 Q8BIF9 Q8BIG2 Q8BIG4 Q8BIH0 Q8BII1 Q8BIJ6 Q8BIL2 Q8BIP0
Q8BIQ3 Q8BIQ5 Q8BIQ8 Q8BIR2 Q8BIW1 Q8BIW9 Q8BIY1 Q8BIY3 Q8BIZ6 Q8BJ05 Q8BJ25
Q8BJ90 Q8BJA3 Q8BJH1 Q8BJI4 Q8BJL0 Q8BJL1 Q8BJM3 Q8BJM7 Q8BJS7 Q8BJS8 Q8BJU9
Q8BJW5 Q8BJW7 Q8BK06 Q8BK26 Q8BK58 Q8BK64 Q8BK75 Q8BK84 Q8BKA3 Q8BKE5 Q8BKE9
Q8BKF1 Q8BKI2 Q8BKK5 Q8BKN5 Q8BKR5 Q8BKT2 Q8BKT3 Q8BKW4 Q8BKX6 Q8BKX7 Q8BKY8
Q8BKZ9 Q8BL06 Q8BL48 Q8BL65 Q8BL74 Q8BL95 Q8BL97 Q8BLA1 Q8BLB0 Q8BLB7 Q8BLB8
Q8BLD6 Q8BLF2 Q8BLH7 Q8BLM0 Q8BLQ0 Q8BLR9 Q8BLU2 Q8BM65 Q8BM85 Q8BMA5 Q8BMB0
Q8BMB3 Q8BMC4 Q8BMD7 Q8BMF3 Q8BMG1 Q8BMI4 Q8BMJ2 Q8BMJ3 Q8BMJ8 Q8BMK0 Q8BMK1
Q8BMK5 Q8BML9 Q8BMQ2 Q8BMQ3 Q8BMQ8 Q8BMU0 Q8BMZ5 Q8BND3 Q8BND4 Q8BNL5 Q8BNN1
Q8BNU0 Q8BNV1 Q8BNW9 Q8BP01 Q8BP27 Q8BP47 Q8BP48 Q8BP56 Q8BP71 Q8BP74 Q8BP78
Q8BP86 Q8BP99 Q8BPA8 Q8BPC6 Q8BPI1 Q8BPK2 Q8BPP0 Q8BPP1 Q8BPW0 Q8BPY9 Q8BPZ8
Q8BQ30 Q8BQ89 Q8BQB6 Q8BQC8 Q8BQH4 Q8BQJ6 Q8BQM4 Q8BQM8 Q8BQM9 Q8BQN5 Q8BQN6
Q8BQR7 Q8BQS4 Q8BQU3 Q8BQY8 Q8BR27 Q8BR63 Q8BR65 Q8BR90 Q8BRB7 Q8BRC6 Q8BRG6
Q8BRH4 Q8BRJ4 Q8BRM2 Q8BRM6 Q8BRV5 Q8BS39 Q8BS45 Q8BS90 Q8BSA9 Q8BSD5 Q8BSF5
Q8BSI6 Q8BSN3 Q8BSN5 Q8BSN9 Q8BSP2 Q8BSQ9 Q8BSV3 Q8BT07 Q8BT51 Q8BT88 Q8BTE0
Q8BTF7 Q8BTF8 Q8BTH6 Q8BTI7 Q8BTI8 Q8BTK5 Q8BTN6 Q8BTQ0 Q8BTR5 Q8BTT6 Q8BTU1
Q8BTV2 Q8BTW3 Q8BTW8 Q8BTY1 Q8BTZ4 Q8BTZ7 Q8BU00 Q8BU03 Q8BU04 Q8BU27 Q8BU30
Q8BUG5 Q8BUH1 Q8BUH8 Q8BUK6 Q8BUR3 Q8BUR9 Q8BUV6 Q8BUY8 Q8BUY9 Q8BUZ3 Q8BV13
Q8BV42 Q8BV49 Q8BV66 Q8BV79 Q8BV84 Q8BVA4 Q8BVE8 Q8BVF2 Q8BVF4 Q8BVF9 Q8BVG4
Q8BVH9 Q8BVI4 Q8BVM7 Q8BVN0 Q8BVN4 Q8BVN8 Q8BVN9 Q8BVP1 Q8BVQ5 Q8BVR0 Q8BVT6
Q8BVV7 Q8BVW0 Q8BVY0 Q8BW00 Q8BW10 Q8BW22 Q8BW49 Q8BW56 Q8BW72 Q8BW74 Q8BW86
Q8BW88 Q8BW94 Q8BWA5 Q8BWA8 Q8BWD2 Q8BWD8 Q8BWF0 Q8BWG4 Q8BWL5 Q8BWQ4 Q8BWR2
Q8BWU3 Q8BWU8 Q8BWW4 Q8BWY3 Q8BWY4 Q8BWZ3 Q8BX05 Q8BX07 Q8BX09 Q8BX13 Q8BX17
Q8BX32 Q8BX46 Q8BX65 Q8BX80 Q8BXC6 Q8BXF8 Q8BXG3 Q8BXJ2 Q8BXJ8 Q8BXK8 Q8BXN7
Q8BXP5 Q8BXQ8 Q8BXR6 Q8BXX2 Q8BXX9 Q8BY46 Q8BY71 Q8BY98 Q8BYG0 Q8BYH3 Q8BYH7
Q8BYK5 Q8BYK6 Q8BYK8 Q8BYL4 Q8BYM7 Q8BYM8 Q8BYN5 Q8BYR1 Q8BYZ7 Q8BZ20 Q8BZ21
Q8BZ25 Q8BZ34 Q8BZ47 Q8BZ52 Q8BZ89 Q8BZ94 Q8BZ97 Q8BZB2 Q8BZH4 Q8BZI0 Q8BZI6
Q8BZJ7 Q8BZJ8 Q8BZL8 Q8BZM1 Q8BZR9 Q8BZS9 Q8BZT2 Q8BZT9 Q8BZW2 Q8BZW4 Q8BZW8
Q8BZX4 Q8C006 Q8C008 Q8C042 Q8C052 Q8C092 Q8C0C0 Q8C0C4 Q8C0C7 Q8C0D0 Q8C0D5
Q8C0D9 Q8C0E3 Q8C0I4 Q8C0J6 Q8C0L9 Q8C0M8 Q8C0M9 Q8C0P5 Q8C0P7 Q8C0Q2 Q8C0Q3
Q8C0R7 Q8C0S4 Q8C0T1 Q8C0W1 Q8C0X0 Q8C0Y0 Q8C0Y1 Q8C120 Q8C152 Q8C156 Q8C159
Q8C176 Q8C181 Q8C190 Q8C1A3 Q8C1B1 Q8C1B7 Q8C1C8 Q8C1D8 Q8C1F5 Q8C1I6 Q8C1M2
Q8C1R3 Q8C1S0 Q8C1Z8 Q8C208 Q8C263 Q8C267 Q8C2A2 Q8C2E4 Q8C2P3 Q8C2S5 Q8C3F7
Q8C3I8 Q8C3L1 Q8C3M9 Q8C3R1 Q8C3W1 Q8C436 Q8C494 Q8C4H2 Q8C4J0 Q8C4J7 Q8C4P0
Q8C4S8 Q8C4V4 Q8C4X1 Q8C525 Q8C551 Q8C552 Q8C569 Q8C570 Q8C5H8 Q8C5K5 Q8C5L7
Q8C5N3 Q8C5N5 Q8C5P5 Q8C5Q4 Q8C5R2 Q8C5S3 Q8C5S7 Q8C5T4 Q8C5T8 Q8C5U9 Q8C5V0
Q8C5V5 Q8C5V8 Q8C5W3 Q8C5W4 Q8C627 Q8C636 Q8C650 Q8C669 Q8C6A8 Q8C6B9 Q8C6C7
Q8C6D4 Q8C6E0 Q8C6I2 Q8C6P8 Q8C6S9 Q8C726 Q8C735 Q8C739 Q8C753 Q8C761 Q8C779

Q8C790 Q8C796 Q8C7B6 Q8C7B8 Q8C7H1 Q8C7Q4 Q8C7R4 Q8C7R7 Q8C7U1 Q8C7V3 Q8C804
Q8C827 Q8C828 Q8C854 Q8C878 Q8C8B0 Q8C8H8 Q8C8J0 Q8C8K3 Q8C8M1 Q8C8T8 Q8C8V1
Q8C8Y5 Q8C966 Q8C9A2 Q8C9H6 Q8C9J3 Q8C9M2 Q8C9R9 Q8C9S4 Q8CA72 Q8CAA7 Q8CAE2
Q8CAF4 Q8CAI1 Q8CAK1 Q8CAS9 Q8CAT8 Q8CAV0 Q8CB49 Q8CB59 Q8CB62 Q8CB77 Q8CBA2
Q8CBE3 Q8CBW3 Q8CBW4 Q8CBX9 Q8CBY0 Q8CBY1 Q8CBY3 Q8CBY8 Q8CC14 Q8CC70 Q8CC86
Q8CC88 Q8CC96 Q8CCA0 Q8CCC3 Q8CCE9 Q8CCG1 Q8CCG4 Q8CCI5 Q8CCJ9 Q8CCK0 Q8CCN5
Q8CCS6 Q8CCT4 Q8CCT7 Q8CCX5 Q8CD15 Q8CD60 Q8CD92 Q8CD94 Q8CDC0 Q8CDC7 Q8CDD8
Q8CDE2 Q8CDF7 Q8CDI6 Q8CDI7 Q8CDK2 Q8CDK3 Q8CDL9 Q8CDM1 Q8CDM4 Q8CDM8 Q8CDN6
Q8CDN8 Q8CDN9 Q8CDP0 Q8CDR2 Q8CDS7 Q8CDT7 Q8CDU4 Q8CDU5 Q8CDU6 Q8CDV0 Q8CDV6
Q8CDW1 Q8CDY7 Q8CDZ2 Q8CE13 Q8CE22 Q8CE46 Q8CE64 Q8CE96 Q8CEC0 Q8CEC6 Q8CEF1
Q8CEG5 Q8CEI1 Q8CEL2 Q8CES0 Q8CEZ0 Q8CEZ4 Q8CF02 Q8CF25 Q8CF36 Q8CF60 Q8CF89
Q8CFA2 Q8CFA7 Q8CFC1 Q8CFC2 Q8CFC7 Q8CFD1 Q8CFD5 Q8CFE2 Q8CFE3 Q8CFF0 Q8CFG4
Q8CFI2 Q8CFI5 Q8CFI7 Q8CFK2 Q8CFL8 Q8CFQ3 Q8CFT2 Q8CFV2 Q8CFV9 Q8CFX8 Q8CG27
Q8CG46 Q8CG47 Q8CG48 Q8CG79 Q8CG80 Q8CGA0 Q8CGA4 Q8CGC4 Q8CGC6 Q8CGF6 Q8CGI1
Q8CGN4 Q8CGP0 Q8CGP1 Q8CGP2 Q8CGP5 Q8CGP6 Q8CGP7 Q8CGS5 Q8CGS6 Q8CGT6 Q8CGU9
Q8CGV2 Q8CGV7 Q8CGW9 Q8CGY6 Q8CH18 Q8CH19 Q8CH20 Q8CH25 Q8CH27 Q8CH40 Q8CH62
Q8CH72 Q8CH84 Q8CHC8 Q8CHG5 Q8CHH5 Q8CHI8 Q8CHP0 Q8CHP5 Q8CHP6 Q8CHQ0 Q8CHR6
Q8CHR9 Q8CHT0 Q8CHT6 Q8CHV6 Q8CHW4 Q8CHW5 Q8CHY6 Q8CHY7 Q8CHZ8 Q8CHZ9 Q8CI03
Q8CI08 Q8CI11 Q8CI12 Q8CI17 Q8CI33 Q8CI43 Q8CI70 Q8CI75 Q8CI78 Q8CI94 Q8CIB9
Q8CID0 Q8CIE2 Q8CIE4 Q8CIG3 Q8CIG9 Q8CIH9 Q8CII0 Q8CII2 Q8CIK8 Q8CIL4 Q8CIM3
Q8CIM8 Q8CIN6 Q8CIS9 Q8CIV8 Q8CIW5 Q8CIZ7 Q8CJ78 Q8CJC5 Q8CJF8 Q8CJF9 Q8CJG0
Q8CJG1 Q8CJH6 Q8CJM2 Q8CJM9 Q8CJP7 Q8CJQ6 Q8CJQ8 Q8CJR5 Q8CJT1 Q8CJT5 Q8CJV9
Q8CJX0 Q8CK04 Q8CK11 Q8CK24 Q8CK28 Q8CK51 Q8CKC6 Q8CKU7 Q8CMH6 Q8CMH7 Q8CMK6
Q8CML1 Q8CML2 Q8CML6 Q8CML7 Q8CMM0 Q8CMM1 Q8CMM6 Q8CMN4 Q8CMN5 Q8CMN6 Q8CMN7
Q8CMP5 Q8CMQ1 Q8CMQ2 Q8CMQ4 Q8CMQ7 Q8CMQ8 Q8CMS3 Q8CMT1 Q8CMT5 Q8CMT6 Q8CMT7
Q8CMT8 Q8CMV4 Q8CMW1 Q8CMW2 Q8CMX3 Q8CMX5 Q8CMY2 Q8CMY4 Q8CMY5 Q8CMZ0 Q8CMZ4
Q8CN01 Q8CN04 Q8CN17 Q8CN22 Q8CN23 Q8CN24 Q8CN27 Q8CN30 Q8CN40 Q8CN61 Q8CN77
Q8CN91 Q8CNA3 Q8CNB8 Q8CNC5 Q8CNC6 Q8CNC7 Q8CNC8 Q8CNC9 Q8CND0 Q8CND1 Q8CNE0
Q8CNE1 Q8CNE3 Q8CNE5 Q8CNE6 Q8CNF4 Q8CNF5 Q8CNF8 Q8CNG0 Q8CNG6 Q8CNH0 Q8CNH7
Q8CNH8 Q8CNH9 Q8CNI2 Q8CNI3 Q8CNI5 Q8CNI7 Q8CNI9 Q8CNK0 Q8CNK1 Q8CNK2 Q8CNK7
Q8CNK9 Q8CNL1 Q8CNL2 Q8CNL3 Q8CNL6 Q8CNL9 Q8CNN7 Q8CNP1 Q8CNP3 Q8CNP5 Q8CNQ1
Q8CNQ7 Q8CNR1 Q8CNS0 Q8CNS1 Q8CNT5 Q8CNU2 Q8CNU3 Q8CNU8 Q8CNV2 Q8CNV8 Q8CNV9
Q8CNW7 Q8CNW8 Q8CNW9 Q8CNX0 Q8CNX4 Q8CNY0 Q8CNY4 Q8CNY5 Q8CNY6 Q8CNY8 Q8CNZ0
Q8CNZ1 Q8CNZ4 Q8CNZ7 Q8CNZ8 Q8CP01 Q8CP08 Q8CP09 Q8CP15 Q8CP16 Q8CP17 Q8CP18
Q8CP22 Q8CP23 Q8CP24 Q8CP25 Q8CP34 Q8CP35 Q8CP37 Q8CP38 Q8CP39 Q8CP40 Q8CP47
Q8CP51 Q8CP62 Q8CP67 Q8CP68 Q8CP70 Q8CP71 Q8CP75 Q8CP83 Q8CP85 Q8CP87 Q8CP90
Q8CP95 Q8CP96 Q8CPA8 Q8CPA9 Q8CPB0 Q8CPB1 Q8CPB2 Q8CPB5 Q8CPB7 Q8CPC2 Q8CPC5
Q8CPC6 Q8CPC7 Q8CPC9 Q8CPD0 Q8CPE6 Q8CPE9 Q8CPF0 Q8CPF4 Q8CPG2 Q8CPG6 Q8CPG7
Q8CPG8 Q8CPG9 Q8CPH0 Q8CPH1 Q8CPH2 Q8CPH4 Q8CPH5 Q8CPH6 Q8CPH7 Q8CPI1 Q8CPI9
Q8CPJ3 Q8CPJ4 Q8CPJ5 Q8CPJ6 Q8CPJ7 Q8CPJ9 Q8CPL0 Q8CPL4 Q8CPL5 Q8CPL6 Q8CPL8
Q8CPN2 Q8CPN3 Q8CPN4 Q8CPN9 Q8CPP0 Q8CPP1 Q8CPP2 Q8CPP4 Q8CPQ4 Q8CPQ5 Q8CPR1
Q8CPR2 Q8CPS8 Q8CPT5 Q8CPT6 Q8CPT8 Q8CPT9 Q8CPU0 Q8CPU3 Q8CPW3 Q8CPW4 Q8CPW8
Q8CPX1 Q8CPX9 Q8CPY4 Q8CPY5 Q8CPY7 Q8CPY8 Q8CPY9 Q8CPZ0 Q8CPZ1 Q8CPZ6 Q8CPZ7
Q8CQ03 Q8CQ25 Q8CQ32 Q8CQ56 Q8CQ62 Q8CQ65 Q8CQ69 Q8CQ77 Q8CQ78 Q8CQ79 Q8CQ81

Q8CQ82 Q8CQ83 Q8CQ84 Q8CQ85 Q8CQ88 Q8CQ89 Q8CQ91 Q8CQ92 Q8CQ93 Q8CQ94 Q8CQ95
Q8CQ96 Q8CQB3 Q8CQB9 Q8CQC2 Q8CQD2 Q8CQD8 Q8CQE8 Q8CQF5 Q8CQF6 Q8CQG5 Q8CQK1
Q8CQK4 Q8CQK5 Q8CQK6 Q8CQK7 Q8CQL3 Q8CQL4 Q8CQN7 Q8CQP4 Q8CQP6 Q8CQQ7 Q8CQQ9
Q8CQR0 Q8CQT5 Q8CQT7 Q8CQT9 Q8CQU1 Q8CQU3 Q8CQU5 Q8CQU6 Q8CQU7 Q8CQU8 Q8CQU9
Q8CQV3 Q8CQV4 Q8CQV5 Q8CQV7 Q8CQV8 Q8CR03 Q8CR04 Q8CR12 Q8CR20 Q8CR21 Q8CR47
Q8CR71 Q8CR97 Q8CRB6 Q8CRC1 Q8CRC6 Q8CRC8 Q8CRD9 Q8CRF7 Q8CRG0 Q8CRG1 Q8CRG2
Q8CRG3 Q8CRG4 Q8CRG5 Q8CRG6 Q8CRG7 Q8CRG8 Q8CRG9 Q8CRH1 Q8CRH2 Q8CRH3 Q8CRH4
Q8CRH5 Q8CRH6 Q8CRH7 Q8CRH8 Q8CRH9 Q8CRI0 Q8CRI1 Q8CRI2 Q8CRI3 Q8CRI4 Q8CRI5
Q8CRI9 Q8CRJ0 Q8CRJ1 Q8CRJ2 Q8CRJ3 Q8CRJ4 Q8CRJ7 Q8CRK4 Q8CRK5 Q8CRL1 Q8CRM3
Q8CRM5 Q8CRM7 Q8CRM8 Q8CRM9 Q8CRN2 Q8CRN3 Q8CRN4 Q8CRN6 Q8CRN9 Q8CRP5 Q8CRP6
Q8CRQ0 Q8CRQ1 Q8CRQ3 Q8CRQ6 Q8CRQ8 Q8CRT1 Q8CRT6 Q8CRT9 Q8CRU0 Q8CRU2 Q8CRU3
Q8CRU4 Q8CRU6 Q8CRU9 Q8CRV4 Q8CRV8 Q8CRV9 Q8CRW4 Q8CRW7 Q8CRY0 Q8CS21 Q8CS25
Q8CS41 Q8CS43 Q8CS46 Q8CS47 Q8CS48 Q8CS54 Q8CS58 Q8CS60 Q8CS61 Q8CS62 Q8CS68
Q8CS69 Q8CS70 Q8CS73 Q8CS74 Q8CS75 Q8CS76 Q8CS77 Q8CS80 Q8CS83 Q8CS87 Q8CS89
Q8CS90 Q8CS91 Q8CS92 Q8CS95 Q8CS97 Q8CS98 Q8CS99 Q8CSA5 Q8CSA7 Q8CSA8 Q8CSA9
Q8CSB0 Q8CSB2 Q8CSB3 Q8CSB6 Q8CSC1 Q8CSC6 Q8CSC7 Q8CSD3 Q8CSD5 Q8CSD7 Q8CSD9
Q8CSE3 Q8CSF2 Q8CSF4 Q8CSF9 Q8CSG0 Q8CSG1 Q8CSG4 Q8CSG7 Q8CSH0 Q8CSH5 Q8CSH9
Q8CSI0 Q8CSI1 Q8CSI3 Q8CSI9 Q8CSJ2 Q8CSJ3 Q8CSK2 Q8CSK6 Q8CSK7 Q8CSL5 Q8CSL9
Q8CSM6 Q8CSM7 Q8CSN0 Q8CSN5 Q8CSN6 Q8CSN9 Q8CSP0 Q8CSP4 Q8CSP8 Q8CSP9 Q8CSQ2
Q8CSR8 Q8CSR9 Q8CSS0 Q8CSS3 Q8CST0 Q8CST1 Q8CST2 Q8CST4 Q8CST6 Q8CST7 Q8CSU0
Q8CSU1 Q8CSU2 Q8CSU3 Q8CSU9 Q8CSV0 Q8CSV3 Q8CSV4 Q8CSV6 Q8CSV8 Q8CSW0 Q8CSW1
Q8CSW3 Q8CSW7 Q8CSX1 Q8CSX5 Q8CSX6 Q8CSX7 Q8CSX8 Q8CSX9 Q8CSY0 Q8CSY5 Q8CSY8
Q8CSY9 Q8CSZ2 Q8CSZ4 Q8CSZ5 Q8CSZ8 Q8CT04 Q8CT08 Q8CT12 Q8CT13 Q8CT15 Q8CT16
Q8CT26 Q8CT27 Q8CT28 Q8CT29 Q8CT30 Q8CT54 Q8CT62 Q8CT64 Q8CT67 Q8CT68 Q8CT69
Q8CT80 Q8CT81 Q8CT82 Q8CT84 Q8CT91 Q8CT93 Q8CTA3 Q8CTA4 Q8CTA8 Q8CTB6 Q8CTD5
Q8CTD6 Q8CTE0 Q8CTE1 Q8CTE3 Q8CTE7 Q8CTF2 Q8CTG5 Q8CTG7 Q8CTG8 Q8CTH8 Q8CTJ9
Q8CTK9 Q8CTN9 Q8CTP4 Q8CTP9 Q8CTQ2 Q8CTQ7 Q8CTQ9 Q8CTR0 Q8CTR3 Q8CTR6 Q8CTR9
Q8CTS8 Q8CTT1 Q8CTT2 Q8CTT4 Q8CTT5 Q8CTT6 Q8CTT9 Q8CTU1 Q8CTU2 Q8CTU3 Q8CTU5
Q8CTV0 Q8CTV1 Q8CTV2 Q8CTV8 Q8CTX1 Q8CTX5 Q8CTX6 Q8CTZ3 Q8CU07 Q8CU16 Q8CU41
Q8CU42 Q8CU52 Q8CU83 Q8CU90 Q8CU95 Q8CUG0 Q8CUG1 Q8CUH1 Q8CUH5 Q8CUK1 Q8CUL8
Q8CUM9 Q8CUN0 Q8CUN1 Q8CUN2 Q8CUP6 Q8CUQ4 Q8CUQ5 Q8CUS7 Q8CUT2 Q8CUT7 Q8CUT9
Q8CUU4 Q8CUW0 Q8CV11 Q8CV19 Q8CV56 Q8CV58 Q8CV64 Q8CV87 Q8CVE9 Q8CVK0 Q8CW10
Q8CW65 Q8CWF6 Q8CWG0 Q8CWG1 Q8CWI5 Q8CWI9 Q8CWJ0 Q8CWJ6 Q8CWK1 Q8CWK3 Q8CWK5
Q8CWK6 Q8CWK7 Q8CWK9 Q8CWL1 Q8CWL2 Q8CWL6 Q8CWL9 Q8CWM0 Q8CWM2 Q8CWM3 Q8CWM5
Q8CWM6 Q8CWM7 Q8CWM8 Q8CWN3 Q8CWN4 Q8CWN5 Q8CWN7 Q8CWN8 Q8CWN9 Q8CWQ0 Q8CWQ1
Q8CWQ3 Q8CWQ4 Q8CWQ5 Q8CWQ7 Q8CWQ9 Q8CWR0 Q8CWR2 Q8CWR5 Q8CWR8 Q8CWR9 Q8CWS0
Q8CWS4 Q8CWS5 Q8CWS7 Q8CWS9 Q8CWT2 Q8CWT3 Q8CWT4 Q8CWU4 Q8CWU6 Q8CWU7 Q8CWU8
Q8CWV0 Q8CWV1 Q8CWV2 Q8CWV3 Q8CWV4 Q8CWV5 Q8CWV6 Q8CWV7 Q8CWV8 Q8CWW0 Q8CWW2
Q8CWW3 Q8CWW4 Q8CWW5 Q8CWW6 Q8CWX0 Q8CWX1 Q8CWX2 Q8CWX4 Q8CWX6 Q8CWX7 Q8CWX9
Q8CWY0 Q8CWY4 Q8CWY5 Q8CWY6 Q8CWZ0 Q8CWZ3 Q8CWZ6 Q8CX00 Q8CX07 Q8CX09 Q8CX13
Q8CX20 Q8CX22 Q8CX26 Q8CX28 Q8CX30 Q8CX31 Q8CX32 Q8CX33 Q8CX40 Q8CX42 Q8CX44
Q8CX45 Q8CX46 Q8CX48 Q8CX49 Q8CX50 Q8CX51 Q8CX52 Q8CX54 Q8CX55 Q8CX59 Q8CX61
Q8CX68 Q8CX76 Q8CXA5 Q8CXB4 Q8CXB7 Q8CXB8 Q8CXC0 Q8CXC4 Q8CXC7 Q8CXC9 Q8CXD2
Q8CXD3 Q8CXD4 Q8CXD9 Q8CXE0 Q8CXE1 Q8CXE2 Q8CXE5 Q8CXF5 Q8CXF9 Q8CXG5 Q8CXG6

Q8CXG7 Q8CXG9 Q8CXH1 Q8CXH2 Q8CXH3 Q8CXH8 Q8CXH9 Q8CXK5 Q8CXK6 Q8CXK7 Q8CXK8
Q8CXL3 Q8CXL4 Q8CXL7 Q8CXM7 Q8CXM8 Q8CXN0 Q8CXN4 Q8CXP1 Q8CXP5 Q8CXP8 Q8CXQ1
Q8CXQ2 Q8CXQ3 Q8CXQ5 Q8CXQ6 Q8CXQ7 Q8CXR2 Q8CXS1 Q8CXS3 Q8CXS6 Q8CXS7 Q8CXS8
Q8CXT5 Q8CXT8 Q8CXU1 Q8CXV3 Q8CXW3 Q8CXW5 Q8CXX5 Q8CXX6 Q8CXZ8 Q8CY19 Q8CY22
Q8CY26 Q8CY27 Q8CY28 Q8CY30 Q8CY34 Q8CY35 Q8CY36 Q8CY38 Q8CY40 Q8CY42 Q8CY45
Q8CY46 Q8CY47 Q8CY49 Q8CY56 Q8CY87 Q8CYD7 Q8CZ70 Q8CZ89 Q8CZD8 Q8CZE3 Q8CZE4
Q8CZE6 Q8CZE8 Q8CZS4 Q8D035 Q8D049 Q8D079 Q8D0D7 Q8D0D8 Q8D0M6 Q8D0U3 Q8D1G8
Q8D1H3 Q8D1I3 Q8D1K0 Q8D1L4 Q8D1U5 Q8D1V0 Q8D1V2 Q8D1V8 Q8D1V9 Q8D1W3 Q8D1W6
Q8D1X1 Q8D1X2 Q8D1X3 Q8D1X5 Q8D1X8 Q8D1X9 Q8D1Y2 Q8D1Y3 Q8D1Y5 Q8D1Y6 Q8D1Y7
Q8D1Y8 Q8D1Y9 Q8D1Z0 Q8D1Z1 Q8D1Z3 Q8D1Z4 Q8D1Z5 Q8D1Z6 Q8D1Z8 Q8D1Z9 Q8D200
Q8D201 Q8D202 Q8D203 Q8D204 Q8D205 Q8D206 Q8D207 Q8D208 Q8D209 Q8D210 Q8D211
Q8D212 Q8D213 Q8D218 Q8D219 Q8D223 Q8D224 Q8D226 Q8D227 Q8D228 Q8D229 Q8D232
Q8D233 Q8D234 Q8D235 Q8D236 Q8D237 Q8D240 Q8D243 Q8D244 Q8D245 Q8D247 Q8D249
Q8D251 Q8D252 Q8D253 Q8D254 Q8D257 Q8D258 Q8D259 Q8D265 Q8D268 Q8D274 Q8D275
Q8D277 Q8D278 Q8D279 Q8D283 Q8D284 Q8D291 Q8D293 Q8D294 Q8D295 Q8D298 Q8D2A1
Q8D2A5 Q8D2A6 Q8D2A7 Q8D2B3 Q8D2B6 Q8D2B9 Q8D2C5 Q8D2E8 Q8D2E9 Q8D2F0 Q8D2F1
Q8D2F3 Q8D2F5 Q8D2G0 Q8D2G2 Q8D2G3 Q8D2G4 Q8D2G5 Q8D2G6 Q8D2H8 Q8D2H9 Q8D2I1
Q8D2J0 Q8D2J1 Q8D2J7 Q8D2J8 Q8D2K0 Q8D2K4 Q8D2K6 Q8D2K8 Q8D2K9 Q8D2L3 Q8D2L6
Q8D2L7 Q8D2L8 Q8D2L9 Q8D2M3 Q8D2M4 Q8D2M5 Q8D2N4 Q8D2N7 Q8D2N8 Q8D2P5 Q8D2P6
Q8D2P7 Q8D2P9 Q8D2Q3 Q8D2Q5 Q8D2Q6 Q8D2Q8 Q8D2R0 Q8D2R2 Q8D2R3 Q8D2R5 Q8D2R7
Q8D2S2 Q8D2S5 Q8D2S6 Q8D2S7 Q8D2T0 Q8D2T4 Q8D2U1 Q8D2U2 Q8D2U3 Q8D2U4 Q8D2V0
Q8D2V1 Q8D2V2 Q8D2V3 Q8D2V5 Q8D2V9 Q8D2W0 Q8D2W3 Q8D2W4 Q8D2W5 Q8D2W7 Q8D2W8
Q8D2W9 Q8D2X0 Q8D2X3 Q8D2X6 Q8D2X7 Q8D2X8 Q8D2X9 Q8D2Y0 Q8D2Y4 Q8D2Y5 Q8D2Y8
Q8D2Z1 Q8D2Z4 Q8D2Z7 Q8D304 Q8D305 Q8D308 Q8D309 Q8D317 Q8D318 Q8D320 Q8D322
Q8D325 Q8D326 Q8D330 Q8D333 Q8D335 Q8D336 Q8D337 Q8D338 Q8D343 Q8D346 Q8D347
Q8D357 Q8D361 Q8D362 Q8D368 Q8D372 Q8D375 Q8D382 Q8D387 Q8D391 Q8D392 Q8D397
Q8D3A7 Q8D3B2 Q8D3B5 Q8D3B6 Q8D3B7 Q8D3B8 Q8D3B9 Q8D3C0 Q8D3C3 Q8D3C5 Q8D3C7
Q8D3C8 Q8D3D3 Q8D3D4 Q8D3D6 Q8D3H1 Q8D3H2 Q8D3H3 Q8D3H4 Q8D3H6 Q8D3H7 Q8D3I0
Q8D3I1 Q8D3I2 Q8D3I7 Q8D3I9 Q8D3J0 Q8D3K0 Q8D3K2 Q8D3K3 Q8D3K4 Q8D3M4 Q8D3M6
Q8D3Q3 Q8D3T7 Q8D3T9 Q8D3Z2 Q8D442 Q8D456 Q8D461 Q8D477 Q8D485 Q8D4B4 Q8D4F7
Q8D4F8 Q8D4H3 Q8D4N5 Q8D4N8 Q8D4P0 Q8D4P3 Q8D4Q0 Q8D4Q2 Q8D4S0 Q8D4T9 Q8D4V5
Q8D4X5 Q8D4Z4 Q8D510 Q8D556 Q8D562 Q8D5H3 Q8D5N2 Q8D5P5 Q8D5Q4 Q8D5R2 Q8D5Z0
Q8D5Z2 Q8D612 Q8D613 Q8D632 Q8D6H8 Q8D6H9 Q8D6I7 Q8D6Q8 Q8D6S1 Q8D737 Q8D757
Q8D781 Q8D788 Q8D7D8 Q8D7E0 Q8D7G5 Q8D7G6 Q8D7G7 Q8D7K4 Q8D7K6 Q8D7M9 Q8D7T8
Q8D7V1 Q8D7Y9 Q8D820 Q8D849 Q8D866 Q8D867 Q8D882 Q8D891 Q8D894 Q8D896 Q8D8B1
Q8D8B2 Q8D8B4 Q8D8C0 Q8D8C1 Q8D8C2 Q8D8C7 Q8D8D2 Q8D8E0 Q8D8F2 Q8D8F4 Q8D8G1
Q8D8G2 Q8D8G4 Q8D8H5 Q8D8I6 Q8D8J1 Q8D8J3 Q8D8J6 Q8D8M6 Q8D8M9 Q8D8N0 Q8D8N2
Q8D8P9 Q8D8Q0 Q8D8Q1 Q8D8Q2 Q8D8Q3 Q8D8Q4 Q8D8Q5 Q8D8Q6 Q8D8Q7 Q8D8R1 Q8D8R2
Q8D8V3 Q8D8W6 Q8D8X2 Q8D900 Q8D911 Q8D915 Q8D921 Q8D926 Q8D935 Q8D937 Q8D951
Q8D985 Q8D992 Q8D994 Q8D9E8 Q8D9F3 Q8D9F8 Q8D9G6 Q8D9H5 Q8D9J0 Q8D9J9 Q8D9M4
Q8D9M7 Q8D9N5 Q8DA11 Q8DA13 Q8DA18 Q8DA19 Q8DA20 Q8DA21 Q8DA31 Q8DA32 Q8DA33
Q8DA35 Q8DA38 Q8DA39 Q8DA40 Q8DA54 Q8DA68 Q8DAF0 Q8DAG7 Q8DAH1 Q8DAH8 Q8DAI5
Q8DAM0 Q8DAN4 Q8DAN6 Q8DAN7 Q8DAN9 Q8DAP0 Q8DAP8 Q8DAP9 Q8DAQ0 Q8DAQ4 Q8DAQ5
Q8DAR0 Q8DAR1 Q8DAR7 Q8DAS0 Q8DAS3 Q8DAU5 Q8DAV0 Q8DB06 Q8DB14 Q8DB22 Q8DB23

Q8DB25 Q8DB36 Q8DB38 Q8DB42 Q8DB44 Q8DBA6 Q8DBB0 Q8DBC2 Q8DBD5 Q8DBD8 Q8DBE1
Q8DBE4 Q8DBE7 Q8DBF5 Q8DBF9 Q8DBG0 Q8DBG1 Q8DBG3 Q8DBG4 Q8DBG5 Q8DBH0 Q8DBH7
Q8DBH9 Q8DBI1 Q8DBI7 Q8DBJ6 Q8DBJ8 Q8DBM6 Q8DBN9 Q8DBS9 Q8DBT0 Q8DBT1 Q8DBT2
Q8DBT6 Q8DBU9 Q8DBV5 Q8DBV8 Q8DBV9 Q8DBW0 Q8DBW2 Q8DBW4 Q8DBW7 Q8DBW9 Q8DBX3
Q8DBX7 Q8DBX8 Q8DBY0 Q8DBZ8 Q8DC06 Q8DC08 Q8DC11 Q8DC12 Q8DC30 Q8DC31 Q8DC32
Q8DC33 Q8DC38 Q8DC47 Q8DC49 Q8DC50 Q8DC51 Q8DC53 Q8DC56 Q8DC57 Q8DC58 Q8DC59
Q8DC60 Q8DC63 Q8DC67 Q8DC73 Q8DC74 Q8DC76 Q8DC84 Q8DC85 Q8DC90 Q8DC93 Q8DC96
Q8DCA0 Q8DCA1 Q8DCA2 Q8DCA3 Q8DCA5 Q8DCA9 Q8DCB0 Q8DCB1 Q8DCB7 Q8DCB9 Q8DCC2
Q8DCC3 Q8DCC5 Q8DCD5 Q8DCE5 Q8DCE7 Q8DCF4 Q8DCF5 Q8DCF6 Q8DCF7 Q8DCG0 Q8DCG1
Q8DCG8 Q8DCJ0 Q8DCJ3 Q8DCJ7 Q8DCK1 Q8DCK2 Q8DCK7 Q8DCL0 Q8DCL3 Q8DCL5 Q8DCL6
Q8DCM0 Q8DCM1 Q8DCM9 Q8DCN0 Q8DCN1 Q8DCN2 Q8DCN7 Q8DCN9 Q8DCP3 Q8DCP4 Q8DCP6
Q8DCP7 Q8DCQ3 Q8DCQ7 Q8DCQ8 Q8DCQ9 Q8DCR0 Q8DCR2 Q8DCR3 Q8DCR7 Q8DCS6 Q8DCS9
Q8DCT6 Q8DCT7 Q8DCT8 Q8DCU1 Q8DCU2 Q8DCU4 Q8DCU9 Q8DCV0 Q8DCV6 Q8DCV7 Q8DCW1
Q8DCW3 Q8DCX0 Q8DCX6 Q8DCY1 Q8DCZ9 Q8DD00 Q8DD03 Q8DD06 Q8DD07 Q8DD14 Q8DD17
Q8DD19 Q8DD20 Q8DD21 Q8DD22 Q8DD23 Q8DD24 Q8DD25 Q8DD27 Q8DD30 Q8DD33 Q8DD39
Q8DD43 Q8DD46 Q8DD50 Q8DD53 Q8DD54 Q8DD81 Q8DD83 Q8DD85 Q8DD95 Q8DDC8 Q8DDD4
Q8DDD5 Q8DDD6 Q8DDD7 Q8DDD8 Q8DDE0 Q8DDE3 Q8DDE4 Q8DDE5 Q8DDG1 Q8DDH8 Q8DDH9
Q8DDI1 Q8DDI3 Q8DDI4 Q8DDI9 Q8DDJ1 Q8DDJ7 Q8DDJ8 Q8DDK1 Q8DDL4 Q8DDL8 Q8DDM6
Q8DDN5 Q8DDN6 Q8DDP9 Q8DDQ8 Q8DDR1 Q8DDS3 Q8DDS6 Q8DDS7 Q8DDU2 Q8DDU4 Q8DDU7
Q8DDV5 Q8DDV6 Q8DDX4 Q8DDX5 Q8DDX7 Q8DDX8 Q8DDY0 Q8DDY1 Q8DDY2 Q8DDY4 Q8DDY6
Q8DE39 Q8DE40 Q8DE41 Q8DE42 Q8DE43 Q8DE44 Q8DE45 Q8DE46 Q8DE47 Q8DE48 Q8DE49
Q8DE50 Q8DE51 Q8DE53 Q8DE54 Q8DE55 Q8DE56 Q8DE57 Q8DE58 Q8DE59 Q8DE61 Q8DE62
Q8DE63 Q8DE64 Q8DE72 Q8DE73 Q8DE75 Q8DE84 Q8DE89 Q8DE90 Q8DE96 Q8DEA0 Q8DEB6
Q8DEC1 Q8DEC2 Q8DEC4 Q8DEC5 Q8DEC6 Q8DED0 Q8DED1 Q8DED2 Q8DED3 Q8DED8 Q8DED9
Q8DEE0 Q8DEE1 Q8DEF3 Q8DEF5 Q8DEG4 Q8DEI1 Q8DEI9 Q8DEJ0 Q8DEJ8 Q8DEK2 Q8DEK5
Q8DEK8 Q8DEL1 Q8DEM0 Q8DEM1 Q8DEM2 Q8DEM9 Q8DEP0 Q8DEP3 Q8DEP5 Q8DEP6 Q8DEP7
Q8DEQ1 Q8DEQ3 Q8DEQ9 Q8DER4 Q8DER5 Q8DER9 Q8DES4 Q8DES7 Q8DET0 Q8DET7 Q8DEU5
Q8DEU6 Q8DEV0 Q8DEV2 Q8DEV7 Q8DEW2 Q8DEX2 Q8DEX9 Q8DEY0 Q8DEY6 Q8DEY7 Q8DEZ0
Q8DEZ1 Q8DEZ4 Q8DEZ5 Q8DEZ6 Q8DEZ8 Q8DEZ9 Q8DF02 Q8DF05 Q8DF07 Q8DF11 Q8DF53
Q8DF55 Q8DF58 Q8DF59 Q8DF66 Q8DF67 Q8DF81 Q8DF91 Q8DF92 Q8DF93 Q8DF94 Q8DF95
Q8DF99 Q8DFA0 Q8DFA3 Q8DFA5 Q8DFA8 Q8DFB2 Q8DFB4 Q8DFB8 Q8DFC9 Q8DFD1 Q8DFD2
Q8DFD3 Q8DFD8 Q8DFE2 Q8DFE6 Q8DFE8 Q8DFF4 Q8DFF5 Q8DFF6 Q8DFF8 Q8DFF9 Q8DFH5
Q8DFK5 Q8DFM0 Q8DFM1 Q8DFM2 Q8DFQ1 Q8DFR1 Q8DFS2 Q8DFS4 Q8DFS5 Q8DG12 Q8DG25
Q8DG26 Q8DG27 Q8DG35 Q8DG49 Q8DG51 Q8DG62 Q8DG71 Q8DG73 Q8DG79 Q8DG98 Q8DGA5
Q8DGC2 Q8DGC3 Q8DGC4 Q8DGC6 Q8DGC8 Q8DGE4 Q8DGG4 Q8DGH0 Q8DGH2 Q8DGH7 Q8DGH9
Q8DGI6 Q8DGI7 Q8DGI8 Q8DGJ7 Q8DGL0 Q8DGM0 Q8DGM2 Q8DGP3 Q8DGP7 Q8DGQ5 Q8DGQ8
Q8DGR0 Q8DGR1 Q8DGR2 Q8DGS4 Q8DGS9 Q8DGU6 Q8DGV8 Q8DGW0 Q8DGW9 Q8DGX5 Q8DH01
Q8DH02 Q8DH04 Q8DH10 Q8DH26 Q8DH31 Q8DH33 Q8DH42 Q8DH56 Q8DH57 Q8DH61 Q8DH70
Q8DH87 Q8DH98 Q8DH99 Q8DHA4 Q8DHB4 Q8DHC4 Q8DHC6 Q8DHE1 Q8DHE3 Q8DHG3 Q8DHG5
Q8DHH2 Q8DHH6 Q8DHK2 Q8DHL1 Q8DHM0 Q8DHN1 Q8DHN4 Q8DHN8 Q8DHP7 Q8DHR2 Q8DHS1
Q8DHT4 Q8DHV7 Q8DHW5 Q8DHW6 Q8DHW7 Q8DHW8 Q8DHY2 Q8DHY6 Q8DHZ6 Q8DI06 Q8DI20
Q8DI26 Q8DI27 Q8DI29 Q8DI39 Q8DI41 Q8DI42 Q8DI43 Q8DI44 Q8DI46 Q8DI53 Q8DI54
Q8DI58 Q8DI59 Q8DI68 Q8DI74 Q8DI76 Q8DI83 Q8DI86 Q8DI91 Q8DI93 Q8DIA2 Q8DIA3
Q8DIA7 Q8DIB2 Q8DIB4 Q8DIE4 Q8DIF1 Q8DIF6 Q8DIF9 Q8DIG8 Q8DIH2 Q8DII3 Q8DII8

Q8DIK0 Q8DIL8 Q8DIM4 Q8DIN0 Q8DIN5 Q8DIP5 Q8DIP7 Q8DIQ7 Q8DIR6 Q8DIS5 Q8DIS6
Q8DIS7 Q8DIS8 Q8DIT5 Q8DIU1 Q8DJ05 Q8DJ26 Q8DJ32 Q8DJ40 Q8DJ45 Q8DJ55 Q8DJ62
Q8DJ63 Q8DJ87 Q8DJ97 Q8DJB1 Q8DJB2 Q8DJB3 Q8DJB8 Q8DJC4 Q8DJC5 Q8DJD3 Q8DJE4
Q8DJE6 Q8DJE9 Q8DJF2 Q8DJI4 Q8DJI6 Q8DJK4 Q8DJK7 Q8DJK9 Q8DJL5 Q8DJN4 Q8DJN6
Q8DJN7 Q8DJP8 Q8DJQ1 Q8DJQ7 Q8DJS3 Q8DJS8 Q8DJT5 Q8DJW4 Q8DJY0 Q8DJY2 Q8DJY3
Q8DJZ1 Q8DJZ9 Q8DK04 Q8DK05 Q8DK09 Q8DK13 Q8DK23 Q8DK29 Q8DK30 Q8DK36 Q8DK65
Q8DK70 Q8DK79 Q8DK85 Q8DK88 Q8DKA0 Q8DKB2 Q8DKB3 Q8DKB4 Q8DKB5 Q8DKB7 Q8DKD7
Q8DKE4 Q8DKF0 Q8DKF1 Q8DKG1 Q8DKH2 Q8DKH7 Q8DKI1 Q8DKJ8 Q8DKK1 Q8DKK2 Q8DKM1
Q8DKM7 Q8DKN4 Q8DKP1 Q8DKQ1 Q8DKQ5 Q8DKR4 Q8DKR6 Q8DKR7 Q8DKS0 Q8DKS3 Q8DKS7
Q8DKT0 Q8DKT7 Q8DKT8 Q8DKU1 Q8DKU2 Q8DKU4 Q8DKU5 Q8DKV6 Q8DKW0 Q8DKX6 Q8DKY2
Q8DKY7 Q8DL09 Q8DL15 Q8DL36 Q8DL37 Q8DL38 Q8DL55 Q8DL56 Q8DL57 Q8DL74 Q8DL83
Q8DL91 Q8DL93 Q8DL94 Q8DLA4 Q8DLA6 Q8DLA9 Q8DLB8 Q8DLC2 Q8DLF2 Q8DLF7 Q8DLF8
Q8DLG1 Q8DLG2 Q8DLG3 Q8DLH0 Q8DLI1 Q8DLI2 Q8DLI3 Q8DLI5 Q8DLI9 Q8DLJ1 Q8DLJ7
Q8DLK3 Q8DLK8 Q8DLL7 Q8DLM1 Q8DLN9 Q8DLQ0 Q8DLS5 Q8DLT3 Q8DLV5 Q8DLV6 Q8DLW0
Q8DLY3 Q8DLZ2 Q8DLZ5 Q8DLZ6 Q8DM00 Q8DM24 Q8DM25 Q8DM26 Q8DM27 Q8DM28 Q8DM42
Q8DM44 Q8DM56 Q8DM63 Q8DM64 Q8DM88 Q8DM90 Q8DMA3 Q8DMA5 Q8DMA8 Q8DMA9 Q8DMD4
Q8DMD8 Q8DMD9 Q8DME3 Q8DMF1 Q8DMF2 Q8DMG9 Q8DMI3 Q8DMI6 Q8DMK6 Q8DMK7 Q8DMK8
Q8DMK9 Q8DML0 Q8DML1 Q8DML2 Q8DML3 Q8DML4 Q8DML6 Q8DML7 Q8DML8 Q8DML9 Q8DMM0
Q8DMM1 Q8DMM2 Q8DMM3 Q8DMM4 Q8DMM5 Q8DMM6 Q8DMM7 Q8DMM8 Q8DMM9 Q8DMN0 Q8DMN1
Q8DMN8 Q8DMP4 Q8DMP6 Q8DMQ4 Q8DMQ7 Q8DMR1 Q8DMS1 Q8DMS3 Q8DMU0 Q8DMV6 Q8DMX3
Q8DN24 Q8DN46 Q8DN52 Q8DN54 Q8DN55 Q8DN69 Q8DN74 Q8DN92 Q8DNC9 Q8DND3 Q8DNE6
Q8DNE8 Q8DNF0 Q8DNF1 Q8DNH5 Q8DNH6 Q8DNI9 Q8DNK7 Q8DNK8 Q8DNL1 Q8DNL8 Q8DNM6
Q8DNM7 Q8DNM8 Q8DNM9 Q8DNP4 Q8DNP8 Q8DNR1 Q8DNR5 Q8DNR7 Q8DNR9 Q8DNV5 Q8DNW1
Q8DNW4 Q8DNX2 Q8DNX8 Q8DNY9 Q8DP01 Q8DP05 Q8DP10 Q8DP12 Q8DP13 Q8DP16 Q8DP23
Q8DP31 Q8DP40 Q8DP42 Q8DP65 Q8DP66 Q8DP67 Q8DP71 Q8DP72 Q8DP79 Q8DPA0 Q8DPA1
Q8DPA2 Q8DPA8 Q8DPC7 Q8DPD1 Q8DPD3 Q8DPE0 Q8DPF0 Q8DPH3 Q8DPH9 Q8DPI2 Q8DPI3
Q8DPI7 Q8DPJ4 Q8DPJ7 Q8DPJ8 Q8DPK7 Q8DPL5 Q8DPM2 Q8DPM9 Q8DPN9 Q8DPP0 Q8DPP1
Q8DPP2 Q8DPP6 Q8DPQ3 Q8DPQ4 Q8DPQ5 Q8DPR3 Q8DPR6 Q8DPR7 Q8DPS3 Q8DPS5 Q8DPS6
Q8DPS9 Q8DPU5 Q8DPU7 Q8DPV4 Q8DPV8 Q8DPW5 Q8DPY7 Q8DPY9 Q8DPZ0 Q8DPZ3 Q8DPZ4
Q8DPZ6 Q8DPZ8 Q8DPZ9 Q8DQ04 Q8DQ05 Q8DQ29 Q8DQ32 Q8DQ33 Q8DQ35 Q8DQ36 Q8DQ37
Q8DQ38 Q8DQ47 Q8DQ49 Q8DQ50 Q8DQ51 Q8DQ56 Q8DQ58 Q8DQ60 Q8DQ61 Q8DQ68 Q8DQ78
Q8DQ85 Q8DQ87 Q8DQ92 Q8DQC4 Q8DQC7 Q8DQD0 Q8DQD1 Q8DQD2 Q8DQD3 Q8DQG3 Q8DQG4
Q8DQG7 Q8DQG9 Q8DQH0 Q8DQI2 Q8DQI3 Q8DQJ9 Q8DQK4 Q8DQK5 Q8DQL5 Q8DQL6 Q8DQM2
Q8DQN1 Q8DQN6 Q8DQQ9 Q8DQS7 Q8DQS8 Q8DQT0 Q8DQT2 Q8DQT6 Q8DQT8 Q8DQU7 Q8DQV1
Q8DQV2 Q8DQV6 Q8DQV8 Q8DQW9 Q8DQX8 Q8DQY0 Q8DQZ1 Q8DQZ7 Q8DR03 Q8DR09 Q8DR10
Q8DR11 Q8DR22 Q8DR26 Q8DR29 Q8DR31 Q8DR48 Q8DR57 Q8DR58 Q8DR77 Q8DRA5 Q8DRA8
Q8DRB0 Q8DRB3 Q8DRB6 Q8DRD2 Q8DRD4 Q8DRE1 Q8DRE2 Q8DRE8 Q8DRF4 Q8DRH2 Q8DRH6
Q8DRH8 Q8DRI4 Q8DRI5 Q8DRM0 Q8DRM1 Q8DRM4 Q8DRM7 Q8DRN4 Q8DRP0 Q8DRP8 Q8DRP9
Q8DRQ2 Q8DRQ7 Q8DRR1 Q8DRR3 Q8DRS4 Q8DRS6 Q8DRS8 Q8DRT7 Q8DRV7 Q8DRV9 Q8DRW2
Q8DRW6 Q8DRW8 Q8DRX0 Q8DRX2 Q8DRX7 Q8DRX8 Q8DRX9 Q8DRY6 Q8DS02 Q8DS08 Q8DS11
Q8DS12 Q8DS16 Q8DS19 Q8DS20 Q8DS21 Q8DS22 Q8DS23 Q8DS24 Q8DS25 Q8DS26 Q8DS27
Q8DS28 Q8DS29 Q8DS30 Q8DS31 Q8DS33 Q8DS34 Q8DS35 Q8DS36 Q8DS37 Q8DS40 Q8DS44
Q8DS46 Q8DS47 Q8DS57 Q8DS85 Q8DS88 Q8DS90 Q8DSB7 Q8DSC8 Q8DSD1 Q8DSD7 Q8DSD8
Q8DSE7 Q8DSE9 Q8DSF4 Q8DSG3 Q8DSG4 Q8DSG5 Q8DSG6 Q8DSI6 Q8DSJ2 Q8DSJ4 Q8DSJ7

Q8DSJ9 Q8DSK1 Q8DSK2 Q8DSK3 Q8DSL1 Q8DSP4 Q8DSP7 Q8DSQ5 Q8DSQ6 Q8DSQ9 Q8DST2
Q8DST6 Q8DST7 Q8DST9 Q8DSU6 Q8DSU9 Q8DSV3 Q8DSX9 Q8DSY0 Q8DSY1 Q8DSY2 Q8DSY4
Q8DSY5 Q8DSY7 Q8DSZ0 Q8DSZ3 Q8DSZ5 Q8DT12 Q8DT23 Q8DT24 Q8DT49 Q8DT52 Q8DT53
Q8DT57 Q8DT66 Q8DT75 Q8DT76 Q8DT87 Q8DT90 Q8DT95 Q8DT96 Q8DTC6 Q8DTC7 Q8DTE3
Q8DTE5 Q8DTG1 Q8DTG3 Q8DTG4 Q8DTG5 Q8DTM1 Q8DTM2 Q8DTM7 Q8DTM9 Q8DTN5 Q8DTN6
Q8DTN7 Q8DTN8 Q8DTP5 Q8DTQ4 Q8DTQ5 Q8DTQ6 Q8DTQ7 Q8DTQ9 Q8DTR1 Q8DTR2 Q8DTR3
Q8DTR4 Q8DTT8 Q8DTT9 Q8DTU0 Q8DTU9 Q8DTV1 Q8DTV2 Q8DTV8 Q8DTX6 Q8DTY0 Q8DU15
Q8DU17 Q8DU30 Q8DU31 Q8DU34 Q8DU62 Q8DU63 Q8DU64 Q8DU67 Q8DU81 Q8DU91 Q8DU94
Q8DU98 Q8DUC3 Q8DUC9 Q8DUE1 Q8DUE5 Q8DUF8 Q8DUG2 Q8DUG4 Q8DUG9 Q8DUH2 Q8DUH9
Q8DUI0 Q8DUI4 Q8DUI9 Q8DUK3 Q8DUK7 Q8DUL0 Q8DUL8 Q8DUL9 Q8DUN5 Q8DUN6 Q8DUN7
Q8DUN9 Q8DUP3 Q8DUP4 Q8DUP5 Q8DUP7 Q8DUQ4 Q8DUQ6 Q8DUR1 Q8DUU2 Q8DUV4 Q8DUV8
Q8DUV9 Q8DUW1 Q8DUW2 Q8DUW3 Q8DUW4 Q8DUW8 Q8DUX9 Q8DV10 Q8DV20 Q8DV21 Q8DV22
Q8DV23 Q8DV44 Q8DV45 Q8DV70 Q8DV72 Q8DV91 Q8DV99 Q8DVB4 Q8DVB5 Q8DVC1 Q8DVC9
Q8DVD3 Q8DVD7 Q8DVE3 Q8DVF2 Q8DVF3 Q8DVF4 Q8DVF5 Q8DVF6 Q8DVG8 Q8DVH2 Q8DVK4
Q8DVK5 Q8DVK6 Q8DVL1 Q8DVL2 Q8DVL3 Q8DVM7 Q8DVM9 Q8DVN0 Q8DVN2 Q8DVN4 Q8DVP8
Q8DVP9 Q8DVQ3 Q8DVQ4 Q8DVR7 Q8DVS2 Q8DVS3 Q8DVT0 Q8DVU6 Q8DVV2 Q8DVV4 Q8DVV5
Q8DVW1 Q8DVX0 Q8DVX4 Q8DVX5 Q8DVX9 Q8DVY6 Q8DVY7 Q8DVZ3 Q8DW14 Q8DW15 Q8DW17
Q8DW19 Q8DW33 Q8DW43 Q8DW97 Q8DWA9 Q8DWB2 Q8DWB3 Q8DWC2 Q8DWE0 Q8DWE1 Q8DWE5
Q8DWG1 Q8DWG2 Q8DWG8 Q8DWH1 Q8DWH2 Q8DWH7 Q8DWH8 Q8DWI4 Q8DWK8 Q8DWL1 Q8DWL6
Q8DWM0 Q8DWM2 Q8DWM8 Q8DWM9 Q8DWN5 Q8DWN9 Q8DWP7 Q8DWQ2 Q8DWQ8 Q8DWS3 Q8DWS6
Q8DWV3 Q8DWV5 Q8DWV9 Q8DWW1 Q8DWW6 Q8DWX0 Q8DWX2 Q8DWY0 Q8DWZ0 Q8DWZ4 Q8DX85
Q8DXC0 Q8DXC1 Q8DXC2 Q8DXC3 Q8DXD5 Q8DXD8 Q8DXF6 Q8DXI7 Q8DXL8 Q8DXM9 Q8DXP1
Q8DXQ6 Q8DXR8 Q8DXR9 Q8DXS5 Q8DXS6 Q8DXS7 Q8DXT0 Q8DXT6 Q8DXT9 Q8DXU2 Q8DXW0
Q8DXW9 Q8DXX3 Q8DXX6 Q8DXY3 Q8DY10 Q8DY22 Q8DY24 Q8DY25 Q8DY26 Q8DY32 Q8DY37
Q8DY64 Q8DY70 Q8DY73 Q8DY77 Q8DY80 Q8DY92 Q8DY93 Q8DY96 Q8DYA5 Q8DYA6 Q8DYB3
Q8DYB5 Q8DYG1 Q8DYG8 Q8DYH1 Q8DYH2 Q8DYJ2 Q8DYJ6 Q8DYL0 Q8DYL6 Q8DYQ7 Q8DYS6
Q8DYT2 Q8DYT4 Q8DYT8 Q8DYU0 Q8DYU4 Q8DYU5 Q8DYU6 Q8DYU7 Q8DYV1 Q8DYV3 Q8DYV5
Q8DYV9 Q8DYW8 Q8DYW9 Q8DYY1 Q8DZ07 Q8DZ10 Q8DZ11 Q8DZ20 Q8DZ21 Q8DZ90 Q8DZ99
Q8DZA4 Q8DZC6 Q8DZG5 Q8DZH0 Q8DZI2 Q8DZI5 Q8DZK0 Q8DZK1 Q8DZK4 Q8DZK7 Q8DZL4
Q8DZL5 Q8DZM3 Q8DZM4 Q8DZM7 Q8DZP5 Q8DZQ2 Q8DZQ5 Q8DZQ7 Q8DZR0 Q8DZT5 Q8DZV0
Q8DZV1 Q8DZX5 Q8DZX7 Q8DZY3 Q8DZZ2 Q8DZZ7 Q8E026 Q8E031 Q8E049 Q8E061 Q8E062
Q8E064 Q8E066 Q8E069 Q8E079 Q8E080 Q8E081 Q8E084 Q8E091 Q8E092 Q8E093 Q8E0A3
Q8E0A8 Q8E0C4 Q8E0C9 Q8E0D7 Q8E0F5 Q8E0G1 Q8E0G6 Q8E0G7 Q8E0G9 Q8E0H0 Q8E0H1
Q8E0H2 Q8E0I1 Q8E0I2 Q8E0J5 Q8E0J8 Q8E0K7 Q8E0L9 Q8E0M8 Q8E0M9 Q8E0N5 Q8E0T7
Q8E0U0 Q8E0V1 Q8E0W1 Q8E0Z9 Q8E121 Q8E130 Q8E146 Q8E155 Q8E166 Q8E168 Q8E177
Q8E181 Q8E186 Q8E197 Q8E1A3 Q8E1A6 Q8E1B0 Q8E1B5 Q8E1E4 Q8E1E5 Q8E1E7 Q8E1F9
Q8E1H3 Q8E1Q1 Q8E1Q3 Q8E1Q4 Q8E1Q7 Q8E1R8 Q8E1R9 Q8E1T0 Q8E1T2 Q8E1T3 Q8E1T4
Q8E1T5 Q8E1X1 Q8E1Y6 Q8E1Z4 Q8E1Z8 Q8E1Z9 Q8E220 Q8E231 Q8E238 Q8E239 Q8E241
Q8E245 Q8E259 Q8E268 Q8E271 Q8E272 Q8E280 Q8E284 Q8E290 Q8E291 Q8E292 Q8E294
Q8E298 Q8E299 Q8E2A0 Q8E2B0 Q8E2B1 Q8E2B5 Q8E2B6 Q8E2B7 Q8E2B8 Q8E2B9 Q8E2C0
Q8E2C1 Q8E2C2 Q8E2C3 Q8E2C4 Q8E2C5 Q8E2C6 Q8E2C7 Q8E2C8 Q8E2D0 Q8E2D1 Q8E2D9
Q8E2G1 Q8E2G4 Q8E2G8 Q8E2H0 Q8E2H3 Q8E2H4 Q8E2I1 Q8E2I7 Q8E2J5 Q8E2K0 Q8E2K7
Q8E2M3 Q8E2M6 Q8E2Q7 Q8E2R1 Q8E2R3 Q8E2R5 Q8E2R7 Q8E2R9 Q8E2S2 Q8E2S4 Q8E2T1
Q8E2U1 Q8E307 Q8E344 Q8E345 Q8E346 Q8E347 Q8E359 Q8E362 Q8E372 Q8E378 Q8E388

Q8E399 Q8E3B1 Q8E3C5 Q8E3D7 Q8E3D9 Q8E3E5 Q8E3E6 Q8E3E7 Q8E3F0 Q8E3F6 Q8E3F9
Q8E3G1 Q8E3H8 Q8E3I8 Q8E3J2 Q8E3J5 Q8E3K2 Q8E3M9 Q8E3P1 Q8E3P3 Q8E3P4 Q8E3P5
Q8E3Q6 Q8E3T0 Q8E3T6 Q8E3T9 Q8E3U2 Q8E3U5 Q8E3V7 Q8E3V8 Q8E3W1 Q8E3X0 Q8E3X1
Q8E3X8 Q8E3Y0 Q8E3Y3 Q8E424 Q8E431 Q8E434 Q8E435 Q8E449 Q8E465 Q8E471 Q8E4B4
Q8E4D4 Q8E4E0 Q8E4E2 Q8E4E6 Q8E4E8 Q8E4F2 Q8E4F3 Q8E4F4 Q8E4F5 Q8E4F9 Q8E4G1
Q8E4G3 Q8E4G7 Q8E4H6 Q8E4H7 Q8E4H8 Q8E4J0 Q8E4L6 Q8E4L8 Q8E4L9 Q8E4M6 Q8E4M7
Q8E4R5 Q8E4R6 Q8E4R7 Q8E4R8 Q8E4S2 Q8E4V8 Q8E4W1 Q8E4W5 Q8E4Y6 Q8E531 Q8E536
Q8E546 Q8E549 Q8E564 Q8E565 Q8E568 Q8E571 Q8E578 Q8E5B5 Q8E5C2 Q8E5C3 Q8E5C6
Q8E5E3 Q8E5F0 Q8E5F3 Q8E5F5 Q8E5I2 Q8E5J8 Q8E5J9 Q8E5M6 Q8E5M8 Q8E5N4 Q8E5P3
Q8E5P8 Q8E5Q9 Q8E5R4 Q8E5S6 Q8E5T7 Q8E5T9 Q8E5U0 Q8E5U1 Q8E5U3 Q8E5U6 Q8E5V5
Q8E5V7 Q8E5V8 Q8E5W1 Q8E5W8 Q8E5W9 Q8E5X0 Q8E5Y0 Q8E5Y5 Q8E600 Q8E611 Q8E629
Q8E635 Q8E640 Q8E641 Q8E643 Q8E644 Q8E645 Q8E647 Q8E656 Q8E657 Q8E671 Q8E680
Q8E693 Q8E6A2 Q8E6A3 Q8E6A9 Q8E6F5 Q8E6F8 Q8E6G8 Q8E6H6 Q8E6I5 Q8E6K1 Q8E6L0
Q8E6M1 Q8E6M3 Q8E6N1 Q8E6N5 Q8E6P1 Q8E6Q1 Q8E6Q7 Q8E6R1 Q8E6R5 Q8E6S0 Q8E6W1
Q8E6W2 Q8E6W5 Q8E6X6 Q8E766 Q8E767 Q8E768 Q8E771 Q8E782 Q8E783 Q8E794 Q8E796
Q8E797 Q8E7E4 Q8E7F2 Q8E7F6 Q8E7F7 Q8E7H6 Q8E7I7 Q8E7J7 Q8E7J8 Q8E7K0 Q8E7K5
Q8E7L9 Q8E7M7 Q8E7N0 Q8E7N1 Q8E7N8 Q8E7P2 Q8E7P8 Q8E7P9 Q8E7Q0 Q8E7Q2 Q8E7Q5
Q8E7Q7 Q8E7Q8 Q8E7Q9 Q8E7R8 Q8E7S2 Q8E7S3 Q8E7S4 Q8E7S5 Q8E7S6 Q8E7S7 Q8E7S8
Q8E7S9 Q8E7T0 Q8E7T1 Q8E7T2 Q8E7T3 Q8E7T4 Q8E7T5 Q8E7T7 Q8E7T8 Q8E7U6 Q8E7W9
Q8E7X2 Q8E7X6 Q8E7X8 Q8E7Y1 Q8E7Y2 Q8E7Y8 Q8E7Z4 Q8E8A9 Q8E8B0 Q8E8D4 Q8E8D7
Q8E8D9 Q8E8E3 Q8E8I0 Q8E8J1 Q8E8J2 Q8E8J7 Q8E8K6 Q8E8N6 Q8E8P2 Q8E8T4 Q8E927
Q8E942 Q8E988 Q8E990 Q8E9D5 Q8E9D9 Q8E9H0 Q8E9H2 Q8E9H5 Q8E9K5 Q8E9L4 Q8E9L5
Q8E9L6 Q8E9L9 Q8E9M0 Q8E9M2 Q8E9M3 Q8E9M4 Q8E9N2 Q8E9N3 Q8E9N4 Q8E9N5 Q8E9N7
Q8E9N9 Q8E9P0 Q8E9P3 Q8E9P6 Q8E9P8 Q8E9R7 Q8E9R9 Q8E9U9 Q8E9V0 Q8E9Y7 Q8E9Z0
Q8EA14 Q8EA30 Q8EA37 Q8EA43 Q8EA52 Q8EA81 Q8EAB6 Q8EAC4 Q8EAE2 Q8EAF7 Q8EAG2
Q8EAG3 Q8EAG5 Q8EAG8 Q8EAH2 Q8EAH3 Q8EAH4 Q8EAH5 Q8EAI6 Q8EAI9 Q8EAJ0 Q8EAL4
Q8EAN7 Q8EAP2 Q8EAQ9 Q8EAR0 Q8EAR2 Q8EAR3 Q8EAR4 Q8EAS7 Q8EAZ2 Q8EAZ9 Q8EB00
Q8EB01 Q8EB09 Q8EB10 Q8EB13 Q8EB40 Q8EB78 Q8EB80 Q8EB81 Q8EB83 Q8EB91 Q8EB92
Q8EB93 Q8EB94 Q8EBB7 Q8EBE6 Q8EBF9 Q8EBH1 Q8EBH2 Q8EBH6 Q8EBH9 Q8EBI4 Q8EBI7
Q8EBN8 Q8EBN9 Q8EBP3 Q8EBP4 Q8EBQ3 Q8EBQ9 Q8EBR2 Q8EBR3 Q8EBR4 Q8EBR5 Q8EBR6
Q8EBR9 Q8EBS0 Q8EBS1 Q8EBS3 Q8EBS5 Q8EBU8 Q8EBX6 Q8EBX8 Q8EBY0 Q8EBY7 Q8EBY9
Q8EBZ7 Q8EBZ8 Q8EBZ9 Q8EC29 Q8EC32 Q8EC33 Q8EC36 Q8EC50 Q8EC52 Q8EC57 Q8ECH6
Q8ECH7 Q8ECI8 Q8ECK0 Q8ECM2 Q8ECM3 Q8ECQ4 Q8ECR0 Q8ECR2 Q8ECU9 Q8ECV0 Q8ECV2
Q8ED55 Q8ED70 Q8ED71 Q8ED72 Q8EDD1 Q8EDD7 Q8EDF2 Q8EDG1 Q8EDG7 Q8EDG9 Q8EDI8
Q8EDI9 Q8EDJ2 Q8EDK3 Q8EDK6 Q8EDK8 Q8EDL3 Q8EDM4 Q8EDN7 Q8EDU8 Q8EDW4 Q8EDW6
Q8EDW7 Q8EDW8 Q8EDX2 Q8EDX4 Q8EDX5 Q8EDX8 Q8EDY2 Q8EDY8 Q8EDZ9 Q8EE14 Q8EE15
Q8EE17 Q8EE23 Q8EE27 Q8EE30 Q8EE60 Q8EE85 Q8EEA2 Q8EEA3 Q8EEB6 Q8EED7 Q8EEE1
Q8EEE6 Q8EEE7 Q8EEE9 Q8EEF0 Q8EEF1 Q8EEF2 Q8EEF3 Q8EEG9 Q8EEH2 Q8EEH8 Q8EEH9
Q8EEI1 Q8EEI4 Q8EEM4 Q8EEN5 Q8EEN9 Q8EEP2 Q8EEP8 Q8EEP9 Q8EEQ1 Q8EEQ9 Q8EER6
Q8EER7 Q8EER8 Q8EER9 Q8EES8 Q8EEU0 Q8EEU3 Q8EEU5 Q8EEU6 Q8EEU9 Q8EEW0 Q8EEW2
Q8EEZ1 Q8EEZ2 Q8EF25 Q8EF26 Q8EF62 Q8EF74 Q8EF96 Q8EF98 Q8EF99 Q8EFA0 Q8EFB0
Q8EFB1 Q8EFB2 Q8EFB3 Q8EFB4 Q8EFB5 Q8EFB6 Q8EFF2 Q8EFF4 Q8EFF5 Q8EFF7 Q8EFF8
Q8EFF9 Q8EFG1 Q8EFG5 Q8EFG7 Q8EFN2 Q8EFN8 Q8EFP8 Q8EFT7 Q8EFU5 Q8EFU8 Q8EFV4
Q8EFW4 Q8EG18 Q8EG19 Q8EG20 Q8EG21 Q8EG22 Q8EG26 Q8EG63 Q8EG64 Q8EG65 Q8EG66

Q8EGC8 Q8EGD7 Q8EGF2 Q8EGF9 Q8EGG1 Q8EGG9 Q8EGH2 Q8EGH3 Q8EGH4 Q8EGH5 Q8EGH8
Q8EGH9 Q8EGJ4 Q8EGP3 Q8EGP9 Q8EGQ9 Q8EGR4 Q8EGR9 Q8EGS0 Q8EGS1 Q8EGU3 Q8EGU7
Q8EGW6 Q8EGW8 Q8EH28 Q8EH62 Q8EH67 Q8EH68 Q8EH70 Q8EH71 Q8EH72 Q8EH78 Q8EH81
Q8EH90 Q8EH94 Q8EH98 Q8EH99 Q8EHA7 Q8EHB3 Q8EHB5 Q8EHC4 Q8EHC8 Q8EHD6 Q8EHD7
Q8EHJ0 Q8EHK0 Q8EHK2 Q8EHK3 Q8EHK4 Q8EHL1 Q8EHL3 Q8EHL4 Q8EHL5 Q8EHL7 Q8EHL9
Q8EHM0 Q8EHM3 Q8EHN5 Q8EHN9 Q8EHP4 Q8EHP8 Q8EHQ4 Q8EHQ5 Q8EHQ6 Q8EHR7 Q8EHR8
Q8EHS6 Q8EHS7 Q8EHT6 Q8EHT7 Q8EHU1 Q8EHU2 Q8EHU9 Q8EHW5 Q8EHZ1 Q8EHZ2 Q8EI15
Q8EI18 Q8EI58 Q8EI67 Q8EI85 Q8EI95 Q8EIB1 Q8EIB2 Q8EIB4 Q8EIC6 Q8EIF2 Q8EIG4
Q8EIG6 Q8EIG9 Q8EIH0 Q8EIJ7 Q8EIK5 Q8EIK8 Q8EIL1 Q8EIM0 Q8EIM7 Q8EIQ6 Q8EIQ7
Q8EIQ8 Q8EIR6 Q8EIV3 Q8EIX3 Q8EIX4 Q8EJ54 Q8EJ69 Q8EJ70 Q8EJ78 Q8EJ79 Q8EJ86
Q8EJ93 Q8EJA7 Q8EJC1 Q8EJF5 Q8EJM1 Q8EJM8 Q8EJP9 Q8EJQ0 Q8EJQ1 Q8EJQ5 Q8EJR7
Q8EJR8 Q8EJR9 Q8EJU6 Q8EJU7 Q8EJW1 Q8EJW2 Q8EJW3 Q8EJW4 Q8EJX1 Q8EJX6 Q8EJZ5
Q8EK07 Q8EK08 Q8EK12 Q8EK19 Q8EK20 Q8EK27 Q8EK28 Q8EK29 Q8EK30 Q8EK32 Q8EK46
Q8EK47 Q8EK48 Q8EK49 Q8EK51 Q8EK52 Q8EK53 Q8EK54 Q8EK55 Q8EK56 Q8EK57 Q8EK58
Q8EK59 Q8EK60 Q8EK61 Q8EK62 Q8EK63 Q8EK64 Q8EK65 Q8EK66 Q8EK67 Q8EK68 Q8EK69
Q8EK70 Q8EK71 Q8EK72 Q8EK73 Q8EK74 Q8EK75 Q8EK76 Q8EK77 Q8EK78 Q8EK81 Q8EK83
Q8EK85 Q8EK90 Q8EK99 Q8EKA0 Q8EKD2 Q8EKD3 Q8EKF2 Q8EKF9 Q8EKG2 Q8EKI6 Q8EKI8
Q8EKJ4 Q8EKJ5 Q8EKJ7 Q8EKK0 Q8EKK8 Q8EKP0 Q8EKQ0 Q8EKQ2 Q8EKQ3 Q8EKQ5 Q8EKQ8
Q8EKQ9 Q8EKR0 Q8EKR8 Q8EKS1 Q8EKS5 Q8EKS6 Q8EKT0 Q8EKT3 Q8EKT4 Q8EKT9 Q8EKU0
Q8EKU3 Q8EKU4 Q8EKU5 Q8EKV4 Q8EKV5 Q8EKX1 Q8EKX9 Q8EL00 Q8EL04 Q8EL19 Q8EL21
Q8EL49 Q8EL52 Q8EL95 Q8ELF0 Q8ELF4 Q8ELH4 Q8ELI2 Q8ELK1 Q8ELL1 Q8ELR0 Q8ELR9
Q8ELT8 Q8ELT9 Q8ELU7 Q8ELW7 Q8EM07 Q8EM37 Q8EM39 Q8EM45 Q8EM52 Q8EM53 Q8EM55
Q8EM58 Q8EM59 Q8EM60 Q8EM72 Q8EM73 Q8EM74 Q8EM86 Q8EMI3 Q8EMJ9 Q8EMK1 Q8EMM9
Q8EMP4 Q8EMP6 Q8EMT9 Q8EMV4 Q8EMZ1 Q8EN03 Q8EN04 Q8EN05 Q8EN08 Q8EN13 Q8EN63
Q8EN66 Q8EN67 Q8EN68 Q8EN69 Q8EN70 Q8EN78 Q8EN87 Q8ENB2 Q8ENE4 Q8ENH9 Q8ENI0
Q8ENI4 Q8ENJ5 Q8ENJ6 Q8ENK1 Q8ENK3 Q8ENK7 Q8ENL3 Q8ENL5 Q8ENM5 Q8ENM6 Q8ENP3
Q8ENP4 Q8ENQ2 Q8ENQ5 Q8ENT9 Q8ENX2 Q8ENX4 Q8ENY0 Q8ENY8 Q8ENZ7 Q8EP04 Q8EP05
Q8EP12 Q8EP25 Q8EP32 Q8EP46 Q8EP79 Q8EP82 Q8EP84 Q8EPA3 Q8EPB3 Q8EPB4 Q8EPB7
Q8EPB9 Q8EPC4 Q8EPD6 Q8EPE2 Q8EPE6 Q8EPE7 Q8EPF0 Q8EPF3 Q8EPF5 Q8EPF6 Q8EPF7
Q8EPH2 Q8EPH4 Q8EPH5 Q8EPH6 Q8EPH7 Q8EPI1 Q8EPJ4 Q8EPJ5 Q8EPJ6 Q8EPM2 Q8EPM7
Q8EPN2 Q8EPP3 Q8EPP6 Q8EPP8 Q8EPQ0 Q8EPQ4 Q8EPQ5 Q8EPQ6 Q8EPQ8 Q8EPR5 Q8EPR6
Q8EPR9 Q8EPS0 Q8EPS9 Q8EPT0 Q8EPT1 Q8EPT2 Q8EPT5 Q8EPT6 Q8EPT8 Q8EPU9 Q8EPV1
Q8EPW0 Q8EPW3 Q8EPW4 Q8EPW5 Q8EPW8 Q8EPW9 Q8EPX8 Q8EPY1 Q8EPY2 Q8EPY4 Q8EPZ2
Q8EQ04 Q8EQ23 Q8EQ28 Q8EQ42 Q8EQ43 Q8EQ44 Q8EQ45 Q8EQ52 Q8EQ56 Q8EQ58 Q8EQ67
Q8EQ73 Q8EQ83 Q8EQ97 Q8EQA8 Q8EQB3 Q8EQB4 Q8EQB6 Q8EQB7 Q8EQB9 Q8EQC1 Q8EQC3
Q8EQD3 Q8EQD4 Q8EQD6 Q8EQD8 Q8EQE2 Q8EQF0 Q8EQG0 Q8EQG4 Q8EQJ1 Q8EQK9 Q8EQM4
Q8EQN3 Q8EQN5 Q8EQP0 Q8EQP2 Q8EQQ9 Q8EQR4 Q8EQR8 Q8EQT7 Q8EQT8 Q8EQT9 Q8EQU1
Q8EQU5 Q8EQU6 Q8EQU9 Q8EQV0 Q8EQV1 Q8EQV2 Q8EQV3 Q8EQV8 Q8EQY4 Q8EQZ4 Q8EQZ7
Q8EQZ8 Q8ER00 Q8ER01 Q8ER04 Q8ER05 Q8ER17 Q8ER21 Q8ER25 Q8ER27 Q8ER28 Q8ER29
Q8ER35 Q8ER36 Q8ER38 Q8ER41 Q8ER44 Q8ER50 Q8ER53 Q8ER54 Q8ER55 Q8ER57 Q8ER59
Q8ER61 Q8ER64 Q8ER66 Q8ER84 Q8ER92 Q8ER96 Q8ER97 Q8ERA3 Q8ERA4 Q8ERE2 Q8ERF4
Q8ERJ2 Q8ERJ6 Q8ERL8 Q8ERL9 Q8ERS1 Q8ERS7 Q8ERS9 Q8ERT3 Q8ERT7 Q8ERU2 Q8ERU3
Q8ERW5 Q8ERW6 Q8ERX9 Q8ERY0 Q8ES27 Q8ES61 Q8ES75 Q8ES77 Q8ES78 Q8ES79 Q8ES91
Q8ES94 Q8ES96 Q8ES97 Q8ES99 Q8ESI2 Q8ESI6 Q8ESJ1 Q8ESK0 Q8ESL3 Q8ESL6 Q8ESP9

Q8ESQ5 Q8ESR4 Q8ESR7 Q8ESR8 Q8ESR9 Q8ESS0 Q8ESS1 Q8ESS2 Q8ESS3 Q8EST0 Q8ESU1

Q8ESU2 Q8ESU3 Q8ESU4 Q8ESU5 Q8ESU7 Q8ESU8 Q8ESU9 Q8ESW9 Q8ESX0 Q8ESX1 Q8ESX3

Q8ESZ2 Q8ESZ3 Q8ET01 Q8ET05 Q8ET11 Q8ET26 Q8ET41 Q8ET50 Q8ET54 Q8ET56 Q8ET75

Q8ET90 Q8ET91 Q8ET92 Q8ET99 Q8ETB7 Q8ETE9 Q8ETG4 Q8ETG9 Q8ETH0 Q8ETM5 Q8ETM7

Q8ETS1 Q8ETS4 Q8ETV3 Q8ETV4 Q8ETV8 Q8ETV9 Q8ETW0 Q8ETW1 Q8ETW2 Q8ETW3 Q8ETW5

Q8ETW6 Q8ETW7 Q8ETW8 Q8ETW9 Q8ETX0 Q8ETX1 Q8ETX2 Q8ETX3 Q8ETX4 Q8ETX5 Q8ETX6

Q8ETX7 Q8ETX8 Q8ETX9 Q8ETY0 Q8ETY1 Q8ETY2 Q8ETY3 Q8ETY4 Q8ETY5 Q8ETY6 Q8ETY7

Q8ETY8 Q8ETZ0 Q8ETZ1 Q8ETZ2 Q8ETZ3 Q8ETZ6 Q8ETZ9 Q8EU02 Q8EU05 Q8EU06 Q8EU10

Q8EU15 Q8EU18 Q8EU33 Q8EU34 Q8EU35 Q8EU37 Q8EU46 Q8EU48 Q8EU53 Q8EU58 Q8EU59

Q8EU77 Q8EU85 Q8EU88 Q8EU89 Q8EU90 Q8EU92 Q8EU94 Q8EU99 Q8EUA1 Q8EUA8 Q8EUB0

Q8EUB2 Q8EUB3 Q8EUB4 Q8EUB6 Q8EUB7 Q8EUB8 Q8EUB9 Q8EUC1 Q8EUC2 Q8EUC3 Q8EUC4

Q8EUC5 Q8EUC6 Q8EUC7 Q8EUC8 Q8EUC9 Q8EUD0 Q8EUD1 Q8EUD5 Q8EUD6 Q8EUD7 Q8EUD8

Q8EUG1 Q8EUG2 Q8EUG3 Q8EUG8 Q8EUG9 Q8EUH0 Q8EUH7 Q8EUI6 Q8EUJ8 Q8EUJ9 Q8EUK2

Q8EUK4 Q8EUK7 Q8EUL1 Q8EUL6 Q8EUM4 Q8EUM5 Q8EUN2 Q8EUN9 Q8EUQ1 Q8EUQ9 Q8EUR6

Q8EUR7 Q8EUS8 Q8EUV2 Q8EUV4 Q8EUV6 Q8EUW1 Q8EUW6 Q8EUW7 Q8EUW9 Q8EUX2 Q8EUX4

Q8EUX9 Q8EUY2 Q8EUY3 Q8EUY4 Q8EV12 Q8EV76 Q8EV77 Q8EV80 Q8EV81 Q8EVC7 Q8EVD0

Q8EVD7 Q8EVF2 Q8EVF5 Q8EVH2 Q8EVH8 Q8EVI1 Q8EVJ0 Q8EVJ1 Q8EVJ8 Q8EVJ9 Q8EVK0

Q8EVK2 Q8EVN6 Q8EVP2 Q8EVU1 Q8EVV1 Q8EVW5 Q8EVW7 Q8EVW9 Q8EW01 Q8EW03 Q8EW04

Q8EW17 Q8EW18 Q8EW19 Q8EW28 Q8EW33 Q8EW34 Q8EW52 Q8EW64 Q8EW68 Q8EW81 Q8EWA5

Q8EWB6 Q8EWB7 Q8EWB8 Q8EWC5 Q8EWD1 Q8EWH6 Q8EWJ2 Q8EWK9 Q8EWL0 Q8EWL4 Q8EWL9

Q8EWM7 Q8EWM9 Q8EWN6 Q8EWP5 Q8EWQ5 Q8EWQ6 Q8EWQ7 Q8EWQ8 Q8EWR4 Q8EWR9 Q8EWS6

Q8EWS9 Q8EWT2 Q8EWT4 Q8EWT5 Q8EWT6 Q8EWT7 Q8EWT9 Q8EWU0 Q8EWU9 Q8EWV0 Q8EWW3

Q8EWW8 Q8EWW9 Q8EWX0 Q8EWX1 Q8EWX7 Q8EWX9 Q8EWY5 Q8EX00 Q8EX02 Q8EX08 Q8EX09

Q8EX11 Q8EX12 Q8EX18 Q8EX19 Q8EX20 Q8EX21 Q8EX24 Q8EX25 Q8EX28 Q8EX97 Q8EXA3

Q8EXA4 Q8EXB9 Q8EXC0 Q8EXC1 Q8EXJ3 Q8EXN2 Q8EXP6 Q8EXP7 Q8EXQ4 Q8EXQ7 Q8EXT3

Q8EXV1 Q8EXX1 Q8EXX3 Q8EXX8 Q8EY44 Q8EY59 Q8EY69 Q8EY78 Q8EY84 Q8EY85 Q8EYA9

Q8EYB0 Q8EYF0 Q8EYG1 Q8EYG2 Q8EYH2 Q8EYK9 Q8EYM3 Q8EYP5 Q8EYP6 Q8EYP7 Q8EYR3

Q8EYT0 Q8EYU4 Q8EYV8 Q8EZ21 Q8EZ24 Q8EZ26 Q8EZ61 Q8EZ81 Q8EZ98 Q8EZC5 Q8EZD6

Q8EZG6 Q8EZH9 Q8EZJ0 Q8EZM8 Q8EZN0 Q8EZN1 Q8EZQ1 Q8EZQ3 Q8EZT4 Q8EZT7 Q8EZT8

Q8EZY7 Q8F031 Q8F052 Q8F0A5 Q8F0E4 Q8F0I0 Q8F0I2 Q8F0Q1 Q8F0R8 Q8F0R9 Q8F0S0

Q8F0S1 Q8F0S2 Q8F0S3 Q8F0S5 Q8F0S9 Q8F0T3 Q8F0T4 Q8F0T5 Q8F0X6 Q8F132 Q8F133

Q8F140 Q8F141 Q8F142 Q8F143 Q8F144 Q8F146 Q8F148 Q8F149 Q8F150 Q8F153 Q8F1F4

Q8F1F5 Q8F1H5 Q8F1S7 Q8F219 Q8F242 Q8F2A8 Q8F2D1 Q8F312 Q8F351 Q8F352 Q8F353

Q8F393 Q8F394 Q8F3A0 Q8F3A1 Q8F3H5 Q8F3I3 Q8F3J3 Q8F3J9 Q8F3K6 Q8F3L1 Q8F3L2

Q8F3L3 Q8F3L4 Q8F3M0 Q8F3P1 Q8F3P4 Q8F3P5 Q8F3Q1 Q8F3Q2 Q8F3Q4 Q8F3Q5 Q8F3V7

Q8F3W3 Q8F3W6 Q8F3X5 Q8F3Z6 Q8F411 Q8F418 Q8F421 Q8F435 Q8F445 Q8F479 Q8F483

Q8F495 Q8F496 Q8F498 Q8F4A0 Q8F4A2 Q8F4D5 Q8F4D8 Q8F4E5 Q8F4E6 Q8F4E9 Q8F4F4

Q8F4F6 Q8F4G5 Q8F4I0 Q8F4I2 Q8F4J4 Q8F4J6 Q8F4P5 Q8F4Q4 Q8F4R6 Q8F509 Q8F522

Q8F523 Q8F525 Q8F5D8 Q8F5E6 Q8F5F1 Q8F5H8 Q8F5I5 Q8F5J2 Q8F5J3 Q8F5J7 Q8F5K1

Q8F5K3 Q8F5K4 Q8F5K6 Q8F5Z7 Q8F625 Q8F661 Q8F6A0 Q8F6B7 Q8F6C0 Q8F6I1 Q8F6I3

Q8F6K1 Q8F6P0 Q8F6P1 Q8F6P4 Q8F6P5 Q8F6Q7 Q8F6Q8 Q8F6Q9 Q8F6R1 Q8F6R2 Q8F6S7

Q8F6W9 Q8F6X4 Q8F6Y3 Q8F708 Q8F710 Q8F729 Q8F737 Q8F738 Q8F743 Q8F746 Q8F7A0

Q8F7C6 Q8F7G5 Q8F7H3 Q8F7J8 Q8F7J9 Q8F7K0 Q8F7K1 Q8F7K3 Q8F7L5 Q8F7N5 Q8F7T2

Q8F7T9 Q8F7U0 Q8F7U1 Q8F7U3 Q8F7V1 Q8F7V2 Q8F7V4 Q8F7V9 Q8F7W8 Q8F7Y2 Q8F7Y6

Q8F812 Q8F814 Q8F832 Q8F839 Q8F874 Q8F8A9 Q8F8C0 Q8F8D9 Q8F8K4 Q8F8N2 Q8F8Q2
Q8F8T4 Q8F8T8 Q8F8T9 Q8F930 Q8F935 Q8F936 Q8F937 Q8F952 Q8F964 Q8F983 Q8F987
Q8F9A3 Q8F9D9 Q8F9E4 Q8F9L0 Q8F9L6 Q8F9N4 Q8F9P6 Q8F9P7 Q8F9Q1 Q8F9R5 Q8F9R6
Q8F9S6 Q8F9T2 Q8F9U1 Q8F9V5 Q8F9W5 Q8F9Y3 Q8FA20 Q8FA32 Q8FA40 Q8FA41 Q8FA42
Q8FA49 Q8FA51 Q8FA52 Q8FA64 Q8FA72 Q8FA88 Q8FAA2 Q8FAD1 Q8FAD6 Q8FAD9 Q8FAE1
Q8FAF3 Q8FAG0 Q8FAG4 Q8FAH0 Q8FAI6 Q8FAI7 Q8FAI9 Q8FAJ4 Q8FAK9 Q8FAL3 Q8FAM7
Q8FAS0 Q8FAT5 Q8FAY8 Q8FB02 Q8FB20 Q8FB22 Q8FB30 Q8FB32 Q8FB35 Q8FB44 Q8FB47
Q8FB55 Q8FB61 Q8FB64 Q8FB68 Q8FB69 Q8FB74 Q8FB75 Q8FB76 Q8FB77 Q8FB78 Q8FB81
Q8FB83 Q8FB88 Q8FB89 Q8FB93 Q8FB96 Q8FB97 Q8FB98 Q8FBA7 Q8FBA9 Q8FBC0 Q8FBC3
Q8FBD0 Q8FBD7 Q8FBD9 Q8FBE0 Q8FBE1 Q8FBE2 Q8FBE5 Q8FBE8 Q8FBE9 Q8FBG9 Q8FBH7
Q8FBI1 Q8FBI4 Q8FBJ0 Q8FBM1 Q8FBN6 Q8FBN9 Q8FBP0 Q8FBP1 Q8FBR0 Q8FBR2 Q8FBR5
Q8FBS4 Q8FBS5 Q8FBS6 Q8FBT4 Q8FBU9 Q8FBV2 Q8FBV3 Q8FBV5 Q8FBX7 Q8FBY0 Q8FBZ5
Q8FC83 Q8FC86 Q8FC87 Q8FC88 Q8FCA2 Q8FCA6 Q8FCB6 Q8FCB7 Q8FCC1 Q8FCD6 Q8FCE3
Q8FCI4 Q8FCI6 Q8FCI7 Q8FCK2 Q8FCL1 Q8FCR6 Q8FCR7 Q8FCR8 Q8FCS8 Q8FCT3 Q8FCT4
Q8FCT6 Q8FCU4 Q8FCV7 Q8FCW0 Q8FCW8 Q8FCY1 Q8FCY2 Q8FCY3 Q8FD03 Q8FD12 Q8FD13
Q8FD15 Q8FD17 Q8FD18 Q8FD47 Q8FD52 Q8FD57 Q8FD58 Q8FD60 Q8FD82 Q8FD84 Q8FD87
Q8FD94 Q8FD95 Q8FD98 Q8FDA0 Q8FDB3 Q8FDB4 Q8FDB5 Q8FDB8 Q8FDC6 Q8FDE5 Q8FDF5
Q8FDG5 Q8FDG6 Q8FDG8 Q8FDH0 Q8FDH9 Q8FDK9 Q8FDM1 Q8FDN6 Q8FDQ2 Q8FDS2 Q8FDU7
Q8FDU8 Q8FE19 Q8FE22 Q8FE27 Q8FE28 Q8FE30 Q8FE31 Q8FE34 Q8FE36 Q8FE65 Q8FE66
Q8FE67 Q8FE69 Q8FE70 Q8FE75 Q8FE90 Q8FE91 Q8FE97 Q8FEA9 Q8FEE5 Q8FEE7 Q8FEF0
Q8FEF2 Q8FEF7 Q8FEF9 Q8FEG6 Q8FEI7 Q8FEI8 Q8FEI9 Q8FEJ0 Q8FEJ1 Q8FEJ2 Q8FEJ5
Q8FEJ6 Q8FEJ7 Q8FEJ8 Q8FEL3 Q8FEL9 Q8FEM8 Q8FEN4 Q8FEN7 Q8FES6 Q8FEY4 Q8FEY5
Q8FEY7 Q8FEY9 Q8FF08 Q8FF10 Q8FF14 Q8FF18 Q8FF24 Q8FF26 Q8FF30 Q8FF44 Q8FF46
Q8FF47 Q8FF53 Q8FF55 Q8FF59 Q8FF64 Q8FF70 Q8FF72 Q8FF82 Q8FF87 Q8FFA1 Q8FFA3
Q8FFB0 Q8FFB5 Q8FFC1 Q8FFC9 Q8FFD4 Q8FFE8 Q8FFF4 Q8FFH0 Q8FFH2 Q8FFJ5 Q8FFK8
Q8FFL1 Q8FFL2 Q8FFL4 Q8FFM5 Q8FFM8 Q8FFN0 Q8FFP0 Q8FFQ5 Q8FFQ7 Q8FFR9 Q8FFS1
Q8FFT8 Q8FFU0 Q8FFU3 Q8FFV3 Q8FFV5 Q8FFX8 Q8FFY0 Q8FFY2 Q8FFY7 Q8FFY8 Q8FG00
Q8FG47 Q8FG48 Q8FG49 Q8FG50 Q8FG51 Q8FG52 Q8FG53 Q8FGA5 Q8FGG0 Q8FGL1 Q8FGN6
Q8FGN7 Q8FGQ0 Q8FGQ2 Q8FGQ5 Q8FGQ6 Q8FGQ9 Q8FGR3 Q8FGS9 Q8FGT3 Q8FGU5 Q8FGZ9
Q8FH00 Q8FH01 Q8FH02 Q8FH03 Q8FH05 Q8FH06 Q8FH10 Q8FH13 Q8FH30 Q8FH32 Q8FH42
Q8FH54 Q8FH55 Q8FH72 Q8FH85 Q8FH87 Q8FH88 Q8FH89 Q8FH99 Q8FHA7 Q8FHA8 Q8FHB1
Q8FHC7 Q8FHD1 Q8FHD2 Q8FHF2 Q8FHF3 Q8FHF4 Q8FHF8 Q8FHG0 Q8FHG5 Q8FHH1 Q8FHK7
Q8FHN0 Q8FHP6 Q8FHT6 Q8FHT8 Q8FHU2 Q8FHU5 Q8FHV4 Q8FHV9 Q8FHW0 Q8FHW7 Q8FHX4
Q8FHX8 Q8FI03 Q8FI04 Q8FI27 Q8FI28 Q8FI31 Q8FI35 Q8FI39 Q8FIB6 Q8FIJ3 Q8FIM4
Q8FIM5 Q8FIN2 Q8FIN3 Q8FIN9 Q8FIP6 Q8FIP7 Q8FIP8 Q8FIR0 Q8FIR2 Q8FIR3 Q8FIR7
Q8FJ41 Q8FJ42 Q8FJ43 Q8FJ44 Q8FJ45 Q8FJ50 Q8FJ51 Q8FJ52 Q8FJ69 Q8FJ70 Q8FJ71
Q8FJ74 Q8FJ79 Q8FJ81 Q8FJ88 Q8FJ93 Q8FJ98 Q8FJA2 Q8FJA3 Q8FJA4 Q8FJB6 Q8FJB7
Q8FJD1 Q8FJE0 Q8FJE7 Q8FJF0 Q8FJJ0 Q8FJK5 Q8FJL3 Q8FJM0 Q8FJM4 Q8FJM6 Q8FJN1
Q8FJP8 Q8FJQ2 Q8FJQ3 Q8FJR2 Q8FJS1 Q8FJS7 Q8FJU5 Q8FJU6 Q8FJW4 Q8FJX7 Q8FJY3
Q8FJY8 Q8FJY9 Q8FJZ1 Q8FK03 Q8FK04 Q8FK07 Q8FK30 Q8FK41 Q8FK44 Q8FK51 Q8FK60
Q8FK65 Q8FK66 Q8FK67 Q8FK74 Q8FK76 Q8FK82 Q8FK83 Q8FK84 Q8FKA2 Q8FKA4 Q8FKA7
Q8FKB7 Q8FKB9 Q8FKC5 Q8FKD5 Q8FKE1 Q8FKE3 Q8FKG1 Q8FKG2 Q8FKG6 Q8FKI7 Q8FKI8
Q8FKL1 Q8FKL2 Q8FKM3 Q8FKM4 Q8FKM5 Q8FKW2 Q8FKY7 Q8FKZ4 Q8FKZ8 Q8FL00 Q8FL05
Q8FL06 Q8FL11 Q8FL13 Q8FL16 Q8FL24 Q8FL25 Q8FL30 Q8FL31 Q8FL39 Q8FL40 Q8FL55

Q8FL56 Q8FL57 Q8FL63 Q8FL65 Q8FL67 Q8FL68 Q8FL75 Q8FL76 Q8FL79 Q8FL80 Q8FL88
Q8FL89 Q8FL92 Q8FL93 Q8FL95 Q8FL96 Q8FL98 Q8FLA3 Q8FLA4 Q8FLA5 Q8FLA6 Q8FLA7
Q8FLB0 Q8FLB1 Q8FLB4 Q8FLB7 Q8FLC5 Q8FLD1 Q8FLD3 Q8FLD8 Q8FLJ5 Q8FLJ6 Q8FLJ8
Q8FLM0 Q8FLN6 Q8FLN8 Q8FLP8 Q8FLP9 Q8FLQ0 Q8FLU6 Q8FLV6 Q8FLY5 Q8FLY8 Q8FM11
Q8FM16 Q8FM44 Q8FM78 Q8FM79 Q8FM80 Q8FM94 Q8FMB0 Q8FMB3 Q8FMB7 Q8FMD0 Q8FMD3
Q8FMF8 Q8FMG0 Q8FMG3 Q8FMH9 Q8FMI3 Q8FMI4 Q8FMI6 Q8FMI8 Q8FML6 Q8FMM2 Q8FMM3
Q8FMN2 Q8FMN5 Q8FMR0 Q8FMR1 Q8FMS2 Q8FMT9 Q8FMU6 Q8FMU8 Q8FMU9 Q8FMX2 Q8FMX4
Q8FMX9 Q8FMY8 Q8FN35 Q8FN36 Q8FN37 Q8FN57 Q8FN62 Q8FN65 Q8FN70 Q8FN76 Q8FN77
Q8FN81 Q8FN82 Q8FN87 Q8FN90 Q8FNA1 Q8FNB3 Q8FND7 Q8FNF4 Q8FNF5 Q8FNF8 Q8FNG2
Q8FNG6 Q8FNH1 Q8FNH2 Q8FNP4 Q8FNP5 Q8FNP8 Q8FNQ2 Q8FNT1 Q8FNT2 Q8FNT5 Q8FNT8
Q8FNU1 Q8FNV0 Q8FNZ0 Q8FNZ1 Q8FNZ2 Q8FNZ6 Q8FNZ7 Q8FNZ9 Q8FP00 Q8FP04 Q8FP16
Q8FP17 Q8FP19 Q8FP30 Q8FP33 Q8FP52 Q8FP55 Q8FP56 Q8FP63 Q8FP66 Q8FP70 Q8FP71
Q8FP74 Q8FP75 Q8FP78 Q8FP80 Q8FP82 Q8FP85 Q8FP91 Q8FP92 Q8FP96 Q8FP97 Q8FPA4
Q8FPA7 Q8FPA8 Q8FPB3 Q8FPB6 Q8FPB7 Q8FPD3 Q8FPD4 Q8FPD5 Q8FPE0 Q8FPE1 Q8FPF8
Q8FPG6 Q8FPG8 Q8FPH9 Q8FPI1 Q8FPI2 Q8FPJ3 Q8FPJ9 Q8FPK2 Q8FPK3 Q8FPK4 Q8FPK5
Q8FPL0 Q8FPL5 Q8FPN2 Q8FPP9 Q8FPQ4 Q8FPQ9 Q8FPR2 Q8FPR3 Q8FPS0 Q8FPT3 Q8FPU9
Q8FPV5 Q8FPX1 Q8FPX6 Q8FPY5 Q8FPZ0 Q8FPZ1 Q8FPZ7 Q8FQ01 Q8FQ12 Q8FQ17 Q8FQ31
Q8FQ44 Q8FQ57 Q8FQ76 Q8FQB2 Q8FQE4 Q8FQG1 Q8FQH8 Q8FQP0 Q8FQP1 Q8FQP8 Q8FQQ3
Q8FQR1 Q8FQR2 Q8FQR4 Q8FQS2 Q8FQV2 Q8FQV5 Q8FQV6 Q8FQW8 Q8FQX0 Q8FQX1 Q8FQX2
Q8FQX3 Q8FQX4 Q8FQY0 Q8FQZ4 Q8FQZ5 Q8FR00 Q8FR20 Q8FR21 Q8FR24 Q8FR25 Q8FR26
Q8FR29 Q8FR39 Q8FR47 Q8FRE7 Q8FRI2 Q8FRJ4 Q8FRM5 Q8FRQ5 Q8FRR3 Q8FRT0 Q8FRU9
Q8FRV5 Q8FRX6 Q8FRZ3 Q8FS07 Q8FS13 Q8FS18 Q8FS19 Q8FS20 Q8FS31 Q8FS32 Q8FS33
Q8FS34 Q8FS35 Q8FS36 Q8FS39 Q8FS51 Q8FS52 Q8FS53 Q8FS54 Q8FS55 Q8FS66 Q8FS69
Q8FS70 Q8FS71 Q8FS73 Q8FS74 Q8FS75 Q8FS76 Q8FS77 Q8FS78 Q8FS79 Q8FS80 Q8FS84
Q8FS85 Q8FS86 Q8FS96 Q8FS97 Q8FSA0 Q8FSA1 Q8FSA5 Q8FSA7 Q8FSB3 Q8FSB6 Q8FSD4
Q8FSD6 Q8FSF0 Q8FSF1 Q8FSF7 Q8FSF8 Q8FSH0 Q8FSH1 Q8FSJ0 Q8FSU7 Q8FSU8 Q8FSV1
Q8FSW7 Q8FT06 Q8FT12 Q8FT19 Q8FT23 Q8FT24 Q8FT29 Q8FT30 Q8FT31 Q8FT32 Q8FT34
Q8FT35 Q8FT38 Q8FT39 Q8FT41 Q8FT42 Q8FT43 Q8FT45 Q8FT46 Q8FT48 Q8FT51 Q8FT52
Q8FT58 Q8FT60 Q8FT61 Q8FT63 Q8FT66 Q8FT67 Q8FT98 Q8FTA2 Q8FTA6 Q8FTA8 Q8FTB1
Q8FTD0 Q8FTD4 Q8FTD5 Q8FTE3 Q8FTE6 Q8FTH7 Q8FTH8 Q8FTJ5 Q8FTK5 Q8FTK6 Q8FTL2
Q8FTL6 Q8FTM1 Q8FTM8 Q8FTM9 Q8FTN0 Q8FTN1 Q8FTN2 Q8FTN3 Q8FTN4 Q8FTN5 Q8FTP0
Q8FTP1 Q8FTQ0 Q8FTQ1 Q8FTQ2 Q8FU05 Q8FU10 Q8FU11 Q8FU18 Q8FU23 Q8FU28 Q8FU35
Q8FU86 Q8FUA2 Q8FUA5 Q8FUA6 Q8FUC8 Q8FUD1 Q8FUI4 Q8FUI5 Q8FUL3 Q8FUL4 Q8FUL7
Q8FUQ2 Q8FUS9 Q8FUT3 Q8FUZ0 Q8FUZ1 Q8FUZ2 Q8FUZ3 Q8FV07 Q8FV09 Q8FV22 Q8FV25
Q8FV30 Q8FV99 Q8FVB2 Q8FVB4 Q8FVB5 Q8FVC0 Q8FVC1 Q8FVD4 Q8FVE4 Q8FVF2 Q8FVF3
Q8FVG9 Q8FVH2 Q8FVM2 Q8FVM4 Q8FVP6 Q8FVR1 Q8FVU9 Q8FVW4 Q8FVY1 Q8FW12 Q8FW50
Q8FW65 Q8FW66 Q8FW67 Q8FW72 Q8FW94 Q8FW95 Q8FWG3 Q8FWG5 Q8FWG9 Q8FWK8 Q8FWN5
Q8FWR0 Q8FWT4 Q8FWU0 Q8FWV5 Q8FWX2 Q8FX38 Q8FX81 Q8FX87 Q8FX90 Q8FX92 Q8FX93
Q8FXF9 Q8FXJ1 Q8FXR6 Q8FXS9 Q8FXT0 Q8FXT1 Q8FXT2 Q8FXT6 Q8FXU4 Q8FXW6 Q8FXW9
Q8FXX1 Q8FXX2 Q8FXX4 Q8FXX5 Q8FXY4 Q8FXY6 Q8FXZ7 Q8FY05 Q8FY07 Q8FY08 Q8FY09
Q8FY11 Q8FY12 Q8FY19 Q8FY21 Q8FY22 Q8FY24 Q8FY28 Q8FY29 Q8FY59 Q8FY98 Q8FYA4
Q8FYA6 Q8FYB3 Q8FYE2 Q8FYE7 Q8FYF0 Q8FYF4 Q8FYF7 Q8FYG9 Q8FYI5 Q8FYI6 Q8FYK0
Q8FYK3 Q8FYL7 Q8FYL8 Q8FYM2 Q8FYM3 Q8FYM9 Q8FYQ3 Q8FYQ7 Q8FYQ9 Q8FYS0 Q8FYT2
Q8FYV6 Q8FYW3 Q8FYX8 Q8FYZ5 Q8FZ01 Q8FZ02 Q8FZ11 Q8FZ13 Q8FZ45 Q8FZ50 Q8FZC4

Q8FZE8 Q8FZF0 Q8FZJ3 Q8FZJ8 Q8FZK4 Q8FZM5 Q8FZP0 Q8FZP2 Q8FZP4 Q8FZP5 Q8FZQ3
Q8FZQ6 Q8FZR2 Q8FZT6 Q8FZU1 Q8FZV8 Q8FZV9 Q8FZW0 Q8FZW2 Q8FZW3 Q8FZW4 Q8G005
Q8G018 Q8G020 Q8G034 Q8G044 Q8G065 Q8G066 Q8G067 Q8G069 Q8G070 Q8G074 Q8G075
Q8G076 Q8G077 Q8G078 Q8G079 Q8G080 Q8G081 Q8G082 Q8G083 Q8G084 Q8G085 Q8G086
Q8G087 Q8G088 Q8G089 Q8G090 Q8G092 Q8G094 Q8G095 Q8G0B6 Q8G0C9 Q8G0D6 Q8G0D7
Q8G0D8 Q8G0D9 Q8G0E8 Q8G0F3 Q8G0F5 Q8G0F7 Q8G0G1 Q8G0H4 Q8G0I4 Q8G0I5 Q8G0I7
Q8G0I9 Q8G0J1 Q8G0K1 Q8G0L3 Q8G0L4 Q8G0L8 Q8G0P0 Q8G0S1 Q8G0S7 Q8G0U3 Q8G0X4
Q8G0X5 Q8G0Y3 Q8G0Z4 Q8G104 Q8G105 Q8G120 Q8G129 Q8G139 Q8G146 Q8G155 Q8G178
Q8G180 Q8G183 Q8G189 Q8G197 Q8G1C8 Q8G1C9 Q8G1E7 Q8G1E8 Q8G1F1 Q8G1K5 Q8G1L5
Q8G1L9 Q8G1M4 Q8G1N0 Q8G1N5 Q8G1P8 Q8G1Q6 Q8G1R0 Q8G1Z7 Q8G1Z8 Q8G1Z9 Q8G204
Q8G228 Q8G242 Q8G253 Q8G256 Q8G257 Q8G274 Q8G276 Q8G290 Q8G292 Q8G2A9 Q8G2B1
Q8G2C1 Q8G2D0 Q8G2E8 Q8G2I3 Q8G2J0 Q8G2L5 Q8G2L7 Q8G2M8 Q8G2N3 Q8G2P5 Q8G2P6
Q8G2P7 Q8G2P8 Q8G2P9 Q8G2Q0 Q8G2Q1 Q8G2R2 Q8G2R5 Q8G2R6 Q8G2U9 Q8G2W6 Q8G2Y5
Q8G2Y6 Q8G304 Q8G310 Q8G312 Q8G340 Q8G353 Q8G374 Q8G376 Q8G381 Q8G3B7 Q8G3C3
Q8G3C4 Q8G3E5 Q8G3E7 Q8G3E9 Q8G3G3 Q8G3G6 Q8G3H1 Q8G3H2 Q8G3H3 Q8G3H4 Q8G3I2
Q8G3M4 Q8G3N0 Q8G3N5 Q8G3N6 Q8G3P4 Q8G3P5 Q8G3P6 Q8G3Q1 Q8G3S1 Q8G3S4 Q8G3T4
Q8G3T9 Q8G3W7 Q8G3X9 Q8G3Y4 Q8G3Y5 Q8G3Z2 Q8G3Z3 Q8G3Z5 Q8G3Z6 Q8G3Z7 Q8G3Z8
Q8G400 Q8G401 Q8G402 Q8G403 Q8G404 Q8G405 Q8G406 Q8G407 Q8G408 Q8G409 Q8G411
Q8G412 Q8G413 Q8G414 Q8G415 Q8G416 Q8G417 Q8G418 Q8G423 Q8G424 Q8G435 Q8G443
Q8G444 Q8G447 Q8G448 Q8G478 Q8G481 Q8G483 Q8G484 Q8G485 Q8G486 Q8G487 Q8G4D1
Q8G4D8 Q8G4E8 Q8G4G1 Q8G4G9 Q8G4H4 Q8G4I8 Q8G4L1 Q8G4L2 Q8G4L4 Q8G4L8 Q8G4M3
Q8G4N5 Q8G4Q2 Q8G4Q4 Q8G4Q6 Q8G4R0 Q8G4R1 Q8G4R4 Q8G4S5 Q8G4S6 Q8G4S7 Q8G4S8
Q8G4S9 Q8G4T0 Q8G4T3 Q8G4T4 Q8G4T9 Q8G4U0 Q8G4U1 Q8G4U2 Q8G4U8 Q8G4V2 Q8G4W1
Q8G4W2 Q8G4X1 Q8G4X3 Q8G4X4 Q8G4Z7 Q8G500 Q8G509 Q8G510 Q8G514 Q8G515 Q8G526
Q8G527 Q8G533 Q8G534 Q8G540 Q8G558 Q8G572 Q8G574 Q8G575 Q8G595 Q8G5A9 Q8G5B0
Q8G5B5 Q8G5B6 Q8G5B7 Q8G5E7 Q8G5E8 Q8G5F0 Q8G5F1 Q8G5F2 Q8G5F3 Q8G5F6 Q8G5G1
Q8G5G8 Q8G5H9 Q8G5I6 Q8G5J9 Q8G5K1 Q8G5L0 Q8G5L2 Q8G5L6 Q8G5L8 Q8G5L9 Q8G5P2
Q8G5P4 Q8G5Q7 Q8G5Q9 Q8G5R0 Q8G5R1 Q8G5R9 Q8G5W1 Q8G5W9 Q8G5X0 Q8G5X3 Q8G5X4
Q8G5X7 Q8G5Y5 Q8G5Z0 Q8G5Z8 Q8G602 Q8G651 Q8G655 Q8G656 Q8G661 Q8G689 Q8G691
Q8G694 Q8G695 Q8G6A8 Q8G6B1 Q8G6B5 Q8G6B7 Q8G6B8 Q8G6B9 Q8G6C0 Q8G6C2 Q8G6C6
Q8G6C7 Q8G6D5 Q8G6D6 Q8G6D7 Q8G6D8 Q8G6E0 Q8G6F6 Q8G6F7 Q8G6I3 Q8G6I4 Q8G6J4
Q8G6J8 Q8G6J9 Q8G6K0 Q8G6K4 Q8G6P9 Q8G6S6 Q8G6T9 Q8G6V1 Q8G6V2 Q8G6W1 Q8G6W2
Q8G6Y1 Q8G6Z6 Q8G6Z9 Q8G700 Q8G709 Q8G756 Q8G757 Q8G758 Q8G759 Q8G767 Q8G768
Q8G769 Q8G777 Q8G784 Q8G7B5 Q8G7C4 Q8G7D7 Q8G7E2 Q8G7F9 Q8G7G1 Q8G7G5 Q8G7H1
Q8G7H2 Q8G7H5 Q8G7I0 Q8G7I1 Q8G7I6 Q8G7J3 Q8G7W6 Q8G7W9 Q8G7X0 Q8G7X1 Q8G7Y3
Q8G7Y4 Q8G7Y6 Q8G7Y7 Q8G7Z8 Q8G813 Q8G815 Q8G816 Q8G817 Q8G818 Q8G830 Q8G863
Q8G864 Q8G879 Q8G8B6 Q8G8D1 Q8G8Y2 Q8G907 Q8G969 Q8G997 Q8G998 Q8G999 Q8G9C6
Q8G9F9 Q8G9G3 Q8G9G4 Q8G9M2 Q8G9P0 Q8GAA0 Q8GAA6 Q8GAH9 Q8GAI3 Q8GAI8 Q8GAJ0
Q8GAK6 Q8GAK7 Q8GAL4 Q8GAN5 Q8GAU3 Q8GAU4 Q8GAV9 Q8GAW0 Q8GAX4 Q8GAZ4 Q8GB16
Q8GB17 Q8GB18 Q8GB19 Q8GB20 Q8GB99 Q8GBB2 Q8GBB4 Q8GBB7 Q8GBD2 Q8GBD3 Q8GBW6
Q8GC81 Q8GC87 Q8GCB0 Q8GCJ0 Q8GCR3 Q8GCR4 Q8GCR5 Q8GCR6 Q8GCV0 Q8GCY1 Q8GDC4
Q8GDE1 Q8GDP4 Q8GDQ6 Q8GDU2 Q8GDU5 Q8GDY3 Q8GE08 Q8GEA9 Q8GED7 Q8GED8 Q8GED9
Q8GEI4 Q8GEK8 Q8GF09 Q8GF77 Q8GF87 Q8GG79 Q8GGJ6 Q8GGL1 Q8GGL2 Q8GGL7 Q8GH23
Q8GH79 Q8GH80 Q8GH81 Q8GH95 Q8GH98 Q8GHA0 Q8GHB1 Q8GHB2 Q8GHK8 Q8GHL1 Q8GHV6

Q8GI14 Q8GI16 Q8GIR7 Q8GIS1 Q8GIT6 Q8GIX7 Q8GIZ5 Q8GJ00 Q8GJ01 Q8GJ74 Q8GJM0
Q8GJN0 Q8GJN6 Q8GJP4 Q8GJP6 Q8GKS1 Q8GKZ1 Q8GLC6 Q8GLE4 Q8GLE5 Q8GLE6 Q8GLE9
Q8GLH5 Q8GLK5 Q8GLK9 Q8GLP8 Q8GM55 Q8GM56 Q8GM57 Q8GMC0 Q8GME2 Q8GMG0 Q8GMG6
Q8GMH2 Q8GMH4 Q8GMJ0 Q8GMR7 Q8GMT0 Q8GN32 Q8GN33 Q8GN53 Q8GNC1 Q8GND0 Q8GND4
Q8GNM0 Q8GNT2 Q8GP72 Q8GPD3 Q8GPG2 Q8GQC5 Q8GQN9 Q8GQS5 Q8GQU1 Q8GR45 Q8GRC3
Q8GRC8 Q8GRF5 Q8GRL4 Q8GRL7 Q8GRN0 Q8GS60 Q8GS71 Q8GSJ1 Q8GSL4 Q8GSL5 Q8GSM7
Q8GSN1 Q8GSN8 Q8GT06 Q8GT20 Q8GT21 Q8GT39 Q8GT73 Q8GTE3 Q8GTF4 Q8GTF5 Q8GTL5
Q8GTQ9 Q8GTR4 Q8GTS0 Q8GTS1 Q8GTS2 Q8GU20 Q8GU25 Q8GU95 Q8GUD7 Q8GUE4 Q8GUG7
Q8GUG9 Q8GUH1 Q8GUL2 Q8GUN5 Q8GUP1 Q8GUP3 Q8GUW4 Q8GV05 Q8GV94 Q8GVA0 Q8GVB5
Q8GVC7 Q8GVE8 Q8GVF9 Q8GVP6 Q8GVZ8 Q8GW05 Q8GW10 Q8GW29 Q8GW31 Q8GW44 Q8GW46
Q8GW57 Q8GW64 Q8GW78 Q8GW80 Q8GWA2 Q8GWA9 Q8GWB2 Q8GWE0 Q8GWF1 Q8GWF4 Q8GWG2
Q8GWI2 Q8GWI5 Q8GWJ4 Q8GWK1 Q8GWK6 Q8GWK7 Q8GWP0 Q8GWP5 Q8GWQ2 Q8GWQ6 Q8GWR1
Q8GWR2 Q8GWR3 Q8GWS0 Q8GWT4 Q8GWU0 Q8GWU1 Q8GWU7 Q8GWV5 Q8GWW7 Q8GX05 Q8GX29
Q8GX46 Q8GX77 Q8GX84 Q8GXB1 Q8GXB7 Q8GXC2 Q8GXC7 Q8GXD6 Q8GXD9 Q8GXE2 Q8GXE9
Q8GXF0 Q8GXF6 Q8GXF8 Q8GXG1 Q8GXH3 Q8GXI9 Q8GXL7 Q8GXM7 Q8GXN9 Q8GXP4 Q8GXQ7
Q8GXR5 Q8GXS3 Q8GXS8 Q8GXT3 Q8GXU5 Q8GXU9 Q8GXV2 Q8GXW1 Q8GXW5 Q8GXW6 Q8GXX0
Q8GXX4 Q8GXX7 Q8GY11 Q8GY23 Q8GY25 Q8GY31 Q8GY42 Q8GY54 Q8GY79 Q8GY84 Q8GY87
Q8GY88 Q8GY89 Q8GY91 Q8GY96 Q8GYA3 Q8GYA6 Q8GYA8 Q8GYB1 Q8GYD2 Q8GYD9 Q8GYE8
Q8GYH7 Q8GYJ2 Q8GYJ3 Q8GYL3 Q8GYL7 Q8GYL9 Q8GYM1 Q8GYM2 Q8GYM3 Q8GYM4 Q8GYM6
Q8GYN9 Q8GYP3 Q8GYP5 Q8GYP6 Q8GYR4 Q8GYU3 Q8GYW8 Q8GYX2 Q8GYX3 Q8GYX8 Q8GYX9
Q8GYY0 Q8GZ13 Q8GZ26 Q8GZ29 Q8GZ38 Q8GZ42 Q8GZ43 Q8GZ45 Q8GZ53 Q8GZ63 Q8GZ84
Q8GZ92 Q8GZA6 Q8GZA8 Q8GZB6 Q8GZM7 Q8GZQ3 Q8H063 Q8H0D2 Q8H0D3 Q8H0F2 Q8H0F6
Q8H0G1 Q8H0G2 Q8H0J6 Q8H0P8 Q8H0S9 Q8H0T4 Q8H0T5 Q8H0T9 Q8H0U8 Q8H0V1 Q8H0V3
Q8H0V4 Q8H0V6 Q8H0W0 Q8H0W2 Q8H0W3 Q8H0X0 Q8H0Y8 Q8H0Z4 Q8H0Z6 Q8H102 Q8H103
Q8H104 Q8H107 Q8H110 Q8H111 Q8H113 Q8H115 Q8H125 Q8H136 Q8H156 Q8H159 Q8H173
Q8H174 Q8H183 Q8H185 Q8H1D2 Q8H1D4 Q8H1D7 Q8H1E7 Q8H1E8 Q8H1F0 Q8H1F6 Q8H1F7
Q8H1F9 Q8H1G0 Q8H1M0 Q8H1P9 Q8H1Q2 Q8H1Q7 Q8H1R7 Q8H1S0 Q8H1S1 Q8H1S4 Q8H1S6
Q8H1T3 Q8H1U4 Q8H1U5 Q8H1Y0 Q8H215 Q8H252 Q8H273 Q8H2B4 Q8H2B5 Q8H2C8 Q8H2C9
Q8H2D0 Q8H2D1 Q8H2P8 Q8H2T0 Q8H2V6 Q8H2X8 Q8H339 Q8H383 Q8H398 Q8H3Q1 Q8H485
Q8H4D4 Q8H4L3 Q8H4S4 Q8H4S6 Q8H506 Q8H507 Q8H594 Q8H5F8 Q8H6A5 Q8H6B1 Q8H7M1
Q8H7U1 Q8H7U8 Q8H7Y8 Q8H859 Q8H8N3 Q8H8T0 Q8H8U5 Q8H936 Q8H991 Q8H9A8 Q8H9B6
Q8H9D2 Q8HA43 Q8HAD9 Q8HAE6 Q8HQQ2 Q8HQQ3 Q8HQQ4 Q8HQQ6 Q8HQQ8 Q8HQR0 Q8HQR1
Q8HQS2 Q8HQS3 Q8HQS7 Q8HQS8 Q8HQT0 Q8HQT1 Q8HQT2 Q8HQT4 Q8HQT5 Q8HQT6 Q8HQT9
Q8HSW1 Q8HTL1 Q8HTL2 Q8HTL3 Q8HTL6 Q8HTL7 Q8HUG7 Q8HUG9 Q8HUH0 Q8HUH1 Q8HUH2
Q8HV04 Q8HVA1 Q8HVY3 Q8HVY4 Q8HVY5 Q8HW57 Q8HWS3 Q8HX86 Q8HXB8 Q8HXB9 Q8HXD5
Q8HXE4 Q8HXH0 Q8HXH6 Q8HXK7 Q8HXL3 Q8HXM1 Q8HXN9 Q8HXP0 Q8HXP1 Q8HXP2 Q8HXP3
Q8HXP4 Q8HXP5 Q8HXP6 Q8HXP7 Q8HXW4 Q8HXW9 Q8HXX1 Q8HXX2 Q8HXX5 Q8HXX8 Q8HXX9
Q8HXY0 Q8HXY4 Q8HXY5 Q8HXY9 Q8HY31 Q8HY33 Q8HY34 Q8HY67 Q8HY68 Q8HY87 Q8HYB8
Q8HYI9 Q8HYW0 Q8HYY4 Q8HZ00 Q8HZ57 Q8HZ58 Q8HZ59 Q8HZ60 Q8HZP9 Q8HZR5 Q8HZV4
Q8HZY8 Q8I009 Q8I030 Q8I0F4 Q8I0H7 Q8I0T3 Q8I0Y4 Q8I145 Q8I1C8 Q8I1E5 Q8I1F4
Q8I1G0 Q8I1G8 Q8I1H7 Q8I1N0 Q8I1N6 Q8I1T1 Q8I1U7 Q8I236 Q8I295 Q8I2S0 Q8I384
Q8I3W2 Q8I3X4 Q8I3Y6 Q8I3Z5 Q8I467 Q8I4F7 Q8I4S1 Q8I4V2 Q8I4V5 Q8I4V8 Q8I4X0
Q8I566 Q8I589 Q8I5G1 Q8I5R7 Q8I615 Q8I659 Q8I7C3 Q8I7D5 Q8I7D6 Q8I7M8 Q8I7N7
Q8I7P9 Q8I7Q0 Q8I7U0 Q8I7X7 Q8I7Z8 Q8I8A2 Q8I8U6 Q8I8U7 Q8I8V0 Q8I8W7 Q8I9B4

Q8IAW3 Q8IAY6 Q8IBB8 Q8IBG1 Q8IBH7 Q8IBP3 Q8IC19 Q8IC32 Q8ID39 Q8ID92 Q8ID94
Q8IDF6 Q8IE51 Q8IE95 Q8IED2 Q8IED5 Q8IEF6 Q8IER7 Q8IFW4 Q8IG39 Q8IG42 Q8IH00
Q8IH18 Q8IH24 Q8IHI1 Q8IHS4 Q8IID4 Q8IIK4 Q8IIS0 Q8IIS5 Q8IIU6 Q8IJP2 Q8IJV6
Q8IKG4 Q8IKN4 Q8IKU0 Q8IL06 Q8IL26 Q8IL84 Q8ILC1 Q8ILC2 Q8ILI6 Q8ILQ7 Q8ILT4
Q8ILW9 Q8IM00 Q8IM04 Q8IM30 Q8IMM9 Q8IMP6 Q8IMU4 Q8IN81 Q8IN94 Q8INF7 Q8INK9
Q8INM3 Q8INR6 Q8INT5 Q8INX3 Q8IPJ3 Q8IPX7 Q8IQ05 Q8IQ13 Q8IQF1 Q8IQQ4 Q8IQV9
Q8IR79 Q8IRB5 Q8IRC7 Q8IRE4 Q8IRG6 Q8IRH5 Q8IRJ8 Q8IRR5 Q8IRW8 Q8IRY7 Q8IS98
Q8ISF8 Q8ISN9 Q8ISP0 Q8ISQ3 Q8IT89 Q8IT98 Q8ITC3 Q8ITI5 Q8ITY4 Q8IU29 Q8IU53
Q8IUB9 Q8IUC0 Q8IUC1 Q8IUC2 Q8IUC3 Q8IUD6 Q8IUE0 Q8IUE1 Q8IUF1 Q8IUG1 Q8IUH3
Q8IUI4 Q8IUM7 Q8IUR0 Q8IUR6 Q8IUW3 Q8IUX4 Q8IUZ0 Q8IUZ5 Q8IV13 Q8IV20 Q8IV32
Q8IV33 Q8IV35 Q8IV38 Q8IV42 Q8IV56 Q8IV76 Q8IVA1 Q8IVB5 Q8IVC4 Q8IVE0 Q8IVF1
Q8IVF4 Q8IVF6 Q8IVG5 Q8IVH4 Q8IVM0 Q8IVP9 Q8IVS8 Q8IVU3 Q8IVU9 Q8IVV2 Q8IVV7
Q8IVW4 Q8IW03 Q8IW19 Q8IW36 Q8IW40 Q8IWA6 Q8IWB6 Q8IWD4 Q8IWF7 Q8IWF9 Q8IWG1
Q8IWL3 Q8IWL8 Q8IWN6 Q8IWP9 Q8IWT0 Q8IWT3 Q8IWU9 Q8IWX7 Q8IWY8 Q8IWZ4 Q8IWZ5
Q8IX04 Q8IX06 Q8IX18 Q8IX29 Q8IXF0 Q8IXL9 Q8IXM2 Q8IXM7 Q8IXN7 Q8IXP5 Q8IXQ8
Q8IXQ9 Q8IXR5 Q8IXR9 Q8IXS0 Q8IXS2 Q8IXT1 Q8IXT2 Q8IXV7 Q8IXW0 Q8IXY8 Q8IXZ3
Q8IY21 Q8IY42 Q8IY51 Q8IY82 Q8IY85 Q8IYA2 Q8IYA7 Q8IYA8 Q8IYB0 Q8IYB1 Q8IYB8
Q8IYB9 Q8IYD1 Q8IYD2 Q8IYD8 Q8IYE0 Q8IYE1 Q8IYF1 Q8IYF3 Q8IYG6 Q8IYI0 Q8IYI8
Q8IYJ2 Q8IYK2 Q8IYL2 Q8IYM0 Q8IYM1 Q8IYM2 Q8IYN0 Q8IYN2 Q8IYN6 Q8IYQ7 Q8IYR0
Q8IYR2 Q8IYS1 Q8IYS4 Q8IYS8 Q8IYT1 Q8IYT2 Q8IYT3 Q8IYT4 Q8IYU4 Q8IYW2 Q8IYX0
Q8IYX1 Q8IYX3 Q8IYX4 Q8IYX7 Q8IYX8 Q8IZ02 Q8IZ13 Q8IZ16 Q8IZ20 Q8IZ41 Q8IZ63
Q8IZC7 Q8IZD0 Q8IZL2 Q8IZM0 Q8IZM8 Q8IZP2 Q8IZP6 Q8IZQ5 Q8IZQ8 Q8IZS5 Q8IZT9
Q8IZU0 Q8IZU1 Q8IZU2 Q8IZY5 Q8J093 Q8J0B2 Q8J0E6 Q8J0E9 Q8J0F0 Q8J0F2 Q8J0F5
Q8J0F7 Q8J0F8 Q8J0F9 Q8J0G0 Q8J0I4 Q8J0L6 Q8J0N2 Q8J0N3 Q8J0N6 Q8J0U0 Q8J0U2
Q8J112 Q8J130 Q8J140 Q8J1F7 Q8J1F8 Q8J1G1 Q8J1G4 Q8J1G7 Q8J1H3 Q8J1K2 Q8J1L3
Q8J1R3 Q8J1R4 Q8J1U9 Q8J1X7 Q8J254 Q8J256 Q8J257 Q8J269 Q8J294 Q8J2J3 Q8J2J8
Q8J2Q6 Q8J2R0 Q8J2R3 Q8J2V8 Q8J2Y1 Q8J308 Q8JFF3 Q8JFP1 Q8JFQ2 Q8JFQ4 Q8JFQ6
Q8JFR1 Q8JFV4 Q8JFW1 Q8JFW4 Q8JFY9 Q8JFZ2 Q8JG30 Q8JGN5 Q8JGR6 Q8JGR7 Q8JGS0
Q8JGS1 Q8JGT5 Q8JH47 Q8JH55 Q8JH70 Q8JH71 Q8JHH5 Q8JHI6 Q8JHJ2 Q8JHU0 Q8JHV9
Q8JI10 Q8JIR6 Q8JIS3 Q8JIS6 Q8JIT5 Q8JIT6 Q8JIY8 Q8JJ04 Q8JJ26 Q8JJ48 Q8JJ64
Q8JJC0 Q8JJE2 Q8JJY5 Q8JL69 Q8JL72 Q8JL75 Q8JL78 Q8JL90 Q8JL91 Q8JL96 Q8JLF5
Q8JLG4 Q8JLI4 Q8JMT1 Q8JMY4 Q8JMY7 Q8JMZ7 Q8JN08 Q8JN59 Q8JN60 Q8JN65 Q8JN66
Q8JN67 Q8JN69 Q8JP02 Q8JPQ9 Q8JPR0 Q8JPR2 Q8JPX5 Q8JPX6 Q8JPY0 Q8JPY1 Q8JSP3
Q8JSP6 Q8JTG9 Q8JTH3 Q8JU52 Q8JU53 Q8JU54 Q8JU55 Q8JU58 Q8JU59 Q8JU60 Q8JU61
Q8JU62 Q8JVB9 Q8JVC0 Q8JVC1 Q8JVC2 Q8JXF6 Q8JYK0 Q8JYK1 Q8JZ13 Q8JZL0 Q8JZL3
Q8JZM0 Q8JZM2 Q8JZM6 Q8JZP3 Q8JZP9 Q8JZQ9 Q8JZS6 Q8JZU0 Q8JZU6 Q8JZV4 Q8JZV7
Q8JZW5 Q8JZW8 Q8JZX3 Q8JZX4 Q8JZY2 Q8JZY4 Q8JZZ6 Q8K003 Q8K004 Q8K010 Q8K013
Q8K019 Q8K023 Q8K039 Q8K057 Q8K070 Q8K088 Q8K0C1 Q8K0C9 Q8K0D0 Q8K0D5 Q8K0E1
Q8K0G7 Q8K0G8 Q8K0H5 Q8K0S0 Q8K0S2 Q8K0S9 Q8K0T2 Q8K0T4 Q8K0U4 Q8K0V4 Q8K0W3
Q8K0W9 Q8K0Y2 Q8K0Y7 Q8K0Z7 Q8K114 Q8K124 Q8K126 Q8K157 Q8K183 Q8K194 Q8K199
Q8K1A0 Q8K1C0 Q8K1C9 Q8K1E6 Q8K1H1 Q8K1I7 Q8K1J6 Q8K1K9 Q8K1L0 Q8K1L2 Q8K1L5
Q8K1L6 Q8K1N4 Q8K1P7 Q8K1Q4 Q8K1R7 Q8K1S5 Q8K1X1 Q8K202 Q8K203 Q8K207 Q8K214
Q8K215 Q8K224 Q8K243 Q8K248 Q8K259 Q8K274 Q8K284 Q8K288 Q8K298 Q8K2A7 Q8K2D0
Q8K2D3 Q8K2D6 Q8K2F0 Q8K2F8 Q8K2H6 Q8K2I2 Q8K2I9 Q8K2J4 Q8K2L8 Q8K2M3 Q8K2P6

Q8K2Q0 Q8K2Q2 Q8K2Q5 Q8K2R5 Q8K2T1 Q8K2U2 Q8K2X2 Q8K2X8 Q8K2Y0 Q8K2Y9 Q8K2Z2
Q8K2Z8 Q8K301 Q8K327 Q8K330 Q8K349 Q8K352 Q8K354 Q8K363 Q8K368 Q8K370 Q8K375
Q8K396 Q8K3A0 Q8K3A2 Q8K3A4 Q8K3A9 Q8K3C3 Q8K3D3 Q8K3E7 Q8K3F3 Q8K3F4 Q8K3F7
Q8K3H5 Q8K3I9 Q8K3K9 Q8K3M5 Q8K3P4 Q8K3P5 Q8K3P7 Q8K3Q3 Q8K3U7 Q8K3V1 Q8K3W0
Q8K3W2 Q8K3X2 Q8K3X3 Q8K3X4 Q8K3X8 Q8K402 Q8K409 Q8K411 Q8K419 Q8K430 Q8K450
Q8K485 Q8K4F6 Q8K4G5 Q8K4H1 Q8K4I6 Q8K4J2 Q8K4K1 Q8K4K3 Q8K4K4 Q8K4L0 Q8K4L4
Q8K4P0 Q8K4Q6 Q8K4R9 Q8K4T3 Q8K4T4 Q8K4T5 Q8K4Z5 Q8K557 Q8K572 Q8K581 Q8K582
Q8K585 Q8K586 Q8K589 Q8K5B8 Q8K5W8 Q8K7S6 Q8K900 Q8K901 Q8K904 Q8K905 Q8K906
Q8K907 Q8K909 Q8K910 Q8K912 Q8K913 Q8K916 Q8K917 Q8K918 Q8K919 Q8K920 Q8K921
Q8K922 Q8K923 Q8K924 Q8K925 Q8K926 Q8K927 Q8K928 Q8K929 Q8K930 Q8K931 Q8K932
Q8K933 Q8K934 Q8K935 Q8K936 Q8K937 Q8K938 Q8K939 Q8K940 Q8K941 Q8K943 Q8K944
Q8K945 Q8K946 Q8K947 Q8K948 Q8K949 Q8K950 Q8K951 Q8K952 Q8K953 Q8K954 Q8K955
Q8K956 Q8K957 Q8K958 Q8K959 Q8K960 Q8K961 Q8K962 Q8K963 Q8K964 Q8K965 Q8K966
Q8K967 Q8K968 Q8K970 Q8K971 Q8K972 Q8K973 Q8K974 Q8K975 Q8K976 Q8K977 Q8K978
Q8K979 Q8K980 Q8K981 Q8K982 Q8K983 Q8K986 Q8K988 Q8K989 Q8K990 Q8K991 Q8K992
Q8K998 Q8K9A0 Q8K9A1 Q8K9A2 Q8K9A3 Q8K9A4 Q8K9A5 Q8K9A6 Q8K9A8 Q8K9A9 Q8K9B0
Q8K9B2 Q8K9B3 Q8K9B5 Q8K9B6 Q8K9B7 Q8K9B8 Q8K9B9 Q8K9C0 Q8K9C1 Q8K9C2 Q8K9C3
Q8K9C4 Q8K9C5 Q8K9C6 Q8K9C7 Q8K9C8 Q8K9C9 Q8K9D0 Q8K9D2 Q8K9D3 Q8K9D4 Q8K9D6
Q8K9D7 Q8K9D8 Q8K9D9 Q8K9E1 Q8K9E2 Q8K9E3 Q8K9E4 Q8K9E5 Q8K9E6 Q8K9E7 Q8K9E9
Q8K9F0 Q8K9F1 Q8K9F2 Q8K9F3 Q8K9F4 Q8K9F5 Q8K9F6 Q8K9F8 Q8K9F9 Q8K9G0 Q8K9G1
Q8K9G2 Q8K9G3 Q8K9G4 Q8K9G5 Q8K9G6 Q8K9G7 Q8K9H0 Q8K9H1 Q8K9H2 Q8K9H3 Q8K9H4
Q8K9H5 Q8K9H6 Q8K9H7 Q8K9H8 Q8K9H9 Q8K9I0 Q8K9I1 Q8K9I2 Q8K9I3 Q8K9I4 Q8K9I6
Q8K9I7 Q8K9I9 Q8K9J1 Q8K9J2 Q8K9J3 Q8K9J7 Q8K9J8 Q8K9K1 Q8K9K4 Q8K9K5 Q8K9K6
Q8K9K7 Q8K9K8 Q8K9K9 Q8K9L1 Q8K9L2 Q8K9L5 Q8K9L6 Q8K9L8 Q8K9L9 Q8K9M2 Q8K9M3
Q8K9M8 Q8K9N0 Q8K9N1 Q8K9N2 Q8K9N3 Q8K9N4 Q8K9N5 Q8K9N6 Q8K9N9 Q8K9P0 Q8K9P1
Q8K9P2 Q8K9P3 Q8K9P4 Q8K9P5 Q8K9P6 Q8K9P7 Q8K9Q0 Q8K9Q1 Q8K9Q2 Q8K9Q3 Q8K9Q5
Q8K9Q7 Q8K9Q8 Q8K9R1 Q8K9R4 Q8K9R5 Q8K9R6 Q8K9R7 Q8K9R9 Q8K9S2 Q8K9S3 Q8K9S7
Q8K9S8 Q8K9S9 Q8K9T0 Q8K9T1 Q8K9T2 Q8K9T5 Q8K9T6 Q8K9T7 Q8K9T8 Q8K9T9 Q8K9U0
Q8K9U1 Q8K9U2 Q8K9U4 Q8K9U5 Q8K9U6 Q8K9U7 Q8K9U8 Q8K9U9 Q8K9V0 Q8K9V1 Q8K9V2
Q8K9V3 Q8K9V4 Q8K9V5 Q8K9V6 Q8K9V8 Q8K9V9 Q8K9W0 Q8K9W1 Q8K9W2 Q8K9W3 Q8K9W4
Q8K9W5 Q8K9W6 Q8K9W7 Q8K9W8 Q8K9W9 Q8K9X0 Q8K9X1 Q8K9X2 Q8K9X3 Q8K9Y8 Q8K9Y9
Q8K9Z0 Q8K9Z1 Q8K9Z2 Q8K9Z4 Q8K9Z5 Q8K9Z6 Q8K9Z7 Q8K9Z8 Q8K9Z9 Q8KA00 Q8KA03
Q8KA05 Q8KA06 Q8KA07 Q8KA09 Q8KA10 Q8KA11 Q8KA13 Q8KA14 Q8KA15 Q8KA16 Q8KA22
Q8KA23 Q8KA24 Q8KA25 Q8KA26 Q8KA27 Q8KA29 Q8KA30 Q8KA31 Q8KA33 Q8KA34 Q8KA40
Q8KA43 Q8KA46 Q8KA47 Q8KA48 Q8KA53 Q8KA54 Q8KA55 Q8KA56 Q8KA57 Q8KA58 Q8KA59
Q8KA60 Q8KA61 Q8KA62 Q8KA63 Q8KA65 Q8KA66 Q8KA67 Q8KA68 Q8KA69 Q8KA70 Q8KA71
Q8KA72 Q8KA73 Q8KA75 Q8KA76 Q8KA78 Q8KA79 Q8KA80 Q8KA81 Q8KA85 Q8KA87 Q8KAA4
Q8KAA5 Q8KAB1 Q8KAD1 Q8KAE1 Q8KAE4 Q8KAE8 Q8KAE9 Q8KAF0 Q8KAG8 Q8KAG9 Q8KAH0
Q8KAH1 Q8KAH2 Q8KAH3 Q8KAH4 Q8KAH5 Q8KAH6 Q8KAH7 Q8KAH8 Q8KAH9 Q8KAI0 Q8KAI1
Q8KAI2 Q8KAI3 Q8KAI4 Q8KAI5 Q8KAI6 Q8KAI7 Q8KAI8 Q8KAI9 Q8KAJ0 Q8KAJ3 Q8KAJ4
Q8KAJ5 Q8KAJ6 Q8KAJ8 Q8KAJ9 Q8KAK0 Q8KAK2 Q8KAK5 Q8KAK6 Q8KAM1 Q8KAM2 Q8KAM3
Q8KAM4 Q8KAM6 Q8KAM7 Q8KAM8 Q8KAM9 Q8KAN0 Q8KAN3 Q8KAN7 Q8KAP4 Q8KAQ7 Q8KAR6
Q8KAS1 Q8KAW2 Q8KAW3 Q8KAW4 Q8KAW7 Q8KAX2 Q8KAX3 Q8KAX9 Q8KAY6 Q8KAY7 Q8KAZ6
Q8KB05 Q8KB10 Q8KB21 Q8KB47 Q8KB57 Q8KB67 Q8KB70 Q8KB71 Q8KBD3 Q8KBD5 Q8KBD8

Q8KBE8 Q8KBF4 Q8KBF5 Q8KBF6 Q8KBF9 Q8KBH1 Q8KBH8 Q8KBI0 Q8KBI7 Q8KBI8 Q8KBJ9
Q8KBK3 Q8KBK4 Q8KBK5 Q8KBK6 Q8KBK7 Q8KBK8 Q8KBK9 Q8KBL4 Q8KBN8 Q8KBP9 Q8KBS4
Q8KBS6 Q8KBT4 Q8KBV8 Q8KBW1 Q8KBW7 Q8KBW8 Q8KBY0 Q8KBY2 Q8KBY3 Q8KBY5 Q8KBZ5
Q8KBZ7 Q8KC00 Q8KC04 Q8KC05 Q8KC06 Q8KC15 Q8KC17 Q8KC25 Q8KC36 Q8KC49 Q8KC50
Q8KC52 Q8KC68 Q8KC69 Q8KC73 Q8KC74 Q8KC79 Q8KC85 Q8KC92 Q8KCA0 Q8KCB0 Q8KCB2
Q8KCB7 Q8KCB8 Q8KCC0 Q8KCC7 Q8KCD1 Q8KCD5 Q8KCD6 Q8KCD7 Q8KCD8 Q8KCE9 Q8KCG4
Q8KCG7 Q8KCG8 Q8KCH5 Q8KCH7 Q8KCH9 Q8KCI9 Q8KCJ0 Q8KCJ3 Q8KCJ4 Q8KCK7 Q8KCK9
Q8KCL0 Q8KCL7 Q8KCM7 Q8KCN1 Q8KCP0 Q8KCQ1 Q8KCQ2 Q8KCR8 Q8KCS2 Q8KCS5 Q8KCT7
Q8KCU0 Q8KCU3 Q8KCU7 Q8KCW2 Q8KCW4 Q8KCY0 Q8KCZ6 Q8KCZ7 Q8KD01 Q8KD05 Q8KD17
Q8KD41 Q8KD46 Q8KD62 Q8KD63 Q8KD64 Q8KD69 Q8KD71 Q8KD74 Q8KD79 Q8KD88 Q8KD89
Q8KD90 Q8KD91 Q8KDC4 Q8KDC5 Q8KDE0 Q8KDE1 Q8KDE2 Q8KDE3 Q8KDH2 Q8KDJ0 Q8KDJ5
Q8KDK5 Q8KDQ7 Q8KDR7 Q8KDS1 Q8KDS2 Q8KDS8 Q8KDS9 Q8KDT4 Q8KDU4 Q8KDU8 Q8KDV6
Q8KE30 Q8KE49 Q8KE61 Q8KE84 Q8KE85 Q8KE87 Q8KEA4 Q8KEA8 Q8KED5 Q8KEF4 Q8KEF9
Q8KEG7 Q8KEG8 Q8KEH1 Q8KEJ1 Q8KEM4 Q8KEP3 Q8KEP5 Q8KER4 Q8KER7 Q8KES1 Q8KES3
Q8KES6 Q8KES7 Q8KET0 Q8KEW2 Q8KEW8 Q8KEW9 Q8KEX0 Q8KEX2 Q8KEX7 Q8KEY6 Q8KEY7
Q8KEZ7 Q8KF00 Q8KF02 Q8KF03 Q8KF11 Q8KF55 Q8KF56 Q8KF60 Q8KF68 Q8KF69 Q8KF74
Q8KFC3 Q8KFC8 Q8KFE7 Q8KFG8 Q8KFI0 Q8KFI4 Q8KFI9 Q8KFJ5 Q8KFJ6 Q8KFK3 Q8KFK6
Q8KFK8 Q8KFL0 Q8KFL5 Q8KFM9 Q8KFN6 Q8KFN8 Q8KFN9 Q8KFQ4 Q8KFQ5 Q8KFR0 Q8KFR1
Q8KFR5 Q8KFS0 Q8KFS5 Q8KFS7 Q8KFS9 Q8KFT0 Q8KFT1 Q8KFT3 Q8KFT6 Q8KFW8 Q8KFZ5
Q8KG04 Q8KG11 Q8KG14 Q8KG15 Q8KG16 Q8KG17 Q8KG18 Q8KG19 Q8KG23 Q8KG38 Q8KG42
Q8KG43 Q8KG45 Q8KG70 Q8KG79 Q8KG83 Q8KG86 Q8KGA2 Q8KGA4 Q8KGB6 Q8KGC0 Q8KGC5
Q8KGC6 Q8KGC9 Q8KGD2 Q8KGD5 Q8KGE0 Q8KGF2 Q8KGF3 Q8KGF7 Q8KGF8 Q8KGF9 Q8KGG4
Q8KGG5 Q8KGG6 Q8KH12 Q8KHE4 Q8KHV6 Q8KHX9 Q8KHZ8 Q8KI76 Q8KIX0 Q8KIX1 Q8KIX2
Q8KIX3 Q8KIX4 Q8KIY0 Q8KIY4 Q8KJ09 Q8KJ13 Q8KJ15 Q8KJ16 Q8KJ17 Q8KJ18 Q8KJ19
Q8KJ20 Q8KJ21 Q8KJ24 Q8KJ25 Q8KKS1 Q8KL44 Q8KLB6 Q8KLI6 Q8KLK5 Q8KLK7 Q8KLM3
Q8KLQ0 Q8KLS9 Q8KLU8 Q8KLV8 Q8KM02 Q8KM30 Q8KM31 Q8KMJ5 Q8KML6 Q8KML7 Q8KML8
Q8KN28 Q8KN33 Q8KN38 Q8KNP2 Q8KNP3 Q8KNX8 Q8KP10 Q8KP37 Q8KPQ7 Q8KPR0 Q8KPR4
Q8KPS9 Q8KPU0 Q8KPU2 Q8KPU9 Q8KQB3 Q8KQC4 Q8KQF5 Q8KQL2 Q8KQT8 Q8KR41 Q8KR59
Q8KR69 Q8KRA0 Q8KRC9 Q8KRD1 Q8KRD3 Q8KRG6 Q8KRM2 Q8KRR5 Q8KRT5 Q8KRZ2 Q8KSB4
Q8KSB6 Q8KSC8 Q8KT95 Q8KT97 Q8KT99 Q8KTA1 Q8KTA3 Q8KTA6 Q8KTA8 Q8KTB0 Q8KTB2
Q8KTB4 Q8KTB6 Q8KTB7 Q8KTB8 Q8KTB9 Q8KTC1 Q8KTE1 Q8KTH8 Q8KTI4 Q8KTP6 Q8KTP9
Q8KTQ9 Q8KTR0 Q8KTR3 Q8KTR6 Q8KTR9 Q8KTS0 Q8KTS4 Q8KTS9 Q8KU07 Q8KU55 Q8KU58
Q8KU93 Q8KUU5 Q8KUV2 Q8KWS8 Q8KWS9 Q8KWT0 Q8KWT1 Q8KWT3 Q8KWT4 Q8KWT5 Q8KWT6
Q8KWX2 Q8KWX4 Q8KX22 Q8KX32 Q8KY27 Q8KYT4 Q8KZ92 Q8KZ93 Q8KZ94 Q8KZA0 Q8KZM7
Q8KZM8 Q8KZM9 Q8KZP5 Q8KZT5 Q8L0V4 Q8L0X4 Q8L0Z4 Q8L163 Q8L164 Q8L1A7 Q8L1A8
Q8L1B1 Q8L1E5 Q8L1H5 Q8L1H9 Q8L1I4 Q8L1Z5 Q8L1Z7 Q8L1Z8 Q8L1Z9 Q8L208 Q8L2A6
Q8L2F3 Q8L2F8 Q8L2I5 Q8L2J6 Q8L2S1 Q8L373 Q8L376 Q8L395 Q8L397 Q8L399 Q8L3A0
Q8L3A9 Q8L3B0 Q8L3B1 Q8L3B2 Q8L3B3 Q8L3B9 Q8L3D3 Q8L3R2 Q8L3R3 Q8L3U3 Q8L3W1
Q8L3X8 Q8L3Z8 Q8L440 Q8L467 Q8L471 Q8L493 Q8L4A7 Q8L4E7 Q8L4E9 Q8L4J2 Q8L4N1
Q8L4Y5 Q8L539 Q8L548 Q8L556 Q8L557 Q8L5A7 Q8L5D8 Q8L5E0 Q8L5K3 Q8L5K4 Q8L5R3
Q8L5T5 Q8L5V0 Q8L5W7 Q8L5W8 Q8L5Y9 Q8L5Z4 Q8L607 Q8L611 Q8L612 Q8L627 Q8L633
Q8L649 Q8L6J1 Q8L6J2 Q8L6J3 Q8L6J4 Q8L6J5 Q8L6Y3 Q8L6Y4 Q8L6Y7 Q8L6Z4 Q8L6Z7
Q8L716 Q8L727 Q8L730 Q8L733 Q8L735 Q8L736 Q8L746 Q8L748 Q8L762 Q8L765 Q8L770
Q8L773 Q8L774 Q8L785 Q8L793 Q8L794 Q8L799 Q8L7B5 Q8L7C2 Q8L7F7 Q8L7G0 Q8L7G4

Q8L7H1 Q8L7I1 Q8L7K0 Q8L7K4 Q8L7K9 Q8L7M4 Q8L7N0 Q8L7N3 Q8L7N8 Q8L7R2 Q8L7S0
Q8L7S3 Q8L7S4 Q8L7S8 Q8L7S9 Q8L7T3 Q8L7T9 Q8L7W2 Q8L7W7 Q8L7Y8 Q8L7Z3 Q8L803
Q8L829 Q8L831 Q8L836 Q8L840 Q8L870 Q8L8A5 Q8L8B8 Q8L8C0 Q8L8L9 Q8L8Q3 Q8L8S2
Q8L8T2 Q8L8T7 Q8L8W6 Q8L8Y0 Q8L8Z8 Q8L910 Q8L925 Q8L953 Q8L960 Q8L968 Q8L981
Q8L983 Q8L9J9 Q8L9S1 Q8L9S3 Q8L9V6 Q8L9X2 Q8L9Y2 Q8LA13 Q8LA16 Q8LA53 Q8LAC4
Q8LAD2 Q8LAH8 Q8LAL2 Q8LAP8 Q8LAS8 Q8LAT0 Q8LAV5 Q8LAW2 Q8LAX0 Q8LAZ7 Q8LB01
Q8LB10 Q8LB33 Q8LB54 Q8LB60 Q8LB65 Q8LB72 Q8LBB7 Q8LBC0 Q8LBK6 Q8LBL5 Q8LBM4
Q8LBN7 Q8LBP6 Q8LBS4 Q8LBU2 Q8LBV4 Q8LBW3 Q8LC03 Q8LC59 Q8LC79 Q8LC83 Q8LCE1
Q8LCE9 Q8LCG7 Q8LCH5 Q8LCH9 Q8LCL3 Q8LCQ3 Q8LCT3 Q8LCV1 Q8LCW9 Q8LCY2 Q8LD03
Q8LD26 Q8LD27 Q8LD46 Q8LD49 Q8LDF9 Q8LDI5 Q8LDM2 Q8LDN8 Q8LDR0 Q8LDS4 Q8LDS5
Q8LDU5 Q8LDW8 Q8LE10 Q8LE42 Q8LE43 Q8LE52 Q8LE94 Q8LEA2 Q8LEB2 Q8LEC0 Q8LED9
Q8LEE8 Q8LEF3 Q8LEG1 Q8LEG3 Q8LEK4 Q8LEM6 Q8LEM8 Q8LEQ0 Q8LEV3 Q8LEV7 Q8LEZ4
Q8LF09 Q8LF41 Q8LF89 Q8LF97 Q8LFA9 Q8LFD3 Q8LFG1 Q8LFH6 Q8LFH7 Q8LFJ5 Q8LFJ6
Q8LFJ8 Q8LFL8 Q8LFQ6 Q8LFS6 Q8LFU0 Q8LFV1 Q8LFV3 Q8LFW5 Q8LG03 Q8LG10 Q8LG70
Q8LG76 Q8LG77 Q8LG95 Q8LGA1 Q8LGD1 Q8LGD5 Q8LGE3 Q8LGG0 Q8LGI3 Q8LGI8 Q8LGJ5
Q8LGM7 Q8LGS1 Q8LGU6 Q8LGU7 Q8LGZ9 Q8LH82 Q8LHA8 Q8LHH9 Q8LHP0 Q8LI30 Q8LI34
Q8LIL0 Q8LJA9 Q8LJS2 Q8LJT8 Q8LJU5 Q8LJW1 Q8LJW3 Q8LK56 Q8LK61 Q8LK93 Q8LKP9
Q8LKW0 Q8LL04 Q8LL05 Q8LL17 Q8LL69 Q8LL87 Q8LLE5 Q8LLR0 Q8LLR1 Q8LLR2 Q8LLV8
Q8LLW7 Q8LMI4 Q8LMR2 Q8LMR9 Q8LN68 Q8LNN8 Q8LNU5 Q8LNZ2 Q8LNZ3 Q8LP17 Q8LPC4
Q8LPF0 Q8LPF1 Q8LPH6 Q8LPI9 Q8LPJ5 Q8LPJ7 Q8LPL5 Q8LPL7 Q8LPN3 Q8LPN7 Q8LPQ3
Q8LPQ9 Q8LPR5 Q8LPS6 Q8LPT9 Q8LPU4 Q8LPU5 Q8LPW2 Q8LQ74 Q8LQH4 Q8LQM5 Q8LR34
Q8LR50 Q8LR63 Q8LR86 Q8LRK8 Q8LRK9 Q8LRM8 Q8LSC2 Q8LSC3 Q8LSN2 Q8LSN3 Q8LSZ4
Q8LT05 Q8LT07 Q8LTE2 Q8LU58 Q8LUX2 Q8LVH2 Q8LW11 Q8LW53 Q8M355 Q8M920 Q8M924
Q8M930 Q8M986 Q8M998 Q8M9A9 Q8M9H0 Q8M9L8 Q8M9T4 Q8M9U3 Q8M9U4 Q8M9U6 Q8M9U7
Q8M9U8 Q8M9U9 Q8M9V0 Q8M9V1 Q8M9V2 Q8M9V3 Q8M9V4 Q8M9V5 Q8M9V6 Q8M9V7 Q8M9W2
Q8M9W7 Q8M9Y1 Q8M9Y2 Q8M9Y6 Q8M9Y7 Q8M9Y8 Q8M9Y9 Q8MA01 Q8MA02 Q8MA03 Q8MA04
Q8MA09 Q8MA10 Q8MA11 Q8MA12 Q8MA18 Q8MA20 Q8MA31 Q8MA72 Q8MAI6 Q8MAI8 Q8MAI9
Q8MAJ0 Q8MC55 Q8MC94 Q8MC99 Q8MCA0 Q8MCA1 Q8MCA4 Q8MCA5 Q8MCJ4 Q8MCJ5 Q8MCK8
Q8MCL7 Q8MCL8 Q8MCL9 Q8MCM0 Q8MCM1 Q8MCM2 Q8MCM3 Q8MCM4 Q8MCM7 Q8MCM8 Q8MCM9
Q8MCN0 Q8MCN1 Q8MCN2 Q8MCN3 Q8MCN7 Q8MCN8 Q8MCN9 Q8MCP4 Q8MCP5 Q8MCP6 Q8MCR5
Q8MCR6 Q8MCR7 Q8MCR8 Q8MCR9 Q8MCS1 Q8MCV5 Q8MD24 Q8MD32 Q8MD78 Q8MDD1 Q8MDT1
Q8MDU0 Q8MEC7 Q8MEC8 Q8MEC9 Q8MED5 Q8MEE6 Q8MEH7 Q8MET5 Q8MEW7 Q8MEW9 Q8MEX2
Q8MEX3 Q8MEX4 Q8MEX5 Q8MEX7 Q8MEX8 Q8MEY0 Q8MEY1 Q8MEY2 Q8MEY4 Q8MFA1 Q8MFF0
Q8MHZ2 Q8MI03 Q8MI06 Q8MI68 Q8MIB5 Q8MIB7 Q8MIB8 Q8MIB9 Q8MIC2 Q8MID1 Q8MID5
Q8MID6 Q8MID8 Q8MIE6 Q8MIE7 Q8MIE9 Q8MIF3 Q8MIF7 Q8MIH1 Q8MIH8 Q8MIK9 Q8MIM3
Q8MIP0 Q8MIP2 Q8MIR0 Q8MJ26 Q8MJ30 Q8MJ36 Q8MJ46 Q8MJ47 Q8MJ87 Q8MJ97 Q8MJ98
Q8MJ99 Q8MJA0 Q8MJC6 Q8MJC8 Q8MJD0 Q8MJD5 Q8MJD6 Q8MJI9 Q8MJJ1 Q8MJK1 Q8MJS8
Q8MJT0 Q8MJU9 Q8MJV0 Q8MJV1 Q8MKD5 Q8MKE8 Q8MKF1 Q8MKH6 Q8MKI3 Q8MKL1 Q8MKW7
Q8MM62 Q8MM75 Q8MMC4 Q8MMN1 Q8MN46 Q8MN49 Q8MNT9 Q8MNU7 Q8MNU8 Q8MNV7 Q8MP20
Q8MPA5 Q8MPA7 Q8MPD6 Q8MPF7 Q8MPM1 Q8MQ70 Q8MQA7 Q8MQH7 Q8MQI6 Q8MQJ9 Q8MQT8
Q8MQU6 Q8MR37 Q8MR62 Q8MS27 Q8MSG8 Q8MSU4 Q8MSV2 Q8MSW4 Q8MSX1 Q8MSX2 Q8MT06
Q8MT08 Q8MT36 Q8MTJ7 Q8MTV8 Q8MUA4 Q8MUC1 Q8MUF6 Q8MUK3 Q8MUR2 Q8MUR4 Q8MX88
Q8MXU4 Q8MYE8 Q8MYF0 Q8MYF1 Q8MYP1 Q8MYQ1 Q8N0N9 Q8N0P0 Q8N0S2 Q8N0T1 Q8N0U4
Q8N0U6 Q8N0U7 Q8N0V1 Q8N0V3 Q8N0W3 Q8N0W5 Q8N0X2 Q8N0X4 Q8N0Y7 Q8N123 Q8N136

Q8N140 Q8N141 Q8N142 Q8N143 Q8N159 Q8N165 Q8N184 Q8N187 Q8N196 Q8N1A0 Q8N1A6
Q8N1B3 Q8N1D0 Q8N1D5 Q8N1E6 Q8N1H7 Q8N1I8 Q8N1L1 Q8N1L9 Q8N1N4 Q8N1P7 Q8N1V2
Q8N1V8 Q8N1W2 Q8N1X5 Q8N228 Q8N239 Q8N257 Q8N268 Q8N283 Q8N2A0 Q8N2B8 Q8N2C9
Q8N2H9 Q8N2I2 Q8N2I9 Q8N2N9 Q8N2R0 Q8N2R8 Q8N2W9 Q8N2Y8 Q8N309 Q8N319 Q8N328
Q8N336 Q8N339 Q8N344 Q8N365 Q8N371 Q8N377 Q8N393 Q8N398 Q8N3A8 Q8N3F0 Q8N3J2
Q8N3J3 Q8N3J5 Q8N3J9 Q8N3L3 Q8N3R3 Q8N3U1 Q8N3X6 Q8N3Y1 Q8N3Z3 Q8N3Z6 Q8N402
Q8N412 Q8N414 Q8N427 Q8N443 Q8N446 Q8N461 Q8N465 Q8N485 Q8N4B4 Q8N4B5 Q8N4E4
Q8N4E7 Q8N4H0 Q8N4J0 Q8N4L8 Q8N4M7 Q8N4N3 Q8N4P2 Q8N4P3 Q8N4P6 Q8N4Q1 Q8N4W9
Q8N4Y2 Q8N531 Q8N535 Q8N554 Q8N584 Q8N587 Q8N5C7 Q8N5I3 Q8N5J4 Q8N5L8 Q8N5M1
Q8N5M4 Q8N5N4 Q8N5Q1 Q8N5S3 Q8N5T2 Q8N5U0 Q8N5X7 Q8N5Z0 Q8N5Z5 Q8N612 Q8N616
Q8N635 Q8N660 Q8N680 Q8N6C7 Q8N6D5 Q8N6F8 Q8N6G2 Q8N6I1 Q8N6K4 Q8N6K7 Q8N6M5
Q8N6M8 Q8N6M9 Q8N6N2 Q8N6N6 Q8N6N7 Q8N6Q8 Q8N6R0 Q8N6U2 Q8N6V4 Q8N6V9 Q8N6W0
Q8N6Y0 Q8N715 Q8N769 Q8N782 Q8N7A1 Q8N7B1 Q8N7B6 Q8N7B9 Q8N7C3 Q8N7E2 Q8N7F7
Q8N7G0 Q8N7H1 Q8N7K0 Q8N7L0 Q8N7M0 Q8N7M2 Q8N7N1 Q8N7P7 Q8N7Q3 Q8N7R0 Q8N7R1
Q8N7U6 Q8N7U7 Q8N7U9 Q8N7X0 Q8N7X1 Q8N7X2 Q8N7X4 Q8N7Y1 Q8N7Z2 Q8N7Z5 Q8N801
Q8N806 Q8N812 Q8N813 Q8N814 Q8N815 Q8N819 Q8N823 Q8N831 Q8N841 Q8N859 Q8N865
Q8N883 Q8N888 Q8N8A2 Q8N8A8 Q8N8B7 Q8N8C0 Q8N8D1 Q8N8D9 Q8N8E1 Q8N8G2 Q8N8G6
Q8N8H1 Q8N8I6 Q8N8J6 Q8N8L2 Q8N8L6 Q8N8M0 Q8N8P6 Q8N8Q3 Q8N8R5 Q8N8U2 Q8N8U3
Q8N8X9 Q8N8Y5 Q8N8Z3 Q8N8Z8 Q8N910 Q8N945 Q8N954 Q8N957 Q8N961 Q8N972 Q8N976
Q8N987 Q8N988 Q8N998 Q8N999 Q8N9B4 Q8N9C0 Q8N9E0 Q8N9F8 Q8N9G6 Q8N9H8 Q8N9H9
Q8N9I9 Q8N9K5 Q8N9L1 Q8N9L7 Q8N9P0 Q8N9P6 Q8N9R0 Q8N9R6 Q8N9S3 Q8N9S9 Q8N9T2
Q8N9U9 Q8N9V2 Q8N9V3 Q8N9V6 Q8N9V7 Q8N9W4 Q8N9W5 Q8N9W6 Q8N9W8 Q8N9X3 Q8N9Y4
Q8N9Z0 Q8N9Z9 Q8NA03 Q8NA19 Q8NA23 Q8NA47 Q8NA54 Q8NA56 Q8NA57 Q8NA61 Q8NA66
Q8NA69 Q8NA70 Q8NA75 Q8NA77 Q8NA92 Q8NA96 Q8NA97 Q8NAA5 Q8NAA6 Q8NAB2 Q8NAE3
Q8NAJ2 Q8NAM6 Q8NAP3 Q8NAP8 Q8NAQ8 Q8NAT2 Q8NAV2 Q8NB12 Q8NB15 Q8NB25 Q8NB42
Q8NB50 Q8NB91 Q8NBA8 Q8NBB2 Q8NBB4 Q8NBC4 Q8NBE8 Q8NBF2 Q8NBF4 Q8NBH2 Q8NBR9
Q8NBT0 Q8NBZ0 Q8NBZ9 Q8NC26 Q8NC38 Q8NC74 Q8NCA5 Q8NCA9 Q8NCE0 Q8NCI6 Q8NCJ5
Q8NCK3 Q8NCN2 Q8NCN5 Q8NCQ2 Q8NCQ5 Q8NCQ7 Q8NCT3 Q8NCU1 Q8NCU4 Q8NCV1 Q8NCX0
Q8NCY6 Q8ND07 Q8ND71 Q8ND90 Q8NDA8 Q8NDB2 Q8NDC4 Q8NDG6 Q8NDH3 Q8NDH6 Q8NDL9
Q8NDM7 Q8NDN9 Q8NDP4 Q8NDV3 Q8NDW4 Q8NDW8 Q8NDY3 Q8NDY4 Q8NDY6 Q8NDZ0 Q8NDZ2
Q8NE18 Q8NE28 Q8NE31 Q8NE65 Q8NEA4 Q8NEC7 Q8NEE0 Q8NEE6 Q8NEE8 Q8NEF3 Q8NEG0
Q8NEG2 Q8NEG5 Q8NEG7 Q8NEH6 Q8NEK5 Q8NEK8 Q8NEM0 Q8NEM1 Q8NEM8 Q8NEN0 Q8NEP7
Q8NEP9 Q8NEQ6 Q8NET4 Q8NEV1 Q8NEV4 Q8NEV8 Q8NEY3 Q8NEZ5 Q8NF64 Q8NF67 Q8NFD2
Q8NFD4 Q8NFH8 Q8NFI4 Q8NFJ8 Q8NFR7 Q8NFU3 Q8NFV5 Q8NFW5 Q8NFW9 Q8NFY9 Q8NFZ0
Q8NFZ5 Q8NG06 Q8NG48 Q8NG50 Q8NG68 Q8NHH1 Q8NHP1 Q8NHP7 Q8NHS0 Q8NHS2 Q8NHS4
Q8NHS9 Q8NHU0 Q8NHU2 Q8NHW5 Q8NHX4 Q8NHY5 Q8NHY6 Q8NHZ7 Q8NI29 Q8NI36 Q8NI37
Q8NI38 Q8NI51 Q8NIC1 Q8NIC8 Q8NID7 Q8NIG3 Q8NIG9 Q8NIH0 Q8NIH3 Q8NIH6 Q8NIL3
Q8NIL4 Q8NIQ8 Q8NIV6 Q8NIW7 Q8NIZ4 Q8NJ52 Q8NJ72 Q8NJA1 Q8NJJ1 Q8NJJ9 Q8NJM2
Q8NJN1 Q8NJN2 Q8NJN3 Q8NJQ3 Q8NJR0 Q8NJR2 Q8NJR4 Q8NJR5 Q8NJR6 Q8NJT4 Q8NJU8
Q8NK47 Q8NK50 Q8NK72 Q8NK83 Q8NKB8 Q8NKC1 Q8NKC2 Q8NKF4 Q8NKF5 Q8NKN9 Q8NKQ0
Q8NKQ2 Q8NKR7 Q8NKR9 Q8NKW8 Q8NKW9 Q8NKX0 Q8NKX3 Q8NKY0 Q8NL02 Q8NL05 Q8NL21
Q8NL22 Q8NL51 Q8NL53 Q8NL65 Q8NL92 Q8NLB6 Q8NLC4 Q8NLE4 Q8NLF0 Q8NLF9 Q8NLG0
Q8NLG1 Q8NLL5 Q8NLN0 Q8NLP6 Q8NLP9 Q8NLS3 Q8NLT9 Q8NLY6 Q8NLY8 Q8NM11 Q8NM16
Q8NM23 Q8NM41 Q8NM44 Q8NM64 Q8NM65 Q8NM79 Q8NM84 Q8NMB0 Q8NMB2 Q8NMB8 Q8NMB9

Q8NMD0 Q8NMD4 Q8NMD7 Q8NMH3 Q8NMH6 Q8NMI4 Q8NMI5 Q8NMJ4 Q8NMJ7 Q8NMK7 Q8NML3
Q8NML5 Q8NMM7 Q8NMN7 Q8NMQ1 Q8NMR4 Q8NMR5 Q8NMT1 Q8NMT3 Q8NMU1 Q8NMU4 Q8NMZ9
Q8NN01 Q8NN02 Q8NN26 Q8NN33 Q8NN37 Q8NN43 Q8NN50 Q8NN51 Q8NN57 Q8NN66 Q8NN99
Q8NNB3 Q8NNB4 Q8NNB7 Q8NNC1 Q8NNC6 Q8NND1 Q8NND3 Q8NNF6 Q8NNH6 Q8NNJ0 Q8NNJ1
Q8NNJ2 Q8NNJ4 Q8NNJ8 Q8NNM6 Q8NNM7 Q8NNN0 Q8NNP0 Q8NNP4 Q8NNT5 Q8NNT9 Q8NNV6
Q8NNV7 Q8NNW4 Q8NNX3 Q8NNX5 Q8NNX7 Q8NNY6 Q8NNZ0 Q8NNZ4 Q8NNZ6 Q8NNZ9 Q8NP01
Q8NP02 Q8NP03 Q8NP06 Q8NP10 Q8NP12 Q8NP26 Q8NP30 Q8NP31 Q8NP37 Q8NP40 Q8NP41
Q8NP46 Q8NP49 Q8NP50 Q8NP66 Q8NP67 Q8NP72 Q8NP73 Q8NP88 Q8NP95 Q8NP96 Q8NPA9
Q8NPB2 Q8NPZ0 Q8NPZ8 Q8NQ07 Q8NQ19 Q8NQ22 Q8NQ23 Q8NQ33 Q8NQ38 Q8NQ39 Q8NQ40
Q8NQ42 Q8NQ43 Q8NQ46 Q8NQ47 Q8NQ52 Q8NQ53 Q8NQ55 Q8NQ56 Q8NQ58 Q8NQ93 Q8NQ94
Q8NQ97 Q8NQ98 Q8NQA1 Q8NQC4 Q8NQD8 Q8NQE1 Q8NQH0 Q8NQH1 Q8NQJ3 Q8NQJ6 Q8NQK6
Q8NQK7 Q8NQL5 Q8NQL7 Q8NQM1 Q8NQM8 Q8NQN6 Q8NQN7 Q8NQP6 Q8NQP7 Q8NQP8 Q8NQU5
Q8NQU9 Q8NQV0 Q8NQV2 Q8NQV7 Q8NQW7 Q8NQX9 Q8NQZ9 Q8NR05 Q8NR09 Q8NR14 Q8NR17
Q8NR18 Q8NR20 Q8NR24 Q8NR39 Q8NR40 Q8NR49 Q8NR57 Q8NR60 Q8NR92 Q8NR94 Q8NRB3
Q8NRC3 Q8NRD4 Q8NRE3 Q8NRE9 Q8NRG3 Q8NRI0 Q8NRM2 Q8NRM3 Q8NRN8 Q8NRP4 Q8NRQ2
Q8NRR3 Q8NRU9 Q8NRV2 Q8NRV3 Q8NRV6 Q8NRW6 Q8NRX4 Q8NRY0 Q8NRY1 Q8NRZ4 Q8NS11
Q8NS12 Q8NS15 Q8NS16 Q8NS17 Q8NS21 Q8NS31 Q8NS38 Q8NS75 Q8NS78 Q8NS80 Q8NS90
Q8NS92 Q8NSC4 Q8NSH0 Q8NSH7 Q8NSH8 Q8NSH9 Q8NSJ4 Q8NSL1 Q8NSL2 Q8NSL3 Q8NSM0
Q8NSM4 Q8NSR1 Q8NSS0 Q8NSS1 Q8NSS3 Q8NSS4 Q8NST4 Q8NST5 Q8NST6 Q8NSV0 Q8NSV1
Q8NSV2 Q8NSV4 Q8NSV5 Q8NSV6 Q8NSV7 Q8NSX3 Q8NSX5 Q8NSX6 Q8NSX7 Q8NSX8 Q8NSZ2
Q8NSZ3 Q8NSZ4 Q8NSZ7 Q8NSZ9 Q8NT00 Q8NT01 Q8NT02 Q8NT05 Q8NT06 Q8NT08 Q8NT09
Q8NT19 Q8NT20 Q8NT21 Q8NT25 Q8NT26 Q8NT28 Q8NT29 Q8NT39 Q8NT57 Q8NT73 Q8NT75
Q8NT90 Q8NTA5 Q8NTA6 Q8NTB0 Q8NTC4 Q8NTD8 Q8NTE1 Q8NTF4 Q8NTQ8 Q8NTQ9 Q8NTR2
Q8NTR6 Q8NTT4 Q8NTU1 Q8NTY7 Q8NU33 Q8NU97 Q8NU98 Q8NUD4 Q8NUD8 Q8NUF9 Q8NUH2
Q8NUH3 Q8NUI2 Q8NUI3 Q8NUI4 Q8NUK7 Q8NUK8 Q8NUK9 Q8NUL1 Q8NUM0 Q8NUM4 Q8NUM5
Q8NUM9 Q8NUN2 Q8NUN3 Q8NUP5 Q8NUQ3 Q8NUQ5 Q8NUQ6 Q8NUR1 Q8NUR2 Q8NUR7 Q8NUT2
Q8NUT7 Q8NUV9 Q8NV07 Q8NV32 Q8NV36 Q8NV45 Q8NV55 Q8NV63 Q8NV65 Q8NV80 Q8NV88
Q8NV89 Q8NV90 Q8NV95 Q8NV99 Q8NVA0 Q8NVA1 Q8NVA3 Q8NVA4 Q8NVA6 Q8NVB4 Q8NVB7
Q8NVC8 Q8NVC9 Q8NVD1 Q8NVE3 Q8NVE6 Q8NVF5 Q8NVF6 Q8NVG0 Q8NVG3 Q8NVG4 Q8NVG5
Q8NVG8 Q8NVH3 Q8NVH4 Q8NVH5 Q8NVH8 Q8NVI5 Q8NVI8 Q8NVI9 Q8NVJ0 Q8NVJ5 Q8NVL7
Q8NVN2 Q8NVS2 Q8NVT1 Q8NVT3 Q8NVT4 Q8NVU3 Q8NVU6 Q8NVV1 Q8NVZ4 Q8NVZ8 Q8NVZ9
Q8NW08 Q8NW14 Q8NW17 Q8NW23 Q8NW24 Q8NW26 Q8NW30 Q8NW33 Q8NW34 Q8NW37 Q8NW49
Q8NW51 Q8NW53 Q8NW54 Q8NW55 Q8NW56 Q8NW58 Q8NW61 Q8NW68 Q8NW72 Q8NW73 Q8NW74
Q8NW75 Q8NW77 Q8NW84 Q8NW87 Q8NW95 Q8NWA1 Q8NWA8 Q8NWA9 Q8NWB0 Q8NWB4 Q8NWB5
Q8NWB7 Q8NWB9 Q8NWC8 Q8NWC9 Q8NWD0 Q8NWE6 Q8NWF2 Q8NWM8 Q8NWN1 Q8NWN3 Q8NWN4
Q8NWN5 Q8NWP0 Q8NWP2 Q8NWP3 Q8NWQ3 Q8NWQ4 Q8NWQ9 Q8NWR6 Q8NWR7 Q8NWS1 Q8NWS3
Q8NWS4 Q8NWS5 Q8NWS9 Q8NWU0 Q8NWU1 Q8NWU2 Q8NWU3 Q8NWU6 Q8NWV1 Q8NWV2 Q8NWV5
Q8NWV8 Q8NWX7 Q8NWX9 Q8NWY9 Q8NWZ0 Q8NWZ1 Q8NWZ3 Q8NWZ5 Q8NWZ6 Q8NWZ7 Q8NWZ8
Q8NWZ9 Q8NX01 Q8NX02 Q8NX05 Q8NX06 Q8NX11 Q8NX12 Q8NX16 Q8NX18 Q8NX19 Q8NX22
Q8NX25 Q8NX29 Q8NX32 Q8NX35 Q8NX38 Q8NX44 Q8NX55 Q8NX56 Q8NX59 Q8NX60 Q8NX62
Q8NX72 Q8NX76 Q8NX78 Q8NX79 Q8NX87 Q8NX88 Q8NX89 Q8NX90 Q8NX91 Q8NX92 Q8NX93
Q8NX94 Q8NX95 Q8NXA0 Q8NXA2 Q8NXC0 Q8NXC2 Q8NXC8 Q8NXD0 Q8NXE7 Q8NXE8 Q8NXE9
Q8NXF0 Q8NXF1 Q8NXF2 Q8NXF3 Q8NXG7 Q8NXH0 Q8NXH7 Q8NXI0 Q8NXI1 Q8NXL2 Q8NXL5
Q8NXL7 Q8NXL9 Q8NXM0 Q8NXM6 Q8NXN2 Q8NXN3 Q8NXQ5 Q8NXS4 Q8NXT8 Q8NXU1 Q8NXW4

Q8NXW5 Q8NXX1 Q8NXX2 Q8NXY1 Q8NXY2 Q8NXY3 Q8NXY7 Q8NXY8 Q8NXY9 Q8NXZ0 Q8NXZ2
Q8NXZ3 Q8NXZ4 Q8NXZ6 Q8NXZ8 Q8NY00 Q8NY02 Q8NY05 Q8NY07 Q8NY69 Q8NY70 Q8NY72
Q8NY74 Q8NY77 Q8NY94 Q8NY95 Q8NYB7 Q8NYC4 Q8NYC7 Q8NYD0 Q8NYH4 Q8NYI0 Q8NYK7
Q8NYL9 Q8NYM5 Q8NYM6 Q8NYM7 Q8NYR1 Q8NYT6 Q8NYV4 Q8NYX2 Q8NYX6 Q8NYY2 Q8NYY3
Q8NYY7 Q8NYZ4 Q8NZ00 Q8NZ02 Q8NZ04 Q8NZ19 Q8NZ20 Q8NZ22 Q8NZ24 Q8NZ25 Q8NZ27
Q8NZ29 Q8NZ30 Q8NZ37 Q8NZ43 Q8NZ45 Q8NZ46 Q8NZ49 Q8NZ53 Q8NZ56 Q8NZ57 Q8NZ98
Q8NZA2 Q8NZB2 Q8NZB4 Q8NZB5 Q8NZB7 Q8NZC6 Q8NZD1 Q8NZD8 Q8NZD9 Q8NZE1 Q8NZF6
Q8NZF7 Q8NZF8 Q8NZG3 Q8NZG7 Q8NZH6 Q8NZI1 Q8NZI5 Q8NZJ0 Q8NZK4 Q8NZL9 Q8NZM0
Q8NZM1 Q8NZM6 Q8NZM7 Q8NZN7 Q8NZU4 Q8NZU7 Q8NZU8 Q8NZV6 Q8NZV7 Q8NZW6 Q8NZW7
Q8NZW9 Q8NZX0 Q8NZX4 Q8NZX8 Q8NZX9 Q8NZY8 Q8NZZ1 Q8P003 Q8P016 Q8P017 Q8P031
Q8P036 Q8P040 Q8P046 Q8P051 Q8P058 Q8P063 Q8P066 Q8P068 Q8P075 Q8P0C0 Q8P0C4
Q8P0C7 Q8P0C9 Q8P0D9 Q8P0E0 Q8P0E6 Q8P0F8 Q8P0G6 Q8P0G7 Q8P0H1 Q8P0H4 Q8P0H7
Q8P0I6 Q8P0J7 Q8P0J9 Q8P0S5 Q8P0S6 Q8P0S7 Q8P0T7 Q8P0V9 Q8P0X2 Q8P0X5 Q8P0Y8
Q8P0Z2 Q8P0Z5 Q8P102 Q8P106 Q8P109 Q8P116 Q8P120 Q8P122 Q8P125 Q8P126 Q8P129
Q8P137 Q8P150 Q8P151 Q8P152 Q8P159 Q8P161 Q8P163 Q8P172 Q8P179 Q8P1A6 Q8P1B6
Q8P1C0 Q8P1C4 Q8P1C5 Q8P1C8 Q8P1D1 Q8P1E3 Q8P1F1 Q8P1F2 Q8P1F3 Q8P1F4 Q8P1G1
Q8P1G9 Q8P1H4 Q8P1H5 Q8P1H7 Q8P1H8 Q8P1H9 Q8P1J2 Q8P1J3 Q8P1K0 Q8P1K1 Q8P1L1
Q8P1M4 Q8P1N3 Q8P1T6 Q8P1T7 Q8P1T8 Q8P1W3 Q8P1W4 Q8P1W7 Q8P1X6 Q8P1Y0 Q8P234
Q8P243 Q8P244 Q8P250 Q8P272 Q8P274 Q8P298 Q8P2A6 Q8P2A7 Q8P2B1 Q8P2B4 Q8P2C9
Q8P2D0 Q8P2D2 Q8P2D3 Q8P2E1 Q8P2H3 Q8P2K1 Q8P2L2 Q8P2N7 Q8P2N8 Q8P2Q5 Q8P2R3
Q8P2R7 Q8P2T2 Q8P2U1 Q8P2W5 Q8P2X6 Q8P2Y2 Q8P2Y3 Q8P2Y5 Q8P2Z4 Q8P302 Q8P308
Q8P310 Q8P312 Q8P315 Q8P322 Q8P327 Q8P336 Q8P337 Q8P340 Q8P3B0 Q8P3B7 Q8P3C4
Q8P3C8 Q8P3D5 Q8P3D7 Q8P3F4 Q8P3H1 Q8P3H2 Q8P3H3 Q8P3H6 Q8P3I4 Q8P3I5 Q8P3J4
Q8P3K1 Q8P3K2 Q8P3K4 Q8P3L1 Q8P3N0 Q8P3Q0 Q8P3S2 Q8P3W6 Q8P3X4 Q8P3Y1 Q8P3Y6
Q8P3Z3 Q8P3Z4 Q8P447 Q8P448 Q8P455 Q8P456 Q8P458 Q8P469 Q8P473 Q8P475 Q8P494
Q8P4C5 Q8P4D0 Q8P4D4 Q8P4D7 Q8P4F2 Q8P4F6 Q8P4F9 Q8P4G0 Q8P4G1 Q8P4H1 Q8P4H2
Q8P4R0 Q8P4S5 Q8P4U3 Q8P4U6 Q8P4V1 Q8P536 Q8P548 Q8P550 Q8P551 Q8P552 Q8P558
Q8P588 Q8P589 Q8P590 Q8P5D4 Q8P5D8 Q8P5E4 Q8P5F3 Q8P5H9 Q8P5K9 Q8P5L1 Q8P5L5
Q8P5L9 Q8P5M4 Q8P5P5 Q8P5Q4 Q8P5R4 Q8P5R7 Q8P5S1 Q8P5S6 Q8P5S9 Q8P5T3 Q8P5T5
Q8P5T7 Q8P5U0 Q8P5U9 Q8P5Z4 Q8P5Z7 Q8P5Z8 Q8P607 Q8P608 Q8P610 Q8P631 Q8P632
Q8P665 Q8P676 Q8P682 Q8P698 Q8P6A0 Q8P6B1 Q8P6E0 Q8P6E3 Q8P6E4 Q8P6E5 Q8P6E7
Q8P6F6 Q8P6M7 Q8P6M8 Q8P6M9 Q8P6N0 Q8P6P1 Q8P6Q4 Q8P6T8 Q8P6T9 Q8P6V6 Q8P6V8
Q8P6X1 Q8P6X3 Q8P6X4 Q8P6X5 Q8P708 Q8P709 Q8P719 Q8P725 Q8P741 Q8P765 Q8P766
Q8P767 Q8P775 Q8P778 Q8P7A0 Q8P7A1 Q8P7C5 Q8P7H4 Q8P7J1 Q8P7J8 Q8P7K1 Q8P7K6
Q8P7L2 Q8P7L3 Q8P7P1 Q8P7P7 Q8P7P8 Q8P7Q8 Q8P7R0 Q8P7R5 Q8P7R6 Q8P7R8 Q8P7S0
Q8P7S2 Q8P7T0 Q8P7U5 Q8P7U7 Q8P7U8 Q8P7U9 Q8P7V0 Q8P7V1 Q8P7X1 Q8P7Z2 Q8P7Z3
Q8P7Z4 Q8P7Z5 Q8P7Z6 Q8P815 Q8P829 Q8P833 Q8P855 Q8P857 Q8P867 Q8P868 Q8P888
Q8P8B2 Q8P8B5 Q8P8C7 Q8P8D9 Q8P8E0 Q8P8E4 Q8P8E9 Q8P8F4 Q8P8F8 Q8P8F9 Q8P8G9
Q8P8H2 Q8P8H3 Q8P8I0 Q8P8J0 Q8P8J2 Q8P8J4 Q8P8J5 Q8P8J6 Q8P8J8 Q8P8J9 Q8P8K2
Q8P8K3 Q8P8L2 Q8P8M6 Q8P8P4 Q8P8P6 Q8P8Q4 Q8P8Q6 Q8P8S1 Q8P8S4 Q8P8U7 Q8P8V6
Q8P8W9 Q8P8Z2 Q8P8Z8 Q8P8Z9 Q8P979 Q8P984 Q8P996 Q8P997 Q8P999 Q8P9A1 Q8P9J6
Q8P9L3 Q8P9M3 Q8P9N3 Q8P9N4 Q8P9N5 Q8P9N8 Q8P9N9 Q8P9P0 Q8P9P1 Q8P9P2 Q8P9P3
Q8P9P5 Q8P9Q0 Q8P9R1 Q8P9R7 Q8P9S7 Q8P9S8 Q8P9S9 Q8P9T0 Q8P9T3 Q8P9T9 Q8P9V1
Q8P9V6 Q8P9V9 Q8P9W9 Q8P9X0 Q8P9X1 Q8P9X7 Q8P9X8 Q8P9Y1 Q8P9Y6 Q8P9Y7 Q8P9Y9

Q8P9Z0 Q8P9Z1 Q8P9Z6 Q8PA92 Q8PA95 Q8PA97 Q8PAA6 Q8PAA7 Q8PAB2 Q8PAB5 Q8PAC0
Q8PAC1 Q8PAC2 Q8PAC4 Q8PAD0 Q8PAD2 Q8PAD3 Q8PAD9 Q8PAE0 Q8PAG3 Q8PAG5 Q8PAH9
Q8PAK8 Q8PAK9 Q8PAL0 Q8PAL1 Q8PAL7 Q8PAT3 Q8PAU0 Q8PAU4 Q8PAV2 Q8PAV3 Q8PAV6
Q8PAV9 Q8PAW7 Q8PAY7 Q8PB40 Q8PB42 Q8PB48 Q8PB50 Q8PB52 Q8PBB3 Q8PBB5 Q8PBB7
Q8PBC0 Q8PBC1 Q8PBC2 Q8PBC3 Q8PBG4 Q8PBG6 Q8PBG9 Q8PBH0 Q8PBH1 Q8PBH2 Q8PBH3
Q8PBH8 Q8PBI2 Q8PBJ4 Q8PBK7 Q8PBL8 Q8PBP3 Q8PBR6 Q8PBV2 Q8PBV4 Q8PBW4 Q8PBW5
Q8PBW9 Q8PBX1 Q8PBX8 Q8PBY0 Q8PBY5 Q8PBY6 Q8PBY7 Q8PC25 Q8PC34 Q8PC35 Q8PC36
Q8PC37 Q8PC38 Q8PC39 Q8PC40 Q8PC41 Q8PC42 Q8PC43 Q8PC44 Q8PC45 Q8PC46 Q8PC47
Q8PC48 Q8PC49 Q8PC50 Q8PC51 Q8PC52 Q8PC53 Q8PC54 Q8PC55 Q8PC56 Q8PC57 Q8PC59
Q8PC61 Q8PC62 Q8PC63 Q8PC64 Q8PC67 Q8PC68 Q8PC73 Q8PC99 Q8PCA4 Q8PCA6 Q8PCB3
Q8PCB8 Q8PCE2 Q8PCE3 Q8PCE4 Q8PCE5 Q8PCE7 Q8PCH1 Q8PCH3 Q8PCH5 Q8PCJ8 Q8PCJ9
Q8PCK4 Q8PCK7 Q8PCK8 Q8PCL4 Q8PCM6 Q8PCM7 Q8PCN3 Q8PCN4 Q8PCN7 Q8PCP3 Q8PCQ7
Q8PCQ9 Q8PCR4 Q8PCR7 Q8PCS4 Q8PCW4 Q8PCW8 Q8PCY1 Q8PD23 Q8PD27 Q8PD33 Q8PD37
Q8PD47 Q8PD48 Q8PD57 Q8PD65 Q8PD66 Q8PD70 Q8PD71 Q8PD87 Q8PDA2 Q8PDA8 Q8PDD1
Q8PDD2 Q8PDF0 Q8PDF1 Q8PDF3 Q8PDG1 Q8PDI0 Q8PDJ3 Q8PDM9 Q8PDQ6 Q8PDR2 Q8PDW3
Q8PDW5 Q8PDW6 Q8PDW8 Q8PDX2 Q8PDY1 Q8PE32 Q8PE48 Q8PE50 Q8PE85 Q8PEB1 Q8PEG8
Q8PEH3 Q8PEH6 Q8PEH9 Q8PEP3 Q8PEP8 Q8PEQ4 Q8PEQ8 Q8PER2 Q8PER4 Q8PET9 Q8PEW5
Q8PEW6 Q8PEW7 Q8PEX0 Q8PEX9 Q8PF09 Q8PF23 Q8PF76 Q8PF98 Q8PFE9 Q8PFF8 Q8PFG5
Q8PFG7 Q8PFH4 Q8PFH5 Q8PFQ4 Q8PFQ5 Q8PFR2 Q8PFR3 Q8PFR5 Q8PFS5 Q8PFS9 Q8PFT1
Q8PFV1 Q8PFX6 Q8PFZ0 Q8PFZ3 Q8PFZ6 Q8PG13 Q8PG17 Q8PG20 Q8PG21 Q8PG22 Q8PG32
Q8PG33 Q8PGC9 Q8PGD0 Q8PGE6 Q8PGH9 Q8PGJ4 Q8PGK0 Q8PGP9 Q8PGQ7 Q8PGR0 Q8PGR5
Q8PGR7 Q8PGU5 Q8PGV2 Q8PGW9 Q8PH04 Q8PH05 Q8PH09 Q8PH13 Q8PH18 Q8PH23 Q8PH31
Q8PH40 Q8PH45 Q8PH49 Q8PH54 Q8PH57 Q8PH61 Q8PH63 Q8PH65 Q8PH70 Q8PHB2 Q8PHB5
Q8PHB6 Q8PHC7 Q8PHC8 Q8PHD0 Q8PHF4 Q8PHF5 Q8PHK1 Q8PHK7 Q8PHL5 Q8PHN2 Q8PHP7
Q8PHQ4 Q8PHR1 Q8PHR3 Q8PHU5 Q8PHU8 Q8PHU9 Q8PHV0 Q8PHV2 Q8PHW1 Q8PHY1 Q8PHY2
Q8PHY3 Q8PHY4 Q8PHZ5 Q8PI07 Q8PI37 Q8PI38 Q8PI56 Q8PI85 Q8PI87 Q8PI88 Q8PI89
Q8PIC2 Q8PIC3 Q8PID3 Q8PID9 Q8PIF6 Q8PIH9 Q8PII0 Q8PII1 Q8PII9 Q8PIJ2 Q8PIM0
Q8PIM1 Q8PIP8 Q8PIT8 Q8PIW4 Q8PIX0 Q8PIX4 Q8PIX9 Q8PIY4 Q8PIY5 Q8PJ05 Q8PJ10
Q8PJ11 Q8PJ20 Q8PJ25 Q8PJ26 Q8PJ28 Q8PJ29 Q8PJ31 Q8PJ38 Q8PJ53 Q8PJ55 Q8PJ56
Q8PJ57 Q8PJ58 Q8PJ59 Q8PJB1 Q8PJE1 Q8PJE2 Q8PJE3 Q8PJE4 Q8PJE5 Q8PJG7 Q8PJH7
Q8PJK3 Q8PJK5 Q8PJL6 Q8PJL7 Q8PJP3 Q8PJR6 Q8PJR9 Q8PJT1 Q8PJW7 Q8PJX2 Q8PJX7
Q8PJY5 Q8PJY6 Q8PJZ7 Q8PK00 Q8PK11 Q8PK23 Q8PK25 Q8PK26 Q8PK29 Q8PK31 Q8PK32
Q8PK34 Q8PK35 Q8PK44 Q8PK58 Q8PK76 Q8PK78 Q8PK86 Q8PK88 Q8PKQ2 Q8PKQ3 Q8PKR0
Q8PKS0 Q8PKT3 Q8PKU2 Q8PKW5 Q8PKW8 Q8PKY6 Q8PKZ1 Q8PL03 Q8PL04 Q8PL06 Q8PL08
Q8PLB3 Q8PLC6 Q8PLD9 Q8PLE0 Q8PLF1 Q8PLG0 Q8PLG1 Q8PLG2 Q8PLG5 Q8PLG6 Q8PLG7
Q8PLG8 Q8PLG9 Q8PLH0 Q8PLH2 Q8PLJ3 Q8PLJ8 Q8PLK9 Q8PLL0 Q8PLL1 Q8PLL4 Q8PLL9
Q8PLN0 Q8PLN5 Q8PLP9 Q8PLQ0 Q8PLQ4 Q8PLQ5 Q8PLQ8 Q8PLR3 Q8PLR4 Q8PLR6 Q8PLR7
Q8PLR8 Q8PLS3 Q8PLY2 Q8PLY5 Q8PLY7 Q8PLZ7 Q8PLZ8 Q8PM04 Q8PM07 Q8PM12 Q8PM13
Q8PM14 Q8PM16 Q8PM31 Q8PM33 Q8PM34 Q8PM39 Q8PM40 Q8PM62 Q8PM64 Q8PM80 Q8PMA9
Q8PMB0 Q8PMB1 Q8PMB2 Q8PMB9 Q8PMI8 Q8PMJ5 Q8PMJ8 Q8PMK5 Q8PMK6 Q8PMK8 Q8PML1
Q8PML9 Q8PMP0 Q8PMU1 Q8PMU6 Q8PMU8 Q8PMV0 Q8PMX2 Q8PMX4 Q8PMY0 Q8PMY1 Q8PMY2
Q8PMY3 Q8PN17 Q8PN19 Q8PN22 Q8PN23 Q8PN24 Q8PN25 Q8PN26 Q8PN34 Q8PN36 Q8PN47
Q8PN49 Q8PN59 Q8PN74 Q8PNB5 Q8PNE9 Q8PNF1 Q8PNG0 Q8PNG1 Q8PNH0 Q8PNH1 Q8PNH8
Q8PNI0 Q8PNI4 Q8PNI5 Q8PNI6 Q8PNN2 Q8PNP8 Q8PNQ7 Q8PNQ9 Q8PNR0 Q8PNR1 Q8PNR2

Q8PNR3 Q8PNR4 Q8PNR5 Q8PNR6 Q8PNR7 Q8PNR8 Q8PNR9 Q8PNS0 Q8PNS1 Q8PNS2 Q8PNS3
Q8PNS4 Q8PNS5 Q8PNS6 Q8PNS7 Q8PNS8 Q8PNS9 Q8PNT0 Q8PNT1 Q8PNT2 Q8PNT3 Q8PNT4
Q8PNT8 Q8PNT9 Q8PNU0 Q8PNU1 Q8PNU4 Q8PNU5 Q8PNV0 Q8PNY6 Q8PNY8 Q8PNZ5 Q8PNZ9
Q8PP24 Q8PP25 Q8PP26 Q8PP27 Q8PP38 Q8PP46 Q8PP72 Q8PP75 Q8PP82 Q8PP84 Q8PPA6
Q8PPA7 Q8PPB2 Q8PPB5 Q8PPB6 Q8PPC2 Q8PPD5 Q8PPD6 Q8PPE2 Q8PPE3 Q8PPF1 Q8PPG1
Q8PPG7 Q8PPG8 Q8PPG9 Q8PPL8 Q8PPL9 Q8PPM0 Q8PPP2 Q8PPP7 Q8PPP8 Q8PPP9 Q8PPQ1
Q8PPR3 Q8PPW9 Q8PPX2 Q8PPX9 Q8PPZ1 Q8PPZ9 Q8PQ06 Q8PQ10 Q8PQ19 Q8PQ21 Q8PQ32
Q8PQ40 Q8PQ41 Q8PQ44 Q8PQ47 Q8PQ48 Q8PQ58 Q8PQ74 Q8PQ80 Q8PQA2 Q8PQA3 Q8PQD7
Q8PQD8 Q8PQE0 Q8PQE8 Q8PQG7 Q8PQI0 Q8PQK3 Q8PQN3 Q8PQV0 Q8PR01 Q8PR13 Q8PR42
Q8PR66 Q8PRF1 Q8PRG0 Q8PRG2 Q8PRQ5 Q8PRX4 Q8PRY4 Q8PRZ8 Q8PRZ9 Q8PS49 Q8PS50
Q8PS69 Q8PS70 Q8PS92 Q8PSJ3 Q8PSK5 Q8PST3 Q8PSV9 Q8PSW0 Q8PT10 Q8PT12 Q8PT13
Q8PT42 Q8PT95 Q8PT96 Q8PT97 Q8PT98 Q8PTA3 Q8PTA5 Q8PTB0 Q8PTC3 Q8PTC5 Q8PTD1
Q8PTD3 Q8PTK0 Q8PTK1 Q8PTN6 Q8PTN8 Q8PTQ6 Q8PTR0 Q8PTR5 Q8PTT5 Q8PTT7 Q8PTT8
Q8PTT9 Q8PTU1 Q8PTU2 Q8PTU3 Q8PTU5 Q8PTX6 Q8PTZ6 Q8PU08 Q8PU18 Q8PU21 Q8PU23
Q8PU28 Q8PU39 Q8PU42 Q8PU52 Q8PU53 Q8PU54 Q8PU55 Q8PU66 Q8PU74 Q8PU75 Q8PU76
Q8PU77 Q8PU78 Q8PU79 Q8PUA4 Q8PUA5 Q8PUA7 Q8PUA8 Q8PUB7 Q8PUB8 Q8PUD6 Q8PUG1
Q8PUG2 Q8PUH2 Q8PUH7 Q8PUH9 Q8PUJ8 Q8PUL7 Q8PUM6 Q8PUM7 Q8PUN1 Q8PUN2 Q8PUP4
Q8PUQ1 Q8PUQ3 Q8PUQ4 Q8PUQ8 Q8PUR2 Q8PUR3 Q8PUR5 Q8PUR6 Q8PUR7 Q8PUR8 Q8PUV4
Q8PUX2 Q8PUX3 Q8PUY4 Q8PUY5 Q8PUZ4 Q8PV16 Q8PV17 Q8PV18 Q8PV19 Q8PV20 Q8PV21
Q8PV26 Q8PV28 Q8PV29 Q8PV30 Q8PV31 Q8PV33 Q8PV34 Q8PV35 Q8PV36 Q8PV37 Q8PV38
Q8PV39 Q8PV40 Q8PV41 Q8PV42 Q8PV43 Q8PV44 Q8PV45 Q8PV46 Q8PV47 Q8PV48 Q8PV49
Q8PV50 Q8PV84 Q8PV85 Q8PV87 Q8PV88 Q8PV89 Q8PV94 Q8PV98 Q8PV99 Q8PVA5 Q8PVB0
Q8PVC6 Q8PVD0 Q8PVD5 Q8PVE6 Q8PVF0 Q8PVF1 Q8PVF4 Q8PVJ4 Q8PVJ7 Q8PVK0 Q8PVP6
Q8PVQ1 Q8PVQ2 Q8PVR7 Q8PVS9 Q8PVT0 Q8PVT4 Q8PVT6 Q8PVV3 Q8PVY4 Q8PVZ0 Q8PVZ4
Q8PW06 Q8PW07 Q8PW37 Q8PW40 Q8PW41 Q8PW42 Q8PW43 Q8PW44 Q8PW45 Q8PW46 Q8PW52
Q8PW55 Q8PW56 Q8PW57 Q8PW58 Q8PW60 Q8PW84 Q8PW86 Q8PW87 Q8PW88 Q8PW90 Q8PW92
Q8PW97 Q8PWA0 Q8PWA7 Q8PWA9 Q8PWF3 Q8PWF5 Q8PWG1 Q8PWH0 Q8PWM8 Q8PWN9 Q8PWQ2
Q8PWS1 Q8PWS2 Q8PWS3 Q8PWS4 Q8PWT1 Q8PWT3 Q8PWT6 Q8PWU0 Q8PWV0 Q8PWV1 Q8PWV5
Q8PWV6 Q8PWX8 Q8PWY1 Q8PWY3 Q8PWY5 Q8PWZ4 Q8PX01 Q8PX04 Q8PX06 Q8PX16 Q8PX17
Q8PX25 Q8PX26 Q8PX29 Q8PX31 Q8PX37 Q8PX39 Q8PX44 Q8PX74 Q8PX75 Q8PXA0 Q8PXA1
Q8PXA5 Q8PXA8 Q8PXA9 Q8PXC3 Q8PXC4 Q8PXC6 Q8PXD4 Q8PXE1 Q8PXE2 Q8PXE6 Q8PXE7
Q8PXE8 Q8PXF0 Q8PXG9 Q8PXH2 Q8PXI0 Q8PXI1 Q8PXJ0 Q8PXJ1 Q8PXJ6 Q8PXJ8 Q8PXK0
Q8PXK5 Q8PXK6 Q8PXL6 Q8PXL7 Q8PXL8 Q8PXV2 Q8PXV5 Q8PXV8 Q8PXW5 Q8PXY1 Q8PY36
Q8PY37 Q8PY39 Q8PY44 Q8PY51 Q8PY52 Q8PY53 Q8PY57 Q8PY58 Q8PY64 Q8PY75 Q8PY83
Q8PY84 Q8PY99 Q8PYC8 Q8PYC9 Q8PYE0 Q8PYF0 Q8PYF6 Q8PYF8 Q8PYH6 Q8PYJ0 Q8PYJ4
Q8PYJ6 Q8PYK1 Q8PYK3 Q8PYK6 Q8PYL9 Q8PYN9 Q8PYP0 Q8PYP2 Q8PYP3 Q8PYP6 Q8PYQ1
Q8PYQ2 Q8PYQ3 Q8PYQ4 Q8PYQ5 Q8PYQ7 Q8PYQ9 Q8PYS1 Q8PYU8 Q8PYW7 Q8PYY0 Q8PYZ0
Q8PYZ1 Q8PZ04 Q8PZ06 Q8PZ09 Q8PZ11 Q8PZ16 Q8PZ21 Q8PZ26 Q8PZ28 Q8PZ49 Q8PZ66
Q8PZ68 Q8PZ69 Q8PZ76 Q8PZ91 Q8PZ92 Q8PZ94 Q8PZ95 Q8PZ96 Q8PZA0 Q8PZA5 Q8PZB2
Q8PZH6 Q8PZI1 Q8PZK7 Q8PZK8 Q8PZL6 Q8PZL8 Q8PZM3 Q8PZN4 Q8PZN5 Q8PZN6 Q8PZP6
Q8PZP7 Q8PZP8 Q8PZQ0 Q8PZQ1 Q8PZR7 Q8PZR8 Q8PZT3 Q8PZU1 Q8PZU2 Q8PZZ8 Q8PZZ9
Q8Q001 Q8Q012 Q8Q024 Q8Q032 Q8Q037 Q8Q038 Q8Q051 Q8Q054 Q8Q078 Q8Q081 Q8Q082
Q8Q083 Q8Q084 Q8Q093 Q8Q095 Q8Q0A4 Q8Q0B5 Q8Q0D6 Q8Q0F2 Q8Q0F4 Q8Q0F9 Q8Q0G3
Q8Q0J1 Q8Q0J2 Q8Q0J4 Q8Q0K3 Q8Q0L5 Q8Q0L8 Q8Q0M2 Q8Q0M4 Q8Q0M5 Q8Q0P3 Q8Q0P9

Q8Q0Q0 Q8Q0Q4 Q8Q0Q5 Q8Q0R2 Q8Q0R3 Q8Q0T0 Q8Q0T5 Q8Q0U0 Q8Q0U4 Q8Q0U5 Q8QFP8
Q8QFR2 Q8QG60 Q8QG78 Q8QG92 Q8QGC4 Q8QGQ7 Q8QGU6 Q8QGV6 Q8QGX4 Q8QHA5 Q8QHF0
Q8QHI2 Q8QHK8 Q8QHL5 Q8QHM9 Q8QHP8 Q8QL13 Q8QL15 Q8QL16 Q8QL17 Q8QL18 Q8QL19
Q8QL20 Q8QL21 Q8QL23 Q8QL24 Q8QL25 Q8QL26 Q8QL27 Q8QL28 Q8QL29 Q8QL31 Q8QL33
Q8QL34 Q8QL35 Q8QL36 Q8QL37 Q8QL38 Q8QL39 Q8QL40 Q8QL41 Q8QL42 Q8QL43 Q8QL44
Q8QL46 Q8QL48 Q8QL49 Q8QL50 Q8QL51 Q8QL85 Q8QME4 Q8QME6 Q8QMN3 Q8QMQ2 Q8QMQ7
Q8QMT5 Q8QMV9 Q8QMY0 Q8QMY9 Q8QMZ9 Q8QN36 Q8QN38 Q8QNT0 Q8QQ16 Q8QQ21 Q8QQ34
Q8QQ67 Q8QV02 Q8QV05 Q8QVL3 Q8QVL4 Q8QVL5 Q8QVL6 Q8QVL7 Q8QVL8 Q8QVL9 Q8QVM0
Q8QVM1 Q8QVU9 Q8QZQ2 Q8QZQ7 Q8QZQ8 Q8QZQ9 Q8QZR1 Q8QZR5 Q8QZR7 Q8QZR8 Q8QZS1
Q8QZT2 Q8QZV6 Q8QZW2 Q8QZX5 Q8QZY1 Q8QZY2 Q8QZY3 Q8QZY9 Q8QZZ7 Q8R003 Q8R010
Q8R016 Q8R023 Q8R033 Q8R034 Q8R040 Q8R050 Q8R059 Q8R071 Q8R080 Q8R086 Q8R089
Q8R0A0 Q8R0A2 Q8R0A5 Q8R0A7 Q8R0C0 Q8R0E5 Q8R0F5 Q8R0F6 Q8R0J8 Q8R0K2 Q8R0K4
Q8R0K9 Q8R0L9 Q8R0P4 Q8R0S1 Q8R0T2 Q8R0V5 Q8R0W1 Q8R0X2 Q8R0Y6 Q8R124 Q8R149
Q8R179 Q8R184 Q8R1B4 Q8R1B8 Q8R1C6 Q8R1D1 Q8R1F0 Q8R1F5 Q8R1F9 Q8R1G1 Q8R1G2
Q8R1G6 Q8R1H0 Q8R1H8 Q8R1K4 Q8R1M0 Q8R1M2 Q8R1N0 Q8R1N4 Q8R1P4 Q8R1Q9 Q8R1Y2
Q8R1Z4 Q8R205 Q8R2H1 Q8R2H4 Q8R2I0 Q8R2K4 Q8R2K7 Q8R2M0 Q8R2M2 Q8R2N0 Q8R2N2
Q8R2Q0 Q8R2Q4 Q8R2R3 Q8R2R5 Q8R2S1 Q8R2S9 Q8R2T8 Q8R2U4 Q8R2U7 Q8R2V2 Q8R2V3
Q8R2W9 Q8R2Y8 Q8R2Y9 Q8R323 Q8R326 Q8R344 Q8R349 Q8R370 Q8R395 Q8R3A2 Q8R3B7
Q8R3C0 Q8R3C1 Q8R3C6 Q8R3F9 Q8R3H9 Q8R3J4 Q8R3J5 Q8R3K3 Q8R3L2 Q8R3L8 Q8R3N1
Q8R3P0 Q8R3P6 Q8R3P9 Q8R3Q6 Q8R3V6 Q8R3W5 Q8R3Y5 Q8R3Y8 Q8R418 Q8R478 Q8R496
Q8R4C2 Q8R4E0 Q8R4E4 Q8R4E6 Q8R4E9 Q8R4G8 Q8R4H7 Q8R4I1 Q8R4I4 Q8R4N0 Q8R4T2
Q8R4U6 Q8R4U7 Q8R4X3 Q8R512 Q8R515 Q8R516 Q8R519 Q8R536 Q8R565 Q8R574 Q8R5A6
Q8R5C5 Q8R5F3 Q8R5F7 Q8R5G2 Q8R5H8 Q8R5I7 Q8R5K4 Q8R5K5 Q8R5M0 Q8R5M5 Q8R5N7
Q8R5N8 Q8R5Q4 Q8R5R4 Q8R5S2 Q8R5S3 Q8R5S5 Q8R5S9 Q8R5T0 Q8R5T7 Q8R5T8 Q8R5U4
Q8R5U7 Q8R5W0 Q8R5W2 Q8R5X3 Q8R5X4 Q8R5X5 Q8R5X6 Q8R5X7 Q8R5X8 Q8R5Y3 Q8R5Y4
Q8R5Z1 Q8R5Z2 Q8R5Z5 Q8R5Z6 Q8R5Z8 Q8R5Z9 Q8R600 Q8R601 Q8R602 Q8R603 Q8R605
Q8R616 Q8R619 Q8R620 Q8R622 Q8R634 Q8R635 Q8R639 Q8R659 Q8R678 Q8R679 Q8R680
Q8R682 Q8R685 Q8R687 Q8R689 Q8R6A2 Q8R6A7 Q8R6B1 Q8R6C8 Q8R6C9 Q8R6E7 Q8R6F5
Q8R6G5 Q8R6G7 Q8R6G8 Q8R6H2 Q8R6I2 Q8R6I7 Q8R6I9 Q8R6K3 Q8R6K4 Q8R6K8 Q8R6K9
Q8R6L0 Q8R6M1 Q8R6M2 Q8R6M3 Q8R6M7 Q8R6M9 Q8R6T8 Q8R6V0 Q8R6V3 Q8R6V4 Q8R6V5
Q8R708 Q8R711 Q8R720 Q8R749 Q8R751 Q8R753 Q8R757 Q8R765 Q8R778 Q8R786 Q8R7B8
Q8R7B9 Q8R7C0 Q8R7C1 Q8R7C2 Q8R7C3 Q8R7C6 Q8R7C8 Q8R7F0 Q8R7I7 Q8R7K0 Q8R7K9
Q8R7L3 Q8R7M2 Q8R7N0 Q8R7N1 Q8R7N2 Q8R7N6 Q8R7Q8 Q8R7S0 Q8R7S6 Q8R7S8 Q8R7S9
Q8R7T1 Q8R7T3 Q8R7T8 Q8R7T9 Q8R7U2 Q8R7U3 Q8R7U4 Q8R7U5 Q8R7U6 Q8R7U7 Q8R7U8
Q8R7U9 Q8R7V0 Q8R7V1 Q8R7V2 Q8R7V3 Q8R7V4 Q8R7V5 Q8R7V6 Q8R7V7 Q8R7V8 Q8R7V9
Q8R7W0 Q8R7W1 Q8R7W2 Q8R7W3 Q8R7W4 Q8R7W5 Q8R7W6 Q8R7W7 Q8R7W8 Q8R7W9 Q8R7X0
Q8R7X1 Q8R7X2 Q8R7X4 Q8R7X7 Q8R7X8 Q8R7X9 Q8R7Y0 Q8R7Y1 Q8R7Y2 Q8R7Y3 Q8R7Y7
Q8R7Y8 Q8R7Y9 Q8R806 Q8R807 Q8R821 Q8R828 Q8R832 Q8R840 Q8R841 Q8R844 Q8R860
Q8R880 Q8R881 Q8R882 Q8R883 Q8R884 Q8R885 Q8R886 Q8R887 Q8R890 Q8R8J4 Q8R8L3
Q8R8M3 Q8R8M4 Q8R8M9 Q8R8N1 Q8R8N2 Q8R8N9 Q8R8Q6 Q8R8R6 Q8R8R7 Q8R8S6 Q8R8W3
Q8R8W4 Q8R8Y7 Q8R8Z6 Q8R8Z8 Q8R909 Q8R914 Q8R918 Q8R922 Q8R924 Q8R933 Q8R943
Q8R961 Q8R965 Q8R966 Q8R973 Q8R977 Q8R979 Q8R984 Q8R986 Q8R9A4 Q8R9A6 Q8R9C2
Q8R9C3 Q8R9C4 Q8R9C6 Q8R9C7 Q8R9D0 Q8R9E0 Q8R9E1 Q8R9F0 Q8R9F8 Q8R9F9 Q8R9G2
Q8R9G4 Q8R9G7 Q8R9H6 Q8R9J1 Q8R9J6 Q8R9J9 Q8R9K5 Q8R9K8 Q8R9L3 Q8R9L4 Q8R9L7

Q8R9M6 Q8R9M7 Q8R9M8 Q8R9M9 Q8R9N0 Q8R9N6 Q8R9P1 Q8R9P3 Q8R9R3 Q8R9R5 Q8R9R6
Q8R9R7 Q8R9S4 Q8R9S6 Q8R9S7 Q8R9T0 Q8R9T1 Q8R9T4 Q8R9T6 Q8R9T7 Q8R9U1 Q8R9U9
Q8R9V2 Q8R9V4 Q8R9W3 Q8R9X1 Q8R9X2 Q8R9X3 Q8R9X4 Q8R9X8 Q8R9X9 Q8R9Y2 Q8R9Y3
Q8R9Y4 Q8RA16 Q8RA21 Q8RA22 Q8RA23 Q8RA24 Q8RA26 Q8RA28 Q8RA30 Q8RA32 Q8RA33
Q8RA37 Q8RA38 Q8RA40 Q8RA42 Q8RA43 Q8RA46 Q8RA52 Q8RA54 Q8RA55 Q8RA56 Q8RA66
Q8RA67 Q8RA69 Q8RA70 Q8RA71 Q8RA72 Q8RA74 Q8RA76 Q8RA78 Q8RA90 Q8RA94 Q8RAA0
Q8RAA1 Q8RAA8 Q8RAB1 Q8RAC2 Q8RAC3 Q8RAC5 Q8RAC8 Q8RAC9 Q8RAD0 Q8RAD1 Q8RAE2
Q8RAE4 Q8RAE5 Q8RAE6 Q8RAE8 Q8RAG2 Q8RAH3 Q8RAH4 Q8RAH7 Q8RAI7 Q8RAI8 Q8RAJ0
Q8RAK6 Q8RAL7 Q8RAL9 Q8RAM9 Q8RAN0 Q8RAN2 Q8RAN3 Q8RAN4 Q8RAR7 Q8RAS5 Q8RAT8
Q8RB11 Q8RB14 Q8RB27 Q8RB30 Q8RB39 Q8RB44 Q8RB46 Q8RB48 Q8RB49 Q8RB52 Q8RB59
Q8RB66 Q8RB67 Q8RB68 Q8RB69 Q8RB70 Q8RB77 Q8RB79 Q8RB93 Q8RBA4 Q8RBA5 Q8RBA7
Q8RBA9 Q8RBB6 Q8RBB8 Q8RBC0 Q8RBC5 Q8RBC6 Q8RBE5 Q8RBG0 Q8RBH4 Q8RBI5 Q8RBI6
Q8RBI7 Q8RBJ3 Q8RBJ4 Q8RBK0 Q8RBK1 Q8RBK5 Q8RBK6 Q8RBL8 Q8RBN5 Q8RBR1 Q8RBR5
Q8RBX8 Q8RC22 Q8RC24 Q8RC25 Q8RC26 Q8RC29 Q8RC30 Q8RC39 Q8RC40 Q8RC41 Q8RC42
Q8RC52 Q8RC54 Q8RC55 Q8RC57 Q8RC58 Q8RC63 Q8RC93 Q8RC98 Q8RCA4 Q8RCD5 Q8RCE4
Q8RCF9 Q8RCG6 Q8RCH7 Q8RCH9 Q8RCM2 Q8RCP3 Q8RCP4 Q8RCS0 Q8RCS8 Q8RCT2 Q8RCT3
Q8RCT6 Q8RCV0 Q8RCV7 Q8RCV9 Q8RCW0 Q8RCW1 Q8RCW2 Q8RCY5 Q8RCY6 Q8RCZ0 Q8RCZ1
Q8RD30 Q8RD35 Q8RD36 Q8RD38 Q8RD44 Q8RD88 Q8RD94 Q8RD98 Q8RDA1 Q8RDA2 Q8RDB2
Q8RDC8 Q8RDD1 Q8RDE6 Q8RDF0 Q8RDI4 Q8RDI5 Q8RDJ5 Q8RDJ9 Q8RDK0 Q8RDK1 Q8RDK2
Q8RDK3 Q8RDK4 Q8RDL3 Q8RDL6 Q8RDM3 Q8RDM4 Q8RDM5 Q8RDM9 Q8RDN0 Q8RDN5 Q8RDN6
Q8RDP7 Q8RDQ1 Q8RDQ2 Q8RDQ3 Q8RDQ4 Q8RDR4 Q8RDR9 Q8RDT4 Q8RDU4 Q8RDV3 Q8RDV8
Q8RDW6 Q8RDX3 Q8RDX7 Q8RDX9 Q8RDZ8 Q8RE08 Q8RE26 Q8RE31 Q8RE41 Q8RE42 Q8RE43
Q8RE44 Q8RE45 Q8RE57 Q8RE97 Q8REA7 Q8REB0 Q8REC3 Q8RED1 Q8RED2 Q8RED7 Q8RED8
Q8REE6 Q8REF0 Q8REF9 Q8REG1 Q8REH1 Q8REH4 Q8REI1 Q8REI2 Q8REI3 Q8REI5 Q8REJ7
Q8REL0 Q8REL1 Q8REM9 Q8REN3 Q8RER7 Q8RET9 Q8REU0 Q8REV1 Q8REV3 Q8REV6 Q8REV7
Q8REW8 Q8RF14 Q8RF15 Q8RF47 Q8RF62 Q8RF63 Q8RF65 Q8RF94 Q8RFB5 Q8RFC2 Q8RFE4
Q8RFE6 Q8RFG1 Q8RFG6 Q8RFG9 Q8RFJ7 Q8RFJ8 Q8RFK2 Q8RFL1 Q8RFN0 Q8RFN7 Q8RFP5
Q8RFS6 Q8RFS8 Q8RFX6 Q8RFX8 Q8RFX9 Q8RFY0 Q8RFY7 Q8RFZ9 Q8RG14 Q8RG35 Q8RG36
Q8RG40 Q8RG42 Q8RG43 Q8RG52 Q8RG55 Q8RG65 Q8RG75 Q8RG80 Q8RG83 Q8RG85 Q8RG86
Q8RG87 Q8RG88 Q8RG89 Q8RG90 Q8RG96 Q8RG97 Q8RG98 Q8RGA2 Q8RGA3 Q8RGE5 Q8RGF0
Q8RGG8 Q8RGH1 Q8RGH2 Q8RGH3 Q8RGH4 Q8RGH8 Q8RGJ4 Q8RGJ5 Q8RGK8 Q8RGM4 Q8RGP6
Q8RGR0 Q8RGR2 Q8RGS0 Q8RGV2 Q8RGV7 Q8RGW9 Q8RGX1 Q8RGX3 Q8RH03 Q8RH05 Q8RH07
Q8RH08 Q8RH11 Q8RH35 Q8RH43 Q8RH44 Q8RH45 Q8RH47 Q8RH57 Q8RH64 Q8RH70 Q8RH71
Q8RH88 Q8RHA1 Q8RHA2 Q8RHA6 Q8RHB4 Q8RHB5 Q8RHC9 Q8RHD0 Q8RHF4 Q8RHH5 Q8RHH9
Q8RHI2 Q8RHI3 Q8RHI4 Q8RHI5 Q8RHI6 Q8RHI7 Q8RHI9 Q8RHJ4 Q8RHJ7 Q8RHJ8 Q8RHJ9
Q8RHK0 Q8RHK1 Q8RHK3 Q8RHM2 Q8RHM6 Q8RHN4 Q8RHN5 Q8RHQ2 Q8RHQ8 Q8RHR7 Q8RHR9
Q8RHS8 Q8RHT8 Q8RHW1 Q8RHX1 Q8RHX2 Q8RHX3 Q8RHX4 Q8RHX7 Q8RHX8 Q8RHY5 Q8RHY7
Q8RHZ9 Q8RI10 Q8RI52 Q8RI57 Q8RI59 Q8RI60 Q8RI63 Q8RI83 Q8RI88 Q8RI89 Q8RI94
Q8RIA1 Q8RIA5 Q8RIE0 Q8RIE2 Q8RIE3 Q8RIE4 Q8RIF4 Q8RIF5 Q8RIF6 Q8RIF7 Q8RIF8
Q8RIF9 Q8RIG0 Q8RIG1 Q8RIG2 Q8RIG3 Q8RIG4 Q8RIG5 Q8RIG6 Q8RIG8 Q8RIG9 Q8RIH0
Q8RIH1 Q8RIH2 Q8RIH3 Q8RIH4 Q8RIH5 Q8RIH7 Q8RII1 Q8RII8 Q8RIJ5 Q8RIK8 Q8RIK9
Q8RIL0 Q8RIL3 Q8RIL9 Q8RIM0 Q8RIQ1 Q8RIQ3 Q8RIR4 Q8RIS8 Q8RIT7 Q8RJB2 Q8RKH2
Q8RKH3 Q8RKI6 Q8RKI9 Q8RKJ0 Q8RL85 Q8RLD7 Q8RLD8 Q8RLD9 Q8RLE0 Q8RLU1 Q8RLV9
Q8RLW0 Q8RLY6 Q8RM02 Q8RM03 Q8RM04 Q8RMD1 Q8RMD4 Q8RMG1 Q8RN03 Q8RN04 Q8RN05

Q8RNM2 Q8RNP4 Q8RP83 Q8RP98 Q8RPQ1 Q8RPQ2 Q8RPX2 Q8RPX7 Q8RPX8 Q8RPX9 Q8RPY7
Q8RPY8 Q8RPZ9 Q8RQ00 Q8RQ01 Q8RQD1 Q8RQD2 Q8RQE7 Q8RQE8 Q8RQE9 Q8RQL3 Q8RQL9
Q8RQM5 Q8RQM6 Q8RQM8 Q8RQM9 Q8RQN0 Q8RQN1 Q8RQN5 Q8RQN6 Q8RQP4 Q8RQP5 Q8RQP9
Q8RQQ0 Q8RR32 Q8RR57 Q8RSQ2 Q8RSQ3 Q8RSQ4 Q8RSS3 Q8RSS4 Q8RSU9 Q8RSX4 Q8RT64
Q8RT65 Q8RT66 Q8RT67 Q8RTI2 Q8RTJ3 Q8RTJ4 Q8RTJ5 Q8RU00 Q8RU01 Q8RU31 Q8RU60
Q8RU74 Q8RU95 Q8RUC1 Q8RUC6 Q8RUD6 Q8RUF8 Q8RUK6 Q8RUN2 Q8RV06 Q8RV79 Q8RV83
Q8RVB2 Q8RVK9 Q8RVL4 Q8RVM0 Q8RVQ9 Q8RW90 Q8RW98 Q8RW99 Q8RWA2 Q8RWB8 Q8RWD0
Q8RWD6 Q8RWD7 Q8RWE8 Q8RWF0 Q8RWG8 Q8RWH3 Q8RWH9 Q8RWJ7 Q8RWL0 Q8RWL4 Q8RWL6
Q8RWM2 Q8RWM3 Q8RWN0 Q8RWN9 Q8RWQ8 Q8RWR2 Q8RWS0 Q8RWS4 Q8RWS8 Q8RWT8 Q8RWU5
Q8RWV0 Q8RWV3 Q8RWV8 Q8RWX7 Q8RWY1 Q8RWY3 Q8RWY4 Q8RWY6 Q8RWY7 Q8RX01 Q8RX22
Q8RX26 Q8RX28 Q8RX32 Q8RX33 Q8RX37 Q8RX66 Q8RX78 Q8RX87 Q8RX97 Q8RXA5 Q8RXA8
Q8RXC8 Q8RXD3 Q8RXD6 Q8RXD9 Q8RXE9 Q8RXF1 Q8RXH2 Q8RXH8 Q8RXK2 Q8RXK6 Q8RXK8
Q8RXM6 Q8RXN4 Q8RXN5 Q8RXN8 Q8RXR6 Q8RXS6 Q8RXT4 Q8RXU4 Q8RXU5 Q8RXU6 Q8RXX5
Q8RXY0 Q8RXY6 Q8RXZ4 Q8RY18 Q8RY22 Q8RY24 Q8RY71 Q8RY73 Q8RY79 Q8RY82 Q8RY97
Q8RY99 Q8RYC2 Q8RYC6 Q8RYD1 Q8RYD3 Q8RYD6 Q8RYD9 Q8RYE9 Q8RYJ8 Q8RYJ9 Q8RYK0
Q8RZA1 Q8RZB5 Q8RZX9 Q8S091 Q8S0G4 Q8S0S6 Q8S151 Q8S1Z0 Q8S1Z1 Q8S2E5 Q8S2E6
Q8S2G0 Q8S2G5 Q8S2T0 Q8S3D1 Q8S3D2 Q8S3D5 Q8S3M3 Q8S3N1 Q8S3N2 Q8S3Q9 Q8S3X0
Q8S4P4 Q8S4P5 Q8S4P6 Q8S4P9 Q8S4W7 Q8S528 Q8S5M8 Q8S625 Q8S7W9 Q8S857 Q8S895
Q8S897 Q8S8C6 Q8S8C7 Q8S8D6 Q8S8D8 Q8S8E8 Q8S8F2 Q8S8F5 Q8S8F7 Q8S8F8 Q8S8F9
Q8S8I1 Q8S8I2 Q8S8M5 Q8S8M9 Q8S8P6 Q8S8Q9 Q8S8S7 Q8S8U0 Q8S8U1 Q8S8U3 Q8S8U9
Q8S8V2 Q8S8V3 Q8S8V4 Q8S8V5 Q8S8V6 Q8S8V7 Q8S8V8 Q8S8V9 Q8S8W2 Q8S8X2 Q8S8X6
Q8S8X9 Q8S8Y0 Q8S8Y1 Q8S8Y5 Q8S8Y6 Q8S904 Q8S905 Q8S911 Q8S919 Q8S920 Q8S929
Q8S932 Q8S949 Q8S983 Q8S985 Q8S9D1 Q8S9G6 Q8S9H7 Q8S9J1 Q8S9J2 Q8S9K3 Q8S9L5
Q8S9N6 Q8S9Z2 Q8S9Z5 Q8SA63 Q8SAE6 Q8SAG3 Q8SAS8 Q8SAY0 Q8SCY1 Q8SDA8 Q8SDC3
Q8SE90 Q8SEL8 Q8SKV2 Q8SKY1 Q8SL86 Q8SL88 Q8SL90 Q8SL92 Q8SL94 Q8SM04 Q8SM14
Q8SM20 Q8SM90 Q8SN66 Q8SPI2 Q8SPQ1 Q8SPQ2 Q8SPS9 Q8SPX7 Q8SQ03 Q8SQ24 Q8SQ26
Q8SQ27 Q8SQH7 Q8SQI1 Q8SQI3 Q8SQI6 Q8SQI7 Q8SQI8 Q8SQI9 Q8SQK0 Q8SQK1 Q8SQK7
Q8SQK8 Q8SQK9 Q8SQM0 Q8SQM3 Q8SQM5 Q8SQN2 Q8SQN7 Q8SQP0 Q8SQP2 Q8SQP3 Q8SQP4
Q8SQP8 Q8SQP9 Q8SQQ3 Q8SQQ7 Q8SQR2 Q8SQR3 Q8SQR8 Q8SQS1 Q8SQS2 Q8SQS4 Q8SQS5
Q8SQS8 Q8SQT7 Q8SQU1 Q8SQU8 Q8SQV6 Q8SQW0 Q8SQW2 Q8SQW5 Q8SQX3 Q8SQY3 Q8SQY7
Q8SQZ4 Q8SQZ6 Q8SQZ7 Q8SR01 Q8SR09 Q8SR10 Q8SR11 Q8SR13 Q8SR18 Q8SR30 Q8SR37
Q8SR40 Q8SR42 Q8SR45 Q8SR49 Q8SR53 Q8SR54 Q8SR63 Q8SR65 Q8SR66 Q8SR69 Q8SR75
Q8SR76 Q8SR83 Q8SR84 Q8SR85 Q8SR86 Q8SR89 Q8SR90 Q8SR96 Q8SR99 Q8SRA7 Q8SRB0
Q8SRB2 Q8SRB3 Q8SRB8 Q8SRB9 Q8SRD2 Q8SRD8 Q8SRE0 Q8SRE1 Q8SRE2 Q8SRF1 Q8SRF5
Q8SRG2 Q8SRG3 Q8SRG9 Q8SRH0 Q8SRH2 Q8SRH5 Q8SRH6 Q8SRI2 Q8SRI3 Q8SRI6 Q8SRJ7
Q8SRK0 Q8SRK1 Q8SRK6 Q8SRK8 Q8SRL5 Q8SRM1 Q8SRM2 Q8SRM7 Q8SRN2 Q8SRP1 Q8SRP2
Q8SRP3 Q8SRP6 Q8SRP9 Q8SRQ8 Q8SRR1 Q8SRR2 Q8SRR3 Q8SRS0 Q8SRS6 Q8SRS8 Q8SRT7
Q8SRU0 Q8SRU2 Q8SRU3 Q8SRU7 Q8SRV0 Q8SRV7 Q8SRW6 Q8SRW8 Q8SRY0 Q8SRY1 Q8SRY4
Q8SRY5 Q8SRZ0 Q8SS01 Q8SS05 Q8SS11 Q8SS18 Q8SS24 Q8SS27 Q8SS29 Q8SS33 Q8SS35
Q8SS40 Q8SS47 Q8SS48 Q8SS49 Q8SS56 Q8SS60 Q8SS62 Q8SS65 Q8SS72 Q8SS73 Q8SS77
Q8SS83 Q8SS85 Q8SS91 Q8SS93 Q8SS96 Q8SS99 Q8SSA2 Q8SSA6 Q8SSA8 Q8SSB1 Q8SSC4
Q8SSC5 Q8SSC8 Q8SSC9 Q8SSD7 Q8SSE4 Q8SSE7 Q8SSF6 Q8SSF9 Q8SSG1 Q8SSG3 Q8SSH0
Q8SSH1 Q8SSH3 Q8SSH4 Q8SSH5 Q8SSI8 Q8SSI9 Q8SSJ5 Q8SSK1 Q8SSK6 Q8SSL0 Q8SSL2
Q8SSL7 Q8SSM2 Q8SSM6 Q8SSM8 Q8SSN5 Q8SSU2 Q8SSV3 Q8SSY6 Q8ST30 Q8ST35 Q8ST42

Q8ST44 Q8ST53 Q8ST62 Q8ST65 Q8ST67 Q8ST79 Q8ST80 Q8ST81 Q8ST83 Q8ST89 Q8ST90
Q8ST93 Q8ST94 Q8ST97 Q8STA9 Q8STB2 Q8STB3 Q8STB5 Q8STD6 Q8STD9 Q8STE0 Q8STE6
Q8STE8 Q8STF0 Q8STF1 Q8STF2 Q8STF6 Q8STG1 Q8STG4 Q8STG8 Q8STI4 Q8STJ1 Q8STJ2
Q8STJ4 Q8STK1 Q8STL0 Q8STL3 Q8STN5 Q8STN7 Q8STP2 Q8STR6 Q8STR8 Q8STS7 Q8STS8
Q8STY8 Q8STZ0 Q8STZ1 Q8SU25 Q8SU27 Q8SU52 Q8SU65 Q8SU76 Q8SU90 Q8SU91 Q8SU92
Q8SU95 Q8SU96 Q8SUB7 Q8SUD4 Q8SUE9 Q8SUH6 Q8SUH8 Q8SUH9 Q8SUI0 Q8SUL0 Q8SUM8
Q8SUN2 Q8SUR9 Q8SUT1 Q8SUT4 Q8SUU2 Q8SUY5 Q8SUZ5 Q8SV03 Q8SV19 Q8SV33 Q8SV35
Q8SV54 Q8SV55 Q8SV56 Q8SV57 Q8SV58 Q8SV60 Q8SV63 Q8SV64 Q8SV71 Q8SV74 Q8SV95
Q8SVB5 Q8SVC0 Q8SVC2 Q8SVD3 Q8SVD9 Q8SVF2 Q8SVF3 Q8SVF4 Q8SVF5 Q8SVF7 Q8SVF8
Q8SVI7 Q8SVJ2 Q8SVJ8 Q8SVL5 Q8SVM5 Q8SVN9 Q8SVP2 Q8SVP3 Q8SVP4 Q8SVP8 Q8SVQ8
Q8SVR6 Q8SVS9 Q8SVT3 Q8SVT6 Q8SVV3 Q8SVY5 Q8SVY6 Q8SVY8 Q8SVY9 Q8SW17 Q8SW18
Q8SW19 Q8SW23 Q8SW28 Q8SW34 Q8SW42 Q8SW43 Q8SW50 Q8SW51 Q8SW70 Q8SW74 Q8SW75
Q8SW76 Q8SW78 Q8SW79 Q8SW83 Q8SW84 Q8SW85 Q8SW86 Q8SW92 Q8SWA6 Q8SWD4 Q8SWE5
Q8SWF6 Q8SWH2 Q8SWH3 Q8SWH4 Q8SWH5 Q8SWH6 Q8SWH7 Q8SWI3 Q8SWL2 Q8SWM6 Q8SWN5
Q8SWN8 Q8SWP1 Q8SWQ4 Q8SWR2 Q8SWR8 Q8SWU7 Q8SWX0 Q8SX68 Q8SX83 Q8SX86 Q8SXD4
Q8SXX4 Q8SY33 Q8SY41 Q8SY96 Q8SYG2 Q8SYK5 Q8SYS7 Q8SZ16 Q8SZZ8 Q8T053 Q8T079
Q8T088 Q8T0D4 Q8T126 Q8T133 Q8T135 Q8T137 Q8T154 Q8T158 Q8T199 Q8T1C6 Q8T1D4
Q8T1E3 Q8T1P1 Q8T1V7 Q8T1V8 Q8T295 Q8T2A4 Q8T2F3 Q8T2H7 Q8T2I8 Q8T2J9 Q8T2K9
Q8T2Q8 Q8T2T7 Q8T305 Q8T380 Q8T3B9 Q8T3G2 Q8T3X9 Q8T3Y0 Q8T4F7 Q8T5C6 Q8T5T1
Q8T5Z4 Q8T626 Q8T697 Q8T6A5 Q8T6B3 Q8T6B4 Q8T6B9 Q8T6C4 Q8T6L5 Q8T6T2 Q8T6Z0
Q8T6Z1 Q8T773 Q8T784 Q8T7J8 Q8T7J9 Q8T806 Q8T880 Q8T882 Q8T888 Q8T8B9 Q8T8C0
Q8T8E6 Q8T8E7 Q8T8M2 Q8T8R1 Q8T938 Q8T940 Q8T9Z7 Q8TA03 Q8TA52 Q8TA81 Q8TA83
Q8TA94 Q8TAA3 Q8TAB5 Q8TAB7 Q8TAC1 Q8TAD7 Q8TAF5 Q8TAF7 Q8TAI1 Q8TAK5 Q8TAK6
Q8TAL5 Q8TAM1 Q8TAM6 Q8TAP4 Q8TAQ5 Q8TAT5 Q8TAU0 Q8TAU3 Q8TAV0 Q8TAW3 Q8TAY7
Q8TB03 Q8TB05 Q8TB33 Q8TB52 Q8TB69 Q8TBB0 Q8TBC4 Q8TBC5 Q8TBE9 Q8TBF8 Q8TBG4
Q8TBJ5 Q8TBK2 Q8TBK6 Q8TBP0 Q8TBR4 Q8TBR5 Q8TBY0 Q8TBY8 Q8TBY9 Q8TBZ0 Q8TBZ5
Q8TBZ6 Q8TBZ8 Q8TBZ9 Q8TC05 Q8TC20 Q8TC21 Q8TC44 Q8TC59 Q8TC90 Q8TC94 Q8TC99
Q8TCB0 Q8TCB7 Q8TCD6 Q8TCE6 Q8TCE9 Q8TCF1 Q8TCH9 Q8TCI5 Q8TCJ0 Q8TCN5 Q8TCX1
Q8TCY9 Q8TCZ7 Q8TD10 Q8TD17 Q8TD23 Q8TD26 Q8TD30 Q8TD31 Q8TD35 Q8TD55 Q8TD57
Q8TD86 Q8TD90 Q8TD91 Q8TD94 Q8TDC0 Q8TDD2 Q8TDG2 Q8TDG4 Q8TDM0 Q8TDN4 Q8TDP1
Q8TDQ7 Q8TDR2 Q8TDR4 Q8TDX5 Q8TDY3 Q8TE02 Q8TE12 Q8TE69 Q8TE73 Q8TE76 Q8TE77
Q8TE85 Q8TE96 Q8TEA1 Q8TEA7 Q8TEB1 Q8TEC5 Q8TED0 Q8TEE9 Q8TEJ3 Q8TET4 Q8TEV8
Q8TF09 Q8TF20 Q8TF21 Q8TF27 Q8TF32 Q8TF45 Q8TF47 Q8TF50 Q8TF61 Q8TF63 Q8TF64
Q8TF65 Q8TF74 Q8TF77 Q8TFD5 Q8TFF3 Q8TFF6 Q8TFF9 Q8TFG7 Q8TFH0 Q8TFH4 Q8TFH5
Q8TFI0 Q8TFJ2 Q8TFK3 Q8TFK8 Q8TFL3 Q8TFM8 Q8TFM9 Q8TFN0 Q8TFN1 Q8TFU8 Q8TFZ1
Q8TG11 Q8TG12 Q8TG13 Q8TG23 Q8TG24 Q8TG40 Q8TG90 Q8TGA0 Q8TGA1 Q8TGE0 Q8TGJ0
Q8TGJ1 Q8TGJ2 Q8TGJ4 Q8TGJ7 Q8TGJ8 Q8TGK0 Q8TGK1 Q8TGK4 Q8TGK6 Q8TGK7 Q8TGK8
Q8TGK9 Q8TGL0 Q8TGL1 Q8TGL2 Q8TGL3 Q8TGL4 Q8TGL5 Q8TGL6 Q8TGL7 Q8TGL8 Q8TGL9
Q8TGM0 Q8TGM1 Q8TGM2 Q8TGM3 Q8TGM4 Q8TGM5 Q8TGM6 Q8TGM7 Q8TGM8 Q8TGN0 Q8TGN1
Q8TGN2 Q8TGN3 Q8TGN4 Q8TGN5 Q8TGN6 Q8TGN8 Q8TGP0 Q8TGP1 Q8TGP2 Q8TGP3 Q8TGP4
Q8TGP5 Q8TGP6 Q8TGP7 Q8TGP8 Q8TGQ1 Q8TGQ3 Q8TGQ4 Q8TGQ5 Q8TGQ6 Q8TGQ8 Q8TGQ9
Q8TGR0 Q8TGR1 Q8TGR4 Q8TGR5 Q8TGR6 Q8TGR8 Q8TGR9 Q8TGS2 Q8TGS6 Q8TGS8 Q8TGS9
Q8TGT0 Q8TGT3 Q8TGT5 Q8TGT6 Q8TGT7 Q8TGT8 Q8TGU1 Q8TGU3 Q8TGU4 Q8TGU5 Q8TGU6
Q8TGU8 Q8TGV0 Q8TGW7 Q8TGX1 Q8TGX2 Q8TGX5 Q8TGX6 Q8TGX7 Q8TGX8 Q8TGX9 Q8TGY1

Q8TGY4 Q8TGY5 Q8TGY6 Q8TGY9 Q8TGZ5 Q8TGZ7 Q8TH04 Q8TH08 Q8TH18 Q8TH22 Q8TH23
Q8TH24 Q8TH25 Q8TH44 Q8TH63 Q8TH84 Q8TH85 Q8TH87 Q8TH90 Q8THA2 Q8THA5 Q8THA9
Q8THB0 Q8THC3 Q8THC4 Q8THC5 Q8THC7 Q8THG2 Q8THG5 Q8THG6 Q8THH1 Q8THH3 Q8THH4
Q8THJ0 Q8THJ1 Q8THJ7 Q8THJ9 Q8THK1 Q8THK3 Q8THL2 Q8THL3 Q8THP0 Q8THP1 Q8THP2
Q8THS6 Q8THS9 Q8THT2 Q8THU2 Q8THW2 Q8TI26 Q8TI28 Q8TI34 Q8TI80 Q8TI81 Q8TI82
Q8TI87 Q8TI88 Q8TI93 Q8TIB0 Q8TIB8 Q8TIB9 Q8TIF3 Q8TIF5 Q8TIM8 Q8TIN0 Q8TIN2
Q8TIN3 Q8TIN4 Q8TIN5 Q8TIN6 Q8TIP4 Q8TIP5 Q8TIR7 Q8TIS9 Q8TIT3 Q8TIT6 Q8TIU3
Q8TIU5 Q8TIV0 Q8TIV9 Q8TIW3 Q8TIY2 Q8TIY5 Q8TIZ2 Q8TJ03 Q8TJ19 Q8TJA1 Q8TJA2
Q8TJB5 Q8TJB7 Q8TJC2 Q8TJC4 Q8TJC5 Q8TJC6 Q8TJD0 Q8TJD5 Q8TJJ1 Q8TJJ4 Q8TJK1
Q8TJM9 Q8TJP7 Q8TJP9 Q8TJQ0 Q8TJQ7 Q8TJS1 Q8TJS2 Q8TJT1 Q8TJT2 Q8TJT3 Q8TJT7
Q8TJU2 Q8TJV8 Q8TJZ5 Q8TJZ6 Q8TJZ8 Q8TJZ9 Q8TK00 Q8TK19 Q8TK32 Q8TK33 Q8TK53
Q8TK55 Q8TK70 Q8TK71 Q8TK72 Q8TK88 Q8TK89 Q8TK90 Q8TK94 Q8TK95 Q8TKB4 Q8TKG2
Q8TKI9 Q8TKK3 Q8TKL5 Q8TKM0 Q8TKM8 Q8TKQ5 Q8TKQ6 Q8TKS2 Q8TKS3 Q8TKT0 Q8TKT5
Q8TKT6 Q8TKU6 Q8TKU7 Q8TKW2 Q8TKW5 Q8TKX2 Q8TKX4 Q8TKX5 Q8TKZ3 Q8TL01 Q8TL02
Q8TL08 Q8TL39 Q8TL41 Q8TL44 Q8TL47 Q8TL48 Q8TLB0 Q8TLB1 Q8TLB2 Q8TLB3 Q8TLB8
Q8TLC0 Q8TLF1 Q8TLH2 Q8TLI4 Q8TLK5 Q8TLL2 Q8TLL3 Q8TLM3 Q8TLM4 Q8TLP3 Q8TLP4
Q8TLP5 Q8TLP6 Q8TLX7 Q8TM02 Q8TM08 Q8TM19 Q8TM54 Q8TMD5 Q8TMD6 Q8TME2 Q8TME4
Q8TMG9 Q8TMI3 Q8TMI6 Q8TMI8 Q8TMK1 Q8TMP0 Q8TMT1 Q8TMW6 Q8TMY6 Q8TN62 Q8TN68
Q8TN69 Q8TN93 Q8TNH5 Q8TNI1 Q8TNN7 Q8TNY3 Q8TNY4 Q8TNY5 Q8TP90 Q8TPA0 Q8TPD4
Q8TPD6 Q8TPF0 Q8TPF4 Q8TPF7 Q8TPJ2 Q8TPT3 Q8TPT4 Q8TPT5 Q8TPT7 Q8TPU4 Q8TPU8
Q8TPU9 Q8TPV0 Q8TPV2 Q8TPW4 Q8TPX0 Q8TPX2 Q8TPX3 Q8TPX5 Q8TPX7 Q8TQ11 Q8TQ37
Q8TQ42 Q8TQ61 Q8TQ68 Q8TQ69 Q8TQD0 Q8TQD3 Q8TQD4 Q8TQF5 Q8TQF7 Q8TQF8 Q8TQH6
Q8TQH7 Q8TQL4 Q8TQL5 Q8TQL6 Q8TQL7 Q8TQL8 Q8TQL9 Q8TQM6 Q8TQP9 Q8TQQ0 Q8TQQ1
Q8TQQ2 Q8TQR1 Q8TQR2 Q8TQR3 Q8TQR9 Q8TQS4 Q8TQU1 Q8TQU3 Q8TQU8 Q8TQV6 Q8TQX8
Q8TQZ1 Q8TQZ3 Q8TR06 Q8TR12 Q8TR14 Q8TR31 Q8TR33 Q8TR65 Q8TR66 Q8TR73 Q8TR74
Q8TR85 Q8TR92 Q8TRA4 Q8TRA5 Q8TRB4 Q8TRB8 Q8TRB9 Q8TRC1 Q8TRC2 Q8TRC3 Q8TRC4
Q8TRF7 Q8TRK7 Q8TRK8 Q8TRL1 Q8TRL2 Q8TRN7 Q8TRR0 Q8TRR1 Q8TRR2 Q8TRR5 Q8TRR6
Q8TRS3 Q8TRS5 Q8TRS6 Q8TRS7 Q8TRS8 Q8TRT0 Q8TRT1 Q8TRT2 Q8TRT3 Q8TRT4 Q8TRT5
Q8TRT6 Q8TRT7 Q8TRT8 Q8TRT9 Q8TRU0 Q8TRU1 Q8TRU2 Q8TRU3 Q8TRU4 Q8TRU5 Q8TRU6
Q8TRU7 Q8TRZ3 Q8TRZ4 Q8TRZ5 Q8TRZ6 Q8TRZ8 Q8TS00 Q8TS07 Q8TS34 Q8TS37 Q8TS38
Q8TS42 Q8TS46 Q8TS47 Q8TS56 Q8TS62 Q8TS71 Q8TS72 Q8TS73 Q8TS74 Q8TS82 Q8TS91
Q8TS96 Q8TSA1 Q8TSA3 Q8TSA6 Q8TSH7 Q8TSI1 Q8TSN5 Q8TSP6 Q8TSP7 Q8TSR7 Q8TSS3
Q8TSS4 Q8TSS8 Q8TST1 Q8TSV7 Q8TSX5 Q8TSZ1 Q8TSZ5 Q8TT00 Q8TT29 Q8TT35 Q8TT38
Q8TT39 Q8TT40 Q8TT41 Q8TT42 Q8TT43 Q8TT44 Q8TT52 Q8TT55 Q8TT56 Q8TT57 Q8TT60
Q8TT87 Q8TT89 Q8TT90 Q8TT93 Q8TT94 Q8TT96 Q8TTA0 Q8TTA3 Q8TTA5 Q8TTA8 Q8TTA9
Q8TTB0 Q8TTB4 Q8TTB6 Q8TTC6 Q8TTH3 Q8TTI0 Q8TTI1 Q8TTK2 Q8TTQ4 Q8TTT4 Q8TTY5
Q8TTZ7 Q8TU06 Q8TU10 Q8TU30 Q8TU55 Q8TU56 Q8TU57 Q8TU65 Q8TU67 Q8TU71 Q8TU79
Q8TU90 Q8TU91 Q8TU94 Q8TUA1 Q8TUA2 Q8TUB6 Q8TUB8 Q8TUB9 Q8TUC0 Q8TUC1 Q8TUD3
Q8TUD7 Q8TUE8 Q8TUE9 Q8TUF7 Q8TUF8 Q8TUG2 Q8TUG3 Q8TUG6 Q8TUH5 Q8TUH7 Q8TUI2
Q8TUJ1 Q8TUM3 Q8TUM4 Q8TUQ4 Q8TUQ5 Q8TUQ9 Q8TUR2 Q8TUS2 Q8TUS5 Q8TUS7 Q8TUT6
Q8TUT7 Q8TUT9 Q8TUU1 Q8TUU3 Q8TUV3 Q8TUV7 Q8TUX2 Q8TUY2 Q8TUY3 Q8TUY4 Q8TUY5
Q8TUY6 Q8TUY7 Q8TUZ3 Q8TUZ4 Q8TUZ5 Q8TUZ9 Q8TV03 Q8TV04 Q8TV06 Q8TV07 Q8TV08
Q8TV10 Q8TV15 Q8TV23 Q8TV34 Q8TV53 Q8TV55 Q8TV58 Q8TV60 Q8TV61 Q8TV83 Q8TV84
Q8TV85 Q8TV89 Q8TV90 Q8TV92 Q8TVA3 Q8TVA6 Q8TVB1 Q8TVB2 Q8TVB4 Q8TVB5 Q8TVB6

Q8TVB7 Q8TVB8 Q8TVB9 Q8TVC0 Q8TVC1 Q8TVD1 Q8TVD2 Q8TVD4 Q8TVD7 Q8TVD8 Q8TVD9
Q8TVE2 Q8TVE5 Q8TVE6 Q8TVE7 Q8TVE8 Q8TVF2 Q8TVG3 Q8TVG5 Q8TVG7 Q8TVH1 Q8TVH5
Q8TVI1 Q8TVJ1 Q8TVJ5 Q8TVJ8 Q8TVL0 Q8TVL8 Q8TVL9 Q8TVM0 Q8TVM4 Q8TVS6 Q8TVS8
Q8TW01 Q8TW09 Q8TW10 Q8TW12 Q8TW13 Q8TW14 Q8TW15 Q8TW16 Q8TW17 Q8TW18 Q8TW19
Q8TW20 Q8TW21 Q8TW23 Q8TW24 Q8TW28 Q8TW29 Q8TW31 Q8TW40 Q8TWD0 Q8TWE6 Q8TWF8
Q8TWG0 Q8TWG3 Q8TWG4 Q8TWG5 Q8TWG9 Q8TWH2 Q8TWH4 Q8TWK0 Q8TWK3 Q8TWL5 Q8TWM7
Q8TWM9 Q8TWN3 Q8TWP2 Q8TWP6 Q8TWQ5 Q8TWR1 Q8TWR2 Q8TWR5 Q8TWR9 Q8TWS2 Q8TWS7
Q8TWU0 Q8TWU1 Q8TWU6 Q8TWU7 Q8TWU9 Q8TWX0 Q8TWY1 Q8TWY2 Q8TWY3 Q8TWY4 Q8TWY8
Q8TWZ3 Q8TX03 Q8TX14 Q8TX15 Q8TX21 Q8TX28 Q8TX33 Q8TX34 Q8TX35 Q8TX36 Q8TX38
Q8TX43 Q8TX44 Q8TX47 Q8TX49 Q8TX50 Q8TX51 Q8TX52 Q8TX56 Q8TX58 Q8TX60 Q8TX67
Q8TX72 Q8TX79 Q8TX83 Q8TX85 Q8TX89 Q8TX91 Q8TX94 Q8TX99 Q8TXA7 Q8TXB4 Q8TXB5
Q8TXB6 Q8TXB7 Q8TXC6 Q8TXC7 Q8TXC8 Q8TXD2 Q8TXD4 Q8TXD5 Q8TXD7 Q8TXF1 Q8TXF2
Q8TXF4 Q8TXF5 Q8TXF6 Q8TXF7 Q8TXF9 Q8TXG3 Q8TXG4 Q8TXI3 Q8TXI4 Q8TXI5 Q8TXJ0
Q8TXJ2 Q8TXJ3 Q8TXJ4 Q8TXJ5 Q8TXJ7 Q8TXK0 Q8TXK6 Q8TXM1 Q8TXM4 Q8TXN1 Q8TXN4
Q8TXN5 Q8TXN9 Q8TXP1 Q8TXP3 Q8TXQ0 Q8TXQ3 Q8TXS2 Q8TXS3 Q8TXS4 Q8TXT2 Q8TXT7
Q8TXT8 Q8TXU4 Q8TXU6 Q8TXU9 Q8TXV5 Q8TXV8 Q8TXW1 Q8TXW5 Q8TXW8 Q8TXX3 Q8TXX9
Q8TXY4 Q8TXZ2 Q8TXZ3 Q8TXZ9 Q8TY00 Q8TY02 Q8TY07 Q8TY09 Q8TY11 Q8TY15 Q8TY18
Q8TY21 Q8TY66 Q8TY70 Q8TY71 Q8TY72 Q8TY76 Q8TY77 Q8TY85 Q8TY86 Q8TY87 Q8TY90
Q8TY91 Q8TY92 Q8TY93 Q8TYA2 Q8TYB1 Q8TYB3 Q8TYB5 Q8TYB6 Q8TYB7 Q8TYC0 Q8TYC1
Q8TYC2 Q8TYC3 Q8TYC4 Q8TYC5 Q8TYC7 Q8TYD4 Q8TYD5 Q8TYD6 Q8TYD7 Q8TYF0 Q8TYF7
Q8TYG5 Q8TYG6 Q8TYK5 Q8TYL4 Q8TYL5 Q8TYL6 Q8TYM1 Q8TYM3 Q8TYM5 Q8TYN8 Q8TYP2
Q8TYP4 Q8TYP6 Q8TYP7 Q8TYR3 Q8TYR4 Q8TYR8 Q8TYS1 Q8TYS9 Q8TYT5 Q8TYT6 Q8TYT7
Q8TYT9 Q8TYV1 Q8TYV3 Q8TYV5 Q8TYW8 Q8TYY6 Q8TYY7 Q8TYY9 Q8TYZ5 Q8TZ08 Q8TZ11
Q8TZ12 Q8TZ14 Q8TZ19 Q8TZ24 Q8TZ33 Q8TZ38 Q8TZ45 Q8TZ52 Q8TZ61 Q8TZ66 Q8TZ81
Q8TZ82 Q8TZ97 Q8TZ98 Q8TZA5 Q8TZA6 Q8TZA7 Q8TZA8 Q8TZB0 Q8TZB2 Q8TZB3 Q8TZB4
Q8TZC1 Q8TZC4 Q8TZC5 Q8TZC9 Q8TZD7 Q8TZD9 Q8TZE1 Q8TZE2 Q8TZE8 Q8TZF3 Q8TZG0
Q8TZH9 Q8TZI2 Q8TZI3 Q8TZI4 Q8TZI8 Q8TZJ2 Q8TZJ3 Q8TZJ8 Q8TZJ9 Q8TZK0 Q8TZK1
Q8TZL1 Q8TZL3 Q8TZL4 Q8TZL5 Q8TZM5 Q8TZQ5 Q8TZQ6 Q8TZR3 Q8TZS0 Q8TZV1 Q8TZV2
Q8TZV6 Q8TZV9 Q8TZW1 Q8TZW4 Q8TZW5 Q8TZX1 Q8TZX4 Q8TZY2 Q8TZY6 Q8TZY8 Q8TZZ2
Q8TZZ8 Q8TZZ9 Q8U000 Q8U001 Q8U002 Q8U003 Q8U004 Q8U005 Q8U006 Q8U007 Q8U008
Q8U009 Q8U010 Q8U011 Q8U012 Q8U013 Q8U014 Q8U015 Q8U016 Q8U017 Q8U018 Q8U020
Q8U036 Q8U039 Q8U040 Q8U052 Q8U053 Q8U061 Q8U064 Q8U067 Q8U071 Q8U076 Q8U077
Q8U082 Q8U085 Q8U086 Q8U088 Q8U089 Q8U092 Q8U093 Q8U094 Q8U0A0 Q8U0A5 Q8U0A6
Q8U0A7 Q8U0A8 Q8U0B3 Q8U0B4 Q8U0B5 Q8U0B6 Q8U0B9 Q8U0C0 Q8U0D4 Q8U0E2 Q8U0E3
Q8U0E4 Q8U0E5 Q8U0E6 Q8U0E7 Q8U0E8 Q8U0F0 Q8U0F2 Q8U0G2 Q8U0G6 Q8U0H4 Q8U0H6
Q8U0I2 Q8U0I8 Q8U0J4 Q8U0J5 Q8U0K5 Q8U0K8 Q8U0K9 Q8U0L6 Q8U0L8 Q8U0L9 Q8U0M0
Q8U0M2 Q8U0M5 Q8U0M6 Q8U0M8 Q8U0N0 Q8U0N1 Q8U0N4 Q8U0N7 Q8U0P1 Q8U0P3 Q8U0P4
Q8U0P5 Q8U0P6 Q8U0P7 Q8U0P9 Q8U0Q4 Q8U0Q5 Q8U0Q6 Q8U0R7 Q8U0R8 Q8U0R9 Q8U0S7
Q8U0U0 Q8U0U1 Q8U0U4 Q8U0U6 Q8U0V6 Q8U0W9 Q8U0X0 Q8U111 Q8U122 Q8U125 Q8U126
Q8U148 Q8U149 Q8U150 Q8U151 Q8U152 Q8U158 Q8U159 Q8U160 Q8U176 Q8U178 Q8U182
Q8U185 Q8U191 Q8U192 Q8U194 Q8U195 Q8U1B3 Q8U1C1 Q8U1C4 Q8U1D7 Q8U1E4 Q8U1E6
Q8U1F0 Q8U1G7 Q8U1H7 Q8U1I5 Q8U1K2 Q8U1K8 Q8U1K9 Q8U1L3 Q8U1L4 Q8U1M0 Q8U1N4
Q8U1N8 Q8U1N9 Q8U1P0 Q8U1P6 Q8U1P9 Q8U1Q1 Q8U1R0 Q8U1R2 Q8U1R4 Q8U1R5 Q8U1R6
Q8U1R8 Q8U1S1 Q8U1S4 Q8U1S5 Q8U1S6 Q8U1S7 Q8U1S9 Q8U1T0 Q8U1T1 Q8U1T8 Q8U1T9

Q8U1U1 Q8U1V2 Q8U1V7 Q8U1W7 Q8U1X0 Q8U1Y4 Q8U1Y5 Q8U1Z9 Q8U203 Q8U207 Q8U217
Q8U218 Q8U221 Q8U227 Q8U232 Q8U248 Q8U259 Q8U260 Q8U261 Q8U262 Q8U263 Q8U297
Q8U2A0 Q8U2A1 Q8U2A2 Q8U2A3 Q8U2A8 Q8U2A9 Q8U2C1 Q8U2D5 Q8U2E4 Q8U2E5 Q8U2E6
Q8U2F9 Q8U2G6 Q8U2H1 Q8U2H3 Q8U2H4 Q8U2H5 Q8U2H9 Q8U2I1 Q8U2K7 Q8U2L3 Q8U2L4
Q8U2L5 Q8U2Q8 Q8U308 Q8U319 Q8U336 Q8U352 Q8U354 Q8U357 Q8U373 Q8U374 Q8U377
Q8U378 Q8U380 Q8U381 Q8U394 Q8U3C9 Q8U3D3 Q8U3D6 Q8U3H1 Q8U3H5 Q8U3H8 Q8U3I5
Q8U3J0 Q8U3J1 Q8U3J2 Q8U3K2 Q8U3K5 Q8U3K8 Q8U3L0 Q8U3L4 Q8U3M2 Q8U3M6 Q8U3M7
Q8U3N5 Q8U3R3 Q8U3R5 Q8U3S2 Q8U3S3 Q8U3S6 Q8U3S7 Q8U3S8 Q8U3S9 Q8U3T0 Q8U3T3
Q8U3V0 Q8U3V3 Q8U3X9 Q8U3Y2 Q8U3Z2 Q8U3Z6 Q8U3Z8 Q8U407 Q8U409 Q8U410 Q8U414
Q8U425 Q8U429 Q8U431 Q8U438 Q8U440 Q8U441 Q8U442 Q8U443 Q8U445 Q8U446 Q8U453
Q8U455 Q8U458 Q8U474 Q8U475 Q8U476 Q8U483 Q8U484 Q8U491 Q8U492 Q8U4A0 Q8U4B6
Q8U4C9 Q8U4D1 Q8U4D3 Q8U4D5 Q8U4E6 Q8U4F3 Q8U4G1 Q8U4G4 Q8U4G5 Q8U4G6 Q8U4H3
Q8U4H6 Q8U4H7 Q8U4I9 Q8U4J3 Q8U4J5 Q8U4K2 Q8U4K6 Q8U4K7 Q8U4L8 Q8U4M2 Q8U4M7
Q8U4N1 Q8U4N3 Q8U4Q3 Q8U4Q7 Q8U4Q8 Q8U4R1 Q8U4R3 Q8U544 Q8U642 Q8U671 Q8U6T2
Q8U705 Q8U7B8 Q8U7F8 Q8U7G6 Q8U7H5 Q8U7H6 Q8U7I5 Q8U7Y1 Q8U7Y5 Q8U822 Q8U8B8
Q8U8I2 Q8U8I9 Q8U8J3 Q8U8J4 Q8U8L4 Q8U8L5 Q8U8M6 Q8U8Q0 Q8U8R2 Q8U8T9 Q8U8Z6
Q8U8Z7 Q8U8Z9 Q8U919 Q8U920 Q8U940 Q8U9F7 Q8U9H0 Q8U9I5 Q8U9I9 Q8U9K1 Q8U9K4
Q8U9K5 Q8U9K6 Q8U9L7 Q8U9L9 Q8U9M5 Q8U9N0 Q8U9Q9 Q8U9T8 Q8U9U6 Q8U9V0 Q8U9V8
Q8U9W1 Q8U9W3 Q8U9X6 Q8U9Z2 Q8UA46 Q8UA56 Q8UA66 Q8UA91 Q8UA92 Q8UAA5 Q8UAA8
Q8UAA9 Q8UAE8 Q8UAM7 Q8UAN8 Q8UAS8 Q8UAS9 Q8UAW0 Q8UAW8 Q8UAX0 Q8UAY9 Q8UB75
Q8UB77 Q8UB93 Q8UBL6 Q8UBM0 Q8UBM1 Q8UBM8 Q8UBP3 Q8UBQ6 Q8UBQ8 Q8UBR0 Q8UBR5
Q8UBR6 Q8UBS0 Q8UBS1 Q8UBS2 Q8UBS4 Q8UBS8 Q8UBV2 Q8UBV5 Q8UBY9 Q8UBZ5 Q8UBZ7
Q8UBZ8 Q8UBZ9 Q8UC03 Q8UC22 Q8UC31 Q8UC37 Q8UC38 Q8UC46 Q8UC47 Q8UC60 Q8UC70
Q8UC79 Q8UC86 Q8UCA5 Q8UCC9 Q8UCD2 Q8UCE7 Q8UCK6 Q8UCM8 Q8UCN6 Q8UCS6 Q8UCS8
Q8UCT2 Q8UCT5 Q8UD04 Q8UD28 Q8UD54 Q8UD57 Q8UD63 Q8UD80 Q8UD91 Q8UD95 Q8UD97
Q8UDA0 Q8UDA9 Q8UDE5 Q8UDE9 Q8UDF7 Q8UDM3 Q8UDM6 Q8UDM9 Q8UDN0 Q8UDN3 Q8UDP9
Q8UDS3 Q8UDU5 Q8UDV0 Q8UE04 Q8UE05 Q8UE06 Q8UE07 Q8UE08 Q8UE09 Q8UE13 Q8UE14
Q8UE15 Q8UE16 Q8UE17 Q8UE18 Q8UE19 Q8UE20 Q8UE21 Q8UE22 Q8UE23 Q8UE24 Q8UE25
Q8UE26 Q8UE27 Q8UE28 Q8UE29 Q8UE30 Q8UE32 Q8UE33 Q8UE34 Q8UE35 Q8UE36 Q8UE38
Q8UE39 Q8UE40 Q8UE41 Q8UE43 Q8UE66 Q8UE71 Q8UE87 Q8UE93 Q8UEA0 Q8UEA1 Q8UEB0
Q8UEB7 Q8UED2 Q8UED3 Q8UEH1 Q8UEK7 Q8UEK8 Q8UEL1 Q8UEM4 Q8UEN6 Q8UEP3 Q8UEP4
Q8UEQ2 Q8UEQ3 Q8UEQ7 Q8UER0 Q8UER6 Q8UER8 Q8UES4 Q8UES7 Q8UES8 Q8UET6 Q8UEU0
Q8UEU8 Q8UEX6 Q8UEY3 Q8UEY5 Q8UEY7 Q8UEY9 Q8UEZ0 Q8UEZ6 Q8UF44 Q8UF49 Q8UF86
Q8UF87 Q8UFA0 Q8UFA2 Q8UFC0 Q8UFD5 Q8UFD6 Q8UFE8 Q8UFF4 Q8UFG1 Q8UFI8 Q8UFL9
Q8UFM0 Q8UFM1 Q8UFM2 Q8UFM3 Q8UFN4 Q8UFR5 Q8UFR8 Q8UFS3 Q8UFS6 Q8UFS8 Q8UFS9
Q8UFT1 Q8UFT9 Q8UFU8 Q8UFV3 Q8UFV9 Q8UFY5 Q8UFY6 Q8UFZ7 Q8UFZ8 Q8UG00 Q8UG34
Q8UG61 Q8UG69 Q8UG70 Q8UG74 Q8UG75 Q8UG81 Q8UG87 Q8UG98 Q8UGA9 Q8UGB6 Q8UGC8
Q8UGD4 Q8UGD5 Q8UGD7 Q8UGE7 Q8UGE8 Q8UGF0 Q8UGG4 Q8UGJ8 Q8UGK2 Q8UGK8 Q8UGL2
Q8UGL3 Q8UGU0 Q8UGV3 Q8UGX7 Q8UH15 Q8UH55 Q8UH56 Q8UH57 Q8UH67 Q8UH69 Q8UH73
Q8UHA1 Q8UHA2 Q8UHA7 Q8UHA8 Q8UHC2 Q8UHD5 Q8UHD7 Q8UHE7 Q8UHG2 Q8UHI3 Q8UHJ6
Q8UHK1 Q8UHK2 Q8UHK4 Q8UHN3 Q8UHP1 Q8UHP8 Q8UHR0 Q8UHU0 Q8UHV1 Q8UHW9 Q8UHX1
Q8UHX2 Q8UHX4 Q8UHX5 Q8UI68 Q8UI70 Q8UI71 Q8UI94 Q8UI98 Q8UI99 Q8UIB2 Q8UIB4
Q8UIB7 Q8UIC6 Q8UIC9 Q8UID0 Q8UID1 Q8UID4 Q8UID9 Q8UIE8 Q8UIF1 Q8UIF2 Q8UIF9
Q8UIG8 Q8UIG9 Q8UIH1 Q8UIH2 Q8UIH4 Q8UIH5 Q8UII1 Q8UII5 Q8UIJ2 Q8UIL2 Q8UIM4

Q8UIN4 Q8UIN5 Q8UIN7 Q8UIN8 Q8UIR1 Q8UIS9 Q8UIV6 Q8UIV7 Q8UIV8 Q8UJ04 Q8UJ05
Q8UJ06 Q8UJ08 Q8UJ09 Q8UJ19 Q8UJ26 Q8UJ43 Q8UJ45 Q8UJ48 Q8UJ51 Q8UJ52 Q8UJ53
Q8UJ54 Q8UJ65 Q8UJ69 Q8UJ85 Q8UJ87 Q8UJ88 Q8UJ89 Q8UJ91 Q8UJ92 Q8UJ94 Q8UJ99
Q8UJA9 Q8UJB0 Q8UJB1 Q8UJC2 Q8UJC4 Q8UJC5 Q8UJC6 Q8UJC7 Q8UJK7 Q8UJW6 Q8UK38
Q8UKK2 Q8UUP2 Q8UUR3 Q8UUZ1 Q8UVK2 Q8UVQ4 Q8UVR3 Q8UVR5 Q8UVV7 Q8UVX0 Q8UVY1
Q8UVY2 Q8UW00 Q8UW19 Q8UW64 Q8UW72 Q8UW74 Q8UW76 Q8UZB3 Q8UZB5 Q8UZB6 Q8UZL8
Q8V2K9 Q8V2L6 Q8V2N1 Q8V2N2 Q8V2N7 Q8V2R6 Q8V2R8 Q8V2V1 Q8V2W4 Q8V2Y0 Q8V2Z3
Q8V336 Q8V3L7 Q8V3R4 Q8V3T6 Q8V3T8 Q8V3U1 Q8V3U2 Q8V3U3 Q8V3U4 Q8V431 Q8V434
Q8V435 Q8V438 Q8V4S4 Q8V4T3 Q8V4T7 Q8V4V3 Q8V4V4 Q8V507 Q8V518 Q8V566 Q8V571
Q8V6W3 Q8V6W4 Q8V736 Q8V7G3 Q8V7G4 Q8V7G5 Q8V7J0 Q8V7J1 Q8V7J2 Q8V987 Q8V9C3
Q8V9C4 Q8V9P2 Q8V9R9 Q8V9S2 Q8V9S7 Q8V9S9 Q8V9U4 Q8VA41 Q8VA42 Q8VA43 Q8VAK2
Q8VBE2 Q8VBT1 Q8VBU8 Q8VBV3 Q8VBV7 Q8VBW5 Q8VBW8 Q8VC03 Q8VC10 Q8VC12 Q8VC16
Q8VC23 Q8VC28 Q8VC30 Q8VC31 Q8VC33 Q8VC34 Q8VC51 Q8VC52 Q8VC56 Q8VC57 Q8VC70
Q8VC85 Q8VC97 Q8VCA8 Q8VCB2 Q8VCB3 Q8VCC6 Q8VCD5 Q8VCD7 Q8VCE1 Q8VCE2 Q8VCE4
Q8VCE6 Q8VCG3 Q8VCI7 Q8VCK3 Q8VCK5 Q8VCM3 Q8VCM4 Q8VCN9 Q8VCP8 Q8VCR7 Q8VCR8
Q8VCS6 Q8VCT9 Q8VCU0 Q8VCW2 Q8VCY6 Q8VCZ7 Q8VCZ8 Q8VD24 Q8VD62 Q8VD63 Q8VD76
Q8VDA5 Q8VDA6 Q8VDC0 Q8VDD9 Q8VDG3 Q8VDG5 Q8VDG6 Q8VDH1 Q8VDI1 Q8VDK1 Q8VDK3
Q8VDL9 Q8VDM4 Q8VDM6 Q8VDP3 Q8VDP4 Q8VDQ9 Q8VDR7 Q8VDS3 Q8VDS4 Q8VDS7 Q8VDW0
Q8VDW4 Q8VDY9 Q8VE08 Q8VE09 Q8VE18 Q8VE33 Q8VE42 Q8VE62 Q8VE65 Q8VE73 Q8VE80
Q8VE88 Q8VE92 Q8VE94 Q8VE95 Q8VE97 Q8VEA4 Q8VEB6 Q8VED5 Q8VED8 Q8VED9 Q8VEE1
Q8VEE4 Q8VEG0 Q8VEG6 Q8VEH6 Q8VEJ1 Q8VEJ4 Q8VEK3 Q8VEK6 Q8VEM9 Q8VH31 Q8VH37
Q8VH51 Q8VHD8 Q8VHE6 Q8VHF5 Q8VHH5 Q8VHI4 Q8VHI6 Q8VHI7 Q8VHJ7 Q8VHL1 Q8VHL5
Q8VHN8 Q8VHP7 Q8VHR5 Q8VHT6 Q8VHT7 Q8VHT8 Q8VHU4 Q8VHV7 Q8VHX2 Q8VHZ4 Q8VHZ7
Q8VI04 Q8VI24 Q8VI33 Q8VI40 Q8VI67 Q8VI84 Q8VIB5 Q8VID5 Q8VIG0 Q8VIG1 Q8VIG2
Q8VIG3 Q8VIH1 Q8VIJ4 Q8VIK1 Q8VIL2 Q8VIM9 Q8VIV2 Q8VJ53 Q8VJU4 Q8VKK0 Q8VL04
Q8VL13 Q8VL14 Q8VL42 Q8VL64 Q8VLR6 Q8VM67 Q8VMM4 Q8VMM5 Q8VMR3 Q8VMU1 Q8VMU2
Q8VMW4 Q8VMY3 Q8VN02 Q8VNN2 Q8VP65 Q8VPF1 Q8VPF2 Q8VPF3 Q8VQ15 Q8VQ84 Q8VQD7
Q8VQF4 Q8VQF5 Q8VQF6 Q8VQP8 Q8VQR8 Q8VQX6 Q8VQZ3 Q8VR31 Q8VRG9 Q8VS06 Q8VS12
Q8VS16 Q8VS50 Q8VS52 Q8VS53 Q8VS60 Q8VS61 Q8VS63 Q8VS69 Q8VS92 Q8VSR0 Q8VT58
Q8VT59 Q8VTT5 Q8VU09 Q8VUP4 Q8VUR8 Q8VUS5 Q8VUW1 Q8VV01 Q8VV84 Q8VV85 Q8VVB5
Q8VVB6 Q8VVB7 Q8VVC1 Q8VVE2 Q8VVE3 Q8VVE4 Q8VVF0 Q8VVH1 Q8VVL0 Q8VW54 Q8VW55
Q8VW57 Q8VW58 Q8VW75 Q8VW76 Q8VW94 Q8VWE6 Q8VWF5 Q8VWF8 Q8VWG4 Q8VWG7 Q8VWI7
Q8VWJ2 Q8VWJ4 Q8VWJ6 Q8VWK0 Q8VWK4 Q8VWM8 Q8VWN6 Q8VWQ4 Q8VWQ5 Q8VWV6 Q8VWY6
Q8VWY7 Q8VX04 Q8VXG7 Q8VXP3 Q8VXQ2 Q8VXQ7 Q8VXQ8 Q8VXQ9 Q8VXU6 Q8VXX4 Q8VXY4
Q8VXZ0 Q8VXZ9 Q8VY05 Q8VY06 Q8VY09 Q8VY64 Q8VY74 Q8VY76 Q8VY78 Q8VY80 Q8VY81
Q8VY84 Q8VY86 Q8VY89 Q8VYA1 Q8VYB1 Q8VYB2 Q8VYB7 Q8VYB9 Q8VYD2 Q8VYD3 Q8VYD6
Q8VYD9 Q8VYF1 Q8VYF5 Q8VYF7 Q8VYG2 Q8VYH0 Q8VYI0 Q8VYI2 Q8VYI5 Q8VYJ7 Q8VYK4
Q8VYK6 Q8VYN6 Q8VYN9 Q8VYP2 Q8VYP6 Q8VYR2 Q8VYR5 Q8VYS2 Q8VYT5 Q8VYU3 Q8VYU4
Q8VYU6 Q8VYV7 Q8VYW2 Q8VYZ0 Q8VYZ5 Q8VZ02 Q8VZ17 Q8VZ19 Q8VZ22 Q8VZ42 Q8VZ49
Q8VZ50 Q8VZ55 Q8VZ59 Q8VZ67 Q8VZ74 Q8VZ79 Q8VZA0 Q8VZB9 Q8VZC3 Q8VZC9 Q8VZE4
Q8VZE7 Q8VZE9 Q8VZF3 Q8VZG1 Q8VZI7 Q8VZI9 Q8VZJ1 Q8VZJ9 Q8VZK0 Q8VZL6 Q8VZM9
Q8VZN1 Q8VZP1 Q8VZP4 Q8VZQ0 Q8VZQ2 Q8VZQ4 Q8VZQ9 Q8VZR0 Q8VZS3 Q8VZS6 Q8VZS9
Q8VZT6 Q8VZT8 Q8VZW3 Q8VZY6 Q8VZY7 Q8W013 Q8W032 Q8W035 Q8W0E4 Q8W0F1 Q8W0W3
Q8W0W4 Q8W0Y9 Q8W0Z7 Q8W0Z9 Q8W104 Q8W108 Q8W117 Q8W119 Q8W127 Q8W171 Q8W1E3

Q8W1K8 Q8W1W6 Q8W1W9 Q8W1X2 Q8W1Y3 Q8W207 Q8W250 Q8W2B7 Q8W2B8 Q8W2F1 Q8W2F2
Q8W2F3 Q8W2X5 Q8W3D9 Q8W3K0 Q8W3K3 Q8W3L9 Q8W3M0 Q8W3M6 Q8W3M7 Q8W3P8 Q8W3U4
Q8W3Y4 Q8W420 Q8W425 Q8W463 Q8W468 Q8W474 Q8W475 Q8W485 Q8W487 Q8W489 Q8W491
Q8W4A5 Q8W4B1 Q8W4C2 Q8W4C3 Q8W4D0 Q8W4D8 Q8W4E1 Q8W4F0 Q8W4F3 Q8W4H1 Q8W4H7
Q8W4I6 Q8W4I9 Q8W4K1 Q8W4K5 Q8W4L0 Q8W4L5 Q8W4M7 Q8W4P1 Q8W4R3 Q8W4R8 Q8W4T9
Q8W519 Q8W566 Q8W575 Q8W586 Q8W593 Q8W595 Q8W5R1 Q8W5S1 Q8W7A1 Q8W8E6 Q8W8J2
Q8W8R9 Q8W8V0 Q8W8W0 Q8WH60 Q8WHJ9 Q8WHK8 Q8WHL4 Q8WHL6 Q8WHL7 Q8WHM0 Q8WHM4
Q8WHM6 Q8WHM7 Q8WHM8 Q8WHW9 Q8WHX2 Q8WHY0 Q8WHY1 Q8WHY2 Q8WHY3 Q8WHY4 Q8WHY5
Q8WHY6 Q8WHY7 Q8WHY8 Q8WHY9 Q8WHZ0 Q8WHZ1 Q8WHZ7 Q8WHZ8 Q8WHZ9 Q8WI00 Q8WI18
Q8WI24 Q8WI25 Q8WI26 Q8WI27 Q8WI35 Q8WIJ9 Q8WIT2 Q8WIU9 Q8WIV4 Q8WIV8 Q8WJ18
Q8WJ26 Q8WJ31 Q8WJ32 Q8WJ37 Q8WJM5 Q8WJN0 Q8WJN2 Q8WJN5 Q8WJN9 Q8WJP1 Q8WJP2
Q8WJP3 Q8WJP5 Q8WJQ6 Q8WJQ9 Q8WJR3 Q8WJR4 Q8WJR5 Q8WJR9 Q8WJV2 Q8WKA5 Q8WKC9
Q8WKE6 Q8WKE7 Q8WKK9 Q8WKL8 Q8WKM1 Q8WKM2 Q8WKM6 Q8WKN0 Q8WKP4 Q8WMC7 Q8WMD3
Q8WME4 Q8WML3 Q8WMS6 Q8WMX6 Q8WMX7 Q8WMX8 Q8WN22 Q8WN55 Q8WN69 Q8WN70 Q8WND5
Q8WNR4 Q8WNR9 Q8WNU4 Q8WNV2 Q8WNV3 Q8WNV5 Q8WNY7 Q8WNY8 Q8WNY9 Q8WNZ0 Q8WNZ1
Q8WNZ2 Q8WNZ3 Q8WNZ4 Q8WNZ5 Q8WNZ6 Q8WNZ7 Q8WNZ8 Q8WNZ9 Q8WP00 Q8WP13 Q8WP14
Q8WP21 Q8WP22 Q8WP23 Q8WP28 Q8WP32 Q8WP33 Q8WPW2 Q8WQ47 Q8WQ86 Q8WQI3 Q8WQI5
Q8WQI7 Q8WQL7 Q8WQX5 Q8WQX6 Q8WR63 Q8WRF3 Q8WSF1 Q8WSW3 Q8WTC1 Q8WTJ4 Q8WTM6
Q8WTP9 Q8WTQ4 Q8WTR2 Q8WTR7 Q8WTS6 Q8WTU0 Q8WTV1 Q8WTY4 Q8WTZ4 Q8WU10 Q8WU43
Q8WU49 Q8WU58 Q8WUB2 Q8WUD4 Q8WUE5 Q8WUH1 Q8WUN7 Q8WUP2 Q8WUQ7 Q8WUU4 Q8WUU5
Q8WUX2 Q8WUY3 Q8WV16 Q8WV37 Q8WV60 Q8WV74 Q8WVB3 Q8WVD3 Q8WVE0 Q8WVF5 Q8WVJ2
Q8WVK2 Q8WVL7 Q8WVM0 Q8WVN8 Q8WVP5 Q8WVS4 Q8WVV4 Q8WVY7 Q8WW01 Q8WW14 Q8WW18
Q8WW24 Q8WW27 Q8WW32 Q8WW33 Q8WW35 Q8WW36 Q8WW59 Q8WWB3 Q8WWB5 Q8WWF6 Q8WWF8
Q8WWH4 Q8WWM1 Q8WWM9 Q8WWR9 Q8WWY6 Q8WX94 Q8WXA3 Q8WXA9 Q8WXB1 Q8WXB4 Q8WXC6
Q8WXE1 Q8WXF0 Q8WXF5 Q8WXG8 Q8WXK4 Q8WXQ3 Q8WXR4 Q8WXT5 Q8WXX0 Q8WY41 Q8WY50
Q8WY54 Q8WYB5 Q8WYK2 Q8WYN3 Q8WYQ4 Q8WYQ9 Q8WYR4 Q8WZ26 Q8WZ33 Q8WZ42 Q8WZ60
Q8WZ69 Q8WZ82 Q8WZA0 Q8WZA8 Q8WZA9 Q8WZB0 Q8WZE0 Q8WZJ7 Q8WZJ8 Q8WZK0 Q8WZK1
Q8WZK3 Q8WZK4 Q8WZK5 Q8WZL4 Q8WZL6 Q8WZN0 Q8WZS3 Q8WZS6 Q8WZW7 Q8WZX5 Q8WZY4
Q8X034 Q8X066 Q8X071 Q8X077 Q8X082 Q8X0F1 Q8X0S4 Q8X0S9 Q8X0V0 Q8X0X0 Q8X0X6
Q8X119 Q8X132 Q8X167 Q8X169 Q8X178 Q8X182 Q8X187 Q8X191 Q8X195 Q8X1D8 Q8X1E6
Q8X1N3 Q8X1P0 Q8X1S6 Q8X1T0 Q8X1T3 Q8X1T6 Q8X1X3 Q8X225 Q8X229 Q8X239 Q8X251
Q8X260 Q8X261 Q8X2F6 Q8X2T9 Q8X2V7 Q8X367 Q8X3T5 Q8X3W8 Q8X444 Q8X485 Q8X487
Q8X4B4 Q8X4C9 Q8X4G0 Q8X4I9 Q8X4L9 Q8X4M5 Q8X4P8 Q8X4Q8 Q8X4R2 Q8X4S5 Q8X4S6
Q8X4T6 Q8X4V5 Q8X4W4 Q8X4Z7 Q8X4Z9 Q8X505 Q8X510 Q8X529 Q8X560 Q8X574 Q8X580
Q8X583 Q8X598 Q8X5C4 Q8X5I5 Q8X5J3 Q8X5J4 Q8X5J5 Q8X5J8 Q8X5J9 Q8X5K0 Q8X5P3
Q8X5P4 Q8X5S3 Q8X5T5 Q8X5U4 Q8X5U9 Q8X5V2 Q8X5V3 Q8X5V6 Q8X5W3 Q8X5W4 Q8X5X6
Q8X5X7 Q8X5Y4 Q8X607 Q8X610 Q8X611 Q8X612 Q8X621 Q8X635 Q8X643 Q8X644 Q8X645
Q8X647 Q8X648 Q8X649 Q8X661 Q8X680 Q8X685 Q8X693 Q8X6C0 Q8X6C3 Q8X6C4 Q8X6C5
Q8X6C7 Q8X6E3 Q8X6I1 Q8X6I8 Q8X6J0 Q8X6J4 Q8X6K2 Q8X6L0 Q8X6Q5 Q8X6R3 Q8X6R5
Q8X6R8 Q8X6R9 Q8X6S9 Q8X6T6 Q8X6X5 Q8X6X6 Q8X6X7 Q8X6X8 Q8X6X9 Q8X6Y0 Q8X6Z5
Q8X701 Q8X702 Q8X707 Q8X711 Q8X715 Q8X716 Q8X723 Q8X724 Q8X727 Q8X730 Q8X732
Q8X733 Q8X735 Q8X742 Q8X743 Q8X769 Q8X7B3 Q8X7B5 Q8X7B6 Q8X7B7 Q8X7D6 Q8X7E7
Q8X7G2 Q8X7G3 Q8X7H0 Q8X7H4 Q8X7I0 Q8X7J8 Q8X7N4 Q8X7N6 Q8X7N7 Q8X7P0 Q8X7P1
Q8X7Q1 Q8X7R5 Q8X7S0 Q8X7U1 Q8X7U2 Q8X7U3 Q8X7U4 Q8X7W6 Q8X7W8 Q8X7X4 Q8X7X7

Q8X7X9 Q8X7Y4 Q8X7Y6 Q8X7Z7 Q8X7Z8 Q8X821 Q8X823 Q8X824 Q8X825 Q8X835 Q8X839
Q8X840 Q8X844 Q8X850 Q8X852 Q8X883 Q8X884 Q8X885 Q8X897 Q8X899 Q8X8A1 Q8X8A2
Q8X8B5 Q8X8B9 Q8X8C1 Q8X8D5 Q8X8E0 Q8X8E1 Q8X8F1 Q8X8F6 Q8X8F8 Q8X8F9 Q8X8G6
Q8X8G9 Q8X8H0 Q8X8H6 Q8X8H9 Q8X8I1 Q8X8J3 Q8X8L4 Q8X8L5 Q8X8L6 Q8X8M9 Q8X8N8
Q8X8P2 Q8X8Q3 Q8X8Q5 Q8X8T3 Q8X8T4 Q8X8T5 Q8X8U4 Q8X8V0 Q8X8V5 Q8X8V7 Q8X8V8
Q8X8W2 Q8X8W8 Q8X8X5 Q8X8X6 Q8X8Y1 Q8X8Y6 Q8X8Y7 Q8X8Y9 Q8X900 Q8X909 Q8X926
Q8X929 Q8X930 Q8X954 Q8X958 Q8X963 Q8X984 Q8X987 Q8X988 Q8X9C6 Q8X9C8 Q8X9F0
Q8X9G9 Q8X9H0 Q8X9H8 Q8X9J6 Q8X9J9 Q8X9K7 Q8X9L2 Q8X9M0 Q8X9M2 Q8X9M3 Q8X9S9
Q8X9T8 Q8X9U4 Q8X9U5 Q8X9U6 Q8X9W5 Q8X9X8 Q8X9Y6 Q8X9Y9 Q8X9Z2 Q8X9Z4 Q8X9Z8
Q8X9Z9 Q8XA00 Q8XA01 Q8XA02 Q8XA10 Q8XA14 Q8XA15 Q8XA22 Q8XA23 Q8XA27 Q8XA32
Q8XA34 Q8XA35 Q8XA36 Q8XA41 Q8XA44 Q8XA46 Q8XA49 Q8XA55 Q8XA65 Q8XA70 Q8XA71
Q8XA72 Q8XA73 Q8XA76 Q8XA82 Q8XA83 Q8XA85 Q8XA87 Q8XA92 Q8XA94 Q8XAA3 Q8XAA4
Q8XAB0 Q8XAC2 Q8XAC3 Q8XAC5 Q8XAD0 Q8XAE3 Q8XAF1 Q8XAF6 Q8XAG0 Q8XAG1 Q8XAG2
Q8XAG3 Q8XAI7 Q8XAK8 Q8XAN1 Q8XAN4 Q8XAP0 Q8XAP1 Q8XAP3 Q8XAS9 Q8XAT0 Q8XAT4
Q8XAT5 Q8XAU1 Q8XAU3 Q8XAU5 Q8XAU7 Q8XAU9 Q8XAV0 Q8XAV1 Q8XAW1 Q8XAW8 Q8XAX1
Q8XAX4 Q8XAY0 Q8XAY5 Q8XAY6 Q8XAZ1 Q8XAZ2 Q8XB14 Q8XB23 Q8XB34 Q8XB35 Q8XB36
Q8XB41 Q8XB43 Q8XB55 Q8XB78 Q8XB99 Q8XBA4 Q8XBB1 Q8XBC7 Q8XBE0 Q8XBE7 Q8XBI4
Q8XBI9 Q8XBJ2 Q8XBJ3 Q8XBJ4 Q8XBK1 Q8XBK3 Q8XBK5 Q8XBK7 Q8XBK9 Q8XBL0 Q8XBL4
Q8XBL5 Q8XBL8 Q8XBN2 Q8XBN8 Q8XBP0 Q8XBP4 Q8XBR7 Q8XBS4 Q8XBS7 Q8XBT3 Q8XBT6
Q8XBW0 Q8XBW1 Q8XBX1 Q8XBZ3 Q8XBZ8 Q8XC04 Q8XCB3 Q8XCB8 Q8XCE7 Q8XCE8 Q8XCF0
Q8XCG3 Q8XCH2 Q8XCH4 Q8XCH5 Q8XCH6 Q8XCH9 Q8XCI6 Q8XCI7 Q8XCJ6 Q8XCJ9 Q8XCK5
Q8XCL9 Q8XCM4 Q8XCQ2 Q8XCQ7 Q8XCR0 Q8XCT6 Q8XCT9 Q8XCU5 Q8XCU6 Q8XCV0 Q8XCV1
Q8XCV3 Q8XCV5 Q8XCW1 Q8XCW6 Q8XCX9 Q8XCY9 Q8XCZ0 Q8XCZ2 Q8XD03 Q8XD09 Q8XD23
Q8XD32 Q8XD33 Q8XD38 Q8XD39 Q8XD41 Q8XD48 Q8XD57 Q8XD58 Q8XD64 Q8XD75 Q8XD79
Q8XD82 Q8XD85 Q8XD86 Q8XD88 Q8XD89 Q8XD99 Q8XDB2 Q8XDB3 Q8XDD9 Q8XDE8 Q8XDE9
Q8XDG0 Q8XDG1 Q8XDG2 Q8XDG3 Q8XDG9 Q8XDI4 Q8XDI5 Q8XDI7 Q8XDJ6 Q8XDJ8 Q8XDL7
Q8XDN0 Q8XDN3 Q8XDN4 Q8XDP0 Q8XDP1 Q8XDS0 Q8XDT8 Q8XDX8 Q8XDX9 Q8XDZ0 Q8XDZ6
Q8XDZ7 Q8XDZ9 Q8XE00 Q8XE07 Q8XE09 Q8XE10 Q8XE27 Q8XE29 Q8XE32 Q8XE45 Q8XE50
Q8XE52 Q8XE68 Q8XE69 Q8XE72 Q8XE74 Q8XE76 Q8XE87 Q8XE94 Q8XEA1 Q8XEA7 Q8XEC1
Q8XEC3 Q8XEC6 Q8XED8 Q8XEE5 Q8XEG2 Q8XEG9 Q8XEH3 Q8XEJ1 Q8XEL4 Q8XEL6 Q8XEM5
Q8XEN6 Q8XEQ0 Q8XES2 Q8XET6 Q8XEU3 Q8XEY8 Q8XEY9 Q8XEZ7 Q8XF03 Q8XF09 Q8XF25
Q8XF41 Q8XF46 Q8XF56 Q8XF77 Q8XF78 Q8XF81 Q8XFC1 Q8XFE0 Q8XFE7 Q8XFF3 Q8XFF4
Q8XFG2 Q8XFI3 Q8XFL2 Q8XFP8 Q8XFQ8 Q8XFV2 Q8XFX8 Q8XFZ8 Q8XG12 Q8XG49 Q8XG51
Q8XG64 Q8XG70 Q8XG78 Q8XG87 Q8XG96 Q8XGA0 Q8XGA4 Q8XGE0 Q8XGF9 Q8XGG5 Q8XGJ8
Q8XGM6 Q8XGN1 Q8XGP4 Q8XGQ8 Q8XGS6 Q8XGT4 Q8XGW7 Q8XGW8 Q8XGY4 Q8XGZ0 Q8XGZ1
Q8XGZ9 Q8XH11 Q8XH25 Q8XH26 Q8XH30 Q8XH31 Q8XH32 Q8XH43 Q8XH44 Q8XH45 Q8XH49
Q8XH63 Q8XH79 Q8XH85 Q8XH97 Q8XHA1 Q8XHA6 Q8XHA7 Q8XHA9 Q8XHB2 Q8XHB3 Q8XHB7
Q8XHC9 Q8XHD3 Q8XHE4 Q8XHG1 Q8XHG8 Q8XHH8 Q8XHI3 Q8XHJ0 Q8XHJ2 Q8XHJ4 Q8XHJ8
Q8XHL1 Q8XHL4 Q8XHL5 Q8XHL7 Q8XHL8 Q8XHL9 Q8XHM4 Q8XHP0 Q8XHP8 Q8XHQ3 Q8XHQ4
Q8XHQ5 Q8XHQ7 Q8XHR1 Q8XHR4 Q8XHR5 Q8XHR6 Q8XHR7 Q8XHR9 Q8XHS0 Q8XHS1 Q8XHS2
Q8XHS3 Q8XHS4 Q8XHS5 Q8XHS6 Q8XHS7 Q8XHS8 Q8XHS9 Q8XHT0 Q8XHT1 Q8XHT2 Q8XHT3
Q8XHT4 Q8XHT5 Q8XHT6 Q8XHT7 Q8XHT8 Q8XHT9 Q8XHU0 Q8XHU1 Q8XHU2 Q8XHU4 Q8XHU6
Q8XHU7 Q8XHU8 Q8XHU9 Q8XHV0 Q8XHV1 Q8XHV5 Q8XHV6 Q8XHV7 Q8XHW7 Q8XHZ5 Q8XHZ7
Q8XI02 Q8XI08 Q8XI22 Q8XI46 Q8XI54 Q8XI55 Q8XI62 Q8XI67 Q8XI68 Q8XI79 Q8XI89

Q8XIA9 Q8XIB3 Q8XIB6 Q8XIB9 Q8XIC4 Q8XID7 Q8XIG8 Q8XIH4 Q8XIH5 Q8XII0 Q8XII2
Q8XII9 Q8XIJ1 Q8XIJ2 Q8XIJ4 Q8XIK2 Q8XIK5 Q8XIN2 Q8XIP3 Q8XIQ9 Q8XIR2 Q8XIR5
Q8XIS2 Q8XIS3 Q8XIS9 Q8XIT0 Q8XIT1 Q8XIT6 Q8XIU0 Q8XIU5 Q8XIU9 Q8XIV4 Q8XIV9
Q8XIW8 Q8XIX0 Q8XIY5 Q8XIY6 Q8XIZ0 Q8XIZ1 Q8XJ13 Q8XJ14 Q8XJ15 Q8XJ16 Q8XJ22
Q8XJ24 Q8XJ27 Q8XJ28 Q8XJ32 Q8XJ38 Q8XJ42 Q8XJ52 Q8XJ55 Q8XJ56 Q8XJ60 Q8XJ67
Q8XJ68 Q8XJ69 Q8XJ75 Q8XJ76 Q8XJ80 Q8XJ87 Q8XJ96 Q8XJ99 Q8XJA8 Q8XJB2 Q8XJC5
Q8XJD6 Q8XJD7 Q8XJD8 Q8XJD9 Q8XJE1 Q8XJE3 Q8XJF4 Q8XJH3 Q8XJH6 Q8XJH7 Q8XJH8
Q8XJH9 Q8XJI6 Q8XJJ7 Q8XJK1 Q8XJK7 Q8XJK8 Q8XJK9 Q8XJL2 Q8XJL3 Q8XJL6 Q8XJL8
Q8XJL9 Q8XJM2 Q8XJM7 Q8XJN0 Q8XJN2 Q8XJN4 Q8XJN8 Q8XJP4 Q8XJP5 Q8XJP6 Q8XJP7
Q8XJP8 Q8XJQ0 Q8XJQ1 Q8XJQ5 Q8XJQ6 Q8XJQ7 Q8XJQ8 Q8XJQ9 Q8XJR1 Q8XJR3 Q8XJR4
Q8XJR8 Q8XJR9 Q8XJS1 Q8XJS3 Q8XJS4 Q8XJS9 Q8XJU2 Q8XJX0 Q8XJX2 Q8XJY5 Q8XJZ6
Q8XK02 Q8XK15 Q8XK30 Q8XK37 Q8XK59 Q8XK60 Q8XKA3 Q8XKA4 Q8XKF6 Q8XKG4 Q8XKG8
Q8XKH1 Q8XKI1 Q8XKI3 Q8XKK0 Q8XKK1 Q8XKK2 Q8XKK5 Q8XKN1 Q8XKP8 Q8XKP9 Q8XKQ7
Q8XKQ8 Q8XKR1 Q8XKS9 Q8XKU0 Q8XKU1 Q8XKU2 Q8XKU7 Q8XKV0 Q8XKX4 Q8XL16 Q8XL46
Q8XL57 Q8XL59 Q8XL62 Q8XL63 Q8XL65 Q8XL84 Q8XL85 Q8XL86 Q8XL87 Q8XL95 Q8XLE8
Q8XLE9 Q8XLF7 Q8XLJ6 Q8XLK7 Q8XLM2 Q8XLN2 Q8XLN7 Q8XLN8 Q8XLP3 Q8XLQ0 Q8XLZ0
Q8XLZ1 Q8XM08 Q8XM22 Q8XM71 Q8XM84 Q8XM99 Q8XMB2 Q8XMI6 Q8XMI8 Q8XMJ2 Q8XMJ3
Q8XMJ6 Q8XMJ7 Q8XMJ8 Q8XMK2 Q8XMK4 Q8XMK6 Q8XMK9 Q8XML8 Q8XMN6 Q8XMP3 Q8XMQ3
Q8XMQ5 Q8XMU5 Q8XMU7 Q8XMW9 Q8XMY2 Q8XMY8 Q8XMZ0 Q8XNB4 Q8XNC2 Q8XNE2 Q8XNE3
Q8XNE7 Q8XNG6 Q8XNH2 Q8XNH7 Q8XNH9 Q8XNI0 Q8XNI1 Q8XNI5 Q8XNJ8 Q8XNK5 Q8XNK6
Q8XNK7 Q8XNL5 Q8XNR7 Q8XNS6 Q8XNV1 Q8XNW5 Q8XNZ3 Q8XP14 Q8XP15 Q8XP22 Q8XP33
Q8XP62 Q8XP74 Q8XP76 Q8XP78 Q8XP97 Q8XPA1 Q8XPA2 Q8XPB8 Q8XPB9 Q8XPD8 Q8XPE2
Q8XPE3 Q8XPE9 Q8XPF9 Q8XPG2 Q8XPH8 Q8XPV7 Q8XQ73 Q8XQ82 Q8XQ89 Q8XQE8 Q8XQG4
Q8XQG9 Q8XQN1 Q8XQR9 Q8XQS6 Q8XR10 Q8XR33 Q8XRC2 Q8XRF9 Q8XRG0 Q8XRG1 Q8XRK5
Q8XRL7 Q8XRM7 Q8XRN1 Q8XRV9 Q8XRY7 Q8XRZ0 Q8XS00 Q8XS01 Q8XS03 Q8XS05 Q8XS35
Q8XS59 Q8XSQ0 Q8XSS8 Q8XSZ5 Q8XT56 Q8XT73 Q8XT76 Q8XTL6 Q8XTQ1 Q8XTV4 Q8XU24
Q8XU28 Q8XU30 Q8XU38 Q8XU39 Q8XU65 Q8XU66 Q8XU90 Q8XU95 Q8XU98 Q8XU99 Q8XUA0
Q8XUB7 Q8XUC6 Q8XUR0 Q8XUY1 Q8XUZ4 Q8XUZ5 Q8XUZ6 Q8XUZ7 Q8XUZ8 Q8XUZ9 Q8XV08
Q8XV09 Q8XV10 Q8XV11 Q8XV12 Q8XV13 Q8XV14 Q8XV15 Q8XV16 Q8XV17 Q8XV18 Q8XV19
Q8XV20 Q8XV21 Q8XV22 Q8XV23 Q8XV24 Q8XV25 Q8XV26 Q8XV27 Q8XV28 Q8XV29 Q8XV30
Q8XV31 Q8XV33 Q8XV34 Q8XV35 Q8XV36 Q8XV37 Q8XV38 Q8XV39 Q8XV43 Q8XV54 Q8XV61
Q8XV62 Q8XV63 Q8XV78 Q8XV79 Q8XV80 Q8XV81 Q8XV83 Q8XV84 Q8XV85 Q8XV86 Q8XV87
Q8XVA0 Q8XVD0 Q8XVD1 Q8XVE4 Q8XVE6 Q8XVE7 Q8XVF3 Q8XVH8 Q8XVH9 Q8XVI2 Q8XVI5
Q8XVI8 Q8XVI9 Q8XVJ7 Q8XVK0 Q8XVK1 Q8XVK2 Q8XVK8 Q8XVK9 Q8XVL0 Q8XVL1 Q8XVL3
Q8XVL4 Q8XVP0 Q8XVP2 Q8XVP5 Q8XVQ0 Q8XVS8 Q8XVT2 Q8XVT3 Q8XVT6 Q8XVU6 Q8XVV4
Q8XVW4 Q8XVW5 Q8XW25 Q8XW26 Q8XW28 Q8XW29 Q8XW30 Q8XW31 Q8XW32 Q8XW36 Q8XW40
Q8XW41 Q8XW45 Q8XW52 Q8XWB0 Q8XWB1 Q8XWB8 Q8XWC0 Q8XWC1 Q8XWC2 Q8XWC4 Q8XWC5
Q8XWD0 Q8XWE1 Q8XWE3 Q8XWE7 Q8XWK9 Q8XWL1 Q8XWM5 Q8XWM7 Q8XWM8 Q8XWM9 Q8XWN0
Q8XWN8 Q8XWQ8 Q8XWR1 Q8XWS3 Q8XWT0 Q8XWT3 Q8XWT9 Q8XWV4 Q8XWV7 Q8XWW2 Q8XWW3
Q8XWX1 Q8XWY3 Q8XX54 Q8XX77 Q8XX80 Q8XX83 Q8XX93 Q8XX95 Q8XX96 Q8XX97 Q8XXA6
Q8XXB0 Q8XXC0 Q8XXC3 Q8XXC4 Q8XXC5 Q8XXC7 Q8XXD0 Q8XXE5 Q8XXE9 Q8XXF4 Q8XXF5
Q8XXF9 Q8XXG0 Q8XXI7 Q8XXJ7 Q8XXL5 Q8XXN8 Q8XXP1 Q8XXP4 Q8XXP6 Q8XXP9 Q8XXS7
Q8XXS8 Q8XXT0 Q8XXT1 Q8XXT2 Q8XXT3 Q8XXT4 Q8XXV4 Q8XXW5 Q8XXX1 Q8XXX3 Q8XXX4
Q8XXX5 Q8XXX8 Q8XXX9 Q8XXY0 Q8XXY2 Q8XY07 Q8XY11 Q8XYH4 Q8XYL0 Q8XYL1 Q8XYN6

Q8XYN9 Q8XYP6 Q8XYP7 Q8XYP8 Q8XYP9 Q8XYW2 Q8XYW3 Q8XYX0 Q8XYX1 Q8XYX8 Q8XYY9
Q8XYZ0 Q8XZ10 Q8XZ19 Q8XZ23 Q8XZ24 Q8XZ25 Q8XZ26 Q8XZ27 Q8XZ28 Q8XZ29 Q8XZ40
Q8XZ76 Q8XZ79 Q8XZ82 Q8XZ83 Q8XZ85 Q8XZ91 Q8XZC2 Q8XZC3 Q8XZG4 Q8XZG9 Q8XZH1
Q8XZH5 Q8XZH7 Q8XZI5 Q8XZI7 Q8XZI8 Q8XZI9 Q8XZJ0 Q8XZJ1 Q8XZJ3 Q8XZJ6 Q8XZJ7
Q8XZK2 Q8XZK5 Q8XZK8 Q8XZM4 Q8XZP1 Q8XZQ6 Q8XZR0 Q8XZR3 Q8XZT5 Q8XZT6 Q8XZT8
Q8XZV4 Q8XZV5 Q8XZV6 Q8XZV8 Q8XZZ5 Q8XZZ9 Q8Y010 Q8Y013 Q8Y014 Q8Y015 Q8Y016
Q8Y019 Q8Y020 Q8Y025 Q8Y026 Q8Y029 Q8Y032 Q8Y033 Q8Y035 Q8Y039 Q8Y040 Q8Y050
Q8Y052 Q8Y062 Q8Y065 Q8Y074 Q8Y077 Q8Y093 Q8Y099 Q8Y0A1 Q8Y0A5 Q8Y0B8 Q8Y0H3
Q8Y0H5 Q8Y0H8 Q8Y0H9 Q8Y0I1 Q8Y0J6 Q8Y0K4 Q8Y0K5 Q8Y0L2 Q8Y0L5 Q8Y0L9 Q8Y0M0
Q8Y0N2 Q8Y0N9 Q8Y0Q3 Q8Y0U6 Q8Y0U9 Q8Y0V1 Q8Y0V3 Q8Y0V7 Q8Y0V8 Q8Y0V9 Q8Y0W0
Q8Y0W7 Q8Y0X8 Q8Y0Y3 Q8Y0Y5 Q8Y0Y6 Q8Y0Y8 Q8Y0Z0 Q8Y0Z5 Q8Y183 Q8Y193 Q8Y199
Q8Y1B3 Q8Y1B4 Q8Y1C3 Q8Y1C6 Q8Y1D0 Q8Y1D2 Q8Y1F2 Q8Y1G1 Q8Y1G2 Q8Y1H8 Q8Y1H9
Q8Y1I7 Q8Y1K9 Q8Y1L2 Q8Y1L3 Q8Y1L4 Q8Y1L5 Q8Y1M4 Q8Y1P8 Q8Y1P9 Q8Y1S1 Q8Y1W3
Q8Y1W6 Q8Y1X7 Q8Y1Y3 Q8Y1Y5 Q8Y1Y6 Q8Y206 Q8Y208 Q8Y211 Q8Y213 Q8Y219 Q8Y231
Q8Y232 Q8Y233 Q8Y235 Q8Y236 Q8Y239 Q8Y240 Q8Y241 Q8Y242 Q8Y245 Q8Y246 Q8Y249
Q8Y270 Q8Y278 Q8Y288 Q8Y2B4 Q8Y2B9 Q8Y2D1 Q8Y2D3 Q8Y2D7 Q8Y2E0 Q8Y2E1 Q8Y2E2
Q8Y2E3 Q8Y2E6 Q8Y2I0 Q8Y2I3 Q8Y2K1 Q8Y2K9 Q8Y2L2 Q8Y2L3 Q8Y2M4 Q8Y2P8 Q8Y2Q7
Q8Y2R5 Q8Y2R9 Q8Y2V6 Q8Y2W3 Q8Y2X7 Q8Y303 Q8Y306 Q8Y339 Q8Y342 Q8Y344 Q8Y347
Q8Y368 Q8Y372 Q8Y387 Q8Y395 Q8Y3A8 Q8Y3B0 Q8Y3C2 Q8Y3C3 Q8Y3C6 Q8Y3C8 Q8Y3D7
Q8Y3D8 Q8Y3E7 Q8Y3F3 Q8Y3G3 Q8Y3H2 Q8Y3H5 Q8Y3H9 Q8Y3I1 Q8Y3I6 Q8Y3I7 Q8Y3I8
Q8Y3I9 Q8Y3L0 Q8Y3M4 Q8Y3M5 Q8Y3N3 Q8Y3N4 Q8Y3P1 Q8Y3P9 Q8Y3R3 Q8Y3R9 Q8Y3S3
Q8Y3T4 Q8Y3T8 Q8Y3U2 Q8Y3X6 Q8Y3X7 Q8Y3Y6 Q8Y420 Q8Y421 Q8Y440 Q8Y441 Q8Y443
Q8Y444 Q8Y445 Q8Y446 Q8Y447 Q8Y449 Q8Y450 Q8Y457 Q8Y458 Q8Y459 Q8Y471 Q8Y491
Q8Y493 Q8Y494 Q8Y495 Q8Y4A2 Q8Y4A6 Q8Y4A7 Q8Y4A8 Q8Y4A9 Q8Y4B2 Q8Y4B3 Q8Y4C4
Q8Y4D8 Q8Y4F5 Q8Y4F6 Q8Y4G1 Q8Y4G3 Q8Y4G9 Q8Y4H0 Q8Y4H1 Q8Y4I2 Q8Y4I3 Q8Y4I4
Q8Y4K4 Q8Y4L2 Q8Y4P3 Q8Y4P5 Q8Y4R7 Q8Y4X1 Q8Y510 Q8Y511 Q8Y551 Q8Y553 Q8Y556
Q8Y563 Q8Y564 Q8Y565 Q8Y570 Q8Y571 Q8Y577 Q8Y5E6 Q8Y5F1 Q8Y5G1 Q8Y5G2 Q8Y5H1
Q8Y5H2 Q8Y5I7 Q8Y5K7 Q8Y5L0 Q8Y5L2 Q8Y5L6 Q8Y5L7 Q8Y5L9 Q8Y5M1 Q8Y5M7 Q8Y5N2
Q8Y5N4 Q8Y5N8 Q8Y5N9 Q8Y5R1 Q8Y5R6 Q8Y5R7 Q8Y5R8 Q8Y5R9 Q8Y5S0 Q8Y5S2 Q8Y5S8
Q8Y5T0 Q8Y5T8 Q8Y5U4 Q8Y5V0 Q8Y5V1 Q8Y5V4 Q8Y5V5 Q8Y5W6 Q8Y5W8 Q8Y5X1 Q8Y5X4
Q8Y5X5 Q8Y5X6 Q8Y5X8 Q8Y5Y0 Q8Y5Y2 Q8Y5Z6 Q8Y5Z7 Q8Y5Z8 Q8Y601 Q8Y602 Q8Y603
Q8Y611 Q8Y613 Q8Y614 Q8Y617 Q8Y620 Q8Y623 Q8Y624 Q8Y626 Q8Y634 Q8Y640 Q8Y641
Q8Y643 Q8Y644 Q8Y660 Q8Y662 Q8Y663 Q8Y664 Q8Y665 Q8Y666 Q8Y667 Q8Y668 Q8Y672
Q8Y673 Q8Y676 Q8Y680 Q8Y691 Q8Y699 Q8Y6A2 Q8Y6A3 Q8Y6B1 Q8Y6B9 Q8Y6C0 Q8Y6C1
Q8Y6C3 Q8Y6C5 Q8Y6C6 Q8Y6D0 Q8Y6D2 Q8Y6D3 Q8Y6D6 Q8Y6E5 Q8Y6H7 Q8Y6I1 Q8Y6I2
Q8Y6J1 Q8Y6J7 Q8Y6J9 Q8Y6K3 Q8Y6K9 Q8Y6M0 Q8Y6M4 Q8Y6M6 Q8Y6M7 Q8Y6Q3 Q8Y6Q4
Q8Y6Q5 Q8Y6Q6 Q8Y6Q7 Q8Y6R2 Q8Y6R8 Q8Y6S5 Q8Y6S8 Q8Y6T4 Q8Y6T6 Q8Y6U0 Q8Y6U1
Q8Y6U2 Q8Y6U3 Q8Y6U4 Q8Y6U5 Q8Y6U8 Q8Y6V0 Q8Y6V4 Q8Y6W0 Q8Y6W7 Q8Y6W8 Q8Y6W9
Q8Y6X2 Q8Y6X3 Q8Y6X4 Q8Y6X5 Q8Y6X8 Q8Y6X9 Q8Y6Y2 Q8Y6Y6 Q8Y6Y9 Q8Y6Z2 Q8Y6Z3
Q8Y6Z5 Q8Y6Z6 Q8Y6Z7 Q8Y6Z8 Q8Y6Z9 Q8Y700 Q8Y708 Q8Y709 Q8Y714 Q8Y722 Q8Y723
Q8Y727 Q8Y729 Q8Y735 Q8Y747 Q8Y752 Q8Y753 Q8Y754 Q8Y755 Q8Y756 Q8Y757 Q8Y766
Q8Y776 Q8Y788 Q8Y789 Q8Y793 Q8Y7A4 Q8Y7A5 Q8Y7A7 Q8Y7B6 Q8Y7B9 Q8Y7C1 Q8Y7C3
Q8Y7C4 Q8Y7C5 Q8Y7C6 Q8Y7D3 Q8Y7D4 Q8Y7D5 Q8Y7F0 Q8Y7F1 Q8Y7F2 Q8Y7F3 Q8Y7F4
Q8Y7F6 Q8Y7G0 Q8Y7G1 Q8Y7G2 Q8Y7G4 Q8Y7G6 Q8Y7G7 Q8Y7H5 Q8Y7H6 Q8Y7I3 Q8Y7J7

Q8Y7J8 Q8Y7J9 Q8Y7K0 Q8Y7K1 Q8Y7K4 Q8Y7K9 Q8Y7L0 Q8Y7M8 Q8Y7N5 Q8Y7N6 Q8Y7N7
Q8Y7P0 Q8Y7P1 Q8Y7P5 Q8Y7P6 Q8Y7Q1 Q8Y7Q2 Q8Y7R3 Q8Y7S7 Q8Y7T0 Q8Y7T9 Q8Y7U4
Q8Y7V1 Q8Y822 Q8Y825 Q8Y832 Q8Y848 Q8Y853 Q8Y866 Q8Y870 Q8Y871 Q8Y8A0 Q8Y8A1
Q8Y8C0 Q8Y8D3 Q8Y8D4 Q8Y8D7 Q8Y8E0 Q8Y8E7 Q8Y8G1 Q8Y8H0 Q8Y8I0 Q8Y8K2 Q8Y8K6
Q8Y8N0 Q8Y8P1 Q8Y8R5 Q8Y8V6 Q8Y914 Q8Y915 Q8Y930 Q8Y9C1 Q8Y9D6 Q8Y9F9 Q8Y9G0
Q8Y9G1 Q8Y9G2 Q8Y9G3 Q8Y9G4 Q8Y9G5 Q8Y9G6 Q8Y9G7 Q8Y9I9 Q8Y9J3 Q8Y9J4 Q8Y9L8
Q8Y9N4 Q8Y9N5 Q8Y9N9 Q8Y9P2 Q8Y9X7 Q8Y9Y1 Q8Y9Y2 Q8Y9Y3 Q8Y9Y4 Q8Y9Z8 Q8Y9Z9
Q8YA20 Q8YA44 Q8YA46 Q8YA66 Q8YA96 Q8YAA3 Q8YAA4 Q8YAB0 Q8YAB1 Q8YAB3 Q8YAB4
Q8YAB5 Q8YAB8 Q8YAC4 Q8YAC5 Q8YAC7 Q8YAD1 Q8YAD3 Q8YAD5 Q8YAE0 Q8YAE1 Q8YAE2
Q8YAE3 Q8YAF2 Q8YAG0 Q8YAG2 Q8YAR1 Q8YAR2 Q8YAR8 Q8YAR9 Q8YAS0 Q8YAS3 Q8YAS5
Q8YAS7 Q8YAT8 Q8YAV8 Q8YB10 Q8YB45 Q8YB47 Q8YB50 Q8YB53 Q8YB57 Q8YBA4 Q8YBH3
Q8YBL2 Q8YBL8 Q8YBP2 Q8YBR2 Q8YBV5 Q8YBV9 Q8YBW1 Q8YC29 Q8YC47 Q8YC54 Q8YC57
Q8YC59 Q8YC60 Q8YC76 Q8YCA5 Q8YCD5 Q8YCE3 Q8YCF2 Q8YCK1 Q8YCL8 Q8YCM8 Q8YCP2
Q8YCQ2 Q8YCQ4 Q8YCV3 Q8YCV6 Q8YCX4 Q8YCX5 Q8YCY1 Q8YCZ1 Q8YD03 Q8YD04 Q8YD09
Q8YD12 Q8YD27 Q8YDA2 Q8YDA5 Q8YDA8 Q8YDB0 Q8YDB2 Q8YDC6 Q8YDC8 Q8YDC9 Q8YDE4
Q8YDE5 Q8YDE6 Q8YDE7 Q8YDL2 Q8YDL6 Q8YDP7 Q8YDY9 Q8YDZ4 Q8YDZ5 Q8YE11 Q8YE19
Q8YE20 Q8YE23 Q8YE30 Q8YE31 Q8YE35 Q8YE36 Q8YE37 Q8YE41 Q8YE49 Q8YE59 Q8YE60
Q8YE73 Q8YE74 Q8YE76 Q8YE77 Q8YE79 Q8YE82 Q8YEA2 Q8YEA9 Q8YEB3 Q8YEB4 Q8YEB5
Q8YEB6 Q8YEB7 Q8YED2 Q8YED5 Q8YED7 Q8YEG1 Q8YEG2 Q8YEG8 Q8YEK2 Q8YEK8 Q8YEL1
Q8YEN0 Q8YEP2 Q8YES3 Q8YES6 Q8YET3 Q8YEV0 Q8YEV2 Q8YEX2 Q8YEZ2 Q8YF55 Q8YF59
Q8YF70 Q8YF71 Q8YF72 Q8YF73 Q8YF86 Q8YFA2 Q8YFA4 Q8YFC9 Q8YFD6 Q8YFH2 Q8YFI3
Q8YFJ0 Q8YFK1 Q8YFK3 Q8YFM2 Q8YFM4 Q8YFN7 Q8YFP0 Q8YFR1 Q8YFR2 Q8YFR6 Q8YFS6
Q8YFU0 Q8YFU1 Q8YFX5 Q8YFY0 Q8YFY2 Q8YFY3 Q8YG60 Q8YG64 Q8YG65 Q8YG76 Q8YG89
Q8YG94 Q8YG99 Q8YGA4 Q8YGA9 Q8YGB9 Q8YGG7 Q8YGH2 Q8YGH3 Q8YGI8 Q8YGJ0 Q8YGJ1
Q8YGL8 Q8YGM5 Q8YGM6 Q8YGN1 Q8YGN4 Q8YGN6 Q8YGQ9 Q8YGR9 Q8YGS6 Q8YGT8 Q8YGU6
Q8YGV8 Q8YGV9 Q8YGW9 Q8YGX8 Q8YGY5 Q8YGY6 Q8YH15 Q8YH17 Q8YH30 Q8YH36 Q8YH68
Q8YH89 Q8YH94 Q8YH95 Q8YHB2 Q8YHC2 Q8YHC4 Q8YHC6 Q8YHC7 Q8YHC8 Q8YHD8 Q8YHF2
Q8YHF5 Q8YHF7 Q8YHF9 Q8YHG4 Q8YHH3 Q8YHH4 Q8YHH6 Q8YHI3 Q8YHJ4 Q8YHL5 Q8YHL6
Q8YHL9 Q8YHM1 Q8YHM2 Q8YHM4 Q8YHM5 Q8YHM6 Q8YHM8 Q8YHM9 Q8YHN0 Q8YHN1 Q8YHN3
Q8YHN4 Q8YHN5 Q8YHN7 Q8YHN8 Q8YHN9 Q8YHP0 Q8YHP3 Q8YHP4 Q8YHP7 Q8YHP8 Q8YHP9
Q8YHQ0 Q8YHQ1 Q8YHR9 Q8YHS6 Q8YHU1 Q8YHU3 Q8YHV6 Q8YHZ6 Q8YHZ7 Q8YHZ8 Q8YI00
Q8YI01 Q8YI02 Q8YI21 Q8YI24 Q8YI29 Q8YI47 Q8YI53 Q8YI56 Q8YI63 Q8YI64 Q8YI68
Q8YI71 Q8YI91 Q8YIB0 Q8YIB8 Q8YIC2 Q8YIG1 Q8YIG3 Q8YIJ3 Q8YIL6 Q8YIR3 Q8YIR8
Q8YIU8 Q8YIV0 Q8YIV5 Q8YIV7 Q8YIW2 Q8YIY0 Q8YIZ8 Q8YJ17 Q8YJ29 Q8YJ42 Q8YJ44
Q8YJ48 Q8YJ71 Q8YJ77 Q8YJ78 Q8YJ80 Q8YJ84 Q8YJ85 Q8YJ91 Q8YJ95 Q8YJ98 Q8YJB0
Q8YJB1 Q8YJC9 Q8YJD7 Q8YJD9 Q8YJE4 Q8YJE7 Q8YJF0 Q8YJF4 Q8YJF9 Q8YJI8 Q8YJJ5
Q8YJJ7 Q8YJK3 Q8YJP2 Q8YJS4 Q8YJS5 Q8YJS6 Q8YJT1 Q8YK41 Q8YKE7 Q8YKG3 Q8YLF9
Q8YLG0 Q8YLJ5 Q8YLJ6 Q8YLJ7 Q8YLJ8 Q8YLK1 Q8YLK6 Q8YLL0 Q8YLL1 Q8YLN2 Q8YLP4
Q8YLP6 Q8YLX9 Q8YLZ0 Q8YM38 Q8YM41 Q8YM45 Q8YM52 Q8YM56 Q8YM62 Q8YM64 Q8YM70
Q8YM73 Q8YM74 Q8YM75 Q8YM77 Q8YM83 Q8YMB5 Q8YMC2 Q8YMD4 Q8YMF7 Q8YMG7 Q8YMH5
Q8YMM6 Q8YMT3 Q8YMT4 Q8YMW6 Q8YMW8 Q8YMX2 Q8YMX6 Q8YMY2 Q8YMY3 Q8YMY9 Q8YMZ0
Q8YN19 Q8YN49 Q8YN62 Q8YN70 Q8YN91 Q8YN94 Q8YN97 Q8YNA5 Q8YNB4 Q8YNB9 Q8YNE3
Q8YNF7 Q8YNF8 Q8YNF9 Q8YNH4 Q8YNI3 Q8YNJ4 Q8YNK2 Q8YNL6 Q8YNL7 Q8YNP4 Q8YNQ6
Q8YNV9 Q8YNW0 Q8YNW5 Q8YNW7 Q8YP11 Q8YP17 Q8YP23 Q8YP43 Q8YP49 Q8YP60 Q8YP61

Q8YP62 Q8YP63 Q8YP64 Q8YP68 Q8YP69 Q8YP73 Q8YP78 Q8YP80 Q8YPC2 Q8YPF9 Q8YPH9
Q8YPI0 Q8YPI1 Q8YPI2 Q8YPI3 Q8YPI4 Q8YPI5 Q8YPI6 Q8YPI7 Q8YPI8 Q8YPI9 Q8YPJ0
Q8YPJ1 Q8YPJ2 Q8YPJ3 Q8YPJ4 Q8YPJ5 Q8YPJ6 Q8YPJ8 Q8YPJ9 Q8YPK0 Q8YPK1 Q8YPK2
Q8YPK3 Q8YPK4 Q8YPK5 Q8YPK6 Q8YPK7 Q8YPK8 Q8YPK9 Q8YPL2 Q8YPL9 Q8YPM0 Q8YPR1
Q8YPR3 Q8YPS9 Q8YPT0 Q8YPT4 Q8YPT9 Q8YPV5 Q8YPW9 Q8YPZ6 Q8YQ06 Q8YQ43 Q8YQ59
Q8YQ66 Q8YQ68 Q8YQ82 Q8YQB1 Q8YQC1 Q8YQC4 Q8YQD7 Q8YQD8 Q8YQE2 Q8YQE6 Q8YQF0
Q8YQF1 Q8YQG6 Q8YQJ1 Q8YQJ5 Q8YQL2 Q8YQL3 Q8YQL7 Q8YQM6 Q8YQR8 Q8YQT0 Q8YQU9
Q8YQX7 Q8YQX8 Q8YQX9 Q8YQY1 Q8YQY6 Q8YQZ0 Q8YQZ2 Q8YQZ3 Q8YQZ4 Q8YQZ8 Q8YQZ9
Q8YR05 Q8YR06 Q8YR77 Q8YR79 Q8YR82 Q8YR87 Q8YRB1 Q8YRC9 Q8YRD8 Q8YRI1 Q8YRI6
Q8YRI9 Q8YRJ1 Q8YRK5 Q8YRL4 Q8YRL8 Q8YRL9 Q8YRN1 Q8YRN2 Q8YRP2 Q8YRP3 Q8YRR1
Q8YRR9 Q8YRT9 Q8YRU9 Q8YRZ2 Q8YS09 Q8YS26 Q8YS28 Q8YS61 Q8YS62 Q8YS75 Q8YS86
Q8YS92 Q8YS95 Q8YSA4 Q8YSA7 Q8YSC1 Q8YSE9 Q8YSI2 Q8YSI5 Q8YSJ2 Q8YSK0 Q8YSK6
Q8YSM8 Q8YSM9 Q8YSU9 Q8YSY4 Q8YSZ1 Q8YSZ3 Q8YSZ5 Q8YT12 Q8YT31 Q8YT32 Q8YT40
Q8YT41 Q8YT42 Q8YT50 Q8YT79 Q8YT80 Q8YTB1 Q8YTC2 Q8YTE0 Q8YTE6 Q8YTI0 Q8YTI2
Q8YTI9 Q8YTJ3 Q8YTT3 Q8YTZ5 Q8YTZ6 Q8YU00 Q8YU19 Q8YU66 Q8YU79 Q8YU89 Q8YU96
Q8YUA3 Q8YUA5 Q8YUA6 Q8YUA7 Q8YUB6 Q8YUC0 Q8YUD4 Q8YUG9 Q8YUH8 Q8YUJ5 Q8YUL9
Q8YUM5 Q8YUR7 Q8YV15 Q8YV57 Q8YV76 Q8YV89 Q8YV98 Q8YVA8 Q8YVB5 Q8YVC1 Q8YVD0
Q8YVG9 Q8YVH1 Q8YVJ2 Q8YVL0 Q8YVM0 Q8YVM2 Q8YVQ0 Q8YVQ3 Q8YVR1 Q8YVS3 Q8YVS4
Q8YVS7 Q8YVS8 Q8YVT3 Q8YVT9 Q8YVU4 Q8YVU5 Q8YVU6 Q8YVX4 Q8YW05 Q8YW18 Q8YW20
Q8YW42 Q8YW43 Q8YW56 Q8YW62 Q8YW74 Q8YW98 Q8YWC1 Q8YWD2 Q8YWD8 Q8YWF0 Q8YWH5
Q8YWL3 Q8YWL4 Q8YWL7 Q8YWR1 Q8YWS8 Q8YWW1 Q8YX02 Q8YX03 Q8YX05 Q8YX22 Q8YX49
Q8YX56 Q8YX57 Q8YX58 Q8YX72 Q8YX96 Q8YX97 Q8YXA2 Q8YXA3 Q8YXB6 Q8YXD1 Q8YXE4
Q8YXF0 Q8YXG5 Q8YXG8 Q8YXG9 Q8YXH5 Q8YXJ1 Q8YXJ6 Q8YXK4 Q8YXK5 Q8YXQ0 Q8YXQ7
Q8YXQ9 Q8YXS7 Q8YXS9 Q8YXW5 Q8YXY3 Q8YXZ4 Q8YY02 Q8YY05 Q8YY12 Q8YY69 Q8YY71
Q8YY90 Q8YYB8 Q8YYG2 Q8YYH3 Q8YYI2 Q8YYI3 Q8YYK4 Q8YYK5 Q8YYP9 Q8YYU7 Q8YYV9
Q8YYW0 Q8YYX9 Q8YZ37 Q8YZ40 Q8YZ70 Q8YZ80 Q8YZ99 Q8YZA0 Q8YZA1 Q8YZH3 Q8YZH7
Q8YZP7 Q8YZP8 Q8YZQ3 Q8YZS3 Q8YZT2 Q8YZT3 Q8YZV0 Q8YZV9 Q8YZZ2 Q8Z006 Q8Z023
Q8Z032 Q8Z064 Q8Z068 Q8Z074 Q8Z080 Q8Z086 Q8Z098 Q8Z0C0 Q8Z0C1 Q8Z0C4 Q8Z0C8
Q8Z0F0 Q8Z0F1 Q8Z0I6 Q8Z0I7 Q8Z0I8 Q8Z0M2 Q8Z0M3 Q8Z0M8 Q8Z0Q9 Q8Z0T4 Q8Z0U0
Q8Z0U2 Q8Z0U3 Q8Z0U8 Q8Z0V2 Q8Z0W8 Q8Z0Y7 Q8Z116 Q8Z118 Q8Z123 Q8Z125 Q8Z127
Q8Z130 Q8Z138 Q8Z146 Q8Z150 Q8Z157 Q8Z163 Q8Z164 Q8Z167 Q8Z168 Q8Z169 Q8Z182
Q8Z183 Q8Z186 Q8Z187 Q8Z193 Q8Z197 Q8Z1D0 Q8Z1E1 Q8Z1R0 Q8Z1T1 Q8Z1T7 Q8Z1U7
Q8Z1V3 Q8Z1V8 Q8Z1W1 Q8Z1W5 Q8Z1W6 Q8Z1W9 Q8Z1X0 Q8Z1X1 Q8Z1X4 Q8Z1X5 Q8Z1X6
Q8Z1X8 Q8Z1Y1 Q8Z1Y2 Q8Z1Y3 Q8Z1Y5 Q8Z1Z3 Q8Z201 Q8Z202 Q8Z205 Q8Z214 Q8Z216
Q8Z217 Q8Z221 Q8Z222 Q8Z223 Q8Z227 Q8Z232 Q8Z233 Q8Z234 Q8Z235 Q8Z259 Q8Z268
Q8Z270 Q8Z277 Q8Z292 Q8Z2B3 Q8Z2B4 Q8Z2C5 Q8Z2D8 Q8Z2E0 Q8Z2E1 Q8Z2F0 Q8Z2F4
Q8Z2H1 Q8Z2H2 Q8Z2H4 Q8Z2H5 Q8Z2H9 Q8Z2L4 Q8Z2L8 Q8Z2M3 Q8Z2N4 Q8Z2N6 Q8Z2N8
Q8Z2P0 Q8Z2Q2 Q8Z2Q7 Q8Z2Q9 Q8Z2R0 Q8Z2R1 Q8Z2R3 Q8Z2R9 Q8Z2S0 Q8Z2T5 Q8Z2U4
Q8Z2U7 Q8Z2V3 Q8Z2V4 Q8Z2V5 Q8Z2V9 Q8Z2X3 Q8Z2X4 Q8Z2X9 Q8Z2Y0 Q8Z2Y2 Q8Z2Y6
Q8Z2Y8 Q8Z2Z0 Q8Z2Z4 Q8Z2Z9 Q8Z300 Q8Z303 Q8Z305 Q8Z307 Q8Z308 Q8Z309 Q8Z311
Q8Z313 Q8Z315 Q8Z316 Q8Z318 Q8Z320 Q8Z322 Q8Z325 Q8Z326 Q8Z327 Q8Z328 Q8Z329
Q8Z334 Q8Z335 Q8Z365 Q8Z377 Q8Z381 Q8Z3A4 Q8Z3A8 Q8Z3B6 Q8Z3B8 Q8Z3E0 Q8Z3F0
Q8Z3F1 Q8Z3G1 Q8Z3H1 Q8Z3H3 Q8Z3H5 Q8Z3H7 Q8Z3H8 Q8Z3H9 Q8Z3I0 Q8Z3I7 Q8Z3I8
Q8Z3K5 Q8Z3K8 Q8Z3L8 Q8Z3L9 Q8Z3M6 Q8Z3M9 Q8Z3R7 Q8Z3U1 Q8Z3U6 Q8Z3U7 Q8Z3V1

Q8Z3V2 Q8Z3W5 Q8Z3X0 Q8Z3X3 Q8Z3X4 Q8Z3X8 Q8Z3X9 Q8Z3Z9 Q8Z410 Q8Z412 Q8Z421
Q8Z427 Q8Z428 Q8Z429 Q8Z433 Q8Z437 Q8Z439 Q8Z446 Q8Z458 Q8Z459 Q8Z460 Q8Z470
Q8Z471 Q8Z472 Q8Z473 Q8Z474 Q8Z495 Q8Z497 Q8Z4A6 Q8Z4C4 Q8Z4C5 Q8Z4C7 Q8Z4D4
Q8Z4D5 Q8Z4D6 Q8Z4E6 Q8Z4G0 Q8Z4I5 Q8Z4J9 Q8Z4K0 Q8Z4K4 Q8Z4K6 Q8Z4L2 Q8Z4L6
Q8Z4M3 Q8Z4N0 Q8Z4N1 Q8Z4N2 Q8Z4N3 Q8Z4P2 Q8Z4P4 Q8Z4P6 Q8Z4Q1 Q8Z4Q3 Q8Z4R0
Q8Z4R2 Q8Z4R8 Q8Z4S2 Q8Z4T0 Q8Z4T3 Q8Z4U3 Q8Z4U8 Q8Z4W1 Q8Z4W4 Q8Z4Z3 Q8Z500
Q8Z524 Q8Z533 Q8Z534 Q8Z544 Q8Z548 Q8Z549 Q8Z560 Q8Z593 Q8Z5A6 Q8Z5A8 Q8Z5B2
Q8Z5C3 Q8Z5C8 Q8Z5G5 Q8Z5H2 Q8Z5J6 Q8Z5J7 Q8Z5J8 Q8Z5J9 Q8Z5K0 Q8Z5K1 Q8Z5K2
Q8Z5M8 Q8Z5M9 Q8Z5N6 Q8Z5N9 Q8Z5Q0 Q8Z5R1 Q8Z5S1 Q8Z5S9 Q8Z5T3 Q8Z5U4 Q8Z5V7
Q8Z5V9 Q8Z5W0 Q8Z5W1 Q8Z5X3 Q8Z662 Q8Z681 Q8Z682 Q8Z688 Q8Z699 Q8Z6E7 Q8Z6F4
Q8Z6F5 Q8Z6F6 Q8Z6F9 Q8Z6G0 Q8Z6G1 Q8Z6G2 Q8Z6G3 Q8Z6G5 Q8Z6G6 Q8Z6H0 Q8Z6H2
Q8Z6I2 Q8Z6I3 Q8Z6I4 Q8Z6I7 Q8Z6I8 Q8Z6I9 Q8Z6K0 Q8Z6K2 Q8Z6L0 Q8Z6P1 Q8Z6Q0
Q8Z6Q2 Q8Z6Q3 Q8Z6R2 Q8Z6R6 Q8Z6R7 Q8Z6X9 Q8Z728 Q8Z747 Q8Z783 Q8Z787 Q8Z788
Q8Z7A8 Q8Z7C9 Q8Z7D5 Q8Z7D9 Q8Z7E0 Q8Z7E5 Q8Z7G6 Q8Z7G9 Q8Z7H6 Q8Z7H9 Q8Z7I6
Q8Z7I7 Q8Z7J7 Q8Z7J8 Q8Z7L2 Q8Z7L5 Q8Z7N9 Q8Z7P9 Q8Z7Q5 Q8Z7R2 Q8Z7R6 Q8Z7S1
Q8Z7S6 Q8Z7Y9 Q8Z7Z5 Q8Z7Z6 Q8Z7Z7 Q8Z803 Q8Z829 Q8Z841 Q8Z861 Q8Z887 Q8Z889
Q8Z890 Q8Z892 Q8Z893 Q8Z896 Q8Z897 Q8Z899 Q8Z8A0 Q8Z8A2 Q8Z8B0 Q8Z8B2 Q8Z8B8
Q8Z8D2 Q8Z8D9 Q8Z8F8 Q8Z8G0 Q8Z8G5 Q8Z8G6 Q8Z8H0 Q8Z8H5 Q8Z8H7 Q8Z8H8 Q8Z8I2
Q8Z8I3 Q8Z8J2 Q8Z8M0 Q8Z8P6 Q8Z8R2 Q8Z8R3 Q8Z8R4 Q8Z8S3 Q8Z8T1 Q8Z8T2 Q8Z8T4
Q8Z8U7 Q8Z8V1 Q8Z8W2 Q8Z8W3 Q8Z8W5 Q8Z8W6 Q8Z8X1 Q8Z8X3 Q8Z8X6 Q8Z8Y0 Q8Z8Y1
Q8Z8Y9 Q8Z903 Q8Z932 Q8Z933 Q8Z983 Q8Z985 Q8Z988 Q8Z996 Q8Z9A6 Q8Z9A7 Q8Z9A8
Q8Z9B1 Q8Z9B4 Q8Z9C1 Q8Z9C2 Q8Z9C3 Q8Z9D2 Q8Z9D3 Q8Z9E2 Q8Z9E3 Q8Z9E5 Q8Z9F9
Q8Z9G7 Q8Z9G8 Q8Z9H0 Q8Z9H3 Q8Z9H4 Q8Z9H5 Q8Z9H9 Q8Z9I0 Q8Z9I1 Q8Z9I2 Q8Z9I3
Q8Z9I4 Q8Z9J4 Q8Z9J5 Q8Z9J7 Q8Z9J8 Q8Z9K8 Q8Z9K9 Q8Z9L0 Q8Z9L2 Q8Z9L3 Q8Z9L4
Q8Z9L5 Q8Z9L6 Q8Z9L7 Q8Z9L8 Q8Z9L9 Q8Z9M0 Q8Z9M9 Q8Z9N2 Q8Z9R1 Q8Z9R5 Q8Z9R8
Q8Z9R9 Q8Z9S8 Q8Z9U2 Q8Z9U4 Q8Z9U5 Q8Z9U9 Q8Z9V5 Q8Z9V6 Q8Z9W6 Q8Z9W7 Q8Z9X0
Q8Z9X7 Q8Z9Y0 Q8Z9Y1 Q8Z9Z1 Q8ZA09 Q8ZA10 Q8ZA46 Q8ZA49 Q8ZA75 Q8ZA76 Q8ZA77
Q8ZA78 Q8ZA84 Q8ZA85 Q8ZA86 Q8ZA87 Q8ZA88 Q8ZA97 Q8ZAA0 Q8ZAA2 Q8ZAA7 Q8ZAB3
Q8ZAC2 Q8ZAD8 Q8ZAG3 Q8ZAI0 Q8ZAL3 Q8ZAL4 Q8ZAL8 Q8ZAN4 Q8ZAN6 Q8ZAN8 Q8ZAP0
Q8ZAP1 Q8ZAP2 Q8ZAP3 Q8ZAP4 Q8ZAP5 Q8ZAP9 Q8ZAQ1 Q8ZAQ2 Q8ZAQ5 Q8ZAQ7 Q8ZAQ8
Q8ZAR2 Q8ZAR3 Q8ZAR4 Q8ZAR7 Q8ZAS2 Q8ZAU5 Q8ZAX1 Q8ZAX6 Q8ZAX7 Q8ZAX8 Q8ZB04
Q8ZB38 Q8ZB39 Q8ZB41 Q8ZB56 Q8ZB62 Q8ZB74 Q8ZB75 Q8ZB81 Q8ZB82 Q8ZB83 Q8ZB84
Q8ZB88 Q8ZB94 Q8ZB98 Q8ZBA2 Q8ZBA7 Q8ZBB3 Q8ZBB5 Q8ZBB8 Q8ZBC0 Q8ZBC2 Q8ZBC3
Q8ZBC4 Q8ZBC5 Q8ZBE1 Q8ZBE9 Q8ZBG8 Q8ZBH1 Q8ZBH3 Q8ZBH8 Q8ZBH9 Q8ZBI0 Q8ZBI2
Q8ZBJ5 Q8ZBJ7 Q8ZBJ8 Q8ZBK6 Q8ZBK7 Q8ZBK8 Q8ZBL1 Q8ZBL9 Q8ZBM5 Q8ZBN1 Q8ZBN6
Q8ZBN7 Q8ZBP1 Q8ZBP2 Q8ZBP3 Q8ZBP6 Q8ZBP7 Q8ZBP8 Q8ZBP9 Q8ZBQ0 Q8ZBQ3 Q8ZBT8
Q8ZBU2 Q8ZBU7 Q8ZBU8 Q8ZBU9 Q8ZBV0 Q8ZBV2 Q8ZBV7 Q8ZBZ0 Q8ZBZ9 Q8ZC08 Q8ZC09
Q8ZC15 Q8ZC17 Q8ZC18 Q8ZC32 Q8ZC33 Q8ZC39 Q8ZC41 Q8ZC42 Q8ZC45 Q8ZC47 Q8ZC49
Q8ZC51 Q8ZC52 Q8ZC64 Q8ZC65 Q8ZC66 Q8ZC72 Q8ZC86 Q8ZC94 Q8ZC96 Q8ZC97 Q8ZC98
Q8ZCA8 Q8ZCC0 Q8ZCC1 Q8ZCD0 Q8ZCD2 Q8ZCF8 Q8ZCF9 Q8ZCG0 Q8ZCG8 Q8ZCG9 Q8ZCK0
Q8ZCP4 Q8ZCP8 Q8ZCQ2 Q8ZCR0 Q8ZCR1 Q8ZCS3 Q8ZCS4 Q8ZCS5 Q8ZCT2 Q8ZCT6 Q8ZCT9
Q8ZCU2 Q8ZCU4 Q8ZCV6 Q8ZCX2 Q8ZCX8 Q8ZCX9 Q8ZD12 Q8ZD27 Q8ZD36 Q8ZD41 Q8ZD43
Q8ZD70 Q8ZD72 Q8ZD80 Q8ZD84 Q8ZD85 Q8ZD86 Q8ZD89 Q8ZD91 Q8ZDA6 Q8ZDC6 Q8ZDD9
Q8ZDE1 Q8ZDF0 Q8ZDF8 Q8ZDG1 Q8ZDG3 Q8ZDG8 Q8ZDG9 Q8ZDH0 Q8ZDJ6 Q8ZDJ9 Q8ZDK3

Q8ZDM8 Q8ZDP7 Q8ZDV4 Q8ZDW5 Q8ZDW6 Q8ZDW7 Q8ZDW8 Q8ZDX0 Q8ZDX1 Q8ZDX2 Q8ZDY2
Q8ZDY4 Q8ZE08 Q8ZE19 Q8ZE20 Q8ZE42 Q8ZE60 Q8ZE81 Q8ZE82 Q8ZEB6 Q8ZED8 Q8ZEF0
Q8ZEG0 Q8ZEG6 Q8ZEG7 Q8ZEG8 Q8ZEG9 Q8ZEH0 Q8ZEI3 Q8ZEJ1 Q8ZEJ5 Q8ZEK1 Q8ZEK7
Q8ZEL2 Q8ZES0 Q8ZES5 Q8ZEU5 Q8ZEU6 Q8ZEU7 Q8ZEU8 Q8ZEV0 Q8ZEV5 Q8ZEV7 Q8ZEW6
Q8ZEX8 Q8ZEX9 Q8ZEY1 Q8ZEY2 Q8ZEY4 Q8ZF52 Q8ZF61 Q8ZF73 Q8ZF85 Q8ZFD6 Q8ZFE0
Q8ZFH8 Q8ZFJ9 Q8ZFN8 Q8ZFQ3 Q8ZFQ5 Q8ZFR0 Q8ZFR1 Q8ZFS2 Q8ZFS3 Q8ZFT9 Q8ZFU1
Q8ZFU4 Q8ZFV7 Q8ZFW5 Q8ZFX4 Q8ZFX5 Q8ZFX6 Q8ZFX7 Q8ZFX8 Q8ZFX9 Q8ZFY0 Q8ZFY1
Q8ZFZ8 Q8ZFZ9 Q8ZG01 Q8ZG08 Q8ZG09 Q8ZG15 Q8ZG65 Q8ZG71 Q8ZG78 Q8ZG93 Q8ZG99
Q8ZGA1 Q8ZGB1 Q8ZGB3 Q8ZGB4 Q8ZGC4 Q8ZGD2 Q8ZGD5 Q8ZGE1 Q8ZGG1 Q8ZGH4 Q8ZGJ1
Q8ZGJ9 Q8ZGK7 Q8ZGK8 Q8ZGM2 Q8ZGM4 Q8ZGM5 Q8ZGM6 Q8ZGR6 Q8ZGV2 Q8ZGV8 Q8ZGV9
Q8ZGW2 Q8ZGW4 Q8ZGW5 Q8ZGW7 Q8ZGW9 Q8ZGX4 Q8ZGY3 Q8ZGY5 Q8ZGY8 Q8ZH00 Q8ZH09
Q8ZH12 Q8ZH14 Q8ZH30 Q8ZH34 Q8ZH36 Q8ZH47 Q8ZH50 Q8ZH54 Q8ZH62 Q8ZH63 Q8ZH64
Q8ZH65 Q8ZH66 Q8ZH68 Q8ZH69 Q8ZH72 Q8ZH75 Q8ZH78 Q8ZH86 Q8ZH96 Q8ZHE7 Q8ZHE9
Q8ZHF1 Q8ZHF4 Q8ZHF5 Q8ZHG0 Q8ZHG2 Q8ZHG3 Q8ZHG6 Q8ZHG7 Q8ZHG8 Q8ZHH3 Q8ZHH8
Q8ZHI2 Q8ZHI6 Q8ZHI7 Q8ZHI8 Q8ZHJ5 Q8ZHK1 Q8ZHK4 Q8ZHK5 Q8ZHU5 Q8ZHU8 Q8ZHV1
Q8ZI40 Q8ZI56 Q8ZI64 Q8ZIC5 Q8ZIC6 Q8ZIE7 Q8ZIE8 Q8ZIF1 Q8ZIF4 Q8ZIF7 Q8ZIF8
Q8ZIG8 Q8ZIG9 Q8ZIH0 Q8ZIH1 Q8ZII0 Q8ZIK1 Q8ZIK5 Q8ZIK6 Q8ZIK7 Q8ZIL4 Q8ZIL5
Q8ZIL6 Q8ZIM0 Q8ZIM3 Q8ZIM6 Q8ZIM7 Q8ZIN2 Q8ZIN3 Q8ZIN5 Q8ZIP0 Q8ZIP1 Q8ZIQ2
Q8ZIQ3 Q8ZIQ4 Q8ZIR0 Q8ZIV4 Q8ZIV7 Q8ZIW2 Q8ZIW3 Q8ZIW4 Q8ZIW9 Q8ZIX0 Q8ZIX3
Q8ZIY0 Q8ZIY3 Q8ZIY4 Q8ZIZ9 Q8ZJ00 Q8ZJ02 Q8ZJ03 Q8ZJ06 Q8ZJ07 Q8ZJ10 Q8ZJ14
Q8ZJ20 Q8ZJ74 Q8ZJ77 Q8ZJ79 Q8ZJ80 Q8ZJ81 Q8ZJ87 Q8ZJ88 Q8ZJ89 Q8ZJ90 Q8ZJ91
Q8ZJ93 Q8ZJ95 Q8ZJ96 Q8ZJ98 Q8ZJ99 Q8ZJA0 Q8ZJA1 Q8ZJA3 Q8ZJA4 Q8ZJA5 Q8ZJA6
Q8ZJA7 Q8ZJA8 Q8ZJA9 Q8ZJB2 Q8ZJB3 Q8ZJB4 Q8ZJB5 Q8ZJB6 Q8ZJB7 Q8ZJB8 Q8ZJC1
Q8ZJD4 Q8ZJF2 Q8ZJF3 Q8ZJF6 Q8ZJF7 Q8ZJG8 Q8ZJG9 Q8ZJH2 Q8ZJI0 Q8ZJI2 Q8ZJI3
Q8ZJI8 Q8ZJJ4 Q8ZJJ5 Q8ZJJ7 Q8ZJK9 Q8ZJL6 Q8ZJM7 Q8ZJN0 Q8ZJN2 Q8ZJN9 Q8ZJP0
Q8ZJP1 Q8ZJP2 Q8ZJP3 Q8ZJP5 Q8ZJP6 Q8ZJP7 Q8ZJP8 Q8ZJQ2 Q8ZJQ3 Q8ZJR9 Q8ZJS0
Q8ZJS4 Q8ZJT1 Q8ZJT2 Q8ZJT3 Q8ZJU8 Q8ZJV1 Q8ZJV7 Q8ZJV8 Q8ZJW4 Q8ZJW6 Q8ZJY9
Q8ZK08 Q8ZK29 Q8ZK31 Q8ZK33 Q8ZK35 Q8ZK57 Q8ZK58 Q8ZK67 Q8ZK71 Q8ZK76 Q8ZK80
Q8ZK81 Q8ZK82 Q8ZK85 Q8ZK86 Q8ZK87 Q8ZKA2 Q8ZKA3 Q8ZKA8 Q8ZKB0 Q8ZKB8 Q8ZKE6
Q8ZKF6 Q8ZKH4 Q8ZKI0 Q8ZKI4 Q8ZKL1 Q8ZKL6 Q8ZKL8 Q8ZKL9 Q8ZKM0 Q8ZKM8 Q8ZKP1
Q8ZKP3 Q8ZKP7 Q8ZKP9 Q8ZKQ0 Q8ZKQ5 Q8ZKQ6 Q8ZKR0 Q8ZKR2 Q8ZKS1 Q8ZKS7 Q8ZKT0
Q8ZKT7 Q8ZKU8 Q8ZKU9 Q8ZKW0 Q8ZKW2 Q8ZKW3 Q8ZKW4 Q8ZKW6 Q8ZKX1 Q8ZKY0 Q8ZKY3
Q8ZKZ1 Q8ZKZ8 Q8ZL03 Q8ZL08 Q8ZL42 Q8ZL46 Q8ZL48 Q8ZL52 Q8ZL56 Q8ZL58 Q8ZL66
Q8ZL67 Q8ZL69 Q8ZL81 Q8ZL83 Q8ZL90 Q8ZL95 Q8ZL96 Q8ZLB5 Q8ZLC8 Q8ZLD7 Q8ZLE2
Q8ZLG5 Q8ZLG6 Q8ZLI0 Q8ZLI3 Q8ZLI7 Q8ZLI8 Q8ZLI9 Q8ZLJ2 Q8ZLJ4 Q8ZLK5 Q8ZLL5
Q8ZLL7 Q8ZLL8 Q8ZLL9 Q8ZLM1 Q8ZLM5 Q8ZLM6 Q8ZLM7 Q8ZLN0 Q8ZLN1 Q8ZLQ6 Q8ZLQ7
Q8ZLQ8 Q8ZLR8 Q8ZLS5 Q8ZLS7 Q8ZLT0 Q8ZLT1 Q8ZLT2 Q8ZLT3 Q8ZLU2 Q8ZLU3 Q8ZLW5
Q8ZLX5 Q8ZLX7 Q8ZLY4 Q8ZM03 Q8ZM06 Q8ZM23 Q8ZM27 Q8ZM36 Q8ZM40 Q8ZM45 Q8ZM46
Q8ZM50 Q8ZM51 Q8ZM71 Q8ZM75 Q8ZM76 Q8ZM79 Q8ZM80 Q8ZM82 Q8ZM86 Q8ZM98 Q8ZMA8
Q8ZMA9 Q8ZMB8 Q8ZMC4 Q8ZMC5 Q8ZMC6 Q8ZMC9 Q8ZMD3 Q8ZMD5 Q8ZME1 Q8ZMF5 Q8ZMF6
Q8ZMF7 Q8ZMF8 Q8ZMF9 Q8ZMG5 Q8ZMH3 Q8ZMJ6 Q8ZMJ7 Q8ZMJ9 Q8ZMK5 Q8ZMK6 Q8ZMS4
Q8ZMX2 Q8ZMX8 Q8ZMX9 Q8ZN19 Q8ZN25 Q8ZN40 Q8ZN41 Q8ZN42 Q8ZN52 Q8ZN58 Q8ZN60
Q8ZN71 Q8ZN74 Q8ZN75 Q8ZN83 Q8ZN89 Q8ZNB2 Q8ZNB9 Q8ZNC4 Q8ZND6 Q8ZNE0 Q8ZNE8
Q8ZNE9 Q8ZNF5 Q8ZNF9 Q8ZNG0 Q8ZNK6 Q8ZNL7 Q8ZNL8 Q8ZNM0 Q8ZNM4 Q8ZNN2 Q8ZNN4

Q8ZNN5 Q8ZNR5 Q8ZNR6 Q8ZNS5 Q8ZNT7 Q8ZNV0 Q8ZNV1 Q8ZNV2 Q8ZNW0 Q8ZNW5 Q8ZNY9
Q8ZNZ2 Q8ZP11 Q8ZP12 Q8ZP19 Q8ZP43 Q8ZP48 Q8ZP51 Q8ZP52 Q8ZP54 Q8ZP65 Q8ZP84
Q8ZP88 Q8ZPB6 Q8ZPC9 Q8ZPD3 Q8ZPE8 Q8ZPI2 Q8ZPK6 Q8ZPL7 Q8ZPL9 Q8ZPM8 Q8ZPM9
Q8ZPN1 Q8ZPQ1 Q8ZPR4 Q8ZPS4 Q8ZPS5 Q8ZPS6 Q8ZPS9 Q8ZPU0 Q8ZPU1 Q8ZPU5 Q8ZPU6
Q8ZPU8 Q8ZPU9 Q8ZPV0 Q8ZPV1 Q8ZPV2 Q8ZPV5 Q8ZPV9 Q8ZPW3 Q8ZPX9 Q8ZPZ1 Q8ZPZ4
Q8ZPZ9 Q8ZQ06 Q8ZQ07 Q8ZQ16 Q8ZQ23 Q8ZQ40 Q8ZQ47 Q8ZQ51 Q8ZQ64 Q8ZQ68 Q8ZQ73
Q8ZQ75 Q8ZQB6 Q8ZQB7 Q8ZQC4 Q8ZQE8 Q8ZQJ5 Q8ZQM1 Q8ZQN4 Q8ZQN7 Q8ZQQ1 Q8ZQQ4
Q8ZQQ5 Q8ZQQ6 Q8ZQQ7 Q8ZQQ9 Q8ZQR1 Q8ZQR2 Q8ZQR4 Q8ZQS2 Q8ZQV6 Q8ZQX1 Q8ZQX5
Q8ZQX7 Q8ZQY4 Q8ZQZ6 Q8ZQZ8 Q8ZR03 Q8ZR04 Q8ZR14 Q8ZR29 Q8ZR41 Q8ZR68 Q8ZR86
Q8ZR87 Q8ZR88 Q8ZRA1 Q8ZRA2 Q8ZRB6 Q8ZRC0 Q8ZRC9 Q8ZRD1 Q8ZRD3 Q8ZRD8 Q8ZRD9
Q8ZRE7 Q8ZRF7 Q8ZRI9 Q8ZRJ0 Q8ZRM2 Q8ZRM4 Q8ZRM7 Q8ZRN5 Q8ZRP3 Q8ZRP4 Q8ZRQ6
Q8ZRQ7 Q8ZRQ8 Q8ZRR0 Q8ZRR1 Q8ZRS3 Q8ZRS4 Q8ZRS5 Q8ZRS6 Q8ZRS8 Q8ZRT5 Q8ZRU1
Q8ZRU2 Q8ZRU4 Q8ZRU7 Q8ZRU8 Q8ZRU9 Q8ZRV8 Q8ZRV9 Q8ZRW8 Q8ZRW9 Q8ZRX0 Q8ZRX2
Q8ZRX3 Q8ZRX4 Q8ZRX5 Q8ZRX6 Q8ZRX8 Q8ZRX9 Q8ZRY7 Q8ZRZ0 Q8ZS17 Q8ZSD2 Q8ZSM3
Q8ZSP2 Q8ZST4 Q8ZST5 Q8ZSU2 Q8ZSU3 Q8ZSU7 Q8ZSV4 Q8ZSV6 Q8ZSY4 Q8ZSY7 Q8ZT00
Q8ZT04 Q8ZT09 Q8ZT11 Q8ZT21 Q8ZT50 Q8ZT61 Q8ZT64 Q8ZT69 Q8ZT70 Q8ZT87 Q8ZT92
Q8ZTA5 Q8ZTA6 Q8ZTA7 Q8ZTA8 Q8ZTB0 Q8ZTB2 Q8ZTB5 Q8ZTB8 Q8ZTC3 Q8ZTD3 Q8ZTD4
Q8ZTE1 Q8ZTE3 Q8ZTE6 Q8ZTE8 Q8ZTF1 Q8ZTF9 Q8ZTG2 Q8ZTG3 Q8ZTH0 Q8ZTI9 Q8ZTJ1
Q8ZTJ2 Q8ZTJ3 Q8ZTJ4 Q8ZTJ7 Q8ZTM9 Q8ZTP3 Q8ZTP4 Q8ZTQ5 Q8ZTQ7 Q8ZTR0 Q8ZTR2
Q8ZTS6 Q8ZTT3 Q8ZTT4 Q8ZTT5 Q8ZTT9 Q8ZTU0 Q8ZTU1 Q8ZTU2 Q8ZTU5 Q8ZTV0 Q8ZTV1
Q8ZTX6 Q8ZTX7 Q8ZTY0 Q8ZTY1 Q8ZTY4 Q8ZTY5 Q8ZTY6 Q8ZTZ0 Q8ZU20 Q8ZU24 Q8ZU30
Q8ZU33 Q8ZU41 Q8ZU54 Q8ZU56 Q8ZU75 Q8ZU79 Q8ZU80 Q8ZU81 Q8ZU95 Q8ZU97 Q8ZU99
Q8ZUA0 Q8ZUE9 Q8ZUF0 Q8ZUF6 Q8ZUG2 Q8ZUJ3 Q8ZUR4 Q8ZUR9 Q8ZV04 Q8ZV05 Q8ZV07
Q8ZV18 Q8ZV44 Q8ZV45 Q8ZV53 Q8ZV65 Q8ZV66 Q8ZV67 Q8ZVA7 Q8ZVB2 Q8ZVB5 Q8ZVE0
Q8ZVE4 Q8ZVF3 Q8ZVG2 Q8ZVI1 Q8ZVI9 Q8ZVK6 Q8ZVL3 Q8ZVM0 Q8ZVM1 Q8ZVM8 Q8ZVM9
Q8ZVN0 Q8ZVN1 Q8ZVN2 Q8ZVN4 Q8ZVP1 Q8ZVP4 Q8ZVQ3 Q8ZVQ9 Q8ZVR0 Q8ZVR4 Q8ZVR7
Q8ZVU9 Q8ZVV7 Q8ZVV9 Q8ZVW0 Q8ZVW1 Q8ZVX7 Q8ZW23 Q8ZW35 Q8ZW36 Q8ZW37 Q8ZW40
Q8ZW41 Q8ZW45 Q8ZW46 Q8ZW50 Q8ZW51 Q8ZW52 Q8ZW59 Q8ZW77 Q8ZW80 Q8ZW89 Q8ZW90
Q8ZWC4 Q8ZWD6 Q8ZWD8 Q8ZWG7 Q8ZWH7 Q8ZWI0 Q8ZWI1 Q8ZWK3 Q8ZWK4 Q8ZWK6 Q8ZWK7
Q8ZWL3 Q8ZWL4 Q8ZWP7 Q8ZWP9 Q8ZWQ7 Q8ZWS2 Q8ZWS3 Q8ZWT2 Q8ZWT4 Q8ZWT5 Q8ZWT6
Q8ZWU4 Q8ZWV0 Q8ZWV9 Q8ZWW5 Q8ZWY4 Q8ZWZ1 Q8ZX17 Q8ZX20 Q8ZX25 Q8ZX28 Q8ZX33
Q8ZX61 Q8ZX62 Q8ZXA6 Q8ZXD8 Q8ZXE0 Q8ZXK6 Q8ZXK7 Q8ZXL4 Q8ZXL6 Q8ZXM4 Q8ZXN8
Q8ZXN9 Q8ZXR6 Q8ZXR9 Q8ZXT3 Q8ZXT5 Q8ZXT6 Q8ZXV9 Q8ZY04 Q8ZY09 Q8ZY14 Q8ZY15
Q8ZY16 Q8ZY17 Q8ZY18 Q8ZY35 Q8ZY36 Q8ZY39 Q8ZY40 Q8ZY44 Q8ZY48 Q8ZY49 Q8ZY96
Q8ZYA5 Q8ZYB3 Q8ZYB4 Q8ZYD2 Q8ZYD8 Q8ZYE0 Q8ZYE1 Q8ZYE2 Q8ZYE5 Q8ZYF2 Q8ZYF4
Q8ZYF5 Q8ZYF6 Q8ZYF8 Q8ZYF9 Q8ZYH0 Q8ZYH2 Q8ZYH7 Q8ZYJ4 Q8ZYK3 Q8ZYK4 Q8ZYK5
Q8ZYL6 Q8ZYM4 Q8ZYM8 Q8ZYN0 Q8ZYN2 Q8ZYN5 Q8ZYP2 Q8ZYP4 Q8ZYP7 Q8ZYP9 Q8ZYQ0
Q8ZYQ1 Q8ZYQ2 Q8ZYQ3 Q8ZYQ4 Q8ZYQ6 Q8ZYQ7 Q8ZYR0 Q8ZYR9 Q8ZYS2 Q8ZYT7 Q8ZYU6
Q8ZYV1 Q8ZYV6 Q8ZYW1 Q8ZYW7 Q8ZYX2 Q8ZZ93 Q8ZZA9 Q8ZZB0 Q8ZZC0 Q8ZZC1 Q8ZZC7
Q8ZZJ7 Q8ZZJ9 Q8ZZK1 Q8ZZK5 Q8ZZL8 Q8ZZL9 Q8ZZM5 Q8ZZP2 Q8ZZP3 Q8ZZU1 Q8ZZU2
Q8ZZW3 Q8ZZX3 Q8ZZX6 Q8ZZZ0 Q90029 Q90030 Q90033 Q90038 Q90054 Q90055 Q90056
Q90162 Q90165 Q90173 Q90176 Q90177 Q90178 Q90179 Q90185 Q90186 Q90229 Q90233
Q90257 Q90259 Q90260 Q90267 Q90268 Q90270 Q90271 Q90303 Q90321 Q90327 Q90339
Q90341 Q90342 Q90346 Q90368 Q90370 Q90382 Q90384 Q90416 Q90417 Q90421 Q90423

Q90435 Q90459 Q90469 Q90470 Q90471 Q90472 Q90476 Q90477 Q90481 Q90485 Q90486
Q90487 Q90497 Q90511 Q90512 Q90595 Q90596 Q90625 Q90635 Q90640 Q90641 Q90643
Q90655 Q90660 Q90667 Q90683 Q90688 Q90690 Q90691 Q90701 Q90705 Q90718 Q90732
Q90733 Q90788 Q90837 Q90850 Q90871 Q90875 Q90876 Q90879 Q90881 Q90888 Q90891
Q90932 Q90941 Q90963 Q90964 Q90966 Q90977 Q90988 Q90VZ3 Q90VZ9 Q90W04 Q90W33
Q90W58 Q90W95 Q90WD0 Q90WE3 Q90WG1 Q90WG2 Q90WG6 Q90WM5 Q90WN4 Q90WN5 Q90WN7
Q90WR8 Q90WU3 Q90X25 Q90X38 Q90X89 Q90X97 Q90X99 Q90XB3 Q90XC2 Q90XD2 Q90XE6
Q90XG0 Q90XH3 Q90XN9 Q90XP0 Q90XV6 Q90XV7 Q90XV8 Q90XV9 Q90XW5 Q90XW6 Q90Y35
Q90YB1 Q90YC0 Q90YG6 Q90YI3 Q90YI8 Q90YJ9 Q90YK7 Q90YK8 Q90YK9 Q90YL3 Q90YP9
Q90YQ0 Q90YQ1 Q90YQ2 Q90YQ4 Q90YQ5 Q90YQ6 Q90YQ7 Q90YQ8 Q90YR4 Q90YR6 Q90YR7
Q90YR8 Q90YS0 Q90YS7 Q90YS9 Q90YT0 Q90YT1 Q90YT2 Q90YT3 Q90YT4 Q90YT5 Q90YT6
Q90YT7 Q90YU3 Q90YU5 Q90YU6 Q90YU8 Q90YU9 Q90YV0 Q90YV2 Q90YV8 Q90YW0 Q90YW1
Q90YW2 Q90YX1 Q90YY5 Q90Z12 Q90Z72 Q90Z80 Q90ZA1 Q90ZE2 Q90ZH7 Q90ZH8 Q90ZH9
Q90ZK2 Q90ZL1 Q90ZL4 Q90ZS6 Q90ZT7 Q90ZY4 Q90ZZ5 Q90ZZ6 Q91012 Q91060 Q91061
Q910F1 Q91154 Q91169 Q91185 Q91191 Q91192 Q91193 Q91196 Q91197 Q91205 Q91206
Q91233 Q91240 Q91241 Q91242 Q91250 Q91279 Q91284 Q91289 Q91291 Q91293 Q91309
Q91311 Q91312 Q91314 Q91315 Q91316 Q91317 Q91318 Q91320 Q91321 Q91348 Q91365
Q91372 Q91375 Q91379 Q91392 Q91398 Q91399 Q913A7 Q91400 Q91424 Q91428 Q91431
Q91437 Q91473 Q91482 Q91483 Q91487 Q914F9 Q914G0 Q914G1 Q914G2 Q914G3 Q914G4
Q914G5 Q914G6 Q914G7 Q914G8 Q914G9 Q914H0 Q914H1 Q914H2 Q914H3 Q914H4 Q914H8
Q914I2 Q914I4 Q914I5 Q914I6 Q914I7 Q914I8 Q914J0 Q914J1 Q914J2 Q914J4 Q914J5
Q914J6 Q914J7 Q914J8 Q914J9 Q914K0 Q914K1 Q914K2 Q914K3 Q914K4 Q914K5 Q914K6
Q914K7 Q914K8 Q914K9 Q914L0 Q914L1 Q914L2 Q914L3 Q914L4 Q914L5 Q914L6 Q914L7
Q914L8 Q914L9 Q914M1 Q914M2 Q914M3 Q914M4 Q914M5 Q914M6 Q914M7 Q914M8 Q914M9
Q914N0 Q914N2 Q914N3 Q91517 Q91518 Q91553 Q91554 Q91555 Q91560 Q91573 Q91574
Q91575 Q91579 Q91584 Q91605 Q91617 Q91622 Q91628 Q91636 Q91637 Q91642 Q91660
Q91661 Q91677 Q91678 Q91684 Q91685 Q91689 Q91690 Q91695 Q91712 Q91715 Q91724
Q91727 Q91731 Q91754 Q91765 Q91770 Q91771 Q91775 Q91783 Q91784 Q91785 Q91790
Q91819 Q91820 Q91822 Q91830 Q91836 Q91837 Q91853 Q91854 Q91857 Q91858 Q91876
Q91879 Q91880 Q91896 Q91898 Q918W2 Q918W3 Q91901 Q91903 Q91904 Q91905 Q91907
Q91924 Q91926 Q91955 Q91957 Q91974 Q91975 Q91981 Q91989 Q91994 Q91998 Q91A21
Q91A24 Q91A25 Q91A27 Q91B74 Q91BI6 Q91C36 Q91CD5 Q91D83 Q91DD4 Q91DD6 Q91DE0
Q91DE1 Q91DM0 Q91DR9 Q91DS2 Q91DS3 Q91E95 Q91E96 Q91EK1 Q91EK2 Q91EK4 Q91F58
Q91F59 Q91F66 Q91F72 Q91F73 Q91F75 Q91F84 Q91F86 Q91F87 Q91F88 Q91F89 Q91F97
Q91F99 Q91FA2 Q91FA3 Q91FA5 Q91FB0 Q91FB1 Q91FB6 Q91FC1 Q91FC9 Q91FD1 Q91FD3
Q91FD5 Q91FD6 Q91FD9 Q91FE0 Q91FE4 Q91FE6 Q91FE8 Q91FE9 Q91FF0 Q91FF5 Q91FG5
Q91FG6 Q91FG9 Q91FH4 Q91FH5 Q91FH6 Q91FH7 Q91FH8 Q91FI1 Q91FI2 Q91FI4 Q91FI9
Q91FJ2 Q91FJ5 Q91FK2 Q91FK7 Q91FK9 Q91FL1 Q91FL5 Q91FL8 Q91FM2 Q91FM3 Q91FN1
Q91FN5 Q91FN7 Q91FN9 Q91FP0 Q91FP2 Q91FQ0 Q91FQ1 Q91FQ5 Q91FR7 Q91FR8 Q91FR9
Q91FS1 Q91FS2 Q91FS5 Q91FS6 Q91FS8 Q91FT0 Q91FT1 Q91FT2 Q91FT4 Q91FT6 Q91FU3
Q91FU4 Q91FU6 Q91FV0 Q91FV2 Q91FV3 Q91FV9 Q91FW0 Q91FW4 Q91FW5 Q91FW9 Q91FX0
Q91FX2 Q91FX3 Q91G49 Q91G54 Q91G55 Q91G56 Q91G57 Q91G65 Q91G67 Q91G70 Q91G76
Q91G88 Q91GM8 Q91HJ9 Q91HK4 Q91HK5 Q91ID2 Q91ID3 Q91ID4 Q91ID5 Q91ID6 Q91ID7
Q91ID8 Q91ID9 Q91IE0 Q91IE1 Q91J26 Q91J29 Q91M93 Q91M95 Q91M96 Q91MF7 Q91MG1

Q91MU3 Q91MX5 Q91PB2 Q91PK4 Q91PQ1 Q91PQ2 Q91PQ3 Q91QN9 Q91QZ1 Q91QZ2 Q91QZ3
Q91QZ8 Q91RA4 Q91RA6 Q91TW8 Q91TW9 Q91UM2 Q91UN2 Q91V16 Q91V27 Q91V51 Q91V64
Q91V76 Q91V81 Q91V83 Q91VB3 Q91VB4 Q91VC0 Q91VC3 Q91VC7 Q91VE1 Q91VE6 Q91VF2
Q91VI7 Q91VJ1 Q91VJ5 Q91VK1 Q91VN1 Q91VN2 Q91VN6 Q91VR5 Q91VT1 Q91VU6 Q91VU7
Q91VV4 Q91VW9 Q91VX2 Q91VY5 Q91VY6 Q91VY9 Q91W18 Q91W40 Q91W45 Q91W59 Q91W61
Q91W63 Q91W67 Q91W78 Q91W82 Q91W96 Q91WA1 Q91WB7 Q91WC0 Q91WC1 Q91WD1 Q91WD4
Q91WE2 Q91WE3 Q91WG3 Q91WJ7 Q91WJ8 Q91WK0 Q91WK1 Q91WK2 Q91WK5 Q91WM1 Q91WQ5
Q91WQ9 Q91WS4 Q91WT4 Q91WT8 Q91WU5 Q91X20 Q91X44 Q91X45 Q91X83 Q91X91 Q91XA5
Q91XA8 Q91XC8 Q91XE0 Q91XF0 Q91XI1 Q91XJ0 Q91XQ0 Q91XW8 Q91Y26 Q91YD3 Q91YD6
Q91YE3 Q91YE5 Q91YE7 Q91YE9 Q91YJ5 Q91YK0 Q91YK2 Q91YL3 Q91YM4 Q91YN0 Q91YP1
Q91YP3 Q91YQ7 Q91YR5 Q91YR7 Q91YU3 Q91YU8 Q91YY4 Q91Z31 Q91Z38 Q91Z46 Q91Z49
Q91Z53 Q91Z58 Q91Z63 Q91Z83 Q91Z96 Q91ZD1 Q91ZD6 Q91ZE0 Q91ZE9 Q91ZF0 Q91ZF7
Q91ZG2 Q91ZJ0 Q91ZJ5 Q91ZK4 Q91ZR3 Q91ZR4 Q91ZS8 Q91ZW1 Q91ZW3 Q91ZW6 Q91ZY6
Q91ZZ3 Q92005 Q92012 Q92020 Q92021 Q92037 Q92044 Q92047 Q92051 Q92055 Q92060
Q92076 Q92091 Q920B9 Q920D2 Q920F6 Q920I9 Q920J3 Q920J4 Q920M2 Q920M5 Q920N2
Q920P5 Q920Q8 Q920R4 Q920S3 Q92117 Q92122 Q92142 Q92146 Q92155 Q92161 Q92162
Q92172 Q92192 Q92193 Q92195 Q921A3 Q921A4 Q921B4 Q921C3 Q921D4 Q921E6 Q921F4
Q921G6 Q921G8 Q921H9 Q921I2 Q921J4 Q921K9 Q921L7 Q921M3 Q921P9 Q921U8 Q921W4
Q921X6 Q921Y2 Q921Z5 Q92203 Q92209 Q92210 Q92236 Q92241 Q92243 Q92247 Q92258
Q92259 Q92260 Q92263 Q92268 Q92269 Q922B2 Q922C1 Q922D8 Q922E6 Q922G2 Q922G7
Q922H1 Q922H2 Q922H4 Q922H7 Q922L6 Q922M5 Q922M7 Q922P9 Q922Q2 Q922Q4 Q922R5
Q922U1 Q922X9 Q922Y1 Q92317 Q92325 Q92335 Q92342 Q92345 Q92346 Q92347 Q92348
Q92350 Q92351 Q92352 Q92355 Q92358 Q92359 Q92360 Q92361 Q92363 Q92368 Q92370
Q92372 Q92373 Q92374 Q92375 Q92376 Q92378 Q92379 Q92380 Q92383 Q92392 Q92396
Q92399 Q923B0 Q923B1 Q923B3 Q923D4 Q923G2 Q923J6 Q923K4 Q923K9 Q923M1 Q923S8
Q923T7 Q923V4 Q923Z3 Q923Z4 Q92403 Q92407 Q92410 Q92411 Q92412 Q92413 Q92429
Q92441 Q92445 Q92447 Q92448 Q92450 Q92481 Q924C5 Q924H7 Q924H9 Q924K1 Q924K2
Q924M5 Q924M7 Q924R9 Q924S9 Q924W5 Q924W6 Q924W9 Q924X7 Q92519 Q92526 Q92527
Q92551 Q92561 Q92564 Q925E7 Q925F0 Q925F4 Q925G0 Q925G1 Q925H2 Q925H3 Q925H6
Q925H7 Q925I0 Q925Q8 Q925T0 Q925W4 Q925W5 Q925W7 Q925X2 Q925X3 Q925X7 Q925Y6
Q925Z2 Q92618 Q92623 Q92628 Q92664 Q92670 Q92698 Q926A5 Q926A6 Q926A7 Q926B0
Q926D8 Q926Q4 Q926R0 Q926R1 Q926R2 Q926R3 Q926T3 Q926U7 Q926U8 Q926V6 Q926V7
Q926W4 Q926X0 Q926X7 Q926Y3 Q926Y8 Q926Z9 Q92753 Q92759 Q92764 Q92771 Q92782
Q92784 Q92786 Q927A3 Q927A7 Q927D8 Q927D9 Q927E4 Q927E5 Q927E6 Q927E8 Q927I2
Q927I5 Q927I6 Q927J4 Q927K7 Q927K9 Q927L2 Q927L4 Q927L6 Q927L7 Q927L8 Q927L9
Q927M2 Q927M3 Q927M4 Q927M5 Q927M6 Q927M8 Q927N4 Q927P1 Q927P2 Q927P3 Q927P4
Q927Q9 Q927S9 Q927T2 Q927T3 Q927T4 Q927U1 Q927U8 Q927U9 Q927V0 Q927V4 Q927V5
Q927W7 Q927X3 Q927Y4 Q92802 Q92826 Q92851 Q92858 Q92871 Q92886 Q928A4 Q928A5
Q928B0 Q928B2 Q928B5 Q928B9 Q928C0 Q928C1 Q928C2 Q928C4 Q928D5 Q928I0 Q928I1
Q928I2 Q928K4 Q928L3 Q928P1 Q928P3 Q928R6 Q928R9 Q928S6 Q928V0 Q928X3 Q928X8
Q92904 Q92947 Q92949 Q92966 Q92979 Q92988 Q92989 Q92990 Q929A8 Q929A9 Q929E8
Q929F0 Q929F3 Q929G1 Q929G2 Q929G7 Q929G8 Q929H5 Q929I1 Q929L4 Q929Q1 Q929Q6
Q929R8 Q929R9 Q929S8 Q929S9 Q929U3 Q929U9 Q929V0 Q929W5 Q929W8 Q929X1 Q929X5
Q929X6 Q929X9 Q929Y1 Q929Y7 Q929Z0 Q929Z2 Q929Z6 Q929Z7 Q92A19 Q92A25 Q92A26

Q92A27 Q92A28 Q92A29 Q92A32 Q92A38 Q92A40 Q92A41 Q92A47 Q92A53 Q92A54 Q92A57
Q92A58 Q92A69 Q92A71 Q92A75 Q92A79 Q92A80 Q92A81 Q92A83 Q92A85 Q92A87 Q92AA1
Q92AA2 Q92AA3 Q92AA6 Q92AA7 Q92AA8 Q92AB2 Q92AB7 Q92AB9 Q92AC0 Q92AC4 Q92AC7
Q92AD1 Q92AD2 Q92AD4 Q92AE2 Q92AE8 Q92AE9 Q92AF1 Q92AF2 Q92AG8 Q92AH0 Q92AH1
Q92AH2 Q92AH3 Q92AH4 Q92AH5 Q92AH6 Q92AH7 Q92AI1 Q92AI2 Q92AI5 Q92AI9 Q92AK3
Q92AL1 Q92AL5 Q92AL6 Q92AM1 Q92AM8 Q92AN6 Q92AN8 Q92AN9 Q92AP1 Q92AP3 Q92AP4
Q92AQ0 Q92AQ2 Q92AQ3 Q92AQ4 Q92AQ7 Q92AR6 Q92AT0 Q92AV0 Q92AV4 Q92AV8 Q92AW7
Q92AX3 Q92AX5 Q92AX9 Q92AY5 Q92AY8 Q92AZ5 Q92AZ9 Q92B01 Q92B02 Q92B78 Q92B79
Q92B80 Q92B81 Q92B82 Q92B90 Q92B94 Q92BA1 Q92BA4 Q92BB1 Q92BB2 Q92BB6 Q92BB7
Q92BB8 Q92BB9 Q92BC0 Q92BC1 Q92BC5 Q92BD4 Q92BD8 Q92BE4 Q92BF1 Q92BF2 Q92BF3
Q92BF5 Q92BF6 Q92BF7 Q92BF8 Q92BG1 Q92BG2 Q92BG5 Q92BG9 Q92BH2 Q92BH6 Q92BH7
Q92BH9 Q92BI0 Q92BI1 Q92BI2 Q92BI3 Q92BI4 Q92BJ1 Q92BJ3 Q92BJ4 Q92BK1 Q92BK9
Q92BL1 Q92BL2 Q92BL6 Q92BL9 Q92BM5 Q92BN6 Q92BN7 Q92BN8 Q92BN9 Q92BP0 Q92BP5
Q92BQ0 Q92BQ1 Q92BQ2 Q92BQ4 Q92BQ5 Q92BQ6 Q92BQ8 Q92BR0 Q92BR1 Q92BR6 Q92BS0
Q92BT5 Q92BV2 Q92BV3 Q92BV7 Q92BV8 Q92BX1 Q92BX2 Q92BX4 Q92BY5 Q92BY8 Q92BZ0
Q92BZ2 Q92BZ3 Q92BZ4 Q92BZ5 Q92C04 Q92C05 Q92C06 Q92C22 Q92C23 Q92C24 Q92C26
Q92C27 Q92C29 Q92C33 Q92C34 Q92C35 Q92C37 Q92C39 Q92C40 Q92C49 Q92C50 Q92C59
Q92C72 Q92C73 Q92C74 Q92C75 Q92C76 Q92C79 Q92C84 Q92C85 Q92CE4 Q92CE5 Q92CH0
Q92CH1 Q92CH2 Q92CH5 Q92CH6 Q92CI0 Q92CI1 Q92CI6 Q92CI7 Q92CK0 Q92CL4 Q92CL7
Q92CN1 Q92CN9 Q92CU0 Q92CU3 Q92CV0 Q92CW3 Q92CW8 Q92CX8 Q92CY2 Q92CY3 Q92CZ4
Q92D10 Q92D11 Q92D33 Q92D46 Q92D47 Q92D53 Q92D57 Q92D64 Q92D89 Q92D94 Q92DB7
Q92DC2 Q92DC9 Q92DF6 Q92DG5 Q92DI8 Q92DM1 Q92DN4 Q92DS8 Q92DU2 Q92DX9 Q92E41
Q92E58 Q92E82 Q92E83 Q92E84 Q92E85 Q92E86 Q92E87 Q92E88 Q92E89 Q92E90 Q92EB7
Q92EC2 Q92EC3 Q92EF1 Q92EG6 Q92EG7 Q92EH1 Q92EH3 Q92EH6 Q92EK7 Q92EQ0 Q92EQ4
Q92EQ5 Q92EQ6 Q92EQ7 Q92ES0 Q92ES1 Q92ES6 Q92ET8 Q92ET9 Q92EU2 Q92EU3 Q92EU4
Q92EU5 Q92EU6 Q92EW5 Q92EW7 Q92EY7 Q92F22 Q92F24 Q92F26 Q92F36 Q92F38 Q92F39
Q92F40 Q92F41 Q92F42 Q92F44 Q92F47 Q92F53 Q92F54 Q92F56 Q92F59 Q92F62 Q92F64
Q92F65 Q92F66 Q92F68 Q92F70 Q92F71 Q92F76 Q92F77 Q92F79 Q92F90 Q92F97 Q92F99
Q92FK7 Q92FQ5 Q92FQ7 Q92FR5 Q92FR6 Q92FR7 Q92FS9 Q92FU8 Q92FV2 Q92FV3 Q92FV4
Q92FV8 Q92FW6 Q92FW7 Q92FW8 Q92FX0 Q92FY0 Q92FZ0 Q92FZ5 Q92FZ7 Q92FZ8 Q92FZ9
Q92G00 Q92G10 Q92G11 Q92G13 Q92G19 Q92G20 Q92G23 Q92G24 Q92G30 Q92G31 Q92G39
Q92G41 Q92G55 Q92G57 Q92G58 Q92G77 Q92G90 Q92GA7 Q92GB1 Q92GB2 Q92GB7 Q92GC0
Q92GC8 Q92GC9 Q92GE1 Q92GE8 Q92GF9 Q92GG0 Q92GG1 Q92GH4 Q92GH5 Q92GH6 Q92GH7
Q92GH8 Q92GI1 Q92GI2 Q92GL1 Q92GL2 Q92GL5 Q92GM6 Q92GM7 Q92GN5 Q92GQ2 Q92GQ4
Q92GR5 Q92GR9 Q92GS5 Q92GS6 Q92GT4 Q92GT5 Q92GU2 Q92GU6 Q92GV0 Q92GV1 Q92GV5
Q92GW0 Q92GW4 Q92GW5 Q92GW6 Q92GW7 Q92GW8 Q92GW9 Q92GX0 Q92GX1 Q92GX2 Q92GX3
Q92GX4 Q92GX5 Q92GX6 Q92GX7 Q92GX8 Q92GX9 Q92GY0 Q92GY1 Q92GY2 Q92GY3 Q92GY4
Q92GY5 Q92GY7 Q92GY8 Q92GY9 Q92GZ0 Q92GZ1 Q92GZ2 Q92GZ5 Q92GZ7 Q92H03 Q92H04
Q92H06 Q92H07 Q92H16 Q92H19 Q92H23 Q92H35 Q92H36 Q92H37 Q92H38 Q92H40 Q92H41
Q92H48 Q92H49 Q92H56 Q92H58 Q92H59 Q92H60 Q92H62 Q92H74 Q92H76 Q92H87 Q92H88
Q92H89 Q92H94 Q92HA7 Q92HB1 Q92HB3 Q92HB4 Q92HC9 Q92HD0 Q92HD1 Q92HD2 Q92HD3
Q92HD4 Q92HD5 Q92HE8 Q92HF3 Q92HF4 Q92HF5 Q92HF9 Q92HG1 Q92HG4 Q92HG8 Q92HH1
Q92HI5 Q92HI8 Q92HJ3 Q92HK6 Q92HK9 Q92HL2 Q92HL4 Q92HL7 Q92HM3 Q92HM4 Q92HM5
Q92HM7 Q92HP1 Q92HR1 Q92HR5 Q92HS0 Q92HT9 Q92HU0 Q92HU7 Q92HU9 Q92HV3 Q92HV5

Q92HV6 Q92HV7 Q92HW4 Q92HY3 Q92HY4 Q92HZ1 Q92HZ6 Q92I02 Q92I08 Q92I11 Q92I14
Q92I21 Q92I22 Q92I24 Q92I25 Q92I26 Q92I30 Q92I35 Q92I37 Q92I38 Q92I39 Q92I41
Q92I60 Q92I70 Q92I71 Q92I74 Q92I77 Q92I78 Q92I83 Q92I87 Q92I89 Q92I92 Q92I93
Q92I97 Q92IA0 Q92IB5 Q92IB8 Q92IC9 Q92IE0 Q92IE5 Q92IE7 Q92IF2 Q92IF9 Q92IG3
Q92IG5 Q92IH1 Q92IH2 Q92IH7 Q92II6 Q92II7 Q92IJ0 Q92IK8 Q92IL0 Q92IL5 Q92IL8
Q92IL9 Q92IM3 Q92IM4 Q92IQ2 Q92IQ7 Q92IR7 Q92IS2 Q92IS6 Q92IT7 Q92IT8 Q92IT9
Q92IU6 Q92IU8 Q92IV0 Q92IV4 Q92IV5 Q92IV6 Q92IW1 Q92IX4 Q92IX6 Q92IX7 Q92IX8
Q92IY7 Q92IZ1 Q92IZ6 Q92J02 Q92J03 Q92J05 Q92J06 Q92J07 Q92J08 Q92J26 Q92J36
Q92J37 Q92J40 Q92J42 Q92J43 Q92J44 Q92J57 Q92J59 Q92J60 Q92J64 Q92J65 Q92J71
Q92J72 Q92J74 Q92J75 Q92J76 Q92J79 Q92J81 Q92J82 Q92J85 Q92J87 Q92J88 Q92J89
Q92J90 Q92J91 Q92J93 Q92J94 Q92J95 Q92JA0 Q92JA4 Q92JA8 Q92JB0 Q92JB4 Q92JB5
Q92JB6 Q92JC9 Q92JD0 Q92JD1 Q92JD2 Q92JD3 Q92JD4 Q92JD5 Q92JD9 Q92JE7 Q92JF1
Q92JF2 Q92JF4 Q92JF5 Q92JF6 Q92JF9 Q92JG5 Q92JG7 Q92JG8 Q92JH2 Q92JH3 Q92JH6
Q92JI0 Q92JI2 Q92JI3 Q92JI4 Q92JI7 Q92JI8 Q92JJ7 Q92JJ8 Q92JK0 Q92JK1 Q92JK2
Q92JK3 Q92JK6 Q92JK7 Q92JK8 Q92JK9 Q92JL0 Q92JL4 Q92JL5 Q92JL7 Q92JL8 Q92JL9
Q92JM0 Q92JM4 Q92JM5 Q92JM6 Q92JM7 Q92JM8 Q92JN5 Q92JP4 Q92JP5 Q92JQ1 Q92JQ4
Q92JQ5 Q92JQ6 Q92JR0 Q92JR5 Q92JR6 Q92JY3 Q92KF8 Q92KH5 Q92KL6 Q92KT8 Q92KW1
Q92KW2 Q92KW3 Q92KW8 Q92KX2 Q92KX6 Q92KY8 Q92KZ3 Q92L19 Q92L21 Q92L25 Q92L39
Q92L41 Q92L42 Q92L43 Q92L46 Q92L66 Q92L68 Q92L73 Q92L76 Q92L94 Q92L99 Q92LA1
Q92LA6 Q92LA8 Q92LB0 Q92LB1 Q92LB2 Q92LB3 Q92LB4 Q92LB7 Q92LB8 Q92LF6 Q92LF7
Q92LH1 Q92LH7 Q92LH8 Q92LI5 Q92LK1 Q92LK3 Q92LP6 Q92LU5 Q92LW9 Q92LX6 Q92M03
Q92M13 Q92M14 Q92M28 Q92M52 Q92M59 Q92M65 Q92M71 Q92M79 Q92M88 Q92M90 Q92M91
Q92M92 Q92M99 Q92MA5 Q92MB5 Q92ME3 Q92ME6 Q92ME7 Q92MF4 Q92MF8 Q92MG0 Q92MH1
Q92MJ1 Q92MK1 Q92MK5 Q92MK7 Q92MQ3 Q92MQ8 Q92MT4 Q92MY3 Q92MY4 Q92MY5 Q92N33
Q92N38 Q92N42 Q92N45 Q92N62 Q92N66 Q92N67 Q92N68 Q92N73 Q92N95 Q92NF5 Q92NH8
Q92NI1 Q92NL3 Q92NL4 Q92NL6 Q92NM0 Q92NM1 Q92NM4 Q92NM8 Q92NN0 Q92NN1 Q92NN5
Q92NQ5 Q92NR2 Q92NT6 Q92P31 Q92P48 Q92P98 Q92PB2 Q92PB4 Q92PB6 Q92PB8 Q92PC8
Q92PD8 Q92PG8 Q92PG9 Q92PH0 Q92PH1 Q92PH7 Q92PI1 Q92PI3 Q92PK5 Q92PK6 Q92PL6
Q92PR8 Q92PR9 Q92PS0 Q92PS4 Q92PU3 Q92PW7 Q92PX0 Q92PY4 Q92PY5 Q92PY8 Q92PZ4
Q92Q10 Q92Q11 Q92Q12 Q92Q15 Q92Q22 Q92Q31 Q92Q32 Q92Q37 Q92Q51 Q92Q52 Q92Q53
Q92Q54 Q92Q55 Q92Q63 Q92Q84 Q92Q90 Q92Q94 Q92QA0 Q92QA1 Q92QB0 Q92QB4 Q92QB5
Q92QD0 Q92QD3 Q92QD5 Q92QD8 Q92QF1 Q92QF2 Q92QF3 Q92QF4 Q92QF5 Q92QF6 Q92QF7
Q92QF8 Q92QF9 Q92QG0 Q92QG1 Q92QG2 Q92QG3 Q92QG4 Q92QG5 Q92QG7 Q92QG8 Q92QG9
Q92QH0 Q92QH1 Q92QH2 Q92QH3 Q92QH4 Q92QH6 Q92QH7 Q92QH8 Q92QH9 Q92QI0 Q92QI1
Q92QI7 Q92QJ0 Q92QJ9 Q92QK2 Q92QK5 Q92QK6 Q92QK7 Q92QK8 Q92QK9 Q92QL0 Q92QL5
Q92QM4 Q92QM8 Q92QN2 Q92QQ2 Q92QR4 Q92QR5 Q92QR7 Q92QT3 Q92QT9 Q92QU0 Q92QU6
Q92QV4 Q92QW2 Q92QX2 Q92QX5 Q92QX9 Q92QY7 Q92QZ0 Q92QZ1 Q92QZ2 Q92QZ7 Q92QZ8
Q92QZ9 Q92R07 Q92R20 Q92R32 Q92R33 Q92R44 Q92R45 Q92R47 Q92R52 Q92R54 Q92R55
Q92R94 Q92RA4 Q92RB2 Q92RC5 Q92RF8 Q92RG0 Q92RG1 Q92RG2 Q92RH3 Q92RH8 Q92RI9
Q92RJ1 Q92RJ3 Q92RL0 Q92RL2 Q92RL5 Q92RM1 Q92RM4 Q92RN6 Q92RN7 Q92RN8 Q92RN9
Q92RP0 Q92RP4 Q92RR6 Q92RR7 Q92RT9 Q92RY3 Q92S04 Q92S07 Q92S14 Q92S21 Q92S22
Q92S23 Q92S25 Q92S26 Q92S27 Q92S49 Q92S97 Q92S99 Q92SA0 Q92SA6 Q92SC4 Q92SC6
Q92SC7 Q92SF3 Q92SF4 Q92SF6 Q92SG0 Q92SG5 Q92SH1 Q92SH4 Q92SH5 Q92SH6 Q92SI3
Q92SI5 Q92SI7 Q92SI9 Q92SJ7 Q92SK0 Q92SK1 Q92SK2 Q92SK4 Q92SK7 Q92SM6 Q92SN3

Q92SN4 Q92SN8 Q92SQ3 Q92SR7 Q92SS9 Q92ST0 Q92ST1 Q92ST2 Q92ST3 Q92ST6 Q92SV4
Q92SV5 Q92SW0 Q92SW1 Q92SW2 Q92SW3 Q92SW4 Q92SW6 Q92SW9 Q92SX1 Q92SY7 Q92SZ1
Q92T07 Q92T25 Q92T27 Q92T28 Q92T47 Q92T48 Q92T49 Q92T50 Q92T74 Q92T88 Q92TA4
Q92TA7 Q92TA9 Q92TB0 Q92TB1 Q92TB2 Q92TB3 Q92TB4 Q92TB5 Q92TC1 Q92TC8 Q92TC9
Q92TD0 Q92TE7 Q92TE9 Q92TF0 Q92TF1 Q92TF2 Q92TI2 Q92TI9 Q92TK9 Q92TP3 Q92TT1
Q92TW1 Q92TY6 Q92TY9 Q92TZ1 Q92UG4 Q92UG5 Q92UI1 Q92UK6 Q92UN9 Q92UV7 Q92UV9
Q92UW6 Q92V04 Q92V09 Q92V27 Q92V80 Q92VB9 Q92VC1 Q92VC6 Q92VM6 Q92VT5 Q92W37
Q92WB5 Q92WP0 Q92WR9 Q92WS1 Q92WX6 Q92X16 Q92XG8 Q92XH8 Q92XP4 Q92XP5 Q92XS8
Q92XV9 Q92Y27 Q92Y45 Q92Y46 Q92YD2 Q92YE0 Q92YE2 Q92YG5 Q92YS6 Q92YY3 Q92ZA4
Q92ZJ6 Q92ZK3 Q92ZL2 Q92ZM6 Q92ZQ3 Q92ZQ4 Q92ZT0 Q92ZT1 Q92ZT2 Q92ZU4 Q93015
Q93034 Q93073 Q93075 Q93079 Q93083 Q93088 Q93099 Q930I4 Q930J0 Q930L2 Q930T2
Q930V2 Q930X9 Q930Y0 Q93104 Q93112 Q93113 Q93129 Q93130 Q93131 Q93132 Q93134
Q93140 Q93169 Q93170 Q931D0 Q931D7 Q931E1 Q931E6 Q931F6 Q931G1 Q931G2 Q931G4
Q931G5 Q931G8 Q931H5 Q931I4 Q931K4 Q931P5 Q931P6 Q931P7 Q931Q1 Q931Q3 Q931Q4
Q931Q6 Q931R3 Q931R7 Q931R8 Q931S8 Q931T1 Q931T2 Q931T8 Q931U1 Q931V3 Q93244
Q93246 Q932A8 Q932F6 Q932F8 Q932G0 Q932G1 Q932G3 Q932I8 Q932K5 Q932K9 Q932L5
Q932M0 Q932V0 Q93341 Q93348 Q93352 Q93353 Q93356 Q93364 Q93367 Q93379 Q933N0
Q93408 Q93425 Q93442 Q93454 Q93459 Q934T0 Q934T6 Q93538 Q93560 Q93571 Q93573
Q93575 Q93591 Q93594 Q935S7 Q935Y2 Q935Y4 Q935Y7 Q935Z3 Q93615 Q93618 Q93635
Q93650 Q936D6 Q936E0 Q936F1 Q936S7 Q936T0 Q936V7 Q936X2 Q936X3 Q93705 Q93712
Q93727 Q93761 Q93794 Q937F6 Q937H2 Q937H8 Q937L4 Q937N6 Q937N7 Q937N8 Q937N9
Q937P0 Q937U7 Q937W6 Q937W7 Q93833 Q93847 Q93899 Q938B4 Q938C8 Q938C9 Q938W0
Q938W7 Q93903 Q939D2 Q939I2 Q939J8 Q939R0 Q939X3 Q93A90 Q93AD7 Q93AG0 Q93AJ8
Q93AK9 Q93AL8 Q93AL9 Q93AM0 Q93AM1 Q93AU4 Q93C64 Q93CC3 Q93CE8 Q93CE9 Q93CH6
Q93CM1 Q93CP9 Q93CX7 Q93D95 Q93DQ9 Q93DW6 Q93E15 Q93E92 Q93ED4 Q93EJ3 Q93EP0
Q93EU6 Q93EU7 Q93FD2 Q93FE6 Q93FQ6 Q93FU7 Q93G07 Q93G08 Q93GB7 Q93GF1 Q93GF2
Q93GG2 Q93GI5 Q93GP6 Q93GP7 Q93GT6 Q93GT7 Q93GT8 Q93GT9 Q93GW2 Q93GX3 Q93H19
Q93HB4 Q93HC3 Q93HD3 Q93HF3 Q93HP9 Q93HT9 Q93HU3 Q93HX6 Q93IE8 Q93IM5 Q93IZ8
Q93J59 Q93JF1 Q93JG0 Q93JH3 Q93JQ8 Q93JQ9 Q93JR5 Q93JT2 Q93JW0 Q93JX5 Q93JY3
Q93K00 Q93K57 Q93K67 Q93K97 Q93KD0 Q93KD1 Q93KF3 Q93L23 Q93L51 Q93LE1 Q93LE2
Q93LE9 Q93LL2 Q93LL7 Q93LP2 Q93LQ8 Q93LR4 Q93M78 Q93MD2 Q93MD5 Q93MH1 Q93MH2
Q93MH3 Q93MI1 Q93MI2 Q93MK7 Q93MK8 Q93MK9 Q93ML3 Q93MM1 Q93MM7 Q93N34 Q93N35
Q93NB3 Q93NC7 Q93NE2 Q93NF6 Q93NG1 Q93NG3 Q93NG6 Q93NH4 Q93PJ0 Q93PJ4 Q93PJ5
Q93PM7 Q93PP8 Q93PP9 Q93PS3 Q93PS7 Q93PU6 Q93PU7 Q93PU8 Q93Q55 Q93Q56 Q93Q64
Q93Q66 Q93QC6 Q93QI2 Q93QX0 Q93R26 Q93R27 Q93R28 Q93R29 Q93R37 Q93R93 Q93RD9
Q93RG0 Q93RJ9 Q93RV9 Q93RW0 Q93RW1 Q93RW2 Q93RX5 Q93RZ5 Q93S00 Q93S23 Q93S39
Q93S40 Q93SF3 Q93SG0 Q93SN7 Q93SN8 Q93ST4 Q93SU8 Q93SW0 Q93SW1 Q93SX7 Q93T05
Q93T12 Q93T20 Q93T81 Q93TF4 Q93TJ5 Q93TL5 Q93TL6 Q93TM7 Q93TM8 Q93TM9 Q93TN0
Q93TU6 Q93TV7 Q93TX3 Q93UJ2 Q93UV4 Q93UZ0 Q93V43 Q93V51 Q93V78 Q93V84 Q93V88
Q93VB8 Q93VC3 Q93VD5 Q93VG5 Q93VG8 Q93VH2 Q93VH6 Q93VH9 Q93VI3 Q93VJ4 Q93VK5
Q93VK9 Q93VM9 Q93VP3 Q93VQ9 Q93VR3 Q93VR4 Q93W20 Q93W22 Q93W28 Q93W34 Q93W77
Q93W88 Q93W93 Q93WC4 Q93WC5 Q93WE4 Q93WF6 Q93WI0 Q93WI2 Q93WJ9 Q93WL3 Q93WN0
Q93WT0 Q93WU2 Q93WU3 Q93WU4 Q93WU6 Q93WU7 Q93WU8 Q93WU9 Q93WV0 Q93WV4 Q93WV5
Q93WV6 Q93WV7 Q93WX6 Q93WY4 Q93X23 Q93XC4 Q93XE1 Q93XE6 Q93XH4 Q93XI4 Q93XK2

Q93XN8 Q93XP7 Q93XQ4 Q93XQ5 Q93XW5 Q93XW7 Q93XX2 Q93XX8 Q93Y00 Q93Y16 Q93Y23
Q93Y32 Q93Y35 Q93Y39 Q93Y52 Q93Y92 Q93Y94 Q93YA3 Q93YF5 Q93YG7 Q93YI9 Q93YN9
Q93YP0 Q93YQ1 Q93YQ3 Q93YR3 Q93YR9 Q93YS6 Q93YU2 Q93YU6 Q93YV0 Q93YV5 Q93YV9
Q93YZ7 Q93Z01 Q93Z17 Q93Z27 Q93Z51 Q93Z53 Q93Z66 Q93Z70 Q93Z96 Q93ZA5 Q93ZB1
Q93ZB6 Q93ZB7 Q93ZC9 Q93ZE2 Q93ZE5 Q93ZF6 Q93ZG3 Q93ZG7 Q93ZG9 Q93ZH1 Q93ZH2
Q93ZJ3 Q93ZJ9 Q93ZK9 Q93ZL5 Q93ZM7 Q93ZN2 Q93ZR1 Q93ZR3 Q93ZS1 Q93ZS9 Q93ZT5
Q93ZT6 Q93ZV7 Q93ZW0 Q93ZX1 Q93ZX6 Q93ZY7 Q94045 Q94053 Q940A2 Q940A6 Q940A7
Q940B0 Q940B8 Q940G8 Q940H7 Q940H8 Q940J1 Q940K0 Q940L5 Q940P8 Q940Q2 Q940T5
Q940T9 Q940V3 Q940X7 Q940X9 Q940Y3 Q940Y8 Q940Z1 Q940Z2 Q940Z5 Q94126 Q94131
Q94165 Q94166 Q941A4 Q941B6 Q941D1 Q941D5 Q941D6 Q941H7 Q941I0 Q941I2 Q941I6
Q941P9 Q941Q2 Q941Q3 Q941R6 Q941T1 Q941T9 Q941Y8 Q941Z0 Q94230 Q94234 Q94250
Q94251 Q94261 Q942F8 Q942P9 Q94361 Q94398 Q943E6 Q943E7 Q943E9 Q943F3 Q943I6
Q943L2 Q943Q3 Q94402 Q94407 Q94425 Q94460 Q94469 Q94480 Q94481 Q94490 Q94497
Q944B0 Q944B6 Q944I4 Q944K3 Q944L7 Q944N1 Q944R7 Q944S1 Q944S2 Q944S3 Q944S9
Q944T2 Q944U4 Q944W6 Q944X7 Q944Z5 Q944Z9 Q94517 Q94521 Q94522 Q94524 Q94529
Q94535 Q94545 Q94546 Q94547 Q94550 Q94554 Q94555 Q94570 Q94571 Q94572 Q94587
Q94596 Q945B4 Q945B5 Q945B6 Q945B7 Q945F4 Q945L0 Q945M9 Q945P0 Q945P1 Q945P8
Q945S8 Q945U1 Q94613 Q94624 Q94636 Q94655 Q94660 Q94688 Q946J2 Q946J8 Q94702
Q94726 Q94738 Q94743 Q94748 Q94757 Q94758 Q94776 Q947C4 Q947C5 Q947G8 Q94818
Q94819 Q94890 Q94899 Q948J9 Q948L5 Q948P5 Q948P6 Q948R0 Q948T6 Q948X8 Q948Z0
Q94901 Q94913 Q94915 Q94972 Q94999 Q949G5 Q949H0 Q949J1 Q949M8 Q949P3 Q949R4
Q949S5 Q949S9 Q949T8 Q949U2 Q949U7 Q949V6 Q949W8 Q949X7 Q94A08 Q94A09 Q94A16
Q94A21 Q94A38 Q94A41 Q94A43 Q94A51 Q94A52 Q94A64 Q94A73 Q94A78 Q94A79 Q94A84
Q94A88 Q94A97 Q94AA4 Q94AB5 Q94AC1 Q94AD1 Q94AD8 Q94AD9 Q94AF0 Q94AF2 Q94AF7
Q94AH6 Q94AH8 Q94AH9 Q94AI1 Q94AI4 Q94AI7 Q94AL9 Q94AM1 Q94AM9 Q94AP0 Q94AQ6
Q94AQ9 Q94AR4 Q94AR6 Q94AR8 Q94AT1 Q94AT3 Q94AU9 Q94AX2 Q94AY1 Q94AY3 Q94B40
Q94B46 Q94B55 Q94B59 Q94B71 Q94B74 Q94BM7 Q94BN2 Q94BP7 Q94BQ2 Q94BR4 Q94BR5
Q94BS8 Q94BT6 Q94BU1 Q94BU8 Q94BV2 Q94BV4 Q94BV5 Q94BY1 Q94BY9 Q94BZ7 Q94C11
Q94C27 Q94C32 Q94C33 Q94C48 Q94C57 Q94C60 Q94C69 Q94C74 Q94C75 Q94C98 Q94CD1
Q94CD5 Q94CE3 Q94CJ8 Q94CK4 Q94CK6 Q94CK9 Q94CL6 Q94CL9 Q94CT7 Q94CU5 Q94CV1
Q94DE2 Q94DH3 Q94DM8 Q94DR2 Q94DT9 Q94DW6 Q94E75 Q94E96 Q94EE9 Q94EH8 Q94EJ2
Q94EY1 Q94EY2 Q94F47 Q94F49 Q94F58 Q94F87 Q94F88 Q94FA4 Q94FA5 Q94FA6 Q94FL5
Q94FL7 Q94FN2 Q94FT1 Q94FT2 Q94FT3 Q94FT4 Q94FT7 Q94FT8 Q94FV7 Q94FY2 Q94FY8
Q94G00 Q94G53 Q94G66 Q94GF1 Q94GL5 Q94H98 Q94HF1 Q94HV8 Q94HW2 Q94IB1 Q94IC4
Q94ID1 Q94ID2 Q94ID3 Q94ID7 Q94IK2 Q94IN5 Q94IR2 Q94JE1 Q94JF2 Q94JI5 Q94JJ4
Q94JJ7 Q94JL3 Q94JM2 Q94JN2 Q94JN3 Q94JQ4 Q94JS8 Q94JV4 Q94JV5 Q94JW0 Q94JX6
Q94JY0 Q94JZ8 Q94K01 Q94K34 Q94K41 Q94K43 Q94K71 Q94K73 Q94K75 Q94KD0 Q94KE2
Q94KL5 Q94KL7 Q94KL8 Q94KR7 Q94KR8 Q94KR9 Q94KU1 Q94KU2 Q94L32 Q94LA4 Q94LP4
Q94LU9 Q94LW3 Q94LW4 Q94LW7 Q94LX1 Q94MV8 Q94PL3 Q94Q55 Q94Q76 Q95000 Q95001
Q95005 Q95008 Q95017 Q95028 Q95032 Q95043 Q95044 Q95046 Q95058 Q95079 Q95081
Q95083 Q95085 Q95086 Q95087 Q95088 Q95095 Q95140 Q95171 Q95175 Q95190 Q95192
Q95202 Q95208 Q95212 Q95216 Q95238 Q95267 Q95274 Q95307 Q95313 Q95332 Q95337
Q95661 Q95666 Q95694 Q95747 Q95749 Q95869 Q95BX6 Q95BX9 Q95BY0 Q95BZ7 Q95CA6
Q95DN9 Q95DP2 Q95DP4 Q95DQ2 Q95DQ5 Q95DQ8 Q95DR2 Q95DR3 Q95E84 Q95EA5 Q95EC9

Q95ED6 Q95ED9 Q95EE0 Q95EE5 Q95EF3 Q95EF8 Q95F35 Q95G88 Q95GJ0 Q95GM2 Q95GM3
Q95GM4 Q95GM5 Q95GM6 Q95GM7 Q95GM9 Q95GN3 Q95GN4 Q95GN5 Q95GN7 Q95GN8 Q95GN9
Q95GQ8 Q95GT2 Q95GT7 Q95H45 Q95H48 Q95H49 Q95H50 Q95H51 Q95H52 Q95H53 Q95H54
Q95H55 Q95H56 Q95H61 Q95H62 Q95H63 Q95J40 Q95J53 Q95JA5 Q95JC8 Q95JD5 Q95JI3
Q95JI7 Q95JI9 Q95JJ2 Q95JJ5 Q95JK1 Q95JK3 Q95JK8 Q95JL1 Q95JL5 Q95JL6 Q95JM8
Q95JN2 Q95JP0 Q95JP6 Q95JQ1 Q95JR0 Q95JR6 Q95JS9 Q95JT3 Q95JU3 Q95JV3 Q95JV5
Q95JW1 Q95JY0 Q95JY5 Q95JY7 Q95K09 Q95K21 Q95K25 Q95K27 Q95K30 Q95K40 Q95K48
Q95K49 Q95K50 Q95K52 Q95K60 Q95K63 Q95K79 Q95K90 Q95KB0 Q95KD0 Q95KD7 Q95KF9
Q95KI0 Q95KI1 Q95KJ0 Q95KJ2 Q95KK4 Q95KL4 Q95KM0 Q95KU9 Q95KV1 Q95L05 Q95L46
Q95L54 Q95LG8 Q95LI3 Q95LL2 Q95LL3 Q95LL8 Q95LM1 Q95LP0 Q95LP5 Q95LR0 Q95LR6
Q95LS7 Q95LT1 Q95LU0 Q95LV5 Q95LY3 Q95LY5 Q95LZ5 Q95M71 Q95M72 Q95MH5 Q95MN7
Q95MN8 Q95MN9 Q95MP0 Q95MP1 Q95MP7 Q95NI4 Q95NN1 Q95NR9 Q95NZ0 Q95P04 Q95P51
Q95PM9 Q95PU1 Q95PX3 Q95PZ0 Q95Q00 Q95Q11 Q95Q17 Q95Q18 Q95Q27 Q95Q34 Q95Q62
Q95Q89 Q95Q98 Q95QC4 Q95QD7 Q95QG8 Q95QI7 Q95QJ7 Q95QK3 Q95QQ2 Q95QT8 Q95QU0
Q95QW0 Q95QY7 Q95R14 Q95RC0 Q95RN0 Q95RR8 Q95RV2 Q95RW8 Q95SK3 Q95SX7 Q95T19
Q95TJ9 Q95TN1 Q95TS5 Q95U54 Q95UN8 Q95V31 Q95V32 Q95V34 Q95V35 Q95V39 Q95V58
Q95V77 Q95V84 Q95VA8 Q95VF7 Q95VY2 Q95W85 Q95W86 Q95WA0 Q95WA2 Q95WY0 Q95X01
Q95X91 Q95X94 Q95XD3 Q95XG7 Q95XN2 Q95XP9 Q95XQ1 Q95XQ8 Q95XR2 Q95XR4 Q95XU6
Q95XX1 Q95XZ5 Q95Y12 Q95Y36 Q95Y41 Q95Y71 Q95Y72 Q95Y80 Q95Y83 Q95Y84 Q95YE9
Q95YF0 Q95YF3 Q95YH0 Q95YJ5 Q95YL7 Q95YM8 Q95ZE8 Q95ZE9 Q95ZG3 Q95ZG4 Q95ZI6
Q95ZK7 Q95ZL9 Q95ZS0 Q96008 Q96033 Q960A1 Q960C5 Q960E8 Q960N3 Q960T2 Q960V3
Q960X4 Q960Z0 Q96255 Q96261 Q96266 Q96270 Q96276 Q96286 Q96291 Q96292 Q96293
Q962A2 Q962P8 Q962Q5 Q962Q6 Q962Q7 Q962Q8 Q962R1 Q962R2 Q962R6 Q962R9 Q962S0
Q962S4 Q962S5 Q962S7 Q962T1 Q962T2 Q962T5 Q962U0 Q962Y8 Q96305 Q96319 Q96323
Q96324 Q96330 Q96331 Q96353 Q96355 Q96356 Q96372 Q96376 Q96386 Q96398 Q963B6
Q963B7 Q96412 Q96418 Q96419 Q96449 Q96460 Q96468 Q96471 Q96474 Q96480 Q96481
Q96482 Q96483 Q96484 Q96493 Q96499 Q964D8 Q964D9 Q964E0 Q964E1 Q964E2 Q964E3
Q96502 Q96529 Q96533 Q96539 Q96540 Q96542 Q96551 Q96552 Q96553 Q96555 Q96556
Q96557 Q96558 Q96559 Q96562 Q96563 Q96564 Q96565 Q96581 Q965H3 Q965S0 Q965S8
Q965T6 Q965W1 Q965W2 Q965X6 Q96623 Q96625 Q96626 Q96629 Q96631 Q96639 Q96676
Q96678 Q96679 Q96680 Q96684 Q96685 Q96687 Q96688 Q96689 Q96690 Q966I0 Q966L3
Q966L5 Q966L9 Q966M5 Q966M6 Q966Q9 Q966R0 Q966T6 Q966X9 Q966Y3 Q96702 Q96704
Q96705 Q96711 Q96713 Q96724 Q96725 Q967G1 Q967R6 Q967S7 Q967U1 Q967W0 Q967Y7
Q967Y8 Q967Y9 Q967Z0 Q96818 Q96895 Q96896 Q968X7 Q969A8 Q969F9 Q969G2 Q969G6
Q969H9 Q969I3 Q969J2 Q969L4 Q969M7 Q969P6 Q969Q6 Q969S8 Q969S9 Q969T4 Q969T7
Q969T9 Q969U7 Q969V4 Q969W3 Q969W8 Q969X6 Q969Y2 Q969Z0 Q96A04 Q96A08 Q96A09
Q96A22 Q96A37 Q96A56 Q96A61 Q96A72 Q96A73 Q96AA8 Q96AB3 Q96AB6 Q96AC6 Q96AH0
Q96AK3 Q96AQ1 Q96AQ9 Q96AT9 Q96AZ1 Q96AZ6 Q96B02 Q96B18 Q96B54 Q96B70 Q96BD6
Q96BF6 Q96BJ8 Q96BM1 Q96BN2 Q96BN6 Q96BP3 Q96BQ3 Q96BQ5 Q96BR5 Q96BR6 Q96BT1
Q96BT7 Q96BU1 Q96BV0 Q96BW5 Q96C00 Q96C01 Q96C11 Q96C23 Q96C28 Q96C45 Q96C55
Q96C74 Q96CB5 Q96CB9 Q96CD0 Q96CG3 Q96CM3 Q96CM4 Q96CN4 Q96CN7 Q96CS4 Q96CT2
Q96CX3 Q96D09 Q96D70 Q96D98 Q96DB5 Q96DE0 Q96DE5 Q96DE9 Q96DG6 Q96DI7 Q96DM1
Q96DN5 Q96DN6 Q96DP5 Q96DT5 Q96DT6 Q96DU9 Q96DX7 Q96DY2 Q96E11 Q96E40 Q96E66
Q96EB1 Q96EF6 Q96EF9 Q96EG3 Q96EH3 Q96EH8 Q96EI5 Q96EK2 Q96EK4 Q96EK5 Q96EL1

Q96EM0 Q96EN9 Q96EP1 Q96EP5 Q96EQ9 Q96ES7 Q96EW2 Q96EX3 Q96EY4 Q96EY8 Q96EY9
Q96EZ4 Q96F10 Q96F44 Q96F45 Q96FA3 Q96FA7 Q96FC9 Q96FF7 Q96FI4 Q96FK6 Q96FN5
Q96FN9 Q96FQ7 Q96FV0 Q96FV2 Q96FX2 Q96FZ2 Q96G04 Q96G21 Q96G25 Q96G27 Q96G42
Q96GC6 Q96GD3 Q96GE5 Q96GE6 Q96GG9 Q96GI7 Q96GJ1 Q96GK7 Q96GT9 Q96GU1 Q96GW9
Q96GX2 Q96H35 Q96H40 Q96H86 Q96HA8 Q96HJ3 Q96HM7 Q96HQ0 Q96HQ2 Q96HT8 Q96HW7
Q96HY7 Q96HZ4 Q96HZ7 Q96I27 Q96I59 Q96I76 Q96IC2 Q96IG2 Q96IJ6 Q96IK1 Q96IM9
Q96IP4 Q96IQ9 Q96IR2 Q96IR3 Q96IR7 Q96IT6 Q96IU4 Q96IW2 Q96IX9 Q96IZ5 Q96IZ6
Q96J01 Q96J77 Q96J87 Q96J88 Q96JB1 Q96JB3 Q96JC4 Q96JD6 Q96JF6 Q96JG8 Q96JG9
Q96JK9 Q96JL9 Q96JN8 Q96JP0 Q96JP2 Q96JP5 Q96JY0 Q96K17 Q96K31 Q96K62 Q96K75
Q96K80 Q96KD3 Q96KF2 Q96KN3 Q96KN4 Q96KP4 Q96KP6 Q96KS9 Q96KT0 Q96KT6 Q96KV7
Q96KW2 Q96KX1 Q96L03 Q96L14 Q96L96 Q96LD4 Q96LD8 Q96LI5 Q96LI6 Q96LI9 Q96LJ8
Q96LK8 Q96LL4 Q96LM1 Q96LM5 Q96LM6 Q96LP2 Q96LP6 Q96LQ0 Q96LR7 Q96LS8 Q96LT6
Q96LW1 Q96LW2 Q96LW9 Q96LX7 Q96LX8 Q96LY2 Q96LZ2 Q96M15 Q96M20 Q96M27 Q96M29
Q96M32 Q96M34 Q96M42 Q96M43 Q96M53 Q96M60 Q96M61 Q96M63 Q96M66 Q96M69 Q96M78
Q96M83 Q96M85 Q96M89 Q96M91 Q96M93 Q96M94 Q96M95 Q96M98 Q96MA1 Q96MA6 Q96MC2
Q96MC4 Q96MC5 Q96MC9 Q96ME1 Q96ME7 Q96MF0 Q96MF4 Q96MH7 Q96MI6 Q96MK2 Q96MM3
Q96MM6 Q96MN5 Q96MP5 Q96MR6 Q96MR7 Q96MR9 Q96MT0 Q96MT4 Q96MU6 Q96MW7 Q96MX6
Q96MY1 Q96MY7 Q96MZ0 Q96MZ4 Q96N11 Q96N20 Q96N22 Q96N23 Q96N38 Q96N53 Q96N58
Q96N76 Q96N95 Q96NB1 Q96NC0 Q96ND8 Q96NG3 Q96NG5 Q96NG8 Q96NI8 Q96NJ1 Q96NJ3
Q96NJ5 Q96NJ6 Q96NL0 Q96NL3 Q96NR7 Q96NS1 Q96NS8 Q96NT1 Q96NT3 Q96NU1 Q96NU7
Q96NX9 Q96NZ1 Q96P09 Q96P15 Q96P16 Q96P26 Q96P47 Q96P50 Q96P64 Q96PC2 Q96PE6
Q96PE7 Q96PF1 Q96PP4 Q96PQ6 Q96PQ7 Q96PS1 Q96PU4 Q96PU9 Q96PV4 Q96PV6 Q96PV7
Q96PX6 Q96PY0 Q96PY6 Q96PZ2 Q96Q07 Q96Q35 Q96Q40 Q96Q77 Q96Q83 Q96QA6 Q96QE5
Q96QF7 Q96QP1 Q96QS6 Q96QU4 Q96QU6 Q96R05 Q96RE9 Q96RJ6 Q96RK1 Q96RP9 Q96RQ3
Q96RR1 Q96RS6 Q96RT8 Q96RU8 Q96S07 Q96S19 Q96S53 Q96S65 Q96S95 Q96SC8 Q96SE7
Q96SF2 Q96SF7 Q96SK3 Q96SL8 Q96SQ5 Q96SR6 Q96SY0 Q96SZ5 Q96SZ6 Q96T17 Q96T59
Q96T66 Q96T68 Q96T75 Q96T92 Q96TJ5 Q96TP5 Q96TU8 Q96U48 Q96U60 Q96UB1 Q96UF1
Q96UF2 Q96UG7 Q96UH7 Q96UI1 Q96UJ9 Q96UK2 Q96UL8 Q96UP2 Q96UP3 Q96UP5 Q96UP6
Q96UQ9 Q96US8 Q96UV5 Q96UW0 Q96UX5 Q96V34 Q96V44 Q96V52 Q96V77 Q96VA4 Q96VB9
Q96VC7 Q96VG0 Q96VG3 Q96VG5 Q96VG8 Q96VH2 Q96VH4 Q96VJ7 Q96VK3 Q96VK7 Q96VL0
Q96VN3 Q96VN5 Q96VP4 Q96VP6 Q96VT4 Q96VU9 Q96W30 Q96W53 Q96W54 Q96W55 Q96WI0
Q96WK6 Q96WL3 Q96WN6 Q96WP5 Q96WP7 Q96WP8 Q96WS6 Q96WT2 Q96WT9 Q96WU9 Q96WV0
Q96WV1 Q96WV9 Q96WW0 Q96WW2 Q96WW3 Q96WX8 Q96WZ1 Q96WZ5 Q96X22 Q96X30 Q96X43
Q96X45 Q96X46 Q96X49 Q96X56 Q96XE0 Q96XN9 Q96XS5 Q96XT2 Q96XT4 Q96XY5 Q96Y23
Q96Y44 Q96Y49 Q96Y60 Q96Y66 Q96Y68 Q96Y77 Q96Y90 Q96Y91 Q96Y94 Q96Y95 Q96Y96
Q96YA4 Q96YA7 Q96YC2 Q96YC3 Q96YD0 Q96YE5 Q96YF6 Q96YI2 Q96YK0 Q96YK1 Q96YK7
Q96YL0 Q96YL5 Q96YL7 Q96YM5 Q96YM9 Q96YN6 Q96YN9 Q96YP3 Q96YP4 Q96YP8 Q96YQ2
Q96YR4 Q96YR5 Q96YR6 Q96YR7 Q96YS2 Q96YU9 Q96YV2 Q96YV3 Q96YV7 Q96YV8 Q96YV9
Q96YW0 Q96YW1 Q96YW2 Q96YW3 Q96YW4 Q96YW5 Q96YW6 Q96YW9 Q96YZ3 Q96YZ9 Q96Z89
Q96ZD6 Q96ZD7 Q96ZD9 Q96ZH0 Q96ZK5 Q96ZL3 Q96ZL5 Q96ZP3 Q96ZT9 Q970N1 Q970Q8
Q970S6 Q970U6 Q970U7 Q970U8 Q970U9 Q970V0 Q970V2 Q970V6 Q970V9 Q970X0 Q970X1
Q970X2 Q970X3 Q970X5 Q970Y4 Q970Y6 Q970Y7 Q970Y8 Q970Y9 Q970Z0 Q970Z1 Q970Z2
Q970Z3 Q970Z4 Q970Z6 Q971A0 Q971A9 Q971B2 Q971B3 Q971B5 Q971C1 Q971C2 Q971C9
Q971D0 Q971D5 Q971D7 Q971D8 Q971E0 Q971G9 Q971H1 Q971I0 Q971I1 Q971I2 Q971I4

Q971I5 Q971I6 Q971J0 Q971J1 Q971J2 Q971J4 Q971K1 Q971K2 Q971K4 Q971K8 Q971K9
Q971S5 Q971S6 Q971T0 Q971T1 Q971T2 Q971T3 Q971T6 Q971T7 Q971T8 Q971U4 Q971U6
Q971U9 Q971V1 Q971V3 Q971V9 Q971W0 Q971W2 Q971W5 Q971W7 Q971X1 Q971Y3 Q971Z4
Q971Z5 Q971Z6 Q971Z7 Q972A1 Q972A2 Q972A3 Q972A7 Q972A8 Q972B1 Q972C0 Q972C1
Q972C3 Q972D1 Q972D2 Q972G5 Q972I2 Q972K6 Q972Q3 Q972S7 Q972V9 Q972W0 Q972W3
Q972W5 Q973C7 Q973C8 Q973E7 Q973E9 Q973F0 Q973F1 Q973F3 Q973F5 Q973F6 Q973F9
Q973G0 Q973G1 Q973J1 Q973R6 Q973T6 Q973W5 Q974F5 Q974L1 Q974M6 Q974N4 Q974N6
Q974Q9 Q974R0 Q974S1 Q974S8 Q974T1 Q974T4 Q974T8 Q974U1 Q974X3 Q974Y0 Q974Y8
Q974Z3 Q974Z6 Q974Z9 Q975A7 Q975C8 Q975D1 Q975D3 Q975D4 Q975D5 Q975D6 Q975E6
Q975E9 Q975F2 Q975F8 Q975F9 Q975G1 Q975G4 Q975G5 Q975G7 Q975G8 Q975G9 Q975H0
Q975H1 Q975H2 Q975H5 Q975H9 Q975I1 Q975I2 Q975I3 Q975I4 Q975I5 Q975I6 Q975I7
Q975I8 Q975I9 Q975J0 Q975J1 Q975J2 Q975J3 Q975J4 Q975J5 Q975J6 Q975J7 Q975J9
Q975K0 Q975K1 Q975K4 Q975K6 Q975K7 Q975L0 Q975L5 Q975L7 Q975M2 Q975M4 Q975M5
Q975N0 Q975N2 Q975N4 Q975N5 Q975N6 Q975N7 Q975N8 Q975P2 Q975P3 Q975P6 Q975P8
Q975Q7 Q975Q8 Q975Q9 Q975R0 Q975R3 Q975R9 Q975S0 Q975S1 Q975S5 Q975S6 Q975T0
Q975T8 Q975T9 Q975U2 Q975U5 Q975U8 Q975U9 Q975V4 Q975W6 Q975X3 Q975X4 Q975Y1
Q975Y9 Q975Z6 Q975Z7 Q975Z8 Q975Z9 Q976A0 Q976A1 Q976A3 Q976A6 Q976A8 Q976B0
Q976B1 Q976B2 Q976C3 Q976E1 Q976E4 Q976E9 Q976F3 Q976G0 Q976G3 Q976G4 Q976H1
Q976H2 Q976H4 Q976H6 Q976H8 Q976I0 Q976I1 Q976I3 Q976I5 Q976J5 Q976J6 Q976J7
Q976J8 Q976J9 Q976K0 Q976K1 Q976K2 Q976K3 Q976L5 Q976M1 Q976V6 Q976X8 Q977Q3
Q977U7 Q977U8 Q977U9 Q977V0 Q977V1 Q977X3 Q977X4 Q977X5 Q977X6 Q977Y1 Q977Z3
Q977Z9 Q978F3 Q978F5 Q978J0 Q978J5 Q978K9 Q978N1 Q978N2 Q978N3 Q978N4 Q978N8
Q978S3 Q978S5 Q978S6 Q978S7 Q978T4 Q978U3 Q978U9 Q978V1 Q978V5 Q978W0 Q978W6
Q978W7 Q978W8 Q978Y8 Q978Z2 Q978Z5 Q979A1 Q979A3 Q979A8 Q979B2 Q979B7 Q979B8
Q979B9 Q979C2 Q979C3 Q979C4 Q979D0 Q979E4 Q979E6 Q979F2 Q979F5 Q979F7 Q979G5
Q979G6 Q979H8 Q979K0 Q979K1 Q979K2 Q979K3 Q979L4 Q979L5 Q979L8 Q979L9 Q979N7
Q979N9 Q979P0 Q979P2 Q979P6 Q979P7 Q979P8 Q979Q0 Q979Q3 Q979Q5 Q979R3 Q979R4
Q979S2 Q979S5 Q979T0 Q979T1 Q979T2 Q979T3 Q979U4 Q979U7 Q979V5 Q979V6 Q979V9
Q979W3 Q979W4 Q979X5 Q979Y3 Q979Y4 Q979Z1 Q979Z3 Q979Z4 Q979Z8 Q97A09 Q97A25
Q97A49 Q97A51 Q97A55 Q97A63 Q97A64 Q97A76 Q97A95 Q97AE6 Q97AF1 Q97AG7 Q97AH7
Q97AI2 Q97AJ3 Q97AJ4 Q97AJ6 Q97AK6 Q97AL5 Q97AN7 Q97AN8 Q97AP1 Q97AR1 Q97AR2
Q97AR9 Q97AU0 Q97AU2 Q97AV6 Q97AY8 Q97AZ4 Q97AZ7 Q97B05 Q97B13 Q97B19 Q97B25
Q97B26 Q97B28 Q97B29 Q97B34 Q97B38 Q97B41 Q97B53 Q97B58 Q97B59 Q97B60 Q97B68
Q97B72 Q97B73 Q97B83 Q97B93 Q97B94 Q97B95 Q97B96 Q97B98 Q97B99 Q97BA1 Q97BA2
Q97BB9 Q97BC1 Q97BC2 Q97BC5 Q97BC6 Q97BD5 Q97BD7 Q97BE8 Q97BG7 Q97BG8 Q97BG9
Q97BH2 Q97BH3 Q97BH4 Q97BJ3 Q97BJ8 Q97BJ9 Q97BK4 Q97BK5 Q97BK6 Q97BK7 Q97BK8
Q97BN1 Q97BN2 Q97BN3 Q97BN6 Q97BN7 Q97BP1 Q97BP2 Q97BP4 Q97BP8 Q97BQ0 Q97BQ3
Q97BQ4 Q97BU5 Q97BU6 Q97BU9 Q97BV0 Q97BV2 Q97BV4 Q97BV5 Q97BV6 Q97BV7 Q97BV9
Q97BW0 Q97BW1 Q97BW2 Q97BW3 Q97BW4 Q97BW5 Q97BW6 Q97BW7 Q97BW8 Q97BW9 Q97BX0
Q97BX1 Q97BX2 Q97BX3 Q97BX4 Q97BX5 Q97BX6 Q97BX7 Q97BZ1 Q97BZ3 Q97BZ4 Q97BZ5
Q97BZ6 Q97BZ8 Q97C04 Q97C05 Q97C09 Q97C11 Q97C13 Q97C14 Q97C15 Q97C22 Q97C23
Q97C26 Q97C32 Q97C35 Q97C36 Q97C45 Q97C60 Q97C75 Q97CA5 Q97CA6 Q97CA8 Q97CB4
Q97CC2 Q97CC6 Q97CC8 Q97CD7 Q97CD8 Q97CD9 Q97CE5 Q97CE6 Q97CE8 Q97CG6 Q97CG9
Q97CI3 Q97CI4 Q97CK6 Q97CL3 Q97CL5 Q97CM5 Q97CM6 Q97CP1 Q97CP5 Q97CQ5 Q97CR4

Q97CR6 Q97CR8 Q97CR9 Q97CS3 Q97CT1 Q97CT6 Q97CT9 Q97CU0 Q97CU1 Q97CU2 Q97CU3
Q97CU4 Q97CV7 Q97CV8 Q97CW2 Q97CW3 Q97CW4 Q97CX2 Q97CX3 Q97CX4 Q97CX9 Q97D53
Q97D54 Q97D55 Q97D80 Q97D82 Q97D83 Q97D87 Q97D94 Q97DC6 Q97DD9 Q97DE2 Q97DK4
Q97DP4 Q97DP6 Q97DQ1 Q97DU2 Q97DY9 Q97DZ9 Q97E05 Q97E14 Q97E56 Q97E62 Q97E63
Q97E66 Q97E82 Q97E86 Q97E89 Q97E91 Q97E93 Q97E97 Q97EB0 Q97EB3 Q97EB4 Q97EB6
Q97EB7 Q97EB8 Q97EB9 Q97EC3 Q97EC9 Q97ED5 Q97ED6 Q97EE0 Q97EE1 Q97EE2 Q97EE3
Q97EE7 Q97EF2 Q97EF3 Q97EF4 Q97EF5 Q97EF6 Q97EG2 Q97EG5 Q97EG6 Q97EG7 Q97EG8
Q97EG9 Q97EH0 Q97EH1 Q97EH2 Q97EH3 Q97EH4 Q97EH5 Q97EH7 Q97EH8 Q97EH9 Q97EI0
Q97EI1 Q97EI2 Q97EI3 Q97EI4 Q97EI5 Q97EI6 Q97EI7 Q97EI8 Q97EI9 Q97EJ0 Q97EJ1
Q97EJ2 Q97EJ3 Q97EJ4 Q97EJ5 Q97EJ6 Q97EJ7 Q97EJ9 Q97EK1 Q97EK2 Q97EK3 Q97EK4
Q97EK5 Q97EK6 Q97EK7 Q97EL1 Q97EL2 Q97EL3 Q97EL4 Q97EQ4 Q97EQ5 Q97ES0 Q97ES6
Q97ET6 Q97ET8 Q97EU2 Q97EV1 Q97EV3 Q97EW5 Q97EW9 Q97EX0 Q97EX7 Q97EX8 Q97EX9
Q97EY4 Q97EY6 Q97EZ2 Q97EZ4 Q97EZ6 Q97F01 Q97F02 Q97F38 Q97F39 Q97F51 Q97F58
Q97F61 Q97F64 Q97F65 Q97F67 Q97F68 Q97F73 Q97F79 Q97F85 Q97FB0 Q97FB5 Q97FC1
Q97FE0 Q97FI7 Q97FJ6 Q97FJ7 Q97FK1 Q97FL0 Q97FL9 Q97FM3 Q97FM9 Q97FP8 Q97FQ5
Q97FQ7 Q97FQ8 Q97FR2 Q97FS3 Q97FS4 Q97FS5 Q97FS6 Q97FT2 Q97FT3 Q97FT6 Q97FT7
Q97FT9 Q97FU0 Q97FU2 Q97FU9 Q97FV2 Q97FX3 Q97FX4 Q97FZ9 Q97G95 Q97GB9 Q97GC0
Q97GF7 Q97GG8 Q97GH6 Q97GH7 Q97GH8 Q97GH9 Q97GI6 Q97GI8 Q97GI9 Q97GJ6 Q97GK4
Q97GK5 Q97GK6 Q97GK7 Q97GK9 Q97GL0 Q97GM6 Q97GQ9 Q97GS1 Q97GT0 Q97GT1 Q97GT2
Q97GT3 Q97GU0 Q97GU2 Q97GU6 Q97GV1 Q97GV2 Q97GV3 Q97GX6 Q97GX8 Q97GY2 Q97GZ2
Q97H05 Q97H06 Q97H19 Q97H50 Q97H75 Q97H80 Q97H81 Q97H84 Q97H93 Q97HA0 Q97HB8
Q97HC8 Q97HC9 Q97HD0 Q97HD1 Q97HD5 Q97HD7 Q97HD8 Q97HE5 Q97HE6 Q97HF0 Q97HF1
Q97HL8 Q97HL9 Q97HT8 Q97I08 Q97I09 Q97I18 Q97I19 Q97I20 Q97I21 Q97I29 Q97I31
Q97I40 Q97I45 Q97I46 Q97I48 Q97I50 Q97I51 Q97I55 Q97I56 Q97I58 Q97I62 Q97I63
Q97I64 Q97I65 Q97I66 Q97I67 Q97I89 Q97I90 Q97I93 Q97I94 Q97I95 Q97I96 Q97I97
Q97IA4 Q97IA7 Q97IA9 Q97IB2 Q97IB7 Q97IC1 Q97IC2 Q97IC4 Q97IC9 Q97ID0 Q97ID1
Q97ID7 Q97IG1 Q97IG2 Q97IG3 Q97IN6 Q97IP6 Q97IQ6 Q97IR8 Q97IU0 Q97IU5 Q97IV1
Q97IZ9 Q97J00 Q97J08 Q97J51 Q97J61 Q97J66 Q97J90 Q97J91 Q97J93 Q97J95 Q97JA9
Q97JB1 Q97JB2 Q97JB4 Q97JC5 Q97JD9 Q97JE0 Q97JE3 Q97JE4 Q97JF4 Q97JG4 Q97JI0
Q97JI4 Q97JI7 Q97JJ2 Q97JJ9 Q97JK0 Q97JL2 Q97JL4 Q97JL5 Q97JL7 Q97JM5 Q97JN2
Q97JN3 Q97JN8 Q97JN9 Q97JQ5 Q97JS8 Q97K22 Q97K30 Q97K32 Q97K78 Q97K92 Q97K94
Q97K95 Q97KC9 Q97KE5 Q97KE6 Q97KG0 Q97KH4 Q97KH6 Q97KH7 Q97KH8 Q97KH9 Q97KI0
Q97KI1 Q97KI2 Q97KI3 Q97KI4 Q97KL6 Q97KL8 Q97KM2 Q97KM3 Q97KN0 Q97KP4 Q97KY6
Q97KZ2 Q97L49 Q97L53 Q97L63 Q97L67 Q97L70 Q97LB6 Q97LC5 Q97LD6 Q97LF4 Q97LF7
Q97LG6 Q97LG7 Q97LG8 Q97LI1 Q97LM4 Q97LN4 Q97LN7 Q97LP1 Q97LP2 Q97LP3 Q97LP4
Q97LP5 Q97LP6 Q97LQ1 Q97LQ2 Q97LQ9 Q97LR2 Q97LS0 Q97LS5 Q97LS6 Q97LS8 Q97LV8
Q97LX0 Q97M37 Q97M82 Q97MB3 Q97MB4 Q97MC5 Q97MD1 Q97ME5 Q97MI6 Q97MK4 Q97MK5
Q97MK9 Q97ML3 Q97MM1 Q97MN6 Q97MP8 Q97MR4 Q97MR5 Q97MT8 Q97MU2 Q97MU4 Q97MU6
Q97MV0 Q97MV3 Q97MX1 Q97N11 Q97N13 Q97N17 Q97N21 Q97N32 Q97N35 Q97N44 Q97N56
Q97N97 Q97NA4 Q97NB7 Q97NC9 Q97ND6 Q97NE3 Q97NE6 Q97NE7 Q97NG0 Q97NG1 Q97NG9
Q97NI5 Q97NL4 Q97NN5 Q97NQ0 Q97NQ2 Q97NQ4 Q97NQ7 Q97NQ8 Q97NV3 Q97NV6 Q97NV7
Q97NV8 Q97NX5 Q97NX6 Q97NZ6 Q97NZ7 Q97P00 Q97P15 Q97P30 Q97P31 Q97P32 Q97P33
Q97P39 Q97P44 Q97P62 Q97P67 Q97P97 Q97P99 Q97PA3 Q97PA6 Q97PH3 Q97PI4 Q97PI9
Q97PK0 Q97PK6 Q97PL0 Q97PM1 Q97PM9 Q97PN7 Q97PN8 Q97PN9 Q97PP0 Q97PP4 Q97PP9

Q97PQ6 Q97PR0 Q97PR7 Q97PS1 Q97PS8 Q97PT2 Q97PW5 Q97PW6 Q97PW7 Q97PX2 Q97PX3
Q97Q15 Q97Q16 Q97Q18 Q97Q26 Q97Q48 Q97Q56 Q97Q59 Q97Q68 Q97Q95 Q97QC6 Q97QD4
Q97QE2 Q97QE4 Q97QE5 Q97QF9 Q97QG5 Q97QG6 Q97QI2 Q97QJ3 Q97QJ8 Q97QL0 Q97QL1
Q97QL2 Q97QL3 Q97QL9 Q97QN4 Q97QP2 Q97QP5 Q97QP9 Q97QQ0 Q97QS5 Q97QS7 Q97QS8
Q97QT2 Q97QU4 Q97QV8 Q97QW0 Q97QW6 Q97QW8 Q97QX6 Q97QY9 Q97QZ0 Q97QZ1 Q97QZ2
Q97R12 Q97R16 Q97R20 Q97R22 Q97R24 Q97R25 Q97R30 Q97R31 Q97R56 Q97R59 Q97R60
Q97R61 Q97R63 Q97R65 Q97R66 Q97R80 Q97R82 Q97R83 Q97R84 Q97R89 Q97R91 Q97R94
Q97R95 Q97RA4 Q97RB5 Q97RC6 Q97RC8 Q97RE1 Q97RH2 Q97RH6 Q97RH7 Q97RI6 Q97RI7
Q97RI9 Q97RM4 Q97RM5 Q97RN5 Q97RN9 Q97RQ1 Q97RQ3 Q97RR9 Q97RS5 Q97RS6 Q97RS9
Q97RU8 Q97RW2 Q97RW5 Q97RX2 Q97RZ6 Q97S07 Q97S25 Q97S28 Q97S31 Q97S34 Q97S36
Q97S45 Q97S56 Q97S57 Q97S61 Q97S73 Q97S89 Q97S93 Q97SA9 Q97SC7 Q97SD7 Q97SE4
Q97SE5 Q97SE6 Q97SE7 Q97SG0 Q97SG5 Q97SG9 Q97SH1 Q97SH8 Q97SI9 Q97SJ0 Q97SJ8
Q97SN4 Q97SQ2 Q97SQ4 Q97SQ9 Q97SR1 Q97SR4 Q97SR6 Q97SS0 Q97SS2 Q97ST4 Q97ST6
Q97SU1 Q97SU3 Q97SU4 Q97SU6 Q97SU7 Q97SV1 Q97SV2 Q97SV3 Q97SV4 Q97SV5 Q97SX0
Q97SX1 Q97SY4 Q97T03 Q97T27 Q97T34 Q97T36 Q97T38 Q97T98 Q97T99 Q97TA2 Q97TA7
Q97TC4 Q97TC5 Q97TD1 Q97TN4 Q97TR7 Q97TX6 Q97TX8 Q97TX9 Q97TY2 Q97TZ8 Q97U03
Q97U19 Q97U21 Q97U27 Q97U28 Q97U29 Q97U30 Q97U63 Q97U94 Q97U96 Q97UA0 Q97UA1
Q97UA2 Q97UB2 Q97UH6 Q97UU4 Q97UV1 Q97V15 Q97VE1 Q97VF1 Q97VM6 Q97VM8 Q97VN4
Q97VP5 Q97VT7 Q97VW7 Q97VW8 Q97VX5 Q97VY2 Q97VY3 Q97VY9 Q97VZ6 Q97VZ9 Q97W02
Q97W04 Q97W05 Q97W08 Q97W36 Q97W44 Q97W51 Q97W56 Q97W59 Q97W62 Q97W70 Q97W73
Q97W92 Q97W94 Q97W96 Q97WC6 Q97WC7 Q97WE5 Q97WE6 Q97WF1 Q97WG4 Q97WG8 Q97WG9
Q97WH0 Q97WH1 Q97WI2 Q97WI6 Q97WJ0 Q97WJ1 Q97WJ5 Q97WJ8 Q97WK3 Q97WK8 Q97WM6
Q97WM8 Q97WN9 Q97WP0 Q97WV8 Q97WW0 Q97WW2 Q97WW9 Q97WX0 Q97WX1 Q97WX2 Q97WX3
Q97WX4 Q97WX5 Q97WY4 Q97X13 Q97X14 Q97X43 Q97X93 Q97XK8 Q97XW1 Q97Y16 Q97Y45
Q97Y84 Q97Y85 Q97Y88 Q97Y90 Q97Y91 Q97Y92 Q97Y96 Q97YC1 Q97YC2 Q97YC5 Q97YC6
Q97YC7 Q97YC8 Q97YD2 Q97YD4 Q97YD5 Q97YJ8 Q97YJ9 Q97YR6 Q97YU3 Q97YX6 Q97Z34
Q97Z43 Q97Z69 Q97Z78 Q97Z79 Q97Z80 Q97Z83 Q97Z84 Q97Z86 Q97Z88 Q97Z89 Q97Z90
Q97Z92 Q97ZC4 Q97ZC5 Q97ZE0 Q97ZE2 Q97ZE3 Q97ZE8 Q97ZF0 Q97ZF1 Q97ZF4 Q97ZF5
Q97ZG0 Q97ZG1 Q97ZG3 Q97ZH0 Q97ZH1 Q97ZH5 Q97ZH6 Q97ZI6 Q97ZI8 Q97ZI9 Q97ZJ9
Q97ZK2 Q97ZK9 Q97ZL0 Q97ZM1 Q97ZN1 Q97ZQ1 Q97ZQ3 Q97ZQ4 Q97ZQ5 Q97ZQ6 Q97ZW3
Q97ZW4 Q97ZX0 Q97ZX2 Q97ZX7 Q97ZY3 Q97ZY5 Q97ZY6 Q97ZY7 Q97ZY8 Q97ZZ0 Q97ZZ6
Q97ZZ8 Q980A0 Q980A1 Q980A3 Q980A5 Q980A6 Q980A7 Q980A8 Q980A9 Q980B1 Q980B5
Q980B6 Q980B8 Q980C1 Q980C3 Q980D1 Q980D9 Q980E8 Q980F0 Q980F8 Q980F9 Q980G0
Q980G6 Q980H6 Q980I4 Q980I5 Q980I7 Q980I8 Q980I9 Q980J7 Q980J9 Q980K0 Q980K2
Q980K7 Q980L3 Q980L4 Q980L5 Q980L7 Q980M1 Q980M3 Q980M4 Q980M5 Q980N4 Q980N5
Q980P7 Q980Q3 Q980Q4 Q980Q5 Q980Q6 Q980Q7 Q980Q8 Q980Q9 Q980R1 Q980R2 Q980R3
Q980R9 Q980S6 Q980S9 Q980T4 Q980T7 Q980T8 Q980U4 Q980U5 Q980U7 Q980U8 Q980V1
Q980V2 Q980V3 Q980V5 Q980W3 Q980W4 Q980W5 Q980W8 Q980W9 Q980X0 Q980X1 Q980Z8
Q98176 Q98185 Q98199 Q981B9 Q981C9 Q981F7 Q981G1 Q981H7 Q981J9 Q981K0 Q981V0
Q981X2 Q98218 Q98224 Q98229 Q98234 Q98262 Q98267 Q98277 Q98298 Q982F2 Q982G0
Q982V5 Q982V6 Q982W8 Q982W9 Q982X3 Q982Z7 Q98314 Q98325 Q983A9 Q983B0 Q983B3
Q983B6 Q983E4 Q983E9 Q983F8 Q983H8 Q983I0 Q983I1 Q983J4 Q983J7 Q983K4 Q983L2
Q983M5 Q983N4 Q983S3 Q983S4 Q983T9 Q983U5 Q98496 Q984K6 Q984N8 Q984P0 Q984P3
Q984S1 Q984S6 Q984S7 Q984S9 Q984T8 Q984T9 Q984U0 Q984X8 Q98514 Q98544 Q985A3

Q985A6 Q985B1 Q985B3 Q985B9 Q985C8 Q985D2 Q985M2 Q985M5 Q985M6 Q985P2 Q985P3
Q985P4 Q985Q5 Q985V6 Q985W1 Q985W2 Q985W3 Q985X9 Q985Y3 Q985Y6 Q985Z4 Q98630
Q98631 Q98632 Q98663 Q986A0 Q986A5 Q986B5 Q986B6 Q986C6 Q986D0 Q986D9 Q986N2
Q986N6 Q986Q6 Q986Q8 Q986R4 Q98745 Q98747 Q98776 Q987C6 Q987C8 Q987N5 Q987R3
Q987S0 Q987S8 Q987S9 Q987T0 Q987T6 Q987V3 Q987X5 Q98829 Q98867 Q98874 Q98875
Q98876 Q98877 Q98878 Q98879 Q988B7 Q988B8 Q988B9 Q988C8 Q988D0 Q988D4 Q988H0
Q988I1 Q988P7 Q988V7 Q98905 Q98924 Q98925 Q98937 Q98948 Q98972 Q989A3 Q989A7
Q989E5 Q989E7 Q989E8 Q989F0 Q989F1 Q989F2 Q989G8 Q989L5 Q989S7 Q989T9 Q98A05
Q98A07 Q98A73 Q98A80 Q98A81 Q98AM3 Q98AM7 Q98AN6 Q98AN7 Q98AP3 Q98AP4 Q98AP7
Q98AT3 Q98AV7 Q98AV8 Q98AX8 Q98AX9 Q98AZ0 Q98AZ3 Q98AZ6 Q98B00 Q98B65 Q98B72
Q98B75 Q98BB4 Q98BB6 Q98BB7 Q98BC8 Q98BG5 Q98BG6 Q98BG8 Q98BG9 Q98BH1 Q98BI3
Q98BI4 Q98BI6 Q98BI7 Q98BI8 Q98BJ1 Q98BJ3 Q98BK1 Q98BK3 Q98BM4 Q98BM7 Q98BV5
Q98BZ8 Q98C03 Q98C10 Q98C21 Q98C27 Q98C31 Q98CB9 Q98CC1 Q98CC2 Q98CH8 Q98CI7
Q98CJ0 Q98CL7 Q98CM3 Q98CN6 Q98CN7 Q98CN8 Q98CP5 Q98CP7 Q98CP8 Q98CP9 Q98CQ1
Q98CR3 Q98CR8 Q98CS9 Q98CT0 Q98CT1 Q98CT3 Q98CT5 Q98CT7 Q98CT8 Q98CU1 Q98CY2
Q98CY3 Q98CY6 Q98CY9 Q98CZ2 Q98CZ3 Q98CZ5 Q98D10 Q98D24 Q98D52 Q98D53 Q98D55
Q98D57 Q98D61 Q98D72 Q98D76 Q98D85 Q98D89 Q98D90 Q98DD1 Q98DD2 Q98DD5 Q98DE8
Q98DI0 Q98DK4 Q98DM0 Q98DM2 Q98DM8 Q98DM9 Q98DN4 Q98DN6 Q98DN7 Q98DR6 Q98DT4
Q98DU8 Q98DV6 Q98DY2 Q98DY3 Q98DY4 Q98DY5 Q98DY6 Q98DZ0 Q98DZ1 Q98DZ2 Q98E33
Q98E34 Q98E51 Q98E57 Q98E70 Q98E81 Q98E86 Q98E92 Q98EB6 Q98EC4 Q98EC5 Q98ED9
Q98EE0 Q98EE1 Q98EF1 Q98EG3 Q98EI6 Q98EI7 Q98EP2 Q98ER2 Q98ES7 Q98ET8 Q98EU7
Q98EV0 Q98EW1 Q98EY9 Q98EZ0 Q98EZ3 Q98EZ4 Q98EZ5 Q98EZ6 Q98F00 Q98F04 Q98F05
Q98F08 Q98F28 Q98F44 Q98F50 Q98F59 Q98F60 Q98F72 Q98F75 Q98F76 Q98F87 Q98F91
Q98F97 Q98FE3 Q98FF1 Q98FG0 Q98FG9 Q98FJ1 Q98FS9 Q98FW0 Q98FX8 Q98FY0 Q98FY1
Q98FZ7 Q98G02 Q98G09 Q98G10 Q98G36 Q98G72 Q98G87 Q98G93 Q98G94 Q98G96 Q98GC6
Q98GG1 Q98GP9 Q98GQ5 Q98GR6 Q98GR7 Q98GS0 Q98GV1 Q98GV2 Q98GV3 Q98GV5 Q98GW1
Q98GY4 Q98H61 Q98H64 Q98H78 Q98HB7 Q98HD8 Q98HE3 Q98HL0 Q98HN3 Q98HQ4 Q98HR4
Q98HS2 Q98HV0 Q98HV6 Q98HV7 Q98HW3 Q98I01 Q98I23 Q98I49 Q98I52 Q98I54 Q98I56
Q98I87 Q98IA7 Q98IF8 Q98IG7 Q98IH6 Q98IH9 Q98II0 Q98IV4 Q98IV5 Q98J52 Q98JB6
Q98JG8 Q98JM4 Q98JM5 Q98JS0 Q98JV5 Q98K73 Q98KA5 Q98KA8 Q98KB1 Q98KB4 Q98KB5
Q98KB6 Q98KC4 Q98KD0 Q98KH9 Q98KJ4 Q98KK0 Q98KL6 Q98KM7 Q98KN6 Q98KN9 Q98KP1
Q98KS5 Q98KT0 Q98KT9 Q98L33 Q98L96 Q98LB1 Q98LB3 Q98LB9 Q98LC0 Q98LC8 Q98LC9
Q98LE1 Q98LE8 Q98LQ5 Q98LQ6 Q98LR2 Q98LT6 Q98LT7 Q98LU3 Q98LU4 Q98LV1 Q98LW4
Q98LX5 Q98LZ1 Q98M06 Q98M36 Q98M38 Q98M41 Q98M51 Q98M55 Q98M57 Q98M73 Q98M86
Q98M90 Q98M91 Q98M93 Q98M95 Q98MA3 Q98MB2 Q98MB3 Q98MB7 Q98MB8 Q98MB9 Q98MD0
Q98ME2 Q98ME3 Q98ME4 Q98ME7 Q98MF0 Q98MK2 Q98MK6 Q98ML8 Q98ML9 Q98MN9 Q98MV4
Q98MV8 Q98MX2 Q98MX9 Q98MY2 Q98N18 Q98N32 Q98N33 Q98N34 Q98N35 Q98N36 Q98N38
Q98N39 Q98N40 Q98N41 Q98N42 Q98N43 Q98N44 Q98N45 Q98N46 Q98N47 Q98N48 Q98N49
Q98N50 Q98N51 Q98N52 Q98N53 Q98N54 Q98N55 Q98N56 Q98N57 Q98N58 Q98N59 Q98N60
Q98N61 Q98N65 Q98N66 Q98N67 Q98N68 Q98N69 Q98N70 Q98NA6 Q98NB7 Q98NC1 Q98NC4
Q98NC9 Q98ND0 Q98ND3 Q98ND4 Q98NF9 Q98NM8 Q98NN0 Q98NN7 Q98NP4 Q98NP5 Q98NP9
Q98NQ5 Q98NQ6 Q98NQ8 Q98NS5 Q98NW7 Q98P28 Q98PA0 Q98PE2 Q98PE3 Q98PE5 Q98PF2
Q98PG4 Q98PH2 Q98PH6 Q98PH7 Q98PI9 Q98PK2 Q98PK4 Q98PK6 Q98PK9 Q98PL1 Q98PL2
Q98PL5 Q98PM0 Q98PM6 Q98PN3 Q98PN5 Q98PN8 Q98PP7 Q98PQ2 Q98PQ3 Q98PR1 Q98PR2

Q98PR3 Q98PU5 Q98PU6 Q98PU8 Q98PV4 Q98PV8 Q98PV9 Q98PW0 Q98PW8 Q98PX9 Q98PY0
Q98PY2 Q98PY4 Q98PY5 Q98PY6 Q98PY8 Q98PY9 Q98PZ0 Q98PZ1 Q98PZ2 Q98PZ3 Q98PZ4
Q98PZ5 Q98PZ6 Q98PZ7 Q98PZ8 Q98PZ9 Q98Q00 Q98Q04 Q98Q05 Q98Q06 Q98Q07 Q98Q08
Q98Q09 Q98Q11 Q98Q12 Q98Q13 Q98Q16 Q98Q19 Q98Q23 Q98Q24 Q98Q29 Q98Q31 Q98Q32
Q98Q36 Q98Q37 Q98Q52 Q98Q55 Q98Q72 Q98Q73 Q98Q74 Q98Q75 Q98Q77 Q98Q81 Q98Q96
Q98Q97 Q98Q98 Q98Q99 Q98QA7 Q98QA8 Q98QB0 Q98QB3 Q98QD6 Q98QD7 Q98QD8 Q98QG1
Q98QG4 Q98QH1 Q98QH2 Q98QI3 Q98QJ8 Q98QJ9 Q98QK8 Q98QK9 Q98QL0 Q98QL4 Q98QM2
Q98QM8 Q98QN1 Q98QN2 Q98QN3 Q98QN7 Q98QN9 Q98QP0 Q98QP6 Q98QP7 Q98QP9 Q98QQ1
Q98QQ2 Q98QS1 Q98QS2 Q98QT4 Q98QV0 Q98QV1 Q98QV2 Q98QV3 Q98QV4 Q98QV7 Q98QV8
Q98QW4 Q98QX1 Q98QX2 Q98QY7 Q98QY9 Q98QZ0 Q98QZ1 Q98R01 Q98R05 Q98R09 Q98R27
Q98R30 Q98R33 Q98R44 Q98R56 Q98R57 Q98R67 Q98R68 Q98R70 Q98R79 Q98R82 Q98R83
Q98R92 Q98R94 Q98R95 Q98RA5 Q98RB3 Q98RB6 Q98RC0 Q98RC1 Q98RC3 Q98RF5 Q98RF7
Q98RF8 Q98RF9 Q98RG4 Q98RH5 Q98RJ3 Q98RJ5 Q98RJ8 Q98RJ9 Q98RK6 Q98RM8 Q98RY4
Q98SC7 Q98SH9 Q98SI1 Q98SJ2 Q98SJ7 Q98SK0 Q98SK9 Q98SL0 Q98SL1 Q98SL2 Q98SM3
Q98SP8 Q98T94 Q98TA5 Q98TR3 Q98TR7 Q98TS0 Q98TT6 Q98TX2 Q98U05 Q98U06 Q98UK4
Q98UK5 Q98VP9 Q99002 Q99020 Q99023 Q99031 Q99039 Q99040 Q99042 Q99069 Q99070
Q99074 Q99078 Q99089 Q99090 Q99091 Q99105 Q99107 Q99133 Q99142 Q99145 Q99148
Q99152 Q99153 Q99160 Q99164 Q99166 Q99177 Q99181 Q99188 Q99189 Q99207 Q99208
Q99210 Q99216 Q99219 Q99247 Q99248 Q99258 Q99278 Q99281 Q99282 Q99283 Q99284
Q99285 Q99296 Q992I2 Q992I9 Q992J0 Q99303 Q99312 Q99314 Q99318 Q99321 Q99326
Q99334 Q99335 Q99338 Q99339 Q99341 Q99344 Q99345 Q99359 Q99369 Q99373 Q99383
Q99388 Q99390 Q99394 Q99395 Q993M1 Q993T1 Q993T3 Q993T4 Q99417 Q99426 Q99440
Q99456 Q99457 Q99471 Q99525 Q99551 Q99581 Q99583 Q99592 Q99598 Q99608 Q99611
Q99613 Q99622 Q99626 Q99633 Q99684 Q99687 Q99707 Q99742 Q99743 Q99750 Q99797
Q997F0 Q997F1 Q997F2 Q99801 Q99811 Q99832 Q99853 Q99865 Q99873 Q99877 Q99879
Q99880 Q99909 Q99929 Q99956 Q99967 Q99973 Q99990 Q99FX5 Q99FX7 Q99FY3 Q99H66
Q99HR5 Q99HS4 Q99J09 Q99J10 Q99J25 Q99J36 Q99J38 Q99J55 Q99J62 Q99J64 Q99J72
Q99J77 Q99J78 Q99J79 Q99J87 Q99J95 Q99JA8 Q99JD2 Q99JG7 Q99JH1 Q99JI4 Q99JP4
Q99JR6 Q99JR8 Q99JT1 Q99JT5 Q99JU7 Q99JW2 Q99JX1 Q99JX4 Q99JY4 Q99K23 Q99K48
Q99K74 Q99K85 Q99K95 Q99K99 Q99KB8 Q99KC8 Q99KE8 Q99KF0 Q99KG3 Q99KI0 Q99KJ5
Q99KK7 Q99KK9 Q99KN2 Q99KP3 Q99KR3 Q99KS6 Q99KW3 Q99KX1 Q99KZ6 Q99L02 Q99L13
Q99L20 Q99L27 Q99L28 Q99L45 Q99L47 Q99L85 Q99LB6 Q99LB7 Q99LC2 Q99LC5 Q99LD8
Q99LD9 Q99LH1 Q99LH4 Q99LI5 Q99LI7 Q99LI9 Q99LJ7 Q99LL5 Q99LM3 Q99LM9 Q99LN9
Q99LQ1 Q99LS1 Q99LS3 Q99LT0 Q99LU8 Q99LW0 Q99LW6 Q99LX5 Q99LZ3 Q99M01 Q99M03
Q99M04 Q99M07 Q99M28 Q99M54 Q99M63 Q99M73 Q99M74 Q99M76 Q99M85 Q99MA0 Q99MA9
Q99MB4 Q99MB5 Q99MC0 Q99MD6 Q99ME3 Q99ME7 Q99MH5 Q99MI7 Q99MI9 Q99MM4 Q99MN9
Q99MR0 Q99MR1 Q99MR6 Q99MT2 Q99MU0 Q99MV1 Q99MV2 Q99MV5 Q99MV7 Q99MW1 Q99MW3
Q99MX0 Q99MX2 Q99MY0 Q99MY8 Q99MZ6 Q99N20 Q99N34 Q99N42 Q99NA5 Q99NA9 Q99NB9
Q99NC0 Q99ND9 Q99NE9 Q99NG0 Q99NG9 Q99NI3 Q99P21 Q99P25 Q99P31 Q99P39 Q99P77
Q99P96 Q99PD2 Q99PF5 Q99PF6 Q99PJ2 Q99PJ7 Q99PK0 Q99PM3 Q99PN3 Q99PP2 Q99PP6
Q99PP7 Q99PQ1 Q99PQ2 Q99PR8 Q99PS0 Q99PT0 Q99PT3 Q99PT9 Q99PU8 Q99PV0 Q99PV5
Q99PY0 Q99PZ1 Q99QM0 Q99R06 Q99R54 Q99R57 Q99R60 Q99R73 Q99R75 Q99R76 Q99R79
Q99R88 Q99RB2 Q99RB9 Q99RD5 Q99RD6 Q99RF5 Q99RK7 Q99RN7 Q99RP3 Q99RQ5 Q99RR4
Q99RW8 Q99RX9 Q99RY0 Q99RY1 Q99S05 Q99S17 Q99S23 Q99S26 Q99S28 Q99S30 Q99S31

Q99S33 Q99S36 Q99S37 Q99S46 Q99S51 Q99S65 Q99S95 Q99SC3 Q99SC8 Q99SD6 Q99SE3
Q99SE4 Q99SG3 Q99SH6 Q99SI6 Q99SJ1 Q99SK3 Q99SQ9 Q99SW6 Q99SX9 Q99SY3 Q99SY9
Q99SZ3 Q99T01 Q99T12 Q99T15 Q99T18 Q99T33 Q99T92 Q99TA0 Q99TA4 Q99TC1 Q99TD2
Q99TD5 Q99TE8 Q99TF3 Q99TF5 Q99TG5 Q99TK6 Q99TK9 Q99TM4 Q99TN4 Q99TQ0 Q99TR3
Q99TX9 Q99TZ0 Q99U14 Q99U17 Q99U32 Q99U34 Q99U41 Q99U42 Q99U50 Q99U59 Q99U60
Q99U66 Q99U75 Q99U78 Q99U83 Q99U87 Q99U93 Q99UA7 Q99UB6 Q99UC2 Q99UC6 Q99UD0
Q99UD1 Q99UE2 Q99UH2 Q99UI1 Q99UJ7 Q99UJ8 Q99UJ9 Q99UK9 Q99UM1 Q99UP2 Q99UQ0
Q99US7 Q99US8 Q99UT3 Q99UU0 Q99UW5 Q99UW8 Q99UX8 Q99UZ0 Q99V08 Q99V09 Q99V32
Q99V34 Q99V47 Q99V48 Q99V50 Q99V77 Q99V86 Q99VB5 Q99VC0 Q99VC2 Q99VC3 Q99VC4
Q99VD0 Q99VD4 Q99VE2 Q99VE8 Q99VF6 Q99VF9 Q99VG5 Q99VH8 Q99VK8 Q99VL4 Q99VM1
Q99VM3 Q99VP5 Q99VR1 Q99VT5 Q99VV3 Q99VX6 Q99W07 Q99W19 Q99W24 Q99W32 Q99W39
Q99W43 Q99W49 Q99W51 Q99W56 Q99W72 Q99W78 Q99W81 Q99W83 Q99W94 Q99WA0 Q99WA2
Q99WA5 Q99WB7 Q99WC3 Q99WJ1 Q99WJ2 Q99WJ6 Q99WM3 Q99WV6 Q99WW4 Q99WW8 Q99WZ6
Q99WZ7 Q99X26 Q99X30 Q99X54 Q99X56 Q99X99 Q99XA5 Q99XB1 Q99XE5 Q99XH0 Q99XH4
Q99XI8 Q99XK8 Q99XK9 Q99XL0 Q99XL5 Q99XL6 Q99XL8 Q99XN7 Q99XP1 Q99XP4 Q99XP5
Q99XQ1 Q99XQ7 Q99XQ9 Q99XR2 Q99XR4 Q99XS1 Q99XV5 Q99XW8 Q99XX2 Q99XY1 Q99XY4
Q99XZ0 Q99XZ5 Q99XZ9 Q99Y12 Q99Y13 Q99Y14 Q99Y18 Q99Y31 Q99Y43 Q99Y46 Q99Y51
Q99Y58 Q99Y66 Q99Y73 Q99Y88 Q99Y98 Q99YC0 Q99YC1 Q99YC7 Q99YE2 Q99YG1 Q99YH1
Q99YH2 Q99YI3 Q99YI4 Q99YI7 Q99YI8 Q99YJ7 Q99YJ8 Q99YK9 Q99YL2 Q99YL3 Q99YL4
Q99YM5 Q99YM7 Q99YP1 Q99YP2 Q99YP3 Q99YR3 Q99YS1 Q99YT1 Q99YT5 Q99YU0 Q99YU5
Q99YW3 Q99YX2 Q99Z28 Q99Z29 Q99Z34 Q99Z49 Q99Z50 Q99Z57 Q99Z77 Q99Z78 Q99Z83
Q99Z86 Q99Z89 Q99Z94 Q99ZA5 Q99ZA6 Q99ZB6 Q99ZD0 Q99ZD3 Q99ZE5 Q99ZE6 Q99ZH1
Q99ZI6 Q99ZI9 Q99ZK9 Q99ZL9 Q99ZM2 Q99ZM6 Q99ZN9 Q99ZP1 Q99ZP5 Q99ZP6 Q99ZQ0
Q99ZQ1 Q99ZR0 Q99ZS9 Q99ZU0 Q99ZU3 Q99ZV6 Q99ZV9 Q99ZW2 Q99ZW8 Q9A059 Q9A071
Q9A072 Q9A076 Q9A077 Q9A088 Q9A0A5 Q9A0B5 Q9A0B6 Q9A0C6 Q9A0D0 Q9A0D6 Q9A0D8
Q9A0E3 Q9A0E4 Q9A0F0 Q9A0F4 Q9A0H3 Q9A0H4 Q9A0I0 Q9A0I1 Q9A0J5 Q9A0L0 Q9A0M0
Q9A0R6 Q9A0R7 Q9A0T3 Q9A0U7 Q9A0V7 Q9A0W7 Q9A115 Q9A126 Q9A131 Q9A148 Q9A149
Q9A176 Q9A178 Q9A186 Q9A187 Q9A190 Q9A196 Q9A1B2 Q9A1B3 Q9A1B6 Q9A1B7 Q9A1C7
Q9A1D7 Q9A1E8 Q9A1F2 Q9A1G2 Q9A1H9 Q9A1I0 Q9A1J4 Q9A1K1 Q9A1K3 Q9A1L1 Q9A1L6
Q9A1P0 Q9A1P8 Q9A1U1 Q9A1U3 Q9A1V0 Q9A1V5 Q9A1V6 Q9A1V8 Q9A1V9 Q9A1W2 Q9A1W3
Q9A1W4 Q9A1W5 Q9A1W7 Q9A1X2 Q9A1X4 Q9A1Y1 Q9A1Y7 Q9A1Z0 Q9A1Z3 Q9A201 Q9A206
Q9A217 Q9A224 Q9A228 Q9A229 Q9A230 Q9A231 Q9A232 Q9A238 Q9A239 Q9A253 Q9A258
Q9A259 Q9A268 Q9A280 Q9A2B1 Q9A2B4 Q9A2C5 Q9A2E3 Q9A2F1 Q9A2H2 Q9A2H3 Q9A2H5
Q9A2I0 Q9A2L1 Q9A2M1 Q9A2M2 Q9A2N5 Q9A2P3 Q9A2P5 Q9A2S6 Q9A2W6 Q9A2X3 Q9A2X6
Q9A311 Q9A321 Q9A341 Q9A342 Q9A345 Q9A347 Q9A352 Q9A353 Q9A354 Q9A383 Q9A385
Q9A3A7 Q9A3C8 Q9A3C9 Q9A3D5 Q9A3E9 Q9A3F5 Q9A3G1 Q9A3G2 Q9A3G6 Q9A3G7 Q9A3G8
Q9A3H7 Q9A3I3 Q9A3J1 Q9A3J2 Q9A3J3 Q9A3K2 Q9A3K3 Q9A3K4 Q9A3M3 Q9A3N5 Q9A3P2
Q9A3P3 Q9A3P8 Q9A3Q9 Q9A3S1 Q9A3U9 Q9A3Y4 Q9A434 Q9A435 Q9A437 Q9A458 Q9A459
Q9A478 Q9A4C3 Q9A4C4 Q9A4D0 Q9A4D6 Q9A4E8 Q9A4F3 Q9A4F4 Q9A4F5 Q9A4I3 Q9A4J7
Q9A4L8 Q9A4N2 Q9A4N3 Q9A4N4 Q9A4N5 Q9A4S4 Q9A4Z6 Q9A515 Q9A545 Q9A566 Q9A594
Q9A595 Q9A597 Q9A5A6 Q9A5A7 Q9A5A9 Q9A5B6 Q9A5B8 Q9A5C1 Q9A5D6 Q9A5D7 Q9A5D9
Q9A5F0 Q9A5F3 Q9A5F9 Q9A5I0 Q9A5I1 Q9A5I8 Q9A5K4 Q9A5K8 Q9A5K9 Q9A5L0 Q9A5L1
Q9A5S0 Q9A5T7 Q9A5U7 Q9A5V1 Q9A5V2 Q9A5V4 Q9A5V5 Q9A5V7 Q9A5Z2 Q9A5Z4 Q9A602
Q9A627 Q9A634 Q9A649 Q9A652 Q9A653 Q9A655 Q9A671 Q9A683 Q9A6B1 Q9A6B8 Q9A6C8

Q9A6H0 Q9A6H4 Q9A6I5 Q9A6J5 Q9A6M3 Q9A6M5 Q9A6N1 Q9A6N3 Q9A6Q5 Q9A6S5 Q9A6T4
Q9A6T5 Q9A6T6 Q9A6Z1 Q9A6Z5 Q9A700 Q9A703 Q9A704 Q9A705 Q9A706 Q9A707 Q9A709
Q9A721 Q9A727 Q9A728 Q9A733 Q9A734 Q9A741 Q9A744 Q9A746 Q9A752 Q9A756 Q9A757
Q9A777 Q9A7H8 Q9A7I5 Q9A7I8 Q9A7K2 Q9A7K3 Q9A7M2 Q9A7M9 Q9A7N4 Q9A7N5 Q9A7N9
Q9A7Q1 Q9A7Q2 Q9A7Q3 Q9A7Q9 Q9A7R9 Q9A7U9 Q9A7V4 Q9A7V7 Q9A7W1 Q9A7W2 Q9A7Y1
Q9A7Y2 Q9A7Z1 Q9A7Z2 Q9A7Z5 Q9A802 Q9A805 Q9A808 Q9A810 Q9A823 Q9A838 Q9A842
Q9A871 Q9A873 Q9A874 Q9A881 Q9A884 Q9A894 Q9A8G9 Q9A8H5 Q9A8H6 Q9A8I3 Q9A8J2
Q9A8J3 Q9A8J4 Q9A8J5 Q9A8J6 Q9A8L2 Q9A8L7 Q9A8M7 Q9A8N3 Q9A8S8 Q9A8S9 Q9A8T0
Q9A8T1 Q9A8T2 Q9A8T4 Q9A8T5 Q9A8T6 Q9A8T7 Q9A8T8 Q9A8T9 Q9A8U0 Q9A8U1 Q9A8U2
Q9A8U3 Q9A8U4 Q9A8U5 Q9A8U6 Q9A8U7 Q9A8U8 Q9A8U9 Q9A8V0 Q9A8V1 Q9A8V2 Q9A8V3
Q9A8V4 Q9A8Y0 Q9A8Y2 Q9A8Z4 Q9A8Z9 Q9A900 Q9A919 Q9A9A7 Q9A9C8 Q9A9D4 Q9A9D9
Q9A9E2 Q9A9E3 Q9A9E4 Q9A9E5 Q9A9E9 Q9A9I6 Q9A9M1 Q9A9M3 Q9A9N8 Q9A9Q6 Q9A9S4
Q9A9T4 Q9A9T7 Q9A9T8 Q9A9V5 Q9A9W0 Q9A9X1 Q9A9Z0 Q9A9Z1 Q9A9Z2 Q9AA08 Q9AA09
Q9AA17 Q9AA39 Q9AA58 Q9AA85 Q9AAA5 Q9AAE1 Q9AAF9 Q9AAG0 Q9AAL3 Q9AAL4 Q9AAL5
Q9AAR4 Q9AAS1 Q9AAT8 Q9AAU1 Q9AAU2 Q9AAU7 Q9AAV6 Q9AAV8 Q9AAV9 Q9AAW1 Q9AAX8
Q9AAY2 Q9AAY3 Q9AAY5 Q9AB72 Q9AB94 Q9ABB2 Q9ABB3 Q9ABB6 Q9ABD2 Q9ABE5 Q9ABE7
Q9ABE9 Q9ABF0 Q9ABF2 Q9ABF3 Q9ABF5 Q9ABF8 Q9ABF9 Q9ABG6 Q9ABH0 Q9ABI2 Q9ABK1
Q9ABK5 Q9ABM9 Q9ABN0 Q9ABN1 Q9ABN3 Q9ABS4 Q9ABS7 Q9ABS9 Q9ABT6 Q9ABU1 Q9ABV2
Q9ABV3 Q9ABW2 Q9ABW5 Q9ABY4 Q9ABY6 Q9ABZ8 Q9ABZ9 Q9AC05 Q9AC08 Q9AC14 Q9AC20
Q9AC22 Q9AC25 Q9AC30 Q9AC31 Q9AC32 Q9AC39 Q9AC48 Q9AC50 Q9AC52 Q9AC53 Q9AC54
Q9AC57 Q9AC58 Q9AC59 Q9ACJ8 Q9AD90 Q9ADA4 Q9ADA7 Q9ADB2 Q9ADG3 Q9ADK0 Q9ADK2
Q9ADP7 Q9ADT1 Q9AE09 Q9AE10 Q9AE11 Q9AE55 Q9AE63 Q9AEM1 Q9AEM9 Q9AEN2 Q9AEP7
Q9AEP8 Q9AEQ5 Q9AES0 Q9AET7 Q9AET8 Q9AET9 Q9AEU1 Q9AEU2 Q9AEV8 Q9AF21 Q9AFA5
Q9AFA6 Q9AFC5 Q9AFG9 Q9AFI5 Q9AG20 Q9AG29 Q9AGA6 Q9AGC0 Q9AGE6 Q9AGE7 Q9AGG3
Q9AGJ6 Q9AGP0 Q9AGP1 Q9AGP2 Q9AGP3 Q9AGP6 Q9AGP8 Q9AGR3 Q9AGS1 Q9AGS7 Q9AGT0
Q9AGU4 Q9AGW0 Q9AGY5 Q9AGY6 Q9AGY7 Q9AGY8 Q9AH77 Q9AHD4 Q9AHF0 Q9AHT6 Q9AI34
Q9AI36 Q9AI65 Q9AIF4 Q9AIF9 Q9AIG8 Q9AIG9 Q9AIK2 Q9AIM2 Q9AIM3 Q9AIN8 Q9AIU0
Q9AIU3 Q9AIU7 Q9AIX6 Q9AIX7 Q9AIX9 Q9AJ37 Q9AJ63 Q9AJ64 Q9AJ75 Q9AJ77 Q9AJ79
Q9AJ80 Q9AJ81 Q9AJ82 Q9AJ83 Q9AJ99 Q9AJC6 Q9AJD6 Q9AJD7 Q9AJE3 Q9AJE4 Q9AJF7
Q9AJG9 Q9AJH1 Q9AJN6 Q9AJQ8 Q9AJS8 Q9AJU0 Q9AJU1 Q9AJU2 Q9AK25 Q9AK76 Q9AK82
Q9AK84 Q9AKA7 Q9AKC4 Q9AKD5 Q9AKE1 Q9AKE3 Q9AKE4 Q9AKE5 Q9AKF3 Q9AKH1 Q9AKI4
Q9AKI7 Q9AKJ0 Q9AKJ8 Q9AKJ9 Q9AKK7 Q9AKM6 Q9AKP0 Q9AKP1 Q9AKP3 Q9AKQ2 Q9AKQ3
Q9AKR8 Q9AKT2 Q9AKT4 Q9AKT8 Q9AL01 Q9AL19 Q9AL83 Q9AL94 Q9AL95 Q9AL99 Q9ALA9
Q9ALJ1 Q9ALJ2 Q9ALJ4 Q9ALN5 Q9ALN6 Q9ALN8 Q9ALN9 Q9ALP0 Q9ALZ7 Q9ALZ8 Q9AM27
Q9AMC5 Q9AME2 Q9AME7 Q9AME8 Q9AMJ8 Q9AMJ9 Q9AMK8 Q9AMN9 Q9AMR9 Q9AMS0 Q9AMS4
Q9AMS6 Q9AMS7 Q9AMV8 Q9AMW4 Q9AMZ0 Q9ANL0 Q9ANL1 Q9ANP1 Q9ANP8 Q9ANR7 Q9ANR9
Q9ANS0 Q9AP01 Q9AP06 Q9AP31 Q9AP33 Q9AP34 Q9AP35 Q9APB2 Q9APC2 Q9APJ2 Q9APM5
Q9APY4 Q9APZ2 Q9AQ50 Q9AQ96 Q9AQ97 Q9AQ98 Q9AQ99 Q9AQA0 Q9AQC6 Q9AQC8 Q9AQI0
Q9AQI4 Q9AQM4 Q9AQT0 Q9AQT2 Q9AQT6 Q9AQU1 Q9AR04 Q9AR19 Q9AR22 Q9AR27 Q9AR38
Q9AR73 Q9AR86 Q9ARF9 Q9ARI6 Q9ARZ9 Q9AS33 Q9AS36 Q9AS75 Q9ASJ1 Q9ASP6 Q9ASR0
Q9ASR4 Q9AST1 Q9ASU7 Q9ASU8 Q9ASV6 Q9ASW8 Q9ASX9 Q9ASY9 Q9ASZ1 Q9ASZ8 Q9ASZ9
Q9AT34 Q9AT35 Q9AT54 Q9AT55 Q9AT56 Q9AT61 Q9AT76 Q9ATB4 Q9ATE5 Q9ATF5 Q9ATY5
Q9AU09 Q9AU10 Q9AU11 Q9AU12 Q9AUV6 Q9AV28 Q9AV47 Q9AV50 Q9AV81 Q9AV88 Q9AV97
Q9AVJ9 Q9AVK0 Q9AVK1 Q9AVK4 Q9AVL4 Q9AVL5 Q9AVL9 Q9AVQ1 Q9AW48 Q9AW90 Q9AWA5

Q9AWB2 Q9AWK2 Q9AWM8 Q9AWM9 Q9AWS0 Q9AWS7 Q9AWZ7 Q9AWZ8 Q9AXA6 Q9AXB0 Q9AXE3
Q9AXH0 Q9AXJ4 Q9AXK1 Q9AXK2 Q9AXM6 Q9AXQ3 Q9AXQ4 Q9AXQ5 Q9AXQ6 Q9AXQ7 Q9AXQ8
Q9AXQ9 Q9AXR0 Q9AXT8 Q9AXU3 Q9AXU4 Q9AY76 Q9AYE4 Q9AYT5 Q9AYT7 Q9AYU0 Q9AYZ1
Q9AZ38 Q9B133 Q9B1A1 Q9B1H9 Q9B1K6 Q9B1K8 Q9B1U9 Q9BA08 Q9BA49 Q9BAB9 Q9BAC0
Q9BAC2 Q9BAC3 Q9BAC5 Q9BBN0 Q9BBN1 Q9BBN7 Q9BBP5 Q9BBP7 Q9BBP8 Q9BBP9 Q9BBQ0
Q9BBQ1 Q9BBQ2 Q9BBQ3 Q9BBQ4 Q9BBQ9 Q9BBR0 Q9BBR1 Q9BBR2 Q9BBS6 Q9BBS7 Q9BBS8
Q9BBS9 Q9BBT3 Q9BBT5 Q9BBU1 Q9BBU2 Q9BCJ9 Q9BDB7 Q9BDJ7 Q9BDY9 Q9BE01 Q9BE11
Q9BE24 Q9BE27 Q9BE39 Q9BE40 Q9BE41 Q9BE52 Q9BE57 Q9BE64 Q9BE70 Q9BE73 Q9BEA9
Q9BEC7 Q9BG88 Q9BG90 Q9BGI0 Q9BGI3 Q9BGM5 Q9BGN8 Q9BGQ5 Q9BGT1 Q9BGW0 Q9BGW4
Q9BGX5 Q9BH04 Q9BH05 Q9BH08 Q9BH80 Q9BHM6 Q9BHU1 Q9BI06 Q9BI14 Q9BI40 Q9BI74
Q9BI88 Q9BIA4 Q9BIB3 Q9BIB4 Q9BIW5 Q9BJK5 Q9BJZ5 Q9BKQ8 Q9BKR0 Q9BKU8 Q9BKZ9
Q9BL02 Q9BLB6 Q9BLC4 Q9BLG0 Q9BM89 Q9BM90 Q9BM91 Q9BM92 Q9BM93 Q9BM94 Q9BM95
Q9BM96 Q9BM97 Q9BM98 Q9BM99 Q9BMK3 Q9BMM0 Q9BMM1 Q9BMM3 Q9BMM8 Q9BMP6 Q9BMQ6
Q9BMX0 Q9BMX3 Q9BMX5 Q9BPL7 Q9BPU3 Q9BPW5 Q9BPX7 Q9BPY8 Q9BPZ2 Q9BPZ3 Q9BQ13
Q9BQ15 Q9BQ24 Q9BQ50 Q9BQ61 Q9BQ65 Q9BQ66 Q9BQ83 Q9BQ87 Q9BQ89 Q9BQ90 Q9BQA5
Q9BQD1 Q9BQE6 Q9BQI3 Q9BQI6 Q9BQI9 Q9BQM9 Q9BQP7 Q9BQW3 Q9BQY9 Q9BR11 Q9BR46
Q9BR84 Q9BRA0 Q9BRA2 Q9BRH9 Q9BRJ7 Q9BRP4 Q9BRP7 Q9BRP9 Q9BRQ3 Q9BRQ4 Q9BRR0
Q9BRT6 Q9BRT8 Q9BRU2 Q9BRV8 Q9BRX5 Q9BRZ2 Q9BS18 Q9BS31 Q9BS40 Q9BS92 Q9BSC4
Q9BSD3 Q9BSE5 Q9BSG1 Q9BSH3 Q9BSH4 Q9BSJ1 Q9BSK1 Q9BSK4 Q9BSQ5 Q9BSU3 Q9BSV6
Q9BSY4 Q9BT43 Q9BT73 Q9BT81 Q9BTA0 Q9BTD1 Q9BTE0 Q9BTE6 Q9BTF0 Q9BTK2 Q9BTK6
Q9BTL4 Q9BTM1 Q9BTM9 Q9BTT4 Q9BTV5 Q9BTV6 Q9BU02 Q9BU19 Q9BU68 Q9BU70 Q9BU89
Q9BUA6 Q9BUB4 Q9BUE0 Q9BUE6 Q9BUG6 Q9BUK0 Q9BUU2 Q9BUV0 Q9BUX1 Q9BUY5 Q9BUY7
Q9BV19 Q9BV29 Q9BV36 Q9BV38 Q9BV44 Q9BV86 Q9BV90 Q9BV97 Q9BV99 Q9BVG3 Q9BVJ7
Q9BVL4 Q9BVM2 Q9BVM4 Q9BVQ7 Q9BVR0 Q9BVS5 Q9BVV2 Q9BVV6 Q9BW11 Q9BW30 Q9BW62
Q9BW66 Q9BWD3 Q9BWJ2 Q9BWJ5 Q9BWK5 Q9BWM5 Q9BWN1 Q9BWT6 Q9BWU1 Q9BWV7 Q9BWW7
Q9BWX5 Q9BX10 Q9BX26 Q9BX46 Q9BX51 Q9BX69 Q9BX70 Q9BX82 Q9BXB7 Q9BXD5 Q9BXG8
Q9BXI3 Q9BXL5 Q9BXR3 Q9BXT4 Q9BXT5 Q9BXT6 Q9BXT8 Q9BXU0 Q9BXU1 Q9BXU2 Q9BXU3
Q9BXU8 Q9BXV9 Q9BXW3 Q9BXX2 Q9BXY5 Q9BY12 Q9BY27 Q9BY31 Q9BY41 Q9BY42 Q9BY66
Q9BY84 Q9BYB4 Q9BYC2 Q9BYD5 Q9BYD9 Q9BYE0 Q9BYE3 Q9BYG7 Q9BYJ4 Q9BYN0 Q9BYN7
Q9BYP8 Q9BYP9 Q9BYQ0 Q9BYQ2 Q9BYQ3 Q9BYQ4 Q9BYQ5 Q9BYQ6 Q9BYQ7 Q9BYQ8 Q9BYQ9
Q9BYR0 Q9BYR2 Q9BYR3 Q9BYR4 Q9BYR5 Q9BYR6 Q9BYR7 Q9BYR8 Q9BYR9 Q9BYS1 Q9BYT5
Q9BYU1 Q9BYU5 Q9BYV1 Q9BYX7 Q9BYZ2 Q9BZ19 Q9BZ81 Q9BZ98 Q9BZA0 Q9BZA5 Q9BZC1
Q9BZE2 Q9BZE3 Q9BZE7 Q9BZF9 Q9BZG8 Q9BZJ0 Q9BZK8 Q9BZL1 Q9BZM3 Q9BZP3 Q9BZQ2
Q9BZR9 Q9BZS9 Q9BZW7 Q9BZY9 Q9C002 Q9C005 Q9C009 Q9C010 Q9C019 Q9C029 Q9C030
Q9C035 Q9C040 Q9C056 Q9C075 Q9C086 Q9C093 Q9C098 Q9C099 Q9C0A1 Q9C0B2 Q9C0D3
Q9C0D4 Q9C0D7 Q9C0F1 Q9C0F3 Q9C0G6 Q9C0H6 Q9C0H9 Q9C0I3 Q9C0I9 Q9C0J9 Q9C0K7
Q9C0N4 Q9C0T1 Q9C0T9 Q9C0U3 Q9C0U6 Q9C0V4 Q9C0V7 Q9C0V9 Q9C0W0 Q9C0W1 Q9C0W2
Q9C0W4 Q9C0X0 Q9C0X2 Q9C0X3 Q9C0X6 Q9C0Y6 Q9C0Y8 Q9C0Z3 Q9C0Z8 Q9C0Z9 Q9C102
Q9C103 Q9C104 Q9C106 Q9C107 Q9C108 Q9C109 Q9C110 Q9C111 Q9C114 Q9C115 Q9C116
Q9C117 Q9C118 Q9C124 Q9C131 Q9C136 Q9C141 Q9C150 Q9C167 Q9C196 Q9C1A4 Q9C1B5
Q9C1B7 Q9C1C2 Q9C1D4 Q9C1J2 Q9C1J4 Q9C1K0 Q9C1K6 Q9C1W6 Q9C1W9 Q9C1X0 Q9C1X1
Q9C1X2 Q9C1X4 Q9C1X5 Q9C224 Q9C239 Q9C245 Q9C270 Q9C285 Q9C291 Q9C2B9 Q9C2H9
Q9C2I5 Q9C2N1 Q9C2P2 Q9C2R2 Q9C2U0 Q9C2U9 Q9C2Y6 Q9C3Z4 Q9C3Z5 Q9C3Z6 Q9C401
Q9C413 Q9C448 Q9C451 Q9C467 Q9C469 Q9C4B7 Q9C4M3 Q9C4M5 Q9C4Z0 Q9C4Z1 Q9C4Z3

Q9C4Z4 Q9C501 Q9C507 Q9C514 Q9C518 Q9C519 Q9C524 Q9C534 Q9C544 Q9C545 Q9C550
Q9C551 Q9C552 Q9C557 Q9C560 Q9C566 Q9C578 Q9C587 Q9C593 Q9C598 Q9C5A4 Q9C5B9
Q9C5C4 Q9C5C5 Q9C5C8 Q9C5D2 Q9C5D7 Q9C5E5 Q9C5E7 Q9C5G0 Q9C5G7 Q9C5H9 Q9C5I1
Q9C5J4 Q9C5J6 Q9C5J7 Q9C5J9 Q9C5K7 Q9C5M1 Q9C5P0 Q9C5P1 Q9C5P4 Q9C5Q2 Q9C5Q8
Q9C5Q9 Q9C5S1 Q9C5T4 Q9C5U3 Q9C5U8 Q9C5V6 Q9C5W3 Q9C5W9 Q9C5X0 Q9C5X2 Q9C5X8
Q9C5X9 Q9C5Y4 Q9C5Z1 Q9C5Z2 Q9C613 Q9C616 Q9C626 Q9C627 Q9C629 Q9C633 Q9C638
Q9C641 Q9C642 Q9C646 Q9C652 Q9C658 Q9C667 Q9C670 Q9C671 Q9C681 Q9C682 Q9C685
Q9C689 Q9C690 Q9C691 Q9C698 Q9C6A9 Q9C6C3 Q9C6C8 Q9C6D2 Q9C6F5 Q9C6F7 Q9C6G0
Q9C6H2 Q9C6H3 Q9C6H4 Q9C6H5 Q9C6I2 Q9C6I4 Q9C6I6 Q9C6I7 Q9C6I8 Q9C6J2 Q9C6J3
Q9C6K3 Q9C6K5 Q9C6L1 Q9C6M5 Q9C6M9 Q9C6N2 Q9C6S2 Q9C6S6 Q9C6T2 Q9C6U1 Q9C6V3
Q9C6V4 Q9C6W6 Q9C6W9 Q9C6X0 Q9C6X9 Q9C6Y3 Q9C6Z0 Q9C6Z2 Q9C6Z3 Q9C701 Q9C707
Q9C710 Q9C713 Q9C717 Q9C718 Q9C723 Q9C725 Q9C748 Q9C758 Q9C768 Q9C774 Q9C777
Q9C778 Q9C793 Q9C7A8 Q9C7B1 Q9C7C3 Q9C7C4 Q9C7E6 Q9C7E7 Q9C7E8 Q9C7F1 Q9C7F5
Q9C7G0 Q9C7G4 Q9C7I3 Q9C7I7 Q9C7I9 Q9C7J9 Q9C7K0 Q9C7M1 Q9C7N3 Q9C7P1 Q9C7P2
Q9C7P4 Q9C7P6 Q9C7R6 Q9C7T4 Q9C7U7 Q9C7V5 Q9C7W7 Q9C7X5 Q9C7X7 Q9C7X8 Q9C7Y2
Q9C7Y4 Q9C7Z1 Q9C800 Q9C801 Q9C810 Q9C813 Q9C829 Q9C866 Q9C869 Q9C870 Q9C876
Q9C878 Q9C895 Q9C899 Q9C8B8 Q9C8D1 Q9C8E1 Q9C8E3 Q9C8F1 Q9C8F7 Q9C8J1 Q9C8J2
Q9C8J4 Q9C8J7 Q9C8K0 Q9C8K1 Q9C8K7 Q9C8L2 Q9C8L4 Q9C8L6 Q9C8M2 Q9C8M3 Q9C8M4
Q9C8N5 Q9C8N7 Q9C8N9 Q9C8P0 Q9C8P7 Q9C8P8 Q9C8R8 Q9C8S9 Q9C8T7 Q9C8T8 Q9C8T9
Q9C8U0 Q9C8U2 Q9C8U9 Q9C8V2 Q9C8V8 Q9C8X7 Q9C8Y1 Q9C8Y4 Q9C8Y5 Q9C8Y6 Q9C8Y7
Q9C8Y8 Q9C8Z4 Q9C8Z9 Q9C912 Q9C918 Q9C923 Q9C928 Q9C932 Q9C944 Q9C948 Q9C950
Q9C951 Q9C955 Q9C966 Q9C968 Q9C969 Q9C971 Q9C977 Q9C983 Q9C991 Q9C998 Q9C9A2
Q9C9A6 Q9C9A9 Q9C9B0 Q9C9B2 Q9C9C4 Q9C9C5 Q9C9C6 Q9C9C8 Q9C9C9 Q9C9D0 Q9C9F0
Q9C9F4 Q9C9F5 Q9C9G7 Q9C9H1 Q9C9H9 Q9C9K2 Q9C9K7 Q9C9L0 Q9C9L2 Q9C9L4 Q9C9M0
Q9C9M2 Q9C9M3 Q9C9M4 Q9C9M6 Q9C9M7 Q9C9N3 Q9C9N6 Q9C9R3 Q9C9R6 Q9C9S2 Q9C9S4
Q9C9T2 Q9C9T3 Q9C9U0 Q9C9U2 Q9C9U5 Q9C9U8 Q9C9V6 Q9C9W3 Q9C9W4 Q9C9W9 Q9C9X0
Q9C9X3 Q9C9X6 Q9C9X7 Q9C9X9 Q9C9Y4 Q9C9Y7 Q9C9Z0 Q9C9Z7 Q9CA02 Q9CA04 Q9CA19
Q9CA23 Q9CA28 Q9CA30 Q9CA36 Q9CA40 Q9CA45 Q9CA51 Q9CA54 Q9CA57 Q9CA58 Q9CA63
Q9CA64 Q9CA66 Q9CA73 Q9CA74 Q9CA78 Q9CA83 Q9CA90 Q9CAA4 Q9CAA5 Q9CAA7 Q9CAA9
Q9CAB6 Q9CAC3 Q9CAD0 Q9CAE3 Q9CAE4 Q9CAE7 Q9CAE9 Q9CAF2 Q9CAF6 Q9CAF9 Q9CAG5
Q9CAG8 Q9CAH0 Q9CAH1 Q9CAI1 Q9CAI3 Q9CAI7 Q9CAJ9 Q9CAK8 Q9CAM1 Q9CAM7 Q9CAM8
Q9CAN0 Q9CAN3 Q9CAN4 Q9CAN5 Q9CAN6 Q9CAP3 Q9CAP9 Q9CAQ2 Q9CAR0 Q9CAR4 Q9CAS1
Q9CAT7 Q9CAU7 Q9CAV0 Q9CAV7 Q9CAX6 Q9CB17 Q9CB23 Q9CB26 Q9CB28 Q9CB36 Q9CB42
Q9CB44 Q9CB45 Q9CB48 Q9CB76 Q9CB81 Q9CB92 Q9CBA8 Q9CBB0 Q9CBB2 Q9CBC6 Q9CBC9
Q9CBD6 Q9CBG0 Q9CBH9 Q9CBI2 Q9CBJ5 Q9CBJ6 Q9CBJ8 Q9CBK0 Q9CBK1 Q9CBK2 Q9CBK7
Q9CBM2 Q9CBM3 Q9CBM9 Q9CBP9 Q9CBQ0 Q9CBQ2 Q9CBS0 Q9CBS3 Q9CBS6 Q9CBS9 Q9CBT0
Q9CBT5 Q9CBU0 Q9CBU3 Q9CBU5 Q9CBW0 Q9CBW1 Q9CBW3 Q9CBW4 Q9CBW6 Q9CBX1 Q9CBX4
Q9CBX6 Q9CBY1 Q9CBY2 Q9CBY3 Q9CBY4 Q9CBY6 Q9CBY7 Q9CBZ0 Q9CBZ2 Q9CBZ3 Q9CBZ4
Q9CBZ5 Q9CBZ6 Q9CBZ7 Q9CBZ8 Q9CC09 Q9CC10 Q9CC11 Q9CC12 Q9CC13 Q9CC14 Q9CC15
Q9CC16 Q9CC17 Q9CC20 Q9CC21 Q9CC22 Q9CC24 Q9CC53 Q9CC54 Q9CC56 Q9CC57 Q9CC59
Q9CC85 Q9CC91 Q9CC97 Q9CCA7 Q9CCA8 Q9CCB5 Q9CCB8 Q9CCE1 Q9CCE2 Q9CCE3 Q9CCF8
Q9CCG2 Q9CCG3 Q9CCH9 Q9CCI3 Q9CCJ4 Q9CCK1 Q9CCK2 Q9CCL9 Q9CCN0 Q9CCN2 Q9CCN8
Q9CCN9 Q9CCP0 Q9CCP1 Q9CCP3 Q9CCP9 Q9CCQ0 Q9CCQ3 Q9CCQ4 Q9CCQ6 Q9CCQ7 Q9CCR1
Q9CCR2 Q9CCR3 Q9CCR4 Q9CCR5 Q9CCR9 Q9CCS0 Q9CCS3 Q9CCS4 Q9CCS5 Q9CCS6 Q9CCS9

Q9CCT0 Q9CCT5 Q9CCW2 Q9CCW4 Q9CCW5 Q9CCW6 Q9CCZ3 Q9CCZ4 Q9CCZ9 Q9CD45 Q9CD48
Q9CD49 Q9CD51 Q9CD52 Q9CD55 Q9CD56 Q9CD57 Q9CD69 Q9CD72 Q9CD75 Q9CD83 Q9CD99
Q9CDC1 Q9CDC4 Q9CDD2 Q9CDE9 Q9CDF4 Q9CDF9 Q9CDG0 Q9CDG1 Q9CDG3 Q9CDG7 Q9CDH8
Q9CDI7 Q9CDK9 Q9CDL1 Q9CDL2 Q9CDL3 Q9CDL9 Q9CDM8 Q9CDM9 Q9CDN0 Q9CDN7 Q9CDP0
Q9CDP6 Q9CDQ6 Q9CDR4 Q9CDR5 Q9CDS1 Q9CDS7 Q9CDT1 Q9CDT4 Q9CDT7 Q9CDV5 Q9CDV7
Q9CDW1 Q9CDW2 Q9CDW3 Q9CDW5 Q9CDW6 Q9CDW7 Q9CDW8 Q9CDW9 Q9CDX0 Q9CDX1 Q9CDX2
Q9CDX3 Q9CDX4 Q9CDX5 Q9CDX7 Q9CDX8 Q9CDX9 Q9CDY0 Q9CDY1 Q9CDY2 Q9CDY3 Q9CDY4
Q9CDZ7 Q9CE00 Q9CE01 Q9CE10 Q9CE11 Q9CE12 Q9CE21 Q9CE35 Q9CE36 Q9CE38 Q9CE39
Q9CE42 Q9CE43 Q9CE46 Q9CE47 Q9CE63 Q9CE70 Q9CE78 Q9CE80 Q9CE86 Q9CE93 Q9CEB4
Q9CEB5 Q9CEB7 Q9CEC5 Q9CEC9 Q9CED2 Q9CEE0 Q9CEE3 Q9CEE4 Q9CEE9 Q9CEF5 Q9CEF6
Q9CEG3 Q9CEH4 Q9CEH8 Q9CEI0 Q9CEI4 Q9CEI6 Q9CEJ0 Q9CEJ1 Q9CEJ4 Q9CEJ7 Q9CEJ8
Q9CEJ9 Q9CEK9 Q9CEL6 Q9CEM5 Q9CEM7 Q9CEN1 Q9CEN6 Q9CEN7 Q9CEP1 Q9CES7 Q9CES8
Q9CET0 Q9CET2 Q9CET9 Q9CEU0 Q9CEU1 Q9CEU2 Q9CEU5 Q9CEV7 Q9CEW0 Q9CEW2 Q9CEX2
Q9CEY1 Q9CEY4 Q9CEY6 Q9CF21 Q9CF34 Q9CF36 Q9CF39 Q9CF41 Q9CF42 Q9CF43 Q9CF50
Q9CF53 Q9CF61 Q9CF72 Q9CF73 Q9CF77 Q9CF79 Q9CF80 Q9CF85 Q9CF91 Q9CF94 Q9CFA6
Q9CFA8 Q9CFB2 Q9CFB4 Q9CFB6 Q9CFB7 Q9CFC0 Q9CFC2 Q9CFC3 Q9CFC8 Q9CFC9 Q9CFE7
Q9CFE8 Q9CFF3 Q9CFF4 Q9CFG0 Q9CFG7 Q9CFG8 Q9CFH4 Q9CFJ0 Q9CFM3 Q9CFM7 Q9CFV1
Q9CFV2 Q9CFW7 Q9CFW8 Q9CFW9 Q9CFX1 Q9CFY8 Q9CFZ0 Q9CG17 Q9CG20 Q9CG28 Q9CG29
Q9CG32 Q9CG33 Q9CG34 Q9CG41 Q9CG42 Q9CG46 Q9CG48 Q9CG49 Q9CG55 Q9CG64 Q9CG65
Q9CG73 Q9CG78 Q9CG79 Q9CG80 Q9CG88 Q9CGA9 Q9CGB9 Q9CGC2 Q9CGD5 Q9CGD7 Q9CGD8
Q9CGE1 Q9CGE3 Q9CGE5 Q9CGE6 Q9CGF0 Q9CGF1 Q9CGG6 Q9CGG8 Q9CGH0 Q9CGJ4 Q9CGJ7
Q9CGJ9 Q9CGK0 Q9CGK1 Q9CGL0 Q9CGL5 Q9CGL7 Q9CGM3 Q9CGM7 Q9CGM8 Q9CGN6 Q9CGS5
Q9CGT5 Q9CGT6 Q9CGU6 Q9CGW5 Q9CGW6 Q9CGX1 Q9CGX3 Q9CGX9 Q9CGY0 Q9CGY1 Q9CGY9
Q9CGZ0 Q9CGZ5 Q9CGZ6 Q9CH00 Q9CH07 Q9CH08 Q9CH10 Q9CH12 Q9CH51 Q9CH65 Q9CH73
Q9CH83 Q9CH84 Q9CH85 Q9CH94 Q9CH98 Q9CHA1 Q9CHB6 Q9CHB8 Q9CHD0 Q9CHD1 Q9CHD2
Q9CHD3 Q9CHD4 Q9CHD5 Q9CHE0 Q9CHE9 Q9CHG3 Q9CHG8 Q9CHH6 Q9CHI1 Q9CHI2 Q9CHI6
Q9CHK0 Q9CHK3 Q9CHM9 Q9CHN1 Q9CHN4 Q9CHN6 Q9CHN8 Q9CHN9 Q9CHQ3 Q9CHQ8 Q9CHT2
Q9CHT4 Q9CHT5 Q9CHT8 Q9CHU2 Q9CHU6 Q9CHV0 Q9CHW2 Q9CHW3 Q9CHW5 Q9CHW7 Q9CHX0
Q9CHX3 Q9CHX5 Q9CHZ4 Q9CI06 Q9CI09 Q9CI15 Q9CI17 Q9CI19 Q9CI20 Q9CI21 Q9CI22
Q9CI63 Q9CI64 Q9CI70 Q9CI75 Q9CI77 Q9CI80 Q9CID5 Q9CID9 Q9CIE4 Q9CIF5 Q9CIG1
Q9CIG4 Q9CIH5 Q9CIH9 Q9CII1 Q9CII4 Q9CII7 Q9CIJ4 Q9CIJ9 Q9CIK4 Q9CIK7 Q9CIL5
Q9CIL6 Q9CIM0 Q9CIP4 Q9CIR1 Q9CIR5 Q9CIR9 Q9CIS0 Q9CIS2 Q9CIS4 Q9CIS5 Q9CIU7
Q9CIU9 Q9CIV4 Q9CIV6 Q9CIV7 Q9CIV8 Q9CIV9 Q9CIW0 Q9CIW1 Q9CIW6 Q9CIX2 Q9CIX4
Q9CIX6 Q9CJ14 Q9CJ17 Q9CJ27 Q9CJ30 Q9CJ41 Q9CJ43 Q9CJ44 Q9CJ45 Q9CJ54 Q9CJ62
Q9CJ63 Q9CJ64 Q9CJ70 Q9CJ73 Q9CJ76 Q9CJ80 Q9CJ86 Q9CJ92 Q9CJ97 Q9CJA6 Q9CJD1
Q9CJD9 Q9CJH1 Q9CJH4 Q9CJI1 Q9CJI9 Q9CJJ0 Q9CJJ1 Q9CJJ2 Q9CJJ8 Q9CJK1 Q9CJL5
Q9CJM2 Q9CJN5 Q9CJN6 Q9CJN7 Q9CJN8 Q9CJN9 Q9CJP3 Q9CJP4 Q9CJR1 Q9CJR4 Q9CJR9
Q9CJS1 Q9CJS9 Q9CJT7 Q9CJU0 Q9CJU5 Q9CJV6 Q9CJV7 Q9CJW1 Q9CJW4 Q9CJW5 Q9CJW9
Q9CJX0 Q9CJX1 Q9CJX3 Q9CJZ9 Q9CK01 Q9CK02 Q9CK03 Q9CK07 Q9CK20 Q9CK24 Q9CK31
Q9CK32 Q9CK40 Q9CK44 Q9CK54 Q9CK55 Q9CK59 Q9CK61 Q9CK62 Q9CK63 Q9CK66 Q9CK84
Q9CK85 Q9CK86 Q9CK89 Q9CK90 Q9CK91 Q9CK92 Q9CK94 Q9CKA2 Q9CKA3 Q9CKA6 Q9CKA9
Q9CKB0 Q9CKB3 Q9CKC0 Q9CKC2 Q9CKD6 Q9CKD7 Q9CKE9 Q9CKF3 Q9CKF5 Q9CKF7 Q9CKG0
Q9CKG2 Q9CKG3 Q9CKG4 Q9CKG8 Q9CKH9 Q9CKJ2 Q9CKJ5 Q9CKJ8 Q9CKK0 Q9CKK4 Q9CKK7
Q9CKL0 Q9CKM2 Q9CKN6 Q9CKN9 Q9CKP2 Q9CKP7 Q9CKP9 Q9CKQ9 Q9CKR4 Q9CKU9 Q9CKV0

Q9CKV2 Q9CKV3 Q9CKV5 Q9CKV9 Q9CKW8 Q9CKY6 Q9CL02 Q9CL12 Q9CL16 Q9CL21 Q9CL27
Q9CL29 Q9CL32 Q9CL33 Q9CL34 Q9CL35 Q9CL36 Q9CL37 Q9CL38 Q9CL39 Q9CL40 Q9CL41
Q9CL42 Q9CL43 Q9CL44 Q9CL45 Q9CL46 Q9CL47 Q9CL48 Q9CL49 Q9CL51 Q9CL52 Q9CL53
Q9CL54 Q9CL60 Q9CL72 Q9CL85 Q9CL86 Q9CL88 Q9CL89 Q9CL92 Q9CLA3 Q9CLB1 Q9CLB9
Q9CLC2 Q9CLC6 Q9CLD4 Q9CLE0 Q9CLE1 Q9CLE2 Q9CLE6 Q9CLE7 Q9CLE9 Q9CLF1 Q9CLF7
Q9CLG4 Q9CLG6 Q9CLG7 Q9CLH5 Q9CLI9 Q9CLJ0 Q9CLJ5 Q9CLJ6 Q9CLL5 Q9CLL7 Q9CLL8
Q9CLL9 Q9CLM0 Q9CLM1 Q9CLM2 Q9CLM3 Q9CLM4 Q9CLN8 Q9CLN9 Q9CLP0 Q9CLP2 Q9CLP9
Q9CLQ1 Q9CLQ4 Q9CLQ6 Q9CLQ8 Q9CLQ9 Q9CLR1 Q9CLR7 Q9CLT2 Q9CLT7 Q9CLT8 Q9CLT9
Q9CLU1 Q9CLU4 Q9CLV5 Q9CLV7 Q9CLW2 Q9CLW3 Q9CLW4 Q9CLW7 Q9CLX3 Q9CLY2 Q9CLY7
Q9CLZ5 Q9CLZ7 Q9CM00 Q9CM13 Q9CM16 Q9CM17 Q9CM18 Q9CM20 Q9CM22 Q9CM49 Q9CM50
Q9CM51 Q9CM57 Q9CM61 Q9CM79 Q9CM85 Q9CM94 Q9CM98 Q9CMA4 Q9CMB1 Q9CMB2 Q9CMB4
Q9CMB5 Q9CMC6 Q9CMC7 Q9CMD1 Q9CMF1 Q9CMF4 Q9CMG9 Q9CMI6 Q9CMI7 Q9CMI8 Q9CMJ6
Q9CMJ8 Q9CMJ9 Q9CMK1 Q9CMK4 Q9CMK5 Q9CMK8 Q9CMK9 Q9CML8 Q9CMM1 Q9CMM8 Q9CMM9
Q9CMN5 Q9CMQ4 Q9CMQ6 Q9CMQ7 Q9CMQ8 Q9CMR2 Q9CMR5 Q9CMS2 Q9CMW8 Q9CMX5 Q9CMY2
Q9CMY9 Q9CMZ2 Q9CMZ3 Q9CN04 Q9CN06 Q9CN08 Q9CN10 Q9CN18 Q9CN20 Q9CN21 Q9CN22
Q9CN24 Q9CN30 Q9CN31 Q9CN36 Q9CN37 Q9CN38 Q9CN39 Q9CN40 Q9CN41 Q9CN42 Q9CN56
Q9CN71 Q9CN81 Q9CN82 Q9CN86 Q9CN88 Q9CN89 Q9CN90 Q9CN91 Q9CN92 Q9CN94 Q9CN96
Q9CN99 Q9CNA0 Q9CNA2 Q9CNA6 Q9CNB0 Q9CNB1 Q9CNB2 Q9CNB3 Q9CNC4 Q9CNC7 Q9CNC8
Q9CND0 Q9CND4 Q9CND8 Q9CND9 Q9CNE0 Q9CNE1 Q9CNE3 Q9CNF0 Q9CNF2 Q9CNF3 Q9CNF6
Q9CNG4 Q9CNG9 Q9CNH1 Q9CNH3 Q9CNH4 Q9CNH7 Q9CNI7 Q9CNJ0 Q9CNJ3 Q9CNL2 Q9CNL5
Q9CNL6 Q9CNM2 Q9CNN1 Q9CNN3 Q9CNN6 Q9CNN7 Q9CNP8 Q9CNS4 Q9CNS7 Q9CNS8 Q9CNT1
Q9CNT5 Q9CNT7 Q9CNU1 Q9CNU2 Q9CNU9 Q9CNV2 Q9CNV3 Q9CNV8 Q9CNV9 Q9CNW9 Q9CNX1
Q9CNX7 Q9CNX8 Q9CNY1 Q9CNY2 Q9CNY5 Q9CNZ0 Q9CP02 Q9CP05 Q9CP11 Q9CP15 Q9CP21
Q9CP22 Q9CP41 Q9CP49 Q9CP53 Q9CP62 Q9CP66 Q9CP69 Q9CP72 Q9CP74 Q9CP85 Q9CP86
Q9CP88 Q9CP90 Q9CP97 Q9CPA2 Q9CPB0 Q9CPB4 Q9CPB5 Q9CPC8 Q9CPD1 Q9CPD3 Q9CPD5
Q9CPD6 Q9CPD7 Q9CPD8 Q9CPE0 Q9CPE8 Q9CPF0 Q9CPF4 Q9CPF5 Q9CPH0 Q9CPH1 Q9CPH2
Q9CPH5 Q9CPH6 Q9CPI4 Q9CPI5 Q9CPK3 Q9CPL6 Q9CPL8 Q9CPN4 Q9CPN6 Q9CPP0 Q9CPR1
Q9CPR7 Q9CPR8 Q9CPS7 Q9CPT0 Q9CPT3 Q9CPV2 Q9CPV4 Q9CPW2 Q9CPW4 Q9CPW7 Q9CPW9
Q9CPX9 Q9CPY0 Q9CPY3 Q9CPY4 Q9CPY6 Q9CPY7 Q9CPZ1 Q9CPZ8 Q9CQ04 Q9CQ07 Q9CQ18
Q9CQ19 Q9CQ21 Q9CQ24 Q9CQ25 Q9CQ28 Q9CQ29 Q9CQ36 Q9CQ37 Q9CQ39 Q9CQ43 Q9CQ46
Q9CQ48 Q9CQ49 Q9CQ60 Q9CQ65 Q9CQ66 Q9CQ70 Q9CQ71 Q9CQ79 Q9CQ84 Q9CQ90 Q9CQA1
Q9CQA5 Q9CQA9 Q9CQB2 Q9CQB7 Q9CQC6 Q9CQE0 Q9CQE1 Q9CQE6 Q9CQE8 Q9CQE9 Q9CQF3
Q9CQF4 Q9CQF6 Q9CQG1 Q9CQG2 Q9CQH7 Q9CQH8 Q9CQI3 Q9CQI4 Q9CQI7 Q9CQI9 Q9CQJ2
Q9CQJ4 Q9CQJ5 Q9CQJ6 Q9CQK1 Q9CQK3 Q9CQK7 Q9CQL0 Q9CQL1 Q9CQL2 Q9CQL7 Q9CQM0
Q9CQM5 Q9CQM6 Q9CQP3 Q9CQQ4 Q9CQQ8 Q9CQR5 Q9CQR6 Q9CQS2 Q9CQS7 Q9CQT1 Q9CQT2
Q9CQT6 Q9CQV3 Q9CQW5 Q9CQX8 Q9CQY2 Q9CQZ1 Q9CQZ7 Q9CR00 Q9CR02 Q9CR04 Q9CR08
Q9CR09 Q9CR10 Q9CR24 Q9CR25 Q9CR29 Q9CR34 Q9CR37 Q9CR40 Q9CR41 Q9CR55 Q9CR70
Q9CR73 Q9CR78 Q9CR80 Q9CR81 Q9CR83 Q9CR86 Q9CR92 Q9CR98 Q9CRA8 Q9CRA9 Q9CRB2
Q9CRB3 Q9CRB6 Q9CRC3 Q9CRC9 Q9CRD0 Q9CRD4 Q9CRT8 Q9CRW3 Q9CS00 Q9CS42 Q9CS74
Q9CSH3 Q9CSN1 Q9CSP9 Q9CSU0 Q9CT10 Q9CTH6 Q9CTN5 Q9CU24 Q9CU62 Q9CU65 Q9CUL5
Q9CUU3 Q9CUX1 Q9CVI2 Q9CVN6 Q9CVW4 Q9CW07 Q9CWB5 Q9CWB7 Q9CWD0 Q9CWD3 Q9CWE0
Q9CWE6 Q9CWG8 Q9CWH0 Q9CWH1 Q9CWH4 Q9CWH5 Q9CWH6 Q9CWI3 Q9CWK3 Q9CWM4 Q9CWN7
Q9CWP8 Q9CWP9 Q9CWQ0 Q9CWQ2 Q9CWR1 Q9CWR2 Q9CWR8 Q9CWS0 Q9CWS1 Q9CWS4 Q9CWT2
Q9CWU0 Q9CWU4 Q9CWU5 Q9CWV0 Q9CWV1 Q9CWV4 Q9CWV6 Q9CWV7 Q9CWW6 Q9CWW7 Q9CWX4

Q9CWX9 Q9CWY3 Q9CWY4 Q9CWY8 Q9CWY9 Q9CWZ3 Q9CX11 Q9CX25 Q9CX34 Q9CX48 Q9CX53
Q9CX56 Q9CX60 Q9CX62 Q9CX66 Q9CX80 Q9CX86 Q9CX97 Q9CXA2 Q9CXB8 Q9CXC3 Q9CXE0
Q9CXE2 Q9CXE6 Q9CXF0 Q9CXF4 Q9CXG3 Q9CXG9 Q9CXJ1 Q9CXK8 Q9CXK9 Q9CXL3 Q9CXP9
Q9CXT6 Q9CXU0 Q9CXU1 Q9CXU9 Q9CXV0 Q9CXV3 Q9CXV9 Q9CXW3 Q9CXY6 Q9CY00 Q9CY02
Q9CY21 Q9CY28 Q9CY34 Q9CY45 Q9CY52 Q9CY57 Q9CY62 Q9CY64 Q9CY66 Q9CY86 Q9CY94
Q9CY97 Q9CYA6 Q9CYC6 Q9CYI0 Q9CYI4 Q9CYP7 Q9CYQ7 Q9CYR0 Q9CYR6 Q9CYS6 Q9CYU6
Q9CYX7 Q9CYY7 Q9CYZ2 Q9CYZ6 Q9CYZ8 Q9CZ00 Q9CZ04 Q9CZ05 Q9CZ15 Q9CZ42 Q9CZ49
Q9CZ57 Q9CZ82 Q9CZ91 Q9CZ96 Q9CZA5 Q9CZB3 Q9CZB6 Q9CZD5 Q9CZG3 Q9CZH3 Q9CZH8
Q9CZJ0 Q9CZJ1 Q9CZJ2 Q9CZJ6 Q9CZK6 Q9CZL5 Q9CZN8 Q9CZP0 Q9CZP7 Q9CZR8 Q9CZS3
Q9CZT4 Q9CZT6 Q9CZU3 Q9CZU6 Q9CZV2 Q9CZV5 Q9CZX0 Q9CZX5 Q9CZY2 Q9D009 Q9D020
Q9D051 Q9D061 Q9D067 Q9D071 Q9D0A0 Q9D0A3 Q9D0B0 Q9D0B1 Q9D0B5 Q9D0B6 Q9D0B8
Q9D0C4 Q9D0D4 Q9D0D5 Q9D0E1 Q9D0E3 Q9D0E5 Q9D0F4 Q9D0F6 Q9D0F9 Q9D0I6 Q9D0I8
Q9D0I9 Q9D0J8 Q9D0K0 Q9D0M0 Q9D0M1 Q9D0N7 Q9D0P7 Q9D0R2 Q9D0R4 Q9D0R8 Q9D0R9
Q9D0T2 Q9D0U1 Q9D0V8 Q9D0W5 Q9D110 Q9D112 Q9D113 Q9D114 Q9D119 Q9D131 Q9D142
Q9D168 Q9D172 Q9D174 Q9D180 Q9D187 Q9D198 Q9D1A0 Q9D1C2 Q9D1C9 Q9D1E4 Q9D1E6
Q9D1F3 Q9D1F4 Q9D1I5 Q9D1J3 Q9D1K7 Q9D1L0 Q9D1P2 Q9D1R2 Q9D1T5 Q9D1U0 Q9D1Z2
Q9D211 Q9D226 Q9D240 Q9D258 Q9D273 Q9D275 Q9D281 Q9D287 Q9D289 Q9D2C2 Q9D2C9
Q9D2D7 Q9D2D9 Q9D2H2 Q9D2H5 Q9D2H6 Q9D2H8 Q9D2I5 Q9D2J7 Q9D2K4 Q9D2L9 Q9D2M8
Q9D2P4 Q9D2Q2 Q9D2Q3 Q9D2Q8 Q9D2U5 Q9D2U9 Q9D2V5 Q9D2X0 Q9D2X5 Q9D2X8 Q9D2Y6
Q9D2Z4 Q9D2Z8 Q9D305 Q9D312 Q9D361 Q9D384 Q9D3D0 Q9D3E6 Q9D3I6 Q9D3J5 Q9D3N2
Q9D3N8 Q9D3P1 Q9D3R6 Q9D3R9 Q9D3S3 Q9D3S5 Q9D3V1 Q9D3V5 Q9D3W1 Q9D3W4 Q9D3W5
Q9D3X5 Q9D417 Q9D424 Q9D439 Q9D454 Q9D478 Q9D479 Q9D483 Q9D485 Q9D486 Q9D489
Q9D495 Q9D498 Q9D4A5 Q9D4B2 Q9D4C1 Q9D4D4 Q9D4F8 Q9D4G2 Q9D4G9 Q9D4H7 Q9D4H8
Q9D4H9 Q9D4I2 Q9D4K4 Q9D4K5 Q9D4K7 Q9D4P7 Q9D4Q1 Q9D4V3 Q9D4V4 Q9D504 Q9D513
Q9D516 Q9D518 Q9D529 Q9D531 Q9D534 Q9D541 Q9D552 Q9D565 Q9D572 Q9D581 Q9D5D8
Q9D5E4 Q9D5H4 Q9D5I4 Q9D5J6 Q9D5K4 Q9D5L7 Q9D5N8 Q9D5P4 Q9D5Q8 Q9D5R2 Q9D5R4
Q9D5S1 Q9D5S7 Q9D5U5 Q9D5U8 Q9D5V2 Q9D5W4 Q9D5W8 Q9D5Y0 Q9D5Y1 Q9D5Z5 Q9D5Z7
Q9D600 Q9D618 Q9D638 Q9D646 Q9D650 Q9D695 Q9D6D8 Q9D6E4 Q9D6K7 Q9D6L8 Q9D6N5
Q9D6P8 Q9D6R2 Q9D6S7 Q9D6T1 Q9D6V8 Q9D6Y9 Q9D6Z1 Q9D706 Q9D711 Q9D722 Q9D727
Q9D735 Q9D738 Q9D757 Q9D772 Q9D783 Q9D799 Q9D7B1 Q9D7B6 Q9D7E3 Q9D7E4 Q9D7G0
Q9D7G4 Q9D7G9 Q9D7H2 Q9D7H3 Q9D7H4 Q9D7I5 Q9D7I9 Q9D7K2 Q9D7M1 Q9D7M8 Q9D7N2
Q9D7P0 Q9D7P9 Q9D7V1 Q9D7V2 Q9D7W5 Q9D7X1 Q9D7X3 Q9D7X8 Q9D7Y9 Q9D7Z3 Q9D805
Q9D819 Q9D820 Q9D842 Q9D845 Q9D864 Q9D883 Q9D898 Q9D8C6 Q9D8C8 Q9D8F1 Q9D8I3
Q9D8L0 Q9D8M4 Q9D8M7 Q9D8N0 Q9D8N2 Q9D8N6 Q9D8P7 Q9D8Q6 Q9D8S4 Q9D8S9 Q9D8T7
Q9D8U7 Q9D8W5 Q9D8X1 Q9D8X5 Q9D8Y7 Q9D8Y8 Q9D902 Q9D903 Q9D924 Q9D937 Q9D939
Q9D945 Q9D967 Q9D975 Q9D979 Q9D994 Q9D995 Q9D9A7 Q9D9B4 Q9D9B7 Q9D9C6 Q9D9C7
Q9D9D3 Q9D9D9 Q9D9F8 Q9D9H3 Q9D9H8 Q9D9I1 Q9D9J2 Q9D9J3 Q9D9K8 Q9D9L5 Q9D9M4
Q9D9M5 Q9D9N5 Q9D9P2 Q9D9R7 Q9D9R9 Q9D9S1 Q9D9S3 Q9D9T0 Q9D9T6 Q9D9T8 Q9D9U4
Q9D9U9 Q9D9V3 Q9D9V4 Q9D9V7 Q9D9W0 Q9D9W6 Q9D9X6 Q9D9Z5 Q9D9Z7 Q9DA08 Q9DA15
Q9DA17 Q9DA42 Q9DA44 Q9DA60 Q9DA64 Q9DA69 Q9DA73 Q9DA80 Q9DA97 Q9DAA3 Q9DAA6
Q9DAA7 Q9DAC7 Q9DAC9 Q9DAE3 Q9DAE7 Q9DAE8 Q9DAF3 Q9DAF8 Q9DAG4 Q9DAG5 Q9DAH1
Q9DAI2 Q9DAI4 Q9DAI9 Q9DAJ2 Q9DAJ4 Q9DAJ5 Q9DAK2 Q9DAK8 Q9DAL3 Q9DAL7 Q9DAM1
Q9DAM9 Q9DAN6 Q9DAN9 Q9DAP0 Q9DAP7 Q9DAQ4 Q9DAQ5 Q9DAR0 Q9DAS2 Q9DAT2 Q9DAT5
Q9DAU9 Q9DAW6 Q9DAW9 Q9DAZ9 Q9DB00 Q9DB26 Q9DB27 Q9DB28 Q9DB30 Q9DB32 Q9DB38

Q9DB40 Q9DB42 Q9DB52 Q9DB54 Q9DB90 Q9DB91 Q9DB94 Q9DB96 Q9DB98 Q9DBA6 Q9DBA8
Q9DBA9 Q9DBB1 Q9DBB4 Q9DBB5 Q9DBB8 Q9DBC0 Q9DBE0 Q9DBE9 Q9DBF7 Q9DBI2 Q9DBJ1
Q9DBL2 Q9DBM1 Q9DBN4 Q9DBN9 Q9DBP5 Q9DBQ9 Q9DBR1 Q9DBR2 Q9DBR4 Q9DBS5 Q9DBS8
Q9DBT3 Q9DBT5 Q9DBT9 Q9DBU3 Q9DBU6 Q9DBV3 Q9DBW3 Q9DBX1 Q9DBX2 Q9DBX7 Q9DBY0
Q9DBY5 Q9DBZ5 Q9DC07 Q9DC22 Q9DC33 Q9DC48 Q9DC63 Q9DC77 Q9DCA5 Q9DCB1 Q9DCB3
Q9DCB4 Q9DCB8 Q9DCC4 Q9DCC7 Q9DCG9 Q9DCH6 Q9DCJ9 Q9DCK4 Q9DCL2 Q9DCL4 Q9DCL9
Q9DCM0 Q9DCM4 Q9DCM7 Q9DCP9 Q9DCQ2 Q9DCR4 Q9DCS2 Q9DCT1 Q9DCT6 Q9DCT8 Q9DCU9
Q9DCV4 Q9DCV7 Q9DCW4 Q9DCY0 Q9DCZ1 Q9DD16 Q9DD18 Q9DD24 Q9DD33 Q9DD48 Q9DDB0
Q9DDD7 Q9DDF1 Q9DDT5 Q9DDU0 Q9DDU1 Q9DDU2 Q9DDY9 Q9DE09 Q9DE13 Q9DE14 Q9DE24
Q9DE25 Q9DE26 Q9DE27 Q9DE46 Q9DEA3 Q9DEB6 Q9DED4 Q9DED5 Q9DED6 Q9DEE9 Q9DEI1
Q9DEN3 Q9DEN4 Q9DEN8 Q9DEN9 Q9DEP0 Q9DEP1 Q9DEQ9 Q9DET5 Q9DEX1 Q9DF25 Q9DF35
Q9DF36 Q9DF42 Q9DF50 Q9DF60 Q9DFG8 Q9DFI0 Q9DFJ9 Q9DFQ7 Q9DFR4 Q9DFR5 Q9DFR6
Q9DFS6 Q9DFZ3 Q9DFZ4 Q9DG02 Q9DG42 Q9DG45 Q9DG67 Q9DG68 Q9DG96 Q9DG97 Q9DG98
Q9DG99 Q9DGA0 Q9DGA2 Q9DGA3 Q9DGA4 Q9DGA5 Q9DGD3 Q9DGI8 Q9DGI9 Q9DGJ0 Q9DGJ1
Q9DGJ2 Q9DGK4 Q9DGN0 Q9DGN1 Q9DGP9 Q9DGQ7 Q9DGS3 Q9DGT6 Q9DGV6 Q9DH17 Q9DH21
Q9DH22 Q9DH37 Q9DH77 Q9DH85 Q9DH90 Q9DHA8 Q9DHF5 Q9DHK3 Q9DHP6 Q9DHQ8 Q9DHR2
Q9DHS8 Q9DHT0 Q9DHU2 Q9DIC5 Q9DK04 Q9DKT8 Q9DKU0 Q9DLD3 Q9DLD6 Q9DRA0 Q9DRA1
Q9DSN8 Q9DSN9 Q9DSQ0 Q9DUB6 Q9DUB7 Q9DUB8 Q9DUB9 Q9DUC0 Q9DUC1 Q9DUC2 Q9DUC3
Q9DUC4 Q9DUC5 Q9DUC9 Q9DUD0 Q9DUD8 Q9DUE0 Q9DUE1 Q9DUE2 Q9DUE3 Q9DUN3 Q9DUT0
Q9DUT1 Q9DUT2 Q9DXE5 Q9DXE7 Q9DXE9 Q9DYE0 Q9E005 Q9E1W2 Q9E6L4 Q9E6L7 Q9E6L8
Q9E6L9 Q9E6M0 Q9E6M1 Q9E6M2 Q9E6M4 Q9E6M5 Q9E6M6 Q9E6M7 Q9E6N0 Q9E6N1 Q9E6N2
Q9E6N4 Q9E6N6 Q9E6N7 Q9E6N9 Q9E6P0 Q9E6P2 Q9E6P3 Q9E6P4 Q9E6P5 Q9E6P7 Q9E6P8
Q9E6P9 Q9E6Q0 Q9E6Q1 Q9E6Q3 Q9E6Q4 Q9E6Q5 Q9E6Q7 Q9E6Q8 Q9E6Q9 Q9E6R0 Q9E6R1
Q9E6R2 Q9E6R3 Q9E6R5 Q9E6S5 Q9E784 Q9E786 Q9E787 Q9E7N7 Q9E7N8 Q9E8F1 Q9E8F2
Q9EMA9 Q9EMI7 Q9EMX1 Q9ENK7 Q9ENK8 Q9ENK9 Q9ENL0 Q9ENL1 Q9ENL2 Q9ENL4 Q9EPB5
Q9EPC5 Q9EPC6 Q9EPH8 Q9EPI8 Q9EPJ0 Q9EPJ3 Q9EPL8 Q9EPM5 Q9EPU0 Q9EPU2 Q9EPU7
Q9EPV5 Q9EPV8 Q9EPX0 Q9EPX5 Q9EPY0 Q9EQ00 Q9EQ10 Q9EQ15 Q9EQ20 Q9EQ28 Q9EQ61
Q9EQ80 Q9EQB9 Q9EQC5 Q9EQC9 Q9EQF5 Q9EQF6 Q9EQH5 Q9EQJ4 Q9EQM3 Q9EQM6 Q9EQN3
Q9EQP0 Q9EQQ0 Q9EQS3 Q9EQW6 Q9EQZ1 Q9ER24 Q9ER34 Q9ER35 Q9ER42 Q9ER56 Q9ER72
Q9ER73 Q9ER74 Q9ER75 Q9ER97 Q9ERA0 Q9ERA5 Q9ERC3 Q9ERD9 Q9ERE2 Q9ERE6 Q9ERE8
Q9ERF3 Q9ERH3 Q9ERH4 Q9ERL7 Q9ERP3 Q9ERQ3 Q9ERQ8 Q9ERR2 Q9ERR8 Q9ERS8 Q9ERU2
Q9ERU3 Q9ERV1 Q9ERZ0 Q9ERZ6 Q9ES00 Q9ES03 Q9ES18 Q9ES34 Q9ES70 Q9ES74 Q9ESB5
Q9ESC8 Q9ESG2 Q9ESJ1 Q9ESL0 Q9ESM0 Q9ESN2 Q9ESN6 Q9ESP5 Q9ESS0 Q9EST2 Q9EST4
Q9ESV0 Q9ESV1 Q9ESV6 Q9ESW0 Q9ESW8 Q9ESX2 Q9ESX4 Q9ESX5 Q9ESZ0 Q9ET01 Q9ET47
Q9ET58 Q9EUE4 Q9EUQ2 Q9EUQ7 Q9EUR1 Q9EUR2 Q9EUR3 Q9EUT5 Q9EUT6 Q9EUT9 Q9EUU8
Q9EUU9 Q9EV05 Q9EV38 Q9EV50 Q9EV79 Q9EV94 Q9EV99 Q9EVE0 Q9EVE1 Q9EVE3 Q9EVE4
Q9EVG2 Q9EVG3 Q9EVG4 Q9EVG5 Q9EVG6 Q9EVG8 Q9EVG9 Q9EVH0 Q9EVH4 Q9EVH5 Q9EVH6
Q9EVH7 Q9EVH8 Q9EVI0 Q9EVI1 Q9EVI3 Q9EVI4 Q9EVI6 Q9EVI7 Q9EVI8 Q9EVR0 Q9EVV4
Q9EVV7 Q9EVX1 Q9EWF7 Q9EWI9 Q9EWJ1 Q9EWJ9 Q9EWK0 Q9EWS0 Q9EWV6 Q9EWW1 Q9EWW3
Q9EWW6 Q9EWW8 Q9EWX7 Q9EX12 Q9EX73 Q9EXC9 Q9EXD3 Q9EXD4 Q9EXD5 Q9EXD6 Q9EXI9
Q9EXL7 Q9EXP1 Q9EXP2 Q9EXQ1 Q9EXU8 Q9EXU9 Q9EXV0 Q9EXV1 Q9EXV2 Q9EXV4 Q9EY50
Q9EY76 Q9EY88 Q9EYG9 Q9EYH3 Q9EYI6 Q9EYI7 Q9EYJ6 Q9EYN8 Q9EYP3 Q9EYT4 Q9EYU0
Q9EYU1 Q9EYU2 Q9EYV3 Q9EYV5 Q9EYW9 Q9EYY5 Q9EYY6 Q9EYZ5 Q9EZ02 Q9EZ08 Q9EZ10
Q9EZ11 Q9EZ12 Q9EZ97 Q9EZJ8 Q9EZK4 Q9EZV1 Q9EZV2 Q9F0C9 Q9F0D4 Q9F0I7 Q9F0J6

Q9F0N8 Q9F0P1 Q9F0Q6 Q9F0R1 Q9F0R4 Q9F0V0 Q9F131 Q9F173 Q9F187 Q9F1K0 Q9F1M9
Q9F1N2 Q9F1N8 Q9F1N9 Q9F1Q3 Q9F1R2 Q9F1S9 Q9F1T4 Q9F1T6 Q9F1V2 Q9F1W4 Q9F1X7
Q9F1Y5 Q9F1Y6 Q9F234 Q9F282 Q9F297 Q9F2F0 Q9F2F9 Q9F2I9 Q9F2S0 Q9F2U7 Q9F305
Q9F309 Q9F314 Q9F315 Q9F316 Q9F323 Q9F326 Q9F344 Q9F364 Q9F3C7 Q9F3E6 Q9F3T4
Q9F3X8 Q9F407 Q9F410 Q9F411 Q9F414 Q9F415 Q9F416 Q9F417 Q9F464 Q9F466 Q9F4E4
Q9F4E6 Q9F4E8 Q9F4F0 Q9F4F2 Q9F4F4 Q9F4F7 Q9F4G3 Q9F4I7 Q9F4I8 Q9F4K4 Q9F5K5
Q9F5K6 Q9F5M1 Q9F5P3 Q9F5P4 Q9F5X1 Q9F5Y0 Q9F5Y3 Q9F618 Q9F664 Q9F665 Q9F672
Q9F6B2 Q9F6X3 Q9F6X5 Q9F6X6 Q9F712 Q9F714 Q9F715 Q9F716 Q9F719 Q9F723 Q9F724
Q9F746 Q9F771 Q9F7A2 Q9F7B2 Q9F7D4 Q9F7E0 Q9F7L5 Q9F7L6 Q9F7Q9 Q9F7R5 Q9F7S4
Q9F7V5 Q9F812 Q9F837 Q9F854 Q9F855 Q9F8I0 Q9F8Q3 Q9F8R8 Q9F8T9 Q9F8X1 Q9F984
Q9F9K6 Q9F9L1 Q9F9U0 Q9F9U1 Q9F9U5 Q9FA03 Q9FA04 Q9FA05 Q9FA11 Q9FA32 Q9FA38
Q9FAF8 Q9FAS2 Q9FAS3 Q9FAS4 Q9FAS7 Q9FAS8 Q9FAW5 Q9FAX1 Q9FAX2 Q9FB57 Q9FBG4
Q9FBM1 Q9FBM3 Q9FBM4 Q9FBN0 Q9FBN6 Q9FBT8 Q9FBV0 Q9FBX1 Q9FC33 Q9FC38 Q9FC99
Q9FCA2 Q9FCA6 Q9FCA7 Q9FCB1 Q9FCC1 Q9FCC2 Q9FCC3 Q9FCC9 Q9FCD1 Q9FCD2 Q9FCD3
Q9FCD5 Q9FCD6 Q9FCG5 Q9FCP2 Q9FCV2 Q9FD28 Q9FD29 Q9FD32 Q9FDA3 Q9FDC7 Q9FDL4
Q9FDL5 Q9FDL9 Q9FDN6 Q9FDN7 Q9FDS1 Q9FDX1 Q9FDX6 Q9FDX8 Q9FDY4 Q9FDZ9 Q9FE01
Q9FE17 Q9FE25 Q9FE58 Q9FE62 Q9FE63 Q9FE64 Q9FE65 Q9FE68 Q9FE83 Q9FE91 Q9FEA2
Q9FEB5 Q9FEC4 Q9FEE2 Q9FEF8 Q9FEP8 Q9FEW2 Q9FEX0 Q9FF09 Q9FF12 Q9FF23 Q9FF30
Q9FF41 Q9FF52 Q9FF58 Q9FF61 Q9FF66 Q9FF79 Q9FF80 Q9FF86 Q9FF90 Q9FFA0 Q9FFA1
Q9FFA2 Q9FFA5 Q9FFA7 Q9FFB6 Q9FFC0 Q9FFC7 Q9FFD2 Q9FFD7 Q9FFD8 Q9FFE0 Q9FFE3
Q9FFF1 Q9FFF4 Q9FFF5 Q9FFF6 Q9FFF9 Q9FFG1 Q9FFG9 Q9FFH0 Q9FFH1 Q9FFI5 Q9FFJ8
Q9FFK0 Q9FFK6 Q9FFL3 Q9FFM6 Q9FFN1 Q9FFN7 Q9FFN9 Q9FFP1 Q9FFP2 Q9FFQ1 Q9FFQ4
Q9FFQ5 Q9FFQ7 Q9FFR6 Q9FFR7 Q9FFS3 Q9FFS4 Q9FFS8 Q9FFS9 Q9FFT0 Q9FFT4 Q9FFT9
Q9FFU1 Q9FFU7 Q9FFV4 Q9FFV5 Q9FFW2 Q9FFW4 Q9FFW6 Q9FFW7 Q9FFX1 Q9FFX2 Q9FFX4
Q9FFY6 Q9FFY9 Q9FFZ2 Q9FFZ6 Q9FG01 Q9FG05 Q9FG13 Q9FG16 Q9FG23 Q9FG29 Q9FG30
Q9FG31 Q9FG36 Q9FG59 Q9FG67 Q9FG68 Q9FG77 Q9FG85 Q9FG90 Q9FG91 Q9FG94 Q9FGB0
Q9FGB8 Q9FGC4 Q9FGC5 Q9FGD1 Q9FGD2 Q9FGD6 Q9FGD7 Q9FGE6 Q9FGE7 Q9FGF0 Q9FGI1
Q9FGJ3 Q9FGL1 Q9FGL9 Q9FGM9 Q9FGN3 Q9FGN4 Q9FGN7 Q9FGN8 Q9FGP6 Q9FGP7 Q9FGP8
Q9FGQ3 Q9FGQ6 Q9FGQ7 Q9FGQ9 Q9FGR2 Q9FGR7 Q9FGR8 Q9FGS0 Q9FGS3 Q9FGS4 Q9FGT1
Q9FGV1 Q9FGW5 Q9FGW9 Q9FGX2 Q9FGY3 Q9FGY4 Q9FGY9 Q9FGZ4 Q9FGZ9 Q9FH04 Q9FH09
Q9FH13 Q9FH22 Q9FH37 Q9FH40 Q9FH50 Q9FH51 Q9FH54 Q9FH59 Q9FH61 Q9FH73 Q9FH77
Q9FH80 Q9FH87 Q9FH99 Q9FHA2 Q9FHA3 Q9FHA7 Q9FHB0 Q9FHB2 Q9FHB3 Q9FHB5 Q9FHE1
Q9FHE4 Q9FHF0 Q9FHF9 Q9FHG2 Q9FHH1 Q9FHH2 Q9FHH8 Q9FHI0 Q9FHI7 Q9FHJ2 Q9FHJ6
Q9FHK0 Q9FHK1 Q9FHK9 Q9FHL4 Q9FHM4 Q9FHN8 Q9FHN9 Q9FHP3 Q9FHQ6 Q9FHR7 Q9FHS6
Q9FHW2 Q9FHW4 Q9FHW7 Q9FHX8 Q9FHY1 Q9FHZ1 Q9FHZ2 Q9FHZ3 Q9FI13 Q9FI16 Q9FI27
Q9FI40 Q9FI48 Q9FI53 Q9FI55 Q9FI61 Q9FI70 Q9FI71 Q9FI72 Q9FI80 Q9FI94 Q9FI96
Q9FI97 Q9FI98 Q9FI99 Q9FIA0 Q9FIA7 Q9FIB2 Q9FIB4 Q9FIB6 Q9FIE9 Q9FIF3 Q9FIF7
Q9FIG8 Q9FIH1 Q9FIH2 Q9FIH4 Q9FIH8 Q9FIH9 Q9FII1 Q9FII2 Q9FII7 Q9FIJ4 Q9FIJ7
Q9FIK0 Q9FIK1 Q9FIK2 Q9FIK7 Q9FIL7 Q9FIM0 Q9FIM4 Q9FIN1 Q9FIP9 Q9FIR9 Q9FIS1
Q9FIT3 Q9FIT4 Q9FIT7 Q9FIT9 Q9FIU6 Q9FIV6 Q9FIW5 Q9FIW9 Q9FIX0 Q9FIX2 Q9FIX3
Q9FIX5 Q9FIX6 Q9FIX8 Q9FIY7 Q9FIZ7 Q9FJ00 Q9FJ10 Q9FJ17 Q9FJ30 Q9FJ32 Q9FJ70
Q9FJ79 Q9FJ80 Q9FJ82 Q9FJ98 Q9FJA2 Q9FJA6 Q9FJB5 Q9FJC0 Q9FJC1 Q9FJE6 Q9FJE8
Q9FJF4 Q9FJF8 Q9FJG2 Q9FJH6 Q9FJH9 Q9FJI1 Q9FJJ3 Q9FJJ4 Q9FJJ5 Q9FJK3 Q9FJK6
Q9FJK7 Q9FJK8 Q9FJL0 Q9FJL3 Q9FJL4 Q9FJL8 Q9FJM5 Q9FJP2 Q9FJP3 Q9FJQ5 Q9FJR0

Q9FJR9 Q9FJT1 Q9FJT2 Q9FJT6 Q9FJT7 Q9FJT8 Q9FJT9 Q9FJU2 Q9FJU3 Q9FJU4 Q9FJV1
Q9FJV2 Q9FJV5 Q9FJW5 Q9FJW6 Q9FJX0 Q9FJX2 Q9FJX5 Q9FJX8 Q9FJX9 Q9FJY7 Q9FK02
Q9FK07 Q9FK14 Q9FK15 Q9FK23 Q9FK25 Q9FK27 Q9FK33 Q9FK35 Q9FK44 Q9FK51 Q9FK54
Q9FK59 Q9FK61 Q9FK81 Q9FK88 Q9FK91 Q9FK93 Q9FKA7 Q9FKA9 Q9FKB3 Q9FKB5 Q9FKB6
Q9FKC0 Q9FKC3 Q9FKC5 Q9FKC6 Q9FKC8 Q9FKD0 Q9FKD5 Q9FKD6 Q9FKD7 Q9FKD9 Q9FKE6
Q9FKE7 Q9FKF0 Q9FKF6 Q9FKF7 Q9FKF9 Q9FKG3 Q9FKG5 Q9FKG6 Q9FKH1 Q9FKI0 Q9FKJ0
Q9FKJ1 Q9FKJ9 Q9FKK2 Q9FKK6 Q9FKK7 Q9FKM5 Q9FKM7 Q9FKN4 Q9FKP0 Q9FKP8 Q9FKQ3
Q9FKQ6 Q9FKR0 Q9FKR3 Q9FKR9 Q9FKS0 Q9FKS4 Q9FKT0 Q9FKT3 Q9FKT4 Q9FKT5 Q9FKT9
Q9FKV5 Q9FKV8 Q9FKW0 Q9FKW2 Q9FKW3 Q9FKW8 Q9FKZ0 Q9FKZ1 Q9FKZ2 Q9FKZ3 Q9FKZ7
Q9FL03 Q9FL12 Q9FL17 Q9FL26 Q9FL33 Q9FL40 Q9FL48 Q9FL62 Q9FL82 Q9FL99 Q9FLA1
Q9FLA2 Q9FLA3 Q9FLB0 Q9FLB3 Q9FLB4 Q9FLC2 Q9FLD8 Q9FLE2 Q9FLE8 Q9FLE9 Q9FLF4
Q9FLF7 Q9FLH5 Q9FLH8 Q9FLI0 Q9FLI1 Q9FLI3 Q9FLJ2 Q9FLJ4 Q9FLJ6 Q9FLK4 Q9FLL0
Q9FLL1 Q9FLL3 Q9FLM1 Q9FLM6 Q9FLM8 Q9FLN4 Q9FLN8 Q9FLP0 Q9FLP5 Q9FLP6 Q9FLP7
Q9FLP9 Q9FLQ3 Q9FLQ4 Q9FLR0 Q9FLR1 Q9FLR3 Q9FLR5 Q9FLS0 Q9FLS9 Q9FLT2 Q9FLU1
Q9FLU7 Q9FLV0 Q9FLV7 Q9FLV8 Q9FLX3 Q9FLX9 Q9FLY0 Q9FLY7 Q9FLY8 Q9FLZ1 Q9FLZ8
Q9FM01 Q9FM03 Q9FM14 Q9FM41 Q9FM54 Q9FM55 Q9FM68 Q9FM80 Q9FM87 Q9FM88 Q9FM89
Q9FM90 Q9FM91 Q9FM92 Q9FM93 Q9FM94 Q9FMA1 Q9FMB4 Q9FMB6 Q9FMC1 Q9FMC2 Q9FMC8
Q9FMD3 Q9FMD9 Q9FME0 Q9FME2 Q9FME3 Q9FMF4 Q9FMF6 Q9FMG1 Q9FMG4 Q9FMH4 Q9FMH5
Q9FML2 Q9FML4 Q9FML6 Q9FMM0 Q9FMM3 Q9FMM4 Q9FMM7 Q9FMN1 Q9FMN4 Q9FMP0 Q9FMP3
Q9FMP4 Q9FMQ1 Q9FMR3 Q9FMR9 Q9FMS4 Q9FMS5 Q9FMS6 Q9FMS7 Q9FMT1 Q9FMT4 Q9FMT5
Q9FMU2 Q9FMU5 Q9FMV0 Q9FMV4 Q9FMW4 Q9FMW5 Q9FMW6 Q9FMW8 Q9FMX2 Q9FMX6 Q9FMX7
Q9FMX8 Q9FMY5 Q9FMY7 Q9FMZ4 Q9FN02 Q9FN07 Q9FN09 Q9FN11 Q9FN19 Q9FN26 Q9FN28
Q9FN29 Q9FN30 Q9FN32 Q9FN41 Q9FN52 Q9FN67 Q9FN69 Q9FN89 Q9FNA2 Q9FNA4 Q9FNB3
Q9FNC7 Q9FND4 Q9FND6 Q9FND7 Q9FND8 Q9FND9 Q9FNE2 Q9FNE9 Q9FNF2 Q9FNF8 Q9FNG3
Q9FNG6 Q9FNG8 Q9FNH4 Q9FNI1 Q9FNI3 Q9FNI6 Q9FNI8 Q9FNI9 Q9FNJ1 Q9FNJ4 Q9FNJ5
Q9FNJ8 Q9FNJ9 Q9FNK0 Q9FNK4 Q9FNK5 Q9FNL2 Q9FNM2 Q9FNM7 Q9FNN2 Q9FNN3 Q9FNN6
Q9FNN7 Q9FNP0 Q9FNP8 Q9FNP9 Q9FNQ0 Q9FNQ1 Q9FNS4 Q9FNV8 Q9FNV9 Q9FNY0 Q9FNY2
Q9FNY3 Q9FNY4 Q9FNY7 Q9FNZ1 Q9FNZ2 Q9FNZ4 Q9FP29 Q9FPE8 Q9FPH0 Q9FPH3 Q9FPJ8
Q9FPK5 Q9FPL6 Q9FPM6 Q9FPN7 Q9FPQ8 Q9FPW3 Q9FPW4 Q9FPW6 Q9FQ02 Q9FQ03 Q9FQ04
Q9FQ08 Q9FQ09 Q9FQ11 Q9FQ19 Q9FQ20 Q9FQ24 Q9FQ25 Q9FQ27 Q9FQA3 Q9FQY8 Q9FR35
Q9FR39 Q9FR62 Q9FR97 Q9FR99 Q9FRA7 Q9FRH4 Q9FRI0 Q9FRS4 Q9FRV4 Q9FRX6 Q9FS29
Q9FS87 Q9FS88 Q9FS90 Q9FSA4 Q9FSB7 Q9FSB8 Q9FSB9 Q9FSC0 Q9FSC1 Q9FSC2 Q9FSD9
Q9FSE4 Q9FSF0 Q9FSV7 Q9FT08 Q9FT36 Q9FT37 Q9FT50 Q9FT69 Q9FT70 Q9FT72 Q9FT73
Q9FT74 Q9FT81 Q9FT92 Q9FTA0 Q9FTA2 Q9FTC7 Q9FTY0 Q9FU27 Q9FU53 Q9FUA4 Q9FUB7
Q9FUB8 Q9FUD1 Q9FUH9 Q9FUJ1 Q9FUL4 Q9FUM1 Q9FUM7 Q9FUM8 Q9FUM9 Q9FUN0 Q9FUR2
Q9FUS6 Q9FUS8 Q9FUS9 Q9FUT0 Q9FUT1 Q9FUW6 Q9FUY2 Q9FUY6 Q9FUZ0 Q9FUZ1 Q9FUZ2
Q9FUZ7 Q9FV02 Q9FV46 Q9FV49 Q9FV50 Q9FV51 Q9FV52 Q9FV53 Q9FV54 Q9FV65 Q9FV66
Q9FV70 Q9FV71 Q9FV81 Q9FVC1 Q9FVC8 Q9FVE6 Q9FVG7 Q9FVG8 Q9FVI1 Q9FVI2 Q9FVL0
Q9FVQ0 Q9FVQ1 Q9FVQ6 Q9FVQ9 Q9FVR6 Q9FVS1 Q9FVS2 Q9FVT2 Q9FVT4 Q9FVU4 Q9FVU9
Q9FVV4 Q9FVV7 Q9FVV8 Q9FVW4 Q9FVX0 Q9FVX1 Q9FVX2 Q9FW03 Q9FW44 Q9FW70 Q9FWA6
Q9FWK4 Q9FWQ2 Q9FWQ5 Q9FWS3 Q9FWS4 Q9FWS6 Q9FWS9 Q9FWT2 Q9FWW3 Q9FWW5 Q9FWW6
Q9FWW7 Q9FWW8 Q9FWW9 Q9FWX2 Q9FWZ1 Q9FX02 Q9FX08 Q9FX09 Q9FX24 Q9FX25 Q9FX35
Q9FX36 Q9FX45 Q9FX53 Q9FX55 Q9FX56 Q9FX60 Q9FX64 Q9FX67 Q9FX77 Q9FX84 Q9FX89
Q9FX92 Q9FX93 Q9FX94 Q9FXA2 Q9FXA3 Q9FXA4 Q9FXA8 Q9FXB2 Q9FXC7 Q9FXE2 Q9FXE3

Q9FXE4 Q9FXG2 Q9FXG8 Q9FXH7 Q9FXI7 Q9FXM3 Q9FXM4 Q9FXM5 Q9FXY7 Q9FY42 Q9FY48
Q9FY49 Q9FY50 Q9FY54 Q9FY60 Q9FY65 Q9FY66 Q9FY69 Q9FY82 Q9FY93 Q9FYA2 Q9FYA6
Q9FYB5 Q9FYB7 Q9FYC4 Q9FYC8 Q9FYE2 Q9FYE4 Q9FYE6 Q9FYF4 Q9FYF9 Q9FYG7 Q9FYH1
Q9FYI3 Q9FYJ1 Q9FYJ2 Q9FYJ6 Q9FYK2 Q9FYN7 Q9FYR6 Q9FYR7 Q9FYS3 Q9FYT6 Q9FYZ9
Q9FZ06 Q9FZ19 Q9FZ21 Q9FZ24 Q9FZ32 Q9FZ48 Q9FZ52 Q9FZ53 Q9FZ57 Q9FZ62 Q9FZ70
Q9FZ75 Q9FZ80 Q9FZ87 Q9FZ89 Q9FZ91 Q9FZ92 Q9FZ99 Q9FZA4 Q9FZD1 Q9FZD4 Q9FZD5
Q9FZD6 Q9FZD9 Q9FZE1 Q9FZE7 Q9FZF3 Q9FZF8 Q9FZG4 Q9FZH0 Q9FZH5 Q9FZI7 Q9FZJ3
Q9FZK1 Q9FZK4 Q9FZN8 Q9FZW7 Q9FZX0 Q9G050 Q9G051 Q9G052 Q9G053 Q9G054 Q9G055
Q9G056 Q9G057 Q9G058 Q9G059 Q9G0B5 Q9G0R3 Q9G1H4 Q9G4F5 Q9GCB9 Q9GE26 Q9GF29
Q9GF32 Q9GF34 Q9GF35 Q9GF36 Q9GF37 Q9GF39 Q9GF40 Q9GF45 Q9GF47 Q9GF50 Q9GF51
Q9GF55 Q9GF59 Q9GF61 Q9GF62 Q9GF63 Q9GFK0 Q9GFK3 Q9GFK8 Q9GFL3 Q9GFL8 Q9GFM3
Q9GFM6 Q9GFM9 Q9GFN2 Q9GFN5 Q9GFN8 Q9GGE2 Q9GGJ6 Q9GGJ8 Q9GGX6 Q9GHB1 Q9GHB2
Q9GHB8 Q9GHC3 Q9GHE1 Q9GHE2 Q9GHE4 Q9GHE5 Q9GHG8 Q9GHG9 Q9GHM0 Q9GHP8 Q9GHR6
Q9GHV0 Q9GI85 Q9GIG3 Q9GIH9 Q9GJR3 Q9GJS7 Q9GJT2 Q9GJU4 Q9GK08 Q9GK13 Q9GK38
Q9GKK4 Q9GKK5 Q9GKM0 Q9GKM1 Q9GKQ1 Q9GKQ2 Q9GKQ3 Q9GKQ4 Q9GKQ5 Q9GKR6 Q9GKR7
Q9GKS9 Q9GKT6 Q9GKT8 Q9GKU1 Q9GKU5 Q9GKV7 Q9GKW3 Q9GKX6 Q9GL21 Q9GL23 Q9GL27
Q9GL32 Q9GL72 Q9GL73 Q9GLL9 Q9GLP8 Q9GLP9 Q9GLQ0 Q9GLQ1 Q9GLQ2 Q9GLQ3 Q9GLQ4
Q9GLQ5 Q9GLQ6 Q9GLQ7 Q9GLQ8 Q9GLQ9 Q9GLX4 Q9GLZ9 Q9GM03 Q9GM44 Q9GMU3 Q9GMV6
Q9GN74 Q9GN94 Q9GNE2 Q9GNF0 Q9GNL2 Q9GNN7 Q9GNV2 Q9GP16 Q9GP32 Q9GP34 Q9GP38
Q9GPH3 Q9GPM4 Q9GPR0 Q9GPS6 Q9GPT6 Q9GPU2 Q9GPZ7 Q9GQ38 Q9GQ60 Q9GQ63 Q9GQN5
Q9GR61 Q9GR88 Q9GRC0 Q9GRF8 Q9GRJ1 Q9GRJ3 Q9GRT1 Q9GRU1 Q9GRW7 Q9GRX4 Q9GRZ3
Q9GS00 Q9GSF8 Q9GSR1 Q9GSR5 Q9GSU4 Q9GSU7 Q9GSX9 Q9GT24 Q9GT49 Q9GTD4 Q9GTW8
Q9GTW9 Q9GTX8 Q9GU37 Q9GU68 Q9GUI6 Q9GUM7 Q9GV27 Q9GV36 Q9GYD9 Q9GYF1 Q9GYG4
Q9GYI0 Q9GYS1 Q9GYS9 Q9GYU7 Q9GYV9 Q9GYX1 Q9GYZ0 Q9GYZ9 Q9GZ01 Q9GZ14 Q9GZ28
Q9GZ69 Q9GZ70 Q9GZ71 Q9GZH3 Q9GZL7 Q9GZL8 Q9GZM3 Q9GZN1 Q9GZN2 Q9GZN7 Q9GZN8
Q9GZP4 Q9GZQ3 Q9GZS3 Q9GZT8 Q9GZU0 Q9GZU2 Q9GZU7 Q9GZU8 Q9GZW5 Q9GZX5 Q9GZY0
Q9GZY1 Q9GZZ0 Q9H000 Q9H008 Q9H069 Q9H074 Q9H091 Q9H094 Q9H095 Q9H098 Q9H0A6
Q9H0A9 Q9H0B3 Q9H0C1 Q9H0C5 Q9H0D2 Q9H0H9 Q9H0I3 Q9H0I9 Q9H0K4 Q9H0K6 Q9H0M4
Q9H0M5 Q9H0N5 Q9H0P0 Q9H0P7 Q9H0R4 Q9H0R6 Q9H0U9 Q9H0W7 Q9H0W9 Q9H0Y0 Q9H116
Q9H161 Q9H171 Q9H175 Q9H1A7 Q9H1B4 Q9H1D9 Q9H1H1 Q9H1K6 Q9H1L0 Q9H1M0 Q9H1P6
Q9H1Q7 Q9H1R3 Q9H1X1 Q9H213 Q9H2A2 Q9H2A3 Q9H2C0 Q9H2C1 Q9H2H0 Q9H2H8 Q9H2I8
Q9H2J1 Q9H2K0 Q9H2M3 Q9H2P9 Q9H2S5 Q9H2S9 Q9H2U6 Q9H2Z4 Q9H320 Q9H321 Q9H322
Q9H334 Q9H347 Q9H361 Q9H379 Q9H3C7 Q9H3H3 Q9H3H9 Q9H3J6 Q9H3L0 Q9H3R0 Q9H3S4
Q9H3Y8 Q9H469 Q9H477 Q9H478 Q9H479 Q9H489 Q9H496 Q9H4B0 Q9H4G8 Q9H4I0 Q9H4I3
Q9H4I8 Q9H4K1 Q9H4M3 Q9H4P4 Q9H4Q3 Q9H4Q4 Q9H4R4 Q9H4S2 Q9H4T2 Q9H4W6 Q9H4Y5
Q9H4Z2 Q9H503 Q9H511 Q9H521 Q9H560 Q9H568 Q9H579 Q9H5F2 Q9H5L6 Q9H5Q4 Q9H5U6
Q9H5X1 Q9H5Z1 Q9H5Z6 Q9H606 Q9H609 Q9H628 Q9H633 Q9H649 Q9H693 Q9H6A0 Q9H6B1
Q9H6E5 Q9H6J7 Q9H6N6 Q9H6P5 Q9H6R0 Q9H6S1 Q9H6X5 Q9H6Y5 Q9H707 Q9H773 Q9H777
Q9H790 Q9H7B4 Q9H7C9 Q9H7H0 Q9H7J1 Q9H7M6 Q9H7R0 Q9H7R5 Q9H7S9 Q9H7T3 Q9H7T9
Q9H7U1 Q9H7X3 Q9H7X7 Q9H7Z3 Q9H7Z6 Q9H808 Q9H816 Q9H825 Q9H832 Q9H840 Q9H857
Q9H867 Q9H869 Q9H871 Q9H875 Q9H892 Q9H898 Q9H8E8 Q9H8G1 Q9H8H0 Q9H8Q6 Q9H8S5
Q9H8U3 Q9H8V8 Q9H8W2 Q9H8W3 Q9H8W5 Q9H8X2 Q9H8X3 Q9H930 Q9H944 Q9H963 Q9H967
Q9H977 Q9H993 Q9H999 Q9H9A7 Q9H9D4 Q9H9F9 Q9H9L3 Q9H9P5 Q9H9R9 Q9H9V9 Q9H9Y2
Q9H9Y6 Q9HA47 Q9HA64 Q9HA77 Q9HA90 Q9HA92 Q9HAA7 Q9HAB8 Q9HAC7 Q9HAC8 Q9HAE3

Q9HAH1 Q9HAH7 Q9HAK2 Q9HAQ2 Q9HAT0 Q9HAT8 Q9HAU6 Q9HAU8 Q9HAW0 Q9HAY2 Q9HAZ1
Q9HB07 Q9HB31 Q9HB65 Q9HBF5 Q9HBH1 Q9HBI5 Q9HBK9 Q9HBL8 Q9HBT7 Q9HBT8 Q9HBU1
Q9HBX3 Q9HBZ2 Q9HC16 Q9HC36 Q9HC38 Q9HCC0 Q9HCC6 Q9HCE0 Q9HCE5 Q9HCG1 Q9HCI6
Q9HCI7 Q9HCJ3 Q9HCJ5 Q9HCK0 Q9HCK1 Q9HCL3 Q9HCM9 Q9HCN2 Q9HCU8 Q9HCY8 Q9HCZ1
Q9HD34 Q9HD40 Q9HD64 Q9HD90 Q9HDF5 Q9HDF6 Q9HDN1 Q9HDQ1 Q9HDQ5 Q9HDS5 Q9HDS6
Q9HDT3 Q9HDT8 Q9HDU2 Q9HDU3 Q9HDU4 Q9HDU5 Q9HDU6 Q9HDU7 Q9HDV0 Q9HDV4 Q9HDV5
Q9HDV9 Q9HDW0 Q9HDW3 Q9HDW4 Q9HDW8 Q9HDX3 Q9HDX6 Q9HDX7 Q9HDX9 Q9HDY1 Q9HDY2
Q9HDY4 Q9HDY5 Q9HDY7 Q9HDZ0 Q9HDZ5 Q9HDZ6 Q9HDZ9 Q9HE01 Q9HE02 Q9HE05 Q9HE06
Q9HE07 Q9HE08 Q9HE09 Q9HE11 Q9HE15 Q9HE16 Q9HE25 Q9HE26 Q9HE63 Q9HE74 Q9HE76
Q9HE87 Q9HEC4 Q9HED8 Q9HEG2 Q9HEG3 Q9HEH1 Q9HEP7 Q9HEQ8 Q9HEQ9 Q9HES8 Q9HET9
Q9HEU0 Q9HEU2 Q9HEV5 Q9HEY7 Q9HF03 Q9HF53 Q9HF61 Q9HF68 Q9HF78 Q9HF97 Q9HFB3
Q9HFC8 Q9HFC9 Q9HFD0 Q9HFE6 Q9HFE7 Q9HFE8 Q9HFF2 Q9HFF5 Q9HFN9 Q9HFQ2 Q9HFQ3
Q9HFQ4 Q9HFQ5 Q9HFQ6 Q9HFQ7 Q9HFR0 Q9HFR6 Q9HFR7 Q9HFS2 Q9HFU4 Q9HFU8 Q9HFU9
Q9HFV5 Q9HFV8 Q9HFW3 Q9HFW8 Q9HFX0 Q9HFX1 Q9HFY6 Q9HG12 Q9HG13 Q9HGE0 Q9HGF8
Q9HGI2 Q9HGI4 Q9HGI6 Q9HGI7 Q9HGI8 Q9HGK2 Q9HGK9 Q9HGL0 Q9HGL2 Q9HGL3 Q9HGL4
Q9HGL5 Q9HGL6 Q9HGL8 Q9HGL9 Q9HGM5 Q9HGM8 Q9HGM9 Q9HGN1 Q9HGN2 Q9HGN4 Q9HGN6
Q9HGN8 Q9HGP0 Q9HGP4 Q9HGP5 Q9HGP9 Q9HGQ0 Q9HGQ2 Q9HGT6 Q9HGU3 Q9HGU4 Q9HGV0
Q9HGX4 Q9HGY1 Q9HGY2 Q9HGY7 Q9HGY8 Q9HGY9 Q9HGZ0 Q9HGZ1 Q9HGZ2 Q9HH00 Q9HH05
Q9HH06 Q9HH07 Q9HH09 Q9HH12 Q9HH21 Q9HH35 Q9HH76 Q9HH78 Q9HH80 Q9HH82 Q9HH83
Q9HH84 Q9HH99 Q9HHA0 Q9HHA1 Q9HHA2 Q9HHB6 Q9HHB8 Q9HHB9 Q9HHC2 Q9HHC4 Q9HHD0
Q9HHE4 Q9HHE9 Q9HHH8 Q9HHJ7 Q9HHK5 Q9HHM7 Q9HHN2 Q9HHN3 Q9HHN4 Q9HHN8 Q9HHR1
Q9HHS8 Q9HHS9 Q9HHT2 Q9HHT6 Q9HHT7 Q9HHU0 Q9HHX2 Q9HHY2 Q9HHY8 Q9HHZ9 Q9HI16
Q9HI21 Q9HI24 Q9HI27 Q9HI35 Q9HI37 Q9HI38 Q9HI47 Q9HI54 Q9HI67 Q9HI69 Q9HI70
Q9HI87 Q9HI94 Q9HI95 Q9HIA9 Q9HIB2 Q9HIC2 Q9HIC5 Q9HIH2 Q9HIH3 Q9HIH4 Q9HIH6
Q9HIH8 Q9HII0 Q9HII1 Q9HII2 Q9HII9 Q9HIJ2 Q9HIK9 Q9HIM0 Q9HIM1 Q9HIN8 Q9HIN9
Q9HIP0 Q9HIP1 Q9HIP2 Q9HIP3 Q9HIP7 Q9HIQ4 Q9HIQ9 Q9HIR0 Q9HIR1 Q9HIR2 Q9HIR3
Q9HIR4 Q9HIR5 Q9HIR8 Q9HIR9 Q9HIS1 Q9HIS2 Q9HIS3 Q9HIS4 Q9HIS6 Q9HIS7 Q9HIS8
Q9HIS9 Q9HIT1 Q9HIT3 Q9HIT4 Q9HIT7 Q9HIW5 Q9HIW7 Q9HIX1 Q9HIX2 Q9HIX3 Q9HIX4
Q9HIX6 Q9HIY0 Q9HIY2 Q9HIY5 Q9HJ08 Q9HJ11 Q9HJ12 Q9HJ14 Q9HJ15 Q9HJ17 Q9HJ19
Q9HJ26 Q9HJ27 Q9HJ28 Q9HJ34 Q9HJ35 Q9HJ36 Q9HJ42 Q9HJ48 Q9HJ56 Q9HJ58 Q9HJ59
Q9HJ60 Q9HJ67 Q9HJ68 Q9HJ69 Q9HJ72 Q9HJ74 Q9HJ78 Q9HJ79 Q9HJ80 Q9HJ83 Q9HJ84
Q9HJ94 Q9HJ95 Q9HJA4 Q9HJA6 Q9HJB1 Q9HJB3 Q9HJD0 Q9HJD1 Q9HJD3 Q9HJD4 Q9HJD6
Q9HJD7 Q9HJD8 Q9HJD9 Q9HJH0 Q9HJJ2 Q9HJJ5 Q9HJJ6 Q9HJL3 Q9HJL5 Q9HJL6 Q9HJL8
Q9HJM1 Q9HJM2 Q9HJM3 Q9HJM5 Q9HJM7 Q9HJN0 Q9HJN5 Q9HJN6 Q9HJP3 Q9HJQ0 Q9HJQ5
Q9HJR8 Q9HJS3 Q9HJT0 Q9HJT3 Q9HJT4 Q9HJT5 Q9HJV9 Q9HJW4 Q9HJW8 Q9HJX7 Q9HJY7
Q9HK01 Q9HK02 Q9HK06 Q9HK14 Q9HK16 Q9HK17 Q9HK20 Q9HK21 Q9HK30 Q9HK31 Q9HK34
Q9HK40 Q9HK42 Q9HK58 Q9HKB7 Q9HKC1 Q9HKD2 Q9HKE4 Q9HKE5 Q9HKF1 Q9HKG3 Q9HKH7
Q9HKI3 Q9HKJ3 Q9HKJ8 Q9HKK0 Q9HKK8 Q9HKL4 Q9HKM2 Q9HKM3 Q9HKM5 Q9HKM6 Q9HKM7
Q9HKN1 Q9HKQ0 Q9HKQ2 Q9HKR0 Q9HKR2 Q9HKR6 Q9HKS5 Q9HKS7 Q9HKT1 Q9HKT2 Q9HKT3
Q9HKT9 Q9HKV1 Q9HKV6 Q9HKV7 Q9HKX0 Q9HKX4 Q9HKX5 Q9HKY3 Q9HKZ2 Q9HKZ7 Q9HL06
Q9HL07 Q9HL08 Q9HL09 Q9HL17 Q9HL27 Q9HL35 Q9HL36 Q9HL57 Q9HL70 Q9HL71 Q9HL74
Q9HL75 Q9HL98 Q9HL99 Q9HLA4 Q9HLA5 Q9HLA6 Q9HLA7 Q9HLB6 Q9HLB8 Q9HLB9 Q9HLC0
Q9HLD5 Q9HLE2 Q9HLE3 Q9HLE4 Q9HLE5 Q9HLE6 Q9HLE7 Q9HLI6 Q9HLI9 Q9HLJ0 Q9HLJ1
Q9HLJ2 Q9HLK5 Q9HLK8 Q9HLL3 Q9HLM8 Q9HLN1 Q9HLP5 Q9HLP8 Q9HLQ1 Q9HLQ2 Q9HLR7

Q9HLR8 Q9HLT5 Q9HLU2 Q9HLV2 Q9HLV3 Q9HLV6 Q9HLV9 Q9HLW6 Q9HLX1 Q9HLX2 Q9HLX5
Q9HLX6 Q9HLX7 Q9HLY1 Q9HLY2 Q9HLY3 Q9HLZ2 Q9HLZ5 Q9HLZ8 Q9HM08 Q9HM12 Q9HM14
Q9HM15 Q9HM16 Q9HM17 Q9HM18 Q9HM19 Q9HM21 Q9HM27 Q9HM28 Q9HM29 Q9HM33 Q9HM46
Q9HM47 Q9HM55 Q9HM60 Q9HM85 Q9HM89 Q9HMA9 Q9HMC3 Q9HMC7 Q9HMD2 Q9HMD4 Q9HME0
Q9HME1 Q9HME3 Q9HMF3 Q9HMG9 Q9HMH2 Q9HMH9 Q9HMJ5 Q9HMK3 Q9HMK4 Q9HMM4 Q9HMM6
Q9HMM9 Q9HMN0 Q9HMN2 Q9HMP0 Q9HMP7 Q9HMQ2 Q9HMQ3 Q9HMQ5 Q9HMR7 Q9HMS3 Q9HMT9
Q9HMU5 Q9HMV2 Q9HMV4 Q9HMV6 Q9HMV8 Q9HMV9 Q9HMX8 Q9HMY5 Q9HMY8 Q9HN05 Q9HN13
Q9HN14 Q9HN20 Q9HN21 Q9HN23 Q9HN24 Q9HN27 Q9HN31 Q9HN33 Q9HN34 Q9HN40 Q9HN43
Q9HN45 Q9HN47 Q9HN50 Q9HN51 Q9HN53 Q9HN56 Q9HN60 Q9HN62 Q9HN64 Q9HN66 Q9HN70
Q9HN72 Q9HN74 Q9HN78 Q9HN82 Q9HN83 Q9HN88 Q9HN97 Q9HNA2 Q9HNF0 Q9HNF4 Q9HNG2
Q9HNG7 Q9HNI0 Q9HNI2 Q9HNI6 Q9HNJ5 Q9HNJ6 Q9HNJ8 Q9HNK9 Q9HNL4 Q9HNL5 Q9HNL6
Q9HNM7 Q9HNN4 Q9HNN7 Q9HNN8 Q9HNP1 Q9HNP3 Q9HNP5 Q9HNP6 Q9HNP9 Q9HNQ0 Q9HNQ2
Q9HNQ6 Q9HNR1 Q9HNR3 Q9HNR6 Q9HNS3 Q9HNS5 Q9HNT1 Q9HNU2 Q9HNU7 Q9HNU8 Q9HNU9
Q9HNV0 Q9HNV4 Q9HNX6 Q9HNX7 Q9HNY7 Q9HNZ0 Q9HNZ1 Q9HP08 Q9HP20 Q9HP27 Q9HP32
Q9HP33 Q9HP40 Q9HP42 Q9HP43 Q9HP55 Q9HP57 Q9HP65 Q9HP68 Q9HP72 Q9HP73 Q9HP78
Q9HP80 Q9HP87 Q9HPA4 Q9HPA5 Q9HPB3 Q9HPB4 Q9HPB6 Q9HPB7 Q9HPB8 Q9HPB9 Q9HPC0
Q9HPC2 Q9HPC3 Q9HPD1 Q9HPD3 Q9HPD4 Q9HPD7 Q9HPE9 Q9HPF0 Q9HPF2 Q9HPG3 Q9HPG4
Q9HPG5 Q9HPG6 Q9HPI4 Q9HPJ7 Q9HPJ8 Q9HPJ9 Q9HPK0 Q9HPK9 Q9HPL5 Q9HPL7 Q9HPL8
Q9HPN4 Q9HPP1 Q9HPQ9 Q9HPR5 Q9HPS3 Q9HPT6 Q9HPU3 Q9HPU6 Q9HPW0 Q9HPW5 Q9HPX1
Q9HPX2 Q9HPX4 Q9HPY4 Q9HPY5 Q9HPZ2 Q9HPZ6 Q9HPZ9 Q9HQ27 Q9HQ29 Q9HQ32 Q9HQ46
Q9HQ52 Q9HQ72 Q9HQ73 Q9HQ86 Q9HQ89 Q9HQ95 Q9HQ97 Q9HQB0 Q9HQB2 Q9HQC1 Q9HQC2
Q9HQC7 Q9HQC9 Q9HQD1 Q9HQD5 Q9HQD6 Q9HQD7 Q9HQD8 Q9HQE1 Q9HQG0 Q9HQG3 Q9HQG6
Q9HQG8 Q9HQH1 Q9HQH8 Q9HQI1 Q9HQI4 Q9HQJ0 Q9HQJ1 Q9HQJ2 Q9HQJ3 Q9HQJ4 Q9HQJ5
Q9HQJ6 Q9HQK6 Q9HQK7 Q9HQL9 Q9HQM3 Q9HQM6 Q9HQN3 Q9HQR5 Q9HQS0 Q9HQS1 Q9HQS4
Q9HQS9 Q9HQT4 Q9HQT7 Q9HQU4 Q9HR31 Q9HR32 Q9HR35 Q9HR36 Q9HR37 Q9HR43 Q9HR44
Q9HR45 Q9HR49 Q9HR52 Q9HR53 Q9HR71 Q9HR76 Q9HRA2 Q9HRA5 Q9HRB2 Q9HRC7 Q9HRC8
Q9HRD6 Q9HRE6 Q9HRE8 Q9HRF0 Q9HRF9 Q9HRG2 Q9HRI1 Q9HRI2 Q9HRM5 Q9HRM7 Q9HRP7
Q9HRT1 Q9HRT6 Q9HRT7 Q9HRT8 Q9HRT9 Q9HRW3 Q9HRW4 Q9HRW6 Q9HRY0 Q9HRY2 Q9HRY3
Q9HS16 Q9HS17 Q9HS19 Q9HS36 Q9HS44 Q9HS52 Q9HS66 Q9HS68 Q9HS70 Q9HS81 Q9HS93
Q9HSA3 Q9HSA4 Q9HSB3 Q9HSB4 Q9HSB6 Q9HSB8 Q9HSB9 Q9HSC0 Q9HSC1 Q9HSC4 Q9HSF3
Q9HSF6 Q9HSF7 Q9HSG3 Q9HSG5 Q9HSG8 Q9HSG9 Q9HSH0 Q9HSH4 Q9HSJ8 Q9HSK7 Q9HSL2
Q9HSL6 Q9HSL7 Q9HSM2 Q9HSM6 Q9HSN7 Q9HSP6 Q9HSS4 Q9HSS7 Q9HST4 Q9HSU8 Q9HSW6
Q9HT00 Q9HT05 Q9HT07 Q9HT09 Q9HT10 Q9HT25 Q9HT35 Q9HT57 Q9HT80 Q9HT81 Q9HT95
Q9HTB6 Q9HTD7 Q9HTE3 Q9HTE9 Q9HTF1 Q9HTH5 Q9HTH8 Q9HTI4 Q9HTJ0 Q9HTJ1 Q9HTK1
Q9HTK7 Q9HTK8 Q9HTK9 Q9HTL0 Q9HTM1 Q9HTM2 Q9HTN2 Q9HTN3 Q9HTN5 Q9HTN8 Q9HTN9
Q9HTQ2 Q9HTQ6 Q9HTS5 Q9HTU7 Q9HTV1 Q9HTW3 Q9HTW5 Q9HTX5 Q9HTX6 Q9HTX7 Q9HTY6
Q9HTZ1 Q9HTZ2 Q9HTZ6 Q9HTZ7 Q9HU36 Q9HU41 Q9HU42 Q9HU43 Q9HU44 Q9HU53 Q9HU55
Q9HU56 Q9HU66 Q9HU72 Q9HU73 Q9HU77 Q9HU78 Q9HU85 Q9HU91 Q9HU92 Q9HUA4 Q9HUB0
Q9HUB6 Q9HUB7 Q9HUC0 Q9HUC5 Q9HUC6 Q9HUC8 Q9HUD0 Q9HUE6 Q9HUF1 Q9HUH4 Q9HUI2
Q9HUI8 Q9HUI9 Q9HUJ2 Q9HUJ8 Q9HUK7 Q9HUK9 Q9HUL3 Q9HUL4 Q9HUL8 Q9HUL9 Q9HUM0
Q9HUM5 Q9HUM6 Q9HUM8 Q9HUM9 Q9HUN0 Q9HUN2 Q9HUN4 Q9HUN6 Q9HUP3 Q9HUP4 Q9HUR2
Q9HUS0 Q9HUS1 Q9HUS2 Q9HUU1 Q9HUU5 Q9HUU6 Q9HUU7 Q9HUU8 Q9HUU9 Q9HUV8 Q9HUV9
Q9HUW0 Q9HUW1 Q9HUW3 Q9HUW9 Q9HUX1 Q9HUX4 Q9HUY1 Q9HV00 Q9HV01 Q9HV11 Q9HV30
Q9HV34 Q9HV35 Q9HV40 Q9HV42 Q9HV43 Q9HV44 Q9HV46 Q9HV50 Q9HV51 Q9HV53 Q9HV55

Q9HV56 Q9HV58 Q9HV59 Q9HV61 Q9HV66 Q9HV67 Q9HV68 Q9HV69 Q9HV70 Q9HV71 Q9HV75
Q9HV77 Q9HV78 Q9HVA2 Q9HVC3 Q9HVC4 Q9HVC5 Q9HVC8 Q9HVD0 Q9HVD7 Q9HVE6 Q9HVF1
Q9HVG4 Q9HVH4 Q9HVI1 Q9HVI7 Q9HVJ9 Q9HVL6 Q9HVL7 Q9HVL8 Q9HVL9 Q9HVM1 Q9HVM4
Q9HVM7 Q9HVN5 Q9HVP7 Q9HVP8 Q9HVQ7 Q9HVR0 Q9HVT7 Q9HVT8 Q9HVT9 Q9HVU3 Q9HVU8
Q9HVV3 Q9HVW7 Q9HVW8 Q9HVW9 Q9HVX0 Q9HVX2 Q9HVX6 Q9HVY2 Q9HVY3 Q9HVZ1 Q9HVZ3
Q9HVZ4 Q9HVZ5 Q9HVZ9 Q9HW02 Q9HW04 Q9HW09 Q9HW11 Q9HW26 Q9HW38 Q9HWB8 Q9HWC1
Q9HWC4 Q9HWC5 Q9HWC6 Q9HWC7 Q9HWC8 Q9HWC9 Q9HWD0 Q9HWD1 Q9HWD2 Q9HWD4 Q9HWD5
Q9HWD6 Q9HWD7 Q9HWD8 Q9HWD9 Q9HWE0 Q9HWE1 Q9HWE2 Q9HWE3 Q9HWE4 Q9HWE5 Q9HWE6
Q9HWE7 Q9HWE8 Q9HWE9 Q9HWF0 Q9HWF1 Q9HWF2 Q9HWF3 Q9HWF4 Q9HWF6 Q9HWF7 Q9HWF8
Q9HWF9 Q9HWG0 Q9HWG3 Q9HWG4 Q9HWG9 Q9HWH2 Q9HWH8 Q9HWH9 Q9HWI0 Q9HWP3 Q9HWS0
Q9HWS2 Q9HWT6 Q9HWT7 Q9HWX1 Q9HWX4 Q9HWX5 Q9HWX6 Q9HWY1 Q9HWY5 Q9HWZ6 Q9HX02
Q9HX08 Q9HX17 Q9HX20 Q9HX21 Q9HX23 Q9HX25 Q9HX33 Q9HX37 Q9HX40 Q9HX48 Q9HX72
Q9HX95 Q9HX97 Q9HXB0 Q9HXD2 Q9HXD6 Q9HXE2 Q9HXE3 Q9HXE4 Q9HXE5 Q9HXF2 Q9HXF7
Q9HXG4 Q9HXH0 Q9HXH8 Q9HXH9 Q9HXI8 Q9HXJ1 Q9HXJ4 Q9HXJ5 Q9HXJ8 Q9HXK5 Q9HXL8
Q9HXM1 Q9HXM6 Q9HXN2 Q9HXP3 Q9HXP9 Q9HXQ0 Q9HXQ1 Q9HXQ2 Q9HXQ6 Q9HXT5 Q9HXU0
Q9HXU7 Q9HXV3 Q9HXV4 Q9HXW0 Q9HXX3 Q9HXY9 Q9HXZ1 Q9HXZ3 Q9HXZ4 Q9HY04 Q9HY05
Q9HY07 Q9HY08 Q9HY25 Q9HY26 Q9HY41 Q9HY42 Q9HY72 Q9HY82 Q9HY84 Q9HYA4 Q9HYC7
Q9HYC9 Q9HYD5 Q9HYE3 Q9HYF0 Q9HYF1 Q9HYF3 Q9HYF4 Q9HYG2 Q9HYG8 Q9HYH1 Q9HYH5
Q9HYI6 Q9HYN9 Q9HYR3 Q9HYR9 Q9HYT6 Q9HYV6 Q9HYW0 Q9HYX7 Q9HYZ3 Q9HYZ7 Q9HZ00
Q9HZ17 Q9HZ46 Q9HZ55 Q9HZ59 Q9HZ62 Q9HZ63 Q9HZ65 Q9HZ66 Q9HZ67 Q9HZ68 Q9HZ70
Q9HZ71 Q9HZ95 Q9HZA3 Q9HZA4 Q9HZC0 Q9HZE0 Q9HZG0 Q9HZG5 Q9HZJ5 Q9HZJ8 Q9HZK0
Q9HZK1 Q9HZK6 Q9HZM4 Q9HZM7 Q9HZM9 Q9HZN2 Q9HZN4 Q9HZN8 Q9HZP6 Q9HZP7 Q9HZP9
Q9HZQ2 Q9HZQ3 Q9HZT9 Q9HZU0 Q9HZU2 Q9HZU3 Q9HZY5 Q9HZZ1 Q9HZZ2 Q9I011 Q9I016
Q9I037 Q9I047 Q9I048 Q9I073 Q9I099 Q9I0A0 Q9I0A1 Q9I0A2 Q9I0A3 Q9I0A4 Q9I0D9
Q9I0H4 Q9I0I1 Q9I0K9 Q9I0L2 Q9I0L7 Q9I0M0 Q9I0M1 Q9I0M6 Q9I0M7 Q9I0N2 Q9I0N3
Q9I0N5 Q9I0Q1 Q9I0Q7 Q9I0Q8 Q9I0S1 Q9I0X4 Q9I136 Q9I137 Q9I138 Q9I154 Q9I163
Q9I1C2 Q9I1F6 Q9I1K2 Q9I1L4 Q9I1L5 Q9I1M0 Q9I1M1 Q9I1M2 Q9I1M3 Q9I1P2 Q9I1Q5
Q9I1T4 Q9I1V0 Q9I1W2 Q9I1W4 Q9I1W5 Q9I1W8 Q9I1X7 Q9I1Z9 Q9I203 Q9I234 Q9I244
Q9I262 Q9I263 Q9I291 Q9I2C0 Q9I2C1 Q9I2C2 Q9I2C3 Q9I2D2 Q9I2E2 Q9I2H6 Q9I2N0
Q9I2P8 Q9I2Q2 Q9I2Q6 Q9I2S7 Q9I2S9 Q9I2T1 Q9I2U0 Q9I2U1 Q9I2U2 Q9I2U6 Q9I2U7
Q9I2U8 Q9I2V5 Q9I2W4 Q9I2W7 Q9I2X0 Q9I2Y2 Q9I315 Q9I321 Q9I340 Q9I341 Q9I342
Q9I344 Q9I347 Q9I351 Q9I352 Q9I382 Q9I383 Q9I387 Q9I3A4 Q9I3C3 Q9I3C5 Q9I3D1
Q9I3D2 Q9I3E3 Q9I3F3 Q9I3F4 Q9I3F5 Q9I3H5 Q9I3H7 Q9I3H9 Q9I3I0 Q9I3I4 Q9I3J5
Q9I3J8 Q9I3J9 Q9I3K1 Q9I3K7 Q9I3S3 Q9I3T5 Q9I3W8 Q9I3W9 Q9I433 Q9I434 Q9I445
Q9I449 Q9I452 Q9I465 Q9I466 Q9I467 Q9I468 Q9I471 Q9I472 Q9I476 Q9I489 Q9I492
Q9I4C8 Q9I4D3 Q9I4E1 Q9I4E3 Q9I4E6 Q9I4E9 Q9I4H5 Q9I4K0 Q9I4K6 Q9I4L0 Q9I4L1
Q9I4N1 Q9I4N6 Q9I4P4 Q9I4S5 Q9I4T5 Q9I4U3 Q9I4V0 Q9I4W0 Q9I4W3 Q9I4W9 Q9I4Z2
Q9I4Z9 Q9I502 Q9I504 Q9I509 Q9I513 Q9I525 Q9I526 Q9I534 Q9I541 Q9I553 Q9I558
Q9I576 Q9I583 Q9I587 Q9I5A4 Q9I5A5 Q9I5F3 Q9I5G3 Q9I5G4 Q9I5G5 Q9I5G9 Q9I5H9
Q9I5I6 Q9I5J6 Q9I5L6 Q9I5Q2 Q9I5Q3 Q9I5Q5 Q9I5Q6 Q9I5Q9 Q9I5R7 Q9I5U0 Q9I5U1
Q9I5U4 Q9I5U5 Q9I5U6 Q9I5U7 Q9I5U8 Q9I5V0 Q9I5V3 Q9I5V7 Q9I5V8 Q9I5W0 Q9I5Y1
Q9I5Y4 Q9I5Z0 Q9I614 Q9I617 Q9I618 Q9I626 Q9I636 Q9I644 Q9I676 Q9I685 Q9I697
Q9I699 Q9I6A8 Q9I6B3 Q9I6B4 Q9I6C8 Q9I6D1 Q9I6E0 Q9I6F1 Q9I6F6 Q9I6G1 Q9I6G2
Q9I6H0 Q9I6J2 Q9I6J9 Q9I6K2 Q9I6M4 Q9I6M5 Q9I6N5 Q9I6P3 Q9I6P6 Q9I6Q8 Q9I6R0

Q9I6S5 Q9I6S8 Q9I6S9 Q9I6Y4 Q9I6Z0 Q9I6Z1 Q9I6Z2 Q9I6Z9 Q9I700 Q9I701 Q9I702
Q9I732 Q9I733 Q9I738 Q9I740 Q9I742 Q9I744 Q9I745 Q9I746 Q9I748 Q9I749 Q9I750
Q9I753 Q9I767 Q9I778 Q9I7A5 Q9I7A8 Q9I7A9 Q9I7B1 Q9I7B7 Q9I7B8 Q9I7C2 Q9I7C3
Q9I7C4 Q9I7H9 Q9I7I7 Q9I7K0 Q9I7K6 Q9I7M2 Q9I7S8 Q9I7T7 Q9I7U4 Q9I7W5 Q9I7X6
Q9I869 Q9I8A3 Q9I8A9 Q9I8B0 Q9I8B3 Q9I8F4 Q9I8G4 Q9I8G9 Q9I8H0 Q9I8J6 Q9I8K7
Q9I8M6 Q9I8S2 Q9I8V0 Q9I8W6 Q9I8X5 Q9I922 Q9I923 Q9I954 Q9I955 Q9I969 Q9I980
Q9I9A7 Q9I9H2 Q9I9H8 Q9I9I0 Q9I9I3 Q9I9J6 Q9I9K0 Q9I9K2 Q9I9K7 Q9I9L0 Q9I9M2
Q9I9R0 Q9IA00 Q9IA11 Q9IA12 Q9IA13 Q9IA14 Q9IA15 Q9IA16 Q9IA17 Q9IA18 Q9IA19
Q9IA20 Q9IA21 Q9IA22 Q9IA23 Q9IA24 Q9IA25 Q9IA26 Q9IA27 Q9IA31 Q9IA76 Q9IA78
Q9IAB2 Q9IAJ2 Q9IAK8 Q9IAL1 Q9IAL2 Q9IAL9 Q9IAM7 Q9IAV3 Q9IAX9 Q9IB50 Q9IB83
Q9IB84 Q9IB89 Q9IBC8 Q9IBD0 Q9IBD1 Q9IBG0 Q9IBG1 Q9IBI4 Q9IBM3 Q9IC37 Q9ICB2
Q9ICC1 Q9ICF7 Q9ICF9 Q9ICH6 Q9IFX1 Q9IG41 Q9IG42 Q9IG43 Q9IG44 Q9IGY6 Q9IIH4
Q9IJX4 Q9IK91 Q9IK92 Q9IKC5 Q9IKC9 Q9IKD0 Q9IKD3 Q9IMM3 Q9IMM4 Q9IMP3 Q9IMP6
Q9INI1 Q9INI2 Q9INI3 Q9INI4 Q9INI5 Q9INJ1 Q9IPJ5 Q9IR52 Q9IVZ9 Q9IW99 Q9IWA0
Q9IWW8 Q9IZU5 Q9IZU6 Q9IZU7 Q9J3N2 Q9J3N3 Q9J3N4 Q9J3N5 Q9J3N6 Q9J3N7 Q9J4Z4
Q9J4Z5 Q9J4Z6 Q9J4Z7 Q9J4Z8 Q9J4Z9 Q9J502 Q9J503 Q9J504 Q9J505 Q9J507 Q9J508
Q9J509 Q9J511 Q9J512 Q9J513 Q9J515 Q9J516 Q9J517 Q9J519 Q9J523 Q9J531 Q9J535
Q9J538 Q9J539 Q9J540 Q9J544 Q9J546 Q9J549 Q9J550 Q9J553 Q9J559 Q9J562 Q9J563
Q9J566 Q9J569 Q9J579 Q9J584 Q9J589 Q9J592 Q9J593 Q9J5A4 Q9J5A5 Q9J5A7 Q9J5B1
Q9J5B5 Q9J5B7 Q9J5B9 Q9J5C0 Q9J5C4 Q9J5C7 Q9J5C8 Q9J5C9 Q9J5D0 Q9J5F7 Q9J5F9
Q9J5G1 Q9J5G3 Q9J5G4 Q9J5G5 Q9J5H2 Q9J5H5 Q9J5H7 Q9J5H8 Q9J5H9 Q9J5I3 Q9J5I7
Q9J5I9 Q9J5J0 Q9J5J1 Q9J5S2 Q9J5U7 Q9J7Z2 Q9J818 Q9J942 Q9JAF2 Q9JF79 Q9JF82
Q9JF93 Q9JFA3 Q9JFA5 Q9JFA8 Q9JFB7 Q9JFC3 Q9JFD9 Q9JFE5 Q9JFF2 Q9JFG1 Q9JFM9
Q9JFS4 Q9JG99 Q9JGF1 Q9JGT6 Q9JGT7 Q9JGT8 Q9JGT9 Q9JGU0 Q9JGU1 Q9JGV3 Q9JH48
Q9JH51 Q9JH65 Q9JH68 Q9JHB4 Q9JHB5 Q9JHC0 Q9JHC9 Q9JHD1 Q9JHD2 Q9JHE6 Q9JHE7
Q9JHG2 Q9JHG6 Q9JHI2 Q9JHI7 Q9JHK4 Q9JHR9 Q9JHS1 Q9JHS9 Q9JHT5 Q9JHU0 Q9JHW0
Q9JHW2 Q9JHW6 Q9JHX2 Q9JHX6 Q9JHY8 Q9JI13 Q9JI20 Q9JI38 Q9JI44 Q9JI48 Q9JI55
Q9JI69 Q9JI70 Q9JI74 Q9JI75 Q9JIB3 Q9JIB4 Q9JIB6 Q9JIC3 Q9JII5 Q9JII7 Q9JIK5
Q9JIL5 Q9JIL8 Q9JIM0 Q9JIM3 Q9JIM4 Q9JIT1 Q9JIW4 Q9JIX0 Q9JIX9 Q9JIY2 Q9JJ25
Q9JJ26 Q9JJ28 Q9JJ43 Q9JJ48 Q9JJ54 Q9JJ78 Q9JJ89 Q9JJ93 Q9JJA4 Q9JJD0 Q9JJF0
Q9JJF3 Q9JJF6 Q9JJH5 Q9JJL0 Q9JJL8 Q9JJN0 Q9JJN2 Q9JJN4 Q9JJN6 Q9JJP6 Q9JJR2
Q9JJR5 Q9JJR6 Q9JJR7 Q9JJR9 Q9JJT0 Q9JJU9 Q9JJV0 Q9JJV1 Q9JJV2 Q9JJW5 Q9JJW6
Q9JJX7 Q9JJY4 Q9JJZ6 Q9JK23 Q9JK24 Q9JK30 Q9JK37 Q9JK42 Q9JK54 Q9JK72 Q9JK81
Q9JK92 Q9JKB3 Q9JKB5 Q9JKD8 Q9JKD9 Q9JKK0 Q9JKL7 Q9JKL8 Q9JKN5 Q9JKP5 Q9JKP7
Q9JKP8 Q9JKQ4 Q9JKS4 Q9JKU3 Q9JKU6 Q9JKU8 Q9JKX4 Q9JKX6 Q9JKY0 Q9JKY1 Q9JKY3
Q9JL10 Q9JL16 Q9JL35 Q9JL60 Q9JLC3 Q9JLC6 Q9JLC8 Q9JLG8 Q9JLG9 Q9JLI7 Q9JLI8
Q9JLJ3 Q9JLM4 Q9JLR5 Q9JLT4 Q9JLV2 Q9JLV6 Q9JLY7 Q9JLZ3 Q9JLZ6 Q9JLZ9 Q9JM00
Q9JM01 Q9JM04 Q9JM05 Q9JM14 Q9JM55 Q9JM73 Q9JM76 Q9JM83 Q9JM93 Q9JM98 Q9JMB1
Q9JMC2 Q9JMD0 Q9JMD1 Q9JMD2 Q9JMG6 Q9JMH6 Q9JMJ2 Q9JMQ1 Q9JMR2 Q9JMR3 Q9JMR6
Q9JMR7 Q9JMR8 Q9JMR9 Q9JMS0 Q9JMS1 Q9JMS2 Q9JMS6 Q9JMS8 Q9JMS9 Q9JMT1 Q9JMT3
Q9JMT7 Q9JMT8 Q9JMT9 Q9JN05 Q9JN41 Q9JN46 Q9JN65 Q9JN69 Q9JP87 Q9JP88 Q9JP92
Q9JPA2 Q9JPA3 Q9JPA5 Q9JPB5 Q9JPB9 Q9JPD1 Q9JQL7 Q9JQM8 Q9JQQ6 Q9JQT2 Q9JQT4
Q9JQX5 Q9JQY4 Q9JR00 Q9JR26 Q9JR30 Q9JR76 Q9JRD8 Q9JRE9 Q9JRF6 Q9JRG1 Q9JRM6
Q9JRP9 Q9JRS8 Q9JRT1 Q9JRW7 Q9JRW9 Q9JRZ6 Q9JRZ9 Q9JS04 Q9JS33 Q9JS45 Q9JS61

Q9JSM1 Q9JSM2 Q9JSM5 Q9JSN0 Q9JSN4 Q9JSQ9 Q9JSR7 Q9JSS3 Q9JSS4 Q9JSS6 Q9JSS7
Q9JST7 Q9JST9 Q9JSU1 Q9JSU8 Q9JSW1 Q9JSX4 Q9JSX6 Q9JSY8 Q9JSY9 Q9JSZ0 Q9JSZ5
Q9JSZ8 Q9JSZ9 Q9JT07 Q9JT15 Q9JT23 Q9JT25 Q9JT57 Q9JT70 Q9JT71 Q9JT75 Q9JT77
Q9JT78 Q9JT81 Q9JT82 Q9JT86 Q9JT95 Q9JT96 Q9JTA1 Q9JTA3 Q9JTA4 Q9JTA6 Q9JTB0
Q9JTB5 Q9JTD9 Q9JTE9 Q9JTF2 Q9JTG4 Q9JTH8 Q9JTH9 Q9JTI3 Q9JTI4 Q9JTI8 Q9JTJ6
Q9JTJ9 Q9JTK1 Q9JTM0 Q9JTM3 Q9JTM4 Q9JTM5 Q9JTM7 Q9JTM8 Q9JTN2 Q9JTP0 Q9JTP4
Q9JTP5 Q9JTP9 Q9JTQ0 Q9JTR0 Q9JTR9 Q9JTS0 Q9JTS2 Q9JTT3 Q9JTT4 Q9JTT7 Q9JTU3
Q9JTW1 Q9JTW6 Q9JTX0 Q9JTX5 Q9JTX7 Q9JTX8 Q9JTX9 Q9JTY8 Q9JTZ1 Q9JTZ3 Q9JTZ5
Q9JTZ6 Q9JU03 Q9JU09 Q9JU11 Q9JU12 Q9JU19 Q9JU21 Q9JU24 Q9JU25 Q9JU26 Q9JU32
Q9JU33 Q9JU34 Q9JU49 Q9JU79 Q9JU81 Q9JU82 Q9JU84 Q9JU97 Q9JU99 Q9JUA2 Q9JUA5
Q9JUB0 Q9JUB2 Q9JUB6 Q9JUC4 Q9JUC7 Q9JUC8 Q9JUC9 Q9JUD5 Q9JUD8 Q9JUD9 Q9JUE0
Q9JUE9 Q9JUF2 Q9JUF4 Q9JUK2 Q9JUK6 Q9JUK7 Q9JUK8 Q9JUP6 Q9JUQ8 Q9JUS4 Q9JUT0
Q9JUT6 Q9JUT8 Q9JUT9 Q9JUU2 Q9JUU5 Q9JUU8 Q9JUU9 Q9JUV1 Q9JUV9 Q9JUW1 Q9JUX7
Q9JUX8 Q9JUY0 Q9JUZ9 Q9JV01 Q9JV02 Q9JV11 Q9JV14 Q9JV18 Q9JV25 Q9JV26 Q9JV28
Q9JV35 Q9JV42 Q9JV52 Q9JV58 Q9JV69 Q9JV73 Q9JV76 Q9JV83 Q9JV90 Q9JV91 Q9JV95
Q9JVA0 Q9JVA1 Q9JVA3 Q9JVA4 Q9JVA5 Q9JVB6 Q9JVC0 Q9JVC2 Q9JVD0 Q9JVD1 Q9JVD3
Q9JVD5 Q9JVD6 Q9JVE0 Q9JVE7 Q9JVF4 Q9JVG3 Q9JVG5 Q9JVG7 Q9JVH3 Q9JVH4 Q9JVH5
Q9JVH6 Q9JVH7 Q9JVI4 Q9JVJ9 Q9JVK3 Q9JVK9 Q9JVL0 Q9JVL1 Q9JVP1 Q9JVP2 Q9JVP8
Q9JVQ9 Q9JVS4 Q9JVS5 Q9JVT3 Q9JVT8 Q9JVU6 Q9JVU8 Q9JVU9 Q9JVV6 Q9JVW1 Q9JVW5
Q9JVW7 Q9JVX8 Q9JVY4 Q9JVY6 Q9JVY8 Q9JVZ6 Q9JW02 Q9JW05 Q9JW08 Q9JW09 Q9JW10
Q9JW13 Q9JW14 Q9JW21 Q9JW31 Q9JW39 Q9JW42 Q9JW45 Q9JW46 Q9JW50 Q9JW52 Q9JW60
Q9JW67 Q9JW77 Q9JW78 Q9JWA3 Q9JWA6 Q9JWB7 Q9JWC5 Q9JWC8 Q9JWD6 Q9JWE5 Q9JWE6
Q9JWF1 Q9JWF3 Q9JWG3 Q9JWG6 Q9JWH1 Q9JWH4 Q9JWI2 Q9JWI4 Q9JWI7 Q9JWI9 Q9JWJ3
Q9JWJ7 Q9JWJ8 Q9JWK3 Q9JWM1 Q9JWM8 Q9JWN9 Q9JWP0 Q9JWQ2 Q9JWQ5 Q9JWS7 Q9JWS8
Q9JWT4 Q9JWT7 Q9JWU6 Q9JWV1 Q9JWV4 Q9JWW8 Q9JWY6 Q9JWY9 Q9JX02 Q9JX03 Q9JX07
Q9JX11 Q9JX12 Q9JX13 Q9JX14 Q9JX15 Q9JX16 Q9JX17 Q9JX18 Q9JX22 Q9JX24 Q9JX33
Q9JX35 Q9JX38 Q9JX40 Q9JX41 Q9JX42 Q9JX48 Q9JX50 Q9JX94 Q9JXA3 Q9JXA4 Q9JXB0
Q9JXB3 Q9JXC1 Q9JXD7 Q9JXE2 Q9JXE3 Q9JXE5 Q9JXE6 Q9JXF1 Q9JXF5 Q9JXF7 Q9JXG3
Q9JXG5 Q9JXG9 Q9JXH2 Q9JXI0 Q9JXI2 Q9JXI7 Q9JXK1 Q9JXK4 Q9JXK5 Q9JXL4 Q9JXM2
Q9JXM4 Q9JXP4 Q9JXP5 Q9JXQ5 Q9JXR2 Q9JXS0 Q9JXS2 Q9JXS6 Q9JXS9 Q9JXT8 Q9JXU9
Q9JXV6 Q9JXV7 Q9JXV8 Q9JXW0 Q9JXW1 Q9JXW2 Q9JXW4 Q9JXW8 Q9JXX4 Q9JXY0 Q9JXY2
Q9JXY5 Q9JXY7 Q9JXZ9 Q9JY01 Q9JY03 Q9JY06 Q9JY09 Q9JY15 Q9JY16 Q9JY18 Q9JY72
Q9JY89 Q9JY93 Q9JY95 Q9JY96 Q9JY99 Q9JYA0 Q9JYB4 Q9JYB5 Q9JYC0 Q9JYC2 Q9JYC3
Q9JYC4 Q9JYC7 Q9JYD2 Q9JYE5 Q9JYF4 Q9JYF7 Q9JYG6 Q9JYH7 Q9JYH8 Q9JYI2 Q9JYJ3
Q9JYJ6 Q9JYJ8 Q9JYL2 Q9JYM1 Q9JYM4 Q9JYM5 Q9JYM6 Q9JYM8 Q9JYM9 Q9JYN5 Q9JYP8
Q9JYQ3 Q9JYQ8 Q9JYQ9 Q9JYS8 Q9JYT0 Q9JYT1 Q9JYT2 Q9JYU1 Q9JYU3 Q9JYU6 Q9JYV1
Q9JYX3 Q9JYX6 Q9JYY0 Q9JYY1 Q9JYY2 Q9JYY3 Q9JYY4 Q9JYY8 Q9JYZ2 Q9JYZ5 Q9JYZ7
Q9JYZ9 Q9JZ00 Q9JZ14 Q9JZ16 Q9JZ17 Q9JZ24 Q9JZ26 Q9JZ29 Q9JZ30 Q9JZ31 Q9JZ38
Q9JZ40 Q9JZ42 Q9JZ56 Q9JZ77 Q9JZ78 Q9JZ80 Q9JZ86 Q9JZ88 Q9JZ92 Q9JZA1 Q9JZA4
Q9JZA5 Q9JZB4 Q9JZF7 Q9JZG1 Q9JZG2 Q9JZG3 Q9JZH1 Q9JZI3 Q9JZI5 Q9JZI6 Q9JZI9
Q9JZL5 Q9JZM7 Q9JZP1 Q9JZP4 Q9JZQ4 Q9JZQ5 Q9JZQ8 Q9JZR0 Q9JZR3 Q9JZR4 Q9JZR6
Q9JZU5 Q9JZU7 Q9JZW3 Q9JZW4 Q9JZW6 Q9JZX9 Q9JZY1 Q9JZY2 Q9JZZ2 Q9JZZ5 Q9K005
Q9K012 Q9K013 Q9K016 Q9K023 Q9K029 Q9K041 Q9K048 Q9K060 Q9K063 Q9K068 Q9K075
Q9K080 Q9K081 Q9K085 Q9K089 Q9K092 Q9K093 Q9K095 Q9K096 Q9K097 Q9K0B0 Q9K0B5

Q9K0B7 Q9K0C5 Q9K0C6 Q9K0C8 Q9K0D0 Q9K0D1 Q9K0D4 Q9K0D9 Q9K0E3 Q9K0G4 Q9K0G6
Q9K0G8 Q9K0H2 Q9K0H3 Q9K0H4 Q9K0H5 Q9K0H6 Q9K0I1 Q9K0J4 Q9K0J8 Q9K0K3 Q9K0K4
Q9K0L7 Q9K0L8 Q9K0M3 Q9K0P6 Q9K0P7 Q9K0Q4 Q9K0Q5 Q9K0Q9 Q9K0U0 Q9K0U3 Q9K0U5
Q9K0V4 Q9K0V9 Q9K0W0 Q9K0X1 Q9K0Y0 Q9K0Y1 Q9K0Y4 Q9K0Y9 Q9K0Z0 Q9K0Z1 Q9K105
Q9K107 Q9K118 Q9K132 Q9K139 Q9K141 Q9K143 Q9K152 Q9K153 Q9K155 Q9K161 Q9K168
Q9K188 Q9K192 Q9K197 Q9K1A2 Q9K1A4 Q9K1A5 Q9K1E9 Q9K1F1 Q9K1F9 Q9K1G0 Q9K1G1
Q9K1G3 Q9K1G6 Q9K1G8 Q9K1H7 Q9K1I0 Q9K1I1 Q9K1I2 Q9K1I4 Q9K1I5 Q9K1J1 Q9K1J3
Q9K1K6 Q9K1K9 Q9K1N8 Q9K1P9 Q9K1Q0 Q9K1Q9 Q9K1R0 Q9K1R6 Q9K1Y4 Q9K2B6 Q9K333
Q9K3D6 Q9K3D7 Q9K3E4 Q9K3J3 Q9K3J6 Q9K3L9 Q9K3R5 Q9K3R6 Q9K3S8 Q9K3T2 Q9K3T8
Q9K3T9 Q9K3U0 Q9K3V1 Q9K3V7 Q9K3Z0 Q9K470 Q9K487 Q9K492 Q9K498 Q9K499 Q9K4A0
Q9K4A7 Q9K4B1 Q9K4C3 Q9K4E3 Q9K4E4 Q9K4E5 Q9K4F4 Q9K4F6 Q9K4F9 Q9K4Q3 Q9K4U5
Q9K4U6 Q9K4V1 Q9K4Y8 Q9K4Y9 Q9K4Z0 Q9K4Z1 Q9K4Z2 Q9K4Z3 Q9K4Z4 Q9K4Z5 Q9K4Z6
Q9K4Z7 Q9K575 Q9K576 Q9K5E4 Q9K5J6 Q9K5J7 Q9K5M8 Q9K5M9 Q9K5N0 Q9K5N8 Q9K5N9
Q9K5P0 Q9K5P5 Q9K5Q8 Q9K5R0 Q9K5T1 Q9K5W7 Q9K5Z2 Q9K5Z5 Q9K5Z6 Q9K5Z8 Q9K623
Q9K625 Q9K656 Q9K672 Q9K681 Q9K682 Q9K6A4 Q9K6A7 Q9K6A8 Q9K6B5 Q9K6B8 Q9K6B9
Q9K6C1 Q9K6D6 Q9K6D7 Q9K6E4 Q9K6E5 Q9K6E9 Q9K6F0 Q9K6F4 Q9K6G3 Q9K6G4 Q9K6G5
Q9K6I0 Q9K6K0 Q9K6K1 Q9K6K2 Q9K6U4 Q9K6V7 Q9K6V8 Q9K6W2 Q9K6X9 Q9K6Y0 Q9K6Y4
Q9K6Y7 Q9K6Z0 Q9K6Z1 Q9K6Z2 Q9K6Z3 Q9K6Z4 Q9K6Z5 Q9K6Z6 Q9K6Z7 Q9K6Z9 Q9K705
Q9K706 Q9K707 Q9K708 Q9K709 Q9K714 Q9K715 Q9K716 Q9K721 Q9K786 Q9K7A0 Q9K7B5
Q9K7C9 Q9K7E3 Q9K7E4 Q9K7E8 Q9K7F0 Q9K7F3 Q9K7G2 Q9K7H6 Q9K7K5 Q9K7L2 Q9K7L8
Q9K7N6 Q9K7Q7 Q9K7Q9 Q9K7R6 Q9K7S8 Q9K7U8 Q9K7V0 Q9K7V3 Q9K7V7 Q9K7W1 Q9K7X9
Q9K7Z8 Q9K801 Q9K804 Q9K808 Q9K813 Q9K815 Q9K820 Q9K821 Q9K829 Q9K838 Q9K843
Q9K849 Q9K855 Q9K857 Q9K859 Q9K861 Q9K864 Q9K866 Q9K867 Q9K868 Q9K869 Q9K888
Q9K893 Q9K895 Q9K896 Q9K897 Q9K8A0 Q9K8A9 Q9K8D8 Q9K8D9 Q9K8E4 Q9K8E7 Q9K8E8
Q9K8E9 Q9K8F0 Q9K8F1 Q9K8F3 Q9K8F4 Q9K8F7 Q9K8F8 Q9K8G0 Q9K8G2 Q9K8G3 Q9K8G8
Q9K8H3 Q9K8H4 Q9K8H8 Q9K8I3 Q9K8I4 Q9K8I7 Q9K8I9 Q9K8J5 Q9K8J7 Q9K8K8 Q9K8U9
Q9K8V2 Q9K8V3 Q9K8V4 Q9K8V5 Q9K8V6 Q9K8V7 Q9K8V8 Q9K8X1 Q9K8X7 Q9K8Y0 Q9K8Y2
Q9K8Z1 Q9K8Z3 Q9K8Z7 Q9K902 Q9K904 Q9K907 Q9K934 Q9K935 Q9K936 Q9K938 Q9K943
Q9K949 Q9K951 Q9K965 Q9K966 Q9K967 Q9K968 Q9K971 Q9K973 Q9K974 Q9K991 Q9K993
Q9K9A8 Q9K9B2 Q9K9C6 Q9K9D1 Q9K9D5 Q9K9G8 Q9K9G9 Q9K9H0 Q9K9H8 Q9K9H9 Q9K9I0
Q9K9I6 Q9K9I9 Q9K9K5 Q9K9K7 Q9K9K8 Q9K9L8 Q9K9L9 Q9K9P8 Q9K9Q1 Q9K9Q6 Q9K9R0
Q9K9R2 Q9K9R5 Q9K9R8 Q9K9R9 Q9K9S0 Q9K9S4 Q9K9S8 Q9K9T1 Q9K9U6 Q9K9V0 Q9K9V4
Q9K9V6 Q9K9V7 Q9K9V8 Q9K9V9 Q9K9W0 Q9K9W1 Q9K9W2 Q9K9W3 Q9K9W8 Q9K9W9 Q9K9Y1
Q9K9Y2 Q9K9Y3 Q9K9Y6 Q9K9Y8 Q9K9Z1 Q9K9Z4 Q9KA05 Q9KA06 Q9KA11 Q9KA12 Q9KA14
Q9KA15 Q9KA16 Q9KA20 Q9KA23 Q9KA24 Q9KA25 Q9KA26 Q9KA27 Q9KA28 Q9KA31 Q9KA57
Q9KA63 Q9KA64 Q9KA65 Q9KA66 Q9KA67 Q9KA69 Q9KA71 Q9KA72 Q9KA73 Q9KA74 Q9KA77
Q9KA79 Q9KA80 Q9KA82 Q9KA83 Q9KA91 Q9KAA3 Q9KAA5 Q9KAA7 Q9KAB7 Q9KAC0 Q9KAC1
Q9KAC3 Q9KAC4 Q9KAD6 Q9KAD7 Q9KAG7 Q9KAG8 Q9KAG9 Q9KAH0 Q9KAH1 Q9KAH2 Q9KAH3
Q9KAH5 Q9KAH8 Q9KAI6 Q9KAI7 Q9KAJ2 Q9KAK0 Q9KAK2 Q9KAK7 Q9KAK9 Q9KAN8 Q9KAP3
Q9KAP6 Q9KAT1 Q9KAV6 Q9KAX0 Q9KB03 Q9KB30 Q9KB75 Q9KB79 Q9KBE3 Q9KBE4 Q9KBE5
Q9KBE6 Q9KBE7 Q9KBG1 Q9KBH7 Q9KBI9 Q9KBJ4 Q9KBM8 Q9KBN7 Q9KBQ0 Q9KBQ2 Q9KBQ3
Q9KBQ4 Q9KBR4 Q9KBU0 Q9KBU1 Q9KBU2 Q9KBV3 Q9KBZ6 Q9KC05 Q9KC06 Q9KC12 Q9KC26
Q9KC32 Q9KC58 Q9KC61 Q9KC63 Q9KC68 Q9KC71 Q9KC78 Q9KC82 Q9KC85 Q9KC86 Q9KC87
Q9KC89 Q9KC92 Q9KC93 Q9KCA3 Q9KCA6 Q9KCA8 Q9KCA9 Q9KCB0 Q9KCB1 Q9KCB2 Q9KCB3

Q9KCB6 Q9KCB7 Q9KCB9 Q9KCC2 Q9KCC4 Q9KCC6 Q9KCC7 Q9KCD4 Q9KCF0 Q9KCF1 Q9KCF9
Q9KCG7 Q9KCI0 Q9KCL0 Q9KCL1 Q9KCL4 Q9KCL9 Q9KCM0 Q9KCM5 Q9KCN2 Q9KCN9 Q9KCP0
Q9KCQ5 Q9KCR4 Q9KCR5 Q9KCT0 Q9KCT3 Q9KCT8 Q9KCU7 Q9KCX2 Q9KCX4 Q9KCY0 Q9KCY6
Q9KCY8 Q9KCZ4 Q9KD10 Q9KD18 Q9KD20 Q9KD27 Q9KD33 Q9KD37 Q9KD39 Q9KD44 Q9KD45
Q9KD46 Q9KD48 Q9KD49 Q9KD50 Q9KD53 Q9KD56 Q9KD65 Q9KD67 Q9KD70 Q9KD71 Q9KD72
Q9KD73 Q9KD74 Q9KD78 Q9KD79 Q9KD89 Q9KD90 Q9KD91 Q9KD93 Q9KDA5 Q9KDD4 Q9KDD7
Q9KDD8 Q9KDE3 Q9KDE4 Q9KDE5 Q9KDE6 Q9KDF2 Q9KDG1 Q9KDG2 Q9KDH0 Q9KDH2 Q9KDI5
Q9KDI6 Q9KDI8 Q9KDI9 Q9KDJ3 Q9KDJ5 Q9KDJ6 Q9KDJ9 Q9KDK0 Q9KDK9 Q9KDL0 Q9KDM4
Q9KDM7 Q9KDN6 Q9KDQ1 Q9KDR2 Q9KDV0 Q9KDV2 Q9KDV3 Q9KDV4 Q9KDW4 Q9KDW8 Q9KDX4
Q9KDX6 Q9KDZ8 Q9KE16 Q9KE27 Q9KE51 Q9KE60 Q9KEB0 Q9KEB3 Q9KEC6 Q9KEC9 Q9KED3
Q9KED4 Q9KED5 Q9KED9 Q9KEE5 Q9KEE6 Q9KEF5 Q9KEI9 Q9KEL8 Q9KER1 Q9KER6 Q9KET9
Q9KEZ3 Q9KEZ4 Q9KF10 Q9KF29 Q9KF37 Q9KF43 Q9KF46 Q9KF49 Q9KF52 Q9KF53 Q9KF55
Q9KF57 Q9KF58 Q9KF60 Q9KF75 Q9KF78 Q9KFD3 Q9KFF1 Q9KFF9 Q9KFH7 Q9KFH9 Q9KFI7
Q9KFP1 Q9KFQ8 Q9KFR8 Q9KFV3 Q9KFX8 Q9KG31 Q9KG32 Q9KG38 Q9KG45 Q9KG46 Q9KG51
Q9KG52 Q9KG53 Q9KG55 Q9KG56 Q9KG58 Q9KG59 Q9KG60 Q9KG61 Q9KG85 Q9KGD4 Q9KGD5
Q9KGE3 Q9KGE4 Q9KGE5 Q9KGE6 Q9KGE7 Q9KGE9 Q9KGF4 Q9KGF6 Q9KGF7 Q9KGF8 Q9KGG1
Q9KGG3 Q9KGG4 Q9KGH4 Q9KGH5 Q9KGH8 Q9KGI8 Q9KGJ3 Q9KGJ4 Q9KGJ5 Q9KGJ7 Q9KGK0
Q9KGK4 Q9KGK8 Q9KGL2 Q9KGL3 Q9KGL5 Q9KGL9 Q9KGM3 Q9KGN4 Q9KGN5 Q9KGN6 Q9KGN8
Q9KGU2 Q9KGU6 Q9KGU7 Q9KGY4 Q9KH06 Q9KH11 Q9KH12 Q9KH19 Q9KH33 Q9KH71 Q9KHA1
Q9KHE2 Q9KHR3 Q9KHS2 Q9KHS6 Q9KHU6 Q9KHU8 Q9KHV6 Q9KI19 Q9KI21 Q9KI47 Q9KI57
Q9KI71 Q9KID9 Q9KIG4 Q9KIP8 Q9KIP9 Q9KIQ6 Q9KIT1 Q9KIZ4 Q9KJ20 Q9KJ21 Q9KJ22
Q9KJ23 Q9KJ24 Q9KJC4 Q9KJE8 Q9KJE9 Q9KJF0 Q9KJF6 Q9KJG4 Q9KJH8 Q9KJL4 Q9KJM5
Q9KJN4 Q9KJT8 Q9KJU0 Q9KJU3 Q9KJU4 Q9KJV4 Q9KK59 Q9KK72 Q9KK75 Q9KKF1 Q9KKJ5
Q9KKL4 Q9KKM4 Q9KKP2 Q9KKP3 Q9KKQ6 Q9KKS5 Q9KKW4 Q9KKY0 Q9KKY1 Q9KKZ9 Q9KL03
Q9KL16 Q9KL18 Q9KL20 Q9KL24 Q9KL40 Q9KL41 Q9KL54 Q9KL62 Q9KL72 Q9KL96 Q9KLB8
Q9KLB9 Q9KLC1 Q9KLC2 Q9KLC3 Q9KLC6 Q9KLC7 Q9KLD1 Q9KLE1 Q9KLG8 Q9KLJ9 Q9KLK2
Q9KLK3 Q9KLK4 Q9KLL6 Q9KLN2 Q9KLP4 Q9KLR1 Q9KLT6 Q9KLW7 Q9KLW8 Q9KLX5 Q9KLX6
Q9KLX7 Q9KLY5 Q9KLY7 Q9KM09 Q9KM17 Q9KM20 Q9KM67 Q9KM69 Q9KM71 Q9KM76 Q9KMA5
Q9KMA6 Q9KMG3 Q9KMG4 Q9KMG5 Q9KMJ4 Q9KMM3 Q9KMN7 Q9KMP4 Q9KMP5 Q9KMT5 Q9KMW1
Q9KMW9 Q9KMX5 Q9KMY3 Q9KMY6 Q9KN00 Q9KN05 Q9KN16 Q9KN38 Q9KN48 Q9KN67 Q9KN74
Q9KNA8 Q9KNB2 Q9KNC3 Q9KNE6 Q9KNE8 Q9KNF3 Q9KNF8 Q9KNG4 Q9KNG5 Q9KNG7 Q9KNI7
Q9KNJ7 Q9KNK0 Q9KNK2 Q9KNL2 Q9KNL4 Q9KNL7 Q9KNM2 Q9KNM3 Q9KNM4 Q9KNP2 Q9KNP5
Q9KNP9 Q9KNQ2 Q9KNQ6 Q9KNQ7 Q9KNQ9 Q9KNR1 Q9KNR6 Q9KNR7 Q9KNS1 Q9KNS6 Q9KNS8
Q9KNT4 Q9KNT5 Q9KNT6 Q9KNT7 Q9KNT8 Q9KNT9 Q9KNV1 Q9KNV2 Q9KNV6 Q9KNV7 Q9KNW0
Q9KNW2 Q9KNW4 Q9KNW8 Q9KNX6 Q9KNX8 Q9KNY1 Q9KNY2 Q9KNY3 Q9KNY4 Q9KNY5 Q9KNY6
Q9KNY7 Q9KNY8 Q9KNY9 Q9KNZ0 Q9KNZ1 Q9KNZ2 Q9KNZ3 Q9KNZ4 Q9KNZ5 Q9KNZ6 Q9KNZ7
Q9KNZ8 Q9KNZ9 Q9KP00 Q9KP01 Q9KP02 Q9KP03 Q9KP05 Q9KP07 Q9KP08 Q9KP09 Q9KP19
Q9KP20 Q9KP21 Q9KP30 Q9KP33 Q9KP34 Q9KP35 Q9KP38 Q9KP43 Q9KP46 Q9KP47 Q9KP62
Q9KP65 Q9KP66 Q9KP68 Q9KP70 Q9KP71 Q9KP73 Q9KP80 Q9KP81 Q9KP82 Q9KP83 Q9KP93
Q9KP97 Q9KPA1 Q9KPA2 Q9KPA4 Q9KPB2 Q9KPB4 Q9KPB5 Q9KPC1 Q9KPC4 Q9KPC6 Q9KPE1
Q9KPE2 Q9KPE3 Q9KPE9 Q9KPF6 Q9KPF9 Q9KPG5 Q9KPG8 Q9KPH7 Q9KPH8 Q9KPH9 Q9KPI4
Q9KPI8 Q9KPK1 Q9KPK4 Q9KPK6 Q9KPK7 Q9KPK8 Q9KPL2 Q9KPL7 Q9KPL8 Q9KPL9 Q9KPM0
Q9KPM2 Q9KPM5 Q9KPN7 Q9KPQ0 Q9KPQ7 Q9KPQ9 Q9KPR9 Q9KPS0 Q9KPS5 Q9KPT1 Q9KPT6
Q9KPT7 Q9KPT8 Q9KPT9 Q9KPU0 Q9KPU4 Q9KPU5 Q9KPU8 Q9KPV0 Q9KPV2 Q9KPV3 Q9KPV4

Q9KPV5 Q9KPV8 Q9KPX0 Q9KPX2 Q9KPX4 Q9KPX6 Q9KPX8 Q9KPY6 Q9KPY7 Q9KQ15 Q9KQ21
Q9KQ22 Q9KQ23 Q9KQ24 Q9KQ25 Q9KQ26 Q9KQ28 Q9KQ47 Q9KQ52 Q9KQ65 Q9KQ82 Q9KQ83
Q9KQ85 Q9KQ91 Q9KQ92 Q9KQA4 Q9KQA7 Q9KQB5 Q9KQC6 Q9KQF5 Q9KQF8 Q9KQF9 Q9KQG2
Q9KQH0 Q9KQH1 Q9KQH3 Q9KQI0 Q9KQI2 Q9KQK0 Q9KQL9 Q9KQM0 Q9KQM3 Q9KQM6 Q9KQN9
Q9KQP1 Q9KQQ6 Q9KQR7 Q9KQS5 Q9KQS6 Q9KQS7 Q9KQS9 Q9KQT2 Q9KQT4 Q9KQT7 Q9KQV6
Q9KQX1 Q9KQX4 Q9KQX6 Q9KQZ5 Q9KQZ9 Q9KR00 Q9KR01 Q9KR02 Q9KR14 Q9KR33 Q9KR57
Q9KR61 Q9KR62 Q9KR70 Q9KRA6 Q9KRA7 Q9KRB0 Q9KRB2 Q9KRB5 Q9KRB6 Q9KRC1 Q9KRC2
Q9KRC6 Q9KRC7 Q9KRC8 Q9KRD1 Q9KRD8 Q9KRK0 Q9KRK5 Q9KRL3 Q9KRL4 Q9KRM2 Q9KRM5
Q9KRP1 Q9KRP7 Q9KRR0 Q9KRS6 Q9KRU5 Q9KRV2 Q9KRV8 Q9KRX4 Q9KRX5 Q9KRY1 Q9KRY6
Q9KRZ3 Q9KRZ5 Q9KS02 Q9KS29 Q9KS67 Q9KS71 Q9KS93 Q9KSA4 Q9KSA5 Q9KSB2 Q9KSB4
Q9KSC2 Q9KSF9 Q9KSH1 Q9KSJ3 Q9KSJ7 Q9KSJ9 Q9KSL4 Q9KSL9 Q9KSM5 Q9KSM8 Q9KSN0
Q9KSN4 Q9KSN6 Q9KSN7 Q9KSP2 Q9KSP7 Q9KSP8 Q9KSQ1 Q9KSQ2 Q9KSQ3 Q9KSQ4 Q9KSR6
Q9KSR8 Q9KSS4 Q9KSS7 Q9KST2 Q9KST3 Q9KST4 Q9KST5 Q9KST6 Q9KST7 Q9KST9 Q9KSU0
Q9KSU2 Q9KSU3 Q9KSU7 Q9KSW0 Q9KSW3 Q9KSW4 Q9KSW7 Q9KSW8 Q9KSW9 Q9KSX0 Q9KSX1
Q9KSX2 Q9KSX4 Q9KSX8 Q9KSZ1 Q9KSZ3 Q9KSZ4 Q9KSZ6 Q9KT00 Q9KT07 Q9KT08 Q9KT47
Q9KT48 Q9KT49 Q9KT50 Q9KT52 Q9KT67 Q9KT69 Q9KT77 Q9KT79 Q9KT81 Q9KT82 Q9KT93
Q9KT97 Q9KTA4 Q9KTA6 Q9KTB6 Q9KTB7 Q9KTD1 Q9KTE0 Q9KTE2 Q9KTE6 Q9KTF1 Q9KTF7
Q9KTF8 Q9KTF9 Q9KTG1 Q9KTK0 Q9KTK4 Q9KTK8 Q9KTL1 Q9KTL3 Q9KTL4 Q9KTM1 Q9KTM3
Q9KTM4 Q9KTM7 Q9KTN0 Q9KTN2 Q9KTP8 Q9KTT7 Q9KTT8 Q9KTU1 Q9KTW2 Q9KTW4 Q9KTW7
Q9KTX0 Q9KTX1 Q9KTX3 Q9KTX4 Q9KTX5 Q9KTX8 Q9KTX9 Q9KTY2 Q9KTY3 Q9KTY9 Q9KTZ1
Q9KU00 Q9KU07 Q9KU12 Q9KU18 Q9KU20 Q9KU27 Q9KU28 Q9KU36 Q9KU37 Q9KU38 Q9KU39
Q9KU44 Q9KU47 Q9KU53 Q9KU56 Q9KU60 Q9KU62 Q9KU64 Q9KU76 Q9KU77 Q9KU78 Q9KU79
Q9KU80 Q9KU82 Q9KU84 Q9KU87 Q9KU89 Q9KU92 Q9KU96 Q9KU97 Q9KUA0 Q9KUA9 Q9KUB7
Q9KUC5 Q9KUC7 Q9KUD0 Q9KUD1 Q9KUD9 Q9KUE1 Q9KUF0 Q9KUF1 Q9KUF7 Q9KUF8 Q9KUF9
Q9KUG0 Q9KUG5 Q9KUH3 Q9KUI6 Q9KUI8 Q9KUI9 Q9KUJ0 Q9KUJ1 Q9KUJ2 Q9KUK8 Q9KUM8
Q9KUN3 Q9KUP2 Q9KUP3 Q9KUP4 Q9KUP7 Q9KUP8 Q9KUP9 Q9KUQ0 Q9KUQ4 Q9KUQ7 Q9KUQ9
Q9KUR1 Q9KUR2 Q9KUR4 Q9KUS1 Q9KUS2 Q9KUS3 Q9KUS4 Q9KUS8 Q9KUS9 Q9KUT0 Q9KUT3
Q9KUT4 Q9KUU2 Q9KUU7 Q9KUW2 Q9KUW5 Q9KUW9 Q9KUX2 Q9KUX3 Q9KUX4 Q9KUX9 Q9KUY4
Q9KUY6 Q9KUY9 Q9KUZ0 Q9KUZ1 Q9KUZ2 Q9KUZ6 Q9KUZ7 Q9KUZ8 Q9KUZ9 Q9KV01 Q9KV02
Q9KV06 Q9KV11 Q9KV12 Q9KV13 Q9KV16 Q9KV17 Q9KV18 Q9KV22 Q9KV26 Q9KV27 Q9KV29
Q9KV30 Q9KV31 Q9KV32 Q9KV33 Q9KV34 Q9KV35 Q9KV37 Q9KV38 Q9KV40 Q9KV52 Q9KV53
Q9KV59 Q9KV60 Q9KV64 Q9KV66 Q9KV80 Q9KV81 Q9KV83 Q9KVC4 Q9KVC5 Q9KVC7 Q9KVC8
Q9KVC9 Q9KVD1 Q9KVD2 Q9KVD5 Q9KVD6 Q9KVE0 Q9KVG5 Q9KVG6 Q9KVG9 Q9KVI4 Q9KVI7
Q9KVJ0 Q9KVK6 Q9KVL2 Q9KVL4 Q9KVL6 Q9KVL7 Q9KVL8 Q9KVM1 Q9KVM7 Q9KVN0 Q9KVP1
Q9KVP6 Q9KVQ6 Q9KVQ7 Q9KVR6 Q9KVS4 Q9KVS8 Q9KVT3 Q9KVT4 Q9KVT5 Q9KVT7 Q9KVT8
Q9KVU0 Q9KVU3 Q9KVU4 Q9KVU5 Q9KVW0 Q9KVW4 Q9KVW7 Q9KVW8 Q9KVX3 Q9KVX4 Q9KVX5
Q9KVY1 Q9KVY2 Q9KVY5 Q9KVY6 Q9KVZ7 Q9KVZ8 Q9KW13 Q9KW14 Q9KW19 Q9KW52 Q9KWA3
Q9KWD1 Q9KWE4 Q9KWE5 Q9KWF6 Q9KWG2 Q9KWG3 Q9KWH3 Q9KWI8 Q9KWJ8 Q9KWK3 Q9KWK4
Q9KWL3 Q9KWN1 Q9KWR0 Q9KWS0 Q9KWS1 Q9KWS2 Q9KWS3 Q9KWS4 Q9KWS5 Q9KWS6 Q9KWS7
Q9KWS8 Q9KWU4 Q9KWU6 Q9KWU7 Q9KWU8 Q9KWW7 Q9KWW9 Q9KWX0 Q9KWZ1 Q9KWZ4 Q9KX08
Q9KX40 Q9KX58 Q9KX62 Q9KX67 Q9KX72 Q9KXN0 Q9KXN1 Q9KXP2 Q9KXP5 Q9KXP9 Q9KXQ0
Q9KXQ4 Q9KXQ5 Q9KXQ6 Q9KXQ9 Q9KXR0 Q9KXR1 Q9KXR2 Q9KXR3 Q9KXR5 Q9KXR6 Q9KXR8
Q9KXS0 Q9KXS1 Q9KXT4 Q9KXU5 Q9KY00 Q9KY04 Q9KY07 Q9KY50 Q9KY56 Q9KYI9 Q9KYP1
Q9KYR2 Q9KYR6 Q9KYR9 Q9KYS1 Q9KYV7 Q9KZ75 Q9KZ78 Q9KZA7 Q9KZC7 Q9KZF5 Q9KZJ5

Q9KZK8 Q9KZK9 Q9KZM1 Q9KZV0 Q9KZV6 Q9KZX7 Q9KZZ7 Q9L012 Q9L016 Q9L082 Q9L0B8
Q9L0C8 Q9L0C9 Q9L0D0 Q9L0D1 Q9L0D2 Q9L0D3 Q9L0D4 Q9L0D5 Q9L0D6 Q9L0D7 Q9L0D9
Q9L0E0 Q9L0H2 Q9L0K4 Q9L0L0 Q9L0L6 Q9L0L7 Q9L0M4 Q9L0Q6 Q9L0Q7 Q9L0Q8 Q9L0T8
Q9L0Y3 Q9L0Y6 Q9L0Z5 Q9L127 Q9L132 Q9L1A3 Q9L1A4 Q9L1A5 Q9L1B1 Q9L1I0 Q9L1I1
Q9L1I2 Q9L1I4 Q9L1K2 Q9L1L8 Q9L1S9 Q9L1U6 Q9L213 Q9L214 Q9L233 Q9L278 Q9L286
Q9L287 Q9L288 Q9L289 Q9L290 Q9L291 Q9L298 Q9L2E9 Q9L2H3 Q9L2H9 Q9L2L4 Q9L385
Q9L3B0 Q9L3B1 Q9L3B2 Q9L3B3 Q9L3B4 Q9L3G6 Q9L3I7 Q9L3Q5 Q9L473 Q9L4D3 Q9L4E1
Q9L4F0 Q9L4G1 Q9L4K0 Q9L4K1 Q9L4K2 Q9L4M7 Q9L4M8 Q9L4N5 Q9L4N6 Q9L4N8 Q9L4P3
Q9L4P4 Q9L4Q7 Q9L4Q8 Q9L4R7 Q9L4S1 Q9L4S2 Q9L4T8 Q9L4U5 Q9L4V5 Q9L516 Q9L543
Q9L555 Q9L558 Q9L560 Q9L5H8 Q9L5N1 Q9L5W4 Q9L5W5 Q9L637 Q9L638 Q9L639 Q9L640
Q9L641 Q9L642 Q9L644 Q9L645 Q9L646 Q9L690 Q9L691 Q9L699 Q9L6A2 Q9L6B4 Q9L6B5
Q9L6B7 Q9L6B8 Q9L6G1 Q9L6I1 Q9L6I2 Q9L6K8 Q9L6L4 Q9L6L9 Q9L6M2 Q9L6M5 Q9L6M9
Q9L6N1 Q9L6Q3 Q9L6S1 Q9L6T3 Q9L6V3 Q9L6W2 Q9L6W9 Q9L6X9 Q9L790 Q9L7A3 Q9L7A5
Q9L7A7 Q9L7B2 Q9L7L3 Q9L7L4 Q9L7L5 Q9L7L6 Q9L7L7 Q9L7L8 Q9L7L9 Q9L7M3 Q9L7P5
Q9L7Q5 Q9L7R9 Q9L7S0 Q9L7T2 Q9L7V3 Q9L7X5 Q9L7X6 Q9L7Z1 Q9L7Z3 Q9L867 Q9L870
Q9L888 Q9L8F6 Q9L9C1 Q9L9C8 Q9L9D7 Q9L9E3 Q9L9E5 Q9L9E6 Q9L9E7 Q9L9E8 Q9L9E9
Q9L9F0 Q9L9F1 Q9L9F2 Q9L9F3 Q9L9F4 Q9L9F5 Q9L9F6 Q9L9F7 Q9L9F8 Q9L9F9 Q9L9G0
Q9L9G1 Q9L9G2 Q9L9G3 Q9L9I4 Q9L9I5 Q9L9I6 Q9L9I7 Q9L9I8 Q9L9J1 Q9L9K3 Q9LA02
Q9LA24 Q9LA96 Q9LAE5 Q9LAG8 Q9LAI0 Q9LAI1 Q9LAI2 Q9LAK5 Q9LAK6 Q9LAM9 Q9LBE7
Q9LBG2 Q9LBQ3 Q9LBV3 Q9LBW4 Q9LBW5 Q9LBW6 Q9LCB4 Q9LCC0 Q9LCC1 Q9LCK0 Q9LCK1
Q9LCQ4 Q9LCQ5 Q9LCQ6 Q9LCR5 Q9LCS5 Q9LCS6 Q9LCS7 Q9LCT0 Q9LCT6 Q9LCU3 Q9LCV9
Q9LCX2 Q9LCX3 Q9LCX4 Q9LCY2 Q9LCY4 Q9LD00 Q9LD12 Q9LD13 Q9LD14 Q9LD28 Q9LD44
Q9LD47 Q9LD55 Q9LD61 Q9LD75 Q9LD90 Q9LD95 Q9LDA7 Q9LDA9 Q9LDD1 Q9LDD4 Q9LDD5
Q9LDE1 Q9LDE2 Q9LDE3 Q9LDJ9 Q9LDM2 Q9LDM4 Q9LDN0 Q9LDN2 Q9LDQ7 Q9LDT3 Q9LDV4
Q9LDW3 Q9LDX1 Q9LDY2 Q9LDY8 Q9LDY9 Q9LDZ1 Q9LE06 Q9LE16 Q9LE22 Q9LE33 Q9LE38
Q9LE42 Q9LE44 Q9LE54 Q9LE73 Q9LE86 Q9LE95 Q9LEB3 Q9LEB4 Q9LEC8 Q9LEI7 Q9LEI8
Q9LEI9 Q9LEJ0 Q9LEK8 Q9LEL5 Q9LEL6 Q9LEM8 Q9LEQ7 Q9LER0 Q9LES2 Q9LES4 Q9LES9
Q9LET0 Q9LEU4 Q9LEU8 Q9LEV0 Q9LEW8 Q9LEX0 Q9LEX5 Q9LEX6 Q9LEX8 Q9LEY4 Q9LEY9
Q9LEZ3 Q9LEZ4 Q9LEZ9 Q9LF10 Q9LF14 Q9LF20 Q9LF22 Q9LF24 Q9LF27 Q9LF30 Q9LF33
Q9LF38 Q9LF41 Q9LF46 Q9LF53 Q9LF54 Q9LF55 Q9LF66 Q9LF68 Q9LF88 Q9LF92 Q9LFA4
Q9LFA5 Q9LFC0 Q9LFC5 Q9LFE0 Q9LFE4 Q9LFF1 Q9LFF6 Q9LFF9 Q9LFG2 Q9LFH3 Q9LFH5
Q9LFJ8 Q9LFJ9 Q9LFK0 Q9LFM6 Q9LFN6 Q9LFP0 Q9LFQ3 Q9LFQ4 Q9LFQ9 Q9LFR7 Q9LFS8
Q9LFT6 Q9LFT8 Q9LFU0 Q9LFU1 Q9LFU5 Q9LFV0 Q9LFV5 Q9LFV6 Q9LFV9 Q9LFW0 Q9LFY0
Q9LG02 Q9LG05 Q9LG11 Q9LG15 Q9LG23 Q9LG26 Q9LG41 Q9LG86 Q9LG97 Q9LGE6 Q9LGH4
Q9LGH8 Q9LGI2 Q9LGQ6 Q9LGU6 Q9LGZ2 Q9LH31 Q9LH42 Q9LH85 Q9LH88 Q9LHB1 Q9LHD3
Q9LHD8 Q9LHE4 Q9LHE9 Q9LHF0 Q9LHG8 Q9LHH5 Q9LHH7 Q9LHI7 Q9LHJ2 Q9LHJ8 Q9LHK1
Q9LHL3 Q9LHL5 Q9LHN5 Q9LHN6 Q9LHN9 Q9LHP1 Q9LHP2 Q9LHQ0 Q9LHQ1 Q9LHQ5 Q9LHQ7
Q9LHR4 Q9LHS8 Q9LHS9 Q9LHY9 Q9LI00 Q9LI77 Q9LI84 Q9LI89 Q9LIA0 Q9LIA1 Q9LIA4
Q9LIA6 Q9LIA8 Q9LIB2 Q9LIB4 Q9LIB5 Q9LID0 Q9LID1 Q9LIE6 Q9LIF1 Q9LIF8 Q9LIG0
Q9LIG6 Q9LIH5 Q9LIH6 Q9LIH9 Q9LII8 Q9LIJ0 Q9LIJ4 Q9LIK3 Q9LIK5 Q9LIK9 Q9LIL1
Q9LIL5 Q9LIN3 Q9LIP2 Q9LIP9 Q9LIQ4 Q9LIQ6 Q9LIQ7 Q9LIQ9 Q9LIR0 Q9LIR1 Q9LIR2
Q9LIR3 Q9LIR4 Q9LIR6 Q9LIR8 Q9LJ39 Q9LJ44 Q9LJ45 Q9LJ48 Q9LJ68 Q9LJ69 Q9LJ74
Q9LJ87 Q9LJ98 Q9LJA0 Q9LJA6 Q9LJB4 Q9LJB5 Q9LJB7 Q9LJB9 Q9LJC0 Q9LJE2 Q9LJE4
Q9LJE5 Q9LJF5 Q9LJF8 Q9LJF9 Q9LJG6 Q9LJH8 Q9LJL3 Q9LJL4 Q9LJN8 Q9LJP0 Q9LJQ9

Q9LJR3 Q9LJW3 Q9LJW5 Q9LJW6 Q9LJX1 Q9LJX4 Q9LJY2 Q9LJZ5 Q9LJZ8 Q9LK08 Q9LK18
Q9LK21 Q9LK22 Q9LK24 Q9LK27 Q9LK32 Q9LK34 Q9LK36 Q9LK38 Q9LK48 Q9LK53 Q9LK57
Q9LK58 Q9LK61 Q9LK70 Q9LK73 Q9LK86 Q9LK95 Q9LKA3 Q9LKA4 Q9LKA5 Q9LKB9 Q9LKC0
Q9LKC3 Q9LKG7 Q9LKI7 Q9LKI8 Q9LKJ1 Q9LKJ3 Q9LKN0 Q9LKP7 Q9LKR4 Q9LKR8 Q9LKU8
Q9LKX9 Q9LKZ3 Q9LL41 Q9LL45 Q9LL84 Q9LLA7 Q9LLQ2 Q9LLQ3 Q9LLR9 Q9LM13 Q9LM18
Q9LM20 Q9LM46 Q9LM50 Q9LM55 Q9LM59 Q9LM63 Q9LM64 Q9LM71 Q9LM73 Q9LM74 Q9LM75
Q9LM83 Q9LM84 Q9LM89 Q9LM90 Q9LM91 Q9LM92 Q9LMA1 Q9LMB0 Q9LMC2 Q9LME2 Q9LME6
Q9LME8 Q9LMF0 Q9LMF1 Q9LMH0 Q9LMH5 Q9LMH6 Q9LMI0 Q9LMK0 Q9LMK5 Q9LMK7 Q9LML2
Q9LML6 Q9LML7 Q9LMM8 Q9LMN4 Q9LMP6 Q9LMR3 Q9LMR4 Q9LMR5 Q9LMT0 Q9LMT1 Q9LMT2
Q9LMU0 Q9LMX8 Q9LMY5 Q9LN01 Q9LN05 Q9LN22 Q9LN63 Q9LN69 Q9LN77 Q9LN95 Q9LNA5
Q9LNB9 Q9LNC5 Q9LNC9 Q9LND0 Q9LND4 Q9LND7 Q9LNE3 Q9LNE4 Q9LNE6 Q9LNE7 Q9LNF4
Q9LNF6 Q9LNG5 Q9LNI1 Q9LNI2 Q9LNI5 Q9LNJ5 Q9LNJ9 Q9LNN1 Q9LNP2 Q9LNP6 Q9LNT9
Q9LNU6 Q9LNV5 Q9LNX4 Q9LP07 Q9LP21 Q9LP37 Q9LP45 Q9LP46 Q9LP56 Q9LP59 Q9LPC4
Q9LPC6 Q9LPC7 Q9LPD2 Q9LPD9 Q9LPH0 Q9LPI7 Q9LPJ2 Q9LPJ7 Q9LPK5 Q9LPL3 Q9LPL5
Q9LPM2 Q9LPM8 Q9LPM9 Q9LPP4 Q9LPR4 Q9LPR8 Q9LPS8 Q9LPS9 Q9LPU6 Q9LPV3 Q9LPV8
Q9LPV9 Q9LPW0 Q9LPW2 Q9LPW3 Q9LPW4 Q9LPW7 Q9LPX2 Q9LPZ1 Q9LQ08 Q9LQ10 Q9LQ12
Q9LQ14 Q9LQ15 Q9LQ16 Q9LQ27 Q9LQ31 Q9LQ46 Q9LQ47 Q9LQ48 Q9LQ54 Q9LQ65 Q9LQ68
Q9LQ81 Q9LQ89 Q9LQ92 Q9LQ95 Q9LQ98 Q9LQB0 Q9LQC1 Q9LQE3 Q9LQE8 Q9LQF0 Q9LQI7
Q9LQI9 Q9LQK0 Q9LQK9 Q9LQL0 Q9LQL4 Q9LQL5 Q9LQM1 Q9LQM3 Q9LQM8 Q9LQN4 Q9LQN6
Q9LQQ1 Q9LQQ2 Q9LQQ4 Q9LQR0 Q9LQU5 Q9LQU7 Q9LQV2 Q9LQW3 Q9LQX2 Q9LQX4 Q9LQX5
Q9LQZ2 Q9LQZ5 Q9LQZ7 Q9LR02 Q9LR09 Q9LR29 Q9LR33 Q9LR44 Q9LR65 Q9LR67 Q9LR73
Q9LR74 Q9LR75 Q9LR97 Q9LRA7 Q9LRB8 Q9LRE5 Q9LRE6 Q9LRH6 Q9LRJ4 Q9LRM4 Q9LRN6
Q9LRN8 Q9LRP6 Q9LRQ1 Q9LRQ7 Q9LRQ8 Q9LRR0 Q9LRR2 Q9LRR4 Q9LRR5 Q9LRR9 Q9LRS0
Q9LRW1 Q9LRW3 Q9LRW4 Q9LRW5 Q9LRW6 Q9LRX6 Q9LRX8 Q9LRY0 Q9LRZ0 Q9LRZ2 Q9LRZ3
Q9LRZ4 Q9LRZ6 Q9LS00 Q9LS04 Q9LS08 Q9LS09 Q9LS10 Q9LS16 Q9LS21 Q9LS24 Q9LS25
Q9LS28 Q9LS39 Q9LS48 Q9LS54 Q9LS56 Q9LS57 Q9LS58 Q9LS72 Q9LS76 Q9LS86 Q9LS88
Q9LS90 Q9LSA5 Q9LSB2 Q9LSB8 Q9LSD0 Q9LSD2 Q9LSD5 Q9LSD6 Q9LSE2 Q9LSF5 Q9LSF7
Q9LSH2 Q9LSI7 Q9LSJ3 Q9LSK8 Q9LSL1 Q9LSL6 Q9LSL8 Q9LSL9 Q9LSM0 Q9LSM5 Q9LSN7
Q9LSP6 Q9LSQ2 Q9LSQ3 Q9LSQ6 Q9LSS3 Q9LSS5 Q9LSS7 Q9LSS8 Q9LST3 Q9LST6 Q9LST7
Q9LSU0 Q9LSU1 Q9LSU3 Q9LSU6 Q9LSU8 Q9LSV0 Q9LSV6 Q9LSW2 Q9LSW6 Q9LSW7 Q9LSW8
Q9LSX8 Q9LSX9 Q9LSY0 Q9LSY1 Q9LSY4 Q9LSY5 Q9LSY6 Q9LSY8 Q9LSY9 Q9LSZ0 Q9LSZ3
Q9LT08 Q9LT10 Q9LT17 Q9LT21 Q9LT23 Q9LT24 Q9LT25 Q9LT27 Q9LT31 Q9LT35 Q9LT45
Q9LT47 Q9LT48 Q9LT57 Q9LT67 Q9LT69 Q9LT72 Q9LT76 Q9LT79 Q9LT81 Q9LT82 Q9LT89
Q9LT93 Q9LTA2 Q9LTA3 Q9LTA6 Q9LTA7 Q9LTA8 Q9LTB2 Q9LTC3 Q9LTC4 Q9LTC7 Q9LTF2
Q9LTF4 Q9LTF5 Q9LTF6 Q9LTF7 Q9LTH2 Q9LTH3 Q9LTH4 Q9LTH7 Q9LTH8 Q9LTI5 Q9LTJ1
Q9LTJ2 Q9LTJ6 Q9LTR9 Q9LTS4 Q9LTS7 Q9LTT3 Q9LTT8 Q9LTT9 Q9LTV2 Q9LTV4 Q9LTV8
Q9LTW4 Q9LTX2 Q9LTX3 Q9LTX9 Q9LTY1 Q9LTY2 Q9LTY6 Q9LU10 Q9LU24 Q9LU36 Q9LU43
Q9LU46 Q9LU47 Q9LU49 Q9LU60 Q9LU68 Q9LU73 Q9LU89 Q9LU90 Q9LU91 Q9LU92 Q9LU93
Q9LUA9 Q9LUB6 Q9LUB7 Q9LUB9 Q9LUC2 Q9LUC4 Q9LUD4 Q9LUD6 Q9LUD9 Q9LUE0 Q9LUE1
Q9LUE2 Q9LUG5 Q9LUG7 Q9LUG9 Q9LUI8 Q9LUI9 Q9LUJ0 Q9LUJ1 Q9LUJ2 Q9LUJ3 Q9LUJ4
Q9LUJ5 Q9LUJ6 Q9LUK1 Q9LUK2 Q9LUK7 Q9LUL5 Q9LUM4 Q9LUM6 Q9LUN5 Q9LUP1 Q9LUP2
Q9LUP4 Q9LUP5 Q9LUP7 Q9LUP8 Q9LUP9 Q9LUQ3 Q9LUQ5 Q9LUQ6 Q9LUQ9 Q9LUR0 Q9LUR2
Q9LUR3 Q9LUS3 Q9LUS5 Q9LUS6 Q9LUS8 Q9LUS9 Q9LUT1 Q9LUT9 Q9LUU3 Q9LUU4 Q9LUU5
Q9LUU7 Q9LUV4 Q9LUV7 Q9LUW5 Q9LUW6 Q9LUX9 Q9LUZ4 Q9LV01 Q9LV03 Q9LV12 Q9LV17

Q9LV21 Q9LV26 Q9LV32 Q9LV35 Q9LV41 Q9LV45 Q9LV63 Q9LV66 Q9LV75 Q9LV77 Q9LV91
Q9LV93 Q9LVA2 Q9LVB8 Q9LVC3 Q9LVC6 Q9LVC9 Q9LVD3 Q9LVE2 Q9LVF0 Q9LVG1 Q9LVG3
Q9LVG8 Q9LVG9 Q9LVI9 Q9LVJ0 Q9LVJ3 Q9LVJ7 Q9LVL4 Q9LVL9 Q9LVN7 Q9LVP1 Q9LVP7
Q9LVQ4 Q9LVQ5 Q9LVQ8 Q9LVR0 Q9LVR1 Q9LVR6 Q9LVS0 Q9LVS3 Q9LVS9 Q9LVT1 Q9LVT3
Q9LVT4 Q9LVU3 Q9LVU8 Q9LVV5 Q9LVW3 Q9LVW7 Q9LVW9 Q9LVX0 Q9LVX1 Q9LVX6 Q9LVY1
Q9LVZ5 Q9LW00 Q9LW09 Q9LW12 Q9LW29 Q9LW32 Q9LW55 Q9LW60 Q9LW64 Q9LW84 Q9LW88
Q9LWB5 Q9LWU9 Q9LWV3 Q9LX07 Q9LX08 Q9LX16 Q9LX26 Q9LX30 Q9LX33 Q9LX45 Q9LX46
Q9LX47 Q9LX48 Q9LX51 Q9LX54 Q9LX55 Q9LX56 Q9LX74 Q9LX85 Q9LX87 Q9LX88 Q9LX89
Q9LXA7 Q9LXA9 Q9LXB4 Q9LXB5 Q9LXC5 Q9LXD1 Q9LXD7 Q9LXD8 Q9LXE1 Q9LXE3 Q9LXE5
Q9LXE8 Q9LXF2 Q9LXF4 Q9LXG0 Q9LXG5 Q9LXG8 Q9LXG9 Q9LXH5 Q9LXI8 Q9LXJ1 Q9LXJ6
Q9LXJ7 Q9LXK1 Q9LXL5 Q9LXM2 Q9LXM4 Q9LXM8 Q9LXN4 Q9LXN8 Q9LXQ0 Q9LXQ1 Q9LXQ3
Q9LXQ4 Q9LXQ6 Q9LXQ7 Q9LXQ8 Q9LXR0 Q9LXR1 Q9LXR4 Q9LXR6 Q9LXR7 Q9LXS0 Q9LXS6
Q9LXS7 Q9LXT3 Q9LXT4 Q9LXU8 Q9LXU9 Q9LXV0 Q9LXV2 Q9LXV4 Q9LXV5 Q9LXW7 Q9LXX3
Q9LXY5 Q9LXZ3 Q9LXZ8 Q9LY16 Q9LY29 Q9LY43 Q9LY48 Q9LY56 Q9LY61 Q9LY75 Q9LY82
Q9LYA9 Q9LYB9 Q9LYC2 Q9LYC5 Q9LYC6 Q9LYC8 Q9LYE3 Q9LYE4 Q9LYE5 Q9LYJ9 Q9LYK6
Q9LYK7 Q9LYK8 Q9LYK9 Q9LYL3 Q9LYL6 Q9LYL9 Q9LYN2 Q9LYN7 Q9LYP4 Q9LYT2 Q9LYT7
Q9LYU0 Q9LYU8 Q9LYU9 Q9LYV3 Q9LYV6 Q9LYV8 Q9LYW0 Q9LYW3 Q9LYY3 Q9LYY5 Q9LYY6
Q9LYY7 Q9LYZ2 Q9LYZ4 Q9LYZ7 Q9LYZ9 Q9LZ00 Q9LZ03 Q9LZ08 Q9LZ15 Q9LZ25 Q9LZ41
Q9LZ46 Q9LZ56 Q9LZ60 Q9LZ78 Q9LZ86 Q9LZ98 Q9LZA5 Q9LZD7 Q9LZD8 Q9LZG7 Q9LZH9
Q9LZI9 Q9LZK4 Q9LZM0 Q9LZM3 Q9LZP3 Q9LZP6 Q9LZP7 Q9LZP9 Q9LZQ9 Q9LZR0 Q9LZS0
Q9LZS3 Q9LZT0 Q9LZT3 Q9LZU5 Q9LZV6 Q9LZW3 Q9LZW9 Q9LZX5 Q9LZX6 Q9LZX7 Q9LZY0
Q9LZY4 Q9LZZ2 Q9LZZ7 Q9M000 Q9M015 Q9M022 Q9M039 Q9M040 Q9M041 Q9M051 Q9M052
Q9M056 Q9M060 Q9M063 Q9M086 Q9M096 Q9M0A5 Q9M0A6 Q9M0B9 Q9M0C5 Q9M0D0 Q9M0E0
Q9M0E1 Q9M0E6 Q9M0F8 Q9M0G4 Q9M0J5 Q9M0K4 Q9M0M7 Q9M0P3 Q9M0Q9 Q9M0R0 Q9M0S0
Q9M0S5 Q9M0S8 Q9M0T3 Q9M0U1 Q9M0U2 Q9M0U6 Q9M0U7 Q9M0U8 Q9M0U9 Q9M0V0 Q9M0V4
Q9M0V6 Q9M0X8 Q9M0Y5 Q9M0Y6 Q9M0Y9 Q9M0Z5 Q9M117 Q9M122 Q9M128 Q9M156 Q9M157
Q9M160 Q9M161 Q9M190 Q9M1A6 Q9M1A7 Q9M1A8 Q9M1A9 Q9M1B0 Q9M1B9 Q9M1C2 Q9M1D1
Q9M1D3 Q9M1D8 Q9M1E6 Q9M1E8 Q9M1G5 Q9M1G8 Q9M1H3 Q9M1H8 Q9M1I1 Q9M1I7 Q9M1J2
Q9M1K0 Q9M1K1 Q9M1K9 Q9M1N8 Q9M1Q1 Q9M1R1 Q9M1R2 Q9M1R4 Q9M1S3 Q9M1S9 Q9M1T1
Q9M1T7 Q9M1U2 Q9M1U4 Q9M1V1 Q9M1V2 Q9M1V8 Q9M1W4 Q9M1W7 Q9M1X0 Q9M1X2 Q9M1X4
Q9M1X5 Q9M1X9 Q9M1Y0 Q9M1Y1 Q9M1Y3 Q9M1Z3 Q9M202 Q9M205 Q9M209 Q9M219 Q9M223
Q9M237 Q9M268 Q9M274 Q9M276 Q9M280 Q9M290 Q9M2A1 Q9M2B0 Q9M2B4 Q9M2B5 Q9M2C4
Q9M2C8 Q9M2C9 Q9M2D0 Q9M2D9 Q9M2E0 Q9M2E2 Q9M2F1 Q9M2F8 Q9M2F9 Q9M2G8 Q9M2H2
Q9M2H5 Q9M2H6 Q9M2H7 Q9M2H8 Q9M2H9 Q9M2I0 Q9M2I2 Q9M2I9 Q9M2J0 Q9M2J1 Q9M2J2
Q9M2J5 Q9M2J6 Q9M2J7 Q9M2K0 Q9M2K4 Q9M2K8 Q9M2N5 Q9M2P4 Q9M2P8 Q9M2Q4 Q9M2S0
Q9M2S3 Q9M2S7 Q9M2T9 Q9M2U1 Q9M2U4 Q9M2V1 Q9M2V9 Q9M2W2 Q9M2W8 Q9M2X3 Q9M2Y4
Q9M2Y7 Q9M2Y8 Q9M2Y9 Q9M2Z2 Q9M2Z3 Q9M2Z5 Q9M302 Q9M310 Q9M315 Q9M316 Q9M322
Q9M336 Q9M337 Q9M338 Q9M339 Q9M347 Q9M349 Q9M350 Q9M352 Q9M358 Q9M364 Q9M365
Q9M367 Q9M384 Q9M385 Q9M3A3 Q9M3A8 Q9M3B4 Q9M3B6 Q9M3B7 Q9M3C3 Q9M3D2 Q9M3I4
Q9M3K5 Q9M3K7 Q9M3N0 Q9M3V8 Q9M401 Q9M424 Q9M427 Q9M439 Q9M462 Q9M4A1 Q9M4A2
Q9M4B0 Q9M4B5 Q9M4B8 Q9M4B9 Q9M4C0 Q9M4C1 Q9M4C2 Q9M4C3 Q9M4C4 Q9M4D8 Q9M4G4
Q9M4G5 Q9M4P3 Q9M4S8 Q9M4T3 Q9M4T8 Q9M4U4 Q9M4U5 Q9M4W3 Q9M4Y3 Q9M531 Q9M547
Q9M548 Q9M565 Q9M568 Q9M573 Q9M593 Q9M5B3 Q9M5G3 Q9M5I8 Q9M5K2 Q9M5K3 Q9M5L0
Q9M5L6 Q9M5M0 Q9M5M1 Q9M5M6 Q9M5M7 Q9M5P4 Q9M5P5 Q9M5P6 Q9M5W4 Q9M5W9 Q9M5Z9

Q9M612 Q9M630 Q9M667 Q9M682 Q9M6E2 Q9M6F0 Q9M6K1 Q9M7C9 Q9M7D0 Q9M7D1 Q9M7I7
Q9M7J4 Q9M7K8 Q9M7M8 Q9M7M9 Q9M7N0 Q9M7N6 Q9M7P6 Q9M7Q3 Q9M7Q7 Q9M7R0 Q9M7T0
Q9M7X7 Q9M811 Q9M876 Q9M884 Q9M885 Q9M886 Q9M888 Q9M891 Q9M8D3 Q9M8J3 Q9M8J5
Q9M8J9 Q9M8K2 Q9M8K6 Q9M8L2 Q9M8L3 Q9M8L5 Q9M8M3 Q9M8M6 Q9M8M7 Q9M8N2 Q9M8R4
Q9M8R7 Q9M8R9 Q9M8T3 Q9M8T5 Q9M8T6 Q9M8U1 Q9M8U4 Q9M8W9 Q9M8X2 Q9M8X9 Q9M8Y0
Q9M907 Q9M9B3 Q9M9C6 Q9M9E7 Q9M9E8 Q9M9F0 Q9M9F1 Q9M9F5 Q9M9F9 Q9M9G6 Q9M9G7
Q9M9H3 Q9M9H4 Q9M9K1 Q9M9L0 Q9M9L6 Q9M9Q4 Q9M9R4 Q9M9S0 Q9M9T0 Q9M9U9 Q9M9V6
Q9M9W1 Q9M9W3 Q9M9X4 Q9M9X9 Q9M9Y4 Q9M9Y9 Q9MA17 Q9MA32 Q9MA49 Q9MA79 Q9MA92
Q9MA95 Q9MA98 Q9MAA2 Q9MAB3 Q9MAB6 Q9MAC5 Q9MAC8 Q9MAC9 Q9MAG5 Q9MAG8 Q9MAH0
Q9MAH1 Q9MAH3 Q9MAH8 Q9MAK9 Q9MAL0 Q9MAL9 Q9MAN1 Q9MAP1 Q9MAP2 Q9MAP3 Q9MAQ0
Q9MAT0 Q9MAT5 Q9MAT6 Q9MAU3 Q9MAV7 Q9MAY3 Q9MB14 Q9MB33 Q9MB35 Q9MB36 Q9MB37
Q9MB38 Q9MB39 Q9MB40 Q9MB41 Q9MB73 Q9MB94 Q9MB95 Q9MBA3 Q9MBB1 Q9MBB3 Q9MBC1
Q9MBC2 Q9MBF6 Q9MBF8 Q9MBG5 Q9MBH1 Q9MCC1 Q9MCD0 Q9MCD1 Q9MCN8 Q9MDA1 Q9MDK3
Q9MDU0 Q9MDX7 Q9ME90 Q9MEF2 Q9MF82 Q9MS97 Q9MS98 Q9MS99 Q9MSA0 Q9MSA1 Q9MSA7
Q9MSP1 Q9MSQ3 Q9MSR8 Q9MSS0 Q9MSS1 Q9MSS3 Q9MSS8 Q9MST4 Q9MSV2 Q9MSV4 Q9MSV6
Q9MT28 Q9MT47 Q9MTD3 Q9MTD5 Q9MTE5 Q9MTE7 Q9MTI3 Q9MTI5 Q9MTI6 Q9MTI7 Q9MTI8
Q9MTI9 Q9MTJ0 Q9MTJ1 Q9MTJ2 Q9MTJ3 Q9MTJ8 Q9MTK0 Q9MTK1 Q9MTK2 Q9MTM2 Q9MTM3
Q9MTM4 Q9MTM5 Q9MTN6 Q9MTP1 Q9MTP9 Q9MTQ0 Q9MTQ1 Q9MTQ6 Q9MUK9 Q9MUL5 Q9MUL6
Q9MUL7 Q9MUL8 Q9MUM1 Q9MUM2 Q9MUM5 Q9MUM6 Q9MUM7 Q9MUN0 Q9MUN1 Q9MUN2 Q9MUN5
Q9MUN9 Q9MUP0 Q9MUP1 Q9MUP2 Q9MUP3 Q9MUP5 Q9MUP6 Q9MUP7 Q9MUP8 Q9MUQ5 Q9MUQ8
Q9MUR2 Q9MUR3 Q9MUR4 Q9MUR5 Q9MUR6 Q9MUR9 Q9MUS5 Q9MUS6 Q9MUS7 Q9MUS8 Q9MUT3
Q9MUT6 Q9MUT8 Q9MUT9 Q9MUU0 Q9MUU1 Q9MUU2 Q9MUU3 Q9MUU4 Q9MUU5 Q9MUU6 Q9MUU7
Q9MUU8 Q9MUU9 Q9MUV0 Q9MUV1 Q9MUV8 Q9MUW8 Q9MUX3 Q9MUY0 Q9MUY2 Q9MUY3 Q9MUY9
Q9MUZ1 Q9MUZ2 Q9MUZ3 Q9MUZ4 Q9MUZ5 Q9MUZ6 Q9MUZ7 Q9MV08 Q9MV14 Q9MV46 Q9MV47
Q9MV48 Q9MV49 Q9MV51 Q9MV56 Q9MV59 Q9MV60 Q9MVD8 Q9MVD9 Q9MVE5 Q9MVP1 Q9MVV7
Q9MVV8 Q9MVV9 Q9MVW4 Q9MVW5 Q9MYN5 Q9MYY8 Q9MZG3 Q9MZT6 Q9MZT7 Q9MZT8 Q9MZT9
Q9MZU0 Q9N003 Q9N010 Q9N012 Q9N077 Q9N085 Q9N0C5 Q9N0C8 Q9N0E7 Q9N0F1 Q9N0F3
Q9N0J3 Q9N0M2 Q9N0N3 Q9N0T5 Q9N0V7 Q9N0W8 Q9N0X0 Q9N121 Q9N1F5 Q9N1Q0 Q9N1U2
Q9N1U3 Q9N287 Q9N2I8 Q9N2K7 Q9N2L7 Q9N2M8 Q9N2N6 Q9N2P6 Q9N2R3 Q9N2U6 Q9N2V2
Q9N303 Q9N337 Q9N358 Q9N359 Q9N361 Q9N363 Q9N373 Q9N390 Q9N392 Q9N3F0 Q9N3F1
Q9N3P1 Q9N3Q4 Q9N3S2 Q9N3S4 Q9N3T2 Q9N425 Q9N428 Q9N4B8 Q9N4D6 Q9N4D9 Q9N4F2
Q9N4G4 Q9N4G9 Q9N4H4 Q9N4I0 Q9N4J8 Q9N4U5 Q9N4V4 Q9N4Z0 Q9N593 Q9N599 Q9N5A1
Q9N5A8 Q9N5K2 Q9N5K3 Q9N5L4 Q9N5M2 Q9N5M6 Q9N5R9 Q9N5S3 Q9N5U5 Q9N5X6 Q9N655
Q9N658 Q9N9V4 Q9N9W8 Q9N9X3 Q9NA26 Q9NA51 Q9NA55 Q9NA56 Q9NA72 Q9NAA5 Q9NAA7
Q9NAE2 Q9NAH2 Q9NAH4 Q9NAH6 Q9NAL4 Q9NAM7 Q9NAN1 Q9NAP9 Q9NAR7 Q9NAS5 Q9NAX8
Q9NB21 Q9NB32 Q9NB33 Q9NB34 Q9NB51 Q9NBJ7 Q9NBK4 Q9NBL2 Q9NBX4 Q9NBX5 Q9NCL7
Q9NCL8 Q9NCP1 Q9NDA1 Q9NDA2 Q9NDC9 Q9NDJ2 Q9NDS0 Q9NE83 Q9NEL2 Q9NES8 Q9NEU2
Q9NEU3 Q9NEU5 Q9NEW6 Q9NEX2 Q9NF14 Q9NFN6 Q9NFP5 Q9NFS4 Q9NFT7 Q9NFT9 Q9NFZ4
Q9NFZ5 Q9NFZ6 Q9NFZ7 Q9NG40 Q9NG42 Q9NG56 Q9NG98 Q9NGA0 Q9NGB4 Q9NGP4 Q9NGP6
Q9NGW9 Q9NH48 Q9NH49 Q9NH52 Q9NH76 Q9NHC6 Q9NHD5 Q9NHP5 Q9NHP7 Q9NHZ4 Q9NHZ5
Q9NHZ6 Q9NI62 Q9NIF3 Q9NIV2 Q9NJA9 Q9NJB5 Q9NJD8 Q9NJE9 Q9NJG9 Q9NJH0 Q9NJN5
Q9NJV4 Q9NK54 Q9NK57 Q9NKV0 Q9NL40 Q9NL98 Q9NLA3 Q9NLA6 Q9NLC2 Q9NNW7 Q9NNX9
Q9NNZ6 Q9NP08 Q9NP50 Q9NP77 Q9NP81 Q9NP98 Q9NPA8 Q9NPB1 Q9NPB8 Q9NPC3 Q9NPC6
Q9NPC8 Q9NPD8 Q9NPE3 Q9NPI7 Q9NPI8 Q9NPJ1 Q9NPJ4 Q9NPJ8 Q9NQ03 Q9NQ31 Q9NQ32

Q9NQ33 Q9NQ35 Q9NQ39 Q9NQ55 Q9NQ69 Q9NQ86 Q9NQ89 Q9NQC1 Q9NQE9 Q9NQF3 Q9NQI0
Q9NQL9 Q9NQM4 Q9NQR7 Q9NQT5 Q9NQV5 Q9NQV7 Q9NQV8 Q9NQW5 Q9NQW7 Q9NQX0 Q9NQX1
Q9NQX6 Q9NQZ3 Q9NQZ8 Q9NR11 Q9NR21 Q9NR45 Q9NR48 Q9NR50 Q9NR64 Q9NR90 Q9NRD1
Q9NRE2 Q9NRG1 Q9NRG7 Q9NRH1 Q9NRH3 Q9NRI5 Q9NRJ2 Q9NRJ4 Q9NRJ5 Q9NRM2 Q9NRN9
Q9NRP2 Q9NRP4 Q9NRP7 Q9NRV9 Q9NRW3 Q9NRX1 Q9NRX3 Q9NRY2 Q9NS37 Q9NS39 Q9NSA3
Q9NSB2 Q9NSB4 Q9NSD4 Q9NSE4 Q9NSJ1 Q9NTI7 Q9NTJ4 Q9NTM9 Q9NTW7 Q9NTX5 Q9NTX9
Q9NU02 Q9NU23 Q9NU39 Q9NU63 Q9NUA8 Q9NUC0 Q9NUD5 Q9NUD7 Q9NUG4 Q9NUG6 Q9NUJ1
Q9NUK0 Q9NUQ6 Q9NUX5 Q9NV06 Q9NV31 Q9NV35 Q9NV39 Q9NV66 Q9NV72 Q9NV88 Q9NVA2
Q9NVD3 Q9NVE4 Q9NVK5 Q9NVL1 Q9NVL8 Q9NVM4 Q9NVQ4 Q9NVR7 Q9NVS9 Q9NVV2 Q9NVW2
Q9NVX2 Q9NVX7 Q9NW08 Q9NW13 Q9NW61 Q9NW64 Q9NWA0 Q9NWB1 Q9NWB7 Q9NWD9 Q9NWF9
Q9NWG9 Q9NWK9 Q9NWL6 Q9NWM3 Q9NWN3 Q9NWQ9 Q9NWS0 Q9NWS1 Q9NWS8 Q9NWT6 Q9NWU2
Q9NWV4 Q9NWW6 Q9NWW7 Q9NWY4 Q9NX01 Q9NX04 Q9NX07 Q9NX08 Q9NX18 Q9NX31 Q9NX38
Q9NX45 Q9NX65 Q9NX70 Q9NXA8 Q9NXF7 Q9NXG0 Q9NXH3 Q9NXJ5 Q9NXL2 Q9NXP7 Q9NXR5
Q9NXR8 Q9NXS3 Q9NXT0 Q9NXU5 Q9NXW9 Q9NXZ1 Q9NXZ2 Q9NY12 Q9NY30 Q9NY33 Q9NY43
Q9NY74 Q9NYC9 Q9NYG5 Q9NYH9 Q9NYJ1 Q9NYK6 Q9NYP3 Q9NYT6 Q9NYW8 Q9NYY8 Q9NZ32
Q9NZ38 Q9NZ81 Q9NZB8 Q9NZC4 Q9NZD4 Q9NZF1 Q9NZI7 Q9NZM4 Q9NZN9 Q9NZU5 Q9NZV6
Q9NZV7 Q9NZY2 Q9P000 Q9P021 Q9P031 Q9P086 Q9P0J1 Q9P0L1 Q9P0M6 Q9P0N9 Q9P0R6
Q9P0T4 Q9P0V9 Q9P0W2 Q9P0W8 Q9P127 Q9P1A2 Q9P1D8 Q9P1F3 Q9P1G2 Q9P1J3 Q9P1U0
Q9P1U1 Q9P1V8 Q9P1W9 Q9P1Z9 Q9P215 Q9P217 Q9P225 Q9P242 Q9P243 Q9P255 Q9P267
Q9P272 Q9P281 Q9P2D7 Q9P2F5 Q9P2G4 Q9P2G9 Q9P2H0 Q9P2H3 Q9P2J3 Q9P2J5 Q9P2J8
Q9P2J9 Q9P2K6 Q9P2N7 Q9P2Q2 Q9P2R7 Q9P2S5 Q9P2S6 Q9P2T0 Q9P2T1 Q9P2W1 Q9P2W6
Q9P2X3 Q9P2X7 Q9P2X8 Q9P2Z0 Q9P305 Q9P326 Q9P370 Q9P374 Q9P376 Q9P383 Q9P3A8
Q9P3A9 Q9P3B0 Q9P3E7 Q9P3I3 Q9P3K9 Q9P3T4 Q9P3T8 Q9P3U1 Q9P3U2 Q9P3U4 Q9P3U8
Q9P3U9 Q9P3V6 Q9P3W1 Q9P3W4 Q9P3W5 Q9P3Y0 Q9P413 Q9P427 Q9P430 Q9P445 Q9P446
Q9P4B5 Q9P4B6 Q9P4B8 Q9P4B9 Q9P4C1 Q9P4C2 Q9P4D1 Q9P4D3 Q9P4D8 Q9P4E9 Q9P4P9
Q9P4R4 Q9P4R5 Q9P4S5 Q9P4T6 Q9P4T7 Q9P4V2 Q9P4W7 Q9P4X3 Q9P4Z1 Q9P534 Q9P546
Q9P567 Q9P5K9 Q9P5M9 Q9P5N3 Q9P5N7 Q9P5P0 Q9P5T3 Q9P5X8 Q9P5Z8 Q9P6B0 Q9P6C8
Q9P6H9 Q9P6I0 Q9P6I1 Q9P6I2 Q9P6I5 Q9P6I6 Q9P6I7 Q9P6I8 Q9P6I9 Q9P6J1 Q9P6J3
Q9P6J4 Q9P6J6 Q9P6J8 Q9P6K0 Q9P6K3 Q9P6K5 Q9P6K7 Q9P6K8 Q9P6K9 Q9P6L2 Q9P6L6
Q9P6L8 Q9P6L9 Q9P6M0 Q9P6M2 Q9P6M3 Q9P6M8 Q9P6M9 Q9P6N2 Q9P6N7 Q9P6N8 Q9P6P0
Q9P6P2 Q9P6P3 Q9P6P6 Q9P6P7 Q9P6P9 Q9P6Q0 Q9P6Q1 Q9P6Q2 Q9P6Q5 Q9P6Q6 Q9P6R0
Q9P6R1 Q9P6R2 Q9P6R4 Q9P6R7 Q9P6R8 Q9P6R9 Q9P6S3 Q9P6S4 Q9P6S5 Q9P6U9 Q9P6V8
Q9P720 Q9P727 Q9P735 Q9P737 Q9P748 Q9P749 Q9P772 Q9P774 Q9P775 Q9P776 Q9P777
Q9P782 Q9P783 Q9P784 Q9P785 Q9P786 Q9P789 Q9P790 Q9P791 Q9P792 Q9P793 Q9P794
Q9P795 Q9P797 Q9P799 Q9P7A0 Q9P7A1 Q9P7B0 Q9P7B4 Q9P7B5 Q9P7B7 Q9P7B8 Q9P7C0
Q9P7C5 Q9P7C6 Q9P7C7 Q9P7C8 Q9P7C9 Q9P7D1 Q9P7D2 Q9P7D3 Q9P7D4 Q9P7D5 Q9P7D9
Q9P7E3 Q9P7F2 Q9P7F4 Q9P7F5 Q9P7G0 Q9P7G6 Q9P7G9 Q9P7H0 Q9P7H1 Q9P7H2 Q9P7H4
Q9P7H6 Q9P7H8 Q9P7H9 Q9P7I0 Q9P7I2 Q9P7I6 Q9P7I7 Q9P7I8 Q9P7J0 Q9P7J1 Q9P7J2
Q9P7J3 Q9P7J4 Q9P7J7 Q9P7K1 Q9P7K2 Q9P7K3 Q9P7K6 Q9P7K8 Q9P7K9 Q9P7L1 Q9P7L3
Q9P7L4 Q9P7L5 Q9P7L6 Q9P7L8 Q9P7L9 Q9P7M1 Q9P7M3 Q9P7M4 Q9P7M5 Q9P7M7 Q9P7M8
Q9P7M9 Q9P7N0 Q9P7N1 Q9P7N3 Q9P7N4 Q9P7N5 Q9P7N9 Q9P7P1 Q9P7P2 Q9P7P6 Q9P7P7
Q9P7P8 Q9P7P9 Q9P7Q1 Q9P7R0 Q9P7R3 Q9P7R4 Q9P7R5 Q9P7R6 Q9P7R9 Q9P7S2 Q9P7S4
Q9P7S6 Q9P7S7 Q9P7S9 Q9P7T0 Q9P7T1 Q9P7T4 Q9P7T6 Q9P7T7 Q9P7T8 Q9P7U1 Q9P7U5
Q9P7U7 Q9P7V3 Q9P7V4 Q9P7V6 Q9P7W1 Q9P7W2 Q9P7W4 Q9P7W5 Q9P7W6 Q9P7W7 Q9P7W9

Q9P7X1 Q9P7X2 Q9P7X3 Q9P7X5 Q9P7X6 Q9P7X7 Q9P7X8 Q9P7Y2 Q9P7Y4 Q9P7Y7 Q9P7Y8
Q9P7Z3 Q9P7Z4 Q9P7Z6 Q9P7Z7 Q9P7Z8 Q9P7Z9 Q9P801 Q9P802 Q9P805 Q9P836 Q9P840
Q9P843 Q9P844 Q9P885 Q9P889 Q9P8C0 Q9P8F8 Q9P8G9 Q9P8I0 Q9P8N0 Q9P8P7 Q9P8R5
Q9P8V3 Q9P8W3 Q9P8X9 Q9P8Z9 Q9P900 Q9P903 Q9P926 Q9P931 Q9P940 Q9P974 Q9P975
Q9P992 Q9P995 Q9P996 Q9P999 Q9P9H1 Q9P9H8 Q9P9I1 Q9P9I3 Q9P9I7 Q9P9I8 Q9P9I9
Q9P9K9 Q9P9L2 Q9P9L3 Q9P9L4 Q9P9L5 Q9P9M0 Q9P9M3 Q9P9M9 Q9P9N0 Q9P9Q9 Q9P9T1
Q9P9T2 Q9P9T3 Q9P9T5 Q9P9T6 Q9P9U0 Q9P9U3 Q9PA21 Q9PA24 Q9PA28 Q9PA69 Q9PA71
Q9PA72 Q9PA75 Q9PA76 Q9PA77 Q9PA78 Q9PA81 Q9PA82 Q9PA83 Q9PA84 Q9PA85 Q9PA86
Q9PA87 Q9PA89 Q9PA90 Q9PA99 Q9PAA9 Q9PAB4 Q9PAB5 Q9PAC7 Q9PAD9 Q9PAE3 Q9PAE6
Q9PAF5 Q9PAF8 Q9PAF9 Q9PAG2 Q9PAG5 Q9PAH1 Q9PAI2 Q9PAL9 Q9PAM0 Q9PAM5 Q9PAM6
Q9PAN0 Q9PAP2 Q9PAP4 Q9PAQ5 Q9PAQ6 Q9PAQ8 Q9PAR4 Q9PAR6 Q9PAR9 Q9PAS1 Q9PAS2
Q9PAS3 Q9PAS7 Q9PAW2 Q9PAX0 Q9PAX1 Q9PAX3 Q9PAY7 Q9PAZ0 Q9PAZ2 Q9PAZ7 Q9PAZ9
Q9PB02 Q9PB03 Q9PB04 Q9PB05 Q9PB06 Q9PB13 Q9PB19 Q9PB21 Q9PB34 Q9PB43 Q9PB58
Q9PB63 Q9PB72 Q9PB76 Q9PB89 Q9PB95 Q9PB96 Q9PB98 Q9PBB6 Q9PBB8 Q9PBC0 Q9PBC2
Q9PBC4 Q9PBC5 Q9PBC6 Q9PBC7 Q9PBC8 Q9PBC9 Q9PBD0 Q9PBD1 Q9PBD3 Q9PBD4 Q9PBD5
Q9PBE1 Q9PBE3 Q9PBG5 Q9PBG8 Q9PBI4 Q9PBI6 Q9PBJ4 Q9PBJ5 Q9PBJ6 Q9PBJ8 Q9PBN4
Q9PBP4 Q9PBQ0 Q9PBR9 Q9PBX0 Q9PBX5 Q9PBY7 Q9PBZ3 Q9PC00 Q9PC02 Q9PC09 Q9PC10
Q9PC24 Q9PC25 Q9PC26 Q9PC29 Q9PC45 Q9PC50 Q9PC57 Q9PC58 Q9PC61 Q9PC73 Q9PC75
Q9PC76 Q9PC77 Q9PC79 Q9PC88 Q9PC89 Q9PC93 Q9PCC3 Q9PCC5 Q9PCF9 Q9PCG3 Q9PCG5
Q9PCH1 Q9PCH2 Q9PCH8 Q9PCI4 Q9PCQ3 Q9PCR2 Q9PCT9 Q9PCX7 Q9PCZ7 Q9PD25 Q9PD40
Q9PD42 Q9PD43 Q9PD45 Q9PD67 Q9PD69 Q9PD73 Q9PD74 Q9PD76 Q9PD77 Q9PD78 Q9PD79
Q9PD81 Q9PD82 Q9PD85 Q9PD92 Q9PD94 Q9PD96 Q9PD98 Q9PDA6 Q9PDB1 Q9PDB9 Q9PDD3
Q9PDD4 Q9PDD7 Q9PDD9 Q9PDE9 Q9PDF2 Q9PDF4 Q9PDF6 Q9PDG4 Q9PDH3 Q9PDH4 Q9PDI7
Q9PDJ4 Q9PDK3 Q9PDK4 Q9PDK5 Q9PDK6 Q9PDL0 Q9PDL1 Q9PDL7 Q9PDM2 Q9PDM3 Q9PDM9
Q9PDN8 Q9PDP1 Q9PDP2 Q9PDP4 Q9PDP5 Q9PDP7 Q9PDQ7 Q9PDR5 Q9PDT5 Q9PDT6 Q9PDU1
Q9PDV8 Q9PDV9 Q9PDW0 Q9PDW9 Q9PDX2 Q9PE17 Q9PE38 Q9PE40 Q9PE41 Q9PE42 Q9PE51
Q9PE53 Q9PE54 Q9PE55 Q9PE57 Q9PE58 Q9PE59 Q9PE60 Q9PE61 Q9PE62 Q9PE63 Q9PE64
Q9PE65 Q9PE66 Q9PE67 Q9PE68 Q9PE69 Q9PE70 Q9PE71 Q9PE72 Q9PE73 Q9PE74 Q9PE75
Q9PE76 Q9PE98 Q9PEA4 Q9PEB0 Q9PEB6 Q9PEC0 Q9PEC1 Q9PEC2 Q9PEC3 Q9PED1 Q9PEF4
Q9PEG4 Q9PEG5 Q9PEH0 Q9PEH7 Q9PEI0 Q9PEI7 Q9PEJ1 Q9PEL0 Q9PEM3 Q9PEM4 Q9PEM5
Q9PEM6 Q9PEM7 Q9PEM9 Q9PEN0 Q9PEN3 Q9PEP8 Q9PEQ0 Q9PER1 Q9PER5 Q9PES3 Q9PES4
Q9PET0 Q9PET2 Q9PET6 Q9PET8 Q9PET9 Q9PEV0 Q9PEV1 Q9PEV5 Q9PEV8 Q9PEV9 Q9PEX2
Q9PEX5 Q9PF20 Q9PF21 Q9PF26 Q9PF27 Q9PF29 Q9PF39 Q9PF46 Q9PF52 Q9PF55 Q9PF56
Q9PF79 Q9PF80 Q9PF85 Q9PF88 Q9PF89 Q9PF94 Q9PF95 Q9PFB0 Q9PFB1 Q9PFB6 Q9PFB8
Q9PFC3 Q9PFD0 Q9PFD5 Q9PFD6 Q9PFD7 Q9PFD8 Q9PFE1 Q9PFE2 Q9PFJ7 Q9PFJ8 Q9PFN3
Q9PFP2 Q9PFQ9 Q9PFS0 Q9PFS7 Q9PFT8 Q9PFT9 Q9PFU1 Q9PFV1 Q9PFV3 Q9PFV5 Q9PFV8
Q9PG37 Q9PG43 Q9PG44 Q9PG47 Q9PG57 Q9PG67 Q9PG68 Q9PG74 Q9PG89 Q9PGA9 Q9PGB0
Q9PGB5 Q9PGC1 Q9PGC4 Q9PGC5 Q9PGE6 Q9PGG4 Q9PGJ7 Q9PGM3 Q9PGQ9 Q9PGR0 Q9PGR1
Q9PGR2 Q9PGR3 Q9PGR5 Q9PGR6 Q9PGR7 Q9PGR8 Q9PGR9 Q9PGS5 Q9PGT5 Q9PGT6 Q9PGU3
Q9PGU4 Q9PGW5 Q9PGW7 Q9PGX4 Q9PGX7 Q9PGY0 Q9PGZ3 Q9PGZ6 Q9PGZ8 Q9PGZ9 Q9PH02
Q9PH03 Q9PH05 Q9PH08 Q9PH12 Q9PH22 Q9PH23 Q9PH30 Q9PH32 Q9PH36 Q9PH37 Q9PH38
Q9PH39 Q9PH47 Q9PH52 Q9PH56 Q9PH57 Q9PH80 Q9PH84 Q9PH91 Q9PH97 Q9PHB0 Q9PHC7
Q9PHE1 Q9PHE3 Q9PHE7 Q9PHE9 Q9PHK2 Q9PHK6 Q9PHK7 Q9PHK8 Q9PHK9 Q9PHL0 Q9PHL7
Q9PHM5 Q9PHM6 Q9PHM7 Q9PHM8 Q9PHM9 Q9PHN0 Q9PHN2 Q9PHN5 Q9PHP0 Q9PHT5 Q9PHT9

Q9PHU4 Q9PHW3 Q9PHW9 Q9PHX1 Q9PHX2 Q9PHY1 Q9PHY2 Q9PHY8 Q9PHZ3 Q9PHZ7 Q9PHZ9
Q9PI02 Q9PI04 Q9PI05 Q9PI08 Q9PI11 Q9PI16 Q9PI17 Q9PI18 Q9PI30 Q9PI32 Q9PI33
Q9PI34 Q9PI35 Q9PI36 Q9PI38 Q9PI55 Q9PI58 Q9PI71 Q9PI73 Q9PIA5 Q9PIB3 Q9PIB4
Q9PIB6 Q9PIB9 Q9PIC0 Q9PIC1 Q9PIC8 Q9PID2 Q9PIE2 Q9PIF1 Q9PIF2 Q9PIF3 Q9PIG7
Q9PIG9 Q9PIH3 Q9PIH4 Q9PIH5 Q9PIH8 Q9PII2 Q9PII5 Q9PII7 Q9PII8 Q9PIJ1 Q9PIK1
Q9PIK2 Q9PIK3 Q9PIK6 Q9PIK9 Q9PIL5 Q9PIL7 Q9PIM0 Q9PIM5 Q9PIP2 Q9PIP3 Q9PIQ0
Q9PIQ1 Q9PIQ3 Q9PIR0 Q9PIR1 Q9PIR7 Q9PIR8 Q9PIS0 Q9PIS2 Q9PIS3 Q9PIT2 Q9PIT8
Q9PIW2 Q9PIW6 Q9PIX2 Q9PIX9 Q9PIY9 Q9PIZ0 Q9PIZ1 Q9PIZ3 Q9PJ01 Q9PJ06 Q9PJ25
Q9PJ28 Q9PJ29 Q9PJ31 Q9PJ40 Q9PJ53 Q9PJ54 Q9PJ66 Q9PJ84 Q9PJ85 Q9PJ98 Q9PJA1
Q9PJB0 Q9PJB1 Q9PJB6 Q9PJB7 Q9PJB8 Q9PJC4 Q9PJC5 Q9PJC6 Q9PJD2 Q9PJD4 Q9PJF4
Q9PJF5 Q9PJF6 Q9PJF7 Q9PJF8 Q9PJF9 Q9PJG1 Q9PJG5 Q9PJI2 Q9PJI3 Q9PJJ6 Q9PJJ7
Q9PJJ9 Q9PJK0 Q9PJK3 Q9PJK4 Q9PJK7 Q9PJL2 Q9PJL4 Q9PJL5 Q9PJL6 Q9PJL7 Q9PJL9
Q9PJM0 Q9PJM1 Q9PJM2 Q9PJM3 Q9PJM4 Q9PJM5 Q9PJM6 Q9PJM7 Q9PJM8 Q9PJM9 Q9PJN0
Q9PJN2 Q9PJN3 Q9PJN4 Q9PJN5 Q9PJN6 Q9PJN7 Q9PJN9 Q9PJP0 Q9PJP1 Q9PJP3 Q9PJP9
Q9PJQ6 Q9PJR8 Q9PJR9 Q9PJS3 Q9PJT0 Q9PJT1 Q9PJT5 Q9PJT7 Q9PJT8 Q9PJU0 Q9PJU2
Q9PJU6 Q9PJU7 Q9PJV5 Q9PJV6 Q9PJV8 Q9PJW0 Q9PJW1 Q9PJW2 Q9PJW3 Q9PJW4 Q9PJW8
Q9PJX1 Q9PJX5 Q9PJX7 Q9PJZ0 Q9PJZ1 Q9PJZ7 Q9PK00 Q9PK01 Q9PK02 Q9PK04 Q9PK06
Q9PK08 Q9PK09 Q9PK16 Q9PK17 Q9PK18 Q9PK21 Q9PK25 Q9PK26 Q9PK27 Q9PK28 Q9PK30
Q9PK32 Q9PK33 Q9PK40 Q9PK41 Q9PK45 Q9PK47 Q9PK48 Q9PK50 Q9PK53 Q9PK58 Q9PK60
Q9PK61 Q9PK62 Q9PK64 Q9PK65 Q9PK66 Q9PK67 Q9PK73 Q9PK75 Q9PK76 Q9PK77 Q9PK78
Q9PK79 Q9PK80 Q9PK84 Q9PK91 Q9PK92 Q9PK95 Q9PK96 Q9PK97 Q9PKA0 Q9PKA2 Q9PKA6
Q9PKA7 Q9PKA8 Q9PKB2 Q9PKB9 Q9PKC1 Q9PKC2 Q9PKC6 Q9PKE4 Q9PKE6 Q9PKF4 Q9PKH2
Q9PKH4 Q9PKH8 Q9PKI0 Q9PKI2 Q9PKI3 Q9PKI4 Q9PKJ0 Q9PKJ5 Q9PKK0 Q9PKK3 Q9PKK4
Q9PKK5 Q9PKK7 Q9PKK8 Q9PKK9 Q9PKL0 Q9PKM3 Q9PKN7 Q9PKP2 Q9PKP3 Q9PKR0 Q9PKR2
Q9PKR3 Q9PKR6 Q9PKR7 Q9PKS5 Q9PKS8 Q9PKT7 Q9PKU0 Q9PKU1 Q9PKU2 Q9PKU9 Q9PKV0
Q9PKV2 Q9PKV8 Q9PKW1 Q9PKW3 Q9PKW4 Q9PKW5 Q9PKW8 Q9PKX7 Q9PKX8 Q9PKY3 Q9PKZ4
Q9PKZ7 Q9PKZ8 Q9PKZ9 Q9PL03 Q9PL07 Q9PL08 Q9PL09 Q9PL10 Q9PL11 Q9PL12 Q9PL13
Q9PL16 Q9PL17 Q9PL20 Q9PL32 Q9PL33 Q9PL36 Q9PL37 Q9PL38 Q9PL40 Q9PL51 Q9PL53
Q9PL59 Q9PL60 Q9PL64 Q9PL68 Q9PL77 Q9PL79 Q9PL83 Q9PL84 Q9PL86 Q9PL88 Q9PL89
Q9PL91 Q9PL92 Q9PL93 Q9PL95 Q9PL99 Q9PLA4 Q9PLA5 Q9PLB2 Q9PLC0 Q9PLC1 Q9PLC2
Q9PLC3 Q9PLC6 Q9PLC9 Q9PLD0 Q9PLD1 Q9PLD4 Q9PLD6 Q9PLD7 Q9PLE0 Q9PLE1 Q9PLE5
Q9PLF1 Q9PLF7 Q9PLG1 Q9PLG5 Q9PLH1 Q9PLH5 Q9PLH7 Q9PLI1 Q9PLJ0 Q9PLJ4 Q9PLJ5
Q9PLJ6 Q9PLJ7 Q9PLK4 Q9PLL6 Q9PLL9 Q9PLM0 Q9PLM1 Q9PLM3 Q9PLM9 Q9PLN1 Q9PLN4
Q9PLN6 Q9PLN9 Q9PLP0 Q9PLP9 Q9PLQ3 Q9PLQ8 Q9PLQ9 Q9PLR1 Q9PLR2 Q9PLR5 Q9PLS6
Q9PLT8 Q9PLU1 Q9PLU2 Q9PLU3 Q9PLU4 Q9PLU8 Q9PLV2 Q9PLV4 Q9PLV9 Q9PLW0 Q9PLW1
Q9PLW2 Q9PLW7 Q9PLX0 Q9PLX1 Q9PLX2 Q9PLX4 Q9PLX5 Q9PLX6 Q9PLX7 Q9PLX8 Q9PLX9
Q9PLY0 Q9PLY2 Q9PLY3 Q9PLY4 Q9PLY5 Q9PLY7 Q9PLY8 Q9PLZ2 Q9PM01 Q9PM24 Q9PM34
Q9PM37 Q9PM39 Q9PM40 Q9PM41 Q9PM63 Q9PM64 Q9PM68 Q9PM71 Q9PM72 Q9PM74 Q9PM75
Q9PM76 Q9PM77 Q9PM78 Q9PM80 Q9PM81 Q9PM82 Q9PM83 Q9PM84 Q9PM85 Q9PMC8 Q9PMD2
Q9PMD4 Q9PMD8 Q9PMD9 Q9PMF9 Q9PMG4 Q9PMG8 Q9PMI3 Q9PMJ7 Q9PMK3 Q9PMK5 Q9PMK7
Q9PML1 Q9PMN3 Q9PMQ3 Q9PMQ5 Q9PMQ6 Q9PMR0 Q9PMS2 Q9PMS6 Q9PMT4 Q9PMT7 Q9PMU0
Q9PMU9 Q9PMV3 Q9PN07 Q9PN13 Q9PN20 Q9PN24 Q9PN25 Q9PN27 Q9PN30 Q9PN35 Q9PN47
Q9PN49 Q9PN51 Q9PN54 Q9PN58 Q9PN59 Q9PN78 Q9PN90 Q9PN94 Q9PN98 Q9PNA7 Q9PNB3
Q9PNB4 Q9PNB8 Q9PNC0 Q9PNC3 Q9PNH7 Q9PNI5 Q9PNI7 Q9PNI9 Q9PNJ0 Q9PNJ2 Q9PNJ6

Q9PNK3 Q9PNL3 Q9PNN2 Q9PNN7 Q9PNP6 Q9PNQ6 Q9PNT0 Q9PNT2 Q9PNU3 Q9PNU6 Q9PNX4
Q9PNX5 Q9PNX9 Q9PNY0 Q9PNY2 Q9PP01 Q9PP04 Q9PP06 Q9PP15 Q9PP26 Q9PP34 Q9PP35
Q9PP65 Q9PP70 Q9PP77 Q9PP78 Q9PP85 Q9PP92 Q9PP96 Q9PP99 Q9PPB0 Q9PPB4 Q9PPB8
Q9PPC1 Q9PPC2 Q9PPE0 Q9PPE4 Q9PPF1 Q9PPF2 Q9PPF3 Q9PPF4 Q9PPG2 Q9PPI9 Q9PPJ3
Q9PPJ4 Q9PPJ5 Q9PPJ7 Q9PPK2 Q9PPK3 Q9PPL4 Q9PPL8 Q9PPM1 Q9PPM7 Q9PPN5 Q9PPN6
Q9PPN8 Q9PPN9 Q9PPP0 Q9PPP2 Q9PPP3 Q9PPP5 Q9PPQ4 Q9PPR2 Q9PPR3 Q9PPS1 Q9PPS2
Q9PPS3 Q9PPT6 Q9PPT7 Q9PPT8 Q9PPT9 Q9PPU2 Q9PPU6 Q9PPU9 Q9PPV0 Q9PPV4 Q9PPV6
Q9PPV7 Q9PPW5 Q9PPW6 Q9PPW7 Q9PPX5 Q9PPX6 Q9PPX7 Q9PPZ2 Q9PPZ6 Q9PPZ7 Q9PQ00
Q9PQ02 Q9PQ21 Q9PQ25 Q9PQ27 Q9PQ29 Q9PQ32 Q9PQ33 Q9PQ36 Q9PQ38 Q9PQ41 Q9PQ42
Q9PQ53 Q9PQ61 Q9PQ68 Q9PQ71 Q9PQ73 Q9PQ74 Q9PQ75 Q9PQ76 Q9PQ78 Q9PQ79 Q9PQ82
Q9PQ83 Q9PQ88 Q9PQ93 Q9PQA4 Q9PQA5 Q9PQA7 Q9PQA9 Q9PQB4 Q9PQB5 Q9PQC0 Q9PQC1
Q9PQC2 Q9PQC6 Q9PQC8 Q9PQD7 Q9PQE0 Q9PQE4 Q9PQE5 Q9PQE6 Q9PQE9 Q9PQF2 Q9PQF8
Q9PQF9 Q9PQH0 Q9PQH1 Q9PQH6 Q9PQI1 Q9PQI5 Q9PQI7 Q9PQJ3 Q9PQJ4 Q9PQJ6 Q9PQK2
Q9PQK5 Q9PQK6 Q9PQK7 Q9PQK9 Q9PQL2 Q9PQL3 Q9PQL4 Q9PQM2 Q9PQM4 Q9PQM5 Q9PQN3
Q9PQN4 Q9PQN5 Q9PQN6 Q9PQN7 Q9PQN8 Q9PQP2 Q9PQP3 Q9PQP4 Q9PQP6 Q9PQP7 Q9PQP8
Q9PQP9 Q9PQQ0 Q9PQQ1 Q9PQQ2 Q9PQQ3 Q9PQQ4 Q9PQQ5 Q9PQQ6 Q9PQQ7 Q9PQQ8 Q9PQQ9
Q9PQR0 Q9PQR1 Q9PQR2 Q9PQR3 Q9PQR4 Q9PQS0 Q9PQS1 Q9PQS5 Q9PQS9 Q9PQT0 Q9PQT2
Q9PQT3 Q9PQT8 Q9PQT9 Q9PQU6 Q9PQV0 Q9PQV5 Q9PQV6 Q9PQV8 Q9PQW1 Q9PQW2 Q9PQW3
Q9PQW4 Q9PQW6 Q9PQW7 Q9PQW8 Q9PQX0 Q9PQX2 Q9PQX3 Q9PQX5 Q9PQX6 Q9PQX7 Q9PQX9
Q9PQY0 Q9PQY2 Q9PQY3 Q9PQY7 Q9PQY8 Q9PQY9 Q9PQZ0 Q9PQZ1 Q9PQZ2 Q9PQZ3 Q9PQZ4
Q9PQZ5 Q9PQZ6 Q9PQZ7 Q9PQZ8 Q9PR00 Q9PR05 Q9PR21 Q9PR23 Q9PR24 Q9PR27 Q9PR28
Q9PR30 Q9PR32 Q9PR33 Q9PR38 Q9PR42 Q9PR43 Q9PR55 Q9PR56 Q9PR57 Q9PR61 Q9PR63
Q9PR67 Q9PR68 Q9PR69 Q9PR71 Q9PR82 Q9PR83 Q9PR87 Q9PR88 Q9PR94 Q9PR95 Q9PR96
Q9PRA5 Q9PRA6 Q9PRB0 Q9PRB1 Q9PRB3 Q9PRB9 Q9PRC5 Q9PRC7 Q9PRC8 Q9PRD4 Q9PRD5
Q9PRE0 Q9PRE1 Q9PRE2 Q9PRI5 Q9PRJ8 Q9PRL8 Q9PRL9 Q9PS27 Q9PSI9 Q9PSJ0 Q9PSV3
Q9PSW9 Q9PSY4 Q9PT26 Q9PT42 Q9PT43 Q9PT61 Q9PT63 Q9PT67 Q9PT68 Q9PT92 Q9PTG8
Q9PTG9 Q9PTJ3 Q9PTJ6 Q9PTK2 Q9PTN2 Q9PTP1 Q9PTQ7 Q9PTS2 Q9PTU0 Q9PTU1 Q9PTU3
Q9PTU5 Q9PTW9 Q9PTX2 Q9PU13 Q9PU20 Q9PU47 Q9PU58 Q9PU65 Q9PU85 Q9PUA8 Q9PUE0
Q9PUK5 Q9PUL2 Q9PUQ1 Q9PV94 Q9PVE2 Q9PVE4 Q9PVG3 Q9PVK2 Q9PVK4 Q9PVK5 Q9PVM0
Q9PVM1 Q9PVM2 Q9PVM3 Q9PVM4 Q9PVN2 Q9PVQ1 Q9PVQ9 Q9PVR2 Q9PVS4 Q9PVU6 Q9PVV4
Q9PVY6 Q9PVY8 Q9PVY9 Q9PVZ3 Q9PVZ9 Q9PW04 Q9PW05 Q9PW06 Q9PW07 Q9PW28 Q9PW58
Q9PW61 Q9PW73 Q9PW80 Q9PWD2 Q9PWD3 Q9PWD4 Q9PWD5 Q9PWD6 Q9PWE8 Q9PWL5 Q9PWL6
Q9PWL7 Q9PWM2 Q9PWM3 Q9PWM5 Q9PWM6 Q9PWX1 Q9PX43 Q9PX53 Q9PX62 Q9PX69 Q9PX71
Q9PY82 Q9PY83 Q9PY93 Q9PY94 Q9PY95 Q9Q1T6 Q9Q1T8 Q9Q2P0 Q9Q2Q3 Q9Q3G5 Q9Q6Q5
Q9Q6X7 Q9Q8J2 Q9Q8K4 Q9Q8L4 Q9Q8L9 Q9Q8N4 Q9Q8N5 Q9Q8N6 Q9Q8P6 Q9Q8Q1 Q9Q8Q2
Q9Q8R9 Q9Q8X0 Q9Q8Y2 Q9Q8Z2 Q9Q907 Q9Q908 Q9Q909 Q9Q921 Q9Q926 Q9Q927 Q9Q944
Q9QAB8 Q9QAQ4 Q9QAQ5 Q9QAQ7 Q9QAR0 Q9QAR4 Q9QAR7 Q9QAS0 Q9QAS1 Q9QAS3 Q9QAW8
Q9QAZ7 Q9QAZ8 Q9QB84 Q9QB92 Q9QB93 Q9QB97 Q9QBB4 Q9QBF1 Q9QBG0 Q9QBU0 Q9QBU1
Q9QBU2 Q9QBU3 Q9QC07 Q9QCE9 Q9QDI8 Q9QEE3 Q9QEE6 Q9QEE7 Q9QEJ4 Q9QEJ5 Q9QJ12
Q9QJ13 Q9QJ16 Q9QJ18 Q9QJ19 Q9QJ20 Q9QJ21 Q9QJ22 Q9QJ23 Q9QJ24 Q9QJ25 Q9QJ26
Q9QJ27 Q9QJ28 Q9QJ32 Q9QJ33 Q9QJ34 Q9QJ38 Q9QJ39 Q9QJ40 Q9QJ47 Q9QJ49 Q9QJ50
Q9QJ52 Q9QJ53 Q9QJ55 Q9QJ56 Q9QJ58 Q9QJT4 Q9QJT5 Q9QJT8 Q9QJT9 Q9QL89 Q9QM72
Q9QM81 Q9QMI1 Q9QNA5 Q9QNA8 Q9QNA9 Q9QNB1 Q9QNB2 Q9QNB3 Q9QNF7 Q9QP77 Q9QPN4
Q9QSK1 Q9QSK2 Q9QSK3 Q9QSK4 Q9QSK5 Q9QSP0 Q9QSP4 Q9QTF2 Q9QU29 Q9QU30 Q9QU31

Q9QUB8 Q9QUG2 Q9QUK0 Q9QUK4 Q9QUK5 Q9QUM7 Q9QUM9 Q9QUR2 Q9QUR6 Q9QUR7 Q9QUS1

Q9QUS6 Q9QUZ8 Q9QVC8 Q9QVN7 Q9QVP4 Q9QVY8 Q9QW08 Q9QWF0 Q9QWG7 Q9QWH1 Q9QWI6

Q9QWL7 Q9QWT9 Q9QX22 Q9QX27 Q9QX47 Q9QX60 Q9QX66 Q9QX67 Q9QX96 Q9QX98 Q9QXA7

Q9QXE2 Q9QXE4 Q9QXF8 Q9QXJ0 Q9QXK2 Q9QXL1 Q9QXL7 Q9QXL8 Q9QXN1 Q9QXN4 Q9QXN5

Q9QXP3 Q9QXP4 Q9QXP6 Q9QXT9 Q9QXV1 Q9QXV3 Q9QXW0 Q9QXY2 Q9QXZ7 Q9QY02 Q9QY14

Q9QY24 Q9QY31 Q9QY36 Q9QY61 Q9QY66 Q9QY83 Q9QY84 Q9QY93 Q9QYB2 Q9QYE3 Q9QYI3

Q9QYI7 Q9QYI8 Q9QYJ3 Q9QYJ7 Q9QYK2 Q9QYK9 Q9QYL0 Q9QYL7 Q9QYP6 Q9QYR2 Q9QYR6

Q9QYU2 Q9QYU4 Q9QYV8 Q9QYW0 Q9QZ04 Q9QZ05 Q9QZ08 Q9QZ10 Q9QZ23 Q9QZ28 Q9QZ41

Q9QZ47 Q9QZ48 Q9QZ57 Q9QZ58 Q9QZ73 Q9QZ76 Q9QZ86 Q9QZA0 Q9QZA1 Q9QZB7 Q9QZD9

Q9QZE7 Q9QZH2 Q9QZH3 Q9QZH7 Q9QZI7 Q9QZM2 Q9QZM5 Q9QZM8 Q9QZM9 Q9QZN0 Q9QZN4

Q9QZP1 Q9QZP2 Q9QZQ0 Q9QZR0 Q9QZR6 Q9QZS2 Q9QZU5 Q9QZU7 Q9QZU9 Q9QZV9 Q9QZW9

Q9QZZ4 Q9R000 Q9R002 Q9R011 Q9R016 Q9R020 Q9R022 Q9R050 Q9R059 Q9R061 Q9R062

Q9R095 Q9R099 Q9R0A3 Q9R0A5 Q9R0A9 Q9R0B7 Q9R0D8 Q9R0G7 Q9R0H5 Q9R0I6 Q9R0I7

Q9R0L4 Q9R0M5 Q9R0N0 Q9R0P3 Q9R0P4 Q9R0U0 Q9R0X0 Q9R0X4 Q9R0Z7 Q9R111 Q9R123

Q9R144 Q9R1A0 Q9R1A9 Q9R1C0 Q9R1C7 Q9R1D1 Q9R1K7 Q9R1M3 Q9R1P1 Q9R1P3 Q9R1P4

Q9R1Q2 Q9R1Q8 Q9R1R0 Q9R1R2 Q9R1R4 Q9R1S8 Q9R1T2 Q9R1T4 Q9R1T5 Q9R1X4 Q9R1Z7

Q9R257 Q9R2E9 Q9R342 Q9R381 Q9R3D1 Q9R4A1 Q9R4E4 Q9R4G7 Q9R4G8 Q9R4J4 Q9R4N6

Q9R4N7 Q9R4N8 Q9R4N9 Q9R4P0 Q9R4P1 Q9R4P2 Q9R4P3 Q9R4P4 Q9R4P5 Q9R4P6 Q9R4P7

Q9R4P8 Q9R5C6 Q9R5K5 Q9R5V5 Q9R5V7 Q9R5V8 Q9R5V9 Q9R6P8 Q9R6P9 Q9R6S7 Q9R6T3

Q9R6X4 Q9R708 Q9R7D5 Q9R7D7 Q9R9H8 Q9R9I9 Q9R9J0 Q9R9J1 Q9R9M8 Q9R9N4 Q9R9T9

Q9R9U0 Q9R9U2 Q9RA05 Q9RA11 Q9RA13 Q9RA36 Q9RA51 Q9RA54 Q9RA57 Q9RA63 Q9RA96

Q9RAA9 Q9RAE4 Q9RAE6 Q9RAE7 Q9RAH6 Q9RAJ1 Q9RAJ5 Q9RAM6 Q9RAM7 Q9RAN9 Q9RB09

Q9RB36 Q9RBG4 Q9RBG5 Q9RBJ0 Q9RBJ3 Q9RBN6 Q9RBP4 Q9RBP5 Q9RBU1 Q9RBW7 Q9RBY6

Q9RC23 Q9RC40 Q9RC41 Q9RC57 Q9RC80 Q9RC82 Q9RC88 Q9RC92 Q9RC95 Q9RC99 Q9RCA1

Q9RCA2 Q9RCA3 Q9RCA4 Q9RCA6 Q9RCA7 Q9RCA8 Q9RCE7 Q9RCE8 Q9RCG0 Q9RCI2 Q9RCT1

Q9RD25 Q9RD27 Q9RDA3 Q9RDD6 Q9RDD7 Q9RDE2 Q9RDE4 Q9RDE5 Q9RDE8 Q9RDE9 Q9RDH8

Q9RDJ9 Q9RDK1 Q9RDK7 Q9RDL5 Q9RDM3 Q9RDS6 Q9RDT9 Q9RDU0 Q9RDV2 Q9RDV3 Q9RDV4

Q9RDV6 Q9RDV7 Q9RDV8 Q9RDV9 Q9RDW6 Q9RDX2 Q9RDX3 Q9RDY2 Q9RE01 Q9RE09 Q9RED7

Q9RED9 Q9REF2 Q9REF4 Q9REJ6 Q9REM0 Q9REQ0 Q9REQ2 Q9REQ3 Q9REQ6 Q9REQ9 Q9RER4

Q9REU4 Q9REV5 Q9REV6 Q9RF06 Q9RF07 Q9RF52 Q9RFD3 Q9RFD4 Q9RFD5 Q9RFD6 Q9RFF5

Q9RFJ6 Q9RFM4 Q9RFN0 Q9RFQ2 Q9RFR0 Q9RFS6 Q9RFW1 Q9RGC3 Q9RGM3 Q9RGQ1 Q9RGR0

Q9RGR9 Q9RGS1 Q9RGS5 Q9RGS6 Q9RGW9 Q9RGX0 Q9RGX2 Q9RGY8 Q9RH04 Q9RH05 Q9RH11

Q9RH29 Q9RH40 Q9RH41 Q9RH43 Q9RH67 Q9RH70 Q9RH74 Q9RH76 Q9RHF4 Q9RHG2 Q9RHG4

Q9RHM3 Q9RHM6 Q9RHM8 Q9RHN1 Q9RHN2 Q9RHS9 Q9RHV9 Q9RHW9 Q9RHX0 Q9RHX2 Q9RHX4

Q9RHX5 Q9RHX6 Q9RI00 Q9RI02 Q9RI18 Q9RJ16 Q9RJ20 Q9RJ47 Q9RJ50 Q9RJ58 Q9RJ61

Q9RJ79 Q9RJ87 Q9RJD5 Q9RJH9 Q9RJM2 Q9RJM5 Q9RJN1 Q9RJS6 Q9RJZ6 Q9RK00 Q9RKA8

Q9RKB5 Q9RKB9 Q9RKK5 Q9RKK6 Q9RKL1 Q9RKL4 Q9RKS2 Q9RKU4 Q9RKU5 Q9RKX1 Q9RKY0

Q9RL17 Q9RL35 Q9RL45 Q9RL48 Q9RL50 Q9RL68 Q9RL77 Q9RL78 Q9RL97 Q9RL99 Q9RLA0

Q9RLB6 Q9RLJ3 Q9RLM3 Q9RLM4 Q9RLM5 Q9RLS9 Q9RLT3 Q9RLT6 Q9RLT9 Q9RLU5 Q9RM66

Q9RMC1 Q9RME2 Q9RME4 Q9RME6 Q9RMG3 Q9RML6 Q9RMM3 Q9RMP7 Q9RMV2 Q9RMV3 Q9RMV4

Q9RMV5 Q9RMV7 Q9RMW0 Q9RMW2 Q9RMW6 Q9RMW7 Q9RMX1 Q9RMX2 Q9RMX3 Q9RMX9 Q9RMY0

Q9RMY4 Q9RMY7 Q9RMY8 Q9RMZ2 Q9RMZ4 Q9RMZ8 Q9RMZ9 Q9RN01 Q9RN04 Q9RN13 Q9RN17

Q9RN22 Q9RN25 Q9RN26 Q9RN28 Q9RN59 Q9RN77 Q9RND6 Q9RND7 Q9RNH6 Q9RNH7 Q9RNL6

Q9RNM2 Q9RNM7 Q9RNM8 Q9RNM9 Q9RNN5 Q9RNN8 Q9RNP3 Q9RNU9 Q9RNY0 Q9RNY7 Q9RNZ1

Q9RNZ5 Q9RNZ9 Q9RP19 Q9RP48 Q9RPD4 Q9RPK1 Q9RPQ3 Q9RPQ4 Q9RPQ5 Q9RPS7 Q9RPT0
Q9RPU2 Q9RPU5 Q9RPV0 Q9RPW2 Q9RPW3 Q9RPW4 Q9RPW5 Q9RPW6 Q9RPW7 Q9RPW8 Q9RQ01
Q9RQ04 Q9RQ13 Q9RQ15 Q9RQ16 Q9RQ17 Q9RQ33 Q9RQ35 Q9RQ47 Q9RQ48 Q9RQ51 Q9RQ52
Q9RQ55 Q9RQ56 Q9RQ65 Q9RQ82 Q9RQ83 Q9RQ85 Q9RQ86 Q9RQ88 Q9RQ89 Q9RQF8 Q9RQI5
Q9RQI6 Q9RQJ6 Q9RQJ9 Q9RQL3 Q9RQQ0 Q9RQV9 Q9RQZ7 Q9RR27 Q9RR28 Q9RR29 Q9RR30
Q9RR31 Q9RR32 Q9RR63 Q9RR70 Q9RR81 Q9RR89 Q9RR90 Q9RR91 Q9RR92 Q9RR97 Q9RRA9
Q9RRB5 Q9RRB7 Q9RRC1 Q9RRC4 Q9RRD7 Q9RRE4 Q9RRE5 Q9RRE9 Q9RRF8 Q9RRG8 Q9RRI2
Q9RRJ1 Q9RRJ4 Q9RRJ5 Q9RRJ6 Q9RRK5 Q9RRL7 Q9RRM7 Q9RRP1 Q9RRP6 Q9RRQ3 Q9RRQ4
Q9RRR1 Q9RRS9 Q9RRT0 Q9RRT8 Q9RRU5 Q9RRW3 Q9RRX5 Q9RRX8 Q9RRX9 Q9RRY0 Q9RS06
Q9RS16 Q9RS19 Q9RS27 Q9RS37 Q9RS38 Q9RS39 Q9RS45 Q9RS64 Q9RSA0 Q9RSC5 Q9RSE6
Q9RSF0 Q9RSG0 Q9RSH2 Q9RSH3 Q9RSI4 Q9RSI9 Q9RSJ5 Q9RSJ6 Q9RSJ7 Q9RSJ8 Q9RSJ9
Q9RSK0 Q9RSK1 Q9RSK7 Q9RSK9 Q9RSL0 Q9RSL1 Q9RSL2 Q9RSL3 Q9RSL4 Q9RSL5 Q9RSN2
Q9RSN7 Q9RSP3 Q9RSQ0 Q9RSQ3 Q9RSQ5 Q9RSQ7 Q9RSR1 Q9RSR5 Q9RSS1 Q9RSS4 Q9RSS6
Q9RSS7 Q9RSS8 Q9RSS9 Q9RST0 Q9RSW0 Q9RSW1 Q9RSW6 Q9RSW7 Q9RSX8 Q9RSY2 Q9RSY5
Q9RSZ6 Q9RSZ7 Q9RT03 Q9RT21 Q9RT38 Q9RT44 Q9RT57 Q9RT63 Q9RT73 Q9RT91 Q9RT99
Q9RTA0 Q9RTA9 Q9RTB6 Q9RTB7 Q9RTD8 Q9RTD9 Q9RTE6 Q9RTF3 Q9RTF5 Q9RTF6 Q9RTG5
Q9RTG8 Q9RTH9 Q9RTI0 Q9RTI6 Q9RTJ5 Q9RTK1 Q9RTK2 Q9RTK7 Q9RTL8 Q9RTN2 Q9RTN7
Q9RTP4 Q9RTQ2 Q9RTR0 Q9RTR6 Q9RTR7 Q9RTR9 Q9RTS6 Q9RTT6 Q9RTU4 Q9RTY2 Q9RTY5
Q9RTY9 Q9RU10 Q9RU11 Q9RU19 Q9RU23 Q9RU32 Q9RU68 Q9RU72 Q9RU74 Q9RU79 Q9RU80
Q9RU81 Q9RU82 Q9RU84 Q9RUA9 Q9RUB5 Q9RUC1 Q9RUD2 Q9RUE2 Q9RUE3 Q9RUF1 Q9RUF2
Q9RUF3 Q9RUF5 Q9RUG0 Q9RUG2 Q9RUG6 Q9RUH3 Q9RUI6 Q9RUK6 Q9RUL6 Q9RUL7 Q9RUL8
Q9RUN0 Q9RUN5 Q9RUN7 Q9RUP2 Q9RUP5 Q9RUP6 Q9RUP8 Q9RUQ2 Q9RUQ9 Q9RUS2 Q9RUS3
Q9RUT7 Q9RUV0 Q9RUV2 Q9RUV5 Q9RUV6 Q9RUV7 Q9RUW4 Q9RUW8 Q9RUY3 Q9RUZ0 Q9RV23
Q9RV24 Q9RV25 Q9RV39 Q9RV46 Q9RV58 Q9RV62 Q9RV66 Q9RV69 Q9RV76 Q9RV79 Q9RV96
Q9RV97 Q9RV98 Q9RVA6 Q9RVA9 Q9RVB9 Q9RVC0 Q9RVC3 Q9RVC4 Q9RVC9 Q9RVD3 Q9RVD6
Q9RVE0 Q9RVE2 Q9RVE3 Q9RVE7 Q9RVF9 Q9RVH4 Q9RVI3 Q9RVK2 Q9RVK3 Q9RVL1 Q9RVM1
Q9RVM9 Q9RVP2 Q9RVP6 Q9RVQ9 Q9RVT0 Q9RVT1 Q9RVT3 Q9RVT6 Q9RVU3 Q9RVV9 Q9RVW0
Q9RVX0 Q9RVY1 Q9RVY3 Q9RVZ8 Q9RW01 Q9RW02 Q9RW10 Q9RW24 Q9RW28 Q9RW44 Q9RW45
Q9RW55 Q9RW68 Q9RW75 Q9RW91 Q9RW93 Q9RW94 Q9RWA4 Q9RWA5 Q9RWB2 Q9RWB4 Q9RWC7
Q9RWD6 Q9RWD7 Q9RWF1 Q9RWG3 Q9RWH9 Q9RWI4 Q9RWJ0 Q9RWJ4 Q9RWK0 Q9RWM1 Q9RWM4
Q9RWM6 Q9RWN1 Q9RWN8 Q9RWN9 Q9RWP7 Q9RWQ9 Q9RWR0 Q9RWR5 Q9RWS1 Q9RWS9 Q9RWV7
Q9RWW0 Q9RX08 Q9RX20 Q9RX22 Q9RX25 Q9RX30 Q9RX32 Q9RX35 Q9RX45 Q9RX51 Q9RX54
Q9RX68 Q9RX75 Q9RX88 Q9RX89 Q9RX91 Q9RX92 Q9RX93 Q9RXC1 Q9RXC9 Q9RXE1 Q9RXF1
Q9RXG4 Q9RXH2 Q9RXI7 Q9RXI8 Q9RXJ0 Q9RXJ1 Q9RXJ2 Q9RXJ3 Q9RXJ4 Q9RXJ5 Q9RXJ6
Q9RXJ7 Q9RXJ8 Q9RXJ9 Q9RXK0 Q9RXK1 Q9RXK2 Q9RXK3 Q9RXK5 Q9RXK6 Q9RXK7 Q9RXL3
Q9RXP0 Q9RXP1 Q9RXR2 Q9RXR4 Q9RXS6 Q9RXS8 Q9RXT0 Q9RXT3 Q9RXT4 Q9RXT9 Q9RXU7
Q9RXV7 Q9RXW9 Q9RXX6 Q9RXY0 Q9RXY1 Q9RXY7 Q9RXZ5 Q9RXZ8 Q9RY06 Q9RY23 Q9RY24
Q9RY28 Q9RY32 Q9RY40 Q9RY41 Q9RY49 Q9RY50 Q9RY51 Q9RY52 Q9RY63 Q9RY64 Q9RY65
Q9RY66 Q9RY70 Q9RY71 Q9RY72 Q9RY77 Q9RY80 Q9RYA1 Q9RYA6 Q9RYB2 Q9RYB4 Q9RYB5
Q9RYC3 Q9RYD4 Q9RYD6 Q9RYD8 Q9RYE3 Q9RYE6 Q9RYG9 Q9RYH5 Q9RYJ3 Q9RYJ4 Q9RYJ6
Q9RYJ8 Q9RYK0 Q9RYK1 Q9RYL9 Q9RYM7 Q9RYM9 Q9RYP0 Q9RYR5 Q9RYR8 Q9RYU4 Q9RYU5
Q9RYV5 Q9RYW7 Q9RYX0 Q9RYX4 Q9RYX8 Q9RYX9 Q9RYY1 Q9RYY6 Q9RYZ2 Q9RZ02 Q9RZ05
Q9RZ06 Q9RZ89 Q9RZE7 Q9RZL8 Q9RZN1 Q9RZS4 Q9RZU9 Q9RZV8 Q9S0K8 Q9S0K9 Q9S0L2
Q9S0Q7 Q9S0Q8 Q9S0Q9 Q9S0R0 Q9S0R1 Q9S0X3 Q9S0X4 Q9S152 Q9S153 Q9S156 Q9S157

Q9S158 Q9S170 Q9S1B7 Q9S1F9 Q9S1G2 Q9S1G8 Q9S1H5 Q9S1M6 Q9S1N4 Q9S1U6 Q9S208
Q9S219 Q9S220 Q9S224 Q9S266 Q9S281 Q9S2B5 Q9S2C0 Q9S2F4 Q9S2G5 Q9S2H7 Q9S2H9
Q9S2K7 Q9S2L4 Q9S2N0 Q9S2N2 Q9S2N4 Q9S2P2 Q9S2Q7 Q9S2R1 Q9S2T7 Q9S2U1 Q9S2V2
Q9S2W4 Q9S2W7 Q9S2W9 Q9S2X2 Q9S2X5 Q9S306 Q9S358 Q9S375 Q9S3K3 Q9S3M0 Q9S3Q0
Q9S3Q5 Q9S3S0 Q9S3S1 Q9S3U4 Q9S3U6 Q9S3U8 Q9S3U9 Q9S3V0 Q9S3V1 Q9S3W0 Q9S3W8
Q9S3Z4 Q9S400 Q9S426 Q9S449 Q9S467 Q9S468 Q9S469 Q9S470 Q9S493 Q9S498 Q9S4D5
Q9S4H7 Q9S4H8 Q9S4K6 Q9S4K9 Q9S4M7 Q9S4T0 Q9S4T3 Q9S4W2 Q9S4W7 Q9S4W9 Q9S4X0
Q9S4X1 Q9S4X2 Q9S4X3 Q9S4X4 Q9S4X5 Q9S4Y6 Q9S4Y7 Q9S4Y8 Q9S515 Q9S586 Q9S597
Q9S5E2 Q9S5G3 Q9S5G4 Q9S5G5 Q9S5G6 Q9S5V6 Q9S5X1 Q9S5Z2 Q9S660 Q9S6P8 Q9S6S1
Q9S6S2 Q9S6S3 Q9S6Z7 Q9S702 Q9S709 Q9S725 Q9S733 Q9S736 Q9S744 Q9S750 Q9S756
Q9S757 Q9S762 Q9S763 Q9S773 Q9S777 Q9S785 Q9S789 Q9S793 Q9S795 Q9S7A0 Q9S7A3
Q9S7B2 Q9S7C0 Q9S7C9 Q9S7D8 Q9S7E4 Q9S7E6 Q9S7F3 Q9S7G6 Q9S7H3 Q9S7H4 Q9S7H8
Q9S7I0 Q9S7I8 Q9S7L8 Q9S7N2 Q9S7P3 Q9S7P8 Q9S7Q2 Q9S7R5 Q9S7S2 Q9S7T1 Q9S7T5
Q9S7T7 Q9S7T9 Q9S7W5 Q9S7X6 Q9S7Y1 Q9S816 Q9S818 Q9S823 Q9S825 Q9S834 Q9S847
Q9S851 Q9S8T0 Q9S8T1 Q9S926 Q9S9E3 Q9S9K9 Q9S9M8 Q9S9N1 Q9S9N9 Q9S9P1 Q9S9P2
Q9S9P3 Q9S9P6 Q9S9P8 Q9S9T0 Q9S9T6 Q9S9T7 Q9S9U6 Q9S9V1 Q9S9V8 Q9S9V9 Q9S9X4
Q9SA01 Q9SA02 Q9SA03 Q9SA07 Q9SA14 Q9SA16 Q9SA18 Q9SA22 Q9SA27 Q9SA32 Q9SA34
Q9SA41 Q9SA42 Q9SA49 Q9SA51 Q9SA60 Q9SA68 Q9SA69 Q9SA76 Q9SA80 Q9SA82 Q9SA94
Q9SA96 Q9SA98 Q9SAA2 Q9SAA6 Q9SAB1 Q9SAB3 Q9SAB4 Q9SAB5 Q9SAC4 Q9SAC6 Q9SAD3
Q9SAD9 Q9SAF1 Q9SAF4 Q9SAG4 Q9SAG8 Q9SAH2 Q9SAH7 Q9SAH9 Q9SAI1 Q9SAI2 Q9SAI5
Q9SAJ5 Q9SAJ6 Q9SAK0 Q9SAK2 Q9SAK4 Q9SAK5 Q9SAL0 Q9SAR0 Q9SAR1 Q9SAV0 Q9SAV1
Q9SAV3 Q9SAZ0 Q9SAZ1 Q9SB00 Q9SB26 Q9SB36 Q9SB60 Q9SB61 Q9SB63 Q9SB64 Q9SB68
Q9SB79 Q9SB80 Q9SB89 Q9SB92 Q9SBF3 Q9SBJ1 Q9SBK9 Q9SBL3 Q9SBL4 Q9SBL5 Q9SBL6
Q9SBL7 Q9SBN4 Q9SBQ5 Q9SBQ7 Q9SBV6 Q9SBV9 Q9SC12 Q9SC13 Q9SC14 Q9SC80 Q9SC88
Q9SCC7 Q9SCC8 Q9SCJ4 Q9SCJ8 Q9SCK1 Q9SCK2 Q9SCK7 Q9SCL2 Q9SCM3 Q9SCM4 Q9SCP4
Q9SCP5 Q9SCP6 Q9SCP7 Q9SCQ2 Q9SCQ5 Q9SCQ7 Q9SCR0 Q9SCR2 Q9SCS3 Q9SCS4 Q9SCT2
Q9SCT6 Q9SCU0 Q9SCU7 Q9SCX3 Q9SCY2 Q9SD02 Q9SD20 Q9SD24 Q9SD25 Q9SD33 Q9SD34
Q9SD40 Q9SD58 Q9SD61 Q9SD70 Q9SD71 Q9SD76 Q9SD82 Q9SDA8 Q9SDD6 Q9SDG6 Q9SDG8
Q9SDK9 Q9SDL7 Q9SDL8 Q9SDM8 Q9SDM9 Q9SDN0 Q9SDS8 Q9SDT1 Q9SDW0 Q9SDX3 Q9SDY5
Q9SE20 Q9SE33 Q9SE37 Q9SE42 Q9SE45 Q9SE60 Q9SE93 Q9SE94 Q9SE95 Q9SE96 Q9SEA0
Q9SEA4 Q9SEC2 Q9SEE4 Q9SEE5 Q9SEH4 Q9SEH5 Q9SEH7 Q9SEI0 Q9SEI2 Q9SEI3 Q9SEI4
Q9SEL7 Q9SEP2 Q9SEP4 Q9SEV0 Q9SEV2 Q9SEW4 Q9SEX2 Q9SEZ1 Q9SEZ3 Q9SEZ4 Q9SEZ6
Q9SF15 Q9SF16 Q9SF21 Q9SF23 Q9SF35 Q9SF37 Q9SF38 Q9SF40 Q9SF41 Q9SF53 Q9SF55
Q9SFC7 Q9SFD5 Q9SFF4 Q9SFG6 Q9SFH9 Q9SFU1 Q9SFV2 Q9SFV6 Q9SFV7 Q9SFV9 Q9SFW6
Q9SFW8 Q9SFX6 Q9SFZ3 Q9SG02 Q9SG85 Q9SG92 Q9SGA0 Q9SGA2 Q9SGA6 Q9SGA8 Q9SGC1
Q9SGD6 Q9SGE0 Q9SGE2 Q9SGE3 Q9SGE9 Q9SGI8 Q9SGP6 Q9SGP8 Q9SGQ6 Q9SGS2 Q9SGS4
Q9SGS5 Q9SGT1 Q9SGT2 Q9SGU3 Q9SGU9 Q9SGZ8 Q9SH13 Q9SH22 Q9SH25 Q9SH26 Q9SH52
Q9SH60 Q9SH88 Q9SHB5 Q9SHD0 Q9SHD3 Q9SHE9 Q9SHF2 Q9SHF3 Q9SHG0 Q9SHH6 Q9SHH7
Q9SHH8 Q9SHI1 Q9SHJ0 Q9SHJ8 Q9SHK2 Q9SHL7 Q9SHP0 Q9SHQ9 Q9SHS7 Q9SHS8 Q9SHT2
Q9SHT3 Q9SHV7 Q9SHX9 Q9SHY1 Q9SHY8 Q9SHZ1 Q9SHZ5 Q9SHZ6 Q9SI02 Q9SI15 Q9SI19
Q9SI20 Q9SI34 Q9SI38 Q9SI52 Q9SI53 Q9SI54 Q9SI61 Q9SI64 Q9SI75 Q9SI78 Q9SI85
Q9SI87 Q9SI96 Q9SIB4 Q9SIC3 Q9SIC4 Q9SIC7 Q9SID3 Q9SIE0 Q9SIE1 Q9SIG8 Q9SIH0
Q9SIH1 Q9SIH5 Q9SII0 Q9SII8 Q9SII9 Q9SIJ0 Q9SIJ3 Q9SIK1 Q9SIK2 Q9SIL5 Q9SIM4
Q9SIM9 Q9SIP7 Q9SIR9 Q9SIS3 Q9SIS4 Q9SIT9 Q9SIU0 Q9SIU9 Q9SIV0 Q9SIV3 Q9SIV7

Q9SIW1 Q9SIW5 Q9SIW9 Q9SIX0 Q9SIY3 Q9SIZ2 Q9SJ04 Q9SJ05 Q9SJ06 Q9SJ07 Q9SJ09
Q9SJ10 Q9SJ20 Q9SJ25 Q9SJ29 Q9SJ32 Q9SJ36 Q9SJ41 Q9SJ44 Q9SJ45 Q9SJ56 Q9SJ66
Q9SJ78 Q9SJ84 Q9SJ85 Q9SJ89 Q9SJ96 Q9SJ98 Q9SJA6 Q9SJA7 Q9SJA8 Q9SJC4 Q9SJC6
Q9SJF3 Q9SJG6 Q9SJG8 Q9SJH0 Q9SJH7 Q9SJI8 Q9SJI9 Q9SJJ3 Q9SJJ5 Q9SJK3 Q9SJK7
Q9SJL0 Q9SJL7 Q9SJL8 Q9SJM1 Q9SJM4 Q9SJM6 Q9SJM7 Q9SJN0 Q9SJN2 Q9SJN6 Q9SJQ6
Q9SJT1 Q9SJT8 Q9SJU0 Q9SJU4 Q9SJU5 Q9SJU8 Q9SJW0 Q9SJW5 Q9SJW6 Q9SJW9 Q9SJX1
Q9SJX7 Q9SJZ3 Q9SJZ5 Q9SJZ6 Q9SK02 Q9SK08 Q9SK22 Q9SK32 Q9SK33 Q9SK47 Q9SK49
Q9SK53 Q9SK55 Q9SK64 Q9SK71 Q9SK74 Q9SK82 Q9SK86 Q9SK87 Q9SK91 Q9SKB3 Q9SKB6
Q9SKB7 Q9SKC1 Q9SKC2 Q9SKC3 Q9SKC4 Q9SKC5 Q9SKD0 Q9SKD9 Q9SKH2 Q9SKI2 Q9SKI4
Q9SKK0 Q9SKK4 Q9SKN2 Q9SKN5 Q9SKN6 Q9SKN7 Q9SKP0 Q9SKP6 Q9SKQ4 Q9SKT1 Q9SKU6
Q9SKX1 Q9SKX4 Q9SKX8 Q9SKY8 Q9SKY9 Q9SKZ1 Q9SKZ2 Q9SKZ3 Q9SL02 Q9SL41 Q9SL42
Q9SL49 Q9SL65 Q9SL67 Q9SL70 Q9SL79 Q9SL92 Q9SL93 Q9SL94 Q9SLA1 Q9SLB6 Q9SLB7
Q9SLD4 Q9SLE0 Q9SLE4 Q9SLF7 Q9SLG0 Q9SLH0 Q9SLH9 Q9SLI2 Q9SLI8 Q9SLL7 Q9SLM6
Q9SLN1 Q9SLN5 Q9SLN8 Q9SLP8 Q9SLV1 Q9SLX9 Q9SLZ0 Q9SLZ4 Q9SM27 Q9SM41 Q9SM59
Q9SM60 Q9SM64 Q9SMB1 Q9SMB8 Q9SMC2 Q9SMG1 Q9SMH3 Q9SMH4 Q9SMI2 Q9SMI3 Q9SMJ3
Q9SMK9 Q9SML4 Q9SML8 Q9SMM0 Q9SMM9 Q9SMN0 Q9SMR2 Q9SMT4 Q9SMT7 Q9SMT9 Q9SMU0
Q9SMU1 Q9SMV7 Q9SMW7 Q9SMX1 Q9SMX5 Q9SMY8 Q9SMZ2 Q9SMZ3 Q9SN09 Q9SN10 Q9SN11
Q9SN20 Q9SN21 Q9SN22 Q9SN23 Q9SN39 Q9SN45 Q9SN47 Q9SN58 Q9SN73 Q9SN74 Q9SN78
Q9SN85 Q9SN89 Q9SN90 Q9SNB0 Q9SNB1 Q9SNB6 Q9SNB8 Q9SNB9 Q9SNC0 Q9SND4 Q9SND9
Q9SNN0 Q9SNQ2 Q9SNW5 Q9SNW6 Q9SNW7 Q9SNW8 Q9SNX2 Q9SNX8 Q9SNY3 Q9SP08 Q9SP32
Q9SP37 Q9SPB1 Q9SPB3 Q9SPG2 Q9SPG3 Q9SPG5 Q9SPG9 Q9SPK5 Q9SPL2 Q9SPN0 Q9SPN1
Q9SPN3 Q9SPP9 Q9SPU3 Q9SPU7 Q9SQ56 Q9SQ64 Q9SQ67 Q9SQ68 Q9SQ69 Q9SQ70 Q9SQ80
Q9SQF4 Q9SQI8 Q9SQI9 Q9SQK3 Q9SQL0 Q9SQL5 Q9SQQ6 Q9SQR2 Q9SQR4 Q9SQT8 Q9SQU4
Q9SQU6 Q9SQV1 Q9SQZ1 Q9SQZ9 Q9SR00 Q9SR01 Q9SR02 Q9SR06 Q9SR07 Q9SR08 Q9SR13
Q9SR15 Q9SR17 Q9SR19 Q9SR24 Q9SR25 Q9SR34 Q9SR36 Q9SR43 Q9SR52 Q9SR66 Q9SR71
Q9SR82 Q9SR86 Q9SR97 Q9SRD0 Q9SRD4 Q9SRE0 Q9SRE5 Q9SRE6 Q9SRE7 Q9SRH9 Q9SRI1
Q9SRK5 Q9SRK6 Q9SRL4 Q9SRL5 Q9SRL8 Q9SRM3 Q9SRM4 Q9SRN4 Q9SRP4 Q9SRP5 Q9SRP7
Q9SRQ2 Q9SRQ5 Q9SRR1 Q9SRR7 Q9SRT2 Q9SRU7 Q9SRV1 Q9SRV2 Q9SRV3 Q9SRW7 Q9SRX1
Q9SRX2 Q9SRX6 Q9SRX7 Q9SRY5 Q9SRY6 Q9SRZ0 Q9SRZ4 Q9SS04 Q9SS17 Q9SS31 Q9SS35
Q9SS38 Q9SS44 Q9SS45 Q9SS47 Q9SS81 Q9SS87 Q9SS97 Q9SSA1 Q9SSB4 Q9SSB5 Q9SSD2
Q9SSE5 Q9SSF1 Q9SSF9 Q9SSK1 Q9SSK2 Q9SSK5 Q9SSK7 Q9SSK9 Q9SSM3 Q9SSM9 Q9SSP5
Q9SSQ2 Q9SSQ4 Q9SSQ8 Q9SSR1 Q9SSR4 Q9SSR6 Q9SSR7 Q9SSR8 Q9SSS5 Q9SSU8 Q9SSW0
Q9SSW1 Q9SSW2 Q9ST02 Q9ST03 Q9ST43 Q9ST48 Q9ST57 Q9ST58 Q9ST59 Q9ST69 Q9ST98
Q9ST99 Q9STB6 Q9STE3 Q9STE5 Q9STE6 Q9STE7 Q9STF1 Q9STF9 Q9STG3 Q9STG6 Q9STJ6
Q9STJ7 Q9STJ9 Q9STK5 Q9STL8 Q9STM4 Q9STN3 Q9STN5 Q9STN7 Q9STQ0 Q9STQ6 Q9STR1
Q9STR4 Q9STS3 Q9STS6 Q9STT1 Q9STU1 Q9STW6 Q9STX0 Q9STX2 Q9STX3 Q9STY4 Q9STY6
Q9SU00 Q9SU25 Q9SU29 Q9SU30 Q9SU38 Q9SU39 Q9SU63 Q9SU75 Q9SU79 Q9SU86 Q9SU87
Q9SU92 Q9SU94 Q9SUA9 Q9SUB0 Q9SUB1 Q9SUC0 Q9SUD8 Q9SUE7 Q9SUF4 Q9SUF5 Q9SUF9
Q9SUG4 Q9SUH5 Q9SUK9 Q9SUM2 Q9SUM5 Q9SUP6 Q9SUP7 Q9SUR6 Q9SUR8 Q9SUR9 Q9SUS1
Q9SUS4 Q9SUS5 Q9SUV9 Q9SUW0 Q9SUW2 Q9SUX7 Q9SUY0 Q9SV01 Q9SV09 Q9SV26 Q9SV36
Q9SV46 Q9SV54 Q9SV55 Q9SV82 Q9SV87 Q9SV96 Q9SV98 Q9SVA0 Q9SVA1 Q9SVA3 Q9SVA6
Q9SVB7 Q9SVC5 Q9SVC6 Q9SVD0 Q9SVD5 Q9SVD7 Q9SVF1 Q9SVG9 Q9SVH3 Q9SVJ9 Q9SVL0
Q9SVM0 Q9SVM4 Q9SVM9 Q9SVN5 Q9SVP6 Q9SVP7 Q9SVQ1 Q9SVR0 Q9SVS0 Q9SVS8 Q9SVU0
Q9SVU6 Q9SVU7 Q9SVV9 Q9SVX4 Q9SVX5 Q9SVY0 Q9SVY6 Q9SVZ6 Q9SW09 Q9SW11 Q9SW33

Q9SW44 Q9SW48 Q9SW70 Q9SW75 Q9SW80 Q9SW95 Q9SW96 Q9SWA6 Q9SWB4 Q9SWB6 Q9SWB8
Q9SWC2 Q9SWF5 Q9SWF9 Q9SWG0 Q9SWG3 Q9SWI6 Q9SWR3 Q9SWW5 Q9SX03 Q9SX04 Q9SX25
Q9SX27 Q9SX29 Q9SX38 Q9SX41 Q9SX45 Q9SX54 Q9SX68 Q9SX78 Q9SX79 Q9SX80 Q9SX97
Q9SX99 Q9SXA0 Q9SXC8 Q9SXD1 Q9SXD5 Q9SXD8 Q9SXD9 Q9SXG8 Q9SXJ6 Q9SXN6 Q9SXP2
Q9SXP7 Q9SXQ6 Q9SXS7 Q9SXU1 Q9SXY0 Q9SXY1 Q9SY00 Q9SY03 Q9SY06 Q9SY07 Q9SY09
Q9SY12 Q9SY17 Q9SY20 Q9SY65 Q9SY66 Q9SY73 Q9SY74 Q9SY75 Q9SY84 Q9SY88 Q9SY96
Q9SYA6 Q9SYD0 Q9SYD4 Q9SYD8 Q9SYG1 Q9SYG4 Q9SYG7 Q9SYH1 Q9SYH2 Q9SYJ4 Q9SYK1
Q9SYK4 Q9SYK9 Q9SYL0 Q9SYL8 Q9SYL9 Q9SYM2 Q9SYP1 Q9SYP3 Q9SYQ0 Q9SYZ9 Q9SZ05
Q9SZ10 Q9SZ20 Q9SZ30 Q9SZ44 Q9SZ52 Q9SZ59 Q9SZ63 Q9SZ69 Q9SZ70 Q9SZ75 Q9SZ83
Q9SZ95 Q9SZA2 Q9SZA4 Q9SZA6 Q9SZA7 Q9SZB4 Q9SZB6 Q9SZD1 Q9SZE1 Q9SZE8 Q9SZF7
Q9SZG3 Q9SZH4 Q9SZI3 Q9SZJ2 Q9SZJ6 Q9SZK1 Q9SZL5 Q9SZL6 Q9SZL7 Q9SZL8 Q9SZP1
Q9SZR0 Q9SZR3 Q9SZR7 Q9SZT8 Q9SZT9 Q9SZV0 Q9SZX3 Q9SZX9 Q9SZZ9 Q9T003 Q9T010
Q9T012 Q9T014 Q9T030 Q9T031 Q9T035 Q9T041 Q9T043 Q9T048 Q9T065 Q9T072 Q9T074
Q9T075 Q9T079 Q9T080 Q9T081 Q9T096 Q9T0A7 Q9T0B0 Q9T0C6 Q9T0C7 Q9T0D6 Q9T0D7
Q9T0E2 Q9T0E4 Q9T0F1 Q9T0F2 Q9T0F3 Q9T0F4 Q9T0G7 Q9T0H2 Q9T0H7 Q9T0I1 Q9T0J4
Q9T0J5 Q9T0J6 Q9T0J9 Q9T0K1 Q9T0K7 Q9T0M8 Q9T0P4 Q9T0R9 Q9T0T0 Q9T0V1 Q9T154
Q9T1L9 Q9T1P4 Q9T1P5 Q9T1P6 Q9T1P7 Q9T1P8 Q9T1P9 Q9T1Q1 Q9T1Q3 Q9T1Q4 Q9T1Q7
Q9T1Q8 Q9T1Q9 Q9T1R1 Q9T1R2 Q9T1R3 Q9T1R4 Q9T1R5 Q9T1R6 Q9T1R7 Q9T1R9 Q9T1S1
Q9T1S2 Q9T1S3 Q9T1S4 Q9T1S6 Q9T1S7 Q9T1S8 Q9T1S9 Q9T1T1 Q9T1T2 Q9T1T3 Q9T1T4
Q9T1T6 Q9T1T8 Q9T1T9 Q9T1U0 Q9T1U1 Q9T1U2 Q9T1U3 Q9T1U4 Q9T1U6 Q9T1U9 Q9T1V0
Q9T1V1 Q9T1V2 Q9T1V3 Q9T1V6 Q9T1V8 Q9T1V9 Q9T1W0 Q9T1W1 Q9T1W5 Q9T1W6 Q9T1W7
Q9T1W8 Q9T1X0 Q9T1X4 Q9T1X7 Q9T216 Q9T390 Q9T399 Q9T3P6 Q9T467 Q9T4F2 Q9T4F6
Q9TEM3 Q9TGM3 Q9TGM4 Q9THV5 Q9THX6 Q9TI97 Q9TIA9 Q9TIB3 Q9TIB6 Q9TIC1 Q9TIQ4
Q9TIQ5 Q9TIR3 Q9TIR8 Q9TIR9 Q9TIS6 Q9TJN6 Q9TJQ5 Q9TJQ6 Q9TJQ7 Q9TJQ8 Q9TJQ9
Q9TJR0 Q9TJR1 Q9TJR5 Q9TK08 Q9TK50 Q9TK52 Q9TKB6 Q9TKB8 Q9TKB9 Q9TKC0 Q9TKD1
Q9TKD3 Q9TKD7 Q9TKI5 Q9TKP0 Q9TKP6 Q9TKP8 Q9TKP9 Q9TKR9 Q9TKS1 Q9TKS4 Q9TKS6
Q9TKU9 Q9TKX3 Q9TKX4 Q9TKX5 Q9TKX7 Q9TKX8 Q9TKZ4 Q9TKZ5 Q9TKZ6 Q9TKZ7 Q9TL03
Q9TL04 Q9TL05 Q9TL06 Q9TL08 Q9TL09 Q9TL10 Q9TL17 Q9TL18 Q9TL19 Q9TL20 Q9TL21
Q9TL22 Q9TL23 Q9TL24 Q9TL25 Q9TL26 Q9TL27 Q9TL28 Q9TL29 Q9TL30 Q9TL38 Q9TLD8
Q9TLQ7 Q9TLQ8 Q9TLR8 Q9TLR9 Q9TLS0 Q9TLS1 Q9TLS3 Q9TLT0 Q9TLT1 Q9TLT2 Q9TLT3
Q9TLT4 Q9TLT5 Q9TLT6 Q9TLT7 Q9TLT8 Q9TLT9 Q9TLU1 Q9TLU2 Q9TLU3 Q9TLU4 Q9TLU5
Q9TLU7 Q9TLU8 Q9TLU9 Q9TLV0 Q9TLV1 Q9TLV2 Q9TLV4 Q9TLV5 Q9TLV6 Q9TLV7 Q9TLV8
Q9TLV9 Q9TLW0 Q9TLW8 Q9TLW9 Q9TLX1 Q9TLX2 Q9TLX6 Q9TLX7 Q9TLX8 Q9TLY1 Q9TLY2
Q9TLY7 Q9TLY9 Q9TLZ0 Q9TLZ1 Q9TLZ9 Q9TM00 Q9TM01 Q9TM05 Q9TM06 Q9TM07 Q9TM10
Q9TM25 Q9TM32 Q9TM33 Q9TM34 Q9TM35 Q9TM37 Q9TM38 Q9TM39 Q9TM55 Q9TMB2 Q9TMC8
Q9TMM9 Q9TMN0 Q9TN87 Q9TN89 Q9TNA1 Q9TNA5 Q9TNB1 Q9TNB2 Q9TQS6 Q9TQX6 Q9TRY0
Q9TS34 Q9TS35 Q9TS53 Q9TS87 Q9TSN7 Q9TSN8 Q9TSN9 Q9TSP1 Q9TSV3 Q9TSV4 Q9TSV5
Q9TSV7 Q9TSW0 Q9TSX5 Q9TSZ2 Q9TSZ7 Q9TT01 Q9TT33 Q9TT34 Q9TT89 Q9TT91 Q9TT99
Q9TTA5 Q9TTD8 Q9TTE5 Q9TTF0 Q9TTI7 Q9TTJ5 Q9TTJ6 Q9TTK0 Q9TTK4 Q9TTR7 Q9TTR8
Q9TTU2 Q9TTW3 Q9TTY4 Q9TU15 Q9TU19 Q9TU47 Q9TU66 Q9TU67 Q9TU71 Q9TUB3 Q9TUC2
Q9TUC3 Q9TUC4 Q9TUG3 Q9TUI5 Q9TUJ2 Q9TV61 Q9TV62 Q9TV63 Q9TV68 Q9TV69 Q9TV70
Q9TVA3 Q9TVL3 Q9TVP3 Q9TVQ5 Q9TVW2 Q9TVW5 Q9TW89 Q9TX44 Q9TXD9 Q9TXH9 Q9TXI0
Q9TXJ1 Q9TXJ8 Q9TXQ1 Q9TXV7 Q9TY03 Q9TY83 Q9TY84 Q9TYK1 Q9TYM6 Q9TYN3 Q9TYP2
Q9TYT0 Q9TYU5 Q9TYU9 Q9TYV5 Q9TZ70 Q9TZ93 Q9TZ99 Q9TZH6 Q9TZL0 Q9TZL8 Q9TZM3

Q9TZN1 Q9TZP5 Q9TZQ3 Q9U0E6 Q9U178 Q9U1I1 Q9U1N0 Q9U1N6 Q9U1Q4 Q9U1R1 Q9U1W2
Q9U1Y5 Q9U213 Q9U221 Q9U226 Q9U239 Q9U263 Q9U281 Q9U297 Q9U299 Q9U2A9 Q9U2C9
Q9U2D6 Q9U2D9 Q9U2G0 Q9U2H1 Q9U2H9 Q9U2M1 Q9U2M4 Q9U2M5 Q9U2M7 Q9U2R0 Q9U2R6
Q9U2T3 Q9U2Y2 Q9U2Z0 Q9U389 Q9U3B6 Q9U3C8 Q9U3D6 Q9U3F4 Q9U3G6 Q9U3L0 Q9U3U0
Q9U3V5 Q9U3W6 Q9U3Z7 Q9U408 Q9U489 Q9U4X2 Q9U4X3 Q9U4X4 Q9U4X5 Q9U501 Q9U518
Q9U568 Q9U5M4 Q9U5P2 Q9U5V6 Q9U5W9 Q9U5Z8 Q9U615 Q9U616 Q9U620 Q9U639 Q9U6A9
Q9U6D2 Q9U6M1 Q9U6M2 Q9U6P6 Q9U6V6 Q9U6V7 Q9U6Y3 Q9U6Y4 Q9U6Y5 Q9U6Y6 Q9U6Y7
Q9U6Y8 Q9U6Y9 Q9U757 Q9U761 Q9U762 Q9U793 Q9U794 Q9U7D0 Q9U7D1 Q9U7E0 Q9U8B8
Q9U8D3 Q9U8F1 Q9U8G7 Q9U8Q4 Q9U8S9 Q9U8U5 Q9U903 Q9U915 Q9U9A3 Q9U9I5 Q9U9J4
Q9U9L1 Q9U9L2 Q9U9Q4 Q9UAE6 Q9UAG7 Q9UAS6 Q9UAV5 Q9UB37 Q9UBC0 Q9UBC1 Q9UBC9
Q9UBD0 Q9UBD5 Q9UBE0 Q9UBF1 Q9UBI1 Q9UBK7 Q9UBK9 Q9UBN7 Q9UBP6 Q9UBP8 Q9UBQ7
Q9UBR1 Q9UBR4 Q9UBT6 Q9UBX0 Q9UBY9 Q9UC06 Q9UC07 Q9UD57 Q9UDT6 Q9UDV6 Q9UDV7
Q9UDW3 Q9UET6 Q9UEU5 Q9UF83 Q9UFB7 Q9UFD9 Q9UFE4 Q9UFF9 Q9UFH2 Q9UFV1 Q9UFV3
Q9UG01 Q9UGB7 Q9UGC7 Q9UGL1 Q9UGL9 Q9UGP5 Q9UGY1 Q9UH03 Q9UH17 Q9UH36 Q9UH64
Q9UH73 Q9UH77 Q9UH90 Q9UHA2 Q9UHA3 Q9UHB4 Q9UHC1 Q9UHC7 Q9UHD4 Q9UHJ3 Q9UHJ6
Q9UHL3 Q9UHN1 Q9UHP6 Q9UHP9 Q9UHQ1 Q9UHQ7 Q9UHR6 Q9UHT4 Q9UHU1 Q9UHV2 Q9UHV8
Q9UHV9 Q9UHY7 Q9UI15 Q9UI17 Q9UI25 Q9UI32 Q9UI36 Q9UI43 Q9UI46 Q9UI47 Q9UI54
Q9UI72 Q9UIC8 Q9UIE0 Q9UIF3 Q9UIF7 Q9UIG5 Q9UII4 Q9UII5 Q9UII6 Q9UIJ7 Q9UIL4
Q9UIM3 Q9UIV8 Q9UIW0 Q9UIY3 Q9UJ04 Q9UJ55 Q9UJ72 Q9UJ98 Q9UJA3 Q9UJA5 Q9UJJ7
Q9UJL9 Q9UJN7 Q9UJQ7 Q9UJT0 Q9UJT1 Q9UJT2 Q9UJT9 Q9UJU3 Q9UJU5 Q9UJV3 Q9UJV8
Q9UJW0 Q9UJW3 Q9UJW7 Q9UJW9 Q9UJX0 Q9UJX3 Q9UJY1 Q9UK10 Q9UK11 Q9UK12 Q9UK13
Q9UK45 Q9UK53 Q9UK73 Q9UK96 Q9UK99 Q9UKA1 Q9UKA2 Q9UKA8 Q9UKB3 Q9UKD2 Q9UKK6
Q9UKN5 Q9UKN7 Q9UKT6 Q9UKT7 Q9UKT8 Q9UKT9 Q9UKW6 Q9UKX2 Q9UKX3 Q9UKZ1 Q9UL16
Q9UL17 Q9UL18 Q9UL36 Q9UL40 Q9UL41 Q9UL42 Q9UL49 Q9UL58 Q9UL59 Q9UL68 Q9ULA0
Q9ULC4 Q9ULC6 Q9ULD0 Q9ULD6 Q9ULE3 Q9ULE4 Q9ULG3 Q9ULI1 Q9ULI2 Q9ULJ1 Q9ULK2
Q9ULK4 Q9ULL0 Q9ULM0 Q9ULM2 Q9ULN7 Q9ULQ0 Q9ULR5 Q9ULW3 Q9ULW8 Q9ULX9 Q9ULZ0
Q9UM07 Q9UM13 Q9UMQ3 Q9UMR3 Q9UMS0 Q9UMY1 Q9UN30 Q9UN81 Q9UNA4 Q9UNH5 Q9UNI6
Q9UNK9 Q9UNN4 Q9UNP9 Q9UNQ2 Q9UNT1 Q9UNZ5 Q9UPM6 Q9UPQ3 Q9UPQ7 Q9UPR3 Q9UPR6
Q9UPS8 Q9UPU9 Q9UPW6 Q9UPY6 Q9UPY8 Q9UQ10 Q9UQ88 Q9UQB9 Q9UQG0 Q9UQK1 Q9UQW6
Q9UQW9 Q9UQX0 Q9UQY0 Q9UQY2 Q9UR05 Q9UR06 Q9UR07 Q9UR08 Q9UR21 Q9UR24 Q9UR27
Q9UR38 Q9UR39 Q9UR66 Q9URM2 Q9URQ3 Q9URQ5 Q9URS0 Q9URS1 Q9URS8 Q9URT2 Q9URT4
Q9URT8 Q9URU2 Q9URU9 Q9URV0 Q9URV2 Q9URV3 Q9URV4 Q9URV8 Q9URV9 Q9URW3 Q9URW6
Q9URW9 Q9URX0 Q9URX3 Q9URX7 Q9URX9 Q9URY0 Q9URY2 Q9URY3 Q9URY5 Q9URY7 Q9URY9
Q9URZ1 Q9URZ2 Q9URZ7 Q9US01 Q9US02 Q9US03 Q9US05 Q9US06 Q9US07 Q9US12 Q9US13
Q9US14 Q9US15 Q9US25 Q9US26 Q9US28 Q9US30 Q9US33 Q9US34 Q9US35 Q9US36 Q9US38
Q9US41 Q9US42 Q9US43 Q9US45 Q9US46 Q9US47 Q9US49 Q9US51 Q9US52 Q9US53 Q9US54
Q9USG9 Q9USH1 Q9USH2 Q9USH5 Q9USH6 Q9USH9 Q9USI0 Q9USI4 Q9USI5 Q9USI6 Q9USJ1
Q9USJ5 Q9USJ6 Q9USJ8 Q9USJ9 Q9USK0 Q9USK1 Q9USK2 Q9USK4 Q9USK7 Q9USL1 Q9USL2
Q9USL3 Q9USL5 Q9USL6 Q9USL7 Q9USM2 Q9USM4 Q9USN2 Q9USN3 Q9USN5 Q9USN6 Q9USN7
Q9USN8 Q9USP0 Q9USP1 Q9USP3 Q9USP4 Q9USP7 Q9USP8 Q9USP9 Q9USQ1 Q9USQ2 Q9USQ4
Q9USQ5 Q9USQ7 Q9USQ8 Q9USQ9 Q9USR0 Q9USR1 Q9USR2 Q9USR4 Q9USR5 Q9USR6 Q9USR9
Q9USS0 Q9USS1 Q9USS2 Q9USS4 Q9USS6 Q9USS7 Q9USS8 Q9UST1 Q9UST4 Q9UST5 Q9UST6
Q9UST7 Q9UST8 Q9UST9 Q9USU2 Q9USU3 Q9USU7 Q9USU8 Q9USU9 Q9USV1 Q9USV2 Q9USV3
Q9USV4 Q9USW0 Q9USW1 Q9USW8 Q9USW9 Q9USX1 Q9USX2 Q9USX3 Q9USX4 Q9USX8 Q9USX9

Q9USY0 Q9USY1 Q9USY2 Q9USY3 Q9USY4 Q9USY8 Q9USZ1 Q9USZ3 Q9USZ4 Q9USZ5 Q9USZ6
Q9UT06 Q9UT08 Q9UT09 Q9UT10 Q9UT12 Q9UT16 Q9UT17 Q9UT19 Q9UT21 Q9UT22 Q9UT23
Q9UT24 Q9UT25 Q9UT28 Q9UT31 Q9UT32 Q9UT33 Q9UT36 Q9UT38 Q9UT39 Q9UT40 Q9UT44
Q9UT46 Q9UT48 Q9UT49 Q9UT50 Q9UT52 Q9UT53 Q9UT54 Q9UT55 Q9UT56 Q9UT57 Q9UT59
Q9UT60 Q9UT63 Q9UT66 Q9UT68 Q9UT70 Q9UT72 Q9UT73 Q9UT74 Q9UT75 Q9UT76 Q9UT77
Q9UT79 Q9UT82 Q9UT85 Q9UT86 Q9UT87 Q9UT93 Q9UT94 Q9UT96 Q9UT97 Q9UT98 Q9UTA1
Q9UTA2 Q9UTA4 Q9UTA5 Q9UTA6 Q9UTA7 Q9UTA8 Q9UTA9 Q9UTB0 Q9UTB2 Q9UTB3 Q9UTB6
Q9UTB8 Q9UTC0 Q9UTC2 Q9UTC3 Q9UTC5 Q9UTC6 Q9UTC7 Q9UTC9 Q9UTD0 Q9UTD1 Q9UTD4
Q9UTD7 Q9UTE1 Q9UTE2 Q9UTE3 Q9UTE4 Q9UTE5 Q9UTE8 Q9UTF0 Q9UTG1 Q9UTG3 Q9UTG5
Q9UTG6 Q9UTG8 Q9UTH1 Q9UTH2 Q9UTH8 Q9UTH9 Q9UTI0 Q9UTI1 Q9UTI2 Q9UTI3 Q9UTI4
Q9UTI7 Q9UTJ2 Q9UTJ3 Q9UTJ4 Q9UTJ8 Q9UTK0 Q9UTK2 Q9UTK3 Q9UTK5 Q9UTK6 Q9UTK9
Q9UTL0 Q9UTL5 Q9UTL6 Q9UTL7 Q9UTL9 Q9UTM0 Q9UTM1 Q9UTM4 Q9UTM5 Q9UTM7 Q9UTM8
Q9UTN0 Q9UTN2 Q9UTN3 Q9UTN4 Q9UTN8 Q9UTN9 Q9UTP0 Q9UTP3 Q9UTP4 Q9UTP9 Q9UTQ0
Q9UTQ1 Q9UTQ4 Q9UTQ5 Q9UTQ8 Q9UTR1 Q9UTR4 Q9UTR7 Q9UTR9 Q9UTS5 Q9UTS6 Q9UTT2
Q9UU76 Q9UU77 Q9UU78 Q9UU80 Q9UU84 Q9UU85 Q9UU88 Q9UU89 Q9UU90 Q9UU93 Q9UU94
Q9UU95 Q9UU96 Q9UUA7 Q9UUA8 Q9UUA9 Q9UUB0 Q9UUB1 Q9UUB3 Q9UUB4 Q9UUB5 Q9UUB7
Q9UUB8 Q9UUC2 Q9UUC6 Q9UUC7 Q9UUC8 Q9UUD2 Q9UUD8 Q9UUD9 Q9UUE0 Q9UUE1 Q9UUE3
Q9UUE4 Q9UUE5 Q9UUE6 Q9UUE7 Q9UUF1 Q9UUF2 Q9UUF3 Q9UUG1 Q9UUG3 Q9UUG5 Q9UUG6
Q9UUG8 Q9UUH2 Q9UUH3 Q9UUH5 Q9UUH6 Q9UUH7 Q9UUI1 Q9UUI2 Q9UUI3 Q9UUI4 Q9UUI7
Q9UUI9 Q9UUJ1 Q9UUJ4 Q9UUJ5 Q9UUJ7 Q9UUJ9 Q9UUK0 Q9UUK1 Q9UUK4 Q9UUK7 Q9UUL2
Q9UUL4 Q9UUM2 Q9UUN3 Q9UUN6 Q9UUN9 Q9UUS2 Q9UUY8 Q9UUZ6 Q9UUZ9 Q9UV10 Q9UV33
Q9UV63 Q9UV64 Q9UV72 Q9UVB8 Q9UVC0 Q9UVE1 Q9UVF3 Q9UVF4 Q9UVF5 Q9UVF8 Q9UVJ4
Q9UVL1 Q9UVN9 Q9UVR2 Q9UVT8 Q9UVW9 Q9UVX3 Q9UVX4 Q9UVY5 Q9UVZ5 Q9UVZ8 Q9UW02
Q9UW12 Q9UW15 Q9UW18 Q9UW19 Q9UW22 Q9UW23 Q9UW26 Q9UW83 Q9UW86 Q9UW96 Q9UWE4
Q9UWF0 Q9UWF8 Q9UWG2 Q9UWG6 Q9UWN5 Q9UWN6 Q9UWN7 Q9UWP5 Q9UWQ4 Q9UWQ5 Q9UWR2
Q9UWR5 Q9UWR8 Q9UWR9 Q9UWT5 Q9UWU0 Q9UWU1 Q9UWV6 Q9UWW1 Q9UWW2 Q9UWX2 Q9UWX3
Q9UWX4 Q9UWX9 Q9UWY8 Q9UX04 Q9UX05 Q9UX07 Q9UX08 Q9UX09 Q9UX10 Q9UX16 Q9UX20
Q9UX21 Q9UX24 Q9UX29 Q9UX31 Q9UX32 Q9UX49 Q9UX78 Q9UX79 Q9UX81 Q9UX83 Q9UX85
Q9UX86 Q9UX87 Q9UX88 Q9UX89 Q9UX90 Q9UX91 Q9UX92 Q9UX93 Q9UX94 Q9UX95 Q9UX97
Q9UX98 Q9UXA0 Q9UXA2 Q9UXA3 Q9UXA5 Q9UXA6 Q9UXA8 Q9UXB1 Q9UXB2 Q9UXB8 Q9UXB9
Q9UXC0 Q9UXC2 Q9UXC4 Q9UXC6 Q9UXC7 Q9UXD0 Q9UXD4 Q9UXE0 Q9UXE4 Q9UXF3 Q9UXF5
Q9UXF6 Q9UXF7 Q9UXF8 Q9UXG1 Q9UXG7 Q9UXG8 Q9UXN8 Q9UXP0 Q9UXR0 Q9UXR8 Q9UXR9
Q9UXS5 Q9UXS6 Q9UXS8 Q9UXT0 Q9UXT1 Q9UXT3 Q9UXT7 Q9UXV5 Q9UXV6 Q9UXW3 Q9UXW5
Q9UXW6 Q9UXX0 Q9UXX2 Q9UXX7 Q9UXX8 Q9UXX9 Q9UXY2 Q9UXZ1 Q9UXZ2 Q9UXZ3 Q9UXZ4
Q9UY08 Q9UY10 Q9UY11 Q9UY16 Q9UY20 Q9UY21 Q9UY24 Q9UY31 Q9UY33 Q9UY36 Q9UY47
Q9UY49 Q9UY53 Q9UY55 Q9UY56 Q9UY67 Q9UY68 Q9UY71 Q9UY84 Q9UY93 Q9UY98 Q9UY99
Q9UYB2 Q9UYB3 Q9UYB5 Q9UYB6 Q9UYB9 Q9UYD1 Q9UYD4 Q9UYF7 Q9UYI4 Q9UYI5 Q9UYI6
Q9UYI7 Q9UYI8 Q9UYJ4 Q9UYJ5 Q9UYK5 Q9UYL3 Q9UYL4 Q9UYQ3 Q9UYQ9 Q9UYR1 Q9UYR5
Q9UYR6 Q9UYR9 Q9UYS3 Q9UYS6 Q9UYS8 Q9UYT0 Q9UYT3 Q9UYT4 Q9UYU5 Q9UYU9 Q9UYV2
Q9UYV8 Q9UYW0 Q9UYX0 Q9UYX2 Q9UYX3 Q9UYX4 Q9UYX5 Q9UYX8 Q9UYY8 Q9UYY9 Q9UYZ0
Q9UYZ1 Q9UYZ2 Q9UYZ3 Q9UYZ4 Q9UYZ5 Q9UZ03 Q9UZ06 Q9UZ07 Q9UZ08 Q9UZ09 Q9UZ13
Q9UZ14 Q9UZ16 Q9UZ20 Q9UZ21 Q9UZ31 Q9UZ35 Q9UZ37 Q9UZ51 Q9UZ54 Q9UZ56 Q9UZ57
Q9UZ62 Q9UZ78 Q9UZ81 Q9UZ84 Q9UZ86 Q9UZ88 Q9UZ94 Q9UZ99 Q9UZA3 Q9UZA8 Q9UZC0
Q9UZC1 Q9UZC8 Q9UZC9 Q9UZD5 Q9UZD7 Q9UZE7 Q9UZE8 Q9UZF0 Q9UZJ5 Q9UZJ6 Q9UZJ7

Q9UZK1 Q9UZK4 Q9UZK7 Q9UZL4 Q9UZM5 Q9UZN0 Q9UZN1 Q9UZP1 Q9UZP4 Q9UZP8 Q9UZQ4
Q9UZQ5 Q9UZR7 Q9UZS9 Q9UZT4 Q9UZT5 Q9UZT8 Q9UZV4 Q9UZV7 Q9UZW0 Q9UZY0 Q9UZZ3
Q9V005 Q9V011 Q9V018 Q9V022 Q9V023 Q9V024 Q9V025 Q9V027 Q9V029 Q9V030 Q9V035
Q9V036 Q9V050 Q9V072 Q9V074 Q9V076 Q9V081 Q9V085 Q9V098 Q9V099 Q9V0A9 Q9V0B5
Q9V0B8 Q9V0B9 Q9V0D0 Q9V0D2 Q9V0D5 Q9V0D9 Q9V0E1 Q9V0E3 Q9V0E4 Q9V0F5 Q9V0F7
Q9V0G0 Q9V0G1 Q9V0G8 Q9V0H5 Q9V0I6 Q9V0I7 Q9V0I8 Q9V0J6 Q9V0J7 Q9V0J8 Q9V0K1
Q9V0L2 Q9V0L6 Q9V0M0 Q9V0M2 Q9V0N0 Q9V0N5 Q9V0N8 Q9V0N9 Q9V0P2 Q9V0P9 Q9V0Q0
Q9V0Q4 Q9V0T9 Q9V0U0 Q9V0V2 Q9V0V3 Q9V0V4 Q9V0V5 Q9V0V6 Q9V0V7 Q9V0W3 Q9V0X3
Q9V0X6 Q9V0X8 Q9V0Y3 Q9V0Y5 Q9V0Y6 Q9V0Y8 Q9V0Y9 Q9V0Z2 Q9V0Z6 Q9V103 Q9V105
Q9V106 Q9V107 Q9V108 Q9V109 Q9V112 Q9V113 Q9V116 Q9V118 Q9V119 Q9V120 Q9V121
Q9V122 Q9V126 Q9V134 Q9V135 Q9V138 Q9V148 Q9V150 Q9V151 Q9V157 Q9V159 Q9V163
Q9V166 Q9V168 Q9V173 Q9V176 Q9V189 Q9V191 Q9V195 Q9V196 Q9V197 Q9V198 Q9V1A0
Q9V1A5 Q9V1A6 Q9V1A9 Q9V1B2 Q9V1B3 Q9V1C6 Q9V1C7 Q9V1D7 Q9V1E3 Q9V1E5 Q9V1E9
Q9V1F2 Q9V1F3 Q9V1F5 Q9V1F8 Q9V1F9 Q9V1G0 Q9V1G3 Q9V1G4 Q9V1G7 Q9V1G8 Q9V1G9
Q9V1H0 Q9V1H1 Q9V1H6 Q9V1H7 Q9V1H8 Q9V1H9 Q9V1I3 Q9V1I4 Q9V1I5 Q9V1I6 Q9V1I9
Q9V1J0 Q9V1J1 Q9V1J7 Q9V1M3 Q9V1N3 Q9V1N4 Q9V1N7 Q9V1N8 Q9V1P1 Q9V1P2 Q9V1P3
Q9V1P5 Q9V1P7 Q9V1P8 Q9V1Q1 Q9V1Q7 Q9V1R0 Q9V1R3 Q9V1S2 Q9V1S9 Q9V1T5 Q9V1T6
Q9V1T7 Q9V1T8 Q9V1T9 Q9V1U0 Q9V1U1 Q9V1U2 Q9V1U3 Q9V1U4 Q9V1U5 Q9V1U6 Q9V1U7
Q9V1U8 Q9V1U9 Q9V1V0 Q9V1V1 Q9V1V2 Q9V1V3 Q9V1V4 Q9V1V5 Q9V1V6 Q9V1V7 Q9V1V9
Q9V1Y5 Q9V1Z8 Q9V1Z9 Q9V200 Q9V201 Q9V202 Q9V207 Q9V216 Q9V225 Q9V233 Q9V242
Q9V247 Q9V249 Q9V251 Q9V253 Q9V254 Q9V262 Q9V270 Q9V277 Q9V280 Q9V281 Q9V287
Q9V288 Q9V290 Q9V291 Q9V292 Q9V298 Q9V2A7 Q9V2A9 Q9V2B7 Q9V2C5 Q9V2C6 Q9V2D7
Q9V2E8 Q9V2F1 Q9V2F2 Q9V2F3 Q9V2F4 Q9V2F5 Q9V2F6 Q9V2F9 Q9V2G1 Q9V2G2 Q9V2G3
Q9V2G4 Q9V2H1 Q9V2H7 Q9V2I3 Q9V2I6 Q9V2J6 Q9V2K6 Q9V2K7 Q9V2K9 Q9V2L2 Q9V2L5
Q9V2M0 Q9V2M4 Q9V2M6 Q9V2N7 Q9V2P6 Q9V2Q7 Q9V2Q9 Q9V2R0 Q9V2R2 Q9V2R5 Q9V2R8
Q9V2S3 Q9V2S9 Q9V2T3 Q9V2T4 Q9V2T5 Q9V2T7 Q9V2T8 Q9V2V5 Q9V2V6 Q9V2W0 Q9V2W4
Q9V2W7 Q9V2W8 Q9V2W9 Q9V2X0 Q9V2X1 Q9V2Z4 Q9V2Z6 Q9V2Z7 Q9V302 Q9V333 Q9V345
Q9V359 Q9V3B4 Q9V3B6 Q9V3C0 Q9V3C1 Q9V3D2 Q9V3D5 Q9V3F3 Q9V3G3 Q9V3G6 Q9V3G7
Q9V3H2 Q9V3H8 Q9V3I5 Q9V3I8 Q9V3J8 Q9V3K3 Q9V3L6 Q9V3N8 Q9V3P0 Q9V3P6 Q9V3R6
Q9V3U2 Q9V3U9 Q9V3V7 Q9V3W0 Q9V3Y2 Q9V411 Q9V429 Q9V431 Q9V439 Q9V444 Q9V452
Q9V460 Q9V461 Q9V464 Q9V468 Q9V4F9 Q9V4M2 Q9V4Q8 Q9V4S8 Q9V4W1 Q9V4Z9 Q9V535
Q9V574 Q9V579 Q9V595 Q9V5C6 Q9V5M3 Q9V5M6 Q9V5P6 Q9V5Q4 Q9V629 Q9V637 Q9V677
Q9V6K1 Q9V6L0 Q9V6L9 Q9V6Q2 Q9V6U8 Q9V6Y3 Q9V727 Q9V780 Q9V7A7 Q9V7P1 Q9V813
Q9V831 Q9V877 Q9V895 Q9V8F3 Q9V8I2 Q9V8M5 Q9V998 Q9V9A7 Q9V9E3 Q9V9K7 Q9V9V9
Q9V9X4 Q9V9Y9 Q9V9Z1 Q9V9Z9 Q9VA12 Q9VA37 Q9VAC8 Q9VAF7 Q9VAH4 Q9VAH9 Q9VAI0
Q9VAI1 Q9VAM9 Q9VAN0 Q9VAN6 Q9VAP2 Q9VAQ1 Q9VAQ5 Q9VAQ7 Q9VAT2 Q9VAY7 Q9VB08
Q9VB14 Q9VB25 Q9VB62 Q9VB74 Q9VBB3 Q9VBG6 Q9VBP3 Q9VBP9 Q9VBQ9 Q9VBU8 Q9VBW6
Q9VBW9 Q9VBX2 Q9VBZ5 Q9VC07 Q9VC45 Q9VC49 Q9VC61 Q9VC99 Q9VCA5 Q9VCB1 Q9VCC0
Q9VCC3 Q9VCD0 Q9VCE1 Q9VCE6 Q9VCE8 Q9VCE9 Q9VCF0 Q9VCI0 Q9VCK0 Q9VCN1 Q9VCP0
Q9VCP1 Q9VCU7 Q9VCU9 Q9VCW3 Q9VCX3 Q9VCY3 Q9VCZ8 Q9VD09 Q9VD25 Q9VD28 Q9VD44
Q9VD51 Q9VD92 Q9VDD9 Q9VDE5 Q9VDL1 Q9VDM6 Q9VDN2 Q9VDQ7 Q9VDR1 Q9VDS6 Q9VDT6
Q9VDY1 Q9VDZ3 Q9VDZ4 Q9VE04 Q9VE09 Q9VE11 Q9VE34 Q9VE61 Q9VE86 Q9VE87 Q9VEA5
Q9VEB3 Q9VEB4 Q9VEB5 Q9VEC1 Q9VEC3 Q9VEI9 Q9VEJ0 Q9VEJ2 Q9VEP1 Q9VEW2 Q9VEX9
Q9VEZ3 Q9VF26 Q9VF30 Q9VF36 Q9VF59 Q9VF71 Q9VF89 Q9VFB2 Q9VFB7 Q9VFC0 Q9VFC8

Q9VFD5 Q9VFE6 Q9VFH6 Q9VFJ2 Q9VFK3 Q9VFP2 Q9VFR0 Q9VFR1 Q9VFS2 Q9VFS8 Q9VG73
Q9VG74 Q9VG76 Q9VG93 Q9VG94 Q9VG95 Q9VG96 Q9VG97 Q9VG98 Q9VGC7 Q9VGE3 Q9VGG6
Q9VGJ5 Q9VGK7 Q9VGN9 Q9VGP0 Q9VGP1 Q9VGP6 Q9VGQ9 Q9VGR2 Q9VGR7 Q9VGU5 Q9VGW6
Q9VGY5 Q9VGZ1 Q9VGZ5 Q9VGZ8 Q9VH07 Q9VH08 Q9VH20 Q9VH36 Q9VH38 Q9VH39 Q9VH70
Q9VH89 Q9VHA0 Q9VHB0 Q9VHB3 Q9VHB9 Q9VHD2 Q9VHD3 Q9VHF2 Q9VHH7 Q9VHH9 Q9VHI4
Q9VHI7 Q9VHJ6 Q9VHM2 Q9VHM6 Q9VHN3 Q9VHN6 Q9VHN9 Q9VHP0 Q9VHS8 Q9VHT9 Q9VHU1
Q9VHV1 Q9VHY5 Q9VI10 Q9VI25 Q9VI58 Q9VI60 Q9VI74 Q9VI82 Q9VI85 Q9VI93 Q9VIE6
Q9VIF0 Q9VIF1 Q9VIF5 Q9VIF6 Q9VIK9 Q9VIM5 Q9VIP2 Q9VIP8 Q9VIQ9 Q9VIT0 Q9VIV3
Q9VIW7 Q9VIZ0 Q9VJ07 Q9VJ29 Q9VJ33 Q9VJ34 Q9VJ38 Q9VJ62 Q9VJ87 Q9VJ97 Q9VJC7
Q9VJD2 Q9VJE4 Q9VJE5 Q9VJG0 Q9VJI9 Q9VJK8 Q9VJL6 Q9VJQ4 Q9VJQ5 Q9VJS8 Q9VJV8
Q9VJY8 Q9VJZ7 Q9VK13 Q9VK33 Q9VK41 Q9VK68 Q9VK71 Q9VK89 Q9VKB4 Q9VKD3 Q9VKD6
Q9VKJ0 Q9VKJ1 Q9VKJ3 Q9VKK1 Q9VKM6 Q9VKN7 Q9VKQ6 Q9VKQ9 Q9VKU7 Q9VKV5 Q9VKV8
Q9VKW2 Q9VKX4 Q9VKY2 Q9VKY8 Q9VL06 Q9VL14 Q9VL18 Q9VL25 Q9VL31 Q9VL52 Q9VL59
Q9VL63 Q9VL72 Q9VL76 Q9VL78 Q9VL96 Q9VLA2 Q9VLC0 Q9VLD6 Q9VLF6 Q9VLJ0 Q9VLJ8
Q9VLJ9 Q9VLK2 Q9VLM6 Q9VLM8 Q9VLN1 Q9VLP4 Q9VLP7 Q9VLR5 Q9VLS9 Q9VLT1 Q9VLT5
Q9VLU5 Q9VLU6 Q9VLV5 Q9VLW2 Q9VM17 Q9VM21 Q9VM35 Q9VM46 Q9VM60 Q9VM71 Q9VM75
Q9VM90 Q9VM91 Q9VM92 Q9VM95 Q9VMA0 Q9VMA2 Q9VMA3 Q9VMC8 Q9VMC9 Q9VMD0 Q9VMD6
Q9VMH7 Q9VMJ7 Q9VML1 Q9VMP9 Q9VMQ7 Q9VMU4 Q9VMU5 Q9VMW7 Q9VMW9 Q9VMX0 Q9VMX1
Q9VMX6 Q9VMY1 Q9VMY9 Q9VN03 Q9VN25 Q9VN27 Q9VN41 Q9VN50 Q9VN95 Q9VNA5 Q9VNA8
Q9VNE1 Q9VNE2 Q9VNG0 Q9VNG1 Q9VNH1 Q9VNI3 Q9VNL4 Q9VNP5 Q9VNS0 Q9VNT5 Q9VNV3
Q9VNX1 Q9VNX8 Q9VP22 Q9VP47 Q9VP65 Q9VPD2 Q9VPD5 Q9VPH2 Q9VPH7 Q9VPH8 Q9VPK7
Q9VPL3 Q9VPM5 Q9VPP5 Q9VPQ6 Q9VPR4 Q9VPU8 Q9VPW1 Q9VPW8 Q9VPY0 Q9VPY8 Q9VQ30
Q9VQ56 Q9VQ79 Q9VQ91 Q9VQC4 Q9VQD6 Q9VQE5 Q9VQF7 Q9VQK0 Q9VQN8 Q9VQS5 Q9VQS6
Q9VQS7 Q9VQU9 Q9VQX4 Q9VQY9 Q9VQZ6 Q9VR05 Q9VR17 Q9VR19 Q9VR35 Q9VR52 Q9VR56
Q9VR59 Q9VR81 Q9VR89 Q9VR93 Q9VR99 Q9VRA2 Q9VRJ0 Q9VRJ1 Q9VRL3 Q9VRN3 Q9VRN5
Q9VRP9 Q9VRQ7 Q9VRV7 Q9VRX3 Q9VS05 Q9VS36 Q9VS38 Q9VS48 Q9VS50 Q9VSA4 Q9VSD7
Q9VSE2 Q9VSF2 Q9VSF3 Q9VSF4 Q9VSH0 Q9VSH4 Q9VSK8 Q9VSK9 Q9VSL3 Q9VSS7 Q9VSY4
Q9VSY6 Q9VSZ6 Q9VT28 Q9VT41 Q9VT57 Q9VT67 Q9VT70 Q9VTC2 Q9VTC4 Q9VTE0 Q9VTE2
Q9VTE5 Q9VTE9 Q9VTF9 Q9VTM5 Q9VTT9 Q9VTU0 Q9VTU4 Q9VTV1 Q9VTY4 Q9VTY6 Q9VTZ6
Q9VU02 Q9VU31 Q9VU41 Q9VU65 Q9VU68 Q9VU70 Q9VU95 Q9VUA0 Q9VUC6 Q9VUJ0 Q9VUK8
Q9VUM0 Q9VUN3 Q9VUQ5 Q9VUQ8 Q9VUR0 Q9VUR3 Q9VUR7 Q9VUU5 Q9VUV9 Q9VUX2 Q9VUY9
Q9VUZ1 Q9VV41 Q9VV43 Q9VV73 Q9VVA0 Q9VVA6 Q9VVB4 Q9VVE2 Q9VVE5 Q9VVI2 Q9VVK8
Q9VVL6 Q9VVN2 Q9VVN4 Q9VVR0 Q9VVR2 Q9VVR6 Q9VVS4 Q9VVU5 Q9VVW8 Q9VVX0 Q9VVY3
Q9VW09 Q9VW14 Q9VW15 Q9VW26 Q9VW27 Q9VW47 Q9VW51 Q9VW56 Q9VW97 Q9VWA2 Q9VWB0
Q9VWD4 Q9VWD6 Q9VWE6 Q9VWF2 Q9VWH4 Q9VWI4 Q9VWN4 Q9VWN9 Q9VWP4 Q9VWQ2 Q9VWV8
Q9VWY6 Q9VX09 Q9VX10 Q9VX25 Q9VX88 Q9VX91 Q9VX98 Q9VXA8 Q9VXB0 Q9VXB3 Q9VXB5
Q9VXE0 Q9VXF1 Q9VXG1 Q9VXG4 Q9VXG8 Q9VXH6 Q9VXK6 Q9VXN0 Q9VXN4 Q9VXP4 Q9VXT5
Q9VXT7 Q9VXY4 Q9VY01 Q9VY28 Q9VY77 Q9VY93 Q9VY99 Q9VYA0 Q9VYA8 Q9VYD1 Q9VYG1
Q9VYH3 Q9VYI3 Q9VYQ0 Q9VYR0 Q9VYS3 Q9VYS4 Q9VYY3 Q9VYY9 Q9VZ64 Q9VZD2 Q9VZD5
Q9VZD8 Q9VZE6 Q9VZF4 Q9VZH1 Q9VZI3 Q9VZM7 Q9VZN4 Q9VZQ1 Q9VZS3 Q9VZV5 Q9VZX9
Q9W020 Q9W021 Q9W032 Q9W040 Q9W058 Q9W062 Q9W074 Q9W088 Q9W0E8 Q9W0G6 Q9W0G7
Q9W0K0 Q9W0K4 Q9W0K7 Q9W0M7 Q9W0P2 Q9W0P3 Q9W0P5 Q9W0P8 Q9W0S7 Q9W0S8 Q9W0T1
Q9W0Y1 Q9W0Y2 Q9W107 Q9W148 Q9W157 Q9W197 Q9W1A2 Q9W1B0 Q9W1G0 Q9W1H4 Q9W1I2
Q9W1I9 Q9W1J3 Q9W1K5 Q9W1R4 Q9W1R5 Q9W1U4 Q9W1U6 Q9W1V1 Q9W1V3 Q9W1V7 Q9W1X7

Q9W1X8 Q9W252 Q9W256 Q9W261 Q9W265 Q9W278 Q9W2D9 Q9W2E7 Q9W2F2 Q9W2H9 Q9W2P0
Q9W2Q7 Q9W2S3 Q9W2U4 Q9W2X7 Q9W2Y3 Q9W328 Q9W351 Q9W352 Q9W369 Q9W3C0 Q9W3C1
Q9W3C2 Q9W3E1 Q9W3J5 Q9W3J8 Q9W3K5 Q9W3L5 Q9W3M2 Q9W3M3 Q9W3M9 Q9W3U1 Q9W3Y0
Q9W3Y4 Q9W3Y5 Q9W3Z3 Q9W401 Q9W415 Q9W420 Q9W445 Q9W484 Q9W485 Q9W4D2 Q9W4I5
Q9W4J5 Q9W4L1 Q9W4M8 Q9W4M9 Q9W4N2 Q9W4S1 Q9W4X7 Q9W517 Q9W539 Q9W590 Q9W596
Q9W5B6 Q9W5B9 Q9W5D0 Q9W5E0 Q9W5E4 Q9W5N0 Q9W5P1 Q9W5X9 Q9W5Z3 Q9W625 Q9W636
Q9W638 Q9W644 Q9W669 Q9W676 Q9W690 Q9W698 Q9W6B3 Q9W6D8 Q9W6G7 Q9W6I9 Q9W6K1
Q9W6K2 Q9W6M2 Q9W6N4 Q9W6S5 Q9W6S8 Q9W6U8 Q9W6X3 Q9W6X9 Q9W6Y0 Q9W6Z2 Q9W704
Q9W706 Q9W707 Q9W719 Q9W739 Q9W747 Q9W757 Q9W758 Q9W769 Q9W785 Q9W790 Q9W7C2
Q9W7E8 Q9W7I5 Q9W7K5 Q9W7L0 Q9W7L1 Q9W7L2 Q9W7L3 Q9W7L4 Q9W7L5 Q9W7M6 Q9W7M8
Q9W7R5 Q9W7R6 Q9WAB1 Q9WAB2 Q9WAL9 Q9WBI3 Q9WC28 Q9WC80 Q9WDG3 Q9WDG4 Q9WDG5
Q9WDG7 Q9WF19 Q9WFZ0 Q9WGY9 Q9WGZ0 Q9WGZ1 Q9WI42 Q9WIB9 Q9WIJ4 Q9WIJ6 Q9WIJ8
Q9WIJ9 Q9WIK1 Q9WJR5 Q9WMB3 Q9WMB4 Q9WMB5 Q9WQ76 Q9WQ77 Q9WRL6 Q9WRL7 Q9WS38
Q9WS39 Q9WS40 Q9WSV7 Q9WSV8 Q9WSZ6 Q9WT06 Q9WT32 Q9WT35 Q9WT39 Q9WT45 Q9WT54
Q9WTJ2 Q9WTJ4 Q9WTJ6 Q9WTJ8 Q9WTK2 Q9WTK8 Q9WTM4 Q9WTP3 Q9WTP7 Q9WTP9 Q9WTQ7
Q9WTT6 Q9WTT7 Q9WTU0 Q9WTV5 Q9WTV7 Q9WTW4 Q9WTX2 Q9WTX7 Q9WTY1 Q9WTY8 Q9WTZ0
Q9WU06 Q9WU11 Q9WU12 Q9WU20 Q9WU28 Q9WU49 Q9WU56 Q9WU84 Q9WUA2 Q9WUB3 Q9WUC4
Q9WUC8 Q9WUD8 Q9WUF2 Q9WUF3 Q9WUH2 Q9WUH5 Q9WUI0 Q9WUJ3 Q9WUJ8 Q9WUK2 Q9WUK4
Q9WUK6 Q9WUL0 Q9WUL8 Q9WUN8 Q9WUQ3 Q9WUQ7 Q9WUR9 Q9WUS0 Q9WUU6 Q9WUV0 Q9WUW8
Q9WUW9 Q9WUZ5 Q9WUZ7 Q9WV04 Q9WV23 Q9WV25 Q9WV32 Q9WV35 Q9WV50 Q9WV59 Q9WV70
Q9WV85 Q9WV93 Q9WVA4 Q9WVB6 Q9WVB7 Q9WVG9 Q9WVH0 Q9WVH3 Q9WVJ2 Q9WVJ5 Q9WVL0
Q9WVL5 Q9WVM3 Q9WVM7 Q9WVM8 Q9WVP6 Q9WVQ0 Q9WVQ5 Q9WVR7 Q9WVR8 Q9WVX8 Q9WW00
Q9WW19 Q9WW25 Q9WW32 Q9WW67 Q9WWG9 Q9WWL3 Q9WWL4 Q9WWN1 Q9WWU5 Q9WWW0 Q9WWW4
Q9WWW5 Q9WX14 Q9WX15 Q9WX16 Q9WX29 Q9WX37 Q9WX39 Q9WX53 Q9WX57 Q9WX64 Q9WX69
Q9WX76 Q9WX94 Q9WX96 Q9WXA9 Q9WXB4 Q9WXB5 Q9WXB7 Q9WXD8 Q9WXE6 Q9WXE8 Q9WXF2
Q9WXG7 Q9WXK7 Q9WXL5 Q9WXM1 Q9WXM9 Q9WXP3 Q9WXP6 Q9WXR9 Q9WXS3 Q9WXS4 Q9WXT2
Q9WXT6 Q9WXU2 Q9WXV2 Q9WXV3 Q9WXV5 Q9WXW3 Q9WXZ2 Q9WXZ3 Q9WXZ5 Q9WXZ7 Q9WY06
Q9WY12 Q9WY14 Q9WY15 Q9WY16 Q9WY22 Q9WY28 Q9WY40 Q9WY43 Q9WY51 Q9WY52 Q9WY54
Q9WY55 Q9WY56 Q9WY57 Q9WY59 Q9WY60 Q9WY68 Q9WY73 Q9WY76 Q9WY79 Q9WY82 Q9WY93
Q9WY94 Q9WY95 Q9WY96 Q9WYA1 Q9WYA3 Q9WYA4 Q9WYB1 Q9WYB3 Q9WYC5 Q9WYC7 Q9WYC8
Q9WYC9 Q9WYD0 Q9WYD1 Q9WYG0 Q9WYG1 Q9WYG6 Q9WYG7 Q9WYG8 Q9WYG9 Q9WYH0 Q9WYH2
Q9WYH8 Q9WYI0 Q9WYI1 Q9WYI2 Q9WYI3 Q9WYI4 Q9WYJ4 Q9WYJ7 Q9WYJ9 Q9WYK0 Q9WYK4
Q9WYK6 Q9WYK8 Q9WYM8 Q9WYM9 Q9WYN2 Q9WYP3 Q9WYP4 Q9WYP5 Q9WYP7 Q9WYQ4 Q9WYS1
Q9WYS7 Q9WYT0 Q9WYT7 Q9WYU3 Q9WYU4 Q9WYU8 Q9WYV0 Q9WYV8 Q9WYW0 Q9WYW2 Q9WYX5
Q9WYX6 Q9WYX8 Q9WYY1 Q9WYY4 Q9WYZ0 Q9WYZ1 Q9WYZ2 Q9WYZ5 Q9WYZ6 Q9WYZ8 Q9WZ09
Q9WZ15 Q9WZ20 Q9WZ21 Q9WZ22 Q9WZ23 Q9WZ24 Q9WZ25 Q9WZ26 Q9WZ27 Q9WZ28 Q9WZ35
Q9WZ44 Q9WZ45 Q9WZ48 Q9WZ56 Q9WZ72 Q9WZ73 Q9WZ74 Q9WZB3 Q9WZB5 Q9WZC1 Q9WZC2
Q9WZC3 Q9WZC6 Q9WZF3 Q9WZF8 Q9WZF9 Q9WZG6 Q9WZG8 Q9WZH8 Q9WZH9 Q9WZI0 Q9WZI1
Q9WZI9 Q9WZJ0 Q9WZJ3 Q9WZJ9 Q9WZK0 Q9WZL2 Q9WZM1 Q9WZM2 Q9WZM6 Q9WZN3 Q9WZP2
Q9WZP3 Q9WZP4 Q9WZP5 Q9WZP7 Q9WZP8 Q9WZQ4 Q9WZR0 Q9WZR1 Q9WZR4 Q9WZS8 Q9WZS9
Q9WZT7 Q9WZU8 Q9WZV3 Q9WZV4 Q9WZV5 Q9WZV9 Q9WZW0 Q9WZW7 Q9WZX6 Q9WZX7 Q9WZY3
Q9WZY4 Q9WZY5 Q9WZZ1 Q9WZZ3 Q9WZZ7 Q9X005 Q9X006 Q9X013 Q9X015 Q9X016 Q9X017
Q9X024 Q9X027 Q9X034 Q9X037 Q9X049 Q9X054 Q9X059 Q9X078 Q9X081 Q9X095 Q9X0C4
Q9X0C5 Q9X0C6 Q9X0C7 Q9X0C8 Q9X0C9 Q9X0D0 Q9X0D1 Q9X0D2 Q9X0D3 Q9X0D7 Q9X0E6

Q9X0G6 Q9X0G8 Q9X0H0 Q9X0H1 Q9X0H5 Q9X0H8 Q9X0H9 Q9X0I0 Q9X0I3 Q9X0I6 Q9X0L3
Q9X0L4 Q9X0N8 Q9X0N9 Q9X0P5 Q9X0Q4 Q9X0R7 Q9X0V7 Q9X0X0 Q9X0X1 Q9X0X2 Q9X0X3
Q9X0X6 Q9X0X7 Q9X0Y0 Q9X0Y1 Q9X0Y2 Q9X0Z6 Q9X0Z7 Q9X0Z9 Q9X100 Q9X105 Q9X108
Q9X112 Q9X113 Q9X137 Q9X139 Q9X145 Q9X165 Q9X166 Q9X172 Q9X183 Q9X186 Q9X191
Q9X192 Q9X199 Q9X1A3 Q9X1A4 Q9X1A5 Q9X1A7 Q9X1B3 Q9X1B4 Q9X1B6 Q9X1B8 Q9X1B9
Q9X1C0 Q9X1C1 Q9X1D0 Q9X1D7 Q9X1E1 Q9X1E4 Q9X1F1 Q9X1F4 Q9X1F5 Q9X1F6 Q9X1F7
Q9X1F8 Q9X1G3 Q9X1G4 Q9X1G5 Q9X1G7 Q9X1G9 Q9X1H4 Q9X1H5 Q9X1H7 Q9X1I0 Q9X1I1
Q9X1I2 Q9X1I3 Q9X1I4 Q9X1I5 Q9X1I6 Q9X1I8 Q9X1J0 Q9X1J1 Q9X1J2 Q9X1J3 Q9X1J7
Q9X1K1 Q9X1K5 Q9X1K7 Q9X1K8 Q9X1K9 Q9X1L0 Q9X1N1 Q9X1N7 Q9X1P2 Q9X1P3 Q9X1P5
Q9X1P7 Q9X1Q0 Q9X1Q2 Q9X1Q3 Q9X1Q4 Q9X1Q5 Q9X1Q6 Q9X1Q7 Q9X1R0 Q9X1R5 Q9X1S1
Q9X1S6 Q9X1S7 Q9X1S8 Q9X1T3 Q9X1T8 Q9X1U0 Q9X1U1 Q9X1V3 Q9X1W1 Q9X1W2 Q9X1W3
Q9X1X1 Q9X1X4 Q9X1X6 Q9X1X7 Q9X1Y4 Q9X1Y7 Q9X1Y8 Q9X1Z5 Q9X214 Q9X215 Q9X216
Q9X220 Q9X231 Q9X232 Q9X233 Q9X234 Q9X235 Q9X239 Q9X240 Q9X242 Q9X251 Q9X255
Q9X276 Q9X278 Q9X282 Q9X284 Q9X286 Q9X287 Q9X288 Q9X289 Q9X290 Q9X291 Q9X295
Q9X299 Q9X2A1 Q9X2A2 Q9X2A3 Q9X2A4 Q9X2A5 Q9X2B6 Q9X2B7 Q9X2D7 Q9X2E0 Q9X2E1
Q9X2E5 Q9X2E6 Q9X2F4 Q9X2H3 Q9X2H4 Q9X2H6 Q9X2H9 Q9X2I8 Q9X2M2 Q9X2N2 Q9X2N4
Q9X2T1 Q9X2V5 Q9X2V8 Q9X2V9 Q9X315 Q9X3W1 Q9X3W2 Q9X3W5 Q9X3X1 Q9X3X2 Q9X3X5
Q9X3X7 Q9X3Y3 Q9X3Y5 Q9X3Y6 Q9X404 Q9X405 Q9X406 Q9X448 Q9X449 Q9X498 Q9X4A0
Q9X4A7 Q9X4D0 Q9X4E0 Q9X4E2 Q9X4E3 Q9X4E5 Q9X4E8 Q9X4F5 Q9X4F7 Q9X4G0 Q9X4H1
Q9X4H2 Q9X4H5 Q9X4I6 Q9X4K2 Q9X4K8 Q9X4M1 Q9X4M8 Q9X4N0 Q9X4P2 Q9X4Q2 Q9X4R2
Q9X4V6 Q9X4W3 Q9X4Y0 Q9X519 Q9X520 Q9X521 Q9X525 Q9X528 Q9X576 Q9X597 Q9X5C9
Q9X5D0 Q9X5D1 Q9X5D2 Q9X5D8 Q9X5E7 Q9X5E8 Q9X5E9 Q9X5F0 Q9X5F1 Q9X5F2 Q9X5F8
Q9X5G1 Q9X5M0 Q9X5M1 Q9X5N0 Q9X5N1 Q9X5P2 Q9X5P5 Q9X5T6 Q9X5U0 Q9X5U8 Q9X5U9
Q9X5V1 Q9X5V4 Q9X5W3 Q9X5W6 Q9X5X4 Q9X5Y0 Q9X5Z9 Q9X600 Q9X601 Q9X670 Q9X682
Q9X696 Q9X697 Q9X698 Q9X6C6 Q9X6C9 Q9X6E7 Q9X6E8 Q9X6G2 Q9X6J4 Q9X6J5 Q9X6J6
Q9X6M5 Q9X6N4 Q9X6R0 Q9X6U2 Q9X6V8 Q9X6V9 Q9X6W6 Q9X6W7 Q9X6W8 Q9X6W9 Q9X6X4
Q9X6Y1 Q9X6Y2 Q9X6Y8 Q9X6Y9 Q9X706 Q9X712 Q9X713 Q9X719 Q9X725 Q9X758 Q9X764
Q9X765 Q9X781 Q9X783 Q9X784 Q9X794 Q9X795 Q9X796 Q9X797 Q9X798 Q9X799 Q9X7A0
Q9X7A1 Q9X7A2 Q9X7B8 Q9X7B9 Q9X7C0 Q9X7C2 Q9X7C3 Q9X7C5 Q9X7C7 Q9X7D6 Q9X7E5
Q9X7F0 Q9X7F1 Q9X7F4 Q9X7F6 Q9X7G7 Q9X7K0 Q9X7L1 Q9X7L2 Q9X7L3 Q9X7L4 Q9X7M4
Q9X7N2 Q9X7Q6 Q9X7R6 Q9X7R9 Q9X7S0 Q9X7T2 Q9X7U5 Q9X7W2 Q9X7W3 Q9X827 Q9X839
Q9X844 Q9X895 Q9X8H0 Q9X8H1 Q9X8H2 Q9X8I0 Q9X8I3 Q9X8I6 Q9X8I9 Q9X8J2 Q9X8K4
Q9X8K6 Q9X8K7 Q9X8K8 Q9X8K9 Q9X8N6 Q9X8N8 Q9X8P6 Q9X8R6 Q9X8R7 Q9X8U2 Q9X8U3
Q9X8U5 Q9X8W0 Q9X909 Q9X928 Q9X980 Q9X981 Q9X9D5 Q9X9E0 Q9X9F2 Q9X9F3 Q9X9F6
Q9X9J3 Q9X9Q0 Q9X9Q6 Q9X9Q7 Q9X9S0 Q9X9U0 Q9X9V6 Q9X9W0 Q9X9X7 Q9X9Z8 Q9XA16
Q9XA76 Q9XAB7 Q9XAC0 Q9XAD5 Q9XAI3 Q9XAI4 Q9XAK6 Q9XAP2 Q9XAP8 Q9XAQ2 Q9XAT2
Q9XAU7 Q9XAY7 Q9XAZ0 Q9XAZ1 Q9XB01 Q9XB14 Q9XB20 Q9XB21 Q9XB22 Q9XB42 Q9XB53
Q9XB58 Q9XB59 Q9XB60 Q9XB61 Q9XBC1 Q9XBG6 Q9XBH4 Q9XBI3 Q9XBK3 Q9XBL1 Q9XBL8
Q9XBM3 Q9XBM6 Q9XBQ0 Q9XBQ3 Q9XBQ8 Q9XBQ9 Q9XBR5 Q9XBR7 Q9XBS9 Q9XC07 Q9XC11
Q9XC19 Q9XC20 Q9XC21 Q9XC67 Q9XC89 Q9XCA6 Q9XCA9 Q9XCB0 Q9XCB1 Q9XCD6 Q9XCI8
Q9XCL6 Q9XCM5 Q9XCV1 Q9XCX7 Q9XCX9 Q9XD08 Q9XD09 Q9XD10 Q9XD11 Q9XD12 Q9XD13
Q9XD14 Q9XD15 Q9XD17 Q9XD18 Q9XD19 Q9XD20 Q9XD21 Q9XD22 Q9XD23 Q9XD24 Q9XD25
Q9XD26 Q9XD27 Q9XD28 Q9XD29 Q9XD30 Q9XD31 Q9XD32 Q9XD33 Q9XD34 Q9XD35 Q9XD36
Q9XD37 Q9XD38 Q9XD64 Q9XD73 Q9XD74 Q9XD77 Q9XDB4 Q9XDH5 Q9XDH6 Q9XDL1 Q9XDM0

Q9XDM1 Q9XDM3 Q9XDM4 Q9XDM6 Q9XDM8 Q9XDN0 Q9XDN3 Q9XDN4 Q9XDN5 Q9XDN6 Q9XDP3
Q9XDU4 Q9XDZ7 Q9XE33 Q9XE58 Q9XE98 Q9XEA0 Q9XEA6 Q9XEA8 Q9XEC1 Q9XEC3 Q9XED7
Q9XED8 Q9XEE6 Q9XEE8 Q9XEF0 Q9XEG7 Q9XEH9 Q9XEI0 Q9XEK6 Q9XEK7 Q9XEK8 Q9XEK9
Q9XER9 Q9XES0 Q9XES5 Q9XET4 Q9XEX4 Q9XF14 Q9XF19 Q9XF37 Q9XF38 Q9XF39 Q9XF40
Q9XF41 Q9XF42 Q9XF47 Q9XF62 Q9XF97 Q9XFA2 Q9XFB0 Q9XFB1 Q9XFB6 Q9XFB7 Q9XFG3
Q9XFH3 Q9XFH8 Q9XFH9 Q9XFI1 Q9XFM0 Q9XFR9 Q9XFS9 Q9XFW3 Q9XFY6 Q9XG56 Q9XG57
Q9XG77 Q9XG83 Q9XG85 Q9XG98 Q9XG99 Q9XGA0 Q9XGA1 Q9XGC9 Q9XGD5 Q9XGD6 Q9XGH7
Q9XGI7 Q9XGI9 Q9XGJ4 Q9XGL4 Q9XGM2 Q9XGN1 Q9XGN3 Q9XGN4 Q9XGP7 Q9XGR4 Q9XGS0
Q9XGV9 Q9XGW0 Q9XGW1 Q9XGX1 Q9XGX2 Q9XGX4 Q9XGX5 Q9XGZ0 Q9XGZ9 Q9XH06 Q9XH32
Q9XH35 Q9XH36 Q9XH37 Q9XH38 Q9XH42 Q9XH43 Q9XH44 Q9XH45 Q9XHE4 Q9XHG2 Q9XHH2
Q9XHH3 Q9XHL7 Q9XHM1 Q9XHM5 Q9XHR2 Q9XHS0 Q9XHV0 Q9XHY7 Q9XHZ3 Q9XHZ8 Q9XI00
Q9XI05 Q9XI07 Q9XI17 Q9XI21 Q9XI36 Q9XI37 Q9XI47 Q9XI84 Q9XI90 Q9XIA2 Q9XIB4
Q9XID4 Q9XIE1 Q9XIE4 Q9XIE8 Q9XIE9 Q9XIF0 Q9XIF2 Q9XIF4 Q9XIF8 Q9XIJ1 Q9XIK0
Q9XIK3 Q9XIK4 Q9XIK5 Q9XIK6 Q9XIL5 Q9XIM0 Q9XIM8 Q9XIN8 Q9XIP5 Q9XIP8 Q9XIQ4
Q9XIQ5 Q9XIR1 Q9XIR4 Q9XJ27 Q9XJ29 Q9XJ32 Q9XJ35 Q9XJ36 Q9XJ54 Q9XJ56 Q9XJ57
Q9XJ58 Q9XJ60 Q9XJ66 Q9XJG4 Q9XJP3 Q9XJQ2 Q9XJQ3 Q9XJQ4 Q9XPI7 Q9XPI8 Q9XPN6
Q9XPP0 Q9XPP1 Q9XPP4 Q9XPP8 Q9XPR5 Q9XPS5 Q9XPS7 Q9XPS8 Q9XPS9 Q9XQ94 Q9XQE1
Q9XQJ6 Q9XQN8 Q9XQQ6 Q9XQQ7 Q9XR19 Q9XS37 Q9XS39 Q9XS41 Q9XS96 Q9XS97 Q9XSB5
Q9XSB7 Q9XSC6 Q9XSE9 Q9XSG3 Q9XSH5 Q9XSI2 Q9XSK0 Q9XSK1 Q9XSK8 Q9XSN2 Q9XSN3
Q9XSR1 Q9XSR3 Q9XST5 Q9XSU0 Q9XSU3 Q9XSU4 Q9XSW2 Q9XT06 Q9XT11 Q9XT28 Q9XT60
Q9XT62 Q9XT73 Q9XTA2 Q9XTF3 Q9XTI0 Q9XTJ4 Q9XTL9 Q9XTQ5 Q9XTR1 Q9XTT8 Q9XTT9
Q9XTU6 Q9XUB2 Q9XUB3 Q9XUC2 Q9XUE6 Q9XUE7 Q9XUK2 Q9XUK7 Q9XUM6 Q9XUN9 Q9XUP3
Q9XUS2 Q9XUV0 Q9XV46 Q9XV51 Q9XV52 Q9XVA4 Q9XVD5 Q9XVH6 Q9XVI9 Q9XVJ2 Q9XVN3
Q9XVR8 Q9XVS1 Q9XVS4 Q9XVS7 Q9XVS8 Q9XVT0 Q9XVV3 Q9XVZ8 Q9XW10 Q9XW12 Q9XW13
Q9XW15 Q9XW16 Q9XW17 Q9XW26 Q9XW42 Q9XW78 Q9XW79 Q9XW88 Q9XWB9 Q9XWE6 Q9XWF4
Q9XWF6 Q9XWG3 Q9XWH0 Q9XWI6 Q9XWJ1 Q9XWN6 Q9XWN9 Q9XWP6 Q9XWQ1 Q9XWQ6 Q9XWR1
Q9XWS0 Q9XWV3 Q9XWV5 Q9XX11 Q9XX14 Q9XX15 Q9XX47 Q9XXD4 Q9XXJ0 Q9XXL3 Q9XXM8
Q9XY07 Q9XY92 Q9XYB7 Q9XYC2 Q9XYD3 Q9XYL0 Q9XYN7 Q9XYP3 Q9XYP7 Q9XYP8 Q9XYU0
Q9XYU1 Q9XYU2 Q9XYZ5 Q9XZ34 Q9XZ56 Q9XZ58 Q9XZ71 Q9XZD6 Q9XZE5 Q9XZF0 Q9XZG5
Q9XZJ3 Q9XZJ4 Q9XZP2 Q9XZS3 Q9XZT1 Q9XZT6 Q9XZT7 Q9Y041 Q9Y091 Q9Y0A7 Q9Y0B7
Q9Y0H3 Q9Y0H7 Q9Y0I1 Q9Y0T3 Q9Y0T4 Q9Y0Y2 Q9Y103 Q9Y105 Q9Y106 Q9Y113 Q9Y118
Q9Y149 Q9Y1A9 Q9Y1B0 Q9Y1C1 Q9Y221 Q9Y222 Q9Y224 Q9Y235 Q9Y236 Q9Y238 Q9Y242
Q9Y247 Q9Y253 Q9Y2A4 Q9Y2B5 Q9Y2B9 Q9Y2D0 Q9Y2D1 Q9Y2E4 Q9Y2F9 Q9Y2G2 Q9Y2G7
Q9Y2G9 Q9Y2H8 Q9Y2I8 Q9Y2J8 Q9Y2L8 Q9Y2L9 Q9Y2M2 Q9Y2M5 Q9Y2P0 Q9Y2P7 Q9Y2P8
Q9Y2Q1 Q9Y2S2 Q9Y2S6 Q9Y2T3 Q9Y2T4 Q9Y2T7 Q9Y2V0 Q9Y2W6 Q9Y2X8 Q9Y2Y1 Q9Y2Y4
Q9Y2Y9 Q9Y2Z2 Q9Y2Z4 Q9Y303 Q9Y305 Q9Y312 Q9Y315 Q9Y324 Q9Y333 Q9Y366 Q9Y3A5
Q9Y3B1 Q9Y3B4 Q9Y3B8 Q9Y3C4 Q9Y3C7 Q9Y3C8 Q9Y3D0 Q9Y3D2 Q9Y3F1 Q9Y3F4 Q9Y3I1
Q9Y3M2 Q9Y3M9 Q9Y3P9 Q9Y3T6 Q9Y3V2 Q9Y421 Q9Y458 Q9Y462 Q9Y466 Q9Y468 Q9Y483
Q9Y4A0 Q9Y4A8 Q9Y4F4 Q9Y4I5 Q9Y4M8 Q9Y4R7 Q9Y4X0 Q9Y4X5 Q9Y4Y9 Q9Y4Z2 Q9Y543
Q9Y547 Q9Y573 Q9Y577 Q9Y586 Q9Y5A6 Q9Y5B8 Q9Y5K1 Q9Y5L5 Q9Y5M1 Q9Y5N5 Q9Y5P2
Q9Y5P3 Q9Y5P6 Q9Y5P8 Q9Y5Q3 Q9Y5Q8 Q9Y5R4 Q9Y5R5 Q9Y5R8 Q9Y5V0 Q9Y5Y2 Q9Y600
Q9Y603 Q9Y605 Q9Y614 Q9Y615 Q9Y617 Q9Y620 Q9Y623 Q9Y651 Q9Y675 Q9Y692 Q9Y697
Q9Y6A4 Q9Y6B2 Q9Y6C7 Q9Y6F7 Q9Y6F8 Q9Y6H1 Q9Y6I0 Q9Y6J9 Q9Y6K8 Q9Y6Q3 Q9Y6R6
Q9Y6S9 Q9Y6V7 Q9Y6X3 Q9Y6X8 Q9Y6Z2 Q9Y6Z4 Q9Y6Z5 Q9Y6Z8 Q9Y6Z9 Q9Y700 Q9Y701

Q9Y702 Q9Y703 Q9Y705 Q9Y707 Q9Y708 Q9Y713 Q9Y720 Q9Y725 Q9Y726 Q9Y738 Q9Y767
Q9Y783 Q9Y794 Q9Y796 Q9Y7B5 Q9Y7B6 Q9Y7C4 Q9Y7C9 Q9Y7D0 Q9Y7D1 Q9Y7D3 Q9Y7D5
Q9Y7F0 Q9Y7G2 Q9Y7G5 Q9Y7I1 Q9Y7I3 Q9Y7I8 Q9Y7J3 Q9Y7J4 Q9Y7J5 Q9Y7J8 Q9Y7K0
Q9Y7K1 Q9Y7K4 Q9Y7K5 Q9Y7K6 Q9Y7K9 Q9Y7L1 Q9Y7L2 Q9Y7L4 Q9Y7L9 Q9Y7M4 Q9Y7M5
Q9Y7M6 Q9Y7M8 Q9Y7N0 Q9Y7N1 Q9Y7N2 Q9Y7N4 Q9Y7N7 Q9Y7P0 Q9Y7P4 Q9Y7P6 Q9Y7P9
Q9Y7Q0 Q9Y7Q1 Q9Y7Q2 Q9Y7Q3 Q9Y7Q6 Q9Y7R1 Q9Y7R3 Q9Y7R4 Q9Y7R7 Q9Y7R8 Q9Y7S0
Q9Y7S2 Q9Y7S3 Q9Y7S6 Q9Y7S8 Q9Y7T1 Q9Y7T2 Q9Y7T3 Q9Y7T7 Q9Y7T8 Q9Y7U4 Q9Y7U7
Q9Y7U8 Q9Y7U9 Q9Y7V1 Q9Y7V2 Q9Y7W4 Q9Y7X2 Q9Y7X5 Q9Y7X8 Q9Y7X9 Q9Y7Y1 Q9Y7Y2
Q9Y7Y3 Q9Y7Y5 Q9Y7Y8 Q9Y7Z1 Q9Y7Z3 Q9Y7Z6 Q9Y7Z7 Q9Y802 Q9Y803 Q9Y804 Q9Y805
Q9Y807 Q9Y808 Q9Y809 Q9Y812 Q9Y813 Q9Y814 Q9Y815 Q9Y818 Q9Y820 Q9Y821 Q9Y822
Q9Y823 Q9Y825 Q9Y826 Q9Y829 Q9Y872 Q9Y875 Q9Y882 Q9Y885 Q9Y896 Q9Y897 Q9Y898
Q9Y899 Q9Y8A5 Q9Y8B5 Q9Y8C1 Q9Y8C2 Q9Y8C7 Q9Y8D1 Q9Y8D3 Q9Y8D9 Q9Y8G3 Q9Y8G8
Q9Y8H3 Q9Y8H7 Q9Y8H8 Q9Y8I1 Q9Y8I2 Q9Y8J2 Q9Y8J3 Q9Y8J4 Q9Y8K8 Q9Y8K9 Q9Y8N2
Q9Y8Q3 Q9Y8T0 Q9Y8T2 Q9Y8T3 Q9Y8T5 Q9Y8T6 Q9Y8T7 Q9Y904 Q9Y914 Q9Y924 Q9Y933
Q9Y935 Q9Y942 Q9Y948 Q9Y9A6 Q9Y9A7 Q9Y9B0 Q9Y9B3 Q9Y9B6 Q9Y9C1 Q9Y9D8 Q9Y9D9
Q9Y9E3 Q9Y9E6 Q9Y9F0 Q9Y9F2 Q9Y9G0 Q9Y9G4 Q9Y9H1 Q9Y9H2 Q9Y9I0 Q9Y9I3 Q9Y9I6
Q9Y9I9 Q9Y9J0 Q9Y9J2 Q9Y9L0 Q9Y9P1 Q9Y9T6 Q9Y9T8 Q9Y9T9 Q9Y9U3 Q9Y9U7 Q9Y9V7
Q9Y9W3 Q9Y9W5 Q9Y9W6 Q9Y9W8 Q9Y9W9 Q9Y9X3 Q9Y9Z0 Q9YA11 Q9YA13 Q9YA15 Q9YA18
Q9YA27 Q9YA42 Q9YA47 Q9YA48 Q9YA49 Q9YA53 Q9YA60 Q9YA64 Q9YA66 Q9YA67 Q9YA72
Q9YA75 Q9YA91 Q9YAB0 Q9YAB6 Q9YAC2 Q9YAC7 Q9YAD3 Q9YAD5 Q9YAD8 Q9YAE7 Q9YAF1
Q9YAF3 Q9YAF9 Q9YAG3 Q9YAH7 Q9YAI1 Q9YAK0 Q9YAN6 Q9YAN8 Q9YAQ8 Q9YAS0 Q9YAS4
Q9YAT1 Q9YAT3 Q9YAU2 Q9YAU3 Q9YAU5 Q9YAU8 Q9YAV0 Q9YAV2 Q9YAV8 Q9YAW0 Q9YAW1
Q9YAW4 Q9YAX2 Q9YAX7 Q9YAZ0 Q9YAZ1 Q9YAZ5 Q9YAZ7 Q9YB01 Q9YB03 Q9YB04 Q9YB13
Q9YB24 Q9YB30 Q9YB37 Q9YB39 Q9YB42 Q9YB44 Q9YB45 Q9YB47 Q9YB48 Q9YB50 Q9YB51
Q9YB53 Q9YB55 Q9YB58 Q9YB60 Q9YB79 Q9YB80 Q9YBA2 Q9YBA5 Q9YBD4 Q9YBD5 Q9YBD6
Q9YBE9 Q9YBF0 Q9YBF1 Q9YBF8 Q9YBG9 Q9YBI2 Q9YBJ4 Q9YBK2 Q9YBK6 Q9YBK7 Q9YBL2
Q9YBM7 Q9YBN4 Q9YBQ2 Q9YBQ6 Q9YBR1 Q9YBR6 Q9YBS5 Q9YBS7 Q9YBU4 Q9YBW7 Q9YBX7
Q9YBX8 Q9YBY0 Q9YBY3 Q9YBY6 Q9YBY8 Q9YBY9 Q9YBZ8 Q9YC00 Q9YC01 Q9YC02 Q9YC03
Q9YC05 Q9YC06 Q9YC08 Q9YC11 Q9YC19 Q9YC65 Q9YC66 Q9YC75 Q9YC88 Q9YC89 Q9YCA4
Q9YCB6 Q9YCI2 Q9YCL9 Q9YCQ6 Q9YCR4 Q9YCR6 Q9YCU5 Q9YCV1 Q9YCV8 Q9YCX3 Q9YCX7
Q9YCY0 Q9YCY3 Q9YCZ5 Q9YD08 Q9YD10 Q9YD18 Q9YD20 Q9YD22 Q9YD25 Q9YD27 Q9YD28
Q9YD56 Q9YD97 Q9YDA1 Q9YDC3 Q9YDC5 Q9YDD2 Q9YDD5 Q9YDD7 Q9YDF6 Q9YDG2 Q9YDH4
Q9YDI2 Q9YDJ9 Q9YDK6 Q9YDM3 Q9YDM5 Q9YDM7 Q9YDP0 Q9YDP3 Q9YDV3 Q9YDW0 Q9YDW8
Q9YDX2 Q9YDY0 Q9YDY1 Q9YDY7 Q9YDY8 Q9YDZ1 Q9YE01 Q9YE02 Q9YE03 Q9YE60 Q9YE64
Q9YE67 Q9YE72 Q9YE84 Q9YE97 Q9YEA1 Q9YEA9 Q9YEB2 Q9YEC5 Q9YED6 Q9YEF2 Q9YEJ9
Q9YEK1 Q9YEK4 Q9YEK6 Q9YEK9 Q9YEL4 Q9YEM5 Q9YEN2 Q9YEN3 Q9YEN9 Q9YEQ4 Q9YEQ5
Q9YER0 Q9YER5 Q9YER8 Q9YES0 Q9YES4 Q9YES5 Q9YES9 Q9YET1 Q9YET5 Q9YEU2 Q9YEV3
Q9YEV5 Q9YEV6 Q9YEV9 Q9YEW8 Q9YEX9 Q9YEY2 Q9YEY5 Q9YEZ1 Q9YEZ5 Q9YEZ8 Q9YF00
Q9YF01 Q9YF02 Q9YF40 Q9YF67 Q9YF74 Q9YF76 Q9YF78 Q9YF79 Q9YF81 Q9YF82 Q9YF83
Q9YF85 Q9YF87 Q9YF89 Q9YF91 Q9YF92 Q9YF93 Q9YF94 Q9YF95 Q9YF96 Q9YF98 Q9YFA8
Q9YFG3 Q9YFI5 Q9YFI6 Q9YFJ2 Q9YFK2 Q9YFK4 Q9YFM0 Q9YFM1 Q9YFM2 Q9YFM7 Q9YFN1
Q9YFP2 Q9YFP5 Q9YFQ1 Q9YFQ8 Q9YFR7 Q9YFR9 Q9YFS7 Q9YFS9 Q9YFT8 Q9YFT9 Q9YFU8
Q9YFV2 Q9YFV3 Q9YFV8 Q9YFY3 Q9YFY5 Q9YFY7 Q9YFY8 Q9YFZ1 Q9YG22 Q9YG32 Q9YG68
Q9YG88 Q9YG90 Q9YGA3 Q9YGA5 Q9YGA7 Q9YGA9 Q9YGB0 Q9YGB1 Q9YGB2 Q9YGB3 Q9YGB4

Q9YGB5 Q9YGD2 Q9YGF2 Q9YGK8 Q9YGL1 Q9YGL2 Q9YGL3 Q9YGL7 Q9YGN1 Q9YGN6 Q9YGQ1
Q9YGS0 Q9YGS6 Q9YGT0 Q9YGT4 Q9YGT5 Q9YGT6 Q9YGV8 Q9YGW1 Q9YGW2 Q9YGW7 Q9YGX5
Q9YGY3 Q9YGY4 Q9YH12 Q9YH13 Q9YH16 Q9YH32 Q9YH33 Q9YH58 Q9YH71 Q9YH95 Q9YH98
Q9YH99 Q9YHA0 Q9YHA1 Q9YHB5 Q9YHC3 Q9YHC5 Q9YHE8 Q9YHT3 Q9YHU0 Q9YHZ5 Q9YHZ6
Q9YHZ7 Q9YI05 Q9YI36 Q9YI37 Q9YI97 Q9YI98 Q9YIB7 Q9YIC0 Q9YIC2 Q9YIT9 Q9YJJ8
Q9YJW2 Q9YJW3 Q9YK98 Q9YKD6 Q9YKK9 Q9YKL8 Q9YLR1 Q9YML3 Q9YMS9 Q9YMV2 Q9YNE2
Q9YNE3 Q9YPD4 Q9YPD5 Q9YPD7 Q9YPJ6 Q9YPK0 Q9YPS1 Q9YPS2 Q9YPS4 Q9YPS6 Q9YPV8
Q9YQ08 Q9YRB2 Q9YRB3 Q9YTR7 Q9YTR8 Q9YTU3 Q9YUC8 Q9YUC9 Q9YUD0 Q9YUD1 Q9YUD2
Q9YUD4 Q9YUS2 Q9YUS3 Q9YVX9 Q9YW06 Q9YW20 Q9YW29 Q9YW39 Q9YWQ0 Q9YWQ1 Q9YWQ2
Q9YX37 Q9YX38 Q9YX47 Q9YX48 Q9YX78 Q9YYS0 Q9YZN9 Q9YZR5 Q9Z035 Q9Z0C9 Q9Z0D0
Q9Z0D1 Q9Z0D2 Q9Z0F6 Q9Z0G7 Q9Z0G8 Q9Z0H1 Q9Z0H8 Q9Z0I6 Q9Z0I7 Q9Z0J5 Q9Z0N1
Q9Z0N2 Q9Z0S1 Q9Z0T0 Q9Z103 Q9Z104 Q9Z111 Q9Z118 Q9Z120 Q9Z129 Q9Z130 Q9Z144
Q9Z148 Q9Z176 Q9Z179 Q9Z180 Q9Z183 Q9Z184 Q9Z185 Q9Z191 Q9Z199 Q9Z1A9 Q9Z1B2
Q9Z1B8 Q9Z1D1 Q9Z1D7 Q9Z1D8 Q9Z1D9 Q9Z1E4 Q9Z1F9 Q9Z1J2 Q9Z1J3 Q9Z1K5 Q9Z1K6
Q9Z1L3 Q9Z1M8 Q9Z1M9 Q9Z1N2 Q9Z1N4 Q9Z1N5 Q9Z1N7 Q9Z1P7 Q9Z1Q1 Q9Z1Q9 Q9Z1R4
Q9Z1X9 Q9Z1Z2 Q9Z1Z6 Q9Z1Z9 Q9Z220 Q9Z221 Q9Z223 Q9Z224 Q9Z238 Q9Z244 Q9Z248
Q9Z255 Q9Z275 Q9Z277 Q9Z287 Q9Z2A5 Q9Z2D6 Q9Z2D7 Q9Z2D8 Q9Z2E1 Q9Z2E4 Q9Z2E5
Q9Z2G0 Q9Z2G1 Q9Z2G9 Q9Z2I9 Q9Z2J9 Q9Z2K0 Q9Z2K1 Q9Z2K3 Q9Z2K5 Q9Z2K8 Q9Z2K9
Q9Z2M2 Q9Z2M7 Q9Z2N6 Q9Z2P4 Q9Z2Q2 Q9Z2Q4 Q9Z2R9 Q9Z2S7 Q9Z2T6 Q9Z2U0 Q9Z2U1
Q9Z2U2 Q9Z2U4 Q9Z2V5 Q9Z2V9 Q9Z2W0 Q9Z2X1 Q9Z2Y0 Q9Z2Y1 Q9Z2Y8 Q9Z2Z9 Q9Z304
Q9Z315 Q9Z320 Q9Z321 Q9Z327 Q9Z331 Q9Z364 Q9Z377 Q9Z395 Q9Z3A0 Q9Z3B7 Q9Z3E6
Q9Z3F0 Q9Z3H5 Q9Z3H6 Q9Z3Q1 Q9Z3Q6 Q9Z3Q7 Q9Z3Q8 Q9Z3Q9 Q9Z3R0 Q9Z3R1 Q9Z3R2
Q9Z3R4 Q9Z3R8 Q9Z3S0 Q9Z3S1 Q9Z3S2 Q9Z3S3 Q9Z3S4 Q9Z3S6 Q9Z3T0 Q9Z3Z6 Q9Z408
Q9Z427 Q9Z431 Q9Z439 Q9Z441 Q9Z448 Q9Z451 Q9Z452 Q9Z462 Q9Z463 Q9Z470 Q9Z471
Q9Z472 Q9Z4P6 Q9Z4Q7 Q9Z4S1 Q9Z4S2 Q9Z4S3 Q9Z4V7 Q9Z4V8 Q9Z4W7 Q9Z4W9 Q9Z4X0
Q9Z4Z5 Q9Z4Z7 Q9Z507 Q9Z512 Q9Z513 Q9Z515 Q9Z518 Q9Z519 Q9Z520 Q9Z523 Q9Z527
Q9Z528 Q9Z565 Q9Z578 Q9Z580 Q9Z581 Q9Z585 Q9Z596 Q9Z598 Q9Z5B5 Q9Z5B6 Q9Z5D3
Q9Z5D7 Q9Z5D9 Q9Z5E9 Q9Z5H5 Q9Z5H8 Q9Z5I4 Q9Z5I7 Q9Z5I8 Q9Z5I9 Q9Z5J1 Q9Z5J2
Q9Z5K8 Q9Z5K9 Q9Z5U2 Q9Z5U7 Q9Z5V0 Q9Z5X0 Q9Z5X5 Q9Z5Y0 Q9Z5Z2 Q9Z613 Q9Z614
Q9Z615 Q9Z616 Q9Z617 Q9Z618 Q9Z660 Q9Z661 Q9Z671 Q9Z6B9 Q9Z6C3 Q9Z6D1 Q9Z6D2
Q9Z6E4 Q9Z6H4 Q9Z6J1 Q9Z6J2 Q9Z6J6 Q9Z6J7 Q9Z6J9 Q9Z6K0 Q9Z6K9 Q9Z6L0 Q9Z6L2
Q9Z6L3 Q9Z6L5 Q9Z6L7 Q9Z6M0 Q9Z6M1 Q9Z6M2 Q9Z6M3 Q9Z6M4 Q9Z6N1 Q9Z6N3 Q9Z6N4
Q9Z6N5 Q9Z6P2 Q9Z6P3 Q9Z6P6 Q9Z6Q0 Q9Z6Q9 Q9Z6R0 Q9Z6R2 Q9Z6R6 Q9Z6R7 Q9Z6R8
Q9Z6R9 Q9Z6S0 Q9Z6S1 Q9Z6S3 Q9Z6S4 Q9Z6S5 Q9Z6S7 Q9Z6T6 Q9Z6U0 Q9Z6U1 Q9Z6U7
Q9Z6V3 Q9Z6V4 Q9Z6V5 Q9Z6V6 Q9Z6V7 Q9Z6V8 Q9Z6W0 Q9Z6W4 Q9Z6W5 Q9Z6W6 Q9Z6W9
Q9Z6X0 Q9Z6X1 Q9Z6X2 Q9Z6X4 Q9Z6X5 Q9Z6Y2 Q9Z6Y8 Q9Z6Z6 Q9Z701 Q9Z705 Q9Z711
Q9Z714 Q9Z716 Q9Z720 Q9Z721 Q9Z729 Q9Z733 Q9Z734 Q9Z735 Q9Z736 Q9Z743 Q9Z755
Q9Z758 Q9Z759 Q9Z760 Q9Z762 Q9Z768 Q9Z773 Q9Z774 Q9Z794 Q9Z798 Q9Z7A0 Q9Z7A1
Q9Z7A2 Q9Z7A3 Q9Z7A4 Q9Z7A5 Q9Z7C9 Q9Z7D1 Q9Z7D3 Q9Z7E4 Q9Z7E8 Q9Z7E9 Q9Z7F0
Q9Z7F1 Q9Z7F2 Q9Z7F9 Q9Z7G1 Q9Z7G2 Q9Z7G3 Q9Z7G4 Q9Z7G7 Q9Z7H0 Q9Z7H1 Q9Z7H2
Q9Z7H3 Q9Z7I5 Q9Z7J1 Q9Z7J2 Q9Z7J4 Q9Z7J5 Q9Z7K0 Q9Z7K4 Q9Z7K6 Q9Z7K7 Q9Z7K8
Q9Z7K9 Q9Z7L5 Q9Z7M0 Q9Z7M3 Q9Z7M5 Q9Z7M9 Q9Z7N8 Q9Z7P1 Q9Z7P2 Q9Z7P5 Q9Z7P6
Q9Z7Q0 Q9Z7Q5 Q9Z7Q7 Q9Z7Q8 Q9Z7Q9 Q9Z7R0 Q9Z7R1 Q9Z7R2 Q9Z7R3 Q9Z7R4 Q9Z7R5
Q9Z7R6 Q9Z7R7 Q9Z7R8 Q9Z7R9 Q9Z7S0 Q9Z7S1 Q9Z7S2 Q9Z7S3 Q9Z7S4 Q9Z7S6 Q9Z7S7

Q9Z7S8 Q9Z7S9 Q9Z7T0 Q9Z7T1 Q9Z7T3 Q9Z7T4 Q9Z7T5 Q9Z7T7 Q9Z7U3 Q9Z7U6 Q9Z7V1
Q9Z7W0 Q9Z7W5 Q9Z7X4 Q9Z7X5 Q9Z7Y0 Q9Z7Y1 Q9Z7Y2 Q9Z7Y3 Q9Z7Y5 Q9Z7Y7 Q9Z7Z2
Q9Z7Z3 Q9Z800 Q9Z801 Q9Z802 Q9Z803 Q9Z805 Q9Z806 Q9Z807 Q9Z808 Q9Z819 Q9Z820
Q9Z826 Q9Z831 Q9Z832 Q9Z833 Q9Z834 Q9Z840 Q9Z843 Q9Z847 Q9Z848 Q9Z849 Q9Z850
Q9Z851 Q9Z852 Q9Z854 Q9Z856 Q9Z862 Q9Z863 Q9Z868 Q9Z873 Q9Z874 Q9Z876 Q9Z8A2
Q9Z8A5 Q9Z8A6 Q9Z8B0 Q9Z8B9 Q9Z8C1 Q9Z8C2 Q9Z8C5 Q9Z8C7 Q9Z8F1 Q9Z8F3 Q9Z8F4
Q9Z8F7 Q9Z8F8 Q9Z8F9 Q9Z8H0 Q9Z8I2 Q9Z8I3 Q9Z8J8 Q9Z8K0 Q9Z8K1 Q9Z8K3 Q9Z8K5
Q9Z8L0 Q9Z8L1 Q9Z8L2 Q9Z8L9 Q9Z8M0 Q9Z8M1 Q9Z8M3 Q9Z8M4 Q9Z8M9 Q9Z8N1 Q9Z8N9
Q9Z8Q7 Q9Z8R3 Q9Z8R4 Q9Z8R5 Q9Z8S4 Q9Z8S6 Q9Z8T3 Q9Z8T4 Q9Z8T7 Q9Z8T8 Q9Z8U0
Q9Z8U5 Q9Z8U6 Q9Z8U7 Q9Z8U8 Q9Z8W5 Q9Z8X8 Q9Z8Z0 Q9Z8Z1 Q9Z8Z4 Q9Z8Z9 Q9Z900
Q9Z904 Q9Z913 Q9Z930 Q9Z934 Q9Z942 Q9Z944 Q9Z946 Q9Z953 Q9Z959 Q9Z960 Q9Z961
Q9Z962 Q9Z963 Q9Z964 Q9Z965 Q9Z968 Q9Z969 Q9Z972 Q9Z984 Q9Z985 Q9Z986 Q9Z987
Q9Z994 Q9Z998 Q9Z999 Q9Z9A0 Q9Z9A1 Q9Z9A2 Q9Z9A3 Q9Z9A4 Q9Z9A5 Q9Z9A7 Q9Z9A8
Q9Z9B6 Q9Z9C2 Q9Z9C4 Q9Z9C7 Q9Z9C8 Q9Z9C9 Q9Z9D4 Q9Z9E2 Q9Z9E9 Q9Z9F0 Q9Z9F4
Q9Z9F6 Q9Z9F7 Q9Z9F9 Q9Z9G6 Q9Z9G7 Q9Z9G8 Q9Z9H0 Q9Z9H5 Q9Z9H6 Q9Z9J0 Q9Z9J4
Q9Z9J6 Q9Z9J7 Q9Z9J8 Q9Z9J9 Q9Z9K0 Q9Z9K1 Q9Z9K2 Q9Z9K3 Q9Z9K4 Q9Z9K5 Q9Z9K6
Q9Z9K7 Q9Z9K8 Q9Z9K9 Q9Z9L0 Q9Z9L1 Q9Z9L2 Q9Z9L3 Q9Z9L4 Q9Z9L5 Q9Z9L6 Q9Z9L7
Q9Z9L8 Q9Z9L9 Q9Z9M0 Q9Z9M1 Q9Z9M2 Q9Z9N7 Q9Z9P2 Q9Z9P6 Q9Z9Q7 Q9Z9S0 Q9Z9T7
Q9Z9U1 Q9Z9W1 Q9Z9W9 Q9Z9X0 Q9ZA11 Q9ZA32 Q9ZA33 Q9ZA56 Q9ZA86 Q9ZA87 Q9ZA98
Q9ZAA0 Q9ZAA1 Q9ZAA7 Q9ZAB0 Q9ZAB8 Q9ZAE1 Q9ZAE3 Q9ZAE4 Q9ZAE5 Q9ZAE6 Q9ZAE7
Q9ZAE8 Q9ZAE9 Q9ZAG3 Q9ZAH3 Q9ZAH5 Q9ZAI9 Q9ZAQ6 Q9ZAQ8 Q9ZAU0 Q9ZAU1 Q9ZAU2
Q9ZAU3 Q9ZAZ5 Q9ZAZ6 Q9ZB06 Q9ZB07 Q9ZB09 Q9ZB10 Q9ZB11 Q9ZB21 Q9ZB26 Q9ZB35
Q9ZB62 Q9ZB63 Q9ZB70 Q9ZB76 Q9ZB77 Q9ZB78 Q9ZB82 Q9ZB91 Q9ZBA9 Q9ZBD8 Q9ZBG3
Q9ZBH0 Q9ZBH3 Q9ZBH5 Q9ZBI5 Q9ZBL0 Q9ZBL1 Q9ZBL2 Q9ZBL5 Q9ZBQ3 Q9ZBQ6 Q9ZBQ7
Q9ZBQ8 Q9ZBR1 Q9ZBR5 Q9ZBR6 Q9ZBR9 Q9ZBS2 Q9ZBT9 Q9ZBW0 Q9ZBX1 Q9ZBX9 Q9ZBY8
Q9ZC49 Q9ZC67 Q9ZC68 Q9ZC70 Q9ZC74 Q9ZC83 Q9ZC84 Q9ZC86 Q9ZC88 Q9ZC89 Q9ZC90
Q9ZC91 Q9ZC93 Q9ZC94 Q9ZCA0 Q9ZCA3 Q9ZCA4 Q9ZCB0 Q9ZCB1 Q9ZCB3 Q9ZCB6 Q9ZCB7
Q9ZCB8 Q9ZCB9 Q9ZCC0 Q9ZCC2 Q9ZCC3 Q9ZCC6 Q9ZCC8 Q9ZCD1 Q9ZCD3 Q9ZCD6 Q9ZCE0
Q9ZCE2 Q9ZCE3 Q9ZCE4 Q9ZCE8 Q9ZCF4 Q9ZCG5 Q9ZCH0 Q9ZCH6 Q9ZCH7 Q9ZCI1 Q9ZCI4
Q9ZCI7 Q9ZCI8 Q9ZCI9 Q9ZCJ2 Q9ZCJ3 Q9ZCJ9 Q9ZCK0 Q9ZCK3 Q9ZCK6 Q9ZCK8 Q9ZCK9
Q9ZCL3 Q9ZCL4 Q9ZCL5 Q9ZCL6 Q9ZCL7 Q9ZCM9 Q9ZCN1 Q9ZCN2 Q9ZCN3 Q9ZCN6 Q9ZCN9
Q9ZCP0 Q9ZCP3 Q9ZCP6 Q9ZCP8 Q9ZCP9 Q9ZCQ4 Q9ZCQ6 Q9ZCQ7 Q9ZCQ8 Q9ZCQ9 Q9ZCR0
Q9ZCR1 Q9ZCR2 Q9ZCR3 Q9ZCR4 Q9ZCR5 Q9ZCR6 Q9ZCR7 Q9ZCR8 Q9ZCR9 Q9ZCS0 Q9ZCS1
Q9ZCS2 Q9ZCS3 Q9ZCS4 Q9ZCS6 Q9ZCS7 Q9ZCS8 Q9ZCS9 Q9ZCT0 Q9ZCT1 Q9ZCT4 Q9ZCT5
Q9ZCT6 Q9ZCT7 Q9ZCT8 Q9ZCT9 Q9ZCU4 Q9ZCU6 Q9ZCU8 Q9ZCU9 Q9ZCV0 Q9ZCV1 Q9ZCV2
Q9ZCV3 Q9ZCV4 Q9ZCV8 Q9ZCW2 Q9ZCW5 Q9ZCW7 Q9ZCX2 Q9ZCX3 Q9ZCX4 Q9ZCX8 Q9ZCX9
Q9ZCY1 Q9ZCY2 Q9ZCZ4 Q9ZCZ6 Q9ZCZ7 Q9ZCZ8 Q9ZCZ9 Q9ZD00 Q9ZD01 Q9ZD02 Q9ZD03
Q9ZD04 Q9ZD05 Q9ZD06 Q9ZD08 Q9ZD09 Q9ZD10 Q9ZD12 Q9ZD15 Q9ZD19 Q9ZD21 Q9ZD24
Q9ZD26 Q9ZD27 Q9ZD28 Q9ZD29 Q9ZD32 Q9ZD33 Q9ZD36 Q9ZD37 Q9ZD40 Q9ZD42 Q9ZD43
Q9ZD44 Q9ZD45 Q9ZD46 Q9ZD49 Q9ZD50 Q9ZD53 Q9ZD54 Q9ZD55 Q9ZD60 Q9ZD61 Q9ZD62
Q9ZD70 Q9ZD73 Q9ZD74 Q9ZD76 Q9ZD77 Q9ZD84 Q9ZD85 Q9ZD87 Q9ZD88 Q9ZD90 Q9ZD92
Q9ZD94 Q9ZD95 Q9ZD97 Q9ZDA2 Q9ZDA3 Q9ZDA4 Q9ZDA7 Q9ZDB1 Q9ZDB3 Q9ZDB4 Q9ZDB5
Q9ZDB6 Q9ZDB7 Q9ZDC2 Q9ZDC3 Q9ZDC7 Q9ZDC8 Q9ZDC9 Q9ZDD0 Q9ZDD2 Q9ZDD5 Q9ZDD6
Q9ZDE1 Q9ZDE5 Q9ZDE6 Q9ZDE7 Q9ZDE8 Q9ZDF1 Q9ZDF3 Q9ZDF6 Q9ZDF8 Q9ZDG2 Q9ZDG5

Q9ZDG6 Q9ZDG7 Q9ZDG8 Q9ZDG9 Q9ZDH6 Q9ZDH7 Q9ZDH8 Q9ZDI0 Q9ZDI3 Q9ZDI9 Q9ZDJ5
Q9ZDJ7 Q9ZDJ8 Q9ZDK2 Q9ZDK3 Q9ZDK5 Q9ZDK8 Q9ZDK9 Q9ZDL1 Q9ZDL2 Q9ZDL9 Q9ZDM1
Q9ZDM6 Q9ZDM8 Q9ZDM9 Q9ZDP8 Q9ZDQ0 Q9ZDQ2 Q9ZDQ3 Q9ZDQ6 Q9ZDR0 Q9ZDR3 Q9ZDR7
Q9ZDS6 Q9ZDS7 Q9ZDS8 Q9ZDT1 Q9ZDT5 Q9ZDT7 Q9ZDT9 Q9ZDU0 Q9ZDU1 Q9ZDU2 Q9ZDU7
Q9ZDU9 Q9ZDV2 Q9ZDV4 Q9ZDV6 Q9ZDV8 Q9ZDV9 Q9ZDW1 Q9ZDW2 Q9ZDW3 Q9ZDW4 Q9ZDW5
Q9ZDW6 Q9ZDX0 Q9ZDX5 Q9ZDX6 Q9ZDX7 Q9ZDX9 Q9ZDY0 Q9ZDY2 Q9ZDY3 Q9ZDY4 Q9ZDY5
Q9ZDZ2 Q9ZDZ5 Q9ZDZ6 Q9ZDZ7 Q9ZDZ8 Q9ZDZ9 Q9ZE00 Q9ZE01 Q9ZE04 Q9ZE07 Q9ZE08
Q9ZE09 Q9ZE10 Q9ZE11 Q9ZE14 Q9ZE16 Q9ZE17 Q9ZE18 Q9ZE20 Q9ZE21 Q9ZE22 Q9ZE23
Q9ZE24 Q9ZE27 Q9ZE28 Q9ZE29 Q9ZE31 Q9ZE35 Q9ZE36 Q9ZE37 Q9ZE38 Q9ZE39 Q9ZE46
Q9ZE47 Q9ZE48 Q9ZE52 Q9ZE55 Q9ZE56 Q9ZE60 Q9ZE61 Q9ZE62 Q9ZE63 Q9ZE65 Q9ZE67
Q9ZE74 Q9ZE76 Q9ZE77 Q9ZE81 Q9ZE82 Q9ZE83 Q9ZE85 Q9ZE87 Q9ZE89 Q9ZE90 Q9ZE91
Q9ZE93 Q9ZEA1 Q9ZEA3 Q9ZEA4 Q9ZEA5 Q9ZEA6 Q9ZEA8 Q9ZEA9 Q9ZEB1 Q9ZEB3 Q9ZEB4
Q9ZEB5 Q9ZEB6 Q9ZEB7 Q9ZEC6 Q9ZED0 Q9ZED2 Q9ZED6 Q9ZED7 Q9ZEE0 Q9ZEE1 Q9ZEE3
Q9ZEH3 Q9ZEH5 Q9ZEJ0 Q9ZEJ6 Q9ZEJ7 Q9ZEL7 Q9ZEM7 Q9ZEN7 Q9ZEP7 Q9ZEP8 Q9ZEQ8
Q9ZER0 Q9ZER9 Q9ZES2 Q9ZET3 Q9ZET4 Q9ZET5 Q9ZET6 Q9ZET8 Q9ZEU3 Q9ZEU4 Q9ZEU5
Q9ZEU6 Q9ZEU7 Q9ZEU8 Q9ZEY8 Q9ZEY9 Q9ZEZ5 Q9ZF22 Q9ZF25 Q9ZF28 Q9ZF31 Q9ZF44
Q9ZF99 Q9ZFA3 Q9ZFA6 Q9ZFA7 Q9ZFA8 Q9ZFC5 Q9ZFC6 Q9ZFC7 Q9ZFD8 Q9ZFD9 Q9ZFE0
Q9ZFE1 Q9ZFE7 Q9ZFF6 Q9ZFF7 Q9ZFK1 Q9ZFL7 Q9ZFL8 Q9ZFM2 Q9ZFN4 Q9ZFR5 Q9ZFR7
Q9ZFR8 Q9ZFR9 Q9ZFU6 Q9ZFU7 Q9ZFU8 Q9ZFV2 Q9ZFV4 Q9ZFV5 Q9ZFV7 Q9ZFV8 Q9ZFY8
Q9ZG11 Q9ZG89 Q9ZG98 Q9ZGE2 Q9ZGE6 Q9ZGE8 Q9ZGE9 Q9ZGF0 Q9ZGG6 Q9ZGH1 Q9ZGH4
Q9ZGH6 Q9ZGH7 Q9ZGH8 Q9ZGI1 Q9ZGI2 Q9ZGI3 Q9ZGI4 Q9ZGI5 Q9ZGM0 Q9ZGM1 Q9ZGM4
Q9ZGN0 Q9ZGR8 Q9ZGR9 Q9ZGS5 Q9ZGW3 Q9ZGW6 Q9ZH20 Q9ZH21 Q9ZH58 Q9ZH75 Q9ZH76
Q9ZH77 Q9ZH99 Q9ZHA2 Q9ZHA4 Q9ZHD6 Q9ZHD7 Q9ZHD8 Q9ZHD9 Q9ZHE0 Q9ZHE1 Q9ZHE2
Q9ZHE3 Q9ZHE4 Q9ZHE5 Q9ZHE6 Q9ZHE7 Q9ZHE8 Q9ZHE9 Q9ZHF0 Q9ZHF6 Q9ZHG4 Q9ZHI0
Q9ZHI1 Q9ZHI2 Q9ZHI4 Q9ZHI5 Q9ZHQ1 Q9ZHQ4 Q9ZHR1 Q9ZHW3 Q9ZHY2 Q9ZHY3 Q9ZHZ1
Q9ZHZ4 Q9ZI33 Q9ZI34 Q9ZI37 Q9ZI38 Q9ZI39 Q9ZI40 Q9ZI41 Q9ZI42 Q9ZI43 Q9ZI44
Q9ZI45 Q9ZI46 Q9ZI47 Q9ZI48 Q9ZI49 Q9ZI50 Q9ZI51 Q9ZI54 Q9ZI56 Q9ZI57 Q9ZI76
Q9ZIB7 Q9ZIE4 Q9ZIG2 Q9ZIM1 Q9ZIM6 Q9ZIM7 Q9ZIM9 Q9ZIN2 Q9ZIN3 Q9ZIQ0 Q9ZIU5
Q9ZIV1 Q9ZIX6 Q9ZIX8 Q9ZIY0 Q9ZJ10 Q9ZJ12 Q9ZJ14 Q9ZJ19 Q9ZJ24 Q9ZJ25 Q9ZJ27
Q9ZJ28 Q9ZJ29 Q9ZJ42 Q9ZJ57 Q9ZJ63 Q9ZJ70 Q9ZJ71 Q9ZJ74 Q9ZJ75 Q9ZJ80 Q9ZJ81
Q9ZJ93 Q9ZJ94 Q9ZJ96 Q9ZJA1 Q9ZJA6 Q9ZJA7 Q9ZJC3 Q9ZJC4 Q9ZJC5 Q9ZJC6 Q9ZJD7
Q9ZJD9 Q9ZJE3 Q9ZJE4 Q9ZJE5 Q9ZJE9 Q9ZJF1 Q9ZJF9 Q9ZJG0 Q9ZJG6 Q9ZJH0 Q9ZJH2
Q9ZJH3 Q9ZJH4 Q9ZJH5 Q9ZJI4 Q9ZJI5 Q9ZJI6 Q9ZJI7 Q9ZJJ0 Q9ZJJ1 Q9ZJJ3 Q9ZJJ4
Q9ZJJ7 Q9ZJJ9 Q9ZJK8 Q9ZJL3 Q9ZJL5 Q9ZJL8 Q9ZJM4 Q9ZJM5 Q9ZJN1 Q9ZJN2 Q9ZJN9
Q9ZJP1 Q9ZJP7 Q9ZJP8 Q9ZJQ0 Q9ZJQ2 Q9ZJQ9 Q9ZJR1 Q9ZJR4 Q9ZJR5 Q9ZJR6 Q9ZJR7
Q9ZJR8 Q9ZJR9 Q9ZJS0 Q9ZJS1 Q9ZJS2 Q9ZJS3 Q9ZJS4 Q9ZJS5 Q9ZJS6 Q9ZJS7 Q9ZJS8
Q9ZJT0 Q9ZJT1 Q9ZJT2 Q9ZJT3 Q9ZJT4 Q9ZJT5 Q9ZJT6 Q9ZJU5 Q9ZJU6 Q9ZJU7 Q9ZJU8
Q9ZJU9 Q9ZJV0 Q9ZJW8 Q9ZJX0 Q9ZJX1 Q9ZJX4 Q9ZJX8 Q9ZJX9 Q9ZJY0 Q9ZJY1 Q9ZJY2
Q9ZJY5 Q9ZJY6 Q9ZJY8 Q9ZJY9 Q9ZJZ7 Q9ZJZ8 Q9ZK05 Q9ZK07 Q9ZK11 Q9ZK14 Q9ZK19
Q9ZK20 Q9ZK21 Q9ZK22 Q9ZK23 Q9ZK24 Q9ZK27 Q9ZK28 Q9ZK37 Q9ZK38 Q9ZK49 Q9ZK53
Q9ZK54 Q9ZK56 Q9ZK60 Q9ZK61 Q9ZK63 Q9ZK65 Q9ZK66 Q9ZK67 Q9ZK72 Q9ZK75 Q9ZK89
Q9ZKA0 Q9ZKA2 Q9ZKB0 Q9ZKB1 Q9ZKB2 Q9ZKB3 Q9ZKC0 Q9ZKC2 Q9ZKC3 Q9ZKD0 Q9ZKD2
Q9ZKD4 Q9ZKD8 Q9ZKE4 Q9ZKE5 Q9ZKE6 Q9ZKE7 Q9ZKF2 Q9ZKF3 Q9ZKF6 Q9ZKF7 Q9ZKF8
Q9ZKF9 Q9ZKG4 Q9ZKG9 Q9ZKH1 Q9ZKH6 Q9ZKM5 Q9ZKM6 Q9ZKM7 Q9ZKM9 Q9ZKP1 Q9ZKP2

Q9ZKP5 Q9ZKP7 Q9ZKQ2 Q9ZKQ6 Q9ZKQ9 Q9ZKR8 Q9ZKS0 Q9ZKS2 Q9ZKS5 Q9ZKS6 Q9ZKS7
Q9ZKT0 Q9ZKT2 Q9ZKU4 Q9ZKU5 Q9ZKU8 Q9ZKW1 Q9ZKW6 Q9ZKW9 Q9ZKX3 Q9ZKX5 Q9ZKY0
Q9ZKY3 Q9ZKY5 Q9ZKY6 Q9ZKZ2 Q9ZKZ9 Q9ZL00 Q9ZL01 Q9ZL08 Q9ZL09 Q9ZL11 Q9ZL12
Q9ZL13 Q9ZL14 Q9ZL18 Q9ZL19 Q9ZL20 Q9ZL21 Q9ZL40 Q9ZL42 Q9ZL44 Q9ZL45 Q9ZL46
Q9ZL49 Q9ZL50 Q9ZL51 Q9ZL64 Q9ZL66 Q9ZL69 Q9ZL74 Q9ZL75 Q9ZL96 Q9ZL98 Q9ZLA1
Q9ZLA5 Q9ZLA7 Q9ZLA9 Q9ZLB5 Q9ZLB6 Q9ZLB9 Q9ZLC8 Q9ZLD4 Q9ZLD6 Q9ZLD9 Q9ZLE3
Q9ZLE4 Q9ZLF9 Q9ZLH0 Q9ZLH1 Q9ZLH2 Q9ZLH3 Q9ZLH6 Q9ZLI1 Q9ZLI5 Q9ZLI6 Q9ZLJ1
Q9ZLJ7 Q9ZLL0 Q9ZLL2 Q9ZLL4 Q9ZLL5 Q9ZLL8 Q9ZLL9 Q9ZLM1 Q9ZLM8 Q9ZLN3 Q9ZLP5
Q9ZLQ0 Q9ZLQ4 Q9ZLQ9 Q9ZLR1 Q9ZLR5 Q9ZLR9 Q9ZLS8 Q9ZLS9 Q9ZLT0 Q9ZLU4 Q9ZLW1
Q9ZLW2 Q9ZLW3 Q9ZLW5 Q9ZLW6 Q9ZLW8 Q9ZLX3 Q9ZLX4 Q9ZLX9 Q9ZLY0 Q9ZLZ3 Q9ZLZ7
Q9ZM10 Q9ZM13 Q9ZM19 Q9ZM26 Q9ZM35 Q9ZM37 Q9ZM39 Q9ZM43 Q9ZM44 Q9ZM45 Q9ZM46
Q9ZM48 Q9ZM51 Q9ZM56 Q9ZM57 Q9ZM60 Q9ZM61 Q9ZM65 Q9ZM67 Q9ZM70 Q9ZM81 Q9ZM94
Q9ZM99 Q9ZMA1 Q9ZMA5 Q9ZMA7 Q9ZMA9 Q9ZMB0 Q9ZMB7 Q9ZMB9 Q9ZMD0 Q9ZMD3 Q9ZMD8
Q9ZMD9 Q9ZME1 Q9ZME5 Q9ZMF0 Q9ZMF2 Q9ZMF4 Q9ZMG1 Q9ZMG6 Q9ZMG9 Q9ZMH1 Q9ZMH7
Q9ZMI1 Q9ZMJ1 Q9ZMJ5 Q9ZMJ6 Q9ZMJ7 Q9ZMK8 Q9ZMK9 Q9ZML2 Q9ZML6 Q9ZML9 Q9ZMM0
Q9ZMM2 Q9ZMM5 Q9ZMN2 Q9ZMN4 Q9ZMN5 Q9ZMN8 Q9ZMP1 Q9ZMP7 Q9ZMP8 Q9ZMQ5 Q9ZMQ6
Q9ZMR1 Q9ZMR8 Q9ZMR9 Q9ZMS3 Q9ZMS7 Q9ZMU4 Q9ZMU7 Q9ZMV1 Q9ZMV2 Q9ZMV3 Q9ZMV4
Q9ZMV6 Q9ZMV7 Q9ZMW2 Q9ZMW3 Q9ZMW4 Q9ZMW6 Q9ZMW7 Q9ZMX5 Q9ZMY2 Q9ZMY3 Q9ZMY6
Q9ZMZ0 Q9ZMZ1 Q9ZMZ2 Q9ZMZ4 Q9ZMZ5 Q9ZMZ6 Q9ZMZ7 Q9ZMZ8 Q9ZN29 Q9ZN30 Q9ZN32
Q9ZN34 Q9ZN36 Q9ZN37 Q9ZN49 Q9ZN50 Q9ZN52 Q9ZN53 Q9ZN54 Q9ZN56 Q9ZN57 Q9ZN78
Q9ZN79 Q9ZN99 Q9ZNA2 Q9ZNA5 Q9ZNA8 Q9ZND4 Q9ZND5 Q9ZND9 Q9ZNE0 Q9ZNH4 Q9ZNJ6
Q9ZNJ8 Q9ZNJ9 Q9ZNK0 Q9ZNL9 Q9ZNM0 Q9ZNN8 Q9ZNQ3 Q9ZNR6 Q9ZNS1 Q9ZNT3 Q9ZNU2
Q9ZNU9 Q9ZNV5 Q9ZNW0 Q9ZNX6 Q9ZNX9 Q9ZP06 Q9ZP19 Q9ZP20 Q9ZP21 Q9ZP27 Q9ZP54
Q9ZP56 Q9ZPB7 Q9ZPC0 Q9ZPE4 Q9ZPF5 Q9ZPI1 Q9ZPN7 Q9ZPN8 Q9ZPN9 Q9ZPP0 Q9ZPQ3
Q9ZPQ5 Q9ZPQ8 Q9ZPR5 Q9ZPS0 Q9ZPS1 Q9ZPS3 Q9ZPS4 Q9ZPW2 Q9ZPW3 Q9ZPW7 Q9ZPX0
Q9ZPX8 Q9ZPX9 Q9ZPY0 Q9ZPY1 Q9ZPY6 Q9ZPY8 Q9ZQ19 Q9ZQ34 Q9ZQ46 Q9ZQ60 Q9ZQ70
Q9ZQ80 Q9ZQ85 Q9ZQ91 Q9ZQ95 Q9ZQ96 Q9ZQ97 Q9ZQ98 Q9ZQ99 Q9ZQA1 Q9ZQA8 Q9ZQC5
Q9ZQE6 Q9ZQF0 Q9ZQF1 Q9ZQF9 Q9ZQG4 Q9ZQG8 Q9ZQH0 Q9ZQH1 Q9ZQI7 Q9ZQM4 Q9ZQR2
Q9ZQR6 Q9ZQV3 Q9ZQV4 Q9ZQV6 Q9ZQV7 Q9ZQV8 Q9ZQV9 Q9ZQW8 Q9ZQX6 Q9ZQZ1 Q9ZQZ6
Q9ZR02 Q9ZR04 Q9ZR07 Q9ZR09 Q9ZR12 Q9ZR14 Q9ZR15 Q9ZR37 Q9ZR39 Q9ZR40 Q9ZR41
Q9ZRA2 Q9ZRA8 Q9ZRA9 Q9ZRB0 Q9ZRB1 Q9ZRB2 Q9ZRB7 Q9ZRF1 Q9ZRI7 Q9ZRI8 Q9ZRJ4
Q9ZRR5 Q9ZRR8 Q9ZRS4 Q9ZRS8 Q9ZRT1 Q9ZRU9 Q9ZRV4 Q9ZRX0 Q9ZRZ8 Q9ZS21 Q9ZS34
Q9ZS40 Q9ZS41 Q9ZS45 Q9ZS91 Q9ZS96 Q9ZSA7 Q9ZSA8 Q9ZSD6 Q9ZSE4 Q9ZSH7 Q9ZSI7
Q9ZSK1 Q9ZSK2 Q9ZSK5 Q9ZSL6 Q9ZSQ4 Q9ZSR8 Q9ZSS6 Q9ZST0 Q9ZST1 Q9ZSV1 Q9ZSW1
Q9ZSW9 Q9ZSY8 Q9ZSY9 Q9ZSZ8 Q9ZT00 Q9ZT42 Q9ZT50 Q9ZT56 Q9ZT71 Q9ZT84 Q9ZT87
Q9ZT91 Q9ZT92 Q9ZT94 Q9ZT95 Q9ZT96 Q9ZTA1 Q9ZTA2 Q9ZTA3 Q9ZTC3 Q9ZTK5 Q9ZTP4
Q9ZTR0 Q9ZTR1 Q9ZTS1 Q9ZTS2 Q9ZTT5 Q9ZTX8 Q9ZTX9 Q9ZU06 Q9ZU13 Q9ZU23 Q9ZU26
Q9ZU27 Q9ZU29 Q9ZU31 Q9ZU32 Q9ZU34 Q9ZU38 Q9ZU51 Q9ZU52 Q9ZU65 Q9ZU67 Q9ZU70
Q9ZU72 Q9ZU75 Q9ZU77 Q9ZU88 Q9ZU90 Q9ZU95 Q9ZUA2 Q9ZUA3 Q9ZUA7 Q9ZUA9 Q9ZUB3
Q9ZUB8 Q9ZUB9 Q9ZUC1 Q9ZUD9 Q9ZUE9 Q9ZUF1 Q9ZUG4 Q9ZUG9 Q9ZUH0 Q9ZUH1 Q9ZUH4
Q9ZUH5 Q9ZUI9 Q9ZUL3 Q9ZUM0 Q9ZUM9 Q9ZUN0 Q9ZUN1 Q9ZUN4 Q9ZUN8 Q9ZUP0 Q9ZUP2
Q9ZUR7 Q9ZUS0 Q9ZUT5 Q9ZUT9 Q9ZUU0 Q9ZUU1 Q9ZUU2 Q9ZUU7 Q9ZUV0 Q9ZUX7 Q9ZUY1
Q9ZUY3 Q9ZUY6 Q9ZUZ6 Q9ZV05 Q9ZV06 Q9ZV19 Q9ZV25 Q9ZV27 Q9ZV31 Q9ZV33 Q9ZV39
Q9ZV48 Q9ZV69 Q9ZV78 Q9ZV88 Q9ZV91 Q9ZV93 Q9ZV94 Q9ZVA3 Q9ZVB5 Q9ZVB9 Q9ZVC1

Q9ZVC2 Q9ZVC9 Q9ZVD0 Q9ZVE0 Q9ZVF2 Q9ZVF3 Q9ZVF4 Q9ZVF5 Q9ZVG8 Q9ZVH0 Q9ZVH3
Q9ZVH4 Q9ZVH8 Q9ZVI1 Q9ZVI3 Q9ZVJ2 Q9ZVJ3 Q9ZVL3 Q9ZVM9 Q9ZVN4 Q9ZVN9 Q9ZVP2
Q9ZVP8 Q9ZVQ3 Q9ZVQ4 Q9ZVQ6 Q9ZVQ8 Q9ZVQ9 Q9ZVR0 Q9ZVR1 Q9ZVR3 Q9ZVR5 Q9ZVT1
Q9ZVT7 Q9ZVT8 Q9ZVW2 Q9ZVX1 Q9ZVX2 Q9ZVX4 Q9ZVX5 Q9ZVY5 Q9ZW02 Q9ZW03 Q9ZW04
Q9ZW06 Q9ZW07 Q9ZW12 Q9ZW13 Q9ZW14 Q9ZW15 Q9ZW16 Q9ZW19 Q9ZW20 Q9ZW21 Q9ZW24
Q9ZW26 Q9ZW27 Q9ZW28 Q9ZW29 Q9ZW30 Q9ZW31 Q9ZW34 Q9ZW35 Q9ZW36 Q9ZW38 Q9ZW74
Q9ZW81 Q9ZW82 Q9ZW84 Q9ZW85 Q9ZW88 Q9ZW96 Q9ZWA1 Q9ZWB4 Q9ZWB7 Q9ZWB8 Q9ZWC5
Q9ZWC6 Q9ZWH8 Q9ZWJ3 Q9ZWM9 Q9ZWQ9 Q9ZWR1 Q9ZWR8 Q9ZWT1 Q9ZWX5 Q9ZX29 Q9ZX92
Q9ZXA5 Q9ZXB2 Q9ZZV8 Q9ZZW1 Q9ZZW2 Q9ZZW3 Q9ZZW4 Q9ZZX0 Q9ZZX7 Q9ZZX8 Q9ZZX9
R1CW23 R1DS73 R1ENF4 R1ERP2 R1F493 R1F9H0 R1G6M4 R4GER2 R4HYA4 R4HZ96 R4JGL8
R4JJJ1 R4JND9 R4JQS2 R4JQT5 R4WH05 R4X244 R4X247 R4X4V6 R4X5P0 R4YVJ8 R4YXW8
R4YZC3 R4ZGN4 R8NBN0 R9QMQ9 R9QMR0 R9QMR1 R9QMR2 R9QMR3 R9QMR4 R9QMR5 R9QMR6
R9QMW1 R9QMW2 R9QMW3 R9QMW4 R9QMW5 R9QMW6 R9QMW7 R9QMW8 R9QMY7 R9QMY8 R9QMY9
R9QMZ0 R9RZK8 R9T6W8 R9UTQ8 S0ASL9 S0DL59 S0DLN5 S0DLN7 S0DLP1 S0DLP2 S0DPL8
S0DQ98 S0DQI7 S0DQN6 S0DQQ0 S0DRI1 S0DRI9 S0DRT4 S0DRW4 S0DRW9 S0DRX0 S0DRX3
S0DRY0 S0DS11 S0DS17 S0DS59 S0DS79 S0DTP6 S0DUX2 S0DUX5 S0DV66 S0DVT6 S0DW94
S0DX56 S0DXJ2 S0DXU0 S0DZI6 S0DZM7 S0E028 S0E2U7 S0E2Y4 S0E3H0 S0E608 S0E627
S0E9A6 S0EA85 S0EBQ6 S0EBV2 S0ECT9 S0EE80 S0EES1 S0EET5 S0EEY3 S0EFU6 S0EG57
S0EGG0 S0EGZ9 S0EH60 S0EHD0 S0EHD6 S0EJ18 S0EJF2 S0EMV0 S0EN43 S0ENH1 S0ENM8
S0ENQ2 S0EX56 S0F332 S2DJ52 S2KHP1 S2L5R8 S3D784 S3D9F1 S3D9F8 S3D9F9 S3DB70
S3DB74 S3DB78 S3DQP3 S3DQP8 S3DQQ3 S3E7P4 S3E7P8 S3E7Q2 S4R2K0 S4S3E3 S4TDL2
S4TF94 S4W172 S4W177 S4W4F3 S4W775 S4W780 S4W887 S5DQS6 S5DWK8 S5FMM4 S5FT07
S5FU55 S5M744 S5N020 S6F527 S7Z6W7 S7Z800 S7Z806 S7Z906 S7ZC75 S7ZEI0 S7ZEI5
S7ZFK6 S7ZFL4 S7ZFL7 S7ZK59 S8AKF3 S8ASK6 S8ASK9 S8AWN1 S8AWN7 S8AWP5 S8AWQ1
S8B3H6 S8B3I4 S8GIB3 T0CUB8 T0D7A2 T1RR71 T1RR72 T1RRI5 T1RRI8 T1RRJ4 T1RRJ6
T1RRL8 T1RRR9 T1RRS0 T1SFR8 T1SH39 T1T504 T2G6Z9 T2G899 T2KLZ8 T2KM09 T2KM13
T2KMF9 T2KN75 T2KN80 T2KN98 T2KNC8 U2PSH1 U2UMQ6 U2ZU49 U3H040 U3H041 U3H042
U3H0A9 U3KRF2 U3KRF8 U3KRG0 U3KYL2 U3LVL5 U3LVZ7 U3LW50 U3MMQ4 U3N9N6 U3NEE3
U3PVA8 U3THC0 U3U715 U3U992 U3UA11 U4PCM1 U5L396 U5L397 U5L3J5 U5L3J7 U5L3J9
U5L3K1 U5L3M6 U5L3M7 U5L3M8 U5L3X0 U5L3X2 U5L406 U5L408 U5L409 U5NH37 U5PZT6
U5S003 U6A492 V5IPE4 V5IVB1 V5LLZ9 V5NBV4 V5QPS4 V5QRX7 V5T8H7 V5TD18 V5TES5
V5TF65 V5XVW4 V5XWI9 V5YW53 V5YXI5 V6CJ04 V6CLA2 V6CLJ5 V7BSD0 V9GXG1 V9GXK1
V9HVX0 V9P9L8 V9P9S4 W0FFD7 W0LYS5 W0T3G1 W0T4A7 W0TA05 W0TAA4 W1QCN1 W3X7R6
W3X7S0 W3X7U2 W3X9K7 W3XA95 W3XAA0 W3XC32 W3XEE6 W4JX79 W5AWH5 W5BGD1 W5DYE3
W5EP13 W5QJZ5 W5U2K2 W5X2N3 W6CW81 W6HUT3 W6JHZ8 W6JQK2 W6PQG8 W6Q0S4 W6Q1E9
W6Q2D7 W6Q4Q9 W6Q9S9 W6QAE7 W6QB10 W6QB19 W6QIM8 W6QIT2 W6QJS4 W6QL00 W6QL41
W6QL47 W6QM20 W6QP10 W6QRI9 W6QV39 W6R4H4 W6RJ11 W7L9E5 W7L9F0 W7LAD1 W7LC91
W7LFK5 W7LKW1 W7LKX1 W7LKX6 W7LL82 W7LUF3 W7LYY0 W7MAD3 W7MC44 W7MGR9 W7MLD5
W7MLD7 W7MLD8 W7MMJ0 W7MPI5 W7MS01 W7MS09 W7MS18 W7MSI0 W7MSI4 W7MSJ7 W7MT28
W7MT31 W7MT41 W7MTI6 W7MTJ1 W7MVT8 W7MWX4 W7MWX7 W7N293 W7N2B2 W7N2C1 W7N2P0
W7N2P6 W7N2Q9 W7N463 W7N466 W7N6M8 W7N6P0 W7NCN7 W7NCP1 W7NDP3 W7NDP8 W7X6T2
W8CV45 W8E7I1 W8JCF0 W8JDE0 W8JMV4 W8JWV8 W8JWW7 W8P570 X1WGV5 X2BTY5 X5CFH4
X5CWH9 X5CY81 X5HYT8 X6R8D5
